# Effects of nano urea on growth and gene expression of *Arabidopsis thaliana* in hydroponics

**DOI:** 10.1101/2024.03.20.585664

**Authors:** Neelam Jangir, Debankona Marik, Devanshu Verma, Arpan Dey, Rajveer Singh Shekhawat, Deep Patel, Pankaj Yadav, Kirti Sankhala, Ayan Sadhukhan

## Abstract

**Introduction:** Hydroponics enables precise control over nutrient delivery, optimized water requirements and growing conditions. The combination of nanotechnology and hydroponics paves the way towards sustainable agriculture with less environmental footprints. We investigated the effects of nano urea on the model plant *Arabidopsis thaliana* in hydroponics.

**Methods:** A growth experiment in a nitrogen-free hydroponic medium compared the effects of a liquid nano urea formulation (NUF) marketed by Indian Farmers Fertilizer Cooperative (IFFCO) to an equimolar bulk urea. Transcriptome analysis identified the molecular mechanisms of growth enhancement. Dynamic light scattering and transmission electron microscopy confirmed NUF’s negative surface charge and sub-100 nm size, correlating its uptake and distribution in the plant.

**Results and discussion:** A two-week growth in the hydroponic medium with 70 μM NUF led to a 20% higher biomass and 16% higher chlorophyll content than a medium with 70 μM urea. Higher doses of NUF inhibited growth, whereas higher equivalent urea doses did not. NUF led to the differential expression of more genes than urea at 12 h to seven days of treatment. Nitrogen assimilation, growth, photosynthesis, and stress tolerance genes showed higher transcript levels in NUF than in urea. On the other hand, NUF led to greater suppression of many negative growth-regulating genes. After seven days of treatment, chlorophyll biosynthesis genes got up-regulated, while chlorophyll catabolism genes got down-regulated at higher levels by NUF than by urea, correlating with the higher chlorophyll content of NUF-treated seedlings. In conclusion, NUF outperformed equimolar urea for the growth promotion of *A. thaliana* at a low concentration in hydroponics, leading to a greater regulation of genes for nitrogen metabolism and chlorophyll biosynthesis. Our results suggest a potential use of NUF as a nitrogen fertilizer for hydroponic agriculture.

## Introduction

The United Nations Department of Economic and Social Affairs (UNDESA) has predicted the global human population to reach 9.8 billion by 2050. The exponential increase in human population demands measures for enhanced food production (Kumar et al., 2020). Sustainable agriculture is the way forward for ensuring food security for a burgeoning population from limited parcel of land. Technological interventions are therefore needed to increase crop productivity to meet the food demands of the world. During late 1960s, Indian farmers adopted high yielding varieties of cereals, irrigation, chemical fertilizers, and pesticides to achieve higher crop yields also termed as the Green Revolution. Over a period, Green Revolution practices have resulted into higher consumption of chemical fertilizers, mainly N, P, and K supplements, which contributed to improved production and self-reliance in Indian agriculture (Khush, 2001). As chemical fertilizers contribute to higher production, they have become popular and are thus overused to attain higher production. Although extensive use of chemical fertilizers has resulted in higher productivity, it portends ecological and environmental consequences like soil, water, and air pollution (Tiwari et al., 2022). The low use efficiency of conventional chemical fertilizers, such as urea, leads to nearly fifty percent of the applied nutrients getting lost through leaching, run-off, and volatilization. The environmental consequence of chemical fertilizers needs to be urgently addressed to achieve sustainable agriculture’s goals and mitigate the impact of climate change. The solution lies in curtailing overuse of conventional chemical fertilizers without penalties on crop productivity. Organic fertilizers alone cannot meet crop nutrition demands unlike chemical fertilizers due to slower release of nutrients (Timsina, 2018; Mirakhorli et al., 2021; Zhai et al., 2022). Alternative fertilizers like nano fertilizers have been under continuous focus during the last decade. The advantages of nano-fertilizer usage include enhanced plant nutrient uptake efficiency, reduced greenhouse gas emissions, minimized leaching, and more precise timing of nutrient release aligned with crop needs (Seleiman et al., 2020).

Indian Farmers Fertilizer Cooperative (IFFCO) has recently developed and patented a nano urea formulation (NUF) for sustainable crop production (Indian patent number 508189). Precise and targeted application of the NUF has shown bio-efficacy in different crops but the molecular mechanism of its action is not yet known. Nano fertilizers reportedly regulate the gene expression of nutrient transporters and other genes related to growth and stress response in a manner or degree different from bulk fertilizers (Khalid et al., 2022). On the other hand, the efficacy of nano fertilizers is dependent on the mode of its application, viz., foliar spray, soil, and hydroponics (Rizwan et al., 2021). In soil applications, nano carbon, and nano calcium carbonate fertilizer synergists in combination with urea induced higher expression levels of wheat nitrate transporter genes, *NRT 2.2* and *NRT 2.3*, and a nitrogen assimilation gene encoding glutamine synthase, than urea alone (Yang et al., 2023). The expression of an aquaporin *AQP2* gene dramatically increased in tomato roots exposed to nano carbon compared to roots from plants grown on a medium with conventional carbon sources, in tissue culture medium (Khodakovskaya et al., 2010).

Numerous studies have revealed that hydroponically cultivated plants show greater growth and phytochemical content than soil grown plants and exhibit improved internalization of nanoparticles (Gashgari et al., 2018; Verdoliva et al., 2021; Fatima et al., 2023). Nano-elicitation promotes plant metabolic activities, stimulating plant productivity (Jadoon et al., 2023). For example, in cucumber, a hydroponic application of a nanoformulation of urea induced higher expression levels of the high-affinity urea transporter gene *CsDUR3* than conventional urea. In addition, *CsDUR3* induction was observed over a more extended period in NUF-treated plants than in urea, leading to an extended uptake (Feil et al., 2021). Hydroponic application of a nano copper fertilizer in wheat led to a lower up-regulation of *NRT* genes and genes encoding auxin efflux proteins than bulk copper fertilizers. This led to lower root growth inhibition under nano copper fertilization (Zhang et al., 2018).

In the current study, NUF marketed by IFFCO was investigated for growth promotion and effectiveness with equimolar urea concentrations on *Arabidopsis thaliana* under hydroponics. The model plant *A. thaliana* represents the physiological system of higher plants, and its well-established genetics makes it suitable to investigate the mechanisms of growth promotion by NUF in hydroponics. Our study sheds light on the difference between the mode of action of NUF and urea. Hitherto unknown molecular mechanisms involved in growth promotion of *A. thaliana* upon NUF, and urea treatment were unraveled through transcriptome and real-time expression analysis of differentially expressed genes. Further, we correlated the physicochemical properties of NUF with the uptake and growth regulation in *A. thaliana*.

## Materials and Methods

### Physicochemical characterization of nano urea

A liquid nano urea formulation marketed by IFFCO (batch no. IPNU291440224, manufactured on 13-07-2023) was procured from the market, and here onwards, referred to as “NUF.” Malvern Zetasizer Ultra Red Label (Malvern Panalytical Ltd., Worcestershire, UK) was used for dynamic light scattering (DLS) and Zeta potential measurements of aqueous solutions of NUF. The observations were taken at 25 ℃. After thoroughly washing the cuvettes with distilled water and ethanol and removing the impurities, the cuvettes were filled with 1 mL of sample and characterized. Each measurement consisted of three one-minute runs to produce an average and standard deviation. Transmission electron microscopy was conducted on the NUF solution with the LVEM-5 Benchtop Electron Microscope (Delong instruments, Brno, Czech Republic). Fourier Transform Infrared (FTIR) was used to investigate the functional groups in urea and NUF, using an IRSpirit spectrophotometer (Shimadzu, Kyoto, Japan) in the wavenumber range of 500—4000 cm^-1^. Lyophilization was performed to extract water from the NUF using a Delvac mini lyodel apparatus (Delvac Pumps Pvt. Ltd., India.). First, NUF was frozen at −20°C and lyophilized at a temperature of −50℃, under a low pressure of 0.185 mbar for 10 hours. Lyophilization eliminated the water molecules from the NUF matrix while maintaining its composition and structural integrity. A field emission scanning electron microscope (FESEM) Apreo 2S (Thermo Fisher Scientific, New Delhi, India) was used to investigate the surface morphology of solid urea and lyophilized nanourea. The dried-powdered samples were placed on carbon tape-affixed aluminum stubs and were gold-coated using a SC7620 mini 374 sputter coater (Quorum Technologies Ltd., Laughton, UK).

### Plant materials and growth conditions

The Columbia ecotype (Col-0) of *A. thaliana* was used for all experiments. The seeds were stored at 4°C under low humidity until usage. These seeds were imbibed for three days with distilled water at 4°C for synchronized and uniform germination, cautiously pipetted on N50 size nylon meshes mounted on 2” × 2” photographic film slide frames, and floated into a tray containing 5 L of hydroponic solution with the help of polystyrene floaters. One hydroponic growth medium had nitrogen, whereas the other had no nitrogen. The pH of both media was maintained at 5.8 by adding NaOH/HCl. The nitrogen-containing solution was a modified Hoagland’s growth medium (Hoagland and Arnon 1938), consisting of 0.24 mM NH_4_H_2_PO_4_, 1.5 mM KNO_3_, 1 mM Ca(NO_3_)_2_.4H_2_O, 2 mM MgSO_4_.7H_2_O, 2.886 μM H_3_BO_3_, 0.571 μM MnCl_2_.4H_2_O, 0.048 μM ZnSO_4_.7H_2_O, 0.020 μM CuSO_4_.5H_2_O, 0.008 μM H_2_MoO_4_, and 0.02 μM C_4_H_4_FeO_6_. The solutions were replaced every three days. After 12 days of growth in the nitrogen-containing nutrient medium, the meshes were shifted to a nitrogen-free solution fabricated from Hoagland’s and MGRL media (Hoagland and Arnon 1938; Fujiwara et al., 1992) for testing fertilization with urea and NUF. This nitrogen-free solution contained 0.24 mM NaH_2_PO_4_.2H_2_O–Na_2_HPO_4_.12H_2_O buffer, 1 mM CaCl_2_.2H_2_O, 2 mM MgSO_4_.7H_2_O, 2.886 μM H_3_BO_3_, 0.571 μM MnCl_2_.4H_2_O, 0.048 μM ZnSO_4_.7H_2_O, 0.020 μM CuSO_4_.5H_2_O, 0.008 μM H_2_MoO_4_, 0.02 μM C_4_H_4_FeO_6_, and 0.75 mM KH_2_PO_4_. For the entire growth regime, a photoperiod of 12 h light (80 μmoles m^-2^ s^-1^ of photosynthetically active radiation) and 12 h darkness was imposed in a controlled growth chamber at a temperature of 22 ± 2°C and relative humidity of 60 ± 5%.

### Fertilization with urea or NUF

An aqueous urea solution of 1.5 M was prepared to match the NUF formulation of 1.5 M (IFFCO certificate of analysis; Supplementary Table S1). These were used as primary stocks diluted for different doses. After 12 d hydroponic growth in nitrogen-containing media, then transferred to the nitrogen-free hydroponic media supplemented with different doses, viz., 0.5, 5, 40, 50, 70, 80, and 100 μM urea and NUF, and were grown further for seven days. On the seventh day, seedlings were carefully removed from the meshes using forceps, placed on solidified agar, and photographed. For long-term hydroponic culture, the seedlings were carefully removed using forceps, and the roots were inserted in holes made in polystyrene foams, floated on the same hydroponic solutions, and grown for another 15 d. On the 15^th^ day, the seedlings were photographed and harvested to determine morphological and biochemical parameters.

### Phenotyping *A. thaliana*

The morphological characteristics of root length, rosette diameter, and biomass were evaluated systematically to compare the growth characteristics of seedlings grown under control, NUF, and urea containing media. Plants were photographed with a scale. From the photographs, the root lengths and rosette diameters were measured using an in-house image-analysis program (Sadhukhan et al., 2017). Shoots and roots were separated with a scalpel and weighed in a precision digital weighing machine (Mettler Toledo India Pvt. Ltd., Mumbai, India) to obtain the biomass.

### Measurement of chlorophyll content

Chlorophyll levels were measured in the leaves collected from all fertilizer-treated and control plants following the methodology outlined by Arnon (1949). One gram of plant leaf was ground in a mortar and mixed with 20 mL of 80% acetone. After centrifugation at 10,000 rpm for 5 minutes, chlorophyll extraction was carried out. A UV-spectrophotometer (Shimadzu Corporation, New Delhi, India) measured the absorbances at 663 and 645 nm, respectively. The measurement of chlorophyll a, chlorophyll b, and total chlorophyll levels were estimated according to the equations: Chlorophyll a (mg/g) = (12.7 × A_663_) – (2.59 × A_645_), Chlorophyll b (mg/g) = (22.9 × A_645_) – (4.7 × A_663_), and Total Chlorophyll (mg/g) = (8.2 × A_663_) + (20.2 × A_645_). The chlorophyll contents were expressed as mg/g tissue fresh weight.

### Measurement of total tissue nitrogen content

Leaves and roots were collected from the exposed plants to determine the total nitrogen content, following the methodology outlined by Koistinen et al., 2020. The process involved homogenizing 1 g of the tissue in a mortar with 2 mL of deionized water. The homogenate was centrifuged at 10,000 rpm for 5 minutes. Then, 5 mL of an oxidizing agent containing 10 g/L K_2_S_2_O_8_ and 6 g/L H_3_BO_3_ in 75 mM NaOH was added. Immediately after that, 3 mL of total nitrogen (TN) mix, prepared by mixing 40 mL of 1% sulfanilamide (dissolved in 2.4 N HCl), 40 mL of 0.07% N-(1-naphthyl)-ethylenediamine dihydrochloride, and 200 mL of 0.5% VCl_3_ (dissolved in 1.2 N HCl), was introduced to the sample. The sample was then incubated at 45°C in a water bath for 30 minutes, and the absorbance was finally measured at 545 nm. The nitrogen content was estimated from a standard curve prepared with Na_2_EDTA.

### Measurement of tissue amino acid content

The exposed plants’ leaves and roots were collected to measure the tissue amino acid content following the methodology outlined by Doi et al., 1981. The tissues were macerated in a mortar using a 2 mL solution containing methanol, chloroform, and deionized water in a 1:1:2 ratio. The final homogenate was centrifuged at 5000 g for 10 minutes at 4°C. The upper phase was carefully aspirated to a new tube, and 4 mL of deionized water was added. One milliliter of ninhydrin reagent, prepared by mixing 1.6 g/L SnCl_2_.2H_2_O in 0.2 M citrate buffer (pH 5.0) with 4% ninhydrin in 2-methoxyethanol, was added to the sample. After that, the sample was left to incubate for 15 minutes at 90°C, and then it was allowed to cool at room temperature. The absorbance of the sample was noted at 570 nm after adding 1 mL of ethanol. The total amino acids were quantified from a standard curve prepared with pure L-leucine and expressed as mg/g tissue fresh weight.

### Transcriptome analysis

Col-0 seeds were grown hydroponically in nitrogen-containing hydroponics media for 12 d and then acclimatized in nitrogen-free hydroponic media for two days. After that, the plants were transferred to urea and NUF-supplemented nitrogen-free hydroponic media for 12 h and seven days in preparation for transcriptome profiling. The snap-freezing of these seedlings in liquid nitrogen allowed for the quick preservation of RNA profiles. Then, the RNA was extracted using the CTAB-LiCl protocol. Its fidelity was then evaluated utilizing the Agilent 4150 TapeStation equipment (Agilent Technologies, California, USA) and assessed with Thermo Fisher’s Qubit 4 fluorometer. RNA-Seq libraries were prepared by NebNext Ultra II RNA library preparation kit (New England Biolabs, Massachusetts, USA) and sequenced using Illumina’s state-of-the-art NovaSeq 6000 V1.5 platform (Illumina, Gurgaon, India). This produced 150 base pair long paired-end reads. The GC content and library complexity of the unprocessed sequence reads were evaluated using the FastQC software (Andrews’ 2010). AdapterRemoval ver. 2.3.2 was implemented for the adapter trimming at Q30 (Schubert, Lindgreen, and Orlando 2016). The sequence reads were aligned to the reference *A. thaliana* genome (GCF000001735.4) using hisat2 version 2.2.1 (Zhang et al., 2021). Using featureCounts v2.0.3, expression quantification was done using the procedures described by Liao, Smyth, and Shi (2014). The R framework was then used to perform differential gene expression (DEG) analysis using the edgeR, DEseq2, and ggplot2 packages, according to Love, Huber, and Anders (2014), Chen, Lun, and Smyth (2016), and Wickham (2016). The gene functional enrichment analysis was executed using the BAR classification superviewer web tool (https://bar.utoronto.ca/).

### Real-time quantitative PCR analysis

Quantitative real-time PCR (qPCR) was adopted to validate the expression levels of genes associated with nitrogen metabolism. Akin to the conditions of the transcriptome experiment, *A. thaliana* seedlings were treated with urea and NUF in a hydroponic environment. The samples were then instantaneously preserved in liquid nitrogen. The tissue samples were ground up through mortars and pestles to extract RNA using Promega’s SV total RNA isolation kit. Following this, the isolated RNA was reverse-transcribed into complementary DNA (cDNA) through reverse transcription using the PrimeScript™ IV 1st strand cDNA Synthesis Mix manufactured by DSS Takara Bio India Pvt. Ltd., located in New Delhi, India. QPCR was conducted utilizing Brilliant II SYBR master mix from Agilent Technologies

India Pvt Ltd (Chandigarh, India) in an Mx3000P qPCR System (Agilent). The standard curve approach was used to determine the relative expression levels of NUF treated compared to N-free unfertilized control seedling samples. The *AtUBQ1* gene was employed as an internal standard for normalizing the cDNA samples. All oligonucleotide primers employed in the qPCR analysis were designed using the Primer3web program version 4.1.0 (Koressaar et al., 2018) and are listed in Supplementary Table S2.

### Statistical analysis

All experiments were independently repeated thrice with complete randomization in the design to nullify environmental or chance effects. Ten biological replicates were used for each treatment in an experiment. Student’s *t*-test at a significance level of *P* < 0.05 was used for pairwise comparisons. One-way ANOVA followed by Tukey’s honestly significant difference (HSD) test was used to compare multiple treatments. The transcriptome data consisted of three replicates, each of ten plants, statistically treated with Fisher’s exact test (Fisher 1945) and Benjamini-Hochberg *false discovery rate* (*FDR*) correction (Benjamini and Hochberg 1995).

## Results

### Physicochemical properties of NUF

The efficiency of nano fertilizer uptake by plants depends on two essential characteristics: particle size and surface charge. A hydrodynamic diameter of 37.56 nm was estimated for 70 μM aqueous solution of NUF (in a hydroponic growth medium background, described later), using DLS (Figure 1A). Zeta potential represents the magnitude of repulsion between neighboring particles, like-charged and unlike-charged, within a dispersion. The zeta potential for 70 μM NUF was −47.8 mV (Figure 1B). The negative zeta potential confirms that NUF carries anionic properties in aqueous solution. The transmission electron microscopy images additionally confirmed the size of NUF in the 30—50 nm range (Figure 1C). We compared the FTIR spectra of NUF and urea (Figure 1D). In urea, prominent triple absorption peaks emerged at 3430.4 cm^-1^, 3330.2 cm^-1^, and 3253.1 cm^-1^, attributed to the asymmetric and symmetric stretching of N-H bonds. Furthermore, the N-H bond deformation frequency was obtained at 1588.5 cm^-1^. The peak at 1675.5 cm^-1^ corresponded to the C=O stretching vibration, and the C-N vibration absorption peak was observed at 1084 cm^-1^ (Onija et al. 2012). For NUF, the sharp double absorption peaks of N-H developed at 3331.9 cm^-1^ and 3241.4 cm^-1^; the higher and lower wavenumbers were attributed to the asymmetric and symmetric stretching absorption peaks of amino groups in urea. The C=O, C-N, and C-O stretching vibrations were observed at 1633.2 cm^-1^, 1449.7 cm^-1^, and 1084.3 cm^-1^, respectively. The FTIR spectra indicated similar molecular signatures of urea and NUF (Figure 1C, Supplementary Table S3). Scanning electron micrographs of lyophilized NUF and urea samples showed their amorphous and crystalline morphology, respectively (Figure 1E).

**Figure 1.**
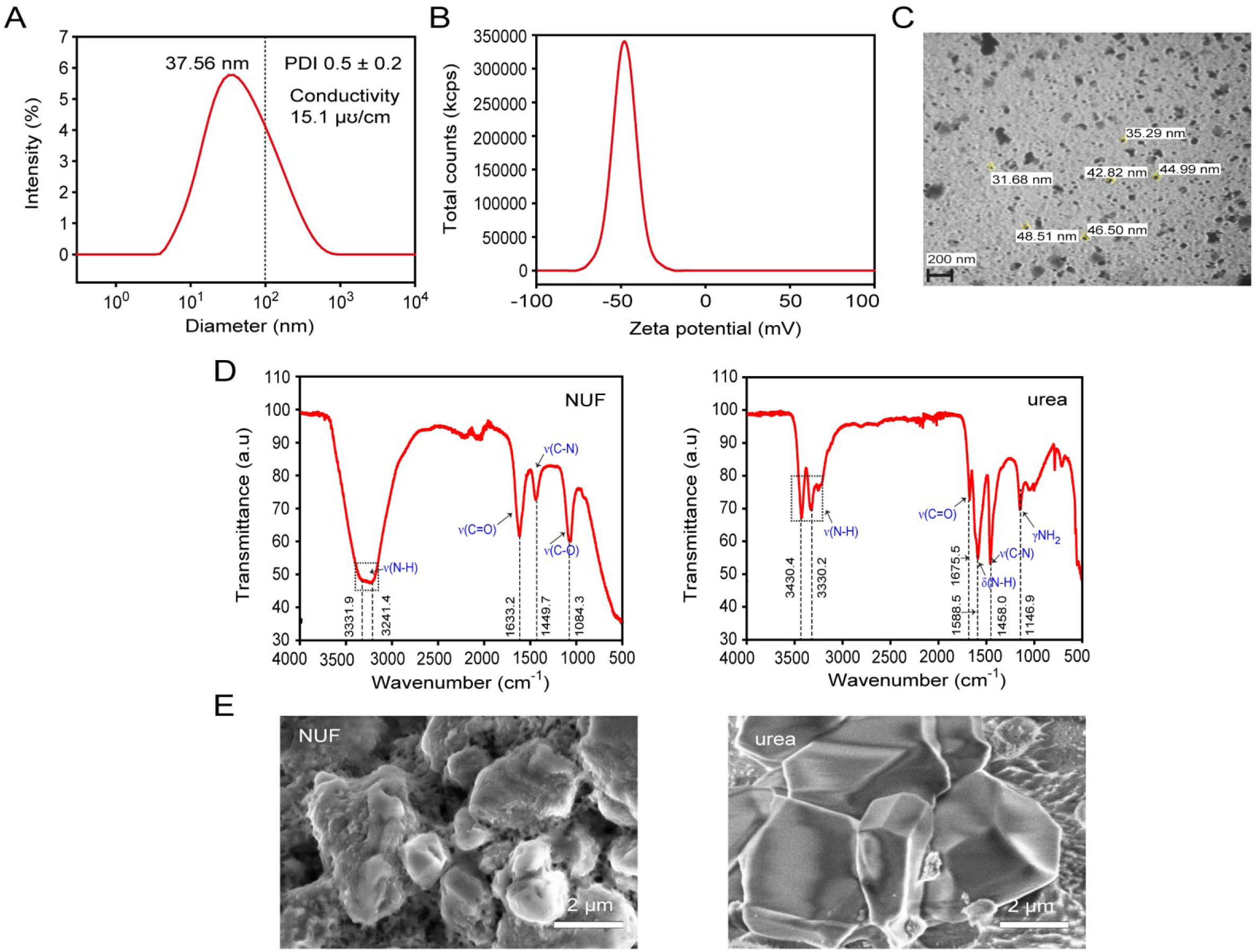
Physicochemical characterization of nano urea. Particle size distribution by dynamic light scattering (DLS) (**A**) and zeta potential (**B**) were measured for the aqueous solution of 70 μM nano urea formulation (NUF) in the same nitrogen-free growth medium background as in the hydroponic fertilization experiment (presented in Figure 2). The percentage intensity of scattered light is plotted against particle size in nm. The dotted line indicates 100 nm. Polydispersity indices (PDI) and ionic conductivities of the solutions (in μ℧/cm) are mentioned in the inset. The total count of scattered photons in kilo counts per second (kcps) was plotted against the zeta potential in mV. (**C**) Transmission electron micrograph of 70 μM solution of NUF is shown. Black scale bar indicates 200 nm. The diameters of six randomly chosen particles are shown. (**D**) Fourier transform infrared spectroscopy (FTIR) spectra of NUF and urea, are shown as plots of transmittance in arbitrary units versus wavenumber in cm^-1^. (**E**) Field-emission scanning electron micrographs (FESEM) show the amorphous and crystalline morphologies of lyophilized NUF and urea. Scale bars indicate 2 μm.

### NUF promotes root growth, chlorophyll, and nitrogen content under short-term hydroponic culture

We investigated the effects of NUF on the growth of *A. thaliana* seedlings in hydroponic culture while keeping equivalent urea concentrations as controls. For uniformity, seedlings were germinated and grown for 12 days in nitrogen-containing hydroponic solution, then transferred to nitrogen-free media alone or supplemented with NUF or urea. To optimize the dose of NUF, we started from a much lower concentration of 0.5 μM, than the recommended dose, and gradually increased the concentration by 10 times, viz., 5 μM and 50 μM. The relative root length, i.e., length in NUF/length in nitrogen-free media (RRL; %), increased at 50 μM. After seven days, the highest RRL was observed at 70 μM, deemed the optimal dose for NUF. However, when we increased the dose, the root growth was inhibited at and above 80 μM. On the other hand, the root growth increased at equivalent higher doses of urea, viz., 50 —100 μM (Figure 1A). The relative shoot (rosette) diameter i.e., diameter in NUF/diameter in nitrogen-free media (RSD; %), increased slightly after 7 d treatment with 50—100 μM NUF, but a more significant increase was obtained in higher 80—100 μM doses of urea (Figure 1A). At the optimal 70 μM concentration, the chlorophyll content of the shoots was significantly higher in the NUF treatment than in the control (Figure 1B), and higher than in urea-containing medium (Figure 1C). The total nitrogen contents in the shoots (Figure 1C) and roots (Figure 1D) increased significantly by NUF and urea compared to control medium. Still, the increase was higher in urea than in NUF at this stage. These results suggested that NUF was a suitable nitrogen fertilizer for *A. thaliana* in hydroponic culture, promoting plant growth at an optimum dose. Meanwhile, the greater enhancement of chlorophyll than urea treatment at the optimal dose and toxicity at higher doses of NUF indicated nitrogen excess in the NUF treatment, possibly due to the higher uptake rate of urea in the nano form.

### NUF enhances biomass, chlorophyll, total tissue nitrogen, and amino acid content greater than urea under long-term hydroponic culture

After the preliminary observations after a week of treatment, we were intrigued to observe the growth effects in *A. thaliana* seedlings after long-term exposure to NUF and urea. After an exposure of 15 d (Figure 1E), the total chlorophyll content remained highest in the NUF treatment, followed by urea and nitrogen-free control (Figure 1F). The shoot and root biomass (Figure 1G), as well as the total tissue nitrogen (Figure 1H) and amino acid content (Figure 1I) in the shoots, was now higher in NUF than urea. Compared to urea, an increase of 20 ± 5% in shoot biomass, 16 ± 2.5% in chlorophyll, 6.5 ± 0.5 % in total shoot nitrogen and 30 ± 5% increase in shoot amino acid content was observed in NUF-treated seedlings. The root nitrogen content in NUF-treated seedlings was similar to that of the urea treated ones but higher than that of the control. These results suggested that NUF outperformed an equivalent dose of urea in the growth enhancement upon long-term treatment.

### NUF alters global gene expression patterns of *A. thaliana* greater than urea

To gain insights into the molecular mechanisms of growth promotion by NUF and urea, we analyzed the differentially expressed genes (DEGs) in NUF versus control and urea versus control at 12 h and 7 d of treatment. At an early 12 h exposure, 589 genes were up-regulated (log_2_fold-change > 1, *FDR* < 0.05), and 409 genes down-regulated under NUF treatment, compared to the nitrogen-free control (log_2_FC < −1, *FDR* < 0.05) (Figure 4A). In contrast, urea treatment induced 279 genes and repressed 187 genes relative to the control (log_2_FC > 1, *FDR* < 0.05) (Figure 4B; Supplementary Table S3). Of the 589 genes up-regulated by NUF, 151 were induced by urea (log_2_FC > 1). 52 genes were down-regulated by urea and NUF (Figure 4C). After 7 d of treatment, the number of differentially expressed genes (−1 < log_2_FC > 1, *FDR* < 0.05) increased in both treatments. Urea treatment induced 977 genes and repressed 573 genes relative to the control (Figure 4A). An even greater number of 1636 genes were up-regulated and 1691 down-regulated under NUF treatment, compared to the nitrogen-free control (Figure 4B). At 7 d, 889 genes were up-regulated by both urea and NUF (log_2_FC > 1), while both down-regulated 490 genes (log_2_FC < −1) (Figure 3C).

**Figure 2.**
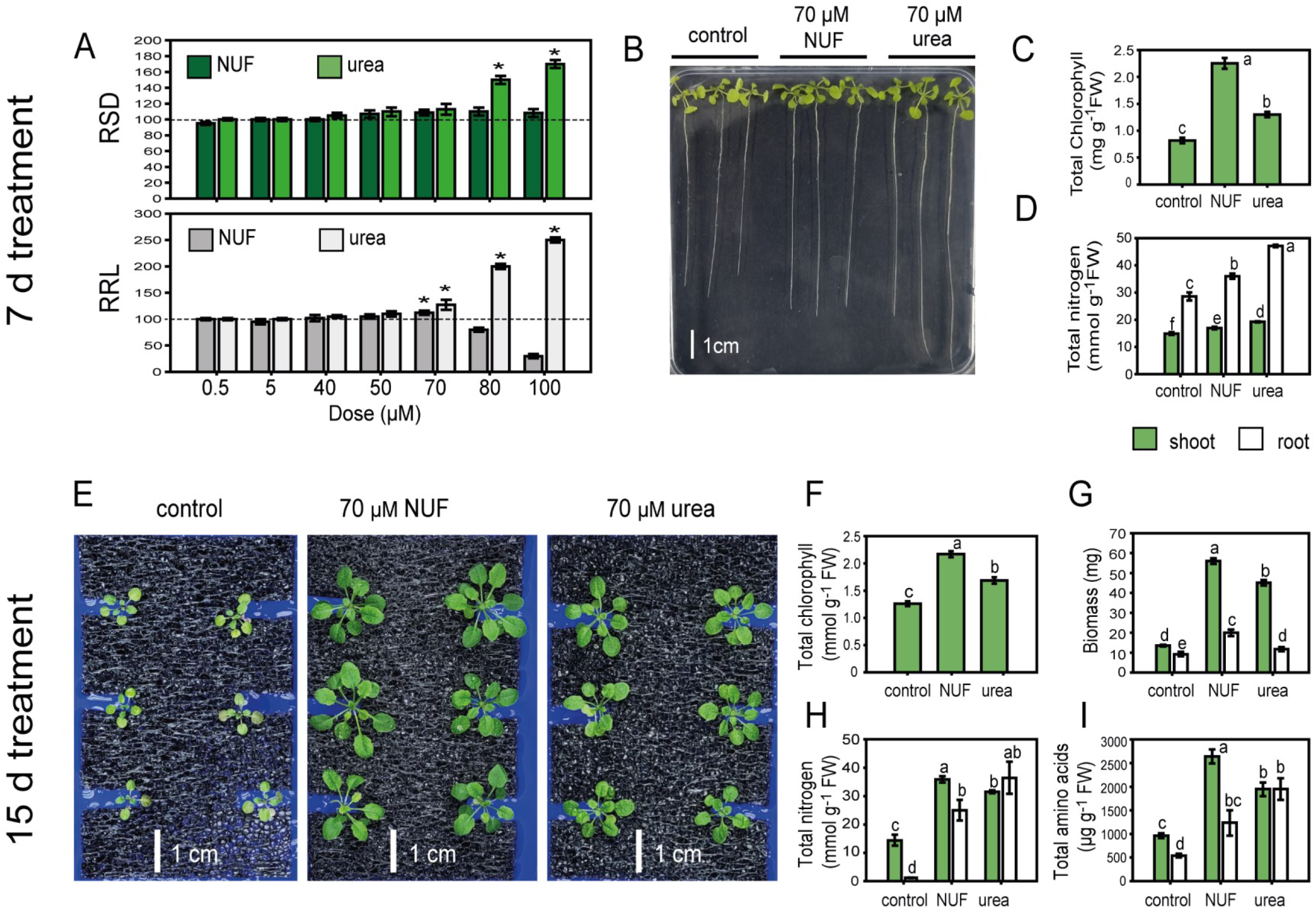
Effect of nano urea on Arabidopsis growth in hydroponics. *Arabidopsis thaliana* plants were grown in hydroponics in nitrogen-containing media for 12 d from germination and, after that, transferred to nitrogen-free media alone (control) or nitrogen-free media supplemented with different doses, 0.5, 5, 40, 50, 70, 80, or 100 μM nano urea formulation (NUF) or urea for seven days (**A**-**D**) or 15 days (**E**-**I**). The seedlings were picked up after seven days, placed carefully on solidified agar plates, and photographed. (**A)** The seedlings’ relative root length (RRL; length of treated root/length of control root) and relative shoot (rosette) diameter (RSD; diameter of treated shoot/diameter of control shoot) in media with different doses of NUF or urea versus control are expressed as percentages. Bars indicate the average of ten seedlings, with standard error. Asterisks indicate significant increases in RRL or RSD from the control (Student’s *t*-test, *P* < 0.05, *N* = 10). (**B)** A representative image of 12-d nitrogen-containing media-grown *A. thaliana* seedlings is shown after further 7 d growth in the nitrogen-less control medium, 70 μM NUF, or 70 μM urea, pickup from hydroponic media and placement on a solidified agar plate. (**C)** The shoots were excised, crushed in mortar and pestle, extracted with acetone, and the chlorophyll contents were measured spectrophotometrically (see Methods). The total chlorophyll contents of seedling shoots in milligrams per gram of tissue fresh weight are shown. (**D)** The total tissue nitrogen contents in shoots and roots are presented after growth in control, urea, or NUF. The shoots and roots were separated and homogenized using a mortar and pestle, and nitrogen contents were measured spectrophotometrically by a biochemical test (see Methods). (**E**) A representative image of 12-d nitrogen-containing media-grown *A. thaliana* seedlings is shown after a further 15 d growth in control, 70 μM NUF, or 70 μM urea. (**F**) The total chlorophyll contents of seedling shoots in milligrams per gram of fresh tissue weight after 15 d growth in control, NUF, and urea are shown. Bars indicate the average of three replicates, each of ten seedlings, with standard error. (**G**) The shoot and root fresh weights were measured after 15 d growth in control, NUF, and urea and presented as biomass in milligrams. Bars indicate the average of ten seedlings, with standard error. The total tissue nitrogen contents (**H**) and amino acids (**I**) in shoots and roots are presented after growth in control, urea, or NUF. The shoots and roots were separated and homogenized using a mortar and pestle, and nitrogen and amino acid contents were measured spectrophotometrically by biochemical tests (see Methods). Bars indicate the average of three replicates, each of ten seedlings, with standard error. Different alphabets indicate significant differences (Tukey’s HSD test, *P* < 0.05, *N* = 3).

**Figure 3.**
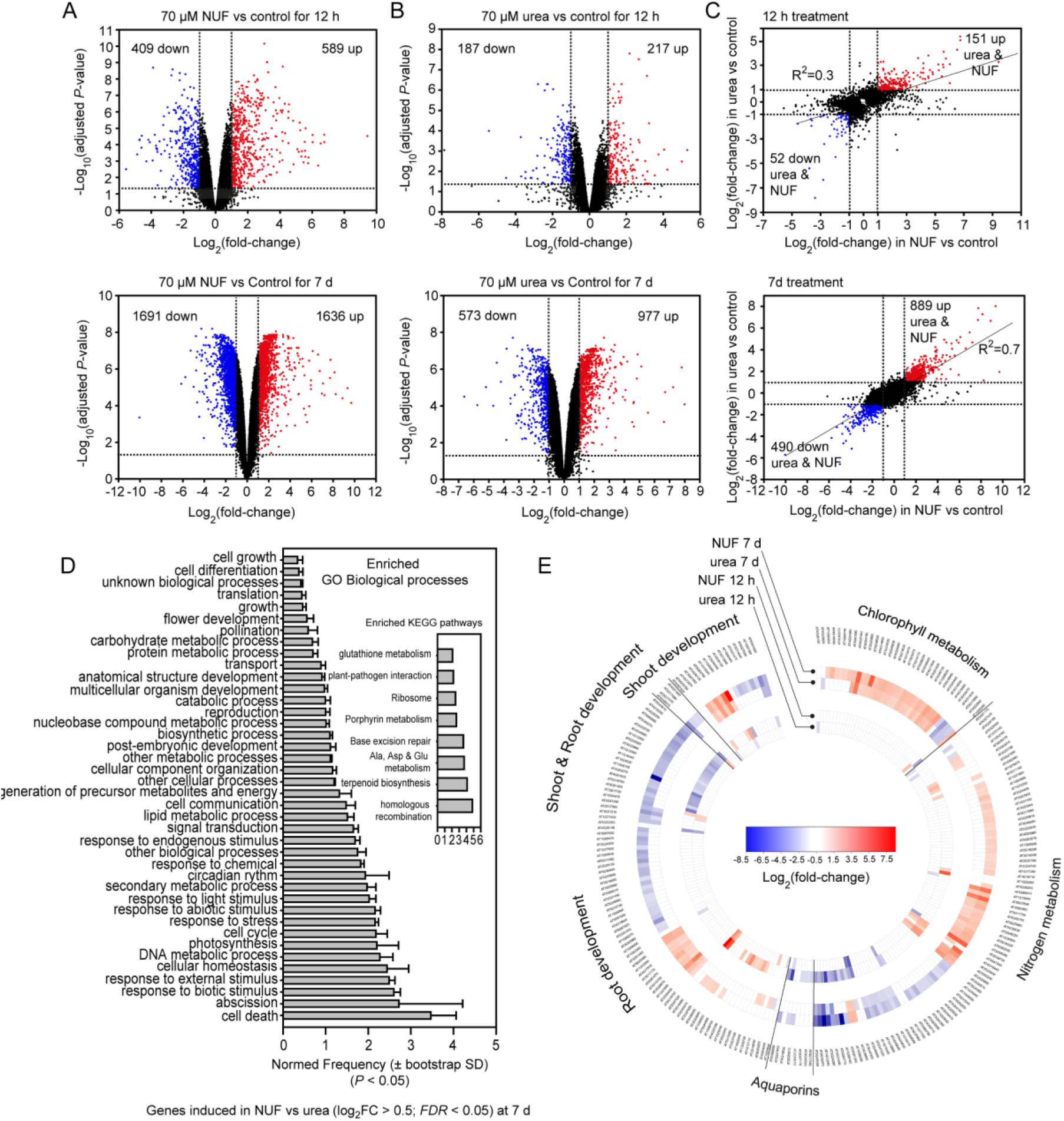
Differential gene expression of *Arabidopsis thaliana* fertilized with nano urea and urea. *A. thaliana* plants were grown for 12 days in nitrogen-containing hydroponic growth media, followed by acclimatization for two days in a nitrogen-free growth medium. After that, the plants were treated separately with 70 μM urea, 70 μM nano urea formulation (NUF), and control (without nitrogen fertilizer) in the nitrogen-free background solution. Seedlings were snap-frozen in liquid nitrogen after 12 h and 7 d post-treatment, total RNA was isolated, and cDNA libraries were prepared and sequenced by Illumina NovaSeq, followed by identifying differentially expressed genes (see Methods). Volcano plots of −log_10_(adjusted *P*-value) versus log_2_FC for both NUF (**A**) and urea (**B**) at 12 h and 7 d of treatment are represented in the figure. Red dots show up-regulated genes with log_2_FC > 1, adjusted *P*-value < 0.05, and down-regulated genes with log_2_FC < −1, adjusted *P*-value < 0.05 by blue dots. Adjusted *P*-value refers to Benjamini-Hochberg’s *false discovery rate* (*FDR*) correction. A correlation plot is shown between genes differentially expressed (adjusted *P*-value < 0.05) in NUF versus urea both for 12 h and 7 d of treatment, each against control (**C**). Enriched Gene Ontology (GO) biological process categories of genes induced in NUF against urea (log_2_FC > 0.5, *P* < 0.05) are plotted against their frequencies of occurrence in the gene set (1057 genes) versus *A. thaliana* genome-wide genes (normed frequencies) together with bootstrap standard deviation (*P* < 0.05), calculated using the BAR Classification SuperViewer tool (**D**). Enriched Kyoto Encyclopedia of Genes and Genomes (KEGG) pathways are shown in the inset. A circular heat map showing clustering of genes in the functional categories “shoot and root development,” “nitrogen metabolism,” “chlorophyll metabolism,” and aquaporins, which were either up-regulated (log_2_FC > 1, *P* < 0.05) or down-regulated (log_2_FC < −1, *P* < 0.05) in NUF treatment for 12h or 7 d are shown. The color scale indicates up-regulation in shades of red and down-regulation in shades of blue according to log_2_FC values (**E**).

**Figure 4.**
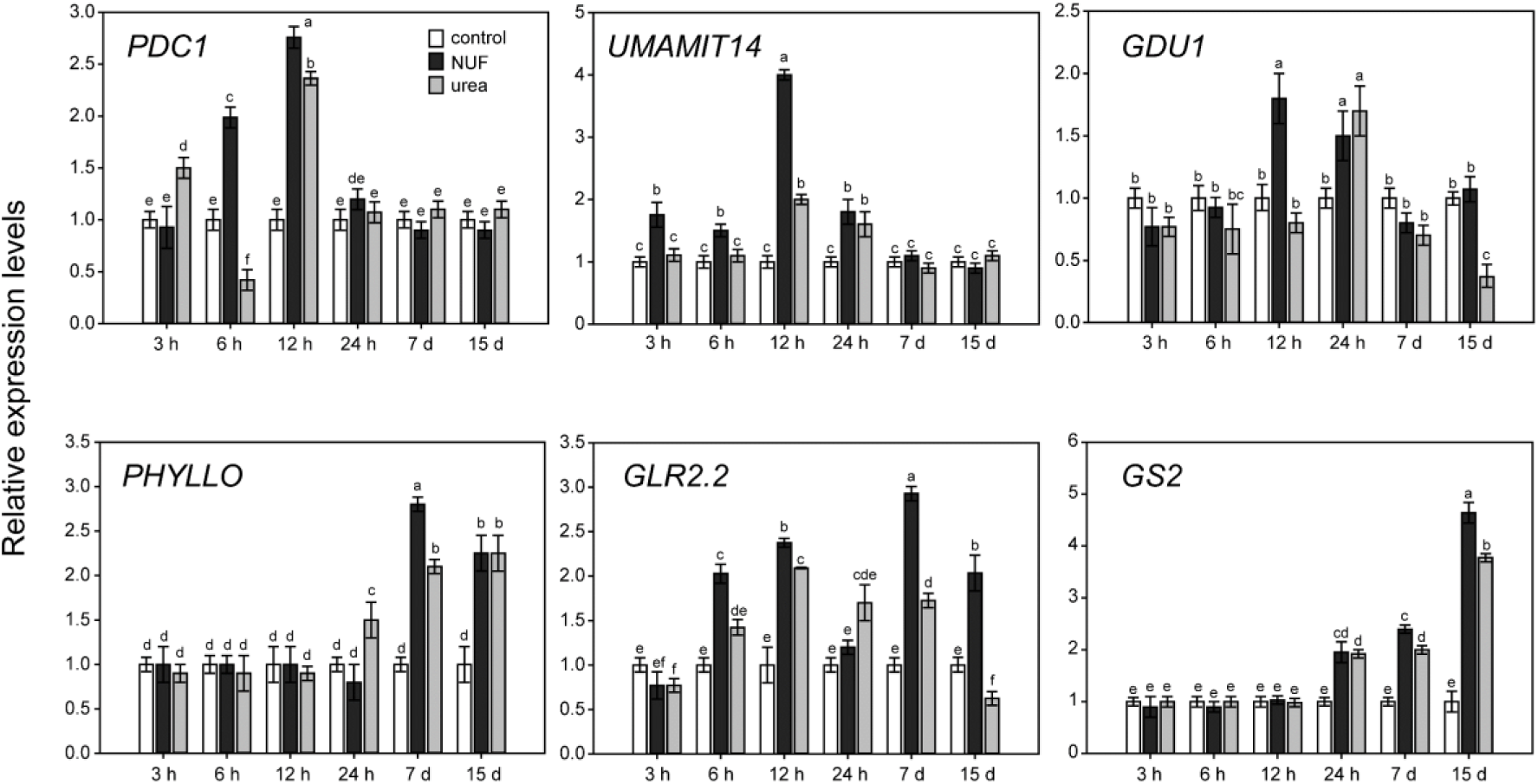
Real-time expression analysis of *Arabidopsis thaliana* genes induced by nano urea. 12-day-old hydroponically grown *Arabidopsis thaliana* seedlings were transferred to nitrogen-free media for two days and then to 70 μM urea, 70 μM nano urea formulation (NUF), and control (without nitrogen) in the nitrogen-free background solution for different time points, viz., 3 h, 6 h, 12 h, 24 h, 7 d, and 15 d. Afterward, the seedlings were harvested to isolate RNA and convert it to cDNA to estimate relative expression levels using real-time PCR (see Methods). The relative fold-change in expression levels of the six highly induced genes in the transcriptome analysis (Table 1), viz., *PDC1*, *UMAMIT14*, *GDU1*, *PHYLLO*, *GLR2.2,* and *GS2*, between control, NUF and urea-treated *A. thaliana* seedling samples are shown in the graph. The expression levels in each sample were further normalized using *AtUBQ1* as the housekeeping gene internal standard due to its steady transcript levels across all samples in the transcriptome data. The bars indicate an average of three biological replicates comprising six seedlings with standard error. Different smaller case alphabets above the error bars indicate significant differences in gene expression between samples (*P* < 0.05, Tukey’s HSD test, *N* = 3 replicates).

**Table 1.**
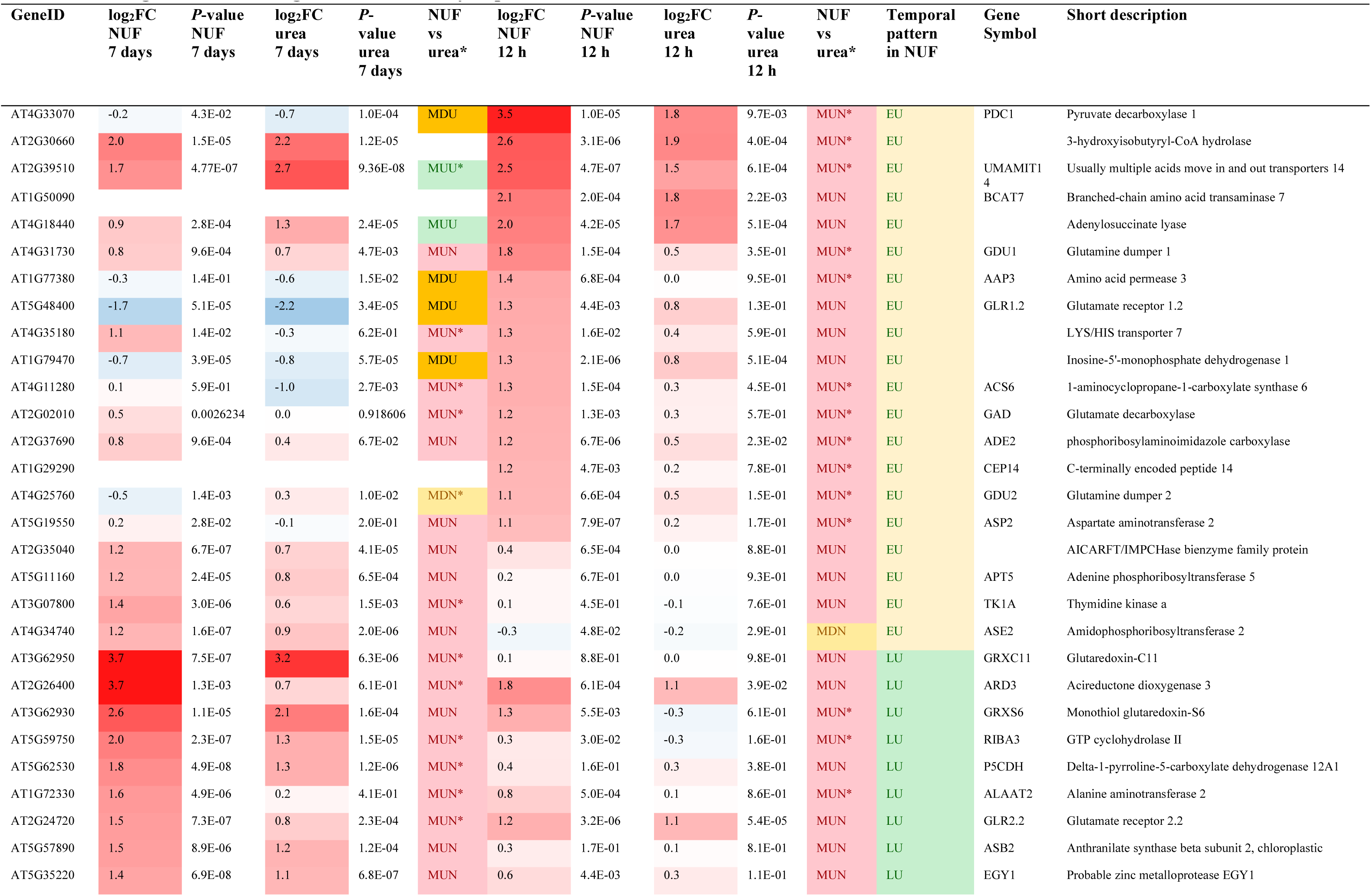
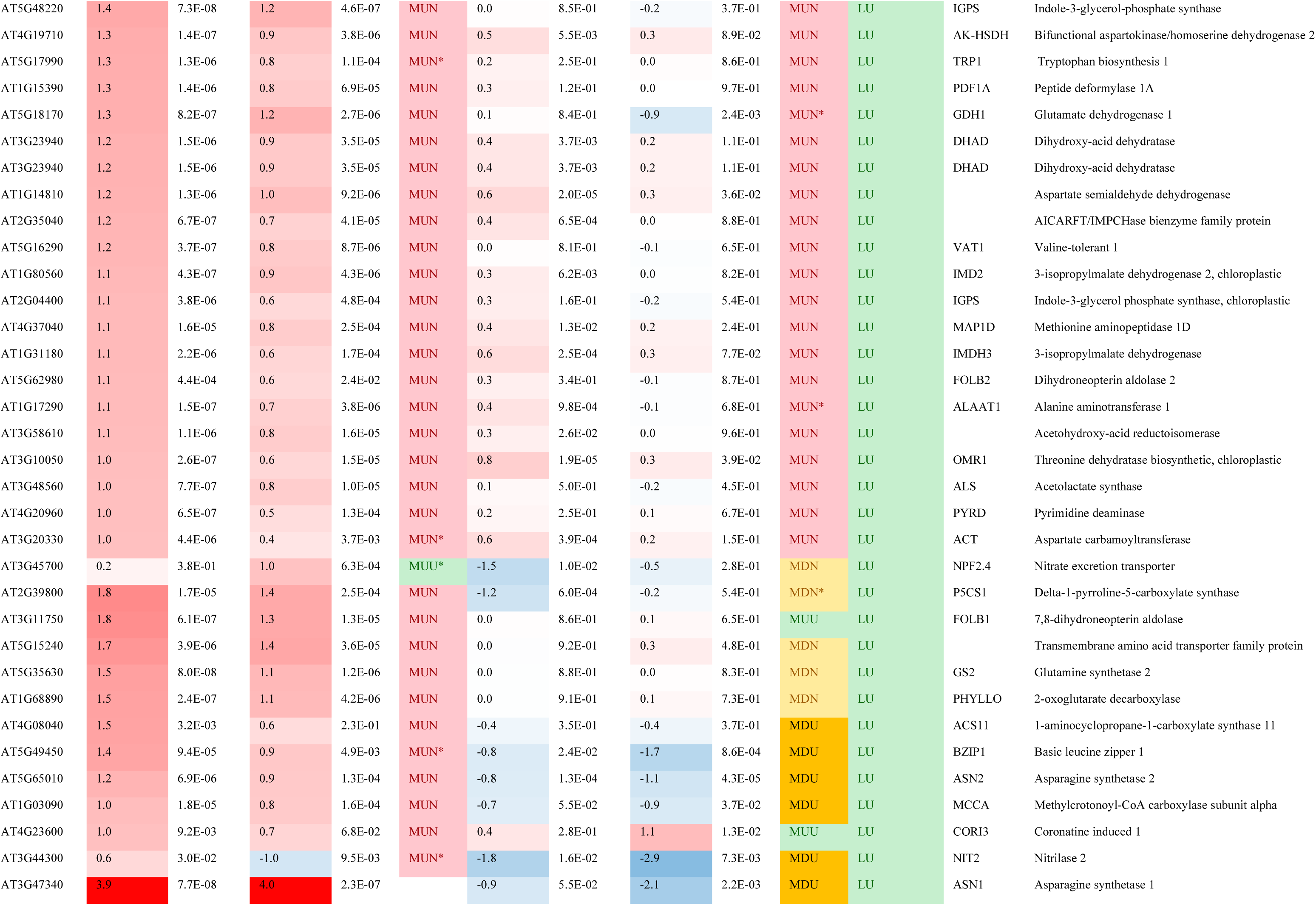
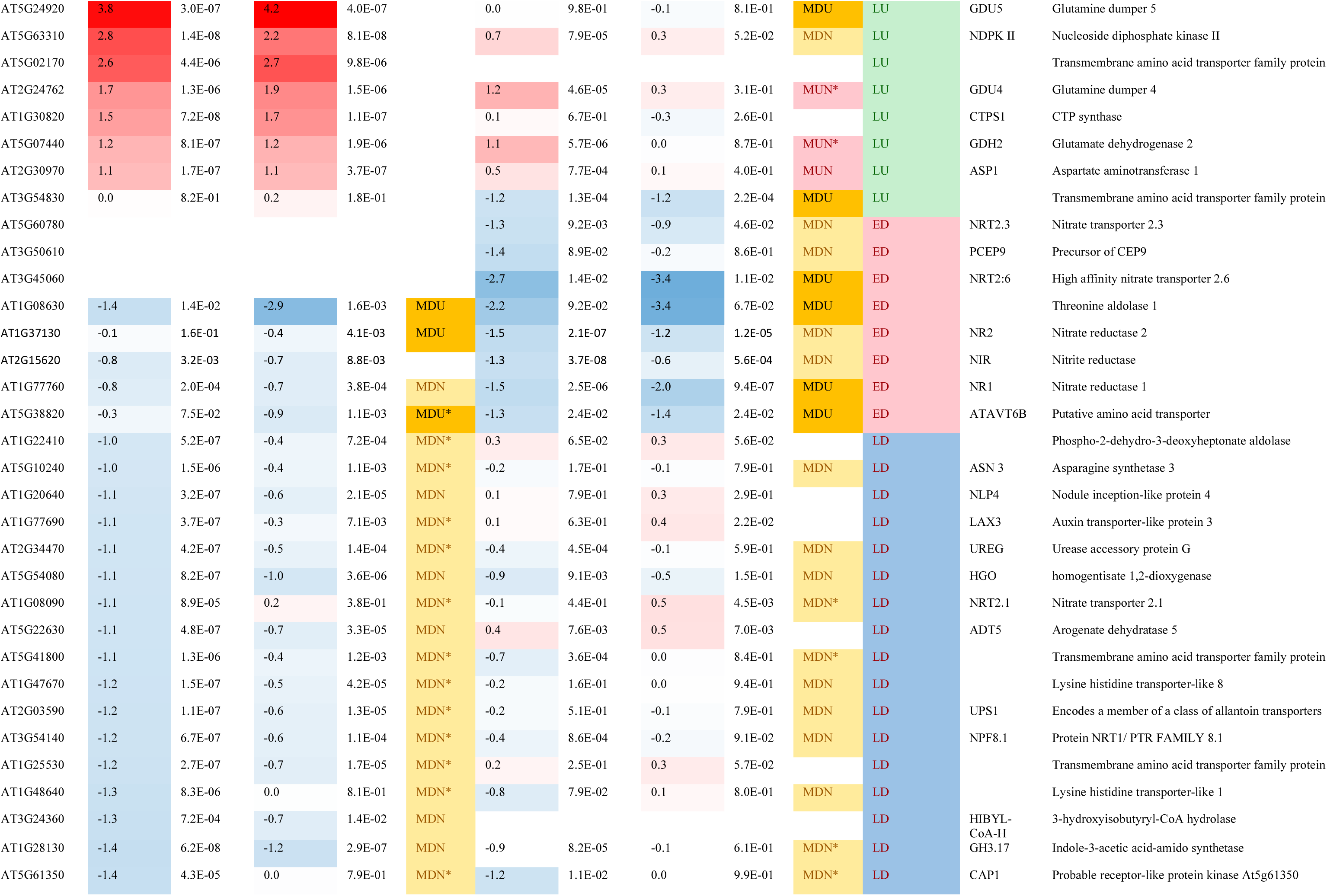
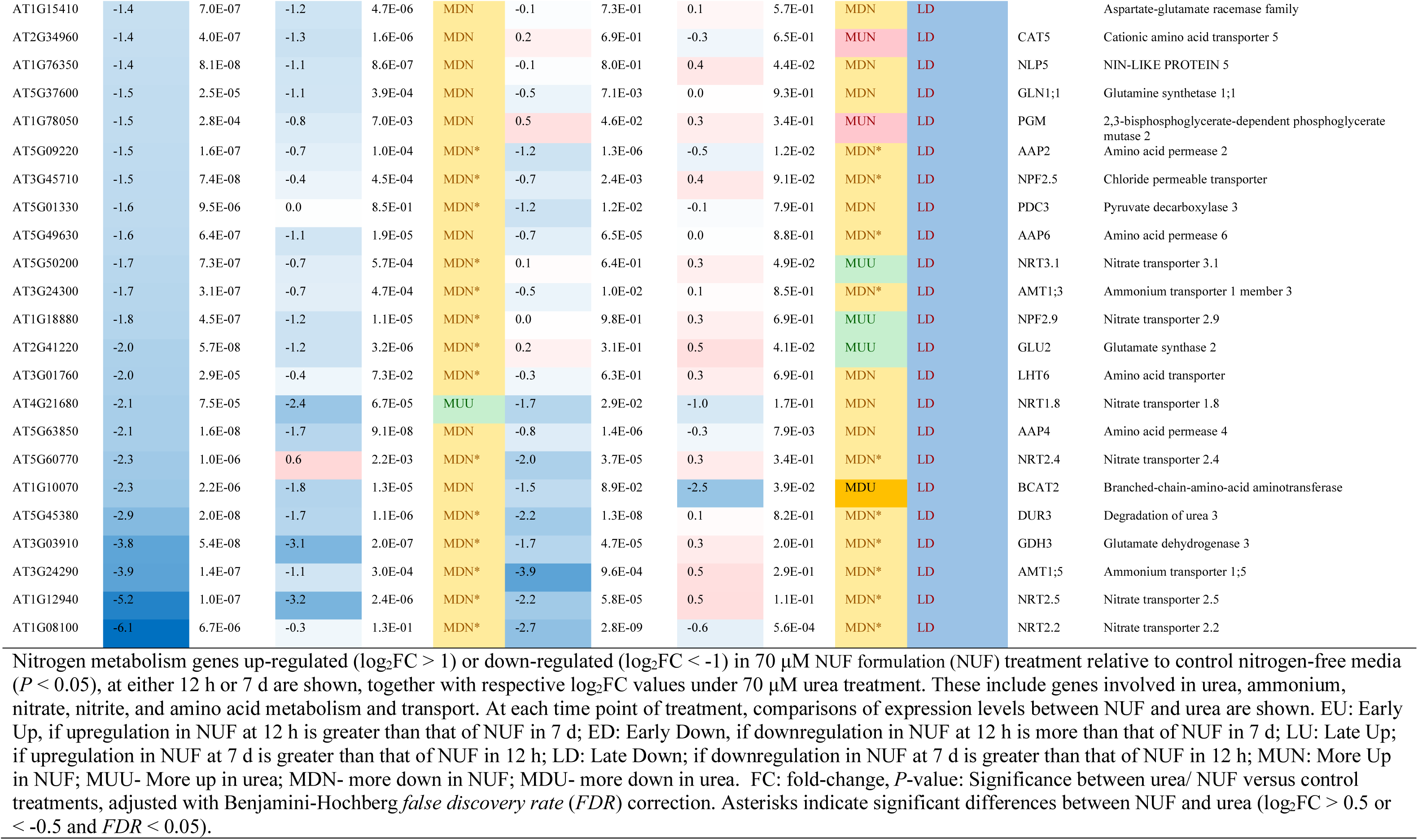
Nitrogen metabolism genes differentially expressed under nano urea and urea treatment.

**Table 2.** Aquaporin genes differentially expressed due to nano urea and urea treatment.

| GeneID | log<sub>2</sub>FC<br>NUF<br>7 days | P-value<br>NUF<br>7 days | log<sub>2</sub>FC<br>urea<br>7 days | P-value<br>urea<br>7 days | Comparison<br>NUF vs urea | log<sub>2</sub>FC<br>NUF<br>12 h | P-value<br>NUF<br>12 h | log<sub>2</sub>FC<br>urea<br>12 h | P-value<br>urea<br>12 h | NUF vs<br>urea | Temporal<br>pattern<br>in NUF | Gene<br>Symbol | Short description |
| --- | --- | --- | --- | --- | --- | --- | --- | --- | --- | --- | --- | --- | --- |
| AT4G17340 | -0.1 | 3.8E-01 | 0.5 | 8.2E-04 | MDN* | -1.2 | 5.4E-07 | -0.3 | 1.1E-01 | MDN* | ED | TIP2;2 | Tonoplast intrinsic protein 2;2 |
| AT3G61430 | -0.1 | 1.1E-01 | 0.3 | 3.8E-03 | MDN | -1.0 | 1.5E-06 | -0.4 | 1.0E-02 | MDN* | ED | PIP1;1 | Plasma membrane intrinsic protein 1;1 |
| AT2G37170 | -0.6 | 1.2E-04 | 0.2 | 1.7E-01 | MDN* | -1.0 | 3.2E-07 | -0.2 | 1.3E-01 | MDN* | ED | PIP2;2 | Plasma membrane intrinsic protein 2;2 |
| AT5G47450 | -0.8 | 9.6E-04 | 0.4 | 2.8E-02 | MDN* | -3.9 | 2.2E-09 | -1.1 | 4.3E-05 | MDN* | ED | TIP2;3 | Tonoplast intrinsic protein 2;3 |
| AT5G60660 | -1.8 | 3.7E-07 | -0.1 | 6.8E-01 | MDN* | -2.1 | 1.0E-07 | -0.8 | 1.0E-03 | MDN* | ED | PIP2;4 | Plasma membrane intrinsic protein 2;4 |
| AT4G19030 | -1.2 | 1.0E-06 | 0.2 | 7.0E-02 | MDN* | -0.8 | 9.2E-04 | -0.3 | 2.6E-01 | MDN* | LD | NIP1;1 | Nod26-like major intrinsic protein 1 |
| AT2G25810 | -1.3 | 4.3E-06 | 0.0 | 9.4E-01 | MDN* | -0.2 | 3.8E-01 | 0.3 | 2.8E-01 | MDN | LD | TIP4-1 | Tonoplast intrinsic protein 4;1 |
| AT3G26520 | 0.4 | 4.1E-03 | 0.9 | 3.5E-05 | MUU* | -1.0 | 1.2E-05 | -0.9 | 2.1E-04 | MDN | LU | TIP1;2 | Tonoplast intrinsic protein 2 |
| AT2G39010 | 0.6 | 3.6E-05 | 0.5 | 2.8E-04 | MUN | -1.1 | 1.0E-06 | -0.6 | 8.7E-04 | MDN | LU | PIP2E | Plasma membrane intrinsic protein 2e |

Aquaporin genes up-regulated ( $\log_2\text{FC} > 1$ ) or down-regulated ( $\log_2\text{FC} < -1$ ) in 70 $\mu\text{M}$ nano urea formulation (NUF) treatment for 12 h or 7 d relative to control nitrogen-free media ( $P < 0.05$ ), are shown, together with respective $\log_2\text{FC}$ values under 70 $\mu\text{M}$ urea treatment. At each time point of treatment, comparisons of expression levels between NUF and urea are shown. EU: Early Up, if upregulation in NUF at 12 h is greater than that of NUF in 7 d; ED: Early Down, if downregulation in NUF at 12 h is more than that of NUF in 7 d; LU: Late Up; if upregulation in NUF at 7 d is greater than that of NUF in 12 h; LD: Late Down; if downregulation in NUF at 7 d is greater than that of NUF in 12 h; MUN: More Up in NUF; MUU- More up in urea; MDN- more down in NUF; MDU- more down in urea. FC: fold-change, $P$ -value: Significance between urea/ NUF versus control treatments, calculated by Fisher's exact test, adjusted with Benjamini-Hochberg *false discovery rate* ( $FDR$ ) correction. Asterisks indicate significant differences between NUF and urea ( $\log_2\text{FC} > 0.5$ or $< -0.5$ and $FDR < 0.05$ ).

To identify further the differences in gene regulation between NUF and urea, we conducted a DEG analysis between NUF and urea. 1449 and 2919 genes showed differential expression (−0.5 < log_2_FC > 0.5, *FDR* < 0.05) between NUF and urea at 12 h and 7 d of treatment (Supplementary Table S4). Gene Ontology (GO) and Kyoto Encyclopedia of Genes and Genomes (KEGG) pathway enrichment analysis was conducted for the genes up-regulated and down-regulated by NUF versus urea (Supplementary Table S5, S6). The results indicated that genes related to growth and development, stress response, cell cycle, translation, metabolism of amino acids, nucleobases, and porphyrins, stress response, photosynthesis, transport, and cell death were enriched in the genes induced in NUF versus urea (Figure 3D). Meanwhile, other subsets of genes related to growth, stress response, and metabolism were down-regulated. Notably, genes for photosynthesis were down-regulated at 12 h but up-regulated at 7 d (Supplementary Table S5).

To gain deeper insights into the important GO functional categories of genes perturbed by NUF, viz. chlorophyll metabolism, nitrogen metabolism, and growth and development, we analyzed their expression patterns in NUF and urea at 12 h and 7 d, relative to control (Figure 3E). We included another category, “aquaporins,” due to their reported roles in urea transport (Li and Wang 2014). This revealed different clusters of early (12 h) or late (7 d) responsive genes, with some gene clusters responding similarly in both urea and NUF, while some up-regulated or down-regulated under NUF but not under urea, with possible functional consequences to the phenotype. Further categorization involved early up-regulated (EU) genes (log_2_FC under NUF at 12 h > 7 d), early down-regulated (ED) genes (log_2_FC NUF at 12 h < 7 d), late up-regulated (LU) genes (log_2_FC NUF at 7 d > 12 h), and late down-regulated (LD) genes (log_2_FC NUF at 7 d < 12 h) along with the patterns: more up-regulated in NUF (MUN), more up-regulated in urea (MUU), more down-regulated in NUF (MDN), and more down-regulated in urea (MDU). Genes having significant differences between NUF and urea (0.5 < log_2_FC > 0.5, *FDR* < 0.05) were identified and marked by asterisks (Figure 3E, Tables 1-4, Supplementary Table S4).

**Table 3.**
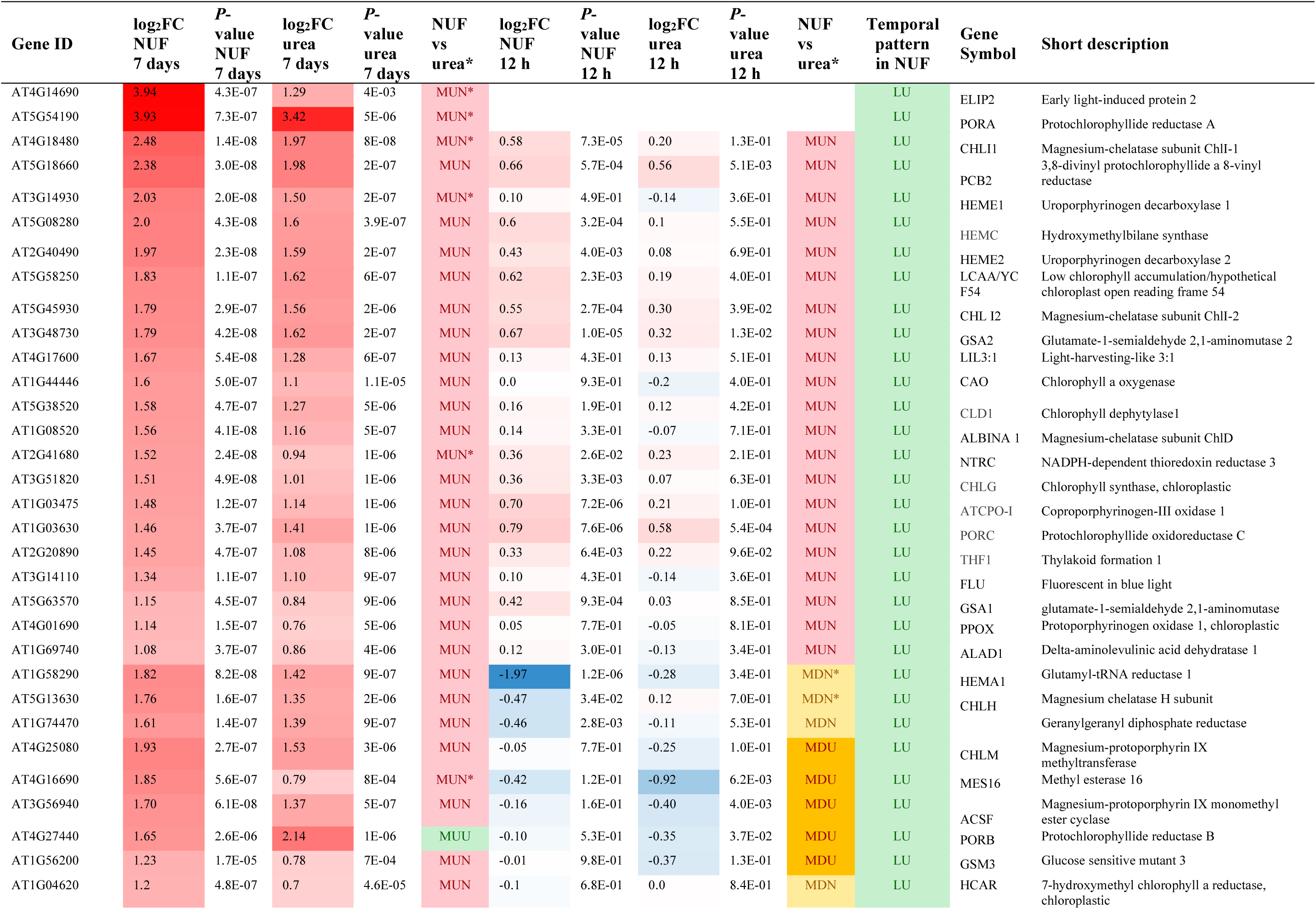
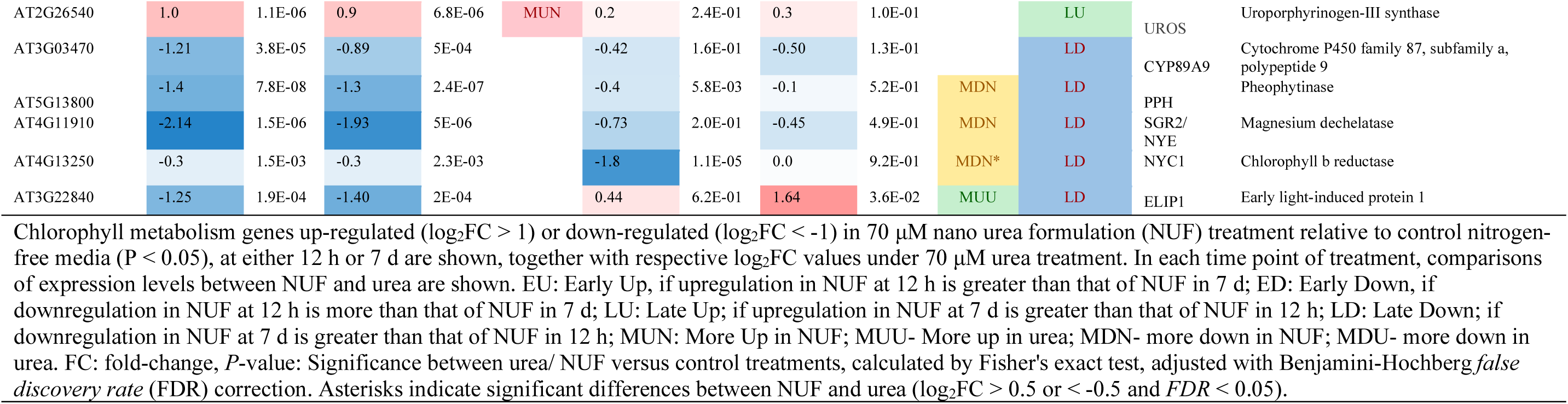
Chlorophyll metabolism genes differentially expressed under nano urea and urea treatment.

**Table 4.**
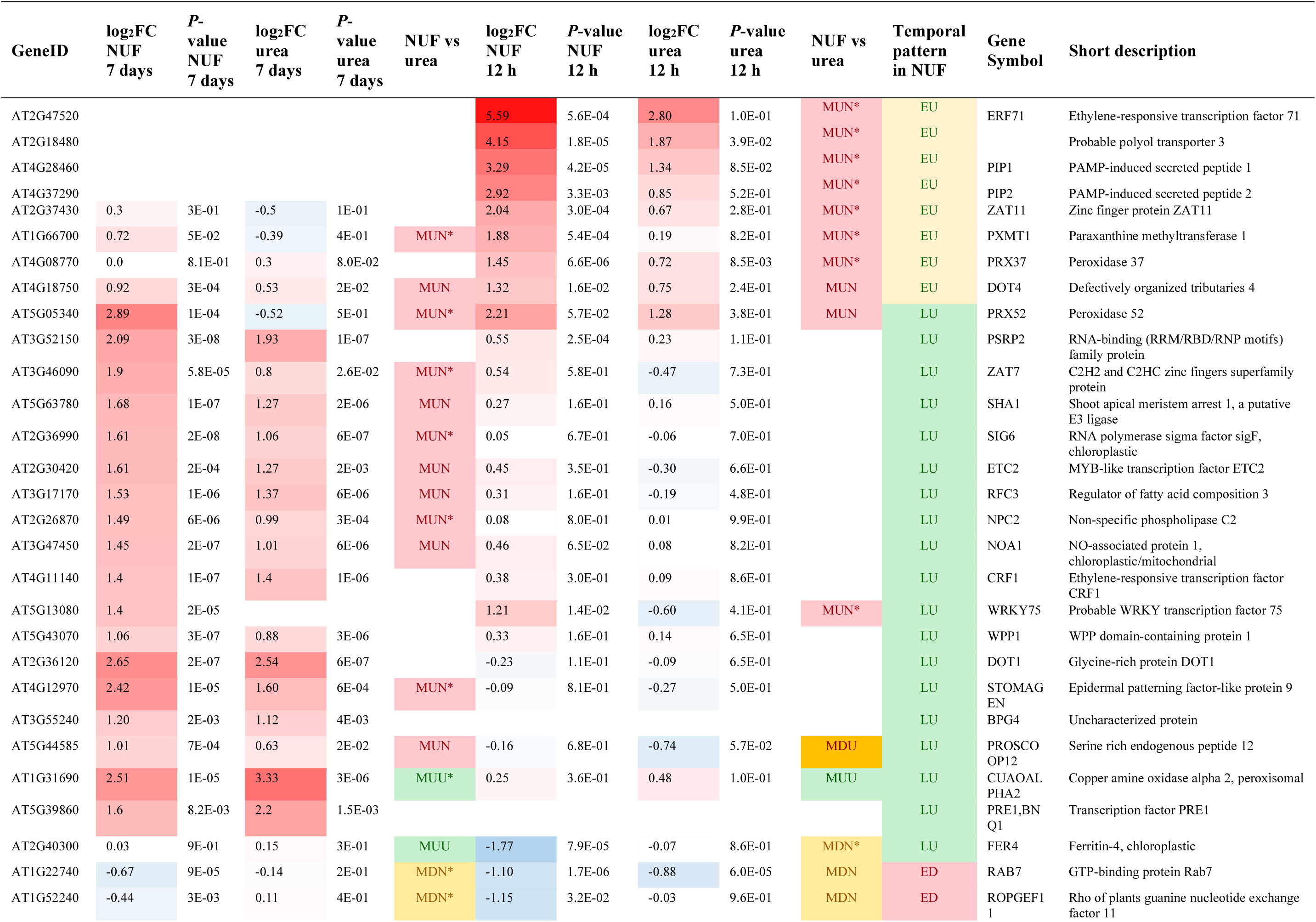
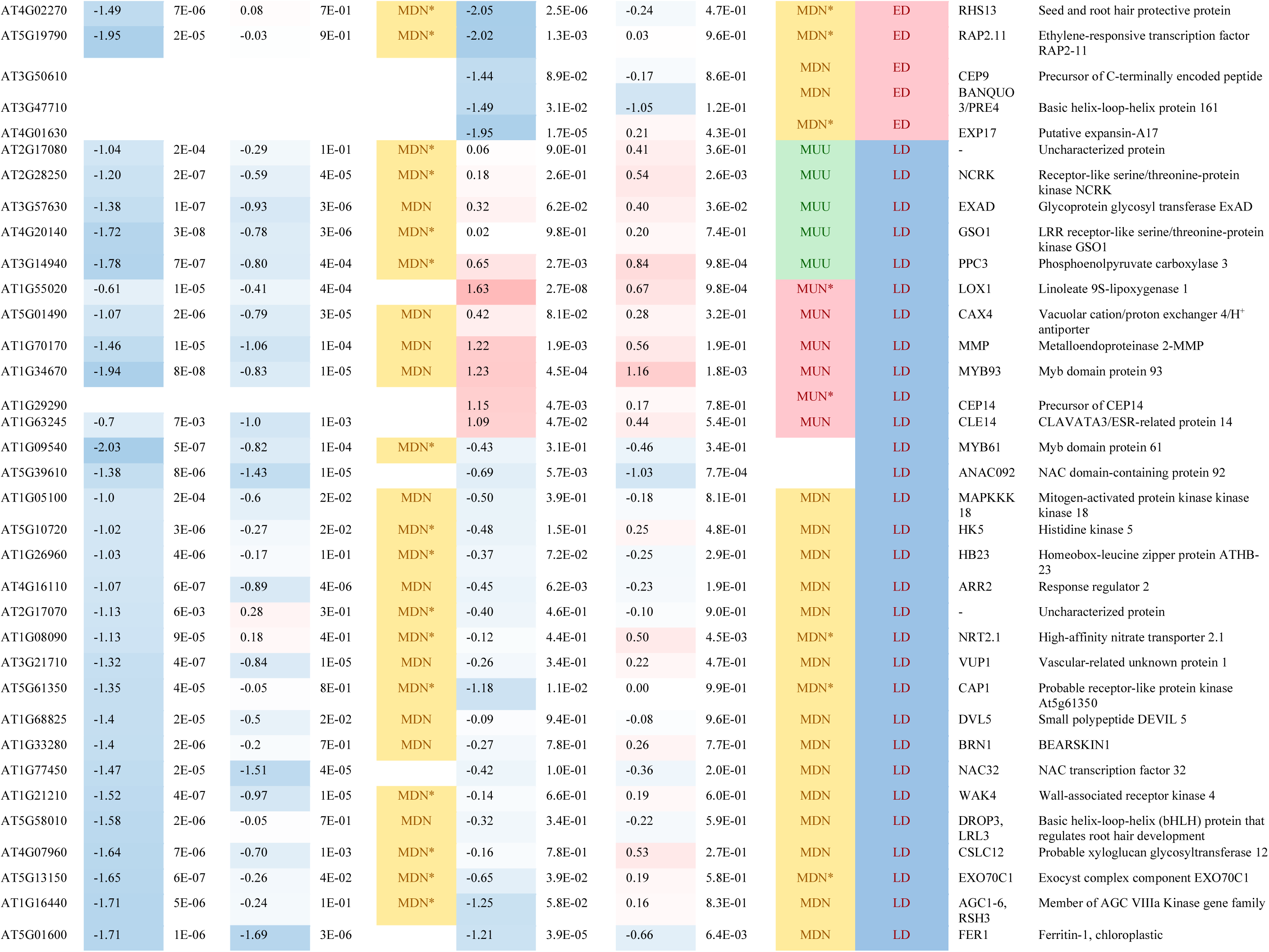
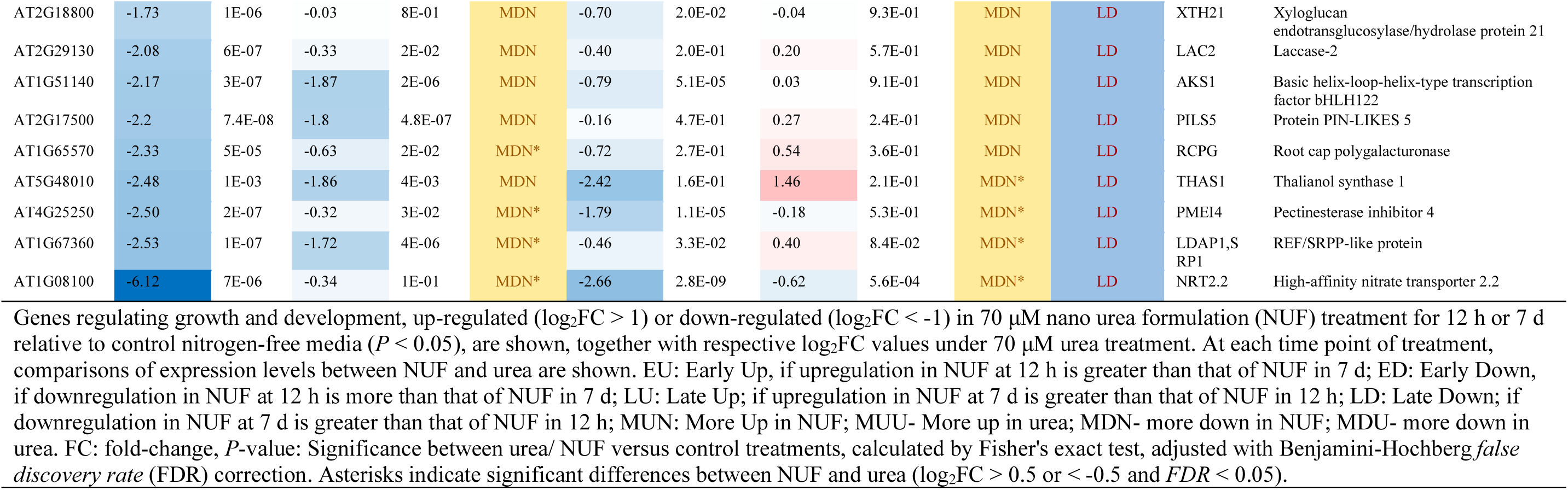
Genes regulating growth and development differentially expressed due to nano urea and urea treatment.

### NUF affects the expression of nitrogen metabolism genes

Two hundred and fifty genes involved in nitrogen metabolism, filtered based on their GO annotations, were differentially expressed under NUF treatment (Supplementary Table S4). Out of the stringently filtered nitrogen metabolism-related DEGs under NUF treatment (−1 < log_2_FC > 1; *FDR* < 0.05), a majority of 51 genes were LU, 21 genes were EU, 8 genes were ED, and 40 genes were LD under NUF. We verified the expression patterns in random DEGs related to nitrogen metabolism by qPCR and found that the expression patterns observed at 7 d persisted till 15 d of treatment (Figure 4). Genes in the nitrogen assimilation pathway were mostly up-regulated by urea and NUF. Among these, *GDH1* (AT5G18170) and *GDH2* (AT5G07440) were significantly induced more, except *GDH3* (AT3G03910), which was more down-regulated under NUF. A critical gene, *aspartate aminotransferase* (*ASP2*; AT5G19550), involved in the biosynthesis of aspartate and aspartate-derived amino acids, was significantly induced more under NUF. All the genes involved in regenerating glutamate from proline, asparagine, and glutamine were up-regulated, while *Delta-1-pyrroline-5-carboxylate dehydrogenase 12A1* (*P5CDH*; AT5G62530) was significantly MUN. *Indole-3-acetic acid-amido synthetase* (*GH3.17*; AT1G28130), catalyzing the production of an auxin-aspartate conjugate, was significantly MDN. *Pyruvate decarboxylase 1* (*PDC1*; AT4G33070) was significantly MDN.

Almost all of the genes related to biosynthesis pathways of amino acids and *de novo* biosynthesis of purines were up-regulated under NUF and urea. Out of these, *phosphoribosylaminoimidazole carboxylase* (*ADE2*; AT2G37690) was significantly MUN. Furthermore, genes involved in various salvage pathways were also up-regulated. Among these genes, *thymidine kinase a* (*TK1A*: AT3G07800), which catalyzes the conversion of thymidine into thymidine monophosphate (TMP), showed a significant MUN pattern. *RIBA3* (AT5G59750), catalyzing the first step in synthesizing the vitamin riboflavin from GTP, was significantly more induced under NUF. Genes involved in valine and asparagine biosynthesis pathways were induced by both urea and NUF, while one gene of the proline biosynthesis pathway, *delta-1-pyrroline-5-carboxylate synthase* (*P5CS1*; AT2G39800), showed EU pattern and showed MUN pattern at 12 h. In addition, two genes, *aspartate aminotransferase* (*ASP2*; AT5G19550) and *acireductone dioxygenase* (*ARD3*; AT2G26400), involved in the methionine salvage pathway via the 5’-methylthioadenosine (MTA) cycle were also significantly MUN. Genes involved in amino acid import or export were mostly up-regulated. Among these, *glutamine dumper* paralogs, *GDU1* (AT4G31730), *GDU2* (AT4G25760), and *GDU4* (AT2G24762), and a multiple amino acid transporter critical for amino acid transport across tissues, *UMAMIT14* (AT2G39510) showed MUN pattern, whereas one amino acid importer gene, *AAP6* (AT5G49630) was MDN. These patterns of expression of the nitrogen metabolism genes, particularly the amino acid and nucleotide biosynthesis genes, could be correlated with the enhanced growth, nitrogen, and amino acid content of the NUF-treated seedlings.

All the genes involved in nitrate uptake showed down-regulation under both urea and NUF. Among these, six *nitrate transporter* genes, *NRT2.1* (AT1G08090), *NRT2.4* (AT5G60770), *NRT2.5* (AT1G12940), *NRT2.2* (AT1G08100), *NRT3.1* (AT5G50200), *NRT2.*1 (AT1G08090), three nitrate and short peptide transporters, *NPF2.4* (AT3G45700), *NPF2.5* (AT3G45710), *NPF8.1* (AT3G54140), as well as two ammonium transporter genes *AMT1;5* (AT3G24290) and *AMT1;3* (AT3G24300) significantly followed the MDN pattern. However, *CEP14* (AT1G29290), a gene that positively regulates nitrate transporters, was MUN. Two paralogs of nitrate reductases converting nitrate (NO_3_ ^-^) to nitrite (NO ^-^), *NR1* (AT1G77760) and *NR2* (AT1G37130) were equally down-regulated by urea and NUF. However, two redox regulators of nitrate reductases, *GRXS6* (AT3G62930) and *GRXC11* (AT3G62950), were significantly MUN.

We also examined the expression of the urea transporter genes under urea and NUF treatment. The high-affinity urea transporter gene *degradation of urea 3* (*DUR3*; AT5G45380) was down-regulated under urea and NUF and showed a significant MDN pattern. Out of the stringently filtered DEGs (−1 < log_2_FC > 1; *FDR* < 0.05), with functional categories “aquaporins” with GO functional categories “water transport” and under NUF treatment, five genes were ED, two genes were LD and two genes were LU. Most were down-regulated under both urea and NUF, with three genes, namely, *NIP1;1* (AT4G19030), *PIP2;2* (AT2G37170), *PIP1;1* (AT3G61430). Only the *plasma membrane intrinsic protein 2E* (*PIP2E*: AT2G39010) was slightly up-regulated under urea and NUF.

### NUF regulates the expression of chlorophyll metabolism genes

Analysis of the DEGs under NUF and urea treatment revealed 62 genes involved in chlorophyll metabolism (Supplementary Table S4). Out of the stringently filtered DEGs under NUF treatment (−1 < log_2_FC > 1; *FDR* < 0.05), a majority of 33 genes were LU, whereas 5 genes were LD, and only one gene *chlorophyll b reductase* (*NYC1*; AT4G13250) showed ED pattern (Table 4). Interestingly, most chlorophyll biosynthesis genes showed up-regulation under NUF and urea, while the chlorophyll catabolism genes showed down-regulation. Out of the genes of the chlorophyll biosynthesis pathway, (*ELIP2*; AT4G14690), *protochlorophyllide reductase A* (*PORA*; AT5G54190), CHLI subunit of magnesium chelatase (*CHLI1*; AT4G18480), *uroporphyrinogen decarboxylase 1* (*HEME1*; AT3G14930), *NADPH-dependent thioredoxin reductase 3* (*NTRC*; AT2G41680), were significantly MUN. All these genes encode enzymes of the porphyrin biosynthesis pathway, except *NTRC*, a positive redox regulator of multiple chlorophyll biosynthesis genes (Brzezowski et al., 2015), and *ELIP2*, a negative regulator of chlorophyll synthesis (Tzvetkova-Chevolleau et al., 2007). *Glutamyl-tRNA reductase 1* (*HEMA1*; AT1G58290), another positive regulator of chlorophyll biosynthesis, showed a significant MDN pattern at 12 h but the opposite MUN pattern at 7 d. On the other hand, the chlorophyll *b*-degrading enzyme *NYC1* showed a significant MDN pattern. *ELIP1* (AT3G22840), an inhibitor of the entire chlorophyll biosynthesis pathway (Rizza et al., 2011), was up-regulated under urea treatment early but was down-regulated by both urea and NUF at 7 d. *Methyl esterase 16* (*MES16;* AT4G16690), a chlorophyll catabolism gene, was down-regulated at 12 h but induced at 7 d. The patterns of expression of most of the chlorophyll metabolism genes could be directly correlated with the enhanced chlorophyll content of the NUF-treated seedlings.

### NUF regulates the expression of genes for growth and development

Four hundred and forty-three genes involved in different GO biological processes involving growth and vegetative development were differentially expressed under NUF treatment (Supplementary Table S4). Out of the stringently filtered DEGs under NUF (−1 < log_2_FC > 1; *FDR* < 0.05), 15 early-responsive genes were related to root development, whereas 58 late-responsive genes regulated both leaf and root development (Table 4). We found 27 up-regulated genes in response to NUF, of which 11 were EU and 18 were LU. Similarly, the down-regulation of 44 genes included seven early-responsive and 39 late-responsive genes (Table 4). Eight EU genes showed a MUN pattern, except for a negative growth regulator *PRX37* (AT4G08770). Five of 19 LU genes were significantly MUN, and a leaf development gene *FER4* (AT2G40300) was MDN at 12 h. Seven genes could be grouped under the ED and MDN category. In comparison, 39 genes were grouped under the LD and MDN pattern category except for a nitrate import and root development gene, *CEP14* (AT1G29290), which was MUN at 12 h. The developmental genes differentially regulated by NUF belong to various functional categories, including transcription factors (TFs), RNA regulators, transporters, enzymes, ribosomal proteins, phytohormone signaling, and redox proteins.

## Discussion

In the current study, we used a hydroponic culture platform to study the effect of NUF on the growth of *A. thaliana*, wherein plant roots were exposed to NUF. The charge and size determine the uptake route of nanoparticles in plants. It was demonstrated that while positively charged nanoparticles cannot enter the root, neutral and negatively charged nanoparticles within the size window 20—100 nm can penetrate the plant roots, and nanoparticles of size ≅ 20 nm get effectively translocated via the xylem (Parkinson et al., 2022). The DLS and TEM analyses confirmed that the NUF size was 30—50 nm, and the zeta potential was negative at the 70 μM concentration used for the growth assay. The clear phenotypic gains observed in the shoots treated with NUF rather than equimolar urea in long-term hydroponic culture (Figure 2E-I) indicate that both NUF and urea were translocated to the shoot. To investigate the effectiveness of NUF as a nitrogen source, we eliminated all other nitrogen sources in the background growth medium. NUF increased the root length compared to seedlings in the nitrogen-free control medium in a dose-dependent manner (Figure 2A). The root and shoot growth inhibition at higher NUF doses, but not urea, suggested a higher uptake rate of urea in the hydroponically grown seedlings. After seven days of treatment, NUF-treated seedlings showed increased chlorophyll content than those treated with urea. After a long-term treatment of 15 days, the biomass of both shoots and roots was higher in NUF, concurrent with the higher tissue nitrogen, chlorophyll and amino acid contents. To understand the molecular mechanisms of the observed phenotypes, we compared the global gene expression patterns between *A. thaliana* seedlings treated with NUF, urea, and nitrogen-free control. In a previous study, expression profiles of hydroponically grown *A. thaliana* were compared between NH_4_NO_3_ as a nitrogen source and urea using a microarray. While many of the genes differentially expressed under urea and NUF treatment in our study were similar to those obtained previously (Mérigout et al., 2008), others differed in expression patterns due to a different, nitrogen-starved control in our case, in contrast to NH_4_NO_3_ used previously. At first glance at the expression patterns, it could be observed that NUF affected the expression of a greater number of genes, induced and suppressed, than equimolar urea concentration at both early 12 h and late 7 d of treatment (Figure 2A, 2B). Moreover, the expression patterns between NUF and urea showed less correlation at 12 h (Figure 2C) but became more correlated with urea after 7 d of treatment. This suggested that NUF caused a higher initial tissue uptake, leading to a transcriptional response different from urea in some of the genes. However, the expression behavior of genes became similar to urea with further long-term exposure, but higher fold-changes of up- or down-regulation were maintained under NUF (Tables 1— 4, S4). The higher fold-change of late-induced genes under NUF was maintained up to 15 d exposure, contributing to the phenotypic advantage observed after 15 d (Figure 4).

We observed an enrichment of the functional categories, amino acid, nucleobase, and DNA metabolism in the genes significantly induced higher in NUF than urea. Genes encoding all the enzymes of the *de novo* purine biosynthesis pathway were induced by both urea and NUF (Moffatt and Ashihara 2002) (Figure 5). Out of these, *ADE2* catalyzing the conversion of phosphoribosylamine to N-formylglycinamidine ribonucleotide (Donovan et al., 2001), and *TK1A* of the pyrimidine salvage pathway catalyzing the conversion of thymidine to its monophosphate form (dTMP) in the cytosol, providing dTTP for DNA repair processes (García et al., 2018; Bitter et al., 2020), were significantly induced more under NUF. Glutamate, glutamine, and asparagine are the amino acids formed from the assimilation of inorganic nitrogen and are essential nitrogen-transporting molecules in plants (Lea and Miflin, 2003). Most of the genes for nitrogen assimilation into amino acids in the chloroplast and mitochondria were induced by both urea and NUF. Two paralogs of *GDH*, *GDH1* and *GDH2*, participating in several steps of nitrogen assimilation (Miflin and Habash, 2002), were induced higher by NUF than urea. GDH activities were significantly impacted by nitrogen availability, and they tended to rise as nitrogen levels increased (Iqbal et al., 2020). The upregulation of *PDC1* under both urea and NUF could be explained by an ammonium excess, which reportedly induces *PDC1* gene expression (Hachiya et al., 2012). PDC1 also activates signaling pathways influencing plant growth and development under anoxia (Kürsteiner et al., 2003). Two genes of the 5’-methylthioadenosine (MTA) cycle, *ASP2*, and *ARD3,* catalyzing the production of methionine and polyamines necessary for growth and cell division (Miyazaki and Yang, 1987; Sauter et al., 2005; Pommerrenig et al., 2011), were significantly induced more under NUF. *P5CDH* (AT5G62530), regenerating glutamate from proline (Deuschle et al., 2004) was again significantly induced more under NUF. γ-aminobutyric acid (GABA) plays an essential part in plant growth and development through the GABA shunt, which is catalyzed by glutamate decarboxylase (GAD) in the cytosol (Li et al., 2021), which was strongly induced by NUF. Most of the amino acid transporters, including four paralogs of *GDU* and *AAP3,* were induced more under NUF. Still, its three other paralogs, *AAP2*, *AAP4*, and *AAP6,* were suppressed more due to their complex regulation under nitrogen starvation and re-supply (Pratelli and Pilot 2014). *UMAMIT14*, a bidirectional cell-to-cell and phloem-mediated long-distance multiple amino acid transporter (Besnard et al., 2016; Besnard et al., 2021), was strongly induced by NUF. Some nitrogen assimilation genes were suppressed early but induced late under NUF treatment. These included *Nitrilase 2* (*NIT2*; AT3G44300), which catalyzes the release of NH_4_^+^ from nitrile compounds generated from different metabolic pathways for subsequent assimilation into amino acids (Howden and Preston, 2009), and *P5CS1*, which catalyzes the rate-limiting step in the biosynthesis of proline (Baumgartner et al., 2005). The temporal regulation of these genes points to the need-based synthesis of these enzymes later for growth processes or due to oscillation under light/dark cycles (Hayashi et al., 2000). *GLR1.2*, functioning in signaling processes, was induced early under NUF, possibly as a defense response, but was suppressed later to counter its negative growth regulation (Hernández-Coronado et al., 2022). The negative regulatory gene *GH3.17* catalyzing the formation of IAA-asp conjugate, leading to degradation of the important growth hormone auxin (Fu et al., 2019), was suppressed at higher levels under NUF, possibly preserving auxin levels for root growth. On the other hand, the high-affinity urea transporter *DUR3*, the paralogs of the NH_4_^+^ transporter *AMT1*, and the nitrate transporter *NRT* were downregulated in NUF treatment, suggesting an ample supply of nitrogen to the plants at the time points of analyzing the transcriptome. Earlier, the expression of these transporter genes was suppressed, coinciding with a higher nitrate accumulation in the cell (Muratore, Espen, and Prinsi 2021). This pattern of expression of these genes could also be attributed to the nitrogen-free background growth medium used in our experiments. In earlier reports, these genes were upregulated in nitrogen-deprived conditions, and their upregulation was reduced upon resupply of nitrogen, suggesting their involvement in regulating plant nitrogen nutrition (Hole et al. 1990; Liu et al., 2003; Wang et al., 2011; Beier et al., 2019). Furthermore, these nitrogen transporter genes were significantly more downregulated in NUF treatment, correlating with the higher nitrogen levels in NUF-treated plants than urea-treated plants (Figure 2). Another sentinel gene involved in nitrogen remobilization and regulating plants’ nitrogen starvation response, *GDH3* (Lezhneva et al., 2014; Safi et al., 2021), showed similar behavior (Table 1, Figure 5). On the other hand, *CEP14* belonging to a family of peptides that induce nitrate transporter genes, resulting in nitrate acquisition in roots (Qiu et al., 2021), playing distinct roles in root and shoot development (Delay et al., 2013), was significantly induced at higher levels by NUF, and can contribute to the observed growth promotion.

**Figure 5.**
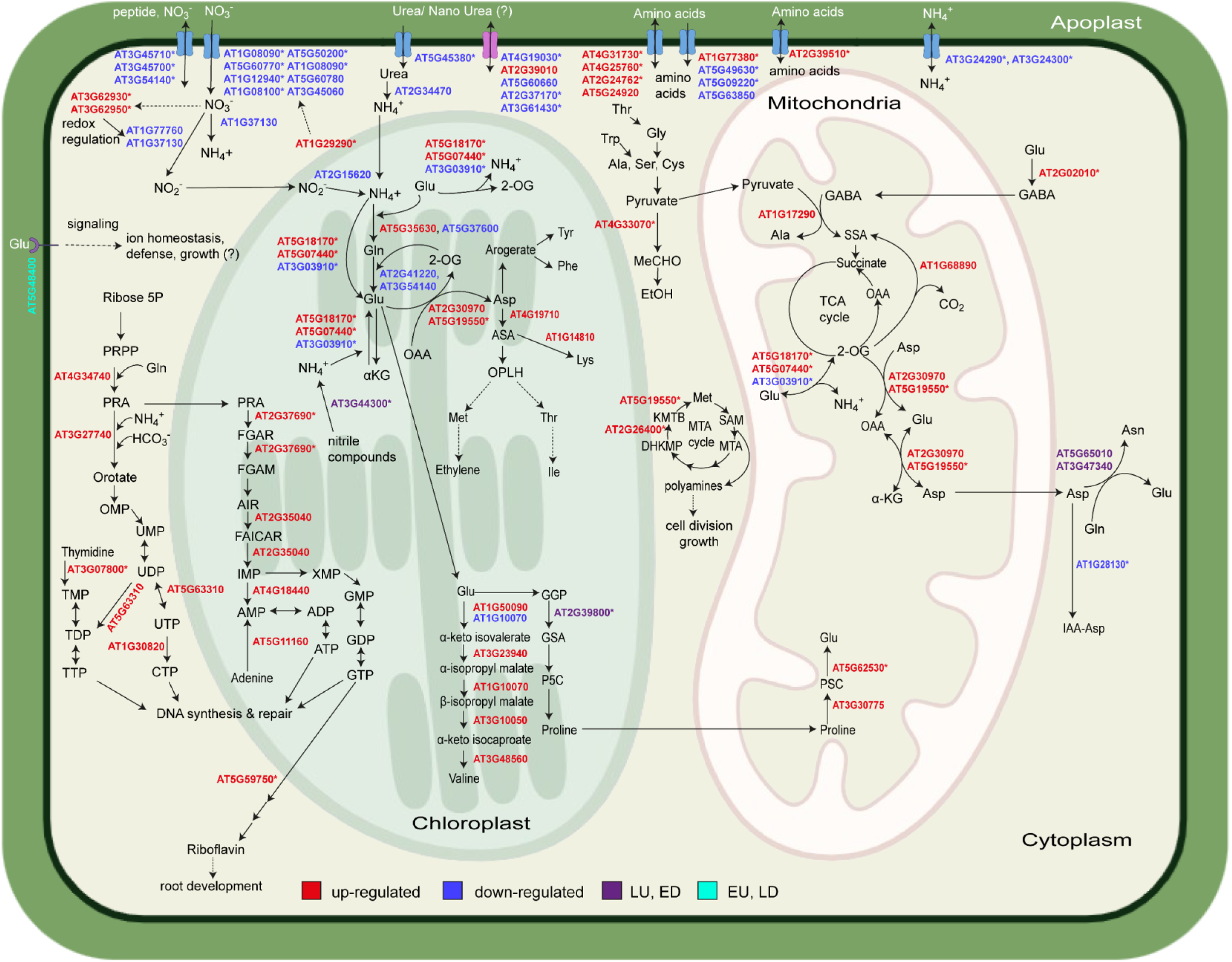
Regulation of nitrogen metabolism genes of *Arabidopsis thaliana* by nano urea. The involvement of genes, differentially regulated under nano urea formulation (NUF) treatment, in different biochemical pathways from urea uptake to nitrogen assimilation in amino acids and nucleotides are shown (see Table 1). Urea is assimilated into plant cells via urea transporter (AT5G45380), converted into ammonia by urease (AT2G34470), and transported to the chloroplast for nitrogen assimilation. Within the chloroplast, ammonia combines with glutamate to form glutamine via glutamine synthetase (AT5G35630, AT5G37600), which can be reversed to form glutamate through GOGAT (AT2G41220, AT3G54140). Aspartate serves as a precursor for several amino acids like tyrosine, phenylalanine, lysine, etc., catalyzed by enzymes including aspartate kinase (AT4G19710) and aspartate semialdehyde dehydrogenase (AT1G14810). Additionally, in the chloroplast, certain amino acids such as proline and valine are synthesized from glutamate with the aid of specific enzymes (AT1G50090, AT3G23940, AT1G10070, AT3G10050, AT3G48560, and AT2G39800). Nucleotide synthesis occurs in the cytoplasm where ribose-5-phosphate is converted into PRA, further processed into IMP, a precursor for ATP and GTP synthesis. In the cytoplasm, PRA converts into orotate and subsequently into UDP, a CTP and TTP synthesis precursor. Glutamate is transported to the mitochondria, where it leads to the formation of GABA. GABA, in turn, combines with pyruvate to produce alanine in the presence of alanine aminotransferase (AT1G17290). Within mitochondria, glutamate and OAA form aspartate, alpha-ketoglutarate, and 2-oxoglutarate via aspartate aminotransferase enzymes (AT2G30970, AT5G19550). Aspartate is then transported to the cytoplasm and converted into asparagine by asparagine synthetases (AT5G65010, AT3G47340). Additionally, proline degradation occurs in the mitochondria, where it is converted into glutamate. In the cytoplasm, the MTP cycle generates methionine and polyamines, which help in cell division. Various transporters are shown on the cell membrane, like those transporting urea, ammonium, nitrate, and amino acid. Genes up-regulated (log_2_FC > 1, *FDR* < 0.05) under NUF at 12 h and/ or 7 d of treatment are shown in red, genes down-regulated (log_2_FC < −1, *FDR* < 0.05) under NUF at 12 h and/ or 7 d of treatment, are shown in blue, genes up-regulated under NUF at 7 d (Late Up; LU) but down-regulated under NUF at 12 h (Early Down; ED) are shown in purple, and genes up-regulated under NUF at 12 h (Early Up; EU) but down-regulated under NUF at 7 d (Early Down; ED) are shown in cyan. Asterisks indicate significant differences in gene expression between urea and NUF (−0.5 < log_2_FC > 0.5, *FDR* < 0.05). ADP, adenosine diphosphate; AIR, aminoimidazole ribonucleotide; αKG, α-ketoglutarate; ASA, argininosuccinic acid; ATP, adenosine triphosphate; CTP, cytidine triphosphate; DHKMP, 1,2-dihydro-3-keto-5-methylthiopentene; FAICAR, 5-formamido-4-imidazolecarboxamide ribonucleotide; FGAM, N-formylglycinamidine ribonucleotide; FGAR, N-formylglycinamide ribonucleotide; GABA, γ-aminobutyric acid; GDP, guanosine diphosphate; GGP, γ-glutamyl phosphate; GMP, guanosine monophosphate; GSA, glutamate-5-semialdehyde; GTP, guanosine triphosphate; IMP, inosine monophosphate; KMTB, α-keto-γ-methylthiobutyric acid; MeCHO, acetaldehyde; MTA, 5’-methylthioadenosine; OAA, oxaloacetic acid; OMP, orotidine monophosphate; OPLH, o-phospho-l-homoserine; P5C, pyrroline-5-carboxylate; PRA, phosphoribosylamine; PRPP, phosphoribosyl diphosphate; SAM, s-adenosylmethionine; SSA, succinic semialdehyde; TCA, tricarboxylic acid; TDP, thymidine diphosphate; TMP, thymidine monophosphate; TTP, thymidine triphosphate; UDP, uridine diphosphate; UMP, uridine monophosphate; UTP, uridine triphosphate; XMP, xanthosine monophosphate.

The advantage of NUF for the enhanced biomass yield of *A. thaliana* can be explained by the observed increased chlorophyll content of the NUF-treated seedlings (Figure 2C, 2F), also evident in the GO and KEGG pathway enrichment analysis where “photosynthesis” and “porphyrin metabolism” were enriched among genes significantly induced more under NUF compared to urea (Figure 3D). Chlorophyll, the light-harvesting pigment of plants, is essential for photosynthesis, the primary biochemical process for growth. The main component of chlorophyll, porphyrin, is a nitrogenous compound. Hence, it is evident that a major portion of the nitrogen supplied through NUF was mobilized into porphyrin biosynthesis. Chlorophyll biosynthesis starts from glutamate, converted into glutamyl-tRNA, and to 5-aminolevulinic acid (ALA), the precursor of tetrapyrrole biosynthesis (Kim et al., 2013). ALA is converted into a four-ring structure uroporphyrinogen III, which, after decarboxylation and oxidation, is combined with magnesium into the tetrapyrrole and ultimately converted to chlorophyll *a* in several steps. We observed the up-regulation of the genes encoding all the enzymes of chlorophyll *a* biosynthesis after 7 d of treatment with both urea and NUF (Figure 6). Again, three genes *PORA*, *CHLI1*, and *HEME1* showed significant higher fold changes under NUF compared to urea. PORA catalyzes the conversion of protochlorophyllide into chlorophyllide through a light-driven reduction (Garrone et al., 2015), forming complexes with PORB. These complexes are essential in regulating specific light-harvesting mechanisms necessary for chlorophyll synthesis. In greening, they additionally offer photoprotection to etiolated seedlings (Reinbothe et al., 2019). CHLI1 is a subunit of the heterotrimeric Mg-chelatase, a key regulatory enzyme in the chlorophyll biosynthesis pathway that catalyzes the conversion of protoporphyrin IX to magnesium protoporphyrin IX in the presence of Mg^2+^ and ATP. (Tran et al., 2023). ChlI stabilizes and activates the heterometric Mg-chelatase by folding and stabilizing the ChlD subunit (Walker and Willows, 1997; Gibson et al., 1999; Gräfe et al., 1999; Rissler et al., 2002). HEME1 is a crucial enzyme that catalyzes the decarboxylation of uroporphyrinogen III to coproporphyrinogen III (Mock et al., 1997), vital for coordinating chlorophyll accumulation during seedling de-etiolation (McCormac et al., 2001). *HEMA1*, encoding the enzyme that reduces the tRNA-bound glutamate to glutamate-l-semialdehyde in chlorophyll biosynthesis (Tanaka et al., 2007; Fang et al., 2016), was suppressed early but induced late by NUF, at higher levels than urea, possibly due to complex negative feedback regulation by other genes including *FLUORESCENT IN BLUE LIGHT* (*FLU*) (Fang et al., 2016). NTRC, an NADPH-dependent redox regulator, is essential in coordinating the activation of many enzymes involved in chlorophyll biosynthesis, which plays a major role in the complex processes involved in plant growth and development (Ojeda et al., 2017). The redox state of several key components in the tetrapyrrole biosynthesis pathway, which includes CHLI1, glutamyl-tRNA reductase (GluTR), magnesium protoporphyrin IX methyltransferase (CHLM), and POR, is regulated by NTRC maintaining a steady flux of intermediates within the tetrapyrrole biosynthesis pathway (Pérez-Ruiz et al., 2006; Brzezowski et al., 2015). *NTRC* knockout mutants have reduced development and pale leaves, indicating the crucial role of this gene in plant growth and development (Pérez-Ruiz et al., 2006; Lepistö et al., 2009). The differential regulation patterns of two negative chlorophyll biosynthesis regulators, *ELIP1* and *ELIP2,* are intriguing. *ELIP1* was induced at 12 h but suppressed after 7 d. *ELIP2* was not detected at 12 h in either urea or NUF but strongly up-regulated at 7 d with significantly higher fold-change under NUF. ELIP proteins inhibit chlorophyll biosynthesis at the early time of synthesis in young leaves, protecting the photocenters from reactive oxygen species (ROS) generated due to excessive energy accumulation when other energy dissipation mechanisms are not fully functional (Montané & Kloppstech, 2000; Hutin et al., 2003; Tzvetkova-Chevolleau et al., 2007). Knocking out of both the *ELIP* paralogs simultaneously does not alter the photooxidative stress response (Casazza et al., 2005). Hence, the induction of *ELIP1* at the early time point and *ELIP2* at the late time point indicate the complex strategies of plants for photoprotection under urea and NUF treatment. In contrast, suppressing *ELIP1* at a late time allowed the continuation of chlorophyll biosynthesis. Chlorophyll *a* is inter-converted to chlorophyll *b* by the chlorophyll cycle, whose genes were mostly induced under urea and NUF. Chlorophyll *b* has significant effects on the efficacy of photosystems, particularly photosystem II (PSII) and photosystem I (PSI), playing a key part in the photosynthesis process (Tanaka and Tanaka, 2011). Chlorophyll degradation starts with its conversion into pheophorbide *a*, which is converted into fluorescent chlorophyll catabolites (FCCs) and transported to the endoplasmic reticulum and cytoplasm, where it further catabolizes into other chlorophyll catabolites. The chlorophyll catabolism gene *NYC1* (AT4G13250) was more downregulated by NUF early, potentially resulting in the maintenance of higher chlorophyll levels in plants, essential for photosynthesis and overall growth. Overexpression of *ZjNYC1* decreased the chlorophyll contents, promoted senescence, and overproduced ROS (Teng et al., 2021). Hence, the downregulation of *NYC1* is crucial to bypassing chlorophyll catabolism.

**Figure 6.**
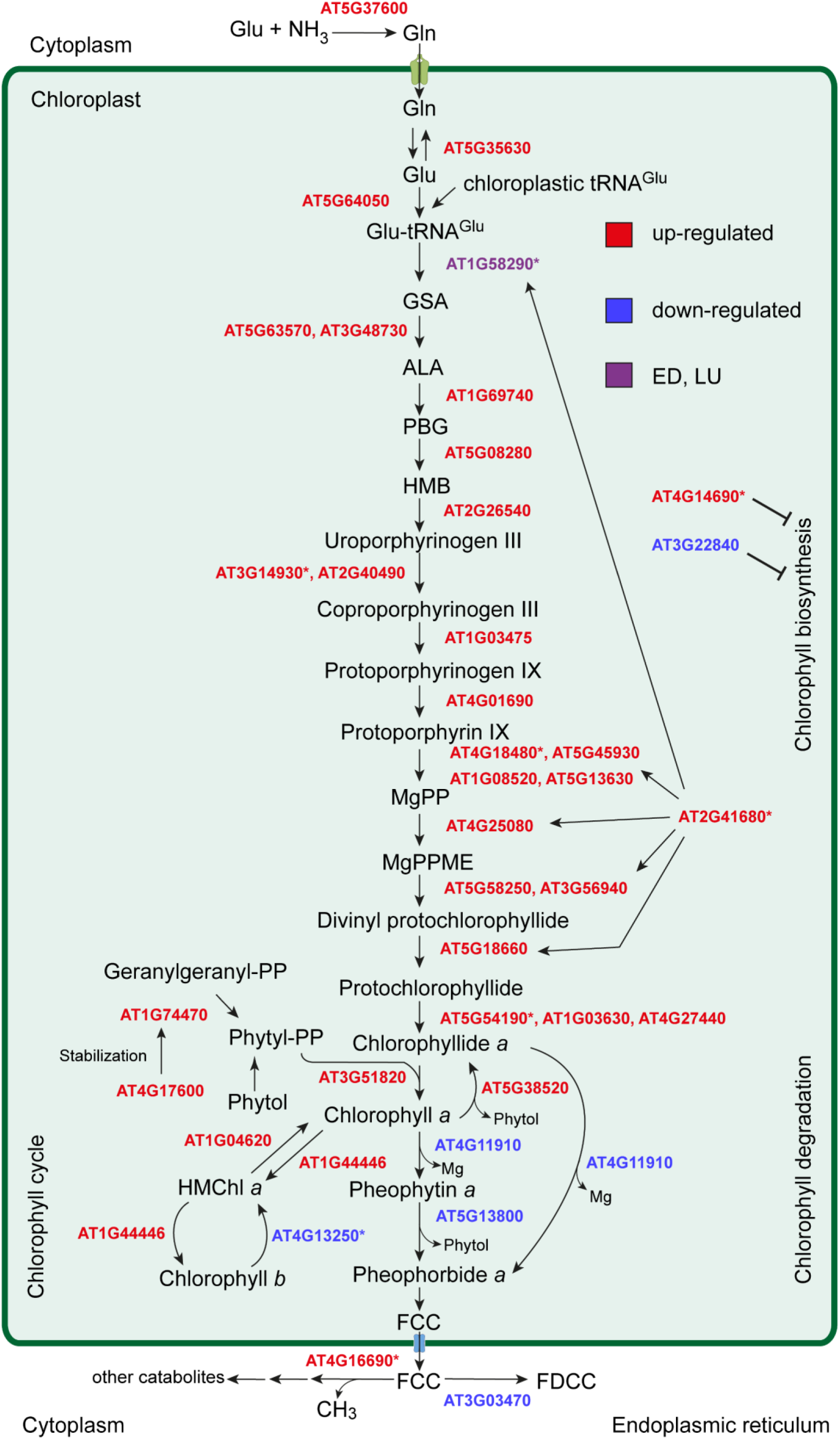
Regulation of chlorophyll metabolism genes of *Arabidopsis thaliana* by nano urea. The involvement of genes differentially regulated under nano urea formulation (NUF) treatment in different biochemical pathways for Chlorophyll biosynthesis and catabolism are shown (see Table 3). Chlorophyll synthesis involves the tetrapyrrole biosynthesis pathway, and 5-aminolevulinic acid (ALA) is the universal precursor for all tetrapyrroles. In the cytoplasm, Glu combines with ammonia to form glutamine (Gln) by glutamine synthetase (AT5G37600), and Gln is transported to the chloroplast by Gln transporters. Gln is converted to Glu by glutamate synthase, but it can also reversibly form Gln by glutamine synthetase 2 (AT5G35630). Glu is first conjugated with chloroplastic tRNA^Glu^ and forms Glu-tRNA^Glu^ by Glu-tRNA synthetase (AT5G64050). After that, Glutamyl-tRNA reductase (AT1G58290) reduces tRNA-bound Glu to glutamate-1-semialdehyde (GSA). Glutamate-1-semialdehyde aminotransferases (AT5G63570, AT3G48370) transaminate GSA to ALA. Two molecules of ALA and four pyrroles combine to form the first closed tetrapyrrole, i.e., uroporphyrinogen III involving 5-aminolevulinic acid dehydratase (AT1G69740), porphobilinogen deaminase (AT5G08280), and uroporphyrinogen III synthase (AT2G26540). The decarboxylation of uroporphyrinogen III forms coproporphyrinogen III, which then undergoes oxidative decarboxylation to form protoporphyrinogen IX, catalyzed by uroporphyrinogen decarboxylases (AT3G14930, AT2G40490) and coproporphyrinogen III oxidase (AT1G03475). Protoporphyrinogen IX is oxidized to Protoporphyrin IX by protoporphyrinogen oxidase (AT4G01690). Now, Mg^2+^ is inserted into protoporphyrin IX forming Mg-protoporphyrin IX (MgPP), catalyzed by Mg-chelatases (AT4G18480, AT5G45930, AT1G08520, AT5G13630), which is converted to Mg-protoporphyrin IX monomethyl ester (Mg-PPME) by Mg-protoporphyrin IX methyltransferase (AT4G25080). Divinyl protochlorophyllide is formed by Mg-protoporphyrin IX monomethyl ester cyclases (AT5G58250, AT3G56940) using Mg-PPME as a substrate. Chlorophyllide *a* is synthesized by divinyl protochlorophyllide reductase (AT5G18660) and protochlorophyllide oxidoreductases (AT5G54190, AT1G03630, AT4G27440) from divinyl protochlorophyllide. Chlorophyllide *a* produces chlorophyll *a* (Chl *a*), catalyzed by chlorophyll synthase (AT3G51820), and a back-conversion to chlorophyllide *a* is catalyzed by chlorophyllase (AT5G38520). In the chlorophyll cycle, Chl *a* is converted to Chl *b* by chlorophyll *a* oxygenase (AT1G4446), via hydroxymethyl chlorophyll (HMChl), and Chl *b* can be reconverted to Chl *a* by Chl *b* reductase (AT4G13250) and 7-hydroxymethyl Chl *a* reductase (AT1G04620). Another way possible to synthesize Chl *a* is by geranylgeranyl reductase (AT1G74470), which reduces the geranylgeranyl pyrophosphate (geranylgeranyl-PP) to phytyl pyrophosphate (phytyl-PP), which along with chlorophyllide *a* synthesize chlorophyll *a* by chlorophyll synthase (AT3G51820). Geranylgeranyl reductase is stabilized by LIL3 (AT4G17600). NTRC (AT2G41680) is a positive regulator of the biosynthetic pathway, and ELIP1 and ELIP2 (AT3G22840, AT4G14690) are negative regulators of chlorophyll biosynthesis. Chl *a* is catabolized by magnesium dechelatase (AT4G11910), pheophytinase (AT5G13800), pheophorbide *a* oxygenase and RCC reductase into fluorescent chlorophyll catabolite (FCC). FCC is transported to the endoplasmic reticulum and converted to fluorescent 1,19-dioxobilin-type chlorophyll catabolite (FDCC) by CYP89A9 (AT3G03470). The Arabidopsis Genome Initiative (AGI) codes of the genes of the above pathway up-regulated (log_2_FC > 1, adjusted *P*-value < 0.05) under NUF at 12 h and/ or 7 d of treatment are shown in red, genes down-regulated (log_2_FC < −1, adjusted *P*-value < 0.05) under NUF at 12 h and/ or 7 d of treatment, are shown in blue, and genes up-regulated under NUF at 7 d (Late Up; LU) but down-regulated under NUF at 12 h (Early Down; ED) are shown in purple. Asterisks indicate significant differences in gene expression between urea and NUF (−0.5 < log_2_FC > 0.5, *FDR* < 0.05).

Growth and developmental EU and MUN category genes encoded TFs, transporters, or RNA-binding regulator proteins (Figure 3E, Table 4). Two TFs, *HRE2* (AT2G47520) and *ZAT11* (AT2G37430) promote root development by regulating cell expansion (Liu et al., 2014; Lee et al., 2015). A carbohydrate transporter gene (AT2G18480) transports sugars through the phloem (Hedhly et al., 2016) and plays a key role in root development (Wintermans et al., 2016). *PIP1* (AT4G28460), *PIP2* (AT4G37290), and *PXMT1* (AT1G66700) play significant roles in lateral root formation (Ghorbani et al., 2015; Chung et al., 2016). Higher up-regulation of these genes under NUF can explain denser lateral roots under NUF treatment (data not presented), leading to higher root biomass (Figure 2G). *DOT4* (AT4G18750), a chloroplast RNA-regulating pentatricopeptide repeat protein induced at higher levels by NUF treatment, has significant roles in leaf development, and its knockout led to deformed leaves and fragmented veins (Petricka et al., 2008; Sun et al., 2015). Among the LU and MUN genes, *PRX52* (AT5G05340), a ROS signaling gene, plays a key role in cell wall synthesis (Zhou et al., 2013; Kwasniewski et al., 2016). *PROSCOOP12* (AT5G44585) regulates root elongation through ROS homeostasis (Guillou et al., 2022). *NOS1* (AT3G47450) encodes a nitric oxide synthase, which regulates plant growth through hormonal signaling (Fang-Qing Guo et al., 2003). *WRKY75* (AT5G13080) has a significant role in lateral root growth (Rosado et al., 2022). *NPC2* (AT2G26870), a phospholipase C, plays a role in root development through the production of phosphocholine (Ngo et al., 2019). Another up-regulated gene, *PRE1* (AT5G39860), regulates plant development using gibberellic acid-mediated pathways (Lee et al., 2006). Another highly induced gene σ*-FACTOR6* (*SIG6*; AT2G36990), regulates thylakoid formation and is key for shoot development (Fenhong et al., 2015). Mesophyll tissue expressed *STOMAGEN* (AT4G12970) increases the production of stomata (Wang et al., 2021). *CRF1* (AT4G11140) promotes cell proliferation during leaf development through cytokinin signaling (Sabaghian et al., 2015), and *AT5G43070* (*WPP1*) is involved in developmental processes (Patel et al., 2004). *RFC3* (AT3G17170), a ribosomal protein, regulates lateral root and shoot development through plastid gene translation (Horiguchi et al., 2003).

The higher down-regulation of several negative growth-regulating genes by NUF than urea at early and late hours can explain the growth advantage of the NUF-treated plants over urea-treated ones (Figure 3E, Table 4). For example, a root apical meristem-localized negative regulator of growth *PRX37* (AT4G08770), whose overexpression led to delayed growth (Pedreira et al., 2011; Tognetti et al., 2017) was up-regulated early at 12 h by both urea and NUF and down-regulated later under NUF treatment but continued to be induced under urea at 7 days, may explain the added advantage of NUF. When overexpressed, the highly down-regulated gene *DOT1* (AT2G36120), encoding a glycine-rich protein, resulted in reduced rosette size and deformed leaves (Petricka et al., 2008). Two genes of the family of C-terminally encoded peptides involved in nitrate import and root and shoot development, *CEP9* (AT3G50610) and *CEP14* (AT1G29290), were suppressed more in NUF under early and late time points, respectively. *CEP9* overexpression caused root swelling and inhibition of lateral root formation (Imin et al., 2013). A negative regulator of bHLH TFs, *PRE4* (AT3G47710), suppressed more under NUF, was found to be up-regulated under the deprivation of the growth hormone gibberellic acid and plays a key role in growth regulation under elevated CO_2_ levels (Ribeiro et al., 2013). MDN genes at 7 d, which correlate with the growth advantage of NUF treatment, include *CLE14* (AT1G63245), a signature protein for short root phenotype in *A. thaliana*, which reduces cell division in the root apical meristem (Meng et al., 2010). *MYB61* (AT1G09540), expressed specifically in guard cells, when knocked out, caused more open stomata (Liang et al., 2005). *MYB93* (AT1G34670) negatively regulates lateral root formation (Gibbs et al., 2014). *NRT2.1* (AT1G08100), a NO_3_^−^ transporter, antagonizes auxin transport and root growth depending on NO_3_^−^ levels and showed drastic suppression by NUF treatment (Lee et al., 2021, Wang et al., 2023). The TF gene *ARR2* (AT4G16110) of the cytokinin signaling pathway, negatively regulating lateral root formation, was found to be down-regulated by both urea and NUF (Mason et al., 2005; Park et al., 2021). *THAS1* (AT5G48010) encoding a thalianol biosynthesis enzyme, whose overexpression led to dwarfed shoots (Field and Osbourn 2008), was up-regulated by urea but down-regulated by NUF at 12 h and down-regulated by both at 7 d. Two leaf senescence-regulating genes, *ANAC092* (AT5G39610) and *ANAC032* (AT1G77450), were suppressed similarly by urea and NUF at 7 d. In addition, *ANAC092* is a negative regulator of root growth (Mahmood et al., 2016; D’Incà et al., 2021). Hence, the advantage of NUF over urea could be explained by the synergistic effect of higher up-regulation of various positive and down-regulation of several negative genes regulating the growth and metabolism of *A. thaliana*.

## Conclusion

In hydroponics we investigated the physiological and molecular responses of the laboratory model plant *A. thaliana* on exposure to NUF or bulk urea equimolar concentrations as the only nitrogen source. Even at a very low concentration (70 μM), NUF treatment for two weeks yielded on average a 20% higher biomass and 16% higher chlorophyll content than urea-treated plants. Gene expression studies showed that NUF follows the expression patterns of urea, but the fold-changes in gene expression were higher in NUF. More chlorophyll and amino acid content in the case of NUF-treated plants can be explained by the enhanced expression of the genes related to nitrogen assimilation, growth, and development. Higher expression of genes related to chlorophyll, amino acid, and nucleobase biosynthesis and greater suppression of catabolism genes further explain the phenotypes. The molecular evidence provided clarifies the regulatory effect of NUF on genes for plant growth, metabolism, and defense. Hydroponics in combination with nano fertilizers has huge potential to improve crop productivity while reducing the environmental impact. Current findings suggest a potential application of NUF at very low concentrations in hydroponic agriculture to increase the biomass yield of plants.

## Data availability statement

The transcriptome raw data are available in the NCBI SRA database with BioProject accession numbers PRJNA1051813 and PRJNA1079889.

## Author contributions

NJ, DM, DV, and AD: investigation, data collection, and writing of the original draft. RSS, DP, and PY: transcriptome data analysis. AS: funding acquisition, design, supervision of biological experiments, data curation, writing, review, and editing of the original draft. KS: funding acquisition and supervision of physicochemical characterizations of NUF.

## Acknowledgments

The authors thank the Department of Bioscience and Bioengineering and the Center for Emerging Technologies for Sustainable Development, IIT Jodhpur, for providing laboratory facilities for the study. The Center for Research & Development of Scientific Instruments, the Center for Ayurtech at IIT Jodhpur, and the Nano Biotechnology Research Center, Kalol, are acknowledged for providing instrumentation facilities for the physicochemical characterization of NUF. The guidance of Prof. Prabuddha Ganguli, Advisor and Adjunct Faculty, School of Management and Entrepreneurship, and Prof. Mitali Mukerji, Head, Department of Bioscience and Bioengineering, IIT Jodhpur are gratefully acknowledged.

## References

Andrews S. (2010). FastQC. Babraham Bioinformatics. http://www.bioinformatics.babraham.ac.uk/

Arnon, D. I. (1949). Copper enzymes in isolated chloroplasts. Polyphenoloxidase in *Beta vulgaris*. Plant Physiol. 24, 1–15. doi: 10.1104/pp.24.1.1

Baumgartner, M. R., Rabier, D., Nassogne, M.-C., Dufier, J.-L., Padovani, J.-P., Kamoun, P., et al. (2005). Δ1-pyrroline-5-carboxylate synthase deficiency: neurodegeneration, cataracts and connective tissue manifestations combined with hyperammonaemia and reduced ornithine, citrulline, arginine and proline. Eur J Pediatr. 164, 31–36. doi: 10.1007/s00431-004-1545-3

Beier, M. P., Fujita, T., Sasaki, K., Kanno, K., Ohashi, M., Tamura, W., et al. (2019). The urea transporter DUR3 contributes to rice production under nitrogen-deficient and field conditions. Physiol. Plant. 167, 75–89. doi: 10.1111/ppl.12872

Benjamini, Y., and Hochberg, Y. (1995). Controlling the False Discovery Rate: A Practical and Powerful Approach to Multiple Testing. J. R. Stat. Soc. Ser. B 57, 289–300. doi: 10.1111/j.2517-6161.1995.tb02031.x

Besnard, J., Pratelli, R., Zhao, C., Sonawala, U., Collakova, E., Pilot, G., et al. (2016). UMAMIT14 is an amino acid exporter involved in phloem unloading in Arabidopsis roots. EXBOTJ. 67, 6385–6397. doi: 10.1093/jxb/erw412

Besnard, J., Sonawala, U., Maharjan, B., Collakova, E., Finlayson, S. A., Pilot, G., et al. (2021). Increased Expression of UMAMIT Amino Acid Transporters Results in Activation of Salicylic Acid Dependent Stress Response. Front. Plant Sci. 11. doi: 10.3389/fpls.2020.606386

Bitter, E. E., Townsend, M. H., Erickson, R., Allen, C., and O’Neill, K. L. (2020). Thymidine kinase 1 through the ages: a comprehensive review. Cell Biosci. 10. doi: 10.1186/s13578-020-00493-1

Brzezowski, P., Richter, A. S., and Grimm, B. (2015). Regulation and function of tetrapyrrole biosynthesis in plants and algae. BBA Bioenergetics. 1847, 968–985. doi: 10.1016/j.bbabio.2015.05.007

Chen, Y., Lun, A.T.L., and Smyth, G.K. (2016). From reads to genes to pathways: Differential expression analysis of RNA-Seq experiments using Rsubread and the edgeR quasi-likelihood pipeline. F1000Res. 5, 1438 doi: 10.12688/F1000RESEARCH.8987.2.

Christ, B., Schelbert, S., Aubry, S., Süssenbacher, I., Müller, T., Kräutler, B., et al., (2011). MES16, a member of the methylesterase protein family, specifically demethylates fluorescent chlorophyll catabolites during chlorophyll breakdown in Arabidopsis. Plant Physiol. 158, 628–641. doi: 10.1104/pp.111.188870

Chung, P. J., Park, B. S., Wang, H., Liu, J., Jang, I.-C., and Chua, N.-H. (2016). Light-inducible MiR163 targets *PXMT1* transcripts to promote seed germination and primary root elongation in *Arabidopsis*. Plant Physiol. 170, 1772–1782. doi: 10.1104/pp.15.01188

Ciftci-Yilmaz, S., Morsy, M. R., Song, L., Coutu, A., Krizek, B. A., Lewis, M. W., et al. (2007). The EAR-motif of the Cys2/His2-type zinc finger protein ZAT7 plays a key role in the defense response of Arabidopsis to salinity stress. J. Biol. Chem. 282, 9260–9268. doi: 10.1074/jbc.m611093200

Casazza, A.P., Rossini, S., Rosso, M.G. et al. (2005). Mutational and expression analysis of ELIP1 and ELIP2 in *Arabidopsis thaliana*. Plant Mol Biol 58, 41–51 doi: 10.1007/s11103-005-4090-1

D’Incà, E., Cazzaniga, S., Foresti, C., Vitulo, N., Bertini, E., Galli, M., et al. (2021). VviNAC33 promotes organ de-greening and represses vegetative growth during the vegetative-to-mature phase transition in grapevine. New Phytol. 231, 726–746. doi: 10.1111/nph.17263

Lally, D., Ingmire, P., Tong, H.-Y., and He, Z.-H. (2001). Antisense expression of a cell wall–associated protein kinase, WAK4, inhibits cell elongation and alters morphology. Plant Cell 13, 1317–1332. doi: 10.1105/tpc.010075

Delay, C., Imin, N., and Djordjevic, M. A. (2013). CEP genes regulate root and shoot development in response to environmental cues and are specific to seed plants. J. Exp. Bot. 64, 5383–5394. doi: 10.1093/jxb/ert332

Deuschle, K., Funck, D., Forlani, G., Stransky, H., Biehl, A., Leister, D., et al. (2004). The Role of Δ1-pyrroline-5-carboxylate dehydrogenase in proline degradation. Plant Cell 16, 3413–3425. doi: 10.1105/tpc.104.023622

Doi, E., Shibata, D., and Matoba, T. (1981). Modified colorimetric ninhydrin methods for peptidase assay. Anal. Biochem. 118, 173–184. doi: 10.1016/0003-2697(81)90175-5

Donovan, M., Schumuke, J. J., Fonzi, W. A., Bonar, S. L., Gheesling-Mullis, K., Jacob, G. S., et al. (2001). Virulence of a phosphoribosylaminoimidazole carboxylase-deficient *Candida albicans* strain in an immunosuppressed murine model of systemic Candidiasis. Infect Immun. 69, 2542–2548. doi: 10.1128/iai.69.4.2542-2548.2001

Fang, Y., Zhao, S., Zhang, F., Zhao, A., Zhang, W., Zhang, M., et al., (2016). The *Arabidopsis* glutamyl-tRNA reductase (GluTR) forms a ternary complex with FLU and GluTR-binding protein. Sci Rep 6. doi: 10.1038/srep19756

Guo, F.-Q., Okamoto, M., and Crawford, N. M. (2003). Identification of a Plant Nitric Oxide Synthase Gene Involved in Hormonal Signaling. Science 302, 100–103. doi: 10.1126/science.1086770

Fatima, H., Hamdani, S. D. A., Ahmed, M., Rajput, T. A., Gul, A., Amir, R., et al. (2023). Anti-MRSA potential of biogenic silver nanoparticles synthesized from hydroponically grown *Foeniculum vulgare*. Phytomed. Plus 3, 100415. doi: 10.1016/j.phyplu.2023.100415

Feil, S. B., Rodegher, G., Gaiotti, F., Alzate Zuluaga, M. Y., Carmona, F. J., Masciocchi, N., et al. (2021). Physiological and molecular investigation of urea uptake dynamics in *Cucumis sativus* L. plants fertilized with urea-doped amorphous calcium phosphate nanoparticles. Front. Plant Sci. 12. doi: 10.3389/fpls.2021.745581

Hu, F., Zhu, Y., Wu, W., Xie, Y., and Huang, J. (2015). Leaf variegation of thylakoid formation 1 is suppressed by mutations of specific σ-Factors in *Arabidopsis*. Plant Physiol. 168, 1066–1075. doi: 10.1104/pp.15.00549

Field, B., and Osbourn, A. E. (2008). Metabolic diversification-independent assembly of operon-like gene clusters in different plants. Science 320, 543–547. doi: 10.1126/science.1154990

Fisher, R. A. (1945). Statistical Method for Research Workers. J. R. Stat. Soc. 108, 235. doi: 10.2307/2981200

Frank, M., and Husted, S. (2023). Is India’s largest fertilizer manufacturer misleading farmers and society using dubious plant and soil science? Plant Soil. doi: 10.1007/s11104-023-06191-4

Fu, X., Shi, Z., Jiang, Y., Jiang, L., Qi, M., Xu, T., et al. (2019). A family of auxin conjugate hydrolases from *Solanum lycopersicum* and analysis of their roles in flower pedicel abscission. BMC Plant Biol. 19. doi: 10.1186/s12870-019-1840-9

Fujiwara, T., Hirai, M. Y., Chino, M., Komeda, Y., and Naito, S. (1992). Effects of sulfur nutrition on expression of the soybean seed storage protein genes in transgenic *Petunia*. Plant physiol. 99, 263–268. doi: 10.1104/pp.99.1.263

Garrone, A., Archipowa, N., Zipfel, P. F., Hermann, G., and Dietzek, B. (2015). Plant protochlorophyllide oxidoreductases A and B. J. Biol. Chem. 290, 28530–28539. doi: 10.1074/jbc.m115.663161

Gashgari, R., Alharbi, K., Mughrbil, K., Jan, A., and Glolam, A. (2018). Comparison between Growing Plants in Hydroponic System and Soil Based System. Proc. World Congr. Mech. Chem. Mater. Eng. doi: 10.11159/icmie18.131

Ghorbani, S., Lin, Y.-C., Parizot, B., et al. (2015). Expanding the repertoire of secretory peptides controlling root development with comparative genome analysis and functional assays. J. Exp. Bot. 66, 5257–5269. doi: 10.1093/jxb/erv346

Gibbs, D. J., Voß, U., Harding, S. A., et al. (2014). *AtMYB93* is a novel negative regulator of lateral root development in *Arabidopsis*. New Phytol. 203, 1194–1207. doi: 10.1111/nph.12879

Gibson, L.C.D., Jensen, P.E., and Hunter, C.N. (1999). Magnesium chelatase from Rhodobacter sphaeroides: initial characterization of the enzyme using purified subunits and evidence for a BchI– BchD complex. Biochem. J. 337, 243–251. doi: 10.1042/bj3370243

Gräfe, S., Saluz, H.-P., Grimm, B., and Hänel, F. (1999). Mg-chelatase of tobacco: The role of the subunit CHL D in the chelation step of protoporphyrin IX. Proc. Natl. Acad. Sci. U.S.A. 96, 1941–1946. doi: 10.1073/pnas.96.5.1941

Guillou, M.-C., Balliau, T., Vergne, E., Canut, H., Chourré, J., Herrera-León, C., et al. (2022). The PROSCOOP10 Gene Encodes Two Extracellular Hydroxylated Peptides and Impacts Flowering Time in Arabidopsis. Plants 11, 3554. doi: 10.3390/plants11243554

Hachiya, T., Watanabe, C. K., Fujimoto, M., Ishikawa, T., Takahara, K., Kawai-Yamada, M., et al. (2012). Nitrate addition alleviates ammonium toxicity without lessening ammonium accumulation, organic acid depletion and inorganic cation depletion in *Arabidopsis thaliana* shoots. Plant Cell Physiol 53, 577–591. doi: 10.1093/pcp/pcs012

Hayashi, F., Ichino, T., Osanai, M., and Wada, K. (2000). Oscillation and Regulation of Proline Content by P5CS and ProDH Gene Expressions in the Light/Dark Cycles in *Arabidopsis thaliana* L. PCP. 41, 1096–1101. doi: 10.1093/pcp/pcd036

Hedhly, A., Vogler, H., Schmid, M. W., Pazmino, D., Gagliardini, V., Santelia, D., et al. (2016). Starch turnover and metabolism during flower and early embryo development. Plant Physiol. 172, 2388–2402. doi: 10.1104/pp.16.00916

Hernández-Coronado, M., Dias Araujo, P. C., Ip, P.-L., Nunes, C. O., Rahni, R., Wudick, M. M., et al. (2022). Plant glutamate receptors mediate a bet-hedging strategy between regeneration and defense. Dev.Cell 57, 451–465.e6. doi: 10.1016/j.devcel.2022.01.013

Hoagland, D.R., and Arnon, D.I. (1938). The water culture method for growing plants without soil. California Agricultural Experiment Station Circulation, 347, 32.

Hole, D. J., Emran, A. M., Fares, Y., and Drew, M. C. (1990). Induction of nitrate transport in maize roots, and kinetics of influx, measured with nitrogen-13. Plant Physiol. 93, 642–647. doi: 10.1104/pp.93.2.642

Horiguchi, G., Kodama, H., and Iba, K. (2003). Mutations in a gene for plastid ribosomal protein S6-like protein reveal a novel developmental process required for the correct organization of lateral root meristem in Arabidopsis. Plant J. 33, 521–529. doi: 10.1046/j.1365-313x.2003.01651.x

Hou, S., Wang, X., Chen, D., Yang, X., Wang, M., Turrà, D., et al. (2014). The secreted peptide PIP1 amplifies immunity through Receptor-Like Kinase 7. PLoS Pathog. 10, e1004331. doi: 10.1371/journal.ppat.1004331

Howden, A. J. M., and Preston, G. M. (2009). Nitrilase enzymes and their role in plant–microbe interactions. Microb. Biotechnol. 2, 441–451. doi: 10.1111/j.1751-7915.2009.00111.x

Hutin, C., Nussaume, L., Moise, N., Moya, I., Kloppstech, K., and Havaux, M. (2003). Early light-induced proteins protect Arabidopsis from photooxidative stress. Proc. Natl. Acad. Sci. U.S.A. 100, 4921–4926. doi: 10.1073/pnas.0736939100

Imin, N., Mohd-Radzman, N. A., Ogilvie, H. A., and Djordjevic, M. A. (2013). The peptide-encoding CEP1 gene modulates lateral root and nodule numbers in *Medicago truncatula*. J. Exp. Bot. 64, 5395– 5409. doi: 10.1093/jxb/ert369

Hwang, I. S., and Hwang, B. K. (2009). The Pepper 9-lipoxygenase gene *CaLOX1* functions in defense and cell death responses to microbial pathogens. Plant Physiol. 152, 948–967. doi: 10.1104/pp.109.147827

Iqbal, A., Dong, Q., Wang, X., Gui, H., Zhang, H., Zhang, X., et al. (2020). Variations in nitrogen metabolism are closely linked with nitrogen uptake and utilization efficiency in cotton genotypes under various nitrogen supplies. Plants 9, 250. doi: 10.3390/plants9020250

Jadoon, L., Gul, A., Fatima, H., and Babar, M. M. (2024). Nano-elicitation and hydroponics: a synergism to enhance plant productivity and secondary metabolism. Planta 259. doi: 10.1007/s00425-024-04353-x

Mahmood, K., El-Kereamy, A., Kim, S.-H., Nambara, E., and Rothstein, S. J. (2016). ANAC032 positively regulates age-dependent and stress-induced senescence in *Arabidopsis thaliana*. Plant Cell Physiol. 57, 2029–2046. doi: 10.1093/pcp/pcw120

Khalid, M. F., Iqbal Khan, R., Jawaid, M. Z., Shafqat, W., Hussain, S., Ahmed, T., et al. (2022). Nanoparticles: the plant saviour under abiotic stresses. Nanomaterials 12, 3915. doi: 10.3390/nano12213915

Khodakovskaya, M. V., de Silva, K., Nedosekin, D. A., Dervishi, E., Biris, A. S., Shashkov, E. V., et al. (2010). Complex genetic, photothermal, and photoacoustic analysis of nanoparticle-plant interactions. Proc. Natl. Acad. Sci. U.S.A. 108, 1028–1033. doi: 10.1073/pnas.1008856108

Khush, G. S. (2001). Green revolution: the way forward. Nat Rev Genet. 2, 815–822. doi: 10.1038/35093585

Kim, M. J., Ruzicka, D., Shin, R., and Schachtman, D. P. (2012). The Arabidopsis AP2/ERF Transcription Factor RAP2.11 modulates plant response to low-potassium conditions. Mol. Plant. 5, 1042–1057. doi: 10.1093/mp/sss003

Kim, S., Schlicke, H., Van Ree, K., Karvonen, K., Subramaniam, A., Richter, A., et al., (2013). Arabidopsis chlorophyll biosynthesis: an essential balance between the methylerythritol phosphate and tetrapyrrole pathways. Plant Cell 25, 4984–4993. doi: 10.1105/tpc.113.119172

Klepek, Y.-S., Geiger, D., Stadler, R., Klebl, F., Landouar-Arsivaud, L., Lemoine, R., et al. (2004). *Arabidopsis* POLYOL TRANSPORTER5, a new member of the monosaccharide transporter-like superfamily, mediates H^+^-symport of numerous substrates, including myo-inositol, glycerol, and ribose. Plant Cell 17, 204–218. doi: 10.1105/tpc.104.026641

Koistinen, J., Sjöblom, M., and Spilling, K. (2017). Determining inorganic and organic phosphorus. Methods Mol. Biol. 87–94. doi: 10.1007/7651_2017_104

Kõressaar, T., Lepamets, M., Kaplinski, L., Raime, K., Andreson, R., and Remm, M. (2018). Primer3_masker: integrating masking of template sequence with primer design software. Bioinformatics 34, 1937–1938. doi: 10.1093/bioinformatics/bty036

Kumar, Y., Tiwari, K.N., Nayak, R.K., Rai, A.S.P., Singh S.P.A.N., Singh, A.N., Kumar, Y., Tomar, H., Singh, T., and Raliya, R. (2020). Nanofertilizers for increasing nutrient use efficiency, yield and economic returns in important winter season crops of Uttar Pradesh. Indian J Fertilisers 16 (8): 772–786

Kürsteiner, O., Dupuis, I., and Kuhlemeier, C. (2003). The *pyruvate decarboxylase1* gene of Arabidopsis is required during anoxia but not other environmental stresses. Plant Physiol. 132, 968– 978. doi: 10.1104/pp.102.016907

Kwasniewski, M., Daszkowska-Golec, A., Janiak, A., Chwialkowska, K., Nowakowska, U., Sablok, G., et al. (2016). Transcriptome analysis reveals the role of the root hairs as environmental sensors to maintain plant functions under water-deficiency conditions. J. Exp. Bot. 67, 1079–1094. doi: 10.1093/jxb/erv498

Kwon, S. I., Cho, H. J., Jung, J. H., Yoshimoto, K., Shirasu, K., and Park, O. K. (2010). The Rab GTPase RabG3b functions in autophagy and contributes to tracheary element differentiation in *Arabidopsis*. Plant J. no-no. doi: 10.1111/j.1365-313x.2010.04315.x

Lea, P. J., and Miflin, B. J. (2003). Glutamate synthase and the synthesis of glutamate in plants. Plant Physiol. Biochem. 41, 555–564. doi: 10.1016/s0981-9428(03)00060-3

Lee, S.-Y., Hwang, E. Y., Seok, H.-Y., Tarte, V. N., Jeong, M. S., Jang, S. B., et al., (2015). *Arabidopsis* AtERF71/HRE2 functions as transcriptional activator via cis-acting GCC box or DRE/CRT element and is involved in root development through regulation of root cell expansion. Plant Cell Rep. 34, 223–231. doi: 10.1007/s00299-014-1701-9

Lee, Y. J., Lee, W. J., Le, Q. T., Hong, S.-W., and Lee, H. (2021). Growth performance can be increased under high nitrate and high salt stress through enhanced nitrate reductase activity in *Arabidopsis* anthocyanin over-producing mutant plants. Front. Plant Sci. 12. doi: 10.3389/fpls.2021.644455

Lepistö, A., Kangasjärvi, S., Luomala, E.-M., Brader, G., Sipari, N., Keränen, M., et al. (2009). Chloroplast NADPH-thioredoxin reductase interacts with photoperiodic development in *Arabidopsis*. Plant Physiol. 149, 1261–1276. doi: 10.1104/pp.108.133777

Lezhneva, L., Kiba, T., Feria-Bourrellier, A., Lafouge, F., Boutet-Mercey, S., Zoufan, P., et al. (2014). The Arabidopsis nitrate transporter NRT2.5 plays a role in nitrate acquisition and remobilization in nitrogen-starved plants. Plant J. 80, 230–241. doi: 10.1111/tpj.12626

Li, C., and Wang, W. (2014). Urea Transport Mediated by Aquaporin Water Channel Proteins. Subcellular Biochemistry, 227–265. doi: 10.1007/978-94-017-9343-8_14

Liang, Y.-K., Dubos, C., Dodd, I. C., Holroyd, G. H., Hetherington, A. M., and Campbell, M. M. (2005). AtMYB61, an R2R3-MYB transcription factor controlling stomatal aperture in *Arabidopsis thaliana*. Curr. Biol. 15, 1201–1206. doi: 10.1016/j.cub.2005.06.041

Liao, Y., Smyth, G.K., and Shi, W. (2014). FeatureCounts: An efficient general purpose program for assigning sequence reads to genomic features. Bioinformatics 30, 923–930. doi: 10.1093/bioinformatics/btt656.

Li, L., Dou, N., Zhang, H., and Wu, C. (2021). The versatile GABA in plants. Plant Signal Behav 16, 1862565. doi: 10.1080/15592324.2020.1862565

Liu, X.-M., An, J., Han, H. J., Kim, S. H., Lim, C. O., Yun, D.-J., et al. (2014). ZAT11, a zinc finger transcription factor, is a negative regulator of nickel ion tolerance in Arabidopsis. Plant Cell Rep. 33, 2015–2021. doi: 10.1007/s00299-014-1675-7

Liu, L.-H., Ludewig, U., Frommer, W. B., and von Wirén, N. (2003). *AtDUR3* Encodes a New Type of High-Affinity Urea/H^+^ Symporter in Arabidopsis. Plant Cell 15, 790–800. doi: 10.1105/tpc.007120

Love, M.I., Huber, W., and Anders, S. (2014). Moderated estimation of fold change and dispersion for RNA-seq data with DESeq2. Genome Biol. 15, 550. doi: 10.1186/s13059-014-0550-8.

Luo, Z., Wang, J., Li, F., Lu, Y., Fang, Z., Fu, M., et al. (2022). The small peptide CEP1 and the NIN-like protein NLP1 regulate NRT2.1 to mediate root nodule formation across nitrate concentrations. Plant cell. 35, 776–794. doi: 10.1093/plcell/koac340

Mason, M. G., Mathews, D. E., Argyros, D. A., et al. (2005) Multiple type-B response regulators mediate cytokinin signal transduction in Arabidopsis. Plant Cell 17(11):3007–18. doi: 10.1105/tpc.105.035451.

McCormac, A. C., Fischer, A., Kumar, A. M., Söll, D., and Terry, M. J. (2001). Regulation of *HEMA1* expression by phytochrome and a plastid signal during de-etiolation in *Arabidopsis thaliana*. Plant J 25, 549–561. doi: 10.1046/j.1365-313x.2001.00986.x

Mei, H., Cheng, N. H., Zhao, J., Park, S., Escareno, R. A., Pittman, J. K., et al. (2009). Root development under metal stress in *Arabidopsis thaliana* requires the H^+^/cation antiporter CAX4. New Phytol 183, 95–105. doi: 10.1111/j.1469-8137.2009.02831.x

Meng, L., and Feldman, L. J. (2010). CLE14/CLE20 peptides may interact with CLAVATA2/CORYNE receptor-like kinases to irreversibly inhibit cell division in the root meristem of Arabidopsis. Planta. 232, 1061–1074. doi: 10.1007/s00425-010-1236-4

Mérigout, P., Lelandais, M., Bitton, F., et al. (2008). Physiological and transcriptomic aspects of urea uptake and assimilation in Arabidopsis plants. Plant Physiol. 147, 1225–1238. doi: 10.1104/pp.108.119339

Miflin, B. J., and Habash, D. Z. (2002). The role of glutamine synthetase and glutamate dehydrogenase in nitrogen assimilation and possibilities for improvement in the nitrogen utilization of crops. J. Exp. Bot. 53, 979–987. doi: 10.1093/jexbot/53.370.979

Mirakhorli, T., Ardebili, Z. O., Ladan-Moghadam, A., and Danaee, E. (2021). Bulk and nanoparticles of zinc oxide exerted their beneficial effects by conferring modifications in transcription factors, histone deacetylase, carbon and nitrogen assimilation, antioxidant biomarkers, and secondary metabolism in soybean. PLoS ONE 16, e0256905. doi: 10.1371/journal.pone.0256905

Miyazaki, J. H., and Yang, S. F. (1987). The methionine salvage pathway in relation to ethylene and polyamine biosynthesis. Physiol Plant 69, 366–370. doi: 10.1111/j.1399-3054.1987.tb04302.x

Mock, H. P., and Grimm, B. (1997). Reduction of uroporphyrinogen decarboxylase by antisense RNA expression affects activities of other enzymes involved in tetrapyrrole biosynthesis and leads to light-dependent necrosis. Plant Physiol. 113, 1101–1112. doi: 10.1104/pp.113.4.1101

Moffatt, B. A., and Ashihara, H. (2002). Purine and pyrimidine nucleotide synthesis and metabolism. The Arabidopsis Book 1, e0018. doi: 10.1199/tab.0018

Montané, M.-H., and Kloppstech, K. (2000). The family of light-harvesting-related proteins (LHCs, ELIPs, HLIPs): was the harvesting of light their primary function? Gene 258, 1–8. doi: 10.1016/s0378-1119(00)00413-3

Moser, J. (2001). V-shaped structure of glutamyl-tRNA reductase, the first enzyme of tRNA-dependent tetrapyrrole biosynthesis. EMBO J 20, 6583–6590. doi: 10.1093/emboj/20.23.6583

Muratore, C., Espen, L., and Prinsi, B. (2021). Nitrogen uptake in plants: the plasma membrane root transport systems from a physiological and proteomic perspective. Plants, 10, 681. doi: 10.3390/plants10040681

Ngo, A. H., Kanehara, K., and Nakamura, Y. (2019). Non-specific phospholipases C, NPC2 and NPC6, are required for root growth in *Arabidopsis*. Plant J. 100, 825–835. doi: 10.1111/tpj.14494

Ohkubo, Y., Tanaka, M., Tabata, R., Ogawa-Ohnishi, M., and Matsubayashi, Y. (2017). Shoot-to-root mobile polypeptides involved in systemic regulation of nitrogen acquisition. Nat. Plants. 3. doi: 10.1038/nplants.2017.29

Ojeda, V., Pérez-Ruiz, J. M., González, M., Nájera, V. A., Sahrawy, M., Serrato, A. J., et al. (2017). NADPH thioredoxin reductase C and thioredoxins act concertedly in seedling development. Plant Physiol. 174, 1436–1448. doi: 10.1104/pp.17.00481

Onija, O., Borodi, Gh., Kacso, I., Pop, M. N., Dadarlat, D., Bratu, I., et al. (2012). Preparation and characterization of urea-oxalic acid solid form. AIP Conf. Proc. doi: 10.1063/1.3681960

Park, J., Lee, S., Park, G., Cho, H., Choi, D., Umeda, M., et al. (2021). CYTOKININ-RESPONSIVE GROWTH REGULATOR regulates cell expansion and cytokinin-mediated cell cycle progression. Plant Physiol. 186, 1734–1746. doi: 10.1093/plphys/kiab180

Parkinson, S. J., Tungsirisurp, S., Joshi, C., Richmond, B. L., Gifford, M. L., Sikder, A., et al. (2022). Polymer nanoparticles pass the plant interface. Nat Commun. 13. doi: 10.1038/s41467-022-35066-y

Patel, S., Rose, A., Meulia, T., Dixit, R., Cyr, R. J., and Meier, I. (2004). Arabidopsis WPP-domain proteins are developmentally associated with the nuclear envelope and promote cell division. Plant cell. 16, 3260–3273. doi: 10.1105/tpc.104.026740

Pedreira, J., Herrera, M. T., Zarra, I., and Revilla, G. (2011). The overexpression of *AtPrx37*, an apoplastic peroxidase, reduces growth in *Arabidopsis*. Physiol Plant. 141, 177–187. doi: 10.1111/j.1399-3054.2010.01427.x

Pedroza-García, J., Nájera-Martínez, M., Mazubert, C., Aguilera-Alvarado, P., Drouin-Wahbi, J., Sánchez-Nieto, S., et al. (2018). Role of pyrimidine salvage pathway in the maintenance of organellar and nuclear genome integrity. Plant J. 97, 430–446. doi: 10.1111/tpj.14128

Pérez-Ruiz, J. M., Spínola, M. C., Kirchsteiger, K., Moreno, J., Sahrawy, M., and Cejudo, F. J. (2006). Rice NTRC Is a High-Efficiency Redox System for Chloroplast Protection against Oxidative Damage. Plant Cell, 18, 2356–2368. doi: 10.1105/tpc.106.041541

Petricka, J. J., Clay, N. K., and Nelson, T. M. (2008). Vein patterning screens and the defectively organized tributaries mutants in Arabidopsis thaliana. Plant J. 56, 251–263. doi: 10.1111/j.1365-313x.2008.03595.x

Pommerrenig, B., Feussner, K., Zierer, W., Rabinovych, V., Klebl, F., Feussner, I., et al. (2011). Phloem-Specific expression of Yang cycle genes and identification of novel Yang cycle enzymes in Plantago and Arabidopsis. Plant Cell, 23, 1904–1919. doi: 10.1105/tpc.110.079657

Poulev, A., O’Neal, J. M., Logendra, S., Pouleva, R. B., Timeva, V., Garvey, A. S., et al. (2003). Elicitation, a New Window into Plant Chemodiversity and Phytochemical Drug Discovery. J. Med. Chem. 46, 2542–2547. doi: 10.1021/jm020359t

Pratelli, R., and Pilot, G. (2014). Regulation of amino acid metabolic enzymes and transporters in plants. Journal of Experimental Botany 65, 5535–5556. doi: 10.1093/jxb/eru320

Provart, N., and Zhu, T. (2003). A browser-based functional classification superviewer for Arabidopsis genomics. Curr. Comput. Mol. Biol. 2003, 271–272.

Qiu, Z., Zhuang, K., Liu, Y., Ge, X., Chen, C., Hu, S., et al. (2021). Functional characterization of C-TERMINALLY ENCODED PEPTIDE (CEP) family in *Brassica rapa* L. Plant Signal Behav. 17. doi: 10.1080/15592324.2021.2021365

Reinbothe, S., Bartsch, S., Rossig, C., Davis, M. Y., Yuan, S., Reinbothe, C., et al. (2019). A Protochlorophyllide (Pchlide) a Oxygenase for Plant Viability. Front. Plant Sci. 10. doi: 10.3389/fpls.2019.00593

Reinders, A., Panshyshyn, J. A., and Ward, J. M. (2005). Analysis of Transport Activity of Arabidopsis Sugar Alcohol Permease Homolog AtPLT5. J Biol Chem. 280, 1594–1602. doi: 10.1074/jbc.m410831200

Ribeiro, D. M., Mueller-Roeber, B., and Schippers, J. H. M. (2013). Promotion of growth by elevated carbon dioxide is coordinated through a flexible transcriptional network in Arabidopsis. Plant Signal Behav. 8, e23356. doi: 10.4161/psb.23356

Rissler, H. M., Collakova, E., DellaPenna, D., Whelan, J., and Pogson, B. J. (2002). Chlorophyll Biosynthesis. Expression of a SecondChl IGene of Magnesium Chelatase in *Arabidopsis* Supports Only Limited Chlorophyll Synthesis. Plant Physiol. 128, 770–779. doi: 10.1104/pp.010625

Rizwan, M., Ali, S., Rehman, M. Z. ur Riaz, M., Adrees, M., Hussain, A., et al. (2021). Effects of nanoparticles on trace element uptake and toxicity in plants: A review. Ecotoxicol. Environ. Saf. 221, 112437. doi: 10.1016/j.ecoenv.2021.112437

Rizza, A., Boccaccini, A., Lopez-Vidriero, I., Costantino, P., and Vittorioso, P. (2011). Inactivation of the ELIP1 and ELIP2 genes affects *Arabidopsis* seed germination. New Phytol. 190, 896–905. doi: 10.1111/j.1469-8137.2010.03637.x

Rosado, D., Ackermann, A., Spassibojko, O., Rossi, M., and Pedmale, U. V. (2022). WRKY transcription factors and ethylene signaling modify root growth during the shade-avoidance response. Plant Physiol. 188, 1294–1311. doi: 10.1093/plphys/kiab493

Sabaghian, E., Drebert, Z., Inzé, D., and Saeys, Y. (2015). An integrated network of Arabidopsis growth regulators and its use for gene prioritization. Sci Rep 5. doi: 10.1038/srep17617

Sadhukhan, A., Kobayashi, Y., Nakano, Y., Iuchi, S., Kobayashi, M., Sahoo, L., et al. (2017). Genome-wide association study reveals that the Aquaporin NIP1;1 contributes to variation in hydrogen peroxide sensitivity in *Arabidopsis thaliana*. Mol Plant 10, 1082–1094. doi: 10.1016/j.molp.2017.07.003

Safi, A., Medici, A., Szponarski, W., Martin, F., Clément-Vidal, A., Marshall-Colon, A., et al. (2021). GARP transcription factors repress Arabidopsis nitrogen starvation response via ROS-dependent and - independent pathways. J Exp Bot 72, 3881–3901. doi: 10.1093/jxb/erab114

Sauter, M., Lorbiecke, R., OuYang, B., Pochapsky, T. C., and Rzewuski, G. (2005). The immediate-early ethylene response gene OsARD1 encodes an acireductone dioxygenase involved in recycling of the ethylene precursor S-adenosylmethionine. Plant J. 44, 718–729. doi: 10.1111/j.1365-313x.2005.02564.x

Schubert, M., Lindgreen, S., and Orlando, L. (2016). AdapterRemoval v2: rapid adapter trimming, identification, and read merging. BMC Res Notes 9. doi: 10.1186/s13104-016-1900-2

Seleiman, M. F., Almutairi, K. F., Alotaibi, M., Shami, A., Alhammad, B. A., and Battaglia, M. L. (2020). Nano-Fertilization as an Emerging Fertilization Technique: Why Can Modern Agriculture Benefit from Its Use? Plants 10, 2. doi: 10.3390/plants10010002

Lee, S., Lee, S., Yang, K.-Y., Kim, Y.-M., Park, S.-Y., Kim, S. Y., et al. (2006). Overexpression of PRE1 and its Homologous Genes Activates Gibberellin-dependent Responses in Arabidopsis thaliana. Plant Cell Physiol. 47, 591–600. doi: 10.1093/pcp/pcj026

Sun, L., Rodriguez, G. R., Clevenger, J. P., Illa-Berenguer, E., Lin, J., Blakeslee, J. J., et al. (2015). Candidate gene selection and detailed morphological evaluations offs8.1, a quantitative trait locus controlling tomato fruit shape. J Exp Bot. 66, 6471–6482. doi: 10.1093/jxb/erv361

Tabata, R., Sumida, K., Yoshii, T., Ohyama, K., Shinohara, H., and Matsubayashi, Y. (2014). Perception of root-derived peptides by shoot LRR-RKs mediates systemic N-demand signaling. Science 346, 343–346. doi: 10.1126/science.1257800

Tanaka, R., and Tanaka, A. (2007). Tetrapyrrole Biosynthesis in Higher Plants. Annu. Rev. Plant Biol. 58, 321–346. doi: 10.1146/annurev.arplant.57.032905.105448

Tanaka, R., and Tanaka, A. (2011). Chlorophyll cycle regulates the construction and destruction of the light-harvesting complexes. Biochim Biophys Acta. 1807, 968–976. doi: 10.1016/j.bbabio.2011.01.002

Teng, K., Tan, P., Guan, J., Dong, D., Liu, L., Guo, Y., et al. (2021). Functional characterization of the chlorophyll b reductase gene NYC1 associated with chlorophyll degradation and photosynthesis in *Zoysia japonica*. EEB 191, 104607. doi: 10.1016/j.envexpbot.2021.104607

Timsina, J. (2018). Can Organic Sources of Nutrients Increase Crop Yields to Meet Global Food Demand? Agronomy 8, 214. doi: 10.3390/agronomy8100214

Tiwari, K.N. (2022). Nano technology based P fertilizers for higher efficiency and agriculture sustainability. APSR 24, 198–207. doi: 10.47815/apsr.2022.10149

Tognetti, V. B., Bielach, A., and Hrtyan, M. (2017). Redox regulation at the site of primary growth: auxin, cytokinin and ROS crosstalk. Plant Cell Environ. 40, 2586–2605. doi: 10.1111/pce.13021

Tran, L. H., Kim, J.-G., and Jung, S. (2023). Expression of the Arabidopsis Mg-chelatase H subunit alleviates iron deficiency-induced stress in transgenic rice. Front. Plant Sci. 14. doi: 10.3389/fpls.2023.1098808

Tzvetkova-Chevolleau, T., Franck, F., Alawady, A. E., Dall’Osto, L., Carrière, F., Bassi, R., et al. (2007). The light stress-induced protein ELIP2 is a regulator of chlorophyll synthesis in *Arabidopsis thaliana*. Plant J. 50, 795–809. doi: 10.1111/j.1365-313x.2007.03090.x

UNDESA. World Population Projected to Reach 9.8 Billion by 2050. United Nations Department of Economic and Social Affairs, New York, USA. https://www.un.org/en/desa/world-population-projected-reach-98-billion-2050-and-112-billion-2100.

Vega-Arroy, JD., Herrera-Estrella, A., Ovando-Vázquez, C. et al., (2024). Inferring co-expression networks of *Arabidopsis thaliana* genes during their interaction with *Trichoderma spp*. Sci Rep. 14, 2466 doi: 10.1038/s41598-023-48332-w

Verdoliva, S. G., Gwyn-Jones, D., Detheridge, A., and Robson, P. (2021). Controlled comparisons between soil and hydroponic systems reveal increased water use efficiency and higher lycopene and β-carotene contents in hydroponically grown tomatoes. Sci Hortic. 279, 109896. doi: 10.1016/j.scienta.2021.109896

Walker, J. C., and Willows, D. R. (1997). Mechanism and regulation of Mg-chelatase. Biochem J. 327, 321–333. doi: 10.1042/bj3270321

Wang, D., Wei, L., Liu, T., Ma, J., Huang, K., Guo, H., et al. (2023). Suppression of ETI by PTI priming to balance plant growth and defense through an MPK3/MPK6-WRKYs-PP2Cs module. Mol Plant 16, 903–918. doi: 10.1016/j.molp.2023.04.004

Wang, S., Zhou, Z., Rahiman, R., Lee, G. S. Y., Yeo, Y. K., Yang, X., et al. (2021). Light regulates stomatal development by modulating paracrine signaling from inner tissues. Nat Commun 12. doi: 10.1038/s41467-021-23728-2

Wang, W., Köhler, B., Cao, F., Liu, G., Gong, Y., Sheng, S., et al. (2011). Rice DUR3 mediates high-affinity urea transport and plays an effective role in improvement of urea acquisition and utilization when expressed in Arabidopsis. New Phytol. 193, 432–444. doi: 10.1111/j.1469-8137.2011.03929.x

Wang, Y., Yuan, Z., Wang, J., Xiao, H., Wan, L., Li, L., et al. (2023). The nitrate transporter NRT2.1 directly antagonizes PIN7-mediated auxin transport for root growth adaptation. Proc. Natl. Acad. Sci. U.S.A. 120. doi: 10.1073/pnas.2221313120

Wickham, H. (2016). Package ‘ggplot2’: Elegant graphics for data analysis. Springer-Verlag New York.

Wintermans, P. C. A., Bakker, P. A. H. M., and Pieterse, C. M. J. (2016). Natural genetic variation in Arabidopsis for responsiveness to plant growth-promoting rhizobacteria. Plant Mol Biol. 90, 623–634. doi: 10.1007/s11103-016-0442-2

Yang, M., Dong, C., and Shi, Y. (2023). Nano fertilizer synergist effects on nitrogen utilization and related gene expression in wheat. BMC Plant Biol. 23. doi: 10.1186/s12870-023-04046-9

You, Q., Dong, N., Yang, H., Feng, F., Xu, Y., Wang, C., et al. (2022). The Arabidopsis Receptor-like Kinase CAP1 promotes shoot growth under ammonium stress. Biology 11, 1452. doi: 10.3390/biology11101452

Zhai, L., Wang, Z., Zhai, Y., Zhang, L., Zheng, M., Yao, H., et al. (2022). Partial substitution of chemical fertilizer by organic fertilizer benefits grain yield, water use efficiency, and economic return of summer maize. Soil Tillage Res. 217, 105287. doi: 10.1016/j.still.2021.105287

Zhang, Y., Park, C., Bennett, C. et al. (2021). Rapid and accurate alignment of nucleotide conversion sequencing reads with HISAT-3N. Genome Res 31, 1290–1295. doi: 10.1101/gr.275193.120.

Zhang, Z., Ke, M., Qu, Q., Peijnenburg, W., Lu, T., Zhang, Q., Ye, Y., Xu, P., Du, B., Sun, L., Qian, H. (2018) Impact of copper nanoparticles and ionic copper exposure on wheat (*Triticum aestivum* L.) root morphology and antioxidant response. Environ Pollut. 239, 689–697. doi:10.1016/j.envpol.2018.04.066

Zhou, X.-Y., Song, L., and Xue, H.-W. (2013). Brassinosteroids regulate the differential growth of arabidopsis hypocotyls through auxin signaling components IAA19 and ARF7. Mol Plant 6, 887–904. doi: 10.1093/mp/sss123

