## Supplemental files for "Effects of nano urea on growth and gene expression of *Arabidopsis thaliana* in hydroponics"

ATOMS INTO ASPIRATIONS

### IFFCO-NANOVENTIONS PVT LTD
##### JV of IFFCO, COIMBATORE

#### NANO UREA
##### Certificate of Analysis

**DOM: 25.08.2023**

| Sl.no | Parameters | Unit | Results |
|---|---|---|---|
| 1 | pH | - | 4.4 ± 0.5 (@25$^0$C) |
| 2 | Size | nm | 37.08 |
| 3 | Zeta | ±mV | -30.2 |
| 4 | Viscosity | cPs | 8.548 |
| 5 | Density | gm/cc | 1.004 |
| 6 | Microbial Contamination | CFU | Nil |
| 7 | Total N | % (w/v) | 4.9 |
| 8 | Purity | % | 95 |
| 9 | Physical Appearance | - | Transparent Liquid |

***1.5 molar of urea is present in the formulation.***

**Expiry Date : 24 months**

Soumiya. P
Chemist
IFFCO-NVPL

IFFCO-Nanoventions Private Limited
CBE-109

**Table S2 List of oligonucleotide primers used for real time PCR**

| S/N | Primer name | Sequence (5'-->3') | AGI Code |
|---|---|---|---|
| 1 | PDC1_RT_Fw | TCACAAGGATGATACGAGCAAAGA | AT4G33070 |
| 2 | PDC1_RT_Rv | ACTGAGGATTGGGAGGACGG | |
| 3 | UMAMIT14_RT_Fw | TGATCTCGAGGCCTAACACCA | AT2G39510 |
| 4 | UMAMIT14_RT_Rv | AATAAGCAACGATCGATCCGGT | |
| 5 | GDU1_RT_Fw | ACCGGCGAGACAACTACTACA | AT4G31730 |
| 6 | GDU1_RT_Rv | GGAATCATGACGAGGACAATTCTCC | |
| 7 | GLR2.2_RT_Fw | AGACACCGACTCGTACATCAAC | AT2G24720 |
| 8 | GLR2.2_RT_Rv | CCGTAGTTTGTCTGGTTTGTTGC | |
| 9 | PHYLLO_RT_Fw | ACCGCGAGATGAGTAAAAGCA | AT1G68890 |
| 10 | PHYLLO_RT_Rv | AGATGTACAGCGTGACCAGC | |
| 11 | GS2_RT_Fw | ACCTCACTTTTGGCAGAGACC | AT5G35630 |
| 12 | GS2_RT_Rv | TGAGAGACCACATAGACACGTGT | |
| 13 | UBQ1 _RT_Fw | TCGTAAGTACAATCAGGATAAGATG | AT3G52590 |
| 14 | UBQ1 _RT_Rv | CACTGAAACAAGAAAAACAAACCCT | |

**Table S3 FTIR peaks and their corresponding bonds of urea and NUF**

| Peak ($cm^{-1}$) | Reference peak ($cm^{-1}$) | Bonds | Reference |
|---|---|---|---|
| | | **Urea** | |
| 3430.4 | 3455 | $\nu_{as}$ (N-H) | Manivannan and Rajendran 2011 |
| | 3426 | | Channab et al. 2023 |
| 3330.2 | 3326 | $\nu_s$ (N-H) | Manivannan and Rajendran 2011 |
| | | | Channab et al. 2023 |
| 3253.1 | 3252 | ν (N-H) | Bayrak and Bayari 2010 |
| | | | Channab et al. 2023 |
| 1675.5 | 1661 | ν (C=O) | Channab et al. 2023 |
| | 1673 | | Onija et al. 2011 |
| | 1679 | | Gangopadhyay et al. 2016 |
| | 1683 | | Kang et al. 2021 |
| 1588.5 | 1588 | ν (N-H) | Channab et al. 2023 |
| | 1629 | ν (C=O) + δ(N-H) | Onija et al. 2011 |
| 1458 | 1444 | ν (C-N) | Channab et al. 2023 |
| | 1453 | | Manivannan and Rajendran 2011 |
| | 1471 | | Gangopadhyay et al. 2016 |
| | 1463 | | Kang et al. 2021 |
| 1146.9 | 1153 | γ ($NH_2$) | Onija et al. 2011 |
| | 1154 | γ ($NH_2$) | Gangopadhyay et al. 2016 |
| | | **NUF** | |
| 3331.9 | 3326 | $\nu_{as}$ (N-H) | Manivannan and Rajendran 2011 |
| | | | Bayrak and Bayari 2010 |
| | | | Channab et al. 2023 |
| 3241.4 | 3252 | $\nu_s$ (N-H) | Manivannan and Rajendran 2011 |
| | | | Bayrak and Bayari 2010 |
| | | | Channab et al. 2023 |
| 1633.2 | 1661 | ν (C=O) | Channab et al. 2023 |
| | 1668 | | Gangopadhyay et al. 2016 |
| | 1673 | | Onija et al. 2011 |
| | 1683 | | Kang et al. 2021 |
| | 1625 | ν (N-H) | Manivannan and Rajendran 2011 |
| 1449.7 | 1444 | ν (C-N) | Channab et al. 2023 |
| | 1453 | | Manivannan and Rajendran 2011 |
| | 1471 | | Gangopadhyay et al. 2016 |
| | 1463 | | Kang et al. 2021 |
| 1084.3 | 1057 | ν (C-O) | Saravanan et al. 2018 |
| | 1055 | δ ($NH_2$) | Onija et al. 2011 |

The different types of vibrations around chemical bonds are denoted by $\nu_{as}$ for asymmetric stretching, $\nu_s$ for symmetric stretching, ν for stretching, δ for deformation, and γ for rocking.

**Supplementary References**


1. Bayrak, C., and Bayari, S. H. (2010). Vibrational and DFT studies of creatinine and its metal complexes. Hacettepe *J. Biol. Chem.* 38(2), 107-118.

2. Channab, B.-E., Elhassani, C. E., Essamlali, Y., Marrane, S. E., Chakir, A., and Zahouily, M. (2023). Urea-Loaded Hydroxyapatite–Carboxylated Cellulose Composites as Slow-Release N Fertilizer Pellets for Efficient Delivery of Nitrogen. *Ind. Eng. Chem. Res.* 62, 14853–14865. doi: 10.1021/acs.iecr.3c00737

3. Fadeeva, I., Deyneko, D., Barbaro, K., et al. (2022). Influence of Synthesis Conditions on Gadolinium-Substituted Tricalcium Phosphate Ceramics and Its Physicochemical, Biological, and Antibacterial Properties. *Nanomaterials (Basel).* 12, 852. doi: 10.3390/nano12050852

4. Gangopadhyay, D., Singh, S. K., Sharma, P., et al. (2016). Spectroscopic and structural study of the newly synthesized heteroligand complex of copper with creatinine and urea. *Spectrochim Acta A Mol Biomol Spectrosc.*154, 200–206. doi: 10.1016/j.saa.2015.10.028

5. Kang, H., An, K., Guan, L., and Tian, D. (2021). Preparation and evaluation of magnetic molecularly imprinted polymers based on konjac glucomannan for urea. *J IRAN CHEM SOC.* doi: 10.1007/s13738-021-02166-3

6. Manivannan, M., & Rajendran, S. (2011). Investigation of inhibitive action of urea-Zn2+ system in the corrosion control of carbon steel in sea water. *Int. J. Eng. Sci. Technol.* 3(11), 8048-8060.

7. Saravanan, N., Ravisankar, V., and Chithambaram, V. (2017). Growth and characterization of novel semi organic nonlinear optical urea lead acetate single crystal by solution growth technique. *J. Mater. Sci.: Mater. Electron.* 29, 5009–5013. doi: 10.1007/s10854-017-8462-5

8. Urnukhsaikhan, E., Bold, B.-E., Gunbileg, A., Sukhbaatar, N., and Mishig-Ochir, T. (2021). Antibacterial activity and characteristics of silver nanoparticles biosynthesized from Carduus crispus. *Sci. Rep.* 11. doi: 10.1038/s41598-021-00520-2

**Table S4 Genes differentially expressed under NUF and urea treatment**

| GeneID | log2FC NUF vs Control 7 days | FDR NUF vs Control 7 days | log2FC Urea vs Control 7 days | FDR Urea vs Control 7 days | log2FC NUF vs Urea 7 days | FDR NUF vs Urea 7 days | log2FC NUF vs Control 12 h | FDR NUF vs Control 12 h | log2FC Urea vs Control 12 h | FDR Urea vs Control 12 h | log2FC NUF vs Urea 12 h | FDR NUF vs Urea 12 h | Protein names |
|---|---|---|---|---|---|---|---|---|---|---|---|---|---|
| AT5G19470 | -10.0 | 4.61E-04 | -5.7 | 9.37E-05 | -4.3 | 9.79E-02 | 1.4 | 5.28E-01 | 0.4 | 8.95E-01 | | | Nudix hydrolase homolog 24 |
| AT1G56600 | -5.7 | 2.90E-06 | -4.2 | 5.05E-06 | -1.6 | 5.30E-02 | -1.0 | 2.19E-02 | 0.0 | 9.76E-01 | -1.0 | 2.14E-02 | Galactinol synthase 2 |
| AT3G59580 | -4.8 | 1.70E-08 | -3.8 | 8.06E-08 | -0.9 | 4.59E-03 | -1.1 | 2.26E-02 | 0.1 | 9.22E-01 | -1.1 | 2.96E-02 | Protein NLP9 |
| AT2G40250 | -4.7 | 1.25E-06 | -2.8 | 9.43E-06 | -1.9 | 6.33E-03 | | | | | | | GDSL esterase |
| AT5G06905 | -4.7 | 2.43E-07 | -0.6 | 1.69E-03 | -4.1 | 1.39E-05 | | | | | | | Cytochrome P450, family 712, subfamily A, polypeptide 2 |
| AT5G04120 | -4.6 | 1.18E-07 | -2.2 | 4.34E-06 | -2.4 | 7.47E-05 | -5.6 | 3.27E-03 | -0.5 | 4.20E-01 | -5.0 | 1.75E-02 | Metal-independent phosphoserine phosphatase |
| AT5G54060 | -4.4 | 4.17E-04 | -3.2 | 9.23E-04 | -1.3 | 2.73E-01 | 1.2 | 2.15E-01 | 2.0 | 3.89E-02 | -0.8 | 1.81E-01 | Anthocyanidin 3-O-glucoside 2'''-O-xylosyltransferase |
| AT5G36150 | -4.4 | 1.77E-05 | -2.1 | 2.31E-04 | -2.4 | 7.22E-03 | | | | | | | Tirucalladienol synthase |
| AT2G19900 | -4.4 | 1.66E-07 | -4.5 | 4.61E-07 | 0.0 | 9.36E-01 | | | | | | | Malic enzyme |
| AT5G06510 | -4.2 | 6.73E-09 | -3.4 | 2.05E-08 | -0.9 | 3.13E-04 | -1.0 | 1.36E-01 | 0.7 | 2.55E-01 | -1.6 | 1.59E-02 | Nuclear transcription factor Y subunit |
| AT4G25010 | -4.2 | 9.91E-08 | -2.2 | 9.11E-07 | -2.0 | 1.84E-04 | -1.3 | 5.19E-02 | 0.6 | 2.86E-01 | -1.9 | 1.10E-02 | Bidirectional sugar transporter SWEET14 |
| AT5G06900 | -4.2 | 2.07E-07 | -0.5 | 2.35E-03 | -3.7 | 1.32E-05 | -1.5 | 3.75E-02 | 0.7 | 2.01E-01 | -2.2 | 1.77E-02 | Cytochrome P450 |
| AT3G52970 | -4.2 | 1.69E-07 | -1.2 | 1.99E-05 | -3.0 | 2.95E-05 | | | | | | | Cytochrome P450, family 76, subfamily G, polypeptide 1 |
| AT1G66390 | -4.1 | 1.81E-05 | -3.5 | 4.03E-05 | -0.6 | 4.46E-01 | 1.0 | 5.49E-01 | 3.1 | 4.13E-02 | | | Transcription factor MYB90 |
| AT1G34060 | -4.0 | 5.22E-08 | -2.8 | 2.77E-07 | -1.3 | 9.38E-04 | -0.8 | 2.45E-02 | -0.3 | 5.00E-01 | -0.6 | 2.07E-01 | Pyridoxal phosphate -dependent transferases superfamily protein |
| AT5G36130 | -4.0 | 2.38E-05 | -1.3 | 1.56E-03 | -2.7 | 2.03E-03 | | | | | | | Uncharacterized protein |
| AT3G24290 | -3.9 | 1.37E-07 | -1.1 | 2.95E-04 | -2.8 | 1.84E-05 | -3.9 | 9.56E-04 | 0.5 | 2.95E-01 | -4.4 | 4.12E-03 | Putative ammonium transporter 1 member 5 |
| AT5G36140 | -3.8 | 8.65E-06 | -1.5 | 2.86E-04 | -2.3 | 1.91E-03 | | | | | | | Cytochrome P450 |
| AT5G16570 | -3.8 | 1.99E-08 | -2.2 | 1.11E-06 | -1.6 | 2.85E-05 | -1.7 | 1.53E-07 | -0.1 | 5.34E-01 | -1.6 | 3.13E-05 | glutamine synthetase |
| AT1G68040 | -3.7 | 6.93E-07 | -0.4 | 1.34E-02 | -3.3 | 2.85E-05 | | | | | | | hypothetical protein |
| AT4G39000 | -3.7 | 3.90E-06 | -0.6 | 1.17E-02 | -3.1 | 1.43E-04 | | | | | | | Endoglucanase 23 |
| AT5G42800 | -3.7 | 6.22E-05 | -2.9 | 3.63E-04 | -0.8 | 2.12E-01 | 0.5 | 2.67E-01 | 1.5 | 1.27E-03 | -1.0 | 1.62E-02 | Dihydroflavonol 4-reductase |
| AT4G19975 | -3.6 | 5.27E-08 | -1.9 | 7.02E-07 | -1.7 | 7.65E-05 | 1.2 | 1.31E-03 | 0.8 | 3.32E-02 | 0.4 | 2.71E-01 | Glycosyltransferase |
| AT1G56650 | -3.6 | 1.60E-04 | -3.1 | 3.63E-04 | -0.5 | 5.72E-01 | -0.2 | 8.94E-01 | 3.2 | 1.05E-02 | -3.4 | 1.17E-02 | Transcription factor MYB75 |
| AT1G19630 | -3.6 | 5.08E-05 | -2.8 | 1.76E-04 | -0.8 | 3.11E-01 | -0.1 | 8.68E-01 | 0.2 | 8.01E-01 | -0.3 | 7.20E-01 | F14P1.4 protein |
| AT2G11810 | -3.5 | 2.67E-06 | -2.1 | 1.82E-05 | -1.4 | 8.71E-03 | | | | | | | monogalactosyldiacylglycerol synthase |
| AT5G61430 | -3.5 | 6.86E-08 | -2.8 | 2.59E-07 | -0.7 | 2.54E-02 | 0.1 | 8.34E-01 | -0.4 | 4.90E-01 | 0.6 | 4.34E-01 | NAC domain containing protein 100 |
| AT1G05650 | -3.4 | 5.55E-07 | -0.4 | 2.09E-02 | -3.0 | 2.63E-05 | -1.9 | 7.94E-05 | -0.1 | 7.70E-01 | -1.8 | 1.73E-03 | F3F20.10 protein |
| AT4G14090 | -3.4 | 3.14E-04 | -2.8 | 8.08E-04 | -0.5 | 5.32E-01 | 0.7 | 1.78E-01 | 1.5 | 9.38E-03 | -0.8 | 9.24E-02 | UDP-glycosyltransferase 75C1 |
| AT4G35190 | -3.3 | 2.21E-07 | -2.9 | 7.18E-07 | -0.4 | 2.35E-01 | 1.2 | 1.85E-02 | 1.1 | 4.37E-02 | 0.1 | 8.69E-01 | Cytokinin riboside 5'-monophosphate phosphoribohydrolase |
| AT4G32940 | -3.2 | 6.73E-09 | -2.4 | 5.40E-08 | -0.8 | 2.69E-05 | -1.1 | 2.35E-07 | -0.5 | 1.79E-03 | -0.6 | 3.63E-03 | Vacuolar-processing enzyme gamma-isozyme |
| AT1G07430 | -3.2 | 4.08E-04 | -2.6 | 1.07E-03 | -0.6 | 5.04E-01 | -2.0 | 2.26E-02 | 0.3 | 6.82E-01 | -2.4 | 4.38E-02 | Protein phosphatase 2C 3 |
| AT4G22880 | -3.2 | 1.81E-04 | -2.4 | 1.08E-03 | -0.8 | 2.41E-01 | 1.5 | 3.80E-02 | 2.3 | 5.63E-03 | -0.7 | 2.52E-01 | Mutant protein of leucoanthocyanidin dioxygenase |
| AT5G51500 | -3.2 | 2.05E-06 | -2.0 | 1.76E-05 | -1.1 | 1.64E-02 | | | | | | | Probable pectinesterase |
| AT5G07680 | -3.2 | 7.33E-07 | -3.0 | 2.23E-06 | -0.2 | 6.44E-01 | -0.4 | 2.85E-01 | -0.3 | 4.25E-01 | -0.1 | 9.30E-01 | NAC domain-containing protein 79 |
| AT3G53980 | -3.2 | 1.89E-08 | -2.8 | 9.13E-08 | -0.4 | 2.59E-02 | -2.0 | 4.23E-05 | -0.4 | 2.60E-01 | -1.6 | 3.42E-03 | Bifunctional inhibitor |
| AT4G37950 | -3.2 | 2.11E-06 | -0.8 | 1.07E-03 | -2.3 | 1.98E-04 | | | | | | | rhamnogalacturonan endolyase |
| AT3G48520 | -3.1 | 8.04E-06 | -2.0 | 7.85E-05 | -1.2 | 2.96E-02 | -0.3 | 8.69E-01 | 1.1 | 3.82E-01 | | | Cytochrome P450 94B3 |
| AT3G61400 | -3.0 | 7.03E-08 | -1.8 | 1.17E-06 | -1.2 | 2.62E-04 | 1.2 | 1.10E-02 | 1.6 | 2.71E-03 | -0.4 | 2.82E-01 | 2-oxoglutarate and Fe-dependent oxygenase superfamily protein |
| AT5G17220 | -3.0 | 3.79E-04 | -2.6 | 9.66E-04 | -0.4 | 6.23E-01 | 1.0 | 1.39E-01 | 1.9 | 8.75E-03 | -0.9 | 9.03E-02 | Glutathione S-transferase F12 |
| AT3G05690 | -3.0 | 1.47E-08 | -2.3 | 1.00E-07 | -0.7 | 3.42E-04 | -0.1 | 5.22E-01 | 0.4 | 7.10E-02 | -0.5 | 3.99E-02 | Nuclear transcription factor Y subunit |
| AT1G15540 | -2.9 | 6.22E-06 | -0.4 | 4.49E-02 | -2.5 | 1.70E-04 | | | | | | | Fe2OG dioxygenase domain-containing protein |
| AT4G12290 | -2.9 | 1.63E-07 | -2.2 | 1.75E-06 | -0.7 | 6.23E-03 | 0.5 | 2.41E-03 | 0.6 | 3.08E-03 | 0.0 | 8.66E-01 | Amine oxidase |
| AT5G37690 | -2.9 | 5.43E-08 | -1.4 | 5.64E-06 | -1.5 | 3.57E-05 | 1.7 | 6.05E-07 | 1.7 | 2.37E-06 | 0.0 | 8.39E-01 | SGNH hydrolase-type esterase superfamily protein |
| AT3G14225 | -2.9 | 1.58E-06 | -1.7 | 1.99E-05 | -1.1 | 7.60E-03 | 1.4 | 4.04E-02 | 0.7 | 4.24E-01 | 0.7 | 3.80E-01 | GDSL-motif lipase 4 |
| AT1G60450 | -2.8 | 5.08E-05 | -0.1 | 7.42E-01 | -2.8 | 3.82E-04 | | | | | | | Galactinol synthase 7 |
| AT2G39370 | -2.8 | 3.97E-07 | -1.8 | 4.59E-06 | -1.0 | 3.95E-03 | 0.5 | 2.65E-01 | 0.4 | 4.14E-01 | 0.1 | 9.11E-01 | Probable membrane-associated kinase regulator 4 |
| AT5G04840 | -2.8 | 1.25E-04 | -1.9 | 8.00E-04 | -0.9 | 1.49E-01 | 0.1 | 8.84E-01 | 0.4 | 1.64E-01 | -0.4 | 3.19E-01 | At5g04840 |
| AT5G10625 | -2.8 | 2.25E-08 | -1.5 | 5.99E-07 | -1.3 | 3.93E-05 | -0.3 | 2.91E-01 | -0.1 | 7.62E-01 | -0.2 | 6.04E-01 | Uncharacterized protein F12B17_10 |
| AT4G38340 | -2.8 | 1.99E-07 | -1.5 | 1.55E-05 | -1.3 | 2.03E-04 | -1.5 | 1.84E-04 | 0.0 | 9.28E-01 | -1.4 | 2.72E-03 | Plant regulator RWP-RK family protein |
| AT1G60470 | -2.8 | 1.14E-06 | -3.3 | 2.17E-06 | 0.5 | 1.70E-01 | -0.5 | 2.07E-01 | 0.2 | 6.35E-01 | -0.7 | 9.33E-02 | Galactinol synthase 4 |
| AT1G51090 | -2.8 | 1.04E-04 | -2.0 | 5.83E-04 | -0.8 | 1.77E-01 | -1.7 | 1.41E-02 | 0.5 | 3.50E-01 | -2.2 | 1.65E-02 | Heavy metal transport |
| AT1G73000 | -2.8 | 1.49E-05 | -1.2 | 7.66E-04 | -1.6 | 4.85E-03 | | | | | | | Abscisic acid receptor PYL3 |
| AT3G16150 | -2.7 | 2.76E-07 | -1.2 | 4.17E-05 | -1.6 | 1.35E-04 | -1.2 | 2.52E-03 | -0.3 | 3.76E-01 | -0.9 | 3.80E-02 | Probable isoaspartyl peptidase |
| AT2G14760 | -2.7 | 2.26E-06 | -0.8 | 9.06E-04 | -1.9 | 2.79E-04 | -2.8 | 6.24E-04 | -0.2 | 6.81E-01 | -2.6 | 1.06E-02 | Basic helix-loop-helix DNA-binding superfamily protein |
| AT1G33090 | -2.7 | 8.45E-07 | -1.3 | 3.80E-05 | -1.4 | 7.42E-04 | | | | | | | Protein DETOXIFICATION 22 |
| AT3G44540 | -2.7 | 2.19E-08 | -2.0 | 2.13E-07 | -0.7 | 6.91E-04 | 1.0 | 2.36E-03 | 0.9 | 7.16E-03 | 0.1 | 8.87E-01 | Fatty acyl-CoA reductase |
| AT1G72830 | -2.7 | 2.19E-08 | -2.0 | 2.47E-07 | -0.7 | 3.98E-04 | -0.2 | 5.78E-01 | 0.1 | 7.81E-01 | -0.3 | 4.73E-01 | Nuclear transcription factor Y subunit |
| AT3G21340 | -2.6 | 4.68E-08 | -0.7 | 6.43E-05 | -1.9 | 1.09E-05 | -0.8 | 5.40E-02 | 0.0 | 9.94E-01 | -0.8 | 8.55E-02 | Leucine-rich repeat protein kinase family protein |
| AT1G18870 | -2.6 | 2.50E-05 | -1.2 | 1.66E-03 | -1.4 | 6.86E-03 | 0.1 | 8.14E-01 | 0.7 | 1.53E-01 | -0.6 | 2.04E-01 | isochorismate synthase |
| AT1G68360 | -2.6 | 5.37E-06 | -2.0 | 3.11E-05 | -0.6 | 1.31E-01 | 0.1 | 8.84E-01 | 0.2 | 5.87E-01 | -0.2 | 7.46E-01 | Zinc finger protein GIS3 |
| AT5G18910 | -2.6 | 2.10E-06 | -0.9 | 6.79E-04 | -1.7 | 4.01E-04 | | | | | | | At5g18910 |
| AT5G07990 | -2.6 | 1.32E-03 | -2.2 | 3.16E-03 | -0.4 | 6.47E-01 | 1.0 | 5.55E-02 | 2.2 | 4.92E-04 | -1.2 | 2.38E-02 | Mutant flavonoid 3'hydroxylase |
| AT1G19230 | -2.6 | 1.93E-08 | -1.4 | 4.89E-07 | -1.2 | 3.33E-05 | 0.5 | 2.76E-01 | 0.8 | 7.18E-02 | -0.4 | 5.11E-01 | Respiratory burst oxidase homolog protein E |
| AT5G42610 | -2.6 | 6.57E-07 | -2.0 | 4.04E-06 | -0.6 | 5.02E-02 | -1.9 | 2.07E-05 | 0.0 | 9.39E-01 | -1.9 | 3.36E-04 | Calcium uniporter protein 4, mitochondrial |
| AT4G20240 | -2.6 | 6.14E-07 | -1.3 | 2.76E-05 | -1.2 | 7.77E-04 | -0.8 | 2.49E-03 | 0.1 | 7.54E-01 | -0.9 | 5.23E-03 | Cytochrome P450, family 71, subfamily A, polypeptide 27 |
| AT1G04220 | -2.5 | 1.41E-07 | -1.3 | 6.58E-06 | -1.2 | 1.88E-04 | 0.4 | 1.08E-01 | 0.4 | 1.15E-01 | 0.0 | 9.28E-01 | 3-ketoacyl-CoA synthase 2 |
| AT5G53380 | -2.5 | 2.79E-06 | -0.8 | 1.42E-03 | -1.8 | 3.51E-04 | | | | | | | Wax ester synthase |
| AT4G39795 | -2.5 | 7.93E-08 | -1.3 | 6.81E-06 | -1.3 | 7.64E-05 | -2.3 | 5.54E-07 | -0.8 | 2.97E-03 | -1.6 | 4.75E-04 | FCS-Like Zinc finger 7 |
| AT5G36110 | -2.5 | 1.04E-07 | -1.6 | 1.32E-06 | -1.0 | 8.82E-04 | | | | | | | Cytochrome P450, family 716, subfamily A, polypeptide 1 |
| AT1G23200 | -2.5 | 8.24E-07 | -1.8 | 7.15E-06 | -0.8 | 1.71E-02 | -0.6 | 2.06E-01 | 0.4 | 3.82E-01 | -1.0 | 3.98E-02 | Probable pectinesterase |
| AT1G35240 | -2.5 | 1.70E-06 | -1.6 | 2.64E-05 | -0.9 | 1.43E-02 | | | | | | | Auxin response factor |
| AT4G26220 | -2.5 | 1.59E-08 | -1.2 | 1.06E-06 | -1.3 | 1.02E-05 | -0.4 | 3.77E-02 | 0.3 | 1.25E-01 | -0.7 | 2.16E-03 | Probable caffeoyl-CoA O-methyltransferase At4g26220 |
| AT4G25250 | -2.5 | 2.35E-07 | -0.3 | 2.76E-02 | -2.2 | 1.39E-05 | -1.8 | 1.13E-05 | -0.2 | 5.26E-01 | -1.6 | 9.16E-04 | Pectinesterase inhibitor 4 |
| AT2G03370 | -2.5 | 7.00E-07 | -1.0 | 1.26E-04 | -1.5 | 2.52E-04 | -1.3 | 3.49E-02 | -0.1 | 8.86E-01 | -1.2 | 1.00E-01 | O-linked-mannose beta-1,4-N-acetylglucosaminyltransferase-like protein |
| AT2G40130 | -2.5 | 1.96E-06 | -2.0 | 1.48E-05 | -0.5 | 8.47E-02 | 0.1 | 7.00E-01 | 0.8 | 2.53E-03 | -0.7 | 9.58E-03 | Double Clp-N motif-containing P-loop nucleoside triphosphate hydrolases superfamily protein |
| AT1G72220 | -2.5 | 1.67E-06 | -1.5 | 2.64E-05 | -1.0 | 6.65E-03 | -0.4 | 3.47E-01 | -0.7 | 1.23E-01 | 0.3 | 5.97E-01 | RING-H2 finger protein ATL54 |

| | | | | | | | | | | | | | |
|---|---|---|---|---|---|---|---|---|---|---|---|---|---|
| AT3G48340 | -2.5 | 4.13E-08 | -0.5 | 2.54E-03 | -2.0 | 6.99E-06 | -1.4 | 1.23E-05 | 0.1 | 8.81E-01 | -1.4 | 4.40E-05 | KDEL-tailed cysteine endopeptidase CEP2 |
| AT5G48010 | -2.5 | 1.13E-03 | -1.9 | 4.21E-03 | -0.6 | 4.44E-01 | -2.4 | 1.56E-01 | 1.5 | 2.15E-01 | -3.9 | 3.53E-02 | Thalianol synthase 1 |
| AT5G08250 | -2.5 | 2.02E-06 | -1.3 | 6.77E-05 | -1.2 | 3.38E-03 | 1.2 | 1.52E-03 | 0.9 | 1.64E-02 | 0.3 | 5.12E-01 | Cytochrome P450 superfamily protein |
| AT2G29750 | -2.5 | 4.59E-08 | -0.7 | 3.27E-04 | -1.7 | 1.02E-05 | -1.9 | 4.76E-08 | 0.2 | 2.88E-01 | -2.2 | 4.02E-06 | Flavonol 3-O-glucosyltransferase UGT71C1 |
| AT3G10710 | -2.5 | 6.77E-07 | -0.8 | 8.98E-04 | -1.7 | 9.04E-05 | -1.7 | 3.53E-03 | 0.0 | 9.80E-01 | -1.7 | 1.01E-02 | Putative pectinesterase |
| AT5G15180 | -2.5 | 1.44E-08 | -1.0 | 1.92E-06 | -1.5 | 6.99E-06 | -1.7 | 1.15E-06 | -0.3 | 1.14E-01 | -1.3 | 2.10E-04 | Peroxidase 56 |
| AT1G29020 | -2.5 | 6.10E-08 | -1.3 | 4.62E-06 | -1.1 | 6.65E-05 | 0.1 | 5.90E-01 | 0.5 | 2.58E-02 | -0.4 | 1.14E-01 | Calcium-binding EF-hand family protein |
| AT1G69480 | -2.5 | 4.08E-04 | -3.0 | 3.71E-04 | 0.5 | 4.89E-01 | | | | | | | Phosphate transporter PHO1 homolog 10 |
| AT1G29000 | -2.5 | 9.98E-06 | -1.2 | 4.52E-04 | -1.3 | 5.69E-03 | | | | | | | HMA domain-containing protein |
| AT4G13620 | -2.4 | 3.81E-06 | -1.0 | 4.60E-04 | -1.5 | 1.30E-03 | -0.3 | 5.03E-01 | -0.1 | 8.94E-01 | -0.2 | 7.14E-01 | Ethylene-responsive transcription factor ERF062 |
| AT1G07460 | -2.4 | 2.03E-05 | -0.4 | 7.24E-02 | -2.0 | 5.73E-04 | | | | | | | non-specific serine |
| AT5G47530 | -2.4 | 1.65E-07 | -1.1 | 1.17E-05 | -1.3 | 1.42E-04 | 0.4 | 3.90E-01 | 0.1 | 8.52E-01 | 0.2 | 6.38E-01 | Cytochrome b561 and DOMON domain-containing protein At5g47530 |
| AT4G09820 | -2.4 | 6.47E-04 | -2.3 | 1.28E-03 | -0.1 | 8.97E-01 | 0.3 | 7.32E-01 | 0.9 | 2.41E-01 | | | Transcription factor TT8 |
| AT5G41730 | -2.4 | 2.68E-05 | -0.8 | 7.53E-03 | -1.6 | 2.95E-03 | | | | | | | Protein kinase family protein |
| AT1G78340 | -2.4 | 1.44E-08 | -1.7 | 9.13E-08 | -0.7 | 1.77E-04 | 2.9 | 7.74E-08 | 1.4 | 8.33E-04 | 1.5 | 2.36E-05 | Glutathione S-transferase U22 |
| AT2G35380 | -2.4 | 2.79E-08 | -1.0 | 4.82E-06 | -1.4 | 1.80E-05 | 1.6 | 1.09E-06 | 1.4 | 4.05E-05 | 0.2 | 2.56E-01 | peroxidase |
| AT2G33380 | -2.4 | 1.51E-04 | -1.8 | 7.71E-04 | -0.6 | 2.51E-01 | -1.2 | 7.54E-02 | 0.9 | 9.84E-02 | -2.1 | 1.81E-02 | Probable peroxygenase 3 |
| AT1G22160 | -2.4 | 4.49E-08 | -1.5 | 1.16E-06 | -0.9 | 1.70E-04 | -0.7 | 6.06E-04 | 0.0 | 9.83E-01 | -0.7 | 7.54E-03 | FCS-Like Zinc finger 5 |
| AT1G06120 | -2.4 | 2.87E-07 | -0.5 | 2.53E-03 | -1.9 | 2.69E-05 | -1.7 | 1.34E-03 | -0.4 | 3.41E-01 | -1.3 | 4.07E-02 | Delta-9 desaturase-like 3 protein |
| AT1G52660 | -2.4 | 4.15E-07 | -0.6 | 8.00E-04 | -1.8 | 4.74E-05 | -1.1 | 1.93E-02 | -0.1 | 9.03E-01 | -1.0 | 3.44E-02 | NB-ARC domain-containing protein |
| AT1G79320 | -2.4 | 3.24E-08 | -1.8 | 2.95E-07 | -0.6 | 1.82E-03 | -1.3 | 9.15E-05 | 0.7 | 6.81E-03 | -2.0 | 1.01E-05 | Metacaspase-6 |
| AT1G53270 | -2.4 | 4.22E-07 | -1.1 | 2.66E-05 | -1.3 | 3.61E-04 | 0.5 | 3.06E-01 | 0.8 | 1.61E-01 | -0.2 | 7.11E-01 | ABC transporter G family member 10 |
| AT5G23190 | -2.4 | 3.24E-08 | -1.4 | 6.98E-07 | -0.9 | 1.35E-04 | 1.5 | 1.64E-06 | 0.9 | 6.32E-04 | 0.5 | 1.87E-02 | Cytochrome P450 86B1 |
| AT1G73810 | -2.4 | 4.96E-08 | -1.9 | 2.97E-07 | -0.4 | 1.57E-02 | 0.7 | 3.66E-02 | 0.0 | 9.42E-01 | 0.6 | 1.22E-01 | Core-2 |
| AT5G19410 | -2.4 | 8.45E-08 | -1.2 | 4.25E-06 | -1.2 | 1.08E-04 | 1.1 | 9.77E-03 | 0.8 | 5.88E-02 | 0.2 | 6.43E-01 | ABC-2 type transporter family protein |
| AT3G47770 | -2.4 | 4.13E-06 | -0.4 | 4.94E-02 | -2.0 | 1.38E-04 | | | | | | | ABC2 homolog 5 |
| AT2G43500 | -2.4 | 1.97E-08 | -1.7 | 2.01E-07 | -0.6 | 4.10E-04 | 0.9 | 2.48E-03 | 0.7 | 2.33E-02 | 0.2 | 5.67E-01 | Plant regulator RWP-RK family protein |
| AT1G10070 | -2.3 | 2.18E-06 | -1.8 | 1.27E-05 | -0.5 | 8.04E-02 | -1.5 | 8.91E-02 | -2.5 | 3.86E-02 | 1.0 | 4.31E-01 | Branched-chain-amino-acid aminotransferase |
| AT4G09020 | -2.3 | 2.16E-06 | -2.1 | 9.35E-06 | -0.2 | 3.56E-01 | -0.2 | 3.14E-01 | 0.3 | 7.90E-02 | -0.5 | 2.14E-02 | Isoamylase 3, chloroplastic |
| AT1G65570 | -2.3 | 4.87E-05 | -0.6 | 2.40E-02 | -1.7 | 2.53E-03 | | | | | | | F5I14.10 |
| AT5G51270 | -2.3 | 1.59E-06 | -0.7 | 1.79E-03 | -1.7 | 1.77E-04 | -1.1 | 7.26E-02 | -0.2 | 8.07E-01 | -0.9 | 2.52E-01 | U-box domain-containing protein kinase family protein |
| AT3G60280 | -2.3 | 2.80E-05 | -0.3 | 1.76E-01 | -2.0 | 4.95E-04 | -2.7 | 1.51E-04 | -0.2 | 6.06E-01 | -2.5 | 3.70E-03 | Uclacyanin-3 |
| AT5G13930 | -2.3 | 1.09E-04 | -2.0 | 4.68E-04 | -0.3 | 4.92E-01 | 1.4 | 7.51E-06 | 2.7 | 3.12E-08 | -1.3 | 3.90E-04 | Chalcone synthase |
| AT5G13580 | -2.3 | 2.19E-08 | -1.3 | 6.81E-07 | -1.0 | 4.58E-05 | 1.5 | 4.97E-05 | 0.9 | 1.36E-02 | 0.7 | 5.34E-02 | ABC transporter G family member 6 |
| AT2G15350 | -2.3 | 3.52E-06 | -0.4 | 5.09E-02 | -2.0 | 1.26E-04 | | | | | | | Putative fucosyltransferase 10 |
| AT5G58360 | -2.3 | 2.86E-04 | -0.8 | 2.42E-02 | -1.5 | 1.80E-02 | | | | | | | Transcription repressor OFP3 |
| AT1G53830 | -2.3 | 7.93E-07 | -0.6 | 2.57E-03 | -1.7 | 7.42E-05 | -0.8 | 4.33E-02 | 0.0 | 9.34E-01 | -0.8 | 9.33E-02 | Pectinesterase 2 |
| AT1G26700 | -2.3 | 4.67E-07 | -0.3 | 2.07E-02 | -2.0 | 2.69E-05 | -1.0 | 3.51E-02 | 0.7 | 7.33E-02 | -1.7 | 1.85E-03 | MLO-like protein 14 |
| AT1G74460 | -2.3 | 7.03E-08 | -1.0 | 3.02E-05 | -1.3 | 2.69E-05 | 1.5 | 4.07E-08 | 1.1 | 4.36E-06 | 0.4 | 1.94E-02 | GDSL esterase |
| AT4G24140 | -2.3 | 1.97E-08 | -1.1 | 1.92E-06 | -1.2 | 1.47E-05 | 1.2 | 6.60E-05 | 0.9 | 2.20E-03 | 0.3 | 2.74E-01 | Probable lysophospholipase BODYGUARD 3 |
| AT5G65040 | -2.3 | 3.73E-07 | -1.5 | 6.00E-06 | -0.8 | 4.32E-03 | 0.0 | 9.64E-01 | 0.1 | 7.35E-01 | -0.1 | 8.13E-01 | Protein INCREASED RESISTANCE TO MYZUS PERSICAE 1 |
| AT2G31990 | -2.3 | 7.44E-08 | -1.5 | 9.64E-07 | -0.8 | 1.17E-03 | -0.4 | 5.85E-02 | -0.4 | 9.45E-02 | 0.0 | 9.63E-01 | Exostosin family protein |
| AT2G23540 | -2.3 | 1.44E-08 | -1.0 | 1.94E-06 | -1.3 | 6.99E-06 | 2.1 | 1.54E-08 | 1.6 | 1.21E-06 | 0.5 | 4.03E-02 | GDSL esterase |
| AT5G07130 | -2.3 | 1.48E-07 | -1.4 | 3.68E-06 | -0.9 | 8.81E-04 | 1.3 | 3.82E-03 | 0.8 | 9.11E-02 | 0.5 | 2.82E-01 | Laccase |
| AT2G22240 | -2.2 | 3.25E-07 | -1.8 | 3.85E-06 | -0.5 | 1.87E-02 | -0.9 | 7.79E-04 | -0.1 | 7.92E-01 | -0.8 | 7.50E-03 | inositol-3-phosphate synthase |
| AT5G56840 | -2.2 | 1.92E-07 | -1.4 | 5.71E-06 | -0.9 | 9.48E-04 | 0.4 | 1.92E-01 | 0.7 | 3.51E-02 | -0.3 | 4.75E-01 | hypothetical protein |
| AT4G17770 | -2.2 | 3.53E-07 | -1.8 | 3.96E-06 | -0.5 | 1.70E-02 | -0.2 | 3.15E-01 | 0.3 | 6.67E-02 | -0.5 | 2.04E-02 | Alpha,alpha-trehalose-phosphate synthase 5 |
| AT3G11430 | -2.2 | 3.14E-08 | -1.1 | 1.92E-06 | -1.1 | 3.67E-05 | 1.7 | 1.18E-06 | 1.1 | 5.07E-04 | 0.6 | 5.96E-03 | Glycerol-3-phosphate acyltransferase 5 |
| AT5G57530 | -2.2 | 1.06E-06 | -0.5 | 5.25E-03 | -1.7 | 7.80E-05 | -2.0 | 1.27E-04 | 0.2 | 4.82E-01 | -2.3 | 9.63E-04 | Probable xyloglucan endotransglucosylase |
| AT5G41570 | -2.2 | 3.51E-07 | -1.2 | 1.06E-05 | -1.0 | 9.23E-04 | 2.3 | 7.01E-05 | 1.4 | 1.26E-02 | 0.9 | 3.32E-02 | Probable WRKY transcription factor 24 |
| AT2G48130 | -2.2 | 1.08E-07 | -0.9 | 5.16E-05 | -1.3 | 3.48E-05 | 0.8 | 1.86E-04 | 0.6 | 4.20E-03 | 0.2 | 3.56E-01 | Non-specific lipid transfer protein GPI-anchored 15 |
| AT3G49760 | -2.2 | 3.81E-07 | -1.0 | 4.04E-05 | -1.2 | 2.44E-04 | -1.3 | 2.33E-02 | -0.3 | 5.57E-01 | -1.0 | 1.83E-01 | BZIP transcription factor-like protein |
| AT5G57540 | -2.2 | 1.76E-05 | -1.2 | 6.67E-04 | -1.1 | 1.50E-02 | -0.3 | 7.54E-01 | 0.0 | 9.69E-01 | | | Putative xyloglucan endotransglucosylase |
| AT5G52020 | -2.2 | 1.74E-07 | -0.7 | 1.37E-04 | -1.5 | 3.82E-05 | -0.4 | 3.37E-01 | 0.0 | 9.68E-01 | -0.4 | 4.01E-01 | Ethylene-responsive transcription factor ERF025 |
| AT1G61270 | -2.2 | 3.11E-06 | -0.8 | 7.11E-04 | -1.4 | 8.23E-04 | | | | | | | Lysine histidine transporter-like 3 |
| AT2G40370 | -2.2 | 7.33E-08 | -1.1 | 4.59E-06 | -1.1 | 9.04E-05 | 2.5 | 6.24E-06 | 1.7 | 1.53E-03 | 0.9 | 6.19E-03 | Laccase-5 |
| AT3G32030 | -2.2 | 1.68E-07 | -0.6 | 5.08E-04 | -1.6 | 2.69E-05 | -1.4 | 4.45E-04 | 0.1 | 7.39E-01 | -1.5 | 4.14E-04 | Terpenoid synthase 30 |
| AT2G17500 | -2.2 | 7.44E-08 | -1.8 | 4.77E-07 | -0.3 | 2.53E-02 | -0.2 | 4.68E-01 | 0.3 | 2.45E-01 | -0.4 | 1.17E-01 | Protein PIN-LIKES 5 |
| AT2G04680 | -2.2 | 2.10E-07 | -0.4 | 1.90E-03 | -1.7 | 2.46E-05 | -0.9 | 2.62E-01 | -0.2 | 8.62E-01 | -0.7 | 4.21E-01 | Cysteine |
| AT5G58860 | -2.2 | 2.38E-08 | -0.9 | 1.34E-05 | -1.3 | 1.09E-05 | 1.6 | 2.82E-07 | 1.4 | 4.36E-06 | 0.1 | 4.92E-01 | Cytochrome P450 86A1 |
| AT4G31250 | -2.2 | 2.19E-06 | -0.8 | 8.34E-04 | -1.4 | 5.02E-04 | | | | | | | Probable LRR receptor-like serine |
| AT1G73040 | -2.2 | 1.96E-07 | -2.1 | 6.46E-07 | -0.1 | 6.72E-01 | -0.1 | 7.40E-01 | 0.3 | 4.40E-01 | | | Jacalin-related lectin 19 |
| AT1G51140 | -2.2 | 2.98E-07 | -1.9 | 1.57E-06 | -0.3 | 1.39E-01 | -0.8 | 5.08E-05 | 0.0 | 9.13E-01 | -0.8 | 7.58E-04 | Transcription factor bHLH122 |
| AT5G36270 | -2.2 | 2.76E-05 | -0.3 | 2.03E-01 | -1.9 | 4.86E-04 | -3.0 | 1.11E-04 | 0.0 | 9.99E-01 | -3.0 | 1.80E-03 | Probable glutathione S-transferase DHAR4 |
| AT1G79250 | -2.2 | 9.29E-06 | -0.9 | 1.70E-03 | -1.3 | 2.38E-03 | | | | | | | Serine |
| AT5G66690 | -2.2 | 8.36E-08 | -1.7 | 1.08E-06 | -0.5 | 4.35E-03 | -1.0 | 3.44E-06 | 0.1 | 4.92E-01 | -1.1 | 1.54E-05 | UDP-glycosyltransferase 72E2 |
| AT2G26420 | -2.2 | 2.53E-07 | -0.5 | 6.21E-04 | -1.6 | 3.30E-05 | -0.7 | 1.90E-01 | 0.6 | 2.61E-01 | -1.3 | 1.87E-02 | Phosphatidylinositol 4-phosphate 5-kinase |
| AT3G15250 | -2.2 | 8.55E-06 | -1.1 | 3.97E-04 | -1.1 | 6.68E-03 | -0.9 | 5.49E-02 | -0.8 | 1.18E-01 | -0.1 | 8.96E-01 | Membrane-associated kinase regulator |
| AT1G63450 | -2.1 | 1.74E-07 | -0.5 | 1.46E-03 | -1.6 | 2.37E-05 | -1.4 | 9.28E-05 | -0.1 | 8.66E-01 | -1.4 | 4.08E-04 | hypothetical protein |
| AT4G11910 | -2.1 | 1.45E-06 | -1.9 | 4.72E-06 | -0.2 | 4.54E-01 | -0.7 | 2.02E-01 | -0.4 | 4.94E-01 | -0.3 | 7.52E-01 | Magnesium dechelatase SGR2, chloroplastic |
| AT3G24310 | -2.1 | 3.19E-07 | -2.0 | 1.11E-06 | -0.1 | 5.25E-01 | | | | | | | MYB-like DNA-binding domain protein |
| AT5G13900 | -2.1 | 2.59E-07 | -1.1 | 1.19E-05 | -1.0 | 4.27E-04 | 0.9 | 1.41E-02 | 0.7 | 1.04E-01 | 0.2 | 4.56E-01 | Non-specific lipid transfer protein GPI-anchored 30 |
| AT1G60095 | -2.1 | 1.70E-08 | -1.4 | 2.28E-07 | -0.7 | 1.32E-04 | 0.2 | 4.92E-01 | 0.7 | 3.72E-03 | -0.5 | 3.71E-02 | Mannose-binding lectin superfamily protein |
| AT3G21180 | -2.1 | 3.80E-07 | -0.4 | 1.08E-02 | -1.8 | 2.69E-05 | -1.1 | 3.73E-02 | 0.3 | 5.00E-01 | -1.4 | 1.29E-02 | Calcium-transporting ATPase 9, plasma membrane-type -ATPase isoform 9) |
| AT4G40070 | -2.1 | 1.15E-06 | -1.3 | 4.38E-05 | -0.8 | 3.52E-03 | -0.7 | 2.47E-04 | -0.1 | 6.34E-01 | -0.7 | 6.19E-03 | RING-H2 finger protein ATL32 |
| AT3G09220 | -2.1 | 2.04E-08 | -1.8 | 1.04E-07 | -0.3 | 2.45E-02 | 1.7 | 1.89E-05 | 1.2 | 1.43E-03 | 0.5 | 1.34E-01 | Laccase |
| AT3G12580 | -2.1 | 2.97E-08 | -2.1 | 9.13E-08 | 0.0 | 9.11E-01 | 0.4 | 8.09E-02 | 1.1 | 3.10E-04 | -0.7 | 1.36E-02 | Heat shock 70 kDa protein 4 |
| AT2G46860 | -2.1 | 1.53E-06 | -0.4 | 1.88E-02 | -1.7 | 7.55E-05 | -1.5 | 1.71E-03 | 0.1 | 8.07E-01 | -1.6 | 3.80E-03 | Soluble inorganic pyrophosphatase 3 |
| AT1G01250 | -2.1 | 5.28E-04 | -1.8 | 1.50E-03 | -0.3 | 6.88E-01 | -1.9 | 1.36E-03 | 0.4 | 2.69E-01 | -2.4 | 2.85E-04 | hypothetical protein |
| AT4G30830 | -2.1 | 6.63E-06 | -1.2 | 1.65E-04 | -0.9 | 1.52E-02 | | | | | | | GTD-binding domain-containing protein |
| AT5G56300 | -2.1 | 2.82E-06 | -2.2 | 5.52E-06 | 0.1 | 7.34E-01 | | | | | | | Gibberellic acid methyltransferase 2 carboxyl methyltransferase) |
| AT2G29130 | -2.1 | 6.10E-07 | -0.3 | 2.05E-02 | -1.8 | 3.30E-05 | -0.4 | 2.04E-01 | 0.2 | 5.67E-01 | -0.6 | 8.56E-02 | Laccase-2 |
| AT3G09240 | -2.1 | 5.03E-07 | -0.9 | 5.83E-05 | -1.1 | 3.53E-04 | -1.7 | 5.77E-03 | 0.0 | 9.81E-01 | -1.7 | 5.37E-03 | Serine |

| | | | | | | | | | | | | | |
|---|---|---|---|---|---|---|---|---|---|---|---|---|---|
| AT3G03650 | -2.1 | 6.74E-08 | -1.3 | 1.28E-06 | -0.8 | 3.82E-04 | -0.4 | 3.96E-01 | 0.0 | 9.79E-01 | -0.3 | 5.12E-01 | AT3G03650 protein |
| AT4G15290 | -2.1 | 2.02E-07 | -0.3 | 1.42E-02 | -1.7 | 1.47E-05 | -1.4 | 1.55E-03 | -0.3 | 4.73E-01 | -1.1 | 2.14E-02 | Cellulose synthase-like protein B5 |
| AT5G24090 | -2.1 | 2.26E-06 | -2.0 | 6.57E-06 | -0.1 | 7.04E-01 | 0.8 | 9.94E-02 | -0.6 | 3.40E-01 | 1.4 | 3.87E-02 | Acidic endochitinase |
| AT5G63850 | -2.1 | 1.59E-08 | -1.7 | 9.13E-08 | -0.4 | 2.45E-03 | -0.8 | 1.37E-06 | -0.3 | 7.92E-03 | -0.4 | 8.36E-03 | Amino acid permease 4 |
| AT3G50400 | -2.1 | 1.63E-07 | -1.0 | 1.66E-05 | -1.1 | 1.38E-04 | 1.8 | 4.02E-06 | 1.4 | 2.59E-04 | 0.4 | 9.84E-02 | GDSL esterase |
| AT3G18170 | -2.1 | 3.81E-08 | -0.4 | 9.94E-04 | -1.6 | 6.99E-06 | -0.9 | 1.56E-05 | 0.1 | 6.68E-01 | -1.0 | 2.32E-04 | Glycosyltransferase family 61 protein |
| AT4G33790 | -2.1 | 1.56E-07 | -0.5 | 5.09E-04 | -1.5 | 2.60E-05 | -1.0 | 2.02E-02 | -0.1 | 8.17E-01 | -0.8 | 4.54E-02 | Fatty acyl-CoA reductase |
| AT1G68850 | -2.1 | 3.55E-08 | -1.0 | 2.92E-06 | -1.1 | 3.37E-05 | 1.8 | 1.87E-07 | 0.8 | 2.04E-03 | 1.0 | 2.73E-04 | Peroxidase 11 |
| AT1G01453 | -2.1 | 4.85E-06 | -0.4 | 2.07E-02 | -1.6 | 2.34E-04 | -2.1 | 1.75E-04 | -0.1 | 7.71E-01 | -2.0 | 1.48E-03 | F22L4.1 protein |
| AT5G51520 | -2.1 | 5.38E-05 | 0.1 | 7.55E-01 | -2.1 | 2.71E-04 | -1.7 | 2.73E-04 | 0.1 | 8.51E-01 | -1.8 | 2.97E-03 | hypothetical protein |
| AT2G04090 | -2.1 | 6.80E-07 | -1.4 | 8.90E-06 | -0.6 | 1.34E-02 | -1.2 | 2.30E-02 | -0.2 | 7.47E-01 | -1.0 | 1.45E-01 | Protein DETOXIFICATION 5 |
| AT1G52820 | -2.0 | 5.28E-07 | 0.3 | 1.96E-02 | -2.4 | 8.63E-06 | -1.0 | 1.34E-02 | 0.3 | 3.29E-01 | -1.4 | 5.81E-03 | hypothetical protein |
| AT3G51560 | -2.0 | 1.02E-04 | -1.5 | 7.61E-04 | -0.6 | 2.22E-01 | | | | | | | Disease resistance protein family |
| AT3G10320 | -2.0 | 2.06E-07 | -1.6 | 1.43E-06 | -0.4 | 2.66E-02 | 0.2 | 4.40E-01 | 0.2 | 6.27E-01 | 0.1 | 9.21E-01 | Xylan glycosyltransferase MUCI21 |
| AT5G10770 | -2.0 | 2.22E-08 | -1.4 | 4.42E-07 | -0.6 | 1.74E-04 | -0.1 | 5.52E-01 | 0.1 | 3.76E-01 | -0.2 | 2.01E-01 | Aspartyl protease family protein At5g10770 |
| AT1G09540 | -2.0 | 5.29E-07 | -0.8 | 1.11E-04 | -1.2 | 2.30E-04 | -0.4 | 3.14E-01 | -0.5 | 3.36E-01 | 0.0 | 9.71E-01 | Transcription factor MYB61 |
| AT2G32300 | -2.0 | 1.79E-07 | -0.3 | 3.33E-02 | -1.8 | 1.13E-05 | -0.8 | 1.16E-02 | -0.1 | 8.01E-01 | -0.7 | 2.97E-02 | Uclacyanin 1 |
| AT5G59530 | -2.0 | 5.79E-08 | -1.4 | 6.67E-07 | -0.6 | 1.58E-03 | -0.3 | 1.89E-01 | 0.2 | 5.18E-01 | -0.5 | 1.05E-01 | 1-aminocyclopropane-1-carboxylate oxidase homolog 11 |
| AT5G57200 | -2.0 | 1.85E-04 | -0.4 | 1.45E-01 | -1.6 | 4.32E-03 | | | | | | | Putative clathrin assembly protein At5g57200 |
| AT4G25480 | -2.0 | 1.85E-04 | -1.4 | 1.36E-03 | -0.6 | 2.16E-01 | | | | | | | Dehydration-responsive element-binding protein 1A |
| AT5G60760 | -2.0 | 1.48E-07 | -0.9 | 3.28E-05 | -1.1 | 7.65E-05 | 0.1 | 6.55E-01 | 0.2 | 3.81E-01 | -0.1 | 7.46E-01 | P-loop containing nucleoside triphosphate hydrolases superfamily protein |
| AT1G12950 | -2.0 | 1.01E-06 | -0.6 | 2.91E-03 | -1.4 | 9.67E-05 | -1.5 | 8.75E-04 | 0.2 | 6.84E-01 | -1.7 | 2.53E-03 | Protein DETOXIFICATION 31 |
| AT1G73860 | -2.0 | 7.27E-07 | -0.8 | 2.10E-04 | -1.2 | 2.33E-04 | -1.3 | 1.04E-01 | 0.6 | 3.58E-01 | -1.9 | 3.98E-02 | P-loop containing nucleoside triphosphate hydrolases superfamily protein |
| AT3G15700 | -2.0 | 5.54E-06 | -0.6 | 5.43E-03 | -1.4 | 5.48E-04 | -1.8 | 1.14E-02 | 0.1 | 9.20E-01 | -1.9 | 3.66E-02 | P-loop containing nucleoside triphosphate hydrolases superfamily protein |
| AT5G49140 | -2.0 | 1.34E-05 | -1.2 | 2.75E-04 | -0.8 | 3.61E-02 | | | | | | | ADP-ribosyl cyclase |
| AT2G03200 | -2.0 | 2.25E-08 | -0.7 | 1.64E-05 | -1.3 | 1.02E-05 | 1.3 | 3.83E-06 | 0.9 | 7.59E-04 | 0.4 | 2.78E-02 | At2g03200 |
| AT5G42590 | -2.0 | 1.23E-04 | -0.7 | 1.69E-02 | -1.3 | 1.06E-02 | | | | | | | Cytochrome P450 71A16 |
| AT3G49960 | -2.0 | 2.42E-06 | -0.6 | 5.69E-03 | -1.4 | 1.98E-04 | -2.4 | 7.30E-05 | 0.2 | 6.46E-01 | -2.6 | 9.48E-04 | Peroxidase 35 |
| AT1G64000 | -2.0 | 1.37E-05 | -1.1 | 3.97E-04 | -0.9 | 1.87E-02 | 2.2 | 8.78E-04 | 1.8 | 8.73E-03 | | | Probable WRKY transcription factor 56 |
| AT5G48410 | -2.0 | 6.57E-07 | -2.6 | 7.82E-07 | 0.6 | 4.54E-02 | 1.8 | 6.48E-04 | 1.0 | 6.95E-02 | 0.8 | 8.34E-02 | Glutamate receptor |
| AT5G03640 | -2.0 | 2.49E-07 | -0.7 | 9.87E-05 | -1.3 | 7.70E-05 | -1.3 | 1.10E-02 | 0.1 | 9.32E-01 | -1.3 | 1.10E-02 | Protein kinase superfamily protein |
| AT1G55740 | -2.0 | 2.25E-08 | -1.4 | 3.00E-07 | -0.6 | 3.96E-04 | -1.1 | 1.15E-05 | -0.1 | 4.31E-01 | -0.9 | 8.52E-04 | Probable galactinol--sucrose galactosyltransferase 1 |
| AT4G00700 | -2.0 | 1.90E-06 | -1.5 | 2.11E-05 | -0.5 | 4.34E-02 | -0.8 | 9.36E-03 | -0.2 | 4.34E-01 | -0.5 | 1.61E-01 | Multiple C2 domain and transmembrane region protein 9 |
| AT1G62710 | -2.0 | 1.30E-06 | -1.7 | 6.65E-06 | -0.3 | 2.11E-01 | 0.3 | 6.01E-01 | 0.5 | 3.17E-01 | -0.2 | 6.74E-01 | Vacuolar-processing enzyme beta-isozyme |
| AT2G03720 | -2.0 | 5.03E-07 | -0.2 | 9.61E-02 | -1.8 | 2.44E-05 | -1.9 | 2.78E-05 | -0.2 | 5.53E-01 | -1.7 | 1.11E-03 | Adenine nucleotide alpha hydrolases-like superfamily protein |
| AT3G10910 | -2.0 | 2.25E-08 | -1.0 | 1.64E-06 | -1.0 | 2.81E-05 | 0.0 | 9.10E-01 | 0.2 | 5.36E-01 | -0.2 | 6.83E-01 | RING-H2 finger protein ATL72 |
| AT1G13300 | -2.0 | 3.62E-03 | -1.1 | 3.53E-02 | -0.9 | 2.14E-01 | -0.1 | 8.21E-01 | -0.5 | 1.49E-01 | 0.4 | 2.80E-01 | HTH myb-type domain-containing protein |
| AT1G01380 | -2.0 | 1.30E-04 | -1.3 | 1.25E-03 | -0.7 | 1.41E-01 | | | | | | | Homeodomain-like superfamily protein |
| AT5G24100 | -2.0 | 2.92E-07 | -0.6 | 7.96E-04 | -1.4 | 3.83E-05 | -0.7 | 5.67E-02 | 0.1 | 8.47E-01 | -0.8 | 7.78E-02 | Inactive leucine-rich repeat receptor-like serine |
| AT4G29180 | -2.0 | 8.02E-07 | -0.3 | 8.28E-02 | -1.7 | 3.04E-05 | -2.5 | 5.15E-04 | -0.1 | 9.19E-01 | -2.4 | 1.66E-03 | Root hair specific 16 |
| AT5G07475 | -2.0 | 7.72E-07 | -0.7 | 5.85E-04 | -1.3 | 1.58E-04 | 1.4 | 1.57E-05 | 1.0 | 9.19E-04 | 0.4 | 9.70E-02 | Phytocyanin domain-containing protein |
| AT5G14340 | -2.0 | 5.60E-07 | -0.4 | 1.02E-02 | -1.6 | 3.57E-05 | -1.7 | 9.26E-05 | -0.1 | 8.53E-01 | -1.6 | 1.15E-03 | MYB transcription factor |
| AT1G27140 | -2.0 | 6.50E-07 | -0.3 | 6.42E-02 | -1.7 | 2.81E-05 | -0.7 | 6.89E-03 | -0.2 | 4.88E-01 | -0.5 | 1.16E-01 | Glutathione S-transferase U14 |
| AT2G41380 | -2.0 | 7.70E-07 | -2.8 | 6.68E-07 | 0.8 | 1.08E-02 | 1.7 | 6.65E-06 | -0.3 | 5.20E-01 | 1.9 | 8.05E-05 | hypothetical protein |
| AT1G05250 | -2.0 | 4.32E-06 | 0.0 | 9.98E-01 | -2.0 | 5.23E-05 | | | | | | | Peroxidase 2 |
| AT1G69240 | -2.0 | 3.11E-06 | -0.3 | 8.22E-02 | -1.7 | 9.50E-05 | -1.3 | 3.75E-03 | -0.1 | 8.94E-01 | -1.3 | 2.31E-02 | Putative methylesterase 15, chloroplastic |
| AT4G33450 | -2.0 | 5.98E-06 | -1.6 | 2.89E-05 | -0.3 | 2.62E-01 | | | | | | | MYB transcription factor |
| AT4G17280 | -2.0 | 2.02E-07 | -0.7 | 2.30E-04 | -1.3 | 4.15E-05 | 0.4 | 7.85E-02 | 0.3 | 3.24E-01 | 0.1 | 6.24E-01 | Cytochrome b561 and DOMON domain-containing protein |
| AT4G05170 | -2.0 | 1.39E-05 | -0.6 | 7.73E-03 | -1.4 | 1.40E-03 | | | | | | | Basic helix-loop-helix DNA-binding superfamily protein |
| AT1G59640 | -2.0 | 4.50E-07 | -1.5 | 3.39E-06 | -0.4 | 4.63E-02 | -0.2 | 4.59E-01 | 0.5 | 1.03E-01 | -0.8 | 3.38E-02 | Transcription factor BPE |
| AT2G41220 | -2.0 | 5.72E-08 | -1.2 | 3.18E-06 | -0.7 | 1.56E-04 | 0.2 | 3.10E-01 | 0.5 | 4.06E-02 | -0.3 | 3.90E-01 | Ferredoxin-dependent glutamate synthase 2, chloroplastic |
| AT1G05240 | -2.0 | 8.56E-04 | 0.2 | 4.81E-01 | -2.2 | 1.72E-03 | | | | | | | Peroxidase |
| AT5G19790 | -1.9 | 2.17E-05 | 0.0 | 8.76E-01 | -1.9 | 1.76E-04 | -2.0 | 1.34E-03 | 0.0 | 9.61E-01 | -2.1 | 7.40E-03 | Ethylene-responsive transcription factor RAP2-11 |
| AT3G47950 | -1.9 | 3.65E-07 | -1.2 | 8.25E-06 | -0.7 | 2.80E-03 | 0.5 | 3.20E-01 | 0.5 | 4.27E-01 | 0.0 | 9.69E-01 | ATPase 4, plasma membrane-type |
| AT4G33330 | -1.9 | 1.95E-07 | -0.5 | 1.17E-03 | -1.5 | 2.69E-05 | -0.7 | 1.24E-01 | 0.2 | 7.78E-01 | -0.9 | 6.41E-02 | Hexosyltransferase |
| AT3G47790 | -1.9 | 2.19E-08 | -2.1 | 8.06E-08 | 0.1 | 3.38E-01 | | | | | | | ABC2 homolog 7 |
| AT1G34670 | -1.9 | 7.93E-08 | -0.8 | 1.41E-05 | -1.1 | 4.81E-05 | 1.2 | 4.55E-04 | 1.2 | 1.83E-03 | 0.1 | 8.48E-01 | Transcription factor MYB93 |
| AT3G47110 | -1.9 | 2.36E-05 | -1.8 | 6.93E-05 | -0.1 | 7.51E-01 | | | | | | | Putative receptor-like protein kinase At3g47110 |
| AT4G25000 | -1.9 | 1.74E-02 | -1.1 | 8.89E-02 | -0.8 | 3.93E-01 | | | | | | | Alpha-amylase 1 |
| AT1G20450 | -1.9 | 3.07E-06 | -1.3 | 1.11E-04 | -0.6 | 1.19E-02 | -0.6 | 1.49E-02 | 0.1 | 8.43E-01 | -0.7 | 5.19E-02 | Dehydrin family protein |
| AT1G09350 | -1.9 | 2.34E-04 | -1.6 | 8.48E-04 | -0.3 | 5.35E-01 | | | | | | | Hexosyltransferase |
| AT2G17770 | -1.9 | 6.63E-07 | -1.2 | 1.36E-05 | -0.7 | 5.72E-03 | | | | | | | bZIP transcription factor 27 |
| AT2G21880 | -1.9 | 9.75E-07 | -0.4 | 1.61E-02 | -1.5 | 5.32E-05 | -1.1 | 1.75E-03 | -0.1 | 8.93E-01 | -1.0 | 1.97E-03 | RAB GTPase homolog 7A |
| AT4G15380 | -1.9 | 4.34E-08 | -1.4 | 4.74E-07 | -0.5 | 1.48E-03 | -0.3 | 2.40E-01 | 0.1 | 8.54E-01 | -0.4 | 2.41E-01 | Cytochrome P450, family 705, subfamily A, polypeptide 4 |
| AT3G46340 | -1.9 | 8.23E-06 | -0.3 | 9.18E-02 | -1.6 | 2.26E-04 | -1.8 | 2.20E-03 | 0.4 | 3.21E-01 | -2.2 | 3.73E-03 | Putative receptor-like protein kinase At3g46340 |
| AT5G59930 | -1.9 | 8.14E-06 | -0.6 | 6.63E-03 | -1.3 | 7.77E-04 | -0.2 | 5.43E-01 | -0.6 | 1.57E-01 | 0.4 | 5.17E-01 | Cysteine |
| AT1G79860 | -1.9 | 2.89E-06 | -0.3 | 2.88E-02 | -1.6 | 1.36E-04 | | | | | | | Rop guanine nucleotide exchange factor 12 |
| AT1G54940 | -1.9 | 7.21E-06 | -0.4 | 4.43E-02 | -1.5 | 2.76E-04 | -1.6 | 9.47E-03 | 0.1 | 9.24E-01 | -1.6 | 1.57E-02 | Putative UDP-glucuronate:xylan alpha-glucuronosyltransferase 4 |
| AT2G35890 | -1.9 | 4.36E-06 | -0.8 | 8.41E-04 | -1.1 | 1.54E-03 | | | | | | | Calcium-dependent protein kinase 25 |
| AT4G09990 | -1.9 | 3.55E-07 | -0.3 | 1.59E-02 | -1.6 | 2.60E-05 | -0.6 | 4.05E-02 | 0.1 | 7.72E-01 | -0.7 | 3.62E-02 | Glucuronoxylan 4-O-methyltransferase 2 |
| AT1G66470 | -1.9 | 1.50E-07 | -0.4 | 1.48E-03 | -1.5 | 1.92E-05 | -0.4 | 1.49E-01 | 0.1 | 6.71E-01 | -0.5 | 9.63E-02 | Transcription factor RHD6 |
| AT1G64930 | -1.9 | 2.25E-05 | -0.5 | 3.34E-02 | -1.4 | 1.16E-03 | | | | | | | Cytochrome P450, family 87, subfamily A, polypeptide 7 |
| AT3G13784 | -1.9 | 1.34E-03 | -1.0 | 2.80E-02 | -0.9 | 1.23E-01 | | | | | | | hypothetical protein |
| AT1G11920 | -1.9 | 7.25E-04 | -0.3 | 2.80E-01 | -1.5 | 9.35E-03 | -1.1 | 2.29E-02 | 0.3 | 5.19E-01 | -1.4 | 2.30E-02 | Pectate lyase |
| AT4G31940 | -1.9 | 1.87E-04 | -2.1 | 1.76E-04 | 0.2 | 6.53E-01 | -3.4 | 3.97E-04 | -7.8 | 9.39E-04 | 4.4 | 8.71E-02 | Xanthotoxin 5-hydroxylase CYP82C4 |
| AT5G23530 | -1.9 | 9.61E-08 | -1.1 | 7.01E-06 | -0.8 | 1.42E-04 | 0.1 | 4.04E-01 | 0.1 | 5.03E-01 | 0.0 | 9.81E-01 | Probable carboxylesterase 18 |
| AT1G48930 | -1.9 | 4.41E-06 | -0.3 | 1.17E-01 | -1.6 | 1.11E-04 | -1.1 | 5.30E-03 | 0.1 | 8.55E-01 | -1.2 | 9.09E-03 | Endoglucanase 5 |
| AT3G62590 | -1.9 | 5.61E-08 | -1.5 | 3.32E-07 | -0.3 | 2.32E-02 | 0.0 | 9.86E-01 | 0.3 | 6.65E-01 | -0.3 | 7.06E-01 | Phospholipase A1 PLIP3, chloroplastic |
| AT1G55290 | -1.9 | 1.59E-05 | -0.5 | 2.78E-02 | -1.4 | 8.52E-04 | | | | | | | Feruloyl CoA ortho-hydroxylase 2 |
| AT5G66400 | -1.9 | 8.58E-06 | -1.8 | 1.90E-05 | 0.0 | 9.35E-01 | -2.4 | 1.45E-02 | -0.6 | 4.19E-01 | -1.8 | 1.89E-01 | Dehydrin family protein |
| AT2G41970 | -1.9 | 2.52E-06 | -0.4 | 2.15E-02 | -1.5 | 1.30E-04 | -1.3 | 7.76E-03 | -0.1 | 8.23E-01 | -1.2 | 3.55E-02 | Protein kinase superfamily protein |
| AT2G18640 | -1.9 | 1.45E-06 | -0.3 | 4.01E-02 | -1.6 | 6.82E-05 | | | | | | | Geranylgeranyl pyrophosphate synthase 4 -farnesyl diphosphate synthase 4) |
| AT2G43880 | -1.9 | 1.50E-06 | 0.0 | 9.36E-01 | -1.8 | 2.81E-05 | -0.6 | 1.23E-01 | 0.3 | 5.40E-01 | -0.9 | 5.01E-02 | Pectin lyase-like superfamily protein |

| | | | | | | | | | | | | | |
|---|---|---|---|---|---|---|---|---|---|---|---|---|---|
| AT5G67430 | -1.9 | 1.90E-06 | -1.8 | 5.32E-06 | -0.1 | 8.42E-01 | 0.5 | 2.14E-01 | 1.0 | 1.89E-02 | -0.5 | 2.92E-01 | Acyl-CoA N-acyltransferases superfamily protein |
| AT1G05990 | -1.9 | 3.31E-06 | -0.3 | 3.11E-02 | -1.5 | 1.61E-04 | -2.6 | 8.46E-03 | -0.2 | 8.23E-01 | | | Calmodulin-like protein 7 |
| AT3G60330 | -1.9 | 1.04E-06 | -0.4 | 1.48E-02 | -1.4 | 5.45E-05 | -0.1 | 8.80E-01 | -0.1 | 7.13E-01 | 0.1 | 8.63E-01 | Plasma membrane ATPase |
| AT1G65060 | -1.8 | 1.79E-03 | -1.5 | 6.77E-03 | -0.4 | 5.24E-01 | 0.0 | 9.17E-01 | 1.0 | 5.56E-04 | -1.0 | 3.05E-03 | 4-coumarate:CoA ligase 3 |
| AT2G39110 | -1.8 | 2.49E-07 | -0.6 | 3.68E-04 | -1.3 | 4.45E-05 | 0.6 | 6.73E-03 | 0.6 | 1.67E-02 | 0.0 | 9.19E-01 | Protein kinase superfamily protein |
| AT1G49475 | -1.8 | 5.33E-07 | -1.0 | 3.02E-05 | -0.9 | 1.10E-03 | -0.1 | 8.19E-01 | 0.1 | 7.74E-01 | -0.2 | 5.98E-01 | B3 domain-containing protein At1g49475 |
| AT1G65310 | -1.8 | 7.85E-07 | -0.5 | 3.75E-03 | -1.3 | 7.27E-05 | -0.9 | 7.61E-03 | 0.0 | 9.27E-01 | -0.9 | 1.81E-02 | Xyloglucan endotransglucosylase |
| AT1G74830 | -1.8 | 2.41E-05 | -0.1 | 5.38E-01 | -1.7 | 2.85E-04 | -1.9 | 1.95E-03 | -0.3 | 6.08E-01 | -1.6 | 2.33E-02 | hypothetical protein |
| AT1G13635 | -1.8 | 5.71E-06 | -0.4 | 2.89E-02 | -1.5 | 2.73E-04 | 0.4 | 4.82E-01 | 0.5 | 4.42E-01 | | | hypothetical protein |
| AT4G39800 | -1.8 | 7.08E-07 | -1.2 | 3.03E-05 | -0.6 | 3.52E-03 | -0.8 | 1.42E-05 | 0.1 | 6.62E-01 | -0.9 | 4.93E-04 | Inositol-3-phosphate synthase isozyme 1 |
| AT3G02610 | -1.8 | 4.69E-07 | -1.5 | 3.00E-06 | -0.3 | 7.96E-02 | -0.9 | 6.56E-03 | -0.3 | 2.66E-01 | -0.5 | 1.42E-01 | stearoyl- 9-desaturase |
| AT2G18370 | -1.8 | 1.74E-07 | -1.2 | 5.03E-06 | -0.6 | 1.10E-03 | 0.4 | 9.21E-02 | 0.4 | 9.92E-02 | 0.0 | 9.40E-01 | Non-specific lipid-transfer protein 8 |
| AT3G54870 | -1.8 | 6.25E-06 | -0.4 | 2.20E-02 | -1.4 | 3.28E-04 | -1.1 | 5.34E-02 | 0.1 | 9.35E-01 | -1.1 | 9.24E-02 | Kinesin-like protein |
| AT4G01890 | -1.8 | 2.43E-05 | -0.1 | 6.89E-01 | -1.8 | 2.60E-04 | | | | | | | Pectin lyase-like superfamily protein |
| AT2G30040 | -1.8 | 1.29E-03 | -2.1 | 9.44E-04 | 0.3 | 6.03E-01 | 0.6 | 3.03E-02 | 0.0 | 9.29E-01 | 0.7 | 6.78E-02 | Mitogen-activated protein kinase kinase kinase 14 |
| AT5G14130 | -1.8 | 1.79E-07 | -1.2 | 2.67E-06 | -0.6 | 3.58E-03 | 0.9 | 3.33E-02 | 0.4 | 4.07E-01 | 0.5 | 2.48E-01 | Peroxidase |
| AT2G32510 | -1.8 | 6.46E-07 | -0.8 | 1.29E-04 | -1.1 | 3.07E-04 | -0.2 | 5.95E-01 | 0.6 | 3.44E-02 | -0.8 | 2.21E-02 | Mitogen-activated protein kinase kinase kinase 17 |
| AT3G07900 | -1.8 | 1.47E-06 | -0.6 | 9.46E-04 | -1.2 | 2.73E-04 | -2.5 | 2.17E-02 | 0.1 | 9.22E-01 | -2.6 | 7.22E-02 | O-fucosyltransferase 24 |
| AT3G61680 | -1.8 | 7.44E-08 | -0.6 | 5.94E-05 | -1.2 | 2.63E-05 | 0.5 | 7.98E-02 | 0.6 | 3.44E-02 | -0.1 | 6.16E-01 | Phospholipase A1 PLIP1, chloroplastic |
| AT2G18980 | -1.8 | 2.50E-07 | -0.1 | 6.01E-01 | -1.8 | 1.02E-05 | -0.7 | 4.15E-03 | 0.1 | 8.20E-01 | -0.7 | 4.03E-03 | Peroxidase 16 |
| AT3G32040 | -1.8 | 1.56E-07 | -0.8 | 3.60E-05 | -1.0 | 8.64E-05 | -0.7 | 1.63E-02 | 0.2 | 5.79E-01 | -0.9 | 3.22E-03 | Geranylgeranyl pyrophosphate synthase 12, chloroplastic -farnesyl diphosphate synthase 12) |
| AT1G51880 | -1.8 | 2.52E-06 | -0.5 | 6.62E-03 | -1.3 | 2.06E-04 | -0.7 | 5.56E-02 | -0.1 | 7.86E-01 | -0.6 | 1.98E-01 | Root hair specific 6 |
| AT3G47740 | -1.8 | 1.55E-06 | -0.3 | 6.15E-02 | -1.6 | 6.08E-05 | -0.4 | 4.17E-01 | 0.1 | 8.87E-01 | | | ABC transporter A family member 3 |
| AT2G26290 | -1.8 | 2.26E-07 | -1.1 | 4.95E-06 | -0.7 | 1.99E-03 | 1.0 | 8.67E-03 | 0.7 | 1.16E-01 | 0.4 | 3.31E-01 | Probable serine |
| AT3G18400 | -1.8 | 3.75E-08 | -1.1 | 9.21E-07 | -0.7 | 1.45E-04 | 0.9 | 2.13E-02 | 0.4 | 4.06E-01 | 0.5 | 1.20E-01 | hypothetical protein |
| AT5G12420 | -1.8 | 7.19E-08 | -0.8 | 1.50E-05 | -1.0 | 4.04E-05 | 0.8 | 4.53E-04 | 0.4 | 6.24E-02 | 0.4 | 6.59E-02 | O-acyltransferase family protein |
| AT1G29025 | -1.8 | 1.45E-07 | -0.5 | 6.26E-04 | -1.3 | 2.37E-05 | -0.7 | 3.19E-03 | 0.2 | 4.06E-01 | -0.9 | 1.65E-03 | F1K23.2 |
| AT1G26360 | -1.8 | 5.11E-05 | -0.2 | 3.49E-01 | -1.6 | 7.14E-04 | -2.8 | 4.43E-05 | -0.3 | 4.22E-01 | -2.5 | 2.27E-03 | Putative methylesterase 13, chloroplastic |
| AT3G21420 | -1.8 | 4.79E-08 | -1.5 | 2.47E-07 | -0.3 | 3.97E-02 | -0.2 | 2.66E-01 | 0.2 | 1.80E-01 | -0.4 | 4.22E-02 | Fe2OG dioxygenase domain-containing protein |
| AT3G45440 | -1.8 | 1.19E-06 | -0.9 | 1.04E-04 | -0.9 | 1.27E-03 | -0.3 | 5.19E-01 | 0.1 | 8.72E-01 | -0.4 | 5.14E-01 | Concanavalin A-like lectin protein kinase family protein |
| AT3G44550 | -1.8 | 8.79E-08 | -1.0 | 3.96E-06 | -0.7 | 2.54E-04 | 1.6 | 3.37E-06 | 1.2 | 2.27E-04 | 0.4 | 8.76E-02 | Fatty acyl-CoA reductase |
| AT1G67810 | -1.8 | 1.79E-06 | -1.6 | 7.18E-06 | -0.2 | 3.60E-01 | 0.6 | 1.39E-02 | 0.4 | 7.84E-02 | 0.1 | 7.04E-01 | SufE-like protein 2, chloroplastic |
| AT3G07490 | -1.8 | 3.47E-05 | -0.1 | 7.00E-01 | -1.7 | 3.16E-04 | -3.2 | 5.02E-05 | -0.2 | 6.06E-01 | -3.0 | 1.15E-03 | Calmodulin-like protein 3 |
| AT5G58910 | -1.8 | 1.06E-05 | -1.1 | 2.66E-04 | -0.7 | 3.12E-02 | 0.7 | 2.55E-01 | 0.3 | 7.72E-01 | | | Laccase |
| AT1G61950 | -1.8 | 5.82E-05 | 0.0 | 8.95E-01 | -1.7 | 4.27E-04 | -1.7 | 2.53E-02 | -0.3 | 7.28E-01 | -1.4 | 1.54E-01 | Calcium-dependent protein kinase 19 |
| AT3G51570 | -1.8 | 1.50E-04 | -0.3 | 1.73E-01 | -1.4 | 3.24E-03 | | | | | | | ADP-ribosyl cyclase |
| AT5G62310 | -1.8 | 2.83E-06 | -0.5 | 6.73E-03 | -1.3 | 2.40E-04 | | | | | | | Probable serine |
| AT5G59490 | -1.8 | 1.41E-05 | -0.8 | 1.79E-03 | -1.0 | 4.77E-03 | 0.3 | 5.58E-01 | 1.3 | 6.67E-03 | -1.0 | 8.96E-03 | Ripening-related protein-like; hydrolase-like |
| AT3G49190 | -1.8 | 3.90E-06 | -0.2 | 1.87E-01 | -1.6 | 9.26E-05 | -0.9 | 2.52E-02 | -0.1 | 8.52E-01 | -0.8 | 9.93E-02 | Wax ester synthase |
| AT4G22120 | -1.8 | 1.97E-08 | -1.1 | 6.67E-07 | -0.7 | 4.08E-05 | -0.3 | 1.37E-01 | 0.1 | 7.19E-01 | -0.5 | 1.17E-01 | Calcium permeable stress-gated cation channel 1 |
| AT2G30210 | -1.8 | 1.90E-07 | -0.7 | 2.72E-04 | -1.1 | 3.70E-05 | 0.1 | 6.56E-01 | 0.3 | 1.49E-01 | -0.2 | 4.12E-01 | Laccase |
| AT5G44610 | -1.8 | 1.03E-07 | -0.5 | 1.35E-03 | -1.3 | 1.39E-05 | -1.2 | 2.68E-04 | -0.4 | 2.15E-01 | -0.8 | 1.10E-02 | Plasma membrane-associated cation-binding protein 2 |
| AT1G55940 | -1.8 | 7.70E-07 | -1.2 | 1.23E-05 | -0.6 | 1.34E-02 | | | | | | | Cytochrome P450 family protein |
| AT4G21340 | -1.8 | 3.47E-05 | -0.4 | 8.76E-02 | -1.4 | 1.16E-03 | | | | | | | Transcription factor bHLH103 |
| AT2G22560 | -1.8 | 5.92E-08 | -0.7 | 1.52E-05 | -1.0 | 3.16E-05 | -0.2 | 4.84E-01 | 0.2 | 3.94E-01 | -0.4 | 1.99E-01 | Protein NETWORKED 2D |
| AT2G15370 | -1.7 | 7.04E-07 | -0.3 | 1.90E-02 | -1.4 | 3.94E-05 | -0.5 | 9.90E-02 | 0.4 | 1.44E-01 | -0.9 | 1.01E-02 | Probable fucosyltransferase 5 |
| AT4G24010 | -1.7 | 1.14E-03 | -1.3 | 6.90E-03 | -0.5 | 3.74E-01 | 0.3 | 6.37E-01 | 1.0 | 5.08E-02 | -0.7 | 2.11E-01 | Uncharacterized protein |
| AT4G22080 | -1.7 | 5.05E-06 | -0.3 | 1.18E-01 | -1.5 | 1.38E-04 | -2.2 | 1.47E-04 | 0.0 | 9.63E-01 | -2.2 | 1.18E-03 | Probable pectate lyase 16 |
| AT5G19560 | -1.7 | 2.06E-06 | -0.5 | 3.59E-03 | -1.3 | 2.24E-04 | -1.1 | 1.72E-02 | -0.2 | 6.39E-01 | -0.9 | 9.64E-02 | ROP uanine nucleotide exchange factor 10 |
| AT1G52890 | -1.7 | 3.87E-03 | -2.1 | 2.77E-03 | 0.3 | 6.50E-01 | -1.7 | 2.12E-01 | -0.6 | 6.68E-01 | -1.1 | 6.31E-01 | NAC domain-containing protein 19 |
| AT3G24300 | -1.7 | 3.13E-07 | -0.7 | 4.68E-04 | -1.1 | 6.46E-05 | -0.5 | 9.97E-03 | 0.1 | 8.46E-01 | -0.5 | 1.19E-02 | Ammonium transporter 1 member 3 |
| AT1G62570 | -1.7 | 2.71E-07 | -1.3 | 4.74E-06 | -0.5 | 4.78E-03 | 0.1 | 7.58E-01 | 0.7 | 2.92E-03 | -0.6 | 8.31E-03 | Flavin-containing monooxygenase FMO GS-OX4 |
| AT2G19060 | -1.7 | 1.51E-07 | -0.5 | 2.83E-04 | -1.2 | 2.95E-05 | -0.3 | 2.91E-01 | 0.5 | 9.82E-02 | -0.8 | 3.20E-03 | GDSL-like Lipase |
| AT2G43050 | -1.7 | 7.84E-07 | -0.5 | 1.47E-03 | -1.2 | 1.05E-04 | -0.1 | 6.54E-01 | -0.1 | 7.86E-01 | 0.0 | 9.49E-01 | Probable pectinesterase |
| AT1G66800 | -1.7 | 2.77E-04 | -0.9 | 9.06E-03 | -0.9 | 6.25E-02 | -3.4 | 4.07E-03 | 0.3 | 7.48E-01 | -3.6 | 1.03E-02 | NAD-binding Rossmann-fold superfamily protein |
| AT4G23980 | -1.7 | 9.91E-08 | -1.3 | 1.11E-06 | -0.4 | 5.39E-03 | -0.8 | 3.36E-05 | -0.3 | 6.45E-02 | -0.5 | 1.26E-02 | Auxin response factor 9 |
| AT2G22950 | -1.7 | 7.27E-05 | 0.0 | 8.27E-01 | -1.7 | 5.89E-04 | | | | | | | Putative calcium-transporting ATPase 7, plasma membrane-type -ATPase isoform 7) |
| AT1G20823 | -1.7 | 3.74E-07 | -1.1 | 1.09E-05 | -0.6 | 2.53E-03 | -1.2 | 4.00E-06 | 0.0 | 9.67E-01 | -1.2 | 3.23E-04 | RING-H2 finger protein ATL80 |
| AT2G14210 | -1.7 | 1.09E-04 | -1.4 | 5.58E-04 | -0.3 | 4.24E-01 | | | | | | | AGAMOUS-like 44 |
| AT3G28345 | -1.7 | 9.21E-07 | -1.5 | 4.79E-06 | -0.2 | 2.25E-01 | -0.3 | 4.17E-01 | -0.3 | 3.51E-01 | 0.1 | 8.88E-01 | ABC transporter B family member 15 |
| AT3G07070 | -1.7 | 9.86E-06 | -0.3 | 7.69E-02 | -1.4 | 3.09E-04 | -1.8 | 1.04E-03 | -0.2 | 6.55E-01 | -1.6 | 1.20E-02 | Probable serine |
| AT2G18800 | -1.7 | 1.19E-06 | 0.0 | 8.16E-01 | -1.7 | 2.69E-05 | -0.7 | 2.02E-02 | 0.0 | 9.32E-01 | -0.7 | 4.11E-02 | Probable xyloglucan endotransglucosylase |
| AT1G16140 | -1.7 | 9.44E-06 | -1.9 | 1.61E-05 | 0.1 | 6.71E-01 | | | | | | | Wall-associated receptor kinase-like 3 |
| AT5G03960 | -1.7 | 4.14E-07 | -0.5 | 1.09E-03 | -1.2 | 5.87E-05 | -0.5 | 9.29E-02 | 0.1 | 9.00E-01 | -0.5 | 7.95E-02 | Protein IQ-DOMAIN 12 |
| AT3G57540 | -1.7 | 3.11E-07 | -1.7 | 8.46E-07 | 0.0 | 8.90E-01 | -0.5 | 7.58E-02 | 0.0 | 9.48E-01 | -0.4 | 2.04E-01 | Remorin 4.1 |
| AT1G73700 | -1.7 | 1.33E-05 | -1.4 | 6.92E-05 | -0.3 | 2.69E-01 | | | | | | | Protein DETOXIFICATION 17 |
| AT1G51470 | -1.7 | 3.05E-07 | -0.3 | 1.81E-02 | -1.4 | 2.10E-05 | -0.7 | 1.42E-02 | -0.1 | 8.96E-01 | -0.7 | 5.01E-02 | Myrosinase 5 |
| AT1G53680 | -1.7 | 2.54E-07 | -0.3 | 2.34E-02 | -1.5 | 1.80E-05 | -0.9 | 2.17E-03 | 0.2 | 5.27E-01 | -1.1 | 3.22E-03 | Glutathione S-transferase U28 |
| AT1G24620 | -1.7 | 6.66E-06 | -0.1 | 6.46E-01 | -1.7 | 9.04E-05 | -1.1 | 7.12E-03 | -0.1 | 8.68E-01 | -1.0 | 1.10E-02 | Probable calcium-binding protein CML25 |
| AT4G20140 | -1.7 | 2.74E-08 | -0.8 | 3.46E-06 | -0.9 | 2.63E-05 | 0.0 | 9.80E-01 | 0.2 | 7.43E-01 | -0.2 | 8.00E-01 | LRR receptor-like serine |
| AT4G15393 | -1.7 | 7.35E-08 | -1.0 | 8.49E-06 | -0.8 | 8.64E-05 | -0.9 | 2.42E-05 | 0.3 | 1.46E-01 | -1.2 | 5.95E-05 | Cytochrome P450, family 702, subfamily A, polypeptide 5 |
| AT4G17550 | -1.7 | 2.78E-07 | -0.6 | 4.63E-04 | -1.1 | 4.64E-05 | -1.1 | 1.22E-04 | -0.1 | 7.73E-01 | -1.0 | 1.27E-03 | Putative glycerol-3-phosphate transporter 4 |
| AT2G02250 | -1.7 | 4.73E-06 | -0.3 | 7.67E-02 | -1.4 | 1.77E-04 | | | | | | | Putative F-box protein PP2-B2 |
| AT3G23630 | -1.7 | 2.25E-06 | -0.9 | 1.31E-04 | -0.8 | 4.14E-03 | 0.7 | 2.28E-01 | 0.3 | 6.85E-01 | 0.4 | 6.17E-01 | Adenylate isopentenyltransferase 7, mitochondrial |
| AT5G01600 | -1.7 | 1.08E-06 | -1.7 | 2.74E-06 | 0.0 | 9.09E-01 | -1.2 | 3.88E-05 | -0.7 | 6.36E-03 | -0.5 | 9.66E-02 | Ferritin-1, chloroplastic |
| AT5G01520 | -1.7 | 1.50E-07 | -1.0 | 8.49E-06 | -0.7 | 3.45E-04 | -1.4 | 1.26E-04 | -0.5 | 1.02E-01 | -1.0 | 8.23E-03 | E3 ubiquitin-protein ligase AIRP2 |
| AT3G25290 | -1.7 | 5.26E-07 | -0.9 | 6.86E-05 | -0.8 | 4.81E-04 | -0.7 | 2.74E-04 | 0.1 | 6.80E-01 | -0.8 | 1.49E-03 | Cytochrome b561 and DOMON domain-containing protein At3g25290 |
| AT1G16440 | -1.7 | 4.77E-06 | -0.2 | 1.15E-01 | -1.5 | 1.41E-04 | -1.3 | 5.83E-02 | 0.2 | 8.28E-01 | -1.4 | 6.91E-02 | Serine |
| AT3G02620 | -1.7 | 2.49E-04 | -1.4 | 8.74E-04 | -0.3 | 5.58E-01 | -2.1 | 1.23E-04 | 0.5 | 1.38E-01 | -2.5 | 4.74E-04 | stearoyl- 9-desaturase |
| AT5G42180 | -1.7 | 1.29E-07 | -0.3 | 1.43E-02 | -1.4 | 1.09E-05 | -0.4 | 2.43E-01 | 0.2 | 5.18E-01 | -0.6 | 5.68E-02 | peroxidase |
| AT1G73580 | -1.7 | 6.47E-06 | 0.1 | 6.67E-01 | -1.8 | 5.69E-05 | -0.9 | 8.09E-02 | 0.0 | 9.42E-01 | -0.8 | 1.31E-01 | Protein C2-DOMAIN ABA-RELATED 3 |
| AT4G26010 | -1.7 | 9.39E-06 | -0.3 | 9.33E-02 | -1.4 | 2.55E-04 | -1.9 | 1.29E-02 | -0.2 | 7.45E-01 | -1.7 | 9.22E-02 | Peroxidase |
| AT1G80730 | -1.7 | 1.68E-06 | -0.8 | 1.75E-04 | -0.9 | 1.55E-03 | | | | | | | Zinc finger protein 1 |

| | | | | | | | | | | | | | |
|---|---|---|---|---|---|---|---|---|---|---|---|---|---|
| AT2G38390 | -1.7 | 2.16E-06 | -0.7 | 1.71E-03 | -1.0 | 4.45E-04 | -0.4 | 8.17E-02 | 0.1 | 7.75E-01 | -0.5 | 7.40E-02 | Peroxidase 23 |
| AT1G33700 | -1.7 | 7.29E-07 | -0.5 | 4.13E-03 | -1.2 | 6.15E-05 | -1.4 | 8.51E-05 | 0.0 | 9.08E-01 | -1.3 | 2.29E-03 | Non-lysosomal glucosylceramidase |
| AT2G18230 | -1.7 | 1.31E-06 | -1.4 | 1.06E-05 | -0.3 | 8.61E-02 | -0.4 | 1.87E-02 | 0.3 | 5.63E-02 | -0.7 | 3.70E-03 | Soluble inorganic pyrophosphatase 2 |
| AT5G48400 | -1.7 | 5.05E-05 | -2.2 | 3.43E-05 | 0.5 | 2.03E-01 | 1.3 | 4.43E-03 | 0.8 | 1.35E-01 | 0.6 | 1.39E-01 | Glutamate receptor 1.2 |
| AT2G44110 | -1.7 | 6.30E-06 | -0.4 | 3.16E-02 | -1.3 | 2.91E-04 | -1.1 | 1.78E-02 | 0.1 | 8.96E-01 | -1.1 | 3.74E-02 | MLO-like protein |
| AT2G38760 | -1.7 | 5.81E-07 | -0.7 | 1.04E-04 | -0.9 | 3.51E-04 | -0.5 | 1.49E-01 | 0.1 | 7.77E-01 | -0.6 | 1.16E-01 | Annexin D3 |
| AT5G43780 | -1.7 | 5.22E-08 | -1.0 | 4.77E-06 | -0.7 | 7.42E-05 | -0.9 | 2.95E-06 | -0.2 | 2.59E-01 | -0.7 | 1.11E-03 | ATP sulfurylase 4, chloroplastic |
| AT1G01360 | -1.7 | 1.62E-07 | -1.4 | 1.11E-06 | -0.3 | 4.15E-02 | -0.6 | 3.24E-04 | -0.2 | 2.15E-01 | -0.4 | 4.23E-02 | Abscisic acid receptor PYL9 |
| AT4G15396 | -1.7 | 3.68E-06 | -1.0 | 9.77E-05 | -0.7 | 1.23E-02 | | | | | | | At4g15396 |
| AT1G33800 | -1.7 | 7.29E-08 | -0.4 | 2.50E-03 | -1.3 | 1.02E-05 | -0.2 | 4.63E-01 | 0.2 | 4.24E-01 | -0.4 | 8.76E-02 | Glucuronoxylan 4-O-methyltransferase 3 |
| AT1G09170 | -1.7 | 3.99E-07 | -0.6 | 3.37E-04 | -1.1 | 9.47E-05 | -0.2 | 5.37E-01 | 0.3 | 4.92E-01 | -0.5 | 2.53E-01 | P-loop nucleoside triphosphate hydrolases superfamily protein with CH domain-containing protein |
| AT5G37990 | -1.7 | 1.80E-07 | -1.2 | 2.78E-06 | -0.4 | 4.67E-03 | -0.5 | 1.81E-01 | 0.4 | 3.75E-01 | -0.9 | 8.26E-02 | Probable S-adenosylmethionine-dependent methyltransferase At5g37990 |
| AT4G28850 | -1.7 | 9.56E-06 | -0.6 | 4.24E-03 | -1.0 | 1.57E-03 | -2.2 | 1.73E-04 | 0.1 | 8.01E-01 | -2.3 | 2.02E-03 | Probable xyloglucan endotransglucosylase |
| AT1G62320 | -1.7 | 2.23E-06 | -0.4 | 1.42E-02 | -1.3 | 1.42E-04 | -0.3 | 3.79E-01 | 0.2 | 5.81E-01 | -0.4 | 2.07E-01 | ERD family protein |
| AT4G00680 | -1.7 | 7.01E-07 | -0.1 | 4.48E-01 | -1.6 | 2.10E-05 | -1.3 | 5.80E-04 | 0.1 | 7.79E-01 | -1.4 | 2.13E-03 | ADF-H domain-containing protein |
| AT2G27510 | -1.7 | 4.41E-06 | -0.7 | 2.24E-03 | -0.9 | 9.58E-04 | 0.1 | 5.42E-01 | 0.3 | 9.73E-02 | -0.2 | 3.40E-01 | Ferredoxin-3, chloroplastic |
| AT2G27010 | -1.7 | 4.01E-07 | -0.5 | 1.35E-03 | -1.2 | 5.11E-05 | -0.9 | 3.20E-03 | 0.1 | 8.05E-01 | -1.0 | 7.80E-03 | Cytochrome P450, family 705, subfamily A, polypeptide 9 |
| AT4G10390 | -1.7 | 2.70E-06 | -0.8 | 2.58E-04 | -0.8 | 2.25E-03 | -0.3 | 3.05E-01 | 0.0 | 9.92E-01 | -0.3 | 3.46E-01 | Probable receptor-like protein kinase At4g10390 |
| AT2G45890 | -1.7 | 7.04E-07 | -0.3 | 1.53E-02 | -1.3 | 4.27E-05 | -0.3 | 1.70E-01 | 0.2 | 4.32E-01 | -0.5 | 8.42E-02 | Rop guanine nucleotide exchange factor 4 |
| AT1G04360 | -1.7 | 1.55E-06 | -0.5 | 2.05E-03 | -1.1 | 2.20E-04 | 0.8 | 8.01E-03 | 0.5 | 1.46E-01 | 0.3 | 2.46E-01 | RING-H2 finger protein ATL1 |
| AT3G16440 | -1.6 | 1.54E-06 | -0.3 | 4.32E-02 | -1.3 | 5.90E-05 | -0.8 | 1.11E-02 | 0.0 | 9.57E-01 | -0.8 | 1.55E-02 | Myrosinase-binding protein-like protein-300B |
| AT3G43890 | -1.6 | 2.02E-06 | -0.6 | 1.30E-03 | -1.1 | 4.16E-04 | -1.6 | 2.21E-03 | -0.4 | 4.24E-01 | -1.3 | 5.80E-02 | Cysteine |
| AT5G04960 | -1.6 | 1.00E-05 | -0.3 | 1.95E-01 | -1.4 | 1.77E-04 | -2.2 | 2.71E-06 | -0.2 | 5.70E-01 | -2.0 | 1.49E-04 | Probable pectinesterase |
| AT5G16600 | -1.6 | 3.78E-07 | -0.7 | 6.50E-05 | -0.9 | 2.61E-04 | -0.1 | 6.87E-01 | 0.4 | 2.26E-01 | -0.5 | 7.74E-02 | MYB transcription factor |
| AT5G01200 | -1.6 | 5.96E-06 | -1.3 | 4.46E-05 | -0.4 | 1.64E-01 | -0.8 | 1.01E-02 | -0.2 | 5.31E-01 | -0.6 | 1.27E-01 | hypothetical protein |
| AT2G19810 | -1.6 | 1.73E-06 | -1.3 | 1.37E-05 | -0.3 | 9.65E-02 | -0.8 | 4.05E-05 | -0.7 | 2.11E-04 | 0.0 | 8.55E-01 | Zinc finger CCCH domain-containing protein 20 |
| AT5G18350 | -1.6 | 6.79E-06 | -1.1 | 1.07E-04 | -0.5 | 4.67E-02 | 0.6 | 1.41E-01 | 0.4 | 3.55E-01 | 0.1 | 7.65E-01 | Disease resistance protein family |
| AT1G04180 | -1.6 | 4.71E-06 | -1.0 | 1.15E-04 | -0.6 | 1.81E-02 | 0.0 | 9.23E-01 | 0.1 | 8.73E-01 | -0.1 | 8.23E-01 | Probable indole-3-pyruvate monooxygenase YUCCA9 |
| AT3G07000 | -1.6 | 8.67E-06 | -0.5 | 6.56E-03 | -1.1 | 9.78E-04 | -0.5 | 1.52E-01 | 0.0 | 9.19E-01 | -0.5 | 1.67E-01 | Zinc finger PHD-type domain-containing protein |
| AT1G74010 | -1.6 | 4.15E-05 | -0.9 | 1.77E-03 | -0.8 | 2.12E-02 | 0.0 | 9.86E-01 | 0.7 | 7.71E-02 | -0.7 | 8.36E-02 | At1g74010 |
| AT3G01990 | -1.6 | 8.05E-07 | -1.4 | 3.54E-06 | -0.2 | 2.95E-01 | | | | | | | ACT domain-containing protein ACR |
| AT4G10370 | -1.6 | 2.69E-05 | -0.5 | 2.41E-02 | -1.2 | 1.85E-03 | -1.0 | 1.03E-01 | 0.2 | 8.01E-01 | -1.2 | 9.54E-02 | Uncharacterized protein AT4g10370 |
| AT4G07960 | -1.6 | 6.59E-06 | -0.7 | 1.24E-03 | -0.9 | 2.24E-03 | -0.2 | 7.76E-01 | 0.5 | 2.69E-01 | -0.7 | 2.11E-01 | Probable xyloglucan glycosyltransferase 12 |
| AT5G06090 | -1.6 | 6.65E-06 | -0.9 | 3.88E-04 | -0.8 | 8.09E-03 | 1.7 | 3.90E-05 | 0.8 | 3.01E-02 | 0.9 | 7.81E-03 | Glycerol-3-phosphate acyltransferase 7 |
| AT5G63590 | -1.6 | 4.45E-07 | -0.8 | 7.19E-05 | -0.9 | 3.37E-04 | 0.2 | 4.79E-01 | 0.5 | 4.43E-02 | -0.3 | 2.69E-01 | Flavonol synthase 3 |
| AT5G55590 | -1.6 | 5.47E-06 | -0.7 | 1.40E-03 | -1.0 | 1.69E-03 | 1.7 | 1.61E-03 | 1.7 | 3.57E-03 | 0.0 | 9.81E-01 | Pectin lyase-like superfamily protein |
| AT3G23090 | -1.6 | 7.03E-08 | -0.6 | 4.08E-05 | -1.0 | 2.69E-05 | -0.5 | 7.14E-03 | 0.1 | 8.27E-01 | -0.6 | 1.18E-02 | TPX2 protein family |
| AT3G58810 | -1.6 | 2.14E-05 | -1.5 | 7.02E-05 | -0.1 | 5.99E-01 | -0.8 | 1.62E-03 | -1.5 | 6.54E-05 | 0.6 | 1.21E-02 | Metal tolerance protein A2 |
| AT2G37090 | -1.6 | 1.09E-06 | -0.4 | 7.46E-03 | -1.3 | 8.71E-05 | -0.5 | 2.01E-01 | -0.1 | 8.46E-01 | -0.4 | 3.24E-01 | Beta-1,4-xylosyltransferase IRX9 |
| AT1G27030 | -1.6 | 1.52E-07 | -1.1 | 3.53E-06 | -0.5 | 1.68E-03 | -0.6 | 1.64E-03 | 0.2 | 3.39E-01 | -0.8 | 7.92E-04 | 2-oxoadipate dioxygenase |
| AT1G30870 | -1.6 | 6.29E-06 | -0.2 | 3.09E-01 | -1.4 | 9.66E-05 | -1.0 | 1.36E-03 | 0.0 | 9.23E-01 | -1.1 | 7.11E-03 | Peroxidase 7 |
| AT2G22290 | -1.6 | 1.74E-04 | 0.0 | 9.64E-01 | -1.6 | 1.02E-03 | | | | | | | Ras-related protein RABH1d |
| AT4G16640 | -1.6 | 3.10E-06 | -0.9 | 1.53E-04 | -0.7 | 5.65E-03 | 0.3 | 4.91E-01 | 0.4 | 4.52E-01 | -0.1 | 9.22E-01 | Peptidase metallopeptidase domain-containing protein |
| AT5G47990 | -1.6 | 1.28E-03 | -1.3 | 4.76E-03 | -0.4 | 4.98E-01 | -0.7 | 2.88E-01 | 1.1 | 4.52E-02 | -1.8 | 2.44E-02 | Cytochrome P450 705A5 |
| AT4G23990 | -1.6 | 5.40E-05 | -1.3 | 2.82E-04 | -0.3 | 3.70E-01 | 0.3 | 5.75E-01 | 1.3 | 4.02E-03 | -1.0 | 1.33E-02 | Cellulose synthase like G3 |
| AT4G25810 | -1.6 | 2.13E-04 | -0.8 | 7.58E-03 | -0.8 | 4.42E-02 | -1.3 | 1.73E-03 | -0.5 | 1.25E-01 | -0.7 | 1.58E-01 | XTR6 |
| AT3G10470 | -1.6 | 7.59E-06 | -0.7 | 1.29E-03 | -0.9 | 3.07E-03 | | | | | | | C2H2-type zinc finger family protein |
| AT2G35270 | -1.6 | 3.36E-06 | -0.9 | 1.92E-04 | -0.7 | 5.35E-03 | -0.3 | 1.98E-01 | 0.1 | 8.64E-01 | -0.4 | 4.80E-02 | AT-hook motif nuclear-localized protein 21 |
| AT3G17070 | -1.6 | 1.04E-05 | -1.0 | 2.13E-04 | -0.6 | 3.50E-02 | -0.6 | 9.97E-03 | 0.0 | 8.78E-01 | -0.7 | 2.69E-02 | peroxidase |
| AT1G77280 | -1.6 | 2.87E-07 | -1.2 | 3.95E-06 | -0.4 | 9.60E-03 | -0.1 | 5.88E-01 | -0.1 | 6.95E-01 | 0.0 | 9.66E-01 | Kinase with adenine nucleotide alpha hydrolases-like domain-containing protein |
| AT1G65180 | -1.6 | 4.15E-06 | -0.7 | 1.07E-03 | -0.9 | 1.51E-03 | -0.5 | 2.18E-01 | 0.3 | 4.54E-01 | -0.8 | 9.04E-02 | Cysteine |
| AT5G26220 | -1.6 | 6.07E-05 | -0.8 | 3.07E-03 | -0.8 | 2.36E-02 | 1.6 | 7.30E-03 | 0.9 | 1.66E-01 | 0.7 | 2.42E-01 | Gamma-glutamylcyclotransferase |
| AT1G47600 | -1.6 | 3.50E-07 | -0.3 | 1.87E-02 | -1.3 | 2.43E-05 | -0.6 | 3.86E-02 | -0.2 | 5.17E-01 | -0.4 | 2.85E-01 | Beta glucosidase 34 |
| AT4G05100 | -1.6 | 7.06E-06 | -0.7 | 9.10E-04 | -0.9 | 3.71E-03 | | | | | | | Transcription factor MYB74 |
| AT5G49630 | -1.6 | 6.41E-07 | -1.1 | 1.86E-05 | -0.5 | 4.48E-03 | -0.7 | 6.53E-05 | 0.0 | 8.76E-01 | -0.6 | 1.83E-03 | Amino acid permease 6 |
| AT5G42580 | -1.6 | 1.36E-04 | -1.3 | 5.06E-04 | -0.3 | 4.94E-01 | | | | | | | Cytochrome P450 705A12 |
| AT1G72125 | -1.6 | 5.15E-07 | -0.6 | 4.12E-04 | -1.0 | 1.24E-04 | -0.5 | 1.48E-01 | -0.1 | 8.63E-01 | -0.4 | 3.37E-01 | Protein NRT1 |
| AT5G49870 | -1.6 | 3.44E-07 | -0.2 | 6.39E-02 | -1.4 | 1.89E-05 | -0.7 | 9.19E-03 | 0.1 | 8.52E-01 | -0.8 | 1.65E-02 | Jacalin-related lectin 48 |
| AT1G73340 | -1.6 | 5.22E-06 | -0.4 | 1.12E-02 | -1.2 | 4.32E-04 | -1.3 | 2.45E-02 | -0.2 | 7.95E-01 | -1.1 | 8.76E-02 | Cytochrome P450 superfamily protein |
| AT5G58010 | -1.6 | 2.16E-06 | 0.0 | 7.34E-01 | -1.5 | 3.57E-05 | -0.3 | 3.45E-01 | -0.2 | 5.88E-01 | -0.1 | 8.48E-01 | Transcription factor LRL3 |
| AT4G17220 | -1.6 | 3.51E-07 | -0.3 | 6.35E-03 | -1.3 | 2.95E-05 | -0.7 | 2.54E-02 | 0.0 | 9.44E-01 | -0.7 | 9.08E-02 | Microtubule-associated protein 70-5 |
| AT3G13840 | -1.6 | 5.21E-06 | -0.4 | 1.23E-02 | -1.2 | 4.47E-04 | | | | | | | Scarecrow-like protein 29 |
| AT5G47980 | -1.6 | 1.42E-04 | -1.4 | 4.15E-04 | -0.2 | 6.19E-01 | 0.0 | 9.73E-01 | 1.8 | 1.01E-02 | -1.7 | 6.04E-03 | BAHD acyltransferase At5g47980 |
| AT4G33880 | -1.6 | 2.62E-06 | -0.3 | 6.69E-02 | -1.3 | 9.53E-05 | -1.0 | 1.31E-03 | 0.2 | 4.92E-01 | -1.2 | 1.71E-03 | Transcription factor RSL2 |
| AT1G16120 | -1.6 | 3.40E-06 | -1.3 | 1.98E-05 | -0.3 | 2.04E-01 | | | | | | | Wall-associated receptor kinase-like 1 |
| AT3G46710 | -1.6 | 1.79E-05 | -0.7 | 1.90E-03 | -0.8 | 7.50E-03 | | | | | | | Putative disease resistance RPP13-like protein 2 |
| AT3G06460 | -1.6 | 5.91E-05 | -0.1 | 6.47E-01 | -1.5 | 5.10E-04 | -0.6 | 8.86E-02 | 0.2 | 6.49E-01 | -0.8 | 5.97E-02 | F24P17.4 protein |
| AT5G65230 | -1.6 | 3.92E-06 | -0.9 | 1.65E-04 | -0.7 | 8.17E-03 | 0.7 | 2.05E-01 | 0.9 | 1.17E-01 | -0.2 | 7.59E-01 | Transcription factor MYB53 |
| AT5G36260 | -1.6 | 8.42E-06 | -0.3 | 5.67E-02 | -1.3 | 3.39E-04 | -0.6 | 3.65E-01 | -0.1 | 9.31E-01 | -0.5 | 5.94E-01 | Aspartic proteinase 36 |
| AT2G38090 | -1.6 | 1.15E-06 | -1.1 | 1.34E-05 | -0.4 | 3.28E-02 | -0.4 | 5.26E-02 | -0.1 | 7.47E-01 | -0.3 | 2.14E-01 | Putative MYB family transcription factor |
| AT3G46400 | -1.6 | 1.71E-06 | -0.1 | 2.30E-01 | -1.4 | 4.65E-05 | -0.9 | 8.59E-02 | 0.0 | 9.57E-01 | -0.8 | 1.25E-01 | Leucine-rich repeat protein kinase family protein |
| AT4G01480 | -1.6 | 2.48E-07 | -0.6 | 2.12E-04 | -0.9 | 6.32E-05 | -0.6 | 1.35E-03 | 0.1 | 6.38E-01 | -0.7 | 3.17E-03 | Soluble inorganic pyrophosphatase 5 |
| AT5G67520 | -1.6 | 8.14E-07 | -0.9 | 3.04E-05 | -0.6 | 3.15E-03 | -0.4 | 3.07E-01 | 0.3 | 4.69E-01 | -0.7 | 1.19E-01 | Adenylyl-sulfate kinase |
| AT1G26680 | -1.6 | 4.71E-05 | -0.5 | 2.33E-02 | -1.1 | 3.75E-03 | | | | | | | Transcriptional factor B3 family protein |
| AT1G57560 | -1.6 | 2.46E-05 | -1.0 | 5.82E-04 | -0.6 | 4.43E-02 | -0.3 | 4.80E-01 | -0.3 | 5.57E-01 | 0.0 | 9.90E-01 | At1g57560 |
| AT1G66020 | -1.6 | 2.76E-06 | -1.6 | 6.27E-06 | 0.0 | 9.28E-01 | -0.1 | 8.95E-01 | 0.1 | 9.28E-01 | | | Uncharacterized protein |
| AT5G56200 | -1.6 | 8.64E-06 | -0.9 | 4.35E-04 | -0.7 | 1.28E-02 | | | | | | | C2H2 type zinc finger transcription factor family |
| AT4G09130 | -1.6 | 1.08E-04 | -0.5 | 4.87E-02 | -1.1 | 6.20E-03 | | | | | | | Putative RING-H2 finger protein ATL37 |
| AT5G01330 | -1.6 | 9.51E-06 | 0.0 | 8.45E-01 | -1.5 | 1.01E-04 | -1.2 | 1.15E-02 | -0.1 | 7.87E-01 | -1.1 | 5.48E-02 | Pyruvate decarboxylase 3 |
| AT2G45290 | -1.5 | 1.10E-07 | -0.8 | 1.93E-05 | -0.8 | 7.81E-05 | 0.1 | 6.87E-01 | 0.3 | 1.93E-01 | -0.2 | 4.55E-01 | Transketolase-2, chloroplastic |
| AT3G01190 | -1.5 | 5.54E-08 | -0.1 | 2.71E-01 | -1.4 | 6.99E-06 | -0.6 | 7.57E-03 | 0.1 | 7.59E-01 | -0.7 | 3.34E-03 | Peroxidase 27 |
| AT3G51860 | -1.5 | 8.66E-05 | -1.8 | 7.29E-05 | 0.3 | 4.35E-01 | -0.1 | 7.84E-01 | 0.0 | 9.78E-01 | -0.1 | 8.38E-01 | Vacuolar cation |
| AT5G67080 | -1.5 | 1.35E-02 | -0.9 | 8.09E-02 | -0.6 | 3.93E-01 | | | | | | | Mitogen-activated protein kinase kinase kinase 19 |

| | | | | | | | | | | | | | |
|---|---|---|---|---|---|---|---|---|---|---|---|---|---|
| AT1G05530 | -1.5 | 2.44E-05 | -0.1 | 4.64E-01 | -1.4 | 3.42E-04 | -0.4 | 3.24E-01 | 0.0 | 9.68E-01 | -0.4 | 4.57E-01 | UDP-glycosyltransferase 75B2 -beta-D-glucosyl transferase 2) |
| AT5G13910 | -1.5 | 1.43E-05 | -1.0 | 2.01E-04 | -0.5 | 6.49E-02 | 0.4 | 3.26E-01 | 0.4 | 4.15E-01 | 0.0 | 9.80E-01 | Ethylene-responsive transcription factor LEP |
| AT5G13700 | -1.5 | 5.37E-06 | -1.1 | 6.64E-05 | -0.4 | 6.79E-02 | | | | | | | Polyamine oxidase 1 -acetylpolyamine oxidase) |
| AT3G45710 | -1.5 | 7.40E-08 | -0.4 | 4.50E-04 | -1.1 | 1.23E-05 | -0.7 | 2.38E-03 | 0.4 | 9.10E-02 | -1.1 | 1.25E-03 | Protein NRT1 |
| AT2G38995 | -1.5 | 1.12E-03 | -0.6 | 6.58E-02 | -0.9 | 5.85E-02 | 0.0 | 9.11E-01 | 0.2 | 6.03E-01 | -0.1 | 6.91E-01 | O-acyltransferase family protein |
| AT3G57380 | -1.5 | 1.83E-07 | -1.4 | 7.56E-07 | -0.1 | 3.65E-01 | 0.7 | 2.21E-02 | 0.1 | 8.39E-01 | 0.6 | 8.53E-02 | Glycosyltransferase |
| AT3G21240 | -1.5 | 7.03E-08 | -0.7 | 1.89E-05 | -0.8 | 3.57E-05 | -0.4 | 3.84E-02 | 0.1 | 6.46E-01 | -0.5 | 3.78E-02 | 4-coumarate:CoA ligase 2 |
| AT4G34131 | -1.5 | 5.07E-06 | -1.0 | 1.23E-04 | -0.5 | 2.05E-02 | -0.3 | 2.65E-01 | 0.1 | 8.56E-01 | -0.4 | 1.27E-01 | UDP-glycosyltransferase 73B3 |
| AT5G09220 | -1.5 | 1.58E-07 | -0.7 | 1.05E-04 | -0.9 | 5.63E-05 | -1.2 | 1.32E-06 | -0.5 | 1.23E-02 | -0.7 | 5.96E-03 | Amino acid permease 2 |
| AT1G51680 | -1.5 | 8.62E-08 | -0.7 | 3.69E-05 | -0.8 | 4.04E-05 | 0.0 | 7.52E-01 | 0.1 | 3.74E-01 | -0.2 | 3.30E-01 | 4-coumarate:CoA ligase 1 |
| AT5G57190 | -1.5 | 2.01E-05 | -1.2 | 1.13E-04 | -0.3 | 2.89E-01 | 0.2 | 5.99E-01 | 0.3 | 3.90E-01 | -0.1 | 7.67E-01 | phosphatidylserine decarboxylase |
| AT5G19100 | -1.5 | 1.29E-04 | -0.5 | 3.07E-02 | -1.0 | 1.11E-02 | -0.9 | 7.31E-02 | -0.2 | 7.32E-01 | -0.7 | 2.31E-01 | Peptidase A1 domain-containing protein |
| AT3G57010 | -1.5 | 7.07E-06 | -0.7 | 9.29E-04 | -0.8 | 3.43E-03 | 0.0 | 8.76E-01 | 0.4 | 1.01E-01 | -0.3 | 1.74E-01 | Protein STRICTOSIDINE SYNTHASE-LIKE 8 |
| AT1G21210 | -1.5 | 3.58E-07 | -1.0 | 1.24E-05 | -0.6 | 1.97E-03 | -0.1 | 6.58E-01 | 0.2 | 5.98E-01 | -0.3 | 3.87E-01 | Wall-associated receptor kinase 4 |
| AT4G39210 | -1.5 | 8.43E-06 | -1.3 | 4.11E-05 | -0.2 | 3.27E-01 | 0.2 | 4.17E-01 | 0.7 | 1.67E-03 | -0.5 | 1.34E-02 | Glucose-1-phosphate adenylyltransferase |
| AT2G37290 | -1.5 | 3.52E-06 | -0.8 | 2.96E-04 | -0.8 | 3.47E-03 | | | | | | | Ypt |
| AT4G35270 | -1.5 | 4.87E-07 | -1.1 | 1.02E-05 | -0.4 | 6.50E-03 | 0.0 | 8.89E-01 | 0.5 | 4.22E-02 | -0.5 | 7.85E-02 | Protein NLP2 |
| AT4G25160 | -1.5 | 7.88E-05 | -0.4 | 8.35E-02 | -1.2 | 2.82E-03 | -0.4 | 5.29E-01 | 0.4 | 4.39E-01 | -0.8 | 1.39E-01 | U-box domain-containing protein 35 |
| AT3G63010 | -1.5 | 1.32E-06 | -1.4 | 4.08E-06 | -0.1 | 6.72E-01 | -0.3 | 1.21E-01 | 0.0 | 8.49E-01 | -0.2 | 3.43E-01 | Gibberellin receptor GID1B |
| AT1G64480 | -1.5 | 1.80E-05 | -0.7 | 2.58E-03 | -0.9 | 5.64E-03 | | | | | | | Calcineurin B-like protein |
| AT1G11300 | -1.5 | 2.27E-05 | -1.5 | 4.27E-05 | 0.0 | 9.46E-01 | 0.7 | 5.53E-02 | 0.7 | 5.40E-02 | -0.1 | 9.14E-01 | Receptor-like serine |
| AT4G20450 | -1.5 | 3.75E-05 | 0.0 | 8.50E-01 | -1.5 | 2.21E-04 | -2.0 | 6.77E-04 | 0.1 | 7.85E-01 | -2.1 | 3.36E-03 | Leucine-rich repeat protein kinase family protein |
| AT1G20440 | -1.5 | 1.95E-07 | -0.7 | 1.44E-04 | -0.8 | 6.48E-05 | -0.7 | 2.42E-04 | -0.1 | 5.70E-01 | -0.6 | 7.05E-03 | Dehydrin COR47 |
| AT1G72230 | -1.5 | 7.29E-08 | -0.6 | 2.95E-05 | -0.9 | 3.16E-05 | 0.3 | 1.24E-01 | 0.4 | 5.07E-02 | -0.1 | 6.17E-01 | hypothetical protein |
| AT2G24430 | -1.5 | 1.73E-06 | -1.3 | 7.65E-06 | -0.2 | 3.49E-01 | | | | | | | hypothetical protein |
| AT5G22410 | -1.5 | 4.15E-05 | -0.3 | 1.38E-01 | -1.2 | 1.14E-03 | -2.0 | 4.07E-03 | -0.2 | 7.50E-01 | -1.8 | 4.48E-02 | Peroxidase 60 |
| AT5G14640 | -1.5 | 5.14E-07 | -1.1 | 7.18E-06 | -0.4 | 1.52E-02 | -0.1 | 6.70E-01 | 0.1 | 4.23E-01 | -0.2 | 3.09E-01 | Shaggy-related protein kinase epsilon |
| AT3G14440 | -1.5 | 1.41E-05 | -1.0 | 2.71E-04 | -0.5 | 4.68E-02 | -1.1 | 9.67E-03 | -0.1 | 8.76E-01 | -1.0 | 4.13E-02 | 9-cis-epoxycarotenoid dioxygenase NCED3, chloroplastic |
| AT1G28470 | -1.5 | 1.10E-06 | -0.9 | 2.90E-05 | -0.6 | 6.57E-03 | 0.7 | 7.02E-02 | 0.2 | 6.40E-01 | 0.5 | 2.75E-01 | NAC domain-containing protein 10 |
| AT4G25820 | -1.5 | 1.09E-05 | 0.0 | 9.51E-01 | -1.5 | 7.31E-05 | -2.0 | 1.22E-06 | -0.1 | 7.47E-01 | -1.9 | 1.26E-04 | Xyloglucan endotransglucosylase |
| AT3G14020 | -1.5 | 2.57E-07 | -1.4 | 9.44E-07 | -0.1 | 4.31E-01 | 0.0 | 9.93E-01 | 0.1 | 8.20E-01 | -0.1 | 8.56E-01 | Nuclear transcription factor Y subunit A-6 |
| AT2G29000 | -1.5 | 2.84E-04 | 0.0 | 9.22E-01 | -1.5 | 1.28E-03 | | | | | | | Leucine-rich repeat protein kinase family protein |
| AT1G07550 | -1.5 | 1.97E-06 | -1.1 | 3.07E-05 | -0.4 | 2.70E-02 | -0.8 | 3.57E-02 | 0.2 | 5.46E-01 | -1.0 | 2.82E-02 | Probable LRR receptor-like serine |
| AT3G21230 | -1.5 | 2.97E-08 | -0.9 | 1.12E-06 | -0.6 | 8.32E-05 | -0.3 | 2.67E-01 | 0.3 | 3.05E-01 | -0.6 | 9.03E-02 | 4-coumarate:CoA ligase 5 |
| AT3G49930 | -1.5 | 2.44E-06 | -0.2 | 1.65E-01 | -1.3 | 7.29E-05 | -0.3 | 3.41E-01 | 0.0 | 9.42E-01 | -0.3 | 5.29E-01 | At3g49930 |
| AT2G21590 | -1.5 | 2.64E-03 | -1.5 | 3.37E-03 | 0.0 | 9.44E-01 | 0.1 | 7.51E-01 | 0.5 | 2.10E-01 | | | Probable glucose-1-phosphate adenylyltransferase large subunit, chloroplastic |
| AT4G20235 | -1.5 | 9.44E-06 | -1.2 | 5.38E-05 | -0.3 | 2.60E-01 | | | | | | | Cytochrome P450, family 71, subfamily A, polypeptide 28 |
| AT1G03920 | -1.5 | 2.70E-06 | -0.6 | 8.52E-04 | -0.9 | 9.61E-04 | -0.5 | 1.88E-01 | 0.5 | 2.27E-01 | -1.0 | 3.64E-02 | Protein kinase family protein |
| AT5G26310 | -1.5 | 1.88E-06 | -0.7 | 3.68E-04 | -0.8 | 1.09E-03 | 0.2 | 4.71E-01 | 0.6 | 3.95E-02 | -0.4 | 1.93E-01 | UDP-glycosyltransferase 72E3 |
| AT2G39040 | -1.5 | 3.66E-05 | -0.1 | 4.85E-01 | -1.4 | 4.47E-04 | -0.3 | 6.43E-01 | 0.1 | 8.85E-01 | -0.4 | 5.88E-01 | Peroxidase 24 |
| AT2G46620 | -1.5 | 2.03E-05 | -1.2 | 1.57E-04 | -0.3 | 1.92E-01 | 0.2 | 2.59E-01 | -0.1 | 7.70E-01 | 0.3 | 1.72E-01 | AAA-ATPase At2g46620 |
| AT4G28720 | -1.5 | 1.16E-05 | -0.9 | 3.23E-04 | -0.6 | 2.89E-02 | -0.5 | 2.99E-01 | 0.3 | 5.95E-01 | | | Probable indole-3-pyruvate monooxygenase YUCCA8 |
| AT5G46040 | -1.5 | 9.13E-05 | -0.1 | 5.91E-01 | -1.4 | 9.78E-04 | | | | | | | Protein NRT1 |
| AT3G21510 | -1.5 | 4.52E-07 | -0.7 | 1.01E-04 | -0.8 | 2.59E-04 | -0.1 | 6.36E-01 | 0.5 | 7.87E-02 | -0.6 | 2.14E-02 | Histidine-containing phosphotransfer protein 1 |
| AT1G19190 | -1.5 | 5.61E-06 | -0.4 | 1.41E-02 | -1.1 | 4.03E-04 | 0.1 | 7.98E-01 | 0.4 | 2.43E-01 | -0.3 | 4.37E-01 | Probable carboxylesterase 1 |
| AT5G45810 | -1.5 | 5.82E-05 | -0.3 | 1.08E-01 | -1.2 | 1.76E-03 | -0.2 | 7.00E-01 | 0.6 | 9.61E-02 | -0.7 | 6.48E-02 | non-specific serine |
| AT5G65530 | -1.5 | 6.93E-07 | -0.5 | 1.06E-03 | -1.0 | 1.21E-04 | -0.6 | 2.21E-02 | -0.1 | 7.14E-01 | -0.5 | 9.44E-02 | Similarity to lectin-like protein kinase |
| AT4G29230 | -1.5 | 6.77E-07 | -0.9 | 1.59E-05 | -0.5 | 5.46E-03 | -0.1 | 8.47E-01 | 0.1 | 8.20E-01 | -0.2 | 7.23E-01 | NAC domain containing protein 75 |
| AT4G21850 | -1.5 | 3.34E-06 | -0.4 | 4.59E-02 | -1.1 | 1.29E-04 | -0.1 | 3.89E-01 | 0.2 | 2.43E-01 | -0.4 | 6.41E-02 | Peptide methionine sulfoxide reductase B9 -S-oxide reductase) |
| AT5G05160 | -1.5 | 8.73E-08 | -1.1 | 1.11E-06 | -0.4 | 3.56E-03 | 0.0 | 9.27E-01 | 0.3 | 1.27E-01 | -0.3 | 1.17E-01 | Probable leucine-rich repeat receptor-like protein kinase At5g05160 |
| AT5G02560 | -1.5 | 2.92E-07 | -0.8 | 3.28E-05 | -0.7 | 3.28E-04 | -0.3 | 2.08E-01 | 0.2 | 5.41E-01 | -0.4 | 8.23E-02 | Histone H2A |
| AT3G29410 | -1.5 | 7.33E-07 | -0.3 | 2.56E-02 | -1.2 | 3.53E-05 | -1.1 | 2.84E-04 | -0.1 | 8.38E-01 | -1.0 | 2.17E-03 | Terpenoid cyclases |
| AT1G78050 | -1.5 | 2.85E-04 | -0.8 | 6.98E-03 | -0.6 | 8.50E-02 | 0.5 | 4.59E-02 | 0.3 | 3.36E-01 | 0.2 | 5.24E-01 | 2,3-bisphosphoglycerate-dependent phosphoglycerate mutase 2 |
| AT1G73010 | -1.5 | 5.56E-05 | -0.9 | 1.06E-03 | -0.5 | 7.64E-02 | -0.2 | 6.84E-01 | 0.2 | 7.44E-01 | -0.4 | 4.03E-01 | PS2 |
| AT1G02390 | -1.5 | 2.85E-04 | -0.9 | 5.73E-03 | -0.6 | 1.01E-01 | -0.5 | 5.40E-02 | 0.1 | 8.17E-01 | -0.6 | 5.11E-02 | Probable glycerol-3-phosphate acyltransferase 2 |
| AT1G77450 | -1.5 | 2.49E-05 | -1.5 | 4.33E-05 | 0.0 | 8.84E-01 | -0.4 | 1.00E-01 | -0.4 | 1.99E-01 | -0.1 | 8.99E-01 | NAC transcription factor 32 |
| AT5G24070 | -1.5 | 5.05E-04 | 0.0 | 8.68E-01 | -1.5 | 1.81E-03 | 0.0 | 9.95E-01 | 0.6 | 1.08E-01 | -0.6 | 1.49E-01 | Probable peroxidase 61 |
| AT2G24130 | -1.5 | 5.35E-05 | -0.9 | 1.30E-03 | -0.6 | 6.22E-02 | | | | | | | Leucine-rich receptor-like protein kinase family protein |
| AT4G36990 | -1.5 | 1.33E-07 | -0.9 | 1.04E-05 | -0.6 | 2.38E-04 | -0.6 | 1.48E-02 | 0.0 | 9.15E-01 | -0.5 | 8.67E-02 | Truncated heat shock factor B1 |
| AT1G70170 | -1.5 | 1.21E-05 | -1.1 | 1.42E-04 | -0.4 | 9.80E-02 | 1.2 | 1.87E-03 | 0.6 | 1.87E-01 | 0.7 | 9.10E-02 | Metalloendoproteinase 2-MMP |
| AT5G37600 | -1.5 | 2.50E-05 | -1.1 | 3.90E-04 | -0.4 | 9.86E-02 | -0.5 | 7.07E-03 | 0.0 | 9.26E-01 | -0.5 | 3.80E-02 | Glutamine synthetase cytosolic isozyme 1-1 |
| AT4G38390 | -1.5 | 3.58E-06 | -0.3 | 4.55E-02 | -1.2 | 1.58E-04 | -1.5 | 1.17E-02 | -0.9 | 9.61E-02 | -0.6 | 5.11E-01 | Protein ROOT HAIR SPECIFIC 17 |
| AT5G17420 | -1.5 | 2.33E-06 | -0.4 | 6.11E-03 | -1.0 | 2.29E-04 | -0.3 | 2.35E-01 | 0.2 | 3.31E-01 | -0.5 | 6.58E-02 | Cellulose synthase A catalytic subunit 7 |
| AT2G17060 | -1.5 | 4.70E-07 | -1.0 | 7.10E-06 | -0.4 | 1.21E-02 | 0.7 | 2.58E-02 | 0.5 | 1.10E-01 | 0.1 | 7.31E-01 | ADP-ribosyl cyclase |
| AT1G34510 | -1.5 | 4.90E-03 | 0.1 | 8.15E-01 | -1.5 | 1.11E-02 | -4.2 | 5.23E-05 | -1.0 | 3.42E-02 | -3.2 | 1.02E-02 | Peroxidase 8 |
| AT5G24240 | -1.5 | 6.80E-06 | -0.7 | 1.02E-03 | -0.8 | 3.42E-03 | -1.1 | 3.05E-02 | -0.1 | 8.41E-01 | -1.0 | 1.46E-01 | Phosphatidylinositol 4-kinase gamma 3 |
| AT4G12050 | -1.5 | 9.61E-08 | -0.4 | 2.55E-04 | -1.0 | 2.37E-05 | -0.6 | 1.75E-02 | 0.0 | 8.98E-01 | -0.6 | 3.49E-02 | AT-hook motif nuclear-localized protein 26 |
| AT2G44450 | -1.5 | 1.85E-06 | -0.5 | 1.90E-03 | -0.9 | 3.05E-04 | -0.4 | 1.65E-01 | 0.2 | 5.89E-01 | -0.5 | 1.27E-01 | Beta glucosidase 15 |
| AT4G16750 | -1.4 | 1.97E-04 | -1.1 | 1.50E-03 | -0.4 | 2.56E-01 | 0.1 | 7.99E-01 | 0.4 | 1.66E-01 | -0.3 | 2.55E-01 | Ethylene-responsive transcription factor ERF039 |
| AT1G72200 | -1.4 | 2.05E-05 | -0.4 | 4.58E-02 | -1.1 | 9.40E-04 | -1.1 | 1.31E-04 | -0.4 | 5.63E-02 | -0.6 | 2.25E-02 | RING-H2 finger protein ATL11 |
| AT2G15390 | -1.4 | 4.10E-06 | -0.7 | 5.57E-04 | -0.7 | 2.49E-03 | 0.3 | 9.90E-02 | -0.1 | 7.03E-01 | 0.4 | 1.15E-01 | Probable fucosyltransferase 4 |
| AT5G47640 | -1.4 | 1.14E-06 | -1.3 | 5.56E-06 | -0.2 | 2.76E-01 | -0.8 | 2.26E-05 | -0.2 | 1.88E-01 | -0.6 | 4.82E-03 | Nuclear transcription factor Y subunit B-2 |
| AT3G07720 | -1.4 | 3.72E-05 | -1.2 | 3.41E-04 | -0.3 | 1.97E-01 | -1.0 | 3.95E-05 | -1.4 | 5.81E-06 | 0.4 | 7.46E-02 | AT3g07720 |
| AT3G05390 | -1.4 | 4.90E-06 | -0.1 | 2.96E-01 | -1.3 | 1.05E-04 | -0.6 | 6.77E-02 | 0.1 | 8.25E-01 | -0.7 | 1.06E-01 | hypothetical protein |
| AT2G45600 | -1.4 | 4.86E-05 | -0.7 | 4.99E-03 | -0.8 | 1.30E-02 | -0.3 | 1.14E-01 | 0.2 | 3.77E-01 | -0.5 | 4.59E-02 | Probable carboxylesterase 8 |
| AT3G04010 | -1.4 | 2.06E-07 | -0.6 | 6.33E-05 | -0.8 | 1.08E-04 | -0.6 | 4.13E-02 | -0.2 | 5.61E-01 | -0.4 | 2.07E-01 | glucan endo-1,3-beta-D-glucosidase |
| AT2G36380 | -1.4 | 7.15E-08 | -1.1 | 7.18E-07 | -0.3 | 5.49E-03 | -0.5 | 2.45E-02 | -0.3 | 3.29E-01 | -0.3 | 4.34E-01 | ABC transporter G family member 34 |
| AT3G23870 | -1.4 | 5.37E-05 | -0.4 | 4.33E-02 | -1.0 | 2.71E-03 | -1.4 | 2.07E-03 | 0.1 | 8.80E-01 | -1.5 | 8.08E-03 | Probable magnesium transporter |
| AT3G62740 | -1.4 | 7.04E-06 | -1.1 | 7.08E-05 | -0.4 | 9.95E-02 | -0.7 | 9.02E-02 | 0.8 | 3.33E-02 | -1.5 | 5.23E-03 | Beta glucosidase 7 |
| AT3G53510 | -1.4 | 2.87E-07 | -0.8 | 2.42E-05 | -0.6 | 4.52E-04 | 0.6 | 2.94E-02 | 0.4 | 1.70E-01 | 0.2 | 6.65E-01 | ABC transporter G family member 20 |
| AT4G11050 | -1.4 | 4.34E-05 | -0.3 | 9.57E-02 | -1.1 | 1.40E-03 | 0.4 | 1.88E-01 | 0.4 | 2.63E-01 | 0.0 | 9.80E-01 | Endoglucanase |
| AT4G25090 | -1.4 | 1.91E-06 | -0.7 | 3.92E-04 | -0.7 | 1.05E-03 | 0.0 | 9.89E-01 | 0.5 | 1.42E-01 | -0.5 | 1.62E-01 | Riboflavin synthase-like superfamily protein |
| AT1G73220 | -1.4 | 3.07E-06 | -1.4 | 7.40E-06 | 0.0 | 9.28E-01 | 3.1 | 2.09E-05 | 2.4 | 1.16E-03 | 0.8 | 4.48E-02 | Organic cation |

| | | | | | | | | | | | | | |
|---|---|---|---|---|---|---|---|---|---|---|---|---|---|
| AT1G76350 | -1.4 | 8.09E-08 | -1.1 | 8.63E-07 | -0.3 | 5.89E-03 | -0.1 | 8.02E-01 | 0.4 | 4.42E-02 | -0.4 | 5.73E-02 | Protein NLP5 |
| AT5G61550 | -1.4 | 4.90E-06 | -0.1 | 3.10E-01 | -1.3 | 9.89E-05 | -0.9 | 4.58E-02 | 0.1 | 9.23E-01 | -1.0 | 1.03E-01 | U-box domain-containing protein kinase family protein |
| AT5G48290 | -1.4 | 1.05E-06 | -0.5 | 7.60E-04 | -0.9 | 2.50E-04 | -0.2 | 3.01E-01 | -0.2 | 3.68E-01 | 0.0 | 9.93E-01 | Heavy metal transport |
| AT4G31710 | -1.4 | 7.43E-06 | -0.6 | 1.17E-03 | -0.8 | 3.67E-03 | -0.3 | 6.47E-01 | 0.4 | 5.86E-01 | -0.7 | 3.93E-01 | Glutamate receptor |
| AT4G30290 | -1.4 | 1.31E-05 | -0.1 | 4.17E-01 | -1.3 | 1.82E-04 | -0.9 | 5.22E-03 | 0.1 | 8.52E-01 | -1.0 | 1.72E-02 | Xyloglucan endotransglucosylase |
| AT1G62540 | -1.4 | 1.40E-04 | -0.9 | 2.15E-03 | -0.5 | 1.33E-01 | 0.2 | 6.03E-01 | 1.2 | 4.07E-03 | -0.9 | 1.84E-02 | Flavin-containing monooxygenase |
| AT1G73160 | -1.4 | 6.22E-05 | 0.1 | 3.91E-01 | -1.6 | 2.23E-04 | -0.3 | 5.95E-01 | 0.5 | 2.53E-01 | -0.8 | 8.64E-02 | F3N23.36 protein |
| AT4G22810 | -1.4 | 4.70E-07 | -0.5 | 4.35E-04 | -0.9 | 1.09E-04 | -0.4 | 1.17E-01 | 0.1 | 7.78E-01 | -0.5 | 1.11E-01 | AT-hook motif nuclear-localized protein 24 |
| AT5G46650 | -1.4 | 2.12E-05 | -0.6 | 4.49E-03 | -0.8 | 5.13E-03 | -0.5 | 2.78E-01 | 0.2 | 8.04E-01 | -0.7 | 2.43E-01 | RING-H2 finger protein ATL30 |
| AT5G38970 | -1.4 | 7.39E-05 | 0.1 | 5.48E-01 | -1.5 | 3.09E-04 | -0.5 | 1.73E-01 | -0.2 | 6.71E-01 | -0.3 | 5.84E-01 | Cytochrome P450 85A1 |
| AT4G17670 | -1.4 | 5.13E-06 | -1.4 | 1.12E-05 | 0.0 | 9.87E-01 | -0.3 | 2.88E-01 | 0.2 | 5.02E-01 | -0.5 | 1.05E-01 | FCS-Like Zinc finger 2 |
| AT1G07150 | -1.4 | 1.82E-02 | -1.5 | 1.78E-02 | 0.1 | 9.02E-01 | 0.0 | 9.23E-01 | 0.2 | 7.07E-01 | -0.1 | 8.13E-01 | F10K1.14 protein |
| AT1G11130 | -1.4 | 6.92E-08 | -1.1 | 5.00E-07 | -0.3 | 1.61E-02 | 0.1 | 8.33E-01 | 0.2 | 4.40E-01 | -0.2 | 6.69E-01 | Leucine-rich repeat protein kinase family protein |
| AT1G69080 | -1.4 | 2.02E-06 | -0.7 | 3.00E-04 | -0.7 | 1.45E-03 | 0.0 | 8.88E-01 | 0.2 | 5.02E-01 | -0.2 | 4.89E-01 | Adenine nucleotide alpha hydrolases-like superfamily protein |
| AT4G18780 | -1.4 | 1.91E-06 | -0.4 | 6.95E-03 | -1.0 | 1.77E-04 | -0.3 | 3.53E-01 | -0.1 | 8.78E-01 | -0.3 | 5.15E-01 | Cellulose synthase A catalytic subunit 8 |
| AT4G39955 | -1.4 | 2.53E-06 | -1.3 | 7.96E-06 | -0.1 | 6.44E-01 | -0.1 | 7.78E-01 | 0.2 | 3.43E-01 | -0.3 | 2.48E-01 | Alpha |
| AT4G19230 | -1.4 | 3.40E-05 | -1.0 | 3.70E-04 | -0.4 | 1.25E-01 | -0.2 | 4.83E-01 | 0.3 | 3.00E-01 | -0.6 | 1.16E-01 | Cytochrome P450, family 707, subfamily A, polypeptide 1 |
| AT4G27290 | -1.4 | 1.67E-06 | -0.8 | 6.80E-05 | -0.6 | 5.38E-03 | -0.5 | 4.08E-01 | 0.4 | 5.33E-01 | -0.8 | 2.25E-01 | Receptor-like serine |
| AT2G47670 | -1.4 | 1.12E-05 | -0.5 | 4.00E-03 | -0.9 | 2.37E-03 | 0.2 | 6.39E-01 | 0.4 | 4.02E-01 | -0.2 | 6.76E-01 | Pectinesterase inhibitor 6 |
| AT1G52800 | -1.4 | 3.96E-06 | -0.1 | 6.59E-01 | -1.3 | 6.16E-05 | -0.6 | 1.06E-01 | 0.6 | 9.08E-02 | -1.2 | 1.33E-02 | hypothetical protein |
| AT1G15125 | -1.4 | 1.06E-04 | -0.7 | 1.15E-02 | -0.7 | 1.81E-02 | -0.2 | 6.41E-01 | -1.3 | 5.16E-03 | 1.1 | 6.10E-03 | hypothetical protein |
| AT1G53163 | -1.4 | 3.19E-05 | -0.3 | 7.86E-02 | -1.1 | 1.35E-03 | 0.3 | 5.40E-01 | 0.0 | 9.83E-01 | 0.3 | 6.70E-01 | F12M16.3 |
| AT5G62020 | -1.4 | 3.23E-05 | -1.2 | 1.23E-04 | -0.2 | 5.08E-01 | -0.2 | 2.88E-01 | -0.3 | 2.07E-01 | 0.1 | 8.41E-01 | Truncated heat shock factor B2a |
| AT3G46810 | -1.4 | 2.01E-05 | -0.6 | 5.22E-03 | -0.8 | 4.35E-03 | -0.5 | 3.47E-01 | 0.3 | 6.53E-01 | -0.8 | 1.85E-01 | Cysteine |
| AT2G28110 | -1.4 | 3.28E-07 | -0.7 | 7.29E-05 | -0.7 | 2.15E-04 | -0.3 | 1.92E-01 | -0.5 | 3.23E-02 | 0.2 | 3.46E-01 | Probable glucuronoxylan glucuronosyltransferase IRX7 |
| AT5G12970 | -1.4 | 1.28E-06 | -0.7 | 1.09E-04 | -0.6 | 1.82E-03 | -0.1 | 8.33E-01 | 0.2 | 6.82E-01 | -0.2 | 5.63E-01 | Multiple C2 domain and transmembrane region protein 5 |
| AT5G45070 | -1.4 | 1.34E-06 | -1.4 | 3.19E-06 | 0.0 | 9.73E-01 | 0.7 | 2.42E-03 | 0.5 | 4.98E-02 | 0.2 | 2.25E-01 | Protein PHLOEM PROTEIN 2-LIKE A8 |
| AT1G72700 | -1.4 | 1.66E-07 | -1.0 | 3.03E-06 | -0.4 | 3.15E-03 | 0.0 | 9.82E-01 | 0.2 | 3.25E-01 | -0.2 | 4.21E-01 | Probable phospholipid-transporting ATPase 5 |
| AT3G23790 | -1.4 | 8.42E-07 | -1.2 | 3.92E-06 | -0.2 | 2.89E-01 | -0.1 | 8.20E-01 | 0.0 | 9.68E-01 | -0.1 | 8.07E-01 | Probable acyl-activating enzyme 16, chloroplastic |
| AT5G03760 | -1.4 | 1.01E-06 | -0.8 | 9.44E-05 | -0.6 | 1.39E-03 | -0.3 | 2.09E-02 | 0.3 | 6.78E-02 | -0.6 | 1.17E-03 | Glucomannan 4-beta-mannosyltransferase 9 |
| AT3G45330 | -1.4 | 1.30E-06 | -0.9 | 2.79E-05 | -0.5 | 1.39E-02 | 0.2 | 7.11E-01 | 0.4 | 3.41E-01 | -0.2 | 6.50E-01 | Putative L-type lectin-domain containing receptor kinase I.1 |
| AT2G40120 | -1.4 | 1.07E-06 | -0.8 | 5.38E-05 | -0.6 | 3.04E-03 | -0.1 | 8.20E-01 | 0.1 | 7.57E-01 | -0.1 | 6.53E-01 | Hypothetical Ser-Thr protein kinase |
| AT1G28130 | -1.4 | 6.15E-08 | -1.2 | 2.87E-07 | -0.1 | 1.17E-01 | -0.9 | 8.24E-05 | -0.1 | 6.11E-01 | -0.8 | 2.31E-03 | Indole-3-acetic acid-amido synthetase GH3.17 |
| AT5G67400 | -1.4 | 1.47E-05 | 0.3 | 1.95E-01 | -1.6 | 4.08E-05 | -1.7 | 9.43E-06 | -0.1 | 7.29E-01 | -1.5 | 3.64E-04 | Peroxidase 73 |
| AT3G57630 | -1.4 | 1.38E-07 | -0.9 | 3.37E-06 | -0.5 | 1.30E-03 | 0.3 | 6.23E-02 | 0.4 | 3.60E-02 | -0.1 | 7.93E-01 | At3g57630 |
| AT1G69490 | -1.4 | 1.46E-04 | -1.5 | 1.60E-04 | 0.1 | 7.72E-01 | -2.1 | 9.25E-04 | -1.6 | 4.78E-03 | -0.5 | 6.25E-01 | NAC transcription factor 29 |
| AT5G39610 | -1.4 | 7.93E-06 | -1.4 | 1.36E-05 | 0.0 | 8.25E-01 | -0.7 | 5.67E-03 | -1.0 | 7.74E-04 | 0.3 | 3.47E-01 | NAC domain-containing protein 92 |
| AT1G08990 | -1.4 | 7.33E-08 | -0.7 | 8.07E-06 | -0.7 | 8.64E-05 | -0.7 | 1.38E-02 | 0.0 | 9.44E-01 | -0.8 | 1.81E-02 | Putative UDP-glucuronate:xylan alpha-glucuronosyltransferase 5 |
| AT3G20935 | -1.4 | 7.76E-06 | -0.6 | 1.43E-03 | -0.8 | 3.42E-03 | -1.3 | 2.19E-02 | -0.2 | 7.48E-01 | -1.1 | 9.93E-02 | Cytochrome P450, family 705, subfamily A, polypeptide 28 |
| AT3G46420 | -1.4 | 1.37E-03 | 0.0 | 8.74E-01 | -1.3 | 6.57E-03 | | | | | | | Protein kinase domain-containing protein |
| AT2G34960 | -1.4 | 3.97E-07 | -1.3 | 1.59E-06 | -0.1 | 4.42E-01 | 0.2 | 6.90E-01 | -0.3 | 6.47E-01 | 0.6 | 3.65E-01 | Cationic amino acid transporter 5 |
| AT5G37540 | -1.4 | 1.37E-06 | -0.4 | 4.56E-03 | -1.0 | 1.38E-04 | -0.9 | 6.47E-05 | -0.2 | 4.42E-01 | -0.8 | 5.53E-03 | Peptidase A1 domain-containing protein |
| AT4G25240 | -1.4 | 9.53E-08 | -0.7 | 1.62E-05 | -0.7 | 7.62E-05 | 0.7 | 7.86E-03 | 0.8 | 4.78E-03 | -0.1 | 7.50E-01 | Monocopper oxidase-like protein SKS1 |
| AT1G55775 | -1.4 | 9.08E-05 | -1.5 | 9.51E-05 | 0.2 | 6.47E-01 | 0.7 | 1.03E-01 | 0.2 | 7.42E-01 | 0.5 | 3.28E-01 | Heavy metal-associated domain protein |
| AT4G38780 | -1.4 | 1.48E-05 | -1.0 | 1.47E-04 | -0.4 | 1.36E-01 | | | | | | | Pre-mRNA-processing-splicing factor 8B |
| AT2G45220 | -1.4 | 6.66E-06 | -0.3 | 1.00E-01 | -1.1 | 1.70E-04 | -1.4 | 3.23E-02 | -0.4 | 5.98E-01 | -1.0 | 7.40E-02 | Probable pectinesterase |
| AT1G66930 | -1.4 | 1.51E-05 | -0.7 | 7.52E-04 | -0.6 | 1.48E-02 | -0.3 | 2.97E-01 | 0.3 | 3.24E-01 | -0.7 | 9.33E-02 | Protein kinase superfamily protein |
| AT1G06090 | -1.4 | 1.35E-05 | -0.2 | 3.15E-01 | -1.2 | 2.03E-04 | -0.6 | 1.26E-02 | 0.1 | 8.48E-01 | -0.7 | 3.34E-02 | Delta-9 desaturase-like 1 protein |
| AT1G47480 | -1.4 | 2.11E-07 | -0.5 | 1.33E-04 | -0.8 | 6.96E-05 | -0.3 | 1.73E-01 | 0.0 | 9.04E-01 | -0.4 | 2.40E-01 | Probable carboxylesterase 2 |
| AT5G15970 | -1.4 | 4.08E-04 | -1.4 | 5.56E-04 | 0.0 | 9.49E-01 | -1.5 | 6.00E-05 | 0.2 | 4.98E-01 | -1.7 | 4.83E-04 | Stress-induced protein KIN2 |
| AT5G61350 | -1.4 | 4.33E-05 | 0.0 | 7.87E-01 | -1.3 | 3.68E-04 | -1.2 | 1.11E-02 | 0.0 | 9.93E-01 | -1.2 | 4.70E-02 | Probable receptor-like protein kinase At5g61350 |
| AT4G40010 | -1.4 | 4.25E-08 | -0.7 | 5.47E-06 | -0.7 | 3.91E-05 | -1.2 | 4.89E-04 | 0.1 | 6.48E-01 | -1.4 | 2.30E-03 | Serine |
| AT5G19740 | -1.4 | 4.76E-07 | -0.9 | 1.89E-05 | -0.5 | 2.20E-03 | -0.8 | 4.03E-03 | -0.4 | 1.02E-01 | -0.3 | 2.71E-01 | Probable glutamate carboxypeptidase LAMP1 |
| AT5G51310 | -1.4 | 3.19E-05 | -0.2 | 2.00E-01 | -1.1 | 6.91E-04 | 0.2 | 3.97E-01 | 0.1 | 8.35E-01 | 0.1 | 6.18E-01 | Gibberellin 20-oxidase-like protein |
| AT3G59010 | -1.4 | 6.59E-05 | -0.5 | 2.71E-02 | -0.9 | 5.07E-03 | -0.4 | 9.61E-02 | 0.2 | 3.68E-01 | -0.6 | 2.38E-02 | Probable pectinesterase |
| AT1G08630 | -1.4 | 1.41E-02 | -2.9 | 1.64E-03 | 1.5 | 8.78E-02 | -2.2 | 9.19E-02 | -3.4 | 6.70E-02 | 1.1 | 5.33E-01 | Threonine aldolase 1 |
| AT1G66400 | -1.4 | 2.90E-04 | -1.2 | 1.00E-03 | -0.2 | 5.82E-01 | -0.2 | 6.38E-01 | 0.3 | 5.22E-01 | -0.5 | 2.41E-01 | Probable calcium-binding protein CML23 |
| AT5G41460 | -1.3 | 1.95E-04 | -1.1 | 1.16E-03 | -0.3 | 3.53E-01 | -0.4 | 8.63E-02 | 0.4 | 1.12E-01 | -0.8 | 1.03E-02 | At5g41460 ) |
| AT1G07870 | -1.3 | 7.01E-07 | -1.0 | 9.39E-06 | -0.4 | 2.31E-02 | -0.4 | 3.90E-02 | -0.1 | 8.62E-01 | -0.4 | 1.75E-01 | Protein kinase superfamily protein |
| AT4G21960 | -1.3 | 4.33E-07 | -0.8 | 4.90E-05 | -0.6 | 6.64E-04 | -0.4 | 6.81E-04 | 0.0 | 9.98E-01 | -0.4 | 1.37E-02 | Peroxidase 42 |
| AT1G04160 | -1.3 | 3.21E-06 | -0.5 | 2.74E-03 | -0.9 | 5.63E-04 | -0.8 | 6.77E-03 | 0.3 | 2.24E-01 | -1.1 | 4.36E-03 | Myosin XI B |
| AT2G28900 | -1.3 | 2.81E-05 | -0.9 | 9.94E-04 | -0.5 | 3.43E-02 | -0.4 | 9.36E-03 | 0.1 | 7.68E-01 | -0.5 | 2.08E-02 | Outer envelope pore protein 16-1, chloroplastic |
| AT1G30500 | -1.3 | 4.07E-06 | -1.2 | 1.57E-05 | -0.1 | 4.67E-01 | -0.3 | 2.43E-01 | -0.3 | 1.62E-01 | 0.1 | 7.95E-01 | Nuclear transcription factor Y subunit |
| AT4G13130 | -1.3 | 1.05E-06 | -0.7 | 7.23E-05 | -0.6 | 2.02E-03 | -1.4 | 4.12E-04 | -0.2 | 5.48E-01 | -1.2 | 5.38E-03 | Cysteine |
| AT3G16920 | -1.3 | 5.78E-07 | 0.0 | 7.44E-01 | -1.4 | 1.23E-05 | -0.3 | 6.48E-02 | -0.1 | 6.39E-01 | -0.2 | 2.83E-01 | Chitinase-like protein |
| AT1G70460 | -1.3 | 9.29E-06 | -0.1 | 4.61E-01 | -1.2 | 1.25E-04 | -1.3 | 1.01E-04 | 0.0 | 9.41E-01 | -1.3 | 2.17E-03 | non-specific serine |
| AT1G27440 | -1.3 | 3.65E-07 | -0.5 | 4.27E-04 | -0.9 | 8.45E-05 | 0.0 | 9.93E-01 | 0.2 | 3.00E-01 | -0.2 | 3.89E-01 | Probable beta-1,4-xylosyltransferase IRX10 |
| AT4G35380 | -1.3 | 1.30E-06 | -0.7 | 2.04E-04 | -0.7 | 9.48E-04 | 0.1 | 5.97E-01 | 0.2 | 4.39E-01 | -0.1 | 8.28E-01 | SEC7-like guanine nucleotide exchange family protein |
| AT3G22600 | -1.3 | 3.86E-07 | -0.4 | 1.43E-03 | -0.9 | 4.65E-05 | 0.8 | 2.96E-04 | 0.3 | 2.19E-01 | 0.5 | 9.48E-03 | Non-specific lipid transfer protein GPI-anchored 5 |
| AT3G29035 | -1.3 | 3.29E-07 | -1.0 | 4.55E-06 | -0.4 | 1.24E-02 | -0.6 | 1.45E-02 | -0.6 | 1.57E-02 | 0.0 | 9.34E-01 | NAC domain-containing protein 59 |
| AT1G05280 | -1.3 | 1.16E-04 | -0.3 | 1.40E-01 | -1.1 | 3.25E-03 | -0.1 | 8.67E-01 | 0.8 | 8.23E-02 | -0.9 | 1.17E-01 | ERV-F 1 provirus ancestral Env polyprotein, putative |
| AT5G39000 | -1.3 | 9.68E-06 | -0.7 | 6.62E-04 | -0.6 | 9.25E-03 | -0.4 | 3.34E-01 | 0.1 | 7.67E-01 | -0.5 | 2.79E-01 | Putative receptor-like protein kinase At5g39000 |
| AT2G25940 | -1.3 | 1.53E-06 | -1.1 | 1.27E-05 | -0.3 | 9.39E-02 | -0.7 | 1.17E-01 | -0.8 | 1.02E-01 | 0.1 | 9.04E-01 | Vacuolar-processing enzyme alpha-isozyme |
| AT1G22990 | -1.3 | 8.09E-07 | -1.1 | 4.95E-06 | -0.2 | 1.42E-01 | -0.8 | 1.59E-01 | -0.2 | 8.40E-01 | -0.7 | 2.52E-01 | Heavy metal transport |
| AT5G24910 | -1.3 | 7.26E-06 | -0.4 | 9.79E-03 | -0.9 | 7.01E-04 | -1.0 | 1.04E-02 | -0.2 | 6.76E-01 | -0.8 | 7.61E-02 | Cytochrome P450, family 714, subfamily A, polypeptide 1 |
| AT4G29800 | -1.3 | 2.70E-06 | -0.3 | 2.29E-02 | -1.0 | 1.58E-04 | 0.0 | 9.62E-01 | 0.4 | 1.58E-01 | -0.4 | 1.78E-01 | Patatin-like protein 8 |
| AT1G67030 | -1.3 | 4.74E-07 | -1.0 | 5.64E-06 | -0.3 | 2.34E-02 | -0.4 | 2.12E-01 | 0.3 | 4.31E-01 | -0.6 | 3.53E-02 | Zinc finger protein 6 |
| AT5G66390 | -1.3 | 1.92E-07 | -0.2 | 5.88E-02 | -1.1 | 1.09E-05 | 0.7 | 2.32E-03 | 0.5 | 4.29E-02 | 0.2 | 3.73E-01 | Peroxidase 72 |
| AT1G17590 | -1.3 | 3.81E-07 | -1.1 | 2.39E-06 | -0.2 | 1.03E-01 | -0.2 | 6.21E-01 | 0.1 | 8.44E-01 | -0.3 | 5.49E-01 | Nuclear transcription factor Y subunit A-8 |
| AT4G23690 | -1.3 | 7.41E-07 | -0.4 | 2.65E-03 | -0.9 | 8.22E-05 | 0.5 | 7.02E-02 | 0.2 | 5.94E-01 | 0.3 | 2.52E-01 | Dirigent protein 6 |
| AT4G30560 | -1.3 | 1.33E-06 | -0.5 | 1.12E-03 | -0.8 | 2.87E-04 | -0.5 | 1.76E-02 | 0.1 | 8.20E-01 | -0.6 | 4.57E-02 | hypothetical protein |
| AT3G07960 | -1.3 | 3.82E-06 | -0.6 | 5.01E-04 | -0.7 | 3.04E-03 | -0.1 | 7.98E-01 | 0.2 | 5.99E-01 | -0.4 | 4.49E-01 | 1-phosphatidylinositol-4-phosphate 5-kinase |
| AT1G04700 | -1.3 | 1.48E-05 | -0.1 | 3.65E-01 | -1.2 | 2.51E-04 | -0.5 | 2.50E-01 | 0.0 | 9.86E-01 | -0.5 | 3.25E-01 | Protein kinase domain-containing protein |

| | | | | | | | | | | | | | |
|---|---|---|---|---|---|---|---|---|---|---|---|---|---|
| AT3G62830 | -1.3 | 6.05E-07 | -0.4 | 2.15E-03 | -0.9 | 7.06E-05 | -0.2 | 1.13E-01 | 0.1 | 3.12E-01 | -0.3 | 3.44E-02 | UDP-glucuronic acid decarboxylase 2 |
| AT1G11160 | -1.3 | 4.01E-06 | -0.3 | 3.58E-02 | -1.0 | 2.00E-04 | -0.8 | 6.16E-02 | 0.1 | 7.83E-01 | -1.0 | 7.26E-02 | Katanin p80 WD40 repeat-containing subunit B1 homolog |
| AT3G20250 | -1.3 | 1.09E-07 | -1.2 | 4.42E-07 | -0.1 | 3.02E-01 | -0.6 | 1.54E-03 | -0.3 | 8.69E-02 | -0.3 | 2.64E-01 | Pumilio 5 |
| AT3G14720 | -1.3 | 2.26E-06 | -1.1 | 1.82E-05 | -0.2 | 1.07E-01 | -0.5 | 1.37E-02 | 0.1 | 5.06E-01 | -0.6 | 2.01E-02 | mitogen-activated protein kinase |
| AT3G23000 | -1.3 | 6.86E-08 | -1.0 | 8.55E-07 | -0.3 | 3.75E-03 | -0.4 | 1.06E-02 | -0.3 | 8.19E-02 | -0.1 | 6.76E-01 | CBL-interacting serine |
| AT4G14580 | -1.3 | 1.74E-07 | -1.3 | 4.61E-07 | 0.0 | 9.73E-01 | 0.0 | 9.08E-01 | 0.2 | 2.20E-01 | -0.2 | 3.46E-01 | CBL-interacting serine |
| AT5G45340 | -1.3 | 1.10E-03 | -1.3 | 1.84E-03 | 0.0 | 9.26E-01 | 0.3 | 2.06E-01 | 0.4 | 9.47E-02 | -0.1 | 7.13E-01 | Abscisic acid 8'-hydroxylase 3 |
| AT2G30490 | -1.3 | 2.49E-07 | -0.5 | 2.96E-04 | -0.8 | 6.08E-05 | -0.1 | 2.37E-01 | 0.1 | 5.94E-01 | -0.2 | 1.78E-01 | Trans-cinnamate 4-monooxygenase |
| AT1G14730 | -1.3 | 2.61E-07 | -0.9 | 4.02E-06 | -0.4 | 6.98E-03 | -0.2 | 2.52E-01 | -0.1 | 6.03E-01 | -0.1 | 7.46E-01 | Probable transmembrane ascorbate ferrireductase 3 |
| AT3G24360 | -1.3 | 7.17E-04 | -0.7 | 1.44E-02 | -0.6 | 1.25E-01 | | | | | | | 3-hydroxyisobutyryl-CoA hydrolase |
| AT2G32430 | -1.3 | 2.91E-06 | -0.4 | 3.99E-03 | -0.9 | 3.88E-04 | 0.2 | 2.97E-01 | 0.6 | 5.01E-03 | -0.4 | 8.52E-02 | Hexosyltransferase |
| AT2G45120 | -1.3 | 7.91E-05 | -0.7 | 3.14E-03 | -0.6 | 4.18E-02 | -0.5 | 1.16E-01 | 0.2 | 5.43E-01 | -0.8 | 7.88E-02 | Zinc finger protein ZAT4 |
| AT5G35580 | -1.3 | 2.47E-07 | -0.7 | 2.59E-05 | -0.6 | 3.42E-04 | 0.8 | 2.22E-04 | 0.5 | 2.05E-02 | 0.3 | 1.16E-01 | Protein kinase superfamily protein |
| AT2G46530 | -1.3 | 3.44E-06 | -0.6 | 6.20E-04 | -0.7 | 1.99E-03 | -0.6 | 3.83E-03 | 0.2 | 3.03E-01 | -0.8 | 1.12E-03 | Auxin response factor 11 |
| AT1G72540 | -1.3 | 4.52E-04 | -1.6 | 3.17E-04 | 0.3 | 4.50E-01 | 1.2 | 3.11E-02 | 0.3 | 7.07E-01 | 0.9 | 1.25E-01 | hypothetical protein |
| AT2G37580 | -1.3 | 7.21E-06 | -1.1 | 4.43E-05 | -0.2 | 2.29E-01 | -0.1 | 5.21E-01 | 0.3 | 2.32E-01 | -0.4 | 1.16E-01 | RING-H2 finger protein ATL33 |
| AT5G10410 | -1.3 | 3.73E-07 | -1.0 | 4.88E-06 | -0.3 | 1.54E-02 | 0.6 | 1.01E-02 | 0.2 | 3.74E-01 | 0.3 | 1.50E-01 | Putative clathrin assembly protein At5g10410 |
| AT2G32990 | -1.3 | 1.16E-06 | -1.0 | 1.39E-05 | -0.3 | 3.58E-02 | -0.3 | 5.78E-02 | 0.6 | 1.36E-03 | -1.0 | 6.62E-04 | Endoglucanase 11 |
| AT1G61860 | -1.3 | 5.59E-05 | -0.4 | 4.91E-02 | -0.9 | 3.12E-03 | | | | | | | Protein kinase domain-containing protein |
| AT1G33260 | -1.3 | 2.29E-06 | -0.7 | 2.26E-04 | -0.6 | 2.60E-03 | 0.0 | 9.01E-01 | 0.2 | 4.66E-01 | -0.2 | 5.99E-01 | Probable receptor-like protein kinase At1g33260 |
| AT5G37790 | -1.3 | 1.31E-07 | -0.5 | 1.33E-04 | -0.8 | 3.37E-05 | -0.3 | 8.15E-02 | 0.2 | 3.84E-01 | -0.4 | 2.76E-02 | Protein kinase domain-containing protein |
| AT4G15093 | -1.3 | 8.69E-08 | -0.7 | 6.12E-06 | -0.6 | 1.66E-04 | -0.2 | 3.03E-01 | 0.0 | 9.61E-01 | -0.2 | 3.30E-01 | Extradiol ring-cleavage dioxygenase |
| AT2G17700 | -1.3 | 8.97E-08 | -1.0 | 7.87E-07 | -0.3 | 1.21E-02 | -0.1 | 6.98E-01 | 0.2 | 4.44E-01 | -0.3 | 3.38E-01 | Serine |
| AT2G34500 | -1.3 | 2.64E-04 | 0.0 | 8.43E-01 | -1.2 | 1.58E-03 | -0.5 | 9.00E-02 | -0.3 | 3.58E-01 | -0.2 | 6.58E-01 | Cytochrome P450 710A1 |
| AT1G08500 | -1.3 | 1.16E-06 | 0.1 | 4.74E-01 | -1.3 | 2.00E-05 | -0.5 | 1.58E-01 | 0.0 | 9.70E-01 | -0.5 | 8.50E-02 | Early nodulin-like protein 18 |
| AT4G10150 | -1.3 | 5.36E-05 | 0.0 | 9.76E-01 | -1.3 | 3.88E-04 | -1.1 | 5.18E-03 | 0.3 | 3.60E-01 | -1.4 | 3.39E-03 | RING-H2 finger protein ATL7 |
| AT2G45750 | -1.3 | 6.52E-07 | -0.3 | 4.81E-03 | -0.9 | 6.36E-05 | -0.2 | 4.46E-01 | 0.0 | 9.67E-01 | -0.2 | 5.87E-01 | Probable methyltransferase PMT16 |
| AT5G58320 | -1.3 | 5.60E-07 | -0.4 | 1.83E-03 | -0.9 | 6.92E-05 | -0.1 | 7.94E-01 | 0.0 | 9.46E-01 | 0.0 | 9.12E-01 | Protein NETWORKED 4A |
| AT1G15310 | -1.3 | 8.03E-04 | -1.1 | 2.47E-03 | -0.2 | 6.12E-01 | 0.2 | 6.49E-01 | 0.5 | 2.68E-01 | -0.3 | 5.70E-01 | Signal recognition particle subunit SRP54 1 |
| AT1G48640 | -1.3 | 8.28E-06 | 0.0 | 8.11E-01 | -1.2 | 9.94E-05 | -0.8 | 7.86E-02 | 0.1 | 8.04E-01 | -0.9 | 9.02E-02 | Lysine histidine transporter-like 1 |
| AT4G15330 | -1.3 | 3.92E-06 | -0.1 | 6.16E-01 | -1.2 | 6.46E-05 | -0.9 | 1.70E-03 | -0.1 | 8.35E-01 | -0.8 | 6.68E-03 | Cytochrome P450, family 705, subfamily A, polypeptide 1 |
| AT5G08640 | -1.3 | 9.21E-04 | -0.5 | 9.18E-02 | -0.8 | 3.18E-02 | 0.8 | 4.60E-04 | 1.8 | 5.32E-07 | -1.0 | 2.73E-04 | Flavonol synthase |
| AT4G04955 | -1.3 | 5.43E-07 | -0.6 | 2.19E-04 | -0.7 | 2.03E-04 | -0.8 | 9.12E-06 | -0.1 | 3.05E-01 | -0.6 | 2.12E-03 | allantoinase |
| AT4G36830 | -1.3 | 1.97E-05 | -0.5 | 3.65E-03 | -0.7 | 5.47E-03 | -0.3 | 3.99E-01 | 0.5 | 8.73E-02 | -0.8 | 2.98E-02 | GNS1 |
| AT5G04970 | -1.3 | 3.01E-05 | 0.0 | 9.17E-01 | -1.3 | 1.93E-04 | 0.1 | 8.06E-01 | 0.3 | 4.34E-01 | -0.2 | 6.59E-01 | Probable pectinesterase |
| AT1G17950 | -1.3 | 1.20E-05 | -0.8 | 2.62E-04 | -0.4 | 4.31E-02 | 1.8 | 1.03E-03 | 1.1 | 5.80E-02 | 0.7 | 9.16E-02 | Transcription factor MYB52 |
| AT1G21360 | -1.3 | 8.44E-05 | -0.7 | 2.52E-03 | -0.5 | 5.78E-02 | -0.1 | 9.12E-01 | 0.2 | 7.67E-01 | -0.3 | 6.25E-01 | Glycolipid transfer protein 2 |
| AT1G27200 | -1.3 | 6.05E-07 | -1.2 | 2.33E-06 | -0.1 | 4.63E-01 | -0.3 | 9.19E-02 | -0.2 | 4.39E-01 | -0.1 | 6.02E-01 | Glycosyltransferase family 92 protein At1g27200 |
| AT3G25110 | -1.3 | 5.14E-07 | -0.6 | 1.94E-04 | -0.7 | 2.16E-04 | 0.4 | 3.27E-03 | 0.4 | 9.52E-03 | 0.0 | 9.24E-01 | Oleoyl-acyl carrier protein thioesterase 1, chloroplastic |
| AT4G29190 | -1.3 | 9.75E-06 | -1.1 | 5.52E-05 | -0.2 | 3.01E-01 | -0.4 | 1.43E-03 | -0.5 | 8.06E-04 | 0.1 | 6.53E-01 | Zinc finger CCCH domain-containing protein 49 |
| AT2G35730 | -1.3 | 6.03E-05 | -0.4 | 2.81E-02 | -0.8 | 5.04E-03 | | | | | | | Heavy metal transport |
| AT4G22770 | -1.3 | 1.72E-05 | -1.0 | 9.91E-05 | -0.2 | 2.86E-01 | -0.4 | 9.15E-02 | -0.1 | 8.57E-01 | -0.4 | 2.47E-01 | AT-hook motif nuclear-localized protein |
| AT1G22530 | -1.3 | 2.92E-05 | -0.3 | 8.00E-02 | -0.9 | 9.56E-04 | -0.9 | 3.25E-05 | -0.6 | 4.05E-03 | -0.3 | 1.58E-01 | PATELLIN 2 |
| AT2G45550 | -1.3 | 2.25E-06 | -1.0 | 1.48E-05 | -0.2 | 1.71E-01 | 0.4 | 4.63E-02 | 0.7 | 6.44E-03 | -0.2 | 4.12E-01 | Cytochrome P450 76C4 |
| AT5G01100 | -1.3 | 1.39E-05 | -1.0 | 7.71E-05 | -0.2 | 2.72E-01 | 0.6 | 1.46E-02 | 0.3 | 3.51E-01 | 0.3 | 2.44E-01 | Protein FRIABLE 1 |
| AT3G46730 | -1.3 | 1.37E-04 | -0.8 | 2.36E-03 | -0.5 | 1.19E-01 | -0.1 | 7.44E-01 | 0.2 | 7.08E-01 | -0.3 | 4.79E-01 | Putative disease resistance RPP13-like protein 3 |
| AT5G04200 | -1.3 | 3.23E-06 | -0.3 | 3.15E-02 | -1.0 | 1.66E-04 | -0.2 | 5.25E-01 | -0.1 | 8.47E-01 | -0.1 | 8.06E-01 | Metacaspase-9 |
| AT3G54950 | -1.2 | 1.51E-06 | -1.0 | 1.34E-05 | -0.3 | 8.61E-02 | 0.4 | 3.31E-02 | 0.7 | 2.41E-03 | -0.3 | 2.03E-01 | Patatin-like protein 7 |
| AT2G35000 | -1.2 | 1.36E-05 | -0.6 | 2.59E-03 | -0.7 | 4.31E-03 | -0.4 | 1.11E-01 | 0.2 | 5.01E-01 | -0.5 | 2.95E-02 | E3 ubiquitin-protein ligase ATL9 |
| AT3G13782 | -1.2 | 1.77E-05 | -0.3 | 5.30E-02 | -1.0 | 8.22E-04 | -0.9 | 8.87E-02 | -0.1 | 8.63E-01 | -0.8 | 1.95E-01 | Nucleosome assembly protein 1;4 |
| AT3G53590 | -1.2 | 3.85E-07 | -0.6 | 6.05E-05 | -0.6 | 3.73E-04 | 0.8 | 4.73E-03 | 0.1 | 8.59E-01 | 0.7 | 2.96E-02 | Putative leucine-rich repeat receptor-like serine |
| AT5G07920 | -1.2 | 1.13E-07 | -1.0 | 1.02E-06 | -0.3 | 1.46E-02 | 0.0 | 8.76E-01 | 0.0 | 9.84E-01 | 0.0 | 9.25E-01 | Diacylglycerol kinase |
| AT1G49100 | -1.2 | 7.34E-04 | 0.4 | 8.49E-02 | -1.6 | 5.87E-04 | | | | | | | Leucine-rich repeat protein kinase family protein |
| AT4G21550 | -1.2 | 8.86E-05 | -0.6 | 5.98E-03 | -0.6 | 2.91E-02 | -0.3 | 3.70E-01 | 0.1 | 8.08E-01 | -0.5 | 2.86E-01 | VP1 |
| AT4G34500 | -1.2 | 2.97E-06 | -0.6 | 4.13E-04 | -0.6 | 2.24E-03 | -0.3 | 7.70E-02 | -0.1 | 8.32E-01 | -0.3 | 2.01E-01 | Probable receptor-like serine |
| AT5G61260 | -1.2 | 4.98E-06 | -0.5 | 2.44E-03 | -0.8 | 1.11E-03 | -0.7 | 8.99E-02 | -0.2 | 7.60E-01 | -0.5 | 2.45E-01 | Calmodulin-binding domain-containing protein |
| AT5G54670 | -1.2 | 2.50E-07 | -0.5 | 1.38E-04 | -0.7 | 8.59E-05 | -0.3 | 3.63E-02 | 0.1 | 6.92E-01 | -0.4 | 5.10E-02 | Kinesin-like protein |
| AT5G59700 | -1.2 | 2.02E-07 | -1.1 | 8.42E-07 | -0.1 | 3.04E-01 | 0.0 | 8.94E-01 | 0.0 | 9.90E-01 | 0.0 | 9.13E-01 | Probable receptor-like protein kinase At5g59700 |
| AT1G14080 | -1.2 | 1.13E-06 | -0.3 | 9.71E-03 | -0.9 | 8.53E-05 | 0.1 | 7.44E-01 | 0.5 | 1.75E-01 | -0.4 | 3.32E-01 | Fucosyltransferase |
| AT5G25820 | -1.2 | 7.04E-07 | -0.6 | 2.18E-04 | -0.7 | 3.28E-04 | 0.0 | 9.31E-01 | 0.4 | 2.06E-01 | -0.3 | 2.86E-01 | Exostosin family protein |
| AT2G38110 | -1.2 | 1.02E-06 | -0.5 | 6.29E-04 | -0.7 | 2.70E-04 | 1.0 | 9.59E-05 | 0.6 | 1.13E-02 | 0.4 | 1.34E-01 | GPAT6 |
| AT3G03500 | -1.2 | 1.71E-05 | -0.4 | 1.70E-02 | -0.9 | 1.43E-03 | -0.9 | 1.82E-02 | 0.0 | 9.87E-01 | -0.9 | 6.90E-02 | hypothetical protein |
| AT3G11410 | -1.2 | 3.26E-06 | -1.3 | 6.11E-06 | 0.0 | 8.42E-01 | -1.1 | 2.60E-05 | -0.1 | 8.43E-01 | -1.0 | 1.43E-03 | Protein phosphatase 2C 37 |
| AT1G15110 | -1.2 | 5.29E-07 | -0.7 | 5.00E-05 | -0.6 | 8.58E-04 | -0.4 | 5.89E-02 | -0.1 | 6.99E-01 | -0.3 | 2.60E-01 | CDP-diacylglycerol--serine O-phosphatidyltransferase |
| AT3G51350 | -1.2 | 4.46E-06 | -0.3 | 8.48E-02 | -1.0 | 1.29E-04 | -0.1 | 5.12E-01 | 0.3 | 1.66E-01 | -0.5 | 9.23E-02 | Eukaryotic aspartyl protease family protein |
| AT1G08340 | -1.2 | 1.60E-06 | 0.1 | 5.79E-01 | -1.3 | 2.47E-05 | -0.5 | 6.19E-02 | -0.3 | 3.31E-01 | -0.2 | 5.18E-01 | Rho GTPase-activating protein 5 |
| AT4G08160 | -1.2 | 4.53E-06 | 0.0 | 9.01E-01 | -1.2 | 5.85E-05 | -0.3 | 3.07E-01 | -0.2 | 6.12E-01 | -0.1 | 7.74E-01 | Glycosyl hydrolase family 10 protein |
| AT1G51440 | -1.2 | 6.36E-05 | -0.8 | 9.40E-04 | -0.4 | 1.31E-01 | -0.5 | 1.58E-01 | 0.1 | 8.19E-01 | -0.6 | 1.43E-01 | Phospholipase A1-Igamma3, chloroplastic |
| AT1G65610 | -1.2 | 1.10E-06 | -0.2 | 3.54E-02 | -1.0 | 4.90E-05 | 0.6 | 5.22E-02 | 0.5 | 8.42E-02 | 0.0 | 9.82E-01 | Endoglucanase 7 |
| AT2G14960 | -1.2 | 5.58E-05 | -0.2 | 1.35E-01 | -1.0 | 1.72E-03 | 0.2 | 7.44E-01 | 0.1 | 9.38E-01 | 0.1 | 8.81E-01 | Probable indole-3-acetic acid-amido synthetase GH3.1 |
| AT3G14050 | -1.2 | 4.32E-07 | -1.0 | 5.49E-06 | -0.3 | 1.86E-02 | 1.0 | 2.91E-05 | -0.2 | 3.31E-01 | 1.2 | 2.66E-04 | Putative GTP pyrophosphokinase |
| AT5G25110 | -1.2 | 7.08E-07 | -1.4 | 8.82E-07 | 0.2 | 2.06E-01 | -0.1 | 7.98E-01 | 0.5 | 7.71E-02 | -0.6 | 3.63E-02 | CBL-interacting serine |
| AT5G13820 | -1.2 | 4.80E-07 | -0.8 | 1.41E-05 | -0.4 | 3.61E-03 | -0.3 | 2.81E-01 | 0.0 | 9.27E-01 | -0.4 | 2.45E-01 | Telomere repeat-binding protein 4 |
| AT5G67180 | -1.2 | 1.45E-04 | -0.9 | 1.55E-03 | -0.4 | 1.84E-01 | -0.2 | 6.10E-01 | 0.5 | 2.88E-01 | -0.7 | 1.24E-01 | Target of early activation tagged 3 |
| AT4G01840 | -1.2 | 1.01E-06 | -0.8 | 2.09E-05 | -0.4 | 1.22E-02 | -0.6 | 2.11E-03 | -0.1 | 5.79E-01 | -0.5 | 3.80E-02 | Two-pore potassium channel 5 |
| AT5G58784 | -1.2 | 3.17E-04 | 0.0 | 9.15E-01 | -1.2 | 1.82E-03 | -0.3 | 5.59E-01 | 0.2 | 7.59E-01 | -0.5 | 4.02E-01 | Dehydrodolichyl diphosphate synthase 5 |
| AT2G38060 | -1.2 | 1.34E-05 | -0.3 | 2.83E-02 | -0.9 | 8.84E-04 | 0.3 | 4.22E-01 | 0.7 | 6.47E-02 | -0.4 | 3.49E-01 | Phosphate transporter 42 |
| AT3G29430 | -1.2 | 1.03E-06 | 0.1 | 3.13E-01 | -1.3 | 1.47E-05 | -0.9 | 7.61E-03 | 0.0 | 9.91E-01 | -0.9 | 2.31E-02 | Geranylgeranyl pyrophosphate synthase 11, chloroplastic -farnesyl diphosphate synthase 11) |
| AT1G04610 | -1.2 | 3.43E-05 | -0.3 | 4.80E-02 | -0.9 | 1.85E-03 | 0.1 | 7.88E-01 | 0.3 | 4.72E-01 | -0.2 | 7.28E-01 | Probable indole-3-pyruvate monooxygenase YUCCA3 |
| AT1G78440 | -1.2 | 3.63E-05 | -0.9 | 5.11E-04 | -0.4 | 1.30E-01 | | | | | | | Gibberellin 2-beta-dioxygenase 1 |
| AT1G76180 | -1.2 | 6.40E-07 | -0.6 | 1.55E-04 | -0.6 | 4.23E-04 | -0.5 | 1.53E-02 | -0.2 | 4.58E-01 | -0.3 | 2.21E-01 | Dehydrin ERD14 |
| AT4G28890 | -1.2 | 3.02E-07 | -0.6 | 3.83E-05 | -0.6 | 3.52E-04 | -0.2 | 6.75E-01 | 0.2 | 6.60E-01 | -0.3 | 1.87E-01 | E3 ubiquitin-protein ligase ATL42 |
| AT1G27170 | -1.2 | 1.65E-06 | -0.9 | 1.70E-05 | -0.3 | 7.27E-02 | -0.4 | 1.59E-01 | -0.1 | 7.22E-01 | -0.3 | 4.84E-01 | Transmembrane receptors |

| | | | | | | | | | | | | | |
|---|---|---|---|---|---|---|---|---|---|---|---|---|---|
| AT4G35390 | -1.2 | 8.49E-04 | -0.6 | 2.28E-02 | -0.6 | 1.10E-01 | 0.0 | 9.34E-01 | 0.0 | 9.70E-01 | -0.1 | 9.14E-01 | AT-hook motif nuclear-localized protein 25 |
| AT5G49850 | -1.2 | 6.68E-06 | -1.1 | 2.91E-05 | -0.1 | 4.11E-01 | 1.2 | 3.46E-04 | 1.5 | 1.39E-04 | -0.3 | 3.65E-01 | Jacalin-related lectin 46 |
| AT3G03470 | -1.2 | 3.84E-05 | -0.9 | 4.86E-04 | -0.3 | 1.17E-01 | -0.4 | 1.61E-01 | -0.5 | 1.35E-01 | 0.1 | 8.96E-01 | Cytochrome P450 89A9 |
| AT2G42350 | -1.2 | 1.01E-06 | -0.8 | 2.86E-05 | -0.4 | 6.77E-03 | 0.1 | 6.52E-01 | 0.3 | 7.29E-02 | -0.2 | 1.87E-01 | RING-H2 finger protein ATL40 |
| AT1G79620 | -1.2 | 4.67E-07 | -0.7 | 3.40E-05 | -0.5 | 1.07E-03 | -0.3 | 2.30E-01 | 0.1 | 6.67E-01 | -0.4 | 1.53E-01 | Leucine-rich repeat protein kinase family protein |
| AT2G42980 | -1.2 | 1.27E-05 | -1.0 | 8.70E-05 | -0.2 | 2.68E-01 | | | | | | | Eukaryotic aspartyl protease family protein |
| AT1G71960 | -1.2 | 1.52E-07 | -0.9 | 2.10E-06 | -0.3 | 5.93E-03 | -0.5 | 1.82E-03 | -0.3 | 6.56E-02 | -0.2 | 3.18E-01 | ABC transporter G family member 25 |
| AT2G23410 | -1.2 | 8.54E-04 | 0.0 | 9.93E-01 | -1.2 | 3.41E-03 | -0.5 | 2.34E-01 | 0.2 | 5.96E-01 | -0.7 | 1.47E-01 | Dehydrodolichyl diphosphate synthase 1 -farnesyl diphosphate specific) 1) |
| AT2G14100 | -1.2 | 1.92E-05 | -0.2 | 1.29E-01 | -1.0 | 5.48E-04 | -0.5 | 4.81E-02 | -0.1 | 8.64E-01 | -0.5 | 1.29E-01 | Cytochrome P450, family 705, subfamily A, polypeptide 13 |
| AT3G20940 | -1.2 | 1.67E-04 | 0.0 | 9.02E-01 | -1.2 | 1.00E-03 | -0.6 | 1.71E-02 | 0.1 | 7.18E-01 | -0.7 | 1.87E-02 | Cytochrome P450, family 705, subfamily A, polypeptide 30 |
| AT1G21140 | -1.2 | 2.99E-04 | -0.4 | 9.60E-02 | -0.8 | 1.03E-02 | -2.0 | 2.05E-05 | -1.0 | 2.22E-03 | -1.0 | 4.42E-02 | Vacuolar iron transporter homolog 1 |
| AT1G45249 | -1.2 | 1.07E-07 | -1.0 | 7.34E-07 | -0.2 | 3.44E-02 | -0.2 | 3.03E-01 | 0.0 | 9.73E-01 | -0.2 | 4.86E-01 | Abscisic acid responsive elements-binding factor 2 |
| AT4G37340 | -1.2 | 9.77E-07 | -0.6 | 1.32E-04 | -0.6 | 1.02E-03 | -0.2 | 6.22E-01 | 0.1 | 7.30E-01 | -0.3 | 3.25E-01 | Cytochrome P450, family 81, subfamily D, polypeptide 3 |
| AT2G37260 | -1.2 | 2.05E-03 | -0.8 | 1.96E-02 | -0.4 | 2.88E-01 | -0.2 | 6.09E-01 | 0.3 | 4.59E-01 | -0.5 | 1.11E-01 | WRKY family transcription factor family protein |
| AT1G12420 | -1.2 | 1.61E-05 | -1.0 | 1.02E-04 | -0.2 | 2.60E-01 | -0.3 | 1.21E-01 | 0.0 | 9.96E-01 | -0.3 | 7.76E-02 | ACT domain-containing protein ACR |
| AT2G28250 | -1.2 | 2.30E-07 | -0.6 | 4.08E-05 | -0.6 | 1.87E-04 | 0.2 | 2.63E-01 | 0.5 | 2.62E-03 | -0.4 | 4.69E-02 | Receptor-like serine |
| AT2G16230 | -1.2 | 2.46E-03 | 0.2 | 5.60E-01 | -1.4 | 4.26E-03 | -0.2 | 7.16E-01 | 0.5 | 1.65E-01 | -0.7 | 9.15E-02 | glucan endo-1,3-beta-D-glucosidase |
| AT4G35060 | -1.2 | 2.42E-04 | -0.5 | 2.14E-02 | -0.7 | 2.77E-02 | -0.8 | 1.34E-02 | 0.0 | 9.80E-01 | -0.8 | 1.51E-02 | Heavy metal-associated isoprenylated plant protein 25 |
| AT1G01520 | -1.2 | 1.64E-02 | -1.7 | 6.72E-03 | 0.5 | 4.89E-01 | 4.2 | 9.11E-04 | -1.0 | 6.05E-01 | | | Homeodomain-like superfamily protein |
| AT2G23300 | -1.2 | 2.57E-07 | -0.6 | 3.05E-05 | -0.6 | 3.17E-04 | -0.3 | 1.20E-01 | 0.3 | 2.48E-01 | -0.6 | 3.11E-02 | Leucine-rich repeat protein kinase family protein |
| AT3G61410 | -1.2 | 4.26E-06 | -0.9 | 4.75E-05 | -0.3 | 7.19E-02 | 0.0 | 9.61E-01 | -0.3 | 4.31E-01 | 0.3 | 5.58E-01 | U-box kinase family protein |
| AT4G27480 | -1.2 | 1.62E-05 | -0.6 | 2.09E-03 | -0.6 | 6.66E-03 | -0.7 | 1.26E-01 | 0.1 | 8.23E-01 | -0.9 | 9.01E-02 | Uncharacterized protein |
| AT2G01430 | -1.2 | 1.50E-06 | -1.3 | 2.97E-06 | 0.1 | 6.84E-01 | 0.6 | 1.26E-01 | 0.7 | 8.99E-02 | -0.1 | 8.34E-01 | Homeobox-leucine zipper protein 17 |
| AT2G34720 | -1.2 | 3.61E-07 | -1.2 | 8.37E-07 | 0.0 | 9.01E-01 | 0.0 | 9.43E-01 | 0.0 | 9.38E-01 | -0.1 | 8.97E-01 | Nuclear transcription factor Y subunit |
| AT3G54140 | -1.2 | 6.68E-07 | -0.6 | 1.12E-04 | -0.6 | 5.06E-04 | -0.4 | 8.62E-04 | -0.2 | 9.08E-02 | -0.2 | 1.64E-01 | Protein NRT1 |
| AT4G05200 | -1.2 | 7.02E-06 | -0.9 | 7.72E-05 | -0.3 | 1.16E-01 | 0.5 | 1.85E-01 | 0.4 | 3.44E-01 | 0.1 | 8.89E-01 | Cysteine-rich RLK 25 |
| AT5G48780 | -1.2 | 2.58E-05 | -0.5 | 6.93E-03 | -0.7 | 5.37E-03 | -0.8 | 1.73E-02 | 0.0 | 9.70E-01 | -0.8 | 5.22E-02 | Disease resistance protein |
| AT1G61890 | -1.2 | 1.33E-06 | -1.0 | 7.92E-06 | -0.2 | 1.88E-01 | -0.1 | 6.47E-01 | 0.4 | 7.07E-02 | -0.4 | 3.64E-02 | Protein DETOXIFICATION |
| AT4G33080 | -1.2 | 1.57E-07 | -0.8 | 4.32E-06 | -0.4 | 1.34E-03 | -0.4 | 1.68E-03 | 0.0 | 8.46E-01 | -0.3 | 3.38E-02 | AGC kinase family protein |
| AT4G35900 | -1.2 | 3.28E-06 | -0.3 | 2.44E-02 | -0.9 | 1.92E-04 | -0.2 | 4.26E-01 | -0.1 | 8.27E-01 | -0.1 | 7.31E-01 | Basic-leucine zipper transcription factor family protein |
| AT5G01820 | -1.2 | 2.95E-07 | -0.8 | 1.62E-05 | -0.4 | 1.00E-03 | -0.6 | 6.43E-04 | 0.2 | 2.62E-01 | -0.8 | 4.99E-04 | CBL-interacting serine |
| AT3G12240 | -1.2 | 3.29E-05 | 0.0 | 7.64E-01 | -1.1 | 3.20E-04 | 0.0 | 9.51E-01 | 0.3 | 5.79E-01 | -0.3 | 5.46E-01 | Serine carboxypeptidase-like 15 |
| AT3G63280 | -1.2 | 8.22E-08 | -0.7 | 4.35E-06 | -0.5 | 2.58E-04 | -0.3 | 1.12E-01 | 0.0 | 9.13E-01 | -0.3 | 2.47E-01 | NIMA-related kinase 4 |
| AT1G16370 | -1.2 | 3.08E-05 | -0.6 | 3.02E-03 | -0.6 | 1.18E-02 | -0.4 | 1.82E-01 | -0.3 | 4.35E-01 | -0.1 | 7.96E-01 | Organic cation |
| AT5G11100 | -1.2 | 1.91E-06 | -0.7 | 5.74E-05 | -0.4 | 1.03E-02 | -0.3 | 3.07E-01 | 0.1 | 8.05E-01 | -0.4 | 2.68E-01 | Synaptotagmin-4 |
| AT5G54840 | -1.2 | 6.71E-05 | -0.9 | 5.63E-04 | -0.3 | 2.13E-01 | -0.1 | 7.43E-01 | -0.2 | 5.17E-01 | 0.1 | 7.89E-01 | At5g54840 |
| AT2G17330 | -1.2 | 1.59E-03 | 0.2 | 3.76E-01 | -1.4 | 2.26E-03 | -1.5 | 2.55E-02 | -0.2 | 8.13E-01 | -1.3 | 2.52E-02 | Putative sterol 14-demethylase-like protein |
| AT5G38000 | -1.2 | 2.14E-05 | -0.6 | 2.05E-03 | -0.6 | 1.11E-02 | -0.2 | 7.55E-01 | 0.6 | 2.23E-01 | | | At5g38000 |
| AT1G28570 | -1.2 | 2.92E-06 | -0.9 | 2.30E-05 | -0.2 | 1.29E-01 | -0.4 | 5.15E-02 | 0.2 | 3.00E-01 | -0.6 | 2.69E-02 | SGNH hydrolase-type esterase superfamily protein |
| AT2G27000 | -1.2 | 1.49E-06 | -0.3 | 2.01E-02 | -0.9 | 8.82E-05 | -0.3 | 2.27E-01 | 0.1 | 8.09E-01 | -0.4 | 1.99E-01 | Cytochrome P450, family 705, subfamily A, polypeptide 8 |
| AT3G24540 | -1.2 | 9.61E-05 | -0.4 | 2.85E-02 | -0.8 | 8.35E-03 | | | | | | | Proline-rich receptor-like protein kinase PERK3 |
| AT1G32900 | -1.2 | 6.75E-05 | -1.9 | 5.37E-06 | 0.7 | 4.90E-03 | 0.0 | 9.81E-01 | 0.7 | 6.25E-02 | -0.7 | 1.06E-01 | Granule-bound starch synthase 1, chloroplastic |
| AT2G25260 | -1.2 | 1.04E-06 | -0.3 | 2.10E-02 | -0.9 | 5.54E-05 | 0.0 | 9.11E-01 | 0.2 | 4.28E-01 | -0.2 | 4.41E-01 | Hydroxyproline O-arabinosyltransferase 2 |
| AT3G17130 | -1.2 | 2.14E-04 | -0.8 | 3.39E-03 | -0.4 | 1.56E-01 | -0.6 | 2.57E-01 | 0.4 | 4.24E-01 | | | hypothetical protein |
| AT3G55950 | -1.2 | 1.23E-06 | -0.6 | 1.36E-04 | -0.6 | 1.47E-03 | 0.0 | 9.37E-01 | -0.1 | 6.82E-01 | 0.2 | 6.91E-01 | Putative serine |
| AT1G78320 | -1.2 | 1.78E-06 | -0.9 | 1.73E-05 | -0.3 | 7.88E-02 | -0.5 | 2.50E-01 | -0.2 | 6.60E-01 | -0.3 | 6.42E-01 | Glutathione S-transferase |
| AT3G55880 | -1.2 | 3.05E-07 | -0.8 | 6.66E-06 | -0.4 | 4.48E-03 | 0.4 | 3.23E-02 | 0.3 | 2.33E-01 | 0.2 | 5.42E-01 | Alpha |
| AT2G12190 | -1.2 | 2.45E-04 | -0.6 | 1.43E-02 | -0.6 | 4.45E-02 | -0.2 | 7.13E-01 | -0.6 | 2.51E-01 | | | Cytochrome P450 superfamily protein |
| AT5G57620 | -1.2 | 2.56E-05 | -0.2 | 2.24E-01 | -1.0 | 5.62E-04 | -0.3 | 4.11E-01 | 0.4 | 3.72E-01 | -0.7 | 1.28E-01 | Transcription factor MYB36 |
| AT3G09290 | -1.2 | 2.35E-05 | -0.6 | 2.39E-03 | -0.6 | 1.05E-02 | 0.7 | 3.66E-02 | 0.5 | 1.82E-01 | 0.2 | 6.18E-01 | Transcriptional regulator TAC1 |
| AT5G18860 | -1.2 | 6.77E-07 | -0.5 | 3.22E-04 | -0.7 | 2.30E-04 | -0.1 | 7.82E-01 | 0.3 | 2.67E-01 | -0.4 | 2.55E-01 | Nucleoside hydrolase 3 |
| AT4G34590 | -1.2 | 1.93E-05 | -1.0 | 1.28E-04 | -0.2 | 2.49E-01 | 0.4 | 9.56E-03 | 0.5 | 6.42E-03 | -0.1 | 7.53E-01 | bZIP transcription factor 11 |
| AT2G01520 | -1.2 | 1.76E-05 | -0.5 | 1.02E-02 | -0.7 | 2.12E-03 | -1.7 | 1.15E-05 | -0.1 | 6.90E-01 | -1.5 | 1.14E-03 | MLP-like protein 328 |
| AT3G05920 | -1.2 | 3.00E-06 | 0.1 | 6.21E-01 | -1.2 | 3.16E-05 | -0.2 | 4.75E-01 | 0.2 | 5.48E-01 | -0.5 | 2.14E-01 | Heavy metal transport |
| AT3G50260 | -1.2 | 1.62E-05 | -0.9 | 1.66E-04 | -0.3 | 1.30E-01 | 0.5 | 6.11E-02 | 0.1 | 8.07E-01 | 0.4 | 1.00E-01 | AP2 |
| AT4G18640 | -1.2 | 6.85E-06 | -0.3 | 2.97E-02 | -0.9 | 3.53E-04 | -0.6 | 7.15E-03 | 0.0 | 9.46E-01 | -0.6 | 1.81E-02 | Protein MALE DISCOVERER 2 |
| AT5G17050 | -1.2 | 2.32E-03 | -1.2 | 2.17E-03 | 0.1 | 8.37E-01 | -0.1 | 8.07E-01 | 0.6 | 2.23E-02 | -0.7 | 2.32E-02 | Flavonol 3-O-glucosyltransferase |
| AT2G39420 | -1.2 | 7.71E-08 | -0.6 | 1.44E-05 | -0.6 | 5.68E-05 | -0.2 | 2.65E-01 | 0.2 | 2.82E-01 | -0.4 | 4.02E-02 | Alpha |
| AT2G23630 | -1.2 | 3.27E-04 | -0.2 | 3.91E-01 | -1.0 | 3.85E-03 | -0.5 | 8.79E-02 | 0.3 | 3.43E-01 | -0.8 | 3.72E-02 | Putative pectinesterase |
| AT5G67230 | -1.2 | 1.47E-05 | -0.6 | 1.88E-03 | -0.6 | 7.30E-03 | -0.5 | 1.39E-01 | -0.5 | 1.33E-01 | 0.0 | 9.36E-01 | Probable beta-1,4-xylosyltransferase IRX14H |
| AT1G03410 | -1.2 | 9.95E-06 | -0.5 | 1.38E-03 | -0.6 | 5.37E-03 | -0.9 | 5.46E-02 | 0.1 | 8.59E-01 | -1.0 | 7.81E-02 | 2-oxoglutarate and Fe-dependent oxygenase superfamily protein |
| AT1G13600 | -1.2 | 1.27E-03 | -0.8 | 9.80E-03 | -0.4 | 3.31E-01 | | | | | | | Basic leucine-zipper 58 |
| AT4G19030 | -1.2 | 1.03E-06 | 0.2 | 6.99E-02 | -1.3 | 1.09E-05 | -0.8 | 9.24E-04 | -0.3 | 2.62E-01 | -0.5 | 1.74E-02 | NOD26-like major intrinsic protein 1 |
| AT3G61880 | -1.2 | 8.19E-06 | -0.2 | 8.24E-02 | -0.9 | 2.70E-04 | -0.9 | 4.21E-03 | 0.5 | 8.41E-02 | -1.3 | 4.74E-04 | Cytochrome p450 78a9 |
| AT4G11320 | -1.2 | 2.99E-06 | -0.6 | 4.58E-04 | -0.5 | 2.22E-03 | -0.8 | 4.57E-05 | -0.2 | 4.19E-01 | -0.7 | 6.93E-03 | Probable cysteine protease RDL5 |
| AT4G34860 | -1.2 | 5.93E-07 | -0.6 | 7.69E-05 | -0.6 | 6.94E-04 | 0.0 | 9.52E-01 | 0.2 | 4.70E-01 | -0.2 | 5.46E-01 | Probable alkaline |
| AT1G07750 | -1.2 | 1.54E-06 | -0.1 | 2.73E-01 | -1.0 | 3.35E-05 | -0.6 | 2.30E-04 | -0.1 | 4.66E-01 | -0.5 | 5.38E-03 | At1g07750 |
| AT1G58070 | -1.2 | 3.28E-06 | -0.1 | 2.90E-01 | -1.0 | 7.81E-05 | -0.1 | 8.82E-01 | -0.1 | 8.92E-01 | 0.0 | 9.99E-01 | At1g58070 |
| AT1G18140 | -1.2 | 1.80E-05 | -0.4 | 9.01E-03 | -0.7 | 2.64E-03 | 2.5 | 4.33E-07 | 1.3 | 1.99E-03 | 1.3 | 1.52E-04 | Laccase |
| AT3G25820 | -1.2 | 1.23E-02 | -0.9 | 3.88E-02 | -0.3 | 6.30E-01 | | | | | | | Terpene synthase-like sequence-1,8-cineole |
| AT1G47670 | -1.2 | 1.49E-07 | -0.5 | 4.19E-05 | -0.6 | 7.92E-05 | -0.2 | 1.64E-01 | 0.0 | 9.38E-01 | -0.2 | 3.86E-01 | Lysine histidine transporter-like 8 |
| AT5G22740 | -1.2 | 6.09E-07 | -0.7 | 3.55E-05 | -0.4 | 1.67E-03 | -0.3 | 8.28E-03 | 0.1 | 5.35E-01 | -0.4 | 1.40E-02 | Glucomannan 4-beta-mannosyltransferase 2 |
| AT5G10280 | -1.2 | 2.45E-07 | -0.5 | 6.86E-05 | -0.6 | 1.54E-04 | -0.2 | 4.08E-01 | 0.0 | 9.29E-01 | -0.2 | 4.22E-01 | Transcription factor MYB92 |
| AT3G45640 | -1.2 | 4.55E-07 | -1.2 | 9.19E-07 | 0.0 | 6.37E-01 | 0.1 | 4.62E-01 | -0.1 | 5.50E-01 | 0.2 | 2.80E-01 | Mitogen-activated protein kinase 3 |
| AT3G19620 | -1.2 | 2.49E-03 | -0.7 | 3.50E-02 | -0.5 | 2.24E-01 | -0.7 | 3.12E-01 | 0.1 | 9.19E-01 | -0.8 | 3.79E-01 | Probable beta-D-xylosidase 5 |
| AT3G12730 | -1.2 | 6.57E-07 | -0.7 | 1.87E-05 | -0.4 | 5.14E-03 | -0.1 | 6.92E-01 | 0.1 | 8.35E-01 | -0.2 | 5.78E-01 | Homeodomain-like superfamily protein |
| AT1G54160 | -1.1 | 1.02E-04 | -1.0 | 3.47E-04 | -0.1 | 6.98E-01 | | | | | | | Nuclear transcription factor Y subunit A-5 |
| AT5G41800 | -1.1 | 1.30E-06 | -0.4 | 1.24E-03 | -0.7 | 2.59E-04 | -0.7 | 3.56E-04 | 0.0 | 8.35E-01 | -0.6 | 4.63E-03 | Probable GABA transporter 2 |
| AT4G28650 | -1.1 | 8.55E-05 | 0.2 | 2.43E-01 | -1.3 | 2.06E-04 | -0.2 | 6.13E-01 | 0.2 | 5.52E-01 | -0.4 | 2.56E-01 | MDIS1-interacting receptor like kinase 1 |
| AT4G36380 | -1.1 | 1.81E-06 | -0.9 | 2.33E-05 | -0.3 | 4.69E-02 | -0.1 | 5.04E-01 | -0.1 | 6.06E-01 | 0.0 | 9.75E-01 | Cytochrome P450 superfamily protein |
| AT3G14590 | -1.1 | 1.61E-06 | -1.0 | 8.09E-06 | -0.1 | 2.98E-01 | -0.2 | 3.44E-01 | 0.2 | 4.63E-01 | -0.4 | 1.39E-01 | Calcium-dependent lipid-binding family protein |
| AT1G68390 | -1.1 | 4.69E-05 | -0.5 | 5.10E-03 | -0.6 | 1.55E-02 | 0.9 | 1.68E-02 | 0.5 | 2.68E-01 | 0.4 | 2.44E-01 | Core-2 |
| AT1G18710 | -1.1 | 2.42E-03 | -1.0 | 7.56E-03 | -0.2 | 7.00E-01 | -0.5 | 1.11E-02 | 0.0 | 9.87E-01 | -0.5 | 4.09E-02 | MYB transcription factor |

| | | | | | | | | | | | | | |
|---|---|---|---|---|---|---|---|---|---|---|---|---|---|
| AT5G22630 | -1.1 | 4.81E-07 | -0.7 | 3.29E-05 | -0.4 | 1.19E-03 | 0.4 | 7.60E-03 | 0.5 | 7.01E-03 | -0.1 | 8.28E-01 | Arogenate dehydratase 5, chloroplastic |
| AT3G09710 | -1.1 | 2.11E-06 | -0.5 | 7.44E-04 | -0.6 | 8.45E-04 | -0.8 | 3.49E-04 | -0.1 | 5.19E-01 | -0.7 | 1.60E-02 | IQ-domain 1 |
| AT4G00940 | -1.1 | 1.97E-05 | -1.0 | 7.02E-05 | -0.1 | 5.85E-01 | 0.2 | 3.47E-01 | 0.0 | 9.76E-01 | 0.2 | 4.59E-01 | Dof zinc finger protein DOF4.1 |
| AT1G16310 | -1.1 | 3.57E-06 | -0.8 | 5.89E-05 | -0.3 | 4.16E-02 | -0.4 | 2.48E-01 | 0.1 | 8.01E-01 | -0.5 | 2.17E-01 | Cation efflux protein cytoplasmic domain-containing protein |
| AT1G75450 | -1.1 | 7.99E-06 | -1.2 | 1.18E-05 | 0.1 | 6.40E-01 | | | | | | | cytokinin dehydrogenase |
| AT4G11230 | -1.1 | 2.41E-06 | -0.7 | 1.65E-04 | -0.5 | 4.39E-03 | 0.1 | 7.37E-01 | 0.2 | 4.58E-01 | -0.1 | 7.23E-01 | Probable respiratory burst oxidase homolog protein I |
| AT4G08570 | -1.1 | 8.95E-04 | -1.2 | 9.70E-04 | 0.1 | 8.72E-01 | 0.5 | 2.65E-01 | 0.5 | 3.08E-01 | 0.0 | 9.89E-01 | Heavy metal-associated isoprenylated plant protein 24 |
| AT4G00460 | -1.1 | 1.16E-05 | -0.1 | 3.77E-01 | -1.0 | 2.21E-04 | 0.0 | 9.51E-01 | 0.3 | 6.83E-01 | -0.2 | 7.19E-01 | RHO guanyl-nucleotide exchange factor 3 |
| AT5G41315 | -1.1 | 5.34E-05 | -0.2 | 1.52E-01 | -0.9 | 1.51E-03 | -0.1 | 9.04E-01 | 0.3 | 5.54E-01 | -0.3 | 4.57E-01 | Basic helix-loop-helix DNA-binding superfamily protein |
| AT3G24670 | -1.1 | 1.14E-06 | -0.1 | 3.61E-01 | -1.0 | 2.85E-05 | -0.3 | 7.57E-02 | 0.0 | 9.76E-01 | -0.3 | 1.67E-01 | Probable pectate lyase 10 |
| AT1G55430 | -1.1 | 6.26E-05 | -0.2 | 3.10E-01 | -1.0 | 1.18E-03 | 0.0 | 9.21E-01 | 0.5 | 1.47E-01 | -0.5 | 2.17E-01 | T5A14.15 protein |
| AT2G17050 | -1.1 | 1.93E-06 | -0.9 | 1.28E-05 | -0.2 | 1.78E-01 | 0.3 | 3.68E-01 | 0.3 | 5.60E-01 | 0.1 | 8.95E-01 | Disease resistance protein |
| AT4G13440 | -1.1 | 4.96E-05 | 0.0 | 9.69E-01 | -1.1 | 3.43E-04 | -0.1 | 8.35E-01 | 0.5 | 2.26E-01 | -0.6 | 6.07E-02 | Calcium-binding EF-hand family protein |
| AT5G54080 | -1.1 | 8.20E-07 | -1.0 | 3.59E-06 | -0.1 | 3.63E-01 | -0.9 | 9.07E-03 | -0.5 | 1.47E-01 | -0.4 | 3.90E-01 | homogentisate 1,2-dioxygenase |
| AT3G20830 | -1.1 | 2.38E-05 | -0.8 | 5.01E-04 | -0.4 | 6.15E-02 | -0.4 | 2.61E-02 | -0.2 | 5.02E-01 | -0.3 | 2.35E-01 | Serine |
| AT5G62580 | -1.1 | 1.71E-07 | -0.6 | 2.51E-05 | -0.5 | 1.69E-04 | 0.3 | 1.02E-01 | 0.4 | 5.66E-02 | -0.1 | 7.57E-01 | TORTIFOLIA1-like protein 3 |
| AT5G07780 | -1.1 | 1.53E-06 | -0.4 | 2.26E-03 | -0.7 | 2.54E-04 | -0.3 | 1.72E-01 | 0.1 | 7.45E-01 | -0.4 | 1.30E-01 | Formin-like protein 19 |
| AT3G27200 | -1.1 | 4.25E-06 | -0.1 | 3.37E-01 | -1.0 | 8.64E-05 | -0.3 | 1.18E-01 | 0.0 | 9.75E-01 | -0.3 | 1.60E-01 | Phytocyanin domain-containing protein |
| AT4G02180 | -1.1 | 1.29E-05 | -0.2 | 1.55E-01 | -0.9 | 3.82E-04 | -0.3 | 5.94E-01 | 0.1 | 9.33E-01 | -0.3 | 6.30E-01 | DC1 domain-containing protein |
| AT4G20210 | -1.1 | 3.85E-07 | -0.3 | 6.97E-03 | -0.9 | 3.35E-05 | -0.4 | 9.51E-02 | 0.1 | 6.70E-01 | -0.5 | 2.78E-02 | Terpenoid synthase 8 |
| AT5G64530 | -1.1 | 1.03E-04 | -1.0 | 4.01E-04 | -0.1 | 5.75E-01 | 0.6 | 2.55E-01 | 0.2 | 7.63E-01 | 0.4 | 5.27E-01 | Xylem NAC domain 1 |
| AT5G57050 | -1.1 | 4.51E-05 | -1.1 | 7.31E-05 | 0.0 | 9.06E-01 | -0.8 | 4.63E-03 | 0.0 | 9.64E-01 | -0.9 | 1.66E-02 | protein-serine |
| AT4G01770 | -1.1 | 7.39E-05 | 0.1 | 5.01E-01 | -1.2 | 2.71E-04 | -0.4 | 1.59E-01 | 0.1 | 6.79E-01 | -0.5 | 1.11E-01 | Glycosyltransferase |
| AT4G17800 | -1.1 | 1.49E-06 | -0.5 | 1.04E-03 | -0.6 | 3.26E-04 | -0.3 | 1.03E-01 | -0.1 | 6.55E-01 | -0.2 | 3.49E-01 | AT-hook motif nuclear-localized protein 23 |
| AT5G44030 | -1.1 | 5.38E-07 | -0.2 | 6.86E-02 | -1.0 | 2.69E-05 | -0.4 | 6.36E-02 | 0.0 | 8.77E-01 | -0.3 | 1.92E-01 | Cellulose synthase |
| AT3G61830 | -1.1 | 1.32E-06 | -0.5 | 2.59E-04 | -0.6 | 9.61E-04 | -0.4 | 7.33E-02 | -0.3 | 2.17E-01 | -0.1 | 8.18E-01 | Auxin response factor |
| AT5G17850 | -1.1 | 1.65E-06 | -0.8 | 2.38E-05 | -0.3 | 3.37E-02 | -0.5 | 2.36E-02 | -0.2 | 4.16E-01 | -0.3 | 2.29E-01 | Cation |
| AT1G65850 | -1.1 | 1.06E-04 | -0.8 | 1.16E-03 | -0.3 | 2.21E-01 | | | | | | | ADP-ribosyl cyclase |
| AT3G53230 | -1.1 | 1.17E-06 | -1.4 | 7.73E-07 | 0.3 | 3.11E-02 | 0.6 | 1.32E-02 | 0.6 | 2.43E-02 | 0.0 | 9.76E-01 | Cell division control protein 48 homolog D |
| AT2G45040 | -1.1 | 1.13E-04 | -0.4 | 5.27E-02 | -0.8 | 7.35E-03 | 0.3 | 4.08E-01 | 0.1 | 8.96E-01 | 0.2 | 6.66E-01 | Metalloendoproteinase 4-MMP |
| AT2G34470 | -1.1 | 4.24E-07 | -0.5 | 1.38E-04 | -0.6 | 1.99E-04 | -0.4 | 4.55E-04 | -0.1 | 5.94E-01 | -0.4 | 2.55E-02 | Urease accessory protein G |
| AT2G22620 | -1.1 | 2.29E-06 | -0.8 | 4.70E-05 | -0.4 | 2.38E-02 | 0.1 | 7.56E-01 | 0.2 | 6.07E-01 | -0.1 | 8.55E-01 | rhamnogalacturonan endolyase |
| AT1G50560 | -1.1 | 2.17E-05 | -0.4 | 6.84E-03 | -0.7 | 3.97E-03 | -0.9 | 1.72E-03 | -0.3 | 3.10E-01 | -0.7 | 2.49E-02 | Cytochrome P450, family 705, subfamily A, polypeptide 25 |
| AT3G55890 | -1.1 | 3.29E-04 | -1.5 | 1.32E-04 | 0.4 | 2.55E-01 | | | | | | | Protein yippee-like |
| AT4G16190 | -1.1 | 4.34E-07 | -1.1 | 1.11E-06 | 0.0 | 9.58E-01 | -0.7 | 5.24E-06 | -0.3 | 2.08E-02 | -0.4 | 1.66E-02 | Probable cysteine protease RD19C |
| AT1G77690 | -1.1 | 3.67E-07 | -0.3 | 7.14E-03 | -0.8 | 2.86E-05 | 0.1 | 6.30E-01 | 0.4 | 2.22E-02 | -0.3 | 3.86E-02 | Auxin transporter-like protein 3 |
| AT3G20130 | -1.1 | 7.85E-07 | -0.4 | 1.90E-03 | -0.7 | 1.21E-04 | -0.6 | 5.00E-03 | -0.2 | 4.31E-01 | -0.5 | 1.09E-01 | Cytochrome P450, family 705, subfamily A, polypeptide 22 |
| AT2G40840 | -1.1 | 1.91E-05 | -1.0 | 8.30E-05 | -0.1 | 4.96E-01 | -0.1 | 7.57E-01 | 0.2 | 4.11E-01 | -0.2 | 3.44E-01 | 4-alpha-glucanotransferase DPE2 |
| AT1G72120 | -1.1 | 1.36E-05 | -0.4 | 5.52E-03 | -0.7 | 2.68E-03 | -0.5 | 9.70E-02 | 0.2 | 5.72E-01 | -0.7 | 6.47E-02 | Protein NRT1 |
| AT2G43040 | -1.1 | 1.21E-06 | -0.2 | 2.63E-02 | -0.9 | 7.08E-05 | -0.6 | 1.56E-02 | 0.0 | 9.66E-01 | -0.6 | 6.92E-02 | Protein NPG1 |
| AT5G66420 | -1.1 | 1.14E-06 | -0.7 | 4.80E-05 | -0.4 | 4.33E-03 | -0.3 | 2.56E-02 | 0.2 | 2.21E-01 | -0.5 | 1.44E-02 | AT5G66420 protein |
| AT1G44970 | -1.1 | 1.24E-05 | 0.0 | 8.34E-01 | -1.1 | 9.80E-05 | -0.5 | 1.04E-01 | -0.1 | 7.81E-01 | -0.4 | 2.53E-01 | Peroxidase 9 |
| AT5G06570 | -1.1 | 8.35E-06 | -0.6 | 5.48E-04 | -0.5 | 9.39E-03 | 0.0 | 9.94E-01 | 0.1 | 5.49E-01 | -0.1 | 6.17E-01 | Alpha |
| AT3G07340 | -1.1 | 6.45E-07 | -0.7 | 2.64E-05 | -0.4 | 3.15E-03 | -0.2 | 2.01E-01 | -0.2 | 4.14E-01 | -0.1 | 8.36E-01 | Basic helix-loop-helix DNA-binding superfamily protein |
| AT4G34000 | -1.1 | 5.13E-07 | -1.0 | 2.01E-06 | -0.1 | 4.38E-01 | -0.5 | 1.66E-02 | 0.0 | 9.66E-01 | -0.5 | 4.73E-02 | BZIP domain-containing protein |
| AT5G24410 | -1.1 | 1.92E-06 | 0.2 | 6.08E-02 | -1.3 | 1.47E-05 | -1.5 | 7.61E-06 | -0.4 | 6.86E-02 | -1.1 | 1.63E-03 | Probable 6-phosphogluconolactonase 4 |
| AT5G65090 | -1.1 | 1.44E-05 | -0.4 | 1.21E-02 | -0.7 | 1.63E-03 | -0.4 | 3.88E-01 | 0.3 | 5.63E-01 | -0.7 | 2.25E-01 | DNAse I-like superfamily protein |
| AT1G76090 | -1.1 | 1.23E-05 | -0.1 | 5.09E-01 | -1.0 | 1.34E-04 | -1.0 | 1.89E-04 | -0.3 | 1.53E-01 | -0.6 | 2.21E-02 | 24-methylenesterol C-methyltransferase 3 |
| AT5G35370 | -1.1 | 4.06E-06 | -0.8 | 7.10E-05 | -0.3 | 3.81E-02 | -0.2 | 4.53E-01 | -0.1 | 6.60E-01 | -0.1 | 8.89E-01 | G-type lectin S-receptor-like serine |
| AT5G63970 | -1.1 | 5.13E-07 | -0.7 | 2.03E-05 | -0.4 | 2.65E-03 | -0.3 | 3.83E-01 | 0.2 | 6.15E-01 | -0.4 | 1.53E-01 | E3 ubiquitin-protein ligase RGLG3 |
| AT4G28500 | -1.1 | 2.54E-04 | -0.2 | 2.12E-01 | -0.9 | 5.71E-03 | | | | | | | NAC domain-containing protein 73 |
| AT1G77920 | -1.1 | 8.35E-07 | -1.2 | 1.11E-06 | 0.1 | 2.82E-01 | 0.0 | 9.92E-01 | 0.4 | 7.71E-02 | -0.4 | 1.10E-01 | Transcription factor TGA7 |
| AT4G11610 | -1.1 | 1.32E-04 | -0.1 | 6.52E-01 | -1.0 | 1.18E-03 | 0.0 | 9.34E-01 | 0.5 | 2.26E-01 | -0.5 | 2.33E-01 | C2 calcium |
| AT5G11540 | -1.1 | 5.58E-05 | -0.1 | 5.03E-01 | -1.0 | 7.02E-04 | -0.5 | 2.10E-01 | 0.2 | 7.26E-01 | -0.6 | 1.92E-01 | L-gulonolactone oxidase 3 |
| AT1G04100 | -1.1 | 3.35E-05 | -0.7 | 9.67E-04 | -0.4 | 5.25E-02 | -0.2 | 6.56E-01 | -0.1 | 8.13E-01 | -0.1 | 9.26E-01 | Auxin-responsive protein IAA10 |
| AT4G08450 | -1.1 | 2.98E-04 | -0.7 | 4.56E-03 | -0.4 | 1.70E-01 | | | | | | | ADP-ribosyl cyclase |
| AT1G23440 | -1.1 | 2.62E-07 | -0.7 | 1.10E-05 | -0.4 | 1.16E-03 | -0.3 | 1.07E-01 | -0.1 | 6.96E-01 | -0.2 | 4.80E-01 | Peptidase C15, pyroglutamyl peptidase I-like protein |
| AT1G67510 | -1.1 | 6.05E-06 | -0.5 | 8.72E-04 | -0.5 | 3.77E-03 | 0.0 | 8.87E-01 | 0.1 | 4.30E-01 | -0.2 | 4.25E-01 | Leucine-rich repeat protein kinase family protein |
| AT1G67820 | -1.1 | 2.61E-05 | -1.0 | 6.32E-05 | 0.0 | 8.83E-01 | -0.2 | 6.69E-01 | 0.2 | 6.07E-01 | -0.3 | 3.93E-01 | protein-serine |
| AT3G55120 | -1.1 | 9.61E-04 | -0.7 | 1.07E-02 | -0.4 | 2.36E-01 | 0.3 | 2.72E-01 | 0.8 | 6.44E-03 | -0.5 | 8.53E-02 | Chalcone-flavonone isomerase family protein |
| AT5G10830 | -1.1 | 1.01E-06 | -0.3 | 4.16E-03 | -0.8 | 1.08E-04 | -0.1 | 7.89E-01 | 0.2 | 4.38E-01 | -0.2 | 3.55E-01 | hypothetical protein |
| AT3G06990 | -1.1 | 2.00E-04 | 0.0 | 9.59E-01 | -1.1 | 1.19E-03 | -0.6 | 8.20E-02 | 0.0 | 9.84E-01 | -0.6 | 9.49E-02 | Cysteine |
| AT5G57110 | -1.1 | 2.14E-07 | -0.4 | 4.80E-04 | -0.7 | 3.65E-05 | -0.2 | 1.82E-01 | 0.1 | 7.99E-01 | -0.2 | 1.94E-01 | Calcium-transporting ATPase 8, plasma membrane-type -ATPase isoform 8) |
| AT3G15890 | -1.1 | 3.53E-06 | -0.4 | 1.73E-03 | -0.6 | 9.78E-04 | -0.3 | 2.00E-01 | 0.1 | 7.67E-01 | -0.4 | 1.26E-01 | PTI1-like tyrosine-protein kinase At3g15890 |
| AT2G15310 | -1.1 | 3.00E-06 | -0.7 | 8.08E-05 | -0.4 | 1.87E-02 | -0.1 | 8.47E-01 | -0.4 | 3.92E-01 | 0.3 | 5.77E-01 | Probable ADP-ribosylation factor At2g15310 |
| AT5G44670 | -1.1 | 5.13E-07 | -0.7 | 1.62E-05 | -0.4 | 3.75E-03 | -0.4 | 2.52E-02 | -0.1 | 8.47E-01 | -0.4 | 1.09E-01 | Galactan beta-1,4-galactosyltransferase GALS2 |
| AT5G45240 | -1.1 | 3.57E-04 | -0.4 | 7.09E-02 | -0.7 | 1.95E-02 | 1.1 | 8.73E-03 | 0.1 | 8.25E-01 | 0.9 | 2.43E-02 | Disease resistance protein |
| AT5G01490 | -1.1 | 2.31E-06 | -0.8 | 3.06E-05 | -0.3 | 4.72E-02 | 0.4 | 8.09E-02 | 0.3 | 3.25E-01 | 0.1 | 6.87E-01 | Vacuolar cation |
| AT4G16110 | -1.1 | 5.56E-07 | -0.9 | 4.18E-06 | -0.2 | 8.26E-02 | -0.5 | 6.21E-03 | -0.2 | 1.87E-01 | -0.2 | 3.17E-01 | Two-component response regulator ARR2 |
| AT5G43370 | -1.1 | 2.46E-04 | -0.1 | 4.54E-01 | -0.9 | 2.83E-03 | -0.3 | 3.69E-01 | 0.0 | 9.66E-01 | -0.3 | 5.49E-01 | Probable inorganic phosphate transporter 1-2 |
| AT4G15300 | -1.1 | 5.65E-05 | 0.1 | 6.17E-01 | -1.1 | 2.35E-04 | 0.3 | 3.54E-01 | 0.9 | 2.02E-02 | -0.5 | 1.27E-01 | Cytochrome P450 like protein |
| AT2G30840 | -1.1 | 3.67E-07 | -0.7 | 1.07E-05 | -0.4 | 3.46E-03 | 0.3 | 2.75E-01 | 0.2 | 4.24E-01 | 0.0 | 9.29E-01 | 2-oxoglutarate and Fe-dependent oxygenase superfamily protein |
| AT3G14610 | -1.1 | 1.03E-06 | -0.9 | 8.61E-06 | -0.2 | 8.86E-02 | -0.2 | 3.82E-01 | -0.1 | 6.27E-01 | -0.1 | 8.73E-01 | Cytochrome P450 |
| AT1G20640 | -1.1 | 3.17E-07 | -0.6 | 2.08E-05 | -0.4 | 7.18E-04 | 0.1 | 7.91E-01 | 0.3 | 2.95E-01 | -0.3 | 5.27E-01 | Protein NLP4 |
| AT3G01080 | -1.1 | 1.69E-04 | -0.9 | 8.93E-04 | -0.2 | 4.14E-01 | -0.5 | 6.87E-02 | -0.3 | 4.06E-01 | -0.3 | 4.66E-01 | WRKY DNA-binding protein 58 |
| AT4G14465 | -1.1 | 6.57E-07 | -0.7 | 1.71E-05 | -0.3 | 5.66E-03 | 0.5 | 1.12E-02 | 0.5 | 3.44E-02 | 0.1 | 8.41E-01 | AT-hook motif nuclear-localized protein 20 |
| AT1G30510 | -1.1 | 1.35E-05 | -0.4 | 1.87E-02 | -0.7 | 9.54E-04 | 0.1 | 5.26E-01 | 0.1 | 4.00E-01 | 0.0 | 8.14E-01 | Ferredoxin--NADP reductase, root isozyme 2, chloroplastic |
| AT3G16390 | -1.1 | 6.93E-07 | 0.1 | 1.82E-01 | -1.2 | 1.05E-05 | -1.4 | 2.79E-05 | -0.3 | 2.50E-01 | -1.1 | 3.10E-04 | Thiohydroximate-O-sulfate sulfur |
| AT4G18430 | -1.1 | 6.86E-06 | -0.1 | 2.72E-01 | -0.9 | 1.38E-04 | 0.5 | 8.66E-02 | 0.3 | 4.30E-01 | 0.2 | 5.28E-01 | Ras-related protein RABA1e |
| AT5G06839 | -1.1 | 2.15E-06 | -0.8 | 2.09E-05 | -0.2 | 8.58E-02 | 0.6 | 3.03E-03 | 0.4 | 5.96E-02 | 0.2 | 3.53E-01 | BZIP transcription factor family protein |
| AT1G02880 | -1.1 | 1.00E-06 | -0.8 | 1.00E-05 | -0.2 | 5.87E-02 | -0.2 | 2.62E-01 | 0.1 | 7.77E-01 | -0.3 | 2.30E-01 | thiamine diphosphokinase |
| AT5G49770 | -1.1 | 2.83E-06 | -0.1 | 6.40E-01 | -1.0 | 3.78E-05 | 0.2 | 3.77E-01 | 0.4 | 6.11E-02 | -0.2 | 3.60E-01 | Probable leucine-rich repeat receptor-like protein kinase At5g49770 |
| AT2G44790 | -1.1 | 7.04E-06 | 0.1 | 5.42E-01 | -1.1 | 3.87E-05 | 0.9 | 8.20E-05 | 0.4 | 8.41E-02 | 0.5 | 9.89E-03 | Uclacyanin-2 |

| | | | | | | | | | | | | | |
|---|---|---|---|---|---|---|---|---|---|---|---|---|---|
| AT5G24960 | -1.1 | 2.17E-05 | -0.2 | 1.52E-01 | -0.9 | 6.21E-04 | -0.6 | 3.77E-02 | 0.1 | 7.83E-01 | -0.7 | 6.02E-02 | Cytochrome P450, family 71, subfamily A, polypeptide 14 |
| AT3G21720 | -1.1 | 1.22E-03 | 0.1 | 7.64E-01 | -1.1 | 3.59E-03 | -0.6 | 4.80E-01 | 0.3 | 7.24E-01 | | | Isocitrate lyase |
| AT4G24120 | -1.1 | 3.25E-06 | -0.9 | 2.61E-05 | -0.2 | 1.30E-01 | -1.2 | 1.45E-04 | -0.4 | 1.13E-01 | -0.8 | 4.77E-03 | Metal-nicotianamine transporter YSL1 |
| AT3G14370 | -1.1 | 2.98E-04 | -0.1 | 4.65E-01 | -0.9 | 3.27E-03 | 0.2 | 5.69E-01 | 0.4 | 1.87E-01 | -0.2 | 5.29E-01 | Serine |
| AT4G24170 | -1.1 | 4.69E-04 | 0.2 | 2.38E-01 | -1.3 | 7.16E-04 | 0.2 | 7.61E-01 | -0.3 | 7.02E-01 | 0.5 | 5.22E-01 | Kinesin-like protein KIN-7I |
| AT5G51490 | -1.1 | 5.80E-04 | 0.3 | 1.02E-01 | -1.4 | 4.95E-04 | -0.1 | 8.95E-01 | 0.6 | 8.08E-02 | -0.7 | 9.23E-02 | Probable pectinesterase |
| AT5G51060 | -1.1 | 5.08E-06 | -0.2 | 1.56E-01 | -0.9 | 1.19E-04 | 0.2 | 6.38E-01 | 0.4 | 2.94E-01 | -0.2 | 5.90E-01 | Respiratory burst oxidase homolog protein C |
| AT4G32250 | -1.1 | 1.28E-05 | -0.8 | 1.42E-04 | -0.3 | 1.23E-01 | -0.5 | 1.97E-02 | -0.2 | 4.19E-01 | -0.3 | 2.74E-01 | Protein KINASE OF THE OUTER CHLOROPLAST MEMBRANE 1 |
| AT1G30690 | -1.1 | 8.45E-07 | -0.4 | 2.49E-03 | -0.7 | 9.67E-05 | -0.4 | 5.83E-03 | -0.1 | 7.66E-01 | -0.3 | 6.35E-02 | Patellin-4 |
| AT2G34660 | -1.1 | 1.17E-06 | -1.1 | 2.54E-06 | 0.0 | 8.71E-01 | 0.0 | 9.92E-01 | 0.4 | 2.13E-01 | -0.4 | 3.11E-01 | ABC-type xenobiotic transporter |
| AT4G02940 | -1.0 | 7.40E-07 | -0.7 | 2.95E-05 | -0.4 | 3.25E-03 | 0.1 | 4.53E-01 | 0.2 | 2.05E-01 | -0.1 | 6.96E-01 | RNA demethylase ALKBH10B |
| AT4G22780 | -1.0 | 2.26E-06 | -0.9 | 1.28E-05 | -0.1 | 2.41E-01 | -0.6 | 4.00E-04 | -0.3 | 8.09E-02 | -0.3 | 1.04E-01 | ACT domain-containing protein ACR7 |
| AT3G47400 | -1.0 | 1.50E-05 | -0.3 | 3.21E-02 | -0.8 | 9.61E-04 | -0.3 | 3.65E-01 | 0.3 | 4.58E-01 | -0.6 | 1.49E-01 | Probable pectinesterase |
| AT1G44170 | -1.0 | 1.84E-07 | -0.6 | 1.34E-05 | -0.4 | 3.74E-04 | 0.1 | 5.55E-01 | 0.1 | 2.81E-01 | -0.1 | 7.11E-01 | Aldehyde dehydrogenase family 3 member H1 |
| AT5G19040 | -1.0 | 3.00E-06 | -0.5 | 9.56E-04 | -0.6 | 1.20E-03 | -0.1 | 5.80E-01 | 0.2 | 3.51E-01 | -0.4 | 1.89E-01 | Adenylate isopentenyltransferase 5, chloroplastic |
| AT3G20570 | -1.0 | 2.10E-06 | -0.4 | 1.50E-03 | -0.6 | 4.24E-04 | 0.1 | 5.95E-01 | 0.2 | 2.64E-01 | -0.1 | 6.27E-01 | Early nodulin-like protein 9 |
| AT5G39380 | -1.0 | 2.75E-06 | -0.6 | 2.21E-04 | -0.5 | 4.21E-03 | -0.1 | 6.38E-01 | 0.2 | 6.27E-01 | -0.3 | 2.86E-01 | Calmodulin-binding domain-containing protein |
| AT1G12480 | -1.0 | 3.41E-04 | -0.5 | 1.51E-02 | -0.5 | 6.82E-02 | | | | | | | Guard cell S-type anion channel SLAC1 |
| AT5G43040 | -1.0 | 4.45E-05 | 0.0 | 9.81E-01 | -1.0 | 3.05E-04 | -0.4 | 1.71E-01 | 0.3 | 4.11E-01 | -0.7 | 7.33E-02 | CHP-rich zinc finger protein-like |
| AT1G76690 | -1.0 | 2.36E-06 | -0.5 | 8.61E-04 | -0.6 | 8.81E-04 | 0.5 | 5.09E-02 | 0.6 | 1.17E-02 | -0.2 | 3.60E-01 | 12-oxophytodienoate reductase 2 |
| AT1G79110 | -1.0 | 2.33E-05 | -1.2 | 1.92E-05 | 0.1 | 3.90E-01 | -0.4 | 1.70E-03 | 0.1 | 4.17E-01 | -0.5 | 3.90E-03 | RING-type domain-containing protein |
| AT3G62820 | -1.0 | 2.87E-04 | -0.2 | 2.80E-01 | -0.8 | 4.63E-03 | -0.3 | 8.65E-02 | 0.1 | 5.30E-01 | -0.4 | 6.76E-02 | Pectinesterase inhibitor domain-containing protein |
| AT1G18910 | -1.0 | 6.87E-07 | -0.9 | 4.77E-06 | -0.2 | 1.15E-01 | -0.1 | 5.62E-01 | 0.1 | 8.52E-01 | -0.2 | 5.01E-01 | Zinc finger protein BRUTUS-like At1g18910 |
| AT3G05580 | -1.0 | 3.20E-06 | -0.4 | 1.47E-03 | -0.6 | 8.82E-04 | -0.3 | 3.23E-02 | -0.2 | 3.74E-01 | -0.2 | 3.90E-01 | Serine |
| AT3G29250 | -1.0 | 1.54E-04 | -0.2 | 2.99E-01 | -0.9 | 2.05E-03 | 0.0 | 9.38E-01 | 0.2 | 2.75E-01 | -0.2 | 1.82E-01 | Short-chain dehydrogenase reductase 4 |
| AT1G14120 | -1.0 | 5.59E-05 | -1.1 | 6.56E-05 | 0.1 | 6.98E-01 | 0.5 | 3.65E-01 | 0.5 | 3.80E-01 | 0.0 | 9.62E-01 | Fe2OG dioxygenase domain-containing protein |
| AT4G18260 | -1.0 | 1.13E-05 | -0.3 | 2.62E-02 | -0.8 | 8.24E-04 | -0.6 | 9.36E-02 | -0.3 | 3.78E-01 | -0.2 | 6.54E-01 | Cytochrome b561 domain-containing protein At4g18260 |
| AT5G27000 | -1.0 | 5.41E-06 | -0.9 | 3.65E-05 | -0.2 | 2.10E-01 | -0.2 | 6.16E-01 | 0.2 | 5.15E-01 | -0.4 | 2.36E-01 | Kinesin-like protein KIN-14G |
| AT1G70800 | -1.0 | 2.27E-03 | -0.8 | 1.23E-02 | -0.3 | 4.28E-01 | 0.2 | 5.13E-01 | 0.5 | 1.67E-01 | -0.3 | 4.40E-01 | Protein C2-DOMAIN ABA-RELATED 6 |
| AT4G13990 | -1.0 | 3.08E-05 | -0.7 | 3.98E-04 | -0.3 | 1.26E-01 | 0.2 | 3.76E-01 | 0.3 | 1.63E-01 | -0.1 | 6.73E-01 | Probable xyloglucan galactosyltransferase GT14 |
| AT1G68470 | -1.0 | 7.92E-06 | -1.1 | 1.26E-05 | 0.1 | 7.56E-01 | -0.1 | 5.57E-01 | 0.2 | 3.16E-01 | -0.3 | 1.60E-01 | Probable xyloglucan galactosyltransferase GT17 |
| AT1G22150 | -1.0 | 1.50E-03 | 0.1 | 6.15E-01 | -1.1 | 2.98E-03 | 1.9 | 1.28E-03 | 1.9 | 2.84E-03 | 0.0 | 9.97E-01 | Sulfate transporter 13 |
| AT2G39410 | -1.0 | 3.48E-06 | -0.3 | 4.76E-03 | -0.7 | 4.87E-04 | -0.6 | 1.29E-02 | 0.0 | 8.78E-01 | -0.6 | 3.52E-02 | Alpha |
| AT1G26960 | -1.0 | 3.86E-06 | -0.2 | 9.94E-02 | -0.9 | 1.35E-04 | -0.4 | 7.18E-02 | -0.2 | 2.91E-01 | -0.1 | 6.71E-01 | Homeobox-leucine zipper protein ATHB-23 |
| AT5G11230 | -1.0 | 3.30E-07 | -0.2 | 1.54E-02 | -0.8 | 2.63E-05 | -0.2 | 3.24E-01 | 0.1 | 7.52E-01 | -0.3 | 2.08E-01 | Probable sugar phosphate |
| AT1G51860 | -1.0 | 4.33E-05 | -0.3 | 3.63E-02 | -0.7 | 2.43E-03 | -0.6 | 1.44E-02 | -0.1 | 6.31E-01 | -0.4 | 1.61E-01 | Leucine-rich repeat protein kinase family protein |
| AT5G10240 | -1.0 | 1.48E-06 | -0.4 | 1.05E-03 | -0.6 | 3.33E-04 | -0.2 | 1.70E-01 | -0.1 | 7.85E-01 | -0.1 | 4.86E-01 | Asparagine synthetase |
| AT5G20860 | -1.0 | 3.06E-05 | -0.6 | 9.54E-04 | -0.4 | 4.68E-02 | 1.6 | 5.94E-05 | 1.1 | 3.92E-03 | 0.5 | 1.25E-01 | Pectinesterase |
| AT5G05110 | -1.0 | 4.38E-07 | -1.0 | 9.57E-07 | 0.0 | 8.30E-01 | -0.6 | 3.27E-04 | -0.4 | 2.43E-02 | -0.3 | 2.05E-01 | Cysteine proteinase inhibitor 7 |
| AT5G25890 | -1.0 | 1.48E-06 | -0.6 | 7.42E-05 | -0.4 | 4.02E-03 | -0.1 | 8.47E-01 | 0.3 | 3.52E-01 | -0.3 | 2.57E-01 | Auxin-responsive protein IAA28 |
| AT3G51360 | -1.0 | 1.20E-06 | -0.4 | 6.12E-04 | -0.6 | 4.30E-04 | 0.0 | 9.00E-01 | 0.1 | 8.71E-01 | 0.0 | 9.67E-01 | Eukaryotic aspartyl protease family protein |
| AT5G02400 | -1.0 | 2.56E-06 | -0.6 | 2.19E-04 | -0.5 | 3.71E-03 | 0.0 | 8.72E-01 | 0.3 | 2.06E-01 | -0.4 | 1.70E-01 | Probable protein phosphatase 2C 66 |
| AT1G49390 | -1.0 | 4.85E-05 | -0.5 | 7.26E-03 | -0.6 | 1.16E-02 | -0.4 | 4.08E-01 | 0.0 | 9.89E-01 | -0.4 | 4.44E-01 | 2-oxoglutarate and Fe-dependent oxygenase superfamily protein |
| AT2G41210 | -1.0 | 1.37E-05 | -1.0 | 2.65E-05 | 0.0 | 9.44E-01 | 0.1 | 8.94E-01 | 0.2 | 6.27E-01 | -0.2 | 7.70E-01 | Phosphatidylinositol 4-phosphate 5-kinase 5 P-5-kinase 5) |
| AT5G17700 | -1.0 | 8.27E-06 | -0.5 | 2.37E-03 | -0.6 | 2.40E-03 | 0.0 | 9.50E-01 | 0.4 | 9.29E-02 | -0.4 | 1.21E-01 | MATE efflux family protein |
| AT1G64330 | -1.0 | 2.43E-07 | -0.5 | 5.03E-05 | -0.5 | 1.66E-04 | -0.4 | 2.19E-03 | -0.1 | 4.35E-01 | -0.3 | 8.50E-02 | At1g64330 |
| AT5G49890 | -1.0 | 6.57E-07 | -0.9 | 2.63E-06 | -0.1 | 3.94E-01 | -0.2 | 3.06E-01 | 0.1 | 6.05E-01 | -0.2 | 2.32E-01 | Chloride channel protein CLC-c |
| AT1G22410 | -1.0 | 5.16E-07 | -0.4 | 7.24E-04 | -0.6 | 1.06E-04 | 0.3 | 6.54E-02 | 0.3 | 5.63E-02 | -0.1 | 8.69E-01 | Phospho-2-dehydro-3-deoxyheptonate aldolase |
| AT1G14350 | -1.0 | 1.46E-07 | -0.8 | 2.16E-06 | -0.3 | 4.74E-03 | -0.1 | 4.24E-01 | 0.0 | 9.50E-01 | -0.1 | 6.39E-01 | Duplicated homeodomain-like superfamily protein |
| AT2G18890 | -1.0 | 1.26E-05 | -0.5 | 2.51E-03 | -0.6 | 4.41E-03 | -0.5 | 1.24E-02 | -0.5 | 1.46E-02 | 0.0 | 9.48E-01 | Protein kinase superfamily protein |
| AT5G45580 | -1.0 | 4.08E-04 | -0.1 | 4.24E-01 | -0.9 | 5.10E-03 | 0.0 | 9.77E-01 | 0.5 | 2.67E-01 | -0.5 | 2.65E-01 | Homeodomain-like superfamily protein |
| AT4G22820 | -1.0 | 1.79E-05 | -0.6 | 6.22E-04 | -0.4 | 2.66E-02 | 0.1 | 7.68E-01 | 0.1 | 6.02E-01 | -0.1 | 8.38E-01 | Zinc finger A20 and AN1 domain-containing stress-associated protein 9 |
| AT1G66270 | -1.0 | 1.06E-05 | 0.3 | 7.40E-02 | -1.3 | 2.85E-05 | -0.2 | 2.78E-01 | 0.1 | 5.49E-01 | -0.3 | 1.26E-01 | Beta-glucosidase 21 |
| AT4G01240 | -1.0 | 3.29E-04 | 0.2 | 2.29E-01 | -1.2 | 5.18E-04 | -0.2 | 6.17E-01 | 0.3 | 5.46E-01 | -0.5 | 2.89E-01 | S-adenosyl-L-methionine-dependent methyltransferases superfamily protein |
| AT1G16060 | -1.0 | 8.08E-06 | -0.4 | 2.63E-03 | -0.6 | 2.41E-03 | -0.3 | 3.34E-01 | 0.1 | 8.12E-01 | -0.4 | 3.25E-01 | ARIA-interacting double AP2 domain protein |
| AT3G02875 | -1.0 | 2.70E-06 | -0.6 | 1.55E-04 | -0.4 | 6.17E-03 | 0.0 | 8.48E-01 | 0.1 | 6.26E-01 | -0.1 | 8.23E-01 | Peptidase M20 |
| AT1G73880 | -1.0 | 5.46E-06 | -0.8 | 3.93E-05 | -0.2 | 1.85E-01 | -0.4 | 6.06E-02 | -0.2 | 2.93E-01 | -0.1 | 6.56E-01 | Flavonol 3-O-glucosyltransferase UGT89B1 |
| AT1G74740 | -1.0 | 2.49E-07 | -0.5 | 5.58E-05 | -0.5 | 1.78E-04 | -0.2 | 2.78E-01 | -0.2 | 2.39E-01 | 0.0 | 9.05E-01 | Calcium-dependent protein kinase 30 |
| AT2G33710 | -1.0 | 3.08E-05 | 0.0 | 8.73E-01 | -1.0 | 2.57E-04 | 0.9 | 3.81E-02 | 0.5 | 3.30E-01 | 0.4 | 3.74E-01 | Integrase-type DNA-binding superfamily protein |
| AT3G52800 | -1.0 | 1.96E-06 | -0.7 | 5.05E-05 | -0.3 | 1.55E-02 | -0.3 | 3.93E-02 | -0.2 | 9.83E-02 | 0.0 | 9.08E-01 | Zinc finger A20 and AN1 domain-containing stress-associated protein 6 |
| AT5G10720 | -1.0 | 3.41E-06 | -0.3 | 1.51E-02 | -0.7 | 2.61E-04 | -0.5 | 1.51E-01 | 0.3 | 4.81E-01 | -0.7 | 6.75E-02 | Histidine kinase 5 |
| AT2G43290 | -1.0 | 9.66E-06 | -0.8 | 8.01E-05 | -0.2 | 1.54E-01 | -0.5 | 8.61E-03 | -0.1 | 5.62E-01 | -0.4 | 1.02E-01 | Calmodulin-like protein 5 |
| AT5G66160 | -1.0 | 6.44E-07 | -0.8 | 6.19E-06 | -0.2 | 4.76E-02 | -0.4 | 2.75E-02 | -0.3 | 7.90E-02 | -0.1 | 8.59E-01 | Receptor homology region transmembrane domain ring H2 motif protein 1 |
| AT5G03190 | -1.0 | 8.80E-06 | -0.7 | 2.33E-04 | -0.3 | 2.91E-02 | -0.9 | 2.61E-04 | 0.3 | 8.78E-02 | -1.2 | 2.49E-04 | hypothetical protein |
| AT3G04570 | -1.0 | 5.67E-06 | -0.9 | 3.41E-05 | -0.2 | 2.32E-01 | -0.1 | 6.78E-01 | 0.3 | 1.01E-01 | -0.3 | 6.40E-02 | AT-hook motif nuclear-localized protein 19 |
| AT1G24320 | -1.0 | 6.83E-06 | -0.5 | 1.46E-03 | -0.5 | 2.82E-03 | -0.5 | 4.12E-02 | 0.4 | 1.47E-01 | -0.9 | 8.62E-03 | Mannosyl-oligosaccharide glucosidase |
| AT2G15300 | -1.0 | 3.81E-06 | -0.2 | 3.82E-02 | -0.8 | 1.99E-04 | 0.0 | 9.40E-01 | 0.1 | 8.90E-01 | 0.0 | 9.41E-01 | Leucine-rich repeat protein kinase family protein |
| AT3G21190 | -1.0 | 1.20E-06 | -0.5 | 3.82E-04 | -0.5 | 5.10E-04 | -0.2 | 2.07E-01 | 0.2 | 3.41E-01 | -0.4 | 8.44E-02 | Protein MANNAN SYNTHESIS-RELATED 1 |
| AT1G12740 | -1.0 | 4.89E-05 | -0.4 | 9.24E-03 | -0.6 | 9.32E-03 | -0.2 | 7.07E-01 | 0.4 | 4.58E-01 | -0.6 | 2.44E-01 | Cytochrome P450, family 87, subfamily A, polypeptide 2 |
| AT4G11220 | -1.0 | 8.69E-07 | -0.6 | 5.64E-05 | -0.4 | 2.14E-03 | -0.1 | 4.60E-01 | 0.0 | 9.47E-01 | -0.1 | 5.32E-01 | Reticulon-like protein B2 |
| AT4G26560 | -1.0 | 6.80E-06 | -0.4 | 4.22E-03 | -0.6 | 1.36E-03 | -0.1 | 8.54E-01 | 0.5 | 9.64E-02 | -0.5 | 8.39E-02 | Calcineurin B-like protein 7 |
| AT5G59450 | -1.0 | 2.69E-06 | -0.8 | 1.69E-05 | -0.2 | 2.03E-01 | 0.3 | 1.73E-01 | 0.2 | 4.48E-01 | 0.1 | 7.68E-01 | Scarecrow-like protein 11 |
| AT2G44160 | -1.0 | 1.09E-06 | -0.4 | 9.68E-04 | -0.6 | 2.38E-04 | -0.2 | 4.26E-02 | 0.0 | 9.79E-01 | -0.2 | 1.68E-01 | Methylenetetrahydrofolate reductase 2 |
| AT2G03450 | -1.0 | 3.72E-07 | -0.5 | 6.19E-05 | -0.5 | 3.77E-04 | 0.1 | 6.34E-01 | 0.1 | 6.06E-01 | 0.0 | 9.51E-01 | Probable inactive purple acid phosphatase 9 |
| AT5G01700 | -1.0 | 6.75E-05 | -0.1 | 6.39E-01 | -0.9 | 7.14E-04 | -1.2 | 5.48E-04 | -0.6 | 3.06E-02 | -0.5 | 1.85E-01 | Probable protein phosphatase 2C 65 |
| AT4G10350 | -1.0 | 1.27E-04 | -0.1 | 5.48E-01 | -0.9 | 1.49E-03 | -0.1 | 8.24E-01 | 0.4 | 3.48E-01 | -0.5 | 2.71E-01 | NAC domain containing protein 70 |
| AT4G30120 | -1.0 | 1.45E-04 | -1.1 | 1.57E-04 | 0.1 | 6.88E-01 | -0.8 | 1.78E-02 | -0.4 | 2.44E-01 | -0.4 | 2.58E-01 | Heavy metal atpase 3 |
| AT3G26330 | -1.0 | 6.77E-07 | -0.8 | 8.10E-06 | -0.2 | 3.35E-02 | -0.1 | 8.09E-01 | -0.1 | 8.52E-01 | 0.0 | 9.97E-01 | Cytochrome P450 71B37 |
| AT5G40760 | -1.0 | 4.22E-07 | -0.6 | 2.72E-05 | -0.4 | 1.07E-03 | 0.1 | 5.02E-01 | 0.1 | 5.25E-01 | 0.0 | 9.70E-01 | Glucose-6-phosphate 1-dehydrogenase 6, cytoplasmic |
| AT1G12290 | -1.0 | 2.14E-06 | -0.9 | 1.18E-05 | -0.1 | 2.57E-01 | -0.4 | 1.82E-01 | -0.1 | 8.36E-01 | -0.3 | 3.66E-01 | Disease resistance protein family |
| AT5G45210 | -1.0 | 6.26E-05 | 0.0 | 9.66E-01 | -1.0 | 4.45E-04 | -0.8 | 1.94E-03 | 0.3 | 2.35E-01 | -1.0 | 2.32E-03 | Disease resistance protein family |
| AT3G55370 | -1.0 | 2.42E-06 | -0.4 | 9.34E-04 | -0.6 | 9.23E-04 | 0.0 | 9.37E-01 | 0.3 | 1.46E-01 | -0.3 | 2.74E-01 | Dof zinc finger protein |
| AT5G52060 | -1.0 | 1.93E-06 | -0.4 | 1.03E-03 | -0.5 | 5.20E-04 | -0.5 | 9.52E-04 | 0.1 | 6.98E-01 | -0.6 | 2.51E-03 | BAG family molecular chaperone regulator 1 |

| | | | | | | | | | | | | | |
|---|---|---|---|---|---|---|---|---|---|---|---|---|---|
| AT3G43960 | -1.0 | 5.00E-05 | 0.2 | 1.89E-01 | -1.2 | 1.09E-04 | -0.5 | 2.33E-02 | 0.0 | 9.02E-01 | -0.4 | 8.81E-02 | Probable cysteine protease RDL3 |
| AT4G04500 | -1.0 | 2.25E-01 | -1.7 | 9.74E-02 | 0.7 | 5.92E-01 | | | | | | | Cysteine-rich RLK 37 |
| AT1G09480 | -1.0 | 2.36E-04 | -0.6 | 4.34E-03 | -0.4 | 1.26E-01 | -0.2 | 5.62E-01 | -0.7 | 1.49E-02 | 0.6 | 1.12E-01 | NAD-binding Rossmann-fold superfamily protein |
| AT2G22660 | -1.0 | 4.60E-07 | -0.6 | 2.80E-05 | -0.4 | 1.43E-03 | -0.6 | 3.35E-05 | -0.2 | 1.14E-01 | -0.4 | 1.71E-02 | DNA-binding protein, putative |
| AT4G33490 | -1.0 | 4.98E-05 | -0.5 | 4.61E-03 | -0.5 | 1.69E-02 | -0.6 | 5.33E-04 | 0.2 | 3.63E-01 | -0.8 | 7.58E-04 | Putative nucellin |
| AT2G46660 | -1.0 | 7.25E-05 | -0.2 | 1.01E-01 | -0.7 | 2.68E-03 | -0.7 | 1.05E-02 | 0.0 | 9.57E-01 | -0.8 | 9.81E-03 | Cytochrome P450 78A6 |
| AT5G51070 | -1.0 | 1.04E-06 | -1.1 | 1.11E-06 | 0.1 | 1.43E-01 | -0.4 | 4.68E-02 | -0.4 | 4.48E-02 | 0.0 | 8.82E-01 | Chaperone protein ClpD, chloroplastic |
| AT1G30640 | -1.0 | 5.06E-07 | -0.9 | 2.19E-06 | -0.1 | 3.63E-01 | 0.1 | 6.06E-01 | 0.3 | 3.41E-01 | -0.1 | 7.41E-01 | T5I8.9 protein |
| AT4G24450 | -1.0 | 1.32E-04 | -1.0 | 2.08E-04 | 0.0 | 9.57E-01 | -0.2 | 6.90E-01 | 0.4 | 4.09E-01 | -0.6 | 2.46E-01 | Alpha-glucan water dikinase 2 |
| AT4G05020 | -1.0 | 1.29E-04 | -0.9 | 4.47E-04 | -0.1 | 5.72E-01 | 1.9 | 1.45E-07 | 0.8 | 2.02E-03 | 1.1 | 2.10E-04 | NADH:ubiquinone reductase |
| AT1G79460 | -1.0 | 2.80E-05 | -0.6 | 1.47E-03 | -0.4 | 2.34E-02 | -0.5 | 1.13E-01 | 0.0 | 9.95E-01 | -0.5 | 1.34E-01 | Ent-kaur-16-ene synthase, chloroplastic |
| AT4G02710 | -1.0 | 1.05E-06 | -0.7 | 1.43E-05 | -0.3 | 3.19E-02 | 0.0 | 9.54E-01 | 0.3 | 3.16E-01 | -0.3 | 4.44E-01 | Protein NETWORKED 1C |
| AT1G06620 | -1.0 | 1.48E-06 | -0.4 | 7.93E-04 | -0.6 | 4.71E-04 | -0.4 | 4.18E-02 | 0.2 | 4.12E-01 | -0.6 | 2.73E-02 | 2-oxoglutarate and Fe-dependent oxygenase superfamily protein |
| AT3G02630 | -1.0 | 1.62E-06 | -0.3 | 6.08E-03 | -0.7 | 1.38E-04 | 0.1 | 5.65E-01 | 0.2 | 1.16E-01 | -0.1 | 4.15E-01 | Stearoyl- 9-desaturase 5, chloroplastic |
| AT3G46260 | -1.0 | 4.34E-04 | -0.1 | 5.77E-01 | -0.9 | 4.34E-03 | | | | | | | Kinase-like protein |
| AT3G09540 | -1.0 | 1.22E-05 | -0.6 | 3.90E-04 | -0.4 | 2.75E-02 | -0.4 | 2.77E-02 | 0.4 | 3.53E-02 | -0.8 | 2.36E-03 | Pectate lyase |
| AT1G33890 | -1.0 | 6.56E-05 | -0.1 | 6.04E-01 | -0.9 | 6.93E-04 | -0.3 | 5.51E-01 | 0.2 | 7.13E-01 | -0.4 | 3.42E-01 | Immune-associated nucleotide-binding protein 3 |
| AT2G34830 | -1.0 | 2.12E-05 | -0.3 | 2.30E-02 | -0.7 | 1.77E-03 | -0.6 | 1.69E-02 | -0.2 | 4.05E-01 | -0.4 | 2.49E-01 | WRKY DNA-binding protein 35 |
| AT2G17150 | -1.0 | 9.17E-06 | -0.7 | 1.78E-04 | -0.3 | 4.92E-02 | -0.2 | 6.52E-01 | -0.1 | 8.37E-01 | -0.1 | 9.14E-01 | Plant regulator RWP-RK family protein |
| AT5G58500 | -1.0 | 1.48E-05 | -0.7 | 2.32E-04 | -0.3 | 8.16E-02 | 0.5 | 1.84E-01 | 0.3 | 5.35E-01 | 0.2 | 6.60E-01 | Protein LIGHT-DEPENDENT SHORT HYPOCOTYLS 5 |
| AT1G52760 | -1.0 | 2.56E-07 | -0.1 | 6.01E-02 | -0.8 | 1.39E-05 | -0.4 | 6.73E-03 | 0.0 | 8.46E-01 | -0.4 | 5.86E-02 | Caffeoylshikimate esterase |
| AT5G45920 | -1.0 | 8.72E-04 | -0.1 | 7.01E-01 | -0.9 | 5.41E-03 | -0.5 | 3.88E-02 | 0.0 | 9.96E-01 | -0.5 | 9.04E-02 | SGNH hydrolase-type esterase superfamily protein |
| AT4G21230 | -1.0 | 6.11E-05 | -0.6 | 1.67E-03 | -0.4 | 7.22E-02 | -0.4 | 4.35E-01 | 0.5 | 3.00E-01 | -0.9 | 5.73E-02 | Cysteine-rich receptor-like protein kinase 27 |
| AT1G75780 | -1.0 | 1.78E-05 | -0.2 | 1.68E-01 | -0.8 | 3.97E-04 | -0.9 | 1.73E-01 | 0.1 | 8.80E-01 | -1.1 | 2.53E-01 | Tubulin beta-1 chain |
| AT4G39030 | -1.0 | 6.69E-05 | -0.5 | 3.97E-03 | -0.4 | 2.76E-02 | 0.1 | 7.75E-01 | 0.4 | 8.08E-02 | -0.4 | 1.92E-01 | Protein DETOXIFICATION 47, chloroplastic |
| AT1G63800 | -1.0 | 2.52E-06 | -0.7 | 6.85E-05 | -0.3 | 1.52E-02 | -0.6 | 5.46E-04 | -0.5 | 1.38E-02 | -0.2 | 4.49E-01 | Ubiquitin-conjugating enzyme 5 |
| AT3G03050 | -1.0 | 1.74E-06 | -0.6 | 1.66E-04 | -0.4 | 2.14E-03 | -0.2 | 3.96E-01 | 0.1 | 8.57E-01 | -0.3 | 4.18E-01 | Cellulose synthase-like protein D3 |
| AT3G09960 | -1.0 | 2.93E-05 | -0.6 | 1.30E-03 | -0.4 | 2.91E-02 | -0.1 | 9.09E-01 | 0.2 | 7.26E-01 | -0.2 | 6.98E-01 | Tyrosine-protein phosphatase At3g09960 |
| AT2G19130 | -1.0 | 1.52E-06 | -1.0 | 3.20E-06 | 0.0 | 8.99E-01 | -0.1 | 6.23E-01 | 0.0 | 8.71E-01 | -0.2 | 5.85E-01 | G-type lectin S-receptor-like serine |
| AT2G21840 | -1.0 | 4.21E-06 | -0.5 | 7.98E-04 | -0.5 | 2.29E-03 | -0.3 | 9.42E-02 | 0.0 | 9.14E-01 | -0.3 | 1.43E-01 | Cysteine |
| AT2G32120 | -1.0 | 3.34E-05 | -0.5 | 1.66E-03 | -0.4 | 2.75E-02 | -0.4 | 2.59E-01 | 0.0 | 9.87E-01 | -0.4 | 4.17E-01 | Heat shock 70 kDa protein 8 |
| AT4G09110 | -1.0 | 4.37E-05 | -0.9 | 1.24E-04 | -0.1 | 7.12E-01 | 0.4 | 1.99E-01 | -0.1 | 7.68E-01 | 0.6 | 3.41E-02 | Putative RING-H2 finger protein ATL35 |
| AT5G06850 | -1.0 | 6.95E-07 | -0.6 | 1.93E-05 | -0.3 | 6.51E-03 | -0.1 | 4.53E-01 | 0.0 | 9.71E-01 | -0.1 | 5.91E-01 | FT-interacting protein 1 |
| AT4G31920 | -1.0 | 1.20E-06 | -0.5 | 2.15E-04 | -0.5 | 8.49E-04 | -0.1 | 6.01E-01 | 0.3 | 5.44E-02 | -0.4 | 3.91E-02 | Two-component response regulator ARR10 |
| AT1G01720 | -1.0 | 2.04E-06 | -1.0 | 3.31E-06 | 0.1 | 6.21E-01 | -0.4 | 2.28E-02 | -0.3 | 4.97E-02 | 0.0 | 9.28E-01 | NAC domain-containing protein 2 |
| AT3G56760 | -1.0 | 6.47E-07 | -0.4 | 2.01E-04 | -0.5 | 3.43E-04 | -0.2 | 2.29E-01 | 0.1 | 6.80E-01 | -0.3 | 1.86E-01 | CDPK-related kinase 7 |
| AT3G16360 | -1.0 | 2.09E-03 | -0.9 | 3.71E-03 | 0.0 | 9.60E-01 | 1.1 | 1.87E-03 | 0.5 | 1.65E-01 | 0.5 | 1.17E-01 | Histidine-containing phosphotransfer protein |
| AT2G37280 | -1.0 | 6.25E-06 | -0.8 | 3.22E-05 | -0.1 | 3.24E-01 | -0.2 | 6.80E-01 | 0.3 | 6.10E-01 | -0.5 | 3.35E-01 | Pleiotropic drug resistance 5 |
| AT4G35790 | -1.0 | 2.99E-07 | -0.6 | 1.34E-05 | -0.3 | 1.39E-03 | -0.3 | 1.03E-02 | -0.1 | 4.32E-01 | -0.2 | 2.57E-01 | Phospholipase D |
| AT5G06250 | -1.0 | 3.02E-04 | -0.1 | 7.07E-01 | -0.9 | 2.49E-03 | 0.0 | 9.78E-01 | 0.9 | 2.07E-02 | -0.9 | 2.89E-02 | AP2 |
| AT1G52750 | -1.0 | 3.71E-05 | 0.1 | 5.25E-01 | -1.0 | 1.65E-04 | -0.6 | 7.17E-02 | -0.2 | 5.28E-01 | -0.4 | 3.29E-01 | Uncharacterized protein F14G24.2 |
| AT4G19720 | -1.0 | 6.57E-07 | -0.5 | 1.72E-04 | -0.5 | 4.37E-04 | 1.8 | 8.37E-07 | 1.4 | 9.39E-05 | 0.4 | 3.89E-02 | GH18 domain-containing protein |
| AT1G12260 | -1.0 | 1.37E-04 | -0.3 | 5.72E-02 | -0.7 | 8.68E-03 | -0.1 | 6.94E-01 | -0.1 | 8.95E-01 | -0.1 | 8.89E-01 | NAC 007 |
| AT5G22500 | -1.0 | 1.83E-05 | -1.0 | 2.76E-05 | 0.0 | 7.89E-01 | 0.3 | 4.51E-01 | 0.0 | 9.91E-01 | 0.3 | 5.64E-01 | Fatty acyl-CoA reductase |
| AT4G38830 | -1.0 | 1.53E-05 | -0.5 | 2.11E-03 | -0.5 | 7.43E-03 | -0.7 | 6.95E-02 | -0.4 | 2.63E-01 | -0.2 | 7.09E-01 | Cysteine-rich receptor-like protein kinase 26 |
| AT4G18940 | -1.0 | 1.92E-04 | 0.4 | 8.25E-03 | -1.4 | 9.80E-05 | -0.7 | 1.61E-02 | -0.2 | 5.06E-01 | -0.5 | 1.72E-01 | RNA ligase |
| AT4G21830 | -1.0 | 1.31E-04 | 0.1 | 3.38E-01 | -1.1 | 2.53E-04 | -0.5 | 1.34E-02 | 0.2 | 2.63E-01 | -0.7 | 6.27E-03 | Peptide methionine sulfoxide reductase B7 -S-oxide reductase) |
| AT3G10660 | -1.0 | 1.01E-06 | -0.2 | 3.95E-02 | -0.8 | 5.11E-05 | -0.4 | 4.46E-02 | 0.2 | 2.86E-01 | -0.7 | 2.21E-02 | Calcium-dependent protein kinase 2 |
| AT5G58300 | -1.0 | 1.98E-07 | -0.5 | 1.97E-05 | -0.4 | 2.76E-04 | 0.0 | 9.85E-01 | 0.1 | 8.41E-01 | -0.1 | 8.47E-01 | Protein kinase domain-containing protein |
| AT3G51160 | -1.0 | 1.69E-06 | -0.2 | 2.02E-02 | -0.7 | 8.91E-05 | -0.1 | 3.31E-01 | 0.1 | 5.70E-01 | -0.2 | 2.25E-01 | GDP-mannose 4,6 dehydratase 2 |
| AT4G30280 | -1.0 | 1.72E-04 | 0.0 | 8.98E-01 | -1.0 | 7.01E-04 | -0.5 | 9.18E-02 | 0.2 | 4.58E-01 | -0.8 | 1.83E-02 | Probable xyloglucan endotransglucosylase |
| AT2G22500 | -1.0 | 6.29E-04 | -1.0 | 8.38E-04 | 0.0 | 9.47E-01 | 0.0 | 8.84E-01 | 0.0 | 9.56E-01 | 0.1 | 8.71E-01 | Mitochondrial uncoupling protein 5 |
| AT3G44840 | -1.0 | 1.84E-04 | -1.3 | 7.64E-05 | 0.3 | 2.32E-01 | 0.3 | 5.58E-01 | 0.8 | 9.31E-02 | -0.5 | 2.13E-01 | Proteinkinase AtPP-like protein |
| AT3G16280 | -1.0 | 7.82E-05 | -0.5 | 3.78E-03 | -0.4 | 3.82E-02 | -0.2 | 6.30E-01 | 0.1 | 8.71E-01 | -0.3 | 4.40E-01 | Integrase-type DNA-binding superfamily protein |
| AT2G27035 | -1.0 | 9.74E-05 | -0.4 | 1.18E-02 | -0.5 | 2.08E-02 | | | | | | | hypothetical protein |
| AT5G65320 | -1.0 | 2.09E-04 | -0.7 | 2.72E-03 | -0.3 | 1.99E-01 | 0.2 | 6.58E-01 | 0.8 | 1.43E-01 | -0.5 | 3.22E-01 | Basic helix-loop-helix DNA-binding superfamily protein |
| AT5G38030 | -1.0 | 2.75E-06 | -0.8 | 1.37E-05 | -0.1 | 3.21E-01 | 0.5 | 6.50E-02 | 0.8 | 6.67E-03 | -0.3 | 3.41E-01 | Protein DETOXIFICATION 30 |
| AT2G36270 | -1.0 | 5.15E-03 | -1.2 | 2.74E-03 | 0.2 | 6.11E-01 | 0.0 | 9.35E-01 | -0.3 | 4.83E-01 | 0.3 | 5.87E-01 | Basic-leucine zipper transcription factor family protein |
| AT1G76680 | -1.0 | 5.81E-05 | -0.3 | 3.53E-02 | -0.6 | 3.33E-03 | 0.8 | 3.72E-04 | 0.2 | 4.80E-01 | 0.6 | 1.40E-02 | 12-oxophytodienoate reductase 1 |
| AT1G61480 | -1.0 | 1.17E-04 | -0.8 | 4.35E-04 | -0.1 | 5.40E-01 | 0.6 | 6.75E-02 | 0.3 | 5.25E-01 | 0.3 | 4.31E-01 | G-type lectin S-receptor-like serine |
| AT3G01470 | -1.0 | 8.53E-07 | -0.7 | 1.96E-05 | -0.3 | 1.01E-02 | -0.4 | 1.61E-03 | -0.3 | 5.17E-02 | -0.2 | 3.79E-01 | Homeobox-leucine zipper protein HAT5 |
| AT3G05290 | -0.9 | 6.59E-07 | -0.6 | 1.86E-05 | -0.3 | 5.60E-03 | -0.6 | 3.19E-04 | -0.4 | 1.51E-02 | -0.2 | 2.56E-01 | Peroxisomal adenine nucleotide carrier 1 |
| AT1G17550 | -0.9 | 1.07E-05 | -0.9 | 3.55E-05 | -0.1 | 5.92E-01 | -0.2 | 2.07E-01 | 0.3 | 3.30E-02 | -0.5 | 6.38E-03 | Protein phosphatase 2C 7 |
| AT4G11820 | -0.9 | 1.81E-06 | -0.3 | 1.32E-02 | -0.7 | 1.11E-04 | -0.2 | 1.84E-01 | 0.1 | 5.58E-01 | -0.3 | 1.44E-01 | Hydroxymethylglutaryl-CoA synthase |
| AT4G25800 | -0.9 | 1.25E-05 | -0.9 | 2.95E-05 | 0.0 | 9.11E-01 | -0.3 | 1.67E-01 | -0.3 | 2.59E-01 | 0.0 | 9.44E-01 | Calmodulin-binding protein 60 D |
| AT3G46410 | -0.9 | 1.50E-02 | 0.9 | 5.27E-03 | -1.8 | 8.72E-04 | | | | | | | Protein kinase superfamily protein |
| AT2G03500 | -0.9 | 1.30E-05 | -0.3 | 7.84E-03 | -0.6 | 2.02E-03 | -0.5 | 6.64E-02 | -0.1 | 6.82E-01 | -0.3 | 3.08E-01 | Myb family transcription factor EFM |
| AT4G00820 | -0.9 | 2.24E-06 | -0.2 | 1.42E-02 | -0.7 | 1.59E-04 | -0.2 | 3.28E-01 | 0.1 | 8.09E-01 | -0.3 | 3.36E-01 | Protein IQ-DOMAIN 17 |
| AT1G73630 | -0.9 | 1.87E-06 | -0.3 | 6.09E-03 | -0.7 | 1.99E-04 | -0.6 | 2.49E-02 | -0.3 | 2.98E-01 | -0.3 | 3.89E-01 | Probable calcium-binding protein CML26 |
| AT2G43220 | -0.9 | 9.35E-04 | 0.0 | 8.94E-01 | -1.0 | 3.56E-03 | -1.6 | 1.75E-03 | -0.2 | 7.10E-01 | -1.4 | 2.31E-02 | Zinc finger PHD-type domain-containing protein |
| AT5G04980 | -0.9 | 4.41E-07 | -0.7 | 5.71E-06 | -0.2 | 2.12E-02 | 0.1 | 6.61E-01 | 0.2 | 5.21E-01 | -0.1 | 8.61E-01 | DNAse I-like superfamily protein |
| AT1G06640 | -0.9 | 1.26E-06 | -0.4 | 9.98E-04 | -0.5 | 2.80E-04 | -0.4 | 2.08E-02 | 0.3 | 1.21E-01 | -0.6 | 6.31E-03 | 2-oxoglutarate and Fe-dependent oxygenase superfamily protein |
| AT4G14640 | -0.9 | 1.69E-04 | 0.0 | 8.06E-01 | -0.9 | 1.37E-03 | 0.1 | 8.76E-01 | 0.4 | 1.79E-01 | -0.4 | 2.12E-01 | Calmodulin 8 |
| AT3G24240 | -0.9 | 2.02E-05 | -0.3 | 2.86E-02 | -0.7 | 1.34E-03 | 0.1 | 8.04E-01 | 0.4 | 1.54E-01 | -0.3 | 2.00E-01 | LRR receptor-like serine |
| AT5G20490 | -0.9 | 4.82E-06 | -0.7 | 6.62E-05 | -0.2 | 6.01E-02 | -0.2 | 1.48E-01 | 0.1 | 6.84E-01 | -0.3 | 1.53E-01 | Myosin family protein with Dil |
| AT5G61980 | -0.9 | 1.88E-05 | -0.3 | 1.22E-02 | -0.6 | 2.53E-03 | -0.7 | 1.11E-01 | -0.1 | 9.32E-01 | -0.7 | 1.95E-01 | ADP-ribosylation factor GTPase-activating protein |
| AT3G14680 | -0.9 | 5.59E-06 | -0.8 | 3.03E-05 | -0.1 | 2.81E-01 | -0.3 | 3.22E-01 | 0.0 | 9.93E-01 | -0.3 | 5.04E-01 | Cytochrome P450 72A14 |
| AT2G03060 | -0.9 | 2.60E-05 | -0.4 | 8.53E-03 | -0.6 | 5.04E-03 | 0.0 | 9.57E-01 | 0.3 | 4.04E-01 | -0.3 | 4.35E-01 | AGAMOUS-like 30 |
| AT2G01910 | -0.9 | 9.50E-07 | -0.5 | 2.49E-04 | -0.5 | 4.82E-04 | -0.5 | 2.13E-04 | -0.1 | 4.04E-01 | -0.4 | 1.61E-02 | 65-kDa microtubule-associated protein 6 |
| AT1G27180 | -0.9 | 9.14E-06 | -0.8 | 3.65E-05 | -0.1 | 4.89E-01 | 0.0 | 9.92E-01 | 0.2 | 6.29E-01 | -0.2 | 6.76E-01 | T7N9.24 |
| AT1G50520 | -0.9 | 5.20E-06 | -0.2 | 3.72E-02 | -0.7 | 2.70E-04 | -0.5 | 3.34E-02 | 0.0 | 9.92E-01 | -0.5 | 7.57E-02 | Cytochrome P450, family 705, subfamily A, polypeptide 27 |
| AT5G47060 | -0.9 | 7.79E-04 | -0.9 | 1.77E-03 | -0.1 | 7.93E-01 | 0.7 | 7.98E-03 | 0.5 | 9.59E-02 | 0.2 | 3.80E-01 | FCS-Like Zinc finger 1 |
| AT1G22550 | -0.9 | 8.62E-06 | -0.1 | 1.79E-01 | -0.8 | 1.96E-04 | -0.8 | 5.62E-03 | -0.3 | 2.62E-01 | -0.5 | 1.54E-01 | Protein NRT1 |

| | | | | | | | | | | | | | |
|---|---|---|---|---|---|---|---|---|---|---|---|---|---|
| AT1G04120 | -0.9 | 5.16E-07 | -0.6 | 4.13E-05 | -0.4 | 1.05E-03 | 0.0 | 8.99E-01 | 0.3 | 1.54E-01 | -0.3 | 2.08E-01 | ABC-type xenobiotic transporter |
| AT2G25790 | -0.9 | 6.99E-06 | -0.9 | 2.54E-05 | -0.1 | 5.72E-01 | -0.2 | 1.99E-01 | 0.0 | 8.98E-01 | -0.2 | 4.18E-01 | Leucine-rich repeat receptor-like serine |
| AT1G02460 | -0.9 | 2.20E-03 | -0.2 | 2.89E-01 | -0.7 | 3.25E-02 | 0.3 | 4.86E-01 | 0.1 | 8.60E-01 | 0.2 | 7.12E-01 | At1g02460 |
| AT1G15210 | -0.9 | 6.73E-06 | -0.3 | 2.39E-02 | -0.7 | 3.57E-04 | -0.3 | 3.28E-01 | 0.2 | 6.63E-01 | -0.5 | 2.70E-01 | ABC transporter G family member 35 |
| AT5G51630 | -0.9 | 8.43E-06 | -0.6 | 1.71E-04 | -0.3 | 4.77E-02 | 0.2 | 3.56E-01 | 0.1 | 6.85E-01 | 0.1 | 8.16E-01 | ADP-ribosyl cyclase |
| AT5G17330 | -0.9 | 1.78E-06 | -0.3 | 1.52E-02 | -0.7 | 1.09E-04 | 0.3 | 3.66E-02 | 0.3 | 6.39E-02 | 0.0 | 9.98E-01 | Glutamate decarboxylase |
| AT3G07570 | -0.9 | 1.23E-06 | -0.2 | 2.50E-02 | -0.7 | 5.97E-05 | -0.3 | 4.27E-02 | 0.0 | 8.89E-01 | -0.3 | 7.40E-02 | Cytochrome b561 and DOMON domain-containing protein At3g07570 |
| AT2G35700 | -0.9 | 8.33E-05 | -0.6 | 1.41E-03 | -0.3 | 1.25E-01 | -0.2 | 6.38E-01 | 0.3 | 3.99E-01 | -0.4 | 2.38E-01 | Ethylene-responsive transcription factor ERF038 |
| AT4G10380 | -0.9 | 7.55E-06 | -0.4 | 7.05E-03 | -0.6 | 9.23E-04 | 0.2 | 4.21E-01 | 0.4 | 1.31E-01 | -0.2 | 4.23E-01 | Probable aquaporin NIP5-1 |
| AT3G19390 | -0.9 | 7.34E-06 | 0.2 | 1.35E-01 | -1.1 | 2.82E-05 | -0.3 | 1.70E-01 | 0.1 | 6.00E-01 | -0.4 | 1.12E-01 | Probable cysteine protease RD21C |
| AT2G17290 | -0.9 | 3.37E-07 | -0.6 | 1.61E-05 | -0.3 | 1.25E-03 | 0.1 | 6.67E-01 | 0.1 | 7.86E-01 | 0.0 | 9.71E-01 | Calcium-dependent protein kinase 6 |
| AT4G15440 | -0.9 | 4.37E-03 | -0.5 | 5.56E-02 | -0.4 | 2.73E-01 | -0.1 | 8.69E-01 | 1.0 | 7.01E-03 | -1.1 | 1.90E-02 | Uncharacterized protein |
| AT4G15490 | -0.9 | 5.04E-04 | -0.8 | 2.34E-03 | -0.2 | 4.63E-01 | -0.2 | 4.05E-01 | 0.6 | 2.29E-02 | -0.8 | 1.66E-02 | UDP-glycosyltransferase 84A3 |
| AT2G38080 | -0.9 | 5.42E-06 | -0.1 | 5.19E-01 | -0.9 | 7.81E-05 | -0.2 | 2.66E-01 | -0.1 | 7.31E-01 | -0.1 | 6.16E-01 | Laccase-4 |
| AT1G34040 | -0.9 | 1.55E-06 | -0.2 | 1.37E-02 | -0.7 | 1.11E-04 | -0.5 | 6.76E-02 | 0.3 | 2.24E-01 | -0.8 | 4.59E-03 | Pyridoxal phosphate -dependent transferases superfamily protein |
| AT1G54100 | -0.9 | 2.66E-05 | -0.9 | 4.68E-05 | 0.0 | 9.43E-01 | -0.3 | 6.53E-02 | -0.6 | 8.31E-03 | 0.2 | 4.12E-01 | Aldehyde dehydrogenase family 7 member B4 |
| AT5G43350 | -0.9 | 2.38E-05 | -0.4 | 1.46E-02 | -0.6 | 2.34E-03 | 0.4 | 5.28E-03 | 0.4 | 1.09E-02 | 0.0 | 9.98E-01 | Inorganic phosphate transporter 1-1 |
| AT2G29660 | -0.9 | 2.35E-06 | -0.3 | 7.22E-03 | -0.6 | 1.98E-04 | -0.4 | 2.37E-02 | 0.0 | 9.08E-01 | -0.4 | 6.13E-02 | At2g29660 family protein) |
| AT3G01220 | -0.9 | 1.02E-04 | 0.0 | 9.85E-01 | -0.9 | 5.63E-04 | -0.1 | 6.93E-01 | 0.2 | 3.75E-01 | -0.4 | 2.19E-01 | Homeobox-leucine zipper protein |
| AT4G24480 | -0.9 | 6.58E-05 | -0.7 | 5.12E-04 | -0.2 | 2.59E-01 | 0.0 | 9.43E-01 | 0.2 | 3.31E-01 | -0.2 | 3.83E-01 | Protein kinase superfamily protein |
| AT5G16910 | -0.9 | 1.01E-06 | -0.6 | 3.15E-05 | -0.3 | 6.15E-03 | 0.0 | 9.87E-01 | 0.1 | 8.21E-01 | -0.1 | 8.47E-01 | Cellulose synthase-like protein D2 |
| AT5G03570 | -0.9 | 1.07E-04 | -0.8 | 3.27E-04 | -0.1 | 6.67E-01 | -1.6 | 3.07E-05 | -1.3 | 2.09E-04 | -0.2 | 5.64E-01 | Solute carrier family 40 member |
| AT5G55250 | -0.9 | 3.20E-04 | -0.7 | 2.58E-03 | -0.2 | 2.99E-01 | -0.7 | 1.54E-02 | -0.1 | 6.99E-01 | -0.6 | 8.51E-02 | Indole-3-acetate O-methyltransferase 1 acetate carboxylmethyltransferase 1) |
| AT5G60300 | -0.9 | 5.18E-06 | -0.5 | 3.94E-04 | -0.4 | 6.47E-03 | -0.7 | 6.55E-02 | -0.3 | 5.64E-01 | -0.4 | 3.81E-01 | non-specific serine |
| AT5G17820 | -0.9 | 1.06E-05 | 0.2 | 1.64E-01 | -1.1 | 3.33E-05 | -0.5 | 1.66E-02 | 0.0 | 9.68E-01 | -0.5 | 2.33E-02 | Peroxidase |
| AT3G25250 | -0.9 | 6.50E-04 | -1.1 | 3.94E-04 | 0.2 | 5.16E-01 | 2.5 | 1.91E-06 | 0.8 | 8.12E-02 | 1.7 | 9.02E-05 | Serine |
| AT1G33750 | -0.9 | 2.73E-04 | 0.1 | 4.63E-01 | -1.0 | 6.31E-04 | -0.2 | 3.62E-01 | 0.3 | 1.49E-01 | -0.5 | 3.99E-02 | Terpenoid synthase 22 |
| AT2G46370 | -0.9 | 5.40E-07 | -0.7 | 6.23E-06 | -0.2 | 3.40E-02 | 0.0 | 8.93E-01 | 0.3 | 4.96E-02 | -0.3 | 1.53E-01 | Auxin-responsive GH3 family protein |
| AT1G70750 | -0.9 | 3.73E-07 | -0.6 | 9.31E-06 | -0.3 | 4.92E-03 | 0.0 | 9.64E-01 | 0.0 | 8.95E-01 | 0.0 | 9.38E-01 | Myosin-binding protein 2 |
| AT2G31960 | -0.9 | 1.34E-06 | -0.8 | 6.58E-06 | -0.1 | 3.08E-01 | -0.3 | 1.91E-01 | 0.0 | 8.93E-01 | -0.2 | 4.11E-01 | 1,3-beta-glucan synthase |
| AT4G29820 | -0.9 | 1.67E-06 | -0.5 | 1.68E-04 | -0.4 | 2.42E-03 | 0.0 | 9.12E-01 | -0.1 | 5.70E-01 | 0.2 | 4.71E-01 | Pre-mRNA cleavage factor Im 25 kDa subunit 1 |
| AT5G17800 | -0.9 | 5.63E-06 | -0.5 | 4.31E-04 | -0.4 | 7.30E-03 | 0.7 | 7.57E-02 | 1.0 | 1.81E-02 | -0.3 | 5.19E-01 | Transcription factor MYB56 |
| AT4G27410 | -0.9 | 1.77E-04 | -0.6 | 2.46E-03 | -0.3 | 1.63E-01 | -1.2 | 3.73E-03 | -0.9 | 2.85E-02 | -0.3 | 6.17E-01 | NAC domain transcriptional regulator superfamily protein |
| AT1G38131 | -0.9 | 1.15E-03 | -0.6 | 1.94E-02 | -0.4 | 2.01E-01 | | | | | | | O-fucosyltransferase 11 |
| AT4G01020 | -0.9 | 1.41E-06 | -0.8 | 7.86E-06 | -0.1 | 2.41E-01 | -0.4 | 8.38E-02 | -0.1 | 7.14E-01 | -0.3 | 3.33E-01 | ATP-dependent RNA helicase DEAH11, chloroplastic |
| AT1G01070 | -0.9 | 1.76E-05 | -0.7 | 1.44E-04 | -0.2 | 2.02E-01 | -0.5 | 2.53E-02 | 0.0 | 9.54E-01 | -0.5 | 7.34E-02 | WAT1-related protein At1g01070 |
| AT5G58940 | -0.9 | 1.45E-04 | -0.7 | 7.72E-04 | -0.2 | 4.02E-01 | -0.1 | 7.08E-01 | -0.2 | 5.67E-01 | 0.1 | 8.83E-01 | Calmodulin-binding receptor-like cytoplasmic kinase 1 |
| AT1G09180 | -0.9 | 2.07E-03 | -0.9 | 2.50E-03 | 0.0 | 9.28E-01 | | | | | | | Putative GTP-binding protein, SAR1B |
| AT1G01780 | -0.9 | 4.21E-05 | 0.0 | 8.75E-01 | -0.9 | 2.09E-04 | -0.7 | 8.97E-04 | -0.2 | 5.01E-01 | -0.6 | 2.81E-02 | LIM domain-containing protein PLIM2b |
| AT1G77420 | -0.9 | 1.19E-06 | -0.3 | 1.31E-03 | -0.6 | 2.59E-04 | -0.1 | 8.36E-01 | 0.1 | 7.68E-01 | -0.1 | 6.66E-01 | Alpha |
| AT4G16350 | -0.9 | 4.08E-05 | -0.1 | 5.27E-01 | -0.8 | 4.87E-04 | -0.7 | 5.09E-03 | 0.1 | 8.57E-01 | -0.8 | 2.21E-02 | Calcineurin B-like protein |
| AT1G17020 | -0.9 | 3.48E-04 | -0.8 | 1.20E-03 | -0.1 | 6.25E-01 | | | | | | | Protein SRG1 |
| AT3G06300 | -0.9 | 2.37E-06 | -0.1 | 2.39E-01 | -0.8 | 4.93E-05 | 0.0 | 7.95E-01 | 0.1 | 5.56E-01 | -0.2 | 4.42E-01 | procollagen-proline 4-dioxygenase |
| AT3G23920 | -0.9 | 1.47E-06 | -0.9 | 2.80E-06 | 0.0 | 7.20E-01 | -0.4 | 9.29E-03 | -0.1 | 4.20E-01 | -0.2 | 2.02E-01 | Beta-amylase 1, chloroplastic |
| AT3G14530 | -0.9 | 1.81E-03 | 0.3 | 9.81E-02 | -1.2 | 1.05E-03 | -1.1 | 5.93E-02 | -0.4 | 5.35E-01 | -0.7 | 3.80E-01 | Geranylgeranyl pyrophosphate synthase 9, chloroplastic -farnesyl diphosphate synthase 9) |
| AT4G09420 | -0.9 | 1.12E-05 | -0.6 | 2.31E-04 | -0.3 | 4.83E-02 | -0.7 | 1.38E-03 | -0.4 | 3.65E-02 | -0.3 | 2.69E-01 | Disease resistance protein |
| AT5G02350 | -0.9 | 5.47E-05 | -0.2 | 2.47E-01 | -0.8 | 1.12E-03 | -0.9 | 5.24E-03 | 0.1 | 6.96E-01 | -1.0 | 1.29E-02 | Cysteine |
| AT1G75390 | -0.9 | 7.65E-07 | -0.5 | 4.81E-05 | -0.4 | 2.21E-03 | 0.2 | 2.65E-01 | 0.4 | 6.72E-02 | -0.2 | 4.70E-01 | Basic leucine-zipper 44 |
| AT2G25410 | -0.9 | 2.04E-04 | -0.4 | 2.53E-02 | -0.5 | 2.53E-02 | -0.2 | 7.34E-01 | 0.2 | 7.89E-01 | -0.4 | 5.70E-01 | RING-H2 finger protein ATL22 |
| AT1G51870 | -0.9 | 1.26E-04 | 0.2 | 8.29E-02 | -1.2 | 1.57E-04 | -1.4 | 3.55E-03 | 0.0 | 9.56E-01 | -1.4 | 1.87E-02 | Protein kinase family protein |
| AT1G52070 | -0.9 | 1.29E-03 | 0.0 | 8.56E-01 | -0.9 | 3.15E-03 | -0.4 | 3.30E-01 | 0.0 | 9.76E-01 | -0.4 | 4.37E-01 | Mannose-binding lectin superfamily protein |
| AT1G02500 | -0.9 | 1.39E-06 | -0.4 | 1.79E-03 | -0.5 | 2.35E-04 | -0.3 | 3.10E-03 | -0.2 | 4.79E-02 | -0.1 | 5.58E-01 | S-adenosylmethionine synthase 1 |
| AT4G04460 | -0.9 | 1.69E-05 | -0.3 | 2.52E-02 | -0.6 | 1.31E-03 | -0.9 | 7.65E-03 | -0.2 | 5.30E-01 | -0.7 | 8.76E-02 | Putative aspartic protease |
| AT4G37320 | -0.9 | 6.65E-06 | -0.6 | 1.57E-04 | -0.3 | 3.20E-02 | 0.3 | 1.06E-01 | 0.4 | 3.62E-02 | -0.1 | 6.32E-01 | Cytochrome P450, family 81, subfamily D, polypeptide 5 |
| AT3G61760 | -0.9 | 2.39E-05 | -0.2 | 8.61E-02 | -0.7 | 1.01E-03 | -0.4 | 2.63E-01 | -0.2 | 6.93E-01 | -0.2 | 5.09E-01 | DYNAMIN-like 1B |
| AT4G37640 | -0.9 | 1.96E-06 | -0.6 | 7.61E-05 | -0.3 | 7.78E-03 | 0.0 | 7.69E-01 | 0.1 | 7.64E-01 | -0.1 | 6.36E-01 | Calcium-transporting ATPase 2, plasma membrane-type -ATPase isoform 2) |
| AT3G14070 | -0.9 | 1.37E-04 | -0.5 | 7.22E-03 | -0.4 | 4.21E-02 | 0.1 | 7.81E-01 | 0.2 | 7.03E-01 | -0.1 | 9.24E-01 | Cation |
| AT4G27260 | -0.9 | 7.87E-06 | -0.8 | 3.87E-05 | -0.1 | 3.46E-01 | -0.6 | 3.48E-03 | -0.5 | 2.66E-02 | -0.1 | 5.33E-01 | Indole-3-acetic acid-amido synthetase GH3.5 |
| AT5G10650 | -0.9 | 5.72E-06 | -0.7 | 3.69E-05 | -0.2 | 2.29E-01 | -0.5 | 5.93E-02 | -0.1 | 6.24E-01 | -0.3 | 3.40E-01 | RING-type E3 ubiquitin transferase |
| AT1G25320 | -0.9 | 5.16E-06 | -0.4 | 2.06E-03 | -0.5 | 1.56E-03 | 0.0 | 9.72E-01 | 0.1 | 7.15E-01 | -0.1 | 7.79E-01 | At1g25320 |
| AT5G45840 | -0.9 | 2.50E-05 | -0.7 | 2.26E-04 | -0.2 | 1.86E-01 | -0.5 | 4.02E-02 | 0.0 | 9.28E-01 | -0.5 | 8.08E-02 | Leucine-rich repeat protein kinase family protein |
| AT5G15130 | -0.9 | 1.89E-05 | -0.6 | 3.35E-04 | -0.3 | 6.88E-02 | 0.8 | 3.94E-04 | 0.7 | 6.31E-03 | 0.2 | 4.06E-01 | WRKY DNA-binding protein 72 |
| AT3G07940 | -0.9 | 2.03E-06 | -0.7 | 2.58E-05 | -0.2 | 4.78E-02 | -0.3 | 5.31E-02 | 0.0 | 8.85E-01 | -0.3 | 1.66E-01 | Calcium-dependent ARF-type GTPase activating protein family |
| AT5G47960 | -0.9 | 6.19E-05 | -0.5 | 4.25E-03 | -0.4 | 2.55E-02 | -0.3 | 5.36E-01 | -0.3 | 6.05E-01 | 0.0 | 9.93E-01 | Ras-related protein RABA4c |
| AT2G20650 | -0.9 | 6.41E-06 | -0.3 | 5.37E-03 | -0.6 | 1.06E-03 | 0.0 | 9.86E-01 | 0.0 | 9.85E-01 | 0.0 | 1.00E+00 | Transmembrane E3 ubiquitin-protein ligase FLY2 |
| AT3G02230 | -0.9 | 4.82E-05 | -0.2 | 2.82E-01 | -0.7 | 6.93E-04 | 0.0 | 7.81E-01 | 0.3 | 2.43E-02 | -0.3 | 9.07E-02 | UDP-arabinopyranose mutase 1 |
| AT5G22300 | -0.9 | 9.66E-05 | -1.0 | 6.37E-05 | 0.1 | 3.80E-01 | -0.4 | 3.88E-01 | -0.2 | 7.03E-01 | -0.2 | 7.24E-01 | Nitrilase 4 |
| AT2G41540 | -0.9 | 1.44E-06 | -0.5 | 1.72E-04 | -0.4 | 1.52E-03 | 0.6 | 1.21E-02 | 0.5 | 2.86E-02 | 0.0 | 9.34E-01 | Glycerol-3-phosphate dehydrogenase GPDHC1, cytosolic |
| AT4G01350 | -0.9 | 2.04E-05 | -0.3 | 3.32E-02 | -0.6 | 1.39E-03 | -0.3 | 3.28E-01 | 0.0 | 9.65E-01 | -0.3 | 3.90E-01 | Cysteine |
| AT2G01950 | -0.9 | 1.14E-06 | -0.4 | 4.08E-04 | -0.5 | 4.52E-04 | -0.2 | 1.66E-01 | 0.0 | 9.29E-01 | -0.2 | 3.89E-01 | Serine |
| AT5G56270 | -0.9 | 4.16E-06 | -1.0 | 3.94E-06 | 0.1 | 2.41E-01 | -0.5 | 9.91E-03 | -0.1 | 6.76E-01 | -0.4 | 1.12E-01 | Probable WRKY transcription factor 2 |
| AT2G28850 | -0.9 | 1.39E-03 | 0.6 | 8.90E-03 | -1.5 | 2.13E-04 | -0.1 | 7.63E-01 | 0.8 | 1.70E-03 | -0.9 | 4.25E-03 | Cytochrome P450 710A3 |
| AT3G14470 | -0.9 | 1.72E-05 | -0.7 | 1.63E-04 | -0.2 | 1.62E-01 | 0.5 | 1.42E-01 | 0.0 | 9.90E-01 | 0.5 | 2.48E-01 | Putative disease resistance RPP13-like protein 1 |
| AT4G39350 | -0.9 | 1.04E-06 | -0.7 | 1.94E-05 | -0.2 | 1.82E-02 | -0.3 | 4.37E-03 | 0.0 | 8.70E-01 | -0.3 | 5.70E-02 | Cellulose synthase A catalytic subunit 2 |
| AT1G47128 | -0.9 | 1.69E-06 | -0.9 | 4.52E-06 | 0.0 | 9.39E-01 | -0.5 | 8.81E-04 | -0.1 | 4.46E-01 | -0.4 | 5.05E-02 | Cysteine proteinase RD21A |
| AT5G54160 | -0.9 | 1.23E-06 | -0.4 | 1.36E-03 | -0.5 | 2.34E-04 | -0.1 | 3.05E-01 | 0.3 | 1.85E-02 | -0.4 | 1.15E-02 | Flavone 3'-O-methyltransferase 1 |
| AT1G67940 | -0.9 | 1.71E-06 | -0.5 | 2.39E-04 | -0.4 | 1.51E-03 | -0.2 | 4.37E-01 | 0.0 | 9.13E-01 | -0.1 | 6.37E-01 | ABC transporter I family member 17 |
| AT3G62260 | -0.9 | 4.17E-06 | -0.7 | 2.56E-05 | -0.1 | 2.39E-01 | -0.9 | 3.06E-04 | -0.1 | 7.81E-01 | -0.8 | 4.39E-03 | protein-serine |
| AT1G55610 | -0.9 | 6.13E-06 | -0.7 | 5.01E-05 | -0.2 | 1.59E-01 | 0.4 | 8.27E-02 | 0.5 | 4.39E-02 | -0.1 | 7.52E-01 | Serine |
| AT2G23000 | -0.9 | 5.78E-02 | -1.2 | 2.63E-02 | 0.3 | 6.33E-01 | 0.6 | 3.25E-01 | 0.6 | 4.15E-01 | 0.0 | 9.80E-01 | Serine carboxypeptidase-like 10 |
| AT4G38230 | -0.9 | 4.13E-06 | -0.3 | 2.81E-03 | -0.5 | 9.23E-04 | 0.0 | 9.98E-01 | 0.2 | 6.71E-01 | -0.2 | 6.96E-01 | Calcium-dependent protein kinase 26 |
| AT2G34020 | -0.9 | 5.91E-04 | 0.0 | 9.80E-01 | -0.9 | 2.49E-03 | -0.2 | 3.51E-01 | 0.3 | 1.66E-01 | -0.5 | 3.12E-02 | Calcium-binding EF-hand family protein |
| AT1G58400 | -0.9 | 2.61E-05 | -0.7 | 2.12E-04 | -0.2 | 2.23E-01 | 0.1 | 7.49E-01 | 0.4 | 1.82E-01 | -0.3 | 3.72E-01 | Putative disease resistance protein At1g58400 |

| | | | | | | | | | | | | | |
|---|---|---|---|---|---|---|---|---|---|---|---|---|---|
| AT3G52060 | -0.9 | 3.25E-06 | -0.2 | 1.26E-01 | -0.7 | 8.25E-05 | -0.3 | 3.15E-02 | -0.1 | 6.30E-01 | -0.2 | 2.93E-01 | At3g52060 |
| AT4G20270 | -0.9 | 3.21E-07 | -0.7 | 5.13E-06 | -0.2 | 8.50E-03 | -0.1 | 7.20E-01 | 0.1 | 7.12E-01 | -0.1 | 5.53E-01 | Leucine-rich repeat receptor-like serine |
| AT2G48080 | -0.9 | 2.22E-02 | 0.3 | 3.37E-01 | -1.2 | 1.14E-02 | 1.8 | 1.61E-05 | 0.6 | 9.29E-02 | 1.2 | 5.38E-04 | hypothetical protein |
| AT2G38750 | -0.9 | 1.88E-04 | -0.2 | 2.91E-01 | -0.7 | 3.15E-03 | -0.1 | 8.31E-01 | 0.5 | 1.96E-01 | -0.6 | 1.78E-01 | AT2G38750 protein |
| AT5G46080 | -0.9 | 1.79E-04 | -0.6 | 1.77E-03 | -0.2 | 2.20E-01 | 0.6 | 1.34E-02 | 0.3 | 3.68E-01 | 0.4 | 2.29E-01 | hypothetical protein |
| AT5G66460 | -0.9 | 7.66E-05 | -0.6 | 1.71E-03 | -0.3 | 8.30E-02 | -0.1 | 6.79E-01 | 0.5 | 8.22E-03 | -0.6 | 1.02E-02 | Mannan endo-1,4-beta-mannosidase 7 |
| AT1G78260 | -0.9 | 3.86E-05 | 0.2 | 7.22E-02 | -1.1 | 6.46E-05 | -0.5 | 9.76E-02 | -0.2 | 6.23E-01 | -0.3 | 3.28E-01 | RNA-binding family protein |
| AT1G69810 | -0.9 | 2.11E-06 | -0.3 | 2.57E-03 | -0.6 | 3.52E-04 | 0.4 | 1.13E-01 | 0.4 | 1.32E-01 | 0.0 | 9.66E-01 | WRKY DNA-binding protein 36 |
| AT3G03520 | -0.9 | 3.90E-07 | -0.4 | 2.23E-04 | -0.5 | 1.34E-04 | -0.2 | 3.42E-01 | 0.1 | 4.57E-01 | -0.3 | 1.01E-01 | Non-specific phospholipase C3 |
| AT2G43840 | -0.9 | 6.81E-05 | -0.7 | 4.09E-04 | -0.2 | 3.88E-01 | | | | | | | Glycosyltransferase |
| AT4G25310 | -0.9 | 9.76E-05 | -0.2 | 2.55E-01 | -0.7 | 1.47E-03 | 0.0 | 9.82E-01 | 0.2 | 3.52E-01 | -0.2 | 3.01E-01 | 2-oxoglutarate and Fe-dependent oxygenase superfamily protein |
| AT1G71340 | -0.9 | 2.36E-06 | -0.7 | 1.69E-05 | -0.1 | 1.50E-01 | -0.7 | 7.82E-04 | -0.3 | 8.84E-02 | -0.3 | 1.59E-01 | Glycerophosphodiester phosphodiesterase GDPD4 |
| AT2G22900 | -0.9 | 2.29E-05 | -0.2 | 5.43E-02 | -0.6 | 9.34E-04 | -0.1 | 4.56E-01 | 0.2 | 1.95E-01 | -0.3 | 7.65E-02 | Putative glycosyltransferase 7 |
| AT2G32620 | -0.9 | 8.18E-04 | 0.1 | 4.34E-01 | -1.0 | 1.50E-03 | -0.2 | 7.17E-01 | 0.7 | 9.07E-02 | -0.9 | 8.74E-02 | Cellulose synthase-like B |
| AT4G01610 | -0.9 | 3.65E-07 | -0.7 | 3.07E-06 | -0.1 | 6.23E-02 | -0.4 | 6.07E-03 | -0.1 | 6.60E-01 | -0.3 | 9.22E-02 | Peptidase C1A papain C-terminal domain-containing protein |
| AT1G45191 | -0.9 | 5.45E-03 | -0.8 | 1.21E-02 | -0.1 | 7.87E-01 | 0.0 | 9.59E-01 | 0.9 | 5.56E-02 | -0.8 | 1.02E-01 | Glycosyl hydrolase superfamily protein |
| AT1G31950 | -0.9 | 8.11E-04 | 0.0 | 8.53E-01 | -0.9 | 2.59E-03 | -0.3 | 3.25E-01 | -0.1 | 7.62E-01 | -0.2 | 6.91E-01 | Terpenoid cyclases |
| AT5G62130 | -0.9 | 7.30E-07 | -0.8 | 2.96E-06 | -0.1 | 4.38E-01 | -0.8 | 8.31E-04 | -0.3 | 1.33E-01 | -0.5 | 8.56E-02 | Post-GPI attachment to proteins factor 3 |
| AT5G49780 | -0.9 | 4.72E-06 | -0.3 | 9.12E-03 | -0.6 | 5.03E-04 | 0.6 | 2.47E-03 | 0.6 | 4.38E-03 | 0.0 | 9.81E-01 | Leucine-rich repeat protein kinase family protein |
| AT5G01190 | -0.9 | 4.12E-04 | 0.1 | 6.91E-01 | -0.9 | 1.43E-03 | -0.7 | 1.14E-01 | 0.0 | 1.00E+00 | -0.7 | 2.36E-01 | Laccase |
| AT1G62975 | -0.9 | 6.63E-04 | -0.6 | 4.83E-03 | -0.2 | 3.40E-01 | -0.1 | 8.05E-01 | 0.6 | 2.62E-01 | -0.7 | 2.33E-01 | Basic helix-loop-helix DNA-binding superfamily protein |
| AT1G63000 | -0.9 | 2.41E-06 | -0.3 | 3.54E-03 | -0.5 | 3.12E-04 | -0.1 | 3.26E-01 | 0.3 | 6.63E-02 | -0.4 | 1.71E-02 | Bifunctional dTDP-4-dehydrorhamnose 3,5-epimerase |
| AT1G64300 | -0.9 | 1.06E-03 | -0.6 | 8.87E-03 | -0.3 | 3.26E-01 | 0.5 | 4.34E-01 | 0.2 | 8.66E-01 | 0.4 | 6.86E-01 | At1g64300 |
| AT4G14940 | -0.9 | 2.31E-05 | -0.2 | 7.55E-02 | -0.7 | 1.05E-03 | 0.1 | 7.65E-01 | 0.4 | 8.42E-02 | -0.4 | 2.28E-01 | Primary amine oxidase 1 |
| AT5G24870 | -0.9 | 8.20E-06 | -0.9 | 1.76E-05 | 0.0 | 9.79E-01 | -0.4 | 3.23E-02 | 0.0 | 9.34E-01 | -0.4 | 8.41E-02 | RING-type E3 ubiquitin transferase |
| AT1G05700 | -0.9 | 2.99E-05 | -0.3 | 2.01E-02 | -0.6 | 3.01E-03 | 1.1 | 4.17E-04 | 0.5 | 6.77E-02 | 0.5 | 5.32E-02 | Leucine-rich repeat transmembrane protein kinase protein |
| AT3G16490 | -0.9 | 3.89E-04 | -0.1 | 4.44E-01 | -0.7 | 4.81E-03 | -0.4 | 2.06E-01 | -0.1 | 8.58E-01 | -0.3 | 4.34E-01 | Protein IQ-domain 26 |
| AT3G59530 | -0.9 | 5.97E-05 | -0.8 | 1.68E-04 | -0.1 | 7.29E-01 | -0.6 | 1.30E-01 | 0.2 | 6.20E-01 | -0.9 | 7.76E-02 | Protein STRICTOSIDINE SYNTHASE-LIKE 13 |
| AT4G24100 | -0.9 | 2.93E-06 | -0.7 | 1.97E-05 | -0.1 | 1.74E-01 | -0.4 | 3.42E-02 | 0.2 | 4.92E-01 | -0.6 | 1.77E-02 | Protein kinase domain-containing protein |
| AT3G48260 | -0.9 | 2.63E-05 | 0.1 | 6.04E-01 | -0.9 | 1.54E-04 | -0.4 | 1.85E-01 | -0.2 | 5.27E-01 | -0.2 | 7.09E-01 | non-specific serine |
| AT3G49160 | -0.9 | 2.99E-05 | -0.7 | 1.74E-04 | -0.1 | 3.34E-01 | -3.7 | 1.69E-04 | -1.4 | 9.28E-03 | -2.3 | 4.11E-02 | pyruvate kinase |
| AT4G29210 | -0.9 | 2.94E-06 | -0.6 | 4.27E-05 | -0.2 | 4.16E-02 | -0.2 | 3.21E-01 | 0.2 | 3.46E-01 | -0.3 | 1.12E-01 | Glutathione hydrolase 3 |
| AT3G15353 | -0.9 | 4.24E-06 | -0.3 | 8.60E-03 | -0.6 | 3.91E-04 | -0.5 | 4.81E-02 | -0.3 | 2.17E-01 | -0.1 | 7.41E-01 | Metallothionein 3 |
| AT3G20640 | -0.9 | 1.50E-04 | -0.4 | 1.38E-02 | -0.5 | 2.69E-02 | 0.1 | 8.73E-01 | 0.2 | 5.54E-01 | -0.1 | 6.57E-01 | Basic helix-loop-helix DNA-binding superfamily protein |
| AT2G28460 | -0.9 | 3.49E-05 | -0.5 | 2.20E-03 | -0.4 | 2.57E-02 | -1.0 | 1.10E-02 | -0.4 | 3.43E-01 | -0.7 | 2.00E-01 | Uncharacterized protein At2g28460 |
| AT4G37690 | -0.9 | 6.90E-04 | -0.5 | 2.39E-02 | -0.4 | 1.06E-01 | 0.0 | 9.98E-01 | 0.6 | 1.90E-01 | -0.6 | 2.28E-01 | Glycosyltransferase 6 |
| AT5G35450 | -0.9 | 7.47E-07 | -0.6 | 1.13E-05 | -0.2 | 2.20E-02 | 0.0 | 9.95E-01 | 0.2 | 5.64E-01 | -0.2 | 5.87E-01 | Disease resistance RPP8-like protein 3 |
| AT5G60360 | -0.9 | 1.56E-06 | -0.8 | 4.55E-06 | 0.0 | 8.51E-01 | -0.3 | 4.98E-03 | 0.0 | 8.96E-01 | -0.3 | 6.40E-02 | Aleurain-like protease |
| AT1G30080 | -0.9 | 6.34E-05 | -0.2 | 1.03E-01 | -0.7 | 2.37E-03 | -0.2 | 4.98E-01 | 0.7 | 1.96E-02 | -0.9 | 6.92E-03 | glucan endo-1,3-beta-D-glucosidase |
| AT3G56000 | -0.9 | 4.75E-04 | 0.2 | 1.48E-01 | -1.1 | 4.84E-04 | -0.8 | 6.14E-03 | -0.3 | 4.14E-01 | -0.6 | 6.40E-02 | Probable glucomannan 4-beta-mannosyltransferase 14 |
| AT3G04060 | -0.9 | 1.11E-05 | -0.6 | 2.22E-04 | -0.3 | 4.99E-02 | 0.2 | 1.83E-01 | 0.5 | 1.34E-02 | -0.2 | 2.11E-01 | NAC domain-containing protein 46 |
| AT1G19940 | -0.9 | 4.36E-04 | -0.2 | 2.76E-01 | -0.7 | 8.11E-03 | 0.1 | 9.00E-01 | 0.4 | 3.38E-01 | | | cellulase |
| AT5G19080 | -0.9 | 6.18E-07 | -0.7 | 5.71E-06 | -0.2 | 5.93E-02 | -0.3 | 1.24E-01 | -0.4 | 4.30E-02 | 0.1 | 6.51E-01 | Probable E3 ubiquitin-protein ligase LUL3 |
| AT1G30760 | -0.9 | 4.57E-06 | -0.4 | 7.75E-04 | -0.4 | 2.91E-03 | 0.8 | 1.62E-04 | 0.6 | 8.67E-03 | 0.3 | 1.90E-01 | FAD-binding Berberine family protein |
| AT1G76580 | -0.9 | 1.87E-05 | -0.8 | 6.43E-05 | -0.1 | 6.21E-01 | -0.3 | 9.75E-02 | -0.1 | 6.59E-01 | -0.2 | 4.42E-01 | SBP-type domain-containing protein |
| AT5G45730 | -0.9 | 1.43E-04 | -0.4 | 2.32E-02 | -0.5 | 1.87E-02 | | | | | | | Cysteine |
| AT4G37160 | -0.9 | 1.88E-03 | 0.0 | 8.83E-01 | -0.8 | 7.44E-03 | -0.4 | 2.27E-01 | 0.3 | 4.81E-01 | -0.7 | 1.12E-01 | Pectinesterase like protein |
| AT5G04340 | -0.9 | 7.48E-04 | -1.0 | 3.68E-04 | 0.2 | 4.62E-01 | 0.5 | 1.04E-01 | -0.1 | 7.78E-01 | 0.6 | 9.29E-02 | Zinc finger protein ZAT6 |
| AT1G79270 | -0.9 | 1.33E-06 | -0.7 | 1.24E-05 | -0.2 | 7.69E-02 | 0.0 | 9.85E-01 | 0.2 | 3.29E-01 | -0.2 | 3.31E-01 | YTH domain-containing family protein |
| AT4G26890 | -0.9 | 3.99E-03 | -0.6 | 3.14E-02 | -0.3 | 3.72E-01 | -0.7 | 1.05E-01 | -0.4 | 3.71E-01 | | | Mitogen-activated protein kinase kinase kinase 16 |
| AT1G32170 | -0.9 | 2.37E-06 | -0.1 | 1.12E-01 | -0.7 | 7.45E-05 | -0.1 | 5.51E-01 | -0.2 | 4.41E-01 | 0.1 | 8.80E-01 | Probable xyloglucan endotransglucosylase |
| AT1G69690 | -0.9 | 1.03E-05 | -0.5 | 5.16E-04 | -0.3 | 1.63E-02 | -0.4 | 5.19E-02 | 0.0 | 9.70E-01 | -0.4 | 1.27E-01 | Transcription factor TCP15 |
| AT5G02270 | -0.8 | 3.26E-05 | 0.0 | 7.44E-01 | -0.9 | 1.34E-04 | 0.0 | 9.25E-01 | 0.6 | 2.53E-03 | -0.7 | 2.81E-03 | Non-intrinsic ABC protein 9 |
| AT5G25810 | -0.8 | 3.73E-06 | 0.0 | 8.79E-01 | -0.9 | 3.57E-05 | -0.5 | 2.29E-02 | 0.3 | 2.18E-01 | -0.8 | 1.54E-03 | Ethylene-responsive transcription factor TINY |
| AT3G13672 | -0.8 | 1.46E-04 | -0.4 | 8.73E-03 | -0.4 | 4.03E-02 | | | | | | | TRAF-like superfamily protein |
| AT3G56950 | -0.8 | 2.12E-06 | -0.4 | 4.38E-04 | -0.4 | 1.18E-03 | 0.2 | 1.98E-01 | 0.2 | 1.63E-01 | 0.0 | 8.69E-01 | Small and basic intrinsic protein 21 |
| AT5G07030 | -0.8 | 1.15E-04 | -0.5 | 4.08E-03 | -0.3 | 5.59E-02 | 0.3 | 4.10E-02 | 0.3 | 1.44E-02 | -0.1 | 6.40E-01 | hypothetical protein |
| AT3G08720 | -0.8 | 3.15E-06 | -0.8 | 1.42E-05 | -0.1 | 3.73E-01 | -0.1 | 6.01E-01 | -0.1 | 8.33E-01 | -0.1 | 8.82E-01 | Serine |
| AT5G62480 | -0.8 | 6.74E-04 | -1.3 | 7.28E-05 | 0.5 | 5.32E-02 | 2.6 | 1.25E-08 | 0.4 | 1.32E-01 | 2.2 | 1.58E-06 | Glutathione S-transferase |
| AT3G53480 | -0.8 | 4.00E-05 | -0.7 | 2.11E-04 | -0.1 | 3.59E-01 | 0.3 | 1.29E-01 | -0.2 | 3.58E-01 | 0.5 | 5.32E-02 | ABC transporter G family member 37 |
| AT3G06860 | -0.8 | 8.75E-07 | -0.6 | 2.29E-05 | -0.3 | 7.61E-03 | 0.1 | 7.18E-01 | 0.2 | 3.37E-01 | -0.1 | 6.34E-01 | Fatty acid multifunctional protein |
| AT1G53070 | -0.8 | 1.25E-04 | -0.2 | 1.29E-01 | -0.6 | 3.80E-03 | 0.3 | 1.88E-01 | 0.6 | 6.35E-03 | -0.3 | 1.51E-01 | Lectin-like protein At1g53070 |
| AT1G16030 | -0.8 | 1.79E-05 | -0.7 | 8.59E-05 | -0.1 | 4.58E-01 | -0.8 | 7.42E-03 | -0.5 | 9.88E-02 | -0.3 | 4.42E-01 | Heat shock 70 kDa protein 5 |
| AT5G58350 | -0.8 | 6.09E-06 | -0.7 | 2.80E-05 | -0.1 | 3.68E-01 | -0.5 | 5.54E-02 | -0.1 | 6.57E-01 | -0.3 | 2.89E-01 | Probable serine |
| AT5G62570 | -0.8 | 5.80E-06 | -0.9 | 6.11E-06 | 0.1 | 3.73E-01 | -0.3 | 1.35E-01 | -0.2 | 3.92E-01 | -0.1 | 7.74E-01 | Calmodulin binding protein-like protein |
| AT5G11920 | -0.8 | 1.07E-05 | -0.9 | 1.92E-05 | 0.0 | 9.12E-01 | 1.4 | 7.15E-08 | 1.3 | 5.32E-07 | 0.1 | 7.42E-01 | 6-&1-fructan exohydrolase |
| AT2G28990 | -0.8 | 1.17E-05 | -0.4 | 2.07E-03 | -0.4 | 4.73E-03 | -0.3 | 3.53E-01 | 0.2 | 5.44E-01 | -0.5 | 1.53E-01 | Probable leucine-rich repeat receptor-like protein kinase At2g28990 |
| AT4G04610 | -0.8 | 1.96E-05 | -0.2 | 6.61E-02 | -0.6 | 6.71E-04 | 0.7 | 7.97E-04 | 0.7 | 2.28E-03 | 0.0 | 9.83E-01 | 5'-adenylylsulfate reductase 1, chloroplastic |
| AT5G58580 | -0.8 | 5.08E-04 | -0.4 | 3.39E-02 | -0.5 | 5.21E-02 | -0.2 | 3.29E-01 | -0.3 | 3.13E-01 | 0.0 | 9.48E-01 | RING-H2 finger protein ATL63 |
| AT1G03080 | -0.8 | 4.55E-07 | -0.6 | 7.65E-06 | -0.2 | 1.14E-02 | 0.2 | 2.47E-01 | 0.1 | 6.08E-01 | 0.1 | 7.85E-01 | Protein NETWORKED 1D |
| AT5G03260 | -0.8 | 3.28E-06 | 0.0 | 8.47E-01 | -0.8 | 4.30E-05 | 0.0 | 8.78E-01 | 0.1 | 5.66E-01 | -0.1 | 7.30E-01 | Laccase |
| AT5G14530 | -0.8 | 6.27E-06 | -0.6 | 1.17E-04 | -0.3 | 4.81E-02 | -0.6 | 8.12E-03 | -0.6 | 2.01E-02 | -0.1 | 8.89E-01 | Protein ANTHESIS POMOTING FACTOR 1 |
| AT5G01060 | -0.8 | 3.45E-04 | -0.1 | 4.64E-01 | -0.7 | 4.21E-03 | -0.7 | 1.00E-01 | -0.1 | 8.08E-01 | -0.6 | 2.44E-01 | Serine |
| AT1G54790 | -0.8 | 4.78E-04 | -0.3 | 1.17E-01 | -0.6 | 1.89E-02 | -0.1 | 8.70E-01 | -0.1 | 8.52E-01 | 0.0 | 9.71E-01 | GDSL esterase |
| AT2G23950 | -0.8 | 1.05E-06 | -0.6 | 3.21E-05 | -0.3 | 7.14E-03 | -0.3 | 7.70E-02 | -0.1 | 6.26E-01 | -0.2 | 4.39E-01 | Leucine-rich repeat protein kinase family protein |
| AT5G41990 | -0.8 | 1.12E-06 | -0.7 | 7.03E-06 | -0.1 | 1.84E-01 | -0.1 | 6.34E-01 | 0.0 | 9.41E-01 | -0.1 | 7.02E-01 | Serine |
| AT1G07230 | -0.8 | 1.09E-05 | -0.7 | 6.36E-05 | -0.1 | 2.76E-01 | -0.4 | 1.68E-02 | -0.1 | 4.61E-01 | -0.2 | 2.82E-01 | Non-specific phospholipase C1 |
| AT2G23560 | -0.8 | 1.77E-04 | -0.3 | 3.50E-02 | -0.5 | 1.61E-02 | -0.3 | 2.55E-01 | 0.1 | 6.46E-01 | -0.4 | 1.43E-01 | Methylesterase 7 |
| AT3G56170 | -0.8 | 1.10E-06 | -0.6 | 1.33E-05 | -0.2 | 4.09E-02 | -0.1 | 4.90E-01 | 0.0 | 9.76E-01 | -0.1 | 6.66E-01 | Staphylococcal-like nuclease CAN1 -dependent nuclease 1) |
| AT4G27180 | -0.8 | 4.01E-06 | -0.6 | 1.08E-04 | -0.3 | 1.88E-02 | -0.2 | 1.39E-01 | 0.3 | 1.30E-01 | -0.5 | 2.71E-02 | Kinesin-like protein KIN-14M |
| AT4G26970 | -0.8 | 1.33E-05 | -0.3 | 1.75E-02 | -0.5 | 1.04E-03 | 0.4 | 1.69E-02 | 0.3 | 8.48E-02 | 0.1 | 7.84E-01 | Aconitate hydratase 2, mitochondrial |
| AT2G13820 | -0.8 | 1.50E-05 | 0.0 | 9.99E-01 | -0.8 | 1.05E-04 | -0.2 | 2.16E-01 | 0.0 | 8.77E-01 | -0.3 | 2.45E-01 | Bifunctional inhibitor |
| AT1G30560 | -0.8 | 1.17E-03 | 0.3 | 1.47E-01 | -1.1 | 9.26E-04 | -0.5 | 1.59E-01 | -0.2 | 6.72E-01 | -0.3 | 5.18E-01 | Putative glycerol-3-phosphate transporter 3 |

| | | | | | | | | | | | | | |
|---|---|---|---|---|---|---|---|---|---|---|---|---|---|
| AT4G14780 | -0.8 | 1.33E-05 | -0.8 | 3.86E-05 | -0.1 | 6.94E-01 | -1.9 | 1.22E-05 | 0.1 | 8.41E-01 | -2.0 | 7.98E-05 | Protein kinase domain-containing protein |
| AT2G01880 | -0.8 | 1.16E-04 | -0.3 | 3.00E-02 | -0.5 | 1.21E-02 | 0.3 | 5.00E-01 | 0.2 | 7.81E-01 | 0.1 | 8.18E-01 | Purple acid phosphatase 7 |
| AT4G32410 | -0.8 | 9.74E-06 | -0.7 | 8.01E-05 | -0.1 | 1.92E-01 | -0.4 | 4.44E-02 | -0.1 | 6.46E-01 | -0.3 | 3.35E-01 | Cellulose synthase A catalytic subunit 1 |
| AT2G18730 | -0.8 | 6.56E-07 | -0.7 | 3.22E-06 | -0.1 | 2.43E-01 | -0.4 | 1.30E-03 | -0.1 | 7.05E-01 | -0.4 | 2.45E-02 | Diacylglycerol kinase 3 |
| AT2G38740 | -0.8 | 2.15E-05 | -0.6 | 4.11E-04 | -0.2 | 6.13E-02 | -0.2 | 2.97E-01 | 0.1 | 8.55E-01 | -0.3 | 2.78E-01 | Haloacid dehalogenase-like hydrolase domain-containing protein Sgpp |
| AT2G15620 | -0.8 | 3.24E-03 | -0.7 | 8.80E-03 | -0.1 | 7.22E-01 | -1.3 | 3.74E-08 | -0.6 | 5.56E-04 | -0.7 | 8.65E-04 | Ferredoxin--nitrite reductase, chloroplastic |
| AT5G18900 | -0.8 | 6.16E-06 | -0.5 | 3.15E-04 | -0.3 | 1.01E-02 | -0.3 | 5.51E-02 | -0.1 | 5.52E-01 | -0.2 | 3.70E-01 | Probable prolyl 4-hydroxylase 4 |
| AT1G77330 | -0.8 | 1.87E-05 | 0.1 | 2.89E-01 | -0.9 | 6.15E-05 | 0.4 | 6.86E-02 | 0.6 | 1.04E-02 | -0.2 | 2.95E-01 | 1-aminocyclopropane-1-carboxylate oxidase 5 |
| AT2G25530 | -0.8 | 8.99E-04 | -0.6 | 7.88E-03 | -0.2 | 2.96E-01 | 0.5 | 1.43E-01 | 0.6 | 1.34E-01 | -0.1 | 9.05E-01 | AFG1-like ATPase family protein |
| AT2G42660 | -0.8 | 1.03E-03 | -0.2 | 3.33E-01 | -0.7 | 1.58E-02 | 0.0 | 9.78E-01 | 0.4 | 4.13E-01 | -0.4 | 4.15E-01 | hypothetical protein |
| AT5G20410 | -0.8 | 1.13E-05 | -0.4 | 2.24E-03 | -0.4 | 4.48E-03 | -0.5 | 6.44E-02 | -0.1 | 8.35E-01 | -0.4 | 2.01E-01 | Monogalactosyldiacylglycerol synthase 2, chloroplastic |
| AT1G70450 | -0.8 | 2.86E-03 | 0.4 | 6.68E-02 | -1.2 | 1.25E-03 | | | | | | | non-specific serine |
| AT3G15940 | -0.8 | 2.06E-06 | -0.5 | 9.41E-05 | -0.3 | 6.26E-03 | -0.1 | 5.43E-01 | 0.2 | 4.70E-01 | -0.3 | 2.19E-01 | Glycosyl transferase family 1 domain-containing protein |
| AT2G16570 | -0.8 | 1.15E-04 | -0.4 | 8.39E-03 | -0.4 | 2.98E-02 | 0.6 | 1.49E-02 | 0.2 | 4.20E-01 | 0.4 | 1.78E-01 | Amidophosphoribosyltransferase 1, chloroplastic |
| AT2G17260 | -0.8 | 8.30E-05 | -0.6 | 1.28E-03 | -0.3 | 1.41E-01 | 0.1 | 7.71E-01 | 0.1 | 8.08E-01 | 0.0 | 9.99E-01 | Glutamate receptor 3.1 |
| AT3G14570 | -0.8 | 5.63E-06 | -0.8 | 1.34E-05 | 0.0 | 9.16E-01 | -0.4 | 2.76E-01 | 0.2 | 5.52E-01 | -0.6 | 1.35E-01 | 1,3-beta-glucan synthase |
| AT1G29230 | -0.8 | 1.04E-04 | 0.4 | 3.21E-03 | -1.2 | 5.13E-05 | -1.4 | 7.97E-03 | 0.0 | 9.93E-01 | -1.4 | 1.67E-02 | CBL-interacting serine |
| AT3G63210 | -0.8 | 1.84E-05 | -0.3 | 7.15E-03 | -0.5 | 2.91E-03 | -0.5 | 1.48E-01 | 0.0 | 9.33E-01 | -0.5 | 2.29E-01 | Protein MARD1 |
| AT4G37940 | -0.8 | 5.21E-06 | -0.5 | 2.35E-04 | -0.3 | 1.39E-02 | -0.3 | 5.06E-01 | -0.2 | 6.95E-01 | -0.1 | 8.84E-01 | AGAMOUS-like 21 |
| AT2G46270 | -0.8 | 2.64E-05 | -0.6 | 2.63E-04 | -0.2 | 1.66E-01 | -0.3 | 2.90E-01 | -0.6 | 3.94E-02 | 0.3 | 3.59E-01 | BZIP domain-containing protein |
| AT4G15400 | -0.8 | 1.41E-05 | -0.4 | 4.02E-03 | -0.5 | 3.63E-03 | 0.1 | 8.26E-01 | 0.3 | 1.22E-01 | -0.3 | 2.50E-01 | BAHD acyltransferase BIA1 |
| AT5G64740 | -0.8 | 2.17E-06 | -0.5 | 2.05E-04 | -0.3 | 3.06E-03 | -0.5 | 1.51E-03 | -0.2 | 3.34E-01 | -0.3 | 1.03E-01 | Cellulose synthase A catalytic subunit 6 |
| AT3G50770 | -0.8 | 1.38E-02 | -1.1 | 4.98E-03 | 0.2 | 5.51E-01 | 0.4 | 4.08E-01 | -0.8 | 1.93E-01 | 1.1 | 6.88E-02 | Probable calcium-binding protein CML41 |
| AT1G69790 | -0.8 | 1.46E-04 | -1.3 | 2.01E-05 | 0.5 | 2.72E-02 | 0.2 | 6.73E-01 | -0.3 | 5.57E-01 | 0.5 | 2.61E-01 | Protein kinase superfamily protein |
| AT3G02880 | -0.8 | 1.93E-06 | -0.4 | 8.70E-04 | -0.4 | 5.77E-04 | -0.1 | 3.24E-01 | -0.1 | 6.39E-01 | -0.1 | 8.31E-01 | Probable inactive receptor kinase At3g02880 |
| AT1G28370 | -0.8 | 1.90E-03 | -1.4 | 1.91E-04 | 0.6 | 7.06E-02 | 1.0 | 2.08E-02 | 0.1 | 8.95E-01 | 0.9 | 3.09E-02 | ERF domain protein 11 |
| AT5G45720 | -0.8 | 2.37E-03 | -0.4 | 5.35E-02 | -0.4 | 1.50E-01 | 0.2 | 3.96E-01 | 0.5 | 5.54E-02 | -0.3 | 3.71E-01 | AAA-type ATPase family protein |
| AT4G35500 | -0.8 | 7.29E-06 | -0.4 | 1.90E-03 | -0.4 | 2.79E-03 | -0.5 | 4.82E-02 | -0.2 | 6.18E-01 | -0.4 | 2.72E-01 | Protein kinase superfamily protein |
| AT3G09270 | -0.8 | 6.09E-06 | -0.8 | 1.91E-05 | -0.1 | 6.39E-01 | 2.2 | 2.82E-07 | 1.0 | 3.44E-03 | 1.2 | 1.75E-04 | Glutathione S-transferase U8 |
| AT2G04240 | -0.8 | 1.85E-06 | -0.4 | 4.91E-04 | -0.4 | 9.61E-04 | -0.6 | 4.37E-04 | -0.2 | 1.73E-01 | -0.4 | 6.16E-02 | Probable E3 ubiquitin-protein ligase XERICO |
| AT2G46740 | -0.8 | 5.36E-05 | 0.2 | 3.96E-02 | -1.1 | 7.31E-05 | -1.0 | 3.81E-03 | -0.3 | 4.14E-01 | -0.7 | 5.04E-02 | L-gulonolactone oxidase 5 |
| AT1G08800 | -0.8 | 1.06E-05 | -0.5 | 4.36E-04 | -0.3 | 1.94E-02 | -0.1 | 7.23E-01 | 0.0 | 9.03E-01 | -0.1 | 6.95E-01 | Myosin-binding protein 1 |
| AT4G20170 | -0.8 | 1.09E-04 | -1.1 | 2.05E-05 | 0.3 | 5.09E-02 | -0.3 | 5.88E-02 | 0.2 | 4.12E-01 | -0.5 | 3.17E-02 | Galactan beta-1,4-galactosyltransferase GALS3 |
| AT1G02340 | -0.8 | 7.48E-03 | -0.4 | 1.54E-01 | -0.4 | 1.82E-01 | | | | | | | Basic helix-loop-helix DNA-binding superfamily protein |
| AT4G22920 | -0.8 | 3.51E-05 | -1.2 | 6.25E-06 | 0.4 | 1.94E-02 | -0.3 | 1.98E-01 | -0.2 | 4.96E-01 | -0.1 | 7.75E-01 | Magnesium dechelatase SGR1, chloroplastic |
| AT5G56460 | -0.8 | 4.75E-06 | -0.8 | 1.16E-05 | 0.0 | 9.17E-01 | 0.3 | 1.39E-01 | 0.1 | 6.36E-01 | 0.2 | 5.14E-01 | Probable serine |
| AT3G51440 | -0.8 | 1.14E-04 | -0.4 | 7.43E-03 | -0.4 | 3.21E-02 | -0.5 | 3.45E-02 | -0.2 | 5.16E-01 | -0.3 | 3.07E-01 | Protein STRICTOSIDINE SYNTHASE-LIKE 6 |
| AT3G01650 | -0.8 | 2.05E-06 | -0.5 | 1.79E-04 | -0.4 | 3.16E-03 | -0.2 | 2.61E-01 | -0.1 | 6.73E-01 | -0.1 | 7.24E-01 | E3 ubiquitin-protein ligase RGLG1 |
| AT5G52320 | -0.8 | 9.42E-03 | -0.3 | 1.93E-01 | -0.5 | 1.63E-01 | -0.1 | 8.93E-01 | 0.9 | 6.00E-03 | -1.0 | 9.21E-03 | Cytochrome P450 |
| AT1G79310 | -0.8 | 5.05E-05 | -0.7 | 2.19E-04 | -0.1 | 4.93E-01 | -0.9 | 3.91E-03 | 0.3 | 1.96E-01 | -1.2 | 1.87E-03 | Metacaspase-7 |
| AT3G25710 | -0.8 | 2.00E-05 | -0.4 | 4.39E-03 | -0.4 | 5.56E-03 | 0.0 | 9.20E-01 | 0.1 | 7.72E-01 | -0.1 | 7.27E-01 | Transcription factor AIG1 |
| AT4G21760 | -0.8 | 4.08E-04 | -0.5 | 6.20E-03 | -0.3 | 1.85E-01 | | | | | | | Beta-glucosidase 47 |
| AT1G06830 | -0.8 | 2.36E-03 | -0.6 | 1.14E-02 | -0.2 | 4.93E-01 | -0.6 | 4.04E-02 | -0.5 | 1.52E-01 | -0.2 | 7.71E-01 | Monothiol glutaredoxin-S11 |
| AT1G10210 | -0.8 | 2.59E-05 | -0.6 | 3.51E-04 | -0.2 | 1.20E-01 | -0.5 | 1.02E-02 | -0.4 | 9.66E-02 | -0.2 | 5.02E-01 | Mitogen-activated protein kinase 1 |
| AT3G18000 | -0.8 | 5.39E-03 | 0.4 | 9.16E-02 | -1.3 | 1.15E-03 | 0.4 | 1.07E-01 | 1.5 | 3.47E-05 | -1.1 | 2.24E-03 | Phosphoethanolamine N-methyltransferase 1 |
| AT3G44970 | -0.8 | 3.27E-02 | -0.5 | 1.87E-01 | -0.3 | 4.27E-01 | 0.3 | 4.46E-01 | 0.5 | 3.10E-01 | -0.1 | 8.18E-01 | Cytochrome P450 superfamily protein |
| AT3G18990 | -0.8 | 8.09E-07 | -0.4 | 9.78E-05 | -0.4 | 9.62E-04 | -0.1 | 3.39E-01 | -0.1 | 7.45E-01 | -0.1 | 7.40E-01 | AP2 |
| AT3G19450 | -0.8 | 2.00E-06 | -0.2 | 5.96E-02 | -0.6 | 7.31E-05 | -0.1 | 2.60E-01 | 0.2 | 1.13E-01 | -0.4 | 2.95E-02 | Cinnamyl alcohol dehydrogenase 4 |
| AT5G01880 | -0.8 | 2.30E-05 | -1.1 | 8.53E-06 | 0.3 | 6.99E-02 | -1.1 | 4.51E-03 | -0.8 | 3.18E-02 | -0.3 | 5.83E-01 | RING-H2 finger protein ATL74 |
| AT4G11920 | -0.8 | 9.85E-06 | -0.4 | 1.08E-03 | -0.4 | 6.87E-03 | -0.2 | 3.62E-01 | -0.3 | 2.73E-01 | 0.1 | 8.62E-01 | Protein FIZZY-RELATED 1 |
| AT2G28810 | -0.8 | 5.21E-06 | -0.5 | 1.72E-04 | -0.3 | 1.80E-02 | -0.1 | 5.93E-01 | 0.0 | 9.93E-01 | -0.1 | 6.90E-01 | Dof zinc finger protein |
| AT1G49860 | -0.8 | 4.48E-05 | 0.0 | 8.79E-01 | -0.8 | 1.92E-04 | -0.7 | 2.13E-04 | -0.4 | 2.00E-02 | -0.3 | 1.66E-01 | Glutathione S-transferase F14 |
| AT5G02880 | -0.8 | 1.25E-05 | -0.9 | 9.35E-06 | 0.1 | 2.53E-01 | -0.2 | 1.14E-01 | -0.2 | 3.30E-01 | -0.1 | 7.93E-01 | E3 ubiquitin-protein ligase |
| AT1G07380 | -0.8 | 3.92E-06 | -0.7 | 1.42E-05 | -0.1 | 5.51E-01 | -0.4 | 6.35E-02 | -0.1 | 7.26E-01 | -0.3 | 2.39E-01 | Neutral ceramidase |
| AT1G26970 | -0.8 | 3.40E-04 | 0.0 | 9.00E-01 | -0.8 | 2.12E-03 | 0.3 | 4.33E-01 | 0.4 | 4.04E-01 | -0.1 | 9.24E-01 | Protein kinase superfamily protein |
| AT1G28610 | -0.8 | 3.04E-03 | -0.7 | 9.30E-03 | -0.1 | 6.65E-01 | -0.1 | 8.45E-01 | 0.8 | 2.04E-02 | -0.9 | 3.12E-02 | GDSL-like Lipase |
| AT1G05260 | -0.8 | 1.25E-06 | 0.4 | 4.39E-04 | -1.2 | 6.99E-06 | -0.1 | 6.58E-01 | 0.3 | 7.41E-02 | -0.4 | 1.40E-02 | Peroxidase 3 |
| AT1G01200 | -0.8 | 2.96E-04 | -0.5 | 6.33E-03 | -0.3 | 1.25E-01 | -0.3 | 3.99E-01 | 0.0 | 9.94E-01 | -0.3 | 5.48E-01 | RABA3 |
| AT5G35670 | -0.8 | 2.30E-05 | -0.5 | 6.88E-04 | -0.3 | 4.30E-02 | -0.1 | 6.92E-01 | 0.4 | 1.73E-01 | -0.5 | 1.27E-01 | Protein IQ-DOMAIN 33 |
| AT4G23050 | -0.8 | 7.76E-07 | -1.0 | 5.40E-07 | 0.2 | 3.02E-02 | -0.4 | 1.05E-01 | 0.0 | 9.32E-01 | -0.4 | 1.87E-01 | PAS domain-containing protein tyrosine kinase family protein |
| AT1G75680 | -0.8 | 5.38E-06 | -0.4 | 1.34E-03 | -0.4 | 2.03E-03 | -0.3 | 1.97E-02 | 0.0 | 8.49E-01 | -0.4 | 6.07E-02 | Endoglucanase 10 |
| AT1G20620 | -0.8 | 1.73E-05 | -0.4 | 2.29E-03 | -0.4 | 7.21E-03 | -1.1 | 8.16E-05 | -0.6 | 2.52E-02 | -0.5 | 3.90E-02 | catalase |
| AT1G72300 | -0.8 | 2.37E-05 | -0.6 | 4.18E-04 | -0.2 | 7.10E-02 | -0.4 | 2.05E-03 | -0.1 | 3.49E-01 | -0.3 | 8.16E-02 | Tyrosine-sulfated glycopeptide receptor 1 |
| AT5G57740 | -0.8 | 6.58E-06 | -0.4 | 1.07E-03 | -0.4 | 3.61E-03 | -0.4 | 6.17E-02 | 0.2 | 5.02E-01 | -0.6 | 4.40E-02 | Uncharacterized protein At5g57740 |
| AT4G00900 | -0.8 | 1.68E-05 | -0.8 | 4.02E-05 | 0.0 | 8.34E-01 | 0.0 | 9.99E-01 | 0.2 | 5.37E-01 | -0.2 | 6.10E-01 | Calcium-transporting ATPase 2, endoplasmic reticulum-type |
| AT4G26470 | -0.8 | 2.32E-06 | -0.5 | 6.06E-05 | -0.3 | 1.79E-02 | 0.2 | 3.22E-01 | 0.2 | 3.45E-01 | 0.0 | 9.61E-01 | Calcium-binding EF-hand family protein |
| AT5G45200 | -0.8 | 5.78E-04 | 0.1 | 6.79E-01 | -0.9 | 1.80E-03 | | | | | | | ADP-ribosyl cyclase |
| AT1G59970 | -0.8 | 1.96E-05 | -0.6 | 1.95E-04 | -0.2 | 1.60E-01 | 0.0 | 9.16E-01 | 0.0 | 9.08E-01 | 0.0 | 9.80E-01 | Metalloendoproteinase 5-MMP |
| AT1G53340 | -0.8 | 4.24E-03 | -0.3 | 1.56E-01 | -0.5 | 1.07E-01 | -0.5 | 2.59E-01 | 0.2 | 7.95E-01 | -0.7 | 2.36E-01 | Cysteine |
| AT5G20960 | -0.8 | 3.29E-06 | -0.4 | 5.86E-04 | -0.4 | 1.93E-03 | -0.1 | 7.34E-01 | 0.0 | 9.38E-01 | -0.1 | 8.84E-01 | Indole-3-acetaldehyde oxidase |
| AT4G28640 | -0.8 | 7.78E-06 | -0.5 | 2.67E-04 | -0.3 | 2.36E-02 | -0.6 | 1.45E-02 | -0.2 | 3.82E-01 | -0.4 | 2.38E-01 | Auxin-responsive protein |
| AT5G63930 | -0.8 | 1.73E-06 | -0.7 | 1.34E-05 | -0.1 | 1.31E-01 | 0.1 | 8.05E-01 | 0.1 | 7.80E-01 | 0.0 | 9.58E-01 | Protein kinase domain-containing protein |
| AT5G40720 | -0.8 | 4.21E-04 | -0.6 | 2.50E-03 | -0.2 | 3.84E-01 | -0.1 | 7.63E-01 | -0.3 | 2.69E-01 | 0.2 | 4.74E-01 | Glycosyltransferase family 92 protein |
| AT2G28960 | -0.8 | 2.04E-05 | -0.4 | 1.84E-03 | -0.4 | 1.11E-02 | 1.0 | 5.58E-05 | 0.7 | 5.25E-03 | 0.3 | 6.34E-02 | Leucine-rich repeat protein kinase family protein |
| AT4G16600 | -0.8 | 2.35E-04 | 0.0 | 8.76E-01 | -0.8 | 1.10E-03 | 0.7 | 3.21E-02 | 0.9 | 1.69E-02 | -0.2 | 6.53E-01 | Putative glucuronosyltransferase PGSIP7 |
| AT2G03140 | -0.8 | 2.10E-05 | -0.9 | 2.34E-05 | 0.1 | 5.85E-01 | -0.1 | 5.59E-01 | 0.2 | 4.04E-01 | -0.3 | 2.21E-01 | Alpha |
| AT4G01850 | -0.8 | 1.87E-05 | -0.2 | 7.25E-02 | -0.6 | 6.31E-04 | -0.2 | 3.88E-02 | 0.0 | 7.67E-01 | -0.2 | 2.57E-01 | S-adenosylmethionine synthase 2 |
| AT3G16340 | -0.8 | 1.61E-05 | -0.6 | 2.53E-04 | -0.2 | 7.19E-02 | 0.1 | 7.37E-01 | 0.1 | 6.86E-01 | 0.0 | 9.24E-01 | Pleiotropic drug resistance 1 |
| AT2G40890 | -0.8 | 2.46E-06 | -0.2 | 1.99E-02 | -0.6 | 1.38E-04 | -0.3 | 3.73E-02 | -0.1 | 6.82E-01 | -0.2 | 2.91E-01 | Cytochrome P450 98A3 |
| AT1G31885 | -0.8 | 1.28E-04 | -0.3 | 4.49E-02 | -0.5 | 8.56E-03 | -0.2 | 3.66E-01 | 0.2 | 5.16E-01 | -0.4 | 1.57E-01 | NOD26-like intrinsic protein 31 |
| AT4G25660 | -0.8 | 1.92E-06 | -0.5 | 9.29E-05 | -0.3 | 5.89E-03 | -0.3 | 5.79E-02 | 0.0 | 8.29E-01 | -0.2 | 1.60E-01 | At4g25660 |
| AT4G39090 | -0.8 | 1.26E-05 | -0.4 | 4.47E-03 | -0.4 | 2.57E-03 | -0.5 | 2.11E-04 | -0.3 | 8.07E-03 | -0.1 | 3.80E-01 | Uncharacterized protein |
| AT1G12200 | -0.8 | 3.58E-06 | -1.0 | 1.43E-06 | 0.3 | 2.12E-02 | 1.4 | 1.52E-08 | 0.5 | 2.87E-03 | 1.0 | 8.05E-05 | Flavin-containing monooxygenase |

| | | | | | | | | | | | | | |
|---|---|---|---|---|---|---|---|---|---|---|---|---|---|
| AT3G12040 | -0.8 | 7.25E-05 | -0.3 | 2.25E-02 | -0.5 | 7.51E-03 | -0.1 | 7.03E-01 | -0.1 | 6.73E-01 | 0.0 | 9.49E-01 | DNA-3-methyladenine glycosylase II |
| AT1G29160 | -0.8 | 6.89E-05 | -0.3 | 2.17E-02 | -0.5 | 7.42E-03 | -0.1 | 5.80E-01 | -0.1 | 8.21E-01 | -0.1 | 8.62E-01 | Dof zinc finger protein DOF1.5 |
| AT2G30020 | -0.8 | 7.62E-04 | -0.7 | 3.04E-03 | -0.1 | 5.43E-01 | 0.0 | 8.75E-01 | 0.0 | 9.55E-01 | 0.0 | 8.55E-01 | Probable protein phosphatase 2C 25 |
| AT1G67750 | -0.8 | 1.42E-04 | -0.3 | 4.12E-02 | -0.5 | 8.90E-03 | -0.3 | 4.32E-02 | 0.1 | 5.41E-01 | -0.4 | 4.27E-02 | Probable pectate lyase 5 |
| AT5G43600 | -0.8 | 3.61E-06 | -0.3 | 3.28E-03 | -0.5 | 5.30E-04 | -0.4 | 9.79E-03 | 0.0 | 9.50E-01 | -0.4 | 4.72E-02 | Ureidoglycolate hydrolase |
| AT2G39220 | -0.8 | 4.21E-05 | -0.7 | 1.37E-04 | -0.1 | 6.46E-01 | -0.3 | 8.99E-02 | 0.2 | 3.09E-01 | -0.6 | 2.28E-02 | Patatin-like protein 6 |
| AT2G28200 | -0.8 | 9.83E-04 | -0.6 | 7.69E-03 | -0.2 | 3.37E-01 | -0.5 | 4.21E-02 | -0.5 | 7.00E-02 | 0.0 | 9.73E-01 | Zinc finger protein ZAT5 |
| AT3G18490 | -0.8 | 1.31E-04 | -0.6 | 8.60E-04 | -0.1 | 3.48E-01 | -0.2 | 6.63E-02 | 0.1 | 5.60E-01 | -0.3 | 6.60E-02 | Protein ASPARTIC PROTEASE IN GUARD CELL 1 |
| AT2G44430 | -0.8 | 3.70E-06 | -0.6 | 4.30E-05 | -0.2 | 7.41E-02 | -0.2 | 2.81E-01 | -0.1 | 6.45E-01 | -0.1 | 7.61E-01 | At2g44430 |
| AT3G17000 | -0.8 | 2.07E-06 | -0.8 | 6.09E-06 | 0.0 | 7.34E-01 | -0.5 | 2.98E-04 | -0.3 | 1.66E-02 | -0.2 | 2.23E-01 | Ubiquitin-conjugating enzyme E2 32 |
| AT2G34410 | -0.8 | 1.82E-05 | -0.3 | 7.33E-03 | -0.5 | 3.10E-03 | 0.0 | 8.76E-01 | 0.1 | 6.50E-01 | -0.2 | 5.98E-01 | Protein REDUCED WALL ACETYLATION 3 |
| AT1G01650 | -0.8 | 5.12E-06 | -0.5 | 2.87E-04 | -0.3 | 8.05E-03 | -0.3 | 4.06E-02 | 0.0 | 9.97E-01 | -0.3 | 1.48E-01 | SIGNAL PEPTIDE PEPTIDASE-LIKE 4 |
| AT1G19390 | -0.8 | 4.95E-04 | -0.8 | 8.64E-04 | 0.0 | 9.65E-01 | | | | | | | Putative wall-associated receptor kinase-like 11 |
| AT5G55600 | -0.8 | 3.41E-06 | -0.7 | 1.22E-05 | -0.1 | 5.54E-01 | -0.3 | 5.92E-02 | -0.1 | 4.79E-01 | -0.2 | 4.93E-01 | BAH domain-containing protein |
| AT5G47450 | -0.8 | 9.58E-04 | 0.4 | 2.76E-02 | -1.2 | 2.36E-04 | -3.9 | 2.15E-09 | -1.1 | 4.33E-05 | -2.8 | 4.02E-06 | Aquaporin TIP2-3 |
| AT1G78360 | -0.8 | 3.00E-03 | -0.7 | 7.84E-03 | -0.1 | 7.42E-01 | | | | | | | Glutathione S-transferase |
| AT5G52830 | -0.8 | 1.13E-05 | -0.5 | 3.76E-04 | -0.3 | 2.84E-02 | -0.1 | 6.97E-01 | -0.2 | 4.52E-01 | 0.1 | 7.75E-01 | Probable WRKY transcription factor 27 |
| AT4G12080 | -0.8 | 3.79E-06 | -0.4 | 8.84E-04 | -0.4 | 1.74E-03 | -0.2 | 4.13E-01 | -0.2 | 2.69E-01 | 0.1 | 8.13E-01 | AT-hook motif nuclear-localized protein 1 |
| AT3G18660 | -0.8 | 9.11E-05 | 0.0 | 9.88E-01 | -0.8 | 6.42E-04 | -0.4 | 3.57E-01 | -0.4 | 3.80E-01 | 0.0 | 9.78E-01 | Hexosyltransferase |
| AT5G66210 | -0.8 | 2.73E-04 | -0.8 | 2.62E-04 | 0.1 | 7.36E-01 | -0.1 | 3.14E-01 | -0.2 | 1.38E-01 | 0.1 | 7.15E-01 | AT5G66210 protein |
| AT1G05570 | -0.8 | 2.58E-05 | -0.5 | 9.25E-04 | -0.3 | 3.17E-02 | -0.3 | 1.38E-01 | -0.2 | 4.57E-01 | -0.1 | 7.24E-01 | 1,3-beta-glucan synthase |
| AT3G01100 | -0.8 | 4.49E-06 | -0.6 | 3.87E-05 | -0.2 | 1.30E-01 | -0.1 | 5.97E-01 | 0.2 | 2.94E-01 | -0.3 | 1.63E-01 | CSC1-like protein HYP1 |
| AT1G56010 | -0.8 | 2.12E-06 | -0.6 | 1.94E-05 | -0.2 | 9.19E-02 | -0.7 | 3.54E-03 | -0.3 | 1.42E-01 | -0.3 | 1.77E-01 | NAC domain-containing protein 21 |
| AT4G00080 | -0.8 | 7.17E-03 | 0.2 | 4.63E-01 | -0.9 | 6.91E-03 | 0.0 | 9.90E-01 | 0.1 | 9.02E-01 | -0.1 | 9.07E-01 | hypothetical protein |
| AT5G60280 | -0.8 | 3.13E-05 | -0.7 | 7.75E-05 | 0.0 | 8.24E-01 | -0.2 | 3.44E-01 | -0.2 | 4.02E-01 | 0.0 | 9.99E-01 | L-type lectin-domain containing receptor kinase I.8 |
| AT5G56660 | -0.8 | 8.86E-06 | -0.5 | 1.78E-04 | -0.2 | 4.59E-02 | -0.2 | 3.20E-01 | 0.0 | 9.13E-01 | -0.2 | 5.72E-01 | IAA-amino acid hydrolase ILR1-like 2 |
| AT4G12270 | -0.8 | 3.30E-04 | -0.5 | 3.59E-03 | -0.2 | 2.51E-01 | | | | | | | Amine oxidase |
| AT4G36360 | -0.8 | 3.24E-05 | -0.5 | 1.89E-03 | -0.3 | 2.14E-02 | -0.2 | 1.41E-01 | 0.3 | 2.00E-02 | -0.5 | 7.26E-03 | Beta-galactosidase like protein |
| AT3G15650 | -0.8 | 1.66E-04 | -0.6 | 1.29E-03 | -0.2 | 2.97E-01 | 0.5 | 5.34E-02 | 0.6 | 6.87E-02 | 0.0 | 9.74E-01 | Alpha |
| AT2G01460 | -0.8 | 2.35E-05 | -0.9 | 1.86E-05 | 0.1 | 3.60E-01 | -0.2 | 5.98E-01 | 0.1 | 8.17E-01 | -0.3 | 4.32E-01 | P-loop containing nucleoside triphosphate hydrolases superfamily protein |
| AT2G35940 | -0.8 | 3.46E-06 | -0.8 | 4.42E-06 | 0.1 | 4.73E-01 | -0.1 | 5.56E-01 | 0.4 | 4.32E-02 | -0.5 | 2.96E-02 | Homeobox domain-containing protein |
| AT3G48700 | -0.8 | 1.15E-02 | -0.9 | 6.12E-03 | 0.2 | 6.49E-01 | | | | | | | Probable carboxylesterase 13 |
| AT1G08510 | -0.8 | 4.92E-06 | -0.6 | 8.08E-05 | -0.2 | 5.20E-02 | -0.5 | 1.43E-03 | -0.3 | 1.12E-01 | -0.3 | 2.27E-01 | Palmitoyl-acyl carrier protein thioesterase, chloroplastic |
| AT5G65683 | -0.8 | 2.04E-05 | -0.3 | 1.13E-02 | -0.5 | 2.35E-03 | -0.3 | 5.03E-02 | 0.2 | 2.42E-01 | -0.4 | 2.12E-02 | Probable E3 ubiquitin-protein ligase WAVH2 |
| AT2G32610 | -0.8 | 1.31E-04 | -0.1 | 6.24E-01 | -0.7 | 1.36E-03 | -0.6 | 9.94E-02 | -0.3 | 4.77E-01 | -0.3 | 5.51E-01 | Cellulose synthase-like protein B1 |
| AT4G39270 | -0.8 | 2.06E-06 | -0.5 | 7.97E-05 | -0.3 | 7.74E-03 | 0.1 | 5.05E-01 | 0.2 | 4.38E-01 | 0.0 | 9.01E-01 | Leucine-rich repeat protein kinase family protein |
| AT5G25620 | -0.8 | 6.79E-05 | -0.4 | 4.50E-03 | -0.4 | 2.81E-02 | 0.3 | 9.65E-02 | 0.2 | 3.06E-01 | 0.1 | 7.19E-01 | Flavin-containing monooxygenase |
| AT5G10520 | -0.8 | 3.11E-05 | -0.5 | 1.07E-03 | -0.3 | 4.12E-02 | 0.3 | 5.98E-02 | 0.0 | 9.87E-01 | 0.3 | 1.34E-01 | ROP binding protein kinases 1 |
| AT4G23000 | -0.8 | 2.52E-04 | -0.6 | 1.36E-03 | -0.1 | 4.56E-01 | -0.5 | 2.05E-01 | 0.1 | 9.00E-01 | -0.6 | 2.77E-01 | Uncharacterized protein AT4g23000 |
| AT1G25510 | -0.8 | 2.22E-04 | -0.3 | 3.90E-02 | -0.5 | 1.80E-02 | -0.2 | 3.75E-01 | 0.1 | 7.43E-01 | -0.3 | 2.40E-01 | Eukaryotic aspartyl protease family protein |
| AT5G08170 | -0.8 | 7.46E-07 | -0.5 | 4.03E-05 | -0.3 | 2.23E-03 | -0.2 | 1.09E-01 | 0.2 | 3.12E-01 | -0.4 | 5.33E-02 | Porphyromonas-type peptidyl-arginine deiminase family protein |
| AT2G43430 | -0.8 | 1.16E-06 | -0.6 | 1.31E-05 | -0.2 | 4.99E-02 | -0.5 | 1.29E-03 | -0.3 | 2.17E-02 | -0.1 | 4.93E-01 | Glyoxalase 2-1 |
| AT4G21534 | -0.8 | 1.34E-05 | -0.8 | 2.76E-05 | 0.0 | 9.80E-01 | -0.5 | 2.38E-02 | -0.5 | 2.60E-02 | 0.0 | 9.36E-01 | Sphingosine kinase 2 |
| AT5G56630 | -0.8 | 2.53E-06 | -0.4 | 3.33E-04 | -0.3 | 2.14E-03 | 0.6 | 1.51E-03 | 0.4 | 3.06E-02 | 0.2 | 4.68E-01 | ATP-dependent 6-phosphofructokinase 7 |
| AT5G52560 | -0.8 | 2.24E-06 | -0.4 | 5.02E-04 | -0.4 | 1.15E-03 | -0.1 | 2.51E-01 | 0.1 | 4.85E-01 | -0.3 | 1.43E-01 | UDP-sugar pyrophosphorylase |
| AT2G26310 | -0.8 | 8.48E-05 | -0.6 | 9.64E-04 | -0.2 | 2.00E-01 | -0.1 | 7.87E-01 | 0.0 | 9.07E-01 | -0.1 | 7.50E-01 | Chalcone-flavanone isomerase family protein |
| AT5G10370 | -0.8 | 5.31E-06 | -0.8 | 6.65E-06 | 0.1 | 4.82E-01 | -0.3 | 9.91E-02 | -0.2 | 3.68E-01 | -0.1 | 7.07E-01 | ATP-dependent RNA helicase DEAH12, chloroplastic |
| AT5G03630 | -0.8 | 3.81E-06 | -0.2 | 6.46E-02 | -0.6 | 1.30E-04 | 0.3 | 2.55E-02 | 0.3 | 1.11E-01 | 0.1 | 7.71E-01 | Monodehydroascorbate reductase 2 |
| AT4G14790 | -0.8 | 1.39E-05 | -0.8 | 2.46E-05 | 0.0 | 9.20E-01 | 0.4 | 2.09E-01 | 0.5 | 1.45E-01 | -0.1 | 8.17E-01 | DExH-box ATP-dependent RNA helicase DExH16, mitochondrial |
| AT1G07040 | -0.8 | 1.34E-06 | -0.6 | 2.30E-05 | -0.2 | 2.41E-02 | -0.3 | 5.14E-02 | -0.4 | 1.96E-02 | 0.1 | 6.85E-01 | 2-oxoadipate dioxygenase |
| AT5G16900 | -0.8 | 2.18E-05 | 0.1 | 2.24E-01 | -0.9 | 7.65E-05 | -1.2 | 6.08E-03 | 0.1 | 7.91E-01 | -1.3 | 7.35E-03 | Leucine-rich repeat protein kinase family protein |
| AT5G35460 | -0.8 | 1.62E-06 | -0.7 | 5.59E-06 | 0.0 | 6.47E-01 | -0.4 | 6.12E-03 | 0.1 | 7.25E-01 | -0.5 | 9.30E-03 | Glycerophosphocholine acyltransferase 1 |
| AT4G31590 | -0.8 | 7.19E-06 | -0.5 | 1.33E-04 | -0.2 | 4.37E-02 | -0.1 | 5.00E-01 | 0.2 | 1.50E-01 | -0.4 | 7.40E-02 | Probable xyloglucan glycosyltransferase 5 |
| AT1G47960 | -0.8 | 3.16E-04 | -0.5 | 5.21E-03 | -0.3 | 1.52E-01 | 0.0 | 9.24E-01 | 0.1 | 8.35E-01 | -0.1 | 8.07E-01 | Pectinesterase inhibitor domain-containing protein |
| AT3G45310 | -0.8 | 1.11E-06 | -0.1 | 3.48E-01 | -0.7 | 2.72E-05 | -0.2 | 1.52E-01 | 0.0 | 8.77E-01 | -0.2 | 3.87E-01 | Cysteine proteinases superfamily protein |
| AT5G45550 | -0.8 | 6.41E-06 | -0.4 | 9.93E-04 | -0.4 | 3.35E-03 | -0.3 | 1.49E-01 | -0.3 | 2.13E-01 | 0.0 | 9.94E-01 | MOB kinase activator-like 1A |
| AT5G07220 | -0.8 | 1.53E-06 | -0.4 | 3.68E-04 | -0.4 | 8.53E-04 | -0.3 | 1.12E-02 | -0.2 | 1.18E-01 | -0.1 | 5.74E-01 | BAG family molecular chaperone regulator 3 |
| AT2G47800 | -0.8 | 3.12E-04 | -1.0 | 9.47E-05 | 0.2 | 1.95E-01 | 0.0 | 9.41E-01 | 0.3 | 2.00E-01 | -0.3 | 2.74E-01 | ABC transporter C family member 4 |
| AT1G80760 | -0.8 | 1.77E-05 | -0.7 | 7.43E-05 | -0.1 | 4.74E-01 | -1.0 | 1.15E-03 | -0.1 | 8.14E-01 | -0.9 | 1.38E-02 | NLM7 |
| AT2G40530 | -0.8 | 8.02E-04 | -0.1 | 7.25E-01 | -0.7 | 5.74E-03 | -0.6 | 2.26E-02 | -0.1 | 8.63E-01 | -0.6 | 7.71E-02 | Precursor of CEP15 |
| AT5G66630 | -0.8 | 3.57E-05 | -0.9 | 3.18E-05 | 0.1 | 4.80E-01 | -0.8 | 4.12E-02 | -0.5 | 1.66E-01 | -0.2 | 7.22E-01 | Protein DA1-related 5 |
| AT5G35200 | -0.8 | 4.64E-07 | -0.5 | 1.86E-05 | -0.3 | 2.38E-03 | 0.0 | 9.94E-01 | -0.1 | 3.96E-01 | 0.1 | 5.17E-01 | Putative clathrin assembly protein At5g35200 |
| AT1G67650 | -0.8 | 2.51E-05 | -1.0 | 1.24E-05 | 0.2 | 1.26E-01 | 0.5 | 2.31E-01 | 0.6 | 1.47E-01 | -0.1 | 7.79E-01 | Signal recognition particle subunit SRP72 |
| AT1G48490 | -0.8 | 7.40E-06 | -0.8 | 1.13E-05 | 0.0 | 7.29E-01 | 0.0 | 9.91E-01 | 0.0 | 8.68E-01 | 0.0 | 8.90E-01 | Protein kinase superfamily protein |
| AT1G04250 | -0.8 | 7.17E-06 | -0.1 | 1.55E-01 | -0.6 | 1.56E-04 | -0.1 | 5.99E-01 | 0.1 | 7.50E-01 | -0.3 | 3.56E-01 | Auxin-responsive protein IAA17 |
| AT3G32047 | -0.8 | 2.30E-03 | -0.3 | 1.44E-01 | -0.5 | 7.11E-02 | | | | | | | Cytochrome P450-like protein |
| AT5G56150 | -0.8 | 1.97E-05 | -0.4 | 2.61E-03 | -0.4 | 7.12E-03 | 0.1 | 6.95E-01 | 0.2 | 4.11E-01 | -0.1 | 7.16E-01 | Ubiquitin-conjugating enzyme 30 |
| AT1G78530 | -0.8 | 2.57E-03 | -0.3 | 1.79E-01 | -0.5 | 6.42E-02 | 0.7 | 1.30E-01 | 0.8 | 8.16E-02 | -0.2 | 7.49E-01 | Protein kinase domain-containing protein |
| AT5G06800 | -0.8 | 5.05E-05 | -0.1 | 5.11E-01 | -0.7 | 6.11E-04 | -0.2 | 5.15E-01 | 0.1 | 8.38E-01 | -0.3 | 3.95E-01 | Myb-like HTH transcriptional regulator family protein |
| AT1G27980 | -0.8 | 5.95E-06 | -0.7 | 1.42E-05 | 0.0 | 8.83E-01 | -0.2 | 4.09E-01 | -0.4 | 1.21E-01 | 0.2 | 4.92E-01 | Sphingosine-1-phosphate lyase |
| AT4G15960 | -0.8 | 1.91E-05 | -0.4 | 2.49E-03 | -0.4 | 8.67E-03 | -0.4 | 2.20E-01 | 0.0 | 9.65E-01 | -0.4 | 4.10E-01 | Alpha |
| AT5G05170 | -0.8 | 8.34E-06 | -0.6 | 1.01E-04 | -0.2 | 1.04E-01 | -0.4 | 3.44E-03 | -0.2 | 3.36E-01 | -0.3 | 1.67E-01 | Cellulose synthase A catalytic subunit 3 |
| AT3G62020 | -0.8 | 7.62E-05 | 0.1 | 2.56E-01 | -0.9 | 1.84E-04 | 0.0 | 9.17E-01 | 0.1 | 5.57E-01 | -0.1 | 6.89E-01 | Germin-like protein |
| AT1G77760 | -0.8 | 2.04E-04 | -0.7 | 3.84E-04 | 0.0 | 9.06E-01 | -1.5 | 2.51E-06 | -2.0 | 9.39E-07 | 0.5 | 4.65E-02 | Nitrate reductase 1 |
| AT3G14510 | -0.8 | 5.26E-03 | 0.4 | 6.31E-02 | -1.1 | 1.70E-03 | -0.4 | 4.31E-01 | -0.3 | 6.56E-01 | -0.1 | 8.49E-01 | Putative geranylgeranyl pyrophosphate synthase 8, chloroplastic -farnesyl diphosphate synthase 8) |
| AT2G18050 | -0.8 | 2.11E-02 | -1.4 | 2.31E-03 | 0.6 | 1.66E-01 | -0.9 | 2.21E-02 | -0.5 | 1.96E-01 | -0.4 | 4.71E-01 | Histone H1-3 |
| AT5G38280 | -0.8 | 7.69E-05 | -0.6 | 7.11E-04 | -0.2 | 2.28E-01 | 0.2 | 4.80E-01 | 0.4 | 1.39E-01 | -0.2 | 4.85E-01 | PR5-like receptor kinase |
| AT5G05790 | -0.8 | 7.02E-06 | -0.2 | 1.31E-02 | -0.5 | 6.05E-04 | 0.3 | 9.70E-02 | 0.4 | 5.48E-02 | -0.1 | 7.30E-01 | At5g05790 |
| AT1G48260 | -0.8 | 1.51E-04 | -0.4 | 6.24E-03 | -0.3 | 5.89E-02 | 0.0 | 8.92E-01 | 0.6 | 2.12E-02 | -0.6 | 6.33E-02 | CBL-interacting protein kinase 17 |
| AT2G40620 | -0.8 | 2.51E-05 | 0.0 | 9.81E-01 | -0.8 | 1.64E-04 | -0.5 | 2.62E-03 | -0.2 | 2.36E-01 | -0.3 | 1.58E-01 | bZIP transcription factor 18 |
| AT5G46250 | -0.8 | 5.16E-06 | -0.4 | 3.98E-04 | -0.3 | 6.15E-03 | -0.2 | 1.61E-01 | -0.1 | 7.95E-01 | -0.2 | 4.97E-01 | La-related protein 6A |
| AT5G59650 | -0.8 | 3.39E-04 | 0.1 | 3.97E-01 | -0.9 | 5.92E-04 | -1.3 | 1.38E-04 | -0.3 | 2.70E-01 | -1.0 | 1.09E-02 | Leucine-rich repeat protein kinase family protein |
| AT5G22850 | -0.8 | 3.97E-05 | -0.4 | 2.53E-03 | -0.3 | 2.12E-02 | -0.1 | 5.06E-01 | 0.0 | 8.76E-01 | -0.1 | 7.77E-01 | hypothetical protein |

| | | | | | | | | | | | | | |
|---|---|---|---|---|---|---|---|---|---|---|---|---|---|
| AT3G51330 | -0.7 | 1.07E-04 | -0.2 | 1.67E-01 | -0.6 | 2.14E-03 | -0.1 | 6.00E-01 | -0.1 | 5.70E-01 | 0.0 | 9.44E-01 | Eukaryotic aspartyl protease family protein |
| AT4G29450 | -0.7 | 1.20E-02 | 0.8 | 3.97E-03 | -1.5 | 6.17E-04 | -2.0 | 2.60E-03 | -0.3 | 5.94E-01 | -1.7 | 2.49E-02 | Leucine-rich repeat protein kinase |
| AT3G17810 | -0.7 | 8.06E-06 | -0.3 | 3.20E-03 | -0.4 | 1.73E-03 | -0.2 | 1.01E-01 | 0.1 | 4.34E-01 | -0.4 | 6.56E-02 | Dihydropyrimidine dehydrogenase ), chloroplastic |
| AT5G57035 | -0.7 | 1.06E-05 | -0.5 | 2.00E-04 | -0.2 | 5.16E-02 | 0.1 | 6.01E-01 | 0.0 | 9.50E-01 | 0.1 | 7.77E-01 | U-box domain-containing protein kinase family protein |
| AT1G78570 | -0.7 | 1.73E-05 | -0.6 | 2.92E-04 | -0.2 | 7.63E-02 | 0.0 | 8.79E-01 | 0.5 | 2.29E-02 | -0.4 | 8.78E-02 | Trifunctional UDP-glucose 4,6-dehydratase |
| AT5G57830 | -0.7 | 9.89E-06 | -0.2 | 2.51E-02 | -0.5 | 7.27E-04 | -0.5 | 1.02E-02 | -0.3 | 1.29E-01 | -0.2 | 3.63E-01 | hypothetical protein |
| AT2G24300 | -0.7 | 2.65E-05 | -0.1 | 5.16E-01 | -0.7 | 3.39E-04 | -0.3 | 1.84E-01 | 0.0 | 9.32E-01 | -0.4 | 1.60E-01 | Calmodulin-binding protein |
| AT1G51170 | -0.7 | 1.44E-04 | -0.6 | 5.98E-04 | -0.1 | 5.21E-01 | -0.2 | 2.69E-01 | 0.0 | 9.58E-01 | -0.2 | 4.59E-01 | Serine |
| AT3G63380 | -0.7 | 1.42E-03 | -1.0 | 3.13E-04 | 0.3 | 2.19E-01 | 1.4 | 1.19E-05 | -0.1 | 8.24E-01 | 1.5 | 2.61E-04 | Calcium-transporting ATPase 12, plasma membrane-type -ATPase isoform 12) |
| AT1G52060 | -0.7 | 3.57E-03 | 0.4 | 6.56E-02 | -1.1 | 8.14E-04 | -0.1 | 7.38E-01 | 0.4 | 3.88E-01 | -0.5 | 2.14E-01 | Jacalin-type lectin domain-containing protein |
| AT5G59940 | -0.7 | 4.35E-04 | 0.1 | 3.34E-01 | -0.9 | 8.23E-04 | -0.1 | 8.47E-01 | 0.1 | 9.13E-01 | -0.2 | 7.72E-01 | Cysteine |
| AT5G10740 | -0.7 | 5.42E-06 | -0.6 | 4.75E-05 | -0.1 | 1.43E-01 | -0.1 | 5.81E-01 | -0.2 | 2.67E-01 | 0.1 | 6.68E-01 | Probable protein phosphatase 2C 69 |
| AT5G13180 | -0.7 | 2.70E-05 | -0.6 | 1.74E-04 | -0.1 | 3.03E-01 | -0.2 | 3.12E-01 | -0.3 | 8.69E-02 | 0.2 | 5.70E-01 | NAC domain-containing protein 83 |
| AT3G43670 | -0.7 | 1.53E-05 | -0.4 | 2.37E-03 | -0.4 | 5.23E-03 | -1.9 | 1.09E-05 | 0.1 | 8.45E-01 | -2.0 | 1.25E-04 | Amine oxidase gamma 2 |
| AT4G36920 | -0.7 | 1.33E-06 | -0.4 | 1.09E-04 | -0.3 | 2.12E-03 | -0.1 | 4.72E-01 | 0.0 | 9.27E-01 | -0.2 | 5.25E-01 | Floral homeotic protein APETALA 2 |
| AT3G46330 | -0.7 | 6.86E-05 | -0.9 | 4.33E-05 | 0.1 | 3.56E-01 | -0.1 | 7.78E-01 | 0.2 | 6.45E-01 | -0.3 | 5.10E-01 | Leucine-rich repeat protein kinase family protein |
| AT1G17450 | -0.7 | 2.44E-06 | -0.6 | 1.33E-05 | -0.1 | 2.88E-01 | 0.0 | 8.43E-01 | 0.2 | 3.65E-01 | -0.2 | 5.74E-01 | B-block binding subunit of TFIIIC |
| AT4G19690 | -0.7 | 1.92E-02 | -1.0 | 4.05E-03 | 0.3 | 3.32E-01 | -0.9 | 1.46E-03 | -2.4 | 1.06E-06 | 1.5 | 8.05E-05 | Fe transport protein 1 transport protein 1) |
| AT3G12560 | -0.7 | 1.20E-06 | -0.4 | 9.66E-05 | -0.3 | 2.13E-03 | -0.4 | 2.15E-02 | 0.1 | 5.67E-01 | -0.5 | 3.07E-02 | Telomere repeat-binding protein 3 |
| AT2G23760 | -0.7 | 3.23E-05 | -0.7 | 1.42E-04 | -0.1 | 4.57E-01 | -0.5 | 1.70E-02 | -0.2 | 3.02E-01 | -0.2 | 3.90E-01 | BEL1-like homeodomain 4 |
| AT2G45670 | -0.7 | 3.38E-06 | -0.4 | 1.10E-03 | -0.4 | 1.15E-03 | -0.4 | 1.31E-03 | -0.2 | 8.51E-02 | -0.2 | 2.56E-01 | Lysophospholipid acyltransferase LPEAT2 |
| AT4G13930 | -0.7 | 7.25E-06 | -0.3 | 1.10E-02 | -0.5 | 6.62E-04 | -0.1 | 5.54E-01 | 0.3 | 4.50E-02 | -0.4 | 3.93E-02 | Serine hydroxymethyltransferase 4 |
| AT3G46930 | -0.7 | 1.02E-04 | -0.6 | 8.15E-04 | -0.2 | 2.70E-01 | 0.4 | 2.26E-01 | 0.3 | 4.14E-01 | 0.1 | 8.78E-01 | Protein kinase superfamily protein |
| AT4G37470 | -0.7 | 1.28E-06 | -0.5 | 5.89E-05 | -0.3 | 4.66E-03 | -0.6 | 1.88E-04 | -0.1 | 4.48E-01 | -0.5 | 7.59E-03 | Probable esterase KAI2 |
| AT2G45660 | -0.7 | 4.45E-05 | -0.5 | 5.36E-04 | -0.2 | 1.38E-01 | -0.1 | 4.15E-01 | 0.0 | 8.55E-01 | -0.1 | 7.04E-01 | AGAMOUS-like 20 |
| AT1G22070 | -0.7 | 1.94E-05 | -0.6 | 2.80E-04 | -0.2 | 9.52E-02 | -0.1 | 4.27E-01 | 0.1 | 4.59E-01 | -0.2 | 1.66E-01 | Uncharacterized protein |
| AT1G11960 | -0.7 | 5.73E-06 | -0.6 | 6.68E-05 | -0.2 | 9.65E-02 | 0.0 | 9.41E-01 | 0.3 | 1.63E-01 | -0.3 | 2.57E-01 | ERD family protein |
| AT2G22630 | -0.7 | 1.71E-03 | -0.1 | 4.14E-01 | -0.6 | 1.99E-02 | 0.4 | 2.79E-01 | 0.3 | 4.79E-01 | 0.1 | 8.65E-01 | Agamous-like MADS-box protein AGL17 |
| AT1G55730 | -0.7 | 1.55E-06 | -0.5 | 2.38E-05 | -0.2 | 3.15E-02 | 0.0 | 9.23E-01 | 0.1 | 5.74E-01 | -0.1 | 5.81E-01 | Vacuolar cation |
| AT4G22950 | -0.7 | 2.80E-03 | -0.5 | 2.54E-02 | -0.3 | 3.15E-01 | 0.1 | 7.75E-01 | 0.2 | 6.05E-01 | -0.1 | 8.44E-01 | AGAMOUS-like 19 |
| AT1G22370 | -0.7 | 1.57E-03 | -0.5 | 2.04E-02 | -0.3 | 2.30E-01 | -0.3 | 4.70E-02 | -0.2 | 3.40E-01 | -0.1 | 5.64E-01 | UDP-glycosyltransferase 85A5 |
| AT5G59320 | -0.7 | 2.52E-02 | -1.1 | 4.72E-03 | 0.4 | 3.36E-01 | -0.7 | 6.35E-01 | 0.2 | 9.15E-01 | -0.9 | 6.66E-01 | Non-specific lipid-transfer protein 3 |
| AT4G34990 | -0.7 | 3.41E-03 | -0.9 | 2.06E-03 | 0.1 | 6.46E-01 | 0.2 | 4.84E-01 | 0.4 | 1.46E-01 | -0.2 | 5.43E-01 | Transcription factor MYB32 |
| AT1G74210 | -0.7 | 7.19E-06 | -0.4 | 8.98E-04 | -0.3 | 4.62E-03 | 0.3 | 9.51E-02 | 0.6 | 9.94E-03 | -0.2 | 3.49E-01 | Glycerophosphodiester phosphodiesterase GDPD5 |
| AT1G14190 | -0.7 | 2.51E-04 | -0.2 | 1.78E-01 | -0.6 | 7.10E-03 | 0.2 | 6.39E-01 | 0.2 | 7.07E-01 | 0.0 | 9.93E-01 | hypothetical protein |
| AT3G08040 | -0.7 | 1.01E-04 | -1.0 | 4.18E-05 | 0.2 | 1.84E-01 | -0.6 | 1.98E-01 | -0.1 | 9.23E-01 | | | MATE efflux family protein |
| AT2G21130 | -0.7 | 6.89E-04 | -0.3 | 6.68E-02 | -0.4 | 3.81E-02 | -0.2 | 2.75E-01 | -0.2 | 4.34E-01 | 0.0 | 9.26E-01 | Peptidyl-prolyl cis-trans isomerase CYP19-2 |
| AT1G09575 | -0.7 | 5.74E-04 | -0.2 | 1.21E-01 | -0.5 | 2.14E-02 | -0.5 | 7.94E-02 | -0.5 | 1.11E-01 | 0.0 | 9.86E-01 | Calcium uniporter protein 6, mitochondrial |
| AT1G02900 | -0.7 | 2.34E-05 | -0.1 | 1.43E-01 | -0.6 | 5.22E-04 | 0.7 | 1.24E-03 | 0.3 | 1.41E-01 | 0.4 | 4.87E-02 | Protein RALF-like 1 |
| AT5G18360 | -0.7 | 2.73E-04 | -0.6 | 2.10E-03 | -0.2 | 3.14E-01 | -0.4 | 3.16E-01 | -0.4 | 3.44E-01 | 0.0 | 9.75E-01 | TIR domain-containing protein |
| AT1G79470 | -0.7 | 3.90E-05 | -0.8 | 5.74E-05 | 0.0 | 8.56E-01 | 1.3 | 2.13E-06 | 0.8 | 5.07E-04 | 0.4 | 5.48E-02 | Inosine-5'-monophosphate dehydrogenase 1 |
| AT3G01720 | -0.7 | 2.37E-05 | -0.2 | 1.38E-01 | -0.6 | 5.21E-04 | -0.2 | 2.14E-01 | 0.2 | 3.99E-01 | -0.4 | 8.95E-02 | Peptidyl serine alpha-galactosyltransferase |
| AT4G04450 | -0.7 | 4.26E-03 | -0.1 | 7.78E-01 | -0.7 | 2.11E-02 | | | | | | | WRKY family transcription factor |
| AT1G43910 | -0.7 | 3.78E-04 | -1.4 | 1.23E-05 | 0.7 | 4.39E-03 | 0.9 | 1.47E-03 | 0.1 | 6.73E-01 | 0.7 | 1.24E-02 | AAA-ATPase At1g43910 |
| AT5G07460 | -0.7 | 3.00E-05 | 0.1 | 2.56E-01 | -0.9 | 7.25E-05 | -0.2 | 1.62E-01 | 0.4 | 3.57E-03 | -0.6 | 1.56E-03 | Peptide methionine sulfoxide reductase A2 -S-oxide reductase) reductase) |
| AT1G24764 | -0.7 | 1.59E-06 | -0.3 | 8.06E-04 | -0.4 | 4.95E-04 | -0.1 | 5.92E-01 | 0.1 | 8.02E-01 | -0.2 | 5.05E-01 | Microtubule-associated proteins 70-2 |
| AT3G03780 | -0.7 | 8.87E-05 | -0.3 | 5.44E-02 | -0.5 | 4.23E-03 | 0.1 | 5.05E-01 | 0.8 | 1.05E-04 | -0.7 | 1.63E-03 | 5-methyltetrahydropteroyltriglutamate--homocysteine methyltransferase 2 |
| AT5G47370 | -0.7 | 3.94E-05 | -0.4 | 4.85E-03 | -0.4 | 1.17E-02 | -0.1 | 3.72E-01 | 0.3 | 9.91E-02 | -0.4 | 4.74E-02 | Homeobox-leucine zipper protein HAT2 |
| AT1G62990 | -0.7 | 3.92E-06 | -0.5 | 1.26E-04 | -0.3 | 1.80E-02 | -0.3 | 2.99E-01 | 0.0 | 9.65E-01 | -0.3 | 4.39E-01 | Uncharacterized protein |
| AT4G09180 | -0.7 | 1.90E-05 | -0.7 | 3.26E-05 | 0.0 | 9.16E-01 | -0.2 | 2.66E-01 | 0.0 | 9.41E-01 | -0.3 | 3.51E-01 | Transcription factor bHLH81 |
| AT5G51770 | -0.7 | 2.14E-05 | -0.6 | 1.32E-04 | -0.1 | 2.97E-01 | -0.4 | 2.02E-03 | -0.1 | 5.45E-01 | -0.4 | 4.52E-02 | Protein kinase superfamily protein |
| AT2G01600 | -0.7 | 6.25E-06 | -0.5 | 1.27E-04 | -0.2 | 3.83E-02 | -0.1 | 7.99E-01 | -0.1 | 7.36E-01 | 0.0 | 9.26E-01 | Putative clathrin assembly protein At2g01600 |
| AT2G38290 | -0.7 | 5.24E-05 | -0.5 | 1.03E-03 | -0.2 | 7.85E-02 | 0.4 | 3.65E-02 | -0.2 | 4.24E-01 | 0.5 | 2.46E-02 | Ammonium transporter 2 |
| AT1G16160 | -0.7 | 1.20E-03 | -0.1 | 7.26E-01 | -0.7 | 7.96E-03 | 0.5 | 2.90E-01 | 0.9 | 7.43E-02 | -0.4 | 5.20E-01 | Wall-associated receptor kinase-like 5 |
| AT5G16720 | -0.7 | 4.64E-06 | -0.1 | 3.53E-01 | -0.7 | 9.50E-05 | 0.1 | 6.34E-01 | -0.1 | 7.70E-01 | 0.1 | 5.05E-01 | Myosin-binding protein 3 |
| AT2G20080 | -0.7 | 1.09E-03 | -0.2 | 1.64E-01 | -0.5 | 3.15E-02 | | | | | | | TIE2 |
| AT5G26230 | -0.7 | 4.34E-05 | -0.1 | 3.77E-01 | -0.6 | 6.83E-04 | -0.2 | 4.64E-01 | 0.3 | 2.12E-01 | -0.5 | 1.06E-01 | Probable membrane-associated kinase regulator 1 |
| AT5G10230 | -0.7 | 6.21E-03 | -1.0 | 1.46E-03 | 0.3 | 3.47E-01 | 1.3 | 8.36E-03 | 1.1 | 3.41E-02 | 0.2 | 7.13E-01 | Annexin D7 |
| AT1G77290 | -0.7 | 2.73E-04 | -0.4 | 1.13E-02 | -0.3 | 7.10E-02 | -0.3 | 2.80E-01 | -0.4 | 1.59E-01 | 0.1 | 7.82E-01 | Glutathione S-transferase TCHQD |
| AT2G29490 | -0.7 | 6.50E-05 | -0.9 | 2.82E-05 | 0.2 | 1.86E-01 | 1.7 | 4.55E-04 | 1.0 | 4.44E-02 | 0.7 | 1.62E-02 | Glutathione S-transferase U1 |
| AT2G16385 | -0.7 | 5.08E-03 | -0.8 | 5.57E-03 | 0.0 | 8.87E-01 | 0.2 | 8.04E-01 | 0.5 | 4.83E-01 | -0.3 | 7.34E-01 | Protein CASPARIAN STRIP INTEGRITY FACTOR 1 |
| AT5G57910 | -0.7 | 2.14E-04 | -0.7 | 4.94E-04 | 0.0 | 7.89E-01 | -0.1 | 6.84E-01 | -0.2 | 3.83E-01 | 0.1 | 7.34E-01 | At5g57910 |
| AT5G54960 | -0.7 | 6.72E-05 | -0.5 | 8.31E-04 | -0.2 | 1.50E-01 | 0.5 | 2.77E-03 | 0.4 | 2.25E-02 | 0.1 | 7.12E-01 | Pyruvate decarboxylase 2 |
| AT4G22070 | -0.7 | 1.76E-03 | -0.4 | 2.94E-02 | -0.3 | 1.99E-01 | 2.2 | 3.34E-04 | 0.8 | 2.06E-01 | 1.3 | 1.74E-02 | Probable WRKY transcription factor 31 |
| AT5G01050 | -0.7 | 1.01E-04 | 0.1 | 5.16E-01 | -0.8 | 3.33E-04 | -0.1 | 9.03E-01 | 0.4 | 2.64E-01 | -0.5 | 1.76E-01 | Laccase-9 |
| AT1G60810 | -0.7 | 3.17E-06 | 0.0 | 7.72E-01 | -0.7 | 4.04E-05 | -0.1 | 3.75E-01 | 0.0 | 9.11E-01 | -0.2 | 4.28E-01 | ATP-citrate synthase alpha chain protein 2 |
| AT2G21550 | -0.7 | 4.59E-06 | -0.4 | 5.56E-04 | -0.3 | 4.25E-03 | 0.3 | 2.18E-01 | 0.5 | 1.11E-01 | -0.1 | 7.01E-01 | Bifunctional dihydrofolate reductase-thymidylate synthase |
| AT3G49880 | -0.7 | 9.85E-06 | -0.3 | 3.52E-03 | -0.4 | 2.37E-03 | -0.3 | 7.16E-02 | 0.1 | 4.71E-01 | -0.4 | 3.63E-02 | Glycosyl hydrolase family protein 43 |
| AT5G63940 | -0.7 | 1.42E-06 | -0.5 | 4.70E-05 | -0.2 | 7.61E-03 | -0.4 | 7.54E-03 | -0.1 | 8.01E-01 | -0.3 | 7.72E-02 | At5g63940 |
| AT5G10030 | -0.7 | 5.40E-06 | -0.2 | 7.95E-03 | -0.5 | 6.25E-04 | -0.2 | 1.60E-01 | -0.3 | 2.22E-02 | 0.2 | 4.20E-01 | Transcription factor TGA4 |
| AT1G54450 | -0.7 | 5.37E-05 | 0.1 | 4.71E-01 | -0.8 | 2.03E-04 | -0.2 | 6.19E-01 | 0.2 | 5.70E-01 | -0.3 | 2.75E-01 | hypothetical protein |
| AT1G23980 | -0.7 | 5.20E-04 | -0.2 | 1.16E-01 | -0.5 | 2.09E-02 | -0.1 | 7.92E-01 | 0.4 | 2.74E-01 | -0.5 | 2.23E-01 | RING-H2 finger protein ATL47 |
| AT4G31650 | -0.7 | 1.48E-04 | -0.6 | 1.04E-03 | -0.2 | 3.29E-01 | -0.3 | 2.18E-01 | 0.1 | 7.48E-01 | -0.3 | 2.05E-01 | Transcriptional factor B3 family protein |
| AT1G51070 | -0.7 | 1.11E-05 | -0.3 | 2.91E-03 | -0.4 | 3.01E-03 | 0.0 | 9.27E-01 | 0.2 | 1.01E-01 | -0.2 | 1.64E-01 | Basic helix-loop-helix DNA-binding superfamily protein |
| AT5G58540 | -0.7 | 8.42E-06 | -0.4 | 5.63E-04 | -0.3 | 9.70E-03 | -0.4 | 6.17E-02 | -0.1 | 7.48E-01 | -0.3 | 2.59E-01 | Protein kinase superfamily protein |
| AT2G33990 | -0.7 | 3.38E-06 | 0.0 | 8.27E-01 | -0.7 | 4.46E-05 | -0.7 | 3.60E-05 | -0.5 | 1.79E-03 | -0.2 | 2.78E-01 | Protein IQ-DOMAIN 9 |
| AT1G67310 | -0.7 | 2.51E-05 | -0.7 | 7.17E-05 | 0.0 | 7.19E-01 | -0.3 | 7.87E-02 | -0.1 | 7.57E-01 | -0.2 | 3.68E-01 | Calmodulin-binding transcription activator 4 |
| AT3G14350 | -0.7 | 3.63E-05 | -0.5 | 6.43E-04 | -0.2 | 8.38E-02 | -0.8 | 2.62E-02 | -0.2 | 5.83E-01 | -0.6 | 2.56E-01 | STRUBBELIG-receptor family 7 |
| AT3G15500 | -0.7 | 6.88E-03 | -0.9 | 3.60E-03 | 0.2 | 6.20E-01 | -0.7 | 2.22E-01 | -1.7 | 2.11E-02 | 1.0 | 2.44E-01 | NAC domain-containing protein 55 |
| AT1G01580 | -0.7 | 1.03E-02 | -1.0 | 2.50E-03 | 0.2 | 3.46E-01 | -1.5 | 2.39E-05 | -2.8 | 5.32E-07 | 1.3 | 6.16E-04 | Ferric reduction oxidase 2 |
| AT4G39850 | -0.7 | 3.03E-06 | -0.7 | 5.84E-06 | 0.0 | 8.80E-01 | -0.1 | 6.13E-01 | 0.1 | 7.85E-01 | -0.2 | 5.44E-01 | Peroxisomal ABC transporter 1 |
| AT1G79530 | -0.7 | 2.25E-05 | -0.3 | 2.07E-02 | -0.5 | 1.68E-03 | 0.3 | 1.22E-01 | 0.4 | 8.86E-02 | -0.1 | 8.39E-01 | Glyceraldehyde-3-phosphate dehydrogenase GAPCP1, chloroplastic |
| AT1G12000 | -0.7 | 1.09E-04 | -0.1 | 4.61E-01 | -0.6 | 1.00E-03 | 0.1 | 5.41E-01 | 0.3 | 5.39E-02 | -0.2 | 3.08E-01 | Pyrophosphate--fructose 6-phosphate 1-phosphotransferase subunit beta 1 |

| | | | | | | | | | | | | | |
|---|---|---|---|---|---|---|---|---|---|---|---|---|---|
| AT5G38540 | -0.7 | 4.18E-04 | -0.6 | 1.38E-03 | -0.1 | 6.77E-01 | 0.6 | 8.24E-02 | 0.9 | 1.29E-02 | -0.3 | 3.40E-01 | Mannose-binding lectin superfamily protein |
| AT5G40390 | -0.7 | 3.72E-05 | -0.4 | 5.76E-03 | -0.4 | 8.16E-03 | 0.1 | 7.08E-01 | 0.4 | 3.16E-02 | -0.4 | 9.04E-02 | Probable galactinol--sucrose galactosyltransferase 5 |
| AT3G07320 | -0.7 | 2.84E-05 | 0.0 | 7.09E-01 | -0.7 | 2.18E-04 | 0.3 | 5.60E-02 | 0.5 | 5.96E-03 | -0.2 | 3.54E-01 | glucan endo-1,3-beta-D-glucosidase |
| AT1G73080 | -0.7 | 4.78E-06 | -0.4 | 2.45E-04 | -0.3 | 9.23E-03 | -0.4 | 1.93E-02 | -0.1 | 8.35E-01 | -0.4 | 1.20E-01 | Leucine-rich repeat receptor-like protein kinase PEPR1 |
| AT3G05020 | -0.7 | 9.11E-04 | 0.0 | 8.41E-01 | -0.7 | 2.02E-03 | 0.1 | 5.79E-01 | 0.3 | 2.25E-01 | -0.2 | 5.40E-01 | Acyl carrier protein 1, chloroplastic |
| AT5G18370 | -0.7 | 1.27E-04 | -0.6 | 4.90E-04 | -0.1 | 5.74E-01 | 0.1 | 7.84E-01 | -0.4 | 2.89E-01 | 0.5 | 2.40E-01 | Disease resistance-like protein DSC2 |
| AT1G08650 | -0.7 | 1.83E-03 | -0.1 | 5.11E-01 | -0.6 | 1.31E-02 | -0.8 | 1.13E-03 | -0.4 | 1.06E-01 | -0.4 | 1.04E-01 | Phosphoenolpyruvate carboxylase kinase 1 |
| AT3G21700 | -0.7 | 4.06E-04 | -0.5 | 5.61E-03 | -0.2 | 1.90E-01 | -0.4 | 8.12E-02 | 0.0 | 8.89E-01 | -0.3 | 1.27E-01 | Ras-related small GTP-binding family protein |
| AT4G19520 | -0.7 | 1.78E-04 | -0.4 | 4.61E-03 | -0.3 | 8.35E-02 | 0.1 | 3.84E-01 | 0.1 | 5.24E-01 | 0.0 | 9.63E-01 | Disease resistance protein family |
| AT1G70140 | -0.7 | 1.02E-03 | -0.5 | 8.83E-03 | -0.2 | 3.14E-01 | 0.2 | 4.65E-01 | 0.3 | 2.86E-01 | -0.1 | 7.57E-01 | Formin-like protein 8 |
| AT3G23150 | -0.7 | 5.19E-05 | -0.5 | 6.28E-04 | -0.2 | 1.55E-01 | 0.8 | 1.27E-02 | 0.5 | 1.62E-01 | 0.3 | 4.27E-01 | Signal transduction histidine kinase, hybrid-type, ethylene sensor |
| AT4G20780 | -0.7 | 5.00E-05 | -0.2 | 8.65E-02 | -0.5 | 1.96E-03 | 0.5 | 4.13E-02 | 0.2 | 4.63E-01 | 0.3 | 3.80E-01 | Calcium-binding protein CML42 |
| AT5G37450 | -0.7 | 1.59E-04 | -0.7 | 3.06E-04 | 0.0 | 9.04E-01 | -0.2 | 4.04E-01 | 0.2 | 3.74E-01 | -0.5 | 1.50E-01 | Leucine-rich repeat protein kinase family protein |
| AT3G23840 | -0.7 | 3.24E-03 | -0.2 | 2.84E-01 | -0.5 | 4.52E-02 | -0.4 | 4.77E-02 | 0.2 | 5.02E-01 | -0.6 | 1.96E-02 | Protein ECERIFERUM 26-like |
| AT3G44720 | -0.7 | 2.40E-05 | -0.7 | 7.05E-05 | 0.0 | 6.88E-01 | 0.0 | 8.85E-01 | 0.0 | 8.21E-01 | 0.0 | 9.40E-01 | Arogenate dehydratase 4, chloroplastic |
| AT1G06550 | -0.7 | 2.25E-05 | -0.3 | 4.98E-03 | -0.4 | 4.81E-03 | 0.4 | 3.32E-03 | 0.5 | 1.81E-03 | -0.1 | 6.76E-01 | 3-hydroxyisobutyryl-CoA hydrolase-like protein 5 |
| AT4G36760 | -0.7 | 2.72E-06 | -0.5 | 5.89E-05 | -0.2 | 2.20E-02 | -0.4 | 1.15E-02 | -0.3 | 1.24E-01 | -0.1 | 5.61E-01 | Aminopeptidase P1 |
| AT1G13900 | -0.7 | 8.05E-06 | -0.3 | 2.60E-03 | -0.4 | 2.04E-03 | -0.1 | 4.38E-01 | 0.0 | 8.73E-01 | -0.2 | 4.63E-01 | Probable inactive purple acid phosphatase 2 |
| AT1G72210 | -0.7 | 2.58E-03 | -0.2 | 1.74E-01 | -0.5 | 6.60E-02 | -0.4 | 1.84E-01 | 0.1 | 8.05E-01 | -0.5 | 1.15E-01 | Basic helix-loop-helix DNA-binding superfamily protein |
| AT5G63710 | -0.7 | 3.37E-05 | -0.6 | 2.55E-04 | -0.1 | 2.42E-01 | 0.3 | 3.65E-02 | 0.2 | 1.14E-01 | 0.0 | 8.45E-01 | Protein kinase domain-containing protein |
| AT1G72160 | -0.7 | 7.80E-06 | -0.1 | 1.13E-01 | -0.6 | 1.93E-04 | -0.6 | 1.38E-04 | -0.3 | 1.69E-02 | -0.2 | 1.86E-01 | Patellin-3 |
| AT5G64370 | -0.7 | 1.52E-06 | -0.5 | 2.43E-05 | -0.2 | 2.72E-02 | 0.1 | 2.57E-01 | 0.4 | 1.19E-02 | -0.2 | 2.29E-01 | Beta-ureidopropionase |
| AT3G50210 | -0.7 | 6.06E-06 | -0.6 | 6.13E-05 | -0.2 | 1.09E-01 | -0.3 | 5.19E-02 | -0.1 | 4.20E-01 | -0.1 | 5.13E-01 | 2-oxoglutarate and Fe-dependent oxygenase superfamily protein |
| AT5G53760 | -0.7 | 5.35E-05 | -0.6 | 3.01E-04 | -0.1 | 3.75E-01 | 0.2 | 5.51E-01 | 0.7 | 1.24E-02 | -0.5 | 9.82E-02 | MLO-like protein 11 |
| AT1G60110 | -0.7 | 3.88E-04 | -0.2 | 1.93E-01 | -0.5 | 8.95E-03 | -0.6 | 4.51E-02 | 0.2 | 5.09E-01 | -0.8 | 3.56E-02 | Mannose-binding lectin superfamily protein |
| AT2G44300 | -0.7 | 1.18E-03 | -0.3 | 6.85E-02 | -0.4 | 6.46E-02 | 0.2 | 2.33E-01 | 0.3 | 1.04E-01 | -0.1 | 6.66E-01 | Non-specific lipid transfer protein GPI-anchored 14 |
| AT1G01600 | -0.7 | 2.01E-04 | -0.5 | 3.72E-03 | -0.2 | 1.34E-01 | 0.7 | 5.15E-03 | 0.6 | 2.43E-02 | 0.1 | 7.61E-01 | Cytochrome P450 86A4 |
| AT2G33570 | -0.7 | 6.44E-06 | -0.4 | 8.98E-04 | -0.3 | 4.09E-03 | -0.4 | 1.33E-02 | -0.1 | 4.79E-01 | -0.3 | 2.21E-01 | Galactan beta-1,4-galactosyltransferase GALS1 |
| AT1G76950 | -0.7 | 6.41E-06 | -0.6 | 3.01E-05 | -0.1 | 3.77E-01 | -0.2 | 1.70E-01 | 0.0 | 8.76E-01 | -0.2 | 4.15E-01 | PH, RCC1 and FYVE domains-containing protein 1 |
| AT5G48560 | -0.7 | 2.49E-04 | -0.6 | 1.21E-03 | -0.1 | 4.45E-01 | 0.4 | 3.28E-02 | 0.5 | 6.24E-03 | -0.2 | 4.54E-01 | Basic helix-loop-helix DNA-binding superfamily protein |
| AT4G14760 | -0.7 | 4.03E-05 | -0.8 | 3.14E-05 | 0.1 | 4.14E-01 | -0.4 | 4.50E-01 | 0.1 | 8.84E-01 | -0.6 | 4.14E-01 | Protein NETWORKED 1B |
| AT5G43060 | -0.7 | 2.12E-05 | 0.0 | 7.51E-01 | -0.7 | 9.80E-05 | 0.0 | 9.81E-01 | 0.0 | 9.75E-01 | 0.0 | 9.91E-01 | Probable cysteine protease RD21B |
| AT3G15510 | -0.7 | 3.88E-04 | -0.7 | 5.32E-04 | 0.0 | 9.35E-01 | -0.3 | 2.07E-01 | -0.5 | 1.21E-01 | 0.1 | 7.55E-01 | NAC transcription factor 56 |
| AT3G03900 | -0.7 | 1.27E-03 | -0.4 | 4.11E-02 | -0.3 | 1.14E-01 | 0.0 | 9.92E-01 | 0.3 | 2.09E-01 | -0.4 | 2.78E-01 | Adenylyl-sulfate kinase |
| AT4G25230 | -0.7 | 2.21E-06 | -0.5 | 3.19E-05 | -0.2 | 3.82E-02 | 0.0 | 8.69E-01 | 0.1 | 6.30E-01 | -0.1 | 5.86E-01 | E3 ubiquitin protein ligase RIN2 |
| AT3G30260 | -0.7 | 5.29E-03 | 0.5 | 2.02E-02 | -1.2 | 9.18E-04 | -0.4 | 3.76E-01 | 0.2 | 6.55E-01 | -0.6 | 1.65E-01 | AGAMOUS-like 79 |
| AT4G02600 | -0.7 | 1.37E-05 | -0.5 | 3.10E-04 | -0.2 | 4.32E-02 | -0.2 | 6.18E-02 | 0.0 | 8.24E-01 | -0.2 | 2.88E-01 | AT4G02600 protein |
| AT5G65970 | -0.7 | 8.33E-05 | -0.4 | 4.31E-03 | -0.3 | 3.77E-02 | -0.3 | 1.41E-01 | -0.2 | 4.23E-01 | -0.1 | 7.52E-01 | MLO-like protein |
| AT1G41830 | -0.7 | 1.69E-05 | -0.2 | 1.00E-01 | -0.5 | 4.48E-04 | -0.1 | 4.86E-01 | 0.3 | 8.76E-02 | -0.4 | 6.35E-02 | Putative pectinesterase |
| AT2G01540 | -0.7 | 7.03E-04 | -0.2 | 1.66E-01 | -0.5 | 1.67E-02 | -0.2 | 2.98E-01 | 0.0 | 9.38E-01 | -0.3 | 3.55E-01 | Protein C2-DOMAIN ABA-RELATED 10 |
| AT5G38020 | -0.7 | 1.13E-04 | -0.7 | 2.01E-04 | 0.0 | 9.69E-01 | -0.1 | 8.68E-01 | 0.7 | 9.26E-02 | -0.7 | 8.59E-02 | S-adenosyl-L-methionine:salicylic acid carboxyl methyltransferase-like |
| AT5G59780 | -0.7 | 3.05E-05 | -0.6 | 1.07E-04 | -0.1 | 5.74E-01 | -0.6 | 1.19E-02 | -0.5 | 4.79E-02 | -0.1 | 8.02E-01 | Transcription factor MYB59 |
| AT3G19000 | -0.7 | 8.58E-06 | -0.6 | 6.32E-05 | -0.1 | 2.16E-01 | 0.2 | 4.76E-01 | 0.3 | 3.41E-01 | -0.1 | 8.42E-01 | 2-oxoglutarate and Fe-dependent oxygenase superfamily protein |
| AT3G03120 | -0.7 | 1.05E-05 | -0.5 | 1.94E-04 | -0.2 | 6.10E-02 | 0.0 | 9.53E-01 | 0.0 | 9.50E-01 | 0.0 | 9.22E-01 | ADP-ribosylation factor |
| AT4G17680 | -0.7 | 9.52E-04 | -0.5 | 5.81E-03 | -0.2 | 4.01E-01 | -0.3 | 1.30E-01 | -0.1 | 5.94E-01 | -0.2 | 4.94E-01 | SBP family protein |
| AT5G65590 | -0.7 | 1.29E-03 | -0.3 | 1.25E-01 | -0.4 | 4.72E-02 | 0.4 | 2.41E-01 | 0.3 | 4.26E-01 | 0.1 | 8.85E-01 | Dof zinc finger protein DOF5.7 |
| AT2G19860 | -0.7 | 2.87E-04 | -0.5 | 3.57E-03 | -0.2 | 1.83E-01 | 0.1 | 3.28E-01 | 0.2 | 1.00E-01 | -0.1 | 5.45E-01 | Phosphotransferase |
| AT5G24590 | -0.7 | 1.70E-06 | -0.6 | 1.09E-05 | -0.1 | 1.85E-01 | -0.4 | 1.91E-03 | -0.2 | 1.44E-01 | -0.2 | 2.07E-01 | NAC domain-containing protein 91 |
| AT3G23600 | -0.7 | 6.94E-06 | -0.3 | 6.34E-03 | -0.4 | 8.97E-04 | -0.2 | 9.95E-02 | 0.0 | 9.66E-01 | -0.2 | 2.57E-01 | Alpha |
| AT5G52510 | -0.7 | 1.07E-05 | -0.5 | 1.86E-04 | -0.2 | 5.77E-02 | -0.4 | 3.42E-03 | -0.1 | 4.78E-01 | -0.3 | 9.04E-02 | Scarecrow-like protein 8 |
| AT3G07330 | -0.7 | 5.99E-06 | -0.3 | 1.45E-03 | -0.4 | 2.05E-03 | -0.2 | 1.22E-01 | -0.1 | 3.88E-01 | -0.1 | 7.46E-01 | Probable xyloglucan glycosyltransferase 6 |
| AT5G56350 | -0.7 | 3.51E-05 | -0.5 | 5.46E-04 | -0.2 | 1.05E-01 | 0.7 | 5.08E-05 | 0.2 | 1.63E-01 | 0.5 | 9.21E-03 | Pyruvate kinase |
| AT1G72950 | -0.7 | 1.10E-03 | -0.6 | 2.55E-03 | -0.1 | 8.24E-01 | | | | | | | TIR domain-containing protein |
| AT3G20015 | -0.7 | 4.01E-05 | -0.2 | 9.12E-02 | -0.5 | 1.23E-03 | 0.0 | 8.18E-01 | 0.2 | 3.44E-01 | -0.1 | 5.20E-01 | Protein ASPARTIC PROTEASE IN GUARD CELL 2 |
| AT1G43800 | -0.7 | 1.14E-02 | -0.5 | 4.66E-02 | -0.2 | 5.47E-01 | 1.1 | 5.63E-02 | 0.2 | 8.00E-01 | 0.9 | 2.19E-01 | Stearoyl- 9-desaturase 6, chloroplastic |
| AT5G60450 | -0.7 | 3.84E-05 | -0.9 | 1.53E-05 | 0.2 | 1.28E-01 | 0.2 | 1.86E-01 | 0.3 | 2.06E-01 | 0.0 | 9.62E-01 | Auxin response factor |
| AT4G22840 | -0.7 | 2.26E-05 | -0.5 | 4.42E-04 | -0.2 | 7.23E-02 | -0.2 | 4.59E-01 | 0.2 | 3.52E-01 | -0.4 | 1.56E-01 | Probable sodium |
| AT1G54570 | -0.7 | 2.45E-04 | -0.7 | 4.45E-04 | 0.0 | 9.46E-01 | 0.0 | 8.31E-01 | 0.3 | 5.14E-02 | -0.3 | 1.59E-01 | Phytyl ester synthase 1, chloroplastic |
| AT2G20030 | -0.7 | 1.90E-04 | -0.5 | 1.68E-03 | -0.2 | 2.77E-01 | 0.3 | 3.51E-01 | 0.4 | 3.59E-01 | 0.0 | 9.50E-01 | Putative RING-H2 finger protein ATL12 |
| AT3G25780 | -0.7 | 2.68E-04 | -0.3 | 5.40E-02 | -0.4 | 1.60E-02 | 0.2 | 3.99E-01 | 0.3 | 3.94E-01 | 0.0 | 9.15E-01 | Allene oxide cyclase 3, chloroplastic |
| AT1G14540 | -0.7 | 9.23E-02 | 0.3 | 3.35E-01 | -1.0 | 3.91E-02 | 2.7 | 1.80E-06 | 0.6 | 2.83E-01 | 2.2 | 1.42E-04 | Peroxidase 4 |
| AT5G15150 | -0.7 | 9.77E-05 | 0.1 | 2.65E-01 | -0.8 | 2.20E-04 | -0.4 | 2.00E-01 | 0.0 | 9.83E-01 | -0.4 | 3.64E-01 | Homeobox-leucine zipper protein |
| AT1G25480 | -0.7 | 7.82E-05 | -0.6 | 4.67E-04 | -0.1 | 3.53E-01 | -0.4 | 7.52E-02 | -0.4 | 6.18E-02 | 0.1 | 8.64E-01 | Aluminum-activated malate transporter 4 |
| AT3G45420 | -0.7 | 2.92E-04 | -0.4 | 1.02E-02 | -0.3 | 8.69E-02 | -0.3 | 3.64E-01 | -0.1 | 8.23E-01 | -0.2 | 6.27E-01 | Putative L-type lectin-domain containing receptor kinase I.4 |
| AT4G27780 | -0.7 | 4.79E-06 | -0.5 | 9.37E-05 | -0.2 | 3.72E-02 | -0.3 | 1.04E-01 | 0.0 | 9.82E-01 | -0.3 | 2.10E-01 | Acyl-CoA binding protein 2 |
| AT5G09870 | -0.7 | 7.71E-05 | -0.6 | 2.19E-04 | 0.0 | 7.44E-01 | -0.5 | 1.04E-03 | 0.1 | 6.17E-01 | -0.6 | 5.23E-03 | Cellulose synthase |
| AT1G63430 | -0.7 | 3.68E-06 | -0.3 | 9.84E-04 | -0.4 | 1.43E-03 | -0.2 | 1.31E-01 | -0.2 | 2.33E-01 | 0.0 | 9.40E-01 | Leucine-rich repeat protein kinase family protein |
| AT5G40850 | -0.7 | 9.95E-03 | -0.3 | 2.23E-01 | -0.4 | 1.35E-01 | -1.5 | 1.71E-07 | -0.9 | 7.72E-05 | -0.5 | 1.65E-02 | Urophorphyrin methylase 1 |
| AT5G54680 | -0.7 | 2.54E-05 | -0.4 | 9.40E-04 | -0.2 | 3.22E-02 | -0.4 | 8.65E-04 | -0.3 | 2.09E-02 | -0.1 | 4.52E-01 | Transcription factor ILR3 |
| AT2G39180 | -0.7 | 5.15E-04 | 0.0 | 7.96E-01 | -0.6 | 3.15E-03 | -0.3 | 2.98E-01 | 0.0 | 9.58E-01 | -0.3 | 3.97E-01 | Serine |
| AT3G20120 | -0.7 | 4.00E-05 | -0.5 | 4.22E-04 | -0.2 | 1.74E-01 | -0.2 | 2.47E-01 | -0.1 | 6.87E-01 | -0.1 | 5.71E-01 | Cytochrome P450, family 705, subfamily A, polypeptide 21 |
| AT1G06840 | -0.7 | 1.59E-06 | -0.4 | 7.43E-05 | -0.3 | 5.35E-03 | 0.4 | 1.12E-02 | 0.1 | 6.98E-01 | 0.4 | 1.25E-01 | Probable LRR receptor-like serine |
| AT1G77200 | -0.7 | 2.03E-03 | -0.3 | 6.15E-02 | -0.3 | 1.29E-01 | -0.6 | 5.67E-02 | -0.3 | 3.90E-01 | -0.3 | 4.71E-01 | Ethylene-responsive transcription factor ERF037 |
| AT2G41300 | -0.7 | 1.23E-04 | -0.2 | 9.49E-02 | -0.5 | 5.34E-03 | 0.1 | 7.12E-01 | 0.3 | 1.93E-01 | -0.2 | 4.11E-01 | Protein STRICTOSIDINE SYNTHASE-LIKE 1 |
| AT5G38240 | -0.7 | 1.21E-02 | -0.6 | 2.43E-02 | -0.1 | 8.36E-01 | -0.4 | 4.01E-01 | -0.1 | 8.66E-01 | | | Protein kinase family protein |
| AT4G12570 | -0.7 | 5.27E-05 | -0.7 | 1.03E-04 | 0.0 | 9.59E-01 | -0.2 | 4.61E-01 | -0.3 | 2.03E-01 | 0.1 | 6.80E-01 | E3 ubiquitin-protein ligase UPL5 |
| AT4G20230 | -0.7 | 4.69E-05 | -0.2 | 3.40E-02 | -0.5 | 3.96E-03 | 0.1 | 8.15E-01 | 0.1 | 8.42E-01 | 0.0 | 9.97E-01 | Terpenoid synthase superfamily protein |
| AT3G10300 | -0.7 | 1.68E-05 | -0.2 | 9.57E-02 | -0.5 | 4.52E-04 | -0.2 | 1.75E-01 | 0.0 | 9.29E-01 | -0.2 | 3.74E-01 | Calcium-binding EF-hand family protein |
| AT3G59700 | -0.7 | 2.59E-04 | -0.7 | 3.15E-04 | 0.0 | 8.77E-01 | 0.4 | 2.04E-01 | 0.0 | 9.69E-01 | 0.4 | 3.72E-01 | L-type lectin-domain containing receptor kinase V.5 |
| AT1G53700 | -0.7 | 1.05E-03 | -0.6 | 4.63E-03 | -0.1 | 5.35E-01 | 0.0 | 9.58E-01 | 0.1 | 8.95E-01 | -0.1 | 8.69E-01 | Serine |
| AT3G47040 | -0.7 | 1.77E-04 | -0.4 | 6.84E-03 | -0.3 | 6.12E-02 | -0.5 | 5.41E-02 | 0.1 | 7.39E-01 | -0.6 | 5.86E-02 | beta-glucosidase |
| AT5G66920 | -0.7 | 4.09E-05 | 0.1 | 1.54E-01 | -0.8 | 7.42E-05 | -0.3 | 1.76E-01 | -0.1 | 8.31E-01 | -0.2 | 4.77E-01 | Pectinesterase like protein |
| AT1G14240 | -0.7 | 1.89E-03 | 0.1 | 3.59E-01 | -0.8 | 2.44E-03 | -0.6 | 6.39E-02 | 0.1 | 7.20E-01 | -0.7 | 6.84E-02 | apyrase |

| | | | | | | | | | | | | | |
|---|---|---|---|---|---|---|---|---|---|---|---|---|---|
| AT2G21620 | -0.7 | 8.62E-06 | -0.5 | 2.97E-04 | -0.2 | 2.22E-02 | -0.2 | 2.32E-01 | -0.2 | 3.67E-01 | 0.0 | 9.51E-01 | Adenine nucleotide alpha hydrolases-like superfamily protein |
| AT1G12140 | -0.7 | 5.83E-06 | -0.4 | 2.36E-04 | -0.2 | 1.39E-02 | 0.0 | 9.27E-01 | 0.3 | 3.80E-02 | -0.3 | 1.09E-01 | Flavin-containing monooxygenase FMO GS-OX5 |
| AT5G57610 | -0.7 | 1.86E-05 | -0.7 | 2.32E-05 | 0.0 | 6.65E-01 | -0.4 | 7.53E-02 | 0.3 | 1.91E-01 | -0.6 | 2.08E-02 | Protein kinase domain-containing protein |
| AT2G39830 | -0.7 | 6.91E-05 | -0.2 | 6.10E-02 | -0.5 | 4.02E-03 | 0.0 | 1.00E+00 | 0.1 | 8.33E-01 | -0.1 | 8.44E-01 | DA1-related protein 2 |
| AT3G17820 | -0.7 | 1.41E-05 | -0.3 | 2.53E-03 | -0.3 | 4.67E-03 | -0.3 | 2.55E-02 | 0.0 | 9.66E-01 | -0.3 | 8.04E-02 | Glutamine synthetase cytosolic isozyme 1-3 |
| AT1G31850 | -0.7 | 1.01E-05 | -0.4 | 5.24E-04 | -0.3 | 1.33E-02 | -0.3 | 2.94E-02 | 0.0 | 8.25E-01 | -0.4 | 8.15E-02 | Probable methyltransferase PMT20 |
| AT5G66790 | -0.7 | 1.48E-04 | -0.6 | 5.55E-04 | -0.1 | 5.68E-01 | -0.1 | 5.62E-01 | -0.5 | 3.89E-02 | 0.3 | 2.13E-01 | Wall-associated receptor kinase-like 21 |
| AT4G35350 | -0.7 | 3.34E-05 | 0.3 | 6.33E-03 | -1.0 | 2.85E-05 | 0.0 | 9.18E-01 | -0.1 | 7.26E-01 | 0.1 | 6.67E-01 | Xylem cysteine peptidase 1 |
| AT5G22290 | -0.7 | 2.25E-04 | -0.8 | 1.06E-04 | 0.2 | 3.36E-01 | 0.0 | 9.51E-01 | -0.1 | 8.49E-01 | 0.0 | 9.12E-01 | NAC domain containing protein 89 |
| AT5G27520 | -0.7 | 1.36E-05 | -0.5 | 2.45E-04 | -0.2 | 6.60E-02 | -0.1 | 3.72E-01 | -0.1 | 7.20E-01 | -0.1 | 7.87E-01 | AT5G27520 protein |
| AT2G32260 | -0.7 | 1.62E-06 | -0.3 | 5.63E-04 | -0.4 | 6.21E-04 | -0.3 | 2.74E-02 | -0.2 | 1.07E-01 | 0.0 | 8.13E-01 | choline-phosphate cytidylyltransferase |
| AT4G13030 | -0.7 | 7.58E-05 | -0.7 | 1.20E-04 | 0.0 | 9.38E-01 | -0.5 | 5.34E-02 | -0.4 | 1.26E-01 | -0.1 | 8.76E-01 | At4g13030 |
| AT3G50740 | -0.7 | 8.00E-05 | -0.6 | 5.57E-04 | -0.1 | 3.16E-01 | -0.2 | 9.78E-02 | -0.6 | 7.36E-05 | 0.5 | 7.48E-03 | UDP-glycosyltransferase 72E1 |
| AT3G46830 | -0.7 | 1.57E-06 | -0.1 | 3.59E-01 | -0.6 | 3.28E-05 | -0.1 | 5.59E-01 | 0.0 | 8.78E-01 | -0.1 | 8.03E-01 | Ras-related protein RABA2c |
| AT4G22130 | -0.7 | 1.99E-05 | -0.4 | 8.36E-04 | -0.3 | 2.44E-02 | -0.2 | 3.30E-01 | 0.1 | 7.00E-01 | -0.3 | 2.40E-01 | STRUBBELIG-receptor family 8 |
| AT1G64390 | -0.7 | 1.02E-03 | -0.3 | 1.04E-01 | -0.4 | 3.27E-02 | 0.0 | 8.05E-01 | 0.7 | 1.76E-04 | -0.6 | 2.36E-03 | Endoglucanase 6 |
| AT5G03520 | -0.7 | 3.19E-06 | -0.3 | 1.06E-03 | -0.4 | 1.10E-03 | -0.4 | 2.51E-03 | -0.2 | 7.82E-02 | -0.2 | 3.43E-01 | Ras-related protein RABE1d |
| AT5G13500 | -0.7 | 2.17E-05 | -0.4 | 1.12E-03 | -0.3 | 1.94E-02 | -0.2 | 4.16E-01 | 0.1 | 6.53E-01 | -0.3 | 2.16E-01 | Hydroxyproline O-arabinosyltransferase 3 |
| AT1G33970 | -0.7 | 1.45E-05 | -0.6 | 4.45E-05 | 0.0 | 6.56E-01 | -0.2 | 1.31E-01 | -0.1 | 5.72E-01 | -0.1 | 5.81E-01 | Immune-associated nucleotide-binding protein 9 |
| AT2G41740 | -0.7 | 8.67E-06 | -0.3 | 3.02E-03 | -0.4 | 1.99E-03 | -0.3 | 4.52E-02 | -0.1 | 4.49E-01 | -0.2 | 4.85E-01 | Villin-2 |
| AT1G09100 | -0.7 | 5.28E-06 | -0.6 | 3.19E-05 | -0.1 | 2.47E-01 | 0.2 | 3.49E-01 | 0.1 | 8.49E-01 | 0.1 | 6.60E-01 | 26S proteasome regulatory subunit 6A homolog B |
| AT1G69580 | -0.7 | 1.50E-05 | -0.4 | 7.70E-04 | -0.3 | 2.02E-02 | -0.3 | 2.48E-01 | -0.1 | 8.63E-01 | -0.2 | 4.68E-01 | Myb family transcription factor PHL8 |
| AT1G61590 | -0.7 | 7.33E-05 | 0.0 | 9.33E-01 | -0.7 | 3.85E-04 | -0.2 | 2.74E-01 | -0.1 | 8.12E-01 | -0.2 | 5.87E-01 | Probable serine |
| AT4G31340 | -0.7 | 9.97E-06 | -0.2 | 8.42E-02 | -0.5 | 2.87E-04 | -0.2 | 1.79E-01 | 0.1 | 4.55E-01 | -0.3 | 1.16E-01 | AT4G31340 protein |
| AT3G22400 | -0.7 | 2.69E-04 | -0.3 | 2.41E-02 | -0.4 | 3.30E-02 | -0.4 | 4.98E-02 | 0.0 | 9.35E-01 | -0.4 | 1.07E-01 | Linoleate 9S-lipoxygenase 5 |
| AT3G25560 | -0.7 | 7.42E-05 | -0.5 | 8.83E-04 | -0.2 | 1.62E-01 | -0.2 | 3.65E-01 | 0.0 | 9.28E-01 | -0.2 | 4.50E-01 | NSP-interacting kinase 2 |
| AT3G46970 | -0.7 | 3.47E-04 | -0.6 | 7.13E-04 | 0.0 | 8.94E-01 | -0.4 | 4.22E-03 | -0.1 | 4.79E-01 | -0.3 | 1.17E-01 | Alpha-glucan phosphorylase 2, cytosolic |
| AT3G45040 | -0.7 | 5.72E-05 | -0.5 | 4.89E-04 | -0.1 | 2.31E-01 | -0.2 | 3.18E-01 | -0.2 | 1.95E-01 | 0.1 | 8.07E-01 | dolichol kinase |
| AT1G22740 | -0.7 | 8.87E-05 | -0.1 | 2.04E-01 | -0.5 | 1.91E-03 | -1.1 | 1.66E-06 | -0.9 | 5.99E-05 | -0.2 | 3.25E-01 | Ras-related protein RABG3b |
| AT3G04420 | -0.7 | 1.28E-03 | -1.1 | 1.17E-04 | 0.4 | 5.74E-02 | 0.9 | 7.13E-02 | 0.8 | 1.79E-01 | 0.1 | 8.31E-01 | AT3g04420 |
| AT1G76490 | -0.7 | 4.57E-06 | -0.4 | 5.16E-04 | -0.3 | 4.11E-03 | -0.5 | 3.49E-03 | -0.5 | 6.14E-03 | 0.0 | 9.62E-01 | hydroxymethylglutaryl-CoA reductase |
| AT1G29400 | -0.7 | 5.05E-05 | -0.6 | 2.88E-04 | -0.1 | 3.53E-01 | -0.1 | 2.18E-01 | 0.0 | 9.95E-01 | -0.1 | 4.07E-01 | Protein MEI2-like 5 |
| AT5G02780 | -0.7 | 9.94E-03 | -1.1 | 1.34E-03 | 0.4 | 2.06E-01 | 0.4 | 1.86E-01 | -0.8 | 4.64E-02 | 1.2 | 3.25E-03 | Glutathione S-transferase L1 |
| AT5G57970 | -0.7 | 2.18E-05 | -0.3 | 1.09E-02 | -0.4 | 3.35E-03 | -0.1 | 6.04E-01 | 0.2 | 5.41E-01 | -0.3 | 3.03E-01 | hypothetical protein |
| AT3G26700 | -0.7 | 2.18E-05 | -0.1 | 5.53E-01 | -0.6 | 2.80E-04 | -0.4 | 4.94E-02 | -0.1 | 7.14E-01 | -0.3 | 2.17E-01 | non-specific serine |
| AT5G24352 | -0.7 | 1.32E-03 | -0.3 | 8.79E-02 | -0.4 | 6.48E-02 | -0.3 | 4.04E-01 | 0.0 | 9.54E-01 | -0.3 | 5.51E-01 | Serine |
| AT3G60180 | -0.7 | 4.78E-05 | -0.7 | 6.39E-05 | 0.0 | 7.86E-01 | -0.3 | 3.45E-01 | -0.1 | 8.35E-01 | -0.2 | 6.06E-01 | Probable UMP-CMP kinase 1 |
| AT3G20500 | -0.7 | 5.73E-06 | -0.6 | 3.82E-05 | -0.1 | 2.16E-01 | -0.2 | 1.28E-01 | 0.0 | 9.00E-01 | -0.2 | 1.85E-01 | Purple acid phosphatase 18 |
| AT4G01120 | -0.7 | 3.71E-06 | -0.3 | 1.07E-03 | -0.4 | 1.39E-03 | -0.6 | 1.04E-04 | -0.6 | 2.31E-04 | 0.0 | 9.43E-01 | G-box binding bZip transcription factor GBF2 |
| AT5G22940 | -0.7 | 1.23E-04 | 0.1 | 5.89E-01 | -0.7 | 3.59E-04 | -0.5 | 3.12E-03 | -0.2 | 1.47E-01 | -0.2 | 2.58E-01 | Glucuronoxylan glucuronosyltransferase, putative |
| AT1G74000 | -0.7 | 1.98E-03 | -0.1 | 4.92E-01 | -0.6 | 1.96E-02 | 0.2 | 5.93E-01 | 0.8 | 4.72E-02 | -0.5 | 1.49E-01 | Protein STRICTOSIDINE SYNTHASE-LIKE 11 |
| AT1G44100 | -0.7 | 1.96E-05 | -0.6 | 1.02E-04 | -0.1 | 3.47E-01 | 0.6 | 2.33E-04 | 0.5 | 6.30E-03 | 0.2 | 4.00E-01 | Amino acid permease 5 |
| AT1G48220 | -0.7 | 5.33E-03 | 0.1 | 4.82E-01 | -0.8 | 7.02E-03 | | | | | | | Protein kinase domain-containing protein |
| AT1G02305 | -0.7 | 7.60E-06 | -0.4 | 4.92E-04 | -0.3 | 9.39E-03 | -0.5 | 7.52E-05 | -0.4 | 4.89E-03 | -0.2 | 2.53E-01 | Cathepsin B-like protease 2 |
| AT3G54820 | -0.7 | 1.46E-04 | -0.6 | 3.91E-04 | 0.0 | 7.46E-01 | -0.4 | 8.97E-02 | 0.0 | 9.04E-01 | -0.3 | 1.99E-01 | Uncharacterized protein |
| AT1G72870 | -0.7 | 3.27E-03 | -0.2 | 2.08E-01 | -0.4 | 7.00E-02 | | | | | | | Disease resistance protein |
| AT2G36570 | -0.7 | 3.14E-05 | -0.4 | 2.65E-03 | -0.3 | 1.36E-02 | -0.1 | 3.93E-01 | 0.2 | 1.88E-01 | -0.3 | 8.10E-02 | Leucine-rich repeat receptor-like protein kinase PXC1 |
| AT5G03340 | -0.7 | 1.72E-05 | -0.7 | 3.15E-05 | 0.0 | 9.65E-01 | -0.1 | 7.91E-01 | 0.2 | 7.75E-01 | -0.3 | 5.87E-01 | Cell division control protein 48 homolog E |
| AT3G13760 | -0.7 | 1.87E-03 | 0.0 | 9.29E-01 | -0.7 | 5.48E-03 | 0.1 | 8.02E-01 | 0.4 | 2.81E-01 | -0.3 | 5.20E-01 | Cysteine |
| AT5G55180 | -0.7 | 2.18E-05 | -0.3 | 4.98E-03 | -0.3 | 4.63E-03 | 0.8 | 1.51E-05 | 0.5 | 1.72E-03 | 0.2 | 1.32E-01 | glucan endo-1,3-beta-D-glucosidase |
| AT2G45050 | -0.7 | 1.22E-03 | 0.4 | 1.12E-02 | -1.1 | 2.31E-04 | -0.5 | 1.55E-02 | -0.1 | 5.29E-01 | -0.3 | 1.79E-01 | GATA transcription factor 2 |
| AT1G77120 | -0.7 | 7.76E-05 | -0.7 | 7.69E-05 | 0.1 | 6.37E-01 | 1.3 | 1.64E-04 | 0.3 | 3.70E-01 | 1.0 | 3.82E-04 | Alcohol dehydrogenase class-P |
| AT5G12200 | -0.7 | 1.80E-05 | -0.5 | 3.66E-04 | -0.2 | 5.47E-02 | -0.2 | 7.69E-02 | -0.1 | 6.73E-01 | -0.1 | 4.18E-01 | Putative dihydropyrimidinase |
| AT2G30350 | -0.7 | 4.07E-03 | -0.3 | 8.37E-02 | -0.3 | 1.77E-01 | -0.3 | 2.16E-01 | -0.3 | 2.38E-01 | 0.0 | 9.73E-01 | Excinuclease ABC, C subunit, N-terminal |
| AT3G20080 | -0.7 | 2.21E-05 | -0.4 | 6.09E-04 | -0.2 | 4.92E-02 | -0.1 | 7.85E-01 | 0.1 | 9.01E-01 | -0.1 | 6.60E-01 | Cytochrome P450, family 705, subfamily A, polypeptide 15 |
| AT2G35880 | -0.7 | 1.25E-05 | -0.1 | 2.61E-01 | -0.6 | 1.99E-04 | -0.7 | 2.79E-05 | -0.5 | 1.52E-03 | -0.2 | 3.00E-01 | Protein WVD2-like 4 |
| AT3G54270 | -0.7 | 2.98E-04 | 0.0 | 7.42E-01 | -0.7 | 1.06E-03 | -0.3 | 2.88E-01 | 0.0 | 8.92E-01 | -0.2 | 4.50E-01 | Probable sucrose-phosphatase 3a |
| AT2G28350 | -0.7 | 5.80E-05 | -0.6 | 1.75E-04 | -0.1 | 6.85E-01 | -0.2 | 1.62E-01 | 0.0 | 9.50E-01 | -0.2 | 2.52E-01 | Auxin response factor |
| AT1G73640 | -0.7 | 2.78E-03 | -0.5 | 1.90E-02 | -0.2 | 4.03E-01 | -0.1 | 7.37E-01 | -0.1 | 8.87E-01 | -0.1 | 9.22E-01 | Ras-related protein RABA6a |
| AT2G23340 | -0.7 | 4.28E-04 | -0.3 | 2.51E-02 | -0.3 | 4.88E-02 | -0.1 | 3.61E-01 | 0.3 | 1.31E-01 | -0.4 | 4.85E-02 | DEAR3 |
| AT5G54500 | -0.7 | 1.90E-04 | -0.3 | 2.16E-02 | -0.3 | 2.15E-02 | 0.4 | 6.77E-04 | -0.1 | 3.82E-01 | 0.6 | 1.92E-03 | NADH dehydrogenase |
| AT5G62165 | -0.7 | 2.58E-04 | -0.3 | 4.04E-02 | -0.4 | 2.15E-02 | 0.0 | 9.95E-01 | 0.3 | 3.52E-01 | -0.3 | 4.28E-01 | AGAMOUS-like 42 |
| AT2G22860 | -0.7 | 1.17E-02 | -0.2 | 3.43E-01 | -0.4 | 1.24E-01 | -0.1 | 8.03E-01 | -0.1 | 8.52E-01 | 0.0 | 9.91E-01 | Phytosulfokines 2 |
| AT3G62210 | -0.7 | 2.15E-03 | 0.0 | 8.57E-01 | -0.6 | 9.51E-03 | 0.3 | 2.99E-01 | 0.4 | 1.29E-01 | -0.1 | 6.85E-01 | Endonuclease or glycosyl hydrolase |
| AT1G02335 | -0.7 | 1.87E-04 | 0.3 | 2.33E-02 | -0.9 | 1.11E-04 | 0.1 | 7.26E-01 | 0.0 | 9.71E-01 | 0.1 | 8.14E-01 | Germin-like protein subfamily 2 member 2 |
| AT3G07010 | -0.7 | 4.25E-05 | 0.2 | 7.03E-02 | -0.8 | 5.76E-05 | -0.8 | 3.03E-05 | -0.3 | 4.48E-02 | -0.5 | 1.75E-02 | Pectate lyase |
| AT1G14390 | -0.7 | 1.32E-02 | 0.0 | 9.71E-01 | -0.7 | 3.36E-02 | | | | | | | Probable LRR receptor-like serine |
| AT2G22430 | -0.7 | 4.73E-05 | -0.4 | 1.45E-03 | -0.2 | 5.15E-02 | -0.6 | 3.35E-05 | -0.3 | 2.55E-02 | -0.3 | 5.98E-02 | Homeobox-leucine zipper protein ATHB-6 |
| AT1G71190 | -0.7 | 4.33E-05 | -0.5 | 7.60E-04 | -0.2 | 9.94E-02 | -0.2 | 1.40E-01 | -0.3 | 1.62E-01 | 0.0 | 9.73E-01 | At1g71190 |
| AT3G57530 | -0.7 | 9.63E-06 | -0.5 | 7.77E-05 | -0.1 | 1.68E-01 | -0.3 | 8.51E-03 | -0.2 | 1.82E-01 | -0.1 | 3.97E-01 | Calcium-dependent protein kinase 32 |
| AT3G56800 | -0.7 | 3.01E-05 | -0.3 | 1.14E-02 | -0.4 | 3.89E-03 | 0.0 | 8.60E-01 | 0.1 | 8.44E-01 | 0.0 | 9.70E-01 | Calmodulin-3 |
| AT5G65500 | -0.7 | 5.04E-03 | -0.8 | 2.21E-03 | 0.2 | 5.20E-01 | | | | | | | U-box domain-containing protein kinase family protein |
| AT5G27320 | -0.7 | 5.45E-05 | -0.4 | 1.14E-03 | -0.2 | 8.09E-02 | -0.4 | 2.64E-02 | -0.3 | 1.52E-01 | -0.1 | 6.80E-01 | Gibberellin receptor GID1C |
| AT5G59260 | -0.6 | 7.31E-04 | -0.5 | 4.16E-03 | -0.1 | 4.24E-01 | -1.1 | 4.59E-04 | -0.5 | 5.63E-02 | -0.6 | 6.53E-02 | Probable L-type lectin-domain containing receptor kinase II.1 |
| AT5G65020 | -0.6 | 1.76E-05 | 0.2 | 6.21E-02 | -0.8 | 3.30E-05 | -0.4 | 1.33E-02 | 0.3 | 1.07E-01 | -0.6 | 3.25E-03 | Annexin |
| AT5G61570 | -0.6 | 5.37E-04 | -0.4 | 6.72E-03 | -0.2 | 2.16E-01 | -0.1 | 6.49E-01 | 0.3 | 1.81E-01 | -0.4 | 8.16E-02 | Disease resistance protein kinase Pto-like protein |
| AT2G23790 | -0.6 | 3.58E-05 | -0.6 | 1.25E-04 | -0.1 | 5.78E-01 | -0.1 | 5.13E-01 | 0.1 | 8.15E-01 | -0.2 | 4.58E-01 | Calcium uniporter protein 2, mitochondrial |
| AT1G47380 | -0.6 | 4.61E-05 | -0.2 | 1.21E-01 | -0.5 | 1.41E-03 | -0.5 | 2.15E-02 | -0.1 | 8.05E-01 | -0.4 | 1.37E-01 | Probable protein phosphatase 2C 12 |
| AT1G17060 | -0.6 | 3.87E-03 | -0.1 | 6.59E-01 | -0.6 | 2.56E-02 | -0.3 | 4.35E-01 | -0.2 | 7.48E-01 | -0.1 | 8.05E-01 | SOB7 |
| AT5G49660 | -0.6 | 4.33E-05 | -0.4 | 1.50E-03 | -0.2 | 4.03E-02 | -0.1 | 7.62E-01 | 0.1 | 5.21E-01 | -0.2 | 4.03E-01 | Protein kinase domain-containing protein |
| AT2G29065 | -0.6 | 6.11E-04 | -0.6 | 1.31E-03 | 0.0 | 8.56E-01 | 0.4 | 7.64E-02 | 0.3 | 3.41E-01 | 0.2 | 6.32E-01 | Scarecrow-like protein 34 |
| AT3G04480 | -0.6 | 7.99E-06 | -0.7 | 1.12E-05 | 0.0 | 6.64E-01 | 0.6 | 5.30E-04 | 0.4 | 1.25E-02 | 0.2 | 4.30E-01 | Diphthine--ammonia ligase |
| AT3G16940 | -0.6 | 6.84E-06 | -0.7 | 1.32E-05 | 0.0 | 9.47E-01 | -0.4 | 3.41E-03 | -0.1 | 6.79E-01 | -0.3 | 6.23E-02 | Calmodulin-binding transcription activator |

| | | | | | | | | | | | | | |
|---|---|---|---|---|---|---|---|---|---|---|---|---|---|
| AT5G05270 | -0.6 | 5.42E-02 | -0.4 | 1.83E-01 | -0.2 | 5.75E-01 | 0.6 | 4.43E-02 | 1.2 | 1.79E-03 | -0.5 | 1.66E-01 | Chalcone-flavonone isomerase family protein |
| AT1G78670 | -0.6 | 3.42E-06 | -0.4 | 3.41E-04 | -0.3 | 3.41E-03 | -0.2 | 7.80E-02 | 0.1 | 6.38E-01 | -0.3 | 9.79E-02 | Probable gamma-glutamyl hydrolase 3 |
| AT1G53500 | -0.6 | 4.17E-04 | -0.2 | 1.57E-01 | -0.5 | 1.04E-02 | -0.1 | 5.26E-01 | 0.1 | 8.03E-01 | -0.2 | 4.53E-01 | Trifunctional UDP-glucose 4,6-dehydratase |
| AT5G20550 | -0.6 | 2.79E-04 | 0.0 | 8.61E-01 | -0.6 | 1.75E-03 | -0.3 | 2.92E-01 | 0.3 | 2.30E-01 | -0.6 | 5.77E-02 | hypothetical protein |
| AT3G16180 | -0.6 | 3.63E-05 | 0.1 | 1.45E-01 | -0.8 | 7.99E-05 | -0.3 | 1.07E-01 | 0.0 | 9.83E-01 | -0.3 | 2.28E-01 | Protein NRT1 |
| AT4G11880 | -0.6 | 2.10E-03 | -0.4 | 1.90E-02 | -0.2 | 3.14E-01 | 0.0 | 9.84E-01 | 0.2 | 5.37E-01 | -0.2 | 5.74E-01 | AGAMOUS-like 14 |
| AT3G16030 | -0.6 | 7.96E-05 | -1.0 | 9.09E-06 | 0.4 | 1.54E-02 | 0.0 | 9.94E-01 | -0.1 | 8.79E-01 | 0.1 | 9.04E-01 | Receptor-like serine |
| AT4G35480 | -0.6 | 1.31E-03 | -0.1 | 6.59E-01 | -0.6 | 9.29E-03 | -0.2 | 5.29E-01 | -0.3 | 3.96E-01 | 0.1 | 8.55E-01 | Probable E3 ubiquitin-protein ligase ATL45 |
| AT2G38610 | -0.6 | 1.11E-05 | -0.6 | 3.32E-05 | 0.0 | 6.77E-01 | -0.5 | 3.45E-03 | -0.5 | 6.42E-03 | 0.0 | 9.83E-01 | KH domain-containing protein At2g38610 |
| AT5G44790 | -0.6 | 1.59E-05 | -0.4 | 4.60E-04 | -0.2 | 3.64E-02 | -0.2 | 1.08E-01 | -0.2 | 3.21E-01 | -0.1 | 8.15E-01 | Copper-transporting ATPase RAN1 |
| AT1G13195 | -0.6 | 1.28E-04 | -0.6 | 2.18E-04 | 0.0 | 9.88E-01 | -0.4 | 9.52E-02 | 0.0 | 9.27E-01 | -0.3 | 1.68E-01 | RING-type domain-containing protein |
| AT1G30900 | -0.6 | 3.13E-05 | -0.1 | 2.94E-01 | -0.5 | 4.42E-04 | -0.1 | 5.94E-01 | 0.2 | 3.71E-01 | -0.3 | 2.45E-01 | Vacuolar-sorting receptor 6 |
| AT5G51350 | -0.6 | 5.29E-05 | -0.3 | 1.42E-02 | -0.4 | 7.77E-03 | -0.5 | 3.27E-02 | 0.0 | 9.30E-01 | -0.5 | 1.18E-01 | Leucine-rich repeat receptor-like protein kinase |
| AT4G22580 | -0.6 | 2.86E-05 | -0.6 | 1.15E-04 | -0.1 | 5.26E-01 | -0.2 | 3.41E-01 | 0.1 | 6.40E-01 | -0.3 | 1.95E-01 | Probable xyloglucan galactosyltransferase GT19 |
| AT5G07270 | -0.6 | 4.41E-06 | -0.6 | 1.76E-05 | -0.1 | 4.61E-01 | 0.0 | 9.04E-01 | 0.0 | 8.63E-01 | -0.1 | 8.17E-01 | XBAT33 |
| AT1G29800 | -0.6 | 3.27E-06 | -0.7 | 4.71E-06 | 0.0 | 5.64E-01 | -0.4 | 1.29E-02 | -0.1 | 4.06E-01 | -0.3 | 2.21E-01 | RING |
| AT1G34260 | -0.6 | 7.76E-06 | -0.8 | 4.71E-06 | 0.1 | 1.30E-01 | 0.0 | 9.23E-01 | 0.3 | 3.79E-01 | -0.3 | 4.12E-01 | 1-phosphatidylinositol-3-phosphate 5-kinase |
| AT1G76760 | -0.6 | 2.53E-03 | -0.3 | 1.08E-01 | -0.4 | 9.87E-02 | 0.0 | 9.85E-01 | 0.3 | 4.55E-01 | -0.3 | 4.39E-01 | Thioredoxin Y1, chloroplastic |
| AT2G26830 | -0.6 | 9.85E-06 | -0.4 | 3.19E-04 | -0.2 | 2.67E-02 | -0.4 | 2.70E-02 | 0.0 | 9.68E-01 | -0.4 | 7.15E-02 | Probable ethanolamine kinase |
| AT5G07070 | -0.6 | 8.71E-05 | -0.3 | 2.06E-02 | -0.4 | 1.09E-02 | -0.6 | 1.47E-02 | -0.1 | 7.79E-01 | -0.5 | 9.90E-02 | CBL-interacting serine |
| AT1G30110 | -0.6 | 5.76E-05 | -0.4 | 1.99E-03 | -0.2 | 4.91E-02 | -0.2 | 2.01E-01 | -0.1 | 8.19E-01 | -0.2 | 4.71E-01 | Nudix hydrolase 25 -tetraphosphatase ) |
| AT1G74940 | -0.6 | 9.71E-06 | -0.3 | 2.23E-03 | -0.3 | 3.00E-03 | -0.6 | 1.04E-03 | -0.3 | 6.13E-02 | -0.3 | 1.84E-01 | FCS-Like Zinc finger 13 |
| AT1G12640 | -0.6 | 3.07E-06 | -0.5 | 5.18E-05 | -0.2 | 3.48E-02 | -0.6 | 8.50E-04 | -0.1 | 5.76E-01 | -0.5 | 2.81E-02 | Lysophospholipid acyltransferase 1 |
| AT3G12110 | -0.6 | 3.35E-05 | 0.0 | 9.57E-01 | -0.6 | 1.77E-04 | -0.7 | 5.81E-05 | -0.1 | 7.08E-01 | -0.7 | 2.81E-03 | Actin-11 |
| AT1G08670 | -0.6 | 1.73E-03 | 0.3 | 3.71E-02 | -1.0 | 6.21E-04 | 0.6 | 2.43E-01 | 0.6 | 2.98E-01 | 0.0 | 9.99E-01 | ENTH |
| AT3G08690 | -0.6 | 1.11E-05 | -0.7 | 1.70E-05 | 0.0 | 8.02E-01 | 0.0 | 9.55E-01 | -0.1 | 8.20E-01 | 0.1 | 8.94E-01 | Ubiquitin-conjugating enzyme E2 11 |
| AT5G62000 | -0.6 | 3.22E-05 | -0.7 | 4.46E-05 | 0.0 | 7.36E-01 | -0.1 | 4.64E-01 | -0.1 | 6.73E-01 | 0.0 | 9.22E-01 | Auxin response factor |
| AT3G06130 | -0.6 | 4.02E-05 | -0.2 | 9.37E-02 | -0.5 | 1.18E-03 | -0.6 | 3.10E-03 | -0.2 | 4.50E-01 | -0.4 | 8.08E-02 | Heavy metal transport |
| AT4G33950 | -0.6 | 3.76E-04 | -0.6 | 1.07E-03 | -0.1 | 6.97E-01 | -0.5 | 1.27E-02 | -0.4 | 9.47E-02 | -0.2 | 5.27E-01 | Protein kinase superfamily protein |
| AT5G08130 | -0.6 | 1.21E-05 | -0.3 | 5.46E-03 | -0.4 | 1.92E-03 | 0.1 | 6.88E-01 | -0.1 | 8.40E-01 | 0.1 | 6.40E-01 | Basic helix-loop-helix DNA-binding superfamily protein |
| AT4G02640 | -0.6 | 3.05E-06 | -0.5 | 1.76E-05 | -0.1 | 2.32E-01 | -0.1 | 3.96E-01 | 0.0 | 9.15E-01 | -0.2 | 4.15E-01 | hypothetical protein |
| AT1G11545 | -0.6 | 7.36E-05 | 0.1 | 1.59E-01 | -0.8 | 1.08E-04 | -0.4 | 6.57E-03 | -0.2 | 3.61E-01 | -0.3 | 1.62E-01 | Probable xyloglucan endotransglucosylase |
| AT3G23340 | -0.6 | 8.62E-04 | -0.6 | 2.60E-03 | -0.1 | 6.77E-01 | -0.2 | 4.19E-01 | 0.0 | 9.37E-01 | -0.2 | 4.41E-01 | Putative casein kinase I |
| AT4G30270 | -0.6 | 6.44E-02 | -0.6 | 1.03E-01 | -0.1 | 8.75E-01 | -0.4 | 5.78E-02 | -0.7 | 6.31E-03 | 0.3 | 2.87E-01 | Xyloglucan endotransglucosylase |
| AT5G54800 | -0.6 | 2.48E-05 | -0.1 | 2.24E-01 | -0.5 | 4.09E-04 | 0.2 | 1.71E-01 | 0.2 | 1.90E-01 | 0.0 | 9.38E-01 | Glucose-6-phosphate |
| AT2G36880 | -0.6 | 3.10E-05 | -0.3 | 9.52E-03 | -0.3 | 4.69E-03 | 0.0 | 9.83E-01 | 0.3 | 3.70E-02 | -0.3 | 1.10E-01 | S-adenosylmethionine synthase 3 |
| AT3G24810 | -0.6 | 3.17E-04 | -0.2 | 1.17E-01 | -0.4 | 1.24E-02 | 0.3 | 3.95E-01 | 0.3 | 3.77E-01 | 0.0 | 9.37E-01 | Cyclin-dependent kinase inhibitor 5 |
| AT3G50650 | -0.6 | 7.33E-05 | -0.4 | 1.44E-03 | -0.2 | 9.79E-02 | -0.2 | 2.35E-01 | -0.3 | 5.18E-02 | 0.1 | 4.54E-01 | Scarecrow-like protein 7 |
| AT1G06850 | -0.6 | 1.52E-05 | -0.1 | 9.07E-02 | -0.5 | 5.85E-04 | -0.4 | 4.12E-02 | -0.1 | 5.79E-01 | -0.3 | 2.88E-01 | F4H5.7 protein |
| AT3G54030 | -0.6 | 1.49E-05 | -0.4 | 1.02E-03 | -0.3 | 1.29E-02 | -0.3 | 1.29E-01 | 0.2 | 3.48E-01 | -0.5 | 4.40E-02 | Serine |
| AT1G10090 | -0.6 | 3.70E-04 | -0.4 | 7.63E-03 | -0.2 | 1.21E-01 | -0.2 | 4.60E-01 | 0.1 | 6.19E-01 | -0.3 | 3.06E-01 | CSC1-like protein At1g10090 |
| AT4G23010 | -0.6 | 1.60E-05 | -0.5 | 1.46E-04 | -0.1 | 1.59E-01 | 0.0 | 9.57E-01 | 0.1 | 7.80E-01 | -0.1 | 8.54E-01 | Uncharacterized protein |
| AT3G54390 | -0.6 | 6.35E-04 | -0.5 | 2.05E-03 | -0.1 | 6.46E-01 | -0.2 | 3.13E-01 | 0.0 | 9.48E-01 | -0.2 | 4.83E-01 | At3g54390 |
| AT3G57650 | -0.6 | 1.37E-05 | -0.3 | 4.36E-03 | -0.3 | 2.70E-03 | 0.0 | 8.04E-01 | 0.2 | 2.03E-01 | -0.2 | 4.06E-01 | DNA-directed RNA polymerase subunit |
| AT1G18860 | -0.6 | 4.85E-04 | -0.5 | 1.73E-03 | -0.1 | 5.97E-01 | 0.3 | 5.20E-01 | 0.4 | 3.49E-01 | -0.2 | 7.82E-01 | WRKY DNA-binding protein 61 |
| AT2G42490 | -0.6 | 1.07E-05 | -0.3 | 3.51E-03 | -0.3 | 2.41E-03 | -0.3 | 4.33E-02 | 0.0 | 9.68E-01 | -0.3 | 1.21E-01 | Amine oxidase zeta, peroxisomal |
| AT3G26380 | -0.6 | 1.29E-05 | -0.5 | 1.66E-04 | -0.2 | 1.07E-01 | -0.2 | 3.06E-01 | 0.2 | 4.43E-01 | -0.4 | 6.45E-02 | Alpha-galactosidase |
| AT2G16740 | -0.6 | 6.88E-05 | -0.3 | 4.27E-03 | -0.3 | 2.71E-02 | 0.1 | 4.93E-01 | 0.1 | 6.01E-01 | 0.0 | 9.78E-01 | Ubiquitin-conjugating enzyme E2 29 |
| AT4G27745 | -0.6 | 3.92E-03 | -0.6 | 7.15E-03 | 0.0 | 8.65E-01 | -0.6 | 4.68E-03 | -0.4 | 6.24E-02 | -0.2 | 4.85E-01 | Protein yippee-like At4g27745 |
| AT4G05390 | -0.6 | 4.97E-03 | -0.3 | 1.59E-01 | -0.4 | 1.09E-01 | 0.4 | 5.24E-03 | 0.0 | 8.94E-01 | 0.4 | 2.18E-02 | Ferredoxin--NADP reductase, chloroplastic |
| AT5G62340 | -0.6 | 1.63E-02 | 0.6 | 1.86E-02 | -1.2 | 1.06E-03 | -0.1 | 5.61E-01 | 0.2 | 4.57E-01 | -0.3 | 2.71E-01 | hypothetical protein |
| AT2G26760 | -0.6 | 3.11E-04 | 0.0 | 8.80E-01 | -0.6 | 1.73E-03 | -0.1 | 6.80E-01 | 0.3 | 5.77E-02 | -0.4 | 5.93E-02 | Cyclin-B1-4 |
| AT1G74410 | -0.6 | 1.13E-04 | -0.5 | 1.28E-03 | -0.2 | 1.95E-01 | -0.4 | 1.24E-01 | 0.0 | 9.18E-01 | -0.4 | 2.30E-01 | NEP1-interacting protein-like 2 |
| AT1G63650 | -0.6 | 1.56E-03 | -0.1 | 6.54E-01 | -0.6 | 1.10E-02 | -0.1 | 8.10E-01 | 0.4 | 2.99E-01 | -0.4 | 2.86E-01 | Transcription factor EGL1 |
| AT4G02630 | -0.6 | 4.75E-06 | -0.6 | 9.20E-06 | 0.0 | 8.81E-01 | -0.6 | 1.24E-04 | -0.1 | 3.80E-01 | -0.5 | 1.04E-02 | Protein kinase domain-containing protein |
| AT1G68710 | -0.6 | 2.06E-05 | -0.4 | 3.98E-04 | -0.2 | 6.25E-02 | -0.2 | 2.06E-01 | 0.3 | 1.34E-01 | -0.6 | 2.93E-02 | Phospholipid-transporting ATPase |
| AT2G04350 | -0.6 | 1.04E-05 | -0.8 | 5.27E-06 | 0.1 | 8.77E-02 | -0.3 | 1.93E-01 | 0.1 | 5.50E-01 | -0.4 | 1.48E-01 | Long chain acyl-CoA synthetase 8 |
| AT1G54340 | -0.6 | 1.55E-06 | -0.5 | 9.53E-06 | -0.1 | 1.99E-01 | -0.1 | 4.09E-01 | 0.0 | 9.88E-01 | -0.1 | 6.04E-01 | NADP specific isocitrate dehydrogenase |
| AT1G05380 | -0.6 | 4.74E-06 | -0.4 | 1.55E-04 | -0.2 | 1.81E-02 | 0.1 | 6.23E-01 | 0.0 | 9.76E-01 | 0.1 | 7.54E-01 | Acyl-CoA N-acyltransferase with RING |
| AT1G24280 | -0.6 | 2.07E-02 | -0.3 | 1.99E-01 | -0.3 | 2.94E-01 | 0.0 | 8.47E-01 | -0.2 | 2.02E-01 | 0.2 | 3.92E-01 | Glucose-6-phosphate 1-dehydrogenase 3, chloroplastic |
| AT3G08850 | -0.6 | 8.33E-05 | -0.6 | 1.67E-04 | 0.0 | 9.20E-01 | -0.3 | 2.06E-01 | -0.1 | 7.67E-01 | -0.2 | 5.84E-01 | Regulatory-associated protein of TOR 1 |
| AT5G56040 | -0.6 | 1.23E-05 | -0.3 | 1.63E-03 | -0.3 | 5.38E-03 | -0.4 | 7.32E-04 | -0.2 | 1.49E-01 | -0.3 | 1.28E-01 | Leucine-rich repeat receptor-like serine |
| AT1G69270 | -0.6 | 3.70E-02 | -0.5 | 8.80E-02 | -0.1 | 7.31E-01 | -0.2 | 5.05E-01 | -0.4 | 2.11E-01 | 0.2 | 5.81E-01 | Probable LRR receptor-like serine |
| AT1G76670 | -0.6 | 3.21E-05 | -0.5 | 1.71E-04 | -0.1 | 3.86E-01 | -0.4 | 7.71E-04 | 0.0 | 7.31E-01 | -0.3 | 2.72E-02 | UDP-rhamnose |
| AT1G47330 | -0.6 | 1.75E-05 | -0.4 | 3.30E-04 | -0.2 | 6.83E-02 | -0.1 | 5.66E-01 | 0.0 | 9.50E-01 | -0.1 | 7.62E-01 | Methyltransferase, putative |
| AT5G04720 | -0.6 | 1.04E-04 | -0.9 | 1.18E-05 | 0.3 | 1.71E-02 | -0.1 | 6.96E-01 | -0.4 | 1.36E-02 | 0.3 | 1.01E-01 | Probable disease resistance protein At5g04720 |
| AT3G57410 | -0.6 | 6.26E-05 | -0.4 | 1.10E-03 | -0.2 | 9.98E-02 | -0.4 | 5.66E-02 | -0.4 | 1.05E-01 | 0.0 | 9.68E-01 | Uncharacterized protein At3g57410 |
| AT4G02410 | -0.6 | 1.82E-05 | -0.7 | 1.29E-05 | 0.1 | 2.73E-01 | 0.0 | 8.39E-01 | -0.2 | 3.06E-01 | 0.2 | 4.73E-01 | L-type lectin-domain containing receptor kinase IV.3 |
| AT5G17450 | -0.6 | 1.49E-02 | -0.4 | 6.60E-02 | -0.2 | 5.19E-01 | -0.2 | 5.56E-01 | 0.1 | 7.80E-01 | -0.3 | 4.23E-01 | HIPP21 |
| AT3G53000 | -0.6 | 5.78E-05 | -0.4 | 9.54E-04 | -0.2 | 1.14E-01 | -0.2 | 4.81E-01 | -0.2 | 4.52E-01 | 0.0 | 9.31E-01 | F-box protein PP2-A15 |
| AT2G45430 | -0.6 | 2.31E-05 | 0.0 | 8.22E-01 | -0.6 | 1.76E-04 | -0.1 | 6.75E-01 | 0.2 | 3.49E-01 | -0.3 | 2.41E-01 | AT-hook motif nuclear-localized protein 22 |
| AT2G27200 | -0.6 | 4.33E-05 | -0.4 | 1.18E-03 | -0.2 | 6.03E-02 | -0.2 | 2.86E-01 | -0.2 | 4.55E-01 | 0.0 | 9.16E-01 | GTPase LSG1-1 |
| AT3G15260 | -0.6 | 3.73E-05 | -0.3 | 2.93E-03 | -0.3 | 1.74E-02 | 0.1 | 6.02E-01 | -0.2 | 5.38E-01 | 0.3 | 3.54E-01 | Probable protein phosphatase 2C 39 |
| AT2G18170 | -0.6 | 1.32E-04 | -0.4 | 2.15E-03 | -0.2 | 1.32E-01 | 0.1 | 3.16E-01 | 0.1 | 4.25E-01 | 0.0 | 9.75E-01 | Mitogen-activated protein kinase 7 |
| AT5G47770 | -0.6 | 3.08E-05 | 0.0 | 7.35E-01 | -0.6 | 1.27E-04 | -0.4 | 1.85E-03 | -0.1 | 5.01E-01 | -0.3 | 7.29E-02 | Farnesyl pyrophosphate synthase 1, mitochondrial -farnesyl diphosphate synthase 1) |
| AT1G66500 | -0.6 | 1.67E-02 | -1.0 | 2.70E-03 | 0.3 | 2.77E-01 | 0.0 | 9.89E-01 | -0.3 | 6.32E-01 | 0.3 | 7.11E-01 | Polyadenylation and cleavage factor homolog 1 |
| AT3G15210 | -0.6 | 1.62E-05 | -0.4 | 6.28E-04 | -0.2 | 2.40E-02 | 0.2 | 2.07E-01 | 0.0 | 8.74E-01 | 0.2 | 2.75E-01 | Ethylene-responsive transcription factor 4 |
| AT2G40820 | -0.6 | 2.01E-04 | -0.2 | 6.89E-02 | -0.4 | 9.34E-03 | -0.2 | 1.76E-01 | 0.1 | 6.55E-01 | -0.3 | 1.19E-01 | Stomatal closure actin-binding-like protein |
| AT5G03530 | -0.6 | 2.05E-03 | -0.3 | 4.39E-02 | -0.3 | 1.58E-01 | -0.5 | 4.22E-02 | -0.1 | 7.38E-01 | -0.4 | 1.99E-01 | Ras-related protein RABC2a |
| AT3G07930 | -0.6 | 5.26E-05 | -0.5 | 2.18E-04 | -0.1 | 5.19E-01 | -0.3 | 1.63E-01 | -0.1 | 6.96E-01 | -0.2 | 5.51E-01 | Methyl-CpG-binding domain protein 4-like protein |
| AT5G47200 | -0.6 | 3.62E-05 | -0.3 | 9.23E-03 | -0.3 | 5.62E-03 | -0.2 | 8.27E-02 | 0.0 | 9.10E-01 | -0.2 | 1.91E-01 | Ras-related protein RABD2b |
| AT5G29000 | -0.6 | 1.14E-04 | -0.6 | 1.81E-04 | 0.0 | 9.63E-01 | -0.2 | 2.64E-01 | -0.1 | 6.47E-01 | -0.1 | 7.41E-01 | PHL1 |
| AT3G04030 | -0.6 | 7.18E-04 | -0.5 | 5.18E-03 | -0.2 | 3.55E-01 | 0.2 | 3.40E-01 | -0.2 | 6.57E-01 | 0.4 | 2.25E-01 | Homeodomain-like superfamily protein |

| | | | | | | | | | | | | | |
|---|---|---|---|---|---|---|---|---|---|---|---|---|---|
| AT2G38940 | -0.6 | 8.20E-05 | 0.0 | 6.76E-01 | -0.7 | 3.16E-04 | 0.3 | 3.11E-01 | 0.4 | 1.84E-01 | -0.1 | 7.86E-01 | Phosphate transporter |
| AT3G11670 | -0.6 | 3.31E-04 | -0.8 | 8.03E-05 | 0.2 | 1.39E-01 | 0.5 | 2.34E-02 | 0.7 | 2.04E-03 | -0.3 | 2.59E-01 | Digalactosyldiacylglycerol synthase 1, chloroplastic |
| AT2G03480 | -0.6 | 2.67E-04 | -0.3 | 3.34E-02 | -0.3 | 2.49E-02 | -0.1 | 5.92E-01 | 0.1 | 5.88E-01 | -0.3 | 2.87E-01 | Methyltransferase |
| AT1G19360 | -0.6 | 2.14E-05 | -0.2 | 3.15E-02 | -0.4 | 1.34E-03 | 0.0 | 9.57E-01 | 0.2 | 7.52E-01 | -0.2 | 7.46E-01 | Arabinosyltransferase RRA3 |
| AT3G46130 | -0.6 | 3.22E-04 | -0.6 | 4.49E-04 | 0.0 | 9.38E-01 | -1.3 | 4.72E-06 | -1.0 | 2.11E-04 | -0.4 | 2.20E-01 | Myb domain protein 48 |
| AT3G47730 | -0.6 | 5.16E-05 | -0.6 | 8.47E-05 | 0.0 | 9.47E-01 | 0.7 | 1.69E-02 | 0.4 | 3.16E-01 | 0.4 | 3.59E-01 | ABC transporter A family member 2 |
| AT1G27660 | -0.6 | 1.94E-04 | -0.2 | 9.16E-02 | -0.4 | 8.48E-03 | -0.3 | 1.35E-01 | 0.0 | 9.44E-01 | -0.3 | 2.44E-01 | Transcription factor bHLH110 |
| AT1G74500 | -0.6 | 7.41E-03 | 0.5 | 1.31E-02 | -1.1 | 7.30E-04 | -0.1 | 8.33E-01 | 0.5 | 1.08E-01 | -0.6 | 7.90E-02 | Transcription factor PRE3 |
| AT2G37170 | -0.6 | 1.20E-04 | 0.2 | 1.72E-01 | -0.8 | 1.39E-04 | -1.0 | 3.21E-07 | -0.2 | 1.27E-01 | -0.8 | 1.83E-04 | Plasma membrane intrinsic protein 2 |
| AT1G20850 | -0.6 | 1.29E-05 | 0.2 | 1.14E-02 | -0.8 | 2.37E-05 | -0.2 | 1.03E-01 | -0.1 | 5.10E-01 | -0.1 | 5.12E-01 | Cysteine protease XCP2 |
| AT1G64970 | -0.6 | 1.60E-04 | -0.6 | 2.03E-04 | 0.0 | 8.64E-01 | -0.6 | 3.91E-03 | 0.1 | 6.64E-01 | -0.7 | 1.11E-02 | Tocopherol O-methyltransferase, chloroplastic |
| AT1G32740 | -0.6 | 6.63E-04 | -0.4 | 6.77E-03 | -0.2 | 2.70E-01 | -1.0 | 4.88E-03 | -0.4 | 2.58E-01 | -0.6 | 1.09E-01 | SBP family protein |
| AT5G41700 | -0.6 | 3.46E-05 | -0.4 | 5.52E-04 | -0.2 | 1.05E-01 | -0.8 | 1.89E-05 | -0.5 | 1.57E-03 | -0.2 | 2.54E-01 | AT5G41700 protein |
| AT1G76340 | -0.6 | 1.23E-04 | -0.3 | 8.66E-03 | -0.3 | 3.10E-02 | -0.2 | 1.64E-01 | 0.1 | 6.44E-01 | -0.4 | 1.04E-01 | GDP-mannose transporter GONST3 |
| AT3G43230 | -0.6 | 4.74E-06 | -0.6 | 6.90E-06 | 0.0 | 6.43E-01 | -0.3 | 2.22E-02 | -0.1 | 5.41E-01 | -0.2 | 2.46E-01 | FYVE-type domain-containing protein |
| AT5G04170 | -0.6 | 1.25E-05 | -0.3 | 4.69E-03 | -0.3 | 2.30E-03 | 0.0 | 7.65E-01 | 0.0 | 8.59E-01 | 0.0 | 9.70E-01 | Probable calcium-binding protein CML50 |
| AT1G55020 | -0.6 | 1.38E-05 | -0.4 | 3.79E-04 | -0.2 | 3.36E-02 | 1.6 | 2.68E-08 | 0.7 | 9.82E-04 | 1.0 | 1.65E-04 | Linoleate 9S-lipoxygenase 1 |
| AT3G13060 | -0.6 | 2.64E-04 | -0.6 | 3.47E-04 | 0.0 | 9.13E-01 | -0.2 | 8.01E-01 | 0.1 | 8.95E-01 | -0.3 | 7.65E-01 | YTH domain-containing family protein |
| AT4G32040 | -0.6 | 1.01E-05 | -0.6 | 1.98E-05 | 0.0 | 9.97E-01 | -0.2 | 3.52E-02 | -0.2 | 3.31E-02 | 0.0 | 8.88E-01 | Homeobox protein knotted-1-like 5 |
| AT3G56310 | -0.6 | 1.12E-05 | -0.4 | 2.93E-04 | -0.2 | 3.55E-02 | -0.2 | 7.24E-02 | 0.0 | 8.72E-01 | -0.2 | 3.12E-01 | Alpha-galactosidase |
| AT5G41120 | -0.6 | 5.73E-04 | -0.5 | 2.11E-03 | -0.1 | 6.10E-01 | 0.2 | 5.27E-01 | 0.6 | 8.95E-02 | -0.4 | 3.79E-01 | Esterase |
| AT1G07810 | -0.6 | 3.49E-05 | -0.5 | 1.79E-04 | -0.1 | 3.75E-01 | 0.0 | 9.37E-01 | -0.1 | 4.89E-01 | 0.2 | 4.90E-01 | Calcium-transporting ATPase 1, endoplasmic reticulum-type |
| AT2G21940 | -0.6 | 6.00E-05 | -0.6 | 1.30E-04 | 0.0 | 8.96E-01 | -0.1 | 6.76E-01 | 0.1 | 5.85E-01 | -0.2 | 3.83E-01 | shikimate kinase |
| AT1G80900 | -0.6 | 2.81E-05 | -0.3 | 2.67E-03 | -0.3 | 1.52E-02 | -0.3 | 1.03E-01 | 0.0 | 9.23E-01 | -0.3 | 1.82E-01 | Magnesium transporter MRS2-10 |
| AT4G33240 | -0.6 | 7.64E-05 | -0.5 | 3.95E-04 | -0.1 | 3.92E-01 | -0.3 | 1.25E-01 | 0.0 | 9.32E-01 | -0.3 | 1.94E-01 | 1-phosphatidylinositol-3-phosphate 5-kinase |
| AT4G21980 | -0.6 | 1.44E-04 | -0.6 | 2.17E-04 | 0.0 | 9.54E-01 | 0.0 | 9.10E-01 | 0.1 | 8.12E-01 | 0.0 | 9.11E-01 | Autophagy-related protein |
| AT3G51960 | -0.6 | 5.63E-03 | -0.3 | 1.26E-01 | -0.3 | 1.68E-01 | -1.0 | 1.41E-02 | -0.5 | 2.09E-01 | -0.5 | 3.25E-01 | Basic leucine zipper 24 |
| AT5G45440 | -0.6 | 2.39E-04 | 0.1 | 6.10E-01 | -0.7 | 7.83E-04 | -0.4 | 1.75E-01 | -0.2 | 5.39E-01 | -0.2 | 6.52E-01 | Probable disease resistance protein At5g45440 |
| AT5G05320 | -0.6 | 1.73E-03 | -0.5 | 9.54E-03 | -0.1 | 4.45E-01 | -0.9 | 7.61E-03 | -1.3 | 1.86E-03 | 0.4 | 5.05E-01 | Monooxygenase 3 |
| AT4G26830 | -0.6 | 4.66E-03 | 0.3 | 5.24E-02 | -0.9 | 1.39E-03 | 0.2 | 7.97E-01 | -0.7 | 2.88E-01 | | | glucan endo-1,3-beta-D-glucosidase |
| AT5G63990 | -0.6 | 2.57E-05 | -0.2 | 7.87E-02 | -0.4 | 9.66E-04 | 0.2 | 2.51E-01 | 0.6 | 3.27E-03 | -0.4 | 6.62E-02 | 3',5'-bisphosphate nucleotidase |
| AT1G10120 | -0.6 | 1.63E-05 | -0.3 | 5.97E-03 | -0.3 | 3.35E-03 | -0.3 | 2.39E-02 | 0.0 | 8.52E-01 | -0.3 | 9.22E-02 | Transcription factor bHLH74 |
| AT2G03730 | -0.6 | 1.01E-05 | -0.5 | 5.49E-05 | -0.1 | 3.04E-01 | -0.5 | 4.85E-03 | -0.2 | 3.73E-01 | -0.4 | 1.56E-01 | ACT domain-containing protein ACR5 |
| AT5G34940 | -0.6 | 2.25E-04 | 0.0 | 9.86E-01 | -0.6 | 1.17E-03 | 0.2 | 4.87E-01 | 0.3 | 2.16E-01 | -0.1 | 5.76E-01 | Glucuronidase 3 |
| AT4G18880 | -0.6 | 1.57E-03 | -0.9 | 1.52E-04 | 0.3 | 7.43E-02 | 0.0 | 9.77E-01 | -0.2 | 4.41E-01 | 0.2 | 4.57E-01 | Heat stress transcription factor A-4a |
| AT1G35190 | -0.6 | 9.84E-06 | -0.6 | 1.64E-05 | 0.0 | 8.59E-01 | 0.2 | 1.49E-01 | 0.2 | 2.30E-01 | 0.0 | 9.73E-01 | 2-oxoglutarate and Fe-dependent oxygenase superfamily protein |
| AT4G31860 | -0.6 | 6.33E-06 | -0.7 | 7.57E-06 | 0.1 | 5.01E-01 | -0.3 | 8.29E-02 | -0.2 | 3.09E-01 | -0.1 | 7.45E-01 | protein-serine |
| AT5G64250 | -0.6 | 3.68E-06 | -0.5 | 2.01E-05 | -0.1 | 2.68E-01 | 0.2 | 1.80E-01 | 0.0 | 9.08E-01 | 0.2 | 2.62E-01 | 2-nitropropane dioxygenase-like protein |
| AT3G62700 | -0.6 | 7.22E-05 | -0.6 | 1.55E-04 | 0.0 | 9.05E-01 | -0.1 | 6.26E-01 | 0.1 | 7.00E-01 | -0.2 | 4.29E-01 | ABC transporter C family member 14 |
| AT4G13290 | -0.6 | 6.16E-04 | -0.4 | 8.09E-03 | -0.2 | 2.22E-01 | 0.2 | 6.42E-01 | 0.9 | 6.40E-03 | -0.7 | 2.79E-02 | Cytochrome P450 71A19 |
| AT1G07560 | -0.6 | 7.27E-04 | -0.4 | 1.60E-02 | -0.2 | 1.40E-01 | -0.1 | 9.06E-01 | 0.3 | 5.01E-01 | -0.3 | 4.72E-01 | Probable LRR receptor-like serine |
| AT1G75540 | -0.6 | 1.55E-05 | -0.2 | 3.91E-02 | -0.4 | 7.08E-04 | -0.2 | 1.24E-01 | -0.2 | 1.96E-01 | 0.0 | 9.68E-01 | B-box zinc finger protein 21 |
| AT2G34790 | -0.6 | 3.18E-04 | -0.1 | 2.07E-01 | -0.5 | 7.52E-03 | -0.1 | 7.96E-01 | -0.5 | 3.33E-01 | 0.3 | 5.76E-01 | Monolignol oxidoreductase AtBBE-like 15 |
| AT3G52250 | -0.6 | 2.08E-04 | -0.5 | 5.63E-04 | 0.0 | 7.14E-01 | -0.1 | 7.95E-01 | -0.1 | 7.95E-01 | 0.0 | 9.77E-01 | Duplicated homeodomain-like superfamily protein |
| AT4G11370 | -0.6 | 2.02E-02 | -0.6 | 1.69E-02 | 0.0 | 8.97E-01 | 0.1 | 8.43E-01 | 0.3 | 6.90E-01 | -0.1 | 8.56E-01 | Probable E3 ubiquitin-protein ligase RHA1A |
| AT1G72310 | -0.6 | 4.77E-05 | -0.1 | 3.73E-01 | -0.5 | 7.02E-04 | 0.0 | 8.87E-01 | 0.0 | 9.66E-01 | 0.0 | 8.78E-01 | RING-H2 finger protein ATL3 |
| AT1G71692 | -0.6 | 4.33E-03 | -0.8 | 1.24E-03 | 0.2 | 3.82E-01 | 0.0 | 9.59E-01 | 0.4 | 2.49E-01 | -0.4 | 2.17E-01 | Agamous-like MADS-box protein AGL12 |
| AT2G37150 | -0.6 | 1.99E-05 | -0.6 | 4.43E-05 | 0.0 | 9.15E-01 | -0.5 | 1.16E-02 | -0.2 | 3.43E-01 | -0.3 | 2.28E-01 | RING-type E3 ubiquitin transferase |
| AT2G17650 | -0.6 | 7.73E-05 | -0.5 | 5.01E-04 | -0.1 | 3.36E-01 | 0.0 | 9.35E-01 | 0.1 | 7.08E-01 | -0.1 | 6.83E-01 | Isovalerate--CoA ligase AAE2 |
| AT2G22125 | -0.6 | 2.03E-04 | -0.5 | 1.30E-03 | -0.1 | 3.51E-01 | -0.3 | 1.41E-01 | -0.2 | 3.53E-01 | -0.1 | 8.39E-01 | Protein CELLULOSE SYNTHASE INTERACTIVE 1 |
| AT5G01800 | -0.6 | 1.09E-05 | -0.4 | 6.33E-04 | -0.2 | 1.28E-02 | -0.3 | 5.18E-02 | -0.1 | 7.34E-01 | -0.2 | 3.03E-01 | Saposin B-type domain-containing protein |
| AT5G18200 | -0.6 | 2.99E-05 | -0.4 | 3.50E-04 | -0.1 | 1.52E-01 | 0.1 | 6.44E-01 | 0.1 | 4.30E-01 | -0.1 | 8.07E-01 | ADP-glucose phosphorylase |
| AT3G03310 | -0.6 | 4.10E-05 | -0.6 | 6.01E-05 | 0.0 | 8.39E-01 | -0.3 | 6.14E-02 | -0.2 | 2.65E-01 | -0.1 | 6.99E-01 | Phospholipase A LCAT3 |
| AT2G35050 | -0.6 | 1.38E-04 | -0.6 | 2.11E-04 | 0.0 | 9.16E-01 | -0.3 | 1.08E-01 | -0.2 | 2.65E-01 | -0.1 | 8.78E-01 | Kinase superfamily with octicosapeptide |
| AT4G37250 | -0.6 | 4.38E-05 | -0.2 | 9.40E-02 | -0.4 | 1.47E-03 | -0.6 | 1.28E-04 | -0.4 | 3.09E-03 | -0.1 | 4.31E-01 | Probable LRR receptor-like serine |
| AT1G35560 | -0.6 | 1.82E-02 | -0.9 | 3.51E-03 | 0.3 | 3.24E-01 | 0.1 | 6.59E-01 | 0.1 | 4.69E-01 | -0.1 | 8.33E-01 | Transcription factor TCP23 |
| AT4G29380 | -0.6 | 2.92E-05 | -0.7 | 2.58E-05 | 0.1 | 4.62E-01 | -0.4 | 1.49E-01 | -0.3 | 2.90E-01 | -0.1 | 9.01E-01 | non-specific serine |
| AT1G02810 | -0.6 | 2.16E-03 | 0.0 | 9.50E-01 | -0.6 | 6.56E-03 | 0.1 | 5.97E-01 | 0.3 | 2.63E-01 | -0.2 | 5.85E-01 | Probable pectinesterase |
| AT1G12550 | -0.6 | 7.69E-05 | -0.5 | 4.82E-04 | -0.1 | 3.46E-01 | -0.3 | 1.53E-01 | 0.0 | 9.60E-01 | -0.3 | 2.25E-01 | Glyoxylate |
| AT5G25880 | -0.6 | 6.22E-03 | 0.3 | 1.16E-01 | -0.9 | 2.65E-03 | -0.5 | 2.91E-01 | 0.0 | 9.58E-01 | -0.4 | 4.89E-01 | Malic enzyme |
| AT1G07670 | -0.6 | 1.43E-05 | -0.2 | 9.22E-03 | -0.4 | 1.66E-03 | 0.1 | 6.36E-01 | 0.3 | 1.04E-01 | -0.2 | 3.84E-01 | Calcium-transporting ATPase 4, endoplasmic reticulum-type |
| AT2G17270 | -0.6 | 8.41E-04 | -0.3 | 1.92E-02 | -0.2 | 1.37E-01 | 0.3 | 3.54E-01 | 0.2 | 4.73E-01 | 0.0 | 9.54E-01 | Mitochondrial phosphate carrier protein 1, mitochondrial |
| AT5G04740 | -0.6 | 1.03E-05 | -0.2 | 3.38E-02 | -0.4 | 4.92E-04 | 0.1 | 4.94E-01 | 0.3 | 1.05E-02 | -0.2 | 1.00E-01 | ACT domain-containing protein ACR |
| AT4G29270 | -0.6 | 1.96E-03 | -0.1 | 4.69E-01 | -0.5 | 1.56E-02 | 0.0 | 9.85E-01 | 0.4 | 8.26E-02 | -0.4 | 1.06E-01 | Acid phosphatase |
| AT5G18830 | -0.6 | 4.47E-05 | -0.5 | 1.61E-04 | -0.1 | 5.67E-01 | -0.2 | 2.54E-01 | 0.1 | 5.03E-01 | -0.3 | 1.35E-01 | Squamosa promoter binding protein-like 7 |
| AT5G06720 | -0.6 | 2.06E-02 | -1.2 | 1.08E-03 | 0.6 | 8.34E-02 | | | | | | | Peroxidase 53 |
| AT1G12160 | -0.6 | 7.25E-05 | -0.1 | 3.05E-01 | -0.5 | 1.27E-03 | -0.5 | 1.53E-01 | 0.0 | 9.96E-01 | -0.5 | 1.66E-01 | Flavin-containing monooxygenase FMO GS-OX-like 1 |
| AT3G20960 | -0.6 | 2.34E-04 | -0.3 | 2.22E-02 | -0.3 | 3.00E-02 | -0.3 | 1.67E-01 | 0.0 | 9.55E-01 | -0.3 | 3.72E-01 | Cytochrome P450, family 705, subfamily A, polypeptide 33 |
| AT1G74800 | -0.6 | 3.43E-05 | -0.3 | 3.47E-03 | -0.3 | 1.41E-02 | -0.1 | 7.85E-01 | 0.2 | 5.80E-01 | -0.2 | 4.73E-01 | Hydroxyproline O-galactosyltransferase GALT5 |
| AT3G54720 | -0.6 | 1.73E-04 | -0.3 | 2.87E-02 | -0.3 | 1.60E-02 | -0.1 | 8.85E-01 | 0.3 | 4.04E-01 | -0.3 | 2.35E-01 | Probable glutamate carboxypeptidase AMP1 |
| AT5G41870 | -0.6 | 1.07E-03 | -0.6 | 1.36E-03 | 0.0 | 9.23E-01 | -0.2 | 7.57E-01 | -0.3 | 7.64E-01 | 0.0 | 9.80E-01 | At5g41870 |
| AT3G62690 | -0.6 | 4.57E-04 | -0.4 | 8.31E-03 | -0.2 | 1.53E-01 | -0.1 | 8.16E-01 | 0.3 | 3.67E-01 | -0.4 | 2.71E-01 | RING-H2 finger protein ATL5 |
| AT3G25400 | -0.6 | 1.33E-03 | -0.4 | 1.01E-02 | -0.2 | 3.53E-01 | 0.1 | 7.54E-01 | 0.2 | 6.54E-01 | -0.1 | 9.02E-01 | dCTP pyrophosphatase 1 |
| AT1G12430 | -0.6 | 3.73E-05 | -0.5 | 2.85E-04 | -0.1 | 2.51E-01 | 0.1 | 6.16E-01 | 0.0 | 9.76E-01 | 0.1 | 7.65E-01 | Kinesin-like protein |
| AT1G07410 | -0.6 | 1.39E-02 | 0.0 | 9.14E-01 | -0.6 | 3.02E-02 | -0.2 | 6.26E-01 | 0.1 | 8.44E-01 | -0.3 | 5.66E-01 | Ras-related protein RABA2b |
| AT2G42790 | -0.6 | 2.06E-05 | -0.7 | 1.34E-05 | 0.1 | 2.48E-01 | -0.4 | 1.42E-02 | -0.4 | 2.14E-02 | 0.0 | 9.77E-01 | Citrate synthase 3, peroxisomal |
| AT1G16670 | -0.6 | 7.56E-04 | -0.5 | 1.74E-03 | 0.0 | 7.82E-01 | 0.4 | 1.58E-02 | 0.1 | 5.34E-01 | 0.3 | 1.59E-01 | Cold-responsive protein kinase 1 |
| AT2G41640 | -0.6 | 6.45E-03 | -0.9 | 8.42E-04 | 0.3 | 1.80E-01 | 0.0 | 9.22E-01 | 0.0 | 9.70E-01 | 0.0 | 9.71E-01 | Glycosyltransferase family 61 protein |
| AT5G15650 | -0.6 | 3.80E-04 | -0.1 | 5.71E-01 | -0.5 | 2.53E-03 | -0.2 | 1.96E-01 | 0.0 | 9.13E-01 | -0.2 | 3.09E-01 | UDP-arabinopyranose mutase 2 |
| AT3G05970 | -0.6 | 1.34E-05 | -0.5 | 1.52E-04 | -0.1 | 1.14E-01 | 0.1 | 2.98E-01 | 0.1 | 4.23E-01 | 0.0 | 9.72E-01 | Long chain acyl-CoA synthetase 6, peroxisomal |
| AT5G03610 | -0.6 | 2.16E-05 | -0.3 | 6.04E-03 | -0.3 | 3.95E-03 | 0.1 | 3.30E-01 | -0.1 | 7.87E-01 | 0.2 | 3.41E-01 | GDSL esterase |
| AT4G28490 | -0.6 | 1.16E-02 | -0.8 | 2.21E-03 | 0.2 | 3.01E-01 | 0.1 | 6.76E-01 | -0.2 | 3.10E-01 | 0.3 | 2.23E-01 | Receptor-like protein kinase 5 |
| AT4G08180 | -0.6 | 1.74E-05 | -0.5 | 1.71E-04 | -0.1 | 1.47E-01 | -0.1 | 6.95E-01 | 0.1 | 8.59E-01 | -0.1 | 6.73E-01 | hypothetical protein |

| | | | | | | | | | | | | | |
|---|---|---|---|---|---|---|---|---|---|---|---|---|---|
| AT5G63410 | -0.6 | 1.86E-05 | -0.5 | 1.12E-04 | -0.1 | 2.96E-01 | -0.2 | 1.98E-01 | 0.0 | 8.08E-01 | -0.1 | 4.98E-01 | Similarity to receptor-like protein kinase |
| AT5G12270 | -0.6 | 1.71E-03 | 0.1 | 3.47E-01 | -0.7 | 2.08E-03 | | | | | | | hypothetical protein |
| AT5G61250 | -0.6 | 4.42E-04 | -0.7 | 2.84E-04 | 0.1 | 5.48E-01 | -0.4 | 4.19E-02 | -0.3 | 1.79E-01 | -0.1 | 7.10E-01 | Heparanase-like protein 2 |
| AT4G12720 | -0.6 | 2.10E-03 | -0.8 | 5.23E-04 | 0.2 | 2.68E-01 | 0.5 | 1.23E-02 | 0.0 | 9.68E-01 | 0.5 | 6.83E-02 | Nudix hydrolase |
| AT3G05670 | -0.6 | 3.80E-06 | -0.4 | 4.99E-05 | -0.1 | 6.08E-02 | 0.1 | 6.01E-01 | 0.1 | 6.06E-01 | 0.0 | 9.62E-01 | hypothetical protein |
| AT4G31670 | -0.6 | 1.36E-04 | -0.5 | 1.04E-03 | -0.1 | 3.01E-01 | -0.2 | 4.43E-01 | 0.1 | 6.48E-01 | -0.3 | 3.27E-01 | Ubiquitin carboxyl-terminal hydrolase 18 |
| AT5G02600 | -0.6 | 2.39E-05 | -0.2 | 2.51E-02 | -0.4 | 2.03E-03 | -0.2 | 2.98E-01 | 0.0 | 9.07E-01 | -0.2 | 3.82E-01 | HMA domain-containing protein |
| AT5G57150 | -0.6 | 3.61E-04 | -0.5 | 2.37E-03 | -0.1 | 3.51E-01 | -0.3 | 1.34E-01 | 0.0 | 9.76E-01 | -0.3 | 2.63E-01 | Basic helix-loop-helix DNA-binding superfamily protein |
| AT1G78940 | -0.6 | 1.33E-03 | -0.1 | 7.13E-01 | -0.5 | 9.28E-03 | -0.2 | 7.23E-01 | -0.1 | 9.22E-01 | | | Kinase with adenine nucleotide alpha hydrolases-like domain-containing protein |
| AT3G14270 | -0.6 | 3.09E-05 | -0.7 | 1.70E-05 | 0.1 | 2.16E-01 | -0.3 | 3.58E-02 | 0.1 | 8.22E-01 | -0.4 | 7.79E-02 | 1-phosphatidylinositol-3-phosphate 5-kinase FAB1B |
| AT2G34300 | -0.6 | 2.02E-05 | -0.2 | 2.83E-02 | -0.4 | 1.18E-03 | -0.2 | 3.05E-01 | 0.2 | 2.11E-01 | -0.4 | 7.40E-02 | Probable methyltransferase PMT25 |
| AT4G39780 | -0.6 | 1.93E-03 | -0.4 | 2.26E-02 | -0.2 | 2.57E-01 | -0.4 | 6.94E-02 | -0.5 | 9.47E-02 | 0.0 | 9.97E-01 | Ethylene-responsive transcription factor ERF060 |
| AT4G26270 | -0.6 | 1.15E-03 | -0.8 | 2.70E-04 | 0.2 | 2.33E-01 | 0.8 | 3.44E-03 | 0.5 | 7.97E-02 | 0.3 | 3.09E-01 | ATP-dependent 6-phosphofructokinase 3 |
| AT4G21910 | -0.6 | 3.05E-04 | -0.4 | 3.56E-03 | -0.2 | 1.95E-01 | 0.1 | 6.90E-01 | 0.6 | 1.16E-02 | -0.5 | 8.36E-02 | Protein DETOXIFICATION |
| AT3G48970 | -0.6 | 3.65E-03 | 0.5 | 6.33E-03 | -1.0 | 3.79E-04 | -0.1 | 7.75E-01 | -0.1 | 8.98E-01 | -0.1 | 9.40E-01 | Heavy metal transport |
| AT1G11910 | -0.6 | 2.02E-05 | -0.5 | 1.20E-04 | -0.1 | 3.38E-01 | -0.3 | 1.19E-02 | -0.1 | 6.71E-01 | -0.2 | 1.75E-01 | Putative aspartic proteinase |
| AT1G02850 | -0.6 | 9.93E-02 | -3.1 | 3.17E-04 | 2.5 | 3.20E-03 | -0.1 | 8.85E-01 | -0.9 | 2.18E-02 | 0.8 | 6.26E-02 | Similar to beta-glucosidases |
| AT2G21060 | -0.6 | 4.57E-04 | -0.1 | 3.81E-01 | -0.5 | 4.45E-03 | -0.3 | 8.41E-03 | -0.3 | 5.55E-03 | 0.1 | 7.72E-01 | Cold shock domain-containing protein 4 |
| AT5G12250 | -0.6 | 1.75E-05 | -0.2 | 2.03E-02 | -0.4 | 1.31E-03 | -0.4 | 5.17E-04 | -0.2 | 5.24E-02 | -0.2 | 2.16E-01 | Tubulin beta-6 chain |
| AT5G22890 | -0.6 | 7.60E-04 | -0.9 | 1.11E-04 | 0.3 | 9.63E-02 | 2.1 | 1.56E-04 | 0.9 | 1.00E-01 | 1.2 | 2.50E-03 | Protein SENSITIVE TO PROTON RHIZOTOXICITY 2 |
| AT3G09980 | -0.6 | 8.45E-06 | -0.3 | 3.76E-03 | -0.3 | 1.62E-03 | -0.1 | 4.41E-01 | 0.0 | 9.79E-01 | -0.1 | 5.70E-01 | RAB6-interacting golgin |
| AT1G03970 | -0.6 | 7.64E-05 | -0.5 | 2.52E-04 | -0.1 | 6.22E-01 | -0.8 | 6.63E-04 | -0.5 | 2.86E-02 | -0.3 | 2.61E-01 | G-box-binding factor 4 |
| AT2G22690 | -0.6 | 2.79E-05 | -0.4 | 4.94E-04 | -0.2 | 9.17E-02 | -0.2 | 4.25E-01 | -0.1 | 8.20E-01 | -0.1 | 7.56E-01 | At2g22690 |
| AT2G36670 | -0.6 | 2.87E-05 | -0.4 | 3.82E-04 | -0.2 | 1.28E-01 | -0.3 | 5.39E-02 | -0.2 | 1.68E-01 | -0.1 | 8.08E-01 | Eukaryotic aspartyl protease family protein |
| AT2G40420 | -0.6 | 2.21E-04 | -0.2 | 4.31E-02 | -0.3 | 1.70E-02 | -0.5 | 9.76E-02 | -0.3 | 3.12E-01 | -0.2 | 7.03E-01 | Amino acid transporter AVT6D |
| AT2G44100 | -0.6 | 2.84E-05 | -0.3 | 3.52E-03 | -0.3 | 9.81E-03 | -0.1 | 5.36E-01 | 0.0 | 9.15E-01 | -0.1 | 6.08E-01 | Guanosine nucleotide diphosphate dissociation inhibitor 1 |
| AT2G22680 | -0.6 | 1.84E-03 | -0.4 | 2.59E-02 | -0.2 | 2.28E-01 | -0.2 | 2.93E-01 | -0.2 | 4.96E-01 | -0.1 | 8.81E-01 | E3 ubiquitin-protein ligase WAVH1 |
| AT1G08290 | -0.6 | 1.75E-03 | -0.7 | 8.83E-04 | 0.1 | 5.30E-01 | -0.1 | 8.05E-01 | 0.1 | 7.23E-01 | -0.2 | 5.53E-01 | Zinc finger protein WIP3 |
| AT3G46060 | -0.6 | 4.06E-05 | -0.2 | 9.04E-02 | -0.4 | 1.23E-03 | -0.2 | 2.37E-01 | -0.1 | 7.73E-01 | -0.1 | 6.29E-01 | Ras-related protein RABE1c |
| AT3G21770 | -0.6 | 2.64E-05 | -0.2 | 3.74E-02 | -0.4 | 1.37E-03 | -0.1 | 6.91E-01 | 0.1 | 4.37E-01 | -0.2 | 2.10E-01 | Peroxidase 30 |
| AT1G04010 | -0.6 | 9.53E-05 | -0.4 | 1.36E-03 | -0.2 | 1.50E-01 | -0.5 | 2.57E-02 | -0.2 | 5.38E-01 | -0.3 | 2.46E-01 | Phospholipid--sterol O-acyltransferase |
| AT2G44940 | -0.6 | 2.31E-03 | -0.2 | 2.00E-01 | -0.4 | 4.07E-02 | -0.5 | 6.46E-03 | 0.3 | 7.82E-02 | -0.9 | 2.17E-03 | Ethylene-responsive transcription factor ERF034 |
| AT1G31050 | -0.6 | 7.62E-04 | -0.5 | 1.51E-03 | 0.0 | 8.57E-01 | -0.4 | 1.74E-01 | 0.2 | 4.93E-01 | -0.6 | 8.21E-02 | Basic helix-loop-helix DNA-binding superfamily protein |
| AT3G08840 | -0.6 | 7.09E-05 | -0.5 | 1.98E-04 | 0.0 | 7.33E-01 | -0.1 | 4.76E-01 | 0.0 | 9.08E-01 | -0.1 | 7.24E-01 | D-alanine-D-alanine ligase family |
| AT5G23860 | -0.6 | 7.93E-06 | -0.1 | 2.19E-01 | -0.5 | 1.43E-04 | -0.1 | 2.46E-01 | 0.0 | 8.73E-01 | -0.1 | 5.38E-01 | Tubulin beta-8 chain |
| AT1G78680 | -0.6 | 6.96E-04 | -0.3 | 2.75E-02 | -0.3 | 7.18E-02 | -0.5 | 2.62E-03 | 0.2 | 2.08E-01 | -0.7 | 2.31E-03 | folate gamma-glutamyl hydrolase |
| AT3G13320 | -0.6 | 4.30E-06 | -0.4 | 6.17E-05 | -0.1 | 5.58E-02 | 0.0 | 8.96E-01 | 0.0 | 9.62E-01 | 0.0 | 9.71E-01 | Vacuolar cation |
| AT4G30910 | -0.6 | 9.10E-05 | -0.5 | 4.44E-04 | -0.1 | 4.23E-01 | -0.4 | 5.21E-02 | -0.7 | 2.50E-03 | 0.3 | 2.22E-01 | AT4G30910 protein |
| AT3G49860 | -0.6 | 1.43E-03 | 0.2 | 1.67E-01 | -0.8 | 9.49E-04 | 0.0 | 9.76E-01 | 0.0 | 9.74E-01 | 0.0 | 9.54E-01 | ADP-ribosylation factor-like protein 8d |
| AT3G10490 | -0.6 | 2.63E-05 | -0.5 | 1.23E-04 | -0.1 | 4.27E-01 | 0.0 | 8.03E-01 | -0.1 | 6.55E-01 | 0.0 | 8.73E-01 | NAC052 |
| AT1G64640 | -0.6 | 2.11E-04 | 0.0 | 7.67E-01 | -0.6 | 5.97E-04 | -0.5 | 1.76E-02 | -0.2 | 4.84E-01 | -0.3 | 2.35E-01 | Early nodulin-like protein 8 |
| AT3G45070 | -0.6 | 5.90E-04 | 0.2 | 1.48E-01 | -0.7 | 5.10E-04 | -0.7 | 9.69E-03 | 0.1 | 7.82E-01 | -0.8 | 7.35E-03 | Sulfotransferase |
| AT1G19700 | -0.6 | 1.51E-04 | -0.7 | 7.41E-05 | 0.1 | 3.13E-01 | -0.1 | 6.26E-01 | 0.3 | 2.20E-01 | -0.4 | 8.42E-02 | BEL1-like homeodomain protein 10 |
| AT3G18370 | -0.6 | 1.00E-04 | -0.6 | 1.94E-04 | 0.0 | 9.32E-01 | 0.2 | 4.18E-01 | 0.1 | 7.00E-01 | 0.1 | 8.64E-01 | AT3g18370 |
| AT3G60740 | -0.6 | 2.65E-05 | -0.3 | 1.52E-03 | -0.2 | 2.26E-02 | 0.1 | 7.84E-01 | 0.0 | 9.40E-01 | 0.0 | 9.16E-01 | ARM repeat superfamily protein |
| AT5G40330 | -0.6 | 3.68E-03 | 0.2 | 3.02E-01 | -0.7 | 3.40E-03 | 0.1 | 7.74E-01 | 0.3 | 3.66E-01 | -0.2 | 5.80E-01 | Transcription factor MYB23 |
| AT2G16530 | -0.6 | 1.40E-04 | -0.1 | 1.39E-01 | -0.4 | 3.82E-03 | -0.2 | 2.81E-01 | 0.0 | 9.83E-01 | -0.2 | 3.90E-01 | Polyprenol reductase |
| AT1G42470 | -0.6 | 1.26E-04 | -0.4 | 3.64E-03 | -0.2 | 7.05E-02 | -0.5 | 5.61E-02 | -0.2 | 4.72E-01 | -0.3 | 4.40E-01 | Patched family protein |
| AT2G20440 | -0.6 | 3.93E-04 | -0.3 | 3.17E-02 | -0.3 | 4.26E-02 | 0.0 | 8.51E-01 | 0.2 | 3.37E-01 | -0.2 | 4.52E-01 | At2g20440 |
| AT1G73500 | -0.6 | 3.82E-03 | -0.9 | 4.35E-04 | 0.3 | 1.29E-01 | 0.2 | 2.84E-01 | -0.1 | 8.04E-01 | 0.3 | 3.04E-01 | Mitogen-activated protein kinase kinase 9 |
| AT1G53590 | -0.6 | 1.80E-05 | -0.3 | 1.10E-03 | -0.2 | 1.51E-02 | -0.3 | 8.52E-02 | -0.1 | 7.25E-01 | -0.2 | 2.87E-01 | C2 domain-containing protein At1g53590 |
| AT1G76430 | -0.6 | 2.96E-03 | -0.6 | 4.84E-03 | 0.0 | 9.45E-01 | 1.2 | 3.20E-03 | 1.0 | 2.62E-02 | 0.2 | 5.64E-01 | Probable inorganic phosphate transporter 1-9 |
| AT5G65240 | -0.6 | 9.04E-05 | -0.6 | 1.65E-04 | 0.0 | 9.71E-01 | -0.1 | 7.46E-01 | 0.0 | 9.43E-01 | 0.0 | 8.83E-01 | Leucine-rich repeat protein kinase family protein |
| AT4G29780 | -0.6 | 5.03E-02 | -0.9 | 7.64E-03 | 0.4 | 3.05E-01 | -0.3 | 2.65E-01 | -0.4 | 1.41E-01 | 0.1 | 7.35E-01 | Uncharacterized protein AT4g29780 |
| AT4G00200 | -0.6 | 1.09E-04 | -0.6 | 2.15E-04 | 0.0 | 9.27E-01 | -0.2 | 3.42E-01 | -0.1 | 7.96E-01 | -0.1 | 6.84E-01 | AT-hook motif nuclear-localized protein 7 |
| AT1G28310 | -0.6 | 7.68E-05 | -0.3 | 5.48E-03 | -0.3 | 2.88E-02 | -0.4 | 9.32E-02 | 0.0 | 8.76E-01 | -0.4 | 1.53E-01 | Dof zinc finger protein |
| AT4G16830 | -0.6 | 1.50E-05 | -0.4 | 1.82E-04 | -0.1 | 1.20E-01 | -0.3 | 7.24E-03 | -0.2 | 7.58E-02 | -0.1 | 6.11E-01 | RGG repeats nuclear RNA binding protein A |
| AT3G07700 | -0.6 | 6.22E-05 | -0.9 | 6.78E-06 | 0.3 | 1.01E-02 | -0.4 | 6.55E-03 | 0.0 | 8.71E-01 | -0.4 | 3.87E-02 | Protein kinase superfamily protein |
| AT1G64660 | -0.6 | 2.16E-03 | -0.9 | 1.82E-04 | 0.3 | 6.99E-02 | -0.8 | 5.54E-03 | -1.0 | 2.32E-03 | 0.2 | 6.26E-01 | Methionine gamma-lyase |
| AT1G60610 | -0.6 | 9.40E-05 | -0.6 | 1.43E-04 | 0.0 | 9.17E-01 | 0.0 | 9.56E-01 | 0.1 | 6.54E-01 | -0.1 | 6.63E-01 | At1g60610 family protein) |
| AT5G16590 | -0.6 | 1.79E-05 | 0.0 | 7.94E-01 | -0.6 | 9.22E-05 | -0.4 | 4.47E-02 | 0.0 | 9.60E-01 | -0.4 | 1.68E-01 | Probable inactive receptor kinase At5g16590 |
| AT1G76500 | -0.6 | 3.73E-03 | -0.1 | 4.26E-01 | -0.4 | 3.69E-02 | -0.2 | 5.06E-01 | -0.3 | 3.78E-01 | 0.1 | 8.64E-01 | AT-hook motif nuclear-localized protein 29 |
| AT4G28940 | -0.6 | 1.89E-04 | 0.1 | 1.42E-01 | -0.7 | 2.18E-04 | 1.9 | 3.44E-06 | 1.6 | 7.45E-05 | 0.2 | 3.48E-01 | At4g28940 |
| AT3G61890 | -0.6 | 4.21E-03 | -0.6 | 5.40E-03 | 0.0 | 9.90E-01 | -0.1 | 7.06E-01 | 0.0 | 9.56E-01 | -0.1 | 8.45E-01 | Homeobox-leucine zipper protein ATHB-12 |
| AT5G10960 | -0.6 | 4.83E-06 | -0.3 | 1.33E-03 | -0.3 | 1.78E-03 | -0.3 | 1.70E-02 | -0.4 | 1.89E-02 | 0.0 | 9.04E-01 | Probable CCR4-associated factor 1 homolog 10 |
| AT1G13860 | -0.6 | 2.18E-04 | -0.5 | 4.47E-04 | 0.0 | 8.84E-01 | -0.1 | 7.94E-01 | 0.2 | 3.58E-01 | -0.3 | 2.77E-01 | Probable methyltransferase PMT4 |
| AT2G40260 | -0.6 | 6.16E-04 | -0.2 | 8.34E-02 | -0.3 | 3.01E-02 | 0.0 | 9.32E-01 | 0.1 | 8.59E-01 | -0.1 | 8.18E-01 | HTH myb-type domain-containing protein |
| AT5G13460 | -0.6 | 8.88E-04 | -0.2 | 1.88E-01 | -0.4 | 2.04E-02 | -0.6 | 3.80E-03 | -0.2 | 2.84E-01 | -0.4 | 1.54E-01 | Protein IQ-DOMAIN 11 |
| AT5G66300 | -0.6 | 7.28E-03 | -0.3 | 7.44E-02 | -0.2 | 3.07E-01 | -0.3 | 4.38E-01 | -0.6 | 1.88E-01 | 0.3 | 6.34E-01 | NAC domain-containing protein 105 |
| AT5G04060 | -0.6 | 1.00E-04 | -0.3 | 7.36E-03 | -0.3 | 2.93E-02 | -0.6 | 8.12E-03 | 0.2 | 5.22E-01 | -0.8 | 1.05E-02 | Probable methyltransferase PMT7 |
| AT1G52880 | -0.6 | 2.02E-03 | -0.8 | 4.38E-04 | 0.2 | 2.35E-01 | -0.2 | 3.40E-01 | 0.1 | 8.49E-01 | -0.3 | 3.05E-01 | NAC domain-containing protein 18 |
| AT2G38700 | -0.6 | 2.27E-05 | -0.1 | 1.06E-01 | -0.4 | 5.92E-04 | -0.2 | 5.86E-02 | 0.0 | 9.22E-01 | -0.3 | 1.58E-01 | Diphosphomevalonate decarboxylase MVD1, peroxisomal |
| AT2G36240 | -0.6 | 2.91E-05 | -0.1 | 9.29E-02 | -0.4 | 1.06E-03 | 0.2 | 1.04E-01 | 0.1 | 5.08E-01 | 0.1 | 5.90E-01 | Pentatricopeptide repeat-containing protein At2g36240 |
| AT1G36160 | -0.6 | 8.72E-05 | -0.5 | 2.19E-04 | 0.0 | 8.09E-01 | -0.2 | 3.83E-01 | 0.1 | 8.29E-01 | -0.3 | 4.14E-01 | Acetyl-CoA carboxylase 1 |
| AT3G44200 | -0.6 | 9.44E-05 | -0.2 | 2.41E-02 | -0.3 | 8.59E-03 | -0.2 | 4.19E-01 | 0.1 | 8.52E-01 | -0.3 | 3.75E-01 | Serine |
| AT4G10955 | -0.6 | 1.33E-03 | -0.2 | 1.56E-01 | -0.4 | 3.65E-02 | -0.3 | 2.18E-01 | 0.2 | 4.04E-01 | -0.5 | 4.89E-02 | GDSL esterase |
| AT5G22250 | -0.6 | 6.86E-03 | -0.8 | 1.27E-03 | 0.2 | 2.79E-01 | 0.8 | 7.23E-03 | 0.0 | 9.57E-01 | 0.8 | 1.37E-02 | Probable CCR4-associated factor 1 homolog 11 |
| AT1G74020 | -0.6 | 8.35E-05 | -0.2 | 5.91E-02 | -0.4 | 3.78E-03 | -0.1 | 6.31E-01 | 0.2 | 2.31E-01 | -0.2 | 1.28E-01 | Protein STRICTOSIDINE SYNTHASE-LIKE 12 |
| AT5G55400 | -0.6 | 8.79E-04 | -0.2 | 7.88E-02 | -0.3 | 4.32E-02 | -0.4 | 1.11E-01 | 0.0 | 9.22E-01 | -0.4 | 2.87E-01 | Fimbrin-3 |
| AT3G24190 | -0.6 | 1.63E-03 | -0.9 | 1.14E-04 | 0.3 | 4.17E-02 | -0.1 | 7.39E-01 | 0.1 | 6.78E-01 | -0.2 | 5.27E-01 | Protein kinase domain-containing protein |
| AT4G34390 | -0.6 | 1.50E-04 | -0.7 | 6.61E-05 | 0.1 | 2.69E-01 | -0.1 | 7.60E-01 | -0.2 | 4.05E-01 | 0.1 | 7.17E-01 | Extra-large GTP-binding protein 2 |
| AT4G16780 | -0.6 | 1.48E-03 | -0.3 | 3.29E-02 | -0.2 | 1.50E-01 | 0.0 | 9.35E-01 | 0.2 | 6.34E-01 | -0.1 | 7.49E-01 | Homeobox-leucine zipper protein HAT4 |
| AT5G49720 | -0.6 | 3.88E-05 | -0.4 | 6.21E-04 | -0.1 | 1.07E-01 | -0.4 | 9.70E-04 | -0.3 | 7.91E-03 | -0.1 | 7.49E-01 | Endoglucanase 25 |

| | | | | | | | | | | | | | |
|---|---|---|---|---|---|---|---|---|---|---|---|---|---|
| AT5G50160 | -0.6 | 6.03E-04 | -0.6 | 9.36E-04 | 0.0 | 9.72E-01 | 0.0 | 8.44E-01 | 0.2 | 2.91E-01 | -0.2 | 4.94E-01 | Ferric reduction oxidase 8 |
| AT5G13110 | -0.6 | 1.77E-03 | -0.4 | 2.44E-02 | -0.2 | 2.10E-01 | -0.1 | 5.73E-01 | -0.1 | 4.96E-01 | 0.0 | 9.07E-01 | Glucose-6-phosphate 1-dehydrogenase 2, chloroplastic |
| AT3G44730 | -0.6 | 3.40E-04 | -0.5 | 1.26E-03 | -0.1 | 5.87E-01 | 0.3 | 3.58E-01 | 0.5 | 2.09E-01 | -0.1 | 7.36E-01 | Kinesin-like protein 1 |
| AT1G03840 | -0.6 | 2.23E-04 | 0.0 | 8.71E-01 | -0.5 | 1.43E-03 | 0.2 | 4.58E-01 | 0.5 | 1.02E-01 | -0.3 | 4.70E-01 | Zinc finger protein MAGPIE |
| AT4G30080 | -0.6 | 6.29E-05 | -0.4 | 8.30E-04 | -0.2 | 1.43E-01 | -0.4 | 1.31E-02 | -0.1 | 8.09E-01 | -0.4 | 9.78E-02 | Auxin response factor 16 |
| AT1G78815 | -0.6 | 2.85E-03 | -0.2 | 2.43E-01 | -0.4 | 5.41E-02 | -0.2 | 7.29E-01 | 0.1 | 9.08E-01 | -0.2 | 6.69E-01 | Protein LIGHT-DEPENDENT SHORT HYPOCOTYLS 7 |
| AT5G48810 | -0.6 | 5.01E-05 | -0.2 | 8.63E-02 | -0.4 | 1.60E-03 | -0.5 | 6.80E-03 | -0.4 | 4.51E-02 | -0.1 | 7.19E-01 | Cytochrome B5 isoform D |
| AT5G50450 | -0.6 | 1.68E-03 | 0.1 | 2.96E-01 | -0.7 | 1.58E-03 | -0.4 | 2.02E-02 | -0.5 | 7.92E-03 | 0.1 | 6.66E-01 | F-box protein At5g50450 |
| AT3G05490 | -0.6 | 4.78E-03 | 0.1 | 7.04E-01 | -0.6 | 5.81E-03 | -0.5 | 1.49E-03 | -0.2 | 3.31E-01 | -0.3 | 5.57E-02 | Protein RALF-like 22 |
| AT4G38580 | -0.6 | 7.02E-05 | -0.5 | 1.32E-04 | 0.0 | 9.52E-01 | -0.2 | 2.28E-01 | -0.2 | 2.36E-01 | 0.0 | 9.51E-01 | Heavy metal-associated isoprenylated plant protein 26 |
| AT2G42880 | -0.6 | 5.30E-05 | -0.3 | 1.99E-03 | -0.2 | 4.09E-02 | -0.4 | 1.48E-01 | 0.1 | 8.52E-01 | -0.4 | 1.90E-01 | mitogen-activated protein kinase |
| AT1G17500 | -0.6 | 2.99E-05 | -0.5 | 2.00E-04 | -0.1 | 2.68E-01 | 0.4 | 8.67E-03 | 0.2 | 2.41E-01 | 0.2 | 3.47E-01 | Probable phospholipid-transporting ATPase 4 |
| AT2G42960 | -0.6 | 2.77E-04 | -0.2 | 4.39E-02 | -0.3 | 2.21E-02 | 0.1 | 5.11E-01 | 0.4 | 1.17E-01 | -0.2 | 4.53E-01 | Protein kinase superfamily protein |
| AT3G61250 | -0.6 | 4.97E-02 | -0.7 | 2.01E-02 | 0.2 | 6.02E-01 | 0.1 | 8.49E-01 | 0.0 | 1.00E+00 | 0.1 | 8.99E-01 | Transcription factor MYB17 |
| AT1G55580 | -0.6 | 1.53E-02 | -0.2 | 2.60E-01 | -0.3 | 2.07E-01 | -0.2 | 6.57E-01 | -0.2 | 7.19E-01 | 0.0 | 9.97E-01 | Scarecrow-like protein 18 |
| AT5G51590 | -0.6 | 2.36E-04 | -0.2 | 4.74E-02 | -0.3 | 1.67E-02 | 0.2 | 3.14E-01 | 0.3 | 1.67E-01 | -0.1 | 7.51E-01 | AT-hook motif nuclear-localized protein |
| AT2G36580 | -0.6 | 1.06E-03 | -0.3 | 6.27E-02 | -0.3 | 5.51E-02 | 0.5 | 1.16E-03 | 0.1 | 6.76E-01 | 0.4 | 3.44E-02 | Pyruvate kinase |
| AT3G43270 | -0.6 | 3.40E-04 | -0.4 | 6.61E-03 | -0.2 | 1.39E-01 | -0.3 | 1.62E-01 | -0.1 | 8.01E-01 | -0.2 | 4.19E-01 | Probable pectinesterase |
| AT1G08200 | -0.6 | 7.58E-05 | -0.3 | 7.31E-03 | -0.3 | 1.87E-02 | -0.3 | 4.98E-03 | 0.1 | 6.76E-01 | -0.3 | 2.21E-02 | UDP-D-apiose |
| AT2G41480 | -0.6 | 8.27E-04 | -0.2 | 6.37E-02 | -0.3 | 4.41E-02 | 0.0 | 8.23E-01 | 0.1 | 7.52E-01 | -0.1 | 5.87E-01 | Peroxidase 25 |
| AT3G45090 | -0.6 | 3.43E-05 | -0.2 | 2.92E-02 | -0.4 | 2.32E-03 | -0.2 | 2.22E-01 | 0.1 | 5.58E-01 | -0.3 | 1.53E-01 | P-loop containing nucleoside triphosphate hydrolases superfamily protein |
| AT1G07260 | -0.6 | 3.07E-04 | -0.3 | 8.77E-03 | -0.2 | 9.58E-02 | 0.1 | 6.38E-01 | 0.0 | 9.69E-01 | 0.1 | 7.07E-01 | UDP-glycosyltransferase 71C3 |
| AT5G67540 | -0.6 | 2.86E-05 | -0.4 | 8.80E-04 | -0.2 | 4.36E-02 | 0.0 | 9.77E-01 | 0.0 | 9.16E-01 | 0.0 | 9.09E-01 | Arabinanase |
| AT3G26690 | -0.6 | 4.86E-04 | -0.2 | 6.82E-02 | -0.3 | 2.71E-02 | -0.4 | 1.43E-01 | -0.5 | 1.65E-01 | 0.0 | 9.77E-01 | Nudix hydrolase 13, mitochondrial |
| AT3G23280 | -0.6 | 2.46E-05 | -0.5 | 5.52E-05 | 0.0 | 9.14E-01 | 0.0 | 7.61E-01 | 0.1 | 2.63E-01 | -0.2 | 2.58E-01 | Putative E3 ubiquitin-protein ligase XBAT35 |
| AT4G24250 | -0.6 | 1.92E-03 | -0.2 | 1.17E-01 | -0.3 | 7.33E-02 | | | | | | | MLO-like protein |
| AT1G66410 | -0.6 | 5.39E-05 | -0.2 | 1.63E-02 | -0.3 | 5.96E-03 | 0.0 | 9.11E-01 | 0.0 | 9.82E-01 | 0.0 | 9.23E-01 | Calmodulin 4 |
| AT1G49720 | -0.5 | 7.74E-05 | -0.3 | 1.01E-02 | -0.3 | 1.47E-02 | -0.3 | 8.83E-02 | -0.2 | 2.13E-01 | -0.1 | 8.55E-01 | Abscisic acid responsive element-binding factor 1 |
| AT4G38380 | -0.5 | 1.74E-04 | -0.6 | 1.11E-04 | 0.1 | 4.49E-01 | 0.9 | 6.56E-02 | 0.2 | 7.86E-01 | 0.7 | 2.05E-01 | Protein DETOXIFICATION |
| AT5G40540 | -0.5 | 2.58E-03 | -0.6 | 2.84E-03 | 0.0 | 9.21E-01 | 0.3 | 2.10E-01 | 0.4 | 1.25E-01 | -0.1 | 7.70E-01 | Protein kinase ATN1 |
| AT4G38530 | -0.5 | 8.92E-04 | -0.5 | 1.98E-03 | 0.0 | 8.29E-01 | -0.1 | 8.55E-01 | 0.1 | 9.22E-01 | -0.2 | 8.22E-01 | Phosphoinositide phospholipase C 3 |
| AT5G10930 | -0.5 | 3.51E-04 | -0.2 | 3.87E-02 | -0.3 | 2.84E-02 | -0.1 | 7.09E-01 | 0.3 | 2.24E-01 | -0.4 | 1.65E-01 | CBL-interacting serine |
| AT1G09090 | -0.5 | 1.46E-03 | -0.1 | 3.90E-01 | -0.4 | 1.75E-02 | -0.8 | 4.52E-02 | -0.3 | 4.64E-01 | -0.5 | 3.98E-01 | Respiratory burst oxidase-like protein |
| AT2G40110 | -0.5 | 3.80E-03 | -0.4 | 2.24E-02 | -0.1 | 4.38E-01 | -0.5 | 9.81E-02 | -0.3 | 3.12E-01 | -0.1 | 7.57E-01 | Protein yippee-like |
| AT3G56200 | -0.5 | 5.68E-04 | 0.0 | 9.43E-01 | -0.5 | 2.58E-03 | 0.0 | 9.21E-01 | -0.2 | 6.50E-01 | 0.1 | 7.67E-01 | Amino acid transporter AVT6C |
| AT5G20730 | -0.5 | 6.70E-05 | -0.5 | 1.17E-04 | 0.0 | 9.93E-01 | -0.3 | 3.12E-01 | -0.2 | 5.88E-01 | -0.1 | 8.36E-01 | Auxin response factor |
| AT1G22540 | -0.5 | 1.55E-04 | -0.2 | 8.55E-02 | -0.4 | 6.20E-03 | -0.3 | 2.03E-01 | 0.1 | 5.50E-01 | -0.4 | 1.08E-01 | Protein NRT1 |
| AT1G73270 | -0.5 | 3.08E-04 | -0.1 | 3.91E-01 | -0.5 | 4.25E-03 | -0.2 | 6.11E-01 | 0.1 | 8.47E-01 | -0.2 | 4.39E-01 | Serine carboxypeptidase-like 6 |
| AT4G12040 | -0.5 | 1.35E-05 | -0.2 | 3.91E-02 | -0.4 | 6.07E-04 | -0.2 | 2.08E-01 | -0.2 | 2.17E-01 | 0.0 | 9.40E-01 | Zinc finger A20 and AN1 domain-containing stress-associated protein 7 |
| AT4G00480 | -0.5 | 1.55E-03 | 0.1 | 4.13E-01 | -0.7 | 2.06E-03 | 0.0 | 8.62E-01 | 0.3 | 2.36E-01 | -0.3 | 1.72E-01 | Basic helix-loop-helix DNA-binding superfamily protein |
| AT2G43490 | -0.5 | 9.07E-06 | -0.2 | 8.20E-03 | -0.3 | 1.21E-03 | 0.1 | 8.02E-01 | 0.3 | 1.37E-01 | -0.3 | 3.06E-01 | Ypt |
| AT1G72880 | -0.5 | 4.38E-05 | -0.2 | 5.29E-02 | -0.4 | 2.03E-03 | -0.1 | 5.29E-01 | 0.0 | 9.75E-01 | -0.1 | 6.23E-01 | AT1G72880 protein |
| AT1G36370 | -0.5 | 2.42E-05 | -0.5 | 1.20E-04 | -0.1 | 4.00E-01 | 0.6 | 1.88E-03 | 0.4 | 3.06E-02 | 0.2 | 4.56E-01 | Serine hydroxymethyltransferase |
| AT2G48010 | -0.5 | 3.56E-04 | -0.6 | 3.93E-04 | 0.0 | 8.23E-01 | -0.1 | 6.23E-01 | -0.1 | 6.40E-01 | 0.0 | 9.81E-01 | Probable LRR receptor-like serine |
| AT4G27830 | -0.5 | 7.86E-04 | -0.7 | 1.65E-04 | 0.2 | 1.68E-01 | 0.0 | 9.55E-01 | -0.1 | 6.50E-01 | 0.1 | 6.24E-01 | Beta glucosidase 10 |
| AT5G11950 | -0.5 | 2.37E-04 | -0.4 | 5.17E-03 | -0.2 | 1.01E-01 | 0.1 | 4.86E-01 | 0.3 | 6.28E-02 | -0.2 | 3.16E-01 | Cytokinin riboside 5'-monophosphate phosphoribohydrolase |
| AT5G42720 | -0.5 | 2.98E-04 | -0.3 | 1.10E-02 | -0.2 | 6.59E-02 | 0.1 | 6.25E-01 | 0.3 | 1.80E-01 | -0.2 | 4.84E-01 | glucan endo-1,3-beta-D-glucosidase |
| AT2G20900 | -0.5 | 1.73E-05 | -0.4 | 1.42E-04 | -0.1 | 1.87E-01 | 0.3 | 2.02E-02 | 0.0 | 9.03E-01 | 0.4 | 4.74E-02 | Diacylglycerol kinase 5 |
| AT4G34880 | -0.5 | 1.56E-03 | -0.2 | 1.14E-01 | -0.3 | 6.30E-02 | -0.4 | 2.61E-01 | 0.2 | 7.09E-01 | -0.6 | 1.92E-01 | Probable amidase At4g34880 |
| AT2G20370 | -0.5 | 1.72E-04 | -0.3 | 6.18E-03 | -0.2 | 5.78E-02 | 0.0 | 8.57E-01 | 0.2 | 3.58E-01 | -0.1 | 5.64E-01 | Xyloglucan galactosyltransferase MUR3 |
| AT3G19580 | -0.5 | 9.80E-04 | -0.8 | 1.41E-04 | 0.3 | 1.16E-01 | 0.4 | 1.16E-01 | 0.0 | 9.60E-01 | 0.4 | 1.14E-01 | Zinc finger protein AZF2 |
| AT1G04990 | -0.5 | 7.47E-04 | -0.5 | 1.39E-03 | 0.0 | 8.93E-01 | -0.4 | 3.12E-02 | 0.1 | 6.07E-01 | -0.6 | 2.20E-02 | Zinc finger protein 2 |
| AT1G24180 | -0.5 | 1.66E-04 | 0.0 | 9.43E-01 | -0.6 | 5.73E-04 | 0.1 | 2.16E-01 | 0.1 | 3.71E-01 | 0.0 | 9.35E-01 | Pyruvate dehydrogenase E1 component subunit alpha-2, mitochondrial |
| AT1G66350 | -0.5 | 8.40E-04 | 0.0 | 8.41E-01 | -0.5 | 4.32E-03 | -0.6 | 2.30E-03 | 0.2 | 3.01E-01 | -0.8 | 1.15E-03 | DELLA protein RGL1 |
| AT1G68760 | -0.5 | 6.51E-03 | -0.5 | 1.01E-02 | 0.0 | 9.21E-01 | 0.3 | 2.04E-01 | 0.0 | 8.84E-01 | 0.2 | 4.45E-01 | Nudix hydrolase 1 |
| AT5G40100 | -0.5 | 3.16E-03 | 0.0 | 9.23E-01 | -0.5 | 1.27E-02 | -0.3 | 5.78E-01 | -0.1 | 9.11E-01 | -0.2 | 7.71E-01 | TIR domain-containing protein |
| AT4G19045 | -0.5 | 2.65E-03 | -0.2 | 1.28E-01 | -0.3 | 8.42E-02 | -0.1 | 7.82E-01 | 0.4 | 1.07E-01 | -0.5 | 9.22E-02 | MOB kinase activator-like 1B |
| AT2G32720 | -0.5 | 3.40E-04 | 0.0 | 7.40E-01 | -0.5 | 1.83E-03 | 0.2 | 9.28E-02 | 0.3 | 3.55E-02 | -0.1 | 6.58E-01 | Cytochrome B5 isoform B |
| AT5G02750 | -0.5 | 1.51E-03 | -0.7 | 4.11E-04 | 0.2 | 2.89E-01 | 0.2 | 4.77E-01 | 0.0 | 9.54E-01 | 0.2 | 5.39E-01 | E3 ubiquitin-protein ligase SGR9, amyloplastic |
| AT1G19220 | -0.5 | 6.11E-04 | -0.2 | 7.11E-02 | -0.3 | 3.14E-02 | 0.1 | 6.09E-01 | 0.1 | 7.36E-01 | 0.0 | 9.62E-01 | Auxin response factor 19 |
| AT3G15020 | -0.5 | 9.83E-04 | 0.2 | 2.05E-01 | -0.7 | 6.45E-04 | -0.1 | 4.81E-01 | 0.1 | 6.62E-01 | -0.3 | 3.03E-01 | Malate dehydrogenase |
| AT5G62220 | -0.5 | 3.34E-04 | -0.3 | 1.19E-02 | -0.2 | 7.08E-02 | -0.4 | 4.80E-02 | 0.1 | 6.68E-01 | -0.5 | 3.26E-02 | Xyloglucan galactosyltransferase XLT2 |
| AT1G05170 | -0.5 | 9.86E-04 | -0.4 | 4.11E-03 | -0.1 | 5.43E-01 | 0.5 | 1.33E-02 | 0.6 | 8.98E-03 | -0.1 | 7.57E-01 | Hexosyltransferase |
| AT4G02590 | -0.5 | 1.21E-05 | -0.4 | 9.54E-05 | -0.1 | 1.85E-01 | -0.1 | 3.53E-01 | -0.1 | 3.64E-01 | 0.0 | 9.57E-01 | Transcription factor UNE12 |
| AT5G40090 | -0.5 | 3.40E-03 | -0.1 | 5.09E-01 | -0.4 | 3.05E-02 | -0.6 | 1.13E-01 | -0.2 | 6.48E-01 | -0.4 | 4.75E-01 | Disease resistance protein CHL1 |
| AT1G76990 | -0.5 | 4.57E-05 | -0.1 | 1.25E-01 | -0.4 | 1.16E-03 | -0.5 | 9.49E-05 | -0.4 | 2.71E-03 | -0.1 | 4.63E-01 | ACT domain-containing protein ACR3 |
| AT2G18960 | -0.5 | 1.30E-04 | -0.5 | 3.62E-04 | 0.0 | 7.51E-01 | -0.2 | 3.83E-01 | 0.1 | 6.64E-01 | -0.3 | 2.94E-01 | Plasma membrane ATPase |
| AT1G34110 | -0.5 | 4.19E-05 | -0.5 | 8.25E-05 | 0.0 | 9.70E-01 | 0.0 | 8.32E-01 | -0.2 | 4.43E-01 | 0.1 | 6.80E-01 | Leucine-rich receptor-like protein kinase family protein |
| AT5G59010 | -0.5 | 2.31E-04 | -0.1 | 4.88E-01 | -0.5 | 2.49E-03 | -0.1 | 5.22E-01 | 0.0 | 9.54E-01 | -0.1 | 7.19E-01 | Serine |
| AT5G58620 | -0.5 | 2.10E-04 | -0.7 | 7.85E-05 | 0.1 | 2.42E-01 | 0.0 | 9.81E-01 | -0.2 | 3.93E-01 | 0.1 | 4.41E-01 | Zinc finger CCCH domain-containing protein 66 |
| AT5G19420 | -0.5 | 2.46E-05 | -0.6 | 1.42E-05 | 0.1 | 2.16E-01 | 0.0 | 8.97E-01 | 0.0 | 9.46E-01 | -0.1 | 8.64E-01 | Regulator of chromosome condensation family with FYVE zinc finger domain-containing protein |
| AT5G38350 | -0.5 | 6.68E-04 | -0.8 | 1.39E-04 | 0.2 | 1.58E-01 | 0.7 | 7.75E-02 | 0.4 | 3.54E-01 | 0.3 | 5.91E-01 | ADP-ribosyl cyclase |
| AT2G16750 | -0.5 | 2.75E-03 | -0.4 | 2.07E-02 | -0.2 | 3.63E-01 | -0.1 | 7.84E-01 | 0.0 | 9.32E-01 | -0.1 | 7.49E-01 | Protein kinase STUNTED |
| AT1G07640 | -0.5 | 8.90E-05 | -0.5 | 3.13E-04 | -0.1 | 5.81E-01 | 0.0 | 9.49E-01 | 0.2 | 1.50E-01 | -0.2 | 2.37E-01 | Dof zinc finger protein |
| AT4G34890 | -0.5 | 7.72E-05 | -0.7 | 3.35E-05 | 0.1 | 2.08E-01 | -0.1 | 6.82E-01 | -0.1 | 5.90E-01 | 0.0 | 9.05E-01 | Xanthine dehydrogenase 1 |
| AT4G32180 | -0.5 | 8.27E-06 | -0.5 | 3.51E-05 | -0.1 | 4.25E-01 | -0.2 | 5.41E-01 | -0.2 | 6.00E-01 | 0.0 | 9.96E-01 | Pantothenate kinase 2 |
| AT5G44110 | -0.5 | 5.01E-04 | 0.2 | 6.89E-02 | -0.7 | 2.76E-04 | 0.1 | 7.40E-01 | 0.8 | 4.31E-03 | -0.7 | 1.44E-02 | P-loop containing nucleoside triphosphate hydrolases superfamily protein |
| AT5G18525 | -0.5 | 3.54E-05 | -0.5 | 5.85E-05 | 0.0 | 9.13E-01 | -0.3 | 1.93E-01 | -0.1 | 5.70E-01 | -0.1 | 7.11E-01 | Protein GFS12 |
| AT5G47240 | -0.5 | 1.10E-02 | 0.1 | 7.38E-01 | -0.6 | 1.27E-02 | -0.3 | 4.24E-01 | 0.3 | 4.95E-01 | -0.6 | 2.01E-01 | Nudix hydrolase |
| AT4G31100 | -0.5 | 5.36E-04 | -0.3 | 1.62E-02 | -0.2 | 1.11E-01 | -0.1 | 7.01E-01 | 0.3 | 2.95E-01 | -0.4 | 1.85E-01 | Wall-associated receptor kinase-like |
| AT4G22730 | -0.5 | 1.18E-03 | -0.3 | 2.78E-02 | -0.2 | 1.52E-01 | 0.0 | 9.36E-01 | 0.2 | 4.63E-01 | -0.2 | 5.95E-01 | At4g22730 |
| AT4G23030 | -0.5 | 1.90E-02 | -0.9 | 1.97E-03 | 0.4 | 2.00E-01 | 0.4 | 2.45E-01 | 0.1 | 8.39E-01 | 0.3 | 4.71E-01 | Protein DETOXIFICATION 49 |
| AT1G11680 | -0.5 | 8.64E-05 | -0.1 | 2.92E-01 | -0.4 | 1.16E-03 | -0.3 | 1.22E-01 | -0.1 | 6.92E-01 | -0.2 | 5.22E-01 | Sterol 14-demethylase |

| | | | | | | | | | | | | | |
|---|---|---|---|---|---|---|---|---|---|---|---|---|---|
| AT2G37740 | -0.5 | 8.91E-03 | -0.3 | 6.63E-02 | -0.2 | 4.04E-01 | | | | | | | Zinc finger protein 10 |
| AT1G73980 | -0.5 | 1.66E-05 | -0.6 | 1.44E-05 | 0.1 | 3.99E-01 | -0.3 | 8.66E-02 | -0.2 | 2.79E-01 | -0.1 | 7.78E-01 | Inorganic pyrophosphatase TTM1 |
| AT1G21730 | -0.5 | 4.20E-05 | -0.2 | 4.45E-02 | -0.4 | 2.44E-03 | -0.2 | 4.02E-01 | 0.0 | 9.65E-01 | -0.2 | 5.85E-01 | Kinesin-like protein KIN-7C, mitochondrial |
| AT3G02580 | -0.5 | 1.51E-04 | -0.2 | 6.65E-02 | -0.3 | 6.20E-03 | -0.1 | 3.40E-01 | 0.0 | 8.86E-01 | -0.2 | 3.90E-01 | Fatty acid hydroxylase domain-containing protein |
| AT5G43900 | -0.5 | 4.05E-05 | -0.5 | 2.08E-04 | -0.1 | 3.99E-01 | 0.1 | 2.35E-01 | 0.2 | 1.63E-01 | 0.0 | 8.23E-01 | Myosin 2 |
| AT5G54650 | -0.5 | 1.28E-04 | -0.3 | 1.09E-02 | -0.3 | 2.85E-02 | 0.2 | 4.79E-01 | 0.2 | 4.59E-01 | 0.0 | 9.38E-01 | Formin-like protein 5 |
| AT3G24450 | -0.5 | 1.33E-03 | -0.3 | 2.39E-02 | -0.2 | 1.91E-01 | -0.3 | 3.87E-01 | -0.2 | 5.76E-01 | -0.1 | 9.05E-01 | Heavy metal transport |
| AT1G52540 | -0.5 | 6.07E-05 | -0.2 | 2.16E-02 | -0.3 | 6.56E-03 | -0.2 | 2.90E-01 | -0.1 | 8.40E-01 | -0.1 | 5.76E-01 | F6D8.24 protein |
| AT3G03690 | -0.5 | 1.92E-03 | -0.2 | 1.49E-01 | -0.3 | 5.51E-02 | -0.4 | 1.20E-01 | -0.1 | 6.46E-01 | -0.2 | 4.18E-01 | Core-2 |
| AT3G17730 | -0.5 | 6.81E-03 | -0.1 | 6.06E-01 | -0.4 | 4.49E-02 | 0.2 | 6.16E-01 | 0.1 | 8.49E-01 | 0.1 | 8.73E-01 | NAC057 |
| AT2G45910 | -0.5 | 5.48E-04 | -0.7 | 1.94E-04 | 0.1 | 2.97E-01 | -0.1 | 7.04E-01 | -0.2 | 5.18E-01 | 0.1 | 8.50E-01 | U-box domain-containing protein 33 |
| AT3G15240 | -0.5 | 5.87E-03 | 0.1 | 5.52E-01 | -0.6 | 6.92E-03 | -0.2 | 3.97E-01 | 0.1 | 6.83E-01 | -0.3 | 2.87E-01 | Uncharacterized protein At3g15240 |
| AT5G56650 | -0.5 | 1.81E-05 | -0.4 | 3.20E-04 | -0.2 | 7.10E-02 | -0.2 | 1.80E-01 | 0.1 | 7.99E-01 | -0.2 | 1.70E-01 | IAA-leucine resistant -like 1 |
| AT2G25520 | -0.5 | 1.51E-04 | -0.2 | 5.54E-02 | -0.3 | 7.38E-03 | 0.0 | 9.11E-01 | 0.0 | 9.19E-01 | 0.0 | 8.67E-01 | Probable sugar phosphate |
| AT2G22800 | -0.5 | 5.92E-05 | -0.3 | 7.87E-03 | -0.3 | 1.54E-02 | 0.2 | 5.38E-01 | -0.2 | 5.18E-01 | 0.4 | 2.44E-01 | Homeobox-leucine zipper protein HAT9 |
| AT5G18070 | -0.5 | 5.08E-04 | -0.1 | 3.94E-01 | -0.4 | 6.11E-03 | -0.2 | 2.37E-01 | -0.2 | 3.39E-01 | 0.0 | 9.53E-01 | Phosphoacetylglucosamine mutase |
| AT3G44400 | -0.5 | 6.03E-05 | -0.6 | 7.26E-05 | 0.0 | 7.40E-01 | -0.1 | 7.74E-01 | -0.2 | 5.14E-01 | 0.1 | 7.47E-01 | ADP-ribosyl cyclase |
| AT4G00720 | -0.5 | 3.55E-05 | -0.2 | 4.50E-02 | -0.4 | 1.68E-03 | -0.4 | 2.81E-02 | -0.3 | 2.19E-01 | -0.2 | 5.57E-01 | Shaggy-related protein kinase theta |
| AT5G28910 | -0.5 | 1.95E-02 | -0.3 | 1.25E-01 | -0.2 | 4.28E-01 | | | | | | | Alpha--fucosyltransferase |
| AT5G20910 | -0.5 | 6.16E-04 | -0.4 | 2.73E-03 | -0.1 | 5.05E-01 | 0.5 | 2.21E-02 | 0.3 | 1.30E-01 | 0.1 | 6.25E-01 | E3 ubiquitin-protein ligase AIP2 |
| AT3G49310 | -0.5 | 2.12E-05 | -0.3 | 2.70E-03 | -0.3 | 8.55E-03 | -0.3 | 3.23E-02 | -0.2 | 2.33E-01 | -0.1 | 5.82E-01 | Molybdate-anion transporter |
| AT2G18750 | -0.5 | 1.73E-04 | -0.4 | 1.79E-03 | -0.1 | 2.12E-01 | -0.2 | 5.55E-02 | -0.2 | 6.22E-02 | 0.0 | 9.32E-01 | Calmodulin-binding protein 60 C |
| AT3G51990 | -0.5 | 1.22E-04 | -0.3 | 1.15E-02 | -0.3 | 2.59E-02 | -0.4 | 2.67E-02 | -0.1 | 7.36E-01 | -0.3 | 1.77E-01 | Serine |
| AT2G29995 | -0.5 | 3.20E-03 | -0.5 | 4.46E-03 | 0.0 | 9.78E-01 | -0.3 | 1.68E-01 | 0.0 | 9.79E-01 | -0.3 | 2.80E-01 | Protein PSY3 |
| AT1G10760 | -0.5 | 1.65E-03 | -0.4 | 1.27E-02 | -0.1 | 3.47E-01 | -0.3 | 2.14E-01 | 0.2 | 5.62E-01 | -0.4 | 1.59E-01 | Alpha-glucan water dikinase 1, chloroplastic |
| AT5G53870 | -0.5 | 7.63E-03 | -0.6 | 3.47E-03 | 0.1 | 5.95E-01 | -0.5 | 1.21E-01 | -0.3 | 3.81E-01 | -0.2 | 7.04E-01 | Early nodulin-like protein 1 |
| AT1G18210 | -0.5 | 1.09E-03 | -0.2 | 1.52E-01 | -0.3 | 2.49E-02 | 0.1 | 6.11E-01 | -0.2 | 2.41E-01 | 0.2 | 1.74E-01 | Probable calcium-binding protein CML27 |
| AT1G62810 | -0.5 | 2.78E-05 | -0.4 | 2.10E-04 | -0.1 | 2.29E-01 | 0.1 | 7.33E-01 | -0.1 | 5.18E-01 | 0.2 | 3.89E-01 | Amine oxidase gamma 1 |
| AT5G15870 | -0.5 | 1.56E-04 | -0.4 | 1.12E-03 | -0.1 | 2.98E-01 | 0.0 | 8.79E-01 | -0.3 | 1.46E-01 | 0.2 | 2.80E-01 | glucan endo-1,3-beta-D-glucosidase |
| AT3G03490 | -0.5 | 5.92E-04 | -0.2 | 6.40E-02 | -0.3 | 3.52E-02 | -0.1 | 8.04E-01 | -0.3 | 1.79E-01 | 0.2 | 3.93E-01 | Peroxisome biogenesis protein 19-1 |
| AT5G06560 | -0.5 | 8.94E-06 | -0.3 | 6.73E-04 | -0.2 | 8.81E-03 | -0.3 | 4.42E-02 | -0.4 | 3.52E-03 | 0.2 | 2.76E-01 | Myosin-binding protein 7 |
| AT4G19640 | -0.5 | 4.88E-05 | -0.4 | 7.01E-04 | -0.1 | 1.20E-01 | -0.1 | 4.97E-01 | -0.3 | 1.27E-01 | 0.2 | 4.87E-01 | Ras-related small GTP-binding family protein |
| AT2G25160 | -0.5 | 1.94E-03 | 0.3 | 1.76E-02 | -0.9 | 3.67E-04 | -0.7 | 3.11E-01 | 1.2 | 2.54E-02 | -1.9 | 1.08E-02 | At2g25160 |
| AT2G35120 | -0.5 | 3.11E-03 | 0.2 | 1.68E-01 | -0.7 | 1.17E-03 | -0.1 | 7.60E-01 | 0.1 | 6.17E-01 | -0.2 | 4.21E-01 | Glycine cleavage system H protein 2, mitochondrial |
| AT1G30650 | -0.5 | 2.35E-04 | -0.2 | 4.03E-02 | -0.3 | 1.90E-02 | 0.2 | 4.65E-01 | 0.6 | 2.87E-02 | -0.4 | 1.55E-01 | Probable WRKY transcription factor 14 |
| AT2G34940 | -0.5 | 1.19E-03 | -0.5 | 2.44E-03 | 0.0 | 8.42E-01 | -0.3 | 3.86E-01 | 0.1 | 8.01E-01 | -0.4 | 3.31E-01 | Vacuolar-sorting receptor 5 |
| AT5G23430 | -0.5 | 1.54E-05 | -0.3 | 2.55E-03 | -0.3 | 5.63E-03 | -0.3 | 1.54E-01 | -0.1 | 7.99E-01 | -0.2 | 4.23E-01 | Katanin p80 WD40 repeat-containing subunit B1 homolog KTN80.4 |
| AT1G75410 | -0.5 | 2.21E-04 | -0.8 | 2.56E-05 | 0.3 | 3.27E-02 | 0.0 | 8.90E-01 | 0.1 | 6.47E-01 | -0.1 | 8.13E-01 | BEL1-like homeodomain 3 |
| AT3G50660 | -0.5 | 2.24E-03 | 0.0 | 9.53E-01 | -0.5 | 6.92E-03 | -0.3 | 7.22E-02 | -0.4 | 2.39E-02 | 0.1 | 6.42E-01 | Cytochrome P450 superfamily protein |
| AT4G38730 | -0.5 | 1.06E-04 | -0.6 | 1.03E-04 | 0.1 | 6.47E-01 | -0.4 | 4.12E-02 | -0.3 | 1.66E-01 | -0.1 | 7.15E-01 | Probable magnesium transporter |
| AT1G74490 | -0.5 | 2.89E-02 | 0.5 | 2.97E-02 | -1.0 | 3.01E-03 | -0.1 | 8.96E-01 | 0.6 | 1.66E-01 | -0.7 | 1.40E-01 | At1g74490 |
| AT3G01090 | -0.5 | 1.35E-05 | -0.4 | 1.48E-04 | -0.1 | 1.19E-01 | -0.2 | 6.05E-02 | 0.0 | 7.39E-01 | -0.2 | 3.05E-01 | SNF1-related protein kinase catalytic subunit alpha KIN10 |
| AT5G45130 | -0.5 | 1.51E-05 | -0.3 | 5.56E-04 | -0.2 | 2.62E-02 | -0.3 | 1.19E-02 | -0.2 | 6.79E-02 | -0.1 | 7.56E-01 | Ras small GTP-binding family protein |
| AT2G22200 | -0.5 | 2.10E-01 | 0.0 | 9.76E-01 | -0.5 | 3.34E-01 | -0.4 | 2.79E-01 | -0.2 | 6.35E-01 | -0.2 | 7.60E-01 | Ethylene-responsive transcription factor ERF056 |
| AT3G63340 | -0.5 | 7.71E-04 | -0.5 | 1.25E-03 | 0.0 | 9.46E-01 | -0.1 | 6.31E-01 | 0.1 | 4.39E-01 | -0.2 | 2.70E-01 | AT3G63340 protein |
| AT5G09890 | -0.5 | 2.51E-05 | -0.2 | 3.44E-02 | -0.4 | 1.36E-03 | -0.1 | 3.70E-01 | 0.0 | 8.40E-01 | -0.2 | 4.22E-01 | Protein kinase |
| AT1G72770 | -0.5 | 9.41E-04 | -0.8 | 9.35E-05 | 0.3 | 6.07E-02 | -0.3 | 7.97E-02 | -0.1 | 7.69E-01 | -0.3 | 3.19E-01 | Protein phosphatase 2C 16 |
| AT2G34140 | -0.5 | 3.42E-03 | -0.2 | 1.29E-01 | -0.3 | 1.11E-01 | 0.2 | 3.93E-01 | 0.0 | 9.90E-01 | 0.2 | 5.03E-01 | Cyclic dof factor 4 |
| AT3G11490 | -0.5 | 2.15E-05 | -0.1 | 1.05E-01 | -0.4 | 6.52E-04 | -0.4 | 9.80E-03 | -0.3 | 8.12E-02 | -0.1 | 5.73E-01 | Rho GTPase-activating protein 4 |
| AT1G28600 | -0.5 | 1.94E-02 | -1.0 | 5.92E-04 | 0.5 | 4.25E-02 | 0.0 | 8.03E-01 | 0.1 | 7.67E-01 | -0.1 | 6.42E-01 | GDSL esterase |
| AT2G46680 | -0.5 | 2.38E-03 | -0.8 | 2.01E-04 | 0.3 | 7.54E-02 | -0.3 | 4.65E-01 | 0.3 | 4.80E-01 | -0.6 | 1.93E-01 | Homeobox-leucine zipper protein |
| AT4G37010 | -0.5 | 8.71E-03 | -0.2 | 2.72E-01 | -0.3 | 1.17E-01 | -0.1 | 7.82E-01 | -0.3 | 5.30E-01 | 0.2 | 7.75E-01 | Centrin 2 |
| AT1G80210 | -0.5 | 8.81E-05 | -0.4 | 3.96E-04 | -0.1 | 4.89E-01 | -0.3 | 1.24E-01 | -0.2 | 3.95E-01 | -0.1 | 7.06E-01 | At1g80210 |
| AT1G07510 | -0.5 | 1.67E-05 | -0.4 | 1.13E-04 | -0.1 | 2.41E-01 | 0.1 | 4.24E-01 | 0.1 | 7.73E-01 | 0.1 | 8.17E-01 | ATP-dependent zinc metalloprotease FTSH 10, mitochondrial |
| AT1G55180 | -0.5 | 1.79E-04 | -0.1 | 2.22E-01 | -0.4 | 4.15E-03 | -0.1 | 7.09E-01 | 0.0 | 9.91E-01 | -0.1 | 7.74E-01 | Phospholipase D |
| AT5G01270 | -0.5 | 4.74E-05 | -0.3 | 2.73E-03 | -0.2 | 2.72E-02 | 0.2 | 1.13E-01 | 0.3 | 5.53E-02 | -0.1 | 7.51E-01 | protein-serine |
| AT2G34560 | -0.5 | 1.21E-04 | 0.0 | 7.02E-01 | -0.5 | 7.33E-04 | -0.3 | 1.98E-02 | 0.1 | 7.20E-01 | -0.3 | 3.63E-02 | AT2G34560 protein |
| AT5G03240 | -0.5 | 7.08E-05 | -0.3 | 7.89E-03 | -0.2 | 1.60E-02 | -0.3 | 3.84E-03 | -0.3 | 1.57E-02 | 0.0 | 8.69E-01 | Polyubiquitin 3 |
| AT3G54020 | -0.5 | 5.76E-05 | -0.4 | 4.92E-04 | -0.1 | 2.43E-01 | -0.2 | 5.06E-01 | 0.3 | 2.44E-01 | -0.4 | 1.01E-01 | Phosphatidylinositol:ceramide inositolphosphotransferase 1 |
| AT5G03080 | -0.5 | 2.21E-05 | -0.2 | 5.57E-03 | -0.3 | 4.66E-03 | -0.2 | 6.94E-02 | 0.0 | 9.75E-01 | -0.2 | 1.87E-01 | Lipid phosphate phosphatase gamma |
| AT1G78420 | -0.5 | 1.66E-05 | -0.6 | 1.10E-05 | 0.1 | 2.33E-01 | 0.0 | 9.12E-01 | 0.0 | 9.41E-01 | 0.0 | 9.93E-01 | E3 ubiquitin-protein ligase DA2 |
| AT5G51050 | -0.5 | 2.27E-04 | -0.4 | 1.81E-03 | -0.1 | 3.03E-01 | -0.1 | 6.90E-01 | 0.2 | 3.64E-01 | -0.4 | 1.99E-01 | Calcium-dependent mitochondrial ATP-magnesium |
| AT5G26780 | -0.5 | 5.19E-05 | -0.2 | 6.70E-02 | -0.4 | 1.93E-03 | 0.1 | 4.84E-01 | 0.3 | 4.49E-02 | -0.2 | 3.08E-01 | Serine hydroxymethyltransferase 2, mitochondrial |
| AT5G40150 | -0.5 | 3.74E-03 | -0.1 | 6.04E-01 | -0.4 | 2.43E-02 | -0.3 | 1.77E-01 | -0.1 | 6.16E-01 | -0.1 | 6.43E-01 | Peroxidase 63 |
| AT5G18930 | -0.5 | 1.38E-03 | -0.3 | 2.86E-02 | -0.2 | 1.81E-01 | 0.0 | 9.49E-01 | 0.0 | 9.60E-01 | 0.1 | 9.26E-01 | S-adenosylmethionine decarboxylase proenzyme 4 |
| AT5G18550 | -0.5 | 6.26E-04 | -0.4 | 2.83E-03 | -0.1 | 5.08E-01 | 0.0 | 8.89E-01 | 0.1 | 5.08E-01 | -0.2 | 4.31E-01 | Zinc finger C-x8-C-x5-C-x3-H type family protein |
| AT5G58670 | -0.5 | 2.65E-04 | -0.3 | 4.08E-03 | -0.2 | 1.65E-01 | -0.4 | 4.25E-02 | 0.0 | 9.87E-01 | -0.4 | 1.41E-01 | Phosphoinositide phospholipase C 1 |
| AT3G55430 | -0.5 | 6.11E-05 | -0.3 | 1.85E-03 | -0.2 | 5.21E-02 | -0.6 | 6.19E-05 | -0.5 | 4.77E-04 | -0.1 | 8.04E-01 | glucan endo-1,3-beta-D-glucosidase |
| AT5G56140 | -0.5 | 1.32E-05 | -0.4 | 1.99E-04 | -0.1 | 7.69E-02 | -0.7 | 1.37E-04 | -0.6 | 9.48E-04 | -0.1 | 8.11E-01 | KH domain-containing protein At5g56140 |
| AT5G66950 | -0.5 | 5.41E-06 | -0.5 | 2.06E-05 | 0.0 | 4.90E-01 | -0.1 | 5.10E-01 | 0.1 | 7.44E-01 | -0.2 | 4.52E-01 | Pyridoxal phosphate -dependent transferases superfamily protein |
| AT1G51630 | -0.5 | 7.10E-05 | -0.2 | 3.33E-02 | -0.3 | 4.50E-03 | -0.1 | 2.81E-01 | 0.1 | 6.34E-01 | -0.2 | 1.32E-01 | Protein MANNAN SYNTHESIS-RELATED 2 |
| AT2G40770 | -0.5 | 2.83E-05 | -0.6 | 3.05E-05 | 0.0 | 6.02E-01 | -0.3 | 2.73E-01 | -0.3 | 3.58E-01 | 0.0 | 9.72E-01 | RING-finger, DEAD-like helicase, PHD and SNF2 domain-containing protein |
| AT2G25090 | -0.5 | 1.46E-04 | 0.1 | 1.27E-01 | -0.6 | 2.03E-04 | -1.8 | 4.77E-04 | -1.0 | 1.20E-02 | -0.7 | 2.51E-01 | SnRK3.18 |
| AT1G71010 | -0.5 | 7.41E-05 | -0.2 | 1.59E-02 | -0.3 | 8.56E-03 | -0.1 | 6.80E-01 | 0.2 | 5.86E-01 | -0.3 | 4.30E-01 | Putative 1-phosphatidylinositol-3-phosphate 5-kinase FAB1C |
| AT2G03890 | -0.5 | 1.74E-04 | -0.4 | 9.28E-04 | -0.1 | 3.97E-01 | -0.5 | 1.31E-03 | -0.6 | 4.08E-04 | 0.1 | 4.64E-01 | 1-phosphatidylinositol 4-kinase |
| AT4G01550 | -0.5 | 4.84E-04 | -0.7 | 7.64E-05 | 0.2 | 9.22E-02 | -0.2 | 4.81E-01 | 0.1 | 7.08E-01 | -0.3 | 3.23E-01 | NAC domain-containing protein 69 |
| AT1G67500 | -0.5 | 1.06E-04 | -0.6 | 8.41E-05 | 0.1 | 5.31E-01 | 0.1 | 6.12E-01 | 0.0 | 9.19E-01 | 0.1 | 8.08E-01 | DNA polymerase |
| AT3G24800 | -0.5 | 9.57E-05 | -0.2 | 1.28E-02 | -0.3 | 1.77E-02 | -0.3 | 1.72E-01 | -0.1 | 7.30E-01 | -0.2 | 5.52E-01 | PRT1 |
| AT5G43100 | -0.5 | 1.52E-05 | -0.4 | 1.00E-04 | -0.1 | 2.51E-01 | 0.0 | 7.69E-01 | -0.1 | 7.25E-01 | 0.1 | 5.97E-01 | Eukaryotic aspartyl protease family protein |
| AT5G62620 | -0.5 | 6.11E-04 | -0.6 | 2.09E-04 | 0.1 | 3.00E-01 | -0.2 | 3.59E-01 | -0.1 | 8.00E-01 | -0.1 | 7.08E-01 | Galactosyltransferase family protein |
| AT5G67470 | -0.5 | 1.41E-05 | -0.5 | 2.59E-05 | 0.0 | 9.59E-01 | 0.1 | 8.03E-01 | 0.2 | 5.34E-01 | -0.1 | 7.67E-01 | Formin-like protein 6 |
| AT4G00550 | -0.5 | 5.30E-05 | -0.5 | 6.43E-05 | 0.0 | 7.33E-01 | 0.0 | 9.42E-01 | 0.1 | 7.17E-01 | -0.1 | 6.76E-01 | Digalactosyldiacylglycerol synthase 2, chloroplastic |
| AT1G01340 | -0.5 | 4.65E-04 | -0.5 | 1.12E-03 | 0.0 | 7.54E-01 | 0.3 | 1.59E-01 | 0.1 | 8.16E-01 | 0.2 | 4.71E-01 | Probable cyclic nucleotide-gated ion channel 10 |

| | | | | | | | | | | | | | |
|---|---|---|---|---|---|---|---|---|---|---|---|---|---|
| AT3G07880 | -0.5 | 1.45E-05 | -0.1 | 2.77E-01 | -0.4 | 2.19E-04 | -0.5 | 8.53E-04 | -0.1 | 5.29E-01 | -0.4 | 3.80E-02 | Rho GDP-dissociation inhibitor 1 |
| AT1G68150 | -0.5 | 1.89E-04 | -0.2 | 1.00E-01 | -0.4 | 7.21E-03 | 0.1 | 5.74E-01 | 0.4 | 8.44E-02 | -0.3 | 2.87E-01 | Probable WRKY transcription factor 9 |
| AT1G78060 | -0.5 | 6.27E-04 | -0.2 | 4.76E-02 | -0.3 | 4.05E-02 | -0.1 | 2.24E-01 | 0.1 | 2.79E-01 | -0.3 | 8.45E-02 | Probable beta-D-xylosidase 7 |
| AT4G09950 | -0.5 | 5.72E-03 | -0.5 | 1.10E-02 | 0.0 | 8.47E-01 | 0.0 | 9.25E-01 | 0.1 | 7.54E-01 | -0.1 | 8.48E-01 | P-loop containing nucleoside triphosphate hydrolases superfamily protein |
| AT2G19880 | -0.5 | 1.28E-05 | -0.2 | 5.78E-03 | -0.3 | 2.10E-03 | -0.1 | 3.45E-01 | 0.1 | 7.69E-01 | -0.2 | 2.07E-01 | ceramide glucosyltransferase |
| AT4G19110 | -0.5 | 2.25E-05 | -0.3 | 9.51E-04 | -0.2 | 2.74E-02 | 0.0 | 9.55E-01 | 0.1 | 6.81E-01 | -0.1 | 7.12E-01 | Protein kinase superfamily protein |
| AT1G63500 | -0.5 | 2.66E-05 | -0.3 | 1.85E-03 | -0.2 | 1.56E-02 | 0.0 | 9.90E-01 | 0.0 | 9.42E-01 | 0.0 | 9.40E-01 | Serine |
| AT2G28240 | -0.5 | 1.84E-03 | -0.5 | 2.97E-03 | 0.0 | 9.44E-01 | -0.3 | 5.07E-01 | 0.1 | 9.27E-01 | -0.3 | 5.17E-01 | ATP-dependent helicase family protein |
| AT5G13530 | -0.5 | 2.40E-05 | -0.5 | 6.39E-05 | 0.0 | 7.58E-01 | 0.0 | 9.22E-01 | -0.2 | 5.16E-01 | 0.1 | 6.92E-01 | RING-type E3 ubiquitin transferase |
| AT5G07620 | -0.5 | 2.41E-02 | -0.8 | 4.71E-03 | 0.3 | 3.48E-01 | | | | | | | Protein kinase domain-containing protein |
| AT1G71870 | -0.5 | 1.10E-03 | 0.3 | 1.41E-02 | -0.8 | 2.21E-04 | -0.1 | 4.86E-01 | 0.1 | 7.39E-01 | -0.2 | 3.27E-01 | Protein DETOXIFICATION |
| AT3G47050 | -0.5 | 6.82E-03 | -0.3 | 5.42E-02 | -0.2 | 3.74E-01 | -0.1 | 8.05E-01 | 0.6 | 1.10E-01 | -0.7 | 4.37E-02 | Beta-D-glucan exohydrolase-like protein |
| AT4G17880 | -0.5 | 3.61E-05 | -0.3 | 1.39E-03 | -0.2 | 3.48E-02 | -0.1 | 7.19E-01 | 0.3 | 5.56E-02 | -0.3 | 6.99E-02 | Transcription factor MYC4 |
| AT5G41690 | -0.5 | 3.34E-03 | -0.5 | 6.79E-03 | 0.0 | 8.64E-01 | 0.0 | 9.73E-01 | 0.1 | 8.99E-01 | -0.1 | 9.37E-01 | RNA-binding family protein |
| AT5G43420 | -0.5 | 1.01E-03 | -0.3 | 1.10E-02 | -0.2 | 2.55E-01 | 0.0 | 8.88E-01 | 0.1 | 7.95E-01 | 0.0 | 9.11E-01 | RING-H2 finger protein ATL16 |
| AT3G51480 | -0.5 | 1.03E-03 | -0.4 | 3.16E-03 | -0.1 | 6.81E-01 | -0.3 | 3.09E-01 | 0.0 | 9.21E-01 | -0.3 | 3.74E-01 | Glutamate receptor |
| AT5G38550 | -0.5 | 1.26E-03 | -0.3 | 2.77E-02 | -0.2 | 1.50E-01 | -0.3 | 4.48E-01 | 0.1 | 7.19E-01 | -0.4 | 2.57E-01 | Mannose-binding lectin superfamily protein |
| AT5G66620 | -0.5 | 1.85E-03 | -0.7 | 4.20E-04 | 0.2 | 2.38E-01 | -1.3 | 3.89E-03 | -0.3 | 5.45E-01 | -1.0 | 6.27E-02 | DA1-related protein 6 |
| AT3G55320 | -0.5 | 6.20E-05 | -0.5 | 9.88E-05 | 0.0 | 9.30E-01 | -0.1 | 7.62E-01 | 0.0 | 8.99E-01 | -0.1 | 7.34E-01 | ABC transporter B family member 20 |
| AT4G31450 | -0.5 | 2.19E-05 | -0.3 | 5.72E-04 | -0.2 | 4.39E-02 | -0.4 | 1.63E-02 | 0.1 | 7.73E-01 | -0.4 | 2.81E-02 | RING-type E3 ubiquitin transferase |
| AT2G19760 | -0.5 | 1.85E-04 | 0.2 | 8.75E-02 | -0.7 | 1.33E-04 | -0.4 | 1.27E-02 | 0.0 | 8.25E-01 | -0.3 | 7.92E-02 | Profilin-1 |
| AT2G17370 | -0.5 | 6.95E-05 | -0.1 | 1.53E-01 | -0.4 | 1.85E-03 | -0.1 | 5.73E-01 | 0.1 | 6.59E-01 | -0.2 | 4.08E-01 | 3-hydroxy-3-methylglutaryl-coenzyme A reductase 2 |
| AT1G07240 | -0.5 | 6.77E-05 | -0.2 | 1.73E-02 | -0.3 | 7.21E-03 | -0.1 | 4.58E-01 | -0.4 | 1.39E-02 | 0.3 | 1.40E-01 | UDP-glycosyltransferase 71C5 |
| AT2G39000 | -0.5 | 6.12E-05 | -0.4 | 6.72E-04 | -0.1 | 1.70E-01 | -0.7 | 8.07E-05 | -0.2 | 1.02E-01 | -0.4 | 2.78E-02 | Acyl-CoA N-acyltransferases superfamily protein |
| AT3G09035 | -0.5 | 9.01E-05 | -0.2 | 1.85E-02 | -0.3 | 1.07E-02 | -0.4 | 2.32E-02 | -0.3 | 1.03E-01 | -0.1 | 7.71E-01 | Legume lectin domain-containing protein |
| AT4G16960 | -0.5 | 1.60E-02 | -0.7 | 3.54E-03 | 0.2 | 3.76E-01 | -0.1 | 9.25E-01 | 0.0 | 9.65E-01 | -0.1 | 9.14E-01 | Disease resistance protein |
| AT3G51520 | -0.5 | 4.17E-05 | -0.2 | 7.01E-03 | -0.3 | 8.32E-03 | -0.4 | 3.80E-03 | -0.2 | 1.30E-01 | -0.2 | 3.43E-01 | Diacylglycerol O-acyltransferase 2 |
| AT1G65080 | -0.5 | 1.42E-03 | -0.4 | 6.71E-03 | -0.1 | 5.21E-01 | 0.5 | 1.37E-01 | 0.1 | 9.29E-01 | 0.5 | 2.88E-01 | ALBINO3-like protein 2, chloroplastic |
| AT2G28260 | -0.5 | 1.68E-04 | -0.3 | 1.08E-02 | -0.2 | 4.30E-02 | -0.3 | 1.64E-01 | -0.1 | 8.25E-01 | -0.2 | 4.04E-01 | Cyclic nucleotide-gated channel 15 |
| AT4G30520 | -0.5 | 3.33E-04 | -0.4 | 2.43E-03 | -0.1 | 3.40E-01 | -0.3 | 7.24E-02 | 0.0 | 9.69E-01 | -0.3 | 1.61E-01 | Probable LRR receptor-like serine |
| AT5G08139 | -0.5 | 3.85E-04 | -0.3 | 5.83E-03 | -0.2 | 1.76E-01 | -0.2 | 1.48E-01 | -0.4 | 3.43E-02 | 0.2 | 5.23E-01 | Uncharacterized protein T22D6_80 |
| AT2G33550 | -0.5 | 4.41E-03 | -0.5 | 5.68E-03 | 0.0 | 1.00E+00 | 0.7 | 1.03E-03 | 0.2 | 4.71E-01 | 0.5 | 1.24E-02 | Trihelix transcription factor ASR3 |
| AT4G36520 | -0.5 | 2.74E-04 | -0.1 | 2.36E-01 | -0.4 | 4.19E-03 | -0.1 | 6.80E-01 | 0.0 | 9.90E-01 | -0.1 | 7.82E-01 | Chaperone DnaJ-domain superfamily protein |
| AT5G40250 | -0.5 | 2.26E-04 | -0.4 | 2.06E-03 | -0.1 | 2.59E-01 | -0.3 | 1.28E-02 | -0.1 | 2.86E-01 | -0.2 | 3.30E-01 | RING-H2 finger protein ATL46 |
| AT3G29200 | -0.5 | 3.60E-05 | -0.2 | 1.35E-02 | -0.3 | 4.12E-03 | -0.2 | 1.31E-01 | 0.1 | 6.06E-01 | -0.3 | 1.20E-01 | Chorismate mutase 1, chloroplastic |
| AT1G67710 | -0.5 | 7.79E-05 | -0.6 | 5.96E-05 | 0.1 | 4.77E-01 | -0.7 | 6.22E-03 | -0.6 | 1.59E-02 | -0.1 | 8.74E-01 | Response regulator 11 |
| AT4G23490 | -0.5 | 1.58E-03 | -0.4 | 4.10E-03 | -0.1 | 7.51E-01 | -0.2 | 2.47E-01 | 0.4 | 5.08E-02 | -0.7 | 1.43E-02 | At4g23490 |
| AT1G17420 | -0.5 | 1.26E-03 | -0.5 | 1.08E-03 | 0.0 | 7.66E-01 | 0.1 | 6.91E-01 | 0.2 | 4.38E-01 | -0.1 | 7.61E-01 | Lipoxygenase |
| AT5G51170 | -0.5 | 9.40E-04 | -0.2 | 6.95E-02 | -0.3 | 5.65E-02 | -0.2 | 2.95E-01 | -0.3 | 2.30E-01 | 0.1 | 8.86E-01 | U6 snRNA phosphodiesterase 1 |
| AT5G23580 | -0.5 | 2.30E-05 | -0.2 | 8.33E-03 | -0.3 | 4.21E-03 | 0.2 | 1.04E-01 | 0.2 | 1.24E-01 | 0.0 | 9.68E-01 | Calcium-dependent protein kinase 12 |
| AT5G16800 | -0.5 | 9.28E-05 | -0.4 | 7.34E-04 | -0.1 | 2.63E-01 | -0.4 | 1.78E-02 | -0.4 | 1.45E-02 | 0.1 | 8.52E-01 | histone acetyltransferase |
| AT5G27420 | -0.5 | 1.63E-02 | -1.2 | 1.42E-04 | 0.7 | 7.97E-03 | 0.4 | 1.76E-01 | -0.5 | 1.26E-01 | 0.8 | 2.10E-02 | E3 ubiquitin-protein ligase ATL31 |
| AT4G17170 | -0.5 | 2.13E-05 | -0.3 | 1.81E-03 | -0.2 | 1.19E-02 | -0.5 | 1.95E-04 | -0.3 | 7.31E-03 | -0.1 | 3.97E-01 | Ras-related protein RABB1c |
| AT5G25160 | -0.5 | 8.27E-04 | -0.3 | 1.23E-02 | -0.2 | 2.00E-01 | 0.2 | 5.95E-01 | 0.1 | 7.31E-01 | 0.0 | 9.56E-01 | Zinc finger protein 3 |
| AT3G16857 | -0.5 | 1.61E-04 | -0.3 | 5.69E-03 | -0.2 | 5.83E-02 | -0.4 | 1.41E-01 | -0.4 | 1.32E-01 | 0.1 | 9.06E-01 | Two-component response regulator |
| AT1G21980 | -0.5 | 1.23E-04 | -0.3 | 4.40E-03 | -0.2 | 5.55E-02 | 0.1 | 5.59E-01 | 0.2 | 3.60E-01 | -0.1 | 8.00E-01 | Phosphatidylinositol 4-phosphate 5-kinase 1 P-5-kinase 1) |
| AT2G15570 | -0.5 | 1.18E-05 | -0.2 | 2.01E-03 | -0.2 | 4.68E-03 | -0.3 | 5.10E-02 | -0.3 | 5.39E-02 | 0.0 | 9.22E-01 | Thioredoxin superfamily protein |
| AT5G64340 | -0.5 | 2.85E-04 | -0.6 | 2.04E-04 | 0.1 | 5.61E-01 | -0.2 | 2.04E-01 | 0.0 | 9.58E-01 | -0.2 | 3.42E-01 | Transcription factor SAC51 |
| AT3G62360 | -0.5 | 1.95E-05 | -0.2 | 1.26E-02 | -0.3 | 1.97E-03 | 0.1 | 5.65E-01 | 0.1 | 5.83E-01 | 0.0 | 9.69E-01 | Carbohydrate-binding-like fold |
| AT2G32390 | -0.5 | 6.22E-04 | -0.1 | 2.38E-01 | -0.4 | 1.16E-02 | 0.0 | 9.29E-01 | 0.2 | 5.29E-01 | -0.3 | 4.76E-01 | Glutamate receptor |
| AT1G27730 | -0.5 | 1.89E-01 | -1.4 | 5.13E-03 | 0.9 | 6.37E-02 | 0.8 | 3.30E-04 | -0.2 | 5.10E-01 | 1.0 | 1.06E-03 | Zinc finger protein ZAT10 |
| AT3G45840 | -0.5 | 3.03E-03 | -0.1 | 6.70E-01 | -0.4 | 1.90E-02 | 0.2 | 4.76E-01 | 0.1 | 8.19E-01 | 0.1 | 7.87E-01 | Cysteine |
| AT3G53400 | -0.5 | 3.55E-03 | -0.3 | 2.54E-02 | -0.1 | 3.67E-01 | 0.0 | 8.16E-01 | 0.4 | 7.42E-03 | -0.4 | 4.25E-02 | Peptide upstream protein |
| AT5G46760 | -0.5 | 2.50E-05 | -0.5 | 6.27E-05 | 0.0 | 8.49E-01 | -0.4 | 8.36E-04 | -0.3 | 1.33E-02 | -0.1 | 5.79E-01 | Transcription factor MYC3 |
| AT4G09160 | -0.5 | 5.37E-05 | -0.1 | 3.07E-01 | -0.4 | 8.25E-04 | -0.3 | 1.74E-02 | -0.3 | 2.59E-02 | 0.0 | 9.86E-01 | SEC14 cytosolic factor family protein |
| AT4G24220 | -0.5 | 6.48E-05 | -0.4 | 2.78E-04 | -0.1 | 5.16E-01 | -0.5 | 5.05E-05 | -0.3 | 1.31E-02 | -0.2 | 9.61E-02 | 3-oxo-Delta-steroid 5-beta-reductase -3-oxosteroid 5-beta-reductase) |
| AT3G22425 | -0.5 | 1.31E-05 | -0.3 | 5.78E-04 | -0.2 | 1.99E-02 | -0.1 | 5.04E-01 | 0.1 | 7.03E-01 | -0.2 | 3.71E-01 | Imidazoleglycerol-phosphate dehydratase |
| AT3G61960 | -0.5 | 4.87E-05 | -0.6 | 3.09E-05 | 0.1 | 3.34E-01 | -0.1 | 7.21E-01 | 0.3 | 1.01E-01 | -0.4 | 9.19E-02 | Protein kinase superfamily protein |
| AT1G68810 | -0.5 | 3.10E-04 | -0.2 | 4.33E-02 | -0.3 | 2.26E-02 | -0.1 | 7.60E-01 | -0.1 | 6.27E-01 | 0.0 | 8.88E-01 | Transcription factor bHLH30 |
| AT1G60890 | -0.5 | 4.55E-04 | -0.5 | 3.76E-04 | 0.1 | 6.77E-01 | 0.2 | 4.35E-01 | 0.3 | 1.51E-01 | -0.2 | 6.10E-01 | Phosphatidylinositol 4-phosphate 5-kinase |
| AT4G32710 | -0.5 | 2.56E-04 | -0.3 | 1.16E-02 | -0.2 | 6.29E-02 | -0.3 | 7.78E-02 | 0.0 | 9.79E-01 | -0.3 | 1.55E-01 | Proline-rich receptor-like protein kinase PERK14 |
| AT1G06490 | -0.5 | 1.05E-03 | -0.5 | 1.32E-03 | 0.0 | 9.21E-01 | 0.1 | 7.37E-01 | -0.1 | 7.95E-01 | 0.3 | 6.11E-01 | 1,3-beta-glucan synthase |
| AT5G24400 | -0.5 | 4.81E-04 | -0.1 | 5.53E-01 | -0.4 | 3.30E-03 | 0.0 | 8.97E-01 | 0.0 | 8.95E-01 | 0.0 | 8.37E-01 | 6-phosphogluconolactonase 3, chloroplastic |
| AT4G17830 | -0.5 | 1.07E-05 | -0.5 | 2.51E-05 | 0.0 | 8.73E-01 | 0.0 | 9.54E-01 | 0.0 | 8.81E-01 | 0.0 | 8.67E-01 | Peptidase M20 |
| AT5G27690 | -0.5 | 1.38E-04 | -0.2 | 2.27E-02 | -0.3 | 1.80E-02 | -0.2 | 4.93E-01 | 0.1 | 8.34E-01 | -0.2 | 4.74E-01 | Heavy metal-associated isoprenylated plant protein 36 |
| AT1G43160 | -0.5 | 6.34E-02 | 0.3 | 2.11E-01 | -0.8 | 1.85E-02 | 0.0 | 9.92E-01 | 0.6 | 2.60E-01 | -0.6 | 2.37E-01 | Ethylene-responsive transcription factor RAP2-6 |
| AT1G17760 | -0.5 | 2.76E-05 | -0.4 | 1.01E-04 | 0.0 | 5.80E-01 | -0.2 | 4.56E-01 | 0.1 | 8.18E-01 | -0.3 | 3.80E-01 | Cleavage stimulation factor subunit 77 |
| AT1G11100 | -0.5 | 3.47E-03 | -0.7 | 6.60E-04 | 0.2 | 2.39E-01 | 0.2 | 6.62E-01 | 0.3 | 5.90E-01 | -0.1 | 9.15E-01 | SNF2 domain-containing protein |
| AT1G68910 | -0.5 | 6.91E-05 | -0.5 | 6.32E-05 | 0.1 | 5.74E-01 | -0.3 | 1.68E-01 | -0.1 | 8.18E-01 | -0.2 | 4.77E-01 | WPP domain-interacting tail-anchored protein 2 |
| AT3G10540 | -0.5 | 3.26E-05 | -0.2 | 1.01E-02 | -0.3 | 5.32E-03 | -0.1 | 7.32E-01 | 0.0 | 8.54E-01 | -0.1 | 6.65E-01 | 3-phosphoinositide-dependent protein kinase 2 |
| AT3G17410 | -0.5 | 8.08E-05 | -0.4 | 3.30E-04 | -0.1 | 5.32E-01 | -0.1 | 4.34E-01 | -0.2 | 4.29E-02 | 0.1 | 3.31E-01 | Receptor-like cytoplasmic kinase 1 |
| AT1G74840 | -0.5 | 4.32E-04 | -0.1 | 3.23E-01 | -0.4 | 4.88E-03 | -0.9 | 1.56E-05 | -0.3 | 4.16E-02 | -0.5 | 9.82E-03 | AT1G74840 protein |
| AT1G50890 | -0.5 | 3.02E-04 | 0.0 | 9.78E-01 | -0.5 | 1.43E-03 | -0.2 | 4.39E-01 | 0.2 | 4.81E-01 | -0.4 | 1.76E-01 | TORTIFOLIA1-like protein 1 |
| AT4G29080 | -0.5 | 5.38E-05 | -0.3 | 2.34E-03 | -0.2 | 3.48E-02 | 0.0 | 9.38E-01 | 0.2 | 2.58E-01 | -0.3 | 3.19E-01 | Auxin-responsive protein IAA27 |
| AT5G65790 | -0.5 | 4.22E-05 | -0.4 | 4.40E-04 | -0.1 | 1.87E-01 | 0.8 | 1.54E-04 | 0.5 | 1.10E-02 | 0.3 | 1.48E-01 | At5g65790 |
| AT1G03040 | -0.5 | 3.27E-05 | -0.4 | 1.10E-04 | 0.0 | 5.96E-01 | 0.0 | 9.81E-01 | -0.1 | 5.08E-01 | 0.1 | 5.62E-01 | Basic helix-loop-helix DNA-binding superfamily protein |
| AT1G64060 | -0.5 | 1.03E-03 | -0.5 | 1.22E-03 | 0.0 | 9.20E-01 | -0.1 | 6.12E-01 | 0.1 | 6.16E-01 | -0.2 | 3.62E-01 | Respiratory burst oxidase homolog protein F |
| AT5G42780 | -0.5 | 6.19E-03 | -0.1 | 6.79E-01 | -0.4 | 3.38E-02 | 0.0 | 9.35E-01 | -0.3 | 3.81E-01 | 0.3 | 5.12E-01 | Zinc-finger homeodomain protein 13 |
| AT2G24640 | -0.5 | 7.22E-04 | -0.2 | 5.89E-02 | -0.3 | 4.56E-02 | 0.1 | 8.04E-01 | 0.2 | 4.80E-01 | -0.1 | 7.09E-01 | AT2G24640 protein |
| AT5G27150 | -0.5 | 5.22E-05 | -0.4 | 3.15E-04 | -0.1 | 3.30E-01 | -0.3 | 2.37E-02 | -0.1 | 5.47E-01 | -0.2 | 2.91E-01 | Sodium |
| AT4G36890 | -0.5 | 2.18E-05 | -0.3 | 1.52E-03 | -0.2 | 1.51E-02 | -0.2 | 3.78E-01 | 0.0 | 9.90E-01 | -0.2 | 4.19E-01 | Beta-1,4-xylosyltransferase IRX14 |
| AT2G38300 | -0.5 | 1.65E-03 | -0.1 | 2.94E-01 | -0.4 | 2.57E-02 | 0.0 | 9.72E-01 | 0.4 | 1.36E-01 | -0.4 | 1.63E-01 | HTH myb-type domain-containing protein |
| AT4G20110 | -0.5 | 4.72E-04 | -0.4 | 3.28E-03 | -0.1 | 3.35E-01 | -0.3 | 1.54E-01 | -0.2 | 3.77E-01 | -0.1 | 8.31E-01 | VACUOLAR SORTING RECEPTOR 7 |

| | | | | | | | | | | | | | |
|---|---|---|---|---|---|---|---|---|---|---|---|---|---|
| AT2G17720 | -0.5 | 1.96E-03 | 0.1 | 6.96E-01 | -0.5 | 2.81E-03 | 0.4 | 3.08E-03 | 0.2 | 1.38E-01 | 0.2 | 2.51E-01 | Prolyl 4-hydroxylase 5 |
| AT4G37530 | -0.5 | 3.76E-04 | 0.4 | 3.56E-03 | -0.8 | 5.75E-05 | 0.1 | 5.96E-01 | 0.1 | 5.59E-01 | 0.0 | 9.31E-01 | Peroxidase |
| AT3G20770 | -0.5 | 6.80E-05 | -0.5 | 1.93E-04 | 0.0 | 7.53E-01 | -0.2 | 9.90E-02 | -0.2 | 1.58E-01 | 0.0 | 9.86E-01 | Protein ETHYLENE INSENSITIVE 3 |
| AT5G59550 | -0.5 | 8.51E-04 | -0.6 | 5.18E-04 | 0.1 | 5.80E-01 | -0.5 | 1.20E-03 | -0.5 | 1.46E-03 | 0.0 | 8.48E-01 | RING-type E3 ubiquitin transferase |
| AT4G36930 | -0.5 | 1.20E-03 | -0.1 | 5.58E-01 | -0.4 | 1.11E-02 | -0.1 | 6.17E-01 | -0.1 | 8.20E-01 | -0.1 | 8.68E-01 | BHLH domain-containing protein |
| AT5G20870 | -0.5 | 4.52E-03 | -0.2 | 2.70E-01 | -0.3 | 7.40E-02 | -0.3 | 5.15E-01 | -0.2 | 7.04E-01 | -0.1 | 9.11E-01 | glucan endo-1,3-beta-D-glucosidase |
| AT1G62310 | -0.5 | 4.40E-03 | -0.5 | 4.33E-03 | 0.0 | 8.86E-01 | -0.1 | 6.47E-01 | 0.2 | 4.42E-01 | -0.3 | 3.09E-01 | Lysine-specific demethylase JMJ29 |
| AT5G16080 | -0.5 | 9.38E-03 | -0.3 | 6.11E-02 | -0.2 | 4.27E-01 | -0.2 | 6.11E-01 | -0.2 | 7.92E-01 | -0.1 | 9.17E-01 | Probable carboxylesterase 17 |
| AT3G48060 | -0.5 | 7.51E-05 | -0.4 | 2.25E-04 | 0.0 | 6.74E-01 | -0.2 | 3.80E-01 | -0.1 | 7.24E-01 | -0.1 | 7.97E-01 | BAH and TFIIS domain-containing protein |
| AT2G42270 | -0.5 | 2.08E-03 | -0.5 | 2.49E-03 | 0.0 | 9.44E-01 | 0.1 | 7.65E-01 | 0.2 | 5.24E-01 | -0.1 | 8.07E-01 | DExH-box ATP-dependent RNA helicase DExH13 |
| AT1G60060 | -0.5 | 3.99E-02 | -0.9 | 3.19E-03 | 0.4 | 1.61E-01 | -0.9 | 5.68E-03 | -1.0 | 3.72E-03 | 0.1 | 7.79E-01 | Serine |
| AT2G22010 | -0.5 | 7.05E-04 | -0.7 | 7.95E-05 | 0.2 | 6.38E-02 | 0.1 | 8.40E-01 | 0.3 | 6.50E-01 | -0.2 | 8.40E-01 | RING-type E3 ubiquitin transferase |
| AT4G35030 | -0.5 | 2.41E-03 | -0.2 | 1.85E-01 | -0.3 | 5.48E-02 | -0.6 | 1.79E-02 | -0.2 | 5.33E-01 | -0.4 | 9.46E-02 | Protein kinase superfamily protein |
| AT3G07130 | -0.5 | 3.77E-04 | -0.2 | 6.73E-02 | -0.3 | 2.10E-02 | -0.1 | 7.07E-01 | 0.2 | 4.46E-01 | -0.3 | 2.47E-01 | Purple acid phosphatase |
| AT5G39660 | -0.5 | 4.46E-04 | -0.4 | 2.24E-03 | -0.1 | 4.63E-01 | -0.8 | 7.09E-02 | -0.6 | 2.05E-01 | -0.2 | 7.79E-01 | Cyclic dof factor 2 |
| AT1G50030 | -0.5 | 1.68E-04 | -0.6 | 1.04E-04 | 0.1 | 4.45E-01 | -0.2 | 4.21E-01 | -0.2 | 5.66E-01 | 0.0 | 9.52E-01 | non-specific serine |
| AT1G26640 | -0.5 | 2.51E-05 | -0.2 | 9.97E-03 | -0.3 | 3.60E-03 | -0.3 | 5.70E-02 | 0.1 | 7.41E-01 | -0.4 | 7.79E-02 | Isopentenyl phosphate kinase |
| AT1G13960 | -0.5 | 3.27E-05 | -0.6 | 2.11E-05 | 0.1 | 3.02E-01 | -0.3 | 8.55E-02 | -0.3 | 1.26E-01 | 0.0 | 9.94E-01 | WRKY DNA-binding protein 4 |
| AT1G67560 | -0.5 | 1.65E-04 | -0.2 | 2.83E-02 | -0.3 | 1.41E-02 | 0.0 | 9.54E-01 | 0.1 | 5.91E-01 | -0.1 | 6.33E-01 | Lipoxygenase 6, chloroplastic |
| AT1G62130 | -0.5 | 2.21E-03 | -0.5 | 2.08E-03 | 0.0 | 8.21E-01 | 0.0 | 9.25E-01 | 0.4 | 4.11E-01 | -0.3 | 5.64E-01 | AAA-type ATPase family protein |
| AT1G58370 | -0.5 | 1.46E-03 | 0.2 | 5.62E-02 | -0.7 | 5.54E-04 | 0.1 | 6.52E-01 | 0.1 | 8.47E-01 | 0.1 | 8.94E-01 | Endo-1,4-beta-xylanase 1 |
| AT1G52050 | -0.5 | 4.41E-02 | 0.7 | 9.32E-03 | -1.1 | 1.16E-03 | -0.3 | 4.43E-01 | 0.2 | 6.51E-01 | -0.5 | 2.30E-01 | Jacalin-related lectin 8 |
| AT2G40140 | -0.5 | 7.54E-03 | -0.8 | 3.69E-04 | 0.4 | 5.40E-02 | 0.5 | 1.19E-03 | 0.0 | 9.56E-01 | 0.5 | 6.12E-03 | Zinc finger CCCH domain-containing protein 29 |
| AT4G37400 | -0.5 | 3.51E-03 | 0.3 | 3.62E-02 | -0.8 | 7.44E-04 | -0.3 | 8.63E-02 | -0.3 | 1.26E-01 | 0.0 | 9.83E-01 | Cytochrome P450 81F3 |
| AT5G35750 | -0.5 | 1.29E-03 | -0.2 | 1.73E-01 | -0.3 | 2.84E-02 | -0.3 | 1.09E-01 | -0.3 | 1.88E-01 | 0.0 | 9.47E-01 | Histidine kinase 2 |
| AT1G20510 | -0.5 | 5.53E-04 | -0.6 | 1.56E-04 | 0.1 | 2.13E-01 | 0.2 | 4.73E-01 | 0.4 | 1.38E-01 | -0.2 | 5.41E-01 | OPC-8:0 CoA ligase1 |
| AT3G63240 | -0.5 | 9.20E-04 | 0.0 | 6.88E-01 | -0.4 | 5.98E-03 | 0.1 | 7.45E-01 | 0.0 | 8.59E-01 | 0.0 | 9.56E-01 | Type I inositol polyphosphate 5-phosphatase 4 |
| AT4G25390 | -0.5 | 6.52E-04 | -0.7 | 1.25E-04 | 0.2 | 1.45E-01 | 0.2 | 2.46E-01 | -0.2 | 3.63E-01 | 0.5 | 6.86E-02 | Receptor-like serine |
| AT4G35860 | -0.5 | 1.80E-05 | -0.3 | 1.18E-03 | -0.2 | 1.42E-02 | -0.3 | 1.80E-02 | -0.2 | 1.51E-01 | -0.1 | 5.80E-01 | hypothetical protein |
| AT2G31390 | -0.5 | 2.32E-03 | 0.1 | 4.54E-01 | -0.6 | 2.03E-03 | 0.2 | 3.79E-01 | 0.2 | 2.37E-01 | -0.1 | 7.54E-01 | Probable fructokinase-1 |
| AT4G38220 | -0.5 | 3.24E-05 | -0.2 | 5.42E-03 | -0.2 | 7.65E-03 | -0.2 | 1.52E-01 | 0.0 | 9.18E-01 | -0.2 | 2.59E-01 | N-acyl-aliphatic-L-amino acid amidohydrolase |
| AT5G59270 | -0.5 | 1.31E-02 | -0.4 | 3.96E-02 | -0.1 | 6.97E-01 | | | | | | | non-specific serine |
| AT5G01710 | -0.5 | 7.23E-05 | -0.3 | 1.29E-03 | -0.1 | 1.02E-01 | -0.2 | 7.02E-02 | -0.1 | 5.79E-01 | -0.1 | 4.60E-01 | Methyltransferase |
| AT5G01040 | -0.5 | 4.02E-03 | 0.2 | 9.04E-02 | -0.7 | 1.14E-03 | -0.2 | 4.34E-01 | 0.2 | 4.14E-01 | -0.5 | 1.77E-01 | Laccase-8 |
| AT1G07530 | -0.5 | 2.13E-05 | -0.5 | 2.85E-05 | 0.0 | 7.51E-01 | -0.5 | 2.51E-03 | -0.5 | 5.87E-03 | 0.0 | 9.79E-01 | Scarecrow-like protein 14 |
| AT2G45510 | -0.5 | 3.59E-04 | -0.5 | 3.37E-04 | 0.0 | 7.39E-01 | -0.3 | 7.33E-02 | -0.1 | 6.56E-01 | -0.2 | 3.80E-01 | At2g45510 |
| AT3G52990 | -0.5 | 2.42E-04 | -0.2 | 2.70E-02 | -0.3 | 2.34E-02 | 0.1 | 2.75E-01 | 0.2 | 1.75E-01 | 0.0 | 8.14E-01 | Pyruvate kinase |
| AT1G80110 | -0.5 | 2.38E-03 | -0.5 | 2.07E-03 | 0.1 | 7.74E-01 | -0.6 | 9.04E-02 | -0.2 | 6.71E-01 | -0.4 | 3.98E-01 | F-box protein PP2-B11 |
| AT3G59350 | -0.5 | 1.93E-04 | 0.2 | 4.54E-02 | -0.7 | 1.36E-04 | -0.5 | 2.28E-03 | -0.7 | 5.77E-04 | 0.2 | 4.60E-01 | Protein kinase superfamily protein |
| AT4G36750 | -0.5 | 2.51E-04 | -0.2 | 1.97E-02 | -0.2 | 3.09E-02 | -0.3 | 5.45E-02 | -0.1 | 6.76E-01 | -0.2 | 3.27E-01 | NADH dehydrogenase |
| AT4G05050 | -0.5 | 3.41E-04 | -0.4 | 1.92E-03 | -0.1 | 4.18E-01 | -0.3 | 1.70E-02 | 0.1 | 3.93E-01 | -0.4 | 1.34E-02 | Polyubiquitin |
| AT5G19110 | -0.5 | 1.69E-01 | -0.4 | 2.97E-01 | -0.1 | 7.93E-01 | -1.3 | 1.30E-02 | -0.2 | 6.90E-01 | -1.1 | 8.90E-02 | Eukaryotic aspartyl protease family protein |
| AT5G25830 | -0.5 | 4.56E-04 | 0.1 | 1.83E-01 | -0.6 | 4.97E-04 | 0.2 | 2.39E-01 | 0.0 | 9.22E-01 | 0.3 | 2.94E-01 | GATA transcription factor 12 |
| AT1G07420 | -0.5 | 5.66E-04 | 0.0 | 6.42E-01 | -0.4 | 3.37E-03 | -0.1 | 2.51E-01 | 0.0 | 8.09E-01 | -0.1 | 5.74E-01 | Methylsterol monooxygenase 2-2 |
| AT2G22250 | -0.5 | 3.18E-05 | -0.3 | 5.49E-04 | -0.1 | 8.43E-02 | 0.2 | 2.98E-01 | 0.2 | 3.86E-01 | 0.0 | 9.92E-01 | Bifunctional aspartate aminotransferase and glutamate |
| AT4G30790 | -0.5 | 1.53E-04 | -0.5 | 3.31E-04 | 0.0 | 8.43E-01 | -0.3 | 1.89E-01 | -0.2 | 4.51E-01 | -0.1 | 8.07E-01 | Autophagy-related protein 11 |
| AT1G50940 | -0.5 | 1.16E-04 | -0.3 | 4.40E-03 | -0.2 | 5.83E-02 | -0.3 | 5.06E-02 | -0.2 | 2.32E-01 | -0.1 | 6.99E-01 | Electron transfer flavoprotein subunit alpha, mitochondrial |
| AT3G05510 | -0.5 | 1.01E-03 | -0.4 | 6.53E-03 | -0.1 | 3.91E-01 | 0.0 | 9.31E-01 | 0.0 | 9.08E-01 | 0.0 | 9.73E-01 | Tafazzin family protein |
| AT2G43980 | -0.5 | 1.78E-04 | -0.2 | 8.41E-02 | -0.3 | 7.87E-03 | 0.0 | 9.60E-01 | 0.2 | 4.38E-01 | -0.2 | 5.07E-01 | inositol-1,3,4-trisphosphate 5 |
| AT1G50360 | -0.5 | 4.37E-05 | -0.3 | 1.26E-03 | -0.2 | 5.34E-02 | 0.1 | 6.52E-01 | 0.0 | 9.46E-01 | 0.0 | 8.19E-01 | Myosin-3 |
| AT3G53970 | -0.5 | 7.35E-05 | -0.3 | 1.23E-03 | -0.1 | 1.08E-01 | 0.1 | 6.50E-01 | 0.0 | 7.99E-01 | 0.0 | 9.49E-01 | PI31 proteasome regulator N-terminal domain-containing protein |
| AT1G23190 | -0.5 | 2.17E-04 | -0.3 | 1.34E-02 | -0.2 | 3.87E-02 | 0.1 | 3.12E-01 | 0.4 | 1.97E-02 | -0.2 | 2.73E-01 | Probable phosphoglucomutase, cytoplasmic 1 |
| AT4G25300 | -0.5 | 1.56E-03 | -0.4 | 3.36E-03 | 0.0 | 8.17E-01 | -0.4 | 7.38E-02 | -0.5 | 3.67E-02 | 0.1 | 7.79E-01 | 2-oxoglutarate and Fe-dependent oxygenase superfamily protein |
| AT3G10570 | -0.5 | 2.39E-02 | -0.4 | 4.79E-02 | -0.1 | 8.31E-01 | 0.1 | 6.08E-01 | 0.3 | 2.17E-01 | -0.2 | 4.91E-01 | At3g10570 |
| AT1G55550 | -0.5 | 1.94E-02 | -0.5 | 2.21E-02 | 0.0 | 9.66E-01 | 0.3 | 4.21E-01 | 0.6 | 1.55E-01 | -0.3 | 5.14E-01 | Kinesin-like protein KIN-14T |
| AT2G35680 | -0.5 | 6.60E-05 | -0.4 | 4.06E-04 | -0.1 | 3.47E-01 | -0.4 | 2.07E-03 | -0.2 | 2.10E-01 | -0.2 | 1.89E-01 | Phosphatidylglycerophosphate phosphatase PTPMT1 |
| AT3G11800 | -0.5 | 6.85E-05 | -0.1 | 9.41E-02 | -0.3 | 2.05E-03 | -0.3 | 5.23E-02 | -0.1 | 4.30E-01 | -0.1 | 5.12E-01 | hypothetical protein |
| AT2G45300 | -0.5 | 3.91E-05 | -0.2 | 1.79E-02 | -0.3 | 3.64E-03 | 0.5 | 3.12E-04 | 0.2 | 1.35E-01 | 0.3 | 6.93E-02 | 3-phosphoshikimate 1-carboxyvinyltransferase |
| AT1G16710 | -0.5 | 7.94E-05 | -0.5 | 4.97E-05 | 0.1 | 3.73E-01 | -0.1 | 4.29E-01 | 0.1 | 8.17E-01 | -0.2 | 4.23E-01 | histone acetyltransferase |
| AT1G68410 | -0.5 | 8.29E-05 | -0.4 | 2.19E-04 | 0.0 | 7.47E-01 | -0.4 | 4.31E-02 | -0.1 | 7.13E-01 | -0.3 | 1.95E-01 | Probable protein phosphatase 2C 15 |
| AT4G28630 | -0.5 | 2.63E-04 | -0.1 | 3.23E-01 | -0.4 | 4.07E-03 | -0.1 | 6.06E-01 | -0.3 | 2.66E-01 | 0.1 | 6.39E-01 | ABC transporter B family member 23, mitochondrial |
| AT4G04960 | -0.5 | 6.57E-04 | -0.5 | 7.88E-04 | 0.0 | 9.06E-01 | -0.3 | 5.16E-02 | -0.5 | 4.41E-03 | 0.2 | 3.42E-01 | Concanavalin A-like lectin protein kinase family protein |
| AT3G06540 | -0.5 | 7.27E-05 | -0.4 | 4.08E-04 | -0.1 | 3.71E-01 | -0.1 | 5.35E-01 | 0.0 | 8.71E-01 | -0.1 | 8.01E-01 | Rab escort protein 1 |
| AT2G23980 | -0.5 | 2.64E-05 | -0.5 | 5.93E-05 | 0.0 | 8.82E-01 | -0.3 | 3.92E-02 | 0.0 | 8.40E-01 | -0.3 | 1.77E-01 | Cyclic nucleotide-gated channel 6 |
| AT3G27000 | -0.5 | 5.06E-04 | -0.4 | 3.36E-03 | -0.1 | 3.71E-01 | 0.0 | 9.44E-01 | 0.1 | 6.63E-01 | -0.1 | 6.43E-01 | Actin-related protein 2 |
| AT2G17040 | -0.5 | 4.87E-02 | -1.2 | 5.99E-04 | 0.8 | 2.18E-02 | 0.0 | 8.99E-01 | -0.2 | 4.83E-01 | 0.2 | 5.63E-01 | At2g17040 |
| AT2G46700 | -0.5 | 4.43E-04 | -0.5 | 6.72E-04 | 0.0 | 9.93E-01 | -0.2 | 1.98E-01 | -0.2 | 3.50E-01 | 0.0 | 9.12E-01 | CDPK-related kinase 3 |
| AT5G20650 | -0.5 | 7.19E-04 | 0.0 | 8.35E-01 | -0.5 | 1.80E-03 | -0.4 | 1.61E-01 | -0.2 | 6.55E-01 | -0.2 | 5.26E-01 | Copper transporter 5 |
| AT1G33900 | -0.5 | 4.06E-03 | 0.4 | 4.81E-03 | -0.9 | 2.71E-04 | -0.5 | 2.87E-02 | 0.2 | 4.98E-01 | -0.7 | 2.53E-02 | Immune-associated nucleotide-binding protein 4 |
| AT1G19490 | -0.5 | 9.16E-03 | -0.4 | 3.68E-02 | -0.1 | 5.93E-01 | | | | | | | BZIP domain-containing protein |
| AT5G53350 | -0.5 | 1.88E-04 | -0.6 | 4.03E-05 | 0.2 | 8.19E-02 | 0.0 | 9.13E-01 | 0.1 | 2.74E-01 | -0.2 | 3.43E-01 | CLP protease regulatory subunit CLPX1, mitochondrial |
| AT2G26600 | -0.5 | 2.61E-05 | -0.5 | 6.05E-05 | 0.0 | 8.62E-01 | -0.2 | 1.88E-01 | 0.0 | 9.46E-01 | -0.2 | 2.91E-01 | glucan endo-1,3-beta-D-glucosidase |
| AT3G47670 | -0.5 | 1.19E-04 | -0.1 | 4.86E-01 | -0.4 | 1.27E-03 | -0.2 | 1.99E-01 | 0.0 | 9.92E-01 | -0.2 | 3.31E-01 | hypothetical protein |
| AT2G20100 | -0.5 | 6.08E-04 | -0.2 | 1.37E-01 | -0.3 | 2.01E-02 | 0.2 | 3.10E-01 | 0.3 | 2.00E-01 | -0.1 | 7.99E-01 | Basic helix-loop-helix DNA-binding superfamily protein |
| AT5G66880 | -0.5 | 6.81E-05 | -0.3 | 1.00E-03 | -0.1 | 1.25E-01 | -0.4 | 2.75E-02 | -0.3 | 1.43E-01 | -0.1 | 6.91E-01 | Serine |
| AT1G64170 | -0.5 | 1.01E-02 | -0.5 | 8.93E-03 | 0.0 | 8.80E-01 | 0.1 | 8.04E-01 | 0.1 | 7.39E-01 | 0.0 | 9.26E-01 | Cation |
| AT1G76550 | -0.5 | 2.14E-04 | 0.0 | 9.93E-01 | -0.5 | 8.64E-04 | 0.3 | 2.06E-02 | 0.1 | 3.73E-01 | 0.2 | 3.01E-01 | Pyrophosphate--fructose 6-phosphate 1-phosphotransferase subunit alpha 2 |
| AT1G19835 | -0.5 | 1.43E-04 | -0.2 | 5.74E-02 | -0.3 | 6.51E-03 | -0.5 | 3.18E-02 | -0.3 | 2.08E-01 | -0.2 | 6.44E-01 | Filament-like plant protein 4 |
| AT1G75640 | -0.5 | 1.68E-04 | -0.2 | 4.34E-02 | -0.3 | 1.32E-02 | -0.3 | 2.02E-01 | -0.1 | 8.20E-01 | -0.2 | 5.00E-01 | F10A5.16 |
| AT3G44320 | -0.5 | 6.22E-04 | 0.2 | 8.75E-02 | -0.6 | 3.68E-04 | 0.1 | 5.66E-01 | 0.3 | 7.72E-02 | -0.2 | 3.19E-01 | Nitrilase 3 |
| AT3G61850 | -0.5 | 9.28E-04 | -0.4 | 2.94E-03 | -0.1 | 6.52E-01 | 0.1 | 5.50E-01 | 0.1 | 6.94E-01 | 0.0 | 9.44E-01 | Dof zinc finger protein |
| AT2G31350 | -0.5 | 1.62E-04 | -0.3 | 1.61E-03 | -0.1 | 2.11E-01 | -0.3 | 7.08E-02 | -0.4 | 4.06E-02 | 0.1 | 7.80E-01 | Glyoxalase 2-5 |
| AT3G55450 | -0.5 | 1.86E-04 | -0.3 | 3.42E-03 | -0.2 | 1.17E-01 | -0.2 | 2.30E-01 | -0.2 | 1.32E-01 | 0.1 | 7.85E-01 | PBS1-like 1 |

| | | | | | | | | | | | | | |
|---|---|---|---|---|---|---|---|---|---|---|---|---|---|
| AT3G49690 | -0.5 | 7.62E-03 | -0.3 | 9.44E-02 | -0.2 | 2.86E-01 | 0.0 | 9.31E-01 | 0.3 | 5.23E-01 | -0.4 | 5.12E-01 | Transcription factor RAX3 |
| AT1G61560 | -0.5 | 3.13E-03 | -0.8 | 1.74E-04 | 0.4 | 4.70E-02 | 1.1 | 4.69E-03 | 0.2 | 6.38E-01 | 0.9 | 3.59E-02 | MLO-like protein |
| AT5G58730 | -0.5 | 1.04E-04 | 0.0 | 7.73E-01 | -0.5 | 3.73E-04 | 0.1 | 6.06E-01 | 0.1 | 4.87E-01 | 0.0 | 8.76E-01 | Inositol 3-kinase |
| AT1G29170 | -0.5 | 4.62E-04 | -0.3 | 6.94E-03 | -0.2 | 1.85E-01 | 0.3 | 4.34E-01 | 0.1 | 7.73E-01 | 0.1 | 7.71E-01 | Protein SCAR |
| AT5G66850 | -0.5 | 9.64E-05 | -0.4 | 2.41E-04 | 0.0 | 7.63E-01 | -0.1 | 4.63E-01 | 0.1 | 6.31E-01 | -0.2 | 2.87E-01 | Mitogen-activated protein kinase kinase kinase 5 |
| AT1G59910 | -0.5 | 2.33E-04 | -0.6 | 1.25E-04 | 0.1 | 3.97E-01 | -0.4 | 7.74E-02 | -0.2 | 4.87E-01 | -0.2 | 5.51E-01 | Formin-like protein 7 |
| AT2G36910 | -0.5 | 1.42E-04 | -0.6 | 6.26E-05 | 0.1 | 2.34E-01 | -0.2 | 4.04E-01 | 0.1 | 8.40E-01 | -0.2 | 4.37E-01 | ABC transporter B family member 1 |
| AT4G31140 | -0.5 | 5.57E-05 | -0.3 | 6.54E-04 | -0.1 | 1.70E-01 | -0.5 | 5.27E-03 | -0.3 | 1.08E-01 | -0.2 | 3.42E-01 | Glucan endo-1,3-beta-glucosidase 5 -beta-glucan endohydrolase 5) -beta-glucanase 5) |
| AT4G13840 | -0.5 | 7.57E-04 | -0.2 | 3.40E-02 | -0.2 | 6.83E-02 | 0.0 | 8.13E-01 | 0.1 | 7.46E-01 | 0.0 | 9.24E-01 | Protein ECERIFERUM 26 |
| AT1G24625 | -0.5 | 2.23E-04 | -0.4 | 1.73E-03 | -0.1 | 3.02E-01 | -0.1 | 6.60E-01 | 0.2 | 3.68E-01 | -0.3 | 2.51E-01 | Zinc finger protein 7 |
| AT4G03230 | -0.5 | 7.69E-04 | -0.5 | 1.07E-03 | 0.0 | 9.81E-01 | -0.1 | 6.67E-01 | 0.3 | 3.28E-01 | -0.4 | 2.30E-01 | non-specific serine |
| AT2G39480 | -0.5 | 3.35E-05 | -0.3 | 3.97E-04 | -0.1 | 1.38E-01 | -0.1 | 2.96E-01 | -0.3 | 3.61E-02 | 0.2 | 3.17E-01 | p-glycoprotein 6 |
| AT2G18650 | -0.5 | 2.03E-03 | -0.1 | 2.16E-01 | -0.3 | 4.52E-02 | -0.1 | 7.97E-01 | 0.2 | 5.82E-01 | -0.3 | 4.43E-01 | Putative RING-H2 finger protein ATL49 |
| AT4G32000 | -0.5 | 1.49E-03 | -0.2 | 1.03E-01 | -0.3 | 6.21E-02 | -0.3 | 1.37E-01 | 0.0 | 9.55E-01 | -0.3 | 2.33E-01 | Serine |
| AT5G42200 | -0.5 | 4.14E-03 | -0.5 | 2.29E-03 | 0.1 | 6.50E-01 | -0.8 | 1.66E-03 | -0.7 | 1.04E-02 | -0.2 | 6.85E-01 | E3 ubiquitin-protein ligase ATL23 |
| AT2G14510 | -0.5 | 5.25E-04 | -0.2 | 3.77E-02 | -0.2 | 4.50E-02 | 0.0 | 9.72E-01 | 0.1 | 7.11E-01 | -0.1 | 6.75E-01 | Leucine-rich repeat protein kinase family protein |
| AT4G29740 | -0.5 | 9.96E-03 | -0.5 | 6.26E-03 | 0.1 | 7.49E-01 | 0.4 | 3.28E-01 | 0.6 | 9.98E-02 | -0.3 | 5.22E-01 | cytokinin dehydrogenase |
| AT1G14370 | -0.5 | 6.34E-04 | -0.6 | 1.47E-04 | 0.2 | 1.83E-01 | 0.0 | 9.02E-01 | 0.0 | 8.71E-01 | 0.1 | 8.22E-01 | Probable serine |
| AT4G40060 | -0.5 | 2.68E-05 | -0.2 | 8.78E-03 | -0.3 | 3.94E-03 | -0.6 | 4.74E-05 | -0.4 | 1.19E-02 | -0.3 | 7.02E-02 | Homeobox-leucine zipper protein ATHB-16 |
| AT3G10480 | -0.5 | 2.42E-05 | -0.5 | 5.12E-05 | 0.0 | 9.25E-01 | -0.1 | 6.56E-01 | -0.1 | 6.67E-01 | 0.0 | 9.80E-01 | NAC domain containing protein 50 |
| AT2G47060 | -0.5 | 4.05E-03 | -0.3 | 2.08E-02 | -0.1 | 4.72E-01 | -0.6 | 3.07E-04 | -0.3 | 4.37E-02 | -0.3 | 1.49E-01 | Protein kinase superfamily protein |
| AT1G73960 | -0.5 | 6.07E-05 | -0.4 | 5.16E-04 | -0.1 | 2.28E-01 | 0.1 | 4.12E-01 | 0.0 | 9.56E-01 | 0.1 | 5.14E-01 | Transcription initiation factor TFIID subunit 2 |
| AT5G16510 | -0.5 | 4.38E-04 | 0.0 | 6.40E-01 | -0.5 | 8.26E-04 | -0.1 | 7.44E-01 | 0.1 | 7.41E-01 | -0.1 | 5.24E-01 | Probable UDP-arabinopyranose mutase 5 |
| AT5G47140 | -0.5 | 8.44E-04 | -0.4 | 3.71E-03 | -0.1 | 5.15E-01 | 0.2 | 2.88E-01 | 0.2 | 1.89E-01 | -0.1 | 7.94E-01 | GATA27 |
| AT2G28970 | -0.5 | 2.04E-03 | -0.1 | 4.18E-01 | -0.4 | 2.22E-02 | 0.3 | 1.14E-01 | 0.2 | 5.46E-01 | 0.2 | 5.55E-01 | Leucine-rich repeat protein kinase family protein |
| AT2G17760 | -0.5 | 6.59E-05 | -0.5 | 3.60E-05 | 0.1 | 2.95E-01 | 0.1 | 2.90E-01 | 0.2 | 3.02E-01 | 0.0 | 9.55E-01 | Aspartyl protease family protein 1 |
| AT4G18520 | -0.5 | 9.60E-04 | -0.1 | 5.66E-01 | -0.4 | 7.87E-03 | -0.1 | 7.69E-01 | 0.0 | 9.43E-01 | -0.1 | 7.49E-01 | Pentatricopeptide repeat-containing protein At4g18520, chloroplastic |
| AT5G40440 | -0.5 | 2.31E-03 | -0.5 | 2.75E-03 | 0.0 | 9.44E-01 | 0.0 | 9.68E-01 | -0.1 | 8.47E-01 | 0.1 | 8.38E-01 | Mitogen-activated protein kinase kinase 3 |
| AT4G11850 | -0.5 | 1.93E-04 | -0.3 | 4.38E-03 | -0.2 | 9.80E-02 | 0.7 | 5.86E-04 | 0.3 | 1.12E-01 | 0.4 | 1.26E-01 | Phospholipase D gamma 1 |
| AT1G68620 | -0.5 | 8.29E-02 | -0.9 | 8.68E-03 | 0.4 | 2.45E-01 | 0.0 | 9.17E-01 | -0.8 | 1.97E-02 | 0.9 | 3.10E-02 | Probable carboxylesterase 6 |
| AT2G25220 | -0.5 | 7.21E-04 | -0.1 | 4.15E-01 | -0.4 | 9.40E-03 | -0.3 | 2.38E-01 | -0.1 | 8.37E-01 | -0.2 | 4.58E-01 | Protein kinase superfamily protein |
| AT3G21360 | -0.5 | 7.49E-03 | -0.5 | 4.27E-03 | 0.1 | 6.84E-01 | -0.3 | 7.52E-02 | 0.0 | 9.55E-01 | -0.3 | 1.51E-01 | Clavaminate synthase-like protein At3g21360 |
| AT2G39770 | -0.5 | 1.16E-04 | -0.2 | 3.49E-02 | -0.3 | 7.82E-03 | -0.1 | 5.84E-01 | 0.0 | 8.17E-01 | -0.1 | 5.58E-01 | Mannose-1-phosphate guanylyltransferase 1 |
| AT2G43800 | -0.5 | 1.15E-04 | -0.4 | 5.17E-04 | -0.1 | 4.69E-01 | 0.1 | 5.59E-01 | 0.2 | 1.35E-01 | -0.2 | 4.85E-01 | Formin-like protein 2 |
| AT4G02860 | -0.5 | 2.04E-05 | -0.5 | 1.77E-05 | 0.1 | 4.24E-01 | -0.2 | 2.73E-01 | -0.1 | 6.95E-01 | -0.1 | 7.27E-01 | At4g02860 |
| AT1G76040 | -0.5 | 1.49E-02 | -0.5 | 7.46E-03 | 0.1 | 6.65E-01 | 0.0 | 9.61E-01 | 0.1 | 7.57E-01 | -0.1 | 8.18E-01 | Calcium-dependent protein kinase 29 |
| AT2G31680 | -0.5 | 3.62E-05 | -0.1 | 3.28E-01 | -0.4 | 5.88E-04 | -0.4 | 5.08E-02 | -0.1 | 7.19E-01 | -0.3 | 2.75E-01 | Ras-related protein RABA5d |
| AT3G20860 | -0.5 | 1.32E-03 | 0.1 | 6.46E-01 | -0.5 | 2.68E-03 | 0.5 | 1.00E-02 | 0.6 | 1.25E-02 | 0.0 | 9.30E-01 | Serine |
| AT3G27560 | -0.5 | 1.05E-04 | -0.4 | 2.27E-04 | 0.0 | 8.64E-01 | 0.1 | 2.92E-01 | 0.1 | 7.87E-01 | 0.1 | 6.63E-01 | AT3g27560 |
| AT3G11910 | -0.5 | 3.06E-05 | -0.3 | 4.44E-04 | -0.1 | 1.08E-01 | -0.1 | 5.35E-01 | -0.1 | 4.79E-01 | 0.0 | 9.23E-01 | ubiquitinyl hydrolase 1 |
| AT4G28530 | -0.5 | 6.69E-02 | -0.7 | 1.34E-02 | 0.3 | 4.08E-01 | -1.5 | 6.66E-03 | -0.1 | 8.53E-01 | -1.4 | 2.78E-02 | At4g28530 |
| AT3G28690 | -0.5 | 1.01E-04 | -0.4 | 6.88E-04 | -0.1 | 3.01E-01 | -0.1 | 3.51E-01 | 0.0 | 8.89E-01 | -0.1 | 5.90E-01 | Protein kinase superfamily protein |
| AT2G06050 | -0.5 | 2.54E-03 | -0.3 | 3.45E-02 | -0.2 | 2.14E-01 | 0.0 | 9.47E-01 | 0.3 | 2.37E-01 | -0.3 | 2.89E-01 | 12-oxophytodienoate reductase 3 |
| AT3G25620 | -0.5 | 3.55E-02 | -0.2 | 3.74E-01 | -0.3 | 2.76E-01 | 0.2 | 7.59E-01 | 0.3 | 6.56E-01 | -0.1 | 9.07E-01 | ABC transporter G family member 21 |
| AT3G10800 | -0.5 | 1.79E-04 | -0.4 | 3.94E-04 | 0.0 | 8.24E-01 | -0.2 | 1.99E-01 | -0.2 | 4.14E-01 | -0.1 | 8.48E-01 | bZIP transcription factor 28 |
| AT5G13870 | -0.5 | 3.28E-02 | 0.1 | 6.96E-01 | -0.5 | 3.54E-02 | -0.2 | 4.25E-01 | 0.3 | 3.79E-01 | -0.5 | 1.34E-01 | Probable xyloglucan endotransglucosylase |
| AT5G10260 | -0.5 | 5.01E-02 | 0.0 | 9.43E-01 | -0.5 | 9.11E-02 | -0.5 | 2.26E-01 | 0.3 | 4.92E-01 | -0.9 | 9.19E-02 | Ras-related protein RABH1e |
| AT1G45688 | -0.5 | 9.84E-05 | -0.3 | 1.38E-03 | -0.1 | 1.45E-01 | -0.3 | 2.98E-03 | -0.1 | 5.13E-01 | -0.3 | 9.29E-02 | Transmembrane protein |
| AT4G08320 | -0.5 | 3.61E-04 | -0.4 | 6.69E-04 | 0.0 | 9.27E-01 | 0.0 | 9.25E-01 | 0.0 | 9.43E-01 | -0.1 | 8.85E-01 | Tetratricopeptide repeat -like superfamily protein |
| AT5G27100 | -0.5 | 2.52E-03 | -0.3 | 5.12E-02 | -0.2 | 1.78E-01 | -0.2 | 5.83E-01 | 0.1 | 7.09E-01 | -0.3 | 4.29E-01 | Glutamate receptor 2.1 |
| AT5G48180 | -0.5 | 5.65E-05 | -0.5 | 2.96E-05 | 0.1 | 2.34E-01 | -0.2 | 2.06E-01 | -0.4 | 4.55E-02 | 0.2 | 5.15E-01 | Thiohydroximate-O-sulfate sulfur |
| AT5G67450 | -0.5 | 2.04E-03 | -0.3 | 2.48E-02 | -0.2 | 2.53E-01 | 0.1 | 8.22E-01 | 0.1 | 7.92E-01 | 0.0 | 9.51E-01 | Zinc finger protein AZF1 |
| AT1G78580 | -0.5 | 1.34E-04 | -0.3 | 2.41E-03 | -0.1 | 1.12E-01 | -0.3 | 3.73E-01 | 0.1 | 7.83E-01 | -0.4 | 2.91E-01 | Alpha,alpha-trehalose-phosphate synthase 1 |
| AT1G08970 | -0.5 | 1.21E-04 | -0.4 | 1.64E-03 | -0.1 | 1.45E-01 | -0.1 | 7.31E-01 | 0.0 | 9.48E-01 | -0.1 | 7.56E-01 | Nuclear transcription factor Y subunit C-9 |
| AT4G04860 | -0.5 | 2.35E-04 | -0.3 | 7.41E-03 | -0.2 | 7.73E-02 | -0.4 | 5.54E-03 | -0.2 | 1.52E-01 | -0.2 | 3.18E-01 | Derlin-2.2 |
| AT3G58760 | -0.5 | 1.00E-04 | -0.2 | 2.85E-02 | -0.3 | 9.12E-03 | -0.1 | 4.37E-01 | -0.1 | 8.10E-01 | -0.1 | 7.66E-01 | AT3g58760 |
| AT2G47980 | -0.5 | 3.53E-05 | -0.4 | 1.98E-04 | -0.1 | 3.40E-01 | -0.3 | 9.56E-02 | -0.2 | 3.25E-01 | -0.1 | 7.72E-01 | Sister-chromatid cohesion protein 3 |
| AT1G79840 | -0.5 | 6.29E-04 | 0.1 | 5.69E-01 | -0.5 | 1.37E-03 | 0.0 | 9.43E-01 | -0.1 | 8.35E-01 | 0.1 | 8.26E-01 | HD-ZIP IV family of homeobox-leucine zipper protein with lipid-binding START domain-containing protein |
| AT1G59780 | -0.5 | 2.26E-03 | 0.0 | 7.43E-01 | -0.4 | 1.29E-02 | 0.3 | 9.76E-02 | 0.3 | 2.65E-01 | 0.1 | 8.07E-01 | Uncharacterized protein |
| AT5G65270 | -0.5 | 3.76E-04 | -0.2 | 8.12E-02 | -0.3 | 1.44E-02 | 0.1 | 7.27E-01 | 0.2 | 3.55E-01 | -0.1 | 6.46E-01 | Ras-related protein RABA4a |
| AT1G48120 | -0.5 | 8.61E-04 | -0.6 | 3.33E-04 | 0.1 | 3.83E-01 | 0.6 | 2.02E-01 | 0.5 | 3.45E-01 | 0.1 | 8.89E-01 | Serine |
| AT1G71697 | -0.5 | 7.28E-04 | -0.5 | 9.61E-04 | 0.0 | 9.59E-01 | 0.0 | 8.70E-01 | 0.0 | 9.98E-01 | 0.0 | 9.18E-01 | Probable choline kinase 1 |
| AT5G48060 | -0.5 | 4.50E-03 | -0.4 | 7.74E-03 | 0.0 | 9.00E-01 | 0.6 | 2.52E-01 | 0.3 | 6.23E-01 | 0.3 | 7.32E-01 | Multiple C2 domain and transmembrane region protein 2 |
| AT1G77130 | -0.5 | 1.29E-04 | -0.1 | 2.47E-01 | -0.4 | 1.93E-03 | -0.3 | 3.66E-02 | 0.1 | 6.06E-01 | -0.3 | 4.27E-02 | Putative UDP-glucuronate:xylan alpha-glucuronosyltransferase 3 |
| AT5G14500 | -0.5 | 8.29E-04 | -0.2 | 2.71E-02 | -0.2 | 9.99E-02 | 0.0 | 8.45E-01 | 0.0 | 9.19E-01 | -0.1 | 8.15E-01 | glucose-6-phosphate 1-epimerase |
| AT5G07470 | -0.5 | 4.65E-04 | -0.2 | 5.54E-02 | -0.3 | 2.50E-02 | -0.3 | 1.06E-02 | -0.1 | 4.96E-01 | -0.2 | 1.88E-01 | Peptide methionine sulfoxide reductase A3 -S-oxide reductase) reductase) |
| AT4G00040 | -0.4 | 1.33E-04 | -0.6 | 2.31E-05 | 0.2 | 5.55E-02 | -0.2 | 3.03E-01 | 0.2 | 4.81E-01 | -0.4 | 1.43E-01 | Type III polyketide synthase C |
| AT2G41880 | -0.4 | 3.44E-04 | -0.4 | 1.99E-03 | -0.1 | 4.08E-01 | 0.0 | 9.13E-01 | -0.4 | 8.66E-02 | 0.4 | 9.47E-02 | guanylate kinase |
| AT3G61070 | -0.4 | 2.25E-04 | -0.2 | 7.23E-02 | -0.3 | 9.57E-03 | -0.4 | 1.79E-02 | 0.1 | 7.87E-01 | -0.5 | 3.10E-02 | Peroxisomal membrane protein 11E |
| AT3G22930 | -0.4 | 6.65E-04 | -0.3 | 4.91E-03 | -0.1 | 3.42E-01 | -0.5 | 1.66E-02 | 0.3 | 1.21E-01 | -0.8 | 4.96E-03 | Calmodulin-like 11 |
| AT4G17615 | -0.4 | 3.69E-04 | -0.3 | 9.47E-03 | -0.2 | 1.12E-01 | 0.0 | 9.62E-01 | 0.2 | 4.04E-01 | -0.2 | 5.13E-01 | Calcineurin B-like protein |
| AT1G76360 | -0.4 | 4.13E-04 | -0.6 | 1.54E-04 | 0.1 | 2.94E-01 | 0.1 | 7.82E-01 | 0.1 | 7.31E-01 | 0.0 | 9.30E-01 | Uncharacterized protein F15M4.14 |
| AT1G45130 | -0.4 | 1.53E-04 | -0.1 | 4.13E-01 | -0.4 | 1.51E-03 | -0.2 | 9.68E-02 | 0.0 | 8.89E-01 | -0.2 | 1.59E-01 | Beta-galactosidase |
| AT5G41260 | -0.4 | 9.44E-05 | -0.4 | 3.26E-04 | 0.0 | 5.86E-01 | -0.2 | 6.66E-02 | -0.2 | 1.48E-01 | 0.0 | 9.08E-01 | Serine |
| AT2G28890 | -0.4 | 4.09E-03 | -0.7 | 5.52E-04 | 0.2 | 1.53E-01 | 0.2 | 2.14E-01 | 0.1 | 7.51E-01 | 0.1 | 6.00E-01 | Probable protein phosphatase 2C 23 |
| AT1G11410 | -0.4 | 7.45E-03 | -0.5 | 7.29E-03 | 0.0 | 8.96E-01 | -0.5 | 7.50E-02 | 0.1 | 6.58E-01 | -0.6 | 8.17E-02 | Receptor-like serine |
| AT4G36900 | -0.4 | 2.15E-04 | -0.4 | 3.44E-04 | 0.0 | 9.98E-01 | -0.4 | 2.84E-02 | -0.5 | 1.14E-02 | 0.1 | 6.77E-01 | Ethylene-responsive transcription factor RAP2-10 |
| AT1G68920 | -0.4 | 1.18E-04 | -0.3 | 9.47E-04 | -0.1 | 2.76E-01 | -0.1 | 7.46E-01 | 0.2 | 3.94E-01 | -0.3 | 3.22E-01 | AT1G68920 protein |
| AT5G41560 | -0.4 | 4.01E-04 | -0.2 | 3.65E-02 | -0.2 | 3.79E-02 | -0.5 | 1.21E-02 | -0.2 | 2.88E-01 | -0.3 | 2.91E-01 | DET1- and DDB1-associated protein 1 |
| AT1G10050 | -0.4 | 2.30E-04 | -0.4 | 1.14E-03 | -0.1 | 4.38E-01 | -0.3 | 1.28E-01 | -0.1 | 6.04E-01 | -0.2 | 4.84E-01 | Glycosyl hydrolase family 10 protein |
| AT2G01210 | -0.4 | 1.04E-02 | -0.3 | 3.73E-02 | -0.1 | 6.29E-01 | 0.2 | 7.82E-01 | 0.3 | 7.82E-01 | 0.0 | 9.76E-01 | Receptor protein kinase-like protein ZAR1 |
| AT3G24530 | -0.4 | 7.55E-04 | -0.3 | 1.54E-02 | -0.2 | 1.42E-01 | -0.4 | 2.80E-02 | 0.0 | 8.49E-01 | -0.3 | 1.34E-01 | hypothetical protein |
| AT1G12470 | -0.4 | 8.56E-05 | -0.4 | 2.65E-04 | 0.0 | 6.46E-01 | -0.1 | 6.23E-01 | 0.0 | 9.90E-01 | -0.1 | 7.57E-01 | Vacuolar sorting protein 18 |

| | | | | | | | | | | | | | |
|---|---|---|---|---|---|---|---|---|---|---|---|---|---|
| AT1G09910 | -0.4 | 5.98E-04 | -0.5 | 3.20E-04 | 0.1 | 4.80E-01 | -0.2 | 3.09E-01 | -0.1 | 6.96E-01 | -0.1 | 7.38E-01 | rhamnogalacturonan endolyase |
| AT1G78700 | -0.4 | 1.10E-03 | -0.3 | 1.54E-02 | -0.2 | 2.19E-01 | -0.9 | 1.95E-01 | -0.2 | 8.56E-01 | -0.7 | 3.96E-01 | BES1 |
| AT5G38610 | -0.4 | 6.36E-03 | 0.2 | 1.37E-01 | -0.6 | 2.58E-03 | -0.1 | 7.08E-01 | 0.2 | 6.58E-01 | -0.3 | 3.02E-01 | At5g38610 |
| AT3G10420 | -0.4 | 3.83E-03 | -0.8 | 1.46E-04 | 0.3 | 2.58E-02 | -0.6 | 2.69E-02 | 0.1 | 7.56E-01 | -0.7 | 4.25E-02 | Protein SEEDLING PLASTID DEVELOPMENT 1 |
| AT2G23780 | -0.4 | 1.65E-04 | -0.2 | 1.91E-02 | -0.2 | 2.05E-02 | -0.2 | 3.68E-01 | -0.1 | 7.88E-01 | -0.1 | 7.23E-01 | E3 ubiquitin-protein ligase RMA |
| AT2G19160 | -0.4 | 1.37E-04 | -0.1 | 8.32E-02 | -0.3 | 5.47E-03 | -0.1 | 8.18E-01 | 0.1 | 8.01E-01 | -0.2 | 6.82E-01 | Uncharacterized protein At2g19160 |
| AT1G51340 | -0.4 | 3.46E-04 | -0.4 | 8.92E-04 | 0.0 | 7.51E-01 | -0.3 | 4.08E-02 | 0.0 | 9.96E-01 | -0.3 | 1.15E-01 | MATE efflux family protein |
| AT5G59660 | -0.4 | 1.22E-02 | -0.1 | 5.02E-01 | -0.3 | 9.20E-02 | | | | | | | Leucine-rich repeat protein kinase family protein |
| AT2G40970 | -0.4 | 7.10E-05 | 0.0 | 4.82E-01 | -0.4 | 7.75E-04 | -0.4 | 2.53E-02 | -0.2 | 1.95E-01 | -0.1 | 5.11E-01 | Transcription factor MYBC1 |
| AT1G15100 | -0.4 | 1.65E-03 | -0.1 | 4.18E-01 | -0.4 | 1.49E-02 | -0.3 | 1.07E-01 | 0.0 | 8.86E-01 | -0.4 | 1.66E-01 | E3 ubiquitin-protein ligase RHA2A |
| AT2G42010 | -0.4 | 1.91E-04 | -0.3 | 2.38E-03 | -0.1 | 1.81E-01 | -0.1 | 6.57E-01 | 0.1 | 8.46E-01 | -0.2 | 5.97E-01 | phospholipase D |
| AT1G01010 | -0.4 | 1.03E-03 | -1.0 | 1.02E-05 | 0.6 | 1.63E-03 | 0.2 | 2.39E-01 | 0.1 | 8.67E-01 | 0.2 | 4.71E-01 | NAC domain-containing protein 1 |
| AT5G63770 | -0.4 | 5.61E-04 | -0.2 | 4.10E-02 | -0.2 | 4.67E-02 | 0.6 | 4.55E-03 | 0.2 | 3.30E-01 | 0.4 | 1.34E-01 | Diacylglycerol kinase 2 |
| AT3G10815 | -0.4 | 2.49E-01 | -1.0 | 3.33E-02 | 0.6 | 3.04E-01 | | | | | | | RING-type E3 ubiquitin transferase |
| AT5G05820 | -0.4 | 1.18E-04 | -0.2 | 7.87E-03 | -0.2 | 3.23E-02 | 0.0 | 8.59E-01 | 0.2 | 2.62E-01 | -0.3 | 2.36E-01 | Probable sugar phosphate |
| AT3G49370 | -0.4 | 3.45E-04 | -0.5 | 1.98E-04 | 0.1 | 4.68E-01 | -0.2 | 4.86E-01 | -0.2 | 5.09E-01 | 0.0 | 9.81E-01 | CDPK-related kinase 6 |
| AT5G19630 | -0.4 | 4.13E-04 | -0.2 | 3.26E-02 | -0.2 | 4.32E-02 | 0.0 | 9.35E-01 | 0.0 | 9.95E-01 | 0.0 | 9.61E-01 | Xaa-Pro dipeptidyl-peptidase-like domain-containing protein |
| AT5G15540 | -0.4 | 1.12E-04 | -0.6 | 2.66E-05 | 0.2 | 9.15E-02 | -0.2 | 2.74E-01 | 0.1 | 8.21E-01 | -0.3 | 2.59E-01 | Sister chromatid cohesion protein SCC2 |
| AT5G16050 | -0.4 | 9.75E-04 | -0.3 | 1.56E-02 | -0.2 | 1.77E-01 | 0.1 | 2.98E-01 | 0.1 | 2.51E-01 | 0.0 | 8.91E-01 | 14-3-3-like protein GF14 upsilon |
| AT1G54510 | -0.4 | 8.96E-05 | -0.4 | 3.17E-04 | 0.0 | 6.01E-01 | -0.1 | 5.15E-01 | -0.2 | 4.40E-01 | 0.0 | 8.97E-01 | NIMA-related serine |
| AT4G02500 | -0.4 | 7.19E-04 | 0.0 | 9.83E-01 | -0.4 | 2.02E-03 | -0.2 | 1.88E-01 | 0.1 | 7.59E-01 | -0.2 | 1.70E-01 | Xyloglucan 6-xylosyltransferase 2 |
| AT2G02040 | -0.4 | 4.61E-05 | -0.3 | 8.46E-04 | -0.1 | 8.54E-02 | -0.4 | 1.13E-02 | -0.2 | 2.60E-01 | -0.2 | 3.33E-01 | Protein NRT1 |
| AT5G65470 | -0.4 | 6.84E-04 | -0.3 | 8.98E-03 | -0.1 | 2.01E-01 | -0.6 | 4.47E-04 | -0.2 | 3.78E-01 | -0.5 | 2.83E-02 | O-fucosyltransferase 39 |
| AT3G26280 | -0.4 | 5.12E-03 | -0.1 | 3.31E-01 | -0.3 | 5.34E-02 | -0.2 | 2.63E-01 | -0.1 | 5.01E-01 | 0.0 | 8.64E-01 | Cytochrome P450, family 71, subfamily B, polypeptide 4 |
| AT1G34300 | -0.4 | 1.88E-03 | -0.3 | 2.09E-02 | -0.1 | 2.61E-01 | -0.2 | 3.02E-01 | -0.5 | 4.57E-02 | 0.3 | 4.00E-01 | G-type lectin S-receptor-like serine |
| AT1G25520 | -0.4 | 4.64E-04 | -0.2 | 6.01E-02 | -0.3 | 2.31E-02 | 0.1 | 6.80E-01 | -0.1 | 7.85E-01 | 0.1 | 5.08E-01 | GDT1-like protein 4 |
| AT2G46420 | -0.4 | 1.79E-03 | -0.5 | 8.77E-04 | 0.1 | 5.35E-01 | 0.1 | 6.27E-01 | 0.2 | 3.73E-01 | -0.1 | 7.41E-01 | Helicase with zinc finger protein |
| AT3G59050 | -0.4 | 8.29E-04 | -0.2 | 4.79E-02 | -0.2 | 5.77E-02 | -0.3 | 9.15E-02 | 0.1 | 7.92E-01 | -0.3 | 1.22E-01 | Polyamine oxidase 3 |
| AT2G41770 | -0.4 | 1.37E-04 | -0.2 | 1.10E-02 | -0.2 | 2.85E-02 | 0.0 | 8.78E-01 | 0.1 | 6.02E-01 | -0.2 | 5.52E-01 | Probable glycosyltransferase STELLO1 |
| AT4G14350 | -0.4 | 8.27E-05 | -0.3 | 6.92E-04 | -0.1 | 2.36E-01 | -0.1 | 5.69E-01 | 0.1 | 6.98E-01 | -0.1 | 4.12E-01 | AGC kinase family protein |
| AT3G45410 | -0.4 | 5.01E-04 | -0.5 | 2.83E-04 | 0.1 | 4.98E-01 | 0.1 | 8.25E-01 | 0.1 | 8.27E-01 | 0.0 | 9.80E-01 | L-type lectin-domain containing receptor kinase I.3 |
| AT1G48790 | -0.4 | 2.14E-04 | -0.3 | 2.44E-03 | -0.1 | 1.90E-01 | -0.1 | 6.28E-01 | 0.1 | 7.67E-01 | -0.1 | 4.86E-01 | AMSH-like ubiquitin thioesterase 1 |
| AT1G13590 | -0.4 | 1.68E-01 | -0.6 | 6.66E-02 | 0.2 | 6.27E-01 | 0.9 | 1.34E-01 | 0.5 | 5.18E-01 | 0.4 | 4.67E-01 | Phytosulfokine |
| AT3G53010 | -0.4 | 1.02E-02 | 0.0 | 8.06E-01 | -0.5 | 2.07E-02 | -0.4 | 2.85E-01 | 0.1 | 8.95E-01 | -0.5 | 3.18E-01 | Uncharacterized protein F8J2_180 |
| AT3G18100 | -0.4 | 7.54E-04 | -0.6 | 2.60E-04 | 0.1 | 3.15E-01 | 0.0 | 9.27E-01 | -0.2 | 7.69E-01 | 0.1 | 8.71E-01 | Myb domain protein 4r1 |
| AT5G24010 | -0.4 | 1.63E-03 | -0.5 | 1.07E-03 | 0.1 | 6.66E-01 | 0.0 | 9.31E-01 | -0.1 | 8.29E-01 | 0.1 | 7.87E-01 | Probable receptor-like protein kinase At5g24010 |
| AT2G44140 | -0.4 | 5.31E-05 | -0.4 | 2.09E-04 | -0.1 | 5.24E-01 | -0.4 | 5.45E-02 | -0.2 | 4.31E-01 | -0.2 | 4.90E-01 | Cysteine protease |
| AT4G12460 | -0.4 | 3.75E-04 | -0.3 | 6.51E-03 | -0.2 | 1.46E-01 | -0.2 | 2.81E-01 | 0.0 | 9.77E-01 | -0.2 | 4.40E-01 | OSBP-related protein 2B |
| AT1G15780 | -0.4 | 1.25E-04 | -0.5 | 7.38E-05 | 0.1 | 3.84E-01 | 0.2 | 3.04E-01 | 0.1 | 5.59E-01 | 0.1 | 8.69E-01 | Mediator of RNA polymerase II transcription subunit 15a-like protein |
| AT2G43060 | -0.4 | 3.43E-03 | -0.3 | 2.91E-02 | -0.1 | 3.39E-01 | -0.1 | 4.61E-01 | -0.3 | 8.24E-02 | 0.2 | 4.06E-01 | Transcription factor IBH1 |
| AT3G07780 | -0.4 | 1.58E-04 | -0.3 | 1.42E-03 | -0.1 | 2.46E-01 | -0.2 | 1.34E-01 | -0.4 | 8.33E-03 | 0.2 | 2.78E-01 | Protein OBERON |
| AT4G29920 | -0.4 | 5.85E-04 | -0.4 | 9.73E-04 | 0.0 | 9.51E-01 | 0.0 | 8.78E-01 | 0.3 | 2.89E-01 | -0.3 | 2.44E-01 | Double Clp-N motif-containing P-loop nucleoside triphosphate hydrolases superfamily protein |
| AT5G47720 | -0.4 | 2.01E-04 | -0.1 | 9.53E-02 | -0.3 | 7.10E-03 | -0.3 | 1.04E-01 | 0.1 | 8.36E-01 | -0.4 | 8.84E-02 | Thiolase family protein |
| AT2G36890 | -0.4 | 1.17E-02 | -0.4 | 2.73E-02 | -0.1 | 7.60E-01 | -0.3 | 2.42E-01 | 0.2 | 5.02E-01 | -0.5 | 8.56E-02 | Duplicated homeodomain-like superfamily protein |
| AT3G51340 | -0.4 | 7.91E-03 | 0.5 | 2.14E-03 | -1.0 | 2.06E-04 | -0.9 | 2.53E-03 | -0.4 | 1.54E-01 | -0.5 | 1.71E-01 | Uncharacterized protein F24M12.380 |
| AT1G52240 | -0.4 | 3.00E-03 | 0.1 | 3.51E-01 | -0.5 | 3.16E-03 | -1.1 | 3.24E-02 | 0.0 | 9.65E-01 | -1.1 | 7.05E-02 | Rop guanine nucleotide exchange factor 11 |
| AT1G20490 | -0.4 | 8.01E-03 | -0.2 | 1.29E-01 | -0.2 | 2.18E-01 | 0.1 | 8.17E-01 | -0.2 | 6.22E-01 | 0.2 | 4.93E-01 | AMP-dependent synthetase and ligase family protein |
| AT5G66700 | -0.4 | 1.66E-02 | -0.8 | 9.87E-04 | 0.4 | 1.05E-01 | -0.4 | 4.02E-01 | 0.4 | 4.19E-01 | -0.7 | 1.37E-01 | Homeobox-leucine zipper protein ATHB-53 |
| AT4G04910 | -0.4 | 1.99E-04 | -0.1 | 2.05E-01 | -0.3 | 3.44E-03 | 0.0 | 9.36E-01 | 0.0 | 9.17E-01 | 0.0 | 8.88E-01 | Vesicle-fusing ATPase |
| AT2G30740 | -0.4 | 5.90E-05 | -0.1 | 2.96E-01 | -0.4 | 8.22E-04 | 0.0 | 9.75E-01 | 0.1 | 6.91E-01 | -0.1 | 7.75E-01 | PTI1-like tyrosine-protein kinase 2 |
| AT1G71170 | -0.4 | 3.48E-03 | -0.4 | 1.24E-02 | -0.1 | 5.94E-01 | -0.1 | 6.38E-01 | -0.1 | 6.27E-01 | 0.0 | 9.58E-01 | Probable 3-hydroxyisobutyrate dehydrogenase-like 2, mitochondrial |
| AT5G48230 | -0.4 | 8.51E-05 | 0.1 | 1.92E-01 | -0.5 | 1.17E-04 | -0.2 | 1.64E-01 | 0.0 | 9.47E-01 | -0.2 | 3.85E-01 | Uncharacterized protein |
| AT5G25220 | -0.4 | 7.41E-04 | -0.7 | 7.51E-05 | 0.2 | 5.37E-02 | -0.2 | 1.22E-01 | 0.1 | 3.81E-01 | -0.3 | 6.54E-02 | Homeobox protein knotted-1-like 3 |
| AT5G22220 | -0.4 | 1.15E-03 | -0.2 | 4.15E-02 | -0.2 | 1.07E-01 | -0.1 | 8.16E-01 | 0.1 | 7.78E-01 | -0.2 | 6.38E-01 | Transcription factor E2FB |
| AT5G62260 | -0.4 | 5.22E-04 | -0.1 | 5.40E-01 | -0.4 | 4.74E-03 | 0.2 | 4.59E-01 | 0.1 | 8.30E-01 | 0.1 | 7.67E-01 | AT-hook motif nuclear-localized protein |
| AT5G26920 | -0.4 | 4.84E-02 | -0.8 | 2.87E-03 | 0.4 | 1.25E-01 | 0.4 | 1.55E-02 | -0.1 | 6.94E-01 | 0.5 | 3.29E-02 | Cam-binding protein 60-like G |
| AT4G35040 | -0.4 | 2.62E-04 | -0.2 | 4.76E-02 | -0.3 | 1.53E-02 | -0.4 | 9.71E-04 | -0.3 | 2.06E-02 | -0.1 | 4.55E-01 | BZIP domain-containing protein |
| AT4G37260 | -0.4 | 2.96E-02 | -0.5 | 2.51E-02 | 0.0 | 8.82E-01 | -0.7 | 1.24E-03 | -0.8 | 8.38E-04 | 0.1 | 6.33E-01 | Transcription factor MYB73 |
| AT1G72730 | -0.4 | 6.97E-03 | 0.2 | 1.70E-01 | -0.6 | 2.34E-03 | 0.0 | 8.87E-01 | 0.2 | 2.17E-01 | -0.2 | 3.74E-01 | Eukaryotic initiation factor 4A-3 |
| AT1G13280 | -0.4 | 3.38E-03 | 0.0 | 8.21E-01 | -0.4 | 1.15E-02 | -0.2 | 2.82E-01 | 0.2 | 3.74E-01 | -0.3 | 1.14E-01 | allene-oxide cyclase |
| AT1G67730 | -0.4 | 2.39E-04 | -0.2 | 2.43E-02 | -0.2 | 2.47E-02 | 0.3 | 2.08E-02 | 0.3 | 2.64E-02 | 0.0 | 9.28E-01 | Very-long-chain 3-oxoacyl-CoA reductase 1 |
| AT1G12060 | -0.4 | 2.41E-02 | 0.0 | 7.80E-01 | -0.5 | 3.50E-02 | 0.0 | 9.96E-01 | 0.1 | 6.67E-01 | -0.1 | 6.66E-01 | BAG family molecular chaperone regulator 5, mitochondrial |
| AT4G03340 | -0.4 | 3.46E-03 | -0.2 | 8.31E-02 | -0.2 | 1.65E-01 | 0.1 | 7.85E-01 | 0.0 | 9.38E-01 | 0.1 | 9.14E-01 | Uncharacterized protein |
| AT2G25290 | -0.4 | 2.03E-02 | -0.6 | 4.53E-03 | 0.2 | 3.79E-01 | 0.1 | 6.81E-01 | 0.5 | 9.82E-02 | -0.3 | 2.09E-01 | Protein PHOX1 |
| AT3G23690 | -0.4 | 2.64E-04 | -0.3 | 3.11E-03 | -0.1 | 1.92E-01 | -0.4 | 7.01E-03 | 0.0 | 9.79E-01 | -0.4 | 2.94E-02 | Transcription factor bHLH77 |
| AT1G73590 | -0.4 | 3.23E-04 | -0.3 | 5.10E-03 | -0.1 | 1.65E-01 | 0.2 | 2.50E-01 | 0.2 | 4.58E-01 | 0.0 | 8.63E-01 | Auxin efflux carrier component 1 |
| AT2G39750 | -0.4 | 3.19E-04 | -0.3 | 2.43E-03 | -0.1 | 3.04E-01 | -0.2 | 3.88E-01 | 0.1 | 7.60E-01 | -0.3 | 3.27E-01 | Probable methyltransferase PMT11 |
| AT3G30841 | -0.4 | 6.59E-04 | -0.1 | 1.20E-01 | -0.3 | 2.35E-02 | -0.3 | 1.07E-01 | -0.2 | 2.40E-01 | -0.1 | 8.48E-01 | Metalloenzyme domain-containing protein |
| AT2G02360 | -0.4 | 5.07E-05 | -0.4 | 8.97E-05 | 0.0 | 9.93E-01 | 0.0 | 9.14E-01 | 0.0 | 8.91E-01 | -0.1 | 8.25E-01 | F-box protein PP2-B10 |
| AT1G35670 | -0.4 | 5.22E-04 | 0.1 | 3.97E-01 | -0.5 | 6.73E-04 | -0.2 | 8.79E-02 | -0.2 | 1.84E-01 | 0.0 | 9.05E-01 | Calcium-dependent protein kinase |
| AT1G33870 | -0.4 | 1.28E-02 | -0.1 | 3.37E-01 | -0.3 | 1.41E-01 | -0.1 | 8.07E-01 | 0.3 | 4.42E-01 | -0.4 | 2.74E-01 | AIG1-like protein; 48352-49494 |
| AT2G20400 | -0.4 | 9.26E-04 | -0.3 | 5.92E-03 | -0.1 | 4.09E-01 | -0.2 | 3.36E-01 | -0.5 | 4.86E-02 | 0.3 | 3.73E-01 | Myb-like HTH transcriptional regulator family protein |
| AT5G04760 | -0.4 | 7.03E-03 | -0.8 | 3.04E-04 | 0.4 | 5.30E-02 | -0.2 | 5.50E-01 | -0.1 | 8.70E-01 | -0.1 | 8.03E-01 | Duplicated homeodomain-like superfamily protein |
| AT3G03550 | -0.4 | 1.64E-03 | -0.1 | 2.81E-01 | -0.3 | 2.35E-02 | -0.2 | 2.19E-01 | -0.2 | 3.82E-01 | 0.0 | 9.10E-01 | RING-H2 finger protein ATL51 |
| AT4G32290 | -0.4 | 2.93E-04 | 0.3 | 1.23E-03 | -0.8 | 4.27E-05 | 0.0 | 8.49E-01 | 0.1 | 5.97E-01 | -0.1 | 7.93E-01 | hypothetical protein |
| AT5G01450 | -0.4 | 1.45E-04 | -0.3 | 2.85E-03 | -0.1 | 1.13E-01 | 0.0 | 8.83E-01 | -0.1 | 8.28E-01 | 0.1 | 7.40E-01 | E3 ubiquitin-protein ligase APD2 |
| AT3G28030 | -0.4 | 1.86E-04 | -0.5 | 2.12E-04 | 0.0 | 8.02E-01 | 0.0 | 8.55E-01 | 0.2 | 3.60E-01 | -0.1 | 5.45E-01 | 5'-3' exonuclease family protein |
| AT4G19120 | -0.4 | 5.60E-04 | -0.1 | 2.75E-01 | -0.3 | 7.30E-03 | -0.3 | 1.18E-02 | 0.3 | 2.00E-02 | -0.7 | 1.68E-03 | Methyltransferase |
| AT5G27450 | -0.4 | 1.27E-04 | 0.2 | 3.41E-02 | -0.6 | 9.04E-05 | -0.4 | 2.19E-02 | 0.2 | 3.78E-01 | -0.5 | 1.31E-02 | Mevalonate kinase |
| AT1G12880 | -0.4 | 1.03E-02 | -0.4 | 1.19E-02 | 0.0 | 9.65E-01 | -0.3 | 3.60E-01 | -0.1 | 8.14E-01 | -0.2 | 6.63E-01 | Nudix hydrolase 12, mitochondrial |
| AT4G11830 | -0.4 | 1.32E-03 | -0.4 | 2.10E-03 | 0.0 | 9.37E-01 | -0.3 | 3.96E-02 | -0.1 | 5.85E-01 | -0.2 | 2.91E-01 | phospholipase D |
| AT4G20910 | -0.4 | 6.95E-04 | -0.3 | 1.06E-02 | -0.1 | 1.96E-01 | 0.1 | 7.84E-01 | 0.2 | 6.02E-01 | -0.1 | 8.14E-01 | Small RNA 2'-O-methyltransferase |
| AT2G30580 | -0.4 | 1.04E-03 | -0.5 | 7.91E-04 | 0.0 | 7.21E-01 | -0.3 | 2.14E-01 | 0.1 | 8.75E-01 | -0.4 | 2.87E-01 | DREB2A-interacting protein 2 |

| | | | | | | | | | | | | | |
|---|---|---|---|---|---|---|---|---|---|---|---|---|---|
| AT4G18820 | -0.4 | 4.12E-04 | -0.1 | 1.20E-01 | -0.3 | 1.30E-02 | 0.1 | 7.02E-01 | 0.2 | 2.10E-01 | -0.2 | 4.89E-01 | Protein STICHEL-like 3 |
| AT5G39500 | -0.4 | 1.89E-04 | -0.2 | 9.59E-03 | -0.2 | 4.57E-02 | -0.1 | 3.65E-01 | 0.1 | 7.19E-01 | -0.2 | 3.20E-01 | ARF guanine-nucleotide exchange factor GNL1 |
| AT1G11670 | -0.4 | 1.34E-04 | 0.2 | 8.79E-03 | -0.6 | 5.23E-05 | -0.1 | 6.78E-01 | 0.5 | 7.21E-03 | -0.6 | 3.87E-03 | Protein DETOXIFICATION 36 |
| AT3G05010 | -0.4 | 4.19E-04 | -0.2 | 1.36E-02 | -0.2 | 9.45E-02 | -0.4 | 3.19E-02 | -0.2 | 4.84E-01 | -0.3 | 3.04E-01 | Protein CANDIDATE G-PROTEIN COUPLED RECEPTOR 2 |
| AT4G31110 | -0.4 | 1.02E-03 | -0.4 | 1.19E-03 | 0.0 | 8.96E-01 | -0.8 | 1.94E-02 | 0.0 | 9.10E-01 | -0.8 | 2.34E-02 | Wall-associated receptor kinase-like |
| AT1G70000 | -0.4 | 2.49E-04 | -0.6 | 5.60E-05 | 0.1 | 1.20E-01 | 0.0 | 8.47E-01 | 0.2 | 2.73E-01 | -0.3 | 2.56E-01 | At1g70000 |
| AT4G26080 | -0.4 | 1.13E-03 | -0.6 | 2.49E-04 | 0.1 | 1.87E-01 | -0.4 | 3.19E-03 | -0.1 | 3.86E-01 | -0.3 | 1.20E-01 | Protein phosphatase 2C 56 |
| AT1G17260 | -0.4 | 2.83E-02 | -0.6 | 8.02E-03 | 0.2 | 4.78E-01 | | | | | | | ATPase 10, plasma membrane-type |
| AT5G67460 | -0.4 | 4.77E-03 | -0.2 | 1.20E-01 | -0.2 | 1.55E-01 | 0.1 | 7.33E-01 | 0.4 | 1.69E-01 | -0.3 | 4.02E-01 | glucan endo-1,3-beta-D-glucosidase |
| AT2G27170 | -0.4 | 1.38E-04 | -0.5 | 9.57E-05 | 0.1 | 4.84E-01 | 0.0 | 8.03E-01 | 0.0 | 9.23E-01 | 0.0 | 9.44E-01 | Structural maintenance of chromosomes family protein |
| AT1G78610 | -0.4 | 2.18E-03 | -0.3 | 3.05E-02 | -0.2 | 2.25E-01 | -0.2 | 2.92E-01 | 0.0 | 9.55E-01 | -0.2 | 4.95E-01 | Mechanosensitive ion channel protein 6 |
| AT5G54590 | -0.4 | 1.26E-03 | -0.2 | 1.09E-01 | -0.3 | 4.86E-02 | -0.1 | 6.58E-01 | 0.2 | 3.96E-01 | -0.3 | 2.60E-01 | Calcium |
| AT2G37790 | -0.4 | 1.18E-02 | -0.2 | 2.29E-01 | -0.2 | 1.73E-01 | -0.4 | 3.77E-02 | -0.1 | 4.77E-01 | -0.2 | 3.42E-01 | Aldo-keto reductase family 4 member C10 |
| AT4G11360 | -0.4 | 3.97E-04 | 0.0 | 9.10E-01 | -0.4 | 1.77E-03 | 0.1 | 8.13E-01 | 0.0 | 9.88E-01 | 0.1 | 8.55E-01 | E3 ubiquitin-protein ligase RHA1B |
| AT1G26100 | -0.4 | 1.09E-02 | -0.1 | 4.21E-01 | -0.3 | 9.83E-02 | -0.7 | 3.87E-02 | -0.2 | 6.73E-01 | -0.5 | 1.89E-01 | Probable transmembrane ascorbate ferrireductase 4 |
| AT1G56310 | -0.4 | 1.64E-02 | -0.3 | 4.52E-02 | -0.1 | 7.03E-01 | 0.0 | 8.96E-01 | -0.1 | 8.63E-01 | 0.0 | 9.70E-01 | Polynucleotidyl transferase, ribonuclease H-like superfamily protein |
| AT3G50630 | -0.4 | 1.86E-03 | 0.0 | 7.55E-01 | -0.5 | 4.02E-03 | -0.4 | 5.15E-02 | 0.0 | 9.44E-01 | -0.4 | 1.53E-01 | KIP-related protein 2 |
| AT2G22300 | -0.4 | 1.80E-03 | -0.6 | 2.91E-04 | 0.2 | 1.54E-01 | 0.0 | 9.40E-01 | -0.1 | 4.63E-01 | 0.2 | 5.21E-01 | Signal responsive 1 |
| AT4G35870 | -0.4 | 2.43E-04 | -0.3 | 5.02E-03 | -0.1 | 1.27E-01 | -0.3 | 4.31E-01 | -0.2 | 7.17E-01 | -0.1 | 8.39E-01 | CSC1-like protein At4g35870 |
| AT2G46070 | -0.4 | 8.57E-03 | -0.3 | 5.20E-02 | -0.1 | 4.46E-01 | -0.5 | 1.25E-02 | -0.1 | 6.59E-01 | -0.4 | 1.13E-01 | Mitogen-activated protein kinase 12 |
| AT5G23170 | -0.4 | 4.66E-03 | -0.2 | 2.24E-01 | -0.3 | 8.92E-02 | -0.3 | 3.84E-01 | 0.0 | 9.13E-01 | -0.2 | 6.17E-01 | Serine |
| AT4G39700 | -0.4 | 8.58E-02 | -0.1 | 7.28E-01 | -0.3 | 2.51E-01 | -0.2 | 7.91E-01 | 0.8 | 1.34E-01 | -1.0 | 1.24E-01 | Heavy metal-associated isoprenylated plant protein 23 |
| AT5G37780 | -0.4 | 3.05E-04 | -0.1 | 2.04E-01 | -0.3 | 5.32E-03 | 0.0 | 9.86E-01 | 0.1 | 6.67E-01 | -0.1 | 7.17E-01 | Calmodulin 1 |
| AT5G59840 | -0.4 | 5.89E-04 | -0.1 | 1.53E-01 | -0.3 | 1.54E-02 | 0.0 | 8.78E-01 | 0.0 | 9.26E-01 | 0.0 | 9.81E-01 | Putative GTP-binding protein ara-3 |
| AT2G26820 | -0.4 | 6.33E-03 | 0.1 | 2.66E-01 | -0.6 | 3.13E-03 | 1.2 | 6.18E-06 | 1.0 | 1.58E-04 | 0.2 | 3.98E-01 | Phloem protein 2-A3 |
| AT1G27960 | -0.4 | 4.46E-04 | -0.1 | 1.60E-01 | -0.3 | 1.30E-02 | 0.0 | 9.00E-01 | -0.1 | 6.45E-01 | 0.1 | 7.77E-01 | YTH domain-containing family protein |
| AT4G08790 | -0.4 | 3.09E-04 | -0.3 | 2.73E-03 | -0.1 | 2.87E-01 | 0.2 | 4.23E-01 | 0.2 | 4.83E-01 | 0.0 | 1.00E+00 | CN hydrolase domain-containing protein |
| AT5G54760 | -0.4 | 1.74E-04 | -0.2 | 2.85E-02 | -0.2 | 1.52E-02 | 0.1 | 3.00E-01 | 0.1 | 4.68E-01 | 0.0 | 9.34E-01 | At5g54760 |
| AT1G29330 | -0.4 | 2.62E-03 | 0.1 | 2.50E-01 | -0.5 | 1.88E-03 | 0.0 | 8.47E-01 | 0.1 | 8.43E-01 | 0.0 | 9.77E-01 | ER lumen protein-retaining receptor |
| AT5G01890 | -0.4 | 1.18E-03 | -0.3 | 1.67E-02 | -0.1 | 2.04E-01 | -0.2 | 4.84E-01 | 0.0 | 9.87E-01 | -0.2 | 6.54E-01 | Leucine-rich repeat receptor-like protein kinase PXC2 |
| AT2G39900 | -0.4 | 4.97E-04 | 0.2 | 3.05E-02 | -0.6 | 1.66E-04 | -0.3 | 1.82E-02 | 0.0 | 8.25E-01 | -0.3 | 1.23E-01 | LIM domain-containing protein WLIM2a |
| AT4G25840 | -0.4 | 4.92E-03 | -0.2 | 8.95E-02 | -0.2 | 1.82E-01 | 0.1 | 5.25E-01 | 0.2 | 3.57E-01 | -0.1 | 7.99E-01 | -glycerol-3-phosphatase 1, mitochondrial uracil phosphatase) -pyrimidinedione 5'-phosphate phosphatase) |
| AT1G02130 | -0.4 | 7.85E-05 | -0.2 | 1.00E-02 | -0.2 | 1.38E-02 | -0.2 | 5.29E-02 | -0.2 | 1.84E-01 | 0.0 | 8.07E-01 | Ras-related protein RABD2a |
| AT3G18780 | -0.4 | 5.87E-04 | 0.0 | 9.70E-01 | -0.4 | 1.91E-03 | -0.2 | 3.45E-02 | -0.2 | 4.73E-02 | 0.0 | 9.65E-01 | Actin 2 |
| AT4G19170 | -0.4 | 5.08E-02 | -0.2 | 4.32E-01 | -0.3 | 2.68E-01 | 0.0 | 9.38E-01 | -0.1 | 4.02E-01 | 0.1 | 4.41E-01 | Probable carotenoid cleavage dioxygenase 4, chloroplastic |
| AT2G17520 | -0.4 | 9.45E-04 | -0.6 | 1.20E-04 | 0.2 | 8.84E-02 | -0.5 | 1.43E-02 | -0.4 | 6.28E-02 | -0.1 | 7.94E-01 | Serine |
| AT1G19310 | -0.4 | 8.60E-04 | -0.3 | 1.20E-02 | -0.1 | 1.97E-01 | -0.4 | 1.67E-02 | -0.4 | 2.36E-02 | 0.0 | 9.85E-01 | E3 ubiquitin-protein ligase RMA |
| AT1G70770 | -0.4 | 1.42E-04 | -0.1 | 4.20E-01 | -0.4 | 1.39E-03 | 0.1 | 6.60E-01 | 0.1 | 5.01E-01 | 0.0 | 8.49E-01 | AT1G70770 protein |
| AT3G48050 | -0.4 | 5.73E-04 | -0.5 | 2.71E-04 | 0.1 | 4.10E-01 | 0.0 | 9.85E-01 | -0.1 | 8.49E-01 | 0.0 | 8.96E-01 | Protein SUO |
| AT3G05210 | -0.4 | 8.00E-05 | -0.3 | 1.27E-03 | -0.1 | 1.28E-01 | -0.2 | 4.26E-01 | -0.2 | 3.07E-01 | 0.1 | 8.25E-01 | Nucleotide repair protein |
| AT4G09760 | -0.4 | 5.31E-03 | -0.6 | 1.30E-03 | 0.1 | 3.32E-01 | -0.6 | 1.13E-02 | -0.4 | 7.76E-02 | -0.2 | 6.49E-01 | Protein kinase superfamily protein |
| AT4G40040 | -0.4 | 2.45E-04 | -0.3 | 3.28E-03 | -0.1 | 1.80E-01 | -0.4 | 1.86E-02 | -0.4 | 2.15E-02 | 0.0 | 9.15E-01 | Histone H3.3 |
| AT4G16150 | -0.4 | 1.45E-04 | -0.5 | 1.36E-04 | 0.0 | 6.37E-01 | -0.3 | 1.81E-02 | -0.3 | 2.01E-02 | 0.0 | 9.12E-01 | Calmodulin-binding transcription activator 5 |
| AT3G58570 | -0.4 | 7.89E-04 | -0.3 | 7.60E-03 | -0.1 | 2.63E-01 | -0.1 | 5.20E-01 | -0.1 | 6.00E-01 | 0.0 | 9.95E-01 | DEAD-box ATP-dependent RNA helicase 52 |
| AT1G21630 | -0.4 | 5.74E-04 | -0.4 | 1.22E-03 | 0.0 | 8.38E-01 | -0.2 | 1.97E-01 | 0.0 | 9.50E-01 | -0.2 | 3.77E-01 | Calcium-binding EF hand family protein |
| AT5G62180 | -0.4 | 5.75E-03 | 0.2 | 8.32E-02 | -0.6 | 1.64E-03 | -0.1 | 5.19E-01 | 0.0 | 9.91E-01 | -0.1 | 6.01E-01 | Probable carboxylesterase 120 |
| AT5G11350 | -0.4 | 9.10E-04 | -0.4 | 1.73E-03 | 0.0 | 8.88E-01 | -0.4 | 1.44E-01 | -0.2 | 4.96E-01 | -0.2 | 6.54E-01 | DNAse I-like superfamily protein |
| AT1G63440 | -0.4 | 1.12E-03 | -0.7 | 5.89E-05 | 0.3 | 2.48E-02 | 0.1 | 6.32E-01 | 0.4 | 8.56E-02 | -0.3 | 2.71E-01 | Probable copper-transporting ATPase HMA5 |
| AT5G53430 | -0.4 | 1.77E-03 | -0.5 | 8.75E-04 | 0.1 | 5.35E-01 | -0.2 | 6.42E-01 | 0.2 | 6.99E-01 | -0.4 | 4.24E-01 | Histone-lysine N-methyltransferase ATX5 |
| AT2G04100 | -0.4 | 2.91E-03 | -0.1 | 2.89E-01 | -0.3 | 4.37E-02 | 0.0 | 8.70E-01 | 0.1 | 7.49E-01 | -0.1 | 6.43E-01 | Protein DETOXIFICATION |
| AT1G03905 | -0.4 | 3.52E-03 | -0.3 | 2.84E-02 | -0.1 | 3.67E-01 | 0.4 | 1.23E-01 | 0.2 | 4.64E-01 | 0.2 | 6.54E-01 | P-loop containing nucleoside triphosphate hydrolases superfamily protein |
| AT1G58030 | -0.4 | 3.98E-03 | -0.5 | 1.28E-03 | 0.1 | 3.98E-01 | -0.2 | 1.79E-01 | -0.2 | 2.49E-01 | 0.0 | 9.98E-01 | Cationic amino acid transporter 2, vacuolar |
| AT3G20040 | -0.4 | 4.44E-03 | -0.1 | 2.67E-01 | -0.3 | 7.16E-02 | -0.1 | 7.65E-01 | -0.1 | 8.80E-01 | 0.0 | 9.46E-01 | Hexokinase-4 |
| AT1G28960 | -0.4 | 5.71E-04 | -0.3 | 3.77E-03 | -0.1 | 3.76E-01 | -0.4 | 8.75E-03 | -0.3 | 9.67E-02 | -0.1 | 5.29E-01 | Nudix hydrolase domain-containing protein |
| AT3G14890 | -0.4 | 1.12E-03 | -0.4 | 3.63E-03 | -0.1 | 6.50E-01 | 0.4 | 9.95E-02 | 0.6 | 3.57E-02 | -0.2 | 6.37E-01 | PARP-type domain-containing protein |
| AT5G07820 | -0.4 | 1.10E-03 | -0.2 | 8.39E-02 | -0.2 | 5.22E-02 | -0.2 | 3.41E-01 | -0.2 | 3.53E-01 | 0.0 | 9.62E-01 | hypothetical protein |
| AT5G16680 | -0.4 | 1.48E-04 | -0.6 | 3.26E-05 | 0.1 | 9.14E-02 | 0.0 | 8.47E-01 | 0.0 | 8.68E-01 | 0.0 | 9.96E-01 | PHD-type domain-containing protein |
| AT1G13980 | -0.4 | 3.88E-04 | -0.2 | 6.72E-02 | -0.3 | 1.78E-02 | -0.3 | 7.99E-02 | -0.2 | 2.04E-01 | 0.0 | 8.60E-01 | ARF guanine-nucleotide exchange factor GNOM |
| AT5G45900 | -0.4 | 1.95E-04 | -0.3 | 2.83E-03 | -0.1 | 1.67E-01 | -0.2 | 4.77E-01 | 0.0 | 9.48E-01 | -0.2 | 5.99E-01 | Ubiquitin-like modifier-activating enzyme atg7 |
| AT1G69910 | -0.4 | 1.27E-03 | -0.3 | 1.36E-02 | -0.1 | 2.67E-01 | 0.0 | 8.56E-01 | -0.1 | 7.26E-01 | 0.0 | 8.87E-01 | LEAF RUST 10 DISEASE-RESISTANCE LOCUS RECEPTOR-LIKE PROTEIN KINASE-like 1.5 |
| AT5G03280 | -0.4 | 2.66E-04 | -0.4 | 1.05E-03 | -0.1 | 5.56E-01 | -0.2 | 3.24E-01 | -0.1 | 7.32E-01 | -0.1 | 7.53E-01 | Ethylene-insensitive protein 2 |
| AT1G61180 | -0.4 | 9.74E-04 | -0.4 | 1.07E-03 | 0.0 | 8.82E-01 | -0.2 | 1.48E-01 | -0.1 | 4.53E-01 | -0.1 | 7.34E-01 | LRR and NB-ARC domains-containing disease resistance protein |
| AT3G24200 | -0.4 | 1.21E-04 | -0.4 | 2.79E-04 | 0.0 | 8.17E-01 | 0.0 | 8.68E-01 | 0.0 | 9.41E-01 | 0.0 | 9.66E-01 | Ubiquinone biosynthesis monooxygenase COQ6, mitochondrial |
| AT2G38280 | -0.4 | 1.55E-04 | -0.3 | 1.89E-03 | -0.1 | 1.72E-01 | 0.4 | 3.20E-03 | 0.4 | 9.74E-03 | 0.0 | 9.23E-01 | AMP deaminase |
| AT2G25740 | -0.4 | 9.11E-04 | -0.4 | 2.58E-03 | 0.0 | 7.09E-01 | 0.0 | 9.79E-01 | 0.2 | 4.56E-01 | -0.2 | 5.15E-01 | Protein cereblon |
| AT4G17530 | -0.4 | 1.27E-04 | -0.3 | 1.24E-03 | -0.1 | 2.25E-01 | -0.2 | 1.48E-01 | 0.0 | 9.76E-01 | -0.2 | 3.33E-01 | Uncharacterized protein |
| AT5G61840 | -0.4 | 1.25E-04 | -0.1 | 2.06E-01 | -0.3 | 2.20E-03 | 0.0 | 7.82E-01 | 0.0 | 8.22E-01 | 0.0 | 9.98E-01 | Probable beta-1,4-xylosyltransferase IRX10L |
| AT1G55250 | -0.4 | 9.01E-05 | -0.4 | 2.32E-04 | 0.0 | 7.59E-01 | -0.1 | 4.71E-01 | 0.1 | 7.79E-01 | -0.2 | 3.93E-01 | E3 ubiquitin protein ligase |
| AT3G20510 | -0.4 | 8.40E-04 | 0.0 | 8.58E-01 | -0.4 | 2.02E-03 | 0.0 | 9.13E-01 | 0.2 | 3.57E-01 | -0.2 | 5.14E-01 | Protein FATTY ACID EXPORT 6 |
| AT3G53490 | -0.4 | 2.07E-03 | -0.1 | 3.96E-01 | -0.3 | 2.35E-02 | 0.1 | 8.18E-01 | -0.1 | 7.59E-01 | 0.1 | 6.02E-01 | At3g53490 |
| AT5G52660 | -0.4 | 1.32E-03 | -0.5 | 9.91E-04 | 0.0 | 7.11E-01 | -0.4 | 8.07E-02 | -0.1 | 6.94E-01 | -0.3 | 2.98E-01 | Protein REVEILLE 6 |
| AT5G35700 | -0.4 | 1.74E-04 | -0.1 | 2.53E-01 | -0.3 | 2.87E-03 | 0.0 | 9.81E-01 | 0.2 | 3.29E-01 | -0.2 | 3.86E-01 | Fimbrin-5 |
| AT4G21410 | -0.4 | 3.75E-04 | -0.1 | 1.52E-01 | -0.3 | 9.16E-03 | 0.1 | 4.55E-01 | -0.1 | 5.27E-01 | 0.2 | 2.70E-01 | Cysteine-rich RLK 29 |
| AT3G07420 | -0.4 | 5.33E-04 | -0.3 | 2.74E-03 | -0.1 | 4.57E-01 | 0.2 | 2.38E-01 | 0.3 | 1.42E-01 | -0.1 | 7.92E-01 | Asparaginyl-tRNA synthetase 2 |
| AT4G32810 | -0.4 | 7.95E-02 | -0.8 | 7.57E-03 | 0.4 | 2.05E-01 | | | | | | | Carotenoid cleavage dioxygenase 8 |
| AT3G57120 | -0.4 | 2.47E-04 | -0.4 | 8.15E-04 | 0.0 | 6.30E-01 | -0.2 | 2.53E-01 | -0.2 | 2.30E-01 | 0.0 | 9.26E-01 | Uncharacterized protein F24I3.200 |
| AT5G47750 | -0.4 | 1.42E-04 | -0.3 | 1.27E-03 | -0.1 | 2.32E-01 | -0.2 | 8.27E-02 | -0.1 | 3.85E-01 | -0.1 | 6.56E-01 | Serine |
| AT5G36220 | -0.4 | 1.05E-02 | -0.7 | 9.93E-04 | 0.3 | 1.49E-01 | 0.5 | 1.91E-01 | 0.3 | 4.36E-01 | 0.1 | 8.22E-01 | Cytochrome p450 81d1 |
| AT1G73670 | -0.4 | 2.93E-04 | -0.3 | 1.52E-03 | -0.1 | 4.25E-01 | 0.0 | 8.50E-01 | 0.1 | 7.11E-01 | -0.1 | 6.38E-01 | Mitogen-activated protein kinase 15 |
| AT5G65730 | -0.4 | 5.11E-02 | 0.2 | 3.79E-01 | -0.6 | 1.96E-02 | -0.9 | 3.92E-04 | -0.6 | 1.23E-02 | -0.3 | 3.97E-01 | Probable xyloglucan endotransglucosylase |
| AT3G07030 | -0.4 | 4.08E-03 | -0.3 | 3.97E-02 | -0.1 | 2.92E-01 | -0.1 | 7.19E-01 | -0.2 | 2.84E-01 | 0.1 | 5.79E-01 | Alba DNA |
| AT4G32850 | -0.4 | 2.29E-04 | -0.3 | 4.61E-03 | -0.1 | 1.16E-01 | -0.2 | 1.53E-01 | 0.0 | 8.03E-01 | -0.1 | 4.99E-01 | Poly polymerase |
| AT1G59820 | -0.4 | 5.53E-04 | -0.5 | 2.96E-04 | 0.1 | 4.80E-01 | -0.1 | 6.68E-01 | -0.2 | 3.77E-01 | 0.1 | 7.43E-01 | Phospholipid-transporting ATPase 3 |

| | | | | | | | | | | | | | |
|---|---|---|---|---|---|---|---|---|---|---|---|---|---|
| AT4G39670 | -0.4 | 5.62E-02 | -0.4 | 4.79E-02 | 0.0 | 9.15E-01 | 0.3 | 2.03E-01 | 0.1 | 9.00E-01 | 0.3 | 2.88E-01 | ACD11 homolog protein |
| AT5G06260 | -0.4 | 6.01E-05 | -0.2 | 1.39E-02 | -0.2 | 7.90E-03 | -0.2 | 1.15E-01 | -0.1 | 7.30E-01 | -0.2 | 4.50E-01 | Similarity to unknown protein |
| AT1G11480 | -0.4 | 5.74E-05 | -0.2 | 6.26E-03 | -0.2 | 1.43E-02 | -0.3 | 1.32E-01 | -0.2 | 3.86E-01 | -0.1 | 7.96E-01 | Eukaryotic translation initiation factor-like protein |
| AT1G17980 | -0.4 | 1.16E-04 | -0.3 | 2.03E-03 | -0.1 | 1.14E-01 | -0.2 | 2.49E-01 | 0.0 | 8.87E-01 | -0.1 | 5.28E-01 | Nuclear poly polymerase 1 ) polymerase I) |
| AT2G43850 | -0.4 | 2.24E-04 | -0.4 | 2.28E-04 | 0.0 | 7.51E-01 | -0.1 | 7.83E-01 | -0.1 | 8.56E-01 | 0.0 | 9.76E-01 | Integrin-linked protein kinase family |
| AT1G68690 | -0.4 | 5.90E-03 | -0.7 | 4.70E-04 | 0.3 | 1.08E-01 | 0.5 | 5.92E-02 | 0.2 | 5.93E-01 | 0.3 | 3.59E-01 | non-specific serine |
| AT5G47820 | -0.4 | 6.08E-04 | -0.3 | 6.11E-03 | -0.1 | 2.46E-01 | -0.3 | 1.03E-01 | -0.1 | 6.08E-01 | -0.2 | 5.27E-01 | Kinesin-like protein KIN-4A |
| AT1G76800 | -0.4 | 9.22E-03 | -0.1 | 2.83E-01 | -0.3 | 1.19E-01 | -1.2 | 2.64E-04 | -1.0 | 1.57E-03 | -0.2 | 6.54E-01 | Vacuolar iron transporter homolog 2 |
| AT4G30160 | -0.4 | 9.44E-04 | -0.2 | 3.63E-02 | -0.2 | 7.75E-02 | -0.1 | 8.31E-01 | 0.2 | 5.94E-01 | -0.2 | 5.13E-01 | Villin-4 |
| AT3G60030 | -0.4 | 7.25E-04 | -0.4 | 6.20E-04 | 0.0 | 7.21E-01 | -0.6 | 2.04E-04 | -0.4 | 6.11E-03 | -0.2 | 4.01E-01 | Squamosa promoter-binding-like protein 12 |
| AT4G34490 | -0.4 | 4.07E-04 | -0.1 | 1.82E-01 | -0.3 | 7.62E-03 | 0.0 | 8.26E-01 | 0.1 | 4.23E-01 | -0.2 | 3.27E-01 | Adenylyl cyclase-associated protein |
| AT1G30360 | -0.4 | 4.73E-04 | -0.2 | 1.91E-02 | -0.2 | 7.00E-02 | -0.5 | 2.43E-02 | -0.4 | 1.62E-01 | -0.2 | 6.61E-01 | ERD4 |
| AT5G11670 | -0.4 | 1.28E-03 | -0.2 | 4.92E-02 | -0.2 | 8.51E-02 | 0.4 | 5.06E-04 | 0.1 | 3.73E-01 | 0.3 | 4.77E-02 | NADP-dependent malic enzyme 2 |
| AT5G56500 | -0.4 | 3.01E-04 | -0.2 | 2.69E-02 | -0.2 | 2.91E-02 | 0.4 | 5.07E-02 | 0.3 | 1.66E-01 | 0.1 | 8.28E-01 | Chaperonin 60 subunit beta 3, chloroplastic |
| AT1G20900 | -0.4 | 8.86E-04 | -0.2 | 2.51E-02 | -0.2 | 1.10E-01 | -0.1 | 8.16E-01 | 0.1 | 8.43E-01 | -0.1 | 7.12E-01 | AT-hook motif nuclear-localized protein 27 |
| AT1G49240 | -0.4 | 8.33E-04 | 0.0 | 7.69E-01 | -0.4 | 1.64E-03 | -0.7 | 1.55E-05 | -0.4 | 4.31E-03 | -0.3 | 1.07E-01 | Actin-8 |
| AT1G04970 | -0.4 | 4.15E-04 | -0.5 | 2.93E-04 | 0.1 | 5.99E-01 | -0.4 | 1.02E-02 | -0.1 | 5.12E-01 | -0.3 | 1.63E-01 | Lipid-binding serum glycoprotein C-terminal domain-containing protein |
| AT5G03200 | -0.4 | 5.41E-04 | -0.4 | 4.19E-04 | 0.0 | 6.65E-01 | -0.2 | 3.18E-01 | -0.2 | 5.05E-01 | 0.0 | 9.02E-01 | Probable E3 ubiquitin-protein ligase LUL1 |
| AT1G75080 | -0.4 | 5.09E-04 | -0.4 | 5.30E-04 | 0.0 | 8.16E-01 | -0.6 | 8.37E-04 | -0.5 | 8.41E-03 | -0.1 | 5.28E-01 | Protein BRASSINAZOLE-RESISTANT 1 |
| AT2G38800 | -0.4 | 4.05E-03 | -0.5 | 2.47E-03 | 0.1 | 6.49E-01 | -0.5 | 7.66E-04 | -0.3 | 5.08E-02 | -0.2 | 2.56E-01 | hypothetical protein |
| AT4G11490 | -0.4 | 1.60E-01 | 1.1 | 8.02E-04 | -1.5 | 9.80E-04 | | | | | | | Putative cysteine-rich receptor-like protein kinase 33 |
| AT4G08500 | -0.4 | 1.35E-03 | -0.5 | 2.99E-04 | 0.1 | 2.22E-01 | -0.1 | 5.15E-01 | -0.2 | 1.46E-01 | 0.1 | 5.22E-01 | Mitogen-activated protein kinase kinase kinase 1 |
| AT3G06590 | -0.4 | 3.47E-04 | -0.1 | 9.31E-02 | -0.3 | 1.22E-02 | -0.6 | 6.54E-04 | -0.3 | 3.32E-02 | -0.2 | 2.99E-01 | Transcription factor bHLH148 |
| AT3G53410 | -0.4 | 3.72E-04 | -0.2 | 2.94E-02 | -0.2 | 4.12E-02 | -0.4 | 1.85E-02 | -0.5 | 1.89E-02 | 0.0 | 9.22E-01 | Probable E3 ubiquitin-protein ligase LUL2 |
| AT1G21590 | -0.4 | 3.07E-02 | -0.3 | 9.36E-02 | -0.1 | 6.30E-01 | 0.2 | 2.73E-01 | 0.2 | 2.27E-01 | 0.0 | 8.85E-01 | Protein kinase domain-containing protein |
| AT3G26550 | -0.4 | 8.06E-03 | 0.0 | 9.17E-01 | -0.4 | 2.86E-02 | -0.4 | 2.66E-01 | -0.4 | 4.03E-01 | -0.1 | 9.23E-01 | CHP-rich zinc finger protein-like |
| AT4G36180 | -0.4 | 1.92E-03 | -0.1 | 4.47E-01 | -0.3 | 1.67E-02 | -0.1 | 7.69E-01 | -0.1 | 8.60E-01 | 0.0 | 9.69E-01 | Probable LRR receptor-like serine |
| AT3G19640 | -0.4 | 7.51E-05 | -0.3 | 4.09E-04 | -0.1 | 3.83E-01 | -0.1 | 4.63E-01 | -0.1 | 6.12E-01 | 0.0 | 9.49E-01 | Magnesium transporter MRS2-3 |
| AT1G62400 | -0.4 | 1.80E-02 | 0.2 | 2.94E-01 | -0.6 | 1.01E-02 | -0.2 | 7.01E-01 | -0.5 | 2.63E-01 | 0.4 | 5.52E-01 | Serine |
| AT5G51780 | -0.4 | 4.50E-03 | 0.3 | 2.22E-02 | -0.7 | 6.58E-04 | -0.7 | 3.76E-02 | 0.1 | 7.98E-01 | -0.8 | 6.29E-02 | Basic helix-loop-helix DNA-binding superfamily protein |
| AT5G10270 | -0.4 | 9.05E-04 | -0.3 | 3.94E-03 | -0.1 | 5.28E-01 | -0.1 | 8.04E-01 | 0.1 | 7.50E-01 | -0.2 | 6.43E-01 | Cyclin-dependent kinase C-1 |
| AT1G77800 | -0.4 | 6.59E-04 | -0.5 | 1.59E-04 | 0.1 | 1.90E-01 | -0.2 | 2.35E-01 | 0.0 | 9.88E-01 | -0.2 | 3.79E-01 | PHD finger family protein |
| AT5G05410 | -0.4 | 6.27E-03 | -0.1 | 5.58E-01 | -0.3 | 4.35E-02 | 0.2 | 6.72E-01 | 0.2 | 7.19E-01 | 0.0 | 9.98E-01 | DRE-binding protein 2A |
| AT1G70080 | -0.4 | 1.52E-01 | -0.4 | 1.67E-01 | 0.0 | 9.93E-01 | -0.8 | 1.38E-01 | -0.3 | 5.83E-01 | -0.5 | 5.20E-01 | Dolabella-3,7-dien-18-ol synthase TPS06 |
| AT2G38840 | -0.4 | 1.81E-04 | -0.1 | 1.17E-01 | -0.3 | 5.53E-03 | -0.1 | 3.99E-01 | 0.2 | 2.94E-01 | -0.3 | 1.17E-01 | At2g38850 |
| AT3G15220 | -0.4 | 1.49E-03 | -0.3 | 8.18E-03 | -0.1 | 4.60E-01 | -0.2 | 4.59E-01 | 0.1 | 8.47E-01 | -0.2 | 4.03E-01 | Protein kinase domain-containing protein |
| AT1G64780 | -0.4 | 6.74E-04 | -0.3 | 1.24E-02 | -0.1 | 1.51E-01 | -0.2 | 3.37E-01 | 0.4 | 4.34E-02 | -0.5 | 1.36E-02 | Ammonium transporter 1 member 2 |
| AT5G43590 | -0.4 | 5.71E-02 | 0.9 | 3.16E-04 | -1.3 | 2.27E-04 | -4.7 | 5.48E-05 | -0.4 | 3.60E-01 | -4.3 | 5.53E-04 | Patatin |
| AT5G49500 | -0.4 | 2.44E-03 | -0.3 | 1.39E-02 | -0.1 | 4.60E-01 | -0.2 | 3.79E-01 | 0.3 | 3.38E-01 | -0.5 | 1.08E-01 | Signal recognition particle, SRP54 subunit protein |
| AT4G01220 | -0.4 | 5.27E-03 | -0.1 | 5.66E-01 | -0.3 | 3.48E-02 | 0.1 | 6.91E-01 | 0.2 | 3.18E-01 | -0.1 | 6.43E-01 | Multifunctional fusion protein |
| AT4G12300 | -0.4 | 1.93E-04 | -0.6 | 2.57E-05 | 0.2 | 4.26E-02 | -0.1 | 4.17E-01 | 0.0 | 9.06E-01 | -0.1 | 6.03E-01 | Cytochrome P450, family 706, subfamily A, polypeptide 4 |
| AT1G58410 | -0.4 | 1.30E-03 | -0.2 | 3.63E-02 | -0.2 | 1.35E-01 | 0.4 | 1.61E-01 | 0.2 | 7.02E-01 | 0.3 | 4.05E-01 | Disease resistance protein family |
| AT2G45620 | -0.4 | 2.63E-04 | -0.3 | 1.13E-03 | -0.1 | 4.93E-01 | -0.2 | 1.17E-01 | -0.4 | 4.06E-02 | 0.1 | 6.24E-01 | UTP:RNA uridylyltransferase 1 |
| AT1G02100 | -0.4 | 1.56E-04 | -0.3 | 2.61E-03 | -0.1 | 1.24E-01 | -0.3 | 3.29E-02 | -0.2 | 9.72E-02 | 0.0 | 8.64E-01 | Leucine carboxyl methyltransferase 1 homolog |
| AT1G03190 | -0.4 | 2.06E-03 | -0.4 | 1.66E-03 | 0.0 | 7.74E-01 | -0.1 | 7.34E-01 | 0.0 | 9.08E-01 | -0.1 | 7.01E-01 | General transcription and DNA repair factor IIH helicase subunit XPD |
| AT2G21970 | -0.4 | 7.56E-05 | -0.3 | 5.66E-04 | -0.1 | 2.81E-01 | 0.2 | 3.22E-01 | 0.2 | 1.79E-01 | -0.1 | 7.16E-01 | Stress enhanced protein 2, chloroplastic |
| AT1G53780 | -0.4 | 3.85E-04 | -0.7 | 1.30E-05 | 0.3 | 5.24E-03 | -0.3 | 9.02E-02 | -0.2 | 3.22E-01 | -0.1 | 7.35E-01 | 26S proteasome regulatory complex ATPase |
| AT1G75840 | -0.4 | 2.62E-04 | -0.1 | 1.15E-01 | -0.3 | 7.22E-03 | -0.1 | 4.50E-01 | -0.1 | 7.43E-01 | -0.1 | 8.45E-01 | Uncharacterized protein |
| AT1G01820 | -0.4 | 1.34E-04 | -0.3 | 2.72E-03 | -0.1 | 9.91E-02 | -0.2 | 8.78E-02 | -0.3 | 2.79E-02 | 0.1 | 6.43E-01 | Peroxisomal membrane protein 11C |
| AT5G41220 | -0.4 | 3.06E-04 | -0.3 | 1.71E-03 | -0.1 | 4.19E-01 | -0.3 | 6.67E-02 | -0.1 | 6.94E-01 | -0.2 | 3.27E-01 | Glutathione S-transferase THETA 3 |
| AT5G26040 | -0.4 | 2.40E-03 | -0.3 | 1.93E-02 | -0.1 | 3.54E-01 | -0.1 | 6.48E-01 | -0.1 | 7.20E-01 | 0.0 | 9.88E-01 | hypothetical protein |
| AT4G14480 | -0.4 | 9.61E-02 | -0.2 | 3.17E-01 | -0.2 | 5.91E-01 | 0.1 | 7.34E-01 | 0.1 | 7.10E-01 | 0.0 | 9.62E-01 | Serine |
| AT2G24650 | -0.4 | 4.67E-03 | -0.4 | 6.93E-03 | 0.0 | 9.52E-01 | -0.2 | 3.73E-01 | 0.0 | 8.90E-01 | -0.2 | 6.02E-01 | B3 domain-containing protein REM13 |
| AT5G13300 | -0.4 | 5.99E-04 | -0.4 | 4.81E-04 | 0.0 | 6.94E-01 | -0.2 | 3.27E-01 | 0.0 | 9.63E-01 | -0.2 | 4.59E-01 | ADP-ribosylation factor GTPase-activating protein AGD3 |
| AT4G00270 | -0.4 | 1.90E-04 | -0.3 | 7.61E-04 | -0.1 | 5.31E-01 | -0.2 | 8.04E-02 | -0.1 | 5.04E-01 | -0.1 | 5.69E-01 | GLABROUS1 enhancer-binding protein |
| AT3G11130 | -0.4 | 2.05E-04 | -0.2 | 3.60E-02 | -0.2 | 1.52E-02 | 0.1 | 5.85E-01 | 0.0 | 8.91E-01 | 0.1 | 6.36E-01 | Clathrin heavy chain 1 |
| AT1G15215 | -0.4 | 2.24E-02 | -0.1 | 4.06E-01 | -0.3 | 1.73E-01 | -0.2 | 3.90E-01 | -0.2 | 6.45E-01 | -0.1 | 8.50E-01 | SAWADEE HOMEODOMAIN protein |
| AT5G66180 | -0.4 | 2.57E-04 | -0.4 | 5.92E-04 | 0.0 | 8.20E-01 | -0.3 | 1.35E-01 | -0.3 | 2.10E-01 | 0.0 | 9.59E-01 | S-adenosyl-L-methionine-dependent methyltransferases superfamily protein |
| AT3G61200 | -0.4 | 5.16E-04 | -0.3 | 5.57E-03 | -0.1 | 2.27E-01 | -0.4 | 1.12E-02 | -0.2 | 3.48E-01 | -0.2 | 2.39E-01 | Thioesterase domain-containing protein |
| AT2G37940 | -0.4 | 5.16E-04 | -0.3 | 9.46E-03 | -0.1 | 1.42E-01 | 0.0 | 7.94E-01 | -0.1 | 6.73E-01 | 0.1 | 5.59E-01 | Phosphatidylinositol:ceramide inositolphosphotransferase 2 |
| AT1G47240 | -0.4 | 2.05E-03 | -0.4 | 4.79E-03 | 0.0 | 7.76E-01 | -0.1 | 7.09E-01 | 0.1 | 6.91E-01 | -0.2 | 5.15E-01 | NRAMP2 |
| AT2G33835 | -0.4 | 5.47E-03 | -0.1 | 4.64E-01 | -0.3 | 4.65E-02 | -0.3 | 1.82E-01 | -0.1 | 6.72E-01 | -0.2 | 5.78E-01 | Protein FRIGIDA-ESSENTIAL 1 |
| AT2G34490 | -0.4 | 3.95E-03 | -0.3 | 2.19E-02 | -0.1 | 4.48E-01 | 0.1 | 5.90E-01 | 0.4 | 1.11E-02 | -0.3 | 7.66E-02 | Cytochrome P450 710A2 |
| AT4G10930 | -0.4 | 9.65E-04 | -0.5 | 3.79E-04 | 0.1 | 3.86E-01 | -0.1 | 5.40E-01 | 0.1 | 5.76E-01 | -0.2 | 2.59E-01 | RING |
| AT5G60160 | -0.4 | 3.21E-04 | -0.2 | 1.54E-02 | -0.2 | 5.26E-02 | -0.2 | 4.55E-02 | -0.1 | 4.58E-01 | -0.1 | 3.79E-01 | AT5g60160 |
| AT1G12050 | -0.4 | 1.52E-03 | -0.4 | 1.11E-03 | 0.0 | 7.08E-01 | -0.1 | 7.65E-01 | 0.1 | 6.77E-01 | -0.2 | 5.30E-01 | Fumarylacetoacetase |
| AT3G18820 | -0.4 | 3.31E-04 | -0.1 | 1.05E-01 | -0.3 | 1.00E-02 | -0.4 | 5.33E-03 | -0.2 | 1.82E-01 | -0.2 | 3.00E-01 | Ras-related protein RABG3f |
| AT1G32810 | -0.4 | 5.10E-04 | -0.5 | 2.96E-04 | 0.1 | 5.16E-01 | 0.0 | 7.85E-01 | -0.2 | 3.19E-01 | 0.1 | 5.35E-01 | Zinc finger PHD-type domain-containing protein |
| AT1G65960 | -0.4 | 2.10E-04 | -0.2 | 3.63E-02 | -0.2 | 1.56E-02 | -0.2 | 6.34E-02 | -0.1 | 3.64E-01 | -0.1 | 6.29E-01 | Glutamate decarboxylase |
| AT5G44410 | -0.4 | 2.80E-03 | 0.0 | 7.17E-01 | -0.3 | 1.39E-02 | 0.1 | 5.58E-01 | 0.4 | 1.88E-02 | -0.3 | 1.03E-01 | Berberine bridge enzyme-like 27 |
| AT2G38600 | -0.4 | 1.18E-01 | 1.2 | 1.42E-04 | -1.6 | 1.92E-04 | -0.9 | 1.58E-02 | -0.1 | 8.37E-01 | -0.8 | 4.85E-02 | HAD superfamily, subfamily IIIB acid phosphatase |
| AT1G08600 | -0.4 | 1.58E-04 | -0.3 | 3.05E-03 | -0.1 | 1.12E-01 | 0.0 | 9.89E-01 | 0.2 | 3.97E-01 | -0.2 | 5.16E-01 | Protein CHROMATIN REMODELING 20 |
| AT3G29270 | -0.4 | 3.78E-04 | -0.4 | 3.14E-04 | 0.0 | 6.80E-01 | -0.1 | 4.43E-01 | 0.0 | 8.77E-01 | -0.1 | 7.08E-01 | RING zinc finger protein-like |
| AT5G54430 | -0.4 | 3.69E-04 | -0.3 | 3.71E-03 | -0.1 | 2.41E-01 | -0.1 | 4.62E-01 | -0.1 | 4.99E-01 | 0.0 | 9.78E-01 | Adenine nucleotide alpha hydrolases-like superfamily protein |
| AT1G47840 | -0.4 | 6.53E-04 | 0.2 | 1.99E-02 | -0.6 | 1.67E-04 | 0.0 | 9.52E-01 | 0.4 | 2.58E-02 | -0.3 | 6.99E-02 | Phosphotransferase |
| AT1G62330 | -0.4 | 5.36E-04 | -0.2 | 7.97E-02 | -0.2 | 2.26E-02 | -0.1 | 3.50E-01 | 0.1 | 5.34E-01 | -0.3 | 1.99E-01 | O-fucosyltransferase 15 |
| AT1G52420 | -0.4 | 1.01E-04 | -0.3 | 7.87E-04 | -0.1 | 2.65E-01 | 0.0 | 7.58E-01 | 0.0 | 9.96E-01 | 0.0 | 8.45E-01 | hypothetical protein |
| AT2G41710 | -0.4 | 4.29E-04 | -0.3 | 1.29E-03 | 0.0 | 6.81E-01 | -0.1 | 4.60E-01 | -0.3 | 1.55E-01 | 0.1 | 5.27E-01 | Integrase-type DNA-binding superfamily protein |
| AT1G51830 | -0.4 | 2.39E-02 | -0.8 | 1.03E-03 | 0.4 | 7.92E-02 | 3.0 | 1.91E-07 | 1.2 | 4.85E-03 | 1.7 | 5.52E-05 | Leucine-rich repeat protein kinase family protein |
| AT3G59290 | -0.4 | 1.79E-03 | -0.3 | 1.90E-02 | -0.1 | 2.82E-01 | 0.2 | 4.53E-01 | 0.1 | 7.61E-01 | 0.1 | 8.18E-01 | Clathrin interactor EPSIN 3 |
| AT2G28290 | -0.4 | 2.79E-04 | -0.5 | 9.87E-05 | 0.1 | 2.35E-01 | -0.1 | 5.81E-01 | -0.1 | 6.35E-01 | 0.0 | 9.95E-01 | P-loop containing nucleoside triphosphate hydrolases superfamily protein |
| AT5G45080 | -0.4 | 4.88E-03 | -0.5 | 1.88E-03 | 0.1 | 4.98E-01 | 0.3 | 9.90E-02 | 0.4 | 3.64E-02 | -0.1 | 5.66E-01 | Protein PHLOEM PROTEIN 2-LIKE A6 |
| AT4G30100 | -0.4 | 3.04E-04 | -0.6 | 2.33E-05 | 0.2 | 1.89E-02 | -0.2 | 5.03E-01 | 0.1 | 7.19E-01 | -0.3 | 4.13E-01 | P-loop containing nucleoside triphosphate hydrolases superfamily protein |

| | | | | | | | | | | | | | |
|---|---|---|---|---|---|---|---|---|---|---|---|---|---|
| AT1G05840 | -0.4 | 5.03E-04 | -0.3 | 2.26E-03 | -0.1 | 5.01E-01 | -0.5 | 1.09E-02 | -0.5 | 2.47E-02 | 0.0 | 9.39E-01 | Eukaryotic aspartyl protease family protein |
| AT3G53960 | -0.4 | 4.79E-02 | -0.2 | 3.60E-01 | -0.2 | 3.35E-01 | -0.1 | 6.54E-01 | 0.0 | 8.51E-01 | -0.1 | 6.15E-01 | Protein NRT1 |
| AT3G57640 | -0.4 | 6.72E-02 | -0.8 | 5.37E-03 | 0.4 | 1.76E-01 | 0.0 | 9.50E-01 | -0.4 | 3.37E-01 | 0.4 | 3.71E-01 | Non-functional pseudokinase ZRK15 |
| AT2G16720 | -0.4 | 9.30E-03 | -0.2 | 9.57E-02 | -0.2 | 3.00E-01 | 0.3 | 2.65E-01 | 0.1 | 8.19E-01 | 0.2 | 4.48E-01 | Transcription factor MYB7 |
| AT5G55560 | -0.4 | 3.84E-03 | -0.4 | 6.16E-03 | 0.0 | 9.17E-01 | -0.1 | 7.84E-01 | 0.0 | 9.64E-01 | -0.1 | 8.84E-01 | hypothetical protein |
| AT5G41180 | -0.4 | 2.30E-03 | -0.4 | 5.15E-03 | 0.0 | 7.96E-01 | 0.2 | 3.01E-01 | 0.0 | 9.91E-01 | 0.2 | 4.02E-01 | Leucine-rich repeat receptor-like protein kinase |
| AT3G09820 | -0.4 | 1.83E-03 | 0.0 | 9.68E-01 | -0.4 | 4.85E-03 | 0.1 | 1.99E-01 | 0.2 | 1.33E-01 | 0.0 | 8.07E-01 | Adenosine kinase 1 |
| AT5G42810 | -0.4 | 7.25E-03 | -0.3 | 4.22E-02 | -0.1 | 4.48E-01 | -0.2 | 4.02E-01 | 0.0 | 9.00E-01 | -0.3 | 3.82E-01 | Inositol-pentakisphosphate 2-kinase P5 2-kinase) |
| AT1G23010 | -0.4 | 2.26E-03 | -0.5 | 4.18E-04 | 0.2 | 2.11E-01 | 0.2 | 6.07E-01 | 0.8 | 1.87E-02 | -0.6 | 8.47E-02 | Multicopper oxidase LPR1 |
| AT3G27580 | -0.4 | 8.04E-04 | -0.3 | 3.82E-03 | -0.1 | 5.02E-01 | 0.5 | 3.07E-02 | 0.2 | 4.45E-01 | 0.3 | 3.46E-01 | Serine |
| AT1G33270 | -0.4 | 8.09E-04 | -0.4 | 1.33E-03 | 0.0 | 9.48E-01 | -0.2 | 4.23E-01 | 0.1 | 6.73E-01 | -0.3 | 2.86E-01 | Patatin |
| AT1G34120 | -0.4 | 2.18E-04 | -0.3 | 3.33E-03 | -0.1 | 1.66E-01 | -0.1 | 5.56E-01 | -0.1 | 6.85E-01 | 0.0 | 9.60E-01 | Inositol polyphosphate 5-phosphatase I |
| AT5G12840 | -0.4 | 5.89E-04 | -0.6 | 5.25E-05 | 0.2 | 4.18E-02 | -0.3 | 2.30E-01 | -0.2 | 3.64E-01 | 0.0 | 9.33E-01 | Nuclear transcription factor Y subunit |
| AT3G53100 | -0.4 | 2.33E-03 | -0.2 | 4.55E-02 | -0.2 | 1.79E-01 | -0.2 | 2.79E-01 | -0.1 | 7.74E-01 | -0.2 | 6.45E-01 | GDSL esterase |
| AT4G33430 | -0.4 | 3.86E-04 | -0.3 | 5.39E-03 | -0.1 | 1.76E-01 | -0.1 | 5.24E-01 | -0.1 | 6.73E-01 | 0.0 | 9.57E-01 | BRI1-associated receptor kinase |
| AT5G41360 | -0.4 | 3.61E-03 | -0.3 | 1.27E-02 | -0.1 | 6.18E-01 | 0.5 | 1.23E-01 | 0.4 | 2.35E-01 | 0.1 | 9.14E-01 | DNA helicase |
| AT4G15180 | -0.4 | 2.22E-03 | -0.6 | 1.90E-04 | 0.2 | 7.38E-02 | 0.1 | 8.97E-01 | 0.3 | 3.70E-01 | -0.3 | 5.43E-01 | Histone-lysine N-methyltransferase ATXR3 |
| AT1G14690 | -0.4 | 1.75E-03 | -0.3 | 1.49E-02 | -0.1 | 3.11E-01 | -0.3 | 3.00E-02 | -0.3 | 1.03E-01 | -0.1 | 8.32E-01 | 65-kDa microtubule-associated protein 7 |
| AT4G30600 | -0.4 | 1.21E-04 | -0.4 | 1.12E-04 | 0.0 | 5.91E-01 | -0.3 | 7.80E-02 | -0.1 | 6.26E-01 | -0.2 | 4.59E-01 | SRP54-type proteins GTP-binding domain-containing protein |
| AT5G44000 | -0.4 | 1.23E-03 | -0.1 | 3.34E-01 | -0.3 | 1.54E-02 | 0.2 | 3.22E-01 | 0.1 | 6.86E-01 | 0.1 | 7.65E-01 | GST C-terminal domain-containing protein |
| AT4G15500 | -0.4 | 7.60E-03 | 0.2 | 4.75E-02 | -0.6 | 1.36E-03 | 0.2 | 3.88E-01 | 0.6 | 5.48E-03 | -0.4 | 4.02E-02 | UDP-glycosyltransferase 84A4 |
| AT3G43600 | -0.4 | 4.37E-04 | -0.3 | 4.08E-03 | -0.1 | 2.54E-01 | -0.2 | 3.49E-01 | 0.0 | 9.21E-01 | -0.2 | 6.01E-01 | Indole-3-acetaldehyde oxidase |
| AT3G16620 | -0.4 | 8.44E-04 | -0.3 | 3.08E-03 | -0.1 | 6.00E-01 | -0.1 | 5.34E-01 | 0.0 | 9.99E-01 | -0.1 | 6.76E-01 | Translocase of chloroplast 120, chloroplastic |
| AT5G42630 | -0.4 | 5.79E-02 | 0.2 | 2.23E-01 | -0.6 | 1.97E-02 | -0.2 | 4.82E-01 | -0.1 | 7.28E-01 | -0.1 | 8.64E-01 | Probable transcription factor KAN4 |
| AT1G08420 | -0.4 | 6.68E-04 | -0.4 | 4.06E-04 | 0.1 | 5.46E-01 | -0.1 | 5.07E-01 | -0.2 | 3.37E-01 | 0.1 | 8.10E-01 | Serine |
| AT3G30875 | -0.4 | 2.88E-03 | -0.1 | 3.75E-01 | -0.3 | 3.37E-02 | -0.6 | 5.53E-02 | -0.1 | 7.01E-01 | -0.5 | 2.75E-01 | Putative ABC transporter B family member 8 |
| AT2G47500 | -0.4 | 5.87E-04 | -0.1 | 2.96E-01 | -0.3 | 7.44E-03 | -0.2 | 6.31E-01 | 0.3 | 4.11E-01 | -0.5 | 2.45E-01 | P-loop nucleoside triphosphate hydrolases superfamily protein with CH domain-containing protein |
| AT2G01270 | -0.4 | 1.76E-04 | -0.1 | 9.51E-02 | -0.3 | 5.96E-03 | -0.1 | 2.87E-01 | -0.2 | 1.75E-01 | 0.1 | 8.02E-01 | Sulfhydryl oxidase |
| AT2G01470 | -0.4 | 6.09E-05 | -0.3 | 5.33E-04 | -0.1 | 2.15E-01 | -0.2 | 1.45E-01 | 0.0 | 8.49E-01 | -0.1 | 4.21E-01 | SEC12-like protein 2 |
| AT2G28720 | -0.4 | 1.46E-03 | -0.2 | 4.82E-02 | -0.2 | 9.60E-02 | 0.0 | 9.21E-01 | 0.0 | 9.21E-01 | 0.0 | 9.89E-01 | Histone H2B.3 |
| AT5G44240 | -0.4 | 3.59E-04 | -0.3 | 1.40E-03 | -0.1 | 5.32E-01 | 0.0 | 9.52E-01 | 0.2 | 5.33E-01 | -0.2 | 5.06E-01 | Phospholipid-transporting ATPase |
| AT3G09940 | -0.4 | 3.73E-02 | 0.3 | 1.12E-01 | -0.7 | 5.23E-03 | -0.3 | 7.58E-02 | 0.0 | 9.20E-01 | -0.4 | 6.40E-02 | Monodehydroascorbate reductase |
| AT3G17880 | -0.4 | 6.85E-04 | -0.4 | 1.16E-03 | 0.0 | 9.28E-01 | 0.0 | 9.98E-01 | 0.0 | 9.83E-01 | 0.0 | 9.84E-01 | TPR repeat-containing thioredoxin TDX |
| AT5G65450 | -0.4 | 1.21E-02 | 0.0 | 9.64E-01 | -0.4 | 3.35E-02 | 0.1 | 7.73E-01 | 0.1 | 8.49E-01 | 0.0 | 9.76E-01 | Ubiquitin-specific protease 17 |
| AT5G41790 | -0.4 | 1.29E-04 | -0.3 | 1.59E-03 | -0.1 | 1.77E-01 | 0.1 | 6.70E-01 | -0.1 | 4.41E-01 | 0.2 | 3.31E-01 | COP1-interactive protein 1 |
| AT5G24430 | -0.4 | 2.45E-04 | -0.4 | 1.97E-04 | 0.0 | 6.25E-01 | -0.4 | 3.45E-03 | -0.5 | 3.36E-03 | 0.0 | 8.31E-01 | CDPK-related kinase 4 |
| AT3G29310 | -0.4 | 1.67E-04 | -0.2 | 3.59E-03 | -0.1 | 1.04E-01 | -0.1 | 2.37E-01 | -0.1 | 4.23E-01 | 0.0 | 8.96E-01 | BAG family molecular chaperone regulator 8, chloroplastic |
| AT2G29900 | -0.4 | 8.48E-04 | 0.1 | 5.09E-01 | -0.4 | 1.47E-03 | -0.1 | 6.50E-01 | 0.0 | 9.75E-01 | -0.1 | 7.79E-01 | Presenilin-like protein At2g29900 |
| AT1G12680 | -0.4 | 3.29E-04 | -0.3 | 2.78E-03 | -0.1 | 2.88E-01 | -0.3 | 6.79E-02 | -0.2 | 3.00E-01 | -0.1 | 6.98E-01 | Serine |
| AT3G25570 | -0.4 | 3.43E-02 | -0.2 | 2.40E-01 | -0.2 | 3.70E-01 | 0.3 | 1.06E-01 | 0.4 | 2.87E-02 | -0.1 | 5.90E-01 | S-adenosylmethionine decarboxylase proenzyme 3 |
| AT2G02800 | -0.4 | 5.71E-04 | -0.2 | 2.06E-02 | -0.2 | 7.62E-02 | 0.0 | 9.28E-01 | 0.1 | 4.40E-01 | -0.1 | 6.25E-01 | Probable serine |
| AT3G29090 | -0.4 | 9.14E-04 | -0.3 | 5.08E-03 | -0.1 | 4.34E-01 | 0.0 | 9.72E-01 | -0.3 | 1.98E-01 | 0.3 | 2.83E-01 | Pectinesterase 31 |
| AT5G49215 | -0.4 | 5.07E-03 | 0.0 | 9.86E-01 | -0.4 | 1.51E-02 | -0.7 | 5.55E-05 | -0.4 | 1.57E-02 | -0.4 | 7.38E-02 | Pectin lyase-like superfamily protein |
| AT5G66760 | -0.4 | 9.22E-04 | -0.2 | 6.20E-02 | -0.2 | 4.83E-02 | 0.2 | 7.38E-02 | 0.3 | 2.89E-02 | -0.1 | 6.66E-01 | Succinate dehydrogenase flavoprotein subunit 1, mitochondrial |
| AT2G40730 | -0.4 | 1.92E-04 | -0.3 | 5.19E-04 | 0.0 | 7.19E-01 | 0.0 | 8.78E-01 | 0.1 | 6.67E-01 | -0.1 | 8.23E-01 | CTEXP |
| AT5G14550 | -0.4 | 5.96E-04 | -0.2 | 4.64E-02 | -0.2 | 4.76E-02 | -0.3 | 1.94E-01 | 0.0 | 9.98E-01 | -0.3 | 2.47E-01 | Core-2 |
| AT4G03190 | -0.4 | 3.07E-03 | 0.1 | 3.72E-01 | -0.5 | 2.66E-03 | -0.6 | 1.80E-01 | -0.2 | 7.82E-01 | -0.5 | 4.93E-01 | GRR1-like protein 1 |
| AT1G48430 | -0.4 | 3.96E-04 | -0.5 | 1.39E-04 | 0.1 | 2.70E-01 | -0.2 | 2.45E-01 | -0.2 | 3.86E-01 | 0.0 | 9.37E-01 | Similar to dihydroxyacetone kinase |
| AT1G27190 | -0.4 | 4.95E-04 | -0.1 | 7.55E-02 | -0.2 | 2.07E-02 | -0.3 | 1.45E-02 | -0.2 | 3.14E-01 | -0.2 | 3.80E-01 | Probable inactive receptor kinase At1g27190 |
| AT1G10280 | -0.4 | 5.55E-03 | -0.4 | 3.92E-03 | 0.0 | 7.52E-01 | 0.3 | 2.76E-01 | 0.3 | 4.27E-01 | 0.1 | 9.24E-01 | hypothetical protein |
| AT1G77740 | -0.4 | 9.63E-04 | -0.2 | 2.22E-02 | -0.2 | 1.25E-01 | -0.1 | 5.81E-01 | -0.1 | 6.60E-01 | 0.0 | 9.95E-01 | Phosphatidylinositol 4-phosphate 5-kinase |
| AT5G49460 | -0.4 | 2.34E-04 | 0.0 | 5.40E-01 | -0.4 | 4.30E-04 | 0.0 | 9.05E-01 | 0.1 | 8.14E-01 | -0.1 | 7.75E-01 | ATP-citrate synthase beta chain protein 2 |
| AT5G12210 | -0.4 | 2.20E-04 | -0.2 | 4.83E-03 | -0.1 | 1.02E-01 | -0.1 | 3.75E-01 | -0.1 | 3.22E-01 | 0.0 | 9.05E-01 | Geranylgeranyl transferase type-2 subunit beta |
| AT1G29790 | -0.4 | 4.85E-04 | 0.0 | 6.19E-01 | -0.3 | 3.19E-03 | -0.1 | 5.59E-01 | 0.0 | 9.45E-01 | -0.1 | 7.58E-01 | Probable methyltransferase At1g29790 |
| AT3G56480 | -0.4 | 1.34E-03 | -0.1 | 4.75E-01 | -0.3 | 1.17E-02 | -0.3 | 1.88E-01 | -0.1 | 7.91E-01 | -0.2 | 5.37E-01 | Stomatal closure-related actin-binding protein 3 |
| AT4G25470 | -0.4 | 2.30E-01 | -0.5 | 1.42E-01 | 0.1 | 7.85E-01 | | | | | | | Dehydration-responsive element-binding protein 1C |
| AT3G27310 | -0.4 | 2.58E-03 | -0.2 | 1.36E-01 | -0.2 | 7.03E-02 | 0.0 | 9.13E-01 | -0.1 | 8.14E-01 | 0.0 | 9.12E-01 | PUX1 |
| AT1G10650 | -0.4 | 6.76E-03 | -0.5 | 1.44E-03 | 0.1 | 3.14E-01 | 0.1 | 6.51E-01 | 0.4 | 7.09E-02 | -0.3 | 2.57E-01 | SBP family protein |
| AT1G31480 | -0.4 | 1.30E-02 | -0.5 | 5.92E-03 | 0.1 | 6.21E-01 | -0.4 | 3.56E-01 | -0.2 | 6.75E-01 | -0.2 | 7.91E-01 | Phospholipase SGR2 |
| AT3G16410 | -0.4 | 3.39E-03 | 0.1 | 3.14E-01 | -0.5 | 2.60E-03 | -0.4 | 3.44E-02 | 0.0 | 8.89E-01 | -0.4 | 8.41E-02 | Thiohydroximate-O-sulfate sulfur |
| AT5G18610 | -0.4 | 1.13E-03 | -0.3 | 6.69E-03 | -0.1 | 4.13E-01 | -0.3 | 1.20E-02 | -0.4 | 1.26E-02 | 0.0 | 9.06E-01 | Serine |
| AT2G26170 | -0.4 | 7.34E-04 | -0.7 | 2.88E-05 | 0.3 | 1.27E-02 | 0.4 | 1.64E-02 | 0.3 | 5.18E-02 | 0.0 | 8.63E-01 | Cytochrome P450 711A1 |
| AT3G52200 | -0.4 | 2.15E-04 | 0.0 | 7.24E-01 | -0.4 | 5.23E-04 | 0.0 | 9.91E-01 | 0.0 | 8.88E-01 | 0.0 | 9.09E-01 | Dihydrolipoamide acetyltransferase component of pyruvate dehydrogenase complex |
| AT2G43790 | -0.4 | 1.95E-04 | -0.1 | 6.62E-02 | -0.2 | 8.22E-03 | -0.1 | 4.81E-01 | -0.1 | 2.62E-01 | 0.1 | 7.35E-01 | Mitogen-activated protein kinase 6 |
| AT5G55090 | -0.4 | 1.59E-02 | -0.2 | 2.51E-01 | -0.2 | 2.23E-01 | 0.8 | 3.32E-02 | 0.4 | 3.18E-01 | 0.3 | 3.36E-01 | hypothetical protein |
| AT4G36870 | -0.4 | 1.96E-03 | -0.4 | 2.04E-03 | 0.0 | 8.86E-01 | -0.3 | 1.28E-01 | -0.1 | 6.16E-01 | -0.2 | 5.52E-01 | BEL1-like homeodomain 2 |
| AT1G55320 | -0.4 | 1.26E-03 | -0.3 | 9.71E-03 | -0.1 | 3.52E-01 | -0.2 | 5.54E-01 | 0.2 | 4.29E-01 | -0.4 | 1.73E-01 | Probable acyl-activating enzyme 18, peroxisomal |
| AT5G58980 | -0.4 | 2.74E-03 | -0.5 | 1.04E-03 | 0.1 | 4.57E-01 | -0.4 | 1.32E-01 | 0.0 | 9.76E-01 | -0.4 | 2.85E-01 | Neutral ceramidase 3 |
| AT5G63190 | -0.4 | 3.84E-03 | -0.2 | 6.62E-02 | -0.2 | 2.00E-01 | -0.5 | 4.81E-04 | -0.7 | 9.86E-05 | 0.2 | 2.75E-01 | MA3 DOMAIN-CONTAINING TRANSLATION REGULATORY FACTOR 1 |
| AT5G05400 | -0.4 | 1.86E-03 | 0.3 | 3.14E-03 | -0.7 | 1.61E-04 | -0.2 | 4.96E-01 | -0.1 | 8.80E-01 | -0.1 | 6.93E-01 | LRR and NB-ARC domains-containing disease resistance protein |
| AT3G49210 | -0.4 | 1.97E-03 | -0.4 | 2.02E-03 | 0.0 | 8.76E-01 | -0.2 | 2.87E-01 | -0.1 | 8.40E-01 | -0.2 | 5.64E-01 | O-acyltransferase family protein |
| AT5G47910 | -0.4 | 7.02E-04 | -0.7 | 1.40E-05 | 0.3 | 3.19E-03 | 0.6 | 3.22E-04 | 0.4 | 1.85E-02 | 0.2 | 2.93E-01 | Respiratory burst oxidase homolog protein D |
| AT1G64650 | -0.4 | 6.85E-04 | 0.0 | 8.27E-01 | -0.4 | 2.68E-03 | 0.0 | 9.99E-01 | 0.2 | 1.80E-01 | -0.2 | 2.45E-01 | Molybdate-anion transporter |
| AT2G39970 | -0.4 | 4.37E-04 | -0.2 | 9.01E-03 | -0.1 | 1.21E-01 | 0.0 | 8.41E-01 | -0.1 | 7.44E-01 | 0.1 | 6.54E-01 | Peroxisomal nicotinamide adenine dinucleotide carrier ) |
| AT3G15580 | -0.4 | 2.59E-04 | -0.4 | 1.54E-04 | 0.1 | 4.56E-01 | -0.5 | 8.42E-03 | -0.2 | 3.57E-01 | -0.3 | 2.47E-01 | Autophagy-related protein 8i |
| AT2G47090 | -0.4 | 1.04E-02 | 0.0 | 7.99E-01 | -0.3 | 3.99E-02 | -0.1 | 6.84E-01 | 0.1 | 8.56E-01 | -0.2 | 5.90E-01 | RING-type E3 ubiquitin transferase |
| AT3G01310 | -0.4 | 8.14E-04 | -0.2 | 1.70E-02 | -0.1 | 1.34E-01 | -0.6 | 9.49E-05 | -0.4 | 4.28E-03 | -0.2 | 3.16E-01 | Inositol hexakisphosphate and diphosphoinositol-pentakisphosphate kinase |
| AT3G16720 | -0.4 | 8.34E-03 | -0.7 | 3.81E-04 | 0.3 | 5.65E-02 | -0.1 | 2.81E-01 | -0.1 | 3.65E-01 | 0.0 | 9.90E-01 | RING-H2 finger protein ATL2 |
| AT3G27870 | -0.4 | 3.07E-03 | -0.5 | 8.52E-04 | 0.1 | 3.53E-01 | -0.2 | 5.67E-01 | 0.0 | 9.27E-01 | -0.1 | 7.51E-01 | Phospholipid-transporting ATPase |
| AT1G03740 | -0.4 | 1.14E-03 | -0.5 | 1.45E-04 | 0.2 | 9.17E-02 | 0.2 | 5.62E-01 | -0.1 | 8.89E-01 | 0.2 | 5.91E-01 | F21B7.34 |
| AT5G19670 | -0.4 | 3.40E-03 | -0.4 | 6.31E-03 | 0.0 | 8.86E-01 | -0.1 | 8.34E-01 | 0.1 | 9.03E-01 | -0.2 | 7.70E-01 | Exostosin GT47 domain-containing protein |
| AT1G49340 | -0.4 | 5.96E-04 | -0.2 | 5.78E-02 | -0.2 | 3.15E-02 | -0.1 | 5.29E-01 | 0.2 | 4.05E-01 | -0.3 | 2.42E-01 | 1-phosphatidylinositol 4-kinase |
| AT1G73740 | -0.4 | 1.50E-03 | -0.5 | 4.32E-04 | 0.1 | 3.08E-01 | 0.0 | 9.65E-01 | 0.0 | 8.45E-01 | 0.0 | 8.96E-01 | Glycosyltransferase -pyrophosphoryl-undecaprenol N-acetylglucosamine transferase) -pyrophosphoryl-undecaprenol N-acetylglucosamine transferase; 62395-63952) |

| | | | | | | | | | | | | | |
|---|---|---|---|---|---|---|---|---|---|---|---|---|---|
| AT4G26940 | -0.4 | 2.63E-04 | -0.2 | 2.22E-02 | -0.2 | 2.94E-02 | -0.1 | 6.03E-01 | 0.2 | 3.85E-01 | -0.2 | 2.35E-01 | Probable beta-1,3-galactosyltransferase 4 |
| AT1G21580 | -0.4 | 1.34E-04 | -0.3 | 3.97E-04 | 0.0 | 6.69E-01 | 0.1 | 5.66E-01 | -0.1 | 8.01E-01 | 0.2 | 4.79E-01 | Zinc finger C-x8-C-x5-C-x3-H type family protein |
| AT1G74380 | -0.4 | 2.10E-04 | -0.2 | 2.18E-02 | -0.2 | 2.30E-02 | -0.1 | 3.77E-01 | 0.1 | 4.93E-01 | -0.2 | 2.22E-01 | Probable xyloglucan 6-xylosyltransferase 5 |
| AT1G12820 | -0.4 | 1.59E-02 | -0.5 | 5.25E-03 | 0.1 | 5.01E-01 | -0.6 | 3.52E-01 | -0.2 | 8.11E-01 | -0.4 | 6.82E-01 | Protein AUXIN SIGNALING F-BOX 3 |
| AT3G05050 | -0.4 | 1.80E-04 | -0.3 | 9.44E-04 | -0.1 | 4.04E-01 | 0.4 | 3.73E-02 | 0.2 | 3.82E-01 | 0.2 | 4.61E-01 | AT3g05050 |
| AT2G43420 | -0.4 | 2.56E-04 | -0.2 | 1.21E-02 | -0.2 | 5.73E-02 | 0.1 | 6.36E-01 | 0.1 | 4.25E-01 | -0.1 | 8.02E-01 | 3beta-hydroxysteroid-dehydrogenase |
| AT3G08590 | -0.4 | 2.87E-03 | -0.6 | 2.59E-04 | 0.2 | 7.79E-02 | 1.1 | 1.15E-06 | 0.2 | 2.00E-01 | 0.9 | 1.73E-04 | Probable 2,3-bisphosphoglycerate-independent phosphoglycerate mutase 2 |
| AT2G46020 | -0.4 | 2.81E-04 | -0.5 | 1.03E-04 | 0.1 | 2.47E-01 | -0.2 | 2.47E-01 | -0.1 | 6.57E-01 | -0.1 | 7.35E-01 | Brahma variant |
| AT5G12400 | -0.4 | 4.58E-04 | -0.4 | 9.37E-04 | 0.0 | 8.42E-01 | -0.2 | 2.90E-01 | -0.2 | 4.82E-01 | 0.0 | 9.07E-01 | PHD-finger and DNA binding domain-containing protein |
| AT2G33700 | -0.4 | 6.63E-04 | -0.4 | 3.27E-04 | 0.1 | 4.59E-01 | -0.5 | 4.41E-03 | -0.3 | 7.35E-02 | -0.2 | 4.34E-01 | protein-serine |
| AT5G16240 | -0.4 | 6.64E-03 | 0.0 | 7.13E-01 | -0.3 | 3.11E-02 | 0.3 | 9.41E-02 | -0.1 | 7.00E-01 | 0.4 | 9.48E-02 | Stearoyl- 9-desaturase 1, chloroplastic |
| AT5G09460 | -0.4 | 3.81E-04 | -0.4 | 2.32E-04 | 0.1 | 5.04E-01 | -0.2 | 1.91E-01 | -0.1 | 7.15E-01 | -0.1 | 6.35E-01 | Transcription factor bHLH143 |
| AT3G25500 | -0.4 | 9.21E-04 | -0.4 | 1.74E-03 | 0.0 | 8.65E-01 | -0.2 | 4.06E-01 | -0.1 | 7.36E-01 | -0.1 | 8.23E-01 | FH2 domain-containing protein |
| AT2G20680 | -0.4 | 1.33E-02 | 0.2 | 1.80E-01 | -0.5 | 5.09E-03 | -0.1 | 6.33E-01 | 0.0 | 9.17E-01 | -0.2 | 6.17E-01 | Mannan endo-1,4-beta-mannosidase 2 |
| AT3G24660 | -0.4 | 1.16E-03 | -0.1 | 5.44E-01 | -0.3 | 9.41E-03 | 0.1 | 5.82E-01 | 0.0 | 8.46E-01 | 0.0 | 8.73E-01 | Putative kinase-like protein TMKL1 |
| AT5G05980 | -0.4 | 7.80E-05 | -0.3 | 9.97E-04 | -0.1 | 1.49E-01 | -0.1 | 5.37E-01 | -0.1 | 7.20E-01 | 0.0 | 9.27E-01 | Folylpolyglutamate synthase |
| AT1G73450 | -0.4 | 5.47E-04 | -0.3 | 1.60E-03 | 0.0 | 6.88E-01 | 0.1 | 8.96E-01 | 0.3 | 5.01E-01 | -0.2 | 6.72E-01 | Protein kinase superfamily protein |
| AT5G64600 | -0.4 | 1.04E-02 | -0.1 | 2.98E-01 | -0.2 | 1.25E-01 | -0.2 | 4.30E-01 | -0.1 | 8.20E-01 | -0.1 | 7.41E-01 | O-fucosyltransferase 38 |
| AT4G24620 | -0.4 | 4.03E-04 | 0.0 | 9.48E-01 | -0.4 | 1.19E-03 | 0.0 | 9.72E-01 | 0.2 | 1.49E-01 | -0.2 | 2.51E-01 | Glucose-6-phosphate isomerase 1, chloroplastic |
| AT5G26667 | -0.4 | 1.48E-03 | 0.0 | 7.19E-01 | -0.4 | 2.41E-03 | 0.0 | 9.61E-01 | 0.1 | 8.57E-01 | 0.0 | 9.08E-01 | UMP-CMP kinase |
| AT3G60270 | -0.4 | 1.60E-02 | 0.3 | 6.13E-02 | -0.6 | 1.93E-03 | 1.0 | 4.17E-04 | 0.5 | 1.16E-01 | 0.6 | 2.19E-02 | Cupredoxin superfamily protein -like protein) |
| AT4G11600 | -0.4 | 4.23E-03 | -0.1 | 3.50E-01 | -0.3 | 4.03E-02 | 0.2 | 1.84E-01 | 0.1 | 4.83E-01 | 0.1 | 7.55E-01 | Glutathione peroxidase |
| AT5G20510 | -0.4 | 1.44E-02 | -0.4 | 1.28E-02 | 0.0 | 8.83E-01 | -0.2 | 4.87E-01 | 0.0 | 9.22E-01 | -0.2 | 5.43E-01 | PHD finger protein ALFIN-LIKE |
| AT5G56340 | -0.4 | 8.89E-05 | -0.2 | 2.39E-03 | -0.1 | 6.99E-02 | -0.3 | 4.12E-02 | -0.2 | 2.30E-01 | -0.1 | 6.32E-01 | RING-type E3 ubiquitin transferase |
| AT3G59950 | -0.4 | 2.12E-04 | -0.4 | 1.21E-04 | 0.1 | 4.24E-01 | -0.3 | 1.16E-01 | -0.1 | 7.84E-01 | -0.2 | 3.70E-01 | Cysteine protease |
| AT3G09190 | -0.4 | 7.28E-03 | -0.3 | 1.65E-02 | 0.0 | 7.84E-01 | 0.5 | 1.08E-01 | 0.2 | 6.42E-01 | 0.3 | 3.94E-01 | Concanavalin A-like lectin family protein |
| AT3G27300 | -0.4 | 3.35E-04 | -0.1 | 6.51E-02 | -0.2 | 1.53E-02 | 0.0 | 8.32E-01 | 0.1 | 5.02E-01 | -0.2 | 4.75E-01 | Glucose-6-phosphate 1-dehydrogenase |
| AT2G16920 | -0.4 | 2.27E-04 | -0.2 | 1.01E-02 | -0.2 | 5.61E-02 | -0.1 | 5.68E-01 | -0.1 | 6.98E-01 | 0.0 | 9.59E-01 | Probable ubiquitin-conjugating enzyme E2 23 |
| AT1G20650 | -0.4 | 1.23E-02 | -0.3 | 3.83E-02 | -0.1 | 6.42E-01 | -1.0 | 3.34E-05 | -0.7 | 1.36E-03 | -0.3 | 2.68E-01 | Probable serine |
| AT3G49500 | -0.4 | 5.73E-04 | -0.2 | 1.50E-02 | -0.1 | 1.13E-01 | 0.1 | 3.94E-01 | 0.2 | 3.81E-01 | 0.0 | 9.41E-01 | RNA-dependent RNA polymerase 6 |
| AT2G47830 | -0.4 | 6.65E-04 | -0.3 | 3.06E-03 | -0.1 | 5.03E-01 | -0.1 | 7.57E-01 | 0.0 | 9.50E-01 | -0.1 | 7.63E-01 | Cation efflux protein cytoplasmic domain-containing protein |
| AT3G60530 | -0.4 | 2.90E-03 | -0.1 | 6.22E-01 | -0.3 | 1.57E-02 | -1.1 | 7.91E-07 | -1.1 | 5.81E-06 | 0.0 | 9.27E-01 | GATA transcription factor 4 |
| AT5G13490 | -0.4 | 6.72E-02 | -0.2 | 2.74E-01 | -0.2 | 5.11E-01 | 0.4 | 4.38E-02 | 0.3 | 1.70E-01 | 0.1 | 7.30E-01 | ADP,ATP carrier protein 2, mitochondrial |
| AT5G07830 | -0.4 | 2.02E-03 | -0.5 | 6.12E-04 | 0.1 | 3.45E-01 | 0.0 | 8.16E-01 | -0.2 | 4.38E-01 | 0.2 | 3.81E-01 | Heparanase-like protein 1 |
| AT5G54280 | -0.4 | 2.06E-03 | -0.3 | 8.38E-03 | -0.1 | 5.54E-01 | -0.6 | 1.94E-02 | -0.5 | 5.33E-02 | -0.1 | 8.83E-01 | Myosin 2 |
| AT5G07720 | -0.4 | 3.21E-02 | 0.1 | 7.40E-01 | -0.4 | 4.30E-02 | 0.0 | 9.20E-01 | 0.5 | 1.96E-01 | -0.5 | 1.55E-01 | Probable xyloglucan 6-xylosyltransferase 3 |
| AT1G75020 | -0.4 | 1.15E-03 | -0.3 | 2.44E-03 | 0.0 | 8.27E-01 | -0.1 | 8.16E-01 | -0.2 | 4.53E-01 | 0.1 | 6.97E-01 | Probable 1-acyl-sn-glycerol-3-phosphate acyltransferase 4 |
| AT1G48480 | -0.4 | 8.56E-04 | -0.1 | 1.33E-01 | -0.2 | 2.20E-02 | -0.2 | 9.80E-02 | -0.3 | 1.35E-01 | 0.0 | 9.90E-01 | Probable inactive receptor kinase At1g48480 |
| AT4G34480 | -0.4 | 1.18E-03 | -0.5 | 1.66E-04 | 0.2 | 1.07E-01 | 0.0 | 8.48E-01 | -0.1 | 7.90E-01 | 0.0 | 9.42E-01 | glucan endo-1,3-beta-D-glucosidase |
| AT5G43380 | -0.4 | 2.95E-03 | 0.0 | 8.45E-01 | -0.3 | 1.40E-02 | 0.1 | 8.68E-01 | 0.2 | 5.76E-01 | -0.2 | 7.50E-01 | Serine |
| AT2G32930 | -0.4 | 1.46E-02 | -0.3 | 4.37E-02 | -0.1 | 6.82E-01 | -0.3 | 2.44E-01 | 0.2 | 4.46E-01 | -0.5 | 7.31E-02 | Zinc finger nuclease 2 |
| AT4G35310 | -0.4 | 1.66E-04 | -0.3 | 7.18E-04 | 0.0 | 5.18E-01 | 0.1 | 2.43E-01 | -0.1 | 3.14E-01 | 0.3 | 9.55E-02 | Calcium-dependent protein kinase 5 |
| AT4G29100 | -0.4 | 4.93E-04 | -0.3 | 1.80E-03 | -0.1 | 5.62E-01 | 0.1 | 4.69E-01 | 0.2 | 8.15E-02 | -0.1 | 4.01E-01 | Basic helix-loop-helix DNA-binding superfamily protein |
| AT3G50050 | -0.4 | 2.36E-03 | -0.2 | 1.17E-01 | -0.2 | 8.64E-02 | -0.1 | 7.34E-01 | -0.1 | 8.91E-01 | -0.1 | 9.20E-01 | At3g50050 |
| AT5G53140 | -0.4 | 4.22E-04 | -0.2 | 1.62E-02 | -0.2 | 7.14E-02 | -0.1 | 4.13E-01 | -0.1 | 3.85E-01 | 0.0 | 9.27E-01 | Probable protein phosphatase 2C 76 |
| AT3G12250 | -0.4 | 4.36E-03 | -0.2 | 4.06E-02 | -0.1 | 3.33E-01 | -0.6 | 3.09E-02 | -0.4 | 1.40E-01 | -0.2 | 6.93E-01 | BZip transcription factor TGA6 |
| AT5G65560 | -0.4 | 2.12E-03 | -0.3 | 1.34E-02 | -0.1 | 4.29E-01 | 0.0 | 9.35E-01 | -0.3 | 3.76E-01 | 0.3 | 5.00E-01 | Pentatricopeptide repeat-containing protein At5g65560 |
| AT4G28540 | -0.4 | 2.18E-04 | -0.2 | 1.57E-02 | -0.2 | 3.23E-02 | 0.0 | 8.61E-01 | 0.2 | 3.52E-01 | -0.1 | 5.62E-01 | Casein kinase 1-like protein 6 |
| AT4G33140 | -0.4 | 2.22E-03 | -0.4 | 2.26E-03 | 0.0 | 8.80E-01 | -0.3 | 5.42E-02 | 0.2 | 2.61E-01 | -0.4 | 2.04E-02 | hypothetical protein |
| AT1G62800 | -0.4 | 1.47E-03 | 0.1 | 4.73E-01 | -0.4 | 1.96E-03 | -0.2 | 3.13E-01 | 0.3 | 2.01E-01 | -0.5 | 5.96E-02 | Aspartate aminotransferase |
| AT1G32190 | -0.4 | 3.30E-02 | 0.3 | 1.02E-01 | -0.6 | 5.84E-03 | 0.0 | 9.29E-01 | -0.2 | 5.98E-01 | 0.2 | 5.86E-01 | Alpha |
| AT2G30400 | -0.4 | 3.79E-02 | 0.3 | 4.87E-02 | -0.7 | 4.82E-03 | -0.4 | 3.53E-01 | -0.2 | 5.99E-01 | -0.1 | 8.62E-01 | Transcription repressor OFP2 |
| AT1G27210 | -0.4 | 2.30E-03 | -0.1 | 2.27E-01 | -0.2 | 3.55E-02 | -0.3 | 7.95E-03 | -0.1 | 5.15E-01 | -0.2 | 1.27E-01 | TORTIFOLIA1-like protein 4 |
| AT1G35515 | -0.4 | 1.98E-02 | -0.6 | 1.88E-03 | 0.3 | 1.81E-01 | | | | | | | Transcription factor MYB8 |
| AT1G08370 | -0.4 | 1.73E-03 | -0.3 | 8.87E-03 | -0.1 | 4.54E-01 | -0.1 | 5.60E-01 | 0.0 | 9.26E-01 | -0.1 | 6.00E-01 | mRNA-decapping enzyme-like protein |
| AT1G20760 | -0.4 | 1.19E-04 | -0.3 | 1.20E-03 | -0.1 | 2.00E-01 | -0.2 | 2.13E-01 | -0.2 | 2.61E-01 | 0.0 | 9.83E-01 | Calcium-binding EF hand family protein |
| AT1G20110 | -0.4 | 6.70E-04 | -0.3 | 2.64E-03 | -0.1 | 5.60E-01 | -0.2 | 6.15E-02 | -0.2 | 2.08E-01 | 0.0 | 8.09E-01 | Protein FREE1 |
| AT1G78280 | -0.4 | 4.37E-03 | -0.4 | 2.20E-03 | 0.1 | 6.01E-01 | 0.0 | 9.27E-01 | 0.0 | 8.73E-01 | 0.0 | 9.57E-01 | Lysine-specific demethylase JMJ21 |
| AT1G05910 | -0.4 | 3.53E-04 | -0.3 | 9.12E-04 | 0.0 | 7.40E-01 | 0.3 | 7.68E-02 | 0.3 | 1.79E-01 | 0.1 | 8.85E-01 | ATPase family AAA domain-containing protein At1g05910 |
| AT5G43330 | -0.4 | 3.07E-03 | -0.3 | 1.21E-02 | -0.1 | 5.71E-01 | 0.4 | 2.00E-02 | 0.1 | 5.75E-01 | 0.3 | 1.50E-01 | Malate dehydrogenase 2, cytoplasmic |
| AT3G46920 | -0.4 | 4.62E-03 | -0.4 | 3.27E-03 | 0.0 | 7.57E-01 | 0.3 | 3.33E-01 | 0.2 | 6.49E-01 | 0.1 | 8.06E-01 | Kinase superfamily with octicosapeptide |
| AT1G54830 | -0.4 | 8.62E-04 | -0.3 | 4.89E-03 | -0.1 | 4.36E-01 | -0.3 | 1.94E-01 | -0.2 | 2.92E-01 | 0.0 | 9.59E-01 | Nuclear transcription factor Y subunit C-3 |
| AT5G51830 | -0.4 | 1.49E-02 | -0.6 | 8.22E-04 | 0.3 | 8.72E-02 | 0.6 | 4.14E-03 | -0.1 | 8.05E-01 | 0.6 | 1.88E-02 | Probable fructokinase-7 |
| AT4G11560 | -0.4 | 1.55E-04 | -0.3 | 6.37E-04 | 0.0 | 5.13E-01 | 0.0 | 7.93E-01 | 0.1 | 7.33E-01 | -0.1 | 5.85E-01 | Bromo-adjacent homology domain-containing protein |
| AT2G21850 | -0.4 | 4.83E-03 | -0.3 | 1.02E-02 | 0.0 | 8.10E-01 | -0.2 | 5.10E-01 | 0.0 | 9.81E-01 | -0.1 | 6.56E-01 | Cysteine |
| AT1G19430 | -0.4 | 2.52E-04 | -0.3 | 1.06E-03 | -0.1 | 5.13E-01 | -0.2 | 3.65E-01 | 0.0 | 8.59E-01 | -0.1 | 6.59E-01 | Methyltransferase |
| AT3G54840 | -0.4 | 3.52E-04 | -0.2 | 1.76E-02 | -0.2 | 5.12E-02 | -0.2 | 1.24E-01 | -0.1 | 3.29E-01 | 0.0 | 8.17E-01 | AT3G54840 protein |
| AT1G79340 | -0.4 | 4.86E-04 | 0.1 | 4.20E-01 | -0.4 | 5.98E-04 | -0.4 | 2.19E-03 | -0.1 | 2.81E-01 | -0.2 | 1.42E-01 | Metacaspase-4 |
| AT3G25610 | -0.4 | 6.47E-03 | -0.8 | 6.67E-05 | 0.4 | 6.38E-03 | 1.0 | 1.22E-05 | 0.1 | 6.01E-01 | 0.9 | 9.75E-04 | Phospholipid-transporting ATPase 10 |
| AT3G04730 | -0.4 | 3.77E-04 | -0.1 | 4.36E-01 | -0.3 | 3.29E-03 | -0.2 | 1.12E-01 | -0.5 | 8.72E-03 | 0.2 | 2.93E-01 | Auxin-responsive protein IAA16 |
| AT5G62500 | -0.4 | 4.49E-03 | 0.1 | 1.47E-01 | -0.5 | 2.08E-03 | -0.1 | 5.80E-01 | 0.2 | 4.34E-01 | -0.3 | 2.70E-01 | Uncharacterized protein |
| AT4G39010 | -0.4 | 4.38E-02 | -0.1 | 6.33E-01 | -0.3 | 1.87E-01 | -0.3 | 4.46E-01 | 0.0 | 9.56E-01 | -0.2 | 6.25E-01 | cellulase |
| AT3G52190 | -0.4 | 5.28E-04 | -0.2 | 5.92E-02 | -0.2 | 3.19E-02 | 0.6 | 1.32E-03 | 0.4 | 4.33E-02 | 0.2 | 3.29E-01 | SEC12-like protein 1 |
| AT5G49900 | -0.4 | 2.77E-03 | -0.4 | 3.05E-03 | 0.0 | 9.25E-01 | 0.1 | 8.60E-01 | 0.3 | 2.73E-01 | -0.3 | 3.76E-01 | Non-lysosomal glucosylceramidase |
| AT4G38620 | -0.4 | 6.94E-03 | -0.4 | 5.85E-03 | 0.0 | 8.49E-01 | 0.6 | 1.27E-04 | 0.3 | 1.23E-02 | 0.2 | 1.78E-01 | Transcription repressor MYB4 |
| AT2G03620 | -0.4 | 4.76E-04 | -0.3 | 1.34E-03 | 0.0 | 7.10E-01 | -0.4 | 3.07E-02 | -0.2 | 4.82E-01 | -0.3 | 3.17E-01 | Magnesium transporter MRS2-5 |
| AT1G22170 | -0.4 | 1.88E-02 | 0.2 | 1.51E-01 | -0.5 | 5.38E-03 | 0.3 | 4.39E-02 | 0.3 | 5.56E-02 | 0.0 | 9.70E-01 | 2,3-bisphosphoglycerate-dependent phosphoglycerate mutase 1 |
| AT4G01820 | -0.4 | 7.96E-02 | 0.4 | 4.94E-02 | -0.7 | 7.59E-03 | -1.8 | 5.13E-03 | -0.2 | 6.95E-01 | -1.6 | 1.74E-02 | p-glycoprotein 3 |
| AT2G19450 | -0.4 | 4.18E-04 | -0.1 | 2.88E-01 | -0.3 | 6.49E-03 | 0.4 | 7.74E-03 | 0.2 | 1.42E-01 | 0.1 | 4.34E-01 | Diacylglycerol O-acyltransferase 1 |
| AT3G54320 | -0.4 | 5.34E-02 | 0.2 | 3.26E-01 | -0.5 | 2.31E-02 | -0.4 | 6.80E-02 | 0.1 | 6.80E-01 | -0.5 | 7.38E-02 | Integrase-type DNA-binding superfamily protein |
| AT5G50375 | -0.4 | 5.81E-03 | 0.2 | 1.70E-01 | -0.5 | 2.05E-03 | -0.2 | 1.77E-01 | 0.1 | 4.02E-01 | -0.3 | 7.15E-02 | Cyclopropyl isomerase |
| AT1G18720 | -0.4 | 5.77E-04 | -0.2 | 5.14E-02 | -0.2 | 3.43E-02 | -0.1 | 2.07E-01 | 0.1 | 7.13E-01 | -0.2 | 2.09E-01 | At1g18720 |
| AT1G69450 | -0.4 | 3.28E-03 | -0.5 | 8.80E-04 | 0.1 | 3.34E-01 | -0.1 | 3.87E-01 | 0.3 | 7.67E-02 | -0.4 | 4.40E-02 | Early-responsive to dehydration stress protein |

| | | | | | | | | | | | | | |
|---|---|---|---|---|---|---|---|---|---|---|---|---|---|
| AT1G32750 | -0.4 | 1.35E-04 | -0.3 | 3.02E-04 | 0.0 | 8.20E-01 | 0.0 | 9.14E-01 | 0.0 | 9.56E-01 | 0.0 | 9.82E-01 | Transcription initiation factor TFIID subunit 1 |
| AT5G63610 | -0.4 | 3.21E-04 | -0.3 | 1.72E-03 | -0.1 | 4.19E-01 | -0.3 | 1.98E-02 | -0.2 | 1.36E-01 | -0.1 | 6.39E-01 | Cyclin-dependent kinase E-1 |
| AT1G47655 | -0.4 | 2.94E-02 | -0.1 | 4.62E-01 | -0.2 | 2.01E-01 | -0.7 | 1.76E-02 | -0.4 | 1.15E-01 | -0.2 | 5.95E-01 | Dof zinc finger protein |
| AT5G36290 | -0.4 | 4.99E-04 | 0.0 | 8.60E-01 | -0.3 | 1.96E-03 | -0.1 | 2.88E-01 | 0.0 | 9.70E-01 | -0.1 | 4.76E-01 | GDT1-like protein 3 |
| AT4G22756 | -0.4 | 2.49E-03 | 0.0 | 9.74E-01 | -0.4 | 7.41E-03 | -0.3 | 1.52E-01 | 0.0 | 9.79E-01 | -0.3 | 2.57E-01 | Methylsterol monooxygenase 1-2 |
| AT1G16130 | -0.4 | 1.75E-02 | -0.9 | 1.39E-04 | 0.6 | 8.30E-03 | -1.0 | 1.72E-02 | -1.1 | 1.65E-02 | 0.1 | 9.10E-01 | Wall-associated receptor kinase-like 2 |
| AT5G08160 | -0.4 | 5.05E-04 | -0.2 | 1.29E-02 | -0.1 | 1.04E-01 | 0.0 | 9.35E-01 | 0.0 | 8.35E-01 | 0.0 | 9.15E-01 | AT5g08160 |
| AT2G21045 | -0.4 | 3.35E-04 | 0.1 | 4.17E-01 | -0.4 | 4.52E-04 | 0.2 | 3.30E-01 | 0.1 | 6.36E-01 | 0.1 | 7.83E-01 | Protein HIGH ARSENIC CONTENT 1, mitochondrial |
| AT2G25430 | -0.4 | 5.18E-04 | -0.3 | 2.56E-03 | -0.1 | 4.70E-01 | -0.3 | 9.62E-02 | -0.4 | 2.16E-02 | 0.2 | 5.48E-01 | Putative clathrin assembly protein At2g25430 |
| AT3G15070 | -0.4 | 1.39E-02 | -0.3 | 4.85E-02 | -0.1 | 6.24E-01 | -0.3 | 3.41E-01 | -0.2 | 5.57E-01 | -0.1 | 8.92E-01 | RING-type E3 ubiquitin transferase |
| AT1G17070 | -0.4 | 1.80E-03 | -0.2 | 4.04E-02 | -0.2 | 1.56E-01 | 0.1 | 4.60E-01 | 0.1 | 8.08E-01 | 0.1 | 7.73E-01 | Septin and tuftelin-interacting protein 1 homolog 1 |
| AT3G17205 | -0.4 | 5.91E-04 | -0.4 | 6.77E-04 | 0.0 | 8.84E-01 | 0.0 | 9.38E-01 | 0.0 | 9.53E-01 | 0.0 | 9.16E-01 | HECT-type E3 ubiquitin transferase |
| AT3G32980 | -0.4 | 2.96E-03 | 0.1 | 4.49E-01 | -0.4 | 2.44E-03 | -0.1 | 4.62E-01 | 0.2 | 2.24E-01 | -0.3 | 3.59E-02 | Peroxidase 32 |
| AT4G20890 | -0.4 | 4.14E-04 | 0.0 | 9.85E-01 | -0.4 | 1.30E-03 | 0.0 | 6.65E-01 | 0.0 | 8.03E-01 | -0.1 | 6.05E-01 | Tubulin beta-9 chain |
| AT1G71410 | -0.4 | 4.01E-03 | -0.3 | 1.90E-02 | -0.1 | 4.89E-01 | -0.2 | 4.97E-01 | -0.2 | 5.18E-01 | 0.0 | 9.71E-01 | SCY1-like protein 2 B |
| AT3G50970 | -0.4 | 1.25E-01 | -0.7 | 8.93E-03 | 0.4 | 1.64E-01 | 0.5 | 2.40E-01 | 0.4 | 3.46E-01 | 0.0 | 9.58E-01 | Dehydrin Xero 2 |
| AT1G58520 | -0.4 | 8.68E-02 | -0.5 | 3.14E-02 | 0.1 | 5.81E-01 | 0.1 | 6.91E-01 | 0.6 | 3.33E-02 | -0.5 | 9.51E-02 | GDSL-like lipase |
| AT2G41820 | -0.4 | 5.36E-03 | -0.3 | 7.55E-03 | 0.0 | 9.60E-01 | -0.4 | 5.30E-02 | 0.1 | 8.22E-01 | -0.5 | 9.58E-02 | Leucine-rich repeat receptor-like tyrosine-protein kinase PXC3 |
| AT3G62760 | -0.4 | 1.66E-01 | 0.6 | 2.35E-02 | -0.9 | 8.06E-03 | -0.2 | 5.08E-01 | 0.5 | 1.13E-01 | -0.7 | 5.79E-02 | Glutathione S-transferase F13 |
| AT5G67560 | -0.4 | 8.39E-05 | 0.0 | 5.60E-01 | -0.3 | 6.73E-04 | -0.3 | 1.14E-01 | -0.2 | 3.53E-01 | -0.1 | 7.63E-01 | ADP-ribosylation factor-like protein 8b |
| AT5G15070 | -0.4 | 5.07E-03 | -0.2 | 5.83E-02 | -0.1 | 2.88E-01 | 0.0 | 9.71E-01 | -0.1 | 8.89E-01 | 0.1 | 8.69E-01 | Inositol hexakisphosphate and diphosphoinositol-pentakisphosphate kinase |
| AT1G49570 | -0.3 | 1.47E-01 | -0.2 | 3.41E-01 | -0.1 | 7.03E-01 | 2.1 | 5.43E-06 | 1.1 | 6.70E-03 | 1.0 | 4.04E-03 | Peroxidase 10 |
| AT4G01010 | -0.3 | 1.95E-02 | -1.0 | 7.87E-05 | 0.7 | 3.88E-03 | 0.0 | 9.43E-01 | -0.5 | 6.52E-02 | 0.5 | 1.10E-01 | Putative cyclic nucleotide-gated ion channel 13 |
| AT1G67880 | -0.3 | 2.06E-03 | -0.3 | 6.91E-03 | -0.1 | 6.39E-01 | -0.2 | 1.34E-01 | 0.0 | 9.85E-01 | -0.2 | 2.01E-01 | Beta-1,4-N-acetylglucosaminyltransferase family protein |
| AT4G30400 | -0.3 | 2.93E-03 | -0.2 | 2.80E-02 | -0.1 | 3.10E-01 | 0.1 | 5.11E-01 | 0.4 | 3.16E-02 | -0.3 | 1.94E-01 | RING-H2 finger protein ATL13 |
| AT3G11240 | -0.3 | 7.07E-03 | -0.1 | 4.32E-01 | -0.3 | 6.37E-02 | -0.1 | 7.25E-01 | -0.2 | 6.45E-01 | 0.1 | 9.10E-01 | Arginyl-tRNA--protein transferase |
| AT5G61140 | -0.3 | 1.97E-03 | -0.4 | 7.33E-04 | 0.1 | 4.22E-01 | -0.1 | 7.07E-01 | 0.1 | 5.44E-01 | -0.2 | 3.97E-01 | U5 small nuclear ribonucleoprotein helicase |
| AT5G03650 | -0.3 | 9.03E-03 | -0.5 | 2.17E-03 | 0.1 | 3.39E-01 | 0.4 | 2.12E-02 | 0.5 | 1.01E-02 | -0.1 | 6.98E-01 | 1,4-alpha-glucan-branching enzyme 2-2, chloroplastic |
| AT5G15530 | -0.3 | 1.69E-02 | 0.3 | 4.75E-02 | -0.6 | 1.68E-03 | -0.1 | 7.00E-01 | 0.2 | 2.20E-01 | -0.2 | 1.87E-01 | Biotin carboxyl carrier protein of acetyl-CoA carboxylase 2, chloroplastic |
| AT1G50700 | -0.3 | 1.38E-02 | -0.2 | 8.35E-02 | -0.1 | 4.61E-01 | 0.0 | 9.24E-01 | -0.1 | 7.72E-01 | 0.1 | 7.29E-01 | Calcium-dependent protein kinase 33 |
| AT1G18550 | -0.3 | 1.81E-02 | -0.3 | 4.35E-02 | -0.1 | 7.58E-01 | 0.1 | 8.40E-01 | 0.3 | 3.82E-01 | -0.2 | 5.93E-01 | Kinesin-like protein KIN-8A |
| AT3G19400 | -0.3 | 9.71E-04 | -0.1 | 2.32E-01 | -0.2 | 1.69E-02 | -0.2 | 1.07E-01 | -0.2 | 3.30E-01 | -0.1 | 7.31E-01 | Cysteine proteinases superfamily protein |
| AT5G22760 | -0.3 | 2.93E-04 | -0.4 | 1.43E-04 | 0.1 | 3.80E-01 | 0.1 | 6.80E-01 | 0.3 | 3.86E-01 | -0.1 | 6.58E-01 | hypothetical protein |
| AT4G36400 | -0.3 | 1.82E-03 | -0.4 | 1.24E-03 | 0.0 | 6.78E-01 | -0.1 | 7.34E-01 | 0.0 | 9.82E-01 | -0.1 | 7.99E-01 | hypothetical protein |
| AT2G28510 | -0.3 | 9.75E-04 | -0.1 | 1.11E-01 | -0.2 | 3.33E-02 | 0.4 | 6.86E-02 | 0.5 | 2.00E-02 | -0.1 | 5.28E-01 | Dof zinc finger protein |
| AT5G64560 | -0.3 | 7.01E-03 | -0.2 | 1.05E-01 | -0.2 | 2.21E-01 | -0.3 | 2.45E-02 | -0.2 | 1.13E-01 | -0.1 | 7.49E-01 | Magnesium transporter MRS2-2 |
| AT1G75250 | -0.3 | 1.59E-01 | 0.2 | 3.89E-01 | -0.5 | 7.13E-02 | -0.2 | 6.64E-01 | -0.1 | 8.78E-01 | -0.1 | 8.82E-01 | Protein RADIALIS-like 6 |
| AT1G53910 | -0.3 | 5.05E-04 | -0.2 | 1.44E-02 | -0.1 | 9.80E-02 | -0.7 | 5.50E-06 | -0.1 | 7.32E-01 | -0.6 | 5.90E-04 | Related to AP2 12 |
| AT3G14130 | -0.3 | 1.28E-02 | -0.6 | 7.45E-04 | 0.3 | 9.52E-02 | -0.4 | 1.04E-01 | -0.2 | 3.96E-01 | -0.1 | 6.20E-01 | #NAME? |
| AT4G15920 | -0.3 | 4.00E-04 | 0.0 | 5.42E-01 | -0.4 | 7.17E-04 | -0.5 | 5.71E-03 | -0.1 | 6.00E-01 | -0.4 | 8.13E-02 | Bidirectional sugar transporter SWEET17 |
| AT3G26210 | -0.3 | 4.32E-02 | -0.7 | 1.70E-03 | 0.3 | 7.60E-02 | 0.1 | 7.65E-01 | -0.4 | 2.33E-01 | 0.5 | 2.28E-01 | Cytochrome P450 71B23 |
| AT4G02010 | -0.3 | 6.32E-04 | -0.3 | 2.80E-03 | -0.1 | 4.98E-01 | 0.2 | 2.67E-01 | 0.1 | 6.47E-01 | 0.1 | 7.35E-01 | AT4g02010 |
| AT4G18390 | -0.3 | 1.72E-02 | -0.3 | 2.40E-02 | 0.0 | 9.38E-01 | -0.1 | 7.68E-01 | 0.2 | 4.70E-01 | -0.2 | 3.84E-01 | Transcription factor TCP2 |
| AT1G01740 | -0.3 | 8.33E-03 | -0.5 | 1.15E-03 | 0.2 | 2.18E-01 | 0.4 | 2.29E-01 | 0.1 | 8.21E-01 | 0.3 | 4.79E-01 | Serine |
| AT1G66180 | -0.3 | 1.05E-03 | -0.4 | 8.79E-04 | 0.0 | 7.26E-01 | -0.3 | 8.51E-03 | -0.5 | 1.64E-03 | 0.1 | 4.29E-01 | Eukaryotic aspartyl protease family protein |
| AT1G45160 | -0.3 | 3.31E-04 | -0.4 | 1.91E-04 | 0.1 | 4.73E-01 | -0.1 | 6.42E-01 | 0.0 | 9.27E-01 | -0.1 | 6.51E-01 | Protein kinase superfamily protein |
| AT5G65710 | -0.3 | 1.23E-02 | -0.3 | 4.06E-02 | -0.1 | 6.38E-01 | -0.2 | 1.76E-01 | 0.0 | 8.46E-01 | -0.2 | 4.44E-01 | LRR receptor-like serine |
| AT5G10760 | -0.3 | 4.15E-01 | -0.8 | 7.40E-02 | 0.5 | 3.12E-01 | -0.1 | 8.58E-01 | -0.5 | 1.76E-01 | 0.5 | 3.48E-01 | Aspartyl protease AED1 |
| AT5G64220 | -0.3 | 2.41E-03 | -0.4 | 1.25E-03 | 0.1 | 5.75E-01 | 0.0 | 8.74E-01 | 0.0 | 9.58E-01 | 0.0 | 9.62E-01 | Calmodulin-binding transcription activator 2 |
| AT3G52930 | -0.3 | 2.78E-02 | 0.3 | 8.18E-02 | -0.6 | 3.08E-03 | 1.1 | 2.45E-06 | 0.5 | 8.19E-03 | 0.6 | 6.16E-03 | Fructose-bisphosphate aldolase 8, cytosolic |
| AT5G14000 | -0.3 | 1.36E-03 | 0.0 | 9.80E-01 | -0.3 | 5.09E-03 | -0.2 | 4.39E-01 | 0.1 | 6.95E-01 | -0.3 | 2.47E-01 | NAC domain-containing protein |
| AT1G07200 | -0.3 | 6.64E-03 | -0.3 | 2.46E-02 | -0.1 | 5.84E-01 | -0.4 | 8.70E-03 | -0.1 | 6.01E-01 | -0.3 | 1.17E-01 | At1g07200 |
| AT1G02740 | -0.3 | 1.23E-02 | -0.4 | 5.02E-03 | 0.1 | 5.84E-01 | -0.1 | 7.24E-01 | -0.3 | 2.06E-01 | 0.2 | 4.74E-01 | Protein MRG2 |
| AT1G58360 | -0.3 | 1.87E-03 | -0.5 | 2.47E-04 | 0.2 | 1.27E-01 | -0.6 | 3.21E-03 | -0.1 | 5.39E-01 | -0.4 | 8.76E-02 | Amino acid permease 1 |
| AT2G38420 | -0.3 | 2.67E-02 | -0.3 | 8.11E-02 | -0.1 | 6.70E-01 | 0.2 | 5.37E-01 | 0.0 | 9.14E-01 | 0.1 | 6.96E-01 | Pentatricopeptide repeat-containing protein At2g38420, mitochondrial |
| AT5G17250 | -0.3 | 1.92E-02 | -0.2 | 9.58E-02 | -0.1 | 5.00E-01 | 0.2 | 6.81E-01 | 0.2 | 6.12E-01 | -0.1 | 9.11E-01 | GPI ethanolamine phosphate transferase 3 |
| AT4G01950 | -0.3 | 2.30E-02 | -0.4 | 1.13E-02 | 0.1 | 6.66E-01 | -0.7 | 4.22E-04 | -0.1 | 8.04E-01 | -0.7 | 9.89E-03 | Glycerol-3-phosphate acyltransferase 3 |
| AT1G16270 | -0.3 | 7.71E-04 | -0.5 | 1.87E-04 | 0.1 | 2.11E-01 | 0.0 | 9.74E-01 | 0.2 | 7.74E-01 | -0.2 | 7.62E-01 | Protein kinase domain-containing protein |
| AT5G45250 | -0.3 | 4.09E-03 | -0.4 | 1.75E-03 | 0.1 | 5.34E-01 | -0.1 | 5.20E-01 | -0.3 | 5.45E-02 | 0.2 | 2.82E-01 | Disease resistance protein RPS4 hydrolase RPS4) |
| AT4G18700 | -0.3 | 7.89E-04 | -0.3 | 2.26E-03 | 0.0 | 6.94E-01 | -0.3 | 6.25E-03 | -0.4 | 3.25E-03 | 0.1 | 6.80E-01 | CBL-interacting serine |
| AT3G43300 | -0.3 | 9.62E-04 | -0.3 | 7.61E-03 | -0.1 | 3.23E-01 | -0.1 | 7.85E-01 | -0.1 | 8.36E-01 | 0.0 | 9.97E-01 | Brefeldin A-inhibited guanine nucleotide-exchange protein 5 |
| AT4G37880 | -0.3 | 3.16E-04 | -0.2 | 1.14E-02 | -0.1 | 6.82E-02 | -0.2 | 9.73E-02 | -0.2 | 1.66E-01 | 0.0 | 9.66E-01 | Protein RMD5 homolog |
| AT2G17265 | -0.3 | 3.91E-03 | 0.0 | 8.75E-01 | -0.4 | 6.76E-03 | -0.3 | 1.42E-02 | -0.1 | 7.89E-01 | -0.3 | 1.16E-01 | Homoserine kinase |
| AT1G52150 | -0.3 | 6.56E-03 | -0.3 | 7.79E-03 | 0.0 | 9.84E-01 | 0.0 | 8.06E-01 | 0.1 | 7.37E-01 | -0.1 | 6.02E-01 | Homeobox-leucine zipper family protein |
| AT5G23450 | -0.3 | 9.19E-04 | -0.3 | 2.34E-03 | 0.0 | 7.49E-01 | -0.3 | 1.43E-02 | -0.2 | 1.56E-01 | -0.1 | 5.69E-01 | Long-chain base kinase 1 |
| AT2G27690 | -0.3 | 3.15E-02 | -0.4 | 1.36E-02 | 0.1 | 6.38E-01 | -0.2 | 7.25E-01 | 0.8 | 2.59E-02 | -1.0 | 1.40E-02 | Cytochrome P450 94C1 |
| AT1G21550 | -0.3 | 1.21E-01 | -0.5 | 5.11E-02 | 0.1 | 6.44E-01 | -0.4 | 3.77E-01 | -0.2 | 6.95E-01 | -0.2 | 7.79E-01 | Probable calcium-binding protein CML44 |
| AT5G09690 | -0.3 | 1.27E-02 | -0.3 | 2.40E-02 | 0.0 | 8.41E-01 | -0.3 | 2.03E-01 | -0.1 | 6.33E-01 | -0.2 | 6.62E-01 | Magnesium transporter |
| AT5G34930 | -0.3 | 4.04E-03 | -0.2 | 4.17E-02 | -0.1 | 3.01E-01 | 0.1 | 4.25E-01 | -0.1 | 6.75E-01 | 0.2 | 2.63E-01 | Arogenate dehydrogenase |
| AT3G54670 | -0.3 | 3.08E-04 | -0.5 | 6.43E-05 | 0.1 | 1.16E-01 | 0.1 | 2.78E-01 | 0.0 | 9.57E-01 | 0.1 | 4.85E-01 | Structural maintenance of chromosomes protein 1 |
| AT1G01190 | -0.3 | 1.48E-02 | 0.2 | 1.91E-01 | -0.5 | 4.20E-03 | 0.4 | 2.15E-03 | 1.1 | 9.39E-07 | -0.7 | 8.47E-04 | Cytochrome P450, family 78, subfamily A, polypeptide 8 |
| AT3G20290 | -0.3 | 5.65E-04 | -0.2 | 1.27E-02 | -0.1 | 1.17E-01 | -0.3 | 3.57E-03 | -0.2 | 1.46E-01 | -0.2 | 3.28E-01 | EH domain-containing protein 1 |
| AT3G13530 | -0.3 | 1.19E-03 | -0.3 | 3.64E-03 | 0.0 | 6.56E-01 | 0.0 | 9.51E-01 | -0.1 | 5.78E-01 | 0.1 | 7.06E-01 | MAP3K epsilon protein kinase 1 |
| AT4G15530 | -0.3 | 8.68E-03 | -0.9 | 1.91E-05 | 0.6 | 8.02E-04 | -0.3 | 2.46E-01 | -0.5 | 1.45E-01 | 0.1 | 7.71E-01 | Pyruvate, phosphate dikinase |
| AT3G19980 | -0.3 | 2.54E-04 | -0.3 | 1.29E-03 | -0.1 | 4.29E-01 | 0.0 | 9.55E-01 | 0.1 | 4.07E-01 | -0.2 | 4.36E-01 | Phytochrome-associated serine |
| AT1G19270 | -0.3 | 1.48E-03 | -0.3 | 4.16E-03 | 0.0 | 7.01E-01 | 0.2 | 2.09E-01 | 0.1 | 8.10E-01 | 0.2 | 4.54E-01 | Protein DA1 |
| AT1G61310 | -0.3 | 3.09E-02 | -0.3 | 4.10E-02 | 0.0 | 9.60E-01 | -0.2 | 6.12E-01 | 0.0 | 9.40E-01 | -0.2 | 5.85E-01 | Probable disease resistance protein At1g61310 |
| AT1G19480 | -0.3 | 8.38E-03 | -0.1 | 3.44E-01 | -0.2 | 9.27E-02 | -0.3 | 9.91E-02 | 0.0 | 9.35E-01 | -0.3 | 1.89E-01 | F18O14.25 |
| AT4G35540 | -0.3 | 2.01E-02 | -0.1 | 4.81E-01 | -0.2 | 1.35E-01 | -0.2 | 5.46E-01 | -0.3 | 3.87E-01 | 0.1 | 8.01E-01 | Plant-specific TFIIB-related protein PTF2 |
| AT4G35970 | -0.3 | 9.95E-03 | -0.2 | 1.05E-01 | -0.1 | 3.00E-01 | -0.1 | 6.37E-01 | -0.1 | 7.76E-01 | 0.0 | 9.43E-01 | L-ascorbate peroxidase |
| AT4G19010 | -0.3 | 3.33E-02 | -0.4 | 2.35E-02 | 0.0 | 8.23E-01 | -0.2 | 4.66E-01 | -0.1 | 6.73E-01 | -0.1 | 9.05E-01 | 4-coumarate--CoA ligase-like 6 |
| AT4G11380 | -0.3 | 4.48E-04 | -0.1 | 5.14E-02 | -0.2 | 2.60E-02 | 0.1 | 6.51E-01 | 0.1 | 4.42E-01 | 0.0 | 8.12E-01 | Beta-adaptin-like protein |
| AT2G01140 | -0.3 | 8.20E-03 | 0.0 | 8.75E-01 | -0.3 | 2.10E-02 | 0.0 | 8.85E-01 | 0.0 | 8.07E-01 | 0.0 | 9.31E-01 | Fructose-bisphosphate aldolase 3, chloroplastic |

| | | | | | | | | | | | | | |
|---|---|---|---|---|---|---|---|---|---|---|---|---|---|
| AT1G01260 | -0.3 | 1.16E-03 | -0.2 | 2.18E-02 | -0.1 | 1.70E-01 | -0.2 | 4.18E-01 | -0.1 | 7.74E-01 | -0.1 | 7.93E-01 | Transcription factor bHLH13 |
| AT5G44380 | -0.3 | 9.99E-04 | 0.3 | 4.95E-03 | -0.6 | 9.94E-05 | 0.6 | 2.65E-03 | 0.4 | 4.35E-02 | 0.2 | 4.90E-01 | FAD-binding PCMH-type domain-containing protein |
| AT5G06710 | -0.3 | 2.17E-02 | -0.1 | 4.64E-01 | -0.2 | 1.48E-01 | -0.1 | 7.74E-01 | 0.2 | 4.02E-01 | -0.3 | 3.09E-01 | Homeobox leucine zipper protein |
| AT3G04880 | -0.3 | 2.35E-03 | -0.2 | 1.60E-02 | -0.1 | 3.74E-01 | -0.3 | 8.63E-03 | -0.1 | 2.89E-01 | -0.2 | 3.04E-01 | DNA damage-repair |
| AT1G71830 | -0.3 | 1.71E-02 | -0.1 | 6.46E-01 | -0.3 | 7.99E-02 | 0.1 | 6.90E-01 | 0.2 | 2.73E-01 | -0.1 | 5.59E-01 | Somatic embryogenesis receptor kinase 1 |
| AT3G05200 | -0.3 | 1.43E-02 | -0.4 | 5.95E-03 | 0.1 | 5.86E-01 | -0.2 | 2.42E-01 | -0.6 | 2.39E-03 | 0.4 | 6.25E-02 | E3 ubiquitin-protein ligase ATL6 |
| AT5G06450 | -0.3 | 5.52E-04 | -0.3 | 3.58E-03 | -0.1 | 3.87E-01 | -0.2 | 4.29E-01 | -0.3 | 3.32E-01 | 0.1 | 8.78E-01 | Protein RISC-INTERACTING CLEARING 3'-5' EXORIBONUCLEASE 2 |
| AT3G09410 | -0.3 | 7.21E-04 | -0.2 | 4.63E-02 | -0.2 | 5.10E-02 | 0.0 | 9.82E-01 | 0.3 | 1.35E-01 | -0.3 | 1.90E-01 | Pectin acetylesterase 5 |
| AT1G73360 | -0.3 | 5.12E-02 | 0.3 | 1.09E-01 | -0.6 | 9.75E-03 | 0.4 | 1.19E-01 | 0.1 | 7.20E-01 | 0.3 | 4.04E-01 | HDG11 |
| AT1G43770 | -0.3 | 1.06E-02 | -0.5 | 1.34E-03 | 0.2 | 2.13E-01 | -0.5 | 8.20E-02 | -0.2 | 4.43E-01 | -0.2 | 4.84E-01 | RING |
| AT3G27190 | -0.3 | 1.66E-03 | -0.3 | 9.52E-03 | -0.1 | 4.17E-01 | 0.1 | 4.45E-01 | 0.2 | 2.94E-01 | -0.1 | 8.07E-01 | Uridine |
| AT3G06440 | -0.3 | 1.45E-03 | 0.0 | 7.80E-01 | -0.3 | 7.43E-03 | -0.4 | 8.37E-02 | -0.3 | 3.00E-01 | -0.1 | 7.27E-01 | Galectin domain-containing protein |
| AT2G24360 | -0.3 | 7.11E-04 | -0.5 | 8.60E-05 | 0.2 | 6.28E-02 | 0.0 | 9.85E-01 | 0.0 | 8.84E-01 | 0.0 | 9.01E-01 | Serine |
| AT1G18040 | -0.3 | 1.09E-02 | -0.1 | 4.74E-01 | -0.3 | 8.28E-02 | 0.5 | 9.00E-02 | 0.4 | 2.74E-01 | 0.1 | 7.49E-01 | Cyclin-dependent kinase D-3 |
| AT3G55730 | -0.3 | 8.77E-03 | -0.1 | 5.65E-01 | -0.3 | 5.47E-02 | 0.7 | 1.34E-03 | 0.6 | 8.90E-03 | 0.1 | 6.99E-01 | MYB transcription factor-like protein |
| AT5G04940 | -0.3 | 1.03E-03 | -0.2 | 4.37E-02 | -0.2 | 8.41E-02 | 0.0 | 9.87E-01 | 0.1 | 6.51E-01 | -0.1 | 7.20E-01 | Histone-lysine N-methyltransferase, H3 lysine-9 specific SUVH1 3-9 homolog protein 1) |
| AT1G69330 | -0.3 | 3.49E-03 | -0.1 | 2.11E-01 | -0.2 | 6.95E-02 | -0.2 | 5.13E-01 | -0.2 | 6.03E-01 | 0.0 | 9.81E-01 | At1g69330 |
| AT1G76650 | -0.3 | 2.24E-01 | -0.6 | 7.38E-02 | 0.2 | 5.54E-01 | 0.5 | 2.85E-01 | -0.2 | 7.68E-01 | 0.8 | 1.76E-01 | Calcium-binding protein CML38 |
| AT1G17120 | -0.3 | 2.49E-03 | -0.3 | 1.29E-02 | -0.1 | 4.65E-01 | -0.4 | 3.27E-03 | -0.3 | 6.72E-02 | -0.2 | 4.27E-01 | Cationic amino acid transporter 8, vacuolar |
| AT1G05810 | -0.3 | 8.75E-03 | 0.1 | 2.54E-01 | -0.5 | 4.20E-03 | -0.6 | 2.42E-03 | -0.5 | 2.06E-02 | -0.1 | 5.54E-01 | Rab GTPase-like A5A protein |
| AT2G34150 | -0.3 | 8.54E-03 | -0.1 | 4.18E-01 | -0.2 | 7.59E-02 | 0.1 | 8.24E-01 | 0.4 | 2.36E-01 | -0.3 | 3.86E-01 | Protein SCAR |
| AT4G02290 | -0.3 | 7.75E-03 | 0.1 | 3.88E-01 | -0.4 | 4.37E-03 | -0.3 | 3.40E-02 | 0.2 | 1.52E-01 | -0.6 | 5.33E-03 | Endoglucanase |
| AT5G46180 | -0.3 | 1.23E-03 | -0.5 | 1.55E-04 | 0.2 | 1.02E-01 | -0.2 | 1.86E-01 | -0.2 | 2.59E-01 | 0.0 | 9.90E-01 | Ornithine aminotransferase, mitochondrial |
| AT1G27770 | -0.3 | 9.66E-03 | -0.6 | 2.83E-04 | 0.3 | 2.98E-02 | 0.0 | 8.85E-01 | 0.0 | 9.22E-01 | 0.0 | 9.97E-01 | Calcium-transporting ATPase |
| AT4G17870 | -0.3 | 7.36E-04 | 0.1 | 1.54E-01 | -0.4 | 4.08E-04 | -0.2 | 3.95E-01 | -0.2 | 4.93E-01 | 0.0 | 9.88E-01 | Abscisic acid receptor PYR1 |
| AT5G11610 | -0.3 | 1.20E-03 | -0.4 | 2.90E-04 | 0.1 | 2.41E-01 | -0.1 | 5.67E-01 | -0.1 | 7.09E-01 | 0.0 | 9.46E-01 | Exostosin family protein |
| AT1G56540 | -0.3 | 2.47E-02 | -0.5 | 3.57E-03 | 0.2 | 2.72E-01 | 0.0 | 9.74E-01 | -0.2 | 7.38E-01 | 0.1 | 7.56E-01 | Disease resistance protein family |
| AT4G12880 | -0.3 | 2.33E-02 | 0.2 | 1.35E-01 | -0.5 | 3.92E-03 | 0.2 | 4.88E-01 | 0.4 | 1.57E-01 | -0.2 | 4.96E-01 | Early nodulin-like protein 19 |
| AT4G29990 | -0.3 | 1.60E-02 | -0.5 | 4.37E-03 | 0.1 | 4.44E-01 | 0.7 | 7.63E-02 | 0.2 | 6.44E-01 | 0.4 | 3.31E-01 | Leucine-rich repeat transmembrane protein kinase protein |
| AT1G75760 | -0.3 | 4.67E-04 | 0.0 | 6.91E-01 | -0.3 | 3.00E-03 | -0.1 | 6.71E-01 | -0.2 | 1.64E-01 | 0.1 | 4.45E-01 | ER lumen protein retaining receptor family protein |
| AT4G02340 | -0.3 | 2.18E-03 | -0.4 | 6.27E-04 | 0.1 | 3.31E-01 | -0.4 | 6.88E-02 | -0.2 | 4.74E-01 | -0.2 | 5.29E-01 | AB hydrolase-1 domain-containing protein |
| AT1G28070 | -0.3 | 1.03E-01 | -0.3 | 1.81E-01 | -0.1 | 8.38E-01 | -0.7 | 3.85E-02 | -0.4 | 2.40E-01 | -0.3 | 5.95E-01 | Protein CURLY FLAG LEAF 2 |
| AT4G08150 | -0.3 | 6.46E-03 | -0.5 | 5.40E-04 | 0.2 | 1.20E-01 | 0.2 | 5.04E-01 | 0.2 | 4.82E-01 | 0.0 | 9.39E-01 | KNAT1 |
| AT2G46500 | -0.3 | 1.07E-03 | -0.4 | 5.36E-04 | 0.1 | 4.80E-01 | 0.1 | 4.50E-01 | 0.1 | 7.18E-01 | 0.0 | 8.62E-01 | 1-phosphatidylinositol 4-kinase |
| AT5G64640 | -0.3 | 2.13E-02 | 0.2 | 1.58E-01 | -0.5 | 5.91E-03 | -0.1 | 4.49E-01 | 0.1 | 6.34E-01 | -0.3 | 2.38E-01 | Probable pectinesterase |
| AT3G16630 | -0.3 | 3.47E-04 | -0.2 | 2.12E-02 | -0.2 | 4.35E-02 | -0.1 | 2.15E-01 | 0.0 | 9.09E-01 | -0.2 | 3.40E-01 | Kinesin-like protein KIN-13A |
| AT3G16090 | -0.3 | 2.74E-04 | -0.3 | 1.37E-03 | -0.1 | 4.48E-01 | -0.2 | 1.31E-01 | 0.0 | 8.23E-01 | -0.2 | 3.42E-01 | ERAD-associated E3 ubiquitin-protein ligase HRD1A |
| AT5G63370 | -0.3 | 3.18E-04 | -0.4 | 1.06E-04 | 0.1 | 2.26E-01 | -0.1 | 7.00E-01 | 0.0 | 9.34E-01 | -0.1 | 7.35E-01 | Protein kinase superfamily protein |
| AT1G54990 | -0.3 | 1.34E-03 | -0.1 | 1.73E-01 | -0.2 | 2.75E-02 | 0.0 | 8.36E-01 | 0.0 | 8.05E-01 | -0.1 | 6.77E-01 | Protein AUXIN RESPONSE 4 |
| AT1G16970 | -0.3 | 7.55E-04 | -0.2 | 1.99E-02 | -0.1 | 1.20E-01 | 0.2 | 1.65E-01 | 0.3 | 1.08E-01 | -0.1 | 8.13E-01 | KU70 |
| AT5G58520 | -0.3 | 6.60E-03 | -0.1 | 4.32E-01 | -0.2 | 5.27E-02 | -0.4 | 6.90E-03 | -0.3 | 5.45E-02 | -0.1 | 6.32E-01 | Protein kinase domain-containing protein |
| AT1G61140 | -0.3 | 1.20E-03 | -0.5 | 7.23E-05 | 0.2 | 3.29E-02 | 0.0 | 9.17E-01 | -0.1 | 6.73E-01 | 0.1 | 8.07E-01 | SNF2 domain-containing protein |
| AT1G70490 | -0.3 | 1.85E-03 | 0.0 | 8.43E-01 | -0.3 | 6.13E-03 | -0.1 | 3.04E-01 | -0.1 | 6.47E-01 | -0.1 | 7.84E-01 | ADP-ribosylation factor 2-B |
| AT1G67340 | -0.3 | 1.72E-02 | -0.4 | 1.19E-02 | 0.0 | 7.96E-01 | 0.3 | 1.46E-01 | 0.1 | 5.93E-01 | 0.1 | 6.11E-01 | F-box protein At1g67340 |
| AT4G29610 | -0.3 | 2.65E-01 | 0.0 | 8.87E-01 | -0.4 | 3.02E-01 | -0.3 | 4.63E-01 | 0.6 | 1.25E-01 | -1.0 | 6.33E-02 | Cytidine deaminase 6 |
| AT2G33510 | -0.3 | 1.26E-02 | -0.2 | 8.84E-02 | -0.1 | 4.01E-01 | -0.7 | 4.93E-03 | -0.7 | 8.09E-03 | 0.0 | 9.88E-01 | ATP-dependent RNA helicase |
| AT4G18465 | -0.3 | 9.45E-03 | -0.4 | 2.74E-03 | 0.1 | 4.45E-01 | -0.2 | 3.72E-01 | -0.1 | 7.84E-01 | -0.1 | 7.37E-01 | Probable pre-mRNA-splicing factor ATP-dependent RNA helicase DEAH9 |
| AT5G47520 | -0.3 | 4.73E-04 | -0.1 | 7.31E-02 | -0.2 | 2.42E-02 | -0.3 | 5.70E-02 | -0.3 | 1.52E-01 | -0.1 | 8.68E-01 | Ras-related protein RABA5a |
| AT5G51740 | -0.3 | 1.61E-03 | -0.4 | 1.10E-03 | 0.0 | 6.82E-01 | -0.2 | 3.57E-01 | -0.1 | 5.76E-01 | 0.0 | 8.89E-01 | Peptidase M48 domain-containing protein |
| AT1G31360 | -0.3 | 1.11E-02 | -0.1 | 3.06E-01 | -0.2 | 1.32E-01 | -0.1 | 8.46E-01 | -0.1 | 8.30E-01 | 0.0 | 9.68E-01 | ATP-dependent DNA helicase |
| AT3G12120 | -0.3 | 9.20E-04 | 0.0 | 9.15E-01 | -0.3 | 2.23E-03 | -0.6 | 4.24E-05 | -0.3 | 1.64E-02 | -0.3 | 7.40E-02 | Delta-fatty-acid desaturase |
| AT1G43890 | -0.3 | 7.37E-03 | -0.2 | 1.21E-01 | -0.2 | 1.85E-01 | -0.2 | 5.04E-01 | -0.1 | 6.23E-01 | 0.0 | 9.75E-01 | hypothetical protein |
| AT5G50820 | -0.3 | 7.14E-03 | 0.0 | 9.68E-01 | -0.3 | 2.15E-02 | 0.0 | 9.06E-01 | 0.1 | 8.37E-01 | -0.1 | 7.70E-01 | NAC domain containing protein 97 |
| AT5G64070 | -0.3 | 5.90E-04 | -0.3 | 9.49E-04 | 0.0 | 9.59E-01 | -0.1 | 4.77E-01 | -0.1 | 6.94E-01 | 0.0 | 9.09E-01 | Uncharacterized protein |
| AT2G27580 | -0.3 | 5.26E-03 | -0.3 | 1.43E-02 | 0.0 | 6.95E-01 | -0.1 | 6.68E-01 | -0.1 | 7.59E-01 | 0.0 | 9.80E-01 | Zinc finger A20 and AN1 domain-containing stress-associated protein 3 |
| AT5G17920 | -0.3 | 2.90E-03 | -0.2 | 1.08E-01 | -0.2 | 9.35E-02 | 0.0 | 8.62E-01 | 0.3 | 7.17E-02 | -0.2 | 2.28E-01 | 5-methyltetrahydropteroyltriglutamate--homocysteine methyltransferase 1 |
| AT4G22910 | -0.3 | 6.22E-03 | 0.1 | 3.19E-01 | -0.4 | 4.31E-03 | 0.0 | 9.16E-01 | 0.4 | 1.20E-01 | -0.4 | 2.29E-01 | Protein FIZZY-RELATED 2 |
| AT5G33280 | -0.3 | 4.27E-03 | -0.2 | 2.54E-02 | -0.1 | 4.28E-01 | -0.1 | 7.74E-01 | 0.0 | 8.86E-01 | -0.1 | 7.42E-01 | Putative chloride channel-like protein CLC-g |
| AT2G32850 | -0.3 | 5.39E-04 | -0.2 | 1.64E-02 | -0.1 | 9.09E-02 | 0.1 | 7.48E-01 | 0.1 | 8.31E-01 | 0.0 | 9.80E-01 | Protein kinase superfamily protein |
| AT1G69170 | -0.3 | 5.45E-03 | -0.2 | 7.98E-02 | -0.1 | 2.36E-01 | -0.5 | 9.54E-02 | 0.0 | 9.66E-01 | -0.5 | 2.28E-01 | Squamosa promoter-binding-like protein 6 |
| AT3G56240 | -0.3 | 6.15E-04 | 0.0 | 6.30E-01 | -0.3 | 3.46E-03 | 0.2 | 3.99E-01 | 0.0 | 9.88E-01 | 0.2 | 5.66E-01 | HMA domain-containing protein |
| AT5G58560 | -0.3 | 1.93E-03 | -0.1 | 2.30E-01 | -0.2 | 2.97E-02 | -0.6 | 2.59E-04 | 0.0 | 9.38E-01 | -0.5 | 5.68E-03 | Farnesol kinase, chloroplastic |
| AT1G54490 | -0.3 | 2.48E-04 | -0.3 | 1.21E-03 | -0.1 | 4.38E-01 | -0.2 | 2.22E-01 | -0.1 | 7.12E-01 | -0.1 | 6.43E-01 | 5'-3' exoribonuclease |
| AT1G28660 | -0.3 | 6.60E-02 | -0.2 | 1.66E-01 | -0.1 | 7.00E-01 | 0.0 | 9.77E-01 | -0.3 | 7.99E-02 | 0.3 | 1.62E-01 | GDSL esterase |
| AT1G09970 | -0.3 | 2.41E-03 | -0.5 | 2.29E-04 | 0.2 | 7.78E-02 | 0.4 | 1.31E-03 | 0.0 | 7.70E-01 | 0.3 | 3.34E-02 | Protein kinase domain-containing protein |
| AT4G16210 | -0.3 | 1.34E-03 | -0.3 | 1.52E-03 | 0.0 | 9.15E-01 | -0.2 | 3.35E-01 | 0.1 | 8.14E-01 | -0.2 | 3.09E-01 | Enoyl-CoA hydratase like protein |
| AT5G46920 | -0.3 | 1.29E-03 | -0.2 | 2.06E-02 | -0.1 | 2.02E-01 | 0.2 | 1.64E-01 | 0.1 | 7.12E-01 | 0.1 | 5.59E-01 | Nuclear intron maturase 2, mitochondrial |
| AT1G48420 | -0.3 | 9.83E-04 | -0.1 | 6.88E-02 | -0.2 | 4.62E-02 | -0.5 | 1.25E-03 | -0.3 | 8.02E-02 | -0.2 | 2.15E-01 | Bifunctional D-cysteine desulfhydrase |
| AT1G12280 | -0.3 | 2.62E-02 | -0.3 | 2.51E-02 | 0.0 | 9.29E-01 | 0.2 | 2.38E-01 | 0.0 | 8.78E-01 | 0.2 | 4.77E-01 | LRR and NB-ARC domains-containing disease resistance protein |
| AT3G09830 | -0.3 | 1.50E-02 | -0.7 | 2.19E-04 | 0.4 | 1.67E-02 | 0.3 | 4.94E-02 | 0.2 | 2.54E-01 | 0.1 | 6.76E-01 | Serine |
| AT4G21600 | -0.3 | 4.79E-03 | 0.4 | 2.29E-03 | -0.7 | 1.78E-04 | -0.2 | 2.97E-01 | 0.0 | 9.79E-01 | -0.2 | 3.89E-01 | Endonuclease 5 |
| AT4G30935 | -0.3 | 1.95E-03 | -0.4 | 3.90E-04 | 0.1 | 2.22E-01 | -0.1 | 6.91E-01 | -0.2 | 3.41E-01 | 0.1 | 6.77E-01 | Probable WRKY transcription factor 32 |
| AT2G04880 | -0.3 | 2.69E-04 | -0.5 | 3.82E-05 | 0.1 | 5.93E-02 | -0.1 | 7.31E-01 | 0.2 | 4.78E-01 | -0.3 | 3.80E-01 | WRKY domain-containing protein |
| AT1G72520 | -0.3 | 1.39E-01 | -0.7 | 1.34E-02 | 0.3 | 2.35E-01 | 0.0 | 9.93E-01 | 0.0 | 9.35E-01 | 0.0 | 9.23E-01 | Lipoxygenase 4, chloroplastic |
| AT2G01530 | -0.3 | 3.92E-03 | 0.0 | 8.85E-01 | -0.3 | 1.08E-02 | -2.1 | 6.87E-07 | -0.3 | 2.64E-01 | -1.8 | 2.49E-04 | MLP-like protein 329 |
| AT2G46590 | -0.3 | 8.75E-03 | -0.6 | 4.22E-04 | 0.3 | 6.59E-02 | 0.2 | 4.77E-01 | 0.2 | 3.51E-01 | -0.1 | 8.43E-01 | Dof zinc finger protein |
| AT1G03750 | -0.3 | 1.35E-02 | -0.5 | 1.84E-03 | 0.2 | 2.42E-01 | -0.4 | 2.59E-01 | -0.1 | 7.65E-01 | -0.3 | 5.14E-01 | Switch 2 |
| AT4G28570 | -0.3 | 4.11E-04 | -0.4 | 1.11E-04 | 0.1 | 1.94E-01 | 0.1 | 7.54E-01 | 0.1 | 5.44E-01 | -0.1 | 8.05E-01 | Long-chain-alcohol oxidase FAO4B |
| AT3G52820 | -0.3 | 1.04E-01 | -0.4 | 4.42E-02 | 0.1 | 6.62E-01 | -0.2 | 7.71E-01 | 0.2 | 8.20E-01 | -0.4 | 6.32E-01 | Purple acid phosphatase 22 |
| AT1G59870 | -0.3 | 7.57E-03 | -0.3 | 1.46E-02 | 0.0 | 8.50E-01 | 0.2 | 2.13E-01 | -0.2 | 3.66E-01 | 0.4 | 9.21E-02 | ABC transporter G family member 36 |
| AT3G20090 | -0.3 | 3.29E-03 | -0.2 | 4.46E-02 | -0.1 | 2.45E-01 | -0.3 | 2.81E-01 | -0.1 | 7.85E-01 | -0.2 | 6.20E-01 | Cytochrome P450, family 705, subfamily A, polypeptide 18 |
| AT4G11660 | -0.3 | 1.42E-02 | -0.5 | 2.89E-03 | 0.1 | 3.52E-01 | -0.5 | 2.33E-01 | -0.1 | 8.35E-01 | -0.4 | 4.74E-01 | Heat stress transcription factor B-2b |
| AT1G60140 | -0.3 | 8.65E-03 | -0.1 | 4.10E-01 | -0.2 | 6.46E-02 | -0.8 | 1.61E-03 | -0.7 | 6.67E-03 | -0.1 | 7.99E-01 | Trehalose phosphate synthase |

| | | | | | | | | | | | | | |
|---|---|---|---|---|---|---|---|---|---|---|---|---|---|
| AT5G63160 | -0.3 | 8.99E-02 | -0.1 | 4.73E-01 | -0.2 | 3.89E-01 | -1.8 | 1.08E-03 | -2.6 | 2.38E-04 | 0.9 | 2.05E-01 | BTB and TAZ domain protein 1 |
| AT3G25510 | -0.3 | 2.86E-02 | -0.8 | 3.91E-04 | 0.5 | 2.26E-02 | 0.4 | 3.47E-01 | 0.0 | 9.97E-01 | 0.4 | 4.52E-01 | ADP-ribosyl cyclase |
| AT5G52230 | -0.3 | 2.91E-02 | -0.5 | 6.56E-03 | 0.1 | 4.09E-01 | 0.0 | 9.25E-01 | 0.1 | 7.91E-01 | -0.2 | 7.16E-01 | Methyl-CpG-binding domain-containing protein 13 |
| AT5G16630 | -0.3 | 2.74E-03 | -0.5 | 4.66E-04 | 0.1 | 2.04E-01 | -0.4 | 4.42E-02 | -0.2 | 3.75E-01 | -0.2 | 4.57E-01 | DNA repair protein Rad4 family |
| AT3G55050 | -0.3 | 2.29E-03 | -0.2 | 1.65E-02 | -0.1 | 3.71E-01 | -0.3 | 2.49E-02 | -0.1 | 8.00E-01 | -0.3 | 1.60E-01 | Probable protein phosphatase 2C 48 |
| AT3G47500 | -0.3 | 1.26E-02 | -0.3 | 2.41E-02 | 0.0 | 8.42E-01 | -0.5 | 1.47E-01 | -0.2 | 6.10E-01 | -0.3 | 5.78E-01 | Cyclic dof factor 3 |
| AT1G27130 | -0.3 | 8.36E-03 | 0.2 | 9.36E-02 | -0.5 | 1.48E-03 | 0.3 | 8.16E-02 | 0.1 | 6.92E-01 | 0.2 | 3.47E-01 | Glutathione S-transferase U13 |
| AT3G11280 | -0.3 | 7.10E-04 | -0.1 | 9.77E-02 | -0.2 | 2.47E-02 | 0.2 | 8.22E-02 | 0.1 | 7.98E-01 | 0.2 | 3.05E-01 | Duplicated homeodomain-like superfamily protein ) |
| AT3G02460 | -0.3 | 1.32E-03 | -0.2 | 5.89E-02 | -0.2 | 7.89E-02 | -0.1 | 5.06E-01 | 0.0 | 8.47E-01 | -0.1 | 4.67E-01 | Ypt |
| AT1G72840 | -0.3 | 5.63E-03 | -0.2 | 4.35E-02 | -0.1 | 3.81E-01 | -0.3 | 4.60E-01 | 0.0 | 9.65E-01 | -0.2 | 6.20E-01 | Disease resistance protein |
| AT3G54810 | -0.3 | 1.09E-01 | -0.6 | 1.44E-02 | 0.2 | 2.76E-01 | -0.4 | 1.12E-02 | -0.1 | 7.11E-01 | -0.3 | 9.61E-02 | GATA transcription factor 8 |
| AT3G07980 | -0.3 | 7.55E-04 | -0.5 | 7.34E-05 | 0.2 | 5.89E-02 | 0.2 | 5.85E-01 | 0.0 | 9.75E-01 | 0.2 | 6.08E-01 | Mitogen-activated protein kinase kinase kinase 6 |
| AT1G17170 | -0.3 | 2.28E-01 | -1.4 | 8.72E-04 | 1.1 | 9.40E-03 | 4.0 | 1.53E-07 | 2.0 | 7.81E-04 | 2.0 | 5.09E-05 | Glutathione S-transferase U24 |
| AT5G07140 | -0.3 | 5.97E-02 | 0.2 | 2.00E-01 | -0.5 | 1.76E-02 | 0.1 | 8.13E-01 | 0.0 | 9.96E-01 | 0.1 | 8.56E-01 | Protein kinase superfamily protein |
| AT4G38240 | -0.3 | 5.14E-04 | -0.2 | 1.23E-02 | -0.1 | 1.17E-01 | -0.2 | 1.96E-01 | -0.1 | 7.57E-01 | -0.1 | 5.40E-01 | Alpha-1,3-mannosyl-glycoprotein 2-beta-N-acetylglucosaminyltransferase |
| AT4G13940 | -0.3 | 8.00E-03 | -0.1 | 6.19E-01 | -0.3 | 3.59E-02 | -0.1 | 6.84E-01 | 0.1 | 7.09E-01 | -0.1 | 5.44E-01 | adenosylhomocysteinase |
| AT1G09710 | -0.3 | 2.66E-02 | -0.2 | 7.97E-02 | -0.1 | 6.82E-01 | -0.2 | 3.68E-01 | -0.4 | 1.69E-01 | 0.2 | 7.06E-01 | Homeodomain-like superfamily protein |
| AT4G25350 | -0.3 | 5.31E-03 | -0.3 | 4.47E-03 | 0.0 | 8.43E-01 | -0.2 | 6.17E-01 | -0.1 | 7.36E-01 | 0.0 | 9.55E-01 | Phosphate transporter PHO1 homolog 4 |
| AT2G36260 | -0.3 | 2.04E-01 | 0.8 | 4.36E-03 | -1.1 | 3.35E-03 | | | | | | | Iron-sulfur assembly protein IscA-like 3, mitochondrial |
| AT3G21215 | -0.3 | 1.00E-03 | -0.3 | 2.06E-03 | 0.0 | 8.26E-01 | 0.0 | 9.15E-01 | -0.1 | 8.21E-01 | 0.1 | 7.52E-01 | AT3G21215 protein family protein) |
| AT1G80290 | -0.3 | 5.24E-03 | -0.1 | 1.96E-01 | -0.2 | 1.06E-01 | 0.0 | 8.08E-01 | 0.0 | 9.86E-01 | 0.1 | 8.60E-01 | Glycosyltransferase family protein 64 C3 |
| AT3G08530 | -0.3 | 5.61E-04 | -0.2 | 6.97E-03 | -0.1 | 2.13E-01 | 0.0 | 8.57E-01 | -0.2 | 3.74E-01 | 0.1 | 5.93E-01 | Clathrin heavy chain 2 |
| AT3G14980 | -0.3 | 9.39E-04 | -0.1 | 1.55E-01 | -0.2 | 2.62E-02 | -0.2 | 2.43E-01 | -0.3 | 2.30E-01 | 0.0 | 9.26E-01 | Acyl-CoA N-acyltransferase with RING |
| AT4G29490 | -0.3 | 5.15E-04 | -0.4 | 1.86E-04 | 0.1 | 3.08E-01 | -0.1 | 4.49E-01 | -0.1 | 4.83E-01 | 0.0 | 9.83E-01 | Aminopeptidase P N-terminal domain-containing protein |
| AT1G64460 | -0.3 | 1.33E-03 | -0.2 | 1.67E-02 | -0.1 | 2.33E-01 | -0.1 | 6.32E-01 | 0.0 | 9.41E-01 | -0.1 | 8.16E-01 | Phosphatidylinositol 4-kinase gamma 2 |
| AT3G56690 | -0.3 | 6.81E-03 | -0.4 | 2.00E-03 | 0.1 | 4.24E-01 | 0.0 | 9.68E-01 | -0.3 | 3.70E-01 | 0.3 | 3.75E-01 | Calmodulin-interacting protein 111 |
| AT2G41870 | -0.3 | 1.93E-02 | -0.1 | 3.24E-01 | -0.2 | 1.68E-01 | -0.5 | 2.94E-02 | -0.1 | 7.17E-01 | -0.4 | 1.56E-01 | Remorin 4.2 |
| AT2G46030 | -0.3 | 1.63E-03 | -0.2 | 4.64E-02 | -0.1 | 1.13E-01 | -0.4 | 1.20E-02 | -0.1 | 5.85E-01 | -0.3 | 1.86E-01 | Ubiquitin-conjugating enzyme 6 |
| AT3G59100 | -0.3 | 3.48E-03 | -0.3 | 5.69E-03 | 0.0 | 9.23E-01 | 0.1 | 7.85E-01 | 0.0 | 9.13E-01 | 0.0 | 9.34E-01 | 1,3-beta-glucan synthase |
| AT1G64530 | -0.3 | 1.36E-03 | -0.2 | 6.39E-02 | -0.2 | 7.01E-02 | -0.4 | 4.95E-02 | -0.3 | 2.41E-01 | -0.1 | 6.47E-01 | Plant regulator RWP-RK family protein |
| AT3G11040 | -0.3 | 5.75E-02 | -0.2 | 2.03E-01 | -0.1 | 5.95E-01 | -0.1 | 6.84E-01 | -0.3 | 3.48E-01 | 0.2 | 6.97E-01 | mannosyl-glycoprotein endo-beta-N-acetylglucosaminidase |
| AT3G49530 | -0.3 | 7.76E-03 | -0.6 | 3.73E-04 | 0.2 | 5.96E-02 | 0.1 | 6.39E-01 | -0.1 | 7.91E-01 | 0.1 | 5.45E-01 | NAC domain containing protein 62 |
| AT1G09020 | -0.3 | 1.17E-03 | -0.3 | 5.22E-03 | -0.1 | 4.96E-01 | -0.1 | 8.26E-01 | 0.1 | 8.17E-01 | -0.1 | 7.15E-01 | Sucrose nonfermenting 4-like protein |
| AT4G35580 | -0.3 | 9.50E-04 | -0.5 | 6.64E-05 | 0.2 | 3.60E-02 | -0.2 | 9.56E-02 | 0.0 | 9.53E-01 | -0.2 | 2.06E-01 | NAC transcription factor-like 9 |
| AT3G56370 | -0.3 | 1.79E-03 | -0.1 | 1.52E-01 | -0.2 | 4.14E-02 | -0.3 | 3.76E-02 | 0.0 | 8.79E-01 | -0.3 | 1.79E-01 | Probable LRR receptor-like serine |
| AT2G28940 | -0.3 | 5.06E-03 | -0.4 | 9.77E-04 | 0.1 | 2.74E-01 | -0.2 | 3.43E-01 | -0.3 | 1.21E-01 | 0.1 | 6.51E-01 | At2g28940 |
| AT5G52210 | -0.3 | 5.27E-03 | -0.2 | 2.37E-02 | -0.1 | 5.31E-01 | -0.2 | 2.45E-01 | -0.1 | 8.18E-01 | -0.1 | 5.73E-01 | At5g52210 |
| AT4G32285 | -0.3 | 6.52E-04 | -0.2 | 4.11E-02 | -0.2 | 4.80E-02 | -0.2 | 3.29E-01 | -0.2 | 2.95E-01 | 0.0 | 9.08E-01 | hypothetical protein |
| AT1G77610 | -0.3 | 2.50E-04 | -0.2 | 2.23E-02 | -0.2 | 2.84E-02 | 0.1 | 4.22E-01 | 0.3 | 5.09E-02 | -0.2 | 3.50E-01 | UDP-galactose transporter 1 |
| AT2G39250 | -0.3 | 2.10E-01 | 0.1 | 6.80E-01 | -0.4 | 1.58E-01 | -1.3 | 2.29E-03 | -0.5 | 2.01E-01 | -0.8 | 1.49E-01 | Integrase-type DNA-binding superfamily protein |
| AT5G04160 | -0.3 | 9.30E-03 | 0.1 | 3.89E-01 | -0.4 | 6.34E-03 | 0.4 | 2.93E-02 | 0.5 | 9.47E-03 | -0.1 | 6.08E-01 | UDP-URONIC ACID TRANSPORTER 1 |
| AT3G27110 | -0.3 | 3.75E-04 | -0.6 | 1.24E-05 | 0.3 | 5.38E-03 | -0.3 | 1.57E-01 | -0.1 | 7.65E-01 | -0.2 | 4.94E-01 | Peptidase family M48 family protein |
| AT3G26840 | -0.3 | 1.44E-02 | -1.0 | 3.39E-05 | 0.6 | 1.65E-03 | -0.1 | 8.54E-01 | 0.0 | 9.55E-01 | 0.0 | 9.50E-01 | Phytyl ester synthase 2, chloroplastic |
| AT5G23870 | -0.3 | 3.65E-03 | 0.1 | 4.61E-01 | -0.4 | 4.20E-03 | -0.3 | 1.94E-01 | -0.2 | 4.10E-01 | -0.1 | 8.32E-01 | Pectin acetylesterase |
| AT4G23850 | -0.3 | 8.91E-04 | -0.3 | 7.10E-04 | 0.0 | 6.73E-01 | -0.1 | 5.69E-01 | -0.1 | 3.81E-01 | 0.1 | 8.22E-01 | Long chain acyl-CoA synthetase 4 |
| AT5G48940 | -0.3 | 1.00E-01 | 0.2 | 3.99E-01 | -0.5 | 4.70E-02 | -0.4 | 1.04E-01 | 0.0 | 9.14E-01 | -0.4 | 1.59E-01 | LRR receptor-like serine |
| AT2G16250 | -0.3 | 8.86E-03 | -0.3 | 8.02E-03 | 0.0 | 8.66E-01 | -0.7 | 4.41E-01 | 0.0 | 9.95E-01 | -0.7 | 5.75E-01 | Leucine-rich repeat protein kinase family protein |
| AT1G09210 | -0.3 | 2.91E-02 | 0.1 | 5.90E-01 | -0.4 | 1.97E-02 | 0.1 | 2.39E-01 | 0.2 | 6.78E-02 | -0.1 | 5.80E-01 | Calreticulin-2 |
| AT3G04850 | -0.3 | 2.33E-02 | -0.2 | 9.76E-02 | -0.1 | 5.75E-01 | 0.5 | 7.70E-02 | 0.4 | 2.84E-01 | 0.2 | 6.63E-01 | Protein tesmin |
| AT4G39910 | -0.3 | 6.53E-04 | -0.2 | 9.58E-03 | -0.1 | 1.79E-01 | -0.2 | 7.43E-02 | -0.3 | 2.60E-02 | 0.1 | 6.44E-01 | hypothetical protein |
| AT4G10590 | -0.3 | 1.28E-03 | -0.4 | 4.80E-04 | 0.1 | 3.96E-01 | 0.0 | 8.35E-01 | -0.1 | 6.96E-01 | 0.1 | 5.55E-01 | Ubiquitin carboxyl-terminal hydrolase 9 |
| AT2G26730 | -0.3 | 1.04E-03 | -0.2 | 2.94E-02 | -0.1 | 1.06E-01 | -0.2 | 3.30E-01 | 0.0 | 9.27E-01 | -0.2 | 4.34E-01 | Probable inactive receptor kinase At2g26730 |
| AT1G76380 | -0.3 | 1.10E-02 | -0.2 | 7.09E-02 | -0.1 | 4.23E-01 | -0.3 | 1.97E-01 | 0.1 | 7.79E-01 | -0.4 | 1.86E-01 | DNA-binding bromodomain-containing protein |
| AT5G42980 | -0.3 | 2.06E-03 | -0.1 | 4.48E-01 | -0.2 | 1.60E-02 | 0.0 | 9.97E-01 | 0.1 | 7.01E-01 | -0.1 | 7.53E-01 | Thioredoxin H3 |
| AT5G41340 | -0.3 | 3.77E-02 | -0.2 | 1.54E-01 | -0.1 | 5.26E-01 | -0.3 | 1.26E-01 | -0.2 | 4.80E-01 | -0.1 | 6.80E-01 | Ubiquitin conjugating enzyme 4 |
| AT5G17570 | -0.3 | 1.27E-01 | -0.2 | 2.70E-01 | -0.1 | 7.54E-01 | 0.3 | 4.15E-01 | 0.4 | 3.83E-01 | -0.1 | 9.18E-01 | hypothetical protein |
| AT2G47190 | -0.3 | 3.53E-01 | -0.5 | 1.93E-01 | 0.2 | 7.33E-01 | | | | | | | ATMYB2 ) |
| AT4G18610 | -0.3 | 4.40E-03 | -0.1 | 3.20E-01 | -0.2 | 5.88E-02 | -0.7 | 2.56E-02 | -0.1 | 8.73E-01 | -0.6 | 6.82E-02 | Protein LIGHT-DEPENDENT SHORT HYPOCOTYLS 9 |
| AT1G08280 | -0.3 | 1.28E-02 | 0.1 | 3.05E-01 | -0.4 | 7.24E-03 | 0.4 | 9.65E-02 | 0.1 | 5.89E-01 | 0.2 | 5.08E-01 | Beta-1,6-galactosyltransferase GALT29A |
| AT1G32790 | -0.3 | 6.67E-04 | -0.2 | 6.81E-03 | -0.1 | 2.56E-01 | -0.1 | 1.79E-01 | -0.1 | 2.57E-01 | 0.0 | 9.92E-01 | CTC-interacting domain 11 |
| AT5G48110 | -0.3 | 2.15E-01 | 0.4 | 1.12E-01 | -0.7 | 2.89E-02 | -0.4 | 1.32E-01 | -0.1 | 7.45E-01 | -0.3 | 4.46E-01 | Inactive terpenoid synthase 20, chloroplastic |
| AT5G50400 | -0.3 | 8.30E-03 | -0.2 | 4.84E-02 | -0.1 | 4.46E-01 | 0.1 | 5.69E-01 | 0.2 | 4.04E-01 | -0.1 | 8.13E-01 | Purple acid phosphatase |
| AT5G45800 | -0.3 | 6.65E-03 | -0.1 | 1.56E-01 | -0.2 | 1.35E-01 | 0.4 | 2.11E-02 | 0.2 | 2.28E-01 | 0.1 | 5.31E-01 | Calmodulin-binding receptor kinase CaMRLK |
| AT3G63150 | -0.3 | 9.19E-04 | -0.2 | 1.11E-02 | -0.1 | 2.22E-01 | 0.0 | 7.91E-01 | 0.0 | 9.64E-01 | 0.0 | 8.32E-01 | Mitochondrial Rho GTPase 2 |
| AT5G55910 | -0.3 | 4.52E-04 | -0.2 | 3.41E-03 | -0.1 | 3.14E-01 | -0.5 | 2.13E-04 | -0.4 | 5.91E-03 | -0.1 | 4.45E-01 | Serine |
| AT5G07010 | -0.3 | 1.89E-02 | -0.7 | 2.40E-04 | 0.4 | 1.77E-02 | -0.1 | 9.38E-01 | -0.1 | 9.30E-01 | 0.0 | 9.85E-01 | Cytosolic sulfotransferase 15 |
| AT3G26170 | -0.3 | 1.70E-01 | -0.4 | 1.23E-01 | 0.1 | 8.47E-01 | 0.1 | 8.45E-01 | -0.1 | 8.63E-01 | 0.2 | 7.56E-01 | Cytochrome P450 71B19 |
| AT5G05700 | -0.3 | 5.83E-04 | -0.3 | 1.36E-03 | 0.0 | 7.82E-01 | -0.2 | 1.20E-01 | 0.0 | 9.49E-01 | -0.2 | 2.32E-01 | Arginyl-tRNA--protein transferase |
| AT5G26210 | -0.3 | 7.03E-04 | -0.2 | 1.10E-02 | -0.1 | 1.70E-01 | -0.3 | 2.73E-02 | -0.3 | 7.17E-02 | 0.0 | 9.09E-01 | PHD finger protein ALFIN-LIKE |
| AT3G51730 | -0.3 | 2.36E-03 | -0.3 | 3.42E-03 | 0.0 | 9.95E-01 | -0.2 | 9.53E-02 | -0.2 | 1.76E-01 | 0.0 | 9.55E-01 | AT3g51730 |
| AT5G58510 | -0.3 | 1.20E-03 | -0.3 | 2.52E-03 | 0.0 | 8.25E-01 | -0.2 | 1.93E-01 | -0.3 | 8.54E-02 | 0.1 | 6.74E-01 | Rab3 GTPase-activating protein catalytic subunit |
| AT3G27890 | -0.3 | 1.59E-03 | 0.0 | 7.05E-01 | -0.3 | 6.93E-03 | 0.0 | 8.57E-01 | 0.1 | 6.53E-01 | -0.1 | 5.57E-01 | NADPH:quinone oxidoreductase |
| AT1G14830 | -0.3 | 8.33E-04 | -0.2 | 1.77E-02 | -0.1 | 1.33E-01 | -0.1 | 3.27E-01 | 0.1 | 8.05E-01 | -0.2 | 3.59E-01 | Phragmoplastin DRP1C |
| AT3G44880 | -0.3 | 2.12E-03 | -0.5 | 7.55E-05 | 0.2 | 1.81E-02 | -0.2 | 2.84E-01 | -0.1 | 7.79E-01 | -0.1 | 6.43E-01 | Pheophorbide a oxygenase, chloroplastic |
| AT1G19100 | -0.3 | 1.33E-03 | -0.3 | 4.47E-03 | 0.0 | 6.39E-01 | 0.1 | 8.68E-01 | 0.0 | 9.22E-01 | 0.1 | 8.19E-01 | MORC6 |
| AT4G24560 | -0.3 | 1.66E-03 | -0.3 | 1.09E-03 | 0.0 | 6.66E-01 | -0.1 | 3.87E-01 | -0.1 | 7.46E-01 | -0.1 | 7.94E-01 | Ubiquitin carboxyl-terminal hydrolase 16 |
| AT4G25880 | -0.3 | 1.01E-03 | -0.5 | 7.21E-05 | 0.2 | 3.46E-02 | -0.1 | 3.03E-01 | -0.1 | 3.66E-01 | 0.0 | 9.95E-01 | Pumilio homolog 6, chloroplastic |
| AT5G40270 | -0.3 | 4.91E-03 | -0.3 | 4.65E-03 | 0.0 | 8.81E-01 | -0.3 | 2.36E-01 | -0.1 | 8.38E-01 | -0.2 | 3.69E-01 | AT5g40270 |
| AT4G14710 | -0.3 | 8.82E-04 | -0.3 | 9.62E-04 | 0.0 | 8.65E-01 | -0.1 | 2.81E-01 | -0.2 | 2.36E-01 | 0.0 | 8.85E-01 | Acireductone dioxygenase -requiring)) -requiring)) |
| AT3G26810 | -0.3 | 9.70E-04 | -0.1 | 1.57E-01 | -0.2 | 2.14E-02 | -0.3 | 3.43E-02 | -0.3 | 3.24E-02 | 0.0 | 8.87E-01 | Protein AUXIN SIGNALING F-BOX 2 |
| AT4G32010 | -0.3 | 6.40E-03 | -0.4 | 3.55E-03 | 0.0 | 6.68E-01 | 0.0 | 9.94E-01 | 0.1 | 5.14E-01 | -0.1 | 5.91E-01 | B3 domain-containing transcription repressor VAL2 |
| AT4G18030 | -0.3 | 3.21E-03 | -0.3 | 6.80E-03 | 0.0 | 8.25E-01 | -0.1 | 3.15E-01 | 0.1 | 5.99E-01 | -0.2 | 1.82E-01 | Probable methyltransferase PMT14 |
| AT3G05190 | -0.3 | 1.37E-01 | -0.3 | 1.20E-01 | 0.0 | 9.01E-01 | 0.2 | 6.55E-01 | 0.5 | 2.23E-01 | -0.3 | 4.54E-01 | Branched-chain-amino-acid aminotransferase-like protein 1 |

| | | | | | | | | | | | | | |
|---|---|---|---|---|---|---|---|---|---|---|---|---|---|
| AT5G67190 | -0.3 | 1.03E-02 | 0.0 | 6.46E-01 | -0.3 | 5.19E-02 | -0.6 | 2.44E-03 | -0.6 | 4.23E-03 | 0.0 | 9.91E-01 | Ethylene-responsive transcription factor ERF010 |
| AT5G04930 | -0.3 | 4.80E-03 | -0.4 | 1.50E-03 | 0.1 | 3.91E-01 | 0.5 | 1.52E-02 | 0.1 | 8.47E-01 | 0.4 | 9.22E-02 | Phospholipid-transporting ATPase 1 |
| AT1G80940 | -0.3 | 2.24E-03 | -0.2 | 1.22E-02 | -0.1 | 4.52E-01 | -0.2 | 2.23E-01 | -0.2 | 2.39E-01 | 0.0 | 9.48E-01 | Snf1 kinase interactor-like protein |
| AT1G18150 | -0.3 | 7.88E-04 | -0.2 | 9.64E-03 | -0.1 | 2.17E-01 | -0.2 | 6.88E-02 | 0.2 | 1.44E-01 | -0.4 | 1.31E-02 | Mitogen-activated protein kinase 8 |
| AT4G20860 | -0.3 | 1.33E-01 | -0.5 | 3.26E-02 | 0.2 | 4.42E-01 | 0.7 | 2.55E-04 | 0.1 | 8.04E-01 | 0.7 | 4.60E-03 | Berberine bridge enzyme-like 22 |
| AT4G18360 | -0.3 | 5.89E-03 | -0.2 | 9.93E-02 | -0.1 | 2.01E-01 | 1.0 | 6.10E-04 | 0.7 | 1.06E-02 | 0.2 | 2.57E-01 | Aldolase-type TIM barrel family protein |
| AT1G33560 | -0.3 | 9.06E-03 | -0.8 | 3.65E-05 | 0.5 | 2.42E-03 | 0.0 | 9.40E-01 | -0.2 | 6.06E-01 | 0.1 | 7.24E-01 | Disease resistance protein ADR1 |
| AT1G08620 | -0.3 | 2.15E-03 | -0.5 | 9.91E-05 | 0.2 | 3.04E-02 | 0.0 | 9.27E-01 | 0.1 | 8.46E-01 | 0.0 | 9.30E-01 | Transcription factor jumonji family protein |
| AT3G46500 | -0.3 | 2.28E-02 | -0.2 | 1.12E-01 | -0.1 | 5.18E-01 | -0.5 | 1.98E-01 | 0.1 | 8.41E-01 | -0.6 | 1.99E-01 | 2-oxoglutarate and Fe-dependent oxygenase superfamily protein |
| AT1G77680 | -0.3 | 4.02E-03 | -0.2 | 5.29E-02 | -0.1 | 2.38E-01 | -0.2 | 5.85E-02 | -0.1 | 6.31E-01 | -0.2 | 4.00E-01 | Inactive exonuclease DIS3L2 |
| AT3G61740 | -0.3 | 5.59E-03 | -0.5 | 5.25E-04 | 0.2 | 1.28E-01 | -0.2 | 6.07E-01 | 0.3 | 4.44E-01 | -0.5 | 2.48E-01 | Histone-lysine N-methyltransferase ATX3 |
| AT2G14720 | -0.3 | 8.25E-04 | -0.4 | 4.64E-04 | 0.1 | 5.00E-01 | 0.1 | 3.73E-01 | 0.1 | 3.97E-01 | 0.0 | 9.65E-01 | Vacuolar-sorting receptor 4 |
| AT1G15750 | -0.3 | 3.17E-03 | -0.3 | 3.85E-03 | 0.0 | 9.46E-01 | 0.0 | 7.96E-01 | -0.2 | 3.45E-01 | 0.1 | 5.97E-01 | Protein TOPLESS |
| AT1G45231 | -0.3 | 8.78E-03 | -0.6 | 3.41E-04 | 0.3 | 5.30E-02 | 0.2 | 4.06E-01 | 0.4 | 1.64E-01 | -0.2 | 6.10E-01 | Trimethylguanosine synthase |
| AT5G65930 | -0.3 | 2.87E-03 | 0.0 | 8.53E-01 | -0.3 | 9.29E-03 | -0.1 | 3.53E-01 | -0.1 | 6.79E-01 | -0.1 | 8.07E-01 | Kinesin-like calmodulin-binding protein |
| AT5G67160 | -0.3 | 2.53E-02 | -0.1 | 4.41E-01 | -0.2 | 1.72E-01 | 0.0 | 9.41E-01 | 0.5 | 1.99E-02 | -0.5 | 5.34E-02 | Protein ENHANCED PSEUDOMONAS SUSCEPTIBILITY 1 |
| AT1G78540 | -0.3 | 2.19E-02 | -0.1 | 4.10E-01 | -0.2 | 1.73E-01 | -0.1 | 8.73E-01 | -0.2 | 6.15E-01 | 0.1 | 7.59E-01 | SH2 domain-containing protein B |
| AT1G10130 | -0.3 | 5.16E-04 | -0.2 | 1.50E-02 | -0.1 | 9.35E-02 | -0.1 | 5.37E-01 | 0.0 | 9.88E-01 | -0.1 | 6.64E-01 | Calcium-transporting ATPase 3, endoplasmic reticulum-type |
| AT5G43430 | -0.3 | 6.46E-03 | -0.1 | 2.41E-01 | -0.2 | 9.76E-02 | -0.2 | 1.92E-01 | 0.0 | 9.38E-01 | -0.2 | 3.06E-01 | Electron transfer flavoprotein subunit beta |
| AT4G38200 | -0.3 | 1.13E-03 | -0.2 | 2.13E-02 | -0.1 | 1.53E-01 | 0.0 | 8.07E-01 | 0.0 | 9.76E-01 | -0.1 | 8.54E-01 | SEC7-like guanine nucleotide exchange family protein |
| AT1G18300 | -0.3 | 8.37E-02 | -0.3 | 8.12E-02 | 0.0 | 9.56E-01 | -0.2 | 5.84E-01 | -0.5 | 1.33E-01 | 0.3 | 4.52E-01 | Nudix hydrolase 4 |
| AT5G13010 | -0.3 | 2.83E-03 | -0.4 | 3.93E-04 | 0.1 | 1.48E-01 | -0.1 | 4.11E-01 | -0.1 | 5.57E-01 | 0.0 | 9.59E-01 | Pre-mRNA-splicing factor ATP-dependent RNA helicase DEAH7 |
| AT3G01300 | -0.3 | 5.79E-04 | -0.2 | 6.56E-03 | -0.1 | 2.34E-01 | -0.4 | 4.75E-03 | -0.3 | 3.23E-02 | -0.1 | 6.98E-01 | Serine |
| AT1G08130 | -0.3 | 1.56E-03 | -0.2 | 1.61E-02 | -0.1 | 2.77E-01 | 0.6 | 4.87E-03 | 0.5 | 2.49E-02 | 0.1 | 7.49E-01 | DNA ligase |
| AT1G68000 | -0.3 | 4.89E-04 | -0.1 | 2.26E-01 | -0.2 | 7.94E-03 | -0.3 | 8.23E-02 | -0.2 | 3.13E-01 | -0.1 | 7.42E-01 | CDP-diacylglycerol--inositol 3-phosphatidyltransferase 1 |
| AT3G54250 | -0.3 | 1.05E-02 | -0.1 | 2.46E-01 | -0.2 | 1.51E-01 | 0.0 | 9.27E-01 | 0.0 | 8.76E-01 | -0.1 | 8.45E-01 | Diphosphomevalonate decarboxylase |
| AT4G38790 | -0.3 | 7.76E-03 | -0.1 | 1.74E-01 | -0.2 | 1.42E-01 | 0.1 | 2.51E-01 | 0.1 | 4.41E-01 | 0.0 | 9.09E-01 | ER lumen protein retaining receptor family protein |
| AT2G01890 | -0.3 | 5.14E-02 | -0.5 | 5.42E-03 | 0.2 | 2.25E-01 | -0.2 | 4.41E-01 | 0.2 | 4.86E-01 | -0.5 | 5.86E-02 | acid phosphatase |
| AT4G13020 | -0.3 | 1.22E-03 | -0.2 | 2.66E-02 | -0.1 | 1.35E-01 | -0.4 | 4.05E-02 | 0.1 | 8.21E-01 | -0.5 | 5.10E-02 | Serine |
| AT3G25730 | -0.3 | 6.33E-03 | -0.4 | 2.44E-03 | 0.1 | 5.25E-01 | 1.3 | 1.37E-03 | 0.9 | 3.55E-02 | 0.4 | 2.15E-01 | AP2 |
| AT4G27630 | -0.3 | 1.07E-02 | -0.2 | 7.72E-02 | -0.1 | 4.04E-01 | 0.0 | 8.76E-01 | 0.0 | 8.91E-01 | -0.1 | 8.01E-01 | GPCR-type G protein 2 |
| AT2G05320 | -0.3 | 4.63E-03 | -0.2 | 1.00E-01 | -0.1 | 1.59E-01 | 0.1 | 7.72E-01 | 0.1 | 7.10E-01 | 0.0 | 9.27E-01 | Alpha-1,6-mannosyl-glycoprotein 2-beta-N-acetylglucosaminyltransferase |
| AT4G14680 | -0.3 | 9.02E-03 | 0.0 | 8.08E-01 | -0.3 | 2.90E-02 | 0.6 | 3.92E-03 | 0.5 | 1.98E-02 | 0.1 | 7.64E-01 | sulfate adenylyltransferase |
| AT1G61690 | -0.3 | 9.75E-04 | -0.3 | 6.56E-04 | 0.0 | 6.40E-01 | -0.2 | 4.18E-01 | -0.2 | 5.23E-01 | 0.0 | 9.78E-01 | At1g61690 |
| AT5G04250 | -0.3 | 3.33E-02 | -0.3 | 4.31E-02 | 0.0 | 9.80E-01 | 0.0 | 9.73E-01 | -0.2 | 6.16E-01 | 0.2 | 6.96E-01 | OVARIAN TUMOR DOMAIN-containing deubiquitinating enzyme 9 |
| AT3G49360 | -0.3 | 7.10E-03 | -0.3 | 1.23E-02 | 0.0 | 8.80E-01 | -0.3 | 4.38E-02 | -0.2 | 1.24E-01 | 0.0 | 8.52E-01 | Probable 6-phosphogluconolactonase 2 |
| AT5G20700 | -0.3 | 3.50E-03 | -0.1 | 4.61E-01 | -0.2 | 2.47E-02 | 0.0 | 6.70E-01 | 0.3 | 4.97E-03 | -0.4 | 1.32E-02 | FCS-Like Zinc finger 14 |
| AT5G55060 | -0.3 | 3.45E-03 | -0.3 | 1.10E-02 | 0.0 | 6.47E-01 | -0.1 | 5.59E-01 | 0.0 | 8.70E-01 | -0.1 | 8.28E-01 | Rab3 GTPase-activating protein catalytic subunit |
| AT4G21430 | -0.3 | 1.44E-02 | -0.1 | 2.34E-01 | -0.2 | 2.08E-01 | -0.1 | 6.79E-01 | 0.0 | 9.33E-01 | -0.1 | 8.39E-01 | Lysine-specific demethylase JMJ28 |
| AT3G27950 | -0.3 | 9.39E-02 | 0.8 | 2.83E-04 | -1.1 | 2.59E-04 | 0.4 | 1.23E-01 | 0.8 | 1.12E-02 | -0.4 | 2.73E-01 | Uncharacterized protein |
| AT4G33940 | -0.3 | 4.97E-04 | -0.3 | 5.16E-04 | 0.0 | 8.21E-01 | -0.3 | 1.23E-02 | -0.5 | 1.80E-03 | 0.2 | 3.80E-01 | Uncharacterized protein At4g33940 |
| AT3G54440 | -0.3 | 4.88E-03 | -0.3 | 7.00E-03 | 0.0 | 9.56E-01 | -0.1 | 7.71E-01 | 0.0 | 9.28E-01 | -0.1 | 7.58E-01 | beta-galactosidase |
| AT1G71800 | -0.3 | 2.82E-03 | -0.2 | 1.67E-02 | -0.1 | 4.46E-01 | 0.0 | 8.56E-01 | 0.3 | 1.92E-01 | -0.3 | 3.74E-01 | Cleavage stimulating factor 64 |
| AT5G38460 | -0.3 | 1.18E-02 | 0.0 | 6.98E-01 | -0.3 | 1.63E-02 | 0.2 | 4.85E-01 | 0.3 | 3.23E-01 | -0.1 | 8.18E-01 | Probable dolichyl pyrophosphate Man9GlcNAc2 alpha-1,3-glucosyltransferase GlcNAc-PP-Dol alpha-1,3-glucosyltransferase) |
| AT3G57810 | -0.3 | 2.46E-02 | -0.2 | 2.11E-01 | -0.1 | 3.32E-01 | -0.2 | 5.06E-01 | -0.3 | 2.94E-01 | 0.1 | 7.51E-01 | OVARIAN TUMOR DOMAIN-containing deubiquitinating enzyme 4 |
| AT3G05120 | -0.3 | 7.63E-04 | -0.2 | 2.42E-02 | -0.1 | 9.78E-02 | 0.0 | 8.59E-01 | -0.2 | 3.95E-01 | 0.2 | 3.80E-01 | Gibberellin receptor GID1A |
| AT5G40830 | -0.3 | 1.15E-03 | 0.0 | 9.70E-01 | -0.3 | 3.95E-03 | 0.2 | 4.12E-01 | 0.2 | 2.51E-01 | -0.1 | 7.39E-01 | At5g40830 |
| AT5G07390 | -0.3 | 1.31E-02 | -0.5 | 5.84E-04 | 0.2 | 7.30E-02 | 0.7 | 6.56E-02 | 0.4 | 3.34E-01 | 0.3 | 5.99E-01 | Respiratory burst oxidase homolog A |
| AT2G17480 | -0.3 | 8.75E-04 | -0.3 | 1.33E-03 | 0.0 | 9.69E-01 | 0.0 | 1.00E+00 | 0.1 | 8.35E-01 | -0.1 | 8.63E-01 | MLO-like protein 8 |
| AT1G18160 | -0.3 | 2.41E-03 | -0.4 | 3.90E-04 | 0.1 | 1.70E-01 | -0.2 | 2.90E-01 | -0.2 | 4.15E-01 | 0.0 | 9.59E-01 | Protein kinase superfamily protein |
| AT5G56900 | -0.3 | 1.26E-03 | -0.3 | 1.02E-03 | 0.0 | 7.48E-01 | -0.4 | 1.20E-02 | -0.3 | 4.70E-02 | -0.1 | 8.09E-01 | CwfJ-like family protein |
| AT5G45160 | -0.3 | 1.23E-03 | -0.4 | 1.65E-04 | 0.1 | 1.13E-01 | -0.2 | 5.96E-01 | 0.0 | 9.53E-01 | -0.2 | 6.59E-01 | Protein ROOT HAIR DEFECTIVE 3 homolog 2 |
| AT1G04130 | -0.3 | 4.83E-02 | -0.2 | 1.58E-01 | -0.1 | 6.14E-01 | 0.1 | 6.48E-01 | 0.1 | 8.12E-01 | 0.0 | 9.19E-01 | At1g04130 |
| AT1G68060 | -0.3 | 9.01E-04 | 0.0 | 6.83E-01 | -0.3 | 4.41E-03 | 0.0 | 9.41E-01 | 0.1 | 3.74E-01 | -0.1 | 4.91E-01 | Microtubule-associated protein 70-1 |
| AT1G23380 | -0.3 | 3.34E-02 | -0.4 | 1.26E-02 | 0.1 | 5.79E-01 | 0.1 | 6.94E-01 | 0.1 | 8.67E-01 | 0.0 | 9.03E-01 | Homeobox protein knotted-1-like 6 |
| AT2G36350 | -0.3 | 9.11E-03 | -0.3 | 2.29E-02 | 0.0 | 7.43E-01 | -0.6 | 1.95E-04 | -0.2 | 1.66E-01 | -0.4 | 3.28E-02 | Serine |
| AT2G41140 | -0.3 | 2.06E-03 | -0.4 | 3.78E-04 | 0.1 | 1.97E-01 | 0.0 | 9.91E-01 | 0.1 | 5.01E-01 | -0.1 | 5.89E-01 | CDPK-related kinase 1 |
| AT1G10630 | -0.3 | 2.53E-02 | 0.1 | 2.90E-01 | -0.4 | 7.89E-03 | -0.1 | 5.62E-01 | -0.1 | 5.94E-01 | 0.0 | 9.79E-01 | ADP-ribosylation factor |
| AT1G67720 | -0.3 | 8.46E-03 | -0.6 | 2.21E-04 | 0.3 | 3.00E-02 | -0.2 | 5.79E-01 | 0.1 | 6.94E-01 | -0.3 | 4.15E-01 | Probable LRR receptor-like serine |
| AT5G11060 | -0.3 | 2.29E-03 | -0.6 | 5.07E-05 | 0.3 | 9.29E-03 | -0.2 | 2.67E-01 | 0.1 | 8.39E-01 | -0.2 | 3.04E-01 | Homeobox protein knotted-1-like 3 |
| AT1G12580 | -0.3 | 4.86E-03 | -0.2 | 2.34E-02 | -0.1 | 4.85E-01 | -0.3 | 3.83E-02 | -0.4 | 4.60E-02 | 0.0 | 9.44E-01 | At1g12580 |
| AT1G30400 | -0.3 | 1.19E-03 | -0.4 | 1.11E-04 | 0.2 | 5.46E-02 | 0.2 | 2.57E-01 | 0.1 | 4.94E-01 | 0.1 | 8.79E-01 | ABC transporter C family member 1 |
| AT5G46520 | -0.3 | 1.70E-02 | -0.3 | 9.76E-03 | 0.0 | 7.33E-01 | 0.6 | 1.44E-02 | 0.2 | 5.65E-01 | 0.4 | 1.09E-01 | ADP-ribosyl cyclase |
| AT1G20980 | -0.3 | 8.63E-04 | -0.4 | 7.43E-05 | 0.2 | 4.33E-02 | 0.2 | 1.48E-01 | 0.1 | 6.46E-01 | 0.1 | 5.81E-01 | Squamosa promoter-binding-like protein 14 |
| AT4G16745 | -0.3 | 9.07E-03 | -0.3 | 1.54E-02 | 0.0 | 9.05E-01 | -0.7 | 8.71E-02 | -0.3 | 4.90E-01 | -0.4 | 5.10E-01 | Exostosin family protein |
| AT3G03300 | -0.3 | 1.19E-02 | -0.5 | 1.33E-03 | 0.2 | 1.89E-01 | 0.0 | 9.86E-01 | 0.3 | 4.56E-01 | -0.2 | 4.99E-01 | Endoribonuclease Dicer homolog 2 |
| AT4G02880 | -0.3 | 2.39E-03 | -0.3 | 5.33E-03 | 0.0 | 7.99E-01 | -0.2 | 1.80E-01 | -0.3 | 4.54E-02 | 0.1 | 5.87E-01 | ELKS |
| AT4G09560 | -0.3 | 1.46E-02 | -0.1 | 2.80E-01 | -0.2 | 1.69E-01 | 0.1 | 8.53E-01 | 0.3 | 3.67E-01 | -0.2 | 5.72E-01 | Receptor homology region, transmembrane domain- and RING domain-containing protein 4 |
| AT5G06530 | -0.3 | 1.06E-02 | -0.8 | 2.58E-05 | 0.5 | 1.10E-03 | -0.7 | 5.48E-04 | -0.6 | 8.70E-03 | -0.2 | 5.32E-01 | ABC-2 type transporter family protein |
| AT2G25850 | -0.3 | 1.45E-03 | -0.2 | 1.55E-02 | -0.1 | 2.61E-01 | -0.1 | 6.38E-01 | 0.0 | 9.97E-01 | -0.1 | 7.63E-01 | polynucleotide adenylyltransferase |
| AT3G27380 | -0.3 | 5.90E-03 | 0.0 | 6.56E-01 | -0.3 | 6.09E-03 | 0.0 | 8.76E-01 | 0.3 | 2.66E-01 | -0.3 | 2.98E-01 | AT3G27380 protein |
| AT3G57420 | -0.3 | 9.89E-04 | -0.2 | 3.84E-02 | -0.1 | 8.53E-02 | 0.0 | 9.01E-01 | -0.1 | 6.63E-01 | 0.1 | 8.13E-01 | Probable glycosyltransferase STELLO2 |
| AT4G02080 | -0.3 | 5.94E-04 | -0.1 | 4.00E-02 | -0.1 | 4.45E-02 | -0.1 | 3.54E-01 | -0.2 | 2.64E-01 | 0.0 | 8.58E-01 | GTP-binding protein SAR1A |
| AT1G27630 | -0.3 | 1.56E-03 | -0.1 | 8.66E-02 | -0.2 | 6.51E-02 | 0.0 | 8.04E-01 | 0.1 | 4.90E-01 | -0.2 | 3.87E-01 | Cyclin T 13 |
| AT5G11650 | -0.3 | 3.34E-03 | -0.4 | 4.03E-04 | 0.1 | 1.39E-01 | 0.2 | 2.52E-01 | 0.1 | 8.01E-01 | 0.1 | 5.90E-01 | Alpha |
| AT1G61210 | -0.3 | 5.71E-03 | -0.5 | 2.67E-04 | 0.2 | 5.15E-02 | 0.0 | 9.82E-01 | 0.2 | 4.54E-01 | -0.2 | 5.24E-01 | At1g61220 |
| AT5G42190 | -0.3 | 3.99E-03 | -0.1 | 4.71E-01 | -0.2 | 2.67E-02 | -0.4 | 3.74E-03 | -0.3 | 2.83E-02 | -0.1 | 7.27E-01 | SKP1-like protein 1B |
| AT2G20190 | -0.3 | 4.48E-03 | -0.3 | 2.28E-03 | 0.1 | 5.96E-01 | -0.1 | 7.46E-01 | 0.0 | 9.34E-01 | 0.0 | 8.97E-01 | CLIP-associated protein |
| AT3G54100 | -0.3 | 1.14E-02 | -0.2 | 3.83E-02 | -0.1 | 6.20E-01 | 0.0 | 9.75E-01 | -0.1 | 6.61E-01 | 0.1 | 7.56E-01 | O-fucosyltransferase 28 |
| AT3G29110 | -0.3 | 2.67E-02 | 0.6 | 2.30E-04 | -0.9 | 1.25E-04 | -0.2 | 5.30E-01 | 0.4 | 1.51E-01 | -0.6 | 8.00E-02 | Uncharacterized protein |
| AT4G00660 | -0.3 | 6.92E-04 | -0.1 | 4.87E-02 | -0.2 | 4.45E-02 | -0.1 | 6.16E-01 | 0.0 | 9.57E-01 | -0.1 | 7.44E-01 | DEAD-box ATP-dependent RNA helicase 8 |
| AT4G23200 | -0.3 | 1.22E-01 | -0.5 | 1.91E-02 | 0.2 | 3.47E-01 | 0.5 | 3.69E-01 | -0.1 | 8.51E-01 | 0.6 | 3.44E-01 | Cysteine-rich RLK 12 |
| AT1G51760 | -0.3 | 1.10E-02 | -0.1 | 4.42E-01 | -0.2 | 7.38E-02 | 0.0 | 9.23E-01 | 0.0 | 9.40E-01 | 0.0 | 1.00E+00 | IAA-amino acid hydrolase ILR1-like 4 |

| | | | | | | | | | | | | | |
|---|---|---|---|---|---|---|---|---|---|---|---|---|---|
| AT4G00240 | -0.3 | 2.13E-02 | -0.3 | 2.29E-02 | 0.0 | 9.71E-01 | 0.2 | 5.55E-01 | 0.4 | 2.03E-01 | -0.2 | 5.73E-01 | Phospholipase D |
| AT3G19510 | -0.3 | 9.52E-03 | -0.3 | 2.11E-02 | 0.0 | 7.86E-01 | 0.3 | 1.49E-01 | 0.1 | 8.44E-01 | 0.2 | 3.54E-01 | Homeobox protein HAT3.1 |
| AT5G67280 | -0.3 | 3.70E-02 | -0.1 | 3.62E-01 | -0.2 | 2.50E-01 | 0.0 | 9.53E-01 | 0.4 | 3.54E-02 | -0.4 | 7.18E-02 | RLK |
| AT5G55500 | -0.3 | 2.24E-03 | -0.3 | 2.79E-03 | 0.0 | 9.71E-01 | -0.2 | 1.42E-01 | -0.3 | 7.72E-02 | 0.1 | 7.80E-01 | Beta-1,2-xylosyltransferase |
| AT5G04410 | -0.3 | 9.79E-04 | -0.4 | 1.76E-04 | 0.1 | 1.37E-01 | -0.1 | 3.48E-01 | -0.1 | 7.19E-01 | 0.0 | 7.88E-01 | NAC domain-containing protein 78 |
| AT3G02570 | -0.3 | 7.19E-03 | -0.1 | 2.19E-01 | -0.2 | 1.05E-01 | 0.1 | 5.66E-01 | 0.0 | 9.23E-01 | 0.1 | 6.33E-01 | Mannose-6-phosphate isomerase 1 |
| AT5G16820 | -0.3 | 7.45E-03 | -0.2 | 4.28E-02 | -0.1 | 4.49E-01 | -0.2 | 4.71E-01 | -0.1 | 8.22E-01 | -0.1 | 7.61E-01 | Heat stress transcription factor A-1b |
| AT2G25620 | -0.3 | 2.60E-02 | 0.1 | 6.07E-01 | -0.3 | 2.23E-02 | 0.4 | 1.13E-02 | 0.3 | 6.32E-02 | 0.1 | 7.27E-01 | Probable protein phosphatase 2C 22 |
| AT2G42200 | -0.3 | 7.93E-02 | -0.4 | 3.46E-02 | 0.1 | 6.55E-01 | 0.1 | 9.16E-01 | 0.5 | 2.58E-01 | -0.4 | 4.05E-01 | SPL9 |
| AT3G23430 | -0.3 | 1.26E-02 | -0.1 | 1.70E-01 | -0.1 | 2.20E-01 | -0.6 | 1.02E-04 | -0.4 | 1.63E-02 | -0.3 | 1.54E-01 | Phosphate transporter PHO1 |
| AT1G70980 | -0.3 | 1.54E-03 | -0.4 | 3.25E-04 | 0.1 | 2.26E-01 | -0.1 | 5.09E-01 | -0.3 | 1.33E-01 | 0.2 | 5.00E-01 | Asparagine--tRNA ligase, cytoplasmic 3 |
| AT4G17180 | -0.3 | 1.73E-02 | 0.2 | 9.04E-02 | -0.5 | 3.89E-03 | -0.1 | 8.58E-01 | -0.1 | 7.87E-01 | 0.0 | 9.31E-01 | glucan endo-1,3-beta-D-glucosidase |
| AT1G59610 | -0.3 | 1.27E-03 | -0.4 | 4.91E-04 | 0.1 | 3.75E-01 | -0.1 | 4.38E-01 | 0.0 | 9.76E-01 | -0.1 | 6.20E-01 | Dynamin-2B |
| AT5G64440 | -0.3 | 9.97E-04 | -0.3 | 6.25E-04 | 0.0 | 5.95E-01 | 0.1 | 5.57E-01 | 0.2 | 3.18E-01 | -0.1 | 7.49E-01 | Fatty acid amide hydrolase |
| AT1G24450 | -0.3 | 7.09E-03 | 0.0 | 9.63E-01 | -0.3 | 1.74E-02 | -0.1 | 4.96E-01 | -0.1 | 8.14E-01 | -0.1 | 8.26E-01 | Protein NUCLEAR FUSION DEFECTIVE 2 |
| AT2G31300 | -0.3 | 1.66E-02 | -0.2 | 6.38E-02 | -0.1 | 5.98E-01 | 0.2 | 6.15E-01 | -0.1 | 8.54E-01 | 0.3 | 5.52E-01 | Actin-related protein 2 |
| AT2G27990 | -0.3 | 9.51E-02 | -0.4 | 5.61E-02 | 0.1 | 7.55E-01 | | | | | | | BEL1-like homeodomain 8 |
| AT2G05710 | -0.3 | 7.47E-03 | -0.3 | 8.07E-03 | 0.0 | 9.23E-01 | 0.1 | 6.02E-01 | 0.4 | 2.76E-02 | -0.3 | 1.90E-01 | Aconitate hydratase 3, mitochondrial |
| AT1G53190 | -0.3 | 8.19E-03 | -0.1 | 2.01E-01 | -0.2 | 1.42E-01 | -0.4 | 1.06E-02 | -0.4 | 9.28E-03 | 0.0 | 8.62E-01 | RING-type E3 ubiquitin transferase |
| AT1G15860 | -0.3 | 1.25E-03 | -0.2 | 2.60E-02 | -0.1 | 1.48E-01 | 0.2 | 2.62E-01 | 0.1 | 4.92E-01 | 0.0 | 8.58E-01 | Defective in cullin neddylation protein |
| AT5G10870 | -0.3 | 2.79E-02 | 0.1 | 4.93E-01 | -0.4 | 2.09E-02 | -0.1 | 6.33E-01 | -0.2 | 4.67E-01 | 0.1 | 8.50E-01 | Chorismate mutase 2 |
| AT3G14100 | -0.3 | 2.20E-03 | -0.2 | 4.47E-02 | -0.1 | 1.53E-01 | -0.1 | 2.50E-01 | 0.0 | 7.67E-01 | -0.1 | 5.95E-01 | Oligouridylate-binding protein 1C -binding protein UBP1C) |
| AT3G05910 | -0.3 | 6.57E-03 | -0.1 | 4.43E-01 | -0.2 | 4.53E-02 | -0.3 | 3.20E-02 | -0.2 | 1.83E-01 | -0.1 | 6.52E-01 | Pectin acetylesterase |
| AT5G55390 | -0.3 | 3.15E-03 | -0.3 | 1.48E-03 | 0.1 | 5.55E-01 | 0.0 | 9.54E-01 | 0.1 | 6.82E-01 | -0.1 | 7.67E-01 | Protein ENHANCED DOWNY MILDEW 2 |
| AT4G23740 | -0.3 | 1.66E-02 | -0.1 | 5.83E-01 | -0.2 | 7.70E-02 | -0.3 | 1.94E-01 | 0.1 | 7.60E-01 | -0.3 | 1.89E-01 | Probable inactive receptor kinase At4g23740 |
| AT3G20630 | -0.3 | 2.16E-03 | -0.3 | 1.68E-03 | 0.0 | 7.62E-01 | 0.0 | 9.73E-01 | 0.0 | 8.51E-01 | 0.0 | 8.62E-01 | Ubiquitin carboxyl-terminal hydrolase 14 |
| AT5G55480 | -0.3 | 4.06E-03 | 0.1 | 4.66E-01 | -0.3 | 3.45E-03 | 0.0 | 8.72E-01 | 0.1 | 6.84E-01 | -0.1 | 8.47E-01 | glycerophosphodiester phosphodiesterase |
| AT3G19770 | -0.3 | 1.21E-03 | -0.2 | 2.84E-02 | -0.1 | 1.37E-01 | -0.1 | 7.24E-01 | -0.1 | 4.94E-01 | 0.1 | 8.07E-01 | Vacuolar sorting protein 9 domain-containing protein |
| AT4G23650 | -0.3 | 2.29E-03 | 0.0 | 6.66E-01 | -0.2 | 1.05E-02 | -0.2 | 6.23E-02 | 0.0 | 9.00E-01 | -0.2 | 2.70E-01 | Calcium-dependent protein kinase 3 |
| AT2G44150 | -0.3 | 3.48E-03 | -0.1 | 1.59E-01 | -0.2 | 8.62E-02 | -0.5 | 8.33E-03 | -0.4 | 7.48E-02 | -0.2 | 5.15E-01 | Histone-lysine N-methyltransferase ASHH3 |
| AT4G02460 | -0.3 | 3.92E-02 | -0.6 | 1.92E-03 | 0.3 | 1.10E-01 | 0.4 | 3.28E-01 | 0.4 | 4.04E-01 | 0.0 | 9.91E-01 | DNA mismatch repair protein |
| AT5G62680 | -0.3 | 9.55E-03 | -0.2 | 4.09E-02 | -0.1 | 5.32E-01 | -0.1 | 6.69E-01 | 0.3 | 8.33E-02 | -0.3 | 9.61E-02 | Protein NRT1 |
| AT1G74030 | -0.3 | 9.51E-03 | 0.2 | 9.19E-02 | -0.4 | 1.72E-03 | 0.2 | 4.45E-01 | 0.4 | 1.68E-01 | -0.2 | 6.40E-01 | Enolase 1, chloroplastic |
| AT3G27010 | -0.3 | 1.27E-02 | -0.4 | 3.53E-03 | 0.1 | 4.41E-01 | -0.4 | 2.95E-02 | -0.3 | 1.07E-01 | -0.1 | 7.77E-01 | Transcription factor TCP20 |
| AT1G02080 | -0.3 | 4.03E-03 | -0.5 | 1.86E-04 | 0.2 | 3.75E-02 | -0.2 | 2.80E-01 | -0.1 | 8.35E-01 | -0.2 | 6.08E-01 | Transcription regulator |
| AT1G51850 | -0.3 | 9.83E-02 | -0.2 | 2.96E-01 | -0.1 | 6.04E-01 | -0.1 | 6.28E-01 | 0.0 | 9.44E-01 | -0.2 | 6.47E-01 | Leucine-rich repeat protein kinase family protein |
| AT1G36060 | -0.3 | 7.65E-02 | 0.3 | 4.36E-02 | -0.6 | 5.61E-03 | -0.7 | 1.68E-02 | -0.2 | 6.04E-01 | -0.5 | 1.04E-01 | Ethylene-responsive transcription factor ERF055 |
| AT4G37520 | -0.3 | 1.80E-02 | -0.2 | 1.04E-01 | -0.1 | 4.21E-01 | 0.2 | 5.32E-01 | -0.4 | 1.96E-01 | 0.6 | 2.94E-02 | Peroxidase |
| AT1G22985 | -0.3 | 1.48E-02 | -0.5 | 1.48E-03 | 0.2 | 1.86E-01 | -0.4 | 2.23E-01 | -0.3 | 4.48E-01 | -0.1 | 8.19E-01 | Ethylene-responsive transcription factor ERF069 |
| AT2G44350 | -0.3 | 7.09E-03 | 0.1 | 5.89E-01 | -0.3 | 6.16E-03 | 0.5 | 2.42E-04 | 0.3 | 1.29E-02 | 0.2 | 3.15E-01 | Citrate synthase |
| AT1G77380 | -0.3 | 1.37E-01 | -0.6 | 1.46E-02 | 0.3 | 2.59E-01 | 1.4 | 6.83E-04 | 0.0 | 9.53E-01 | 1.3 | 2.39E-03 | Amino acid permease 3 |
| AT4G27300 | -0.3 | 1.25E-02 | -0.5 | 8.20E-04 | 0.2 | 9.88E-02 | 0.7 | 1.06E-03 | 0.1 | 8.25E-01 | 0.6 | 1.03E-02 | G-type lectin S-receptor-like serine |
| AT1G28420 | -0.3 | 6.94E-04 | -0.3 | 6.39E-04 | 0.0 | 7.79E-01 | 0.2 | 1.27E-01 | 0.2 | 2.29E-01 | 0.0 | 9.49E-01 | Homeobox-DDT domain protein RLT1 |
| AT5G54140 | -0.3 | 4.55E-03 | -0.2 | 1.28E-02 | 0.0 | 7.04E-01 | 0.1 | 7.82E-01 | 0.3 | 1.16E-01 | -0.3 | 2.36E-01 | IAA-amino acid hydrolase ILR1-like 3 |
| AT3G62290 | -0.3 | 2.98E-03 | -0.1 | 5.19E-01 | -0.2 | 1.88E-02 | -0.6 | 1.13E-04 | -0.5 | 1.98E-03 | -0.1 | 5.82E-01 | ADP-ribosylation factor |
| AT3G19290 | -0.3 | 1.79E-02 | -0.3 | 7.33E-03 | 0.1 | 5.90E-01 | -0.4 | 7.16E-03 | -0.5 | 4.57E-03 | 0.1 | 7.76E-01 | ABRE binding factor 4 |
| AT4G17260 | -0.3 | 5.61E-03 | -0.1 | 4.57E-01 | -0.2 | 4.24E-02 | 0.9 | 4.52E-05 | 0.1 | 6.77E-01 | 0.8 | 1.29E-03 | L-lactate dehydrogenase |
| AT4G15430 | -0.3 | 3.38E-02 | -0.3 | 1.81E-02 | 0.1 | 7.15E-01 | 0.4 | 2.83E-01 | 0.2 | 5.61E-01 | 0.1 | 8.26E-01 | Hyperosmolality-gated Ca2+ permeable channel 1.6 |
| AT1G56560 | -0.3 | 9.87E-04 | -0.4 | 2.06E-04 | 0.1 | 1.91E-01 | 0.1 | 6.40E-01 | 0.1 | 8.03E-01 | 0.0 | 9.31E-01 | Alkaline |
| AT5G23630 | -0.3 | 5.98E-03 | -0.2 | 3.33E-02 | -0.1 | 4.40E-01 | 0.2 | 2.27E-01 | 0.2 | 2.93E-01 | 0.0 | 9.97E-01 | Probable manganese-transporting ATPase PDR2 |
| AT2G28070 | -0.3 | 2.48E-03 | -0.2 | 5.23E-02 | -0.1 | 1.66E-01 | -0.1 | 3.18E-01 | 0.0 | 9.07E-01 | -0.1 | 5.70E-01 | ABC transporter G family member 3 |
| AT2G35100 | -0.3 | 2.69E-03 | -0.2 | 7.46E-03 | 0.0 | 7.08E-01 | -0.3 | 1.30E-01 | -0.1 | 7.38E-01 | -0.2 | 3.82E-01 | Probable arabinosyltransferase ARAD1 |
| AT1G13210 | -0.3 | 3.21E-03 | -0.3 | 1.27E-03 | 0.1 | 4.61E-01 | 0.3 | 5.16E-02 | 0.0 | 9.68E-01 | 0.3 | 1.49E-01 | Probable phospholipid-transporting ATPase 11 |
| AT4G30780 | -0.3 | 1.43E-02 | -0.5 | 7.21E-04 | 0.2 | 8.40E-02 | -0.6 | 3.99E-01 | -0.4 | 6.72E-01 | -0.2 | 8.45E-01 | ATP-dependent DNA helicase |
| AT4G28710 | -0.3 | 4.99E-03 | -0.4 | 1.52E-03 | 0.1 | 4.11E-01 | 0.2 | 3.47E-01 | 0.1 | 7.37E-01 | 0.1 | 7.77E-01 | Myosin-14 |
| AT4G30060 | -0.3 | 6.09E-03 | 0.0 | 8.24E-01 | -0.3 | 1.03E-02 | 0.2 | 3.30E-01 | 0.3 | 1.21E-01 | -0.1 | 5.87E-01 | hypothetical protein |
| AT2G33580 | -0.3 | 6.18E-02 | -0.7 | 9.50E-04 | 0.4 | 3.48E-02 | 0.4 | 1.35E-01 | 0.5 | 1.75E-01 | 0.0 | 9.90E-01 | Protein LYK5 |
| AT3G46200 | -0.3 | 1.24E-02 | 0.1 | 3.27E-01 | -0.4 | 7.89E-03 | -0.4 | 9.80E-02 | -0.2 | 4.44E-01 | -0.2 | 5.90E-01 | Nudix hydrolase homolog 9 |
| AT2G31970 | -0.3 | 2.19E-03 | -0.5 | 8.80E-05 | 0.2 | 2.69E-02 | 0.4 | 1.70E-01 | 0.1 | 7.60E-01 | 0.3 | 5.12E-01 | DNA repair protein RAD50 |
| AT3G23300 | -0.3 | 2.00E-03 | -0.3 | 4.27E-03 | 0.0 | 8.08E-01 | -0.2 | 1.21E-01 | 0.1 | 3.96E-01 | -0.3 | 7.31E-02 | Probable methyltransferase PMT1 |
| AT5G43670 | -0.3 | 5.71E-03 | -0.3 | 4.34E-03 | 0.0 | 7.92E-01 | 0.0 | 9.37E-01 | 0.0 | 9.81E-01 | 0.0 | 9.41E-01 | Protein transport protein SEC23 G |
| AT1G11190 | -0.3 | 6.74E-02 | -0.5 | 9.25E-03 | 0.2 | 3.08E-01 | -0.4 | 3.83E-01 | 0.5 | 2.19E-01 | -0.8 | 9.20E-02 | Endonuclease 1 |
| AT1G17040 | -0.3 | 6.62E-02 | -0.4 | 1.11E-02 | 0.2 | 3.48E-01 | -0.5 | 7.04E-02 | -0.2 | 5.08E-01 | -0.3 | 4.45E-01 | SH2 domain-containing protein A |
| AT1G53165 | -0.3 | 2.91E-02 | -0.3 | 2.69E-02 | 0.0 | 9.21E-01 | -0.3 | 1.12E-01 | -0.3 | 9.31E-02 | 0.0 | 8.88E-01 | Protein kinase superfamily protein |
| AT4G35780 | -0.3 | 1.46E-02 | -0.2 | 3.66E-02 | 0.0 | 7.23E-01 | -0.4 | 1.01E-02 | -0.4 | 1.22E-02 | 0.0 | 9.33E-01 | STY17 |
| AT1G19290 | -0.3 | 2.96E-02 | -0.4 | 7.88E-03 | 0.1 | 4.53E-01 | -0.3 | 4.92E-01 | 0.0 | 9.71E-01 | -0.3 | 6.18E-01 | Putative pentatricopeptide repeat-containing protein At1g19290 |
| AT1G21480 | -0.3 | 1.97E-02 | 0.1 | 4.03E-01 | -0.4 | 1.27E-02 | -0.2 | 1.96E-01 | -0.2 | 2.13E-01 | 0.0 | 9.62E-01 | Exostosin family protein |
| AT3G63260 | -0.3 | 1.09E-02 | -0.4 | 2.26E-03 | 0.1 | 3.30E-01 | -0.3 | 5.29E-01 | -0.1 | 8.35E-01 | -0.2 | 8.31E-01 | ATMRK1 |
| AT1G16890 | -0.3 | 9.38E-04 | -0.2 | 3.44E-03 | 0.0 | 5.73E-01 | -0.1 | 7.56E-01 | 0.1 | 8.02E-01 | -0.1 | 6.36E-01 | Ubiquitin-conjugating enzyme 36 |
| AT2G21190 | -0.3 | 1.24E-03 | -0.1 | 3.82E-02 | -0.1 | 1.02E-01 | 0.1 | 4.72E-01 | 0.0 | 8.88E-01 | 0.1 | 7.53E-01 | ER lumen protein retaining receptor family protein |
| AT2G41460 | -0.3 | 2.40E-02 | -0.2 | 1.19E-01 | -0.1 | 4.96E-01 | -0.1 | 7.47E-01 | -0.1 | 8.19E-01 | 0.0 | 9.80E-01 | DNA- endonuclease |
| AT5G49950 | -0.3 | 8.43E-03 | -0.4 | 1.37E-03 | 0.1 | 2.56E-01 | 0.2 | 2.95E-01 | 0.2 | 3.30E-01 | 0.0 | 9.77E-01 | Alpha |
| AT2G32010 | -0.3 | 8.42E-02 | 0.0 | 9.56E-01 | -0.3 | 1.56E-01 | -0.3 | 1.48E-01 | 0.0 | 9.34E-01 | -0.2 | 2.83E-01 | Type IV inositol polyphosphate 5-phosphatase 7 |
| AT4G39990 | -0.3 | 2.48E-03 | 0.0 | 9.64E-01 | -0.3 | 6.63E-03 | -0.2 | 1.18E-01 | -0.1 | 3.90E-01 | -0.1 | 7.35E-01 | Ras-related protein RABA4b |
| AT3G23560 | -0.3 | 3.40E-03 | -0.3 | 2.21E-03 | 0.0 | 7.06E-01 | 0.6 | 4.57E-04 | 0.3 | 1.32E-01 | 0.4 | 5.81E-02 | MATE efflux family protein |
| AT1G75880 | -0.3 | 7.52E-02 | 0.2 | 2.90E-01 | -0.4 | 2.96E-02 | 0.7 | 9.18E-02 | 0.3 | 5.13E-01 | 0.3 | 4.85E-01 | hypothetical protein |
| AT2G22090 | -0.3 | 3.94E-02 | -0.3 | 1.97E-02 | 0.1 | 7.00E-01 | -0.4 | 4.05E-01 | -0.2 | 7.75E-01 | -0.2 | 7.24E-01 | RNA-binding family protein |
| AT5G65810 | -0.3 | 9.72E-03 | 0.2 | 1.08E-01 | -0.4 | 2.04E-03 | -0.1 | 3.51E-01 | 0.2 | 2.49E-01 | -0.3 | 1.12E-01 | Probable pectin methylesterase CGR3 |
| AT3G19100 | -0.3 | 7.05E-03 | -0.1 | 3.20E-01 | -0.2 | 7.86E-02 | 0.0 | 9.27E-01 | 0.1 | 7.41E-01 | -0.1 | 7.30E-01 | CDPK-related kinase 2 |
| AT4G35220 | -0.3 | 1.71E-03 | -0.1 | 2.35E-01 | -0.2 | 2.83E-02 | 0.0 | 7.40E-01 | -0.1 | 7.86E-01 | 0.0 | 9.99E-01 | Cyclase-like protein 2 |
| AT1G72990 | -0.3 | 1.38E-02 | -0.3 | 4.77E-03 | 0.1 | 5.23E-01 | 0.0 | 8.96E-01 | 0.3 | 3.16E-01 | -0.2 | 4.48E-01 | Beta-galactosidase |
| AT1G71070 | -0.3 | 1.97E-03 | -0.1 | 6.00E-02 | -0.1 | 1.10E-01 | -0.1 | 4.77E-01 | 0.1 | 4.53E-01 | -0.3 | 2.24E-01 | At1g71070 |
| AT5G26330 | -0.3 | 4.79E-02 | 0.1 | 5.82E-01 | -0.3 | 3.95E-02 | 0.0 | 8.86E-01 | 0.1 | 7.01E-01 | -0.2 | 6.40E-01 | hypothetical protein |

| | | | | | | | | | | | | | |
|---|---|---|---|---|---|---|---|---|---|---|---|---|---|
| AT3G54610 | -0.3 | 4.08E-03 | -0.3 | 3.00E-03 | 0.0 | 7.69E-01 | 0.2 | 2.81E-01 | 0.2 | 4.14E-01 | 0.0 | 9.42E-01 | Histone acetyltransferase GCN5 |
| AT1G60490 | -0.3 | 2.25E-03 | -0.6 | 1.16E-05 | 0.4 | 1.00E-03 | 0.0 | 8.05E-01 | 0.1 | 3.18E-01 | -0.1 | 5.69E-01 | Phosphatidylinositol 3-kinase VPS34 |
| AT3G53370 | -0.3 | 8.50E-02 | 0.3 | 3.96E-02 | -0.6 | 6.35E-03 | 0.1 | 5.52E-01 | 0.3 | 1.79E-01 | -0.2 | 5.11E-01 | S1FA-like DNA-binding protein |
| AT5G38860 | -0.3 | 4.00E-02 | -0.1 | 3.17E-01 | -0.1 | 3.43E-01 | -0.6 | 2.46E-02 | -0.5 | 1.27E-01 | -0.2 | 6.75E-01 | BES1-interacting Myc-like protein 3 |
| AT1G08110 | -0.3 | 1.71E-03 | -0.2 | 2.32E-02 | -0.1 | 2.14E-01 | -0.3 | 1.30E-02 | -0.2 | 2.18E-01 | -0.1 | 4.59E-01 | Lactoylglutathione lyase |
| AT1G72280 | -0.3 | 6.55E-02 | -0.3 | 9.17E-02 | 0.0 | 9.27E-01 | 0.2 | 3.27E-01 | 0.0 | 9.07E-01 | 0.2 | 5.30E-01 | hypothetical protein |
| AT5G04480 | -0.3 | 2.01E-02 | -0.3 | 1.44E-02 | 0.0 | 8.10E-01 | 0.0 | 9.80E-01 | 0.1 | 6.60E-01 | -0.1 | 7.21E-01 | AT5g04480 |
| AT5G26030 | -0.3 | 7.13E-03 | -0.3 | 2.38E-03 | 0.1 | 4.56E-01 | 0.3 | 4.53E-03 | 0.3 | 2.94E-02 | 0.1 | 7.48E-01 | Ferrochelatase-1, chloroplastic |
| AT1G60860 | -0.3 | 8.92E-03 | -0.2 | 3.73E-02 | -0.1 | 5.58E-01 | 0.0 | 9.75E-01 | 0.2 | 5.70E-01 | -0.2 | 6.25E-01 | ADP-ribosylation factor GTPase-activating protein |
| AT3G56100 | -0.3 | 1.13E-01 | 0.2 | 3.03E-01 | -0.4 | 4.18E-02 | 0.2 | 5.73E-01 | 0.3 | 5.41E-01 | 0.0 | 9.41E-01 | Meristematic receptor-like kinase |
| AT5G02410 | -0.3 | 1.19E-02 | 0.0 | 7.28E-01 | -0.2 | 5.00E-02 | 0.0 | 8.40E-01 | 0.3 | 1.23E-01 | -0.2 | 2.87E-01 | Dol-P-Glc:GlcManGlcNAc-PP-Dol alpha-1,2-glucosyltransferase |
| AT1G19710 | -0.3 | 3.01E-03 | 0.0 | 7.22E-01 | -0.2 | 1.57E-02 | 0.3 | 1.04E-01 | 0.3 | 1.94E-01 | 0.0 | 9.26E-01 | hypothetical protein |
| AT2G44950 | -0.3 | 7.84E-03 | -0.3 | 5.00E-03 | 0.0 | 7.35E-01 | -0.1 | 5.27E-01 | -0.1 | 4.37E-01 | 0.0 | 9.04E-01 | E3 ubiquitin protein ligase |
| AT5G24800 | -0.3 | 3.31E-03 | -0.1 | 1.89E-01 | -0.2 | 6.21E-02 | -0.9 | 2.30E-04 | -0.7 | 3.21E-03 | -0.2 | 4.76E-01 | Basic leucine zipper 9 |
| AT4G24780 | -0.3 | 5.42E-02 | 0.2 | 2.54E-01 | -0.4 | 1.27E-02 | -0.3 | 1.13E-02 | 0.1 | 5.53E-01 | -0.4 | 1.74E-02 | Probable pectate lyase 18 |
| AT5G20620 | -0.3 | 8.52E-04 | -0.3 | 1.40E-03 | 0.0 | 9.48E-01 | 0.0 | 9.32E-01 | 0.1 | 3.03E-01 | -0.1 | 4.57E-01 | Polyubiquitin 4 |
| AT3G01640 | -0.3 | 3.68E-03 | 0.1 | 4.00E-01 | -0.3 | 3.37E-03 | -0.5 | 3.04E-03 | -0.5 | 5.74E-03 | 0.0 | 9.97E-01 | Glucuronokinase G |
| AT2G47180 | -0.3 | 2.92E-02 | -0.4 | 7.16E-03 | 0.1 | 4.24E-01 | -0.5 | 2.87E-01 | -0.3 | 5.78E-01 | -0.2 | 8.02E-01 | Galactinol synthase 1 |
| AT5G61460 | -0.3 | 5.79E-02 | -0.3 | 6.27E-02 | 0.0 | 9.73E-01 | 0.6 | 1.71E-01 | 0.4 | 4.49E-01 | 0.2 | 7.66E-01 | Structural maintenance of chromosomes protein 6B |
| AT1G79570 | -0.3 | 2.25E-03 | -0.4 | 2.17E-04 | 0.1 | 8.99E-02 | 0.0 | 9.28E-01 | 0.0 | 9.80E-01 | 0.0 | 9.37E-01 | Protein kinase domain-containing protein |
| AT5G13570 | -0.3 | 2.32E-03 | -0.4 | 3.50E-04 | 0.1 | 1.67E-01 | -0.3 | 1.03E-01 | -0.4 | 3.95E-02 | 0.1 | 6.81E-01 | mRNA-decapping enzyme subunit 2 GpppN-mRNA hydrolase DCP2) |
| AT2G40060 | -0.3 | 3.24E-03 | 0.0 | 9.36E-01 | -0.3 | 8.21E-03 | -0.2 | 9.92E-02 | -0.1 | 6.45E-01 | -0.1 | 5.14E-01 | Clathrin light chain 2 |
| AT1G08680 | -0.3 | 1.55E-02 | -0.3 | 1.05E-02 | 0.0 | 7.83E-01 | -0.2 | 3.76E-01 | -0.1 | 8.01E-01 | -0.1 | 7.45E-01 | ARF GAP-like zinc finger-containing protein ZIGA4 |
| AT4G12610 | -0.3 | 4.29E-03 | -0.2 | 6.34E-02 | -0.1 | 2.14E-01 | 0.1 | 5.73E-01 | 0.1 | 4.34E-01 | -0.1 | 8.60E-01 | Transcription initiation factor IIF subunit alpha |
| AT1G65920 | -0.3 | 7.43E-02 | -0.2 | 1.87E-01 | -0.1 | 7.12E-01 | -0.1 | 8.42E-01 | 0.1 | 8.40E-01 | -0.2 | 6.90E-01 | F12P19.9 protein family with FYVE zinc finger domain-containing protein) |
| AT4G19040 | -0.3 | 5.08E-03 | -0.4 | 3.37E-04 | 0.2 | 7.64E-02 | -0.1 | 7.32E-01 | 0.2 | 2.28E-01 | -0.3 | 1.82E-01 | Protein ENHANCED DISEASE RESISTANCE 2-like |
| AT3G26670 | -0.3 | 6.64E-03 | -0.3 | 2.33E-03 | 0.1 | 4.84E-01 | 0.1 | 4.24E-01 | 0.0 | 9.53E-01 | 0.1 | 6.20E-01 | Probable magnesium transporter NIPA8 |
| AT3G24503 | -0.3 | 6.14E-03 | -0.1 | 2.92E-01 | -0.2 | 6.91E-02 | 0.5 | 3.46E-04 | 0.1 | 4.59E-01 | 0.4 | 2.58E-02 | Aldehyde dehydrogenase family 2 member C4 |
| AT2G39450 | -0.3 | 1.56E-02 | -0.1 | 1.88E-01 | -0.1 | 2.36E-01 | 0.0 | 8.85E-01 | 0.1 | 7.04E-01 | -0.1 | 6.08E-01 | Metal tolerance protein 11 |
| AT3G17250 | -0.3 | 2.52E-02 | -0.4 | 2.66E-03 | 0.2 | 2.16E-01 | -0.4 | 3.49E-02 | -0.2 | 4.59E-01 | -0.3 | 2.96E-01 | Probable protein phosphatase 2C 43 |
| AT1G33910 | -0.3 | 1.12E-01 | 0.2 | 2.83E-01 | -0.4 | 3.96E-02 | 0.5 | 1.70E-01 | 0.6 | 1.10E-01 | -0.1 | 8.07E-01 | Immune-associated nucleotide-binding protein 5 |
| AT2G41110 | -0.3 | 3.50E-02 | 0.0 | 9.19E-01 | -0.3 | 5.16E-02 | -0.2 | 4.08E-01 | -0.2 | 5.75E-01 | 0.0 | 9.36E-01 | Calmodulin 2 |
| AT5G09810 | -0.3 | 4.56E-03 | -0.1 | 5.18E-01 | -0.2 | 2.73E-02 | 0.0 | 9.18E-01 | 0.0 | 7.53E-01 | 0.0 | 8.69E-01 | Actin-7 |
| AT4G22790 | -0.3 | 3.74E-02 | 0.1 | 4.07E-01 | -0.4 | 2.25E-02 | -0.2 | 4.60E-01 | 0.2 | 6.17E-01 | -0.4 | 2.04E-01 | Protein DETOXIFICATION 56 |
| AT4G08940 | -0.3 | 1.63E-01 | 0.1 | 6.82E-01 | -0.3 | 1.41E-01 | 0.3 | 4.20E-01 | 0.2 | 6.23E-01 | 0.1 | 9.05E-01 | At4g08940 |
| AT1G07630 | -0.3 | 1.13E-03 | -0.5 | 1.75E-05 | 0.3 | 3.61E-03 | 0.1 | 7.40E-01 | -0.2 | 3.75E-01 | 0.2 | 2.32E-01 | Probable protein phosphatase 2C 4 |
| AT5G27840 | -0.3 | 1.20E-02 | -0.3 | 1.27E-02 | 0.0 | 9.52E-01 | -0.2 | 1.28E-01 | -0.1 | 5.34E-01 | -0.1 | 6.41E-01 | Serine |
| AT3G51895 | -0.3 | 2.86E-02 | -0.2 | 1.42E-01 | -0.1 | 4.74E-01 | 0.1 | 4.77E-01 | 0.4 | 9.34E-02 | -0.2 | 3.74E-01 | Sulfate transporter 31 |
| AT4G13770 | -0.3 | 3.81E-02 | 0.1 | 5.40E-01 | -0.3 | 2.19E-02 | -0.6 | 5.02E-04 | 0.5 | 1.03E-02 | -1.1 | 2.22E-04 | AT4g13770 |
| AT5G02380 | -0.3 | 3.59E-02 | 0.1 | 6.77E-01 | -0.3 | 2.84E-02 | -0.2 | 3.57E-01 | -0.1 | 7.42E-01 | -0.1 | 7.82E-01 | Metallothionein-like protein 2B |
| AT5G41240 | -0.3 | 1.30E-02 | -0.3 | 1.55E-02 | 0.0 | 9.88E-01 | -0.3 | 9.29E-02 | -0.1 | 6.28E-01 | -0.2 | 4.72E-01 | Glutathione S-transferase T2 |
| AT5G02800 | -0.3 | 2.62E-03 | -0.3 | 1.12E-03 | 0.1 | 5.01E-01 | -0.3 | 5.98E-02 | -0.3 | 6.24E-02 | 0.0 | 9.34E-01 | Probable serine |
| AT1G09780 | -0.3 | 3.35E-02 | 0.0 | 9.51E-01 | -0.3 | 6.00E-02 | 0.1 | 5.86E-01 | 0.1 | 4.83E-01 | 0.0 | 8.91E-01 | 2,3-bisphosphoglycerate-independent phosphoglycerate mutase 1 |
| AT3G62800 | -0.3 | 5.91E-03 | -0.3 | 3.62E-03 | 0.0 | 7.06E-01 | 0.0 | 8.81E-01 | 0.0 | 8.99E-01 | 0.0 | 9.98E-01 | Double-stranded RNA-binding protein 4 |
| AT5G45710 | -0.3 | 1.21E-02 | -0.2 | 2.92E-02 | 0.0 | 7.48E-01 | -0.2 | 4.17E-01 | -0.3 | 2.10E-01 | 0.1 | 7.44E-01 | Winged-helix DNA-binding transcription factor family protein |
| AT4G19650 | -0.3 | 3.71E-02 | -0.3 | 2.53E-02 | 0.0 | 8.05E-01 | 0.3 | 5.57E-01 | 0.2 | 7.03E-01 | 0.1 | 9.28E-01 | Mitochondrial transcription termination factor family protein |
| AT2G14835 | -0.3 | 4.58E-03 | 0.0 | 9.83E-01 | -0.3 | 1.05E-02 | -0.1 | 6.74E-01 | -0.1 | 3.79E-01 | 0.1 | 7.46E-01 | Expressed protein |
| AT1G50670 | -0.3 | 6.27E-03 | -0.1 | 1.64E-01 | -0.1 | 1.28E-01 | 0.1 | 7.86E-01 | 0.0 | 8.47E-01 | 0.0 | 9.92E-01 | OVARIAN TUMOR DOMAIN-containing deubiquitinating enzyme 2 |
| AT3G21750 | -0.3 | 4.60E-02 | -0.4 | 6.37E-03 | 0.2 | 2.80E-01 | -0.5 | 3.18E-04 | 0.1 | 4.06E-01 | -0.6 | 9.32E-04 | UDP-glycosyltransferase 71B1 |
| AT4G02730 | -0.3 | 1.46E-02 | -0.1 | 1.65E-01 | -0.1 | 2.84E-01 | 0.2 | 5.89E-01 | 0.1 | 6.70E-01 | 0.0 | 9.92E-01 | COMPASS-like H3K4 histone methylase component WDR5B |
| AT4G10070 | -0.3 | 1.74E-02 | -0.4 | 2.80E-03 | 0.1 | 2.93E-01 | 0.2 | 5.00E-01 | -0.1 | 6.76E-01 | 0.3 | 3.52E-01 | KH domain-containing protein |
| AT1G06290 | -0.3 | 3.33E-03 | -0.2 | 1.03E-02 | 0.0 | 6.57E-01 | 0.0 | 7.92E-01 | 0.2 | 2.33E-01 | -0.2 | 4.68E-01 | Acyl-coenzyme A oxidase 3, peroxisomal |
| AT2G40095 | -0.3 | 2.75E-02 | -0.3 | 1.89E-02 | 0.0 | 8.16E-01 | 0.8 | 6.04E-03 | 0.1 | 8.35E-01 | 0.7 | 2.46E-02 | Alpha |
| AT1G77300 | -0.3 | 2.45E-02 | -0.4 | 2.57E-03 | 0.2 | 2.00E-01 | 0.0 | 9.32E-01 | 0.1 | 8.46E-01 | -0.2 | 8.11E-01 | Histone-lysine N-methyltransferase |
| AT4G08170 | -0.3 | 9.83E-03 | -0.3 | 1.05E-02 | 0.0 | 9.48E-01 | 0.0 | 9.08E-01 | 0.1 | 6.84E-01 | -0.1 | 8.00E-01 | Inositol-tetrakisphosphate 1-kinase |
| AT1G77480 | -0.3 | 6.91E-03 | -0.2 | 6.58E-02 | -0.1 | 3.26E-01 | 0.1 | 6.08E-01 | -0.2 | 1.70E-01 | 0.3 | 1.05E-01 | Eukaryotic aspartyl protease family protein |
| AT1G15220 | -0.3 | 4.26E-02 | -0.2 | 1.71E-01 | -0.1 | 5.57E-01 | -0.3 | 1.22E-01 | -0.5 | 1.42E-02 | 0.2 | 4.05E-01 | Cytochrome c-type biogenesis CcmH-like mitochondrial protein |
| AT4G38960 | -0.3 | 1.57E-02 | -0.4 | 1.12E-03 | 0.2 | 1.32E-01 | -0.9 | 8.02E-02 | -0.3 | 6.44E-01 | -0.6 | 3.14E-01 | B-box type zinc finger family protein |
| AT3G58630 | -0.3 | 2.88E-03 | -0.2 | 1.23E-02 | -0.1 | 5.55E-01 | 0.0 | 8.74E-01 | -0.1 | 6.07E-01 | 0.1 | 7.93E-01 | Sequence-specific DNA binding transcription factor |
| AT3G17430 | -0.3 | 1.00E-02 | 0.0 | 6.40E-01 | -0.2 | 4.63E-02 | 0.0 | 9.45E-01 | 0.2 | 3.83E-01 | -0.2 | 3.95E-01 | Probable sugar phosphate |
| AT2G29140 | -0.3 | 1.13E-03 | -0.4 | 9.97E-05 | 0.1 | 5.86E-02 | 0.4 | 5.32E-03 | 0.0 | 8.62E-01 | 0.4 | 3.66E-02 | Pumilio homolog 3 |
| AT5G07580 | -0.3 | 1.54E-02 | -0.2 | 5.95E-02 | -0.1 | 5.57E-01 | 0.0 | 8.91E-01 | -0.2 | 2.38E-01 | 0.2 | 4.22E-01 | Ethylene-responsive transcription factor ERF106 |
| AT4G16340 | -0.3 | 3.44E-03 | -0.2 | 4.24E-02 | -0.1 | 2.36E-01 | 0.1 | 7.40E-01 | 0.1 | 5.06E-01 | -0.1 | 7.92E-01 | SPK1 |
| AT3G09840 | -0.3 | 7.81E-03 | -0.4 | 1.74E-03 | 0.1 | 3.07E-01 | -0.1 | 7.62E-01 | -0.1 | 7.79E-01 | 0.0 | 9.83E-01 | Cell division control protein 48 homolog A |
| AT3G04530 | -0.3 | 4.30E-01 | -0.1 | 7.72E-01 | -0.2 | 7.01E-01 | 1.7 | 5.02E-05 | 0.9 | 2.12E-02 | 0.8 | 2.33E-02 | Phosphoenolpyruvate carboxylase kinase 2 |
| AT5G61390 | -0.3 | 4.82E-02 | -0.4 | 1.16E-02 | 0.1 | 4.53E-01 | 0.0 | 9.63E-01 | 0.1 | 8.32E-01 | -0.1 | 7.84E-01 | Protein NEN2 |
| AT2G18000 | -0.3 | 3.28E-02 | -0.2 | 1.48E-01 | -0.1 | 5.32E-01 | 0.6 | 1.14E-01 | 0.3 | 4.62E-01 | 0.2 | 5.79E-01 | Transcription initiation factor TFIID subunit 14 |
| AT2G39260 | -0.3 | 1.63E-03 | -0.3 | 5.31E-04 | 0.1 | 3.48E-01 | 0.0 | 1.00E+00 | 0.1 | 7.06E-01 | -0.1 | 7.56E-01 | Regulator of nonsense transcripts UPF2 |
| AT1G05620 | -0.3 | 3.73E-02 | 0.3 | 2.41E-02 | -0.5 | 2.10E-03 | -0.1 | 4.80E-01 | 0.1 | 6.03E-01 | -0.2 | 2.79E-01 | Probable uridine nucleosidase 2 |
| AT2G30920 | -0.3 | 7.43E-03 | 0.0 | 7.97E-01 | -0.2 | 2.96E-02 | 0.0 | 9.50E-01 | 0.0 | 9.12E-01 | 0.0 | 8.85E-01 | Ubiquinone biosynthesis O-methyltransferase, mitochondrial |
| AT3G17630 | -0.3 | 6.52E-02 | -0.5 | 3.96E-03 | 0.3 | 1.47E-01 | -0.3 | 6.24E-01 | -0.1 | 9.27E-01 | | | Cation |
| AT5G63650 | -0.3 | 6.31E-02 | 0.1 | 5.30E-01 | -0.3 | 4.54E-02 | -0.2 | 4.70E-01 | -0.3 | 4.92E-01 | 0.0 | 9.75E-01 | AT5G63650 protein |
| AT1G21540 | -0.3 | 2.90E-02 | 0.0 | 8.89E-01 | -0.2 | 7.67E-02 | 0.0 | 9.17E-01 | -0.1 | 7.87E-01 | 0.1 | 8.91E-01 | Probable acyl-activating enzyme 9 |
| AT4G24805 | -0.3 | 1.68E-02 | -0.2 | 8.64E-02 | -0.1 | 4.88E-01 | -0.3 | 4.27E-02 | 0.0 | 9.87E-01 | -0.3 | 1.03E-01 | hypothetical protein |
| AT2G02680 | -0.3 | 9.36E-02 | 0.2 | 1.34E-01 | -0.5 | 1.59E-02 | -0.7 | 4.13E-02 | 0.1 | 7.99E-01 | -0.8 | 6.02E-02 | Cysteine |
| AT1G18390 | -0.3 | 1.31E-02 | -0.5 | 2.31E-04 | 0.3 | 2.13E-02 | 0.3 | 1.85E-01 | -0.1 | 8.64E-01 | 0.3 | 2.22E-01 | LEAF RUST 10 DISEASE-RESISTANCE LOCUS RECEPTOR-LIKE PROTEIN KINASE-like 1.2 |
| AT1G65450 | -0.3 | 2.38E-01 | 0.1 | 7.68E-01 | -0.3 | 2.27E-01 | 0.3 | 4.79E-01 | 0.4 | 3.78E-01 | -0.1 | 8.21E-01 | Acyltransferase GLAUCE |
| AT4G12020 | -0.3 | 2.39E-02 | -0.5 | 1.23E-03 | 0.2 | 9.83E-02 | -0.3 | 1.78E-01 | -0.1 | 8.21E-01 | -0.3 | 4.58E-01 | Protein kinase family protein |
| AT2G36310 | -0.3 | 2.27E-03 | 0.1 | 9.98E-02 | -0.4 | 6.90E-04 | -0.3 | 5.51E-03 | 0.1 | 4.51E-01 | -0.4 | 1.06E-02 | Uridine nucleosidase 1 |
| AT5G04090 | -0.3 | 5.31E-03 | -0.4 | 7.76E-04 | 0.1 | 2.08E-01 | 0.1 | 5.47E-01 | 0.1 | 7.64E-01 | 0.0 | 8.98E-01 | Histidine-tRNA ligase |
| AT2G21010 | -0.3 | 6.46E-02 | 0.6 | 5.63E-04 | -0.8 | 3.17E-04 | -0.9 | 5.84E-02 | -0.1 | 8.36E-01 | -0.7 | 2.03E-01 | Calcium-dependent lipid-binding family protein |
| AT3G62280 | -0.3 | 2.72E-01 | 0.6 | 1.68E-02 | -0.8 | 8.79E-03 | 0.6 | 3.24E-02 | 0.8 | 1.87E-02 | -0.1 | 7.56E-01 | GDSL-like Lipase |
| AT2G33620 | -0.3 | 2.78E-02 | -0.2 | 3.64E-02 | 0.0 | 9.57E-01 | -0.3 | 5.07E-02 | 0.0 | 8.37E-01 | -0.4 | 7.97E-02 | AT-hook motif nuclear-localized protein 10 |

| | | | | | | | | | | | | | |
|---|---|---|---|---|---|---|---|---|---|---|---|---|---|
| AT5G19430 | -0.3 | 2.45E-02 | -0.5 | 1.49E-03 | 0.2 | 1.19E-01 | -0.5 | 5.75E-03 | -0.3 | 7.78E-02 | -0.2 | 4.76E-01 | AT5g19430 |
| AT1G67420 | -0.3 | 2.31E-02 | -0.3 | 1.53E-02 | 0.0 | 7.88E-01 | -0.3 | 3.88E-01 | 0.0 | 9.38E-01 | -0.2 | 5.37E-01 | Zn-dependent exopeptidases superfamily protein |
| AT5G40670 | -0.3 | 1.38E-03 | -0.3 | 1.08E-03 | 0.0 | 7.43E-01 | 0.1 | 2.95E-01 | 0.1 | 3.41E-01 | 0.0 | 9.81E-01 | Cystinosin homolog |
| AT1G56670 | -0.3 | 4.11E-02 | -0.3 | 1.93E-02 | 0.1 | 6.71E-01 | -0.3 | 4.31E-01 | 0.4 | 3.01E-01 | -0.8 | 1.10E-01 | GDSL-like Lipase |
| AT3G13610 | -0.3 | 1.19E-02 | 0.2 | 9.09E-02 | -0.4 | 1.82E-03 | 0.7 | 6.54E-04 | 0.4 | 8.65E-02 | 0.4 | 7.45E-02 | Feruloyl CoA ortho-hydroxylase 1 |
| AT1G63690 | -0.3 | 1.35E-02 | -0.1 | 2.54E-01 | -0.1 | 1.57E-01 | -0.3 | 5.42E-03 | -0.2 | 8.55E-02 | -0.1 | 4.59E-01 | Uncharacterized protein At1g63690 |
| AT1G02520 | -0.3 | 5.80E-02 | 0.0 | 7.15E-01 | -0.2 | 1.77E-01 | 2.0 | 1.06E-07 | 0.3 | 2.36E-01 | 1.7 | 2.94E-05 | ABC transporter B family member 11 |
| AT1G20090 | -0.3 | 8.92E-03 | 0.1 | 5.17E-01 | -0.3 | 6.56E-03 | -0.4 | 1.05E-02 | -0.5 | 1.02E-02 | 0.1 | 8.69E-01 | Rac-like GTP-binding protein ARAC4 |
| AT2G39270 | -0.3 | 1.04E-02 | -0.4 | 2.05E-03 | 0.1 | 3.20E-01 | -0.4 | 4.69E-02 | 0.1 | 5.73E-01 | -0.5 | 5.65E-02 | Probable adenylate kinase 6, chloroplastic |
| AT3G08760 | -0.3 | 4.78E-03 | -0.4 | 4.52E-04 | 0.1 | 1.19E-01 | 0.2 | 8.89E-02 | -0.1 | 3.28E-01 | 0.4 | 4.71E-02 | ATSIK |
| AT3G20910 | -0.3 | 2.09E-02 | -0.5 | 5.17E-04 | 0.3 | 4.32E-02 | 0.2 | 4.37E-01 | -0.2 | 4.37E-01 | 0.4 | 1.57E-01 | Nuclear transcription factor Y subunit |
| AT5G61600 | -0.3 | 4.18E-01 | -0.7 | 5.77E-02 | 0.4 | 2.71E-01 | -0.1 | 7.22E-01 | -0.6 | 7.88E-03 | 0.6 | 5.23E-02 | Ethylene-responsive transcription factor ERF104 |
| AT2G33730 | -0.3 | 1.81E-03 | -0.3 | 6.43E-04 | 0.1 | 3.71E-01 | 0.1 | 4.45E-01 | 0.0 | 8.49E-01 | 0.1 | 7.72E-01 | DEAD-box ATP-dependent RNA helicase 21 |
| AT4G14147 | -0.2 | 3.76E-02 | 0.0 | 7.06E-01 | -0.2 | 1.33E-01 | -0.1 | 5.48E-01 | -0.1 | 5.88E-01 | 0.0 | 9.92E-01 | Actin-related protein 2 |
| AT2G37060 | -0.2 | 3.45E-02 | -0.1 | 4.36E-01 | -0.2 | 2.29E-01 | 0.0 | 9.60E-01 | -0.1 | 8.40E-01 | 0.1 | 8.27E-01 | Nuclear transcription factor Y subunit B-8 |
| AT1G29280 | -0.2 | 5.77E-03 | 0.1 | 9.28E-02 | -0.4 | 1.18E-03 | -0.1 | 4.26E-01 | 0.0 | 8.73E-01 | -0.1 | 5.89E-01 | WRKY family transcription factor |
| AT3G02130 | -0.2 | 8.41E-03 | -0.4 | 4.89E-04 | 0.2 | 7.88E-02 | -0.3 | 1.09E-01 | -0.2 | 3.93E-01 | -0.1 | 6.98E-01 | Receptor-like protein kinase 2 |
| AT5G54830 | -0.2 | 7.90E-03 | -0.2 | 6.64E-02 | -0.1 | 3.43E-01 | 0.1 | 6.54E-01 | 0.1 | 5.96E-01 | 0.0 | 9.26E-01 | Cytochrome b561, DM13 and DOMON domain-containing protein At5g54830 |
| AT2G35620 | -0.2 | 1.07E-02 | -0.3 | 1.02E-02 | 0.0 | 9.12E-01 | -0.3 | 1.92E-01 | 0.1 | 8.39E-01 | -0.3 | 2.25E-01 | LRR receptor-like serine |
| AT5G06100 | -0.2 | 5.93E-03 | -0.3 | 4.10E-03 | 0.0 | 7.60E-01 | -0.3 | 1.49E-01 | -0.2 | 3.82E-01 | -0.1 | 7.79E-01 | Transcription factor MYB33 |
| AT3G47160 | -0.2 | 7.27E-03 | 0.2 | 3.50E-02 | -0.4 | 8.24E-04 | -0.3 | 4.33E-02 | -0.3 | 1.69E-01 | -0.1 | 7.71E-01 | RING |
| AT5G53010 | -0.2 | 8.17E-03 | -0.4 | 1.37E-03 | 0.1 | 2.73E-01 | 0.0 | 8.94E-01 | 0.0 | 9.36E-01 | -0.1 | 8.52E-01 | Calcium-transporting ATPase |
| AT5G15170 | -0.2 | 8.15E-03 | -0.3 | 5.98E-03 | 0.0 | 7.95E-01 | -0.1 | 5.90E-01 | 0.0 | 9.04E-01 | -0.2 | 5.87E-01 | Tyrosyl-DNA phosphodiesterase 1 |
| AT4G35920 | -0.2 | 3.06E-02 | 0.0 | 6.97E-01 | -0.2 | 9.83E-02 | -0.3 | 3.26E-02 | -0.1 | 6.71E-01 | -0.2 | 2.42E-01 | Protein MID1-COMPLEMENTING ACTIVITY 1 |
| AT3G08650 | -0.2 | 1.83E-02 | -0.2 | 8.05E-02 | -0.1 | 5.20E-01 | -0.2 | 2.69E-02 | 0.0 | 9.32E-01 | -0.2 | 1.27E-01 | hypothetical protein |
| AT2G35350 | -0.2 | 2.59E-02 | -0.2 | 6.25E-02 | 0.0 | 7.45E-01 | 0.4 | 1.59E-01 | 0.6 | 2.75E-02 | -0.2 | 4.12E-01 | Protein phosphatase 2C 29 |
| AT2G38050 | -0.2 | 6.82E-03 | 0.0 | 9.19E-01 | -0.3 | 1.43E-02 | 0.0 | 8.91E-01 | 0.1 | 4.79E-01 | -0.1 | 6.73E-01 | Steroid 5-alpha-reductase DET2 -22,23-dihydroxycampest-4-en-3-one synthase) -22-hydroxy-5alpha-ergostan-3-one synthase) |
| AT3G56960 | -0.2 | 7.99E-02 | -0.5 | 5.24E-03 | 0.3 | 1.67E-01 | 0.3 | 4.50E-01 | 0.1 | 9.00E-01 | 0.2 | 6.68E-01 | Phosphatidylinositol 4-phosphate 5-kinase 4 P-5-kinase 4) |
| AT5G14940 | -0.2 | 2.04E-02 | -0.2 | 9.41E-02 | -0.1 | 5.22E-01 | -0.4 | 2.14E-01 | -0.5 | 1.37E-01 | 0.1 | 7.96E-01 | Protein NRT1 |
| AT1G08450 | -0.2 | 3.82E-01 | -0.3 | 2.93E-01 | 0.1 | 8.51E-01 | -0.1 | 4.89E-01 | -0.4 | 4.57E-02 | 0.3 | 2.76E-01 | Calreticulin like protein |
| AT3G16050 | -0.2 | 1.34E-01 | 0.1 | 5.17E-01 | -0.3 | 8.28E-02 | 0.0 | 8.54E-01 | 0.2 | 3.95E-01 | -0.2 | 3.24E-01 | Pyridoxal 5'-phosphate synthase-like subunit PDX1.2 |
| AT5G11850 | -0.2 | 8.46E-03 | -0.2 | 3.42E-02 | -0.1 | 5.46E-01 | 0.1 | 5.91E-01 | 0.1 | 4.23E-01 | 0.0 | 8.27E-01 | AT5g11850 |
| AT3G16950 | -0.2 | 8.17E-03 | 0.0 | 7.83E-01 | -0.2 | 2.54E-02 | -0.1 | 3.97E-01 | -0.1 | 6.36E-01 | 0.0 | 9.00E-01 | Dihydrolipoyl dehydrogenase |
| AT1G79750 | -0.2 | 4.21E-03 | -0.1 | 1.91E-01 | -0.1 | 7.59E-02 | -0.1 | 1.52E-01 | -0.1 | 5.59E-01 | -0.1 | 6.93E-01 | NADP-dependent malic enzyme 4, chloroplastic |
| AT5G38560 | -0.2 | 3.59E-03 | -0.3 | 2.92E-03 | 0.0 | 8.08E-01 | -0.1 | 5.41E-01 | -0.1 | 8.40E-01 | -0.1 | 8.32E-01 | non-specific serine |
| AT1G06420 | -0.2 | 9.99E-02 | 0.2 | 1.03E-01 | -0.5 | 1.63E-02 | 0.2 | 5.57E-01 | 0.2 | 5.65E-01 | 0.0 | 9.67E-01 | DNA ligase-like protein |
| AT2G14120 | -0.2 | 2.65E-03 | -0.3 | 1.40E-03 | 0.0 | 5.82E-01 | 0.1 | 4.60E-01 | -0.1 | 7.52E-01 | 0.1 | 4.27E-01 | Dynamin related protein |
| AT1G53920 | -0.2 | 7.04E-02 | 0.1 | 4.24E-01 | -0.3 | 3.58E-02 | 0.0 | 9.22E-01 | 0.6 | 3.27E-03 | -0.6 | 1.75E-02 | GLIP5 |
| AT1G53570 | -0.2 | 6.03E-03 | -0.3 | 4.12E-03 | 0.0 | 7.52E-01 | -0.1 | 6.73E-01 | -0.2 | 3.57E-01 | 0.1 | 6.98E-01 | hypothetical protein |
| AT1G14900 | -0.2 | 7.50E-03 | 0.0 | 9.62E-01 | -0.2 | 1.68E-02 | 0.1 | 4.61E-01 | 0.0 | 9.27E-01 | 0.2 | 5.58E-01 | HMG-Y-related protein A |
| AT5G42140 | -0.2 | 4.67E-03 | -0.3 | 1.37E-03 | 0.1 | 4.00E-01 | -0.2 | 4.68E-01 | -0.2 | 4.50E-01 | 0.0 | 9.41E-01 | Regulator of chromosome condensation family with FYVE zinc finger domain-containing protein |
| AT2G18160 | -0.2 | 1.24E-02 | 0.2 | 7.17E-02 | -0.4 | 1.60E-03 | 0.5 | 7.13E-04 | 0.4 | 1.49E-02 | 0.1 | 4.36E-01 | bZIP transcription factor 2 |
| AT4G29940 | -0.2 | 2.23E-02 | -0.2 | 2.90E-02 | 0.0 | 9.69E-01 | 0.1 | 6.56E-01 | -0.2 | 3.07E-01 | 0.3 | 2.11E-01 | Pathogenesis related homeodomain protein A |
| AT5G51890 | -0.2 | 1.07E-02 | 0.3 | 5.00E-03 | -0.5 | 4.30E-04 | -0.4 | 3.65E-02 | -0.1 | 5.74E-01 | -0.3 | 2.85E-01 | Peroxidase 66 |
| AT3G32940 | -0.2 | 2.34E-02 | -0.4 | 4.53E-03 | 0.1 | 3.52E-01 | 0.1 | 7.52E-01 | 0.0 | 9.85E-01 | 0.1 | 8.45E-01 | RNA-binding KH domain-containing protein |
| AT5G16270 | -0.2 | 1.04E-02 | -0.4 | 1.23E-03 | 0.1 | 1.77E-01 | -0.2 | 3.21E-01 | 0.0 | 9.27E-01 | -0.2 | 4.14E-01 | Sister chromatid cohesion 1 protein 4 |
| AT1G71030 | -0.2 | 8.32E-02 | 0.0 | 9.75E-01 | -0.2 | 1.11E-01 | -1.4 | 2.42E-03 | -2.2 | 4.92E-04 | 0.9 | 2.30E-01 | At1g71030 |
| AT1G28760 | -0.2 | 3.44E-02 | -0.1 | 2.25E-01 | -0.1 | 4.09E-01 | 0.2 | 5.24E-01 | 0.1 | 6.73E-01 | 0.0 | 9.44E-01 | F1K23.6 3'-pyrophosphohydrolase, putative ) |
| AT1G16410 | -0.2 | 2.09E-01 | 0.1 | 5.48E-01 | -0.4 | 1.01E-01 | -1.0 | 4.17E-03 | 1.1 | 1.70E-03 | -2.1 | 1.97E-04 | Dihomomethionine N-hydroxylase |
| AT4G22690 | -0.2 | 3.04E-02 | -0.2 | 1.52E-01 | -0.1 | 4.64E-01 | 0.5 | 7.97E-03 | 0.0 | 9.92E-01 | 0.5 | 3.63E-02 | Cytochrome P450 like protein |
| AT3G19820 | -0.2 | 1.59E-02 | 0.1 | 5.77E-01 | -0.3 | 1.18E-02 | -0.3 | 5.70E-03 | -0.1 | 2.86E-01 | -0.2 | 2.78E-01 | Delta-sterol reductase |
| AT1G01610 | -0.2 | 1.23E-02 | -0.1 | 1.72E-01 | -0.1 | 2.12E-01 | 0.4 | 5.73E-03 | 0.0 | 9.60E-01 | 0.4 | 2.79E-02 | Glycerol-3-phosphate 2-O-acyltransferase 4 |
| AT1G79670 | -0.2 | 7.77E-03 | -0.3 | 7.32E-03 | 0.0 | 8.97E-01 | -0.1 | 3.99E-01 | -0.2 | 2.63E-01 | 0.1 | 7.99E-01 | Wall-associated receptor kinase-like |
| AT1G61820 | -0.2 | 2.42E-01 | 0.1 | 4.85E-01 | -0.4 | 1.22E-01 | 0.4 | 1.97E-02 | -0.2 | 2.95E-01 | 0.6 | 8.71E-03 | Beta glucosidase 46 |
| AT5G53370 | -0.2 | 1.78E-02 | -0.5 | 1.87E-04 | 0.3 | 1.02E-02 | -0.3 | 2.37E-02 | -0.1 | 6.71E-01 | -0.2 | 2.42E-01 | Pectinesterase |
| AT1G55830 | -0.2 | 5.35E-02 | -0.1 | 3.46E-01 | -0.1 | 3.74E-01 | -0.3 | 1.75E-01 | -0.4 | 8.34E-02 | 0.1 | 7.41E-01 | Coiled-coil protein |
| AT4G03060 | -0.2 | 4.15E-01 | 0.1 | 7.50E-01 | -0.3 | 3.31E-01 | -1.4 | 1.51E-02 | 0.7 | 8.91E-02 | -2.1 | 6.00E-03 | Probable inactive 2-oxoglutarate-dependent dioxygenase AOP2 |
| AT2G04430 | -0.2 | 4.41E-03 | 0.0 | 8.18E-01 | -0.2 | 1.79E-02 | 1.1 | 1.89E-05 | 0.5 | 2.22E-02 | 0.6 | 1.81E-02 | Nudix hydrolase |
| AT1G30450 | -0.2 | 1.34E-02 | -0.1 | 2.43E-01 | -0.1 | 1.68E-01 | 0.0 | 9.29E-01 | 0.0 | 9.72E-01 | 0.0 | 9.82E-01 | Cation-chloride co-transporter 1 |
| AT2G44610 | -0.2 | 1.54E-02 | -0.1 | 5.81E-01 | -0.2 | 6.82E-02 | -0.2 | 6.68E-02 | -0.1 | 3.55E-01 | -0.1 | 6.77E-01 | Ras-related protein RABH1b |
| AT1G59750 | -0.2 | 5.44E-03 | -0.3 | 1.78E-03 | 0.1 | 4.33E-01 | -0.3 | 3.26E-02 | -0.3 | 8.67E-02 | 0.0 | 9.01E-01 | Auxin response factor |
| AT3G56080 | -0.2 | 2.10E-01 | -0.3 | 1.50E-01 | 0.1 | 8.36E-01 | 0.0 | 9.35E-01 | 0.7 | 3.95E-02 | -0.7 | 4.87E-02 | Methyltransferase |
| AT4G21390 | -0.2 | 4.99E-02 | -0.1 | 3.02E-01 | -0.1 | 4.05E-01 | 0.8 | 5.34E-03 | 0.2 | 5.99E-01 | 0.6 | 6.10E-02 | Receptor-like serine |
| AT3G05420 | -0.2 | 4.35E-03 | -0.1 | 1.87E-01 | -0.1 | 7.83E-02 | 0.1 | 5.58E-01 | 0.2 | 3.66E-01 | -0.1 | 7.96E-01 | Acyl-CoA-binding domain-containing protein 4 |
| AT4G21710 | -0.2 | 6.19E-03 | -0.4 | 7.94E-04 | 0.1 | 1.74E-01 | -0.1 | 5.94E-01 | -0.2 | 3.51E-01 | 0.1 | 7.49E-01 | DNA-directed RNA polymerase II subunit 2 |
| AT5G61480 | -0.2 | 8.31E-03 | -0.2 | 1.64E-02 | 0.0 | 8.18E-01 | -0.1 | 4.09E-01 | 0.0 | 8.41E-01 | -0.2 | 4.33E-01 | Leucine-rich repeat receptor-like protein kinase TDR |
| AT1G62560 | -0.2 | 2.09E-01 | -0.1 | 7.97E-01 | -0.2 | 3.82E-01 | -0.7 | 6.40E-02 | 0.8 | 2.92E-02 | -1.4 | 4.12E-03 | Flavin-containing monooxygenase |
| AT1G25540 | -0.2 | 8.60E-03 | -0.4 | 7.61E-04 | 0.1 | 1.33E-01 | -0.3 | 1.18E-01 | -0.3 | 1.50E-01 | 0.0 | 9.92E-01 | Mediator of RNA polymerase II transcription subunit 25 |
| AT5G14310 | -0.2 | 5.31E-03 | -0.2 | 1.08E-02 | 0.0 | 8.22E-01 | 0.0 | 9.38E-01 | 0.1 | 8.95E-01 | -0.1 | 8.48E-01 | Probable carboxylesterase 16 |
| AT5G43710 | -0.2 | 1.29E-02 | -0.1 | 1.24E-01 | -0.1 | 3.16E-01 | -0.3 | 2.72E-01 | -0.1 | 8.54E-01 | -0.2 | 5.53E-01 | alpha-1,2-Mannosidase |
| AT3G07160 | -0.2 | 5.97E-03 | -0.3 | 1.41E-03 | 0.1 | 3.11E-01 | -0.1 | 4.70E-01 | -0.1 | 6.02E-01 | 0.0 | 9.68E-01 | Putative glucan synthase |
| AT5G13050 | -0.2 | 9.09E-03 | 0.0 | 8.82E-01 | -0.2 | 2.80E-02 | -0.1 | 4.67E-01 | 0.1 | 7.35E-01 | -0.2 | 3.12E-01 | 5-formyltetrahydrofolate cyclo-ligase |
| AT1G05150 | -0.2 | 2.00E-03 | -0.1 | 3.79E-01 | -0.2 | 1.89E-02 | -0.1 | 6.75E-01 | 0.0 | 9.90E-01 | -0.1 | 7.84E-01 | Uncharacterized TPR repeat-containing protein At1g05150 |
| AT1G04830 | -0.2 | 1.24E-02 | -0.3 | 7.49E-03 | 0.0 | 7.31E-01 | -0.2 | 1.88E-01 | -0.2 | 2.85E-01 | 0.0 | 9.61E-01 | Ypt |
| AT1G79430 | -0.2 | 1.55E-02 | -0.1 | 3.77E-01 | -0.2 | 1.36E-01 | -0.1 | 5.75E-01 | -0.1 | 5.92E-01 | 0.0 | 9.81E-01 | WDY |
| AT3G25840 | -0.2 | 7.03E-03 | -0.4 | 8.40E-04 | 0.1 | 1.64E-01 | -0.1 | 6.80E-01 | 0.0 | 9.48E-01 | 0.0 | 8.38E-01 | non-specific serine |
| AT1G63270 | -0.2 | 1.80E-02 | -0.2 | 9.11E-02 | -0.1 | 4.94E-01 | -0.2 | 2.02E-01 | -0.5 | 1.45E-02 | 0.3 | 2.41E-01 | ABC transporter I family member 1 |
| AT3G23810 | -0.2 | 7.13E-02 | -0.1 | 5.93E-01 | -0.2 | 2.42E-01 | 0.2 | 1.37E-01 | 0.6 | 2.34E-04 | -0.5 | 7.26E-03 | Adenosylhomocysteinase 2 |
| AT3G09100 | -0.2 | 5.44E-03 | -0.5 | 1.22E-04 | 0.2 | 1.70E-02 | 0.0 | 8.76E-01 | 0.1 | 6.22E-01 | -0.1 | 5.92E-01 | AT3g09100 |
| AT2G31280 | -0.2 | 5.58E-02 | -0.4 | 7.73E-03 | 0.2 | 2.93E-01 | 0.0 | 9.88E-01 | -0.1 | 7.19E-01 | 0.1 | 7.26E-01 | Transcription factor bHLH155 |
| AT2G03220 | -0.2 | 2.05E-02 | -0.1 | 3.47E-01 | -0.1 | 1.79E-01 | 0.1 | 7.66E-01 | 0.0 | 9.20E-01 | 0.0 | 9.20E-01 | Galactoside 2-alpha-L-fucosyltransferase -fucosyltransferase) |
| AT5G37000 | -0.2 | 6.34E-02 | -0.2 | 1.45E-01 | -0.1 | 7.59E-01 | -0.2 | 6.49E-01 | 0.1 | 9.18E-01 | -0.2 | 6.47E-01 | Exostosin GT47 domain-containing protein |
| AT3G63030 | -0.2 | 2.17E-02 | -0.3 | 4.38E-03 | 0.1 | 3.66E-01 | 0.4 | 1.84E-02 | 0.2 | 1.68E-01 | 0.1 | 5.69E-01 | Methyl-CpG-binding domain-containing protein 4 |

| | | | | | | | | | | | | | |
|---|---|---|---|---|---|---|---|---|---|---|---|---|---|
| AT1G65580 | -0.2 | 3.84E-03 | -0.3 | 1.49E-03 | 0.1 | 4.87E-01 | 0.0 | 9.01E-01 | 0.1 | 8.19E-01 | -0.1 | 7.75E-01 | Endonuclease |
| AT4G01660 | -0.2 | 2.05E-02 | -0.3 | 1.63E-02 | 0.0 | 8.59E-01 | 0.5 | 6.17E-02 | 0.3 | 2.39E-01 | 0.1 | 7.27E-01 | Protein ABC transporter 1, mitochondrial |
| AT1G22880 | -0.2 | 3.16E-02 | 0.0 | 8.70E-01 | -0.3 | 4.77E-02 | -0.2 | 3.11E-01 | -0.2 | 4.31E-01 | 0.0 | 9.48E-01 | cellulase |
| AT3G06620 | -0.2 | 1.64E-02 | -0.4 | 1.24E-03 | 0.2 | 1.33E-01 | -0.2 | 1.80E-01 | -0.1 | 6.82E-01 | -0.1 | 5.83E-01 | PAS domain-containing protein tyrosine kinase family protein |
| AT2G46240 | -0.2 | 2.91E-02 | -0.1 | 3.83E-01 | -0.1 | 2.23E-01 | -0.5 | 4.56E-02 | -0.2 | 3.72E-01 | -0.2 | 4.98E-01 | BAG family molecular chaperone regulator 6 |
| AT1G78600 | -0.2 | 2.61E-02 | -0.1 | 2.37E-01 | -0.1 | 2.86E-01 | 0.2 | 4.09E-01 | -0.1 | 6.56E-01 | 0.3 | 1.59E-01 | Light-regulated zinc finger protein 1 |
| AT2G41010 | -0.2 | 1.10E-02 | -0.4 | 1.06E-03 | 0.1 | 1.62E-01 | 0.3 | 2.58E-01 | -0.1 | 8.20E-01 | 0.3 | 2.54E-01 | Calmodulin-binding protein 25 |
| AT4G26630 | -0.2 | 2.27E-02 | -0.1 | 2.97E-01 | -0.1 | 2.09E-01 | 0.0 | 8.87E-01 | 0.1 | 5.75E-01 | -0.1 | 5.62E-01 | DEK-C domain-containing protein |
| AT5G07180 | -0.2 | 1.34E-01 | -0.3 | 8.91E-02 | 0.1 | 8.01E-01 | -0.1 | 8.68E-01 | 0.0 | 9.89E-01 | -0.1 | 8.83E-01 | ERECTA-like 2 |
| AT2G20760 | -0.2 | 1.54E-02 | 0.0 | 8.20E-01 | -0.3 | 1.87E-02 | 0.1 | 6.55E-01 | 0.0 | 9.19E-01 | 0.0 | 8.62E-01 | Clathrin light chain 1 |
| AT1G52040 | -0.2 | 4.90E-01 | 0.1 | 8.55E-01 | -0.3 | 4.59E-01 | -0.1 | 8.99E-01 | 0.1 | 9.27E-01 | -0.1 | 8.65E-01 | Myrosinase-binding protein 1 |
| AT3G16170 | -0.2 | 4.85E-02 | -0.5 | 1.06E-03 | 0.3 | 4.80E-02 | 0.2 | 4.25E-01 | 0.0 | 9.15E-01 | 0.1 | 6.49E-01 | Malonate--CoA ligase |
| AT3G11270 | -0.2 | 2.24E-02 | -0.3 | 7.23E-03 | 0.1 | 5.19E-01 | 0.5 | 3.81E-03 | 0.2 | 1.77E-01 | 0.2 | 2.14E-01 | Mov34 |
| AT5G65210 | -0.2 | 2.40E-02 | 0.0 | 9.76E-01 | -0.2 | 3.83E-02 | 0.2 | 2.86E-01 | 0.0 | 9.72E-01 | 0.2 | 4.09E-01 | BZIP transcription factor family protein |
| AT3G05530 | -0.2 | 4.52E-03 | -0.2 | 1.75E-02 | 0.0 | 5.80E-01 | -0.1 | 5.22E-01 | -0.2 | 3.33E-01 | 0.1 | 8.01E-01 | 26S proteasome regulatory subunit 6A homolog A |
| AT4G29040 | -0.2 | 1.16E-02 | -0.3 | 3.34E-03 | 0.1 | 4.09E-01 | 0.3 | 1.16E-02 | 0.3 | 2.51E-02 | 0.0 | 9.78E-01 | 26S proteasome regulatory subunit 4 homolog A |
| AT3G23640 | -0.2 | 9.79E-03 | -0.3 | 1.87E-03 | 0.1 | 2.97E-01 | -0.3 | 5.11E-02 | -0.2 | 2.20E-01 | -0.1 | 7.39E-01 | Alpha glucosidase-like protein |
| AT2G30440 | -0.2 | 6.03E-03 | -0.2 | 7.41E-03 | 0.0 | 9.67E-01 | -0.1 | 5.44E-01 | -0.2 | 3.86E-02 | 0.2 | 2.32E-01 | Thylakoidal processing peptidase 1, chloroplastic |
| AT5G42020 | -0.2 | 1.36E-01 | 0.1 | 7.27E-01 | -0.3 | 1.02E-01 | 0.3 | 5.49E-02 | 0.4 | 1.88E-02 | -0.1 | 6.49E-01 | Heat shock protein 70 family protein |
| AT5G49810 | -0.2 | 4.51E-03 | -0.2 | 2.15E-02 | -0.1 | 4.96E-01 | -0.1 | 7.44E-01 | 0.1 | 6.02E-01 | -0.1 | 4.87E-01 | Methionine S-methyltransferase |
| AT2G22770 | -0.2 | 2.29E-02 | -0.1 | 4.33E-01 | -0.2 | 1.59E-01 | -0.2 | 6.45E-01 | 0.4 | 2.18E-01 | -0.6 | 1.43E-01 | Transcription factor NAI1 |
| AT1G07890 | -0.2 | 7.35E-02 | -0.2 | 2.14E-01 | -0.1 | 6.29E-01 | -0.6 | 1.56E-04 | 0.2 | 1.93E-01 | -0.8 | 5.83E-04 | L-ascorbate peroxidase |
| AT3G53280 | -0.2 | 1.02E-01 | 0.0 | 7.56E-01 | -0.3 | 9.32E-02 | 0.0 | 8.95E-01 | -0.1 | 3.97E-01 | 0.1 | 5.69E-01 | AT3g53280 |
| AT2G33630 | -0.2 | 1.09E-02 | -0.2 | 8.18E-02 | -0.1 | 3.54E-01 | -0.2 | 2.49E-01 | -0.2 | 2.38E-01 | 0.0 | 9.27E-01 | 3-beta hydroxysteroid dehydrogenase |
| AT5G42220 | -0.2 | 6.18E-03 | -0.3 | 9.94E-04 | 0.1 | 2.23E-01 | -0.1 | 5.29E-01 | -0.1 | 7.00E-01 | 0.0 | 9.37E-01 | Ubiquitin-like domain-containing protein |
| AT4G38350 | -0.2 | 7.64E-03 | -0.4 | 7.97E-04 | 0.1 | 1.43E-01 | -0.1 | 7.34E-01 | 0.0 | 9.52E-01 | -0.1 | 8.73E-01 | Patched family protein |
| AT5G19450 | -0.2 | 3.33E-03 | -0.3 | 1.00E-03 | 0.1 | 3.77E-01 | -0.3 | 1.05E-02 | -0.3 | 1.75E-02 | 0.0 | 9.81E-01 | EF-hand domain-containing protein |
| AT3G27330 | -0.2 | 5.61E-02 | 0.1 | 6.08E-01 | -0.3 | 4.92E-02 | 0.1 | 4.54E-01 | 0.0 | 8.61E-01 | 0.2 | 3.76E-01 | RING-type domain-containing protein |
| AT4G21060 | -0.2 | 2.09E-02 | -0.2 | 1.95E-02 | 0.0 | 9.21E-01 | -0.1 | 8.55E-01 | 0.3 | 5.53E-01 | -0.3 | 4.52E-01 | Galactosyltransferase family protein |
| AT5G63630 | -0.2 | 4.46E-02 | -0.3 | 1.35E-02 | 0.1 | 5.19E-01 | 0.2 | 3.75E-01 | 0.1 | 6.67E-01 | 0.1 | 8.32E-01 | DEAD-box ATP-dependent RNA helicase 31 |
| AT1G79880 | -0.2 | 3.04E-02 | 0.0 | 7.40E-01 | -0.2 | 1.07E-01 | -0.4 | 7.85E-02 | -0.3 | 1.85E-01 | -0.1 | 8.78E-01 | La protein 2 |
| AT2G39350 | -0.2 | 6.46E-02 | -0.2 | 1.27E-01 | 0.0 | 7.96E-01 | 0.3 | 8.96E-02 | 0.2 | 3.28E-01 | 0.1 | 7.34E-01 | ABC transporter G family member 1 |
| AT1G25470 | -0.2 | 1.38E-02 | -0.1 | 1.59E-01 | -0.1 | 2.78E-01 | -0.1 | 6.54E-01 | -0.2 | 3.77E-01 | 0.1 | 7.49E-01 | AP2 |
| AT5G23280 | -0.2 | 2.56E-02 | -0.1 | 5.46E-01 | -0.2 | 1.23E-01 | -0.3 | 1.17E-01 | -0.1 | 7.46E-01 | -0.2 | 4.41E-01 | Transcription factor TCP7 |
| AT3G60580 | -0.2 | 1.10E-01 | -0.2 | 1.43E-01 | 0.0 | 9.54E-01 | 0.1 | 9.21E-01 | 0.0 | 9.70E-01 | 0.0 | 9.75E-01 | Zinc finger protein ZAT9 |
| AT1G04680 | -0.2 | 4.62E-02 | 0.3 | 4.27E-02 | -0.5 | 2.65E-03 | -0.3 | 6.65E-02 | 0.1 | 7.93E-01 | -0.4 | 1.06E-01 | Probable pectate lyase 1 |
| AT1G31150 | -0.2 | 1.79E-02 | -0.1 | 2.40E-01 | -0.1 | 2.37E-01 | -0.4 | 6.07E-02 | 0.0 | 8.82E-01 | -0.4 | 8.76E-02 | K-box domain-containing protein |
| AT1G69780 | -0.2 | 4.06E-02 | 0.0 | 8.22E-01 | -0.3 | 5.11E-02 | -0.3 | 3.50E-02 | -0.2 | 1.69E-01 | -0.1 | 7.24E-01 | Homeobox-leucine zipper protein ATHB-13 |
| AT3G61970 | -0.2 | 6.22E-02 | -0.1 | 4.39E-01 | -0.1 | 3.45E-01 | -0.3 | 2.01E-01 | -0.3 | 1.55E-01 | 0.1 | 8.71E-01 | B3 domain-containing transcription factor NGA2 |
| AT3G18060 | -0.2 | 9.58E-03 | 0.0 | 5.53E-01 | -0.3 | 7.25E-03 | -0.3 | 5.01E-03 | -0.2 | 5.48E-02 | -0.1 | 6.27E-01 | Actin-interacting protein 1-2 |
| AT3G17390 | -0.2 | 8.23E-02 | 0.1 | 4.80E-01 | -0.3 | 3.66E-02 | 0.0 | 9.04E-01 | 0.1 | 7.30E-01 | 0.0 | 8.45E-01 | S-adenosylmethionine synthase 4 |
| AT1G17160 | -0.2 | 9.46E-03 | -0.2 | 1.31E-02 | 0.0 | 9.51E-01 | -0.2 | 1.88E-01 | 0.0 | 9.32E-01 | -0.2 | 4.11E-01 | Ribokinase |
| AT1G44180 | -0.2 | 1.20E-01 | -0.8 | 8.80E-04 | 0.5 | 1.97E-02 | | | | | | | Aminoacylase, putative |
| AT5G20930 | -0.2 | 6.56E-03 | -0.2 | 6.46E-03 | 0.0 | 9.05E-01 | 0.0 | 9.85E-01 | -0.1 | 8.43E-01 | 0.1 | 8.75E-01 | Serine |
| AT5G10020 | -0.2 | 3.27E-02 | -0.1 | 2.08E-01 | -0.1 | 3.88E-01 | -0.3 | 1.20E-01 | -0.2 | 4.09E-01 | -0.1 | 6.79E-01 | Probable inactive receptor kinase At5g10020 |
| AT5G64813 | -0.2 | 3.50E-03 | -0.3 | 2.04E-03 | 0.0 | 6.56E-01 | -0.1 | 3.77E-01 | -0.1 | 6.53E-01 | -0.1 | 8.62E-01 | Small GTPase LIP1 |
| AT1G11600 | -0.2 | 1.25E-01 | -0.1 | 4.08E-01 | -0.1 | 5.56E-01 | -0.2 | 4.48E-01 | -0.2 | 5.20E-01 | 0.0 | 9.98E-01 | Cytochrome P450 like protein |
| AT5G01780 | -0.2 | 2.77E-02 | -0.4 | 3.61E-03 | 0.1 | 2.60E-01 | 0.0 | 9.51E-01 | 0.0 | 9.31E-01 | 0.0 | 8.87E-01 | 2-oxoglutarate-dependent dioxygenase family protein |
| AT5G57360 | -0.2 | 1.94E-02 | -0.3 | 2.79E-03 | 0.1 | 2.49E-01 | -0.4 | 1.66E-02 | -0.4 | 1.97E-02 | 0.0 | 9.23E-01 | Galactose oxidase |
| AT5G11730 | -0.2 | 6.52E-03 | -0.2 | 8.39E-03 | 0.0 | 9.90E-01 | -0.2 | 2.53E-01 | -0.1 | 7.91E-01 | -0.1 | 5.88E-01 | At5g11730 |
| AT5G01470 | -0.2 | 2.35E-02 | -0.1 | 2.37E-01 | -0.1 | 3.02E-01 | -0.2 | 4.83E-01 | -0.1 | 8.10E-01 | -0.1 | 8.07E-01 | S-adenosyl-L-methionine-dependent methyltransferases superfamily protein |
| AT3G09340 | -0.2 | 3.89E-01 | 1.5 | 5.45E-05 | -1.8 | 1.93E-04 | | | | | | | Amino acid transporter AVT1F |
| AT3G04080 | -0.2 | 4.19E-03 | -0.1 | 2.09E-01 | -0.1 | 7.08E-02 | -0.1 | 5.20E-01 | -0.1 | 6.83E-01 | 0.0 | 9.40E-01 | apyrase |
| AT2G47680 | -0.2 | 1.67E-02 | -0.4 | 9.22E-04 | 0.2 | 1.04E-01 | 0.1 | 7.09E-01 | 0.0 | 9.32E-01 | 0.1 | 8.67E-01 | DExH-box ATP-dependent RNA helicase DExH8 |
| AT2G47270 | -0.2 | 4.30E-01 | 0.4 | 1.76E-01 | -0.6 | 8.71E-02 | 0.0 | 9.86E-01 | 0.3 | 5.59E-01 | -0.4 | 5.16E-01 | Transcription factor UPBEAT1 |
| AT4G31150 | -0.2 | 9.15E-02 | -0.3 | 2.93E-02 | 0.1 | 5.62E-01 | -0.2 | 4.99E-01 | -0.4 | 2.63E-01 | 0.2 | 7.33E-01 | hypothetical protein |
| AT5G12110 | -0.2 | 9.16E-02 | -0.1 | 3.03E-01 | -0.1 | 5.90E-01 | 0.2 | 6.87E-01 | 0.3 | 3.79E-01 | -0.2 | 7.17E-01 | Elongation factor 1-beta 1 |
| AT5G56720 | -0.2 | 1.61E-01 | 0.0 | 9.68E-01 | -0.2 | 2.69E-01 | 0.1 | 8.47E-01 | 0.2 | 7.18E-01 | -0.1 | 8.62E-01 | Malate dehydrogenase |
| AT1G64380 | -0.2 | 4.27E-01 | 0.2 | 5.92E-01 | -0.4 | 2.64E-01 | -0.8 | 1.29E-02 | 0.2 | 6.80E-01 | -1.0 | 2.24E-02 | Ethylene-responsive transcription factor ERF061 |
| AT4G39370 | -0.2 | 1.75E-02 | -0.2 | 2.28E-02 | 0.0 | 9.68E-01 | 0.0 | 8.71E-01 | 0.0 | 9.70E-01 | 0.0 | 9.35E-01 | Ubiquitin-specific protease 27 |
| AT2G37440 | -0.2 | 3.11E-02 | -0.3 | 9.53E-03 | 0.1 | 5.09E-01 | -0.4 | 2.52E-01 | 0.4 | 1.87E-01 | -0.8 | 2.42E-02 | DNAse I-like superfamily protein |
| AT1G16420 | -0.2 | 2.48E-01 | -0.2 | 3.09E-01 | 0.0 | 9.38E-01 | 0.6 | 1.14E-01 | 0.0 | 9.81E-01 | 0.6 | 1.77E-01 | Metacaspase-8 |
| AT5G06950 | -0.2 | 2.07E-02 | -0.3 | 7.29E-03 | 0.1 | 5.46E-01 | -0.1 | 5.63E-01 | -0.1 | 6.75E-01 | 0.0 | 9.65E-01 | Transcription factor TGA2 |
| AT1G13350 | -0.2 | 1.59E-02 | -0.4 | 1.74E-03 | 0.1 | 1.93E-01 | -0.1 | 7.36E-01 | 0.1 | 8.19E-01 | -0.2 | 5.90E-01 | non-specific serine |
| AT2G19110 | -0.2 | 2.88E-02 | -0.2 | 1.01E-01 | -0.1 | 5.80E-01 | -0.2 | 4.53E-01 | 0.0 | 9.92E-01 | -0.2 | 5.87E-01 | Putative cadmium |
| AT2G16485 | -0.2 | 5.69E-03 | -0.3 | 8.47E-04 | 0.1 | 2.09E-01 | 0.2 | 1.07E-01 | 0.1 | 4.86E-01 | 0.1 | 6.27E-01 | GW repeat-and PHD finger-containing protein NERD |
| AT5G24670 | -0.2 | 1.02E-02 | -0.3 | 2.00E-03 | 0.1 | 3.14E-01 | -0.2 | 1.11E-01 | 0.0 | 9.75E-01 | -0.2 | 1.85E-01 | Cytidine |
| AT4G29140 | -0.2 | 2.09E-02 | -0.3 | 1.41E-02 | 0.0 | 7.93E-01 | 0.1 | 4.93E-01 | -0.2 | 2.88E-01 | 0.3 | 1.32E-01 | Protein DETOXIFICATION 51 |
| AT3G19300 | -0.2 | 1.33E-01 | -0.1 | 4.90E-01 | -0.1 | 5.17E-01 | 0.3 | 3.94E-01 | -0.1 | 9.11E-01 | 0.4 | 4.51E-01 | Protein kinase domain-containing protein |
| AT4G32200 | -0.2 | 1.24E-01 | 0.0 | 9.91E-01 | -0.2 | 2.05E-01 | 0.1 | 7.52E-01 | 0.1 | 8.89E-01 | 0.0 | 9.25E-01 | DNA-binding HORMA family protein |
| AT2G38490 | -0.2 | 2.38E-01 | 0.7 | 3.02E-03 | -0.9 | 2.19E-03 | 0.2 | 6.76E-01 | 0.3 | 5.65E-01 | -0.1 | 8.79E-01 | CBL-interacting serine |
| AT4G31550 | -0.2 | 6.02E-03 | -0.2 | 2.00E-02 | 0.0 | 6.23E-01 | 0.1 | 3.55E-01 | 0.1 | 5.10E-01 | 0.0 | 9.47E-01 | WRKY DNA-binding protein 11 |
| AT3G52910 | -0.2 | 1.66E-01 | 0.0 | 9.16E-01 | -0.2 | 2.14E-01 | -0.3 | 2.72E-01 | 0.1 | 6.56E-01 | -0.4 | 1.28E-01 | Growth-regulating factor 4 |
| AT2G39330 | -0.2 | 5.65E-01 | 0.1 | 8.33E-01 | -0.3 | 5.14E-01 | 1.2 | 4.48E-01 | 2.6 | 9.33E-02 | -1.4 | 3.31E-01 | Jacalin-related lectin 23 |
| AT3G15820 | -0.2 | 9.72E-03 | -0.1 | 2.94E-01 | -0.1 | 1.03E-01 | -0.6 | 2.94E-04 | -0.4 | 8.58E-03 | -0.2 | 4.11E-01 | Phosphatidylcholine:diacylglycerol cholinephosphotransferase 1 |
| AT1G44820 | -0.2 | 1.14E-02 | -0.4 | 6.47E-04 | 0.2 | 8.25E-02 | -0.1 | 3.82E-01 | 0.2 | 3.24E-01 | -0.3 | 1.42E-01 | At1g44820 |
| AT4G18340 | -0.2 | 1.15E-01 | 0.4 | 1.77E-02 | -0.6 | 3.92E-03 | 0.6 | 9.37E-03 | -0.1 | 7.30E-01 | 0.7 | 1.75E-02 | glucan endo-1,3-beta-D-glucosidase |
| AT3G10330 | -0.2 | 7.36E-03 | -0.1 | 2.03E-01 | -0.1 | 1.26E-01 | -0.2 | 2.85E-01 | -0.1 | 7.35E-01 | -0.1 | 7.04E-01 | Transcription initiation factor IIB-2 |
| AT1G22910 | -0.2 | 3.63E-02 | 0.0 | 8.55E-01 | -0.2 | 8.80E-02 | -0.1 | 6.56E-01 | 0.0 | 9.52E-01 | -0.1 | 7.89E-01 | At1g22910 |
| AT3G51630 | -0.2 | 9.96E-03 | -0.4 | 2.59E-04 | 0.2 | 2.88E-02 | -0.2 | 2.33E-01 | 0.1 | 6.81E-01 | -0.3 | 2.01E-01 | AT3g51630 |
| AT3G63080 | -0.2 | 3.37E-02 | -0.1 | 4.37E-01 | -0.1 | 1.89E-01 | -0.1 | 3.15E-01 | -0.2 | 2.78E-01 | 0.0 | 9.04E-01 | Glutathione peroxidase |
| AT2G14050 | -0.2 | 1.99E-01 | -0.1 | 6.46E-01 | -0.1 | 5.18E-01 | 0.6 | 1.69E-01 | 0.4 | 4.24E-01 | 0.2 | 7.23E-01 | DNA helicase |
| AT1G02890 | -0.2 | 1.97E-02 | -0.3 | 3.58E-03 | 0.1 | 3.05E-01 | 0.1 | 5.33E-01 | 0.0 | 8.44E-01 | 0.1 | 8.31E-01 | AAA-type ATPase family protein |

| | | | | | | | | | | | | | |
|---|---|---|---|---|---|---|---|---|---|---|---|---|---|
| AT5G59150 | -0.2 | 1.26E-02 | 0.1 | 3.29E-01 | -0.3 | 5.76E-03 | -0.2 | 2.06E-01 | 0.0 | 9.46E-01 | -0.2 | 2.97E-01 | RAB GTPase homolog A2D |
| AT3G17750 | -0.2 | 1.34E-02 | -0.1 | 1.21E-01 | -0.1 | 3.15E-01 | 0.0 | 9.88E-01 | 0.0 | 9.87E-01 | 0.0 | 9.78E-01 | Ser-Thr protein kinase-like protein |
| AT4G01540 | -0.2 | 4.19E-02 | -0.3 | 8.06E-03 | 0.1 | 3.83E-01 | 0.3 | 1.87E-01 | 0.3 | 2.32E-01 | 0.0 | 9.92E-01 | NAC with transmembrane motif1 |
| AT4G23620 | -0.2 | 8.43E-02 | -0.1 | 5.23E-01 | -0.1 | 3.45E-01 | 0.6 | 9.71E-04 | 0.5 | 8.95E-03 | 0.1 | 6.49E-01 | Ribosomal protein L25 |
| AT3G53710 | -0.2 | 1.57E-03 | -0.1 | 2.81E-02 | -0.1 | 1.65E-01 | -0.2 | 1.32E-01 | -0.2 | 1.39E-01 | 0.0 | 9.34E-01 | Probable ADP-ribosylation factor GTPase-activating protein AGD6 |
| AT4G25320 | -0.2 | 5.14E-03 | -0.2 | 8.57E-03 | 0.0 | 8.92E-01 | 0.3 | 1.20E-02 | 0.2 | 1.38E-01 | 0.1 | 5.27E-01 | AT-hook motif nuclear-localized protein 3 |
| AT3G09770 | -0.2 | 1.26E-02 | -0.2 | 4.64E-02 | -0.1 | 5.74E-01 | -0.3 | 1.79E-02 | -0.2 | 6.88E-02 | 0.0 | 8.22E-01 | E3 ubiquitin-protein ligase |
| AT3G50070 | -0.2 | 2.93E-02 | -0.2 | 6.91E-02 | 0.0 | 7.46E-01 | 0.0 | 9.06E-01 | 0.1 | 8.39E-01 | -0.1 | 7.72E-01 | Cyclin-D3-3 |
| AT1G48210 | -0.2 | 4.52E-02 | -0.4 | 4.39E-03 | 0.2 | 2.02E-01 | 0.1 | 7.97E-01 | 0.0 | 9.83E-01 | 0.0 | 8.94E-01 | Protein kinase domain-containing protein |
| AT1G78020 | -0.2 | 1.35E-01 | -0.1 | 7.25E-01 | -0.2 | 3.07E-01 | -0.4 | 3.15E-02 | 0.0 | 9.72E-01 | -0.4 | 1.43E-01 | FCS-Like Zinc finger 6 |
| AT1G64760 | -0.2 | 8.93E-02 | -0.2 | 1.38E-01 | 0.0 | 8.86E-01 | 0.0 | 9.22E-01 | 0.2 | 3.92E-01 | -0.2 | 3.72E-01 | Glucan endo-1,3-beta-glucosidase 8 -beta-glucan endohydrolase 8) -beta-glucanase 8) |
| AT5G39030 | -0.2 | 5.46E-02 | -0.4 | 3.84E-03 | 0.2 | 1.65E-01 | 0.3 | 1.64E-01 | 0.2 | 3.50E-01 | 0.1 | 8.59E-01 | Protein kinase superfamily protein |
| AT5G46370 | -0.2 | 3.02E-01 | 0.7 | 3.08E-03 | -0.9 | 3.00E-03 | -1.4 | 1.85E-02 | -0.7 | 2.49E-01 | | | Two-pore potassium channel 2 |
| AT3G06400 | -0.2 | 5.81E-03 | -0.3 | 1.88E-03 | 0.1 | 4.19E-01 | 0.0 | 9.87E-01 | 0.1 | 7.55E-01 | -0.1 | 7.93E-01 | Chromatin-remodeling protein 11 |
| AT5G42320 | -0.2 | 8.41E-03 | -0.1 | 2.58E-01 | -0.1 | 1.13E-01 | -0.1 | 6.91E-01 | 0.0 | 9.15E-01 | -0.1 | 6.68E-01 | Peptidase M14 carboxypeptidase A domain-containing protein |
| AT1G03520 | -0.2 | 2.86E-02 | -0.1 | 5.62E-01 | -0.2 | 1.45E-01 | -0.2 | 4.60E-01 | 0.2 | 4.70E-01 | -0.3 | 1.99E-01 | Core-2 |
| AT3G02020 | -0.2 | 1.14E-01 | 0.0 | 7.88E-01 | -0.3 | 9.80E-02 | -0.9 | 1.26E-03 | 0.7 | 1.05E-02 | -1.6 | 2.85E-04 | Aspartokinase |
| AT3G62560 | -0.2 | 5.32E-02 | -0.3 | 3.13E-02 | 0.0 | 7.61E-01 | 0.0 | 9.43E-01 | -0.1 | 7.90E-01 | 0.1 | 8.64E-01 | hypothetical protein |
| AT4G18890 | -0.2 | 6.54E-02 | -0.3 | 3.36E-02 | 0.1 | 7.20E-01 | 0.1 | 7.33E-01 | 0.3 | 2.32E-01 | -0.2 | 4.88E-01 | Protein BZR1 homolog |
| AT1G59580 | -0.2 | 2.07E-03 | -0.2 | 6.19E-03 | 0.0 | 6.60E-01 | -0.2 | 2.11E-01 | -0.2 | 1.24E-01 | 0.1 | 7.40E-01 | Mitogen-activated protein kinase 2 |
| AT1G72150 | -0.2 | 1.03E-01 | 0.1 | 4.73E-01 | -0.3 | 4.40E-02 | -0.5 | 2.03E-04 | -0.8 | 2.35E-05 | 0.3 | 1.18E-01 | Patellin-1 |
| AT3G07690 | -0.2 | 1.03E-02 | -0.3 | 1.01E-03 | 0.1 | 1.63E-01 | 0.0 | 9.05E-01 | -0.2 | 5.33E-01 | 0.2 | 5.15E-01 | Glycerol-3-phosphate dehydrogenase |
| AT2G01490 | -0.2 | 3.56E-03 | -0.2 | 2.50E-02 | -0.1 | 3.68E-01 | -0.3 | 1.67E-02 | -0.2 | 5.02E-02 | 0.0 | 8.96E-01 | Phytanoyl-CoA dioxygenase |
| AT4G34680 | -0.2 | 5.83E-02 | -0.2 | 8.13E-02 | 0.0 | 9.33E-01 | 0.3 | 1.37E-01 | 0.0 | 9.03E-01 | 0.4 | 2.13E-01 | GATA transcription factor 3 |
| AT3G13380 | -0.2 | 1.33E-01 | -0.7 | 9.81E-04 | 0.5 | 1.89E-02 | 0.1 | 5.70E-01 | 0.2 | 4.40E-01 | -0.1 | 8.50E-01 | Receptor-like protein kinase BRI1-like 3 |
| AT2G27480 | -0.2 | 6.52E-02 | -0.2 | 1.24E-01 | 0.0 | 8.21E-01 | -0.8 | 4.10E-02 | -0.5 | 2.02E-01 | -0.3 | 5.85E-01 | Calcium-binding EF-hand family protein |
| AT4G10040 | -0.2 | 5.99E-02 | 0.2 | 1.80E-01 | -0.4 | 1.15E-02 | 0.5 | 1.27E-01 | 0.4 | 2.29E-01 | 0.0 | 9.25E-01 | Cytochrome c-2 |
| AT5G35330 | -0.2 | 6.85E-03 | -0.2 | 9.99E-03 | 0.0 | 9.43E-01 | -0.5 | 3.91E-03 | -0.5 | 4.49E-03 | 0.0 | 9.03E-01 | Uncharacterized protein |
| AT5G56730 | -0.2 | 1.99E-02 | -0.3 | 6.43E-03 | 0.1 | 5.13E-01 | 0.0 | 9.56E-01 | 0.1 | 8.59E-01 | -0.1 | 7.99E-01 | Peptidase M16 C-terminal domain-containing protein |
| AT4G27130 | -0.2 | 1.91E-02 | 0.1 | 4.29E-01 | -0.3 | 9.54E-03 | -0.1 | 2.71E-01 | -0.2 | 1.00E-01 | 0.1 | 6.52E-01 | Protein translation factor SUI1 homolog 1 |
| AT4G11530 | -0.2 | 4.95E-02 | -0.3 | 1.87E-02 | 0.1 | 6.03E-01 | -0.3 | 1.98E-01 | -0.2 | 5.16E-01 | -0.1 | 7.13E-01 | Cysteine-rich RLK 34 |
| AT5G24260 | -0.2 | 8.58E-03 | 0.0 | 9.88E-01 | -0.2 | 1.80E-02 | -0.2 | 1.51E-01 | -0.2 | 2.83E-01 | 0.0 | 9.16E-01 | Dipeptidyl peptidase IV-like protein |
| AT5G24850 | -0.2 | 4.33E-02 | -0.1 | 2.08E-01 | -0.1 | 4.77E-01 | 0.1 | 8.77E-01 | 0.1 | 8.71E-01 | 0.0 | 9.82E-01 | Cryptochrome DASH, chloroplastic |
| AT1G67970 | -0.2 | 1.68E-01 | -0.6 | 4.37E-03 | 0.4 | 7.13E-02 | 0.2 | 3.49E-01 | -0.3 | 2.19E-01 | 0.4 | 5.70E-02 | Heat stress transcription factor A-8 |
| AT1G10320 | -0.2 | 1.24E-02 | -0.5 | 2.31E-04 | 0.2 | 2.36E-02 | 0.0 | 9.78E-01 | 0.0 | 9.69E-01 | 0.0 | 9.90E-01 | Zinc finger C-x8-C-x5-C-x3-H type family protein |
| AT2G14750 | -0.2 | 5.69E-02 | 0.0 | 7.15E-01 | -0.3 | 4.54E-02 | 0.0 | 9.87E-01 | 0.5 | 2.14E-02 | -0.6 | 5.13E-02 | Adenylyl-sulfate kinase 1, chloroplastic |
| AT4G33070 | -0.2 | 4.34E-02 | -0.7 | 1.03E-04 | 0.5 | 3.39E-03 | 3.5 | 1.03E-05 | 1.8 | 9.74E-03 | 1.7 | 3.64E-04 | Pyruvate decarboxylase 1 |
| AT2G13560 | -0.2 | 6.91E-03 | -0.3 | 3.87E-03 | 0.0 | 6.37E-01 | 0.1 | 5.06E-01 | 0.1 | 3.76E-01 | 0.0 | 8.53E-01 | NAD-dependent malic enzyme 1, mitochondrial |
| AT1G11660 | -0.2 | 7.85E-03 | 0.0 | 8.67E-01 | -0.2 | 1.27E-02 | -0.2 | 2.93E-01 | 0.0 | 8.76E-01 | -0.1 | 5.96E-01 | Heat shock protein 70 family protein |
| AT1G54170 | -0.2 | 1.86E-02 | -0.3 | 2.21E-03 | 0.1 | 2.17E-01 | -0.1 | 6.22E-01 | 0.0 | 8.73E-01 | -0.2 | 5.85E-01 | Polyadenylate-binding protein-interacting protein 3 -binding protein-interacting protein 3) |
| AT2G27040 | -0.2 | 4.50E-03 | -0.2 | 1.82E-02 | 0.0 | 5.43E-01 | -0.1 | 7.53E-01 | 0.1 | 7.86E-01 | -0.1 | 6.17E-01 | Protein argonaute 4 |
| AT5G60620 | -0.2 | 3.24E-03 | -0.1 | 4.58E-02 | -0.1 | 2.14E-01 | -0.3 | 2.19E-02 | -0.1 | 4.05E-01 | -0.2 | 3.48E-01 | Glycerol-3-phosphate acyltransferase 9 |
| AT1G63490 | -0.2 | 1.03E-01 | -0.4 | 1.26E-02 | 0.2 | 2.79E-01 | 0.2 | 3.69E-01 | 0.4 | 9.80E-02 | -0.2 | 5.13E-01 | Transcription factor jumonji domain-containing protein |
| AT3G49060 | -0.2 | 3.51E-02 | -0.3 | 4.63E-03 | 0.1 | 2.67E-01 | 0.2 | 3.95E-01 | 0.2 | 5.34E-01 | 0.0 | 9.57E-01 | U-box domain-containing protein kinase family protein |
| AT4G18800 | -0.2 | 2.30E-02 | 0.0 | 7.55E-01 | -0.2 | 7.71E-02 | 0.1 | 6.28E-01 | 0.0 | 9.01E-01 | 0.1 | 8.36E-01 | Ras-related protein RABA1d |
| AT3G15790 | -0.2 | 5.87E-02 | 0.0 | 9.32E-01 | -0.2 | 1.10E-01 | 0.0 | 9.29E-01 | 0.2 | 5.72E-01 | -0.1 | 7.13E-01 | Methyl-CpG-binding domain-containing protein 11 |
| AT5G63260 | -0.2 | 8.26E-03 | -0.3 | 1.10E-03 | 0.1 | 2.10E-01 | -0.3 | 1.46E-01 | -0.1 | 7.42E-01 | -0.2 | 4.74E-01 | Zinc finger C-x8-C-x5-C-x3-H type family protein |
| AT5G41480 | -0.2 | 5.52E-02 | -0.1 | 2.14E-01 | -0.1 | 5.49E-01 | -0.2 | 3.29E-01 | 0.0 | 9.29E-01 | -0.2 | 5.39E-01 | Dihydrofolate synthetase |
| AT4G10570 | -0.2 | 1.29E-02 | -0.2 | 5.40E-02 | -0.1 | 5.51E-01 | 0.0 | 9.60E-01 | 0.0 | 9.95E-01 | 0.0 | 9.78E-01 | Ubiquitin carboxyl-terminal hydrolase 10 |
| AT5G48150 | -0.2 | 1.05E-02 | -0.4 | 5.85E-04 | 0.2 | 7.39E-02 | 0.0 | 8.18E-01 | 0.2 | 4.11E-01 | -0.1 | 6.68E-01 | Scarecrow-like transcription factor PAT1 |
| AT5G66910 | -0.2 | 2.67E-02 | -0.4 | 7.72E-04 | 0.2 | 5.40E-02 | 0.5 | 1.01E-02 | 0.5 | 1.22E-02 | 0.0 | 9.26E-01 | Probable disease resistance protein At5g66910 |
| AT5G47650 | -0.2 | 2.82E-02 | -0.3 | 1.34E-02 | 0.0 | 6.69E-01 | -0.5 | 8.52E-02 | -0.1 | 7.72E-01 | -0.4 | 3.50E-01 | Nudix hydrolase |
| AT5G17980 | -0.2 | 3.33E-02 | -0.5 | 6.81E-04 | 0.3 | 4.16E-02 | -0.2 | 5.20E-01 | 0.0 | 9.17E-01 | -0.2 | 5.16E-01 | Multiple C2 domain and transmembrane region protein 16 |
| AT1G20270 | -0.2 | 1.50E-02 | -0.1 | 2.34E-01 | -0.1 | 2.13E-01 | 0.2 | 2.67E-01 | 0.2 | 2.72E-01 | 0.0 | 9.50E-01 | 2-oxoglutarate and Fe-dependent oxygenase superfamily protein |
| AT5G15730 | -0.2 | 1.06E-02 | -0.1 | 5.07E-01 | -0.2 | 6.47E-02 | 0.3 | 2.12E-02 | 0.1 | 6.99E-01 | 0.2 | 1.88E-01 | Protein kinase domain-containing protein |
| AT1G57870 | -0.2 | 5.95E-03 | -0.2 | 3.97E-03 | 0.0 | 7.43E-01 | -0.1 | 5.52E-01 | 0.0 | 8.40E-01 | -0.1 | 4.71E-01 | non-specific serine |
| AT2G25070 | -0.2 | 4.79E-02 | -0.1 | 2.97E-01 | -0.1 | 3.66E-01 | -0.1 | 6.44E-01 | -0.2 | 2.77E-01 | 0.1 | 6.51E-01 | Probable protein phosphatase 2C 21 |
| AT3G59990 | -0.2 | 5.63E-03 | -0.1 | 6.67E-02 | -0.1 | 2.50E-01 | -0.1 | 4.86E-01 | -0.1 | 4.16E-01 | 0.0 | 9.09E-01 | Methionine aminopeptidase 2B |
| AT3G10350 | -0.2 | 4.84E-02 | 0.0 | 8.55E-01 | -0.2 | 1.07E-01 | 0.2 | 5.99E-02 | 0.2 | 2.35E-01 | 0.1 | 7.58E-01 | P-loop containing nucleoside triphosphate hydrolases superfamily protein |
| AT3G46620 | -0.2 | 9.86E-02 | -0.5 | 2.69E-03 | 0.3 | 6.59E-02 | 0.2 | 3.66E-01 | 0.1 | 7.58E-01 | 0.1 | 7.51E-01 | E3 ubiquitin-protein ligase RDUF1 |
| AT3G57710 | -0.2 | 3.35E-02 | 0.0 | 9.17E-01 | -0.2 | 5.89E-02 | 0.2 | 4.06E-01 | 0.1 | 8.18E-01 | 0.1 | 6.78E-01 | Serine |
| AT2G29960 | -0.2 | 8.68E-03 | 0.1 | 1.00E-01 | -0.3 | 1.65E-03 | 0.0 | 8.60E-01 | 0.1 | 6.85E-01 | 0.0 | 8.49E-01 | Peptidyl-prolyl cis-trans isomerase CYP19-4 |
| AT1G63050 | -0.2 | 5.48E-02 | 0.1 | 2.96E-01 | -0.3 | 1.93E-02 | -0.1 | 6.00E-01 | 0.1 | 5.73E-01 | -0.3 | 3.65E-01 | Lysophospholipid acyltransferase 2 |
| AT5G65310 | -0.2 | 1.89E-02 | 0.0 | 9.78E-01 | -0.2 | 3.45E-02 | -0.4 | 7.76E-04 | -0.1 | 6.83E-01 | -0.3 | 2.74E-02 | Homeobox-leucine zipper protein |
| AT1G53320 | -0.2 | 3.81E-03 | -0.1 | 3.29E-02 | -0.1 | 3.14E-01 | -0.4 | 1.31E-02 | -0.2 | 1.64E-01 | -0.1 | 5.11E-01 | Tubby-like F-box protein 7 |
| AT5G10380 | -0.2 | 4.24E-01 | -0.8 | 1.47E-02 | 0.6 | 8.12E-02 | -0.3 | 4.78E-01 | -0.7 | 7.76E-02 | 0.5 | 3.72E-01 | E3 ubiquitin-protein ligase RING1 |
| AT4G30410 | -0.2 | 9.78E-02 | 0.4 | 1.14E-02 | -0.6 | 3.00E-03 | -0.5 | 1.34E-02 | -0.3 | 1.37E-01 | -0.2 | 4.84E-01 | Transcription factor IBH1-like 1 |
| AT5G28080 | -0.2 | 7.93E-02 | 0.2 | 1.27E-01 | -0.4 | 1.53E-02 | -0.3 | 4.65E-01 | 0.3 | 4.68E-01 | -0.5 | 9.19E-02 | non-specific serine |
| AT3G24550 | -0.2 | 2.63E-02 | -0.2 | 4.21E-02 | 0.0 | 8.93E-01 | -0.2 | 8.74E-02 | -0.2 | 1.28E-01 | 0.0 | 9.98E-01 | Proline-rich receptor-like protein kinase PERK1 |
| AT1G76030 | -0.2 | 1.21E-02 | 0.1 | 4.04E-01 | -0.3 | 6.16E-03 | -0.2 | 1.71E-02 | -0.2 | 6.04E-02 | 0.0 | 8.73E-01 | V-type proton ATPase subunit B1 -ATPase subunit B isoform 1) |
| AT2G23100 | -0.2 | 5.76E-02 | 0.0 | 7.95E-01 | -0.2 | 7.44E-02 | -0.5 | 1.64E-01 | -0.3 | 5.27E-01 | -0.2 | 6.77E-01 | Uncharacterized protein At2g23100 |
| AT1G05500 | -0.2 | 9.20E-03 | 0.0 | 9.32E-01 | -0.2 | 2.09E-02 | 0.0 | 9.23E-01 | 0.1 | 5.60E-01 | -0.1 | 5.71E-01 | Calcium-dependent lipid-binding family protein |
| AT3G53720 | -0.2 | 3.89E-02 | -0.2 | 7.72E-02 | 0.0 | 8.06E-01 | 0.0 | 9.13E-01 | -0.1 | 7.92E-01 | 0.1 | 6.85E-01 | Cation |
| AT1G55850 | -0.2 | 3.84E-02 | -0.5 | 9.32E-04 | 0.2 | 4.99E-02 | 0.3 | 2.10E-02 | -0.4 | 3.86E-02 | 0.7 | 1.17E-03 | Cellulose synthase-like protein E1 |
| AT1G70630 | -0.2 | 8.29E-02 | -0.2 | 7.24E-02 | 0.0 | 9.04E-01 | -0.1 | 5.95E-01 | -0.2 | 5.40E-01 | 0.0 | 9.32E-01 | Nucleotide-diphospho-sugar transferase family protein |
| AT5G26600 | -0.2 | 7.24E-03 | 0.0 | 7.12E-01 | -0.2 | 8.54E-03 | -0.1 | 5.80E-01 | 0.0 | 7.86E-01 | -0.1 | 5.08E-01 | Probable L-cysteine desulfhydrase, chloroplastic |
| AT1G11880 | -0.2 | 1.39E-01 | -0.3 | 6.01E-02 | 0.1 | 6.49E-01 | -0.2 | 6.16E-01 | 0.2 | 5.62E-01 | -0.4 | 2.71E-01 | GPI mannosyltransferase 2 |
| AT5G48000 | -0.2 | 5.69E-01 | -0.5 | 2.35E-01 | 0.3 | 5.94E-01 | -0.1 | 9.47E-01 | 1.3 | 1.10E-01 | -1.4 | 1.56E-01 | Cytochrome P450, family 708, subfamily A, polypeptide 2 |
| AT1G75440 | -0.2 | 1.40E-01 | -0.5 | 7.14E-03 | 0.3 | 1.37E-01 | 0.2 | 5.71E-01 | -0.3 | 5.03E-01 | 0.5 | 2.68E-01 | Probable ubiquitin-conjugating enzyme E2 16 |
| AT3G15350 | -0.2 | 1.37E-02 | -0.3 | 5.67E-03 | 0.1 | 5.84E-01 | -0.2 | 1.74E-01 | 0.2 | 3.66E-01 | -0.4 | 3.64E-02 | hypothetical protein |
| AT5G58760 | -0.2 | 5.08E-02 | -0.3 | 1.13E-02 | 0.1 | 4.26E-01 | 0.2 | 5.39E-01 | 0.2 | 5.41E-01 | 0.0 | 9.58E-01 | Damaged DNA binding 2 |
| AT2G36850 | -0.2 | 7.10E-03 | -0.4 | 2.32E-04 | 0.2 | 3.08E-02 | 0.0 | 8.95E-01 | -0.1 | 7.08E-01 | 0.1 | 8.48E-01 | Callose synthase 10 |
| AT1G75210 | -0.2 | 7.94E-02 | 0.0 | 9.18E-01 | -0.2 | 1.46E-01 | 0.0 | 9.57E-01 | 0.1 | 6.76E-01 | -0.1 | 7.75E-01 | Cytosolic IMP-GMP specific 5-nucleotidase, putative |

| | | | | | | | | | | | | | |
|---|---|---|---|---|---|---|---|---|---|---|---|---|---|
| AT5G64280 | -0.2 | 3.16E-02 | -0.3 | 6.52E-03 | 0.1 | 3.79E-01 | -0.2 | 5.36E-01 | 0.0 | 9.98E-01 | -0.2 | 6.32E-01 | Dicarboxylate transporter 2.2, chloroplastic |
| AT1G06570 | -0.2 | 6.14E-02 | -0.6 | 2.42E-04 | 0.4 | 5.74E-03 | -0.8 | 5.07E-03 | -1.2 | 8.69E-04 | 0.4 | 2.66E-01 | 4-hydroxyphenylpyruvate dioxygenase |
| AT3G57330 | -0.2 | 1.17E-02 | -0.2 | 3.39E-02 | 0.0 | 6.77E-01 | 0.4 | 1.74E-02 | 0.2 | 3.91E-01 | 0.2 | 3.12E-01 | Calcium-transporting ATPase |
| AT4G16430 | -0.2 | 1.00E-02 | -0.3 | 1.65E-03 | 0.1 | 2.62E-01 | 0.0 | 9.92E-01 | 0.0 | 9.23E-01 | 0.0 | 9.28E-01 | Transcription factor bHLH3 |
| AT1G53730 | -0.2 | 1.63E-02 | -0.2 | 2.79E-02 | 0.0 | 8.63E-01 | -0.1 | 6.53E-01 | -0.1 | 8.73E-01 | -0.1 | 8.82E-01 | STRUBBELIG-receptor family 6 |
| AT3G12070 | -0.2 | 2.84E-01 | -0.3 | 2.04E-01 | 0.1 | 8.45E-01 | 0.2 | 8.17E-01 | 0.2 | 8.19E-01 | | | Geranylgeranyl transferase type-2 subunit beta |
| AT2G18090 | -0.2 | 2.27E-02 | -0.3 | 3.30E-03 | 0.1 | 2.71E-01 | -0.2 | 1.04E-01 | -0.3 | 5.40E-02 | 0.1 | 7.65E-01 | PHD finger family protei |
| AT4G13350 | -0.2 | 2.21E-02 | -0.3 | 4.76E-03 | 0.1 | 3.57E-01 | -0.1 | 5.99E-01 | -0.1 | 2.94E-01 | 0.1 | 6.99E-01 | At4g13350 -interacting GTPase) |
| AT3G28950 | -0.2 | 1.39E-02 | 0.2 | 5.80E-02 | -0.4 | 1.76E-03 | 0.2 | 5.82E-01 | 0.0 | 9.90E-01 | 0.2 | 6.86E-01 | Protein AIG2 C |
| AT1G63700 | -0.2 | 9.27E-03 | -0.3 | 7.61E-04 | 0.1 | 1.22E-01 | -0.1 | 8.14E-01 | 0.0 | 8.64E-01 | -0.1 | 7.52E-01 | Mitogen-activated protein kinase kinase kinase YODA |
| AT4G37100 | -0.2 | 4.20E-02 | -0.3 | 1.75E-02 | 0.1 | 6.27E-01 | 0.0 | 9.38E-01 | 0.1 | 7.80E-01 | -0.1 | 8.69E-01 | hypothetical protein |
| AT3G13870 | -0.2 | 6.43E-03 | -0.1 | 4.84E-02 | -0.1 | 3.62E-01 | 0.1 | 7.80E-01 | 0.0 | 9.36E-01 | 0.0 | 9.25E-01 | Protein ROOT HAIR DEFECTIVE 3 |
| AT2G32290 | -0.2 | 3.93E-01 | -0.3 | 3.14E-01 | 0.1 | 8.77E-01 | 0.8 | 1.02E-01 | 1.1 | 3.65E-02 | -0.3 | 5.94E-01 | Beta-amylase |
| AT2G25910 | -0.2 | 1.86E-02 | -0.1 | 9.49E-02 | -0.1 | 4.60E-01 | -0.3 | 4.56E-03 | -0.2 | 2.02E-01 | -0.2 | 2.53E-01 | 3'-5' exonuclease domain-containing protein |
| AT1G43700 | -0.2 | 1.94E-02 | -0.3 | 4.34E-03 | 0.1 | 3.80E-01 | -0.3 | 3.57E-02 | -0.4 | 1.46E-02 | 0.1 | 6.81E-01 | Transcription factor VIP1 |
| AT3G51950 | -0.2 | 2.49E-02 | -0.2 | 7.24E-02 | 0.0 | 6.65E-01 | -0.3 | 4.88E-03 | -0.1 | 4.84E-01 | -0.2 | 1.32E-01 | Zinc finger CCCH domain-containing protein 46 |
| AT2G27490 | -0.2 | 6.69E-02 | -0.2 | 1.23E-01 | 0.0 | 8.21E-01 | -0.3 | 2.40E-01 | -0.4 | 1.44E-01 | 0.1 | 8.02E-01 | Dephospho-CoA kinase |
| AT5G39990 | -0.2 | 2.37E-02 | 0.1 | 5.62E-01 | -0.3 | 1.80E-02 | 0.3 | 2.10E-02 | 0.4 | 1.16E-02 | -0.1 | 7.39E-01 | Beta-glucuronosyltransferase GlcAT14A |
| AT1G33360 | -0.2 | 2.79E-02 | -0.1 | 4.03E-01 | -0.1 | 1.96E-01 | -0.2 | 1.75E-01 | 0.0 | 8.68E-01 | -0.3 | 1.79E-01 | CLP protease regulatory subunit CLPX3, mitochondrial |
| AT1G08060 | -0.2 | 2.61E-02 | -0.3 | 8.19E-03 | 0.1 | 5.11E-01 | -0.3 | 1.15E-01 | -0.1 | 6.28E-01 | -0.2 | 5.02E-01 | ATP-dependent helicase family protein |
| AT2G07680 | -0.2 | 1.03E-01 | -0.2 | 7.95E-02 | 0.0 | 8.70E-01 | 0.4 | 2.60E-01 | 0.4 | 3.09E-01 | 0.0 | 9.91E-01 | Multidrug resistance-associated protein 11 |
| AT1G48620 | -0.2 | 2.09E-02 | -0.2 | 5.52E-02 | 0.0 | 7.04E-01 | -0.3 | 2.35E-03 | -0.2 | 8.88E-02 | -0.1 | 3.58E-01 | T1N15.25 |
| AT5G01090 | -0.2 | 1.94E-02 | -0.1 | 2.39E-01 | -0.1 | 2.29E-01 | -0.1 | 3.61E-01 | -0.1 | 6.66E-01 | -0.1 | 8.40E-01 | Legume lectin domain-containing protein |
| AT5G14950 | -0.2 | 2.58E-02 | -0.3 | 1.07E-02 | 0.1 | 6.08E-01 | 0.1 | 6.34E-01 | 0.0 | 8.49E-01 | 0.1 | 5.64E-01 | Alpha-mannosidase 2 |
| AT2G32320 | -0.2 | 6.04E-02 | -0.3 | 2.31E-02 | 0.1 | 6.11E-01 | -0.1 | 7.56E-01 | 0.0 | 8.91E-01 | -0.1 | 7.35E-01 | tRNA guanylyltransferase |
| AT2G42330 | -0.2 | 1.59E-02 | -0.3 | 1.28E-03 | 0.1 | 1.49E-01 | -0.3 | 1.75E-01 | -0.1 | 5.07E-01 | -0.1 | 7.24E-01 | Septin and tuftelin-interacting protein 1 homolog 2 |
| AT1G22870 | -0.2 | 8.72E-03 | -0.5 | 8.49E-05 | 0.3 | 8.33E-03 | -0.3 | 3.34E-01 | -0.2 | 5.88E-01 | -0.1 | 8.29E-01 | Kinase family with ARM repeat domain-containing protein |
| AT2G23520 | -0.2 | 1.16E-02 | -0.2 | 9.46E-03 | 0.0 | 8.49E-01 | 0.3 | 3.79E-02 | 0.1 | 6.99E-01 | 0.2 | 1.82E-01 | Pyridoxal phosphate -dependent transferases superfamily protein |
| AT5G50370 | -0.2 | 1.34E-01 | 0.2 | 1.13E-01 | -0.4 | 1.60E-02 | 0.3 | 1.25E-01 | 0.2 | 2.69E-01 | 0.0 | 8.76E-01 | adenylate kinase |
| AT2G39200 | -0.2 | 3.21E-01 | -0.4 | 5.35E-02 | 0.2 | 3.25E-01 | -0.2 | 6.08E-01 | -0.1 | 7.73E-01 | 0.0 | 9.31E-01 | MLO-like protein |
| AT4G38950 | -0.2 | 5.38E-02 | -0.1 | 2.90E-01 | -0.1 | 4.08E-01 | 0.0 | 8.95E-01 | 0.2 | 5.59E-01 | -0.2 | 5.14E-01 | Kinesin-like protein KIN-7F |
| AT5G15980 | -0.2 | 1.47E-02 | -0.2 | 6.16E-02 | -0.1 | 5.40E-01 | 0.5 | 1.10E-02 | 0.4 | 4.93E-02 | 0.1 | 7.71E-01 | Small ribosomal subunit protein mS81 |
| AT2G24570 | -0.2 | 4.11E-02 | -0.1 | 3.05E-01 | -0.1 | 3.29E-01 | 0.3 | 1.01E-01 | 0.2 | 3.91E-01 | 0.1 | 7.17E-01 | Probable WRKY transcription factor 17 |
| AT1G72710 | -0.2 | 6.53E-03 | -0.2 | 4.09E-03 | 0.0 | 6.83E-01 | -0.2 | 9.29E-02 | -0.1 | 6.20E-01 | -0.1 | 5.04E-01 | Casein kinase 1-like protein 2 |
| AT1G79950 | -0.2 | 6.30E-02 | -0.3 | 1.82E-02 | 0.1 | 5.13E-01 | -0.2 | 3.42E-01 | -0.2 | 5.42E-01 | -0.1 | 8.91E-01 | RAD3-like DNA-binding helicase protein |
| AT1G80170 | -0.2 | 5.37E-02 | 0.1 | 6.23E-01 | -0.3 | 4.93E-02 | 0.0 | 9.61E-01 | 0.0 | 9.92E-01 | 0.0 | 9.67E-01 | Pectin lyase-like superfamily protein |
| AT4G35290 | -0.2 | 1.85E-01 | -0.3 | 1.16E-01 | 0.1 | 7.89E-01 | 0.2 | 6.17E-01 | 0.3 | 5.38E-01 | -0.1 | 9.12E-01 | Glutamate receptor |
| AT2G33150 | -0.2 | 1.66E-02 | -0.1 | 9.78E-02 | -0.1 | 4.28E-01 | -0.1 | 6.21E-01 | 0.0 | 8.56E-01 | 0.0 | 8.89E-01 | 3-ketoacyl-CoA thiolase 2, peroxisomal |
| AT5G15920 | -0.2 | 1.50E-02 | -0.5 | 1.21E-04 | 0.3 | 8.77E-03 | 0.3 | 3.05E-01 | 0.2 | 4.38E-01 | 0.0 | 9.40E-01 | Structural maintenance of chromosomes protein 5 |
| AT3G15640 | -0.2 | 6.54E-02 | 0.2 | 3.99E-02 | -0.5 | 3.23E-03 | 0.1 | 5.67E-01 | 0.2 | 5.01E-01 | 0.0 | 9.11E-01 | Rubredoxin-like superfamily protein |
| AT3G22790 | -0.2 | 5.84E-02 | 0.1 | 1.75E-01 | -0.3 | 1.22E-02 | 0.1 | 6.80E-01 | 0.3 | 2.07E-01 | -0.2 | 4.91E-01 | Protein NETWORKED 1A |
| AT3G18270 | -0.2 | 1.85E-02 | -0.2 | 1.92E-02 | 0.0 | 9.51E-01 | -0.3 | 8.22E-02 | -0.3 | 1.25E-01 | 0.0 | 9.86E-01 | Dipeptide epimerase |
| AT1G66250 | -0.2 | 7.87E-02 | 0.1 | 2.94E-01 | -0.3 | 2.67E-02 | 0.6 | 2.89E-03 | 0.5 | 1.89E-02 | 0.1 | 6.99E-01 | Glucan endo-1,3-beta-glucosidase 2 -beta-glucan endohydrolase 2) -beta-glucanase 2) |
| AT5G63110 | -0.2 | 1.57E-02 | -0.2 | 1.42E-02 | 0.0 | 9.01E-01 | -0.1 | 6.95E-01 | 0.0 | 9.43E-01 | -0.1 | 7.36E-01 | Histone deacetylase |
| AT4G36945 | -0.2 | 8.35E-02 | 0.1 | 1.90E-01 | -0.4 | 2.07E-02 | -0.3 | 1.28E-01 | -0.1 | 7.73E-01 | -0.2 | 4.10E-01 | MAP3K-like protein kinase |
| AT4G27470 | -0.2 | 6.76E-02 | 0.0 | 7.13E-01 | -0.2 | 1.95E-01 | 0.1 | 4.82E-01 | 0.1 | 5.22E-01 | 0.0 | 9.88E-01 | E3 ubiquitin-protein ligase RMA3 |
| AT5G53890 | -0.2 | 2.06E-02 | -0.3 | 4.56E-03 | 0.1 | 3.89E-01 | 0.0 | 9.75E-01 | 0.0 | 9.76E-01 | 0.0 | 9.99E-01 | Phytosulfokine receptor 2 |
| AT5G17380 | -0.2 | 3.76E-02 | -0.2 | 1.05E-01 | 0.0 | 6.63E-01 | -0.1 | 7.24E-01 | -0.1 | 7.54E-01 | 0.0 | 9.89E-01 | 2-hydroxyacyl-CoA lyase |
| AT1G16010 | -0.2 | 3.96E-03 | -0.3 | 2.40E-04 | 0.1 | 6.03E-02 | -0.1 | 7.20E-01 | 0.1 | 7.32E-01 | -0.1 | 5.72E-01 | Magnesium transporter MRS2-1 |
| AT5G61780 | -0.2 | 1.04E-02 | 0.0 | 8.41E-01 | -0.2 | 1.41E-02 | 0.1 | 6.77E-01 | 0.2 | 2.15E-01 | -0.1 | 5.50E-01 | Ribonuclease TUDOR 2 |
| AT2G16880 | -0.2 | 4.25E-02 | -0.2 | 4.49E-02 | 0.0 | 9.68E-01 | 0.0 | 9.87E-01 | -0.1 | 7.91E-01 | 0.1 | 8.32E-01 | Pentatricopeptide repeat-containing protein At2g16880 |
| AT2G40930 | -0.2 | 7.67E-02 | -0.3 | 1.65E-02 | 0.1 | 4.17E-01 | -0.3 | 1.92E-01 | -0.2 | 4.68E-01 | -0.1 | 7.87E-01 | Ubiquitin carboxyl-terminal hydrolase 5 |
| AT3G60080 | -0.2 | 6.40E-03 | -0.2 | 1.88E-02 | 0.0 | 6.69E-01 | -0.2 | 1.77E-01 | -0.1 | 2.97E-01 | 0.0 | 9.37E-01 | RING-type E3 ubiquitin transferase |
| AT2G40460 | -0.2 | 7.66E-02 | -0.5 | 2.26E-03 | 0.3 | 6.99E-02 | 0.2 | 2.37E-01 | 0.6 | 2.53E-03 | -0.4 | 7.20E-02 | Protein NRT1 |
| AT4G15090 | -0.2 | 2.01E-02 | -0.3 | 1.90E-03 | 0.1 | 1.82E-01 | 0.1 | 7.75E-01 | 0.1 | 8.51E-01 | 0.0 | 9.78E-01 | Protein FAR-RED IMPAIRED RESPONSE 1 |
| AT4G36800 | -0.2 | 7.17E-03 | -0.1 | 2.21E-01 | -0.1 | 1.05E-01 | -0.3 | 1.11E-01 | -0.1 | 6.68E-01 | -0.2 | 5.12E-01 | NEDD8-conjugating enzyme Ubc12 |
| AT5G55260 | -0.2 | 3.30E-02 | 0.0 | 9.68E-01 | -0.2 | 7.00E-02 | 0.0 | 9.72E-01 | 0.2 | 2.94E-01 | -0.2 | 3.68E-01 | Serine |
| AT2G42500 | -0.2 | 1.13E-02 | 0.0 | 8.03E-01 | -0.2 | 3.20E-02 | -0.1 | 5.59E-01 | -0.1 | 4.88E-01 | 0.0 | 9.14E-01 | Serine |
| AT5G05660 | -0.2 | 3.28E-02 | 0.0 | 6.08E-01 | -0.2 | 1.34E-01 | -0.3 | 1.92E-02 | -0.1 | 4.58E-01 | -0.2 | 2.67E-01 | NF-X1-type zinc finger protein NFXL2 |
| AT3G60860 | -0.2 | 2.21E-02 | -0.2 | 2.81E-02 | 0.0 | 9.71E-01 | 0.1 | 6.58E-01 | 0.1 | 6.22E-01 | 0.0 | 9.40E-01 | Brefeldin A-inhibited guanine nucleotide-exchange protein 2 |
| AT3G55030 | -0.2 | 2.97E-02 | -0.1 | 1.67E-01 | -0.1 | 4.42E-01 | -0.3 | 7.49E-03 | -0.3 | 5.88E-02 | -0.1 | 6.27E-01 | CDP-diacylglycerol--glycerol-3-phosphate 3-phosphatidyltransferase 2 |
| AT5G05750 | -0.2 | 4.09E-02 | -0.2 | 4.65E-02 | 0.0 | 9.93E-01 | -0.5 | 2.50E-03 | -0.6 | 8.69E-04 | 0.1 | 5.58E-01 | DNAJ heat shock N-terminal domain-containing protein |
| AT3G06650 | -0.2 | 2.69E-02 | -0.2 | 2.76E-02 | 0.0 | 9.38E-01 | -0.3 | 2.56E-01 | 0.0 | 9.46E-01 | -0.2 | 4.69E-01 | ATP-citrate synthase beta chain protein 1 |
| AT1G50620 | -0.2 | 1.44E-02 | -0.2 | 2.25E-02 | 0.0 | 9.12E-01 | 0.0 | 9.42E-01 | -0.1 | 6.01E-01 | 0.1 | 5.97E-01 | PHD-type domain-containing protein |
| AT1G06270 | -0.2 | 1.29E-01 | -0.2 | 2.22E-01 | 0.0 | 8.42E-01 | 0.4 | 1.39E-01 | 0.4 | 2.15E-01 | 0.0 | 9.58E-01 | Pentatricopeptide repeat-containing protein At1g06270 |
| AT5G60980 | -0.2 | 1.24E-02 | 0.0 | 7.88E-01 | -0.2 | 1.48E-02 | -0.1 | 2.43E-01 | -0.2 | 2.31E-02 | 0.1 | 3.76E-01 | Nuclear transport factor 2 |
| AT3G46180 | -0.2 | 1.05E-01 | 0.0 | 8.53E-01 | -0.2 | 2.28E-01 | 0.1 | 8.38E-01 | 0.2 | 4.28E-01 | -0.2 | 6.32E-01 | UDP-galactose transporter 5 |
| AT2G16640 | -0.2 | 5.05E-03 | -0.1 | 9.84E-02 | -0.1 | 1.62E-01 | -0.1 | 3.84E-01 | -0.2 | 2.83E-01 | 0.1 | 8.46E-01 | Translocase of chloroplast 132, chloroplastic |
| AT3G54690 | -0.2 | 1.88E-02 | -0.1 | 2.71E-01 | -0.1 | 2.05E-01 | -0.8 | 3.25E-06 | -0.2 | 8.61E-02 | -0.6 | 2.19E-03 | Probable arabinose 5-phosphate isomerase |
| AT3G08680 | -0.2 | 1.05E-02 | -0.1 | 4.89E-02 | -0.1 | 4.99E-01 | -0.2 | 2.01E-01 | 0.0 | 8.78E-01 | -0.2 | 4.77E-01 | AT3G08680 protein |
| AT4G38600 | -0.2 | 1.83E-02 | -0.5 | 1.03E-04 | 0.3 | 4.53E-03 | 0.0 | 9.45E-01 | 0.0 | 9.90E-01 | 0.0 | 9.78E-01 | E3 ubiquitin-protein ligase |
| AT1G60630 | -0.2 | 1.95E-01 | 0.0 | 9.34E-01 | -0.2 | 3.25E-01 | -0.3 | 3.96E-01 | -0.1 | 7.92E-01 | -0.2 | 7.35E-01 | Inactive leucine-rich repeat receptor-like serine |
| AT1G69960 | -0.2 | 9.98E-03 | -0.1 | 1.01E-01 | -0.1 | 2.79E-01 | 0.1 | 5.94E-01 | 0.0 | 8.58E-01 | 0.0 | 8.67E-01 | Serine |
| AT5G64270 | -0.2 | 1.50E-02 | -0.4 | 6.25E-04 | 0.2 | 5.54E-02 | -0.1 | 5.03E-01 | -0.3 | 1.88E-01 | 0.1 | 6.25E-01 | TOG domain-containing protein |
| AT3G45430 | -0.2 | 1.85E-01 | 0.5 | 4.80E-03 | -0.7 | 2.72E-03 | -0.5 | 1.31E-01 | -0.8 | 4.09E-02 | 0.3 | 5.39E-01 | Probable L-type lectin-domain containing receptor kinase I.5 |
| AT1G29820 | -0.2 | 1.36E-02 | -0.1 | 4.73E-01 | -0.1 | 9.05E-02 | 0.1 | 4.86E-01 | 0.1 | 8.17E-01 | 0.1 | 8.19E-01 | Magnesium transporter CorA-like family protein |
| AT5G42650 | -0.2 | 2.17E-01 | 0.2 | 1.55E-01 | -0.4 | 2.63E-02 | 0.9 | 2.34E-03 | 0.5 | 7.72E-02 | 0.3 | 3.69E-01 | Allene oxide synthase, chloroplastic |
| AT4G10750 | -0.2 | 4.06E-02 | 0.0 | 7.06E-01 | -0.2 | 1.40E-01 | -0.3 | 2.93E-01 | 0.0 | 9.81E-01 | -0.3 | 3.91E-01 | HpcH |
| AT1G10370 | -0.2 | 2.71E-01 | -0.4 | 7.16E-02 | 0.2 | 4.65E-01 | -0.3 | 6.51E-01 | 1.1 | 6.99E-02 | -1.4 | 6.73E-02 | Glutathione S-transferase U17 |
| AT2G18350 | -0.2 | 1.14E-01 | -0.1 | 4.59E-01 | -0.1 | 4.82E-01 | 0.2 | 4.33E-01 | -0.2 | 5.38E-01 | 0.5 | 2.44E-01 | Zinc-finger homeodomain protein 6 |
| AT1G05120 | -0.2 | 1.45E-02 | -0.3 | 9.62E-04 | 0.1 | 1.12E-01 | 0.0 | 9.01E-01 | 0.2 | 3.85E-01 | -0.2 | 4.05E-01 | Helicase protein with RING |
| AT1G17820 | -0.2 | 4.37E-02 | -0.3 | 8.54E-03 | 0.1 | 3.84E-01 | -0.1 | 6.40E-01 | -0.1 | 7.02E-01 | 0.0 | 9.97E-01 | Uncharacterized protein At1g17820 |
| AT2G32800 | -0.2 | 1.36E-01 | -0.3 | 2.42E-02 | 0.1 | 3.62E-01 | -0.4 | 5.06E-03 | -0.5 | 9.83E-04 | 0.1 | 4.17E-01 | Receptor like protein kinase S.2 |

| | | | | | | | | | | | |
|---|---|---|---|---|---|---|---|---|---|---|---|
| AT1G10670 | -0.2 | 2.06E-02 | 0.1 | 2.19E-01 | -0.3 | 5.23E-03 | -0.2 | 1.70E-01 | 0.0 | 9.19E-01 | -0.1 | 4.42E-01 | ATP citrate synthase |
| AT4G19840 | -0.2 | 1.06E-02 | 0.0 | 9.31E-01 | -0.2 | 1.72E-02 | -0.1 | 4.23E-01 | 0.1 | 5.78E-01 | -0.2 | 2.89E-01 | Protein PHLOEM PROTEIN 2-LIKE A1 |
| AT5G43170 | -0.2 | 7.84E-02 | -0.4 | 3.08E-03 | 0.2 | 1.06E-01 | -0.1 | 8.16E-01 | -0.1 | 8.50E-01 | 0.0 | 9.97E-01 | Zinc finger protein AZF3 |
| AT4G19180 | -0.2 | 1.28E-02 | -0.1 | 6.15E-02 | -0.1 | 5.07E-01 | -0.1 | 6.37E-01 | 0.0 | 9.54E-01 | -0.2 | 6.70E-01 | Probable apyrase 7 |
| AT3G25760 | -0.2 | 3.18E-01 | 0.2 | 2.29E-01 | -0.4 | 6.16E-02 | 1.5 | 3.28E-03 | 1.4 | 1.02E-02 | 0.1 | 9.10E-01 | Allene oxide cyclase 1, chloroplastic |
| AT2G44410 | -0.2 | 3.21E-02 | -0.3 | 2.72E-03 | 0.1 | 1.71E-01 | -0.2 | 2.08E-01 | -0.4 | 1.09E-01 | 0.1 | 7.23E-01 | E3 ubiquitin-protein ligase RMA |
| AT1G27020 | -0.2 | 3.77E-01 | 0.2 | 3.42E-01 | -0.4 | 1.12E-01 | 0.5 | 1.23E-01 | -0.7 | 7.17E-02 | 1.3 | 3.42E-03 | 2-oxoadipate dioxygenase |
| AT4G38650 | -0.2 | 1.99E-01 | 0.0 | 7.83E-01 | -0.2 | 4.21E-01 | -0.2 | 5.73E-01 | 0.0 | 9.92E-01 | -0.3 | 5.83E-01 | GH10 domain-containing protein |
| AT3G55740 | -0.2 | 6.67E-02 | -0.5 | 6.65E-04 | 0.3 | 2.11E-02 | 0.2 | 1.29E-01 | 0.4 | 8.23E-03 | -0.2 | 2.68E-01 | Proline transporter 2 |
| AT3G01070 | -0.2 | 2.96E-01 | 0.4 | 4.29E-02 | -0.6 | 2.09E-02 | 0.6 | 1.39E-01 | 0.3 | 5.82E-01 | 0.3 | 4.99E-01 | hypothetical protein |
| AT1G61040 | -0.2 | 8.21E-03 | -0.3 | 1.13E-03 | 0.1 | 2.15E-01 | -0.1 | 6.66E-01 | -0.2 | 3.80E-01 | 0.1 | 7.19E-01 | Protein RTF1 homolog |
| AT1G10870 | -0.2 | 7.03E-02 | -0.3 | 2.92E-02 | 0.1 | 6.36E-01 | -0.2 | 3.71E-01 | 0.0 | 9.02E-01 | -0.2 | 6.37E-01 | ADP-ribosylation factor GTPase-activating protein |
| AT1G20575 | -0.2 | 1.54E-02 | 0.0 | 7.32E-01 | -0.2 | 5.67E-02 | -0.2 | 2.80E-01 | 0.0 | 9.15E-01 | -0.2 | 3.74E-01 | Dolichol-phosphate mannosyltransferase subunit 1 |
| AT5G57000 | -0.2 | 2.76E-01 | -0.3 | 1.28E-01 | 0.1 | 6.77E-01 | 0.0 | 9.44E-01 | 0.0 | 9.96E-01 | 0.0 | 9.61E-01 | At5g57000 |
| AT5G13290 | -0.2 | 1.46E-01 | -0.1 | 2.91E-01 | 0.0 | 7.80E-01 | -0.5 | 1.23E-01 | -0.4 | 2.78E-01 | -0.1 | 8.31E-01 | Inactive leucine-rich repeat receptor-like protein kinase CORYNE |
| AT5G46510 | -0.2 | 3.98E-02 | -0.3 | 8.49E-03 | 0.1 | 4.02E-01 | 0.1 | 5.44E-01 | 0.1 | 8.14E-01 | 0.0 | 8.69E-01 | ADP-ribosyl cyclase |
| AT4G34180 | -0.2 | 3.14E-02 | -0.3 | 9.98E-03 | 0.1 | 4.96E-01 | 0.0 | 8.36E-01 | -0.3 | 1.01E-01 | 0.3 | 2.32E-01 | Cyclase family protein |
| AT4G27100 | -0.2 | 6.24E-02 | -0.1 | 5.46E-01 | -0.1 | 2.69E-01 | -0.4 | 2.06E-02 | -0.3 | 2.02E-01 | -0.2 | 5.09E-01 | Uncharacterized protein At4g27100 |
| AT5G37510 | -0.2 | 1.16E-02 | -0.1 | 1.70E-01 | -0.1 | 2.06E-01 | 0.1 | 2.74E-01 | 0.1 | 3.78E-01 | 0.0 | 9.83E-01 | AT5G37510 protein |
| AT3G46960 | -0.2 | 4.94E-02 | -0.2 | 4.04E-02 | 0.0 | 8.86E-01 | -0.1 | 8.84E-01 | -0.1 | 7.92E-01 | 0.1 | 9.09E-01 | RNA helicase, ATP-dependent, SK12 |
| AT2G39710 | -0.2 | 1.35E-01 | -0.4 | 1.24E-02 | 0.2 | 2.21E-01 | 0.1 | 6.51E-01 | 0.1 | 6.55E-01 | 0.0 | 9.66E-01 | Eukaryotic aspartyl protease family protein |
| AT1G22960 | -0.2 | 6.67E-02 | -0.2 | 1.55E-01 | 0.0 | 7.35E-01 | 0.3 | 2.46E-01 | 0.1 | 7.77E-01 | 0.2 | 5.93E-01 | Pentatricopeptide repeat-containing protein At1g22960, mitochondrial |
| AT1G02145 | -0.2 | 2.36E-02 | -0.1 | 2.44E-01 | -0.1 | 2.78E-01 | -0.1 | 7.43E-01 | 0.1 | 3.68E-01 | -0.2 | 3.13E-01 | Mannosyltransferase |
| AT3G01380 | -0.2 | 7.22E-02 | -0.2 | 8.56E-02 | 0.0 | 9.82E-01 | 0.2 | 3.49E-01 | 0.3 | 2.13E-01 | -0.1 | 7.89E-01 | GPI ethanolamine phosphate transferase 1 |
| AT2G30250 | -0.2 | 1.11E-01 | -0.8 | 7.69E-05 | 0.6 | 1.09E-03 | 0.2 | 1.53E-01 | -0.3 | 1.40E-01 | 0.5 | 2.13E-02 | Probable WRKY transcription factor 25 |
| AT5G04020 | -0.2 | 4.19E-02 | -0.2 | 3.15E-02 | 0.0 | 8.49E-01 | 0.1 | 4.46E-01 | 0.1 | 4.92E-01 | 0.0 | 9.82E-01 | Calmodulin binding protein PICBP |
| AT5G54310 | -0.2 | 1.65E-02 | -0.2 | 5.56E-02 | 0.0 | 6.14E-01 | 0.1 | 7.73E-01 | 0.1 | 5.68E-01 | -0.1 | 8.28E-01 | ADP-ribosylation factor GTPase-activating protein AGD5 |
| AT5G01020 | -0.2 | 1.01E-02 | -0.1 | 2.72E-01 | -0.1 | 1.15E-01 | -0.4 | 7.79E-04 | -0.3 | 3.02E-02 | -0.1 | 3.54E-01 | Probable serine |
| AT4G35880 | -0.2 | 1.53E-01 | 0.0 | 8.42E-01 | -0.2 | 1.68E-01 | 0.1 | 6.99E-01 | 0.2 | 4.39E-01 | -0.1 | 7.43E-01 | Eukaryotic aspartyl protease family protein |
| AT1G58060 | -0.2 | 6.22E-02 | -0.3 | 1.07E-02 | 0.1 | 3.53E-01 | -0.1 | 7.60E-01 | 0.0 | 9.99E-01 | -0.1 | 7.95E-01 | DExH-box ATP-dependent RNA helicase DExH7, chloroplastic |
| AT4G21940 | -0.2 | 2.95E-01 | -0.4 | 5.33E-02 | 0.2 | 3.72E-01 | 0.1 | 7.87E-01 | 0.3 | 6.69E-01 | -0.1 | 8.83E-01 | Calcium-dependent protein kinase 15 |
| AT2G45500 | -0.2 | 4.66E-02 | -0.4 | 1.24E-03 | 0.2 | 6.11E-02 | -0.2 | 3.47E-01 | 0.0 | 8.73E-01 | -0.2 | 3.26E-01 | AAA-type ATPase family protein |
| AT2G40435 | -0.2 | 5.36E-01 | -0.4 | 2.43E-01 | 0.2 | 6.19E-01 | -0.4 | 2.69E-01 | 0.2 | 6.33E-01 | -0.6 | 1.70E-01 | Transcription factor SCREAM-like protein |
| AT1G30810 | -0.2 | 1.58E-01 | -0.4 | 8.41E-03 | 0.2 | 1.36E-01 | 0.0 | 9.64E-01 | 0.2 | 5.36E-01 | -0.2 | 6.45E-01 | Lysine-specific demethylase JMJ18 monodemethylase JMJ18) |
| AT3G27400 | -0.2 | 1.11E-01 | 0.0 | 7.94E-01 | -0.2 | 2.60E-01 | -0.2 | 3.87E-01 | 0.3 | 3.20E-01 | -0.5 | 8.76E-02 | Pectate lyase |
| AT5G58290 | -0.2 | 3.16E-02 | -0.2 | 1.60E-02 | 0.0 | 6.63E-01 | 0.1 | 2.61E-01 | 0.0 | 9.79E-01 | 0.1 | 4.59E-01 | 26S proteasome regulatory subunit 6B homolog |
| AT1G70670 | -0.2 | 1.22E-01 | 0.1 | 5.61E-01 | -0.3 | 7.52E-02 | -0.3 | 1.85E-01 | 0.0 | 9.43E-01 | -0.3 | 2.57E-01 | Probable peroxygenase 4 |
| AT1G20050 | -0.2 | 9.38E-02 | 0.2 | 1.26E-01 | -0.4 | 1.06E-02 | 0.0 | 7.91E-01 | 0.0 | 9.27E-01 | 0.0 | 9.29E-01 | Probable 3-beta-hydroxysteroid-Delta,Delta-isomerase -Delta sterol isomerase) |
| AT4G11570 | -0.2 | 8.74E-02 | -0.5 | 5.52E-04 | 0.4 | 1.05E-02 | -0.1 | 6.35E-01 | 0.2 | 3.14E-01 | -0.2 | 1.88E-01 | 5-amino-6-uracil phosphatase, chloroplastic -pyrimidinedione 5'-phosphate phosphatase) |
| AT1G56145 | -0.2 | 2.31E-02 | -0.2 | 1.97E-02 | 0.0 | 8.86E-01 | 0.1 | 2.75E-01 | -0.2 | 2.09E-01 | 0.3 | 6.50E-02 | Leucine-rich repeat transmembrane protein kinase |
| AT1G62640 | -0.2 | 1.15E-02 | 0.1 | 1.29E-01 | -0.3 | 2.45E-03 | 0.1 | 3.72E-01 | 0.1 | 4.07E-01 | 0.0 | 9.73E-01 | Beta-ketoacyl- synthase III, chloroplastic |
| AT2G26300 | -0.2 | 8.34E-03 | -0.1 | 1.99E-01 | -0.1 | 1.37E-01 | -0.1 | 4.27E-01 | -0.2 | 2.37E-01 | 0.1 | 7.68E-01 | Guanine nucleotide-binding protein alpha-1 subunit |
| AT2G40850 | -0.2 | 1.32E-01 | -0.3 | 3.57E-02 | 0.1 | 5.04E-01 | -0.2 | 7.34E-01 | 0.6 | 2.69E-01 | -0.9 | 1.98E-01 | Phosphatidylinositol 4-kinase gamma 1 |
| AT1G33930 | -0.2 | 1.25E-01 | 1.0 | 1.53E-05 | -1.1 | 3.57E-05 | -1.6 | 5.43E-05 | 0.1 | 7.11E-01 | -1.7 | 4.14E-04 | AIG1-type G domain-containing protein |
| AT4G11800 | -0.2 | 3.89E-02 | -0.3 | 3.66E-03 | 0.1 | 1.97E-01 | -0.1 | 7.34E-01 | 0.1 | 7.21E-01 | -0.2 | 5.71E-01 | hypothetical protein |
| AT4G18050 | -0.2 | 3.46E-01 | -0.2 | 4.31E-01 | 0.0 | 9.36E-01 | -0.1 | 9.27E-01 | 0.7 | 4.65E-01 | -0.8 | 4.53E-01 | p-glycoprotein 9 |
| AT3G56730 | -0.2 | 3.13E-01 | 0.1 | 6.95E-01 | -0.3 | 2.52E-01 | 0.1 | 6.78E-01 | 0.3 | 4.33E-01 | -0.1 | 7.58E-01 | Endonuclease or glycosyl hydrolase |
| AT1G49300 | -0.2 | 9.90E-02 | -0.1 | 2.38E-01 | -0.1 | 6.84E-01 | 0.0 | 8.26E-01 | -0.1 | 6.22E-01 | 0.1 | 8.32E-01 | RABG3E |
| AT2G29390 | -0.2 | 5.86E-02 | 0.1 | 5.44E-01 | -0.3 | 3.62E-02 | -0.1 | 3.80E-01 | 0.0 | 8.72E-01 | -0.2 | 4.34E-01 | SMO2-2 |
| AT3G48440 | -0.2 | 1.05E-01 | -0.2 | 6.71E-02 | 0.0 | 8.02E-01 | 0.3 | 2.66E-01 | 0.6 | 3.79E-02 | -0.3 | 3.72E-01 | Zinc finger CCCH domain-containing protein 43 |
| AT5G56000 | -0.2 | 1.10E-01 | 0.1 | 5.62E-01 | -0.3 | 6.14E-02 | 0.5 | 1.70E-03 | 0.3 | 3.88E-02 | 0.2 | 4.30E-01 | Heat shock protein 90-4 |
| AT5G06680 | -0.2 | 1.06E-02 | -0.2 | 7.46E-03 | 0.0 | 7.90E-01 | 0.0 | 9.92E-01 | 0.0 | 9.82E-01 | 0.0 | 9.79E-01 | Gamma-tubulin complex component 3 |
| AT5G07080 | -0.2 | 7.81E-02 | 0.3 | 1.16E-02 | -0.5 | 1.91E-03 | -0.8 | 3.06E-04 | -0.3 | 8.78E-02 | -0.5 | 5.96E-02 | HXXXD-type acyl-transferase family protein |
| AT2G22480 | -0.2 | 9.07E-03 | -0.2 | 8.22E-03 | 0.0 | 8.89E-01 | 0.1 | 5.02E-01 | 0.3 | 5.04E-02 | -0.2 | 2.97E-01 | ATP-dependent 6-phosphofructokinase 5, chloroplastic |
| AT5G10080 | -0.2 | 2.45E-01 | 0.0 | 9.47E-01 | -0.2 | 3.82E-01 | -0.4 | 5.73E-01 | 0.2 | 8.60E-01 | | | Aspartic proteinase-like protein 1 |
| AT1G35460 | -0.2 | 1.74E-01 | -0.1 | 3.76E-01 | -0.1 | 7.15E-01 | -0.1 | 5.63E-01 | -0.1 | 7.59E-01 | 0.0 | 9.23E-01 | Basic helix-loop-helix DNA-binding superfamily protein |
| AT4G00330 | -0.2 | 1.30E-02 | -0.3 | 1.10E-03 | 0.1 | 1.38E-01 | 0.0 | 9.94E-01 | -0.4 | 4.80E-03 | 0.4 | 1.75E-02 | Calmodulin-binding receptor-like cytoplasmic kinase 2 |
| AT5G48880 | -0.2 | 3.30E-01 | -0.2 | 2.81E-01 | 0.0 | 9.09E-01 | 0.2 | 6.33E-01 | 0.9 | 1.90E-02 | -0.7 | 1.11E-01 | PKT2 |
| AT3G55270 | -0.2 | 3.89E-02 | -0.4 | 4.54E-04 | 0.3 | 1.89E-02 | -0.1 | 5.16E-01 | -0.1 | 4.78E-01 | 0.0 | 9.27E-01 | Protein-tyrosine-phosphatase MKP1 |
| AT1G08720 | -0.2 | 3.34E-02 | -0.3 | 8.03E-03 | 0.1 | 4.31E-01 | -0.5 | 4.31E-03 | -0.2 | 2.50E-01 | -0.3 | 1.96E-01 | Protein kinase superfamily protein |
| AT3G06480 | -0.2 | 1.00E-02 | -0.2 | 3.31E-03 | 0.1 | 4.86E-01 | -0.2 | 1.84E-01 | -0.2 | 3.55E-01 | 0.0 | 8.81E-01 | DEAD box RNA helicase family protein |
| AT3G25140 | -0.2 | 2.01E-02 | -0.1 | 2.36E-01 | -0.1 | 2.35E-01 | 0.0 | 8.71E-01 | 0.1 | 3.73E-01 | -0.2 | 3.95E-01 | Galacturonosyltransferase 8 |
| AT5G62930 | -0.2 | 3.21E-02 | 0.1 | 3.19E-01 | -0.3 | 1.26E-02 | -0.1 | 5.15E-01 | -0.1 | 5.77E-01 | 0.0 | 9.97E-01 | SGNH hydrolase-type esterase superfamily protein |
| AT5G53290 | -0.2 | 9.90E-02 | 0.1 | 1.78E-01 | -0.3 | 2.30E-02 | 0.3 | 2.86E-01 | 0.5 | 1.30E-01 | -0.2 | 6.32E-01 | Ethylene-responsive transcription factor CRF3 |
| AT5G65700 | -0.2 | 5.90E-02 | -0.1 | 2.03E-01 | -0.1 | 5.67E-01 | -0.1 | 2.37E-01 | -0.1 | 4.79E-01 | 0.0 | 8.70E-01 | Leucine-rich repeat receptor-like serine |
| AT4G28880 | -0.2 | 2.43E-02 | -0.2 | 1.35E-02 | 0.0 | 6.95E-01 | -0.3 | 3.04E-02 | -0.3 | 3.66E-02 | 0.0 | 9.36E-01 | Casein kinase 1-like protein 3 |
| AT5G13520 | -0.2 | 8.71E-02 | 0.1 | 2.87E-01 | -0.3 | 2.62E-02 | 0.0 | 9.39E-01 | 0.1 | 6.14E-01 | -0.2 | 6.25E-01 | Leucine aminopeptidase |
| AT5G56610 | -0.2 | 6.83E-02 | 0.0 | 7.56E-01 | -0.2 | 7.86E-02 | 0.1 | 8.01E-01 | 0.3 | 2.80E-01 | -0.2 | 5.04E-01 | Uncharacterized protein |
| AT5G38470 | -0.2 | 2.17E-02 | -0.2 | 5.33E-02 | 0.0 | 7.35E-01 | 0.1 | 4.40E-01 | 0.0 | 8.11E-01 | 0.1 | 8.02E-01 | Rad23 UV excision repair protein family |
| AT1G66090 | -0.2 | 4.79E-01 | -0.4 | 1.63E-01 | 0.2 | 5.38E-01 | 0.2 | 6.28E-01 | -0.5 | 1.68E-01 | 0.7 | 1.18E-01 | Disease resistance protein |
| AT1G77620 | -0.2 | 4.23E-02 | -0.4 | 2.05E-03 | 0.2 | 1.12E-01 | -0.1 | 8.19E-01 | 0.2 | 6.27E-01 | -0.3 | 5.38E-01 | P-loop containing nucleoside triphosphate hydrolases superfamily protein |
| AT5G52450 | -0.2 | 1.64E-02 | -0.1 | 1.05E-01 | -0.1 | 4.03E-01 | 0.5 | 4.02E-03 | 0.0 | 9.50E-01 | 0.5 | 1.06E-02 | Protein DETOXIFICATION 16 |
| AT3G48750 | -0.2 | 1.13E-02 | 0.0 | 4.76E-01 | -0.1 | 6.84E-02 | -0.1 | 2.82E-01 | -0.2 | 2.37E-01 | 0.0 | 8.81E-01 | Protein kinase domain-containing protein |
| AT4G36710 | -0.2 | 2.41E-02 | -0.1 | 9.31E-02 | -0.1 | 5.77E-01 | -0.2 | 1.72E-01 | -0.2 | 2.38E-01 | 0.0 | 9.83E-01 | Scarecrow-like protein 15 |
| AT3G11200 | -0.2 | 1.91E-02 | 0.0 | 6.49E-01 | -0.2 | 7.12E-02 | 0.0 | 8.24E-01 | 0.1 | 5.38E-01 | -0.1 | 5.01E-01 | PHD finger protein ALFIN-LIKE |
| AT1G62660 | -0.2 | 6.40E-02 | 0.5 | 5.16E-04 | -0.7 | 1.66E-04 | -0.3 | 6.00E-02 | 0.2 | 2.44E-01 | -0.5 | 1.46E-02 | Glycosyl hydrolases family 32 protein |
| AT1G29950 | -0.2 | 1.32E-01 | -0.4 | 6.02E-03 | 0.2 | 1.10E-01 | -0.2 | 2.08E-01 | -0.1 | 7.60E-01 | -0.1 | 5.86E-01 | Transcription factor bHLH144 |
| AT3G58500 | -0.2 | 3.60E-02 | -0.1 | 3.42E-01 | -0.1 | 2.60E-01 | -0.2 | 1.70E-01 | -0.2 | 7.96E-02 | 0.1 | 7.25E-01 | Serine |
| AT3G52890 | -0.2 | 1.40E-02 | -0.1 | 9.53E-02 | -0.1 | 3.89E-01 | -0.2 | 3.74E-01 | -0.1 | 7.33E-01 | -0.1 | 7.93E-01 | Serine |
| AT5G60690 | -0.2 | 3.98E-02 | -0.2 | 1.38E-02 | 0.1 | 5.58E-01 | 0.1 | 5.67E-01 | 0.3 | 7.34E-02 | -0.2 | 2.71E-01 | Homeobox-leucine zipper protein REVOLUTA |
| AT1G30570 | -0.2 | 1.66E-02 | -0.1 | 1.19E-01 | -0.1 | 3.86E-01 | -0.4 | 3.64E-02 | -0.1 | 5.54E-01 | -0.3 | 3.29E-01 | Probable receptor-like protein kinase At1g30570 |
| AT3G10360 | -0.2 | 1.14E-01 | -0.3 | 4.72E-02 | 0.1 | 6.33E-01 | -0.2 | 6.69E-01 | 0.2 | 7.48E-01 | -0.3 | 5.19E-01 | Pumilio homolog 4 |
| AT4G24740 | -0.2 | 1.11E-02 | -0.3 | 1.75E-03 | 0.1 | 2.49E-01 | -0.3 | 1.91E-02 | -0.1 | 7.39E-01 | -0.2 | 1.54E-01 | LAMMER-type protein kinase AFC2 |

| | | | | | | | | | | | | | |
|---|---|---|---|---|---|---|---|---|---|---|---|---|---|
| AT3G26400 | -0.2 | 2.01E-02 | -0.2 | 1.61E-02 | 0.0 | 8.61E-01 | 0.0 | 7.66E-01 | -0.2 | 1.24E-01 | 0.2 | 3.55E-01 | Eukaryotic translation initiation factor 4B1 |
| AT4G27270 | -0.2 | 1.06E-01 | 0.4 | 6.54E-03 | -0.5 | 1.83E-03 | 0.0 | 9.92E-01 | 0.1 | 7.91E-01 | -0.1 | 7.95E-01 | NADH dehydrogenase |
| AT5G28490 | -0.2 | 8.67E-02 | 0.0 | 9.52E-01 | -0.2 | 1.60E-01 | 0.1 | 8.62E-01 | 0.0 | 9.45E-01 | 0.0 | 9.56E-01 | Protein LIGHT-DEPENDENT SHORT HYPOCOTYLS 1 |
| AT4G13980 | -0.2 | 2.38E-02 | -0.1 | 1.54E-01 | -0.1 | 4.12E-01 | -0.4 | 4.66E-03 | -0.4 | 9.28E-03 | 0.0 | 9.69E-01 | Heat stress transcription factor A-5 |
| AT1G63740 | -0.2 | 5.22E-02 | -0.2 | 2.30E-02 | 0.1 | 6.59E-01 | 0.1 | 8.02E-01 | -0.3 | 2.98E-01 | 0.4 | 2.48E-01 | Disease resistance protein family |
| AT1G50260 | -0.2 | 1.06E-01 | -0.3 | 3.57E-02 | 0.1 | 5.74E-01 | 0.3 | 3.08E-01 | 0.1 | 8.40E-01 | 0.2 | 5.68E-01 | At1g50260 |
| AT3G58160 | -0.2 | 2.50E-01 | -0.3 | 1.23E-01 | 0.1 | 6.87E-01 | -0.1 | 9.21E-01 | -0.1 | 9.17E-01 | 0.0 | 9.88E-01 | P-loop containing nucleoside triphosphate hydrolases superfamily protein |
| AT3G05320 | -0.2 | 3.64E-01 | -0.2 | 3.68E-01 | 0.0 | 9.82E-01 | -0.3 | 2.27E-01 | 0.0 | 9.91E-01 | -0.3 | 3.72E-01 | O-fucosyltransferase 23 |
| AT1G73460 | -0.2 | 4.88E-02 | -0.4 | 1.53E-03 | 0.2 | 6.82E-02 | 0.0 | 9.60E-01 | 0.0 | 9.17E-01 | -0.1 | 8.99E-01 | Protein kinase superfamily protein |
| AT4G40030 | -0.2 | 4.10E-02 | 0.0 | 9.74E-01 | -0.2 | 6.05E-02 | -0.2 | 8.07E-02 | -0.2 | 1.88E-01 | 0.0 | 9.07E-01 | Histone superfamily protein |
| AT3G09010 | -0.2 | 7.73E-02 | -0.4 | 4.41E-03 | 0.2 | 1.47E-01 | 0.6 | 8.37E-03 | 0.5 | 3.02E-02 | 0.1 | 8.07E-01 | Putative receptor ser |
| AT4G26140 | -0.2 | 1.23E-01 | 0.2 | 9.97E-02 | -0.4 | 1.60E-02 | -0.4 | 2.72E-01 | -0.2 | 6.20E-01 | -0.2 | 7.49E-01 | Beta-galactosidase |
| AT4G08350 | -0.2 | 1.18E-02 | -0.4 | 2.91E-04 | 0.2 | 2.61E-02 | -0.1 | 3.40E-01 | 0.0 | 9.77E-01 | -0.1 | 5.29E-01 | Putative transcription elongation factor SPT5 homolog 1 |
| AT4G37790 | -0.2 | 1.26E-01 | -0.1 | 2.26E-01 | 0.0 | 8.06E-01 | 0.2 | 4.91E-01 | 0.2 | 3.60E-01 | -0.1 | 8.51E-01 | Homeobox-leucine zipper protein HAT22 |
| AT1G79520 | -0.2 | 5.50E-02 | -0.5 | 6.04E-04 | 0.3 | 2.28E-02 | 0.0 | 9.98E-01 | 0.2 | 4.28E-01 | -0.2 | 4.70E-01 | Cation efflux family protein |
| AT5G44180 | -0.2 | 4.29E-02 | -0.4 | 1.80E-03 | 0.2 | 8.34E-02 | -0.2 | 2.00E-01 | -0.1 | 4.34E-01 | 0.0 | 8.50E-01 | Homeodomain-like transcriptional regulator |
| AT4G31970 | -0.2 | 8.85E-01 | 0.9 | 3.83E-01 | -1.1 | 4.17E-01 | 6.5 | 8.75E-05 | 4.2 | 8.59E-03 | 2.3 | 6.99E-04 | Xanthotoxin 5-hydroxylase CYP82C2 |
| AT4G38740 | -0.2 | 4.59E-02 | 0.3 | 8.79E-03 | -0.5 | 9.06E-04 | 0.0 | 9.11E-01 | 0.2 | 1.54E-01 | -0.2 | 2.01E-01 | Peptidyl-prolyl cis-trans isomerase CYP18-3 |
| AT1G51710 | -0.2 | 3.67E-02 | -0.2 | 3.73E-02 | 0.0 | 9.53E-01 | 0.1 | 2.26E-01 | 0.1 | 3.04E-01 | 0.0 | 9.97E-01 | Ubiquitin carboxyl-terminal hydrolase |
| AT4G09520 | -0.2 | 3.35E-02 | -0.1 | 1.48E-01 | -0.1 | 5.18E-01 | 0.0 | 9.45E-01 | 0.1 | 7.99E-01 | -0.1 | 7.71E-01 | Cofactor-independent phosphoglycerate mutase |
| AT5G06600 | -0.2 | 9.59E-03 | -0.1 | 5.84E-02 | -0.1 | 4.20E-01 | 0.1 | 6.33E-01 | 0.0 | 7.89E-01 | 0.1 | 5.85E-01 | Ubiquitin carboxyl-terminal hydrolase |
| AT3G19950 | -0.2 | 1.80E-02 | 0.0 | 5.62E-01 | -0.1 | 8.75E-02 | -0.2 | 4.75E-02 | -0.2 | 1.17E-01 | 0.0 | 8.91E-01 | RING-type E3 ubiquitin transferase |
| AT5G61810 | -0.2 | 8.96E-02 | -0.2 | 1.18E-01 | 0.0 | 9.37E-01 | 0.1 | 6.23E-01 | 0.5 | 8.82E-03 | -0.4 | 7.66E-02 | Mitochondrial substrate carrier family protein |
| AT5G57160 | -0.2 | 3.32E-02 | -0.3 | 6.59E-03 | 0.1 | 3.74E-01 | 0.0 | 9.03E-01 | -0.2 | 3.21E-01 | 0.2 | 3.68E-01 | DNA ligase IV |
| AT4G35800 | -0.2 | 3.97E-02 | -0.3 | 2.83E-03 | 0.1 | 1.35E-01 | 0.2 | 2.50E-01 | 0.1 | 6.79E-01 | 0.1 | 7.20E-01 | DNA-directed RNA polymerase II subunit RPB1 |
| AT2G28470 | -0.2 | 1.55E-01 | 0.2 | 1.27E-01 | -0.4 | 1.67E-02 | 0.0 | 9.11E-01 | 0.4 | 3.61E-02 | -0.4 | 1.13E-01 | Beta-galactosidase |
| AT5G15950 | -0.2 | 2.49E-01 | -0.3 | 6.43E-02 | 0.1 | 4.59E-01 | -0.1 | 8.14E-01 | 0.0 | 9.54E-01 | -0.1 | 8.14E-01 | S-adenosylmethionine decarboxylase proenzyme 2 |
| AT4G33360 | -0.2 | 6.16E-02 | 0.2 | 9.26E-02 | -0.3 | 6.01E-03 | -0.3 | 5.22E-02 | 0.0 | 8.35E-01 | -0.3 | 9.44E-02 | NAD-binding Rossmann-fold superfamily protein |
| AT2G24830 | -0.2 | 6.38E-02 | -0.2 | 5.20E-02 | 0.0 | 8.87E-01 | -0.2 | 3.18E-01 | -0.3 | 3.58E-01 | 0.0 | 9.83E-01 | Zinc finger CCCH domain-containing protein 22 |
| AT3G56580 | -0.2 | 4.62E-02 | -0.2 | 6.70E-02 | 0.0 | 9.15E-01 | 0.1 | 7.79E-01 | 0.2 | 4.74E-01 | -0.1 | 7.53E-01 | E3 ubiquitin-protein ligase RZF1 |
| AT2G40270 | -0.2 | 2.88E-02 | -0.3 | 1.64E-03 | 0.1 | 1.06E-01 | 0.3 | 1.59E-02 | 0.1 | 4.84E-01 | 0.2 | 2.42E-01 | Protein kinase family protein |
| AT4G28320 | -0.2 | 8.50E-02 | -0.1 | 4.55E-01 | -0.1 | 4.11E-01 | 0.0 | 9.74E-01 | 0.3 | 4.94E-01 | -0.3 | 4.68E-01 | Mannan endo-1,4-beta-mannosidase 5 |
| AT1G24470 | -0.2 | 2.55E-01 | -0.1 | 4.29E-01 | 0.0 | 8.15E-01 | | | | | | | Very-long-chain 3-oxoacyl-CoA reductase-like protein At1g24470 |
| AT5G66120 | -0.2 | 1.09E-02 | -0.1 | 5.58E-02 | 0.0 | 4.75E-01 | 0.1 | 2.15E-01 | 0.1 | 4.68E-01 | 0.0 | 8.51E-01 | 3-dehydroquinate synthase, chloroplastic |
| AT5G08590 | -0.2 | 1.36E-02 | -0.1 | 4.96E-02 | 0.0 | 5.88E-01 | -0.1 | 4.83E-01 | 0.0 | 9.00E-01 | -0.1 | 7.53E-01 | SNF1-related protein kinase 2.1 |
| AT1G15890 | -0.2 | 1.02E-01 | -0.1 | 5.61E-01 | -0.1 | 3.74E-01 | -0.4 | 1.17E-01 | -0.6 | 3.36E-02 | 0.2 | 5.99E-01 | Disease resistance protein family |
| AT3G54420 | -0.2 | 1.13E-01 | -0.5 | 1.38E-03 | 0.3 | 3.42E-02 | 0.3 | 3.74E-01 | 0.2 | 6.63E-01 | 0.1 | 8.40E-01 | Endochitinase EP3 |
| AT1G07210 | -0.2 | 3.80E-02 | 0.0 | 5.96E-01 | -0.1 | 1.52E-01 | 0.3 | 1.10E-02 | 0.2 | 2.24E-01 | 0.2 | 3.75E-01 | Small ribosomal subunit protein bS18c |
| AT3G25980 | -0.2 | 1.49E-01 | 0.2 | 5.73E-02 | -0.4 | 1.15E-02 | 0.2 | 2.07E-01 | 0.1 | 4.62E-01 | 0.1 | 8.18E-01 | DNA-binding HORMA family protein |
| AT2G45030 | -0.2 | 3.82E-02 | -0.3 | 6.36E-03 | 0.1 | 3.23E-01 | 0.3 | 1.70E-01 | 0.2 | 4.95E-01 | 0.1 | 7.52E-01 | Elongation factor G-2, mitochondrial |
| AT1G29470 | -0.2 | 1.36E-02 | 0.0 | 9.83E-01 | -0.2 | 2.38E-02 | 0.1 | 4.34E-01 | 0.2 | 2.75E-01 | -0.1 | 8.07E-01 | Probable methyltransferase PMT24 |
| AT3G53620 | -0.2 | 3.05E-02 | 0.0 | 5.61E-01 | -0.2 | 2.21E-02 | -0.5 | 2.30E-03 | -0.1 | 6.42E-01 | -0.4 | 3.35E-02 | Soluble inorganic pyrophosphatase 4 |
| AT5G45490 | -0.2 | 3.82E-02 | 0.2 | 1.14E-02 | -0.4 | 9.61E-04 | -0.2 | 1.69E-01 | 0.0 | 8.13E-01 | -0.1 | 4.73E-01 | Probable disease resistance protein At5g45490 |
| AT1G11840 | -0.2 | 7.12E-02 | 0.0 | 6.83E-01 | -0.2 | 5.18E-02 | 0.1 | 4.72E-01 | 0.4 | 2.25E-02 | -0.3 | 1.64E-01 | Lactoylglutathione lyase |
| AT3G28730 | -0.2 | 1.15E-02 | -0.2 | 6.09E-03 | 0.0 | 6.60E-01 | -0.2 | 5.51E-02 | -0.2 | 1.34E-01 | 0.0 | 9.12E-01 | FACT complex subunit SSRP1 |
| AT1G43580 | -0.2 | 1.21E-02 | 0.1 | 3.45E-01 | -0.2 | 5.93E-03 | -0.4 | 9.10E-04 | -0.3 | 2.22E-02 | -0.1 | 4.01E-01 | Protein PHLOEM UNLOADING MODULATOR |
| AT3G23410 | -0.2 | 1.31E-01 | 0.0 | 7.94E-01 | -0.1 | 2.70E-01 | 0.0 | 8.57E-01 | 0.2 | 5.03E-01 | -0.2 | 4.39E-01 | Long-chain-alcohol oxidase |
| AT4G26570 | -0.2 | 4.89E-02 | -0.1 | 2.06E-01 | -0.1 | 4.95E-01 | -0.2 | 7.71E-02 | -0.1 | 6.73E-01 | -0.1 | 4.07E-01 | Calcineurin B-like protein 3 |
| AT5G11490 | -0.2 | 1.62E-02 | -0.1 | 1.37E-01 | -0.1 | 3.17E-01 | 0.1 | 5.56E-01 | 0.0 | 9.95E-01 | 0.1 | 6.97E-01 | Beta-adaptin-like protein |
| AT2G47070 | -0.2 | 6.54E-02 | -0.4 | 2.81E-03 | 0.2 | 8.72E-02 | -0.3 | 1.33E-02 | -0.2 | 2.36E-01 | -0.1 | 4.46E-01 | Squamosa promoter binding protein-like 1 |
| AT3G03305 | -0.2 | 2.17E-01 | -0.3 | 4.03E-02 | 0.2 | 3.83E-01 | -0.1 | 9.17E-01 | 0.1 | 9.18E-01 | -0.2 | 8.58E-01 | Calcineurin-like phosphoesterase domain-containing protein |
| AT5G15280 | -0.2 | 9.02E-02 | 0.1 | 5.12E-01 | -0.2 | 5.62E-02 | 0.1 | 7.74E-01 | 0.1 | 5.90E-01 | -0.1 | 8.45E-01 | Pentatricopeptide repeat-containing protein At5g15280, mitochondrial |
| AT4G00710 | -0.2 | 1.95E-02 | -0.2 | 1.36E-02 | 0.0 | 7.89E-01 | 0.0 | 7.58E-01 | 0.2 | 1.51E-01 | -0.2 | 1.76E-01 | Serine |
| AT4G15020 | -0.2 | 2.65E-02 | -0.2 | 1.06E-02 | 0.0 | 6.01E-01 | 0.1 | 6.41E-01 | 0.0 | 9.55E-01 | 0.1 | 7.93E-01 | HAT transposon superfamily |
| AT5G55660 | -0.2 | 9.15E-02 | 0.0 | 9.61E-01 | -0.2 | 1.39E-01 | 0.1 | 7.29E-01 | 0.1 | 8.19E-01 | 0.0 | 9.81E-01 | DEK-C domain-containing protein |
| AT5G27950 | -0.2 | 8.08E-02 | -0.2 | 5.34E-02 | 0.0 | 8.09E-01 | -0.3 | 7.74E-02 | -0.1 | 7.11E-01 | -0.2 | 3.40E-01 | P-loop containing nucleoside triphosphate hydrolases superfamily protein |
| AT5G44090 | -0.2 | 1.03E-02 | -0.3 | 7.76E-04 | 0.1 | 1.12E-01 | 0.0 | 9.55E-01 | -0.1 | 5.22E-01 | 0.1 | 5.55E-01 | Calcium-binding EF-hand family protein |
| AT1G01490 | -0.2 | 6.65E-02 | 0.1 | 5.84E-01 | -0.2 | 3.91E-02 | 0.0 | 8.75E-01 | -0.5 | 4.51E-02 | 0.4 | 1.53E-01 | Heavy metal transport |
| AT4G34430 | -0.2 | 1.87E-02 | -0.1 | 5.03E-02 | 0.0 | 6.96E-01 | 0.0 | 8.60E-01 | 0.0 | 9.11E-01 | 0.1 | 8.31E-01 | SWI |
| AT5G07350 | -0.2 | 4.83E-02 | 0.1 | 2.88E-01 | -0.3 | 1.30E-02 | 0.1 | 3.79E-01 | 0.1 | 7.49E-01 | 0.1 | 8.11E-01 | Ribonuclease TUDOR 1 |
| AT3G02650 | -0.2 | 1.21E-01 | 0.0 | 8.41E-01 | -0.2 | 1.29E-01 | 0.4 | 1.54E-02 | 0.4 | 1.54E-02 | 0.0 | 8.78E-01 | Small ribosomal subunit protein mS80 |
| AT1G26110 | -0.2 | 3.72E-02 | -0.2 | 2.97E-02 | 0.0 | 8.47E-01 | 0.0 | 6.96E-01 | 0.0 | 8.73E-01 | 0.1 | 7.12E-01 | Protein decapping 5 |
| AT3G47090 | -0.2 | 3.78E-01 | -0.4 | 9.20E-02 | 0.2 | 4.53E-01 | 0.0 | 9.51E-01 | 0.2 | 7.75E-01 | -0.2 | 8.41E-01 | Leucine-rich repeat protein kinase family protein |
| AT2G18770 | -0.2 | 1.03E-01 | 0.0 | 6.52E-01 | -0.1 | 3.11E-01 | -0.1 | 6.86E-01 | -0.1 | 7.83E-01 | 0.0 | 9.71E-01 | Signal recognition particle receptor subunit beta |
| AT3G48680 | -0.2 | 2.43E-02 | 0.0 | 9.74E-01 | -0.2 | 3.85E-02 | -0.3 | 9.98E-03 | -0.3 | 5.13E-02 | -0.1 | 7.70E-01 | Gamma carbonic anhydrase-like 2, mitochondrial |
| AT3G15380 | -0.2 | 2.98E-02 | -0.2 | 5.08E-02 | 0.0 | 8.64E-01 | -0.2 | 1.90E-01 | -0.3 | 2.16E-01 | 0.0 | 9.72E-01 | Choline transporter protein 1 |
| AT3G21220 | -0.2 | 7.93E-02 | -0.2 | 1.28E-01 | 0.0 | 8.59E-01 | 0.4 | 1.24E-02 | 0.2 | 3.30E-01 | 0.2 | 2.71E-01 | Mitogen-activated protein kinase kinase 5 |
| AT5G53340 | -0.2 | 1.06E-01 | -0.2 | 1.35E-01 | 0.0 | 9.51E-01 | 0.0 | 9.96E-01 | 0.3 | 5.11E-01 | -0.3 | 5.87E-01 | Hydroxyproline O-galactosyltransferase HPGT1 |
| AT2G40030 | -0.2 | 8.61E-02 | -0.2 | 1.39E-01 | 0.0 | 8.69E-01 | 0.1 | 7.71E-01 | 0.1 | 8.06E-01 | 0.0 | 9.98E-01 | DNA-directed RNA polymerase |
| AT3G03140 | -0.2 | 4.56E-02 | -0.3 | 9.78E-03 | 0.1 | 4.13E-01 | -0.1 | 7.91E-01 | 0.0 | 9.75E-01 | -0.1 | 8.80E-01 | T17B22.17 protein |
| AT4G36240 | -0.2 | 1.44E-01 | -0.1 | 2.54E-01 | 0.0 | 8.17E-01 | -0.3 | 4.39E-02 | 0.2 | 2.93E-01 | -0.5 | 2.07E-02 | GATA transcription factor 7 |
| AT5G24360 | -0.2 | 2.64E-02 | -0.3 | 2.70E-03 | 0.1 | 2.04E-01 | -0.1 | 4.57E-01 | 0.1 | 6.34E-01 | -0.3 | 3.14E-01 | AT5g24360 |
| AT2G18150 | -0.2 | 2.20E-01 | -0.3 | 3.21E-02 | 0.2 | 3.15E-01 | 1.3 | 8.75E-05 | 1.0 | 2.19E-03 | 0.3 | 2.40E-01 | Peroxidase 15 |
| AT2G02780 | -0.2 | 4.48E-02 | 0.1 | 2.81E-01 | -0.3 | 1.61E-02 | -0.1 | 7.46E-01 | 0.1 | 7.92E-01 | -0.2 | 6.31E-01 | Leucine-rich repeat protein kinase family protein |
| AT5G20000 | -0.2 | 1.22E-01 | -0.2 | 3.79E-02 | 0.1 | 5.32E-01 | 0.3 | 6.49E-02 | 0.2 | 2.37E-01 | 0.1 | 7.44E-01 | 26S proteasome regulatory subunit 8 homolog B |
| AT3G44610 | -0.2 | 9.85E-02 | 0.0 | 9.93E-01 | -0.2 | 1.60E-01 | -0.5 | 6.53E-02 | -0.2 | 4.34E-01 | -0.2 | 4.90E-01 | At3g44610 |
| AT3G01590 | -0.2 | 2.08E-02 | -0.2 | 1.91E-02 | 0.0 | 9.11E-01 | -0.3 | 3.89E-02 | -0.2 | 1.27E-01 | 0.0 | 8.30E-01 | glucose-6-phosphate 1-epimerase |
| AT2G35690 | -0.2 | 6.88E-02 | -0.1 | 1.27E-01 | 0.0 | 8.22E-01 | -0.3 | 4.15E-02 | -0.1 | 4.63E-01 | -0.2 | 3.65E-01 | Putative peroxisomal acyl-coenzyme A oxidase 1.2 |
| AT1G70710 | -0.2 | 1.16E-01 | 0.0 | 9.64E-01 | -0.2 | 1.74E-01 | -0.3 | 5.28E-02 | -0.2 | 3.85E-01 | -0.1 | 4.85E-01 | Endoglucanase 8 |
| AT3G52290 | -0.2 | 1.95E-01 | 0.1 | 4.52E-01 | -0.3 | 9.22E-02 | -0.5 | 2.34E-02 | -0.1 | 6.83E-01 | -0.4 | 1.80E-01 | Protein IQ-DOMAIN 3 |
| AT1G79730 | -0.2 | 3.40E-02 | -0.2 | 1.73E-02 | 0.0 | 6.98E-01 | 0.0 | 8.35E-01 | 0.0 | 9.08E-01 | 0.1 | 8.07E-01 | Protein PAF1 homolog |
| AT5G56680 | -0.2 | 1.75E-01 | -0.1 | 4.06E-01 | -0.1 | 6.65E-01 | 0.1 | 2.61E-01 | 0.0 | 9.19E-01 | 0.2 | 3.55E-01 | Asparagine--tRNA ligase, cytoplasmic 1 |
| AT4G12750 | -0.2 | 1.87E-01 | -0.2 | 7.94E-02 | 0.1 | 6.56E-01 | 0.4 | 6.65E-02 | 0.1 | 6.32E-01 | 0.3 | 3.53E-01 | Homeodomain-like transcriptional regulator |

| | | | | | | | | | | | | | |
|---|---|---|---|---|---|---|---|---|---|---|---|---|---|
| AT2G37250 | -0.2 | 1.39E-01 | 0.1 | 2.34E-01 | -0.3 | 2.72E-02 | 0.2 | 8.64E-02 | 0.0 | 9.76E-01 | 0.2 | 1.99E-01 | Adenylate kinase 1, chloroplastic |
| AT5G60430 | -0.2 | 2.02E-01 | -0.2 | 1.01E-01 | 0.1 | 7.08E-01 | -0.9 | 2.48E-02 | -0.3 | 3.76E-01 | -0.5 | 2.63E-01 | ribonuclease P |
| AT2G47000 | -0.2 | 3.47E-01 | -0.9 | 4.98E-04 | 0.8 | 3.06E-03 | 1.3 | 3.72E-06 | 0.3 | 1.35E-01 | 1.0 | 1.92E-03 | ATP binding cassette subfamily B4 |
| AT1G54460 | -0.2 | 8.09E-02 | -0.1 | 2.28E-01 | -0.1 | 6.33E-01 | -0.1 | 2.22E-01 | -0.1 | 7.58E-01 | -0.1 | 6.18E-01 | hypothetical protein |
| AT5G18270 | -0.2 | 1.46E-01 | -0.7 | 3.34E-04 | 0.5 | 5.41E-03 | 1.0 | 2.65E-04 | 0.7 | 6.44E-03 | 0.2 | 2.59E-01 | NAC domain containing protein 87 |
| AT5G27210 | -0.2 | 1.06E-01 | -0.1 | 5.82E-01 | -0.1 | 3.78E-01 | 0.1 | 8.53E-01 | 0.0 | 9.48E-01 | 0.1 | 8.25E-01 | Protein CANDIDATE G-PROTEIN COUPLED RECEPTOR 8 |
| AT2G17530 | -0.2 | 2.50E-02 | 0.0 | 8.06E-01 | -0.2 | 3.36E-02 | 0.1 | 6.13E-01 | 0.1 | 5.86E-01 | 0.0 | 9.38E-01 | At2g17530 |
| AT5G49020 | -0.2 | 8.13E-02 | 0.0 | 7.24E-01 | -0.1 | 2.19E-01 | 0.0 | 9.97E-01 | 0.1 | 5.01E-01 | -0.1 | 5.71E-01 | Probable histone-arginine methyltransferase 1.4 |
| AT5G41150 | -0.2 | 3.55E-02 | -0.2 | 1.36E-02 | 0.0 | 5.96E-01 | 0.1 | 6.63E-01 | 0.0 | 8.72E-01 | 0.1 | 8.96E-01 | Restriction endonuclease, type II-like superfamily protein |
| AT5G02460 | -0.2 | 2.21E-01 | -0.2 | 2.52E-01 | 0.0 | 9.91E-01 | 0.1 | 8.78E-01 | 0.2 | 5.79E-01 | -0.2 | 7.60E-01 | Dof zinc finger protein DOF5.1 |
| AT1G15920 | -0.2 | 3.83E-02 | -0.1 | 2.65E-01 | -0.1 | 3.64E-01 | -0.2 | 2.67E-01 | -0.2 | 4.04E-01 | 0.0 | 9.43E-01 | Probable CCR4-associated factor 1 homolog 2 |
| AT1G74360 | -0.2 | 2.32E-01 | -0.6 | 1.65E-03 | 0.5 | 2.11E-02 | 0.6 | 1.70E-02 | -0.2 | 4.92E-01 | 0.9 | 1.75E-02 | Probable LRR receptor-like serine |
| AT4G14370 | -0.2 | 4.83E-01 | -0.8 | 1.20E-02 | 0.6 | 6.97E-02 | 1.1 | 3.03E-04 | 0.4 | 2.03E-01 | 0.7 | 2.74E-02 | Disease resistance protein family |
| AT1G13090 | -0.2 | 4.30E-02 | -0.3 | 3.15E-03 | 0.1 | 1.54E-01 | -0.3 | 3.85E-02 | 0.0 | 8.71E-01 | -0.3 | 5.92E-02 | Cytochrome P450, family 71, subfamily B, polypeptide 28 |
| AT3G27320 | -0.2 | 5.61E-02 | -0.2 | 2.90E-02 | 0.0 | 7.15E-01 | 0.0 | 8.77E-01 | 0.0 | 9.88E-01 | 0.0 | 9.12E-01 | hypothetical protein |
| AT5G26360 | -0.2 | 3.25E-02 | 0.0 | 6.87E-01 | -0.2 | 2.64E-02 | 0.2 | 2.98E-02 | 0.1 | 2.60E-01 | 0.1 | 6.01E-01 | T-complex protein 1 subunit gamma |
| AT1G01710 | -0.2 | 3.75E-02 | -0.1 | 3.63E-01 | -0.1 | 2.64E-01 | -0.1 | 7.29E-01 | 0.0 | 9.17E-01 | 0.0 | 8.85E-01 | Acyl-CoA hydrolase 2 |
| AT5G09630 | -0.2 | 1.45E-01 | -0.3 | 3.64E-02 | 0.1 | 4.79E-01 | 0.1 | 7.58E-01 | -0.1 | 8.89E-01 | 0.2 | 6.51E-01 | hypothetical protein |
| AT5G65110 | -0.2 | 4.10E-02 | -0.2 | 2.38E-02 | 0.0 | 7.47E-01 | 0.1 | 6.11E-01 | -0.3 | 2.42E-01 | 0.4 | 1.84E-01 | Acyl-coenzyme A oxidase |
| AT5G63840 | -0.2 | 2.14E-02 | 0.0 | 6.24E-01 | -0.1 | 8.22E-02 | 0.1 | 3.76E-01 | 0.1 | 4.30E-01 | 0.0 | 9.83E-01 | Glycosyl hydrolases family 31 protein |
| AT1G30490 | -0.2 | 4.34E-02 | -0.4 | 5.01E-04 | 0.2 | 2.09E-02 | -0.2 | 9.04E-02 | 0.0 | 8.97E-01 | -0.2 | 2.25E-01 | Homeobox-leucine zipper protein ATHB-9 |
| AT1G74310 | -0.2 | 1.19E-01 | -0.3 | 2.02E-02 | 0.1 | 3.68E-01 | -0.2 | 4.11E-01 | 0.2 | 4.49E-01 | -0.5 | 1.82E-01 | Heat shock protein 101 |
| AT5G56010 | -0.2 | 2.46E-01 | -0.2 | 1.57E-01 | 0.0 | 7.76E-01 | 0.3 | 1.08E-02 | 0.5 | 1.05E-03 | -0.2 | 2.62E-01 | Heat shock protein 90-3 |
| AT4G33090 | -0.2 | 3.53E-02 | -0.1 | 3.82E-01 | -0.1 | 2.29E-01 | -0.2 | 1.00E-01 | -0.1 | 6.04E-01 | -0.1 | 5.17E-01 | Aminopeptidase M1 |
| AT5G50330 | -0.2 | 3.04E-01 | -0.2 | 3.58E-01 | 0.0 | 9.57E-01 | -0.3 | 2.10E-01 | -0.2 | 5.76E-01 | -0.2 | 6.71E-01 | Protein kinase superfamily protein |
| AT4G11860 | -0.2 | 6.45E-02 | -0.1 | 1.69E-01 | 0.0 | 6.76E-01 | -0.2 | 2.77E-01 | 0.0 | 9.81E-01 | -0.2 | 3.82E-01 | hypothetical protein |
| AT4G17750 | -0.2 | 8.52E-02 | -0.3 | 7.42E-03 | 0.1 | 2.17E-01 | 0.1 | 8.15E-01 | 0.1 | 7.16E-01 | -0.1 | 9.10E-01 | Heat stress transcription factor A-1a |
| AT1G79380 | -0.2 | 1.07E-01 | -0.2 | 1.46E-01 | 0.0 | 9.29E-01 | -0.3 | 5.31E-02 | -0.1 | 8.21E-01 | -0.2 | 2.41E-01 | E3 ubiquitin-protein ligase RGLG4 |
| AT3G27700 | -0.2 | 8.30E-02 | -0.4 | 2.29E-03 | 0.2 | 6.74E-02 | -0.1 | 6.87E-01 | -0.2 | 6.20E-01 | 0.0 | 9.28E-01 | Zinc finger CCCH domain-containing protein 41 |
| AT5G41580 | -0.2 | 7.57E-02 | -0.3 | 7.92E-03 | 0.1 | 2.44E-01 | -0.1 | 7.12E-01 | -0.1 | 8.33E-01 | 0.0 | 9.48E-01 | E4 SUMO-protein ligase PIAL2 |
| AT4G18950 | -0.2 | 2.13E-01 | -0.2 | 2.43E-01 | 0.0 | 9.73E-01 | 0.9 | 2.90E-06 | 0.3 | 2.63E-02 | 0.5 | 2.67E-03 | AT4g18950 |
| AT2G02970 | -0.2 | 7.37E-02 | -0.2 | 2.24E-02 | 0.1 | 5.23E-01 | -0.1 | 3.17E-01 | -0.1 | 4.58E-01 | 0.0 | 9.48E-01 | apyrase |
| AT5G56280 | -0.2 | 6.60E-02 | -0.1 | 2.81E-01 | -0.1 | 4.88E-01 | 0.2 | 2.45E-01 | 0.2 | 2.51E-01 | 0.0 | 9.42E-01 | COP9 signalosome complex subunit 6a |
| AT4G28350 | -0.2 | 3.18E-01 | -0.1 | 4.13E-01 | 0.0 | 9.15E-01 | 0.3 | 1.76E-01 | 0.0 | 8.93E-01 | 0.2 | 3.97E-01 | Probable L-type lectin-domain containing receptor kinase VII.2 |
| AT5G67480 | -0.2 | 4.66E-01 | -0.5 | 5.50E-02 | 0.3 | 2.36E-01 | -0.9 | 1.39E-03 | -1.4 | 9.86E-05 | 0.5 | 1.07E-01 | BTB |
| AT5G63910 | -0.2 | 4.44E-02 | -0.1 | 3.15E-01 | -0.1 | 3.31E-01 | -0.1 | 6.86E-01 | -0.1 | 6.10E-01 | 0.0 | 9.20E-01 | Farnesylcysteine lyase |
| AT5G22030 | -0.2 | 6.40E-02 | -0.1 | 1.37E-01 | 0.0 | 7.58E-01 | -0.1 | 4.39E-01 | -0.2 | 3.72E-01 | 0.0 | 9.05E-01 | UBP8 |
| AT4G28070 | -0.2 | 2.39E-01 | -0.1 | 4.21E-01 | 0.0 | 7.97E-01 | -0.3 | 4.37E-01 | -0.3 | 4.22E-01 | 0.0 | 9.53E-01 | AFG1-like ATPase family protein |
| AT4G09140 | -0.2 | 5.76E-02 | -0.2 | 3.24E-02 | 0.0 | 7.50E-01 | 0.2 | 3.20E-01 | 0.1 | 6.87E-01 | 0.1 | 7.47E-01 | MUTL-homologue 1 |
| AT2G43200 | -0.2 | 1.74E-01 | -0.1 | 3.34E-01 | 0.0 | 7.71E-01 | 0.2 | 3.72E-01 | 0.2 | 4.52E-01 | 0.0 | 9.88E-01 | Probable methyltransferase PMT19 |
| AT1G54115 | -0.2 | 4.86E-02 | -0.2 | 1.89E-02 | 0.0 | 6.03E-01 | -0.2 | 2.23E-01 | -0.2 | 2.50E-01 | 0.0 | 9.67E-01 | Cation |
| AT2G28620 | -0.2 | 5.79E-02 | 0.1 | 2.07E-01 | -0.3 | 1.55E-02 | 0.2 | 3.32E-01 | 0.3 | 1.70E-01 | -0.1 | 7.41E-01 | P-loop containing nucleoside triphosphate hydrolases superfamily protein |
| AT5G61580 | -0.2 | 4.79E-02 | -0.2 | 1.34E-02 | 0.1 | 4.89E-01 | -0.2 | 2.57E-01 | 0.1 | 8.01E-01 | -0.2 | 2.64E-01 | ATP-dependent 6-phosphofructokinase 4, chloroplastic |
| AT1G55090 | -0.2 | 9.61E-02 | -0.4 | 1.54E-03 | 0.2 | 3.72E-02 | -0.1 | 7.36E-01 | 0.0 | 9.66E-01 | -0.1 | 8.52E-01 | Glutamine-dependent NAD synthetase synthase ) synthetase) |
| AT1G26550 | -0.2 | 3.60E-02 | 0.0 | 8.59E-01 | -0.1 | 7.71E-02 | 0.1 | 5.00E-01 | 0.1 | 5.88E-01 | 0.0 | 9.97E-01 | Peptidyl-prolyl cis-trans isomerase |
| AT2G31270 | -0.2 | 2.20E-01 | 0.1 | 5.73E-01 | -0.2 | 1.37E-01 | 0.2 | 2.14E-01 | 0.3 | 2.25E-01 | 0.0 | 9.49E-01 | CDT1-like protein a, chloroplastic |
| AT5G46560 | -0.2 | 1.26E-01 | 0.0 | 6.98E-01 | -0.1 | 3.35E-01 | 0.1 | 7.65E-01 | 0.0 | 9.32E-01 | 0.1 | 7.79E-01 | Man1-Src1p-carboxy-terminal domain protein |
| AT3G21690 | -0.2 | 4.92E-02 | -0.4 | 2.80E-04 | 0.3 | 7.66E-03 | 0.4 | 4.53E-04 | 0.1 | 2.86E-01 | 0.3 | 4.82E-02 | Protein DETOXIFICATION 40 |
| AT1G79930 | -0.2 | 1.08E-01 | 0.0 | 9.62E-01 | -0.2 | 1.35E-01 | 0.2 | 6.49E-02 | 0.1 | 5.99E-01 | 0.1 | 4.51E-01 | Heat shock 70 kDa protein 14 |
| AT4G24530 | -0.2 | 7.06E-02 | -0.2 | 2.85E-02 | 0.1 | 6.31E-01 | -0.3 | 1.49E-01 | -0.3 | 2.61E-01 | 0.0 | 9.11E-01 | O-fucosyltransferase 31 |
| AT3G16640 | -0.2 | 9.50E-02 | -0.1 | 2.78E-01 | -0.1 | 6.11E-01 | -0.5 | 5.15E-03 | -0.5 | 1.19E-02 | 0.0 | 9.81E-01 | Translationally-controlled tumor protein 1 |
| AT1G28440 | -0.2 | 4.55E-02 | -0.1 | 4.12E-01 | -0.1 | 2.57E-01 | -0.1 | 7.09E-01 | 0.1 | 7.98E-01 | -0.1 | 6.22E-01 | Receptor-like protein kinase HSL1 |
| AT3G46220 | -0.2 | 3.38E-02 | -0.3 | 3.97E-03 | 0.1 | 2.41E-01 | 0.2 | 2.06E-01 | 0.0 | 9.12E-01 | 0.2 | 2.99E-01 | E3 UFM1-protein ligase-like protein |
| AT3G12680 | -0.2 | 2.06E-02 | -0.2 | 8.46E-03 | 0.0 | 6.02E-01 | 0.0 | 9.19E-01 | 0.0 | 9.75E-01 | 0.0 | 9.20E-01 | Zinc finger CCCH domain-containing protein 37 |
| AT5G11770 | -0.2 | 1.04E-01 | 0.0 | 7.43E-01 | -0.2 | 8.21E-02 | -0.1 | 2.55E-01 | 0.0 | 9.78E-01 | -0.1 | 4.40E-01 | NADH dehydrogenase iron-sulfur protein 7, mitochondrial |
| AT2G04890 | -0.2 | 6.53E-02 | -0.1 | 1.07E-01 | 0.0 | 8.66E-01 | -0.1 | 7.04E-01 | -0.2 | 5.47E-01 | 0.1 | 8.57E-01 | Scarecrow-like protein 21 |
| AT3G46290 | -0.2 | 1.60E-01 | -0.4 | 4.34E-03 | 0.2 | 6.92E-02 | -0.2 | 7.01E-01 | 0.0 | 9.83E-01 | -0.2 | 8.01E-01 | Receptor-like protein kinase HERK 1 |
| AT3G06410 | -0.2 | 3.73E-02 | -0.3 | 4.58E-03 | 0.1 | 2.48E-01 | -0.2 | 1.74E-01 | 0.2 | 3.64E-01 | -0.4 | 7.76E-02 | Zinc finger CCCH domain-containing protein 34 |
| AT2G01710 | -0.2 | 6.94E-02 | -0.1 | 2.23E-01 | -0.1 | 5.96E-01 | -0.1 | 3.73E-01 | -0.3 | 1.08E-01 | 0.1 | 5.57E-01 | At2g01710 |
| AT1G66140 | -0.2 | 2.84E-01 | 0.0 | 8.09E-01 | -0.1 | 4.97E-01 | 0.0 | 7.83E-01 | -0.2 | 2.30E-01 | 0.2 | 1.72E-01 | Zinc finger protein 4 |
| AT1G32090 | -0.2 | 1.74E-01 | 0.1 | 6.51E-01 | -0.2 | 1.25E-01 | -0.1 | 7.52E-01 | 0.1 | 7.63E-01 | -0.2 | 5.95E-01 | CSC1-like protein At1g32090 |
| AT1G35780 | -0.2 | 5.71E-02 | -0.1 | 2.57E-01 | -0.1 | 4.67E-01 | -0.1 | 6.13E-01 | 0.0 | 9.02E-01 | -0.1 | 5.84E-01 | At1g35780 |
| AT3G13330 | -0.2 | 3.95E-02 | -0.5 | 5.25E-05 | 0.4 | 1.21E-03 | 0.7 | 4.86E-04 | 0.3 | 1.37E-01 | 0.4 | 8.74E-02 | Proteasome activator subunit 4 |
| AT5G20320 | -0.2 | 1.11E-01 | -0.4 | 3.61E-03 | 0.2 | 8.47E-02 | 0.0 | 8.90E-01 | -0.1 | 6.05E-01 | 0.2 | 5.95E-01 | Dicer-like 4 |
| AT2G03240 | -0.2 | 1.29E-01 | -0.4 | 5.52E-03 | 0.2 | 1.11E-01 | 0.2 | 2.75E-01 | 0.3 | 1.33E-01 | -0.1 | 7.24E-01 | Phosphate transporter PHO1 homolog 5 |
| AT3G62200 | -0.2 | 7.70E-02 | -0.2 | 4.99E-02 | 0.0 | 8.07E-01 | 0.0 | 9.29E-01 | 0.1 | 8.41E-01 | -0.1 | 9.10E-01 | Endonuclease or glycosyl hydrolase |
| AT2G42390 | -0.2 | 4.50E-02 | -0.1 | 3.11E-01 | -0.1 | 3.48E-01 | -0.2 | 6.88E-02 | -0.1 | 6.11E-01 | -0.2 | 4.16E-01 | Expressed protein |
| AT1G19000 | -0.2 | 1.60E-01 | -0.2 | 1.89E-01 | 0.0 | 9.75E-01 | -0.2 | 2.62E-02 | -0.2 | 4.36E-02 | 0.0 | 9.93E-01 | hypothetical protein |
| AT3G09900 | -0.2 | 3.45E-02 | -0.2 | 5.77E-03 | 0.1 | 3.22E-01 | -0.4 | 4.57E-03 | -0.3 | 3.65E-02 | -0.1 | 6.38E-01 | RABE1e |
| AT2G38440 | -0.2 | 1.02E-01 | -0.3 | 1.54E-02 | 0.1 | 3.14E-01 | -0.1 | 5.17E-01 | -0.1 | 7.79E-01 | -0.1 | 8.62E-01 | Protein SCAR2 |
| AT4G12670 | -0.2 | 7.74E-02 | -0.2 | 3.28E-02 | 0.0 | 6.56E-01 | 0.2 | 3.90E-01 | -0.1 | 8.62E-01 | 0.3 | 3.48E-01 | At4g12670 |
| AT2G40280 | -0.2 | 6.68E-02 | -0.1 | 9.46E-02 | 0.0 | 9.18E-01 | 0.0 | 9.10E-01 | 0.2 | 4.19E-01 | -0.1 | 6.02E-01 | Probable methyltransferase PMT23 |
| AT1G02270 | -0.2 | 1.44E-01 | -0.2 | 3.74E-02 | 0.1 | 4.77E-01 | 0.0 | 8.33E-01 | -0.1 | 6.95E-01 | 0.0 | 8.86E-01 | Calcium-binding endonuclease |
| AT3G19270 | -0.2 | 3.13E-01 | -0.5 | 1.33E-02 | 0.3 | 1.18E-01 | | | | | | | Cytochrome P450, family 707, subfamily A, polypeptide 4 |
| AT1G79000 | -0.2 | 7.92E-02 | -0.4 | 7.08E-04 | 0.3 | 1.71E-02 | -0.2 | 2.69E-01 | -0.1 | 6.90E-01 | -0.1 | 7.21E-01 | histone acetyltransferase |
| AT1G34190 | -0.2 | 3.94E-02 | -0.2 | 9.97E-03 | 0.1 | 4.32E-01 | -0.1 | 6.52E-01 | -0.2 | 1.79E-01 | 0.1 | 4.91E-01 | NAC domain-containing protein 17 |
| AT5G66770 | -0.2 | 3.53E-02 | 0.0 | 7.82E-01 | -0.1 | 1.02E-01 | 0.3 | 1.93E-02 | 0.1 | 5.62E-01 | 0.2 | 1.79E-01 | Scarecrow-like protein 4 |
| AT5G61190 | -0.2 | 1.02E-01 | -0.2 | 2.16E-02 | 0.1 | 4.17E-01 | 0.1 | 7.16E-01 | 0.1 | 6.33E-01 | 0.0 | 9.10E-01 | Endonuclease or glycosyl hydrolase with C2H2-type zinc finger domain-containing protein |
| AT2G20990 | -0.2 | 6.19E-02 | 0.0 | 6.98E-01 | -0.2 | 4.72E-02 | -0.2 | 1.05E-01 | -0.1 | 5.62E-01 | -0.1 | 6.09E-01 | Synaptotagmin A |
| AT2G29190 | -0.2 | 2.23E-02 | -0.2 | 3.05E-03 | 0.1 | 2.55E-01 | 0.2 | 1.27E-01 | 0.1 | 4.57E-01 | 0.1 | 7.15E-01 | Pumilio homolog 2 |
| AT4G23730 | -0.2 | 7.31E-02 | -0.1 | 1.60E-01 | 0.0 | 7.44E-01 | -0.2 | 2.28E-01 | -0.1 | 3.99E-01 | 0.0 | 9.10E-01 | glucose-6-phosphate 1-epimerase |
| AT1G18640 | -0.2 | 8.49E-02 | -0.1 | 3.84E-01 | -0.1 | 4.53E-01 | -0.2 | 2.51E-01 | -0.2 | 3.57E-01 | 0.0 | 9.70E-01 | phosphoserine phosphatase |
| AT4G17090 | -0.2 | 2.26E-01 | -0.4 | 1.27E-02 | 0.2 | 1.24E-01 | 0.4 | 5.54E-03 | 0.5 | 2.53E-03 | -0.1 | 6.18E-01 | Beta-amylase 3, chloroplastic |

| | | | | | | | | | | | | | |
|---|---|---|---|---|---|---|---|---|---|---|---|---|---|
| AT5G61520 | -0.2 | 2.24E-01 | -0.3 | 2.74E-02 | 0.2 | 2.75E-01 | 0.0 | 9.67E-01 | -0.4 | 1.63E-02 | 0.4 | 5.10E-02 | Major facilitator superfamily protein |
| AT2G21070 | -0.2 | 1.80E-01 | 0.0 | 7.44E-01 | -0.1 | 4.11E-01 | 0.1 | 7.92E-01 | -0.2 | 6.19E-01 | 0.2 | 4.48E-01 | U6 small nuclear RNA -N)-methyltransferase |
| AT2G42530 | -0.2 | 7.12E-01 | 0.4 | 3.44E-01 | -0.5 | 2.51E-01 | -0.9 | 1.42E-02 | 0.8 | 3.31E-02 | -1.6 | 1.34E-03 | Protein COLD-REGULATED 15B, chloroplastic |
| AT3G01260 | -0.2 | 3.39E-01 | 0.5 | 5.26E-03 | -0.7 | 3.59E-03 | -0.7 | 1.67E-03 | -0.1 | 7.84E-01 | -0.7 | 1.57E-02 | Galactose mutarotase-like superfamily protein |
| AT1G60270 | -0.2 | 5.47E-01 | -0.3 | 2.89E-01 | 0.1 | 6.85E-01 | -0.4 | 1.88E-01 | 0.5 | 6.24E-02 | -1.0 | 1.67E-02 | Beta glucosidase 6 |
| AT4G26760 | -0.2 | 1.47E-01 | 0.1 | 2.23E-01 | -0.3 | 3.12E-02 | -0.1 | 6.35E-01 | 0.2 | 4.54E-01 | -0.3 | 2.68E-01 | 65-kDa microtubule-associated protein 2 |
| AT3G51120 | -0.2 | 1.35E-01 | -0.3 | 1.65E-02 | 0.1 | 2.73E-01 | 0.0 | 8.81E-01 | -0.1 | 6.96E-01 | 0.1 | 8.53E-01 | Zinc finger CCCH domain-containing protein 44 |
| AT3G01460 | -0.2 | 6.25E-02 | -0.2 | 3.32E-02 | 0.0 | 7.25E-01 | 0.0 | 8.81E-01 | 0.1 | 7.44E-01 | -0.1 | 6.97E-01 | Methyl-CpG-binding domain-containing protein 9 |
| AT3G49650 | -0.2 | 1.29E-01 | 0.3 | 1.27E-02 | -0.4 | 3.43E-03 | -0.4 | 1.76E-01 | 0.0 | 9.26E-01 | -0.3 | 3.34E-01 | Kinesin-like protein KIN-8B |
| AT1G71230 | -0.2 | 1.95E-01 | 0.1 | 5.97E-01 | -0.2 | 1.36E-01 | 0.0 | 9.47E-01 | 0.2 | 6.83E-01 | -0.2 | 6.80E-01 | COP9-signalosome 5B |
| AT4G26690 | -0.2 | 1.82E-01 | 0.0 | 8.34E-01 | -0.1 | 3.20E-01 | -0.3 | 3.42E-01 | -0.4 | 1.45E-01 | 0.2 | 6.72E-01 | Glycerophosphodiester phosphodiesterase GDPDL3 |
| AT2G28390 | -0.2 | 4.40E-02 | -0.1 | 1.71E-01 | -0.1 | 5.37E-01 | 0.0 | 9.53E-01 | 0.0 | 8.57E-01 | 0.0 | 9.22E-01 | Vacuolar fusion protein MON1 homolog |
| AT4G14340 | -0.2 | 3.51E-02 | -0.1 | 4.20E-02 | 0.0 | 9.84E-01 | -0.3 | 3.98E-02 | -0.1 | 4.44E-01 | -0.2 | 3.68E-01 | Casein kinase I |
| AT3G63270 | -0.2 | 8.38E-02 | -0.2 | 1.49E-02 | 0.1 | 3.72E-01 | -0.3 | 1.83E-01 | -0.3 | 3.20E-01 | 0.0 | 9.14E-01 | Protein ANTAGONIST OF LIKE HETEROCHROMATIN PROTEIN 1 |
| AT5G05870 | -0.2 | 1.57E-01 | -0.3 | 1.60E-02 | 0.1 | 2.44E-01 | -0.3 | 6.65E-02 | -0.2 | 2.35E-01 | -0.1 | 7.41E-01 | UDP-glycosyltransferase 76C1 |
| AT3G10130 | -0.2 | 1.41E-01 | -0.1 | 2.80E-01 | 0.0 | 7.53E-01 | -0.1 | 2.94E-01 | 0.0 | 8.03E-01 | -0.1 | 6.38E-01 | Heme-binding-like protein At3g10130, chloroplastic |
| AT3G03450 | -0.2 | 3.02E-01 | 0.1 | 3.06E-01 | -0.3 | 1.02E-01 | 0.2 | 3.30E-01 | 0.6 | 2.42E-02 | -0.3 | 1.90E-01 | DELLA protein RGL2 |
| AT1G50110 | -0.1 | 3.74E-01 | 0.1 | 5.59E-01 | -0.2 | 2.22E-01 | 0.5 | 9.03E-02 | 0.4 | 1.51E-01 | 0.0 | 9.63E-01 | Branched-chain-amino-acid aminotransferase |
| AT1G13450 | -0.1 | 1.61E-01 | -0.1 | 1.87E-01 | 0.0 | 9.77E-01 | -0.2 | 1.35E-01 | -0.2 | 1.60E-01 | 0.0 | 9.77E-01 | Homeodomain-like superfamily protein |
| AT4G18710 | -0.1 | 5.14E-02 | 0.0 | 6.43E-01 | -0.1 | 1.66E-01 | -0.2 | 1.93E-01 | -0.2 | 1.43E-01 | 0.0 | 8.41E-01 | non-specific serine |
| AT5G35080 | -0.1 | 6.07E-02 | -0.1 | 4.07E-01 | -0.1 | 3.47E-01 | 0.0 | 8.91E-01 | 0.1 | 5.95E-01 | -0.1 | 7.41E-01 | Protein OS-9 homolog |
| AT1G80000 | -0.1 | 5.15E-02 | -0.2 | 3.40E-02 | 0.0 | 8.05E-01 | -0.2 | 1.92E-01 | -0.3 | 1.16E-01 | 0.1 | 8.03E-01 | CASC3 |
| AT1G07090 | -0.1 | 8.11E-02 | 0.0 | 7.74E-01 | -0.1 | 2.04E-01 | -0.1 | 6.86E-01 | -0.2 | 2.60E-01 | 0.1 | 5.16E-01 | Protein LIGHT-DEPENDENT SHORT HYPOCOTYLS 6 |
| AT5G05570 | -0.1 | 5.02E-02 | -0.2 | 4.58E-02 | 0.0 | 9.28E-01 | -0.1 | 5.18E-01 | 0.1 | 6.03E-01 | -0.2 | 3.47E-01 | Transducin family protein |
| AT5G06960 | -0.1 | 1.36E-01 | -0.4 | 1.25E-03 | 0.3 | 2.43E-02 | 0.2 | 2.78E-01 | -0.1 | 7.34E-01 | 0.3 | 2.04E-01 | OCS-element binding factor 5 |
| AT3G11770 | -0.1 | 4.41E-02 | 0.2 | 3.14E-02 | -0.3 | 2.61E-03 | -0.2 | 2.64E-01 | -0.2 | 1.78E-01 | 0.1 | 8.36E-01 | Protein RISC-INTERACTING CLEARING 3'-5' EXORIBONUCLEASE 1 |
| AT3G01910 | -0.1 | 5.92E-02 | -0.1 | 4.39E-01 | -0.1 | 2.95E-01 | -0.1 | 6.09E-01 | 0.0 | 7.68E-01 | 0.0 | 9.47E-01 | Sulfite oxidase |
| AT5G12950 | -0.1 | 1.19E-01 | 0.1 | 3.33E-01 | -0.2 | 3.67E-02 | -0.3 | 4.03E-02 | -0.2 | 2.98E-01 | -0.1 | 5.54E-01 | Proline-tRNA ligase |
| AT5G12480 | -0.1 | 1.06E-01 | -0.2 | 6.44E-02 | 0.0 | 7.75E-01 | 0.0 | 9.20E-01 | 0.0 | 9.84E-01 | 0.0 | 9.34E-01 | Calmodulin-domain protein kinase 7 |
| AT5G19200 | -0.1 | 2.59E-01 | -0.2 | 2.03E-01 | 0.0 | 8.79E-01 | -0.8 | 6.77E-04 | -0.6 | 6.39E-03 | -0.2 | 5.50E-01 | 3-dehydrosphinganine reductase TSC10B |
| AT1G10550 | -0.1 | 4.97E-01 | 0.2 | 2.57E-01 | -0.4 | 1.41E-01 | -1.1 | 1.37E-02 | -0.5 | 2.43E-01 | -0.6 | 2.03E-01 | Probable xyloglucan endotransglucosylase |
| AT3G48530 | -0.1 | 7.87E-02 | 0.2 | 1.15E-02 | -0.4 | 1.51E-03 | -0.6 | 2.67E-04 | -0.8 | 5.41E-05 | 0.2 | 2.48E-01 | SNF1-related protein kinase regulatory subunit gamma-1 |
| AT3G12810 | -0.1 | 8.62E-02 | -0.4 | 1.22E-03 | 0.2 | 3.32E-02 | -0.2 | 7.69E-01 | 0.1 | 9.45E-01 | -0.2 | 7.81E-01 | SNF2 domain-containing protein |
| AT4G31270 | -0.1 | 2.71E-01 | 0.0 | 7.76E-01 | -0.2 | 2.55E-01 | -0.4 | 1.86E-01 | -0.5 | 1.48E-01 | 0.1 | 9.05E-01 | At4g31270 |
| AT1G54260 | -0.1 | 1.64E-01 | -0.2 | 4.76E-02 | 0.1 | 5.20E-01 | 0.0 | 9.21E-01 | 0.2 | 3.22E-01 | -0.2 | 3.46E-01 | Uncharacterized protein At1g54260 |
| AT4G29350 | -0.1 | 1.14E-01 | 0.0 | 1.00E+00 | -0.1 | 1.55E-01 | -0.2 | 7.66E-02 | 0.0 | 9.01E-01 | -0.2 | 2.25E-01 | Profilin-2 |
| AT5G09680 | -0.1 | 5.41E-02 | -0.2 | 2.79E-02 | 0.0 | 7.14E-01 | -0.3 | 1.71E-01 | -0.4 | 5.11E-02 | 0.2 | 6.27E-01 | Reduced lateral root formation |
| AT1G49660 | -0.1 | 8.65E-02 | 0.0 | 9.79E-01 | -0.1 | 1.20E-01 | 0.1 | 4.60E-01 | 0.3 | 1.20E-01 | -0.2 | 5.23E-01 | Probable carboxylesterase 5 |
| AT1G51980 | -0.1 | 5.61E-02 | 0.0 | 7.55E-01 | -0.2 | 4.90E-02 | 0.2 | 2.53E-02 | 0.2 | 6.15E-02 | 0.0 | 9.46E-01 | Insulinase protein |
| AT3G56030 | -0.1 | 1.37E-01 | -0.3 | 7.30E-03 | 0.2 | 1.50E-01 | 0.3 | 1.28E-01 | -0.1 | 8.69E-01 | 0.4 | 1.59E-01 | Pentatricopeptide repeat-containing protein At3g56030, mitochondrial |
| AT2G37550 | -0.1 | 4.21E-02 | 0.1 | 1.82E-01 | -0.2 | 8.38E-03 | -0.1 | 3.53E-01 | 0.0 | 9.76E-01 | -0.1 | 4.80E-01 | ADP-ribosylation factor GTPase-activating protein AGD7 |
| AT4G16900 | -0.1 | 8.06E-02 | -0.3 | 1.03E-02 | 0.1 | 2.91E-01 | -0.1 | 7.34E-01 | -0.2 | 4.94E-01 | 0.1 | 7.98E-01 | Disease resistance protein family |
| AT1G50510 | -0.1 | 6.72E-02 | -0.2 | 2.52E-02 | 0.0 | 6.02E-01 | 0.2 | 4.19E-01 | 0.2 | 4.66E-01 | 0.0 | 9.88E-01 | Pseudouridine-5'-phosphate glycosidase |
| AT1G06070 | -0.1 | 9.60E-02 | -0.1 | 1.65E-01 | 0.0 | 8.37E-01 | 0.0 | 8.98E-01 | 0.0 | 9.83E-01 | 0.0 | 9.40E-01 | At1g06070 transcription factor family protein) |
| AT2G29560 | -0.1 | 3.73E-02 | 0.0 | 9.16E-01 | -0.2 | 5.42E-02 | 0.1 | 5.12E-01 | 0.1 | 5.44E-01 | 0.0 | 9.81E-01 | Cytosolic enolase 3 |
| AT1G20920 | -0.1 | 1.26E-01 | -0.2 | 6.72E-02 | 0.0 | 7.14E-01 | -0.1 | 3.74E-01 | -0.2 | 8.03E-02 | 0.1 | 5.03E-01 | DEAD-box ATP-dependent RNA helicase 42 |
| AT1G01040 | -0.1 | 1.06E-01 | -0.3 | 3.25E-03 | 0.2 | 7.57E-02 | -0.1 | 7.38E-01 | 0.0 | 8.38E-01 | 0.0 | 9.65E-01 | Dicer-like 1 |
| AT4G17840 | -0.1 | 1.68E-01 | -0.6 | 2.15E-04 | 0.5 | 2.34E-03 | -0.6 | 2.10E-02 | -0.3 | 2.86E-01 | -0.3 | 3.75E-01 | hypothetical protein |
| AT2G44830 | -0.1 | 1.72E-01 | -0.2 | 7.35E-02 | 0.1 | 6.48E-01 | -0.1 | 5.62E-01 | -0.1 | 7.07E-01 | 0.0 | 9.48E-01 | Uncharacterized protein At2g44830 |
| AT3G12480 | -0.1 | 1.02E-01 | -0.2 | 5.91E-02 | 0.0 | 7.63E-01 | -0.1 | 7.18E-01 | 0.0 | 9.70E-01 | -0.2 | 7.62E-01 | At3g12480 |
| AT5G08390 | -0.1 | 1.58E-01 | -0.3 | 1.93E-02 | 0.1 | 2.81E-01 | -0.1 | 8.75E-01 | 0.0 | 9.27E-01 | -0.1 | 8.40E-01 | Katanin p80 WD40 repeat-containing subunit B1 homolog |
| AT5G41410 | -0.1 | 1.00E-01 | -0.5 | 3.97E-04 | 0.3 | 7.52E-03 | 0.0 | 9.12E-01 | 0.2 | 3.51E-01 | -0.2 | 3.99E-01 | Homeobox protein BEL1 homolog |
| AT3G42640 | -0.1 | 2.22E-01 | -0.2 | 8.03E-02 | 0.1 | 5.89E-01 | -0.1 | 8.94E-01 | 0.0 | 9.36E-01 | 0.0 | 9.83E-01 | ATPase 8, plasma membrane-type |
| AT4G12110 | -0.1 | 7.56E-02 | 0.0 | 6.35E-01 | -0.1 | 2.41E-01 | 0.0 | 8.44E-01 | 0.0 | 9.06E-01 | 0.0 | 9.82E-01 | Methylsterol monooxygenase 1-1 |
| AT3G23780 | -0.1 | 3.60E-02 | -0.3 | 1.89E-03 | 0.1 | 1.10E-01 | -0.1 | 7.84E-01 | 0.0 | 8.82E-01 | 0.0 | 9.61E-01 | DNA-directed RNA polymerases IV and V subunit 2 |
| AT1G17190 | -0.1 | 1.45E-01 | 0.7 | 2.59E-05 | -0.9 | 4.32E-05 | -0.9 | 4.26E-04 | -0.1 | 6.39E-01 | -0.8 | 5.24E-03 | Glutathione S-transferase U26 |
| AT4G21585 | -0.1 | 2.56E-01 | -0.1 | 7.03E-01 | -0.1 | 5.58E-01 | 0.1 | 6.80E-01 | 0.3 | 2.04E-01 | -0.2 | 4.14E-01 | Aspergillus nuclease S1 |
| AT3G17940 | -0.1 | 1.56E-01 | 0.1 | 2.04E-01 | -0.3 | 2.70E-02 | -0.2 | 2.25E-01 | 0.0 | 9.26E-01 | -0.2 | 3.20E-01 | Aldose 1-epimerase |
| AT1G74120 | -0.1 | 2.60E-01 | -0.2 | 1.56E-01 | 0.0 | 7.73E-01 | 0.3 | 3.44E-01 | -0.1 | 8.22E-01 | 0.4 | 3.35E-01 | Transcription termination factor MTERF15, mitochondrial |
| AT1G80070 | -0.1 | 4.03E-02 | -0.3 | 4.43E-04 | 0.2 | 1.55E-02 | 0.2 | 3.30E-01 | 0.1 | 8.49E-01 | 0.1 | 6.73E-01 | Pre-mRNA-processing-splicing factor 8A |
| AT3G11730 | -0.1 | 2.17E-01 | -0.1 | 4.76E-01 | -0.1 | 6.72E-01 | 0.0 | 8.70E-01 | 0.0 | 9.83E-01 | -0.1 | 9.02E-01 | Ras-related protein RABD1 |
| AT1G75860 | -0.1 | 1.60E-01 | -0.4 | 5.33E-03 | 0.2 | 9.18E-02 | 0.1 | 5.92E-01 | 0.0 | 9.09E-01 | 0.1 | 8.13E-01 | DNA ligase |
| AT1G65660 | -0.1 | 1.09E-01 | -0.2 | 2.70E-02 | 0.1 | 4.59E-01 | -0.2 | 9.41E-02 | -0.2 | 3.20E-01 | -0.1 | 7.77E-01 | Pre-mRNA-splicing factor SLU7-A |
| AT3G45400 | -0.1 | 2.78E-01 | 0.0 | 8.96E-01 | -0.1 | 4.43E-01 | -0.2 | 3.78E-01 | 0.3 | 1.42E-01 | -0.5 | 4.92E-02 | Exostosin GT47 domain-containing protein |
| AT3G19515 | -0.1 | 7.39E-02 | 0.0 | 7.01E-01 | -0.2 | 6.33E-02 | 0.0 | 9.97E-01 | 0.0 | 9.90E-01 | 0.0 | 9.96E-01 | Apoptosis inhibitor |
| AT1G01960 | -0.1 | 9.67E-02 | -0.1 | 1.32E-01 | 0.0 | 9.26E-01 | -0.1 | 7.42E-01 | -0.1 | 7.52E-01 | 0.0 | 9.76E-01 | Brefeldin A-inhibited guanine nucleotide-exchange protein 3 |
| AT1G29720 | -0.1 | 3.70E-01 | 0.0 | 7.82E-01 | -0.2 | 2.95E-01 | 0.3 | 1.77E-01 | 0.2 | 3.34E-01 | 0.0 | 8.68E-01 | Probable LRR receptor-like serine |
| AT5G65050 | -0.1 | 4.26E-02 | 0.0 | 4.85E-01 | -0.1 | 2.07E-01 | -0.1 | 5.63E-01 | 0.0 | 9.15E-01 | -0.1 | 7.84E-01 | AGAMOUS-like 31 |
| AT4G35550 | -0.1 | 1.81E-01 | -0.3 | 1.30E-02 | 0.2 | 1.86E-01 | -0.2 | 3.22E-01 | 0.1 | 8.47E-01 | -0.2 | 3.53E-01 | WUSCHEL-related homeobox 13 |
| AT1G80350 | -0.1 | 8.55E-02 | 0.1 | 2.68E-01 | -0.2 | 2.24E-02 | -0.4 | 3.57E-03 | -0.3 | 1.06E-02 | 0.0 | 9.17E-01 | Katanin p60 ATPase-containing subunit A1 |
| AT1G47750 | -0.1 | 1.89E-01 | -0.1 | 2.36E-01 | 0.0 | 9.46E-01 | -0.2 | 5.61E-01 | -0.1 | 8.64E-01 | -0.1 | 8.28E-01 | Peroxisomal membrane protein 11A |
| AT5G14770 | -0.1 | 3.58E-01 | -0.1 | 6.83E-01 | -0.1 | 7.11E-01 | 0.5 | 1.66E-01 | 0.4 | 2.84E-01 | 0.1 | 9.19E-01 | Pentatricopeptide repeat-containing protein At5g14770, mitochondrial |
| AT2G22450 | -0.1 | 1.52E-01 | 0.1 | 4.64E-01 | -0.2 | 7.41E-02 | -0.2 | 7.54E-02 | -0.3 | 7.03E-02 | 0.0 | 9.01E-01 | Monofunctional riboflavin biosynthesis protein RIBA 2, chloroplastic |
| AT1G34270 | -0.1 | 1.33E-01 | 0.1 | 2.42E-01 | -0.3 | 3.66E-02 | -0.1 | 7.05E-01 | -0.1 | 7.31E-01 | 0.0 | 9.88E-01 | Exostosin family protein |
| AT5G43270 | -0.1 | 2.24E-01 | 0.0 | 9.98E-01 | -0.1 | 2.96E-01 | -0.6 | 3.26E-03 | -0.2 | 2.87E-01 | -0.4 | 1.09E-01 | Squamosa promoter-binding-like protein 2 |
| AT2G36400 | -0.1 | 1.49E-01 | 0.1 | 4.14E-01 | -0.2 | 6.02E-02 | 0.1 | 5.12E-01 | 0.0 | 8.60E-01 | 0.2 | 3.71E-01 | Growth-regulating factor 3 |
| AT2G35630 | -0.1 | 3.67E-02 | -0.2 | 9.81E-03 | 0.1 | 4.53E-01 | -0.1 | 8.14E-01 | -0.1 | 8.71E-01 | 0.0 | 9.87E-01 | ARM repeat superfamily protein |
| AT5G11960 | -0.1 | 8.21E-02 | 0.0 | 7.87E-01 | -0.2 | 7.74E-02 | 0.0 | 8.88E-01 | 0.1 | 8.04E-01 | -0.1 | 7.36E-01 | Probable magnesium transporter NIPA9 |
| AT3G08550 | -0.1 | 4.09E-02 | 0.0 | 9.64E-01 | -0.1 | 7.19E-02 | 0.0 | 8.85E-01 | 0.0 | 9.42E-01 | 0.0 | 9.80E-01 | Glycosyltransferase-like KOBITO 1 |
| AT2G18030 | -0.1 | 2.44E-01 | 0.1 | 5.64E-01 | -0.2 | 1.40E-01 | -0.1 | 5.91E-01 | -0.1 | 4.04E-01 | 0.1 | 8.28E-01 | Peptide methionine sulfoxide reductase A5 -S-oxide reductase) reductase) |
| AT2G27030 | -0.1 | 2.19E-01 | 0.0 | 9.87E-01 | -0.1 | 2.65E-01 | 0.0 | 9.11E-01 | -0.1 | 8.57E-01 | 0.1 | 8.21E-01 | Calmodulin 5 |
| AT5G09330 | -0.1 | 8.58E-02 | -0.4 | 5.59E-04 | 0.3 | 1.23E-02 | 0.0 | 9.72E-01 | 0.0 | 9.10E-01 | 0.0 | 9.46E-01 | NAC domain-containing protein 82 |
| AT4G21160 | -0.1 | 1.93E-01 | -0.2 | 7.08E-02 | 0.1 | 5.86E-01 | 0.0 | 7.96E-01 | -0.1 | 5.13E-01 | 0.1 | 7.53E-01 | ADP-ribosylation factor GTPase-activating protein AGD12 |

| | | | | | | | | | | | | | |
|---|---|---|---|---|---|---|---|---|---|---|---|---|---|
| AT1G34420 | -0.1 | 4.78E-01 | -0.6 | 1.66E-02 | 0.5 | 9.24E-02 | 0.1 | 7.77E-01 | 0.0 | 9.76E-01 | 0.1 | 8.63E-01 | Protein kinase domain-containing protein |
| AT1G29810 | -0.1 | 1.31E-01 | -0.1 | 5.30E-01 | -0.1 | 4.64E-01 | 0.3 | 1.35E-01 | 0.0 | 9.04E-01 | 0.3 | 3.52E-01 | 4a-hydroxytetrahydrobiopterin dehydratase |
| AT5G63800 | -0.1 | 2.24E-01 | 0.1 | 2.74E-01 | -0.3 | 4.84E-02 | -0.5 | 2.79E-03 | -0.3 | 1.46E-01 | -0.3 | 2.60E-01 | beta-galactosidase |
| AT1G37130 | -0.1 | 1.62E-01 | -0.4 | 4.09E-03 | 0.2 | 5.62E-02 | -1.5 | 2.08E-07 | -1.2 | 1.16E-05 | -0.3 | 1.31E-01 | AT1G37130 protein |
| AT1G14910 | -0.1 | 9.29E-02 | -0.2 | 2.92E-02 | 0.1 | 5.37E-01 | -0.3 | 2.64E-01 | -0.2 | 3.93E-01 | 0.0 | 9.41E-01 | ENTH |
| AT2G01440 | -0.1 | 5.05E-02 | -0.4 | 1.93E-04 | 0.3 | 5.89E-03 | 0.0 | 9.93E-01 | 0.0 | 9.83E-01 | 0.0 | 9.94E-01 | ATP-dependent DNA helicase homolog RECG, chloroplastic |
| AT5G18280 | -0.1 | 1.06E-01 | 0.0 | 5.93E-01 | -0.2 | 6.44E-02 | -0.2 | 1.39E-01 | -0.2 | 5.73E-02 | 0.1 | 7.04E-01 | Apyrase 2 |
| AT1G76720 | -0.1 | 2.59E-01 | 0.2 | 8.32E-02 | -0.4 | 2.64E-02 | -0.1 | 7.39E-01 | -0.4 | 1.68E-01 | 0.3 | 4.33E-01 | Eukaryotic translation initiation factor 5B |
| AT5G38340 | -0.1 | 2.38E-01 | -0.4 | 7.57E-03 | 0.3 | 9.53E-02 | 0.1 | 7.54E-01 | 0.1 | 8.33E-01 | 0.0 | 9.72E-01 | ADP-ribosyl cyclase |
| AT3G63250 | -0.1 | 8.76E-02 | 0.1 | 2.74E-01 | -0.2 | 2.36E-02 | -0.3 | 3.43E-02 | -0.2 | 3.86E-01 | -0.2 | 4.06E-01 | Homocysteine methyltransferase 2 |
| AT5G51290 | -0.1 | 1.20E-01 | -0.3 | 6.74E-03 | 0.2 | 1.43E-01 | 0.0 | 9.67E-01 | -0.1 | 6.63E-01 | 0.1 | 6.98E-01 | Diacylglycerol kinase family protein |
| AT4G34640 | -0.1 | 5.18E-02 | -0.1 | 6.45E-02 | 0.0 | 9.73E-01 | -0.2 | 1.44E-01 | 0.0 | 8.71E-01 | -0.2 | 2.37E-01 | Squalene synthase |
| AT3G22220 | -0.1 | 8.13E-02 | -0.2 | 5.37E-02 | 0.0 | 8.09E-01 | -0.1 | 5.53E-01 | -0.1 | 6.69E-01 | 0.0 | 9.70E-01 | BED-type domain-containing protein |
| AT3G51370 | -0.1 | 9.77E-02 | -0.3 | 2.20E-03 | 0.2 | 4.67E-02 | -0.3 | 8.22E-03 | -0.3 | 2.19E-02 | 0.0 | 9.42E-01 | Probable protein phosphatase 2C 46 |
| AT5G57630 | -0.1 | 2.42E-01 | 0.1 | 2.59E-01 | -0.3 | 5.66E-02 | -0.3 | 1.45E-01 | -0.4 | 8.13E-02 | 0.1 | 7.79E-01 | CBL-interacting protein kinase 21 |
| AT4G00990 | -0.1 | 1.05E-01 | -0.2 | 1.28E-02 | 0.1 | 2.76E-01 | -0.1 | 6.87E-01 | 0.0 | 9.37E-01 | -0.1 | 8.19E-01 | Transcription factor jumonji domain-containing protein |
| AT1G31930 | -0.1 | 7.39E-02 | -0.1 | 7.49E-02 | 0.0 | 9.67E-01 | -0.1 | 5.67E-01 | -0.3 | 4.68E-02 | 0.2 | 2.27E-01 | Extra-large guanine nucleotide-binding protein 3 |
| AT1G33080 | -0.1 | 1.59E-01 | 0.0 | 7.91E-01 | -0.1 | 3.39E-01 | -0.2 | 3.17E-01 | 0.0 | 9.64E-01 | -0.2 | 4.46E-01 | Protein DETOXIFICATION |
| AT1G74590 | -0.1 | 3.00E-01 | -0.4 | 1.51E-02 | 0.3 | 1.42E-01 | 0.9 | 9.25E-04 | 0.5 | 7.58E-02 | 0.4 | 1.16E-01 | Glutathione S-transferase U10 |
| AT5G36890 | -0.1 | 1.32E-01 | -0.1 | 4.24E-01 | -0.1 | 5.50E-01 | -0.3 | 5.75E-02 | -0.2 | 1.46E-01 | 0.0 | 8.70E-01 | Beta-glucosidase 42 |
| AT1G09440 | -0.1 | 2.25E-01 | 0.2 | 6.47E-02 | -0.3 | 1.97E-02 | 0.1 | 8.10E-01 | -0.3 | 3.93E-01 | 0.3 | 3.46E-01 | Protein kinase superfamily protein |
| AT1G51790 | -0.1 | 2.98E-01 | -0.2 | 2.59E-01 | 0.0 | 9.18E-01 | 0.1 | 5.42E-01 | 0.0 | 9.77E-01 | 0.1 | 6.51E-01 | Leucine-rich repeat protein kinase family protein |
| AT5G64020 | -0.1 | 6.87E-02 | -0.2 | 6.90E-03 | 0.1 | 2.27E-01 | 0.0 | 9.97E-01 | 0.1 | 6.17E-01 | -0.1 | 7.04E-01 | Protein trichome birefringence-like 14 |
| AT1G01910 | -0.1 | 8.90E-02 | 0.0 | 5.54E-01 | -0.1 | 3.13E-01 | -0.1 | 3.92E-01 | -0.2 | 1.77E-01 | 0.1 | 7.10E-01 | P-loop containing nucleoside triphosphate hydrolases superfamily protein |
| AT1G60130 | -0.1 | 3.85E-01 | 0.5 | 3.11E-03 | -0.7 | 3.30E-03 | -0.5 | 1.79E-01 | 0.4 | 2.50E-01 | -0.9 | 3.65E-02 | Mannose-binding lectin superfamily protein |
| AT4G13710 | -0.1 | 2.22E-01 | 0.2 | 3.71E-02 | -0.4 | 1.27E-02 | -0.3 | 1.49E-01 | -0.1 | 7.87E-01 | -0.2 | 4.42E-01 | Pectate lyase |
| AT1G11940 | -0.1 | 2.90E-01 | -0.1 | 4.82E-01 | 0.0 | 8.10E-01 | -0.2 | 5.67E-01 | 0.0 | 9.77E-01 | -0.2 | 6.36E-01 | Core-2 |
| AT5G46470 | -0.1 | 1.99E-01 | -0.4 | 2.98E-03 | 0.3 | 3.72E-02 | 0.2 | 2.73E-01 | 0.2 | 3.96E-01 | 0.0 | 9.62E-01 | Disease resistance protein RPS6 |
| AT1G10200 | -0.1 | 4.10E-02 | 0.3 | 9.85E-04 | -0.4 | 2.07E-04 | -0.1 | 3.70E-01 | -0.1 | 4.18E-01 | 0.0 | 9.83E-01 | GATA type zinc finger transcription factor family protein |
| AT1G22020 | -0.1 | 6.53E-02 | -0.2 | 4.77E-03 | 0.1 | 1.68E-01 | -0.1 | 4.18E-01 | -0.2 | 4.16E-01 | 0.0 | 9.58E-01 | Serine hydroxymethyltransferase 6 |
| AT1G20480 | -0.1 | 4.10E-01 | 0.1 | 7.06E-01 | -0.2 | 3.26E-01 | | | | | | | 4-coumarate--CoA ligase-like 2 |
| AT4G26400 | -0.1 | 1.95E-01 | -0.2 | 8.65E-02 | 0.1 | 6.60E-01 | -0.3 | 2.67E-02 | -0.4 | 3.64E-03 | 0.1 | 3.89E-01 | RING-type E3 ubiquitin transferase |
| AT5G05310 | -0.1 | 1.30E-01 | -0.3 | 1.31E-02 | 0.1 | 2.47E-01 | -0.2 | 3.77E-01 | -0.3 | 2.41E-01 | 0.1 | 8.00E-01 | ADP,ATP carrier protein |
| AT1G58340 | -0.1 | 2.59E-01 | -0.6 | 1.01E-03 | 0.5 | 1.18E-02 | -0.1 | 8.59E-01 | -0.7 | 1.23E-01 | 0.6 | 1.04E-01 | Protein DETOXIFICATION |
| AT2G31160 | -0.1 | 3.76E-01 | -0.2 | 1.42E-01 | 0.1 | 5.83E-01 | 0.0 | 9.85E-01 | 0.2 | 4.22E-01 | -0.2 | 4.76E-01 | Protein LIGHT-DEPENDENT SHORT HYPOCOTYLS 3 |
| AT1G53440 | -0.1 | 3.43E-01 | -0.3 | 3.11E-02 | 0.2 | 2.02E-01 | -0.1 | 8.90E-01 | 0.0 | 9.81E-01 | -0.1 | 9.08E-01 | Probable LRR receptor-like serine |
| AT5G03300 | -0.1 | 1.34E-01 | 0.1 | 1.48E-01 | -0.3 | 1.64E-02 | 0.2 | 1.31E-01 | 0.2 | 1.42E-01 | 0.0 | 9.37E-01 | Adenosine kinase |
| AT5G44800 | -0.1 | 7.02E-02 | -0.4 | 4.45E-04 | 0.2 | 1.15E-02 | 0.1 | 5.04E-01 | 0.0 | 8.51E-01 | 0.1 | 7.98E-01 | Protein CHROMATIN REMODELING 4 |
| AT5G58950 | -0.1 | 9.48E-02 | -0.2 | 2.38E-02 | 0.1 | 4.36E-01 | -0.3 | 3.28E-02 | -0.1 | 5.45E-01 | -0.2 | 3.34E-01 | hypothetical protein |
| AT2G40940 | -0.1 | 1.09E-01 | -0.2 | 1.91E-02 | 0.1 | 3.44E-01 | 0.1 | 7.50E-01 | -0.1 | 4.57E-01 | 0.2 | 3.34E-01 | Ethylene response sensor 1 |
| AT5G24318 | -0.1 | 3.08E-01 | -0.1 | 2.77E-01 | 0.0 | 9.29E-01 | -0.5 | 4.12E-02 | -0.4 | 1.27E-01 | -0.1 | 8.11E-01 | glucan endo-1,3-beta-D-glucosidase |
| AT4G25870 | -0.1 | 1.38E-01 | 0.1 | 5.07E-01 | -0.2 | 7.13E-02 | 0.0 | 8.46E-01 | 0.0 | 9.28E-01 | 0.0 | 9.73E-01 | At4g25870 |
| AT2G39100 | -0.1 | 2.28E-01 | -0.2 | 5.41E-02 | 0.1 | 4.61E-01 | -0.4 | 1.73E-01 | -0.3 | 2.64E-01 | 0.0 | 9.51E-01 | Putative RING zinc finger protein |
| AT3G58030 | -0.1 | 5.37E-02 | -0.2 | 4.29E-03 | 0.1 | 1.70E-01 | -0.3 | 2.93E-02 | -0.2 | 1.59E-01 | -0.1 | 6.83E-01 | E3 ubiquitin-protein ligase RMA |
| AT2G04570 | -0.1 | 3.33E-01 | 0.0 | 9.40E-01 | -0.1 | 3.72E-01 | 0.0 | 9.38E-01 | 0.5 | 2.46E-01 | -0.5 | 2.75E-01 | GDSL esterase |
| AT1G80550 | -0.1 | 1.18E-01 | 0.0 | 7.41E-01 | -0.1 | 2.99E-01 | -0.2 | 1.59E-01 | -0.2 | 2.37E-01 | 0.0 | 9.77E-01 | Pentatricopeptide repeat-containing protein At1g80550, mitochondrial |
| AT1G50740 | -0.1 | 1.75E-01 | -0.3 | 9.27E-03 | 0.2 | 1.40E-01 | 0.0 | 9.91E-01 | -0.1 | 6.85E-01 | 0.1 | 7.39E-01 | Protein FATTY ACID EXPORT 5 |
| AT3G11830 | -0.1 | 1.79E-01 | 0.1 | 3.30E-01 | -0.2 | 4.70E-02 | 0.0 | 9.31E-01 | 0.0 | 9.18E-01 | 0.0 | 8.84E-01 | T-complex protein 1 subunit eta |
| AT3G55980 | -0.1 | 6.34E-01 | -0.9 | 6.65E-03 | 0.8 | 2.20E-02 | 0.2 | 3.90E-01 | -0.3 | 7.70E-02 | 0.5 | 3.03E-02 | Salt-inducible zinc finger 1 |
| AT3G49870 | -0.1 | 1.45E-01 | -0.1 | 4.20E-01 | -0.1 | 5.74E-01 | 0.1 | 7.77E-01 | -0.2 | 3.44E-01 | 0.2 | 3.25E-01 | ADP-ribosylation factor-like A1C |
| AT4G39080 | -0.1 | 1.16E-01 | 0.1 | 5.50E-01 | -0.2 | 5.88E-02 | -0.1 | 6.05E-01 | -0.1 | 6.72E-01 | 0.0 | 1.00E+00 | V-type proton ATPase subunit a3 -ATPase subunit a isoform 3) |
| AT1G48050 | -0.1 | 1.31E-01 | -0.2 | 9.77E-02 | 0.0 | 8.57E-01 | 0.2 | 2.98E-01 | 0.1 | 8.35E-01 | 0.1 | 6.18E-01 | ATP-dependent DNA helicase 2 subunit KU80 |
| AT1G09300 | -0.1 | 1.85E-01 | -0.2 | 9.29E-02 | 0.0 | 7.15E-01 | 0.4 | 8.34E-02 | 0.2 | 5.63E-01 | 0.2 | 4.27E-01 | Intermediate cleaving peptidase 55, mitochondrial |
| AT1G04050 | -0.1 | 4.00E-01 | 0.1 | 7.49E-01 | -0.2 | 3.39E-01 | 0.1 | 7.57E-01 | 0.3 | 4.11E-01 | -0.2 | 6.36E-01 | Histone-lysine N-methyltransferase SUVR2-like protein |
| AT4G16144 | -0.1 | 1.90E-01 | -0.2 | 3.62E-02 | 0.1 | 3.89E-01 | 0.0 | 9.46E-01 | 0.1 | 6.88E-01 | -0.1 | 7.92E-01 | Associated molecule with the SH3 domain of STAM 3 |
| AT2G02090 | -0.1 | 1.25E-01 | -0.2 | 2.29E-02 | 0.1 | 3.76E-01 | -0.1 | 6.53E-01 | 0.0 | 9.03E-01 | -0.1 | 8.72E-01 | Protein CHROMATIN REMODELING 19 |
| AT2G20580 | -0.1 | 5.50E-02 | -0.2 | 2.43E-02 | 0.0 | 6.26E-01 | 0.1 | 7.25E-01 | 0.1 | 5.51E-01 | -0.1 | 8.52E-01 | 26S proteasome non-ATPase regulatory subunit 2 homolog A |
| AT2G01750 | -0.1 | 3.17E-01 | 0.0 | 9.17E-01 | -0.1 | 4.66E-01 | 0.3 | 1.49E-01 | 0.3 | 2.23E-01 | 0.0 | 9.69E-01 | Microtubule-associated proteins 70-3 |
| AT3G53930 | -0.1 | 1.20E-01 | -0.3 | 9.97E-03 | 0.1 | 2.09E-01 | -0.1 | 7.68E-01 | 0.0 | 9.35E-01 | 0.0 | 8.94E-01 | hypothetical protein |
| AT5G04560 | -0.1 | 8.08E-02 | -0.2 | 8.88E-03 | 0.1 | 2.43E-01 | 0.1 | 8.04E-01 | 0.1 | 5.11E-01 | -0.1 | 7.55E-01 | Transcriptional activator DEMETER |
| AT3G13930 | -0.1 | 1.47E-01 | 0.2 | 4.56E-02 | -0.3 | 6.42E-03 | 0.3 | 9.94E-02 | 0.2 | 1.93E-01 | 0.0 | 9.46E-01 | Acetyltransferase component of pyruvate dehydrogenase complex |
| AT2G42670 | -0.1 | 2.61E-01 | 0.0 | 7.61E-01 | -0.1 | 5.11E-01 | -0.5 | 1.59E-02 | -0.2 | 4.29E-01 | -0.3 | 2.39E-01 | cysteine dioxygenase |
| AT2G33610 | -0.1 | 5.71E-02 | 0.0 | 5.00E-01 | -0.1 | 2.63E-01 | -0.1 | 3.07E-01 | -0.2 | 3.28E-01 | 0.0 | 9.62E-01 | SWI |
| AT3G46370 | -0.1 | 5.08E-01 | 0.4 | 3.92E-02 | -0.6 | 2.54E-02 | -2.1 | 4.65E-04 | -0.2 | 6.51E-01 | -1.9 | 9.43E-03 | Leucine-rich repeat protein kinase family protein |
| AT1G63460 | -0.1 | 4.03E-01 | -0.1 | 4.24E-01 | 0.0 | 9.99E-01 | 0.1 | 6.53E-01 | 0.1 | 8.35E-01 | 0.1 | 9.18E-01 | Glutathione peroxidase |
| AT1G08960 | -0.1 | 1.84E-01 | -0.1 | 2.37E-01 | 0.0 | 9.36E-01 | 0.0 | 8.92E-01 | 0.1 | 7.84E-01 | -0.1 | 7.02E-01 | Cation |
| AT5G02190 | -0.1 | 4.46E-01 | 0.3 | 8.38E-02 | -0.4 | 4.52E-02 | 0.5 | 2.81E-01 | 0.4 | 4.98E-01 | 0.1 | 8.54E-01 | Aspartic proteinase PCS1 |
| AT4G31420 | -0.1 | 5.48E-02 | -0.1 | 1.59E-01 | 0.0 | 6.38E-01 | -0.4 | 5.12E-03 | -0.3 | 4.58E-02 | -0.1 | 5.88E-01 | Zinc finger protein 622 |
| AT1G26270 | -0.1 | 6.21E-02 | 0.0 | 7.76E-01 | -0.1 | 1.47E-01 | 0.0 | 8.76E-01 | -0.1 | 3.31E-01 | 0.1 | 5.31E-01 | Phosphatidylinositol 4-kinase gamma 5 |
| AT5G25610 | -0.1 | 3.47E-01 | -0.1 | 6.28E-01 | -0.1 | 7.21E-01 | -0.3 | 1.22E-01 | -0.1 | 8.44E-01 | -0.2 | 3.77E-01 | BURP domain protein RD22 |
| AT4G33910 | -0.1 | 6.87E-02 | -0.2 | 2.31E-02 | 0.0 | 5.59E-01 | -0.4 | 6.72E-03 | -0.3 | 4.30E-02 | -0.1 | 6.68E-01 | Probable prolyl 4-hydroxylase 9 |
| AT1G32860 | -0.1 | 2.01E-01 | 0.1 | 4.65E-01 | -0.2 | 9.93E-02 | 0.1 | 7.43E-01 | 0.1 | 8.26E-01 | 0.0 | 9.74E-01 | Glucan endo-1,3-beta-glucosidase 11 -beta-glucan endohydrolase 11) -beta-glucanase 11) |
| AT5G12010 | -0.1 | 7.02E-02 | -0.5 | 9.73E-05 | 0.3 | 2.04E-03 | -0.3 | 4.83E-02 | -0.3 | 1.66E-01 | -0.1 | 7.98E-01 | DDE Tnp4 domain-containing protein |
| AT1G79550 | -0.1 | 2.08E-01 | 0.1 | 3.80E-01 | -0.2 | 6.42E-02 | 0.6 | 1.27E-04 | 0.4 | 2.89E-03 | 0.1 | 4.66E-01 | Phosphoglycerate kinase 3, cytosolic |
| AT3G53610 | -0.1 | 5.21E-02 | -0.1 | 2.85E-01 | -0.1 | 4.01E-01 | -0.1 | 2.36E-01 | -0.1 | 6.60E-01 | -0.1 | 7.22E-01 | Ras-related protein RABE1a |
| AT2G27470 | -0.1 | 3.53E-01 | 0.1 | 6.46E-01 | -0.2 | 2.44E-01 | 0.4 | 3.79E-02 | 0.2 | 4.58E-01 | 0.2 | 3.80E-01 | DNA polymerase II subunit B4 |
| AT1G49710 | -0.1 | 1.31E-01 | -0.2 | 4.94E-02 | 0.1 | 6.03E-01 | 0.2 | 3.27E-01 | 0.1 | 5.69E-01 | 0.0 | 8.74E-01 | Putative fucosyltransferase-like protein |
| AT3G12020 | -0.1 | 1.28E-01 | -0.3 | 6.70E-03 | 0.1 | 1.30E-01 | -0.1 | 4.20E-01 | -0.1 | 7.36E-01 | -0.1 | 8.24E-01 | P-loop containing nucleoside triphosphate hydrolases superfamily protein |
| AT5G10610 | -0.1 | 1.18E-01 | -0.2 | 2.97E-02 | 0.1 | 4.76E-01 | 0.0 | 9.83E-01 | -0.1 | 5.29E-01 | 0.1 | 5.81E-01 | Cytochrome P450, family 81, subfamily K, polypeptide 1 |
| AT5G09290 | -0.1 | 2.97E-01 | -0.4 | 5.94E-03 | 0.3 | 6.14E-02 | -0.1 | 8.92E-01 | 0.3 | 3.46E-01 | -0.4 | 3.03E-01 | Probable SAL4 phosphatase ,5'-bisphosphate nucleotidase 4) ,5'-bisphosphonucleoside 3'-phosphohydrolase 4) |
| AT5G09250 | -0.1 | 3.40E-01 | -0.1 | 5.81E-01 | -0.1 | 7.68E-01 | 0.3 | 2.97E-01 | 0.0 | 9.41E-01 | 0.2 | 5.11E-01 | SsDNA-binding transcriptional regulator |
| AT1G15340 | -0.1 | 1.35E-01 | 0.0 | 8.68E-01 | -0.1 | 2.34E-01 | -0.1 | 4.16E-01 | -0.2 | 3.55E-01 | 0.0 | 8.93E-01 | Methyl-CpG-binding domain-containing protein 10 |
| AT5G26340 | -0.1 | 5.10E-01 | -0.2 | 2.41E-01 | 0.1 | 6.24E-01 | -0.3 | 4.68E-01 | -1.3 | 7.42E-03 | 1.0 | 8.19E-02 | Sugar transport protein 13 |

| | | | | | | | | | | | | | |
|---|---|---|---|---|---|---|---|---|---|---|---|---|---|
| AT4G36050 | -0.1 | 3.75E-01 | -0.3 | 7.24E-02 | 0.2 | 3.77E-01 | -1.1 | 1.63E-02 | -0.6 | 2.13E-01 | -0.6 | 2.36E-01 | DNA- endonuclease 2 |
| AT5G11260 | -0.1 | 4.12E-01 | 0.0 | 7.68E-01 | -0.1 | 6.88E-01 | 0.1 | 8.75E-01 | 0.0 | 9.89E-01 | 0.0 | 9.20E-01 | Basic-leucine zipper transcription factor family protein |
| AT5G44080 | -0.1 | 7.07E-02 | -0.1 | 1.06E-01 | 0.0 | 8.94E-01 | -0.5 | 3.42E-03 | -0.4 | 1.70E-02 | -0.1 | 7.84E-01 | Basic leucine zipper transcription factor transcription factor family protein) |
| AT5G36940 | -0.1 | 1.63E-01 | 0.0 | 9.67E-01 | -0.1 | 2.04E-01 | -0.3 | 6.39E-02 | -0.3 | 1.24E-01 | 0.0 | 9.29E-01 | Cationic amino acid transporter 3, mitochondrial |
| AT1G61360 | -0.1 | 2.49E-01 | -0.4 | 7.44E-03 | 0.2 | 8.21E-02 | 0.2 | 3.26E-01 | 0.0 | 8.99E-01 | 0.2 | 3.83E-01 | G-type lectin S-receptor-like serine |
| AT3G04320 | -0.1 | 4.59E-01 | 0.5 | 7.51E-03 | -0.6 | 7.32E-03 | 0.4 | 2.04E-01 | 0.7 | 4.60E-02 | -0.3 | 4.39E-01 | Uncharacterized protein |
| AT3G04810 | -0.1 | 2.70E-01 | 0.0 | 7.89E-01 | -0.1 | 5.00E-01 | -0.7 | 1.31E-01 | -0.2 | 6.95E-01 | -0.5 | 4.21E-01 | Serine |
| AT3G62150 | -0.1 | 4.43E-01 | 0.0 | 9.53E-01 | -0.1 | 5.50E-01 | 0.4 | 9.94E-02 | -0.6 | 7.24E-02 | 1.0 | 7.26E-03 | ABC transporter B family member 21 |
| AT3G17850 | -0.1 | 9.48E-02 | -0.2 | 5.70E-03 | 0.1 | 1.36E-01 | -0.2 | 3.29E-01 | -0.2 | 3.97E-01 | 0.0 | 9.98E-01 | Probable serine |
| AT1G74690 | -0.1 | 7.54E-02 | 0.1 | 5.77E-02 | -0.3 | 4.74E-03 | 0.1 | 3.77E-01 | 0.1 | 2.67E-01 | 0.0 | 8.39E-01 | Protein IQ-DOMAIN 31 |
| AT1G73200 | -0.1 | 5.58E-02 | -0.2 | 4.46E-03 | 0.1 | 1.78E-01 | -0.3 | 1.48E-01 | -0.2 | 3.20E-01 | -0.1 | 8.72E-01 | SMP-LTD domain-containing protein |
| AT3G61050 | -0.1 | 1.29E-01 | 0.1 | 3.33E-01 | -0.2 | 3.51E-02 | -0.2 | 7.16E-02 | -0.1 | 4.65E-01 | -0.1 | 5.63E-01 | hypothetical protein |
| AT4G39890 | -0.1 | 4.93E-01 | -0.2 | 2.32E-01 | 0.1 | 6.56E-01 | -0.2 | 5.02E-01 | -0.1 | 7.12E-01 | -0.1 | 9.02E-01 | Ras-related protein RABH1c |
| AT2G43070 | -0.1 | 6.87E-02 | 0.0 | 5.54E-01 | -0.1 | 2.56E-01 | -0.1 | 4.90E-01 | 0.0 | 8.59E-01 | -0.1 | 5.14E-01 | SIGNAL PEPTIDE PEPTIDASE-LIKE 3 |
| AT2G26330 | -0.1 | 3.21E-01 | -0.1 | 4.39E-01 | 0.0 | 8.80E-01 | -0.2 | 2.16E-01 | -0.1 | 6.35E-01 | -0.1 | 6.96E-01 | LRR receptor-like serine |
| AT1G49820 | -0.1 | 9.80E-02 | -0.1 | 6.88E-02 | 0.0 | 8.34E-01 | -0.2 | 1.35E-01 | -0.2 | 1.46E-01 | 0.0 | 9.48E-01 | Methylthioribose kinase |
| AT5G19980 | -0.1 | 2.06E-01 | 0.0 | 7.04E-01 | -0.1 | 4.46E-01 | -0.1 | 3.27E-01 | -0.1 | 4.66E-01 | 0.0 | 9.57E-01 | GDP-fucose transporter 1 |
| AT2G22310 | -0.1 | 2.08E-01 | -0.2 | 1.07E-01 | 0.0 | 7.26E-01 | -0.6 | 1.75E-02 | -0.3 | 2.09E-01 | -0.3 | 4.40E-01 | Ubiquitin-specific protease 4 |
| AT1G67580 | -0.1 | 1.40E-01 | -0.3 | 8.68E-03 | 0.1 | 1.50E-01 | -0.2 | 2.23E-01 | -0.2 | 1.91E-01 | 0.0 | 8.94E-01 | F12A21.29 |
| AT1G13930 | -0.1 | 4.53E-01 | 0.1 | 4.29E-01 | -0.3 | 1.66E-01 | -0.4 | 4.99E-02 | 0.1 | 5.85E-01 | -0.5 | 5.10E-02 | Nodulin-related protein 2 |
| AT1G12570 | -0.1 | 6.98E-01 | 0.1 | 7.14E-01 | -0.2 | 5.40E-01 | -0.3 | 3.93E-01 | -0.1 | 8.95E-01 | -0.3 | 5.95E-01 | Glucose-methanol-choline oxidoreductase N-terminal domain-containing protein |
| AT4G18160 | -0.1 | 2.60E-01 | -0.1 | 4.20E-01 | 0.0 | 8.24E-01 | -0.4 | 2.53E-02 | -0.3 | 1.73E-01 | -0.2 | 6.03E-01 | Two-pore potassium channel 3 |
| AT5G11460 | -0.1 | 3.47E-01 | 0.0 | 9.81E-01 | -0.1 | 4.59E-01 | 0.3 | 2.67E-01 | 0.1 | 7.35E-01 | 0.2 | 6.44E-01 | FCS-Like Zinc finger 10 |
| AT5G26760 | -0.1 | 1.18E-01 | -0.1 | 1.86E-01 | 0.0 | 8.61E-01 | -0.1 | 5.93E-01 | -0.2 | 1.96E-01 | 0.1 | 5.40E-01 | Putative RNA polymerase II subunit B1 CTD phosphatase RPAP2 homolog |
| AT5G18190 | -0.1 | 1.21E-01 | -0.3 | 3.39E-03 | 0.2 | 6.38E-02 | 0.0 | 9.78E-01 | 0.1 | 7.50E-01 | -0.1 | 7.58E-01 | Casein kinase-like protein |
| AT5G64960 | -0.1 | 9.65E-02 | -0.4 | 5.99E-04 | 0.2 | 1.29E-02 | 0.2 | 7.20E-01 | 0.2 | 8.30E-01 | 0.0 | 9.62E-01 | Cdc2-like protein kinase |
| AT5G63870 | -0.1 | 9.02E-02 | -0.2 | 6.25E-03 | 0.1 | 1.65E-01 | 0.0 | 7.64E-01 | -0.1 | 6.87E-01 | 0.0 | 9.28E-01 | Serine |
| AT1G78550 | -0.1 | 1.49E-01 | -0.2 | 6.76E-02 | 0.0 | 6.82E-01 | 0.1 | 5.85E-01 | 0.1 | 7.44E-01 | 0.0 | 9.37E-01 | Fe2OG dioxygenase domain-containing protein |
| AT5G15080 | -0.1 | 8.76E-02 | -0.2 | 5.02E-03 | 0.1 | 1.40E-01 | -0.1 | 5.70E-01 | -0.2 | 3.04E-01 | 0.1 | 7.13E-01 | Serine |
| AT2G21120 | -0.1 | 1.13E-01 | -0.3 | 2.43E-03 | 0.2 | 5.26E-02 | 0.0 | 7.85E-01 | -0.2 | 2.27E-01 | 0.2 | 1.59E-01 | Probable magnesium transporter NIPA6 |
| AT3G19710 | -0.1 | 4.71E-01 | 0.3 | 1.16E-01 | -0.4 | 4.55E-02 | -0.7 | 7.61E-03 | 0.6 | 2.86E-02 | -1.3 | 1.54E-03 | hypothetical protein |
| AT2G22840 | -0.1 | 1.57E-01 | 0.1 | 1.16E-01 | -0.3 | 1.91E-02 | 0.1 | 8.36E-01 | 0.1 | 8.52E-01 | 0.0 | 9.93E-01 | Growth-regulating factor 1 |
| AT3G10940 | -0.1 | 5.02E-01 | 0.2 | 1.72E-01 | -0.4 | 8.47E-02 | 0.1 | 6.77E-01 | 0.5 | 1.44E-02 | -0.4 | 9.00E-02 | Phosphoglucan phosphatase LSF2, chloroplastic |
| AT4G23160 | -0.1 | 7.46E-01 | -1.3 | 1.18E-02 | 1.2 | 4.14E-02 | | | | | | | Uncharacterized protein |
| AT1G21700 | -0.1 | 6.93E-02 | -0.2 | 7.67E-03 | 0.1 | 2.43E-01 | 0.1 | 7.67E-01 | 0.0 | 9.64E-01 | 0.1 | 8.75E-01 | SWI |
| AT1G05670 | -0.1 | 2.60E-01 | -0.1 | 2.22E-01 | 0.0 | 9.06E-01 | 0.6 | 2.17E-01 | 0.2 | 7.58E-01 | 0.4 | 5.52E-01 | Pentatricopeptide repeat-containing protein At1g05670, mitochondrial |
| AT3G44680 | -0.1 | 3.57E-01 | -0.1 | 4.11E-01 | 0.0 | 9.63E-01 | 0.3 | 1.09E-01 | 0.2 | 2.29E-01 | 0.0 | 9.05E-01 | Histone deacetylase 9 |
| AT4G02480 | -0.1 | 1.32E-01 | -0.2 | 4.93E-02 | 0.0 | 5.70E-01 | -0.2 | 2.74E-01 | -0.2 | 1.66E-01 | 0.1 | 7.95E-01 | AAA+ ATPase domain-containing protein |
| AT4G24150 | -0.1 | 5.25E-01 | 0.1 | 7.31E-01 | -0.2 | 4.23E-01 | -0.2 | 6.00E-01 | 0.0 | 9.64E-01 | -0.3 | 6.32E-01 | Growth-regulating factor |
| AT2G03270 | -0.1 | 2.66E-01 | -0.2 | 1.23E-01 | 0.1 | 6.75E-01 | -0.5 | 1.14E-02 | -0.6 | 1.01E-02 | 0.1 | 8.79E-01 | DNA-binding protein |
| AT5G21105 | -0.1 | 2.04E-01 | -0.2 | 8.34E-02 | 0.1 | 6.09E-01 | 0.0 | 7.87E-01 | 0.1 | 5.80E-01 | 0.0 | 8.33E-01 | L-ascorbate oxidase |
| AT1G77470 | -0.1 | 2.80E-01 | -0.1 | 6.45E-01 | -0.1 | 6.30E-01 | 0.1 | 7.54E-01 | 0.0 | 9.97E-01 | 0.1 | 8.14E-01 | RFC5 |
| AT5G52640 | -0.1 | 5.03E-01 | -0.7 | 3.04E-03 | 0.6 | 1.81E-02 | 0.6 | 5.13E-02 | 1.0 | 5.69E-03 | -0.4 | 1.21E-01 | Heat shock protein 90-1 |
| AT4G22330 | -0.1 | 1.37E-01 | -0.2 | 3.06E-02 | 0.1 | 4.25E-01 | -0.3 | 3.61E-02 | -0.3 | 3.61E-02 | 0.0 | 9.02E-01 | Alkaline ceramidase |
| AT5G01560 | -0.1 | 1.06E-01 | -0.2 | 5.09E-02 | 0.0 | 6.91E-01 | -0.2 | 1.91E-01 | -0.2 | 2.88E-01 | 0.0 | 9.66E-01 | Lectin-domain containing receptor kinase VI.4 |
| AT2G47330 | -0.1 | 8.11E-02 | -0.2 | 4.12E-03 | 0.1 | 1.20E-01 | -0.1 | 7.84E-01 | 0.1 | 6.53E-01 | -0.2 | 5.49E-01 | DEAD-box ATP-dependent RNA helicase 24 |
| AT4G18020 | -0.1 | 2.02E-01 | -0.2 | 6.56E-02 | 0.1 | 5.42E-01 | 0.0 | 9.96E-01 | 0.1 | 6.71E-01 | -0.1 | 6.96E-01 | Two-component response regulator-like APRR2 |
| AT2G21410 | -0.1 | 8.72E-02 | -0.1 | 4.31E-01 | -0.1 | 3.98E-01 | -0.3 | 1.74E-02 | 0.0 | 7.81E-01 | -0.3 | 1.56E-01 | V-type proton ATPase subunit a2 -ATPase subunit a isoform 2) |
| AT2G46800 | -0.1 | 7.70E-02 | 0.1 | 8.99E-02 | -0.2 | 7.21E-03 | -0.6 | 1.13E-03 | -0.4 | 2.07E-02 | -0.2 | 4.63E-01 | Metal tolerance protein 1 |
| AT3G26750 | -0.1 | 2.88E-01 | -0.1 | 4.36E-01 | 0.0 | 8.52E-01 | -0.3 | 2.08E-01 | -0.2 | 6.09E-01 | -0.2 | 6.71E-01 | HECT-like ubiquitin-conjugating enzyme -binding protein |
| AT3G49600 | -0.1 | 1.19E-01 | 0.0 | 9.63E-01 | -0.1 | 1.62E-01 | 0.4 | 5.27E-03 | 0.3 | 4.23E-02 | 0.1 | 6.53E-01 | ubiquitinyl hydrolase 1 |
| AT1G66730 | -0.1 | 3.88E-01 | -0.4 | 1.12E-02 | 0.3 | 8.10E-02 | 0.2 | 3.29E-01 | 0.1 | 7.99E-01 | 0.1 | 6.77E-01 | DNA ligase ATP-dependent C-terminal domain-containing protein |
| AT5G26751 | -0.1 | 1.04E-01 | -0.1 | 1.73E-01 | 0.0 | 8.42E-01 | -0.1 | 7.38E-01 | -0.1 | 4.13E-01 | 0.1 | 7.39E-01 | Shaggy-related protein kinase alpha |
| AT5G47790 | -0.1 | 1.74E-01 | -0.2 | 3.40E-02 | 0.1 | 3.98E-01 | 0.0 | 9.13E-01 | -0.1 | 8.49E-01 | 0.0 | 9.48E-01 | Protein phosphatase 1 regulatory inhibitor subunit PPP1R8 homolog |
| AT2G22910 | -0.1 | 2.14E-01 | -0.1 | 1.99E-01 | 0.0 | 9.45E-01 | 0.0 | 9.03E-01 | 0.1 | 8.17E-01 | 0.0 | 9.20E-01 | Probable amino-acid acetyltransferase NAGS1, chloroplastic |
| AT1G53750 | -0.1 | 1.26E-01 | -0.2 | 5.22E-02 | 0.0 | 6.10E-01 | 0.2 | 1.45E-02 | 0.2 | 1.19E-01 | 0.1 | 6.58E-01 | 26S proteasome regulatory subunit 7 homolog A |
| AT3G50750 | -0.1 | 4.09E-01 | 0.1 | 3.13E-01 | -0.3 | 1.20E-01 | -0.3 | 6.73E-02 | -0.5 | 1.70E-02 | 0.2 | 5.90E-01 | BES1 |
| AT1G25350 | -0.1 | 1.18E-01 | -0.1 | 5.89E-02 | 0.0 | 6.92E-01 | 0.2 | 3.06E-01 | 0.2 | 2.02E-01 | -0.1 | 8.07E-01 | Glutamine-tRNA ligase, putative |
| AT5G25370 | -0.1 | 2.92E-01 | -0.4 | 6.39E-03 | 0.3 | 6.81E-02 | 0.7 | 1.06E-01 | 0.5 | 3.50E-01 | 0.2 | 7.23E-01 | Phospholipase D |
| AT1G01120 | -0.1 | 3.13E-01 | -0.3 | 2.80E-02 | 0.2 | 1.83E-01 | -0.3 | 1.05E-01 | -0.3 | 1.02E-01 | 0.0 | 8.98E-01 | 3-ketoacyl-CoA synthase 1 |
| AT5G47710 | -0.1 | 1.73E-01 | -0.1 | 9.82E-02 | 0.0 | 7.56E-01 | 0.0 | 9.09E-01 | -0.1 | 5.60E-01 | 0.2 | 5.00E-01 | Protein C2-DOMAIN ABA-RELATED 11 |
| AT1G60440 | -0.1 | 1.08E-01 | -0.3 | 1.39E-03 | 0.2 | 3.32E-02 | 0.2 | 3.18E-01 | 0.1 | 7.01E-01 | 0.1 | 7.58E-01 | Pantothenate kinase 1 |
| AT2G25680 | -0.1 | 2.79E-01 | 0.3 | 1.25E-02 | -0.4 | 5.37E-03 | -1.4 | 6.32E-06 | -0.3 | 1.35E-01 | -1.1 | 7.22E-04 | Molybdate transporter 1 |
| AT4G23150 | -0.1 | 8.49E-01 | -1.2 | 1.10E-01 | 1.1 | 2.19E-01 | -1.4 | 1.17E-01 | -1.6 | 1.03E-01 | 0.3 | 8.87E-01 | Cysteine-rich receptor-like protein kinase 7 |
| AT4G04710 | -0.1 | 5.62E-01 | -0.7 | 1.02E-02 | 0.6 | 5.42E-02 | 1.5 | 8.49E-04 | 1.3 | 6.07E-03 | 0.2 | 6.01E-01 | Calcium-dependent kinase-like protein |
| AT2G37840 | -0.1 | 2.75E-01 | -0.2 | 1.46E-01 | 0.0 | 7.16E-01 | -0.3 | 2.92E-01 | -0.1 | 8.54E-01 | -0.2 | 4.89E-01 | Protein kinase superfamily protein |
| AT1G62950 | -0.1 | 5.09E-01 | 0.0 | 8.38E-01 | -0.2 | 4.78E-01 | -0.3 | 4.26E-01 | 0.1 | 9.06E-01 | -0.4 | 4.22E-01 | Protein kinase domain-containing protein |
| AT3G15355 | -0.1 | 1.12E-01 | -0.3 | 1.47E-03 | 0.2 | 3.37E-02 | 0.0 | 7.97E-01 | -0.2 | 1.23E-01 | 0.2 | 3.07E-01 | Probable ubiquitin-conjugating enzyme E2 25 |
| AT1G24140 | -0.1 | 7.11E-01 | -1.0 | 1.53E-02 | 0.9 | 5.26E-02 | 0.6 | 1.01E-01 | 0.1 | 8.08E-01 | 0.5 | 2.76E-01 | Metalloendoproteinase 3-MMP |
| AT1G06400 | -0.1 | 1.14E-01 | -0.1 | 1.61E-01 | 0.0 | 9.01E-01 | -0.2 | 2.69E-01 | -0.4 | 4.16E-02 | 0.2 | 4.16E-01 | Ras-related protein RABA1a |
| AT5G52882 | -0.1 | 2.82E-01 | 0.3 | 3.34E-02 | -0.4 | 9.33E-03 | -0.5 | 1.50E-03 | -0.4 | 2.16E-02 | -0.1 | 5.11E-01 | P-loop containing nucleoside triphosphate hydrolases superfamily protein |
| AT3G02750 | -0.1 | 1.93E-01 | -0.1 | 3.36E-01 | 0.0 | 7.90E-01 | -0.4 | 4.63E-03 | -0.4 | 1.41E-02 | 0.0 | 9.26E-01 | Probable protein phosphatase 2C 33 |
| AT3G11340 | -0.1 | 4.58E-01 | 0.1 | 6.22E-01 | -0.2 | 2.79E-01 | 0.2 | 1.96E-01 | 0.5 | 1.57E-02 | -0.2 | 2.45E-01 | UDP-glycosyltransferase 76B1 |
| AT5G57940 | -0.1 | 1.64E-01 | -0.2 | 4.27E-02 | 0.1 | 4.67E-01 | -0.2 | 3.88E-01 | -0.1 | 6.40E-01 | -0.1 | 8.82E-01 | Probable cyclic nucleotide-gated ion channel 5 |
| AT3G61240 | -0.1 | 9.95E-02 | -0.1 | 1.78E-01 | 0.0 | 8.14E-01 | 0.1 | 2.83E-01 | 0.2 | 2.51E-01 | 0.0 | 8.89E-01 | DEAD-box ATP-dependent RNA helicase 12 |
| AT5G17000 | -0.1 | 3.45E-01 | -0.4 | 1.03E-02 | 0.3 | 7.62E-02 | 0.2 | 3.40E-01 | 0.3 | 1.68E-01 | -0.1 | 7.43E-01 | Zinc-binding dehydrogenase family protein |
| AT2G04850 | -0.1 | 3.82E-01 | -0.2 | 2.04E-01 | 0.1 | 7.15E-01 | 0.7 | 2.63E-02 | 0.5 | 1.29E-01 | 0.2 | 6.90E-01 | Cytochrome b561 and DOMON domain-containing protein At2g04850 |
| AT4G29900 | -0.1 | 2.93E-01 | -0.4 | 1.88E-03 | 0.3 | 1.47E-02 | -0.1 | 3.96E-01 | -0.3 | 5.30E-02 | 0.2 | 3.47E-01 | Calcium-transporting ATPase |
| AT1G56140 | -0.1 | 1.46E-01 | -0.3 | 2.67E-03 | 0.2 | 4.59E-02 | 0.3 | 1.03E-01 | -0.1 | 8.21E-01 | 0.3 | 1.56E-01 | Probable LRR receptor-like serine |
| AT1G27600 | -0.1 | 9.07E-02 | 0.0 | 9.44E-01 | -0.1 | 1.17E-01 | -0.3 | 1.25E-02 | -0.4 | 5.11E-03 | 0.1 | 6.58E-01 | Probable beta-1,4-xylosyltransferase IRX9H |
| AT5G43010 | -0.1 | 2.05E-01 | -0.1 | 1.79E-01 | 0.0 | 9.17E-01 | 0.1 | 3.02E-01 | 0.1 | 5.23E-01 | 0.0 | 8.84E-01 | 26S proteasome regulatory subunit 10B homolog A |
| AT5G26830 | -0.1 | 1.24E-01 | -0.1 | 7.73E-02 | 0.0 | 7.76E-01 | 0.0 | 9.25E-01 | 0.0 | 9.45E-01 | 0.0 | 9.97E-01 | Threonine--tRNA ligase, mitochondrial 1 |
| AT2G24580 | -0.1 | 1.77E-01 | 0.3 | 7.64E-03 | -0.4 | 2.23E-03 | 0.1 | 6.33E-01 | 0.0 | 9.97E-01 | 0.1 | 6.80E-01 | Probable sarcosine oxidase |

| | | | | | | | | | | | | | |
|---|---|---|---|---|---|---|---|---|---|---|---|---|---|
| AT4G31750 | -0.1 | 2.13E-01 | -0.1 | 4.17E-01 | 0.0 | 7.37E-01 | -0.1 | 7.31E-01 | 0.0 | 9.02E-01 | 0.0 | 9.11E-01 | protein-serine |
| AT3G43810 | -0.1 | 1.88E-01 | 0.3 | 3.75E-03 | -0.4 | 1.19E-03 | 0.2 | 3.53E-01 | 0.2 | 4.46E-01 | 0.0 | 9.95E-01 | Z-box binding factor 3 |
| AT2G29730 | -0.1 | 1.78E-01 | 0.3 | 2.43E-03 | -0.4 | 1.19E-03 | -0.2 | 1.01E-01 | -0.1 | 4.35E-01 | -0.1 | 6.40E-01 | UDP-glycosyltransferase 71D1 |
| AT3G15010 | -0.1 | 1.35E-01 | -0.2 | 3.53E-02 | 0.1 | 4.75E-01 | 0.1 | 3.88E-01 | 0.1 | 7.70E-01 | 0.1 | 7.87E-01 | UBP1-associated protein 2C |
| AT1G48850 | -0.1 | 1.62E-01 | 0.1 | 4.11E-01 | -0.2 | 5.51E-02 | 0.4 | 1.46E-03 | 0.3 | 1.34E-02 | 0.1 | 7.08E-01 | Chorismate synthase |
| AT1G05630 | -0.1 | 2.21E-01 | -0.2 | 5.90E-02 | 0.1 | 4.78E-01 | 0.0 | 9.72E-01 | 0.1 | 8.71E-01 | -0.1 | 8.66E-01 | Endonuclease |
| AT2G17430 | -0.1 | 5.40E-01 | -0.6 | 1.38E-02 | 0.5 | 6.97E-02 | | | | | | | MLO-like protein 7 |
| AT5G17600 | -0.1 | 2.54E-01 | -0.1 | 2.45E-01 | 0.0 | 9.60E-01 | -0.4 | 9.36E-02 | -0.1 | 8.07E-01 | -0.3 | 3.31E-01 | RING-H2 finger protein ATL52 |
| AT4G29260 | -0.1 | 1.15E-01 | 0.0 | 8.37E-01 | -0.1 | 1.13E-01 | -0.1 | 6.42E-01 | 0.2 | 2.97E-01 | -0.2 | 2.13E-01 | Acid phosphatase-like protein |
| AT5G15270 | -0.1 | 1.68E-01 | -0.2 | 1.54E-02 | 0.1 | 2.17E-01 | -0.2 | 1.10E-01 | 0.0 | 9.66E-01 | -0.2 | 2.26E-01 | RNA-binding KH domain-containing protein |
| AT1G17750 | -0.1 | 4.17E-01 | 0.0 | 9.59E-01 | -0.1 | 5.57E-01 | 0.2 | 6.87E-01 | 0.6 | 1.98E-01 | -0.4 | 4.59E-01 | Leucine-rich repeat receptor-like protein kinase PEPR2 |
| AT3G60400 | -0.1 | 3.26E-01 | -0.3 | 4.11E-02 | 0.1 | 2.90E-01 | 0.1 | 7.44E-01 | -0.2 | 5.07E-01 | 0.3 | 3.91E-01 | Transcription termination factor MTEF18, mitochondrial |
| AT1G22310 | -0.1 | 2.97E-01 | -0.1 | 2.27E-01 | 0.0 | 8.67E-01 | -0.3 | 7.78E-02 | -0.3 | 2.36E-01 | -0.1 | 7.49E-01 | MBD8 |
| AT5G09740 | -0.1 | 1.36E-01 | -0.3 | 4.80E-03 | 0.2 | 9.81E-02 | -0.2 | 2.95E-01 | -0.2 | 3.66E-01 | 0.0 | 9.97E-01 | Histone acetyltransferase of the MYST family 2 |
| AT4G00150 | -0.1 | 2.57E-01 | -0.3 | 1.71E-02 | 0.2 | 1.62E-01 | -0.5 | 5.12E-03 | -0.2 | 2.50E-01 | -0.3 | 2.22E-01 | Scarecrow-like protein 6 |
| AT4G15410 | -0.1 | 1.42E-01 | -0.2 | 5.03E-02 | 0.0 | 5.82E-01 | 0.1 | 6.57E-01 | 0.0 | 8.63E-01 | 0.0 | 8.87E-01 | Plant UBX domain-containing protein 5 |
| AT5G61970 | -0.1 | 2.03E-01 | 0.1 | 4.93E-01 | -0.2 | 8.97E-02 | -0.1 | 5.21E-01 | 0.0 | 9.14E-01 | -0.1 | 7.79E-01 | Signal recognition particle subunit SRP68 |
| AT1G14290 | -0.1 | 1.74E-01 | 0.4 | 3.76E-04 | -0.5 | 2.89E-04 | -0.4 | 1.17E-02 | -0.5 | 4.91E-03 | 0.1 | 6.31E-01 | Sphingoid base hydroxylase 2 |
| AT5G44750 | -0.1 | 3.25E-01 | -0.4 | 4.38E-03 | 0.3 | 3.46E-02 | 0.1 | 7.24E-01 | 0.0 | 9.59E-01 | 0.1 | 8.59E-01 | DNA repair protein REV1 |
| AT4G38680 | -0.1 | 4.59E-01 | 0.4 | 2.29E-02 | -0.5 | 1.05E-02 | -0.1 | 7.60E-01 | 0.0 | 8.28E-01 | -0.1 | 6.51E-01 | Cold shock protein 2 |
| AT2G20830 | -0.1 | 1.81E-01 | 0.0 | 8.22E-01 | -0.1 | 1.72E-01 | -0.2 | 1.35E-01 | -0.1 | 5.36E-01 | -0.1 | 6.56E-01 | glutamate formimidoyltransferase |
| AT4G39050 | -0.1 | 2.34E-01 | -0.2 | 6.03E-02 | 0.1 | 4.53E-01 | -0.2 | 1.74E-01 | 0.0 | 7.92E-01 | -0.2 | 2.12E-01 | Kinesin-like protein KIN-7D, mitochondrial |
| AT1G68840 | -0.1 | 5.65E-01 | -0.5 | 2.55E-02 | 0.4 | 8.72E-02 | -0.2 | 2.74E-01 | -1.0 | 4.18E-04 | 0.7 | 1.40E-02 | AP2 |
| AT5G04110 | -0.1 | 2.23E-01 | -0.2 | 9.03E-02 | 0.1 | 6.27E-01 | -0.1 | 6.07E-01 | 0.0 | 8.81E-01 | -0.1 | 5.92E-01 | DNA topoisomerase 2 |
| AT2G47730 | -0.1 | 4.18E-01 | 0.2 | 2.02E-01 | -0.3 | 6.85E-02 | 0.1 | 6.36E-01 | 0.1 | 5.57E-01 | 0.0 | 8.96E-01 | glutathione transferase |
| AT4G10130 | -0.1 | 3.32E-01 | 0.2 | 1.34E-01 | -0.3 | 4.98E-02 | -0.2 | 3.08E-01 | -0.4 | 9.78E-02 | 0.2 | 5.56E-01 | DNAJ heat shock N-terminal domain-containing protein |
| AT4G13780 | -0.1 | 1.61E-01 | 0.0 | 9.34E-01 | -0.1 | 1.81E-01 | 0.0 | 9.27E-01 | -0.1 | 6.34E-01 | 0.1 | 7.72E-01 | Methionine--tRNA ligase, cytoplasmic |
| AT2G41040 | -0.1 | 3.70E-01 | -0.2 | 2.18E-01 | 0.0 | 7.52E-01 | 0.0 | 9.32E-01 | 0.1 | 8.44E-01 | -0.1 | 8.14E-01 | Uncharacterized methyltransferase At2g41040, chloroplastic |
| AT5G09600 | -0.1 | 2.29E-01 | -0.1 | 2.07E-01 | 0.0 | 9.35E-01 | 0.2 | 7.62E-01 | 0.4 | 5.47E-01 | -0.2 | 7.78E-01 | Succinate dehydrogenase 3-1 |
| AT1G64400 | -0.1 | 2.57E-01 | 0.1 | 4.88E-01 | -0.2 | 1.08E-01 | -0.3 | 1.86E-02 | 0.0 | 8.59E-01 | -0.3 | 1.16E-01 | Long chain acyl-CoA synthetase 3 |
| AT3G55770 | -0.1 | 1.86E-01 | 0.0 | 9.80E-01 | -0.1 | 2.26E-01 | -0.3 | 1.41E-02 | -0.3 | 3.55E-02 | 0.0 | 9.43E-01 | Transcription factor L2 |
| AT2G25180 | -0.1 | 2.58E-01 | 0.0 | 9.11E-01 | -0.1 | 3.81E-01 | -0.3 | 2.55E-01 | -0.1 | 8.08E-01 | -0.2 | 6.11E-01 | Two-component response regulator ARR12 |
| AT2G02470 | -0.1 | 1.66E-01 | -0.1 | 3.55E-01 | 0.0 | 7.14E-01 | -0.1 | 5.17E-01 | -0.2 | 3.99E-01 | 0.0 | 8.48E-01 | PHD finger protein ALFIN-LIKE |
| AT3G05870 | -0.1 | 4.68E-01 | -0.1 | 4.46E-01 | 0.0 | 9.53E-01 | -0.1 | 8.65E-01 | -0.1 | 7.89E-01 | 0.1 | 9.18E-01 | Anaphase-promoting complex subunit 11 |
| AT4G24390 | -0.1 | 1.41E-01 | 0.0 | 9.10E-01 | -0.1 | 2.50E-01 | -0.1 | 5.10E-01 | -0.1 | 6.18E-01 | 0.0 | 9.74E-01 | F-box protein FBX14 |
| AT1G80460 | -0.1 | 1.79E-01 | 0.1 | 3.25E-01 | -0.2 | 4.64E-02 | 0.1 | 2.31E-01 | 0.1 | 6.71E-01 | 0.1 | 7.15E-01 | glycerol kinase |
| AT4G34215 | -0.1 | 2.72E-01 | -0.2 | 7.16E-02 | 0.1 | 4.64E-01 | -0.3 | 2.85E-02 | -0.1 | 4.08E-01 | -0.2 | 4.11E-01 | Probable carbohydrate esterase At4g34215 |
| AT4G17950 | -0.1 | 1.40E-01 | -0.1 | 5.43E-02 | 0.0 | 6.15E-01 | -0.1 | 6.33E-01 | -0.2 | 4.16E-01 | 0.1 | 7.84E-01 | AT-hook motif nuclear-localized protein |
| AT5G66130 | -0.1 | 4.00E-01 | -0.6 | 2.19E-03 | 0.5 | 1.79E-02 | -0.3 | 3.99E-01 | -0.3 | 4.12E-01 | | | RADIATION SENSITIVE 17 |
| AT1G13050 | -0.1 | 4.79E-01 | -0.2 | 3.11E-01 | 0.1 | 7.86E-01 | 0.0 | 9.27E-01 | 0.1 | 8.17E-01 | -0.1 | 9.10E-01 | Proline-rich receptor-like kinase |
| AT3G13222 | -0.1 | 2.01E-01 | 0.1 | 5.07E-01 | -0.2 | 1.04E-01 | 0.1 | 6.46E-01 | 0.1 | 8.37E-01 | 0.0 | 9.14E-01 | GBF-interacting protein 1 |
| AT1G32530 | -0.1 | 2.10E-01 | -0.1 | 3.47E-01 | 0.0 | 8.19E-01 | -0.1 | 7.81E-01 | -0.4 | 1.19E-01 | 0.3 | 3.09E-01 | MND1-interacting protein 1 |
| AT5G21274 | -0.1 | 3.49E-01 | 0.0 | 7.35E-01 | -0.1 | 6.24E-01 | 0.2 | 3.84E-01 | 0.0 | 8.76E-01 | 0.1 | 6.67E-01 | Calmodulin-6 |
| AT3G57430 | -0.1 | 5.54E-01 | 0.0 | 8.99E-01 | -0.1 | 7.34E-01 | 0.2 | 6.59E-01 | -0.1 | 9.04E-01 | 0.2 | 5.78E-01 | Pentatricopeptide repeat-containing protein At3g57430, chloroplastic |
| AT1G02170 | -0.1 | 1.89E-01 | -0.2 | 7.66E-02 | 0.0 | 6.17E-01 | -0.2 | 1.06E-01 | -0.2 | 3.03E-01 | -0.1 | 8.13E-01 | Metacaspase-1 |
| AT5G51340 | -0.1 | 1.52E-01 | -0.3 | 2.70E-03 | 0.2 | 4.61E-02 | 0.0 | 8.45E-01 | 0.0 | 9.39E-01 | 0.1 | 8.40E-01 | Sister chromatid cohesion protein SCC4 |
| AT3G14960 | -0.1 | 2.86E-01 | 0.0 | 9.19E-01 | -0.1 | 4.26E-01 | 0.2 | 2.54E-01 | -0.2 | 4.12E-01 | 0.3 | 6.91E-02 | Probable beta-1,3-galactosyltransferase 13 |
| AT1G16190 | -0.1 | 3.26E-01 | 0.1 | 3.64E-01 | -0.2 | 1.11E-01 | 0.2 | 1.00E-01 | 0.2 | 2.49E-01 | 0.0 | 8.47E-01 | Ubiquitin receptor RAD23 |
| AT4G22300 | -0.1 | 2.03E-01 | 0.1 | 1.90E-01 | -0.2 | 3.90E-02 | -0.1 | 5.53E-01 | -0.1 | 7.84E-01 | 0.0 | 8.78E-01 | Probable carboxylesterase SOBER1-like |
| AT4G13460 | -0.1 | 2.23E-01 | -0.2 | 3.58E-02 | 0.1 | 3.33E-01 | -0.3 | 1.20E-01 | 0.0 | 9.75E-01 | -0.3 | 2.99E-01 | Histone-lysine N-methyltransferase family member SUVH9 3-9 homolog protein 9) |
| AT1G09810 | -0.1 | 3.13E-01 | -0.2 | 1.16E-01 | 0.1 | 5.93E-01 | 0.2 | 3.53E-01 | -0.1 | 7.33E-01 | 0.3 | 2.62E-01 | YTH domain-containing family protein |
| AT3G57390 | -0.1 | 3.22E-01 | -0.3 | 1.67E-02 | 0.2 | 1.42E-01 | -0.2 | 3.42E-01 | -0.6 | 2.30E-02 | 0.4 | 2.35E-01 | AGAMOUS-like 18 |
| AT4G12010 | -0.1 | 2.07E-01 | -0.3 | 6.73E-03 | 0.2 | 9.27E-02 | -0.1 | 5.90E-01 | -0.3 | 2.37E-01 | 0.1 | 6.36E-01 | Disease resistance-like protein DSC1 |
| AT1G26800 | -0.1 | 5.61E-01 | 0.1 | 4.56E-01 | -0.2 | 2.57E-01 | -0.6 | 4.28E-02 | -0.9 | 7.99E-03 | 0.3 | 4.71E-01 | E3 ubiquitin-protein ligase MPSR1 |
| AT5G42080 | -0.1 | 1.57E-01 | -0.1 | 1.58E-01 | 0.0 | 9.60E-01 | -0.1 | 4.58E-01 | -0.1 | 5.88E-01 | 0.0 | 9.71E-01 | Dynamin-like protein |
| AT2G46600 | -0.1 | 5.03E-01 | -0.6 | 2.83E-03 | 0.5 | 1.33E-02 | 0.1 | 7.32E-01 | -0.5 | 1.98E-01 | 0.6 | 1.64E-01 | hypothetical protein |
| AT4G09680 | -0.1 | 2.28E-01 | -0.4 | 2.20E-03 | 0.3 | 2.71E-02 | 0.0 | 9.85E-01 | 0.1 | 8.73E-01 | -0.1 | 8.79E-01 | CST complex subunit CTC1 |
| AT1G79350 | -0.1 | 1.55E-01 | -0.2 | 7.34E-03 | 0.1 | 1.20E-01 | 0.1 | 7.59E-01 | 0.0 | 9.70E-01 | 0.1 | 8.62E-01 | Uncharacterized protein At1g79350 |
| AT1G68720 | -0.1 | 1.07E-01 | -0.3 | 7.82E-04 | 0.2 | 1.58E-02 | -0.1 | 7.59E-01 | 0.1 | 8.04E-01 | -0.2 | 6.44E-01 | tRNA) deaminase, chloroplastic |
| AT2G47170 | -0.1 | 2.81E-01 | 0.1 | 1.56E-01 | -0.2 | 3.43E-02 | -0.1 | 6.37E-01 | 0.0 | 8.74E-01 | 0.0 | 8.83E-01 | ADP-ribosylation factor 1 |
| AT5G16890 | -0.1 | 1.81E-01 | -0.1 | 3.11E-01 | 0.0 | 8.15E-01 | -0.1 | 8.06E-01 | -0.1 | 8.27E-01 | 0.0 | 9.95E-01 | Exostosin GT47 domain-containing protein |
| AT5G05780 | -0.1 | 3.21E-01 | 0.1 | 2.08E-01 | -0.2 | 5.27E-02 | 0.1 | 6.33E-01 | 0.1 | 3.82E-01 | -0.1 | 7.75E-01 | RP non-ATPase subunit 8A |
| AT1G78800 | -0.1 | 2.04E-01 | -0.1 | 4.69E-01 | 0.0 | 6.64E-01 | -0.2 | 7.54E-02 | -0.4 | 2.00E-02 | 0.1 | 6.00E-01 | Alpha-1,3 |
| AT3G62240 | -0.1 | 1.25E-01 | -0.2 | 1.11E-02 | 0.1 | 2.08E-01 | -0.1 | 5.64E-01 | -0.2 | 3.45E-01 | 0.1 | 7.68E-01 | RING-type E3 ubiquitin transferase |
| AT1G47290 | -0.1 | 2.20E-01 | 0.2 | 3.05E-02 | -0.3 | 8.35E-03 | -0.3 | 6.60E-02 | 0.0 | 9.22E-01 | -0.3 | 2.29E-01 | 3beta-hydroxysteroid-dehydrogenase |
| AT1G79690 | -0.1 | 1.94E-01 | -0.2 | 1.46E-02 | 0.1 | 1.74E-01 | 0.1 | 7.61E-01 | 0.2 | 3.49E-01 | -0.1 | 6.20E-01 | Nudix hydrolase 3 |
| AT2G40900 | -0.1 | 2.34E-01 | 0.0 | 7.01E-01 | -0.1 | 1.65E-01 | 0.9 | 1.17E-04 | 0.3 | 1.30E-01 | 0.6 | 7.38E-03 | WAT1-related protein At2g40900 |
| AT2G32450 | -0.1 | 3.64E-01 | -0.1 | 5.69E-01 | 0.0 | 8.06E-01 | -0.4 | 1.21E-01 | -0.3 | 2.32E-01 | -0.1 | 9.10E-01 | Uncharacterized TPR repeat-containing protein At2g32450 |
| AT5G35210 | -0.1 | 1.88E-01 | -0.3 | 5.09E-03 | 0.2 | 6.56E-02 | -0.2 | 2.86E-01 | -0.3 | 1.42E-01 | 0.1 | 7.18E-01 | PHD type transcription factor with transmembrane domain protein |
| AT3G19130 | -0.1 | 2.82E-01 | 0.2 | 5.24E-02 | -0.3 | 1.38E-02 | 0.2 | 1.04E-01 | 0.1 | 4.65E-01 | 0.1 | 6.54E-01 | Polyadenylate-binding protein RBP47B -binding protein RBP47B) |
| AT5G03680 | -0.1 | 4.02E-01 | 0.0 | 9.09E-01 | -0.1 | 4.40E-01 | -0.6 | 1.15E-01 | 0.0 | 9.94E-01 | -0.6 | 2.05E-01 | Trihelix transcription factor PTL |
| AT2G41700 | -0.1 | 2.93E-01 | -0.2 | 7.69E-02 | 0.1 | 4.58E-01 | 0.3 | 2.52E-01 | 0.1 | 9.03E-01 | 0.3 | 4.94E-01 | ATP-binding cassette A1 |
| AT1G33060 | -0.1 | 3.00E-01 | -0.1 | 3.53E-01 | 0.0 | 9.60E-01 | -0.2 | 1.77E-01 | -0.3 | 7.58E-02 | 0.1 | 7.21E-01 | NAC 014 |
| AT5G44740 | -0.1 | 4.36E-01 | 0.0 | 9.59E-01 | -0.1 | 5.72E-01 | -0.3 | 6.06E-01 | 0.0 | 9.90E-01 | -0.3 | 6.75E-01 | Y-family DNA polymerase H |
| AT5G28850 | -0.1 | 1.30E-01 | -0.1 | 3.04E-01 | 0.0 | 6.92E-01 | 0.1 | 6.72E-01 | 0.1 | 7.86E-01 | 0.0 | 9.70E-01 | Serine |
| AT2G27230 | -0.1 | 3.21E-01 | -0.2 | 5.14E-02 | 0.1 | 3.17E-01 | 0.0 | 9.69E-01 | 0.1 | 7.29E-01 | -0.1 | 8.06E-01 | Transcription factor LHW |
| AT3G46460 | -0.1 | 3.43E-01 | 0.0 | 7.39E-01 | -0.1 | 6.22E-01 | -0.1 | 5.52E-01 | 0.0 | 9.79E-01 | -0.1 | 7.10E-01 | Ubiquitin-conjugating enzyme 13 |
| AT1G20100 | -0.1 | 3.36E-01 | -0.2 | 7.02E-02 | 0.1 | 3.76E-01 | 0.2 | 2.93E-01 | 0.0 | 8.47E-01 | 0.1 | 6.14E-01 | DNA ligase-like protein |
| AT1G18030 | -0.1 | 4.22E-01 | 0.1 | 6.98E-01 | -0.1 | 3.21E-01 | -0.1 | 7.08E-01 | -0.3 | 2.26E-01 | 0.2 | 5.26E-01 | Probable protein phosphatase 2C 8 |
| AT5G66310 | -0.1 | 1.85E-01 | -0.1 | 1.73E-01 | 0.0 | 9.44E-01 | 0.0 | 8.70E-01 | -0.1 | 7.16E-01 | 0.1 | 8.74E-01 | Kinesin-like protein KIN-7H |
| AT1G52000 | -0.1 | 7.71E-01 | -0.1 | 8.85E-01 | 0.0 | 9.18E-01 | -0.1 | 7.15E-01 | 0.8 | 4.09E-02 | -0.9 | 5.31E-02 | Jacalin-related lectin 5 |
| AT3G52280 | -0.1 | 1.89E-01 | 0.0 | 6.02E-01 | -0.1 | 5.17E-01 | 0.0 | 9.37E-01 | 0.0 | 8.49E-01 | -0.1 | 8.27E-01 | General transcription factor group E6 |
| AT5G19440 | -0.1 | 4.40E-01 | -0.2 | 1.89E-01 | 0.1 | 5.95E-01 | 0.1 | 5.24E-01 | 0.0 | 9.91E-01 | 0.1 | 6.79E-01 | At5g19440 -binding Rossmann-fold superfamily protein) |

| | | | | | | | | | | | | | |
|---|---|---|---|---|---|---|---|---|---|---|---|---|---|
| AT3G11520 | -0.1 | 3.45E-01 | 0.1 | 4.32E-01 | -0.2 | 1.54E-01 | -0.1 | 5.81E-01 | 0.2 | 4.68E-01 | -0.3 | 2.24E-01 | CYCLIN B13 |
| AT5G64200 | -0.1 | 3.05E-01 | -0.1 | 5.57E-01 | 0.0 | 7.29E-01 | -0.1 | 6.44E-01 | -0.1 | 2.40E-01 | 0.1 | 6.04E-01 | Serine |
| AT1G77080 | -0.1 | 2.28E-01 | -0.1 | 1.44E-01 | 0.0 | 7.90E-01 | -0.2 | 1.37E-01 | -0.1 | 4.31E-01 | -0.1 | 7.42E-01 | K-box region and MADS-box transcription factor family protein |
| AT5G55990 | -0.1 | 2.07E-01 | 0.0 | 5.91E-01 | -0.1 | 1.20E-01 | 0.0 | 8.29E-01 | 0.1 | 6.90E-01 | 0.0 | 8.75E-01 | Calcineurin B-like protein |
| AT2G18196 | -0.1 | 5.19E-01 | -0.3 | 1.24E-01 | 0.2 | 4.16E-01 | 0.2 | 4.79E-01 | -0.3 | 3.46E-01 | 0.4 | 1.59E-01 | Heavy metal transport |
| AT4G02610 | -0.1 | 2.26E-01 | -0.2 | 6.33E-02 | 0.1 | 5.08E-01 | -0.4 | 1.07E-02 | -0.3 | 6.29E-02 | -0.1 | 6.83E-01 | Tryptophan synthase alpha chain |
| AT5G57565 | -0.1 | 5.39E-01 | -0.2 | 2.42E-01 | 0.1 | 6.26E-01 | -0.3 | 3.24E-01 | -0.2 | 5.74E-01 | -0.1 | 8.62E-01 | At5g57565 |
| AT3G20310 | -0.1 | 2.07E-01 | 0.0 | 5.45E-01 | 0.0 | 5.80E-01 | -0.2 | 3.84E-02 | -0.2 | 2.28E-01 | -0.1 | 6.32E-01 | Ethylene-responsive transcription factor 7 |
| AT3G14790 | -0.1 | 1.25E-01 | 0.1 | 3.55E-01 | -0.2 | 3.94E-02 | -0.1 | 6.37E-01 | 0.1 | 6.56E-01 | -0.2 | 4.38E-01 | Rhamnose biosynthesis 3 |
| AT1G04910 | -0.1 | 2.27E-01 | 0.0 | 6.01E-01 | -0.1 | 5.72E-01 | -0.2 | 1.92E-01 | -0.1 | 5.46E-01 | -0.1 | 7.21E-01 | O-fucosyltransferase 1 |
| AT5G01930 | -0.1 | 3.28E-01 | 0.2 | 4.64E-02 | -0.3 | 1.85E-02 | 0.2 | 3.86E-01 | 0.2 | 3.99E-01 | 0.0 | 9.60E-01 | Mannan endo-1,4-beta-mannosidase 6 |
| AT3G25900 | -0.1 | 4.10E-01 | 0.3 | 9.67E-03 | -0.4 | 7.51E-03 | 0.7 | 4.89E-03 | 0.3 | 2.35E-01 | 0.4 | 1.50E-01 | Homocysteine S-methyltransferase family protein |
| AT3G13050 | -0.1 | 1.70E-01 | -0.2 | 2.05E-02 | 0.1 | 2.77E-01 | -0.3 | 9.76E-02 | -0.3 | 1.42E-01 | 0.0 | 9.84E-01 | Organic cation |
| AT5G45260 | -0.1 | 2.37E-01 | -0.2 | 4.53E-02 | 0.1 | 3.83E-01 | 0.2 | 4.35E-01 | -0.1 | 8.78E-01 | 0.3 | 4.54E-01 | Disease resistance protein RRS1 |
| AT3G10850 | -0.1 | 2.56E-01 | 0.0 | 8.29E-01 | -0.1 | 2.23E-01 | -0.2 | 1.04E-01 | -0.3 | 3.51E-02 | 0.1 | 6.58E-01 | Hydroxyacylglutathione hydrolase cytoplasmic |
| AT2G26740 | -0.1 | 3.00E-01 | -0.3 | 7.19E-03 | 0.2 | 5.67E-02 | -0.2 | 2.85E-01 | -0.2 | 3.32E-01 | 0.0 | 9.78E-01 | ATsEH ) |
| AT4G03390 | -0.1 | 2.00E-01 | -0.1 | 2.10E-01 | 0.0 | 9.93E-01 | 0.0 | 8.22E-01 | 0.1 | 4.92E-01 | -0.1 | 6.74E-01 | Protein STRUBBELIG-RECEPTOR FAMILY 3 |
| AT3G14160 | -0.1 | 3.06E-01 | -0.1 | 1.32E-01 | 0.1 | 6.48E-01 | 0.3 | 1.94E-01 | -0.2 | 6.71E-01 | 0.5 | 1.03E-01 | 2-oxoglutarate-dependent dioxygenase family protein |
| AT5G51690 | -0.1 | 5.85E-01 | -0.4 | 3.78E-02 | 0.3 | 1.43E-01 | -0.7 | 2.52E-01 | -0.3 | 7.09E-01 | -0.5 | 6.36E-01 | 1-amino-cyclopropane-1-carboxylate synthase 12 |
| AT1G45332 | -0.1 | 2.50E-01 | -0.2 | 1.41E-02 | 0.1 | 1.42E-01 | 0.6 | 5.93E-02 | 0.2 | 6.75E-01 | 0.4 | 3.18E-01 | Translation elongation factor EFG |
| AT1G61500 | -0.1 | 3.82E-01 | -0.4 | 8.11E-03 | 0.3 | 6.43E-02 | 1.0 | 4.95E-03 | 0.7 | 5.00E-02 | 0.3 | 4.96E-01 | Receptor-like serine |
| AT3G50530 | -0.1 | 2.90E-01 | -0.1 | 1.71E-01 | 0.0 | 7.53E-01 | 0.1 | 5.93E-01 | 0.2 | 1.08E-01 | -0.1 | 4.27E-01 | CDPK-related kinase 5 |
| AT5G12350 | -0.1 | 1.90E-01 | -0.1 | 5.58E-02 | 0.1 | 5.13E-01 | -0.1 | 6.16E-01 | 0.1 | 8.28E-01 | -0.2 | 5.48E-01 | Regulator of chromosome condensation family with FYVE zinc finger domain-containing protein |
| AT2G35635 | -0.1 | 3.83E-01 | 0.0 | 8.26E-01 | -0.1 | 5.84E-01 | 0.1 | 7.65E-01 | 0.1 | 7.76E-01 | 0.0 | 9.77E-01 | Ubiquitin-NEDD8-like protein RUB2 |
| AT3G53990 | -0.1 | 3.47E-01 | 0.2 | 8.89E-02 | -0.3 | 2.53E-02 | -0.3 | 2.67E-01 | 0.0 | 8.95E-01 | -0.2 | 4.98E-01 | AT3G53990 protein |
| AT4G36260 | -0.1 | 4.39E-01 | -0.2 | 9.39E-02 | 0.1 | 3.95E-01 | -0.6 | 1.20E-01 | 0.2 | 7.23E-01 | -0.8 | 6.40E-02 | Protein SHI RELATED SEQUENCE 2 |
| AT2G18250 | -0.1 | 4.95E-01 | 0.0 | 8.68E-01 | -0.1 | 4.78E-01 | -0.2 | 3.24E-01 | -0.2 | 3.08E-01 | 0.0 | 9.34E-01 | Phosphopantetheine adenylyltransferase |
| AT1G14400 | -0.1 | 2.30E-01 | -0.1 | 1.19E-01 | 0.0 | 6.91E-01 | -0.2 | 6.78E-02 | -0.2 | 1.56E-01 | 0.0 | 9.17E-01 | Ubiquitin-conjugating enzyme E2 1 |
| AT5G66100 | -0.1 | 2.27E-01 | 0.0 | 7.79E-01 | -0.1 | 1.99E-01 | 0.0 | 8.66E-01 | 0.0 | 9.22E-01 | -0.1 | 8.10E-01 | Winged-helix DNA-binding transcription factor family protein |
| AT4G08310 | -0.1 | 2.02E-01 | -0.1 | 2.23E-01 | 0.0 | 9.88E-01 | -0.1 | 5.06E-01 | 0.0 | 8.36E-01 | -0.1 | 8.19E-01 | DNA ligase |
| AT5G41370 | -0.1 | 3.24E-01 | -0.4 | 1.23E-03 | 0.3 | 9.55E-03 | -0.1 | 6.58E-01 | 0.0 | 9.24E-01 | -0.1 | 8.34E-01 | General transcription and DNA repair factor IIH helicase subunit XPB1 |
| AT5G17060 | -0.1 | 2.40E-01 | -0.1 | 4.72E-01 | 0.0 | 7.29E-01 | 0.3 | 1.19E-01 | 0.2 | 1.77E-01 | 0.0 | 9.87E-01 | ADP-ribosylation factor |
| AT2G39550 | -0.1 | 2.62E-01 | -0.1 | 4.39E-01 | 0.0 | 8.00E-01 | -0.3 | 1.55E-01 | -0.1 | 5.67E-01 | -0.1 | 6.49E-01 | Geranylgeranyl transferase type-1 subunit beta |
| AT5G15700 | -0.1 | 3.81E-01 | -0.5 | 2.08E-03 | 0.4 | 1.74E-02 | 0.9 | 1.49E-03 | 0.3 | 3.01E-01 | 0.6 | 6.02E-02 | DNA-directed RNA polymerase |
| AT4G35890 | -0.1 | 1.52E-01 | -0.1 | 2.36E-01 | 0.0 | 8.58E-01 | 0.2 | 1.22E-01 | 0.3 | 6.63E-02 | -0.1 | 7.50E-01 | Winged-helix DNA-binding transcription factor family protein |
| AT2G17220 | -0.1 | 3.86E-01 | -0.4 | 3.43E-03 | 0.3 | 2.33E-02 | 0.2 | 3.56E-01 | -0.1 | 7.99E-01 | 0.2 | 3.80E-01 | Probable serine |
| AT1G50640 | -0.1 | 1.72E-01 | -0.1 | 2.04E-01 | 0.0 | 9.61E-01 | -0.2 | 1.35E-01 | -0.3 | 2.42E-02 | 0.1 | 4.98E-01 | Ethylene-responsive transcription factor 3 |
| AT2G13800 | -0.1 | 2.76E-01 | -0.3 | 2.87E-03 | 0.2 | 3.32E-02 | 0.1 | 6.25E-01 | 0.0 | 9.64E-01 | 0.1 | 7.70E-01 | Somatic embryogenesis receptor-like kinase 5 |
| AT5G60550 | -0.1 | 1.57E-01 | -0.1 | 1.94E-01 | 0.0 | 9.46E-01 | 0.0 | 8.71E-01 | 0.1 | 3.74E-01 | -0.1 | 5.67E-01 | Serine |
| AT4G25260 | -0.1 | 5.35E-01 | 0.4 | 2.57E-02 | -0.5 | 1.75E-02 | -0.4 | 5.53E-03 | -0.1 | 6.51E-01 | -0.4 | 7.31E-02 | Pectinesterase inhibitor 7 |
| AT5G04510 | -0.1 | 2.59E-01 | 0.0 | 9.25E-01 | -0.1 | 2.87E-01 | -0.2 | 3.03E-01 | 0.1 | 8.19E-01 | -0.3 | 2.41E-01 | non-specific serine |
| AT3G02780 | -0.1 | 3.26E-01 | 0.4 | 2.55E-03 | -0.5 | 1.35E-03 | -0.1 | 4.30E-01 | -0.1 | 7.32E-01 | -0.1 | 8.27E-01 | isopentenyl-diphosphate Delta-isomerase |
| AT1G03430 | -0.1 | 3.07E-01 | -0.2 | 1.02E-01 | 0.1 | 5.44E-01 | 0.0 | 9.92E-01 | 0.0 | 9.48E-01 | 0.0 | 9.58E-01 | Histidine-containing phosphotransfer protein |
| AT2G41790 | -0.1 | 1.28E-01 | -0.3 | 1.55E-03 | 0.2 | 2.59E-02 | 0.0 | 9.18E-01 | 0.1 | 7.86E-01 | 0.0 | 8.94E-01 | Insulin-degrading enzyme-like 1, peroxisomal |
| AT3G26590 | -0.1 | 3.61E-01 | -0.3 | 1.08E-02 | 0.2 | 7.38E-02 | 0.3 | 9.70E-02 | 0.2 | 5.13E-01 | 0.2 | 5.72E-01 | MATE efflux family protein |
| AT4G01680 | -0.1 | 5.19E-01 | 0.1 | 6.33E-01 | -0.2 | 3.57E-01 | -1.1 | 1.73E-03 | -0.8 | 1.67E-02 | -0.3 | 4.72E-01 | Myb domain protein 55 |
| AT5G01850 | -0.1 | 3.53E-01 | -0.4 | 1.67E-03 | 0.3 | 1.47E-02 | 0.1 | 8.48E-01 | 0.3 | 2.64E-01 | -0.3 | 4.68E-01 | Protein kinase ATN1-like protein |
| AT4G11110 | -0.1 | 4.35E-01 | -0.3 | 1.36E-02 | 0.3 | 7.70E-02 | 0.0 | 9.70E-01 | 0.1 | 8.08E-01 | -0.1 | 8.23E-01 | Protein SPA1-RELATED 2 |
| AT2G23450 | -0.1 | 3.48E-01 | -0.3 | 1.66E-02 | 0.2 | 1.14E-01 | -0.3 | 1.63E-01 | -0.4 | 8.24E-02 | 0.1 | 7.64E-01 | Wall-associated receptor kinase-like 14 |
| AT4G14070 | -0.1 | 3.12E-01 | -0.2 | 5.51E-02 | 0.1 | 3.42E-01 | 0.0 | 9.98E-01 | 0.1 | 7.00E-01 | -0.1 | 7.64E-01 | Long-chain-fatty-acid-- ligase AEE15, chloroplastic |
| AT3G18430 | -0.1 | 3.76E-01 | -0.1 | 4.06E-01 | 0.0 | 9.89E-01 | 0.2 | 3.79E-01 | 0.1 | 7.15E-01 | 0.1 | 7.98E-01 | hypothetical protein |
| AT3G20420 | -0.1 | 5.26E-01 | -0.5 | 7.04E-03 | 0.4 | 3.88E-02 | 0.2 | 4.35E-01 | 0.0 | 9.99E-01 | 0.2 | 5.50E-01 | Ribonuclease 3-like protein 2 |
| AT5G65940 | -0.1 | 1.85E-01 | 0.0 | 5.98E-01 | -0.1 | 5.00E-01 | -0.2 | 2.09E-01 | -0.1 | 5.27E-01 | -0.1 | 7.71E-01 | 3-hydroxyisobutyryl-CoA hydrolase |
| AT3G60190 | -0.1 | 2.58E-01 | 0.0 | 9.21E-01 | -0.1 | 2.73E-01 | 0.0 | 9.21E-01 | -0.3 | 3.24E-02 | 0.3 | 8.55E-02 | UMP-CMP kinase |
| AT5G61030 | -0.1 | 5.66E-01 | 0.1 | 6.17E-01 | -0.2 | 3.36E-01 | 0.7 | 7.70E-05 | 0.6 | 2.05E-03 | 0.2 | 4.37E-01 | Glycine-rich RNA-binding protein 3, mitochondrial |
| AT4G24670 | -0.1 | 3.52E-01 | 0.2 | 2.13E-02 | -0.3 | 1.08E-02 | -0.4 | 3.97E-02 | -0.1 | 6.26E-01 | -0.3 | 2.67E-01 | Tryptophan aminotransferase-related protein 2 |
| AT2G43000 | -0.1 | 7.56E-01 | -0.8 | 2.74E-02 | 0.7 | 8.38E-02 | 2.1 | 6.82E-05 | 0.8 | 1.11E-01 | 1.3 | 6.04E-03 | NAC domain containing protein 42 |
| AT4G30170 | -0.1 | 2.80E-01 | 0.8 | 1.07E-05 | -0.9 | 2.78E-05 | -0.3 | 1.93E-01 | 0.2 | 5.02E-01 | -0.5 | 1.77E-02 | Peroxidase 45 |
| AT4G24400 | -0.1 | 1.80E-01 | -0.1 | 5.63E-02 | 0.0 | 5.30E-01 | 0.1 | 4.61E-01 | 0.1 | 5.79E-01 | 0.0 | 9.77E-01 | non-specific serine |
| AT1G76270 | -0.1 | 2.57E-01 | 0.1 | 1.02E-01 | -0.2 | 2.87E-02 | 0.2 | 4.49E-01 | 0.0 | 9.84E-01 | 0.2 | 5.71E-01 | O-fucosyltransferase 16 |
| AT2G38660 | -0.1 | 1.92E-01 | -0.1 | 7.04E-02 | 0.0 | 5.94E-01 | -0.1 | 3.95E-01 | 0.0 | 8.45E-01 | -0.2 | 3.76E-01 | Tetrahydrofolate dehydrogenase |
| AT3G04160 | -0.1 | 4.32E-01 | -0.3 | 3.72E-02 | 0.2 | 2.05E-01 | -0.3 | 3.08E-01 | 0.0 | 8.99E-01 | -0.2 | 5.50E-01 | U11 |
| AT5G20350 | -0.1 | 1.95E-01 | -0.2 | 4.24E-02 | 0.1 | 3.95E-01 | -0.2 | 1.04E-01 | -0.2 | 3.00E-01 | -0.1 | 8.28E-01 | Protein S-acyltransferase 24 |
| AT5G19990 | -0.1 | 2.86E-01 | 0.0 | 6.65E-01 | -0.1 | 1.77E-01 | -0.1 | 4.34E-01 | -0.1 | 3.42E-01 | 0.0 | 8.64E-01 | Regulatory particle triple-A ATPase 6A |
| AT4G33650 | -0.1 | 1.84E-01 | -0.2 | 1.37E-02 | 0.1 | 1.80E-01 | 0.2 | 2.86E-01 | 0.0 | 9.79E-01 | 0.2 | 4.34E-01 | Dynamin-related protein 3A |
| AT1G30270 | -0.1 | 2.96E-01 | -0.1 | 4.19E-01 | 0.0 | 8.69E-01 | 0.4 | 3.65E-03 | 0.0 | 8.19E-01 | 0.4 | 5.27E-02 | non-specific serine |
| AT5G46750 | -0.1 | 1.68E-01 | 0.0 | 6.76E-01 | -0.1 | 1.15E-01 | -0.2 | 2.04E-01 | -0.2 | 2.09E-01 | 0.0 | 9.40E-01 | Probable ADP-ribosylation factor GTPase-activating protein AGD9 |
| AT1G21640 | -0.1 | 4.47E-01 | -0.3 | 1.44E-02 | 0.2 | 7.23E-02 | -0.5 | 1.34E-01 | 0.0 | 9.79E-01 | -0.5 | 2.72E-01 | NAD kinase 2 |
| AT1G43900 | -0.1 | 1.64E-01 | -0.1 | 2.72E-01 | 0.0 | 8.26E-01 | -0.1 | 3.22E-01 | 0.0 | 8.39E-01 | -0.1 | 6.66E-01 | Probable protein phosphatase 2C 11 |
| AT2G31010 | -0.1 | 5.03E-01 | 0.2 | 7.33E-02 | -0.3 | 4.39E-02 | -0.7 | 7.38E-03 | -0.7 | 8.75E-03 | 0.0 | 9.39E-01 | Protein kinase domain-containing protein |
| AT4G22710 | -0.1 | 4.42E-01 | -0.2 | 1.12E-01 | 0.1 | 4.45E-01 | 2.6 | 6.81E-07 | 1.3 | 3.27E-03 | 1.4 | 3.84E-04 | Cytochrome P450 - like protein |
| AT5G22770 | -0.1 | 1.85E-01 | -0.1 | 2.12E-01 | 0.0 | 9.72E-01 | 0.0 | 9.27E-01 | 0.0 | 8.73E-01 | 0.1 | 8.42E-01 | AP-2 complex subunit alpha-1 |
| AT1G67325 | -0.1 | 3.67E-01 | 0.0 | 6.72E-01 | -0.1 | 2.43E-01 | -0.1 | 6.65E-01 | -0.1 | 4.92E-01 | 0.0 | 8.39E-01 | Ran BP2 |
| AT1G22190 | -0.1 | 5.72E-01 | -0.2 | 1.42E-01 | 0.2 | 3.93E-01 | -0.5 | 5.53E-03 | -0.5 | 8.87E-03 | 0.0 | 9.64E-01 | Ethylene-responsive transcription factor RAP2-13 |
| AT3G46000 | -0.1 | 1.95E-01 | 0.0 | 4.83E-01 | -0.1 | 7.99E-02 | -0.3 | 1.31E-01 | -0.1 | 5.99E-01 | -0.1 | 5.93E-01 | AT3G46000 protein |
| AT2G41070 | -0.1 | 5.34E-01 | -0.1 | 6.42E-01 | 0.0 | 9.25E-01 | 0.0 | 8.98E-01 | -0.1 | 7.86E-01 | 0.1 | 9.01E-01 | Basic-leucine zipper transcription factor family protein |
| AT4G01000 | -0.1 | 1.65E-01 | -0.1 | 5.68E-02 | 0.0 | 5.46E-01 | -0.3 | 7.94E-03 | -0.4 | 1.36E-03 | 0.1 | 4.05E-01 | Replication stress response regulator SDE2 |
| AT5G56790 | -0.1 | 3.41E-01 | -0.2 | 3.52E-02 | 0.1 | 2.38E-01 | 0.1 | 6.46E-01 | 0.2 | 2.73E-01 | -0.1 | 5.76E-01 | Protein kinase domain-containing protein |
| AT2G31370 | -0.1 | 2.97E-01 | 0.0 | 9.60E-01 | -0.1 | 3.91E-01 | 0.1 | 6.01E-01 | 0.2 | 4.34E-01 | -0.1 | 8.39E-01 | Basic-leucine zipper transcription factor family protein |
| AT1G55460 | -0.1 | 3.91E-01 | 0.0 | 7.38E-01 | 0.0 | 6.78E-01 | -0.2 | 2.34E-01 | -0.2 | 3.74E-01 | 0.0 | 9.40E-01 | KIN17-like protein |
| AT2G26430 | -0.1 | 2.18E-01 | -0.1 | 3.71E-01 | 0.0 | 7.97E-01 | -0.5 | 6.57E-04 | -0.5 | 3.49E-03 | -0.1 | 8.07E-01 | Arginine-rich cyclin 1 |
| AT3G59140 | -0.1 | 3.39E-01 | -0.6 | 1.19E-04 | 0.5 | 1.04E-03 | 0.7 | 3.90E-02 | 0.9 | 1.30E-02 | -0.2 | 6.05E-01 | ABC transporter C family member 10 |
| AT1G51890 | -0.1 | 5.67E-01 | -0.5 | 5.01E-03 | 0.5 | 2.24E-02 | 0.9 | 4.39E-05 | 0.3 | 8.92E-02 | 0.6 | 1.69E-02 | Leucine-rich repeat protein kinase family protein |

| | | | | | | | | | | | | | |
|---|---|---|---|---|---|---|---|---|---|---|---|---|---|
| AT5G08470 | -0.1 | 3.20E-01 | -0.2 | 2.45E-02 | 0.1 | 1.84E-01 | 0.0 | 9.30E-01 | 0.0 | 9.30E-01 | 0.1 | 8.94E-01 | Peroxisomal ATPase PEX1 |
| AT1G25390 | -0.1 | 2.56E-01 | -0.2 | 1.15E-02 | 0.1 | 1.13E-01 | 0.2 | 2.40E-01 | 0.1 | 7.52E-01 | 0.1 | 6.02E-01 | LEAF RUST 10 DISEASE-RESISTANCE LOCUS RECEPTOR-LIKE PROTEIN KINASE-like 1.1 |
| AT4G31350 | -0.1 | 2.25E-01 | -0.1 | 2.35E-01 | 0.0 | 9.90E-01 | 0.0 | 9.60E-01 | 0.2 | 2.64E-01 | -0.2 | 3.43E-01 | hypothetical protein |
| AT4G30360 | -0.1 | 3.26E-01 | -0.3 | 5.85E-03 | 0.2 | 4.87E-02 | -0.2 | 1.75E-01 | -0.1 | 4.29E-01 | -0.1 | 8.16E-01 | Cyclic nucleotide-gated ion channel 17 |
| AT5G16760 | -0.1 | 2.03E-01 | 0.0 | 7.07E-01 | -0.1 | 1.43E-01 | 0.1 | 5.06E-01 | 0.0 | 9.30E-01 | 0.1 | 7.52E-01 | Inositol-tetrakisphosphate 1-kinase 1 P 5 |
| AT1G80300 | -0.1 | 2.98E-01 | -0.1 | 3.00E-01 | 0.0 | 9.70E-01 | 0.0 | 8.17E-01 | 0.0 | 9.23E-01 | 0.0 | 9.56E-01 | ADP,ATP carrier protein |
| AT4G35360 | -0.1 | 3.62E-01 | -0.2 | 2.27E-02 | 0.2 | 1.47E-01 | 0.0 | 8.75E-01 | 0.2 | 2.23E-01 | -0.2 | 3.87E-01 | At4g35360 |
| AT3G44260 | -0.1 | 7.99E-01 | -0.7 | 5.50E-02 | 0.6 | 1.26E-01 | 0.2 | 4.04E-01 | -0.1 | 8.05E-01 | 0.3 | 3.71E-01 | Probable CCR4-associated factor 1 homolog 9 |
| AT4G30200 | -0.1 | 2.50E-01 | 0.0 | 6.47E-01 | -0.1 | 1.53E-01 | -0.3 | 2.65E-02 | -0.3 | 1.09E-01 | -0.1 | 8.04E-01 | VIN3-like protein 2 |
| AT3G54980 | -0.1 | 3.72E-01 | -0.1 | 1.96E-01 | 0.0 | 7.15E-01 | 0.4 | 2.75E-02 | 0.0 | 9.96E-01 | 0.4 | 8.69E-02 | Pentatricopeptide repeat-containing protein At3g54980, mitochondrial |
| AT1G77810 | -0.1 | 5.01E-01 | -0.1 | 4.23E-01 | 0.0 | 9.07E-01 | 0.1 | 8.36E-01 | -0.1 | 6.92E-01 | 0.2 | 5.82E-01 | Hexosyltransferase |
| AT1G14130 | -0.1 | 3.36E-01 | -0.2 | 1.63E-02 | 0.2 | 1.33E-01 | -0.1 | 7.36E-01 | -0.1 | 8.04E-01 | 0.0 | 9.86E-01 | 2-oxoglutarate and Fe-dependent oxygenase superfamily protein |
| AT5G25760 | -0.1 | 3.89E-01 | -0.1 | 2.57E-01 | 0.0 | 7.98E-01 | 0.1 | 8.05E-01 | 0.2 | 4.18E-01 | -0.1 | 6.66E-01 | Protein PEROXIN-4 |
| AT3G24120 | -0.1 | 2.20E-01 | -0.2 | 3.55E-02 | 0.1 | 3.38E-01 | -0.5 | 3.12E-04 | -0.4 | 3.86E-03 | -0.1 | 5.76E-01 | hypothetical protein |
| AT5G14040 | -0.1 | 4.74E-01 | 0.2 | 1.11E-01 | -0.3 | 4.43E-02 | 0.4 | 5.76E-03 | 0.2 | 1.34E-01 | 0.1 | 4.31E-01 | Mitochondrial phosphate carrier protein 3, mitochondrial |
| AT2G23150 | -0.1 | 3.53E-01 | -0.1 | 1.18E-01 | 0.1 | 5.47E-01 | 0.3 | 3.83E-01 | 0.4 | 2.91E-01 | -0.1 | 8.54E-01 | Metal transporter Nramp3 |
| AT3G60220 | -0.1 | 6.76E-01 | 0.4 | 3.60E-02 | -0.5 | 3.84E-02 | -0.3 | 1.38E-01 | -0.1 | 6.92E-01 | -0.2 | 4.45E-01 | E3 ubiquitin-protein ligase ATL4 |
| AT3G57880 | -0.1 | 3.52E-01 | -0.4 | 2.59E-03 | 0.3 | 2.04E-02 | -0.4 | 6.78E-02 | -0.3 | 1.93E-01 | -0.1 | 8.31E-01 | FT-interacting protein 3 |
| AT1G08460 | -0.1 | 3.31E-01 | -0.1 | 1.14E-01 | 0.1 | 5.47E-01 | -0.3 | 9.27E-02 | -0.2 | 2.10E-01 | 0.0 | 8.67E-01 | Histone deacetylase 8 |
| AT1G72010 | -0.1 | 4.24E-01 | -0.2 | 4.33E-02 | 0.1 | 2.00E-01 | 0.1 | 5.26E-01 | 0.0 | 9.26E-01 | 0.1 | 7.61E-01 | Transcription factor TCP22 |
| AT5G14480 | -0.1 | 5.05E-01 | 0.1 | 5.77E-01 | -0.2 | 3.06E-01 | 0.1 | 6.12E-01 | -0.3 | 3.24E-01 | 0.4 | 1.67E-01 | Beta-1,4-N-acetylglucosaminyltransferase family protein |
| AT1G27520 | -0.1 | 2.74E-01 | -0.2 | 2.78E-02 | 0.1 | 2.32E-01 | -0.3 | 1.01E-01 | -0.2 | 2.33E-01 | -0.1 | 8.78E-01 | Alpha-mannosidase I MNS5 |
| AT5G08680 | -0.1 | 3.84E-01 | 0.1 | 5.14E-01 | -0.1 | 1.71E-01 | 0.2 | 1.06E-01 | 0.2 | 2.59E-01 | 0.0 | 8.87E-01 | ATP synthase subunit beta-3, mitochondrial |
| AT4G20840 | -0.1 | 2.60E-01 | 0.1 | 1.00E-01 | -0.2 | 2.65E-02 | 0.0 | 8.02E-01 | 0.2 | 2.84E-01 | -0.1 | 4.86E-01 | Berberine bridge enzyme-like 21 |
| AT5G60120 | -0.1 | 2.49E-01 | -0.1 | 2.03E-01 | 0.0 | 8.93E-01 | -0.1 | 7.24E-01 | 0.1 | 7.17E-01 | -0.2 | 5.12E-01 | Target of early activation tagged 2 |
| AT1G18690 | -0.1 | 3.01E-01 | 0.1 | 9.84E-02 | -0.2 | 3.34E-02 | -0.3 | 1.99E-01 | -0.2 | 4.27E-01 | -0.1 | 8.30E-01 | Xyloglucan 6-xylosyltransferase 5 |
| AT4G17640 | -0.1 | 2.53E-01 | -0.1 | 6.65E-02 | 0.1 | 4.63E-01 | -0.2 | 1.17E-01 | -0.1 | 3.45E-01 | -0.1 | 7.57E-01 | Casein kinase II subunit beta |
| AT1G73250 | -0.1 | 3.29E-01 | 0.1 | 4.02E-01 | -0.2 | 1.24E-01 | -0.2 | 1.41E-01 | -0.2 | 2.39E-01 | 0.0 | 9.50E-01 | GDP-L-fucose synthase 1 |
| AT5G48430 | -0.1 | 4.80E-01 | 0.8 | 6.39E-05 | -0.9 | 1.30E-04 | 1.3 | 8.95E-08 | 0.7 | 1.34E-04 | 0.5 | 2.90E-03 | At5g48430 |
| AT5G43850 | -0.1 | 7.30E-01 | -0.6 | 2.01E-02 | 0.5 | 5.15E-02 | -0.2 | 2.29E-01 | -0.1 | 6.59E-01 | -0.1 | 7.03E-01 | Acireductone dioxygenase 4 -requiring) 4) -requiring) 4) |
| AT4G17570 | -0.1 | 2.67E-01 | -0.4 | 5.38E-04 | 0.3 | 5.06E-03 | 0.2 | 2.22E-01 | 0.3 | 5.35E-02 | -0.1 | 5.38E-01 | GATA transcription factor 26 |
| AT3G56060 | -0.1 | 5.25E-01 | 0.4 | 4.73E-03 | -0.5 | 4.30E-03 | -0.3 | 9.39E-02 | -0.3 | 1.06E-01 | 0.0 | 9.54E-01 | ADHESION OF CALYX EDGES-like protein |
| AT1G56210 | -0.1 | 4.01E-01 | 0.2 | 4.27E-02 | -0.3 | 2.20E-02 | 0.0 | 8.37E-01 | -0.1 | 7.59E-01 | 0.0 | 9.28E-01 | Heavy metal-associated isoprenylated plant protein 35 |
| AT5G19770 | -0.1 | 3.27E-01 | 0.3 | 9.12E-03 | -0.3 | 3.61E-03 | -0.2 | 4.27E-02 | -0.3 | 1.57E-02 | 0.1 | 6.73E-01 | Tubulin alpha-3 chain |
| AT3G58750 | -0.1 | 2.28E-01 | 0.0 | 5.05E-01 | -0.1 | 1.05E-01 | -0.4 | 1.06E-02 | -0.4 | 1.20E-02 | 0.0 | 9.16E-01 | Citrate synthase 2, peroxisomal |
| AT3G17365 | -0.1 | 7.20E-01 | -0.4 | 1.48E-01 | 0.3 | 3.38E-01 | -0.1 | 8.62E-01 | 0.2 | 6.76E-01 | -0.2 | 6.04E-01 | S-adenosyl-L-methionine-dependent methyltransferases superfamily protein |
| AT1G55510 | -0.1 | 5.14E-01 | -0.2 | 7.33E-02 | 0.2 | 2.88E-01 | -0.3 | 1.47E-01 | -0.6 | 2.05E-02 | 0.3 | 3.87E-01 | 2-oxoisovalerate dehydrogenase subunit beta 1, mitochondrial |
| AT4G02980 | -0.1 | 3.34E-01 | 0.0 | 7.66E-01 | -0.1 | 2.84E-01 | 0.2 | 2.35E-01 | 0.2 | 2.36E-01 | 0.0 | 9.41E-01 | Auxin-binding protein 1 |
| AT3G09350 | -0.1 | 5.06E-01 | -0.2 | 8.99E-02 | 0.1 | 3.35E-01 | 0.1 | 6.32E-01 | 0.0 | 9.58E-01 | 0.1 | 7.64E-01 | Fes1A |
| AT1G31730 | -0.1 | 3.25E-01 | -0.1 | 2.63E-01 | 0.0 | 8.81E-01 | 0.0 | 9.86E-01 | 0.0 | 9.05E-01 | 0.1 | 9.07E-01 | AP-4 complex subunit epsilon |
| AT5G49470 | -0.1 | 3.45E-01 | -0.2 | 5.44E-02 | 0.1 | 3.18E-01 | 0.0 | 8.81E-01 | -0.1 | 7.86E-01 | 0.0 | 9.18E-01 | Protein kinase |
| AT5G14720 | -0.1 | 2.73E-01 | -0.2 | 1.94E-02 | 0.1 | 1.57E-01 | -0.2 | 3.28E-01 | -0.1 | 6.77E-01 | -0.1 | 7.93E-01 | Protein kinase domain-containing protein |
| AT5G10630 | -0.1 | 2.92E-01 | -0.1 | 1.76E-01 | 0.0 | 7.68E-01 | 0.0 | 9.96E-01 | 0.1 | 8.32E-01 | -0.1 | 8.48E-01 | Uncharacterized protein At5g10630 |
| AT2G24420 | -0.1 | 3.74E-01 | -0.3 | 1.46E-02 | 0.2 | 8.54E-02 | -0.2 | 6.32E-02 | -0.1 | 2.96E-01 | -0.1 | 7.18E-01 | AT2G24420 protein |
| AT2G21230 | -0.1 | 2.87E-01 | -0.2 | 2.19E-02 | 0.1 | 1.79E-01 | -0.2 | 3.25E-01 | -0.2 | 3.27E-01 | 0.0 | 9.42E-01 | Basic-leucine zipper transcription factor family protein |
| AT3G05380 | -0.1 | 3.64E-01 | -0.6 | 1.44E-04 | 0.5 | 1.12E-03 | 0.1 | 6.50E-01 | 0.0 | 9.97E-01 | 0.1 | 7.29E-01 | Protein ALWAYS EARLY 2 |
| AT5G22780 | -0.1 | 2.45E-01 | 0.0 | 7.59E-01 | -0.1 | 1.89E-01 | 0.2 | 2.13E-01 | -0.1 | 5.76E-01 | 0.3 | 1.03E-01 | AP-2 complex subunit alpha-2 |
| AT1G54290 | -0.1 | 4.05E-01 | -0.1 | 1.55E-01 | 0.1 | 5.80E-01 | -0.1 | 7.49E-01 | 0.1 | 7.94E-01 | -0.1 | 5.77E-01 | Protein translation factor SUI1 homolog 2 |
| AT4G24470 | -0.1 | 3.26E-01 | -0.2 | 1.64E-02 | 0.1 | 1.24E-01 | -0.2 | 1.22E-01 | -0.1 | 5.70E-01 | -0.1 | 5.71E-01 | GATA-type zinc finger protein with TIFY domain-containing protein |
| AT4G23550 | -0.1 | 5.09E-01 | -0.1 | 6.80E-01 | 0.0 | 8.67E-01 | 0.7 | 4.60E-04 | 0.3 | 2.06E-01 | 0.5 | 2.95E-02 | WRKY family transcription factor |
| AT4G25180 | -0.1 | 2.95E-01 | -0.3 | 2.07E-03 | 0.2 | 2.19E-02 | -0.5 | 1.01E-02 | -0.3 | 1.31E-01 | -0.2 | 4.63E-01 | DNA-directed RNA polymerase III subunit RPC4 |
| AT1G34180 | -0.1 | 6.11E-01 | -0.5 | 7.46E-03 | 0.5 | 3.62E-02 | 0.5 | 1.50E-01 | 0.1 | 7.71E-01 | 0.3 | 4.19E-01 | NAC domain containing protein 16 |
| AT2G23080 | -0.1 | 3.43E-01 | -0.1 | 1.71E-01 | 0.0 | 6.90E-01 | 0.0 | 9.42E-01 | 0.2 | 2.81E-01 | -0.2 | 3.31E-01 | Casein kinase II subunit alpha-3 |
| AT5G45510 | -0.1 | 3.48E-01 | 0.0 | 7.73E-01 | -0.1 | 5.89E-01 | 0.2 | 5.50E-02 | 0.2 | 7.46E-02 | 0.0 | 9.64E-01 | Probable disease resistance protein At5g45510 |
| AT3G50500 | -0.1 | 4.52E-01 | 0.0 | 9.07E-01 | -0.1 | 5.90E-01 | -0.1 | 2.90E-01 | -0.3 | 2.39E-02 | 0.2 | 3.31E-01 | SNF1-related protein kinase 2.2 |
| AT3G29160 | -0.1 | 2.26E-01 | 0.0 | 4.44E-01 | 0.0 | 7.22E-01 | 0.1 | 5.08E-01 | 0.2 | 1.50E-01 | -0.1 | 5.32E-01 | SNF1-related protein kinase catalytic subunit alpha KIN11 |
| AT1G48380 | -0.1 | 5.11E-01 | -0.2 | 6.66E-02 | 0.2 | 2.67E-01 | 0.1 | 7.47E-01 | 0.2 | 3.28E-01 | -0.1 | 5.95E-01 | DNA-binding protein RHL1 |
| AT3G29760 | -0.1 | 5.02E-01 | -0.2 | 1.84E-01 | 0.1 | 5.43E-01 | 0.0 | 8.87E-01 | 0.1 | 8.64E-01 | -0.1 | 7.90E-01 | Haloacid dehalogenase-like hydrolase superfamily protein |
| AT1G23730 | -0.1 | 5.73E-01 | -0.1 | 3.32E-01 | 0.1 | 7.24E-01 | -0.3 | 2.51E-01 | -0.4 | 2.16E-01 | 0.1 | 9.04E-01 | Carbonic anhydrase |
| AT2G45280 | -0.1 | 5.33E-01 | 0.0 | 7.77E-01 | -0.1 | 4.58E-01 | 0.5 | 1.57E-01 | 0.4 | 2.51E-01 | 0.0 | 9.51E-01 | RAS associated with diabetes protein 51C |
| AT1G16700 | -0.1 | 2.59E-01 | 0.0 | 8.62E-01 | -0.1 | 2.41E-01 | 0.1 | 5.58E-01 | 0.0 | 8.46E-01 | 0.1 | 8.48E-01 | NADH dehydrogenase iron-sulfur protein 8-B, mitochondrial |
| AT4G04950 | -0.1 | 2.48E-01 | -0.2 | 3.74E-02 | 0.1 | 3.12E-01 | 0.0 | 9.77E-01 | -0.1 | 4.65E-01 | 0.1 | 5.37E-01 | Monothiol glutaredoxin-S17 |
| AT3G21270 | -0.1 | 4.89E-01 | 0.1 | 2.99E-01 | -0.2 | 1.44E-01 | 1.0 | 5.26E-06 | 0.0 | 9.45E-01 | 1.0 | 2.93E-04 | Dof zinc finger protein DOF3.1 |
| AT2G25170 | -0.1 | 2.92E-01 | -0.2 | 4.45E-02 | 0.1 | 2.96E-01 | -0.1 | 6.71E-01 | 0.0 | 9.59E-01 | -0.1 | 7.49E-01 | Chromatin remodeling factor CHD3 |
| AT1G33980 | -0.1 | 2.32E-01 | -0.2 | 1.00E-02 | 0.1 | 1.06E-01 | 0.0 | 1.00E+00 | -0.1 | 7.16E-01 | 0.1 | 7.64E-01 | Regulator of nonsense transcripts UPF3 |
| AT4G37580 | -0.1 | 6.59E-01 | 0.0 | 9.64E-01 | -0.1 | 7.66E-01 | 0.1 | 7.67E-01 | -0.4 | 1.14E-01 | 0.5 | 1.00E-01 | Acyl-CoA N-acyltransferases superfamily protein |
| AT5G23540 | -0.1 | 5.21E-01 | -0.1 | 4.22E-01 | 0.0 | 8.71E-01 | 0.1 | 6.13E-01 | 0.0 | 8.65E-01 | 0.1 | 5.80E-01 | Mov34 |
| AT1G77460 | -0.1 | 4.54E-01 | -0.1 | 3.42E-01 | 0.0 | 8.52E-01 | 0.2 | 3.61E-01 | 0.1 | 8.42E-01 | 0.1 | 6.87E-01 | C2 domain-containing protein |
| AT5G35410 | -0.1 | 5.64E-01 | -0.1 | 5.18E-01 | 0.0 | 9.44E-01 | -0.1 | 8.59E-01 | -0.2 | 5.66E-01 | 0.1 | 7.49E-01 | non-specific serine |
| AT3G10190 | -0.1 | 4.22E-01 | -0.1 | 2.70E-01 | 0.0 | 7.82E-01 | -0.4 | 9.10E-02 | -0.3 | 1.91E-01 | -0.1 | 9.04E-01 | Probable calcium-binding protein CML36 |
| AT4G00570 | -0.1 | 2.12E-01 | -0.1 | 3.42E-02 | 0.1 | 3.14E-01 | 0.2 | 4.21E-01 | 0.1 | 7.55E-01 | 0.1 | 8.32E-01 | Malic enzyme |
| AT3G18830 | -0.1 | 5.52E-01 | -0.1 | 4.10E-01 | 0.0 | 8.22E-01 | -0.3 | 9.97E-03 | -0.3 | 1.35E-02 | 0.0 | 9.41E-01 | Polyol transporter 5 |
| AT5G35735 | -0.1 | 5.95E-01 | -0.1 | 3.13E-01 | 0.1 | 6.54E-01 | 0.6 | 2.85E-05 | 0.0 | 8.61E-01 | 0.6 | 1.41E-03 | Cytochrome b561 and DOMON domain-containing protein At5g35735 |
| AT4G39520 | -0.1 | 3.28E-01 | 0.0 | 5.53E-01 | 0.0 | 7.65E-01 | 0.2 | 5.86E-02 | 0.0 | 8.94E-01 | 0.2 | 2.45E-01 | Developmentally-regulated G-protein 3 |
| AT4G21903 | -0.1 | 5.64E-01 | -0.6 | 9.99E-04 | 0.6 | 5.56E-03 | 1.0 | 4.23E-04 | 0.8 | 4.31E-03 | 0.2 | 5.16E-01 | Protein DETOXIFICATION |
| AT5G65960 | -0.1 | 3.38E-01 | -0.1 | 2.74E-01 | 0.0 | 8.89E-01 | -0.1 | 3.83E-01 | -0.2 | 1.89E-01 | 0.1 | 7.27E-01 | Uncharacterized protein |
| AT5G66730 | -0.1 | 3.79E-01 | -0.4 | 6.87E-04 | 0.3 | 4.41E-03 | -0.2 | 7.34E-02 | -0.3 | 6.70E-02 | 0.0 | 9.00E-01 | Zinc finger protein ENHYDROUS |
| AT3G02250 | -0.1 | 5.74E-01 | -0.1 | 4.01E-01 | 0.0 | 8.01E-01 | -0.1 | 6.35E-01 | 0.1 | 4.63E-01 | -0.2 | 3.04E-01 | Rhamnogalacturonan I rhamnosyltransferase 1 |
| AT3G10760 | -0.1 | 3.13E-01 | -0.1 | 2.11E-01 | 0.0 | 8.08E-01 | 0.0 | 7.60E-01 | 0.0 | 8.99E-01 | -0.1 | 6.87E-01 | hypothetical protein |
| AT5G22400 | -0.1 | 4.26E-01 | 0.1 | 2.03E-01 | -0.2 | 8.70E-02 | 0.1 | 7.69E-01 | 0.0 | 9.00E-01 | 0.0 | 9.39E-01 | Rho GTPase-activating protein 1 |
| AT4G02550 | -0.1 | 5.78E-01 | -0.2 | 2.38E-01 | 0.1 | 5.89E-01 | 0.3 | 1.10E-01 | -0.1 | 7.25E-01 | 0.4 | 9.34E-02 | Myb |
| AT1G10600 | -0.1 | 4.11E-01 | 0.0 | 9.96E-01 | -0.1 | 5.06E-01 | -0.2 | 3.34E-01 | -0.1 | 7.36E-01 | -0.1 | 7.53E-01 | AMSH-like ubiquitin thioesterase 2 |
| AT2G21430 | -0.1 | 5.78E-01 | 0.0 | 8.84E-01 | -0.1 | 5.73E-01 | -0.2 | 6.53E-01 | 0.6 | 2.27E-01 | -0.8 | 9.21E-02 | Probable cysteine protease RD19B |

| | | | | | | | | | | | | | |
|---|---|---|---|---|---|---|---|---|---|---|---|---|---|
| AT3G54130 | -0.1 | 3.45E-01 | 0.1 | 4.70E-01 | -0.1 | 1.63E-01 | -0.1 | 7.56E-01 | 0.1 | 7.44E-01 | -0.2 | 5.87E-01 | ubiquitinyl hydrolase 1 |
| AT4G03550 | -0.1 | 2.77E-01 | -0.1 | 1.94E-01 | 0.0 | 8.24E-01 | 0.1 | 3.32E-01 | 0.0 | 8.73E-01 | 0.2 | 3.94E-01 | Callose synthase 12 |
| AT5G58380 | -0.1 | 3.06E-01 | -0.2 | 4.60E-02 | 0.1 | 3.15E-01 | 0.1 | 7.37E-01 | -0.1 | 7.50E-01 | 0.2 | 5.38E-01 | non-specific serine |
| AT3G49390 | -0.1 | 3.24E-01 | 0.0 | 9.11E-01 | -0.1 | 3.32E-01 | 0.2 | 1.45E-01 | 0.2 | 2.08E-01 | 0.0 | 9.95E-01 | Polyadenylate-binding protein-interacting protein 10 -binding protein-interacting protein 10) |
| AT1G78870 | -0.1 | 2.78E-01 | 0.0 | 4.86E-01 | 0.0 | 7.64E-01 | 0.0 | 9.55E-01 | 0.0 | 9.19E-01 | 0.0 | 9.65E-01 | Ubiquitin-conjugating enzyme 35 |
| AT4G01810 | -0.1 | 3.53E-01 | -0.1 | 8.18E-02 | 0.1 | 4.09E-01 | 0.2 | 2.65E-01 | 0.0 | 9.68E-01 | 0.2 | 4.48E-01 | Protein transport protein SEC23 A |
| AT3G22980 | -0.1 | 4.05E-01 | -0.1 | 1.25E-01 | 0.1 | 5.05E-01 | 0.3 | 8.48E-02 | 0.0 | 8.42E-01 | 0.2 | 2.52E-01 | Tr-type G domain-containing protein |
| AT4G04970 | -0.1 | 4.03E-01 | 0.1 | 3.91E-01 | -0.1 | 1.45E-01 | 0.1 | 4.02E-01 | 0.1 | 6.75E-01 | 0.0 | 8.74E-01 | Callose synthase 11 |
| AT1G10290 | -0.1 | 2.80E-01 | -0.2 | 1.42E-02 | 0.1 | 1.12E-01 | 0.1 | 7.29E-01 | 0.1 | 4.47E-01 | -0.1 | 7.54E-01 | AT1G10290 protein |
| AT3G07520 | -0.1 | 7.44E-01 | -0.5 | 6.08E-02 | 0.4 | 1.61E-01 | -0.2 | 5.68E-01 | -0.3 | 3.02E-01 | 0.2 | 6.89E-01 | Glutamate receptor |
| AT4G01320 | -0.1 | 4.13E-01 | 0.1 | 3.40E-01 | -0.2 | 1.20E-01 | 0.2 | 3.25E-02 | 0.0 | 8.30E-01 | 0.2 | 1.99E-01 | CAAX prenyl protease |
| AT4G15130 | -0.1 | 4.25E-01 | -0.3 | 1.33E-02 | 0.2 | 8.18E-02 | 0.2 | 2.32E-01 | 0.1 | 6.90E-01 | 0.1 | 6.72E-01 | Choline-phosphate cytidylyltransferase 2 |
| AT3G49720 | -0.1 | 3.90E-01 | 0.2 | 4.20E-02 | -0.3 | 1.52E-02 | -0.1 | 3.92E-01 | 0.1 | 4.09E-01 | -0.2 | 1.87E-01 | Probable pectin methylesterase CGR2 |
| AT5G56740 | -0.1 | 5.15E-01 | 0.0 | 7.66E-01 | -0.1 | 4.15E-01 | 0.0 | 9.41E-01 | 0.0 | 8.67E-01 | 0.1 | 8.21E-01 | Histone acetyltransferase type B catalytic subunit |
| AT4G16650 | -0.1 | 2.81E-01 | -0.1 | 7.61E-02 | 0.1 | 4.72E-01 | -0.3 | 1.82E-01 | -0.1 | 7.31E-01 | -0.2 | 5.81E-01 | O-fucosyltransferase 29 |
| AT4G36195 | -0.1 | 2.44E-01 | -0.3 | 6.64E-04 | 0.2 | 5.74E-03 | -0.1 | 7.13E-01 | 0.1 | 6.17E-01 | -0.1 | 4.71E-01 | Prolyl carboxypeptidase like protein |
| AT5G15830 | -0.1 | 6.88E-01 | 0.1 | 7.03E-01 | -0.1 | 5.14E-01 | 0.3 | 4.92E-01 | -0.3 | 5.86E-01 | 0.5 | 2.92E-01 | BZIP3 |
| AT1G73730 | -0.1 | 3.36E-01 | -0.3 | 4.98E-03 | 0.2 | 4.19E-02 | 0.1 | 6.75E-01 | 0.1 | 6.19E-01 | 0.0 | 9.27E-01 | ETHYLENE-INSENSITIVE3-like 3 |
| AT1G73440 | -0.1 | 5.67E-01 | 0.0 | 7.21E-01 | 0.0 | 8.90E-01 | -0.3 | 2.74E-01 | -0.3 | 3.48E-01 | 0.0 | 9.83E-01 | At1g73440 |
| AT4G30480 | -0.1 | 4.24E-01 | -0.1 | 2.82E-01 | 0.0 | 7.96E-01 | -0.1 | 4.59E-01 | 0.0 | 8.92E-01 | -0.2 | 4.86E-01 | AT4G30480 protein -like superfamily protein) |
| AT3G10220 | -0.1 | 4.11E-01 | 0.0 | 6.68E-01 | -0.1 | 2.86E-01 | -0.4 | 1.06E-02 | -0.3 | 6.22E-02 | -0.1 | 6.81E-01 | Tubulin-folding cofactor B |
| AT1G27370 | -0.1 | 3.69E-01 | -0.2 | 1.88E-02 | 0.1 | 1.33E-01 | 0.3 | 2.07E-01 | 0.1 | 6.80E-01 | 0.2 | 6.08E-01 | Squamosa promoter binding protein-like 10 |
| AT5G08780 | -0.1 | 5.37E-01 | 0.0 | 9.87E-01 | -0.1 | 6.16E-01 | 0.1 | 8.13E-01 | 0.0 | 9.64E-01 | 0.1 | 8.30E-01 | At5g08780 |
| AT3G62600 | -0.1 | 5.99E-01 | -0.1 | 6.17E-01 | 0.0 | 9.99E-01 | 0.4 | 4.07E-03 | 0.3 | 3.84E-02 | 0.1 | 6.65E-01 | DnaJ protein ERDJ3B |
| AT2G39720 | -0.1 | 2.77E-01 | -0.2 | 6.39E-03 | 0.1 | 5.51E-02 | -0.4 | 3.34E-03 | -0.3 | 3.38E-02 | -0.1 | 6.51E-01 | RING-type E3 ubiquitin transferase |
| AT1G45976 | -0.1 | 5.84E-01 | -0.3 | 2.18E-02 | 0.3 | 8.74E-02 | -0.7 | 1.87E-03 | -0.7 | 2.92E-03 | 0.0 | 9.63E-01 | SBP1 |
| AT5G13160 | -0.1 | 3.29E-01 | -0.1 | 9.44E-02 | 0.1 | 4.84E-01 | -0.2 | 2.65E-01 | -0.1 | 6.35E-01 | -0.1 | 7.53E-01 | Serine |
| AT1G58470 | -0.1 | 5.41E-01 | -0.4 | 9.32E-03 | 0.3 | 5.06E-02 | -0.2 | 4.99E-01 | -0.2 | 6.83E-01 | -0.1 | 9.12E-01 | RNA-binding protein 1 |
| AT2G26140 | -0.1 | 2.71E-01 | -0.1 | 1.08E-01 | 0.0 | 5.87E-01 | 0.2 | 1.66E-01 | 0.2 | 1.57E-01 | 0.0 | 9.10E-01 | ATP-dependent zinc metalloprotease FTSH 4, mitochondrial |
| AT4G01130 | -0.1 | 7.56E-01 | 0.8 | 2.51E-03 | -0.8 | 5.40E-03 | -0.6 | 1.22E-02 | -0.3 | 3.03E-01 | -0.3 | 2.36E-01 | GDSL-like Lipase |
| AT5G17510 | -0.1 | 2.77E-01 | -0.2 | 2.76E-03 | 0.2 | 2.76E-02 | 0.3 | 8.92E-02 | 0.3 | 9.99E-02 | 0.0 | 9.53E-01 | Glutamine-rich protein |
| AT2G40980 | -0.1 | 3.25E-01 | -0.1 | 4.37E-01 | 0.0 | 8.94E-01 | -0.1 | 4.93E-01 | -0.1 | 5.35E-01 | 0.0 | 9.86E-01 | Protein kinase superfamily protein |
| AT3G15290 | -0.1 | 3.48E-01 | 0.0 | 8.17E-01 | -0.1 | 3.16E-01 | -0.5 | 4.94E-04 | -0.4 | 6.30E-03 | -0.1 | 5.22E-01 | hypothetical protein |
| AT5G43910 | -0.1 | 6.23E-01 | -0.2 | 2.83E-01 | 0.1 | 6.08E-01 | 0.1 | 5.69E-01 | 0.1 | 8.32E-01 | 0.1 | 8.62E-01 | PfkB-like carbohydrate kinase family protein |
| AT1G10940 | -0.1 | 4.82E-01 | -0.2 | 8.69E-02 | 0.1 | 3.08E-01 | 0.0 | 7.95E-01 | 0.0 | 8.25E-01 | 0.0 | 9.97E-01 | Protein kinase superfamily protein |
| AT1G16570 | -0.1 | 4.10E-01 | -0.1 | 5.27E-01 | 0.0 | 9.06E-01 | 0.1 | 5.03E-01 | 0.1 | 6.32E-01 | 0.0 | 9.60E-01 | UDP-glycosyltransferase TURAN |
| AT1G14740 | -0.1 | 3.64E-01 | -0.1 | 6.91E-02 | 0.1 | 3.37E-01 | -0.2 | 1.14E-01 | -0.5 | 4.97E-03 | 0.2 | 2.14E-01 | Protein OBERON 3 |
| AT3G42670 | -0.1 | 5.63E-01 | 0.1 | 5.88E-01 | -0.1 | 3.46E-01 | -0.1 | 7.74E-01 | 0.2 | 6.64E-01 | -0.3 | 5.39E-01 | Chromatin remodeling 38 |
| AT2G33470 | -0.1 | 3.33E-01 | 0.2 | 1.88E-02 | -0.3 | 6.96E-03 | -0.1 | 7.65E-01 | 0.0 | 8.83E-01 | -0.1 | 7.27E-01 | Glycolipid transfer protein 1 |
| AT3G57280 | -0.1 | 3.66E-01 | 0.0 | 7.30E-01 | -0.1 | 2.69E-01 | 0.1 | 3.40E-01 | 0.2 | 2.54E-01 | 0.0 | 8.37E-01 | Protein FATTY ACID EXPORT 1, chloroplastic |
| AT1G04430 | -0.1 | 3.56E-01 | 0.0 | 9.87E-01 | -0.1 | 4.24E-01 | -0.4 | 3.14E-03 | -0.2 | 2.04E-01 | -0.2 | 2.41E-01 | Probable methyltransferase PMT8 |
| AT5G57580 | -0.1 | 3.71E-01 | -0.4 | 5.26E-04 | 0.3 | 3.20E-03 | -0.1 | 3.99E-01 | -0.3 | 2.77E-02 | 0.2 | 2.52E-01 | Calmodulin-binding protein 60 B |
| AT2G02390 | -0.1 | 3.72E-01 | -0.3 | 1.33E-03 | 0.3 | 9.05E-03 | 0.2 | 5.28E-02 | 0.1 | 5.87E-01 | 0.2 | 3.97E-01 | Glutathione S-transferase zeta 1 |
| AT3G21640 | -0.1 | 5.50E-01 | -0.1 | 6.19E-01 | 0.0 | 9.54E-01 | 0.0 | 9.68E-01 | -0.1 | 4.58E-01 | 0.1 | 5.81E-01 | Peptidyl-prolyl cis-trans isomerase FKBP42 |
| AT5G10170 | -0.1 | 5.16E-01 | 0.4 | 1.34E-03 | -0.5 | 1.31E-03 | 0.1 | 5.68E-01 | 0.2 | 2.11E-01 | -0.1 | 5.73E-01 | Probable inositol 3-phosphate synthase isozyme 3 |
| AT2G21240 | -0.1 | 4.48E-01 | -0.2 | 4.92E-02 | 0.1 | 2.20E-01 | -0.5 | 6.44E-04 | -0.5 | 1.52E-03 | 0.0 | 9.80E-01 | Protein BASIC PENTACYSTEINE4 |
| AT1G74680 | -0.1 | 3.89E-01 | -0.1 | 2.99E-01 | 0.0 | 8.64E-01 | -0.1 | 5.25E-01 | -0.1 | 5.12E-01 | 0.0 | 9.55E-01 | hypothetical protein |
| AT1G04040 | -0.1 | 5.05E-01 | 0.3 | 1.32E-02 | -0.4 | 8.28E-03 | -0.2 | 5.70E-01 | 0.0 | 9.49E-01 | -0.2 | 6.51E-01 | At1g04040 |
| AT2G21300 | -0.1 | 3.67E-01 | -0.2 | 2.51E-02 | 0.1 | 1.53E-01 | -0.1 | 4.74E-01 | 0.0 | 9.27E-01 | -0.1 | 7.15E-01 | Kinesin-like protein KIN-7E |
| AT5G03415 | -0.1 | 3.80E-01 | -0.1 | 1.37E-01 | 0.1 | 5.63E-01 | 0.1 | 6.60E-01 | 0.0 | 9.81E-01 | 0.1 | 7.32E-01 | Transcription factor-like protein DPB |
| AT1G75990 | -0.1 | 4.27E-01 | 0.0 | 9.12E-01 | -0.1 | 4.27E-01 | 0.0 | 8.27E-01 | 0.1 | 4.72E-01 | -0.2 | 4.25E-01 | 26S proteasome non-ATPase regulatory subunit 3 homolog B |
| AT1G26370 | -0.1 | 4.15E-01 | -0.1 | 1.37E-01 | 0.1 | 5.21E-01 | 0.1 | 6.61E-01 | 0.0 | 8.96E-01 | 0.1 | 6.46E-01 | Pre-mRNA-splicing factor ATP-dependent RNA helicase DEAH10 |
| AT5G44460 | -0.1 | 5.90E-01 | 0.3 | 4.51E-02 | -0.3 | 3.97E-02 | 0.5 | 1.08E-01 | 0.6 | 3.38E-02 | -0.2 | 5.83E-01 | Probable calcium-binding protein CML43 |
| AT5G13440 | -0.1 | 4.22E-01 | 0.0 | 9.98E-01 | -0.1 | 4.82E-01 | 0.3 | 2.09E-02 | 0.2 | 1.37E-01 | 0.1 | 6.76E-01 | Cytochrome b-c1 complex subunit Rieske-2, mitochondrial |
| AT4G29120 | -0.1 | 4.92E-01 | 0.0 | 8.84E-01 | -0.1 | 4.62E-01 | -0.1 | 5.43E-01 | -0.1 | 6.35E-01 | 0.0 | 9.84E-01 | Probable 3-hydroxyisobutyrate dehydrogenase-like 1, mitochondrial |
| AT2G25970 | -0.1 | 3.92E-01 | -0.2 | 3.07E-02 | 0.1 | 1.58E-01 | 0.1 | 7.94E-01 | 0.1 | 8.56E-01 | 0.0 | 9.89E-01 | F17H15.1 |
| AT5G66320 | -0.1 | 5.31E-01 | -0.2 | 1.22E-01 | 0.1 | 3.84E-01 | 0.0 | 9.49E-01 | -0.3 | 1.24E-01 | 0.3 | 1.92E-01 | GATA transcription factor 5 |
| AT4G24520 | -0.1 | 3.89E-01 | -0.1 | 1.76E-01 | 0.0 | 6.18E-01 | -0.2 | 5.84E-02 | -0.2 | 1.00E-01 | 0.0 | 9.86E-01 | NADPH--cytochrome P450 reductase |
| AT5G59670 | -0.1 | 8.51E-01 | -0.1 | 7.47E-01 | 0.1 | 9.08E-01 | 0.9 | 5.60E-04 | 0.6 | 2.06E-02 | 0.3 | 2.93E-01 | Receptor-like protein kinase At5g59670 |
| AT2G37230 | -0.1 | 5.45E-01 | 0.1 | 3.17E-01 | -0.2 | 1.62E-01 | 0.3 | 1.04E-01 | 0.2 | 4.71E-01 | 0.1 | 6.45E-01 | Large ribosomal subunit protein mL102 |
| AT1G01110 | -0.1 | 7.03E-01 | 0.3 | 8.11E-02 | -0.4 | 7.95E-02 | -0.3 | 1.28E-01 | -0.2 | 3.07E-01 | -0.1 | 8.21E-01 | Protein IQ-DOMAIN 18 |
| AT5G62760 | -0.1 | 4.71E-01 | -0.2 | 6.82E-02 | 0.1 | 2.95E-01 | -0.1 | 7.47E-01 | -0.1 | 7.07E-01 | 0.0 | 9.47E-01 | P-loop containing nucleoside triphosphate hydrolases superfamily protein |
| AT3G02770 | -0.1 | 4.78E-01 | 0.0 | 7.06E-01 | 0.0 | 8.03E-01 | 0.0 | 9.63E-01 | -0.2 | 3.68E-01 | 0.2 | 3.80E-01 | Putative 4-hydroxy-4-methyl-2-oxoglutarate aldolase 1 |
| AT5G03290 | -0.1 | 3.51E-01 | 0.1 | 4.09E-01 | -0.1 | 1.18E-01 | 0.2 | 1.35E-01 | 0.0 | 9.72E-01 | 0.2 | 2.99E-01 | Isocitrate dehydrogenase catalytic subunit 5, mitochondrial -specific ICDH 5) |
| AT4G24730 | -0.1 | 4.49E-01 | 0.1 | 2.61E-01 | -0.2 | 1.20E-01 | -0.4 | 4.04E-02 | -0.2 | 2.17E-01 | -0.1 | 6.49E-01 | Manganese-dependent ADP-ribose |
| AT4G14560 | -0.1 | 6.79E-01 | 0.1 | 4.30E-01 | -0.2 | 3.12E-01 | 0.2 | 4.42E-01 | 0.2 | 4.55E-01 | 0.0 | 9.65E-01 | Auxin-responsive protein IAA1 |
| AT5G41070 | -0.1 | 7.63E-01 | -0.4 | 6.36E-02 | 0.4 | 1.56E-01 | 0.1 | 7.38E-01 | 0.2 | 3.90E-01 | -0.1 | 6.47E-01 | Double-stranded RNA-binding protein 5 |
| AT2G24690 | -0.1 | 5.53E-01 | -0.2 | 9.42E-02 | 0.1 | 3.15E-01 | -0.1 | 7.61E-01 | -0.2 | 4.71E-01 | 0.1 | 7.58E-01 | Transcriptional factor B3 family protein |
| AT4G19020 | -0.1 | 5.72E-01 | 0.0 | 7.45E-01 | 0.0 | 8.65E-01 | -0.2 | 6.32E-01 | 0.1 | 8.38E-01 | -0.2 | 5.30E-01 | DNA -methyltransferase |
| AT5G33320 | -0.1 | 4.51E-01 | 0.1 | 1.11E-01 | -0.2 | 4.16E-02 | 0.3 | 2.42E-03 | 0.4 | 3.98E-03 | 0.0 | 9.37E-01 | Phosphoenolpyruvate |
| AT3G17330 | -0.1 | 4.59E-01 | 0.0 | 7.39E-01 | 0.0 | 7.54E-01 | 0.2 | 4.25E-01 | 0.3 | 2.00E-01 | -0.1 | 6.68E-01 | YTH domain-containing family protein |
| AT2G44980 | -0.1 | 6.19E-01 | -0.4 | 1.51E-02 | 0.3 | 6.27E-02 | -0.1 | 7.52E-01 | -0.2 | 5.17E-01 | 0.1 | 7.85E-01 | SNF2 domain-containing protein |
| AT1G44350 | -0.1 | 8.22E-01 | 0.2 | 3.97E-01 | -0.3 | 3.66E-01 | -0.4 | 4.35E-01 | -0.3 | 6.53E-01 | -0.1 | 9.07E-01 | IAA-amino acid hydrolase ILR1-like 6 |
| AT2G04560 | -0.1 | 6.34E-01 | 0.0 | 7.36E-01 | 0.0 | 9.37E-01 | -0.2 | 5.97E-01 | -0.1 | 8.37E-01 | -0.1 | 8.63E-01 | Probable lipid-A-disaccharide synthase, mitochondrial |
| AT3G58580 | -0.1 | 3.60E-01 | -0.2 | 1.87E-02 | 0.1 | 1.25E-01 | 0.1 | 7.87E-01 | 0.3 | 2.06E-01 | -0.3 | 4.52E-01 | Carbon catabolite repressor protein 4 homolog 2 |
| AT4G29720 | -0.1 | 4.49E-01 | 0.0 | 7.84E-01 | 0.0 | 7.05E-01 | -0.3 | 9.65E-02 | 0.0 | 9.68E-01 | -0.3 | 2.00E-01 | Probable polyamine oxidase 5 |
| AT5G22070 | -0.1 | 4.67E-01 | 0.1 | 1.37E-01 | -0.2 | 6.47E-02 | 0.0 | 8.31E-01 | 0.0 | 9.92E-01 | 0.0 | 8.80E-01 | Core-2 |
| AT4G19880 | -0.1 | 4.60E-01 | 0.1 | 2.52E-01 | -0.2 | 9.66E-02 | 0.4 | 2.49E-03 | 0.0 | 7.98E-01 | 0.5 | 1.17E-02 | Glutathione S-transferase family protein |
| AT4G32890 | -0.1 | 5.44E-01 | 0.5 | 7.98E-04 | -0.5 | 1.37E-03 | -0.4 | 1.21E-01 | 0.0 | 9.70E-01 | -0.4 | 2.27E-01 | GATA transcription factor 9 |
| AT5G62070 | -0.1 | 5.79E-01 | -0.3 | 3.35E-02 | 0.2 | 1.20E-01 | 0.5 | 2.30E-03 | 0.4 | 1.17E-02 | 0.1 | 8.07E-01 | Protein IQ-DOMAIN 23 |
| AT1G70740 | -0.1 | 4.91E-01 | -0.3 | 9.92E-03 | 0.2 | 5.40E-02 | 0.1 | 6.89E-01 | 0.4 | 2.04E-01 | -0.3 | 5.04E-01 | Protein kinase domain-containing protein |
| AT2G18380 | -0.1 | 6.68E-01 | 0.1 | 5.78E-01 | -0.1 | 4.14E-01 | 0.1 | 8.84E-01 | 0.1 | 8.37E-01 | 0.0 | 9.54E-01 | GATA transcription factor 20 |
| AT2G01630 | -0.1 | 4.22E-01 | 0.0 | 9.52E-01 | -0.1 | 4.63E-01 | 0.1 | 3.59E-01 | 0.1 | 7.14E-01 | 0.1 | 7.97E-01 | glucan endo-1,3-beta-D-glucosidase |

| | | | | | | | | | | | | | |
|---|---|---|---|---|---|---|---|---|---|---|---|---|---|
| AT1G73750 | -0.1 | 6.86E-01 | 0.0 | 9.46E-01 | -0.1 | 6.91E-01 | -0.2 | 7.68E-01 | 0.3 | 6.04E-01 | -0.5 | 4.68E-01 | Alpha |
| AT1G73930 | -0.1 | 3.35E-01 | 0.0 | 6.46E-01 | -0.1 | 2.19E-01 | 0.0 | 9.01E-01 | 0.3 | 3.03E-01 | -0.2 | 4.66E-01 | AT1G73930 protein |
| AT1G70570 | -0.1 | 3.04E-01 | -0.1 | 3.31E-01 | 0.0 | 9.87E-01 | 0.0 | 9.11E-01 | 0.1 | 3.49E-01 | -0.1 | 4.03E-01 | Anthranilate phosphoribosyltransferase |
| AT1G54580 | -0.1 | 5.96E-01 | 0.2 | 1.38E-01 | -0.2 | 7.20E-02 | 0.2 | 2.36E-01 | 0.2 | 3.46E-01 | 0.0 | 9.75E-01 | Acyl carrier protein 2, chloroplastic |
| AT2G01610 | -0.1 | 5.73E-01 | 0.1 | 3.64E-01 | -0.2 | 2.16E-01 | -0.4 | 6.26E-02 | 0.0 | 9.24E-01 | -0.4 | 1.89E-01 | Pectinesterase inhibitor domain-containing protein |
| AT5G02860 | -0.1 | 5.70E-01 | 0.1 | 1.82E-01 | -0.2 | 1.09E-01 | 0.2 | 4.08E-01 | 0.0 | 9.91E-01 | 0.2 | 5.28E-01 | Pentatricopeptide repeat-containing protein At5g02860 |
| AT1G29310 | -0.1 | 5.70E-01 | 0.2 | 4.78E-02 | -0.3 | 2.84E-02 | 0.2 | 7.06E-02 | 0.2 | 7.67E-02 | 0.0 | 9.27E-01 | SecY protein transport family protein |
| AT3G47690 | -0.1 | 6.87E-01 | 0.2 | 2.62E-01 | -0.2 | 2.07E-01 | 0.4 | 1.46E-01 | 0.5 | 1.30E-01 | -0.1 | 8.85E-01 | Microtubule-associated protein RP |
| AT5G49665 | -0.1 | 6.01E-01 | 0.0 | 9.14E-01 | 0.0 | 7.45E-01 | 0.3 | 1.62E-01 | 0.3 | 1.69E-01 | 0.0 | 9.40E-01 | E3 ubiquitin-protein ligase WAV3 |
| AT1G53470 | -0.1 | 7.13E-01 | 0.4 | 2.37E-02 | -0.4 | 3.03E-02 | 0.1 | 9.06E-01 | 0.5 | 2.16E-01 | -0.4 | 3.25E-01 | Mechanosensitive ion channel protein 4 |
| AT5G56590 | -0.1 | 6.13E-01 | 0.1 | 2.12E-01 | -0.2 | 1.44E-01 | -0.4 | 1.85E-02 | -0.3 | 6.90E-02 | -0.1 | 8.02E-01 | Glucan endo-1,3-beta-glucosidase 13 -beta-glucan endohydrolase 13) -beta-glucanase 13) |
| AT3G50960 | -0.1 | 6.16E-01 | 0.2 | 6.63E-02 | -0.3 | 5.42E-02 | 0.1 | 8.72E-01 | 0.0 | 9.71E-01 | 0.0 | 9.41E-01 | Thioredoxin domain-containing protein PLP3A |
| AT2G25140 | -0.1 | 4.44E-01 | -0.3 | 2.60E-03 | 0.2 | 1.67E-02 | 0.7 | 2.73E-03 | 0.2 | 4.33E-01 | 0.5 | 2.42E-02 | Chaperone protein ClpB4, mitochondrial |
| AT3G24860 | -0.1 | 6.60E-01 | -0.2 | 1.50E-01 | 0.1 | 3.79E-01 | -0.1 | 6.85E-01 | 0.0 | 9.30E-01 | -0.1 | 8.26E-01 | Homeodomain-like superfamily protein |
| AT1G43850 | -0.1 | 7.48E-01 | -0.6 | 9.96E-03 | 0.5 | 3.11E-02 | -0.7 | 6.28E-01 | -0.2 | 9.03E-01 | -0.5 | 8.29E-01 | Transcriptional corepressor SEUSS |
| AT1G71440 | -0.1 | 4.39E-01 | -0.1 | 7.48E-02 | 0.1 | 3.29E-01 | -0.2 | 5.31E-01 | -0.1 | 7.38E-01 | -0.1 | 8.97E-01 | Tubulin-folding cofactor E |
| AT4G19460 | -0.1 | 5.87E-01 | -0.1 | 5.54E-01 | 0.0 | 9.53E-01 | 0.2 | 6.86E-01 | -0.1 | 8.43E-01 | 0.3 | 6.21E-01 | Glycosyltransferase |
| AT5G36160 | -0.1 | 6.68E-01 | -0.5 | 6.16E-03 | 0.4 | 1.94E-02 | -0.3 | 3.83E-02 | 0.1 | 6.15E-01 | -0.4 | 3.99E-02 | Tyrosine aminotransferase |
| AT5G17410 | -0.1 | 4.80E-01 | -0.1 | 4.28E-01 | 0.0 | 9.28E-01 | -0.2 | 2.31E-01 | -0.1 | 7.19E-01 | -0.1 | 6.35E-01 | Gamma-tubulin complex component |
| AT3G28140 | -0.1 | 5.63E-01 | 0.0 | 7.38E-01 | -0.1 | 4.44E-01 | -0.3 | 3.24E-01 | -0.1 | 6.96E-01 | -0.1 | 7.61E-01 | RNA ligase |
| AT4G28470 | -0.1 | 4.06E-01 | 0.0 | 9.08E-01 | 0.0 | 5.42E-01 | 0.1 | 3.72E-01 | 0.1 | 7.64E-01 | 0.1 | 7.80E-01 | 26S proteasome non-ATPase regulatory subunit 2 homolog B |
| AT5G19280 | -0.1 | 4.01E-01 | -0.1 | 2.33E-01 | 0.0 | 7.42E-01 | -0.1 | 4.60E-01 | -0.2 | 3.11E-01 | 0.1 | 8.19E-01 | protein-serine |
| AT5G58190 | -0.1 | 4.19E-01 | -0.1 | 1.88E-01 | 0.0 | 6.37E-01 | 0.1 | 6.64E-01 | 0.2 | 4.19E-01 | -0.1 | 7.79E-01 | YTH domain-containing family protein |
| AT5G08120 | -0.1 | 5.93E-01 | 0.0 | 8.52E-01 | -0.1 | 5.38E-01 | -0.2 | 2.46E-01 | 0.0 | 8.80E-01 | -0.1 | 4.71E-01 | Protein MICROTUBULE BINDING PROTEIN 2C |
| AT2G40190 | -0.1 | 5.25E-01 | 0.0 | 6.26E-01 | 0.0 | 9.28E-01 | 0.1 | 5.52E-01 | 0.1 | 4.97E-01 | 0.0 | 9.23E-01 | GDP-Man:ManGlcNAc-PP-Dol alpha-1,2-mannosyltransferase |
| AT1G75420 | -0.1 | 5.61E-01 | -0.1 | 4.95E-01 | 0.0 | 9.23E-01 | 0.0 | 9.59E-01 | -0.1 | 7.78E-01 | 0.1 | 7.57E-01 | F1B16.5 protein |
| AT2G24100 | -0.1 | 5.96E-01 | -0.4 | 4.91E-03 | 0.3 | 2.22E-02 | -0.3 | 3.02E-01 | -0.6 | 4.02E-02 | 0.4 | 2.87E-01 | ATP-dependent DNA helicase |
| AT1G23490 | -0.1 | 5.55E-01 | 0.2 | 1.25E-01 | -0.2 | 6.00E-02 | 0.0 | 9.90E-01 | -0.2 | 1.93E-01 | 0.2 | 2.94E-01 | ADP-ribosylation factor 2-A |
| AT2G23093 | -0.1 | 6.32E-01 | -0.1 | 2.39E-01 | 0.1 | 5.50E-01 | -0.1 | 7.79E-01 | 0.2 | 7.06E-01 | -0.3 | 5.50E-01 | Molybdate-anion transporter |
| AT1G56340 | -0.1 | 6.39E-01 | 0.2 | 1.74E-01 | -0.2 | 9.95E-02 | 0.3 | 1.92E-02 | 0.2 | 1.18E-01 | 0.1 | 7.34E-01 | Calreticulin |
| AT5G67320 | -0.1 | 4.39E-01 | 0.0 | 9.49E-01 | -0.1 | 5.45E-01 | 0.0 | 9.72E-01 | 0.0 | 9.40E-01 | 0.0 | 9.37E-01 | WD40 repeat-containing protein HOS15 |
| AT1G57700 | -0.1 | 5.62E-01 | -0.1 | 3.40E-01 | 0.0 | 7.48E-01 | 0.1 | 7.40E-01 | 0.2 | 4.10E-01 | -0.1 | 6.76E-01 | Protein kinase superfamily protein |
| AT1G16900 | -0.1 | 5.30E-01 | 0.0 | 6.24E-01 | -0.1 | 3.33E-01 | -0.1 | 5.02E-01 | 0.0 | 9.76E-01 | -0.2 | 5.67E-01 | Dol-P-Man:ManGlcNAc-PP-Dol alpha-1,2-mannosyltransferase GlcNAc-PP-Dol alpha-1,2-mannosyltransferase) |
| AT2G25980 | -0.1 | 7.16E-01 | 0.6 | 1.37E-03 | -0.7 | 2.25E-03 | 0.2 | 4.15E-01 | 0.3 | 1.48E-01 | -0.2 | 5.59E-01 | Mannose-binding lectin superfamily protein |
| AT3G15800 | -0.1 | 7.04E-01 | -0.1 | 6.61E-01 | 0.0 | 9.47E-01 | -0.1 | 9.05E-01 | 0.4 | 3.99E-01 | -0.5 | 3.20E-01 | glucan endo-1,3-beta-D-glucosidase |
| AT4G17430 | -0.1 | 6.33E-01 | 0.0 | 7.55E-01 | 0.0 | 9.16E-01 | 0.1 | 5.17E-01 | 0.2 | 4.80E-01 | 0.0 | 9.32E-01 | O-fucosyltransferase 30 |
| AT4G36220 | -0.1 | 5.99E-01 | 0.0 | 9.16E-01 | -0.1 | 5.72E-01 | -0.4 | 9.63E-03 | -0.1 | 8.16E-01 | -0.4 | 9.11E-02 | Cytochrome P450 84A1 |
| AT3G10370 | -0.1 | 4.25E-01 | -0.1 | 1.27E-01 | 0.1 | 4.78E-01 | 0.0 | 9.17E-01 | 0.2 | 3.50E-01 | -0.2 | 5.03E-01 | Glycerol-3-phosphate dehydrogenase SDP6, mitochondrial |
| AT1G30825 | -0.1 | 5.37E-01 | 0.0 | 6.47E-01 | -0.1 | 3.63E-01 | -0.2 | 4.14E-01 | -0.3 | 1.14E-01 | 0.2 | 5.26E-01 | Actin-related protein 2 |
| AT5G12850 | -0.1 | 4.03E-01 | -0.2 | 1.97E-02 | 0.1 | 1.12E-01 | -0.1 | 6.88E-01 | 0.0 | 9.32E-01 | -0.1 | 6.86E-01 | Zinc finger CCCH domain-containing protein 56 |
| AT4G20740 | -0.1 | 7.01E-01 | 0.0 | 7.94E-01 | -0.1 | 5.95E-01 | -0.1 | 6.37E-01 | -0.2 | 3.71E-01 | 0.1 | 7.17E-01 | Pentatricopeptide repeat-containing protein At4g20740 |
| AT5G03040 | -0.1 | 5.02E-01 | -0.1 | 3.16E-01 | 0.0 | 7.49E-01 | -0.3 | 3.73E-02 | -0.1 | 4.79E-01 | -0.2 | 3.95E-01 | Protein IQ-DOMAIN 2 |
| AT3G61820 | -0.1 | 6.21E-01 | 0.3 | 2.98E-02 | -0.3 | 1.91E-02 | 0.1 | 6.76E-01 | 0.5 | 2.36E-03 | -0.4 | 2.94E-02 | Eukaryotic aspartyl protease family protein |
| AT1G03310 | -0.1 | 6.53E-01 | -0.2 | 1.45E-01 | 0.1 | 3.38E-01 | 0.1 | 5.72E-01 | 0.6 | 1.47E-02 | -0.4 | 1.13E-01 | Isoamylase 2, chloroplastic |
| AT1G01620 | -0.1 | 5.74E-01 | 0.7 | 4.02E-05 | -0.8 | 8.05E-05 | -0.9 | 1.17E-05 | -0.8 | 1.58E-04 | -0.1 | 7.20E-01 | Plasma membrane intrinsic protein 1C |
| AT4G24570 | -0.1 | 8.92E-01 | -0.5 | 2.00E-01 | 0.5 | 3.12E-01 | -0.2 | 2.53E-01 | -0.4 | 3.26E-02 | 0.2 | 3.80E-01 | Mitochondrial uncoupling protein 4 |
| AT2G31580 | -0.1 | 7.82E-01 | -0.3 | 1.02E-01 | 0.3 | 2.27E-01 | -0.1 | 8.45E-01 | -0.1 | 7.12E-01 | 0.1 | 8.74E-01 | tRNA guanylyltransferase |
| AT4G15880 | -0.1 | 4.10E-01 | 0.0 | 6.31E-01 | 0.0 | 7.99E-01 | 0.1 | 2.67E-01 | 0.1 | 6.76E-01 | 0.1 | 7.44E-01 | Cysteine proteinases superfamily protein |
| AT5G19780 | -0.1 | 5.76E-01 | 0.3 | 2.15E-02 | -0.3 | 1.30E-02 | -0.3 | 1.11E-02 | -0.4 | 6.33E-03 | 0.1 | 7.47E-01 | Tubulin alpha-5 chain |
| AT1G26810 | -0.1 | 6.58E-01 | 0.0 | 8.43E-01 | 0.0 | 8.66E-01 | -0.6 | 7.75E-02 | -0.3 | 3.97E-01 | -0.3 | 5.24E-01 | Beta-1,3-galactosyltransferase GALT1 |
| AT3G26730 | -0.1 | 4.23E-01 | -0.2 | 6.47E-03 | 0.2 | 3.47E-02 | 0.2 | 1.27E-01 | 0.1 | 7.95E-01 | 0.2 | 4.14E-01 | Putative RING zinc finger protein |
| AT3G53570 | -0.1 | 6.09E-01 | -0.1 | 5.75E-01 | 0.0 | 9.55E-01 | 0.0 | 8.80E-01 | 0.2 | 5.87E-01 | -0.1 | 7.68E-01 | Serine |
| AT1G16780 | -0.1 | 6.37E-01 | -0.1 | 5.60E-01 | 0.0 | 9.19E-01 | 0.0 | 9.39E-01 | 0.0 | 9.83E-01 | 0.0 | 9.40E-01 | Pyrophosphate-energized membrane proton pump 3 -PPase 3) |
| AT2G41705 | -0.1 | 5.52E-01 | -0.4 | 2.15E-03 | 0.3 | 1.19E-02 | -0.1 | 7.75E-01 | 0.1 | 8.57E-01 | -0.1 | 6.90E-01 | Camphor resistance CrcB family protein |
| AT2G28550 | -0.1 | 5.96E-01 | -0.2 | 1.27E-01 | 0.1 | 3.28E-01 | -0.3 | 6.70E-02 | 0.4 | 2.29E-02 | -0.6 | 3.25E-03 | Related to AP2.7 |
| AT3G51550 | -0.1 | 4.95E-01 | -0.1 | 8.32E-02 | 0.1 | 2.86E-01 | -0.3 | 2.42E-03 | -0.5 | 8.92E-04 | 0.1 | 5.61E-01 | Receptor-like protein kinase FERONIA |
| AT5G35180 | -0.1 | 4.47E-01 | -0.1 | 2.67E-01 | 0.0 | 7.32E-01 | 0.0 | 8.89E-01 | 0.1 | 6.21E-01 | -0.1 | 6.14E-01 | ENHANCED DISEASE RESISTANCE protein |
| AT3G43800 | -0.1 | 6.00E-01 | 0.3 | 8.13E-03 | -0.4 | 6.22E-03 | 0.1 | 6.06E-01 | 0.2 | 3.52E-01 | -0.1 | 7.27E-01 | Glutathione S-transferase U27 |
| AT1G48280 | -0.1 | 6.10E-01 | -0.1 | 3.75E-01 | 0.0 | 7.45E-01 | -0.3 | 7.99E-02 | -0.2 | 4.29E-01 | -0.2 | 5.54E-01 | INCREASED PETAL GROWTH ANISOTROPY 1-like protein 2 |
| AT2G21830 | -0.1 | 6.61E-01 | 0.2 | 1.55E-01 | -0.2 | 1.16E-01 | -0.5 | 2.04E-02 | -0.1 | 6.03E-01 | -0.4 | 1.46E-01 | Cysteine |
| AT5G19290 | -0.1 | 6.73E-01 | 0.2 | 1.96E-01 | -0.2 | 1.22E-01 | -0.2 | 1.48E-01 | 0.0 | 8.72E-01 | -0.1 | 4.36E-01 | Alpha |
| AT1G52740 | -0.1 | 5.62E-01 | 0.2 | 4.72E-02 | -0.2 | 2.48E-02 | -0.4 | 2.55E-02 | -0.4 | 4.52E-02 | 0.0 | 9.98E-01 | Probable histone H2A variant 3 |
| AT3G61160 | -0.1 | 7.14E-01 | 0.3 | 6.55E-02 | -0.3 | 6.59E-02 | -0.1 | 5.15E-01 | -0.2 | 2.60E-01 | 0.1 | 6.64E-01 | non-specific serine |
| AT4G05320 | -0.1 | 5.81E-01 | -0.1 | 1.68E-01 | 0.1 | 4.23E-01 | -0.2 | 1.43E-02 | -0.1 | 6.17E-01 | -0.2 | 1.98E-01 | Polyubiquitin |
| AT4G35260 | 0.0 | 5.88E-01 | 0.1 | 5.74E-01 | -0.1 | 3.12E-01 | 0.3 | 1.25E-02 | 0.2 | 8.13E-02 | 0.1 | 7.33E-01 | Isocitrate dehydrogenase regulatory subunit 1, mitochondrial -specific ICDH 1) |
| AT5G14420 | 0.0 | 4.99E-01 | -0.1 | 2.75E-01 | 0.0 | 7.05E-01 | 0.0 | 8.99E-01 | -0.1 | 7.11E-01 | 0.1 | 8.36E-01 | E3 ubiquitin-protein ligase RGLG2 |
| AT3G13090 | 0.0 | 7.13E-01 | -0.5 | 5.69E-03 | 0.4 | 2.34E-02 | 1.0 | 8.70E-03 | 0.8 | 5.60E-02 | 0.2 | 6.47E-01 | ABC-type xenobiotic transporter |
| AT2G22720 | 0.0 | 3.92E-01 | -0.1 | 6.62E-02 | 0.1 | 3.14E-01 | -0.3 | 3.50E-02 | -0.2 | 1.10E-01 | 0.0 | 8.51E-01 | At2g22720 |
| AT3G62120 | 0.0 | 6.54E-01 | 0.3 | 1.66E-02 | -0.4 | 1.22E-02 | 0.1 | 3.66E-01 | 0.0 | 9.53E-01 | 0.1 | 5.01E-01 | proline--tRNA ligase |
| AT4G08900 | 0.0 | 4.69E-01 | -0.2 | 1.21E-02 | 0.2 | 6.38E-02 | 0.1 | 6.45E-01 | 0.1 | 6.14E-01 | 0.0 | 9.41E-01 | Arginase 1, mitochondrial |
| AT3G20100 | 0.0 | 6.89E-01 | -0.2 | 1.35E-01 | 0.1 | 3.09E-01 | 0.1 | 7.13E-01 | 0.1 | 8.01E-01 | 0.0 | 9.80E-01 | AT3g20100 |
| AT5G01310 | 0.0 | 5.83E-01 | -0.2 | 3.81E-02 | 0.2 | 1.50E-01 | -0.1 | 8.74E-01 | 0.1 | 7.70E-01 | -0.2 | 6.85E-01 | APRATAXIN-like protein |
| AT3G05250 | 0.0 | 7.07E-01 | -0.1 | 3.56E-01 | 0.1 | 6.45E-01 | -0.4 | 2.62E-02 | -0.5 | 3.23E-02 | 0.0 | 9.63E-01 | At3g05250 |
| AT2G28380 | 0.0 | 5.69E-01 | -0.1 | 2.39E-01 | 0.1 | 5.74E-01 | 0.1 | 5.53E-01 | 0.1 | 6.17E-01 | 0.0 | 9.95E-01 | Double-stranded RNA-binding protein 2 |
| AT3G16270 | 0.0 | 4.51E-01 | 0.0 | 8.50E-01 | 0.0 | 6.41E-01 | 0.0 | 8.96E-01 | -0.1 | 3.36E-01 | 0.1 | 3.80E-01 | Protein MODIFIED TRANSPORT TO THE VACUOLE 1 |
| AT2G43360 | 0.0 | 6.35E-01 | 0.3 | 1.52E-02 | -0.3 | 1.25E-02 | 0.2 | 2.13E-01 | 0.2 | 2.85E-01 | 0.0 | 9.99E-01 | Biotin synthase, mitochondrial |
| AT5G57590 | 0.0 | 7.55E-01 | 0.0 | 9.14E-01 | -0.1 | 7.32E-01 | 0.4 | 1.25E-01 | 0.1 | 7.23E-01 | 0.3 | 4.53E-01 | Bifunctional dethiobiotin synthetase |
| AT1G34340 | 0.0 | 7.02E-01 | -0.3 | 1.53E-02 | 0.3 | 4.73E-02 | 0.1 | 7.03E-01 | 0.0 | 9.90E-01 | 0.1 | 7.49E-01 | Alpha |
| AT2G37630 | 0.0 | 5.25E-01 | -0.2 | 3.03E-02 | 0.1 | 1.23E-01 | -0.3 | 7.74E-02 | -0.1 | 7.60E-01 | -0.2 | 3.35E-01 | Transcription factor AS1 |
| AT1G19570 | 0.0 | 7.44E-01 | 0.2 | 1.81E-01 | -0.2 | 1.27E-01 | 0.2 | 6.21E-01 | 0.6 | 6.13E-02 | -0.5 | 2.56E-01 | Glutathione S-transferase DHAR1, mitochondrial |
| AT3G06810 | 0.0 | 5.60E-01 | -0.2 | 1.68E-02 | 0.2 | 6.95E-02 | -0.1 | 5.98E-01 | 0.0 | 9.06E-01 | -0.1 | 8.27E-01 | Probable acyl-CoA dehydrogenase IBR3 |
| AT3G55110 | 0.0 | 8.30E-01 | -0.1 | 5.89E-01 | 0.1 | 7.86E-01 | 0.7 | 1.98E-02 | 0.7 | 3.40E-02 | 0.0 | 9.81E-01 | ABC transporter G family member 18 |
| AT2G01330 | 0.0 | 6.45E-01 | -0.2 | 5.13E-02 | 0.2 | 1.70E-01 | -0.2 | 7.21E-01 | 0.3 | 4.55E-01 | -0.5 | 2.74E-01 | Actin-interacting protein 1-1 |

| | | | | | | | | | | | | | |
|---|---|---|---|---|---|---|---|---|---|---|---|---|---|
| AT5G16990 | 0.0 | 6.99E-01 | -0.3 | 3.90E-02 | 0.2 | 1.07E-01 | 0.2 | 2.07E-01 | 0.1 | 4.85E-01 | 0.1 | 7.94E-01 | NADP-dependent alkenal double bond reductase P2 |
| AT1G79870 | 0.0 | 5.41E-01 | 0.0 | 8.38E-01 | -0.1 | 4.61E-01 | -0.1 | 4.61E-01 | 0.2 | 2.62E-01 | -0.3 | 1.53E-01 | Glyoxylate |
| AT5G58470 | 0.0 | 5.56E-01 | -0.1 | 7.86E-02 | 0.1 | 2.47E-01 | 0.1 | 4.29E-01 | 0.1 | 7.86E-01 | 0.1 | 8.11E-01 | Transcription initiation factor TFIID subunit 15b |
| AT3G24870 | 0.0 | 8.05E-01 | -0.4 | 4.75E-02 | 0.4 | 1.15E-01 | -0.1 | 9.56E-01 | 0.1 | 9.18E-01 | -0.2 | 8.97E-01 | Helicase |
| AT3G19490 | 0.0 | 5.32E-01 | 0.1 | 4.03E-01 | -0.1 | 1.91E-01 | -0.1 | 6.59E-01 | 0.1 | 7.51E-01 | -0.2 | 4.96E-01 | Sodium |
| AT5G65750 | 0.0 | 5.38E-01 | -0.3 | 2.49E-03 | 0.2 | 1.19E-02 | 0.2 | 2.65E-01 | -0.1 | 8.53E-01 | 0.3 | 2.79E-01 | oxoglutarate dehydrogenase |
| AT4G25720 | 0.0 | 6.35E-01 | 0.1 | 4.89E-01 | -0.1 | 3.26E-01 | 0.2 | 4.55E-01 | 0.0 | 9.35E-01 | 0.2 | 4.81E-01 | AT4G25720 protein |
| AT1G58100 | 0.0 | 5.40E-01 | -0.1 | 8.44E-02 | 0.1 | 2.78E-01 | -0.2 | 1.77E-02 | -0.3 | 3.58E-03 | 0.1 | 5.00E-01 | Transcription factor TCP8 |
| AT5G22010 | 0.0 | 5.07E-01 | -0.1 | 8.41E-02 | 0.1 | 3.05E-01 | 0.1 | 5.83E-01 | 0.0 | 9.66E-01 | 0.1 | 7.56E-01 | Replication factor C subunit 1 |
| AT2G41900 | 0.0 | 6.16E-01 | -0.3 | 7.27E-03 | 0.2 | 2.44E-02 | 0.1 | 4.08E-01 | 0.1 | 6.17E-01 | 0.0 | 9.10E-01 | Zinc finger CCCH domain-containing protein 30 |
| AT1G20260 | 0.0 | 5.49E-01 | 0.2 | 2.24E-02 | -0.2 | 1.27E-02 | -0.1 | 1.47E-01 | 0.0 | 9.76E-01 | -0.1 | 3.43E-01 | Vacuolar-type H+-ATPase subunit B3 |
| AT5G05860 | 0.0 | 6.11E-01 | -0.1 | 1.75E-01 | 0.1 | 4.44E-01 | -0.1 | 8.16E-01 | 0.1 | 6.46E-01 | -0.2 | 5.39E-01 | UDP-glycosyltransferase 76C2 |
| AT2G42640 | 0.0 | 7.89E-01 | -0.5 | 1.35E-02 | 0.5 | 4.27E-02 | 0.2 | 6.96E-01 | 0.3 | 5.99E-01 | -0.1 | 8.96E-01 | Protein kinase domain-containing protein |
| AT4G32330 | 0.0 | 5.36E-01 | 0.0 | 5.36E-01 | -0.1 | 2.58E-01 | 0.0 | 7.41E-01 | 0.1 | 2.69E-01 | -0.2 | 2.45E-01 | TPX2 C-terminal domain-containing protein |
| AT1G75130 | 0.0 | 5.62E-01 | -0.1 | 3.86E-01 | 0.0 | 8.00E-01 | -0.2 | 2.77E-01 | -0.1 | 7.87E-01 | -0.1 | 6.62E-01 | Cytochrome P450, family 721, subfamily A, polypeptide 1 |
| AT3G12640 | 0.0 | 5.40E-01 | -0.2 | 2.57E-02 | 0.1 | 1.12E-01 | -0.1 | 9.22E-01 | 0.0 | 9.92E-01 | -0.1 | 9.57E-01 | RRM domain-containing protein |
| AT4G10890 | 0.0 | 7.30E-01 | 0.0 | 7.29E-01 | 0.0 | 9.88E-01 | 0.1 | 8.16E-01 | 0.1 | 8.73E-01 | 0.0 | 9.86E-01 | Uncharacterized protein AT4g10890 |
| AT1G36990 | 0.0 | 5.86E-01 | -0.1 | 2.48E-01 | 0.1 | 5.83E-01 | 0.0 | 9.98E-01 | 0.0 | 9.73E-01 | 0.0 | 9.73E-01 | C-jun-amino-terminal kinase-interacting protein |
| AT3G12050 | 0.0 | 6.49E-01 | 0.0 | 9.94E-01 | 0.0 | 7.02E-01 | -0.2 | 3.34E-01 | 0.0 | 9.88E-01 | -0.2 | 4.49E-01 | Aha1 domain-containing protein |
| AT2G45960 | 0.0 | 6.40E-01 | 0.3 | 8.10E-03 | -0.3 | 6.45E-03 | -0.5 | 4.94E-05 | -0.4 | 2.13E-03 | -0.1 | 3.88E-01 | AT2G45960 protein |
| AT3G17240 | 0.0 | 6.78E-01 | 0.0 | 7.22E-01 | -0.1 | 4.78E-01 | 0.4 | 5.76E-03 | 0.2 | 9.11E-02 | 0.1 | 5.30E-01 | Dihydrolipoyl dehydrogenase |
| AT1G71790 | 0.0 | 6.06E-01 | 0.3 | 7.44E-03 | -0.3 | 8.43E-03 | -0.4 | 1.97E-02 | -0.3 | 8.75E-02 | -0.1 | 7.51E-01 | Probable F-actin-capping protein subunit beta |
| AT5G12080 | 0.0 | 5.98E-01 | -0.1 | 3.54E-01 | 0.0 | 7.26E-01 | 0.1 | 6.33E-01 | 0.2 | 4.13E-01 | -0.1 | 7.96E-01 | Mechanosensitive ion channel protein 10 |
| AT1G53460 | 0.0 | 6.89E-01 | -0.1 | 3.83E-01 | 0.1 | 6.90E-01 | -0.2 | 4.19E-01 | 0.0 | 9.01E-01 | -0.1 | 6.80E-01 | At1g53460 |
| AT2G20300 | 0.0 | 6.12E-01 | -0.1 | 3.55E-01 | 0.0 | 7.10E-01 | 0.1 | 7.34E-01 | 0.2 | 3.20E-01 | -0.1 | 6.09E-01 | Protein kinase domain-containing protein |
| AT4G17020 | 0.0 | 6.17E-01 | -0.2 | 4.30E-02 | 0.1 | 1.52E-01 | 0.1 | 6.84E-01 | 0.1 | 5.18E-01 | -0.1 | 8.50E-01 | RNA polymerase II transcription factor B subunit 2 |
| AT3G23310 | 0.0 | 5.94E-01 | -0.1 | 1.78E-01 | 0.1 | 4.52E-01 | -0.2 | 2.62E-01 | -0.1 | 4.03E-01 | 0.0 | 9.35E-01 | AGC kinase family protein |
| AT1G01510 | 0.0 | 5.40E-01 | 0.0 | 9.26E-01 | 0.0 | 5.36E-01 | -0.2 | 7.47E-02 | -0.2 | 2.27E-01 | -0.1 | 8.28E-01 | C-terminal binding protein AN |
| AT2G02850 | 0.0 | 8.28E-01 | -0.6 | 8.60E-03 | 0.6 | 2.37E-02 | 0.7 | 4.67E-02 | 0.5 | 2.16E-01 | 0.2 | 6.96E-01 | Basic blue protein |
| AT4G14850 | 0.0 | 7.88E-01 | -0.3 | 5.31E-02 | 0.3 | 1.34E-01 | -0.1 | 8.70E-01 | -0.1 | 9.22E-01 | 0.0 | 9.80E-01 | Pentatricopeptide repeat-containing protein At4g14850 |
| AT2G27450 | 0.0 | 5.87E-01 | 0.2 | 2.71E-02 | -0.2 | 1.83E-02 | 0.0 | 9.86E-01 | 0.0 | 8.49E-01 | 0.0 | 8.89E-01 | Nitrilase-like protein 1 |
| AT3G49400 | 0.0 | 7.10E-01 | -0.2 | 5.20E-02 | 0.2 | 1.55E-01 | 0.2 | 4.32E-01 | 0.1 | 6.90E-01 | 0.1 | 8.61E-01 | Uncharacterized protein F2K15.260 |
| AT5G64240 | 0.0 | 7.25E-01 | 0.0 | 8.26E-01 | 0.0 | 9.28E-01 | -0.2 | 1.79E-01 | -0.2 | 2.62E-01 | 0.0 | 9.70E-01 | Metacaspase 3 |
| AT2G36490 | 0.0 | 7.63E-01 | -0.3 | 7.69E-02 | 0.2 | 1.66E-01 | -0.1 | 3.03E-01 | -0.2 | 1.68E-01 | 0.1 | 7.71E-01 | DNA glycosylase |
| AT1G49950 | 0.0 | 5.08E-01 | -0.2 | 2.93E-02 | 0.1 | 1.23E-01 | 0.0 | 8.88E-01 | -0.1 | 7.40E-01 | 0.1 | 6.97E-01 | Telomere repeat-binding factor 1 |
| AT2G01060 | 0.0 | 5.52E-01 | -0.2 | 4.49E-02 | 0.1 | 1.61E-01 | -0.2 | 1.22E-01 | -0.2 | 3.23E-01 | -0.1 | 8.23E-01 | Myb family transcription factor PHL7 |
| AT5G47630 | 0.0 | 7.30E-01 | 0.3 | 1.06E-02 | -0.4 | 1.53E-02 | 0.3 | 3.27E-01 | 0.1 | 8.98E-01 | 0.3 | 5.93E-01 | Acyl carrier protein 3, mitochondrial |
| AT1G20200 | 0.0 | 6.03E-01 | -0.1 | 1.25E-01 | 0.1 | 3.22E-01 | 0.2 | 1.61E-01 | 0.2 | 1.96E-01 | 0.0 | 9.74E-01 | 26S proteasome non-ATPase regulatory subunit 3 homolog A |
| AT3G22550 | 0.0 | 7.91E-01 | -0.1 | 4.01E-01 | 0.1 | 6.26E-01 | 0.0 | 9.49E-01 | 0.9 | 2.51E-03 | -0.8 | 1.03E-02 | FCS-Like Zinc finger 8 |
| AT2G27210 | 0.0 | 6.03E-01 | -0.2 | 4.80E-02 | 0.1 | 1.53E-01 | -0.1 | 6.77E-01 | 0.0 | 9.06E-01 | -0.1 | 8.66E-01 | Serine |
| AT5G59680 | 0.0 | 7.24E-01 | 0.1 | 2.56E-01 | -0.2 | 1.74E-01 | 0.3 | 9.67E-02 | 0.1 | 6.56E-01 | 0.2 | 3.74E-01 | Probable LRR receptor-like serine |
| AT1G55040 | 0.0 | 7.29E-01 | -0.2 | 1.31E-01 | 0.1 | 3.06E-01 | 0.4 | 2.53E-01 | 0.0 | 9.55E-01 | 0.4 | 4.18E-01 | Uncharacterized protein F14C21.55 family protein) |
| AT5G22330 | 0.0 | 5.93E-01 | 0.0 | 5.32E-01 | -0.1 | 3.09E-01 | 0.1 | 2.50E-01 | 0.0 | 9.36E-01 | 0.2 | 3.70E-01 | RuvB-like protein 1 |
| AT2G43180 | 0.0 | 6.59E-01 | -0.1 | 4.07E-01 | 0.0 | 7.40E-01 | 0.0 | 9.08E-01 | 0.0 | 9.81E-01 | 0.0 | 9.55E-01 | Phosphoenolpyruvate carboxylase family protein |
| AT3G22170 | 0.0 | 5.91E-01 | -0.2 | 6.24E-03 | 0.2 | 2.36E-02 | 0.0 | 9.15E-01 | 0.0 | 9.19E-01 | 0.0 | 9.90E-01 | Protein FAR-RED ELONGATED HYPOCOTYL 3 |
| AT2G24520 | 0.0 | 7.43E-01 | -0.1 | 3.48E-01 | 0.1 | 5.98E-01 | 0.0 | 9.56E-01 | 0.1 | 7.34E-01 | -0.1 | 8.22E-01 | Plasma membrane ATPase |
| AT3G18520 | 0.0 | 5.25E-01 | 0.0 | 6.58E-01 | 0.0 | 8.96E-01 | -0.3 | 5.29E-02 | -0.2 | 1.58E-01 | -0.1 | 8.37E-01 | Histone deacetylase 15 |
| AT1G14185 | 0.0 | 8.41E-01 | 0.2 | 3.92E-01 | -0.2 | 3.79E-01 | -1.4 | 1.46E-02 | -1.1 | 4.63E-02 | -0.3 | 7.19E-01 | At1g14180 |
| AT4G12980 | 0.0 | 8.25E-01 | 0.1 | 6.09E-01 | -0.1 | 5.02E-01 | -0.3 | 6.74E-02 | 0.0 | 9.14E-01 | -0.3 | 7.93E-02 | Cytochrome b561 and DOMON domain-containing protein At4g12980 |
| AT1G05460 | 0.0 | 6.73E-01 | 0.0 | 7.92E-01 | 0.0 | 9.17E-01 | 0.1 | 7.29E-01 | 0.0 | 8.71E-01 | 0.0 | 9.28E-01 | RNA helicase |
| AT2G23770 | 0.0 | 7.24E-01 | 0.0 | 8.70E-01 | 0.0 | 8.96E-01 | 0.9 | 4.39E-04 | 0.6 | 1.44E-02 | 0.3 | 2.08E-01 | LysM domain receptor-like kinase 4 |
| AT1G01220 | 0.0 | 5.32E-01 | -0.1 | 1.98E-01 | 0.0 | 5.45E-01 | -0.1 | 6.48E-01 | 0.1 | 8.20E-01 | -0.2 | 5.23E-01 | L-fucokinase |
| AT1G52620 | 0.0 | 8.56E-01 | -0.2 | 2.69E-01 | 0.2 | 4.29E-01 | -0.2 | 7.43E-01 | 0.1 | 8.74E-01 | -0.3 | 6.69E-01 | Pentatricopeptide repeat-containing protein At1g52620 |
| AT1G47570 | 0.0 | 6.29E-01 | -0.1 | 3.79E-01 | 0.0 | 7.30E-01 | 0.0 | 7.74E-01 | 0.1 | 7.75E-01 | 0.0 | 9.73E-01 | F16N3.15 |
| AT5G43990 | 0.0 | 7.53E-01 | -0.2 | 1.96E-01 | 0.1 | 3.93E-01 | -0.3 | 3.54E-01 | 0.0 | 9.99E-01 | -0.3 | 5.19E-01 | SET-domain containing protein lysine methyltransferase family protein |
| AT3G13445 | 0.0 | 6.42E-01 | 0.0 | 8.28E-01 | -0.1 | 5.55E-01 | 0.1 | 7.75E-01 | 0.0 | 9.19E-01 | 0.1 | 7.82E-01 | TATA binding protein 1 |
| AT1G32330 | 0.0 | 6.87E-01 | -0.1 | 2.03E-01 | 0.1 | 4.45E-01 | 0.0 | 9.45E-01 | 0.1 | 7.07E-01 | -0.2 | 7.21E-01 | Heat shock transcription factor A1D |
| AT2G29590 | 0.0 | 7.71E-01 | 0.1 | 3.63E-01 | -0.2 | 3.12E-01 | -0.3 | 3.72E-01 | -0.3 | 3.46E-01 | 0.1 | 9.26E-01 | Thioesterase domain-containing protein |
| AT1G60220 | 0.0 | 6.76E-01 | -0.2 | 1.01E-01 | 0.1 | 2.56E-01 | -0.2 | 2.28E-01 | 0.0 | 9.80E-01 | -0.2 | 3.49E-01 | Ubiquitin-like-specific protease 1D |
| AT5G53300 | 0.0 | 5.96E-01 | -0.1 | 2.94E-01 | 0.0 | 6.22E-01 | -0.3 | 9.94E-02 | -0.3 | 1.55E-01 | 0.0 | 9.92E-01 | Ubiquitin-conjugating enzyme 10 |
| AT1G18580 | 0.0 | 6.20E-01 | -0.1 | 2.04E-01 | 0.1 | 4.86E-01 | 0.0 | 8.43E-01 | -0.1 | 6.03E-01 | 0.1 | 8.05E-01 | Hexosyltransferase GAUT11 |
| AT3G44110 | 0.0 | 6.67E-01 | -0.2 | 2.28E-02 | 0.2 | 6.31E-02 | 0.1 | 2.03E-01 | 0.1 | 6.19E-01 | 0.1 | 7.32E-01 | DnaJ protein homolog atj3 |
| AT3G05330 | 0.0 | 7.90E-01 | 0.0 | 8.63E-01 | -0.1 | 7.19E-01 | 0.4 | 7.98E-02 | 0.3 | 2.70E-01 | 0.1 | 7.31E-01 | ATTAN |
| AT5G12100 | 0.0 | 6.95E-01 | -0.1 | 4.73E-01 | 0.0 | 7.85E-01 | 0.0 | 8.61E-01 | 0.1 | 8.08E-01 | 0.0 | 9.42E-01 | Pentatricopeptide repeat-containing protein At5g12100, mitochondrial |
| AT1G78920 | 0.0 | 6.74E-01 | -0.2 | 4.09E-02 | 0.2 | 1.07E-01 | 0.0 | 9.40E-01 | 0.0 | 9.47E-01 | 0.0 | 9.97E-01 | Pyrophosphate-energized membrane proton pump 2 -PPase 2) |
| AT1G62010 | 0.0 | 8.25E-01 | 0.1 | 4.72E-01 | -0.1 | 4.38E-01 | 0.3 | 3.95E-01 | 0.1 | 8.67E-01 | 0.2 | 6.52E-01 | Mitochondrial transcription termination factor family protein |
| AT3G02990 | 0.0 | 8.31E-01 | -0.2 | 1.70E-01 | 0.2 | 3.14E-01 | -0.3 | 4.45E-01 | 0.1 | 7.90E-01 | -0.5 | 4.06E-01 | Heat stress transcription factor A-1e |
| AT4G01280 | 0.0 | 6.35E-01 | -0.2 | 1.11E-02 | 0.2 | 4.42E-02 | 0.0 | 9.65E-01 | -0.3 | 1.77E-01 | 0.3 | 3.04E-01 | A_IG002N01.20 protein |
| AT5G56870 | 0.0 | 8.62E-01 | 0.7 | 5.67E-03 | -0.7 | 7.94E-03 | -0.4 | 2.38E-01 | -0.8 | 4.39E-02 | 0.4 | 3.73E-01 | Beta-galactosidase 4 |
| AT4G14960 | 0.0 | 6.86E-01 | 0.2 | 5.86E-02 | -0.2 | 4.00E-02 | -0.1 | 3.60E-01 | 0.0 | 9.94E-01 | -0.1 | 5.05E-01 | Tubulin alpha chain |
| AT5G50180 | 0.0 | 6.66E-01 | 0.2 | 5.96E-02 | -0.2 | 4.49E-02 | 0.0 | 8.81E-01 | 0.2 | 4.55E-01 | -0.2 | 4.15E-01 | At5g50180 |
| AT4G03240 | 0.0 | 6.68E-01 | 0.0 | 6.72E-01 | -0.1 | 4.72E-01 | 0.1 | 8.21E-01 | -0.2 | 5.17E-01 | 0.3 | 4.48E-01 | Frataxin, mitochondrial |
| AT5G59160 | 0.0 | 6.55E-01 | 0.2 | 2.46E-02 | -0.2 | 1.76E-02 | -0.1 | 6.11E-01 | 0.0 | 8.46E-01 | 0.0 | 8.70E-01 | Serine |
| AT5G01840 | 0.0 | 8.09E-01 | 0.1 | 3.77E-01 | -0.2 | 3.38E-01 | -0.1 | 7.93E-01 | 0.3 | 3.46E-01 | -0.4 | 2.83E-01 | Transcription repressor OFP1 |
| AT5G44290 | 0.0 | 7.60E-01 | -0.3 | 1.39E-02 | 0.3 | 3.35E-02 | 0.1 | 5.93E-01 | 0.0 | 8.78E-01 | 0.1 | 8.39E-01 | Protein kinase domain-containing protein |
| AT4G36290 | 0.0 | 6.49E-01 | -0.2 | 4.09E-02 | 0.1 | 1.33E-01 | 0.0 | 9.11E-01 | -0.2 | 4.20E-01 | 0.1 | 5.85E-01 | Protein MICRORCHIDIA 1 |
| AT4G38570 | 0.0 | 7.35E-01 | 0.1 | 4.75E-01 | -0.1 | 3.68E-01 | 0.0 | 8.80E-01 | -0.1 | 6.50E-01 | 0.1 | 8.06E-01 | CDP-diacylglycerol--inositol 3-phosphatidyltransferase |
| AT1G68100 | 0.0 | 6.13E-01 | -0.1 | 2.00E-01 | 0.1 | 4.77E-01 | -0.1 | 6.32E-01 | -0.1 | 4.68E-01 | 0.1 | 8.42E-01 | ZIP metal ion transporter family |
| AT5G61060 | 0.0 | 6.47E-01 | -0.4 | 3.15E-04 | 0.4 | 1.34E-03 | 0.0 | 9.17E-01 | -0.2 | 3.22E-01 | 0.2 | 3.72E-01 | Histone deacetylase 5 |
| AT4G24190 | 0.0 | 8.39E-01 | 0.0 | 8.27E-01 | 0.0 | 9.81E-01 | 0.5 | 7.97E-04 | 0.4 | 1.02E-02 | 0.1 | 6.42E-01 | Chaperone protein htpG family protein |
| AT1G60940 | 0.0 | 6.86E-01 | 0.1 | 5.04E-01 | -0.1 | 3.27E-01 | -0.1 | 6.46E-01 | -0.2 | 2.77E-01 | 0.1 | 6.51E-01 | Serine |
| AT2G33540 | 0.0 | 6.94E-01 | -0.2 | 2.78E-02 | 0.2 | 8.22E-02 | 0.0 | 8.84E-01 | 0.0 | 9.86E-01 | 0.0 | 9.33E-01 | RNA polymerase II C-terminal domain phosphatase-like |
| AT1G53940 | 0.0 | 7.68E-01 | 0.7 | 3.50E-05 | -0.8 | 1.38E-04 | -0.9 | 8.68E-04 | -0.1 | 6.54E-01 | -0.8 | 1.18E-02 | GDSL esterase |

| | | | | | | | | | | | | | |
|---|---|---|---|---|---|---|---|---|---|---|---|---|---|
| AT5G28900 | 0.0 | 6.85E-01 | 0.0 | 5.72E-01 | 0.0 | 8.87E-01 | -0.3 | 5.28E-03 | -0.2 | 8.27E-02 | -0.1 | 4.86E-01 | Probable serine |
| AT2G20420 | 0.0 | 6.96E-01 | 0.2 | 3.43E-02 | -0.2 | 2.49E-02 | 0.3 | 2.60E-02 | 0.2 | 2.35E-01 | 0.1 | 5.82E-01 | Succinate--CoA ligase subunit beta, mitochondrial |
| AT1G77260 | 0.0 | 6.51E-01 | -0.1 | 1.36E-01 | 0.1 | 3.31E-01 | -0.1 | 7.69E-01 | 0.1 | 5.58E-01 | -0.2 | 4.59E-01 | Probable methyltransferase PMT10 |
| AT3G03210 | 0.0 | 7.45E-01 | 0.0 | 9.43E-01 | 0.0 | 7.49E-01 | 0.0 | 9.48E-01 | 0.1 | 9.01E-01 | -0.1 | 8.61E-01 | Protein ALTERED XYLOGLUCAN 9 |
| AT3G04690 | 0.0 | 8.25E-01 | 0.1 | 3.75E-01 | -0.2 | 3.56E-01 | -0.2 | 7.20E-01 | 0.2 | 7.02E-01 | -0.5 | 4.74E-01 | Receptor-like protein kinase ANXUR1 |
| AT1G48360 | 0.0 | 8.33E-01 | 0.1 | 6.54E-01 | -0.1 | 5.78E-01 | -0.1 | 6.87E-01 | -0.2 | 5.46E-01 | 0.1 | 8.74E-01 | Fanconi-associated nuclease |
| AT3G56850 | 0.0 | 7.17E-01 | -0.1 | 5.23E-01 | 0.0 | 8.12E-01 | -0.2 | 1.54E-01 | -0.2 | 1.69E-01 | 0.0 | 9.62E-01 | ABSCISIC ACID-INSENSITIVE 5-like protein 2 |
| AT5G42400 | 0.0 | 7.24E-01 | -0.2 | 3.93E-02 | 0.2 | 1.12E-01 | 0.0 | 9.17E-01 | 0.0 | 9.32E-01 | -0.1 | 8.74E-01 | SET domain protein 25 |
| AT1G30970 | 0.0 | 7.00E-01 | -0.2 | 8.73E-03 | 0.2 | 2.91E-02 | 0.2 | 4.19E-01 | 0.2 | 5.97E-01 | 0.0 | 9.06E-01 | Zinc finger family protein |
| AT5G51300 | 0.0 | 6.86E-01 | -0.3 | 4.32E-03 | 0.2 | 1.39E-02 | 0.2 | 4.93E-02 | 0.1 | 5.58E-01 | 0.2 | 3.71E-01 | Splicing factor-like protein 1 |
| AT2G41060 | 0.0 | 7.08E-01 | -0.2 | 1.92E-02 | 0.2 | 6.05E-02 | 0.0 | 9.93E-01 | -0.1 | 6.59E-01 | 0.1 | 6.98E-01 | UBP1-associated protein 2B |
| AT1G20960 | 0.0 | 6.46E-01 | -0.1 | 7.79E-02 | 0.1 | 1.97E-01 | 0.2 | 2.79E-01 | 0.1 | 7.99E-01 | 0.1 | 6.71E-01 | DExH-box ATP-dependent RNA helicase DExH12 |
| AT4G32690 | 0.0 | 7.37E-01 | 0.0 | 9.06E-01 | 0.0 | 8.74E-01 | -0.7 | 4.85E-03 | -0.1 | 8.14E-01 | -0.6 | 1.74E-02 | Two-on-two hemoglobin-3 |
| AT4G18910 | 0.0 | 7.16E-01 | 0.1 | 1.16E-01 | -0.2 | 9.23E-02 | 0.1 | 3.86E-01 | 0.4 | 1.17E-02 | -0.2 | 1.62E-01 | NOD26-like intrinsic protein 12 |
| AT2G22830 | 0.0 | 8.11E-01 | 0.3 | 4.60E-02 | -0.3 | 5.63E-02 | -0.3 | 1.75E-01 | -0.1 | 8.72E-01 | -0.3 | 3.79E-01 | Squalene monooxygenase |
| AT5G45560 | 0.0 | 7.47E-01 | 0.0 | 6.24E-01 | 0.0 | 8.87E-01 | -0.1 | 7.64E-01 | 0.0 | 9.58E-01 | -0.1 | 8.56E-01 | Pleckstrin homology domain-containing protein |
| AT1G74700 | 0.0 | 8.66E-01 | -0.3 | 1.36E-01 | 0.2 | 2.51E-01 | 0.7 | 1.46E-02 | 0.5 | 1.13E-01 | 0.2 | 5.69E-01 | tRNase Z TRZ1 |
| AT5G17530 | 0.0 | 6.40E-01 | 0.0 | 6.38E-01 | 0.0 | 9.83E-01 | 0.0 | 8.80E-01 | 0.2 | 4.30E-01 | -0.3 | 3.71E-01 | phosphoglucomutase |
| AT1G53290 | 0.0 | 7.07E-01 | 0.3 | 1.28E-03 | -0.3 | 2.25E-03 | -0.2 | 9.21E-02 | 0.0 | 8.08E-01 | -0.3 | 1.16E-01 | Hexosyltransferase |
| AT3G55140 | 0.0 | 7.47E-01 | -0.1 | 4.64E-01 | 0.0 | 7.25E-01 | -0.2 | 6.43E-02 | -0.1 | 4.07E-01 | -0.1 | 5.69E-01 | Pectate lyase |
| AT4G37280 | 0.0 | 7.39E-01 | 0.1 | 4.44E-01 | -0.1 | 3.39E-01 | -0.2 | 3.06E-01 | 0.0 | 9.58E-01 | -0.1 | 4.80E-01 | Protein MRG1 |
| AT1G16540 | 0.0 | 8.16E-01 | -0.4 | 7.30E-03 | 0.4 | 2.25E-02 | 0.4 | 1.40E-01 | 0.2 | 4.61E-01 | 0.1 | 6.74E-01 | Molybdenum cofactor sulfurase |
| AT2G16790 | 0.0 | 9.07E-01 | -0.3 | 2.73E-01 | 0.2 | 4.03E-01 | 0.3 | 4.59E-01 | 0.0 | 9.90E-01 | 0.3 | 6.06E-01 | Gluconokinase |
| AT3G55260 | 0.0 | 7.34E-01 | 0.1 | 3.40E-01 | -0.1 | 2.42E-01 | 0.0 | 9.10E-01 | 0.0 | 7.98E-01 | 0.0 | 8.94E-01 | Beta-hexosaminidase 1 |
| AT3G02740 | 0.0 | 7.35E-01 | -0.1 | 2.21E-01 | 0.1 | 4.23E-01 | 0.1 | 3.73E-01 | 0.0 | 9.97E-01 | 0.1 | 5.33E-01 | Eukaryotic aspartyl protease family protein |
| AT3G10810 | 0.0 | 7.61E-01 | -0.1 | 1.73E-01 | 0.1 | 3.31E-01 | 0.1 | 5.86E-01 | 0.1 | 6.26E-01 | 0.0 | 9.90E-01 | RING-type E3 ubiquitin transferase |
| AT1G43690 | 0.0 | 6.53E-01 | -0.1 | 1.02E-01 | 0.1 | 2.51E-01 | 0.0 | 7.94E-01 | 0.2 | 1.58E-01 | -0.2 | 3.94E-01 | Deubiquitinating enzyme MINDY-3 |
| AT4G25640 | 0.0 | 8.06E-01 | -0.3 | 7.98E-03 | 0.3 | 1.79E-02 | 0.2 | 2.33E-01 | 0.0 | 9.50E-01 | 0.2 | 3.86E-01 | Protein DETOXIFICATION |
| AT3G51880 | 0.0 | 7.04E-01 | 0.0 | 5.12E-01 | -0.1 | 3.42E-01 | -0.1 | 5.01E-01 | -0.3 | 1.06E-01 | 0.2 | 4.54E-01 | High mobility group B1 |
| AT5G40880 | 0.0 | 8.06E-01 | -0.3 | 2.16E-02 | 0.3 | 6.27E-02 | 0.4 | 3.27E-01 | 0.5 | 2.08E-01 | -0.1 | 7.62E-01 | WD-40 repeat family protein |
| AT4G16765 | 0.0 | 7.75E-01 | -0.3 | 1.54E-02 | 0.2 | 4.59E-02 | 0.0 | 9.09E-01 | 0.1 | 6.67E-01 | -0.1 | 6.41E-01 | 2-oxoglutarate and Fe-dependent oxygenase superfamily protein |
| AT4G17890 | 0.0 | 7.11E-01 | 0.2 | 4.75E-02 | -0.2 | 4.00E-02 | 0.1 | 5.90E-01 | 0.1 | 5.62E-01 | 0.0 | 9.41E-01 | Probable ADP-ribosylation factor GTPase-activating protein AGD8 |
| AT5G44510 | 0.0 | 8.38E-01 | -0.3 | 3.87E-02 | 0.3 | 8.47E-02 | 0.3 | 2.43E-01 | 0.4 | 1.18E-01 | -0.1 | 7.07E-01 | ADP-ribosyl cyclase |
| AT2G46430 | 0.0 | 8.47E-01 | -0.6 | 1.02E-03 | 0.6 | 2.83E-03 | 0.7 | 1.01E-03 | 0.3 | 1.80E-01 | 0.4 | 1.04E-01 | Probable cyclic nucleotide-gated ion channel 3 |
| AT5G43960 | 0.0 | 8.17E-01 | 0.1 | 4.14E-01 | -0.1 | 3.34E-01 | 0.4 | 1.87E-03 | 0.2 | 1.45E-01 | 0.2 | 2.36E-01 | Gb\|AAD20086.1 family protein with RNA binding domain-containing protein) |
| AT3G24715 | 0.0 | 9.12E-01 | 0.0 | 9.10E-01 | -0.1 | 8.57E-01 | -0.8 | 2.81E-01 | -0.2 | 8.28E-01 | -0.6 | 5.79E-01 | Kinase superfamily with octicosapeptide |
| AT1G61960 | 0.0 | 8.79E-01 | -0.1 | 7.67E-01 | 0.0 | 9.07E-01 | 0.3 | 2.63E-01 | 0.4 | 2.68E-01 | 0.0 | 9.47E-01 | Mitochondrial transcription termination factor family protein |
| AT3G13860 | 0.0 | 8.53E-01 | 0.1 | 3.49E-01 | -0.2 | 3.17E-01 | 0.5 | 3.57E-04 | 0.4 | 1.06E-02 | 0.2 | 3.70E-01 | Chaperonin CPN60-like 2, mitochondrial |
| AT5G08110 | 0.0 | 8.49E-01 | -0.2 | 1.66E-01 | 0.2 | 3.00E-01 | -0.1 | 7.56E-01 | 0.2 | 6.05E-01 | -0.4 | 4.58E-01 | UBQ, helicase-c and DEAD-like helicase domain-containing protein |
| AT3G28860 | 0.0 | 7.62E-01 | 0.3 | 4.13E-03 | -0.3 | 4.48E-03 | -0.2 | 2.36E-01 | 0.0 | 9.98E-01 | -0.2 | 3.84E-01 | ABC transporter B family member 19 |
| AT1G04820 | 0.0 | 8.18E-01 | 0.3 | 3.08E-02 | -0.3 | 2.92E-02 | 0.0 | 7.71E-01 | 0.1 | 6.95E-01 | -0.1 | 5.61E-01 | Tubulin alpha-4 chain |
| AT1G79500 | 0.0 | 8.25E-01 | 0.3 | 3.17E-02 | -0.3 | 3.93E-02 | -0.1 | 5.81E-01 | 0.1 | 7.82E-01 | -0.2 | 4.71E-01 | 2-dehydro-3-deoxyphosphooctonate aldolase 1 |
| AT3G55850 | 0.0 | 8.15E-01 | -0.1 | 1.88E-01 | 0.1 | 3.38E-01 | -0.1 | 6.86E-01 | 0.0 | 8.93E-01 | -0.1 | 8.87E-01 | Amidohydrolase family |
| AT1G29750 | 0.0 | 8.05E-01 | -0.1 | 2.33E-01 | 0.1 | 4.06E-01 | 0.0 | 8.97E-01 | 0.2 | 6.10E-01 | -0.2 | 5.91E-01 | Receptor-like kinase in flowers 1 |
| AT4G32210 | 0.0 | 8.11E-01 | 0.0 | 7.28E-01 | 0.0 | 9.25E-01 | 0.1 | 8.96E-01 | 0.3 | 4.75E-01 | -0.2 | 6.42E-01 | Succinate dehydrogenase subunit 3-2, mitochondrial |
| AT5G42950 | 0.0 | 7.18E-01 | -0.1 | 1.11E-01 | 0.1 | 2.41E-01 | -0.2 | 4.51E-01 | -0.4 | 7.21E-02 | 0.2 | 3.80E-01 | Protein ESSENTIAL FOR POTEXVIRUS ACCUMULATION 1 |
| AT3G06350 | 0.0 | 7.35E-01 | -0.1 | 2.30E-01 | 0.1 | 4.21E-01 | 0.0 | 7.92E-01 | 0.0 | 8.61E-01 | 0.0 | 9.81E-01 | Bifunctional 3-dehydroquinate dehydratase |
| AT5G63680 | 0.0 | 7.39E-01 | -0.1 | 1.43E-01 | 0.1 | 2.90E-01 | 0.2 | 1.11E-01 | 0.0 | 8.49E-01 | 0.2 | 1.50E-01 | Pyruvate kinase |
| AT2G26710 | 0.0 | 8.92E-01 | 0.0 | 8.22E-01 | -0.1 | 7.65E-01 | -0.2 | 2.98E-01 | 0.2 | 5.45E-01 | -0.4 | 1.20E-01 | Cytochrome P450 734A1 |
| AT1G62305 | 0.0 | 8.65E-01 | 0.0 | 9.45E-01 | 0.0 | 8.48E-01 | 0.2 | 4.36E-01 | 0.1 | 7.26E-01 | 0.1 | 8.38E-01 | At1g62305 |
| AT3G53030 | 0.0 | 7.65E-01 | -0.2 | 5.33E-02 | 0.1 | 1.29E-01 | -0.5 | 6.51E-03 | -0.4 | 1.53E-02 | 0.0 | 9.40E-01 | Ser |
| AT5G07100 | 0.0 | 8.61E-01 | -0.2 | 1.26E-01 | 0.2 | 2.24E-01 | -0.2 | 6.75E-01 | -0.9 | 6.55E-02 | 0.7 | 1.07E-01 | At5g07100 |
| AT2G02760 | 0.0 | 8.61E-01 | -0.1 | 2.59E-01 | 0.1 | 3.74E-01 | -0.3 | 6.00E-02 | -0.2 | 3.08E-01 | -0.1 | 6.98E-01 | Ubiquitin-conjugating enzyme E2 2 |
| AT1G68370 | 0.0 | 7.24E-01 | -0.2 | 2.18E-02 | 0.1 | 5.95E-02 | -0.3 | 3.08E-02 | -0.3 | 4.49E-02 | 0.0 | 9.80E-01 | Chaperone protein dnaJ 15 |
| AT4G36780 | 0.0 | 7.76E-01 | 0.1 | 3.15E-01 | -0.1 | 2.40E-01 | -0.2 | 5.49E-02 | -0.2 | 7.89E-02 | 0.0 | 9.81E-01 | Protein BZR1 homolog |
| AT1G54610 | 0.0 | 7.66E-01 | -0.2 | 2.17E-02 | 0.2 | 5.15E-02 | 0.1 | 7.87E-01 | 0.0 | 8.99E-01 | 0.1 | 7.49E-01 | Probable serine |
| AT2G40830 | 0.0 | 7.61E-01 | -0.3 | 1.26E-03 | 0.3 | 4.02E-03 | 0.1 | 5.22E-01 | 0.1 | 7.01E-01 | 0.0 | 9.35E-01 | Probable E3 ubiquitin-protein ligase RHC1A |
| AT3G14230 | 0.0 | 8.22E-01 | -0.1 | 4.06E-01 | 0.1 | 5.74E-01 | 0.1 | 5.75E-01 | -0.3 | 5.64E-02 | 0.3 | 5.89E-02 | hypothetical protein |
| AT1G70810 | 0.0 | 9.14E-01 | -0.5 | 2.54E-02 | 0.5 | 5.20E-02 | 0.9 | 1.45E-02 | 0.4 | 3.82E-01 | 0.5 | 2.27E-01 | Protein C2-DOMAIN ABA-RELATED 7 |
| AT1G08610 | 0.0 | 8.80E-01 | 0.0 | 8.33E-01 | 0.0 | 7.67E-01 | 0.2 | 3.50E-01 | 0.1 | 6.83E-01 | 0.1 | 7.82E-01 | Pentatricopeptide repeat-containing protein At1g08610 |
| AT4G11170 | 0.0 | 9.30E-01 | -0.6 | 3.97E-02 | 0.5 | 8.51E-02 | 2.1 | 3.66E-06 | 0.5 | 1.99E-01 | 1.6 | 4.52E-04 | Putative disease resistance protein At4g11170 |
| AT1G48110 | 0.0 | 8.40E-01 | -0.1 | 5.77E-01 | 0.0 | 7.55E-01 | 0.1 | 5.61E-01 | 0.1 | 4.72E-01 | 0.0 | 8.94E-01 | YTH domain-containing family protein |
| AT4G12760 | 0.0 | 9.35E-01 | -0.1 | 6.62E-01 | 0.1 | 7.69E-01 | 0.1 | 8.36E-01 | 0.1 | 8.81E-01 | 0.0 | 9.90E-01 | RPA-interacting protein C-terminal domain-containing protein |
| AT1G35580 | 0.0 | 8.31E-01 | 0.2 | 7.09E-02 | -0.2 | 6.33E-02 | -0.3 | 3.15E-02 | -0.3 | 2.72E-02 | 0.0 | 8.61E-01 | Alkaline |
| AT2G41410 | 0.0 | 8.78E-01 | -0.4 | 3.47E-03 | 0.4 | 7.12E-03 | 0.1 | 6.19E-01 | -0.2 | 4.45E-01 | 0.3 | 3.11E-01 | Probable calcium-binding protein CML35 |
| AT4G33300 | 0.0 | 8.28E-01 | -0.3 | 3.34E-03 | 0.3 | 7.52E-03 | -0.1 | 6.42E-01 | -0.3 | 4.69E-02 | 0.2 | 2.34E-01 | Probable disease resistance protein At4g33300 |
| AT1G08750 | 0.0 | 8.57E-01 | 0.0 | 6.91E-01 | -0.1 | 6.23E-01 | -0.2 | 3.96E-01 | -0.1 | 8.07E-01 | -0.1 | 6.91E-01 | Putative GPI-anchor transamidase |
| AT1G63900 | 0.0 | 7.82E-01 | -0.2 | 5.01E-03 | 0.2 | 1.41E-02 | -0.3 | 1.29E-01 | -0.4 | 8.80E-02 | 0.1 | 8.23E-01 | RING-type E3 ubiquitin transferase |
| AT4G20400 | 0.0 | 7.65E-01 | -0.1 | 7.53E-02 | 0.1 | 1.63E-01 | 0.1 | 6.08E-01 | 0.0 | 9.76E-01 | 0.1 | 7.21E-01 | PKDM7B |
| AT1G48900 | 0.0 | 7.87E-01 | 0.0 | 7.41E-01 | 0.0 | 5.81E-01 | 0.0 | 8.64E-01 | 0.0 | 8.28E-01 | 0.1 | 7.70E-01 | Signal recognition particle 54 kDa protein |
| AT5G48655 | 0.0 | 8.44E-01 | -0.2 | 3.75E-02 | 0.2 | 8.13E-02 | -0.2 | 9.04E-02 | -0.3 | 4.86E-02 | 0.1 | 7.79E-01 | Putative RING zinc finger |
| AT5G53320 | 0.0 | 8.47E-01 | -0.2 | 9.93E-02 | 0.1 | 2.01E-01 | 0.7 | 2.00E-02 | 0.5 | 1.69E-01 | 0.2 | 5.12E-01 | Probable inactive receptor kinase At5g53320 |
| AT3G21260 | 0.0 | 9.02E-01 | -0.1 | 4.99E-01 | 0.1 | 6.44E-01 | -0.1 | 8.04E-01 | 0.0 | 8.77E-01 | 0.0 | 9.76E-01 | GLTP3 |
| AT1G78690 | 0.0 | 8.62E-01 | 0.0 | 9.24E-01 | 0.0 | 8.26E-01 | -0.5 | 5.84E-03 | -0.3 | 8.67E-02 | -0.2 | 4.54E-01 | Tafazzin family protein |
| AT5G18580 | 0.0 | 7.73E-01 | 0.0 | 9.64E-01 | 0.0 | 8.43E-01 | -0.2 | 1.40E-01 | -0.1 | 6.19E-01 | -0.1 | 5.98E-01 | Probable serine |
| AT2G03120 | 0.0 | 8.03E-01 | 0.1 | 1.46E-01 | -0.1 | 1.20E-01 | 0.3 | 1.32E-02 | 0.2 | 1.38E-01 | 0.1 | 5.99E-01 | Signal peptide peptidase |
| AT3G04670 | 0.0 | 8.35E-01 | -0.1 | 1.24E-01 | 0.1 | 2.24E-01 | -0.2 | 2.78E-01 | -0.2 | 1.68E-01 | 0.1 | 7.96E-01 | AT3G04670 protein |
| AT3G47890 | 0.0 | 8.39E-01 | 0.0 | 8.01E-01 | 0.0 | 9.62E-01 | 0.1 | 7.01E-01 | 0.1 | 7.56E-01 | 0.0 | 1.00E+00 | hypothetical protein |
| AT1G69010 | 0.0 | 8.49E-01 | 0.0 | 8.80E-01 | 0.0 | 9.82E-01 | -0.1 | 7.54E-01 | -0.1 | 6.33E-01 | 0.0 | 8.99E-01 | Transcription factor BIM2 |
| AT1G04860 | 0.0 | 8.03E-01 | -0.1 | 1.21E-01 | 0.1 | 2.33E-01 | 0.1 | 7.53E-01 | -0.1 | 5.09E-01 | 0.2 | 3.88E-01 | Ubiquitin carboxyl-terminal hydrolase 2 |
| AT5G44400 | 0.0 | 8.38E-01 | 0.0 | 6.42E-01 | -0.1 | 5.44E-01 | 0.6 | 3.73E-03 | 0.5 | 1.34E-02 | 0.1 | 8.73E-01 | Berberine bridge enzyme-like 26 |
| AT5G42990 | 0.0 | 8.61E-01 | -0.1 | 5.48E-01 | 0.0 | 7.21E-01 | -0.2 | 4.12E-01 | -0.1 | 8.51E-01 | -0.2 | 7.25E-01 | UBC18 |
| AT1G60930 | 0.0 | 9.04E-01 | -0.1 | 6.00E-01 | 0.1 | 7.43E-01 | -0.1 | 9.40E-01 | 0.4 | 4.92E-01 | -0.5 | 4.87E-01 | ATP-dependent DNA helicase Q-like 4B |

| | | | | | | | | | | | | | |
|---|---|---|---|---|---|---|---|---|---|---|---|---|---|
| AT3G47520 | 0.0 | 8.78E-01 | 0.1 | 2.11E-01 | -0.2 | 1.92E-01 | 0.4 | 2.66E-03 | 0.1 | 3.90E-01 | 0.2 | 1.05E-01 | Malate dehydrogenase, chloroplastic |
| AT3G12980 | 0.0 | 8.61E-01 | -0.3 | 1.25E-02 | 0.3 | 2.90E-02 | -0.1 | 8.02E-01 | -0.2 | 4.58E-01 | 0.1 | 7.19E-01 | histone acetyltransferase |
| AT1G64040 | 0.0 | 8.09E-01 | 0.0 | 5.59E-01 | 0.0 | 7.57E-01 | -0.2 | 1.03E-01 | -0.3 | 4.59E-02 | 0.1 | 7.17E-01 | Serine |
| AT1G48860 | 0.0 | 8.13E-01 | 0.2 | 1.66E-02 | -0.2 | 1.75E-02 | 0.1 | 4.15E-01 | 0.1 | 5.76E-01 | 0.0 | 9.54E-01 | 3-phosphoshikimate 1-carboxyvinyltransferase |
| AT3G14150 | 0.0 | 8.88E-01 | 0.1 | 4.89E-01 | -0.1 | 4.72E-01 | -0.4 | 2.00E-02 | -0.3 | 1.24E-01 | -0.1 | 6.36E-01 | #NAME? |
| AT3G24040 | 0.0 | 8.30E-01 | 0.2 | 1.15E-02 | -0.2 | 1.49E-02 | -0.2 | 2.38E-01 | 0.1 | 4.84E-01 | -0.3 | 1.45E-01 | Glycosyltransferase |
| AT2G45160 | 0.0 | 8.20E-01 | 0.1 | 4.07E-01 | -0.1 | 3.32E-01 | -0.1 | 6.10E-01 | -0.1 | 7.47E-01 | 0.0 | 9.57E-01 | Scarecrow-like protein 27 |
| AT1G70580 | 0.0 | 9.09E-01 | -0.2 | 2.05E-01 | 0.2 | 2.82E-01 | -0.4 | 1.21E-02 | 0.0 | 9.92E-01 | -0.4 | 6.50E-02 | Glutamate--glyoxylate aminotransferase 2 |
| AT5G56030 | 0.0 | 9.22E-01 | 0.0 | 7.86E-01 | -0.1 | 7.37E-01 | 0.4 | 1.39E-02 | 0.4 | 1.47E-02 | 0.0 | 8.69E-01 | Heat shock protein 81-2 |
| AT5G55670 | 0.0 | 8.40E-01 | -0.2 | 1.72E-02 | 0.2 | 3.83E-02 | 0.0 | 8.51E-01 | 0.0 | 8.70E-01 | -0.1 | 7.88E-01 | RNA-binding family protein |
| AT5G46260 | 0.0 | 9.10E-01 | -0.3 | 3.64E-02 | 0.3 | 7.53E-02 | 0.5 | 3.92E-02 | 0.1 | 8.05E-01 | 0.4 | 1.81E-01 | ADP-ribosyl cyclase |
| AT2G22780 | 0.0 | 8.19E-01 | 0.0 | 6.95E-01 | 0.0 | 5.66E-01 | 0.0 | 8.08E-01 | 0.1 | 6.79E-01 | 0.0 | 8.83E-01 | Malate dehydrogenase 1, peroxisomal |
| AT1G53210 | 0.0 | 8.62E-01 | 0.1 | 1.24E-01 | -0.1 | 1.13E-01 | -0.4 | 2.51E-03 | -0.3 | 1.38E-02 | 0.0 | 8.14E-01 | Sodium |
| AT1G21690 | 0.0 | 9.17E-01 | 0.1 | 3.96E-01 | -0.1 | 4.10E-01 | -0.1 | 7.51E-01 | -0.1 | 8.49E-01 | 0.0 | 9.64E-01 | ATPase family associated with various cellular activities |
| AT4G03510 | 0.0 | 9.40E-01 | 0.3 | 1.24E-01 | -0.3 | 1.54E-01 | -0.3 | 2.08E-01 | -0.9 | 1.01E-03 | 0.6 | 2.39E-02 | E3 ubiquitin-protein ligase RMA1 |
| AT5G13000 | 0.0 | 8.81E-01 | -0.2 | 4.96E-02 | 0.2 | 8.34E-02 | 0.0 | 9.29E-01 | 0.1 | 5.60E-01 | -0.2 | 5.71E-01 | 1,3-beta-glucan synthase |
| AT1G07250 | 0.0 | 8.61E-01 | -0.1 | 4.93E-01 | 0.0 | 6.44E-01 | -0.5 | 6.94E-04 | -0.4 | 8.13E-03 | -0.1 | 6.33E-01 | Flavonol 3-O-glucosyltransferase UGT71C4 |
| AT4G03415 | 0.0 | 8.41E-01 | 0.1 | 4.59E-02 | -0.2 | 4.86E-02 | -0.3 | 1.21E-01 | -0.2 | 3.43E-01 | -0.1 | 7.96E-01 | Probable protein phosphatase 2C 52 |
| AT1G51200 | 0.0 | 8.86E-01 | -0.1 | 2.35E-01 | 0.1 | 3.29E-01 | -0.2 | 2.39E-01 | -0.2 | 2.60E-01 | 0.0 | 9.53E-01 | Zinc finger A20 and AN1 domain-containing stress-associated protein 2 |
| AT2G18760 | 0.0 | 8.67E-01 | -0.3 | 6.91E-03 | 0.3 | 1.92E-02 | 0.1 | 8.43E-01 | 0.2 | 6.55E-01 | -0.1 | 8.28E-01 | Chromatin remodeling 8 |
| AT2G18510 | 0.0 | 8.74E-01 | 0.0 | 5.98E-01 | 0.0 | 7.49E-01 | 0.6 | 1.53E-04 | 0.3 | 6.55E-02 | 0.3 | 5.37E-02 | Putative spliceosome-associated protein family protein) |
| AT2G32710 | 0.0 | 8.86E-01 | 0.0 | 6.33E-01 | -0.1 | 5.93E-01 | -0.5 | 5.41E-03 | -0.5 | 8.59E-03 | 0.0 | 9.80E-01 | Cyclin-dependent kinase inhibitor domain-containing protein |
| AT5G10860 | 0.0 | 9.02E-01 | -0.1 | 5.89E-01 | 0.0 | 7.05E-01 | -0.5 | 7.84E-04 | -0.6 | 1.03E-03 | 0.0 | 8.45E-01 | CBS domain-containing protein CBSX3, mitochondrial |
| AT5G63960 | 0.0 | 8.95E-01 | -0.2 | 1.51E-01 | 0.1 | 2.53E-01 | 0.3 | 1.56E-01 | 0.1 | 6.41E-01 | 0.2 | 5.31E-01 | DNA polymerase |
| AT5G61760 | 0.0 | 8.87E-01 | 0.0 | 6.77E-01 | -0.1 | 6.28E-01 | 0.1 | 3.48E-01 | -0.2 | 3.00E-01 | 0.3 | 1.08E-01 | Inositol polyphosphate multikinase beta |
| AT3G23210 | 0.0 | 8.88E-01 | -0.2 | 7.33E-02 | 0.2 | 1.40E-01 | 0.0 | 9.47E-01 | 0.0 | 9.81E-01 | 0.0 | 9.82E-01 | Basic helix-loop-helix DNA-binding superfamily protein |
| AT2G03150 | 0.0 | 8.68E-01 | 0.0 | 7.00E-01 | 0.0 | 8.47E-01 | 0.3 | 6.22E-02 | 0.2 | 1.21E-01 | 0.0 | 9.56E-01 | ATP |
| AT2G36320 | 0.0 | 9.04E-01 | 0.4 | 2.12E-03 | -0.4 | 3.20E-03 | -0.6 | 3.29E-03 | -0.6 | 5.48E-03 | 0.0 | 9.46E-01 | Zinc finger A20 and AN1 domain-containing stress-associated protein 4 |
| AT1G75000 | 0.0 | 9.36E-01 | -0.2 | 3.49E-01 | 0.1 | 4.73E-01 | 0.5 | 1.31E-01 | 0.4 | 3.68E-01 | 0.1 | 7.16E-01 | F25A4.4 |
| AT3G02090 | 0.0 | 8.71E-01 | 0.0 | 5.59E-01 | -0.1 | 4.89E-01 | 0.1 | 2.31E-01 | 0.1 | 3.22E-01 | 0.0 | 9.94E-01 | Putative mitochondrial processing peptidase |
| AT2G47850 | 0.0 | 8.59E-01 | -0.1 | 2.05E-01 | 0.1 | 3.30E-01 | -0.1 | 3.79E-01 | -0.1 | 6.98E-01 | -0.1 | 8.28E-01 | Zinc finger C-x8-C-x5-C-x3-H type family protein |
| AT5G20300 | 0.0 | 8.89E-01 | 0.0 | 8.80E-01 | 0.0 | 8.01E-01 | -0.2 | 2.71E-01 | -0.2 | 2.51E-01 | 0.0 | 9.11E-01 | Avirulence induced gene family protein |
| AT5G43810 | 0.0 | 8.93E-01 | 0.0 | 9.77E-01 | 0.0 | 8.93E-01 | -0.1 | 8.09E-01 | 0.0 | 9.03E-01 | 0.0 | 9.62E-01 | Protein argonaute 10 |
| AT2G22740 | 0.0 | 9.10E-01 | -0.1 | 4.66E-01 | 0.1 | 5.98E-01 | -0.3 | 2.06E-01 | -0.4 | 1.08E-01 | 0.1 | 7.23E-01 | Histone-lysine N-methyltransferase, H3 lysine-9 specific SUVH6 3-9 homolog protein 6) |
| AT3G02530 | 0.0 | 8.93E-01 | 0.1 | 4.16E-01 | -0.1 | 3.84E-01 | 0.2 | 6.82E-02 | 0.2 | 1.73E-01 | 0.0 | 8.94E-01 | hypothetical protein |
| AT4G26190 | 0.0 | 9.07E-01 | 0.0 | 6.76E-01 | -0.1 | 6.27E-01 | 0.2 | 1.82E-01 | 0.2 | 3.83E-01 | 0.0 | 8.78E-01 | Haloacid dehalogenase-like hydrolase superfamily protein |
| AT5G59740 | 0.0 | 9.14E-01 | 0.1 | 1.96E-01 | -0.2 | 2.37E-01 | 0.3 | 2.53E-01 | 0.3 | 4.80E-01 | 0.1 | 8.46E-01 | UDP-galactose |
| AT1G15280 | 0.0 | 8.90E-01 | -0.3 | 2.37E-03 | 0.3 | 5.31E-03 | 0.1 | 7.29E-01 | 0.0 | 9.97E-01 | 0.1 | 7.82E-01 | CASC3 |
| AT4G04210 | 0.0 | 8.94E-01 | -0.1 | 1.21E-01 | 0.1 | 1.81E-01 | 0.0 | 9.36E-01 | 0.2 | 2.44E-01 | -0.2 | 3.10E-01 | Plant UBX domain-containing protein 4 |
| AT2G36460 | 0.0 | 8.95E-01 | -0.3 | 7.86E-03 | 0.2 | 1.48E-02 | 0.3 | 1.14E-02 | -0.2 | 3.44E-01 | 0.5 | 4.39E-03 | Fructose-bisphosphate aldolase |
| AT3G22200 | 0.0 | 8.88E-01 | 0.0 | 6.84E-01 | 0.0 | 6.14E-01 | 0.1 | 5.89E-01 | 0.4 | 8.82E-02 | -0.3 | 3.32E-01 | Gamma-aminobutyrate transaminase POP2, mitochondrial |
| AT2G06510 | 0.0 | 9.04E-01 | -0.2 | 8.03E-02 | 0.2 | 1.47E-01 | -0.2 | 3.69E-01 | -0.3 | 1.69E-01 | 0.1 | 6.80E-01 | Replication protein A subunit |
| AT4G39100 | 0.0 | 8.76E-01 | -0.1 | 1.22E-01 | 0.1 | 1.95E-01 | -0.5 | 4.09E-03 | -0.3 | 4.82E-02 | -0.1 | 5.91E-01 | PHD finger family protein |
| AT1G11310 | 0.0 | 9.12E-01 | -0.2 | 7.38E-02 | 0.2 | 1.11E-01 | 0.4 | 2.55E-02 | 0.3 | 1.42E-01 | 0.1 | 7.34E-01 | At1g11310 |
| AT2G28100 | 0.0 | 9.16E-01 | 0.0 | 6.59E-01 | -0.1 | 6.41E-01 | -0.1 | 6.27E-01 | 0.2 | 5.37E-01 | -0.3 | 2.98E-01 | alpha-L-fucosidase |
| AT1G51590 | 0.0 | 8.99E-01 | 0.0 | 7.64E-01 | 0.0 | 8.81E-01 | 0.0 | 8.53E-01 | 0.1 | 3.43E-01 | -0.1 | 5.75E-01 | Mannosyl-oligosaccharide 1,2-alpha-mannosidase MNS1 |
| AT4G15810 | 0.0 | 9.21E-01 | 0.0 | 7.67E-01 | 0.0 | 8.70E-01 | 0.3 | 1.94E-01 | 0.3 | 2.25E-01 | 0.0 | 9.78E-01 | P-loop containing nucleoside triphosphate hydrolases superfamily protein |
| AT1G17890 | 0.0 | 9.18E-01 | 0.2 | 3.26E-02 | -0.2 | 4.02E-02 | 0.1 | 4.70E-01 | 0.1 | 4.49E-01 | 0.0 | 9.35E-01 | GDP-L-fucose synthase |
| AT1G52160 | 0.0 | 9.31E-01 | -0.1 | 4.96E-01 | 0.1 | 6.17E-01 | 0.4 | 6.90E-03 | 0.0 | 8.72E-01 | 0.4 | 5.31E-02 | tRNase Z TRZ3, mitochondrial |
| AT2G44670 | 0.0 | 9.45E-01 | 0.4 | 1.59E-02 | -0.4 | 2.09E-02 | 0.2 | 3.05E-01 | 0.1 | 7.12E-01 | 0.1 | 7.37E-01 | FCS-Like Zinc finger 3 |
| AT4G13730 | 0.0 | 8.81E-01 | -0.1 | 1.00E-01 | 0.1 | 1.76E-01 | 0.0 | 8.34E-01 | -0.1 | 7.47E-01 | 0.0 | 9.22E-01 | Uncharacterized protein AT4g13730 |
| AT2G27500 | 0.0 | 9.31E-01 | 0.2 | 1.17E-01 | -0.2 | 1.46E-01 | 0.2 | 3.62E-01 | -0.3 | 1.66E-01 | 0.5 | 6.70E-02 | Glucan endo-1,3-beta-glucosidase 14 -beta-glucan endohydrolase 14) -beta-glucanase 14) |
| AT5G39340 | 0.0 | 9.66E-01 | 0.0 | 9.88E-01 | 0.0 | 9.82E-01 | 0.1 | 5.66E-01 | 0.1 | 6.07E-01 | 0.0 | 9.90E-01 | Histidine-containing phosphotransfer protein |
| AT1G33680 | 0.0 | 9.22E-01 | -0.3 | 6.38E-03 | 0.3 | 1.35E-02 | 0.0 | 9.97E-01 | 0.0 | 9.86E-01 | 0.0 | 9.87E-01 | KH domain-containing protein |
| AT1G62740 | 0.0 | 8.77E-01 | -0.1 | 1.57E-01 | 0.1 | 2.43E-01 | 0.2 | 3.36E-01 | 0.2 | 5.68E-01 | 0.1 | 8.85E-01 | Hsp70-Hsp90 organizing protein 2 |
| AT3G25210 | 0.0 | 9.19E-01 | 0.0 | 9.59E-01 | 0.0 | 9.04E-01 | -0.1 | 3.52E-01 | -0.3 | 8.39E-02 | 0.1 | 5.08E-01 | Pentatricopeptide repeat-containing protein At3g25210, mitochondrial |
| AT5G23690 | 0.0 | 9.53E-01 | -0.1 | 6.26E-01 | 0.1 | 7.23E-01 | 0.0 | 9.22E-01 | 0.4 | 1.17E-01 | -0.4 | 2.33E-01 | Polynucleotide adenylyltransferase family protein |
| AT2G29080 | 0.0 | 8.84E-01 | -0.1 | 4.60E-02 | 0.1 | 7.67E-02 | 0.3 | 3.44E-02 | 0.1 | 3.68E-01 | 0.1 | 5.08E-01 | ATP-dependent zinc metalloprotease FTSH 3, mitochondrial |
| AT4G32880 | 0.0 | 9.19E-01 | -0.2 | 2.02E-02 | 0.2 | 4.16E-02 | 0.1 | 7.81E-01 | 0.0 | 9.12E-01 | 0.0 | 9.37E-01 | Homeobox-leucine zipper protein ATHB-8 |
| AT1G59830 | 0.0 | 8.95E-01 | -0.1 | 2.79E-01 | 0.1 | 3.89E-01 | -0.3 | 6.12E-03 | -0.5 | 1.02E-03 | 0.1 | 3.91E-01 | Serine |
| AT1G19340 | 0.0 | 9.59E-01 | 0.0 | 9.34E-01 | 0.0 | 9.15E-01 | -0.2 | 3.42E-01 | -0.3 | 1.53E-01 | 0.1 | 6.74E-01 | Methyltransferase-like protein 2 |
| AT1G08010 | 0.0 | 9.14E-01 | 0.0 | 5.93E-01 | 0.0 | 5.75E-01 | -0.1 | 7.45E-01 | 0.1 | 6.86E-01 | -0.1 | 5.31E-01 | GATA transcription factor 11 |
| AT5G51980 | 0.0 | 9.23E-01 | 0.0 | 7.61E-01 | 0.0 | 7.29E-01 | 0.1 | 6.81E-01 | 0.0 | 8.66E-01 | 0.0 | 9.09E-01 | C3H1-type domain-containing protein |
| AT1G20950 | 0.0 | 9.27E-01 | 0.1 | 2.22E-01 | -0.1 | 2.24E-01 | 0.2 | 1.33E-01 | 0.3 | 2.06E-02 | -0.1 | 4.40E-01 | Pyrophosphate--fructose 6-phosphate 1-phosphotransferase subunit alpha 1 |
| AT1G14000 | 0.0 | 9.17E-01 | -0.2 | 1.07E-02 | 0.2 | 1.95E-02 | 0.0 | 8.13E-01 | 0.1 | 6.13E-01 | -0.1 | 8.40E-01 | Serine |
| AT3G19740 | 0.0 | 9.15E-01 | 0.0 | 6.89E-01 | 0.0 | 7.98E-01 | 0.0 | 9.85E-01 | 0.1 | 8.46E-01 | -0.1 | 8.39E-01 | P-loop containing nucleoside triphosphate hydrolases superfamily protein |
| AT2G21050 | 0.0 | 9.27E-01 | 0.0 | 6.95E-01 | 0.0 | 6.74E-01 | 0.1 | 6.38E-01 | -0.1 | 5.47E-01 | 0.2 | 3.05E-01 | Auxin transporter-like protein 2 |
| AT4G12420 | 0.0 | 9.38E-01 | 0.4 | 1.34E-03 | -0.4 | 2.23E-03 | -0.3 | 1.23E-02 | -0.3 | 3.89E-02 | 0.0 | 8.93E-01 | Monocopper oxidase-like protein SKU5 |
| AT3G15120 | 0.0 | 9.21E-01 | -0.2 | 4.05E-03 | 0.2 | 9.82E-03 | -0.3 | 1.51E-01 | -0.1 | 8.04E-01 | -0.3 | 4.23E-01 | P-loop containing nucleoside triphosphate hydrolases superfamily protein |
| AT3G24880 | 0.0 | 9.65E-01 | -0.4 | 2.18E-02 | 0.4 | 4.45E-02 | 0.1 | 8.93E-01 | 0.3 | 7.22E-01 | -0.2 | 8.46E-01 | Helicase |
| AT3G63400 | 0.0 | 9.59E-01 | -0.1 | 4.21E-01 | 0.1 | 4.98E-01 | 0.1 | 3.68E-01 | -0.2 | 3.09E-01 | 0.3 | 9.14E-02 | Peptidyl-prolyl cis-trans isomerase CYP63 |
| AT2G46505 | 0.0 | 9.59E-01 | 0.2 | 1.09E-01 | -0.2 | 1.25E-01 | 0.1 | 5.80E-01 | 0.1 | 5.43E-01 | 0.0 | 9.26E-01 | Succinate dehydrogenase subunit 4, mitochondrial |
| AT3G49780 | 0.0 | 9.62E-01 | 0.1 | 6.26E-01 | -0.1 | 6.52E-01 | -0.1 | 7.96E-01 | -0.5 | 9.76E-02 | 0.4 | 2.75E-01 | Phytosulfokines 3 |
| AT4G11350 | 0.0 | 9.79E-01 | -0.2 | 2.72E-01 | 0.2 | 3.71E-01 | 0.4 | 3.21E-01 | 0.3 | 5.46E-01 | | | Transferring glycosyl group transferase |
| AT1G14230 | 0.0 | 9.55E-01 | -0.1 | 2.80E-01 | 0.1 | 3.65E-01 | -0.2 | 3.07E-01 | 0.1 | 6.55E-01 | -0.3 | 2.25E-01 | Probable apyrase 4 |
| AT2G47310 | 0.0 | 9.69E-01 | -0.1 | 3.65E-01 | 0.1 | 4.63E-01 | 0.1 | 8.19E-01 | 0.2 | 5.66E-01 | -0.1 | 7.87E-01 | At2g47310 |
| AT2G21470 | 0.0 | 9.45E-01 | -0.2 | 2.77E-02 | 0.2 | 4.80E-02 | 0.1 | 2.09E-01 | 0.1 | 2.83E-01 | 0.0 | 9.97E-01 | SUMO-activating enzyme subunit |
| AT5G65620 | 0.0 | 9.52E-01 | -0.1 | 1.14E-01 | 0.1 | 1.51E-01 | 0.2 | 5.17E-01 | 0.2 | 6.23E-01 | 0.0 | 9.84E-01 | oligopeptidase A |
| AT1G51450 | 0.0 | 9.54E-01 | 0.1 | 3.61E-01 | -0.1 | 4.05E-01 | -0.1 | 5.55E-01 | 0.0 | 8.79E-01 | -0.1 | 8.15E-01 | Protein TRAUCO |
| AT5G04895 | 0.0 | 9.67E-01 | -0.3 | 3.63E-02 | 0.3 | 7.21E-02 | 0.7 | 2.05E-02 | 0.2 | 5.48E-01 | 0.5 | 2.04E-01 | RNA helicase |
| AT2G19120 | 0.0 | 9.68E-01 | -0.2 | 3.21E-02 | 0.2 | 6.54E-02 | 0.4 | 2.89E-01 | 0.3 | 4.96E-01 | 0.1 | 8.76E-01 | P-loop containing nucleoside triphosphate hydrolases superfamily protein |
| AT5G35930 | 0.0 | 9.63E-01 | 0.0 | 7.23E-01 | 0.0 | 7.98E-01 | -0.2 | 3.31E-01 | -0.1 | 6.65E-01 | -0.1 | 7.67E-01 | AMP-dependent synthetase and ligase family protein |
| AT3G45240 | 0.0 | 9.51E-01 | 0.0 | 6.59E-01 | 0.0 | 7.39E-01 | 0.0 | 7.78E-01 | 0.1 | 5.32E-01 | -0.1 | 4.68E-01 | Geminivirus rep interacting kinase 1 |

| | | | | | | | | | | | | | |
|---|---|---|---|---|---|---|---|---|---|---|---|---|---|
| AT1G78080 | 0.0 | 9.67E-01 | -0.2 | 3.21E-02 | 0.2 | 4.66E-02 | -0.3 | 1.49E-02 | -0.1 | 3.40E-01 | -0.1 | 3.75E-01 | Ethylene-responsive transcription factor RAP2-4 |
| AT4G18930 | 0.0 | 9.59E-01 | 0.0 | 6.13E-01 | 0.0 | 6.16E-01 | -0.3 | 4.09E-02 | -0.2 | 2.76E-01 | -0.1 | 5.81E-01 | Cyclic phosphodiesterase |
| AT3G19280 | 0.0 | 9.75E-01 | -0.1 | 2.19E-01 | 0.1 | 3.04E-01 | -0.1 | 6.58E-01 | -0.1 | 8.80E-01 | -0.1 | 8.52E-01 | Alpha 1, 3 fucosyltransferase-like protein |
| AT5G53310 | 0.0 | 9.64E-01 | 0.1 | 1.66E-01 | -0.1 | 1.93E-01 | -0.2 | 1.89E-01 | -0.2 | 1.36E-01 | 0.0 | 8.46E-01 | TH1 domain-containing protein |
| AT1G24150 | 0.0 | 9.69E-01 | -0.2 | 1.16E-02 | 0.2 | 2.27E-02 | 0.6 | 2.51E-03 | 0.0 | 9.26E-01 | 0.6 | 7.21E-03 | Formin-like protein |
| AT4G16710 | 0.0 | 9.81E-01 | -0.3 | 3.38E-02 | 0.3 | 6.23E-02 | -0.2 | 5.59E-01 | -0.3 | 3.60E-01 | 0.1 | 7.87E-01 | hypothetical protein |
| AT1G01300 | 0.0 | 9.74E-01 | 0.3 | 6.45E-03 | -0.3 | 9.55E-03 | 0.2 | 8.96E-02 | 0.2 | 2.08E-01 | 0.0 | 9.11E-01 | Aspartyl protease family protein 2 |
| AT2G38380 | 0.0 | 9.80E-01 | 0.4 | 4.37E-03 | -0.4 | 6.81E-03 | 0.6 | 5.02E-05 | 0.7 | 4.26E-05 | -0.1 | 3.74E-01 | Peroxidase 22 |
| AT3G13235 | 0.0 | 9.71E-01 | -0.1 | 3.00E-01 | 0.1 | 3.66E-01 | 0.1 | 3.82E-01 | 0.1 | 7.86E-01 | 0.1 | 7.80E-01 | Ubiquitin family protein |
| AT3G49850 | 0.0 | 9.81E-01 | -0.1 | 3.28E-01 | 0.1 | 4.09E-01 | 0.0 | 9.41E-01 | 0.1 | 5.76E-01 | -0.1 | 5.90E-01 | Telomere repeat-binding factor 3 |
| AT1G16350 | 0.0 | 9.86E-01 | 0.2 | 2.75E-02 | -0.2 | 3.98E-02 | 0.2 | 8.46E-02 | 0.2 | 3.45E-01 | 0.1 | 7.24E-01 | Putative inosine-5'-monophosphate dehydrogenase |
| AT4G34040 | 0.0 | 9.86E-01 | -0.3 | 2.30E-03 | 0.3 | 4.79E-03 | -0.2 | 3.10E-01 | -0.1 | 6.39E-01 | -0.1 | 8.16E-01 | E3 ubiquitin-protein ligase MBR2 |
| AT5G08670 | 0.0 | 9.90E-01 | 0.1 | 1.82E-01 | -0.1 | 2.08E-01 | 0.0 | 9.85E-01 | 0.0 | 8.04E-01 | 0.0 | 8.57E-01 | ATP synthase subunit beta-1, mitochondrial |
| AT2G22540 | 0.0 | 9.92E-01 | -0.1 | 1.86E-01 | 0.1 | 2.18E-01 | -0.3 | 1.96E-02 | -0.1 | 6.45E-01 | -0.3 | 2.04E-01 | hypothetical protein |
| AT5G56930 | 0.0 | 9.92E-01 | -0.1 | 3.71E-01 | 0.1 | 4.52E-01 | 0.0 | 8.63E-01 | -0.2 | 5.41E-01 | 0.1 | 7.49E-01 | Zinc finger CCCH domain-containing protein 65 |
| AT4G39400 | 0.0 | 9.93E-01 | 0.0 | 9.57E-01 | 0.0 | 9.55E-01 | -0.3 | 4.67E-02 | -0.2 | 2.47E-01 | -0.1 | 6.84E-01 | Protein BRASSINOSTEROID INSENSITIVE 1 |
| AT5G51280 | 0.0 | 9.92E-01 | -0.1 | 4.42E-01 | 0.1 | 5.02E-01 | 0.1 | 4.06E-01 | -0.1 | 6.90E-01 | 0.2 | 3.06E-01 | DEAD-box ATP-dependent RNA helicase 35 |
| AT4G16580 | 0.0 | 9.93E-01 | 0.0 | 8.96E-01 | 0.0 | 9.18E-01 | -0.1 | 4.52E-01 | -0.2 | 4.31E-01 | 0.0 | 9.37E-01 | Probable protein phosphatase 2C 55 |
| AT4G37120 | 0.0 | 9.93E-01 | -0.1 | 1.46E-01 | 0.1 | 1.76E-01 | 0.1 | 1.89E-01 | 0.0 | 9.23E-01 | 0.1 | 4.53E-01 | Pre-mRNA-splicing factor SLU7-B |
| AT4G33410 | 0.0 | 9.99E-01 | 0.1 | 7.31E-02 | -0.1 | 1.09E-01 | -0.1 | 3.13E-01 | 0.0 | 9.11E-01 | -0.2 | 3.81E-01 | Signal peptide peptidase-like 1 |
| AT1G31770 | 0.0 | 1.00E+00 | 0.4 | 8.51E-03 | -0.4 | 1.59E-02 | 0.6 | 3.06E-04 | 0.4 | 1.46E-02 | 0.2 | 2.58E-01 | ABC transporter G family member 14 |
| AT1G63290 | 0.0 | 9.99E-01 | 0.2 | 1.04E-01 | -0.2 | 1.37E-01 | 0.0 | 9.52E-01 | 0.0 | 9.50E-01 | 0.0 | 9.95E-01 | Ribulose-phosphate 3-epimerase |
| AT5G55300 | 0.0 | 9.96E-01 | -0.1 | 1.78E-01 | 0.1 | 2.15E-01 | -0.1 | 5.43E-01 | 0.0 | 7.66E-01 | 0.0 | 9.11E-01 | DNA topoisomerase |
| AT3G04290 | 0.0 | 9.98E-01 | 0.0 | 8.76E-01 | 0.0 | 8.89E-01 | -0.2 | 1.84E-01 | -0.1 | 6.26E-01 | -0.1 | 6.56E-01 | GDSL esterase |
| AT1G30280 | 0.0 | 9.99E-01 | 0.6 | 2.71E-02 | -0.6 | 5.91E-02 | -0.4 | 4.37E-01 | -0.2 | 7.72E-01 | -0.2 | 8.15E-01 | Chaperone DnaJ-domain superfamily protein |
| AT2G30780 | 0.0 | 9.95E-01 | -0.1 | 4.78E-01 | 0.1 | 5.41E-01 | 0.1 | 5.52E-01 | 0.2 | 3.40E-01 | -0.1 | 7.79E-01 | Pentatricopeptide repeat-containing protein At2g30780 |
| AT1G55520 | 0.0 | 9.93E-01 | 0.1 | 1.35E-01 | -0.1 | 1.87E-01 | 0.0 | 9.39E-01 | 0.0 | 9.48E-01 | 0.0 | 9.98E-01 | TATA binding protein 2 |
| AT3G55410 | 0.0 | 9.94E-01 | 0.0 | 5.54E-01 | 0.0 | 5.90E-01 | 0.5 | 5.79E-03 | 0.2 | 2.04E-01 | 0.2 | 3.22E-01 | oxoglutarate dehydrogenase |
| AT4G19510 | 0.0 | 9.95E-01 | -0.2 | 7.91E-02 | 0.2 | 1.10E-01 | 0.1 | 6.44E-01 | -0.1 | 7.06E-01 | 0.1 | 4.72E-01 | Disease resistance protein |
| AT1G30000 | 0.0 | 9.91E-01 | 0.0 | 8.83E-01 | 0.0 | 9.07E-01 | 0.0 | 9.25E-01 | 0.0 | 9.65E-01 | 0.0 | 9.83E-01 | alpha-1,2-Mannosidase |
| AT3G18210 | 0.0 | 9.91E-01 | 0.1 | 2.92E-01 | -0.1 | 3.67E-01 | 0.3 | 1.80E-01 | 0.0 | 9.23E-01 | 0.2 | 3.52E-01 | 2-oxoglutarate and iron-dependent oxygenase domain-containing protein CP2 |
| AT3G03340 | 0.0 | 9.86E-01 | -0.2 | 4.99E-03 | 0.2 | 9.73E-03 | 0.2 | 2.73E-01 | 0.2 | 3.73E-01 | 0.0 | 9.73E-01 | LUC7 related protein |
| AT5G12040 | 0.0 | 9.86E-01 | 0.0 | 8.38E-01 | 0.0 | 8.45E-01 | 0.2 | 5.17E-02 | 0.3 | 8.62E-03 | -0.1 | 4.58E-01 | hypothetical protein |
| AT1G14170 | 0.0 | 9.85E-01 | 0.3 | 3.32E-03 | -0.3 | 7.06E-03 | 0.0 | 9.81E-01 | 0.0 | 9.68E-01 | 0.0 | 9.63E-01 | AT1G14170 protein |
| AT1G67680 | 0.0 | 9.86E-01 | 0.1 | 1.79E-01 | -0.1 | 2.19E-01 | 0.2 | 9.23E-02 | 0.1 | 3.73E-01 | 0.1 | 7.32E-01 | Signal recognition particle subunit SRP72 |
| AT3G43920 | 0.0 | 9.89E-01 | -0.2 | 2.65E-01 | 0.2 | 3.36E-01 | 0.0 | 9.33E-01 | -0.3 | 5.73E-01 | 0.2 | 6.90E-01 | Dicer-like 3 |
| AT1G50370 | 0.0 | 9.85E-01 | 0.0 | 7.94E-01 | 0.0 | 8.06E-01 | 0.0 | 9.78E-01 | 0.0 | 9.38E-01 | 0.0 | 9.65E-01 | Phytochrome-associated serine |
| AT3G06270 | 0.0 | 9.86E-01 | 0.0 | 7.04E-01 | 0.0 | 7.38E-01 | 0.3 | 2.96E-02 | -0.2 | 3.26E-01 | 0.5 | 1.48E-02 | Probable protein phosphatase 2C 35 |
| AT3G44630 | 0.0 | 9.91E-01 | -0.4 | 2.58E-02 | 0.4 | 4.20E-02 | 0.3 | 1.50E-01 | 0.1 | 5.96E-01 | 0.2 | 5.59E-01 | ADP-ribosyl cyclase |
| AT1G18670 | 0.0 | 9.84E-01 | -0.1 | 6.20E-01 | 0.1 | 6.61E-01 | -0.1 | 7.97E-01 | 0.0 | 9.97E-01 | -0.1 | 8.63E-01 | Protein kinase superfamily protein |
| AT5G45020 | 0.0 | 9.83E-01 | 0.1 | 3.21E-01 | -0.1 | 4.10E-01 | -0.2 | 3.55E-01 | -0.1 | 8.63E-01 | -0.2 | 6.07E-01 | AT5g45020 |
| AT4G14000 | 0.0 | 9.80E-01 | 0.0 | 6.76E-01 | 0.0 | 7.39E-01 | 0.1 | 6.66E-01 | 0.0 | 8.25E-01 | 0.0 | 9.33E-01 | Methyltransferase family protein |
| AT1G03770 | 0.0 | 9.84E-01 | -0.1 | 4.90E-01 | 0.1 | 5.51E-01 | 0.4 | 1.37E-01 | 0.3 | 3.28E-01 | 0.1 | 8.15E-01 | RING 1B |
| AT4G30000 | 0.0 | 9.80E-01 | 0.0 | 7.41E-01 | 0.0 | 7.67E-01 | 0.1 | 7.70E-01 | 0.2 | 6.01E-01 | -0.1 | 8.24E-01 | Putative dihydropteroate synthase |
| AT4G03250 | 0.0 | 9.77E-01 | -0.1 | 3.64E-01 | 0.1 | 4.26E-01 | -0.1 | 6.99E-01 | -0.2 | 5.09E-01 | 0.1 | 8.26E-01 | At4g03250 |
| AT5G62990 | 0.0 | 9.84E-01 | -0.3 | 7.93E-02 | 0.3 | 1.27E-01 | 0.7 | 4.87E-03 | 0.5 | 8.67E-02 | 0.3 | 4.03E-01 | Ubiquitin carboxyl-terminal hydrolase family protein |
| AT1G12500 | 0.0 | 9.78E-01 | 0.2 | 4.32E-02 | -0.2 | 6.26E-02 | -0.2 | 3.70E-01 | 0.0 | 9.82E-01 | -0.2 | 4.87E-01 | Probable sugar phosphate |
| AT3G02890 | 0.0 | 9.73E-01 | -0.1 | 1.70E-01 | 0.1 | 2.21E-01 | 0.0 | 8.47E-01 | 0.0 | 9.92E-01 | 0.0 | 8.87E-01 | Zinc finger PHD-type domain-containing protein |
| AT2G44490 | 0.0 | 9.73E-01 | -0.1 | 2.88E-01 | 0.1 | 3.07E-01 | 0.2 | 5.48E-02 | -0.1 | 5.18E-01 | 0.3 | 5.81E-02 | Beta-glucosidase 26, peroxisomal |
| AT5G58230 | 0.0 | 9.68E-01 | -0.1 | 4.34E-01 | 0.1 | 4.63E-01 | 0.2 | 1.22E-01 | 0.0 | 8.40E-01 | 0.2 | 3.95E-01 | Histone-binding protein MSI1 |
| AT5G26940 | 0.0 | 9.69E-01 | 0.0 | 9.85E-01 | 0.0 | 9.88E-01 | -0.2 | 2.23E-01 | -0.1 | 8.14E-01 | -0.1 | 5.05E-01 | Exonuclease DPD1, chloroplastic |
| AT1G32150 | 0.0 | 9.82E-01 | -0.1 | 3.26E-01 | 0.1 | 3.82E-01 | -0.2 | 3.53E-01 | 0.0 | 9.30E-01 | -0.2 | 5.73E-01 | bZIP transcription factor 68 |
| AT4G00340 | 0.0 | 9.72E-01 | -0.1 | 1.79E-01 | 0.1 | 2.08E-01 | 0.1 | 7.26E-01 | 0.2 | 4.57E-01 | -0.1 | 7.60E-01 | Putative receptor-like protein kinase |
| AT1G17110 | 0.0 | 9.59E-01 | -0.1 | 6.91E-02 | 0.1 | 8.84E-02 | -0.1 | 8.27E-01 | -0.1 | 8.64E-01 | 0.0 | 9.96E-01 | Ubiquitin-specific protease 15 |
| AT3G10572 | 0.0 | 9.80E-01 | 0.1 | 6.29E-01 | -0.1 | 7.04E-01 | -0.2 | 3.96E-01 | -0.3 | 9.80E-02 | 0.2 | 5.17E-01 | 3-phosphoinositide-dependent protein kinase-1 |
| AT4G37310 | 0.0 | 9.77E-01 | -0.3 | 2.95E-02 | 0.3 | 4.39E-02 | -0.2 | 2.24E-01 | -0.2 | 3.79E-01 | 0.0 | 9.16E-01 | Cytochrome P450, family 81, subfamily H, polypeptide 1 |
| AT1G16560 | 0.0 | 9.63E-01 | 0.2 | 2.74E-02 | -0.2 | 4.45E-02 | 0.1 | 3.56E-01 | 0.1 | 4.32E-01 | 0.0 | 9.99E-01 | Post-GPI attachment to proteins factor 3 |
| AT4G30570 | 0.0 | 9.80E-01 | 0.0 | 9.37E-01 | 0.0 | 9.64E-01 | 0.0 | 9.47E-01 | -0.2 | 8.14E-01 | 0.2 | 7.93E-01 | Probable mannose-1-phosphate guanylyltransferase 3 |
| AT5G16450 | 0.0 | 9.51E-01 | 0.0 | 7.59E-01 | 0.0 | 8.35E-01 | 0.1 | 7.18E-01 | 0.0 | 9.73E-01 | 0.1 | 8.19E-01 | Putative 4-hydroxy-4-methyl-2-oxoglutarate aldolase 2 |
| AT2G20140 | 0.0 | 9.59E-01 | 0.2 | 5.01E-02 | -0.2 | 7.70E-02 | 0.1 | 2.70E-01 | 0.0 | 9.63E-01 | 0.1 | 4.98E-01 | 26S proteasome regulatory subunit 4 homolog B |
| AT5G46070 | 0.0 | 9.40E-01 | -0.1 | 3.34E-01 | 0.1 | 3.43E-01 | 0.2 | 2.49E-01 | 0.1 | 5.09E-01 | 0.0 | 8.52E-01 | Guanylate-binding family protein |
| AT5G48900 | 0.0 | 9.62E-01 | 0.4 | 3.20E-03 | -0.4 | 7.51E-03 | -0.5 | 2.63E-02 | 0.0 | 9.85E-01 | -0.5 | 5.18E-02 | Pectate lyase |
| AT1G58684 | 0.0 | 9.95E-01 | 0.4 | 6.62E-01 | -0.4 | 7.48E-01 | | | | | | | XW6 |
| AT5G15450 | 0.0 | 9.67E-01 | -0.6 | 1.59E-03 | 0.6 | 2.68E-03 | 0.3 | 2.00E-01 | 0.6 | 3.52E-02 | -0.3 | 4.53E-01 | Chaperone protein ClpB3, chloroplastic |
| AT1G28390 | 0.0 | 9.61E-01 | 0.2 | 1.31E-01 | -0.2 | 2.03E-01 | -0.3 | 1.86E-01 | -0.2 | 3.28E-01 | 0.0 | 9.14E-01 | Protein kinase superfamily protein |
| AT5G37830 | 0.0 | 9.38E-01 | -0.1 | 1.88E-01 | 0.1 | 1.96E-01 | 0.1 | 8.10E-01 | 0.1 | 7.84E-01 | 0.0 | 9.57E-01 | 5-oxoprolinase 1 |
| AT5G58200 | 0.0 | 9.50E-01 | -0.2 | 1.17E-01 | 0.2 | 1.38E-01 | -0.2 | 2.75E-01 | -0.2 | 1.91E-01 | 0.1 | 7.99E-01 | hypothetical protein |
| AT3G20650 | 0.0 | 9.38E-01 | -0.1 | 3.83E-01 | 0.1 | 4.08E-01 | 0.1 | 6.87E-01 | 0.0 | 9.57E-01 | 0.1 | 8.16E-01 | mRNA cap guanine-N7 methyltransferase |
| AT2G29110 | 0.0 | 9.80E-01 | -0.7 | 3.03E-02 | 0.7 | 5.61E-02 | -0.3 | 5.02E-01 | -0.3 | 6.21E-01 | 0.0 | 9.59E-01 | Glutamate receptor |
| AT5G10790 | 0.0 | 9.36E-01 | 0.0 | 7.65E-01 | 0.0 | 7.45E-01 | 0.1 | 5.37E-01 | 0.1 | 4.19E-01 | 0.0 | 8.70E-01 | Ubiquitin-specific protease 22 |
| AT5G05210 | 0.0 | 9.54E-01 | 0.1 | 3.86E-01 | -0.1 | 4.68E-01 | 0.3 | 3.31E-02 | 0.1 | 5.53E-01 | 0.2 | 3.28E-01 | AT5g05210 |
| AT1G77550 | 0.0 | 9.56E-01 | 0.1 | 5.78E-01 | -0.1 | 6.71E-01 | 0.3 | 8.25E-02 | 0.3 | 1.40E-01 | 0.0 | 9.66E-01 | At1g77550 |
| AT1G14685 | 0.0 | 9.27E-01 | 0.1 | 8.01E-02 | -0.1 | 1.27E-01 | -0.3 | 1.21E-02 | -0.3 | 3.43E-02 | 0.0 | 8.96E-01 | GAGA-binding transcriptional activator |
| AT3G06010 | 0.0 | 9.21E-01 | -0.2 | 4.05E-02 | 0.2 | 4.99E-02 | 0.0 | 9.23E-01 | 0.0 | 8.84E-01 | 0.0 | 9.70E-01 | Probable ATP-dependent DNA helicase CHR12 |
| AT1G51660 | 0.0 | 9.34E-01 | -0.4 | 7.70E-04 | 0.4 | 1.80E-03 | 0.6 | 3.27E-04 | 0.2 | 3.30E-01 | 0.4 | 2.58E-02 | Mitogen-activated protein kinase kinase 4 |
| AT4G18300 | 0.0 | 9.30E-01 | -0.3 | 4.14E-03 | 0.3 | 8.68E-03 | 0.0 | 8.83E-01 | 0.0 | 9.15E-01 | 0.0 | 9.97E-01 | eIF-2B GDP-GTP exchange factor subunit epsilon |
| AT2G22420 | 0.0 | 9.58E-01 | -0.1 | 4.20E-01 | 0.1 | 4.68E-01 | -0.1 | 6.92E-01 | -0.1 | 7.44E-01 | 0.0 | 1.00E+00 | Peroxidase 17 |
| AT1G69640 | 0.0 | 9.10E-01 | 0.0 | 7.50E-01 | 0.0 | 6.97E-01 | 0.1 | 5.25E-01 | 0.0 | 8.43E-01 | 0.1 | 8.24E-01 | Sphinganine C4-monooxygenase 1 |
| AT4G38040 | 0.0 | 9.08E-01 | 0.0 | 7.12E-01 | 0.0 | 6.74E-01 | -0.3 | 9.36E-02 | 0.0 | 9.18E-01 | -0.3 | 1.41E-01 | Exostosin family protein |
| AT4G16660 | 0.0 | 9.31E-01 | -0.1 | 5.66E-01 | 0.1 | 5.43E-01 | 0.6 | 8.25E-05 | 0.5 | 1.99E-03 | 0.1 | 4.95E-01 | Heat shock protein 70 family protein |
| AT1G09250 | 0.0 | 9.30E-01 | 0.2 | 5.05E-02 | -0.2 | 8.63E-02 | -0.7 | 2.74E-04 | -0.4 | 1.14E-02 | -0.2 | 2.30E-01 | Transcription factor bHLH149 |
| AT4G04800 | 0.0 | 9.45E-01 | 0.0 | 8.89E-01 | 0.0 | 8.58E-01 | 0.0 | 9.81E-01 | -0.1 | 4.94E-01 | 0.1 | 5.95E-01 | Peptide methionine sulfoxide reductase B3 -S-oxide reductase) |
| AT3G11900 | 0.0 | 9.07E-01 | 0.0 | 5.08E-01 | 0.1 | 4.84E-01 | 0.0 | 8.87E-01 | 0.3 | 2.64E-01 | -0.3 | 2.57E-01 | Amino acid transporter ANT1 |

| | | | | | | | | | | | | | |
|---|---|---|---|---|---|---|---|---|---|---|---|---|---|
| AT3G19760 | 0.0 | 9.35E-01 | 0.1 | 4.57E-01 | -0.1 | 5.41E-01 | 0.1 | 3.96E-01 | 0.0 | 9.97E-01 | 0.1 | 5.21E-01 | Eukaryotic initiation factor 4A-III homolog |
| AT5G01960 | 0.0 | 8.98E-01 | -0.3 | 2.56E-03 | 0.3 | 4.24E-03 | 0.1 | 5.20E-01 | 0.0 | 8.37E-01 | 0.2 | 4.89E-01 | hypothetical protein |
| AT5G19460 | 0.0 | 9.45E-01 | 0.0 | 7.35E-01 | 0.0 | 8.21E-01 | 0.1 | 5.96E-01 | 0.2 | 3.45E-01 | -0.1 | 7.31E-01 | Nudix hydrolase 20, chloroplastic |
| AT5G44720 | 0.0 | 9.45E-01 | 0.2 | 1.86E-01 | -0.2 | 2.47E-01 | 0.1 | 7.00E-01 | 0.2 | 3.90E-01 | -0.1 | 7.34E-01 | Molybdenum cofactor sulfurase family protein |
| AT1G80360 | 0.0 | 9.19E-01 | -0.1 | 2.27E-01 | 0.1 | 2.23E-01 | -0.6 | 4.05E-05 | -0.2 | 2.25E-01 | -0.5 | 7.27E-03 | Aromatic aminotransferase ISS1 |
| AT2G32070 | 0.0 | 9.27E-01 | -0.3 | 6.30E-03 | 0.3 | 1.02E-02 | 0.1 | 7.17E-01 | -0.1 | 8.35E-01 | 0.1 | 6.55E-01 | Probable CCR4-associated factor 1 homolog 7 |
| AT2G20290 | 0.0 | 9.41E-01 | 0.0 | 7.94E-01 | 0.0 | 8.77E-01 | 0.3 | 4.19E-01 | 0.4 | 2.15E-01 | -0.2 | 7.15E-01 | Myosin-like protein XIG |
| AT4G20310 | 0.0 | 9.39E-01 | -0.1 | 3.81E-01 | 0.1 | 4.14E-01 | -0.5 | 7.84E-02 | 0.0 | 9.44E-01 | -0.6 | 1.57E-01 | Endopeptidase S2P |
| AT4G14410 | 0.0 | 9.39E-01 | -0.2 | 1.90E-01 | 0.2 | 2.05E-01 | -0.1 | 3.58E-01 | -0.1 | 4.31E-01 | 0.0 | 9.98E-01 | BHLH104 |
| AT4G05160 | 0.0 | 8.82E-01 | -0.1 | 1.05E-01 | 0.1 | 1.08E-01 | -0.1 | 4.38E-01 | 0.0 | 9.70E-01 | -0.1 | 5.45E-01 | 4-coumarate--CoA ligase-like 7 |
| AT1G51390 | 0.0 | 9.28E-01 | -0.1 | 2.84E-01 | 0.1 | 3.19E-01 | -0.2 | 5.83E-01 | -0.1 | 6.91E-01 | 0.0 | 9.63E-01 | NifU-like protein 5, mitochondrial |
| AT2G29630 | 0.0 | 9.36E-01 | -0.2 | 7.77E-02 | 0.2 | 8.53E-02 | 0.0 | 8.86E-01 | 0.3 | 1.45E-01 | -0.3 | 1.71E-01 | ThiaminC |
| AT3G44370 | 0.0 | 9.15E-01 | -0.2 | 8.61E-02 | 0.2 | 1.13E-01 | 0.5 | 1.43E-02 | 0.4 | 9.29E-02 | 0.1 | 6.35E-01 | Membrane insertion protein, OxaA |
| AT2G14080 | 0.0 | 9.43E-01 | 0.0 | 8.08E-01 | 0.0 | 7.86E-01 | -0.1 | 4.82E-01 | 0.0 | 9.04E-01 | -0.1 | 7.39E-01 | Disease resistance protein family |
| AT2G32400 | 0.0 | 9.08E-01 | -0.1 | 2.51E-01 | 0.1 | 2.70E-01 | 0.5 | 2.86E-02 | 0.3 | 1.72E-01 | 0.1 | 6.25E-01 | Glutamate receptor 3.7 |
| AT1G58210 | 0.0 | 9.21E-01 | 0.0 | 9.36E-01 | 0.0 | 9.92E-01 | 0.0 | 9.31E-01 | -0.1 | 8.63E-01 | 0.1 | 8.45E-01 | Uncharacterized protein F16M22.3 |
| AT4G20870 | 0.0 | 9.10E-01 | 0.3 | 8.09E-03 | -0.3 | 1.80E-02 | -0.4 | 1.08E-02 | -0.3 | 6.55E-02 | -0.1 | 7.15E-01 | Fatty acid hydroxylase domain-containing protein |
| AT5G59440 | 0.0 | 8.90E-01 | -0.1 | 4.21E-01 | 0.1 | 4.11E-01 | -0.2 | 2.30E-01 | -0.3 | 7.06E-02 | 0.1 | 5.29E-01 | dTMP kinase |
| AT5G16560 | 0.0 | 8.91E-01 | 0.0 | 9.93E-01 | 0.0 | 9.22E-01 | -0.1 | 7.02E-01 | -0.2 | 4.15E-01 | 0.1 | 7.39E-01 | Transcription repressor KAN1 |
| AT1G14610 | 0.0 | 8.72E-01 | -0.1 | 1.76E-01 | 0.1 | 1.60E-01 | 0.2 | 1.77E-01 | 0.1 | 6.23E-01 | 0.1 | 6.81E-01 | Valine--tRNA ligase, mitochondrial 1 |
| AT5G63790 | 0.0 | 9.15E-01 | -0.3 | 2.46E-02 | 0.3 | 3.17E-02 | 0.1 | 5.34E-01 | -0.3 | 1.04E-01 | 0.3 | 6.36E-02 | hypothetical protein |
| AT3G58990 | 0.0 | 9.30E-01 | 0.5 | 4.60E-03 | -0.5 | 8.45E-03 | -0.9 | 2.70E-04 | 0.8 | 9.90E-04 | -1.6 | 6.27E-05 | 3-isopropylmalate dehydratase small subunit 3 alkylmalate dehydratase small subunit 3) |
| AT2G23460 | 0.0 | 8.84E-01 | -0.1 | 2.25E-01 | 0.1 | 2.21E-01 | -0.2 | 2.00E-01 | -0.2 | 4.75E-01 | -0.1 | 8.12E-01 | Extra-large guanine nucleotide-binding protein 1 |
| AT3G54740 | 0.0 | 8.94E-01 | 0.2 | 5.95E-02 | -0.2 | 1.20E-01 | -0.3 | 6.71E-02 | -0.2 | 2.51E-01 | -0.1 | 7.52E-01 | Uncharacterized protein T5N23_100 |
| AT1G69250 | 0.0 | 9.18E-01 | 0.0 | 7.88E-01 | 0.0 | 7.47E-01 | 0.3 | 1.01E-01 | 0.3 | 2.30E-01 | 0.0 | 8.66E-01 | At1g69250 family protein with RNA binding domain-containing protein) |
| AT5G61590 | 0.0 | 9.34E-01 | 0.0 | 9.24E-01 | 0.0 | 8.76E-01 | -0.1 | 6.85E-01 | -1.3 | 1.03E-03 | 1.1 | 5.02E-03 | Ethylene-responsive transcription factor ERF107 |
| AT5G05960 | 0.0 | 9.10E-01 | -0.1 | 5.51E-01 | 0.1 | 5.39E-01 | 0.1 | 4.94E-01 | 0.2 | 5.33E-01 | 0.0 | 9.78E-01 | hypothetical protein |
| AT1G26260 | 0.0 | 9.17E-01 | 0.0 | 9.55E-01 | 0.0 | 9.74E-01 | 0.3 | 3.95E-01 | 0.5 | 8.25E-02 | -0.3 | 3.82E-01 | Cryptochrome-interacting basic-helix-loop-helix 5 |
| AT1G01050 | 0.0 | 8.43E-01 | 0.2 | 2.60E-02 | -0.2 | 5.31E-02 | 0.0 | 9.91E-01 | 0.0 | 9.88E-01 | 0.0 | 9.97E-01 | inorganic diphosphatase |
| AT1G70560 | 0.0 | 8.85E-01 | 0.4 | 1.30E-03 | -0.4 | 4.16E-03 | 0.3 | 7.22E-02 | 0.1 | 6.00E-01 | 0.2 | 4.32E-01 | L-tryptophan--pyruvate aminotransferase 1 |
| AT1G14040 | 0.0 | 8.85E-01 | -0.1 | 3.66E-01 | 0.1 | 3.67E-01 | 1.4 | 1.15E-07 | 0.7 | 4.68E-04 | 0.7 | 1.50E-03 | Phosphate transporter PHO1 homolog 3 |
| AT5G39760 | 0.0 | 8.77E-01 | 0.3 | 6.14E-03 | -0.3 | 1.65E-02 | 0.3 | 7.80E-02 | 0.0 | 8.76E-01 | 0.2 | 2.13E-01 | Zinc-finger homeodomain protein 10 |
| AT4G29810 | 0.0 | 8.98E-01 | -0.2 | 1.19E-01 | 0.2 | 1.22E-01 | 0.5 | 1.64E-03 | 0.3 | 7.01E-02 | 0.2 | 2.60E-01 | AT4G29810 protein |
| AT1G78490 | 0.0 | 8.86E-01 | 0.2 | 7.37E-02 | -0.2 | 1.35E-01 | 0.1 | 8.44E-01 | 0.2 | 3.35E-01 | -0.2 | 5.38E-01 | At1g78490 |
| AT2G24230 | 0.0 | 9.04E-01 | 0.3 | 1.39E-02 | -0.3 | 3.38E-02 | 0.1 | 7.77E-01 | 0.2 | 3.46E-01 | -0.1 | 6.21E-01 | Probable LRR receptor-like serine |
| AT3G17100 | 0.0 | 8.81E-01 | 0.2 | 1.15E-01 | -0.1 | 1.86E-01 | -0.9 | 1.03E-05 | -0.7 | 2.69E-04 | -0.2 | 3.80E-01 | Transcription factor bHLH147 |
| AT3G59420 | 0.0 | 8.96E-01 | 0.0 | 7.68E-01 | 0.0 | 8.92E-01 | 0.0 | 9.17E-01 | 0.2 | 6.76E-01 | -0.2 | 6.39E-01 | hypothetical protein |
| AT1G13110 | 0.0 | 8.71E-01 | 0.1 | 2.57E-01 | -0.1 | 3.63E-01 | 0.3 | 1.16E-02 | 0.2 | 1.70E-01 | 0.1 | 5.08E-01 | Cytochrome P450 71B7 |
| AT4G08480 | 0.0 | 8.91E-01 | -0.3 | 1.46E-02 | 0.3 | 2.33E-02 | 0.0 | 9.10E-01 | -0.3 | 4.58E-01 | 0.3 | 4.54E-01 | Mitogen-activated protein kinase kinase kinase 9 |
| AT5G44340 | 0.0 | 8.64E-01 | 0.2 | 6.27E-02 | -0.2 | 1.06E-01 | -0.3 | 9.56E-03 | -0.2 | 1.11E-01 | -0.1 | 5.95E-01 | Tubulin beta-4 chain |
| AT4G30490 | 0.0 | 8.55E-01 | -0.1 | 1.38E-01 | 0.1 | 1.37E-01 | -0.3 | 8.32E-02 | -0.4 | 2.36E-02 | 0.1 | 6.05E-01 | AFG1-like ATPase family protein |
| AT4G14920 | 0.0 | 8.38E-01 | 0.0 | 5.87E-01 | 0.1 | 5.08E-01 | -0.1 | 6.78E-01 | 0.0 | 9.44E-01 | -0.1 | 7.17E-01 | Acyl-CoA N-acyltransferase with RING |
| AT3G17310 | 0.0 | 8.48E-01 | -0.1 | 2.51E-01 | 0.1 | 2.40E-01 | -0.2 | 1.49E-01 | -0.2 | 2.08E-01 | 0.0 | 9.90E-01 | S-adenosyl-L-methionine-dependent methyltransferases superfamily protein |
| AT3G55830 | 0.0 | 8.49E-01 | -0.2 | 5.27E-02 | 0.2 | 5.81E-02 | 0.1 | 8.68E-01 | 0.0 | 9.48E-01 | 0.0 | 9.60E-01 | Glycosylinositol phosphorylceramide mannosyl transferase 1 |
| AT1G75660 | 0.0 | 8.49E-01 | -0.2 | 1.07E-02 | 0.3 | 1.27E-02 | -0.1 | 7.49E-01 | -0.2 | 5.05E-01 | 0.1 | 7.95E-01 | 5'-3' exoribonuclease |
| AT2G35430 | 0.0 | 8.91E-01 | 0.0 | 9.49E-01 | 0.0 | 9.60E-01 | -0.2 | 4.55E-01 | -0.4 | 1.21E-01 | 0.2 | 4.82E-01 | Zinc finger CCCH domain-containing protein 28 |
| AT3G20410 | 0.0 | 8.15E-01 | 0.2 | 3.46E-02 | -0.1 | 7.20E-02 | 0.2 | 1.70E-01 | -0.2 | 1.48E-01 | 0.4 | 2.96E-02 | Calcium-dependent protein kinase 9 |
| AT4G29130 | 0.0 | 8.42E-01 | 0.1 | 3.05E-01 | -0.1 | 4.46E-01 | 0.0 | 7.84E-01 | 0.1 | 4.46E-01 | -0.1 | 6.72E-01 | Hexokinase-1 |
| AT4G36860 | 0.0 | 7.96E-01 | 0.1 | 3.70E-01 | 0.0 | 5.77E-01 | 0.0 | 9.22E-01 | 0.1 | 5.55E-01 | -0.1 | 6.97E-01 | LIM domain-containing protein |
| AT1G46480 | 0.0 | 9.10E-01 | 0.0 | 9.88E-01 | 0.0 | 9.22E-01 | 0.2 | 6.38E-01 | 0.6 | 1.09E-01 | -0.4 | 2.61E-01 | WUSCHEL related homeobox 4 |
| AT5G23300 | 0.0 | 9.07E-01 | 0.1 | 3.85E-01 | -0.1 | 5.16E-01 | 0.7 | 9.10E-04 | 0.5 | 1.64E-02 | 0.2 | 4.38E-01 | Dihydroorotate dehydrogenase , mitochondrial |
| AT3G04130 | 0.0 | 8.93E-01 | -0.1 | 3.18E-01 | 0.1 | 3.38E-01 | 0.1 | 7.94E-01 | -0.1 | 7.81E-01 | 0.2 | 6.39E-01 | Pentatricopeptide repeat-containing protein At3g04130, mitochondrial |
| AT2G36530 | 0.0 | 9.05E-01 | 0.3 | 7.26E-02 | -0.3 | 1.12E-01 | 0.6 | 2.71E-04 | 0.1 | 4.46E-01 | 0.5 | 9.12E-03 | phosphopyruvate hydratase |
| AT1G56680 | 0.0 | 9.47E-01 | 0.6 | 1.68E-02 | -0.6 | 4.48E-02 | -0.1 | 6.56E-01 | 0.4 | 1.94E-01 | -0.5 | 1.39E-01 | Chitin-binding type-1 domain-containing protein |
| AT3G20540 | 0.0 | 8.50E-01 | -0.2 | 5.00E-02 | 0.2 | 5.83E-02 | 0.2 | 1.63E-01 | 0.0 | 8.93E-01 | 0.2 | 3.99E-01 | Polymerase gamma 1 |
| AT3G53760 | 0.0 | 8.50E-01 | -0.1 | 4.20E-01 | 0.1 | 3.96E-01 | 0.0 | 9.44E-01 | -0.1 | 8.61E-01 | 0.1 | 8.48E-01 | Gamma-tubulin complex component |
| AT5G49800 | 0.0 | 8.93E-01 | 0.5 | 2.02E-03 | -0.5 | 7.75E-03 | -0.2 | 4.26E-01 | 0.1 | 5.76E-01 | -0.3 | 2.40E-01 | Polyketide cyclase |
| AT1G08470 | 0.0 | 8.16E-01 | 0.0 | 6.81E-01 | 0.0 | 8.79E-01 | -0.1 | 3.38E-01 | 0.0 | 8.05E-01 | -0.2 | 3.34E-01 | Protein STRICTOSIDINE SYNTHASE-LIKE 3 |
| AT1G15550 | 0.0 | 8.93E-01 | -0.3 | 4.97E-02 | 0.3 | 6.99E-02 | -0.1 | 8.30E-01 | 0.1 | 8.25E-01 | -0.2 | 7.24E-01 | Gibberellin 3-beta-dioxygenase 1 |
| AT1G06180 | 0.0 | 9.08E-01 | -0.2 | 1.91E-01 | 0.2 | 2.26E-01 | 0.1 | 8.57E-01 | -0.3 | 3.73E-01 | 0.4 | 3.56E-01 | Transcription factor MYB13 |
| AT5G05620 | 0.0 | 8.58E-01 | 0.2 | 6.92E-02 | -0.2 | 1.43E-01 | 0.2 | 1.64E-01 | 0.2 | 3.74E-01 | 0.1 | 8.27E-01 | Tubulin gamma-2 chain |
| AT1G55110 | 0.0 | 7.97E-01 | -0.2 | 7.64E-03 | 0.2 | 1.05E-02 | -0.1 | 4.18E-01 | -0.4 | 5.36E-02 | 0.2 | 3.19E-01 | hypothetical protein |
| AT4G15230 | 0.0 | 8.85E-01 | -0.6 | 1.47E-03 | 0.6 | 3.75E-03 | 0.1 | 9.32E-01 | 0.8 | 1.63E-01 | -0.7 | 2.55E-01 | Pleiotropic drug resistance 2 |
| AT2G03340 | 0.0 | 8.21E-01 | -0.2 | 5.06E-02 | 0.2 | 5.00E-02 | -0.2 | 1.69E-01 | -0.1 | 3.70E-01 | 0.0 | 8.49E-01 | Probable WRKY transcription factor 3 |
| AT2G44750 | 0.0 | 8.70E-01 | -0.4 | 4.54E-03 | 0.4 | 9.11E-03 | 0.0 | 9.10E-01 | -0.4 | 3.20E-01 | 0.4 | 2.45E-01 | thiamine diphosphokinase |
| AT1G73100 | 0.0 | 7.90E-01 | 0.0 | 8.53E-01 | 0.0 | 7.01E-01 | 0.1 | 5.25E-01 | 0.0 | 9.22E-01 | 0.2 | 5.19E-01 | Histone-lysine N-methyltransferase, H3 lysine-9 specific SUVH3 3-9 homolog protein 3) |
| AT5G19010 | 0.0 | 8.46E-01 | 0.0 | 8.09E-01 | 0.0 | 9.62E-01 | -0.2 | 3.28E-01 | 0.0 | 9.66E-01 | -0.3 | 3.90E-01 | Mitogen-activated protein kinase 16 |
| AT3G20010 | 0.0 | 8.33E-01 | 0.0 | 8.12E-01 | 0.0 | 7.07E-01 | 0.2 | 3.79E-01 | 0.3 | 1.42E-01 | -0.1 | 6.32E-01 | SNF2 domain-containing protein |
| AT5G08430 | 0.0 | 8.48E-01 | -0.1 | 5.59E-01 | 0.1 | 4.93E-01 | -0.3 | 1.92E-01 | 0.1 | 8.08E-01 | -0.4 | 1.83E-01 | SWIB |
| AT4G02070 | 0.0 | 8.53E-01 | -0.1 | 4.55E-01 | 0.1 | 4.26E-01 | 0.2 | 3.48E-01 | -0.2 | 5.28E-01 | 0.4 | 1.95E-01 | DNA mismatch repair protein |
| AT3G05520 | 0.0 | 8.40E-01 | 0.0 | 7.49E-01 | 0.0 | 9.19E-01 | -0.4 | 6.84E-03 | -0.2 | 1.61E-01 | -0.2 | 3.62E-01 | Subunits of heterodimeric actin filament capping protein Capz superfamily |
| AT4G27720 | 0.0 | 8.15E-01 | 0.0 | 8.39E-01 | 0.0 | 9.88E-01 | -0.1 | 3.88E-01 | 0.2 | 1.90E-01 | -0.3 | 8.55E-02 | Molybdate-anion transporter |
| AT5G62610 | 0.0 | 7.50E-01 | 0.0 | 5.54E-01 | 0.1 | 4.16E-01 | -0.2 | 5.52E-02 | -0.3 | 4.11E-02 | 0.0 | 8.43E-01 | Transcription factor bHLH79 |
| AT2G30910 | 0.0 | 8.63E-01 | -0.1 | 3.95E-01 | 0.1 | 3.80E-01 | -0.1 | 6.92E-01 | 0.0 | 9.75E-01 | -0.1 | 8.15E-01 | Actin-related protein 2 |
| AT3G20840 | 0.0 | 9.34E-01 | 0.3 | 2.52E-01 | -0.2 | 3.80E-01 | 0.2 | 6.80E-01 | 0.9 | 2.07E-02 | -0.7 | 6.50E-02 | AP2-like ethylene-responsive transcription factor PLT1 |
| AT4G38910 | 0.0 | 9.02E-01 | 0.2 | 2.88E-01 | -0.1 | 4.29E-01 | -0.6 | 3.49E-02 | -0.1 | 6.73E-01 | -0.4 | 1.86E-01 | Protein BASIC PENTACYSTEINE5 |
| AT5G38630 | 0.0 | 8.49E-01 | 0.2 | 5.50E-02 | -0.2 | 1.09E-01 | 0.1 | 4.80E-01 | -0.1 | 3.80E-01 | 0.2 | 1.73E-01 | Transmembrane ascorbate ferrireductase 2 |
| AT2G24390 | 0.0 | 8.03E-01 | 0.0 | 8.84E-01 | 0.0 | 9.45E-01 | 0.5 | 1.80E-02 | 0.1 | 5.45E-01 | 0.3 | 2.08E-01 | Putative gamma-glutamylcyclotransferase |
| AT4G11330 | 0.0 | 9.02E-01 | -0.3 | 5.57E-02 | 0.4 | 6.92E-02 | -0.1 | 7.96E-01 | -0.1 | 4.93E-01 | 0.1 | 7.46E-01 | Mitogen-activated protein kinase |
| AT4G25900 | 0.0 | 8.58E-01 | 0.2 | 6.63E-02 | -0.2 | 1.18E-01 | 0.6 | 2.45E-03 | 0.2 | 4.53E-01 | 0.4 | 6.51E-02 | glucose-6-phosphate 1-epimerase |
| AT2G26100 | 0.0 | 7.93E-01 | -0.1 | 1.93E-01 | 0.1 | 1.75E-01 | 0.0 | 9.46E-01 | 0.0 | 8.90E-01 | 0.0 | 9.40E-01 | Hexosyltransferase |
| AT1G67660 | 0.0 | 8.57E-01 | 0.2 | 1.60E-01 | -0.1 | 2.79E-01 | 0.0 | 9.59E-01 | 0.0 | 9.12E-01 | 0.0 | 9.56E-01 | F12A21.19 |
| AT5G26680 | 0.0 | 8.33E-01 | 0.1 | 1.43E-01 | -0.1 | 2.76E-01 | 0.6 | 7.17E-03 | 0.4 | 1.27E-01 | 0.2 | 3.90E-01 | Flap endonuclease 1 |

| | | | | | | | | | | | | | |
|---|---|---|---|---|---|---|---|---|---|---|---|---|---|
| AT2G17740 | 0.0 | 9.68E-01 | 0.0 | 9.50E-01 | 0.0 | 9.82E-01 | -0.4 | 6.38E-01 | -3.4 | 5.61E-02 | 2.9 | 5.37E-02 | DC1 domain-containing protein |
| AT1G26460 | 0.0 | 8.34E-01 | 0.1 | 5.00E-01 | 0.0 | 6.83E-01 | 0.3 | 3.22E-02 | 0.2 | 3.37E-01 | 0.2 | 4.78E-01 | Pentatricopeptide repeat-containing protein At1g26460, mitochondrial |
| AT2G38950 | 0.0 | 7.37E-01 | -0.2 | 1.52E-02 | 0.2 | 1.54E-02 | 0.1 | 6.61E-01 | -0.1 | 7.38E-01 | 0.1 | 4.71E-01 | Probable inactive lysine-specific demethylase JMJ19 |
| AT2G40950 | 0.0 | 7.62E-01 | -0.3 | 2.80E-03 | 0.3 | 4.00E-03 | 0.2 | 2.12E-01 | 0.1 | 6.86E-01 | 0.1 | 6.39E-01 | bZIP transcription factor 17 |
| AT1G10900 | 0.0 | 8.08E-01 | -0.1 | 3.77E-01 | 0.1 | 3.07E-01 | 0.2 | 3.20E-01 | 0.4 | 3.69E-02 | -0.2 | 3.14E-01 | Phosphatidylinositol 4-phosphate 5-kinase 7 P-5-kinase 7) |
| AT3G15530 | 0.0 | 8.03E-01 | -0.1 | 2.11E-01 | 0.1 | 1.74E-01 | -0.2 | 1.45E-01 | -0.2 | 2.04E-01 | 0.0 | 9.97E-01 | hypothetical protein |
| AT3G57870 | 0.0 | 7.91E-01 | 0.1 | 1.68E-01 | -0.1 | 2.91E-01 | 0.0 | 8.05E-01 | 0.0 | 9.84E-01 | 0.0 | 8.62E-01 | SUMO-conjugating enzyme SCE1 |
| AT1G13170 | 0.0 | 8.10E-01 | 0.1 | 4.17E-01 | -0.1 | 6.17E-01 | -0.1 | 8.27E-01 | 0.2 | 4.58E-01 | -0.3 | 4.20E-01 | OSBP-related protein 1D |
| AT2G21380 | 0.0 | 8.52E-01 | -0.2 | 1.98E-01 | 0.2 | 1.78E-01 | -0.3 | 2.81E-01 | 0.0 | 9.20E-01 | -0.3 | 3.85E-01 | Kinesin-like protein KIN-7M, chloroplastic |
| AT1G11000 | 0.0 | 8.29E-01 | 0.2 | 2.45E-02 | -0.2 | 6.67E-02 | -0.2 | 4.79E-01 | 0.2 | 4.48E-01 | -0.3 | 2.00E-01 | MLO-like protein |
| AT1G79050 | 0.0 | 7.36E-01 | -0.1 | 9.62E-02 | 0.1 | 7.96E-02 | 0.1 | 3.42E-01 | 0.0 | 9.22E-01 | 0.1 | 5.59E-01 | RecA DNA recombination family protein |
| AT2G42680 | 0.0 | 8.29E-01 | -0.1 | 5.13E-01 | 0.1 | 4.19E-01 | 0.1 | 2.48E-01 | 0.2 | 2.45E-01 | 0.0 | 9.24E-01 | Multiprotein-bridging factor 1a |
| AT1G65790 | 0.0 | 9.49E-01 | -0.3 | 4.10E-01 | 0.3 | 4.46E-01 | 1.2 | 2.60E-03 | 0.5 | 3.18E-01 | 0.7 | 6.51E-02 | Receptor kinase 1 |
| AT1G30330 | 0.0 | 8.10E-01 | 0.0 | 9.98E-01 | 0.0 | 8.43E-01 | 0.0 | 9.38E-01 | 0.1 | 4.84E-01 | -0.1 | 6.35E-01 | Auxin response factor |
| AT2G17560 | 0.0 | 8.21E-01 | 0.2 | 2.98E-02 | -0.2 | 5.95E-02 | 0.2 | 2.50E-01 | 0.3 | 3.07E-01 | 0.0 | 9.83E-01 | High mobility group B protein 4 |
| AT1G06770 | 0.0 | 8.48E-01 | -0.2 | 1.45E-01 | 0.2 | 1.59E-01 | -0.3 | 3.23E-01 | -0.2 | 4.54E-01 | 0.0 | 9.35E-01 | DREB2A-interacting protein 1 |
| AT5G06050 | 0.0 | 8.17E-01 | 0.3 | 7.98E-03 | -0.3 | 2.39E-02 | 0.1 | 6.90E-01 | 0.1 | 5.99E-01 | 0.0 | 9.02E-01 | Probable methyltransferase PMT12 |
| AT5G07800 | 0.0 | 8.81E-01 | 0.0 | 8.61E-01 | 0.0 | 9.78E-01 | -0.1 | 7.58E-01 | 0.1 | 8.13E-01 | -0.3 | 6.04E-01 | Flavin-containing monooxygenase FMO GS-OX-like 9 |
| AT2G28590 | 0.0 | 8.40E-01 | -0.2 | 1.62E-01 | 0.2 | 1.75E-01 | 0.1 | 6.50E-01 | 0.3 | 3.88E-01 | -0.1 | 7.36E-01 | Protein kinase superfamily protein |
| AT5G02030 | 0.0 | 8.40E-01 | -0.2 | 5.87E-02 | 0.3 | 6.93E-02 | -0.2 | 3.35E-01 | -0.3 | 1.56E-01 | 0.1 | 7.19E-01 | BEL1-like homeodomain protein 9 |
| AT4G31120 | 0.0 | 7.67E-01 | 0.0 | 5.77E-01 | 0.0 | 8.21E-01 | 0.2 | 7.80E-02 | 0.0 | 9.17E-01 | 0.2 | 1.71E-01 | Protein arginine N-methyltransferase 1.5 |
| AT1G05710 | 0.0 | 8.79E-01 | -0.1 | 5.57E-01 | 0.1 | 5.38E-01 | -0.2 | 4.48E-01 | 0.0 | 9.00E-01 | -0.2 | 4.76E-01 | Basic helix-loop-helix DNA-binding superfamily protein |
| AT1G66340 | 0.0 | 7.74E-01 | -0.1 | 1.42E-01 | 0.2 | 1.18E-01 | 0.0 | 9.51E-01 | 0.0 | 9.38E-01 | 0.0 | 9.11E-01 | Ethylene receptor 1 |
| AT1G06410 | 0.0 | 7.65E-01 | -0.1 | 1.37E-01 | 0.2 | 1.02E-01 | 0.0 | 7.54E-01 | -0.1 | 2.57E-01 | 0.2 | 2.57E-01 | Probable alpha,alpha-trehalose-phosphate synthase 7 |
| AT4G23310 | 0.0 | 9.63E-01 | -1.0 | 1.39E-01 | 1.0 | 2.01E-01 | | | | | | | Cysteine-rich RLK 23 |
| AT4G26720 | 0.0 | 7.67E-01 | 0.1 | 9.87E-02 | -0.1 | 2.20E-01 | 0.0 | 7.41E-01 | -0.1 | 3.15E-01 | 0.1 | 6.31E-01 | Serine |
| AT3G23750 | 0.0 | 7.40E-01 | 0.0 | 9.70E-01 | 0.0 | 7.50E-01 | -0.3 | 3.77E-02 | -0.1 | 5.17E-01 | -0.2 | 3.85E-01 | Putative receptor-like kinase |
| AT2G41890 | 0.0 | 7.72E-01 | -0.1 | 2.78E-01 | 0.1 | 2.31E-01 | 0.2 | 3.24E-01 | 0.0 | 9.36E-01 | 0.1 | 5.53E-01 | G-type lectin S-receptor-like serine |
| AT1G16040 | 0.0 | 8.00E-01 | 0.1 | 3.71E-01 | -0.1 | 5.89E-01 | 0.2 | 4.64E-01 | 0.1 | 8.25E-01 | 0.1 | 7.64E-01 | At1g16040 |
| AT2G18520 | 0.0 | 8.66E-01 | 0.2 | 1.75E-01 | -0.2 | 3.14E-01 | 0.3 | 1.55E-01 | 0.0 | 9.05E-01 | 0.4 | 1.96E-01 | Pentatricopeptide repeat-containing protein At2g18520, mitochondrial |
| AT1G26850 | 0.0 | 6.85E-01 | 0.0 | 7.24E-01 | 0.0 | 9.82E-01 | 0.1 | 2.51E-01 | 0.1 | 3.33E-01 | 0.0 | 9.98E-01 | Methyltransferase |
| AT2G30590 | 0.0 | 6.76E-01 | -0.1 | 9.33E-02 | 0.1 | 7.09E-02 | -0.2 | 1.35E-01 | -0.4 | 3.48E-03 | 0.2 | 1.63E-01 | Probable WRKY transcription factor 21 |
| AT1G07940 | 0.0 | 8.22E-01 | 0.3 | 3.53E-02 | -0.3 | 6.88E-02 | 0.2 | 3.68E-01 | 0.0 | 9.66E-01 | 0.2 | 5.33E-01 | Elongation factor 1-alpha 1 |
| AT1G44760 | 0.0 | 7.96E-01 | 0.0 | 9.23E-01 | 0.0 | 7.71E-01 | -0.3 | 3.75E-02 | -0.2 | 1.65E-01 | -0.1 | 7.15E-01 | Adenine nucleotide alpha hydrolases-like superfamily protein |
| AT2G39360 | 0.0 | 7.17E-01 | 0.0 | 9.95E-01 | 0.0 | 7.74E-01 | -0.2 | 3.04E-01 | -0.1 | 6.28E-01 | -0.1 | 8.03E-01 | Probable receptor-like protein kinase At2g39360 |
| AT5G11530 | 0.0 | 7.32E-01 | -0.2 | 1.73E-02 | 0.3 | 1.96E-02 | 0.0 | 9.74E-01 | -0.3 | 2.74E-01 | 0.3 | 3.62E-01 | Protein EMBRYONIC FLOWER 1 |
| AT2G43900 | 0.0 | 8.05E-01 | 0.2 | 6.97E-02 | -0.2 | 1.64E-01 | -0.6 | 1.95E-01 | -0.2 | 6.67E-01 | -0.4 | 5.63E-01 | Endonuclease |
| AT1G55350 | 0.0 | 7.45E-01 | -0.1 | 1.26E-01 | 0.2 | 9.84E-02 | 0.1 | 7.99E-01 | 0.0 | 9.11E-01 | 0.0 | 9.53E-01 | Calpain-type cysteine protease family |
| AT2G22390 | 0.0 | 8.42E-01 | 0.1 | 4.92E-01 | -0.1 | 6.94E-01 | 0.0 | 9.29E-01 | -0.1 | 8.25E-01 | 0.1 | 9.10E-01 | Putative Ras-related protein RABA4e |
| AT2G40300 | 0.0 | 8.60E-01 | 0.2 | 3.35E-01 | -0.1 | 4.97E-01 | -1.8 | 7.89E-05 | -0.1 | 8.58E-01 | -1.7 | 1.34E-03 | Ferritin-4, chloroplastic |
| AT5G58575 | 0.0 | 7.80E-01 | 0.0 | 9.13E-01 | 0.0 | 9.04E-01 | -0.2 | 3.79E-01 | -0.3 | 1.88E-01 | 0.1 | 6.86E-01 | SAGA-associated factor 11 |
| AT4G04340 | 0.0 | 7.49E-01 | 0.1 | 1.88E-01 | -0.1 | 3.43E-01 | -0.6 | 1.15E-04 | -0.2 | 8.67E-02 | -0.3 | 4.90E-02 | Protein OSCA1 ]i increase 1) |
| AT4G16360 | 0.0 | 6.82E-01 | 0.0 | 8.62E-01 | 0.0 | 6.06E-01 | 0.2 | 8.00E-02 | 0.1 | 4.17E-01 | 0.1 | 6.37E-01 | SNF1-related protein kinase regulatory subunit beta-2 |
| AT5G35570 | 0.0 | 6.59E-01 | -0.1 | 1.63E-01 | 0.1 | 1.12E-01 | -0.1 | 4.68E-01 | -0.3 | 8.42E-02 | 0.2 | 3.37E-01 | O-fucosyltransferase 35 |
| AT2G33840 | 0.0 | 8.03E-01 | 0.1 | 3.25E-01 | -0.1 | 5.16E-01 | 0.3 | 6.87E-02 | 0.3 | 9.47E-02 | 0.0 | 9.81E-01 | Tyrosine--tRNA ligase 1, cytoplasmic |
| AT4G24330 | 0.0 | 7.53E-01 | 0.2 | 1.13E-01 | -0.1 | 2.21E-01 | 0.1 | 5.68E-01 | 0.0 | 9.91E-01 | 0.1 | 7.49E-01 | At4g24330 |
| AT4G05330 | 0.0 | 7.84E-01 | 0.2 | 3.51E-02 | -0.2 | 9.97E-02 | -0.3 | 1.09E-02 | -0.4 | 7.92E-03 | 0.1 | 8.23E-01 | ARF-GAP domain 13 |
| AT2G40660 | 0.0 | 7.90E-01 | 0.2 | 7.03E-02 | -0.2 | 1.47E-01 | 0.2 | 1.34E-01 | 0.3 | 3.51E-02 | -0.1 | 5.50E-01 | Nucleic acid-binding, OB-fold-like protein |
| AT1G45233 | 0.0 | 7.35E-01 | 0.1 | 5.26E-01 | 0.0 | 8.06E-01 | 0.0 | 8.45E-01 | 0.1 | 7.62E-01 | 0.0 | 9.16E-01 | THO complex subunit 5A |
| AT4G39540 | 0.0 | 7.96E-01 | -0.1 | 5.46E-01 | 0.1 | 4.43E-01 | 0.2 | 3.09E-01 | 0.1 | 7.84E-01 | 0.1 | 6.54E-01 | shikimate kinase |
| AT1G76510 | 0.0 | 7.31E-01 | 0.0 | 8.69E-01 | 0.0 | 6.69E-01 | -0.1 | 6.42E-01 | -0.1 | 8.40E-01 | 0.0 | 9.07E-01 | ARID |
| AT3G58970 | 0.0 | 6.13E-01 | -0.2 | 2.10E-02 | 0.2 | 1.60E-02 | -0.1 | 5.68E-01 | 0.0 | 9.75E-01 | -0.1 | 5.93E-01 | Magnesium transporter MRS2-4 |
| AT3G04630 | 0.0 | 7.55E-01 | 0.2 | 2.37E-02 | -0.2 | 6.16E-02 | -0.1 | 6.76E-01 | 0.0 | 7.76E-01 | -0.1 | 5.40E-01 | WVD2-like 1 |
| AT3G44850 | 0.0 | 7.65E-01 | 0.1 | 3.90E-01 | -0.1 | 6.29E-01 | 0.3 | 2.26E-01 | 0.3 | 1.76E-01 | -0.1 | 8.64E-01 | Protein kinase superfamily protein |
| AT1G20630 | 0.0 | 6.54E-01 | 0.1 | 2.53E-01 | -0.1 | 5.21E-01 | -0.5 | 3.39E-03 | -0.1 | 4.68E-01 | -0.4 | 4.53E-02 | Catalase-1 |
| AT2G47760 | 0.0 | 6.57E-01 | 0.1 | 1.78E-01 | -0.1 | 4.19E-01 | -0.2 | 3.06E-01 | -0.2 | 3.16E-01 | 0.0 | 9.61E-01 | Dol-P-Man:ManGlcNAc-PP-Dol alpha-1,3-mannosyltransferase |
| AT5G37350 | 0.0 | 7.20E-01 | -0.1 | 2.08E-01 | 0.1 | 1.56E-01 | 0.2 | 2.30E-01 | 0.0 | 9.35E-01 | 0.2 | 4.26E-01 | Serine |
| AT4G24790 | 0.0 | 7.61E-01 | 0.3 | 2.08E-02 | -0.2 | 6.71E-02 | -0.7 | 2.30E-02 | -0.4 | 1.74E-01 | -0.3 | 5.61E-01 | AAA-type ATPase family protein |
| AT4G29890 | 0.0 | 8.28E-01 | -0.4 | 2.03E-02 | 0.4 | 2.76E-02 | -0.5 | 3.10E-02 | -0.3 | 2.40E-01 | -0.2 | 5.22E-01 | Choline monooxygenase, chloroplastic |
| AT3G54540 | 0.0 | 6.30E-01 | -0.1 | 3.74E-01 | 0.1 | 2.12E-01 | -0.1 | 6.37E-01 | -0.2 | 4.83E-01 | 0.1 | 8.55E-01 | ABC transporter F family member 4 |
| AT3G03950 | 0.0 | 6.65E-01 | -0.2 | 2.34E-02 | 0.2 | 2.05E-02 | 0.3 | 5.38E-02 | 0.0 | 9.74E-01 | 0.3 | 1.49E-01 | YTH domain-containing family protein |
| AT5G17890 | 0.0 | 6.77E-01 | -0.1 | 2.18E-01 | 0.1 | 1.45E-01 | 0.2 | 9.78E-02 | -0.2 | 1.77E-01 | 0.4 | 2.14E-02 | Protein DA1-related 4 |
| AT5G39730 | 0.0 | 7.22E-01 | 0.4 | 2.20E-03 | -0.3 | 7.25E-03 | 0.2 | 3.94E-01 | 0.2 | 4.22E-01 | 0.0 | 9.64E-01 | AIG2-like protein B |
| AT1G49350 | 0.0 | 6.90E-01 | -0.1 | 3.28E-01 | 0.1 | 2.44E-01 | -0.3 | 3.46E-01 | 0.0 | 9.98E-01 | -0.3 | 4.56E-01 | Pseudouridine kinase |
| AT5G62670 | 0.0 | 7.42E-01 | 0.1 | 4.37E-01 | 0.0 | 6.82E-01 | -0.9 | 2.68E-04 | 0.0 | 9.84E-01 | -0.9 | 2.27E-03 | ATPase 11, plasma membrane-type |
| AT5G20070 | 0.0 | 6.49E-01 | -0.1 | 2.16E-01 | 0.1 | 1.27E-01 | 0.2 | 1.21E-01 | 0.3 | 1.97E-02 | -0.1 | 4.26E-01 | NAD diphosphatase |
| AT1G09195 | 0.0 | 8.40E-01 | -0.3 | 8.61E-02 | 0.3 | 1.00E-01 | -0.1 | 8.15E-01 | -0.1 | 7.39E-01 | 0.0 | 9.14E-01 | Ppx-GppA phosphatase |
| AT1G01290 | 0.0 | 6.62E-01 | -0.2 | 3.16E-02 | 0.2 | 2.91E-02 | -0.1 | 4.99E-01 | 0.0 | 8.64E-01 | -0.2 | 5.02E-01 | Cyclic pyranopterin monophosphate synthase, mitochondrial |
| AT3G58510 | 0.0 | 6.46E-01 | 0.0 | 6.11E-01 | 0.1 | 3.78E-01 | 0.2 | 2.82E-01 | 0.1 | 5.47E-01 | 0.1 | 8.48E-01 | DEAD-box ATP-dependent RNA helicase 11 |
| AT1G44800 | 0.0 | 6.59E-01 | 0.8 | 5.99E-06 | -0.7 | 3.37E-05 | 0.8 | 7.70E-07 | 0.3 | 1.40E-02 | 0.5 | 2.81E-03 | WAT1-related protein At1g44800 |
| AT2G47250 | 0.0 | 5.78E-01 | 0.0 | 8.52E-01 | 0.0 | 7.68E-01 | 0.1 | 5.85E-01 | 0.0 | 9.40E-01 | 0.1 | 6.64E-01 | RNA helicase |
| AT5G19690 | 0.0 | 7.18E-01 | 0.2 | 2.42E-02 | -0.2 | 6.04E-02 | 0.3 | 1.33E-02 | 0.1 | 4.41E-01 | 0.2 | 2.86E-01 | dolichyl-diphosphooligosaccharide--protein glycotransferase |
| AT3G27240 | 0.0 | 7.48E-01 | 0.2 | 8.35E-02 | -0.2 | 1.70E-01 | -0.1 | 4.99E-01 | 0.3 | 2.29E-02 | -0.4 | 2.13E-02 | Cytochrome c1 1, heme protein, mitochondrial |
| AT1G78660 | 0.0 | 6.27E-01 | 0.3 | 8.35E-04 | -0.3 | 3.47E-03 | 0.0 | 8.69E-01 | 0.1 | 6.60E-01 | -0.1 | 8.32E-01 | folate gamma-glutamyl hydrolase |
| AT5G08300 | 0.0 | 7.39E-01 | 0.2 | 1.02E-01 | -0.1 | 2.08E-01 | 0.3 | 2.47E-02 | 0.2 | 3.04E-01 | 0.2 | 4.62E-01 | Succinate--CoA ligase subunit alpha-1, mitochondrial |
| AT5G49945 | 0.0 | 6.58E-01 | 0.1 | 1.13E-01 | -0.1 | 2.94E-01 | -0.2 | 1.90E-01 | 0.0 | 9.18E-01 | -0.2 | 2.63E-01 | Uncharacterized protein At5g49945 |
| AT2G42300 | 0.0 | 6.27E-01 | 0.1 | 1.86E-01 | -0.1 | 4.48E-01 | -0.3 | 1.40E-01 | -0.2 | 4.38E-01 | -0.1 | 7.28E-01 | Basic helix-loop-helix DNA-binding superfamily protein |
| AT5G62690 | 0.0 | 7.12E-01 | 0.2 | 1.08E-01 | -0.1 | 2.29E-01 | -0.2 | 3.10E-02 | -0.2 | 2.07E-01 | -0.1 | 6.73E-01 | Tubulin beta-2 chain |
| AT1G15170 | 0.0 | 6.52E-01 | -0.1 | 8.65E-02 | 0.2 | 6.83E-02 | 0.3 | 4.37E-01 | 0.4 | 3.48E-01 | -0.1 | 8.62E-01 | MATE efflux family protein |
| AT3G11850 | 0.0 | 6.87E-01 | -0.1 | 1.44E-01 | 0.2 | 1.07E-01 | -0.1 | 6.02E-01 | -0.1 | 5.70E-01 | 0.0 | 9.48E-01 | F26K24.14 protein |
| AT3G16460 | 0.0 | 7.34E-01 | 0.4 | 3.85E-03 | -0.3 | 1.04E-02 | -0.4 | 1.77E-01 | 0.2 | 4.71E-01 | -0.6 | 9.19E-02 | Jacalin-related lectin 34 |
| AT1G54080 | 0.0 | 6.57E-01 | 0.0 | 8.01E-01 | 0.0 | 8.90E-01 | 0.0 | 8.22E-01 | 0.1 | 4.71E-01 | -0.1 | 4.54E-01 | Oligouridylate-binding protein 1A |
| AT3G19960 | 0.0 | 6.15E-01 | -0.1 | 7.08E-02 | 0.2 | 4.24E-02 | -0.2 | 9.03E-02 | -0.1 | 3.35E-01 | -0.1 | 7.32E-01 | Myosin 1 |

| | | | | | | | | | | | | | |
|---|---|---|---|---|---|---|---|---|---|---|---|---|---|
| AT5G08310 | 0.0 | 8.37E-01 | -0.1 | 4.49E-01 | 0.2 | 4.23E-01 | 0.1 | 9.27E-01 | 0.4 | 4.87E-01 | | | Putative pentatricopeptide repeat-containing protein At5g08310, mitochondrial |
| AT1G59760 | 0.0 | 6.39E-01 | 0.0 | 9.82E-01 | 0.0 | 7.14E-01 | 0.1 | 7.49E-01 | 0.0 | 9.60E-01 | 0.1 | 8.75E-01 | DExH-box ATP-dependent RNA helicase DExH9 |
| AT4G26540 | 0.0 | 7.49E-01 | 0.4 | 2.90E-03 | -0.4 | 1.06E-02 | -0.3 | 7.66E-02 | -0.2 | 2.81E-01 | -0.1 | 7.43E-01 | LRR receptor-like serine |
| AT4G35850 | 0.0 | 8.00E-01 | 0.3 | 8.09E-02 | -0.2 | 1.67E-01 | 0.2 | 2.57E-01 | 0.1 | 6.16E-01 | 0.1 | 7.58E-01 | Pentatricopeptide repeat-containing protein At4g35850, mitochondrial |
| AT2G29580 | 0.0 | 7.33E-01 | 0.2 | 8.99E-02 | -0.1 | 2.26E-01 | 0.3 | 4.07E-01 | 0.2 | 7.33E-01 | 0.1 | 7.90E-01 | Zinc finger CCCH domain-containing protein 25 |
| AT5G01950 | 0.0 | 6.52E-01 | -0.2 | 1.06E-02 | 0.3 | 9.29E-03 | 0.0 | 8.36E-01 | -0.1 | 6.66E-01 | 0.1 | 8.29E-01 | Uncharacterized protein At5g01950 |
| AT1G05470 | 0.0 | 8.07E-01 | 0.3 | 6.77E-02 | -0.2 | 1.57E-01 | -0.1 | 7.65E-01 | -0.1 | 6.06E-01 | 0.1 | 8.46E-01 | Type IV inositol polyphosphate 5-phosphatase 6 |
| AT4G02560 | 0.0 | 6.96E-01 | -0.2 | 1.18E-01 | 0.2 | 9.17E-02 | 0.2 | 5.20E-01 | 0.2 | 5.98E-01 | 0.0 | 9.95E-01 | Homeobox protein LUMINIDEPENDENS |
| AT2G33290 | 0.0 | 6.61E-01 | 0.1 | 2.92E-01 | -0.1 | 5.96E-01 | 0.1 | 7.35E-01 | -0.1 | 7.74E-01 | 0.2 | 5.19E-01 | Uncharacterized protein |
| AT4G14716 | 0.0 | 5.65E-01 | 0.1 | 7.40E-02 | -0.1 | 2.32E-01 | 0.1 | 5.00E-01 | 0.3 | 2.90E-01 | -0.1 | 7.41E-01 | Acireductone dioxygenase -requiring)) -requiring)) |
| AT1G02300 | 0.0 | 6.95E-01 | 0.1 | 4.74E-01 | 0.0 | 7.81E-01 | -0.3 | 3.90E-02 | 0.1 | 6.88E-01 | -0.4 | 5.06E-02 | Peptidase C1A propeptide domain-containing protein |
| AT1G20670 | 0.0 | 7.29E-01 | 0.0 | 9.56E-01 | 0.0 | 8.20E-01 | -0.1 | 7.09E-01 | -0.1 | 7.28E-01 | 0.0 | 9.82E-01 | At1g20670 |
| AT1G79410 | 0.0 | 6.64E-01 | -0.3 | 6.16E-03 | 0.3 | 7.30E-03 | 0.1 | 4.58E-01 | 0.1 | 5.75E-01 | 0.0 | 9.71E-01 | Organic cation |
| AT4G02050 | 0.0 | 6.23E-01 | 0.1 | 7.88E-02 | -0.1 | 2.27E-01 | -0.2 | 1.87E-01 | -0.1 | 7.67E-01 | -0.1 | 5.48E-01 | Sugar transport protein 7 |
| AT2G31650 | 0.0 | 7.52E-01 | 0.1 | 4.71E-01 | 0.0 | 7.38E-01 | 0.4 | 3.35E-01 | 0.5 | 2.79E-01 | -0.1 | 8.63E-01 | Histone H3-lysine N-trimethyltransferase ATX1 |
| AT3G56050 | 0.0 | 6.83E-01 | 0.0 | 6.22E-01 | 0.0 | 9.29E-01 | -0.2 | 3.89E-02 | -0.1 | 3.23E-01 | -0.1 | 5.66E-01 | Probable inactive receptor-like protein kinase At3g56050 |
| AT4G19190 | 0.0 | 6.03E-01 | -0.2 | 1.46E-02 | 0.2 | 1.24E-02 | -0.3 | 8.41E-02 | -0.3 | 6.42E-02 | 0.1 | 8.25E-01 | Uncharacterized zinc finger CCHC domain-containing protein At4g19190 |
| AT4G16845 | 0.0 | 6.45E-01 | -0.3 | 7.95E-03 | 0.3 | 8.39E-03 | -0.1 | 7.96E-01 | 0.1 | 5.95E-01 | -0.2 | 4.84E-01 | VEFS-Box of polycomb protein |
| AT5G59830 | 0.0 | 6.34E-01 | 0.0 | 9.42E-01 | 0.0 | 6.49E-01 | -0.1 | 6.53E-01 | -0.1 | 8.25E-01 | 0.0 | 9.19E-01 | DNA-binding protein |
| AT2G28930 | 0.0 | 6.46E-01 | 0.1 | 1.34E-01 | -0.1 | 3.39E-01 | 0.1 | 5.89E-01 | -0.1 | 6.51E-01 | 0.2 | 3.91E-01 | Uncharacterized protein At2g28930 |
| AT2G41490 | 0.0 | 6.34E-01 | -0.1 | 1.08E-01 | 0.2 | 7.78E-02 | -0.2 | 4.68E-01 | -0.2 | 3.20E-01 | 0.1 | 8.07E-01 | UDP-N-acetylglucosamine--dolichyl-phosphate N-acetylglucosaminephosphotransferase |
| AT1G60710 | 0.0 | 6.18E-01 | 0.1 | 2.12E-01 | -0.1 | 4.77E-01 | -0.1 | 3.68E-01 | -0.1 | 6.43E-01 | 0.0 | 8.66E-01 | Probable aldo-keto reductase 4 |
| AT1G63750 | 0.0 | 7.51E-01 | -0.6 | 8.84E-04 | 0.6 | 1.72E-03 | -0.1 | 7.35E-01 | -0.2 | 4.20E-01 | 0.1 | 6.60E-01 | Disease resistance protein family |
| AT5G06320 | 0.0 | 8.14E-01 | -0.2 | 1.62E-01 | 0.3 | 1.37E-01 | -0.3 | 1.28E-01 | -0.5 | 1.38E-02 | 0.2 | 2.90E-01 | NDR1 |
| AT1G07920 | 0.0 | 7.78E-01 | 0.4 | 2.23E-02 | -0.3 | 4.93E-02 | 0.1 | 3.76E-01 | 0.0 | 9.50E-01 | 0.1 | 4.72E-01 | Elongation factor 1-alpha |
| AT5G47250 | 0.0 | 7.10E-01 | 0.4 | 3.29E-03 | -0.3 | 1.54E-02 | 0.3 | 3.19E-01 | 0.0 | 9.88E-01 | 0.3 | 4.47E-01 | Probable disease resistance protein At5g47250 |
| AT5G57350 | 0.0 | 6.82E-01 | 0.3 | 1.41E-02 | -0.2 | 3.90E-02 | 0.1 | 6.92E-01 | 0.0 | 9.49E-01 | 0.0 | 8.62E-01 | Plasma membrane ATPase |
| AT2G44730 | 0.0 | 6.23E-01 | 0.0 | 7.67E-01 | 0.1 | 4.86E-01 | -0.3 | 1.77E-02 | -0.4 | 3.52E-03 | 0.1 | 4.86E-01 | Alcohol dehydrogenase transcription factor Myb |
| AT1G35510 | 0.0 | 6.39E-01 | 0.0 | 7.01E-01 | 0.1 | 4.57E-01 | -0.1 | 7.12E-01 | 0.0 | 9.91E-01 | -0.1 | 8.21E-01 | O-fucosyltransferase 9 |
| AT5G43020 | 0.0 | 6.88E-01 | 0.3 | 1.45E-02 | -0.2 | 5.13E-02 | -0.1 | 4.08E-01 | 0.0 | 9.86E-01 | -0.1 | 5.21E-01 | Leucine-rich repeat protein kinase family protein |
| AT5G27540 | 0.0 | 5.74E-01 | -0.2 | 4.07E-02 | 0.2 | 2.44E-02 | 0.4 | 2.27E-02 | 0.2 | 3.46E-01 | 0.2 | 4.18E-01 | Mitochondrial Rho GTPase 1 |
| AT2G44660 | 0.0 | 6.57E-01 | 0.2 | 2.63E-02 | -0.2 | 9.81E-02 | -0.3 | 3.33E-01 | 0.0 | 9.64E-01 | -0.2 | 5.32E-01 | Probable dolichyl pyrophosphate Glc1Man9GlcNAc2 alpha-1,3-glucosyltransferase ManGlcNAc-PP-dolichyl alpha-1,3-glucosyltransferase) |
| AT3G09090 | 0.0 | 5.23E-01 | 0.0 | 5.85E-01 | 0.0 | 9.57E-01 | 0.1 | 3.08E-01 | 0.1 | 5.15E-01 | 0.0 | 9.04E-01 | Protein DEFECTIVE IN EXINE FORMATION 1 |
| AT2G05940 | 0.0 | 7.01E-01 | -0.5 | 9.77E-04 | 0.5 | 1.44E-03 | 0.8 | 1.52E-05 | 0.5 | 3.60E-03 | 0.3 | 6.79E-02 | Serine |
| AT1G62600 | 0.0 | 6.20E-01 | -0.1 | 1.50E-01 | 0.2 | 9.85E-02 | -0.3 | 4.99E-02 | -0.1 | 6.59E-01 | -0.2 | 2.83E-01 | Flavin-containing monooxygenase |
| AT1G66260 | 0.0 | 6.43E-01 | -0.1 | 2.56E-01 | 0.1 | 1.45E-01 | -0.1 | 6.93E-01 | 0.0 | 9.09E-01 | -0.1 | 7.15E-01 | THO complex subunit 4C |
| AT3G26290 | 0.0 | 8.05E-01 | -0.3 | 1.35E-01 | 0.3 | 1.11E-01 | 0.0 | 9.24E-01 | 0.3 | 3.21E-01 | -0.4 | 2.25E-01 | Cytochrome P450, family 71, subfamily B, polypeptide 26 |
| AT4G38510 | 0.0 | 5.95E-01 | 0.1 | 1.35E-01 | -0.1 | 3.45E-01 | -0.1 | 4.85E-01 | 0.0 | 8.35E-01 | 0.0 | 8.21E-01 | Vacuolar proton pump subunit B |
| AT5G04590 | 0.0 | 5.80E-01 | 0.2 | 1.87E-02 | -0.2 | 6.35E-02 | -0.7 | 1.58E-04 | 0.1 | 5.95E-01 | -0.8 | 5.38E-04 | Assimilatory sulfite reductase , chloroplastic ) |
| AT5G58787 | 0.0 | 6.33E-01 | 0.2 | 7.87E-02 | -0.1 | 2.35E-01 | 0.2 | 2.89E-01 | -0.1 | 4.75E-01 | 0.3 | 1.53E-01 | RING |
| AT1G51805 | 0.0 | 6.58E-01 | 0.1 | 2.07E-01 | -0.1 | 4.32E-01 | 0.2 | 2.51E-01 | 0.1 | 6.82E-01 | 0.1 | 7.20E-01 | Leucine-rich repeat protein kinase family protein |
| AT5G42380 | 0.0 | 9.30E-01 | -1.1 | 6.77E-02 | 1.2 | 1.05E-01 | 2.3 | 7.42E-05 | 0.7 | 2.90E-01 | 1.6 | 2.96E-04 | Calcium-binding protein CML37 |
| AT1G79920 | 0.0 | 5.21E-01 | -0.2 | 3.52E-02 | 0.2 | 1.76E-02 | 0.3 | 4.96E-01 | 0.2 | 6.24E-01 | 0.0 | 9.70E-01 | Heat shock protein 70 family protein |
| AT4G02450 | 0.0 | 7.36E-01 | 0.2 | 1.07E-01 | -0.2 | 2.17E-01 | 0.3 | 1.50E-02 | 0.2 | 1.95E-01 | 0.1 | 5.12E-01 | Co-chaperone protein p23 |
| AT2G04550 | 0.0 | 7.17E-01 | -0.1 | 5.33E-01 | 0.1 | 3.91E-01 | 0.0 | 8.16E-01 | -0.1 | 4.57E-01 | 0.2 | 4.06E-01 | Protein-tyrosine-phosphatase IBR5 |
| AT1G29030 | 0.0 | 6.38E-01 | -0.1 | 2.12E-01 | 0.2 | 1.49E-01 | 0.8 | 3.77E-03 | 0.4 | 1.96E-01 | 0.4 | 1.22E-01 | Apoptosis inhibitory protein 5 |
| AT4G23660 | 0.0 | 6.62E-01 | -0.1 | 3.77E-01 | 0.1 | 2.65E-01 | 0.1 | 8.19E-01 | 0.0 | 9.28E-01 | 0.1 | 7.88E-01 | 4-hydroxybenzoate polyprenyltransferase, mitochondrial |
| AT3G51710 | 0.0 | 7.32E-01 | 0.4 | 8.76E-03 | -0.3 | 3.77E-02 | -0.1 | 5.62E-01 | -0.3 | 2.84E-01 | 0.1 | 6.66E-01 | hypothetical protein |
| AT4G08770 | 0.0 | 8.11E-01 | 0.3 | 8.05E-02 | -0.3 | 1.50E-01 | 1.5 | 6.60E-06 | 0.7 | 8.52E-03 | 0.7 | 7.91E-03 | Peroxidase 37 |
| AT4G11130 | 0.0 | 6.38E-01 | -0.2 | 1.24E-01 | 0.2 | 9.27E-02 | -0.7 | 8.29E-04 | -0.3 | 7.48E-02 | -0.4 | 1.26E-01 | RNA-dependent RNA polymerase 2 |
| AT1G61380 | 0.0 | 5.85E-01 | -0.3 | 5.30E-03 | 0.3 | 4.74E-03 | -0.1 | 7.79E-01 | -0.4 | 3.47E-01 | 0.3 | 6.05E-01 | S-domain-1 29 |
| AT1G61370 | 0.0 | 7.26E-01 | -0.3 | 6.51E-02 | 0.3 | 6.55E-02 | -0.1 | 8.68E-01 | 0.1 | 8.79E-01 | -0.1 | 7.92E-01 | S-locus lectin protein kinase family protein |
| AT4G32390 | 0.0 | 6.39E-01 | 0.2 | 2.47E-02 | -0.2 | 7.33E-02 | 0.1 | 3.32E-01 | 0.0 | 9.82E-01 | 0.2 | 5.01E-01 | Probable sugar phosphate |
| AT5G18480 | 0.0 | 5.75E-01 | 0.0 | 8.84E-01 | 0.0 | 7.36E-01 | 0.0 | 8.20E-01 | 0.0 | 8.51E-01 | 0.1 | 7.63E-01 | Inositol phosphorylceramide glucuronosyltransferase 1 |
| AT1G71090 | 0.0 | 5.52E-01 | -0.1 | 3.10E-01 | 0.1 | 1.50E-01 | -0.1 | 2.46E-01 | 0.0 | 9.31E-01 | -0.2 | 3.46E-01 | Protein PIN-LIKES 2 |
| AT3G52880 | 0.0 | 5.12E-01 | -0.1 | 1.36E-01 | 0.2 | 5.86E-02 | 0.1 | 3.74E-01 | 0.0 | 9.55E-01 | 0.1 | 6.13E-01 | Monodehydroascorbate reductase 1 |
| AT4G28980 | 0.0 | 5.34E-01 | 0.0 | 5.95E-01 | 0.0 | 9.60E-01 | -0.2 | 2.27E-01 | -0.1 | 3.65E-01 | 0.0 | 9.41E-01 | Cyclin-dependent kinase F-1 |
| AT2G32730 | 0.0 | 5.27E-01 | -0.1 | 4.11E-01 | 0.1 | 1.86E-01 | 0.0 | 7.99E-01 | 0.0 | 9.66E-01 | 0.0 | 9.11E-01 | 26S proteasome non-ATPase regulatory subunit 1 homolog A |
| AT3G01540 | 0.0 | 5.09E-01 | -0.1 | 1.22E-01 | 0.2 | 5.22E-02 | -0.1 | 3.40E-01 | -0.2 | 1.48E-01 | 0.1 | 6.64E-01 | DEAD-box ATP-dependent RNA helicase 14 |
| AT2G28600 | 0.0 | 7.47E-01 | 0.4 | 1.06E-02 | -0.4 | 3.57E-02 | 0.4 | 2.83E-02 | 0.1 | 6.78E-01 | 0.3 | 1.46E-01 | DEAD |
| AT3G46820 | 0.0 | 5.27E-01 | 0.1 | 3.65E-01 | 0.0 | 8.04E-01 | 0.1 | 4.35E-01 | 0.1 | 7.62E-01 | 0.1 | 8.17E-01 | Serine |
| AT3G14180 | 0.0 | 4.85E-01 | -0.1 | 7.88E-02 | 0.2 | 3.83E-02 | -0.3 | 5.63E-02 | -0.3 | 1.55E-01 | -0.1 | 8.40E-01 | Trihelix transcription factor ASIL2 |
| AT2G24630 | 0.0 | 7.26E-01 | 0.0 | 8.23E-01 | 0.1 | 6.32E-01 | 0.0 | 9.08E-01 | 0.0 | 9.08E-01 | 0.1 | 8.56E-01 | Probable xyloglucan glycosyltransferase 8 |
| AT1G16150 | 0.0 | 7.06E-01 | -0.1 | 2.63E-01 | 0.2 | 2.12E-01 | -0.2 | 6.16E-01 | 0.2 | 4.67E-01 | -0.4 | 2.50E-01 | Wall-associated receptor kinase-like 4 |
| AT5G05080 | 0.0 | 4.98E-01 | 0.2 | 7.30E-03 | -0.2 | 3.30E-02 | -0.2 | 5.25E-02 | -0.2 | 1.71E-01 | 0.0 | 8.29E-01 | Ubiquitin-conjugating enzyme E2 22 |
| AT1G15970 | 0.0 | 5.78E-01 | 0.0 | 7.47E-01 | 0.0 | 8.73E-01 | -0.2 | 1.66E-01 | -0.2 | 4.35E-01 | -0.1 | 7.87E-01 | DNA glycosylase superfamily protein |
| AT1G08700 | 0.0 | 5.87E-01 | 0.1 | 1.45E-01 | -0.1 | 3.91E-01 | 0.1 | 4.29E-01 | 0.0 | 9.75E-01 | 0.1 | 6.36E-01 | Presenilin-like protein At1g08700 |
| AT5G43440 | 0.0 | 6.31E-01 | -0.1 | 4.34E-01 | 0.1 | 2.65E-01 | 0.0 | 9.02E-01 | -0.1 | 6.35E-01 | 0.1 | 5.76E-01 | 1-aminocyclopropane-1-carboxylate oxidase homolog 9 |
| AT3G02280 | 0.0 | 5.17E-01 | -0.2 | 1.07E-02 | 0.3 | 8.90E-03 | 0.0 | 9.94E-01 | 0.0 | 8.73E-01 | -0.1 | 8.87E-01 | NADPH-dependent diflavin oxidoreductase 1 |
| AT1G04280 | 0.0 | 6.34E-01 | 0.3 | 1.29E-02 | -0.2 | 4.25E-02 | 0.1 | 5.94E-01 | -0.2 | 9.84E-02 | 0.3 | 9.25E-02 | Calmodulin calcium-dependent NAD kinase |
| AT2G38730 | 0.0 | 7.39E-01 | 0.3 | 3.95E-02 | -0.3 | 1.14E-01 | 0.1 | 5.52E-01 | 0.0 | 9.55E-01 | 0.1 | 7.41E-01 | Peptidyl-prolyl cis-trans isomerase CYP22 |
| AT2G30470 | 0.0 | 5.20E-01 | 0.0 | 8.49E-01 | 0.0 | 7.16E-01 | -0.1 | 2.98E-01 | -0.2 | 1.27E-01 | 0.1 | 7.02E-01 | B3 domain-containing transcription repressor VAL1 |
| AT1G35710 | 0.0 | 8.86E-01 | -0.6 | 6.97E-02 | 0.7 | 7.14E-02 | 0.8 | 4.18E-03 | 0.1 | 8.58E-01 | 0.7 | 4.11E-02 | Probable leucine-rich repeat receptor-like protein kinase At1g35710 |
| AT4G16130 | 0.0 | 6.02E-01 | -0.2 | 3.58E-02 | 0.3 | 2.54E-02 | -0.2 | 2.67E-01 | -0.2 | 3.52E-01 | 0.0 | 9.80E-01 | L-arabinokinase |
| AT5G45190 | 0.0 | 4.54E-01 | 0.0 | 6.93E-01 | 0.0 | 7.88E-01 | -0.1 | 7.44E-01 | -0.1 | 7.79E-01 | 0.0 | 9.95E-01 | Cyclin family protein |
| AT1G18480 | 0.0 | 6.35E-01 | -0.1 | 2.04E-01 | 0.2 | 1.32E-01 | 0.2 | 3.30E-01 | 0.1 | 7.09E-01 | 0.1 | 7.58E-01 | Shewanella-like protein phosphatase 2 |
| AT2G22530 | 0.0 | 6.32E-01 | -0.1 | 4.03E-01 | 0.1 | 2.67E-01 | -0.2 | 6.52E-01 | 0.0 | 9.49E-01 | -0.1 | 8.13E-01 | GPI ethanolamine phosphate transferase 2 |
| AT1G10230 | 0.0 | 7.89E-01 | 0.1 | 4.93E-01 | -0.1 | 7.33E-01 | -0.2 | 3.79E-01 | -0.1 | 5.71E-01 | 0.0 | 9.05E-01 | SKP1-like protein |
| AT1G11650 | 0.0 | 4.82E-01 | 0.2 | 2.38E-02 | -0.1 | 1.00E-01 | 0.0 | 7.11E-01 | -0.1 | 2.20E-01 | 0.2 | 1.98E-01 | Polyadenylate-binding protein RBP45B -binding protein RBP45B) |
| AT5G47010 | 0.0 | 5.04E-01 | -0.2 | 4.98E-03 | 0.3 | 3.32E-03 | 0.3 | 5.59E-02 | 0.2 | 2.65E-01 | 0.1 | 7.27E-01 | Regulator of nonsense transcripts 1 homolog |
| AT4G23250 | 0.0 | 7.98E-01 | -0.4 | 4.15E-02 | 0.5 | 4.56E-02 | 0.7 | 2.87E-04 | 0.7 | 7.40E-04 | 0.0 | 9.94E-01 | Cysteine-rich receptor-like protein kinase 17 |
| AT1G45180 | 0.0 | 5.02E-01 | -0.2 | 4.39E-02 | 0.2 | 2.79E-02 | 0.0 | 9.22E-01 | -0.1 | 7.21E-01 | 0.1 | 6.17E-01 | RING-type E3 ubiquitin transferase |
| AT5G59300 | 0.0 | 5.41E-01 | 0.2 | 5.60E-02 | -0.1 | 2.13E-01 | 0.2 | 4.06E-01 | 0.2 | 5.87E-01 | 0.0 | 9.27E-01 | Ubiquitin-conjugating enzyme E2 7 |

| | | | | | | | | | | | | | |
|---|---|---|---|---|---|---|---|---|---|---|---|---|---|
| AT5G56950 | 0.0 | 5.92E-01 | 0.2 | 3.55E-02 | -0.2 | 1.16E-01 | 0.4 | 1.93E-02 | 0.2 | 3.53E-01 | 0.2 | 3.47E-01 | NFA3 |
| AT4G22260 | 0.0 | 4.87E-01 | -0.2 | 1.93E-02 | 0.2 | 1.11E-02 | 0.0 | 9.11E-01 | -0.1 | 6.69E-01 | 0.1 | 8.10E-01 | Ubiquinol oxidase 4, chloroplastic |
| AT3G61520 | 0.0 | 6.10E-01 | -0.1 | 4.10E-01 | 0.1 | 2.68E-01 | 0.2 | 4.88E-01 | 0.1 | 8.59E-01 | 0.1 | 7.53E-01 | Pentatricopeptide repeat-containing protein At3g61520, mitochondrial |
| AT1G36730 | 0.0 | 4.72E-01 | 0.0 | 9.06E-01 | 0.0 | 6.12E-01 | -0.2 | 1.45E-02 | -0.1 | 2.73E-01 | -0.1 | 4.14E-01 | Probable eukaryotic translation initiation factor 5-1 |
| AT4G04720 | 0.0 | 4.60E-01 | 0.0 | 4.76E-01 | 0.0 | 9.98E-01 | 0.1 | 6.36E-01 | -0.1 | 7.48E-01 | 0.1 | 4.84E-01 | Calcium-dependent protein kinase 21 |
| AT5G62050 | 0.0 | 5.07E-01 | 0.0 | 8.03E-01 | 0.1 | 4.23E-01 | 0.5 | 1.87E-03 | 0.2 | 3.70E-01 | 0.3 | 6.55E-02 | Membrane insertase YidC |
| AT3G47540 | 0.0 | 6.95E-01 | -0.2 | 2.30E-01 | 0.2 | 1.79E-01 | 1.0 | 1.61E-03 | 0.5 | 1.78E-01 | 0.6 | 4.67E-02 | Putative endochitinase |
| AT5G57980 | 0.0 | 8.76E-01 | 0.6 | 4.92E-02 | -0.6 | 1.22E-01 | -0.3 | 4.83E-01 | 0.0 | 9.90E-01 | -0.3 | 6.06E-01 | DNA-directed RNA polymerase subunit 5-like protein 1 |
| AT4G25600 | 0.0 | 7.43E-01 | 0.1 | 5.63E-01 | 0.0 | 8.34E-01 | -0.2 | 6.26E-01 | 0.1 | 9.07E-01 | -0.2 | 5.39E-01 | procollagen-proline 4-dioxygenase |
| AT2G37380 | 0.0 | 7.68E-01 | 0.5 | 6.96E-03 | -0.5 | 2.62E-02 | -0.3 | 2.14E-01 | 0.3 | 1.60E-01 | -0.7 | 3.52E-02 | Probable membrane-associated kinase regulator 3 |
| AT2G17390 | 0.0 | 3.98E-01 | 0.0 | 4.46E-01 | 0.0 | 9.73E-01 | 0.0 | 7.08E-01 | -0.1 | 5.49E-01 | 0.0 | 8.68E-01 | Ankyrin repeat domain-containing protein 2B |
| AT1G68200 | 0.0 | 8.08E-01 | -0.5 | 4.57E-02 | 0.5 | 6.21E-02 | -0.4 | 4.08E-01 | -0.3 | 5.35E-01 | -0.1 | 9.55E-01 | hypothetical protein |
| AT5G06810 | 0.1 | 7.64E-01 | 0.1 | 5.34E-01 | -0.1 | 7.91E-01 | 0.3 | 1.78E-01 | 0.1 | 7.34E-01 | 0.2 | 5.14E-01 | Transcription termination factor family protein |
| AT5G12440 | 0.1 | 5.15E-01 | 0.1 | 3.50E-01 | 0.0 | 7.99E-01 | -0.2 | 1.65E-01 | -0.3 | 5.56E-02 | 0.1 | 6.37E-01 | CCCH-type zinc fingerfamily protein with RNA-binding domain-containing protein |
| AT3G57230 | 0.1 | 6.17E-01 | -0.1 | 2.57E-01 | 0.2 | 1.57E-01 | 0.1 | 4.21E-01 | 0.1 | 6.45E-01 | 0.0 | 8.92E-01 | AGAMOUS-like 16 |
| AT5G34850 | 0.1 | 4.34E-01 | 0.2 | 2.65E-02 | -0.1 | 1.23E-01 | 0.0 | 9.32E-01 | 0.1 | 6.85E-01 | -0.1 | 7.94E-01 | Purple acid phosphatase |
| AT5G64100 | 0.1 | 6.54E-01 | 0.6 | 2.99E-04 | -0.6 | 1.06E-03 | 0.0 | 9.01E-01 | 0.2 | 3.14E-01 | -0.2 | 2.83E-01 | peroxidase |
| AT1G21160 | 0.1 | 5.07E-01 | -0.2 | 4.03E-02 | 0.2 | 2.46E-02 | -0.1 | 3.72E-01 | -0.2 | 2.54E-01 | 0.1 | 8.19E-01 | Eukaryotic translation initiation factor 5B |
| AT3G22950 | 0.1 | 5.54E-01 | 0.1 | 3.99E-01 | 0.0 | 8.24E-01 | 0.3 | 1.25E-01 | 0.0 | 9.76E-01 | 0.3 | 2.42E-01 | ADP-ribosylation factor C1 |
| AT5G14620 | 0.1 | 5.96E-01 | 0.1 | 4.23E-01 | 0.0 | 8.17E-01 | -0.6 | 1.07E-03 | -0.3 | 5.85E-02 | -0.3 | 2.44E-01 | DNA -methyltransferase DRM2 |
| AT1G50410 | 0.1 | 4.54E-01 | -0.2 | 1.62E-02 | 0.2 | 9.62E-03 | 0.1 | 8.10E-01 | 0.1 | 8.05E-01 | 0.0 | 9.75E-01 | SNF2 domain-containing protein |
| AT1G28350 | 0.1 | 5.39E-01 | 0.3 | 6.85E-03 | -0.2 | 3.35E-02 | 0.3 | 4.33E-02 | 0.1 | 4.58E-01 | 0.2 | 4.31E-01 | Nucleotidylyl transferase superfamily protein |
| AT1G67440 | 0.1 | 6.01E-01 | 0.2 | 1.22E-01 | -0.1 | 3.47E-01 | -0.3 | 1.50E-01 | 0.1 | 7.64E-01 | -0.3 | 1.32E-01 | Minichromosome maintenance family protein |
| AT1G11260 | 0.1 | 6.72E-01 | 0.3 | 1.75E-02 | -0.3 | 4.87E-02 | -0.7 | 8.25E-04 | -1.1 | 4.05E-05 | 0.4 | 5.96E-02 | Sugar transport protein 1 |
| AT1G10490 | 0.1 | 6.07E-01 | -0.1 | 3.28E-01 | 0.2 | 1.99E-01 | 0.4 | 3.56E-02 | 0.2 | 4.80E-01 | 0.2 | 3.74E-01 | RNA cytidine acetyltransferase 1 |
| AT5G55200 | 0.1 | 7.16E-01 | -0.2 | 1.37E-01 | 0.3 | 1.13E-01 | 1.3 | 9.19E-07 | 0.3 | 2.38E-01 | 1.0 | 8.05E-05 | GrpE protein homolog 1, mitochondrial |
| AT2G27820 | 0.1 | 5.86E-01 | 0.3 | 1.27E-02 | -0.2 | 4.76E-02 | -0.1 | 5.09E-01 | 0.3 | 4.57E-02 | -0.4 | 3.09E-02 | Arogenate dehydratase 3, chloroplastic |
| AT4G39980 | 0.1 | 6.14E-01 | 0.2 | 1.05E-01 | -0.1 | 2.71E-01 | 0.4 | 5.82E-03 | 0.3 | 3.50E-02 | 0.1 | 7.67E-01 | Phospho-2-dehydro-3-deoxyheptonate aldolase 1, chloroplastic |
| AT4G35600 | 0.1 | 5.94E-01 | -0.2 | 3.83E-02 | 0.3 | 2.78E-02 | -0.1 | 3.99E-01 | -0.3 | 2.23E-02 | 0.2 | 2.35E-01 | Protein kinase superfamily protein |
| AT1G18450 | 0.1 | 3.88E-01 | -0.1 | 2.56E-01 | 0.1 | 8.53E-02 | -0.2 | 1.41E-01 | -0.2 | 1.53E-01 | 0.0 | 9.49E-01 | Actin-related protein 4 |
| AT1G21070 | 0.1 | 6.37E-01 | 0.3 | 3.20E-02 | -0.2 | 1.23E-01 | -0.2 | 3.40E-01 | 0.0 | 9.25E-01 | -0.1 | 5.72E-01 | UDP-rhamnose |
| AT5G20010 | 0.1 | 5.16E-01 | 0.1 | 4.05E-01 | 0.0 | 8.53E-01 | -0.2 | 4.75E-02 | -0.2 | 9.21E-02 | 0.0 | 9.70E-01 | GTP-binding nuclear protein |
| AT1G17130 | 0.1 | 4.84E-01 | 0.0 | 8.75E-01 | 0.0 | 6.59E-01 | 0.2 | 1.15E-01 | 0.1 | 4.19E-01 | 0.1 | 7.13E-01 | Splicing factor YJU2 |
| AT1G04310 | 0.1 | 6.59E-01 | -0.3 | 2.37E-02 | 0.4 | 2.43E-02 | -0.6 | 1.81E-03 | -0.5 | 5.12E-03 | 0.0 | 9.01E-01 | ERS2 |
| AT5G13270 | 0.1 | 6.52E-01 | -0.2 | 2.23E-01 | 0.2 | 1.65E-01 | -0.3 | 3.79E-01 | 0.0 | 9.82E-01 | -0.3 | 4.68E-01 | Pentatricopeptide repeat-containing protein At5g13270, chloroplastic |
| AT1G56720 | 0.1 | 6.09E-01 | 0.4 | 4.03E-03 | -0.3 | 1.94E-02 | -0.6 | 2.85E-03 | -0.5 | 2.23E-02 | -0.1 | 6.49E-01 | Protein kinase domain-containing protein |
| AT4G20380 | 0.1 | 5.12E-01 | 0.0 | 9.57E-01 | 0.0 | 6.19E-01 | -0.2 | 7.01E-02 | -0.4 | 1.27E-02 | 0.1 | 4.71E-01 | LSD1 zinc finger family protein |
| AT1G01140 | 0.1 | 6.25E-01 | 0.1 | 2.94E-01 | -0.1 | 6.08E-01 | -0.2 | 2.41E-01 | -0.1 | 4.70E-01 | 0.0 | 8.56E-01 | CBL-interacting serine |
| AT1G54390 | 0.1 | 4.92E-01 | 0.0 | 9.01E-01 | 0.0 | 6.56E-01 | 0.0 | 9.76E-01 | 0.1 | 8.09E-01 | -0.1 | 8.15E-01 | PHD finger protein ING |
| AT5G47050 | 0.1 | 5.41E-01 | 0.0 | 7.55E-01 | 0.1 | 4.22E-01 | -0.2 | 2.38E-01 | 0.0 | 9.63E-01 | -0.2 | 2.89E-01 | SBP family protein |
| AT1G30460 | 0.1 | 5.02E-01 | 0.0 | 9.87E-01 | 0.1 | 5.92E-01 | 0.1 | 6.30E-01 | 0.1 | 5.90E-01 | 0.0 | 9.42E-01 | Cleavage and polyadenylation specificity factor 30 |
| AT4G27230 | 0.1 | 4.33E-01 | 0.3 | 1.25E-03 | -0.3 | 7.35E-03 | -0.1 | 6.57E-01 | -0.1 | 3.91E-01 | 0.1 | 7.48E-01 | Probable histone H2A.3 |
| AT5G56450 | 0.1 | 6.23E-01 | 0.2 | 6.42E-02 | -0.2 | 2.21E-01 | 0.2 | 4.59E-01 | 0.3 | 3.22E-01 | -0.1 | 7.98E-01 | Probable ADP,ATP carrier protein At5g56450 |
| AT3G22780 | 0.1 | 4.36E-01 | 0.0 | 9.68E-01 | 0.1 | 4.84E-01 | 0.3 | 7.38E-02 | 0.3 | 1.51E-01 | 0.0 | 9.25E-01 | CRC domain-containing protein TSO1 |
| AT1G11930 | 0.1 | 3.58E-01 | -0.1 | 2.80E-01 | 0.1 | 8.10E-02 | -0.3 | 4.32E-02 | -0.2 | 3.23E-01 | -0.1 | 5.80E-01 | Pyridoxal phosphate homeostasis protein |
| AT2G11520 | 0.1 | 4.28E-01 | -0.2 | 1.01E-02 | 0.3 | 5.84E-03 | 0.2 | 7.22E-02 | 0.1 | 7.13E-01 | 0.2 | 3.49E-01 | Calmodulin-binding receptor-like cytoplasmic kinase 3 |
| AT2G02160 | 0.1 | 3.91E-01 | 0.1 | 1.74E-01 | 0.0 | 6.13E-01 | -0.1 | 4.08E-01 | -0.3 | 2.42E-02 | 0.2 | 2.24E-01 | Zinc finger CCCH domain-containing protein 17 |
| AT2G17870 | 0.1 | 5.39E-01 | 0.1 | 1.96E-01 | -0.1 | 5.17E-01 | 0.1 | 5.08E-01 | -0.1 | 7.35E-01 | 0.1 | 4.02E-01 | Cold shock domain-containing protein 3 |
| AT1G68550 | 0.1 | 5.83E-01 | 0.1 | 1.90E-01 | -0.1 | 4.92E-01 | 0.3 | 8.26E-02 | -0.1 | 4.84E-01 | 0.4 | 4.80E-02 | Ethylene-responsive transcription factor ERF118 |
| AT2G46650 | 0.1 | 6.68E-01 | 0.3 | 2.20E-02 | -0.3 | 6.40E-02 | -0.8 | 3.46E-03 | 0.5 | 5.17E-02 | -1.3 | 1.67E-03 | Cytochrome B5 isoform C |
| AT4G24240 | 0.1 | 5.03E-01 | -0.1 | 1.66E-01 | 0.2 | 7.94E-02 | 0.0 | 8.40E-01 | -0.3 | 6.23E-02 | 0.3 | 2.00E-01 | WRKY DNA-binding protein 7 |
| AT4G34650 | 0.1 | 8.24E-01 | -0.1 | 6.87E-01 | 0.2 | 5.99E-01 | -0.5 | 2.23E-01 | 0.2 | 7.48E-01 | -0.6 | 2.07E-01 | Squalene synthase 2 |
| AT5G57270 | 0.1 | 5.63E-01 | 0.0 | 9.53E-01 | 0.0 | 6.78E-01 | 0.0 | 9.57E-01 | 0.1 | 6.65E-01 | -0.1 | 7.61E-01 | Core-2 |
| AT4G01940 | 0.1 | 7.04E-01 | -0.2 | 2.78E-01 | 0.2 | 2.07E-01 | -0.1 | 6.53E-01 | -0.1 | 6.95E-01 | 0.0 | 9.96E-01 | NifU-like protein 1, chloroplastic |
| AT5G50210 | 0.1 | 4.11E-01 | 0.0 | 5.24E-01 | 0.0 | 9.04E-01 | -0.2 | 8.76E-02 | -0.2 | 2.69E-01 | -0.1 | 8.15E-01 | Quinolinate synthase, chloroplastic |
| AT5G38830 | 0.1 | 4.02E-01 | -0.1 | 3.41E-01 | 0.1 | 1.15E-01 | 0.1 | 6.37E-01 | 0.0 | 8.66E-01 | 0.1 | 6.29E-01 | Cysteine--tRNA ligase 2, cytoplasmic |
| AT1G78150 | 0.1 | 4.75E-01 | 0.2 | 5.30E-02 | -0.1 | 2.07E-01 | -0.2 | 9.58E-02 | -0.2 | 3.86E-02 | 0.1 | 6.83E-01 | N-lysine methyltransferase |
| AT4G06599 | 0.1 | 3.78E-01 | 0.0 | 7.61E-01 | 0.0 | 6.48E-01 | 0.2 | 1.79E-01 | 0.2 | 2.07E-01 | 0.0 | 9.70E-01 | Ubiquitin-like domain-containing CTD phosphatase |
| AT5G37770 | 0.1 | 7.56E-01 | -0.2 | 2.13E-01 | 0.3 | 1.70E-01 | -0.2 | 5.49E-01 | -0.2 | 5.83E-01 | 0.0 | 9.82E-01 | Calcium-binding protein CML24 |
| AT1G14920 | 0.1 | 5.58E-01 | 0.4 | 3.76E-03 | -0.3 | 1.60E-02 | -0.3 | 3.23E-02 | -0.2 | 1.14E-01 | -0.1 | 8.35E-01 | DELLA protein GAI |
| AT3G55350 | 0.1 | 5.88E-01 | 0.0 | 8.11E-01 | 0.0 | 8.26E-01 | 0.1 | 7.87E-01 | 0.1 | 8.02E-01 | 0.0 | 9.81E-01 | Protein ALP1-like |
| AT5G54770 | 0.1 | 5.65E-01 | 0.0 | 6.83E-01 | 0.0 | 9.12E-01 | 0.0 | 8.70E-01 | 0.2 | 2.62E-01 | -0.2 | 4.28E-01 | Thiamine thiazole synthase, chloroplastic |
| AT4G02580 | 0.1 | 5.31E-01 | 0.3 | 1.51E-02 | -0.2 | 5.82E-02 | -0.1 | 6.12E-01 | 0.0 | 8.92E-01 | 0.0 | 8.56E-01 | AT4G02580 protein |
| AT1G63250 | 0.1 | 6.41E-01 | -0.1 | 3.92E-01 | 0.2 | 2.72E-01 | 0.7 | 1.25E-02 | 0.4 | 2.20E-01 | 0.3 | 2.78E-01 | Probable DEAD-box ATP-dependent RNA helicase 48 |
| AT5G10900 | 0.1 | 6.46E-01 | -0.1 | 5.61E-01 | 0.1 | 3.89E-01 | -0.2 | 6.32E-01 | -0.1 | 9.20E-01 | -0.1 | 7.96E-01 | Calcineurin-like metallo-phosphoesterase superfamily protein |
| AT5G22840 | 0.1 | 5.44E-01 | 0.1 | 3.77E-01 | 0.0 | 8.09E-01 | 0.3 | 1.21E-01 | -0.2 | 3.91E-01 | 0.5 | 4.69E-02 | Protein kinase domain-containing protein |
| AT4G28220 | 0.1 | 4.77E-01 | 0.4 | 1.50E-03 | -0.3 | 8.65E-03 | 0.0 | 9.49E-01 | -0.1 | 7.65E-01 | 0.1 | 8.48E-01 | NADH:ubiquinone reductase |
| AT1G67890 | 0.1 | 4.57E-01 | -0.1 | 1.31E-01 | 0.2 | 5.81E-02 | -0.2 | 2.94E-01 | 0.1 | 8.06E-01 | -0.3 | 2.80E-01 | PAS domain-containing protein tyrosine kinase family protein |
| AT2G21710 | 0.1 | 6.93E-01 | 0.4 | 1.45E-02 | -0.4 | 5.90E-02 | 0.4 | 1.73E-01 | -0.1 | 8.01E-01 | 0.6 | 1.92E-01 | Mitochondrial transcription termination factor family protein |
| AT5G50870 | 0.1 | 4.01E-01 | 0.0 | 9.44E-01 | 0.1 | 5.17E-01 | 0.0 | 9.97E-01 | -0.1 | 5.94E-01 | 0.1 | 6.80E-01 | E2 ubiquitin-conjugating enzyme |
| AT4G10730 | 0.1 | 5.51E-01 | -0.2 | 8.42E-02 | 0.2 | 4.65E-02 | 0.2 | 4.50E-01 | 0.1 | 7.59E-01 | 0.1 | 7.88E-01 | Protein kinase superfamily protein |
| AT1G15020 | 0.1 | 4.14E-01 | 0.2 | 1.48E-02 | -0.1 | 9.36E-02 | -0.1 | 6.06E-01 | -0.2 | 1.80E-01 | 0.2 | 4.42E-01 | Sulfhydryl oxidase 1 |
| AT3G54080 | 0.1 | 4.55E-01 | 0.0 | 9.78E-01 | 0.1 | 5.64E-01 | -0.1 | 8.70E-01 | 0.1 | 9.09E-01 | -0.1 | 8.13E-01 | Legume lectin domain-containing protein |
| AT4G25550 | 0.1 | 5.54E-01 | 0.1 | 2.03E-01 | -0.1 | 5.26E-01 | 0.1 | 5.65E-01 | -0.1 | 6.69E-01 | 0.1 | 3.90E-01 | Uncharacterized protein |
| AT5G06660 | 0.1 | 6.00E-01 | 0.4 | 8.79E-03 | -0.3 | 3.53E-02 | -0.1 | 7.17E-01 | -0.2 | 4.63E-01 | 0.1 | 7.35E-01 | Calcium load-activated calcium channel |
| AT5G65640 | 0.1 | 4.87E-01 | 0.0 | 6.10E-01 | 0.0 | 9.05E-01 | -0.6 | 4.83E-03 | 0.0 | 8.57E-01 | -0.7 | 1.03E-02 | Transcription factor bHLH93 |
| AT5G64360 | 0.1 | 5.40E-01 | 0.1 | 1.73E-01 | -0.1 | 4.98E-01 | -0.4 | 6.45E-03 | -0.4 | 8.19E-03 | 0.0 | 9.37E-01 | AT5G64360 protein |
| AT3G18035 | 0.1 | 3.43E-01 | 0.1 | 1.51E-01 | 0.0 | 6.17E-01 | -0.5 | 3.70E-04 | -0.4 | 6.35E-03 | -0.1 | 5.50E-01 | At3g18035 |
| AT2G47210 | 0.1 | 4.36E-01 | -0.1 | 3.94E-01 | 0.1 | 1.49E-01 | 0.1 | 6.48E-01 | 0.1 | 4.57E-01 | -0.1 | 8.29E-01 | SWR1-complex protein 4 |
| AT5G54900 | 0.1 | 6.09E-01 | 0.2 | 1.25E-01 | -0.1 | 3.23E-01 | 0.4 | 3.60E-03 | 0.3 | 1.47E-02 | 0.0 | 8.43E-01 | Polyadenylate-binding protein RBP45A -binding protein RBP45A) |
| AT1G62360 | 0.1 | 6.39E-01 | -0.2 | 2.08E-01 | 0.2 | 1.51E-01 | -0.2 | 4.83E-01 | -0.3 | 4.06E-01 | 0.1 | 8.84E-01 | Homeobox protein SHOOT MERISTEMLESS |
| AT5G45430 | 0.1 | 4.20E-01 | 0.0 | 7.08E-01 | 0.1 | 2.83E-01 | 0.1 | 6.00E-01 | 0.2 | 4.11E-01 | -0.1 | 8.11E-01 | Protein kinase superfamily protein -like protein) |
| AT3G51850 | 0.1 | 4.34E-01 | 0.0 | 5.49E-01 | 0.1 | 2.16E-01 | -0.1 | 3.55E-01 | -0.1 | 6.88E-01 | -0.1 | 8.29E-01 | Calcium-dependent protein kinase 13 |

| | | | | | | | | | | | | | |
|---|---|---|---|---|---|---|---|---|---|---|---|---|---|
| AT5G43320 | 0.1 | 4.61E-01 | 0.0 | 8.75E-01 | 0.1 | 4.20E-01 | 0.0 | 8.94E-01 | 0.0 | 8.53E-01 | 0.0 | 8.15E-01 | Casein kinase I-like 8 |
| AT3G18524 | 0.1 | 5.10E-01 | 0.2 | 5.99E-02 | -0.1 | 2.57E-01 | 0.2 | 3.44E-01 | 0.0 | 9.50E-01 | 0.2 | 4.99E-01 | DNA mismatch repair protein MSH2 |
| AT1G60700 | 0.1 | 6.95E-01 | 0.1 | 7.18E-01 | 0.0 | 9.97E-01 | -0.1 | 7.95E-01 | 0.1 | 8.71E-01 | -0.2 | 6.75E-01 | FHA domain-containing protein |
| AT4G14430 | 0.1 | 5.58E-01 | 0.2 | 1.23E-01 | -0.1 | 3.56E-01 | -0.2 | 1.99E-01 | -0.4 | 2.48E-02 | 0.2 | 3.89E-01 | Enoyl-CoA delta isomerase 2, peroxisomal ,Delta-enoyl CoA isomerase 2) |
| AT5G07340 | 0.1 | 4.22E-01 | 0.0 | 5.93E-01 | 0.1 | 2.29E-01 | 0.3 | 6.70E-03 | 0.3 | 2.52E-02 | 0.0 | 8.46E-01 | Calreticulin family protein |
| AT4G30820 | 0.1 | 5.35E-01 | 0.0 | 7.89E-01 | 0.0 | 7.99E-01 | -0.3 | 2.05E-01 | -0.2 | 4.00E-01 | -0.1 | 8.64E-01 | Cyclin-dependent kinase-activating kinase assembly factor-related |
| AT1G28320 | 0.1 | 4.08E-01 | 0.0 | 8.68E-01 | 0.1 | 3.88E-01 | 0.1 | 6.48E-01 | -0.1 | 8.19E-01 | 0.2 | 5.18E-01 | Glyoxysomal processing protease, glyoxysomal |
| AT5G49570 | 0.1 | 4.81E-01 | -0.2 | 4.59E-02 | 0.3 | 2.44E-02 | 0.3 | 3.57E-01 | 0.1 | 9.19E-01 | 0.3 | 5.87E-01 | Peptide-N-asparagine amidase |
| AT1G04400 | 0.1 | 3.78E-01 | 0.0 | 9.74E-01 | 0.1 | 4.18E-01 | -0.4 | 1.33E-03 | -0.2 | 9.62E-02 | -0.2 | 2.16E-01 | Cryptochrome-2 |
| AT1G09770 | 0.1 | 3.68E-01 | 0.0 | 6.82E-01 | 0.1 | 2.36E-01 | 0.0 | 7.78E-01 | -0.2 | 2.95E-01 | 0.1 | 5.78E-01 | Cell division cycle 5-like protein |
| AT4G32130 | 0.1 | 3.96E-01 | 0.1 | 1.27E-01 | -0.1 | 5.06E-01 | 0.1 | 5.59E-01 | 0.0 | 8.33E-01 | 0.1 | 4.76E-01 | ER membrane protein complex subunit 7 homolog |
| AT2G28060 | 0.1 | 6.05E-01 | -0.2 | 1.28E-01 | 0.3 | 8.78E-02 | 0.1 | 6.94E-01 | 0.0 | 9.76E-01 | 0.2 | 7.63E-01 | SNF1-related protein kinase regulatory subunit beta-3 |
| AT3G02720 | 0.1 | 3.84E-01 | 0.1 | 1.85E-01 | 0.0 | 6.54E-01 | -0.3 | 2.77E-02 | -0.1 | 4.11E-01 | -0.2 | 3.68E-01 | Protein DJ-1 homolog D |
| AT5G08580 | 0.1 | 4.54E-01 | 0.1 | 1.36E-01 | -0.1 | 4.88E-01 | 0.2 | 1.26E-01 | 0.2 | 3.98E-01 | 0.1 | 7.24E-01 | Calcium-binding EF hand family protein |
| AT2G02230 | 0.1 | 3.96E-01 | 0.1 | 9.94E-02 | -0.1 | 4.45E-01 | 0.1 | 7.76E-01 | 0.1 | 7.02E-01 | 0.0 | 9.24E-01 | F-box protein PP2-B1 |
| AT3G30180 | 0.1 | 5.36E-01 | 0.5 | 5.87E-04 | -0.4 | 4.02E-03 | -0.8 | 9.75E-05 | -0.9 | 9.86E-05 | 0.1 | 6.99E-01 | Cytochrome P450 85A2 |
| AT3G21160 | 0.1 | 4.09E-01 | 0.1 | 1.10E-01 | -0.1 | 4.55E-01 | 0.1 | 6.09E-01 | 0.0 | 9.37E-01 | 0.1 | 8.09E-01 | alpha-1,2-Mannosidase |
| AT3G06060 | 0.1 | 3.59E-01 | 0.3 | 1.69E-03 | -0.2 | 1.32E-02 | -0.1 | 4.97E-01 | 0.0 | 8.75E-01 | -0.1 | 7.49E-01 | 3-dehydrosphinganine reductase |
| AT2G26580 | 0.1 | 5.76E-01 | 0.2 | 1.20E-01 | -0.1 | 3.60E-01 | 0.0 | 8.34E-01 | 0.0 | 8.57E-01 | -0.1 | 7.21E-01 | Axial regulator YABBY 5 |
| AT5G33370 | 0.1 | 7.94E-01 | 0.4 | 1.20E-01 | -0.3 | 2.56E-01 | 0.4 | 1.67E-01 | -0.1 | 9.17E-01 | 0.5 | 1.80E-01 | GDSL esterase |
| AT2G20770 | 0.1 | 5.15E-01 | 0.2 | 1.36E-01 | -0.1 | 4.54E-01 | 0.1 | 7.94E-01 | 0.1 | 8.53E-01 | 0.0 | 9.88E-01 | LanC-like protein GCL2 |
| AT4G10970 | 0.1 | 5.08E-01 | -0.1 | 1.60E-01 | 0.2 | 7.42E-02 | 0.0 | 8.43E-01 | -0.2 | 3.52E-01 | 0.1 | 5.97E-01 | Ribosome maturation factor |
| AT3G55130 | 0.1 | 7.09E-01 | -0.3 | 1.51E-01 | 0.3 | 1.05E-01 | 0.1 | 7.73E-01 | 0.1 | 6.80E-01 | 0.0 | 9.13E-01 | ABC transporter G family member 19 |
| AT4G04885 | 0.1 | 3.76E-01 | -0.1 | 2.07E-01 | 0.2 | 7.23E-02 | 0.2 | 4.39E-01 | 0.2 | 4.02E-01 | 0.0 | 9.16E-01 | Polyadenylation and cleavage factor homolog 4 |
| AT2G14110 | 0.1 | 6.81E-01 | 0.0 | 8.25E-01 | 0.1 | 6.02E-01 | 0.1 | 7.58E-01 | 0.3 | 4.33E-01 | -0.2 | 7.17E-01 | Haloacid dehalogenase-like hydrolase superfamily protein |
| AT2G27600 | 0.1 | 3.83E-01 | -0.1 | 7.09E-02 | 0.2 | 2.29E-02 | 0.1 | 5.83E-01 | 0.1 | 4.33E-01 | 0.0 | 8.50E-01 | Protein SUPPRESSOR OF K TRANSPORT GROWTH DEFECT 1 |
| AT3G47750 | 0.1 | 7.36E-01 | 0.2 | 2.28E-01 | -0.2 | 4.63E-01 | 0.2 | 6.89E-01 | 0.5 | 3.54E-01 | | | ABC transporter A family member 4 |
| AT4G16760 | 0.1 | 3.52E-01 | -0.3 | 4.14E-03 | 0.3 | 2.04E-03 | 0.0 | 8.04E-01 | -0.2 | 2.85E-01 | 0.1 | 5.32E-01 | Acyl-coenzyme A oxidase |
| AT1G53170 | 0.1 | 5.54E-01 | 0.0 | 6.84E-01 | 0.1 | 3.94E-01 | -0.4 | 9.23E-02 | -0.1 | 6.96E-01 | -0.2 | 4.12E-01 | Ethylene-responsive transcription factor 8 |
| AT1G22800 | 0.1 | 4.80E-01 | 0.1 | 3.57E-01 | 0.0 | 8.46E-01 | 0.0 | 9.96E-01 | 0.0 | 9.55E-01 | 0.0 | 9.65E-01 | Methyltransferase type 11 domain-containing protein |
| AT1G13260 | 0.1 | 7.60E-01 | -0.2 | 2.62E-01 | 0.3 | 1.89E-01 | -0.1 | 6.75E-01 | -0.6 | 2.52E-03 | 0.5 | 1.16E-02 | AP2 |
| AT1G42990 | 0.1 | 5.74E-01 | -0.3 | 3.45E-02 | 0.3 | 2.02E-02 | 0.2 | 3.07E-01 | -0.2 | 5.45E-01 | 0.4 | 1.62E-01 | bZIP transcription factor 60 |
| AT5G06410 | 0.1 | 6.52E-01 | 0.2 | 2.01E-01 | -0.1 | 4.81E-01 | 0.0 | 9.29E-01 | 0.1 | 8.39E-01 | -0.1 | 9.14E-01 | Iron-sulfur cluster co-chaperone protein HscB homolog |
| AT3G43240 | 0.1 | 3.28E-01 | 0.0 | 9.78E-01 | 0.1 | 4.16E-01 | -0.1 | 7.24E-01 | 0.1 | 7.59E-01 | -0.1 | 5.68E-01 | AT-rich interactive domain-containing protein 4 |
| AT2G34250 | 0.1 | 4.11E-01 | 0.1 | 1.61E-01 | -0.1 | 5.57E-01 | 0.2 | 3.97E-02 | 0.1 | 2.54E-01 | 0.1 | 6.80E-01 | AT2G34250 protein |
| AT5G42520 | 0.1 | 3.29E-01 | 0.0 | 7.24E-01 | 0.1 | 2.37E-01 | -0.4 | 2.04E-03 | -0.3 | 2.97E-02 | -0.1 | 5.12E-01 | GAGA-binding transcriptional activator |
| AT5G58090 | 0.1 | 3.26E-01 | 0.2 | 1.41E-02 | -0.1 | 1.06E-01 | 0.0 | 9.50E-01 | -0.2 | 1.40E-01 | 0.2 | 2.24E-01 | Glucan endo-1,3-beta-glucosidase 6 -beta-glucan endohydrolase 6) -beta-glucanase 6) |
| AT3G56860 | 0.1 | 2.72E-01 | 0.0 | 5.35E-01 | 0.1 | 1.26E-01 | 0.0 | 9.83E-01 | 0.0 | 9.12E-01 | 0.0 | 9.32E-01 | UBP1-associated protein 2A |
| AT1G08810 | 0.1 | 5.27E-01 | 0.2 | 3.51E-02 | -0.2 | 1.62E-01 | -0.1 | 7.09E-01 | 0.0 | 9.20E-01 | -0.1 | 8.54E-01 | Transcription factor MYB60 |
| AT4G26300 | 0.1 | 2.48E-01 | 0.0 | 5.48E-01 | 0.0 | 6.51E-01 | 0.2 | 2.04E-01 | 0.2 | 2.67E-01 | 0.0 | 9.97E-01 | arginine--tRNA ligase |
| AT1G50200 | 0.1 | 2.91E-01 | -0.2 | 1.48E-02 | 0.2 | 4.71E-03 | 0.1 | 3.31E-01 | 0.0 | 8.95E-01 | 0.1 | 6.47E-01 | Alanine--tRNA ligase |
| AT1G14510 | 0.1 | 4.39E-01 | 0.2 | 6.92E-02 | -0.1 | 2.93E-01 | -0.1 | 5.01E-01 | -0.2 | 7.56E-02 | 0.2 | 3.36E-01 | PHD finger protein ALFIN-LIKE 7 |
| AT3G61010 | 0.1 | 5.27E-01 | -0.2 | 8.04E-02 | 0.3 | 4.21E-02 | 0.0 | 9.90E-01 | 0.0 | 9.91E-01 | 0.0 | 9.83E-01 | mannosyl-glycoprotein endo-beta-N-acetylglucosaminidase |
| AT1G52500 | 0.1 | 5.56E-01 | -0.1 | 4.14E-01 | 0.2 | 2.46E-01 | 0.1 | 8.84E-01 | 0.3 | 4.77E-01 | -0.3 | 6.53E-01 | MUTM homolog-1 |
| AT2G38550 | 0.1 | 4.56E-01 | 0.2 | 3.01E-02 | -0.2 | 1.45E-01 | 0.2 | 5.13E-02 | 0.1 | 4.29E-01 | 0.1 | 4.99E-01 | Protein FATTY ACID EXPORT 3, chloroplastic |
| AT3G57300 | 0.1 | 5.87E-01 | -0.3 | 3.25E-02 | 0.4 | 2.16E-02 | -0.2 | 3.02E-01 | 0.0 | 9.23E-01 | -0.2 | 5.21E-01 | Chromatin-remodeling ATPase INO80 |
| AT3G61540 | 0.1 | 3.89E-01 | 0.0 | 7.82E-01 | 0.0 | 6.49E-01 | -0.1 | 4.25E-01 | -0.3 | 1.33E-01 | 0.1 | 5.90E-01 | AT3g61540 |
| AT1G49050 | 0.1 | 2.97E-01 | -0.5 | 6.61E-05 | 0.5 | 9.50E-05 | 0.1 | 3.96E-01 | -0.2 | 1.46E-01 | 0.3 | 6.34E-02 | Eukaryotic aspartyl protease family protein |
| AT3G50880 | 0.1 | 3.61E-01 | -0.1 | 1.55E-01 | 0.2 | 5.49E-02 | 0.1 | 7.41E-01 | 0.0 | 8.78E-01 | 0.0 | 9.40E-01 | At3g50880 ) |
| AT2G31230 | 0.1 | 6.50E-01 | -0.1 | 5.34E-01 | 0.2 | 3.48E-01 | 1.0 | 9.74E-04 | 1.0 | 1.99E-03 | 0.0 | 9.78E-01 | Ethylene-responsive transcription factor 15 |
| AT1G18560 | 0.1 | 5.81E-01 | -0.1 | 5.73E-01 | 0.1 | 3.52E-01 | 0.1 | 8.85E-01 | 0.1 | 7.72E-01 | -0.1 | 8.94E-01 | F25I16.10 protein |
| AT5G04220 | 0.1 | 5.58E-01 | 0.3 | 3.55E-02 | -0.2 | 1.53E-01 | 0.1 | 8.73E-01 | 0.1 | 7.51E-01 | -0.1 | 8.80E-01 | Calcium-dependent lipid-binding family protein |
| AT5G40910 | 0.1 | 5.56E-01 | -0.3 | 1.27E-02 | 0.4 | 9.33E-03 | 0.0 | 9.95E-01 | 0.0 | 9.91E-01 | 0.0 | 9.99E-01 | Disease resistance protein family |
| AT1G78240 | 0.1 | 3.72E-01 | 0.1 | 3.13E-01 | 0.0 | 9.01E-01 | -0.2 | 1.52E-01 | 0.0 | 9.91E-01 | -0.2 | 3.16E-01 | Probable pectin methyltransferase QUA2 |
| AT3G51790 | 0.1 | 5.27E-01 | 0.2 | 1.68E-01 | -0.1 | 5.03E-01 | 0.3 | 1.12E-01 | 0.0 | 8.81E-01 | 0.2 | 2.47E-01 | Cytochrome c-type biogenesis protein CcmE homolog, mitochondrial |
| AT2G44710 | 0.1 | 2.55E-01 | -0.1 | 3.81E-02 | 0.2 | 1.04E-02 | 0.0 | 8.59E-01 | -0.1 | 7.33E-01 | 0.1 | 6.58E-01 | At2g44720 |
| AT5G67070 | 0.1 | 4.83E-01 | 0.3 | 4.98E-03 | -0.3 | 2.40E-02 | -0.2 | 2.52E-01 | 0.0 | 8.75E-01 | -0.2 | 3.27E-01 | Protein RALF-like 34 |
| AT1G02360 | 0.1 | 6.77E-01 | 0.3 | 6.95E-02 | -0.2 | 1.88E-01 | 0.8 | 6.93E-05 | 0.4 | 3.12E-02 | 0.4 | 3.27E-02 | At1g02360 |
| AT4G35100 | 0.1 | 4.52E-01 | 0.5 | 2.08E-04 | -0.5 | 1.05E-03 | -0.9 | 1.61E-05 | -0.7 | 2.72E-04 | -0.1 | 6.30E-01 | Aquaporin PIP2-7 |
| AT3G13430 | 0.1 | 4.48E-01 | 0.1 | 2.54E-01 | 0.0 | 7.32E-01 | -0.1 | 6.83E-01 | -0.1 | 6.89E-01 | 0.0 | 9.70E-01 | RING-type E3 ubiquitin transferase |
| AT3G45610 | 0.1 | 5.99E-01 | 0.0 | 9.20E-01 | 0.1 | 7.56E-01 | -0.7 | 1.16E-02 | -0.3 | 3.70E-01 | -0.5 | 1.55E-01 | Dof zinc finger protein DOF3.2 |
| AT5G61790 | 0.1 | 6.38E-01 | 0.1 | 3.86E-01 | -0.1 | 7.10E-01 | 0.7 | 9.40E-06 | 0.6 | 2.00E-04 | 0.1 | 5.58E-01 | Calnexin homolog 1 |
| AT1G52280 | 0.1 | 5.30E-01 | 0.2 | 8.21E-02 | -0.1 | 2.89E-01 | 0.0 | 9.37E-01 | 0.1 | 6.27E-01 | -0.2 | 6.39E-01 | Ras-related protein RABG3d |
| AT5G12140 | 0.1 | 5.31E-01 | 0.2 | 4.53E-02 | -0.2 | 1.57E-01 | 0.0 | 9.86E-01 | 0.1 | 8.43E-01 | -0.1 | 8.52E-01 | Cysteine proteinase inhibitor 1 |
| AT3G20050 | 0.1 | 4.44E-01 | 0.2 | 8.04E-02 | -0.1 | 3.10E-01 | 0.3 | 3.52E-02 | 0.1 | 6.17E-01 | 0.2 | 3.13E-01 | T-complex protein 1 subunit alpha |
| AT4G38900 | 0.1 | 3.28E-01 | 0.0 | 9.16E-01 | 0.1 | 3.48E-01 | -0.2 | 2.88E-01 | -0.1 | 6.82E-01 | -0.1 | 7.27E-01 | bZIP transcription factor 29 |
| AT1G28120 | 0.1 | 3.79E-01 | -0.1 | 2.80E-01 | 0.2 | 9.29E-02 | 0.0 | 9.72E-01 | 0.0 | 9.80E-01 | 0.0 | 9.64E-01 | OVARIAN TUMOR DOMAIN-containing deubiquitinating enzyme 1 |
| AT1G11790 | 0.1 | 4.79E-01 | -0.1 | 3.71E-01 | 0.2 | 1.69E-01 | 0.0 | 9.46E-01 | 0.1 | 6.33E-01 | -0.1 | 7.21E-01 | Arogenate dehydratase |
| AT1G04810 | 0.1 | 3.29E-01 | -0.1 | 1.11E-01 | 0.2 | 3.01E-02 | 0.2 | 5.33E-01 | 0.1 | 8.37E-01 | 0.1 | 8.29E-01 | 26S proteasome non-ATPase regulatory subunit 1 homolog B |
| AT5G18620 | 0.1 | 3.56E-01 | 0.0 | 8.04E-01 | 0.0 | 5.76E-01 | 0.1 | 5.05E-01 | 0.1 | 5.77E-01 | 0.0 | 9.98E-01 | hypothetical protein |
| AT5G46640 | 0.1 | 6.03E-01 | 0.2 | 2.41E-01 | -0.1 | 5.74E-01 | -0.2 | 5.34E-01 | 0.4 | 1.84E-01 | -0.6 | 7.08E-02 | AT-hook motif nuclear-localized protein |
| AT3G25740 | 0.1 | 6.26E-01 | -0.3 | 7.21E-02 | 0.4 | 5.50E-02 | 0.1 | 7.63E-01 | 0.1 | 7.58E-01 | 0.0 | 9.71E-01 | Methionine aminopeptidase |
| AT5G57140 | 0.1 | 5.80E-01 | -0.4 | 8.89E-03 | 0.5 | 9.90E-03 | -0.3 | 6.48E-01 | 0.0 | 9.76E-01 | -0.3 | 7.38E-01 | Probable inactive purple acid phosphatase 28 |
| AT5G47080 | 0.1 | 5.15E-01 | -0.3 | 2.03E-02 | 0.4 | 1.38E-02 | 0.0 | 8.76E-01 | 0.0 | 9.65E-01 | 0.0 | 8.81E-01 | Casein kinase II subunit beta |
| AT3G49730 | 0.1 | 5.41E-01 | 0.1 | 4.20E-01 | 0.0 | 8.65E-01 | 0.3 | 1.61E-01 | -0.1 | 6.98E-01 | 0.4 | 1.25E-01 | Pentatricopeptide repeat-containing protein At3g49730 |
| AT4G37650 | 0.1 | 3.47E-01 | 0.1 | 7.09E-02 | -0.1 | 3.86E-01 | -0.1 | 5.87E-01 | 0.0 | 9.31E-01 | -0.1 | 7.85E-01 | Protein SHORT-ROOT |
| AT1G25580 | 0.1 | 3.90E-01 | -0.1 | 2.26E-01 | 0.2 | 8.32E-02 | 0.2 | 2.53E-01 | 0.1 | 5.38E-01 | 0.1 | 8.13E-01 | NAC domain-containing protein |
| AT3G26932 | 0.1 | 5.91E-01 | 0.3 | 7.69E-02 | -0.2 | 2.60E-01 | -0.1 | 6.32E-01 | -0.3 | 3.02E-01 | 0.1 | 6.40E-01 | DsRNA-binding protein 3 |
| AT3G03270 | 0.1 | 6.81E-01 | -0.2 | 2.58E-01 | 0.3 | 1.81E-01 | 0.7 | 4.33E-03 | 0.2 | 5.22E-01 | 0.5 | 6.02E-02 | AT3G03270 protein |
| AT5G64610 | 0.1 | 2.98E-01 | 0.0 | 6.56E-01 | 0.1 | 2.00E-01 | -0.1 | 7.89E-01 | -0.1 | 5.88E-01 | 0.1 | 8.27E-01 | Histone acetyltransferase of the MYST family 1 |
| AT3G54230 | 0.1 | 2.74E-01 | -0.3 | 3.20E-03 | 0.3 | 1.58E-03 | -0.1 | 7.84E-01 | -0.2 | 4.86E-01 | 0.1 | 7.43E-01 | Suppressor of abi3-5 |
| AT3G62980 | 0.1 | 3.74E-01 | 0.2 | 8.07E-02 | -0.1 | 3.73E-01 | -0.5 | 9.21E-04 | -0.3 | 4.50E-02 | -0.2 | 3.01E-01 | Protein TRANSPORT INHIBITOR RESPONSE 1 |
| AT3G50360 | 0.1 | 5.49E-01 | 0.1 | 3.35E-01 | 0.0 | 7.48E-01 | 0.2 | 2.69E-01 | 0.2 | 5.32E-01 | 0.1 | 8.31E-01 | EF-hand domain-containing protein |

| | | | | | | | | | | | | | |
|---|---|---|---|---|---|---|---|---|---|---|---|---|---|
| AT1G49480 | 0.1 | 3.16E-01 | 0.1 | 3.11E-01 | 0.0 | 9.70E-01 | -0.4 | 2.28E-02 | -0.4 | 3.18E-02 | 0.0 | 9.62E-01 | B3 domain-containing protein REM19 |
| AT3G42170 | 0.1 | 2.99E-01 | 0.1 | 2.63E-01 | 0.0 | 9.21E-01 | -0.1 | 7.24E-01 | 0.0 | 9.12E-01 | -0.1 | 7.34E-01 | BED zinc finger and hAT dimerization domain-containing protein DAYSLEEPER |
| AT1G21450 | 0.1 | 4.08E-01 | -0.1 | 2.71E-01 | 0.2 | 9.76E-02 | 0.0 | 9.31E-01 | -0.3 | 7.21E-02 | 0.3 | 1.20E-01 | Scarecrow-like protein 1 |
| AT4G21470 | 0.1 | 3.71E-01 | -0.2 | 7.21E-02 | 0.2 | 2.70E-02 | 0.2 | 2.66E-01 | 0.3 | 1.27E-01 | -0.1 | 7.29E-01 | Bifunctional riboflavin kinase |
| AT3G62220 | 0.1 | 3.21E-01 | -0.2 | 4.25E-02 | 0.2 | 1.70E-02 | -0.3 | 5.38E-02 | -0.2 | 2.65E-01 | -0.1 | 6.47E-01 | At3g62220 |
| AT5G48160 | 0.1 | 3.23E-01 | 0.0 | 6.05E-01 | 0.1 | 1.76E-01 | -0.4 | 5.45E-03 | -0.6 | 3.18E-04 | 0.3 | 1.56E-01 | Protein OBERON 2 |
| AT3G23730 | 0.1 | 6.38E-01 | 0.3 | 9.35E-02 | -0.2 | 2.51E-01 | -0.7 | 1.40E-04 | -0.8 | 2.61E-04 | 0.0 | 8.85E-01 | Probable xyloglucan endotransglucosylase |
| AT5G17770 | 0.1 | 3.64E-01 | 0.3 | 8.92E-03 | -0.2 | 5.46E-02 | 0.2 | 4.94E-02 | 0.0 | 7.64E-01 | 0.2 | 3.02E-01 | NADH--cytochrome b5 reductase 1 |
| AT5G60790 | 0.1 | 3.76E-01 | 0.0 | 8.91E-01 | 0.1 | 5.22E-01 | 0.3 | 9.52E-03 | 0.0 | 7.74E-01 | 0.2 | 1.13E-01 | ABC transporter F family member 1 |
| AT1G09700 | 0.1 | 4.97E-01 | 0.2 | 1.04E-01 | -0.1 | 3.85E-01 | 0.3 | 4.87E-02 | -0.1 | 6.65E-01 | 0.4 | 3.71E-02 | DsRNA-binding domain-like superfamily protein |
| AT1G21440 | 0.1 | 5.80E-01 | 0.3 | 3.81E-02 | -0.2 | 1.27E-01 | -0.6 | 1.93E-03 | 0.2 | 2.99E-01 | -0.8 | 3.33E-03 | At1g21440 |
| AT1G05590 | 0.1 | 4.43E-01 | 0.1 | 1.55E-01 | -0.1 | 5.35E-01 | 0.2 | 3.85E-01 | 0.4 | 7.41E-02 | -0.2 | 4.03E-01 | Beta-hexosaminidase 2 |
| AT4G02405 | 0.1 | 2.96E-01 | -0.1 | 2.82E-01 | 0.2 | 8.28E-02 | 0.1 | 6.83E-01 | -0.1 | 8.30E-01 | 0.2 | 4.40E-01 | S-adenosyl-L-methionine-dependent methyltransferases superfamily protein |
| AT1G80340 | 0.1 | 5.92E-01 | 0.3 | 6.46E-02 | -0.2 | 2.36E-01 | 1.0 | 8.80E-02 | 1.2 | 8.04E-02 | | | Gibberellin 3-beta-dioxygenase 2 |
| AT5G08370 | 0.1 | 3.78E-01 | 0.1 | 1.07E-01 | -0.1 | 4.85E-01 | -0.2 | 3.78E-01 | 0.2 | 3.53E-01 | -0.4 | 9.70E-02 | Alpha-galactosidase 2 |
| AT3G02680 | 0.1 | 6.26E-01 | -0.1 | 6.84E-01 | 0.1 | 4.63E-01 | -0.1 | 5.97E-01 | 0.0 | 9.20E-01 | -0.1 | 8.00E-01 | Nibrin homolog |
| AT2G39840 | 0.1 | 2.18E-01 | 0.0 | 6.90E-01 | 0.1 | 1.55E-01 | 0.2 | 3.32E-01 | 0.0 | 8.78E-01 | 0.1 | 6.20E-01 | Serine |
| AT1G76810 | 0.1 | 4.47E-01 | 0.0 | 9.66E-01 | 0.1 | 5.36E-01 | 0.2 | 3.29E-02 | 0.0 | 8.91E-01 | 0.2 | 1.91E-01 | Eukaryotic translation initiation factor 5B |
| AT1G20610 | 0.1 | 5.38E-01 | 0.4 | 4.80E-03 | -0.3 | 2.89E-02 | -0.2 | 3.55E-01 | -0.1 | 7.91E-01 | -0.1 | 6.73E-01 | Cyclin-B2-3 |
| AT1G73020 | 0.1 | 4.00E-01 | 0.0 | 8.55E-01 | 0.1 | 6.05E-01 | 0.0 | 9.81E-01 | -0.1 | 7.21E-01 | 0.1 | 7.03E-01 | Anoctamin-like protein At1g73020 |
| AT3G21280 | 0.1 | 4.37E-01 | -0.1 | 3.07E-01 | 0.2 | 1.34E-01 | 0.0 | 9.77E-01 | -0.1 | 7.55E-01 | 0.1 | 7.55E-01 | Ubiquitin carboxyl-terminal hydrolase |
| AT4G39280 | 0.1 | 3.02E-01 | 0.0 | 8.55E-01 | 0.1 | 4.67E-01 | 0.2 | 1.59E-01 | 0.1 | 6.91E-01 | 0.1 | 5.61E-01 | phenylalanine--tRNA ligase |
| AT5G59380 | 0.1 | 4.00E-01 | 0.1 | 1.31E-01 | -0.1 | 5.37E-01 | -0.2 | 2.10E-01 | -0.3 | 1.21E-01 | 0.1 | 7.74E-01 | Methyl-CpG-binding domain-containing protein 6 |
| AT3G25040 | 0.1 | 3.52E-01 | 0.3 | 8.61E-03 | -0.2 | 6.12E-02 | 0.2 | 3.47E-01 | 0.3 | 1.88E-01 | -0.1 | 7.41E-01 | ER lumen protein-retaining receptor B |
| AT5G49970 | 0.1 | 2.93E-01 | 0.0 | 5.76E-01 | 0.0 | 6.93E-01 | -0.1 | 6.01E-01 | 0.0 | 9.63E-01 | -0.1 | 7.01E-01 | NADH-hydrate epimerase HX epimerase) |
| AT4G21800 | 0.1 | 3.64E-01 | -0.1 | 2.04E-01 | 0.2 | 6.65E-02 | 0.4 | 1.21E-02 | 0.1 | 6.59E-01 | 0.3 | 1.41E-01 | GPN-loop GTPase |
| AT1G31710 | 0.1 | 3.74E-01 | 0.4 | 1.65E-03 | -0.3 | 1.15E-02 | 0.1 | 6.16E-01 | 0.0 | 9.23E-01 | 0.1 | 8.07E-01 | Amine oxidase alpha 3, peroxisomal |
| AT5G49060 | 0.1 | 3.43E-01 | 0.0 | 7.82E-01 | 0.1 | 6.02E-01 | 0.0 | 8.92E-01 | -0.2 | 5.24E-01 | 0.2 | 4.73E-01 | Chaperone protein dnaJ 49 |
| AT2G31740 | 0.1 | 3.73E-01 | -0.2 | 8.23E-02 | 0.2 | 3.15E-02 | 0.2 | 2.38E-01 | -0.1 | 6.79E-01 | 0.3 | 2.04E-01 | Methyltransferase type 11 domain-containing protein |
| AT1G69390 | 0.1 | 3.49E-01 | 0.1 | 1.43E-01 | -0.1 | 6.04E-01 | 0.0 | 8.20E-01 | -0.1 | 5.06E-01 | 0.1 | 7.31E-01 | Cell division topological specificity factor homolog, chloroplastic |
| AT3G55920 | 0.1 | 4.42E-01 | 0.3 | 1.79E-02 | -0.2 | 1.07E-01 | -0.2 | 2.34E-01 | -0.1 | 7.05E-01 | -0.1 | 6.36E-01 | Peptidyl-prolyl cis-trans isomerase CYP21-2 |
| AT3G58670 | 0.1 | 3.35E-01 | 0.1 | 3.71E-01 | 0.0 | 9.81E-01 | -0.3 | 2.90E-02 | -0.2 | 1.45E-01 | -0.1 | 7.14E-01 | cysteine dioxygenase |
| AT4G06746 | 0.1 | 7.38E-01 | 0.4 | 9.64E-02 | -0.3 | 2.56E-01 | 1.4 | 1.31E-03 | -0.1 | 9.37E-01 | 1.4 | 6.08E-03 | Ethylene-responsive transcription factor RAP2-9 |
| AT5G66240 | 0.1 | 1.80E-01 | 0.0 | 9.78E-01 | 0.1 | 2.44E-01 | 0.0 | 9.87E-01 | 0.0 | 7.85E-01 | 0.0 | 8.28E-01 | Transducin |
| AT3G09440 | 0.1 | 5.53E-01 | -0.5 | 2.42E-03 | 0.6 | 2.01E-03 | 0.4 | 4.24E-03 | 0.7 | 2.76E-04 | -0.3 | 8.35E-02 | Heat shock 70 kDa protein 3 |
| AT1G18490 | 0.1 | 3.57E-01 | 0.2 | 5.53E-02 | -0.1 | 3.22E-01 | -0.1 | 6.81E-01 | 0.0 | 9.35E-01 | -0.1 | 8.42E-01 | Plant cysteine oxidase 3 |
| AT1G28090 | 0.1 | 3.65E-01 | 0.2 | 1.97E-02 | -0.2 | 1.40E-01 | 0.3 | 1.23E-01 | 0.2 | 4.96E-01 | 0.1 | 6.35E-01 | Polynucleotide adenylyltransferase family protein |
| AT2G45700 | 0.1 | 3.39E-01 | 0.0 | 8.37E-01 | 0.1 | 3.17E-01 | -0.3 | 1.18E-01 | -0.3 | 1.66E-01 | 0.0 | 9.94E-01 | Sterile alpha motif domain-containing protein |
| AT5G62390 | 0.1 | 4.18E-01 | 0.2 | 6.15E-02 | -0.1 | 2.65E-01 | -0.2 | 3.33E-01 | -0.3 | 1.06E-01 | 0.1 | 5.85E-01 | BAG family molecular chaperone regulator 7 |
| AT4G15955 | 0.1 | 3.57E-01 | 0.0 | 7.20E-01 | 0.1 | 2.85E-01 | 0.0 | 8.95E-01 | 0.0 | 9.02E-01 | -0.1 | 7.98E-01 | Alpha |
| AT5G64950 | 0.1 | 5.48E-01 | 0.2 | 2.25E-01 | -0.1 | 5.98E-01 | 0.4 | 1.65E-01 | -0.4 | 3.15E-01 | 0.8 | 3.27E-02 | Emb\|CAB62602.1 |
| AT5G56890 | 0.1 | 3.45E-01 | -0.1 | 3.68E-01 | 0.2 | 1.06E-01 | 0.0 | 8.57E-01 | 0.2 | 4.19E-01 | -0.2 | 3.97E-01 | Protein kinase domain-containing protein |
| AT3G21630 | 0.1 | 2.03E-01 | -0.1 | 3.56E-02 | 0.2 | 7.77E-03 | 0.1 | 2.43E-01 | 0.0 | 8.32E-01 | 0.2 | 2.82E-01 | Protein kinase domain-containing protein |
| AT2G25310 | 0.1 | 4.02E-01 | 0.1 | 5.12E-01 | 0.0 | 9.05E-01 | -0.2 | 1.20E-01 | -0.1 | 3.98E-01 | -0.1 | 7.30E-01 | ER membrane protein complex subunit-like protein |
| AT1G44835 | 0.1 | 3.70E-01 | 0.3 | 1.44E-02 | -0.2 | 1.03E-01 | 0.2 | 2.90E-01 | 0.2 | 3.45E-01 | 0.0 | 9.90E-01 | Uncharacterized protein At1g44835 |
| AT2G26770 | 0.1 | 2.34E-01 | 0.1 | 3.28E-01 | 0.0 | 8.83E-01 | 0.1 | 3.72E-01 | 0.1 | 5.46E-01 | 0.0 | 9.35E-01 | Stomatal closure-related actin-binding protein 1 |
| AT1G15910 | 0.1 | 2.62E-01 | 0.1 | 1.17E-01 | 0.0 | 6.56E-01 | -0.2 | 8.28E-02 | -0.2 | 9.53E-02 | 0.0 | 9.54E-01 | Factor of DNA methylation 1 |
| AT1G45000 | 0.1 | 3.26E-01 | 0.1 | 3.13E-01 | 0.0 | 9.51E-01 | 0.2 | 8.79E-02 | 0.1 | 2.49E-01 | 0.0 | 8.47E-01 | AAA-type ATPase family protein |
| AT5G03140 | 0.1 | 3.40E-01 | 0.1 | 1.08E-01 | -0.1 | 5.18E-01 | 0.0 | 8.62E-01 | 0.0 | 9.84E-01 | 0.0 | 8.90E-01 | L-type lectin-domain containing receptor kinase VIII.2 |
| AT4G25650 | 0.1 | 3.47E-01 | -0.1 | 3.40E-01 | 0.2 | 9.63E-02 | -0.4 | 4.37E-02 | -0.2 | 3.54E-01 | -0.2 | 5.13E-01 | hypothetical protein |
| AT5G08530 | 0.1 | 2.26E-01 | 0.0 | 5.18E-01 | 0.0 | 6.45E-01 | -0.1 | 6.34E-01 | -0.1 | 5.09E-01 | 0.0 | 8.86E-01 | NADH dehydrogenase flavoprotein 1, mitochondrial |
| AT3G18380 | 0.1 | 2.97E-01 | 0.0 | 8.33E-01 | 0.1 | 4.98E-01 | 0.1 | 5.95E-01 | 0.0 | 9.79E-01 | 0.1 | 7.48E-01 | hypothetical protein |
| AT1G53710 | 0.1 | 2.50E-01 | 0.1 | 1.39E-01 | 0.0 | 7.44E-01 | 0.0 | 8.53E-01 | -0.2 | 3.43E-01 | 0.2 | 3.39E-01 | Calcineurin-like metallo-phosphoesterase superfamily protein |
| AT1G13120 | 0.1 | 2.77E-01 | -0.2 | 3.34E-02 | 0.3 | 1.16E-02 | 0.3 | 1.13E-01 | 0.1 | 5.69E-01 | 0.2 | 5.42E-01 | mRNA export factor GLE1 |
| AT5G20080 | 0.1 | 3.26E-01 | 0.3 | 7.59E-03 | -0.2 | 5.50E-02 | 0.4 | 8.45E-03 | 0.3 | 5.01E-02 | 0.1 | 7.50E-01 | NADH-cytochrome b5 reductase-like protein |
| AT3G43610 | 0.1 | 5.10E-01 | 0.1 | 5.31E-01 | 0.0 | 9.98E-01 | 0.0 | 9.25E-01 | 0.4 | 1.09E-01 | -0.4 | 1.54E-01 | Spc97 |
| AT2G17380 | 0.1 | 4.64E-01 | 0.3 | 3.88E-02 | -0.2 | 1.81E-01 | -0.2 | 4.48E-01 | -0.3 | 2.01E-01 | 0.1 | 6.82E-01 | AP complex subunit sigma |
| AT2G20280 | 0.1 | 3.06E-01 | 0.2 | 4.11E-02 | -0.1 | 2.76E-01 | 0.1 | 6.68E-01 | 0.1 | 5.28E-01 | 0.0 | 8.68E-01 | Zinc finger CCCH domain-containing protein 21 |
| AT4G02020 | 0.1 | 2.73E-01 | -0.1 | 1.27E-01 | 0.2 | 3.17E-02 | -0.3 | 7.78E-02 | -0.3 | 1.45E-01 | 0.0 | 9.37E-01 | SET domain-containing protein |
| AT5G20890 | 0.1 | 4.16E-01 | 0.2 | 1.09E-01 | -0.1 | 4.19E-01 | 0.2 | 1.83E-01 | 0.1 | 5.83E-01 | 0.1 | 7.30E-01 | T-complex protein 1 subunit beta |
| AT3G12630 | 0.1 | 5.22E-01 | -0.1 | 6.37E-01 | 0.1 | 3.10E-01 | -0.4 | 4.60E-02 | -0.6 | 1.03E-02 | 0.2 | 5.31E-01 | Zinc finger A20 and AN1 domain-containing stress-associated protein 5 |
| AT5G20500 | 0.1 | 4.50E-01 | 0.3 | 2.31E-02 | -0.2 | 1.15E-01 | 0.0 | 9.99E-01 | -0.1 | 8.25E-01 | 0.1 | 8.52E-01 | Glutaredoxin-C4 |
| AT4G17500 | 0.1 | 6.55E-01 | -0.3 | 1.86E-01 | 0.3 | 1.22E-01 | 0.6 | 2.27E-02 | -0.2 | 4.73E-01 | 0.8 | 1.83E-02 | Ethylene-responsive transcription factor 1A |
| AT5G19150 | 0.1 | 2.57E-01 | 0.1 | 3.10E-01 | 0.0 | 9.54E-01 | -0.2 | 2.37E-01 | -0.1 | 6.25E-01 | -0.1 | 7.23E-01 | ATP-dependent -NADH-hydrate dehydratase HX dehydratase) |
| AT4G35987 | 0.1 | 3.97E-01 | 0.1 | 1.87E-01 | -0.1 | 6.74E-01 | 0.0 | 9.85E-01 | 0.1 | 6.12E-01 | -0.1 | 6.88E-01 | Calmodulin-lysine N-methyltransferase |
| AT3G18190 | 0.1 | 2.97E-01 | 0.2 | 2.05E-02 | -0.1 | 1.48E-01 | -0.1 | 6.58E-01 | 0.0 | 9.63E-01 | -0.1 | 8.22E-01 | T-complex protein 1 subunit delta |
| AT3G03940 | 0.1 | 1.90E-01 | -0.2 | 2.47E-02 | 0.2 | 5.03E-03 | -0.1 | 4.57E-01 | -0.1 | 6.01E-01 | 0.0 | 9.62E-01 | Protein kinase family protein |
| AT2G28660 | 0.1 | 2.81E-01 | 0.2 | 1.07E-02 | -0.2 | 1.13E-01 | 0.0 | 9.05E-01 | -0.2 | 3.57E-01 | 0.2 | 5.05E-01 | HMA domain-containing protein |
| AT5G60900 | 0.1 | 6.59E-01 | -0.6 | 1.34E-02 | 0.7 | 1.29E-02 | 0.8 | 3.42E-03 | 0.4 | 2.27E-01 | 0.5 | 1.38E-01 | G-type lectin S-receptor-like serine |
| AT1G73300 | 0.1 | 6.98E-01 | 0.5 | 2.58E-02 | -0.4 | 9.67E-02 | | | | | | | Serine carboxypeptidase-like 2 |
| AT2G42030 | 0.1 | 2.89E-01 | -0.1 | 4.86E-01 | 0.1 | 1.31E-01 | -0.4 | 2.40E-02 | -0.3 | 6.34E-02 | 0.0 | 8.86E-01 | E3 ubiquitin-protein ligase RMA |
| AT3G07220 | 0.1 | 2.27E-01 | 0.0 | 5.05E-01 | 0.1 | 1.07E-01 | -0.4 | 3.19E-02 | -0.2 | 3.15E-01 | -0.2 | 5.04E-01 | FHA domain-containing protein FHA2 |
| AT5G40740 | 0.1 | 3.65E-01 | 0.0 | 9.82E-01 | 0.1 | 4.36E-01 | -0.4 | 2.74E-02 | -0.3 | 2.02E-01 | -0.1 | 6.05E-01 | AUGMIN subunit 6 |
| AT1G06645 | 0.1 | 3.75E-01 | 0.1 | 3.49E-01 | 0.0 | 9.46E-01 | -0.1 | 8.32E-01 | 0.1 | 8.59E-01 | -0.1 | 7.17E-01 | 2-oxoglutarate and Fe-dependent oxygenase superfamily protein |
| AT1G10610 | 0.1 | 6.52E-01 | -0.1 | 6.22E-01 | 0.2 | 4.31E-01 | -0.8 | 1.57E-02 | -0.2 | 5.60E-01 | -0.6 | 1.68E-01 | Basic helix-loop-helix DNA-binding superfamily protein |
| AT4G25440 | 0.1 | 3.26E-01 | 0.0 | 6.33E-01 | 0.1 | 2.17E-01 | -0.4 | 4.45E-02 | -0.4 | 3.42E-02 | 0.1 | 8.33E-01 | Zinc finger CCCH domain-containing protein 48 |
| AT5G45050 | 0.1 | 3.67E-01 | -0.3 | 1.92E-02 | 0.4 | 9.23E-03 | 0.0 | 9.42E-01 | -0.2 | 5.70E-01 | 0.2 | 5.61E-01 | Disease resistance protein |
| AT5G67380 | 0.1 | 1.71E-01 | 0.1 | 3.89E-01 | 0.0 | 6.74E-01 | -0.1 | 5.15E-01 | -0.2 | 3.25E-01 | 0.1 | 7.82E-01 | Casein kinase II subunit alpha-1 |
| AT4G32660 | 0.1 | 3.08E-01 | -0.1 | 1.41E-01 | 0.2 | 4.13E-02 | -0.1 | 7.63E-01 | 0.1 | 5.82E-01 | -0.2 | 4.11E-01 | Protein kinase superfamily protein |
| AT5G62190 | 0.1 | 2.47E-01 | 0.1 | 2.12E-01 | 0.0 | 9.03E-01 | 0.1 | 5.38E-01 | -0.2 | 1.11E-01 | 0.3 | 8.55E-02 | DEAD-box ATP-dependent RNA helicase 7 |
| AT4G36090 | 0.1 | 4.19E-01 | -0.2 | 1.81E-01 | 0.2 | 7.33E-02 | 0.6 | 1.71E-02 | 0.4 | 1.66E-01 | 0.2 | 4.82E-01 | Oxidoreductase, 2OG-Fe oxygenase family protein |
| AT4G30190 | 0.1 | 3.62E-01 | 0.3 | 5.08E-03 | -0.3 | 3.21E-02 | 0.5 | 3.20E-03 | 0.2 | 2.51E-01 | 0.3 | 2.11E-01 | Plasma membrane ATPase |
| AT1G53840 | 0.1 | 3.47E-01 | 0.4 | 1.91E-03 | -0.3 | 1.27E-02 | -0.1 | 3.55E-01 | 0.0 | 9.60E-01 | -0.1 | 5.98E-01 | Pectinesterase 1 |

| | | | | | | | | | | | | | |
|---|---|---|---|---|---|---|---|---|---|---|---|---|---|
| AT5G42920 | 0.1 | 2.09E-01 | 0.1 | 4.46E-01 | 0.0 | 6.98E-01 | -0.2 | 2.64E-01 | -0.1 | 6.62E-01 | -0.1 | 7.00E-01 | THO complex subunit 5B |
| AT1G65820 | 0.1 | 2.87E-01 | -0.1 | 4.39E-01 | 0.2 | 1.05E-01 | 0.4 | 9.53E-02 | 0.0 | 9.01E-01 | 0.3 | 2.80E-01 | Microsomal glutathione s-transferase |
| AT5G67580 | 0.1 | 2.49E-01 | 0.2 | 5.12E-02 | -0.1 | 4.06E-01 | 0.0 | 9.17E-01 | -0.1 | 5.10E-01 | 0.1 | 5.23E-01 | Telomere repeat-binding factor 2 |
| AT5G48380 | 0.1 | 5.36E-01 | -0.3 | 4.39E-02 | 0.4 | 2.24E-02 | 0.2 | 8.89E-02 | -0.2 | 1.84E-01 | 0.4 | 2.76E-02 | Probably inactive leucine-rich repeat receptor-like protein kinase At5g48380 |
| AT1G54150 | 0.1 | 2.41E-01 | -0.2 | 1.38E-02 | 0.3 | 5.39E-03 | -0.2 | 2.22E-01 | -0.2 | 1.18E-01 | 0.1 | 7.76E-01 | RING-type E3 ubiquitin transferase |
| AT4G01990 | 0.1 | 4.21E-01 | 0.0 | 8.57E-01 | 0.1 | 4.18E-01 | 0.5 | 2.38E-02 | 0.3 | 2.73E-01 | 0.2 | 3.54E-01 | Pentatricopeptide repeat-containing protein At4g01990, mitochondrial |
| AT5G25170 | 0.1 | 4.54E-01 | 0.2 | 1.49E-01 | -0.1 | 5.23E-01 | 0.2 | 4.09E-01 | 0.0 | 8.98E-01 | 0.1 | 6.36E-01 | PPPDE putative thiol peptidase family protein |
| AT5G63700 | 0.1 | 4.36E-01 | 0.0 | 9.23E-01 | 0.1 | 4.66E-01 | -0.1 | 6.55E-01 | -0.4 | 1.59E-01 | 0.3 | 4.62E-01 | Zinc ion binding |
| AT2G06025 | 0.1 | 3.44E-01 | 0.0 | 8.03E-01 | 0.1 | 2.95E-01 | -0.3 | 5.84E-02 | -0.2 | 1.69E-01 | -0.1 | 8.44E-01 | Acyl-CoA N-acyltransferases superfamily protein |
| AT5G13430 | 0.1 | 4.17E-01 | 0.3 | 1.77E-02 | -0.2 | 9.02E-02 | 0.1 | 4.04E-01 | 0.1 | 6.12E-01 | 0.0 | 9.17E-01 | Cytochrome b-c1 complex subunit Rieske-1, mitochondrial |
| AT1G29350 | 0.1 | 3.32E-01 | 0.1 | 2.73E-01 | 0.0 | 8.95E-01 | 0.2 | 2.29E-01 | -0.1 | 6.83E-01 | 0.3 | 1.68E-01 | RNA polymerase II degradation factor-like protein |
| AT2G46110 | 0.1 | 5.14E-01 | 0.2 | 1.48E-01 | -0.1 | 4.80E-01 | -0.1 | 8.56E-01 | 0.1 | 6.63E-01 | -0.2 | 5.67E-01 | 3-methyl-2-oxobutanoate hydroxymethyltransferase 1, mitochondrial |
| AT5G04750 | 0.1 | 5.67E-01 | -0.1 | 5.36E-01 | 0.2 | 2.83E-01 | -0.3 | 1.28E-01 | -0.2 | 5.04E-01 | -0.1 | 6.71E-01 | At5g04750 |
| AT2G01220 | 0.1 | 2.76E-01 | 0.0 | 9.25E-01 | 0.1 | 3.99E-01 | -0.1 | 7.87E-01 | 0.1 | 5.37E-01 | -0.2 | 4.34E-01 | Nucleotidylyl transferase superfamily protein |
| AT3G08020 | 0.1 | 1.85E-01 | -0.2 | 1.87E-02 | 0.3 | 5.34E-03 | -0.2 | 5.69E-01 | -0.1 | 6.71E-01 | 0.0 | 9.79E-01 | PHD finger family protein |
| AT5G06740 | 0.1 | 4.64E-01 | 0.1 | 4.30E-01 | 0.0 | 9.46E-01 | 1.2 | 8.80E-05 | 0.1 | 7.10E-01 | 1.1 | 1.90E-03 | non-specific serine |
| AT3G53180 | 0.1 | 2.09E-01 | 0.1 | 4.42E-01 | 0.0 | 6.93E-01 | 0.0 | 9.76E-01 | -0.1 | 6.45E-01 | 0.1 | 7.50E-01 | Nodulin |
| AT5G65870 | 0.1 | 7.84E-01 | -0.4 | 2.53E-01 | 0.5 | 2.28E-01 | 0.4 | 3.49E-01 | 0.7 | 1.13E-01 | -0.3 | 5.11E-01 | Phytosulfokines 5 |
| AT4G14610 | 0.1 | 2.92E-01 | -0.2 | 3.00E-02 | 0.3 | 1.25E-02 | 0.4 | 1.49E-01 | 0.0 | 9.20E-01 | 0.3 | 3.58E-01 | Probable disease resistance protein At4g14610 |
| AT2G37050 | 0.1 | 3.78E-01 | 0.0 | 9.98E-01 | 0.1 | 4.43E-01 | -0.2 | 2.29E-01 | -0.1 | 5.30E-01 | -0.1 | 8.11E-01 | Leucine-rich repeat protein kinase family protein |
| AT1G11720 | 0.1 | 5.42E-01 | -0.1 | 3.91E-01 | 0.2 | 1.93E-01 | 0.0 | 9.36E-01 | 0.4 | 2.18E-02 | -0.4 | 4.38E-02 | SS3 |
| AT2G43950 | 0.1 | 2.17E-01 | -0.1 | 1.00E-01 | 0.2 | 2.02E-02 | 0.1 | 6.38E-01 | 0.0 | 8.12E-01 | 0.1 | 5.66E-01 | Outer envelope pore protein 37, chloroplastic |
| AT2G35320 | 0.1 | 2.44E-01 | 0.1 | 1.86E-01 | 0.0 | 8.67E-01 | -0.5 | 7.62E-02 | -0.2 | 4.34E-01 | -0.2 | 5.83E-01 | Eyes absent homolog |
| AT5G59540 | 0.1 | 2.49E-01 | -0.1 | 1.86E-01 | 0.2 | 3.60E-02 | 0.4 | 2.45E-03 | 0.2 | 8.78E-02 | 0.2 | 3.34E-01 | 1-aminocyclopropane-1-carboxylate oxidase homolog 12 |
| AT5G25754 | 0.1 | 2.53E-01 | -0.1 | 4.74E-01 | 0.1 | 1.16E-01 | 0.1 | 7.03E-01 | -0.1 | 5.45E-01 | 0.2 | 3.59E-01 | Eukaryotic translation initiation factor 3 subunit L |
| AT3G17450 | 0.1 | 2.42E-01 | -0.2 | 1.30E-02 | 0.3 | 5.07E-03 | -0.2 | 2.35E-01 | -0.2 | 2.94E-01 | 0.0 | 9.95E-01 | hypothetical protein |
| AT3G10030 | 0.1 | 2.35E-01 | -0.1 | 2.12E-01 | 0.2 | 4.68E-02 | -0.3 | 2.85E-02 | -0.4 | 2.50E-02 | 0.0 | 8.63E-01 | UMP kinase |
| AT2G44180 | 0.1 | 2.02E-01 | -0.1 | 4.75E-01 | 0.1 | 8.78E-02 | 0.1 | 5.57E-01 | -0.2 | 1.74E-01 | 0.3 | 9.23E-02 | Methionine aminopeptidase 2 |
| AT2G23070 | 0.1 | 1.46E-01 | 0.1 | 1.73E-01 | 0.0 | 9.74E-01 | -0.1 | 5.70E-01 | -0.1 | 3.72E-01 | 0.1 | 8.07E-01 | Casein kinase II subunit alpha-4, chloroplastic |
| AT1G08210 | 0.1 | 2.26E-01 | 0.0 | 5.67E-01 | 0.0 | 6.17E-01 | -0.4 | 4.46E-02 | -0.3 | 1.52E-01 | -0.1 | 7.88E-01 | Eukaryotic aspartyl protease family protein |
| AT2G32860 | 0.1 | 3.99E-01 | 0.1 | 3.29E-01 | 0.0 | 8.91E-01 | 0.0 | 9.78E-01 | 0.2 | 2.11E-01 | -0.2 | 2.70E-01 | Beta-glucosidase 33 |
| AT1G63480 | 0.1 | 4.25E-01 | 0.2 | 1.20E-01 | -0.1 | 4.89E-01 | 0.0 | 9.37E-01 | 0.0 | 9.69E-01 | 0.0 | 9.32E-01 | AT-hook motif nuclear-localized protein |
| AT1G80230 | 0.1 | 4.05E-01 | 0.4 | 6.45E-03 | -0.3 | 3.66E-02 | 0.1 | 6.03E-01 | 0.1 | 6.41E-01 | 0.0 | 9.77E-01 | Cytochrome c oxidase subunit 5b-2, mitochondrial |
| AT2G26240 | 0.1 | 4.56E-01 | 0.2 | 1.09E-01 | -0.1 | 3.97E-01 | -0.2 | 3.68E-01 | -0.4 | 2.83E-02 | 0.3 | 1.68E-01 | Protein FATTY ACID EXPORT 7 |
| AT1G34470 | 0.1 | 2.37E-01 | -0.1 | 2.50E-01 | 0.2 | 5.98E-02 | 0.0 | 9.48E-01 | -0.1 | 7.99E-01 | 0.1 | 7.89E-01 | Probable magnesium transporter NIPA3 |
| AT1G61660 | 0.1 | 2.81E-01 | -0.1 | 3.79E-01 | 0.2 | 9.65E-02 | -0.2 | 1.53E-01 | -0.2 | 2.60E-01 | 0.0 | 9.39E-01 | Basic helix-loop-helix DNA-binding superfamily protein |
| AT3G15540 | 0.1 | 3.00E-01 | 0.2 | 1.74E-02 | -0.2 | 1.61E-01 | 0.3 | 1.05E-01 | 0.1 | 8.31E-01 | 0.3 | 3.12E-01 | Auxin-responsive protein IAA19 |
| AT4G36680 | 0.1 | 5.31E-01 | 0.1 | 5.37E-01 | 0.0 | 9.88E-01 | 0.7 | 1.12E-03 | 0.4 | 8.73E-02 | 0.3 | 1.47E-01 | Small ribosomal subunit protein mL103 |
| AT1G50010 | 0.1 | 3.21E-01 | 0.4 | 3.77E-03 | -0.3 | 2.67E-02 | -0.3 | 5.20E-02 | -0.1 | 6.54E-01 | -0.2 | 3.70E-01 | Tubulin alpha-2 chain |
| AT4G22010 | 0.1 | 4.39E-01 | 0.4 | 4.55E-03 | -0.3 | 2.37E-02 | 0.0 | 9.11E-01 | 0.2 | 3.86E-01 | -0.1 | 5.66E-01 | At4g22010 |
| AT4G09980 | 0.1 | 2.04E-01 | -0.1 | 2.15E-01 | 0.2 | 4.14E-02 | 0.2 | 4.44E-01 | 0.1 | 7.03E-01 | 0.1 | 8.74E-01 | EMB1691 |
| AT2G43140 | 0.1 | 5.21E-01 | 0.1 | 4.71E-01 | 0.0 | 9.36E-01 | 0.9 | 3.22E-03 | 0.7 | 2.21E-02 | 0.2 | 6.68E-01 | Basic helix-loop-helix DNA-binding superfamily protein |
| AT3G53110 | 0.1 | 1.61E-01 | 0.2 | 3.94E-02 | -0.1 | 4.35E-01 | 0.0 | 9.88E-01 | -0.1 | 2.16E-01 | 0.1 | 3.42E-01 | LOS4 |
| AT5G17440 | 0.1 | 2.85E-01 | -0.1 | 4.96E-01 | 0.2 | 1.33E-01 | -0.2 | 1.92E-01 | -0.2 | 1.44E-01 | 0.0 | 8.56E-01 | AT5g17440 |
| AT1G10430 | 0.1 | 2.00E-01 | 0.2 | 3.84E-02 | -0.1 | 3.87E-01 | -0.1 | 3.65E-01 | -0.2 | 1.44E-01 | 0.1 | 6.79E-01 | Serine |
| AT5G05460 | 0.1 | 3.83E-01 | 0.0 | 9.98E-01 | 0.1 | 4.61E-01 | 0.2 | 2.02E-01 | 0.0 | 9.73E-01 | 0.2 | 3.23E-01 | Cytosolic endo-beta-N-acetylglucosaminidase 1 |
| AT2G25480 | 0.1 | 2.75E-01 | 0.2 | 1.34E-02 | -0.2 | 1.28E-01 | 0.2 | 1.80E-01 | 0.2 | 2.68E-01 | 0.0 | 9.74E-01 | TPX2 C-terminal domain-containing protein |
| AT1G79220 | 0.1 | 5.39E-01 | 0.2 | 1.61E-01 | -0.1 | 4.89E-01 | -0.1 | 7.48E-01 | 0.0 | 9.38E-01 | -0.1 | 8.83E-01 | Mitochondrial transcription termination factor family protein |
| AT3G04380 | 0.1 | 6.25E-01 | 0.1 | 6.74E-01 | 0.0 | 9.81E-01 | 0.4 | 5.66E-01 | 0.9 | 2.18E-01 | -0.5 | 5.28E-01 | SET-domain containing protein lysine methyltransferase family protein |
| AT1G12220 | 0.1 | 4.04E-01 | 0.0 | 7.98E-01 | 0.1 | 6.59E-01 | -0.2 | 4.08E-01 | -0.8 | 1.64E-03 | 0.6 | 2.90E-02 | Disease resistance protein family |
| AT2G29640 | 0.1 | 5.07E-01 | 0.0 | 8.28E-01 | 0.1 | 4.73E-01 | 0.1 | 9.18E-01 | 0.2 | 7.54E-01 | -0.1 | 8.54E-01 | Josephin-like protein |
| AT4G12790 | 0.1 | 1.82E-01 | 0.0 | 8.54E-01 | 0.1 | 3.15E-01 | 0.1 | 3.52E-01 | 0.1 | 7.63E-01 | 0.1 | 7.42E-01 | GPN-loop GTPase 3 |
| AT5G47230 | 0.1 | 5.79E-01 | -0.6 | 5.16E-03 | 0.7 | 5.39E-03 | 0.9 | 8.74E-04 | 0.3 | 3.77E-01 | 0.6 | 2.04E-02 | Ethylene-responsive transcription factor 5 |
| AT1G71900 | 0.1 | 2.14E-01 | 0.1 | 1.77E-01 | 0.0 | 9.01E-01 | -0.1 | 7.63E-01 | 0.0 | 9.59E-01 | -0.1 | 7.44E-01 | Probable magnesium transporter |
| AT2G17975 | 0.1 | 5.29E-01 | -0.1 | 6.73E-01 | 0.2 | 3.84E-01 | -0.1 | 7.86E-01 | -0.1 | 8.01E-01 | 0.0 | 9.86E-01 | At2g17975 family protein) |
| AT5G07710 | 0.1 | 2.76E-01 | -0.3 | 1.26E-02 | 0.4 | 5.92E-03 | -0.2 | 2.85E-01 | 0.0 | 9.60E-01 | -0.2 | 4.79E-01 | Protein NEN1 |
| AT3G16840 | 0.1 | 3.72E-01 | 0.0 | 9.68E-01 | 0.1 | 4.18E-01 | 0.4 | 2.51E-02 | 0.2 | 4.22E-01 | 0.2 | 3.79E-01 | ATP-dependent RNA helicase |
| AT5G02290 | 0.1 | 3.37E-01 | -0.3 | 1.40E-02 | 0.4 | 5.76E-03 | 0.1 | 6.66E-01 | -0.3 | 1.54E-01 | 0.4 | 1.07E-01 | Probable serine |
| AT2G23430 | 0.1 | 3.76E-01 | 0.0 | 8.04E-01 | 0.1 | 3.38E-01 | -0.2 | 3.28E-01 | 0.2 | 5.65E-01 | -0.4 | 2.16E-01 | Cyclin-dependent kinase inhibitor 1 |
| AT2G27350 | 0.1 | 2.13E-01 | 0.0 | 6.70E-01 | 0.1 | 1.42E-01 | -0.1 | 4.00E-01 | -0.2 | 3.10E-01 | 0.0 | 8.50E-01 | OVARIAN TUMOR DOMAIN-containing deubiquitinating enzyme 6 |
| AT5G14080 | 0.1 | 2.87E-01 | 0.1 | 5.50E-01 | 0.0 | 7.32E-01 | -0.3 | 8.29E-02 | -0.2 | 2.26E-01 | -0.1 | 8.31E-01 | Pentatricopeptide repeat-containing protein At5g14080 |
| AT5G41920 | 0.1 | 2.56E-01 | 0.2 | 2.83E-02 | -0.1 | 2.61E-01 | -0.5 | 7.62E-03 | -0.3 | 1.02E-01 | -0.2 | 4.74E-01 | Scarecrow-like protein 23 |
| AT5G16940 | 0.1 | 7.41E-01 | -0.1 | 8.41E-01 | 0.2 | 6.65E-01 | -0.5 | 4.45E-01 | 0.0 | 9.94E-01 | -0.5 | 5.48E-01 | hypothetical protein |
| AT3G48080 | 0.1 | 5.91E-01 | -0.3 | 1.05E-01 | 0.4 | 6.92E-02 | 0.0 | 9.72E-01 | -0.1 | 8.63E-01 | 0.1 | 8.66E-01 | Protein EDS1B |
| AT3G54350 | 0.1 | 1.68E-01 | -0.1 | 2.75E-01 | 0.2 | 4.16E-02 | -0.3 | 5.39E-02 | -0.4 | 3.34E-02 | 0.1 | 8.02E-01 | Uncharacterized protein T12E18_40 |
| AT2G15050 | 0.1 | 5.45E-01 | 0.3 | 4.75E-02 | -0.2 | 1.70E-01 | 0.1 | 5.01E-01 | 0.0 | 9.09E-01 | 0.1 | 7.53E-01 | LTP7 |
| AT2G43400 | 0.1 | 2.45E-01 | -0.1 | 2.96E-01 | 0.2 | 6.88E-02 | -0.2 | 4.11E-01 | -0.3 | 2.03E-01 | 0.1 | 6.66E-01 | Electron transfer flavoprotein-ubiquinone oxidoreductase |
| AT2G03510 | 0.1 | 2.72E-01 | 0.1 | 5.45E-01 | 0.0 | 6.92E-01 | 0.3 | 1.58E-02 | 0.1 | 6.60E-01 | 0.2 | 1.78E-01 | Expressed protein |
| AT1G71310 | 0.1 | 4.17E-01 | -0.1 | 6.82E-01 | 0.2 | 3.08E-01 | 0.3 | 3.43E-01 | 0.1 | 6.96E-01 | 0.1 | 7.79E-01 | RAD52-1B.2 |
| AT5G46790 | 0.1 | 3.08E-01 | 0.1 | 4.89E-01 | 0.0 | 8.07E-01 | 0.0 | 9.51E-01 | 0.0 | 9.27E-01 | 0.0 | 9.05E-01 | Abscisic acid receptor PYL1 |
| AT4G02840 | 0.1 | 3.21E-01 | 0.3 | 8.06E-03 | -0.2 | 6.68E-02 | 0.2 | 2.49E-01 | 0.0 | 9.59E-01 | 0.2 | 4.46E-01 | Small nuclear ribonucleoprotein SmD1b |
| AT5G59880 | 0.1 | 2.54E-01 | 0.2 | 1.59E-02 | -0.1 | 1.36E-01 | 0.0 | 9.55E-01 | -0.1 | 7.79E-01 | 0.1 | 7.92E-01 | Actin depolymerizing factor 3 |
| AT4G26430 | 0.1 | 3.11E-01 | 0.1 | 4.04E-01 | 0.0 | 9.17E-01 | 0.4 | 7.78E-02 | 0.1 | 6.32E-01 | 0.3 | 3.70E-01 | COP9 signalosome complex subunit 6 |
| AT3G18640 | 0.1 | 1.77E-01 | -0.2 | 2.10E-02 | 0.3 | 5.00E-03 | 0.3 | 1.14E-01 | 0.2 | 3.26E-01 | 0.1 | 7.85E-01 | Zinc finger CCCH domain-containing protein 38 |
| AT1G66750 | 0.1 | 2.43E-01 | -0.1 | 1.17E-01 | 0.2 | 3.15E-02 | 0.3 | 1.72E-01 | 0.2 | 4.31E-01 | 0.1 | 7.80E-01 | Cyclin-dependent kinase D-2 |
| AT3G06580 | 0.1 | 1.64E-01 | 0.1 | 3.05E-01 | 0.0 | 7.73E-01 | -0.1 | 4.16E-01 | 0.0 | 9.27E-01 | -0.1 | 6.47E-01 | Galactokinase |
| AT4G26000 | 0.1 | 2.13E-01 | 0.0 | 9.69E-01 | 0.1 | 3.02E-01 | -0.2 | 1.60E-01 | -0.3 | 1.26E-01 | 0.0 | 8.68E-01 | RNA-binding KH domain-containing protein PEPPER |
| AT2G38770 | 0.1 | 2.45E-01 | 0.0 | 9.49E-01 | 0.1 | 2.82E-01 | 0.2 | 1.17E-01 | -0.1 | 7.53E-01 | 0.3 | 1.60E-01 | P-loop containing nucleoside triphosphate hydrolases superfamily protein |
| AT4G30530 | 0.1 | 2.37E-01 | 0.2 | 1.67E-02 | -0.1 | 1.52E-01 | 0.2 | 3.73E-01 | 0.3 | 1.74E-01 | -0.1 | 7.02E-01 | Gamma-glutamyl peptidase 1 |
| AT1G25440 | 0.1 | 3.69E-01 | 0.3 | 4.29E-02 | -0.2 | 2.24E-01 | -0.4 | 5.51E-03 | -0.1 | 5.09E-01 | -0.3 | 1.10E-01 | Zinc finger protein CONSTANS-LIKE 16 |
| AT5G47870 | 0.1 | 2.08E-01 | 0.2 | 9.91E-03 | -0.1 | 1.24E-01 | 0.3 | 4.64E-02 | 0.2 | 2.03E-01 | 0.1 | 7.36E-01 | DNA repair RAD52-like protein 2, chloroplastic |
| AT2G35530 | 0.1 | 1.87E-01 | 0.0 | 9.27E-01 | 0.1 | 2.30E-01 | 0.0 | 9.80E-01 | 0.1 | 8.21E-01 | -0.1 | 8.22E-01 | bZIP transcription factor 16 |
| AT5G65080 | 0.1 | 6.40E-01 | 0.0 | 9.00E-01 | 0.1 | 6.25E-01 | -0.1 | 7.34E-01 | -0.1 | 8.01E-01 | 0.0 | 9.83E-01 | K-box region |

| | | | | | | | | | | | | | |
|---|---|---|---|---|---|---|---|---|---|---|---|---|---|
| AT1G62820 | 0.1 | 4.71E-01 | 0.0 | 7.82E-01 | 0.1 | 7.38E-01 | 0.1 | 8.02E-01 | -0.1 | 8.32E-01 | 0.2 | 6.96E-01 | Probable calcium-binding protein CML14 |
| AT5G19350 | 0.1 | 1.32E-01 | 0.0 | 8.42E-01 | 0.1 | 2.47E-01 | 0.0 | 9.08E-01 | -0.1 | 8.28E-01 | 0.1 | 7.79E-01 | Polyadenylate-binding protein RBP47B' -binding protein RBP47B') |
| AT5G50320 | 0.1 | 1.79E-01 | -0.1 | 2.73E-01 | 0.2 | 4.44E-02 | -0.2 | 7.66E-02 | -0.3 | 1.67E-02 | 0.1 | 5.29E-01 | Elongator complex protein 3 acetyltransferase) |
| AT1G78790 | 0.1 | 2.69E-01 | 0.0 | 7.71E-01 | 0.1 | 2.45E-01 | -0.3 | 1.41E-01 | -0.2 | 3.73E-01 | -0.1 | 7.91E-01 | Centromere protein X |
| AT5G62350 | 0.1 | 3.05E-01 | 0.5 | 6.19E-04 | -0.4 | 4.33E-03 | -0.1 | 7.45E-01 | -0.1 | 7.80E-01 | 0.0 | 9.92E-01 | hypothetical protein |
| AT1G34020 | 0.1 | 2.91E-01 | 0.1 | 4.47E-01 | 0.0 | 8.34E-01 | -0.1 | 7.36E-01 | 0.2 | 3.94E-01 | -0.3 | 3.09E-01 | UDP-rhamnose |
| AT4G04320 | 0.1 | 1.65E-01 | -0.1 | 1.73E-01 | 0.2 | 2.81E-02 | 0.0 | 8.68E-01 | 0.1 | 5.96E-01 | -0.1 | 5.68E-01 | Malonyl-CoA decarboxylase family protein |
| AT1G47230 | 0.1 | 2.59E-01 | 0.1 | 5.56E-01 | 0.0 | 6.81E-01 | -0.1 | 4.64E-01 | 0.0 | 9.95E-01 | -0.1 | 6.08E-01 | Cyclin N-terminal domain-containing protein |
| AT5G22040 | 0.1 | 2.66E-01 | 0.1 | 5.01E-01 | 0.0 | 7.48E-01 | 0.0 | 8.67E-01 | -0.1 | 6.51E-01 | 0.2 | 5.70E-01 | At5g22040 |
| AT5G67200 | 0.1 | 3.31E-01 | 0.0 | 7.03E-01 | 0.1 | 6.55E-01 | 0.4 | 6.32E-02 | 0.3 | 1.22E-01 | 0.0 | 9.39E-01 | Probable inactive receptor kinase At5g67200 |
| AT1G80260 | 0.1 | 2.30E-01 | -0.2 | 3.13E-02 | 0.3 | 8.59E-03 | -0.2 | 4.26E-01 | -0.1 | 8.25E-01 | -0.1 | 7.56E-01 | Gamma-tubulin complex component |
| AT3G03090 | 0.1 | 2.54E-01 | 0.2 | 1.67E-02 | -0.1 | 1.79E-01 | 0.0 | 9.96E-01 | 0.2 | 4.03E-01 | -0.2 | 4.57E-01 | D-xylose-proton symporter-like 1 |
| AT1G21240 | 0.1 | 8.43E-01 | -0.9 | 1.51E-01 | 1.0 | 1.76E-01 | 0.9 | 1.19E-01 | -1.3 | 1.68E-01 | 2.2 | 1.72E-02 | Wall-associated receptor kinase-like |
| AT4G31170 | 0.1 | 1.59E-01 | -0.2 | 6.45E-03 | 0.3 | 1.62E-03 | 0.0 | 8.65E-01 | -0.2 | 2.00E-01 | 0.1 | 4.05E-01 | Protein kinase superfamily protein |
| AT5G11020 | 0.1 | 3.01E-01 | 0.3 | 1.04E-02 | -0.2 | 1.08E-01 | 0.3 | 2.59E-01 | 0.1 | 6.55E-01 | 0.1 | 7.34E-01 | Protein kinase domain-containing protein |
| AT2G42910 | 0.1 | 2.69E-01 | 0.2 | 3.80E-02 | -0.1 | 2.91E-01 | 0.3 | 6.13E-02 | 0.0 | 8.89E-01 | 0.3 | 1.21E-01 | Phosphoribosyltransferase domain-containing protein |
| AT3G13690 | 0.1 | 2.26E-01 | 0.1 | 1.98E-01 | 0.0 | 9.19E-01 | -0.2 | 1.31E-01 | -0.1 | 7.09E-01 | -0.2 | 5.08E-01 | Kinase with adenine nucleotide alpha hydrolases-like domain-containing protein |
| AT4G36410 | 0.1 | 4.68E-01 | 0.8 | 1.83E-04 | -0.7 | 1.46E-03 | 1.0 | 3.88E-05 | 0.6 | 4.23E-03 | 0.3 | 9.22E-02 | Probable ubiquitin-conjugating enzyme E2 17 |
| AT5G04620 | 0.1 | 2.84E-01 | 0.2 | 3.06E-02 | -0.1 | 2.59E-01 | -0.1 | 7.45E-01 | -0.1 | 9.09E-01 | -0.1 | 9.11E-01 | Biotin F |
| AT3G24490 | 0.1 | 1.32E-01 | 0.0 | 9.93E-01 | 0.1 | 1.96E-01 | -0.1 | 6.31E-01 | -0.1 | 6.06E-01 | 0.0 | 9.55E-01 | Alcohol dehydrogenase transcription factor Myb |
| AT5G07400 | 0.1 | 4.70E-01 | -0.1 | 7.27E-01 | 0.2 | 3.86E-01 | 0.7 | 1.42E-01 | 0.8 | 1.34E-01 | -0.1 | 9.09E-01 | FHA domain-containing protein |
| AT1G09430 | 0.1 | 1.28E-01 | 0.1 | 7.23E-02 | 0.0 | 7.32E-01 | 0.0 | 9.14E-01 | -0.3 | 1.03E-01 | 0.2 | 1.97E-01 | ATP-citrate synthase alpha chain protein 3 |
| AT5G64920 | 0.1 | 2.47E-01 | 0.1 | 1.96E-01 | 0.0 | 8.82E-01 | -0.4 | 2.76E-02 | -0.4 | 5.17E-02 | 0.0 | 9.80E-01 | RING-type E3 ubiquitin transferase |
| AT4G21540 | 0.1 | 1.32E-01 | -0.1 | 7.91E-02 | 0.2 | 1.14E-02 | -0.1 | 3.02E-01 | -0.2 | 2.22E-01 | 0.0 | 8.55E-01 | Sphingosine kinase 1 |
| AT5G25480 | 0.1 | 3.64E-01 | -0.1 | 6.49E-01 | 0.2 | 2.62E-01 | 0.1 | 5.88E-01 | -0.1 | 8.96E-01 | 0.2 | 5.94E-01 | tRNA -C)-methyltransferase 2 -methyltransferase-like protein 2) |
| AT5G02630 | 0.1 | 2.74E-01 | -0.1 | 2.02E-01 | 0.2 | 6.09E-02 | 0.2 | 3.94E-01 | -0.2 | 3.45E-01 | 0.4 | 9.47E-02 | Protein CANDIDATE G-PROTEIN COUPLED RECEPTOR 6 |
| AT3G47780 | 0.1 | 2.88E-01 | 0.0 | 9.86E-01 | 0.1 | 3.67E-01 | 2.0 | 3.19E-08 | 1.0 | 1.36E-04 | 0.9 | 7.24E-04 | ABC transporter A family member 7 |
| AT2G20340 | 0.1 | 4.66E-01 | -0.2 | 2.45E-01 | 0.3 | 1.10E-01 | 0.1 | 3.63E-01 | 0.5 | 4.69E-03 | -0.3 | 8.54E-02 | Uncharacterized protein |
| AT4G29360 | 0.1 | 1.81E-01 | 0.3 | 4.60E-03 | -0.2 | 7.82E-02 | -0.2 | 3.09E-01 | -0.1 | 6.76E-01 | -0.1 | 7.27E-01 | glucan endo-1,3-beta-D-glucosidase |
| AT1G63020 | 0.1 | 4.17E-01 | 0.1 | 4.93E-01 | 0.0 | 9.50E-01 | 0.3 | 2.55E-01 | 0.3 | 3.35E-01 | 0.0 | 9.84E-01 | DNA-directed RNA polymerase |
| AT2G36720 | 0.1 | 2.71E-01 | -0.1 | 2.49E-01 | 0.2 | 6.12E-02 | 0.0 | 9.27E-01 | 0.0 | 1.00E+00 | 0.0 | 9.57E-01 | Acyl-CoA N-acyltransferase with RING |
| AT1G70410 | 0.1 | 1.95E-01 | 0.2 | 1.44E-02 | -0.1 | 1.60E-01 | 0.0 | 7.91E-01 | -0.2 | 1.06E-01 | 0.1 | 3.35E-01 | Beta carbonic anhydrase 4 |
| AT3G61610 | 0.1 | 3.08E-01 | 0.2 | 9.04E-02 | -0.1 | 5.17E-01 | 0.3 | 1.47E-01 | 0.3 | 1.49E-01 | 0.0 | 9.39E-01 | glucose-6-phosphate 1-epimerase |
| AT1G11800 | 0.1 | 2.05E-01 | 0.1 | 4.81E-01 | 0.0 | 6.65E-01 | 0.4 | 4.71E-02 | 0.3 | 1.97E-01 | 0.1 | 7.29E-01 | At1g11800 |
| AT3G27230 | 0.1 | 1.35E-01 | 0.1 | 8.81E-02 | 0.0 | 8.08E-01 | 0.3 | 7.32E-02 | 0.3 | 6.02E-02 | 0.0 | 8.54E-01 | At3g27230 |
| AT5G24530 | 0.1 | 6.99E-01 | -0.4 | 1.21E-01 | 0.6 | 8.02E-02 | 0.6 | 7.57E-03 | 0.4 | 6.59E-02 | 0.1 | 6.30E-01 | Protein DOWNY MILDEW RESISTANCE 6 -Fe oxygenase-like protein DMR6) |
| AT5G65060 | 0.1 | 1.22E-01 | 0.0 | 9.59E-01 | 0.1 | 2.06E-01 | 0.0 | 9.73E-01 | -0.1 | 6.35E-01 | 0.2 | 6.67E-01 | At5g65060 |
| AT1G76560 | 0.1 | 3.67E-01 | 0.3 | 1.31E-02 | -0.2 | 1.06E-01 | -0.4 | 5.22E-02 | -0.7 | 3.63E-03 | 0.3 | 3.04E-01 | Calvin cycle protein CP12-3, chloroplastic |
| AT1G79680 | 0.1 | 7.03E-01 | -0.5 | 1.03E-01 | 0.7 | 9.58E-02 | 0.9 | 7.86E-02 | 0.0 | 9.88E-01 | 1.0 | 1.27E-01 | Wall-associated receptor kinase-like 10 |
| AT4G17620 | 0.1 | 1.03E-01 | -0.1 | 3.95E-01 | 0.2 | 4.03E-02 | -0.2 | 2.22E-01 | 0.0 | 8.45E-01 | -0.1 | 5.06E-01 | Decapping nuclease |
| AT5G62940 | 0.1 | 3.58E-01 | 0.2 | 1.94E-01 | -0.1 | 7.33E-01 | 0.0 | 9.46E-01 | 0.0 | 8.70E-01 | 0.1 | 8.47E-01 | Dof zinc finger protein DOF5.6 |
| AT2G26560 | 0.1 | 4.71E-01 | -0.1 | 4.20E-01 | 0.2 | 1.67E-01 | 1.1 | 1.04E-04 | -0.8 | 6.54E-03 | 1.8 | 8.26E-05 | Patatin-like protein 2 |
| AT1G77990 | 0.1 | 3.64E-01 | 0.4 | 4.32E-03 | -0.3 | 4.06E-02 | 0.2 | 3.36E-01 | 0.2 | 3.01E-01 | 0.0 | 9.11E-01 | STAS domain-containing protein |
| AT5G06460 | 0.1 | 1.24E-01 | 0.0 | 7.38E-01 | 0.1 | 1.02E-01 | 0.4 | 1.15E-02 | 0.1 | 3.83E-01 | 0.2 | 1.89E-01 | Ubiquitin-activating enzyme E1 2 |
| AT3G08950 | 0.1 | 3.95E-01 | 0.2 | 1.00E-01 | -0.1 | 4.65E-01 | 0.5 | 1.95E-02 | 0.2 | 4.88E-01 | 0.3 | 2.47E-01 | Protein SCO1 homolog 1, mitochondrial |
| AT3G50890 | 0.1 | 4.67E-01 | 0.4 | 3.03E-02 | -0.2 | 1.73E-01 | -0.2 | 4.64E-01 | -0.2 | 3.57E-01 | 0.1 | 8.64E-01 | Zinc-finger homeodomain protein 7 |
| AT2G21250 | 0.1 | 1.40E-01 | 0.2 | 1.95E-02 | -0.1 | 2.86E-01 | -0.1 | 3.54E-01 | -0.1 | 6.42E-01 | -0.1 | 8.48E-01 | NAD-linked oxidoreductase superfamily protein |
| AT2G18670 | 0.1 | 3.77E-01 | 0.2 | 8.18E-02 | -0.1 | 4.19E-01 | -0.1 | 6.20E-01 | -0.6 | 2.42E-02 | 0.5 | 1.25E-01 | RING-H2 finger protein ATL56 |
| AT4G00830 | 0.1 | 8.43E-02 | 0.0 | 7.85E-01 | 0.1 | 7.68E-02 | 0.1 | 6.16E-01 | 0.0 | 9.47E-01 | 0.0 | 7.99E-01 | Heterogeneous nuclear ribonucleoprotein Q |
| AT2G27050 | 0.1 | 2.29E-01 | -0.1 | 5.28E-01 | 0.2 | 1.07E-01 | -0.4 | 7.94E-02 | -0.3 | 2.18E-01 | -0.1 | 8.51E-01 | ETHYLENE INSENSITIVE 3-like 1 protein |
| AT1G22950 | 0.1 | 1.75E-01 | 0.2 | 1.28E-02 | -0.1 | 1.95E-01 | 0.7 | 5.30E-03 | 0.4 | 8.65E-02 | 0.2 | 4.40E-01 | Fe2OG dioxygenase domain-containing protein |
| AT1G18890 | 0.1 | 2.80E-01 | -0.2 | 5.61E-02 | 0.3 | 1.58E-02 | 0.0 | 8.57E-01 | -0.4 | 6.09E-03 | 0.4 | 2.62E-02 | Calcium-dependent protein kinase 10 |
| AT1G08780 | 0.1 | 3.09E-01 | 0.3 | 1.13E-02 | -0.2 | 1.01E-01 | -0.1 | 6.96E-01 | -0.1 | 7.91E-01 | 0.0 | 9.77E-01 | Prefoldin subunit 4 |
| AT1G04210 | 0.1 | 1.17E-01 | -0.2 | 3.06E-02 | 0.3 | 4.74E-03 | 0.1 | 5.54E-01 | 0.1 | 8.68E-01 | 0.1 | 8.27E-01 | Protein kinase domain-containing protein |
| AT1G18840 | 0.1 | 1.47E-01 | 0.1 | 3.15E-01 | 0.0 | 7.17E-01 | 0.0 | 7.71E-01 | 0.0 | 9.62E-01 | -0.1 | 8.07E-01 | Protein IQ-DOMAIN 30 |
| AT3G47000 | 0.1 | 1.49E-01 | 0.1 | 1.66E-01 | 0.0 | 9.89E-01 | -0.4 | 3.81E-02 | -0.2 | 3.92E-01 | -0.2 | 4.71E-01 | beta-glucosidase |
| AT3G57050 | 0.1 | 1.52E-01 | 0.3 | 6.98E-03 | -0.1 | 1.10E-01 | -0.2 | 1.97E-01 | 0.0 | 9.27E-01 | -0.2 | 3.83E-01 | Cystathionine beta-lyase |
| AT4G36480 | 0.1 | 9.43E-02 | 0.0 | 5.65E-01 | 0.1 | 3.25E-01 | -0.1 | 6.14E-01 | 0.0 | 9.49E-01 | -0.1 | 7.93E-01 | Long-chain base1 |
| AT4G00730 | 0.1 | 1.88E-01 | 0.2 | 7.34E-02 | -0.1 | 5.90E-01 | -0.1 | 6.52E-01 | 0.0 | 8.53E-01 | 0.0 | 9.11E-01 | Homeobox-leucine zipper family protein |
| AT1G31970 | 0.1 | 3.45E-01 | 0.1 | 4.59E-01 | 0.0 | 8.93E-01 | 0.5 | 1.16E-03 | 0.3 | 8.20E-02 | 0.2 | 2.31E-01 | DEAD-box ATP-dependent RNA helicase 5 |
| AT5G60750 | 0.1 | 1.16E-01 | -0.2 | 4.76E-02 | 0.3 | 6.94E-03 | -0.1 | 5.20E-01 | -0.2 | 3.03E-01 | 0.1 | 7.77E-01 | CAAX amino terminal protease family protein |
| AT1G51600 | 0.1 | 1.33E-01 | 0.1 | 2.79E-01 | 0.0 | 7.45E-01 | -0.2 | 2.15E-01 | -0.1 | 4.80E-01 | -0.1 | 7.98E-01 | GATA transcription factor 28 |
| AT4G33770 | 0.1 | 2.47E-01 | 0.2 | 3.64E-02 | -0.1 | 3.33E-01 | 0.1 | 6.79E-01 | 0.2 | 4.44E-01 | -0.1 | 7.87E-01 | Inositol-tetrakisphosphate 1-kinase |
| AT1G71210 | 0.1 | 1.90E-01 | -0.1 | 5.71E-01 | 0.2 | 1.20E-01 | 0.4 | 1.12E-01 | 0.2 | 5.27E-01 | 0.2 | 5.84E-01 | Pentatricopeptide repeat-containing protein At1g71210, mitochondrial |
| AT3G55380 | 0.1 | 2.48E-01 | 0.3 | 1.89E-02 | -0.1 | 1.87E-01 | 0.2 | 8.33E-02 | 0.1 | 5.21E-01 | 0.1 | 5.37E-01 | Ubiquitin-conjugating enzyme 14 |
| AT5G03380 | 0.1 | 3.45E-01 | 0.4 | 1.08E-02 | -0.3 | 6.99E-02 | 0.7 | 6.79E-04 | -0.1 | 5.03E-01 | 0.8 | 2.22E-03 | Heavy metal transport |
| AT1G30910 | 0.1 | 2.10E-01 | -0.1 | 4.51E-01 | 0.2 | 9.32E-02 | 0.0 | 9.31E-01 | 0.1 | 5.44E-01 | -0.1 | 5.70E-01 | MOSC domain-containing protein |
| AT1G63470 | 0.1 | 2.56E-01 | 0.2 | 5.15E-02 | -0.1 | 4.02E-01 | -0.3 | 6.35E-02 | -0.1 | 6.09E-01 | -0.2 | 3.66E-01 | AT-hook motif nuclear-localized protein 5 |
| AT1G01350 | 0.1 | 1.06E-01 | 0.2 | 4.42E-02 | 0.0 | 6.46E-01 | -0.2 | 1.97E-01 | -0.2 | 2.02E-01 | 0.0 | 9.48E-01 | Zinc finger CCCH domain-containing protein 1 |
| AT1G05830 | 0.1 | 1.48E-01 | -0.1 | 1.58E-01 | 0.2 | 2.31E-02 | -0.1 | 7.46E-01 | 0.0 | 1.00E+00 | -0.1 | 8.05E-01 | Histone-lysine N-methyltransferase ATX2 |
| AT2G16390 | 0.1 | 2.71E-01 | 0.0 | 6.79E-01 | 0.1 | 5.95E-01 | -0.2 | 2.88E-01 | -0.1 | 8.56E-01 | -0.2 | 5.85E-01 | Protein CHROMATIN REMODELING 35 |
| AT1G16650 | 0.1 | 2.87E-01 | 0.0 | 7.29E-01 | 0.1 | 5.81E-01 | 0.4 | 2.25E-01 | 0.0 | 9.70E-01 | 0.4 | 2.47E-01 | At1g16650 |
| AT2G46580 | 0.1 | 3.95E-01 | 0.0 | 9.54E-01 | 0.1 | 5.27E-01 | -0.1 | 7.81E-01 | 0.0 | 9.62E-01 | 0.0 | 8.97E-01 | Pyridoxine |
| AT3G57660 | 0.1 | 2.62E-01 | -0.1 | 3.83E-01 | 0.2 | 9.27E-02 | 0.7 | 7.55E-04 | 0.4 | 6.45E-02 | 0.3 | 1.30E-01 | DNA-directed RNA polymerase subunit |
| AT3G62720 | 0.1 | 3.56E-01 | -0.1 | 3.34E-01 | 0.2 | 1.10E-01 | -0.7 | 2.73E-04 | -0.5 | 5.96E-03 | -0.2 | 3.56E-01 | Xyloglucan 6-xylosyltransferase 1 |
| AT4G26640 | 0.1 | 1.17E-01 | -0.1 | 1.82E-01 | 0.2 | 1.97E-02 | 0.0 | 9.94E-01 | 0.1 | 6.90E-01 | -0.1 | 7.50E-01 | WRKY family transcription factor family protein |
| AT3G12800 | 0.1 | 1.51E-01 | -0.1 | 2.66E-01 | 0.2 | 3.20E-02 | -0.3 | 1.01E-01 | 0.0 | 9.53E-01 | -0.3 | 2.48E-01 | Peroxisomal 2,4-dienoyl-CoA reductase |
| AT5G07360 | 0.1 | 2.90E-01 | 0.0 | 7.88E-01 | 0.1 | 5.10E-01 | 0.0 | 8.74E-01 | -0.1 | 7.19E-01 | 0.1 | 8.63E-01 | Amidase family protein |
| AT5G53560 | 0.1 | 3.90E-01 | 0.1 | 4.48E-01 | 0.0 | 9.62E-01 | 0.3 | 1.81E-01 | 0.1 | 6.26E-01 | 0.1 | 6.74E-01 | Cytochrome b5 isoform E |
| AT1G18500 | 0.1 | 8.76E-02 | 0.2 | 2.94E-03 | -0.1 | 7.44E-02 | 0.3 | 1.75E-02 | 0.1 | 3.97E-01 | 0.2 | 3.49E-01 | 2-isopropylmalate synthase 1, chloroplastic |
| AT2G34650 | 0.1 | 4.47E-01 | -0.1 | 7.09E-01 | 0.2 | 3.48E-01 | 0.3 | 1.78E-01 | 0.2 | 3.77E-01 | 0.1 | 8.37E-01 | Protein kinase PINOID |
| AT5G23010 | 0.1 | 3.64E-01 | 0.3 | 2.35E-02 | -0.2 | 1.35E-01 | -0.7 | 6.89E-04 | 0.5 | 1.80E-02 | -1.2 | 3.17E-04 | Methylthioalkylmalate synthase 1 |
| AT3G17700 | 0.1 | 3.46E-01 | -0.3 | 4.72E-02 | 0.4 | 1.81E-02 | 0.1 | 4.23E-01 | 0.0 | 9.76E-01 | 0.2 | 5.27E-01 | Cyclic nucleotide-binding transporter 1 |

| | | | | | | | | | | | | | |
|---|---|---|---|---|---|---|---|---|---|---|---|---|---|
| AT4G00520 | 0.1 | 2.35E-01 | 0.0 | 6.58E-01 | 0.2 | 1.70E-01 | 0.0 | 8.82E-01 | 0.3 | 2.20E-01 | -0.3 | 1.94E-01 | Acyl-CoA thioesterase family protein |
| AT3G52570 | 0.1 | 1.94E-01 | -0.1 | 5.94E-01 | 0.2 | 1.28E-01 | -0.2 | 2.97E-01 | -0.1 | 8.65E-01 | -0.2 | 4.76E-01 | AT3g52570 |
| AT5G54910 | 0.1 | 3.21E-01 | 0.0 | 7.22E-01 | 0.1 | 6.22E-01 | 0.9 | 5.49E-05 | 0.4 | 3.31E-02 | 0.5 | 2.08E-02 | DEAD-box ATP-dependent RNA helicase 32 |
| AT4G32120 | 0.1 | 1.13E-01 | 0.1 | 2.89E-01 | 0.0 | 6.69E-01 | -0.2 | 3.11E-01 | -0.2 | 2.10E-01 | 0.1 | 8.29E-01 | Hexosyltransferase |
| AT5G08415 | 0.1 | 1.44E-01 | 0.1 | 1.21E-01 | 0.0 | 9.07E-01 | 0.0 | 9.93E-01 | 0.1 | 8.01E-01 | -0.1 | 8.16E-01 | Lipoyl synthase, chloroplastic |
| AT5G02680 | 0.1 | 2.11E-01 | 0.1 | 4.02E-01 | 0.0 | 7.65E-01 | -0.1 | 7.60E-01 | 0.0 | 9.92E-01 | -0.1 | 8.12E-01 | Uncharacterized protein |
| AT3G09720 | 0.1 | 3.27E-01 | 0.0 | 9.52E-01 | 0.1 | 4.55E-01 | 0.4 | 6.55E-02 | 0.4 | 4.00E-02 | -0.1 | 7.34E-01 | DEAD-box ATP-dependent RNA helicase 57 |
| AT1G20693 | 0.1 | 1.03E-01 | 0.0 | 7.49E-01 | 0.1 | 2.38E-01 | -0.1 | 7.69E-01 | -0.1 | 5.98E-01 | 0.1 | 8.48E-01 | NFD2 |
| AT5G40010 | 0.1 | 3.73E-01 | -0.6 | 2.63E-03 | 0.7 | 2.43E-03 | 1.0 | 2.75E-03 | 0.8 | 1.92E-02 | 0.2 | 5.39E-01 | AAA+ ATPase domain-containing protein |
| AT3G15590 | 0.1 | 1.65E-01 | -0.1 | 5.68E-01 | 0.2 | 9.68E-02 | 0.5 | 8.71E-03 | 0.1 | 5.19E-01 | 0.3 | 1.67E-01 | Pentatricopeptide repeat-containing protein At3g15590, mitochondrial |
| AT1G58180 | 0.1 | 3.39E-01 | 0.0 | 8.94E-01 | 0.1 | 4.91E-01 | -0.2 | 4.51E-01 | -0.6 | 2.58E-02 | 0.4 | 2.35E-01 | Carbonic anhydrase |
| AT3G49725 | 0.1 | 2.98E-01 | -0.1 | 6.18E-01 | 0.2 | 2.01E-01 | 0.8 | 1.36E-03 | 0.4 | 9.96E-02 | 0.4 | 1.67E-01 | Hflx-type G domain-containing protein |
| AT4G22140 | 0.1 | 1.10E-01 | 0.2 | 1.56E-02 | -0.1 | 3.13E-01 | 0.1 | 5.75E-01 | 0.1 | 7.92E-01 | 0.0 | 9.05E-01 | Uncharacterized protein |
| AT4G11970 | 0.1 | 2.76E-01 | 0.1 | 2.58E-01 | 0.0 | 9.48E-01 | 0.5 | 8.71E-02 | 0.4 | 2.56E-01 | 0.1 | 7.86E-01 | YTH domain-containing family protein |
| AT4G06598 | 0.1 | 3.60E-01 | -0.2 | 2.15E-01 | 0.3 | 8.57E-02 | -0.2 | 7.76E-01 | 0.2 | 8.30E-01 | -0.4 | 6.77E-01 | BZIP domain-containing protein |
| AT2G42590 | 0.1 | 1.20E-01 | 0.2 | 1.61E-02 | -0.1 | 2.69E-01 | -0.1 | 3.76E-01 | -0.2 | 1.77E-01 | 0.1 | 7.20E-01 | General regulatory factor 9 |
| AT3G44600 | 0.1 | 8.49E-02 | 0.0 | 9.30E-01 | 0.1 | 1.10E-01 | 0.2 | 1.11E-01 | 0.1 | 7.67E-01 | 0.2 | 4.23E-01 | Peptidyl-prolyl cis-trans isomerase CYP71 |
| AT3G59110 | 0.1 | 2.01E-01 | 0.0 | 6.10E-01 | 0.1 | 5.29E-01 | -0.5 | 5.33E-02 | -0.4 | 1.68E-01 | -0.1 | 7.96E-01 | Protein kinase domain-containing protein |
| AT1G27360 | 0.1 | 1.35E-01 | 0.1 | 2.52E-01 | 0.0 | 8.01E-01 | -0.1 | 7.00E-01 | 0.4 | 1.50E-01 | -0.5 | 6.45E-02 | SPL11 |
| AT5G60590 | 0.1 | 2.12E-01 | 0.1 | 5.75E-01 | 0.1 | 5.91E-01 | 0.4 | 3.13E-02 | 0.0 | 9.91E-01 | 0.4 | 1.05E-01 | Threonylcarbamoyl-AMP synthase |
| AT1G53050 | 0.1 | 1.58E-01 | 0.0 | 6.65E-01 | 0.2 | 1.10E-01 | 0.1 | 6.84E-01 | 0.1 | 6.00E-01 | 0.0 | 9.12E-01 | F8L10.9 protein |
| AT3G09970 | 0.1 | 1.19E-01 | 0.1 | 1.44E-01 | 0.0 | 9.67E-01 | 0.1 | 3.44E-01 | 0.0 | 9.50E-01 | 0.1 | 5.55E-01 | Calcineurin-like metallo-phosphoesterase superfamily protein |
| AT3G03640 | 0.1 | 7.35E-02 | 0.1 | 2.88E-01 | 0.0 | 5.16E-01 | -0.1 | 5.32E-01 | 0.1 | 8.05E-01 | -0.2 | 4.84E-01 | Probable inactive beta-glucosidase 25 |
| AT2G01930 | 0.1 | 2.61E-01 | 0.1 | 5.26E-01 | 0.0 | 7.10E-01 | -0.4 | 2.43E-02 | -0.5 | 4.82E-03 | 0.2 | 4.87E-01 | Protein BASIC PENTACYSTEINE1 |
| AT3G13790 | 0.1 | 2.16E-01 | 0.2 | 1.35E-01 | 0.0 | 7.72E-01 | 0.1 | 4.39E-01 | -0.1 | 6.53E-01 | 0.2 | 3.27E-01 | Glycosyl hydrolases family 32 protein |
| AT4G35050 | 0.1 | 2.38E-01 | 0.1 | 2.64E-01 | 0.0 | 9.87E-01 | 0.1 | 6.58E-01 | 0.1 | 7.16E-01 | 0.0 | 9.97E-01 | WD-40 repeat-containing protein MSI3 |
| AT3G20870 | 0.1 | 9.19E-02 | 0.1 | 4.01E-01 | 0.1 | 4.60E-01 | 0.1 | 5.39E-01 | 0.1 | 7.87E-01 | 0.0 | 8.74E-01 | ZIP metal ion transporter family |
| AT5G53130 | 0.1 | 1.86E-01 | -0.2 | 2.34E-02 | 0.4 | 5.15E-03 | -0.2 | 4.66E-01 | -0.2 | 4.06E-01 | 0.1 | 8.99E-01 | Cyclic nucleotide-gated ion channel 1 |
| AT1G49970 | 0.1 | 7.54E-02 | 0.0 | 4.96E-01 | 0.1 | 3.11E-01 | 0.1 | 7.06E-01 | 0.1 | 7.23E-01 | 0.0 | 9.76E-01 | ATP-dependent Clp protease proteolytic subunit-related protein 1, chloroplastic |
| AT5G42620 | 0.1 | 1.39E-01 | 0.1 | 2.03E-01 | 0.0 | 8.85E-01 | 0.2 | 1.55E-01 | 0.1 | 4.58E-01 | 0.1 | 7.70E-01 | Metalloendopeptidase |
| AT2G37585 | 0.1 | 3.70E-01 | 0.0 | 9.83E-01 | 0.1 | 4.49E-01 | -0.3 | 1.27E-01 | 0.0 | 9.52E-01 | -0.3 | 2.14E-01 | Putative RING zinc finger protein |
| AT2G39930 | 0.1 | 2.76E-01 | 0.1 | 5.59E-01 | 0.1 | 7.01E-01 | -0.6 | 2.84E-01 | 0.1 | 9.27E-01 | -0.7 | 3.42E-01 | Isoamylase 1 |
| AT4G39940 | 0.1 | 2.93E-01 | 0.3 | 1.56E-02 | -0.2 | 1.17E-01 | 0.0 | 9.81E-01 | 0.5 | 4.60E-02 | -0.4 | 1.14E-01 | Adenylyl-sulfate kinase 2, chloroplastic |
| AT5G03730 | 0.1 | 2.03E-01 | -0.2 | 5.21E-02 | 0.3 | 9.62E-03 | 0.0 | 9.32E-01 | -0.2 | 4.68E-01 | 0.2 | 4.93E-01 | Serine |
| AT3G44670 | 0.1 | 2.07E-01 | -0.4 | 2.34E-03 | 0.5 | 1.03E-03 | 0.0 | 9.23E-01 | -0.1 | 7.13E-01 | 0.1 | 8.15E-01 | ADP-ribosyl cyclase |
| AT4G10760 | 0.1 | 9.72E-02 | 0.0 | 7.25E-01 | 0.2 | 8.68E-02 | 0.1 | 6.90E-01 | 0.1 | 8.56E-01 | 0.1 | 9.22E-01 | N6-adenosine-methyltransferase MT-A70-like |
| AT1G02560 | 0.1 | 9.02E-02 | 0.0 | 5.78E-01 | 0.2 | 5.01E-02 | 0.0 | 9.70E-01 | 0.0 | 9.67E-01 | 0.0 | 9.52E-01 | ATP-dependent Clp protease proteolytic subunit 5, chloroplastic |
| AT1G55630 | 0.1 | 2.72E-01 | 0.1 | 4.55E-01 | 0.0 | 8.09E-01 | 0.3 | 2.44E-01 | 0.4 | 1.77E-01 | -0.1 | 8.42E-01 | Pentatricopeptide repeat-containing protein At1g55630 |
| AT2G32000 | 0.1 | 1.32E-01 | 0.0 | 8.04E-01 | 0.1 | 2.87E-01 | -0.1 | 8.60E-01 | -0.1 | 8.76E-01 | 0.0 | 9.97E-01 | DNA topoisomerase 3-beta |
| AT3G59970 | 0.1 | 1.01E-01 | 0.3 | 2.27E-03 | -0.2 | 4.66E-02 | -0.1 | 7.40E-01 | 0.1 | 7.39E-01 | -0.1 | 5.81E-01 | Methylenetetrahydrofolate reductase |
| AT4G13640 | 0.1 | 1.34E-01 | 0.1 | 2.88E-01 | 0.0 | 7.34E-01 | -0.4 | 5.47E-02 | -0.5 | 1.75E-02 | 0.1 | 6.23E-01 | UNE16 |
| AT3G14310 | 0.1 | 3.70E-01 | 0.6 | 1.32E-03 | -0.5 | 7.93E-03 | 0.2 | 9.41E-02 | 0.0 | 9.37E-01 | 0.2 | 2.87E-01 | Pectinesterase |
| AT3G63200 | 0.1 | 1.33E-01 | 0.2 | 1.12E-02 | -0.1 | 1.84E-01 | -0.4 | 4.26E-03 | 0.0 | 8.23E-01 | -0.4 | 2.21E-02 | Probable inactive patatin-like protein 9 |
| AT1G33390 | 0.1 | 2.76E-01 | -0.1 | 6.72E-01 | 0.2 | 2.01E-01 | 0.7 | 1.95E-04 | 0.1 | 4.63E-01 | 0.6 | 1.11E-02 | ATP-dependent RNA helicase DEAH13 |
| AT3G23900 | 0.1 | 9.53E-02 | -0.1 | 4.55E-01 | 0.2 | 4.30E-02 | -0.1 | 6.89E-01 | -0.2 | 2.10E-01 | 0.1 | 5.11E-01 | RNA recognition motif -containing protein |
| AT4G22285 | 0.1 | 1.22E-01 | -0.1 | 5.09E-01 | 0.2 | 6.66E-02 | 0.4 | 1.85E-02 | 0.1 | 6.83E-01 | 0.3 | 1.68E-01 | Ubiquitin C-terminal hydrolases superfamily protein |
| AT1G05350 | 0.1 | 1.22E-01 | 0.2 | 1.97E-02 | -0.1 | 3.42E-01 | 0.2 | 3.36E-01 | 0.1 | 5.26E-01 | 0.0 | 9.13E-01 | Ubiquitin-like modifier-activating enzyme 5 |
| AT5G14450 | 0.1 | 2.00E-01 | 0.3 | 1.01E-02 | -0.2 | 1.49E-01 | -0.1 | 6.23E-01 | -0.1 | 5.97E-01 | 0.0 | 9.42E-01 | GDSL esterase |
| AT2G05170 | 0.1 | 7.81E-02 | 0.1 | 1.98E-01 | 0.0 | 6.74E-01 | 0.0 | 9.32E-01 | -0.1 | 4.33E-01 | 0.1 | 5.85E-01 | Vacuolar protein-sorting-associated protein 11 homolog |
| AT1G09850 | 0.1 | 1.80E-01 | 0.3 | 5.46E-03 | -0.2 | 8.58E-02 | 0.0 | 8.53E-01 | 0.3 | 3.87E-02 | -0.3 | 1.21E-01 | At1g09850 |
| AT4G11280 | 0.1 | 5.90E-01 | -1.0 | 2.71E-03 | 1.1 | 3.15E-03 | 1.3 | 1.49E-04 | 0.3 | 4.47E-01 | 1.0 | 5.23E-03 | 1-aminocyclopropane-1-carboxylate synthase 6 |
| AT3G60390 | 0.1 | 2.63E-01 | 0.2 | 5.60E-02 | -0.1 | 4.18E-01 | -0.4 | 2.03E-02 | -0.6 | 1.81E-03 | 0.3 | 2.87E-01 | Homeobox-leucine zipper protein HAT3 |
| AT1G22450 | 0.1 | 2.17E-01 | 0.4 | 2.31E-03 | -0.3 | 2.38E-02 | 0.0 | 7.90E-01 | -0.1 | 7.65E-01 | 0.1 | 6.40E-01 | Cytochrome c oxidase subunit 6b-1 |
| AT5G45140 | 0.1 | 1.52E-01 | -0.2 | 7.90E-02 | 0.3 | 1.42E-02 | 0.5 | 7.46E-03 | 0.2 | 3.54E-01 | 0.3 | 1.97E-01 | DNA-directed RNA polymerase |
| AT4G30800 | 0.1 | 5.78E-01 | 0.5 | 4.35E-02 | -0.4 | 1.77E-01 | 0.7 | 7.87E-03 | 0.6 | 3.12E-02 | 0.1 | 7.97E-01 | Small ribosomal subunit protein uS17y |
| AT4G36470 | 0.1 | 3.51E-01 | -0.1 | 3.61E-01 | 0.3 | 1.31E-01 | 0.3 | 7.75E-02 | -0.1 | 8.47E-01 | 0.4 | 9.55E-02 | At4g36470 |
| AT5G16070 | 0.1 | 1.90E-01 | 0.2 | 4.19E-02 | -0.1 | 4.09E-01 | 0.1 | 2.56E-01 | 0.0 | 8.55E-01 | 0.2 | 3.22E-01 | T-complex protein 1 subunit zeta 2 |
| AT4G27960 | 0.1 | 4.58E-01 | -0.1 | 7.47E-01 | 0.2 | 3.34E-01 | 0.1 | 6.31E-01 | 0.1 | 3.57E-01 | -0.1 | 7.52E-01 | Ubiquitin conjugating enzyme 9 |
| AT2G33170 | 0.1 | 2.37E-01 | -0.1 | 2.28E-01 | 0.3 | 5.12E-02 | 0.2 | 2.91E-01 | 0.0 | 8.77E-01 | 0.2 | 4.99E-01 | Probable leucine-rich repeat receptor-like protein kinase At2g33170 |
| AT1G75230 | 0.1 | 1.16E-01 | 0.0 | 8.54E-01 | 0.1 | 2.41E-01 | -0.2 | 2.27E-01 | -0.1 | 5.79E-01 | -0.1 | 7.43E-01 | DNA glycosylase superfamily protein |
| AT2G44530 | 0.1 | 5.94E-02 | 0.0 | 6.67E-01 | 0.2 | 4.57E-02 | 0.1 | 4.01E-01 | 0.0 | 8.16E-01 | 0.2 | 3.89E-01 | ribose-phosphate diphosphokinase |
| AT2G46040 | 0.1 | 2.16E-01 | 0.1 | 2.29E-01 | 0.0 | 9.93E-01 | 0.2 | 5.95E-01 | 0.0 | 9.76E-01 | 0.2 | 7.01E-01 | ARID |
| AT3G04580 | 0.1 | 9.23E-02 | -0.1 | 4.84E-01 | 0.2 | 4.71E-02 | 0.2 | 1.23E-01 | 0.1 | 6.18E-01 | 0.1 | 5.49E-01 | Protein EIN4 |
| AT3G15660 | 0.1 | 1.08E-01 | 0.2 | 9.64E-03 | -0.1 | 2.09E-01 | 0.2 | 3.11E-01 | 0.0 | 9.02E-01 | 0.1 | 5.98E-01 | Monothiol glutaredoxin-S15, mitochondrial |
| AT2G34690 | 0.1 | 4.81E-02 | 0.2 | 3.05E-03 | -0.1 | 1.40E-01 | -0.1 | 5.81E-01 | -0.2 | 2.22E-01 | 0.1 | 5.91E-01 | Accelerated cell death 11 |
| AT2G23320 | 0.1 | 1.83E-01 | -0.3 | 5.88E-03 | 0.5 | 1.95E-03 | -0.2 | 1.49E-01 | -0.8 | 3.77E-04 | 0.6 | 2.25E-02 | WRKY DNA-binding protein 15 |
| AT3G48190 | 0.1 | 1.03E-01 | -0.2 | 3.16E-02 | 0.3 | 5.10E-03 | -0.4 | 4.05E-02 | -0.4 | 1.21E-01 | -0.1 | 8.34E-01 | non-specific serine |
| AT5G44370 | 0.1 | 1.12E-01 | 0.1 | 1.53E-01 | 0.0 | 9.28E-01 | 0.1 | 5.44E-01 | 0.3 | 1.23E-01 | -0.2 | 4.38E-01 | Probable anion transporter 5 |
| AT1G51430 | 0.1 | 2.31E-01 | 0.2 | 1.33E-01 | 0.0 | 7.61E-01 | -0.3 | 9.24E-02 | 0.2 | 3.96E-01 | -0.5 | 3.86E-02 | Uncharacterized protein F5D21.20 |
| AT2G43710 | 0.1 | 8.58E-02 | 0.2 | 1.25E-02 | -0.1 | 2.84E-01 | 0.2 | 6.39E-02 | 0.1 | 4.68E-01 | 0.1 | 5.42E-01 | Acyl- desaturase |
| AT3G12700 | 0.1 | 1.05E-01 | 0.2 | 1.46E-02 | -0.1 | 3.15E-01 | 0.9 | 9.58E-05 | 0.2 | 2.67E-01 | 0.6 | 5.14E-03 | Peptidase A1 domain-containing protein |
| AT2G30980 | 0.1 | 7.17E-02 | 0.1 | 3.69E-01 | 0.1 | 4.03E-01 | 0.0 | 7.29E-01 | -0.1 | 4.87E-01 | 0.1 | 8.03E-01 | Shaggy-related protein kinase zeta |
| AT1G13710 | 0.1 | 3.73E-01 | 0.4 | 2.34E-02 | -0.2 | 1.60E-01 | -0.3 | 2.36E-01 | -0.2 | 4.63E-01 | -0.1 | 8.41E-01 | Cytochrome P450 78A5 |
| AT2G27060 | 0.1 | 2.80E-01 | 0.0 | 7.37E-01 | 0.1 | 5.55E-01 | 0.0 | 9.60E-01 | 0.0 | 9.78E-01 | 0.0 | 9.94E-01 | Leucine-rich repeat receptor-like protein kinase |
| AT2G26670 | 0.1 | 1.27E-01 | -0.1 | 4.51E-01 | 0.2 | 4.91E-02 | -0.2 | 3.54E-02 | -0.2 | 1.91E-01 | -0.1 | 7.20E-01 | Heme oxygenase 1, chloroplastic |
| AT1G27450 | 0.1 | 3.37E-01 | 0.3 | 2.77E-02 | -0.2 | 1.69E-01 | 0.1 | 7.08E-01 | -0.1 | 6.72E-01 | 0.2 | 4.68E-01 | adenine phosphoribosyltransferase |
| AT3G59630 | 0.1 | 1.02E-01 | 0.0 | 9.41E-01 | 0.1 | 1.87E-01 | 0.2 | 4.15E-01 | 0.0 | 9.13E-01 | 0.1 | 6.55E-01 | 2-histidine synthase subunit 2 |
| AT5G66860 | 0.1 | 4.14E-01 | 0.1 | 4.11E-01 | 0.0 | 9.76E-01 | 0.3 | 8.28E-02 | 0.1 | 6.80E-01 | 0.2 | 2.94E-01 | Ribosomal protein L25 |
| AT1G52245 | 0.1 | 4.15E-01 | 1.1 | 4.16E-05 | -0.9 | 4.72E-04 | -0.1 | 7.68E-01 | -0.2 | 6.80E-01 | 0.1 | 9.09E-01 | Dynein light chain |
| AT5G57300 | 0.1 | 1.33E-01 | 0.2 | 8.17E-02 | 0.0 | 7.88E-01 | 0.1 | 6.20E-01 | 0.1 | 8.19E-01 | 0.0 | 9.01E-01 | 2-methoxy-6-polyprenyl-1,4-benzoquinol methylase, mitochondrial |
| AT2G14170 | 0.1 | 1.50E-01 | 0.3 | 6.19E-03 | -0.2 | 9.66E-02 | -0.1 | 5.10E-01 | -0.2 | 5.02E-01 | 0.0 | 9.48E-01 | methylmalonate-semialdehyde dehydrogenase |
| AT4G35740 | 0.1 | 2.91E-01 | -0.1 | 5.54E-01 | 0.2 | 1.74E-01 | -0.3 | 3.53E-01 | -0.1 | 8.97E-01 | -0.2 | 4.61E-01 | hypothetical protein |
| AT3G57340 | 0.1 | 9.57E-02 | -0.1 | 2.84E-01 | 0.2 | 2.68E-02 | -0.3 | 5.36E-02 | -0.4 | 9.62E-03 | 0.1 | 4.76E-01 | AT3G57340 protein ) |

| | | | | | | | | | | | | | |
|---|---|---|---|---|---|---|---|---|---|---|---|---|---|
| AT5G43760 | 0.1 | 6.72E-02 | 0.1 | 1.71E-01 | 0.0 | 6.88E-01 | -0.2 | 2.05E-01 | -0.3 | 3.65E-02 | 0.1 | 4.88E-01 | 3-ketoacyl-CoA synthase 20 |
| AT1G73170 | 0.1 | 3.39E-01 | 0.0 | 8.11E-01 | 0.1 | 5.51E-01 | -0.3 | 5.29E-02 | 0.1 | 5.17E-01 | -0.4 | 3.11E-02 | P-loop containing nucleoside triphosphate hydrolases superfamily protein |
| AT5G60190 | 0.1 | 2.46E-01 | 0.2 | 1.09E-01 | -0.1 | 6.69E-01 | 0.1 | 5.56E-01 | -0.5 | 5.13E-02 | 0.6 | 3.48E-02 | NEDD8-specific protease 1 |
| AT1G01950 | 0.1 | 1.79E-01 | -0.2 | 9.29E-02 | 0.3 | 1.77E-02 | -0.2 | 3.75E-01 | 0.1 | 8.41E-01 | -0.2 | 3.82E-01 | Kinesin-like protein |
| AT1G16400 | 0.1 | 1.93E-01 | 0.5 | 7.79E-04 | -0.3 | 1.01E-02 | -0.5 | 4.15E-03 | 0.3 | 5.63E-02 | -0.8 | 4.95E-04 | Hexahomomethionine N-hydroxylase |
| AT4G17895 | 0.1 | 1.27E-01 | -0.2 | 2.19E-02 | 0.4 | 4.78E-03 | 0.0 | 9.54E-01 | -0.1 | 6.61E-01 | 0.1 | 7.64E-01 | Ubiquitin carboxyl-terminal hydrolase 20 |
| AT5G57390 | 0.1 | 3.28E-01 | 0.1 | 3.84E-01 | 0.0 | 9.65E-01 | 0.7 | 5.50E-02 | 0.4 | 2.90E-01 | 0.2 | 4.80E-01 | AINTEGUMENTA-like 5 |
| AT1G53510 | 0.1 | 5.59E-02 | 0.1 | 1.49E-01 | 0.0 | 6.74E-01 | -0.1 | 7.71E-01 | 0.0 | 8.83E-01 | 0.0 | 9.48E-01 | Mitogen-activated protein kinase 18 |
| AT5G62880 | 0.1 | 8.91E-02 | 0.1 | 8.18E-02 | 0.0 | 9.32E-01 | -0.3 | 9.52E-02 | -0.3 | 1.33E-01 | 0.0 | 9.97E-01 | RAC-like 10 |
| AT5G28050 | 0.1 | 5.67E-02 | 0.2 | 1.79E-02 | -0.1 | 5.10E-01 | -0.2 | 3.53E-01 | -0.1 | 7.67E-01 | -0.1 | 7.66E-01 | Cytidine |
| AT1G55880 | 0.1 | 8.61E-02 | 0.1 | 1.59E-01 | 0.0 | 8.19E-01 | 0.0 | 8.07E-01 | -0.2 | 3.25E-01 | 0.2 | 3.08E-01 | At1g55880 |
| AT5G49840 | 0.1 | 2.01E-01 | 0.1 | 4.77E-01 | 0.1 | 6.59E-01 | -0.1 | 7.05E-01 | 0.0 | 9.04E-01 | -0.1 | 8.78E-01 | CLP protease regulatory subunit CLPX-like |
| AT5G46240 | 0.1 | 4.47E-01 | 0.6 | 8.04E-03 | -0.4 | 5.46E-02 | 0.2 | 4.12E-01 | 0.2 | 5.25E-01 | 0.0 | 9.71E-01 | Potassium channel KAT1 |
| AT3G55440 | 0.1 | 2.30E-01 | 0.3 | 1.24E-02 | -0.2 | 1.20E-01 | 0.1 | 3.06E-01 | 0.0 | 9.24E-01 | 0.1 | 3.80E-01 | Triosephosphate isomerase, cytosolic |
| AT5G67300 | 0.1 | 5.77E-01 | -0.3 | 2.76E-01 | 0.4 | 1.37E-01 | -0.6 | 1.27E-02 | -0.4 | 9.67E-02 | -0.2 | 6.49E-01 | Transcription factor MYB44 |
| AT1G05020 | 0.1 | 4.25E-01 | 0.5 | 6.80E-03 | -0.4 | 5.36E-02 | 0.5 | 2.24E-01 | -0.3 | 6.71E-01 | 0.8 | 1.12E-01 | Clathrin coat assembly protein AP180 |
| AT1G26190 | 0.1 | 1.01E-01 | 0.2 | 6.29E-02 | 0.0 | 7.91E-01 | 0.0 | 9.70E-01 | 0.1 | 7.49E-01 | -0.1 | 8.07E-01 | Phosphoribulokinase |
| AT1G79010 | 0.1 | 4.44E-02 | 0.3 | 2.05E-03 | -0.1 | 8.72E-02 | 0.1 | 7.36E-01 | 0.0 | 8.35E-01 | 0.1 | 6.81E-01 | hypothetical protein |
| AT2G03670 | 0.1 | 2.02E-01 | 0.0 | 9.75E-01 | 0.1 | 3.11E-01 | 0.2 | 5.90E-01 | -0.1 | 7.91E-01 | 0.3 | 5.02E-01 | Cell division cycle 48B |
| AT1G15790 | 0.1 | 5.19E-01 | -0.5 | 5.27E-02 | 0.6 | 3.79E-02 | 0.5 | 6.43E-02 | -0.2 | 6.01E-01 | 0.7 | 3.58E-02 | Mediator complex subunit 15 KIX domain-containing protein |
| AT5G21100 | 0.1 | 1.83E-01 | -0.1 | 2.99E-01 | 0.2 | 4.58E-02 | -0.1 | 4.51E-01 | 0.4 | 3.13E-02 | -0.5 | 2.32E-02 | L-ascorbate oxidase |
| AT5G05200 | 0.1 | 1.44E-01 | -0.1 | 2.52E-01 | 0.2 | 2.90E-02 | -0.1 | 6.31E-01 | 0.1 | 7.82E-01 | -0.2 | 5.48E-01 | Uncharacterized aarF domain-containing protein kinase At5g05200, chloroplastic |
| AT5G04360 | 0.1 | 2.30E-01 | 0.0 | 8.57E-01 | 0.1 | 3.93E-01 | -0.2 | 5.03E-01 | 0.0 | 8.89E-01 | -0.2 | 4.21E-01 | Pullulanase 1, chloroplastic |
| AT1G12270 | 0.1 | 8.32E-02 | 0.2 | 1.70E-02 | -0.1 | 3.93E-01 | 0.1 | 6.28E-01 | 0.0 | 9.71E-01 | 0.1 | 7.95E-01 | Hsp70-Hsp90 organizing protein 1 |
| AT1G09160 | 0.1 | 7.21E-02 | 0.3 | 1.68E-03 | -0.2 | 5.81E-02 | 0.1 | 5.80E-01 | 0.0 | 9.83E-01 | 0.1 | 7.02E-01 | Probable protein phosphatase 2C 5 |
| AT2G16870 | 0.1 | 2.85E-01 | -0.2 | 2.19E-01 | 0.3 | 7.08E-02 | 0.3 | 4.09E-01 | 0.1 | 7.67E-01 | 0.1 | 7.69E-01 | Disease resistance protein family , putative) |
| AT5G17790 | 0.1 | 6.90E-02 | 0.2 | 2.24E-02 | -0.1 | 5.58E-01 | 0.2 | 3.89E-01 | 0.2 | 4.06E-01 | 0.0 | 9.65E-01 | Zinc finger protein VAR3, chloroplastic |
| AT1G66740 | 0.1 | 1.28E-01 | 0.2 | 4.29E-02 | -0.1 | 5.74E-01 | 0.0 | 8.96E-01 | -0.1 | 6.06E-01 | 0.2 | 5.91E-01 | Probable histone chaperone ASF1A |
| AT3G19120 | 0.1 | 1.60E-01 | 0.3 | 1.27E-02 | -0.1 | 2.01E-01 | -0.1 | 6.80E-01 | -0.3 | 7.58E-02 | 0.2 | 2.66E-01 | AT3g19120 |
| AT1G03470 | 0.1 | 2.85E-01 | 0.1 | 4.22E-01 | 0.0 | 8.74E-01 | 1.1 | 1.46E-03 | -0.4 | 3.49E-01 | 1.4 | 8.29E-04 | Protein NETWORKED 3A |
| AT5G47740 | 0.1 | 5.80E-01 | 0.3 | 1.90E-01 | -0.2 | 5.15E-01 | 1.4 | 1.44E-03 | 0.6 | 2.30E-01 | 0.8 | 9.89E-03 | Adenine nucleotide alpha hydrolases-like superfamily protein |
| AT1G07790 | 0.1 | 2.65E-01 | 0.4 | 7.56E-03 | -0.3 | 7.13E-02 | 0.2 | 1.33E-01 | 0.2 | 2.62E-01 | 0.0 | 9.21E-01 | Histone H2B |
| AT1G71990 | 0.1 | 2.94E-01 | 0.3 | 2.13E-02 | -0.2 | 1.99E-01 | 0.0 | 8.44E-01 | -0.4 | 5.33E-02 | 0.4 | 1.56E-01 | Alpha--fucosyltransferase -L-fucosyltransferase) |
| AT5G60850 | 0.1 | 2.10E-01 | 0.2 | 1.52E-01 | 0.0 | 8.47E-01 | -0.8 | 1.89E-04 | -0.7 | 8.24E-04 | -0.1 | 8.61E-01 | Dof zinc finger protein DOF5.4 |
| AT5G63860 | 0.1 | 4.87E-02 | 0.1 | 2.91E-01 | 0.1 | 3.80E-01 | -0.2 | 1.52E-01 | -0.1 | 2.99E-01 | 0.0 | 9.13E-01 | Ultraviolet-B receptor UVR8 |
| AT1G09320 | 0.1 | 1.12E-01 | 0.0 | 8.82E-01 | 0.2 | 1.44E-01 | -0.1 | 7.25E-01 | -0.3 | 8.39E-02 | 0.3 | 2.52E-01 | Protein AGENET DOMAIN -CONTAINING P1 |
| AT3G54360 | 0.1 | 5.27E-02 | 0.0 | 8.52E-01 | 0.1 | 1.09E-01 | -0.2 | 3.07E-02 | -0.1 | 3.08E-01 | -0.1 | 5.01E-01 | Protein NCA1 |
| AT1G03900 | 0.1 | 6.53E-02 | 0.2 | 2.81E-02 | 0.0 | 6.26E-01 | 0.0 | 9.65E-01 | -0.1 | 5.61E-01 | 0.1 | 5.96E-01 | Uncharacterized protein At1g03900 |
| AT1G62120 | 0.1 | 3.93E-01 | 0.3 | 6.97E-02 | -0.2 | 3.65E-01 | 0.0 | 9.89E-01 | -0.4 | 1.73E-01 | 0.4 | 2.63E-01 | Mitochondrial transcription termination factor family protein |
| AT2G43100 | 0.1 | 3.35E-01 | 0.5 | 4.73E-03 | -0.4 | 3.21E-02 | -0.3 | 9.61E-02 | 0.8 | 9.60E-04 | -1.1 | 5.53E-04 | 3-isopropylmalate dehydratase small subunit 2 alkylmalate dehydratase small subunit 2) |
| AT5G24340 | 0.1 | 2.38E-01 | -0.1 | 3.09E-01 | 0.3 | 7.68E-02 | 0.0 | 9.40E-01 | 0.2 | 5.99E-01 | -0.2 | 7.24E-01 | 3'-5' exonuclease domain-containing protein |
| AT1G09130 | 0.1 | 5.46E-02 | 0.1 | 2.47E-01 | 0.1 | 4.72E-01 | 0.0 | 8.59E-01 | 0.0 | 9.95E-01 | 0.0 | 9.02E-01 | ATP-dependent Clp protease proteolytic subunit |
| AT3G59600 | 0.1 | 1.40E-01 | 0.2 | 1.17E-01 | 0.0 | 9.04E-01 | 0.1 | 7.44E-01 | 0.0 | 9.03E-01 | 0.1 | 7.29E-01 | Uncharacterized protein |
| AT5G35910 | 0.1 | 9.74E-02 | 0.0 | 6.47E-01 | 0.2 | 6.69E-02 | 0.4 | 7.11E-03 | 0.0 | 8.06E-01 | 0.3 | 7.37E-02 | Putative nucleolar protein |
| AT5G65720 | 0.1 | 5.40E-02 | -0.1 | 4.77E-01 | 0.2 | 2.48E-02 | 0.2 | 2.36E-02 | 0.1 | 3.20E-01 | 0.1 | 4.51E-01 | AT5G65720 protein -like 1) |
| AT2G31450 | 0.1 | 1.28E-01 | 0.0 | 8.47E-01 | 0.2 | 1.50E-01 | -0.2 | 4.14E-01 | 0.0 | 9.56E-01 | -0.2 | 5.94E-01 | ATNTH1 |
| AT1G06670 | 0.1 | 3.97E-02 | -0.1 | 3.50E-01 | 0.2 | 1.55E-02 | 0.4 | 1.73E-01 | 0.3 | 3.90E-01 | 0.1 | 8.48E-01 | RNA helicase |
| AT1G70190 | 0.1 | 2.28E-01 | 0.3 | 1.75E-02 | -0.2 | 1.93E-01 | 0.0 | 9.81E-01 | 0.0 | 9.29E-01 | 0.0 | 9.20E-01 | F20P5.9 protein |
| AT5G15510 | 0.1 | 3.26E-01 | 0.3 | 5.36E-02 | -0.2 | 3.55E-01 | 0.1 | 8.47E-01 | 0.3 | 3.41E-01 | -0.3 | 5.13E-01 | TPX2 protein family |
| AT1G30210 | 0.1 | 8.31E-02 | 0.3 | 2.36E-03 | -0.2 | 6.89E-02 | -0.1 | 4.35E-01 | 0.1 | 4.81E-01 | -0.2 | 1.63E-01 | Transcription factor TCP24 |
| AT1G48610 | 0.1 | 2.81E-01 | 0.4 | 1.24E-02 | -0.2 | 1.16E-01 | 0.1 | 5.39E-01 | 0.1 | 7.54E-01 | 0.0 | 9.10E-01 | AT hook motif-containing protein |
| AT5G59920 | 0.1 | 1.10E-01 | 0.0 | 6.46E-01 | 0.2 | 8.40E-02 | 0.3 | 1.82E-01 | 0.1 | 6.05E-01 | 0.1 | 6.50E-01 | Cysteine |
| AT5G11710 | 0.1 | 3.73E-02 | 0.1 | 1.95E-01 | 0.1 | 4.37E-01 | 0.0 | 8.56E-01 | 0.0 | 9.65E-01 | 0.0 | 8.74E-01 | Clathrin interactor EPSIN 1 |
| AT2G02060 | 0.1 | 4.69E-01 | -0.5 | 5.91E-02 | 0.6 | 3.60E-02 | -0.6 | 6.29E-02 | -0.7 | 4.55E-02 | 0.1 | 8.07E-01 | Uncharacterized protein At2g02060 |
| AT2G39280 | 0.1 | 3.18E-01 | -0.1 | 6.25E-01 | 0.2 | 2.07E-01 | -0.1 | 8.63E-01 | 0.2 | 5.72E-01 | -0.2 | 4.78E-01 | Ypt |
| AT1G76150 | 0.1 | 4.08E-02 | 0.1 | 3.67E-01 | 0.1 | 2.73E-01 | -0.2 | 2.24E-01 | -0.2 | 1.20E-01 | 0.1 | 7.62E-01 | Enoyl-CoA hydratase 2, peroxisomal |
| AT1G46264 | 0.1 | 1.81E-01 | 0.1 | 6.08E-01 | 0.1 | 5.08E-01 | -0.1 | 6.28E-01 | 0.0 | 9.48E-01 | -0.1 | 6.73E-01 | Heat stress transcription factor B-4 |
| AT5G13030 | 0.1 | 3.37E-02 | -0.1 | 3.40E-01 | 0.2 | 1.17E-02 | 0.2 | 7.85E-02 | 0.1 | 4.48E-01 | 0.1 | 6.13E-01 | Selenoprotein O |
| AT1G19970 | 0.1 | 4.55E-01 | 0.0 | 9.32E-01 | 0.2 | 4.98E-01 | 0.0 | 9.81E-01 | 0.3 | 3.09E-01 | -0.3 | 4.15E-01 | T20H2.24 protein |
| AT5G46880 | 0.1 | 3.65E-01 | 0.1 | 4.72E-01 | 0.0 | 9.15E-01 | -0.1 | 7.83E-01 | -0.1 | 8.10E-01 | 0.0 | 9.97E-01 | Homeobox-leucine zipper protein HDG5 |
| AT2G13540 | 0.1 | 6.73E-02 | 0.1 | 1.25E-01 | 0.0 | 8.07E-01 | 0.0 | 8.18E-01 | 0.1 | 4.12E-01 | -0.1 | 6.46E-01 | Nuclear cap-binding protein subunit 1 |
| AT5G16170 | 0.1 | 2.18E-01 | 0.1 | 4.44E-01 | 0.1 | 7.34E-01 | 1.0 | 1.06E-03 | 0.5 | 8.57E-02 | 0.5 | 1.58E-01 | Uncharacterized protein T21H19_90 |
| AT3G52430 | 0.1 | 2.87E-01 | -0.3 | 5.11E-02 | 0.4 | 1.40E-02 | 0.3 | 1.39E-01 | 0.0 | 8.89E-01 | 0.3 | 3.22E-01 | Lipase-like PAD4 |
| AT3G25860 | 0.1 | 1.10E-01 | 0.4 | 2.39E-03 | -0.2 | 4.68E-02 | 0.1 | 5.81E-01 | 0.0 | 8.76E-01 | 0.0 | 8.56E-01 | Dihydrolipoyllysine-residue acetyltransferase component 4 of pyruvate dehydrogenase complex, chloroplastic |
| AT1G31420 | 0.1 | 7.70E-02 | 0.1 | 3.14E-01 | 0.1 | 4.86E-01 | -0.1 | 5.23E-01 | -0.1 | 4.54E-01 | 0.0 | 9.12E-01 | Leucine-rich repeat protein kinase family protein |
| AT5G63120 | 0.1 | 4.32E-02 | 0.0 | 7.64E-01 | 0.2 | 4.96E-02 | 0.1 | 6.09E-01 | 0.1 | 5.49E-01 | 0.0 | 9.18E-01 | P-loop containing nucleoside triphosphate hydrolases superfamily protein |
| AT1G27900 | 0.1 | 3.89E-02 | -0.1 | 8.45E-02 | 0.3 | 5.23E-03 | 0.4 | 5.21E-02 | 0.3 | 9.47E-02 | 0.0 | 9.61E-01 | RNA helicase family protein |
| AT3G47120 | 0.1 | 1.56E-01 | 0.0 | 7.83E-01 | 0.1 | 3.36E-01 | 0.0 | 8.08E-01 | -0.1 | 5.85E-01 | 0.1 | 8.14E-01 | RNA recognition motif -containing protein |
| AT3G62310 | 0.1 | 6.99E-02 | 0.0 | 5.33E-01 | 0.2 | 3.83E-02 | 0.1 | 6.02E-01 | 0.0 | 9.93E-01 | 0.1 | 7.31E-01 | RNA helicase |
| AT1G36280 | 0.1 | 2.54E-01 | 0.2 | 2.07E-01 | 0.0 | 8.96E-01 | 0.5 | 2.62E-02 | 0.5 | 3.30E-02 | 0.0 | 9.59E-01 | Adenylosuccinate lyase |
| AT1G07660 | 0.1 | 7.42E-02 | 0.2 | 1.38E-02 | -0.1 | 3.85E-01 | 0.1 | 5.49E-01 | -0.1 | 5.96E-01 | 0.3 | 2.90E-01 | Histone H4 |
| AT2G23890 | 0.1 | 1.42E-01 | 0.0 | 8.41E-01 | 0.1 | 2.88E-01 | 0.6 | 2.23E-03 | 0.2 | 4.08E-01 | 0.4 | 4.42E-02 | hypothetical protein |
| AT1G21100 | 0.1 | 1.50E-01 | -0.2 | 8.21E-02 | 0.3 | 1.51E-02 | 1.1 | 8.46E-05 | 0.3 | 3.62E-01 | 0.9 | 5.14E-03 | Indole glucosinolate O-methyltransferase 1 |
| AT2G32810 | 0.1 | 4.25E-02 | 0.0 | 9.49E-01 | 0.1 | 7.74E-02 | -0.2 | 4.49E-01 | 0.0 | 9.71E-01 | -0.2 | 6.54E-01 | beta-galactosidase |
| AT3G04520 | 0.1 | 1.40E-01 | 0.1 | 5.14E-01 | 0.1 | 4.86E-01 | 0.0 | 9.49E-01 | 0.2 | 1.60E-01 | -0.2 | 2.14E-01 | Threonine aldolase 2 |
| AT1G18630 | 0.1 | 2.58E-01 | 0.2 | 1.38E-01 | -0.1 | 7.39E-01 | 0.5 | 7.07E-03 | 0.3 | 1.25E-01 | 0.2 | 3.90E-01 | Small ribosomal subunit protein mS86 |
| AT5G44100 | 0.1 | 2.69E-02 | 0.0 | 4.55E-01 | 0.2 | 1.36E-02 | 0.0 | 9.23E-01 | 0.1 | 5.01E-01 | -0.1 | 5.12E-01 | Casein kinase 1-like protein 7 |
| AT1G64230 | 0.1 | 1.55E-01 | -0.1 | 3.78E-01 | 0.2 | 4.78E-02 | -0.5 | 8.26E-04 | -0.5 | 4.72E-03 | -0.1 | 8.07E-01 | Ubiquitin-conjugating enzyme 28 |
| AT4G04770 | 0.1 | 4.90E-02 | 0.0 | 9.50E-01 | 0.2 | 6.63E-02 | -1.7 | 1.54E-05 | 0.2 | 6.36E-01 | -1.9 | 2.36E-05 | UPF0051 protein ABCI8, chloroplastic |
| AT2G14740 | 0.1 | 8.61E-02 | 0.1 | 3.87E-01 | 0.1 | 4.42E-01 | 0.1 | 7.00E-01 | 0.1 | 5.52E-01 | -0.1 | 8.69E-01 | Vacuolar-sorting receptor 3 |
| AT3G25430 | 0.1 | 2.87E-01 | 0.1 | 4.43E-01 | 0.0 | 8.42E-01 | 0.0 | 9.27E-01 | -0.2 | 6.74E-01 | 0.2 | 6.42E-01 | Polynucleotidyl transferase, ribonuclease H-like superfamily protein |
| AT2G43700 | 0.1 | 2.60E-01 | -0.4 | 1.16E-02 | 0.6 | 5.96E-03 | -0.1 | 7.40E-01 | 0.2 | 7.41E-01 | -0.3 | 5.77E-01 | Concanavalin A-like lectin protein kinase family protein |
| AT5G13330 | 0.1 | 3.66E-01 | -1.0 | 8.12E-04 | 1.1 | 1.06E-03 | 0.2 | 6.77E-01 | -1.5 | 2.77E-02 | 1.7 | 2.08E-02 | AP2 |

| | | | | | | | | | | | | | |
|---|---|---|---|---|---|---|---|---|---|---|---|---|---|
| AT5G62030 | 0.1 | 1.27E-01 | 0.3 | 1.86E-02 | -0.1 | 3.30E-01 | 0.1 | 5.37E-01 | -0.1 | 7.79E-01 | 0.2 | 4.31E-01 | 2-histidine synthase subunit 1 |
| AT5G60890 | 0.1 | 2.29E-01 | 0.2 | 2.03E-01 | 0.0 | 9.28E-01 | -0.2 | 3.14E-01 | 0.4 | 9.43E-02 | -0.6 | 2.92E-02 | Transcription factor MYB34 |
| AT2G23290 | 0.1 | 5.29E-01 | -0.1 | 6.97E-01 | 0.2 | 3.69E-01 | -0.3 | 1.46E-01 | -0.6 | 5.64E-03 | 0.4 | 1.87E-01 | At2g23280 |
| AT4G18010 | 0.1 | 2.60E-01 | -0.4 | 2.10E-02 | 0.5 | 6.12E-03 | -0.2 | 3.19E-01 | -0.6 | 3.06E-02 | 0.4 | 2.82E-01 | Myo-inositol polyphosphate 5-phosphatase 2 |
| AT5G60730 | 0.1 | 5.50E-01 | 0.5 | 4.90E-02 | -0.4 | 2.18E-01 | 1.1 | 2.74E-02 | 0.7 | 2.04E-01 | 0.4 | 5.16E-01 | ATPase GET3C |
| AT1G15530 | 0.1 | 8.97E-02 | 0.2 | 7.87E-02 | 0.0 | 9.17E-01 | -0.4 | 2.07E-02 | -0.6 | 1.64E-03 | 0.2 | 2.66E-01 | L-type lectin-domain containing receptor kinase S.1 |
| AT5G43280 | 0.2 | 4.02E-02 | 0.1 | 9.95E-02 | 0.0 | 7.20E-01 | 0.1 | 3.82E-01 | 0.2 | 2.95E-01 | 0.0 | 8.50E-01 | Delta-Delta-dienoyl-CoA isomerase, peroxisomal |
| AT4G33200 | 0.2 | 1.49E-01 | -0.3 | 6.96E-03 | 0.5 | 1.90E-03 | -0.1 | 6.28E-01 | 0.0 | 9.48E-01 | -0.1 | 6.66E-01 | Myosin |
| AT5G54510 | 0.2 | 1.96E-01 | 0.2 | 8.00E-02 | -0.1 | 6.27E-01 | -0.1 | 8.15E-01 | 0.0 | 9.27E-01 | 0.0 | 9.24E-01 | Indole-3-acetic acid-amido synthetase GH3.6 |
| AT3G06140 | 0.2 | 1.99E-01 | 0.3 | 3.29E-02 | -0.1 | 3.67E-01 | -0.7 | 1.27E-03 | -0.5 | 1.08E-02 | -0.1 | 6.28E-01 | E3 ubiquitin-protein ligase |
| AT2G29400 | 0.2 | 4.28E-02 | 0.1 | 1.96E-01 | 0.1 | 4.77E-01 | -0.1 | 2.45E-01 | -0.2 | 7.82E-02 | 0.1 | 6.21E-01 | Serine |
| AT1G20140 | 0.2 | 1.39E-01 | 0.1 | 5.49E-01 | 0.1 | 4.70E-01 | 0.9 | 5.98E-04 | 0.4 | 8.52E-02 | 0.4 | 7.42E-02 | SKP1-like protein 4 |
| AT3G18850 | 0.2 | 1.07E-01 | 0.4 | 1.44E-03 | -0.2 | 4.18E-02 | -0.4 | 2.24E-01 | -0.3 | 3.47E-01 | -0.1 | 9.11E-01 | 1-acylglycerol-3-phosphate O-acyltransferase |
| AT1G10850 | 0.2 | 9.82E-02 | 0.2 | 1.10E-01 | 0.0 | 9.95E-01 | -0.2 | 2.09E-01 | 0.1 | 5.23E-01 | -0.3 | 1.16E-01 | Protein kinase domain-containing protein |
| AT5G43740 | 0.2 | 2.35E-01 | 0.0 | 9.10E-01 | 0.2 | 2.69E-01 | 0.3 | 1.72E-01 | 0.2 | 3.16E-01 | 0.0 | 9.01E-01 | Probable disease resistance protein At5g43740 |
| AT5G03320 | 0.2 | 5.19E-02 | -0.1 | 2.00E-01 | 0.3 | 1.13E-02 | 0.3 | 4.34E-02 | 0.0 | 8.32E-01 | 0.3 | 9.34E-02 | Serine |
| AT1G72550 | 0.2 | 2.37E-02 | 0.1 | 2.74E-01 | 0.1 | 2.34E-01 | 0.0 | 7.41E-01 | -0.1 | 2.98E-01 | 0.2 | 2.71E-01 | phenylalanine--tRNA ligase |
| AT3G07480 | 0.2 | 8.01E-02 | 0.2 | 1.24E-02 | -0.1 | 2.97E-01 | 0.0 | 9.31E-01 | 0.0 | 9.09E-01 | 0.0 | 9.71E-01 | 2Fe-2S ferredoxin-like superfamily protein |
| AT1G04440 | 0.2 | 7.86E-02 | 0.1 | 3.67E-01 | 0.1 | 4.32E-01 | -0.1 | 6.24E-01 | 0.0 | 9.79E-01 | -0.1 | 7.73E-01 | Casein kinase 1-like protein 13 |
| AT4G40000 | 0.2 | 3.15E-01 | -0.1 | 6.99E-01 | 0.2 | 2.59E-01 | 0.7 | 9.24E-02 | 0.5 | 3.21E-01 | 0.2 | 7.22E-01 | S-adenosyl-L-methionine-dependent methyltransferases superfamily protein |
| AT3G13040 | 0.2 | 5.80E-02 | 0.1 | 3.87E-01 | 0.1 | 3.30E-01 | -0.1 | 4.21E-01 | 0.2 | 2.72E-01 | -0.3 | 1.27E-01 | HTH myb-type domain-containing protein |
| AT4G02120 | 0.2 | 1.24E-01 | -0.1 | 1.81E-01 | 0.3 | 2.19E-02 | -0.3 | 6.94E-02 | 0.0 | 9.87E-01 | -0.3 | 1.59E-01 | CTP synthase |
| AT4G00430 | 0.2 | 4.53E-02 | 0.1 | 1.96E-01 | 0.1 | 4.98E-01 | -0.5 | 8.38E-04 | -0.2 | 2.38E-01 | -0.3 | 7.19E-02 | Plasma membrane intrinsic protein 14 |
| AT5G61160 | 0.2 | 7.42E-01 | 0.6 | 1.86E-01 | -0.4 | 4.09E-01 | 0.6 | 2.10E-01 | -0.8 | 1.52E-01 | 1.4 | 3.99E-02 | Agmatine coumaroyltransferase |
| AT1G24490 | 0.2 | 3.92E-02 | -0.1 | 2.88E-01 | 0.2 | 1.36E-02 | 0.5 | 2.79E-03 | 0.3 | 7.67E-02 | 0.2 | 3.34E-01 | Uncharacterized protein At1g24500 |
| AT2G17410 | 0.2 | 3.69E-02 | 0.0 | 7.31E-01 | 0.2 | 3.49E-02 | -0.1 | 5.84E-01 | -0.1 | 4.37E-01 | 0.0 | 8.62E-01 | ARID |
| AT1G74330 | 0.2 | 1.28E-01 | -0.2 | 3.89E-02 | 0.4 | 7.60E-03 | -0.5 | 3.55E-02 | -0.5 | 4.20E-02 | 0.0 | 9.63E-01 | Protein kinase superfamily protein |
| AT3G52850 | 0.2 | 2.72E-02 | 0.1 | 4.08E-01 | 0.1 | 1.72E-01 | -0.2 | 2.32E-01 | -0.1 | 8.57E-01 | -0.2 | 5.49E-01 | EGF-like calcium-binding domain-containing protein |
| AT5G08100 | 0.2 | 5.64E-02 | 0.0 | 6.64E-01 | 0.2 | 4.31E-02 | -0.3 | 2.61E-02 | -0.1 | 2.93E-01 | -0.1 | 4.62E-01 | N-terminal nucleophile aminohydrolases superfamily protein |
| AT4G31180 | 0.2 | 5.17E-02 | 0.0 | 7.11E-01 | 0.2 | 4.14E-02 | 0.1 | 3.42E-01 | -0.1 | 3.67E-01 | 0.2 | 1.54E-01 | aspartate--tRNA ligase |
| AT5G57020 | 0.2 | 5.26E-02 | 0.3 | 3.56E-03 | -0.1 | 1.38E-01 | -0.2 | 1.01E-01 | -0.3 | 1.81E-02 | 0.1 | 4.61E-01 | Glycylpeptide N-tetradecanoyltransferase 1 |
| AT5G58270 | 0.2 | 2.15E-02 | 0.0 | 7.60E-01 | 0.1 | 6.56E-02 | -0.1 | 3.81E-01 | 0.0 | 8.43E-01 | -0.1 | 6.91E-01 | ABC transporter B family member 25, mitochondrial |
| AT3G45850 | 0.2 | 1.15E-01 | 0.1 | 3.19E-01 | 0.1 | 6.21E-01 | -0.2 | 4.19E-01 | -0.1 | 8.43E-01 | -0.1 | 7.34E-01 | P-loop containing nucleoside triphosphate hydrolases superfamily protein |
| AT5G08550 | 0.2 | 1.23E-01 | -0.1 | 2.45E-01 | 0.3 | 3.29E-02 | -0.2 | 7.75E-01 | -0.3 | 5.92E-01 | 0.2 | 8.39E-01 | Transcriptional repressor ILP1 |
| AT4G26910 | 0.2 | 1.16E-01 | 0.2 | 8.41E-02 | 0.0 | 8.32E-01 | 0.5 | 6.06E-04 | 0.0 | 9.45E-01 | 0.4 | 1.27E-02 | Dihydrolipoyllysine-residue succinyltransferase component of 2-oxoglutarate dehydrogenase complex 2, mitochondrial |
| AT5G20400 | 0.2 | 1.35E-01 | -0.2 | 5.52E-02 | 0.4 | 8.41E-03 | 0.7 | 1.65E-03 | 0.0 | 9.03E-01 | 0.6 | 7.11E-03 | hypothetical protein |
| AT5G18500 | 0.2 | 2.51E-02 | 0.0 | 4.46E-01 | 0.1 | 1.45E-01 | -0.1 | 2.88E-01 | -0.2 | 2.98E-01 | 0.0 | 9.46E-01 | Probable receptor-like protein kinase At5g18500 |
| AT5G09410 | 0.2 | 4.44E-02 | -0.1 | 2.23E-01 | 0.2 | 1.10E-02 | -0.2 | 2.56E-01 | 0.0 | 8.59E-01 | -0.2 | 2.89E-01 | Ethylene induced calmodulin binding protein |
| AT3G18730 | 0.2 | 2.93E-01 | 0.1 | 5.24E-01 | 0.1 | 7.78E-01 | 0.6 | 6.50E-02 | 0.6 | 1.21E-01 | 0.0 | 9.29E-01 | Protein TONSOKU |
| AT5G37055 | 0.2 | 2.17E-01 | 0.0 | 7.59E-01 | 0.1 | 4.67E-01 | -0.1 | 7.77E-01 | -0.1 | 8.73E-01 | 0.0 | 9.62E-01 | SWR1 complex subunit 6 |
| AT2G28880 | 0.2 | 1.89E-01 | 0.0 | 9.02E-01 | 0.2 | 2.29E-01 | 0.0 | 8.65E-01 | 0.2 | 5.34E-01 | -0.2 | 4.48E-01 | Aminodeoxychorismate synthase, chloroplastic |
| AT5G61380 | 0.2 | 2.62E-02 | 0.2 | 2.29E-02 | 0.0 | 9.01E-01 | -0.1 | 5.53E-01 | -0.4 | 3.02E-03 | 0.3 | 4.67E-02 | Two-component response regulator-like APRR1 |
| AT1G64630 | 0.2 | 4.40E-02 | 0.0 | 9.02E-01 | 0.1 | 1.03E-01 | 0.0 | 9.89E-01 | -0.1 | 7.57E-01 | 0.1 | 8.14E-01 | Probable serine |
| AT1G11950 | 0.2 | 1.45E-01 | -0.1 | 3.19E-01 | 0.3 | 5.27E-02 | 0.1 | 6.31E-01 | -0.1 | 5.02E-01 | 0.2 | 3.01E-01 | Transcription factor jumonji domain-containing protein |
| AT1G47260 | 0.2 | 7.48E-02 | 0.4 | 1.81E-03 | -0.2 | 4.76E-02 | 0.2 | 1.63E-02 | 0.2 | 8.07E-02 | 0.0 | 8.01E-01 | Gamma carbonic anhydrase 2, mitochondrial |
| AT3G48470 | 0.2 | 2.27E-01 | -0.1 | 3.91E-01 | 0.3 | 1.03E-01 | 0.1 | 7.74E-01 | 0.1 | 8.82E-01 | 0.0 | 9.47E-01 | Embryo defective 2423 |
| AT3G07610 | 0.2 | 3.39E-02 | -0.2 | 7.68E-03 | 0.4 | 9.12E-04 | 0.5 | 8.19E-03 | 0.2 | 3.97E-01 | 0.3 | 1.97E-01 | Uncharacterized protein At3g07610 |
| AT4G16860 | 0.2 | 2.28E-01 | -0.2 | 1.99E-01 | 0.3 | 3.57E-02 | 0.2 | 2.26E-01 | -0.2 | 2.95E-01 | 0.4 | 6.50E-02 | Disease resistance protein family |
| AT3G03773 | 0.2 | 1.39E-01 | 0.1 | 5.58E-01 | 0.1 | 4.65E-01 | 0.1 | 5.06E-01 | 0.1 | 7.91E-01 | 0.1 | 8.37E-01 | Co-chaperone protein p23 |
| AT3G24440 | 0.2 | 5.22E-02 | 0.1 | 5.29E-01 | 0.1 | 2.35E-01 | 0.1 | 4.20E-01 | 0.2 | 4.09E-01 | 0.0 | 9.45E-01 | Fibronectin type III domain-containing protein |
| AT2G34710 | 0.2 | 1.51E-01 | -0.1 | 4.59E-01 | 0.2 | 6.40E-02 | -0.2 | 3.20E-01 | -0.1 | 7.47E-01 | -0.1 | 7.23E-01 | Homeobox-leucine zipper protein ATHB-14 |
| AT5G49520 | 0.2 | 3.09E-01 | -0.5 | 1.17E-02 | 0.7 | 6.53E-03 | 0.2 | 5.92E-01 | 0.4 | 3.00E-01 | -0.2 | 6.98E-01 | WRKY domain-containing protein |
| AT4G14360 | 0.2 | 2.72E-02 | 0.0 | 7.49E-01 | 0.1 | 7.89E-02 | -0.1 | 7.02E-01 | 0.2 | 3.70E-01 | -0.3 | 2.89E-01 | Probable methyltransferase PMT3 |
| AT1G54270 | 0.2 | 4.49E-02 | 0.1 | 4.69E-01 | 0.1 | 2.21E-01 | 0.2 | 9.90E-02 | 0.0 | 7.56E-01 | 0.1 | 4.11E-01 | Eif4a-2 |
| AT2G23180 | 0.2 | 1.84E-01 | 0.0 | 8.21E-01 | 0.1 | 3.71E-01 | 0.4 | 1.20E-01 | 0.6 | 3.42E-02 | -0.2 | 5.50E-01 | Cytochrome P450, family 96, subfamily A, polypeptide 1 |
| AT3G57220 | 0.2 | 4.46E-02 | 0.3 | 1.82E-03 | -0.1 | 9.90E-02 | 0.0 | 9.91E-01 | -0.3 | 6.88E-02 | 0.3 | 1.27E-01 | UDP-N-acetylglucosamine--dolichyl-phosphate N-acetylglucosaminephosphotransferase |
| AT5G05560 | 0.2 | 6.04E-02 | -0.1 | 2.48E-01 | 0.3 | 1.95E-02 | 0.2 | 3.99E-01 | 0.2 | 5.76E-01 | 0.0 | 9.20E-01 | E3 ubiquitin ligase |
| AT1G69310 | 0.2 | 1.06E-01 | -0.1 | 3.01E-01 | 0.3 | 3.64E-02 | -0.1 | 7.02E-01 | 0.0 | 9.38E-01 | -0.1 | 8.30E-01 | WRKY DNA-binding protein 57 |
| AT1G29740 | 0.2 | 3.41E-01 | 0.8 | 5.17E-04 | -0.6 | 6.58E-03 | 0.0 | 8.93E-01 | -0.2 | 5.72E-01 | 0.2 | 7.52E-01 | Leucine-rich repeat transmembrane protein kinase |
| AT3G24320 | 0.2 | 2.02E-01 | 0.0 | 9.20E-01 | 0.1 | 3.44E-01 | 0.1 | 7.68E-01 | 0.1 | 8.61E-01 | 0.0 | 9.64E-01 | MUTL protein homolog 1 |
| AT5G66060 | 0.2 | 3.43E-02 | 0.2 | 6.69E-03 | -0.1 | 3.74E-01 | -0.2 | 2.02E-01 | -0.3 | 3.03E-02 | 0.1 | 4.48E-01 | 2-oxoglutarate and Fe-dependent oxygenase superfamily protein |
| AT3G51180 | 0.2 | 2.64E-01 | 0.0 | 8.38E-01 | 0.2 | 2.81E-01 | 0.4 | 2.11E-01 | 0.2 | 7.18E-01 | 0.2 | 5.87E-01 | Zinc finger CCCH domain-containing protein 45 |
| AT5G42940 | 0.2 | 2.56E-02 | -0.1 | 1.55E-01 | 0.3 | 5.51E-03 | -0.1 | 7.05E-01 | -0.2 | 5.10E-01 | 0.1 | 8.19E-01 | Probable E3 ubiquitin-protein ligase RHG1A |
| AT3G05840 | 0.2 | 3.97E-02 | -0.1 | 9.15E-02 | 0.3 | 4.18E-03 | 0.2 | 5.86E-02 | 0.1 | 4.00E-01 | 0.1 | 5.99E-01 | Shaggy-related protein kinase gamma |
| AT1G75950 | 0.2 | 8.54E-02 | 0.0 | 6.79E-01 | 0.1 | 2.35E-01 | 0.1 | 6.49E-01 | 0.0 | 8.21E-01 | 0.1 | 5.95E-01 | SKP1-like protein 1A |
| AT5G24120 | 0.2 | 4.08E-01 | -0.4 | 8.10E-02 | 0.5 | 3.05E-02 | 0.0 | 9.46E-01 | 0.6 | 1.88E-01 | -0.6 | 2.57E-01 | RNA polymerase sigma factor sigE, chloroplastic |
| AT3G44735 | 0.2 | 5.27E-01 | -0.2 | 5.47E-01 | 0.3 | 2.99E-01 | -0.5 | 7.27E-02 | 0.1 | 8.35E-01 | -0.6 | 1.15E-01 | Putative phytosulfokines 6 |
| AT1G25380 | 0.2 | 1.15E-01 | 0.2 | 1.37E-01 | 0.0 | 9.67E-01 | 0.2 | 1.07E-01 | 0.2 | 3.06E-01 | 0.1 | 8.11E-01 | Nicotinamide adenine dinucleotide transporter 2, mitochondrial transporter 2) |
| AT2G15820 | 0.2 | 1.17E-01 | 0.0 | 7.71E-01 | 0.1 | 2.82E-01 | 0.3 | 1.83E-01 | 0.3 | 1.63E-01 | 0.0 | 9.05E-01 | Pentatricopeptide repeat-containing protein At2g15820, chloroplastic |
| AT1G07910 | 0.2 | 7.72E-02 | 0.1 | 3.66E-01 | 0.1 | 4.56E-01 | -0.1 | 8.16E-01 | 0.0 | 9.75E-01 | -0.1 | 9.04E-01 | hypothetical protein |
| AT1G74720 | 0.2 | 2.59E-02 | 0.0 | 9.49E-01 | 0.2 | 5.20E-02 | 0.2 | 2.23E-01 | 0.0 | 9.21E-01 | 0.1 | 4.63E-01 | Protein QUIRKY |
| AT4G19420 | 0.2 | 2.73E-01 | 0.2 | 1.83E-01 | 0.0 | 8.09E-01 | -0.1 | 4.22E-01 | -0.5 | 8.19E-03 | 0.4 | 8.35E-02 | Pectin acetylesterase |
| AT2G19470 | 0.2 | 7.90E-02 | -0.1 | 2.76E-01 | 0.3 | 2.24E-02 | -0.1 | 7.17E-01 | -0.1 | 7.98E-01 | 0.0 | 9.81E-01 | Casein kinase 1-like protein 5 |
| AT4G38810 | 0.2 | 8.34E-02 | 0.3 | 4.55E-03 | -0.2 | 1.26E-01 | -0.2 | 5.36E-02 | -0.2 | 2.82E-01 | -0.1 | 6.90E-01 | Calcium-binding EF-hand family protein |
| AT4G00360 | 0.2 | 1.12E-01 | -0.1 | 5.66E-01 | 0.2 | 5.93E-02 | -0.2 | 7.51E-01 | 0.0 | 9.96E-01 | -0.2 | 8.39E-01 | Cytochrome P450 86A2 |
| AT4G13870 | 0.2 | 3.15E-01 | 0.0 | 9.64E-01 | 0.2 | 4.51E-01 | -0.2 | 5.04E-01 | -0.1 | 8.37E-01 | -0.1 | 7.72E-01 | 3'-5' exonuclease |
| AT1G17520 | 0.2 | 9.26E-02 | 0.0 | 7.21E-01 | 0.2 | 8.90E-02 | -0.3 | 1.02E-01 | -0.3 | 1.45E-01 | 0.0 | 9.92E-01 | MYB transcription factor |
| AT3G59080 | 0.2 | 1.57E-01 | -0.4 | 4.54E-03 | 0.6 | 1.96E-03 | 0.5 | 3.46E-02 | 0.0 | 9.66E-01 | 0.5 | 9.20E-02 | AT3g59080 |
| AT5G09590 | 0.2 | 1.12E-01 | -0.1 | 5.76E-01 | 0.2 | 6.20E-02 | 1.0 | 1.37E-05 | 0.5 | 5.60E-03 | 0.4 | 5.11E-02 | Heat shock 70 kDa protein 10, mitochondrial |
| AT5G63550 | 0.2 | 1.14E-01 | 0.2 | 3.33E-02 | -0.1 | 5.18E-01 | 0.3 | 5.21E-02 | 0.2 | 2.33E-01 | 0.1 | 7.31E-01 | DEK-C domain-containing protein |
| AT3G28760 | 0.2 | 2.05E-01 | 0.3 | 3.75E-02 | -0.1 | 3.91E-01 | 0.7 | 6.29E-03 | 0.7 | 8.47E-03 | 0.0 | 9.46E-01 | 3-dehydroquinate synthase |
| AT3G13580 | 0.2 | 2.33E-01 | 0.3 | 5.38E-02 | -0.1 | 4.11E-01 | -0.1 | 5.52E-01 | -0.3 | 1.47E-01 | 0.2 | 4.63E-01 | Large ribosomal subunit protein uL30w |
| AT5G10290 | 0.2 | 4.33E-02 | 0.0 | 8.86E-01 | 0.2 | 5.65E-02 | 0.3 | 1.15E-02 | 0.0 | 9.96E-01 | 0.3 | 6.35E-02 | hypothetical protein |

| | | | | | | | | | | | | | |
|---|---|---|---|---|---|---|---|---|---|---|---|---|---|
| AT5G53000 | 0.2 | 2.59E-02 | 0.0 | 7.42E-01 | 0.2 | 2.65E-02 | 0.1 | 3.47E-01 | 0.1 | 6.07E-01 | 0.0 | 8.61E-01 | PP2A regulatory subunit TAP46 |
| AT4G00970 | 0.2 | 4.15E-01 | 0.3 | 1.16E-01 | -0.2 | 4.68E-01 | -0.1 | 7.51E-01 | -0.2 | 2.50E-01 | 0.1 | 4.70E-01 | Cysteine-rich RLK 41 |
| AT1G03590 | 0.2 | 1.51E-02 | 0.1 | 9.30E-02 | 0.1 | 4.23E-01 | -0.4 | 7.74E-03 | -0.5 | 5.01E-03 | 0.1 | 7.75E-01 | Probable protein phosphatase 2C 1 |
| AT5G67270 | 0.2 | 2.58E-01 | 0.5 | 2.67E-03 | -0.4 | 3.48E-02 | 0.4 | 3.34E-02 | 0.5 | 1.58E-02 | -0.1 | 7.24E-01 | Microtubule-associated protein RP |
| AT5G44070 | 0.2 | 1.05E-01 | -0.1 | 3.37E-01 | 0.3 | 2.95E-02 | 0.3 | 2.75E-02 | 0.2 | 3.37E-01 | 0.2 | 4.75E-01 | glutathione gamma-glutamylcysteinyltransferase |
| AT4G20070 | 0.2 | 2.19E-01 | 0.0 | 7.32E-01 | 0.2 | 1.65E-01 | -0.1 | 5.02E-01 | -0.2 | 2.95E-01 | 0.1 | 7.51E-01 | Allantoate amidohydrolase |
| AT4G03560 | 0.2 | 4.18E-02 | 0.2 | 1.64E-02 | 0.0 | 5.72E-01 | -0.2 | 5.23E-02 | -0.1 | 2.20E-01 | -0.1 | 7.80E-01 | Two pore calcium channel protein 1 |
| AT1G50480 | 0.2 | 4.66E-02 | 0.1 | 1.78E-01 | 0.1 | 5.43E-01 | -0.2 | 2.00E-01 | -0.2 | 3.47E-01 | 0.0 | 9.30E-01 | Formate--tetrahydrofolate ligase |
| AT5G21482 | 0.2 | 2.31E-01 | -0.1 | 4.55E-01 | 0.3 | 1.10E-01 | -0.3 | 1.16E-01 | 0.1 | 7.49E-01 | -0.4 | 1.15E-01 | Cytokinin dehydrogenase 7 |
| AT3G16470 | 0.2 | 4.89E-02 | 0.4 | 8.50E-04 | -0.2 | 2.89E-02 | 0.3 | 4.31E-01 | 0.7 | 8.53E-02 | -0.4 | 4.63E-01 | Jacalin-type lectin domain-containing protein |
| AT1G76710 | 0.2 | 6.06E-02 | 0.1 | 3.69E-01 | 0.1 | 3.90E-01 | -0.3 | 1.64E-01 | -0.3 | 2.79E-01 | 0.0 | 9.20E-01 | SET domain group 26 |
| AT5G63080 | 0.2 | 9.88E-02 | -0.2 | 1.06E-01 | 0.3 | 1.41E-02 | 0.1 | 7.81E-01 | -0.1 | 7.60E-01 | 0.2 | 5.73E-01 | Arginine-specific demethylase JMJ20 monodemethylase JMJ20) |
| AT2G48100 | 0.2 | 4.49E-02 | 0.2 | 4.44E-02 | 0.0 | 9.53E-01 | 0.0 | 8.49E-01 | -0.2 | 1.39E-01 | 0.2 | 2.90E-01 | RNA exonuclease 4 |
| AT1G02660 | 0.2 | 2.09E-01 | 0.0 | 9.50E-01 | 0.2 | 3.25E-01 | -0.3 | 2.22E-01 | -1.0 | 5.43E-03 | 0.6 | 1.13E-01 | Phospholipase A1 PLIP2, chloroplastic |
| AT3G50410 | 0.2 | 2.37E-01 | -0.2 | 2.33E-01 | 0.3 | 6.27E-02 | 0.1 | 6.72E-01 | -0.2 | 6.39E-01 | 0.3 | 3.78E-01 | Dof zinc finger protein DOF3.4 |
| AT1G50920 | 0.2 | 1.48E-01 | 0.2 | 1.44E-01 | 0.0 | 9.47E-01 | 0.5 | 8.74E-05 | 0.2 | 5.61E-02 | 0.3 | 6.52E-02 | Nucleolar GTP-binding protein 1 |
| AT5G39900 | 0.2 | 9.20E-02 | 0.2 | 9.50E-02 | 0.0 | 9.73E-01 | 0.4 | 1.26E-01 | 0.4 | 1.33E-01 | 0.0 | 9.41E-01 | Translation factor GUF1 homolog, mitochondrial |
| AT5G23020 | 0.2 | 5.96E-02 | 0.3 | 3.61E-03 | -0.1 | 1.21E-01 | -0.2 | 3.04E-01 | 0.3 | 8.53E-02 | -0.5 | 8.17E-03 | Methylthioalkylmalate synthase 3, chloroplastic |
| AT4G23060 | 0.2 | 1.28E-01 | -0.3 | 1.29E-02 | 0.5 | 3.17E-03 | -0.5 | 1.65E-02 | -0.5 | 3.40E-02 | 0.0 | 9.41E-01 | Protein IQ-DOMAIN 22 |
| AT4G08590 | 0.2 | 1.11E-01 | 0.0 | 6.84E-01 | 0.1 | 3.36E-01 | -0.1 | 8.34E-01 | -0.1 | 7.63E-01 | 0.1 | 9.28E-01 | E3 ubiquitin-protein ligase ORTHRUS |
| AT1G72180 | 0.2 | 6.05E-02 | 0.1 | 1.17E-01 | 0.0 | 7.93E-01 | 0.0 | 9.46E-01 | -0.1 | 8.16E-01 | 0.0 | 8.98E-01 | Receptor protein-tyrosine kinase CEPR2 |
| AT4G38130 | 0.2 | 3.18E-02 | 0.0 | 7.59E-01 | 0.1 | 8.63E-02 | 0.0 | 7.33E-01 | -0.1 | 6.20E-01 | 0.1 | 5.02E-01 | Histone deacetylase |
| AT3G21810 | 0.2 | 1.12E-01 | -0.2 | 5.39E-02 | 0.4 | 8.67E-03 | 0.3 | 7.83E-02 | 0.0 | 8.57E-01 | 0.3 | 2.07E-01 | Zinc finger C-x8-C-x5-C-x3-H type family protein |
| AT5G16260 | 0.2 | 4.32E-02 | -0.1 | 3.05E-01 | 0.2 | 1.57E-02 | 0.0 | 8.45E-01 | -0.2 | 4.14E-01 | 0.1 | 6.37E-01 | At5g16260 family protein) |
| AT1G55970 | 0.2 | 1.98E-01 | -0.2 | 1.30E-01 | 0.4 | 2.99E-02 | 0.0 | 9.77E-01 | 0.0 | 9.83E-01 | 0.0 | 9.99E-01 | histone acetyltransferase |
| AT2G31880 | 0.2 | 4.45E-01 | -0.7 | 1.23E-02 | 0.8 | 6.27E-03 | -0.1 | 8.78E-01 | -0.3 | 5.42E-01 | 0.2 | 7.28E-01 | Leucine-rich repeat receptor-like serine |
| AT2G15270 | 0.2 | 9.40E-02 | 0.2 | 8.16E-02 | 0.0 | 9.16E-01 | 0.2 | 4.77E-01 | -0.1 | 8.39E-01 | 0.3 | 4.66E-01 | PRKR-interacting protein 1 |
| AT2G27150 | 0.2 | 3.76E-02 | -0.2 | 2.41E-02 | 0.4 | 2.12E-03 | -0.3 | 3.99E-01 | 0.1 | 8.45E-01 | -0.4 | 3.88E-01 | Abscisic aldehyde oxidase 3 |
| AT4G16330 | 0.2 | 3.54E-02 | -0.1 | 1.24E-01 | 0.3 | 6.11E-03 | -0.6 | 6.41E-04 | -0.6 | 1.99E-03 | 0.0 | 9.13E-01 | 2-oxoglutarate and Fe-dependent oxygenase superfamily protein |
| AT2G14260 | 0.2 | 2.42E-02 | -0.1 | 4.67E-01 | 0.2 | 1.36E-02 | -0.3 | 3.09E-02 | -0.3 | 7.24E-02 | 0.0 | 9.29E-01 | Proline iminopeptidase |
| AT2G46160 | 0.2 | 1.28E-01 | -0.1 | 4.96E-01 | 0.2 | 7.44E-02 | -0.2 | 3.88E-01 | -0.2 | 3.64E-01 | 0.0 | 9.40E-01 | RING-H2 finger protein ATL67 |
| AT5G58800 | 0.2 | 1.24E-01 | 0.1 | 4.92E-01 | 0.1 | 4.67E-01 | -0.2 | 2.11E-01 | -0.2 | 2.55E-01 | 0.0 | 9.85E-01 | NADH dehydrogenase |
| AT1G34430 | 0.2 | 5.15E-02 | 0.3 | 1.81E-03 | -0.2 | 6.81E-02 | 0.0 | 9.82E-01 | 0.1 | 7.75E-01 | -0.1 | 7.88E-01 | Dihydrolipoyllysine-residue acetyltransferase component 5 of pyruvate dehydrogenase complex, chloroplastic |
| AT1G09630 | 0.2 | 6.72E-02 | 0.4 | 7.71E-04 | -0.2 | 2.47E-02 | 0.1 | 7.48E-01 | 0.2 | 3.61E-01 | -0.1 | 6.43E-01 | Ras-related protein RABA2a |
| AT1G32440 | 0.2 | 5.50E-02 | -0.1 | 4.37E-01 | 0.2 | 2.44E-02 | 0.3 | 1.20E-01 | 0.3 | 9.78E-02 | 0.0 | 8.78E-01 | Plastidial pyruvate kinase 3, chloroplastic |
| AT2G22100 | 0.2 | 1.82E-01 | 0.2 | 7.80E-02 | -0.1 | 6.58E-01 | 0.2 | 2.79E-01 | 0.0 | 8.83E-01 | 0.2 | 4.74E-01 | UBP1-associated proteins 1B |
| AT2G01820 | 0.2 | 2.12E-02 | 0.0 | 9.49E-01 | 0.2 | 3.20E-02 | -0.1 | 6.63E-01 | -0.1 | 5.91E-01 | 0.0 | 9.24E-01 | Receptor-like kinase TMK3 |
| AT1G59700 | 0.2 | 9.68E-02 | -0.2 | 3.11E-02 | 0.4 | 4.41E-03 | -0.1 | 4.76E-01 | 0.0 | 8.73E-01 | -0.1 | 7.49E-01 | Glutathione S-transferase U16 |
| AT1G65295 | 0.2 | 2.23E-01 | 0.1 | 5.22E-01 | 0.1 | 6.51E-01 | -0.1 | 7.81E-01 | 0.0 | 9.50E-01 | -0.1 | 9.04E-01 | Ubiquitin carboxyl-terminal hydrolase |
| AT3G43520 | 0.2 | 6.72E-02 | 0.2 | 1.05E-01 | 0.0 | 8.82E-01 | 0.2 | 1.20E-01 | 0.1 | 4.58E-01 | 0.1 | 6.85E-01 | Protein FATTY ACID EXPORT 2, chloroplastic |
| AT3G12760 | 0.2 | 1.92E-02 | 0.2 | 1.85E-02 | 0.0 | 9.31E-01 | 0.0 | 8.03E-01 | 0.0 | 7.70E-01 | 0.1 | 6.77E-01 | Defective in cullin neddylation protein |
| AT2G38670 | 0.2 | 2.40E-02 | 0.2 | 6.33E-03 | -0.1 | 4.31E-01 | 0.1 | 3.72E-01 | 0.2 | 1.38E-01 | -0.1 | 6.51E-01 | Ethanolamine-phosphate cytidylyltransferase |
| AT2G02220 | 0.2 | 5.88E-02 | -0.1 | 2.51E-01 | 0.3 | 1.55E-02 | 0.4 | 1.39E-02 | 0.0 | 9.90E-01 | 0.4 | 4.20E-02 | Phytosulfokine receptor 1 |
| AT4G37270 | 0.2 | 6.43E-02 | -0.2 | 2.29E-02 | 0.4 | 2.19E-03 | -0.1 | 8.36E-01 | 0.2 | 5.43E-01 | -0.2 | 4.64E-01 | Probable cadmium |
| AT4G36270 | 0.2 | 3.14E-01 | 0.0 | 8.92E-01 | 0.2 | 3.57E-01 | -0.2 | 6.32E-01 | 0.5 | 1.90E-01 | -0.6 | 6.96E-02 | Protein MICRORCHIDIA 3 |
| AT1G48160 | 0.2 | 1.60E-01 | 0.3 | 3.12E-02 | -0.1 | 4.02E-01 | 0.3 | 1.33E-01 | 0.0 | 9.27E-01 | 0.2 | 2.84E-01 | Signal recognition particle 19 kDa protein |
| AT1G33990 | 0.2 | 9.04E-02 | -0.1 | 5.59E-01 | 0.2 | 5.19E-02 | -0.3 | 5.13E-02 | -0.3 | 3.53E-02 | 0.1 | 8.14E-01 | Putative methylesterase 14, chloroplastic |
| AT3G20740 | 0.2 | 6.38E-02 | 0.1 | 4.37E-01 | 0.1 | 3.37E-01 | -0.1 | 3.95E-01 | 0.0 | 8.96E-01 | -0.1 | 6.66E-01 | Polycomb group protein FERTILIZATION-INDEPENDENT ENDOSPERM |
| AT2G42950 | 0.2 | 9.76E-02 | -0.2 | 5.49E-02 | 0.4 | 8.50E-03 | 0.0 | 9.17E-01 | 0.1 | 7.75E-01 | -0.1 | 8.78E-01 | hypothetical protein |
| AT5G26710 | 0.2 | 4.67E-02 | 0.2 | 2.26E-02 | 0.0 | 6.63E-01 | 0.2 | 6.81E-02 | 0.2 | 2.61E-01 | 0.1 | 7.78E-01 | Glutamyl-tRNA synthetase |
| AT2G42290 | 0.2 | 1.16E-01 | 0.3 | 2.88E-02 | -0.1 | 4.87E-01 | -0.4 | 9.18E-02 | -0.6 | 3.51E-02 | 0.2 | 7.10E-01 | Protein kinase domain-containing protein |
| AT3G21950 | 0.2 | 3.08E-01 | 0.6 | 2.34E-03 | -0.5 | 2.60E-02 | 0.1 | 8.56E-01 | 0.1 | 8.83E-01 | 0.0 | 1.00E+00 | S-adenosyl-L-methionine-dependent methyltransferases superfamily protein |
| AT4G10620 | 0.2 | 1.75E-01 | 0.1 | 5.46E-01 | 0.1 | 5.46E-01 | 0.3 | 2.67E-01 | 0.2 | 5.17E-01 | 0.1 | 8.39E-01 | G domain-containing protein |
| AT4G19600 | 0.2 | 1.99E-02 | 0.0 | 5.05E-01 | 0.1 | 1.12E-01 | 0.0 | 9.47E-01 | -0.2 | 2.00E-01 | 0.2 | 2.60E-01 | Cyclin-T1-4 |
| AT2G41130 | 0.2 | 1.16E-01 | 0.0 | 8.02E-01 | 0.1 | 2.78E-01 | 0.0 | 9.78E-01 | 0.1 | 8.35E-01 | -0.1 | 8.42E-01 | Transcription factor bHLH106 |
| AT3G13100 | 0.2 | 2.37E-01 | -0.4 | 1.97E-02 | 0.6 | 8.02E-03 | 1.1 | 1.28E-02 | 1.0 | 4.71E-02 | 0.2 | 7.79E-01 | ABC-type xenobiotic transporter |
| AT2G37710 | 0.2 | 1.40E-01 | -0.2 | 7.44E-02 | 0.4 | 9.29E-03 | 0.4 | 4.85E-03 | 0.1 | 5.73E-01 | 0.3 | 6.91E-02 | L-type lectin-domain containing receptor kinase IV.1 |
| AT1G44110 | 0.2 | 1.32E-01 | 0.3 | 1.73E-02 | -0.1 | 3.15E-01 | 0.0 | 8.68E-01 | 0.2 | 2.95E-01 | -0.3 | 3.08E-01 | Cyclin-A1-1 |
| AT5G60970 | 0.2 | 7.46E-02 | -0.1 | 6.12E-01 | 0.2 | 6.41E-02 | -0.3 | 3.38E-01 | -0.3 | 2.54E-01 | 0.1 | 8.15E-01 | Transcription factor TCP5 |
| AT5G16440 | 0.2 | 1.64E-01 | 0.4 | 5.94E-03 | -0.2 | 9.09E-02 | 0.0 | 9.18E-01 | -0.1 | 6.59E-01 | 0.1 | 8.04E-01 | isopentenyl-diphosphate Delta-isomerase |
| AT4G27650 | 0.2 | 3.83E-02 | 0.1 | 4.74E-01 | 0.1 | 2.11E-01 | -0.1 | 7.82E-01 | -0.2 | 2.82E-01 | 0.2 | 5.17E-01 | Protein PELOTA 1 |
| AT3G46640 | 0.2 | 2.17E-01 | 0.5 | 2.77E-03 | -0.4 | 4.12E-02 | -0.4 | 1.75E-02 | -0.5 | 1.25E-02 | 0.1 | 7.87E-01 | Homeodomain-like superfamily protein |
| AT5G21160 | 0.2 | 2.52E-02 | 0.1 | 7.17E-02 | 0.0 | 6.64E-01 | 0.0 | 9.06E-01 | -0.4 | 6.82E-03 | 0.4 | 1.59E-02 | LA RNA-binding protein |
| AT5G10510 | 0.2 | 2.16E-01 | -0.1 | 5.33E-01 | 0.3 | 1.32E-01 | 0.2 | 8.29E-01 | 0.4 | 6.68E-01 | -0.2 | 8.55E-01 | AINTEGUMENTA-like 6 |
| AT5G47810 | 0.2 | 6.39E-02 | -0.2 | 2.52E-02 | 0.4 | 3.50E-03 | -0.3 | 1.08E-01 | -0.4 | 1.13E-01 | 0.0 | 9.42E-01 | ATP-dependent 6-phosphofructokinase 2 |
| AT1G53230 | 0.2 | 1.32E-01 | 0.3 | 3.37E-02 | -0.1 | 5.00E-01 | -0.2 | 5.30E-01 | 0.2 | 6.53E-01 | -0.4 | 3.36E-01 | Transcription factor TCP3 |
| AT3G51670 | 0.2 | 4.11E-02 | 0.2 | 4.96E-02 | 0.0 | 9.82E-01 | 0.6 | 2.57E-04 | 0.0 | 9.22E-01 | 0.6 | 5.41E-03 | Patellin-6 |
| AT1G17580 | 0.2 | 6.73E-02 | -0.1 | 3.45E-01 | 0.3 | 2.36E-02 | -0.3 | 1.07E-01 | -0.2 | 3.00E-01 | -0.1 | 8.10E-01 | XI-1 |
| AT1G74100 | 0.2 | 1.11E-01 | 0.3 | 1.77E-02 | -0.1 | 3.13E-01 | 0.3 | 2.52E-01 | -0.1 | 8.05E-01 | 0.4 | 2.57E-01 | Cytosolic sulfotransferase 16 |
| AT1G63350 | 0.2 | 1.95E-01 | -0.3 | 5.71E-02 | 0.5 | 1.57E-02 | 0.0 | 9.88E-01 | -0.7 | 2.12E-02 | 0.7 | 4.40E-02 | Disease resistance protein family |
| AT2G05160 | 0.2 | 2.02E-01 | 0.3 | 2.34E-02 | -0.2 | 2.87E-01 | -0.1 | 8.05E-01 | 0.2 | 4.38E-01 | -0.3 | 3.36E-01 | CCCH-type zinc fingerfamily protein with RNA-binding domain-containing protein |
| AT5G39020 | 0.2 | 1.23E-01 | 0.0 | 7.26E-01 | 0.2 | 1.09E-01 | 0.5 | 4.47E-02 | 0.0 | 9.08E-01 | 0.4 | 7.99E-02 | Probable receptor-like protein kinase At5g39020 |
| AT2G22400 | 0.2 | 5.70E-02 | 0.1 | 1.70E-01 | 0.0 | 6.24E-01 | 0.3 | 3.65E-02 | 0.0 | 8.92E-01 | 0.3 | 9.90E-02 | S-adenosyl-L-methionine-dependent methyltransferases superfamily protein |
| AT2G20980 | 0.2 | 2.29E-01 | 0.2 | 1.06E-01 | -0.1 | 6.88E-01 | 0.4 | 3.02E-01 | 0.5 | 2.41E-01 | -0.1 | 8.75E-01 | Minichromosome maintenance 10 |
| AT4G11160 | 0.2 | 6.17E-02 | 0.0 | 6.47E-01 | 0.2 | 4.42E-02 | 0.1 | 3.65E-01 | 0.1 | 4.88E-01 | 0.0 | 9.73E-01 | At4g11160 |
| AT3G04110 | 0.2 | 6.32E-02 | 0.0 | 6.78E-01 | 0.2 | 5.39E-02 | -0.1 | 7.65E-01 | -0.1 | 6.81E-01 | 0.0 | 9.12E-01 | Glutamate receptor 1.1 |
| AT1G51420 | 0.2 | 9.03E-02 | -0.3 | 2.27E-02 | 0.4 | 3.64E-03 | 1.3 | 7.77E-05 | 0.7 | 2.01E-02 | 0.6 | 2.21E-02 | Sucrose-phosphatase |
| AT3G47910 | 0.2 | 4.34E-02 | -0.2 | 4.34E-02 | 0.4 | 3.04E-03 | 0.4 | 1.34E-01 | 0.2 | 4.72E-01 | 0.1 | 6.76E-01 | Ubiquitin carboxyl-terminal hydrolase-related protein |
| AT1G67840 | 0.2 | 1.10E-01 | 0.0 | 7.97E-01 | 0.2 | 9.93E-02 | -0.2 | 7.78E-02 | 0.0 | 8.49E-01 | -0.2 | 2.98E-01 | Chloroplast sensor kinase, chloroplastic |
| AT4G28700 | 0.2 | 2.30E-01 | 0.3 | 8.69E-02 | -0.1 | 6.06E-01 | | | | | | | Ammonium transporter 1 member 4 |
| AT2G19480 | 0.2 | 8.93E-02 | 0.2 | 6.58E-02 | 0.0 | 8.28E-01 | 0.2 | 4.14E-02 | 0.0 | 9.80E-01 | 0.2 | 1.51E-01 | Nucleosome assembly protein 12 |
| AT3G03960 | 0.2 | 1.24E-01 | 0.3 | 2.69E-02 | -0.1 | 3.96E-01 | 0.3 | 3.99E-02 | 0.2 | 1.68E-01 | 0.1 | 7.82E-01 | T-complex protein 1 subunit theta |
| AT2G17320 | 0.2 | 7.11E-02 | 0.1 | 2.34E-01 | 0.1 | 5.98E-01 | -0.1 | 7.83E-01 | -0.1 | 6.80E-01 | 0.0 | 8.92E-01 | Pantothenate kinase |

| | | | | | | | | | | | | | |
|---|---|---|---|---|---|---|---|---|---|---|---|---|---|
| AT1G02205 | 0.2 | 2.24E-01 | 0.4 | 2.33E-02 | -0.2 | 2.53E-01 | 0.1 | 5.70E-01 | 0.0 | 9.65E-01 | 0.1 | 7.22E-01 | Fatty acid hydroxylase superfamily |
| AT3G60490 | 0.2 | 2.37E-01 | 0.3 | 4.10E-02 | -0.2 | 3.86E-01 | -0.2 | 5.32E-01 | 0.0 | 9.50E-01 | -0.2 | 5.17E-01 | Ethylene-responsive transcription factor ERF035 |
| AT3G59410 | 0.2 | 1.13E-01 | -0.2 | 1.95E-01 | 0.3 | 2.34E-02 | -0.1 | 5.25E-01 | -0.1 | 6.56E-01 | 0.0 | 9.56E-01 | non-specific serine |
| AT1G02050 | 0.2 | 1.51E-01 | 0.1 | 3.30E-01 | 0.1 | 7.31E-01 | -0.2 | 3.73E-01 | -0.2 | 4.73E-01 | 0.0 | 9.66E-01 | Type III polyketide synthase A |
| AT1G65520 | 0.2 | 5.67E-02 | -0.1 | 3.89E-01 | 0.3 | 2.66E-02 | 0.3 | 1.80E-01 | 0.1 | 8.50E-01 | 0.2 | 4.45E-01 | Enoyl-CoA delta isomerase 1, peroxisomal ,Delta-enoyl CoA isomerase 1) |
| AT1G16220 | 0.2 | 1.67E-01 | 0.3 | 3.75E-02 | -0.1 | 4.59E-01 | 0.3 | 2.82E-01 | 0.4 | 1.95E-01 | -0.1 | 8.27E-01 | Probable protein phosphatase 2C 6 |
| AT5G43830 | 0.2 | 8.42E-02 | 0.7 | 9.80E-05 | -0.5 | 1.50E-03 | -0.3 | 4.16E-03 | -0.3 | 9.28E-03 | 0.0 | 9.92E-01 | DUF3700 domain-containing protein |
| AT3G12600 | 0.2 | 1.10E-01 | 0.3 | 1.46E-02 | -0.1 | 2.90E-01 | 0.1 | 7.15E-01 | -0.1 | 5.41E-01 | 0.2 | 4.06E-01 | Nudix hydrolase 16, mitochondrial |
| AT5G48640 | 0.2 | 1.00E-01 | 0.0 | 9.92E-01 | 0.2 | 1.64E-01 | -0.1 | 7.36E-01 | 0.2 | 4.36E-01 | -0.3 | 2.56E-01 | Cyclin family protein |
| AT5G45100 | 0.2 | 8.55E-02 | -0.1 | 2.31E-01 | 0.3 | 2.09E-02 | 0.2 | 4.14E-01 | 0.2 | 5.11E-01 | 0.0 | 9.87E-01 | BOI-related E3 ubiquitin-protein ligase 1 |
| AT5G53970 | 0.2 | 2.48E-01 | -0.4 | 3.87E-02 | 0.5 | 1.11E-02 | -0.4 | 8.18E-02 | -0.7 | 1.44E-02 | 0.3 | 4.66E-01 | Probable aminotransferase TAT2 |
| AT2G27100 | 0.2 | 2.12E-02 | 0.0 | 4.96E-01 | 0.2 | 1.28E-02 | 0.1 | 6.82E-01 | 0.0 | 8.65E-01 | 0.1 | 6.64E-01 | Serrate RNA effector molecule |
| AT3G16810 | 0.2 | 2.33E-01 | 0.0 | 9.74E-01 | 0.2 | 2.92E-01 | 0.7 | 5.84E-04 | 0.3 | 1.04E-01 | 0.4 | 4.77E-02 | Pumilio homolog 24 |
| AT1G59560 | 0.2 | 1.80E-01 | 0.1 | 4.38E-01 | 0.1 | 6.68E-01 | 0.1 | 8.01E-01 | -0.2 | 6.50E-01 | 0.3 | 5.18E-01 | RING-type E3 ubiquitin transferase |
| AT1G50140 | 0.2 | 2.29E-02 | -0.1 | 2.11E-01 | 0.3 | 6.67E-03 | 0.0 | 9.02E-01 | 0.0 | 9.72E-01 | 0.0 | 9.01E-01 | P-loop containing nucleoside triphosphate hydrolases superfamily protein |
| AT2G30720 | 0.2 | 1.05E-01 | -0.1 | 4.74E-01 | 0.3 | 5.75E-02 | 0.0 | 9.29E-01 | 0.2 | 5.48E-01 | -0.2 | 5.38E-01 | At2g30720 |
| AT1G58440 | 0.2 | 4.65E-02 | 0.1 | 2.66E-01 | 0.1 | 4.00E-01 | -0.2 | 1.53E-01 | 0.0 | 9.86E-01 | -0.2 | 3.37E-01 | Squalene epoxidase 1 |
| AT5G40530 | 0.2 | 2.45E-01 | 0.2 | 2.13E-01 | 0.0 | 9.27E-01 | 0.5 | 4.10E-02 | 0.5 | 6.91E-02 | 0.0 | 9.70E-01 | Ribosomal RNA-processing protein 8 |
| AT5G06670 | 0.2 | 1.35E-01 | 0.1 | 3.90E-01 | 0.1 | 6.18E-01 | 0.0 | 8.76E-01 | -0.1 | 6.85E-01 | 0.2 | 5.82E-01 | Kinesin-like protein |
| AT2G40100 | 0.2 | 2.87E-01 | 0.1 | 4.64E-01 | 0.1 | 8.08E-01 | 0.3 | 3.31E-01 | 0.9 | 7.04E-03 | -0.6 | 1.33E-02 | Chlorophyll a-b binding protein, chloroplastic |
| AT2G40700 | 0.2 | 1.37E-01 | 0.0 | 8.38E-01 | 0.2 | 2.84E-01 | 0.5 | 1.24E-02 | 0.2 | 2.86E-01 | 0.3 | 2.77E-01 | ATP-dependent RNA helicase |
| AT5G21170 | 0.2 | 2.42E-01 | 0.7 | 1.41E-03 | -0.5 | 1.61E-02 | -0.6 | 1.22E-02 | -1.0 | 5.46E-04 | 0.4 | 1.32E-01 | 5'-AMP-activated protein kinase beta-2 subunit protein |
| AT5G47100 | 0.2 | 5.91E-02 | 0.4 | 2.30E-03 | -0.2 | 1.01E-01 | -0.2 | 2.71E-01 | -0.2 | 4.28E-01 | 0.0 | 9.12E-01 | Calcineurin B-like protein |
| AT1G75240 | 0.2 | 7.33E-02 | 0.2 | 1.03E-01 | 0.0 | 9.26E-01 | 0.0 | 9.96E-01 | -0.4 | 2.74E-02 | 0.4 | 2.40E-02 | Zinc-finger homeodomain protein 5 |
| AT1G71130 | 0.2 | 6.62E-02 | 0.0 | 8.03E-01 | 0.2 | 1.69E-01 | -0.3 | 4.31E-02 | 0.0 | 9.68E-01 | -0.3 | 9.14E-02 | Ethylene-responsive transcription factor ERF070 |
| AT5G64860 | 0.2 | 1.56E-01 | 0.2 | 1.26E-01 | 0.0 | 8.93E-01 | -0.2 | 1.64E-01 | -0.1 | 6.71E-01 | -0.1 | 5.79E-01 | 4-alpha-glucanotransferase |
| AT2G38680 | 0.2 | 2.62E-01 | 0.1 | 5.72E-01 | 0.1 | 6.76E-01 | -0.1 | 7.96E-01 | 0.0 | 9.60E-01 | -0.1 | 8.05E-01 | 5'-nucleotidase |
| AT3G12130 | 0.2 | 1.20E-02 | 0.1 | 4.05E-01 | 0.1 | 9.80E-02 | 0.1 | 4.59E-01 | 0.0 | 8.00E-01 | 0.1 | 4.35E-01 | KH domain-containing protein |
| AT5G24630 | 0.2 | 2.45E-01 | 0.1 | 4.77E-01 | 0.1 | 7.44E-01 | 0.3 | 4.61E-01 | 0.1 | 8.89E-01 | 0.2 | 6.53E-01 | Double-stranded DNA binding protein |
| AT2G42750 | 0.2 | 1.09E-01 | -0.2 | 1.41E-01 | 0.4 | 1.30E-02 | 2.0 | 1.92E-07 | 0.4 | 1.78E-01 | 1.7 | 1.62E-04 | At2g42750 |
| AT2G36305 | 0.2 | 9.54E-02 | 0.2 | 1.50E-01 | 0.0 | 8.82E-01 | -0.6 | 2.77E-02 | -0.4 | 1.62E-01 | -0.2 | 5.18E-01 | Farnesylated protein-converting enzyme 2 |
| AT2G44020 | 0.2 | 7.75E-02 | -0.1 | 4.27E-01 | 0.3 | 4.25E-02 | 0.2 | 3.48E-01 | 0.0 | 9.04E-01 | 0.2 | 4.02E-01 | Mitochondrial transcription termination factor family protein |
| AT2G32090 | 0.2 | 1.79E-01 | 0.2 | 2.70E-01 | 0.0 | 8.72E-01 | -0.5 | 4.34E-02 | -0.5 | 1.09E-01 | -0.1 | 8.78E-01 | At2g32090 |
| AT4G27910 | 0.2 | 2.50E-02 | -0.1 | 8.78E-02 | 0.3 | 4.41E-03 | 0.1 | 8.55E-01 | 0.1 | 7.79E-01 | 0.0 | 9.26E-01 | Histone-lysine N-methyltransferase ATX4 |
| AT2G37120 | 0.2 | 6.55E-02 | 0.4 | 2.51E-03 | -0.2 | 9.39E-02 | -0.1 | 7.95E-01 | -0.1 | 6.83E-01 | 0.0 | 8.87E-01 | DNA-binding protein S1FA2 |
| AT5G22060 | 0.2 | 4.28E-02 | -0.3 | 9.79E-03 | 0.4 | 9.26E-04 | 0.3 | 2.22E-01 | 0.1 | 8.46E-01 | 0.2 | 5.49E-01 | Chaperone protein dnaJ 2 |
| AT3G26610 | 0.2 | 4.46E-01 | 0.8 | 3.78E-03 | -0.6 | 3.22E-02 | 0.1 | 5.72E-01 | 0.3 | 1.84E-01 | -0.2 | 5.50E-01 | Pectin lyase-like superfamily protein |
| AT5G09350 | 0.2 | 9.31E-02 | 0.0 | 7.56E-01 | 0.1 | 2.51E-01 | -0.1 | 8.69E-01 | 0.0 | 9.43E-01 | -0.1 | 8.45E-01 | Phosphatidylinositol 4-kinase beta 2 |
| AT3G18500 | 0.2 | 7.16E-02 | 0.1 | 2.94E-01 | 0.1 | 4.98E-01 | 0.2 | 1.53E-01 | 0.2 | 3.86E-01 | 0.1 | 8.18E-01 | DNAse I-like superfamily protein |
| AT3G08580 | 0.2 | 1.26E-01 | 0.4 | 6.15E-03 | -0.2 | 1.09E-01 | 0.2 | 6.17E-02 | 0.1 | 2.73E-01 | 0.1 | 7.54E-01 | ADP |
| AT5G66350 | 0.2 | 1.79E-01 | -0.2 | 1.98E-01 | 0.4 | 4.27E-02 | 0.0 | 9.95E-01 | 0.0 | 9.60E-01 | 0.0 | 9.69E-01 | Lateral root primordium protein-like protein |
| AT4G27060 | 0.2 | 2.42E-02 | 0.1 | 2.67E-01 | 0.1 | 2.55E-01 | -0.1 | 5.35E-01 | 0.0 | 9.11E-01 | -0.1 | 7.58E-01 | Microtubule-associated protein TORTIFOLIA1 |
| AT1G48040 | 0.2 | 4.14E-02 | 0.1 | 3.67E-01 | 0.1 | 3.06E-01 | 0.5 | 1.71E-02 | 0.0 | 8.82E-01 | 0.4 | 4.96E-02 | Probable protein phosphatase 2C 13 |
| AT1G75820 | 0.2 | 9.36E-02 | 0.3 | 2.02E-02 | -0.1 | 3.95E-01 | -0.1 | 7.04E-01 | -0.1 | 6.48E-01 | 0.0 | 9.35E-01 | Receptor protein kinase CLAVATA1 |
| AT3G11620 | 0.2 | 7.86E-02 | 0.3 | 1.98E-02 | -0.1 | 4.82E-01 | -0.2 | 3.56E-01 | -0.4 | 1.43E-01 | 0.2 | 6.22E-01 | Alpha |
| AT2G46510 | 0.2 | 2.18E-01 | 0.0 | 8.06E-01 | 0.2 | 2.18E-01 | -0.3 | 2.29E-01 | -0.1 | 8.57E-01 | -0.2 | 5.18E-01 | Transcription factor ABA-INDUCIBLE bHLH-TYPE |
| AT2G24200 | 0.2 | 2.98E-02 | 0.2 | 1.69E-02 | 0.0 | 7.04E-01 | 0.0 | 9.29E-01 | 0.0 | 8.36E-01 | 0.0 | 9.23E-01 | Putative leucine aminopeptidase |
| AT3G51030 | 0.2 | 3.14E-01 | 0.3 | 7.55E-02 | -0.2 | 4.62E-01 | 0.0 | 9.79E-01 | 0.3 | 4.98E-01 | -0.3 | 5.32E-01 | Thioredoxin H1 |
| AT4G12340 | 0.2 | 1.18E-01 | 0.3 | 2.22E-02 | -0.1 | 3.82E-01 | 0.0 | 9.17E-01 | -0.1 | 5.76E-01 | 0.1 | 5.95E-01 | AT4g12340 |
| AT4G09570 | 0.2 | 1.03E-01 | 0.1 | 4.97E-01 | 0.1 | 3.97E-01 | 0.3 | 4.75E-02 | 0.1 | 5.06E-01 | 0.1 | 4.26E-01 | Calcium-dependent protein kinase 4 |
| AT5G26610 | 0.2 | 1.97E-02 | -0.1 | 7.22E-02 | 0.3 | 2.71E-03 | -0.3 | 1.37E-01 | 0.0 | 9.79E-01 | -0.3 | 2.20E-01 | G-patch domain-containing protein |
| AT3G45300 | 0.2 | 1.32E-01 | 0.1 | 4.16E-01 | 0.1 | 5.61E-01 | -0.2 | 2.82E-01 | -0.7 | 3.56E-03 | 0.5 | 5.65E-02 | Isovaleryl-CoA dehydrogenase, mitochondrial |
| AT5G45300 | 0.2 | 2.53E-02 | 0.0 | 9.64E-01 | 0.2 | 5.65E-02 | 0.1 | 6.87E-01 | 0.1 | 4.92E-01 | -0.1 | 8.29E-01 | Beta-amylase |
| AT4G32730 | 0.2 | 1.49E-02 | 0.0 | 6.53E-01 | 0.2 | 6.80E-02 | 0.3 | 1.54E-01 | 0.2 | 2.43E-01 | 0.0 | 9.61E-01 | Homeodomain-like protein |
| AT3G04940 | 0.2 | 5.76E-02 | 0.2 | 3.53E-02 | 0.0 | 7.81E-01 | -0.2 | 1.50E-01 | -0.2 | 1.13E-01 | 0.0 | 8.62E-01 | Bifunctional L-3-cyanoalanine synthase |
| AT3G54280 | 0.2 | 5.20E-02 | 0.0 | 9.63E-01 | 0.2 | 7.88E-02 | 0.3 | 8.41E-02 | 0.2 | 4.06E-01 | 0.1 | 6.30E-01 | ROOT GROWTH DEFECTIVE 3 |
| AT1G31230 | 0.2 | 6.96E-02 | 0.2 | 3.44E-02 | 0.0 | 6.88E-01 | 0.2 | 4.99E-01 | 0.5 | 5.24E-02 | -0.4 | 2.65E-01 | Bifunctional aspartokinase |
| AT5G37020 | 0.2 | 1.03E-01 | -0.1 | 4.25E-01 | 0.3 | 4.39E-02 | 0.2 | 6.03E-01 | 0.2 | 4.81E-01 | -0.1 | 8.76E-01 | Auxin response factor |
| AT3G12920 | 0.2 | 6.38E-02 | 0.3 | 1.56E-02 | -0.1 | 4.53E-01 | 0.1 | 5.43E-01 | -0.1 | 7.80E-01 | 0.1 | 4.53E-01 | Probable BOI-related E3 ubiquitin-protein ligase 3 |
| AT3G18030 | 0.2 | 4.82E-02 | 0.1 | 1.74E-01 | 0.1 | 5.83E-01 | -0.2 | 2.41E-01 | -0.3 | 7.08E-02 | 0.1 | 5.41E-01 | Phosphopantothenoylcysteine decarboxylase |
| AT1G68020 | 0.2 | 2.43E-02 | 0.2 | 6.22E-02 | 0.0 | 7.15E-01 | -0.5 | 1.02E-03 | -0.6 | 5.07E-04 | 0.1 | 5.91E-01 | Alpha,alpha-trehalose-phosphate synthase 6 |
| AT4G22720 | 0.2 | 4.59E-02 | 0.1 | 5.32E-01 | 0.1 | 2.14E-01 | -0.1 | 7.40E-01 | -0.2 | 2.37E-01 | 0.1 | 5.20E-01 | Glycoprotease 2 |
| AT3G24140 | 0.2 | 2.28E-01 | 0.1 | 6.79E-01 | 0.1 | 5.29E-01 | 0.1 | 8.43E-01 | 0.0 | 9.46E-01 | 0.0 | 9.48E-01 | Basic helix-loop-helix DNA-binding superfamily protein |
| AT3G09650 | 0.2 | 3.00E-02 | 0.0 | 6.70E-01 | 0.2 | 1.08E-01 | 0.0 | 9.40E-01 | 0.2 | 4.79E-01 | -0.1 | 6.12E-01 | Pentatricopeptide repeat-containing protein At3g09650, chloroplastic |
| AT1G16280 | 0.2 | 2.75E-02 | 0.1 | 4.80E-01 | 0.1 | 1.73E-01 | 0.5 | 2.15E-02 | 0.3 | 1.34E-01 | 0.1 | 6.02E-01 | DEAD-box ATP-dependent RNA helicase 36 |
| AT1G61620 | 0.2 | 1.09E-02 | 0.1 | 1.39E-01 | 0.1 | 2.35E-01 | -0.2 | 6.68E-02 | -0.3 | 2.29E-02 | 0.1 | 6.48E-01 | E3 ubiquitin-protein ligase CSU1 |
| AT2G36480 | 0.2 | 9.70E-03 | -0.1 | 3.06E-01 | 0.3 | 5.19E-03 | 0.0 | 8.56E-01 | -0.1 | 5.76E-01 | 0.1 | 7.35E-01 | Uncharacterized protein At2g36475 |
| AT4G24020 | 0.2 | 6.02E-02 | 0.0 | 6.66E-01 | 0.1 | 1.81E-01 | -0.2 | 1.07E-01 | -0.1 | 3.43E-01 | -0.1 | 7.84E-01 | NIN like protein 7 |
| AT3G50700 | 0.2 | 7.08E-02 | 0.4 | 2.71E-03 | -0.2 | 1.01E-01 | 0.0 | 9.23E-01 | -0.3 | 1.34E-01 | 0.3 | 2.03E-01 | Zinc finger protein GAI-ASSOCIATED FACTOR 1 |
| AT2G45880 | 0.2 | 4.15E-02 | -0.2 | 3.23E-02 | 0.4 | 2.99E-03 | 0.0 | 8.76E-01 | -0.1 | 6.59E-01 | 0.1 | 8.26E-01 | Beta-amylase 7 |
| AT3G45700 | 0.2 | 3.78E-01 | 1.0 | 6.26E-04 | -0.8 | 7.24E-03 | -1.5 | 1.00E-02 | -0.5 | 2.84E-01 | -0.9 | 2.03E-01 | Protein NRT1 |
| AT2G02765 | 0.2 | 2.99E-01 | -0.2 | 2.46E-01 | 0.4 | 8.22E-02 | -0.2 | 5.88E-01 | -0.2 | 5.14E-01 | 0.1 | 9.09E-01 | Uncharacterized protein At2g02770 |
| AT4G14500 | 0.2 | 2.05E-02 | 0.1 | 3.22E-01 | 0.1 | 1.88E-01 | -0.1 | 6.96E-01 | -0.2 | 6.66E-01 | 0.0 | 9.48E-01 | Polyketide cyclase |
| AT5G66900 | 0.2 | 1.22E-01 | 0.0 | 8.95E-01 | 0.2 | 1.51E-01 | 0.3 | 2.53E-02 | -0.4 | 1.63E-02 | 0.7 | 6.65E-04 | Disease resistance protein family |
| AT1G06900 | 0.2 | 9.57E-03 | 0.2 | 4.13E-03 | 0.0 | 5.88E-01 | 0.0 | 7.90E-01 | 0.0 | 8.78E-01 | 0.0 | 9.67E-01 | Nardilysin-like |
| AT4G28390 | 0.2 | 5.18E-01 | -0.7 | 6.42E-02 | 0.9 | 4.18E-02 | -0.1 | 7.32E-01 | -0.6 | 1.68E-01 | 0.4 | 3.97E-01 | ADP,ATP carrier protein 3, mitochondrial |
| AT5G25265 | 0.2 | 1.99E-02 | 0.4 | 6.20E-04 | -0.2 | 4.39E-02 | 0.0 | 8.88E-01 | -0.3 | 4.32E-03 | 0.3 | 2.83E-02 | Hydroxyproline O-arabinosyltransferase 1 |
| AT1G67120 | 0.2 | 6.94E-02 | -0.2 | 6.29E-02 | 0.4 | 5.94E-03 | 0.7 | 7.50E-04 | 0.0 | 9.81E-01 | 0.7 | 5.86E-03 | Midasin |
| AT1G20696 | 0.2 | 1.31E-01 | 0.1 | 3.24E-01 | 0.1 | 6.59E-01 | 0.1 | 6.69E-01 | -0.1 | 8.75E-01 | 0.2 | 6.59E-01 | HMG box domain-containing protein |
| AT1G52410 | 0.2 | 2.03E-01 | 0.3 | 4.61E-02 | -0.1 | 4.35E-01 | 0.7 | 6.65E-02 | 0.7 | 8.05E-02 | 0.0 | 9.50E-01 | TSK-associating protein 1 |
| AT1G61850 | 0.2 | 2.88E-02 | 0.1 | 3.71E-01 | 0.1 | 2.14E-01 | 0.0 | 9.98E-01 | 0.0 | 9.16E-01 | 0.0 | 9.22E-01 | Phospholipase A I-like protein |
| AT1G69526 | 0.2 | 1.53E-01 | 0.4 | 8.58E-03 | -0.2 | 1.66E-01 | -0.4 | 3.53E-01 | 0.1 | 7.77E-01 | -0.5 | 2.64E-01 | Methyltransferase type 11 domain-containing protein |
| AT1G24510 | 0.2 | 6.10E-02 | 0.2 | 3.09E-02 | 0.0 | 6.78E-01 | 0.1 | 3.97E-01 | 0.0 | 9.11E-01 | 0.1 | 6.87E-01 | T-complex protein 1 subunit epsilon |

| | | | | | | | | | | | | | |
|---|---|---|---|---|---|---|---|---|---|---|---|---|---|
| AT4G09510 | 0.2 | 4.56E-02 | 0.3 | 8.85E-03 | -0.1 | 3.84E-01 | 0.1 | 6.15E-01 | 0.2 | 2.02E-01 | -0.1 | 5.33E-01 | Alkaline |
| AT2G34770 | 0.2 | 1.67E-01 | 0.2 | 1.59E-01 | 0.0 | 9.52E-01 | -0.8 | 6.40E-05 | -0.9 | 8.26E-05 | 0.1 | 6.96E-01 | Dihydroceramide fatty acyl 2-hydroxylase FAH1 |
| AT5G09840 | 0.2 | 1.15E-01 | 0.3 | 2.41E-02 | -0.1 | 4.41E-01 | 0.9 | 2.32E-03 | 0.3 | 3.31E-01 | 0.6 | 6.42E-02 | Emb\|CAB71880.1 |
| AT4G31460 | 0.2 | 1.19E-01 | 0.3 | 2.60E-02 | -0.1 | 4.33E-01 | 0.1 | 4.24E-01 | 0.0 | 8.87E-01 | 0.1 | 7.21E-01 | Large ribosomal subunit protein bL28c |
| AT2G40000 | 0.2 | 4.30E-01 | -0.4 | 1.46E-01 | 0.6 | 5.27E-02 | -0.9 | 6.09E-03 | -1.3 | 1.18E-03 | 0.4 | 4.22E-01 | Nematode resistance protein-like HSPRO2 |
| AT4G27680 | 0.2 | 1.24E-02 | 0.0 | 7.09E-01 | 0.2 | 1.55E-02 | 0.2 | 2.51E-01 | 0.2 | 2.54E-01 | 0.0 | 9.40E-01 | AAA+ ATPase domain-containing protein |
| AT5G66510 | 0.2 | 4.78E-02 | 0.4 | 1.93E-03 | -0.2 | 7.81E-02 | 0.2 | 1.20E-01 | 0.1 | 4.42E-01 | 0.1 | 7.21E-01 | Gamma carbonic anhydrase 3, mitochondrial |
| AT2G33980 | 0.2 | 3.14E-01 | 0.0 | 8.83E-01 | 0.2 | 3.50E-01 | -0.2 | 4.70E-01 | -0.1 | 7.20E-01 | -0.1 | 8.67E-01 | hypothetical protein |
| AT4G28010 | 0.2 | 1.46E-01 | 0.1 | 5.56E-01 | 0.1 | 4.98E-01 | 0.4 | 3.40E-01 | 0.5 | 3.14E-01 | -0.1 | 9.13E-01 | Pentatricopeptide repeat-containing protein At4g28010 |
| AT3G50000 | 0.2 | 8.92E-03 | 0.1 | 1.26E-01 | 0.1 | 2.14E-01 | -0.1 | 3.24E-01 | -0.1 | 2.31E-01 | 0.0 | 8.49E-01 | Casein kinase II subunit alpha-2 |
| AT5G55310 | 0.2 | 2.66E-02 | -0.3 | 4.02E-03 | 0.5 | 4.71E-04 | 0.1 | 7.57E-01 | 0.1 | 4.36E-01 | -0.1 | 7.31E-01 | DNA topoisomerase 1 beta |
| AT4G15940 | 0.2 | 5.71E-02 | 0.2 | 6.84E-02 | 0.0 | 9.79E-01 | 0.5 | 7.16E-03 | 0.2 | 2.43E-01 | 0.2 | 3.01E-01 | Probable acylpyruvase FAHD1, mitochondrial |
| AT5G55070 | 0.2 | 4.38E-02 | 0.1 | 1.82E-01 | 0.1 | 5.10E-01 | 0.6 | 8.39E-05 | 0.3 | 1.17E-02 | 0.2 | 1.51E-01 | Dihydrolipoyllysine-residue succinyltransferase component of 2-oxoglutarate dehydrogenase complex 1, mitochondrial |
| AT5G01180 | 0.2 | 2.57E-01 | 0.3 | 1.06E-01 | -0.1 | 6.32E-01 | -0.3 | 5.83E-01 | -0.8 | 1.77E-01 | 0.5 | 4.75E-01 | Protein NRT1 |
| AT3G12620 | 0.2 | 1.27E-02 | 0.0 | 8.38E-01 | 0.2 | 3.70E-02 | 0.0 | 8.63E-01 | 0.0 | 9.41E-01 | 0.0 | 9.63E-01 | Probable protein phosphatase 2C 38 |
| AT4G33580 | 0.2 | 4.17E-02 | 0.2 | 1.69E-02 | -0.1 | 6.02E-01 | 0.0 | 9.96E-01 | 0.1 | 5.98E-01 | -0.1 | 6.35E-01 | Beta carbonic anhydrase 5, chloroplastic |
| AT3G02030 | 0.2 | 8.69E-02 | 0.1 | 3.20E-01 | 0.1 | 5.69E-01 | 0.0 | 9.93E-01 | -0.1 | 8.54E-01 | | | Transferase |
| AT3G12690 | 0.2 | 7.16E-02 | 0.2 | 4.45E-02 | 0.0 | 7.95E-01 | -0.2 | 6.04E-01 | -0.1 | 8.71E-01 | -0.1 | 8.44E-01 | Serine |
| AT3G03000 | 0.2 | 1.13E-01 | -0.1 | 3.71E-01 | 0.3 | 4.54E-02 | -0.3 | 2.75E-01 | -0.4 | 1.49E-01 | 0.1 | 7.53E-01 | Probable calcium-binding protein CML18 |
| AT3G16060 | 0.2 | 1.39E-02 | 0.1 | 2.82E-01 | 0.1 | 1.47E-01 | -0.2 | 2.52E-01 | 0.0 | 8.83E-01 | -0.1 | 5.35E-01 | Kinesin-like protein KIN-13B |
| AT5G28540 | 0.2 | 2.00E-01 | 0.1 | 5.42E-01 | 0.1 | 5.73E-01 | 0.6 | 2.78E-05 | 0.5 | 8.06E-04 | 0.1 | 4.45E-01 | Heat shock 70 kDa protein BIP1 |
| AT4G28610 | 0.2 | 1.06E-02 | 0.0 | 4.85E-01 | 0.2 | 7.37E-03 | -0.2 | 7.98E-02 | -0.4 | 4.28E-03 | 0.2 | 2.34E-01 | Protein PHOSPHATE STARVATION RESPONSE 1 |
| AT2G01570 | 0.2 | 3.65E-02 | 0.1 | 3.39E-01 | 0.1 | 2.65E-01 | -0.2 | 3.56E-02 | -0.3 | 1.53E-02 | 0.1 | 7.05E-01 | DELLA protein RGA |
| AT5G55380 | 0.2 | 1.17E-01 | -0.1 | 4.96E-01 | 0.3 | 7.10E-02 | -0.5 | 7.43E-02 | -0.1 | 6.40E-01 | -0.3 | 3.59E-01 | Probable long-chain-alcohol O-fatty-acyltransferase 1 |
| AT1G32490 | 0.2 | 5.93E-03 | 0.0 | 9.77E-01 | 0.2 | 1.43E-02 | 0.1 | 4.39E-01 | -0.1 | 8.01E-01 | 0.2 | 3.84E-01 | RNA helicase |
| AT1G09280 | 0.2 | 4.65E-02 | 0.1 | 3.63E-01 | 0.1 | 3.25E-01 | -0.4 | 3.75E-02 | -0.4 | 2.79E-02 | 0.1 | 8.39E-01 | Rhodanese-like domain-containing protein 6 |
| AT5G40650 | 0.2 | 2.23E-02 | 0.2 | 2.38E-02 | 0.0 | 9.66E-01 | -0.3 | 7.80E-02 | 0.1 | 5.44E-01 | -0.4 | 4.84E-02 | Succinate dehydrogenase iron-sulfur subunit, mitochondrial |
| AT5G41900 | 0.2 | 5.43E-01 | -0.2 | 6.29E-01 | 0.4 | 3.49E-01 | -1.0 | 8.53E-04 | -0.3 | 2.65E-01 | -0.7 | 4.59E-02 | Probable lysophospholipase BODYGUARD 2 |
| AT1G62770 | 0.2 | 4.01E-02 | 0.4 | 7.46E-04 | -0.2 | 4.12E-02 | -0.6 | 6.22E-03 | -0.6 | 1.53E-02 | 0.0 | 9.07E-01 | Pectinesterase inhibitor 9 |
| AT5G58003 | 0.2 | 7.00E-02 | 0.0 | 7.55E-01 | 0.2 | 2.00E-01 | 0.1 | 4.27E-01 | 0.1 | 7.79E-01 | 0.1 | 7.87E-01 | RNA polymerase II C-terminal domain phosphatase-like |
| AT1G03960 | 0.2 | 2.22E-02 | -0.2 | 7.10E-02 | 0.4 | 3.50E-03 | 0.1 | 7.90E-01 | 0.3 | 4.00E-01 | -0.2 | 6.61E-01 | Calcium-binding EF hand family protein |
| AT2G22640 | 0.2 | 1.78E-01 | -0.1 | 6.08E-01 | 0.3 | 1.24E-01 | 0.0 | 8.73E-01 | -0.1 | 7.13E-01 | 0.1 | 8.71E-01 | Protein BRICK 1 |
| AT1G16590 | 0.2 | 1.94E-01 | 0.0 | 9.35E-01 | 0.2 | 3.30E-01 | -0.2 | 5.19E-01 | 0.1 | 8.55E-01 | -0.3 | 4.11E-01 | DNA-binding HORMA family protein |
| AT5G04130 | 0.2 | 1.62E-02 | 0.3 | 3.14E-03 | -0.1 | 3.41E-01 | 0.5 | 5.22E-03 | 0.3 | 9.88E-02 | 0.2 | 3.93E-01 | DNA gyrase subunit B, mitochondrial |
| AT5G23760 | 0.2 | 1.26E-02 | 0.0 | 9.20E-01 | 0.2 | 3.46E-02 | 0.3 | 4.34E-01 | 0.1 | 8.31E-01 | 0.2 | 7.57E-01 | HMA domain-containing protein |
| AT1G18800 | 0.2 | 9.94E-02 | 0.2 | 1.60E-01 | 0.0 | 8.54E-01 | 0.5 | 9.85E-03 | 0.2 | 3.72E-01 | 0.3 | 2.25E-01 | NRP2 |
| AT4G30250 | 0.2 | 1.69E-01 | 0.3 | 7.14E-02 | -0.1 | 6.59E-01 | 0.3 | 4.43E-01 | 0.1 | 8.93E-01 | 0.3 | 6.65E-01 | P-loop containing nucleoside triphosphate hydrolases superfamily protein |
| AT5G25900 | 0.2 | 1.31E-02 | 0.1 | 6.66E-02 | 0.1 | 4.80E-01 | -0.1 | 4.82E-01 | -0.3 | 3.95E-02 | 0.2 | 2.56E-01 | Ent-kaurene oxidase, chloroplastic |
| AT4G17490 | 0.2 | 3.20E-01 | -0.6 | 1.72E-02 | 0.8 | 8.56E-03 | 1.3 | 3.07E-05 | 0.1 | 7.60E-01 | 1.2 | 7.94E-04 | Ethylene-responsive transcription factor 6 |
| AT4G20940 | 0.2 | 3.68E-02 | 0.3 | 1.59E-02 | 0.0 | 6.47E-01 | 0.0 | 9.77E-01 | -0.1 | 7.40E-01 | 0.1 | 7.87E-01 | Protein kinase domain-containing protein |
| AT3G14660 | 0.2 | 1.86E-02 | 0.1 | 1.59E-01 | 0.1 | 3.14E-01 | -0.3 | 2.88E-01 | -0.3 | 2.95E-01 | 0.0 | 9.37E-01 | Cytochrome P450, family 72, subfamily A, polypeptide 13 |
| AT5G39960 | 0.2 | 6.35E-02 | 0.0 | 6.73E-01 | 0.2 | 2.20E-01 | 0.2 | 3.79E-01 | 0.1 | 7.91E-01 | 0.1 | 7.07E-01 | GTPase Der |
| AT1G03990 | 0.2 | 2.06E-01 | 0.0 | 9.49E-01 | 0.2 | 3.46E-01 | 1.1 | 9.35E-03 | 0.8 | 8.62E-02 | 0.3 | 4.34E-01 | Long-chain-alcohol oxidase |
| AT1G80510 | 0.2 | 1.61E-02 | 0.0 | 6.69E-01 | 0.2 | 7.20E-02 | 0.1 | 6.28E-01 | 0.0 | 9.52E-01 | 0.1 | 6.52E-01 | Amino acid transporter AVT6E |
| AT1G64620 | 0.2 | 3.39E-02 | 0.1 | 5.50E-01 | 0.1 | 1.81E-01 | 0.0 | 8.58E-01 | 0.1 | 6.19E-01 | -0.1 | 7.85E-01 | Dof zinc finger protein DOF1.8 |
| AT3G14740 | 0.2 | 2.09E-01 | 0.5 | 7.79E-03 | -0.3 | 1.16E-01 | 0.0 | 8.92E-01 | 0.2 | 5.97E-01 | -0.1 | 7.24E-01 | hypothetical protein |
| AT2G38040 | 0.2 | 1.33E-02 | 0.1 | 7.58E-02 | 0.1 | 4.44E-01 | 0.0 | 7.78E-01 | -0.1 | 3.68E-01 | 0.1 | 6.62E-01 | Carboxyltransferase alpha subunit |
| AT2G05210 | 0.2 | 7.35E-02 | 0.2 | 1.04E-01 | 0.0 | 9.25E-01 | -0.2 | 3.48E-01 | 0.0 | 9.87E-01 | -0.2 | 5.01E-01 | Protection of telomeres protein 1a |
| AT1G05160 | 0.2 | 3.40E-02 | 0.0 | 9.03E-01 | 0.2 | 8.88E-02 | -0.5 | 3.20E-02 | -0.3 | 1.62E-01 | -0.2 | 6.73E-01 | Ent-kaurenoic acid oxidase 1 |
| AT5G27620 | 0.2 | 6.54E-02 | 0.2 | 1.26E-01 | 0.0 | 8.09E-01 | 0.5 | 2.07E-02 | 0.0 | 9.65E-01 | 0.5 | 4.65E-02 | Cyclin H1 |
| AT4G37750 | 0.2 | 2.37E-02 | 0.1 | 2.20E-01 | 0.1 | 3.12E-01 | 0.0 | 9.45E-01 | 0.1 | 7.85E-01 | -0.1 | 7.66E-01 | AP2-like ethylene-responsive transcription factor ANT |
| AT1G28210 | 0.2 | 9.92E-02 | -0.1 | 3.42E-01 | 0.3 | 4.46E-02 | 1.1 | 2.87E-03 | 0.4 | 3.91E-01 | 0.7 | 5.71E-02 | DNAJ heat shock family protein |
| AT5G05600 | 0.2 | 1.46E-01 | 0.3 | 5.54E-02 | -0.1 | 6.17E-01 | 0.2 | 6.86E-01 | 0.6 | 2.83E-01 | -0.3 | 6.23E-01 | 2-oxoglutarate and Fe-dependent oxygenase superfamily protein |
| AT4G04700 | 0.2 | 1.36E-02 | 0.3 | 3.03E-03 | -0.1 | 3.83E-01 | 0.7 | 8.65E-04 | 0.6 | 7.21E-03 | 0.1 | 5.51E-01 | Calcium-dependent protein kinase 27 |
| AT1G20330 | 0.2 | 2.42E-02 | 0.2 | 1.42E-02 | 0.0 | 7.09E-01 | -0.2 | 2.06E-01 | -0.3 | 1.06E-01 | 0.1 | 7.53E-01 | 24-methylenesterol C-methyltransferase 2 |
| AT2G25710 | 0.2 | 1.75E-01 | 0.2 | 2.62E-01 | 0.0 | 8.77E-01 | 0.1 | 7.29E-01 | 0.1 | 7.50E-01 | 0.0 | 9.87E-01 | Holocarboxylase synthase 1 |
| AT1G29850 | 0.2 | 1.78E-02 | 0.1 | 1.29E-01 | 0.1 | 3.61E-01 | 0.0 | 9.14E-01 | 0.0 | 9.27E-01 | 0.0 | 9.97E-01 | Double-stranded DNA-binding family protein |
| AT4G09640 | 0.2 | 2.02E-02 | 0.2 | 1.65E-02 | 0.0 | 8.72E-01 | 0.0 | 9.99E-01 | -0.1 | 8.52E-01 | 0.1 | 8.76E-01 | Probable magnesium transporter NIPA5 |
| AT3G42790 | 0.2 | 7.78E-03 | 0.1 | 2.22E-01 | 0.1 | 1.13E-01 | 0.0 | 9.86E-01 | -0.2 | 1.31E-01 | 0.3 | 2.31E-01 | PHD finger protein ALFIN-LIKE 3 |
| AT3G48350 | 0.2 | 8.00E-02 | 0.6 | 4.35E-04 | -0.4 | 1.47E-02 | 0.1 | 7.97E-01 | 0.2 | 5.05E-01 | -0.1 | 6.98E-01 | Cysteine proteinases superfamily protein |
| AT1G72090 | 0.2 | 1.04E-02 | 0.1 | 8.13E-02 | 0.1 | 3.63E-01 | -0.2 | 1.94E-01 | -0.1 | 6.62E-01 | -0.1 | 6.21E-01 | Threonylcarbamoyladenosine tRNA methylthiotransferase A37 methylthiotransferase) |
| AT5G01500 | 0.2 | 3.62E-02 | 0.4 | 1.22E-03 | -0.2 | 7.33E-02 | 0.2 | 1.37E-01 | 0.2 | 3.64E-01 | 0.1 | 8.11E-01 | Thylakoid ADP,ATP carrier protein, chloroplastic |
| AT5G50000 | 0.2 | 2.25E-02 | 0.3 | 1.83E-03 | -0.1 | 1.46E-01 | -0.3 | 2.72E-03 | -0.3 | 2.28E-02 | -0.1 | 6.97E-01 | Protein kinase |
| AT3G53580 | 0.2 | 1.44E-02 | 0.3 | 2.80E-03 | -0.1 | 3.13E-01 | 0.2 | 1.54E-01 | 0.0 | 9.96E-01 | 0.2 | 3.31E-01 | Diaminopimelate epimerase, chloroplastic |
| AT5G48960 | 0.2 | 2.70E-02 | 0.2 | 1.50E-02 | 0.0 | 7.27E-01 | 0.3 | 9.40E-02 | -0.1 | 7.98E-01 | 0.3 | 1.14E-01 | HAD-superfamily hydrolase, subfamily IG, 5'-nucleotidase |
| AT3G24010 | 0.2 | 1.75E-02 | 0.1 | 7.91E-02 | 0.1 | 5.31E-01 | -0.1 | 5.73E-01 | 0.0 | 9.45E-01 | -0.1 | 7.21E-01 | AT3G24010 protein |
| AT3G18110 | 0.2 | 9.78E-02 | 0.4 | 8.89E-03 | -0.2 | 2.25E-01 | 0.4 | 5.39E-02 | 0.5 | 2.41E-02 | -0.1 | 6.83E-01 | Pentatricopeptide repeat-containing protein At3g18110, chloroplastic |
| AT3G57030 | 0.2 | 1.35E-02 | 0.3 | 8.45E-04 | -0.1 | 1.08E-01 | 0.0 | 9.49E-01 | -0.2 | 2.47E-01 | 0.2 | 3.20E-01 | Protein STRICTOSIDINE SYNTHASE-LIKE 10 |
| AT5G01670 | 0.2 | 2.14E-01 | -0.1 | 4.54E-01 | 0.3 | 1.08E-01 | 0.4 | 1.86E-01 | -0.1 | 8.57E-01 | 0.5 | 9.13E-02 | NAD-linked oxidoreductase superfamily protein |
| AT3G05640 | 0.2 | 1.22E-01 | 0.8 | 1.06E-04 | -0.6 | 2.12E-03 | -1.0 | 6.73E-03 | -0.4 | 2.61E-01 | -0.6 | 2.01E-01 | Probable protein phosphatase 2C 34 |
| AT5G46270 | 0.2 | 5.90E-02 | 0.1 | 3.25E-01 | 0.1 | 4.17E-01 | 0.1 | 6.95E-01 | -0.1 | 7.54E-01 | 0.2 | 4.93E-01 | ADP-ribosyl cyclase |
| AT2G43810 | 0.2 | 5.32E-02 | 0.2 | 8.79E-02 | 0.0 | 8.69E-01 | 0.3 | 1.10E-01 | 0.2 | 2.68E-01 | 0.1 | 8.35E-01 | Sm-like protein LSM36B |
| AT1G80270 | 0.2 | 1.14E-01 | 0.2 | 7.77E-02 | 0.0 | 8.26E-01 | 0.8 | 5.60E-04 | 0.5 | 2.57E-02 | 0.3 | 2.83E-01 | Pentatricopeptide repeat-containing protein At1g80270, mitochondrial |
| AT5G10480 | 0.2 | 5.14E-03 | 0.3 | 8.54E-04 | -0.1 | 2.19E-01 | 0.3 | 5.06E-03 | 0.3 | 1.94E-02 | 0.0 | 8.74E-01 | Very-long-chain -3-hydroxyacyl-CoA dehydratase |
| AT4G18460 | 0.2 | 6.20E-02 | 0.3 | 6.83E-03 | -0.1 | 2.69E-01 | -0.1 | 7.83E-01 | 0.0 | 9.40E-01 | -0.1 | 8.89E-01 | D-aminoacyl-tRNA deacylase |
| AT2G45240 | 0.2 | 3.40E-02 | 0.1 | 2.31E-01 | 0.1 | 3.66E-01 | 0.2 | 3.22E-01 | -0.2 | 3.86E-01 | 0.3 | 1.26E-01 | Methionine aminopeptidase |
| AT4G13630 | 0.2 | 1.11E-02 | 0.0 | 8.91E-01 | 0.2 | 3.20E-02 | 0.1 | 7.81E-01 | 0.1 | 6.29E-01 | 0.0 | 8.70E-01 | GTD-binding domain-containing protein |
| AT3G26340 | 0.2 | 4.51E-02 | 0.2 | 1.25E-01 | 0.0 | 6.72E-01 | 0.2 | 1.59E-01 | 0.0 | 9.24E-01 | 0.2 | 3.20E-01 | Proteasome subunit beta type-5-B |
| AT2G01120 | 0.2 | 1.46E-01 | 0.3 | 6.23E-02 | -0.1 | 6.65E-01 | 0.3 | 1.67E-01 | -0.2 | 6.59E-01 | 0.5 | 8.56E-02 | Origin of replication complex subunit 4 |
| AT2G24765 | 0.2 | 1.62E-02 | 0.4 | 8.85E-04 | -0.2 | 9.21E-02 | 0.0 | 9.26E-01 | -0.1 | 7.64E-01 | 0.0 | 8.69E-01 | ADP-ribosylation factor 3 |
| AT1G19510 | 0.2 | 2.35E-01 | 0.5 | 8.68E-03 | -0.3 | 1.18E-01 | 0.5 | 9.79E-02 | -0.1 | 8.05E-01 | 0.6 | 1.22E-01 | Protein RADIALIS-like 5 |
| AT4G25630 | 0.2 | 3.23E-01 | 0.4 | 9.88E-02 | -0.2 | 4.88E-01 | 0.6 | 5.80E-04 | 0.5 | 5.85E-03 | 0.1 | 6.39E-01 | rRNA 2'-O-methyltransferase fibrillarin 2 |
| AT1G52310 | 0.2 | 1.65E-02 | -0.1 | 2.11E-01 | 0.3 | 5.45E-03 | 0.2 | 3.38E-01 | 0.1 | 5.17E-01 | 0.0 | 9.12E-01 | Protein kinase family protein |

| | | | | | | | | | | | | | |
|---|---|---|---|---|---|---|---|---|---|---|---|---|---|
| AT3G01610 | 0.2 | 1.02E-02 | -0.1 | 1.71E-01 | 0.3 | 3.59E-03 | 0.0 | 9.62E-01 | 0.0 | 9.98E-01 | 0.0 | 9.78E-01 | Cell division cycle 48C |
| AT1G72130 | 0.2 | 2.47E-02 | 0.2 | 5.57E-02 | 0.0 | 7.51E-01 | 0.2 | 4.79E-01 | 0.4 | 1.56E-01 | -0.2 | 5.50E-01 | Protein NRT1 |
| AT1G18590 | 0.2 | 1.31E-01 | 0.5 | 2.49E-03 | -0.3 | 4.83E-02 | 0.2 | 4.31E-01 | 0.6 | 1.83E-02 | -0.4 | 1.53E-01 | Cytosolic sulfotransferase 17 |
| AT5G42740 | 0.2 | 1.15E-02 | 0.2 | 7.02E-03 | 0.0 | 7.12E-01 | 0.4 | 6.04E-03 | 0.2 | 1.42E-01 | 0.2 | 3.97E-01 | Glucose-6-phosphate isomerase |
| AT3G47060 | 0.2 | 6.27E-03 | -0.3 | 6.42E-04 | 0.5 | 9.47E-05 | -0.1 | 8.05E-01 | -0.1 | 9.38E-01 | -0.1 | 9.27E-01 | ATP-dependent zinc metalloprotease FTSH 7, chloroplastic |
| AT1G09660 | 0.2 | 1.75E-02 | 0.2 | 4.94E-02 | 0.0 | 6.79E-01 | -0.1 | 3.95E-01 | 0.0 | 8.98E-01 | -0.1 | 6.72E-01 | KH domain-containing protein At1g09660 |
| AT4G28270 | 0.2 | 1.35E-01 | 0.1 | 4.40E-01 | 0.1 | 5.52E-01 | 0.3 | 8.99E-03 | 0.1 | 6.28E-01 | 0.3 | 1.09E-01 | E3 ubiquitin-protein ligase RMA2 |
| AT5G10980 | 0.2 | 1.11E-02 | 0.1 | 7.66E-02 | 0.1 | 3.88E-01 | -0.2 | 3.18E-01 | -0.3 | 1.58E-01 | 0.1 | 7.24E-01 | Histone H3.3 |
| AT1G32930 | 0.2 | 4.47E-02 | 0.2 | 6.92E-02 | 0.0 | 9.05E-01 | 0.1 | 6.80E-01 | 0.2 | 4.23E-01 | -0.1 | 7.48E-01 | Beta-1,6-galactosyltransferase GALT31A |
| AT4G23210 | 0.2 | 5.70E-01 | -0.1 | 8.45E-01 | 0.3 | 5.31E-01 | 0.0 | 9.53E-01 | -1.1 | 6.55E-02 | 1.1 | 6.48E-02 | Cysteine-rich RLK 13 |
| AT5G67630 | 0.2 | 5.94E-02 | 0.1 | 6.46E-01 | 0.2 | 2.02E-01 | 0.3 | 4.65E-02 | 0.0 | 8.35E-01 | 0.3 | 9.07E-02 | RuvB-like helicase |
| AT5G19485 | 0.2 | 6.82E-03 | 0.0 | 6.07E-01 | 0.2 | 8.02E-03 | 0.1 | 5.48E-01 | -0.1 | 5.30E-01 | 0.2 | 3.11E-01 | Transferases |
| AT3G62420 | 0.2 | 7.55E-02 | 0.3 | 1.79E-02 | -0.1 | 4.23E-01 | 0.2 | 1.91E-01 | -0.1 | 6.41E-01 | 0.3 | 1.20E-01 | bZIP transcription factor 53 |
| AT1G61870 | 0.2 | 1.05E-01 | 0.2 | 1.08E-01 | 0.0 | 9.78E-01 | 0.4 | 1.68E-02 | 0.1 | 5.11E-01 | 0.3 | 1.54E-01 | Small ribosomal subunit protein mS86 |
| AT2G36740 | 0.2 | 3.28E-02 | 0.1 | 5.90E-01 | 0.2 | 1.50E-01 | 0.1 | 7.64E-01 | 0.0 | 9.57E-01 | 0.1 | 7.85E-01 | DNA binding protein SWC2 |
| AT4G26870 | 0.2 | 1.02E-02 | 0.1 | 9.13E-02 | 0.1 | 3.12E-01 | 0.0 | 9.29E-01 | 0.0 | 8.13E-01 | 0.1 | 7.99E-01 | Aspartate--tRNA ligase 1, cytoplasmic |
| AT3G56090 | 0.2 | 2.00E-01 | 0.3 | 1.15E-01 | -0.1 | 7.51E-01 | -0.1 | 5.07E-01 | 0.0 | 9.85E-01 | -0.1 | 6.01E-01 | Ferritin |
| AT1G66200 | 0.2 | 8.01E-02 | 0.8 | 6.93E-05 | -0.6 | 1.07E-03 | 0.8 | 1.58E-05 | 0.6 | 2.18E-03 | 0.3 | 8.34E-02 | glutamine synthetase |
| AT4G19210 | 0.2 | 1.73E-02 | 0.0 | 5.99E-01 | 0.2 | 7.30E-02 | -0.2 | 1.92E-01 | 0.0 | 9.61E-01 | -0.2 | 4.00E-01 | ABC transporter E family member 2 |
| AT5G12310 | 0.2 | 6.36E-02 | 0.1 | 6.55E-01 | 0.2 | 2.04E-01 | -0.1 | 4.20E-01 | -0.1 | 4.37E-01 | 0.0 | 9.66E-01 | RING-type domain-containing protein |
| AT4G34160 | 0.2 | 6.69E-03 | 0.1 | 6.64E-02 | 0.1 | 3.08E-01 | -0.3 | 7.06E-02 | 0.0 | 9.27E-01 | -0.2 | 2.29E-01 | Cyclin-D3-1 |
| AT2G40240 | 0.2 | 1.65E-01 | 0.1 | 7.23E-01 | 0.2 | 4.18E-01 | 0.7 | 5.65E-02 | -0.2 | 7.76E-01 | 0.8 | 2.04E-02 | Pentatricopeptide repeat-containing protein At2g40240, mitochondrial |
| AT1G64550 | 0.2 | 3.98E-03 | 0.1 | 1.03E-01 | 0.1 | 1.27E-01 | 0.1 | 5.57E-01 | 0.0 | 8.25E-01 | 0.1 | 5.32E-01 | ABC transporter F family member 3 |
| AT5G26180 | 0.2 | 8.11E-02 | -0.1 | 6.31E-01 | 0.3 | 6.60E-02 | 0.2 | 1.52E-01 | 0.0 | 9.20E-01 | 0.2 | 3.19E-01 | 25S rRNA )-methyltransferase NSUN5 |
| AT5G45420 | 0.2 | 1.32E-02 | 0.3 | 2.44E-03 | -0.1 | 3.11E-01 | 0.1 | 6.55E-01 | 0.1 | 7.14E-01 | 0.0 | 9.98E-01 | Transcription factor MAMYB |
| AT3G11250 | 0.2 | 5.52E-02 | 0.4 | 3.71E-03 | -0.2 | 1.50E-01 | 0.4 | 3.18E-03 | 0.4 | 1.31E-02 | 0.0 | 8.41E-01 | Large ribosomal subunit protein uL10x |
| AT5G22690 | 0.2 | 6.74E-02 | -0.3 | 1.83E-02 | 0.5 | 3.02E-03 | 0.0 | 9.65E-01 | -0.2 | 5.35E-01 | 0.2 | 5.83E-01 | ADP-ribosyl cyclase |
| AT1G29940 | 0.2 | 3.16E-02 | -0.2 | 1.10E-01 | 0.4 | 5.28E-03 | 0.7 | 2.04E-03 | 0.2 | 3.52E-01 | 0.5 | 8.41E-02 | DNA-directed RNA polymerase I subunit 2 |
| AT3G46030 | 0.2 | 4.31E-02 | 0.4 | 4.74E-03 | -0.1 | 2.09E-01 | 0.1 | 8.10E-01 | -0.1 | 7.79E-01 | 0.2 | 6.71E-01 | Histone H2B.7 |
| AT5G51410 | 0.2 | 5.12E-03 | 0.0 | 7.04E-01 | 0.2 | 7.01E-03 | -0.1 | 3.04E-01 | -0.2 | 2.01E-01 | 0.1 | 8.29E-01 | Arginine-aspartate-rich RNA binding protein-like |
| AT4G19060 | 0.2 | 9.96E-02 | 0.1 | 6.46E-01 | 0.2 | 3.18E-01 | -0.3 | 2.15E-01 | 0.0 | 8.73E-01 | -0.2 | 4.36E-01 | Probable disease resistance protein At4g19060 |
| AT3G30300 | 0.2 | 1.51E-02 | 0.0 | 9.83E-01 | 0.2 | 3.10E-02 | -0.1 | 7.16E-01 | 0.0 | 9.36E-01 | -0.1 | 7.46E-01 | O-fucosyltransferase 27 |
| AT2G30280 | 0.2 | 4.59E-02 | 0.1 | 2.45E-01 | 0.1 | 4.59E-01 | -0.2 | 4.18E-01 | -0.3 | 1.41E-01 | 0.2 | 6.06E-01 | RNA-directed DNA methylation 4 |
| AT3G10870 | 0.2 | 6.01E-02 | 0.5 | 7.00E-04 | -0.3 | 3.16E-02 | -0.1 | 8.45E-01 | -0.1 | 7.70E-01 | 0.0 | 9.26E-01 | Methylesterase 17 |
| AT5G04710 | 0.2 | 7.05E-03 | 0.0 | 7.05E-01 | 0.2 | 3.01E-02 | 0.0 | 9.30E-01 | 0.0 | 9.03E-01 | 0.0 | 8.60E-01 | Aspartyl aminopeptidase |
| AT3G27120 | 0.2 | 1.08E-01 | 0.2 | 1.56E-01 | 0.0 | 9.18E-01 | 0.3 | 4.18E-01 | 0.1 | 7.76E-01 | 0.1 | 7.61E-01 | hypothetical protein |
| AT5G11640 | 0.2 | 5.48E-03 | 0.2 | 1.24E-02 | 0.0 | 7.84E-01 | 0.1 | 6.46E-01 | 0.1 | 6.85E-01 | 0.0 | 9.86E-01 | Thioredoxin superfamily protein |
| AT1G13100 | 0.2 | 9.78E-03 | 0.2 | 5.42E-02 | 0.1 | 4.67E-01 | 0.4 | 1.30E-02 | 0.2 | 4.68E-01 | 0.3 | 1.53E-01 | Cytochrome P450 71B29 |
| AT5G08790 | 0.2 | 1.42E-01 | -0.8 | 4.51E-04 | 1.0 | 2.43E-04 | 0.9 | 1.07E-05 | 0.0 | 9.48E-01 | 0.9 | 3.17E-04 | Protein ATAF2 |
| AT1G75520 | 0.2 | 8.62E-02 | -0.4 | 1.16E-02 | 0.6 | 2.90E-03 | 0.9 | 9.18E-02 | 0.7 | 2.36E-01 | 0.2 | 8.09E-01 | Protein SHI RELATED SEQUENCE 5 |
| AT5G45400 | 0.2 | 7.18E-02 | 0.0 | 8.86E-01 | 0.2 | 1.67E-01 | 0.1 | 8.85E-01 | 0.2 | 7.67E-01 | -0.1 | 8.98E-01 | Replication protein A 70 kDa DNA-binding subunit C |
| AT3G11450 | 0.2 | 2.58E-02 | 0.3 | 6.25E-03 | -0.1 | 4.03E-01 | 0.1 | 3.37E-01 | -0.1 | 5.06E-01 | 0.3 | 1.70E-01 | DnaJ and Myb-like DNA-binding domain-containing protein |
| AT5G03560 | 0.2 | 4.26E-02 | 0.0 | 8.81E-01 | 0.2 | 6.51E-02 | 0.0 | 8.38E-01 | -0.1 | 5.65E-01 | 0.2 | 4.97E-01 | AT5g03560 |
| AT1G09620 | 0.2 | 5.13E-03 | 0.1 | 9.37E-02 | 0.1 | 1.74E-01 | 0.1 | 4.61E-01 | 0.0 | 9.21E-01 | 0.2 | 5.28E-01 | leucine--tRNA ligase |
| AT2G33860 | 0.2 | 1.00E-02 | 0.2 | 2.04E-02 | 0.0 | 8.13E-01 | -0.1 | 4.95E-01 | 0.0 | 8.90E-01 | -0.1 | 7.53E-01 | Auxin response factor |
| AT5G17490 | 0.2 | 8.25E-02 | 0.6 | 6.50E-04 | -0.4 | 2.35E-02 | -0.7 | 2.36E-02 | -0.4 | 1.66E-01 | -0.2 | 5.45E-01 | DELLA protein RGL3 |
| AT4G27280 | 0.2 | 5.47E-01 | -0.9 | 3.58E-02 | 1.1 | 2.24E-02 | 0.3 | 2.31E-01 | -0.2 | 5.19E-01 | 0.6 | 1.15E-01 | Calcium-binding protein KRP1 |
| AT5G55760 | 0.2 | 5.64E-02 | 0.1 | 2.01E-01 | 0.1 | 5.91E-01 | 0.3 | 1.55E-01 | 0.2 | 3.66E-01 | 0.1 | 8.18E-01 | NAD-dependent protein deacetylase SRT1 |
| AT3G43210 | 0.2 | 2.24E-02 | -0.1 | 3.72E-01 | 0.3 | 1.30E-02 | 0.0 | 8.46E-01 | 0.1 | 7.43E-01 | -0.1 | 5.56E-01 | Kinesin-like protein KIN-7B |
| AT4G23180 | 0.2 | 2.92E-01 | -0.1 | 6.83E-01 | 0.3 | 1.89E-01 | 0.1 | 3.14E-01 | 0.0 | 9.01E-01 | 0.2 | 3.90E-01 | Uncharacterized protein |
| AT5G16000 | 0.2 | 1.56E-02 | 0.4 | 1.10E-03 | -0.1 | 1.29E-01 | -0.3 | 2.92E-02 | -0.3 | 1.19E-01 | -0.1 | 7.79E-01 | Protein NSP-INTERACTING KINASE 1 |
| AT5G53770 | 0.2 | 5.32E-02 | 0.1 | 6.40E-01 | 0.2 | 2.01E-01 | 0.2 | 2.93E-01 | 0.0 | 9.76E-01 | 0.2 | 4.11E-01 | Nucleotidyltransferase family protein |
| AT5G25800 | 0.2 | 1.73E-02 | 0.1 | 2.75E-01 | 0.1 | 2.17E-01 | 0.2 | 2.81E-01 | -0.1 | 7.38E-01 | 0.3 | 2.43E-01 | Polynucleotidyl transferase, ribonuclease H-like superfamily protein |
| AT1G29730 | 0.2 | 9.08E-02 | 0.4 | 9.08E-03 | -0.2 | 2.54E-01 | -0.3 | 2.19E-01 | 0.0 | 9.85E-01 | -0.3 | 3.68E-01 | Leucine-rich repeat transmembrane protein kinase |
| AT5G55050 | 0.2 | 3.82E-02 | -0.1 | 3.50E-01 | 0.3 | 1.85E-02 | 0.1 | 8.10E-01 | 0.4 | 2.79E-01 | -0.3 | 4.88E-01 | GDSL esterase |
| AT3G55040 | 0.2 | 2.41E-02 | 0.4 | 1.79E-03 | -0.1 | 1.46E-01 | 0.2 | 2.03E-01 | 0.1 | 6.90E-01 | 0.1 | 5.92E-01 | Glutathione S-transferase L2, chloroplastic |
| AT5G16690 | 0.2 | 1.52E-01 | 0.2 | 1.52E-01 | 0.0 | 9.61E-01 | 0.4 | 3.35E-01 | 0.4 | 3.50E-01 | 0.0 | 9.63E-01 | Origin recognition complex subunit 3 |
| AT1G09060 | 0.2 | 2.47E-02 | -0.2 | 3.66E-02 | 0.4 | 1.82E-03 | -0.3 | 1.16E-01 | -0.1 | 5.21E-01 | -0.1 | 6.20E-01 | E3 ubiquitin-protein ligase JMJ24 |
| AT3G22330 | 0.2 | 2.16E-02 | -0.1 | 2.20E-01 | 0.3 | 5.76E-03 | 0.5 | 5.78E-04 | 0.1 | 3.48E-01 | 0.4 | 4.58E-02 | DEAD-box ATP-dependent RNA helicase 53, mitochondrial |
| AT2G45810 | 0.2 | 8.10E-03 | 0.1 | 1.39E-01 | 0.1 | 1.81E-01 | 0.1 | 7.83E-01 | 0.1 | 7.87E-01 | 0.0 | 9.68E-01 | DEAD-box ATP-dependent RNA helicase 6 |
| AT1G76890 | 0.2 | 1.84E-02 | 0.4 | 1.46E-03 | -0.1 | 1.47E-01 | -0.6 | 8.50E-05 | -0.6 | 1.58E-04 | 0.0 | 8.41E-01 | Trihelix transcription factor GT-2 |
| AT5G26690 | 0.2 | 5.29E-01 | -0.6 | 1.61E-01 | 0.8 | 1.05E-01 | -0.2 | 5.44E-01 | 0.8 | 5.48E-03 | -1.0 | 2.78E-03 | Heavy metal-associated isoprenylated plant protein 2 |
| AT3G57940 | 0.2 | 1.16E-01 | 0.1 | 4.51E-01 | 0.1 | 4.97E-01 | 1.1 | 4.68E-04 | 0.6 | 6.49E-02 | 0.5 | 7.83E-02 | RNA cytidine acetyltransferase |
| AT3G19930 | 0.2 | 8.99E-02 | 0.2 | 1.64E-01 | 0.0 | 8.13E-01 | -0.2 | 2.47E-01 | -0.4 | 6.28E-02 | 0.2 | 5.68E-01 | Sugar transport protein 4 |
| AT3G58120 | 0.2 | 1.52E-01 | 0.6 | 1.46E-03 | -0.4 | 2.47E-02 | -0.6 | 1.89E-03 | -0.5 | 1.42E-02 | -0.1 | 6.71E-01 | Basic-leucine zipper transcription factor family protein |
| AT5G20020 | 0.2 | 1.43E-01 | 0.5 | 1.10E-02 | -0.2 | 1.66E-01 | 0.2 | 1.72E-01 | 0.0 | 8.74E-01 | 0.2 | 2.42E-01 | GTP-binding nuclear protein Ran-2 |
| AT5G09230 | 0.2 | 5.15E-02 | 0.0 | 6.76E-01 | 0.3 | 5.11E-02 | -0.3 | 1.17E-01 | -0.1 | 6.77E-01 | -0.2 | 4.42E-01 | Sirtuin 2 |
| AT5G17760 | 0.2 | 8.71E-02 | -0.7 | 7.45E-04 | 0.9 | 3.37E-04 | 0.6 | 6.29E-03 | 0.1 | 7.37E-01 | 0.5 | 4.82E-02 | AAA-ATPase At5g17760 |
| AT2G26970 | 0.2 | 2.09E-02 | 0.1 | 1.19E-01 | 0.1 | 4.59E-01 | -0.1 | 7.01E-01 | -0.2 | 4.82E-01 | 0.1 | 8.05E-01 | AT2G26970 protein |
| AT3G02065 | 0.2 | 1.94E-02 | 0.1 | 1.78E-01 | 0.1 | 3.23E-01 | 0.1 | 7.34E-01 | -0.2 | 2.75E-01 | 0.2 | 2.27E-01 | DEAD-box ATP-dependent RNA helicase 41 |
| AT2G27360 | 0.2 | 1.85E-02 | 0.1 | 4.61E-01 | 0.2 | 1.23E-01 | 0.1 | 6.03E-01 | 0.3 | 2.67E-01 | -0.2 | 6.30E-01 | GDSL esterase |
| AT5G59890 | 0.2 | 2.16E-01 | 0.1 | 7.01E-01 | 0.1 | 4.77E-01 | 0.0 | 8.22E-01 | 0.0 | 9.66E-01 | 0.1 | 8.54E-01 | ADF-H domain-containing protein |
| AT5G62540 | 0.2 | 1.89E-01 | 0.0 | 9.70E-01 | 0.2 | 2.51E-01 | 0.5 | 1.50E-02 | 0.2 | 3.13E-01 | 0.2 | 3.24E-01 | Ubiquitin-conjugating enzyme E2 3 |
| AT1G24400 | 0.2 | 1.65E-01 | 0.2 | 3.45E-01 | 0.1 | 7.54E-01 | -0.4 | 2.24E-01 | 0.0 | 9.65E-01 | -0.4 | 3.83E-01 | Lysine histidine transporter 2 |
| AT1G30960 | 0.2 | 1.63E-01 | 0.2 | 1.61E-01 | 0.0 | 9.65E-01 | 0.8 | 1.49E-02 | 0.5 | 1.77E-01 | 0.3 | 3.87E-01 | GTP-binding protein ERG |
| AT5G22140 | 0.2 | 5.56E-02 | 0.0 | 7.04E-01 | 0.2 | 1.85E-01 | 0.5 | 3.52E-02 | 0.5 | 1.08E-01 | 0.1 | 7.96E-01 | FAD |
| AT2G39990 | 0.2 | 2.08E-02 | 0.2 | 1.58E-02 | 0.0 | 8.21E-01 | 0.1 | 7.00E-01 | -0.1 | 4.34E-01 | 0.2 | 3.22E-01 | Eukaryotic translation initiation factor 3 subunit F |
| AT5G24940 | 0.2 | 1.44E-01 | 0.7 | 1.07E-03 | -0.4 | 2.52E-02 | -0.3 | 4.61E-01 | -0.1 | 8.64E-01 | -0.2 | 7.39E-01 | Probable protein phosphatase 2C 71 |
| AT5G50960 | 0.2 | 3.10E-02 | 0.0 | 7.00E-01 | 0.2 | 9.98E-02 | 0.2 | 2.83E-01 | -0.1 | 5.19E-01 | 0.3 | 1.46E-01 | Cytosolic Fe-S cluster assembly factor NBP35 |
| AT2G48000 | 0.2 | 4.09E-02 | 0.1 | 5.87E-01 | 0.2 | 1.83E-01 | 0.0 | 9.27E-01 | 0.1 | 7.57E-01 | -0.1 | 7.20E-01 | hypothetical protein |
| AT4G15350 | 0.2 | 3.70E-01 | 0.5 | 4.51E-02 | -0.3 | 3.04E-01 | 0.3 | 4.30E-01 | 0.2 | 6.96E-01 | 0.1 | 8.56E-01 | Cytochrome P450 like protein |
| AT4G26610 | 0.2 | 8.22E-03 | 0.0 | 8.50E-01 | 0.2 | 1.43E-02 | -0.4 | 9.49E-02 | -0.5 | 6.63E-02 | 0.1 | 8.32E-01 | Serine |
| AT3G13062 | 0.2 | 2.03E-01 | 0.2 | 2.39E-01 | 0.0 | 9.62E-01 | -0.2 | 2.10E-01 | -0.7 | 1.60E-03 | 0.5 | 4.82E-02 | Polyketide cyclase |

| | | | | | | | | | | | | | |
|---|---|---|---|---|---|---|---|---|---|---|---|---|---|
| AT2G42810 | 0.2 | 2.00E-03 | 0.0 | 9.44E-01 | 0.2 | 5.65E-03 | 0.1 | 5.85E-01 | -0.1 | 4.42E-01 | 0.2 | 2.62E-01 | Serine |
| AT1G48300 | 0.2 | 3.04E-02 | 0.2 | 6.82E-02 | 0.0 | 7.67E-01 | 0.5 | 2.62E-04 | 0.0 | 7.98E-01 | 0.4 | 8.51E-03 | Diacylglycerol O-acyltransferase 3 |
| AT5G14210 | 0.2 | 5.15E-02 | 0.2 | 1.06E-01 | 0.0 | 7.73E-01 | -0.1 | 7.20E-01 | 0.1 | 7.68E-01 | -0.2 | 5.64E-01 | Protein kinase domain-containing protein |
| AT5G57330 | 0.2 | 1.69E-02 | 0.3 | 1.05E-02 | 0.0 | 7.32E-01 | 0.2 | 9.41E-02 | 0.2 | 2.42E-01 | 0.0 | 8.64E-01 | glucose-6-phosphate 1-epimerase |
| AT1G36310 | 0.2 | 5.30E-02 | 0.1 | 3.46E-01 | 0.1 | 3.53E-01 | 0.2 | 7.69E-02 | 0.1 | 6.55E-01 | 0.1 | 4.16E-01 | Methyltransferase type 11 domain-containing protein |
| AT3G23570 | 0.2 | 4.33E-02 | 0.2 | 1.25E-01 | 0.1 | 6.48E-01 | 0.1 | 6.79E-01 | 0.0 | 9.66E-01 | 0.1 | 8.22E-01 | Alpha |
| AT1G30580 | 0.2 | 3.52E-02 | 0.3 | 2.56E-02 | 0.0 | 8.09E-01 | 0.2 | 2.60E-02 | 0.1 | 2.00E-01 | 0.1 | 6.54E-01 | Obg-like ATPase 1 |
| AT3G24170 | 0.2 | 1.38E-02 | 0.0 | 9.51E-01 | 0.2 | 2.25E-02 | 0.5 | 1.36E-04 | 0.4 | 1.02E-03 | 0.0 | 7.82E-01 | Glutathione reductase, cytosolic |
| AT2G36070 | 0.2 | 2.50E-02 | 0.1 | 2.35E-01 | 0.1 | 2.96E-01 | 0.9 | 5.94E-04 | 0.4 | 1.00E-01 | 0.4 | 7.19E-02 | Mitochondrial import inner membrane translocase subunit TIM44-2 |
| AT3G04600 | 0.2 | 7.76E-03 | 0.1 | 3.83E-01 | 0.2 | 6.69E-02 | 0.1 | 6.37E-01 | -0.2 | 1.69E-01 | 0.2 | 1.47E-01 | tryptophan--tRNA ligase |
| AT1G21270 | 0.2 | 4.77E-02 | -0.2 | 7.90E-02 | 0.4 | 4.29E-03 | 0.5 | 5.36E-02 | 0.1 | 8.88E-01 | 0.4 | 2.09E-01 | Wall-associated receptor kinase 2 |
| AT1G45248 | 0.2 | 1.84E-01 | 0.0 | 7.94E-01 | 0.2 | 4.01E-01 | -0.2 | 5.51E-01 | -0.1 | 7.56E-01 | -0.1 | 9.07E-01 | Nucleolar histone methyltransferase-related protein |
| AT1G42440 | 0.2 | 1.70E-02 | -0.1 | 2.76E-01 | 0.3 | 6.18E-03 | 0.4 | 1.36E-02 | 0.1 | 8.23E-01 | 0.3 | 1.12E-01 | At1g42440 |
| AT5G06580 | 0.2 | 9.74E-03 | 0.1 | 4.56E-01 | 0.2 | 7.35E-02 | -0.1 | 7.55E-01 | -0.1 | 3.82E-01 | 0.1 | 6.90E-01 | D-lactate dehydrogenase , mitochondrial |
| AT3G61210 | 0.2 | 2.08E-01 | 0.1 | 6.03E-01 | 0.1 | 5.56E-01 | 0.3 | 2.50E-01 | 0.2 | 3.83E-01 | 0.0 | 9.33E-01 | hypothetical protein |
| AT2G45790 | 0.2 | 6.26E-03 | 0.3 | 2.96E-03 | 0.0 | 5.77E-01 | 0.1 | 4.05E-01 | 0.0 | 9.91E-01 | 0.1 | 6.03E-01 | Phosphomannomutase |
| AT5G41210 | 0.2 | 3.13E-02 | 0.0 | 7.77E-01 | 0.2 | 8.59E-02 | 0.1 | 5.53E-01 | 0.1 | 6.34E-01 | 0.0 | 9.98E-01 | Glutathione S-transferase T1 |
| AT4G20930 | 0.2 | 1.49E-01 | 0.0 | 9.50E-01 | 0.2 | 2.14E-01 | -0.3 | 3.37E-01 | 0.2 | 5.53E-01 | -0.5 | 1.69E-01 | 3-hydroxyisobutyrate dehydrogenase |
| AT2G15430 | 0.2 | 2.88E-02 | 0.2 | 5.44E-02 | 0.0 | 8.23E-01 | 0.3 | 4.22E-02 | 0.0 | 8.52E-01 | 0.3 | 1.44E-01 | DNA-directed RNA polymerase family protein |
| AT1G48140 | 0.2 | 5.38E-02 | 0.3 | 3.03E-02 | 0.0 | 7.56E-01 | 0.2 | 5.12E-01 | 0.2 | 5.24E-01 | 0.0 | 9.70E-01 | Dolichol-phosphate mannosyltransferase subunit 3 |
| AT4G38800 | 0.2 | 1.71E-02 | 0.3 | 4.97E-03 | -0.1 | 4.32E-01 | -0.2 | 8.82E-02 | -0.1 | 3.83E-01 | -0.1 | 7.08E-01 | 5'-methylthioadenosine nucleosidase |
| AT1G22510 | 0.2 | 5.62E-02 | 0.1 | 3.40E-01 | 0.1 | 4.07E-01 | -0.2 | 3.60E-01 | -0.4 | 1.33E-01 | 0.2 | 6.43E-01 | E3 ubiquitin-protein ligase RNF170 |
| AT5G02250 | 0.2 | 3.17E-02 | 0.0 | 6.39E-01 | 0.2 | 1.30E-01 | -0.1 | 7.45E-01 | -0.1 | 7.05E-01 | 0.0 | 9.48E-01 | Ribonuclease II, chloroplastic |
| AT4G19890 | 0.2 | 1.74E-01 | 0.0 | 9.70E-01 | 0.2 | 2.90E-01 | -0.1 | 7.94E-01 | -0.2 | 6.96E-01 | 0.1 | 9.12E-01 | Pentatricopeptide repeat-containing protein At4g19890 |
| AT5G19550 | 0.2 | 2.80E-02 | -0.1 | 1.96E-01 | 0.4 | 5.96E-03 | 1.1 | 7.89E-07 | 0.2 | 1.75E-01 | 0.8 | 3.55E-04 | Aspartate aminotransferase |
| AT1G60260 | 0.2 | 2.47E-01 | 0.1 | 7.65E-01 | 0.2 | 4.93E-01 | -0.1 | 6.06E-01 | 0.1 | 8.14E-01 | -0.2 | 5.16E-01 | Putative beta-glucosidase 5 |
| AT1G79540 | 0.2 | 4.93E-02 | 0.1 | 4.82E-01 | 0.1 | 2.70E-01 | -0.1 | 8.06E-01 | -0.2 | 6.11E-01 | 0.1 | 8.07E-01 | Pentatricopeptide repeat-containing protein At1g79540 |
| AT4G14330 | 0.2 | 1.02E-01 | 0.5 | 3.66E-03 | -0.3 | 1.05E-01 | 0.1 | 7.87E-01 | 0.3 | 3.38E-01 | -0.2 | 5.91E-01 | Kinesin-like protein KIN-10A |
| AT1G27680 | 0.2 | 3.89E-02 | 0.2 | 8.16E-02 | 0.0 | 7.85E-01 | 0.2 | 5.28E-01 | 0.3 | 3.22E-01 | -0.1 | 7.29E-01 | Glucose-1-phosphate adenylyltransferase large subunit 2, chloroplastic |
| AT1G18335 | 0.2 | 8.41E-02 | 0.1 | 6.02E-01 | 0.2 | 3.13E-01 | 0.3 | 4.14E-01 | -0.1 | 8.11E-01 | 0.4 | 3.54E-01 | N-alpha-acetyltransferase 40 |
| AT1G03000 | 0.2 | 2.36E-02 | 0.0 | 7.96E-01 | 0.3 | 3.11E-02 | -0.5 | 1.09E-02 | -0.3 | 2.39E-01 | -0.3 | 3.06E-01 | Peroxisomal ATPase PEX6 |
| AT1G54410 | 0.2 | 1.76E-02 | 0.5 | 2.86E-04 | -0.3 | 1.81E-02 | -0.1 | 6.47E-01 | 0.0 | 9.08E-01 | -0.1 | 6.78E-01 | Dehydrin HIRD11 |
| AT5G13660 | 0.2 | 3.98E-02 | 0.0 | 9.07E-01 | 0.2 | 1.01E-01 | 0.0 | 9.51E-01 | -0.1 | 6.71E-01 | 0.1 | 7.48E-01 | N-lysine methyltransferase |
| AT1G67490 | 0.2 | 8.86E-03 | 0.1 | 1.62E-01 | 0.1 | 1.90E-01 | 0.4 | 7.02E-02 | 0.2 | 3.79E-01 | 0.2 | 6.11E-01 | Mannosyl-oligosaccharide glucosidase |
| AT5G15120 | 0.2 | 1.53E-01 | 0.3 | 6.67E-02 | -0.1 | 6.72E-01 | 0.8 | 2.74E-02 | 0.1 | 8.55E-01 | 0.7 | 6.25E-02 | Plant cysteine oxidase 1 |
| AT1G51060 | 0.2 | 1.60E-02 | 0.4 | 2.25E-03 | -0.1 | 2.21E-01 | 0.2 | 2.91E-01 | 0.2 | 2.78E-01 | 0.0 | 9.16E-01 | Probable histone H2A.1 |
| AT3G27820 | 0.2 | 5.18E-03 | -0.1 | 4.24E-01 | 0.3 | 3.45E-03 | 0.1 | 4.59E-01 | 0.2 | 1.45E-01 | -0.1 | 5.83E-01 | Monodehydroascorbate reductase 4, peroxisomal |
| AT3G46540 | 0.2 | 2.70E-02 | 0.1 | 1.49E-01 | 0.1 | 4.37E-01 | 0.3 | 5.05E-02 | 0.3 | 9.59E-02 | 0.0 | 9.63E-01 | ENTH |
| AT5G03770 | 0.2 | 1.51E-01 | -0.1 | 6.71E-01 | 0.3 | 1.34E-01 | -0.1 | 8.11E-01 | 0.1 | 8.75E-01 | -0.2 | 7.50E-01 | KDO transferase A |
| AT1G06560 | 0.2 | 7.53E-02 | 0.2 | 1.92E-01 | 0.1 | 7.01E-01 | 0.0 | 9.08E-01 | -0.5 | 4.25E-02 | 0.4 | 8.13E-02 | rRNA )-methyltransferase NOP2C |
| AT4G31770 | 0.2 | 2.45E-02 | 0.0 | 8.56E-01 | 0.3 | 4.21E-02 | 0.0 | 9.20E-01 | -0.3 | 1.99E-01 | 0.3 | 2.48E-01 | Debranching enzyme 1 |
| AT1G72740 | 0.2 | 1.49E-02 | 0.1 | 3.13E-01 | 0.1 | 1.56E-01 | -0.2 | 1.80E-01 | -0.2 | 3.86E-01 | -0.1 | 8.49E-01 | MYB transcription factor |
| AT2G31320 | 0.2 | 2.55E-02 | 0.2 | 5.58E-02 | 0.0 | 7.86E-01 | -0.2 | 4.71E-01 | 0.1 | 9.07E-01 | -0.3 | 4.40E-01 | Poly polymerase 1 ADP-ribosyltransferase 1) |
| AT1G02410 | 0.2 | 8.02E-03 | 0.0 | 7.63E-01 | 0.2 | 3.16E-02 | 0.1 | 5.95E-01 | 0.1 | 6.92E-01 | 0.0 | 9.82E-01 | Cytochrome c oxidase assembly protein COX11, mitochondrial |
| AT4G02540 | 0.2 | 5.26E-02 | 0.1 | 5.91E-01 | 0.2 | 2.01E-01 | -0.3 | 4.27E-02 | -0.1 | 5.73E-01 | -0.2 | 3.53E-01 | Cysteine |
| AT5G50860 | 0.2 | 4.33E-03 | 0.1 | 2.17E-01 | 0.1 | 7.61E-02 | -0.1 | 5.97E-01 | 0.0 | 9.65E-01 | -0.1 | 6.91E-01 | Protein kinase domain-containing protein |
| AT5G27120 | 0.2 | 2.12E-01 | 0.2 | 2.39E-01 | 0.0 | 9.83E-01 | 0.6 | 3.94E-03 | 0.3 | 1.72E-01 | 0.3 | 2.12E-01 | Probable nucleolar protein 5-1 |
| AT2G15530 | 0.2 | 9.18E-03 | 0.0 | 9.05E-01 | 0.2 | 1.76E-02 | 0.2 | 2.70E-01 | 0.2 | 3.74E-01 | 0.0 | 9.69E-01 | RING-type E3 ubiquitin transferase |
| AT5G37680 | 0.2 | 1.31E-01 | 0.0 | 9.36E-01 | 0.2 | 1.85E-01 | -0.3 | 2.91E-01 | -0.1 | 8.41E-01 | -0.2 | 6.04E-01 | ADP-ribosylation factor-like A1A |
| AT3G51920 | 0.2 | 1.51E-02 | 0.1 | 3.65E-01 | 0.2 | 1.20E-01 | -0.1 | 7.06E-01 | -0.3 | 1.93E-01 | 0.2 | 4.50E-01 | Calmodulin-like protein 9 |
| AT5G40870 | 0.2 | 1.17E-02 | 0.1 | 1.15E-01 | 0.1 | 3.05E-01 | 0.0 | 9.74E-01 | -0.1 | 7.01E-01 | 0.1 | 7.21E-01 | Uridine |
| AT2G29200 | 0.2 | 6.98E-02 | 0.0 | 9.11E-01 | 0.2 | 1.38E-01 | 0.5 | 1.20E-02 | 0.4 | 1.05E-01 | 0.2 | 5.93E-01 | Pumilio homolog 1 |
| AT1G55530 | 0.2 | 9.48E-03 | 0.0 | 9.37E-01 | 0.2 | 2.36E-02 | -0.2 | 3.28E-02 | -0.5 | 4.54E-04 | 0.3 | 8.21E-02 | RING-type E3 ubiquitin transferase |
| AT1G26340 | 0.2 | 6.08E-02 | 0.4 | 6.91E-03 | -0.2 | 2.66E-01 | 0.2 | 2.36E-01 | 0.2 | 4.96E-01 | 0.1 | 7.89E-01 | Cytochrome b5 isoform A |
| AT2G34040 | 0.2 | 5.42E-03 | 0.2 | 2.91E-02 | 0.1 | 4.53E-01 | 0.1 | 3.99E-01 | 0.0 | 9.01E-01 | 0.1 | 6.77E-01 | Apoptosis inhibitory protein 5 |
| AT5G11720 | 0.2 | 8.04E-03 | 0.2 | 4.04E-02 | 0.1 | 4.75E-01 | -0.3 | 1.58E-02 | -0.2 | 2.52E-01 | -0.1 | 4.55E-01 | AT5g11720 |
| AT3G45260 | 0.2 | 1.94E-02 | -0.3 | 1.49E-02 | 0.5 | 9.28E-04 | 0.0 | 8.12E-01 | -0.3 | 4.06E-02 | 0.3 | 1.43E-01 | Zinc finger protein BALDIBIS |
| AT5G26860 | 0.2 | 2.67E-02 | 0.0 | 7.74E-01 | 0.2 | 7.96E-02 | 1.1 | 4.13E-05 | 0.5 | 3.52E-02 | 0.6 | 2.96E-02 | AT5g26860 |
| AT3G57550 | 0.2 | 8.42E-03 | 0.1 | 1.04E-01 | 0.1 | 2.42E-01 | -0.1 | 5.43E-01 | -0.4 | 7.08E-02 | 0.3 | 3.40E-01 | guanylate kinase |
| AT3G11210 | 0.2 | 4.10E-02 | 0.0 | 7.64E-01 | 0.3 | 5.18E-02 | 0.2 | 5.02E-01 | 0.1 | 6.69E-01 | 0.0 | 9.23E-01 | GDSL esterase |
| AT5G05450 | 0.2 | 6.07E-03 | 0.2 | 1.42E-02 | 0.0 | 7.66E-01 | 0.1 | 7.69E-01 | -0.2 | 3.27E-01 | 0.3 | 3.02E-01 | DEAD-box ATP-dependent RNA helicase 18 |
| AT3G59060 | 0.2 | 7.04E-03 | 0.0 | 6.07E-01 | 0.3 | 6.36E-03 | -0.5 | 6.02E-02 | -0.5 | 6.42E-02 | 0.0 | 9.57E-01 | Transcription factor PIF5 |
| AT1G77850 | 0.2 | 4.51E-02 | 0.0 | 8.70E-01 | 0.3 | 6.80E-02 | 0.1 | 6.42E-01 | 0.3 | 2.91E-01 | -0.2 | 6.21E-01 | Auxin response factor |
| AT3G23580 | 0.2 | 2.33E-02 | 0.0 | 8.29E-01 | 0.3 | 3.51E-02 | 0.2 | 3.99E-01 | 0.0 | 8.93E-01 | 0.2 | 3.43E-01 | Ribonucleoside-diphosphate reductase small chain A |
| AT5G02100 | 0.2 | 2.11E-02 | 0.1 | 5.03E-01 | 0.2 | 1.14E-01 | 0.0 | 8.87E-01 | 0.0 | 8.50E-01 | 0.1 | 7.85E-01 | Oxysterol-binding family protein |
| AT4G37090 | 0.2 | 1.34E-01 | 0.4 | 3.30E-02 | -0.1 | 4.79E-01 | 0.6 | 1.50E-02 | 0.5 | 6.53E-02 | 0.1 | 7.48E-01 | UDP-N-acetylmuramoyl-L-alanyl-D-glutamate-2, 6-diaminopimelate ligase |
| AT1G74710 | 0.2 | 2.36E-01 | -0.3 | 1.21E-01 | 0.6 | 2.72E-02 | 0.4 | 5.93E-02 | 0.7 | 3.62E-03 | -0.3 | 2.67E-01 | Isochorismate synthase 1, chloroplastic |
| AT4G37760 | 0.2 | 7.30E-02 | -0.2 | 2.42E-01 | 0.4 | 1.59E-02 | -0.3 | 1.36E-01 | -0.1 | 8.01E-01 | -0.2 | 3.75E-01 | Squalene epoxidase 3 |
| AT1G48175 | 0.2 | 7.69E-02 | 0.1 | 5.23E-01 | 0.1 | 3.39E-01 | 0.1 | 7.69E-01 | -0.4 | 2.79E-01 | 0.5 | 2.13E-01 | Cytidine |
| AT5G41130 | 0.2 | 4.21E-02 | -0.1 | 3.86E-01 | 0.3 | 2.32E-02 | -0.4 | 6.02E-02 | -0.1 | 7.43E-01 | -0.3 | 2.45E-01 | Esterase |
| AT4G33050 | 0.2 | 8.40E-02 | -0.5 | 5.02E-03 | 0.7 | 9.12E-04 | 0.5 | 5.13E-03 | -0.3 | 1.47E-01 | 0.8 | 3.19E-03 | Calmodulin-binding family protein |
| AT3G06910 | 0.2 | 3.54E-02 | -0.1 | 3.78E-01 | 0.3 | 2.03E-02 | 0.1 | 6.37E-01 | 0.1 | 6.94E-01 | 0.0 | 1.00E+00 | Ubiquitin-like-specific protease 1A |
| AT1G12460 | 0.2 | 9.22E-02 | 0.2 | 2.96E-01 | 0.1 | 6.01E-01 | 0.2 | 5.90E-01 | -0.1 | 8.31E-01 | 0.2 | 5.47E-01 | Leucine-rich repeat protein kinase family protein |
| AT1G10170 | 0.2 | 9.51E-03 | -0.4 | 4.39E-04 | 0.7 | 8.33E-05 | 0.3 | 1.26E-01 | -0.4 | 8.26E-02 | 0.8 | 1.62E-02 | NF-X-like 1 |
| AT1G54030 | 0.2 | 1.24E-02 | 0.3 | 5.09E-03 | -0.1 | 5.61E-01 | -0.1 | 7.66E-01 | 0.2 | 3.45E-01 | -0.2 | 3.18E-01 | GDSL-like Lipase |
| AT3G60160 | 0.2 | 2.35E-01 | -0.5 | 4.77E-02 | 0.7 | 1.70E-02 | 0.8 | 3.55E-03 | 0.7 | 9.14E-03 | 0.0 | 9.26E-01 | ABC-type xenobiotic transporter |
| AT1G55150 | 0.2 | 1.04E-02 | -0.1 | 1.49E-01 | 0.4 | 3.29E-03 | 0.3 | 7.04E-02 | 0.1 | 7.31E-01 | 0.2 | 3.21E-01 | DEA-box RNA helicase family protein |
| AT1G75340 | 0.2 | 7.38E-03 | 0.2 | 1.46E-02 | 0.0 | 8.34E-01 | 0.1 | 6.58E-01 | 0.2 | 5.66E-01 | 0.0 | 9.06E-01 | Zinc finger C-x8-C-x5-C-x3-H type family protein |
| AT1G60770 | 0.2 | 3.93E-02 | 0.3 | 2.58E-02 | 0.0 | 8.02E-01 | 0.4 | 1.80E-02 | 0.3 | 9.82E-02 | 0.1 | 6.98E-01 | Large ribosomal subunit protein mL101 |
| AT3G15080 | 0.2 | 7.15E-02 | 0.2 | 1.45E-01 | 0.0 | 7.88E-01 | 0.4 | 5.37E-02 | 0.0 | 8.76E-01 | 0.3 | 1.68E-01 | RNA exonuclease 4 |
| AT5G26110 | 0.2 | 5.43E-02 | 0.1 | 4.85E-01 | 0.2 | 2.75E-01 | -0.1 | 6.79E-01 | -0.1 | 4.49E-01 | 0.1 | 7.89E-01 | non-specific serine |
| AT1G13870 | 0.2 | 1.18E-02 | 0.2 | 8.07E-02 | 0.1 | 4.09E-01 | -0.1 | 7.15E-01 | -0.1 | 6.33E-01 | 0.0 | 9.05E-01 | Protein KTI12 homolog |
| AT5G16310 | 0.2 | 5.29E-02 | 0.0 | 7.73E-01 | 0.3 | 6.71E-02 | 0.0 | 9.64E-01 | 0.0 | 9.41E-01 | 0.0 | 9.80E-01 | Ubiquitin carboxyl-terminal hydrolase |

| | | | | | | | | | | | | | |
|---|---|---|---|---|---|---|---|---|---|---|---|---|---|
| AT2G20570 | 0.2 | 4.08E-02 | 0.1 | 2.37E-01 | 0.1 | 4.01E-01 | -0.4 | 1.74E-02 | 0.2 | 3.45E-01 | -0.5 | 9.07E-03 | GBF's pro-rich region-interacting factor 1 |
| AT3G53650 | 0.2 | 8.73E-02 | 0.7 | 5.19E-04 | -0.4 | 1.70E-02 | -0.4 | 1.75E-01 | 0.1 | 8.57E-01 | -0.4 | 1.37E-01 | Histone H2B.8 |
| AT2G35840 | 0.2 | 1.54E-02 | -0.3 | 1.39E-02 | 0.5 | 6.36E-04 | -0.2 | 1.68E-01 | 0.1 | 8.00E-01 | -0.2 | 1.74E-01 | Probable sucrose-phosphatase 2 |
| AT4G22980 | 0.2 | 3.95E-02 | -0.1 | 4.01E-01 | 0.3 | 2.44E-02 | 0.1 | 5.66E-01 | -0.1 | 8.48E-01 | 0.2 | 5.06E-01 | Molybdenum cofactor sulfurase-like protein |
| AT5G54580 | 0.2 | 7.04E-02 | 0.3 | 2.97E-02 | -0.1 | 6.47E-01 | 0.4 | 4.23E-02 | 0.0 | 9.00E-01 | 0.4 | 8.56E-02 | Organelle RRM domain-containing protein 2, mitochondrial |
| AT2G46710 | 0.2 | 6.12E-02 | 0.4 | 5.22E-03 | -0.2 | 1.95E-01 | -0.2 | 1.04E-01 | -0.3 | 3.49E-02 | 0.1 | 6.51E-01 | Rho GTPase-activating protein 3 |
| AT4G13950 | 0.2 | 7.73E-02 | 0.2 | 1.31E-01 | 0.0 | 8.59E-01 | 0.2 | 5.96E-01 | 0.1 | 8.70E-01 | 0.1 | 8.43E-01 | Protein RALF-like 31 |
| AT5G58900 | 0.2 | 9.17E-03 | 0.4 | 1.09E-03 | -0.1 | 1.86E-01 | 0.1 | 6.10E-01 | 0.0 | 9.56E-01 | 0.1 | 6.66E-01 | Homeodomain-like transcriptional regulator |
| AT2G35160 | 0.2 | 7.04E-02 | 0.2 | 2.59E-01 | 0.1 | 5.81E-01 | 0.6 | 1.00E-01 | 0.1 | 8.13E-01 | 0.4 | 3.27E-01 | SU3-9 homolog 5 |
| AT4G35410 | 0.2 | 2.91E-02 | 0.3 | 8.93E-03 | -0.1 | 5.31E-01 | 0.4 | 1.68E-01 | 0.4 | 1.71E-01 | 0.0 | 9.39E-01 | AP complex mu |
| AT3G60050 | 0.2 | 2.34E-02 | 0.2 | 5.69E-02 | 0.0 | 7.46E-01 | 0.0 | 8.99E-01 | 0.1 | 6.73E-01 | -0.1 | 8.07E-01 | Pentatricopeptide repeat-containing protein At3g60050 |
| AT1G51700 | 0.2 | 1.99E-01 | -0.2 | 2.80E-01 | 0.5 | 5.10E-02 | 0.1 | 5.78E-01 | 0.0 | 9.97E-01 | 0.1 | 6.80E-01 | Dof zinc finger protein DOF1.7 |
| AT1G68580 | 0.2 | 7.09E-03 | 0.0 | 8.66E-01 | 0.3 | 1.18E-02 | -0.2 | 2.95E-01 | -0.3 | 1.29E-01 | 0.1 | 7.11E-01 | Agenet and bromo-adjacent homology domain-containing protein |
| AT5G51230 | 0.2 | 7.41E-03 | 0.0 | 7.43E-01 | 0.2 | 2.93E-02 | 0.0 | 9.18E-01 | 0.0 | 9.19E-01 | 0.0 | 9.92E-01 | Polycomb protein VEFS-Box domain-containing protein |
| AT1G74090 | 0.2 | 3.47E-02 | 0.4 | 2.44E-03 | -0.2 | 1.30E-01 | -0.7 | 1.72E-04 | 0.3 | 9.55E-02 | -0.9 | 2.77E-04 | Cytosolic sulfotransferase 18 |
| AT2G42520 | 0.2 | 6.21E-03 | 0.0 | 7.94E-01 | 0.2 | 2.12E-02 | 0.0 | 9.31E-01 | 0.0 | 9.07E-01 | -0.1 | 8.70E-01 | RNA helicase |
| AT1G51510 | 0.2 | 3.40E-03 | 0.3 | 3.90E-03 | 0.0 | 9.29E-01 | 0.1 | 3.73E-01 | 0.0 | 9.11E-01 | 0.1 | 6.47E-01 | RNA-binding protein Y14 |
| AT1G71040 | 0.2 | 1.18E-02 | -0.1 | 4.12E-01 | 0.3 | 6.33E-03 | 0.1 | 6.96E-01 | -0.2 | 3.00E-01 | 0.3 | 2.44E-01 | Multicopper oxidase LPR2 |
| AT5G43700 | 0.2 | 5.56E-02 | 0.1 | 4.65E-01 | 0.2 | 2.78E-01 | -0.1 | 7.21E-01 | -0.3 | 2.69E-01 | 0.2 | 5.56E-01 | Auxin-responsive protein IAA4 |
| AT3G51080 | 0.2 | 1.86E-01 | 0.2 | 2.96E-01 | 0.0 | 8.54E-01 | -0.4 | 5.14E-02 | -0.4 | 8.65E-02 | 0.0 | 9.67E-01 | GATA transcription factor 6 |
| AT3G20660 | 0.2 | 8.79E-02 | 0.3 | 8.38E-02 | 0.0 | 9.51E-01 | 0.0 | 8.80E-01 | -0.7 | 3.49E-02 | 0.6 | 8.08E-02 | Organic cation |
| AT3G46010 | 0.2 | 1.55E-02 | 0.2 | 6.23E-02 | 0.1 | 5.57E-01 | 0.1 | 6.33E-01 | 0.1 | 7.98E-01 | 0.0 | 9.44E-01 | Actin depolymerizing factor 1 |
| AT4G01880 | 0.2 | 1.27E-02 | 0.2 | 2.27E-02 | 0.0 | 8.66E-01 | 0.3 | 2.03E-01 | -0.1 | 8.66E-01 | 0.3 | 2.58E-01 | tRNA:mX modification enzyme TRM13 |
| AT3G47930 | 0.2 | 1.14E-02 | 0.2 | 2.27E-02 | 0.0 | 8.20E-01 | 0.4 | 1.32E-01 | 0.3 | 2.06E-01 | 0.0 | 9.71E-01 | L-galactono-1,4-lactone dehydrogenase |
| AT2G24990 | 0.2 | 7.75E-02 | 0.2 | 2.41E-01 | 0.1 | 6.12E-01 | 0.4 | 3.31E-02 | 0.0 | 9.83E-01 | 0.4 | 1.07E-01 | Serine |
| AT1G77840 | 0.2 | 5.82E-03 | 0.3 | 4.36E-03 | 0.0 | 7.80E-01 | 0.4 | 4.93E-03 | 0.0 | 9.98E-01 | 0.4 | 2.00E-02 | Probable eukaryotic translation initiation factor 5-2 |
| AT3G55340 | 0.2 | 5.06E-02 | 0.2 | 9.95E-02 | 0.0 | 8.03E-01 | 0.7 | 3.22E-03 | 0.4 | 6.81E-02 | 0.2 | 3.85E-01 | Phragmoplastin interacting protein 1 |
| AT5G38880 | 0.2 | 5.31E-03 | 0.1 | 2.13E-01 | 0.2 | 8.98E-02 | -0.1 | 5.72E-01 | -0.2 | 4.31E-01 | 0.1 | 8.60E-01 | AUGMIN subunit 5 |
| AT5G08520 | 0.2 | 3.27E-02 | 0.3 | 2.90E-02 | 0.0 | 9.10E-01 | -0.6 | 2.54E-01 | -0.5 | 3.64E-01 | 0.0 | 9.61E-01 | Transcription factor SRM1 |
| AT3G52390 | 0.2 | 9.40E-02 | 0.3 | 3.45E-02 | -0.1 | 6.20E-01 | 0.6 | 2.21E-02 | 0.4 | 1.76E-01 | 0.2 | 4.79E-01 | At3g52390 |
| AT3G24030 | 0.2 | 4.80E-03 | 0.1 | 2.84E-01 | 0.2 | 6.71E-02 | -0.5 | 6.97E-03 | -0.2 | 3.31E-01 | -0.3 | 1.69E-01 | Hydroxyethylthiazole kinase |
| AT3G20150 | 0.2 | 2.74E-02 | 0.2 | 3.51E-02 | 0.0 | 9.73E-01 | 0.2 | 3.03E-01 | 0.2 | 5.82E-01 | 0.1 | 8.03E-01 | Kinesin motor family protein |
| AT1G53390 | 0.2 | 7.36E-03 | 0.1 | 4.07E-01 | 0.2 | 6.13E-02 | 0.3 | 9.78E-02 | 0.2 | 2.04E-01 | 0.0 | 9.08E-01 | ABC transporter G family member 24 |
| AT3G52660 | 0.2 | 1.90E-03 | 0.0 | 7.93E-01 | 0.2 | 8.56E-03 | -0.1 | 5.34E-01 | -0.1 | 7.28E-01 | 0.0 | 9.08E-01 | At3g52660 family protein) |
| AT4G04570 | 0.2 | 1.05E-01 | -0.2 | 1.52E-01 | 0.5 | 1.42E-02 | 0.0 | 8.99E-01 | -0.2 | 2.10E-01 | 0.2 | 3.91E-01 | Cysteine-rich RLK 40 |
| AT1G75120 | 0.2 | 1.11E-01 | 0.1 | 4.18E-01 | 0.1 | 5.29E-01 | 0.2 | 5.32E-01 | 0.0 | 9.70E-01 | 0.2 | 5.14E-01 | Arabinosyltransferase RRA1 |
| AT5G66650 | 0.2 | 2.76E-01 | 0.8 | 2.10E-03 | -0.6 | 3.17E-02 | 1.7 | 3.49E-03 | 0.2 | 7.92E-01 | 1.4 | 3.08E-02 | Calcium uniporter protein 3, mitochondrial |
| AT4G31810 | 0.2 | 6.36E-02 | 0.4 | 1.58E-02 | -0.1 | 4.74E-01 | 0.7 | 3.44E-04 | 0.4 | 3.40E-02 | 0.3 | 1.57E-01 | Small ribosomal subunit protein mS47 |
| AT5G38850 | 0.2 | 5.48E-02 | 0.1 | 5.06E-01 | 0.2 | 2.76E-01 | -0.5 | 3.24E-02 | -0.5 | 4.78E-02 | 0.0 | 9.97E-01 | TIR domain-containing protein |
| AT3G17090 | 0.2 | 4.49E-03 | 0.2 | 9.52E-03 | 0.0 | 8.06E-01 | -0.2 | 3.73E-01 | -0.2 | 3.60E-01 | 0.0 | 9.35E-01 | Probable protein phosphatase 2C 42 |
| AT1G28580 | 0.2 | 8.57E-03 | 0.1 | 4.60E-01 | 0.2 | 5.62E-02 | -0.2 | 1.04E-01 | -0.2 | 8.94E-02 | 0.0 | 8.87E-01 | GDSL esterase |
| AT5G12220 | 0.2 | 7.84E-02 | 0.1 | 6.72E-01 | 0.2 | 2.56E-01 | 0.4 | 5.07E-02 | 0.1 | 8.43E-01 | 0.3 | 2.11E-01 | Las1-like family protein |
| AT1G06150 | 0.2 | 1.43E-02 | 0.0 | 6.84E-01 | 0.2 | 6.04E-02 | 0.1 | 7.94E-01 | -0.1 | 7.93E-01 | 0.1 | 6.04E-01 | Transcription factor EMB1444 |
| AT2G43640 | 0.3 | 3.99E-02 | 0.5 | 1.69E-03 | -0.2 | 1.02E-01 | 0.0 | 9.61E-01 | -0.2 | 5.05E-01 | 0.2 | 4.76E-01 | Signal recognition particle 14 kDa protein |
| AT1G14520 | 0.3 | 3.58E-01 | -0.6 | 8.60E-02 | 0.8 | 3.36E-02 | -0.6 | 1.64E-03 | -0.2 | 2.51E-01 | -0.4 | 9.68E-02 | Inositol oxygenase |
| AT1G59620 | 0.3 | 1.03E-01 | -0.3 | 1.32E-01 | 0.5 | 1.70E-02 | 0.3 | 2.69E-01 | 0.3 | 3.23E-01 | 0.0 | 9.92E-01 | Disease resistance protein family |
| AT4G23270 | 0.3 | 1.27E-02 | 0.0 | 9.22E-01 | 0.3 | 2.05E-02 | 0.2 | 3.38E-01 | -0.1 | 6.46E-01 | 0.3 | 2.08E-01 | Cysteine-rich RLK 19 |
| AT2G31570 | 0.3 | 3.58E-03 | 0.3 | 3.01E-03 | 0.0 | 7.98E-01 | 0.4 | 1.47E-02 | 0.1 | 4.26E-01 | 0.2 | 2.62E-01 | Glutathione peroxidase |
| AT2G26910 | 0.3 | 4.86E-02 | 0.1 | 3.10E-01 | 0.1 | 3.57E-01 | -0.1 | 8.15E-01 | -0.1 | 6.45E-01 | 0.1 | 8.58E-01 | ABC transporter G family member 32 |
| AT5G19090 | 0.3 | 7.52E-03 | 0.2 | 2.80E-02 | 0.1 | 5.79E-01 | -0.3 | 3.04E-01 | 0.1 | 8.52E-01 | -0.3 | 2.56E-01 | HMA domain-containing protein |
| AT3G51740 | 0.3 | 5.71E-02 | 0.4 | 1.30E-02 | -0.1 | 4.34E-01 | 0.1 | 4.51E-01 | 0.1 | 5.01E-01 | 0.0 | 9.84E-01 | Probably inactive leucine-rich repeat receptor-like protein kinase IMK2 |
| AT2G21650 | 0.3 | 1.49E-01 | 0.7 | 1.64E-03 | -0.5 | 2.93E-02 | -0.4 | 1.66E-01 | 0.2 | 4.59E-01 | -0.6 | 4.90E-02 | Protein RADIALIS-like 2 |
| AT1G02060 | 0.3 | 2.78E-01 | -0.4 | 1.98E-01 | 0.6 | 6.60E-02 | 0.0 | 9.88E-01 | -0.2 | 8.51E-01 | 0.2 | 8.39E-01 | Pentatricopeptide repeat-containing protein At1g02060, chloroplastic |
| AT4G26100 | 0.3 | 1.44E-03 | 0.1 | 6.14E-02 | 0.1 | 8.30E-02 | -0.2 | 1.58E-01 | -0.1 | 7.93E-01 | -0.2 | 4.53E-01 | Casein kinase 1-like protein 1 |
| AT5G17630 | 0.3 | 1.06E-02 | 0.2 | 5.53E-02 | 0.1 | 4.79E-01 | 0.3 | 1.40E-01 | 0.5 | 2.36E-02 | -0.2 | 4.64E-01 | Xylulose 5-phosphate |
| AT2G19750 | 0.3 | 7.83E-02 | 0.4 | 1.14E-02 | -0.2 | 2.92E-01 | 0.5 | 3.23E-02 | 0.3 | 2.61E-01 | 0.2 | 5.40E-01 | Small ribosomal subunit protein eS30z |
| AT3G53380 | 0.3 | 2.78E-02 | 0.5 | 1.12E-03 | -0.2 | 8.61E-02 | -0.2 | 2.80E-01 | 0.0 | 8.04E-01 | -0.2 | 2.63E-01 | L-type lectin-domain containing receptor kinase VIII.1 |
| AT5G44020 | 0.3 | 3.70E-02 | 1.0 | 1.86E-05 | -0.7 | 3.53E-04 | -0.4 | 2.55E-02 | -0.7 | 4.22E-03 | 0.2 | 4.36E-01 | HAD superfamily, subfamily IIIB acid phosphatase |
| AT4G32060 | 0.3 | 7.86E-03 | 0.1 | 2.19E-01 | 0.1 | 1.15E-01 | -0.2 | 5.65E-02 | -0.2 | 1.44E-01 | 0.0 | 9.04E-01 | AT4G32060 protein |
| AT5G48820 | 0.3 | 2.06E-01 | 0.3 | 1.62E-01 | 0.0 | 8.82E-01 | 0.3 | 4.37E-01 | 0.2 | 7.82E-01 | 0.2 | 7.90E-01 | Inhibitor |
| AT5G19760 | 0.3 | 1.11E-02 | 0.5 | 2.89E-04 | -0.2 | 2.64E-02 | 0.2 | 6.43E-02 | 0.3 | 2.04E-02 | -0.1 | 6.36E-01 | Mitochondrial dicarboxylate |
| AT1G61010 | 0.3 | 1.54E-03 | 0.1 | 3.34E-02 | 0.1 | 1.46E-01 | 0.1 | 6.62E-01 | 0.0 | 9.65E-01 | 0.1 | 7.50E-01 | Cleavage and polyadenylation specificity factor subunit 3-I |
| AT2G34630 | 0.3 | 6.09E-03 | 0.0 | 6.97E-01 | 0.2 | 2.49E-02 | 0.4 | 2.18E-02 | 0.3 | 6.52E-02 | 0.0 | 8.70E-01 | Solanesyl diphosphate synthase 3, chloroplastic |
| AT1G27880 | 0.3 | 8.76E-02 | 0.3 | 9.62E-02 | 0.0 | 9.81E-01 | 0.3 | 4.02E-01 | 0.8 | 2.25E-02 | -0.5 | 1.95E-01 | DEAD |
| AT2G42120 | 0.3 | 3.78E-02 | 0.1 | 4.56E-01 | 0.2 | 2.35E-01 | -0.1 | 6.91E-01 | 0.0 | 9.11E-01 | -0.1 | 6.32E-01 | DNA polymerase delta small subunit |
| AT1G23000 | 0.3 | 7.01E-02 | 0.4 | 9.46E-03 | -0.2 | 3.16E-01 | -0.2 | 3.16E-01 | -0.1 | 7.15E-01 | -0.1 | 7.35E-01 | Heavy metal transport |
| AT3G58680 | 0.3 | 2.85E-03 | 0.3 | 2.92E-03 | 0.0 | 8.66E-01 | -0.1 | 5.36E-01 | -0.2 | 1.40E-01 | 0.1 | 4.85E-01 | Multiprotein-bridging factor 1b |
| AT3G25910 | 0.3 | 3.64E-03 | 0.0 | 6.54E-01 | 0.2 | 1.75E-02 | -0.3 | 3.31E-02 | -0.6 | 2.53E-03 | 0.2 | 3.01E-01 | AT3g25910 |
| AT5G27680 | 0.3 | 3.80E-02 | 0.1 | 6.73E-01 | 0.2 | 1.58E-01 | 0.0 | 9.66E-01 | 0.2 | 6.51E-01 | -0.2 | 6.74E-01 | ATP-dependent DNA helicase |
| AT3G11470 | 0.3 | 8.87E-02 | 0.1 | 4.51E-01 | 0.1 | 4.33E-01 | 0.0 | 9.54E-01 | -0.1 | 8.63E-01 | 0.1 | 8.55E-01 | 4'-phosphopantetheinyl transferase superfamily |
| AT5G64120 | 0.3 | 6.72E-01 | -0.3 | 6.14E-01 | 0.6 | 3.93E-01 | -0.8 | 3.83E-01 | -3.2 | 2.18E-02 | 2.4 | 5.25E-02 | Peroxidase 71 |
| AT1G12010 | 0.3 | 1.06E-01 | 0.4 | 3.40E-02 | -0.1 | 5.41E-01 | -0.6 | 2.30E-02 | -0.2 | 5.20E-01 | -0.4 | 1.50E-01 | 1-aminocyclopropane-1-carboxylate oxidase 3 |
| AT5G53900 | 0.3 | 2.22E-02 | 0.3 | 1.91E-02 | 0.0 | 8.88E-01 | 0.1 | 4.18E-01 | -0.2 | 3.68E-01 | 0.3 | 1.21E-01 | Serine |
| AT3G17840 | 0.3 | 8.75E-03 | 0.1 | 1.09E-01 | 0.1 | 2.41E-01 | -0.1 | 6.22E-01 | 0.1 | 4.33E-01 | -0.2 | 2.94E-01 | Probable inactive receptor kinase RLK902 |
| AT5G55700 | 0.3 | 2.31E-02 | 0.1 | 5.98E-01 | 0.2 | 1.12E-01 | -0.3 | 1.33E-01 | -0.4 | 8.77E-02 | 0.1 | 8.19E-01 | Beta-amylase |
| AT3G53810 | 0.3 | 2.27E-02 | -0.2 | 1.48E-01 | 0.4 | 5.88E-03 | 0.0 | 9.03E-01 | -0.4 | 7.49E-02 | 0.4 | 7.46E-02 | L-type lectin-domain containing receptor kinase IV.2 |
| AT1G07360 | 0.3 | 4.40E-03 | 0.1 | 3.77E-01 | 0.2 | 3.92E-02 | 0.2 | 4.96E-01 | 0.1 | 6.92E-01 | 0.0 | 9.22E-01 | Zinc finger CCCH domain-containing protein 4 |
| AT1G74320 | 0.3 | 6.81E-03 | 0.2 | 4.98E-02 | 0.1 | 3.95E-01 | -0.2 | 3.90E-01 | 0.0 | 9.30E-01 | -0.2 | 5.37E-01 | Probable choline kinase 2 |
| AT1G09560 | 0.3 | 3.89E-02 | 0.1 | 6.55E-01 | 0.2 | 1.28E-01 | -0.2 | 2.37E-01 | -1.0 | 1.10E-05 | 0.8 | 5.83E-04 | Germin-like protein subfamily 2 member 1 |
| AT3G45780 | 0.3 | 6.98E-02 | 0.2 | 1.95E-01 | 0.1 | 6.49E-01 | -0.6 | 5.62E-04 | -0.2 | 3.72E-01 | -0.4 | 3.75E-02 | Phototropin-1 |
| AT3G22660 | 0.3 | 1.61E-01 | 0.3 | 1.31E-01 | 0.0 | 8.95E-01 | 0.5 | 2.19E-03 | 0.2 | 2.81E-01 | 0.3 | 8.97E-02 | Probable rRNA-processing protein EBP2 homolog |
| AT1G44900 | 0.3 | 8.08E-03 | 0.2 | 3.20E-02 | 0.1 | 5.82E-01 | 0.5 | 7.35E-02 | 0.4 | 1.45E-01 | 0.0 | 9.21E-01 | DNA replication licensing factor MCM2 |
| AT4G25280 | 0.3 | 2.75E-02 | 0.1 | 2.61E-01 | 0.1 | 3.07E-01 | -0.1 | 8.07E-01 | 0.0 | 9.81E-01 | 0.0 | 8.79E-01 | Probable UMP-CMP kinase 2 |

| | | | | | | | | | | | | | |
|---|---|---|---|---|---|---|---|---|---|---|---|---|---|
| AT3G48690 | 0.3 | 1.06E-02 | -0.1 | 2.01E-01 | 0.4 | 3.07E-03 | -0.6 | 2.02E-04 | -0.7 | 2.38E-04 | 0.1 | 7.43E-01 | Probable carboxylesterase 12 |
| AT4G35230 | 0.3 | 1.53E-02 | 0.0 | 7.27E-01 | 0.2 | 5.16E-02 | -0.2 | 9.04E-02 | -0.3 | 1.34E-02 | 0.1 | 4.36E-01 | Serine |
| AT2G17120 | 0.3 | 1.30E-01 | 0.1 | 7.12E-01 | 0.2 | 3.13E-01 | 0.2 | 9.04E-02 | 0.0 | 8.70E-01 | 0.3 | 1.32E-01 | LysM domain-containing GPI-anchored protein 2 |
| AT3G28580 | 0.3 | 2.58E-01 | -1.0 | 4.30E-03 | 1.3 | 3.06E-03 | 2.3 | 8.25E-06 | 0.8 | 7.26E-02 | 1.5 | 1.35E-03 | AAA+ ATPase domain-containing protein |
| AT2G01900 | 0.3 | 2.39E-01 | 0.0 | 9.01E-01 | 0.3 | 2.93E-01 | -0.9 | 3.03E-02 | -0.2 | 6.35E-01 | -0.6 | 1.36E-01 | DNAse I-like superfamily protein |
| AT4G39950 | 0.3 | 6.67E-02 | 0.5 | 2.32E-03 | -0.3 | 7.04E-02 | 0.4 | 2.02E-02 | 0.3 | 9.42E-02 | 0.1 | 7.86E-01 | Cytochrome P450, family 79, subfamily B, polypeptide 2 |
| AT3G57080 | 0.3 | 8.75E-03 | 0.2 | 1.53E-02 | 0.0 | 8.80E-01 | 0.0 | 9.15E-01 | -0.2 | 1.90E-01 | 0.2 | 3.29E-01 | DNA-directed RNA polymerase V subunit 5A |
| AT1G80370 | 0.3 | 9.26E-02 | 0.6 | 1.66E-03 | -0.3 | 5.56E-02 | -0.2 | 5.22E-01 | 0.0 | 9.55E-01 | -0.2 | 5.83E-01 | Cyclin-A2-4 |
| AT3G22460 | 0.3 | 4.04E-02 | -0.1 | 6.35E-01 | 0.3 | 4.38E-02 | -0.7 | 2.69E-02 | -1.0 | 4.94E-03 | 0.3 | 4.53E-01 | Putative inactive cysteine synthase 2 -lyase 4) |
| AT3G45980 | 0.3 | 4.80E-03 | 0.4 | 4.19E-04 | -0.1 | 1.04E-01 | -0.1 | 5.81E-01 | -0.3 | 2.49E-01 | 0.2 | 6.51E-01 | Histone H2B.6 |
| AT2G27760 | 0.3 | 2.13E-02 | 0.0 | 7.76E-01 | 0.3 | 3.42E-02 | -0.3 | 3.78E-01 | -0.1 | 7.82E-01 | -0.2 | 7.03E-01 | tRNA dimethylallyltransferase 2 |
| AT4G20330 | 0.3 | 6.41E-02 | 0.0 | 7.81E-01 | 0.3 | 6.88E-02 | 0.0 | 9.16E-01 | 0.0 | 9.81E-01 | 0.0 | 9.28E-01 | Transcription initiation factor IIE subunit beta |
| AT5G03406 | 0.3 | 4.92E-02 | 0.1 | 3.00E-01 | 0.1 | 4.17E-01 | 0.1 | 7.45E-01 | 0.1 | 8.04E-01 | 0.0 | 9.92E-01 | Class II aaRS and biotin synthetases superfamily protein |
| AT1G70940 | 0.3 | 2.84E-02 | 0.0 | 7.76E-01 | 0.2 | 7.69E-02 | -0.5 | 1.55E-03 | -0.3 | 5.84E-02 | -0.2 | 2.82E-01 | Auxin efflux carrier component 3 |
| AT1G70730 | 0.3 | 5.23E-03 | 0.0 | 7.67E-01 | 0.3 | 6.82E-03 | 0.1 | 7.99E-01 | 0.2 | 3.46E-01 | -0.2 | 5.79E-01 | phosphoglucomutase |
| AT5G06811 | 0.3 | 1.53E-02 | 0.0 | 9.01E-01 | 0.3 | 3.43E-02 | 0.3 | 2.85E-01 | -0.2 | 4.29E-01 | 0.5 | 9.68E-02 | Uncharacterized protein |
| AT5G60990 | 0.3 | 1.71E-02 | 0.2 | 1.13E-01 | 0.1 | 4.12E-01 | 0.4 | 1.53E-02 | 0.0 | 8.47E-01 | 0.3 | 8.97E-02 | DEAD-box ATP-dependent RNA helicase 10 |
| AT3G28180 | 0.3 | 2.09E-02 | 0.1 | 3.44E-01 | 0.2 | 1.67E-01 | -0.2 | 9.86E-02 | -0.2 | 1.31E-01 | 0.0 | 9.83E-01 | Xyloglucan glycosyltransferase 4 |
| AT5G58110 | 0.3 | 1.08E-02 | 0.1 | 3.28E-01 | 0.2 | 1.01E-01 | -0.1 | 5.36E-01 | -0.2 | 2.33E-01 | 0.1 | 6.57E-01 | Activator of Hsp90 ATPase AHSA1-like N-terminal domain-containing protein |
| AT5G39840 | 0.3 | 1.85E-02 | 0.1 | 2.46E-01 | 0.1 | 2.52E-01 | 0.7 | 1.39E-02 | 0.3 | 3.35E-01 | 0.4 | 2.46E-01 | DExH-box ATP-dependent RNA helicase DExH18, mitochondrial |
| AT1G17960 | 0.3 | 3.05E-01 | -1.1 | 6.45E-03 | 1.3 | 4.49E-03 | 1.1 | 1.67E-02 | 0.6 | 2.57E-01 | 0.5 | 2.62E-01 | threonine--tRNA ligase |
| AT1G64850 | 0.3 | 7.24E-03 | 0.3 | 8.20E-03 | 0.0 | 9.70E-01 | -0.2 | 5.17E-01 | 0.0 | 8.93E-01 | -0.1 | 7.80E-01 | hypothetical protein |
| AT2G29570 | 0.3 | 2.73E-02 | 0.4 | 6.62E-03 | -0.1 | 4.40E-01 | 0.2 | 2.67E-01 | 0.0 | 8.86E-01 | 0.1 | 5.15E-01 | Proliferating cell nuclear antigen 2 |
| AT2G37520 | 0.3 | 9.59E-03 | 0.0 | 7.94E-01 | 0.3 | 1.43E-02 | 0.1 | 6.30E-01 | 0.0 | 9.03E-01 | 0.1 | 6.32E-01 | Acyl-CoA N-acyltransferase with RING |
| AT5G62430 | 0.3 | 1.12E-01 | 0.3 | 8.65E-02 | 0.0 | 8.70E-01 | -1.1 | 2.24E-02 | -0.9 | 7.02E-02 | -0.2 | 8.14E-01 | CDF1 |
| AT5G11010 | 0.3 | 2.09E-02 | 0.2 | 1.63E-01 | 0.1 | 3.71E-01 | 0.0 | 9.90E-01 | 0.0 | 9.40E-01 | 0.0 | 9.34E-01 | Polynucleotide 5'-hydroxyl-kinase NOL9 |
| AT5G42250 | 0.3 | 3.09E-03 | 0.1 | 6.98E-02 | 0.1 | 1.44E-01 | -0.6 | 2.02E-03 | -0.2 | 2.87E-01 | -0.4 | 5.70E-02 | Alcohol dehydrogenase-like 7 |
| AT5G38940 | 0.3 | 7.16E-02 | 1.2 | 1.92E-05 | -1.0 | 2.61E-04 | -0.1 | 6.40E-01 | 0.1 | 7.83E-01 | -0.2 | 4.44E-01 | Germin-like protein |
| AT3G17510 | 0.3 | 1.50E-01 | 0.1 | 4.77E-01 | 0.1 | 5.32E-01 | -0.4 | 7.76E-03 | -0.4 | 7.62E-03 | 0.0 | 8.59E-01 | CBL-interacting serine |
| AT5G19310 | 0.3 | 3.92E-03 | 0.0 | 7.41E-01 | 0.2 | 2.02E-02 | -0.1 | 4.39E-01 | -0.3 | 1.39E-01 | 0.1 | 5.85E-01 | Homeotic protein regulator |
| AT5G20950 | 0.3 | 7.07E-02 | 0.6 | 2.09E-03 | -0.3 | 5.87E-02 | -0.1 | 6.15E-01 | -0.2 | 5.09E-02 | 0.2 | 2.64E-01 | Beta-D-glucan exohydrolase-like protein |
| AT1G25450 | 0.3 | 1.55E-02 | 0.4 | 1.80E-03 | -0.1 | 2.06E-01 | 0.0 | 8.87E-01 | -0.1 | 4.80E-01 | 0.2 | 4.72E-01 | 3-ketoacyl-CoA synthase 5 |
| AT1G05200 | 0.3 | 3.46E-02 | -0.1 | 5.34E-01 | 0.3 | 2.38E-02 | -0.4 | 5.60E-01 | 0.0 | 9.70E-01 | -0.4 | 6.05E-01 | Glutamate receptor 3.4 |
| AT4G36150 | 0.3 | 8.34E-02 | -0.3 | 8.60E-02 | 0.6 | 1.07E-02 | 0.6 | 2.25E-02 | 0.3 | 3.86E-01 | 0.3 | 2.80E-01 | ADP-ribosyl cyclase |
| AT5G36880 | 0.3 | 6.34E-03 | 0.2 | 2.89E-02 | 0.1 | 5.17E-01 | 0.1 | 4.82E-01 | 0.1 | 6.87E-01 | 0.0 | 9.26E-01 | Acetyl-coenzyme A synthetase |
| AT5G63510 | 0.3 | 1.46E-03 | 0.4 | 3.12E-04 | -0.1 | 2.02E-01 | -0.1 | 4.15E-01 | 0.0 | 9.76E-01 | -0.1 | 6.25E-01 | Gamma carbonic anhydrase like 1 |
| AT3G26720 | 0.3 | 1.69E-03 | 0.4 | 2.23E-04 | -0.1 | 1.21E-01 | -0.2 | 7.78E-02 | -0.2 | 5.77E-02 | 0.0 | 8.44E-01 | Alpha-mannosidase |
| AT3G61650 | 0.3 | 1.44E-02 | 0.1 | 2.88E-01 | 0.2 | 1.73E-01 | 0.3 | 2.48E-01 | 0.4 | 9.04E-02 | -0.1 | 6.39E-01 | Tubulin gamma-1 chain |
| AT4G39160 | 0.3 | 5.23E-03 | 0.0 | 9.76E-01 | 0.3 | 1.53E-02 | -0.2 | 3.68E-01 | -0.2 | 3.03E-01 | 0.0 | 8.83E-01 | Homeodomain-like superfamily protein |
| AT2G24645 | 0.3 | 7.65E-02 | 0.3 | 9.80E-02 | 0.0 | 9.62E-01 | 0.2 | 5.91E-01 | 0.1 | 8.56E-01 | 0.1 | 8.33E-01 | Transcriptional factor B3 family protein |
| AT4G37330 | 0.3 | 1.62E-02 | 0.0 | 7.10E-01 | 0.3 | 1.76E-02 | -0.2 | 6.71E-02 | -0.1 | 3.75E-01 | -0.1 | 6.36E-01 | At4g37330 |
| AT3G45290 | 0.3 | 7.97E-02 | -0.2 | 2.17E-01 | 0.5 | 2.20E-02 | 0.1 | 6.81E-01 | -0.2 | 5.78E-01 | 0.3 | 4.08E-01 | MLO-like protein 3 |
| AT2G23620 | 0.3 | 1.95E-01 | -0.1 | 5.51E-01 | 0.4 | 1.30E-01 | -1.4 | 8.63E-03 | -0.3 | 6.16E-01 | -1.1 | 4.80E-02 | Methylesterase 1 |
| AT5G09760 | 0.3 | 1.21E-02 | 0.4 | 7.44E-04 | -0.2 | 1.02E-01 | -0.1 | 3.41E-01 | 0.1 | 5.34E-01 | -0.2 | 1.96E-01 | Plant invertase |
| AT5G58710 | 0.3 | 1.72E-02 | 0.3 | 6.09E-03 | -0.1 | 5.09E-01 | 0.4 | 3.67E-03 | 0.1 | 4.28E-01 | 0.3 | 1.15E-01 | Peptidyl-prolyl cis-trans isomerase CYP20-1 |
| AT3G55360 | 0.3 | 9.15E-03 | 0.4 | 1.30E-03 | -0.1 | 2.04E-01 | 0.3 | 3.16E-02 | 0.3 | 1.06E-01 | 0.0 | 8.48E-01 | Very-long-chain enoyl-CoA reductase |
| AT5G47620 | 0.3 | 1.39E-03 | 0.0 | 9.10E-01 | 0.3 | 4.59E-03 | -0.2 | 1.55E-01 | -0.1 | 3.10E-01 | 0.0 | 9.06E-01 | Putative RNA-binding protein family protein) |
| AT1G20910 | 0.3 | 5.50E-03 | 0.3 | 2.17E-03 | -0.1 | 5.32E-01 | 0.1 | 5.72E-01 | -0.1 | 8.59E-01 | 0.2 | 5.16E-01 | Uncharacterized protein |
| AT1G66670 | 0.3 | 6.32E-03 | 0.3 | 8.73E-03 | 0.0 | 9.81E-01 | 0.3 | 1.36E-02 | 0.2 | 1.56E-01 | 0.1 | 5.43E-01 | ATP-dependent Clp protease proteolytic subunit 3, chloroplastic |
| AT1G32360 | 0.3 | 1.34E-02 | 0.1 | 4.83E-01 | 0.2 | 8.59E-02 | -0.1 | 3.78E-01 | -0.1 | 5.66E-01 | 0.0 | 9.27E-01 | Zinc finger CCCH domain-containing protein 12 |
| AT3G56490 | 0.3 | 1.96E-02 | 0.2 | 5.99E-02 | 0.1 | 6.45E-01 | 0.2 | 3.30E-01 | 0.1 | 7.67E-01 | 0.1 | 7.39E-01 | Adenylylsulfatase HINT1 |
| AT5G14780 | 0.3 | 2.43E-02 | -0.1 | 3.92E-01 | 0.4 | 1.03E-02 | -0.5 | 7.67E-02 | -1.0 | 1.56E-03 | 0.6 | 5.01E-02 | Formate dehydrogenase, mitochondrial |
| AT4G32790 | 0.3 | 4.57E-02 | 0.0 | 7.45E-01 | 0.2 | 1.49E-01 | -0.4 | 4.40E-02 | -0.5 | 2.18E-02 | 0.1 | 7.16E-01 | Uncharacterized protein At4g32790 |
| AT2G41290 | 0.3 | 6.02E-02 | 0.3 | 5.14E-02 | 0.0 | 9.01E-01 | -0.3 | 4.09E-01 | 0.1 | 7.81E-01 | -0.4 | 2.28E-01 | Protein STRICTOSIDINE SYNTHASE-LIKE 2 |
| AT1G77030 | 0.3 | 7.74E-02 | 0.2 | 2.24E-01 | 0.1 | 6.29E-01 | 0.5 | 8.50E-03 | 0.1 | 6.17E-01 | 0.4 | 1.05E-01 | RNA helicase |
| AT4G10320 | 0.3 | 1.32E-03 | 0.0 | 7.12E-01 | 0.2 | 5.78E-03 | 0.1 | 2.46E-01 | 0.1 | 7.03E-01 | 0.1 | 7.13E-01 | Isoleucine--tRNA ligase, cytoplasmic |
| AT3G24560 | 0.3 | 4.16E-02 | 0.0 | 8.08E-01 | 0.3 | 6.26E-02 | -0.2 | 6.83E-01 | -0.1 | 8.91E-01 | -0.1 | 8.63E-01 | tRNA-lysidine synthetase |
| AT4G08390 | 0.3 | 2.05E-02 | 0.3 | 1.43E-02 | 0.0 | 7.84E-01 | -0.1 | 5.06E-01 | 0.6 | 2.39E-03 | -0.7 | 1.14E-03 | L-ascorbate peroxidase |
| AT4G14510 | 0.3 | 1.68E-02 | 0.3 | 1.49E-02 | 0.0 | 8.92E-01 | 0.2 | 5.94E-01 | 0.4 | 3.07E-01 | -0.2 | 5.90E-01 | CFM3B |
| AT1G26520 | 0.3 | 4.64E-02 | 0.2 | 2.14E-01 | 0.1 | 5.08E-01 | 0.2 | 4.84E-01 | 0.4 | 1.51E-01 | -0.2 | 5.28E-01 | Uncharacterized protein At1g26520 |
| AT5G19320 | 0.3 | 3.13E-02 | 0.2 | 2.04E-01 | 0.1 | 3.84E-01 | 0.1 | 3.95E-01 | 0.2 | 2.73E-01 | -0.1 | 8.29E-01 | RAN GTPase-activating protein 2 |
| AT3G03770 | 0.3 | 2.98E-02 | -0.2 | 8.11E-02 | 0.5 | 4.01E-03 | 0.1 | 7.92E-01 | 0.4 | 3.95E-02 | -0.4 | 9.79E-02 | Protein kinase domain-containing protein |
| AT2G45970 | 0.3 | 3.14E-03 | 0.1 | 3.82E-01 | 0.2 | 2.91E-02 | 0.1 | 5.82E-01 | -0.1 | 7.31E-01 | 0.2 | 4.68E-01 | Cytochrome P450 86A8 |
| AT1G55120 | 0.3 | 1.74E-03 | 0.3 | 7.94E-04 | -0.1 | 5.11E-01 | 0.5 | 7.07E-03 | 0.5 | 1.47E-02 | 0.0 | 9.71E-01 | Beta-fructofuranosidase 5 |
| AT4G01040 | 0.3 | 2.19E-03 | 0.3 | 6.30E-04 | -0.1 | 3.46E-01 | 0.0 | 9.60E-01 | -0.1 | 7.12E-01 | 0.1 | 7.44E-01 | Glycosyl hydrolase superfamily protein |
| AT2G47510 | 0.3 | 7.35E-03 | 0.3 | 3.22E-03 | -0.1 | 5.40E-01 | 0.2 | 2.20E-02 | 0.0 | 8.07E-01 | 0.2 | 1.85E-01 | Fumarate hydratase 1, mitochondrial |
| AT5G14590 | 0.3 | 2.24E-03 | 0.1 | 2.15E-01 | 0.2 | 3.90E-02 | 0.0 | 8.04E-01 | 0.1 | 8.03E-01 | 0.0 | 9.70E-01 | Isocitrate dehydrogenase , chloroplastic |
| AT3G28940 | 0.3 | 1.60E-02 | 0.3 | 1.58E-02 | 0.0 | 9.11E-01 | 0.3 | 1.68E-01 | 0.0 | 9.08E-01 | 0.3 | 4.19E-01 | Protein AIG2 B |
| AT2G20060 | 0.3 | 1.98E-02 | 0.3 | 2.24E-02 | 0.0 | 9.88E-01 | 0.3 | 2.81E-02 | 0.2 | 3.53E-01 | 0.2 | 3.18E-01 | 50S ribosomal protein L4 |
| AT2G26680 | 0.3 | 7.19E-03 | 0.3 | 4.81E-03 | 0.0 | 7.56E-01 | 0.0 | 9.34E-01 | 0.2 | 1.66E-01 | -0.2 | 2.49E-01 | Expressed protein |
| AT1G67280 | 0.3 | 3.80E-03 | 0.1 | 4.45E-01 | 0.2 | 2.76E-02 | -0.1 | 4.16E-01 | 0.0 | 8.49E-01 | -0.1 | 7.31E-01 | Probable lactoylglutathione lyase, chloroplastic |
| AT3G12490 | 0.3 | 2.21E-02 | 0.1 | 4.99E-01 | 0.2 | 1.15E-01 | -0.3 | 9.73E-02 | -0.1 | 5.72E-01 | -0.2 | 5.31E-01 | Cysteine proteinase inhibitor |
| AT4G35020 | 0.3 | 2.49E-03 | 0.0 | 6.18E-01 | 0.2 | 1.57E-02 | -0.1 | 7.34E-01 | -0.2 | 2.72E-01 | 0.1 | 5.79E-01 | RAC-like 3 |
| AT2G43350 | 0.3 | 2.23E-02 | 0.0 | 9.22E-01 | 0.3 | 5.66E-02 | 0.3 | 1.92E-01 | 0.1 | 5.96E-01 | 0.1 | 6.74E-01 | Glutathione peroxidase |
| AT5G65900 | 0.3 | 2.18E-02 | 0.3 | 1.71E-02 | 0.0 | 8.57E-01 | 0.5 | 3.91E-02 | 0.3 | 3.33E-01 | 0.2 | 5.05E-01 | DEAD-box ATP-dependent RNA helicase 27 |
| AT5G52470 | 0.3 | 4.02E-02 | 0.2 | 1.71E-01 | 0.1 | 5.10E-01 | 0.3 | 2.44E-02 | 0.0 | 8.56E-01 | 0.3 | 6.16E-02 | rRNA 2'-O-methyltransferase fibrillarin 1 |
| AT3G04890 | 0.3 | 8.67E-03 | 0.2 | 2.32E-02 | 0.0 | 7.16E-01 | -0.3 | 1.04E-01 | -0.2 | 2.71E-01 | -0.1 | 8.21E-01 | Adenine phosphoribosyltransferase-like protein, putative |
| AT5G11880 | 0.3 | 6.68E-03 | 0.4 | 6.73E-04 | -0.1 | 1.28E-01 | 0.4 | 2.05E-03 | 0.1 | 4.28E-01 | 0.3 | 9.48E-02 | Diaminopimelate decarboxylase 2, chloroplastic |
| AT5G02500 | 0.3 | 2.49E-02 | 0.1 | 3.89E-01 | 0.2 | 1.71E-01 | 0.1 | 5.55E-01 | -0.1 | 3.36E-01 | 0.2 | 2.36E-01 | Heat shock cognate protein 70-1 |
| AT5G66410 | 0.3 | 2.54E-02 | 0.5 | 9.63E-04 | -0.2 | 7.49E-02 | 0.2 | 2.01E-01 | 0.2 | 2.92E-01 | 0.0 | 9.81E-01 | Thioredoxin domain-containing protein PLP3B |
| AT1G64600 | 0.3 | 3.85E-02 | 0.0 | 8.10E-01 | 0.2 | 1.19E-01 | 0.8 | 1.22E-02 | 0.5 | 1.77E-01 | 0.3 | 3.36E-01 | At1g64600 |
| AT4G29540 | 0.3 | 7.02E-02 | -0.1 | 4.20E-01 | 0.4 | 4.25E-02 | 0.2 | 6.07E-01 | 0.1 | 8.52E-01 | 0.1 | 8.37E-01 | Bacterial transferase hexapeptide repeat-containing protein |
| AT3G22370 | 0.3 | 5.95E-02 | -0.8 | 1.59E-04 | 1.1 | 7.80E-05 | 1.3 | 8.38E-07 | 0.3 | 8.34E-02 | 1.0 | 2.77E-04 | Ubiquinol oxidase 1a, mitochondrial |

| | | | | | | | | | | | | | |
|---|---|---|---|---|---|---|---|---|---|---|---|---|---|
| AT1G45050 | 0.3 | 4.08E-03 | 0.3 | 8.24E-03 | 0.0 | 8.38E-01 | -0.1 | 6.13E-01 | -0.2 | 4.06E-01 | 0.1 | 8.05E-01 | Ubiquitin-conjugating enzyme family protein |
| AT1G79650 | 0.3 | 2.36E-03 | 0.2 | 9.51E-03 | 0.0 | 5.52E-01 | 0.1 | 5.52E-01 | -0.1 | 6.14E-01 | 0.2 | 3.98E-01 | Ubiquitin receptor RAD23 |
| AT2G47550 | 0.3 | 3.44E-02 | 0.7 | 1.05E-04 | -0.5 | 5.04E-03 | 0.6 | 3.29E-02 | 0.7 | 3.24E-02 | -0.1 | 8.90E-01 | Probable pectinesterase |
| AT5G05610 | 0.3 | 1.11E-02 | 0.2 | 9.85E-02 | 0.1 | 3.21E-01 | 0.1 | 4.75E-01 | -0.2 | 1.79E-01 | 0.3 | 9.08E-02 | PHD finger protein ALFIN-LIKE 1 |
| AT4G35620 | 0.3 | 3.04E-02 | 0.6 | 4.89E-04 | -0.3 | 3.63E-02 | 0.2 | 4.66E-01 | 0.0 | 9.86E-01 | 0.2 | 5.32E-01 | Cyclin B22 |
| AT1G67110 | 0.3 | 1.23E-01 | 1.0 | 1.22E-04 | -0.7 | 3.16E-03 | 0.9 | 1.21E-02 | 0.8 | 3.54E-02 | 0.1 | 8.46E-01 | Cytochrome P450, family 735, subfamily A, polypeptide 2 |
| AT5G55130 | 0.3 | 1.26E-02 | 0.0 | 9.87E-01 | 0.3 | 2.81E-02 | 0.2 | 3.18E-01 | 0.1 | 8.49E-01 | 0.1 | 6.03E-01 | Co-factor for nitrate, reductase and xanthine dehydrogenase 5 |
| AT5G44210 | 0.3 | 9.62E-02 | 0.4 | 1.58E-02 | -0.2 | 3.77E-01 | 0.3 | 2.90E-01 | -0.1 | 8.53E-01 | 0.4 | 2.29E-01 | Ethylene-responsive transcription factor 9 |
| AT4G29050 | 0.3 | 2.92E-02 | 0.1 | 3.79E-01 | 0.2 | 2.34E-01 | 0.1 | 6.21E-01 | -0.3 | 2.96E-01 | 0.4 | 1.50E-01 | non-specific serine |
| AT1G65410 | 0.3 | 5.11E-03 | 0.0 | 7.03E-01 | 0.3 | 7.80E-03 | 0.0 | 8.93E-01 | 0.0 | 9.19E-01 | 0.0 | 9.98E-01 | Protein TRIGALACTOSYLDIACYLGLYCEROL 3, chloroplastic |
| AT4G17510 | 0.3 | 4.74E-03 | 0.1 | 1.02E-01 | 0.1 | 1.65E-01 | 0.1 | 6.86E-01 | 0.0 | 8.49E-01 | 0.1 | 6.29E-01 | Ubiquitin carboxyl-terminal hydrolase 3 |
| AT4G22860 | 0.3 | 5.17E-02 | 0.3 | 3.40E-02 | 0.0 | 7.96E-01 | 0.0 | 9.73E-01 | 0.2 | 7.53E-01 | -0.2 | 8.13E-01 | At4g22860 |
| AT3G16850 | 0.3 | 1.27E-02 | 0.4 | 3.43E-03 | -0.1 | 3.96E-01 | -0.4 | 2.51E-03 | -0.4 | 1.22E-02 | 0.0 | 8.28E-01 | Pectin lyase-like superfamily protein |
| AT1G27650 | 0.3 | 5.95E-03 | 0.2 | 5.37E-02 | 0.1 | 3.11E-01 | -0.1 | 4.16E-01 | -0.3 | 9.87E-03 | 0.2 | 1.18E-01 | U2 snRNP auxiliary factor small subunit |
| AT1G55890 | 0.3 | 2.81E-02 | 0.3 | 1.04E-02 | -0.1 | 5.78E-01 | 0.2 | 2.68E-01 | 0.0 | 8.64E-01 | 0.1 | 5.39E-01 | Small ribosomal subunit protein mS78 |
| AT5G51560 | 0.3 | 1.42E-02 | 0.3 | 5.42E-03 | -0.1 | 5.65E-01 | -0.4 | 7.85E-03 | -0.2 | 2.48E-01 | -0.2 | 2.65E-01 | Protein kinase domain-containing protein |
| AT1G29990 | 0.3 | 6.30E-03 | 0.2 | 1.34E-02 | 0.0 | 7.90E-01 | 0.2 | 3.71E-01 | 0.2 | 4.49E-01 | 0.0 | 9.98E-01 | Prefoldin subunit 6 |
| AT5G27600 | 0.3 | 3.27E-02 | -0.5 | 2.55E-03 | 0.8 | 4.71E-04 | 0.3 | 1.23E-01 | 0.0 | 9.76E-01 | 0.3 | 2.50E-01 | Long chain acyl-CoA synthetase 7, peroxisomal |
| AT3G14990 | 0.3 | 1.48E-03 | 0.3 | 1.15E-03 | 0.0 | 6.97E-01 | 0.3 | 9.63E-03 | -0.2 | 1.14E-01 | 0.6 | 5.38E-03 | Protein DJ-1 homolog A |
| AT4G20850 | 0.3 | 2.38E-03 | 0.1 | 1.08E-01 | 0.2 | 7.62E-02 | -0.1 | 2.53E-01 | -0.1 | 4.73E-01 | 0.0 | 8.89E-01 | Tripeptidyl-peptidase 2 |
| AT1G75510 | 0.3 | 1.03E-02 | 0.1 | 5.60E-01 | 0.2 | 5.73E-02 | 0.3 | 9.26E-02 | 0.1 | 4.70E-01 | 0.1 | 5.92E-01 | TFIIF beta subunit HTH domain-containing protein |
| AT1G73660 | 0.3 | 1.11E-02 | 0.0 | 6.91E-01 | 0.2 | 4.21E-02 | 0.0 | 8.63E-01 | -0.4 | 8.16E-02 | 0.3 | 1.92E-01 | Probable serine |
| AT3G19630 | 0.3 | 2.37E-03 | 0.2 | 1.81E-02 | 0.1 | 3.67E-01 | -0.1 | 4.30E-01 | -0.1 | 6.30E-01 | 0.0 | 9.11E-01 | Radical SAM superfamily protein |
| AT5G53470 | 0.3 | 9.99E-04 | 0.1 | 2.18E-01 | 0.2 | 2.03E-02 | 0.1 | 5.55E-01 | 0.1 | 5.73E-01 | 0.0 | 9.72E-01 | Acyl-CoA-binding domain-containing protein 1 |
| AT5G14520 | 0.3 | 3.20E-02 | 0.3 | 1.88E-02 | 0.0 | 7.39E-01 | 0.3 | 1.54E-02 | 0.0 | 8.04E-01 | 0.3 | 1.10E-01 | Pescadillo homolog |
| AT3G48710 | 0.3 | 1.13E-02 | -0.1 | 4.95E-01 | 0.3 | 9.33E-03 | 0.0 | 9.44E-01 | 0.1 | 6.13E-01 | -0.1 | 6.53E-01 | DEK domain-containing chromatin-associated protein 1 |
| AT4G18240 | 0.3 | 2.26E-02 | 0.2 | 9.36E-02 | 0.1 | 5.30E-01 | 0.3 | 1.66E-01 | 0.3 | 2.77E-01 | 0.0 | 9.45E-01 | Probable starch synthase 4, chloroplastic |
| AT3G07410 | 0.3 | 3.39E-02 | 0.2 | 5.95E-02 | 0.0 | 8.57E-01 | 0.0 | 8.36E-01 | -0.1 | 6.90E-01 | 0.0 | 8.85E-01 | Ras-related protein RABA5b |
| AT1G09520 | 0.3 | 7.43E-03 | 0.0 | 7.43E-01 | 0.2 | 3.31E-02 | -0.2 | 7.54E-02 | -0.4 | 6.93E-03 | 0.2 | 3.60E-01 | hypothetical protein |
| AT2G35240 | 0.3 | 9.29E-02 | 0.3 | 1.29E-01 | 0.0 | 9.18E-01 | 0.2 | 1.98E-01 | 0.0 | 8.76E-01 | 0.2 | 4.63E-01 | Multiple organellar RNA editing factor 6, mitochondrial |
| AT1G06700 | 0.3 | 4.91E-03 | 0.3 | 3.20E-03 | 0.0 | 7.14E-01 | -0.1 | 6.06E-01 | -0.1 | 7.00E-01 | 0.0 | 9.84E-01 | PTI1-like tyrosine-protein kinase 1 |
| AT5G51540 | 0.3 | 2.51E-02 | 0.2 | 1.34E-01 | 0.1 | 4.91E-01 | 0.5 | 1.48E-01 | -0.1 | 9.25E-01 | 0.5 | 1.54E-01 | Mitochondrial intermediate peptidase, mitochondrial |
| AT1G12440 | 0.3 | 4.32E-03 | 0.2 | 7.71E-02 | 0.1 | 1.78E-01 | -0.6 | 2.25E-03 | -0.8 | 3.36E-04 | 0.3 | 2.85E-01 | Zinc finger A20 and AN1 domain-containing stress-associated protein 1 |
| AT5G28290 | 0.3 | 1.34E-02 | 0.2 | 1.48E-01 | 0.1 | 2.90E-01 | -0.3 | 1.17E-01 | 0.0 | 8.56E-01 | -0.2 | 3.27E-01 | NIMA-related kinase 3 |
| AT1G61990 | 0.3 | 1.47E-02 | 0.2 | 9.73E-02 | 0.1 | 4.17E-01 | 0.3 | 6.95E-02 | 0.2 | 3.30E-01 | 0.1 | 6.18E-01 | At1g61990 |
| AT1G65650 | 0.3 | 9.48E-03 | 0.2 | 6.23E-02 | 0.1 | 3.97E-01 | 0.1 | 2.98E-01 | 0.0 | 7.92E-01 | 0.1 | 6.90E-01 | Ubiquitin carboxyl-terminal hydrolase 2 |
| AT2G37590 | 0.3 | 5.17E-02 | 0.1 | 3.68E-01 | 0.1 | 3.76E-01 | 0.0 | 9.99E-01 | 0.0 | 9.79E-01 | 0.0 | 9.81E-01 | Dof zinc finger protein DOF2.4 |
| AT1G56180 | 0.3 | 2.96E-02 | 0.0 | 7.97E-01 | 0.2 | 9.21E-02 | 0.0 | 8.89E-01 | -0.2 | 3.48E-01 | 0.2 | 4.83E-01 | ATP-dependent zinc metalloprotease |
| AT5G48360 | 0.3 | 7.87E-03 | 0.3 | 4.09E-03 | 0.0 | 6.57E-01 | 0.0 | 9.61E-01 | 0.1 | 5.49E-01 | -0.1 | 6.02E-01 | Formin-like protein 9 |
| AT3G02760 | 0.3 | 2.56E-03 | 0.2 | 1.46E-02 | 0.1 | 4.37E-01 | 0.4 | 9.55E-04 | 0.2 | 7.01E-02 | 0.2 | 2.57E-01 | Histidine--tRNA ligase, cytoplasmic |
| AT1G76540 | 0.3 | 2.40E-03 | 0.4 | 4.97E-04 | -0.1 | 2.52E-01 | 0.0 | 7.84E-01 | -0.1 | 6.46E-01 | 0.1 | 5.50E-01 | Cyclin-dependent kinase B2-1 |
| AT1G18340 | 0.3 | 1.35E-02 | 0.4 | 1.52E-03 | -0.1 | 2.16E-01 | 0.0 | 9.82E-01 | -0.2 | 5.87E-01 | 0.1 | 6.36E-01 | General transcription and DNA repair factor IIH subunit TFB4 |
| AT1G31812 | 0.3 | 1.82E-02 | 0.3 | 2.02E-02 | 0.0 | 9.55E-01 | 0.0 | 9.29E-01 | -0.1 | 7.85E-01 | 0.0 | 8.80E-01 | Acyl-CoA-binding domain-containing protein 6 |
| AT1G30420 | 0.3 | 4.34E-03 | 0.1 | 4.37E-01 | 0.2 | 3.89E-02 | 0.4 | 4.75E-02 | 0.1 | 5.79E-01 | 0.2 | 3.64E-01 | Multidrug resistance-associated protein 12 |
| AT2G13790 | 0.3 | 8.27E-02 | -0.1 | 7.36E-01 | 0.3 | 7.02E-02 | 0.3 | 1.04E-01 | 0.0 | 9.68E-01 | 0.3 | 2.13E-01 | Somatic embryogenesis receptor kinase 4 |
| AT1G60800 | 0.3 | 4.29E-03 | 0.3 | 3.95E-03 | 0.0 | 8.49E-01 | 0.1 | 4.41E-01 | 0.0 | 9.27E-01 | 0.1 | 5.41E-01 | NSP-interacting kinase 3 |
| AT5G49760 | 0.3 | 4.57E-03 | 0.3 | 1.04E-02 | 0.0 | 7.92E-01 | 0.2 | 3.63E-02 | 0.1 | 4.02E-01 | 0.1 | 4.77E-01 | Leucine-rich repeat receptor protein kinase HPCA1 |
| AT3G48160 | 0.3 | 7.67E-02 | 0.4 | 2.66E-02 | -0.1 | 5.84E-01 | 0.1 | 8.04E-01 | 0.3 | 5.86E-01 | -0.2 | 7.98E-01 | E2F transcription factor-like E2FE |
| AT1G48310 | 0.3 | 3.19E-03 | 0.2 | 6.91E-02 | 0.1 | 1.77E-01 | 0.1 | 6.38E-01 | 0.1 | 8.49E-01 | 0.0 | 8.97E-01 | Chromatin remodeling factor18 |
| AT1G14360 | 0.3 | 7.30E-02 | 0.2 | 2.15E-01 | 0.1 | 6.34E-01 | 0.3 | 2.48E-02 | 0.3 | 2.59E-02 | 0.0 | 9.19E-01 | UDP-galactose |
| AT5G11330 | 0.3 | 4.73E-03 | 0.1 | 1.71E-01 | 0.2 | 9.84E-02 | 0.2 | 1.64E-01 | 0.1 | 3.36E-01 | 0.0 | 8.99E-01 | FAD |
| AT2G41670 | 0.3 | 1.56E-01 | 0.2 | 3.47E-01 | 0.1 | 7.22E-01 | 0.7 | 5.54E-02 | 0.5 | 1.74E-01 | 0.1 | 7.32E-01 | P-loop containing nucleoside triphosphate hydrolases superfamily protein |
| AT1G09830 | 0.3 | 2.55E-02 | 0.4 | 8.84E-03 | -0.1 | 5.48E-01 | 0.6 | 1.41E-03 | 0.2 | 1.79E-01 | 0.3 | 8.62E-02 | Phosphoribosylamine--glycine ligase, chloroplastic |
| AT1G67630 | 0.3 | 2.39E-02 | 0.3 | 1.39E-02 | 0.0 | 7.49E-01 | 0.2 | 4.25E-01 | 0.3 | 2.78E-01 | -0.1 | 8.00E-01 | DNA polymerase alpha subunit B |
| AT3G07050 | 0.3 | 7.59E-02 | 0.3 | 1.01E-01 | 0.0 | 9.38E-01 | 0.5 | 2.42E-04 | 0.2 | 5.84E-02 | 0.3 | 1.03E-01 | Guanine nucleotide-binding protein-like NSN1 |
| AT3G26220 | 0.3 | 1.28E-01 | -0.2 | 3.30E-01 | 0.5 | 4.75E-02 | -0.3 | 2.58E-01 | -0.9 | 1.18E-02 | 0.6 | 1.97E-01 | At3g26230 |
| AT4G32720 | 0.3 | 4.60E-02 | 0.3 | 5.02E-02 | 0.0 | 9.82E-01 | 0.4 | 2.99E-03 | 0.2 | 2.18E-01 | 0.2 | 1.58E-01 | La protein 1 |
| AT4G28860 | 0.3 | 4.16E-03 | 0.0 | 6.17E-01 | 0.2 | 2.08E-02 | 0.0 | 8.02E-01 | 0.0 | 8.76E-01 | 0.1 | 7.39E-01 | Casein kinase 1-like protein 4 |
| AT2G04390 | 0.3 | 6.26E-02 | 0.5 | 8.56E-03 | -0.2 | 2.66E-01 | 0.3 | 1.18E-01 | 0.3 | 1.82E-01 | 0.0 | 9.93E-01 | Small ribosomal subunit protein eS17z |
| AT1G29040 | 0.3 | 4.72E-03 | 0.2 | 6.66E-02 | 0.1 | 2.38E-01 | -0.1 | 7.28E-01 | 0.0 | 9.68E-01 | -0.1 | 7.53E-01 | 50S ribosomal protein L34 |
| AT4G22220 | 0.3 | 2.38E-03 | 0.1 | 4.29E-01 | 0.2 | 1.88E-02 | 0.0 | 8.08E-01 | 0.0 | 8.80E-01 | 0.0 | 9.81E-01 | Iron-sulfur cluster assembly protein 1 |
| AT5G25930 | 0.3 | 3.72E-02 | -0.5 | 5.78E-03 | 0.8 | 9.16E-04 | 0.9 | 7.23E-03 | 0.1 | 8.67E-01 | 0.8 | 5.53E-02 | Kinase family with leucine-rich repeat domain-containing protein |
| AT2G02955 | 0.3 | 2.98E-02 | -0.2 | 1.23E-01 | 0.5 | 7.28E-03 | 0.5 | 5.85E-02 | 0.1 | 8.31E-01 | 0.4 | 1.27E-01 | TATA box-binding protein-associated factor RNA polymerase I subunit B |
| AT1G14020 | 0.3 | 1.39E-03 | 0.2 | 4.83E-03 | 0.0 | 6.24E-01 | 0.3 | 1.32E-01 | 0.4 | 9.83E-02 | -0.1 | 8.58E-01 | Rhamnogalacturonan I rhamnosyltransferase 1 |
| AT5G06420 | 0.3 | 3.16E-02 | -0.1 | 3.85E-01 | 0.4 | 2.19E-02 | -0.2 | 5.39E-01 | -0.3 | 5.73E-01 | 0.0 | 9.86E-01 | Zinc finger CCCH domain-containing protein 51 |
| AT3G18090 | 0.3 | 3.78E-02 | 0.0 | 9.21E-01 | 0.3 | 7.03E-02 | 0.4 | 2.68E-01 | 0.2 | 6.85E-01 | 0.2 | 6.72E-01 | DNA-directed RNA polymerase subunit beta |
| AT5G11470 | 0.3 | 2.99E-02 | -0.1 | 5.60E-01 | 0.4 | 3.11E-02 | 0.0 | 9.28E-01 | 0.1 | 7.79E-01 | -0.1 | 8.75E-01 | Protein ANTI-SILENCING 1 |
| AT1G73180 | 0.3 | 6.71E-03 | 0.2 | 6.45E-02 | 0.1 | 3.10E-01 | 0.3 | 7.41E-02 | 0.1 | 6.21E-01 | 0.2 | 3.99E-01 | Eukaryotic translation initiation factor 2A |
| AT3G44480 | 0.3 | 7.18E-03 | -0.3 | 1.19E-02 | 0.6 | 3.87E-04 | -0.1 | 3.94E-01 | -0.1 | 5.35E-01 | 0.0 | 9.52E-01 | ADP-ribosyl cyclase |
| AT5G45750 | 0.3 | 1.28E-02 | 0.4 | 4.93E-03 | -0.1 | 5.56E-01 | 0.0 | 9.62E-01 | 0.0 | 8.66E-01 | 0.1 | 8.70E-01 | Ras-related protein RABA1c |
| AT3G16010 | 0.3 | 5.16E-03 | 0.2 | 3.20E-02 | 0.1 | 4.47E-01 | -0.2 | 4.18E-01 | 0.0 | 9.75E-01 | -0.2 | 5.96E-01 | Pentatricopeptide repeat-containing protein At3g16010 |
| AT3G42660 | 0.3 | 5.46E-03 | 0.2 | 4.56E-02 | 0.1 | 3.64E-01 | 0.2 | 3.20E-01 | 0.1 | 5.44E-01 | 0.0 | 8.80E-01 | Protein ENHANCER OF LHP1 1 |
| AT1G58848 | 0.3 | 5.97E-01 | 0.6 | 2.82E-01 | -0.3 | 6.54E-01 | 0.4 | 6.16E-01 | 0.5 | 5.99E-01 | -0.1 | 9.56E-01 | Probable disease resistance protein RF9 |
| AT1G23935 | 0.3 | 4.16E-02 | 0.3 | 2.86E-02 | 0.0 | 8.21E-01 | 0.5 | 1.12E-01 | 0.1 | 7.50E-01 | 0.4 | 4.11E-01 | Apoptosis inhibitory protein |
| AT3G18930 | 0.3 | 7.71E-03 | 0.1 | 3.73E-01 | 0.2 | 8.68E-02 | -0.3 | 4.22E-02 | -0.4 | 4.63E-02 | 0.0 | 9.48E-01 | RING-H2 finger protein ATL65 |
| AT5G51700 | 0.3 | 1.78E-02 | 0.1 | 6.24E-01 | 0.2 | 8.75E-02 | 0.3 | 6.82E-02 | 0.1 | 6.74E-01 | 0.2 | 3.70E-01 | Cysteine and histidine-rich domain-containing protein RAR1 |
| AT5G52920 | 0.3 | 6.26E-03 | 0.3 | 1.23E-02 | 0.0 | 8.45E-01 | 0.0 | 8.84E-01 | -0.1 | 5.11E-01 | 0.1 | 4.96E-01 | Plastidial pyruvate kinase 2 |
| AT1G03930 | 0.3 | 9.49E-03 | 0.0 | 7.92E-01 | 0.3 | 1.41E-02 | -0.1 | 2.51E-01 | 0.1 | 6.90E-01 | -0.2 | 2.32E-01 | Casein kinase 1-like protein 9 |
| AT4G03110 | 0.3 | 1.71E-03 | 0.5 | 1.33E-04 | -0.2 | 6.03E-02 | -0.5 | 1.42E-04 | -0.4 | 2.69E-03 | -0.1 | 4.71E-01 | RNA-binding protein BRN1 |
| AT1G51950 | 0.3 | 1.94E-03 | 0.0 | 8.95E-01 | 0.3 | 4.85E-03 | -0.3 | 2.18E-02 | -0.4 | 7.74E-03 | 0.1 | 6.36E-01 | Auxin-responsive protein IAA18 |
| AT3G05060 | 0.3 | 9.98E-02 | 0.4 | 2.49E-02 | -0.1 | 4.44E-01 | 0.5 | 1.40E-02 | 0.3 | 2.01E-01 | 0.2 | 4.75E-01 | Probable nucleolar protein 5-2 |
| AT5G04490 | 0.3 | 1.74E-03 | 0.1 | 2.14E-01 | 0.2 | 3.06E-02 | -0.1 | 7.81E-01 | -0.2 | 4.98E-01 | 0.1 | 7.67E-01 | Phytol kinase 1 VTE5 |
| AT4G38460 | 0.3 | 7.37E-03 | 0.1 | 3.60E-01 | 0.2 | 6.46E-02 | -0.1 | 4.85E-01 | -0.1 | 6.96E-01 | 0.0 | 9.09E-01 | Heterodimeric geranylgeranyl pyrophosphate synthase small subunit, chloroplastic |

| | | | | | | | | | | | | | |
|---|---|---|---|---|---|---|---|---|---|---|---|---|---|
| AT4G16920 | 0.3 | 3.06E-03 | 0.1 | 5.24E-01 | 0.2 | 2.72E-02 | 0.6 | 2.07E-02 | 0.0 | 9.60E-01 | 0.6 | 6.76E-02 | Disease resistance RPP5 like protein family) |
| AT5G38780 | 0.3 | 7.43E-02 | -0.4 | 6.07E-02 | 0.6 | 8.43E-03 | 0.6 | 2.50E-01 | 0.0 | 9.92E-01 | 0.6 | 2.56E-01 | S-adenosyl-L-methionine-dependent methyltransferases superfamily protein |
| AT5G54470 | 0.3 | 1.41E-01 | 0.4 | 9.04E-02 | -0.1 | 8.03E-01 | | | | | | | At5g54470 |
| AT4G02890 | 0.3 | 2.24E-03 | 0.0 | 8.53E-01 | 0.3 | 7.15E-03 | -0.2 | 1.79E-01 | 0.2 | 2.37E-01 | -0.3 | 3.98E-02 | Polyubiquitin 14 |
| AT5G54230 | 0.3 | 1.41E-01 | 0.2 | 2.76E-01 | 0.1 | 7.88E-01 | 0.1 | 7.71E-01 | -0.3 | 4.78E-01 | 0.5 | 2.74E-01 | Myb domain protein 49 |
| AT5G10330 | 0.3 | 7.68E-03 | 0.3 | 1.00E-02 | 0.0 | 9.88E-01 | -0.1 | 8.28E-01 | -0.1 | 8.09E-01 | 0.0 | 9.62E-01 | Imidazole acetol-phosphate transaminase |
| AT5G11590 | 0.3 | 2.48E-01 | 0.4 | 1.33E-01 | -0.1 | 7.34E-01 | 0.0 | 9.37E-01 | 0.7 | 2.05E-02 | -0.7 | 3.96E-02 | Dehydration-responsive element-binding protein 3 |
| AT1G06143 | 0.3 | 4.82E-02 | 0.0 | 9.17E-01 | 0.3 | 1.25E-01 | 0.4 | 2.24E-01 | 0.3 | 3.73E-01 | 0.1 | 9.12E-01 | Pentatricopeptide repeat-containing protein At1g06143 |
| AT1G75330 | 0.3 | 1.31E-02 | 0.4 | 5.12E-03 | -0.1 | 5.40E-01 | 0.2 | 1.24E-01 | -0.1 | 4.26E-01 | 0.3 | 5.32E-02 | Ornithine transcarbamylase, chloroplastic |
| AT1G31540 | 0.3 | 8.82E-03 | 0.0 | 9.88E-01 | 0.3 | 1.85E-02 | 0.4 | 3.75E-02 | -0.1 | 6.99E-01 | 0.5 | 4.66E-02 | ADP-ribosyl cyclase |
| AT1G04350 | 0.3 | 3.37E-02 | -0.2 | 2.51E-01 | 0.4 | 8.95E-03 | 0.1 | 7.40E-01 | 0.1 | 5.63E-01 | -0.1 | 8.33E-01 | 1-aminocyclopropane-1-carboxylate oxidase homolog 6 |
| AT1G15480 | 0.3 | 2.13E-02 | 0.3 | 3.97E-02 | 0.0 | 8.41E-01 | 0.5 | 3.59E-02 | 0.1 | 6.63E-01 | 0.3 | 2.36E-01 | Pentatricopeptide repeat-containing protein At1g15480, mitochondrial |
| AT5G48620 | 0.3 | 1.75E-02 | 0.0 | 7.92E-01 | 0.3 | 2.01E-02 | 0.0 | 8.95E-01 | -0.1 | 5.05E-01 | 0.2 | 4.82E-01 | Probable disease resistance RPP8-like protein 4 |
| AT5G02490 | 0.3 | 3.32E-01 | -0.7 | 5.23E-02 | 1.0 | 1.80E-02 | 0.3 | 1.20E-01 | 0.7 | 7.66E-03 | -0.3 | 2.31E-01 | Heat shock 70 kDa protein 2 |
| AT1G51940 | 0.3 | 1.24E-02 | 0.4 | 4.13E-03 | -0.1 | 5.03E-01 | -0.7 | 4.92E-04 | -0.6 | 2.41E-03 | -0.1 | 8.11E-01 | LYK3 |
| AT4G21670 | 0.3 | 2.57E-03 | -0.3 | 4.62E-03 | 0.6 | 1.76E-04 | 0.2 | 3.95E-01 | 0.2 | 5.70E-01 | 0.0 | 9.32E-01 | RNA polymerase II C-terminal domain phosphatase-like 1 |
| AT1G29960 | 0.3 | 7.71E-02 | 0.1 | 5.49E-01 | 0.2 | 3.38E-01 | 0.0 | 8.85E-01 | -0.1 | 8.62E-01 | 0.1 | 7.54E-01 | Uncharacterized protein T1P2.16 |
| AT1G70150 | 0.3 | 2.50E-02 | 0.2 | 1.78E-01 | 0.1 | 4.10E-01 | -0.2 | 2.95E-01 | -0.6 | 3.45E-02 | 0.3 | 3.43E-01 | MYND-type domain-containing protein |
| AT4G09720 | 0.3 | 3.14E-03 | 0.3 | 9.01E-03 | 0.0 | 6.81E-01 | 0.2 | 7.51E-02 | 0.2 | 3.17E-01 | 0.1 | 7.11E-01 | RAB GTPase homolog G3A |
| AT1G22940 | 0.3 | 3.32E-03 | -0.1 | 4.23E-01 | 0.4 | 3.50E-03 | 0.0 | 9.18E-01 | -0.1 | 5.55E-01 | 0.1 | 6.85E-01 | Thiamine biosynthetic bifunctional enzyme TH1, chloroplastic |
| AT2G27190 | 0.3 | 4.72E-03 | 0.4 | 1.08E-03 | -0.1 | 3.25E-01 | 0.4 | 1.18E-02 | 0.1 | 7.77E-01 | 0.3 | 7.46E-02 | Iron |
| AT5G58220 | 0.3 | 5.29E-03 | 0.1 | 1.18E-01 | 0.1 | 1.49E-01 | -0.1 | 7.99E-01 | -0.3 | 1.90E-01 | 0.2 | 3.83E-01 | Uric acid degradation bifunctional protein TTL |
| AT1G50710 | 0.3 | 4.33E-03 | 0.1 | 1.39E-01 | 0.2 | 1.22E-01 | -0.2 | 2.62E-01 | 0.1 | 7.03E-01 | -0.3 | 2.08E-01 | AUGMIN subunit 4 |
| AT2G03820 | 0.3 | 2.99E-03 | 0.1 | 5.06E-01 | 0.2 | 1.92E-02 | 0.2 | 9.32E-02 | -0.1 | 3.69E-01 | 0.3 | 5.47E-02 | 60S ribosomal export protein NMD3 |
| AT1G68080 | 0.3 | 1.28E-02 | 0.2 | 1.21E-01 | 0.1 | 3.47E-01 | -0.2 | 4.03E-01 | -0.3 | 2.30E-01 | 0.1 | 7.39E-01 | procollagen-proline 3-dioxygenase |
| AT2G45060 | 0.3 | 4.39E-03 | 0.4 | 1.87E-03 | -0.1 | 5.47E-01 | 0.1 | 5.69E-01 | 0.0 | 8.31E-01 | 0.1 | 5.35E-01 | Alanine-tRNA ligase |
| AT5G44450 | 0.3 | 3.43E-03 | 0.2 | 2.87E-02 | 0.1 | 3.54E-01 | 0.1 | 6.56E-01 | -0.2 | 4.25E-01 | 0.2 | 2.92E-01 | Alpha amino-terminal protein methyltransferase |
| AT5G05970 | 0.3 | 2.03E-03 | 0.1 | 3.45E-01 | 0.2 | 2.71E-02 | 0.0 | 8.93E-01 | -0.1 | 5.30E-01 | 0.1 | 6.87E-01 | Protein NEDD1 |
| AT4G27940 | 0.3 | 8.28E-03 | -0.1 | 2.43E-01 | 0.4 | 4.30E-03 | 0.3 | 2.69E-01 | -0.2 | 6.06E-01 | 0.5 | 9.77E-02 | Mitochondrial carrier protein MTM1 |
| AT3G09390 | 0.3 | 1.59E-01 | -0.2 | 2.78E-01 | 0.5 | 3.49E-02 | -0.1 | 8.04E-01 | -0.2 | 4.78E-01 | 0.2 | 7.37E-01 | Metallothionein-like protein 2A |
| AT5G44320 | 0.3 | 2.65E-03 | 0.1 | 1.16E-01 | 0.2 | 7.88E-02 | 0.0 | 9.96E-01 | -0.1 | 2.69E-01 | 0.1 | 3.68E-01 | Eukaryotic translation initiation factor 3 subunit D |
| AT2G30620 | 0.3 | 1.09E-03 | 0.3 | 3.17E-03 | 0.0 | 6.99E-01 | -0.2 | 2.05E-01 | -0.4 | 5.70E-02 | 0.2 | 5.87E-01 | Winged-helix DNA-binding transcription factor family protein |
| AT5G57210 | 0.3 | 1.83E-03 | 0.1 | 1.22E-01 | 0.2 | 6.27E-02 | 0.1 | 6.84E-01 | 0.1 | 7.14E-01 | 0.0 | 9.83E-01 | Ypt |
| AT1G05230 | 0.3 | 1.18E-01 | 0.0 | 8.83E-01 | 0.3 | 1.58E-01 | -0.3 | 7.47E-01 | 0.0 | 9.76E-01 | -0.3 | 8.37E-01 | Homeodomain GLABROUS 2 |
| AT5G45940 | 0.3 | 1.03E-01 | 0.0 | 9.46E-01 | 0.3 | 1.94E-01 | -0.2 | 6.49E-01 | 0.0 | 9.88E-01 | -0.2 | 6.69E-01 | Nudix hydrolase homolog 11 |
| AT1G15960 | 0.3 | 1.76E-01 | 0.0 | 8.53E-01 | 0.3 | 3.45E-01 | -0.1 | 7.97E-01 | 0.0 | 9.98E-01 | -0.1 | 8.44E-01 | At1g15960 |
| AT1G79640 | 0.3 | 2.02E-02 | 0.3 | 2.06E-02 | 0.0 | 9.37E-01 | 0.3 | 3.63E-01 | 0.4 | 2.76E-01 | -0.1 | 8.56E-01 | Protein kinase superfamily protein |
| AT3G63520 | 0.3 | 9.83E-03 | -0.1 | 3.24E-01 | 0.4 | 4.31E-03 | 0.0 | 9.60E-01 | 0.1 | 8.33E-01 | 0.0 | 8.98E-01 | Carotenoid 9,10-cleavage dioxygenase 1 |
| AT4G09900 | 0.3 | 1.13E-01 | -0.3 | 1.17E-01 | 0.6 | 1.55E-02 | -0.2 | 3.40E-01 | 0.0 | 9.58E-01 | -0.2 | 5.40E-01 | Putative methylesterase 12, chloroplastic |
| AT3G13670 | 0.3 | 2.00E-03 | -0.1 | 9.51E-02 | 0.4 | 6.21E-04 | -0.1 | 8.85E-01 | 0.0 | 9.88E-01 | -0.1 | 9.14E-01 | Casein kinase-like protein |
| AT2G46470 | 0.3 | 1.11E-03 | 0.4 | 5.01E-04 | -0.1 | 4.75E-01 | 0.1 | 5.04E-01 | 0.1 | 7.79E-01 | 0.0 | 8.70E-01 | Uncharacterized protein |
| AT4G38300 | 0.3 | 5.83E-02 | -0.1 | 4.97E-01 | 0.4 | 4.53E-02 | | | | | | | Glycosyl hydrolase family 10 protein |
| AT3G48090 | 0.3 | 3.56E-02 | -0.2 | 8.88E-02 | 0.5 | 4.08E-03 | 0.6 | 4.52E-03 | 0.3 | 2.08E-01 | 0.3 | 1.79E-01 | Protein EDS1 |
| AT5G67030 | 0.3 | 6.80E-02 | -0.1 | 5.50E-01 | 0.4 | 3.68E-02 | 0.1 | 7.66E-01 | 0.2 | 5.04E-01 | -0.1 | 7.46E-01 | Zeaxanthin epoxidase, chloroplastic |
| AT4G13360 | 0.3 | 5.23E-03 | 0.1 | 2.36E-01 | 0.2 | 7.89E-02 | 0.0 | 7.71E-01 | 0.0 | 8.86E-01 | 0.0 | 9.55E-01 | 3-hydroxyisobutyryl-CoA hydrolase |
| AT1G56330 | 0.3 | 2.98E-03 | 0.4 | 1.46E-03 | -0.1 | 5.31E-01 | 0.4 | 2.33E-03 | 0.3 | 4.52E-02 | 0.1 | 4.87E-01 | GTP-binding protein SAR1B |
| AT1G22920 | 0.3 | 2.65E-03 | 0.2 | 1.90E-02 | 0.1 | 3.70E-01 | 0.1 | 3.20E-01 | -0.1 | 6.42E-01 | 0.2 | 2.23E-01 | COP9 signalosome complex subunit 5a |
| AT1G51620 | 0.3 | 1.53E-01 | 0.1 | 7.66E-01 | 0.2 | 3.71E-01 | 0.5 | 3.88E-01 | 0.2 | 8.49E-01 | 0.4 | 5.87E-01 | Protein kinase superfamily protein |
| AT4G18250 | 0.3 | 6.89E-02 | -0.2 | 3.53E-01 | 0.5 | 3.02E-02 | 1.1 | 8.92E-05 | 0.0 | 9.66E-01 | 1.1 | 1.79E-03 | Receptor Serine |
| AT5G17880 | 0.3 | 2.33E-02 | 0.2 | 8.96E-02 | 0.1 | 5.76E-01 | -0.1 | 6.51E-01 | -0.3 | 9.96E-02 | 0.2 | 3.29E-01 | ADP-ribosyl cyclase |
| AT2G39660 | 0.3 | 2.93E-03 | 0.0 | 7.35E-01 | 0.3 | 5.06E-03 | 0.0 | 9.07E-01 | -0.3 | 1.57E-01 | 0.3 | 1.78E-01 | Serine |
| AT5G22110 | 0.3 | 3.21E-02 | 0.3 | 4.66E-02 | 0.0 | 9.37E-01 | 0.4 | 2.44E-01 | -0.2 | 7.51E-01 | 0.6 | 1.83E-01 | DNA polymerase epsilon subunit |
| AT1G09640 | 0.3 | 7.37E-03 | 0.2 | 4.53E-02 | 0.1 | 4.23E-01 | 0.1 | 1.90E-01 | -0.1 | 2.06E-01 | 0.3 | 6.51E-02 | AT1G09640 protein |
| AT2G26040 | 0.3 | 1.56E-01 | 0.5 | 2.29E-02 | -0.2 | 3.45E-01 | 0.9 | 8.20E-02 | 1.1 | 6.28E-02 | -0.1 | 8.19E-01 | Abscisic acid receptor PYL2 |
| AT3G26630 | 0.3 | 5.51E-03 | -0.1 | 3.55E-01 | 0.4 | 4.51E-03 | -0.2 | 2.18E-01 | 0.0 | 9.32E-01 | -0.2 | 2.99E-01 | Pentatricopeptide repeat-containing protein At3g26630, chloroplastic |
| AT1G42540 | 0.3 | 2.64E-03 | 0.1 | 4.26E-01 | 0.2 | 2.36E-02 | 0.2 | 4.14E-01 | 0.2 | 4.96E-01 | 0.0 | 9.90E-01 | Glutamate receptor 3.3 |
| AT5G59800 | 0.3 | 7.76E-03 | 0.2 | 1.39E-01 | 0.1 | 2.07E-01 | -0.6 | 1.95E-03 | -0.4 | 3.41E-02 | -0.2 | 3.66E-01 | Methyl-CpG-binding domain-containing protein 7 |
| AT5G20980 | 0.3 | 5.72E-03 | 0.1 | 3.76E-01 | 0.2 | 6.46E-02 | 0.2 | 4.66E-01 | 0.1 | 8.53E-01 | 0.1 | 6.65E-01 | 5-methyltetrahydropteroyltriglutamate--homocysteine methyltransferase 3, chloroplastic |
| AT4G11240 | 0.3 | 7.63E-04 | 0.3 | 1.47E-03 | 0.0 | 8.65E-01 | 0.3 | 5.85E-02 | 0.0 | 9.00E-01 | 0.2 | 2.07E-01 | Serine |
| AT1G10840 | 0.3 | 1.06E-02 | 0.2 | 5.28E-02 | 0.1 | 4.83E-01 | 0.1 | 5.89E-01 | -0.1 | 5.58E-01 | 0.2 | 3.29E-01 | Eukaryotic translation initiation factor 3 subunit H |
| AT5G63030 | 0.3 | 2.57E-03 | 0.1 | 2.78E-01 | 0.2 | 3.61E-02 | -0.2 | 4.46E-01 | -0.2 | 3.82E-01 | 0.0 | 9.07E-01 | Glutaredoxin-C1 |
| AT3G04210 | 0.3 | 2.55E-02 | 0.0 | 7.66E-01 | 0.3 | 2.55E-02 | 1.0 | 6.52E-06 | 0.2 | 3.45E-01 | 0.9 | 4.95E-04 | Putative disease resistance protein |
| AT5G14850 | 0.3 | 6.46E-03 | 0.1 | 1.88E-01 | 0.2 | 1.39E-01 | -0.2 | 4.56E-01 | 0.0 | 9.07E-01 | -0.2 | 5.08E-01 | Mannosyltransferase |
| AT2G23740 | 0.3 | 4.27E-03 | -0.1 | 2.48E-01 | 0.4 | 2.84E-03 | 0.2 | 7.13E-01 | 0.1 | 8.72E-01 | 0.1 | 9.07E-01 | Histone-lysine N-methyltransferase SUVR5 |
| AT5G54640 | 0.3 | 6.60E-04 | 0.1 | 7.24E-02 | 0.2 | 3.20E-02 | 0.0 | 9.45E-01 | -0.1 | 6.08E-01 | 0.2 | 5.79E-01 | Histone H2A |
| AT3G61860 | 0.3 | 1.72E-03 | 0.0 | 5.84E-01 | 0.3 | 1.01E-02 | 0.0 | 9.85E-01 | -0.2 | 1.96E-01 | 0.2 | 3.35E-01 | Serine |
| AT5G66750 | 0.3 | 1.97E-02 | 0.3 | 2.83E-02 | 0.0 | 9.41E-01 | 0.5 | 1.45E-02 | 0.3 | 2.06E-01 | 0.2 | 3.70E-01 | ATP-dependent DNA helicase DDM1 |
| AT4G34910 | 0.3 | 9.94E-03 | 0.2 | 8.72E-02 | 0.1 | 3.47E-01 | 0.5 | 5.28E-03 | 0.1 | 7.92E-01 | 0.5 | 4.83E-02 | ATP-dependent RNA helicase |
| AT4G25940 | 0.3 | 1.38E-02 | 0.1 | 3.11E-01 | 0.2 | 1.42E-01 | 0.3 | 2.72E-01 | 0.1 | 8.94E-01 | 0.2 | 4.87E-01 | Putative clathrin assembly protein At4g25940 |
| AT1G24440 | 0.3 | 2.43E-03 | 0.4 | 1.34E-03 | 0.0 | 6.18E-01 | 0.1 | 4.37E-01 | 0.0 | 9.04E-01 | 0.1 | 4.59E-01 | RING |
| AT3G07200 | 0.3 | 1.37E-02 | 0.2 | 7.75E-02 | 0.1 | 4.74E-01 | -0.1 | 7.82E-01 | -0.2 | 3.70E-01 | 0.2 | 6.25E-01 | Putative RING zinc finger protein |
| AT1G06220 | 0.3 | 7.74E-04 | 0.1 | 1.34E-01 | 0.2 | 2.02E-02 | 0.3 | 2.35E-02 | 0.3 | 1.37E-01 | 0.1 | 7.21E-01 | 110 kDa U5 small nuclear ribonucleoprotein component CLO |
| AT4G38890 | 0.3 | 4.73E-03 | 0.0 | 6.82E-01 | 0.3 | 6.08E-03 | 0.1 | 4.18E-01 | 0.1 | 4.85E-01 | 0.0 | 1.00E+00 | tRNA-dihydrouridine synthase -like |
| AT4G39380 | 0.3 | 3.09E-02 | 0.3 | 5.70E-02 | 0.0 | 8.55E-01 | 0.1 | 8.98E-01 | 0.0 | 9.42E-01 | 0.1 | 8.78E-01 | TSL-kinase interacting-like protein |
| AT5G47970 | 0.3 | 4.51E-03 | 0.0 | 8.68E-01 | 0.3 | 1.03E-02 | -0.1 | 6.17E-01 | -0.1 | 5.20E-01 | 0.0 | 8.99E-01 | tRNA-dihydrouridine synthase |
| AT2G03280 | 0.3 | 8.53E-02 | 0.1 | 7.80E-01 | 0.3 | 2.31E-01 | -0.3 | 2.75E-01 | -0.1 | 6.62E-01 | -0.1 | 6.96E-01 | Rhamnogalacturonan I rhamnosyltransferase 4 |
| AT1G67690 | 0.3 | 3.46E-02 | 0.3 | 3.12E-02 | 0.0 | 9.14E-01 | 0.1 | 8.91E-01 | 0.1 | 7.78E-01 | -0.1 | 8.96E-01 | Probable thimet oligopeptidase |
| AT1G14460 | 0.3 | 1.15E-02 | 0.1 | 5.18E-01 | 0.2 | 7.59E-02 | 0.0 | 9.49E-01 | 0.2 | 4.62E-01 | -0.1 | 6.05E-01 | Protein STICHEL-like 1 |
| AT1G34210 | 0.3 | 2.57E-03 | 0.1 | 1.15E-01 | 0.2 | 9.11E-02 | 0.0 | 8.81E-01 | 0.1 | 6.46E-01 | -0.1 | 8.08E-01 | Somatic embryogenesis receptor kinase 2 |
| AT5G22370 | 0.3 | 1.00E-02 | 0.1 | 3.40E-01 | 0.2 | 1.12E-01 | -0.1 | 5.07E-01 | -0.2 | 2.20E-01 | 0.1 | 6.57E-01 | GPN-loop GTPase QQT1 |
| AT5G24270 | 0.3 | 3.64E-02 | 0.0 | 8.63E-01 | 0.3 | 5.87E-02 | 0.6 | 9.04E-02 | 0.1 | 7.72E-01 | 0.4 | 3.47E-01 | Calcineurin B-like protein |
| AT3G48830 | 0.3 | 1.16E-01 | -0.2 | 2.88E-01 | 0.5 | 4.32E-02 | 0.9 | 2.60E-02 | 0.0 | 9.58E-01 | 0.8 | 8.85E-02 | tRNA nucleotidyltransferase |
| AT3G57800 | 0.3 | 3.18E-02 | 0.3 | 7.67E-02 | 0.1 | 7.35E-01 | -0.8 | 2.29E-04 | -0.6 | 2.34E-03 | -0.1 | 5.74E-01 | hypothetical protein |

| | | | | | | | | | | | | | |
|---|---|---|---|---|---|---|---|---|---|---|---|---|---|
| AT2G31955 | 0.3 | 1.08E-03 | -0.1 | 4.26E-01 | 0.4 | 1.47E-03 | 0.1 | 3.11E-01 | -0.2 | 1.59E-01 | 0.3 | 4.36E-02 | GTP 3',8-cyclase |
| AT2G32415 | 0.3 | 5.13E-03 | 0.1 | 3.19E-01 | 0.2 | 6.99E-02 | -0.2 | 5.52E-01 | -0.4 | 1.88E-01 | 0.2 | 5.73E-01 | Polynucleotidyl transferase, ribonuclease H fold protein with HRDC domain-containing protein |
| AT3G21250 | 0.3 | 1.05E-02 | -0.2 | 1.63E-01 | 0.5 | 3.28E-03 | 0.0 | 9.76E-01 | 0.2 | 6.90E-01 | -0.2 | 7.47E-01 | ABC-type xenobiotic transporter |
| AT1G19485 | 0.3 | 9.47E-04 | 0.0 | 6.49E-01 | 0.3 | 2.08E-03 | 0.1 | 6.80E-01 | 0.0 | 9.54E-01 | 0.1 | 8.31E-01 | hypothetical protein |
| AT4G02930 | 0.3 | 5.80E-02 | 0.4 | 2.00E-02 | -0.1 | 5.38E-01 | 0.8 | 7.28E-06 | 0.5 | 2.79E-03 | 0.3 | 6.44E-02 | Elongation factor Tu, mitochondrial |
| AT1G73820 | 0.3 | 1.62E-02 | 0.0 | 7.04E-01 | 0.3 | 7.33E-02 | -0.1 | 7.57E-01 | 0.0 | 9.28E-01 | -0.1 | 9.08E-01 | RNA polymerase II subunit A C-terminal domain phosphatase SSU72 |
| AT2G17800 | 0.3 | 7.01E-03 | 0.3 | 2.19E-02 | 0.1 | 6.49E-01 | -0.1 | 3.84E-01 | 0.0 | 9.32E-01 | -0.1 | 5.01E-01 | Rac-like GTP-binding protein ARAC1 |
| AT1G21710 | 0.3 | 1.06E-02 | 0.2 | 8.15E-02 | 0.1 | 3.87E-01 | -0.2 | 2.79E-01 | -0.1 | 7.25E-01 | -0.1 | 7.03E-01 | N-glycosylase |
| AT3G01120 | 0.3 | 2.05E-03 | 0.3 | 5.04E-03 | 0.0 | 7.71E-01 | 0.3 | 2.16E-02 | 0.5 | 8.00E-04 | -0.2 | 1.53E-01 | Cystathionine gamma-synthase 1, chloroplastic -lyase) |
| AT2G30500 | 0.3 | 1.26E-03 | 0.2 | 1.56E-02 | 0.1 | 2.30E-01 | -0.3 | 8.64E-03 | -0.3 | 3.88E-02 | -0.1 | 7.84E-01 | Kinase interacting family protein |
| AT2G13840 | 0.3 | 1.29E-02 | 0.1 | 5.52E-01 | 0.2 | 8.44E-02 | -0.3 | 2.13E-01 | -0.3 | 3.09E-01 | 0.0 | 9.65E-01 | At2g13860 |
| AT3G46580 | 0.3 | 8.50E-02 | 0.0 | 9.01E-01 | 0.3 | 1.27E-01 | -0.1 | 8.43E-01 | -0.1 | 6.20E-01 | 0.1 | 8.13E-01 | Methyl-CpG-binding domain-containing protein 5 |
| AT2G24960 | 0.3 | 2.11E-02 | 0.0 | 7.23E-01 | 0.3 | 9.67E-02 | -0.2 | 7.80E-01 | 0.1 | 9.11E-01 | -0.3 | 7.36E-01 | Myb |
| AT4G32800 | 0.3 | 5.14E-02 | 0.1 | 3.95E-01 | 0.2 | 3.36E-01 | -0.3 | 2.19E-01 | 0.1 | 8.20E-01 | -0.4 | 2.60E-01 | Ethylene-responsive transcription factor ERF043 |
| AT2G41530 | 0.3 | 4.57E-04 | 0.2 | 2.92E-03 | 0.1 | 3.72E-01 | -0.1 | 5.43E-01 | -0.1 | 7.72E-01 | 0.0 | 8.98E-01 | S-formylglutathione hydrolase |
| AT3G13160 | 0.3 | 2.70E-02 | 0.3 | 3.16E-02 | 0.0 | 9.97E-01 | 0.4 | 2.08E-02 | 0.2 | 1.93E-01 | 0.1 | 5.55E-01 | Small ribosomal subunit protein mS79 |
| AT1G10500 | 0.3 | 7.06E-03 | -0.1 | 3.65E-01 | 0.4 | 4.41E-03 | -0.1 | 5.82E-01 | -0.1 | 7.11E-01 | 0.0 | 9.64E-01 | Uncharacterized protein At1g10500 |
| AT1G25560 | 0.3 | 5.73E-02 | -0.4 | 2.64E-02 | 0.7 | 2.23E-03 | -0.1 | 5.13E-01 | -0.8 | 4.62E-04 | 0.7 | 9.89E-03 | AP2 |
| AT2G45590 | 0.3 | 4.23E-03 | 0.2 | 7.61E-02 | 0.1 | 1.76E-01 | -0.1 | 7.18E-01 | -0.1 | 7.09E-01 | 0.0 | 9.62E-01 | Receptor-like serine |
| AT4G27950 | 0.3 | 1.23E-01 | 0.4 | 7.02E-02 | -0.1 | 7.81E-01 | 0.3 | 4.71E-01 | 0.3 | 4.20E-01 | -0.1 | 9.00E-01 | Ethylene-responsive transcription factor CRF4 |
| AT4G09460 | 0.3 | 6.74E-03 | 0.3 | 9.88E-03 | 0.0 | 9.46E-01 | -0.3 | 2.50E-01 | -0.6 | 1.76E-02 | 0.3 | 2.65E-01 | Transcription repressor MYB6 |
| AT1G33830 | 0.3 | 3.16E-01 | 1.8 | 8.83E-05 | -1.5 | 1.05E-03 | | | | | | | Immune-associated nucleotide-binding protein 1 |
| AT4G11840 | 0.3 | 3.51E-02 | 0.1 | 7.35E-01 | 0.3 | 1.14E-01 | 0.7 | 2.32E-03 | 0.4 | 1.10E-01 | 0.3 | 1.90E-01 | Phospholipase D |
| AT2G43010 | 0.3 | 2.17E-02 | 0.3 | 2.05E-02 | 0.0 | 9.15E-01 | -0.4 | 4.13E-01 | -0.4 | 4.68E-01 | 0.0 | 9.97E-01 | Phytochrome interacting factor 4 |
| AT1G68990 | 0.3 | 3.90E-03 | -0.1 | 2.11E-01 | 0.4 | 2.26E-03 | 0.8 | 8.20E-05 | 0.3 | 1.77E-01 | 0.5 | 1.01E-02 | At1g68990 |
| AT1G72275 | 0.3 | 3.09E-02 | 0.3 | 6.90E-02 | 0.0 | 7.87E-01 | 0.0 | 9.12E-01 | 0.0 | 9.23E-01 | 0.1 | 8.64E-01 | Uncharacterized protein T9N14.23 |
| AT5G63400 | 0.3 | 1.42E-02 | 0.6 | 7.08E-04 | -0.3 | 6.70E-02 | 0.2 | 8.04E-02 | 0.0 | 9.40E-01 | 0.2 | 2.84E-01 | adenylate kinase |
| AT1G29900 | 0.3 | 2.95E-03 | 0.1 | 1.73E-01 | 0.2 | 5.99E-02 | 0.3 | 4.80E-02 | 0.1 | 8.12E-01 | 0.3 | 2.52E-01 | Carbamoyl-phosphate synthase large chain, chloroplastic |
| AT1G07615 | 0.3 | 1.13E-02 | 0.0 | 8.43E-01 | 0.3 | 2.24E-02 | -0.1 | 8.36E-01 | -0.2 | 3.36E-01 | 0.2 | 5.64E-01 | Probable GTP-binding protein OBGM, mitochondrial |
| AT5G43630 | 0.3 | 7.03E-03 | -0.1 | 6.09E-01 | 0.4 | 6.61E-03 | 0.1 | 6.46E-01 | 0.3 | 1.69E-01 | -0.2 | 4.76E-01 | Uncharacterized protein At5g43630 family protein) |
| AT4G31580 | 0.3 | 8.26E-03 | 0.2 | 1.25E-01 | 0.1 | 2.02E-01 | 0.2 | 7.05E-02 | 0.1 | 4.68E-01 | 0.1 | 5.78E-01 | Serine |
| AT5G53850 | 0.3 | 1.55E-03 | 0.1 | 4.96E-01 | 0.3 | 1.09E-02 | -0.1 | 6.65E-01 | -0.1 | 6.73E-01 | 0.0 | 9.73E-01 | Probable methylthioribulose-1-phosphate dehydratase |
| AT4G31790 | 0.3 | 7.24E-02 | 0.2 | 3.64E-01 | 0.2 | 4.34E-01 | 0.8 | 4.91E-05 | 0.2 | 2.17E-01 | 0.6 | 3.56E-03 | diphthine methyl ester synthase |
| AT3G53420 | 0.3 | 2.77E-03 | 0.4 | 4.63E-04 | -0.1 | 1.72E-01 | -0.7 | 1.37E-04 | -0.4 | 3.52E-02 | -0.3 | 1.19E-01 | PIP2A |
| AT5G40810 | 0.3 | 5.13E-04 | 0.3 | 1.33E-03 | 0.0 | 7.58E-01 | 0.0 | 8.45E-01 | -0.1 | 6.59E-01 | 0.1 | 5.93E-01 | Cytochrome c1 2, heme protein, mitochondrial |
| AT2G34810 | 0.3 | 2.01E-02 | 0.6 | 3.73E-04 | -0.3 | 3.22E-02 | 0.8 | 2.86E-02 | 1.0 | 1.98E-02 | -0.1 | 7.99E-01 | Berberine bridge enzyme-like 16 |
| AT1G08845 | 0.3 | 5.95E-02 | 0.3 | 7.59E-02 | 0.0 | 9.56E-01 | -0.1 | 7.62E-01 | 0.0 | 9.05E-01 | 0.0 | 8.95E-01 | Ribosomal L18p |
| AT4G32520 | 0.3 | 1.76E-03 | 0.1 | 9.01E-02 | 0.2 | 6.71E-02 | 0.4 | 1.26E-03 | 0.1 | 4.71E-01 | 0.3 | 5.91E-02 | Serine hydroxymethyltransferase 3, chloroplastic |
| AT5G20060 | 0.3 | 2.93E-03 | 0.5 | 2.93E-04 | -0.2 | 1.06E-01 | 0.0 | 9.64E-01 | 0.0 | 9.98E-01 | 0.0 | 9.73E-01 | Phospholipase |
| AT3G61950 | 0.3 | 1.17E-01 | 0.3 | 1.04E-01 | 0.0 | 9.07E-01 | -0.2 | 5.34E-01 | 0.3 | 4.72E-01 | -0.5 | 2.03E-01 | Basic helix-loop-helix DNA-binding superfamily protein |
| AT5G67240 | 0.3 | 2.47E-03 | 0.3 | 8.46E-03 | 0.0 | 6.24E-01 | 0.4 | 5.74E-02 | 0.2 | 4.46E-01 | 0.2 | 4.57E-01 | Small RNA degrading nuclease 3 |
| AT5G41880 | 0.3 | 1.11E-02 | 0.2 | 4.29E-02 | 0.1 | 5.96E-01 | 0.6 | 1.93E-02 | 0.4 | 1.28E-01 | 0.2 | 6.32E-01 | DNA primase |
| AT4G21090 | 0.3 | 1.16E-02 | 0.3 | 1.82E-02 | 0.0 | 9.18E-01 | 0.1 | 6.30E-01 | -0.2 | 3.19E-01 | 0.3 | 1.52E-01 | Adrenodoxin-like protein 2, mitochondrial |
| AT5G66610 | 0.3 | 4.11E-02 | -0.1 | 5.45E-01 | 0.4 | 3.82E-02 | -0.8 | 1.37E-01 | -0.2 | 7.51E-01 | -0.6 | 4.35E-01 | DA1-related protein 7 |
| AT4G26780 | 0.3 | 2.54E-02 | 0.3 | 2.48E-02 | 0.0 | 9.40E-01 | 0.9 | 7.62E-05 | 0.5 | 2.41E-02 | 0.4 | 3.44E-02 | GrpE protein homolog 2, mitochondrial |
| AT5G64770 | 0.3 | 1.96E-01 | 0.4 | 1.53E-01 | 0.0 | 8.74E-01 | 0.0 | 8.96E-01 | 0.2 | 4.86E-01 | -0.3 | 4.72E-01 | Protein GOLVEN 2 -related protein CLEL9) |
| AT4G26200 | 0.3 | 3.79E-01 | -0.4 | 3.71E-01 | 0.7 | 1.54E-01 | 0.8 | 2.69E-02 | -0.2 | 6.44E-01 | 1.0 | 2.42E-02 | 1-aminocyclopropane-1-carboxylate synthase 7 |
| AT1G55930 | 0.3 | 1.11E-02 | 0.1 | 4.65E-01 | 0.2 | 8.69E-02 | 0.0 | 9.08E-01 | -0.1 | 7.70E-01 | 0.1 | 7.31E-01 | CBS domain-containing protein |
| AT1G55900 | 0.3 | 9.69E-03 | 0.2 | 6.55E-02 | 0.1 | 4.08E-01 | 0.6 | 3.21E-03 | 0.3 | 2.55E-01 | 0.3 | 1.29E-01 | Mitochondrial import inner membrane translocase subunit TIM50 |
| AT3G13800 | 0.3 | 1.03E-02 | 0.4 | 7.54E-03 | 0.0 | 8.04E-01 | 0.1 | 6.59E-01 | -0.1 | 6.98E-01 | 0.2 | 4.91E-01 | At3g13800 |
| AT5G14610 | 0.3 | 1.31E-03 | 0.4 | 8.37E-04 | 0.0 | 6.47E-01 | 0.0 | 9.00E-01 | -0.1 | 7.34E-01 | 0.1 | 6.76E-01 | DEAD box RNA helicase family protein |
| AT1G64950 | 0.3 | 4.32E-03 | -0.1 | 1.95E-01 | 0.4 | 1.68E-03 | 0.1 | 5.99E-01 | -0.1 | 4.14E-01 | 0.2 | 2.61E-01 | Cytochrome P450, family 89, subfamily A, polypeptide 5 |
| AT1G66910 | 0.3 | 1.10E-02 | -0.1 | 4.59E-01 | 0.4 | 9.39E-03 | 0.2 | 3.68E-01 | 0.0 | 9.90E-01 | 0.2 | 5.05E-01 | Protein kinase superfamily protein |
| AT4G37740 | 0.3 | 2.93E-02 | 0.3 | 3.71E-02 | 0.0 | 9.72E-01 | 0.1 | 8.67E-01 | 0.2 | 5.17E-01 | -0.2 | 6.57E-01 | Growth-regulating factor 2 |
| AT1G11350 | 0.3 | 1.24E-02 | 0.4 | 3.75E-03 | -0.1 | 4.70E-01 | 0.0 | 8.26E-01 | -0.3 | 1.03E-01 | 0.3 | 1.28E-01 | G-type lectin S-receptor-like serine |
| AT2G34970 | 0.3 | 1.40E-03 | 0.1 | 1.51E-01 | 0.2 | 3.61E-02 | 0.4 | 1.59E-02 | 0.2 | 2.88E-01 | 0.2 | 3.36E-01 | eIF-2B GDP-GTP exchange factor subunit epsilon |
| AT3G13460 | 0.3 | 6.81E-04 | 0.1 | 1.64E-01 | 0.2 | 1.49E-02 | 0.1 | 6.50E-01 | -0.2 | 1.21E-01 | 0.2 | 1.03E-01 | YTH domain-containing family protein |
| AT5G22050 | 0.3 | 2.44E-03 | 0.2 | 5.09E-02 | 0.1 | 1.70E-01 | -0.4 | 3.97E-02 | -0.4 | 5.81E-02 | 0.0 | 9.99E-01 | Protein kinase superfamily protein |
| AT1G02840 | 0.3 | 7.71E-04 | 0.1 | 6.58E-02 | 0.2 | 3.83E-02 | -0.2 | 5.31E-02 | -0.2 | 5.64E-02 | 0.0 | 9.15E-01 | Serine |
| AT4G23100 | 0.3 | 8.55E-04 | 0.4 | 3.38E-04 | -0.1 | 3.45E-01 | 0.0 | 6.69E-01 | 0.1 | 2.35E-01 | -0.1 | 5.69E-01 | Glutamate--cysteine ligase |
| AT5G07320 | 0.3 | 8.84E-04 | 0.4 | 6.02E-04 | 0.0 | 6.45E-01 | 0.1 | 6.62E-01 | -0.1 | 8.46E-01 | 0.2 | 6.11E-01 | Calcium-dependent mitochondrial ATP-magnesium |
| AT5G01320 | 0.3 | 7.92E-02 | -1.0 | 1.09E-03 | 1.4 | 5.74E-04 | | | | | | | Pyruvate decarboxylase 4 |
| AT2G26840 | 0.3 | 7.95E-02 | 0.3 | 1.49E-01 | 0.1 | 8.17E-01 | 0.2 | 6.83E-01 | 0.3 | 4.84E-01 | -0.1 | 8.24E-01 | MutS2 |
| AT3G57190 | 0.3 | 4.56E-03 | 0.4 | 1.49E-03 | -0.1 | 4.56E-01 | 0.2 | 2.87E-01 | 0.0 | 9.42E-01 | 0.2 | 4.92E-01 | Peptide chain release factor PrfB3, chloroplastic |
| AT5G45060 | 0.3 | 1.21E-02 | -0.1 | 2.38E-01 | 0.5 | 4.94E-03 | 0.1 | 6.47E-01 | -0.3 | 2.24E-01 | 0.4 | 8.42E-02 | Disease resistance protein RPS4B |
| AT5G47320 | 0.3 | 2.51E-02 | 0.3 | 7.41E-02 | 0.1 | 6.55E-01 | 0.1 | 3.60E-01 | 0.0 | 8.59E-01 | 0.1 | 6.54E-01 | Small ribosomal subunit protein uS19m |
| AT5G12330 | 0.3 | 2.75E-02 | -0.3 | 9.80E-02 | 0.6 | 5.73E-03 | 0.5 | 1.05E-01 | 0.3 | 4.07E-01 | 0.2 | 6.11E-01 | LRP1 |
| AT3G25700 | 0.3 | 1.11E-02 | 0.5 | 1.98E-03 | -0.1 | 3.08E-01 | 0.3 | 1.41E-01 | 0.5 | 1.05E-02 | -0.3 | 2.44E-01 | At3g25700 |
| AT3G57260 | 0.3 | 6.55E-01 | -0.9 | 2.73E-01 | 1.2 | 1.70E-01 | -0.8 | 2.78E-01 | -0.9 | 2.81E-01 | 0.1 | 9.38E-01 | glucan endo-1,3-beta-D-glucosidase |
| AT5G14430 | 0.3 | 9.26E-04 | 0.2 | 4.83E-02 | 0.2 | 6.09E-02 | 0.2 | 1.72E-01 | 0.0 | 9.63E-01 | 0.2 | 3.82E-01 | Methyltransferase |
| AT4G36190 | 0.3 | 3.81E-03 | -0.1 | 1.42E-01 | 0.5 | 1.36E-03 | -0.2 | 1.64E-01 | 0.0 | 9.11E-01 | -0.2 | 2.57E-01 | At4g36190 |
| AT1G11780 | 0.3 | 1.07E-02 | 0.4 | 6.57E-03 | 0.0 | 7.45E-01 | 0.3 | 8.75E-02 | 0.1 | 8.01E-01 | 0.3 | 2.21E-01 | Alpha-ketoglutarate-dependent dioxygenase alkB |
| AT1G73680 | 0.3 | 5.10E-03 | -0.1 | 5.38E-01 | 0.4 | 5.22E-03 | 0.1 | 4.69E-01 | -0.3 | 1.47E-01 | 0.4 | 4.57E-02 | Alpha dioxygenase |
| AT4G01250 | 0.3 | 1.13E-02 | 0.0 | 9.76E-01 | 0.3 | 3.17E-02 | -0.3 | 3.03E-01 | -1.0 | 5.01E-03 | 0.7 | 9.29E-02 | WRKY transcription factor 22 |
| AT2G02070 | 0.3 | 8.87E-04 | 0.2 | 1.56E-02 | 0.1 | 1.59E-01 | -0.3 | 7.58E-02 | -0.3 | 4.90E-02 | 0.1 | 7.94E-01 | Protein indeterminate-domain 5, chloroplastic |
| AT2G27980 | 0.3 | 1.50E-03 | 0.0 | 7.22E-01 | 0.3 | 7.78E-03 | -0.1 | 5.41E-01 | 0.1 | 7.93E-01 | -0.2 | 4.78E-01 | Uncharacterized protein At2g27980 |
| AT3G50920 | 0.3 | 1.97E-02 | 0.2 | 1.43E-01 | 0.1 | 3.90E-01 | 0.0 | 9.28E-01 | -0.2 | 3.60E-01 | 0.2 | 5.03E-01 | Lipid phosphate phosphatase epsilon 1, chloroplastic |
| AT4G12620 | 0.3 | 2.13E-02 | 0.4 | 1.29E-02 | 0.0 | 7.69E-01 | 0.0 | 9.28E-01 | 0.0 | 9.48E-01 | -0.1 | 9.01E-01 | Origin of replication complex subunit 1B |
| AT5G06110 | 0.3 | 1.29E-02 | 0.3 | 3.00E-02 | 0.0 | 7.56E-01 | 0.3 | 1.51E-02 | -0.1 | 5.09E-01 | 0.4 | 1.62E-02 | Cell division related protein-like |
| AT4G24040 | 0.3 | 4.62E-02 | 0.0 | 9.51E-01 | 0.3 | 9.27E-02 | -0.1 | 8.04E-01 | -0.6 | 8.42E-02 | 0.5 | 2.04E-01 | Trehalase |
| AT3G52050 | 0.3 | 5.93E-03 | 0.2 | 3.96E-02 | 0.1 | 4.22E-01 | -0.1 | 7.84E-01 | 0.1 | 7.12E-01 | -0.2 | 5.59E-01 | 5'-3' exonuclease family protein |
| AT4G14910 | 0.3 | 1.59E-03 | 0.2 | 5.75E-02 | 0.2 | 1.02E-01 | 0.2 | 2.51E-01 | 0.2 | 1.77E-01 | -0.1 | 8.39E-01 | Imidazoleglycerol-phosphate dehydratase |
| AT3G01420 | 0.3 | 5.53E-04 | 0.1 | 2.25E-01 | 0.2 | 8.75E-03 | 3.0 | 7.30E-11 | 2.2 | 1.64E-08 | 0.8 | 5.83E-04 | Alpha-dioxygenase 1 |

| | | | | | | | | | | | | | |
|---|---|---|---|---|---|---|---|---|---|---|---|---|---|
| AT2G16780 | 0.3 | 2.71E-03 | 0.2 | 2.17E-02 | 0.1 | 3.64E-01 | 0.1 | 5.75E-01 | -0.1 | 7.90E-01 | 0.2 | 4.95E-01 | WD-40 repeat-containing protein MSI2 |
| AT4G30890 | 0.3 | 6.04E-04 | 0.1 | 1.52E-01 | 0.2 | 1.47E-02 | 0.4 | 6.27E-03 | 0.2 | 3.05E-01 | 0.2 | 2.44E-01 | Ubiquitin carboxyl-terminal hydrolase 24 |
| AT2G04630 | 0.3 | 2.29E-02 | 0.2 | 1.28E-01 | 0.1 | 4.33E-01 | 0.1 | 3.89E-01 | 0.0 | 9.72E-01 | 0.2 | 4.63E-01 | DNA-directed RNA polymerases II and V subunit 6B |
| AT2G28450 | 0.3 | 1.04E-03 | 0.1 | 8.89E-02 | 0.2 | 4.77E-02 | 0.0 | 9.18E-01 | -0.1 | 6.77E-01 | 0.1 | 8.13E-01 | Zinc finger family protein |
| AT5G42540 | 0.3 | 9.40E-04 | 0.1 | 3.47E-01 | 0.3 | 1.02E-02 | 0.1 | 5.20E-01 | -0.1 | 5.75E-01 | 0.2 | 3.51E-01 | 5'-3' exoribonuclease |
| AT4G24510 | 0.3 | 1.66E-02 | 0.3 | 2.39E-02 | 0.0 | 9.37E-01 | -0.3 | 1.79E-01 | -0.2 | 4.87E-01 | -0.1 | 7.34E-01 | Protein ECERIFERUM 2 |
| AT3G06200 | 0.3 | 3.33E-03 | 0.2 | 4.81E-02 | 0.1 | 2.28E-01 | 0.1 | 5.78E-01 | -0.1 | 4.76E-01 | 0.2 | 2.41E-01 | Guanylate kinase 3, chloroplastic |
| AT4G16480 | 0.3 | 1.44E-03 | 0.2 | 1.03E-02 | 0.1 | 3.89E-01 | -0.4 | 2.01E-02 | -0.4 | 2.96E-02 | 0.0 | 9.99E-01 | Inositol transporter 4 |
| AT4G16630 | 0.3 | 4.76E-02 | 0.2 | 1.92E-01 | 0.1 | 5.22E-01 | 0.6 | 1.66E-04 | 0.3 | 9.27E-02 | 0.3 | 4.73E-02 | DEAD-box ATP-dependent RNA helicase 28 |
| AT3G11580 | 0.3 | 1.82E-02 | -0.1 | 3.28E-01 | 0.5 | 1.14E-02 | 0.3 | 2.68E-01 | 0.2 | 4.22E-01 | 0.0 | 9.20E-01 | AP2 |
| AT3G57320 | 0.3 | 5.21E-03 | 0.3 | 7.06E-03 | 0.0 | 9.81E-01 | -0.3 | 1.16E-01 | -0.2 | 3.39E-01 | -0.1 | 7.67E-01 | Threonine-tRNA ligase 2 |
| AT4G38710 | 0.3 | 3.65E-03 | 0.3 | 4.17E-03 | 0.0 | 9.25E-01 | 0.3 | 4.65E-03 | 0.1 | 5.50E-01 | 0.3 | 1.04E-01 | Eukaryotic translation initiation factor 4B3 |
| AT2G25300 | 0.3 | 4.91E-03 | 0.3 | 1.59E-02 | 0.1 | 6.66E-01 | 0.4 | 1.20E-01 | 0.1 | 6.48E-01 | 0.2 | 4.34E-01 | Hexosyltransferase |
| AT3G06470 | 0.3 | 1.36E-03 | 0.1 | 4.93E-01 | 0.3 | 1.05E-02 | -0.1 | 2.87E-01 | -0.5 | 1.81E-03 | 0.4 | 4.04E-02 | F24P17.3 protein |
| AT4G00490 | 0.3 | 2.82E-03 | -0.1 | 3.39E-01 | 0.4 | 1.96E-03 | -0.1 | 7.60E-01 | 0.2 | 3.34E-01 | -0.3 | 2.91E-01 | Beta-amylase 2, chloroplastic |
| AT1G19715 | 0.3 | 8.46E-03 | 0.2 | 1.35E-01 | 0.2 | 1.93E-01 | 0.0 | 9.39E-01 | 0.3 | 2.80E-01 | -0.3 | 3.97E-01 | Jacalin-related lectin 3 |
| AT2G31360 | 0.3 | 6.58E-03 | 0.5 | 9.07E-04 | -0.2 | 1.76E-01 | -0.3 | 3.32E-03 | -0.2 | 1.05E-01 | -0.1 | 3.90E-01 | Delta-9 acyl-lipid desaturase 2 |
| AT4G32300 | 0.3 | 3.45E-04 | 0.1 | 8.58E-02 | 0.2 | 1.39E-02 | 0.0 | 9.38E-01 | -0.3 | 3.50E-02 | 0.3 | 1.08E-01 | G-type lectin S-receptor-like serine |
| AT5G03740 | 0.3 | 4.77E-03 | 0.3 | 1.88E-02 | 0.1 | 5.58E-01 | 0.2 | 1.66E-01 | 0.0 | 8.28E-01 | 0.1 | 4.55E-01 | Histone deacetylase HDT3 |
| AT4G11420 | 0.3 | 3.59E-03 | 0.2 | 8.61E-02 | 0.2 | 1.38E-01 | 0.2 | 2.01E-01 | 0.0 | 8.47E-01 | 0.2 | 4.85E-01 | Eukaryotic translation initiation factor 3 subunit A |
| AT4G09550 | 0.3 | 2.75E-03 | 0.2 | 1.03E-01 | 0.2 | 1.14E-01 | -0.3 | 3.61E-01 | -0.4 | 2.89E-01 | 0.1 | 8.73E-01 | Mitotic-spindle organizing protein 1B |
| AT5G16180 | 0.3 | 1.34E-02 | 0.2 | 1.92E-01 | 0.2 | 2.37E-01 | -0.2 | 3.12E-01 | -0.2 | 2.82E-01 | 0.0 | 9.16E-01 | Chloroplast splicing factor-like protein |
| AT1G50420 | 0.3 | 2.00E-03 | 0.0 | 8.23E-01 | 0.4 | 4.96E-03 | 0.0 | 7.73E-01 | 0.0 | 9.17E-01 | 0.0 | 9.25E-01 | Scarecrow-like protein 3 |
| AT1G54310 | 0.3 | 1.24E-02 | 0.5 | 1.64E-03 | -0.2 | 2.50E-01 | 0.2 | 3.84E-01 | -0.2 | 5.40E-01 | 0.5 | 1.72E-01 | Uncharacterized protein F20D21.13 |
| AT1G46768 | 0.3 | 1.50E-02 | 0.2 | 2.10E-01 | 0.2 | 2.51E-01 | -0.4 | 1.12E-01 | -0.7 | 1.45E-02 | 0.3 | 3.50E-01 | Related to AP2 1 |
| AT2G38010 | 0.3 | 1.88E-03 | 0.2 | 3.12E-02 | 0.1 | 1.89E-01 | 0.3 | 1.43E-01 | 0.4 | 4.46E-02 | -0.1 | 5.72E-01 | Neutral ceramidase |
| AT1G08260 | 0.3 | 1.62E-02 | 0.0 | 9.02E-01 | 0.3 | 5.01E-02 | -0.1 | 9.23E-01 | 0.0 | 9.82E-01 | -0.1 | 9.22E-01 | DNA polymerase epsilon catalytic subunit |
| AT5G50780 | 0.3 | 2.18E-03 | 0.0 | 6.31E-01 | 0.4 | 3.88E-03 | -0.2 | 1.89E-01 | 0.0 | 8.37E-01 | -0.2 | 4.75E-01 | Histidine kinase-, DNA gyrase B-, and HSP90-like ATPase family protein |
| AT2G19520 | 0.3 | 5.93E-04 | 0.3 | 2.55E-03 | 0.1 | 5.17E-01 | 0.3 | 4.55E-02 | 0.1 | 7.17E-01 | 0.2 | 3.01E-01 | WD-40 repeat-containing protein MSI4 |
| AT5G40490 | 0.3 | 2.24E-02 | 0.1 | 5.14E-01 | 0.2 | 1.28E-01 | 0.3 | 1.54E-01 | -0.1 | 8.19E-01 | 0.3 | 1.67E-01 | RNA-binding family protein |
| AT1G19140 | 0.3 | 1.40E-03 | 0.1 | 1.10E-01 | 0.2 | 5.22E-02 | 0.2 | 9.97E-02 | 0.4 | 2.14E-02 | -0.1 | 5.35E-01 | Ubiquinone biosynthesis protein |
| AT1G61980 | 0.3 | 1.60E-02 | 0.2 | 7.58E-02 | 0.1 | 5.27E-01 | 0.2 | 3.81E-01 | 0.0 | 9.47E-01 | 0.2 | 4.40E-01 | At1g61980 |
| AT2G02810 | 0.3 | 1.80E-01 | 0.2 | 5.19E-01 | 0.2 | 5.83E-01 | 0.3 | 1.81E-01 | 0.2 | 3.47E-01 | 0.0 | 8.83E-01 | UDP-galactose |
| AT3G48170 | 0.3 | 1.76E-04 | 0.3 | 2.78E-04 | 0.0 | 9.68E-01 | -0.1 | 3.43E-01 | 0.0 | 8.75E-01 | -0.1 | 6.25E-01 | Aminoaldehyde dehydrogenase ALDH10A9, peroxisomal |
| AT3G23490 | 0.3 | 6.26E-02 | 0.0 | 8.82E-01 | 0.4 | 7.61E-02 | -0.2 | 2.35E-01 | -0.1 | 6.12E-01 | -0.1 | 7.56E-01 | Cyanase |
| AT2G30160 | 0.3 | 2.66E-03 | 0.2 | 2.38E-02 | 0.1 | 3.38E-01 | 0.0 | 9.28E-01 | -0.3 | 1.47E-01 | 0.3 | 2.42E-01 | Mitochondrial substrate carrier family protein |
| AT5G10390 | 0.3 | 1.90E-02 | 0.5 | 2.49E-03 | -0.2 | 2.46E-01 | 0.5 | 3.42E-02 | 0.2 | 5.18E-01 | 0.3 | 2.58E-01 | Histone H3.1 |
| AT2G42070 | 0.3 | 5.63E-03 | 0.4 | 4.18E-03 | 0.0 | 7.88E-01 | 0.1 | 5.64E-01 | -0.2 | 2.11E-01 | 0.3 | 1.35E-01 | Nudix hydrolase homolog 23 |
| AT2G33480 | 0.3 | 7.98E-02 | -0.2 | 2.55E-01 | 0.6 | 2.39E-02 | 0.2 | 4.08E-01 | 0.0 | 9.24E-01 | 0.2 | 4.89E-01 | NAC domain containing protein 41 |
| AT5G57440 | 0.3 | 1.29E-03 | 0.5 | 1.07E-04 | -0.2 | 6.47E-02 | -0.2 | 2.10E-01 | 0.0 | 7.99E-01 | -0.1 | 5.52E-01 | -glycerol-3-phosphatase 2 |
| AT3G57470 | 0.3 | 1.51E-02 | 0.1 | 4.27E-01 | 0.2 | 1.14E-01 | -0.1 | 5.70E-01 | -0.2 | 5.63E-01 | 0.0 | 9.53E-01 | Insulinase family protein |
| AT3G60630 | 0.3 | 1.28E-03 | 0.3 | 3.30E-03 | 0.0 | 7.28E-01 | 0.0 | 8.76E-01 | 0.1 | 8.16E-01 | -0.1 | 7.44E-01 | Scarecrow-like protein 22 |
| AT4G15830 | 0.3 | 4.36E-02 | 0.6 | 2.05E-03 | -0.3 | 1.16E-01 | 0.1 | 5.31E-01 | 0.3 | 3.15E-01 | -0.1 | 7.70E-01 | ARM repeat superfamily protein |
| AT5G07300 | 0.3 | 9.82E-04 | 0.2 | 6.44E-02 | 0.2 | 5.19E-02 | 0.2 | 3.07E-01 | 0.0 | 9.79E-01 | 0.2 | 4.68E-01 | Uncharacterized protein |
| AT4G16950 | 0.3 | 3.63E-03 | 0.0 | 8.69E-01 | 0.4 | 6.23E-03 | 0.1 | 5.21E-01 | -0.2 | 1.28E-01 | 0.3 | 8.36E-02 | ADP-ribosyl cyclase |
| AT5G55190 | 0.3 | 3.65E-03 | 0.3 | 6.27E-03 | 0.0 | 9.15E-01 | 0.1 | 6.22E-01 | 0.0 | 9.64E-01 | 0.1 | 8.12E-01 | GTP-binding nuclear protein |
| AT1G54250 | 0.3 | 1.95E-02 | 0.5 | 2.88E-03 | -0.2 | 2.94E-01 | 0.3 | 1.56E-01 | 0.1 | 8.03E-01 | 0.2 | 4.54E-01 | DNA-directed RNA polymerases II and V subunit 8A |
| AT5G50280 | 0.3 | 4.91E-03 | 0.2 | 6.27E-02 | 0.1 | 2.68E-01 | 0.0 | 9.39E-01 | 0.0 | 8.77E-01 | 0.0 | 9.37E-01 | Pentatricopeptide repeat-containing protein At5g50280, chloroplastic |
| AT1G12990 | 0.3 | 1.08E-02 | 0.1 | 2.88E-01 | 0.2 | 1.26E-01 | -0.1 | 7.54E-01 | -0.2 | 2.35E-01 | 0.2 | 4.41E-01 | F3F19.2 protein |
| AT4G38520 | 0.3 | 5.61E-04 | 0.0 | 6.57E-01 | 0.4 | 1.02E-03 | -0.2 | 1.01E-01 | 0.0 | 8.57E-01 | -0.2 | 3.31E-01 | Probable protein phosphatase 2C 64 |
| AT3G08990 | 0.3 | 1.47E-02 | 0.2 | 1.22E-01 | 0.1 | 3.85E-01 | 0.0 | 9.29E-01 | -0.1 | 9.00E-01 | 0.0 | 9.70E-01 | Protein yippee-like At3g08990 |
| AT1G69520 | 0.3 | 8.62E-02 | -0.3 | 1.51E-01 | 0.7 | 1.97E-02 | 0.5 | 1.31E-01 | 0.1 | 9.22E-01 | 0.5 | 2.70E-01 | Methyltransferase type 11 domain-containing protein |
| AT3G27925 | 0.3 | 1.83E-03 | 0.0 | 6.46E-01 | 0.4 | 2.42E-03 | -0.1 | 3.72E-01 | 0.1 | 5.61E-01 | -0.2 | 1.72E-01 | Protease Do-like 1, chloroplastic |
| AT1G62110 | 0.3 | 3.64E-02 | 0.3 | 9.24E-02 | 0.1 | 7.29E-01 | 0.4 | 5.51E-02 | 0.0 | 9.55E-01 | 0.4 | 1.61E-01 | At1g62110 |
| AT5G50920 | 0.3 | 2.65E-03 | 0.0 | 9.88E-01 | 0.3 | 6.43E-03 | 0.0 | 9.01E-01 | 0.0 | 9.78E-01 | 0.0 | 9.62E-01 | ATP-dependent Clp protease, ATP-binding subunit |
| AT1G72890 | 0.3 | 1.40E-02 | 0.0 | 9.86E-01 | 0.3 | 3.66E-02 | 0.3 | 2.40E-01 | -0.1 | 8.09E-01 | 0.3 | 1.72E-01 | Similar to part of disease resistance protein |
| AT5G20050 | 0.3 | 2.17E-03 | 0.0 | 8.56E-01 | 0.3 | 7.63E-03 | 0.0 | 9.11E-01 | -0.3 | 6.38E-02 | 0.2 | 1.73E-01 | Probable receptor-like protein kinase At5g20050 |
| AT5G46160 | 0.3 | 1.18E-02 | 0.4 | 1.21E-02 | 0.0 | 9.42E-01 | 0.2 | 3.51E-01 | -0.1 | 8.17E-01 | 0.3 | 3.52E-01 | Large ribosomal subunit protein uL14mz |
| AT1G17470 | 0.3 | 2.26E-04 | 0.3 | 8.77E-04 | 0.0 | 5.44E-01 | 0.0 | 9.05E-01 | -0.1 | 5.85E-01 | 0.1 | 5.85E-01 | Developmentally-regulated G-protein 1 |
| AT1G50490 | 0.3 | 1.26E-02 | 0.6 | 9.23E-04 | -0.2 | 1.31E-01 | 0.1 | 4.67E-01 | 0.2 | 4.24E-01 | 0.0 | 9.23E-01 | Ubiquitin-conjugating enzyme E2 20 |
| AT1G65860 | 0.3 | 4.17E-02 | 0.3 | 1.03E-01 | 0.1 | 7.13E-01 | -0.9 | 2.72E-02 | 0.7 | 3.90E-02 | -1.6 | 3.80E-03 | Flavin-containing monooxygenase FMO GS-OX1 |
| AT5G19180 | 0.3 | 1.75E-04 | 0.2 | 6.15E-03 | 0.1 | 6.60E-02 | 0.2 | 1.31E-01 | -0.1 | 7.64E-01 | 0.2 | 1.70E-01 | NEDD8-activating enzyme E1 catalytic subunit |
| AT4G28440 | 0.3 | 5.60E-03 | 0.2 | 4.34E-02 | 0.1 | 3.52E-01 | 0.2 | 2.04E-01 | 0.1 | 6.33E-01 | 0.1 | 6.81E-01 | Uncharacterized protein At4g28440 |
| AT2G05790 | 0.3 | 3.52E-03 | 0.5 | 7.12E-04 | -0.1 | 2.35E-01 | -0.2 | 5.42E-02 | -0.2 | 1.41E-01 | 0.0 | 9.00E-01 | glucan endo-1,3-beta-D-glucosidase |
| AT5G07290 | 0.3 | 2.62E-03 | 0.2 | 3.58E-02 | 0.1 | 2.34E-01 | 0.3 | 5.07E-02 | 0.2 | 3.75E-01 | 0.2 | 5.02E-01 | Protein MEI2-like 4 |
| AT4G33260 | 0.3 | 2.88E-03 | 0.4 | 2.36E-03 | 0.0 | 8.00E-01 | 0.0 | 9.51E-01 | 0.1 | 7.87E-01 | -0.1 | 8.44E-01 | Cell division cycle 20.2, cofactor of APC complex |
| AT3G23100 | 0.3 | 1.48E-02 | 0.4 | 7.23E-03 | -0.1 | 6.66E-01 | 0.4 | 2.12E-02 | 0.0 | 8.52E-01 | 0.3 | 1.32E-01 | Carbamoyl-phosphate synthetase large subunit-like ATP-binding domain-containing protein |
| AT4G03120 | 0.3 | 3.37E-03 | 0.3 | 1.05E-02 | 0.0 | 6.66E-01 | 0.0 | 7.61E-01 | -0.1 | 7.95E-01 | 0.1 | 6.42E-01 | U1 small nuclear ribonucleoprotein C |
| AT5G57960 | 0.3 | 1.02E-02 | -0.1 | 5.58E-01 | 0.4 | 1.01E-02 | 0.2 | 2.03E-01 | 0.2 | 2.79E-01 | 0.0 | 9.82E-01 | GTP binding protein-like |
| AT1G60690 | 0.3 | 3.32E-03 | 0.2 | 4.97E-02 | 0.1 | 2.40E-01 | -0.3 | 2.15E-01 | 0.1 | 8.52E-01 | -0.3 | 2.68E-01 | Probable aldo-keto reductase 3 |
| AT3G16260 | 0.3 | 7.36E-03 | 0.2 | 4.15E-02 | 0.1 | 4.61E-01 | 0.1 | 6.59E-01 | 0.0 | 9.26E-01 | 0.1 | 8.27E-01 | ribonuclease Z |
| AT5G63600 | 0.3 | 6.01E-04 | 0.4 | 1.82E-04 | -0.1 | 2.34E-01 | 0.2 | 8.83E-02 | -0.1 | 7.47E-01 | 0.3 | 7.99E-02 | Flavonol synthase 5 |
| AT5G67330 | 0.3 | 4.40E-03 | 0.0 | 8.84E-01 | 0.4 | 8.12E-03 | 1.8 | 5.94E-08 | 0.1 | 6.28E-01 | 1.6 | 2.65E-05 | Metal transporter Nramp4 |
| AT3G46950 | 0.3 | 3.91E-02 | 0.1 | 3.81E-01 | 0.2 | 3.02E-01 | 0.5 | 4.05E-02 | 0.1 | 8.92E-01 | 0.5 | 1.05E-01 | Mitochondrial transcription termination factor family protein |
| AT5G41190 | 0.3 | 4.47E-03 | 0.3 | 6.99E-03 | 0.0 | 9.22E-01 | 0.3 | 5.71E-03 | 0.0 | 9.83E-01 | 0.3 | 3.46E-02 | RNA-binding NOB1-like protein |
| AT2G36010 | 0.3 | 1.51E-02 | 0.3 | 4.24E-02 | 0.1 | 7.15E-01 | 0.4 | 2.23E-01 | 0.1 | 7.38E-01 | 0.2 | 5.64E-01 | E2F transcription factor 3 |
| AT5G51120 | 0.3 | 8.66E-03 | 0.2 | 4.80E-02 | 0.1 | 4.46E-01 | 0.1 | 5.01E-01 | 0.1 | 3.73E-01 | 0.0 | 8.44E-01 | Polyadenylate-binding protein 1 |
| AT5G42820 | 0.3 | 2.06E-02 | 0.1 | 6.19E-01 | 0.3 | 1.13E-01 | -0.1 | 7.88E-01 | -0.3 | 5.96E-01 | 0.1 | 8.32E-01 | Splicing factor U2af small subunit B |
| AT5G43470 | 0.3 | 4.02E-03 | 0.1 | 4.47E-01 | 0.3 | 2.90E-02 | 0.0 | 7.44E-01 | -0.2 | 1.19E-01 | 0.3 | 1.12E-01 | Disease resistance protein RPP8 |
| AT5G47070 | 0.3 | 5.26E-02 | -0.1 | 5.13E-01 | 0.5 | 3.83E-02 | -0.1 | 2.92E-01 | -0.5 | 2.00E-03 | 0.4 | 5.03E-02 | Probable serine |
| AT1G52710 | 0.3 | 2.50E-03 | 0.3 | 4.09E-03 | 0.0 | 9.26E-01 | -0.3 | 1.70E-01 | -0.3 | 1.19E-01 | 0.1 | 8.52E-01 | Uncharacterized protein |
| AT2G03800 | 0.3 | 1.13E-03 | -0.1 | 2.85E-01 | 0.4 | 1.21E-03 | 0.1 | 3.70E-01 | 0.1 | 4.53E-01 | 0.0 | 9.92E-01 | D-aminoacyl-tRNA deacylase |
| AT5G22830 | 0.3 | 5.91E-03 | 0.2 | 1.42E-01 | 0.2 | 1.41E-01 | -0.1 | 6.07E-01 | -0.1 | 8.24E-01 | -0.1 | 8.87E-01 | Magnesium transporter MRS2-11, chloroplastic |

| | | | | | | | | | | | | | |
|---|---|---|---|---|---|---|---|---|---|---|---|---|---|
| AT1G65800 | 0.3 | 2.61E-02 | -0.1 | 3.59E-01 | 0.5 | 1.07E-02 | 0.8 | 1.00E-03 | 0.5 | 2.32E-02 | 0.2 | 4.15E-01 | Receptor kinase 2 |
| AT2G16710 | 0.3 | 6.44E-04 | 0.1 | 1.97E-01 | 0.2 | 1.57E-02 | 0.2 | 3.22E-01 | -0.1 | 7.86E-01 | 0.2 | 3.22E-01 | Iron-sulfur cluster biosynthesis family protein |
| AT2G34450 | 0.3 | 2.20E-02 | 0.2 | 3.12E-01 | 0.2 | 2.39E-01 | 0.4 | 2.38E-01 | 0.6 | 9.71E-02 | -0.2 | 5.94E-01 | High mobility group B protein 14 |
| AT1G21920 | 0.3 | 1.49E-02 | 0.5 | 1.69E-03 | -0.2 | 2.00E-01 | -0.4 | 2.84E-02 | -0.6 | 3.54E-03 | 0.2 | 3.14E-01 | Histone H3 K4-specific methyltransferase SET7 |
| AT1G71180 | 0.3 | 4.46E-03 | 0.2 | 4.93E-02 | 0.1 | 2.95E-01 | -0.2 | 1.74E-01 | -0.1 | 5.78E-01 | -0.1 | 6.74E-01 | Probable 3-hydroxyisobutyrate dehydrogenase-like 3, mitochondrial |
| AT2G20610 | 0.3 | 5.90E-04 | 0.3 | 1.45E-03 | 0.0 | 7.86E-01 | 0.0 | 7.90E-01 | 0.3 | 7.59E-02 | -0.2 | 2.50E-01 | S-alkyl-thiohydroximate lyase SUR1 |
| AT1G62085 | 0.4 | 4.43E-03 | 0.4 | 4.05E-03 | 0.0 | 8.67E-01 | -0.1 | 5.72E-01 | -0.2 | 4.26E-01 | 0.1 | 8.48E-01 | F19K23.4 protein |
| AT1G75200 | 0.4 | 6.36E-03 | 0.2 | 1.54E-01 | 0.2 | 1.47E-01 | 0.3 | 5.86E-02 | -0.1 | 7.72E-01 | 0.4 | 8.76E-02 | S-adenosyl-L-methionine-dependent tRNA 4-demethylwyosine synthase |
| AT5G67100 | 0.4 | 1.50E-02 | 0.2 | 1.04E-01 | 0.1 | 4.14E-01 | 0.3 | 4.65E-01 | 0.2 | 6.51E-01 | 0.1 | 9.19E-01 | DNA polymerase alpha catalytic subunit |
| AT2G19490 | 0.4 | 4.13E-02 | 0.1 | 6.17E-01 | 0.3 | 1.79E-01 | 1.0 | 4.32E-04 | 0.8 | 5.41E-03 | 0.2 | 5.05E-01 | DNA repair protein recA homolog 3, mitochondrial |
| AT3G56150 | 0.4 | 2.58E-03 | 0.1 | 6.01E-01 | 0.3 | 1.35E-02 | 0.2 | 1.23E-01 | 0.1 | 6.60E-01 | 0.1 | 5.61E-01 | Eukaryotic translation initiation factor 3 subunit C |
| AT2G13290 | 0.4 | 1.62E-03 | 0.0 | 6.62E-01 | 0.3 | 1.07E-02 | 0.1 | 4.10E-01 | -0.2 | 4.17E-01 | 0.3 | 1.33E-01 | hypothetical protein |
| AT5G10910 | 0.4 | 2.13E-03 | 0.3 | 1.53E-02 | 0.1 | 3.94E-01 | 0.2 | 5.16E-01 | 0.4 | 1.04E-01 | -0.3 | 4.01E-01 | MraW methylase family protein |
| AT4G24015 | 0.4 | 1.14E-02 | 0.8 | 8.35E-05 | -0.4 | 8.58E-03 | -0.1 | 7.19E-01 | -0.2 | 3.68E-01 | 0.1 | 6.82E-01 | Probable E3 ubiquitin-protein ligase RHA4A |
| AT5G10070 | 0.4 | 1.77E-02 | -0.2 | 2.22E-01 | 0.5 | 6.86E-03 | 0.6 | 4.37E-03 | 0.2 | 5.09E-01 | 0.4 | 7.74E-02 | 18S rRNA aminocarboxypropyltransferase |
| AT3G63090 | 0.4 | 1.10E-02 | 0.3 | 3.01E-02 | 0.1 | 7.10E-01 | 0.5 | 1.35E-02 | 0.2 | 4.54E-01 | 0.3 | 1.86E-01 | Ubiquitin carboxyl-terminal hydrolase family protein |
| AT1G63880 | 0.4 | 2.50E-04 | 0.1 | 2.67E-01 | 0.3 | 3.58E-03 | 0.0 | 7.73E-01 | -0.3 | 4.22E-02 | 0.3 | 8.65E-02 | Disease resistance protein family |
| AT2G06990 | 0.4 | 5.20E-04 | 0.1 | 2.98E-01 | 0.3 | 7.20E-03 | 0.1 | 4.85E-01 | 0.2 | 4.75E-01 | 0.0 | 9.46E-01 | DExH-box ATP-dependent RNA helicase DExH10 |
| AT5G52950 | 0.4 | 1.50E-02 | 0.4 | 6.96E-03 | -0.1 | 6.46E-01 | 0.5 | 2.66E-01 | 0.1 | 8.49E-01 | 0.4 | 5.40E-01 | LIM domain protein |
| AT5G65380 | 0.4 | 2.76E-03 | 0.0 | 7.19E-01 | 0.3 | 1.16E-02 | -0.1 | 6.80E-01 | -0.4 | 1.89E-02 | 0.4 | 7.65E-02 | MATE efflux family protein |
| AT5G42440 | 0.4 | 2.19E-02 | 0.0 | 8.84E-01 | 0.4 | 3.92E-02 | 0.4 | 9.95E-02 | 0.0 | 9.66E-01 | 0.4 | 2.07E-01 | Protein kinase domain-containing protein |
| AT4G25730 | 0.4 | 1.88E-02 | 0.2 | 2.64E-01 | 0.2 | 2.02E-01 | 0.6 | 9.48E-05 | 0.2 | 1.51E-01 | 0.4 | 2.25E-02 | Putative rRNA methyltransferase |
| AT1G74560 | 0.4 | 3.92E-03 | 0.3 | 1.33E-02 | 0.1 | 6.14E-01 | 0.7 | 2.25E-05 | 0.3 | 2.98E-02 | 0.4 | 2.70E-02 | NAP1-related protein 1 |
| AT2G17580 | 0.4 | 1.88E-03 | 0.2 | 4.77E-02 | 0.2 | 1.53E-01 | 0.1 | 6.98E-01 | 0.1 | 9.00E-01 | 0.1 | 8.85E-01 | Polynucleotide adenylyltransferase family protein |
| AT1G08880 | 0.4 | 3.34E-04 | 0.2 | 1.16E-02 | 0.1 | 7.33E-02 | 0.1 | 3.95E-01 | 0.0 | 8.84E-01 | 0.1 | 7.02E-01 | Probable histone H2AXa |
| AT1G73760 | 0.4 | 2.18E-04 | 0.2 | 3.43E-03 | 0.1 | 1.57E-01 | -0.5 | 1.45E-03 | -0.3 | 5.40E-02 | -0.2 | 3.15E-01 | RING-type E3 ubiquitin transferase |
| AT5G46740 | 0.4 | 6.71E-03 | 0.2 | 1.28E-01 | 0.2 | 1.97E-01 | 0.7 | 4.95E-04 | 0.4 | 2.86E-02 | 0.3 | 2.14E-01 | Ubiquitin-specific protease 21 |
| AT1G03210 | 0.4 | 1.11E-03 | 0.1 | 3.78E-01 | 0.3 | 1.23E-02 | -0.1 | 6.40E-01 | -0.1 | 4.23E-01 | 0.1 | 7.92E-01 | Uncharacterized protein |
| AT4G19540 | 0.4 | 7.97E-03 | 0.4 | 4.20E-03 | -0.1 | 6.83E-01 | 0.0 | 9.95E-01 | 0.2 | 6.18E-01 | -0.2 | 6.39E-01 | Iron-sulfur protein required for NADH dehydrogenase, mitochondrial |
| AT5G09920 | 0.4 | 1.67E-02 | -0.1 | 6.90E-01 | 0.4 | 1.90E-02 | 0.3 | 2.39E-01 | 0.1 | 8.90E-01 | 0.2 | 5.12E-01 | DNA-directed RNA polymerase II subunit 4 |
| AT1G11755 | 0.4 | 5.77E-03 | 0.3 | 1.60E-02 | 0.0 | 7.22E-01 | 0.3 | 1.82E-01 | 0.0 | 8.99E-01 | 0.2 | 3.81E-01 | ditrans,polycis-polyprenyl diphosphate synthase |
| AT1G66530 | 0.4 | 2.23E-04 | 0.2 | 3.90E-02 | 0.2 | 1.82E-02 | 0.0 | 9.52E-01 | 0.2 | 1.92E-01 | -0.2 | 2.68E-01 | Arginine--tRNA ligase, cytoplasmic |
| AT5G40060 | 0.4 | 4.10E-03 | -0.1 | 5.83E-01 | 0.4 | 5.93E-03 | 0.2 | 5.17E-01 | 0.2 | 4.59E-01 | 0.0 | 9.16E-01 | ADP-ribosyl cyclase |
| AT1G60620 | 0.4 | 4.78E-03 | 0.1 | 5.25E-01 | 0.3 | 3.87E-02 | 0.2 | 3.37E-01 | 0.2 | 3.73E-01 | 0.0 | 9.80E-01 | DNA-directed RNA polymerases I and III subunit RPAC1 |
| AT5G01290 | 0.4 | 1.63E-03 | 0.0 | 7.49E-01 | 0.3 | 8.44E-03 | 0.2 | 1.88E-01 | 0.0 | 9.82E-01 | 0.2 | 2.60E-01 | mRNA guanylyltransferase |
| AT3G22310 | 0.4 | 4.78E-02 | 0.2 | 3.81E-01 | 0.2 | 2.95E-01 | 0.8 | 3.31E-04 | 0.4 | 3.46E-02 | 0.3 | 1.69E-01 | Mitochondrial RNA helicase 1 |
| AT4G39640 | 0.4 | 4.64E-03 | 0.5 | 4.97E-04 | -0.2 | 1.30E-01 | -0.3 | 1.69E-02 | -0.2 | 7.90E-02 | -0.1 | 7.87E-01 | Glutathione hydrolase 1 |
| AT1G71050 | 0.4 | 3.71E-02 | 0.2 | 2.40E-01 | 0.2 | 4.17E-01 | -0.2 | 6.06E-01 | -0.3 | 3.68E-01 | 0.1 | 7.49E-01 | Heavy metal-associated isoprenylated plant protein 20 |
| AT4G12310 | 0.4 | 4.71E-03 | 0.0 | 9.68E-01 | 0.4 | 1.16E-02 | 0.3 | 3.40E-02 | 0.3 | 2.03E-02 | -0.1 | 7.49E-01 | At4g12310 |
| AT2G39290 | 0.4 | 1.63E-03 | 0.3 | 2.90E-03 | 0.0 | 8.96E-01 | 0.2 | 1.89E-01 | 0.2 | 2.62E-01 | 0.0 | 9.89E-01 | CDP-diacylglycerol--glycerol-3-phosphate 3-phosphatidyltransferase 1, chloroplastic |
| AT5G08620 | 0.4 | 2.87E-03 | 0.3 | 2.27E-02 | 0.1 | 3.56E-01 | 0.3 | 7.20E-02 | 0.0 | 9.87E-01 | 0.3 | 1.26E-01 | DEAD-box ATP-dependent RNA helicase 25 |
| AT1G11580 | 0.4 | 2.11E-03 | 0.6 | 1.27E-04 | -0.2 | 3.73E-02 | -0.4 | 3.68E-03 | -0.1 | 4.52E-01 | -0.3 | 8.23E-02 | Pectinesterase |
| AT1G56070 | 0.4 | 4.36E-03 | 0.2 | 4.38E-02 | 0.1 | 2.92E-01 | 0.4 | 6.80E-03 | 0.1 | 6.43E-01 | 0.3 | 1.31E-01 | Elongation factor 2 |
| AT3G47620 | 0.4 | 7.36E-03 | 0.1 | 3.31E-01 | 0.2 | 7.15E-02 | -0.1 | 5.03E-01 | -0.4 | 3.44E-02 | 0.3 | 1.67E-01 | TCP domain-containing protein |
| AT3G16570 | 0.4 | 7.79E-02 | 0.0 | 9.21E-01 | 0.4 | 9.90E-02 | -0.6 | 4.52E-03 | -0.5 | 3.67E-02 | -0.1 | 6.83E-01 | Rapid alkalinization factor 23 |
| AT5G01810 | 0.4 | 1.06E-02 | 0.4 | 5.83E-03 | -0.1 | 6.54E-01 | 0.1 | 5.26E-01 | -0.2 | 1.10E-01 | 0.3 | 6.88E-02 | non-specific serine |
| AT5G63450 | 0.4 | 3.96E-02 | 1.0 | 7.59E-05 | -0.7 | 3.39E-03 | | | | | | | Cytochrome P450, family 94, subfamily B, polypeptide 1 |
| AT1G65590 | 0.4 | 7.39E-03 | 0.5 | 1.18E-03 | -0.2 | 2.47E-01 | -0.1 | 3.89E-01 | 0.0 | 8.94E-01 | -0.2 | 4.32E-01 | Beta-hexosaminidase 3 |
| AT5G05550 | 0.4 | 1.41E-02 | 0.3 | 2.34E-02 | 0.0 | 8.95E-01 | -0.4 | 5.16E-02 | -0.6 | 1.36E-02 | 0.2 | 5.69E-01 | Sequence-specific DNA binding transcription factor |
| AT1G10590 | 0.4 | 3.22E-04 | 0.3 | 1.33E-03 | 0.0 | 5.36E-01 | 0.1 | 4.61E-01 | 0.0 | 9.08E-01 | 0.2 | 5.42E-01 | Nucleic acid-binding, OB-fold-like protein |
| AT2G20020 | 0.4 | 3.65E-03 | 0.2 | 7.77E-02 | 0.2 | 1.66E-01 | 0.4 | 1.67E-02 | 0.2 | 4.01E-01 | 0.2 | 2.34E-01 | CRS2-associated factor 1, chloroplastic |
| AT2G40550 | 0.4 | 2.93E-03 | 0.3 | 6.01E-03 | 0.0 | 8.38E-01 | 0.4 | 1.01E-01 | 0.5 | 4.67E-02 | -0.1 | 6.71E-01 | Mini-chromosome maintenance complex-binding protein |
| AT5G43250 | 0.4 | 1.81E-01 | 0.6 | 4.91E-02 | -0.2 | 5.29E-01 | 0.7 | 1.31E-02 | 0.1 | 9.12E-01 | 0.6 | 4.82E-02 | Nuclear factor Y, subunit C13 |
| AT3G22960 | 0.4 | 2.25E-04 | 0.3 | 1.91E-03 | 0.1 | 2.80E-01 | -0.1 | 7.17E-01 | -0.1 | 6.45E-01 | 0.0 | 9.17E-01 | Plastidial pyruvate kinase 1, chloroplastic |
| AT4G02900 | 0.4 | 6.25E-03 | 0.1 | 4.18E-01 | 0.3 | 6.79E-02 | -0.5 | 2.81E-01 | 0.1 | 8.86E-01 | -0.6 | 2.45E-01 | ERD family protein |
| AT3G60910 | 0.4 | 1.46E-02 | 0.2 | 1.20E-01 | 0.1 | 3.63E-01 | 0.0 | 9.49E-01 | 0.0 | 8.91E-01 | 0.0 | 9.44E-01 | Uncharacterized protein T4C21_320 |
| AT4G35830 | 0.4 | 4.89E-04 | 0.3 | 5.09E-03 | 0.1 | 2.48E-01 | 0.1 | 6.00E-01 | -0.1 | 4.49E-01 | 0.2 | 3.04E-01 | Aconitate hydratase |
| AT1G20840 | 0.4 | 1.04E-03 | 0.6 | 6.83E-05 | -0.2 | 2.90E-02 | -0.4 | 4.59E-02 | -0.1 | 6.79E-01 | -0.3 | 2.96E-01 | Monosaccharide-sensing protein 1 |
| AT3G56630 | 0.4 | 5.00E-03 | 0.1 | 4.95E-01 | 0.3 | 3.85E-02 | 0.0 | 8.14E-01 | 0.0 | 8.47E-01 | 0.1 | 6.85E-01 | Cytochrome P450, family 94, subfamily D, polypeptide 2 |
| AT1G63770 | 0.4 | 2.72E-04 | 0.1 | 1.54E-01 | 0.3 | 6.02E-03 | 0.1 | 7.10E-01 | 0.0 | 9.74E-01 | 0.0 | 8.47E-01 | Peptidase M1 family protein |
| AT3G19010 | 0.4 | 5.42E-03 | 0.0 | 9.15E-01 | 0.4 | 9.54E-03 | 0.5 | 1.76E-02 | 0.1 | 8.49E-01 | 0.5 | 1.18E-01 | 2-oxoglutarate and Fe-dependent oxygenase superfamily protein |
| AT3G13960 | 0.4 | 2.09E-02 | 0.0 | 7.97E-01 | 0.3 | 8.19E-02 | -0.3 | 3.49E-01 | -0.4 | 2.56E-01 | 0.1 | 8.38E-01 | Growth-regulating factor |
| AT2G39810 | 0.4 | 4.07E-04 | 0.2 | 5.39E-02 | 0.2 | 2.59E-02 | 0.1 | 4.98E-01 | -0.1 | 6.94E-01 | 0.2 | 3.40E-01 | Ubiquitin-protein ligase |
| AT5G11380 | 0.4 | 2.18E-03 | 0.1 | 2.59E-01 | 0.2 | 3.87E-02 | 0.1 | 7.47E-01 | 0.1 | 8.41E-01 | 0.0 | 9.65E-01 | 1-D-deoxyxylulose 5-phosphate synthase-like protein |
| AT3G60130 | 0.4 | 2.72E-03 | 0.6 | 1.02E-04 | -0.3 | 2.72E-02 | -0.4 | 4.13E-03 | -0.3 | 1.81E-02 | 0.0 | 8.42E-01 | Beta glucosidase 16 |
| AT5G22750 | 0.4 | 4.41E-03 | 0.0 | 9.74E-01 | 0.4 | 1.68E-02 | 0.5 | 1.11E-01 | 0.3 | 3.70E-01 | 0.2 | 6.81E-01 | DNA repair protein RAD5A |
| AT4G31500 | 0.4 | 8.37E-03 | 0.5 | 1.83E-03 | -0.1 | 3.04E-01 | 0.4 | 1.97E-02 | 0.2 | 2.94E-01 | 0.2 | 4.66E-01 | Cytochrome P450 83B1 |
| AT4G29530 | 0.4 | 7.49E-02 | 0.2 | 2.43E-01 | 0.1 | 6.28E-01 | -0.1 | 7.95E-01 | 0.0 | 1.00E+00 | -0.1 | 8.38E-01 | Pyridoxal phosphate phosphatase-related protein |
| AT1G09080 | 0.4 | 6.81E-01 | -0.9 | 4.46E-01 | 1.2 | 3.52E-01 | 2.2 | 7.93E-04 | 2.7 | 2.94E-04 | -0.5 | 2.29E-01 | hypothetical protein |
| AT1G63855 | 0.4 | 9.48E-03 | 0.4 | 6.39E-03 | 0.0 | 7.74E-01 | 0.4 | 1.06E-01 | 0.0 | 9.59E-01 | 0.4 | 2.12E-01 | Methyltransferase family protein |
| AT5G10560 | 0.4 | 1.01E-03 | 0.2 | 1.54E-02 | 0.1 | 1.84E-01 | 0.1 | 4.06E-01 | 0.0 | 9.23E-01 | 0.1 | 5.08E-01 | Probable beta-D-xylosidase 6 |
| AT3G20280 | 0.4 | 3.80E-03 | 0.2 | 1.67E-01 | 0.2 | 1.02E-01 | 0.0 | 9.11E-01 | -0.2 | 6.33E-01 | 0.1 | 7.56E-01 | PHD-type domain-containing protein |
| AT4G35070 | 0.4 | 3.10E-02 | 0.2 | 2.07E-01 | 0.2 | 4.18E-01 | 0.1 | 7.32E-01 | -0.1 | 8.26E-01 | 0.2 | 6.24E-01 | SBP family protein |
| AT1G18270 | 0.4 | 7.48E-04 | 0.5 | 1.85E-04 | -0.1 | 1.91E-01 | -0.4 | 3.73E-02 | -0.1 | 4.80E-01 | -0.2 | 3.55E-01 | F15H18.22 |
| AT4G02060 | 0.4 | 7.92E-04 | 0.2 | 1.66E-02 | 0.1 | 1.51E-01 | 0.4 | 1.58E-02 | 0.1 | 5.60E-01 | 0.3 | 1.84E-01 | DNA replication licensing factor MCM7 |
| AT3G23070 | 0.4 | 1.25E-02 | 0.4 | 7.67E-03 | -0.1 | 7.44E-01 | 0.1 | 5.68E-01 | 0.2 | 5.24E-01 | 0.0 | 9.06E-01 | CRM-domain containing factor CFM3A, chloroplastic |
| AT1G18360 | 0.4 | 2.90E-02 | 0.3 | 4.17E-02 | 0.0 | 9.22E-01 | -0.4 | 1.52E-01 | -0.2 | 5.80E-01 | -0.2 | 6.18E-01 | Alpha |
| AT4G01710 | 0.4 | 4.56E-02 | 0.1 | 7.27E-01 | 0.3 | 1.54E-01 | 0.1 | 8.38E-01 | -0.1 | 8.10E-01 | 0.1 | 6.93E-01 | Actin-related protein 2 |
| AT3G01850 | 0.4 | 4.46E-04 | 0.0 | 8.93E-01 | 0.4 | 1.48E-03 | -0.1 | 5.67E-01 | -0.2 | 1.20E-01 | 0.2 | 4.21E-01 | Ribulose-phosphate 3-epimerase |
| AT5G59770 | 0.4 | 5.05E-04 | 0.2 | 1.98E-02 | 0.2 | 7.44E-02 | 0.0 | 8.76E-01 | 0.1 | 4.41E-01 | -0.2 | 3.99E-01 | Very-long-chain -3-hydroxyacyl-CoA dehydratase |
| AT3G21890 | 0.4 | 2.37E-01 | -0.1 | 7.43E-01 | 0.5 | 2.19E-01 | | | | | | | B-box domain protein 31 |
| AT5G65460 | 0.4 | 1.70E-02 | -0.1 | 4.94E-01 | 0.5 | 1.25E-02 | -0.1 | 6.79E-01 | 0.2 | 4.33E-01 | -0.3 | 3.02E-01 | Kinesin like protein for actin based chloroplast movement 2 |
| AT1G56510 | 0.4 | 3.93E-03 | -0.1 | 3.67E-01 | 0.5 | 3.03E-03 | 0.2 | 2.19E-01 | 0.1 | 7.56E-01 | 0.1 | 5.81E-01 | Disease resistance protein ADR2 |

| | | | | | | | | | | | | | |
|---|---|---|---|---|---|---|---|---|---|---|---|---|---|
| AT4G36570 | 0.4 | 2.95E-02 | 1.1 | 3.93E-05 | -0.7 | 1.65E-03 | 0.6 | 2.18E-02 | 0.2 | 5.96E-01 | 0.5 | 1.68E-01 | Protein RADIALIS-like 3 |
| AT5G52870 | 0.4 | 1.18E-02 | 0.0 | 8.44E-01 | 0.3 | 4.36E-02 | 0.3 | 1.31E-01 | 0.0 | 9.64E-01 | 0.3 | 3.03E-01 | Membrane-associated kinase regulator |
| AT3G58560 | 0.4 | 7.84E-05 | -0.1 | 1.65E-01 | 0.5 | 1.07E-04 | 0.3 | 5.14E-02 | 0.1 | 7.79E-01 | 0.2 | 2.54E-01 | Carbon catabolite repressor protein 4 homolog 1 |
| AT3G10140 | 0.4 | 1.79E-02 | 0.1 | 4.83E-01 | 0.3 | 1.34E-01 | 0.1 | 8.42E-01 | 0.0 | 9.55E-01 | 0.1 | 8.11E-01 | DNA repair protein recA homolog 2, mitochondrial |
| AT1G68130 | 0.4 | 2.07E-03 | 0.1 | 4.62E-01 | 0.3 | 2.14E-02 | -0.2 | 2.29E-01 | 0.1 | 7.37E-01 | -0.3 | 1.56E-01 | Protein indeterminate-domain 14 |
| AT4G15850 | 0.4 | 1.01E-02 | 0.1 | 3.01E-01 | 0.2 | 1.39E-01 | 0.4 | 1.80E-01 | -0.1 | 9.17E-01 | 0.5 | 1.61E-01 | DEAD-box ATP-dependent RNA helicase 1 |
| AT2G46340 | 0.4 | 4.37E-03 | 0.0 | 8.19E-01 | 0.4 | 6.55E-03 | 0.3 | 2.36E-02 | 0.4 | 2.15E-02 | 0.0 | 8.78E-01 | Protein SUPPRESSOR OF PHYA-105 1 |
| AT1G16340 | 0.4 | 1.87E-03 | 0.4 | 1.38E-03 | 0.0 | 7.32E-01 | 0.1 | 6.20E-01 | 0.1 | 6.72E-01 | 0.0 | 9.97E-01 | 3-deoxy-8-phosphooctulonate synthase |
| AT4G29220 | 0.4 | 3.50E-04 | 0.1 | 1.57E-01 | 0.3 | 7.62E-03 | -0.1 | 7.69E-01 | 0.1 | 8.47E-01 | -0.1 | 6.87E-01 | ATP-dependent 6-phosphofructokinase |
| AT2G23380 | 0.4 | 4.41E-04 | 0.2 | 6.38E-02 | 0.2 | 2.66E-02 | -0.1 | 7.63E-01 | -0.1 | 7.72E-01 | 0.0 | 9.76E-01 | SET domain-containing protein |
| AT1G71140 | 0.4 | 1.26E-02 | 0.2 | 2.67E-01 | 0.2 | 1.76E-01 | 0.7 | 1.11E-02 | 0.1 | 8.86E-01 | 0.6 | 5.76E-02 | Protein DETOXIFICATION 14 |
| AT1G54220 | 0.4 | 2.00E-03 | 0.3 | 9.44E-03 | 0.1 | 5.09E-01 | 0.1 | 4.22E-01 | 0.2 | 2.73E-01 | -0.1 | 7.99E-01 | Dihydrolipoyllysine-residue acetyltransferase component 3 of pyruvate dehydrogenase complex, mitochondrial |
| AT1G30545 | 0.4 | 2.80E-02 | -0.1 | 7.47E-01 | 0.4 | 4.31E-02 | -0.3 | 5.17E-01 | -0.4 | 3.91E-01 | 0.1 | 8.59E-01 | Trimethylguanosine synthase |
| AT5G57710 | 0.4 | 4.63E-03 | -0.2 | 9.52E-02 | 0.6 | 1.02E-03 | -0.2 | 1.87E-01 | -0.2 | 3.31E-01 | 0.0 | 9.28E-01 | Protein SUPPRESSOR OF MAX2 1 |
| AT2G18600 | 0.4 | 6.84E-02 | 0.0 | 8.62E-01 | 0.4 | 1.05E-01 | 0.3 | 3.04E-01 | 0.0 | 9.93E-01 | 0.4 | 3.90E-01 | Probable NEDD8-conjugating enzyme Ubc12-like |
| AT1G07880 | 0.4 | 7.40E-02 | 0.5 | 3.60E-02 | -0.1 | 6.97E-01 | 0.3 | 2.78E-01 | 0.4 | 1.56E-01 | -0.1 | 7.53E-01 | Mitogen-activated protein kinase |
| AT1G03830 | 0.4 | 2.31E-03 | 0.3 | 2.59E-02 | 0.1 | 2.87E-01 | 0.1 | 4.60E-01 | 0.1 | 5.49E-01 | 0.0 | 9.88E-01 | F11M21.24 protein |
| AT5G62370 | 0.4 | 4.11E-02 | 0.0 | 8.44E-01 | 0.3 | 1.29E-01 | 0.8 | 5.70E-02 | 0.6 | 2.06E-01 | 0.2 | 7.44E-01 | Pentatricopeptide repeat-containing protein At5g62370 |
| AT1G26150 | 0.4 | 2.73E-03 | -0.1 | 4.18E-01 | 0.5 | 2.12E-03 | 0.0 | 9.88E-01 | 0.2 | 2.59E-01 | -0.3 | 2.80E-01 | non-specific serine |
| AT1G51380 | 0.4 | 7.75E-02 | 0.3 | 2.15E-01 | 0.1 | 6.77E-01 | 1.2 | 5.47E-03 | 0.9 | 4.10E-02 | 0.2 | 5.52E-01 | DEA-box RNA helicase family protein |
| AT2G03667 | 0.4 | 3.46E-03 | 0.0 | 8.19E-01 | 0.3 | 1.70E-02 | 0.0 | 9.36E-01 | 0.1 | 7.87E-01 | -0.1 | 7.48E-01 | Asparagine synthase family protein |
| AT2G43750 | 0.4 | 5.42E-04 | 0.5 | 1.02E-04 | -0.1 | 1.14E-01 | -0.1 | 5.11E-01 | -0.2 | 2.13E-01 | 0.1 | 6.32E-01 | Cysteine synthase, chloroplastic |
| AT1G18660 | 0.4 | 9.93E-04 | -0.1 | 5.23E-01 | 0.4 | 1.36E-03 | 0.0 | 9.75E-01 | 0.1 | 7.48E-01 | -0.1 | 8.11E-01 | Zinc finger family protein |
| AT2G43760 | 0.4 | 5.95E-04 | 0.2 | 4.05E-02 | 0.2 | 5.29E-02 | -0.4 | 2.63E-02 | -0.5 | 3.14E-02 | 0.0 | 9.45E-01 | Molybdopterin synthase catalytic subunit |
| AT3G07270 | 0.4 | 6.46E-04 | 0.4 | 1.38E-03 | 0.0 | 8.29E-01 | -0.2 | 1.79E-01 | -0.2 | 9.79E-02 | 0.1 | 7.72E-01 | GTP cyclohydrolase 1 |
| AT3G27340 | 0.4 | 2.24E-03 | 0.3 | 1.16E-02 | 0.1 | 4.89E-01 | -0.1 | 7.52E-01 | -0.1 | 9.00E-01 | 0.0 | 9.22E-01 | Myb domain protein |
| AT1G48630 | 0.4 | 4.37E-02 | 0.6 | 5.05E-03 | -0.2 | 2.20E-01 | 0.5 | 7.21E-03 | 0.4 | 3.06E-02 | 0.1 | 8.28E-01 | Small ribosomal subunit protein RACK1y |
| AT1G10060 | 0.4 | 4.45E-03 | 0.5 | 8.17E-04 | -0.1 | 2.64E-01 | 0.0 | 8.55E-01 | -0.2 | 5.76E-01 | 0.2 | 4.68E-01 | Branched-chain-amino-acid aminotransferase |
| AT5G63380 | 0.4 | 1.12E-03 | 0.0 | 7.91E-01 | 0.4 | 5.09E-03 | -0.1 | 5.32E-01 | 0.1 | 8.56E-01 | -0.2 | 5.16E-01 | 4-coumarate--CoA ligase-like 9 |
| AT2G43970 | 0.4 | 2.33E-03 | 0.0 | 7.70E-01 | 0.3 | 8.70E-03 | 0.0 | 9.46E-01 | -0.2 | 1.84E-01 | 0.2 | 3.27E-01 | La-related protein 6B |
| AT5G17620 | 0.4 | 1.69E-02 | 0.2 | 2.84E-01 | 0.2 | 2.06E-01 | 0.0 | 8.48E-01 | -0.2 | 4.33E-01 | 0.1 | 6.68E-01 | AUGMIN subunit 7 |
| AT5G46450 | 0.4 | 3.26E-03 | 0.2 | 4.87E-02 | 0.2 | 2.14E-01 | 0.2 | 2.27E-01 | -0.3 | 8.26E-02 | 0.6 | 7.11E-03 | Disease resistance protein family |
| AT3G03420 | 0.4 | 1.85E-03 | 0.1 | 5.52E-01 | 0.3 | 1.56E-02 | 0.4 | 3.56E-02 | 0.1 | 8.39E-01 | 0.4 | 1.18E-01 | Mitochondrial inner membrane protease ATP23 |
| AT1G14770 | 0.4 | 2.76E-03 | 0.1 | 2.07E-01 | 0.2 | 6.45E-02 | 0.0 | 9.53E-01 | 0.0 | 9.89E-01 | 0.0 | 9.77E-01 | Zinc finger PHD-type domain-containing protein |
| AT2G36250 | 0.4 | 1.31E-04 | 0.4 | 2.03E-04 | 0.0 | 9.45E-01 | 0.2 | 1.47E-01 | 0.1 | 5.30E-01 | 0.1 | 7.01E-01 | Tubulin |
| AT3G16480 | 0.4 | 2.54E-04 | 0.4 | 5.30E-04 | 0.0 | 8.49E-01 | 0.1 | 2.37E-01 | 0.2 | 2.72E-01 | 0.0 | 9.69E-01 | AT3G16480 protein |
| AT5G48450 | 0.4 | 1.21E-02 | 0.2 | 1.10E-01 | 0.1 | 3.57E-01 | -0.3 | 5.35E-01 | 0.0 | 9.84E-01 | -0.3 | 5.06E-01 | SKU5 similar 3 |
| AT5G02820 | 0.4 | 4.07E-03 | 0.6 | 2.18E-04 | -0.2 | 6.22E-02 | 0.1 | 5.14E-01 | 0.0 | 9.41E-01 | 0.1 | 6.56E-01 | DNA topoisomerase 6 subunit A |
| AT1G06520 | 0.4 | 2.42E-02 | 0.2 | 1.51E-01 | 0.1 | 4.52E-01 | -0.5 | 2.30E-01 | -0.2 | 6.55E-01 | -0.3 | 6.67E-01 | Glycerol-3-phosphate acyltransferase 1 |
| AT3G04120 | 0.4 | 5.74E-03 | 0.1 | 3.21E-01 | 0.3 | 5.92E-02 | 1.2 | 1.05E-06 | 0.3 | 1.02E-01 | 0.9 | 3.97E-04 | Glyceraldehyde-3-phosphate dehydrogenase GAPC1, cytosolic |
| AT1G69830 | 0.4 | 4.94E-03 | 0.3 | 1.32E-02 | 0.0 | 7.25E-01 | 0.3 | 1.25E-01 | 0.3 | 1.13E-01 | 0.0 | 8.80E-01 | Alpha-amylase 3, chloroplastic |
| AT3G57490 | 0.4 | 8.75E-03 | 0.3 | 2.01E-02 | 0.0 | 7.68E-01 | 0.6 | 9.21E-05 | 0.3 | 2.60E-02 | 0.3 | 8.38E-02 | S5 DRBM domain-containing protein |
| AT1G72040 | 0.4 | 6.22E-04 | 0.2 | 1.13E-02 | 0.1 | 1.52E-01 | 0.5 | 2.00E-02 | 0.2 | 4.37E-01 | 0.3 | 2.41E-01 | Deoxyguanosine kinase, putative |
| AT1G33330 | 0.4 | 5.85E-04 | 0.1 | 1.18E-01 | 0.2 | 2.30E-02 | -0.1 | 8.19E-01 | 0.0 | 9.80E-01 | -0.1 | 8.45E-01 | Class I peptide chain release factor |
| AT5G14200 | 0.4 | 1.13E-02 | 0.6 | 1.73E-03 | -0.2 | 2.24E-01 | -0.3 | 6.28E-02 | 0.6 | 2.18E-03 | -0.9 | 5.83E-04 | Isopropylmalate dehydrogenase 1 |
| AT4G24265 | 0.4 | 4.45E-02 | 0.3 | 9.38E-02 | 0.1 | 8.04E-01 | 0.2 | 5.97E-01 | 0.7 | 1.08E-01 | -0.4 | 2.34E-01 | At4g24265 |
| AT1G04410 | 0.4 | 4.79E-04 | 0.5 | 2.50E-04 | -0.1 | 4.19E-01 | 0.2 | 1.10E-01 | 0.0 | 9.30E-01 | 0.1 | 3.47E-01 | Malate dehydrogenase 1, cytoplasmic |
| AT5G23405 | 0.4 | 2.85E-02 | -0.1 | 7.73E-01 | 0.4 | 4.18E-02 | 0.2 | 5.79E-01 | 0.0 | 9.89E-01 | 0.2 | 7.26E-01 | HMG box domain-containing protein |
| AT1G71850 | 0.4 | 2.00E-02 | 0.2 | 2.26E-01 | 0.2 | 2.81E-01 | 0.6 | 1.15E-02 | 0.3 | 3.64E-01 | 0.4 | 2.24E-01 | F14O23.23 protein |
| AT4G32840 | 0.4 | 1.73E-03 | 0.6 | 2.04E-04 | -0.2 | 1.24E-01 | 0.1 | 7.42E-01 | 0.0 | 9.90E-01 | 0.1 | 7.99E-01 | ATP-dependent 6-phosphofructokinase |
| AT5G60960 | 0.4 | 8.20E-03 | 0.5 | 3.37E-03 | -0.1 | 5.58E-01 | 0.4 | 3.40E-03 | 0.2 | 9.73E-02 | 0.2 | 2.95E-01 | Small ribosomal subunit protein mL104 |
| AT3G62470 | 0.4 | 9.68E-03 | 0.4 | 5.95E-03 | -0.1 | 7.33E-01 | -0.2 | 3.95E-01 | -0.2 | 2.95E-01 | 0.1 | 8.64E-01 | Pentatricopeptide repeat-containing protein At3g62470, mitochondrial |
| AT1G80710 | 0.4 | 9.88E-03 | 0.3 | 3.86E-02 | 0.1 | 6.19E-01 | 0.2 | 7.20E-01 | 0.1 | 9.07E-01 | 0.1 | 8.89E-01 | DROUGHT SENSITIVE 1 |
| AT4G23810 | 0.4 | 7.92E-02 | -0.1 | 5.26E-01 | 0.5 | 4.91E-02 | 0.2 | 2.72E-01 | -0.2 | 4.55E-01 | 0.4 | 1.29E-01 | Probable WRKY transcription factor 53 |
| AT1G63160 | 0.4 | 1.49E-02 | 0.3 | 8.71E-02 | 0.1 | 4.64E-01 | 0.4 | 1.29E-01 | 0.1 | 8.19E-01 | 0.3 | 3.90E-01 | Replication factor C subunit 2 |
| AT3G49220 | 0.4 | 2.15E-04 | 0.4 | 5.42E-04 | 0.0 | 7.88E-01 | -0.1 | 3.79E-01 | -0.1 | 7.41E-01 | -0.1 | 8.09E-01 | Plant invertase |
| AT4G08850 | 0.4 | 5.70E-03 | -0.2 | 5.54E-02 | 0.6 | 8.20E-04 | 0.4 | 5.45E-03 | -0.1 | 4.96E-01 | 0.5 | 1.23E-02 | MDIS1-interacting receptor like kinase 2 |
| AT5G01075 | 0.4 | 2.70E-02 | 0.8 | 6.54E-04 | -0.4 | 5.37E-02 | -0.5 | 5.11E-02 | -0.3 | 3.92E-01 | -0.3 | 4.85E-01 | At5g01075 |
| AT4G37210 | 0.4 | 8.45E-05 | 0.1 | 6.39E-02 | 0.3 | 4.03E-03 | 0.1 | 4.26E-01 | 0.0 | 9.95E-01 | 0.1 | 5.91E-01 | Tetratricopeptide repeat -like superfamily protein |
| AT1G06390 | 0.4 | 7.72E-05 | 0.2 | 6.28E-03 | 0.2 | 2.26E-02 | -0.2 | 1.96E-01 | -0.3 | 8.49E-02 | 0.1 | 6.74E-01 | non-specific serine |
| AT1G32380 | 0.4 | 6.57E-03 | 0.3 | 2.75E-02 | 0.1 | 5.60E-01 | 0.3 | 2.25E-02 | 0.0 | 7.95E-01 | 0.2 | 1.56E-01 | ribose-phosphate diphosphokinase |
| AT3G20060 | 0.4 | 1.10E-04 | 0.4 | 2.82E-04 | 0.0 | 7.54E-01 | -0.2 | 4.14E-02 | -0.3 | 3.42E-02 | 0.0 | 8.62E-01 | Ubiquitin-conjugating enzyme E2 19 |
| AT5G25120 | 0.4 | 5.77E-03 | 0.4 | 1.27E-02 | 0.0 | 7.96E-01 | 0.6 | 7.80E-03 | 0.3 | 1.54E-01 | 0.2 | 3.95E-01 | Cytochrome P450 71B11 |
| AT5G44635 | 0.4 | 2.95E-03 | 0.2 | 1.11E-01 | 0.2 | 1.13E-01 | 0.7 | 3.27E-03 | 0.7 | 4.75E-03 | 0.0 | 9.43E-01 | DNA replication licensing factor MCM6 |
| AT5G04330 | 0.4 | 2.09E-02 | 0.4 | 2.51E-02 | 0.0 | 9.97E-01 | -0.7 | 1.45E-02 | -0.4 | 2.37E-01 | -0.4 | 3.44E-01 | Cytochrome P450 84A4 |
| AT1G71260 | 0.4 | 1.95E-03 | 0.4 | 3.29E-03 | 0.0 | 9.18E-01 | 0.6 | 1.46E-03 | 0.1 | 5.94E-01 | 0.4 | 1.48E-02 | Single-stranded DNA-binding protein WHY2, mitochondrial |
| AT5G02840 | 0.4 | 4.11E-02 | 0.0 | 9.15E-01 | 0.4 | 7.91E-02 | 0.0 | 8.02E-01 | -0.1 | 4.74E-01 | 0.1 | 4.01E-01 | hypothetical protein |
| AT5G22860 | 0.4 | 1.25E-02 | 0.0 | 9.95E-01 | 0.4 | 3.29E-02 | 0.2 | 5.42E-01 | 0.0 | 9.39E-01 | 0.2 | 5.66E-01 | Prolylcarboxypeptidase-like protein |
| AT1G18070 | 0.4 | 1.53E-04 | 0.3 | 2.85E-03 | 0.1 | 1.18E-01 | 0.3 | 3.19E-03 | 0.2 | 1.53E-01 | 0.2 | 3.09E-01 | T10F20.8 protein |
| AT5G08020 | 0.4 | 6.01E-03 | 0.3 | 1.71E-02 | 0.1 | 7.01E-01 | 0.5 | 5.48E-02 | 0.4 | 1.76E-01 | 0.1 | 7.79E-01 | Replication protein A 70 kDa DNA-binding subunit B |
| AT2G17450 | 0.4 | 9.59E-03 | 0.4 | 1.46E-02 | 0.0 | 9.21E-01 | -0.4 | 1.02E-02 | -0.5 | 8.98E-03 | 0.1 | 8.57E-01 | Probable E3 ubiquitin-protein ligase ATL44 |
| AT3G23020 | 0.4 | 3.01E-02 | 0.2 | 3.30E-01 | 0.2 | 3.00E-01 | 0.4 | 3.37E-01 | 0.1 | 8.60E-01 | 0.3 | 5.97E-01 | Pentatricopeptide repeat-containing protein At3g23020 |
| AT3G24180 | 0.4 | 2.33E-04 | 0.2 | 4.99E-02 | 0.2 | 1.32E-02 | 0.5 | 7.31E-03 | 0.3 | 1.82E-01 | 0.2 | 3.72E-01 | Non-lysosomal glucosylceramidase |
| AT1G55870 | 0.4 | 7.33E-04 | 0.0 | 6.44E-01 | 0.3 | 5.92E-03 | 0.5 | 9.34E-02 | 0.2 | 5.37E-01 | 0.3 | 4.96E-01 | Polynucleotidyl transferase, ribonuclease H-like superfamily protein |
| AT1G50300 | 0.4 | 8.80E-04 | 0.1 | 3.39E-01 | 0.3 | 1.24E-02 | 0.0 | 8.03E-01 | -0.3 | 9.79E-02 | 0.2 | 2.26E-01 | TBP-associated factor 15 |
| AT5G10700 | 0.4 | 3.32E-03 | 0.3 | 9.75E-03 | 0.1 | 7.02E-01 | -0.2 | 4.68E-01 | -0.3 | 3.53E-01 | 0.1 | 8.40E-01 | aminoacyl-tRNA hydrolase |
| AT5G64420 | 0.4 | 2.26E-04 | 0.1 | 1.57E-01 | 0.3 | 5.32E-03 | 0.4 | 8.66E-03 | 0.1 | 7.26E-01 | 0.3 | 1.13E-01 | DNA polymerase V family |
| AT3G19170 | 0.4 | 2.05E-03 | 0.2 | 5.49E-02 | 0.2 | 1.23E-01 | 0.3 | 9.36E-03 | 0.1 | 3.83E-01 | 0.2 | 2.35E-01 | Presequence protease 1 |
| AT2G03390 | 0.4 | 2.10E-03 | 0.0 | 6.68E-01 | 0.4 | 3.01E-03 | -0.1 | 2.87E-01 | -0.2 | 3.47E-01 | 0.0 | 9.91E-01 | UvrB |
| AT4G34870 | 0.4 | 1.93E-03 | 0.5 | 5.03E-04 | -0.1 | 2.62E-01 | 0.4 | 1.66E-02 | 0.4 | 6.74E-02 | 0.1 | 8.16E-01 | Peptidyl-prolyl cis-trans isomerase CYP18-4 |
| AT4G37070 | 0.4 | 1.63E-03 | 0.6 | 9.38E-05 | -0.2 | 4.50E-02 | -0.4 | 1.33E-01 | -0.1 | 7.67E-01 | -0.3 | 4.14E-01 | Patatin |
| AT1G26090 | 0.4 | 5.37E-03 | 0.2 | 7.12E-02 | 0.2 | 2.72E-01 | -0.2 | 5.08E-01 | -0.2 | 6.45E-01 | 0.0 | 9.46E-01 | Uncharacterized protein At1g26090, chloroplastic |
| AT4G28990 | 0.4 | 3.87E-03 | 0.2 | 6.35E-02 | 0.2 | 2.20E-01 | 0.1 | 7.67E-01 | 0.0 | 9.08E-01 | 0.1 | 7.30E-01 | RNA-binding protein-like protein |

| | | | | | | | | | | | | | |
|---|---|---|---|---|---|---|---|---|---|---|---|---|---|
| AT3G10070 | 0.4 | 6.74E-04 | 0.0 | 9.01E-01 | 0.4 | 3.66E-03 | 0.3 | 3.52E-01 | 0.0 | 9.95E-01 | 0.3 | 4.39E-01 | Transcription initiation factor TFIID subunit 12 |
| AT1G13640 | 0.4 | 6.32E-03 | 0.3 | 2.58E-02 | 0.1 | 5.46E-01 | 0.0 | 8.20E-01 | -0.3 | 9.15E-02 | 0.3 | 1.14E-01 | Phosphatidylinositol 4-kinase gamma 6 |
| AT3G24495 | 0.4 | 7.91E-03 | 0.1 | 3.65E-01 | 0.3 | 9.28E-02 | 0.5 | 5.25E-02 | 0.3 | 2.41E-01 | 0.2 | 6.76E-01 | MUTS homolog 7 |
| AT5G46030 | 0.4 | 7.54E-04 | 0.2 | 4.23E-02 | 0.2 | 5.75E-02 | 0.1 | 8.26E-01 | 0.0 | 9.66E-01 | 0.0 | 9.24E-01 | Transcription elongation factor 1 homolog |
| AT5G24970 | 0.4 | 1.13E-03 | 0.0 | 8.32E-01 | 0.4 | 6.82E-03 | -0.2 | 3.58E-01 | -0.1 | 6.67E-01 | -0.1 | 8.00E-01 | Protein kinase superfamily protein |
| AT4G22930 | 0.4 | 1.44E-03 | 0.6 | 1.24E-04 | -0.2 | 6.51E-02 | 0.3 | 3.05E-02 | 0.1 | 3.98E-01 | 0.1 | 4.30E-01 | Dihydroorotase, mitochondrial |
| AT2G18390 | 0.4 | 2.07E-03 | 0.3 | 7.42E-03 | 0.1 | 6.19E-01 | 0.0 | 9.52E-01 | -0.1 | 8.58E-01 | 0.0 | 9.25E-01 | ADP-ribosylation factor-like protein 2 |
| AT1G76170 | 0.4 | 1.17E-01 | -0.1 | 8.27E-01 | 0.5 | 1.45E-01 | -0.2 | 6.92E-01 | 0.1 | 8.57E-01 | -0.3 | 6.30E-01 | Cytoplasmic tRNA 2-thiolation protein 1 |
| AT1G07570 | 0.4 | 2.27E-03 | 0.1 | 4.71E-01 | 0.3 | 2.09E-02 | -0.5 | 2.23E-02 | -0.4 | 1.13E-01 | -0.1 | 6.79E-01 | Protein kinase superfamily protein |
| AT3G25770 | 0.4 | 2.75E-02 | 0.7 | 1.23E-03 | -0.4 | 7.42E-02 | 1.9 | 1.52E-05 | 1.5 | 5.46E-04 | 0.4 | 3.96E-01 | Allene oxide cyclase 2, chloroplastic |
| AT2G28305 | 0.4 | 6.01E-02 | 0.2 | 3.24E-01 | 0.2 | 4.27E-01 | -0.2 | 4.54E-01 | -0.2 | 3.21E-01 | 0.1 | 8.32E-01 | Cytokinin riboside 5'-monophosphate phosphoribohydrolase |
| AT5G27670 | 0.4 | 1.61E-04 | 0.5 | 7.56E-05 | -0.1 | 3.14E-01 | -0.2 | 3.73E-01 | -0.1 | 4.83E-01 | 0.0 | 9.80E-01 | Probable histone H2A.5 |
| AT2G38880 | 0.4 | 1.40E-03 | 0.1 | 2.09E-01 | 0.3 | 3.01E-02 | -0.1 | 4.91E-01 | -0.3 | 6.66E-02 | 0.2 | 3.72E-01 | AT2G38880 protein |
| AT3G12900 | 0.4 | 2.08E-01 | 0.5 | 1.67E-01 | -0.1 | 8.95E-01 | -3.8 | 2.60E-05 | -5.4 | 1.05E-04 | 1.7 | 1.10E-01 | Scopoletin 8-hydroxylase |
| AT4G27240 | 0.4 | 8.82E-04 | 0.4 | 1.34E-03 | 0.0 | 9.85E-01 | -0.3 | 1.09E-01 | -0.3 | 2.11E-01 | 0.0 | 9.16E-01 | At4g27240 family protein) |
| AT1G71460 | 0.4 | 4.28E-03 | 0.4 | 1.12E-02 | 0.0 | 7.55E-01 | 0.4 | 1.48E-01 | 0.0 | 9.79E-01 | 0.4 | 2.63E-01 | Pentatricopeptide repeat-containing protein At1g71460, chloroplastic |
| AT3G51000 | 0.4 | 1.06E-03 | 0.0 | 8.56E-01 | 0.4 | 4.43E-03 | -0.3 | 2.12E-01 | -0.4 | 1.06E-01 | 0.1 | 7.39E-01 | AB hydrolase-1 domain-containing protein |
| AT1G27930 | 0.4 | 1.20E-04 | 0.4 | 1.85E-04 | 0.0 | 9.26E-01 | 0.1 | 5.82E-01 | -0.1 | 7.35E-01 | 0.2 | 4.77E-01 | Arabinogalactan O-methyltransferase 1 |
| AT5G57990 | 0.4 | 1.13E-03 | 0.1 | 1.30E-01 | 0.2 | 3.03E-02 | 0.4 | 2.19E-03 | 0.1 | 4.80E-01 | 0.3 | 6.88E-02 | Ubiquitin carboxyl-terminal hydrolase 23 |
| AT1G07930 | 0.4 | 7.66E-03 | 0.6 | 1.33E-03 | -0.2 | 2.37E-01 | 0.4 | 9.99E-04 | 0.1 | 2.63E-01 | 0.3 | 9.02E-02 | Elongation factor 1-alpha |
| AT5G13840 | 0.4 | 7.90E-04 | 0.4 | 1.19E-03 | 0.0 | 9.92E-01 | 0.1 | 6.66E-01 | 0.2 | 5.13E-01 | -0.1 | 8.45E-01 | Protein FIZZY-RELATED 3 |
| AT1G61970 | 0.4 | 2.63E-03 | 0.2 | 1.10E-01 | 0.2 | 1.13E-01 | 0.3 | 2.31E-01 | -0.1 | 7.79E-01 | 0.4 | 1.65E-01 | At1g61970 |
| AT3G49680 | 0.4 | 4.10E-03 | 0.5 | 2.49E-03 | -0.1 | 6.50E-01 | 0.2 | 8.88E-02 | 0.4 | 2.87E-03 | -0.2 | 1.75E-01 | Branched-chain-amino-acid aminotransferase |
| AT5G58390 | 0.4 | 1.29E-03 | 0.1 | 2.34E-01 | 0.3 | 2.31E-02 | 0.4 | 1.38E-01 | 0.1 | 8.17E-01 | 0.3 | 3.39E-01 | Peroxidase 67 |
| AT5G59430 | 0.4 | 2.38E-03 | 0.2 | 4.48E-02 | 0.2 | 1.74E-01 | -0.1 | 5.57E-01 | -0.1 | 7.90E-01 | 0.0 | 8.90E-01 | Telomere repeat-binding protein 1 |
| AT1G07820 | 0.4 | 2.49E-03 | 0.5 | 5.92E-04 | -0.1 | 2.62E-01 | -0.1 | 7.82E-01 | -0.2 | 4.46E-01 | 0.1 | 7.11E-01 | Histone H4 |
| AT5G27470 | 0.4 | 6.50E-04 | 0.2 | 1.62E-02 | 0.2 | 1.13E-01 | 0.1 | 5.44E-01 | 0.0 | 9.93E-01 | 0.1 | 7.20E-01 | Serine--tRNA ligase, cytoplasmic synthetase) |
| AT2G04270 | 0.4 | 1.36E-02 | 0.5 | 3.85E-03 | -0.1 | 4.53E-01 | 0.0 | 9.57E-01 | -0.1 | 9.12E-01 | 0.0 | 9.55E-01 | Putative endoribonuclease E |
| AT1G49580 | 0.4 | 7.65E-04 | 0.3 | 1.40E-02 | 0.1 | 1.66E-01 | 0.1 | 5.25E-01 | 0.2 | 1.14E-01 | -0.1 | 4.46E-01 | CDPK-related kinase 8 |
| AT1G31340 | 0.4 | 1.71E-04 | 0.5 | 8.59E-05 | -0.1 | 3.35E-01 | 0.2 | 3.15E-01 | 0.2 | 4.37E-01 | 0.0 | 9.70E-01 | Ubiquitin-NEDD8-like protein RUB1 |
| AT5G66230 | 0.4 | 3.67E-04 | 0.6 | 4.11E-05 | -0.2 | 5.57E-02 | 0.2 | 3.03E-01 | 0.2 | 2.69E-01 | 0.0 | 9.07E-01 | At5g66230 |
| AT2G20050 | 0.4 | 6.27E-04 | 0.2 | 6.64E-02 | 0.2 | 3.18E-02 | -0.3 | 1.15E-01 | -0.2 | 3.20E-01 | -0.1 | 8.25E-01 | protein-serine |
| AT3G46610 | 0.4 | 1.33E-03 | 0.2 | 1.40E-01 | 0.2 | 4.37E-02 | 0.1 | 6.76E-01 | 0.1 | 8.06E-01 | 0.0 | 9.53E-01 | Protein LOW PHOTOSYNTHETIC EFFICIENCY 1, chloroplastic |
| AT3G28920 | 0.4 | 1.05E-04 | 0.4 | 3.34E-04 | 0.0 | 6.35E-01 | 0.1 | 3.36E-01 | 0.1 | 7.53E-01 | 0.1 | 7.62E-01 | Zinc-finger homeodomain protein 9 |
| AT3G26520 | 0.4 | 4.05E-03 | 0.9 | 3.51E-05 | -0.5 | 3.30E-03 | -1.0 | 1.15E-05 | -0.9 | 2.06E-04 | -0.1 | 6.07E-01 | Aquaporin TIP1-2 |
| AT2G44050 | 0.4 | 2.21E-04 | 0.4 | 2.02E-04 | 0.0 | 6.76E-01 | 0.1 | 4.77E-01 | 0.1 | 8.05E-01 | 0.1 | 8.36E-01 | 6,7-dimethyl-8-ribityllumazine synthase, chloroplastic |
| AT5G11170 | 0.4 | 6.59E-05 | 0.3 | 1.82E-03 | 0.1 | 6.31E-02 | 0.2 | 1.97E-01 | 0.0 | 8.73E-01 | 0.2 | 4.63E-01 | DEAD |
| AT5G46290 | 0.4 | 9.93E-04 | 0.5 | 6.07E-04 | -0.1 | 5.45E-01 | 0.3 | 4.48E-02 | 0.1 | 4.90E-01 | 0.2 | 4.35E-01 | beta-ketoacyl- synthase I |
| AT2G25100 | 0.4 | 5.39E-04 | 0.3 | 5.12E-03 | 0.1 | 2.95E-01 | -0.1 | 6.61E-01 | -0.2 | 4.78E-01 | 0.1 | 8.37E-01 | Ribonuclease |
| AT3G61150 | 0.4 | 5.49E-04 | 0.3 | 5.14E-03 | 0.1 | 2.82E-01 | -0.3 | 1.47E-02 | -0.2 | 5.78E-02 | 0.0 | 8.31E-01 | Homeobox-leucine zipper protein HDG1 |
| AT2G19640 | 0.4 | 3.00E-02 | 0.3 | 1.37E-01 | 0.1 | 5.17E-01 | 0.0 | 8.72E-01 | 0.0 | 9.48E-01 | 0.0 | 9.56E-01 | Histone-lysine N-methyltransferase ASHR2 |
| AT4G16265 | 0.4 | 2.39E-03 | 0.5 | 1.45E-03 | -0.1 | 6.68E-01 | 0.6 | 6.46E-02 | 0.2 | 5.66E-01 | 0.3 | 3.04E-01 | DNA-directed RNA polymerase subunit |
| AT4G23750 | 0.4 | 3.36E-03 | 0.7 | 1.59E-04 | -0.3 | 4.87E-02 | -0.1 | 5.99E-01 | -0.2 | 3.97E-01 | 0.1 | 8.13E-01 | Ethylene-responsive transcription factor CRF2 |
| AT5G27740 | 0.4 | 5.87E-04 | 0.3 | 8.86E-03 | 0.1 | 1.91E-01 | 0.1 | 4.68E-01 | 0.0 | 8.55E-01 | 0.1 | 4.83E-01 | Replication factor C subunit 3 |
| AT1G33110 | 0.4 | 2.06E-04 | -0.2 | 2.96E-02 | 0.6 | 9.92E-05 | 0.5 | 8.91E-04 | 0.2 | 1.26E-01 | 0.3 | 1.42E-01 | MATE efflux family protein |
| AT2G29540 | 0.4 | 2.94E-02 | 0.5 | 1.77E-02 | -0.1 | 7.74E-01 | 0.6 | 1.98E-02 | 0.3 | 2.72E-01 | 0.3 | 3.86E-01 | DNA-directed RNA polymerase 14 kDa subunit ) and III 14 kDa subunit) |
| AT1G17920 | 0.4 | 1.58E-02 | 0.4 | 1.60E-02 | 0.0 | 9.28E-01 | 0.2 | 6.01E-01 | 0.0 | 9.32E-01 | 0.2 | 6.25E-01 | Homeobox-leucine zipper protein HDG12 |
| AT3G55610 | 0.4 | 4.98E-04 | 0.3 | 3.31E-03 | 0.1 | 3.67E-01 | -0.1 | 4.79E-01 | 0.0 | 8.87E-01 | -0.2 | 4.65E-01 | Delta 1-pyrroline-5-carboxylate synthase 2 |
| AT4G13040 | 0.4 | 3.40E-03 | 0.1 | 4.62E-01 | 0.3 | 3.11E-02 | 0.2 | 1.88E-01 | -0.1 | 5.79E-01 | 0.4 | 1.00E-01 | Integrase-type DNA-binding superfamily protein |
| AT5G49940 | 0.4 | 5.70E-04 | 0.2 | 8.44E-02 | 0.2 | 2.23E-02 | 0.0 | 7.94E-01 | -0.1 | 7.20E-01 | 0.0 | 9.31E-01 | NifU-like protein 2, chloroplastic |
| AT5G14760 | 0.4 | 1.89E-03 | -0.2 | 1.45E-01 | 0.6 | 1.22E-03 | 1.1 | 7.62E-05 | 0.3 | 2.20E-01 | 0.7 | 7.52E-03 | L-aspartate oxidase, chloroplastic |
| AT5G51600 | 0.4 | 4.76E-04 | 0.5 | 1.54E-04 | -0.1 | 2.81E-01 | 0.1 | 6.82E-01 | 0.2 | 5.66E-01 | -0.1 | 8.89E-01 | 65-kDa microtubule-associated protein 3 |
| AT5G48070 | 0.4 | 4.16E-03 | 0.2 | 7.06E-02 | 0.2 | 2.14E-01 | 1.1 | 4.59E-04 | 0.8 | 1.01E-02 | 0.3 | 3.09E-01 | Xyloglucan endotransglucosylase |
| AT1G29880 | 0.4 | 2.13E-04 | 0.1 | 2.49E-01 | 0.3 | 3.18E-03 | 0.2 | 3.43E-01 | 0.1 | 8.05E-01 | 0.1 | 7.18E-01 | Glycine--tRNA ligase, mitochondrial 1 |
| AT2G43130 | 0.4 | 3.19E-04 | 0.4 | 1.03E-03 | 0.0 | 6.46E-01 | 0.1 | 6.05E-01 | -0.1 | 6.74E-01 | 0.2 | 4.40E-01 | Uncharacterized protein |
| AT5G20920 | 0.4 | 1.71E-03 | 0.4 | 4.78E-03 | 0.0 | 7.10E-01 | 0.1 | 7.58E-01 | -0.1 | 8.17E-01 | 0.1 | 6.40E-01 | Eukaryotic translation initiation factor 2 subunit beta |
| AT5G47220 | 0.4 | 2.30E-02 | -0.4 | 3.08E-02 | 0.8 | 1.70E-03 | 0.7 | 7.47E-03 | -0.1 | 8.71E-01 | 0.8 | 2.39E-02 | Ethylene-responsive transcription factor 2 |
| AT1G16800 | 0.4 | 9.64E-04 | 0.1 | 5.17E-01 | 0.3 | 9.69E-03 | 0.4 | 1.77E-01 | 0.4 | 2.57E-01 | 0.0 | 9.75E-01 | P-loop containing nucleoside triphosphate hydrolases superfamily protein |
| AT1G22500 | 0.4 | 2.51E-02 | 0.9 | 2.55E-04 | -0.5 | 1.89E-02 | -0.2 | 4.84E-01 | -0.5 | 1.38E-01 | 0.3 | 5.31E-01 | E3 ubiquitin-protein ligase ATL15 |
| AT5G06150 | 0.4 | 1.35E-03 | 0.7 | 8.12E-05 | -0.2 | 4.18E-02 | 0.1 | 7.40E-01 | 0.2 | 5.25E-01 | -0.1 | 8.02E-01 | Cyclin family protein |
| AT1G04020 | 0.4 | 2.05E-03 | 0.3 | 9.14E-03 | 0.1 | 5.53E-01 | 0.7 | 7.36E-02 | 0.5 | 2.31E-01 | 0.2 | 7.81E-01 | Breast cancer associated RING 1 |
| AT5G03430 | 0.4 | 9.86E-04 | 0.1 | 5.69E-01 | 0.4 | 6.38E-03 | 0.1 | 3.88E-01 | 0.1 | 7.05E-01 | 0.0 | 8.28E-01 | Uncharacterized protein At5g03430 |
| AT1G72900 | 0.4 | 4.38E-02 | -0.5 | 3.40E-02 | 0.9 | 3.14E-03 | 1.3 | 1.17E-04 | 0.2 | 5.34E-01 | 1.1 | 2.36E-03 | Similar to part of disease resistance protein domain-containing protein) |
| AT3G14400 | 0.4 | 1.51E-04 | 0.2 | 1.92E-02 | 0.2 | 1.92E-02 | 0.2 | 8.12E-02 | 0.1 | 6.58E-01 | 0.2 | 4.26E-01 | Ubiquitin carboxyl-terminal hydrolase 25 |
| AT4G26070 | 0.4 | 3.25E-02 | 0.2 | 2.84E-01 | 0.2 | 3.07E-01 | 0.3 | 1.52E-02 | 0.0 | 9.30E-01 | 0.3 | 7.68E-02 | MAP kinase |
| AT3G13150 | 0.4 | 5.10E-03 | 0.3 | 4.56E-02 | 0.1 | 3.59E-01 | 0.5 | 1.20E-01 | 0.4 | 3.03E-01 | 0.1 | 8.23E-01 | Pentatricopeptide repeat-containing protein At3g13150 |
| AT5G01920 | 0.4 | 2.47E-03 | 0.4 | 1.91E-03 | 0.0 | 7.52E-01 | -0.3 | 6.28E-02 | -0.1 | 5.74E-01 | -0.2 | 4.02E-01 | Serine |
| AT4G08620 | 0.4 | 1.08E-02 | 0.9 | 8.64E-05 | -0.5 | 6.53E-03 | -0.4 | 5.10E-02 | 0.1 | 6.88E-01 | -0.5 | 4.32E-02 | SULTR1 |
| AT4G19350 | 0.4 | 1.12E-02 | 0.1 | 5.72E-01 | 0.3 | 6.54E-02 | 0.1 | 7.18E-01 | -0.1 | 6.85E-01 | 0.2 | 5.22E-01 | At4g19350 |
| AT4G04695 | 0.4 | 1.98E-03 | 0.4 | 5.28E-03 | 0.0 | 7.30E-01 | 0.5 | 3.53E-03 | 0.4 | 6.19E-02 | 0.2 | 4.18E-01 | Calcium-dependent protein kinase 31 |
| AT1G74260 | 0.4 | 8.82E-05 | 0.2 | 9.12E-03 | 0.2 | 1.82E-02 | 0.3 | 3.20E-02 | 0.1 | 5.83E-01 | 0.2 | 3.31E-01 | Probable phosphoribosylformylglycinamidine synthase, chloroplastic |
| AT2G33740 | 0.4 | 1.00E-03 | 0.0 | 7.27E-01 | 0.4 | 6.05E-03 | -0.1 | 5.30E-01 | -0.2 | 4.41E-01 | 0.0 | 8.98E-01 | Protein CutA, chloroplastic |
| AT5G07240 | 0.4 | 1.68E-03 | 0.1 | 4.86E-01 | 0.3 | 1.35E-02 | 0.1 | 5.62E-01 | 0.1 | 5.00E-01 | 0.0 | 9.07E-01 | IQ-domain 24 |
| AT3G50950 | 0.4 | 5.37E-03 | -0.2 | 2.02E-01 | 0.6 | 1.91E-03 | 0.3 | 8.90E-02 | 0.0 | 8.89E-01 | 0.4 | 1.59E-01 | Disease resistance RPP13-like protein 4 |
| AT5G63420 | 0.4 | 1.02E-03 | 0.3 | 5.07E-03 | 0.1 | 4.93E-01 | 0.6 | 1.33E-02 | 0.3 | 2.78E-01 | 0.3 | 2.85E-01 | Ribonuclease J |
| AT2G20860 | 0.4 | 9.54E-05 | 0.2 | 2.53E-02 | 0.2 | 8.14E-03 | -0.1 | 3.02E-01 | 0.0 | 8.73E-01 | -0.1 | 5.89E-01 | Lipoyl synthase, mitochondrial |
| AT5G05670 | 0.4 | 2.11E-04 | 0.3 | 7.05E-03 | 0.2 | 8.43E-02 | -0.3 | 9.19E-02 | -0.2 | 2.04E-01 | 0.0 | 8.85E-01 | Signal recognition particle receptor subunit beta |
| AT5G18400 | 0.4 | 1.01E-03 | 0.1 | 3.32E-01 | 0.3 | 1.35E-02 | 0.5 | 2.60E-03 | 0.2 | 2.86E-01 | 0.3 | 1.18E-01 | Anamorsin homolog |
| AT4G36670 | 0.4 | 5.48E-02 | 0.2 | 4.75E-01 | 0.3 | 2.66E-01 | -0.8 | 6.40E-02 | -1.8 | 2.68E-03 | 1.0 | 1.07E-01 | Probable polyol transporter 6 |
| AT5G08690 | 0.4 | 9.88E-04 | 0.4 | 3.85E-03 | 0.1 | 5.76E-01 | 0.3 | 2.76E-02 | 0.2 | 1.84E-01 | 0.1 | 6.90E-01 | ATP synthase subunit beta-2, mitochondrial |
| AT1G47830 | 0.4 | 6.77E-03 | 0.2 | 2.36E-01 | 0.2 | 1.07E-01 | 0.2 | 4.29E-01 | 0.0 | 9.32E-01 | 0.1 | 6.58E-01 | AP-2 complex subunit sigma |
| AT3G10010 | 0.4 | 3.87E-03 | -0.2 | 2.02E-01 | 0.6 | 2.52E-03 | 0.1 | 8.72E-01 | -0.4 | 2.69E-01 | 0.4 | 2.88E-01 | DEMETER-like protein 2 |
| AT1G51610 | 0.4 | 6.08E-04 | 0.3 | 1.13E-02 | 0.1 | 1.50E-01 | 0.0 | 9.47E-01 | 0.1 | 5.83E-01 | -0.1 | 6.00E-01 | Metal tolerance protein C4 |

| | | | | | | | | | | | | |
|---|---|---|---|---|---|---|---|---|---|---|---|---|
| AT3G11710 | 0.4 | 3.31E-03 | 0.1 | 2.67E-01 | 0.3 | 4.30E-02 | 0.4 | 3.21E-03 | 0.2 | 2.69E-01 | 0.2 | 1.94E-01 | Lysine--tRNA ligase, cytoplasmic |
| AT3G23890 | 0.4 | 9.15E-04 | 0.5 | 4.91E-04 | -0.1 | 5.34E-01 | -0.1 | 4.56E-01 | 0.0 | 8.56E-01 | -0.1 | 7.58E-01 | DNA topoisomerase 2 |
| AT4G30860 | 0.4 | 5.37E-03 | 0.5 | 2.23E-03 | -0.1 | 5.67E-01 | -0.1 | 8.04E-01 | -0.1 | 8.01E-01 | 0.0 | 9.70E-01 | Histone-lysine N-methyltransferase ASHR3 |
| AT1G05310 | 0.4 | 3.25E-02 | 0.9 | 5.32E-04 | -0.5 | 3.68E-02 | 0.1 | 8.84E-01 | -0.1 | 8.08E-01 | 0.2 | 7.08E-01 | Probable pectinesterase 8 |
| AT1G31550 | 0.4 | 6.45E-04 | 0.1 | 2.02E-01 | 0.3 | 1.61E-02 | 0.5 | 1.89E-01 | 0.5 | 2.06E-01 | 0.0 | 9.62E-01 | GDSL esterase |
| AT1G75270 | 0.4 | 4.29E-04 | 0.2 | 1.82E-02 | 0.2 | 7.23E-02 | 0.8 | 7.30E-04 | 0.5 | 4.67E-02 | 0.3 | 2.08E-01 | Glutathione S-transferase DHAR2 |
| AT2G41630 | 0.4 | 1.19E-04 | 0.1 | 3.39E-01 | 0.3 | 1.52E-03 | 0.0 | 8.45E-01 | -0.2 | 2.62E-01 | 0.1 | 4.80E-01 | Transcription initiation factor IIB-1 |
| AT4G17460 | 0.4 | 7.78E-03 | 0.6 | 6.70E-04 | -0.2 | 1.26E-01 | -0.2 | 3.18E-01 | -0.2 | 3.74E-01 | 0.0 | 9.95E-01 | Homeobox-leucine zipper protein HAT1 |
| AT4G39120 | 0.4 | 1.12E-03 | 0.2 | 8.37E-02 | 0.2 | 5.98E-02 | 0.1 | 7.24E-01 | 0.2 | 4.69E-01 | -0.1 | 7.85E-01 | Bifunctional phosphatase IMPL2, chloroplastic |
| AT2G24500 | 0.4 | 4.73E-04 | 0.2 | 2.42E-02 | 0.2 | 5.69E-02 | 0.3 | 2.86E-02 | 0.0 | 9.76E-01 | 0.3 | 8.10E-02 | Cytoplasmic 60S subunit biogenesis factor REI1 homolog 2 |
| AT1G09000 | 0.4 | 8.88E-04 | 0.1 | 5.52E-01 | 0.4 | 8.04E-03 | -0.1 | 6.14E-01 | 0.1 | 7.59E-01 | -0.2 | 4.85E-01 | NPK1-related protein kinase 1 |
| AT4G29750 | 0.4 | 4.77E-03 | 0.3 | 5.86E-02 | 0.2 | 2.50E-01 | 0.3 | 5.18E-02 | 0.2 | 3.21E-01 | 0.1 | 5.40E-01 | CRM domain-containing protein |
| AT3G07630 | 0.4 | 2.79E-04 | 0.2 | 9.50E-03 | 0.2 | 8.30E-02 | 0.4 | 1.14E-02 | 0.3 | 1.07E-01 | 0.1 | 5.23E-01 | Arogenate dehydratase |
| AT5G19300 | 0.4 | 1.71E-02 | 0.2 | 1.55E-01 | 0.2 | 3.35E-01 | 1.0 | 7.60E-04 | 0.5 | 8.94E-02 | 0.5 | 1.16E-01 | At5g19300 |
| AT5G60540 | 0.4 | 4.05E-03 | 0.1 | 3.40E-01 | 0.3 | 4.70E-02 | 0.3 | 1.14E-02 | 0.0 | 8.47E-01 | 0.3 | 9.58E-02 | Probable pyridoxal 5'-phosphate synthase subunit PDX2 |
| AT1G08410 | 0.4 | 4.20E-03 | 0.2 | 1.80E-01 | 0.2 | 9.24E-02 | 0.4 | 1.87E-02 | 0.2 | 4.38E-01 | 0.2 | 2.44E-01 | GTPase LSG1-2 |
| AT1G67320 | 0.4 | 1.46E-03 | 0.5 | 1.11E-03 | 0.0 | 7.39E-01 | 0.1 | 7.62E-01 | 0.2 | 5.14E-01 | -0.1 | 7.58E-01 | DNA primase large subunit |
| AT3G54150 | 0.4 | 4.40E-02 | -0.2 | 3.01E-01 | 0.6 | 2.24E-02 | 1.2 | 2.87E-03 | -0.1 | 8.37E-01 | 1.3 | 5.20E-03 | Embryonic abundant protein-like |
| AT5G16390 | 0.4 | 3.81E-04 | 0.5 | 2.72E-04 | -0.1 | 5.62E-01 | 0.0 | 8.94E-01 | 0.1 | 7.33E-01 | 0.0 | 8.68E-01 | Biotin carboxyl carrier protein of acetyl-CoA carboxylase |
| AT1G52030 | 0.4 | 3.93E-02 | 0.6 | 6.92E-03 | -0.2 | 3.57E-01 | 0.5 | 6.37E-01 | 0.8 | 4.29E-01 | -0.3 | 8.07E-01 | Jacalin-type lectin domain-containing protein |
| AT3G60960 | 0.4 | 8.40E-03 | 0.2 | 1.51E-01 | 0.2 | 2.09E-01 | 0.0 | 9.46E-01 | -0.1 | 8.17E-01 | 0.0 | 8.87E-01 | Pentatricopeptide repeat-containing protein At3g60960, mitochondrial |
| AT1G09820 | 0.4 | 2.51E-03 | 0.2 | 5.93E-02 | 0.2 | 1.75E-01 | 0.0 | 9.23E-01 | -0.1 | 7.48E-01 | 0.2 | 7.19E-01 | Pentatricopeptide repeat-containing protein At1g09820 |
| AT2G19800 | 0.4 | 2.37E-02 | 0.9 | 2.43E-04 | -0.5 | 1.52E-02 | -0.5 | 1.95E-01 | -0.4 | 3.23E-01 | -0.1 | 9.31E-01 | Inositol oxygenase 2 |
| AT1G79150 | 0.4 | 8.92E-03 | 0.3 | 6.16E-02 | 0.1 | 3.98E-01 | 0.7 | 2.18E-04 | 0.3 | 1.66E-01 | 0.5 | 1.71E-02 | Nucleolar complex-associated protein 3 |
| AT3G45860 | 0.4 | 2.16E-01 | -0.6 | 1.01E-01 | 1.1 | 2.27E-02 | 0.3 | 5.88E-01 | -0.3 | 6.48E-01 | 0.7 | 3.64E-01 | Cysteine-rich receptor-like protein kinase 4 |
| AT2G35920 | 0.4 | 7.94E-04 | 0.1 | 2.92E-01 | 0.3 | 1.15E-02 | 0.0 | 9.73E-01 | -0.1 | 8.01E-01 | 0.1 | 8.07E-01 | RNA helicase |
| AT1G67520 | 0.4 | 3.03E-02 | -0.4 | 8.20E-02 | 0.8 | 5.78E-03 | 0.2 | 7.47E-01 | -0.5 | 5.68E-01 | 0.7 | 3.22E-01 | Receptor-like serine |
| AT4G21400 | 0.4 | 5.69E-04 | 0.0 | 7.54E-01 | 0.5 | 1.81E-03 | 0.5 | 5.31E-02 | -0.1 | 8.27E-01 | 0.5 | 4.89E-02 | Cysteine-rich RLK 28 |
| AT5G15350 | 0.4 | 7.18E-03 | 0.5 | 2.59E-03 | -0.1 | 4.66E-01 | -0.3 | 4.11E-02 | -0.2 | 2.06E-01 | -0.1 | 7.13E-01 | Early nodulin-like protein 17 |
| AT1G73875 | 0.4 | 6.63E-04 | 0.1 | 1.37E-01 | 0.3 | 2.40E-02 | -0.1 | 6.88E-01 | -0.1 | 8.56E-01 | 0.0 | 9.11E-01 | DNAse I-like superfamily protein |
| AT4G31990 | 0.4 | 6.70E-05 | 0.3 | 2.38E-03 | 0.2 | 4.90E-02 | 0.4 | 3.35E-04 | 0.3 | 2.26E-02 | 0.2 | 2.74E-01 | Aspartate aminotransferase |
| AT4G01720 | 0.4 | 2.35E-03 | -0.2 | 2.00E-01 | 0.6 | 1.40E-03 | -0.1 | 6.72E-01 | -0.6 | 7.66E-03 | 0.5 | 4.14E-02 | Probable WRKY transcription factor 47 |
| AT2G15690 | 0.4 | 1.88E-02 | 0.4 | 4.95E-02 | 0.1 | 7.30E-01 | 0.3 | 3.49E-01 | -0.2 | 6.03E-01 | 0.5 | 1.98E-01 | Pentatricopeptide repeat-containing protein At2g15690, mitochondrial |
| AT5G13780 | 0.4 | 8.23E-04 | 0.2 | 3.51E-02 | 0.2 | 7.16E-02 | 0.1 | 5.17E-01 | -0.1 | 3.36E-01 | 0.2 | 1.93E-01 | N-terminal acetyltransferase A complex catalytic subunit NAA10 |
| AT2G40860 | 0.4 | 1.88E-03 | 0.0 | 7.84E-01 | 0.5 | 5.09E-03 | -0.5 | 1.66E-01 | -0.2 | 6.66E-01 | -0.3 | 5.13E-01 | Protein kinase family protein |
| AT3G01660 | 0.4 | 2.40E-03 | 0.3 | 9.71E-03 | 0.1 | 5.86E-01 | 0.5 | 1.02E-01 | 0.5 | 1.06E-01 | 0.0 | 9.31E-01 | hypothetical protein |
| AT3G24090 | 0.4 | 1.25E-03 | 0.1 | 3.95E-01 | 0.3 | 1.71E-02 | 0.4 | 1.12E-01 | 0.6 | 2.25E-02 | -0.2 | 4.84E-01 | Glutamine--fructose-6-phosphate aminotransferase 1 |
| AT5G64620 | 0.4 | 6.77E-03 | 0.8 | 1.02E-04 | -0.4 | 1.40E-02 | -0.4 | 2.12E-02 | -0.2 | 2.60E-01 | -0.2 | 4.72E-01 | Cell wall |
| AT3G52340 | 0.4 | 2.61E-03 | 0.0 | 8.71E-01 | 0.4 | 5.28E-03 | 0.0 | 8.74E-01 | -0.1 | 5.90E-01 | 0.1 | 5.81E-01 | Sucrose-phosphatase |
| AT1G66150 | 0.4 | 3.45E-04 | 0.4 | 1.67E-03 | 0.1 | 4.69E-01 | -0.5 | 5.63E-03 | -0.4 | 2.80E-02 | -0.1 | 7.90E-01 | hypothetical protein |
| AT3G16700 | 0.4 | 5.26E-03 | 0.2 | 9.77E-02 | 0.2 | 2.13E-01 | 0.4 | 9.11E-02 | 0.4 | 1.01E-01 | 0.0 | 9.56E-01 | Fumarylacetoacetate hydrolase family |
| AT1G03160 | 0.4 | 1.40E-03 | 0.3 | 1.27E-02 | 0.1 | 2.96E-01 | 0.1 | 6.80E-01 | 0.3 | 1.48E-01 | -0.2 | 3.79E-01 | FZO-like protein |
| AT3G52100 | 0.4 | 4.92E-04 | 0.3 | 1.28E-02 | 0.2 | 1.22E-01 | -0.1 | 5.05E-01 | -0.1 | 4.81E-01 | 0.0 | 9.49E-01 | RING |
| AT5G19490 | 0.4 | 5.05E-02 | -0.6 | 3.54E-02 | 1.0 | 4.75E-03 | | | | | | | Histone superfamily protein |
| AT3G62370 | 0.4 | 8.92E-04 | 0.3 | 9.30E-03 | 0.1 | 2.70E-01 | 0.0 | 9.89E-01 | -0.2 | 2.89E-01 | 0.2 | 3.86E-01 | Heme binding protein |
| AT1G79440 | 0.4 | 9.12E-05 | 0.5 | 6.44E-05 | -0.1 | 4.61E-01 | -0.1 | 2.88E-01 | -0.2 | 2.39E-01 | 0.0 | 8.84E-01 | Succinate-semialdehyde dehydrogenase, mitochondrial -dependent succinic semialdehyde dehydrogenase) |
| AT5G64840 | 0.4 | 3.63E-03 | 0.0 | 9.95E-01 | 0.4 | 7.98E-03 | 0.1 | 5.76E-01 | 0.1 | 8.38E-01 | 0.1 | 8.71E-01 | ABC transporter F family member 5 |
| AT1G05010 | 0.4 | 4.79E-04 | 0.3 | 1.48E-02 | 0.2 | 9.02E-02 | 0.0 | 9.20E-01 | -0.1 | 7.10E-01 | 0.1 | 7.15E-01 | 1-aminocyclopropane-1-carboxylate oxidase 4 |
| AT5G44930 | 0.4 | 1.09E-02 | 0.0 | 9.53E-01 | 0.4 | 3.52E-02 | 0.2 | 7.35E-01 | 0.1 | 9.18E-01 | 0.1 | 8.94E-01 | Exostosin family protein |
| AT5G55230 | 0.4 | 4.21E-05 | 0.3 | 3.60E-04 | 0.1 | 2.28E-01 | -0.3 | 8.85E-02 | -0.3 | 5.35E-02 | 0.1 | 7.99E-01 | Microtubule-associated proteins 65-1 |
| AT5G60860 | 0.4 | 1.60E-03 | 0.3 | 7.63E-03 | 0.1 | 5.23E-01 | -0.8 | 1.50E-03 | -0.7 | 3.27E-03 | 0.0 | 9.48E-01 | Ras-related protein RABA1f |
| AT3G14390 | 0.4 | 2.61E-03 | 0.5 | 1.03E-03 | -0.1 | 4.73E-01 | 0.4 | 5.36E-03 | 0.1 | 5.02E-01 | 0.3 | 1.29E-01 | Diaminopimelate decarboxylase 1, chloroplastic |
| AT2G25000 | 0.4 | 7.55E-03 | -0.2 | 2.59E-01 | 0.6 | 4.70E-03 | 0.0 | 9.19E-01 | 0.4 | 1.97E-01 | -0.4 | 3.21E-01 | WRKY DNA-binding protein 60 |
| AT1G65930 | 0.4 | 8.77E-04 | 0.6 | 9.98E-05 | -0.2 | 6.44E-02 | -0.1 | 5.64E-01 | -0.3 | 1.70E-02 | 0.2 | 1.62E-01 | Cytosolic isocitrate dehydrogenase |
| AT5G62700 | 0.4 | 3.68E-04 | 0.3 | 5.81E-03 | 0.1 | 1.63E-01 | -0.1 | 5.10E-01 | -0.1 | 6.24E-01 | 0.0 | 9.80E-01 | Tubulin beta-3 chain |
| AT1G34130 | 0.4 | 8.62E-05 | 0.2 | 5.37E-03 | 0.2 | 2.92E-02 | 0.3 | 8.02E-02 | 0.1 | 5.13E-01 | 0.1 | 5.32E-01 | Dolichyl-diphosphooligosaccharide--protein glycosyltransferase subunit STT3B |
| AT5G11520 | 0.4 | 5.89E-05 | 0.4 | 3.08E-04 | 0.1 | 4.04E-01 | 0.3 | 1.33E-02 | 0.1 | 3.92E-01 | 0.2 | 2.86E-01 | Aspartate aminotransferase 3, chloroplastic |
| AT4G35000 | 0.4 | 2.60E-04 | 0.3 | 4.38E-03 | 0.1 | 1.45E-01 | 0.0 | 7.63E-01 | -0.2 | 1.90E-01 | 0.1 | 4.54E-01 | L-ascorbate peroxidase 3 |
| AT3G49660 | 0.4 | 2.86E-03 | 0.2 | 1.19E-01 | 0.2 | 1.08E-01 | 0.2 | 4.55E-01 | 0.1 | 8.59E-01 | 0.2 | 7.23E-01 | COMPASS-like H3K4 histone methylase component WDR5A |
| AT3G08870 | 0.4 | 9.54E-03 | -0.3 | 1.33E-01 | 0.7 | 3.45E-03 | 1.8 | 4.12E-04 | 1.2 | 1.80E-02 | 0.6 | 1.59E-01 | Concanavalin A-like lectin protein kinase family protein |
| AT1G65250 | 0.4 | 6.18E-03 | 0.0 | 9.45E-01 | 0.4 | 2.15E-02 | 0.1 | 8.06E-01 | 0.1 | 7.44E-01 | 0.0 | 9.27E-01 | Probable inactive receptor-like protein kinase At1g65250 |
| AT5G39850 | 0.4 | 7.44E-03 | -0.5 | 7.32E-03 | 0.9 | 4.65E-04 | 1.1 | 1.18E-04 | 0.4 | 1.93E-01 | 0.8 | 6.76E-03 | Small ribosomal subunit protein uS4y |
| AT5G06210 | 0.4 | 3.89E-03 | 0.4 | 1.20E-02 | 0.1 | 6.73E-01 | -0.1 | 5.06E-01 | -0.2 | 3.58E-01 | 0.1 | 8.39E-01 | Small RNA-binding protein 11, chloroplastic |
| AT3G27360 | 0.4 | 6.03E-03 | 0.6 | 5.83E-04 | -0.2 | 1.53E-01 | 0.1 | 7.63E-01 | -0.1 | 7.24E-01 | 0.3 | 5.66E-01 | Histone H3.1 |
| AT1G43860 | 0.4 | 3.99E-05 | 0.3 | 4.35E-04 | 0.1 | 1.68E-01 | 0.3 | 4.88E-02 | 0.1 | 6.82E-01 | 0.2 | 3.11E-01 | At1g43860 |
| AT2G05220 | 0.4 | 5.59E-03 | 0.6 | 1.62E-03 | -0.1 | 3.69E-01 | 0.3 | 7.97E-02 | 0.2 | 4.08E-01 | 0.1 | 6.50E-01 | Small ribosomal subunit protein eS17y |
| AT3G63170 | 0.4 | 2.40E-03 | 0.5 | 1.47E-03 | -0.1 | 6.60E-01 | 0.0 | 9.35E-01 | 0.2 | 1.01E-01 | -0.2 | 2.08E-01 | Fatty-acid-binding protein 1 |
| AT4G38225 | 0.4 | 1.56E-04 | 0.3 | 1.92E-03 | 0.1 | 1.76E-01 | -0.1 | 6.63E-01 | -0.1 | 8.27E-01 | 0.0 | 9.27E-01 | Glycerol kinase |
| AT1G19350 | 0.4 | 2.29E-03 | 0.0 | 9.81E-01 | 0.4 | 5.29E-03 | -0.6 | 4.10E-04 | -0.4 | 1.97E-02 | -0.2 | 3.19E-01 | Protein BZR1 homolog |
| AT2G04520 | 0.4 | 1.00E-03 | 0.4 | 4.37E-03 | 0.1 | 5.20E-01 | 0.4 | 1.22E-01 | 0.1 | 7.60E-01 | 0.3 | 4.55E-01 | Eukaryotic translation initiation factor 4C |
| AT1G09010 | 0.4 | 3.83E-04 | 0.4 | 1.73E-03 | 0.1 | 4.88E-01 | -0.3 | 1.89E-01 | -0.4 | 9.97E-02 | 0.1 | 7.43E-01 | Mannosylglycoprotein endo-beta-mannosidase |
| AT5G42100 | 0.4 | 5.08E-04 | 0.2 | 2.26E-02 | 0.2 | 6.39E-02 | -0.2 | 1.65E-01 | -0.3 | 6.21E-02 | 0.1 | 6.63E-01 | glucan endo-1,3-beta-D-glucosidase |
| AT3G54180 | 0.4 | 2.53E-03 | 0.4 | 2.72E-03 | 0.0 | 9.17E-01 | 0.2 | 3.10E-01 | 0.3 | 1.77E-01 | -0.1 | 7.47E-01 | Cyclin-dependent kinase B1-1 |
| AT2G27970 | 0.4 | 1.04E-03 | 0.5 | 1.08E-03 | 0.0 | 8.65E-01 | 0.4 | 1.15E-01 | 0.3 | 3.07E-01 | 0.1 | 8.13E-01 | Cyclin-dependent kinases regulatory subunit 2 |
| AT5G54260 | 0.4 | 1.15E-03 | 0.0 | 9.84E-01 | 0.4 | 5.19E-03 | 0.1 | 7.76E-01 | -0.1 | 6.23E-01 | 0.2 | 4.24E-01 | Double-strand break repair protein |
| AT2G22230 | 0.4 | 9.72E-04 | 0.7 | 7.02E-05 | -0.2 | 4.13E-02 | 0.1 | 4.11E-01 | 0.2 | 1.30E-01 | -0.1 | 5.94E-01 | 3-hydroxyacyl- dehydratase |
| AT3G17611 | 0.4 | 1.01E-03 | -0.1 | 3.90E-01 | 0.5 | 1.63E-03 | 0.2 | 3.24E-01 | 0.1 | 7.57E-01 | 0.1 | 6.96E-01 | Rhomboid-like protein 14, mitochondrial |
| AT2G07690 | 0.4 | 2.65E-04 | 0.2 | 2.58E-02 | 0.2 | 3.42E-02 | 0.0 | 9.35E-01 | 0.1 | 7.38E-01 | -0.1 | 8.37E-01 | DNA replication licensing factor MCM5 |
| AT2G24120 | 0.4 | 1.79E-03 | 0.3 | 2.71E-02 | 0.2 | 2.18E-01 | 0.7 | 4.07E-03 | 0.2 | 5.03E-01 | 0.5 | 6.42E-02 | DNA-directed RNA polymerase |
| AT1G51500 | 0.4 | 8.34E-03 | 0.1 | 3.92E-01 | 0.3 | 6.52E-02 | -0.5 | 8.78E-03 | -0.3 | 2.04E-01 | -0.2 | 3.84E-01 | ABC transporter G family member 12 |
| AT3G50100 | 0.4 | 8.82E-03 | 0.5 | 8.35E-03 | 0.0 | 8.96E-01 | 0.3 | 3.44E-01 | 0.3 | 3.46E-01 | 0.0 | 9.51E-01 | Small RNA degrading nuclease 1 |
| AT4G30960 | 0.4 | 8.84E-04 | 0.2 | 1.05E-01 | 0.3 | 2.90E-02 | 0.4 | 2.47E-04 | 0.3 | 1.89E-02 | 0.2 | 2.41E-01 | CBL-interacting serine |
| AT4G16970 | 0.4 | 4.72E-03 | 0.4 | 9.42E-03 | 0.0 | 8.56E-01 | 0.2 | 4.60E-01 | 0.0 | 9.96E-01 | 0.2 | 5.81E-01 | Kinase like protein |

| | | | | | | | | | | | | | |
|---|---|---|---|---|---|---|---|---|---|---|---|---|---|
| AT3G51780 | 0.4 | 7.47E-05 | 0.0 | 7.64E-01 | 0.4 | 5.58E-04 | 0.1 | 6.69E-01 | 0.0 | 9.70E-01 | 0.1 | 8.12E-01 | BAG family molecular chaperone regulator 4 |
| AT1G67070 | 0.4 | 9.46E-03 | 0.3 | 7.24E-02 | 0.1 | 4.11E-01 | 0.2 | 6.52E-01 | 0.0 | 9.59E-01 | 0.2 | 6.73E-01 | mannose-6-phosphate isomerase |
| AT4G14440 | 0.4 | 2.66E-03 | 0.3 | 3.20E-02 | 0.2 | 2.45E-01 | 0.2 | 2.83E-01 | 0.0 | 8.97E-01 | 0.1 | 5.49E-01 | Enoyl-CoA delta isomerase 3 ,Delta-enoyl CoA isomerase 3) |
| AT1G11330 | 0.4 | 2.60E-02 | 0.0 | 8.14E-01 | 0.4 | 7.00E-02 | 1.0 | 1.87E-06 | 0.4 | 8.71E-03 | 0.5 | 5.72E-03 | Receptor-like serine |
| AT5G53050 | 0.4 | 1.19E-02 | -0.1 | 7.49E-01 | 0.5 | 1.89E-02 | -0.4 | 5.12E-01 | -0.6 | 3.61E-01 | 0.2 | 8.27E-01 | Alpha |
| AT4G30310 | 0.4 | 7.72E-05 | 0.0 | 8.42E-01 | 0.5 | 2.84E-04 | -0.3 | 2.00E-01 | -0.1 | 7.21E-01 | -0.2 | 5.33E-01 | FGGY family of carbohydrate kinase |
| AT2G29720 | 0.4 | 2.68E-02 | -0.2 | 3.26E-01 | 0.6 | 1.30E-02 | -0.1 | 8.22E-01 | -0.3 | 2.74E-01 | 0.2 | 4.48E-01 | FAD |
| AT5G55830 | 0.4 | 1.03E-02 | 0.3 | 8.89E-02 | 0.2 | 3.83E-01 | 0.1 | 7.69E-01 | 0.2 | 5.18E-01 | -0.1 | 7.68E-01 | Probable L-type lectin-domain containing receptor kinase S.7 |
| AT1G07745 | 0.4 | 8.89E-03 | 0.2 | 2.36E-01 | 0.3 | 1.61E-01 | -0.1 | 7.82E-01 | -0.1 | 8.39E-01 | 0.0 | 9.86E-01 | DNA repair RAD51-like protein |
| AT2G16440 | 0.4 | 1.69E-03 | 0.4 | 7.37E-03 | 0.1 | 5.44E-01 | 0.3 | 9.97E-02 | 0.2 | 3.50E-01 | 0.1 | 7.29E-01 | DNA replication licensing factor MCM4 |
| AT4G26510 | 0.4 | 3.62E-05 | 0.3 | 6.82E-04 | 0.1 | 8.20E-02 | 0.1 | 7.95E-01 | 0.1 | 7.17E-01 | 0.0 | 9.24E-01 | Uridine kinase-like protein 4 |
| AT3G03060 | 0.4 | 1.08E-02 | 0.4 | 3.50E-02 | 0.1 | 6.54E-01 | 1.4 | 7.98E-06 | 0.6 | 1.94E-02 | 0.7 | 6.10E-03 | AAA+ ATPase domain-containing protein |
| AT3G19860 | 0.4 | 6.31E-04 | 0.2 | 4.45E-02 | 0.2 | 4.87E-02 | -0.4 | 2.41E-03 | -0.6 | 4.08E-04 | 0.2 | 3.58E-01 | hypothetical protein |
| AT4G03080 | 0.4 | 5.30E-04 | 0.1 | 4.57E-01 | 0.4 | 4.45E-03 | 0.2 | 4.72E-01 | 0.0 | 9.46E-01 | 0.2 | 6.93E-01 | Serine |
| AT1G18090 | 0.4 | 7.53E-03 | 0.3 | 6.56E-02 | 0.2 | 3.78E-01 | 0.2 | 7.09E-01 | -0.1 | 8.05E-01 | 0.3 | 5.89E-01 | Exonuclease 1 |
| AT5G41750 | 0.4 | 7.20E-02 | -0.6 | 5.42E-02 | 1.0 | 6.62E-03 | 0.5 | 4.27E-02 | -0.2 | 5.03E-01 | 0.8 | 2.52E-02 | Disease resistance protein family |
| AT4G17520 | 0.4 | 6.91E-03 | 0.4 | 1.05E-02 | 0.0 | 9.48E-01 | 0.4 | 1.67E-03 | 0.2 | 1.15E-01 | 0.2 | 2.57E-01 | RGG repeats nuclear RNA binding protein B |
| AT1G21230 | 0.4 | 3.90E-02 | 0.6 | 1.24E-02 | -0.1 | 5.57E-01 | 1.4 | 7.12E-03 | 0.9 | 8.38E-02 | 0.4 | 3.48E-01 | Wall-associated receptor kinase 5 |
| AT3G18080 | 0.4 | 1.66E-03 | 0.7 | 1.99E-04 | -0.2 | 9.16E-02 | -0.4 | 1.16E-02 | -0.1 | 7.13E-01 | -0.3 | 9.29E-02 | Beta-glucosidase 44 |
| AT5G25490 | 0.4 | 5.93E-03 | 0.3 | 4.99E-02 | 0.1 | 3.81E-01 | -0.1 | 7.64E-01 | 0.0 | 9.88E-01 | -0.1 | 7.96E-01 | RanBP2-type domain-containing protein |
| AT4G37870 | 0.4 | 1.16E-04 | 0.3 | 2.49E-03 | 0.1 | 9.74E-02 | 0.2 | 2.06E-01 | -0.1 | 4.46E-01 | 0.3 | 8.98E-02 | phosphoenolpyruvate carboxykinase |
| AT3G13350 | 0.4 | 1.58E-03 | 0.3 | 1.11E-02 | 0.1 | 3.95E-01 | 0.0 | 9.41E-01 | -0.1 | 5.88E-01 | 0.1 | 6.85E-01 | High mobility group B protein 10 |
| AT3G53920 | 0.4 | 3.63E-04 | 0.1 | 4.79E-01 | 0.4 | 3.24E-03 | -0.4 | 1.89E-02 | -0.3 | 1.05E-01 | -0.1 | 7.15E-01 | RNApolymerase sigma-subunit C |
| AT3G48810 | 0.4 | 3.91E-03 | 0.2 | 1.24E-01 | 0.2 | 1.45E-01 | 0.2 | 7.79E-01 | 0.3 | 5.79E-01 | -0.2 | 7.87E-01 | Pentatricopeptide repeat-containing protein At3g48810 |
| AT2G32480 | 0.4 | 1.18E-04 | 0.3 | 1.06E-03 | 0.1 | 2.35E-01 | 0.1 | 3.65E-01 | 0.2 | 8.25E-02 | -0.1 | 5.14E-01 | Chloroplastic membrane metallo proteases |
| AT4G19380 | 0.4 | 1.65E-02 | 0.8 | 6.89E-04 | -0.3 | 7.80E-02 | 0.1 | 7.65E-01 | -0.3 | 2.79E-01 | 0.3 | 2.28E-01 | Long-chain-alcohol oxidase |
| AT4G19150 | 0.4 | 2.11E-04 | 0.4 | 1.61E-03 | 0.1 | 3.23E-01 | -0.1 | 5.90E-01 | -0.2 | 3.43E-01 | 0.1 | 7.67E-01 | Ankyrin repeat family protein |
| AT3G13920 | 0.4 | 1.47E-03 | 0.3 | 1.07E-02 | 0.1 | 3.65E-01 | 0.1 | 3.86E-01 | -0.1 | 2.00E-01 | 0.2 | 1.10E-01 | Eukaryotic translation initiation factor 4A1 |
| AT5G22130 | 0.4 | 1.12E-03 | 0.3 | 3.55E-02 | 0.2 | 1.22E-01 | 0.5 | 1.74E-02 | 0.3 | 2.34E-01 | 0.2 | 4.12E-01 | GPI mannosyltransferase 1 |
| AT1G10920 | 0.4 | 7.84E-03 | -0.1 | 6.80E-01 | 0.5 | 1.31E-02 | 0.0 | 9.12E-01 | -0.2 | 5.33E-01 | 0.2 | 6.44E-01 | NB-ARC domain-containing disease resistance protein |
| AT3G55560 | 0.4 | 1.66E-03 | 0.3 | 3.25E-02 | 0.2 | 1.65E-01 | 0.3 | 3.89E-02 | -0.2 | 2.15E-01 | 0.5 | 1.67E-02 | AT-hook motif nuclear-localized protein 15 |
| AT3G28540 | 0.4 | 5.31E-02 | 0.0 | 8.36E-01 | 0.5 | 5.73E-02 | 0.0 | 9.92E-01 | -0.3 | 7.76E-01 | 0.3 | 8.03E-01 | AAA+ ATPase domain-containing protein |
| AT1G54630 | 0.4 | 3.69E-04 | 0.4 | 1.43E-03 | 0.1 | 5.64E-01 | 0.4 | 1.01E-01 | 0.2 | 4.06E-01 | 0.1 | 7.13E-01 | Acyl carrier protein 3 |
| AT2G17340 | 0.4 | 7.07E-04 | 0.1 | 2.33E-01 | 0.3 | 1.12E-02 | 0.0 | 8.87E-01 | 0.2 | 2.28E-01 | -0.2 | 4.03E-01 | Damage-control phosphatase At2g17340 |
| AT3G18580 | 0.4 | 2.77E-03 | 0.3 | 4.76E-02 | 0.2 | 2.14E-01 | 0.4 | 2.72E-02 | 0.1 | 6.51E-01 | 0.3 | 1.44E-01 | Nucleic acid-binding, OB-fold-like protein |
| AT2G45740 | 0.4 | 6.46E-04 | 0.4 | 3.79E-03 | 0.1 | 4.08E-01 | -0.1 | 3.24E-01 | -0.1 | 5.09E-01 | 0.0 | 9.26E-01 | At2g45740 |
| AT1G05690 | 0.4 | 4.15E-03 | -0.3 | 7.73E-02 | 0.7 | 1.55E-03 | 0.1 | 8.04E-01 | 0.1 | 8.97E-01 | 0.0 | 9.62E-01 | BTB |
| AT4G01370 | 0.4 | 3.03E-05 | 0.2 | 3.10E-02 | 0.3 | 1.89E-03 | 0.3 | 1.19E-02 | 0.1 | 4.36E-01 | 0.2 | 2.08E-01 | Mitogen-activated protein kinase 4 |
| AT1G74340 | 0.4 | 5.20E-04 | 0.4 | 1.65E-03 | 0.0 | 6.60E-01 | 0.5 | 1.70E-02 | 0.3 | 2.11E-01 | 0.2 | 4.59E-01 | Domain X domain-containing protein |
| AT5G19030 | 0.5 | 1.11E-03 | 0.2 | 1.49E-01 | 0.3 | 3.92E-02 | 0.0 | 9.00E-01 | 0.0 | 9.71E-01 | 0.0 | 9.62E-01 | RNA-binding family protein |
| AT3G53700 | 0.5 | 2.25E-03 | 0.5 | 1.47E-03 | -0.1 | 6.81E-01 | 0.2 | 2.29E-01 | 0.2 | 4.95E-01 | 0.1 | 8.17E-01 | Pentatricopeptide repeat-containing protein At3g53700, chloroplastic |
| AT5G11250 | 0.5 | 3.42E-03 | 0.0 | 7.61E-01 | 0.4 | 1.70E-02 | 0.5 | 6.68E-02 | 0.0 | 9.61E-01 | 0.5 | 1.98E-01 | ADP-ribosyl cyclase |
| AT1G58380 | 0.5 | 6.76E-03 | 0.4 | 1.19E-02 | 0.0 | 8.69E-01 | 0.4 | 2.18E-02 | 0.2 | 3.35E-01 | 0.2 | 3.23E-01 | Small ribosomal subunit protein uS5z |
| AT4G02230 | 0.5 | 1.22E-03 | 0.4 | 3.48E-03 | 0.0 | 7.07E-01 | 0.2 | 1.49E-01 | -0.1 | 6.54E-01 | 0.3 | 1.46E-01 | Large ribosomal subunit protein eL19y |
| AT3G23480 | 0.5 | 2.13E-02 | 0.6 | 7.54E-03 | -0.1 | 5.60E-01 | 0.0 | 9.99E-01 | -0.4 | 5.17E-01 | 0.4 | 5.72E-01 | Cyclopropane-fatty-acyl-phospholipid synthase |
| AT3G18600 | 0.5 | 2.95E-02 | 0.3 | 1.23E-01 | 0.1 | 5.28E-01 | 0.7 | 2.95E-04 | 0.3 | 6.13E-02 | 0.3 | 9.86E-02 | DEAD-box ATP-dependent RNA helicase 51 |
| AT4G07950 | 0.5 | 7.78E-03 | 0.4 | 2.01E-02 | 0.1 | 7.36E-01 | 0.1 | 7.49E-01 | -0.3 | 1.98E-01 | 0.4 | 1.01E-01 | DNA-directed RNA polymerase subunit |
| AT1G05600 | 0.5 | 9.57E-04 | 0.4 | 5.54E-03 | 0.1 | 4.59E-01 | -0.3 | 1.98E-01 | -0.4 | 1.47E-01 | 0.1 | 8.65E-01 | Pentatricopeptide repeat-containing protein At1g05600 |
| AT1G06760 | 0.5 | 8.85E-05 | 0.2 | 6.90E-03 | 0.2 | 2.38E-02 | -0.3 | 1.10E-02 | -0.3 | 2.57E-02 | 0.0 | 9.62E-01 | H15 domain-containing protein |
| AT3G17465 | 0.5 | 1.53E-03 | 0.4 | 2.47E-03 | 0.0 | 9.30E-01 | 0.2 | 6.21E-02 | -0.1 | 6.38E-01 | 0.3 | 6.10E-02 | Large ribosomal subunit protein uL3m |
| AT2G33210 | 0.5 | 5.61E-03 | 0.4 | 8.16E-03 | 0.0 | 9.52E-01 | 0.9 | 3.44E-06 | 0.5 | 2.79E-03 | 0.4 | 3.14E-02 | Heat shock protein 60-2 |
| AT5G13360 | 0.5 | 2.45E-03 | 0.3 | 5.63E-02 | 0.2 | 1.64E-01 | 0.0 | 8.71E-01 | 0.0 | 9.81E-01 | 0.1 | 8.64E-01 | Auxin reponsive-like protein |
| AT5G48910 | 0.5 | 1.56E-03 | 0.1 | 2.38E-01 | 0.3 | 2.92E-02 | -0.1 | 5.08E-01 | 0.3 | 6.70E-02 | -0.4 | 5.14E-02 | Pentatricopeptide repeat-containing protein At5g48910 |
| AT1G71710 | 0.5 | 4.58E-03 | 0.1 | 3.10E-01 | 0.3 | 5.85E-02 | -0.1 | 5.71E-01 | 0.1 | 6.04E-01 | -0.3 | 3.74E-01 | Inositol polyphosphate-related phosphatase domain-containing protein |
| AT1G55500 | 0.5 | 2.14E-04 | 0.0 | 7.85E-01 | 0.4 | 1.23E-03 | 0.4 | 3.77E-03 | 0.3 | 4.69E-02 | 0.1 | 5.68E-01 | YTH domain-containing family protein |
| AT3G13940 | 0.5 | 6.37E-03 | 0.1 | 6.83E-01 | 0.4 | 3.21E-02 | 0.9 | 4.37E-05 | 0.3 | 9.42E-02 | 0.5 | 1.04E-02 | DNA binding |
| AT3G61060 | 0.5 | 1.17E-02 | 1.0 | 9.14E-05 | -0.6 | 7.06E-03 | -1.1 | 3.40E-03 | -0.7 | 4.74E-02 | -0.4 | 4.31E-01 | Phloem protein 2-A13 |
| AT1G53030 | 0.5 | 9.88E-02 | -0.1 | 6.83E-01 | 0.6 | 9.77E-02 | 0.0 | 9.15E-01 | -0.1 | 7.83E-01 | 0.2 | 7.54E-01 | Cytochrome c oxidase copper chaperone 2 |
| AT3G12670 | 0.5 | 1.14E-03 | 0.2 | 1.35E-01 | 0.3 | 3.04E-02 | 0.2 | 1.11E-01 | 0.0 | 8.98E-01 | 0.2 | 2.64E-01 | CTP synthase |
| AT5G51880 | 0.5 | 2.48E-04 | 0.2 | 9.44E-02 | 0.3 | 1.02E-02 | -0.1 | 6.12E-01 | -0.2 | 4.25E-01 | 0.1 | 8.17E-01 | Fe2OG dioxygenase domain-containing protein |
| AT4G00955 | 0.5 | 7.30E-03 | 0.0 | 8.90E-01 | 0.4 | 2.53E-02 | 0.4 | 1.32E-01 | 0.3 | 3.05E-01 | 0.1 | 8.07E-01 | Uncharacterized protein At4g00955 |
| AT5G64670 | 0.5 | 1.17E-03 | 0.4 | 7.56E-03 | 0.1 | 3.85E-01 | 0.5 | 3.56E-04 | 0.4 | 8.21E-03 | 0.1 | 4.20E-01 | Ribosomal protein L18e |
| AT3G16780 | 0.5 | 1.82E-02 | 0.7 | 3.31E-03 | -0.2 | 2.97E-01 | 0.4 | 1.36E-02 | 0.4 | 1.36E-02 | 0.0 | 8.75E-01 | Large ribosomal subunit protein eL19z |
| AT1G14440 | 0.5 | 1.85E-03 | 0.2 | 7.21E-02 | 0.2 | 1.09E-01 | -0.1 | 4.12E-01 | -0.2 | 2.49E-01 | 0.1 | 7.93E-01 | Zinc-finger homeodomain protein 4 |
| AT2G46090 | 0.5 | 3.72E-04 | 0.3 | 1.14E-02 | 0.2 | 1.02E-01 | -0.1 | 4.79E-01 | 0.0 | 9.24E-01 | -0.1 | 7.15E-01 | LCBK2 |
| AT3G51800 | 0.5 | 5.18E-03 | 0.5 | 3.87E-03 | -0.1 | 7.49E-01 | 0.5 | 1.22E-04 | 0.3 | 1.96E-02 | 0.2 | 1.53E-01 | ERBB-3 BINDING PROTEIN 1 |
| AT2G31800 | 0.5 | 4.13E-04 | 0.4 | 1.36E-03 | 0.1 | 6.20E-01 | -0.1 | 6.01E-01 | 0.0 | 9.13E-01 | -0.1 | 8.19E-01 | Protein kinase domain-containing protein |
| AT5G20990 | 0.5 | 4.39E-05 | 0.3 | 6.55E-04 | 0.1 | 1.13E-01 | 0.3 | 1.71E-02 | -0.1 | 4.38E-01 | 0.5 | 1.65E-02 | molybdopterin molybdotransferase |
| AT5G65410 | 0.5 | 5.53E-04 | 0.4 | 3.04E-03 | 0.1 | 4.45E-01 | -0.1 | 4.51E-01 | -0.3 | 1.74E-01 | 0.1 | 6.20E-01 | Zinc-finger homeodomain protein 1 |
| AT3G01820 | 0.5 | 2.31E-03 | 0.2 | 6.69E-02 | 0.2 | 1.27E-01 | 0.4 | 5.73E-03 | -0.1 | 6.66E-01 | 0.5 | 9.20E-03 | Probable adenylate kinase 7, mitochondrial |
| AT1G66280 | 0.5 | 2.62E-04 | 0.8 | 1.26E-05 | -0.3 | 6.20E-03 | -0.2 | 1.58E-01 | 0.3 | 2.80E-02 | -0.5 | 4.77E-03 | BGLU22 |
| AT1G58602 | 0.5 | 7.11E-04 | 0.3 | 2.30E-02 | 0.2 | 9.20E-02 | 0.3 | 3.28E-02 | 0.1 | 5.18E-01 | 0.2 | 3.01E-01 | Protein RECOGNITION OF PERONOSPORA PARASITICA 7 |
| AT3G54920 | 0.5 | 9.78E-04 | 0.6 | 3.28E-04 | -0.1 | 3.21E-01 | -0.3 | 1.28E-02 | -0.4 | 4.78E-03 | 0.1 | 6.02E-01 | Probable pectate lyase 13 |
| AT4G09800 | 0.5 | 1.36E-03 | 0.5 | 7.82E-04 | -0.1 | 5.41E-01 | 0.0 | 9.44E-01 | -0.2 | 3.89E-01 | 0.2 | 4.52E-01 | Small ribosomal subunit protein uS13z |
| AT5G28300 | 0.5 | 1.11E-03 | 0.2 | 6.11E-02 | 0.2 | 6.12E-02 | 0.1 | 4.93E-01 | 0.0 | 8.88E-01 | 0.1 | 5.22E-01 | Trihelix transcription factor GTL2 |
| AT5G04900 | 0.5 | 5.69E-04 | 0.3 | 1.21E-02 | 0.2 | 1.50E-01 | 0.0 | 9.30E-01 | 0.2 | 3.66E-01 | -0.3 | 3.90E-01 | Chlorophyll b reductase NOL, chloroplastic |
| AT4G39830 | 0.5 | 3.47E-02 | -0.2 | 4.56E-01 | 0.6 | 2.61E-02 | -0.3 | 2.78E-01 | -0.2 | 3.67E-01 | 0.0 | 9.72E-01 | L-ascorbate oxidase |
| AT4G16890 | 0.5 | 1.40E-03 | 0.0 | 7.59E-01 | 0.4 | 5.62E-03 | 0.1 | 5.20E-01 | -0.2 | 2.34E-01 | 0.3 | 1.08E-01 | ADP-ribosyl cyclase |
| AT4G36280 | 0.5 | 1.31E-02 | 0.3 | 8.64E-02 | 0.1 | 4.56E-01 | 0.1 | 8.43E-01 | 0.0 | 9.76E-01 | 0.0 | 9.12E-01 | Protein MICRORCHIDIA 2 |
| AT5G61560 | 0.5 | 1.50E-03 | 0.1 | 6.06E-01 | 0.4 | 1.22E-02 | -0.4 | 1.62E-01 | -0.6 | 5.63E-02 | 0.2 | 6.32E-01 | U-box domain-containing protein kinase family protein |
| AT5G02010 | 0.5 | 1.63E-03 | 0.4 | 2.83E-03 | 0.0 | 9.15E-01 | -0.4 | 7.59E-02 | -0.2 | 3.58E-01 | -0.2 | 5.90E-01 | Rop guanine nucleotide exchange factor 7 |
| AT4G29000 | 0.5 | 1.42E-03 | 0.2 | 1.83E-01 | 0.3 | 3.78E-02 | -0.1 | 8.16E-01 | -0.2 | 4.06E-01 | 0.1 | 6.51E-01 | Protein tesmin |
| AT4G25110 | 0.5 | 8.64E-03 | 0.3 | 9.76E-02 | 0.2 | 2.81E-01 | 0.0 | 9.90E-01 | -0.3 | 3.11E-01 | 0.3 | 4.20E-01 | Metacaspase-2 |

| | | | | | | | | | | | | | |
|---|---|---|---|---|---|---|---|---|---|---|---|---|---|
| AT2G34060 | 0.5 | 1.49E-03 | 0.5 | 8.75E-04 | -0.1 | 6.17E-01 | -0.1 | 8.18E-01 | 0.2 | 3.46E-01 | -0.3 | 2.94E-01 | Peroxidase 19 |
| AT2G15980 | 0.5 | 3.10E-04 | 0.1 | 2.60E-01 | 0.4 | 7.26E-03 | 0.0 | 9.33E-01 | 0.0 | 9.48E-01 | 0.0 | 9.98E-01 | Pentatricopeptide repeat-containing protein At2g15980 |
| AT2G45850 | 0.5 | 2.36E-03 | 0.4 | 6.71E-03 | 0.1 | 7.05E-01 | -0.6 | 8.03E-03 | -0.4 | 8.84E-02 | -0.2 | 3.59E-01 | AT-hook motif nuclear-localized protein |
| AT1G07370 | 0.5 | 1.19E-02 | 0.5 | 5.87E-03 | -0.1 | 6.55E-01 | 0.2 | 3.38E-01 | 0.3 | 2.73E-01 | -0.1 | 8.72E-01 | Proliferating cellular nuclear antigen 1 |
| AT3G57290 | 0.5 | 1.89E-04 | 0.4 | 1.03E-03 | 0.1 | 4.19E-01 | 0.2 | 5.42E-02 | 0.0 | 9.92E-01 | 0.2 | 1.78E-01 | Eukaryotic translation initiation factor 3 subunit E |
| AT2G01670 | 0.5 | 1.16E-03 | 0.4 | 3.73E-03 | 0.1 | 6.32E-01 | 0.2 | 3.79E-01 | 0.0 | 9.21E-01 | 0.2 | 4.77E-01 | Nudix hydrolase 17, mitochondrial |
| AT2G03050 | 0.5 | 2.74E-03 | 0.5 | 3.13E-03 | 0.0 | 9.24E-01 | 0.2 | 3.54E-01 | 0.2 | 3.79E-01 | 0.0 | 9.75E-01 | Transcription termination factor MTEF1, chloroplastic |
| AT4G26110 | 0.5 | 2.37E-04 | 0.4 | 1.41E-03 | 0.1 | 3.83E-01 | 0.4 | 3.43E-03 | 0.1 | 3.74E-01 | 0.3 | 1.41E-01 | Nucleosome assembly protein11 |
| AT1G48650 | 0.5 | 1.68E-04 | 0.1 | 4.71E-01 | 0.4 | 1.61E-03 | 0.3 | 9.63E-02 | 0.2 | 2.48E-01 | 0.1 | 8.41E-01 | RNA helicase |
| AT1G57610 | 0.5 | 3.43E-04 | 0.5 | 5.16E-04 | 0.0 | 9.79E-01 | 0.1 | 5.06E-01 | 0.0 | 9.84E-01 | 0.1 | 6.73E-01 | Calcium uniporter protein |
| AT2G47990 | 0.5 | 1.50E-02 | 0.3 | 6.43E-02 | 0.1 | 5.54E-01 | 0.4 | 4.51E-02 | 0.2 | 5.28E-01 | 0.2 | 3.84E-01 | Protein SLOW WALKER 1 |
| AT5G58770 | 0.5 | 3.78E-02 | -0.4 | 1.11E-01 | 0.8 | 6.33E-03 | 0.1 | 7.19E-01 | 0.3 | 4.30E-01 | -0.1 | 7.22E-01 | Dehydrodolichyl diphosphate synthase 2 |
| AT3G01790 | 0.5 | 2.01E-03 | 0.5 | 2.51E-03 | 0.0 | 9.75E-01 | 0.0 | 8.62E-01 | -0.1 | 6.60E-01 | 0.1 | 6.14E-01 | At1g01640 |
| AT1G28670 | 0.5 | 3.47E-04 | 0.4 | 1.75E-03 | 0.1 | 4.48E-01 | 0.1 | 5.36E-01 | 0.0 | 9.65E-01 | 0.1 | 7.35E-01 | GDSL esterase |
| AT3G51840 | 0.5 | 2.79E-04 | 0.5 | 5.38E-04 | 0.0 | 9.26E-01 | -0.4 | 8.96E-04 | -0.5 | 7.50E-04 | 0.1 | 7.32E-01 | Short-chain acyl CoA oxidase |
| AT4G09620 | 0.5 | 3.68E-03 | 0.0 | 7.45E-01 | 0.4 | 1.80E-02 | -0.1 | 6.58E-01 | -0.3 | 1.51E-01 | 0.2 | 4.52E-01 | At4g09620 |
| AT4G20320 | 0.5 | 2.01E-04 | 0.4 | 5.26E-04 | 0.0 | 7.44E-01 | 0.0 | 9.74E-01 | 0.1 | 4.73E-01 | -0.1 | 5.73E-01 | CTP synthase |
| AT3G61550 | 0.5 | 3.17E-04 | 0.1 | 2.03E-01 | 0.3 | 7.22E-03 | -0.2 | 4.22E-01 | -0.1 | 6.19E-01 | 0.0 | 9.03E-01 | RING-H2 finger protein ATL68 |
| AT2G18330 | 0.5 | 4.96E-03 | 0.2 | 1.40E-01 | 0.2 | 1.35E-01 | 0.7 | 2.48E-03 | 0.4 | 6.24E-02 | 0.2 | 3.37E-01 | AAA+ ATPase domain-containing protein |
| AT1G11280 | 0.5 | 1.68E-04 | 0.2 | 8.29E-02 | 0.3 | 7.62E-03 | 0.4 | 2.76E-02 | 0.3 | 1.32E-01 | 0.1 | 7.06E-01 | Receptor-like serine |
| AT1G50250 | 0.5 | 1.15E-03 | 0.0 | 8.95E-01 | 0.5 | 3.75E-03 | 0.0 | 9.18E-01 | 0.1 | 8.76E-01 | -0.1 | 8.35E-01 | ATP-dependent zinc metalloprotease FTSH 1, chloroplastic |
| AT5G05540 | 0.5 | 8.02E-04 | 0.3 | 2.34E-02 | 0.2 | 1.21E-01 | 0.0 | 9.45E-01 | -0.1 | 5.54E-01 | 0.1 | 6.58E-01 | Exonuclease domain-containing protein |
| AT4G00020 | 0.5 | 1.18E-03 | 0.4 | 6.02E-03 | 0.1 | 5.02E-01 | 0.2 | 5.22E-01 | 0.2 | 7.15E-01 | 0.1 | 9.09E-01 | BREAST CANCER 2 like 2A |
| AT5G66450 | 0.5 | 9.03E-04 | 0.2 | 1.48E-01 | 0.3 | 3.01E-02 | -0.3 | 1.24E-01 | -0.1 | 4.90E-01 | -0.1 | 5.88E-01 | Phosphatidic acid phosphatase family protein |
| AT3G22880 | 0.5 | 8.87E-03 | 0.5 | 1.04E-02 | 0.0 | 9.68E-01 | 0.2 | 6.52E-01 | 0.2 | 6.73E-01 | 0.0 | 9.80E-01 | Uncharacterized protein |
| AT3G06510 | 0.5 | 1.29E-04 | 0.0 | 9.80E-01 | 0.5 | 5.24E-04 | 0.1 | 3.29E-01 | 0.2 | 2.10E-01 | -0.1 | 8.08E-01 | Galactolipid galactosyltransferase SFR2, chloroplastic |
| AT2G19690 | 0.5 | 5.66E-02 | 0.2 | 4.47E-01 | 0.3 | 3.25E-01 | 0.4 | 1.61E-01 | 0.1 | 6.98E-01 | 0.2 | 5.15E-01 | Phospholipase A2-beta |
| AT1G71980 | 0.5 | 2.82E-03 | 0.4 | 7.46E-03 | 0.1 | 7.32E-01 | 0.1 | 5.09E-01 | 0.0 | 9.95E-01 | 0.1 | 6.62E-01 | hypothetical protein |
| AT1G31800 | 0.5 | 5.66E-04 | 0.3 | 6.68E-03 | 0.1 | 2.17E-01 | 0.2 | 1.01E-01 | 0.3 | 3.06E-02 | -0.1 | 5.47E-01 | Uncharacterized protein At1g31800 |
| AT1G09940 | 0.5 | 6.96E-05 | 0.3 | 5.10E-03 | 0.2 | 2.72E-02 | -0.4 | 3.81E-03 | -0.2 | 9.29E-02 | -0.2 | 3.45E-01 | Glutamyl-tRNA reductase 2, chloroplastic |
| AT4G37670 | 0.5 | 3.56E-04 | 0.1 | 4.21E-01 | 0.4 | 5.39E-03 | 0.6 | 1.24E-01 | 0.3 | 5.91E-01 | 0.3 | 5.39E-01 | N-acetyltransferase domain-containing protein |
| AT1G18370 | 0.5 | 1.04E-03 | 0.6 | 4.25E-04 | -0.1 | 4.27E-01 | 0.3 | 1.01E-01 | 0.4 | 9.27E-02 | 0.0 | 9.05E-01 | Kinesin-like protein KIN-7A |
| AT3G59550 | 0.5 | 2.01E-02 | 0.3 | 8.66E-02 | 0.1 | 5.72E-01 | 0.6 | 7.46E-02 | 0.0 | 9.25E-01 | 0.5 | 2.03E-01 | Sister chromatid cohesion 1 protein 3 |
| AT1G77670 | 0.5 | 2.38E-04 | 0.3 | 3.64E-03 | 0.1 | 1.66E-01 | 0.5 | 1.74E-03 | 0.2 | 3.54E-01 | 0.3 | 9.19E-02 | At1g77670 -dependent transferases superfamily protein) |
| AT5G61420 | 0.5 | 2.85E-04 | 0.5 | 5.63E-04 | 0.0 | 8.97E-01 | -0.2 | 1.97E-01 | 0.3 | 1.45E-01 | -0.5 | 1.49E-02 | Transcription factor MYB28 |
| AT3G04680 | 0.5 | 3.01E-04 | 0.3 | 1.05E-02 | 0.2 | 8.49E-02 | 0.2 | 2.83E-01 | 0.1 | 7.26E-01 | 0.1 | 6.32E-01 | Protein CLP1 homolog |
| AT1G05900 | 0.5 | 3.31E-04 | 0.1 | 4.74E-01 | 0.4 | 3.88E-03 | -0.4 | 6.70E-02 | 0.0 | 8.76E-01 | -0.3 | 1.50E-01 | Endonuclease III homolog 2, chloroplastic lyase 2) |
| AT1G51570 | 0.5 | 2.44E-04 | 0.3 | 1.11E-02 | 0.2 | 5.36E-02 | -0.1 | 5.23E-01 | -0.3 | 1.13E-01 | 0.2 | 4.39E-01 | FT-interacting protein 4 |
| AT5G22650 | 0.5 | 2.41E-03 | 0.4 | 4.96E-03 | 0.0 | 8.47E-01 | 0.7 | 5.50E-06 | 0.4 | 5.98E-03 | 0.3 | 3.42E-02 | Histone deacetylase HDT2 |
| AT5G65740 | 0.5 | 6.25E-03 | 0.4 | 1.51E-02 | 0.0 | 7.85E-01 | -0.2 | 4.51E-01 | 0.2 | 3.99E-01 | -0.4 | 1.53E-01 | AT5G65740 protein |
| AT2G30950 | 0.5 | 2.19E-04 | 0.3 | 1.08E-02 | 0.2 | 4.92E-02 | -0.1 | 8.06E-01 | 0.0 | 9.82E-01 | -0.1 | 8.53E-01 | FtsH extracellular protease family |
| AT1G71750 | 0.5 | 1.27E-03 | 0.2 | 1.95E-01 | 0.3 | 3.37E-02 | 0.6 | 2.16E-02 | 0.4 | 2.18E-01 | 0.2 | 4.71E-01 | Hypoxanthine phosphoribosyltransferase |
| AT3G02060 | 0.5 | 4.18E-04 | 0.1 | 3.01E-01 | 0.4 | 6.73E-03 | 0.4 | 1.93E-02 | -0.1 | 8.33E-01 | 0.5 | 3.87E-02 | ATP-dependent DNA helicase At3g02060, chloroplastic |
| AT5G61020 | 0.5 | 2.67E-04 | 0.4 | 2.17E-03 | 0.1 | 2.90E-01 | 0.5 | 8.07E-04 | 0.1 | 5.11E-01 | 0.4 | 3.32E-02 | YTH domain-containing family protein |
| AT1G47210 | 0.5 | 1.14E-03 | 0.6 | 6.07E-04 | -0.1 | 5.46E-01 | 0.3 | 1.65E-01 | 0.3 | 2.55E-01 | 0.0 | 9.67E-01 | Cyclin-dependent protein kinase 32 |
| AT1G04170 | 0.5 | 6.29E-04 | 0.5 | 1.21E-03 | 0.0 | 9.01E-01 | 0.3 | 1.35E-02 | 0.2 | 2.84E-01 | 0.2 | 3.71E-01 | protein-synthesizing GTPase |
| AT3G10920 | 0.5 | 4.97E-05 | 0.4 | 2.67E-04 | 0.1 | 4.05E-01 | 0.3 | 1.63E-02 | 0.3 | 5.54E-02 | 0.0 | 8.80E-01 | Superoxide dismutase |
| AT1G60660 | 0.5 | 8.97E-04 | 0.6 | 3.22E-04 | -0.1 | 3.74E-01 | 0.4 | 7.58E-02 | 0.0 | 9.18E-01 | 0.4 | 1.19E-01 | Cytochrome B5-like protein |
| AT1G63360 | 0.5 | 3.21E-04 | 0.3 | 8.13E-03 | 0.2 | 1.04E-01 | 0.1 | 7.50E-01 | -0.3 | 1.16E-01 | 0.3 | 1.00E-01 | Disease resistance protein family |
| AT3G58180 | 0.5 | 2.19E-04 | 0.2 | 8.93E-02 | 0.3 | 1.06E-02 | 0.2 | 2.92E-01 | -0.1 | 5.72E-01 | 0.3 | 1.81E-01 | Deoxyhypusine hydroxylase |
| AT1G06730 | 0.5 | 1.06E-04 | 0.5 | 1.98E-04 | 0.0 | 9.65E-01 | 0.0 | 9.64E-01 | 0.0 | 9.81E-01 | 0.0 | 9.65E-01 | At1g06730 |
| AT3G17770 | 0.5 | 3.05E-04 | 0.3 | 9.77E-03 | 0.2 | 8.01E-02 | 1.0 | 1.92E-05 | 0.2 | 3.50E-01 | 0.8 | 2.31E-03 | 3,4-dihydroxy-2-butanone kinase |
| AT3G60880 | 0.5 | 1.11E-03 | 0.2 | 4.95E-02 | 0.2 | 8.48E-02 | 0.2 | 1.63E-01 | 0.1 | 6.80E-01 | 0.1 | 5.66E-01 | 4-hydroxy-tetrahydrodipicolinate synthase 1, chloroplastic |
| AT3G07540 | 0.5 | 3.84E-03 | 0.3 | 3.44E-02 | 0.2 | 3.41E-01 | 0.0 | 8.62E-01 | -0.1 | 4.65E-01 | 0.1 | 6.83E-01 | Formin-like protein 10 |
| AT4G09730 | 0.5 | 1.89E-04 | 0.5 | 2.42E-04 | 0.0 | 8.88E-01 | 0.7 | 4.91E-04 | 0.1 | 6.00E-01 | 0.5 | 1.67E-02 | DEAD-box ATP-dependent RNA helicase 39 |
| AT4G22350 | 0.5 | 4.96E-04 | 0.3 | 7.64E-03 | 0.2 | 2.01E-01 | 0.2 | 4.75E-01 | 0.0 | 9.17E-01 | 0.1 | 6.98E-01 | AT4g22410 |
| AT1G15250 | 0.5 | 1.98E-03 | 0.4 | 1.46E-02 | 0.1 | 3.84E-01 | 0.4 | 1.20E-01 | -0.2 | 6.11E-01 | 0.6 | 9.34E-02 | Large ribosomal subunit protein eL37z |
| AT2G38960 | 0.5 | 1.30E-04 | 0.2 | 1.54E-02 | 0.2 | 2.32E-02 | 0.2 | 1.86E-01 | -0.1 | 7.05E-01 | 0.3 | 1.80E-01 | Endoplasmic reticulum oxidoreductins 2 |
| AT4G35950 | 0.5 | 1.23E-04 | 0.3 | 2.32E-03 | 0.1 | 1.27E-01 | -0.3 | 2.75E-01 | -0.1 | 6.81E-01 | -0.1 | 6.68E-01 | Rac-like GTP-binding protein ARAC6 |
| AT3G49120 | 0.5 | 3.63E-03 | 0.2 | 1.33E-01 | 0.3 | 9.32E-02 | 0.1 | 8.10E-01 | 0.2 | 4.36E-01 | -0.2 | 6.64E-01 | Peroxidase 34 |
| AT2G29550 | 0.5 | 7.02E-04 | 0.4 | 6.11E-03 | 0.1 | 3.00E-01 | -0.1 | 8.19E-01 | 0.1 | 7.59E-01 | -0.2 | 6.66E-01 | Tubulin beta-7 chain |
| AT5G06550 | 0.5 | 5.52E-03 | 0.3 | 8.38E-02 | 0.2 | 2.35E-01 | 0.6 | 1.15E-02 | 0.2 | 4.68E-01 | 0.4 | 1.05E-01 | Arginine-specific demethylase JMJ22 monodemethylase JMJ22) |
| AT5G50340 | 0.5 | 7.17E-05 | -0.1 | 2.32E-01 | 0.6 | 1.69E-04 | 0.2 | 1.85E-01 | 0.1 | 7.81E-01 | 0.2 | 4.80E-01 | DNA repair protein RadA-like protein |
| AT5G08540 | 0.5 | 5.95E-05 | 0.5 | 1.65E-04 | 0.0 | 7.25E-01 | -0.1 | 5.86E-01 | -0.1 | 6.67E-01 | 0.0 | 9.95E-01 | Ribosomal RNA small subunit methyltransferase J |
| AT1G50460 | 0.5 | 8.77E-04 | 0.5 | 6.14E-04 | -0.1 | 6.61E-01 | -0.3 | 1.42E-01 | -0.2 | 3.55E-01 | -0.1 | 8.25E-01 | Phosphotransferase |
| AT1G66880 | 0.5 | 2.99E-02 | -0.3 | 1.35E-01 | 0.8 | 4.82E-03 | 0.6 | 6.85E-03 | 0.0 | 9.44E-01 | 0.6 | 3.31E-02 | Protein kinase superfamily protein |
| AT4G25150 | 0.5 | 1.20E-02 | 0.7 | 1.67E-03 | -0.2 | 2.52E-01 | -0.2 | 6.32E-01 | 0.3 | 3.57E-01 | -0.4 | 2.07E-01 | Acid phosphatase-like protein |
| AT3G26300 | 0.5 | 5.15E-04 | 0.2 | 4.50E-02 | 0.3 | 3.84E-02 | 0.0 | 8.81E-01 | -0.1 | 6.66E-01 | 0.1 | 8.17E-01 | Cytochrome P450 71B34 |
| AT3G25600 | 0.5 | 2.65E-03 | 0.1 | 3.20E-01 | 0.3 | 3.59E-02 | 0.5 | 5.16E-02 | 0.1 | 7.63E-01 | 0.4 | 2.58E-01 | Probable calcium-binding protein CML16 |
| AT4G23320 | 0.5 | 5.61E-02 | -0.5 | 1.16E-01 | 1.0 | 1.29E-02 |  |  |  |  |  |  | Cysteine-rich RLK 24 |
| AT1G15570 | 0.5 | 4.64E-04 | 0.7 | 8.91E-05 | -0.2 | 1.45E-01 | -0.2 | 2.82E-01 | 0.0 | 9.84E-01 | -0.2 | 3.77E-01 | Cyclin-A2-3 |
| AT4G30140 | 0.5 | 1.54E-03 | 0.6 | 7.08E-04 | -0.1 | 4.90E-01 | 0.2 | 2.85E-01 | 0.1 | 7.48E-01 | 0.1 | 6.51E-01 | GDSL esterase |
| AT1G66920 | 0.5 | 1.21E-02 | -0.1 | 5.55E-01 | 0.6 | 1.26E-02 | -0.3 | 4.42E-01 | -0.4 | 3.03E-01 | 0.1 | 8.13E-01 | Protein kinase superfamily protein |
| AT1G49980 | 0.5 | 4.87E-03 | 0.0 | 9.58E-01 | 0.5 | 1.64E-02 | 0.0 | 8.60E-01 | 0.1 | 6.61E-01 | -0.1 | 8.21E-01 | DNA polymerase kappa |
| AT1G26160 | 0.5 | 2.15E-05 | 0.3 | 6.61E-04 | 0.2 | 4.55E-02 | -0.1 | 3.06E-01 | -0.2 | 2.06E-01 | 0.1 | 8.28E-01 | 5'-deoxynucleotidase |
| AT4G22745 | 0.5 | 9.97E-04 | 0.1 | 3.53E-01 | 0.4 | 1.44E-02 | 0.2 | 3.36E-01 | 0.0 | 9.56E-01 | 0.2 | 4.95E-01 | hypothetical protein |
| AT5G57480 | 0.5 | 2.52E-05 | 0.9 | 5.34E-07 | -0.5 | 1.96E-04 | 0.5 | 1.54E-02 | -0.2 | 3.09E-01 | 0.7 | 6.52E-03 | AAA-ATPase At5g57480 |
| AT1G80420 | 0.5 | 2.64E-04 | 0.2 | 2.09E-02 | 0.2 | 4.16E-02 | -0.1 | 5.13E-01 | -0.3 | 1.91E-01 | 0.1 | 6.01E-01 | DNA-repair protein XRCC1 |
| AT5G25757 | 0.5 | 9.68E-05 | 0.5 | 2.26E-04 | 0.0 | 8.25E-01 | 0.4 | 3.55E-02 | 0.1 | 6.98E-01 | 0.3 | 2.43E-01 | Eukaryotic translation initiation factor 3 subunit L |
| AT4G16566 | 0.5 | 4.65E-04 | 0.1 | 5.89E-01 | 0.4 | 4.53E-03 | 0.0 | 9.04E-01 | 0.0 | 9.58E-01 | 0.0 | 9.75E-01 | HIT domain-containing protein |
| AT1G09200 | 0.5 | 6.73E-03 | 0.7 | 8.80E-04 | -0.2 | 1.72E-01 | 0.2 | 1.72E-01 | 0.2 | 2.95E-01 | 0.0 | 9.48E-01 | Histone H3.1 |
| AT1G77250 | 0.5 | 1.92E-03 | 0.2 | 8.53E-02 | 0.3 | 9.82E-02 | 0.0 | 8.80E-01 | -0.1 | 7.99E-01 | 0.0 | 9.19E-01 | At1g77250 |
| AT4G13050 | 0.5 | 7.52E-05 | 0.4 | 7.44E-04 | 0.1 | 2.05E-01 | 0.2 | 2.27E-01 | 0.0 | 8.95E-01 | 0.2 | 2.12E-01 | Oleoyl-acyl carrier protein thioesterase 2, chloroplastic |

| | | | | | | | | | | | | | |
|---|---|---|---|---|---|---|---|---|---|---|---|---|---|
| AT1G49180 | 0.5 | 5.64E-03 | 0.3 | 4.88E-02 | 0.2 | 3.70E-01 | -0.5 | 2.38E-02 | -0.4 | 6.65E-02 | -0.1 | 8.38E-01 | Protein kinase family protein |
| AT1G08830 | 0.5 | 1.13E-04 | 0.4 | 8.63E-04 | 0.1 | 2.92E-01 | 0.2 | 9.40E-02 | 0.1 | 5.70E-01 | 0.1 | 5.60E-01 | superoxide dismutase |
| AT4G31310 | 0.5 | 1.13E-02 | -0.1 | 4.95E-01 | 0.6 | 9.79E-03 | 0.0 | 8.55E-01 | 0.1 | 7.69E-01 | 0.0 | 9.22E-01 | Putative gamma-glutamylcyclotransferase |
| AT4G24800 | 0.5 | 4.82E-05 | 0.5 | 4.64E-05 | -0.1 | 5.47E-01 | -0.1 | 3.66E-01 | -0.2 | 1.32E-01 | 0.1 | 6.35E-01 | MA3 DOMAIN-CONTAINING TRANSLATION REGULATORY FACTOR 3 |
| AT3G10610 | 0.5 | 1.00E-02 | 0.6 | 3.27E-03 | -0.1 | 4.77E-01 | 0.2 | 2.20E-01 | 0.2 | 3.99E-01 | 0.0 | 9.04E-01 | Small ribosomal subunit protein eS17x |
| AT4G27800 | 0.5 | 2.40E-04 | 0.1 | 3.23E-01 | 0.4 | 2.86E-03 | -0.4 | 1.19E-02 | -0.2 | 2.35E-01 | -0.2 | 3.82E-01 | Protein phosphatase 2C 57 |
| AT2G30790 | 0.5 | 2.58E-02 | 0.8 | 1.62E-03 | -0.3 | 1.43E-01 | 0.6 | 1.15E-01 | 0.0 | 9.59E-01 | 0.6 | 1.90E-01 | Putative oxygen-evolving enhancer protein 2-2 |
| AT4G13430 | 0.5 | 1.55E-04 | 0.6 | 7.86E-05 | -0.1 | 2.94E-01 | -0.2 | 8.40E-02 | 0.3 | 1.62E-02 | -0.5 | 3.98E-03 | AT4G13430 protein |
| AT1G04730 | 0.5 | 1.26E-03 | 0.3 | 1.90E-02 | 0.2 | 2.38E-01 | 0.6 | 3.59E-02 | 0.6 | 7.49E-02 | 0.0 | 9.14E-01 | P-loop containing nucleoside triphosphate hydrolases superfamily protein |
| AT1G01060 | 0.5 | 1.28E-03 | -0.8 | 1.21E-04 | 1.3 | 2.81E-05 | | | | | | | Homeodomain-like superfamily protein |
| AT1G78650 | 0.5 | 6.21E-04 | 0.4 | 6.11E-03 | 0.1 | 3.00E-01 | 0.4 | 4.04E-02 | 0.6 | 1.75E-02 | -0.1 | 7.04E-01 | DNA polymerase delta subunit 3 |
| AT3G44150 | 0.5 | 1.01E-03 | 0.5 | 1.09E-03 | 0.0 | 8.85E-01 | 0.1 | 5.86E-01 | 0.2 | 3.02E-01 | -0.1 | 7.17E-01 | Expp1 protein |
| AT3G48040 | 0.5 | 3.38E-04 | 0.6 | 1.96E-04 | -0.1 | 5.05E-01 | 0.1 | 8.90E-01 | 0.0 | 9.99E-01 | 0.1 | 9.27E-01 | Uncharacterized protein |
| AT1G15080 | 0.5 | 7.19E-04 | 0.3 | 9.86E-03 | 0.2 | 2.44E-01 | -0.6 | 1.86E-02 | -0.4 | 1.27E-01 | -0.2 | 5.77E-01 | Lipid phosphate phosphatase 2 |
| AT5G51940 | 0.5 | 3.14E-04 | 0.2 | 5.22E-02 | 0.3 | 1.95E-02 | 0.3 | 1.80E-01 | 0.1 | 6.92E-01 | 0.2 | 5.52E-01 | DNA-directed RNA polymerases II, IV and V subunit 6A |
| AT5G09790 | 0.5 | 2.61E-03 | 0.2 | 1.08E-01 | 0.3 | 1.20E-01 | -0.2 | 4.96E-01 | -0.1 | 7.70E-01 | -0.1 | 8.27E-01 | TRITHORAX-RELATED PROTEIN 5 |
| AT1G48830 | 0.5 | 1.61E-03 | 0.7 | 3.82E-04 | -0.2 | 2.29E-01 | 0.5 | 4.39E-03 | 0.1 | 4.34E-01 | 0.3 | 1.06E-01 | Small ribosomal subunit protein eS7z |
| AT2G16430 | 0.5 | 1.96E-05 | 0.4 | 7.43E-05 | 0.0 | 5.31E-01 | 0.0 | 9.56E-01 | 0.5 | 1.53E-03 | -0.5 | 7.35E-03 | Purple acid phosphatase 10 |
| AT5G46420 | 0.5 | 2.99E-04 | 0.5 | 3.17E-04 | 0.0 | 7.83E-01 | 0.2 | 1.15E-01 | 0.1 | 4.38E-01 | 0.1 | 6.72E-01 | 16S rRNA processing protein RimM family |
| AT2G42380 | 0.5 | 2.11E-03 | 0.9 | 5.41E-05 | -0.4 | 1.61E-02 | -1.0 | 2.38E-03 | -0.6 | 4.72E-02 | -0.4 | 3.36E-01 | Basic-leucine zipper transcription factor family protein |
| AT2G46440 | 0.5 | 3.91E-03 | 0.0 | 9.97E-01 | 0.5 | 9.55E-03 | 1.2 | 5.99E-05 | 0.5 | 4.19E-02 | 0.6 | 2.30E-02 | Cyclic nucleotide-gated channels |
| AT5G35530 | 0.5 | 5.12E-04 | 0.5 | 5.23E-04 | 0.0 | 7.64E-01 | 0.1 | 1.90E-01 | 0.1 | 7.30E-01 | 0.1 | 6.14E-01 | Small ribosomal subunit protein uS3x |
| AT3G04910 | 0.5 | 6.08E-04 | 0.5 | 1.19E-03 | 0.0 | 8.87E-01 | -0.3 | 6.35E-03 | -0.1 | 5.04E-01 | -0.2 | 1.36E-01 | non-specific serine |
| AT1G01030 | 0.5 | 8.18E-03 | 0.2 | 3.01E-01 | 0.3 | 1.21E-01 | -0.2 | 5.27E-01 | 0.0 | 9.73E-01 | -0.2 | 5.54E-01 | AP2 |
| AT5G54250 | 0.5 | 2.95E-04 | 0.2 | 3.05E-02 | 0.3 | 3.30E-02 | -0.3 | 6.19E-02 | 0.1 | 4.31E-01 | -0.4 | 2.98E-02 | Cyclic nucleotide-gated cation channel 4 |
| AT5G65260 | 0.5 | 5.62E-05 | 0.2 | 1.81E-02 | 0.3 | 6.21E-03 | -0.1 | 6.74E-01 | -0.1 | 5.79E-01 | 0.0 | 9.10E-01 | Polyadenylate-binding protein 2 -binding protein 2) -binding protein 2) -binding protein II) |
| AT3G25890 | 0.5 | 1.06E-03 | 0.5 | 2.78E-03 | 0.0 | 7.37E-01 | 0.8 | 5.98E-05 | -0.1 | 5.93E-01 | 0.9 | 3.75E-04 | Ethylene-responsive transcription factor ERF119 |
| AT5G58480 | 0.5 | 1.64E-03 | 0.7 | 2.20E-04 | -0.2 | 1.45E-01 | 0.2 | 3.09E-01 | 0.1 | 8.36E-01 | 0.1 | 5.77E-01 | Glucan endo-1,3-beta-glucosidase 9 -beta-glucan endohydrolase 9) -beta-glucanase 9) |
| AT3G10700 | 0.5 | 3.39E-04 | 0.4 | 4.20E-03 | 0.1 | 2.28E-01 | -0.2 | 2.37E-01 | -0.4 | 7.57E-02 | 0.2 | 6.14E-01 | Galacturonic acid kinase |
| AT5G59910 | 0.5 | 1.38E-04 | 0.6 | 1.09E-04 | -0.1 | 5.10E-01 | 0.3 | 1.26E-01 | 0.1 | 8.41E-01 | 0.2 | 3.88E-01 | Histone H2B.11 |
| AT4G26850 | 0.5 | 1.61E-03 | 0.1 | 5.57E-01 | 0.4 | 9.61E-03 | 0.4 | 3.59E-03 | 0.4 | 6.99E-03 | 0.0 | 9.78E-01 | GDP-L-galactose phosphorylase 1 |
| AT3G25100 | 0.5 | 1.41E-03 | 0.6 | 1.00E-03 | -0.1 | 6.97E-01 | -0.1 | 7.90E-01 | 0.0 | 9.34E-01 | -0.1 | 7.84E-01 | CDC45 -like protein |
| AT2G42610 | 0.5 | 1.51E-03 | 0.5 | 2.61E-03 | 0.0 | 9.03E-01 | 0.2 | 5.39E-01 | 0.1 | 7.73E-01 | 0.1 | 8.75E-01 | Protein LIGHT-DEPENDENT SHORT HYPOCOTYLS 10 |
| AT5G43210 | 0.5 | 1.14E-02 | 0.1 | 4.40E-01 | 0.4 | 1.07E-01 | -0.4 | 3.61E-01 | 0.2 | 7.03E-01 | -0.5 | 2.85E-01 | Emb\|CAB76036.1 |
| AT4G32770 | 0.5 | 4.63E-03 | 0.5 | 8.84E-03 | 0.0 | 8.38E-01 | -0.6 | 2.60E-02 | 0.2 | 4.87E-01 | -0.8 | 2.53E-02 | Tocopherol cyclase, chloroplastic |
| AT1G64740 | 0.5 | 1.54E-03 | 0.2 | 8.06E-02 | 0.3 | 7.96E-02 | -0.3 | 4.98E-01 | -0.3 | 5.52E-01 | 0.0 | 9.97E-01 | Tubulin alpha-1 chain |
| AT3G18040 | 0.5 | 1.23E-04 | 0.3 | 4.27E-03 | 0.2 | 6.73E-02 | -0.3 | 5.86E-02 | -0.1 | 6.04E-01 | -0.2 | 3.50E-01 | MAP kinase 9 |
| AT3G53190 | 0.5 | 2.90E-05 | 0.5 | 9.63E-05 | 0.0 | 6.23E-01 | 0.0 | 7.12E-01 | 0.0 | 9.89E-01 | -0.1 | 8.05E-01 | Probable pectate lyase 12 |
| AT5G66720 | 0.5 | 1.19E-05 | 0.4 | 9.75E-05 | 0.1 | 1.85E-01 | 0.2 | 3.11E-02 | 0.0 | 8.05E-01 | 0.2 | 1.89E-01 | Probable protein phosphatase 2C 80 |
| AT1G79600 | 0.5 | 9.31E-05 | 0.0 | 9.03E-01 | 0.5 | 3.42E-04 | -0.1 | 7.54E-01 | 0.1 | 8.52E-01 | -0.1 | 6.67E-01 | Protein ACTIVITY OF BC1 COMPLEX KINASE 3, chloroplastic |
| AT3G22320 | 0.5 | 1.76E-03 | 0.3 | 4.27E-02 | 0.2 | 1.38E-01 | 0.2 | 3.23E-01 | -0.1 | 7.60E-01 | 0.3 | 3.09E-01 | DNA-directed RNA polymerases II and IV subunit 5A |
| AT2G29890 | 0.5 | 6.41E-04 | 0.3 | 1.49E-02 | 0.2 | 1.45E-01 | -0.4 | 1.38E-01 | 0.0 | 9.96E-01 | -0.4 | 2.29E-01 | Putative villin 1 VLN1 |
| AT3G07770 | 0.5 | 3.64E-05 | 0.4 | 6.27E-04 | 0.1 | 9.21E-02 | 0.6 | 1.04E-03 | 0.4 | 1.19E-02 | 0.1 | 6.04E-01 | Heat shock protein 90-6, mitochondrial |
| AT4G30920 | 0.5 | 3.33E-04 | 0.5 | 9.54E-04 | 0.0 | 6.98E-01 | -0.1 | 4.17E-01 | -0.3 | 5.30E-02 | 0.2 | 3.43E-01 | Leucine aminopeptidase 2, chloroplastic |
| AT3G52090 | 0.5 | 2.51E-05 | 0.4 | 1.72E-04 | 0.1 | 2.68E-01 | 0.2 | 8.61E-02 | 0.3 | 5.85E-02 | -0.1 | 8.26E-01 | DNA-directed RNA polymerase, RBP11-like protein |
| AT5G25080 | 0.5 | 4.77E-04 | 0.4 | 2.95E-03 | 0.1 | 4.03E-01 | 0.2 | 2.75E-01 | 0.0 | 9.87E-01 | 0.2 | 4.31E-01 | Nuclear nucleic acid-binding protein C1D |
| AT5G63980 | 0.5 | 5.65E-04 | 0.5 | 1.12E-03 | 0.0 | 8.60E-01 | 0.2 | 2.78E-01 | 0.4 | 3.79E-02 | -0.2 | 4.15E-01 | 3',5'-bisphosphate nucleotidase |
| AT1G14890 | 0.5 | 2.33E-03 | 0.4 | 1.33E-02 | 0.1 | 4.55E-01 | 0.1 | 6.48E-01 | -0.4 | 5.02E-02 | 0.4 | 6.02E-02 | Plant invertase |
| AT5G42310 | 0.5 | 1.58E-04 | 0.1 | 1.77E-01 | 0.4 | 3.22E-03 | 0.0 | 9.01E-01 | 0.1 | 6.84E-01 | -0.1 | 7.92E-01 | Pentatricopeptide repeat-containing protein At5g42310, chloroplastic |
| AT5G44390 | 0.5 | 7.08E-03 | 0.0 | 9.69E-01 | 0.5 | 2.05E-02 | 1.0 | 3.80E-02 | 0.0 | 9.57E-01 | 1.0 | 4.82E-02 | Berberine bridge enzyme-like 25 |
| AT1G21250 | 0.5 | 6.63E-02 | -0.6 | 3.91E-02 | 1.1 | 3.24E-03 | 0.8 | 4.93E-03 | 0.1 | 8.25E-01 | 0.8 | 5.27E-02 | Wall-associated receptor kinase 1 |
| AT2G16770 | 0.5 | 2.49E-04 | 0.1 | 1.60E-01 | 0.4 | 8.41E-03 | 0.1 | 5.11E-01 | 0.1 | 7.84E-01 | 0.1 | 8.61E-01 | Basic leucine zipper 23 |
| AT5G08570 | 0.5 | 6.52E-05 | 0.2 | 3.08E-02 | 0.3 | 4.50E-03 | 0.0 | 9.91E-01 | 0.0 | 9.70E-01 | 0.0 | 9.85E-01 | Pyruvate kinase |
| AT3G22300 | 0.5 | 3.21E-03 | 0.3 | 3.92E-02 | 0.2 | 2.63E-01 | 0.5 | 1.54E-03 | 0.1 | 5.99E-01 | 0.4 | 4.29E-02 | Small ribosomal subunit protein uS10m |
| AT4G33460 | 0.5 | 2.05E-03 | 0.5 | 2.55E-03 | 0.0 | 9.59E-01 | 0.1 | 5.52E-01 | 0.1 | 4.46E-01 | 0.0 | 8.80E-01 | ABC transporter I family member 10 |
| AT4G14365 | 0.5 | 5.40E-02 | -0.5 | 8.13E-02 | 1.0 | 5.32E-03 | 0.2 | 4.36E-01 | -0.2 | 3.85E-01 | 0.4 | 1.47E-01 | Putative E3 ubiquitin-protein ligase XBAT34 |
| AT1G06160 | 0.5 | 5.67E-02 | 0.3 | 2.32E-01 | 0.2 | 5.38E-01 | 1.5 | 1.02E-04 | 0.3 | 4.52E-01 | 1.2 | 4.12E-03 | Ethylene-responsive transcription factor ERF094 |
| AT3G55580 | 0.5 | 7.63E-03 | 0.5 | 1.15E-02 | 0.0 | 9.71E-01 |  |  |  |  |  |  | Regulator of chromosome condensation family protein |
| AT5G44870 | 0.5 | 7.33E-04 | 0.1 | 3.96E-01 | 0.4 | 8.72E-03 | 0.4 | 2.61E-02 | 0.1 | 7.42E-01 | 0.3 | 1.73E-01 | ADP-ribosyl cyclase |
| AT3G22890 | 0.5 | 1.04E-04 | 0.6 | 8.66E-05 | -0.1 | 5.10E-01 | 0.2 | 2.42E-01 | 0.1 | 5.56E-01 | 0.1 | 8.14E-01 | ATP sulfurylase 1, chloroplastic |
| AT4G12330 | 0.5 | 1.44E-02 | 0.3 | 1.35E-01 | 0.2 | 3.63E-01 | -1.8 | 8.72E-04 | -0.4 | 3.53E-01 | -1.4 | 1.12E-02 | Cytochrome P450, family 706, subfamily A, polypeptide 7 |
| AT3G47460 | 0.5 | 1.20E-03 | 0.1 | 4.42E-01 | 0.4 | 1.50E-02 | 0.4 | 2.25E-01 | 0.5 | 9.50E-02 | -0.2 | 5.82E-01 | Structural maintenance of chromosomes protein |
| AT5G64520 | 0.5 | 1.32E-03 | 0.2 | 1.26E-01 | 0.3 | 5.75E-02 | -0.3 | 3.58E-01 | -0.3 | 5.44E-01 | -0.1 | 9.04E-01 | Homolog of X-ray repair cross complementing 2 |
| AT2G30410 | 0.5 | 1.80E-04 | 0.5 | 3.33E-04 | 0.0 | 9.25E-01 | -0.2 | 2.73E-01 | -0.2 | 4.73E-01 | 0.0 | 8.95E-01 | Tubulin-folding cofactor A |
| AT1G17970 | 0.5 | 1.35E-03 | 0.4 | 1.58E-02 | 0.2 | 2.60E-01 | -0.2 | 3.06E-01 | -0.3 | 9.66E-02 | 0.1 | 6.25E-01 | RING-type E3 ubiquitin transferase |
| AT1G68830 | 0.5 | 5.88E-04 | 0.0 | 6.73E-01 | 0.6 | 1.06E-03 | 0.1 | 4.37E-01 | 0.0 | 8.35E-01 | 0.2 | 3.47E-01 | Serine |
| AT5G60200 | 0.5 | 9.28E-05 | 0.3 | 5.99E-03 | 0.2 | 3.68E-02 | 0.0 | 9.03E-01 | 0.1 | 5.86E-01 | -0.1 | 7.49E-01 | Dof zinc finger protein DOF5.3 |
| AT1G26230 | 0.5 | 7.26E-04 | 0.6 | 4.06E-04 | -0.1 | 5.41E-01 | 0.3 | 4.60E-01 | 0.4 | 3.23E-01 | -0.1 | 7.79E-01 | TCP-1 |
| AT1G18330 | 0.5 | 1.15E-03 | 0.8 | 7.95E-05 | -0.3 | 4.57E-02 | | | | | | | hypothetical protein |
| AT5G48030 | 0.5 | 1.42E-04 | 0.1 | 5.97E-01 | 0.5 | 1.28E-03 | 0.6 | 7.55E-04 | 0.2 | 3.67E-01 | 0.4 | 3.59E-02 | Chaperone protein dnaJ GFA2, mitochondrial |
| AT2G18220 | 0.5 | 2.16E-03 | 0.3 | 2.55E-02 | 0.2 | 2.62E-01 | 0.5 | 1.71E-03 | 0.1 | 5.40E-01 | 0.4 | 4.40E-02 | Nucleolar complex-associated protein 2 |
| AT1G08980 | 0.5 | 7.20E-04 | 0.7 | 1.18E-04 | -0.2 | 1.27E-01 | -0.3 | 2.93E-02 | -0.4 | 1.51E-02 | 0.1 | 7.15E-01 | Amidase 1 |
| AT5G27380 | 0.5 | 6.29E-05 | 0.4 | 3.78E-04 | 0.1 | 3.56E-01 | 0.3 | 3.34E-02 | -0.1 | 3.88E-01 | 0.4 | 2.32E-02 | Glutathione synthetase, chloroplastic |
| AT5G04885 | 0.5 | 1.23E-04 | 0.5 | 4.14E-04 | 0.0 | 6.38E-01 | 0.2 | 3.51E-01 | 0.2 | 3.62E-01 | 0.0 | 9.49E-01 | Uncharacterized protein At5g04885 |
| AT4G04890 | 0.5 | 3.71E-04 | 0.4 | 2.52E-03 | 0.1 | 3.48E-01 | -0.1 | 6.09E-01 | -0.2 | 2.82E-01 | 0.1 | 6.84E-01 | Homeobox-leucine zipper protein PROTODERMAL FACTOR 2 |
| AT1G24040 | 0.5 | 3.25E-04 | 0.2 | 3.73E-02 | 0.3 | 3.36E-02 | 0.0 | 8.94E-01 | -0.1 | 6.66E-01 | 0.1 | 7.84E-01 | N-acetyltransferase domain-containing protein |
| AT5G51020 | 0.5 | 1.61E-04 | 0.1 | 2.18E-01 | 0.4 | 3.21E-03 | -0.2 | 2.57E-01 | 0.0 | 9.92E-01 | -0.2 | 3.97E-01 | Chromophore lyase CRL, chloroplastic |
| AT1G51800 | 0.5 | 1.28E-02 | 0.0 | 9.21E-01 | 0.5 | 2.14E-02 | 0.2 | 1.84E-01 | 0.1 | 6.33E-01 | 0.1 | 6.39E-01 | LRR receptor-like serine |
| AT4G08920 | 0.5 | 8.63E-04 | 0.3 | 3.12E-02 | 0.2 | 8.49E-02 | -0.2 | 6.02E-02 | -0.3 | 2.55E-02 | 0.1 | 7.01E-01 | Cryptochrome-1 |
| AT2G32030 | 0.5 | 1.79E-02 | 0.0 | 9.71E-01 | 0.5 | 5.37E-02 | 0.6 | 7.98E-02 | -0.6 | 1.68E-01 | 1.2 | 4.72E-03 | hypothetical protein |
| AT3G55280 | 0.5 | 1.29E-03 | 0.6 | 5.42E-04 | -0.1 | 4.09E-01 | 0.5 | 1.69E-02 | 0.4 | 7.12E-02 | 0.1 | 8.13E-01 | Ribosomal protein L23AB |
| AT3G13730 | 0.5 | 2.92E-03 | 0.2 | 1.76E-01 | 0.3 | 8.00E-02 | 1.2 | 3.79E-05 | 0.9 | 2.09E-03 | 0.3 | 1.31E-01 | Cytochrome P450, family 90, subfamily D, polypeptide 1 |
| AT1G53430 | 0.5 | 4.95E-04 | 0.1 | 4.25E-01 | 0.4 | 4.53E-03 | 0.3 | 2.33E-01 | -0.2 | 3.77E-01 | 0.5 | 5.62E-02 | Leucine-rich repeat transmembrane protein kinase |

| | | | | | | | | | | | | | |
|---|---|---|---|---|---|---|---|---|---|---|---|---|---|
| AT5G65360 | 0.5 | 7.00E-04 | 0.7 | 1.98E-04 | -0.2 | 2.19E-01 | 0.3 | 3.54E-02 | 0.2 | 1.48E-01 | 0.1 | 7.81E-01 | Histone H3.1 |
| AT5G54380 | 0.5 | 2.04E-03 | 0.4 | 1.32E-02 | 0.1 | 3.91E-01 | -0.5 | 3.02E-03 | -0.4 | 1.70E-02 | -0.1 | 7.63E-01 | Receptor-like protein kinase THESEUS 1 |
| AT3G10090 | 0.5 | 2.31E-03 | 0.4 | 2.32E-02 | 0.2 | 3.05E-01 | 0.4 | 1.39E-01 | 0.2 | 4.00E-01 | 0.1 | 7.67E-01 | Small ribosomal subunit protein eS28z |
| AT5G40610 | 0.5 | 2.11E-03 | 0.8 | 1.19E-04 | -0.3 | 4.57E-02 | 0.3 | 3.20E-02 | 0.0 | 9.79E-01 | 0.3 | 9.43E-02 | Glycerol-3-phosphate dehydrogenase 1, chloroplastic |
| AT3G47390 | 0.5 | 5.26E-05 | 0.1 | 2.67E-01 | 0.4 | 1.12E-03 | 0.1 | 4.56E-01 | 0.1 | 7.11E-01 | 0.0 | 8.57E-01 | 5-amino-6-uracil reductase |
| AT1G08000 | 0.5 | 2.71E-04 | 0.3 | 1.56E-02 | 0.2 | 5.78E-02 | 0.1 | 6.02E-01 | -0.1 | 8.10E-01 | 0.2 | 5.15E-01 | GATA transcription factor 10 |
| AT2G46870 | 0.5 | 3.30E-03 | 0.2 | 1.97E-01 | 0.3 | 7.16E-02 | 0.2 | 1.94E-01 | 0.3 | 8.43E-02 | -0.1 | 6.23E-01 | B3 domain-containing transcription factor NGA1 |
| AT3G14690 | 0.5 | 1.72E-04 | 0.2 | 8.50E-02 | 0.3 | 6.02E-03 | -0.1 | 8.38E-01 | -0.1 | 7.72E-01 | 0.0 | 9.29E-01 | Cytochrome P450, family 72, subfamily A, polypeptide 15 |
| AT3G09735 | 0.5 | 5.34E-04 | 0.4 | 2.62E-03 | 0.1 | 4.57E-01 | 0.1 | 6.28E-01 | 0.0 | 8.98E-01 | 0.2 | 6.55E-01 | DNA-binding protein S1FA3 |
| AT4G02220 | 0.5 | 1.79E-03 | 0.1 | 6.67E-01 | 0.5 | 1.15E-02 | 0.3 | 4.77E-02 | -0.1 | 8.19E-01 | 0.4 | 9.07E-02 | Putative zinc finger protein |
| AT5G17780 | 0.5 | 9.54E-03 | -0.1 | 7.52E-01 | 0.6 | 1.79E-02 | 1.3 | 2.73E-02 | 1.1 | 1.13E-01 | | | Probable lysophospholipase BODYGUARD 4 |
| AT5G39040 | 0.5 | 4.31E-05 | 0.2 | 5.03E-02 | 0.4 | 2.25E-03 | 0.6 | 1.38E-04 | 0.2 | 3.13E-01 | 0.5 | 1.46E-02 | Transporter associated with antigen processing protein 2 |
| AT5G40820 | 0.5 | 1.03E-03 | 0.1 | 6.33E-01 | 0.5 | 8.95E-03 | 0.0 | 9.75E-01 | 0.1 | 7.39E-01 | -0.1 | 7.79E-01 | non-specific serine |
| AT4G17040 | 0.5 | 1.06E-05 | 0.2 | 1.30E-02 | 0.3 | 9.47E-04 | -0.2 | 1.28E-01 | -0.2 | 1.76E-01 | 0.0 | 9.95E-01 | ATP-dependent Clp protease proteolytic subunit-related protein 4, chloroplastic |
| AT3G04590 | 0.5 | 9.82E-06 | 0.1 | 5.54E-02 | 0.4 | 3.94E-04 | 0.0 | 9.17E-01 | -0.1 | 5.47E-01 | 0.1 | 7.09E-01 | AT-hook motif nuclear-localized protein |
| AT2G15890 | 0.5 | 4.68E-03 | 1.0 | 1.00E-04 | -0.5 | 1.44E-02 | -0.5 | 1.70E-02 | -1.1 | 1.38E-04 | 0.6 | 4.12E-02 | CCG-binding protein 1 |
| AT5G14930 | 0.5 | 2.11E-04 | -0.2 | 8.33E-02 | 0.7 | 1.77E-04 | -0.1 | 4.60E-01 | -0.4 | 1.44E-02 | 0.3 | 1.15E-01 | Senescence-associated gene 101 |
| AT5G08340 | 0.5 | 9.51E-04 | 0.3 | 5.35E-02 | 0.3 | 7.90E-02 | -0.1 | 7.30E-01 | -0.3 | 2.05E-01 | 0.2 | 4.10E-01 | FAD synthase |
| AT5G25090 | 0.5 | 2.43E-02 | 0.9 | 1.39E-03 | -0.4 | 1.27E-01 | 0.1 | 7.33E-01 | 0.4 | 2.55E-01 | -0.2 | 5.45E-01 | Early nodulin-like protein 13 |
| AT1G16920 | 0.5 | 4.64E-04 | 0.7 | 1.45E-04 | -0.1 | 2.52E-01 | -0.1 | 3.27E-01 | -0.2 | 2.78E-01 | 0.0 | 9.02E-01 | Ras-related protein RABA1b |
| AT5G38530 | 0.5 | 5.80E-05 | 0.1 | 9.33E-02 | 0.4 | 1.90E-03 | 0.2 | 7.47E-02 | 0.4 | 6.87E-03 | -0.2 | 3.19E-01 | tryptophan synthase |
| AT4G21900 | 0.5 | 1.13E-04 | 0.4 | 2.17E-03 | 0.2 | 1.29E-01 | 0.1 | 7.57E-01 | 0.0 | 9.54E-01 | 0.1 | 8.84E-01 | PRORP3 |
| AT5G51795 | 0.5 | 1.20E-02 | 0.9 | 6.81E-04 | -0.3 | 1.07E-01 | 0.7 | 2.74E-02 | 0.4 | 2.03E-01 | 0.2 | 5.66E-01 | KIN17-like protein KLP |
| AT4G25270 | 0.5 | 2.19E-03 | 0.3 | 4.99E-02 | 0.2 | 1.89E-01 | 0.0 | 9.68E-01 | 0.4 | 4.76E-01 | -0.4 | 5.80E-01 | Pentatricopeptide repeat-containing protein At4g25270, chloroplastic |
| AT5G16150 | 0.5 | 1.92E-04 | 0.4 | 1.38E-03 | 0.1 | 3.23E-01 | -0.5 | 1.39E-02 | -0.4 | 6.85E-02 | -0.1 | 7.86E-01 | Plastidic glucose transporter 4 |
| AT2G21370 | 0.5 | 1.46E-03 | 0.4 | 6.72E-03 | 0.1 | 5.09E-01 | 0.2 | 4.65E-01 | 0.4 | 5.55E-02 | -0.3 | 2.87E-01 | Xylulose kinase-1 |
| AT1G58110 | 0.5 | 1.60E-04 | 0.6 | 1.78E-04 | 0.0 | 7.90E-01 | 0.0 | 9.73E-01 | 0.1 | 7.12E-01 | -0.1 | 7.33E-01 | BZIP domain-containing protein |
| AT4G35570 | 0.5 | 1.40E-03 | 0.3 | 2.37E-02 | 0.2 | 1.82E-01 | 0.3 | 1.37E-01 | 0.2 | 4.14E-01 | 0.1 | 7.58E-01 | High mobility group B protein 5 |
| AT3G23990 | 0.5 | 2.24E-03 | 0.5 | 5.07E-03 | 0.0 | 8.02E-01 | 0.5 | 1.10E-02 | 0.3 | 1.40E-01 | 0.2 | 5.50E-01 | Chaperonin CPN60, mitochondrial |
| AT1G26390 | 0.5 | 4.50E-01 | 0.1 | 8.53E-01 | 0.4 | 6.70E-01 | 1.5 | 2.66E-02 | -0.1 | 9.66E-01 | 1.6 | 5.52E-02 | Berberine bridge enzyme-like 4 |
| AT4G08470 | 0.5 | 9.01E-04 | 0.0 | 9.50E-01 | 0.5 | 2.48E-03 | 0.2 | 3.03E-01 | -0.3 | 2.24E-01 | 0.5 | 7.51E-02 | MAPK |
| AT1G32870 | 0.5 | 1.31E-03 | -0.6 | 1.15E-03 | 1.2 | 8.28E-05 | 1.0 | 5.76E-05 | -0.2 | 5.01E-01 | 1.2 | 3.08E-04 | NAC domain protein 13 |
| AT1G77940 | 0.5 | 5.12E-03 | 0.8 | 6.82E-04 | -0.3 | 1.56E-01 | 0.4 | 3.57E-02 | 0.2 | 3.25E-01 | 0.2 | 5.00E-01 | Large ribosomal subunit protein eL30y |
| AT1G27400 | 0.5 | 7.75E-03 | 0.7 | 2.59E-03 | -0.2 | 4.40E-01 | 0.3 | 3.23E-02 | 0.2 | 3.19E-01 | 0.2 | 4.94E-01 | Large ribosomal subunit protein uL22z |
| AT2G18328 | 0.5 | 1.20E-01 | 0.4 | 2.35E-01 | 0.1 | 7.81E-01 | -0.3 | 2.07E-01 | 0.2 | 4.70E-01 | -0.5 | 1.09E-01 | Protein RADIALIS-like 4 |
| AT3G61310 | 0.5 | 5.09E-05 | 0.2 | 1.50E-02 | 0.3 | 7.35E-03 | 0.0 | 8.86E-01 | -0.1 | 6.92E-01 | 0.1 | 6.61E-01 | AT-hook motif nuclear-localized protein 11 |
| AT1G54690 | 0.5 | 7.03E-03 | 0.5 | 9.46E-03 | 0.0 | 9.67E-01 | 0.3 | 1.55E-01 | 0.3 | 2.64E-01 | 0.0 | 9.38E-01 | Probable histone H2AXb |
| AT5G35360 | 0.5 | 2.03E-05 | 0.5 | 7.31E-05 | 0.0 | 6.20E-01 | 0.0 | 9.41E-01 | 0.0 | 9.66E-01 | 0.0 | 9.35E-01 | Biotin carboxylase |
| AT3G20780 | 0.5 | 7.39E-05 | 0.3 | 5.75E-03 | 0.2 | 3.07E-02 | 0.1 | 8.26E-01 | 0.1 | 6.60E-01 | -0.1 | 8.45E-01 | DNA topoisomerase 6 subunit B |
| AT5G46330 | 0.5 | 1.25E-03 | 0.4 | 5.02E-03 | 0.1 | 5.59E-01 | 0.3 | 5.98E-02 | -0.4 | 2.38E-02 | 0.7 | 4.07E-03 | LRR receptor-like serine |
| AT5G59030 | 0.5 | 4.31E-05 | 0.4 | 9.69E-04 | 0.2 | 6.92E-02 | -0.1 | 4.85E-01 | -0.4 | 2.68E-02 | 0.3 | 2.28E-01 | Copper transport protein |
| AT3G09920 | 0.5 | 2.33E-04 | 0.2 | 3.90E-02 | 0.3 | 1.80E-02 | -0.3 | 2.61E-01 | -0.1 | 7.80E-01 | -0.2 | 6.03E-01 | Phosphatidylinositol 4-phosphate 5-kinase 9 P-5-kinase 9) |
| AT1G20010 | 0.5 | 6.99E-05 | 0.7 | 1.87E-05 | -0.2 | 7.13E-02 | -0.3 | 2.14E-02 | -0.4 | 1.66E-02 | 0.1 | 8.33E-01 | Tubulin beta-5 chain |
| AT2G43690 | 0.5 | 3.29E-03 | 0.3 | 7.14E-02 | 0.2 | 2.05E-01 | -0.2 | 6.00E-01 | 0.1 | 8.65E-01 | -0.3 | 4.56E-01 | Concanavalin A-like lectin protein kinase family protein |
| AT4G05090 | 0.5 | 5.35E-05 | 0.5 | 7.89E-05 | 0.0 | 8.67E-01 | -0.2 | 3.09E-01 | 0.0 | 9.69E-01 | -0.1 | 4.59E-01 | Inositol monophosphatase family protein |
| AT1G55450 | 0.5 | 1.33E-03 | 0.0 | 8.86E-01 | 0.5 | 4.29E-03 | 0.7 | 4.22E-05 | 0.1 | 6.76E-01 | 0.6 | 2.97E-03 | S-adenosyl-L-methionine-dependent methyltransferases superfamily protein |
| AT2G29940 | 0.5 | 4.38E-03 | -0.2 | 3.92E-01 | 0.7 | 5.35E-03 | 0.2 | 7.67E-01 | 0.1 | 8.46E-01 | 0.0 | 9.69E-01 | ABC transporter G family member 31 |
| AT1G19850 | 0.5 | 4.02E-05 | 0.2 | 1.65E-02 | 0.3 | 5.65E-03 | 0.1 | 6.17E-01 | 0.0 | 9.25E-01 | 0.1 | 6.72E-01 | Auxin response factor |
| AT1G74060 | 0.5 | 1.11E-03 | 0.6 | 5.62E-04 | -0.1 | 4.66E-01 | 0.2 | 1.31E-01 | 0.1 | 7.85E-01 | 0.1 | 4.52E-01 | Large ribosomal subunit protein eL6y |
| AT3G22190 | 0.5 | 1.06E-04 | 0.2 | 6.56E-02 | 0.3 | 6.03E-03 | -0.1 | 7.34E-01 | 0.0 | 9.80E-01 | -0.1 | 8.35E-01 | Protein IQ-DOMAIN 5 |
| AT2G35450 | 0.5 | 6.52E-04 | 0.5 | 1.76E-03 | 0.0 | 7.36E-01 | -0.2 | 1.39E-01 | 0.0 | 9.73E-01 | -0.3 | 2.40E-01 | Catalytic |
| AT5G22510 | 0.5 | 6.61E-05 | 0.1 | 2.39E-01 | 0.4 | 1.06E-03 | -0.1 | 5.12E-01 | 0.0 | 9.81E-01 | -0.2 | 6.29E-01 | Alkaline |
| AT2G17900 | 0.5 | 2.32E-04 | 0.3 | 1.46E-02 | 0.2 | 5.58E-02 | 0.0 | 9.82E-01 | 0.4 | 3.42E-01 | -0.4 | 4.60E-01 | SET domain group 37 |
| AT5G49700 | 0.5 | 1.76E-02 | -0.2 | 4.32E-01 | 0.7 | 1.69E-02 | 0.2 | 5.53E-01 | -1.5 | 1.07E-03 | 1.7 | 1.69E-03 | AT-hook motif nuclear-localized protein 17 |
| AT1G78290 | 0.5 | 4.11E-04 | 0.2 | 1.62E-01 | 0.4 | 1.09E-02 | 0.1 | 5.37E-01 | 0.1 | 8.35E-01 | 0.1 | 8.21E-01 | Serine |
| AT3G05600 | 0.5 | 5.17E-04 | 0.5 | 1.06E-03 | 0.0 | 8.76E-01 | 0.4 | 6.79E-02 | -0.1 | 8.00E-01 | 0.4 | 8.08E-02 | AB hydrolase-1 domain-containing protein |
| AT2G26975 | 0.5 | 1.56E-04 | 0.5 | 6.00E-04 | 0.1 | 5.55E-01 | 0.0 | 9.57E-01 | -0.4 | 6.22E-02 | 0.4 | 1.45E-01 | Copper transporter 6 |
| AT3G13490 | 0.5 | 2.02E-05 | 0.2 | 7.21E-03 | 0.3 | 3.57E-03 | 0.3 | 1.08E-02 | 0.1 | 4.13E-01 | 0.2 | 2.26E-01 | Lysine--tRNA ligase, chloroplastic |
| AT5G48840 | 0.5 | 6.92E-04 | 0.4 | 1.29E-02 | 0.2 | 1.75E-01 | 0.5 | 3.56E-03 | 0.1 | 6.22E-01 | 0.4 | 5.34E-02 | Pantoate--beta-alanine ligase |
| AT3G62750 | 0.5 | 2.29E-03 | 0.6 | 1.55E-03 | -0.1 | 7.00E-01 | -0.4 | 2.52E-02 | -0.1 | 7.26E-01 | -0.3 | 1.61E-01 | Beta glucosidase 8 |
| AT4G04880 | 0.5 | 1.32E-04 | 0.4 | 3.46E-03 | 0.2 | 1.01E-01 | 0.2 | 1.93E-01 | 0.1 | 7.51E-01 | 0.2 | 5.52E-01 | Adenosine |
| AT3G13450 | 0.5 | 7.71E-05 | 0.7 | 1.34E-05 | -0.2 | 4.63E-02 | -0.5 | 2.08E-02 | -0.7 | 2.53E-03 | 0.3 | 2.04E-01 | 2-oxoisovalerate dehydrogenase subunit beta 2, mitochondrial |
| AT2G30860 | 0.5 | 4.00E-04 | 0.9 | 2.37E-05 | -0.4 | 1.37E-02 | 0.2 | 4.50E-01 | 0.2 | 4.45E-01 | 0.0 | 9.44E-01 | Glutathione S-transferase F9 |
| AT5G27610 | 0.5 | 1.29E-03 | -0.2 | 3.01E-01 | 0.7 | 1.76E-03 | 0.3 | 4.77E-01 | 0.0 | 9.96E-01 | 0.3 | 5.96E-01 | Protein ALWAYS EARLY 1 |
| AT5G02180 | 0.5 | 1.25E-03 | 0.4 | 1.66E-02 | 0.2 | 2.32E-01 | | | | | | | Amino acid transporter AVT1D |
| AT2G20725 | 0.5 | 3.66E-05 | 0.5 | 8.08E-05 | 0.0 | 8.79E-01 | -0.2 | 9.19E-02 | -0.2 | 7.25E-02 | 0.0 | 8.64E-01 | CAAX amino terminal protease family protein |
| AT2G29120 | 0.5 | 1.70E-03 | -0.3 | 5.59E-02 | 0.8 | 5.55E-04 | 0.3 | 3.23E-01 | -0.5 | 2.97E-01 | 0.8 | 7.77E-02 | Glutamate receptor 2.7 |
| AT3G49670 | 0.5 | 4.82E-05 | 0.5 | 1.05E-04 | 0.0 | 9.10E-01 | 0.0 | 9.75E-01 | -0.1 | 5.25E-01 | 0.1 | 6.30E-01 | Leucine-rich repeat receptor-like serine |
| AT3G54460 | 0.5 | 7.53E-05 | 0.4 | 1.90E-03 | 0.2 | 7.83E-02 | 0.1 | 7.61E-01 | 0.1 | 7.99E-01 | 0.0 | 9.97E-01 | SNF2 domain-containing protein |
| AT3G54110 | 0.5 | 7.80E-05 | 0.6 | 6.32E-05 | -0.1 | 5.13E-01 | 0.3 | 7.55E-03 | 0.1 | 7.37E-01 | 0.3 | 8.76E-02 | Mitochondrial uncoupling protein 1 |
| AT2G02010 | 0.5 | 2.62E-03 | 0.0 | 9.19E-01 | 0.5 | 1.28E-02 | 1.2 | 1.32E-03 | 0.3 | 5.75E-01 | 0.9 | 9.81E-03 | Glutamate decarboxylase |
| AT5G54390 | 0.5 | 2.90E-05 | 0.4 | 7.12E-04 | 0.2 | 5.64E-02 | -0.1 | 3.47E-01 | 0.2 | 1.27E-01 | -0.3 | 6.02E-02 | PAP-specific phosphatase HAL2-like ,5'-bisphosphate nucleotidase) ,5-bisphosphonucleoside 3'-phosphohydrolase) |
| AT2G23420 | 0.5 | 2.86E-03 | -0.2 | 3.21E-01 | 0.7 | 2.26E-03 | 0.2 | 3.04E-01 | -0.1 | 7.19E-01 | 0.2 | 2.52E-01 | Nicotinate phosphoribosyltransferase |
| AT5G47435 | 0.5 | 1.81E-05 | 0.4 | 4.86E-04 | 0.2 | 4.56E-02 | -0.1 | 4.89E-01 | 0.0 | 8.77E-01 | -0.1 | 7.71E-01 | Formyltetrahydrofolate deformylase 1, mitochondrial |
| AT1G72910 | 0.5 | 5.40E-02 | -0.5 | 1.21E-01 | 1.0 | 9.12E-03 | 1.3 | 6.63E-05 | 0.6 | 4.73E-02 | 0.7 | 2.22E-02 | Similar to part of disease resistance protein domain-containing protein) |
| AT5G01550 | 0.5 | 1.01E-01 | -0.4 | 3.48E-01 | 0.9 | 4.73E-02 | 2.0 | 7.25E-04 | 0.4 | 6.42E-01 | 1.6 | 1.90E-03 | Lectin receptor kinase a4.1 |
| AT2G26080 | 0.5 | 2.49E-03 | 0.1 | 6.65E-01 | 0.5 | 1.16E-02 | 0.2 | 4.25E-01 | 0.2 | 4.36E-01 | 0.0 | 9.52E-01 | Glycine dehydrogenase 2, mitochondrial 2) |
| AT5G63920 | 0.5 | 6.38E-04 | 0.2 | 1.75E-01 | 0.4 | 2.12E-02 | 0.2 | 6.32E-01 | 0.3 | 5.46E-01 | -0.1 | 9.12E-01 | DNA topoisomerase 3-alpha |
| AT5G42390 | 0.5 | 2.81E-05 | 0.1 | 4.69E-01 | 0.5 | 2.87E-04 | 0.2 | 2.09E-01 | 0.2 | 3.22E-01 | 0.0 | 9.69E-01 | Insulinase family protein |
| AT2G45340 | 0.5 | 5.42E-04 | 0.4 | 8.72E-03 | 0.2 | 1.67E-01 | -0.5 | 8.62E-03 | -0.2 | 2.46E-01 | -0.2 | 1.26E-01 | Leucine-rich repeat protein kinase family protein |
| AT5G07280 | 0.5 | 1.24E-02 | 0.5 | 2.54E-02 | 0.0 | 8.48E-01 | -0.1 | 8.91E-01 | 0.1 | 7.08E-01 | -0.2 | 6.28E-01 | Leucine-rich repeat receptor protein kinase EMS1 |
| AT1G49650 | 0.5 | 5.25E-04 | 0.9 | 2.81E-05 | -0.3 | 2.10E-02 | -0.2 | 2.88E-01 | -0.3 | 8.54E-02 | 0.1 | 5.93E-01 | Probable carboxylesterase 4, mitochondrial |
| AT2G19385 | 0.5 | 1.15E-03 | 0.4 | 1.37E-02 | 0.2 | 2.61E-01 | 0.6 | 3.34E-03 | 0.2 | 5.55E-01 | 0.5 | 5.95E-02 | hypothetical protein |

| | | | | | | | | | | | | | |
|---|---|---|---|---|---|---|---|---|---|---|---|---|---|
| AT5G60390 | 0.5 | 2.22E-03 | 0.5 | 3.13E-03 | 0.0 | 9.95E-01 | 0.4 | 9.80E-03 | 0.1 | 4.92E-01 | 0.2 | 1.97E-01 | Elongation factor 1-alpha 4 |
| AT1G70530 | 0.5 | 1.57E-03 | 0.1 | 4.73E-01 | 0.4 | 1.28E-02 | 0.3 | 1.22E-01 | -0.2 | 4.92E-01 | 0.4 | 8.26E-02 | Cysteine-rich RLK 3 |
| AT2G27800 | 0.5 | 1.30E-03 | 0.3 | 3.73E-02 | 0.2 | 1.53E-01 | -0.1 | 7.92E-01 | -0.1 | 9.05E-01 | 0.0 | 9.48E-01 | Pentatricopeptide repeat-containing protein At2g27800, mitochondrial |
| AT5G58870 | 0.5 | 4.60E-04 | 0.0 | 6.91E-01 | 0.6 | 9.12E-04 | 0.2 | 4.83E-01 | 0.2 | 4.50E-01 | 0.0 | 9.26E-01 | ATP-dependent zinc metalloprotease FTSH 9, chloroplastic |
| AT5G58240 | 0.5 | 1.59E-04 | 0.3 | 1.35E-02 | 0.3 | 3.43E-02 | 0.1 | 5.24E-01 | -0.1 | 6.90E-01 | 0.2 | 3.90E-01 | FRAGILE HISTIDINE TRIAD |
| AT4G23230 | 0.5 | 2.29E-02 | -0.2 | 4.87E-01 | 0.7 | 2.25E-02 | -0.4 | 4.17E-01 | -0.6 | 2.29E-01 | 0.2 | 7.10E-01 | Cysteine-rich receptor-like protein kinase 15 |
| AT1G75010 | 0.5 | 3.71E-04 | 0.3 | 3.36E-02 | 0.3 | 3.92E-02 | 0.2 | 3.14E-01 | 0.3 | 2.96E-01 | 0.0 | 9.27E-01 | Protein ACCUMULATION AND REPLICATION OF CHLOROPLASTS 3, chloroplastic |
| AT5G58140 | 0.6 | 6.25E-04 | 0.2 | 5.52E-02 | 0.3 | 3.58E-02 | -0.1 | 5.62E-01 | 0.1 | 5.74E-01 | -0.2 | 3.46E-01 | Phototropin 2 |
| AT1G53670 | 0.6 | 1.37E-04 | 0.4 | 1.20E-03 | 0.1 | 2.50E-01 | 0.1 | 4.39E-01 | 0.0 | 8.66E-01 | 0.1 | 7.50E-01 | Peptide-methionine -S-oxide reductase |
| AT3G12080 | 0.6 | 3.38E-05 | 0.6 | 3.33E-05 | -0.1 | 5.74E-01 | -0.1 | 6.80E-01 | -0.3 | 7.07E-02 | 0.3 | 1.78E-01 | GTPase Der |
| AT3G12710 | 0.6 | 1.23E-03 | 0.7 | 5.62E-04 | -0.1 | 4.89E-01 | -0.3 | 9.52E-02 | -0.2 | 2.33E-01 | -0.1 | 8.24E-01 | DNA-3-methyladenine glycosylase I |
| AT4G11010 | 0.6 | 4.72E-05 | 0.7 | 2.82E-05 | -0.1 | 2.62E-01 | 0.5 | 2.80E-04 | 0.4 | 2.84E-03 | 0.1 | 7.01E-01 | Nucleoside diphosphate kinase III, chloroplastic |
| AT1G72360 | 0.6 | 4.50E-03 | 0.4 | 3.67E-02 | 0.2 | 3.75E-01 | 0.3 | 2.41E-01 | -0.3 | 3.48E-01 | 0.6 | 5.37E-02 | Integrase-type DNA-binding superfamily protein |
| AT3G11945 | 0.6 | 4.64E-05 | 0.7 | 2.64E-05 | -0.1 | 2.85E-01 | 0.1 | 3.18E-01 | 0.0 | 7.67E-01 | 0.1 | 7.34E-01 | Homogentisate prenyltransferase |
| AT5G04800 | 0.6 | 7.15E-04 | 0.7 | 2.22E-04 | -0.2 | 2.53E-01 | 0.3 | 1.93E-02 | 0.1 | 6.10E-01 | 0.2 | 2.03E-01 | Small ribosomal subunit protein eS17w |
| AT1G15500 | 0.6 | 2.94E-05 | 0.4 | 6.11E-04 | 0.2 | 7.30E-02 | 0.1 | 7.06E-01 | 0.0 | 8.32E-01 | 0.1 | 6.49E-01 | ADP,ATP carrier protein 2, chloroplastic |
| AT3G16420 | 0.6 | 1.00E-04 | 0.8 | 1.28E-05 | -0.3 | 2.04E-02 | 0.0 | 7.62E-01 | 0.2 | 2.42E-01 | -0.1 | 5.25E-01 | PYK10-binding protein 1 |
| AT3G10080 | 0.6 | 4.71E-03 | 0.5 | 1.12E-02 | 0.1 | 7.81E-01 | 0.0 | 9.78E-01 | 0.0 | 9.14E-01 | 0.0 | 9.36E-01 | Germin-like protein |
| AT1G24360 | 0.6 | 1.38E-04 | 0.5 | 6.87E-04 | 0.1 | 4.53E-01 | 0.4 | 2.07E-03 | 0.2 | 1.65E-01 | 0.2 | 2.21E-01 | 3-oxoacyl- reductase, chloroplastic |
| AT5G56260 | 0.6 | 1.45E-04 | 0.2 | 1.13E-01 | 0.4 | 4.24E-03 | 0.2 | 1.32E-01 | 0.0 | 8.18E-01 | 0.2 | 3.71E-01 | 4-hydroxy-4-methyl-2-oxoglutarate aldolase |
| AT5G62470 | 0.6 | 6.59E-03 | -0.1 | 5.66E-01 | 0.7 | 7.51E-03 | -0.2 | 4.19E-01 | 0.0 | 9.62E-01 | -0.2 | 4.91E-01 | Transcription factor MYB96 |
| AT2G26800 | 0.6 | 1.12E-05 | 0.0 | 8.81E-01 | 0.5 | 9.04E-05 | -0.1 | 5.57E-01 | -0.1 | 6.37E-01 | 0.0 | 9.90E-01 | hydroxymethylglutaryl-CoA lyase |
| AT2G23820 | 0.6 | 6.69E-04 | 0.6 | 7.15E-04 | 0.0 | 8.49E-01 | 0.0 | 8.55E-01 | 0.0 | 9.24E-01 | -0.1 | 8.33E-01 | 5'-deoxynucleotidase |
| AT5G05000 | 0.6 | 2.46E-05 | 0.5 | 1.32E-04 | 0.1 | 3.72E-01 | 0.2 | 1.21E-01 | 0.1 | 7.17E-01 | 0.2 | 4.83E-01 | Translocase of chloroplast 34, chloroplastic |
| AT3G54220 | 0.6 | 2.68E-05 | 0.3 | 3.28E-03 | 0.3 | 1.08E-02 | -0.2 | 1.10E-01 | -0.2 | 2.97E-01 | -0.1 | 8.29E-01 | Protein SCARECROW |
| AT4G01026 | 0.6 | 8.28E-03 | 0.4 | 5.13E-02 | 0.2 | 4.63E-01 | 0.4 | 3.75E-01 | 0.0 | 9.96E-01 | 0.4 | 4.39E-01 | Abscisic acid receptor PYL7 |
| AT3G07650 | 0.6 | 2.20E-03 | 0.6 | 2.03E-03 | 0.0 | 8.24E-01 | -0.4 | 1.83E-03 | -0.7 | 2.25E-04 | 0.2 | 1.78E-01 | Zinc finger protein CONSTANS-LIKE 9 |
| AT3G53730 | 0.6 | 2.67E-04 | 0.6 | 1.94E-04 | -0.1 | 5.37E-01 | -0.1 | 7.40E-01 | -0.1 | 5.97E-01 | 0.1 | 8.78E-01 | Histone H4 |
| AT4G23140 | 0.6 | 1.93E-01 | -0.5 | 2.99E-01 | 1.0 | 5.48E-02 | 0.1 | 6.66E-01 | -0.2 | 6.53E-01 | 0.3 | 3.99E-01 | Cysteine-rich receptor-like protein kinase 6 |
| AT2G17820 | 0.6 | 2.88E-04 | 0.4 | 6.54E-03 | 0.2 | 1.14E-01 | 0.2 | 2.33E-01 | -0.1 | 4.77E-01 | 0.3 | 1.20E-01 | Histidine kinase 1 |
| AT1G78140 | 0.6 | 5.21E-05 | 0.3 | 3.68E-03 | 0.2 | 2.25E-02 | -0.4 | 6.05E-03 | -0.2 | 1.78E-01 | -0.2 | 3.01E-01 | Uncharacterized methyltransferase At1g78140, chloroplastic |
| AT2G35130 | 0.6 | 1.26E-03 | 0.3 | 3.39E-02 | 0.2 | 1.35E-01 | 0.0 | 8.71E-01 | 0.1 | 7.58E-01 | -0.1 | 6.47E-01 | Tetratricopeptide repeat -like superfamily protein |
| AT5G25475 | 0.6 | 3.64E-05 | 0.5 | 1.63E-04 | 0.1 | 5.03E-01 | -0.2 | 3.13E-01 | -0.1 | 5.95E-01 | -0.1 | 8.36E-01 | AP2 |
| AT5G60600 | 0.6 | 3.48E-05 | 0.4 | 7.26E-04 | 0.2 | 7.57E-02 | 0.2 | 1.04E-01 | 0.1 | 4.46E-01 | 0.1 | 6.79E-01 | 4-hydroxy-3-methylbut-2-enyl diphosphate synthase |
| AT1G68400 | 0.6 | 1.76E-04 | 0.7 | 7.89E-05 | -0.1 | 3.11E-01 | -0.1 | 3.25E-01 | -0.3 | 2.65E-02 | 0.2 | 2.59E-01 | Probable leucine-rich repeat receptor-like protein kinase At1g68400 |
| AT1G65070 | 0.6 | 7.87E-04 | 0.1 | 3.01E-01 | 0.4 | 1.28E-02 | 0.0 | 9.96E-01 | 0.3 | 3.06E-01 | -0.3 | 3.32E-01 | DNA mismatch repair protein MutS, type 2 |
| AT2G42740 | 0.6 | 1.31E-03 | 0.8 | 2.26E-04 | -0.2 | 1.79E-01 | 0.6 | 5.11E-03 | 0.0 | 9.92E-01 | 0.6 | 2.33E-02 | Large ribosomal subunit protein uL5z |
| AT5G02960 | 0.6 | 4.79E-03 | 0.7 | 1.70E-03 | -0.2 | 4.32E-01 | 0.2 | 2.52E-01 | 0.0 | 9.01E-01 | 0.2 | 5.06E-01 | Small ribosomal subunit protein uS12y |
| AT1G06790 | 0.6 | 3.12E-04 | 0.4 | 2.28E-03 | 0.1 | 3.69E-01 | 0.1 | 6.38E-01 | -0.3 | 4.31E-01 | 0.4 | 1.87E-01 | DNA-directed RNA polymerase subunit |
| AT2G16700 | 0.6 | 4.54E-04 | 0.1 | 2.89E-01 | 0.4 | 8.07E-03 | -0.1 | 8.22E-01 | -0.3 | 4.04E-01 | 0.2 | 6.28E-01 | Actin depolymerizing factor 5 |
| AT3G17590 | 0.6 | 8.19E-04 | 0.1 | 4.07E-01 | 0.4 | 1.03E-02 | -0.4 | 1.61E-02 | -0.3 | 1.19E-01 | -0.1 | 6.28E-01 | Transcription regulatory protein SNF5, putative |
| AT2G29090 | 0.6 | 1.62E-02 | -0.3 | 2.02E-01 | 0.9 | 7.46E-03 | | | | | | | Cytochrome P450, family 707, subfamily A, polypeptide 2 |
| AT1G22650 | 0.6 | 2.50E-03 | 0.3 | 4.94E-02 | 0.2 | 1.87E-01 | -0.4 | 1.05E-01 | -0.2 | 3.22E-01 | -0.1 | 7.45E-01 | Alkaline |
| AT5G39710 | 0.6 | 4.87E-05 | 0.1 | 4.15E-01 | 0.5 | 8.96E-04 | 0.5 | 3.83E-02 | 0.1 | 8.82E-01 | 0.4 | 1.58E-01 | Pentatricopeptide repeat-containing protein At5g39710 |
| AT1G73230 | 0.6 | 6.32E-04 | 0.7 | 1.84E-04 | -0.2 | 2.18E-01 | 0.3 | 3.07E-02 | 0.1 | 4.07E-01 | 0.2 | 4.53E-01 | Nascent polypeptide-associated complex subunit beta |
| AT1G27120 | 0.6 | 1.39E-03 | 0.3 | 4.59E-02 | 0.3 | 1.17E-01 | -0.3 | 2.19E-01 | 0.1 | 8.59E-01 | -0.4 | 2.56E-01 | Galactosyltransferase family protein |
| AT1G63660 | 0.6 | 9.64E-04 | 0.3 | 7.34E-02 | 0.3 | 4.56E-02 | 0.4 | 1.14E-02 | 0.1 | 6.20E-01 | 0.3 | 1.38E-01 | GMP synthase |
| AT1G15160 | 0.6 | 2.42E-03 | 0.4 | 2.90E-02 | 0.2 | 2.95E-01 | 0.2 | 5.90E-01 | 0.4 | 3.60E-01 | -0.2 | 7.03E-01 | Protein DETOXIFICATION 11 |
| AT2G38270 | 0.6 | 1.31E-04 | 0.6 | 1.43E-04 | 0.0 | 7.56E-01 | 0.3 | 6.77E-02 | 0.0 | 9.40E-01 | 0.3 | 1.91E-01 | Bifunctional monothiol glutaredoxin-S16, chloroplastic |
| AT1G80870 | 0.6 | 2.51E-05 | 0.4 | 7.30E-04 | 0.2 | 5.56E-02 | -0.2 | 4.58E-01 | 0.2 | 4.07E-01 | -0.3 | 1.26E-01 | Putative receptor-like protein kinase At1g80870 |
| AT1G06650 | 0.6 | 8.81E-06 | 0.1 | 2.00E-01 | 0.5 | 1.92E-04 | -0.2 | 1.76E-01 | -0.3 | 2.65E-02 | 0.2 | 4.53E-01 | 1-aminocyclopropane-1-carboxylate oxidase homolog 3 |
| AT5G02790 | 0.6 | 4.67E-05 | 0.3 | 1.20E-02 | 0.3 | 6.35E-03 | 0.0 | 8.28E-01 | 0.1 | 4.24E-01 | -0.1 | 6.55E-01 | GST N-terminal domain-containing protein |
| AT2G26440 | 0.6 | 1.14E-03 | -0.1 | 3.95E-01 | 0.7 | 1.15E-03 | 0.7 | 3.11E-04 | -0.1 | 5.01E-01 | 0.9 | 1.75E-03 | Probable pectinesterase |
| AT3G44620 | 0.6 | 2.96E-05 | 0.4 | 2.71E-04 | 0.1 | 1.91E-01 | 0.3 | 7.20E-02 | 0.3 | 6.05E-02 | 0.0 | 8.63E-01 | acid phosphatase |
| AT5G20380 | 0.6 | 1.68E-04 | 0.2 | 9.85E-02 | 0.4 | 6.13E-03 | 0.0 | 9.60E-01 | -0.1 | 4.96E-01 | 0.1 | 5.90E-01 | Phosphate transporter 45 |
| AT1G18080 | 0.6 | 1.57E-03 | 0.6 | 1.10E-03 | -0.1 | 6.54E-01 | 0.2 | 3.11E-02 | 0.1 | 6.16E-01 | 0.2 | 2.71E-01 | Small ribosomal subunit protein RACK1z |
| AT3G51270 | 0.6 | 4.14E-05 | 0.4 | 6.92E-04 | 0.2 | 9.79E-02 | 0.0 | 7.63E-01 | -0.1 | 6.90E-01 | 0.1 | 5.55E-01 | non-specific serine |
| AT2G44620 | 0.6 | 5.40E-04 | 0.6 | 6.20E-04 | 0.0 | 8.47E-01 | 0.0 | 8.77E-01 | 0.0 | 9.55E-01 | 0.0 | 9.61E-01 | Acyl carrier protein 1, mitochondrial |
| AT1G57720 | 0.6 | 1.38E-04 | 0.5 | 7.23E-04 | 0.1 | 4.37E-01 | 0.1 | 3.31E-01 | -0.1 | 7.35E-01 | 0.2 | 2.78E-01 | Uncharacterized protein At1g09640 |
| AT4G36530 | 0.6 | 2.12E-04 | 0.4 | 2.52E-03 | 0.2 | 1.92E-01 | 0.0 | 8.90E-01 | 0.1 | 5.68E-01 | -0.1 | 5.12E-01 | Alpha |
| AT5G16930 | 0.6 | 2.88E-03 | 0.1 | 6.70E-01 | 0.5 | 1.92E-02 | 0.8 | 2.63E-02 | 0.2 | 6.86E-01 | 0.6 | 1.55E-01 | AAA+ ATPase domain-containing protein |
| AT1G69370 | 0.6 | 1.52E-04 | 0.6 | 2.15E-04 | 0.0 | 9.10E-01 | 0.1 | 7.64E-01 | 0.2 | 4.00E-01 | -0.2 | 6.54E-01 | Chorismate mutase 3, chloroplastic |
| AT5G47040 | 0.6 | 2.44E-04 | 0.4 | 5.59E-03 | 0.2 | 1.04E-01 | -0.5 | 9.43E-03 | -0.3 | 8.91E-02 | -0.1 | 5.63E-01 | LON2 |
| AT4G00370 | 0.6 | 1.57E-04 | 0.4 | 5.02E-03 | 0.2 | 6.77E-02 | 0.5 | 1.28E-02 | 0.4 | 4.19E-02 | 0.1 | 8.62E-01 | Ascorbate transporter, chloroplastic |
| AT1G80050 | 0.6 | 6.97E-05 | 0.5 | 6.04E-04 | 0.1 | 2.60E-01 | -0.1 | 6.80E-01 | 0.0 | 9.30E-01 | -0.1 | 8.37E-01 | adenine phosphoribosyltransferase |
| AT5G09380 | 0.6 | 6.95E-03 | 0.1 | 5.81E-01 | 0.5 | 5.00E-02 | 0.0 | 9.23E-01 | -0.1 | 8.10E-01 | 0.1 | 9.02E-01 | RNA polymerase III RPC4 |
| AT5G35970 | 0.6 | 1.84E-03 | -0.2 | 2.91E-01 | 0.7 | 1.19E-03 | -0.4 | 2.03E-01 | -0.1 | 8.07E-01 | -0.3 | 5.11E-01 | DNA helicase-like |
| AT5G49480 | 0.6 | 1.89E-03 | 0.2 | 2.55E-01 | 0.4 | 2.70E-02 | 1.4 | 8.80E-06 | 0.7 | 6.24E-03 | 0.6 | 3.52E-02 | Calcium-binding protein CP1 |
| AT3G48720 | 0.6 | 1.25E-03 | 1.1 | 2.32E-05 | -0.5 | 5.50E-03 | 0.1 | 6.89E-01 | -0.2 | 1.80E-01 | 0.3 | 1.71E-01 | HXXXD-type acyl-transferase family protein |
| AT5G24160 | 0.6 | 1.50E-02 | 0.3 | 2.63E-01 | 0.3 | 1.91E-01 | 0.1 | 4.71E-01 | 0.2 | 3.11E-01 | -0.1 | 7.90E-01 | Squalene monooxygenase |
| AT2G17970 | 0.6 | 1.87E-03 | 0.1 | 4.35E-01 | 0.4 | 2.19E-02 | 0.0 | 8.87E-01 | 0.3 | 4.19E-01 | -0.2 | 5.85E-01 | 2-oxoglutarate and Fe-dependent oxygenase superfamily protein |
| AT1G62390 | 0.6 | 1.41E-05 | 0.3 | 3.56E-03 | 0.3 | 3.60E-03 | 0.1 | 3.99E-01 | -0.1 | 5.77E-01 | 0.2 | 2.46E-01 | Protein CLMP1 |
| AT2G01320 | 0.6 | 9.06E-05 | 0.1 | 2.53E-01 | 0.5 | 1.62E-03 | -0.2 | 2.95E-01 | -0.1 | 5.95E-01 | -0.1 | 8.07E-01 | ABC-2 type transporter family protein |
| AT5G38480 | 0.6 | 1.11E-05 | 0.4 | 1.69E-04 | 0.1 | 8.18E-02 | 0.2 | 1.46E-01 | 0.2 | 3.26E-01 | 0.0 | 8.78E-01 | 14-3-3-like protein GF14 psi |
| AT1G35720 | 0.6 | 9.55E-05 | 1.1 | 1.98E-06 | -0.6 | 5.10E-04 | 0.2 | 2.62E-01 | 0.5 | 2.37E-03 | -0.4 | 2.96E-02 | Annexin D1 |
| AT1G12410 | 0.6 | 5.21E-06 | 0.4 | 2.67E-04 | 0.2 | 1.06E-02 | -0.2 | 6.25E-02 | -0.1 | 2.25E-01 | 0.0 | 7.98E-01 | ATP-dependent Clp protease proteolytic subunit-related protein 2, chloroplastic |
| AT4G03205 | 0.6 | 5.42E-04 | 0.4 | 5.12E-03 | 0.1 | 3.03E-01 | -0.3 | 8.74E-02 | -0.2 | 3.48E-01 | -0.1 | 6.73E-01 | coproporphyrinogen oxidase |
| AT5G15520 | 0.6 | 1.85E-03 | 0.7 | 5.03E-04 | -0.2 | 3.23E-01 | 0.3 | 1.88E-01 | 0.3 | 3.16E-01 | 0.0 | 9.43E-01 | Small ribosomal subunit protein eS19y |
| AT5G52860 | 0.6 | 8.37E-04 | 0.3 | 3.14E-02 | 0.3 | 1.07E-01 | 0.5 | 7.67E-02 | -0.2 | 7.00E-01 | 0.7 | 5.03E-02 | ABC transporter G family member 8 |
| AT2G19590 | 0.6 | 1.79E-04 | 0.5 | 1.08E-03 | 0.1 | 3.97E-01 | 0.9 | 3.20E-03 | 0.7 | 3.81E-02 | 0.2 | 3.56E-01 | hypothetical protein |
| AT3G07040 | 0.6 | 2.45E-04 | 0.0 | 9.68E-01 | 0.6 | 1.54E-03 | 0.2 | 5.69E-01 | 0.2 | 6.27E-01 | 0.0 | 9.99E-01 | Disease resistance protein RPM1 |
| AT3G55660 | 0.6 | 7.27E-05 | 0.7 | 1.96E-05 | -0.2 | 1.10E-01 | 0.2 | 3.16E-01 | 0.3 | 1.50E-01 | -0.1 | 6.96E-01 | ROP guanine nucleotide exchange factor 6 |
| AT3G51910 | 0.6 | 2.53E-03 | 1.0 | 6.43E-05 | -0.4 | 1.88E-02 | -0.2 | 4.96E-01 | -0.8 | 1.19E-02 | 0.6 | 1.12E-01 | At3g51910 |

| | | | | | | | | | | | | | |
|---|---|---|---|---|---|---|---|---|---|---|---|---|---|
| AT4G01100 | 0.6 | 1.92E-05 | 0.5 | 1.01E-04 | 0.1 | 3.86E-01 | 0.2 | 1.05E-01 | -0.1 | 6.13E-01 | 0.3 | 9.17E-02 | Adenine nucleotide transporter 1 |
| AT2G31610 | 0.6 | 3.09E-04 | 0.6 | 2.57E-04 | -0.1 | 6.13E-01 | 0.3 | 1.12E-02 | 0.2 | 1.58E-01 | 0.1 | 5.09E-01 | Small ribosomal subunit protein uS3z |
| AT2G37420 | 0.6 | 1.22E-04 | 0.3 | 1.97E-02 | 0.3 | 2.04E-02 | 0.7 | 1.79E-02 | 0.5 | 2.02E-01 | 0.3 | 4.01E-01 | Kinesin-like protein KIN-5B |
| AT2G14440 | 0.6 | 3.02E-03 | 0.0 | 8.37E-01 | 0.5 | 1.81E-02 | 0.6 | 2.29E-01 | 0.7 | 1.61E-01 | -0.1 | 7.76E-01 | Leucine-rich repeat protein kinase family protein |
| AT5G40380 | 0.6 | 9.47E-05 | 0.1 | 5.24E-01 | 0.5 | 1.26E-03 | -0.4 | 1.99E-01 | -0.3 | 3.28E-01 | -0.1 | 9.25E-01 | Cysteine-rich RLK 42 |
| AT3G29320 | 0.6 | 6.20E-04 | 0.4 | 7.94E-03 | 0.2 | 2.10E-01 | -0.3 | 1.75E-01 | -0.1 | 6.76E-01 | -0.1 | 6.16E-01 | Alpha-glucan phosphorylase 1 |
| AT4G04490 | 0.6 | 2.46E-02 | -0.8 | 1.54E-02 | 1.4 | 1.54E-03 | 0.0 | 9.31E-01 | -0.6 | 2.07E-01 | 0.6 | 3.26E-01 | Cysteine-rich receptor-like protein kinase 36 |
| AT4G19500 | 0.6 | 1.03E-03 | 0.2 | 1.55E-01 | 0.4 | 2.63E-02 | 0.2 | 1.62E-01 | 0.0 | 9.95E-01 | 0.2 | 2.46E-01 | Nucleoside-triphosphatase |
| AT3G13070 | 0.6 | 1.04E-04 | 0.4 | 9.80E-04 | 0.1 | 2.21E-01 | 0.2 | 1.65E-01 | 0.0 | 8.73E-01 | 0.1 | 4.39E-01 | Putative DUF21 domain-containing protein At3g13070, chloroplastic |
| AT3G48590 | 0.6 | 3.56E-04 | 0.6 | 6.98E-04 | 0.0 | 8.83E-01 | -0.2 | 1.77E-01 | -0.3 | 6.23E-02 | 0.1 | 6.39E-01 | Nuclear transcription factor Y subunit C-1 |
| AT3G56840 | 0.6 | 2.61E-04 | 0.2 | 1.53E-01 | 0.4 | 8.65E-03 | 0.0 | 9.29E-01 | 0.0 | 8.76E-01 | 0.0 | 9.60E-01 | FAD-dependent oxidoreductase family protein |
| AT2G24820 | 0.6 | 1.11E-04 | 0.0 | 9.13E-01 | 0.6 | 4.38E-04 | -0.2 | 6.09E-01 | 0.1 | 8.11E-01 | -0.4 | 4.91E-01 | hypothetical protein |
| AT1G11820 | 0.6 | 3.90E-04 | 0.5 | 2.07E-03 | 0.1 | 4.36E-01 | 0.2 | 4.00E-01 | 0.0 | 9.60E-01 | 0.2 | 5.23E-01 | glucan endo-1,3-beta-D-glucosidase |
| AT3G16565 | 0.6 | 5.02E-04 | 0.1 | 4.77E-01 | 0.5 | 6.33E-03 | 0.2 | 3.57E-01 | 0.0 | 8.76E-01 | 0.1 | 6.25E-01 | Threonyl and alanyl tRNA synthetase second additional domain-containing protein |
| AT5G53460 | 0.6 | 2.56E-04 | 0.9 | 3.50E-05 | -0.3 | 4.51E-02 | 0.7 | 1.27E-05 | 0.3 | 1.87E-02 | 0.4 | 3.12E-02 | Glutamate synthase 1 , chloroplastic |
| AT1G51310 | 0.6 | 5.75E-05 | 0.3 | 7.06E-03 | 0.3 | 1.85E-02 | 0.2 | 4.07E-01 | -0.1 | 8.20E-01 | 0.2 | 3.20E-01 | tRNA-5-taurinomethyluridine 2-sulfurtransferase |
| AT3G61580 | 0.6 | 2.21E-05 | 0.4 | 8.45E-04 | 0.2 | 3.07E-02 | -0.3 | 3.10E-02 | -0.3 | 6.79E-02 | 0.0 | 9.54E-01 | Delta-fatty-acid desaturase 1 -sphingolipid desaturase 1) -desaturase 1) |
| AT4G36250 | 0.6 | 4.43E-05 | 0.3 | 5.40E-03 | 0.3 | 1.20E-02 | 0.1 | 3.26E-01 | 0.3 | 9.30E-02 | -0.1 | 5.71E-01 | Aldehyde dehydrogenase family 3 member F1 |
| AT1G08840 | 0.6 | 6.27E-04 | 0.1 | 2.95E-01 | 0.4 | 1.24E-02 | 0.1 | 7.36E-01 | -0.1 | 8.12E-01 | 0.2 | 5.12E-01 | DNA replication ATP-dependent helicase |
| AT1G47530 | 0.6 | 1.27E-05 | 0.3 | 1.06E-03 | 0.3 | 9.54E-03 | 0.1 | 7.30E-01 | -0.3 | 4.42E-02 | 0.4 | 6.17E-02 | Protein DETOXIFICATION |
| AT1G18570 | 0.6 | 1.53E-02 | 0.1 | 6.51E-01 | 0.5 | 7.05E-02 | 0.7 | 1.73E-03 | -0.3 | 2.69E-01 | 1.0 | 2.92E-03 | Transcription factor MYB51 |
| AT3G47370 | 0.6 | 2.18E-03 | 0.9 | 2.87E-04 | -0.3 | 1.19E-01 | 0.5 | 6.04E-03 | 0.3 | 1.61E-01 | 0.2 | 3.72E-01 | Small ribosomal subunit protein uS10y |
| AT1G07610 | 0.6 | 5.80E-03 | 0.1 | 6.87E-01 | 0.5 | 2.94E-02 | -0.1 | 7.67E-01 | -0.8 | 2.28E-02 | 0.7 | 7.55E-02 | Metallothionein-like protein 1C |
| AT2G02450 | 0.6 | 7.07E-04 | 0.3 | 3.10E-02 | 0.3 | 8.02E-02 | 0.3 | 1.44E-01 | 0.2 | 3.91E-01 | 0.1 | 7.27E-01 | NAC domain-containing protein 35 |
| AT5G64110 | 0.6 | 4.97E-02 | 0.5 | 9.24E-02 | 0.1 | 8.18E-01 | 0.0 | 9.91E-01 | -0.8 | 1.57E-02 | 0.7 | 3.59E-02 | Peroxidase 70 |
| AT4G23260 | 0.6 | 8.15E-03 | -0.1 | 5.45E-01 | 0.7 | 7.64E-03 | 0.4 | 1.55E-01 | -0.2 | 6.94E-01 | 0.6 | 6.40E-02 | Cysteine-rich RLK 18 |
| AT5G65685 | 0.6 | 1.52E-03 | 0.2 | 1.51E-01 | 0.4 | 4.49E-02 | 0.0 | 9.79E-01 | 0.1 | 5.31E-01 | -0.1 | 6.24E-01 | At5g65685 |
| AT2G29450 | 0.6 | 1.55E-03 | 0.4 | 3.03E-02 | 0.2 | 1.55E-01 | -0.2 | 4.91E-01 | 0.2 | 4.08E-01 | -0.4 | 2.25E-01 | Glutathione S-transferase U5 |
| AT2G24180 | 0.6 | 1.25E-05 | 0.1 | 1.93E-01 | 0.5 | 2.51E-04 | 0.4 | 8.84E-04 | -0.1 | 5.57E-01 | 0.5 | 3.42E-03 | Cytochrome P450 71B6 |
| AT3G10270 | 0.6 | 8.35E-06 | 0.3 | 9.23E-04 | 0.3 | 6.38E-03 | 0.4 | 1.77E-01 | 0.2 | 6.95E-01 | 0.2 | 5.39E-01 | DNA gyrase subunit B |
| AT2G29650 | 0.6 | 1.08E-04 | 0.2 | 9.40E-02 | 0.4 | 3.44E-03 | 0.4 | 2.14E-01 | 0.3 | 4.61E-01 | 0.1 | 8.15E-01 | Phosphate transporter 41 |
| AT1G69620 | 0.6 | 2.00E-03 | 0.7 | 9.28E-04 | -0.1 | 4.77E-01 | 0.1 | 6.09E-01 | 0.0 | 9.35E-01 | 0.1 | 8.07E-01 | Large ribosomal subunit protein eL34y |
| AT4G29590 | 0.6 | 9.71E-05 | 0.5 | 5.40E-04 | 0.1 | 3.81E-01 | 0.3 | 8.18E-02 | 0.5 | 1.89E-02 | -0.2 | 5.40E-01 | AT4g29590 |
| AT1G14310 | 0.6 | 1.85E-04 | 0.4 | 4.53E-03 | 0.2 | 1.18E-01 | 0.0 | 9.71E-01 | -0.3 | 2.57E-01 | 0.3 | 2.68E-01 | F14L17.7 protein superfamily protein) |
| AT2G39010 | 0.6 | 3.62E-05 | 0.5 | 2.76E-04 | 0.1 | 2.63E-01 | -1.1 | 9.98E-07 | -0.6 | 8.69E-04 | -0.5 | 3.04E-02 | Plasma membrane intrinsic protein 2E |
| AT3G49640 | 0.6 | 7.71E-04 | 0.3 | 5.23E-02 | 0.3 | 6.50E-02 | 0.1 | 6.67E-01 | 0.0 | 9.98E-01 | 0.1 | 7.71E-01 | Uncharacterized protein T9C5.230 |
| AT3G23830 | 0.6 | 3.20E-03 | 0.7 | 1.15E-03 | -0.1 | 4.57E-01 | 0.5 | 1.68E-02 | 0.2 | 4.31E-01 | 0.3 | 2.53E-01 | Glycine-rich RNA-binding protein 4, mitochondrial |
| AT5G45775 | 0.6 | 1.03E-03 | 0.6 | 1.47E-03 | 0.0 | 9.73E-01 | 0.3 | 5.15E-02 | 0.1 | 5.68E-01 | 0.2 | 3.69E-01 | Large ribosomal subunit protein uL5y |
| AT1G54440 | 0.6 | 4.95E-05 | 0.1 | 2.07E-01 | 0.5 | 1.26E-03 | 0.3 | 1.18E-01 | 0.2 | 3.11E-01 | 0.1 | 7.95E-01 | Polynucleotidyl transferase, ribonuclease H fold protein with HRDC domain-containing protein |
| AT1G66940 | 0.6 | 3.69E-03 | 0.8 | 6.70E-04 | -0.2 | 2.40E-01 | 0.5 | 4.02E-03 | -0.1 | 6.99E-01 | 0.6 | 3.98E-03 | At1g66930 |
| AT1G05970 | 0.6 | 6.41E-05 | 0.3 | 5.33E-03 | 0.3 | 3.07E-02 | -0.4 | 6.72E-02 | -0.4 | 4.35E-02 | 0.1 | 8.19E-01 | ASI1-immunoprecipitated protein 1 |
| AT5G49650 | 0.6 | 1.80E-05 | 0.6 | 5.11E-05 | 0.0 | 7.17E-01 | -0.3 | 5.41E-02 | -0.2 | 2.44E-01 | -0.1 | 7.20E-01 | Xylulose kinase |
| AT1G24610 | 0.6 | 9.21E-05 | 0.4 | 1.21E-03 | 0.2 | 1.66E-01 | -0.1 | 3.20E-01 | 0.0 | 9.12E-01 | -0.2 | 4.25E-01 | SET domain-containing protein |
| AT3G06740 | 0.6 | 4.73E-04 | 0.5 | 1.87E-03 | 0.1 | 5.60E-01 | 0.3 | 2.43E-01 | 0.2 | 4.89E-01 | 0.1 | 8.39E-01 | GATA transcription factor 15 |
| AT4G32700 | 0.6 | 2.64E-04 | 0.0 | 9.57E-01 | 0.6 | 1.58E-03 | -0.3 | 2.97E-01 | 0.0 | 9.37E-01 | -0.3 | 3.13E-01 | Helicase and polymerase-containing protein TEBICHI |
| AT3G18130 | 0.6 | 2.58E-03 | 0.7 | 1.24E-03 | -0.1 | 5.17E-01 | 0.5 | 3.32E-03 | 0.4 | 1.56E-02 | 0.1 | 8.03E-01 | Small ribosomal subunit protein RACK1x |
| AT4G39880 | 0.6 | 1.26E-03 | 0.6 | 2.37E-03 | 0.0 | 8.66E-01 | 0.1 | 8.01E-01 | -0.1 | 8.65E-01 | 0.1 | 7.49E-01 | Uncharacterized protein |
| AT3G22680 | 0.6 | 2.18E-04 | 0.3 | 1.68E-02 | 0.3 | 4.30E-02 | -0.1 | 8.16E-01 | -0.3 | 2.63E-01 | 0.2 | 4.70E-01 | Protein RDM1 |
| AT4G38420 | 0.6 | 1.53E-03 | 0.7 | 7.44E-04 | -0.1 | 5.26E-01 | 0.3 | 2.11E-01 | -0.1 | 8.00E-01 | 0.4 | 1.20E-01 | SKU5 similar 9 |
| AT1G63970 | 0.6 | 6.88E-06 | 0.2 | 4.79E-03 | 0.4 | 1.09E-03 | 0.0 | 8.33E-01 | 0.2 | 3.58E-01 | -0.1 | 5.61E-01 | 2-C-methyl-D-erythritol 2,4-cyclodiphosphate synthase |
| AT4G02800 | 0.6 | 5.15E-04 | 0.8 | 1.02E-04 | -0.2 | 1.59E-01 | 0.4 | 1.11E-01 | 0.2 | 4.28E-01 | 0.1 | 6.74E-01 | At4g02800 |
| AT1G22570 | 0.6 | 7.71E-05 | 0.8 | 1.98E-05 | -0.2 | 9.68E-02 | 0.0 | 8.61E-01 | -0.4 | 1.27E-01 | 0.5 | 1.54E-01 | Major facilitator superfamily protein |
| AT1G63640 | 0.6 | 1.54E-05 | 0.3 | 5.02E-03 | 0.3 | 3.73E-03 | 0.2 | 4.17E-01 | 0.2 | 3.44E-01 | 0.0 | 8.94E-01 | P-loop nucleoside triphosphate hydrolases superfamily protein with CH domain-containing protein |
| AT5G21326 | 0.6 | 2.04E-05 | 0.2 | 2.53E-02 | 0.4 | 1.42E-03 | -0.1 | 6.25E-01 | -0.1 | 3.74E-01 | 0.1 | 7.66E-01 | CBL-interacting serine |
| AT3G62130 | 0.6 | 3.42E-06 | 0.3 | 1.08E-03 | 0.3 | 1.42E-03 | -0.2 | 1.39E-01 | -0.2 | 1.94E-01 | 0.0 | 1.00E+00 | L-cysteine desulfhydrase |
| AT4G20980 | 0.6 | 4.22E-05 | 0.2 | 2.10E-02 | 0.4 | 3.64E-03 | 0.4 | 3.07E-03 | 0.2 | 1.88E-01 | 0.2 | 2.49E-01 | Eukaryotic translation initiation factor 3 subunit D |
| AT2G45560 | 0.6 | 3.03E-04 | 0.5 | 2.28E-03 | 0.1 | 3.13E-01 | 0.0 | 9.18E-01 | 0.0 | 8.72E-01 | 0.0 | 9.56E-01 | Cytochrome P450, family 76, subfamily C, polypeptide 1 |
| AT3G21820 | 0.6 | 9.16E-05 | 0.4 | 2.75E-03 | 0.2 | 8.00E-02 | 0.4 | 9.79E-02 | 0.3 | 2.08E-01 | 0.1 | 8.81E-01 | Histone-lysine N-methyltransferase ATXR2 |
| AT1G14840 | 0.6 | 5.32E-04 | 0.3 | 3.32E-02 | 0.3 | 5.51E-02 | -0.1 | 4.22E-01 | 0.0 | 8.45E-01 | -0.1 | 7.41E-01 | Microtubule-associated proteins 70-4 |
| AT2G26980 | 0.6 | 3.47E-05 | 0.3 | 6.56E-03 | 0.3 | 9.81E-03 | -0.4 | 1.51E-02 | -0.9 | 2.27E-04 | 0.5 | 5.69E-02 | non-specific serine |
| AT1G09795 | 0.6 | 4.83E-05 | 0.5 | 5.41E-04 | 0.1 | 1.61E-01 | 0.1 | 6.10E-01 | 0.0 | 7.91E-01 | 0.0 | 9.32E-01 | ATP phosphoribosyltransferase 2, chloroplastic |
| AT1G02930 | 0.6 | 5.49E-02 | -0.2 | 5.89E-01 | 0.8 | 3.32E-02 | 0.7 | 9.38E-02 | -0.6 | 2.20E-01 | 1.3 | 2.87E-02 | Glutathione S-transferase F6 |
| AT1G52190 | 0.6 | 4.29E-03 | 1.1 | 1.46E-04 | -0.5 | 2.66E-02 | -0.6 | 4.35E-04 | -0.7 | 7.77E-04 | 0.0 | 8.96E-01 | Protein NRT1 |
| AT1G24020 | 0.6 | 9.12E-04 | 0.6 | 1.12E-03 | 0.0 | 9.12E-01 | 0.8 | 1.95E-03 | 0.7 | 8.75E-03 | 0.1 | 8.39E-01 | MLP-like protein 423 |
| AT1G18680 | 0.6 | 9.38E-04 | 0.2 | 2.16E-01 | 0.4 | 2.48E-02 | -0.3 | 1.10E-01 | -0.4 | 9.67E-02 | 0.0 | 9.01E-01 | F6A14.21 protein |
| AT5G22280 | 0.6 | 2.80E-04 | 0.4 | 8.58E-03 | 0.2 | 9.40E-02 | 0.2 | 3.49E-01 | 0.1 | 8.40E-01 | 0.2 | 6.82E-01 | Peptidyl-prolyl cis-trans isomerase G |
| AT3G10740 | 0.6 | 1.33E-03 | 0.6 | 1.46E-03 | 0.0 | 9.00E-01 | -0.3 | 1.46E-02 | -0.4 | 8.59E-03 | 0.1 | 7.50E-01 | non-reducing end alpha-L-arabinofuranosidase |
| AT1G66980 | 0.6 | 3.44E-04 | 0.2 | 1.66E-01 | 0.4 | 7.33E-03 | 0.4 | 6.49E-02 | 0.3 | 1.92E-01 | 0.1 | 8.40E-01 | glycerophosphodiester phosphodiesterase |
| AT4G16500 | 0.6 | 2.65E-04 | 0.8 | 8.86E-05 | -0.2 | 2.36E-01 | 0.1 | 6.44E-01 | -0.1 | 7.63E-01 | 0.2 | 5.33E-01 | Cysteine proteinase inhibitor 4 |
| AT4G15800 | 0.6 | 4.13E-03 | 0.1 | 5.15E-01 | 0.5 | 2.96E-02 | -0.4 | 6.13E-02 | -0.4 | 1.06E-01 | 0.0 | 9.80E-01 | Protein RALF-like 33 |
| AT5G44050 | 0.6 | 1.10E-03 | 0.4 | 1.56E-02 | 0.2 | 2.50E-01 | 0.4 | 3.80E-01 | 0.3 | 5.85E-01 | 0.1 | 9.01E-01 | Uncharacterized protein At5g44050 |
| AT1G12240 | 0.6 | 2.59E-05 | 0.6 | 3.20E-05 | 0.0 | 6.44E-01 | -0.4 | 3.54E-03 | 0.0 | 9.90E-01 | -0.4 | 2.66E-02 | beta-fructofuranosidase |
| AT4G26900 | 0.6 | 1.81E-05 | 0.4 | 7.35E-04 | 0.2 | 2.57E-02 | 0.2 | 4.32E-01 | 0.2 | 4.57E-01 | 0.0 | 9.66E-01 | Imidazole glycerol phosphate synthase hisHF, chloroplastic |
| AT5G23330 | 0.6 | 2.11E-05 | 0.2 | 6.96E-02 | 0.4 | 1.11E-03 | 0.0 | 8.52E-01 | 0.0 | 8.76E-01 | 0.0 | 1.00E+00 | FAD synthase |
| AT2G42280 | 0.6 | 1.90E-04 | 0.4 | 2.20E-03 | 0.2 | 2.27E-01 | -0.1 | 5.56E-01 | -0.3 | 2.09E-01 | 0.2 | 6.05E-01 | Basic helix-loop-helix DNA-binding superfamily protein |
| AT2G41940 | 0.6 | 1.06E-04 | 0.2 | 1.18E-01 | 0.4 | 3.63E-03 | -0.1 | 7.39E-01 | -0.1 | 7.81E-01 | 0.0 | 9.98E-01 | Zinc finger protein 8 |
| AT5G54170 | 0.6 | 6.57E-04 | 0.0 | 7.66E-01 | 0.6 | 3.92E-03 | -0.3 | 6.65E-02 | -0.5 | 6.39E-03 | 0.2 | 3.22E-01 | hypothetical protein |
| AT2G05990 | 0.6 | 3.32E-05 | 0.6 | 8.71E-05 | 0.0 | 8.24E-01 | 0.4 | 9.93E-04 | 0.2 | 4.51E-02 | 0.2 | 3.43E-01 | Enoyl- reductase , chloroplastic |
| AT1G03680 | 0.6 | 2.85E-05 | 0.8 | 9.38E-06 | -0.2 | 5.72E-02 | 0.0 | 8.71E-01 | 0.1 | 6.17E-01 | -0.1 | 7.80E-01 | Thioredoxin M1, chloroplastic |
| AT1G62630 | 0.6 | 2.21E-04 | 0.3 | 3.36E-02 | 0.3 | 2.29E-02 | 0.4 | 2.67E-02 | 0.2 | 3.41E-01 | 0.2 | 3.80E-01 | Probable disease resistance protein At1g62630 |
| AT3G03920 | 0.6 | 4.22E-04 | 0.6 | 7.51E-04 | 0.0 | 9.31E-01 | 0.4 | 1.14E-02 | 0.1 | 7.99E-01 | 0.4 | 8.76E-02 | Putative H |
| AT2G27960 | 0.6 | 2.03E-04 | 0.4 | 6.78E-03 | 0.2 | 7.70E-02 | 0.1 | 7.53E-01 | 0.0 | 9.38E-01 | 0.0 | 8.92E-01 | Cyclin-dependent kinases regulatory subunit 1 |
| AT1G77640 | 0.6 | 2.15E-01 | -0.5 | 3.72E-01 | 1.1 | 1.00E-01 | | | | | | | Ethylene-responsive transcription factor ERF013 |

| | | | | | | | | | | | | | |
|---|---|---|---|---|---|---|---|---|---|---|---|---|---|
| AT1G50840 | 0.6 | 4.23E-05 | 0.1 | 2.62E-01 | 0.5 | 7.33E-04 | 0.3 | 7.19E-02 | 0.0 | 9.97E-01 | 0.3 | 1.50E-01 | Polymerase gamma 2 |
| AT4G31850 | 0.6 | 1.22E-03 | 0.4 | 1.12E-02 | 0.2 | 3.13E-01 | 0.2 | 7.40E-01 | 0.4 | 4.93E-01 | -0.2 | 7.84E-01 | Pentatricopeptide repeat-containing protein At4g31850, chloroplastic |
| AT3G48930 | 0.6 | 1.70E-03 | 0.7 | 1.07E-03 | -0.1 | 6.04E-01 | 0.3 | 8.49E-02 | 0.1 | 5.97E-01 | 0.2 | 4.70E-01 | Small ribosomal subunit protein uS17z |
| AT4G08870 | 0.6 | 7.18E-03 | 0.3 | 1.76E-01 | 0.3 | 1.31E-01 | 1.0 | 5.10E-02 | 1.4 | 1.06E-02 | -0.5 | 5.01E-01 | AT4G08870 protein |
| AT5G64140 | 0.6 | 1.13E-03 | 0.8 | 2.46E-04 | -0.2 | 1.84E-01 | 0.4 | 5.63E-02 | 0.3 | 1.87E-01 | 0.1 | 8.22E-01 | Small ribosomal subunit protein eS28x |
| AT1G17840 | 0.6 | 5.43E-05 | 0.3 | 1.27E-02 | 0.3 | 7.34E-03 | -0.2 | 5.25E-01 | -0.2 | 6.09E-01 | 0.0 | 9.94E-01 | ABC transporter G family member 11 |
| AT2G01760 | 0.6 | 9.38E-04 | 0.5 | 5.94E-03 | 0.1 | 4.41E-01 | -0.7 | 2.58E-02 | -0.7 | 2.80E-02 | 0.0 | 9.36E-01 | Response regulator 14 |
| AT3G57750 | 0.6 | 8.22E-05 | 0.2 | 4.98E-02 | 0.4 | 6.21E-03 | 0.4 | 1.43E-01 | 0.1 | 7.16E-01 | 0.3 | 4.97E-01 | Non-functional pseudokinase ZED1 |
| AT5G21060 | 0.6 | 4.76E-05 | 0.5 | 4.86E-04 | 0.1 | 1.87E-01 | 0.1 | 5.42E-01 | 0.1 | 4.53E-01 | 0.0 | 9.00E-01 | Homoserine dehydrogenase |
| AT1G36180 | 0.6 | 1.30E-04 | -0.1 | 2.86E-01 | 0.7 | 3.10E-04 | -0.1 | 8.03E-01 | 0.2 | 7.15E-01 | -0.3 | 5.62E-01 | Acetyl-CoA carboxylase 2 |
| AT4G01330 | 0.6 | 1.39E-03 | 0.7 | 7.01E-04 | -0.1 | 5.35E-01 | -0.7 | 5.17E-04 | -0.8 | 5.07E-04 | 0.1 | 7.24E-01 | A_IG002N01.22 protein |
| AT1G70610 | 0.6 | 1.02E-03 | 0.2 | 2.84E-01 | 0.5 | 1.48E-02 | 0.0 | 9.53E-01 | 0.2 | 6.01E-01 | -0.2 | 5.96E-01 | Transporter associated with antigen processing protein 1 |
| AT5G23140 | 0.6 | 9.92E-06 | 0.4 | 1.98E-04 | 0.2 | 4.96E-02 | -0.1 | 3.87E-01 | -0.1 | 3.96E-01 | 0.0 | 9.59E-01 | ATP-dependent Clp protease proteolytic subunit 2, mitochondrial |
| AT3G44300 | 0.6 | 2.96E-02 | -1.0 | 9.53E-03 | 1.7 | 1.58E-03 | -1.8 | 1.57E-02 | -2.9 | 7.34E-03 | 1.0 | 5.20E-01 | Nitrilase 2 |
| AT1G49630 | 0.6 | 2.23E-05 | 0.3 | 1.19E-02 | 0.4 | 3.58E-03 | 0.5 | 2.53E-03 | 0.1 | 5.75E-01 | 0.4 | 4.02E-02 | Presequence protease 2 |
| AT1G75670 | 0.6 | 1.28E-03 | 0.3 | 3.95E-02 | 0.3 | 1.29E-01 | 0.7 | 9.65E-03 | 0.2 | 6.14E-01 | 0.6 | 7.79E-02 | DNA-directed RNA polymerase subunit |
| AT2G36230 | 0.6 | 2.85E-04 | 0.6 | 3.88E-04 | 0.0 | 9.25E-01 | 0.3 | 2.22E-01 | 0.1 | 6.38E-01 | 0.1 | 7.14E-01 | 1--5- imidazole-4-carboxamide isomerase, chloroplastic |
| AT2G42210 | 0.6 | 2.10E-05 | 0.6 | 2.90E-05 | 0.0 | 7.17E-01 | 0.0 | 8.82E-01 | 0.0 | 9.88E-01 | 0.0 | 9.14E-01 | Outer envelope pore protein 16-3, chloroplastic |
| AT4G32830 | 0.6 | 2.80E-05 | 0.4 | 7.04E-04 | 0.2 | 6.95E-02 | 0.3 | 1.50E-01 | 0.3 | 1.62E-01 | 0.0 | 9.48E-01 | Serine |
| AT4G04740 | 0.6 | 1.54E-03 | 0.0 | 9.98E-01 | 0.6 | 5.62E-03 | 0.0 | 9.06E-01 | 0.0 | 9.40E-01 | 0.0 | 9.86E-01 | Calcium-dependent protein kinase 23 |
| AT4G24660 | 0.6 | 6.26E-05 | 0.6 | 1.29E-04 | 0.0 | 9.30E-01 | -0.3 | 1.81E-01 | 0.0 | 9.60E-01 | -0.2 | 2.92E-01 | Homeobox protein 22 |
| AT2G47110 | 0.6 | 1.19E-03 | 0.6 | 2.12E-03 | 0.0 | 9.25E-01 | 0.2 | 3.63E-02 | 0.1 | 2.01E-01 | 0.1 | 7.34E-01 | Ubiquitin-ribosomal protein eS31y fusion protein |
| AT1G54000 | 0.6 | 3.36E-04 | 0.9 | 4.25E-05 | -0.3 | 4.54E-02 | 0.3 | 8.45E-02 | 0.3 | 4.64E-02 | -0.1 | 7.64E-01 | GDSL esterase |
| AT3G03460 | 0.6 | 1.74E-03 | 0.4 | 2.07E-02 | 0.2 | 2.93E-01 | 0.5 | 1.06E-01 | 0.4 | 2.73E-01 | 0.1 | 7.99E-01 | Mediator of RNA polymerase II transcription subunit-like protein |
| AT1G02730 | 0.6 | 2.48E-05 | 0.5 | 1.52E-04 | 0.1 | 3.22E-01 | 0.1 | 6.77E-01 | 0.1 | 7.59E-01 | 0.0 | 9.92E-01 | Cellulose synthase-like protein D5 |
| AT5G06340 | 0.6 | 6.98E-06 | 0.4 | 3.10E-04 | 0.2 | 1.47E-02 | -0.1 | 2.65E-01 | -0.2 | 2.05E-01 | 0.0 | 8.62E-01 | Nudix hydrolase 27, chloroplastic |
| AT3G16430 | 0.6 | 1.80E-05 | 0.9 | 2.19E-06 | -0.3 | 5.15E-03 | -0.2 | 2.69E-01 | 0.4 | 4.59E-02 | -0.6 | 6.16E-03 | PBPII |
| AT3G63130 | 0.6 | 9.68E-05 | 0.4 | 2.26E-03 | 0.2 | 8.75E-02 | 0.2 | 1.80E-01 | 0.2 | 2.57E-01 | 0.0 | 9.91E-01 | RAN GTPase-activating protein 1 |
| AT4G04850 | 0.6 | 5.58E-04 | 0.1 | 3.62E-01 | 0.5 | 5.96E-03 | 0.2 | 5.01E-01 | 0.2 | 3.75E-01 | -0.1 | 8.39E-01 | K efflux antiporter 3, chloroplastic |
| AT1G14880 | 0.6 | 3.82E-01 | 0.4 | 5.78E-01 | 0.2 | 8.14E-01 | 0.6 | 2.56E-01 | -0.5 | 4.36E-01 | 1.1 | 1.18E-01 | Protein PLANT CADMIUM RESISTANCE 1 |
| AT5G13960 | 0.6 | 1.69E-04 | 0.2 | 5.55E-02 | 0.4 | 1.23E-02 | 0.1 | 5.59E-01 | 0.0 | 9.85E-01 | 0.1 | 5.63E-01 | Histone-lysine N-methyltransferase, H3 lysine-9 specific SUVH4-like protein |
| AT1G33811 | 0.6 | 1.67E-03 | 0.9 | 2.78E-04 | -0.2 | 1.72E-01 | 0.1 | 3.00E-01 | 0.0 | 9.66E-01 | 0.1 | 4.48E-01 | GDSL esterase |
| AT4G34260 | 0.6 | 2.81E-06 | 0.5 | 1.86E-05 | 0.1 | 1.91E-01 | -0.3 | 1.74E-01 | -0.2 | 2.90E-01 | 0.0 | 9.44E-01 | Alpha-L-fucosidase 2 |
| AT2G04540 | 0.6 | 5.21E-05 | 0.3 | 9.41E-03 | 0.3 | 1.32E-02 | -0.3 | 1.02E-01 | -0.3 | 1.70E-01 | 0.0 | 9.44E-01 | beta-ketoacyl- synthase I |
| AT1G52980 | 0.6 | 1.69E-05 | 0.4 | 7.33E-04 | 0.2 | 2.27E-02 | 0.2 | 5.52E-02 | 0.0 | 7.65E-01 | 0.2 | 3.23E-01 | Nuclear |
| AT5G54130 | 0.6 | 2.22E-04 | 0.3 | 1.60E-02 | 0.3 | 4.54E-02 | -0.2 | 4.31E-01 | -0.3 | 1.51E-01 | 0.2 | 6.02E-01 | Endonuclease |
| AT5G55120 | 0.6 | 4.16E-05 | 0.5 | 4.01E-04 | 0.1 | 1.97E-01 | 0.2 | 4.14E-01 | 0.2 | 4.66E-01 | 0.0 | 9.90E-01 | GDP-L-galactose phosphorylase 2 |
| AT4G33270 | 0.6 | 1.29E-04 | 0.7 | 9.23E-05 | -0.1 | 5.04E-01 | 0.2 | 4.06E-01 | 0.1 | 8.03E-01 | 0.1 | 7.53E-01 | hypothetical protein |
| AT3G07110 | 0.6 | 1.87E-03 | 0.8 | 6.61E-04 | -0.2 | 3.59E-01 | 0.5 | 5.22E-04 | 0.2 | 1.23E-01 | 0.3 | 9.87E-02 | Ribosomal protein L13 family protein |
| AT1G14600 | 0.6 | 2.08E-02 | -0.3 | 3.71E-01 | 0.9 | 1.51E-02 | -0.2 | 7.52E-01 | 0.0 | 9.97E-01 | -0.2 | 8.31E-01 | Putative Myb family transcription factor At1g14600 |
| AT3G03990 | 0.6 | 3.20E-05 | 0.3 | 5.59E-03 | 0.3 | 7.45E-03 | -0.3 | 9.90E-02 | -0.5 | 8.75E-03 | 0.2 | 2.72E-01 | Strigolactone esterase D14 |
| AT3G19830 | 0.6 | 5.49E-05 | 0.3 | 9.70E-03 | 0.3 | 1.24E-02 | 0.0 | 9.78E-01 | 0.0 | 8.63E-01 | 0.0 | 9.05E-01 | Calcium-dependent lipid-binding family protein |
| AT3G21090 | 0.6 | 7.79E-04 | -0.1 | 4.39E-01 | 0.7 | 1.88E-03 | -0.4 | 2.65E-01 | -0.7 | 6.72E-02 | 0.3 | 4.54E-01 | ABC transporter G family member 15 |
| AT4G31840 | 0.6 | 8.71E-04 | 0.7 | 3.60E-04 | -0.1 | 4.15E-01 | 0.1 | 4.06E-01 | 0.2 | 3.39E-01 | 0.0 | 8.86E-01 | Early nodulin-like protein 15 |
| AT5G65070 | 0.6 | 1.31E-02 | 0.1 | 7.77E-01 | 0.5 | 5.73E-02 | 0.1 | 8.86E-01 | -0.1 | 8.45E-01 | 0.2 | 7.78E-01 | K-box region and MADS-box transcription factor family protein |
| AT5G45950 | 0.6 | 3.71E-04 | 0.8 | 8.40E-05 | -0.2 | 1.51E-01 | -0.4 | 2.07E-02 | 0.0 | 8.49E-01 | -0.5 | 4.40E-02 | GDSL esterase |
| AT1G44790 | 0.6 | 5.05E-05 | 0.5 | 7.81E-04 | 0.2 | 1.32E-01 | 0.2 | 3.42E-01 | 0.4 | 8.33E-02 | -0.2 | 4.98E-01 | Gamma-glutamylcyclotransferase |
| AT5G02470 | 0.6 | 5.08E-05 | 0.4 | 2.70E-03 | 0.2 | 3.91E-02 | 0.0 | 9.41E-01 | 0.0 | 9.22E-01 | 0.1 | 8.86E-01 | Transcription factor-like protein DPA |
| AT2G20180 | 0.6 | 1.19E-03 | 0.5 | 3.94E-03 | 0.1 | 6.58E-01 | -0.3 | 4.97E-01 | -0.3 | 5.49E-01 | 0.0 | 9.97E-01 | Phytochrome interacting factor 3-like 5 |
| AT1G20930 | 0.6 | 1.31E-04 | 0.5 | 8.28E-04 | 0.1 | 3.85E-01 | -0.1 | 6.13E-01 | 0.0 | 9.99E-01 | -0.1 | 6.86E-01 | Cyclin-dependent kinase B2-2 |
| AT1G27695 | 0.6 | 5.61E-04 | 0.4 | 6.77E-03 | 0.2 | 2.52E-01 | 0.1 | 7.81E-01 | 0.2 | 4.02E-01 | -0.2 | 6.66E-01 | Protein TRIGALACTOSYLDIACYLGLYCEROL 5, chloroplastic |
| AT2G37470 | 0.6 | 7.30E-04 | 0.6 | 9.77E-04 | 0.0 | 9.67E-01 | 0.0 | 9.86E-01 | -0.2 | 5.79E-01 | 0.2 | 6.68E-01 | Histone H2B.4 |
| AT5G46570 | 0.6 | 3.59E-05 | 0.5 | 1.81E-04 | 0.1 | 4.20E-01 | 0.4 | 3.39E-02 | 0.4 | 3.99E-02 | 0.0 | 9.48E-01 | Serine |
| AT1G06430 | 0.6 | 1.30E-04 | 0.0 | 8.54E-01 | 0.6 | 4.19E-04 | -0.3 | 6.68E-02 | -0.1 | 7.48E-01 | -0.2 | 3.05E-01 | T2D23.13 protein |
| AT2G38170 | 0.6 | 4.24E-04 | 0.0 | 9.15E-01 | 0.6 | 1.56E-03 | -0.8 | 4.55E-04 | -0.9 | 1.16E-03 | 0.0 | 9.71E-01 | Vacuolar cation |
| AT1G15140 | 0.6 | 2.86E-05 | 0.8 | 1.29E-05 | -0.2 | 1.27E-01 | 0.5 | 1.88E-04 | 0.3 | 1.17E-02 | 0.2 | 2.46E-01 | F9L1.8 protein -binding oxidoreductase) |
| AT1G54350 | 0.6 | 3.36E-04 | 0.3 | 2.64E-02 | 0.3 | 3.72E-02 | -0.2 | 4.54E-01 | 0.1 | 8.37E-01 | -0.3 | 4.60E-01 | ABCD2 |
| AT5G19830 | 0.6 | 8.58E-05 | 0.3 | 3.03E-02 | 0.4 | 1.08E-02 | 0.3 | 2.04E-01 | 0.0 | 9.95E-01 | 0.3 | 3.31E-01 | Peptidyl-tRNA hydrolase, mitochondrial |
| AT3G20970 | 0.6 | 2.51E-05 | 0.4 | 1.33E-03 | 0.2 | 2.64E-02 | 0.2 | 3.92E-01 | 0.0 | 9.84E-01 | 0.2 | 5.71E-01 | NFU domain protein 4 |
| AT5G61000 | 0.6 | 2.24E-05 | 0.5 | 2.90E-04 | 0.1 | 1.40E-01 | 0.5 | 1.49E-02 | 0.4 | 1.10E-01 | 0.1 | 5.93E-01 | Replication protein A 70 kDa DNA-binding subunit D |
| AT4G34570 | 0.6 | 2.22E-05 | 0.5 | 3.43E-04 | 0.2 | 1.12E-01 | 0.3 | 1.15E-01 | 0.2 | 5.55E-01 | 0.2 | 5.16E-01 | dihydrofolate reductase |
| AT5G59870 | 0.6 | 4.22E-04 | 0.8 | 1.44E-04 | -0.2 | 2.35E-01 | 0.2 | 1.50E-01 | 0.2 | 3.35E-01 | 0.0 | 8.73E-01 | Probable histone H2A.7 |
| AT3G29350 | 0.6 | 5.64E-04 | 0.4 | 1.19E-02 | 0.2 | 1.41E-01 | 0.0 | 9.44E-01 | -0.1 | 7.55E-01 | 0.1 | 8.41E-01 | Histidine-containing phosphotransfer protein 2 |
| AT2G45990 | 0.6 | 7.72E-05 | 0.4 | 3.00E-03 | 0.2 | 4.86E-02 | -0.1 | 3.14E-01 | -0.2 | 3.05E-01 | 0.0 | 9.33E-01 | AT2G45990 protein |
| AT5G16370 | 0.6 | 3.12E-04 | 0.5 | 2.03E-03 | 0.1 | 3.52E-01 | -0.3 | 1.53E-01 | -0.8 | 1.94E-03 | 0.5 | 8.82E-02 | Probable acyl-activating enzyme 5, peroxisomal |
| AT4G33010 | 0.6 | 4.94E-04 | 0.3 | 7.34E-02 | 0.4 | 2.20E-02 | 0.5 | 6.58E-02 | 0.5 | 8.99E-02 | 0.0 | 9.71E-01 | Glycine cleavage system P protein |
| AT3G06820 | 0.6 | 1.28E-04 | 0.1 | 2.67E-01 | 0.5 | 2.58E-03 | 0.1 | 6.83E-01 | 0.0 | 9.55E-01 | 0.1 | 8.13E-01 | Mov34 |
| AT5G44190 | 0.6 | 6.30E-05 | 0.5 | 3.10E-04 | 0.1 | 4.50E-01 | -0.2 | 3.14E-01 | -0.3 | 2.09E-01 | 0.1 | 7.96E-01 | GOLDEN2-like 2 |
| AT5G16130 | 0.6 | 1.62E-03 | 0.7 | 1.73E-03 | 0.0 | 8.57E-01 | 0.3 | 2.34E-02 | 0.2 | 3.12E-01 | 0.2 | 4.54E-01 | Small ribosomal subunit protein eS7x |
| AT5G24210 | 0.6 | 5.42E-04 | 0.1 | 3.64E-01 | 0.5 | 5.49E-03 | 0.8 | 5.72E-03 | -0.2 | 6.09E-01 | 1.0 | 1.32E-02 | Alpha |
| AT4G00100 | 0.6 | 1.10E-03 | 0.7 | 1.25E-03 | 0.0 | 8.66E-01 | 0.2 | 1.30E-01 | 0.0 | 9.69E-01 | 0.2 | 3.04E-01 | Small ribosomal subunit protein uS15y |
| AT3G53340 | 0.6 | 2.87E-05 | 0.2 | 4.03E-02 | 0.4 | 2.00E-03 | 0.0 | 8.43E-01 | 0.0 | 9.67E-01 | 0.0 | 9.28E-01 | Nuclear factor Y, subunit B10 |
| AT3G53750 | 0.6 | 9.97E-06 | 0.7 | 9.67E-06 | -0.1 | 4.08E-01 | 0.2 | 2.40E-01 | 0.2 | 3.46E-01 | 0.0 | 9.77E-01 | Actin-3 |
| AT4G20830 | 0.6 | 3.89E-02 | 0.3 | 3.15E-01 | 0.3 | 3.48E-01 | 0.7 | 2.37E-01 | 0.1 | 8.90E-01 | 0.6 | 4.26E-01 | Berberine bridge enzyme-like 19 |
| AT4G37150 | 0.6 | 6.81E-03 | 0.4 | 6.79E-02 | 0.2 | 3.35E-01 | 0.3 | 6.03E-01 | 0.7 | 1.60E-01 | -0.4 | 4.75E-01 | Methylesterase 9 |
| AT1G09900 | 0.6 | 7.61E-05 | 0.5 | 6.80E-04 | 0.1 | 2.55E-01 | 0.2 | 3.31E-01 | -0.1 | 6.65E-01 | 0.3 | 2.38E-01 | Pentatricopeptide repeat-containing protein At1g09900 |
| AT1G02780 | 0.6 | 5.69E-04 | 0.6 | 1.20E-03 | 0.0 | 8.61E-01 | 0.3 | 7.02E-02 | 0.1 | 6.11E-01 | 0.2 | 4.52E-01 | Large ribosomal subunit protein eL19x |
| AT3G15030 | 0.6 | 5.25E-06 | 0.4 | 2.98E-04 | 0.2 | 9.49E-03 | 0.0 | 9.80E-01 | 0.3 | 2.51E-01 | -0.3 | 3.53E-01 | Transcription factor TCP4 |
| AT1G31500 | 0.6 | 8.73E-06 | 0.4 | 3.68E-04 | 0.2 | 2.30E-02 | 0.6 | 5.04E-03 | 0.4 | 1.07E-01 | 0.2 | 3.44E-01 | Endonuclease |
| AT3G15352 | 0.6 | 5.01E-04 | 0.4 | 1.61E-02 | 0.2 | 1.22E-01 | 0.9 | 2.59E-02 | 0.2 | 6.57E-01 | 0.6 | 1.03E-01 | Cytochrome c oxidase 17 |
| AT3G13560 | 0.6 | 3.96E-04 | 0.4 | 1.12E-02 | 0.2 | 1.08E-01 | 0.2 | 1.04E-01 | 0.0 | 9.96E-01 | 0.2 | 1.97E-01 | glucan endo-1,3-beta-D-glucosidase |
| AT5G03910 | 0.6 | 1.88E-05 | 0.4 | 1.73E-03 | 0.3 | 1.28E-02 | -0.1 | 6.10E-01 | 0.0 | 9.08E-01 | -0.1 | 8.28E-01 | ABC transporter B family member 29, chloroplastic |
| AT2G13360 | 0.6 | 1.06E-03 | 0.5 | 9.66E-03 | 0.2 | 2.97E-01 | -0.3 | 3.77E-02 | 0.0 | 9.28E-01 | -0.3 | 1.25E-01 | Serine--glyoxylate aminotransferase |

| | | | | | | | | | | | | | |
|---|---|---|---|---|---|---|---|---|---|---|---|---|---|
| AT4G05520 | 0.6 | 3.57E-04 | 0.5 | 2.28E-03 | 0.1 | 4.02E-01 | 0.3 | 1.30E-01 | 0.3 | 1.39E-01 | 0.0 | 9.48E-01 | EH domain-containing protein 2 |
| AT4G34030 | 0.6 | 2.33E-05 | 0.4 | 6.40E-04 | 0.2 | 4.66E-02 | -0.2 | 4.83E-02 | -0.4 | 2.62E-03 | 0.2 | 2.12E-01 | 3-methylcrotonyl-CoA carboxylase non-biotinylated subunit |
| AT3G46530 | 0.6 | 5.14E-04 | 0.1 | 3.55E-01 | 0.5 | 6.65E-03 | 0.3 | 2.55E-01 | 0.0 | 9.90E-01 | 0.3 | 4.00E-01 | Disease resistance protein RPP13 |
| AT5G54180 | 0.6 | 3.66E-04 | 0.8 | 7.86E-05 | -0.2 | 1.50E-01 | 0.4 | 6.13E-02 | 0.5 | 4.91E-02 | -0.1 | 8.37E-01 | Transcription termination factor MTERF8, chloroplastic |
| AT3G08505 | 0.6 | 8.79E-04 | 0.1 | 4.55E-01 | 0.5 | 1.09E-02 | 0.1 | 5.83E-01 | 0.1 | 8.03E-01 | 0.1 | 8.83E-01 | E3 ubiquitin-protein ligase makorin |
| AT4G10920 | 0.6 | 2.05E-03 | 0.3 | 7.99E-02 | 0.3 | 9.63E-02 | 0.2 | 2.80E-01 | 0.1 | 8.50E-01 | 0.2 | 5.70E-01 | RNA polymerase II transcriptional coactivator KELP |
| AT5G15410 | 0.6 | 9.35E-05 | 0.5 | 8.12E-04 | 0.1 | 2.46E-01 | -0.6 | 2.41E-03 | -0.4 | 5.63E-02 | -0.2 | 3.90E-01 | Cyclic nucleotide-regulated ion channel family protein |
| AT3G59890 | 0.6 | 8.05E-03 | 0.8 | 2.50E-03 | -0.2 | 4.76E-01 | 0.7 | 6.29E-03 | 0.3 | 3.23E-01 | 0.4 | 9.27E-02 | 4-hydroxy-tetrahydrodipicolinate reductase 2, chloroplastic |
| AT3G48990 | 0.6 | 4.23E-06 | 0.4 | 1.81E-04 | 0.2 | 1.23E-02 | -0.4 | 1.81E-02 | -0.7 | 9.64E-04 | 0.3 | 1.52E-01 | Oxalate--CoA ligase |
| AT5G18100 | 0.6 | 4.84E-04 | 0.4 | 8.00E-03 | 0.2 | 1.86E-01 | -0.6 | 1.99E-02 | -0.1 | 7.13E-01 | -0.5 | 1.27E-01 | Superoxide dismutase copper |
| AT5G24580 | 0.6 | 1.86E-03 | 0.9 | 2.02E-04 | -0.3 | 1.12E-01 | -0.1 | 8.45E-01 | -0.1 | 7.15E-01 | 0.1 | 8.78E-01 | Heavy metal transport |
| AT2G20850 | 0.6 | 9.99E-06 | 0.2 | 6.72E-02 | 0.5 | 4.87E-04 | 0.0 | 9.26E-01 | -0.1 | 6.64E-01 | 0.1 | 7.51E-01 | STRUBBELIG-receptor family 1 |
| AT1G47490 | 0.6 | 5.52E-06 | 0.3 | 9.03E-04 | 0.3 | 3.11E-03 | 0.2 | 1.30E-01 | 0.2 | 2.96E-01 | 0.0 | 8.58E-01 | Polyadenylate-binding protein RBP47C -binding protein RBP47C) |
| AT5G57220 | 0.6 | 1.01E-02 | 0.0 | 8.82E-01 | 0.6 | 3.36E-02 | 2.6 | 1.30E-08 | 1.0 | 9.83E-04 | 1.6 | 4.95E-05 | Cytochrome P450 81F2 |
| AT1G74960 | 0.6 | 5.21E-06 | 0.3 | 1.35E-03 | 0.3 | 1.96E-03 | -0.1 | 3.70E-01 | -0.1 | 5.56E-01 | 0.0 | 9.22E-01 | 3-oxoacyl- synthase |
| AT5G67590 | 0.6 | 4.63E-05 | 0.5 | 8.61E-04 | 0.2 | 8.91E-02 | 0.2 | 2.92E-01 | 0.1 | 5.38E-01 | 0.1 | 8.78E-01 | NADH dehydrogenase iron-sulfur protein 4, mitochondrial |
| AT4G22530 | 0.6 | 8.05E-04 | -0.1 | 5.69E-01 | 0.7 | 1.78E-03 | 0.9 | 1.33E-03 | 0.5 | 8.27E-02 | 0.4 | 1.53E-01 | hypothetical protein |
| AT4G12320 | 0.6 | 1.29E-04 | 0.3 | 1.00E-02 | 0.3 | 2.70E-02 | 1.0 | 1.49E-06 | 0.8 | 7.72E-05 | 0.2 | 2.31E-01 | At4g12320 |
| AT1G29250 | 0.6 | 7.34E-04 | 0.7 | 8.36E-04 | 0.0 | 8.96E-01 | 0.6 | 2.52E-02 | 0.4 | 2.30E-01 | 0.2 | 5.40E-01 | Alba DNA |
| AT5G08380 | 0.6 | 2.33E-04 | 0.2 | 2.00E-01 | 0.5 | 5.76E-03 | 0.0 | 9.26E-01 | -0.1 | 7.83E-01 | 0.0 | 8.70E-01 | Alpha-galactosidase |
| AT5G64380 | 0.6 | 1.50E-04 | 0.6 | 3.76E-04 | 0.0 | 7.99E-01 | -0.2 | 1.94E-01 | -0.2 | 3.57E-01 | 0.0 | 8.97E-01 | fructose-bisphosphatase |
| AT4G15110 | 0.6 | 6.58E-04 | 0.4 | 1.07E-02 | 0.2 | 1.98E-01 | -0.5 | 3.77E-02 | -0.2 | 4.67E-01 | -0.3 | 3.31E-01 | Cytochrome P450 97B3, chloroplastic |
| AT2G13440 | 0.6 | 4.52E-06 | 0.2 | 7.76E-03 | 0.4 | 5.34E-04 | 0.1 | 5.44E-01 | 0.0 | 9.14E-01 | 0.2 | 5.55E-01 | Glucose-inhibited division family A protein |
| AT2G16600 | 0.6 | 2.15E-05 | 0.6 | 7.33E-05 | 0.0 | 6.56E-01 | 0.4 | 1.39E-01 | 0.1 | 6.59E-01 | 0.2 | 5.50E-01 | Peptidyl-prolyl cis-trans isomerase CYP19-1 |
| AT3G50930 | 0.7 | 8.10E-02 | -1.1 | 2.33E-02 | 1.8 | 4.20E-03 | 1.8 | 1.71E-06 | 0.4 | 1.76E-01 | 1.4 | 2.85E-04 | AAA+ ATPase domain-containing protein |
| AT3G06770 | 0.7 | 3.57E-05 | 0.7 | 5.47E-05 | 0.0 | 8.66E-01 | -0.7 | 1.34E-03 | -0.5 | 3.59E-02 | -0.3 | 3.42E-01 | F3E22.9 protein |
| AT1G11340 | 0.7 | 1.35E-02 | 0.6 | 1.86E-02 | 0.0 | 9.76E-01 | 0.2 | 4.98E-01 | 0.0 | 9.96E-01 | 0.2 | 5.81E-01 | Receptor-like serine |
| AT1G15690 | 0.7 | 6.35E-06 | 0.6 | 3.83E-05 | 0.1 | 2.88E-01 | 0.1 | 5.65E-01 | -0.1 | 4.33E-01 | 0.2 | 2.51E-01 | H-exporting diphosphatase |
| AT3G19360 | 0.7 | 4.48E-04 | 0.5 | 4.07E-03 | 0.2 | 3.04E-01 | 0.3 | 1.06E-01 | 0.0 | 8.72E-01 | 0.3 | 2.74E-01 | Zinc finger CCCH domain-containing protein 39 |
| AT2G22870 | 0.7 | 1.45E-04 | 0.5 | 1.58E-03 | 0.2 | 2.27E-01 | 0.8 | 1.59E-03 | 0.4 | 1.38E-01 | 0.4 | 1.22E-01 | GTP-binding protein At2g22870 |
| AT3G12290 | 0.7 | 3.61E-06 | 0.4 | 1.77E-04 | 0.2 | 9.06E-03 | 0.0 | 9.85E-01 | -0.1 | 5.73E-01 | 0.1 | 6.93E-01 | Bifunctional protein FolD 2 |
| AT3G04070 | 0.7 | 3.91E-03 | 0.3 | 1.49E-01 | 0.4 | 1.12E-01 | 0.0 | 9.57E-01 | 0.1 | 8.87E-01 | -0.1 | 9.40E-01 | NAC domain containing protein 47 ) |
| AT5G01870 | 0.7 | 1.62E-03 | 1.1 | 5.02E-05 | -0.5 | 1.61E-02 | 0.3 | 2.01E-01 | 0.1 | 6.94E-01 | 0.2 | 5.95E-01 | Non-specific lipid-transfer protein |
| AT3G48870 | 0.7 | 2.39E-05 | 0.3 | 5.36E-03 | 0.3 | 5.19E-03 | 0.3 | 5.98E-01 | 0.2 | 8.04E-01 | 0.1 | 9.08E-01 | Clp ATPase |
| AT1G70250 | 0.7 | 1.24E-04 | 0.0 | 7.18E-01 | 0.7 | 4.81E-04 | 0.4 | 4.69E-02 | 0.3 | 3.33E-01 | 0.2 | 5.36E-01 | Protein kinase domain-containing protein |
| AT3G45140 | 0.7 | 2.35E-02 | 0.5 | 6.06E-02 | 0.1 | 7.17E-01 | 0.9 | 5.29E-03 | 1.0 | 4.97E-03 | -0.1 | 8.25E-01 | Lipoxygenase |
| AT3G09405 | 0.7 | 4.39E-03 | 0.2 | 2.60E-01 | 0.4 | 8.40E-02 | 1.2 | 1.46E-03 | -0.3 | 5.59E-01 | 1.5 | 2.27E-03 | Pectin acetylesterase |
| AT2G13610 | 0.7 | 7.50E-05 | 0.4 | 5.66E-03 | 0.3 | 2.98E-02 | -0.6 | 6.10E-04 | -0.7 | 3.36E-04 | 0.1 | 5.88E-01 | ABC transporter G family member 5 |
| AT1G49560 | 0.7 | 6.56E-05 | 0.6 | 2.22E-04 | 0.1 | 6.36E-01 | 0.4 | 2.72E-02 | 0.3 | 1.46E-01 | 0.1 | 6.80E-01 | Homeodomain-like superfamily protein |
| AT2G29350 | 0.7 | 1.82E-01 | -1.1 | 9.41E-02 | 1.8 | 2.64E-02 | -1.6 | 2.26E-01 | -2.9 | 1.12E-01 | 1.3 | 6.06E-01 | Senescence-associated protein 13 |
| AT3G06850 | 0.7 | 5.21E-05 | 0.6 | 1.99E-04 | 0.1 | 5.65E-01 | -0.3 | 6.88E-02 | -0.6 | 5.85E-03 | 0.2 | 2.15E-01 | Lipoamide acyltransferase component of branched-chain alpha-keto acid dehydrogenase complex, mitochondrial transferase) |
| AT2G25210 | 0.7 | 1.44E-03 | 0.9 | 3.36E-04 | -0.2 | 2.18E-01 | 0.2 | 2.56E-01 | 0.2 | 4.67E-01 | 0.1 | 8.87E-01 | Large ribosomal subunit protein eL39z |
| AT3G45930 | 0.7 | 4.98E-04 | 0.6 | 8.41E-04 | 0.0 | 9.37E-01 | 0.4 | 1.51E-01 | 0.1 | 7.51E-01 | 0.2 | 5.01E-01 | Histone H4 |
| AT4G02770 | 0.7 | 7.80E-05 | 1.0 | 8.22E-06 | -0.4 | 1.10E-02 | 0.1 | 6.82E-01 | 0.0 | 8.71E-01 | 0.0 | 9.16E-01 | Photosystem I reaction center subunit II-1, chloroplastic |
| AT3G61110 | 0.7 | 1.80E-04 | 0.7 | 2.76E-04 | 0.0 | 9.36E-01 | 0.5 | 3.59E-02 | 0.2 | 5.00E-01 | 0.3 | 3.92E-01 | Small ribosomal subunit protein eS27y |
| AT3G53870 | 0.7 | 6.96E-05 | 0.6 | 2.91E-04 | 0.1 | 5.30E-01 | 0.4 | 9.28E-04 | 0.2 | 1.23E-01 | 0.2 | 1.75E-01 | Small ribosomal subunit protein uS3y |
| AT3G09480 | 0.7 | 1.13E-02 | 0.7 | 8.42E-03 | -0.1 | 8.36E-01 | -0.1 | 8.82E-01 | 0.0 | 9.58E-01 | -0.1 | 8.66E-01 | Histone H2B.5 |
| AT1G31170 | 0.7 | 1.43E-02 | 0.1 | 8.41E-01 | 0.6 | 4.67E-02 | -0.1 | 7.96E-01 | 0.0 | 9.95E-01 | -0.1 | 8.79E-01 | sulfiredoxin |
| AT3G01490 | 0.7 | 2.34E-04 | 0.9 | 4.68E-05 | -0.2 | 1.04E-01 | -0.2 | 2.19E-01 | -0.6 | 1.07E-03 | 0.4 | 4.03E-02 | hypothetical protein |
| AT5G38660 | 0.7 | 7.71E-05 | 0.5 | 8.12E-04 | 0.2 | 1.93E-01 | 0.1 | 4.79E-01 | -0.1 | 7.40E-01 | 0.2 | 3.78E-01 | Acclimation of photosynthesis to environment |
| AT4G31800 | 0.7 | 1.76E-02 | 0.2 | 5.73E-01 | 0.5 | 9.72E-02 | 0.8 | 8.67E-03 | 0.1 | 7.67E-01 | 0.7 | 4.80E-02 | WRKY DNA-binding protein 18 |
| AT2G31380 | 0.7 | 2.88E-04 | 0.4 | 1.51E-02 | 0.3 | 5.01E-02 | | | | | | | B-box zinc finger protein 25 |
| AT1G22840 | 0.7 | 1.69E-04 | 0.5 | 1.54E-03 | 0.2 | 2.43E-01 | 0.5 | 2.74E-02 | 0.2 | 4.16E-01 | 0.3 | 3.89E-01 | CYTOCHROME C-1 |
| AT2G31070 | 0.7 | 2.02E-05 | 0.1 | 4.00E-01 | 0.6 | 2.76E-04 | 0.1 | 6.29E-01 | 0.1 | 8.72E-01 | 0.1 | 8.64E-01 | Transcription factor TCP10 |
| AT5G66055 | 0.7 | 3.88E-06 | 0.3 | 8.85E-04 | 0.3 | 1.89E-03 | -0.1 | 6.13E-01 | -0.1 | 4.84E-01 | 0.0 | 8.72E-01 | Ankyrin repeat domain-containing protein, chloroplastic |
| AT1G63170 | 0.7 | 1.87E-06 | 0.3 | 7.61E-04 | 0.4 | 7.61E-04 | 0.0 | 9.45E-01 | 0.0 | 9.53E-01 | 0.0 | 9.97E-01 | E3 ubiquitin-protein ligase At1g63170 |
| AT2G38310 | 0.7 | 7.20E-04 | 0.7 | 6.15E-04 | -0.1 | 6.90E-01 | 0.4 | 2.74E-02 | -0.2 | 4.07E-01 | 0.6 | 2.68E-02 | Abscisic acid receptor PYL4 |
| AT4G31210 | 0.7 | 2.39E-04 | 0.3 | 6.61E-02 | 0.4 | 1.47E-02 | 0.8 | 3.26E-03 | 0.4 | 1.41E-01 | 0.4 | 2.21E-01 | DNA topoisomerase |
| AT1G27050 | 0.7 | 3.33E-04 | 0.4 | 9.25E-03 | 0.2 | 1.13E-01 | 0.5 | 5.35E-02 | 0.2 | 5.94E-01 | 0.3 | 3.61E-01 | Uncharacterized protein At1g27050 |
| AT1G76130 | 0.7 | 2.01E-05 | 0.4 | 1.46E-03 | 0.3 | 1.82E-02 | -0.2 | 3.74E-01 | -0.1 | 6.72E-01 | -0.1 | 8.23E-01 | Probable alpha-amylase 2 |
| AT5G11200 | 0.7 | 7.00E-06 | 0.3 | 1.29E-03 | 0.3 | 3.17E-03 | 0.3 | 7.61E-03 | 0.1 | 4.00E-01 | 0.2 | 2.08E-01 | DEAD |
| AT5G19890 | 0.7 | 5.99E-04 | 0.6 | 2.62E-03 | 0.1 | 5.10E-01 | 0.9 | 2.38E-04 | 0.5 | 1.86E-02 | 0.3 | 7.50E-02 | Peroxidase |
| AT5G63890 | 0.7 | 1.66E-06 | 0.5 | 1.50E-05 | 0.1 | 9.20E-02 | 0.3 | 3.77E-02 | 0.1 | 4.36E-01 | 0.2 | 4.26E-01 | Histidinol dehydrogenase, chloroplastic |
| AT3G01970 | 0.7 | 2.70E-02 | -0.2 | 5.02E-01 | 0.9 | 2.61E-02 | 0.5 | 2.02E-01 | -1.0 | 7.86E-02 | 1.5 | 2.14E-02 | Probable WRKY transcription factor 45 |
| AT3G28480 | 0.7 | 3.98E-05 | 0.3 | 2.34E-02 | 0.4 | 3.96E-03 | 0.6 | 3.21E-03 | 0.2 | 3.05E-01 | 0.4 | 1.38E-01 | procollagen-proline 4-dioxygenase |
| AT5G16970 | 0.7 | 1.24E-04 | 0.3 | 1.98E-02 | 0.4 | 1.48E-02 | 0.1 | 5.12E-01 | 0.0 | 9.49E-01 | 0.1 | 7.39E-01 | NADPH-dependent oxidoreductase 2-alkenal reductase |
| AT1G76160 | 0.7 | 1.80E-05 | 0.6 | 8.79E-05 | 0.1 | 4.24E-01 | 0.0 | 9.67E-01 | -0.3 | 3.01E-02 | 0.3 | 9.34E-02 | AT1G76160 protein |
| AT3G05560 | 0.7 | 1.09E-03 | 0.7 | 1.24E-03 | 0.0 | 8.67E-01 | 0.3 | 5.16E-02 | 0.1 | 5.03E-01 | 0.2 | 4.43E-01 | Large ribosomal subunit protein eL22z |
| AT4G18100 | 0.7 | 1.17E-03 | 0.8 | 7.61E-04 | -0.1 | 5.88E-01 | 0.3 | 7.66E-02 | 0.1 | 5.83E-01 | 0.2 | 4.67E-01 | Large ribosomal subunit protein eL32z |
| AT5G53540 | 0.7 | 2.45E-06 | 0.4 | 1.54E-04 | 0.3 | 5.80E-03 | 0.0 | 9.22E-01 | -0.1 | 3.73E-01 | 0.2 | 4.18E-01 | 26S proteasome regulatory particle chain RPT6-like protein |
| AT2G19190 | 0.7 | 1.44E-01 | -0.1 | 8.54E-01 | 0.8 | 1.66E-01 | -0.6 | 5.20E-01 | -2.0 | 7.85E-02 | 1.4 | 1.75E-01 | Senescence-induced receptor-like serine |
| AT4G10100 | 0.7 | 5.35E-05 | 0.2 | 8.94E-02 | 0.5 | 2.54E-03 | -0.3 | 1.20E-01 | -0.4 | 6.97E-02 | 0.1 | 8.06E-01 | Molybdopterin synthase sulfur carrier subunit |
| AT3G06430 | 0.7 | 5.95E-06 | 0.6 | 1.71E-05 | 0.0 | 7.33E-01 | 0.1 | 6.22E-01 | -0.1 | 6.44E-01 | 0.1 | 4.40E-01 | Pentatricopeptide repeat-containing protein At3g06430, chloroplastic |
| AT4G30825 | 0.7 | 3.17E-04 | 0.5 | 3.03E-03 | 0.2 | 2.76E-01 | 0.3 | 1.71E-01 | 0.3 | 1.32E-01 | -0.1 | 8.57E-01 | Pentatricopeptide repeat-containing protein At4g30825, chloroplastic |
| AT5G47210 | 0.7 | 2.97E-04 | 0.5 | 5.28E-03 | 0.2 | 1.40E-01 | 0.4 | 1.89E-03 | 0.2 | 2.15E-01 | 0.2 | 1.73E-01 | RGG repeats nuclear RNA binding protein C |
| AT3G01510 | 0.7 | 2.49E-05 | 0.6 | 1.64E-04 | 0.1 | 3.15E-01 | 0.7 | 1.99E-03 | 0.7 | 3.08E-03 | 0.0 | 9.47E-01 | Like SEX4 1 |
| AT1G19670 | 0.7 | 8.85E-03 | 0.9 | 1.88E-03 | -0.2 | 3.29E-01 | 0.5 | 2.87E-01 | 0.7 | 1.88E-01 | -0.2 | 8.13E-01 | Chlorophyllase-1 |
| AT5G60270 | 0.7 | 5.74E-03 | 0.2 | 4.54E-01 | 0.5 | 5.77E-02 | -0.5 | 2.83E-02 | -0.5 | 3.24E-02 | 0.0 | 9.51E-01 | L-type lectin-domain containing receptor kinase I.7 |
| AT5G22620 | 0.7 | 9.51E-06 | 0.4 | 4.88E-04 | 0.3 | 1.48E-02 | 0.0 | 7.81E-01 | 0.1 | 4.81E-01 | -0.1 | 7.63E-01 | Phosphoglycerate |
| AT2G30200 | 0.7 | 1.70E-05 | 0.6 | 8.49E-05 | 0.1 | 3.86E-01 | 0.4 | 4.38E-03 | 0.3 | 5.69E-02 | 0.1 | 5.40E-01 | S-malonyltransferase |
| AT4G38430 | 0.7 | 5.75E-05 | 0.4 | 1.87E-03 | 0.2 | 5.00E-02 | -0.2 | 1.96E-01 | -0.2 | 4.17E-01 | -0.1 | 8.62E-01 | Rop guanine nucleotide exchange factor 1 |
| AT5G64290 | 0.7 | 2.23E-06 | 0.4 | 1.19E-04 | 0.3 | 6.10E-03 | 0.4 | 1.07E-02 | 0.3 | 7.50E-02 | 0.1 | 6.93E-01 | Dicarboxylate transporter 2.1, chloroplastic |
| AT4G16990 | 0.7 | 3.19E-05 | 0.3 | 5.10E-03 | 0.3 | 8.19E-03 | 0.3 | 1.11E-01 | -0.2 | 2.69E-01 | 0.5 | 2.66E-02 | At4g16990 |

| | | | | | | | | | | | | | |
|---|---|---|---|---|---|---|---|---|---|---|---|---|---|
| AT3G20320 | 0.7 | 9.49E-06 | 0.5 | 1.38E-04 | 0.2 | 8.51E-02 | 0.0 | 8.30E-01 | 0.1 | 7.51E-01 | 0.0 | 9.23E-01 | Protein TRIGALACTOSYLDIACYLGLYCEROL 2, chloroplastic |
| AT5G24020 | 0.7 | 7.20E-06 | 0.6 | 4.22E-05 | 0.1 | 2.94E-01 | 0.2 | 2.10E-01 | -0.2 | 4.51E-01 | 0.4 | 1.07E-01 | Septum site-determining protein minD homolog, chloroplastic |
| AT3G16350 | 0.7 | 4.79E-05 | 0.7 | 6.49E-05 | 0.0 | 8.01E-01 | -0.4 | 1.76E-01 | -0.3 | 4.00E-01 | -0.1 | 8.15E-01 | At3g16350 |
| AT4G37910 | 0.7 | 5.76E-04 | 0.3 | 9.07E-02 | 0.4 | 2.10E-02 | 1.2 | 7.70E-07 | 0.4 | 3.11E-02 | 0.8 | 1.79E-03 | Heat shock 70 kDa protein 9, mitochondrial |
| AT5G53170 | 0.7 | 3.11E-05 | 0.3 | 1.41E-02 | 0.4 | 3.42E-03 | 0.0 | 8.58E-01 | 0.0 | 9.20E-01 | 0.0 | 9.78E-01 | AAA+ ATPase domain-containing protein |
| AT2G45440 | 0.7 | 1.41E-05 | 0.4 | 7.13E-04 | 0.3 | 2.05E-02 | 0.4 | 6.57E-03 | 0.1 | 4.22E-01 | 0.3 | 1.67E-01 | 4-hydroxy-tetrahydrodipicolinate synthase 2, chloroplastic |
| AT3G20480 | 0.7 | 6.75E-04 | 0.3 | 4.38E-02 | 0.3 | 7.51E-02 | -0.1 | 9.11E-01 | -0.1 | 8.67E-01 | 0.0 | 9.46E-01 | tetraacyldisaccharide 4'-kinase |
| AT1G75150 | 0.7 | 3.14E-05 | 0.4 | 2.45E-03 | 0.3 | 2.45E-02 | 0.3 | 4.83E-01 | 0.2 | 6.73E-01 | 0.1 | 9.11E-01 | DNA ligase-like protein |
| AT2G40880 | 0.7 | 2.70E-02 | 0.3 | 3.89E-01 | 0.4 | 1.92E-01 | -0.2 | 3.85E-01 | -0.2 | 4.98E-01 | 0.0 | 9.77E-01 | Cysteine proteinase inhibitor 3 |
| AT1G23310 | 0.7 | 4.16E-05 | 0.4 | 2.66E-03 | 0.3 | 2.25E-02 | 0.0 | 9.43E-01 | 0.0 | 9.19E-01 | 0.0 | 9.69E-01 | Glutamate--glyoxylate aminotransferase 1 |
| AT5G09830 | 0.7 | 2.57E-03 | 0.2 | 3.11E-01 | 0.5 | 3.80E-02 | 0.2 | 4.34E-01 | 0.1 | 8.73E-01 | 0.1 | 7.02E-01 | Protein BOLA2 |
| AT5G01410 | 0.7 | 2.45E-05 | 0.2 | 2.19E-02 | 0.4 | 1.87E-03 | 0.3 | 7.83E-03 | 0.2 | 1.68E-01 | 0.1 | 4.18E-01 | Pyridoxal 5'-phosphate synthase subunit PDX1.3 |
| AT3G05470 | 0.7 | 1.55E-04 | 0.2 | 9.03E-02 | 0.4 | 8.75E-03 | 0.0 | 9.21E-01 | 0.1 | 7.95E-01 | -0.1 | 7.51E-01 | Formin-like protein 11 |
| AT2G41100 | 0.7 | 3.43E-03 | 0.3 | 1.11E-01 | 0.4 | 1.05E-01 | 0.4 | 2.58E-01 | -0.9 | 3.95E-02 | 1.3 | 1.57E-02 | Calcium-binding EF hand family protein |
| AT2G43610 | 0.7 | 7.08E-04 | 0.8 | 3.88E-04 | -0.1 | 4.98E-01 | 0.3 | 9.70E-02 | 0.0 | 9.15E-01 | 0.3 | 1.62E-01 | Endochitinase At2g43610 |
| AT4G28780 | 0.7 | 2.88E-03 | 0.8 | 1.66E-03 | -0.1 | 6.40E-01 | 0.0 | 9.39E-01 | 0.2 | 3.00E-01 | -0.2 | 3.37E-01 | GDSL esterase |
| AT3G48425 | 0.7 | 4.01E-05 | 0.4 | 1.60E-03 | 0.2 | 5.04E-02 | 0.3 | 3.35E-01 | 0.4 | 1.60E-01 | -0.1 | 6.99E-01 | DNA- endonuclease |
| AT1G80830 | 0.7 | 1.66E-05 | 0.4 | 1.03E-03 | 0.3 | 1.53E-02 | -0.1 | 4.80E-01 | -0.1 | 6.04E-01 | 0.0 | 9.71E-01 | Metal transporter Nramp1 |
| AT3G20270 | 0.7 | 6.02E-04 | 0.3 | 8.57E-02 | 0.4 | 3.27E-02 | 0.0 | 9.82E-01 | 0.1 | 6.73E-01 | -0.1 | 7.03E-01 | Lipid-binding serum glycoprotein family protein |
| AT5G46430 | 0.7 | 3.93E-05 | 0.5 | 5.62E-04 | 0.2 | 1.22E-01 | -0.1 | 7.39E-01 | -0.2 | 3.97E-01 | 0.1 | 6.98E-01 | Large ribosomal subunit protein eL32y |
| AT1G01210 | 0.7 | 5.53E-04 | 0.4 | 2.15E-02 | 0.3 | 9.16E-02 | 0.6 | 1.94E-03 | 0.2 | 2.66E-01 | 0.4 | 9.88E-02 | DNA-directed RNA polymerase subunit |
| AT4G21750 | 0.7 | 1.77E-05 | 0.5 | 3.43E-04 | 0.2 | 6.43E-02 | -0.1 | 5.00E-01 | -0.2 | 3.02E-01 | 0.1 | 7.90E-01 | Uncharacterized protein |
| AT1G08490 | 0.7 | 1.20E-06 | 0.2 | 1.67E-02 | 0.5 | 7.54E-05 | -0.1 | 7.33E-01 | -0.1 | 7.13E-01 | 0.0 | 9.53E-01 | Cysteine desulfurase 1, chloroplastic |
| AT3G46210 | 0.7 | 9.32E-05 | 0.3 | 1.65E-02 | 0.4 | 1.61E-02 | 0.3 | 1.38E-01 | 0.0 | 9.80E-01 | 0.3 | 2.52E-01 | Exosome complex exonuclease RRP46 homolog |
| AT1G74050 | 0.7 | 8.34E-05 | 0.7 | 1.28E-04 | 0.0 | 8.77E-01 | 0.0 | 8.26E-01 | -0.2 | 2.63E-01 | 0.1 | 5.12E-01 | Large ribosomal subunit protein eL6x |
| AT3G16400 | 0.7 | 5.62E-06 | 0.6 | 3.63E-05 | 0.1 | 2.53E-01 | -0.1 | 6.36E-01 | 0.2 | 4.41E-01 | -0.3 | 3.28E-01 | Thiohydroximate-O-sulfate sulfur |
| AT2G37980 | 0.7 | 6.03E-05 | 0.9 | 1.34E-05 | -0.2 | 6.40E-02 | 0.1 | 7.15E-01 | 0.3 | 3.48E-01 | -0.2 | 6.47E-01 | O-fucosyltransferase 19 |
| AT1G64820 | 0.7 | 3.48E-03 | 1.1 | 1.87E-04 | -0.4 | 5.96E-02 | | | | | | | Protein DETOXIFICATION 7 |
| AT4G28706 | 0.7 | 6.01E-06 | 0.6 | 3.03E-05 | 0.1 | 3.40E-01 | 0.0 | 7.83E-01 | 0.1 | 4.31E-01 | -0.1 | 7.11E-01 | At4g29227 |
| AT3G62250 | 0.7 | 1.43E-03 | 0.7 | 2.48E-03 | 0.0 | 9.30E-01 | 0.3 | 8.93E-02 | 0.1 | 4.81E-01 | 0.1 | 5.90E-01 | Ubiquitin-ribosomal protein eS31x fusion protein |
| AT1G13080 | 0.7 | 1.23E-04 | 0.6 | 5.34E-04 | 0.1 | 4.86E-01 | -0.1 | 7.34E-01 | 0.4 | 5.62E-03 | -0.5 | 9.95E-03 | Cytochrome P450 71B2 |
| AT3G54710 | 0.7 | 2.86E-04 | 0.5 | 2.62E-03 | 0.2 | 3.12E-01 | 0.7 | 2.07E-02 | 0.5 | 8.95E-02 | 0.1 | 7.38E-01 | CDT1-like protein b |
| AT5G48570 | 0.7 | 3.93E-04 | 0.1 | 6.04E-01 | 0.6 | 4.13E-03 | 0.4 | 2.64E-01 | 0.7 | 4.93E-02 | -0.3 | 3.78E-01 | Peptidyl-prolyl cis-trans isomerase FKBP65 |
| AT4G01700 | 0.7 | 3.18E-04 | 0.6 | 1.17E-03 | 0.1 | 5.86E-01 | 1.2 | 1.17E-05 | 0.6 | 1.97E-02 | 0.7 | 5.38E-03 | Glycoside hydrolase family 19 catalytic domain-containing protein |
| AT4G12830 | 0.7 | 2.11E-04 | 0.9 | 7.33E-05 | -0.2 | 2.28E-01 | 0.6 | 1.09E-03 | 0.5 | 1.00E-02 | 0.1 | 6.17E-01 | AT4g12830 |
| AT4G36130 | 0.7 | 1.80E-03 | 0.6 | 5.62E-03 | 0.1 | 6.68E-01 | 0.2 | 1.92E-01 | 0.1 | 4.05E-01 | 0.0 | 8.78E-01 | Large ribosomal subunit protein uL2x |
| AT1G31160 | 0.7 | 4.79E-05 | 0.6 | 1.66E-04 | 0.1 | 6.29E-01 | 0.1 | 4.89E-01 | 0.2 | 5.02E-01 | 0.0 | 9.62E-01 | HISTIDINE TRIAD NUCLEOTIDE-BINDING 2 |
| AT4G31060 | 0.7 | 3.92E-04 | 0.7 | 5.83E-04 | 0.0 | 9.61E-01 | -0.5 | 1.39E-01 | -0.4 | 1.94E-01 | 0.0 | 9.78E-01 | Ethylene-responsive transcription factor ERF015 |
| AT4G34900 | 0.7 | 5.03E-03 | 1.0 | 5.92E-04 | -0.3 | 1.68E-01 | -0.5 | 1.79E-01 | 0.3 | 3.97E-01 | -0.8 | 7.51E-02 | Xanthine dehydrogenase 2 |
| AT2G35260 | 0.7 | 1.46E-05 | 0.5 | 2.39E-04 | 0.2 | 7.78E-02 | -0.2 | 2.40E-01 | 0.0 | 8.59E-01 | -0.2 | 2.84E-01 | At2g35260 |
| AT4G37730 | 0.7 | 2.38E-05 | 0.8 | 1.05E-05 | -0.2 | 1.33E-01 | 0.4 | 1.44E-01 | 0.3 | 4.25E-01 | 0.2 | 7.08E-01 | BZIP transcription factor-like protein |
| AT2G31200 | 0.7 | 1.62E-03 | 0.3 | 1.32E-01 | 0.4 | 5.19E-02 | 0.5 | 6.48E-02 | 0.3 | 3.57E-01 | 0.2 | 6.27E-01 | Actin-depolymerizing factor 6 |
| AT5G28840 | 0.7 | 3.21E-05 | 0.5 | 3.98E-04 | 0.2 | 1.41E-01 | 0.0 | 8.34E-01 | -0.5 | 3.09E-03 | 0.4 | 1.48E-02 | GDP-mannose 3,5-epimerase |
| AT2G04842 | 0.7 | 2.93E-05 | 0.4 | 3.02E-03 | 0.3 | 1.18E-02 | 0.3 | 4.45E-02 | 0.1 | 6.48E-01 | 0.2 | 3.23E-01 | Threonine--tRNA ligase, chloroplastic |
| AT4G33510 | 0.7 | 7.40E-06 | 0.5 | 1.01E-04 | 0.2 | 8.33E-02 | 0.2 | 2.71E-01 | -0.1 | 7.78E-01 | 0.2 | 2.50E-01 | Phospho-2-dehydro-3-deoxyheptonate aldolase |
| AT4G26700 | 0.7 | 4.80E-05 | 0.2 | 7.11E-02 | 0.5 | 1.91E-03 | 0.2 | 4.71E-01 | 0.1 | 8.07E-01 | 0.1 | 7.99E-01 | AT4G26700 protein |
| AT4G26150 | 0.7 | 1.56E-03 | 0.6 | 3.36E-03 | 0.0 | 8.23E-01 | -0.3 | 2.51E-01 | -1.1 | 1.02E-03 | 0.8 | 3.27E-03 | Putative GATA transcription factor 22 |
| AT4G23700 | 0.7 | 5.67E-02 | 0.6 | 1.04E-01 | 0.1 | 8.22E-01 | -1.6 | 4.39E-04 | -1.0 | 1.11E-02 | -0.6 | 1.78E-01 | Cation |
| AT1G33140 | 0.7 | 3.66E-04 | 0.8 | 3.40E-04 | -0.1 | 6.88E-01 | 0.4 | 1.76E-01 | 0.0 | 9.81E-01 | 0.4 | 3.56E-01 | Large ribosomal subunit protein uL6z |
| AT1G20720 | 0.7 | 1.22E-04 | 0.5 | 1.42E-03 | 0.2 | 2.44E-01 | 0.1 | 8.33E-01 | 0.3 | 6.95E-01 | -0.1 | 8.64E-01 | RAD3-like DNA-binding helicase protein |
| AT4G17390 | 0.7 | 1.18E-03 | 0.8 | 8.13E-04 | -0.1 | 6.19E-01 | 0.3 | 2.56E-02 | 0.1 | 4.70E-01 | 0.2 | 3.23E-01 | Large ribosomal subunit protein eL15y |
| AT2G37130 | 0.7 | 3.10E-04 | 1.0 | 5.80E-05 | -0.3 | 9.25E-02 | -0.5 | 5.66E-03 | -0.8 | 4.22E-04 | 0.3 | 1.41E-01 | peroxidase |
| AT5G10160 | 0.7 | 6.52E-05 | 0.6 | 2.59E-04 | 0.1 | 5.16E-01 | -0.1 | 4.32E-01 | 0.0 | 9.22E-01 | -0.1 | 6.91E-01 | 3-hydroxyacyl- dehydratase |
| AT2G19740 | 0.7 | 6.59E-04 | 0.6 | 1.53E-03 | 0.0 | 7.99E-01 | 0.2 | 2.58E-01 | -0.1 | 6.55E-01 | 0.3 | 2.06E-01 | Large ribosomal subunit protein eL31z |
| AT3G56980 | 0.7 | 1.16E-02 | -0.3 | 3.64E-01 | 1.0 | 1.12E-02 | 3.6 | 2.76E-07 | 0.1 | 8.68E-01 | 3.5 | 2.65E-05 | Transcription factor ORG3 |
| AT3G57720 | 0.7 | 3.74E-04 | 0.1 | 4.23E-01 | 0.6 | 6.56E-03 | -0.2 | 5.95E-01 | 0.1 | 8.49E-01 | -0.3 | 5.29E-01 | Non-functional pseudokinase ZRK2 |
| AT1G15710 | 0.7 | 8.69E-06 | 0.6 | 3.34E-05 | 0.1 | 5.33E-01 | 0.3 | 8.03E-02 | 0.2 | 3.85E-01 | 0.1 | 5.96E-01 | hypothetical protein |
| AT2G45710 | 0.7 | 3.06E-05 | 0.4 | 2.17E-03 | 0.3 | 2.32E-02 | 0.7 | 1.03E-02 | 0.3 | 3.92E-01 | 0.4 | 2.20E-01 | Small ribosomal subunit protein eS27z |
| AT5G59690 | 0.7 | 5.21E-05 | 0.8 | 5.16E-05 | -0.1 | 5.56E-01 | 0.2 | 3.03E-01 | 0.0 | 9.76E-01 | 0.2 | 4.54E-01 | Histone H4 |
| AT1G01560 | 0.7 | 9.25E-02 | -0.9 | 1.04E-01 | 1.6 | 1.93E-02 | 0.8 | 4.95E-02 | -0.1 | 8.16E-01 | 0.9 | 6.60E-02 | Mitogen-activated protein kinase |
| AT5G04310 | 0.7 | 7.04E-04 | 0.4 | 3.84E-02 | 0.3 | 8.57E-02 | 0.0 | 9.88E-01 | -0.5 | 2.03E-01 | 0.5 | 2.55E-01 | Pectate lyase |
| AT2G44310 | 0.7 | 3.14E-04 | 0.5 | 3.13E-03 | 0.2 | 2.36E-01 | 0.2 | 4.62E-01 | -0.1 | 8.78E-01 | 0.3 | 5.02E-01 | hypothetical protein |
| AT3G60510 | 0.7 | 1.22E-04 | 0.5 | 1.56E-03 | 0.2 | 1.93E-01 | 0.0 | 8.77E-01 | 0.0 | 9.95E-01 | 0.0 | 9.14E-01 | 3-hydroxyisobutyryl-CoA hydrolase |
| AT3G53430 | 0.7 | 4.48E-04 | 0.7 | 6.39E-04 | 0.0 | 9.36E-01 | 0.4 | 1.01E-01 | 0.1 | 6.68E-01 | 0.2 | 4.60E-01 | Large ribosomal subunit protein uL11y |
| AT4G36580 | 0.7 | 7.84E-05 | 0.1 | 4.08E-01 | 0.6 | 1.60E-03 | 1.7 | 6.76E-05 | 0.8 | 5.19E-02 | 0.9 | 1.57E-02 | AAA-type ATPase family protein |
| AT3G26744 | 0.7 | 8.07E-06 | 0.4 | 8.36E-04 | 0.3 | 6.23E-03 | -0.3 | 3.34E-02 | -0.3 | 6.13E-02 | 0.0 | 9.75E-01 | Basic helix-loop-helix DNA-binding superfamily protein |
| AT3G57830 | 0.7 | 1.71E-04 | 0.8 | 1.64E-04 | -0.1 | 6.85E-01 | -0.3 | 2.52E-01 | -0.2 | 4.75E-01 | -0.1 | 8.56E-01 | Leucine-rich repeat protein kinase family protein |
| AT4G27820 | 0.7 | 8.15E-04 | 0.5 | 1.03E-02 | 0.2 | 2.16E-01 | 0.3 | 5.93E-02 | 0.3 | 1.13E-01 | 0.0 | 9.44E-01 | Beta glucosidase 9 |
| AT5G08720 | 0.7 | 4.93E-04 | 0.2 | 2.66E-01 | 0.5 | 8.27E-03 | -0.3 | 5.53E-02 | -0.1 | 7.48E-01 | -0.3 | 1.72E-01 | At5g08720 |
| AT3G02920 | 0.7 | 1.64E-05 | 0.4 | 1.88E-03 | 0.3 | 1.18E-02 | 0.3 | 2.22E-01 | 0.3 | 1.27E-01 | -0.1 | 7.83E-01 | Replication protein A, subunit RPA32 |
| AT3G11110 | 0.7 | 2.94E-05 | 0.5 | 5.21E-04 | 0.2 | 1.04E-01 | -0.3 | 1.83E-01 | -0.3 | 2.25E-01 | 0.0 | 9.94E-01 | RING-H2 finger protein ATL66 |
| AT5G16470 | 0.7 | 4.04E-05 | 0.3 | 6.17E-03 | 0.4 | 9.29E-03 | 0.0 | 8.70E-01 | -0.3 | 2.69E-01 | 0.3 | 2.96E-01 | Protein METHYLENE BLUE SENSITIVITY 2 |
| AT2G16370 | 0.7 | 2.54E-04 | 0.5 | 4.96E-03 | 0.2 | 1.38E-01 | 0.1 | 4.15E-01 | 0.1 | 4.67E-01 | 0.0 | 9.92E-01 | Thymidylate synthase 1 |
| AT4G00620 | 0.7 | 9.02E-05 | 0.3 | 4.58E-02 | 0.4 | 6.58E-03 | 0.6 | 5.33E-04 | 0.4 | 4.20E-02 | 0.3 | 1.54E-01 | Bifunctional protein FolD 4, chloroplastic |
| AT5G55280 | 0.7 | 2.56E-05 | 0.7 | 5.00E-05 | 0.0 | 1.00E+00 | 0.6 | 1.91E-03 | 0.2 | 4.02E-01 | 0.4 | 6.50E-02 | Cell division protein FtsZ homolog 1, chloroplastic |
| AT1G79560 | 0.7 | 8.99E-06 | 0.5 | 1.13E-04 | 0.2 | 9.54E-02 | 0.6 | 9.84E-04 | 0.1 | 5.44E-01 | 0.5 | 2.14E-02 | FTSH protease 12 |
| AT4G18590 | 0.7 | 1.06E-04 | 0.4 | 4.18E-03 | 0.3 | 6.27E-02 | 0.0 | 9.43E-01 | -0.3 | 4.12E-01 | 0.3 | 3.78E-01 | Replication protein A 14 kDa subunit B |
| AT2G25060 | 0.7 | 2.65E-04 | 1.0 | 4.22E-05 | -0.3 | 7.98E-02 | 0.1 | 4.84E-01 | 0.2 | 3.41E-01 | -0.1 | 8.23E-01 | hypothetical protein |
| AT4G24830 | 0.7 | 2.08E-04 | 0.6 | 9.70E-04 | 0.1 | 4.76E-01 | 0.4 | 3.29E-02 | 0.3 | 9.79E-02 | 0.1 | 8.76E-01 | argininosuccinate synthase |
| AT5G38110 | 0.7 | 1.46E-05 | 0.6 | 5.56E-05 | 0.1 | 5.71E-01 | 0.1 | 7.82E-01 | -0.1 | 6.30E-01 | 0.2 | 4.87E-01 | Histone chaperone ASF1B |
| AT1G72140 | 0.7 | 1.18E-02 | 1.2 | 4.74E-04 | -0.5 | 6.98E-02 | -0.2 | 7.61E-01 | -0.4 | 5.56E-01 | 0.2 | 8.06E-01 | Protein NRT1 |
| AT3G44160 | 0.7 | 3.78E-05 | 0.4 | 1.95E-03 | 0.3 | 3.48E-02 | 0.2 | 2.51E-01 | 0.1 | 8.02E-01 | 0.2 | 5.11E-01 | Outer envelope protein 39, chloroplastic |
| AT4G11290 | 0.7 | 2.49E-04 | 0.8 | 1.50E-04 | -0.1 | 4.74E-01 | 1.0 | 1.97E-04 | 0.5 | 4.49E-02 | 0.5 | 1.87E-02 | Peroxidase 39 |

| | | | | | | | | | | | | | |
|---|---|---|---|---|---|---|---|---|---|---|---|---|---|
| AT3G07670 | 0.7 | 1.52E-05 | 0.5 | 2.36E-04 | 0.2 | 8.42E-02 | 0.3 | 5.89E-02 | 0.2 | 2.85E-01 | 0.1 | 6.71E-01 | hypothetical protein |
| AT5G02150 | 0.7 | 4.71E-04 | 0.5 | 8.67E-03 | 0.2 | 1.88E-01 | -0.4 | 1.91E-01 | -0.6 | 6.76E-02 | 0.2 | 5.94E-01 | Fes1C |
| AT2G24270 | 0.7 | 4.38E-05 | 0.4 | 2.27E-03 | 0.3 | 2.88E-02 | -0.2 | 1.67E-01 | 0.0 | 8.16E-01 | -0.1 | 4.39E-01 | Aldehyde dehydrogenase 11A3 |
| AT4G33670 | 0.7 | 5.29E-06 | 0.7 | 1.86E-05 | 0.0 | 5.87E-01 | -0.4 | 6.52E-03 | -0.4 | 1.39E-02 | 0.0 | 9.64E-01 | L-galactose dehydrogenase |
| AT1G80600 | 0.7 | 7.34E-06 | 0.4 | 6.02E-04 | 0.3 | 7.30E-03 | 0.2 | 9.36E-02 | 0.3 | 3.67E-02 | -0.1 | 6.81E-01 | Acetylornithine aminotransferase, chloroplastic |
| AT1G65290 | 0.7 | 2.72E-05 | 0.6 | 1.12E-04 | 0.1 | 5.20E-01 | 0.5 | 6.23E-03 | 0.2 | 2.46E-01 | 0.3 | 2.59E-01 | Acyl carrier protein 2, mitochondrial |
| AT1G19860 | 0.7 | 1.48E-06 | 0.2 | 6.02E-03 | 0.5 | 1.71E-04 | 0.1 | 7.28E-01 | 0.1 | 7.00E-01 | 0.0 | 9.45E-01 | Zinc finger C-x8-C-x5-C-x3-H type family protein |
| AT5G49740 | 0.7 | 8.59E-03 | 0.6 | 2.55E-02 | 0.1 | 6.61E-01 | 0.3 | 6.51E-02 | -0.2 | 3.34E-01 | 0.6 | 1.98E-02 | Ferric reduction oxidase 7 |
| AT1G21150 | 0.7 | 2.65E-03 | 0.6 | 7.48E-03 | 0.1 | 7.55E-01 | -0.5 | 1.97E-01 | 0.1 | 9.17E-01 | -0.5 | 1.52E-01 | Eukaryotic translation initiation factor 5B |
| AT4G02260 | 0.7 | 1.79E-05 | 0.3 | 8.43E-03 | 0.4 | 2.45E-03 | 0.0 | 7.65E-01 | 0.1 | 4.14E-01 | -0.1 | 6.76E-01 | GTP diphosphokinase |
| AT5G05440 | 0.7 | 2.89E-04 | 0.6 | 8.54E-04 | 0.1 | 6.78E-01 | 0.3 | 1.22E-01 | -0.4 | 6.56E-02 | 0.6 | 1.29E-02 | Abscisic acid receptor PYL5 |
| AT2G23200 | 0.7 | 1.14E-04 | -0.1 | 5.30E-01 | 0.8 | 3.72E-04 | 0.5 | 3.92E-01 | 0.5 | 5.08E-01 | 0.0 | 9.67E-01 | Probable receptor-like protein kinase At2g23200 |
| AT1G79720 | 0.7 | 3.19E-06 | 0.5 | 1.01E-04 | 0.2 | 1.78E-02 | -0.4 | 1.21E-02 | -0.2 | 2.31E-01 | -0.2 | 3.27E-01 | hypothetical protein |
| AT5G03370 | 0.7 | 2.80E-05 | 0.4 | 2.48E-03 | 0.3 | 1.79E-02 | 0.1 | 5.92E-01 | 0.4 | 1.09E-01 | -0.2 | 3.69E-01 | acylphosphatase |
| AT1G61190 | 0.7 | 1.97E-05 | 0.3 | 1.54E-02 | 0.4 | 1.81E-03 | 0.2 | 2.00E-01 | -0.2 | 2.58E-01 | 0.3 | 6.91E-02 | LRR and NB-ARC domains-containing disease resistance protein |
| AT5G60040 | 0.7 | 2.64E-04 | 0.1 | 6.78E-01 | 0.6 | 2.62E-03 | 0.8 | 5.20E-03 | 0.2 | 5.39E-01 | 0.5 | 4.87E-02 | DNA-directed RNA polymerase subunit |
| AT5G47930 | 0.7 | 3.77E-04 | 0.7 | 5.58E-04 | 0.0 | 9.46E-01 | 0.3 | 1.89E-02 | 0.2 | 2.19E-01 | 0.1 | 5.23E-01 | Small ribosomal subunit protein eS27w |
| AT3G03010 | 0.7 | 8.30E-06 | 0.4 | 8.68E-04 | 0.3 | 8.59E-03 | 0.2 | 1.25E-01 | -0.3 | 1.24E-01 | 0.5 | 1.06E-02 | aminoacyl-tRNA hydrolase |
| AT5G08070 | 0.7 | 1.15E-04 | 0.5 | 3.76E-03 | 0.3 | 8.87E-02 | -0.2 | 7.19E-01 | 0.1 | 9.19E-01 | -0.2 | 7.21E-01 | Transcription factor TCP17 |
| AT1G59124 | 0.7 | 4.31E-03 | 0.4 | 1.04E-01 | 0.3 | 1.81E-01 | | | | | | | PRM1 homolog |
| AT4G00050 | 0.7 | 5.31E-04 | 0.3 | 9.94E-02 | 0.4 | 2.15E-02 | -0.1 | 7.25E-01 | 0.2 | 6.09E-01 | -0.3 | 4.36E-01 | Basic helix-loop-helix DNA-binding superfamily protein |
| AT2G32520 | 0.7 | 2.07E-06 | 0.6 | 2.38E-05 | 0.2 | 6.40E-02 | 0.2 | 2.53E-01 | -0.1 | 5.27E-01 | 0.3 | 1.53E-01 | Alpha |
| AT5G09315 | 0.7 | 8.17E-05 | 0.4 | 8.02E-03 | 0.3 | 2.96E-02 | 0.2 | 2.90E-01 | 0.0 | 9.66E-01 | 0.2 | 3.78E-01 | Uncharacterized protein T5E8_120 |
| AT5G56600 | 0.7 | 2.42E-06 | 0.6 | 2.62E-05 | 0.1 | 7.96E-02 | 0.0 | 8.82E-01 | 0.3 | 6.36E-02 | -0.3 | 1.77E-01 | hypothetical protein |
| AT1G52290 | 0.7 | 2.83E-04 | 0.4 | 1.23E-02 | 0.3 | 6.33E-02 | 0.2 | 2.93E-01 | 0.0 | 9.30E-01 | 0.2 | 5.22E-01 | non-specific serine |
| AT5G60930 | 0.7 | 2.19E-05 | 0.1 | 2.04E-01 | 0.6 | 4.58E-04 | 0.3 | 1.20E-01 | 0.2 | 4.79E-01 | 0.1 | 6.67E-01 | P-loop containing nucleoside triphosphate hydrolases superfamily protein |
| AT3G04870 | 0.7 | 3.83E-06 | 0.2 | 2.85E-02 | 0.5 | 1.87E-04 | 0.0 | 9.43E-01 | 0.0 | 8.96E-01 | 0.0 | 9.52E-01 | Zeta-carotene desaturase, chloroplastic |
| AT1G66700 | 0.7 | 5.16E-02 | -0.4 | 3.64E-01 | 1.1 | 3.43E-02 | 1.9 | 5.41E-04 | 0.2 | 8.17E-01 | 1.7 | 2.94E-03 | Paraxanthine methyltransferase 1 |
| AT3G01180 | 0.7 | 4.09E-05 | 0.4 | 5.57E-03 | 0.4 | 1.05E-02 | 0.1 | 2.85E-01 | 0.3 | 6.70E-02 | -0.1 | 5.30E-01 | Starch synthase 2, chloroplastic |
| AT3G16100 | 0.7 | 9.24E-05 | 0.4 | 4.61E-03 | 0.3 | 4.43E-02 | 0.1 | 6.47E-01 | 0.0 | 9.56E-01 | 0.1 | 8.01E-01 | Ras-related protein RABG3c |
| AT3G56300 | 0.7 | 5.68E-04 | 0.5 | 8.98E-03 | 0.2 | 2.27E-01 | -0.1 | 7.85E-01 | 0.4 | 3.69E-01 | -0.5 | 2.01E-01 | Cysteinyl-tRNA synthetase, class Ia family protein |
| AT3G15360 | 0.7 | 2.59E-05 | 0.4 | 1.30E-03 | 0.3 | 2.42E-02 | 0.1 | 6.77E-01 | 0.1 | 5.21E-01 | 0.0 | 8.50E-01 | Thioredoxin M4, chloroplastic |
| AT1G62960 | 0.7 | 5.88E-05 | 0.1 | 2.39E-01 | 0.6 | 1.04E-03 | 0.1 | 6.50E-01 | 0.0 | 9.81E-01 | 0.1 | 7.19E-01 | Probable aminotransferase ACS10 |
| AT4G24230 | 0.7 | 2.24E-04 | 0.8 | 2.48E-04 | 0.0 | 8.07E-01 | -0.6 | 1.78E-03 | -1.0 | 9.86E-05 | 0.4 | 1.26E-01 | Acyl-CoA-binding domain 3 |
| AT3G63420 | 0.7 | 1.04E-04 | 0.3 | 2.83E-02 | 0.4 | 1.14E-02 | 0.1 | 5.93E-01 | -0.2 | 3.82E-01 | 0.3 | 1.94E-01 | Guanine nucleotide-binding protein subunit gamma 1 |
| AT3G21110 | 0.7 | 6.54E-05 | 0.5 | 2.07E-03 | 0.3 | 6.24E-02 | 0.2 | 2.29E-01 | 0.3 | 1.96E-01 | 0.0 | 8.89E-01 | Phosphoribosylaminoimidazole-succinocarboxamide synthase, chloroplastic |
| AT2G39400 | 0.7 | 1.48E-04 | 0.7 | 3.67E-04 | 0.0 | 7.82E-01 | -0.3 | 4.50E-01 | -1.3 | 2.13E-03 | 1.1 | 2.05E-02 | Alpha |
| AT2G26550 | 0.7 | 1.89E-05 | 0.4 | 2.83E-03 | 0.3 | 8.27E-03 | 0.2 | 3.18E-01 | 0.1 | 8.08E-01 | 0.1 | 6.30E-01 | Heme oxygenase 2 |
| AT4G13170 | 0.7 | 2.08E-04 | 0.8 | 1.85E-04 | -0.1 | 6.51E-01 | 0.6 | 5.81E-05 | 0.3 | 1.07E-02 | 0.2 | 1.14E-01 | Large ribosomal subunit protein uL13x |
| AT5G19510 | 0.7 | 2.32E-05 | 0.6 | 1.44E-04 | 0.1 | 3.27E-01 | 0.1 | 1.58E-01 | -0.1 | 6.90E-01 | 0.2 | 1.96E-01 | Elongation factor 1-beta 2 |
| AT4G37510 | 0.7 | 5.04E-06 | 0.6 | 4.19E-05 | 0.1 | 1.56E-01 | 0.3 | 7.02E-02 | 0.1 | 6.36E-01 | 0.2 | 3.18E-01 | Ribonuclease III domain-containing protein RNC1, chloroplastic |
| AT3G09500 | 0.7 | 2.03E-04 | 0.8 | 2.14E-04 | -0.1 | 7.15E-01 | 0.1 | 5.29E-01 | 0.0 | 9.48E-01 | 0.1 | 7.39E-01 | Large ribosomal subunit protein uL29z |
| AT5G05690 | 0.7 | 8.26E-05 | 0.9 | 4.90E-05 | -0.1 | 3.16E-01 | -0.2 | 1.73E-01 | -0.5 | 2.84E-03 | 0.3 | 1.13E-01 | Cytochrome P450 superfamily protein |
| AT2G25870 | 0.7 | 6.15E-06 | 0.6 | 5.56E-05 | 0.1 | 1.36E-01 | 0.2 | 3.37E-01 | 0.0 | 9.71E-01 | 0.2 | 4.64E-01 | Endoribonuclease YBEY, chloroplastic |
| AT3G16770 | 0.7 | 3.49E-03 | 0.6 | 1.71E-02 | 0.2 | 4.99E-01 | -0.9 | 9.63E-03 | -1.3 | 2.60E-03 | 0.3 | 4.51E-01 | Ethylene-responsive transcription factor RAP2-3 |
| AT5G64300 | 0.7 | 1.37E-06 | 0.3 | 4.72E-04 | 0.4 | 5.92E-04 | 0.2 | 1.04E-01 | -0.1 | 4.56E-01 | 0.3 | 5.53E-02 | GTP cyclohydrolase II |
| AT4G23240 | 0.7 | 3.69E-04 | 0.5 | 9.90E-03 | 0.3 | 1.38E-01 | -0.5 | 2.62E-01 | -0.4 | 4.58E-01 | -0.1 | 8.42E-01 | Cysteine-rich RLK 16 |
| AT3G13065 | 0.7 | 2.31E-03 | 0.5 | 2.91E-02 | 0.2 | 2.94E-01 | | | | | | | At3g13065 |
| AT1G48450 | 0.7 | 3.71E-06 | 0.5 | 1.40E-04 | 0.2 | 1.30E-02 | 0.0 | 9.84E-01 | -0.1 | 5.09E-01 | 0.1 | 5.69E-01 | Alanine-tRNA ligase, putative |
| AT2G28740 | 0.7 | 4.39E-05 | 0.7 | 1.02E-04 | 0.0 | 8.33E-01 | 0.2 | 4.40E-01 | 0.1 | 6.53E-01 | 0.0 | 9.18E-01 | Histone H4 |
| AT5G52520 | 0.7 | 1.45E-06 | 0.5 | 2.97E-05 | 0.2 | 1.84E-02 | 0.4 | 4.88E-02 | 0.3 | 1.10E-01 | 0.0 | 9.16E-01 | Proline--tRNA ligase, chloroplastic |
| AT2G24050 | 0.7 | 1.50E-06 | 0.5 | 6.43E-05 | 0.3 | 6.09E-03 | 0.5 | 2.42E-03 | 0.2 | 1.84E-01 | 0.3 | 1.93E-01 | Eukaryotic translation initiation factor isoform 4G-2 4G-2) |
| AT1G32310 | 0.7 | 1.29E-05 | 0.6 | 1.26E-04 | 0.1 | 1.75E-01 | 0.0 | 9.36E-01 | -0.1 | 6.86E-01 | 0.2 | 6.80E-01 | L-gulonolactone oxidase |
| AT3G30390 | 0.7 | 9.36E-06 | 1.1 | 1.70E-06 | -0.3 | 5.33E-03 | 0.1 | 6.38E-01 | 0.1 | 5.31E-01 | 0.0 | 8.91E-01 | Amino acid transporter AVT6A |
| AT1G23020 | 0.7 | 7.94E-04 | 0.1 | 4.05E-01 | 0.6 | 8.07E-03 | 2.1 | 2.33E-07 | -0.2 | 5.39E-01 | 2.3 | 2.02E-05 | Ferric reduction oxidase 3 |
| AT2G37270 | 0.7 | 2.14E-04 | 0.8 | 2.45E-04 | 0.0 | 7.72E-01 | 0.5 | 2.11E-04 | 0.2 | 1.02E-01 | 0.3 | 6.73E-02 | Small ribosomal subunit protein uS7z |
| AT1G33265 | 0.7 | 7.18E-05 | 0.6 | 8.00E-04 | 0.2 | 2.03E-01 | 0.2 | 4.45E-01 | 0.0 | 9.96E-01 | 0.2 | 5.96E-01 | Protein FATTY ACID EXPORT 4, chloroplastic |
| AT1G63780 | 0.7 | 1.56E-04 | 0.5 | 2.71E-03 | 0.2 | 1.39E-01 | 0.7 | 1.83E-04 | 0.3 | 1.49E-01 | 0.5 | 1.77E-02 | Putative U3 small nucleolar ribonucleoprotein |
| AT3G05350 | 0.7 | 3.25E-06 | 0.3 | 1.28E-03 | 0.4 | 1.06E-03 | 0.0 | 8.78E-01 | 0.0 | 9.83E-01 | 0.0 | 9.04E-01 | Xaa-Pro aminopeptidase P |
| AT3G61080 | 0.7 | 1.14E-04 | 0.7 | 2.51E-04 | 0.0 | 8.63E-01 | -0.4 | 2.28E-02 | 0.0 | 9.84E-01 | -0.4 | 7.79E-02 | protein-ribulosamine 3-kinase |
| AT5G23070 | 0.7 | 6.92E-05 | 0.5 | 2.30E-03 | 0.3 | 6.15E-02 | 0.3 | 3.39E-02 | 0.1 | 7.00E-01 | 0.2 | 2.30E-01 | Thymidine kinase b |
| AT4G12400 | 0.7 | 7.62E-05 | 0.4 | 8.25E-03 | 0.3 | 2.41E-02 | 0.5 | 6.15E-03 | 0.8 | 9.83E-04 | -0.2 | 3.66E-01 | Hsp70-Hsp90 organizing protein 3 |
| AT5G62790 | 0.7 | 1.93E-05 | 0.4 | 1.10E-03 | 0.3 | 1.80E-02 | 0.1 | 3.49E-01 | 0.2 | 9.69E-02 | -0.1 | 5.58E-01 | 1-deoxy-D-xylulose-5-phosphate reductoisomerase |
| AT1G64770 | 0.7 | 1.30E-04 | 0.4 | 4.97E-03 | 0.3 | 5.42E-02 | 0.2 | 1.15E-01 | 0.1 | 2.81E-01 | 0.0 | 8.69E-01 | NDH-dependent cyclic electron flow 1 |
| AT1G17330 | 0.7 | 6.80E-05 | 0.2 | 8.88E-02 | 0.5 | 3.22E-03 | 0.4 | 2.21E-02 | 0.0 | 9.11E-01 | 0.4 | 1.06E-01 | T13M22.1 protein |
| AT3G28930 | 0.7 | 1.83E-03 | 0.6 | 5.97E-03 | 0.1 | 6.27E-01 | 0.5 | 9.59E-02 | 0.0 | 9.57E-01 | 0.5 | 2.04E-01 | Putative gamma-glutamylcyclotransferase |
| AT3G10840 | 0.7 | 3.05E-04 | 0.3 | 3.57E-02 | 0.4 | 2.94E-02 | 0.3 | 1.27E-01 | 0.6 | 9.47E-03 | -0.3 | 2.17E-01 | Alpha |
| AT2G03980 | 0.7 | 3.25E-05 | 0.6 | 3.79E-04 | 0.2 | 1.61E-01 | 0.4 | 7.43E-03 | 0.1 | 5.71E-01 | 0.3 | 9.21E-02 | Uncharacterized protein |
| AT3G52115 | 0.7 | 1.94E-03 | 0.6 | 8.61E-03 | 0.1 | 5.80E-01 | 0.5 | 1.12E-01 | 0.1 | 7.73E-01 | 0.4 | 3.33E-01 | Protein gamma response 1 |
| AT4G34670 | 0.7 | 7.17E-05 | 0.6 | 4.01E-04 | 0.1 | 3.94E-01 | 0.2 | 2.88E-02 | 0.0 | 8.99E-01 | 0.2 | 1.86E-01 | Small ribosomal subunit protein eS1y |
| AT2G31810 | 0.7 | 1.94E-05 | 0.8 | 1.91E-05 | -0.1 | 5.04E-01 | 0.0 | 9.76E-01 | -0.5 | 1.19E-02 | 0.5 | 2.08E-02 | Acetolactate synthase small subunit 1, chloroplastic |
| AT4G39330 | 0.7 | 1.39E-03 | 0.8 | 1.25E-03 | -0.1 | 7.63E-01 | 0.0 | 8.32E-01 | 0.2 | 1.86E-01 | -0.3 | 1.99E-01 | Probable cinnamyl alcohol dehydrogenase 9 |
| AT5G46490 | 0.7 | 8.26E-05 | 0.2 | 1.17E-01 | 0.5 | 3.20E-03 | 0.0 | 9.89E-01 | -0.4 | 4.80E-02 | 0.4 | 9.78E-02 | Disease resistance protein family |
| AT3G60245 | 0.7 | 1.28E-03 | 0.9 | 6.43E-04 | -0.1 | 4.80E-01 | 0.2 | 2.39E-01 | 0.1 | 5.60E-01 | 0.1 | 7.98E-01 | Large ribosomal subunit protein eL43y |
| AT4G19410 | 0.7 | 1.50E-04 | 0.8 | 1.07E-04 | -0.1 | 4.56E-01 | 0.0 | 7.65E-01 | -0.3 | 3.80E-02 | 0.4 | 2.63E-02 | Pectin acetylesterase |
| AT5G57660 | 0.7 | 4.51E-04 | 0.5 | 1.04E-02 | 0.3 | 1.16E-01 | -1.2 | 3.22E-05 | -1.2 | 1.56E-04 | -0.1 | 8.47E-01 | Zinc finger protein CONSTANS-LIKE 5 |
| AT5G27770 | 0.7 | 1.03E-04 | 0.8 | 1.23E-04 | -0.1 | 7.34E-01 | 0.3 | 2.07E-01 | 0.0 | 9.30E-01 | 0.2 | 4.23E-01 | Large ribosomal subunit protein eL22y |
| AT3G51470 | 0.7 | 8.37E-05 | 0.6 | 6.05E-04 | 0.1 | 3.33E-01 | 0.1 | 6.38E-01 | 0.0 | 9.79E-01 | 0.1 | 7.67E-01 | protein-serine |
| AT1G26130 | 0.7 | 2.40E-05 | 0.3 | 2.64E-02 | 0.5 | 2.24E-03 | 0.2 | 3.77E-01 | 0.0 | 9.52E-01 | 0.2 | 5.85E-01 | Phospholipid-transporting ATPase |
| AT2G03710 | 0.7 | 8.22E-04 | 0.2 | 1.85E-01 | 0.5 | 2.48E-02 | 0.1 | 6.13E-01 | -0.1 | 7.47E-01 | 0.2 | 4.68E-01 | Sep-04 |
| AT1G79700 | 0.7 | 4.28E-04 | 0.8 | 4.76E-04 | 0.0 | 8.43E-01 | -0.8 | 2.73E-03 | -1.4 | 9.86E-05 | 0.6 | 6.79E-02 | Integrase-type DNA-binding superfamily protein |
| AT3G23670 | 0.7 | 4.65E-05 | 0.7 | 2.02E-04 | 0.1 | 5.19E-01 | -0.3 | 3.47E-01 | 0.0 | 9.83E-01 | -0.3 | 4.02E-01 | Kinesin-like protein |
| AT5G03690 | 0.7 | 2.14E-05 | 0.5 | 5.57E-04 | 0.2 | 6.08E-02 | 0.1 | 5.53E-01 | 0.0 | 8.62E-01 | 0.1 | 5.63E-01 | Fructose-bisphosphate aldolase |

| | | | | | | | | | | | | | |
|---|---|---|---|---|---|---|---|---|---|---|---|---|---|
| AT3G48310 | 0.7 | 2.30E-05 | 0.0 | 8.08E-01 | 0.7 | 2.03E-04 | 0.0 | 9.70E-01 | -0.2 | 2.72E-01 | 0.2 | 3.49E-01 | Cytochrome P450 71A22 |
| AT3G47490 | 0.7 | 3.19E-06 | 0.5 | 1.59E-04 | 0.3 | 9.92E-03 | -0.2 | 3.80E-01 | -0.1 | 8.68E-01 | -0.1 | 6.66E-01 | At3g47490 |
| AT2G16650 | 0.7 | 1.87E-05 | 0.3 | 2.34E-02 | 0.5 | 2.02E-03 | 0.1 | 7.12E-01 | -0.3 | 1.82E-01 | 0.3 | 1.57E-01 | ribonuclease P |
| AT3G51450 | 0.7 | 1.42E-04 | 0.6 | 1.51E-03 | 0.2 | 2.32E-01 | 0.8 | 2.51E-02 | 0.6 | 1.47E-01 | 0.2 | 6.76E-01 | Protein STRICTOSIDINE SYNTHASE-LIKE 7 |
| AT5G10400 | 0.7 | 1.07E-05 | 0.8 | 1.34E-05 | -0.1 | 6.20E-01 | 0.4 | 7.53E-02 | 0.2 | 4.06E-01 | 0.2 | 5.78E-01 | Histone H3.1 |
| AT4G32900 | 0.7 | 8.88E-06 | 0.6 | 5.16E-05 | 0.1 | 3.27E-01 | 0.1 | 7.11E-01 | -0.2 | 3.86E-01 | 0.3 | 2.68E-01 | aminoacyl-tRNA hydrolase |
| AT2G24190 | 0.7 | 5.73E-04 | 0.1 | 7.32E-01 | 0.7 | 4.81E-03 | 0.0 | 9.84E-01 | 0.6 | 1.76E-01 | -0.6 | 2.31E-01 | Short-chain dehydrogenase |
| AT3G55010 | 0.7 | 4.80E-04 | 0.6 | 3.02E-03 | 0.2 | 3.71E-01 | 0.5 | 2.49E-03 | 0.3 | 1.72E-01 | 0.3 | 1.59E-01 | Phosphoribosylformylglycinamidine cyclo-ligase, chloroplastic |
| AT1G51460 | 0.7 | 2.97E-03 | 0.3 | 1.74E-01 | 0.4 | 9.86E-02 | 0.1 | 9.08E-01 | 0.4 | 4.70E-01 | | | ABC transporter G family member 13 |
| AT4G02420 | 0.7 | 2.50E-03 | 0.2 | 3.22E-01 | 0.5 | 2.89E-02 | 0.4 | 6.97E-03 | 0.3 | 3.96E-02 | 0.1 | 7.54E-01 | L-type lectin-domain containing receptor kinase IV.4 |
| AT5G18820 | 0.7 | 1.56E-04 | 0.5 | 3.64E-03 | 0.2 | 1.17E-01 | 1.1 | 4.46E-04 | 0.3 | 3.63E-01 | 0.8 | 1.31E-02 | TCP-1 |
| AT1G58080 | 0.7 | 8.15E-06 | 0.5 | 2.82E-04 | 0.3 | 2.15E-02 | 0.0 | 8.84E-01 | -0.1 | 6.00E-01 | 0.1 | 7.79E-01 | ATP phosphoribosyltransferase 1, chloroplastic |
| AT1G54780 | 0.7 | 1.38E-05 | 0.8 | 1.55E-05 | -0.1 | 4.93E-01 | 0.3 | 4.33E-02 | 0.3 | 1.02E-01 | 0.0 | 9.28E-01 | UPF0603 protein At1g54780, chloroplastic |
| AT3G52500 | 0.7 | 1.03E-05 | 0.7 | 2.66E-05 | 0.0 | 8.48E-01 | -0.1 | 5.30E-01 | 0.0 | 7.47E-01 | 0.0 | 9.15E-01 | At3g52500 |
| AT5G13120 | 0.7 | 2.91E-06 | 0.7 | 1.34E-05 | 0.1 | 4.06E-01 | 0.3 | 5.31E-02 | 0.2 | 2.06E-01 | 0.1 | 7.87E-01 | Photosynthetic NDH subunit of lumenal location 5, chloroplastic |
| AT1G03030 | 0.7 | 2.68E-05 | 0.2 | 1.17E-01 | 0.6 | 1.11E-03 | 0.2 | 4.41E-01 | 0.0 | 9.67E-01 | 0.1 | 6.36E-01 | hypothetical protein |
| AT4G16250 | 0.7 | 2.63E-05 | 0.5 | 1.38E-03 | 0.3 | 2.99E-02 | 0.3 | 1.00E-01 | -0.2 | 4.07E-01 | 0.5 | 3.22E-02 | Phytochrome D |
| AT3G11940 | 0.7 | 1.58E-04 | 0.6 | 8.52E-04 | 0.1 | 4.24E-01 | 0.1 | 4.91E-01 | -0.1 | 6.99E-01 | 0.2 | 4.02E-01 | Small ribosomal subunit protein uS7y |
| AT2G18020 | 0.7 | 5.26E-04 | 0.7 | 1.46E-03 | 0.1 | 7.28E-01 | 0.3 | 4.30E-03 | 0.2 | 1.89E-01 | 0.2 | 3.21E-01 | Large ribosomal subunit protein uL2z |
| AT5G60340 | 0.7 | 1.20E-05 | 0.3 | 3.92E-03 | 0.4 | 2.76E-03 | -0.1 | 4.46E-01 | -0.4 | 6.54E-02 | 0.2 | 4.06E-01 | Adenylate kinase isoenzyme 6 homolog |
| AT3G61530 | 0.7 | 1.56E-05 | 0.4 | 3.82E-03 | 0.4 | 5.96E-03 | 0.2 | 4.22E-01 | -0.2 | 3.82E-01 | 0.3 | 1.64E-01 | 3-methyl-2-oxobutanoate hydroxymethyltransferase |
| AT2G03550 | 0.8 | 7.14E-04 | 0.6 | 3.94E-03 | 0.1 | 4.37E-01 | 0.1 | 7.28E-01 | -0.1 | 7.18E-01 | 0.1 | 5.66E-01 | Probable carboxylesterase 7 |
| AT3G15680 | 0.8 | 1.56E-05 | 0.5 | 4.22E-04 | 0.2 | 4.46E-02 | 0.1 | 6.54E-01 | 0.0 | 9.11E-01 | 0.1 | 6.67E-01 | hypothetical protein |
| AT3G46940 | 0.8 | 2.50E-05 | 0.6 | 2.82E-04 | 0.2 | 1.63E-01 | 0.4 | 1.09E-01 | 0.2 | 6.08E-01 | 0.3 | 4.87E-01 | Deoxyuridine 5'-triphosphate nucleotidohydrolase |
| AT3G61220 | 0.8 | 1.76E-04 | 0.2 | 9.08E-02 | 0.5 | 6.42E-03 | 0.6 | 1.43E-02 | 0.9 | 2.35E-03 | -0.3 | 3.98E-01 | Short-chain dehydrogenase |
| AT3G08730 | 0.8 | 3.09E-06 | 0.4 | 4.28E-04 | 0.3 | 2.45E-03 | -0.1 | 6.11E-01 | 0.0 | 9.15E-01 | -0.1 | 8.14E-01 | Serine |
| AT2G26930 | 0.8 | 1.84E-06 | 0.7 | 6.57E-06 | 0.0 | 5.86E-01 | 0.4 | 9.01E-03 | 0.3 | 8.85E-02 | 0.1 | 5.54E-01 | 4-diphosphocytidyl-2-C-methyl-D-erythritol kinase, chloroplastic -2-C-methyl-D-erythritol kinase) |
| AT3G14600 | 0.8 | 7.65E-05 | 0.8 | 1.33E-04 | 0.0 | 9.51E-01 | 0.0 | 7.09E-01 | 0.0 | 9.54E-01 | 0.0 | 8.64E-01 | Large ribosomal subunit protein eL20x |
| AT1G23400 | 0.8 | 1.85E-05 | 0.6 | 1.72E-04 | 0.2 | 1.70E-01 | 0.1 | 4.37E-01 | 0.0 | 9.15E-01 | 0.1 | 6.18E-01 | CRS2-associated factor 2, chloroplastic |
| AT2G15910 | 0.8 | 8.12E-05 | 0.6 | 4.18E-04 | 0.1 | 4.23E-01 | 0.2 | 4.78E-01 | -0.3 | 3.78E-01 | 0.5 | 1.93E-01 | CSL zinc finger domain-containing protein |
| AT3G22750 | 0.8 | 8.32E-05 | 0.5 | 3.15E-03 | 0.3 | 6.12E-02 | -0.1 | 5.28E-01 | -0.4 | 2.94E-02 | 0.3 | 2.09E-01 | Kinase-like protein |
| AT1G78010 | 0.8 | 2.90E-05 | 0.5 | 1.40E-03 | 0.3 | 3.39E-02 | -0.2 | 3.99E-01 | -0.1 | 6.38E-01 | 0.0 | 8.45E-01 | TrmE-type G domain-containing protein |
| AT3G49100 | 0.8 | 1.46E-03 | 0.4 | 4.90E-02 | 0.3 | 1.18E-01 | 0.4 | 1.21E-01 | 0.2 | 5.09E-01 | 0.2 | 6.23E-01 | Signal recognition particle 9 kDa protein |
| AT3G56070 | 0.8 | 2.40E-04 | 0.6 | 1.44E-03 | 0.1 | 3.73E-01 | 0.7 | 1.02E-03 | 0.5 | 2.57E-02 | 0.2 | 3.04E-01 | Peptidyl-prolyl cis-trans isomerase CYP19-3 |
| AT3G52750 | 0.8 | 5.00E-06 | 0.8 | 8.20E-06 | 0.0 | 7.56E-01 | 0.4 | 6.90E-03 | 0.5 | 7.56E-03 | 0.0 | 8.85E-01 | Tubulin |
| AT5G20040 | 0.8 | 1.09E-05 | 0.8 | 1.72E-05 | 0.0 | 8.04E-01 | 0.6 | 6.88E-03 | 0.2 | 3.00E-01 | 0.3 | 1.70E-01 | Isopentenyltransferase 9 |
| AT2G37770 | 0.8 | 8.11E-02 | -2.6 | 6.00E-03 | 3.4 | 2.78E-03 | 0.5 | 7.74E-01 | 0.6 | 7.28E-01 | -0.2 | 9.42E-01 | NADPH-dependent aldo-keto reductase, chloroplastic |
| AT3G60840 | 0.8 | 6.64E-05 | 0.6 | 5.15E-04 | 0.1 | 3.01E-01 | 0.1 | 6.44E-01 | 0.2 | 4.00E-01 | -0.1 | 7.79E-01 | 65-kDa microtubule-associated protein 4 |
| AT4G21820 | 0.8 | 4.37E-05 | 0.7 | 1.19E-04 | 0.0 | 8.15E-01 | 0.2 | 6.02E-01 | -0.2 | 7.50E-01 | 0.4 | 4.37E-01 | Binding |
| AT2G16400 | 0.8 | 2.02E-06 | 0.5 | 1.19E-04 | 0.3 | 5.41E-03 | 0.1 | 5.84E-01 | -0.2 | 1.75E-01 | 0.3 | 1.20E-01 | BEL1-like homeodomain protein 7 |
| AT5G66005 | 0.8 | 9.59E-04 | 0.9 | 5.61E-04 | -0.1 | 5.55E-01 | 0.2 | 5.43E-01 | 0.3 | 3.57E-01 | -0.1 | 8.11E-01 | Expressed protein |
| AT4G25960 | 0.8 | 4.14E-05 | 0.5 | 9.06E-04 | 0.2 | 7.03E-02 | -0.1 | 7.74E-01 | -0.2 | 2.74E-01 | 0.2 | 5.15E-01 | ABC transporter B family member 2-like |
| AT5G24300 | 0.8 | 7.83E-06 | 0.4 | 8.41E-04 | 0.3 | 5.64E-03 | 0.5 | 2.01E-02 | 0.5 | 3.36E-02 | 0.0 | 9.89E-01 | Starch synthase 1, chloroplastic |
| AT3G10230 | 0.8 | 3.54E-06 | 0.4 | 8.33E-04 | 0.4 | 1.64E-03 | 0.0 | 9.06E-01 | 0.0 | 8.17E-01 | 0.0 | 9.26E-01 | Lycopene beta cyclase, chloroplastic |
| AT4G05450 | 0.8 | 2.62E-05 | 0.3 | 2.21E-02 | 0.5 | 3.44E-03 | 0.6 | 1.16E-02 | 0.6 | 2.25E-02 | 0.0 | 9.58E-01 | Adrenodoxin-like protein 1, mitochondrial |
| AT4G21860 | 0.8 | 1.82E-06 | 0.6 | 1.49E-05 | 0.1 | 1.23E-01 | -0.1 | 4.61E-01 | 0.0 | 8.92E-01 | -0.1 | 7.19E-01 | Peptide methionine sulfoxide reductase B2, chloroplastic -S-oxide reductasemethionine) |
| AT3G49410 | 0.8 | 2.02E-02 | 0.4 | 2.78E-01 | 0.4 | 2.55E-01 | 0.1 | 8.27E-01 | 0.4 | 5.35E-01 | -0.2 | 6.87E-01 | Transcription factor IIIC, subunit 5 |
| AT3G02470 | 0.8 | 5.47E-06 | 0.7 | 2.85E-05 | 0.1 | 3.55E-01 | 0.2 | 7.70E-02 | 0.0 | 7.92E-01 | 0.2 | 1.55E-01 | S-adenosylmethionine decarboxylase proenzyme 1 |
| AT5G15050 | 0.8 | 4.97E-06 | 0.6 | 3.65E-05 | 0.1 | 2.06E-01 | -0.2 | 4.06E-01 | 0.2 | 4.14E-01 | -0.4 | 1.73E-01 | Beta-glucuronosyltransferase GlcAT14B |
| AT1G12520 | 0.8 | 3.04E-06 | 0.6 | 3.88E-05 | 0.2 | 6.62E-02 | 0.0 | 7.77E-01 | -0.2 | 1.80E-01 | 0.1 | 4.05E-01 | Copper chaperone for superoxide dismutase, chloroplastic |
| AT4G39510 | 0.8 | 1.28E-03 | 0.6 | 6.43E-03 | 0.1 | 4.86E-01 | 0.3 | 6.15E-02 | 0.3 | 1.12E-01 | 0.0 | 9.50E-01 | AT4g39510 |
| AT5G38990 | 0.8 | 7.96E-05 | 0.1 | 2.97E-01 | 0.6 | 1.11E-03 | 0.3 | 7.70E-02 | 0.0 | 9.46E-01 | 0.3 | 2.29E-01 | Probable receptor-like protein kinase At5g38990 |
| AT1G10830 | 0.8 | 1.66E-06 | 0.4 | 2.10E-04 | 0.3 | 2.13E-03 | 0.1 | 5.20E-01 | 0.1 | 6.56E-01 | 0.0 | 9.61E-01 | 15-cis-zeta-carotene isomerase |
| AT1G30700 | 0.8 | 2.24E-03 | 0.2 | 4.24E-01 | 0.6 | 2.51E-02 | 1.7 | 2.13E-07 | 0.5 | 3.36E-02 | 1.2 | 1.14E-04 | Berberine bridge enzyme-like 8 |
| AT1G50170 | 0.8 | 1.12E-05 | 0.4 | 3.03E-03 | 0.4 | 4.01E-03 | 0.5 | 1.27E-02 | 0.2 | 3.97E-01 | 0.3 | 2.35E-01 | Sirohydrochlorin ferrochelatase B |
| AT4G05190 | 0.8 | 1.01E-04 | 0.3 | 2.17E-02 | 0.4 | 1.25E-02 | 0.2 | 5.33E-01 | 0.1 | 7.01E-01 | 0.0 | 9.40E-01 | Kinesin-like protein |
| AT5G16870 | 0.8 | 4.24E-06 | 0.5 | 1.55E-04 | 0.2 | 2.02E-02 | -0.2 | 3.15E-01 | -0.2 | 4.29E-01 | 0.0 | 9.53E-01 | aminoacyl-tRNA hydrolase |
| AT3G46100 | 0.8 | 1.13E-06 | 0.5 | 3.94E-05 | 0.3 | 6.76E-03 | 0.1 | 3.96E-01 | 0.1 | 6.56E-01 | 0.0 | 8.68E-01 | Histidine--tRNA ligase, chloroplastic |
| AT3G17470 | 0.8 | 8.86E-04 | 0.2 | 2.42E-01 | 0.5 | 1.71E-02 | -0.5 | 3.16E-02 | -0.1 | 6.80E-01 | -0.4 | 2.14E-01 | GTP diphosphokinase |
| AT3G05180 | 0.8 | 1.23E-03 | 0.6 | 1.02E-02 | 0.2 | 3.57E-01 | -0.1 | 7.06E-01 | 0.3 | 3.73E-01 | -0.4 | 2.52E-01 | GDSL esterase |
| AT3G16870 | 0.8 | 1.29E-04 | 0.5 | 2.67E-03 | 0.2 | 1.20E-01 | 0.3 | 1.15E-01 | 0.2 | 4.90E-01 | 0.2 | 6.39E-01 | GATA transcription factor 17 |
| AT1G61420 | 0.8 | 8.71E-03 | 0.0 | 9.57E-01 | 0.8 | 3.43E-02 | 0.4 | 5.20E-01 | 0.5 | 4.85E-01 | -0.1 | 9.26E-01 | S-locus lectin protein kinase family protein |
| AT3G23805 | 0.8 | 1.39E-03 | 0.6 | 1.06E-02 | 0.2 | 3.73E-01 | -0.1 | 7.23E-01 | -0.1 | 8.89E-01 | -0.1 | 9.16E-01 | Protein RALF-like 24 |
| AT1G31190 | 0.8 | 1.24E-05 | 0.6 | 1.27E-04 | 0.2 | 1.35E-01 | 0.2 | 8.54E-02 | 0.2 | 1.39E-01 | 0.0 | 9.87E-01 | Phosphatase IMPL1, chloroplastic |
| AT4G34840 | 0.8 | 5.18E-05 | 0.4 | 4.67E-03 | 0.3 | 2.43E-02 | -0.2 | 4.61E-01 | -0.2 | 4.75E-01 | 0.0 | 9.69E-01 | Phosphorylase superfamily protein |
| AT1G73805 | 0.8 | 3.79E-03 | -0.3 | 2.02E-01 | 1.1 | 2.15E-03 | 0.7 | 2.44E-02 | 0.0 | 9.62E-01 | 0.7 | 4.89E-02 | Protein SAR DEFICIENT 1 |
| AT3G44310 | 0.8 | 8.19E-06 | 0.3 | 7.30E-03 | 0.5 | 1.03E-03 | -0.2 | 7.45E-02 | 0.1 | 3.50E-01 | -0.4 | 4.01E-02 | Nitrilase 1 |
| AT3G14650 | 0.8 | 5.07E-05 | 0.2 | 7.32E-02 | 0.5 | 1.83E-03 | 0.1 | 8.43E-01 | 0.1 | 7.59E-01 | -0.1 | 9.20E-01 | Cytochrome P450 72A11 |
| AT2G38460 | 0.8 | 6.17E-05 | 0.6 | 9.55E-04 | 0.2 | 1.40E-01 | -0.1 | 8.33E-01 | 0.2 | 6.75E-01 | -0.3 | 5.00E-01 | Solute carrier family 40 member |
| AT3G61630 | 0.8 | 2.61E-05 | -0.4 | 1.02E-02 | 1.1 | 3.32E-05 | 0.2 | 3.20E-01 | -0.6 | 1.57E-02 | 0.8 | 8.07E-03 | Ethylene-responsive transcription factor CRF6 |
| AT5G27550 | 0.8 | 2.65E-04 | 0.4 | 1.90E-02 | 0.4 | 4.72E-02 | 0.0 | 9.56E-01 | 0.0 | 9.96E-01 | 0.0 | 9.72E-01 | Kinesin-like protein KIN-14S |
| AT5G59820 | 0.8 | 2.08E-03 | -0.4 | 1.40E-01 | 1.1 | 1.38E-03 | 1.5 | 7.60E-04 | 0.0 | 9.87E-01 | 1.5 | 2.41E-03 | Zinc finger protein ZAT12 |
| AT3G62060 | 0.8 | 2.70E-05 | 0.6 | 3.60E-04 | 0.2 | 1.49E-01 | 0.0 | 9.56E-01 | 0.0 | 9.30E-01 | 0.0 | 9.14E-01 | Pectin acetylesterase |
| AT3G05800 | 0.8 | 1.35E-04 | 0.8 | 2.57E-04 | 0.0 | 9.33E-01 | -0.4 | 2.37E-02 | -0.3 | 8.03E-02 | -0.1 | 8.22E-01 | Transcription factor bHLH150 |
| AT3G58690 | 0.8 | 2.79E-05 | 0.5 | 1.71E-03 | 0.3 | 2.91E-02 | -0.4 | 6.46E-02 | -0.3 | 2.67E-01 | -0.1 | 6.87E-01 | Protein kinase superfamily protein |
| AT1G70600 | 0.8 | 4.23E-04 | 0.8 | 6.73E-04 | 0.0 | 9.89E-01 | 0.4 | 1.13E-02 | 0.3 | 1.66E-01 | 0.2 | 4.64E-01 | Large ribosomal subunit protein uL15x |
| AT4G37490 | 0.8 | 3.18E-04 | 0.4 | 2.78E-02 | 0.4 | 4.41E-02 | 0.3 | 1.35E-01 | 0.1 | 5.74E-01 | 0.2 | 5.94E-01 | Cyclin-B1-1 |
| AT1G57820 | 0.8 | 4.54E-05 | 0.6 | 4.65E-04 | 0.2 | 1.88E-01 | 0.1 | 7.04E-01 | 0.1 | 7.95E-01 | 0.0 | 9.81E-01 | E3 ubiquitin-protein ligase ORTHRUS 2 |
| AT3G52590 | 0.8 | 7.23E-05 | 0.7 | 1.87E-04 | 0.0 | 8.07E-01 | 0.3 | 1.37E-01 | 0.1 | 8.08E-01 | 0.2 | 4.36E-01 | Ubiquitin-ribosomal protein eL40y fusion protein |
| AT2G28800 | 0.8 | 6.04E-05 | 0.6 | 4.47E-04 | 0.1 | 2.88E-01 | -0.1 | 4.14E-01 | -0.1 | 3.91E-01 | 0.0 | 9.27E-01 | 63 kDa inner membrane family protein |
| AT4G23220 | 0.8 | 1.71E-02 | 0.3 | 3.17E-01 | 0.5 | 1.80E-01 | -0.1 | 8.36E-01 | -0.3 | 2.62E-01 | 0.3 | 4.52E-01 | Uncharacterized protein |
| AT1G62150 | 0.8 | 6.18E-05 | 0.5 | 1.72E-03 | 0.3 | 8.49E-02 | 0.3 | 1.11E-01 | 0.1 | 8.29E-01 | 0.3 | 3.51E-01 | Mitochondrial transcription termination factor family protein |
| AT2G41840 | 0.8 | 1.72E-05 | 0.7 | 6.47E-05 | 0.1 | 5.83E-01 | 0.2 | 1.05E-01 | 0.0 | 8.55E-01 | 0.2 | 1.99E-01 | Small ribosomal subunit protein uS5x |

| | | | | | | | | | | | | | |
|---|---|---|---|---|---|---|---|---|---|---|---|---|---|
| AT1G11750 | 0.8 | 1.89E-06 | 0.5 | 6.92E-05 | 0.3 | 9.00E-03 | 0.1 | 2.95E-01 | 0.1 | 7.04E-01 | 0.1 | 7.46E-01 | ATP-dependent Clp protease proteolytic subunit |
| AT1G33120 | 0.8 | 4.13E-04 | 0.7 | 9.39E-04 | 0.0 | 8.29E-01 | 0.4 | 1.46E-01 | 0.0 | 9.75E-01 | 0.4 | 2.74E-01 | Large ribosomal subunit protein uL6z |
| AT1G61520 | 0.8 | 3.65E-05 | 1.0 | 1.69E-05 | -0.2 | 1.48E-01 | -0.4 | 1.50E-02 | -0.4 | 2.19E-02 | 0.0 | 9.59E-01 | Photosystem I chlorophyll a |
| AT3G29290 | 0.8 | 1.27E-05 | 0.4 | 5.20E-03 | 0.4 | 3.04E-03 | 0.1 | 7.53E-01 | 0.0 | 8.33E-01 | 0.0 | 9.81E-01 | Pentatricopeptide repeat-containing protein At3g29290 |
| AT5G05730 | 0.8 | 1.48E-05 | 0.6 | 2.04E-04 | 0.2 | 9.51E-02 | 0.1 | 6.59E-01 | -0.2 | 3.54E-01 | 0.4 | 1.78E-01 | anthranilate synthase |
| AT5G41740 | 0.8 | 1.79E-04 | -0.3 | 7.87E-02 | 1.1 | 1.88E-04 | 0.4 | 1.52E-01 | -0.2 | 5.01E-01 | 0.6 | 6.60E-02 | Disease resistance protein family |
| AT5G65990 | 0.8 | 2.41E-05 | 0.2 | 1.06E-01 | 0.6 | 8.53E-04 | -0.2 | 3.67E-01 | -0.1 | 6.90E-01 | -0.1 | 8.22E-01 | Amino acid transporter AVT3A |
| AT3G21150 | 0.8 | 1.40E-03 | 0.0 | 9.35E-01 | 0.8 | 7.66E-03 | | | | | | | B-box zinc finger protein 32 |
| AT4G30760 | 0.8 | 1.76E-04 | 0.3 | 3.43E-02 | 0.4 | 1.49E-02 | -0.1 | 8.13E-01 | -0.3 | 4.07E-01 | 0.2 | 6.63E-01 | Endonuclease or glycosyl hydrolase |
| AT1G55915 | 0.8 | 1.68E-05 | 0.3 | 8.68E-03 | 0.4 | 3.98E-03 | 0.3 | 2.12E-01 | 0.3 | 4.09E-01 | 0.1 | 8.51E-01 | At1g55915 |
| AT4G33680 | 0.8 | 2.89E-06 | 0.7 | 1.62E-05 | 0.1 | 2.79E-01 | 0.5 | 1.07E-03 | 0.3 | 4.75E-02 | 0.2 | 3.13E-01 | LL-diaminopimelate aminotransferase, chloroplastic |
| AT2G37690 | 0.8 | 9.58E-04 | 0.4 | 6.75E-02 | 0.4 | 5.67E-02 | 1.2 | 6.69E-06 | 0.5 | 2.25E-02 | 0.7 | 7.54E-03 | phosphoribosylaminoimidazole carboxylase |
| AT3G30775 | 0.8 | 4.50E-04 | 1.0 | 1.61E-04 | -0.2 | 2.54E-01 | -0.2 | 4.36E-01 | -0.8 | 2.99E-02 | 0.5 | 1.98E-01 | Proline dehydrogenase 1, mitochondrial |
| AT3G21530 | 0.8 | 5.53E-04 | 0.3 | 1.26E-01 | 0.5 | 2.15E-02 | 0.5 | 1.33E-01 | 0.0 | 9.81E-01 | 0.5 | 1.73E-01 | hypothetical protein |
| AT5G43080 | 0.8 | 1.93E-04 | 0.6 | 1.64E-03 | 0.2 | 3.18E-01 | 0.7 | 3.00E-02 | 0.3 | 5.09E-01 | 0.4 | 2.57E-01 | Cyclin A31 |
| AT3G01370 | 0.8 | 2.62E-05 | 0.6 | 4.32E-04 | 0.2 | 8.98E-02 | 0.2 | 5.19E-01 | 0.2 | 4.81E-01 | 0.0 | 9.15E-01 | CRM-domain containing factor CFM2, chloroplastic |
| AT1G09530 | 0.8 | 9.63E-06 | 0.1 | 3.66E-01 | 0.7 | 2.21E-04 | -0.5 | 2.19E-02 | -0.8 | 2.21E-03 | 0.3 | 2.90E-01 | Transcription factor PIF3 |
| AT5G61540 | 0.8 | 2.04E-04 | 0.2 | 1.30E-01 | 0.5 | 8.41E-03 | 0.2 | 2.40E-01 | -0.1 | 8.46E-01 | 0.3 | 2.61E-01 | Isoaspartyl peptidase |
| AT3G54660 | 0.8 | 2.45E-06 | 0.4 | 5.15E-04 | 0.4 | 1.47E-03 | 0.3 | 8.26E-02 | 0.2 | 2.74E-01 | 0.1 | 8.06E-01 | Glutathione reductase, chloroplastic |
| AT3G25470 | 0.8 | 2.43E-06 | 0.6 | 3.16E-05 | 0.2 | 5.37E-02 | 0.2 | 7.99E-02 | 0.0 | 9.15E-01 | 0.2 | 1.63E-01 | Hemolysin-like |
| AT3G50060 | 0.8 | 3.90E-02 | -0.3 | 4.13E-01 | 1.1 | 2.32E-02 | -0.8 | 3.75E-03 | -0.9 | 4.20E-03 | 0.1 | 9.16E-01 | Transcription factor MYB77 |
| AT4G23130 | 0.8 | 2.71E-04 | 0.3 | 1.26E-01 | 0.5 | 1.03E-02 | 0.8 | 9.17E-03 | 0.2 | 5.35E-01 | 0.5 | 9.40E-02 | Uncharacterized protein |
| AT3G14420 | 0.8 | 6.99E-06 | 0.4 | 7.35E-04 | 0.4 | 5.59E-03 | 0.2 | 2.09E-01 | 0.2 | 1.11E-01 | -0.1 | 7.42E-01 | #NAME? |
| AT5G24150 | 0.8 | 1.49E-04 | 0.1 | 7.39E-01 | 0.7 | 1.10E-03 | -0.7 | 4.67E-02 | 0.8 | 1.16E-02 | -1.6 | 8.67E-04 | Squalene monooxygenase |
| AT3G44860 | 0.8 | 1.78E-02 | 0.3 | 2.98E-01 | 0.5 | 1.92E-01 | 0.7 | 6.91E-01 | 1.2 | 5.12E-01 | -0.5 | 8.31E-01 | Farnesoic acid carboxyl-O-methyltransferase |
| AT2G46310 | 0.8 | 4.47E-04 | 0.2 | 1.71E-01 | 0.5 | 1.56E-02 | 0.3 | 1.25E-01 | -0.4 | 1.13E-01 | 0.8 | 9.77E-03 | Ethylene-responsive transcription factor CRF5 |
| AT4G32260 | 0.8 | 1.15E-05 | 0.7 | 3.97E-05 | 0.1 | 6.28E-01 | 0.0 | 8.72E-01 | 0.1 | 7.30E-01 | -0.1 | 6.86E-01 | AT4g32260 |
| AT1G52300 | 0.8 | 3.27E-04 | 0.9 | 3.63E-04 | -0.1 | 7.79E-01 | 0.3 | 6.30E-02 | 0.1 | 7.94E-01 | 0.3 | 3.11E-01 | Large ribosomal subunit protein eL37y |
| AT5G67220 | 0.8 | 1.01E-06 | 0.3 | 1.07E-03 | 0.5 | 2.13E-04 | 0.1 | 1.80E-01 | 0.0 | 7.37E-01 | 0.2 | 2.14E-01 | hypothetical protein |
| AT3G02730 | 0.8 | 1.83E-05 | 0.6 | 3.46E-04 | 0.2 | 6.99E-02 | 0.3 | 3.33E-02 | 0.4 | 3.16E-02 | 0.0 | 8.74E-01 | Thioredoxin F1, chloroplastic |
| AT3G11510 | 0.8 | 9.26E-05 | 0.8 | 1.93E-04 | 0.0 | 9.33E-01 | 0.4 | 6.40E-02 | 0.1 | 7.99E-01 | 0.3 | 2.88E-01 | Small ribosomal subunit protein uS11y |
| AT5G22880 | 0.8 | 5.18E-06 | 0.8 | 1.37E-05 | 0.0 | 8.59E-01 | 0.4 | 4.15E-02 | 0.1 | 6.37E-01 | 0.3 | 3.24E-01 | Histone H2B.10 |
| AT4G29410 | 0.8 | 5.05E-04 | 0.8 | 8.95E-04 | 0.0 | 9.45E-01 | 0.4 | 4.80E-03 | 0.2 | 2.74E-01 | 0.2 | 2.45E-01 | Large ribosomal subunit protein eL28y |
| AT3G02560 | 0.8 | 7.16E-05 | 0.8 | 9.24E-05 | 0.0 | 7.50E-01 | 0.1 | 5.62E-01 | -0.1 | 3.75E-01 | 0.2 | 2.05E-01 | Small ribosomal subunit protein eS7y |
| AT5G11300 | 0.8 | 2.99E-05 | 0.6 | 3.44E-04 | 0.2 | 1.91E-01 | 0.1 | 8.86E-01 | 0.1 | 7.67E-01 | -0.1 | 8.95E-01 | Mitotic-like cyclin 3B from Arabidopsi |
| AT1G69550 | 0.8 | 1.09E-04 | 0.2 | 1.70E-01 | 0.6 | 3.19E-03 | 0.3 | 1.30E-01 | 0.4 | 1.08E-01 | -0.1 | 8.61E-01 | ADP-ribosyl cyclase |
| AT1G29630 | 0.8 | 6.33E-06 | 0.3 | 6.62E-03 | 0.5 | 1.36E-03 | 0.2 | 6.48E-01 | 0.0 | 9.80E-01 | 0.2 | 6.59E-01 | 5'-3' exonuclease family protein |
| AT4G19810 | 0.8 | 4.33E-06 | 0.4 | 7.84E-04 | 0.4 | 3.05E-03 | 1.8 | 1.29E-07 | 0.5 | 3.49E-02 | 1.3 | 1.85E-05 | Class V chitinase |
| AT3G15060 | 0.8 | 1.41E-05 | 0.5 | 9.23E-04 | 0.3 | 1.80E-02 | -0.2 | 2.10E-01 | -0.5 | 3.27E-02 | 0.2 | 3.79E-01 | Ras-related protein RABA1g |
| AT3G09780 | 0.8 | 1.64E-03 | 0.8 | 2.79E-03 | 0.0 | 9.47E-01 | -0.1 | 7.73E-01 | -0.2 | 4.83E-01 | 0.1 | 7.43E-01 | Serine |
| AT5G62710 | 0.8 | 4.38E-06 | 0.2 | 1.80E-02 | 0.6 | 3.57E-04 | 0.0 | 8.69E-01 | -0.1 | 6.65E-01 | 0.1 | 6.26E-01 | Receptor-like protein kinase |
| AT3G29000 | 0.8 | 1.57E-02 | -0.9 | 3.56E-02 | 1.7 | 2.71E-03 | 2.7 | 4.66E-05 | 1.2 | 4.93E-02 | 1.4 | 4.03E-03 | Probable calcium-binding protein CML45 |
| AT2G06925 | 0.8 | 4.76E-06 | 0.8 | 8.90E-06 | 0.0 | 8.69E-01 | -0.2 | 1.40E-01 | -0.3 | 7.84E-02 | 0.1 | 7.91E-01 | Phospholipase A2 family protein |
| AT5G27850 | 0.8 | 2.33E-04 | 0.8 | 5.83E-04 | 0.0 | 7.99E-01 | 0.4 | 5.38E-02 | 0.1 | 7.80E-01 | 0.3 | 2.62E-01 | Ribosomal protein L18e |
| AT5G39740 | 0.8 | 3.86E-05 | 0.7 | 2.72E-04 | 0.1 | 2.90E-01 | 0.5 | 3.85E-04 | 0.3 | 4.35E-02 | 0.2 | 1.99E-01 | Ribosomal protein L5 eukaryotic C-terminal domain-containing protein |
| AT4G10060 | 0.8 | 1.43E-04 | 0.6 | 1.64E-03 | 0.2 | 1.86E-01 | -0.1 | 5.63E-01 | -0.1 | 6.76E-01 | 0.0 | 9.78E-01 | Non-lysosomal glucosylceramidase |
| AT5G53450 | 0.8 | 3.52E-04 | 0.1 | 5.72E-01 | 0.7 | 2.45E-03 | 2.8 | 2.87E-07 | 0.2 | 7.42E-01 | 2.7 | 1.08E-04 | OBP3-responsive protein 1 |
| AT4G28950 | 0.8 | 2.55E-03 | 0.7 | 7.57E-03 | 0.1 | 7.18E-01 | 0.1 | 7.79E-01 | -0.2 | 6.45E-01 | 0.4 | 4.96E-01 | Rac-like GTP-binding protein ARAC7 |
| AT5G53940 | 0.8 | 2.23E-03 | 0.3 | 1.51E-01 | 0.5 | 6.82E-02 | 0.2 | 5.61E-01 | 0.1 | 8.57E-01 | 0.1 | 8.38E-01 | Protein yippee-like |
| AT2G23030 | 0.8 | 8.64E-05 | 0.2 | 2.40E-01 | 0.6 | 2.20E-03 | 0.2 | 3.65E-01 | 0.3 | 2.62E-01 | -0.1 | 8.14E-01 | Serine |
| AT5G62410 | 0.8 | 4.61E-06 | 0.4 | 2.25E-03 | 0.4 | 1.47E-03 | 0.3 | 3.24E-01 | 0.2 | 5.96E-01 | 0.1 | 8.52E-01 | Structural maintenance of chromosomes protein 2-1 |
| AT3G28200 | 0.8 | 1.21E-05 | 0.4 | 2.36E-03 | 0.4 | 4.40E-03 | -0.2 | 3.82E-01 | 0.0 | 9.83E-01 | -0.2 | 5.04E-01 | Peroxidase 31 |
| AT4G01900 | 0.8 | 3.80E-05 | 0.7 | 1.59E-04 | 0.1 | 5.14E-01 | 0.1 | 5.02E-01 | 0.2 | 1.39E-01 | -0.1 | 5.47E-01 | Nitrogen regulatory protein P-II homolog |
| AT1G34030 | 0.8 | 3.13E-05 | 0.8 | 7.02E-05 | 0.0 | 9.30E-01 | 0.6 | 1.01E-03 | 0.2 | 2.75E-01 | 0.4 | 8.55E-02 | Small ribosomal subunit protein uS13z |
| AT1G53090 | 0.8 | 9.30E-05 | 0.2 | 1.14E-01 | 0.6 | 3.78E-03 | 0.2 | 2.58E-01 | 0.2 | 3.74E-01 | 0.0 | 9.62E-01 | SPA1-related 4 |
| AT5G63530 | 0.8 | 7.24E-07 | 0.8 | 1.56E-06 | 0.0 | 7.96E-01 | 0.6 | 5.92E-04 | 0.6 | 3.08E-03 | 0.1 | 8.60E-01 | FP3 |
| AT3G03970 | 0.8 | 3.75E-05 | 0.3 | 2.38E-02 | 0.5 | 4.74E-03 | -0.3 | 9.92E-02 | -0.2 | 2.94E-01 | -0.1 | 7.64E-01 | Protein SINE2 |
| AT5G10540 | 0.8 | 1.77E-06 | 0.7 | 1.34E-05 | 0.1 | 1.44E-01 | -0.2 | 2.92E-01 | -0.1 | 7.46E-01 | -0.1 | 6.97E-01 | Probable cytosolic oligopeptidase A |
| AT3G04840 | 0.8 | 2.93E-04 | 0.8 | 7.23E-04 | 0.0 | 8.01E-01 | 0.4 | 5.07E-03 | 0.2 | 1.88E-01 | 0.2 | 2.99E-01 | Small ribosomal subunit protein eS1z |
| AT3G57560 | 0.8 | 1.36E-05 | 0.7 | 6.85E-05 | 0.1 | 3.70E-01 | 0.3 | 4.84E-02 | 0.1 | 7.65E-01 | 0.2 | 2.27E-01 | Acetylglutamate kinase, chloroplastic |
| AT3G45030 | 0.8 | 2.72E-04 | 0.8 | 3.76E-04 | 0.0 | 8.98E-01 | 0.3 | 8.21E-02 | 0.1 | 8.03E-01 | 0.2 | 3.59E-01 | Small ribosomal subunit protein uS10z |
| AT4G30110 | 0.8 | 1.29E-04 | 1.1 | 3.05E-05 | -0.2 | 1.15E-01 | 0.0 | 9.93E-01 | 0.1 | 8.20E-01 | -0.1 | 8.33E-01 | Cadmium |
| AT1G53350 | 0.8 | 6.94E-06 | 0.4 | 1.29E-03 | 0.4 | 4.84E-03 | 0.2 | 4.40E-01 | 0.1 | 8.72E-01 | 0.1 | 6.98E-01 | Probable disease resistance RPP8-like protein 2 |
| AT3G54640 | 0.8 | 1.26E-04 | 0.6 | 1.07E-03 | 0.2 | 2.59E-01 | 0.8 | 2.98E-04 | 0.2 | 3.28E-01 | 0.6 | 2.82E-02 | Tryptophan synthase alpha chain, chloroplastic |
| AT2G40330 | 0.8 | 5.20E-04 | 0.9 | 3.22E-04 | -0.1 | 5.46E-01 | 0.7 | 2.22E-04 | 0.2 | 2.86E-01 | 0.5 | 1.48E-02 | Abscisic acid receptor PYL6 |
| AT5G54490 | 0.8 | 1.09E-02 | -0.1 | 7.93E-01 | 0.9 | 2.54E-02 | 2.0 | 2.87E-04 | 0.0 | 9.66E-01 | 1.9 | 2.59E-03 | Calcium-binding protein PBP1 |
| AT2G09990 | 0.8 | 4.04E-05 | 0.6 | 4.42E-04 | 0.2 | 1.62E-01 | 0.3 | 8.09E-02 | 0.1 | 7.10E-01 | 0.2 | 4.08E-01 | Small ribosomal subunit protein uS9z |
| AT4G16563 | 0.8 | 4.50E-03 | 0.6 | 3.27E-02 | 0.2 | 3.94E-01 | -0.7 | 4.86E-04 | -0.9 | 1.11E-04 | 0.2 | 3.33E-01 | Probable aspartyl protease At4g16563 |
| AT4G31390 | 0.8 | 1.67E-05 | 0.4 | 4.36E-03 | 0.4 | 4.23E-03 | 0.3 | 9.39E-02 | 0.1 | 7.80E-01 | 0.2 | 3.37E-01 | Protein kinase superfamily protein |
| AT2G29460 | 0.8 | 2.17E-02 | -1.1 | 2.57E-02 | 1.9 | 3.08E-03 | -0.6 | 3.95E-01 | -2.0 | 3.89E-02 | 1.4 | 2.47E-01 | Glutathione S-transferase U4 |
| AT1G28560 | 0.8 | 2.09E-04 | 0.3 | 4.94E-02 | 0.5 | 2.05E-02 | -0.1 | 7.55E-01 | 0.0 | 9.46E-01 | -0.2 | 6.69E-01 | snRNA activating complex family protein |
| AT1G74440 | 0.8 | 1.46E-04 | 0.2 | 1.88E-01 | 0.6 | 4.19E-03 | 0.4 | 2.23E-02 | -0.3 | 1.85E-01 | 0.7 | 5.89E-03 | ER membrane protein, putative |
| AT1G70070 | 0.8 | 5.95E-06 | 0.5 | 7.15E-04 | 0.4 | 4.63E-03 | 0.0 | 9.54E-01 | 0.0 | 9.60E-01 | 0.0 | 9.34E-01 | DExH-box ATP-dependent RNA helicase DExH15 chloroplastic |
| AT2G35340 | 0.8 | 4.77E-05 | 0.3 | 3.41E-02 | 0.5 | 4.58E-03 | 0.4 | 3.94E-01 | 0.3 | 5.74E-01 | 0.1 | 9.08E-01 | RNA helicase |
| AT3G47570 | 0.8 | 8.37E-05 | 0.6 | 1.95E-03 | 0.3 | 1.01E-01 | 0.4 | 1.90E-02 | -0.1 | 6.94E-01 | 0.5 | 3.09E-02 | Probable LRR receptor-like serine |
| AT4G04870 | 0.8 | 4.70E-05 | 0.4 | 1.37E-02 | 0.4 | 8.33E-03 | 0.3 | 9.99E-02 | 0.3 | 1.97E-01 | 0.0 | 9.15E-01 | Cardiolipin synthase , mitochondrial |
| AT1G03130 | 0.8 | 1.93E-05 | 1.1 | 5.79E-06 | -0.3 | 3.76E-02 | -0.6 | 6.94E-04 | -0.1 | 7.41E-01 | -0.5 | 1.75E-02 | Photosystem I reaction center subunit II-2, chloroplastic |
| AT2G16500 | 0.8 | 1.34E-06 | 0.5 | 7.97E-05 | 0.3 | 3.76E-03 | 0.3 | 6.13E-02 | 0.1 | 6.45E-01 | 0.2 | 3.73E-01 | Arginine decarboxylase 1, chloroplastic |
| AT3G47640 | 0.8 | 1.06E-04 | 0.5 | 5.87E-03 | 0.3 | 4.69E-02 | 1.6 | 1.07E-05 | 0.8 | 8.71E-03 | 0.7 | 2.41E-02 | Transcription factor bHLH47 |
| AT1G02640 | 0.8 | 3.39E-04 | 1.1 | 7.41E-05 | -0.3 | 1.43E-01 | -0.4 | 1.33E-02 | -0.9 | 1.95E-04 | 0.4 | 5.34E-02 | Probable beta-D-xylosidase 2 |
| AT2G04080 | 0.8 | 2.95E-05 | 0.4 | 8.49E-03 | 0.4 | 6.44E-03 | 0.3 | 2.66E-01 | 0.1 | 6.32E-01 | 0.1 | 7.51E-01 | Protein DETOXIFICATION |
| AT3G48320 | 0.8 | 1.09E-04 | 0.2 | 1.40E-01 | 0.6 | 3.99E-03 | 0.4 | 1.13E-01 | 0.3 | 3.95E-01 | 0.2 | 6.88E-01 | Cytochrome P450 71A21 |
| AT5G17400 | 0.8 | 1.42E-05 | 0.5 | 1.24E-03 | 0.3 | 1.34E-02 | -0.4 | 3.04E-02 | -0.2 | 4.13E-01 | -0.2 | 3.43E-01 | ADP |

| | | | | | | | | | | | | | |
|---|---|---|---|---|---|---|---|---|---|---|---|---|---|
| AT5G16540 | 0.8 | 5.25E-05 | 0.5 | 1.82E-03 | 0.3 | 4.97E-02 | -0.2 | 4.32E-01 | 0.1 | 7.62E-01 | -0.2 | 3.01E-01 | Zinc finger nuclease 3 |
| AT1G72940 | 0.8 | 1.47E-03 | -0.1 | 7.36E-01 | 0.9 | 3.75E-03 | 0.5 | 2.42E-02 | -0.7 | 1.35E-02 | 1.2 | 1.47E-03 | At1g72940 |
| AT1G05140 | 0.8 | 1.42E-06 | 0.7 | 1.26E-05 | 0.1 | 9.39E-02 | 0.1 | 4.44E-01 | 0.1 | 5.36E-01 | 0.0 | 9.95E-01 | Probable membrane metalloprotease ARASP2, chloroplastic |
| AT2G40750 | 0.8 | 1.52E-03 | 0.1 | 6.65E-01 | 0.7 | 8.81E-03 | 1.6 | 3.94E-06 | 1.4 | 8.61E-05 | 0.2 | 3.73E-01 | Probable WRKY transcription factor 54 |
| AT1G04870 | 0.8 | 4.38E-06 | 0.4 | 7.71E-04 | 0.4 | 2.68E-03 | 0.4 | 1.61E-03 | 0.2 | 1.95E-01 | 0.2 | 1.62E-01 | Protein arginine N-methyltransferase PRMT10 |
| AT2G39460 | 0.8 | 3.82E-05 | 0.8 | 1.38E-04 | 0.1 | 6.16E-01 | 0.6 | 1.02E-03 | 0.3 | 8.91E-02 | 0.3 | 2.28E-01 | Large ribosomal subunit protein uL23z |
| AT4G09320 | 0.8 | 2.58E-04 | 0.9 | 2.99E-04 | -0.1 | 7.94E-01 | 0.3 | 9.56E-02 | 0.1 | 6.25E-01 | 0.2 | 4.59E-01 | Nucleoside diphosphate kinase 1 |
| AT2G19730 | 0.8 | 2.48E-05 | 0.8 | 7.57E-05 | 0.0 | 7.32E-01 | 0.5 | 2.68E-03 | 0.3 | 1.33E-01 | 0.2 | 2.74E-01 | Large ribosomal subunit protein eL28z |
| AT1G78380 | 0.8 | 9.34E-06 | 0.4 | 2.76E-03 | 0.4 | 2.49E-03 | 0.4 | 1.68E-02 | 0.1 | 5.07E-01 | 0.3 | 2.29E-01 | Glutathione S-transferase U19 |
| AT4G04350 | 0.8 | 5.20E-06 | 0.6 | 1.68E-04 | 0.3 | 2.14E-02 | 0.3 | 1.78E-01 | 0.2 | 4.31E-01 | 0.1 | 7.93E-01 | Leucine--tRNA ligase, chloroplastic |
| AT1G72680 | 0.8 | 2.12E-04 | 0.2 | 3.16E-01 | 0.7 | 3.87E-03 | 0.0 | 8.92E-01 | -0.5 | 7.13E-02 | 0.4 | 1.83E-01 | Probable cinnamyl alcohol dehydrogenase 1 |
| AT1G79080 | 0.8 | 2.46E-06 | 0.5 | 2.85E-04 | 0.4 | 3.69E-03 | 0.0 | 9.81E-01 | 0.1 | 4.29E-01 | -0.1 | 5.38E-01 | Pentatricopeptide repeat-containing protein At1g79080, chloroplastic |
| AT4G26860 | 0.8 | 4.53E-06 | 0.3 | 3.55E-03 | 0.5 | 7.76E-04 | 0.3 | 1.53E-02 | 0.2 | 3.43E-01 | 0.2 | 2.80E-01 | Pyridoxal phosphate homeostasis protein |
| AT3G03670 | 0.8 | 1.51E-04 | 0.7 | 1.07E-03 | 0.2 | 3.10E-01 | 1.3 | 2.19E-07 | 0.5 | 2.84E-03 | 0.7 | 3.13E-04 | Peroxidase 28 |
| AT2G41560 | 0.8 | 1.33E-05 | 0.8 | 2.99E-05 | 0.0 | 9.08E-01 | -0.3 | 1.82E-01 | -0.5 | 2.24E-02 | 0.3 | 3.43E-01 | Calcium-transporting ATPase |
| AT1G33170 | 0.8 | 8.15E-06 | 0.7 | 8.52E-05 | 0.2 | 1.20E-01 | 0.1 | 4.10E-01 | 0.1 | 6.38E-01 | 0.0 | 9.07E-01 | Methyltransferase |
| AT4G12390 | 0.8 | 8.32E-06 | 0.8 | 2.37E-05 | 0.0 | 7.22E-01 | 0.2 | 3.22E-01 | 0.1 | 8.21E-01 | 0.1 | 6.87E-01 | hypothetical protein |
| AT1G16110 | 0.8 | 1.97E-05 | 0.3 | 4.45E-02 | 0.6 | 1.37E-03 | 0.0 | 9.73E-01 | -0.3 | 4.61E-01 | 0.3 | 5.55E-01 | Wall-associated receptor kinase-like 6 |
| AT3G16730 | 0.8 | 5.83E-06 | 0.1 | 6.08E-01 | 0.8 | 1.10E-04 | -0.3 | 2.80E-01 | -0.1 | 8.31E-01 | -0.2 | 5.60E-01 | Condensin-2 complex subunit H2 |
| AT3G01060 | 0.8 | 2.87E-06 | 0.6 | 6.74E-05 | 0.2 | 2.34E-02 | 0.7 | 5.89E-03 | 0.6 | 4.83E-02 | 0.2 | 5.36E-01 | Lysine-tRNA ligase |
| AT3G52840 | 0.8 | 3.86E-04 | 1.0 | 2.08E-04 | -0.1 | 4.49E-01 | 0.1 | 7.09E-01 | -0.3 | 2.42E-02 | 0.4 | 2.93E-02 | Beta-galactosidase |
| AT3G06483 | 0.8 | 1.73E-06 | 0.7 | 1.06E-05 | 0.1 | 2.33E-01 | -0.1 | 5.49E-01 | 0.0 | 8.62E-01 | 0.0 | 8.19E-01 | Protein-serine |
| AT3G26085 | 0.8 | 6.14E-05 | 0.2 | 1.11E-01 | 0.6 | 2.34E-03 | 0.0 | 9.35E-01 | -0.1 | 6.70E-01 | 0.1 | 6.68E-01 | CAAX amino terminal protease family protein |
| AT4G33470 | 0.8 | 1.93E-05 | 0.8 | 7.24E-05 | 0.1 | 5.60E-01 | 0.1 | 7.01E-01 | 0.1 | 5.69E-01 | 0.0 | 8.73E-01 | HDA14 |
| AT1G26840 | 0.8 | 2.58E-04 | 0.6 | 5.76E-03 | 0.3 | 1.50E-01 | 0.6 | 1.23E-01 | 0.6 | 2.12E-01 | 0.1 | 9.28E-01 | Origin of replication complex subunit 6 |
| AT4G32980 | 0.8 | 1.26E-04 | 0.5 | 1.00E-02 | 0.4 | 3.03E-02 | 0.5 | 1.80E-02 | 0.7 | 3.49E-03 | -0.2 | 4.60E-01 | Homeobox domain-containing protein |
| AT5G53200 | 0.8 | 4.15E-02 | 1.1 | 1.32E-02 | -0.3 | 5.20E-01 | | | | | | | Transcription factor TRY |
| AT2G45480 | 0.8 | 1.62E-03 | 0.8 | 4.79E-03 | 0.1 | 7.19E-01 | -0.1 | 7.69E-01 | -0.3 | 5.24E-01 | 0.2 | 7.84E-01 | Growth-regulating factor |
| AT5G36930 | 0.8 | 1.03E-04 | 0.5 | 7.73E-03 | 0.4 | 3.27E-02 | 0.6 | 2.53E-03 | 0.1 | 5.52E-01 | 0.5 | 4.41E-02 | Disease resistance protein family |
| AT3G02040 | 0.8 | 6.52E-04 | 0.7 | 3.57E-03 | 0.1 | 4.86E-01 | 1.3 | 1.38E-02 | 0.7 | 2.59E-01 | 0.6 | 2.52E-01 | Glycerophosphodiester phosphodiesterase GDPD1, chloroplastic |
| AT5G39550 | 0.8 | 2.72E-05 | 0.6 | 3.98E-04 | 0.2 | 1.22E-01 | 0.3 | 1.77E-01 | 0.2 | 3.45E-01 | 0.0 | 8.96E-01 | E3 ubiquitin-protein ligase ORTHRUS 1 |
| AT5G08650 | 0.8 | 4.38E-06 | 0.5 | 2.42E-04 | 0.3 | 8.59E-03 | 0.2 | 2.59E-01 | 0.2 | 3.62E-01 | 0.0 | 9.81E-01 | Translation factor GUF1 homolog, mitochondrial |
| AT5G39190 | 0.8 | 9.77E-04 | 0.3 | 1.24E-01 | 0.5 | 4.58E-02 | 0.8 | 6.29E-03 | 0.7 | 4.39E-02 | 0.2 | 5.39E-01 | Germin-like protein |
| AT1G50180 | 0.8 | 6.15E-04 | 0.0 | 8.75E-01 | 0.8 | 4.63E-03 | 1.0 | 7.79E-03 | 0.8 | 5.63E-02 | 0.2 | 5.46E-01 | Putative disease resistance protein At1g50180 |
| AT5G17300 | 0.8 | 6.74E-03 | 0.6 | 5.93E-02 | 0.3 | 3.63E-01 | -0.6 | 4.81E-02 | -0.8 | 2.84E-02 | 0.1 | 7.97E-01 | Homeodomain-like superfamily protein |
| AT1G18485 | 0.8 | 3.88E-05 | 0.8 | 1.62E-04 | 0.1 | 5.45E-01 | 0.0 | 9.87E-01 | -0.1 | 6.39E-01 | 0.1 | 6.56E-01 | Pentatricopeptide repeat-containing protein At1g18485 |
| AT3G24830 | 0.8 | 8.42E-06 | 0.7 | 4.18E-05 | 0.1 | 3.92E-01 | 0.1 | 2.70E-01 | 0.0 | 9.66E-01 | 0.1 | 4.80E-01 | Large ribosomal subunit protein uL13y |
| AT1G03220 | 0.8 | 6.79E-05 | 0.8 | 1.70E-04 | 0.0 | 8.17E-01 | 1.1 | 4.96E-07 | 0.2 | 2.57E-01 | 0.9 | 2.05E-04 | Eukaryotic aspartyl protease family protein |
| AT1G23100 | 0.8 | 3.17E-03 | 0.8 | 4.37E-03 | 0.0 | 9.74E-01 | 1.1 | 3.04E-03 | 0.7 | 6.15E-02 | 0.4 | 3.33E-01 | GroES-like family protein protein) |
| AT5G13280 | 0.8 | 6.63E-06 | 0.5 | 8.24E-04 | 0.4 | 4.73E-03 | 0.2 | 1.22E-01 | 0.0 | 9.97E-01 | 0.2 | 2.17E-01 | Aspartokinase 1, chloroplastic |
| AT4G02510 | 0.8 | 2.70E-06 | 0.5 | 3.53E-04 | 0.4 | 2.45E-03 | 0.3 | 3.07E-01 | 0.0 | 9.87E-01 | 0.3 | 4.49E-01 | Translocase of chloroplast 159, chloroplastic |
| AT1G77720 | 0.9 | 2.68E-05 | 0.3 | 3.28E-02 | 0.6 | 2.35E-03 | -0.1 | 7.63E-01 | 0.1 | 6.89E-01 | -0.2 | 5.55E-01 | Serine |
| AT2G45270 | 0.9 | 2.28E-06 | 0.6 | 4.16E-05 | 0.2 | 3.75E-02 | 0.2 | 1.92E-01 | 0.1 | 8.11E-01 | 0.2 | 5.12E-01 | Probable tRNA N6-adenosine threonylcarbamoyltransferase, mitochondrial A synthase) A37 threonylcarbamoyladenosine biosynthesis protein GCP1) |
| AT2G32440 | 0.9 | 7.39E-05 | 0.4 | 1.10E-02 | 0.4 | 1.58E-02 | -0.1 | 7.36E-01 | -0.2 | 3.49E-01 | 0.1 | 6.51E-01 | Ent-kaurenoic acid oxidase 2 |
| AT3G48420 | 0.9 | 4.36E-05 | 1.1 | 1.91E-05 | -0.2 | 1.57E-01 | 0.2 | 1.25E-01 | 0.4 | 2.40E-02 | -0.2 | 4.66E-01 | CBBY-like protein |
| AT1G14860 | 0.9 | 4.05E-05 | 0.8 | 1.27E-04 | 0.1 | 6.86E-01 | -0.2 | 6.48E-01 | -0.4 | 3.28E-01 | 0.2 | 6.91E-01 | Nudix hydrolase 18, mitochondrial |
| AT3G28040 | 0.9 | 6.44E-06 | 0.6 | 9.19E-05 | 0.2 | 6.92E-02 | 0.1 | 3.95E-01 | 0.0 | 8.95E-01 | 0.2 | 4.07E-01 | Probably inactive leucine-rich repeat receptor-like protein kinase At3g28040 |
| AT5G51910 | 0.9 | 4.23E-04 | 0.7 | 2.09E-03 | 0.1 | 5.09E-01 | 0.0 | 9.62E-01 | -0.6 | 2.39E-01 | 0.5 | 3.18E-01 | Transcription factor TCP19 |
| AT4G33420 | 0.9 | 4.38E-04 | 0.7 | 1.86E-03 | 0.1 | 5.25E-01 | 1.0 | 2.95E-06 | 0.2 | 2.08E-01 | 0.7 | 5.42E-04 | Peroxidase |
| AT4G18440 | 0.9 | 2.83E-04 | 1.3 | 2.44E-05 | -0.5 | 2.46E-02 | 2.0 | 4.17E-05 | 1.7 | 5.07E-04 | 0.3 | 6.49E-01 | Adenylosuccinate lyase |
| AT4G16720 | 0.9 | 1.62E-05 | 0.7 | 1.65E-04 | 0.2 | 1.59E-01 | 0.3 | 2.28E-03 | 0.1 | 3.28E-01 | 0.2 | 1.51E-01 | Large ribosomal subunit protein eL15z |
| AT4G11470 | 0.9 | 1.98E-03 | 0.6 | 2.22E-02 | 0.3 | 2.94E-01 | 0.3 | 5.55E-01 | -0.6 | 2.09E-01 | 0.9 | 3.44E-02 | Cysteine-rich RLK 31 |
| AT5G50740 | 0.9 | 1.25E-05 | 1.2 | 2.58E-06 | -0.3 | 1.41E-02 | -0.3 | 8.91E-02 | 0.0 | 9.98E-01 | -0.3 | 2.22E-01 | At5g50730 |
| AT4G16800 | 0.9 | 3.77E-06 | 0.8 | 1.34E-05 | 0.1 | 6.20E-01 | -0.1 | 7.66E-01 | 0.2 | 4.69E-01 | -0.2 | 4.07E-01 | Probable enoyl-CoA hydratase 2, mitochondrial |
| AT2G36080 | 0.9 | 1.03E-05 | 0.5 | 1.80E-03 | 0.4 | 6.35E-03 | 0.0 | 9.88E-01 | -0.2 | 5.89E-01 | 0.2 | 5.69E-01 | AP2 |
| AT3G23550 | 0.9 | 7.83E-04 | 1.0 | 2.86E-04 | -0.2 | 3.67E-01 | 0.5 | 1.37E-01 | -1.3 | 6.19E-03 | 1.8 | 1.59E-03 | Protein DETOXIFICATION 18 |
| AT4G12740 | 0.9 | 1.13E-03 | 0.7 | 7.94E-03 | 0.2 | 4.16E-01 | 0.3 | 5.11E-01 | 0.3 | 5.22E-01 | 0.0 | 9.70E-01 | Adenine DNA glycosylase |
| AT3G05590 | 0.9 | 1.39E-05 | 0.7 | 1.40E-04 | 0.2 | 1.56E-01 | 0.5 | 4.35E-03 | 0.2 | 2.15E-01 | 0.3 | 2.56E-01 | Ribosomal protein L18e |
| AT1G31650 | 0.9 | 5.97E-06 | 0.5 | 5.31E-04 | 0.4 | 7.35E-03 | -0.1 | 5.66E-01 | -0.2 | 3.58E-01 | 0.1 | 7.63E-01 | RHO guanyl-nucleotide exchange factor 14 |
| AT2G40510 | 0.9 | 1.78E-05 | 0.7 | 1.12E-04 | 0.1 | 2.99E-01 | 0.4 | 2.49E-03 | 0.3 | 5.63E-02 | 0.1 | 4.59E-01 | Small ribosomal subunit protein eS26z |
| AT2G15290 | 0.9 | 7.08E-07 | 0.6 | 1.61E-05 | 0.2 | 9.62E-03 | 0.2 | 1.77E-01 | 0.1 | 7.10E-01 | 0.1 | 5.25E-01 | Protein TIC 21, chloroplastic |
| AT5G38290 | 0.9 | 1.89E-05 | 0.7 | 1.25E-04 | 0.1 | 3.07E-01 | 0.4 | 2.90E-02 | 0.2 | 5.34E-01 | 0.3 | 2.61E-01 | Peptidyl-tRNA hydrolase family protein |
| AT3G46320 | 0.9 | 4.12E-05 | 0.8 | 9.69E-05 | 0.0 | 8.45E-01 | 0.4 | 1.64E-01 | 0.2 | 4.66E-01 | 0.1 | 7.49E-01 | Histone H4 |
| AT5G38250 | 0.9 | 5.23E-03 | -0.2 | 5.41E-01 | 1.1 | 9.75E-03 | -0.7 | 2.23E-01 | -1.5 | 4.27E-02 | 0.7 | 4.58E-01 | Protein kinase domain-containing protein |
| AT1G64355 | 0.9 | 6.94E-06 | 0.7 | 4.58E-05 | 0.1 | 2.44E-01 | -0.2 | 1.97E-01 | -0.2 | 3.58E-01 | 0.0 | 9.12E-01 | 1-acyl-sn-glycerol-3-phosphate acyltransferase |
| AT4G30870 | 0.9 | 1.82E-05 | 0.3 | 4.65E-02 | 0.6 | 1.37E-03 | 0.1 | 7.55E-01 | 0.1 | 7.30E-01 | 0.0 | 9.59E-01 | Crossover junction endonuclease MUS81 |
| AT4G14770 | 0.9 | 1.82E-06 | 0.3 | 3.38E-03 | 0.5 | 3.57E-04 | 0.0 | 9.77E-01 | 0.2 | 6.17E-01 | -0.2 | 5.99E-01 | TESMIN |
| AT4G28680 | 0.9 | 1.55E-03 | 0.7 | 1.20E-02 | 0.2 | 4.17E-01 | 1.0 | 6.37E-02 | 0.2 | 8.08E-01 | | | L-tyrosine decarboxylase |
| AT5G38910 | 0.9 | 1.07E-04 | -0.9 | 6.13E-04 | 1.8 | 3.23E-05 | 4.2 | 1.82E-09 | 3.1 | 2.06E-07 | 1.1 | 4.14E-04 | RmlC-like cupins superfamily protein |
| AT3G52630 | 0.9 | 2.26E-03 | 0.6 | 3.58E-02 | 0.3 | 2.34E-01 | 0.6 | 1.07E-01 | 0.5 | 2.48E-01 | 0.1 | 8.39E-01 | Replication protein A 14 kDa subunit A |
| AT3G47480 | 0.9 | 3.49E-03 | -0.2 | 4.88E-01 | 1.1 | 5.96E-03 | 0.6 | 2.26E-01 | -0.2 | 7.70E-01 | 0.8 | 2.12E-01 | Probable calcium-binding protein CML47 |
| AT3G02870 | 0.9 | 9.44E-06 | 0.9 | 1.89E-05 | 0.0 | 9.93E-01 | 0.4 | 2.68E-02 | 0.4 | 2.28E-02 | 0.0 | 8.79E-01 | Inositol-1-monophosphatase |
| AT1G24340 | 0.9 | 1.19E-05 | 0.5 | 1.55E-03 | 0.4 | 8.67E-03 | 0.0 | 9.35E-01 | 0.0 | 9.92E-01 | 0.0 | 9.51E-01 | FAD |
| AT4G25130 | 0.9 | 2.67E-07 | 0.5 | 3.39E-05 | 0.4 | 3.35E-04 | 0.3 | 5.44E-02 | 0.3 | 1.03E-01 | 0.0 | 9.71E-01 | Peptide methionine sulfoxide reductase A4, chloroplastic -S-oxide reductase) reductase) |
| AT4G31050 | 0.9 | 2.28E-04 | 0.9 | 2.89E-04 | 0.0 | 8.47E-01 | 0.1 | 7.67E-01 | -0.1 | 7.49E-01 | 0.1 | 5.84E-01 | Octanoyltransferase LIP2p, chloroplastic |
| AT3G27740 | 0.9 | 9.40E-06 | 0.6 | 1.45E-04 | 0.2 | 7.11E-02 | 0.3 | 4.30E-03 | 0.0 | 9.27E-01 | 0.3 | 4.47E-02 | Carbamoyl phosphate synthetase A |
| AT2G37390 | 0.9 | 1.31E-03 | 1.0 | 5.59E-04 | -0.2 | 4.44E-01 | -0.2 | 4.07E-01 | 0.2 | 5.04E-01 | -0.4 | 1.73E-01 | Protein SODIUM POTASSIUM ROOT DEFECTIVE 2 |
| AT4G37550 | 0.9 | 5.22E-06 | 0.6 | 1.51E-04 | 0.3 | 2.26E-02 | -0.7 | 1.56E-04 | -0.5 | 3.60E-03 | -0.2 | 3.73E-01 | Acetamidase |
| AT5G57800 | 0.9 | 5.90E-06 | 0.6 | 1.88E-04 | 0.3 | 2.04E-02 | -0.3 | 3.20E-02 | -0.2 | 1.43E-01 | -0.1 | 7.73E-01 | Very-long-chain aldehyde decarbonylase CER3 |
| AT3G50240 | 0.9 | 1.65E-04 | 0.5 | 5.87E-03 | 0.3 | 7.06E-02 | -0.2 | 2.97E-01 | 0.0 | 9.17E-01 | -0.2 | 4.13E-01 | Kinesin-like protein |
| AT1G62380 | 0.9 | 4.95E-05 | 0.6 | 7.50E-04 | 0.2 | 1.21E-01 | 0.4 | 5.22E-03 | 0.0 | 8.21E-01 | 0.3 | 7.65E-02 | 1-aminocyclopropane-1-carboxylate oxidase 2 |
| AT5G14100 | 0.9 | 2.12E-05 | 0.5 | 1.60E-03 | 0.3 | 2.09E-02 | 0.0 | 9.27E-01 | -0.2 | 4.40E-01 | 0.2 | 5.53E-01 | NAP14 |
| AT1G11290 | 0.9 | 3.54E-05 | 0.7 | 5.23E-04 | 0.2 | 1.53E-01 | 0.4 | 1.10E-01 | 0.0 | 9.68E-01 | 0.4 | 2.28E-01 | Pentatricopeptide repeat-containing protein At1g11290, chloroplastic |

| | | | | | | | | | | | | | |
|---|---|---|---|---|---|---|---|---|---|---|---|---|---|
| AT3G54400 | 0.9 | 1.09E-05 | 0.7 | 9.09E-05 | 0.2 | 1.95E-01 | -0.2 | 2.89E-01 | 0.0 | 8.27E-01 | -0.2 | 2.55E-01 | AT3g54400 |
| AT5G20480 | 0.9 | 1.15E-05 | 0.2 | 2.15E-01 | 0.7 | 3.68E-04 | 0.6 | 1.92E-01 | 0.1 | 8.76E-01 | 0.5 | 2.71E-01 | LRR receptor-like serine |
| AT2G07180 | 0.9 | 7.60E-06 | 0.5 | 6.05E-04 | 0.4 | 7.83E-03 | 0.2 | 8.35E-02 | -0.2 | 2.79E-01 | 0.4 | 2.98E-02 | Probable serine |
| AT3G09150 | 0.9 | 1.05E-05 | 0.7 | 9.91E-05 | 0.2 | 1.74E-01 | 0.2 | 4.12E-01 | 0.0 | 8.94E-01 | 0.1 | 6.69E-01 | Phytochromobilin:ferredoxin oxidoreductase, chloroplast |
| AT1G13270 | 0.9 | 4.89E-06 | 0.6 | 1.33E-04 | 0.3 | 2.79E-02 | 0.3 | 2.98E-02 | 0.0 | 9.34E-01 | 0.3 | 8.51E-02 | Methionine aminopeptidase 1B |
| AT2G02500 | 0.9 | 2.51E-05 | 0.8 | 8.01E-05 | 0.1 | 6.92E-01 | 0.2 | 4.67E-01 | 0.1 | 7.92E-01 | 0.1 | 8.07E-01 | 2-C-methyl-D-erythritol 4-phosphate cytidylyltransferase, chloroplastic |
| AT4G00750 | 0.9 | 2.02E-05 | 0.6 | 8.95E-04 | 0.3 | 2.99E-02 | -0.1 | 7.34E-01 | -0.2 | 4.94E-01 | 0.1 | 7.51E-01 | Probable methyltransferase PMT15 |
| AT2G36170 | 0.9 | 4.74E-05 | 0.8 | 1.71E-04 | 0.1 | 6.01E-01 | 0.3 | 7.42E-02 | 0.0 | 9.63E-01 | 0.3 | 2.26E-01 | Ubiquitin-like domain-containing protein |
| AT3G53320 | 0.9 | 6.51E-06 | 0.6 | 3.40E-04 | 0.3 | 1.64E-02 | -0.1 | 6.50E-01 | -0.2 | 3.69E-01 | 0.1 | 6.92E-01 | At3g53320 |
| AT1G74040 | 0.9 | 3.28E-07 | 0.5 | 2.58E-05 | 0.4 | 8.15E-04 | 0.1 | 4.68E-01 | 0.0 | 9.38E-01 | 0.1 | 5.61E-01 | 2-isopropylmalate synthase 1 |
| AT1G66230 | 0.9 | 8.08E-05 | 0.7 | 7.96E-04 | 0.2 | 2.61E-01 | -0.4 | 3.61E-01 | -1.0 | 3.10E-02 | 0.6 | 2.82E-01 | Transcription factor MYB20 |
| AT4G27740 | 0.9 | 6.27E-05 | 0.6 | 1.25E-03 | 0.3 | 1.07E-01 | -0.1 | 7.16E-01 | -0.2 | 5.53E-01 | 0.1 | 8.27E-01 | Protein yippee-like At4g27740 |
| AT1G06820 | 0.9 | 1.82E-05 | 0.3 | 1.55E-02 | 0.5 | 2.01E-03 | 0.2 | 1.19E-01 | 0.1 | 5.96E-01 | 0.1 | 5.64E-01 | Carotenoid isomerase |
| AT3G49910 | 0.9 | 2.51E-05 | 0.8 | 8.79E-05 | 0.1 | 6.39E-01 | 0.3 | 1.65E-01 | 0.1 | 7.46E-01 | 0.2 | 5.29E-01 | Large ribosomal subunit protein uL24z |
| AT5G28450 | 0.9 | 3.29E-04 | 1.2 | 6.64E-05 | -0.3 | 1.25E-01 | | | | | | | Chlorophyll a-b binding protein, chloroplastic |
| AT3G57610 | 0.9 | 2.30E-06 | 0.6 | 5.69E-05 | 0.3 | 1.91E-02 | 0.5 | 7.19E-04 | 0.3 | 2.79E-02 | 0.2 | 3.45E-01 | Adenylosuccinate synthetase, chloroplastic |
| AT1G71695 | 0.9 | 2.87E-05 | 1.0 | 2.11E-05 | -0.1 | 3.08E-01 | 0.1 | 6.01E-01 | 0.1 | 4.21E-01 | -0.1 | 8.13E-01 | Peroxidase 12 |
| AT5G63290 | 0.9 | 5.89E-05 | 0.2 | 2.39E-01 | 0.7 | 1.95E-03 | 0.5 | 9.94E-02 | 0.3 | 3.41E-01 | 0.1 | 6.97E-01 | Radical S-adenosyl methionine domain-containing protein 1, mitochondrial |
| AT2G47260 | 0.9 | 6.63E-06 | 0.3 | 1.09E-02 | 0.6 | 1.09E-03 | 0.2 | 5.96E-01 | 0.0 | 9.38E-01 | 0.1 | 7.24E-01 | WRKY transcription factor 23 |
| AT5G19730 | 0.9 | 4.11E-05 | 0.3 | 5.89E-02 | 0.6 | 2.63E-03 | 0.2 | 4.93E-01 | 0.3 | 3.59E-01 | -0.1 | 8.31E-01 | Pectinesterase |
| AT1G10700 | 0.9 | 2.78E-06 | 0.7 | 5.48E-05 | 0.2 | 3.45E-02 | -0.1 | 3.96E-01 | -0.1 | 3.32E-01 | 0.0 | 8.96E-01 | Phosphoribosyl pyrophosphate synthase 3 |
| AT1G61570 | 0.9 | 9.32E-04 | 0.7 | 6.56E-03 | 0.2 | 3.60E-01 | 0.7 | 6.17E-03 | 0.3 | 2.28E-01 | 0.3 | 1.98E-01 | Mitochondrial import inner membrane translocase subunit TIM13 |
| AT2G36870 | 0.9 | 7.04E-05 | 0.5 | 8.26E-03 | 0.4 | 1.93E-02 | 0.4 | 1.95E-01 | 0.7 | 2.04E-02 | -0.3 | 3.23E-01 | Xyloglucan endotransglucosylase |
| AT2G39390 | 0.9 | 1.07E-04 | 0.8 | 2.70E-04 | 0.0 | 7.98E-01 | 0.5 | 6.89E-03 | 0.3 | 2.06E-01 | 0.2 | 3.27E-01 | Large ribosomal subunit protein uL29y |
| AT1G56520 | 0.9 | 5.13E-05 | 0.4 | 2.05E-02 | 0.5 | 7.06E-03 | 0.3 | 2.10E-01 | 0.0 | 9.27E-01 | 0.3 | 2.61E-01 | Disease resistance protein family |
| AT3G02150 | 0.9 | 2.27E-04 | 0.2 | 3.75E-01 | 0.7 | 3.94E-03 | 0.0 | 9.54E-01 | -0.4 | 1.99E-01 | 0.3 | 2.25E-01 | Transcription factor TCP13 |
| AT1G61300 | 0.9 | 2.14E-05 | 0.3 | 1.40E-02 | 0.5 | 2.39E-03 | 0.3 | 8.40E-02 | 0.0 | 9.21E-01 | 0.3 | 1.70E-01 | LRR and NB-ARC domains-containing disease resistance protein |
| AT3G50790 | 0.9 | 2.60E-06 | 0.7 | 2.76E-05 | 0.2 | 8.19E-02 | -0.4 | 3.65E-02 | -0.4 | 7.80E-02 | 0.0 | 9.32E-01 | Esterase |
| AT3G10310 | 0.9 | 1.45E-05 | 0.5 | 9.81E-04 | 0.3 | 1.89E-02 | 0.3 | 3.70E-01 | 0.3 | 2.98E-01 | -0.1 | 8.61E-01 | P-loop nucleoside triphosphate hydrolases superfamily protein with CH domain-containing protein |
| AT4G14880 | 0.9 | 3.97E-07 | 0.8 | 1.51E-06 | 0.0 | 5.63E-01 | -0.2 | 2.68E-02 | 0.0 | 7.40E-01 | -0.2 | 2.28E-01 | Cysteine synthase |
| AT3G55750 | 0.9 | 4.75E-05 | 0.8 | 2.44E-04 | 0.1 | 4.17E-01 | 0.6 | 7.91E-04 | 0.5 | 1.17E-02 | 0.1 | 5.92E-01 | Large ribosomal subunit protein eL33x |
| AT3G02080 | 0.9 | 1.06E-05 | 0.8 | 4.70E-05 | 0.1 | 4.66E-01 | 0.1 | 5.92E-01 | -0.1 | 7.01E-01 | 0.2 | 4.64E-01 | Small ribosomal subunit protein eS19z |
| AT2G46400 | 0.9 | 1.95E-03 | -0.7 | 2.38E-02 | 1.6 | 5.63E-04 | 0.4 | 3.01E-01 | -0.1 | 9.32E-01 | 0.5 | 3.31E-01 | Probable WRKY transcription factor 46 |
| AT1G26410 | 0.9 | 1.10E-02 | 0.4 | 1.97E-01 | 0.5 | 2.08E-01 | 1.1 | 4.31E-02 | -0.6 | 4.75E-01 | 1.7 | 3.59E-02 | Berberine bridge enzyme-like 6 |
| AT4G15975 | 0.9 | 9.00E-04 | 0.4 | 6.53E-02 | 0.5 | 6.71E-02 | 0.9 | 1.78E-02 | 0.0 | 9.43E-01 | 0.9 | 6.08E-02 | RING-H2 finger protein ATL17 |
| AT1G36390 | 0.9 | 2.25E-05 | 0.8 | 8.46E-05 | 0.1 | 5.74E-01 | 0.4 | 4.01E-02 | 0.2 | 2.48E-01 | 0.1 | 6.11E-01 | GrpE protein homolog |
| AT5G64050 | 0.9 | 3.73E-07 | 0.6 | 7.43E-06 | 0.2 | 7.32E-03 | 0.3 | 1.13E-02 | 0.2 | 8.27E-02 | 0.1 | 6.82E-01 | Glutamate--tRNA ligase, chloroplastic |
| AT5G63470 | 0.9 | 1.51E-05 | 1.0 | 1.32E-05 | -0.1 | 3.96E-01 | -0.1 | 6.58E-01 | -0.3 | 2.84E-01 | 0.2 | 6.35E-01 | Nuclear transcription factor Y subunit C-4 |
| AT4G01037 | 0.9 | 2.75E-05 | 0.5 | 1.85E-03 | 0.4 | 1.97E-02 | 0.9 | 1.90E-04 | 0.1 | 8.35E-01 | 0.8 | 3.93E-03 | Protein WHAT'S THIS FACTOR 1 homolog, chloroplastic |
| AT4G39970 | 0.9 | 8.48E-05 | 0.8 | 2.54E-04 | 0.1 | 6.71E-01 | -0.1 | 5.28E-01 | 0.1 | 5.12E-01 | -0.2 | 2.24E-01 | Haloacid dehalogenase-like hydrolase domain-containing protein At4g39970 |
| AT1G69720 | 0.9 | 6.80E-06 | 0.3 | 1.06E-02 | 0.6 | 9.96E-04 | 0.3 | 5.94E-01 | -0.7 | 3.21E-01 | 1.0 | 2.04E-01 | Heme oxygenase 3, chloroplastic |
| AT4G34970 | 0.9 | 6.12E-04 | 0.8 | 2.50E-03 | 0.1 | 5.98E-01 | -0.5 | 2.91E-01 | -1.1 | 6.30E-02 | 0.6 | 3.42E-01 | Actin-depolymerizing factor 9 |
| AT1G14710 | 0.9 | 1.72E-06 | 0.7 | 1.75E-05 | 0.2 | 7.64E-02 | 0.4 | 2.32E-03 | 0.1 | 6.71E-01 | 0.3 | 5.63E-02 | hypothetical protein |
| AT5G44260 | 0.9 | 1.48E-03 | 0.9 | 2.88E-03 | 0.0 | 8.69E-01 | -0.6 | 4.21E-02 | -1.2 | 3.05E-03 | 0.5 | 2.74E-01 | Zinc finger CCCH domain-containing protein 61 |
| AT3G43720 | 0.9 | 2.72E-07 | 0.8 | 1.23E-06 | 0.1 | 2.81E-01 | -0.3 | 7.64E-02 | -0.1 | 4.16E-01 | -0.1 | 6.12E-01 | Non-specific lipid transfer protein GPI-anchored 2 |
| AT3G02450 | 0.9 | 6.97E-07 | 0.8 | 3.60E-06 | 0.1 | 2.41E-01 | 0.4 | 3.02E-02 | 0.1 | 4.76E-01 | 0.2 | 3.13E-01 | Probable inactive ATP-dependent zinc metalloprotease FTSHI 3, chloroplastic |
| AT2G40690 | 0.9 | 2.79E-04 | 0.8 | 1.52E-03 | 0.1 | 4.53E-01 | 0.4 | 6.82E-02 | 0.2 | 5.15E-01 | 0.2 | 3.90E-01 | Glycerol-3-phosphate dehydrogenase |
| AT1G13750 | 0.9 | 3.96E-05 | 0.6 | 8.13E-04 | 0.3 | 8.96E-02 | 0.6 | 2.69E-03 | 0.4 | 5.14E-02 | 0.2 | 3.52E-01 | Probable inactive purple acid phosphatase 1 |
| AT1G23740 | 0.9 | 3.57E-06 | 0.6 | 7.85E-05 | 0.3 | 2.95E-02 | 0.6 | 3.23E-04 | 0.7 | 5.07E-04 | 0.0 | 8.19E-01 | NADPH-dependent alkenal |
| AT5G16710 | 0.9 | 1.27E-05 | 0.9 | 3.36E-05 | 0.0 | 8.20E-01 | 0.2 | 1.74E-01 | 0.1 | 4.57E-01 | 0.1 | 8.02E-01 | Glutathione S-transferase DHAR3, chloroplastic |
| AT4G36390 | 0.9 | 3.08E-06 | 0.7 | 3.21E-05 | 0.2 | 8.64E-02 | 0.3 | 3.87E-03 | 0.2 | 1.56E-01 | 0.2 | 3.27E-01 | CDK5RAP1-like protein |
| AT2G44370 | 0.9 | 1.09E-02 | 0.5 | 1.78E-01 | 0.4 | 2.10E-01 | 1.5 | 9.98E-07 | 0.6 | 1.12E-02 | 0.9 | 1.56E-03 | At2g44370 |
| AT1G03055 | 0.9 | 1.78E-03 | 0.3 | 2.80E-01 | 0.6 | 3.67E-02 | | | | | | | Beta-carotene isomerase D27-like protein |
| AT3G04340 | 0.9 | 1.50E-06 | 0.6 | 9.82E-05 | 0.4 | 3.47E-03 | 0.3 | 5.04E-02 | 0.3 | 1.80E-01 | 0.1 | 8.04E-01 | Probable inactive ATP-dependent zinc metalloprotease FTSHI 5, chloroplastic |
| AT3G09260 | 0.9 | 3.81E-06 | 1.0 | 3.85E-06 | -0.1 | 2.46E-01 | 0.2 | 3.49E-01 | 0.5 | 4.54E-02 | -0.3 | 3.82E-01 | PYK10 |
| AT1G08540 | 0.9 | 2.31E-06 | 0.9 | 4.98E-06 | 0.0 | 9.05E-01 | 0.4 | 3.76E-02 | 0.3 | 9.44E-02 | 0.0 | 9.14E-01 | RNA polymerase sigma factor sigB |
| AT2G23170 | 0.9 | 5.25E-05 | 0.7 | 4.98E-04 | 0.2 | 2.41E-01 | 0.5 | 2.28E-01 | -0.5 | 3.30E-01 | 1.0 | 6.53E-02 | Indole-3-acetic acid-amido synthetase GH3.3 |
| AT5G04140 | 0.9 | 2.60E-06 | 0.4 | 2.39E-03 | 0.5 | 4.66E-04 | -0.3 | 3.06E-01 | 0.2 | 6.17E-01 | -0.5 | 1.98E-01 | Ferredoxin-dependent glutamate synthase |
| AT5G10920 | 0.9 | 7.87E-06 | 0.6 | 1.66E-04 | 0.3 | 4.34E-02 | 0.4 | 9.85E-03 | 0.2 | 2.30E-01 | 0.2 | 3.33E-01 | Argininosuccinate lyase, chloroplastic |
| AT4G03210 | 0.9 | 8.42E-05 | 0.7 | 1.16E-03 | 0.2 | 1.42E-01 | -0.2 | 3.29E-01 | -0.3 | 1.92E-01 | 0.1 | 7.78E-01 | Xyloglucan endotransglucosylase |
| AT4G00400 | 0.9 | 1.27E-05 | 0.9 | 3.11E-05 | 0.0 | 8.37E-01 | -0.2 | 9.79E-02 | -0.2 | 1.96E-01 | 0.0 | 9.30E-01 | Probable glycerol-3-phosphate acyltransferase 8 |
| AT1G23360 | 0.9 | 2.75E-05 | 0.7 | 4.43E-04 | 0.2 | 1.11E-01 | 0.1 | 7.04E-01 | 0.0 | 8.75E-01 | 0.1 | 6.39E-01 | 2-phytyl-1,4-beta-naphthoquinone methyltransferase, chloroplastic |
| AT4G05180 | 0.9 | 2.33E-05 | 1.1 | 1.11E-05 | -0.2 | 1.17E-01 | -0.3 | 2.00E-02 | -0.1 | 5.81E-01 | -0.2 | 2.56E-01 | Oxygen-evolving enhancer protein 3-2, chloroplastic |
| AT5G59480 | 0.9 | 5.99E-05 | 0.6 | 1.64E-03 | 0.3 | 7.38E-02 | -0.1 | 6.16E-01 | 0.3 | 1.52E-01 | -0.5 | 9.79E-02 | Haloacid dehalogenase-like hydrolase superfamily protein |
| AT4G23190 | 0.9 | 7.73E-04 | -0.1 | 5.49E-01 | 1.0 | 1.59E-03 | 0.2 | 4.12E-01 | -0.3 | 2.60E-01 | 0.4 | 1.20E-01 | Uncharacterized protein |
| AT5G14060 | 0.9 | 8.32E-06 | 0.6 | 1.78E-04 | 0.3 | 4.30E-02 | -0.2 | 2.01E-01 | -0.1 | 4.82E-01 | -0.1 | 7.72E-01 | Aspartokinase |
| AT5G39980 | 0.9 | 1.01E-05 | 0.6 | 2.73E-04 | 0.3 | 4.52E-02 | 0.4 | 1.02E-01 | 0.3 | 2.29E-01 | 0.1 | 8.73E-01 | Pentatricopeptide repeat-containing protein At5g39980, chloroplastic |
| AT5G55460 | 0.9 | 9.66E-03 | 0.4 | 3.09E-01 | 0.6 | 1.46E-01 | 0.0 | 9.31E-01 | -0.3 | 5.82E-01 | 0.4 | 5.01E-01 | At5g55460 |
| AT2G20450 | 0.9 | 2.21E-04 | 0.9 | 4.54E-04 | 0.0 | 8.87E-01 | 0.7 | 2.33E-03 | 0.4 | 5.47E-02 | 0.2 | 4.31E-01 | Large ribosomal subunit protein eL14z |
| AT5G58330 | 0.9 | 5.65E-06 | 0.5 | 7.44E-04 | 0.4 | 3.94E-03 | 0.0 | 7.50E-01 | 0.1 | 6.36E-01 | 0.0 | 8.89E-01 | Malate dehydrogenase |
| AT5G61170 | 0.9 | 1.31E-04 | 0.8 | 5.75E-04 | 0.1 | 5.12E-01 | 0.6 | 2.48E-04 | 0.3 | 3.12E-02 | 0.2 | 1.72E-01 | Small ribosomal subunit protein eS19x |
| AT4G32915 | 0.9 | 3.60E-05 | 0.8 | 1.77E-04 | 0.1 | 4.29E-01 | 0.5 | 4.15E-03 | 0.3 | 1.26E-01 | 0.2 | 3.49E-01 | Glutamyl-tRNA amidotransferase subunit C, chloroplastic |
| AT3G20230 | 0.9 | 2.07E-06 | 0.8 | 1.10E-05 | 0.1 | 3.06E-01 | 0.1 | 5.12E-01 | -0.1 | 8.02E-01 | 0.2 | 4.84E-01 | AT3G20230 protein |
| AT4G25910 | 0.9 | 1.44E-06 | 0.7 | 2.38E-05 | 0.2 | 3.11E-02 | -0.3 | 3.28E-02 | -0.3 | 4.27E-02 | 0.0 | 9.59E-01 | AT4G25910 protein |
| AT2G19380 | 0.9 | 1.18E-05 | 0.5 | 2.44E-03 | 0.4 | 6.33E-03 | 0.2 | 4.72E-01 | 0.3 | 4.86E-01 | 0.0 | 9.71E-01 | UBP1-associated proteins 1C |
| AT1G67740 | 0.9 | 9.97E-06 | 0.9 | 1.88E-05 | 0.0 | 8.96E-01 | 0.2 | 2.29E-01 | 0.2 | 3.41E-01 | 0.0 | 9.73E-01 | Photosystem II core complex proteins psbY, chloroplastic |
| AT1G30680 | 0.9 | 7.84E-06 | 0.5 | 1.07E-03 | 0.4 | 5.60E-03 | 0.9 | 6.04E-05 | 0.4 | 8.30E-02 | 0.6 | 1.61E-02 | SF4 helicase domain-containing protein |
| AT1G35530 | 0.9 | 7.59E-05 | 0.3 | 8.72E-02 | 0.6 | 4.73E-03 | 0.5 | 1.80E-01 | 0.4 | 3.76E-01 | 0.1 | 8.27E-01 | DEAD |
| AT2G29760 | 0.9 | 8.93E-06 | 0.6 | 4.25E-04 | 0.3 | 2.31E-02 | 0.7 | 6.05E-03 | 0.1 | 7.63E-01 | 0.6 | 5.01E-02 | Pentatricopeptide repeat-containing protein At2g29760, chloroplastic |
| AT4G31000 | 0.9 | 4.31E-06 | 0.6 | 1.74E-04 | 0.3 | 1.76E-02 | -0.6 | 2.74E-02 | -0.3 | 2.96E-01 | -0.3 | 3.80E-01 | Calmodulin-binding protein |
| AT2G47240 | 0.9 | 1.39E-06 | 0.8 | 1.14E-05 | 0.2 | 1.22E-01 | 0.1 | 3.03E-01 | -0.1 | 6.57E-01 | 0.2 | 2.33E-01 | Long-chain-fatty-acid--CoA ligase |
| AT1G14250 | 0.9 | 7.68E-03 | 0.9 | 9.03E-03 | 0.0 | 9.81E-01 | 0.5 | 3.75E-02 | 0.8 | 5.89E-03 | -0.3 | 4.11E-01 | Probable apyrase 5 |
| AT5G48300 | 0.9 | 4.83E-06 | 0.5 | 3.83E-04 | 0.4 | 6.13E-03 | 0.2 | 6.56E-02 | 0.4 | 9.81E-03 | -0.2 | 4.10E-01 | ADPG pyrophosphorylase small subunit |

| | | | | | | | | | | | | | |
|---|---|---|---|---|---|---|---|---|---|---|---|---|---|
| AT2G40590 | 0.9 | 3.27E-05 | 0.7 | 5.08E-04 | 0.3 | 1.01E-01 | 0.5 | 4.64E-04 | 0.3 | 2.86E-02 | 0.2 | 2.90E-01 | Small ribosomal subunit protein eS26y |
| AT3G28900 | 0.9 | 2.10E-04 | 1.1 | 1.50E-04 | -0.1 | 5.10E-01 | 0.5 | 1.29E-02 | 0.1 | 6.92E-01 | 0.4 | 1.59E-01 | Large ribosomal subunit protein eL34x |
| AT2G46750 | 0.9 | 3.75E-05 | 0.1 | 4.57E-01 | 0.8 | 6.80E-04 | 1.8 | 7.31E-05 | 0.8 | 5.78E-02 | 1.0 | 5.38E-03 | L-gulonolactone oxidase 2 |
| AT3G61470 | 0.9 | 1.03E-05 | 1.1 | 8.87E-06 | -0.1 | 2.71E-01 | -0.6 | 2.98E-04 | -0.4 | 1.57E-02 | -0.2 | 2.98E-01 | Photosystem I chlorophyll a |
| AT3G26180 | 0.9 | 1.16E-04 | 0.8 | 4.07E-04 | 0.1 | 5.96E-01 | 0.5 | 2.27E-03 | 0.2 | 1.35E-01 | 0.2 | 2.01E-01 | Cytochrome P450, family 71, subfamily B, polypeptide 20 |
| AT3G26230 | 0.9 | 1.28E-04 | 0.2 | 2.12E-01 | 0.7 | 3.26E-03 | 0.3 | 1.36E-01 | 0.1 | 7.80E-01 | 0.2 | 4.36E-01 | Uncharacterized protein |
| AT2G37190 | 0.9 | 3.50E-05 | 0.8 | 1.68E-04 | 0.1 | 4.58E-01 | 0.3 | 1.01E-01 | 0.0 | 9.58E-01 | 0.3 | 2.70E-01 | Large ribosomal subunit protein uL11z |
| AT3G25520 | 0.9 | 3.40E-05 | 0.8 | 2.21E-04 | 0.2 | 3.15E-01 | 0.6 | 3.48E-05 | 0.2 | 1.30E-01 | 0.4 | 2.05E-02 | Ribosomal protein |
| AT4G24710 | 0.9 | 1.02E-04 | 0.6 | 3.70E-03 | 0.3 | 7.57E-02 | 0.4 | 1.84E-01 | 0.2 | 6.84E-01 | 0.3 | 4.23E-01 | Pachytene checkpoint protein 2 homolog |
| AT3G16290 | 0.9 | 2.30E-06 | 0.6 | 9.55E-05 | 0.3 | 8.56E-03 | 0.3 | 1.29E-01 | 0.4 | 7.46E-02 | -0.1 | 7.57E-01 | Probable inactive ATP-dependent zinc metalloprotease FTSHI 2, chloroplastic |
| AT1G60230 | 0.9 | 4.15E-07 | 0.6 | 2.82E-05 | 0.4 | 1.58E-03 | 0.0 | 7.98E-01 | 0.0 | 8.43E-01 | 0.1 | 6.99E-01 | T13D8.12 protein |
| AT1G34380 | 0.9 | 5.19E-05 | 0.3 | 8.45E-02 | 0.6 | 3.13E-03 | 0.9 | 1.71E-03 | 0.1 | 7.02E-01 | 0.8 | 1.65E-02 | 5'-3' exonuclease family protein |
| AT1G53240 | 0.9 | 2.12E-06 | 0.8 | 1.59E-05 | 0.1 | 1.63E-01 | 0.5 | 3.70E-04 | 0.4 | 7.04E-03 | 0.1 | 5.54E-01 | Malate dehydrogenase 1, mitochondrial |
| AT2G04450 | 0.9 | 1.62E-01 | 0.2 | 7.70E-01 | 0.7 | 4.08E-01 | 0.0 | 9.53E-01 | -0.4 | 5.57E-01 | 0.4 | 5.61E-01 | Nudix hydrolase 6 |
| AT5G44680 | 0.9 | 6.03E-05 | 1.0 | 7.10E-05 | -0.1 | 7.41E-01 | -0.3 | 9.55E-02 | -0.5 | 4.78E-03 | 0.3 | 2.39E-01 | DNA-3-methyladenine glycosylase I |
| AT4G01995 | 0.9 | 1.88E-06 | 0.6 | 9.33E-05 | 0.3 | 7.94E-03 | -0.1 | 8.15E-01 | 0.1 | 5.82E-01 | -0.2 | 5.02E-01 | hypothetical protein |
| AT2G07170 | 0.9 | 1.75E-05 | 0.7 | 4.65E-04 | 0.3 | 5.57E-02 | -0.1 | 7.39E-01 | 0.1 | 7.61E-01 | -0.2 | 5.79E-01 | TORTIFOLIA1-like protein 2 |
| AT3G47010 | 0.9 | 1.37E-05 | 0.4 | 1.50E-02 | 0.6 | 1.88E-03 | 0.0 | 9.81E-01 | -0.2 | 4.66E-01 | 0.2 | 5.10E-01 | beta-glucosidase |
| AT1G70520 | 0.9 | 6.35E-06 | 0.4 | 3.94E-03 | 0.5 | 1.23E-03 | 0.3 | 3.97E-02 | 0.0 | 9.56E-01 | 0.3 | 1.08E-01 | Cysteine-rich receptor-like protein kinase 2 |
| AT5G23740 | 0.9 | 1.99E-05 | 0.9 | 4.11E-05 | 0.0 | 9.94E-01 | 0.3 | 6.77E-02 | 0.0 | 9.38E-01 | 0.3 | 2.11E-01 | Small ribosomal subunit protein uS17x |
| AT5G51970 | 0.9 | 4.41E-06 | 0.7 | 1.08E-04 | 0.3 | 2.67E-02 | -0.2 | 1.56E-01 | -0.3 | 1.78E-02 | 0.2 | 3.34E-01 | Sorbitol dehydrogenase |
| AT3G19720 | 0.9 | 4.57E-06 | 0.4 | 1.54E-03 | 0.5 | 1.43E-03 | 0.2 | 2.87E-01 | 0.2 | 4.13E-01 | 0.0 | 9.60E-01 | P-loop containing nucleoside triphosphate hydrolases superfamily protein |
| AT2G28815 | 0.9 | 1.03E-03 | -0.4 | 1.85E-01 | 1.3 | 1.37E-03 | 2.2 | 6.01E-03 | 0.5 | 6.98E-01 | 1.7 | 1.02E-02 | Uncharacterized protein |
| AT5G42270 | 0.9 | 1.19E-05 | 0.5 | 8.67E-04 | 0.4 | 1.10E-02 | 0.1 | 7.33E-01 | 0.2 | 3.20E-01 | -0.1 | 6.09E-01 | AT5G42270 protein |
| AT1G11475 | 0.9 | 4.25E-04 | 0.5 | 3.36E-02 | 0.5 | 4.61E-02 | 0.8 | 6.28E-03 | 0.3 | 3.70E-01 | 0.5 | 1.71E-01 | DNA-directed RNA polymerases II, IV and V subunit 10 |
| AT3G57460 | 0.9 | 9.52E-03 | 0.1 | 8.37E-01 | 0.9 | 4.36E-02 | 0.0 | 9.73E-01 | -0.4 | 5.23E-01 | 0.4 | 4.98E-01 | Uncharacterized protein T8H10.60 |
| AT4G31290 | 0.9 | 1.73E-05 | 0.9 | 3.49E-05 | 0.0 | 9.57E-01 | -0.3 | 3.05E-02 | -0.3 | 2.39E-02 | 0.1 | 8.35E-01 | Gamma-glutamylcyclotransferase 2-2 |
| AT3G17210 | 0.9 | 4.44E-07 | 1.0 | 9.87E-07 | 0.0 | 7.92E-01 | 0.2 | 3.55E-01 | -0.1 | 7.60E-01 | 0.2 | 3.50E-01 | Stress-response A |
| AT2G05100 | 0.9 | 1.41E-05 | 0.9 | 3.96E-05 | 0.0 | 7.88E-01 | -0.9 | 1.15E-02 | -0.6 | 8.75E-02 | -0.3 | 5.76E-01 | Chlorophyll a-b binding protein, chloroplastic |
| AT1G80640 | 0.9 | 1.05E-05 | 0.8 | 8.91E-05 | 0.2 | 1.79E-01 | 0.2 | 2.12E-01 | 0.0 | 8.97E-01 | 0.2 | 2.72E-01 | Probable receptor-like protein kinase At1g80640 |
| AT5G16140 | 0.9 | 1.50E-03 | 0.8 | 6.82E-03 | 0.2 | 5.43E-01 | 0.2 | 4.56E-01 | 0.1 | 6.16E-01 | 0.0 | 9.36E-01 | Chloroplastic group IIB intron splicing facilitator CRS2-B, chloroplastic |
| AT5G53580 | 0.9 | 3.24E-06 | 0.7 | 5.48E-05 | 0.2 | 3.94E-02 | 0.2 | 6.03E-02 | 0.2 | 7.04E-02 | 0.0 | 9.44E-01 | Pyridoxal reductase, chloroplastic |
| AT5G63810 | 0.9 | 7.15E-07 | 1.2 | 3.32E-07 | -0.3 | 6.38E-03 | -0.2 | 2.25E-01 | -0.3 | 1.51E-01 | 0.1 | 8.27E-01 | beta-galactosidase |
| AT1G43670 | 0.9 | 7.66E-06 | 0.8 | 3.46E-05 | 0.1 | 4.45E-01 | 0.3 | 1.91E-02 | 0.1 | 4.21E-01 | 0.2 | 2.88E-01 | Fructose-1,6-bisphosphatase, cytosolic |
| AT1G55960 | 0.9 | 7.64E-04 | 0.4 | 8.09E-02 | 0.6 | 3.17E-02 | -0.3 | 1.50E-01 | -0.2 | 3.29E-01 | -0.1 | 8.52E-01 | Polyketide cyclase |
| AT2G39310 | 0.9 | 9.89E-07 | 0.7 | 2.19E-05 | 0.3 | 1.22E-02 | 0.4 | 2.22E-02 | 0.2 | 2.49E-01 | 0.2 | 5.13E-01 | Jacalin-related lectin 22 |
| AT4G25450 | 0.9 | 4.81E-07 | 0.4 | 4.88E-04 | 0.6 | 1.25E-04 | 0.1 | 7.58E-01 | 0.0 | 9.19E-01 | 0.1 | 9.13E-01 | Non-intrinsic ABC protein 8 |
| AT4G16680 | 0.9 | 1.53E-04 | -0.5 | 4.16E-02 | 1.4 | 1.97E-04 | 1.3 | 2.85E-02 | -0.1 | 9.12E-01 | 1.4 | 6.27E-02 | RNA helicase |
| AT3G19220 | 0.9 | 5.38E-07 | 0.6 | 6.05E-05 | 0.4 | 1.12E-03 | 0.0 | 9.85E-01 | -0.1 | 4.81E-01 | 0.1 | 5.39E-01 | Protein disulfide-isomerase SCO2 |
| AT4G34610 | 0.9 | 1.84E-06 | 0.4 | 8.40E-04 | 0.5 | 7.33E-04 | 0.4 | 5.35E-02 | 0.1 | 6.86E-01 | 0.3 | 1.93E-01 | BEL1-like homeodomain protein 6 |
| AT4G31870 | 0.9 | 9.21E-05 | 0.3 | 1.03E-01 | 0.7 | 4.31E-03 | | | | | | | Glutathione peroxidase 7 |
| AT1G51820 | 0.9 | 4.57E-02 | -0.5 | 3.78E-01 | 1.5 | 2.92E-02 | | | | | | | Leucine-rich repeat protein kinase family protein |
| AT3G17830 | 0.9 | 6.13E-05 | 0.7 | 6.88E-04 | 0.2 | 2.04E-01 | 0.4 | 1.90E-01 | 0.1 | 6.95E-01 | 0.2 | 5.63E-01 | At3g17830 |
| AT2G34010 | 0.9 | 4.28E-04 | 1.0 | 5.77E-04 | 0.0 | 9.46E-01 | | | | | | | Verprolin |
| AT3G57020 | 0.9 | 1.15E-05 | 0.7 | 1.89E-04 | 0.2 | 7.04E-02 | 0.5 | 5.93E-02 | 0.3 | 4.24E-01 | 0.2 | 2.71E-01 | Protein STRICTOSIDINE SYNTHASE-LIKE 9 |
| AT5G16120 | 0.9 | 1.81E-05 | 0.6 | 8.51E-04 | 0.4 | 2.46E-02 | -0.3 | 1.87E-01 | -0.6 | 8.98E-03 | 0.4 | 1.71E-01 | Alpha |
| AT3G49170 | 0.9 | 5.48E-05 | 0.7 | 6.91E-04 | 0.2 | 1.97E-01 | 0.9 | 2.51E-03 | 0.4 | 2.83E-01 | 0.5 | 6.91E-02 | Pentatricopeptide repeat-containing protein At3g49170, chloroplastic |
| AT1G59720 | 0.9 | 1.65E-04 | 0.6 | 5.71E-03 | 0.3 | 9.68E-02 | -0.2 | 7.94E-01 | 0.1 | 9.40E-01 | -0.3 | 7.63E-01 | Pentatricopeptide repeat-containing protein At1g59720, chloroplastic |
| AT5G59970 | 1.0 | 4.09E-04 | 0.7 | 4.20E-03 | 0.2 | 2.52E-01 | 0.4 | 3.99E-02 | 0.1 | 5.90E-01 | 0.3 | 3.09E-01 | Histone H4 |
| AT5G35170 | 1.0 | 1.03E-05 | 0.8 | 7.89E-05 | 0.2 | 2.15E-01 | -0.2 | 6.58E-02 | -0.2 | 1.53E-01 | 0.0 | 9.12E-01 | Adenylate kinase-like protein |
| AT3G59820 | 1.0 | 2.16E-05 | 0.5 | 5.07E-03 | 0.5 | 6.49E-03 | 0.8 | 7.70E-05 | 0.2 | 4.83E-01 | 0.7 | 2.42E-03 | Mitochondrial proton |
| AT5G05740 | 1.0 | 7.14E-06 | 1.0 | 9.14E-06 | -0.1 | 5.73E-01 | 0.4 | 6.88E-03 | 0.2 | 2.26E-01 | 0.2 | 2.91E-01 | Ethylene-dependent gravitropism-deficient and yellow-green-like 2 |
| AT5G39050 | 1.0 | 1.54E-04 | -0.1 | 6.79E-01 | 1.0 | 5.26E-04 | -0.3 | 1.92E-01 | -0.9 | 1.51E-03 | 0.6 | 3.93E-02 | Phenolic glucoside malonyltransferase 1 |
| AT1G20700 | 1.0 | 4.77E-04 | 0.8 | 1.74E-03 | 0.1 | 6.19E-01 | 0.0 | 9.08E-01 | 0.1 | 8.37E-01 | -0.1 | 7.75E-01 | WUSCHEL related homeobox 14 |
| AT5G49030 | 1.0 | 2.34E-07 | 0.7 | 3.05E-06 | 0.2 | 1.23E-02 | 0.3 | 1.73E-01 | 0.2 | 3.75E-01 | 0.1 | 8.56E-01 | isoleucine--tRNA ligase |
| AT5G45170 | 1.0 | 4.60E-07 | 0.6 | 2.76E-05 | 0.4 | 1.56E-03 | 0.0 | 9.97E-01 | 0.1 | 5.91E-01 | -0.1 | 6.81E-01 | Haloacid dehalogenase-like hydrolase superfamily protein |
| AT3G10160 | 1.0 | 7.94E-07 | 0.5 | 9.47E-05 | 0.4 | 1.18E-03 | 0.1 | 3.98E-01 | 0.2 | 2.80E-01 | -0.1 | 8.23E-01 | Folylpolyglutamate synthase |
| AT1G50320 | 1.0 | 5.75E-07 | 0.7 | 1.18E-05 | 0.3 | 1.13E-02 | 0.1 | 6.02E-01 | 0.1 | 4.63E-01 | 0.0 | 8.54E-01 | Thioredoxin X, chloroplastic |
| AT4G32320 | 1.0 | 3.45E-06 | 0.5 | 1.25E-03 | 0.5 | 1.44E-03 | -0.4 | 9.11E-03 | -0.1 | 7.18E-01 | -0.3 | 1.14E-01 | Putative L-ascorbate peroxidase 6 |
| AT5G08610 | 1.0 | 2.33E-06 | 0.9 | 7.37E-06 | 0.0 | 6.67E-01 | 0.7 | 7.64E-04 | 0.1 | 8.63E-01 | 0.7 | 8.43E-03 | DEAD-box ATP-dependent RNA helicase 26 |
| AT3G46900 | 1.0 | 4.52E-05 | 0.3 | 4.30E-02 | 0.6 | 4.46E-03 | -0.7 | 3.75E-02 | -1.5 | 1.94E-03 | 0.7 | 9.25E-02 | Copper transporter 2 |
| AT4G24350 | 1.0 | 1.80E-05 | 0.8 | 1.09E-04 | 0.1 | 2.95E-01 | 0.6 | 2.85E-02 | 0.4 | 2.18E-01 | 0.2 | 5.93E-01 | Uncharacterized protein AT4g24350 |
| AT2G30390 | 1.0 | 1.55E-06 | 0.7 | 2.68E-05 | 0.2 | 2.69E-02 | 0.2 | 2.22E-01 | 0.2 | 2.16E-01 | 0.0 | 9.23E-01 | Ferrochelatase |
| AT1G67470 | 1.0 | 1.40E-04 | 0.3 | 7.94E-02 | 0.6 | 8.23E-03 | 0.0 | 8.96E-01 | -0.2 | 3.68E-01 | 0.3 | 3.79E-01 | Inactive serine |
| AT5G48460 | 1.0 | 4.34E-05 | 0.6 | 1.97E-03 | 0.4 | 3.91E-02 | -0.3 | 8.42E-02 | -0.1 | 7.26E-01 | -0.2 | 3.76E-01 | Fimbrin-2 |
| AT4G18730 | 1.0 | 3.71E-06 | 0.8 | 2.38E-05 | 0.1 | 2.42E-01 | 0.3 | 4.43E-02 | 0.1 | 8.07E-01 | 0.3 | 2.40E-01 | Large ribosomal subunit protein uL5y |
| AT1G76730 | 1.0 | 8.65E-06 | 0.5 | 1.81E-03 | 0.5 | 3.32E-03 | 0.2 | 1.69E-01 | 0.1 | 6.47E-01 | 0.1 | 6.36E-01 | 5-formyltetrahydrofolate cyclo-ligase-like protein COG0212 |
| AT1G33290 | 1.0 | 1.83E-06 | 0.6 | 7.30E-05 | 0.3 | 9.50E-03 | 0.0 | 8.27E-01 | -0.1 | 3.69E-01 | 0.2 | 3.43E-01 | P-loop containing nucleoside triphosphate hydrolases superfamily protein |
| AT5G48730 | 1.0 | 1.50E-06 | 0.8 | 1.39E-05 | 0.2 | 1.06E-01 | 0.3 | 2.70E-01 | 0.1 | 7.12E-01 | 0.2 | 6.40E-01 | Pentatricopeptide repeat-containing protein At5g48730, chloroplastic |
| AT3G26310 | 1.0 | 1.70E-05 | 0.3 | 2.10E-02 | 0.6 | 1.82E-03 | 0.1 | 8.29E-01 | 0.0 | 9.27E-01 | 0.0 | 9.57E-01 | Cytochrome P450 71B35 |
| AT5G25440 | 1.0 | 3.70E-05 | 0.3 | 7.04E-02 | 0.7 | 1.92E-03 | -0.1 | 8.16E-01 | -0.3 | 2.02E-01 | 0.2 | 4.10E-01 | Protein kinase superfamily protein |
| AT1G12860 | 1.0 | 2.90E-05 | 0.7 | 3.31E-04 | 0.2 | 1.53E-01 | -0.1 | 5.43E-01 | -0.4 | 3.55E-02 | 0.3 | 1.52E-01 | Basic helix-loop-helix DNA-binding superfamily protein |
| AT4G39960 | 1.0 | 3.97E-07 | 0.7 | 1.26E-05 | 0.3 | 3.44E-03 | 0.3 | 6.29E-03 | 0.1 | 4.12E-01 | 0.2 | 1.28E-01 | DnaJ-like protein |
| AT1G48520 | 1.0 | 5.86E-06 | 0.8 | 6.14E-05 | 0.2 | 1.10E-01 | 0.2 | 5.81E-02 | 0.1 | 5.96E-01 | 0.1 | 3.44E-01 | Glutamyl-tRNA amidotransferase subunit B, chloroplastic |
| AT5G27240 | 1.0 | 2.17E-05 | 0.2 | 1.45E-01 | 0.7 | 8.80E-04 | 0.4 | 3.93E-01 | 0.2 | 6.96E-01 | 0.2 | 8.23E-01 | A_TM021B04.9 protein |
| AT3G20440 | 1.0 | 1.06E-05 | 0.5 | 1.57E-03 | 0.4 | 7.28E-03 | 0.8 | 2.58E-02 | -0.3 | 5.72E-01 | 1.1 | 1.65E-02 | 1,4-alpha-glucan-branching enzyme 3, chloroplastic |
| AT2G37620 | 1.0 | 1.00E-06 | 0.6 | 4.22E-05 | 0.3 | 5.12E-03 | 0.1 | 7.22E-01 | 0.1 | 6.76E-01 | 0.0 | 9.28E-01 | Actin-1 |
| AT2G38330 | 1.0 | 1.06E-06 | 0.7 | 1.61E-05 | 0.2 | 2.93E-02 | -0.2 | 9.53E-02 | -0.4 | 1.23E-02 | 0.2 | 3.91E-01 | Protein DETOXIFICATION 44, chloroplastic |
| AT5G07900 | 1.0 | 5.34E-05 | 0.9 | 2.16E-04 | 0.1 | 5.72E-01 | 0.3 | 4.37E-01 | 0.3 | 6.08E-01 | 0.1 | 9.22E-01 | hypothetical protein |
| AT5G48760 | 1.0 | 3.34E-06 | 0.7 | 9.91E-05 | 0.3 | 1.68E-02 | 0.3 | 4.73E-02 | 0.1 | 6.23E-01 | 0.2 | 3.58E-01 | Large ribosomal subunit protein uL13w |
| AT4G04840 | 1.0 | 2.30E-04 | 1.7 | 5.02E-06 | -0.8 | 1.88E-03 | 0.7 | 9.85E-03 | 0.3 | 3.32E-01 | 0.4 | 2.30E-01 | Peptide-methionine -S-oxide reductase |
| AT3G14840 | 1.0 | 5.95E-06 | 0.6 | 2.21E-04 | 0.3 | 1.71E-02 | 0.7 | 3.06E-05 | 0.2 | 1.41E-01 | 0.5 | 1.17E-02 | Probable leucine-rich repeat receptor-like serine |
| AT5G58120 | 1.0 | 2.56E-04 | 0.1 | 6.91E-01 | 0.9 | 2.53E-03 | 0.3 | 1.89E-01 | -0.3 | 1.77E-01 | 0.6 | 2.49E-02 | ADP-ribosyl cyclase |

| | | | | | | | | | | | | | |
|---|---|---|---|---|---|---|---|---|---|---|---|---|---|
| AT1G68110 | 1.0 | 2.50E-03 | 1.1 | 1.98E-03 | -0.1 | 7.35E-01 | -0.4 | 1.75E-02 | -0.4 | 3.79E-02 | 0.0 | 9.28E-01 | Putative clathrin assembly protein At1g68110 |
| AT5G09660 | 1.0 | 5.16E-06 | 0.8 | 3.96E-05 | 0.2 | 1.88E-01 | 0.2 | 7.34E-02 | 0.1 | 3.58E-01 | 0.1 | 6.72E-01 | Malate dehydrogenase |
| AT3G49110 | 1.0 | 1.19E-05 | 0.9 | 4.04E-05 | 0.1 | 5.96E-01 | 0.2 | 4.96E-01 | -0.2 | 6.70E-01 | 0.4 | 2.13E-01 | Peroxidase 33 |
| AT1G17050 | 1.0 | 1.01E-05 | 0.2 | 5.54E-02 | 0.7 | 3.82E-04 | 0.2 | 1.19E-01 | 0.3 | 4.27E-02 | -0.1 | 6.51E-01 | Solanesyl diphosphate synthase 2, chloroplastic ) |
| AT5G19700 | 1.0 | 7.62E-03 | 1.0 | 8.50E-03 | 0.0 | 9.45E-01 | -0.5 | 1.62E-01 | -0.9 | 4.17E-02 | 0.3 | 5.74E-01 | Protein DETOXIFICATION 52 |
| AT1G72250 | 1.0 | 1.02E-05 | 0.5 | 3.46E-03 | 0.5 | 3.44E-03 | 0.1 | 7.60E-01 | 0.3 | 3.73E-01 | -0.2 | 6.43E-01 | Di-glucose binding protein with Kinesin motor domain-containing protein |
| AT1G31330 | 1.0 | 7.17E-06 | 1.2 | 4.71E-06 | -0.2 | 1.27E-01 | -0.9 | 6.18E-06 | -0.2 | 1.72E-01 | -0.7 | 2.81E-03 | Photosystem I reaction center subunit III, chloroplastic |
| AT1G70290 | 1.0 | 8.06E-04 | 0.8 | 3.42E-03 | 0.2 | 5.23E-01 | -0.6 | 7.16E-03 | -1.0 | 2.72E-04 | 0.4 | 6.91E-02 | Trehalose-6-phosphatase synthase S8 |
| AT4G29150 | 1.0 | 2.06E-04 | 0.7 | 4.23E-03 | 0.3 | 1.53E-01 | 0.1 | 8.10E-01 | -0.1 | 8.40E-01 | 0.2 | 7.06E-01 | DUF4005 domain-containing protein |
| AT2G35390 | 1.0 | 3.54E-07 | 0.7 | 8.15E-06 | 0.3 | 5.32E-03 | 0.3 | 5.26E-02 | -0.1 | 6.41E-01 | 0.4 | 2.41E-02 | Ribose-phosphate pyrophosphokinase |
| AT2G37500 | 1.0 | 2.33E-07 | 0.6 | 1.16E-05 | 0.3 | 1.15E-03 | 0.4 | 9.68E-02 | 0.1 | 7.99E-01 | 0.3 | 3.09E-01 | Arginine biosynthesis bifunctional protein ArgJ, chloroplastic |
| AT2G22330 | 1.0 | 6.53E-07 | 1.4 | 2.14E-07 | -0.4 | 9.23E-04 | 0.0 | 9.87E-01 | 0.0 | 9.83E-01 | 0.0 | 9.80E-01 | Cytochrome P450, family 79, subfamily B, polypeptide 3 |
| AT4G02990 | 1.0 | 1.28E-06 | 0.5 | 3.99E-04 | 0.5 | 7.13E-04 | 0.7 | 1.22E-04 | 0.2 | 3.62E-01 | 0.5 | 1.10E-02 | Transcription termination factor MTERF4, chloroplastic |
| AT1G59990 | 1.0 | 7.11E-06 | 0.7 | 2.21E-04 | 0.3 | 2.62E-02 | 0.8 | 1.44E-04 | 0.3 | 1.97E-01 | 0.5 | 7.35E-03 | DEA-box RNA helicase family protein |
| AT4G17300 | 1.0 | 3.73E-07 | 0.6 | 1.39E-05 | 0.3 | 2.53E-03 | 0.3 | 1.11E-01 | 0.1 | 7.02E-01 | 0.2 | 4.68E-01 | asparagine--tRNA ligase |
| AT4G27310 | 1.0 | 1.06E-04 | 0.5 | 9.78E-03 | 0.5 | 2.14E-02 | | | | | | | B-box type zinc finger family protein |
| AT3G18440 | 1.0 | 2.74E-07 | 0.7 | 7.12E-06 | 0.3 | 3.50E-03 | 0.4 | 4.17E-03 | 0.2 | 2.66E-01 | 0.2 | 1.72E-01 | Aluminum-activated malate transporter 9 |
| AT5G25140 | 1.0 | 1.90E-05 | 0.3 | 3.40E-02 | 0.6 | 1.77E-03 | -0.3 | 4.45E-01 | -0.8 | 5.18E-02 | 0.5 | 3.23E-01 | Cytochrome P450 71B13 |
| AT4G00950 | 1.0 | 5.50E-04 | 1.2 | 2.16E-04 | -0.2 | 3.67E-01 | -0.5 | 1.77E-02 | -0.8 | 1.22E-03 | 0.3 | 2.48E-01 | Uncharacterized protein At4g00950 |
| AT3G02660 | 1.0 | 3.19E-07 | 0.7 | 5.68E-06 | 0.2 | 1.03E-02 | 0.3 | 1.85E-02 | 0.4 | 8.75E-03 | -0.1 | 6.95E-01 | Tyrosine--tRNA ligase, chloroplastic |
| AT5G63180 | 1.0 | 2.40E-05 | 1.1 | 2.60E-05 | -0.1 | 5.39E-01 | -0.1 | 5.17E-01 | 0.2 | 2.75E-01 | -0.3 | 1.48E-01 | Probable pectate lyase 22 |
| AT4G15233 | 1.0 | 3.25E-04 | 0.4 | 5.19E-02 | 0.5 | 2.58E-02 | 0.3 | 4.11E-01 | -0.8 | 4.65E-02 | 1.1 | 6.38E-03 | ABC-2 and Plant PDR ABC-type transporter family protein |
| AT3G18750 | 1.0 | 1.03E-05 | 0.5 | 2.17E-03 | 0.5 | 4.80E-03 | 0.1 | 7.98E-01 | 0.2 | 4.63E-01 | -0.1 | 6.74E-01 | With no lysine kinase 6 |
| AT1G15660 | 1.0 | 4.41E-06 | 0.6 | 5.06E-04 | 0.4 | 4.67E-03 | 0.1 | 4.28E-01 | 0.1 | 5.96E-01 | 0.0 | 9.40E-01 | Centromere protein C |
| AT3G57180 | 1.0 | 2.66E-06 | 0.7 | 4.50E-05 | 0.2 | 4.53E-02 | 0.5 | 6.13E-02 | 0.2 | 5.75E-01 | 0.3 | 2.76E-01 | GTP-binding protein BRASSINAZOLE INSENSITIVE PALE GREEN 2, chloroplastic |
| AT4G13720 | 1.0 | 1.59E-06 | 0.8 | 2.09E-05 | 0.2 | 5.17E-02 | 0.0 | 8.55E-01 | -0.1 | 8.04E-01 | 0.1 | 7.31E-01 | Inosine triphosphate pyrophosphatase |
| AT5G47110 | 1.0 | 5.94E-07 | 0.7 | 1.19E-05 | 0.3 | 1.19E-02 | 0.0 | 9.16E-01 | 0.0 | 8.07E-01 | 0.0 | 9.14E-01 | Light-harvesting complex-like protein 3 isotype 2, chloroplastic |
| AT2G18450 | 1.0 | 6.37E-05 | 0.7 | 8.84E-04 | 0.3 | 1.41E-01 | -0.2 | 4.56E-01 | 0.0 | 9.86E-01 | -0.2 | 5.18E-01 | Succinate dehydrogenase flavoprotein subunit 2, mitochondrial |
| AT4G29390 | 1.0 | 2.41E-05 | 0.5 | 3.78E-03 | 0.5 | 9.13E-03 | 0.4 | 5.41E-02 | 0.3 | 2.53E-01 | 0.1 | 6.42E-01 | Small ribosomal subunit protein eS30z |
| AT5G02450 | 1.0 | 1.24E-04 | 0.9 | 3.99E-04 | 0.1 | 6.70E-01 | 0.2 | 3.08E-01 | 0.0 | 9.24E-01 | 0.2 | 5.59E-01 | Large ribosomal subunit protein eL36x |
| AT2G36160 | 1.0 | 3.96E-06 | 0.8 | 3.94E-05 | 0.2 | 1.13E-01 | 0.2 | 3.26E-01 | 0.0 | 9.42E-01 | 0.1 | 5.63E-01 | Small ribosomal subunit protein uS11z |
| AT1G80480 | 1.0 | 2.92E-07 | 0.9 | 1.55E-06 | 0.1 | 1.76E-01 | 0.5 | 3.93E-04 | 0.2 | 9.64E-02 | 0.3 | 8.13E-02 | At1g80480 |
| AT5G56860 | 1.0 | 2.97E-06 | 0.8 | 2.58E-05 | 0.2 | 1.28E-01 | -0.6 | 4.63E-03 | -0.7 | 2.41E-03 | 0.1 | 5.59E-01 | GATA transcription factor 21 |
| AT5G20230 | 1.0 | 1.32E-03 | 0.6 | 2.25E-02 | 0.4 | 1.72E-01 | -0.7 | 1.65E-01 | -1.7 | 3.23E-03 | 1.1 | 6.75E-02 | Blue copper protein |
| AT1G31860 | 1.0 | 1.81E-05 | 0.9 | 9.28E-05 | 0.1 | 3.98E-01 | 0.2 | 3.16E-01 | 0.0 | 9.59E-01 | 0.2 | 5.37E-01 | Histidine biosynthesis bifunctional protein |
| AT5G28060 | 1.0 | 4.64E-06 | 0.7 | 9.37E-05 | 0.3 | 3.84E-02 | 0.2 | 4.02E-01 | -0.1 | 6.17E-01 | 0.3 | 2.75E-01 | 40S ribosomal protein S24 |
| AT1G45110 | 1.0 | 7.98E-05 | 0.5 | 1.86E-02 | 0.5 | 1.54E-02 | 0.4 | 4.35E-01 | 0.6 | 2.42E-01 | -0.2 | 6.86E-01 | Tetrapyrrole methylase domain-containing protein |
| AT2G21960 | 1.0 | 7.99E-07 | 0.6 | 3.83E-05 | 0.4 | 3.21E-03 | 0.2 | 3.04E-01 | 0.3 | 1.10E-01 | -0.1 | 6.51E-01 | Expressed protein |
| AT2G19870 | 1.0 | 2.36E-06 | 0.7 | 4.99E-05 | 0.3 | 2.72E-02 | 0.6 | 5.84E-04 | 0.2 | 3.29E-01 | 0.4 | 4.40E-02 | rRNA methyltransferase 1, mitochondrial |
| AT5G10690 | 1.0 | 5.88E-06 | 0.9 | 1.91E-05 | 0.1 | 6.39E-01 | -0.2 | 3.90E-01 | -0.1 | 6.18E-01 | -0.1 | 8.78E-01 | Pentatricopeptide repeat-containing protein At5g10690 |
| AT1G11303 | 1.0 | 2.42E-04 | 0.3 | 1.30E-01 | 0.7 | 1.01E-02 | 0.4 | 4.35E-01 | 0.1 | 9.05E-01 | 0.3 | 6.59E-01 | G-type lectin S-receptor-like serine |
| AT5G47840 | 1.0 | 4.12E-07 | 0.6 | 2.38E-05 | 0.4 | 1.39E-03 | -0.2 | 8.36E-02 | -0.1 | 3.53E-01 | -0.1 | 7.19E-01 | adenylate kinase |
| AT3G10670 | 1.0 | 2.27E-07 | 0.6 | 1.20E-05 | 0.4 | 8.30E-04 | 0.2 | 2.26E-02 | 0.1 | 4.21E-01 | 0.1 | 3.34E-01 | ABC transporter I family member 6, chloroplastic |
| AT5G66270 | 1.0 | 4.26E-05 | 0.5 | 4.98E-03 | 0.4 | 2.14E-02 | 0.3 | 5.26E-01 | 0.1 | 8.58E-01 | 0.2 | 7.80E-01 | Zinc finger CCCH domain-containing protein 68 |
| AT1G63680 | 1.0 | 3.97E-07 | 0.8 | 6.11E-06 | 0.2 | 1.58E-02 | 0.4 | 1.99E-02 | 0.0 | 9.38E-01 | 0.4 | 5.85E-02 | UDP-N-acetylmuramoyl-L-alanyl-D-glutamate--2,6-diaminopimelate ligase MurE homolog, chloroplastic |
| AT4G15560 | 1.0 | 1.52E-05 | 0.8 | 1.95E-04 | 0.2 | 1.13E-01 | -0.1 | 7.02E-01 | 0.0 | 9.92E-01 | -0.1 | 8.03E-01 | 1-deoxy-D-xylulose-5-phosphate synthase |
| AT3G26830 | 1.0 | 1.40E-02 | -0.3 | 5.62E-01 | 1.2 | 1.75E-02 | 0.4 | 2.85E-01 | -1.0 | 5.25E-02 | 1.4 | 2.14E-02 | PAD3 |
| AT4G02850 | 1.0 | 7.40E-05 | 1.3 | 2.31E-05 | -0.3 | 1.25E-01 | -0.1 | 7.17E-01 | 0.0 | 9.57E-01 | -0.1 | 8.15E-01 | At4g02850 |
| AT2G36950 | 1.0 | 7.33E-07 | 0.9 | 2.70E-06 | 0.1 | 5.46E-01 | 0.3 | 2.66E-02 | -0.1 | 5.78E-01 | 0.4 | 2.86E-02 | Heavy metal-associated isoprenylated plant protein 5 |
| AT5G05130 | 1.0 | 1.25E-05 | 0.6 | 1.10E-03 | 0.4 | 1.44E-02 | -0.2 | 4.25E-01 | 0.0 | 9.71E-01 | -0.2 | 5.33E-01 | Putative SWI |
| AT3G20330 | 1.0 | 4.41E-06 | 0.4 | 3.69E-03 | 0.6 | 8.62E-04 | 0.6 | 3.93E-04 | 0.2 | 1.47E-01 | 0.3 | 3.97E-02 | aspartate carbamoyltransferase |
| AT5G48250 | 1.0 | 4.30E-06 | 1.1 | 5.97E-06 | -0.1 | 5.83E-01 | -0.1 | 6.58E-01 | -0.1 | 6.40E-01 | 0.0 | 9.46E-01 | Zinc finger protein CONSTANS-LIKE 10 |
| AT4G36430 | 1.0 | 7.71E-07 | 0.1 | 2.50E-01 | 0.9 | 2.86E-05 | 2.5 | 1.60E-08 | 1.1 | 3.77E-04 | 1.4 | 2.65E-05 | Peroxidase 49 |
| AT3G57600 | 1.0 | 5.22E-04 | 1.0 | 8.83E-04 | 0.0 | 9.62E-01 | 0.6 | 2.30E-01 | 0.4 | 5.06E-01 | 0.2 | 7.97E-01 | Dehydration-responsive element-binding protein 2F |
| AT3G52210 | 1.0 | 2.29E-06 | 0.5 | 1.04E-03 | 0.5 | 1.15E-03 | 0.2 | 2.56E-01 | 0.1 | 6.76E-01 | 0.1 | 6.91E-01 | mRNA cap guanine-N7 methyltransferase 2 )-methyltransferase 2) |
| AT5G65890 | 1.0 | 3.84E-05 | 0.8 | 3.33E-04 | 0.2 | 2.44E-01 | -0.1 | 8.45E-01 | -0.1 | 6.47E-01 | 0.1 | 8.30E-01 | ACT domain-containing protein ACR |
| AT3G19210 | 1.0 | 1.41E-04 | 0.2 | 2.25E-01 | 0.8 | 4.38E-03 | 0.6 | 8.87E-02 | 0.4 | 3.81E-01 | 0.2 | 6.43E-01 | DNA repair |
| AT3G01920 | 1.0 | 2.20E-05 | 0.5 | 3.63E-03 | 0.5 | 9.00E-03 | 0.3 | 2.02E-01 | 0.2 | 5.60E-01 | 0.1 | 6.80E-01 | Threonylcarbamoyl-AMP synthase |
| AT1G29600 | 1.0 | 1.18E-04 | 0.2 | 2.63E-01 | 0.8 | 3.56E-03 | 0.4 | 4.23E-01 | 0.5 | 2.68E-01 | -0.2 | 7.60E-01 | Zinc finger C-x8-C-x5-C-x3-H type family protein |
| AT5G55630 | 1.0 | 6.94E-07 | 0.7 | 2.20E-05 | 0.3 | 5.39E-03 | 0.1 | 5.12E-01 | -0.1 | 3.68E-01 | 0.2 | 2.01E-01 | Two-pore potassium channel 1 |
| AT1G69040 | 1.0 | 4.31E-06 | 1.1 | 4.04E-06 | -0.1 | 2.87E-01 | -0.2 | 2.25E-01 | -0.3 | 7.35E-02 | 0.1 | 5.62E-01 | ACT domain-containing protein ACR |
| AT5G19210 | 1.0 | 1.53E-05 | 0.6 | 9.79E-04 | 0.4 | 2.14E-02 | 0.0 | 8.84E-01 | 0.2 | 4.94E-01 | -0.2 | 6.50E-01 | At5g19210 |
| AT5G56670 | 1.0 | 1.68E-05 | 1.0 | 3.78E-05 | 0.0 | 9.47E-01 | 0.5 | 4.56E-02 | 0.2 | 3.98E-01 | 0.2 | 5.27E-01 | Small ribosomal subunit protein eS30z |
| AT4G13800 | 1.0 | 1.94E-03 | 0.2 | 5.62E-01 | 0.8 | 1.80E-02 | | | | | | | Probable magnesium transporter |
| AT5G16340 | 1.0 | 9.96E-05 | 0.8 | 7.24E-04 | 0.2 | 3.53E-01 | -0.1 | 8.62E-01 | -0.2 | 7.36E-01 | 0.1 | 8.78E-01 | Probable acyl-activating enzyme 6 |
| AT3G13750 | 1.0 | 1.38E-04 | 1.3 | 5.08E-05 | -0.3 | 1.72E-01 | -0.5 | 1.72E-02 | -0.8 | 9.44E-04 | 0.3 | 1.11E-01 | beta-galactosidase |
| AT1G74270 | 1.0 | 2.54E-05 | 0.7 | 6.92E-04 | 0.3 | 4.67E-02 | 0.5 | 1.96E-02 | 0.2 | 4.48E-01 | 0.3 | 3.14E-01 | Large ribosomal subunit protein eL33y |
| AT3G25940 | 1.0 | 4.93E-04 | 0.5 | 3.18E-02 | 0.5 | 6.20E-02 | 0.7 | 2.44E-02 | 0.5 | 1.74E-01 | 0.2 | 4.59E-01 | DNA-directed RNA polymerase subunit |
| AT5G44585 | 1.0 | 6.81E-04 | 0.6 | 1.72E-02 | 0.4 | 1.31E-01 | -0.2 | 6.79E-01 | -0.7 | 5.73E-02 | 0.6 | 2.21E-01 | Serine rich endogenous peptide 12 |
| AT5G58650 | 1.0 | 3.86E-04 | 0.6 | 1.94E-02 | 0.4 | 7.08E-02 | 0.7 | 3.22E-02 | -0.4 | 3.28E-01 | 1.1 | 1.86E-02 | Protein PSY1 |
| AT4G20960 | 1.0 | 6.46E-07 | 0.5 | 1.30E-04 | 0.5 | 6.31E-04 | 0.2 | 2.55E-01 | 0.1 | 6.73E-01 | 0.1 | 7.06E-01 | Riboflavin biosynthesis protein PYRD, chloroplastic |
| AT3G48560 | 1.0 | 7.72E-07 | 0.8 | 1.03E-05 | 0.2 | 3.50E-02 | 0.1 | 5.03E-01 | -0.2 | 4.46E-01 | 0.3 | 2.29E-01 | Acetolactate synthase |
| AT1G78970 | 1.0 | 1.42E-05 | 1.0 | 2.77E-05 | 0.0 | 9.63E-01 | 0.1 | 7.95E-01 | 0.1 | 9.13E-01 | 0.1 | 9.32E-01 | Lupeol synthase 1 -2,3-epoxysqualene synthase) |
| AT3G25120 | 1.0 | 4.04E-07 | 0.7 | 1.46E-05 | 0.3 | 3.78E-03 | 0.6 | 1.60E-02 | 0.1 | 8.94E-01 | 0.5 | 9.14E-02 | At2g45250 |
| AT1G52827 | 1.0 | 4.74E-02 | 0.2 | 6.78E-01 | 0.8 | 1.77E-01 | -0.5 | 3.87E-01 | -0.3 | 7.04E-01 | -0.2 | 8.15E-01 | Protein CYSTEINE-RICH TRANSMEMBRANE MODULE 2 |
| AT3G08940 | 1.0 | 1.06E-05 | 1.1 | 1.78E-05 | 0.0 | 8.07E-01 | -0.6 | 3.03E-04 | -0.7 | 4.92E-04 | 0.1 | 8.24E-01 | Chlorophyll a-b binding protein CP29.2, chloroplastic |
| AT1G48570 | 1.0 | 1.25E-05 | 0.8 | 1.71E-04 | 0.2 | 1.09E-01 | 0.9 | 5.05E-05 | -0.2 | 5.01E-01 | 1.1 | 4.04E-04 | RanBP2-type domain-containing protein |
| AT3G48850 | 1.0 | 2.86E-02 | -0.7 | 2.15E-01 | 1.7 | 1.64E-02 | 3.0 | 9.24E-04 | 1.3 | 1.97E-01 | 1.7 | 2.81E-02 | Mitochondrial phosphate carrier protein 2, mitochondrial |
| AT1G48030 | 1.0 | 1.13E-06 | 0.9 | 8.96E-06 | 0.2 | 1.24E-01 | 0.3 | 2.91E-02 | 0.3 | 1.81E-02 | -0.1 | 7.73E-01 | Dihydrolipoyl dehydrogenase 1, mitochondrial |
| AT1G14720 | 1.0 | 1.83E-06 | 0.8 | 1.69E-05 | 0.2 | 9.27E-02 | 0.2 | 1.20E-01 | -0.2 | 2.36E-01 | 0.4 | 4.27E-02 | Xyloglucan endo-transglycosylase |
| AT1G11870 | 1.0 | 7.83E-07 | 0.8 | 8.31E-06 | 0.2 | 5.81E-02 | 0.4 | 3.43E-02 | 0.1 | 7.31E-01 | 0.3 | 1.80E-01 | serine--tRNA ligase |
| AT5G02610 | 1.0 | 1.24E-05 | 0.8 | 1.34E-04 | 0.2 | 1.26E-01 | 0.6 | 6.24E-04 | 0.3 | 7.69E-02 | 0.3 | 1.74E-01 | Ribosomal L29 family protein |
| AT5G22570 | 1.0 | 9.48E-02 | -0.5 | 5.04E-01 | 1.6 | 8.83E-02 | 1.5 | 1.25E-02 | 0.6 | 4.20E-01 | 0.9 | 9.36E-02 | Probable WRKY transcription factor 38 |

| | | | | | | | | | | | | | |
|---|---|---|---|---|---|---|---|---|---|---|---|---|---|
| AT3G10050 | 1.0 | 2.62E-07 | 0.6 | 1.48E-05 | 0.4 | 8.67E-04 | 0.8 | 1.86E-05 | 0.3 | 3.90E-02 | 0.4 | 1.43E-02 | Threonine dehydratase biosynthetic, chloroplastic |
| AT2G31170 | 1.0 | 1.96E-06 | 0.9 | 1.50E-05 | 0.2 | 1.56E-01 | 0.4 | 1.37E-02 | 0.1 | 6.94E-01 | 0.3 | 1.02E-01 | cysteine--tRNA ligase |
| AT3G04920 | 1.0 | 1.61E-05 | 0.8 | 1.72E-04 | 0.2 | 1.43E-01 | 0.4 | 2.15E-02 | 0.2 | 3.75E-01 | 0.2 | 3.89E-01 | 40S ribosomal protein S24 |
| AT3G22150 | 1.0 | 1.07E-04 | 0.8 | 7.73E-04 | 0.2 | 3.30E-01 | 0.2 | 3.81E-01 | 0.1 | 5.83E-01 | 0.0 | 8.93E-01 | Pentatricopeptide repeat-containing protein At3g22150, chloroplastic |
| AT4G26370 | 1.0 | 1.33E-06 | 1.1 | 2.33E-06 | 0.0 | 6.87E-01 | 0.4 | 2.23E-02 | 0.2 | 4.06E-01 | 0.2 | 2.91E-01 | Antitermination NusB domain-containing protein |
| AT2G38810 | 1.0 | 6.83E-06 | 0.9 | 5.24E-05 | 0.2 | 2.09E-01 | 0.2 | 4.18E-01 | 0.1 | 8.35E-01 | 0.1 | 7.50E-01 | Probable histone H2A variant 2 |
| AT1G18440 | 1.0 | 1.21E-06 | 0.6 | 1.83E-04 | 0.5 | 1.68E-03 | -0.1 | 7.02E-01 | -0.1 | 8.10E-01 | 0.0 | 9.64E-01 | Peptidyl-tRNA hydrolase family protein |
| AT4G30580 | 1.0 | 2.02E-07 | 0.7 | 5.32E-06 | 0.3 | 2.44E-03 | -0.1 | 6.73E-01 | -0.3 | 5.99E-02 | 0.2 | 2.38E-01 | 1-acyl-sn-glycerol-3-phosphate acyltransferase LPAT1, chloroplastic |
| AT1G17480 | 1.0 | 2.97E-06 | 0.5 | 1.35E-03 | 0.5 | 1.50E-03 | 0.1 | 6.65E-01 | 0.1 | 7.06E-01 | 0.0 | 9.94E-01 | Protein IQ-DOMAIN 7 |
| AT1G03090 | 1.0 | 1.78E-05 | 0.8 | 1.59E-04 | 0.2 | 1.93E-01 | -0.7 | 5.51E-02 | -0.9 | 3.68E-02 | 0.2 | 7.71E-01 | Methylcrotonoyl-CoA carboxylase subunit alpha, mitochondrial |
| AT2G30424 | 1.0 | 5.16E-06 | 0.9 | 2.60E-05 | 0.1 | 3.72E-01 | 0.2 | 4.80E-01 | -0.2 | 6.01E-01 | 0.3 | 3.10E-01 | Homeodomain-like superfamily protein |
| AT5G06770 | 1.0 | 3.35E-06 | 0.8 | 5.53E-05 | 0.2 | 5.32E-02 | 0.2 | 1.70E-01 | 0.1 | 7.17E-01 | 0.1 | 5.39E-01 | KH domain-containing protein |
| AT4G35090 | 1.0 | 2.07E-06 | 0.3 | 1.43E-02 | 0.8 | 1.36E-04 | -1.6 | 1.80E-06 | -0.1 | 6.23E-01 | -1.4 | 9.71E-05 | Catalase |
| AT1G53250 | 1.0 | 1.01E-04 | 1.4 | 2.32E-05 | -0.3 | 8.54E-02 | -0.3 | 4.01E-01 | -0.4 | 2.27E-01 | 0.2 | 7.56E-01 | F12M16.15 |
| AT1G23205 | 1.0 | 4.75E-06 | 1.1 | 8.22E-06 | 0.0 | 7.65E-01 | -0.1 | 6.98E-01 | 0.2 | 2.99E-01 | -0.3 | 2.10E-01 | hypothetical protein |
| AT5G20290 | 1.0 | 5.76E-06 | 0.8 | 1.01E-04 | 0.3 | 5.16E-02 | 0.3 | 2.96E-02 | 0.1 | 6.25E-01 | 0.2 | 2.94E-01 | Small ribosomal subunit protein eS8z |
| AT5G38060 | 1.0 | 9.51E-06 | 0.8 | 1.71E-04 | 0.3 | 7.60E-02 | 0.4 | 1.07E-01 | 0.1 | 7.72E-01 | 0.3 | 3.72E-01 | Carboxypeptidase |
| AT5G10470 | 1.0 | 6.20E-06 | 0.4 | 6.90E-03 | 0.7 | 7.16E-04 | 0.3 | 1.68E-01 | 0.3 | 2.61E-01 | 0.0 | 9.71E-01 | Kinesin-like protein KIN-14A |
| AT4G14970 | 1.0 | 9.19E-06 | 0.4 | 7.54E-03 | 0.6 | 2.11E-03 | 0.2 | 6.70E-01 | 0.5 | 2.20E-01 | -0.3 | 4.79E-01 | Fanconi anemia group D2 protein |
| AT3G51420 | 1.0 | 2.51E-06 | 0.8 | 4.41E-05 | 0.3 | 3.28E-02 | 0.0 | 9.33E-01 | -0.2 | 1.97E-01 | 0.2 | 2.46E-01 | Protein STRICTOSIDINE SYNTHASE-LIKE 4 |
| AT1G14410 | 1.0 | 2.11E-06 | 0.8 | 2.94E-05 | 0.2 | 5.60E-02 | 0.4 | 1.95E-02 | 0.1 | 6.56E-01 | 0.3 | 1.73E-01 | Single-stranded DNA-binding protein WHY1, chloroplastic |
| AT1G44920 | 1.0 | 1.99E-07 | 0.8 | 3.85E-06 | 0.3 | 4.50E-03 | -0.1 | 5.12E-01 | 0.1 | 6.47E-01 | -0.2 | 3.18E-01 | T12C22.21 protein |
| AT1G14980 | 1.1 | 4.33E-05 | 0.7 | 9.26E-04 | 0.3 | 7.30E-02 | 0.7 | 5.86E-03 | 0.4 | 1.96E-01 | 0.3 | 3.09E-01 | Chaperonin 10 |
| AT5G41010 | 1.1 | 3.51E-05 | 0.4 | 1.68E-02 | 0.6 | 4.88E-03 | 0.8 | 6.28E-03 | 0.0 | 9.27E-01 | 0.7 | 3.24E-02 | DNA-directed RNA polymerases II, IV and V subunit 12 |
| AT1G33960 | 1.1 | 8.95E-02 | 0.0 | 9.59E-01 | 1.1 | 1.77E-01 | -1.3 | 1.34E-01 | -1.9 | 8.33E-02 | 0.5 | 7.36E-01 | P-loop containing nucleoside triphosphate hydrolases superfamily protein |
| AT5G06730 | 1.1 | 1.77E-05 | 0.2 | 2.31E-01 | 0.9 | 6.55E-04 | 3.3 | 1.45E-07 | 1.2 | 1.18E-02 | 2.1 | 2.94E-05 | Peroxidase 54 |
| AT5G08330 | 1.1 | 1.23E-06 | 1.0 | 3.09E-06 | 0.0 | 9.76E-01 | -0.8 | 4.72E-06 | -0.7 | 6.20E-05 | -0.1 | 6.97E-01 | Transcription factor TCP21 |
| AT3G53680 | 1.1 | 7.02E-05 | 0.9 | 3.14E-04 | 0.1 | 5.13E-01 | 0.6 | 5.98E-02 | 0.0 | 9.48E-01 | 0.5 | 7.85E-02 | Acyl-CoA N-acyltransferase with RING |
| AT1G12770 | 1.1 | 3.27E-07 | 0.7 | 1.15E-05 | 0.3 | 2.43E-03 | 0.2 | 2.20E-01 | -0.2 | 2.18E-01 | 0.3 | 3.85E-02 | ISE1 |
| AT3G26320 | 1.1 | 1.61E-04 | 0.6 | 1.08E-02 | 0.4 | 5.27E-02 | 0.3 | 6.67E-01 | -0.2 | 7.67E-01 | 0.5 | 4.57E-01 | Cytochrome P450 71B36 |
| AT2G18120 | 1.1 | 1.86E-04 | 0.7 | 6.36E-03 | 0.4 | 9.95E-02 | | | | | | | SHI-related sequence 4 |
| AT3G58610 | 1.1 | 1.05E-06 | 0.8 | 1.59E-05 | 0.3 | 2.91E-02 | 0.3 | 2.56E-02 | 0.0 | 9.60E-01 | 0.3 | 9.49E-02 | Acetohydroxy-acid reductoisomerase |
| AT5G45670 | 1.1 | 3.02E-06 | 0.9 | 1.92E-05 | 0.1 | 2.23E-01 | 0.1 | 3.74E-01 | 0.0 | 9.66E-01 | 0.1 | 5.73E-01 | hypothetical protein |
| AT4G33000 | 1.1 | 1.12E-04 | 0.8 | 1.28E-03 | 0.3 | 2.06E-01 | -0.4 | 4.43E-02 | -0.2 | 5.18E-01 | -0.3 | 3.80E-01 | Calcineurin B-like protein |
| AT3G63140 | 1.1 | 2.21E-06 | 1.1 | 3.60E-06 | -0.1 | 5.56E-01 | 0.2 | 1.79E-01 | 0.3 | 9.16E-02 | -0.1 | 7.39E-01 | Chloroplast stem-loop binding protein of 41 kDa a, chloroplastic |
| AT3G22420 | 1.1 | 1.60E-05 | 0.7 | 6.65E-04 | 0.4 | 2.69E-02 | -0.2 | 2.94E-01 | -0.4 | 3.95E-02 | 0.2 | 3.91E-01 | With no lysine kinase 2 |
| AT1G64900 | 1.1 | 5.00E-06 | 0.1 | 3.84E-01 | 1.0 | 7.60E-05 | 0.9 | 6.77E-04 | 0.8 | 2.71E-03 | 0.1 | 8.62E-01 | Cytochrome P450 89A2 |
| AT1G33720 | 1.1 | 2.27E-04 | 0.2 | 3.95E-01 | 0.9 | 3.78E-03 | -0.1 | 8.01E-01 | -0.7 | 1.01E-01 | 0.6 | 2.49E-01 | Cytochrome P450, family 76, subfamily C, polypeptide 6 |
| AT3G20670 | 1.1 | 1.37E-06 | 0.9 | 1.22E-05 | 0.2 | 9.38E-02 | 0.4 | 7.19E-02 | 0.1 | 7.87E-01 | 0.3 | 3.07E-01 | Probable histone H2A.2 |
| AT4G34220 | 1.1 | 1.23E-05 | 1.0 | 2.68E-05 | 0.0 | 9.15E-01 | -0.5 | 6.93E-03 | -0.4 | 8.25E-02 | -0.2 | 4.45E-01 | Receptor protein kinase-like protein At4g34220 |
| AT1G54500 | 1.1 | 3.29E-07 | 1.0 | 1.42E-06 | 0.1 | 3.75E-01 | -0.2 | 2.31E-01 | 0.0 | 8.61E-01 | -0.2 | 2.72E-01 | At1g54500 |
| AT2G30970 | 1.1 | 1.65E-07 | 1.1 | 3.67E-07 | 0.0 | 7.24E-01 | 0.5 | 7.70E-04 | 0.1 | 3.97E-01 | 0.3 | 4.51E-02 | Aspartate aminotransferase |
| AT3G54900 | 1.1 | 1.14E-06 | 0.9 | 9.83E-06 | 0.2 | 1.02E-01 | 0.4 | 3.75E-02 | 0.2 | 5.21E-01 | 0.3 | 3.71E-01 | Monothiol glutaredoxin-S14, chloroplastic |
| AT1G17290 | 1.1 | 1.51E-07 | 0.7 | 3.77E-06 | 0.3 | 2.05E-03 | 0.4 | 9.79E-04 | -0.1 | 6.81E-01 | 0.5 | 5.14E-03 | Alanine aminotransferase 1, mitochondrial |
| AT3G02830 | 1.1 | 9.48E-06 | 0.3 | 5.04E-02 | 0.8 | 3.82E-04 | -0.2 | 2.82E-01 | 0.4 | 3.67E-02 | -0.6 | 7.42E-03 | Zinc finger protein 1 |
| AT1G80380 | 1.1 | 3.20E-06 | 0.9 | 1.71E-05 | 0.1 | 3.18E-01 | 0.0 | 7.33E-01 | 0.0 | 9.45E-01 | 0.1 | 7.67E-01 | P-loop containing nucleoside triphosphate hydrolases superfamily protein |
| AT5G13730 | 1.1 | 3.97E-06 | 0.9 | 3.14E-05 | 0.2 | 1.67E-01 | 0.6 | 2.29E-03 | 0.1 | 5.88E-01 | 0.4 | 5.48E-02 | RNA polymerase sigma factor sigD, chloroplastic |
| AT1G19920 | 1.1 | 2.82E-07 | 0.6 | 2.28E-05 | 0.4 | 5.74E-04 | 0.2 | 1.52E-01 | 0.2 | 1.79E-01 | 0.0 | 9.64E-01 | sulfate adenylyltransferase |
| AT3G10720 | 1.1 | 2.35E-07 | 0.8 | 2.79E-06 | 0.2 | 1.66E-02 | 0.5 | 2.83E-02 | 0.2 | 4.69E-01 | 0.3 | 3.73E-01 | Probable pectinesterase |
| AT4G01800 | 1.1 | 1.35E-06 | 0.6 | 9.29E-05 | 0.4 | 3.09E-03 | -0.1 | 8.78E-01 | 0.0 | 9.92E-01 | -0.1 | 9.08E-01 | chloroplast protein-transporting ATPase |
| AT4G23800 | 1.1 | 9.80E-07 | 0.8 | 1.64E-05 | 0.3 | 2.02E-02 | 0.1 | 7.10E-01 | 0.3 | 1.60E-01 | -0.2 | 4.16E-01 | HMG box protein |
| AT1G21650 | 1.1 | 7.50E-07 | 0.5 | 4.38E-04 | 0.6 | 3.57E-04 | 0.6 | 4.48E-02 | 0.1 | 8.54E-01 | 0.5 | 1.72E-01 | F8K7.7 protein |
| AT5G20140 | 1.1 | 3.40E-06 | 0.7 | 2.27E-04 | 0.4 | 6.48E-03 | 0.3 | 1.83E-01 | 0.2 | 4.81E-01 | 0.1 | 6.99E-01 | SOUL heme-binding family protein |
| AT4G21380 | 1.1 | 2.94E-06 | 0.4 | 3.49E-03 | 0.6 | 5.61E-04 | 1.8 | 3.36E-06 | 0.8 | 1.33E-02 | 1.0 | 4.09E-03 | Receptor kinase 3 |
| AT4G25835 | 1.1 | 1.87E-04 | 1.3 | 7.82E-05 | -0.2 | 2.95E-01 | -0.3 | 3.75E-02 | -0.9 | 9.86E-05 | 0.6 | 1.07E-02 | AAA-ATPase At4g25835 |
| AT2G19400 | 1.1 | 1.07E-04 | 0.8 | 1.47E-03 | 0.3 | 1.86E-01 | -0.2 | 5.81E-01 | -0.1 | 9.00E-01 | -0.1 | 7.96E-01 | AGC kinase family protein |
| AT3G51150 | 1.1 | 1.05E-05 | 0.9 | 1.04E-04 | 0.2 | 1.53E-01 | -0.2 | 2.79E-01 | -0.1 | 5.96E-01 | -0.1 | 7.98E-01 | ATP binding microtubule motor family protein |
| AT3G59480 | 1.1 | 1.92E-03 | 1.3 | 6.72E-04 | -0.2 | 3.82E-01 | -1.0 | 4.28E-02 | -1.2 | 2.18E-02 | 0.3 | 7.23E-01 | Probable fructokinase-4 |
| AT5G62980 | 1.1 | 4.40E-04 | 0.6 | 2.44E-02 | 0.5 | 6.68E-02 | 0.3 | 3.45E-01 | -0.1 | 8.72E-01 | 0.3 | 3.90E-01 | Dihydroneopterin aldolase 2 |
| AT1G69523 | 1.1 | 1.42E-03 | 1.0 | 4.73E-03 | 0.1 | 6.56E-01 | 0.5 | 1.27E-01 | 0.5 | 1.44E-01 | 0.0 | 9.62E-01 | Methyltransferase type 11 domain-containing protein |
| AT3G47430 | 1.1 | 6.42E-05 | 0.4 | 2.84E-02 | 0.6 | 5.94E-03 | -0.2 | 4.08E-01 | -0.1 | 6.37E-01 | -0.1 | 8.59E-01 | Peroxisomal membrane protein 11B |
| AT5G20850 | 1.1 | 8.53E-05 | 0.6 | 1.31E-02 | 0.5 | 2.24E-02 | 0.1 | 8.39E-01 | 0.1 | 8.25E-01 | 0.0 | 9.71E-01 | DNA repair protein RAD51 homolog |
| AT3G55400 | 1.1 | 4.60E-07 | 0.8 | 1.04E-05 | 0.3 | 7.30E-03 | 0.6 | 5.96E-04 | 0.2 | 2.35E-01 | 0.4 | 5.31E-02 | methionine--tRNA ligase |
| AT1G19720 | 1.1 | 7.18E-05 | 0.6 | 3.92E-03 | 0.4 | 3.66E-02 | 0.0 | 9.48E-01 | -0.2 | 5.52E-01 | 0.2 | 5.68E-01 | Pentatricopeptide repeat-containing protein At1g19720 |
| AT1G41880 | 1.1 | 4.41E-06 | 0.8 | 8.79E-05 | 0.3 | 3.76E-02 | 0.8 | 1.37E-04 | 0.5 | 1.02E-02 | 0.3 | 2.57E-01 | Large ribosomal subunit protein eL33z |
| AT1G31180 | 1.1 | 2.18E-06 | 0.6 | 1.65E-04 | 0.4 | 3.78E-03 | 0.6 | 2.50E-04 | 0.3 | 7.71E-02 | 0.3 | 6.50E-02 | 3-isopropylmalate dehydrogenase |
| AT4G37040 | 1.1 | 1.56E-05 | 0.8 | 2.53E-04 | 0.3 | 8.20E-02 | 0.4 | 1.26E-02 | 0.2 | 2.40E-01 | 0.2 | 3.41E-01 | Methionine aminopeptidase 1D, chloroplastic |
| AT4G33520 | 1.1 | 4.50E-07 | 0.9 | 4.77E-06 | 0.2 | 3.48E-02 | 0.2 | 1.72E-01 | 0.0 | 9.14E-01 | 0.1 | 3.71E-01 | P-type ATP-ase 1 |
| AT3G51600 | 1.1 | 3.17E-07 | 0.9 | 2.04E-06 | 0.1 | 1.27E-01 | 0.3 | 1.50E-01 | -0.1 | 8.47E-01 | 0.4 | 2.24E-01 | Non-specific lipid-transfer protein 5 |
| AT2G47590 | 1.1 | 2.73E-07 | 0.8 | 5.61E-06 | 0.3 | 5.39E-03 | 0.6 | 3.30E-04 | 0.5 | 1.81E-03 | 0.0 | 8.57E-01 | Blue-light photoreceptor PHR2 |
| AT4G14490 | 1.1 | 2.08E-05 | 0.7 | 7.80E-04 | 0.4 | 4.34E-02 | -0.1 | 6.61E-01 | -0.1 | 7.60E-01 | 0.0 | 9.77E-01 | FHA domain-containing protein At4g14490 |
| AT3G18290 | 1.1 | 1.79E-05 | 0.5 | 3.79E-03 | 0.6 | 4.70E-03 | 2.9 | 1.06E-07 | 0.8 | 2.86E-02 | 2.1 | 2.32E-04 | Zinc finger protein BRUTUS |
| AT2G04400 | 1.1 | 3.78E-06 | 0.6 | 4.78E-04 | 0.5 | 3.10E-03 | 0.3 | 1.60E-01 | -0.2 | 5.35E-01 | 0.4 | 9.20E-02 | Indole-3-glycerol phosphate synthase, chloroplastic |
| AT5G60150 | 1.1 | 2.57E-06 | 0.7 | 1.01E-04 | 0.3 | 1.50E-02 | 0.6 | 8.94E-02 | 0.4 | 2.57E-01 | 0.1 | 7.80E-01 | Emb\|CAB67639.1 |
| AT1G16390 | 1.1 | 3.65E-05 | 1.2 | 2.89E-05 | -0.1 | 4.31E-01 | -0.2 | 4.98E-01 | -0.5 | 1.08E-01 | 0.3 | 4.60E-01 | Organic cation |
| AT5G57655 | 1.1 | 4.20E-06 | 1.1 | 1.16E-05 | 0.0 | 8.40E-01 | -0.4 | 5.16E-02 | -0.7 | 3.23E-03 | 0.3 | 2.08E-01 | Xylose isomerase |
| AT5G45780 | 1.1 | 3.37E-05 | 0.6 | 4.80E-03 | 0.5 | 1.42E-02 | 0.3 | 2.61E-01 | 0.0 | 9.34E-01 | 0.2 | 4.18E-01 | Leucine-rich repeat protein kinase family protein |
| AT2G25880 | 1.1 | 1.70E-06 | 0.8 | 4.46E-05 | 0.3 | 1.97E-02 | 0.3 | 1.57E-01 | 0.2 | 4.00E-01 | 0.1 | 7.62E-01 | Aurora kinase |
| AT5G63020 | 1.1 | 4.57E-06 | 0.7 | 3.68E-04 | 0.4 | 7.50E-03 | 0.1 | 8.41E-01 | -0.2 | 4.15E-01 | 0.3 | 3.81E-01 | Probable disease resistance protein At5g63020 |
| AT5G48600 | 1.1 | 1.75E-06 | 0.6 | 2.90E-04 | 0.5 | 1.69E-03 | 0.2 | 4.78E-01 | 0.2 | 4.12E-01 | 0.0 | 9.09E-01 | Structural maintenance of chromosomes protein |
| AT4G34350 | 1.1 | 3.50E-07 | 0.7 | 1.07E-05 | 0.3 | 3.46E-03 | -0.1 | 4.92E-01 | -0.2 | 1.12E-01 | 0.1 | 4.59E-01 | 4-hydroxy-3-methylbut-2-enyl diphosphate reductase, chloroplastic |
| AT3G22230 | 1.1 | 4.41E-06 | 0.8 | 7.02E-05 | 0.3 | 5.41E-02 | 0.5 | 3.87E-02 | 0.2 | 5.42E-01 | 0.3 | 3.72E-01 | Large ribosomal subunit protein eL27y |
| AT3G60750 | 1.1 | 1.06E-06 | 0.8 | 2.53E-05 | 0.3 | 1.24E-02 | 0.4 | 2.42E-02 | 0.3 | 1.21E-01 | 0.1 | 7.53E-01 | transketolase |

| | | | | | | | | | | | | | |
|---|---|---|---|---|---|---|---|---|---|---|---|---|---|
| AT5G05940 | 1.1 | 2.38E-05 | 0.8 | 5.06E-04 | 0.3 | 8.72E-02 | 0.3 | 3.43E-01 | 0.2 | 5.51E-01 | 0.1 | 8.88E-01 | ROP guanine nucleotide exchange factor 5 |
| AT5G59250 | 1.1 | 8.82E-07 | 0.7 | 5.61E-05 | 0.4 | 2.37E-03 | -0.3 | 6.32E-02 | -0.2 | 3.45E-01 | -0.1 | 6.48E-01 | D-xylose-proton symporter-like 3, chloroplastic |
| AT5G63950 | 1.1 | 1.56E-05 | 0.3 | 9.71E-02 | 0.8 | 5.37E-04 | 0.1 | 6.26E-01 | 0.1 | 5.90E-01 | 0.0 | 9.31E-01 | Chromatin remodeling 24 |
| AT5G45820 | 1.1 | 5.81E-07 | 0.8 | 1.18E-05 | 0.3 | 1.08E-02 | | | | | | | CBL-interacting serine |
| AT1G02970 | 1.1 | 3.25E-05 | 0.7 | 2.27E-03 | 0.4 | 2.89E-02 | 0.2 | 2.99E-01 | 0.2 | 3.21E-01 | 0.0 | 9.66E-01 | Wee1-like protein kinase |
| AT1G76410 | 1.1 | 4.95E-05 | 1.0 | 1.39E-04 | 0.1 | 7.82E-01 | -0.6 | 2.03E-01 | -1.0 | 8.46E-02 | 0.4 | 6.37E-01 | RING-H2 finger protein ATL8 |
| AT2G36690 | 1.1 | 1.94E-04 | 0.3 | 1.30E-01 | 0.8 | 6.57E-03 | 1.4 | 5.85E-06 | 0.6 | 1.91E-02 | 0.8 | 3.14E-03 | 2-oxoglutarate and Fe-dependent oxygenase superfamily protein |
| AT5G56580 | 1.1 | 1.23E-06 | 0.7 | 8.55E-05 | 0.4 | 4.23E-03 | 0.3 | 1.30E-01 | 0.3 | 2.78E-01 | 0.1 | 8.78E-01 | Mitogen-activated protein kinase kinase 6 |
| AT1G65190 | 1.1 | 1.64E-06 | 0.7 | 9.08E-05 | 0.4 | 4.58E-03 | 0.5 | 7.91E-04 | 0.1 | 5.03E-01 | 0.4 | 4.15E-02 | Protein kinase superfamily protein |
| AT4G24970 | 1.1 | 4.54E-05 | 0.4 | 4.00E-02 | 0.7 | 4.43E-03 | 0.7 | 5.25E-03 | 0.6 | 1.44E-02 | 0.0 | 9.09E-01 | Protein MICRORCHIDIA 7 |
| AT2G44480 | 1.1 | 1.49E-04 | 0.5 | 2.61E-02 | 0.6 | 2.45E-02 | 0.7 | 2.72E-01 | -0.4 | 6.93E-01 | 1.1 | 2.12E-01 | Beta glucosidase 17 |
| AT1G17220 | 1.1 | 5.18E-06 | 0.5 | 1.67E-03 | 0.6 | 1.65E-03 | 0.2 | 3.88E-01 | 0.2 | 2.57E-01 | -0.1 | 7.99E-01 | Translation initiation factor IF-2, chloroplastic |
| AT3G20200 | 1.1 | 2.45E-05 | 0.6 | 4.96E-03 | 0.5 | 1.02E-02 | 0.1 | 8.62E-01 | 0.3 | 5.58E-01 | | | Ser |
| AT4G24810 | 1.1 | 1.02E-05 | 0.9 | 1.01E-04 | 0.2 | 1.61E-01 | 0.1 | 7.94E-01 | 0.2 | 5.15E-01 | -0.1 | 6.69E-01 | At4g24810 |
| AT1G80560 | 1.1 | 4.34E-07 | 0.9 | 4.34E-06 | 0.2 | 3.92E-02 | 0.3 | 6.21E-03 | 0.0 | 8.21E-01 | 0.3 | 2.82E-02 | 3-isopropylmalate dehydrogenase 2, chloroplastic |
| AT5G26000 | 1.1 | 9.05E-06 | 1.2 | 1.39E-05 | -0.1 | 7.03E-01 | -0.2 | 1.88E-01 | 0.0 | 9.50E-01 | -0.2 | 4.21E-01 | Myrosinase 1 |
| AT4G10340 | 1.1 | 4.20E-06 | 1.3 | 3.24E-06 | -0.2 | 1.39E-01 | -0.3 | 5.18E-03 | -0.4 | 2.14E-03 | 0.1 | 6.06E-01 | Chlorophyll a-b binding protein CP26, chloroplastic |
| AT1G69770 | 1.1 | 1.49E-06 | 0.7 | 9.61E-05 | 0.4 | 3.94E-03 | 0.4 | 5.72E-03 | 0.2 | 2.16E-01 | 0.2 | 2.58E-01 | DNA -methyltransferase CMT3 |
| AT1G66520 | 1.1 | 4.91E-05 | 0.8 | 7.41E-04 | 0.3 | 1.51E-01 | 0.4 | 4.42E-01 | 0.3 | 6.35E-01 | 0.1 | 8.93E-01 | Methionyl-tRNA formyltransferase, mitochondrial |
| AT1G10930 | 1.1 | 8.57E-07 | 0.7 | 4.41E-05 | 0.4 | 4.02E-03 | 0.6 | 4.30E-03 | 0.1 | 5.83E-01 | 0.4 | 6.86E-02 | ATP-dependent DNA helicase Q-like 4A |
| AT2G32540 | 1.1 | 1.21E-05 | 0.6 | 2.46E-03 | 0.6 | 4.85E-03 | 0.2 | 2.09E-01 | 0.1 | 6.54E-01 | 0.1 | 6.38E-01 | Cellulose synthase-like protein B4 |
| AT3G48110 | 1.1 | 1.43E-07 | 0.7 | 5.76E-06 | 0.4 | 7.16E-04 | 0.5 | 1.51E-02 | 0.3 | 3.25E-01 | 0.3 | 2.95E-01 | Glycine--tRNA ligase, chloroplastic |
| AT3G27540 | 1.1 | 2.62E-06 | 0.8 | 9.36E-05 | 0.3 | 1.82E-02 | 0.1 | 8.69E-01 | 0.2 | 6.84E-01 | -0.1 | 8.52E-01 | Beta-1,4-N-acetylglucosaminyltransferase family protein |
| AT3G54560 | 1.1 | 2.53E-06 | 0.9 | 2.44E-05 | 0.2 | 9.54E-02 | 0.3 | 1.19E-01 | 0.2 | 3.67E-01 | 0.1 | 7.57E-01 | Histone H2A variant 1 |
| AT1G29660 | 1.1 | 1.81E-07 | 0.8 | 4.71E-06 | 0.3 | 2.42E-03 | -0.4 | 8.36E-03 | -0.3 | 1.41E-01 | -0.2 | 4.65E-01 | GDSL esterase |
| AT5G53210 | 1.1 | 8.64E-06 | 1.0 | 6.39E-05 | 0.2 | 2.61E-01 | 0.2 | 5.90E-01 | 0.0 | 9.37E-01 | 0.3 | 5.00E-01 | Basic helix-loop-helix DNA-binding superfamily protein |
| AT1G54010 | 1.1 | 1.37E-05 | 0.9 | 1.00E-04 | 0.2 | 2.29E-01 | 0.7 | 2.27E-03 | 0.4 | 8.04E-02 | 0.3 | 3.38E-01 | Inactive GDSL esterase |
| AT4G17360 | 1.1 | 1.24E-05 | 0.6 | 2.49E-03 | 0.5 | 5.66E-03 | -0.1 | 7.69E-01 | 0.1 | 6.48E-01 | -0.2 | 4.52E-01 | Formyl transferase |
| AT5G16715 | 1.1 | 4.49E-07 | 0.6 | 4.90E-05 | 0.5 | 7.31E-04 | 0.4 | 9.52E-02 | 0.2 | 4.34E-01 | 0.2 | 6.38E-01 | Valine--tRNA ligase, chloroplastic |
| AT5G16810 | 1.1 | 3.30E-07 | 0.8 | 9.53E-06 | 0.3 | 4.19E-03 | 0.3 | 9.02E-02 | 0.0 | 9.65E-01 | 0.3 | 1.72E-01 | Protein kinase superfamily protein |
| AT1G75100 | 1.1 | 9.99E-07 | 0.5 | 4.41E-04 | 0.6 | 3.86E-04 | 0.2 | 6.15E-01 | -0.2 | 5.57E-01 | 0.4 | 2.90E-01 | J domain-containing protein required for chloroplast accumulation response 1 |
| AT5G60800 | 1.1 | 3.30E-04 | 0.7 | 8.47E-03 | 0.4 | 1.14E-01 | -0.2 | 4.06E-01 | -0.5 | 1.00E-01 | 0.2 | 5.09E-01 | Heavy metal transport |
| AT1G77210 | 1.1 | 2.93E-04 | 1.2 | 2.56E-04 | -0.1 | 6.82E-01 | -0.1 | 4.91E-01 | -0.7 | 3.12E-03 | 0.6 | 3.14E-02 | Sugar transporter 14 |
| AT4G21210 | 1.1 | 1.58E-07 | 0.8 | 2.54E-06 | 0.3 | 5.71E-03 | 0.5 | 1.98E-03 | 0.3 | 4.24E-02 | 0.2 | 4.82E-01 | Uncharacterized protein |
| AT1G61430 | 1.1 | 1.34E-06 | 0.7 | 9.29E-05 | 0.4 | 4.59E-03 | 0.1 | 6.11E-01 | -0.1 | 7.60E-01 | 0.2 | 4.48E-01 | S-locus lectin protein kinase family protein |
| AT5G49160 | 1.1 | 2.31E-06 | 0.7 | 2.24E-04 | 0.5 | 3.16E-03 | 0.4 | 1.37E-01 | 0.3 | 3.17E-01 | 0.1 | 8.58E-01 | DNA -methyltransferase |
| AT1G17600 | 1.1 | 5.68E-04 | 0.7 | 2.41E-02 | 0.5 | 1.04E-01 | 0.6 | 1.64E-01 | 0.2 | 7.51E-01 | 0.4 | 4.39E-01 | Disease resistance protein family |
| AT3G53130 | 1.1 | 8.00E-06 | 1.1 | 1.66E-05 | 0.0 | 9.92E-01 | 0.4 | 2.20E-02 | 0.1 | 7.08E-01 | 0.3 | 1.26E-01 | Carotene epsilon-monooxygenase, chloroplastic |
| AT1G56500 | 1.1 | 5.05E-06 | 0.8 | 1.39E-04 | 0.4 | 2.39E-02 | 0.3 | 9.36E-03 | 0.2 | 7.42E-02 | 0.1 | 6.56E-01 | Protein SUPPRESSOR OF QUENCHING 1, chloroplastic |
| AT5G66190 | 1.1 | 1.42E-06 | 0.9 | 1.21E-05 | 0.2 | 1.15E-01 | 0.3 | 1.52E-02 | 0.2 | 6.33E-02 | 0.0 | 8.17E-01 | Ferredoxin--NADP reductase, chloroplastic |
| AT3G15140 | 1.1 | 2.02E-05 | 0.8 | 5.26E-04 | 0.3 | 6.02E-02 | 1.0 | 4.53E-04 | 0.6 | 2.92E-02 | 0.4 | 1.80E-01 | Polynucleotidyl transferase, ribonuclease H-like superfamily protein |
| AT4G27710 | 1.1 | 7.30E-07 | 0.8 | 2.27E-05 | 0.4 | 5.57E-03 | 0.0 | 8.45E-01 | -0.1 | 7.05E-01 | 0.2 | 5.46E-01 | Cytochrome P450, family 709, subfamily B, polypeptide 3 |
| AT1G03400 | 1.1 | 5.73E-06 | 0.5 | 2.17E-03 | 0.6 | 1.68E-03 | 0.3 | 9.19E-02 | -0.2 | 3.33E-01 | 0.4 | 4.13E-02 | 2-oxoglutarate and Fe-dependent oxygenase superfamily protein |
| AT5G10620 | 1.1 | 2.11E-05 | 0.8 | 6.10E-04 | 0.3 | 6.68E-02 | -0.2 | 6.43E-01 | 0.0 | 9.59E-01 | -0.2 | 6.73E-01 | RNA methyltransferase At5g10620 |
| AT1G15730 | 1.1 | 1.20E-07 | 0.7 | 7.01E-06 | 0.4 | 3.82E-04 | -0.1 | 7.48E-01 | -0.1 | 5.92E-01 | 0.0 | 8.41E-01 | At1g15730 |
| AT1G64150 | 1.1 | 1.23E-06 | 0.9 | 1.75E-05 | 0.3 | 3.83E-02 | -0.3 | 1.43E-01 | 0.0 | 9.07E-01 | -0.2 | 3.53E-01 | Protein PAM71, chloroplastic |
| AT5G60670 | 1.1 | 1.78E-05 | 1.0 | 1.16E-04 | 0.2 | 2.72E-01 | 0.6 | 8.38E-03 | 0.2 | 3.51E-01 | 0.3 | 2.37E-01 | Large ribosomal subunit protein uL11x |
| AT3G14330 | 1.1 | 3.26E-06 | 1.0 | 2.53E-05 | 0.2 | 1.81E-01 | 0.3 | 1.72E-01 | 0.0 | 9.87E-01 | 0.3 | 2.52E-01 | Pentatricopeptide repeat-containing protein At3g14330 |
| AT2G39880 | 1.2 | 1.18E-05 | 1.0 | 5.68E-05 | 0.1 | 4.56E-01 | 0.1 | 7.76E-01 | -0.1 | 7.36E-01 | 0.2 | 5.49E-01 | Transcription factor MYB25 |
| AT1G05820 | 1.2 | 3.37E-05 | 0.6 | 8.56E-03 | 0.6 | 1.01E-02 | 0.4 | 3.30E-01 | 0.5 | 2.69E-01 | -0.1 | 8.59E-01 | SIGNAL PEPTIDE PEPTIDASE-LIKE 5 |
| AT5G62360 | 1.2 | 8.42E-05 | 2.2 | 1.40E-06 | -1.0 | 3.64E-04 | 0.6 | 4.48E-02 | -0.1 | 8.74E-01 | 0.7 | 8.76E-02 | Pectinesterase inhibitor domain-containing protein |
| AT2G18940 | 1.2 | 1.96E-06 | 0.7 | 1.54E-04 | 0.5 | 3.96E-03 | 0.1 | 7.25E-01 | 0.0 | 9.26E-01 | 0.1 | 7.22E-01 | Pentatricopeptide repeat-containing protein At2g18940, chloroplastic |
| AT1G64860 | 1.2 | 3.02E-07 | 0.7 | 2.76E-05 | 0.5 | 5.55E-04 | -1.1 | 4.43E-05 | 0.2 | 4.17E-01 | -1.3 | 1.65E-04 | RNA polymerase sigma factor sigA |
| AT4G23290 | 1.2 | 1.35E-06 | 0.7 | 9.87E-05 | 0.5 | 2.83E-03 | 0.4 | 1.72E-01 | -0.2 | 6.74E-01 | 0.5 | 8.34E-02 | Uncharacterized protein |
| AT5G63570 | 1.2 | 4.49E-07 | 0.8 | 9.08E-06 | 0.3 | 9.45E-03 | 0.4 | 9.25E-04 | 0.0 | 8.47E-01 | 0.4 | 1.49E-02 | glutamate-1-semialdehyde 2,1-aminomutase |
| AT5G16290 | 1.2 | 3.69E-07 | 0.8 | 8.69E-06 | 0.3 | 6.10E-03 | 0.0 | 8.08E-01 | -0.1 | 6.45E-01 | 0.1 | 5.38E-01 | VAT1 |
| AT3G14415 | 1.2 | 3.10E-07 | 0.9 | 3.11E-06 | 0.2 | 3.52E-02 | 0.1 | 2.45E-01 | 0.0 | 9.53E-01 | 0.1 | 3.60E-01 | #NAME? |
| AT4G02790 | 1.2 | 3.81E-06 | 1.0 | 2.41E-05 | 0.2 | 2.19E-01 | 0.6 | 4.87E-04 | 0.2 | 2.58E-01 | 0.4 | 4.98E-02 | DAR GTPase 3, chloroplastic |
| AT1G26560 | 1.2 | 3.56E-06 | 1.0 | 2.37E-05 | 0.2 | 2.28E-01 | -0.2 | 4.98E-01 | 0.5 | 1.34E-01 | -0.7 | 7.74E-02 | Beta-glucosidase 40 |
| AT3G18740 | 1.2 | 6.44E-06 | 0.8 | 2.10E-04 | 0.4 | 2.10E-02 | 0.5 | 5.30E-02 | 0.1 | 8.01E-01 | 0.4 | 2.72E-01 | Large ribosomal subunit protein eL30x |
| AT1G11430 | 1.2 | 2.89E-07 | 1.0 | 1.90E-06 | 0.2 | 1.05E-01 | 0.4 | 4.72E-03 | 0.0 | 9.09E-01 | 0.3 | 4.34E-02 | Multiple organellar RNA editing factor 9, chloroplastic |
| AT2G21900 | 1.2 | 1.87E-04 | -0.6 | 5.11E-02 | 1.8 | 2.93E-04 | 0.7 | 1.78E-02 | 1.3 | 1.76E-04 | -0.7 | 3.62E-02 | Probable WRKY transcription factor 59 |
| AT5G11160 | 1.2 | 2.39E-05 | 0.8 | 6.53E-04 | 0.3 | 6.40E-02 | 0.2 | 6.75E-01 | 0.0 | 9.27E-01 | 0.2 | 6.86E-01 | adenine phosphoribosyltransferase |
| AT5G13650 | 1.2 | 5.93E-07 | 0.6 | 8.78E-05 | 0.5 | 6.57E-04 | 0.3 | 9.51E-02 | 0.1 | 7.16E-01 | 0.2 | 4.18E-01 | Elongation factor family protein |
| AT5G26742 | 1.2 | 4.01E-07 | 0.7 | 1.77E-05 | 0.4 | 2.12E-03 | 0.4 | 1.78E-02 | 0.0 | 9.51E-01 | 0.4 | 6.15E-02 | RNA helicase |
| AT4G24540 | 1.2 | 1.74E-05 | 0.6 | 2.17E-03 | 0.5 | 1.02E-02 | 0.0 | 9.39E-01 | 0.0 | 9.79E-01 | 0.0 | 9.41E-01 | Uncharacterized protein |
| AT1G49730 | 1.2 | 1.96E-06 | 0.8 | 8.36E-05 | 0.4 | 8.82E-03 | -0.3 | 1.32E-01 | -0.4 | 3.62E-02 | 0.2 | 5.37E-01 | Protein kinase superfamily protein |
| AT5G18670 | 1.2 | 3.68E-06 | 1.4 | 2.74E-06 | -0.2 | 1.38E-01 | -0.3 | 1.07E-01 | -0.2 | 2.60E-01 | -0.1 | 8.41E-01 | Inactive beta-amylase 9 |
| AT4G39250 | 1.2 | 1.01E-02 | 1.8 | 7.52E-04 | -0.6 | 1.23E-01 | | | | | | | Protein RADIALIS-like 1 |
| AT1G69200 | 1.2 | 1.17E-06 | 1.0 | 5.97E-06 | 0.1 | 3.03E-01 | 1.0 | 1.86E-04 | 0.1 | 7.12E-01 | 0.9 | 1.43E-03 | Fructokinase-like 2, chloroplastic |
| AT3G58140 | 1.2 | 2.56E-07 | 0.7 | 2.33E-05 | 0.5 | 4.47E-04 | 0.3 | 1.97E-02 | 0.2 | 1.51E-01 | 0.1 | 6.52E-01 | Phenylalanine--tRNA ligase, chloroplastic |
| AT1G30720 | 1.2 | 2.31E-03 | 0.8 | 2.42E-02 | 0.4 | 2.79E-01 | -0.3 | 4.69E-01 | -1.0 | 1.96E-02 | 0.7 | 9.63E-02 | Berberine bridge enzyme-like 10 |
| AT2G39730 | 1.2 | 1.93E-06 | 1.1 | 8.87E-06 | 0.1 | 4.26E-01 | 0.3 | 2.18E-02 | 0.2 | 2.13E-01 | 0.1 | 5.59E-01 | Ribulose bisphosphate carboxylase |
| AT1G80030 | 1.2 | 3.51E-07 | 0.7 | 2.08E-05 | 0.4 | 1.10E-03 | 0.3 | 1.60E-02 | 0.3 | 4.17E-02 | 0.0 | 9.27E-01 | Molecular chaperone Hsp40 |
| AT3G14220 | 1.2 | 1.36E-07 | 1.0 | 1.07E-06 | 0.2 | 2.80E-02 | -0.1 | 5.12E-01 | -0.1 | 7.67E-01 | 0.0 | 8.81E-01 | GDSL esterase |
| AT1G52510 | 1.2 | 2.85E-06 | 0.7 | 1.91E-04 | 0.5 | 5.19E-03 | -0.1 | 7.47E-01 | 0.1 | 8.56E-01 | -0.2 | 6.91E-01 | Uncharacterized protein At1g52510 |
| AT1G75900 | 1.2 | 1.01E-05 | 0.6 | 1.68E-03 | 0.5 | 5.32E-03 | -0.2 | 2.30E-01 | 0.1 | 7.72E-01 | -0.3 | 2.24E-01 | GDSL esterase |
| AT4G34740 | 1.2 | 1.64E-07 | 0.9 | 2.01E-06 | 0.3 | 1.04E-02 | -0.3 | 4.77E-02 | -0.2 | 2.91E-01 | -0.1 | 4.68E-01 | Amidophosphoribosyltransferase 2, chloroplastic |
| AT1G56190 | 1.2 | 4.69E-07 | 1.0 | 3.57E-06 | 0.2 | 1.00E-01 | 0.5 | 1.42E-02 | 0.3 | 1.09E-01 | 0.1 | 6.49E-01 | Phosphoglycerate kinase |
| AT2G45770 | 1.2 | 7.70E-07 | 0.5 | 6.88E-04 | 0.7 | 2.17E-04 | 0.2 | 2.28E-01 | 0.1 | 6.24E-01 | 0.1 | 7.20E-01 | Cell division protein FtsY homolog, chloroplastic |
| AT5G07370 | 1.2 | 6.09E-08 | 0.9 | 1.06E-06 | 0.3 | 1.39E-03 | -0.1 | 6.27E-01 | -0.2 | 1.31E-01 | 0.1 | 4.54E-01 | Inositol polyphosphate multikinase alpha |
| AT5G21940 | 1.2 | 1.14E-04 | 1.4 | 5.02E-05 | -0.3 | 2.48E-01 | -0.8 | 4.61E-03 | -1.7 | 5.41E-05 | 0.8 | 3.78E-02 | Protein OXIDATIVE STRESS 3 LIKE 1 |
| AT4G27370 | 1.2 | 4.10E-05 | 0.7 | 3.92E-03 | 0.5 | 2.37E-02 | 0.1 | 7.62E-01 | 0.3 | 5.51E-01 | -0.1 | 8.03E-01 | P-loop containing nucleoside triphosphate hydrolases superfamily protein |

| | | | | | | | | | | | | | |
|---|---|---|---|---|---|---|---|---|---|---|---|---|---|
| AT4G01460 | 1.2 | 4.94E-05 | 1.0 | 3.14E-04 | 0.2 | 3.55E-01 | -0.5 | 2.27E-03 | -0.5 | 1.32E-02 | -0.1 | 7.35E-01 | Transcription factor bHLH57 |
| AT4G36730 | 1.2 | 8.35E-07 | 0.7 | 5.76E-05 | 0.4 | 2.38E-03 | -0.2 | 1.53E-01 | -0.6 | 2.18E-03 | 0.4 | 7.19E-02 | hypothetical protein |
| AT2G35040 | 1.2 | 6.71E-07 | 0.7 | 4.09E-05 | 0.5 | 1.92E-03 | 0.4 | 6.53E-04 | 0.0 | 8.77E-01 | 0.4 | 1.14E-02 | AICARFT |
| AT3G08770 | 1.2 | 2.72E-06 | 1.1 | 1.19E-05 | 0.1 | 4.28E-01 | 0.8 | 9.12E-05 | 0.4 | 3.41E-02 | 0.4 | 7.19E-02 | Non-specific lipid-transfer protein |
| AT3G44050 | 1.2 | 3.80E-07 | 0.6 | 7.67E-05 | 0.5 | 4.47E-04 | -0.2 | 3.31E-01 | 0.2 | 3.97E-01 | -0.5 | 1.38E-01 | Kinesin-like protein KIN-12E |
| AT1G45474 | 1.2 | 1.33E-06 | 0.8 | 5.15E-05 | 0.4 | 6.33E-03 | 0.0 | 9.29E-01 | 0.0 | 9.82E-01 | 0.0 | 9.40E-01 | Photosystem I chlorophyll a |
| AT1G79790 | 1.2 | 4.84E-07 | 0.8 | 1.47E-05 | 0.4 | 4.59E-03 | -0.2 | 1.42E-01 | -0.3 | 6.96E-02 | 0.1 | 7.49E-01 | Haloacid dehalogenase-like hydrolase superfamily protein |
| AT2G01850 | 1.2 | 3.99E-06 | 0.7 | 1.97E-04 | 0.4 | 9.38E-03 | -0.3 | 9.73E-02 | -1.0 | 3.39E-04 | 0.7 | 2.45E-02 | Probable xyloglucan endotransglucosylase |
| AT3G47800 | 1.2 | 1.01E-05 | 1.4 | 7.73E-06 | -0.2 | 2.26E-01 | 0.1 | 4.65E-01 | -0.6 | 6.81E-03 | 0.7 | 2.12E-03 | Aldose 1-epimerase |
| AT4G19820 | 1.2 | 6.37E-04 | 1.6 | 1.33E-04 | -0.4 | 1.59E-01 | | | | | | | Glycosyl hydrolase family protein with chitinase insertion domain-containing protein |
| AT5G58370 | 1.2 | 9.38E-07 | 0.8 | 4.42E-05 | 0.4 | 5.09E-03 | 0.6 | 3.75E-03 | 0.3 | 2.41E-01 | 0.4 | 1.28E-01 | Similarity to GTP-binding protein CGPA |
| AT3G13180 | 1.2 | 8.20E-07 | 1.0 | 4.95E-06 | 0.1 | 1.94E-01 | 0.5 | 2.29E-03 | 0.1 | 7.07E-01 | 0.4 | 3.29E-02 | NOL1 |
| AT5G48545 | 1.2 | 3.47E-07 | 0.6 | 9.53E-05 | 0.6 | 2.68E-04 | -0.1 | 2.86E-01 | -0.3 | 5.63E-02 | 0.1 | 5.10E-01 | Histidine triad nucleotide-binding 3 |
| AT5G65010 | 1.2 | 6.94E-06 | 0.9 | 1.30E-04 | 0.3 | 4.60E-02 | -0.8 | 1.28E-04 | -1.1 | 4.28E-05 | 0.3 | 1.26E-01 | Asparagine synthetase |
| AT3G16080 | 1.2 | 1.37E-05 | 0.9 | 1.99E-04 | 0.3 | 8.58E-02 | 0.6 | 1.61E-03 | 0.3 | 6.70E-02 | 0.2 | 3.07E-01 | Ribosomal protein L37 |
| AT5G41080 | 1.2 | 2.45E-05 | 1.9 | 1.70E-06 | -0.7 | 1.73E-03 | 0.3 | 1.98E-01 | 0.1 | 7.50E-01 | 0.2 | 5.58E-01 | glycerophosphodiester phosphodiesterase |
| AT1G26380 | 1.2 | 5.83E-04 | 0.6 | 5.51E-02 | 0.6 | 4.14E-02 | 2.0 | 1.10E-07 | 0.4 | 1.31E-01 | 1.6 | 8.05E-05 | Berberine bridge enzyme-like 3 |
| AT1G77060 | 1.2 | 2.32E-06 | 1.1 | 6.98E-06 | 0.0 | 7.48E-01 | -0.2 | 2.74E-01 | -0.1 | 6.03E-01 | -0.1 | 7.80E-01 | Putative carboxyphosphonoenolpyruvate mutase |
| AT1G07110 | 1.2 | 5.73E-06 | 0.7 | 4.32E-04 | 0.5 | 6.89E-03 | 0.0 | 9.09E-01 | -0.1 | 7.20E-01 | 0.1 | 8.43E-01 | 6-phosphofructo-2-kinase |
| AT4G27600 | 1.2 | 2.18E-07 | 0.8 | 6.21E-06 | 0.4 | 2.21E-03 | 0.0 | 9.49E-01 | -0.1 | 7.39E-01 | 0.1 | 7.42E-01 | Carbohydrate kinase-like protein |
| AT3G17040 | 1.2 | 1.45E-06 | 0.8 | 5.62E-05 | 0.4 | 6.75E-03 | -0.2 | 1.71E-01 | -0.1 | 6.38E-01 | -0.1 | 6.11E-01 | High chlorophyll fluorescent 107 |
| AT2G43590 | 1.2 | 3.87E-04 | 0.5 | 7.66E-02 | 0.7 | 1.80E-02 | 0.5 | 7.30E-02 | -0.1 | 7.80E-01 | 0.6 | 8.38E-02 | Endochitinase At2g43590 |
| AT5G25980 | 1.2 | 3.20E-06 | 1.3 | 5.30E-06 | -0.1 | 6.35E-01 | -0.1 | 5.97E-01 | 0.0 | 8.91E-01 | -0.1 | 6.25E-01 | Thioglucosidase |
| AT5G45390 | 1.2 | 7.38E-08 | 0.7 | 3.22E-06 | 0.4 | 3.10E-04 | 0.2 | 1.27E-01 | 0.1 | 3.92E-01 | 0.1 | 7.87E-01 | ATP-dependent Clp protease proteolytic subunit 4, chloroplastic |
| AT5G55740 | 1.2 | 1.09E-05 | 0.9 | 2.06E-04 | 0.3 | 6.61E-02 | 0.4 | 2.21E-02 | 0.2 | 3.02E-01 | 0.2 | 3.86E-01 | Pentatricopeptide repeat-containing protein At5g55740, chloroplastic |
| AT2G41090 | 1.2 | 4.45E-06 | 0.5 | 2.26E-03 | 0.7 | 1.05E-03 | 0.8 | 7.45E-05 | 0.6 | 2.13E-03 | 0.2 | 4.34E-01 | AT2G41090 protein |
| AT1G78370 | 1.2 | 1.38E-04 | 1.3 | 1.66E-04 | -0.1 | 7.57E-01 | 0.6 | 4.46E-02 | 1.1 | 2.52E-03 | -0.5 | 2.16E-01 | Glutathione S-transferase U20 |
| AT3G19050 | 1.2 | 9.80E-06 | 0.6 | 2.46E-03 | 0.6 | 4.09E-03 | 0.1 | 7.52E-01 | 0.2 | 6.55E-01 | -0.1 | 8.98E-01 | Kinesin-like protein KIN-12D |
| AT4G22305 | 1.2 | 1.41E-05 | 0.5 | 7.53E-03 | 0.7 | 3.12E-03 | 0.4 | 6.48E-02 | 0.2 | 3.85E-01 | 0.2 | 5.64E-01 | Alpha |
| AT1G25230 | 1.2 | 1.55E-05 | 1.3 | 1.94E-05 | -0.1 | 6.69E-01 | -0.4 | 4.03E-02 | -0.7 | 2.34E-03 | 0.3 | 1.69E-01 | acid phosphatase |
| AT1G73830 | 1.2 | 2.71E-05 | 0.5 | 1.49E-02 | 0.7 | 4.55E-03 | -0.3 | 3.14E-01 | -0.6 | 8.31E-02 | 0.3 | 5.27E-01 | BHLH domain-containing protein |
| AT4G24280 | 1.2 | 1.09E-07 | 0.9 | 2.05E-06 | 0.3 | 2.79E-03 | 0.7 | 2.07E-03 | 0.3 | 2.65E-01 | 0.4 | 1.15E-01 | Heat shock 70 kDa protein 6, chloroplastic |
| AT1G10360 | 1.2 | 1.01E-06 | 0.9 | 1.87E-05 | 0.3 | 1.91E-02 | -0.2 | 1.75E-01 | -0.2 | 2.24E-01 | 0.0 | 9.93E-01 | Glutathione S-transferase |
| AT1G33340 | 1.2 | 2.05E-05 | 1.8 | 2.01E-06 | -0.6 | 3.42E-03 | -0.1 | 8.89E-01 | 0.4 | 3.32E-01 | -0.5 | 3.19E-01 | Putative clathrin assembly protein At1g33340 |
| AT2G02580 | 1.2 | 2.05E-05 | 1.5 | 9.22E-06 | -0.3 | 1.23E-01 | 0.9 | 1.29E-02 | 1.1 | 3.08E-03 | -0.3 | 4.46E-01 | Cytochrome P450 71B9 |
| AT3G11930 | 1.2 | 7.53E-07 | 1.1 | 4.01E-06 | 0.1 | 2.58E-01 | -0.2 | 3.24E-01 | -0.3 | 1.46E-01 | 0.1 | 7.11E-01 | Adenine nucleotide alpha hydrolases-like superfamily protein |
| AT4G23940 | 1.2 | 1.68E-07 | 0.8 | 7.09E-06 | 0.4 | 9.78E-04 | 0.5 | 8.51E-02 | 0.4 | 2.16E-01 | 0.1 | 8.53E-01 | Probable inactive ATP-dependent zinc metalloprotease FTSHI 1, chloroplastic |
| AT4G37930 | 1.2 | 1.26E-06 | 0.9 | 2.62E-05 | 0.3 | 1.81E-02 | 0.2 | 1.46E-01 | 0.2 | 1.11E-01 | 0.0 | 8.39E-01 | Serine hydroxymethyltransferase 1, mitochondrial |
| AT5G10570 | 1.2 | 2.08E-03 | 1.2 | 2.79E-03 | 0.0 | 9.60E-01 | | | | | | | Transcription factor bHLH61 |
| AT2G35660 | 1.2 | 2.06E-05 | 0.5 | 8.87E-03 | 0.7 | 4.04E-03 | -0.5 | 1.89E-02 | -0.3 | 1.63E-01 | -0.2 | 5.34E-01 | FAD |
| AT5G22800 | 1.2 | 2.41E-07 | 0.6 | 4.85E-05 | 0.6 | 1.87E-04 | 0.4 | 5.88E-02 | 0.3 | 2.03E-01 | 0.1 | 7.93E-01 | Probable alanine--tRNA ligase, chloroplastic |
| AT3G60210 | 1.2 | 1.03E-06 | 0.8 | 4.18E-05 | 0.4 | 5.89E-03 | 0.3 | 1.43E-01 | 0.2 | 4.89E-01 | 0.1 | 7.11E-01 | 10 kDa chaperonin 1, chloroplastic |
| AT1G30410 | 1.2 | 3.05E-06 | 0.5 | 4.17E-03 | 0.7 | 5.61E-04 | 0.1 | 7.98E-01 | -0.1 | 7.80E-01 | 0.2 | 6.39E-01 | ABC-type xenobiotic transporter |
| AT3G23940 | 1.2 | 1.49E-06 | 0.9 | 3.47E-05 | 0.4 | 1.51E-02 | 0.4 | 3.67E-03 | 0.2 | 1.11E-01 | 0.2 | 3.05E-01 | Dihydroxy-acid dehydratase, chloroplastic |
| AT3G17609 | 1.2 | 7.72E-06 | 0.3 | 9.09E-02 | 0.9 | 3.47E-04 | 0.8 | 4.46E-02 | 0.6 | 1.47E-01 | 0.2 | 7.71E-01 | BZip transcription factor AtbZip64 |
| AT4G25120 | 1.2 | 1.62E-06 | 0.5 | 1.93E-03 | 0.7 | 4.37E-04 | 0.2 | 5.17E-01 | 0.1 | 6.92E-01 | 0.0 | 9.33E-01 | ATP-dependent DNA helicase SRS2-like protein At4g25120 |
| AT5G51820 | 1.2 | 2.39E-06 | 1.1 | 1.06E-05 | 0.1 | 4.30E-01 | 0.4 | 2.42E-02 | 0.2 | 3.26E-01 | 0.2 | 3.80E-01 | Phosphoglucomutase, chloroplastic |
| AT3G06730 | 1.2 | 7.25E-07 | 1.1 | 2.81E-06 | 0.1 | 4.75E-01 | 0.6 | 1.31E-03 | 0.1 | 6.45E-01 | 0.5 | 2.96E-02 | Thioredoxin-like protein CITRX, chloroplastic |
| AT5G15210 | 1.2 | 7.68E-07 | 1.0 | 6.36E-06 | 0.2 | 9.76E-02 | 0.2 | 3.82E-01 | -0.1 | 6.76E-01 | 0.3 | 2.83E-01 | Zinc-finger homeodomain protein 8 |
| AT5G18430 | 1.2 | 5.43E-05 | 0.8 | 1.77E-03 | 0.4 | 6.69E-02 | -0.4 | 1.14E-01 | -0.1 | 6.53E-01 | -0.3 | 4.39E-01 | hypothetical protein |
| AT4G18170 | 1.2 | 1.38E-05 | 0.6 | 4.05E-03 | 0.6 | 5.76E-03 | -0.5 | 2.06E-01 | -1.3 | 1.09E-02 | 0.8 | 1.99E-01 | WRKY transcription factor 28 |
| AT1G32080 | 1.2 | 8.63E-07 | 1.1 | 4.35E-06 | 0.1 | 3.22E-01 | 0.3 | 2.84E-02 | 0.3 | 6.95E-02 | 0.0 | 9.37E-01 | Plastidal glycolate |
| AT5G57030 | 1.2 | 7.10E-06 | 1.0 | 6.05E-05 | 0.2 | 1.57E-01 | 0.2 | 2.74E-01 | 0.4 | 4.31E-02 | -0.2 | 4.18E-01 | Lycopene epsilon cyclase, chloroplastic |
| AT5G61950 | 1.2 | 1.28E-05 | 0.5 | 1.14E-02 | 0.7 | 2.37E-03 | | | | | | | C2H2-type domain-containing protein |
| AT3G28460 | 1.2 | 1.76E-06 | 1.3 | 3.54E-06 | 0.0 | 8.21E-01 | 0.4 | 4.06E-02 | 0.0 | 8.89E-01 | 0.4 | 7.78E-02 | rRNA methyltransferase YlbH |
| AT1G54385 | 1.2 | 3.25E-06 | 0.8 | 2.64E-04 | 0.5 | 6.11E-03 | 0.1 | 7.95E-01 | 0.4 | 1.25E-01 | -0.3 | 1.57E-01 | Protein SINE1 |
| AT5G50250 | 1.2 | 1.53E-06 | 0.9 | 4.17E-05 | 0.4 | 1.19E-02 | 0.4 | 8.26E-03 | 0.2 | 3.98E-01 | 0.3 | 1.98E-01 | RNA-binding protein CP31B, chloroplastic |
| AT1G17100 | 1.2 | 7.75E-08 | 0.7 | 6.67E-06 | 0.5 | 1.40E-04 | 0.1 | 4.68E-01 | 0.2 | 2.07E-01 | -0.1 | 6.38E-01 | At1g17100 |
| AT3G17350 | 1.2 | 2.45E-07 | 1.2 | 8.67E-07 | 0.1 | 5.39E-01 | -0.1 | 6.93E-01 | 0.1 | 8.47E-01 | -0.2 | 4.53E-01 | Wall-associated receptor kinase carboxy-terminal protein |
| AT4G39040 | 1.2 | 8.32E-07 | 1.0 | 6.28E-06 | 0.2 | 1.07E-01 | 0.1 | 4.14E-01 | -0.1 | 3.42E-01 | 0.2 | 1.08E-01 | At4g39040 domain protein) |
| AT1G16880 | 1.2 | 1.24E-06 | 0.8 | 4.36E-05 | 0.4 | 7.32E-03 | 0.2 | 1.94E-01 | 0.2 | 1.51E-01 | 0.0 | 8.38E-01 | ACT domain-containing protein ACR |
| AT1G35420 | 1.2 | 1.65E-06 | 0.7 | 1.18E-04 | 0.5 | 3.28E-03 | -0.3 | 9.38E-02 | -0.2 | 3.10E-01 | -0.1 | 7.82E-01 | Carboxymethylenebutenolidase homolog |
| AT5G48630 | 1.2 | 1.25E-05 | 0.8 | 6.25E-04 | 0.4 | 2.12E-02 | 0.2 | 4.26E-01 | -0.1 | 7.81E-01 | 0.3 | 3.31E-01 | Cyclin family protein |
| AT5G55580 | 1.2 | 7.40E-07 | 0.8 | 4.40E-05 | 0.4 | 3.13E-03 | 0.4 | 1.71E-02 | -0.1 | 6.17E-01 | 0.5 | 2.41E-02 | Mitochondrial transcription termination factor family protein |
| AT3G48460 | 1.2 | 6.58E-05 | 0.7 | 4.28E-03 | 0.5 | 2.99E-02 | 0.3 | 2.21E-01 | 0.4 | 1.19E-01 | -0.1 | 7.63E-01 | GDSL esterase |
| AT3G06980 | 1.2 | 4.59E-07 | 1.3 | 8.38E-07 | -0.1 | 5.46E-01 | 0.3 | 2.03E-01 | -0.1 | 5.96E-01 | 0.4 | 1.21E-01 | DEAD-box ATP-dependent RNA helicase 50 |
| AT1G08380 | 1.2 | 8.24E-07 | 1.2 | 2.16E-06 | 0.0 | 9.88E-01 | -0.3 | 8.51E-02 | -0.2 | 2.41E-01 | -0.1 | 8.48E-01 | Photosystem I subunit O |
| AT1G68260 | 1.2 | 1.48E-07 | 0.7 | 1.32E-05 | 0.5 | 3.49E-04 | -0.1 | 3.13E-01 | -0.1 | 3.86E-01 | 0.0 | 9.98E-01 | Acyl-acyl carrier protein thioesterase ATL3, chloroplastic |
| AT5G61410 | 1.2 | 9.61E-08 | 0.9 | 1.28E-06 | 0.3 | 5.49E-03 | -0.5 | 4.23E-04 | 0.1 | 3.83E-01 | -0.7 | 9.33E-04 | Ribulose-5-phosphate-3-epimerase, chloroplastic |
| AT5G14650 | 1.2 | 1.12E-03 | 0.9 | 1.12E-02 | 0.3 | 3.03E-01 | -1.5 | 2.47E-03 | 0.8 | 2.69E-02 | -2.3 | 1.29E-03 | Pectin lyase-like superfamily protein |
| AT2G40340 | 1.2 | 6.30E-05 | 0.7 | 8.51E-03 | 0.6 | 2.04E-02 | 1.9 | 1.80E-03 | 1.0 | 1.61E-01 | 1.0 | 4.27E-02 | Integrase-type DNA-binding superfamily protein |
| AT5G46580 | 1.3 | 3.10E-07 | 1.0 | 3.24E-06 | 0.3 | 2.66E-02 | 0.7 | 1.94E-03 | 0.3 | 1.96E-01 | 0.4 | 1.48E-01 | Pentatricopeptide repeat-containing protein At5g46580, chloroplastic |
| AT5G11480 | 1.3 | 4.25E-08 | 1.1 | 2.77E-07 | 0.2 | 2.37E-02 | 0.5 | 3.26E-04 | 0.1 | 2.64E-01 | 0.3 | 4.06E-02 | AT5g11480 |
| AT2G05070 | 1.3 | 1.76E-06 | 1.2 | 5.51E-06 | 0.0 | 7.66E-01 | -1.0 | 1.21E-03 | -0.7 | 1.70E-02 | -0.3 | 4.55E-01 | Chlorophyll a-b binding protein 2.2, chloroplastic |
| AT4G15360 | 1.3 | 4.13E-05 | 0.8 | 1.57E-03 | 0.4 | 5.33E-02 | 0.8 | 4.85E-02 | 0.5 | 3.06E-01 | 0.3 | 5.69E-01 | Cytochrome P450 like protein |
| AT1G54820 | 1.3 | 1.17E-05 | 1.1 | 4.42E-05 | 0.1 | 5.18E-01 | -0.2 | 2.63E-01 | 0.1 | 7.20E-01 | -0.2 | 2.24E-01 | Protein kinase superfamily protein |
| AT5G12860 | 1.3 | 6.61E-07 | 0.7 | 7.14E-05 | 0.5 | 1.04E-03 | 0.3 | 5.59E-02 | 0.0 | 9.65E-01 | 0.3 | 1.28E-01 | Dicarboxylate transporter 1 |
| AT1G15390 | 1.3 | 1.36E-06 | 0.8 | 6.88E-05 | 0.4 | 5.55E-03 | 0.3 | 1.20E-01 | 0.0 | 9.71E-01 | 0.3 | 2.65E-01 | Peptide deformylase 1A, chloroplastic |
| AT4G31780 | 1.3 | 2.57E-07 | 0.9 | 6.55E-06 | 0.4 | 3.12E-03 | -0.1 | 6.84E-01 | -0.1 | 4.89E-01 | 0.1 | 8.24E-01 | monogalactosyldiacylglycerol synthase |
| AT5G02830 | 1.3 | 3.19E-06 | 0.6 | 7.53E-04 | 0.6 | 1.62E-03 | 0.5 | 2.74E-02 | 0.5 | 4.20E-02 | 0.0 | 9.98E-01 | Pentatricopeptide repeat-containing protein At5g02830, chloroplastic |
| AT3G20110 | 1.3 | 1.27E-04 | 1.6 | 3.21E-05 | -0.3 | 1.25E-01 | 0.6 | 8.90E-02 | 0.9 | 1.22E-02 | -0.3 | 3.48E-01 | Cytochrome P450 705A20 |
| AT4G17810 | 1.3 | 8.82E-07 | 1.0 | 1.30E-05 | 0.3 | 3.17E-02 | -0.3 | 1.04E-01 | -0.5 | 8.44E-03 | 0.2 | 2.63E-01 | At4g17810 |
| AT3G47470 | 1.3 | 1.30E-06 | 1.2 | 4.35E-06 | 0.1 | 6.83E-01 | -0.5 | 2.32E-03 | -0.8 | 1.84E-04 | 0.3 | 1.55E-01 | Chlorophyll a-b binding protein, chloroplastic |

| | | | | | | | | | | | | | |
|---|---|---|---|---|---|---|---|---|---|---|---|---|---|
| AT3G62410 | 1.3 | 5.86E-07 | 1.3 | 1.41E-06 | 0.0 | 8.70E-01 | 0.1 | 3.50E-01 | 0.0 | 8.67E-01 | 0.1 | 6.28E-01 | Calvin cycle protein CP12-2, chloroplastic |
| AT1G50240 | 1.3 | 2.03E-06 | 0.7 | 2.61E-04 | 0.5 | 2.64E-03 | -0.3 | 2.85E-01 | -0.2 | 6.12E-01 | -0.1 | 7.84E-01 | Serine |
| AT3G10690 | 1.3 | 2.62E-07 | 0.8 | 1.69E-05 | 0.5 | 7.32E-04 | 0.9 | 4.44E-05 | 0.3 | 6.28E-02 | 0.5 | 2.71E-02 | DNA gyrase subunit A, chloroplastic |
| AT3G14620 | 1.3 | 4.33E-05 | 0.1 | 4.55E-01 | 1.1 | 5.10E-04 | 0.7 | 3.76E-02 | -0.1 | 7.71E-01 | 0.8 | 5.25E-02 | AT3g14620 |
| AT5G18570 | 1.3 | 6.02E-07 | 1.1 | 4.19E-06 | 0.2 | 1.13E-01 | 0.5 | 6.56E-03 | 0.2 | 3.35E-01 | 0.3 | 1.93E-01 | GTP-binding protein OBGC, chloroplastic |
| AT2G40020 | 1.3 | 2.44E-06 | 1.0 | 2.56E-05 | 0.2 | 9.83E-02 | 0.1 | 6.87E-01 | -0.1 | 5.40E-01 | 0.2 | 3.27E-01 | Nucleolar histone methyltransferase-related protein |
| AT2G30170 | 1.3 | 3.06E-05 | 0.7 | 4.60E-03 | 0.6 | 1.05E-02 | -0.2 | 1.34E-01 | -0.1 | 6.44E-01 | -0.1 | 4.89E-01 | Protein phosphatase |
| AT5G17990 | 1.3 | 1.33E-06 | 0.8 | 1.08E-04 | 0.5 | 2.41E-03 | 0.2 | 2.52E-01 | 0.0 | 8.60E-01 | 0.2 | 5.30E-01 | Anthranilate phosphoribosyltransferase, chloroplastic |
| AT5G47560 | 1.3 | 2.02E-06 | 0.6 | 9.19E-04 | 0.7 | 6.24E-04 | -0.7 | 1.51E-02 | -1.1 | 1.57E-03 | 0.4 | 2.02E-01 | Tonoplast dicarboxylate transporter |
| AT1G74370 | 1.3 | 1.90E-06 | 0.9 | 8.45E-05 | 0.4 | 1.06E-02 | -0.1 | 8.08E-01 | -0.3 | 3.65E-01 | 0.2 | 5.42E-01 | hypothetical protein |
| AT5G49910 | 1.3 | 1.53E-07 | 1.0 | 2.22E-06 | 0.3 | 7.20E-03 | 1.2 | 2.47E-06 | 0.6 | 2.40E-03 | 0.5 | 2.50E-02 | Heat shock 70 kDa protein 7, chloroplastic |
| AT1G23870 | 1.3 | 1.07E-05 | 1.3 | 1.65E-05 | 0.0 | 8.06E-01 | -0.6 | 2.75E-02 | -1.0 | 2.84E-03 | 0.4 | 2.97E-01 | Probable alpha,alpha-trehalose-phosphate synthase 9 |
| AT4G08780 | 1.3 | 3.16E-04 | 0.8 | 8.83E-03 | 0.5 | 1.10E-01 | 2.2 | 4.97E-07 | 1.1 | 2.09E-03 | 1.1 | 2.85E-04 | Peroxidase 38 |
| AT3G17790 | 1.3 | 2.39E-05 | 0.7 | 4.39E-03 | 0.6 | 1.04E-02 | 1.4 | 4.45E-06 | 0.8 | 1.99E-03 | 0.5 | 2.12E-02 | Purple acid phosphatase 17 |
| AT2G47400 | 1.3 | 2.91E-07 | 1.1 | 1.54E-06 | 0.1 | 2.11E-01 | -0.1 | 6.09E-01 | -0.2 | 4.09E-01 | 0.1 | 7.99E-01 | Calvin cycle protein CP12-1, chloroplastic |
| AT5G44580 | 1.3 | 5.34E-05 | 0.5 | 2.90E-02 | 0.8 | 3.73E-03 | 0.6 | 7.82E-03 | 0.3 | 1.84E-01 | 0.3 | 4.11E-01 | Serine rich endogenous peptide 10 |
| AT4G03280 | 1.3 | 3.23E-07 | 1.2 | 1.59E-06 | 0.1 | 2.73E-01 | -0.1 | 3.23E-01 | 0.0 | 8.08E-01 | -0.2 | 3.36E-01 | Cytochrome b6-f complex iron-sulfur subunit, chloroplastic |
| AT4G02330 | 1.3 | 8.47E-05 | 0.4 | 6.85E-02 | 0.9 | 4.02E-03 | 0.2 | 2.58E-01 | 0.1 | 7.83E-01 | 0.2 | 5.48E-01 | Probable pectinesterase |
| AT1G03650 | 1.3 | 2.21E-05 | 0.6 | 1.12E-02 | 0.7 | 4.71E-03 | 0.4 | 1.26E-01 | -0.4 | 2.36E-01 | 0.9 | 2.46E-02 | N-acetyltransferase domain-containing protein |
| AT3G01440 | 1.3 | 1.26E-05 | 1.4 | 1.51E-05 | -0.1 | 5.92E-01 | 0.7 | 8.98E-04 | 0.8 | 1.32E-03 | -0.1 | 8.59E-01 | Photosynthetic NDH subunit of lumenal location 3, chloroplastic |
| AT3G48360 | 1.3 | 1.17E-05 | 0.9 | 5.58E-04 | 0.4 | 2.72E-02 | -1.2 | 1.62E-02 | -2.1 | 7.31E-04 | 0.9 | 2.69E-02 | BTB and TAZ domain protein 2 |
| AT3G17680 | 1.3 | 7.56E-07 | 1.3 | 1.95E-06 | 0.0 | 9.81E-01 | -0.2 | 4.15E-01 | 0.0 | 8.94E-01 | -0.2 | 4.36E-01 | Kinase interacting family protein |
| AT4G19710 | 1.3 | 1.35E-07 | 0.9 | 3.77E-06 | 0.4 | 1.25E-03 | 0.5 | 5.54E-03 | 0.3 | 8.90E-02 | 0.2 | 4.79E-01 | Bifunctional aspartokinase |
| AT5G66530 | 1.3 | 2.09E-07 | 1.3 | 5.25E-07 | 0.0 | 9.50E-01 | 0.2 | 6.81E-02 | 0.2 | 2.05E-01 | 0.0 | 8.39E-01 | glucose-6-phosphate 1-epimerase |
| AT3G47380 | 1.3 | 1.41E-04 | 1.3 | 2.15E-04 | 0.0 | 9.33E-01 | 0.0 | 8.63E-01 | 0.0 | 9.16E-01 | -0.1 | 8.27E-01 | Pectinesterase inhibitor 11 |
| AT1G80840 | 1.3 | 2.35E-03 | -0.2 | 5.76E-01 | 1.5 | 5.79E-03 | -0.2 | 6.23E-01 | -1.0 | 9.35E-03 | 0.8 | 4.44E-02 | Probable WRKY transcription factor 40 |
| AT2G38620 | 1.3 | 5.00E-06 | 0.7 | 7.76E-04 | 0.6 | 4.59E-03 | 0.4 | 1.07E-01 | 0.5 | 9.63E-02 | -0.1 | 9.02E-01 | Cyclin-dependent kinase B12 |
| AT2G04700 | 1.3 | 2.33E-07 | 0.9 | 5.97E-06 | 0.4 | 2.92E-03 | 0.1 | 6.72E-01 | 0.1 | 6.90E-01 | 0.0 | 9.74E-01 | Ferredoxin-thioredoxin reductase catalytic chain, chloroplastic |
| AT4G26230 | 1.3 | 1.30E-05 | 0.8 | 5.41E-04 | 0.5 | 2.12E-02 | 0.5 | 3.43E-03 | 0.3 | 1.91E-01 | 0.3 | 2.58E-01 | Large ribosomal subunit protein eL31y |
| AT1G77540 | 1.3 | 1.39E-05 | 0.7 | 3.06E-03 | 0.6 | 5.39E-03 | 0.0 | 9.56E-01 | 0.1 | 6.94E-01 | -0.1 | 7.88E-01 | Acetyltransferase At1g77540 |
| AT2G04030 | 1.3 | 1.45E-06 | 0.9 | 3.38E-05 | 0.4 | 1.55E-02 | 1.3 | 1.60E-08 | 0.4 | 2.71E-03 | 0.9 | 1.12E-04 | Heat shock protein 90-5, chloroplastic |
| AT4G32590 | 1.3 | 1.39E-05 | 0.8 | 5.23E-04 | 0.5 | 2.50E-02 | -0.2 | 3.29E-01 | -0.1 | 5.76E-01 | -0.1 | 8.40E-01 | 2Fe-2S ferredoxin-like superfamily protein |
| AT1G12790 | 1.3 | 8.87E-05 | 0.9 | 2.24E-03 | 0.4 | 1.07E-01 | 0.1 | 7.99E-01 | -0.2 | 6.50E-01 | 0.3 | 3.81E-01 | Protein PARTING DANCERS |
| AT1G32200 | 1.3 | 4.00E-07 | 1.2 | 1.92E-06 | 0.1 | 2.78E-01 | -0.1 | 7.24E-01 | 0.1 | 5.83E-01 | -0.2 | 3.01E-01 | AT1G32200 protein |
| AT1G59540 | 1.3 | 6.57E-07 | 0.7 | 9.32E-05 | 0.6 | 1.04E-03 | 0.3 | 2.07E-01 | 0.3 | 2.99E-01 | 0.0 | 9.70E-01 | Kinesin-like protein |
| AT1G07010 | 1.3 | 3.81E-07 | 0.4 | 1.10E-03 | 0.9 | 5.85E-05 | 0.6 | 6.79E-03 | 0.4 | 5.43E-02 | 0.1 | 5.84E-01 | Calcineurin-like metallo-phosphoesterase superfamily protein |
| AT5G04150 | 1.3 | 6.11E-04 | 0.5 | 1.54E-01 | 0.8 | 2.39E-02 | 3.6 | 1.90E-06 | 0.0 | 9.87E-01 | 3.6 | 3.85E-05 | Basic helix-loop-helix DNA-binding superfamily protein |
| AT3G10520 | 1.3 | 1.81E-05 | 1.0 | 1.75E-04 | 0.3 | 1.58E-01 | 0.2 | 1.51E-01 | 0.0 | 9.87E-01 | 0.2 | 3.52E-01 | Non-symbiotic hemoglobin 2 |
| AT1G32060 | 1.3 | 1.01E-06 | 1.2 | 4.38E-06 | 0.1 | 4.22E-01 | 0.2 | 1.58E-01 | 0.1 | 4.36E-01 | 0.1 | 8.02E-01 | Phosphoribulokinase, chloroplastic |
| AT1G63570 | 1.3 | 7.69E-06 | 1.5 | 5.16E-06 | -0.2 | 1.65E-01 | -1.0 | 4.87E-04 | -1.6 | 5.41E-05 | 0.6 | 1.52E-01 | Receptor-like protein kinase-related family protein |
| AT4G37610 | 1.3 | 3.34E-04 | 1.8 | 5.35E-05 | -0.5 | 8.57E-02 | -0.2 | 2.98E-01 | -0.9 | 1.53E-03 | 0.7 | 3.52E-02 | BTB |
| AT5G51010 | 1.3 | 6.49E-08 | 0.9 | 1.74E-06 | 0.4 | 5.40E-04 | -0.7 | 2.02E-03 | 0.2 | 3.02E-01 | -0.9 | 1.75E-03 | Rubredoxin-like domain-containing protein |
| AT4G04040 | 1.3 | 5.59E-07 | 1.0 | 7.01E-06 | 0.3 | 2.72E-02 | 0.2 | 2.04E-01 | -0.1 | 4.12E-01 | 0.3 | 7.65E-02 | Pyrophosphate--fructose 6-phosphate 1-phosphotransferase subunit beta 2 |
| AT5G14660 | 1.3 | 1.01E-07 | 1.2 | 5.65E-07 | 0.2 | 7.93E-02 | 0.3 | 3.11E-02 | 0.0 | 8.69E-01 | 0.3 | 1.32E-01 | Peptide deformylase 1B, chloroplastic |
| AT3G18390 | 1.3 | 5.57E-07 | 0.9 | 1.48E-05 | 0.4 | 5.81E-03 | 0.3 | 1.94E-01 | 0.2 | 4.20E-01 | 0.1 | 8.37E-01 | CRS1 |
| AT3G18680 | 1.3 | 2.95E-07 | 1.1 | 2.86E-06 | 0.3 | 3.08E-02 | 0.6 | 4.07E-03 | 0.1 | 7.34E-01 | 0.5 | 2.74E-02 | UMP kinase |
| AT1G13820 | 1.3 | 1.15E-07 | 1.0 | 2.33E-06 | 0.4 | 2.21E-03 | 0.1 | 4.96E-01 | 0.2 | 3.37E-01 | -0.1 | 8.18E-01 | hypothetical protein |
| AT3G25660 | 1.3 | 1.06E-07 | 1.0 | 1.41E-06 | 0.3 | 5.56E-03 | 0.1 | 6.86E-01 | 0.0 | 8.72E-01 | 0.0 | 9.18E-01 | Glutamyl-tRNA amidotransferase subunit A, chloroplastic |
| AT1G04640 | 1.3 | 3.00E-07 | 0.7 | 7.03E-05 | 0.6 | 2.71E-04 | -0.1 | 5.42E-01 | -0.4 | 2.63E-02 | 0.3 | 1.67E-01 | Octanoyltransferase LIP2, mitochondrial |
| AT2G43820 | 1.3 | 3.68E-06 | 0.8 | 4.35E-04 | 0.6 | 3.60E-03 | 0.4 | 8.47E-03 | -0.2 | 1.77E-01 | 0.6 | 6.10E-03 | UDP-glycosyltransferase 74F2 |
| AT2G32230 | 1.3 | 9.92E-08 | 1.0 | 1.78E-06 | 0.4 | 2.41E-03 | 0.2 | 3.75E-01 | 0.2 | 3.37E-01 | 0.0 | 8.89E-01 | Proteinaceous RNase P 1, chloroplastic |
| AT2G25450 | 1.3 | 4.74E-07 | 1.1 | 5.78E-06 | 0.3 | 3.06E-02 | 0.1 | 6.54E-01 | -0.1 | 8.17E-01 | 0.2 | 5.63E-01 | Probable 2-oxoacid dependent dioxygenase |
| AT1G66970 | 1.3 | 1.91E-04 | 0.9 | 2.63E-03 | 0.4 | 1.67E-01 | 0.6 | 2.55E-03 | 0.2 | 3.85E-01 | 0.4 | 9.86E-02 | glycerophosphodiester phosphodiesterase |
| AT5G66570 | 1.3 | 3.28E-07 | 1.1 | 2.33E-06 | 0.2 | 9.68E-02 | -0.2 | 1.51E-01 | -0.2 | 3.11E-01 | 0.0 | 9.02E-01 | Oxygen-evolving enhancer protein 1-1, chloroplastic |
| AT3G56400 | 1.3 | 9.67E-04 | 0.3 | 2.87E-01 | 1.0 | 1.24E-02 | 1.8 | 4.96E-06 | 1.0 | 2.79E-03 | 0.8 | 3.29E-02 | Probable WRKY transcription factor 70 |
| AT4G33220 | 1.3 | 4.10E-08 | 1.1 | 2.97E-07 | 0.2 | 1.41E-02 | 0.1 | 6.80E-01 | -0.2 | 1.53E-01 | 0.3 | 7.10E-02 | Pectinesterase |
| AT1G64680 | 1.3 | 1.37E-07 | 0.9 | 4.01E-06 | 0.4 | 1.18E-03 | 0.3 | 4.36E-02 | 0.1 | 4.17E-01 | 0.1 | 4.59E-01 | Beta-carotene isomerase D27 |
| AT1G73530 | 1.3 | 6.57E-07 | 1.1 | 8.49E-06 | 0.3 | 3.11E-02 | 0.2 | 1.60E-01 | 0.0 | 9.36E-01 | 0.2 | 3.60E-01 | Organelle RRM domain-containing protein 6, chloroplastic |
| AT2G42170 | 1.3 | 8.34E-04 | 1.1 | 5.60E-03 | 0.3 | 4.10E-01 | -1.1 | 9.72E-04 | -0.7 | 2.64E-02 | -0.5 | 3.21E-01 | Actin family protein |
| AT4G11080 | 1.3 | 5.43E-07 | 0.9 | 3.06E-05 | 0.5 | 2.42E-03 | 0.1 | 5.14E-01 | 0.1 | 6.41E-01 | 0.0 | 9.73E-01 | High mobility group B protein 13 |
| AT2G21800 | 1.3 | 8.68E-06 | 0.7 | 3.11E-03 | 0.7 | 3.95E-03 | 0.5 | 2.02E-01 | 0.4 | 3.65E-01 | 0.1 | 8.64E-01 | Essential meiotic endonuclease 1A |
| AT3G59040 | 1.4 | 7.70E-07 | 1.3 | 2.78E-06 | 0.1 | 5.61E-01 | 0.2 | 2.27E-01 | -0.2 | 4.22E-01 | 0.4 | 5.70E-02 | Tetratricopeptide repeat -like superfamily protein |
| AT1G66540 | 1.4 | 5.46E-06 | 0.7 | 1.88E-03 | 0.7 | 2.19E-03 | 0.0 | 9.36E-01 | 0.3 | 5.30E-01 | -0.2 | 6.35E-01 | Cytochrome P450 superfamily protein |
| AT1G02150 | 1.4 | 9.06E-08 | 1.2 | 5.52E-07 | 0.2 | 6.12E-02 | 0.2 | 4.80E-02 | -0.1 | 7.68E-01 | 0.3 | 6.90E-02 | Pentatricopeptide repeat-containing protein At1g02150 |
| AT5G23310 | 1.4 | 3.69E-07 | 1.1 | 3.24E-06 | 0.2 | 5.65E-02 | 1.0 | 8.44E-05 | 0.4 | 7.18E-02 | 0.6 | 1.87E-02 | Superoxide dismutase 3, chloroplastic |
| AT1G49010 | 1.4 | 6.06E-07 | 0.9 | 3.54E-05 | 0.5 | 2.44E-03 | -1.2 | 2.84E-04 | 0.2 | 4.74E-01 | -1.4 | 6.65E-04 | At1g49010 |
| AT5G48220 | 1.4 | 7.29E-08 | 1.2 | 4.61E-07 | 0.2 | 4.34E-02 | 0.0 | 8.53E-01 | -0.2 | 3.71E-01 | 0.2 | 3.07E-01 | indole-3-glycerol-phosphate synthase |
| AT2G38540 | 1.4 | 1.52E-06 | 1.2 | 1.13E-05 | 0.2 | 1.59E-01 | 0.1 | 4.50E-01 | 0.1 | 8.24E-01 | 0.1 | 7.88E-01 | Non-specific lipid-transfer protein 1 |
| AT4G20130 | 1.4 | 8.42E-07 | 1.1 | 1.06E-05 | 0.3 | 3.93E-02 | 0.3 | 6.67E-02 | 0.0 | 9.82E-01 | 0.2 | 1.48E-01 | Plastid transcriptionally active 14 |
| AT4G39620 | 1.4 | 2.31E-07 | 0.9 | 1.44E-05 | 0.5 | 8.94E-04 | 0.4 | 4.42E-02 | 0.1 | 6.71E-01 | 0.3 | 1.95E-01 | Tetratricopeptide repeat -like superfamily protein |
| AT2G32960 | 1.4 | 1.20E-04 | 0.8 | 6.31E-03 | 0.5 | 5.96E-02 | 1.2 | 2.33E-04 | 0.6 | 7.80E-02 | 0.7 | 3.47E-02 | Tyrosine-protein phosphatase DSP2 |
| AT4G27520 | 1.4 | 4.43E-07 | 1.1 | 5.83E-06 | 0.3 | 2.36E-02 | 0.1 | 7.01E-01 | 0.0 | 8.87E-01 | 0.0 | 9.18E-01 | Early nodulin-like protein 2 |
| AT1G02280 | 1.4 | 5.72E-07 | 1.3 | 2.22E-06 | 0.1 | 4.47E-01 | 0.1 | 4.10E-01 | -0.2 | 2.60E-01 | 0.3 | 9.81E-02 | Translocase of chloroplast 33, chloroplastic |
| AT5G07810 | 1.4 | 2.82E-06 | 0.8 | 4.57E-04 | 0.6 | 2.82E-03 | 0.0 | 9.51E-01 | 0.1 | 7.11E-01 | -0.2 | 7.31E-01 | SNF2 domain-containing protein |
| AT2G35800 | 1.4 | 1.24E-07 | 1.0 | 3.07E-06 | 0.4 | 1.48E-03 | 0.3 | 2.69E-02 | -0.1 | 4.77E-01 | 0.4 | 1.87E-02 | Mitochondrial substrate carrier family protein |
| AT1G16260 | 1.4 | 8.25E-06 | 0.7 | 2.17E-03 | 0.7 | 3.43E-03 | 0.6 | 3.16E-02 | 0.2 | 6.57E-01 | 0.4 | 1.92E-01 | Wall-associated receptor kinase-like |
| AT2G25900 | 1.4 | 2.88E-05 | 1.3 | 8.13E-05 | 0.1 | 7.55E-01 | -1.0 | 2.53E-04 | -1.4 | 4.33E-05 | 0.4 | 1.87E-01 | Zinc finger CCCH domain-containing protein 23 |
| AT4G38160 | 1.4 | 2.98E-07 | 1.3 | 1.20E-06 | 0.1 | 3.72E-01 | 0.1 | 6.34E-01 | -0.2 | 2.81E-01 | 0.2 | 1.62E-01 | Mitochondrial transcription termination factor family protein |
| AT5G28770 | 1.4 | 3.88E-06 | 1.5 | 4.77E-06 | -0.1 | 4.66E-01 | -0.1 | 6.25E-01 | -0.5 | 1.87E-02 | 0.4 | 1.01E-01 | BZIP transcription factor family protein |
| AT5G03940 | 1.4 | 1.41E-07 | 1.0 | 2.33E-06 | 0.4 | 3.94E-03 | -0.1 | 8.08E-01 | -0.1 | 7.95E-01 | 0.0 | 9.62E-01 | Signal recognition particle subunit SRP54, chloroplastic |
| AT2G31730 | 1.4 | 2.07E-05 | 1.0 | 4.08E-04 | 0.4 | 8.34E-02 | -0.4 | 9.99E-02 | -0.6 | 5.63E-02 | 0.1 | 8.01E-01 | Transcription factor bHLH154 |
| AT1G55370 | 1.4 | 1.16E-06 | 1.1 | 1.71E-05 | 0.3 | 3.37E-02 | 0.1 | 7.30E-01 | -0.1 | 7.14E-01 | 0.2 | 4.89E-01 | Protein NDH-DEPENDENT CYCLIC ELECTRON FLOW 5 |
| AT1G78860 | 1.4 | 3.20E-06 | 1.2 | 2.01E-05 | 0.2 | 2.24E-01 | 1.2 | 5.39E-06 | 0.8 | 1.40E-03 | 0.5 | 2.57E-02 | EP1-like glycoprotein 4 lectin family protein) |

| | | | | | | | | | | | | | |
|---|---|---|---|---|---|---|---|---|---|---|---|---|---|
| AT1G69730 | 1.4 | 1.24E-06 | 0.5 | 2.46E-03 | 0.9 | 1.83E-04 | 1.0 | 7.67E-06 | 0.5 | 7.60E-03 | 0.5 | 2.38E-02 | Wall-associated receptor kinase-like 9 |
| AT3G56970 | 1.4 | 6.15E-04 | -0.1 | 7.55E-01 | 1.5 | 2.89E-03 | 3.1 | 8.37E-07 | -0.5 | 4.74E-01 | 3.6 | 2.94E-05 | Transcription factor ORG2 |
| AT4G11900 | 1.4 | 5.17E-06 | 0.6 | 3.64E-03 | 0.8 | 1.24E-03 | 0.8 | 3.64E-04 | 0.4 | 4.75E-02 | 0.4 | 1.19E-01 | Receptor-like serine |
| AT5G46800 | 1.4 | 7.94E-07 | 0.7 | 1.46E-04 | 0.7 | 6.67E-04 | 0.1 | 3.32E-01 | 0.0 | 7.76E-01 | 0.1 | 7.26E-01 | Mitochondrial carnitine |
| AT5G33300 | 1.4 | 1.15E-06 | 0.9 | 6.82E-05 | 0.5 | 4.25E-03 | 0.2 | 3.62E-01 | 0.2 | 2.09E-01 | -0.1 | 7.49E-01 | Tesmin |
| AT2G02740 | 1.4 | 7.30E-07 | 0.9 | 4.66E-05 | 0.5 | 2.70E-03 | 0.8 | 2.72E-03 | 0.1 | 8.42E-01 | 0.8 | 1.78E-02 | Single-stranded DNA-binding protein WHY3, chloroplastic |
| AT2G39670 | 1.4 | 9.15E-08 | 0.7 | 1.36E-05 | 0.7 | 9.50E-05 | 0.5 | 2.60E-03 | 0.0 | 9.98E-01 | 0.5 | 1.99E-02 | Radical SAM superfamily protein |
| AT2G36000 | 1.4 | 4.03E-07 | 1.2 | 2.50E-06 | 0.2 | 1.33E-01 | -0.1 | 6.03E-01 | -0.5 | 2.53E-03 | 0.4 | 1.81E-02 | At2g36000 |
| AT4G21070 | 1.4 | 2.06E-06 | 0.7 | 7.94E-04 | 0.7 | 1.25E-03 | 0.4 | 2.73E-01 | 0.0 | 9.71E-01 | 0.4 | 3.23E-01 | Protein BREAST CANCER SUSCEPTIBILITY 1 homolog |
| AT2G21860 | 1.4 | 1.47E-07 | 1.0 | 3.39E-06 | 0.4 | 2.06E-03 | 0.6 | 5.45E-03 | 0.3 | 1.62E-01 | 0.3 | 2.39E-01 | At2g21860 |
| AT4G23300 | 1.4 | 4.16E-06 | 0.7 | 1.41E-03 | 0.7 | 1.68E-03 | 0.0 | 8.87E-01 | 0.0 | 9.70E-01 | 0.1 | 8.78E-01 | Protein kinase domain-containing protein |
| AT4G14540 | 1.4 | 7.09E-07 | 1.1 | 1.26E-05 | 0.3 | 1.78E-02 | 0.1 | 8.61E-01 | -0.1 | 7.93E-01 | 0.1 | 7.33E-01 | Nuclear transcription factor Y subunit B-3 |
| AT1G14150 | 1.4 | 1.54E-05 | 1.0 | 2.48E-04 | 0.4 | 7.44E-02 | 0.1 | 5.01E-01 | 0.3 | 8.91E-02 | -0.2 | 4.00E-01 | Photosynthetic NDH subunit of lumenal location 2, chloroplastic |
| AT1G18265 | 1.4 | 1.73E-05 | 1.1 | 2.08E-04 | 0.3 | 1.37E-01 | 0.3 | 4.58E-01 | 0.4 | 3.66E-01 | -0.1 | 8.46E-01 | Protein FLOURY 1-like |
| AT2G18700 | 1.4 | 1.47E-04 | 1.1 | 9.87E-04 | 0.3 | 3.31E-01 | -1.1 | 4.36E-03 | -1.4 | 1.51E-03 | 0.4 | 4.71E-01 | Probable alpha,alpha-trehalose-phosphate synthase 11 |
| AT3G23700 | 1.4 | 2.43E-07 | 1.1 | 3.56E-06 | 0.3 | 9.29E-03 | 0.0 | 7.78E-01 | -0.1 | 6.81E-01 | 0.1 | 5.47E-01 | AT3g23700 |
| AT1G10522 | 1.4 | 3.78E-06 | 1.3 | 1.27E-05 | 0.1 | 6.42E-01 | 0.8 | 1.71E-03 | 0.2 | 5.89E-01 | 0.6 | 3.63E-02 | Protein PLASTID REDOX INSENSITIVE 2, chloroplastic |
| AT4G04640 | 1.4 | 1.43E-07 | 1.1 | 1.43E-06 | 0.3 | 1.71E-02 | 0.3 | 2.14E-02 | 0.2 | 9.98E-02 | 0.1 | 7.89E-01 | ATP synthase gamma chain 1, chloroplastic |
| AT5G54270 | 1.4 | 8.87E-07 | 1.5 | 1.66E-06 | -0.1 | 5.74E-01 | -0.5 | 2.24E-02 | -0.6 | 1.90E-02 | 0.1 | 8.33E-01 | Chlorophyll a-b binding protein 3, chloroplastic |
| AT3G24430 | 1.4 | 1.16E-07 | 1.0 | 2.26E-06 | 0.4 | 2.44E-03 | 0.2 | 2.63E-01 | 0.2 | 3.09E-01 | 0.0 | 9.73E-01 | Fe-S cluster assembly factor HCF101, chloroplastic |
| AT4G26520 | 1.4 | 3.43E-07 | 1.2 | 2.83E-06 | 0.2 | 5.23E-02 | -0.1 | 6.77E-01 | -0.2 | 1.15E-01 | 0.2 | 3.27E-01 | Fructose-bisphosphate aldolase |
| AT5G21222 | 1.4 | 4.35E-07 | 0.9 | 3.33E-05 | 0.5 | 1.16E-03 | -0.3 | 5.36E-02 | -0.2 | 4.04E-01 | -0.1 | 4.51E-01 | Pentatricopeptide repeat-containing protein At5g21222 |
| AT5G42700 | 1.4 | 3.39E-06 | 0.9 | 2.19E-04 | 0.5 | 9.68E-03 | 0.6 | 8.59E-02 | 0.5 | 1.96E-01 | 0.1 | 8.59E-01 | AP2 |
| AT2G47920 | 1.4 | 1.30E-05 | 1.0 | 4.25E-04 | 0.4 | 4.49E-02 | -0.3 | 4.85E-01 | 0.0 | 9.61E-01 | -0.3 | 6.49E-01 | Protein NETWORKED 3C |
| AT3G07800 | 1.4 | 3.01E-06 | 0.6 | 1.48E-03 | 0.8 | 9.22E-04 | 0.1 | 4.51E-01 | -0.1 | 7.57E-01 | 0.2 | 3.01E-01 | Thymidine kinase a |
| AT1G09750 | 1.4 | 8.42E-07 | 1.5 | 1.70E-06 | -0.1 | 6.51E-01 | -0.8 | 4.22E-04 | -0.6 | 1.22E-02 | -0.2 | 2.80E-01 | Aspartyl protease AED3 |
| AT5G49450 | 1.4 | 9.41E-05 | 0.9 | 4.91E-03 | 0.5 | 4.72E-02 | -0.8 | 2.36E-02 | -1.7 | 8.59E-04 | 0.9 | 1.17E-01 | Basic leucine zipper 1 |
| AT5G35220 | 1.4 | 6.92E-08 | 1.1 | 6.80E-07 | 0.3 | 6.44E-03 | 0.6 | 4.42E-03 | 0.3 | 1.06E-01 | 0.2 | 2.69E-01 | Probable zinc metalloprotease EGY1, chloroplastic |
| AT5G02370 | 1.4 | 2.28E-06 | 0.9 | 1.77E-04 | 0.5 | 4.85E-03 | -0.1 | 8.48E-01 | 0.2 | 6.69E-01 | -0.3 | 5.84E-01 | Kinesin-like protein |
| AT5G22580 | 1.4 | 1.31E-05 | 1.2 | 1.07E-04 | 0.3 | 1.85E-01 | 0.3 | 1.52E-01 | 0.0 | 9.55E-01 | 0.3 | 2.58E-01 | Stress-response A |
| AT1G62750 | 1.4 | 1.29E-07 | 1.0 | 3.86E-06 | 0.5 | 1.22E-03 | 0.5 | 7.79E-03 | 0.3 | 2.05E-01 | 0.3 | 3.33E-01 | Elongation factor G, chloroplastic |
| AT5G09240 | 1.4 | 2.50E-05 | 0.9 | 1.33E-03 | 0.5 | 2.92E-02 | 0.3 | 2.70E-01 | 0.0 | 9.48E-01 | 0.3 | 4.71E-01 | SsDNA-binding transcriptional regulator |
| AT1G48605 | 1.4 | 2.69E-05 | 0.8 | 2.90E-03 | 0.6 | 1.63E-02 | -0.3 | 1.98E-01 | -0.7 | 9.73E-03 | 0.4 | 1.93E-01 | Probable phosphopantothenoylcysteine decarboxylase |
| AT4G16390 | 1.4 | 7.93E-08 | 1.2 | 7.11E-07 | 0.3 | 1.29E-02 | 0.7 | 5.91E-05 | 0.2 | 2.68E-01 | 0.5 | 6.98E-03 | Pentatricopeptide repeat-containing protein At4g16390, chloroplastic |
| AT4G10500 | 1.4 | 6.19E-02 | 0.3 | 6.98E-01 | 1.1 | 2.37E-01 | 1.4 | 1.96E-06 | 1.6 | 2.48E-06 | -0.2 | 4.00E-01 | Protein DMR6-LIKE OXYGENASE 1 -Fe oxygenase-like protein DLO1) |
| AT5G47500 | 1.4 | 2.65E-06 | 1.5 | 4.64E-06 | -0.1 | 7.20E-01 | -0.4 | 1.62E-01 | 0.1 | 8.63E-01 | -0.5 | 1.83E-01 | Pectinesterase |
| AT5G26820 | 1.4 | 2.57E-07 | 1.1 | 4.00E-06 | 0.4 | 8.12E-03 | -0.2 | 2.83E-01 | -0.1 | 5.76E-01 | -0.1 | 7.92E-01 | Solute carrier family 40 member 3, chloroplastic |
| AT5G16400 | 1.4 | 7.25E-07 | 1.3 | 4.33E-06 | 0.2 | 1.97E-01 | -0.1 | 5.53E-01 | -0.1 | 4.89E-01 | 0.0 | 9.15E-01 | Thioredoxin F2, chloroplastic |
| AT5G02760 | 1.4 | 6.71E-04 | 1.6 | 4.29E-04 | -0.2 | 5.80E-01 | 0.0 | 9.79E-01 | -0.2 | 6.72E-01 | 0.1 | 7.41E-01 | Probable protein phosphatase 2C 67 |
| AT2G21530 | 1.4 | 7.44E-08 | 1.0 | 2.23E-06 | 0.5 | 5.74E-04 | 0.3 | 1.26E-02 | 0.4 | 9.57E-03 | -0.1 | 7.96E-01 | At2g21530 |
| AT1G12800 | 1.4 | 1.21E-07 | 1.1 | 1.92E-06 | 0.4 | 4.41E-03 | 0.5 | 4.03E-02 | 0.4 | 1.72E-01 | 0.1 | 7.06E-01 | Nucleic acid-binding, OB-fold-like protein |
| AT1G15520 | 1.4 | 8.45E-03 | -0.2 | 7.40E-01 | 1.7 | 2.43E-02 | -0.8 | 1.76E-01 | -1.2 | 7.14E-02 | 0.4 | 4.57E-01 | Pleiotropic drug resistance 12 |
| AT2G38025 | 1.4 | 1.09E-05 | 0.7 | 3.05E-03 | 0.7 | 3.75E-03 | 0.1 | 6.13E-01 | -0.1 | 4.92E-01 | 0.2 | 3.29E-01 | Ubiquitin thioesterase OTU |
| AT4G23820 | 1.4 | 4.72E-06 | 1.5 | 7.01E-06 | -0.1 | 6.37E-01 | -0.6 | 3.53E-04 | -0.7 | 4.40E-04 | 0.1 | 7.67E-01 | Pectin lyase-like superfamily protein |
| AT3G47450 | 1.5 | 1.84E-07 | 1.0 | 5.58E-06 | 0.4 | 2.04E-03 | 0.5 | 6.50E-02 | 0.1 | 8.19E-01 | 0.4 | 1.86E-01 | NO-associated protein 1, chloroplastic |
| AT4G08040 | 1.5 | 3.17E-03 | 0.6 | 2.33E-01 | 0.9 | 6.81E-02 | -0.4 | 3.50E-01 | -0.4 | 3.66E-01 | 0.0 | 9.70E-01 | 1-aminocyclopropane-1-carboxylate synthase 11 |
| AT5G08030 | 1.5 | 4.86E-04 | 1.6 | 4.80E-04 | -0.1 | 7.15E-01 | | | | | | | glycerophosphodiester phosphodiesterase |
| AT1G33790 | 1.5 | 2.38E-06 | 0.9 | 1.69E-04 | 0.5 | 5.75E-03 | 0.4 | 7.99E-02 | -0.1 | 8.92E-01 | 0.5 | 1.07E-01 | Jacalin lectin family protein |
| AT5G01530 | 1.5 | 3.86E-07 | 1.4 | 1.18E-06 | 0.0 | 8.09E-01 | -0.5 | 5.39E-04 | -0.3 | 7.44E-02 | -0.3 | 1.39E-01 | Chlorophyll a-b binding protein CP29.1, chloroplastic |
| AT1G67830 | 1.5 | 8.27E-07 | 1.1 | 1.30E-05 | 0.3 | 2.72E-02 | -0.2 | 2.89E-01 | -0.1 | 8.32E-01 | -0.2 | 5.18E-01 | Alpha-L-fucosidase 3 |
| AT3G25190 | 1.5 | 2.24E-06 | 1.5 | 3.57E-06 | -0.1 | 6.24E-01 | -1.4 | 1.79E-04 | -0.7 | 1.87E-02 | -0.7 | 4.14E-02 | Vacuolar iron transporter homolog 2.1 |
| AT4G14150 | 1.5 | 1.15E-06 | 0.6 | 2.60E-03 | 0.9 | 2.30E-04 | 0.1 | 7.63E-01 | 0.2 | 5.61E-01 | -0.1 | 8.27E-01 | Kinesin-like protein KIN-12A |
| AT3G63480 | 1.5 | 4.89E-06 | 0.8 | 1.02E-03 | 0.7 | 4.12E-03 | 0.4 | 4.22E-01 | 0.2 | 6.77E-01 | 0.1 | 8.53E-01 | Kinesin-like protein KIN-1 |
| AT5G64580 | 1.5 | 2.67E-07 | 1.1 | 5.54E-06 | 0.4 | 4.75E-03 | 0.6 | 1.17E-03 | 0.5 | 1.85E-02 | 0.2 | 5.09E-01 | EMB3144 |
| AT4G19530 | 1.5 | 5.64E-06 | 1.3 | 3.04E-05 | 0.2 | 2.95E-01 | -0.2 | 4.53E-01 | -0.4 | 2.01E-01 | 0.2 | 6.66E-01 | Probable disease resistance protein At4g19530 |
| AT1G55200 | 1.5 | 2.14E-05 | 1.4 | 8.11E-05 | 0.1 | 6.04E-01 | 0.2 | 6.14E-01 | 0.2 | 7.42E-01 | 0.0 | 9.55E-01 | Kinase with adenine nucleotide alpha hydrolases-like domain-containing protein |
| AT1G74970 | 1.5 | 2.92E-07 | 1.5 | 7.02E-07 | 0.0 | 8.48E-01 | 0.1 | 7.68E-01 | -0.1 | 7.38E-01 | 0.1 | 5.95E-01 | Small ribosomal subunit protein uS9c |
| AT2G41250 | 1.5 | 5.47E-06 | 1.0 | 2.08E-04 | 0.5 | 1.77E-02 | -0.3 | 1.42E-01 | -0.7 | 5.07E-03 | 0.4 | 1.88E-01 | Haloacid dehalogenase-like hydrolase superfamily protein |
| AT1G76100 | 1.5 | 3.65E-07 | 1.3 | 1.90E-06 | 0.1 | 2.52E-01 | -0.3 | 4.73E-02 | -0.4 | 3.12E-02 | 0.1 | 8.05E-01 | Plastocyanin |
| AT1G68890 | 1.5 | 2.37E-07 | 1.1 | 4.21E-06 | 0.4 | 6.17E-03 | 0.0 | 9.12E-01 | 0.1 | 7.30E-01 | -0.2 | 7.11E-01 | 2-oxoglutarate decarboxylase |
| AT5G17170 | 1.5 | 2.83E-06 | 1.1 | 4.42E-05 | 0.4 | 4.18E-02 | -1.9 | 1.89E-04 | 0.4 | 3.88E-01 | -2.3 | 2.32E-04 | Rubredoxin family protein |
| AT4G21840 | 1.5 | 5.23E-03 | 0.1 | 8.10E-01 | 1.3 | 3.63E-02 | 0.4 | 3.31E-01 | -0.2 | 6.55E-01 | | | Peptide methionine sulfoxide reductase B8 -S-oxide reductase) |
| AT1G01370 | 1.5 | 2.89E-07 | 1.0 | 1.51E-05 | 0.5 | 1.54E-03 | 0.4 | 2.29E-01 | 0.1 | 7.87E-01 | 0.2 | 5.61E-01 | HTR12 |
| AT2G24680 | 1.5 | 4.72E-05 | 0.6 | 2.35E-02 | 0.8 | 7.59E-03 | 0.0 | 9.69E-01 | 0.6 | 9.66E-02 | -0.5 | 1.72E-01 | Transcriptional factor B3 family protein |
| AT1G15180 | 1.5 | 6.46E-07 | 1.1 | 9.95E-06 | 0.3 | 2.21E-02 | -0.2 | 4.96E-01 | -0.4 | 1.65E-01 | 0.2 | 4.98E-01 | MATE efflux family protein |
| AT5G09650 | 1.5 | 2.10E-07 | 1.2 | 1.56E-06 | 0.2 | 6.46E-02 | 0.5 | 2.35E-04 | 0.3 | 5.30E-02 | 0.3 | 1.36E-01 | Soluble inorganic pyrophosphatase 6, chloroplastic |
| AT1G03475 | 1.5 | 1.18E-07 | 1.1 | 1.47E-06 | 0.3 | 8.09E-03 | 0.7 | 7.22E-06 | 0.2 | 9.99E-02 | 0.5 | 3.09E-03 | Coproporphyrinogen-III oxidase 1, chloroplastic |
| AT1G04710 | 1.5 | 6.10E-08 | 0.6 | 6.92E-05 | 0.9 | 2.69E-05 | 0.1 | 5.61E-01 | 0.2 | 5.03E-01 | 0.0 | 9.20E-01 | 3-ketoacyl-CoA thiolase 1, peroxisomal |
| AT2G22360 | 1.5 | 5.70E-08 | 0.9 | 2.52E-06 | 0.6 | 1.99E-04 | 0.6 | 5.67E-04 | 0.2 | 1.61E-01 | 0.3 | 8.43E-02 | Chaperone protein dnaJ A6, chloroplastic |
| AT3G06500 | 1.5 | 4.94E-08 | 1.3 | 3.23E-07 | 0.2 | 2.37E-02 | 0.7 | 4.33E-03 | 0.1 | 8.32E-01 | 0.6 | 5.12E-02 | Alkaline |
| AT3G02380 | 1.5 | 7.38E-05 | 1.0 | 2.06E-03 | 0.5 | 7.43E-02 | -0.8 | 1.68E-01 | -0.2 | 7.72E-01 | -0.6 | 4.95E-01 | Zinc finger protein CONSTANS-LIKE 2 |
| AT1G80850 | 1.5 | 1.89E-06 | 1.1 | 5.15E-05 | 0.4 | 1.73E-02 | 0.3 | 3.25E-01 | 0.3 | 3.69E-01 | 0.0 | 9.83E-01 | hypothetical protein |
| AT3G26200 | 1.5 | 7.08E-04 | 1.4 | 2.04E-03 | 0.1 | 7.05E-01 | 0.5 | 1.01E-01 | -1.0 | 2.77E-02 | 1.6 | 2.53E-03 | Cytochrome P450 71B22 |
| AT5G63140 | 1.5 | 8.22E-07 | 1.3 | 4.71E-06 | 0.2 | 2.31E-01 | 0.3 | 8.85E-02 | -0.4 | 1.17E-01 | 0.7 | 6.93E-03 | Probable inactive purple acid phosphatase 29 |
| AT1G29930 | 1.5 | 3.61E-07 | 1.5 | 8.48E-07 | 0.0 | 8.38E-01 | -0.5 | 3.57E-04 | -0.7 | 1.76E-04 | 0.1 | 4.88E-01 | Chlorophyll a-b binding protein 1, chloroplastic |
| AT4G19700 | 1.5 | 2.16E-06 | 0.7 | 1.55E-03 | 0.8 | 6.94E-04 | 0.0 | 8.78E-01 | -0.4 | 7.04E-02 | 0.4 | 1.12E-01 | E3 ubiquitin-protein ligase BOI -like protein) |
| AT3G54890 | 1.5 | 6.15E-07 | 1.5 | 1.69E-06 | 0.0 | 9.59E-01 | -0.4 | 2.42E-03 | -0.5 | 4.65E-03 | 0.0 | 9.66E-01 | Chlorophyll a-b binding protein, chloroplastic |
| AT4G30620 | 1.5 | 1.54E-06 | 1.3 | 1.34E-05 | 0.2 | 1.11E-01 | 0.3 | 9.58E-02 | 0.1 | 7.22E-01 | 0.2 | 3.89E-01 | Nucleoid-associated protein At4g30620, chloroplastic |
| AT5G35630 | 1.5 | 8.04E-08 | 1.1 | 1.18E-06 | 0.4 | 3.66E-03 | 0.0 | 8.76E-01 | 0.0 | 8.32E-01 | -0.1 | 7.54E-01 | glutamine synthetase |
| AT2G44290 | 1.5 | 4.73E-04 | 0.9 | 1.86E-02 | 0.6 | 7.70E-02 | 0.5 | 1.90E-02 | 0.0 | 9.06E-01 | 0.4 | 6.23E-02 | Non-specific lipid transfer protein GPI-anchored 13 |
| AT3G21300 | 1.5 | 2.10E-06 | 1.3 | 1.67E-05 | 0.2 | 1.43E-01 | 0.7 | 1.01E-03 | 0.0 | 9.87E-01 | 0.7 | 6.16E-03 | RNA methyltransferase family protein |
| AT1G52400 | 1.5 | 1.23E-05 | 1.3 | 8.78E-05 | 0.3 | 2.45E-01 | 0.8 | 2.26E-02 | 0.6 | 1.21E-01 | 0.2 | 7.43E-01 | Beta-D-glucopyranosyl abscisate beta-glucosidase |
| AT3G51820 | 1.5 | 4.94E-08 | 1.0 | 1.24E-06 | 0.5 | 4.13E-04 | 0.4 | 3.29E-03 | 0.1 | 6.30E-01 | 0.3 | 6.65E-02 | Chlorophyll synthase, chloroplastic |

| | | | | | | | | | | | | | |
|---|---|---|---|---|---|---|---|---|---|---|---|---|---|
| AT3G26570 | 1.5 | 1.51E-07 | 1.1 | 2.58E-06 | 0.4 | 4.74E-03 | 0.4 | 1.78E-03 | 0.3 | 2.04E-02 | 0.1 | 6.25E-01 | Inorganic phosphate transporter 2-1, chloroplastic |
| AT1G30820 | 1.5 | 7.19E-08 | 1.7 | 1.09E-07 | -0.2 | 5.19E-02 | 0.1 | 6.65E-01 | -0.3 | 2.60E-01 | 0.4 | 1.02E-01 | CTP synthase |
| AT4G26530 | 1.5 | 3.71E-06 | 1.6 | 6.20E-06 | -0.1 | 6.60E-01 | 0.1 | 7.41E-01 | 0.3 | 1.65E-01 | -0.2 | 3.54E-01 | Fructose-bisphosphate aldolase 5, cytosolic |
| AT5G57890 | 1.5 | 8.87E-06 | 1.2 | 1.25E-04 | 0.3 | 8.77E-02 | 0.3 | 1.66E-01 | 0.1 | 8.10E-01 | 0.2 | 4.49E-01 | Anthranilate synthase beta subunit 2, chloroplastic |
| AT1G14270 | 1.5 | 5.65E-08 | 1.2 | 5.12E-07 | 0.3 | 7.23E-03 | 0.2 | 1.71E-01 | 0.2 | 2.11E-01 | 0.0 | 9.84E-01 | CAAX amino terminal protease family protein |
| AT4G38970 | 1.5 | 9.16E-08 | 1.3 | 6.84E-07 | 0.2 | 3.21E-02 | 0.3 | 2.42E-02 | 0.2 | 1.46E-01 | 0.1 | 6.98E-01 | Fructose-bisphosphate aldolase |
| AT2G41680 | 1.5 | 2.39E-08 | 0.9 | 9.57E-07 | 0.6 | 8.87E-05 | 0.4 | 2.63E-02 | 0.2 | 2.06E-01 | 0.1 | 6.04E-01 | NADPH-dependent thioredoxin reductase 3 |
| AT2G22610 | 1.5 | 3.18E-06 | 0.8 | 7.44E-04 | 0.7 | 2.15E-03 | 0.0 | 9.92E-01 | 0.1 | 8.43E-01 | -0.1 | 8.66E-01 | Di-glucose binding protein with Kinesin motor domain-containing protein |
| AT1G27950 | 1.5 | 1.64E-07 | 1.4 | 8.41E-07 | 0.2 | 1.78E-01 | 0.0 | 9.77E-01 | -0.1 | 6.42E-01 | 0.1 | 6.75E-01 | Non-specific lipid transfer protein GPI-anchored 1 |
| AT1G52700 | 1.5 | 2.55E-05 | 1.7 | 2.01E-05 | -0.2 | 3.71E-01 | 0.0 | 9.68E-01 | -0.1 | 8.05E-01 | 0.1 | 7.58E-01 | Alpha |
| AT3G10180 | 1.5 | 3.03E-06 | 0.6 | 5.33E-03 | 0.9 | 5.55E-04 | 0.3 | 4.65E-01 | 0.2 | 7.49E-01 | 0.1 | 8.38E-01 | P-loop containing nucleoside triphosphate hydrolases superfamily protein |
| AT4G01580 | 1.5 | 7.33E-05 | 1.4 | 2.29E-04 | 0.1 | 6.87E-01 | -0.4 | 1.54E-01 | -0.3 | 3.74E-01 | -0.1 | 8.18E-01 | B3 domain-containing protein At4g01580 |
| AT5G50915 | 1.5 | 1.32E-03 | 0.5 | 2.56E-01 | 1.0 | 2.68E-02 | -0.5 | 2.54E-01 | -0.5 | 3.18E-01 | 0.0 | 9.92E-01 | Basic helix-loop-helix DNA-binding superfamily protein |
| AT5G19220 | 1.5 | 1.59E-05 | 0.8 | 2.38E-03 | 0.7 | 7.60E-03 | -0.1 | 9.33E-01 | 0.3 | 6.41E-01 | -0.4 | 6.19E-01 | Glucose-1-phosphate adenylyltransferase large subunit 1, chloroplastic |
| AT1G76110 | 1.5 | 8.74E-06 | 1.2 | 1.09E-04 | 0.3 | 1.01E-01 | -0.1 | 8.33E-01 | -0.1 | 7.16E-01 | 0.1 | 8.72E-01 | High mobility group B protein 9 |
| AT2G28605 | 1.5 | 6.71E-07 | 1.1 | 2.11E-05 | 0.5 | 5.96E-03 | 0.3 | 6.67E-02 | 0.2 | 3.44E-01 | 0.1 | 6.61E-01 | PsbP C-terminal domain-containing protein |
| AT2G28000 | 1.5 | 1.69E-07 | 1.4 | 7.41E-07 | 0.1 | 3.14E-01 | 1.1 | 3.67E-07 | 0.4 | 2.20E-02 | 0.8 | 4.74E-04 | Chaperonin 60 subunit alpha 1, chloroplastic |
| AT5G24155 | 1.5 | 5.73E-06 | 0.6 | 8.18E-03 | 0.9 | 1.02E-03 | 0.3 | 4.83E-01 | 0.6 | 2.35E-01 | -0.3 | 6.85E-01 | Squalene monooxygenase |
| AT1G02220 | 1.5 | 2.87E-07 | 0.0 | 7.42E-01 | 1.5 | 1.47E-05 | 2.2 | 7.69E-06 | 1.5 | 1.29E-03 | 0.7 | 2.96E-02 | NAC domain-containing protein 3 |
| AT4G18253 | 1.5 | 1.18E-04 | 0.8 | 1.86E-02 | 0.8 | 1.77E-02 | 0.3 | 4.31E-01 | -0.9 | 7.86E-02 | 1.2 | 3.67E-02 | Receptor Serine |
| AT2G24720 | 1.5 | 7.27E-07 | 0.8 | 2.26E-04 | 0.8 | 4.87E-04 | 1.2 | 3.25E-06 | 1.1 | 5.41E-05 | 0.1 | 5.98E-01 | Glutamate receptor 2.2 |
| AT4G03100 | 1.6 | 2.69E-07 | 1.0 | 1.67E-05 | 0.6 | 9.98E-04 | -0.2 | 3.69E-01 | 0.1 | 6.38E-01 | -0.3 | 2.45E-01 | Rho GTPase-activating protein 2 |
| AT1G77090 | 1.6 | 8.68E-07 | 1.0 | 4.25E-05 | 0.6 | 3.24E-03 | 0.6 | 1.68E-03 | 0.5 | 1.04E-02 | 0.1 | 7.04E-01 | PsbP domain-containing protein 4, chloroplastic |
| AT4G34710 | 1.6 | 1.67E-08 | 0.5 | 2.11E-05 | 1.0 | 6.99E-06 | 1.6 | 6.97E-07 | 0.6 | 9.38E-03 | 1.0 | 3.50E-03 | Arginine decarboxylase 2 |
| AT4G22970 | 1.6 | 2.35E-07 | 0.6 | 3.80E-04 | 0.9 | 7.49E-05 | 0.4 | 1.75E-01 | 0.2 | 6.23E-01 | 0.2 | 5.64E-01 | separase |
| AT4G18970 | 1.6 | 1.94E-06 | 1.7 | 2.85E-06 | -0.1 | 5.16E-01 | -0.4 | 1.19E-02 | -0.2 | 1.76E-01 | -0.2 | 4.57E-01 | GDSL-like Lipase |
| AT2G46820 | 1.6 | 2.42E-07 | 1.4 | 1.32E-06 | 0.2 | 1.69E-01 | 0.0 | 9.49E-01 | 0.1 | 5.87E-01 | -0.1 | 6.06E-01 | Protein CURVATURE THYLAKOID 1B, chloroplastic |
| AT3G53460 | 1.6 | 2.12E-06 | 1.7 | 2.77E-06 | -0.2 | 3.42E-01 | 0.5 | 4.02E-03 | -0.3 | 1.37E-01 | 0.7 | 4.88E-04 | Chloroplast RNA-binding protein 29 |
| AT4G19985 | 1.6 | 2.54E-07 | 1.0 | 1.90E-05 | 0.6 | 7.30E-04 | 0.3 | 1.48E-01 | 0.3 | 1.14E-01 | -0.1 | 8.72E-01 | hypothetical protein |
| AT3G29230 | 1.6 | 4.24E-08 | 1.4 | 2.59E-07 | 0.2 | 3.01E-02 | 0.4 | 2.69E-02 | 0.3 | 1.28E-01 | 0.1 | 6.79E-01 | Pentatricopeptide repeat-containing protein At3g29230 |
| AT1G08520 | 1.6 | 4.10E-08 | 1.2 | 5.37E-07 | 0.4 | 1.58E-03 | 0.1 | 3.30E-01 | -0.1 | 7.05E-01 | 0.2 | 2.52E-01 | Magnesium-chelatase subunit ChlD, chloroplastic |
| AT4G16155 | 1.6 | 1.68E-07 | 0.9 | 1.46E-05 | 0.7 | 2.97E-04 | 0.2 | 2.22E-01 | 0.2 | 2.95E-01 | 0.0 | 1.00E+00 | Dihydrolipoyl dehydrogenase 2, chloroplastic |
| AT1G10960 | 1.6 | 5.91E-08 | 0.9 | 4.34E-06 | 0.7 | 1.13E-04 | 0.3 | 9.12E-02 | 0.2 | 2.86E-01 | 0.1 | 8.02E-01 | Ferredoxin-1, chloroplastic |
| AT2G37220 | 1.6 | 7.03E-08 | 1.3 | 5.16E-07 | 0.3 | 2.26E-02 | 0.4 | 1.07E-02 | 0.0 | 9.68E-01 | 0.4 | 5.37E-02 | RNA-binding protein CP29B, chloroplastic |
| AT1G20020 | 1.6 | 3.09E-05 | 1.2 | 3.65E-04 | 0.4 | 1.49E-01 | 0.3 | 9.85E-03 | 0.3 | 8.80E-02 | 0.1 | 5.94E-01 | Ferredoxin--NADP reductase, chloroplastic |
| AT2G25840 | 1.6 | 3.47E-07 | 1.2 | 5.54E-06 | 0.4 | 1.23E-02 | 0.1 | 8.06E-01 | -0.1 | 7.21E-01 | 0.1 | 5.40E-01 | tryptophan--tRNA ligase |
| AT1G19250 | 1.6 | 1.35E-05 | 0.6 | 1.48E-02 | 0.9 | 2.01E-03 | 1.3 | 9.71E-06 | 1.7 | 3.43E-06 | -0.4 | 1.15E-01 | Probable flavin-containing monooxygenase 1 |
| AT3G53900 | 1.6 | 5.59E-07 | 1.4 | 2.79E-06 | 0.1 | 2.67E-01 | 0.5 | 5.12E-03 | 0.2 | 3.55E-01 | 0.3 | 1.58E-01 | Uracil phosphoribosyltransferase, chloroplastic |
| AT4G25700 | 1.6 | 1.04E-07 | 0.8 | 3.11E-05 | 0.8 | 6.96E-05 | 1.1 | 1.61E-05 | 0.0 | 8.65E-01 | 1.0 | 1.01E-03 | beta-carotene 3-hydroxylase |
| AT1G44446 | 1.6 | 5.03E-07 | 1.1 | 1.06E-05 | 0.4 | 9.72E-03 | 0.0 | 9.29E-01 | -0.2 | 3.97E-01 | 0.2 | 4.40E-01 | Pheophorbide a oxygenase family protein with Rieske 2Fe-2S domain-containing protein |
| AT5G38520 | 1.6 | 4.71E-07 | 1.3 | 4.77E-06 | 0.3 | 4.33E-02 | 0.2 | 1.89E-01 | 0.1 | 4.21E-01 | 0.0 | 8.49E-01 | Alpha |
| AT3G48280 | 1.6 | 6.58E-07 | 0.6 | 3.26E-03 | 1.0 | 1.35E-04 | 0.4 | 3.42E-01 | 0.4 | 4.68E-01 | 0.0 | 9.51E-01 | Uncharacterized protein |
| AT4G14630 | 1.6 | 2.02E-07 | 1.3 | 2.19E-06 | 0.3 | 1.86E-02 | 0.9 | 2.79E-05 | 0.7 | 9.64E-04 | 0.2 | 3.14E-01 | Germin-like protein subfamily 1 member 8 |
| AT1G03230 | 1.6 | 1.33E-06 | 1.3 | 1.24E-05 | 0.3 | 7.92E-02 | 0.4 | 5.07E-03 | 0.0 | 9.55E-01 | 0.4 | 4.80E-02 | Eukaryotic aspartyl protease family protein |
| AT1G73480 | 1.6 | 2.52E-05 | 1.0 | 1.42E-03 | 0.6 | 2.09E-02 | 0.3 | 4.77E-01 | 0.0 | 9.93E-01 | 0.3 | 5.66E-01 | Alpha |
| AT1G30730 | 1.6 | 1.43E-05 | 0.9 | 1.46E-03 | 0.7 | 1.01E-02 | -0.6 | 5.26E-02 | -1.1 | 2.13E-03 | 0.6 | 1.70E-01 | Berberine bridge enzyme-like 11 |
| AT5G49730 | 1.6 | 1.67E-04 | 1.5 | 3.88E-04 | 0.1 | 8.46E-01 | -0.1 | 5.13E-01 | -0.7 | 8.90E-03 | 0.5 | 6.17E-02 | Ferric reduction oxidase 6 |
| AT3G16370 | 1.6 | 1.68E-07 | 1.6 | 4.67E-07 | 0.0 | 9.93E-01 | -0.2 | 1.64E-01 | -0.2 | 2.55E-01 | 0.0 | 9.77E-01 | GDSL esterase |
| AT2G21330 | 1.6 | 1.74E-07 | 1.3 | 1.92E-06 | 0.3 | 1.84E-02 | 0.7 | 1.05E-04 | 0.4 | 1.63E-02 | 0.3 | 1.04E-01 | Fructose-bisphosphate aldolase |
| AT4G11175 | 1.6 | 1.01E-07 | 1.2 | 1.75E-06 | 0.4 | 2.53E-03 | 0.2 | 2.67E-01 | 0.2 | 3.10E-01 | 0.0 | 9.77E-01 | Translation initiation factor IF-1, chloroplastic |
| AT3G06030 | 1.6 | 8.42E-07 | 0.7 | 8.27E-04 | 0.9 | 2.41E-04 | 0.2 | 3.48E-01 | 0.2 | 4.73E-01 | 0.0 | 9.64E-01 | Mitogen-activated protein kinase kinase kinase 3 |
| AT1G72330 | 1.6 | 4.92E-06 | 0.2 | 4.15E-01 | 1.4 | 1.16E-04 | 0.8 | 5.03E-04 | 0.1 | 8.64E-01 | 0.7 | 4.77E-03 | Alanine aminotransferase 2, mitochondrial |
| AT2G30420 | 1.6 | 2.29E-04 | 1.3 | 2.12E-03 | 0.3 | 2.81E-01 | 0.4 | 3.48E-01 | -0.3 | 6.60E-01 | 0.7 | 1.99E-01 | MYB-like transcription factor ETC2 |
| AT4G37800 | 1.6 | 1.62E-06 | 1.3 | 1.44E-05 | 0.3 | 1.05E-01 | 0.1 | 4.41E-01 | 0.3 | 1.08E-01 | -0.2 | 4.63E-01 | Probable xyloglucan endotransglucosylase |
| AT1G43560 | 1.6 | 2.25E-08 | 1.3 | 2.13E-07 | 0.3 | 3.23E-03 | 0.4 | 1.53E-02 | 0.2 | 3.01E-01 | 0.2 | 3.91E-01 | Thioredoxin Y2, chloroplastic |
| AT3G22760 | 1.6 | 1.12E-06 | 0.9 | 2.20E-04 | 0.7 | 1.11E-03 | -0.1 | 7.13E-01 | 0.2 | 5.66E-01 | -0.3 | 3.88E-01 | Tesmin |
| AT5G35790 | 1.6 | 1.59E-06 | 1.3 | 1.36E-05 | 0.3 | 9.95E-02 | 0.1 | 4.40E-01 | -0.2 | 2.95E-01 | 0.3 | 1.22E-01 | Glucose-6-phosphate 1-dehydrogenase 1, chloroplastic |
| AT1G66430 | 1.6 | 1.59E-08 | 1.2 | 1.90E-07 | 0.5 | 1.71E-04 | 0.2 | 5.49E-02 | 0.1 | 3.18E-01 | 0.1 | 6.60E-01 | Probable fructokinase-6, chloroplastic |
| AT1G74470 | 1.6 | 1.40E-07 | 1.4 | 8.76E-07 | 0.2 | 7.33E-02 | -0.5 | 2.76E-03 | -0.1 | 5.34E-01 | -0.3 | 3.85E-02 | Geranylgeranyl diphosphate reductase, chloroplastic |
| AT1G02920 | 1.6 | 2.42E-05 | 0.6 | 2.19E-02 | 1.0 | 1.76E-03 | 1.5 | 1.29E-04 | 0.1 | 8.49E-01 | 1.4 | 4.91E-03 | Glutathione S-transferase F7 |
| AT2G36990 | 1.6 | 2.38E-08 | 1.1 | 6.42E-07 | 0.5 | 1.59E-04 | 0.1 | 6.75E-01 | -0.1 | 7.04E-01 | 0.1 | 5.03E-01 | RNA polymerase sigma factor sigF, chloroplastic |
| AT1G60950 | 1.6 | 1.04E-07 | 1.6 | 2.90E-07 | 0.0 | 9.29E-01 | -0.7 | 5.02E-04 | 0.2 | 3.53E-01 | -0.9 | 7.94E-04 | Ferredoxin-2, chloroplastic |
| AT2G05620 | 1.6 | 4.22E-05 | 0.7 | 1.09E-02 | 0.9 | 5.78E-03 | 0.4 | 2.85E-02 | 0.1 | 7.18E-01 | 0.3 | 1.92E-01 | Protein PROTON GRADIENT REGULATION 5, chloroplastic |
| AT1G80770 | 1.6 | 3.55E-08 | 1.2 | 5.12E-07 | 0.4 | 8.94E-04 | 0.3 | 4.25E-02 | 0.0 | 9.46E-01 | 0.3 | 1.12E-01 | GTP-binding protein, putative |
| AT5G39120 | 1.6 | 1.07E-06 | 0.5 | 6.43E-03 | 1.1 | 1.77E-04 | 2.7 | 3.25E-06 | 1.1 | 2.36E-02 | 1.6 | 1.29E-04 | Germin-like protein subfamily 1 member 15 |
| AT3G54050 | 1.6 | 1.51E-07 | 1.3 | 1.39E-06 | 0.3 | 2.51E-02 | 0.3 | 2.82E-02 | 0.3 | 1.34E-01 | 0.1 | 7.45E-01 | Fructose-1,6-bisphosphatase 1, chloroplastic |
| AT4G19130 | 1.6 | 1.94E-06 | 0.9 | 5.74E-04 | 0.8 | 1.58E-03 | 0.6 | 1.53E-01 | 0.2 | 7.85E-01 | 0.4 | 4.30E-01 | Replication protein A subunit |
| AT1G75960 | 1.6 | 5.18E-06 | 2.0 | 2.40E-06 | -0.4 | 4.44E-02 | 0.2 | 4.85E-01 | -0.3 | 4.86E-01 | 0.5 | 2.26E-01 | Probable acyl-activating enzyme 8 |
| AT5G42760 | 1.6 | 5.67E-06 | 0.7 | 7.51E-03 | 1.0 | 1.05E-03 | 1.0 | 7.22E-02 | 0.8 | 2.21E-01 | 0.2 | 7.89E-01 | Leucine carboxyl methyltransferase |
| AT3G63110 | 1.6 | 2.83E-07 | 1.5 | 1.38E-06 | 0.1 | 2.38E-01 | 0.4 | 5.94E-02 | -0.4 | 4.52E-02 | 0.8 | 1.95E-03 | Adenylate isopentenyltransferase 3, chloroplastic |
| AT5G17710 | 1.6 | 3.17E-08 | 1.3 | 3.61E-07 | 0.4 | 1.69E-03 | 0.4 | 2.69E-03 | 0.0 | 8.01E-01 | 0.4 | 2.66E-02 | GrpE protein homolog |
| AT1G64720 | 1.6 | 2.68E-06 | 1.4 | 1.82E-05 | 0.2 | 2.05E-01 | -0.2 | 1.72E-01 | -0.4 | 3.14E-02 | 0.2 | 3.89E-01 | Polyketide cyclase |
| AT5G47550 | 1.6 | 2.33E-07 | 1.6 | 5.64E-07 | 0.0 | 9.46E-01 | -0.3 | 1.70E-01 | -0.5 | 6.70E-02 | 0.1 | 5.81E-01 | Cysteine proteinase inhibitor 5 |
| AT5G39860 | 1.6 | 8.22E-03 | 2.2 | 1.53E-03 | -0.6 | 2.57E-01 | | | | | | | Transcription factor PRE1 |
| AT1G01080 | 1.6 | 7.34E-07 | 1.4 | 6.09E-06 | 0.3 | 7.86E-02 | 0.4 | 4.16E-03 | 0.2 | 1.34E-01 | 0.2 | 2.87E-01 | RNA-binding family protein |
| AT1G17180 | 1.6 | 9.89E-06 | -0.4 | 1.70E-01 | 2.0 | 7.77E-05 | 4.5 | 9.93E-07 | 1.9 | 1.16E-02 | 2.6 | 1.33E-05 | Glutathione S-transferase U25 |
| AT5G08760 | 1.6 | 3.99E-06 | 0.6 | 1.31E-02 | 1.1 | 4.32E-04 | 1.5 | 7.42E-05 | 0.6 | 1.12E-01 | 0.9 | 1.02E-02 | Serine rich endogenous peptide 15 |
| AT1G29670 | 1.6 | 2.67E-07 | 1.6 | 7.50E-07 | 0.0 | 9.29E-01 | -0.2 | 4.90E-02 | -0.4 | 1.01E-02 | 0.1 | 5.12E-01 | GDSL-like Lipase |
| AT2G32020 | 1.6 | 2.16E-05 | 0.3 | 2.93E-01 | 1.4 | 6.59E-04 | 4.3 | 3.40E-07 | 1.8 | 5.43E-03 | 2.5 | 9.81E-06 | N-acetyltransferase domain-containing protein |
| AT1G15820 | 1.6 | 1.75E-07 | 1.7 | 3.67E-07 | -0.1 | 4.95E-01 | 0.0 | 9.78E-01 | -0.1 | 6.52E-01 | 0.1 | 6.91E-01 | Chlorophyll a-b binding protein, chloroplastic |
| AT3G63410 | 1.6 | 2.19E-08 | 1.2 | 3.22E-07 | 0.5 | 4.75E-04 | 0.3 | 2.84E-02 | 0.3 | 5.42E-02 | 0.0 | 9.89E-01 | 2-methyl-6-phytyl-1,4-hydroquinone methyltransferase, chloroplastic |
| AT2G44650 | 1.6 | 3.00E-07 | 1.2 | 6.00E-06 | 0.5 | 5.43E-03 | 0.2 | 3.21E-01 | -0.1 | 7.70E-01 | 0.3 | 3.12E-01 | 10 kDa chaperonin 2, chloroplastic |
| AT5G18700 | 1.6 | 6.52E-07 | 0.9 | 1.16E-04 | 0.7 | 7.05E-04 | 0.1 | 6.52E-01 | 0.2 | 2.61E-01 | -0.1 | 5.96E-01 | Serine |

| | | | | | | | | | | | | | |
|---|---|---|---|---|---|---|---|---|---|---|---|---|---|
| AT5G11270 | 1.6 | 4.25E-08 | 1.2 | 8.41E-07 | 0.5 | 6.37E-04 | 0.3 | 1.26E-01 | -0.1 | 6.57E-01 | 0.4 | 1.28E-01 | Protein OVEREXPRESSOR OF CATIONIC PEROXIDASE 3 |
| AT4G27440 | 1.6 | 2.57E-06 | 2.1 | 1.25E-06 | -0.5 | 2.42E-02 | -0.1 | 5.31E-01 | -0.3 | 3.66E-02 | 0.2 | 2.04E-01 | Protochlorophyllide reductase B, chloroplastic |
| AT1G75040 | 1.7 | 1.68E-02 | 0.8 | 2.07E-01 | 0.9 | 2.28E-01 | 0.3 | 2.00E-01 | -0.2 | 3.67E-01 | 0.5 | 9.14E-02 | Pathogenesis-related protein 5 |
| AT3G14210 | 1.7 | 1.72E-07 | 1.2 | 3.23E-06 | 0.5 | 4.46E-03 | -0.1 | 4.89E-01 | 0.4 | 3.50E-02 | -0.5 | 3.12E-02 | GDSL-like lipase |
| AT2G32590 | 1.7 | 3.92E-07 | 0.8 | 2.93E-04 | 0.9 | 1.69E-04 | -0.1 | 7.60E-01 | 0.2 | 5.09E-01 | -0.3 | 3.72E-01 | Condensin complex subunit 2 |
| AT1G72670 | 1.7 | 1.31E-06 | 0.9 | 3.07E-04 | 0.8 | 1.11E-03 | -0.5 | 2.66E-01 | 0.1 | 8.39E-01 | -0.6 | 3.08E-01 | Protein IQ-DOMAIN 8 |
| AT1G49130 | 1.7 | 1.30E-05 | 1.5 | 7.29E-05 | 0.2 | 3.71E-01 | -0.9 | 1.02E-03 | -0.9 | 1.68E-03 | 0.0 | 9.48E-01 | B-box type zinc finger protein with CCT domain-containing protein |
| AT1G60550 | 1.7 | 1.18E-07 | 1.2 | 2.04E-06 | 0.4 | 3.03E-03 | -0.4 | 1.02E-02 | -0.4 | 1.75E-02 | 0.0 | 9.86E-01 | 1,4-dihydroxy-2-naphthoyl-CoA synthase, peroxisomal |
| AT1G32550 | 1.7 | 1.00E-07 | 1.3 | 1.38E-06 | 0.4 | 4.33E-03 | 0.1 | 6.28E-01 | 0.2 | 4.35E-01 | -0.1 | 8.19E-01 | 2Fe-2S ferredoxin-like superfamily protein |
| AT4G15760 | 1.7 | 5.29E-07 | 0.8 | 1.98E-04 | 0.9 | 2.60E-04 | 0.0 | 9.98E-01 | -1.0 | 1.88E-04 | 1.0 | 9.75E-04 | FAD-binding domain-containing protein |
| AT4G17600 | 1.7 | 5.42E-08 | 1.3 | 6.00E-07 | 0.4 | 3.59E-03 | 0.1 | 4.27E-01 | 0.1 | 5.10E-01 | 0.0 | 1.00E+00 | Light-harvesting complex-like protein 3 isotype 1, chloroplastic |
| AT3G16450 | 1.7 | 1.19E-06 | 1.3 | 1.64E-05 | 0.4 | 3.40E-02 | -0.3 | 1.80E-01 | -0.1 | 7.54E-01 | -0.2 | 4.91E-01 | Jacalin-related lectin 33 |
| AT3G13080 | 1.7 | 1.66E-05 | -0.6 | 1.32E-02 | 2.3 | 2.69E-05 | 0.9 | 3.91E-05 | -0.1 | 8.11E-01 | 0.9 | 9.81E-04 | Multidrug resistance-associated protein 3 |
| AT3G57520 | 1.7 | 8.36E-08 | 1.8 | 1.76E-07 | -0.2 | 2.01E-01 | -0.3 | 4.52E-01 | -0.9 | 1.89E-02 | 0.6 | 6.96E-02 | galactinol--sucrose galactosyltransferase |
| AT3G55800 | 1.7 | 9.23E-08 | 1.5 | 4.74E-07 | 0.2 | 1.39E-01 | 0.5 | 3.27E-04 | 0.4 | 6.99E-03 | 0.1 | 4.76E-01 | Sedoheptulose-1,7-bisphosphatase, chloroplastic P2ase) |
| AT1G53520 | 1.7 | 4.03E-06 | 1.6 | 1.50E-05 | 0.1 | 5.46E-01 | -0.2 | 9.57E-02 | -0.3 | 1.02E-01 | 0.0 | 9.44E-01 | Fatty-acid-binding protein 3, chloroplastic |
| AT5G51110 | 1.7 | 3.37E-08 | 1.4 | 2.57E-07 | 0.3 | 1.03E-02 | 0.4 | 1.15E-02 | 0.1 | 7.87E-01 | 0.3 | 1.10E-01 | 4a-hydroxytetrahydrobiopterin dehydratase |
| AT3G13470 | 1.7 | 6.77E-08 | 1.5 | 3.91E-07 | 0.2 | 5.10E-02 | 1.2 | 5.81E-05 | 0.4 | 1.38E-01 | 0.8 | 5.81E-03 | RuBisCO large subunit-binding protein subunit beta, chloroplastic |
| AT1G56120 | 1.7 | 9.32E-04 | 0.4 | 3.83E-01 | 1.3 | 1.27E-02 | 1.4 | 5.60E-04 | 0.2 | 6.83E-01 | 1.2 | 8.31E-03 | Leucine-rich repeat transmembrane protein kinase |
| AT5G57560 | 1.7 | 1.20E-03 | 0.3 | 4.65E-01 | 1.4 | 1.22E-02 | -1.1 | 1.07E-02 | -1.7 | 1.51E-03 | 0.7 | 1.70E-01 | Xyloglucan endotransglucosylase |
| AT4G27030 | 1.7 | 1.74E-07 | 0.8 | 6.05E-05 | 0.9 | 1.08E-04 | 0.5 | 6.86E-02 | 0.5 | 9.01E-02 | 0.0 | 9.81E-01 | Fatty acid desaturase 4, chloroplastic |
| AT3G56940 | 1.7 | 6.10E-08 | 1.4 | 5.35E-07 | 0.3 | 1.09E-02 | -0.2 | 1.59E-01 | -0.4 | 4.01E-03 | 0.2 | 1.43E-01 | magnesium-protoporphyrin IX monomethyl ester cyclase |
| AT1G05190 | 1.7 | 4.53E-08 | 1.5 | 2.77E-07 | 0.2 | 3.98E-02 | 0.2 | 2.47E-01 | 0.0 | 9.02E-01 | 0.2 | 3.47E-01 | Large ribosomal subunit protein uL6c |
| AT3G27060 | 1.7 | 3.05E-07 | 0.9 | 4.14E-05 | 0.8 | 3.42E-04 | 1.5 | 1.75E-06 | 0.2 | 5.53E-01 | 1.4 | 2.10E-04 | Ribonucleoside-diphosphate reductase small chain C |
| AT4G11460 | 1.7 | 3.40E-04 | 1.7 | 5.94E-04 | 0.0 | 9.54E-01 | 1.0 | 3.57E-02 | -0.8 | 2.40E-01 | 1.8 | 8.20E-03 | Cysteine-rich RLK 30 |
| AT4G11890 | 1.7 | 1.45E-03 | -0.1 | 8.93E-01 | 1.8 | 5.22E-03 | -0.5 | 2.12E-01 | -1.2 | 1.99E-02 | 0.7 | 3.03E-01 | Protein kinase superfamily protein |
| AT3G16000 | 1.7 | 7.71E-08 | 1.4 | 7.66E-07 | 0.3 | 1.06E-02 | 0.3 | 2.85E-02 | 0.1 | 3.07E-01 | 0.1 | 4.50E-01 | MAR-binding filament-like protein 1 |
| AT4G21270 | 1.7 | 7.17E-07 | 0.8 | 7.07E-04 | 1.0 | 2.26E-04 | -0.1 | 7.49E-01 | -0.3 | 4.20E-01 | 0.2 | 7.17E-01 | Kinesin-like protein KIN-14C |
| AT2G26180 | 1.7 | 1.33E-06 | 1.1 | 7.47E-05 | 0.6 | 4.77E-03 | 0.0 | 9.93E-01 | 0.2 | 3.73E-01 | -0.2 | 4.19E-01 | Protein IQ-DOMAIN 6 |
| AT1G13700 | 1.7 | 1.65E-05 | 1.1 | 1.07E-03 | 0.6 | 1.95E-02 | -0.9 | 1.09E-02 | -1.1 | 6.36E-03 | 0.2 | 7.38E-01 | Probable 6-phosphogluconolactonase 1 |
| AT1G11860 | 1.7 | 7.78E-08 | 1.3 | 1.08E-06 | 0.4 | 4.25E-03 | 0.3 | 3.63E-02 | 0.3 | 6.37E-02 | 0.0 | 9.98E-01 | Aminomethyltransferase |
| AT2G43570 | 1.7 | 1.61E-04 | 0.4 | 2.82E-01 | 1.3 | 3.47E-03 | 1.9 | 2.82E-07 | 0.6 | 2.50E-02 | 1.3 | 2.85E-04 | Endochitinase CHI |
| AT5G52570 | 1.7 | 5.83E-06 | 1.0 | 6.72E-04 | 0.7 | 5.23E-03 | | | | | | | Beta-carotene 3-hydroxylase 2, chloroplastic |
| AT1G32990 | 1.7 | 7.29E-08 | 1.7 | 2.54E-07 | 0.1 | 5.89E-01 | 0.4 | 1.09E-02 | 0.2 | 1.33E-01 | 0.1 | 5.31E-01 | Large ribosomal subunit protein uL11c |
| AT3G28510 | 1.7 | 4.97E-03 | -0.1 | 9.32E-01 | 1.8 | 1.62E-02 | 2.0 | 2.49E-07 | 1.4 | 5.41E-05 | 0.6 | 3.04E-02 | AAA-ATPase At3g28510 |
| AT2G39510 | 1.7 | 4.77E-07 | 2.7 | 9.36E-08 | -0.9 | 8.64E-05 | 2.5 | 4.72E-07 | 1.5 | 6.11E-04 | 1.1 | 1.87E-03 | WAT1-related protein At2g39510 |
| AT3G27830 | 1.7 | 3.51E-08 | 1.5 | 2.13E-07 | 0.2 | 3.97E-02 | 0.3 | 7.98E-02 | 0.2 | 3.21E-01 | 0.1 | 7.47E-01 | Large ribosomal subunit protein bL12cz |
| AT4G13890 | 1.7 | 8.87E-07 | 1.7 | 2.84E-06 | 0.1 | 6.96E-01 | 1.4 | 3.25E-06 | 1.2 | 9.27E-05 | 0.2 | 3.23E-01 | Serine hydroxymethyltransferase |
| AT4G23280 | 1.7 | 1.37E-05 | 0.6 | 3.40E-02 | 1.1 | 1.20E-03 | 0.3 | 6.92E-01 | -1.8 | 7.22E-02 | 2.1 | 3.80E-02 | Cysteine-rich RLK 20 |
| AT5G20740 | 1.8 | 2.15E-07 | 1.5 | 1.75E-06 | 0.3 | 4.71E-02 | 0.0 | 9.28E-01 | 0.0 | 9.74E-01 | 0.0 | 9.74E-01 | Plant invertase |
| AT3G52720 | 1.8 | 2.31E-07 | 1.7 | 7.87E-07 | 0.1 | 5.55E-01 | 0.5 | 3.96E-02 | 0.7 | 1.24E-02 | -0.2 | 6.21E-01 | Alpha carbonic anhydrase 1, chloroplastic |
| AT3G55330 | 1.8 | 4.77E-08 | 1.6 | 2.38E-07 | 0.2 | 1.07E-01 | 0.0 | 9.33E-01 | 0.0 | 8.21E-01 | 0.1 | 8.06E-01 | PsbP-like protein 1, chloroplastic |
| AT5G64940 | 1.8 | 1.69E-07 | 1.0 | 1.68E-05 | 0.8 | 2.74E-04 | 0.6 | 2.98E-03 | 0.5 | 1.93E-02 | 0.1 | 7.36E-01 | Protein ACTIVITY OF BC1 COMPLEX KINASE 8, chloroplastic |
| AT1G22400 | 1.8 | 7.73E-05 | 1.1 | 2.80E-03 | 0.6 | 5.45E-02 | -0.8 | 1.09E-01 | -1.5 | 1.36E-02 | 0.7 | 1.20E-01 | UDP-glycosyltransferase 85A1 |
| AT5G13630 | 1.8 | 1.60E-07 | 1.4 | 2.09E-06 | 0.4 | 1.01E-02 | -0.5 | 3.38E-02 | 0.1 | 7.02E-01 | -0.6 | 2.14E-02 | magnesium chelatase |
| AT5G25130 | 1.8 | 2.28E-06 | 1.0 | 4.18E-04 | 0.8 | 2.44E-03 | -0.7 | 1.62E-02 | -0.4 | 1.49E-01 | -0.3 | 4.16E-01 | Uncharacterized protein |
| AT5G50950 | 1.8 | 1.30E-06 | 1.8 | 3.20E-06 | 0.0 | 9.82E-01 | -0.5 | 1.06E-02 | -0.3 | 1.96E-01 | -0.2 | 4.45E-01 | fumarate hydratase |
| AT1G71500 | 1.8 | 2.38E-08 | 1.4 | 2.33E-07 | 0.4 | 4.05E-03 | 0.0 | 9.88E-01 | 0.0 | 7.70E-01 | 0.0 | 8.23E-01 | Rieske domain-containing protein |
| AT3G63160 | 1.8 | 2.99E-07 | 1.9 | 6.00E-07 | -0.1 | 5.61E-01 | 0.3 | 1.42E-01 | 0.1 | 7.27E-01 | 0.2 | 5.30E-01 | AT3g63160 |
| AT1G77490 | 1.8 | 1.65E-07 | 1.5 | 1.04E-06 | 0.2 | 8.03E-02 | -0.1 | 6.70E-01 | -0.1 | 7.96E-01 | 0.0 | 9.62E-01 | L-ascorbate peroxidase |
| AT5G24110 | 1.8 | 1.17E-05 | 0.6 | 3.78E-02 | 1.2 | 1.05E-03 | 0.4 | 5.03E-01 | -0.6 | 4.38E-01 | | | Probable WRKY transcription factor 30 |
| AT1G09450 | 1.8 | 1.50E-06 | 1.1 | 1.30E-04 | 0.7 | 3.37E-03 | 0.2 | 3.77E-01 | 0.1 | 6.96E-01 | 0.1 | 8.13E-01 | Serine |
| AT3G11750 | 1.8 | 6.14E-07 | 1.3 | 1.31E-05 | 0.4 | 1.26E-02 | 0.0 | 8.59E-01 | 0.1 | 6.53E-01 | -0.1 | 8.28E-01 | 7,8-dihydroneopterin aldolase |
| AT5G02120 | 1.8 | 3.06E-05 | 1.0 | 2.83E-03 | 0.8 | 1.30E-02 | 0.5 | 2.56E-02 | 0.4 | 1.19E-01 | 0.1 | 7.57E-01 | Light-harvesting complex-like protein OHP1, chloroplastic |
| AT1G67700 | 1.8 | 5.42E-08 | 1.3 | 7.87E-07 | 0.5 | 1.75E-03 | 0.1 | 2.06E-01 | -0.1 | 6.26E-01 | 0.2 | 1.62E-01 | Protein HHL1, chloroplastic |
| AT3G12780 | 1.8 | 2.38E-08 | 1.5 | 1.88E-07 | 0.3 | 1.45E-02 | 0.3 | 9.35E-03 | 0.3 | 2.22E-02 | 0.0 | 9.64E-01 | Phosphoglycerate kinase 1, chloroplastic |
| AT2G24710 | 1.8 | 3.02E-07 | 1.9 | 5.12E-07 | -0.1 | 4.15E-01 | -3.0 | 5.61E-07 | -1.2 | 2.42E-04 | -1.7 | 3.86E-03 | Glutamate receptor |
| AT3G48730 | 1.8 | 4.24E-08 | 1.6 | 2.13E-07 | 0.2 | 1.32E-01 | 0.7 | 1.03E-05 | 0.3 | 1.27E-02 | 0.3 | 2.67E-02 | Glutamate-1-semialdehyde 2,1-aminomutase 2, chloroplastic |
| AT4G30950 | 1.8 | 1.97E-08 | 1.2 | 3.76E-07 | 0.6 | 1.71E-04 | 0.2 | 9.41E-02 | 0.1 | 5.10E-01 | 0.1 | 5.88E-01 | Omega-6 fatty acid desaturase, chloroplastic |
| AT5G45930 | 1.8 | 2.92E-07 | 1.6 | 1.80E-06 | 0.2 | 1.05E-01 | 0.6 | 2.68E-04 | 0.3 | 3.90E-02 | 0.3 | 1.18E-01 | Magnesium-chelatase subunit ChlI-2, chloroplastic |
| AT3G63190 | 1.8 | 5.74E-08 | 1.5 | 4.64E-07 | 0.3 | 1.32E-02 | 0.4 | 4.90E-03 | 0.1 | 5.80E-01 | 0.3 | 8.85E-02 | Ribosome-recycling factor, chloroplastic |
| AT5G51720 | 1.8 | 2.97E-06 | 1.1 | 2.02E-04 | 0.7 | 4.69E-03 | -2.8 | 6.41E-05 | 0.8 | 1.48E-02 | -3.6 | 8.64E-05 | CDGSH iron-sulfur domain-containing protein NEET |
| AT5G62530 | 1.8 | 4.93E-08 | 1.3 | 1.18E-06 | 0.5 | 4.62E-04 | 0.4 | 1.56E-01 | 0.3 | 3.81E-01 | 0.1 | 8.07E-01 | Delta-1-pyrroline-5-carboxylate dehydrogenase 12A1, mitochondrial |
| AT3G46490 | 1.8 | 7.71E-08 | 1.8 | 2.13E-07 | 0.0 | 8.32E-01 | 0.0 | 8.81E-01 | -0.1 | 4.62E-01 | 0.2 | 4.47E-01 | Uncharacterized protein F12A12.10 |
| AT4G38470 | 1.8 | 1.81E-07 | 1.8 | 4.99E-07 | 0.0 | 8.98E-01 | -0.2 | 2.15E-01 | -0.7 | 4.07E-03 | 0.4 | 9.61E-02 | ACT-like protein tyrosine kinase family protein |
| AT1G29910 | 1.8 | 1.53E-07 | 1.8 | 4.74E-07 | 0.0 | 7.82E-01 | 0.2 | 4.54E-01 | -0.3 | 3.54E-01 | 0.4 | 1.68E-01 | Chlorophyll a-b binding protein 3, chloroplastic |
| AT1G58290 | 1.8 | 8.20E-08 | 1.4 | 9.19E-07 | 0.4 | 7.79E-03 | -2.0 | 1.19E-06 | -0.3 | 3.44E-01 | -1.7 | 1.09E-04 | Glutamyl-tRNA reductase 1, chloroplastic |
| AT1G78830 | 1.8 | 9.88E-07 | 1.7 | 3.94E-06 | 0.1 | 4.94E-01 | 0.4 | 3.07E-02 | -0.8 | 5.07E-04 | 1.1 | 8.11E-05 | EP1-like glycoprotein 2 lectin family protein) |
| AT1G62780 | 1.8 | 5.79E-08 | 1.5 | 5.25E-07 | 0.4 | 6.73E-03 | 0.8 | 8.09E-05 | 0.5 | 1.31E-02 | 0.3 | 9.22E-02 | Dimethylallyl, adenosine tRNA methylthiotransferase |
| AT5G20720 | 1.8 | 3.17E-08 | 1.6 | 2.13E-07 | 0.3 | 2.67E-02 | 0.7 | 2.02E-04 | 0.1 | 5.10E-01 | 0.6 | 1.07E-02 | 20 kDa chaperonin, chloroplastic |
| AT1G54960 | 1.8 | 1.03E-06 | 0.8 | 7.46E-04 | 1.0 | 3.66E-04 | 0.3 | 1.28E-01 | 0.3 | 2.50E-01 | 0.0 | 9.11E-01 | Mitogen-activated protein kinase kinase kinase 2 |
| AT5G58250 | 1.8 | 1.06E-07 | 1.6 | 5.64E-07 | 0.2 | 1.19E-01 | 0.6 | 2.29E-03 | 0.2 | 4.01E-01 | 0.4 | 8.01E-02 | YCF54 |
| AT3G27690 | 1.8 | 1.60E-06 | 2.1 | 1.83E-06 | -0.3 | 2.08E-01 | 0.0 | 9.67E-01 | -0.1 | 8.28E-01 | 0.1 | 8.88E-01 | Chlorophyll a-b binding protein, chloroplastic |
| AT2G38530 | 1.8 | 2.43E-07 | 0.8 | 1.43E-04 | 1.0 | 8.64E-05 | 0.1 | 7.43E-01 | -0.2 | 6.84E-01 | 0.3 | 5.20E-01 | Non-specific lipid-transfer protein |
| AT5G21930 | 1.8 | 1.65E-06 | 1.3 | 4.04E-05 | 0.5 | 1.35E-02 | -0.1 | 4.60E-01 | -0.3 | 2.39E-01 | 0.1 | 6.71E-01 | P-type ATPase of Arabidopsis 2 |
| AT4G36540 | 1.8 | 3.29E-06 | 1.7 | 1.21E-05 | 0.1 | 5.61E-01 | -0.3 | 1.65E-02 | -0.3 | 1.76E-02 | 0.0 | 9.00E-01 | Transcription factor BEE 2 |
| AT3G60290 | 1.8 | 3.93E-05 | 2.1 | 2.64E-05 | -0.3 | 2.82E-01 | -0.4 | 2.36E-01 | -0.6 | 1.10E-01 | 0.2 | 6.71E-01 | Fe2OG dioxygenase domain-containing protein |
| AT2G39800 | 1.8 | 1.66E-05 | 1.4 | 2.49E-04 | 0.5 | 7.94E-02 | -1.2 | 5.98E-04 | -0.2 | 5.38E-01 | -1.0 | 2.44E-02 | Delta-1-pyrroline-5-carboxylate synthase |
| AT5G17670 | 1.8 | 1.96E-07 | 1.3 | 6.36E-06 | 0.6 | 1.51E-03 | -0.2 | 1.91E-01 | -0.1 | 5.06E-01 | -0.1 | 7.41E-01 | GPI inositol-deacylase |
| AT1G18400 | 1.8 | 2.42E-06 | 1.2 | 1.53E-04 | 0.6 | 7.30E-03 | 0.1 | 8.91E-01 | 0.4 | 6.04E-01 | -0.3 | 7.66E-01 | BR enhanced expression 1 |
| AT4G16690 | 1.9 | 5.60E-07 | 0.8 | 8.40E-04 | 1.1 | 1.69E-04 | -0.4 | 1.22E-01 | -0.9 | 6.20E-03 | 0.5 | 1.90E-01 | pFDCC methylesterase MES16 |
| AT5G66470 | 1.9 | 3.75E-08 | 1.4 | 4.89E-07 | 0.5 | 1.16E-03 | 0.7 | 3.95E-05 | 0.4 | 1.01E-02 | 0.3 | 5.23E-02 | GTPase ERA-like, chloroplastic |
| AT3G48900 | 1.9 | 1.71E-05 | 0.8 | 1.52E-02 | 1.1 | 2.48E-03 | -0.2 | 5.78E-01 | 0.0 | 9.84E-01 | -0.2 | 7.10E-01 | Single-stranded DNA endonuclease family protein |

| | | | | | | | | | | | | | |
|---|---|---|---|---|---|---|---|---|---|---|---|---|---|
| AT3G55630 | 1.9 | 1.75E-07 | 1.5 | 1.69E-06 | 0.3 | 2.26E-02 | -0.1 | 6.14E-01 | 0.1 | 5.03E-01 | -0.2 | 3.09E-01 | Folylpolyglutamate synthase |
| AT2G28190 | 1.9 | 1.89E-08 | 1.4 | 2.13E-07 | 0.5 | 6.25E-04 | 0.0 | 7.88E-01 | 0.1 | 6.76E-01 | 0.0 | 8.87E-01 | Superoxide dismutase 2, chloroplastic |
| AT3G47650 | 1.9 | 6.77E-08 | 1.3 | 1.25E-06 | 0.5 | 1.15E-03 | 0.4 | 1.25E-01 | 0.3 | 2.86E-01 | 0.1 | 8.91E-01 | Protein BUNDLE SHEATH DEFECTIVE 2, chloroplastic |
| AT1G09390 | 1.9 | 3.90E-07 | 1.2 | 2.86E-05 | 0.7 | 1.15E-03 | 0.5 | 9.36E-02 | 0.0 | 9.19E-01 | 0.5 | 1.46E-01 | GDSL esterase |
| AT5G14910 | 1.9 | 1.67E-08 | 1.6 | 9.13E-08 | 0.2 | 1.71E-02 | 0.1 | 3.20E-01 | -0.1 | 4.26E-01 | 0.2 | 1.61E-01 | AT5g14910 |
| AT5G43450 | 1.9 | 2.31E-06 | 0.0 | 9.40E-01 | 1.9 | 3.16E-05 | 1.6 | 2.82E-07 | -0.5 | 3.42E-02 | 2.1 | 1.58E-06 | 2-oxoglutarate and Fe-dependent oxygenase superfamily protein |
| AT5G55220 | 1.9 | 6.10E-08 | 1.5 | 5.71E-07 | 0.4 | 6.44E-03 | 0.7 | 2.95E-04 | 0.3 | 4.98E-02 | 0.3 | 7.97E-02 | Trigger factor-like protein TIG, Chloroplastic |
| AT5G23910 | 1.9 | 1.53E-06 | 1.0 | 3.31E-04 | 0.9 | 1.20E-03 | 0.3 | 1.17E-01 | 0.2 | 3.60E-01 | 0.1 | 7.63E-01 | Kinesin motor domain-containing protein |
| AT1G60600 | 1.9 | 2.25E-08 | 1.6 | 1.76E-07 | 0.3 | 1.06E-02 | 0.3 | 9.19E-02 | 0.2 | 4.14E-01 | 0.1 | 6.40E-01 | 2-carboxy-1,4-naphthoquinone phytyltransferase, chloroplastic |
| AT5G64810 | 1.9 | 1.20E-05 | 0.0 | 8.89E-01 | 1.9 | 2.38E-04 | 2.1 | 2.56E-05 | 1.2 | 9.23E-03 | 0.9 | 2.67E-03 | Probable WRKY transcription factor 51 |
| AT2G25080 | 1.9 | 9.61E-08 | 1.8 | 3.67E-07 | 0.1 | 4.63E-01 | -0.1 | 5.03E-01 | -0.4 | 1.57E-02 | 0.3 | 1.57E-01 | Phospholipid hydroperoxide glutathione peroxidase 1, chloroplastic |
| AT3G26060 | 1.9 | 7.44E-08 | 1.6 | 5.12E-07 | 0.3 | 3.72E-02 | 0.2 | 2.95E-01 | 0.3 | 1.20E-01 | -0.1 | 6.44E-01 | thioredoxin-dependent peroxiredoxin |
| AT3G46090 | 1.9 | 5.82E-05 | 0.8 | 2.60E-02 | 1.1 | 7.41E-03 | 0.5 | 5.76E-01 | -0.5 | 7.32E-01 | 1.0 | 4.49E-01 | Zinc finger protein ZAT7 |
| AT5G44568 | 1.9 | 3.97E-06 | 1.0 | 1.10E-03 | 1.0 | 1.69E-03 | 1.1 | 4.56E-03 | 0.1 | 9.34E-01 | 1.0 | 3.18E-02 | Serine rich endogenous peptide 4 |
| AT2G24020 | 1.9 | 3.72E-07 | 1.6 | 3.20E-06 | 0.3 | 4.50E-02 | 0.5 | 5.16E-03 | 0.1 | 8.19E-01 | 0.4 | 5.07E-02 | Nucleoid-associated protein At2g24020, chloroplastic |
| AT1G68880 | 1.9 | 3.68E-06 | 2.0 | 6.09E-06 | -0.1 | 7.06E-01 | 4.8 | 3.62E-06 | 2.4 | 8.04E-03 | 2.4 | 3.74E-05 | Basic leucine zipper 8 |
| AT1G26420 | 1.9 | 2.36E-04 | 0.3 | 4.65E-01 | 1.6 | 3.59E-03 | 0.2 | 7.76E-01 | -1.4 | 6.88E-02 | 1.6 | 7.08E-02 | Berberine bridge enzyme-like 7 |
| AT2G20635 | 1.9 | 1.10E-06 | 0.9 | 9.91E-04 | 1.1 | 3.28E-04 | -0.3 | 1.77E-01 | 0.0 | 8.86E-01 | -0.3 | 3.65E-01 | Mitotic checkpoint serine |
| AT3G56650 | 1.9 | 7.54E-08 | 1.6 | 6.33E-07 | 0.3 | 1.44E-02 | 0.1 | 3.70E-01 | -0.2 | 2.35E-01 | 0.3 | 1.00E-01 | PsbP domain-containing protein 6, chloroplastic |
| AT4G29060 | 1.9 | 3.13E-08 | 1.5 | 3.36E-07 | 0.4 | 2.43E-03 | 0.7 | 5.96E-05 | 0.4 | 2.90E-02 | 0.4 | 4.81E-02 | Polyprotein of EF-Ts, chloroplastic |
| AT2G34180 | 1.9 | 9.18E-07 | 2.4 | 4.67E-07 | -0.5 | 8.44E-03 | -0.7 | 5.78E-02 | 0.2 | 6.35E-01 | -0.9 | 4.48E-02 | CBL-interacting serine |
| AT4G25080 | 1.9 | 2.67E-07 | 1.5 | 2.94E-06 | 0.4 | 2.39E-02 | 0.0 | 7.71E-01 | -0.3 | 1.03E-01 | 0.2 | 2.71E-01 | Magnesium-protoporphyrin IX methyltransferase |
| AT2G06850 | 1.9 | 9.58E-07 | 1.8 | 4.18E-06 | 0.2 | 4.23E-01 | -0.3 | 5.50E-02 | -0.7 | 9.48E-04 | 0.4 | 9.68E-02 | Xyloglucan endotransglucosylase |
| AT1G60000 | 2.0 | 1.67E-08 | 1.4 | 2.13E-07 | 0.6 | 1.45E-04 | 0.4 | 1.13E-02 | 0.1 | 6.40E-01 | 0.3 | 1.27E-01 | RNA-binding family protein |
| AT1G73870 | 2.0 | 3.39E-06 | 1.4 | 9.78E-05 | 0.6 | 1.70E-02 | | | | | | | Zinc finger protein CONSTANS-LIKE 7 |
| AT2G36800 | 2.0 | 3.25E-06 | -0.1 | 7.88E-01 | 2.0 | 4.45E-05 | 0.2 | 3.26E-01 | -0.4 | 1.80E-01 | 0.6 | 2.27E-02 | UDP-glycosyltransferase 73C5 |
| AT4G24770 | 2.0 | 1.99E-08 | 1.7 | 1.09E-07 | 0.3 | 2.42E-02 | 0.2 | 1.17E-01 | 0.0 | 8.73E-01 | 0.2 | 1.67E-01 | 31 kDa ribonucleoprotein, chloroplastic |
| AT2G39470 | 2.0 | 2.87E-07 | 1.6 | 3.04E-06 | 0.4 | 2.26E-02 | 0.4 | 1.04E-02 | 0.2 | 1.29E-01 | 0.1 | 4.59E-01 | Photosynthetic NDH subunit of lumenal location 1, chloroplastic |
| AT5G11450 | 2.0 | 4.45E-08 | 1.6 | 3.39E-07 | 0.3 | 9.64E-03 | 0.7 | 2.80E-05 | 0.1 | 4.69E-01 | 0.6 | 1.95E-03 | PsbP domain protein |
| AT1G29920 | 2.0 | 5.32E-08 | 1.8 | 2.59E-07 | 0.2 | 1.18E-01 | -1.2 | 5.71E-03 | -1.4 | 3.90E-03 | 0.2 | 7.70E-01 | Chlorophyll a-b binding protein 2, chloroplastic |
| AT5G65690 | 2.0 | 2.48E-07 | 2.1 | 3.96E-07 | -0.1 | 2.57E-01 | -1.3 | 5.75E-03 | -0.4 | 3.23E-01 | -0.8 | 1.05E-01 | phosphoenolpyruvate carboxykinase |
| AT2G30660 | 2.0 | 1.51E-05 | 2.2 | 1.17E-05 | -0.3 | 2.30E-01 | 2.6 | 3.12E-06 | 1.9 | 3.95E-04 | 0.7 | 2.23E-02 | 3-hydroxyisobutyryl-CoA hydrolase |
| AT5G58310 | 2.0 | 5.29E-07 | 1.6 | 6.26E-06 | 0.4 | 3.28E-02 | 0.9 | 1.29E-02 | 1.1 | 5.57E-03 | -0.2 | 6.10E-01 | Methylesterase 18 |
| AT5G08280 | 2.0 | 4.25E-08 | 1.6 | 3.93E-07 | 0.4 | 5.75E-03 | 0.6 | 3.18E-04 | 0.1 | 5.49E-01 | 0.5 | 1.35E-02 | hydroxymethylbilane synthase |
| AT5G44572 | 2.0 | 1.79E-04 | 0.9 | 4.10E-02 | 1.1 | 1.55E-02 | -0.8 | 5.53E-02 | -1.6 | 3.56E-03 | 0.8 | 1.74E-01 | Serine rich endogenous peptide 6 |
| AT5G27390 | 2.0 | 2.43E-08 | 1.6 | 2.38E-07 | 0.4 | 3.18E-03 | 0.3 | 1.02E-01 | 0.0 | 8.55E-01 | 0.2 | 2.82E-01 | Uncharacterized protein At5g27390 |
| AT5G36925 | 2.0 | 4.57E-04 | 0.7 | 1.28E-01 | 1.2 | 1.43E-02 | 0.5 | 3.87E-01 | 0.1 | 8.78E-01 | 0.3 | 6.41E-01 | Peptide ARACIN 1 |
| AT4G15510 | 2.0 | 7.44E-08 | 1.7 | 5.75E-07 | 0.3 | 1.69E-02 | 0.3 | 3.47E-02 | 0.1 | 6.83E-01 | 0.3 | 2.11E-01 | PsbP domain-containing protein 1, chloroplastic |
| AT3G54090 | 2.0 | 1.56E-07 | 1.5 | 2.38E-06 | 0.5 | 5.39E-03 | 1.1 | 3.88E-05 | 0.2 | 5.49E-01 | 0.9 | 1.41E-03 | Fructokinase-like 1, chloroplastic |
| AT5G59750 | 2.0 | 2.29E-07 | 1.3 | 1.48E-05 | 0.7 | 6.38E-04 | 0.3 | 3.01E-02 | -0.3 | 1.64E-01 | 0.6 | 7.35E-03 | GTP cyclohydrolase II |
| AT2G35410 | 2.0 | 1.44E-08 | 1.5 | 9.13E-08 | 0.5 | 2.64E-04 | 0.4 | 2.01E-03 | 0.1 | 4.55E-01 | 0.3 | 8.15E-02 | Putative chloroplast RNA binding protein family protein) |
| AT5G39580 | 2.0 | 7.92E-03 | 0.0 | 9.80E-01 | 2.0 | 2.81E-02 | 1.2 | 2.17E-03 | -1.2 | 2.36E-02 | 2.3 | 2.85E-04 | peroxidase |
| AT3G23080 | 2.0 | 6.97E-07 | 2.3 | 6.98E-07 | -0.3 | 9.84E-02 | -0.5 | 2.45E-02 | -0.6 | 6.62E-03 | 0.2 | 5.39E-01 | Membrane related protein-like |
| AT4G17560 | 2.0 | 1.89E-08 | 1.8 | 1.04E-07 | 0.3 | 1.66E-02 | 0.4 | 1.55E-02 | 0.0 | 9.04E-01 | 0.4 | 1.12E-01 | Large ribosomal subunit protein bL19cz |
| AT3G15190 | 2.0 | 1.59E-08 | 1.6 | 1.04E-07 | 0.4 | 1.50E-03 | 0.3 | 2.24E-02 | 0.0 | 8.07E-01 | 0.2 | 1.20E-01 | Small ribosomal subunit protein bS20c |
| AT2G38140 | 2.0 | 2.25E-08 | 1.7 | 1.90E-07 | 0.4 | 7.47E-03 | 0.0 | 9.93E-01 | 0.0 | 9.78E-01 | 0.0 | 9.86E-01 | Small ribosomal subunit protein bTHXc |
| AT2G33800 | 2.0 | 2.19E-08 | 1.8 | 1.37E-07 | 0.3 | 1.80E-02 | 0.2 | 1.35E-01 | 0.0 | 8.16E-01 | 0.2 | 1.78E-01 | Small ribosomal subunit protein uS5c |
| AT2G47560 | 2.0 | 1.60E-06 | 1.8 | 1.13E-05 | 0.2 | 1.85E-01 | | | | | | | RING-H2 finger protein ATL64 |
| AT5G61070 | 2.0 | 1.35E-06 | 1.2 | 1.96E-04 | 0.8 | 1.86E-03 | 0.3 | 5.19E-01 | -0.4 | 4.39E-01 | 0.7 | 2.01E-01 | Histone deacetylase 18 |
| AT4G28080 | 2.1 | 1.01E-07 | 1.4 | 2.86E-06 | 0.7 | 1.02E-03 | 0.2 | 2.03E-01 | -0.1 | 6.07E-01 | 0.4 | 9.58E-02 | Protein REDUCED CHLOROPLAST COVERAGE 2 |
| AT2G30750 | 2.1 | 2.10E-05 | 1.1 | 3.04E-03 | 0.9 | 7.30E-03 | 2.6 | 4.76E-08 | 1.4 | 1.38E-04 | 1.2 | 1.97E-03 | hypothetical protein |
| AT1G68670 | 2.1 | 5.05E-07 | 1.7 | 5.06E-06 | 0.4 | 4.28E-02 | 0.2 | 1.70E-01 | 0.0 | 8.94E-01 | 0.2 | 2.53E-01 | Transcription factor HHO2 |
| AT1G73110 | 2.1 | 4.14E-07 | 1.3 | 2.66E-05 | 0.8 | 1.16E-03 | 0.2 | 2.01E-01 | 0.1 | 4.76E-01 | 0.1 | 7.89E-01 | Ribulose bisphosphate carboxylase |
| AT2G36200 | 2.1 | 5.13E-07 | 1.1 | 1.28E-04 | 1.0 | 3.37E-04 | 0.1 | 5.55E-01 | 0.2 | 2.54E-01 | -0.1 | 6.79E-01 | P-loop containing nucleoside triphosphate hydrolases superfamily protein |
| AT1G28010 | 2.1 | 6.78E-08 | 1.4 | 2.10E-06 | 0.6 | 4.52E-04 | 0.2 | 4.56E-01 | -0.3 | 2.02E-01 | 0.5 | 8.37E-02 | ABC transporter B family member 14 |
| AT2G02930 | 2.1 | 1.26E-04 | 0.8 | 5.60E-02 | 1.2 | 1.06E-02 | 0.0 | 9.82E-01 | -1.5 | 6.72E-02 | 1.5 | 1.14E-01 | Glutathione S-transferase F3 |
| AT1G20060 | 2.1 | 1.49E-06 | 1.1 | 3.62E-04 | 1.0 | 1.04E-03 | 0.2 | 5.06E-01 | 0.2 | 4.67E-01 | 0.0 | 9.11E-01 | Kinesin-like protein |
| AT3G50820 | 2.1 | 7.19E-08 | 1.8 | 4.67E-07 | 0.3 | 4.13E-02 | 0.2 | 9.73E-02 | 0.1 | 4.08E-01 | 0.1 | 6.77E-01 | Oxygen-evolving enhancer protein 1-2, chloroplastic |
| AT1G20340 | 2.1 | 2.37E-08 | 2.0 | 9.19E-08 | 0.1 | 6.32E-01 | 0.2 | 2.90E-01 | 0.0 | 8.57E-01 | 0.1 | 6.07E-01 | Plastocyanin major isoform, chloroplastic |
| AT5G18600 | 2.1 | 5.20E-06 | 2.2 | 8.31E-06 | -0.1 | 6.63E-01 | -0.3 | 1.73E-01 | -0.3 | 3.85E-01 | -0.1 | 8.20E-01 | Monothiol glutaredoxin-S2 |
| AT1G42970 | 2.1 | 1.03E-07 | 1.6 | 1.12E-06 | 0.5 | 1.01E-02 | 0.5 | 2.69E-04 | 0.4 | 5.85E-03 | 0.1 | 4.40E-01 | Glyceraldehyde-3-phosphate dehydrogenase GAPB, chloroplastic |
| AT3G52150 | 2.1 | 2.97E-08 | 1.9 | 1.36E-07 | 0.2 | 1.79E-01 | 0.5 | 2.53E-04 | 0.2 | 1.08E-01 | 0.3 | 7.42E-02 | Small ribosomal subunit protein cS22 |
| AT5G20630 | 2.1 | 8.93E-07 | 2.2 | 1.74E-06 | -0.1 | 6.02E-01 | 0.0 | 9.81E-01 | 0.0 | 9.17E-01 | 0.0 | 9.48E-01 | Germin-like protein subfamily 3 member 3 |
| AT2G43030 | 2.1 | 1.59E-08 | 1.7 | 9.13E-08 | 0.4 | 2.24E-03 | 0.3 | 3.27E-02 | 0.0 | 8.61E-01 | 0.3 | 9.17E-02 | Large ribosomal subunit protein uL3c |
| AT4G37180 | 2.1 | 1.97E-08 | 1.8 | 1.36E-07 | 0.4 | 3.47E-03 | 0.9 | 2.17E-04 | 0.0 | 9.90E-01 | 0.9 | 3.98E-03 | hypothetical protein |
| AT4G37370 | 2.1 | 2.86E-06 | -0.4 | 2.00E-01 | 2.5 | 3.35E-05 | 0.8 | 3.56E-03 | -0.9 | 1.05E-02 | 1.6 | 1.73E-04 | Cytochrome P450, family 81, subfamily D, polypeptide 8 |
| AT3G27850 | 2.1 | 1.59E-08 | 1.7 | 1.04E-07 | 0.4 | 1.95E-03 | 0.2 | 1.31E-01 | 0.0 | 9.81E-01 | 0.2 | 2.81E-01 | Large ribosomal subunit protein bL12cx |
| AT3G63540 | 2.1 | 2.75E-08 | 1.4 | 6.75E-07 | 0.7 | 1.99E-04 | 0.1 | 5.28E-01 | 0.0 | 8.97E-01 | 0.1 | 5.73E-01 | Thylakoid lumenal 19 kDa protein, chloroplastic |
| AT3G13120 | 2.1 | 1.67E-08 | 1.8 | 9.13E-08 | 0.3 | 1.47E-02 | 0.4 | 1.05E-02 | 0.0 | 8.50E-01 | 0.4 | 8.69E-02 | Small ribosomal subunit protein uS10c |
| AT1G78630 | 2.1 | 2.25E-08 | 1.9 | 1.12E-07 | 0.2 | 6.38E-02 | 0.0 | 9.23E-01 | -0.1 | 7.06E-01 | 0.1 | 6.85E-01 | Large ribosomal subunit protein uL13c |
| AT5G27290 | 2.1 | 7.38E-08 | 1.8 | 5.40E-07 | 0.3 | 1.97E-02 | 0.1 | 7.29E-01 | 0.1 | 5.20E-01 | -0.1 | 8.02E-01 | Stress regulated protein |
| AT2G44040 | 2.1 | 3.72E-08 | 1.6 | 4.53E-07 | 0.5 | 1.43E-03 | 0.6 | 1.12E-04 | 0.2 | 3.23E-01 | 0.5 | 6.77E-03 | 4-hydroxy-tetrahydrodipicolinate reductase |
| AT1G76450 | 2.2 | 3.76E-06 | 1.4 | 1.71E-04 | 0.8 | 8.62E-03 | 0.4 | 4.42E-03 | 0.2 | 1.46E-01 | 0.2 | 2.88E-01 | PsbP domain-containing protein 3, chloroplastic |
| AT5G51100 | 2.2 | 2.25E-08 | 1.7 | 2.42E-07 | 0.4 | 1.48E-03 | 0.7 | 5.91E-05 | 0.1 | 6.16E-01 | 0.6 | 3.05E-03 | Superoxide dismutase |
| AT1G22890 | 2.2 | 3.05E-07 | 1.1 | 1.17E-04 | 1.1 | 1.66E-04 | -0.4 | 5.16E-02 | -1.2 | 1.65E-04 | 0.7 | 1.81E-02 | Serine rich endogenous peptide 14 |
| AT2G32530 | 2.2 | 1.15E-06 | 0.9 | 2.50E-03 | 1.3 | 2.06E-04 | 1.0 | 1.27E-03 | 0.9 | 3.27E-03 | 0.0 | 9.49E-01 | Cellulose synthase-like protein B3 |
| AT2G37720 | 2.2 | 4.18E-06 | 1.4 | 2.87E-04 | 0.8 | 6.65E-03 | | | | | | | Protein trichome birefringence-like 15 |
| AT4G01310 | 2.2 | 1.44E-08 | 1.9 | 8.06E-08 | 0.3 | 2.05E-02 | 0.3 | 3.81E-02 | 0.0 | 8.98E-01 | 0.2 | 1.32E-01 | Large ribosomal subunit protein uL5c |
| AT4G34190 | 2.2 | 3.44E-06 | 1.3 | 1.95E-04 | 0.9 | 6.46E-03 | 0.4 | 2.18E-02 | 0.2 | 4.53E-01 | 0.3 | 3.15E-01 | Stress enhanced protein 1, chloroplastic |
| AT1G78820 | 2.2 | 1.88E-07 | 2.4 | 3.36E-07 | -0.2 | 3.31E-01 | 0.3 | 7.35E-02 | -0.1 | 4.67E-01 | 0.4 | 5.07E-02 | EP1-like glycoprotein 1 |
| AT4G21280 | 2.2 | 1.70E-08 | 1.8 | 1.04E-07 | 0.4 | 6.61E-03 | 0.1 | 3.06E-01 | 0.1 | 3.82E-01 | 0.0 | 9.97E-01 | PSBQA |
| AT3G23450 | 2.2 | 2.94E-06 | 1.9 | 1.66E-05 | 0.3 | 2.43E-01 | 0.5 | 9.77E-03 | -0.1 | 8.28E-01 | 0.5 | 3.79E-02 | Transmembrane protein |
| AT3G16240 | 2.2 | 1.46E-07 | 2.1 | 5.12E-07 | 0.1 | 5.39E-01 | -1.0 | 3.48E-04 | -1.5 | 2.96E-05 | 0.5 | 7.66E-02 | AT3G16240 protein |

| | | | | | | | | | | | | | |
|---|---|---|---|---|---|---|---|---|---|---|---|---|---|
| AT1G19150 | 2.2 | 1.02E-07 | 1.9 | 6.00E-07 | 0.3 | 7.08E-02 | 0.1 | 4.10E-01 | 0.2 | 2.59E-01 | -0.1 | 7.88E-01 | Photosystem I chlorophyll a |
| AT1G14700 | 2.2 | 4.84E-07 | 2.0 | 2.31E-06 | 0.2 | 2.86E-01 | 0.2 | 2.90E-01 | 0.1 | 6.26E-01 | 0.1 | 7.81E-01 | Purple acid phosphatase |
| AT5G47190 | 2.2 | 1.24E-07 | 1.7 | 1.83E-06 | 0.6 | 4.47E-03 | 0.3 | 1.97E-02 | 0.0 | 8.05E-01 | 0.3 | 1.36E-01 | Large ribosomal subunit protein bL19cy |
| AT2G45490 | 2.2 | 2.88E-06 | 1.3 | 3.40E-04 | 0.9 | 3.16E-03 | -0.2 | 3.77E-01 | -0.2 | 5.61E-01 | 0.0 | 9.12E-01 | Aurora kinase |
| AT5G44575 | 2.2 | 4.64E-03 | -0.1 | 9.48E-01 | 2.3 | 2.06E-02 | | | | | | | Serine rich endogenous peptide 8 |
| AT2G34620 | 2.2 | 1.08E-06 | 1.9 | 9.23E-06 | 0.4 | 7.85E-02 | -0.6 | 4.03E-04 | 0.0 | 8.47E-01 | -0.7 | 2.41E-03 | Mitochondrial transcription termination factor family protein |
| AT2G22140 | 2.2 | 4.27E-06 | 1.4 | 3.98E-04 | 0.8 | 5.14E-03 | -0.3 | 3.94E-01 | -0.1 | 7.56E-01 | -0.1 | 7.77E-01 | Essential meiotic endonuclease 1B |
| AT4G09650 | 2.3 | 1.67E-08 | 2.1 | 8.06E-08 | 0.1 | 2.37E-01 | 0.4 | 4.76E-03 | 0.5 | 4.89E-03 | -0.1 | 8.41E-01 | ATP synthase subunit delta, chloroplastic sector subunit delta) |
| AT5G40950 | 2.3 | 1.44E-08 | 1.8 | 8.06E-08 | 0.4 | 1.02E-03 | 0.4 | 1.29E-02 | 0.1 | 6.67E-01 | 0.3 | 1.57E-01 | Large ribosomal subunit protein bL27c |
| AT3G52380 | 2.3 | 1.70E-08 | 1.8 | 1.12E-07 | 0.4 | 2.45E-03 | 0.9 | 1.40E-05 | 0.2 | 3.03E-01 | 0.7 | 2.74E-03 | RNA-binding protein CP33, chloroplastic |
| AT3G12500 | 2.3 | 7.03E-08 | 1.7 | 9.07E-07 | 0.5 | 2.54E-03 | 1.6 | 2.46E-05 | 1.0 | 4.33E-03 | 0.6 | 3.46E-02 | hypothetical protein |
| AT3G16530 | 2.3 | 2.39E-06 | 1.9 | 2.32E-05 | 0.4 | 8.51E-02 | 0.7 | 9.23E-02 | -0.5 | 2.91E-01 | 1.2 | 2.40E-02 | Lectin-like protein At3g16530 |
| AT4G09010 | 2.3 | 2.04E-08 | 2.1 | 9.13E-08 | 0.2 | 1.79E-01 | 0.0 | 8.20E-01 | 0.0 | 9.12E-01 | 0.0 | 9.63E-01 | Ascorbate peroxidase 4 |
| AT2G01918 | 2.3 | 7.12E-06 | 1.8 | 1.09E-04 | 0.5 | 5.36E-02 | 0.2 | 4.78E-01 | 0.2 | 6.30E-01 | 0.0 | 9.39E-01 | PQL3 |
| AT3G44890 | 2.3 | 1.44E-08 | 1.8 | 9.13E-08 | 0.5 | 1.30E-03 | 0.4 | 1.91E-03 | 0.2 | 1.62E-01 | 0.2 | 1.95E-01 | Large ribosomal subunit protein bL9c |
| AT1G55490 | 2.3 | 2.19E-08 | 2.0 | 1.12E-07 | 0.3 | 3.81E-02 | 0.8 | 1.03E-05 | 0.3 | 1.02E-01 | 0.6 | 2.27E-03 | Chaperonin 60 subunit beta 1, chloroplastic |
| AT2G24090 | 2.3 | 1.65E-08 | 1.9 | 1.04E-07 | 0.4 | 2.23E-03 | 0.3 | 9.95E-02 | 0.0 | 8.89E-01 | 0.3 | 3.16E-01 | Large ribosomal subunit protein bL35c |
| AT1G35680 | 2.3 | 2.25E-08 | 2.0 | 1.59E-07 | 0.3 | 2.14E-02 | 0.4 | 2.63E-02 | 0.0 | 9.36E-01 | 0.3 | 1.27E-01 | Large ribosomal subunit protein bL21c |
| AT1G65500 | 2.3 | 6.58E-04 | 0.2 | 7.35E-01 | 2.1 | 6.27E-03 | 0.5 | 3.02E-01 | -0.3 | 6.73E-01 | 0.8 | 2.46E-01 | Secreted transmembrane peptide 6 |
| AT2G41240 | 2.3 | 3.11E-04 | 0.6 | 3.04E-01 | 1.7 | 6.61E-03 | 4.6 | 3.64E-07 | -0.6 | 5.30E-01 | 5.1 | 3.08E-05 | Transcription factor bHLH100 |
| AT4G20820 | 2.3 | 3.53E-07 | 2.6 | 4.52E-07 | -0.3 | 9.66E-02 | -0.3 | 4.91E-01 | 0.3 | 4.56E-01 | -0.7 | 1.95E-01 | Berberine bridge enzyme-like 18 |
| AT5G39620 | 2.3 | 3.89E-06 | 2.9 | 1.65E-06 | -0.6 | 1.31E-02 | | | | | | | Uncharacterized protein |
| AT1G75690 | 2.3 | 1.44E-08 | 1.9 | 8.81E-08 | 0.4 | 2.08E-03 | 0.3 | 1.28E-01 | 0.2 | 3.92E-01 | 0.1 | 7.76E-01 | Protein disulfide-isomerase LQY1, chloroplastic |
| AT1G56050 | 2.3 | 3.55E-08 | 1.8 | 3.83E-07 | 0.5 | 1.91E-03 | 1.3 | 7.56E-07 | 0.5 | 1.22E-02 | 0.8 | 5.26E-04 | Obg-like ATPase 1 |
| AT2G37070 | 2.3 | 2.06E-06 | 1.1 | 1.33E-03 | 1.2 | 6.93E-04 | -0.5 | 2.33E-01 | -0.2 | 6.66E-01 | -0.3 | 6.73E-01 | Uncharacterized protein At2g37070 |
| AT2G33450 | 2.3 | 3.06E-08 | 1.7 | 4.42E-07 | 0.6 | 6.54E-04 | 0.4 | 5.93E-02 | 0.3 | 1.60E-01 | 0.1 | 8.56E-01 | Large ribosomal subunit protein bL28c |
| AT3G16250 | 2.3 | 1.80E-07 | 2.1 | 8.63E-07 | 0.2 | 1.70E-01 | 0.6 | 6.97E-03 | 0.6 | 1.64E-02 | 0.0 | 9.67E-01 | Photosynthetic NDH subunit of subcomplex B 3, chloroplastic |
| AT3G57240 | 2.4 | 4.16E-03 | 1.0 | 2.03E-01 | 1.4 | 8.66E-02 | 0.3 | 6.40E-01 | -1.7 | 9.82E-02 | 2.0 | 8.66E-02 | glucan endo-1,3-beta-D-glucosidase |
| AT5G18660 | 2.4 | 2.97E-08 | 2.0 | 2.28E-07 | 0.4 | 7.21E-03 | 0.7 | 5.67E-04 | 0.6 | 5.07E-03 | 0.1 | 6.95E-01 | Divinyl chlorophyllide a 8-vinyl-reductase, chloroplastic |
| AT2G03760 | 2.4 | 6.75E-08 | 0.6 | 1.47E-03 | 1.8 | 1.09E-05 | 2.0 | 4.52E-07 | -0.4 | 2.81E-01 | 2.4 | 7.42E-06 | Cytosolic sulfotransferase 12 |
| AT4G25750 | 2.4 | 2.25E-04 | 1.3 | 1.91E-02 | 1.0 | 4.06E-02 | -0.1 | 8.56E-01 | -1.0 | 1.37E-01 | 0.9 | 1.90E-01 | ABC transporter G family member 4 |
| AT3G51870 | 2.4 | 9.53E-08 | 2.2 | 4.77E-07 | 0.2 | 1.08E-01 | 0.7 | 3.45E-03 | 0.1 | 7.31E-01 | 0.6 | 7.75E-03 | Probable envelope ADP,ATP carrier protein, chloroplastic |
| AT1G69710 | 2.4 | 3.60E-06 | 1.3 | 1.00E-03 | 1.1 | 2.00E-03 | 0.3 | 4.74E-01 | 0.5 | 3.07E-01 | -0.2 | 7.93E-01 | Putative regulator of chromosome condensation; 48393-44372 family with FYVE zinc finger domain-containing protein) |
| AT4G34620 | 2.4 | 1.89E-08 | 1.9 | 1.70E-07 | 0.5 | 1.38E-03 | 0.1 | 5.40E-01 | -0.2 | 3.74E-01 | 0.3 | 2.25E-01 | Small ribosomal subunit protein bS16cy |
| AT1G29100 | 2.4 | 6.85E-06 | -0.8 | 8.91E-02 | 3.2 | 6.69E-05 | -1.5 | 1.93E-03 | 0.1 | 8.28E-01 | -1.6 | 3.17E-03 | Heavy metal transport |
| AT2G34430 | 2.4 | 3.68E-06 | 2.6 | 4.84E-06 | -0.2 | 4.40E-01 | -1.8 | 8.37E-07 | -2.3 | 5.32E-07 | 0.5 | 1.90E-01 | Chlorophyll a-b binding protein, chloroplastic |
| AT1G52100 | 2.4 | 5.41E-07 | 1.3 | 2.44E-04 | 1.2 | 2.58E-04 | 0.1 | 8.97E-01 | -0.5 | 2.35E-01 | 0.6 | 8.45E-02 | Mannose-binding lectin superfamily protein |
| AT1G48350 | 2.5 | 1.63E-07 | 1.8 | 2.40E-06 | 0.6 | 5.11E-03 | 0.2 | 2.87E-01 | -0.1 | 7.92E-01 | 0.2 | 2.01E-01 | Large ribosomal subunit protein uL18c |
| AT1G15380 | 2.5 | 1.41E-08 | 2.7 | 2.05E-08 | -0.2 | 4.79E-02 | -0.3 | 6.47E-02 | -0.7 | 1.18E-03 | 0.4 | 8.55E-02 | Glyoxylase I 4 |
| AT3G04720 | 2.5 | 3.86E-07 | 1.7 | 1.26E-05 | 0.8 | 2.45E-03 | 0.9 | 7.30E-03 | -0.4 | 3.28E-01 | 1.3 | 6.72E-03 | Hevein-like preproprotein |
| AT5G48490 | 2.5 | 1.90E-07 | 2.0 | 1.75E-06 | 0.5 | 2.29E-02 | -0.9 | 6.90E-03 | -1.3 | 1.01E-03 | 0.5 | 2.89E-01 | Bifunctional inhibitor |
| AT4G18480 | 2.5 | 1.44E-08 | 2.0 | 8.06E-08 | 0.5 | 5.64E-04 | 0.6 | 7.30E-05 | 0.2 | 1.25E-01 | 0.4 | 2.42E-02 | Magnesium-chelatase subunit ChlI-1, chloroplastic |
| AT1G65240 | 2.5 | 8.57E-07 | 1.3 | 3.02E-04 | 1.1 | 5.11E-04 | | | | | | | Aspartic proteinase 39 |
| AT4G01970 | 2.5 | 1.67E-08 | 2.0 | 1.16E-07 | 0.5 | 7.26E-04 | 0.5 | 1.13E-01 | 0.7 | 1.92E-02 | -0.3 | 4.44E-01 | Stachyose synthase |
| AT1G21910 | 2.5 | 1.63E-04 | 0.5 | 3.59E-01 | 2.0 | 3.01E-03 | -0.7 | 6.97E-02 | -0.7 | 1.05E-01 | 0.0 | 9.71E-01 | Ethylene-responsive transcription factor ERF012 |
| AT1G31690 | 2.5 | 1.21E-05 | 3.3 | 2.93E-06 | -0.8 | 5.11E-03 | 0.2 | 3.56E-01 | 0.5 | 1.02E-01 | -0.2 | 3.69E-01 | Amine oxidase alpha 2, peroxisomal |
| AT5G38420 | 2.5 | 2.25E-08 | 2.5 | 9.13E-08 | 0.0 | 9.43E-01 | 0.6 | 9.25E-04 | 0.4 | 2.31E-02 | 0.2 | 4.39E-01 | Ribulose bisphosphate carboxylase small subunit 2B, chloroplastic |
| AT1G25550 | 2.5 | 8.03E-07 | 1.8 | 2.58E-05 | 0.8 | 5.15E-03 | 0.2 | 2.60E-01 | -0.7 | 2.54E-03 | 0.9 | 2.08E-03 | Myb-like transcription factor family protein |
| AT3G27160 | 2.5 | 1.97E-08 | 2.0 | 1.79E-07 | 0.5 | 2.02E-03 | 0.3 | 6.04E-02 | 0.0 | 8.87E-01 | 0.3 | 2.42E-01 | Ribosomal protein S21 family protein |
| AT4G22755 | 2.6 | 3.77E-07 | 1.9 | 1.01E-05 | 0.7 | 3.06E-03 | 0.1 | 6.84E-01 | 0.0 | 9.50E-01 | 0.2 | 7.34E-01 | Methylsterol monooxygenase 1-3 |
| AT1G66050 | 2.6 | 1.46E-05 | 1.7 | 7.81E-04 | 0.8 | 1.88E-02 | 0.0 | 9.30E-01 | -0.1 | 9.41E-01 | 0.0 | 9.99E-01 | E3 ubiquitin-protein ligase ORTHRUS 5 |
| AT3G01330 | 2.6 | 2.78E-07 | 1.6 | 2.95E-05 | 1.0 | 3.57E-04 | -0.1 | 8.54E-01 | 0.1 | 7.67E-01 | -0.1 | 6.71E-01 | E2F transcription factor-like E2FF |
| AT3G17360 | 2.6 | 2.48E-06 | 1.5 | 3.68E-04 | 1.1 | 1.88E-03 | 0.3 | 2.59E-01 | 0.3 | 3.28E-01 | 0.0 | 9.99E-01 | Phragmoplast orienting kinesin 1 |
| AT3G03630 | 2.6 | 6.53E-08 | 1.7 | 3.37E-06 | 0.9 | 1.39E-04 | 1.3 | 1.37E-05 | 0.3 | 2.01E-01 | 0.9 | 1.92E-03 | S-sulfo-L-cysteine synthase , chloroplastic |
| AT5G03350 | 2.6 | 4.98E-03 | 1.3 | 9.19E-02 | 1.2 | 1.59E-01 | 2.5 | 2.73E-07 | 1.6 | 1.76E-04 | 1.0 | 1.33E-02 | Lectin-like protein |
| AT3G15850 | 2.6 | 1.44E-08 | 2.0 | 9.13E-08 | 0.6 | 7.73E-04 | 0.1 | 2.95E-01 | -0.5 | 8.46E-04 | 0.7 | 6.57E-04 | Palmitoyl-monogalactosyldiacylglycerol delta-7 desaturase, chloroplastic |
| AT1G55810 | 2.6 | 1.67E-08 | 2.5 | 8.06E-08 | 0.1 | 1.88E-01 | 0.0 | 9.00E-01 | -0.1 | 6.48E-01 | 0.2 | 5.67E-01 | Uridine kinase |
| AT3G62930 | 2.6 | 1.10E-05 | 2.1 | 1.64E-04 | 0.6 | 7.03E-02 | 1.3 | 5.47E-03 | -0.3 | 6.05E-01 | 1.6 | 8.18E-03 | Monothiol glutaredoxin-S6 |
| AT3G27730 | 2.6 | 1.99E-06 | 1.4 | 7.71E-04 | 1.2 | 8.50E-04 | | | | | | | Uncharacterized protein |
| AT2G16060 | 2.6 | 6.99E-06 | 1.6 | 9.31E-04 | 1.1 | 4.66E-03 | 1.8 | 3.15E-04 | -0.6 | 3.42E-01 | 2.4 | 6.82E-04 | Non-symbiotic hemoglobin 1 |
| AT2G47750 | 2.6 | 1.97E-06 | 2.4 | 9.08E-06 | 0.2 | 3.57E-01 | 0.4 | 4.30E-01 | 0.3 | 7.14E-01 | 0.2 | 8.15E-01 | Putative indole-3-acetic acid-amido synthetase GH3.9 |
| AT1G63990 | 2.6 | 1.80E-05 | 1.5 | 2.73E-03 | 1.1 | 7.63E-03 | -0.1 | 7.46E-01 | 0.1 | 9.02E-01 | -0.2 | 6.59E-01 | Sporulation 11-2 |
| AT2G43620 | 2.7 | 2.79E-03 | 0.4 | 6.77E-01 | 2.3 | 1.82E-02 | -0.6 | 3.13E-01 | -1.8 | 2.00E-02 | 1.2 | 1.78E-01 | Endochitinase At2g43620 |
| AT4G25050 | 2.7 | 1.70E-08 | 2.3 | 1.04E-07 | 0.4 | 7.80E-03 | 0.2 | 2.62E-01 | 0.0 | 9.92E-01 | 0.2 | 4.42E-01 | Acyl carrier protein 4 |
| AT5G38430 | 2.7 | 1.44E-08 | 2.4 | 8.06E-08 | 0.3 | 2.39E-02 | 0.9 | 1.63E-06 | 0.4 | 6.28E-03 | 0.5 | 6.10E-03 | Ribulose bisphosphate carboxylase small subunit 1B, chloroplastic |
| AT5G14740 | 2.7 | 7.29E-08 | 2.3 | 4.96E-07 | 0.4 | 3.45E-02 | -0.3 | 5.79E-02 | -0.3 | 4.66E-02 | 0.1 | 8.23E-01 | Carbonic anhydrase |
| AT2G26695 | 2.7 | 4.34E-07 | 2.5 | 2.04E-06 | 0.2 | 2.41E-01 | 2.7 | 3.31E-07 | 1.1 | 1.00E-02 | 1.6 | 1.30E-04 | RanBP2-type domain-containing protein |
| AT5G38930 | 2.8 | 1.97E-08 | 3.4 | 5.07E-08 | -0.6 | 3.26E-04 | -0.6 | 8.71E-03 | -0.7 | 6.11E-03 | 0.1 | 7.98E-01 | Germin-like protein subfamily 1 member 10 |
| AT5G25190 | 2.8 | 1.68E-07 | 1.4 | 8.20E-05 | 1.4 | 7.70E-05 | | | | | | | Ethylene-responsive transcription factor ERF003 |
| AT5G20250 | 2.8 | 1.48E-07 | 2.9 | 3.58E-07 | -0.1 | 7.32E-01 | -0.1 | 8.89E-01 | -1.4 | 2.56E-04 | 1.4 | 2.85E-04 | galactinol--sucrose galactosyltransferase |
| AT5G63310 | 2.8 | 1.41E-08 | 2.2 | 8.06E-08 | 0.6 | 1.87E-04 | 0.7 | 7.86E-05 | 0.3 | 5.25E-02 | 0.4 | 3.78E-02 | Nucleoside diphosphate kinase II, chloroplastic |
| AT2G18300 | 2.8 | 7.99E-07 | 2.7 | 2.67E-06 | 0.1 | 6.18E-01 | -0.2 | 3.84E-01 | -0.8 | 1.94E-03 | 0.6 | 2.68E-02 | Basic helix-loop-helix DNA-binding superfamily protein |
| AT3G15356 | 2.8 | 7.19E-08 | 2.3 | 5.16E-07 | 0.5 | 1.64E-02 | 0.6 | 6.46E-02 | -1.1 | 5.12E-03 | 1.6 | 1.12E-03 | Lectin-like protein LEC |
| AT2G36750 | 2.8 | 3.81E-04 | -0.5 | 5.48E-01 | 3.3 | 1.63E-03 | | | | | | | UDP-glycosyltransferase 73C1 |
| AT5G24420 | 2.8 | 7.53E-07 | 2.4 | 6.26E-06 | 0.4 | 5.84E-02 | 0.1 | 8.65E-01 | 0.5 | 5.12E-01 | -0.4 | 7.29E-01 | Probable 6-phosphogluconolactonase 5 |
| AT5G19120 | 2.8 | 4.58E-05 | 2.6 | 1.37E-04 | 0.2 | 6.74E-01 | 0.2 | 5.12E-01 | -1.3 | 1.80E-03 | 1.5 | 4.03E-03 | hypothetical protein |
| AT5G05340 | 2.9 | 1.23E-04 | -0.5 | 5.02E-01 | 3.4 | 1.18E-03 | 2.2 | 5.75E-02 | 1.3 | 3.83E-01 | | | Peroxidase 52 |
| AT2G24850 | 3.0 | 1.81E-02 | 1.5 | 2.76E-01 | 1.4 | 2.78E-01 | 1.2 | 3.20E-01 | 0.6 | 7.40E-01 | 0.6 | 7.11E-01 | Probable aminotransferase TAT3 |
| AT1G49160 | 3.1 | 6.82E-08 | 2.8 | 2.95E-07 | 0.2 | 1.19E-01 | 1.0 | 5.41E-04 | -0.2 | 5.19E-01 | 1.2 | 1.09E-03 | Protein kinase superfamily protein |
| AT1G02620 | 3.2 | 2.59E-05 | 2.1 | 1.51E-03 | 1.1 | 1.40E-02 | -0.4 | 3.81E-01 | -1.3 | 2.34E-02 | 0.9 | 1.99E-01 | Ras-related small GTP-binding family protein |
| AT2G27420 | 3.3 | 1.44E-08 | 2.2 | 2.54E-07 | 1.1 | 1.47E-05 | | | | | | | Cysteine proteinase |
| AT3G01500 | 3.4 | 1.89E-08 | 3.2 | 9.13E-08 | 0.2 | 2.41E-01 | 0.4 | 2.08E-02 | 0.5 | 2.05E-02 | -0.1 | 8.73E-01 | Carbonic anhydrase |
| AT4G16260 | 3.4 | 4.59E-08 | 2.6 | 5.65E-07 | 0.8 | 1.02E-03 | 2.1 | 7.50E-06 | 0.9 | 2.26E-02 | 1.2 | 7.21E-03 | glucan endo-1,3-beta-D-glucosidase |

| | | | | | | | | | | | | | |
|---|---|---|---|---|---|---|---|---|---|---|---|---|---|
| AT1G15040 | 3.5 | 7.95E-05 | 2.9 | 5.02E-04 | 0.5 | 1.91E-01 | 1.0 | 2.57E-01 | -0.3 | 8.58E-01 | | | Putative glutamine amidotransferase GAT1_2.1 |
| AT1G73260 | 3.5 | 9.95E-06 | 2.1 | 7.07E-04 | 1.4 | 5.37E-03 | 0.4 | 1.45E-01 | -0.2 | 7.04E-01 | 0.6 | 8.90E-02 | Uncharacterized protein |
| AT2G04070 | 3.5 | 2.87E-05 | -0.3 | 6.78E-01 | 3.8 | 4.47E-04 | 3.7 | 3.27E-08 | 1.2 | 9.14E-03 | 2.5 | 9.81E-06 | Protein DETOXIFICATION 4 |
| AT2G26400 | 3.7 | 1.32E-03 | 0.7 | 6.15E-01 | 3.0 | 2.40E-02 | 1.8 | 6.06E-04 | 1.1 | 3.88E-02 | 0.7 | 1.61E-01 | Acireductone dioxygenase -requiring)) -requiring)) |
| AT5G13370 | 3.8 | 1.67E-08 | 2.4 | 5.71E-07 | 1.4 | 1.20E-05 | 0.4 | 5.88E-02 | -0.4 | 8.39E-02 | 0.8 | 1.36E-03 | Indole-3-acetic acid-amido synthetase GH3.15 |
| AT4G26260 | 3.8 | 2.35E-07 | 3.8 | 5.71E-07 | 0.0 | 8.83E-01 | -2.0 | 4.56E-02 | -2.2 | 5.24E-02 | | | Inositol oxygenase |
| AT1G66760 | 3.8 | 7.75E-08 | 2.8 | 2.07E-06 | 1.0 | 1.66E-04 | 0.9 | 3.73E-02 | -0.7 | 2.79E-01 | 1.6 | 1.49E-02 | AT1G66760 protein |
| AT2G04040 | 3.8 | 8.03E-06 | 0.6 | 3.23E-01 | 3.2 | 1.82E-04 | 0.4 | 3.00E-01 | -2.4 | 1.80E-03 | 2.8 | 2.16E-03 | Protein DETOXIFICATION 1 |
| AT2G30770 | 3.8 | 3.05E-04 | 1.0 | 3.15E-01 | 2.8 | 6.95E-03 | -1.1 | 3.51E-01 | -3.1 | 7.55E-02 | | | Indoleacetaldoxime dehydratase |
| AT1G78850 | 3.9 | 2.32E-08 | 3.1 | 2.16E-07 | 0.8 | 2.99E-03 | 1.0 | 5.43E-06 | -0.1 | 5.18E-01 | 1.2 | 1.62E-04 | EP1-like glycoprotein 3 lectin family protein) |
| AT5G54190 | 3.9 | 7.27E-07 | 3.4 | 5.15E-06 | 0.5 | 3.92E-02 | | | | | | | Protochlorophyllide reductase A, chloroplastic |
| AT3G47340 | 3.9 | 7.71E-08 | 4.0 | 2.26E-07 | 0.0 | 9.51E-01 | -0.9 | 5.52E-02 | -2.1 | 2.19E-03 | 1.2 | 1.51E-01 | Asparagine synthetase |
| AT3G01600 | 4.0 | 6.07E-07 | 0.0 | 9.80E-01 | 4.0 | 2.76E-05 | 2.8 | 3.58E-03 | 0.4 | 7.69E-01 | 2.3 | 2.38E-02 | NAC domain containing protein 44 |
| AT4G13300 | 4.2 | 1.17E-06 | 3.1 | 3.40E-05 | 1.1 | 2.08E-03 | 0.8 | 7.19E-02 | 0.9 | 6.94E-02 | -0.1 | 9.19E-01 | -gamma-bisabolene synthase 2 |
| AT2G13810 | 4.3 | 3.97E-06 | 1.3 | 4.86E-02 | 3.1 | 1.89E-04 | 2.6 | 1.53E-07 | 1.8 | 4.38E-05 | 0.8 | 5.24E-03 | Aminotransferase ALD1, chloroplastic |
| AT1G66040 | 4.5 | 2.56E-06 | 3.3 | 7.28E-05 | 1.2 | 3.43E-03 | | | | | | | Putative E3 ubiquitin-protein ligase ORTHRUS 4 |
| AT2G18190 | 4.5 | 1.25E-03 | -1.1 | 5.22E-01 | 5.5 | 3.50E-03 | | | | | | | AAA-ATPase At2g18190 |
| AT4G19800 | 4.6 | 7.33E-08 | 4.9 | 1.62E-07 | -0.3 | 9.42E-02 | -2.3 | 2.88E-06 | -0.4 | 1.41E-01 | -1.9 | 4.74E-04 | GH18 domain-containing protein |
| AT4G19760 | 4.7 | 1.46E-07 | 3.0 | 1.27E-05 | 1.7 | 5.17E-05 | 2.3 | 3.11E-03 | 2.2 | 7.66E-03 | 0.1 | 9.42E-01 | hypothetical protein |
| AT2G18193 | 4.8 | 2.02E-07 | 0.7 | 1.09E-01 | 4.1 | 1.39E-05 | 1.7 | 5.52E-07 | 0.2 | 5.03E-01 | 1.5 | 9.28E-05 | AAA-ATPase At2g18193 |
| AT4G13310 | 4.9 | 4.80E-07 | 3.5 | 1.49E-05 | 1.4 | 3.09E-04 | -0.4 | 1.40E-01 | 0.2 | 4.57E-01 | -0.6 | 7.37E-02 | Cytochrome P450, family 71, subfamily A, polypeptide 20 |
| AT4G19750 | 5.2 | 2.83E-07 | 3.3 | 2.02E-05 | 1.8 | 7.55E-05 | 1.7 | 4.56E-04 | 1.8 | 9.83E-04 | 0.0 | 9.42E-01 | Glycosyl hydrolase family protein with chitinase insertion domain-containing protein |
| AT2G38340 | 5.5 | 3.27E-04 | -0.7 | 6.77E-01 | 6.2 | 1.73E-03 | | | | | | | Dehydration-responsive element-binding protein 2E |
| AT2G20800 | 5.8 | 1.12E-04 | 0.0 | 9.80E-01 | 5.7 | 1.36E-03 | 3.5 | 7.15E-03 | 0.6 | 7.79E-01 | | | External alternative NADH-ubiquinone oxidoreductase B4, mitochondrial NDB4) |
| AT3G05950 | 6.6 | 1.68E-07 | 4.1 | 1.42E-05 | 2.5 | 1.40E-05 | -2.9 | 1.20E-06 | -0.8 | 9.74E-03 | -2.1 | 1.15E-03 | Germin-like protein subfamily 1 member 7 |
| AT2G04050 | 6.8 | 1.35E-06 | 0.1 | 9.25E-01 | 6.7 | 4.03E-05 | 4.1 | 8.66E-07 | 0.0 | 9.76E-01 | 4.1 | 3.73E-05 | Protein DETOXIFICATION 3 |
| AT3G21460 | 7.8 | 3.26E-07 | 7.8 | 8.30E-07 | 0.0 | 8.87E-01 | 1.0 | 5.51E-03 | 0.2 | 6.24E-01 | 0.8 | 7.24E-02 | Monothiol glutaredoxin-S10 |
| AT1G05680 | 8.3 | 2.47E-05 | 1.1 | 4.72E-01 | 7.1 | 2.59E-04 | 2.3 | 7.50E-05 | -1.0 | 2.09E-01 | 3.3 | 3.22E-04 | UDP-glycosyltransferase 74E2 |
| AT1G61120 | | | | | | | -3.587503077 | 3.10E-02 | -1.527566278 | 2.42E-01 | -2.1 | 4.19E-01 | #NAME? |
| AT1G06100 | | | | | | | 0.867289328 | 3.10E-02 | 0.662259669 | 1.41E-01 | 0.2 | 6.38E-01 | Delta-9 desaturase-like 2 protein |
| AT1G01480 | | | | | | | 0.8 | 1.70E-01 | -0.2 | 7.83E-01 | 1.0 | 1.62E-01 | 1-aminocyclopropane-1-carboxylate synthase 2 |
| AT5G65800 | | | | | | | -0.7 | 3.31E-01 | 0.8 | 1.83E-01 | | | 1-aminocyclopropane-1-carboxylate synthase 5 |
| AT2G30540 | | | | | | | -0.407210046 | 4.04E-01 | -0.082498029 | 9.04E-01 | -0.3 | 6.58E-01 | Monothiol glutaredoxin-S9 |
| AT2G38240 | | | | | | | 0.924112681 | 6.91E-01 | 0.101978694 | 9.79E-01 | 0.8 | 8.06E-01 | Jasmonate-induced oxygenase 4 |
| AT3G49620 | | | | | | | 0.297140177 | 8.70E-01 | -1.464545091 | 5.33E-01 | 1.8 | 5.21E-01 | Probable 2-oxoglutarate-dependent dioxygenase DIN11 |
| AT3G22450 | | | | | | | 0.6 | 1.03E-01 | -0.1 | 9.29E-01 | 0.7 | 1.41E-01 | Expressed protein |
| AT4G15215 | | | | | | | -0.1 | 9.06E-01 | -0.1 | 9.15E-01 | | | Pleiotropic drug resistance 13 |
| AT3G55090 | | | | | | | 1.2 | 6.42E-04 | 0.5 | 1.24E-01 | 0.6 | 6.13E-02 | ABC transporter G family member 16 |
| AT5G52050 | | | | | | | -2.2 | 1.47E-05 | -1.9 | 9.86E-05 | -0.3 | 6.02E-01 | Protein DETOXIFICATION 50 |
| AT4G29340 | | | | | | | 1.18316417 | 8.15E-02 | 1.143196712 | 1.31E-01 | 0.0 | 9.69E-01 | Profilin-4 |
| AT5G16230 | | | | | | | 0.3 | 5.35E-01 | 0.2 | 6.80E-01 | 0.0 | 9.46E-01 | stearoyl- 9-desaturase |
| AT3G14360 | | | | | | | -0.4 | 4.68E-01 | 0.2 | 6.96E-01 | -0.6 | 3.02E-01 | Triacylglycerol lipase OBL1 |
| AT2G41450 | | | | | | | 0.8 | 7.80E-02 | 1.0 | 5.34E-02 | -0.2 | 7.95E-01 | N-acetyltransferase |
| AT3G12230 | | | | | | | 3.819631276 | 5.09E-03 | 3.553316839 | 1.64E-02 | 0.3 | 8.18E-01 | Serine carboxypeptidase-like 14 |
| AT1G63730 | | | | | | | 0.2 | 7.10E-01 | -0.1 | 8.43E-01 | 0.3 | 6.35E-01 | Disease resistance protein family |
| AT3G04220 | | | | | | | 0.9 | 7.90E-03 | 0.6 | 6.18E-02 | 0.2 | 5.38E-01 | Disease resistance protein family |
| AT1G59218 | | | | | | | -0.3 | 8.22E-01 | -0.6 | 6.94E-01 | | | Probable disease resistance protein RDL6 |
| AT4G25434 | | | | | | | 1.0 | 4.12E-02 | 1.0 | 6.23E-02 | 0.0 | 9.98E-01 | Nudix hydrolase |
| AT1G09500 | | | | | | | 0.7 | 3.42E-01 | 1.0 | 2.07E-01 | | | At1g09500 |
| AT4G24080 | | | | | | | -0.4 | 5.57E-01 | -0.1 | 9.30E-01 | -0.3 | 7.40E-01 | Aldolase like |
| AT1G60750 | | | | | | | 2.1 | 8.93E-03 | 1.4 | 1.15E-01 | | | Probable aldo-keto reductase 6 |
| AT1G63710 | | | | | | | 0.4 | 5.57E-01 | 0.8 | 2.09E-01 | | | Cytochrome P450 86A7 |
| AT1G32350 | | | | | | | 2.3 | 2.18E-03 | -2.7 | 6.18E-02 | 4.9 | 2.97E-03 | Ubiquinol oxidase |
| AT4G15210 | | | | | | | 1.416441047 | 3.93E-01 | 3.264402639 | 4.11E-02 | -1.8 | 2.22E-01 | Beta-amylase |
| AT3G10290 | | | | | | | 0.5 | 2.49E-01 | -0.3 | 6.08E-01 | 0.8 | 1.04E-01 | Probable sugar phosphate |
| AT1G07290 | | | | | | | -0.1 | 8.88E-01 | 0.3 | 7.05E-01 | -0.4 | 5.63E-01 | Golgi nucleotide sugar transporter 2 |
| AT2G34360 | | | | | | | -0.2 | 7.66E-01 | 0.4 | 4.20E-01 | -0.6 | 3.08E-01 | MATE efflux family protein |
| AT5G55340 | | | | | | | 0.6 | 1.87E-01 | 0.1 | 9.30E-01 | 0.6 | 3.79E-01 | Probable long-chain-alcohol O-fatty-acyltransferase 5 |
| AT4G37990 | | | | | | | -1.2 | 6.75E-01 | -1.1 | 7.57E-01 | -0.1 | 9.87E-01 | Cinnamyl alcohol dehydrogenase 8 |
| AT1G62290 | | | | | | | -0.2 | 5.84E-01 | -0.3 | 2.84E-01 | 0.2 | 6.33E-01 | Saposin-like aspartyl protease family protein |
| AT5G40000 | | | | | | | 2.764887818 | 6.05E-04 | 2.368225572 | 4.97E-03 | 0.4 | 4.41E-01 | AAA-ATPase At5g40000 |
| AT3G22910 | | | | | | | 0.8 | 1.70E-01 | -0.3 | 7.12E-01 | 1.1 | 1.53E-01 | Putative calcium-transporting ATPase 13, plasma membrane-type -ATPase isoform 13) |
| AT1G24290 | | | | | | | -0.2 | 6.84E-01 | -0.2 | 8.01E-01 | 0.0 | 9.46E-01 | AAA-type ATPase family protein |
| AT3G05740 | | | | | | | 0.2 | 7.64E-01 | 0.4 | 4.33E-01 | -0.3 | 6.82E-01 | ATP-dependent DNA helicase |
| AT1G54680 | | | | | | | -0.4 | 3.54E-01 | -0.5 | 3.14E-01 | 0.1 | 8.87E-01 | Uncharacterized protein At1g54680 |
| AT1G61550 | | | | | | | 2.0 | 2.16E-03 | 0.9 | 2.22E-01 | 1.1 | 5.41E-02 | G-type lectin S-receptor-like serine |
| AT5G65600 | | | | | | | 2.0 | 5.47E-05 | 0.8 | 1.13E-01 | 1.2 | 3.83E-03 | Protein kinase domain-containing protein |
| AT3G59750 | | | | | | | 1.0 | 1.28E-01 | 1.1 | 1.45E-01 | | | Putative L-type lectin-domain containing receptor kinase V.8 |
| AT2G29220 | | | | | | | 5.2 | 4.69E-06 | 3.0 | 3.21E-03 | 2.2 | 2.07E-04 | Probable inactive L-type lectin-domain containing receptor kinase III.1 |
| AT2G29250 | | | | | | | 6.041807988 | 7.30E-05 | 3.053792066 | 4.17E-02 | 3.0 | 8.65E-04 | Putative inactive L-type lectin-domain containing receptor kinase III.2 |
| AT1G73690 | | | | | | | 0.5 | 1.94E-01 | 0.5 | 2.30E-01 | 0.0 | 9.86E-01 | Cyclin-dependent kinase D-1 |
| AT3G13170 | | | | | | | 0.3 | 6.69E-01 | 0.2 | 8.14E-01 | | | Spo11 |
| AT4G13000 | | | | | | | 0.0 | 9.72E-01 | 0.1 | 9.44E-01 | | | AGC kinase family protein |
| AT5G40030 | | | | | | | -0.2 | 7.18E-01 | 0.4 | 3.78E-01 | -0.5 | 2.07E-01 | Protein kinase |
| AT1G05100 | | | | | | | -0.5 | 3.89E-01 | -0.2 | 8.11E-01 | -0.3 | 7.02E-01 | Mitogen-activated protein kinase kinase kinase 18 |
| AT3G01200 | | | | | | | -0.3 | 3.86E-01 | -0.2 | 7.21E-01 | -0.2 | 7.78E-01 | Pyruvate, phosphate dikinase regulatory protein 2 |
| AT2G26700 | | | | | | | 0.1 | 8.13E-01 | 0.1 | 8.17E-01 | 0.0 | 9.80E-01 | Protein kinase domain-containing protein |
| AT3G57760 | | | | | | | -0.1 | 8.12E-01 | -0.3 | 5.63E-01 | 0.2 | 7.98E-01 | Non-functional pseudokinase ZRK6 |
| AT5G06940 | | | | | | | -0.3 | 5.65E-01 | -0.1 | 8.29E-01 | -0.2 | 8.55E-01 | Probably inactive leucine-rich repeat receptor-like protein kinase At5g06940 |
| AT5G62230 | | | | | | | 0.3 | 4.90E-01 | 0.2 | 7.26E-01 | 0.1 | 8.50E-01 | ERECTA-like 1 |
| AT1G70430 | | | | | | | -0.3 | 4.16E-01 | 0.1 | 8.26E-01 | | | Protein kinase domain-containing protein |
| AT1G51810 | | | | | | | -1.200121848 | 4.14E-02 | 0.013610767 | 9.87E-01 | -1.2 | 6.56E-02 | Probable LRR receptor-like serine |
| AT3G24400 | | | | | | | 1.7 | 9.07E-05 | 0.6 | 1.47E-01 | 1.1 | 3.80E-03 | Proline-rich receptor-like protein kinase PERK2 |

| | | | | | | | |
|---|---|---|---|---|---|---|---|
| AT3G53840 | -0.4 | 4.59E-01 | 0.2 | 6.72E-01 | -0.6 | 2.43E-01 | Wall-associated receptor kinase-like 15 |
| AT3G57770 | -0.4 | 2.49E-01 | -0.3 | 4.54E-01 | -0.1 | 8.67E-01 | Serine |
| AT1G56130 | 0.8 | 4.65E-02 | 0.4 | 3.45E-01 | 0.3 | 4.58E-01 | Leucine-rich repeat transmembrane protein kinase |
| AT3G05140 | -0.3 | 5.94E-01 | -0.2 | 7.58E-01 | -0.1 | 9.14E-01 | ROP binding protein kinases 2 |
| AT4G11480 | 2.358887124 | 4.73E-04 | 1.462728343 | 3.41E-02 | 0.9 | 5.34E-02 | Cysteine-rich RLK 32 |
| AT1G61490 | 0.7 | 9.20E-02 | 0.0 | 9.55E-01 | 0.8 | 1.16E-01 | S-locus lectin protein kinase family protein |
| AT4G09012 | 0.3 | 5.25E-01 | -0.2 | 6.73E-01 | 0.5 | 3.53E-01 | Mitochondrial ribosomal protein L27 |
| AT3G60120 | 6.772965766 | 7.31E-05 | 5.014092659 | 3.38E-03 | 1.8 | 2.72E-03 | Beta-glucosidase 27 |
| AT5G28510 | 2.989171254 | 2.19E-03 | 0.364058215 | 8.19E-01 | | | Beta-glucosidase 24 |
| AT1G50090 | 2.1 | 2.02E-04 | 1.8 | 2.16E-03 | 0.3 | 2.97E-01 | Putative branched-chain-amino-acid aminotransferase 7 |
| AT2G14920 | -0.3 | 6.07E-01 | -0.1 | 9.39E-01 | -0.2 | 6.91E-01 | Cytosolic sulfotransferase 10 |
| AT5G58660 | -0.2 | 7.03E-01 | -0.7 | 1.63E-01 | 0.5 | 3.20E-01 | 2-oxoglutarate and Fe-dependent oxygenase superfamily protein |
| AT1G54530 | -0.4 | 3.84E-01 | 0.3 | 4.63E-01 | -0.7 | 1.81E-01 | EF-hand domain-containing protein |
| AT3G01830 | -0.005833681 | 9.96E-01 | 0.023134102 | 9.86E-01 | | | Probable calcium-binding protein CML40 |
| AT3G24110 | 0.0 | 9.39E-01 | 0.1 | 9.17E-01 | -0.1 | 8.74E-01 | Calcium-binding EF-hand family protein |
| AT5G46830 | 0.0 | 9.69E-01 | 0.6 | 5.26E-01 | | | Transcription factor bHLH28 |
| AT1G52570 | 0.2 | 6.40E-01 | 0.1 | 8.54E-01 | | | Phospholipase D alpha 2 |
| AT4G21490 | 0.5 | 6.90E-01 | 0.3 | 8.74E-01 | | | NADH:ubiquinone reductase |
| AT3G51380 | -0.7 | 1.54E-01 | 0.4 | 3.98E-01 | | | Protein IQ-DOMAIN 20 |
| AT5G28650 | 0.6 | 1.40E-01 | 0.0 | 9.88E-01 | 0.6 | 1.70E-01 | Probable WRKY transcription factor 74 |
| AT1G16905 | -0.2 | 8.23E-01 | 0.1 | 9.30E-01 | | | F6I1.9 protein |
| AT1G53060 | 0.0 | 9.54E-01 | 0.1 | 8.20E-01 | -0.1 | 8.75E-01 | Lectin-like protein At1g53060 |
| AT1G56250 | 2.3 | 8.19E-03 | 0.5 | 6.63E-01 | 1.7 | 5.43E-02 | F-box protein VBF |
| AT1G58160 | 0.1 | 8.82E-01 | -0.2 | 6.79E-01 | 0.3 | 5.07E-01 | Jacalin-related lectin 15 |
| AT2G02310 | -0.3 | 5.52E-01 | 0.0 | 9.65E-01 | -0.4 | 6.11E-01 | Putative F-box protein PP2-B6 |
| AT5G18470 | 1.0 | 1.05E-01 | 0.3 | 7.53E-01 | 0.7 | 2.89E-01 | Bulb-type lectin domain-containing protein |
| AT5G35940 | -1.1 | 9.87E-02 | -0.7 | 2.86E-01 | -0.3 | 7.04E-01 | Jacalin-related lectin 41 |
| AT2G18480 | 4.154471232 | 1.79E-05 | 1.874452259 | 3.93E-02 | 2.3 | 1.25E-03 | Probable polyol transporter 3 |
| AT2G28210 | 0.2 | 4.38E-01 | 0.2 | 5.05E-01 | 0.0 | 9.98E-01 | carbonic anhydrase |
| AT1G02800 | 0.1 | 9.10E-01 | 0.1 | 9.00E-01 | | | Endoglucanase 1 |
| AT2G43600 | -0.2 | 6.55E-01 | 0.4 | 4.57E-01 | -0.6 | 2.78E-01 | Chitinase family protein |
| AT4G15480 | 1.7 | 3.43E-02 | 2.3 | 9.14E-03 | | | UDP-glycosyltransferase 84A1 |
| AT5G05390 | 1.868994307 | 1.46E-03 | 1.509185879 | 1.46E-02 | 0.4 | 4.87E-01 | Laccase-12 |
| AT1G49620 | -0.1 | 8.91E-01 | 0.1 | 8.01E-01 | -0.2 | 7.26E-01 | Cyclin-dependent kinase inhibitor |
| AT2G17620 | 0.2 | 5.54E-01 | 0.6 | 8.19E-02 | | | Cyclin-B2-1 |
| AT5G39890 | 0.3 | 4.78E-01 | 0.2 | 7.82E-01 | | | cysteine dioxygenase |
| AT5G50260 | -0.8 | 2.39E-01 | 0.7 | 3.14E-01 | | | KDEL-tailed cysteine endopeptidase CEP1 |
| AT2G46760 | 0.0 | 9.39E-01 | 0.3 | 6.23E-01 | -0.3 | 6.32E-01 | Probable L-gulonolactone oxidase 6 |
| AT5G56490 | -0.6 | 2.28E-01 | -0.2 | 6.92E-01 | -0.3 | 6.55E-01 | Probable L-gulonolactone oxidase 4 |
| AT1G48980 | 2.496238212 | 2.38E-04 | 1.033438583 | 1.46E-01 | 1.5 | 5.61E-03 | F27J15.23 |
| AT2G18620 | 0.4 | 5.94E-01 | 1.5 | 1.17E-02 | -1.2 | 4.11E-02 | Geranylgeranyl pyrophosphate synthase 7, chloroplastic -farnesyl diphosphate synthase 7) |
| AT4G35810 | 0.8 | 9.94E-02 | 0.0 | 9.59E-01 | 0.7 | 8.84E-02 | Fe2OG dioxygenase domain-containing protein |
| AT1G16640 | -0.1 | 8.97E-01 | -0.2 | 7.00E-01 | 0.2 | 8.22E-01 | B3 domain-containing protein At1g16640 |
| AT1G65910 | -0.7 | 2.16E-01 | -0.1 | 9.41E-01 | -0.6 | 3.09E-01 | NAC028 |
| AT2G24700 | -0.4 | 3.08E-01 | 0.0 | 9.92E-01 | -0.4 | 3.83E-01 | B3 domain-containing protein REM10 |
| AT2G30130 | 0.2 | 7.24E-01 | -0.1 | 8.88E-01 | 0.2 | 6.49E-01 | LOB domain-containing protein 12 |
| AT3G06160 | -0.591435051 | 4.43E-02 | -0.03207038 | 9.39E-01 | -0.6 | 1.24E-01 | AP2 |
| AT3G18960 | -0.5 | 9.22E-02 | -0.8 | 3.97E-02 | 0.2 | 6.78E-01 | AP2 |
| AT4G31620 | -0.3 | 4.35E-01 | 0.0 | 9.90E-01 | -0.3 | 5.30E-01 | TF-B3 domain-containing protein |
| AT4G36160 | -0.1 | 9.07E-01 | 0.3 | 3.94E-01 | -0.4 | 4.10E-01 | NAC domain containing protein 76 |
| AT5G05330 | -0.5 | 1.57E-01 | -0.3 | 5.55E-01 | -0.3 | 6.32E-01 | HMG box domain-containing protein |
| AT1G02230 | 0.5 | 2.87E-01 | 0.2 | 7.70E-01 | 0.3 | 6.23E-01 | NAC domain-containing protein 4 |
| AT1G28360 | -0.1 | 8.38E-01 | 0.0 | 9.70E-01 | -0.1 | 9.22E-01 | Ethylene-responsive transcription factor 12 |
| AT1G51190 | 0.2 | 7.20E-01 | 0.1 | 8.71E-01 | 0.1 | 9.17E-01 | AP2-like ethylene-responsive transcription factor PLT2 |
| AT1G68120 | 0.042540542 | 9.44E-01 | 0.371949842 | 4.99E-01 | -0.3 | 6.08E-01 | GAGA-binding transcriptional activator |
| AT1G71520 | -0.01710607 | 9.89E-01 | -2.164814473 | 1.15E-01 | | | Ethylene-responsive transcription factor ERF020 |
| AT1G75490 | -1.0 | 1.84E-01 | -2.0 | 3.80E-02 | 1.0 | 4.06E-01 | At1g75490 |
| AT3G12910 | 2.3 | 4.47E-04 | 0.7 | 3.81E-01 | 1.7 | 8.79E-03 | NAC domain transcriptional regulator superfamily protein |
| AT3G18010 | -0.4 | 1.08E-01 | -0.5 | 4.65E-02 | 0.1 | 7.02E-01 | WUSCHEL-related homeobox 1 |
| AT4G17785 | 1.8 | 1.66E-03 | 1.1 | 6.40E-02 | 0.7 | 2.06E-01 | Transcription factor MYB39 |
| AT5G11190 | -0.1 | 8.09E-01 | -0.3 | 6.77E-01 | 0.1 | 8.88E-01 | Ethylene-responsive transcription factor SHINE 2 |
| AT5G37260 | 0.0 | 8.48E-01 | -0.7 | 5.46E-04 | 0.7 | 3.00E-03 | Protein REVEILLE 2 |
| AT5G59570 | -0.438780179 | 3.95E-03 | -0.43204822 | 8.04E-03 | 0.0 | 9.81E-01 | Transcription factor BOA |
| AT5G64750 | 0.4 | 6.34E-01 | 0.2 | 8.63E-01 | | | Ethylene-responsive transcription factor ABR1 |
| AT1G14687 | 0.1 | 7.60E-01 | -0.4 | 4.62E-01 | 0.5 | 3.70E-01 | Zinc-finger homeodomain protein 14 |
| AT2G02540 | 0.4 | 2.72E-01 | 0.1 | 8.66E-01 | 0.3 | 5.38E-01 | Zinc-finger homeodomain protein 3 |
| AT2G44840 | 0.3 | 6.39E-01 | 0.0 | 9.89E-01 | 0.3 | 7.26E-01 | Ethylene-responsive transcription factor 13 |
| AT3G23240 | 0.9 | 1.95E-01 | 0.0 | 9.72E-01 | 0.9 | 1.76E-01 | Ethylene-responsive transcription factor 1B |
| AT2G41340 | -0.1 | 7.49E-01 | -0.7 | 3.12E-02 | 0.6 | 9.58E-02 | RNA polymerase II fifth largest subunit, D |
| AT2G15400 | 0.0 | 9.77E-01 | -0.2 | 6.20E-01 | 0.2 | 6.58E-01 | DNA-directed RNA polymerase family protein |
| AT5G52170 | 0.3 | 6.59E-01 | 0.6 | 4.05E-01 | -0.3 | 7.66E-01 | Homeodomain GLABROUS 7 |
| AT5G63690 | 0.2 | 6.04E-01 | -0.4 | 3.16E-01 | 0.5 | 1.57E-01 | Uncharacterized protein |
| AT5G56960 | 2.5 | 1.71E-05 | 1.4 | 1.13E-02 | 1.2 | 1.03E-02 | Basic helix-loop-helix DNA-binding family protein |
| AT4G13570 | 0.8 | 1.59E-01 | 0.8 | 1.79E-01 | 0.0 | 9.66E-01 | Histone H2A |
| AT5G65350 | 0.4 | 2.71E-01 | 0.5 | 1.46E-01 | -0.2 | 7.66E-01 | Histone H3-like 5 |
| AT5G49110 | -0.5 | 2.87E-01 | -0.1 | 9.44E-01 | | | Fanconi anemia group I-like protein |
| AT1G75090 | -0.5 | 3.04E-01 | -0.1 | 9.23E-01 | -0.4 | 5.04E-01 | hypothetical protein |
| AT3G50870 | 0.0 | 9.98E-01 | 0.2 | 6.33E-01 | -0.2 | 6.72E-01 | GATA transcription factor 18 |
| AT1G16070 | 0.6 | 2.64E-01 | 1.3 | 1.44E-02 | | | Tubby like protein 8 |
| AT3G58070 | 0.4 | 4.82E-01 | 1.0 | 1.01E-01 | -0.6 | 4.42E-01 | C2H2-type domain-containing protein |
| AT5G60910 | 0.2 | 7.71E-01 | 0.6 | 2.60E-01 | -0.4 | 4.96E-01 | AGAMOUS-like 8 |
| AT1G47760 | 0.5 | 4.52E-01 | 0.5 | 5.05E-01 | | | AGAMOUS-like 102 |
| AT4G20970 | -1.636825776 | 4.10E-01 | 0.531733644 | 7.93E-01 | -2.2 | 3.76E-01 | Transcription factor bHLH162 |

| | | | | | | | |
|---|---|---|---|---|---|---|---|
| AT3G46080 | 0.5 | 5.10E-01 | -1.0 | 3.01E-01 | | | Zinc finger protein ZAT8 |
| AT3G57670 | -0.1 | 8.97E-01 | 0.2 | 7.44E-01 | -0.3 | 7.01E-01 | Zinc finger protein WIP2 |
| AT2G37430 | 2.0 | 2.95E-04 | 0.7 | 2.78E-01 | 1.4 | 1.57E-02 | Zinc finger protein ZAT11 |
| AT5G13790 | 0.3 | 6.09E-01 | 0.1 | 9.26E-01 | 0.2 | 8.13E-01 | AGAMOUS-like 15 |
| AT4G00870 | -0.879006809 | 3.00E-01 | 0.81754191 | 2.83E-01 | | | Transcription factor bHLH14 |
| AT5G13080 | 1.2 | 1.41E-02 | -0.6 | 4.06E-01 | 1.8 | 9.63E-03 | Probable WRKY transcription factor 75 |
| AT1G29860 | -0.6 | 2.81E-01 | -1.4 | 3.71E-02 | | | WRKY transcription factor |
| AT2G40740 | 0.9 | 1.08E-02 | 0.3 | 5.95E-01 | 0.7 | 1.03E-01 | WRKY DNA-binding protein 55 |
| AT3G22830 | -0.796496589 | 7.49E-02 | 0.156775046 | 7.67E-01 | -1.0 | 9.02E-02 | HSFA6B |
| AT3G49950 | 0.009187931 | 9.85E-01 | -0.239968871 | 5.94E-01 | 0.2 | 6.43E-01 | Scarecrow-like protein 32 |
| AT5G01900 | 2.2 | 3.65E-03 | 1.5 | 8.20E-02 | 0.8 | 2.85E-01 | Probable WRKY transcription factor 62 |
| AT5G03720 | 0.6 | 3.83E-02 | -0.2 | 4.98E-01 | 0.8 | 2.58E-02 | HSFA3 |
| AT2G47520 | 5.6 | 5.58E-04 | 2.8 | 1.04E-01 | 2.8 | 1.15E-03 | Ethylene-responsive transcription factor ERF071 |
| AT1G74080 | 1.1 | 7.51E-03 | 0.5 | 2.77E-01 | 0.6 | 1.28E-01 | Transcription factor MYB122 |
| AT2G31180 | 0.9 | 1.01E-01 | 0.4 | 6.16E-01 | 0.5 | 3.04E-01 | Transcription factor MYB14 |
| AT4G11140 | 0.4 | 3.04E-01 | 0.1 | 8.64E-01 | 0.3 | 4.81E-01 | Ethylene-responsive transcription factor CRF1 |
| AT5G61890 | -0.431963164 | 8.13E-01 | -0.250966897 | 9.16E-01 | -0.2 | 9.53E-01 | Ethylene-responsive transcription factor ERF114 |
| AT4G39070 | -0.3 | 2.97E-01 | -0.2 | 5.30E-01 | -0.1 | 8.40E-01 | B-box zinc finger protein 20 |
| AT1G70920 | 1.7 | 3.49E-03 | 0.9 | 2.10E-01 | 0.9 | 1.08E-01 | Homeobox-leucine zipper protein 18 |
| AT1G78930 | 0.8 | 4.84E-02 | -0.2 | 8.17E-01 | 1.0 | 3.85E-02 | At1g78930 |
| AT3G18870 | -0.5 | 2.03E-01 | -0.3 | 5.33E-01 | -0.2 | 7.16E-01 | At3g18870 |
| AT5G23930 | 0.5 | 2.03E-01 | 0.1 | 8.08E-01 | 0.4 | 4.52E-01 | Mitochondrial transcription termination factor family protein |
| AT5G45040 | -0.1 | 7.76E-01 | -0.1 | 8.59E-01 | 0.0 | 9.70E-01 | Cytochrome c-553 |
| AT1G30010 | -0.7 | 2.66E-01 | 0.1 | 9.03E-01 | -0.9 | 2.91E-01 | Nuclear intron maturase 1, mitochondrial |
| AT5G64710 | 0.4 | 3.26E-01 | 0.8 | 8.04E-02 | -0.4 | 4.40E-01 | Emb\|CAB71880.1 |
| AT5G25500 | 0.2 | 5.84E-01 | -0.2 | 7.86E-01 | 0.4 | 4.81E-01 | Exosome complex exonuclease |
| AT2G37870 | -1.192863085 | 3.90E-01 | -0.126834068 | 9.46E-01 | -1.1 | 5.55E-01 | Bifunctional inhibitor |
| AT4G13260 | -0.3 | 5.22E-01 | 0.1 | 9.38E-01 | | | Flavin-containing monooxygenase |
| AT1G49880 | -0.2 | 5.91E-01 | -0.4 | 3.48E-01 | 0.2 | 7.16E-01 | Sulfhydryl oxidase |
| AT4G16820 | 0.499429874 | 4.77E-01 | -0.44933521 | 6.44E-01 | | | Phospholipase A1-Ibeta2, chloroplastic |
| AT1G22380 | -0.085113632 | 8.14E-01 | 0.235872232 | 5.00E-01 | -0.3 | 4.08E-01 | UDP-glycosyltransferase 85A3 |
| AT1G03980 | 1.180951848 | 1.31E-01 | 0.864392797 | 3.59E-01 | 0.3 | 7.48E-01 | glutathione gamma-glutamylcysteinyltransferase |
| AT1G69920 | 5.606312222 | 1.03E-05 | 4.062615406 | 8.84E-04 | 1.5 | 8.79E-04 | Glutathione S-transferase |
| AT1G69930 | -0.388097054 | 7.07E-01 | -2.234297499 | 8.85E-02 | | | Glutathione S-transferase U11 |
| AT2G29420 | -0.1 | 7.66E-01 | -0.4 | 3.98E-01 | 0.3 | 6.72E-01 | Glutathione S-transferase U7 |
| AT2G29440 | 0.5 | 2.96E-01 | 0.1 | 8.76E-01 | 0.4 | 4.71E-01 | Glutathione S-transferase U6 |
| AT1G75110 | -0.4 | 3.63E-01 | -0.2 | 7.54E-01 | -0.2 | 7.55E-01 | Arabinosyltransferase RRA2 |
| AT4G19970 | 2.0 | 4.81E-04 | 0.1 | 9.42E-01 | 1.9 | 6.65E-04 | Glycosyltransferase |
| AT4G37420 | 2.4 | 6.05E-03 | -0.5 | 7.51E-01 | | | Glycosyltransferase family 92 protein |
| AT5G25970 | 0.586942461 | 2.65E-01 | 0.269164823 | 7.08E-01 | 0.3 | 6.49E-01 | T1N24.19 protein |
| AT3G02240 | 1.0 | 1.30E-02 | 0.4 | 3.55E-01 | 0.5 | 2.21E-01 | Protein GOLVEN 4 -related protein CLEL4) |
| AT2G44690 | 0.3 | 5.94E-01 | 0.7 | 1.48E-01 | -0.4 | 4.50E-01 | hypothetical protein |
| AT1G10300 | 0.0 | 9.86E-01 | -0.1 | 8.65E-01 | 0.1 | 8.63E-01 | Nucleolar GTP-binding protein 1 |
| AT3G17660 | -0.2 | 6.72E-01 | -0.4 | 3.54E-01 | 0.2 | 6.99E-01 | Arf-GAP domain-containing protein |
| AT5G50550 | -0.184383117 | 9.31E-01 | 0.244316723 | 9.19E-01 | -0.4 | 8.64E-01 | SEC12-like protein |
| AT1G01700 | -0.4 | 3.36E-01 | 0.1 | 8.06E-01 | | | RHO guanyl-nucleotide exchange factor 2 |
| AT2G47770 | -3.099456293 | 1.62E-01 | -1.735446442 | 3.66E-01 | -1.4 | 7.46E-01 | Translocator protein homolog |
| AT1G34540 | -4.3 | 1.08E-03 | -0.4 | 5.40E-01 | -3.9 | 2.31E-02 | Cytochrome P450 family protein |
| AT2G45570 | -1.7 | 9.43E-02 | -2.3 | 5.97E-02 | 0.6 | 7.77E-01 | Cytochrome P450 76C2 |
| AT3G28740 | 0.531005633 | 7.92E-01 | -1.082308221 | 6.92E-01 | 1.6 | 5.95E-01 | Uncharacterized protein |
| AT4G15310 | 1.590455412 | 3.62E-02 | 1.987531604 | 1.56E-02 | -0.4 | 5.79E-01 | Cytochrome P450, family 702, subfamily A, polypeptide 3 |
| AT1G14550 | 2.212907455 | 1.45E-03 | -0.568007959 | 5.86E-01 | 2.8 | 2.72E-03 | Peroxidase 5 |
| AT3G55840 | -0.5 | 3.28E-01 | -0.9 | 1.04E-01 | 0.4 | 5.72E-01 | Nematode resistance protein-like HSPRO1 |
| AT1G11730 | -0.3 | 4.37E-01 | -0.3 | 5.05E-01 | 0.0 | 9.92E-01 | Hexosyltransferase |
| AT5G57500 | 0.1 | 7.65E-01 | 0.1 | 7.91E-01 | 0.0 | 9.91E-01 | Hexosyltransferase |
| AT1G29290 | 1.2 | 4.66E-03 | 0.2 | 7.80E-01 | 1.0 | 3.80E-02 | Precursor of CEP14 |
| AT3G50610 | -1.444551791 | 8.90E-02 | -0.171437779 | 8.63E-01 | | | Precursor of CEP9 |
| AT5G66815 | -0.930928657 | 1.43E-01 | 0.169047223 | 8.25E-01 | | | Precursor of CEP5 |
| AT5G38200 | 4.131284708 | 1.11E-06 | 0.965715884 | 2.01E-01 | 3.2 | 1.22E-05 | Gb\|AAF31027.1 |
| AT1G54020 | 0.343913032 | 8.41E-01 | 1.195504404 | 4.40E-01 | -0.9 | 6.65E-01 | GDSL-like Lipase |
| AT4G16230 | 0.009112258 | 9.94E-01 | 0.714525069 | 4.00E-01 | -0.7 | 5.01E-01 | GDSL esterase |
| AT2G23590 | 0.3 | 5.07E-01 | 0.2 | 6.20E-01 | 0.0 | 9.66E-01 | Methylesterase 8 |
| AT3G60140 | -1.9 | 8.90E-02 | -4.5 | 2.18E-02 | 2.6 | 2.33E-01 | Glycosyl hydrolase superfamily protein |
| AT5G57550 | -0.9 | 6.63E-03 | -1.6 | 4.92E-04 | 0.7 | 8.19E-02 | Probable xyloglucan endotransglucosylase |
| AT2G14620 | 0.2 | 7.09E-01 | -0.1 | 9.37E-01 | 0.3 | 6.72E-01 | Xyloglucan endotransglucosylase |
| AT5G06760 | -3.2 | 1.52E-01 | -0.7 | 7.13E-01 | -2.5 | 4.54E-01 | Late embryogenesis abundant protein 46 |
| AT1G05440 | 0.3 | 5.99E-01 | -0.2 | 8.25E-01 | 0.5 | 4.92E-01 | C-8 sterol isomerase |
| AT1G58150 | -1.0 | 1.64E-03 | -0.2 | 5.17E-01 | -0.8 | 2.97E-02 | Phosphoglycerate kinase |
| AT5G66800 | -0.8 | 3.44E-02 | -0.6 | 1.19E-01 | -0.2 | 7.87E-01 | Emb\|CAB62433.1\| |
| AT1G65890 | -0.014232817 | 9.94E-01 | 1.879099709 | 1.21E-01 | -1.9 | 2.00E-01 | Probable acyl-activating enzyme 12, peroxisomal |
| AT1G53990 | 1.4 | 1.17E-03 | 0.5 | 3.32E-01 | 0.9 | 1.75E-02 | GDSL esterase |
| AT5G40990 | 4.179661958 | 8.39E-07 | 2.724826744 | 3.87E-04 | 1.5 | 7.58E-04 | GDSL esterase |
| AT5G59310 | -2.783099852 | 2.99E-01 | -3.057184484 | 3.29E-01 | 0.3 | 9.69E-01 | Non-specific lipid-transfer protein |
| AT5G59330 | 0.3 | 5.80E-01 | -0.2 | 7.78E-01 | 0.6 | 2.11E-01 | Bifunctional inhibitor |
| AT1G36640 | 2.0 | 1.11E-02 | 1.1 | 2.57E-01 | 0.9 | 1.78E-01 | Serine rich endogenous peptide 20 |
| AT5G44567 | -1.253720059 | 1.62E-01 | -2.518140898 | 3.79E-02 | 1.3 | 4.66E-01 | Serine rich endogenous peptide 2 |
| AT5G51465 | 1.1 | 4.40E-02 | -1.0 | 2.04E-01 | 2.1 | 6.93E-03 | Serine rich endogenous peptide 18 |
| AT1G80160 | -1.267823095 | 3.29E-01 | -1.726689779 | 2.58E-01 | 0.5 | 8.64E-01 | F18B13.24 protein |
| AT5G41650 | 0.866115974 | 6.63E-02 | 0.265339496 | 6.92E-01 | 0.6 | 2.57E-01 | Gb\|AAF24581.1 |
| AT3G45590 | 0.0 | 9.37E-01 | -0.1 | 8.42E-01 | 0.2 | 8.03E-01 | tRNA-splicing endonuclease subunit Sen2-1 |
| AT3G15720 | 0.7 | 2.61E-01 | 1.2 | 4.99E-02 | | | Pectin lyase-like superfamily protein |
| AT4G20050 | 0.5 | 3.29E-01 | 0.3 | 6.71E-01 | | | Pectin lyase-like superfamily protein |
| AT5G09710 | 0.4 | 3.40E-01 | 0.0 | 9.80E-01 | 0.4 | 4.59E-01 | Putative magnesium transporter MRS2-9 |

| | | | | | | | | | | | | | |
|---|---|---|---|---|---|---|---|---|---|---|---|---|---|
| AT5G03860 | | | | | | | -0.3 | 5.63E-01 | -1.1 | 5.87E-02 | 0.8 | 2.47E-01 | Malate synthase |
| AT5G39100 | | | | | | | 1.777224002 | 4.52E-03 | 0.996197476 | 1.53E-01 | 0.8 | 1.51E-01 | Germin-like protein subfamily 1 member 13 |
| AT5G39110 | | | | | | | 1.068142303 | 5.74E-02 | 0.200907383 | 8.20E-01 | 0.9 | 1.55E-01 | Germin-like protein subfamily 1 member 14 |
| AT5G39130 | | | | | | | 2.0 | 3.51E-04 | 1.1 | 4.28E-02 | 0.8 | 1.24E-02 | Germin-like protein subfamily 1 member 16 |
| AT5G39150 | | | | | | | 0.131421656 | 8.97E-01 | -0.189627766 | 8.76E-01 | 0.3 | 7.64E-01 | Germin-like protein subfamily 1 member 17 |
| AT5G39160 | | | | | | | 1.0 | 2.88E-02 | 0.5 | 3.29E-01 | 0.5 | 3.38E-01 | Germin-like protein |
| AT5G39180 | | | | | | | 0.720455751 | 4.19E-01 | -0.038441545 | 9.81E-01 | | | Germin-like protein subfamily 1 member 19 |
| AT3G58060 | | | | | | | -0.1 | 9.20E-01 | -1.2 | 1.67E-01 | | | Putative metal tolerance protein C3 |
| AT1G06330 | | | | | | | 1.495728668 | 3.42E-02 | 1.214980467 | 1.28E-01 | | | Heavy metal transport |
| AT1G67180 | | | | | | | 0.7 | 2.08E-01 | 0.7 | 2.89E-01 | | | RING-type E3 ubiquitin transferase BRCA1 |
| AT1G70650 | | | | | | | 0.3 | 5.13E-01 | 0.1 | 8.64E-01 | 0.2 | 7.27E-01 | Ran BP2 |
| AT3G17460 | | | | | | | -0.1 | 7.05E-01 | -0.3 | 4.24E-01 | 0.2 | 7.56E-01 | Uncharacterized protein |
| AT3G27500 | | | | | | | 0.1 | 8.68E-01 | 0.5 | 5.01E-01 | -0.3 | 6.17E-01 | Cysteine |
| AT3G53530 | | | | | | | -0.4 | 3.40E-01 | -0.5 | 2.91E-01 | 0.1 | 9.05E-01 | Protein SODIUM POTASSIUM ROOT DEFECTIVE 3 |
| AT4G10600 | | | | | | | 0.6 | 9.96E-02 | 0.4 | 4.14E-01 | | | RING |
| AT4G16807 | | | | | | | 0.3 | 4.11E-01 | 0.1 | 8.50E-01 | | | Uncharacterized protein AT4g16810 |
| AT4G38410 | | | | | | | 0.91052025 | 1.80E-01 | 0.938101142 | 2.10E-01 | 0.0 | 9.80E-01 | At4g38410 |
| AT5G05365 | | | | | | | 0.7 | 2.56E-01 | 0.3 | 6.92E-01 | | | hypothetical protein |
| AT5G41980 | | | | | | | -0.1 | 7.49E-01 | -0.2 | 5.94E-01 | 0.1 | 8.56E-01 | hypothetical protein |
| AT5G42840 | | | | | | | -1.0 | 3.85E-02 | -0.1 | 8.71E-01 | -0.9 | 1.39E-01 | CHP-rich zinc finger protein-like |
| AT5G49120 | | | | | | | -0.3 | 5.54E-01 | 0.1 | 8.93E-01 | | | FCS-Like Zinc finger 15 |
| AT5G52720 | | | | | | | 2.5 | 7.82E-04 | 1.6 | 4.70E-02 | 1.0 | 6.10E-02 | Copper transport protein family |
| AT1G19640 | | | | | | | -0.2 | 8.12E-01 | 0.9 | 9.46E-02 | -1.1 | 1.20E-01 | Jasmonate O-methyltransferase |
| AT1G07160 | | | | | | | 1.4 | 2.90E-02 | 0.7 | 3.96E-01 | | | Probable protein phosphatase 2C 2 |
| AT2G40180 | | | | | | | 1.2 | 2.30E-02 | -0.2 | 8.59E-01 | | | Probable protein phosphatase 2C 30 |
| AT4G21590 | | | | | | | -0.3 | 5.88E-01 | 0.4 | 4.92E-01 | -0.7 | 2.57E-01 | Aspergillus nuclease S1 |
| AT3G55970 | | | | | | | -3.065697787 | 7.62E-02 | -0.425673703 | 7.92E-01 | -2.6 | 2.78E-01 | Jasmonate-regulated gene 21 |
| AT5G55720 | | | | | | | 0.5 | 3.97E-01 | -0.1 | 8.90E-01 | 0.6 | 2.23E-01 | Putative pectate lyase 21 |
| AT2G24350 | | | | | | | -0.5 | 4.04E-01 | 0.1 | 8.46E-01 | -0.6 | 3.89E-01 | At2g24350 binding protein) family protein) |
| AT5G58610 | | | | | | | 1.1 | 1.41E-01 | 0.3 | 7.91E-01 | | | Gb\|AAC80581.1 |
| AT4G30370 | | | | | | | 0.1 | 8.72E-01 | -0.1 | 8.15E-01 | 0.2 | 7.11E-01 | RING-H2 finger protein ATL14 |
| AT5G05810 | | | | | | | 0.2 | 4.95E-01 | -0.1 | 8.37E-01 | 0.3 | 4.01E-01 | RING-H2 finger protein ATL43 |
| AT5G07040 | | | | | | | -0.5 | 2.66E-01 | -0.4 | 4.38E-01 | -0.1 | 8.79E-01 | Putative RING-H2 finger protein ATL69 |
| AT5G21140 | | | | | | | 0.1 | 8.71E-01 | 0.3 | 6.48E-01 | -0.2 | 7.89E-01 | Non-structural maintenance of chromosomes element 1 homolog |
| AT1G13790 | | | | | | | 0.8 | 4.67E-02 | 0.4 | 3.87E-01 | 0.4 | 4.32E-01 | Factor of DNA methylation 4 |
| AT3G10200 | | | | | | | 0.1 | 7.37E-01 | 0.2 | 7.42E-01 | 0.0 | 9.75E-01 | Methyltransferase |
| AT5G54400 | | | | | | | -0.2 | 7.19E-01 | 0.2 | 6.91E-01 | | | hypothetical protein |
| AT2G38720 | | | | | | | 0.6 | 1.60E-01 | -0.1 | 8.56E-01 | 0.8 | 2.03E-01 | 65-kDa microtubule-associated protein 5 |
| AT2G44190 | | | | | | | 0.0 | 9.89E-01 | 0.4 | 5.06E-01 | -0.4 | 4.95E-01 | Protein ENDOSPERM DEFECTIVE 1 |
| AT3G60660 | | | | | | | 0.4 | 4.34E-01 | 0.1 | 8.81E-01 | 0.3 | 7.08E-01 | Spindle and kinetochore-associated protein 1 homolog |
| AT3G12520 | | | | | | | 0.3 | 4.05E-01 | 0.2 | 6.65E-01 | 0.1 | 8.63E-01 | Sulfate transporter 42 |
| AT3G17690 | | | | | | | 0.6 | 3.84E-01 | 0.3 | 7.44E-01 | 0.3 | 6.52E-01 | Cyclic nucleotide gated channel 19 |
| AT4G21480 | | | | | | | -0.1 | 7.71E-01 | -0.7 | 1.03E-01 | 0.6 | 2.71E-01 | hypothetical protein |
| AT1G09680 | | | | | | | 0.7 | 2.81E-02 | 0.0 | 9.90E-01 | 0.7 | 4.80E-02 | Putative pentatricopeptide repeat-containing protein At1g09680 |
| AT1G16830 | | | | | | | 0.5 | 3.29E-01 | 0.2 | 8.17E-01 | 0.3 | 6.40E-01 | Putative pentatricopeptide repeat-containing protein At1g16830 |
| AT2G15630 | | | | | | | -0.4 | 4.34E-01 | -0.2 | 7.30E-01 | -0.2 | 8.34E-01 | Pentatricopeptide repeat-containing protein At2g15630, mitochondrial |
| AT2G17140 | | | | | | | 0.7 | 1.02E-01 | 0.4 | 4.74E-01 | 0.3 | 5.14E-01 | Pentatricopeptide repeat-containing protein At2g17140 |
| AT2G17525 | | | | | | | 0.0 | 9.85E-01 | -0.1 | 9.64E-01 | 0.1 | 9.49E-01 | Pentatricopeptide repeat-containing protein At2g17525, mitochondrial |
| AT2G39230 | | | | | | | 1.1 | 1.07E-01 | 0.6 | 4.92E-01 | | | Pentatricopeptide repeat-containing protein At2g39230, mitochondrial |
| AT3G07290 | | | | | | | 0.6 | 6.46E-02 | 0.2 | 6.30E-01 | 0.4 | 3.31E-01 | Pentatricopeptide repeat-containing protein At3g07290, mitochondrial |
| AT3G18020 | | | | | | | -0.1 | 9.05E-01 | 0.1 | 9.21E-01 | -0.2 | 8.39E-01 | Pentatricopeptide repeat-containing protein At3g18020 |
| AT4G17915 | | | | | | | 0.3 | 3.68E-01 | -0.1 | 8.11E-01 | 0.5 | 3.44E-01 | Putative pentatricopeptide repeat-containing protein At4g17915 |
| AT5G18475 | | | | | | | 0.0 | 9.49E-01 | -0.2 | 8.00E-01 | 0.2 | 7.74E-01 | Pentatricopeptide repeat-containing protein At5g18475 |
| AT5G38730 | | | | | | | 0.9 | 3.70E-02 | 0.8 | 9.71E-02 | 0.1 | 8.55E-01 | Pentatricopeptide repeat-containing protein At5g38730 |
| AT5G46680 | | | | | | | -0.2 | 6.31E-01 | -0.2 | 6.17E-01 | 0.0 | 9.61E-01 | Pentatricopeptide repeat-containing protein At5g46680 |
| AT5G48650 | | | | | | | -0.2 | 6.87E-01 | 0.0 | 9.68E-01 | -0.1 | 8.07E-01 | Nuclear transport factor 2 family protein with RNA binding domain-containing protein |
| AT5G61370 | | | | | | | 1.1 | 4.43E-03 | 0.5 | 2.34E-01 | 0.6 | 1.47E-01 | Pentatricopeptide repeat-containing protein At5g61370, mitochondrial |
| AT5G64490 | | | | | | | 0.588019134 | 3.27E-01 | -0.21484068 | 8.16E-01 | 0.8 | 2.96E-01 | Pumilio homolog 26 |
| AT2G24140 | | | | | | | -0.3 | 5.30E-01 | 0.0 | 9.60E-01 | | | Expressed protein |
| AT2G37210 | | | | | | | 0.5 | 3.59E-01 | 0.5 | 4.21E-01 | 0.0 | 1.00E+00 | Cytokinin riboside 5'-monophosphate phosphoribohydrolase |
| AT2G19940 | 0.8 | 4.41E-06 | 0.6 | 5.15E-05 | 0.2 | 8.49E-02 | 0.2 | 1.75E-01 | 0.2 | 2.72E-01 | 0.0 | 9.71E-01 | N-acetyl-gamma-glutamyl-phosphate reductase |
| AT1G30540 | 0.1 | 4.71E-01 | -0.1 | 3.54E-01 | 0.1 | 1.62E-01 | 0.1 | 7.42E-01 | 0.0 | 9.31E-01 | 0.0 | 8.93E-01 | N-acetyl-D-glucosamine kinase |
| AT3G58130 | 0.2 | 1.91E-01 | 0.2 | 7.87E-02 | -0.1 | 6.52E-01 | -0.2 | 4.60E-01 | -0.1 | 7.94E-01 | -0.1 | 8.07E-01 | N-acetylglucosaminylphosphatidylinositol deacetylase |
| AT2G27340 | 0.5 | 5.27E-03 | 0.2 | 2.59E-01 | 0.3 | 9.51E-02 | 0.0 | 9.94E-01 | -0.1 | 7.38E-01 | 0.1 | 7.94E-01 | N-acetylglucosaminylphosphatidylinositol deacetylase |
| AT2G39020 | 0.3 | 1.05E-02 | 0.2 | 3.24E-02 | 0.1 | 6.45E-01 | 0.0 | 7.82E-01 | 0.0 | 8.69E-01 | -0.1 | 7.31E-01 | GCN5-related N-acetyltransferase 8 |
| AT1G32070 | 0.4 | 4.03E-03 | 0.2 | 2.10E-01 | 0.3 | 6.76E-02 | 0.0 | 9.58E-01 | 0.0 | 8.70E-01 | 0.0 | 8.62E-01 | NSI |
| AT2G04845 | 0.8 | 2.23E-03 | 0.4 | 6.85E-02 | 0.4 | 1.36E-01 | 0.4 | 1.40E-01 | 0.1 | 8.41E-01 | 0.3 | 3.03E-01 | Acyl-CoA N-acyltransferases superfamily protein |
| AT1G72030 | 0.8 | 1.87E-04 | 0.5 | 4.36E-03 | 0.3 | 1.01E-01 | 0.1 | 7.41E-01 | 0.2 | 4.66E-01 | -0.1 | 7.75E-01 | GCN5-related N-acetyltransferase 10, chloroplastic |
| AT2G39030 | | | | | | | -2.006284293 | 5.58E-01 | -0.067638493 | 9.88E-01 | -1.9 | 7.19E-01 | L-ornithine N5-acetyltransferase NATA1 |
| AT5G11340 | 0.0 | 8.57E-01 | 0.3 | 3.03E-02 | -0.3 | 3.82E-02 | 0.2 | 2.29E-01 | 0.0 | 9.91E-01 | 0.2 | 3.23E-01 | Acyl-CoA N-acyltransferases superfamily protein |
| AT3G05630 | -0.8 | 7.02E-04 | -0.7 | 1.60E-03 | -0.1 | 7.97E-01 | -0.1 | 9.27E-01 | 0.3 | 6.50E-01 | -0.4 | 6.25E-01 | Phospholipase D zeta 2 |
| AT3G15730 | -0.6 | 2.27E-05 | -0.5 | 2.01E-04 | -0.1 | 2.02E-01 | -0.2 | 2.25E-01 | 0.0 | 9.66E-01 | -0.2 | 3.96E-01 | phospholipase D |
| AT3G16785 | -0.1 | 4.24E-02 | -0.2 | 2.04E-02 | 0.0 | 6.81E-01 | 0.0 | 9.46E-01 | -0.1 | 5.30E-01 | 0.2 | 5.42E-01 | Phospholipase |
| AT4G21220 | 0.4 | 7.78E-03 | 0.2 | 1.63E-01 | 0.2 | 2.03E-01 | 0.5 | 1.91E-01 | 0.6 | 9.36E-02 | -0.2 | 6.89E-01 | LpxD2 |
| AT3G19480 | 1.5 | 5.71E-07 | 1.0 | 1.64E-05 | 0.4 | 5.26E-03 | 0.0 | 8.96E-01 | 0.1 | 5.42E-01 | -0.2 | 4.23E-01 | D-3-phosphoglycerate dehydrogenase 3, chloroplastic |
| AT3G25530 | 0.4 | 8.58E-05 | 0.4 | 1.01E-04 | 0.0 | 7.27E-01 | 0.0 | 8.89E-01 | 0.1 | 6.30E-01 | 0.0 | 7.87E-01 | Glyoxylate reductase 1 |
| AT1G16300 | -0.2 | 1.59E-01 | 0.2 | 9.59E-02 | -0.4 | 1.81E-02 | 0.3 | 1.76E-01 | 0.3 | 2.65E-01 | 0.0 | 9.62E-01 | Glyceraldehyde-3-phosphate dehydrogenase |
| AT1G17650 | 2.0 | 5.05E-07 | 1.8 | 3.00E-06 | 0.2 | 1.56E-01 | 0.2 | 3.07E-01 | 0.1 | 5.86E-01 | 0.1 | 8.31E-01 | hypothetical protein |
| AT1G12900 | 2.6 | 1.44E-08 | 2.1 | 9.13E-08 | 0.6 | 5.55E-04 | 0.3 | 1.39E-02 | 0.2 | 1.00E-01 | 0.1 | 6.80E-01 | Glyceraldehyde 3-phosphate dehydrogenase A subunit 2 |
| AT1G14810 | 1.2 | 1.33E-06 | 1.0 | 9.20E-06 | 0.2 | 1.61E-01 | 0.6 | 2.01E-05 | 0.3 | 3.59E-02 | 0.4 | 2.72E-02 | Semialdehyde dehydrogenase family protein |
| AT1G72190 | -0.1 | 1.94E-01 | -0.2 | 1.67E-01 | 0.0 | 9.13E-01 | -0.1 | 6.04E-01 | -0.2 | 4.47E-01 | 0.1 | 8.52E-01 | Phosphoglycerate dehydrogenase, putative; 33424-31403 |
| AT2G45630 | 0.4 | 1.50E-04 | 0.1 | 4.64E-01 | 0.4 | 2.03E-03 | -0.2 | 4.21E-01 | -0.3 | 1.34E-01 | 0.2 | 5.81E-01 | Putative glycerate dehydrogenase |
| AT1G68010 | 1.2 | 6.79E-07 | 0.9 | 1.41E-05 | 0.3 | 1.19E-02 | 0.0 | 8.46E-01 | 0.0 | 9.85E-01 | 0.0 | 8.91E-01 | Hydroxypyruvate reductase |

| | | | | | | | | | | | | | |
|---|---|---|---|---|---|---|---|---|---|---|---|---|---|
| AT1G17745 | -1.7 | 1.51E-06 | -0.9 | 2.44E-04 | -0.8 | 1.06E-03 | 0.0 | 9.98E-01 | 0.4 | 1.43E-01 | -0.4 | 1.98E-01 | D-3-phosphoglycerate dehydrogenase 2, chloroplastic |
| AT5G15490 | -1.3 | 1.69E-06 | -0.5 | 2.37E-03 | -0.8 | 2.30E-04 | -0.2 | 2.46E-01 | 0.3 | 1.45E-01 | -0.4 | 4.12E-02 | UDP-glucose 6-dehydrogenase 3 |
| AT5G39320 | -0.9 | 7.08E-06 | 0.0 | 8.62E-01 | -0.9 | 6.23E-05 | -0.2 | 3.74E-01 | 0.3 | 2.02E-01 | -0.4 | 7.68E-02 | UDP-glucose 6-dehydrogenase 4 |
| AT3G29360 | -0.7 | 9.61E-06 | -0.3 | 3.04E-03 | -0.4 | 2.34E-03 | -0.1 | 5.48E-01 | 0.0 | 8.08E-01 | -0.1 | 5.34E-01 | UDP-glucose 6-dehydrogenase 2 |
| AT1G26570 | 0.3 | 3.32E-02 | 0.3 | 1.62E-02 | -0.1 | 6.91E-01 | 0.1 | 7.82E-01 | 0.0 | 9.66E-01 | 0.1 | 8.14E-01 | UDP-glucose 6-dehydrogenase 1 |
| AT2G27860 | 0.1 | 1.17E-01 | 0.2 | 4.82E-02 | 0.0 | 6.06E-01 | -0.3 | 5.01E-03 | -0.2 | 2.18E-02 | 0.0 | 8.53E-01 | UDP-D-apiose |
| AT1G47370 | 2.3 | 3.61E-05 | 1.3 | 4.20E-03 | 1.0 | 1.02E-02 | 1.2 | 6.29E-02 | 0.2 | 8.30E-01 | 1.0 | 1.72E-01 | Disease resistance protein RBA1 hydrolase RBA1) hydrolase RBA1) |
| AT1G72930 | 0.8 | 5.09E-05 | 0.3 | 3.38E-02 | 0.5 | 3.20E-03 | 1.4 | 5.30E-07 | 0.6 | 5.75E-03 | 0.8 | 1.63E-03 | Toll |
| AT5G08740 | 0.6 | 1.12E-04 | 0.3 | 5.51E-03 | 0.2 | 4.24E-02 | 0.0 | 8.76E-01 | 0.2 | 2.37E-01 | -0.2 | 3.96E-01 | Alternative NADH-ubiquinone oxidoreductase C1, chloroplastic |
| AT2G35510 | -0.3 | 3.79E-04 | -0.3 | 8.10E-04 | 0.0 | 8.24E-01 | 0.0 | 9.07E-01 | 0.0 | 9.60E-01 | 0.0 | 9.07E-01 | Uncharacterized protein At2g35510 |
| AT1G32230 | 0.0 | 7.93E-01 | -0.4 | 8.83E-04 | 0.3 | 2.35E-03 | 0.0 | 7.69E-01 | -0.1 | 5.72E-01 | 0.1 | 5.12E-01 | WWE protein-protein interaction domain protein family |
| AT1G23550 | 0.2 | 1.83E-01 | -0.1 | 5.62E-01 | 0.4 | 1.26E-01 | 0.6 | 1.20E-01 | 0.4 | 3.60E-01 | 0.2 | 7.23E-01 | Probable inactive poly polymerase SRO2 |
| AT1G70440 | | | | | | | 4.6 | 4.77E-03 | 2.6 | 1.74E-01 | | | Probable inactive poly polymerase SRO3 |
| AT4G02390 | 1.3 | 1.92E-07 | 0.4 | 1.81E-03 | 0.9 | 3.16E-05 | 0.0 | 9.84E-01 | 0.1 | 7.35E-01 | -0.1 | 7.85E-01 | Poly polymerase |
| AT2G47650 | -1.0 | 7.72E-07 | -0.4 | 1.05E-03 | -0.6 | 1.39E-04 | -0.1 | 6.86E-01 | 0.0 | 9.83E-01 | -0.1 | 8.05E-01 | UDP-glucuronate decarboxylase |
| AT5G59290 | -0.8 | 3.42E-06 | -0.1 | 1.93E-01 | -0.7 | 7.37E-05 | -0.2 | 1.66E-01 | 0.1 | 7.72E-01 | -0.2 | 1.75E-01 | UDP-glucuronate decarboxylase |
| AT2G28760 | -0.7 | 3.82E-05 | -0.1 | 2.06E-01 | -0.6 | 8.21E-04 | 0.4 | 1.74E-02 | 0.5 | 4.91E-03 | -0.1 | 5.24E-01 | UDP-glucuronate decarboxylase |
| AT3G46440 | -0.4 | 6.03E-04 | -0.2 | 3.10E-02 | -0.2 | 5.55E-02 | -0.1 | 4.21E-01 | 0.1 | 3.09E-01 | -0.3 | 1.39E-01 | UDP-glucuronic acid decarboxylase 5 |
| AT3G53520 | -0.2 | 1.39E-02 | 0.0 | 9.85E-01 | -0.2 | 2.47E-02 | -0.1 | 5.23E-01 | -0.2 | 2.06E-01 | 0.1 | 6.52E-01 | UDP-glucuronate decarboxylase |
| AT1G78590 | -0.1 | 1.29E-01 | -0.1 | 1.30E-01 | 0.0 | 9.67E-01 | -0.5 | 2.63E-02 | -0.4 | 1.19E-01 | -0.1 | 6.77E-01 | NAD kinase 3 |
| AT3G21070 | -1.0 | 2.73E-06 | -0.6 | 7.76E-05 | -0.3 | 1.48E-02 | -0.3 | 3.16E-02 | -0.2 | 2.00E-01 | -0.1 | 5.75E-01 | NAD kinase 1 |
| AT1G07180 | 1.4 | 8.24E-07 | 0.6 | 2.64E-04 | 0.7 | 4.06E-04 | 1.0 | 5.85E-02 | 0.5 | 4.98E-01 | 0.5 | 3.76E-01 | Internal alternative NADH-ubiquinone oxidoreductase A1, mitochondrial H dehydrogenase 1) NDA1) |
| AT2G29990 | -0.2 | 9.22E-02 | -0.6 | 5.24E-04 | 0.4 | 1.16E-02 | 0.0 | 8.93E-01 | -0.1 | 8.37E-01 | 0.1 | 7.65E-01 | Internal alternative NADH-ubiquinone oxidoreductase A2, mitochondrial NDA2) |
| AT1G09420 | | | | | | | 0.0 | 9.77E-01 | -0.1 | 9.41E-01 | 0.1 | 9.65E-01 | Glucose-6-phosphate 1-dehydrogenase |
| AT1G64190 | -1.1 | 5.69E-04 | -0.8 | 4.16E-03 | -0.3 | 3.09E-01 | 0.1 | 6.32E-01 | -0.1 | 6.32E-01 | 0.1 | 4.40E-01 | 6-phosphogluconate dehydrogenase, decarboxylating 1, chloroplastic |
| AT5G41670 | -0.8 | 1.20E-03 | -0.5 | 3.28E-02 | -0.4 | 1.12E-01 | 0.2 | 2.67E-01 | 0.0 | 9.40E-01 | 0.1 | 5.33E-01 | 6-phosphogluconate dehydrogenase, decarboxylating 3, chloroplastic |
| AT3G02360 | -0.7 | 1.02E-05 | -0.4 | 2.46E-03 | -0.4 | 3.12E-03 | 0.0 | 7.61E-01 | 0.0 | 8.22E-01 | 0.0 | 9.96E-01 | 6-phosphogluconate dehydrogenase, decarboxylating 2 |
| AT4G21580 | -0.4 | 1.61E-04 | -0.6 | 2.95E-05 | 0.2 | 6.81E-02 | -0.1 | 5.85E-01 | -0.1 | 4.83E-01 | 0.0 | 8.91E-01 | AT4G21580 protein |
| AT1G05180 | 0.2 | 3.92E-02 | 0.0 | 5.37E-01 | 0.1 | 1.80E-01 | 0.2 | 2.26E-01 | -0.1 | 6.66E-01 | 0.3 | 1.74E-01 | NAD-binding Rossmann-fold superfamily protein |
| AT1G21840 | 0.3 | 1.66E-01 | -0.2 | 4.03E-01 | 0.4 | 7.84E-02 | -0.1 | 8.92E-01 | -0.2 | 6.52E-01 | 0.2 | 7.97E-01 | Urease accessory protein F |
| AT1G67550 | -0.2 | 3.83E-02 | -0.3 | 3.17E-03 | 0.1 | 1.72E-01 | 0.0 | 9.45E-01 | 0.0 | 9.15E-01 | -0.1 | 8.90E-01 | Urease |
| AT1G09240 | -0.8 | 3.05E-04 | -0.5 | 9.64E-03 | -0.3 | 8.32E-02 | | | | | | | Nicotianamine synthase 3 |
| AT5G04950 | 1.0 | 4.54E-04 | 0.9 | 8.69E-04 | 0.0 | 8.86E-01 | -0.4 | 5.84E-02 | -0.6 | 2.07E-02 | 0.2 | 6.49E-01 | Nicotianamine synthase 1 |
| AT5G56080 | 2.2 | 7.76E-05 | 1.6 | 1.57E-03 | 0.6 | 8.72E-02 | | | | | | | Nicotianamine synthase 2 |
| AT1G56430 | 2.5 | 4.25E-08 | 2.1 | 3.32E-07 | 0.4 | 8.04E-03 | 1.9 | 3.19E-08 | 0.5 | 2.79E-02 | 1.5 | 2.55E-05 | Probable nicotianamine synthase 4 |
| AT2G22570 | -0.3 | 1.74E-03 | -0.2 | 6.38E-02 | -0.2 | 1.05E-01 | -0.2 | 4.07E-01 | -0.1 | 7.11E-01 | -0.1 | 8.33E-01 | Isochorismatase-like domain-containing protein |
| AT5G23220 | 3.5 | 1.08E-07 | 2.2 | 9.44E-06 | 1.3 | 8.35E-05 | 1.2 | 2.84E-03 | 0.8 | 4.25E-02 | 0.3 | 3.77E-01 | Nicotinamidase 3 |
| AT5G55810 | -0.3 | 7.32E-02 | -0.3 | 1.00E-01 | 0.0 | 9.40E-01 | -0.3 | 4.20E-01 | -0.2 | 7.03E-01 | -0.1 | 7.99E-01 | Nicotinate |
| AT4G36940 | -0.4 | 8.55E-04 | 0.0 | 6.81E-01 | -0.4 | 1.82E-03 | 0.0 | 9.74E-01 | 0.1 | 4.92E-01 | -0.1 | 5.62E-01 | Nicotinate phosphoribosyltransferase |
| AT5G05880 | -0.6 | 1.31E-03 | 0.0 | 9.18E-01 | -0.6 | 5.51E-03 | 0.2 | 6.14E-01 | 0.5 | 9.88E-02 | -0.4 | 2.98E-01 | UDP-glycosyltransferase 76C4 |
| AT5G05890 | 0.2 | 6.60E-02 | 0.6 | 1.09E-04 | -0.4 | 3.10E-03 | -0.2 | 1.86E-01 | -0.4 | 3.89E-02 | 0.2 | 5.05E-01 | UDP-glycosyltransferase 76C5 |
| AT2G01350 | 0.2 | 9.44E-03 | 0.1 | 1.77E-01 | 0.1 | 1.67E-01 | 0.2 | 1.64E-01 | 0.1 | 7.65E-01 | 0.1 | 5.05E-01 | Nicotinate-nucleotide pyrophosphorylase like protein |
| AT1G08100 | -6.1 | 6.68E-06 | -0.3 | 1.33E-01 | -5.8 | 8.12E-05 | -2.7 | 2.75E-09 | -0.6 | 5.59E-04 | -2.0 | 1.85E-05 | High-affinity nitrate transporter 2.2 |
| AT1G12940 | -5.2 | 1.03E-07 | -3.2 | 2.39E-06 | -2.1 | 2.15E-04 | -2.2 | 5.81E-05 | 0.5 | 1.11E-01 | -2.8 | 1.74E-04 | High affinity nitrate transporter 2.5 |
| AT5G60770 | -2.3 | 1.03E-06 | 0.6 | 2.15E-03 | -2.9 | 8.63E-06 | -2.0 | 3.68E-05 | 0.3 | 3.43E-01 | -2.3 | 3.38E-04 | High affinity nitrate transporter 2.4 |
| AT5G50200 | -1.7 | 7.27E-07 | -0.7 | 5.72E-04 | -0.9 | 1.93E-04 | 0.1 | 6.38E-01 | 0.3 | 4.88E-02 | -0.2 | 2.17E-01 | High-affinity nitrate transporter 3.1 |
| AT1G08090 | -1.1 | 8.87E-05 | 0.2 | 3.76E-01 | -1.3 | 1.71E-04 | -0.1 | 4.35E-01 | 0.5 | 4.55E-03 | -0.6 | 6.20E-03 | High-affinity nitrate transporter 2.1 |
| AT5G14570 | 0.4 | 6.73E-04 | 0.1 | 2.51E-01 | 0.3 | 1.15E-02 | -0.4 | 2.13E-02 | -0.1 | 7.70E-01 | -0.3 | 1.13E-01 | NRT2.7 |
| AT3G45060 | | | | | | | -2.728517893 | 1.39E-02 | -3.449423129 | 1.11E-02 | 0.7 | 7.18E-01 | High affinity nitrate transporter 2.6 |
| AT5G60780 | | | | | | | -1.3 | 9.19E-03 | -0.9 | 4.55E-02 | -0.3 | 5.57E-01 | High affinity nitrate transporter 2.3 |
| AT1G32450 | 0.4 | 2.07E-03 | 0.6 | 2.05E-04 | -0.2 | 7.78E-02 | -1.0 | 3.25E-06 | -0.2 | 2.70E-01 | -0.8 | 8.43E-04 | Protein NRT1 |
| AT3G21670 | 0.3 | 3.84E-02 | -0.4 | 2.85E-02 | 0.7 | 1.90E-03 | 0.1 | 5.97E-01 | 0.5 | 1.57E-02 | -0.4 | 1.00E-01 | Protein NRT1 |
| AT1G12110 | 0.8 | 1.16E-04 | 1.0 | 5.86E-05 | -0.2 | 2.65E-01 | 0.1 | 2.33E-01 | 0.1 | 4.38E-01 | 0.0 | 9.04E-01 | Protein NRT1 |
| AT4G21680 | -2.1 | 7.54E-05 | -2.4 | 6.74E-05 | 0.3 | 5.19E-01 | -1.7 | 2.85E-02 | -1.0 | 1.74E-01 | -0.8 | 3.97E-01 | Protein NRT1 |
| AT1G18880 | -1.8 | 4.45E-07 | -1.2 | 1.06E-05 | -0.6 | 4.32E-03 | 0.0 | 9.81E-01 | 0.3 | 6.92E-01 | -0.2 | 7.55E-01 | Protein NRT1 |
| AT3G45680 | -0.6 | 3.08E-03 | -0.5 | 1.21E-02 | -0.1 | 5.94E-01 | -0.1 | 8.34E-01 | 0.8 | 4.06E-02 | -0.9 | 3.98E-02 | Protein NRT1 |
| AT3G27170 | -2.2 | 2.25E-08 | -1.0 | 4.49E-06 | -1.2 | 1.39E-05 | 0.5 | 2.48E-01 | 1.1 | 1.03E-02 | -0.7 | 1.94E-01 | Chloride channel protein CLC-b |
| AT3G05040 | -0.4 | 2.69E-04 | -0.4 | 2.55E-04 | 0.0 | 7.25E-01 | 0.4 | 1.15E-01 | 0.3 | 4.26E-01 | 0.2 | 6.71E-01 | ARM repeat superfamily protein |
| AT3G04490 | -0.2 | 7.59E-02 | -0.3 | 1.24E-02 | 0.1 | 3.40E-01 | -0.2 | 3.43E-01 | -0.2 | 4.55E-01 | 0.0 | 9.62E-01 | Exportin-4 protein |
| AT5G06120 | -0.5 | 2.35E-04 | -0.4 | 4.71E-04 | 0.0 | 8.81E-01 | -0.1 | 8.49E-01 | 0.1 | 6.96E-01 | -0.2 | 5.81E-01 | ARM repeat superfamily protein |
| AT5G17020 | -0.3 | 6.34E-04 | -0.2 | 2.53E-03 | 0.0 | 5.35E-01 | 0.2 | 1.57E-01 | 0.1 | 3.64E-01 | 0.0 | 8.64E-01 | Protein EXPORTIN 1A |
| AT3G03110 | 0.3 | 8.99E-04 | 0.1 | 2.02E-01 | 0.2 | 2.09E-02 | 0.1 | 7.81E-01 | -0.1 | 8.21E-01 | 0.1 | 6.98E-01 | Protein EXPORTIN 1B |
| AT2G46520 | 0.5 | 8.23E-05 | 0.2 | 5.63E-02 | 0.3 | 4.05E-03 | 0.3 | 1.54E-01 | 0.0 | 8.93E-01 | 0.2 | 4.10E-01 | Exportin-2 |
| AT1G72560 | 0.2 | 5.34E-03 | 0.0 | 4.92E-01 | 0.2 | 4.15E-02 | 0.1 | 5.88E-01 | 0.0 | 9.92E-01 | 0.1 | 7.10E-01 | Exportin-T ) |
| AT4G02150 | 0.1 | 3.48E-01 | -0.1 | 1.03E-01 | 0.2 | 3.50E-02 | 0.0 | 9.81E-01 | 0.2 | 6.64E-01 | -0.2 | 7.11E-01 | ARM repeat superfamily protein |
| AT1G02690 | 0.6 | 7.87E-05 | 0.5 | 5.36E-04 | 0.1 | 3.21E-01 | 0.2 | 1.71E-01 | 0.1 | 7.56E-01 | 0.1 | 5.12E-01 | Importin subunit alpha |
| AT3G06720 | -0.5 | 8.45E-05 | -0.4 | 1.18E-03 | -0.1 | 1.39E-01 | 0.0 | 8.47E-01 | 0.0 | 7.72E-01 | 0.1 | 7.11E-01 | Importin subunit alpha-1 |
| AT1G09270 | -0.3 | 4.42E-04 | -0.1 | 2.17E-01 | -0.2 | 7.13E-03 | -0.1 | 4.77E-01 | 0.0 | 9.62E-01 | -0.1 | 5.98E-01 | Importin subunit alpha-4 |
| AT5G03070 | -0.2 | 1.09E-02 | -0.2 | 4.09E-02 | -0.1 | 5.95E-01 | 0.2 | 1.63E-01 | 0.1 | 5.20E-01 | 0.1 | 6.86E-01 | Importin subunit alpha-9 |
| AT5G19820 | -0.1 | 1.33E-01 | 0.0 | 8.45E-01 | -0.1 | 2.41E-01 | 0.2 | 1.36E-01 | -0.1 | 5.52E-01 | 0.3 | 1.11E-01 | ARM repeat superfamily protein |
| AT3G08947 | 0.6 | 9.05E-05 | 0.2 | 1.00E-01 | 0.4 | 3.98E-03 | 0.2 | 3.52E-01 | 0.1 | 5.63E-01 | 0.0 | 9.03E-01 | ARM repeat superfamily protein |
| AT5G53480 | -0.3 | 6.73E-04 | -0.3 | 1.67E-03 | 0.0 | 7.71E-01 | -0.1 | 4.37E-01 | -0.1 | 6.99E-01 | -0.1 | 8.81E-01 | Importin subunit beta-1 |
| AT2G16950 | -0.2 | 2.24E-02 | -0.1 | 7.04E-02 | 0.0 | 6.33E-01 | 0.3 | 9.29E-02 | 0.0 | 8.63E-01 | 0.2 | 3.37E-01 | Transportin-1 |
| AT5G02200 | -1.5 | 3.72E-04 | -0.8 | 1.18E-02 | -0.7 | 7.61E-02 | | | | | | | Protein FAR-RED-ELONGATED HYPOCOTYL 1-LIKE |
| AT2G37678 | 0.0 | 8.38E-01 | 0.1 | 4.44E-01 | -0.1 | 4.06E-01 | -0.3 | 2.10E-01 | -0.2 | 5.13E-01 | -0.1 | 7.92E-01 | Protein FAR-RED ELONGATED HYPOCOTYL 1 |
| AT1G66080 | 0.1 | 5.71E-01 | -0.1 | 3.25E-01 | 0.2 | 1.98E-01 | 0.2 | 6.87E-01 | 0.4 | 3.64E-01 | -0.2 | 7.00E-01 | At1g66080 |
| AT1G10390 | -0.4 | 2.35E-04 | -0.3 | 5.59E-04 | 0.0 | 8.07E-01 | -0.2 | 1.51E-01 | -0.2 | 1.80E-01 | 0.0 | 9.71E-01 | Nuclear pore complex protein NUP98A |
| AT4G37130 | 0.4 | 3.45E-04 | 0.0 | 9.34E-01 | 0.4 | 1.48E-03 | 0.0 | 9.39E-01 | -0.1 | 8.44E-01 | 0.1 | 9.12E-01 | Nuclear pore complex protein NUP58 |
| AT1G19660 | -0.8 | 6.68E-05 | -0.5 | 1.05E-03 | -0.2 | 1.18E-01 | -0.5 | 1.12E-04 | -0.3 | 1.78E-02 | -0.2 | 1.67E-01 | Wound-responsive family protein |
| AT1G75380 | -0.7 | 1.99E-04 | -0.2 | 9.14E-02 | -0.5 | 6.69E-03 | 0.0 | 9.64E-01 | 0.0 | 7.86E-01 | 0.0 | 8.46E-01 | Bifunctional nuclease 1 |
| AT5G66050 | 0.0 | 6.11E-01 | 0.1 | 2.64E-01 | 0.0 | 5.87E-01 | -0.5 | 3.89E-03 | -0.5 | 6.51E-03 | 0.0 | 9.79E-01 | Wound-responsive family protein |
| AT5G60230 | 0.3 | 3.04E-02 | 0.5 | 5.52E-03 | -0.2 | 3.66E-01 | -0.5 | 7.25E-02 | -0.5 | 8.48E-02 | 0.0 | 9.73E-01 | tRNA-intron lyase |
| AT3G55220 | -0.5 | 1.68E-02 | -0.5 | 2.67E-02 | 0.0 | 9.41E-01 | 0.6 | 4.11E-01 | 0.4 | 6.17E-01 | 0.1 | 8.97E-01 | Spliceosome-associated protein 130 B |

| | | | | | | | | | | | | | |
|---|---|---|---|---|---|---|---|---|---|---|---|---|---|
| AT1G20225 | -0.5 | 7.18E-05 | -0.4 | 2.15E-04 | 0.0 | 6.61E-01 | -0.1 | 7.17E-01 | 0.2 | 2.83E-01 | -0.2 | 2.04E-01 | T20H2.2 protein |
| AT3G57910 | -0.5 | 7.26E-04 | -0.3 | 1.10E-02 | -0.2 | 2.01E-01 | -0.1 | 7.00E-01 | 0.2 | 5.94E-01 | -0.3 | 4.38E-01 | G-patch domain-containing protein |
| AT4G05420 | -0.3 | 2.07E-03 | -0.3 | 1.88E-03 | 0.0 | 8.14E-01 | -0.1 | 4.24E-01 | -0.1 | 5.20E-01 | 0.0 | 9.88E-01 | Damaged DNA binding protein 1A |
| AT1G31280 | -0.2 | 1.87E-02 | -0.8 | 5.21E-06 | 0.6 | 1.38E-04 | -0.2 | 4.63E-01 | 0.0 | 9.32E-01 | -0.2 | 4.80E-01 | Protein argonaute 2 |
| AT1G65440 | -0.2 | 2.71E-02 | -0.3 | 5.10E-03 | 0.1 | 3.18E-01 | -0.1 | 6.66E-01 | -0.1 | 3.97E-01 | 0.1 | 7.76E-01 | Transcription elongation factor spt6 |
| AT3G09850 | -0.2 | 7.02E-03 | -0.3 | 5.53E-04 | 0.1 | 1.00E-01 | -0.4 | 6.08E-02 | -0.2 | 3.23E-01 | -0.1 | 6.57E-01 | G-patch domain-containing protein |
| AT5G51660 | -0.1 | 9.11E-02 | -0.2 | 8.98E-03 | 0.1 | 2.29E-01 | 0.1 | 5.70E-01 | 0.0 | 9.82E-01 | 0.1 | 6.56E-01 | Cleavage and polyadenylation specificity factor 160 |
| AT5G51730 | -0.1 | 3.32E-01 | -0.1 | 4.35E-01 | 0.0 | 9.10E-01 | -0.1 | 6.17E-01 | -0.1 | 8.58E-01 | -0.1 | 8.48E-01 | RRM domain-containing protein |
| AT4G25020 | -0.1 | 5.75E-01 | -0.2 | 2.19E-01 | 0.1 | 5.56E-01 | -0.2 | 6.73E-01 | 0.3 | 5.75E-01 | -0.5 | 3.33E-01 | At4g25020 |
| AT5G05100 | -0.1 | 3.92E-01 | -0.1 | 2.80E-01 | 0.0 | 8.25E-01 | -0.5 | 7.07E-03 | -0.7 | 1.73E-03 | 0.2 | 4.54E-01 | Gb\|AAF19572.1 |
| AT3G54760 | 0.0 | 4.97E-01 | -0.1 | 1.11E-01 | 0.1 | 3.69E-01 | 0.0 | 9.88E-01 | 0.1 | 7.38E-01 | -0.1 | 8.02E-01 | hypothetical protein |
| AT5G08535 | 0.0 | 7.22E-01 | -0.2 | 1.93E-01 | 0.1 | 4.12E-01 | -0.2 | 5.00E-01 | 0.1 | 8.29E-01 | -0.3 | 4.67E-01 | AT5g08540 |
| AT1G48410 | 0.0 | 8.79E-01 | -0.2 | 3.62E-02 | 0.2 | 3.78E-02 | -0.1 | 5.40E-01 | -0.1 | 7.17E-01 | 0.0 | 9.37E-01 | Uncharacterized protein |
| AT3G04620 | 0.1 | 6.98E-01 | 0.5 | 6.02E-03 | -0.5 | 3.04E-02 | 0.2 | 5.69E-01 | -0.1 | 8.47E-01 | 0.4 | 5.05E-01 | hypothetical protein |
| AT3G10770 | 0.2 | 3.71E-02 | 0.1 | 4.68E-01 | 0.1 | 1.90E-01 | -0.3 | 7.49E-03 | -0.5 | 1.43E-03 | 0.1 | 3.96E-01 | Single-stranded nucleic acid binding R3H protein |
| AT4G39680 | 0.2 | 1.64E-02 | 0.1 | 4.48E-01 | 0.1 | 1.02E-01 | 0.1 | 5.29E-01 | 0.1 | 5.57E-01 | 0.0 | 9.73E-01 | SAP domain-containing protein |
| AT1G03250 | 0.2 | 1.03E-02 | 0.1 | 3.26E-01 | 0.1 | 1.04E-01 | 0.2 | 3.17E-01 | 0.0 | 8.88E-01 | 0.2 | 2.23E-01 | R3H domain protein |
| AT5G22120 | 0.2 | 2.46E-03 | 0.1 | 4.07E-01 | 0.2 | 2.46E-02 | 0.1 | 7.84E-01 | -0.3 | 1.95E-01 | 0.4 | 1.11E-01 | Coiled-coil domain-containing protein R3HCC1L |
| AT3G50670 | 0.3 | 6.26E-03 | 0.1 | 3.17E-01 | 0.2 | 6.41E-02 | 0.0 | 9.60E-01 | -0.1 | 3.37E-01 | 0.1 | 3.69E-01 | U1SNRNP |
| AT1G63980 | 0.3 | 3.03E-04 | 0.3 | 5.85E-04 | 0.0 | 9.02E-01 | 0.0 | 8.54E-01 | -0.2 | 1.54E-01 | 0.2 | 1.93E-01 | D111 |
| AT1G20220 | 0.4 | 1.29E-02 | 0.4 | 1.59E-02 | 0.0 | 9.98E-01 | 0.2 | 2.90E-01 | 0.2 | 1.70E-01 | -0.1 | 7.81E-01 | T20H2.2 protein |
| AT1G76010 | 0.5 | 2.46E-05 | 0.4 | 2.16E-04 | 0.1 | 2.08E-01 | 0.1 | 1.43E-01 | -0.1 | 5.22E-01 | 0.2 | 1.20E-01 | DNA |
| AT2G34160 | 0.5 | 2.29E-02 | 0.0 | 8.57E-01 | 0.5 | 6.92E-02 | 0.2 | 5.99E-01 | 0.2 | 5.76E-01 | 0.0 | 9.53E-01 | DNA |
| AT3G56680 | 0.7 | 2.03E-06 | 0.6 | 1.90E-05 | 0.1 | 9.14E-02 | 0.1 | 4.42E-01 | -0.1 | 6.58E-01 | 0.3 | 2.28E-01 | Single-stranded nucleic acid binding R3H protein |
| AT2G40960 | 0.8 | 1.06E-04 | 0.8 | 1.78E-04 | 0.0 | 9.66E-01 | 0.1 | 7.44E-01 | 0.1 | 8.43E-01 | 0.0 | 9.58E-01 | Expressed protein |
| AT5G30510 | 1.7 | 7.44E-08 | 1.2 | 1.77E-06 | 0.5 | 1.04E-03 | 0.3 | 1.38E-02 | 0.1 | 3.51E-01 | 0.2 | 2.99E-01 | AT5G30510 protein |
| AT1G71720 | 1.8 | 3.10E-07 | 1.5 | 2.12E-06 | 0.2 | 9.11E-02 | -0.1 | 7.06E-01 | 0.0 | 9.17E-01 | -0.1 | 8.64E-01 | Nucleic acid-binding proteins superfamily |
| AT5G03495 | | | | | | | 0.1 | 9.31E-01 | -0.3 | 6.86E-01 | | | RNA-binding family protein |
| AT5G53700 | | | | | | | 2.4 | 8.70E-03 | 1.9 | 5.99E-02 | | | RRM domain-containing protein |
| AT3G16310 | 0.8 | 1.52E-05 | 0.6 | 1.25E-04 | 0.1 | 1.98E-01 | 0.3 | 2.77E-02 | 0.0 | 9.10E-01 | 0.3 | 9.11E-02 | Nuclear pore complex protein NUP35 |
| AT1G80780 | 0.2 | 6.44E-03 | 0.0 | 7.36E-01 | 0.2 | 2.28E-02 | -0.4 | 2.66E-03 | -0.4 | 3.05E-03 | 0.0 | 8.62E-01 | poly-specific ribonuclease |
| AT3G13440 | -0.1 | 3.12E-01 | 0.1 | 4.20E-01 | -0.2 | 1.30E-01 | -0.4 | 5.92E-02 | -0.4 | 5.79E-02 | 0.0 | 9.10E-01 | N6-DNA-methyltransferase-like protein |
| AT5G64150 | 0.3 | 4.72E-02 | 0.3 | 2.13E-02 | -0.1 | 6.76E-01 | 0.6 | 1.53E-02 | 0.0 | 9.30E-01 | 0.5 | 6.85E-02 | Methyltransferase small domain-containing protein |
| AT3G58470 | 0.5 | 2.21E-04 | 0.3 | 5.98E-03 | 0.2 | 1.06E-01 | -0.1 | 6.41E-01 | -0.3 | 1.54E-01 | 0.2 | 4.34E-01 | Protein-lysine N-methyltransferase At3g58470 |
| AT3G60040 | -0.4 | 1.90E-02 | -0.4 | 2.63E-02 | 0.0 | 9.68E-01 | -0.4 | 1.11E-01 | -0.2 | 4.13E-01 | -0.2 | 6.49E-01 | F-box family protein |
| AT5G51080 | 0.3 | 7.24E-02 | 0.2 | 1.65E-01 | 0.1 | 7.55E-01 | -0.1 | 6.44E-01 | 0.2 | 6.57E-01 | -0.3 | 2.92E-01 | Similarity to RNase H |
| AT1G24090 | | | | | | | 1.0 | 2.79E-03 | 0.4 | 3.49E-01 | 0.7 | 8.13E-02 | F3I6.1 protein |
| AT4G09490 | -0.3 | 4.60E-02 | -0.2 | 2.75E-01 | -0.1 | 4.31E-01 | -0.5 | 1.72E-01 | 0.1 | 7.68E-01 | -0.6 | 1.38E-01 | Polynucleotidyl transferase, ribonuclease H-like superfamily protein |
| AT3G01410 | 1.2 | 6.57E-07 | 0.8 | 4.18E-05 | 0.5 | 2.37E-03 | 0.3 | 5.12E-02 | 0.4 | 5.05E-02 | 0.0 | 9.12E-01 | Polynucleotidyl transferase, ribonuclease H-like superfamily protein |
| AT4G19670 | 0.0 | 8.41E-01 | -0.3 | 7.90E-03 | 0.3 | 1.01E-02 | 0.2 | 4.38E-01 | 0.3 | 1.90E-01 | -0.1 | 6.69E-01 | RING |
| AT5G60250 | 1.7 | 1.32E-05 | 0.2 | 5.63E-01 | 1.5 | 3.10E-04 | 0.4 | 2.64E-01 | 0.4 | 2.63E-01 | 0.0 | 9.44E-01 | Uncharacterized protein |
| AT4G28830 | 0.4 | 2.76E-03 | 0.2 | 1.94E-01 | 0.3 | 6.83E-02 | -0.3 | 1.39E-01 | -0.3 | 1.85E-01 | 0.0 | 9.98E-01 | AT4G28830 protein |
| AT3G26410 | 0.2 | 2.69E-01 | 0.2 | 2.79E-01 | 0.0 | 9.90E-01 | 0.0 | 9.74E-01 | -0.1 | 6.98E-01 | 0.1 | 7.00E-01 | Uncharacterized protein At3g26410 |
| AT1G27750 | -0.4 | 5.15E-04 | -0.3 | 1.14E-03 | 0.0 | 8.21E-01 | -0.3 | 1.89E-01 | -0.2 | 4.73E-01 | -0.1 | 7.92E-01 | Nucleic acid binding protein |
| AT5G47430 | 0.0 | 6.19E-01 | -0.1 | 3.29E-02 | 0.1 | 1.07E-01 | -0.1 | 3.24E-01 | -0.2 | 1.63E-01 | 0.1 | 7.49E-01 | DWNN domain, a CCHC-type zinc finger |
| AT3G62330 | -0.2 | 7.15E-03 | -0.3 | 1.90E-03 | 0.1 | 3.92E-01 | -0.3 | 7.98E-02 | 0.0 | 8.84E-01 | -0.3 | 2.67E-01 | At3g62330 family protein) |
| AT3G24070 | -0.2 | 1.24E-02 | -0.2 | 1.49E-02 | 0.0 | 9.97E-01 | -0.5 | 9.83E-04 | -0.4 | 3.99E-03 | 0.0 | 8.60E-01 | At3g24060 family protein) |
| AT3G42860 | -0.2 | 2.20E-01 | -0.3 | 5.17E-02 | 0.1 | 4.64E-01 | 0.2 | 7.18E-01 | 0.0 | 9.81E-01 | 0.2 | 7.46E-01 | Uncharacterized protein At3g42860 family protein) |
| AT2G28910 | -0.2 | 2.96E-02 | -0.1 | 1.04E-01 | 0.0 | 5.92E-01 | -0.7 | 1.68E-04 | -0.6 | 1.38E-03 | -0.1 | 7.84E-01 | CAX-interacting protein 4 |
| AT1G74250 | 0.0 | 9.86E-01 | -0.1 | 2.60E-01 | 0.1 | 3.14E-01 | 0.1 | 3.42E-01 | 0.0 | 9.50E-01 | 0.1 | 4.80E-01 | DNAJ protein JJJ1 homolog |
| AT5G26749 | 0.0 | 9.65E-01 | 0.0 | 8.09E-01 | 0.0 | 8.75E-01 | 0.0 | 9.41E-01 | 0.2 | 6.67E-01 | -0.2 | 6.51E-01 | C2H2 and C2HC zinc fingers superfamily protein |
| AT1G49590 | 0.1 | 1.29E-01 | 0.0 | 8.58E-01 | 0.1 | 2.53E-01 | -0.1 | 6.15E-01 | 0.0 | 9.17E-01 | -0.1 | 5.90E-01 | C2H2 and C2HC zinc fingers superfamily protein |
| AT3G02860 | 0.2 | 2.42E-02 | 0.1 | 4.11E-01 | 0.1 | 1.75E-01 | -0.3 | 5.51E-02 | -0.4 | 6.40E-02 | 0.0 | 9.59E-01 | Protein ABA AND ROS SENSITIVE 1 |
| AT3G05760 | 0.2 | 5.23E-02 | 0.3 | 1.01E-02 | -0.1 | 3.90E-01 | -0.1 | 5.45E-01 | -0.1 | 7.92E-01 | 0.0 | 8.69E-01 | C2H2 and C2HC zinc fingers superfamily protein |
| AT2G32600 | 0.2 | 1.48E-02 | 0.2 | 1.59E-02 | 0.0 | 9.61E-01 | 0.0 | 9.45E-01 | -0.2 | 2.30E-01 | 0.2 | 2.28E-01 | Matrin-type domain-containing protein |
| AT5G52380 | 0.2 | 1.25E-01 | 0.4 | 3.62E-02 | -0.1 | 5.43E-01 | 0.5 | 2.94E-02 | 0.1 | 7.08E-01 | 0.4 | 1.33E-01 | CCHC-type domain-containing protein |
| AT1G75560 | 0.3 | 2.34E-03 | 0.3 | 2.61E-03 | 0.0 | 9.29E-01 | -0.1 | 7.22E-01 | 0.0 | 9.00E-01 | -0.1 | 7.11E-01 | F10A5.22 |
| AT4G36020 | 0.3 | 8.86E-03 | 0.2 | 1.15E-01 | 0.2 | 2.32E-01 | 0.4 | 2.58E-02 | 0.2 | 3.85E-01 | 0.2 | 2.49E-01 | Cold shock domain protein 1 |
| AT3G02820 | 0.5 | 1.30E-02 | 0.5 | 1.39E-02 | 0.0 | 9.53E-01 | 0.2 | 5.34E-01 | 0.0 | 9.60E-01 | 0.1 | 7.11E-01 | hypothetical protein |
| AT3G43590 | 0.6 | 1.42E-04 | 0.1 | 3.71E-01 | 0.5 | 2.37E-03 | 0.4 | 4.05E-01 | 0.4 | 4.68E-01 | 0.0 | 9.99E-01 | Uncharacterized protein At3g43590 family protein) |
| AT2G36930 | 0.6 | 2.09E-04 | 0.5 | 1.25E-03 | 0.1 | 3.68E-01 | -0.2 | 2.48E-01 | -0.4 | 3.59E-02 | 0.2 | 3.77E-01 | Expressed protein family protein) |
| AT5G20220 | 1.2 | 3.68E-05 | 0.4 | 7.59E-02 | 0.8 | 2.43E-03 | | | | | | | Zinc knuckle family protein |
| AT5G13920 | | | | | | | -0.3 | 6.77E-01 | -0.3 | 6.63E-01 | 0.1 | 9.61E-01 | Emb\|CAB86673.1 |
| AT5G50300 | -0.8 | 2.91E-04 | 0.2 | 2.46E-01 | -1.0 | 4.47E-04 | -0.2 | 2.77E-01 | 0.3 | 1.91E-01 | -0.5 | 2.44E-02 | Xanthine |
| AT1G17410 | 0.2 | 1.60E-01 | 0.1 | 6.87E-01 | 0.1 | 4.25E-01 | -0.3 | 2.40E-01 | -0.9 | 1.73E-03 | 0.6 | 3.82E-02 | nucleoside-diphosphate kinase |
| AT1G70330 | -0.7 | 4.48E-06 | -0.4 | 8.28E-04 | -0.4 | 2.31E-03 | -0.4 | 1.80E-02 | 0.0 | 9.76E-01 | -0.4 | 8.76E-02 | hypothetical protein |
| AT4G05120 | -0.3 | 1.76E-02 | 0.2 | 1.99E-01 | -0.5 | 6.02E-03 | -0.3 | 2.31E-02 | -0.4 | 1.22E-02 | 0.1 | 7.46E-01 | Equilibrative nucleotide transporter 3 |
| AT4G05130 | | | | | | | -0.1 | 8.02E-01 | -0.3 | 5.14E-01 | 0.2 | 7.42E-01 | Equilibrative nucleotide transporter 4 |
| AT4G29690 | -0.7 | 2.55E-03 | -0.1 | 4.65E-01 | -0.6 | 2.25E-02 | 0.1 | 8.70E-01 | 0.1 | 6.90E-01 | -0.1 | 8.45E-01 | Alkaline-phosphatase-like family protein ) |
| AT4G29680 | -0.3 | 1.02E-03 | 0.0 | 5.25E-01 | -0.3 | 1.38E-03 | 0.1 | 5.81E-01 | 0.3 | 7.88E-02 | -0.2 | 3.56E-01 | Alkaline-phosphatase-like family protein ) |
| AT5G42770 | 0.4 | 2.42E-02 | 0.2 | 2.45E-01 | 0.2 | 3.02E-01 | 0.0 | 9.16E-01 | 0.2 | 5.83E-01 | -0.2 | 5.64E-01 | Maf-like protein |
| AT5G66550 | 1.1 | 6.83E-04 | 0.7 | 1.36E-02 | 0.4 | 1.62E-01 | -0.3 | 1.99E-01 | 0.0 | 8.94E-01 | -0.3 | 3.85E-01 | Maf-like protein |
| AT3G33520 | 0.1 | 2.56E-02 | 0.0 | 5.03E-01 | 0.2 | 1.82E-02 | -0.1 | 7.08E-01 | -0.3 | 1.93E-01 | 0.2 | 4.44E-01 | Actin-related protein 6 |
| AT4G14930 | 0.1 | 2.29E-01 | 0.1 | 5.66E-01 | 0.1 | 6.17E-01 | -0.6 | 1.39E-02 | -0.4 | 1.08E-01 | -0.2 | 5.68E-01 | Uncharacterized protein At4g14930 |
| AT3G28450 | -0.4 | 1.21E-04 | -0.5 | 7.86E-05 | 0.1 | 4.34E-01 | 0.1 | 6.56E-01 | 0.0 | 8.68E-01 | 0.1 | 6.23E-01 | Inactive LRR receptor-like serine |
| AT1G34200 | -0.1 | 1.65E-01 | 0.0 | 7.29E-01 | -0.1 | 3.78E-01 | -0.1 | 7.93E-01 | -0.2 | 2.30E-01 | 0.2 | 4.75E-01 | F23M19.12 protein |
| AT5G62950 | -0.1 | 6.20E-01 | 0.0 | 9.06E-01 | -0.1 | 6.07E-01 | 0.0 | 8.42E-01 | 0.0 | 9.68E-01 | 0.0 | 9.28E-01 | DNA-directed RNA polymerase III subunit RPC9 |
| AT3G12610 | 0.0 | 9.08E-01 | 0.6 | 1.38E-04 | -0.6 | 2.95E-04 | -0.4 | 1.48E-03 | -0.1 | 3.04E-01 | -0.3 | 7.87E-02 | DNA damage-repair |
| AT4G15950 | 0.3 | 1.63E-01 | 0.1 | 6.73E-01 | 0.2 | 4.41E-01 | -0.1 | 6.81E-01 | -0.1 | 7.24E-01 | 0.0 | 9.97E-01 | DNA-directed RNA polymerases IV and V subunit 4 |
| AT3G20790 | 0.4 | 2.89E-04 | 0.4 | 4.25E-04 | 0.0 | 9.72E-01 | -0.2 | 1.18E-01 | 0.0 | 8.49E-01 | -0.2 | 3.44E-01 | NAD-binding Rossmann-fold superfamily protein |
| AT1G66130 | 0.8 | 3.78E-06 | 0.4 | 1.06E-03 | 0.4 | 1.61E-03 | 0.1 | 7.75E-01 | 0.3 | 3.44E-01 | -0.2 | 5.59E-01 | NAD-binding Rossmann-fold superfamily protein |
| AT3G28956 | | | | | | | 0.4 | 2.58E-01 | 0.2 | 6.13E-01 | 0.2 | 7.31E-01 | DNA-directed RNA polymerase III subunit RPC9 |
| AT4G09670 | -0.4 | 1.85E-04 | -0.2 | 1.81E-02 | -0.2 | 2.34E-02 | -0.1 | 4.74E-01 | 0.0 | 7.98E-01 | -0.1 | 4.18E-01 | Uncharacterized oxidoreductase At4g09670 |

| | | | | | | | | | | | | | |
|---|---|---|---|---|---|---|---|---|---|---|---|---|---|
| AT3G03910 | -3.8 | 5.42E-08 | -3.1 | 2.01E-07 | -0.7 | 2.01E-02 | -1.7 | 4.73E-05 | 0.3 | 2.02E-01 | -2.1 | 8.19E-05 | Glutamate dehydrogenase |
| AT5G23250 | 0.4 | 1.56E-03 | 0.5 | 2.92E-04 | -0.1 | 1.96E-01 | 0.4 | 5.54E-03 | 0.2 | 2.18E-01 | 0.2 | 2.71E-01 | Succinate--CoA ligase subunit alpha, mitochondrial |
| AT4G32272 | -0.2 | 5.67E-02 | -0.2 | 1.39E-01 | 0.0 | 7.23E-01 | -0.1 | 5.50E-01 | -0.2 | 5.85E-01 | 0.0 | 9.82E-01 | Nucleotide |
| AT2G04650 | -0.4 | 4.57E-03 | 0.0 | 9.57E-01 | -0.4 | 1.24E-02 | 0.1 | 8.07E-01 | 0.1 | 5.65E-01 | -0.1 | 7.77E-01 | ADP-glucose pyrophosphorylase family protein |
| AT1G52780 | 0.0 | 6.51E-01 | -0.2 | 4.27E-02 | 0.2 | 1.24E-01 | 0.5 | 1.02E-03 | 0.3 | 1.31E-01 | 0.3 | 1.41E-01 | PII, uridylyltransferase |
| AT3G61690 | 0.0 | 7.88E-01 | -0.2 | 3.42E-02 | 0.2 | 7.81E-02 | 0.0 | 9.30E-01 | 0.1 | 6.33E-01 | -0.1 | 7.64E-01 | Uncharacterized protein F15G16.80 |
| AT1G74910 | 0.8 | 3.15E-06 | 0.5 | 2.19E-04 | 0.3 | 5.16E-03 | -0.3 | 5.29E-02 | -0.6 | 4.65E-03 | 0.2 | 2.38E-01 | F25A4.12 protein |
| AT2G40520 | 0.8 | 8.87E-04 | 0.6 | 8.29E-03 | 0.2 | 3.19E-01 | -0.3 | 4.43E-01 | -0.4 | 2.83E-01 | 0.1 | 7.54E-01 | Polymerase nucleotidyl transferase domain-containing protein |
| AT5G20130 | 1.5 | 7.44E-08 | 1.1 | 1.24E-06 | 0.4 | 2.41E-03 | 0.2 | 2.94E-01 | -0.1 | 5.10E-01 | 0.3 | 1.93E-01 | Sulfate adenylyltransferase subunit |
| AT1G22660 | 0.0 | 6.02E-01 | 0.0 | 9.95E-01 | 0.0 | 6.81E-01 | 0.1 | 7.04E-01 | -0.1 | 8.63E-01 | 0.2 | 6.08E-01 | Polynucleotide adenylyltransferase family protein |
| AT2G44270 | 0.0 | 9.83E-01 | -0.1 | 3.72E-01 | 0.1 | 4.29E-01 | 0.1 | 7.94E-01 | 0.0 | 9.87E-01 | 0.1 | 8.54E-01 | Cytoplasmic tRNA 2-thiolation protein 1 |
| AT4G35910 | 0.2 | 4.06E-02 | 0.0 | 9.99E-01 | 0.2 | 7.92E-02 | -0.3 | 2.04E-01 | -0.3 | 2.27E-01 | 0.0 | 9.63E-01 | Cytoplasmic tRNA 2-thiolation protein 2 |
| AT5G24770 | -0.4 | 2.19E-01 | 0.6 | 4.83E-02 | -1.0 | 1.54E-02 | 0.9 | 2.03E-01 | 2.0 | 1.47E-02 | -1.0 | 2.28E-01 | Vegetative storage protein 2 |
| AT5G51260 | -0.2 | 7.34E-02 | -0.1 | 5.72E-01 | -0.1 | 2.88E-01 | 0.3 | 2.24E-01 | 0.3 | 1.76E-01 | -0.1 | 8.62E-01 | HAD superfamily, subfamily IIIB acid phosphatase |
| AT5G58070 | 0.1 | 7.95E-02 | 0.1 | 4.72E-01 | 0.1 | 3.43E-01 | 0.2 | 1.92E-01 | 0.0 | 9.96E-01 | 0.2 | 2.77E-01 | Temperature-induced lipocalin-1 |
| AT5G24780 | 3.9 | 1.38E-07 | 3.8 | 4.67E-07 | 0.1 | 4.22E-01 | 1.7 | 1.99E-01 | 2.3 | 1.08E-01 | -0.6 | 6.79E-01 | Vegetative storage protein 1 |
| AT3G47860 | 1.2 | 2.92E-07 | 1.1 | 1.64E-06 | 0.1 | 1.50E-01 | -0.4 | 8.25E-03 | -0.1 | 8.00E-01 | -0.3 | 7.40E-02 | Chloroplastic lipocalin |
| AT5G01360 | -2.5 | 2.87E-07 | -0.9 | 4.96E-05 | -1.5 | 9.80E-05 | -0.5 | 1.77E-01 | -0.3 | 5.21E-01 | -0.2 | 6.70E-01 | Trichome birefringence-like protein |
| AT1G29050 | -2.0 | 6.75E-08 | -1.0 | 1.20E-05 | -1.0 | 4.04E-05 | -0.3 | 4.70E-02 | 0.2 | 2.19E-01 | -0.4 | 8.87E-03 | TRICHOME BIREFRINGENCE-LIKE 38 |
| AT5G15900 | -1.9 | 3.25E-06 | -0.5 | 5.27E-03 | -1.4 | 3.07E-04 | -0.4 | 5.13E-01 | 0.2 | 8.18E-01 | -0.5 | 4.48E-01 | TRICHOME BIREFRINGENCE-LIKE 19 |
| AT5G15890 | -1.9 | 2.68E-06 | -0.5 | 5.16E-03 | -1.4 | 2.35E-04 | -0.8 | 8.51E-02 | 0.0 | 9.73E-01 | -0.8 | 1.63E-01 | Trichome birefringence-like N-terminal domain-containing protein |
| AT2G38320 | -1.7 | 6.18E-07 | -0.5 | 2.11E-03 | -1.3 | 7.02E-05 | 0.1 | 8.11E-01 | 0.0 | 8.81E-01 | 0.0 | 9.75E-01 | TRICHOME BIREFRINGENCE-LIKE 34 |
| AT5G06230 | -1.5 | 2.01E-06 | -1.3 | 1.07E-05 | -0.2 | 2.78E-01 | | | | | | | TRICHOME BIREFRINGENCE-LIKE 9 |
| AT4G01080 | -1.3 | 1.43E-02 | -1.2 | 2.15E-02 | -0.1 | 9.24E-01 | 0.5 | 6.12E-01 | 2.3 | 9.74E-03 | -1.8 | 4.65E-02 | Protein trichome birefringence-like 26 |
| AT2G31110 | -0.8 | 2.85E-05 | -0.4 | 4.73E-03 | -0.4 | 7.31E-03 | 0.2 | 3.29E-01 | -0.1 | 6.92E-01 | 0.3 | 2.33E-01 | Trichome birefringence-like protein |
| AT2G42570 | -0.7 | 4.23E-04 | 0.2 | 1.04E-01 | -1.0 | 2.72E-04 | -0.6 | 1.26E-04 | 0.0 | 9.91E-01 | -0.6 | 2.97E-03 | Protein trichome birefringence-like 39 |
| AT1G60790 | -0.5 | 4.16E-04 | -0.1 | 1.70E-01 | -0.4 | 1.01E-02 | -0.3 | 5.21E-02 | 0.0 | 9.23E-01 | -0.3 | 1.09E-01 | Protein trichome birefringence-like 2 |
| AT2G40150 | -0.5 | 2.13E-04 | -0.1 | 2.63E-01 | -0.4 | 3.98E-03 | -0.4 | 1.01E-01 | 0.1 | 7.00E-01 | -0.5 | 7.55E-02 | Trichome birefringence-like N-terminal domain-containing protein |
| AT4G23790 | -0.4 | 5.74E-04 | -0.1 | 3.84E-01 | -0.4 | 6.37E-03 | -0.1 | 2.81E-01 | 0.1 | 5.30E-01 | -0.2 | 1.68E-01 | Protein trichome birefringence-like 24 |
| AT4G25360 | -0.4 | 1.23E-04 | -0.2 | 6.68E-03 | -0.2 | 3.75E-02 | -0.2 | 2.12E-01 | 0.1 | 5.59E-01 | -0.3 | 1.51E-01 | Protein trichome birefringence-like 18 |
| AT4G11090 | -0.3 | 7.16E-04 | -0.1 | 1.12E-01 | -0.2 | 2.64E-02 | -0.3 | 3.60E-02 | -0.1 | 4.84E-01 | -0.2 | 3.52E-01 | Protein trichome birefringence-like 23 |
| AT1G48880 | -0.3 | 8.71E-02 | -0.1 | 5.37E-01 | -0.2 | 3.59E-01 | 0.1 | 7.95E-01 | 0.0 | 9.90E-01 | 0.1 | 8.07E-01 | Protein trichome berefringence-like 7 |
| AT5G06700 | -0.2 | 2.73E-02 | 0.0 | 8.21E-01 | -0.2 | 6.64E-02 | -0.3 | 1.24E-02 | -0.1 | 3.72E-01 | -0.2 | 3.08E-01 | Protein trichome birefringence |
| AT2G14530 | -0.2 | 4.25E-02 | 0.0 | 7.22E-01 | -0.2 | 1.39E-01 | -0.2 | 2.08E-01 | -0.2 | 2.79E-01 | 0.0 | 9.83E-01 | Protein trichome birefringence-like 13 |
| AT3G14850 | -0.2 | 1.20E-01 | -0.1 | 4.32E-01 | -0.1 | 5.28E-01 | -0.5 | 7.21E-02 | 0.0 | 9.39E-01 | -0.5 | 1.96E-01 | Protein trichome birefringence-like 41 |
| AT5G19160 | -0.2 | 2.51E-01 | -0.1 | 6.81E-01 | -0.1 | 5.74E-01 | 0.0 | 9.86E-01 | 0.0 | 9.90E-01 | 0.0 | 9.97E-01 | TRICHOME BIREFRINGENCE-LIKE 11 |
| AT3G54260 | -0.2 | 1.06E-01 | 0.2 | 1.09E-01 | -0.3 | 1.23E-02 | -0.3 | 3.58E-02 | 0.0 | 9.74E-01 | -0.3 | 1.01E-01 | Protein trichome birefringence-like 36 |
| AT3G62390 | 0.0 | 9.83E-01 | 0.4 | 7.19E-04 | -0.4 | 2.23E-03 | -0.3 | 2.28E-01 | -0.1 | 8.82E-01 | -0.3 | 4.27E-01 | TRICHOME BIREFRINGENCE-LIKE 6 |
| AT2G30010 | 0.0 | 9.99E-01 | 0.4 | 1.17E-02 | -0.4 | 1.99E-02 | 0.0 | 9.73E-01 | -0.2 | 2.61E-01 | 0.2 | 3.59E-01 | TRICHOME BIREFRINGENCE-LIKE 45 |
| AT3G61020 | 0.0 | 5.86E-01 | 0.1 | 3.92E-01 | 0.0 | 7.90E-01 | -0.1 | 5.03E-01 | -0.2 | 3.21E-01 | 0.1 | 7.95E-01 | Putative truncated protein trichome birefringence-like 46 |
| AT2G34070 | 0.1 | 2.83E-01 | 0.1 | 5.13E-01 | 0.1 | 7.54E-01 | -0.7 | 4.53E-04 | -0.5 | 6.11E-03 | -0.1 | 5.35E-01 | TRICHOME BIREFRINGENCE-LIKE 37 |
| AT1G70230 | 0.1 | 8.02E-02 | 0.0 | 7.64E-01 | 0.1 | 1.97E-01 | 0.0 | 7.41E-01 | 0.1 | 6.24E-01 | -0.1 | 5.11E-01 | Xyloglucan O-acetyltransferase 1 |
| AT5G64470 | 0.2 | 1.43E-01 | 0.2 | 1.55E-01 | 0.0 | 9.97E-01 | 0.1 | 6.22E-01 | 0.1 | 7.74E-01 | 0.0 | 9.34E-01 | Trichome birefringence-like protein |
| AT5G58600 | 0.2 | 2.14E-01 | 0.4 | 9.26E-03 | -0.2 | 1.30E-01 | 0.0 | 9.80E-01 | -0.3 | 3.49E-01 | 0.2 | 3.63E-01 | Protein PMR5 |
| AT5G20590 | 0.3 | 6.33E-03 | 0.2 | 2.64E-02 | 0.1 | 5.51E-01 | 0.0 | 8.73E-01 | 0.1 | 5.96E-01 | -0.1 | 7.24E-01 | Uncharacterized protein At5g20590 |
| AT1G01430 | 0.4 | 3.29E-03 | 0.3 | 6.33E-03 | 0.0 | 8.46E-01 | -0.2 | 1.48E-01 | -0.3 | 8.04E-03 | 0.2 | 2.42E-01 | Protein trichome birefringence-like 25 |
| AT3G06080 | 0.6 | 1.86E-04 | 0.3 | 1.29E-02 | 0.3 | 3.61E-02 | -0.4 | 2.65E-02 | -0.6 | 3.57E-03 | 0.2 | 4.04E-01 | TBL10 |
| AT2G40320 | -1.4 | 1.16E-06 | -0.3 | 8.40E-03 | -1.1 | 9.43E-05 | -0.2 | 3.00E-01 | 0.1 | 7.71E-01 | -0.3 | 1.50E-01 | Protein trichome birefringence-like 33 |
| AT3G55990 | -1.2 | 9.58E-07 | -0.2 | 4.01E-02 | -1.0 | 4.65E-05 | -0.2 | 5.95E-01 | 0.1 | 7.70E-01 | -0.3 | 4.52E-01 | Xylan O-acetyltransferase 1 |
| AT3G11030 | -0.6 | 1.28E-04 | -0.2 | 7.88E-02 | -0.4 | 5.40E-03 | -0.2 | 3.41E-01 | -0.1 | 8.47E-01 | -0.1 | 6.02E-01 | Protein trichome birefringence-like 32 |
| AT5G01620 | -0.3 | 2.04E-02 | 0.1 | 5.21E-01 | -0.3 | 1.61E-02 | -0.1 | 5.92E-01 | 0.0 | 9.00E-01 | -0.1 | 5.83E-01 | Protein trichome birefringence-like 35 |
| AT3G51970 | -0.6 | 5.47E-04 | 0.1 | 2.48E-01 | -0.8 | 7.78E-04 | -0.5 | 2.65E-01 | 0.1 | 8.79E-01 | -0.6 | 2.87E-01 | Acyl-CoA--sterol O-acyltransferase 1 |
| AT4G17910 | -0.1 | 1.93E-01 | 0.1 | 5.66E-01 | -0.2 | 1.22E-01 | 0.4 | 5.15E-02 | 0.4 | 6.45E-02 | 0.0 | 9.66E-01 | Transferases, transferring acyl groups |
| AT5G13640 | 0.0 | 8.74E-01 | -0.2 | 3.35E-02 | 0.3 | 3.55E-02 | -0.1 | 5.20E-01 | 0.0 | 8.47E-01 | -0.2 | 5.11E-01 | Phospholipid:diacylglycerol acyltransferase |
| AT1G80950 | 0.3 | 3.57E-03 | 0.2 | 5.19E-03 | 0.0 | 9.52E-01 | -0.1 | 5.07E-01 | -0.1 | 7.31E-01 | 0.0 | 8.94E-01 | Phospholipid |
| AT1G27480 | 1.6 | 1.63E-07 | 0.9 | 1.42E-05 | 0.7 | 2.92E-04 | 0.2 | 8.22E-02 | 0.2 | 3.08E-01 | 0.1 | 7.11E-01 | Lecithin-cholesterol acyltransferase-like 1 |
| AT4G19860 | -0.3 | 5.70E-03 | -0.1 | 1.88E-01 | -0.2 | 1.00E-01 | -0.6 | 2.34E-03 | -0.3 | 1.07E-01 | -0.3 | 2.62E-01 | Lecithin-cholesterol acyltransferase-like 4 |
| AT1G20380 | 0.5 | 4.11E-03 | 0.3 | 2.38E-02 | 0.1 | 4.77E-01 | 1.1 | 9.39E-04 | 0.5 | 1.58E-01 | 0.6 | 5.90E-02 | Prolyl endopeptidase |
| AT5G24380 | -1.3 | 2.73E-06 | -0.7 | 1.77E-04 | -0.6 | 4.79E-03 | 0.3 | 3.22E-01 | 0.4 | 1.96E-01 | -0.1 | 7.88E-01 | AT5G24380 protein |
| AT1G65730 | -1.2 | 3.40E-04 | -0.6 | 1.44E-02 | -0.6 | 5.84E-02 | -0.3 | 5.27E-01 | -0.3 | 4.99E-01 | 0.0 | 9.49E-01 | Probable metal-nicotianamine transporter YSL7 |
| AT4G10770 | -1.1 | 1.30E-06 | -0.6 | 2.40E-04 | -0.5 | 8.71E-04 | -0.3 | 2.47E-02 | 0.2 | 2.30E-01 | -0.4 | 8.17E-03 | Oligopeptide transporter 7 |
| AT5G64410 | -0.7 | 1.08E-04 | -0.6 | 1.07E-03 | -0.2 | 2.04E-01 | -0.1 | 4.66E-01 | -0.2 | 3.15E-01 | 0.1 | 7.93E-01 | Oligopeptide transporter 4 |
| AT5G41000 | -0.5 | 1.18E-04 | -0.1 | 3.22E-01 | -0.5 | 2.01E-03 | 0.0 | 9.08E-01 | 0.2 | 4.89E-01 | -0.2 | 6.24E-01 | YELLOW STRIPE like 4 |
| AT1G48370 | -0.4 | 3.33E-03 | -0.3 | 2.14E-02 | -0.1 | 4.40E-01 | 0.4 | 4.78E-01 | 0.7 | 2.15E-01 | -0.3 | 6.74E-01 | YELLOW STRIPE like 8 |
| AT3G27020 | -0.3 | 3.85E-03 | -0.2 | 5.75E-02 | -0.1 | 2.02E-01 | -0.2 | 3.36E-01 | 0.0 | 9.28E-01 | -0.2 | 4.45E-01 | Probable metal-nicotianamine transporter YSL6 |
| AT5G55930 | 0.1 | 7.26E-01 | -0.2 | 2.60E-01 | 0.3 | 1.93E-01 | 0.1 | 8.53E-01 | -0.3 | 6.83E-01 | 0.5 | 5.73E-01 | Oligopeptide transporter 1 |
| AT3G17650 | 0.2 | 2.27E-02 | 0.1 | 1.88E-01 | 0.1 | 3.17E-01 | -0.2 | 3.75E-02 | -0.3 | 1.81E-02 | 0.1 | 7.42E-01 | Uncharacterized protein |
| AT4G27730 | 0.3 | 6.55E-02 | 0.2 | 3.45E-01 | 0.2 | 4.35E-01 | -0.2 | 5.44E-01 | -0.4 | 2.06E-01 | 0.2 | 5.59E-01 | Oligopeptide transporter 6 |
| AT5G53550 | 0.4 | 4.96E-03 | 0.2 | 6.76E-02 | 0.2 | 2.30E-01 | 0.0 | 9.34E-01 | -0.4 | 7.41E-02 | 0.4 | 1.82E-01 | YELLOW STRIPE like 3 |
| AT4G16370 | 1.6 | 9.46E-08 | 1.0 | 3.10E-06 | 0.6 | 6.58E-04 | 1.9 | 3.93E-06 | 0.8 | 1.70E-02 | 1.1 | 8.72E-03 | Oligopeptide transporter 3 |
| AT3G29590 | -3.2 | 4.38E-04 | -2.3 | 1.83E-03 | -0.9 | 2.89E-01 | 1.4 | 7.05E-02 | 2.6 | 2.84E-03 | -1.2 | 4.95E-02 | Malonyl-CoA:anthocyanidin 5-O-glucoside-6"-O-malonyltransferase |
| AT3G29670 | -1.9 | 6.10E-08 | -1.2 | 1.04E-06 | -0.7 | 5.58E-04 | 0.2 | 4.02E-01 | -0.1 | 7.67E-01 | 0.3 | 3.43E-01 | Phenolic glucoside malonyltransferase 2 |
| AT4G34050 | -1.4 | 1.06E-07 | -0.5 | 1.82E-04 | -0.9 | 2.81E-05 | 0.0 | 8.30E-01 | 0.3 | 7.57E-02 | -0.2 | 1.90E-01 | S-adenosyl-L-methionine-dependent methyltransferases superfamily protein |
| AT3G62000 | 1.1 | 1.16E-05 | 0.7 | 3.09E-04 | 0.3 | 4.73E-02 | 0.0 | 8.54E-01 | 0.1 | 6.51E-01 | -0.1 | 8.30E-01 | S-adenosyl-L-methionine-dependent methyltransferases superfamily protein |
| AT1G67980 | | | | | | | 0.1 | 8.92E-01 | -3.0 | 9.41E-03 | 3.1 | 1.75E-02 | Caffeoyl-CoA 3-O-methyltransferase |
| AT4G35160 | -1.7 | 1.71E-06 | -1.0 | 7.94E-05 | -0.6 | 4.29E-03 | 0.4 | 1.33E-02 | 0.4 | 4.20E-02 | 0.1 | 8.47E-01 | Uncharacterized protein |
| AT1G21120 | 0.6 | 2.07E-02 | -0.1 | 7.77E-01 | 0.7 | 3.00E-02 | 2.0 | 1.03E-05 | 0.9 | 2.09E-02 | 1.1 | 4.04E-03 | O-methyltransferase family protein |
| AT1G77520 | -2.8 | 1.83E-08 | -2.1 | 1.28E-07 | -0.7 | 6.00E-04 | -0.3 | 3.84E-01 | 0.8 | 1.25E-02 | -1.2 | 7.36E-03 | O-methyltransferase family protein |
| AT1G77530 | -2.1 | 2.25E-08 | -1.6 | 2.19E-07 | -0.5 | 1.25E-03 | -0.6 | 4.82E-02 | 0.1 | 8.09E-01 | -0.7 | 6.05E-02 | O-methyltransferase family protein |
| AT1G21130 | -0.6 | 5.53E-04 | -0.1 | 2.81E-01 | -0.4 | 7.23E-03 | 0.0 | 8.19E-01 | -0.6 | 2.16E-03 | 0.6 | 1.62E-02 | Indole glucosinolate O-methyltransferase 4 |
| AT1G76790 | -0.1 | 4.60E-01 | 0.1 | 4.09E-01 | -0.2 | 1.73E-01 | -0.1 | 8.26E-01 | 0.5 | 9.45E-02 | -0.6 | 1.11E-01 | Indole glucosinolate O-methyltransferase 5 |
| AT1G21110 | 0.3 | 8.45E-02 | -0.4 | 2.08E-02 | 0.7 | 3.73E-03 | 2.9 | 4.33E-07 | 1.9 | 1.76E-04 | 1.0 | 2.84E-03 | Indole glucosinolate O-methyltransferase 3 |
| AT1G33030 | | | | | | | -0.3 | 7.00E-01 | -2.8 | 2.41E-02 | 2.5 | 1.05E-01 | At1g33030 |

| | | | | | | | | | | | | | |
|---|---|---|---|---|---|---|---|---|---|---|---|---|---|
| AT3G61990 | 0.4 | 2.92E-03 | 0.1 | 1.98E-01 | 0.2 | 5.20E-02 | 0.2 | 1.92E-01 | 0.2 | 1.32E-01 | -0.1 | 8.21E-01 | S-adenosyl-L-methionine-dependent methyltransferases superfamily protein |
| AT5G51130 | 0.3 | 4.18E-02 | 0.0 | 9.74E-01 | 0.3 | 9.70E-02 | 0.3 | 2.59E-01 | 0.1 | 8.09E-01 | 0.2 | 4.93E-01 | RNA methyltransferase |
| AT1G24735 | -1.8 | 2.92E-07 | -0.7 | 9.87E-05 | -1.1 | 1.15E-04 | -0.9 | 6.74E-02 | 0.0 | 9.89E-01 | -0.9 | 9.61E-02 | F5A9.20 |
| AT4G35630 | -0.2 | 4.07E-02 | -0.1 | 6.31E-01 | -0.2 | 1.40E-01 | 0.4 | 9.43E-03 | 0.4 | 1.71E-02 | 0.0 | 9.92E-01 | Phosphoserine aminotransferase 1, chloroplastic |
| AT2G17630 | -0.1 | 5.65E-01 | 0.3 | 1.96E-02 | -0.4 | 1.55E-02 | 0.1 | 5.67E-01 | 0.0 | 8.82E-01 | 0.1 | 8.19E-01 | Phosphoserine aminotransferase 2, chloroplastic |
| AT1G61800 | -4.6 | 6.83E-05 | -4.7 | 1.01E-04 | 0.0 | 9.74E-01 | -0.4 | 7.58E-01 | -1.3 | 3.97E-01 | | | Glucose-6-phosphate |
| AT5G46110 | 1.0 | 3.09E-06 | 0.8 | 3.44E-05 | 0.2 | 8.43E-02 | -0.2 | 1.07E-01 | -0.1 | 3.91E-01 | -0.1 | 7.40E-01 | Glucose-6-phosphate |
| AT3G01550 | 1.4 | 2.79E-06 | 1.1 | 3.26E-05 | 0.3 | 7.45E-02 | 0.1 | 7.24E-01 | 0.1 | 8.33E-01 | 0.0 | 9.57E-01 | Phosphoenolpyruvate |
| AT3G54470 | 0.6 | 1.20E-04 | 0.6 | 2.97E-04 | 0.0 | 7.95E-01 | 0.5 | 1.04E-03 | 0.4 | 6.82E-03 | 0.1 | 7.53E-01 | Uridine 5'-monophosphate synthase |
| AT2G29370 | -1.5 | 5.83E-04 | -2.3 | 1.74E-04 | 0.8 | 1.28E-01 | | | | | | | Tropinone reductase homolog At2g29370 |
| AT2G43020 | -1.3 | 9.17E-07 | -0.9 | 2.38E-05 | -0.4 | 7.05E-03 | -0.8 | 3.22E-04 | 0.0 | 9.42E-01 | -0.8 | 3.63E-03 | PAO2 |
| AT5G39410 | -1.1 | 2.03E-06 | -0.6 | 1.33E-04 | -0.4 | 3.82E-03 | -0.4 | 4.02E-03 | 0.0 | 9.48E-01 | -0.4 | 1.28E-02 | Probable mitochondrial saccharopine dehydrogenase-like oxidoreductase At5g39410 |
| AT1G60680 | -0.7 | 9.97E-06 | -0.2 | 4.56E-02 | -0.5 | 3.88E-04 | 0.1 | 5.59E-01 | 0.2 | 2.08E-01 | -0.1 | 5.61E-01 | NAD-linked oxidoreductase superfamily protein |
| AT2G47140 | -0.6 | 8.40E-05 | 0.3 | 5.07E-03 | -1.0 | 4.42E-05 | 0.3 | 2.22E-01 | 0.4 | 2.27E-01 | 0.0 | 9.28E-01 | Short-chain dehydrogenase reductase 3b |
| AT3G08710 | -0.6 | 1.04E-05 | -0.4 | 5.99E-04 | -0.2 | 1.26E-02 | 0.3 | 2.30E-01 | 0.3 | 3.19E-01 | 0.0 | 9.84E-01 | Thioredoxin H9 |
| AT3G44190 | -0.6 | 2.25E-05 | -0.2 | 2.12E-02 | -0.4 | 1.62E-03 | 0.1 | 5.06E-01 | 0.3 | 4.79E-02 | -0.2 | 1.70E-01 | FAD |
| AT5G40942 | -0.6 | 1.94E-03 | -0.1 | 6.49E-01 | -0.5 | 1.40E-02 | 0.1 | 5.80E-01 | 0.3 | 2.64E-01 | -0.1 | 6.43E-01 | Putative ribonucleoside-diphosphate reductase small chain B |
| AT1G75280 | -0.6 | 5.54E-05 | -0.4 | 1.71E-03 | -0.2 | 5.12E-02 | -0.1 | 5.82E-01 | 0.0 | 8.80E-01 | -0.1 | 5.67E-01 | Isoflavone reductase homolog P3 |
| AT4G05530 | -0.6 | 5.65E-05 | -0.4 | 6.05E-04 | -0.1 | 1.67E-01 | 0.1 | 2.81E-01 | 0.2 | 1.33E-01 | -0.1 | 7.06E-01 | Short-chain dehydrogenase |
| AT1G02020 | -0.5 | 3.43E-05 | -0.5 | 1.51E-04 | -0.1 | 4.92E-01 | 0.1 | 7.83E-01 | 0.1 | 5.87E-01 | -0.1 | 8.32E-01 | Nitroreductase family protein |
| AT3G08970 | -0.5 | 2.48E-02 | -0.7 | 8.02E-03 | 0.2 | 4.96E-01 | 0.4 | 2.91E-01 | 0.8 | 6.28E-02 | -0.4 | 4.21E-01 | DnaJ protein ERDJ3A |
| AT1G65840 | -0.4 | 1.50E-03 | -0.2 | 1.24E-01 | -0.3 | 4.13E-02 | 0.2 | 2.22E-01 | 0.7 | 5.07E-04 | -0.5 | 1.34E-02 | Putative polyamine oxidase |
| AT4G30210 | -0.4 | 2.74E-03 | -0.5 | 8.32E-04 | 0.1 | 3.35E-01 | -0.2 | 1.50E-01 | -0.2 | 1.22E-01 | 0.0 | 8.78E-01 | AT4g30210 |
| AT3G47360 | -0.4 | 7.35E-02 | -0.4 | 8.61E-02 | 0.0 | 9.88E-01 | -0.5 | 7.67E-02 | 0.0 | 9.23E-01 | -0.4 | 2.05E-01 | 11-beta-hydroxysteroid dehydrogenase-like 3 |
| AT1G62610 | -0.3 | 6.83E-03 | -0.3 | 2.97E-02 | -0.1 | 5.57E-01 | -0.4 | 2.36E-01 | -0.2 | 6.14E-01 | -0.2 | 6.82E-01 | At1g62610 -binding Rossmann-fold superfamily protein) |
| AT3G03350 | -0.3 | 1.61E-01 | -0.6 | 1.49E-02 | 0.3 | 2.19E-01 | | | | | | | NAD-binding Rossmann-fold superfamily protein |
| AT3G03360 | -0.2 | 2.66E-02 | -0.3 | 4.73E-03 | 0.1 | 3.39E-01 | -0.3 | 1.93E-01 | 0.2 | 4.63E-01 | -0.5 | 9.58E-02 | F-box domain-containing protein |
| AT4G25570 | -0.2 | 1.95E-02 | 0.1 | 5.42E-01 | -0.3 | 1.27E-02 | -0.2 | 7.98E-02 | -0.1 | 7.03E-01 | -0.1 | 4.19E-01 | Cytochrome b561 |
| AT4G16310 | -0.2 | 2.50E-02 | -0.3 | 2.44E-03 | 0.1 | 1.93E-01 | 0.1 | 7.94E-01 | -0.1 | 7.79E-01 | 0.1 | 6.59E-01 | Lysine-specific histone demethylase 1 homolog 3 |
| AT3G55290 | -0.1 | 2.61E-01 | 0.1 | 3.81E-01 | -0.3 | 1.05E-01 | -0.3 | 3.53E-01 | 0.4 | 3.43E-01 | -0.7 | 6.67E-02 | hypothetical protein |
| AT4G13010 | -0.1 | 6.59E-02 | -0.3 | 8.61E-04 | 0.2 | 2.28E-02 | 0.0 | 9.53E-01 | 0.0 | 9.13E-01 | 0.0 | 8.98E-01 | Chloroplast envelope quinone oxidoreductase homolog |
| AT4G15765 | -0.1 | 3.78E-01 | 0.1 | 5.42E-01 | -0.2 | 2.23E-01 | -0.2 | 3.88E-01 | -0.2 | 4.25E-01 | 0.0 | 9.81E-01 | FAD |
| AT3G23510 | -0.1 | 4.01E-01 | 0.0 | 8.99E-01 | -0.1 | 4.23E-01 | -0.3 | 3.76E-01 | -0.2 | 6.98E-01 | -0.1 | 8.05E-01 | Amine oxidase domain-containing protein |
| AT5G61510 | -0.1 | 7.76E-02 | -0.2 | 7.64E-03 | 0.1 | 2.15E-01 | -0.2 | 5.49E-02 | -0.3 | 4.30E-02 | 0.0 | 8.64E-01 | Putative quinone oxidoreductase |
| AT1G62830 | -0.1 | 4.60E-01 | -0.1 | 4.85E-01 | 0.0 | 9.95E-01 | 0.1 | 6.67E-01 | 0.1 | 7.99E-01 | 0.0 | 9.51E-01 | Lysine-specific histone demethylase 1 homolog 1 |
| AT3G53880 | 0.0 | 9.93E-01 | 0.1 | 2.69E-01 | -0.1 | 3.45E-01 | -0.1 | 7.47E-01 | 0.1 | 7.81E-01 | -0.2 | 5.46E-01 | NAD-linked oxidoreductase superfamily protein |
| AT3G15090 | 0.0 | 7.40E-01 | 0.0 | 7.22E-01 | 0.0 | 9.73E-01 | 0.1 | 7.65E-01 | 0.2 | 2.62E-01 | -0.1 | 4.65E-01 | hypothetical protein |
| AT3G13682 | 0.0 | 8.12E-01 | 0.1 | 5.05E-01 | -0.1 | 7.27E-01 | 0.4 | 4.10E-01 | 0.1 | 8.53E-01 | 0.2 | 6.66E-01 | Lysine-specific histone demethylase 1 homolog 2 |
| AT5G10050 | 0.1 | 4.93E-01 | 0.1 | 2.83E-01 | 0.0 | 7.32E-01 | -0.2 | 6.95E-02 | 0.1 | 7.48E-01 | -0.3 | 1.01E-01 | 3-oxoacyl- reductase |
| AT5G63620 | 0.1 | 2.57E-01 | 0.1 | 6.68E-02 | -0.1 | 4.62E-01 | -0.2 | 1.11E-01 | -0.2 | 1.40E-01 | 0.0 | 9.80E-01 | Alcohol dehydrogenase-like protein |
| AT5G23440 | 0.1 | 2.16E-01 | 0.1 | 8.75E-02 | 0.0 | 6.23E-01 | 0.0 | 8.70E-01 | -0.1 | 7.36E-01 | 0.0 | 8.90E-01 | hypothetical protein |
| AT5G67290 | 0.1 | 8.66E-02 | -0.1 | 6.98E-02 | 0.3 | 8.00E-03 | -0.3 | 4.20E-02 | -0.1 | 7.48E-01 | -0.2 | 1.74E-01 | Uncharacterized protein At5g67290 |
| AT2G22890 | 0.1 | 4.04E-01 | 0.5 | 5.00E-03 | -0.4 | 4.52E-02 | 0.0 | 9.46E-01 | -0.2 | 4.08E-01 | 0.2 | 5.12E-01 | Fatty acid desaturase 4-like 2, chloroplastic |
| AT3G10390 | 0.1 | 7.37E-02 | -0.1 | 3.58E-01 | 0.2 | 3.10E-02 | 0.0 | 8.92E-01 | 0.0 | 9.09E-01 | 0.0 | 9.98E-01 | Protein FLOWERING locus D-like protein |
| AT3G03330 | 0.2 | 3.80E-02 | 0.1 | 2.18E-01 | 0.1 | 4.19E-01 | -0.3 | 2.24E-02 | -0.3 | 3.80E-02 | 0.0 | 9.90E-01 | hypothetical protein |
| AT5G18210 | 0.2 | 1.44E-01 | 0.0 | 9.18E-01 | 0.2 | 1.95E-01 | -0.2 | 4.68E-01 | -0.3 | 2.39E-01 | 0.1 | 7.19E-01 | hypothetical protein |
| AT5G49555 | 0.2 | 2.93E-02 | 0.1 | 5.53E-01 | 0.2 | 1.38E-01 | -0.1 | 4.89E-01 | 0.0 | 9.44E-01 | -0.1 | 5.67E-01 | Amine oxidase domain-containing protein |
| AT2G22650 | 0.2 | 2.46E-01 | 0.3 | 1.22E-01 | -0.1 | 7.05E-01 | -0.5 | 7.74E-02 | -0.4 | 1.90E-01 | -0.1 | 8.39E-01 | FAD dependent oxidoreductase domain-containing protein |
| AT5G48440 | 0.3 | 1.14E-03 | 0.2 | 3.62E-02 | 0.1 | 1.09E-01 | 0.0 | 8.88E-01 | -0.1 | 7.94E-01 | 0.0 | 9.17E-01 | At5g48440 |
| AT2G29320 | 0.3 | 6.74E-03 | 0.1 | 4.59E-01 | 0.2 | 4.94E-02 | 0.3 | 1.06E-01 | 0.2 | 3.82E-01 | 0.1 | 6.75E-01 | NAD-binding Rossmann-fold superfamily protein |
| AT5G62630 | 0.3 | 4.17E-03 | 0.5 | 9.37E-04 | -0.1 | 2.93E-01 | 0.8 | 1.28E-06 | 0.4 | 4.97E-03 | 0.4 | 7.54E-03 | HIPL2 protein |
| AT5G65205 | 0.4 | 3.89E-02 | 0.0 | 8.89E-01 | 0.3 | 1.11E-01 | 0.0 | 8.99E-01 | -0.1 | 7.32E-01 | 0.2 | 6.81E-01 | Uncharacterized protein At5g65205 |
| AT4G27760 | 0.4 | 3.97E-04 | 0.4 | 5.92E-04 | 0.0 | 9.92E-01 | -0.2 | 1.12E-01 | -0.2 | 1.19E-01 | 0.0 | 9.42E-01 | Dehydrogenase |
| AT3G45770 | 0.4 | 9.57E-05 | 0.1 | 8.41E-02 | 0.3 | 3.78E-03 | -0.1 | 5.50E-01 | -0.2 | 3.03E-01 | 0.1 | 7.40E-01 | Polyketide synthase, enoylreductase family protein |
| AT1G19540 | 0.4 | 4.37E-02 | 0.4 | 4.86E-02 | 0.0 | 9.92E-01 | -0.6 | 1.25E-01 | -0.8 | 5.04E-02 | 0.2 | 6.40E-01 | NmrA-like domain-containing protein |
| AT5G38900 | 0.5 | 8.04E-02 | -0.3 | 3.34E-01 | 0.8 | 3.79E-02 | 1.6 | 2.79E-04 | -0.4 | 4.57E-01 | 2.0 | 1.54E-03 | Thioredoxin superfamily protein |
| AT2G47130 | 0.5 | 3.65E-03 | -0.2 | 1.55E-01 | 0.8 | 1.62E-03 | 0.7 | 6.14E-03 | 0.2 | 6.53E-01 | 0.6 | 5.76E-02 | Short-chain dehydrogenase reductase 3a |
| AT5G06060 | 0.6 | 5.56E-05 | 0.3 | 1.10E-02 | 0.3 | 8.56E-03 | 0.0 | 8.61E-01 | 0.0 | 9.31E-01 | 0.0 | 9.70E-01 | Tropinone reductase homolog At5g06060 |
| AT5G08410 | 0.8 | 3.17E-04 | 0.6 | 2.36E-03 | 0.2 | 3.10E-01 | -0.6 | 1.13E-02 | 0.2 | 4.51E-01 | -0.8 | 1.32E-02 | Ferredoxin |
| AT1G57770 | 0.8 | 6.07E-05 | 0.6 | 4.52E-04 | 0.1 | 2.69E-01 | 0.5 | 2.70E-02 | 0.4 | 1.76E-01 | 0.2 | 5.95E-01 | FAD |
| AT1G07440 | 0.8 | 2.95E-04 | 0.3 | 8.11E-02 | 0.5 | 1.30E-02 | 0.0 | 7.58E-01 | 0.0 | 8.19E-01 | -0.1 | 6.67E-01 | NAD-binding Rossmann-fold superfamily protein |
| AT5G02240 | 0.8 | 1.50E-06 | 0.7 | 9.67E-06 | 0.1 | 2.08E-01 | -0.1 | 4.47E-01 | 0.0 | 8.07E-01 | -0.1 | 4.53E-01 | NAD-binding domain-containing protein |
| AT2G37760 | 0.9 | 1.25E-04 | -0.4 | 5.38E-02 | 1.3 | 1.54E-04 | 0.4 | 1.03E-01 | 0.4 | 8.28E-02 | -0.1 | 8.64E-01 | NAD-linked oxidoreductase superfamily protein |
| AT2G29360 | 1.0 | 8.42E-05 | 0.3 | 5.96E-02 | 0.6 | 4.11E-03 | -0.1 | 6.53E-01 | 0.0 | 9.57E-01 | -0.1 | 7.88E-01 | Uncharacterized protein |
| AT3G23530 | 1.0 | 5.94E-07 | 1.0 | 1.61E-06 | 0.0 | 9.35E-01 | 0.1 | 3.05E-01 | -0.1 | 3.90E-01 | 0.2 | 1.18E-01 | Amine oxidase domain-containing protein |
| AT1G63940 | 1.1 | 4.60E-07 | 0.9 | 3.88E-06 | 0.2 | 7.33E-02 | 0.2 | 9.90E-02 | -0.1 | 5.35E-01 | 0.3 | 9.78E-02 | Monodehydroascorbate reductase 6 |
| AT2G29330 | 1.1 | 4.82E-06 | 0.7 | 3.95E-04 | 0.4 | 9.22E-03 | 1.0 | 1.03E-03 | 0.6 | 8.01E-02 | 0.5 | 9.42E-02 | Tropinone reductase homolog At2g29330 |
| AT5G07440 | 1.2 | 8.11E-07 | 1.2 | 1.92E-06 | 0.0 | 8.43E-01 | 1.1 | 5.74E-06 | 0.0 | 8.73E-01 | 1.2 | 2.13E-04 | Glutamate dehydrogenase 2 |
| AT2G29170 | 1.2 | 1.37E-04 | 0.8 | 4.21E-03 | 0.4 | 8.02E-02 | 0.3 | 6.07E-01 | 0.3 | 6.77E-01 | | | Tropinone reductase homolog At2g29170 |
| AT4G30720 | 1.2 | 3.28E-07 | 1.0 | 3.92E-06 | 0.3 | 2.24E-02 | 0.0 | 8.65E-01 | -0.2 | 4.63E-01 | 0.2 | 4.16E-01 | FAD |
| AT1G04620 | 1.2 | 4.80E-07 | 0.7 | 4.58E-05 | 0.5 | 1.32E-03 | -0.1 | 6.83E-01 | 0.0 | 8.37E-01 | -0.1 | 5.91E-01 | 7-hydroxymethyl chlorophyll a reductase, chloroplastic |
| AT5G18170 | 1.3 | 8.24E-07 | 1.2 | 2.68E-06 | 0.0 | 6.88E-01 | 0.1 | 8.41E-01 | -0.9 | 2.41E-03 | 1.0 | 3.06E-03 | AT5g18170 |
| AT3G11170 | 1.3 | 1.01E-07 | 1.1 | 7.82E-07 | 0.2 | 3.15E-02 | 0.3 | 2.69E-02 | 0.1 | 3.28E-01 | 0.1 | 4.28E-01 | Omega-3 fatty acid desaturase, chloroplast |
| AT2G29310 | 1.4 | 8.52E-05 | 1.1 | 8.94E-04 | 0.3 | 2.05E-01 | -0.2 | 4.00E-01 | -0.3 | 2.73E-01 | 0.1 | 7.83E-01 | NAD-binding Rossmann-fold superfamily protein |
| AT1G07450 | 1.6 | 4.63E-05 | 0.7 | 2.70E-02 | 0.9 | 4.74E-03 | | | | | | | Tropinone reductase homolog At1g07450 |
| AT3G03980 | 1.7 | 2.67E-07 | 1.1 | 1.13E-05 | 0.5 | 1.79E-03 | 0.1 | 5.71E-01 | 0.1 | 8.17E-01 | 0.1 | 8.79E-01 | NADPH-dependent aldehyde reductase-like protein, chloroplastic |
| AT1G50450 | 1.7 | 3.75E-08 | 1.1 | 1.32E-06 | 0.6 | 1.59E-04 | 0.4 | 1.98E-02 | 0.3 | 9.56E-02 | 0.1 | 7.47E-01 | Saccharopine dehydrogenase NADP binding domain-containing protein |
| AT2G29300 | 1.8 | 1.00E-04 | 1.5 | 9.39E-04 | 0.3 | 2.69E-01 | | | | | | | NAD-binding Rossmann-fold superfamily protein |
| AT2G29340 | 1.9 | 9.61E-08 | 1.1 | 1.34E-05 | 0.9 | 1.10E-04 | 0.1 | 7.11E-01 | -0.6 | 5.75E-02 | 0.7 | 6.51E-02 | Tropinone reductase homolog At2g29340 |
| AT2G29290 | 2.0 | 2.92E-07 | 1.6 | 2.63E-06 | 0.4 | 3.45E-02 | -0.2 | 1.61E-01 | -0.3 | 1.12E-01 | 0.1 | 8.39E-01 | Tropinone reductase homolog At2g29290 |
| AT3G09580 | 2.2 | 4.47E-07 | 1.8 | 4.19E-06 | 0.4 | 4.54E-02 | 0.0 | 9.07E-01 | 0.0 | 9.65E-01 | 0.0 | 8.87E-01 | hypothetical protein |
| AT2G37660 | 2.5 | 1.44E-08 | 1.9 | 9.13E-08 | 0.6 | 3.39E-04 | 0.7 | 9.83E-05 | 0.3 | 5.54E-02 | 0.4 | 3.80E-02 | AT2G37660 protein |
| AT2G30670 | 2.8 | 5.72E-08 | 1.6 | 7.42E-06 | 1.2 | 4.46E-05 | 2.9 | 2.31E-05 | 2.1 | 1.80E-03 | 0.8 | 7.60E-02 | Tropinone reductase homolog At2g30670 |

| | | | | | | | | | | | | | |
|---|---|---|---|---|---|---|---|---|---|---|---|---|---|
| AT2G29150 | | | | | | | 6.765769212 | 2.58E-05 | 5.313088085 | 9.28E-04 | 1.5 | 3.05E-03 | Tropinone reductase homolog At2g29150 |
| AT2G29260 | | | | | | | 0.7 | 1.55E-01 | 0.7 | 2.01E-01 | 0.0 | 9.98E-01 | Tropinone reductase homolog At2g29260, chloroplastic |
| AT3G51680 | | | | | | | -0.4 | 5.46E-01 | 0.6 | 3.39E-01 | -1.0 | 1.02E-01 | Short-chain dehydrogenase reductase 2a |
| AT4G12870 | 1.7 | 2.36E-05 | 1.5 | 1.74E-04 | 0.3 | 2.82E-01 | 0.7 | 2.89E-01 | 0.5 | 5.06E-01 | 0.2 | 8.67E-01 | Gamma interferon responsive lysosomal thiol reductase family protein |
| AT5G16010 | 0.9 | 1.21E-05 | 0.4 | 5.44E-03 | 0.5 | 2.09E-03 | -0.1 | 5.06E-01 | 0.0 | 9.98E-01 | -0.1 | 6.61E-01 | 3-oxo-5-alpha-steroid 4-dehydrogenase family protein |
| AT4G35760 | 0.7 | 1.82E-05 | 0.4 | 1.55E-03 | 0.3 | 1.15E-02 | 0.1 | 5.15E-01 | 0.1 | 4.83E-01 | 0.0 | 9.22E-01 | NADH dehydrogenase s |
| AT4G22110 | -1.1 | 6.35E-06 | -0.3 | 2.07E-02 | -0.8 | 4.45E-04 | -0.2 | 6.36E-01 | 0.6 | 3.56E-02 | -0.8 | 2.34E-02 | GroES-like zinc-binding dehydrogenase family protein |
| AT1G22440 | -0.6 | 2.63E-05 | 0.0 | 7.26E-01 | -0.5 | 2.34E-04 | 0.3 | 3.27E-01 | 0.4 | 1.13E-01 | -0.2 | 4.59E-01 | Uncharacterized protein |
| AT3G56460 | 0.0 | 8.33E-01 | 0.1 | 4.68E-01 | 0.0 | 6.50E-01 | -0.2 | 5.92E-02 | -0.2 | 7.69E-02 | 0.0 | 9.65E-01 | GroES-like zinc-binding alcohol dehydrogenase family protein |
| AT1G64710 | 0.1 | 5.09E-01 | -0.1 | 3.99E-01 | 0.2 | 1.75E-01 | 0.3 | 1.78E-01 | 0.1 | 8.56E-01 | 0.2 | 4.59E-01 | GroES-like zinc-binding alcohol dehydrogenase family protein |
| AT1G49670 | 0.2 | 7.38E-02 | 0.0 | 9.53E-01 | 0.2 | 9.95E-02 | 0.2 | 1.63E-01 | 0.2 | 2.30E-01 | 0.0 | 9.97E-01 | ARP protein ) |
| AT1G22430 | 1.1 | 2.61E-05 | 0.7 | 8.13E-04 | 0.4 | 4.16E-02 | -0.3 | 7.75E-02 | -0.3 | 1.56E-01 | 0.0 | 9.26E-01 | GroES-like zinc-binding dehydrogenase family protein |
| AT1G07080 | -0.6 | 1.86E-05 | -0.3 | 4.10E-03 | -0.3 | 4.55E-03 | -0.4 | 4.33E-03 | -0.2 | 1.64E-01 | -0.2 | 2.98E-01 | F10K1.21 protein |
| AT4G26160 | -0.6 | 2.72E-05 | -0.5 | 1.73E-04 | -0.1 | 3.15E-01 | -0.4 | 3.27E-02 | -0.2 | 3.02E-01 | -0.2 | 4.86E-01 | Thioredoxin-like 2-1, chloroplastic |
| AT1G45145 | -0.2 | 2.34E-01 | 0.2 | 4.21E-01 | -0.4 | 8.28E-02 | 0.1 | 7.43E-01 | 0.1 | 8.39E-01 | 0.0 | 9.71E-01 | Thioredoxin H5 |
| AT4G37200 | 0.2 | 4.30E-02 | 0.0 | 6.22E-01 | 0.1 | 1.52E-01 | -0.1 | 4.56E-01 | 0.1 | 5.86E-01 | -0.2 | 2.59E-01 | Thioredoxin-like protein HCF164, chloroplastic |
| AT4G29670 | 0.2 | 1.64E-02 | 0.4 | 1.02E-03 | -0.2 | 1.02E-01 | 0.3 | 7.90E-02 | 0.2 | 1.89E-01 | 0.0 | 8.96E-01 | Thioredoxin-like 2-2, chloroplastic |
| AT1G31020 | 0.3 | 6.26E-03 | 0.1 | 3.20E-01 | 0.2 | 8.22E-02 | 0.2 | 5.01E-01 | 0.1 | 8.71E-01 | 0.1 | 7.65E-01 | Thioredoxin O2, mitochondrial |
| AT2G35010 | 0.5 | 2.26E-04 | 0.3 | 7.03E-03 | 0.2 | 8.27E-02 | 0.3 | 4.84E-02 | 0.2 | 1.26E-01 | 0.0 | 8.83E-01 | Thioredoxin O1, mitochondrial |
| AT4G12900 | 0.6 | 8.33E-04 | 0.7 | 5.20E-04 | -0.1 | 5.94E-01 | 0.0 | 9.90E-01 | 0.3 | 4.00E-01 | -0.3 | 4.91E-01 | Gamma interferon responsive lysosomal thiol reductase family protein |
| AT1G76080 | 0.9 | 5.05E-07 | 0.7 | 9.70E-06 | 0.2 | 1.19E-02 | 0.2 | 9.91E-02 | 0.2 | 1.78E-01 | 0.0 | 9.64E-01 | Thioredoxin-like protein CDSP32, chloroplastic |
| AT5G39950 | -0.4 | 4.09E-03 | -0.1 | 2.65E-01 | -0.3 | 5.20E-02 | -0.1 | 5.91E-01 | -0.1 | 8.36E-01 | -0.1 | 8.61E-01 | Thioredoxin H2 |
| AT1G72970 | 0.3 | 1.93E-02 | 0.3 | 9.76E-03 | -0.1 | 6.77E-01 | 0.2 | 7.53E-02 | 0.3 | 2.29E-02 | -0.1 | 6.05E-01 | Uncharacterized protein At1g72970 |
| AT4G11960 | -0.4 | 1.48E-03 | -0.5 | 8.33E-04 | 0.1 | 5.78E-01 | 0.0 | 9.77E-01 | 0.2 | 2.18E-01 | -0.2 | 3.53E-01 | PGR5-like B |
| AT5G59400 | 0.3 | 5.74E-03 | -0.1 | 3.74E-01 | 0.3 | 4.74E-03 | -1.9 | 3.56E-04 | -0.3 | 4.00E-01 | -1.6 | 4.53E-03 | PGR5-like A protein |
| AT4G22890 | 0.8 | 4.14E-05 | 0.7 | 2.46E-04 | 0.1 | 3.56E-01 | 0.4 | 2.47E-03 | 0.1 | 3.33E-01 | 0.3 | 1.36E-01 | PGR5-LIKE A |
| AT5G56090 | 0.2 | 1.04E-01 | -0.2 | 1.04E-01 | 0.4 | 1.35E-02 | 0.9 | 4.35E-04 | 0.4 | 9.10E-02 | 0.5 | 2.66E-02 | Cytochrome c oxidase assembly protein COX15 |
| AT4G39230 | -0.1 | 1.56E-01 | -0.1 | 2.41E-01 | 0.0 | 8.67E-01 | 0.6 | 1.13E-02 | 0.4 | 1.28E-01 | 0.2 | 4.63E-01 | Phenylcoumaran benzylic ether reductase 1 |
| AT1G74880 | 0.7 | 4.30E-04 | 0.5 | 4.30E-03 | 0.2 | 2.45E-01 | 0.2 | 5.60E-02 | 0.2 | 1.50E-01 | 0.0 | 8.92E-01 | NADH-quinone oxidoreductase subunit O, chloroplastic H dehydrogenase subunit O) |
| AT5G58260 | 0.8 | 1.92E-05 | 0.7 | 1.39E-04 | 0.1 | 2.40E-01 | -0.1 | 7.06E-01 | 0.3 | 1.05E-01 | -0.3 | 1.03E-01 | NADH-quinone oxidoreductase subunit N, chloroplastic H dehydrogenase subunit N) |
| AT4G37925 | 0.9 | 4.29E-04 | 0.5 | 1.61E-02 | 0.4 | 7.24E-02 | 0.2 | 1.64E-01 | 0.1 | 4.80E-01 | 0.1 | 7.62E-01 | NADH-quinone oxidoreductase subunit M, chloroplastic H dehydrogenase subunit M) |
| AT1G06080 | -0.5 | 5.48E-04 | 0.4 | 1.60E-03 | -1.0 | 7.81E-05 | -1.0 | 1.13E-03 | -0.1 | 6.63E-01 | -0.8 | 1.10E-02 | Delta 9 desaturase 1 |
| AT2G29980 | 0.3 | 3.48E-03 | 0.6 | 9.77E-05 | -0.3 | 1.62E-02 | 0.0 | 7.41E-01 | 0.0 | 8.59E-01 | 0.0 | 9.56E-01 | Fatty acid desaturase 3 |
| AT1G06360 | 0.5 | 7.18E-02 | 0.9 | 4.52E-03 | -0.4 | 1.54E-01 | 0.0 | 8.06E-01 | -0.2 | 2.31E-01 | 0.3 | 1.99E-01 | Fatty acid desaturase family protein |
| AT5G05580 | 1.8 | 1.34E-07 | 1.1 | 6.01E-06 | 0.7 | 5.16E-04 | 0.4 | 1.24E-01 | 0.4 | 1.72E-01 | 0.0 | 9.98E-01 | Fatty acid desaturase 8 |
| AT1G21400 | -0.8 | 1.44E-03 | -1.3 | 1.48E-04 | 0.5 | 7.16E-02 | -0.6 | 1.13E-01 | -1.2 | 1.17E-02 | 0.6 | 3.57E-01 | Thiamin diphosphate-binding fold superfamily protein |
| AT5G09300 | -0.7 | 6.47E-06 | -0.2 | 2.18E-02 | -0.5 | 4.82E-04 | -0.2 | 3.40E-01 | 0.2 | 4.04E-01 | -0.5 | 1.35E-01 | Thiamin diphosphate-binding fold superfamily protein |
| AT1G23800 | -0.6 | 1.61E-05 | -0.2 | 1.96E-02 | -0.4 | 1.43E-03 | -0.4 | 1.15E-01 | 0.0 | 9.18E-01 | -0.4 | 2.09E-01 | Aldehyde dehydrogenase 2B7 |
| AT1G74920 | -0.1 | 1.03E-01 | 0.0 | 9.62E-01 | -0.1 | 1.52E-01 | -0.2 | 2.08E-02 | -0.1 | 7.12E-01 | -0.2 | 1.93E-01 | Aldehyde dehydrogenase 10A8 |
| AT3G66658 | 0.0 | 5.89E-01 | 0.0 | 6.27E-01 | -0.1 | 3.64E-01 | 0.0 | 9.73E-01 | 0.1 | 7.24E-01 | -0.1 | 7.59E-01 | Aldehyde dehydrogenase |
| AT3G48000 | 0.2 | 1.75E-02 | 0.0 | 6.89E-01 | 0.2 | 6.06E-02 | 0.0 | 7.79E-01 | -0.2 | 2.94E-01 | 0.1 | 5.40E-01 | ALDH2B4 |
| AT4G34240 | 0.7 | 1.07E-05 | 0.3 | 3.57E-03 | 0.4 | 2.45E-03 | -0.1 | 7.19E-01 | 0.1 | 6.08E-01 | -0.2 | 4.43E-01 | Aldehyde dehydrogenase 3I1 |
| AT1G13440 | 0.7 | 2.46E-06 | 0.6 | 2.08E-05 | 0.1 | 1.32E-01 | 0.1 | 5.19E-01 | -0.1 | 3.87E-01 | 0.2 | 2.35E-01 | Putative glyceraldehyde-3-phosphate dehydrogenase |
| AT3G26650 | 1.6 | 5.03E-07 | 1.0 | 2.10E-05 | 0.6 | 2.79E-03 | 0.0 | 7.38E-01 | 0.0 | 9.12E-01 | 0.1 | 7.27E-01 | Glyceraldehyde 3-phosphate dehydrogenase A subunit |
| AT2G46890 | -0.6 | 1.89E-04 | 0.0 | 8.58E-01 | -0.7 | 6.93E-04 | -0.1 | 4.64E-01 | 0.2 | 1.68E-01 | -0.3 | 4.44E-02 | Steroid 5-alpha reductase C-terminal domain-containing protein |
| AT1G73650 | 0.6 | 8.16E-06 | 0.1 | 3.72E-01 | 0.6 | 1.31E-04 | 0.2 | 2.07E-01 | 0.0 | 9.28E-01 | 0.2 | 3.18E-01 | 3-oxo-5-alpha-steroid 4-dehydrogenase |
| AT1G18180 | 0.7 | 2.38E-04 | 0.4 | 1.03E-02 | 0.3 | 7.98E-02 | -0.3 | 3.08E-01 | -0.4 | 3.37E-01 | 0.0 | 9.82E-01 | 3-oxo-5-alpha-steroid 4-dehydrogenase |
| AT5G37980 | -3.4 | 5.13E-07 | -2.4 | 2.67E-06 | -1.0 | 1.77E-02 | -0.9 | 5.20E-02 | 0.7 | 6.24E-02 | -1.7 | 7.00E-03 | Enoyl reductase domain-containing protein |
| AT1G50430 | 0.0 | 7.91E-01 | 0.2 | 3.68E-02 | -0.2 | 7.64E-02 | -0.2 | 1.63E-01 | -0.2 | 2.10E-01 | 0.0 | 9.88E-01 | Ergosterol biosynthesis ERG4 |
| AT3G52940 | 0.1 | 5.44E-01 | 0.3 | 1.47E-02 | -0.2 | 6.60E-02 | 0.0 | 8.38E-01 | 0.0 | 8.54E-01 | -0.1 | 7.64E-01 | Ergosterol biosynthesis ERG4 |
| AT1G65560 | 0.2 | 1.36E-01 | -0.1 | 3.65E-01 | 0.3 | 5.86E-02 | 0.1 | 6.34E-01 | 0.0 | 9.32E-01 | 0.1 | 6.58E-01 | At1g65560 |
| AT3G59840 | 0.7 | 1.49E-04 | 0.6 | 1.06E-03 | 0.1 | 3.21E-01 | 0.1 | 5.51E-01 | 0.0 | 9.08E-01 | 0.2 | 5.82E-01 | Allyl alcohol dehydrogenase-like protein |
| AT1G51720 | 0.2 | 8.77E-03 | 0.0 | 8.68E-01 | 0.2 | 1.59E-02 | 0.0 | 9.26E-01 | -0.1 | 8.28E-01 | 0.0 | 9.11E-01 | NADP-specific glutatamate dehydrogenase, putative |
| AT5G62420 | -3.3 | 7.09E-07 | -2.4 | 3.95E-06 | -0.8 | 4.08E-02 | -0.6 | 2.88E-01 | -0.8 | 1.89E-01 | 0.2 | 7.93E-01 | NADP-dependent oxidoreductase domain-containing protein |
| AT5G14700 | -0.8 | 6.22E-03 | -0.7 | 1.82E-02 | -0.1 | 6.72E-01 | 0.4 | 2.05E-01 | 0.3 | 3.82E-01 | 0.1 | 8.84E-01 | NAD-binding Rossmann-fold superfamily protein |
| AT4G29010 | -0.3 | 2.37E-03 | -0.2 | 2.46E-02 | -0.1 | 2.76E-01 | -0.1 | 5.73E-01 | -0.1 | 5.62E-01 | 0.0 | 9.53E-01 | AIM1 protein |
| AT4G37980 | -0.2 | 1.33E-01 | -0.2 | 1.83E-01 | 0.0 | 9.18E-01 | -0.5 | 7.90E-03 | -0.1 | 4.87E-01 | -0.3 | 1.80E-01 | AT4G37980 protein |
| AT1G09490 | 0.0 | 9.83E-01 | 0.2 | 2.70E-01 | -0.2 | 3.45E-01 | -0.1 | 3.45E-01 | -0.1 | 5.26E-01 | 0.0 | 9.24E-01 | NAD-binding Rossmann-fold superfamily protein |
| AT1G52340 | 0.1 | 2.39E-01 | 0.1 | 3.52E-01 | 0.0 | 8.75E-01 | -0.1 | 5.49E-01 | 0.1 | 6.17E-01 | -0.3 | 2.79E-01 | Xanthoxin dehydrogenase |
| AT5G61830 | 0.4 | 3.59E-03 | 0.2 | 1.90E-01 | 0.2 | 8.37E-02 | 0.0 | 9.20E-01 | -0.1 | 7.86E-01 | 0.1 | 7.34E-01 | Short-chain dehydrogenase |
| AT1G10310 | 0.5 | 1.89E-04 | 0.3 | 1.33E-02 | 0.2 | 3.99E-02 | 0.3 | 6.15E-02 | 0.3 | 1.13E-01 | 0.0 | 9.58E-01 | NADPH-dependent pterin aldehyde reductase |
| AT1G59960 | 0.7 | 1.47E-05 | 0.5 | 1.99E-04 | 0.2 | 1.07E-01 | -0.1 | 5.02E-01 | -0.1 | 8.27E-01 | -0.1 | 8.23E-01 | NADP-dependent oxidoreductase domain-containing protein |
| AT2G23910 | | | | | | | 0.3 | 1.63E-01 | 0.5 | 2.93E-02 | -0.2 | 4.11E-01 | At2g23910 -binding Rossmann-fold superfamily protein) |
| AT4G37970 | -1.8 | 5.97E-06 | -1.1 | 1.30E-04 | -0.7 | 2.09E-02 | -1.0 | 2.57E-03 | -0.2 | 4.68E-01 | -0.7 | 5.30E-02 | Cinnamyl alcohol dehydrogenase 6 |
| AT4G34230 | -1.0 | 6.03E-06 | -0.8 | 8.86E-05 | -0.3 | 5.57E-02 | 0.1 | 4.90E-01 | 0.3 | 1.92E-01 | -0.1 | 6.33E-01 | Cinnamyl alcohol dehydrogenase 5 |
| AT1G78950 | -0.7 | 1.47E-04 | -0.2 | 4.91E-02 | -0.5 | 1.10E-02 | 0.9 | 6.65E-02 | 0.4 | 5.55E-01 | 0.5 | 4.00E-01 | Terpene cyclase |
| AT3G45130 | -0.6 | 9.50E-03 | 0.3 | 1.68E-01 | -0.9 | 4.45E-03 | | | | | | | Terpene cyclase |
| AT1G66960 | -0.4 | 4.69E-01 | -0.3 | 6.09E-01 | -0.1 | 8.96E-01 | | | | | | | Terpene cyclase |
| AT2G07050 | 0.2 | 1.03E-02 | 0.1 | 7.34E-02 | 0.1 | 3.76E-01 | -0.1 | 6.63E-01 | 0.0 | 9.01E-01 | -0.1 | 6.84E-01 | Terpene cyclase |
| AT1G78960 | 0.2 | 2.55E-02 | -0.3 | 1.97E-02 | 0.5 | 1.75E-03 | 0.2 | 5.23E-01 | 0.3 | 3.67E-01 | -0.1 | 8.28E-01 | Terpene cyclase |
| AT4G15340 | 1.1 | 6.23E-04 | 0.9 | 3.47E-03 | 0.2 | 4.72E-01 | -0.4 | 2.82E-01 | 0.0 | 9.71E-01 | -0.4 | 4.01E-01 | Pentacyclic triterpene synthase 1 |
| AT4G15370 | 5.2 | 7.49E-07 | 2.2 | 1.67E-03 | 3.0 | 4.27E-05 | 3.0 | 1.63E-02 | 2.5 | 6.70E-02 | 0.5 | 6.79E-01 | Terpene cyclase |
| AT5G14220 | 0.3 | 6.57E-03 | 0.3 | 2.09E-02 | 0.1 | 6.54E-01 | 0.1 | 7.11E-01 | 0.0 | 9.68E-01 | 0.1 | 7.23E-01 | Protoporphyrinogen oxidase |
| AT4G01690 | 1.1 | 1.48E-07 | 0.8 | 4.52E-06 | 0.4 | 1.32E-03 | 0.0 | 7.75E-01 | -0.1 | 8.06E-01 | 0.1 | 6.35E-01 | Protoporphyrinogen oxidase 1, chloroplastic |
| AT4G17470 | -0.4 | 4.89E-01 | 0.3 | 5.48E-01 | -0.7 | 2.86E-01 | 1.1 | 1.56E-01 | 1.7 | 3.38E-02 | -0.6 | 4.75E-01 | Alpha |
| AT3G60340 | 0.0 | 6.27E-01 | 0.0 | 6.16E-01 | 0.0 | 9.76E-01 | -0.3 | 3.22E-02 | -0.2 | 2.94E-01 | -0.1 | 5.27E-01 | AT3G60340 protein |
| AT1G57590 | -1.8 | 1.15E-04 | -1.0 | 4.40E-03 | -0.9 | 3.43E-02 | -0.3 | 2.51E-01 | 0.5 | 6.20E-02 | -0.8 | 1.72E-02 | Pectin acetylesterase 2 |
| AT5G26670 | 0.0 | 7.80E-01 | 0.2 | 1.89E-01 | -0.2 | 1.72E-01 | -0.2 | 2.48E-01 | 0.3 | 1.08E-01 | -0.6 | 2.37E-02 | Pectin acetylesterase 10 |
| AT5G45280 | 0.1 | 4.87E-01 | 0.1 | 3.62E-01 | 0.0 | 8.33E-01 | -0.1 | 2.54E-01 | 0.2 | 6.64E-02 | -0.4 | 2.23E-02 | Pectin acetylesterase 11 |
| AT2G46930 | 0.4 | 4.88E-03 | 0.2 | 1.81E-01 | 0.2 | 1.04E-01 | 0.0 | 8.62E-01 | 0.2 | 3.30E-01 | -0.2 | 5.35E-01 | Pectin acetylesterase 3 |
| AT1G10770 | 2.5 | 3.19E-06 | 2.2 | 1.98E-05 | 0.3 | 2.10E-01 | | | | | | | At1g10770 |
| AT3G49330 | -0.8 | 2.49E-03 | 0.1 | 4.20E-01 | -0.9 | 3.34E-03 | 0.6 | 1.03E-01 | 0.8 | 3.23E-02 | -0.2 | 5.87E-01 | Plant invertase |
| AT2G03360 | | | | | | | 0.2 | 8.06E-01 | 0.9 | 1.46E-01 | -0.7 | 2.88E-01 | Uncharacterized protein At2g03360 |

| | | | | | | | | | | | | | |
|---|---|---|---|---|---|---|---|---|---|---|---|---|---|
| AT2G40570 | 0.0 | 7.11E-01 | 0.0 | 8.64E-01 | -0.1 | 6.56E-01 | 0.1 | 7.40E-01 | -0.1 | 6.94E-01 | 0.2 | 4.77E-01 | Initiator tRNA phosphoribosyl transferase family protein |
| AT1G80690 | -1.0 | 4.26E-06 | -0.3 | 1.24E-02 | -0.7 | 3.59E-04 | -0.5 | 2.60E-03 | -0.2 | 3.97E-01 | -0.4 | 6.85E-02 | hypothetical protein |
| AT1G78880 | -0.6 | 2.69E-06 | -0.4 | 7.41E-05 | -0.2 | 1.52E-02 | -0.3 | 2.83E-02 | -0.1 | 4.17E-01 | -0.2 | 4.03E-01 | F9K20.7 protein |
| AT1G47740 | -0.5 | 1.58E-04 | -0.1 | 3.35E-01 | -0.4 | 1.88E-03 | -0.2 | 3.15E-01 | 0.1 | 7.72E-01 | -0.3 | 3.27E-01 | PPPDE domain-containing protein |
| AT4G25680 | -0.3 | 6.96E-03 | -0.2 | 5.17E-02 | -0.1 | 3.63E-01 | -0.3 | 1.07E-01 | 0.0 | 9.93E-01 | -0.3 | 2.35E-01 | PPPDE domain-containing protein |
| AT3G07090 | -0.3 | 6.51E-03 | -0.3 | 1.52E-02 | 0.0 | 7.50E-01 | 0.2 | 2.33E-01 | 0.5 | 1.70E-02 | -0.3 | 2.06E-01 | PPPDE putative thiol peptidase family protein |
| AT4G30550 | -0.1 | 5.16E-01 | 0.0 | 9.06E-01 | -0.1 | 6.69E-01 | -0.5 | 7.04E-03 | -0.4 | 4.29E-02 | -0.1 | 7.40E-01 | Gamma-glutamyl peptidase 3 |
| AT4G31980 | 0.0 | 6.95E-01 | -0.1 | 3.64E-01 | 0.0 | 6.54E-01 | 0.0 | 7.92E-01 | -0.1 | 5.66E-01 | 0.1 | 7.92E-01 | Uncharacterized protein At4g31980 |
| AT5G47310 | 0.0 | 8.73E-01 | 0.0 | 4.52E-01 | 0.0 | 5.98E-01 | -0.1 | 5.46E-01 | 0.0 | 9.21E-01 | -0.1 | 7.74E-01 | hypothetical protein |
| AT1G73990 | 0.1 | 5.65E-01 | 0.0 | 9.34E-01 | 0.1 | 5.64E-01 | -0.2 | 2.67E-01 | 0.0 | 9.91E-01 | -0.2 | 4.34E-01 | Signal peptide peptidase |
| AT4G17486 | 0.1 | 4.38E-01 | 0.1 | 9.16E-02 | -0.1 | 3.73E-01 | -0.4 | 8.16E-04 | -0.4 | 3.05E-03 | 0.0 | 9.04E-01 | PPPDE putative thiol peptidase family protein |
| AT2G39960 | 0.1 | 4.32E-01 | 0.2 | 2.64E-02 | -0.2 | 1.35E-01 | 0.1 | 6.87E-01 | 0.0 | 8.64E-01 | 0.0 | 9.10E-01 | Signal peptidase complex subunit 2 |
| AT2G25190 | 0.1 | 1.21E-01 | -0.2 | 1.11E-01 | 0.3 | 1.59E-02 | 0.0 | 9.08E-01 | 0.0 | 9.81E-01 | 0.0 | 9.19E-01 | hypothetical protein |
| AT4G04200 | 0.4 | 1.70E-02 | 0.3 | 7.85E-02 | 0.1 | 5.32E-01 | 0.3 | 1.60E-01 | -0.1 | 8.69E-01 | 0.3 | 2.13E-01 | Signal peptidase complex subunit 2 |
| AT1G19740 | 0.4 | 1.54E-04 | 0.3 | 5.72E-03 | 0.2 | 5.90E-02 | -0.4 | 5.51E-03 | 0.0 | 1.00E+00 | -0.4 | 2.75E-02 | ATP-dependent protease La domain protein |
| AT5G67170 | 0.4 | 9.38E-03 | 0.1 | 3.80E-01 | 0.3 | 1.03E-01 | -0.3 | 2.71E-01 | -0.2 | 5.70E-01 | -0.1 | 7.91E-01 | SEC-C motif-containing protein |
| AT4G18370 | 1.3 | 1.09E-04 | 0.9 | 3.38E-03 | 0.5 | 7.28E-02 | 0.1 | 4.34E-01 | 0.1 | 8.49E-01 | 0.1 | 7.35E-01 | Protease HhoA like |
| AT2G38870 | 1.5 | 7.03E-05 | 1.2 | 6.99E-04 | 0.3 | 1.91E-01 | -0.1 | 6.04E-01 | -0.3 | 2.54E-01 | 0.2 | 6.38E-01 | Putative protease inhibitor |
| AT2G26470 | -0.1 | 4.03E-01 | -0.1 | 2.55E-01 | 0.0 | 7.81E-01 | -0.3 | 1.52E-01 | -0.1 | 5.90E-01 | -0.2 | 6.00E-01 | At2g26470 |
| AT4G31985 | 1.3 | 4.33E-06 | 1.0 | 4.58E-05 | 0.3 | 9.84E-02 | 0.6 | 1.63E-02 | 0.3 | 3.88E-01 | 0.4 | 3.38E-01 | PPPDE domain-containing protein |
| AT4G31300 | 0.0 | 8.91E-01 | 0.1 | 4.23E-01 | -0.1 | 5.40E-01 | 0.3 | 4.09E-02 | 0.1 | 3.45E-01 | 0.1 | 5.08E-01 | Proteasome subunit beta |
| AT1G73330 | -1.5 | 1.13E-06 | -0.8 | 3.13E-04 | -0.7 | 6.08E-04 | -1.9 | 5.32E-09 | -0.7 | 5.20E-04 | -1.2 | 7.11E-05 | Kunitz trypsin inhibitor 1 |
| AT3G02980 | -0.4 | 3.87E-02 | -0.2 | 3.43E-01 | -0.2 | 3.24E-01 | -0.1 | 7.93E-01 | 0.1 | 9.10E-01 | -0.2 | 7.37E-01 | histone acetyltransferase |
| AT2G38130 | 0.0 | 9.84E-01 | -0.1 | 5.04E-01 | 0.1 | 5.51E-01 | 0.0 | 9.85E-01 | -0.1 | 8.19E-01 | 0.1 | 8.33E-01 | N-alpha-acetyltransferase MAK3 |
| AT1G03150 | 0.4 | 2.54E-03 | 0.5 | 1.06E-03 | -0.1 | 5.03E-01 | 0.2 | 4.15E-01 | 0.1 | 8.24E-01 | 0.1 | 7.53E-01 | N-terminal acetyltransferase B complex catalytic subunit NAA20 |
| AT1G80080 | 0.8 | 9.37E-05 | 0.6 | 8.15E-04 | 0.2 | 2.70E-01 | -0.2 | 3.09E-01 | -0.1 | 8.24E-01 | -0.1 | 6.05E-01 | Protein TOO MANY MOUTHS |
| AT4G13340 | 1.1 | 1.51E-03 | 0.6 | 4.94E-02 | 0.5 | 9.95E-02 | -0.5 | 1.10E-02 | -0.8 | 5.46E-04 | 0.3 | 1.12E-01 | Leucine-rich repeat extensin-like protein 3 |
| AT1G33612 | 0.3 | 1.64E-01 | 0.4 | 3.78E-02 | -0.2 | 4.77E-01 | 0.3 | 2.81E-01 | 0.8 | 1.24E-02 | -0.5 | 1.46E-01 | Leucine-rich repeat family protein |
| AT2G18660 | 1.8 | 1.52E-03 | 0.3 | 6.07E-01 | 1.6 | 1.15E-02 | 0.1 | 8.65E-01 | -0.6 | 3.26E-01 | 0.7 | 3.01E-01 | EG45-like domain containing protein 2 |
| AT1G65380 | -0.4 | 2.01E-03 | -0.2 | 3.50E-02 | -0.2 | 1.96E-01 | 0.1 | 6.38E-01 | 0.1 | 7.44E-01 | 0.0 | 9.71E-01 | Receptor-like protein CLAVATA2 |
| AT4G04810 | 0.6 | 1.69E-02 | 1.3 | 1.35E-04 | -0.7 | 1.23E-02 | -0.3 | 3.42E-01 | -0.4 | 2.47E-01 | 0.1 | 8.49E-01 | MsrB domain-containing protein |
| AT4G04830 | 1.4 | 3.98E-07 | 1.6 | 3.67E-07 | -0.2 | 4.59E-02 | 0.0 | 1.00E+00 | -0.2 | 3.49E-01 | 0.2 | 4.63E-01 | Methionine sulfoxide reductase B5 |
| AT5G61640 | -1.3 | 4.55E-07 | -1.2 | 2.02E-06 | -0.1 | 3.11E-01 | -0.1 | 8.47E-01 | -0.1 | 6.50E-01 | 0.1 | 8.14E-01 | peptide-methionine -S-oxide reductase -S-oxide reductase) |
| AT1G53770 | -0.1 | 2.06E-01 | -0.2 | 3.45E-02 | 0.1 | 3.60E-01 | -0.4 | 5.85E-02 | -0.2 | 5.50E-01 | -0.3 | 4.12E-01 | O-fucosyltransferase family protein |
| AT5G50420 | 0.0 | 6.50E-01 | 0.2 | 2.69E-02 | -0.2 | 2.49E-02 | -0.1 | 8.34E-01 | 0.2 | 5.63E-01 | -0.3 | 4.59E-01 | O-fucosyltransferase 36 |
| AT1G17270 | 0.2 | 7.03E-02 | 0.0 | 7.96E-01 | 0.2 | 8.49E-02 | -0.1 | 7.97E-01 | -0.2 | 3.84E-01 | 0.1 | 5.92E-01 | O-fucosyltransferase 5 |
| AT1G21880 | 0.1 | 4.96E-01 | 0.3 | 1.70E-02 | -0.2 | 9.13E-02 | 0.2 | 1.72E-01 | 0.1 | 6.87E-01 | 0.1 | 5.66E-01 | LysM domain-containing GPI-anchored protein 1 |
| AT1G77630 | 0.2 | 1.31E-02 | 0.4 | 1.13E-03 | -0.1 | 1.63E-01 | 0.4 | 1.15E-01 | 0.3 | 3.46E-01 | 0.1 | 7.26E-01 | LysM domain-containing GPI-anchored protein 3 |
| AT2G15790 | -0.6 | 2.90E-04 | -0.1 | 3.18E-01 | -0.5 | 4.35E-03 | 0.3 | 1.71E-01 | 0.2 | 2.87E-01 | 0.0 | 9.31E-01 | PPIase cyclophilin-type domain-containing protein |
| AT4G34960 | -0.3 | 4.21E-03 | 0.0 | 8.55E-01 | -0.3 | 1.49E-02 | 0.0 | 7.67E-01 | 0.1 | 7.83E-01 | -0.1 | 5.91E-01 | Peptidyl-prolyl cis-trans isomerase |
| AT2G47320 | -0.3 | 1.94E-02 | 0.2 | 1.28E-01 | -0.4 | 3.47E-03 | -0.3 | 5.03E-02 | -0.1 | 7.87E-01 | -0.2 | 2.01E-01 | Peptidyl-prolyl cis-trans isomerase CYP21-3, mitochondrial |
| AT3G54010 | -0.2 | 4.67E-03 | -0.3 | 5.48E-04 | 0.1 | 1.45E-01 | 0.3 | 3.49E-02 | 0.1 | 5.47E-01 | 0.2 | 3.59E-01 | Peptidyl-prolyl cis-trans isomerase PASTICCINO1 |
| AT4G33060 | -0.1 | 5.67E-01 | -0.3 | 1.60E-02 | 0.2 | 6.80E-02 | -0.1 | 4.37E-01 | 0.0 | 9.09E-01 | -0.1 | 6.81E-01 | Peptidyl-prolyl cis-trans isomerase CYP57 |
| AT1G26940 | 0.0 | 8.72E-01 | 0.3 | 7.64E-03 | -0.3 | 2.12E-02 | 0.0 | 9.03E-01 | 0.2 | 2.67E-01 | -0.2 | 4.21E-01 | Cyclophilin-like peptidyl-prolyl cis-trans isomerase family protein |
| AT3G12340 | 0.0 | 8.25E-01 | 0.1 | 4.18E-01 | -0.1 | 6.25E-01 | 0.7 | 2.00E-02 | 0.1 | 7.54E-01 | 0.5 | 1.13E-01 | peptidylprolyl isomerase |
| AT4G25340 | 0.1 | 7.00E-01 | 0.2 | 2.19E-01 | -0.2 | 4.34E-01 | 0.5 | 1.07E-02 | 0.4 | 6.76E-02 | 0.1 | 7.19E-01 | peptidylprolyl isomerase |
| AT3G25220 | 0.1 | 4.24E-01 | 0.4 | 5.40E-03 | -0.3 | 3.35E-02 | 0.2 | 4.49E-01 | 0.1 | 6.27E-01 | 0.0 | 9.41E-01 | Peptidyl-prolyl cis-trans isomerase FKBP15-1 |
| AT5G64350 | 0.1 | 2.42E-01 | 0.2 | 7.30E-02 | -0.1 | 4.89E-01 | -0.1 | 7.36E-01 | -0.1 | 8.64E-01 | 0.0 | 9.44E-01 | Peptidyl-prolyl cis-trans isomerase FKBP12 |
| AT2G36130 | 0.1 | 2.50E-01 | 0.2 | 9.15E-02 | -0.1 | 5.83E-01 | -0.2 | 4.45E-01 | -0.2 | 3.86E-01 | 0.0 | 8.89E-01 | Peptidyl-prolyl cis-trans isomerase |
| AT1G01940 | 0.2 | 1.79E-01 | 0.2 | 5.27E-02 | -0.1 | 5.34E-01 | 0.5 | 2.26E-03 | 0.3 | 9.42E-02 | 0.2 | 2.58E-01 | Peptidyl-prolyl cis-trans isomerase |
| AT4G17070 | 0.2 | 5.12E-02 | 0.1 | 3.63E-01 | 0.1 | 3.35E-01 | 0.3 | 2.52E-02 | -0.2 | 2.02E-01 | 0.6 | 1.21E-02 | PPIase cyclophilin-type domain-containing protein |
| AT5G48580 | 0.2 | 9.06E-02 | 0.4 | 2.73E-03 | -0.2 | 7.14E-02 | 0.3 | 9.39E-02 | 0.2 | 3.58E-01 | 0.1 | 7.39E-01 | Peptidyl-prolyl cis-trans isomerase FKBP15-2 |
| AT3G66654 | 0.2 | 5.00E-03 | 0.1 | 4.12E-02 | 0.1 | 3.34E-01 | 0.0 | 8.57E-01 | -0.1 | 6.26E-01 | 0.1 | 8.23E-01 | Peptidyl-prolyl cis-trans isomerase CYP21-4 |
| AT4G32420 | 0.2 | 4.39E-03 | -0.2 | 2.59E-02 | 0.4 | 4.95E-04 | 0.1 | 3.66E-01 | 0.2 | 2.84E-01 | 0.0 | 8.46E-01 | PPIase cyclophilin-type domain-containing protein |
| AT3G22942 | 0.3 | 2.77E-02 | 0.1 | 3.88E-01 | 0.2 | 2.25E-01 | 0.3 | 2.47E-01 | -0.1 | 7.85E-01 | 0.4 | 2.27E-01 | Peptidyl-prolyl cis-trans isomerase |
| AT3G55520 | 0.4 | 1.76E-03 | 0.4 | 1.92E-03 | 0.0 | 9.07E-01 | -0.6 | 3.32E-03 | -0.3 | 1.32E-01 | -0.3 | 2.59E-01 | Peptidyl-prolyl cis-trans isomerase FKBP20-1 |
| AT2G18040 | 0.4 | 8.28E-03 | 0.1 | 4.27E-01 | 0.3 | 5.88E-02 | 0.3 | 3.28E-02 | 0.2 | 1.41E-01 | 0.1 | 7.89E-01 | Peptidyl-prolyl cis-trans isomerase Pin1 |
| AT5G13410 | 0.5 | 8.57E-04 | 0.5 | 5.18E-04 | -0.1 | 5.85E-01 | -0.2 | 1.55E-01 | -0.1 | 3.76E-01 | -0.1 | 8.13E-01 | peptidylprolyl isomerase |
| AT1G18170 | 0.9 | 1.23E-06 | 0.7 | 1.76E-05 | 0.2 | 3.60E-02 | 0.1 | 4.05E-01 | 0.1 | 8.04E-01 | 0.1 | 7.63E-01 | Peptidyl-prolyl cis-trans isomerase FKBP17-2, chloroplastic |
| AT5G35100 | 0.9 | 1.03E-06 | 0.6 | 2.66E-05 | 0.3 | 9.75E-03 | -0.1 | 6.70E-01 | -0.1 | 4.77E-01 | 0.1 | 8.30E-01 | Cyclophilin-like peptidyl-prolyl cis-trans isomerase family protein |
| AT5G19370 | 1.0 | 2.61E-07 | 0.7 | 7.93E-06 | 0.3 | 2.35E-03 | 0.2 | 2.96E-01 | 0.1 | 5.94E-01 | 0.1 | 8.28E-01 | Rhodanese-like |
| AT1G20810 | 1.2 | 2.11E-06 | 1.0 | 9.94E-06 | 0.1 | 3.67E-01 | 0.0 | 8.86E-01 | 0.1 | 7.92E-01 | -0.1 | 7.47E-01 | peptidylprolyl isomerase |
| AT4G19830 | 1.4 | 3.78E-07 | 1.2 | 2.68E-06 | 0.2 | 1.08E-01 | -0.3 | 1.31E-01 | -0.1 | 4.41E-01 | -0.1 | 6.90E-01 | Peptidyl-prolyl cis-trans isomerase FKBP17-1, chloroplastic |
| AT4G39710 | 1.5 | 8.87E-07 | 1.3 | 4.59E-06 | 0.2 | 2.55E-01 | 0.0 | 9.82E-01 | 0.1 | 7.34E-01 | -0.1 | 7.59E-01 | peptidylprolyl isomerase |
| AT3G10060 | 1.7 | 6.11E-07 | 1.2 | 1.33E-05 | 0.5 | 8.52E-03 | 0.2 | 1.67E-01 | 0.2 | 2.90E-01 | 0.0 | 9.47E-01 | Peptidyl-prolyl cis-trans isomerase FKBP16-4, chloroplastic |
| AT1G74070 | 1.8 | 1.10E-07 | 1.8 | 3.56E-07 | 0.1 | 6.55E-01 | 0.3 | 5.50E-02 | 0.2 | 2.61E-01 | 0.1 | 6.82E-01 | Cyclophilin-like peptidyl-prolyl cis-trans isomerase family protein |
| AT3G60370 | 1.8 | 5.48E-07 | 1.1 | 6.01E-05 | 0.7 | 9.68E-04 | 0.3 | 1.39E-01 | 0.0 | 8.71E-01 | 0.2 | 2.48E-01 | peptidylprolyl isomerase |
| AT1G73655 | 1.8 | 1.07E-06 | 1.6 | 6.65E-06 | 0.2 | 1.63E-01 | 0.2 | 1.76E-01 | 0.1 | 7.90E-01 | 0.1 | 5.06E-01 | Peptidyl-prolyl cis-trans isomerase FKBP17-3, chloroplastic |
| AT4G26555 | 1.9 | 1.20E-07 | 1.6 | 1.01E-06 | 0.3 | 2.39E-02 | -0.5 | 6.40E-03 | -0.5 | 7.00E-03 | 0.0 | 8.81E-01 | peptidylprolyl isomerase |
| AT2G43560 | 2.0 | 5.84E-07 | 1.5 | 9.53E-06 | 0.5 | 1.44E-02 | 0.2 | 7.97E-02 | 0.0 | 9.84E-01 | 0.2 | 1.97E-01 | peptidylprolyl isomerase |
| AT3G15520 | 2.0 | 5.03E-07 | 1.8 | 3.03E-06 | 0.2 | 1.53E-01 | -0.1 | 4.54E-01 | 0.1 | 6.97E-01 | -0.2 | 3.42E-01 | Cyclophilin-like peptidyl-prolyl cis-trans isomerase family protein |
| AT3G62030 | 2.1 | 4.12E-08 | 2.2 | 1.07E-07 | -0.1 | 6.26E-01 | 0.2 | 1.91E-01 | 0.2 | 3.00E-01 | 0.0 | 9.67E-01 | Peptidyl-prolyl cis-trans isomerase |
| AT3G01480 | 2.1 | 4.77E-08 | 1.8 | 3.32E-07 | 0.3 | 2.10E-02 | 0.4 | 7.13E-03 | 0.1 | 3.73E-01 | 0.2 | 1.74E-01 | Peptidyl-prolyl cis-trans isomerase CYP38, chloroplastic |
| AT5G45680 | 2.4 | 5.00E-08 | 2.1 | 2.90E-07 | 0.3 | 3.17E-02 | 0.5 | 1.50E-02 | 0.0 | 9.09E-01 | 0.4 | 8.36E-02 | Peptidyl-prolyl cis-trans isomerase FKBP13, chloroplastic |
| AT4G08960 | -0.3 | 4.00E-03 | -0.1 | 5.10E-01 | -0.3 | 2.82E-02 | 0.2 | 3.21E-01 | 0.1 | 7.02E-01 | 0.1 | 7.70E-01 | Serine |
| AT3G25230 | 0.0 | 7.56E-01 | 0.1 | 4.77E-01 | -0.1 | 3.47E-01 | 0.4 | 1.18E-03 | 0.4 | 8.95E-03 | 0.1 | 7.35E-01 | peptidylprolyl isomerase |
| AT2G30695 | 1.6 | 2.06E-07 | 1.3 | 2.09E-06 | 0.3 | 2.55E-02 | 0.5 | 3.42E-03 | 0.2 | 1.78E-01 | 0.2 | 2.38E-01 | At2g30700 |
| AT1G53720 | 0.3 | 4.95E-02 | 0.3 | 3.54E-02 | 0.0 | 8.40E-01 | 0.1 | 5.10E-01 | -0.1 | 8.41E-01 | 0.2 | 4.93E-01 | Peptidyl-prolyl cis-trans isomerase |
| AT5G67530 | -0.1 | 2.13E-01 | -0.1 | 1.49E-01 | 0.0 | 8.32E-01 | -0.1 | 5.57E-01 | -0.1 | 5.71E-01 | 0.0 | 9.70E-01 | Peptidyl-prolyl cis-trans isomerase CYP65 |
| AT3G11630 | 2.0 | 2.14E-08 | 1.6 | 1.88E-07 | 0.4 | 3.32E-03 | 0.3 | 7.33E-02 | 0.2 | 1.96E-01 | 0.0 | 8.75E-01 | 2-Cys peroxiredoxin BAS1, chloroplastic |
| AT5G56290 | 0.1 | 1.32E-01 | -0.1 | 4.56E-01 | 0.2 | 5.46E-02 | -0.2 | 2.62E-01 | -0.1 | 4.63E-01 | 0.0 | 8.98E-01 | Peroxisome biogenesis protein 5 |
| AT1G29260 | 0.1 | 7.38E-02 | 0.1 | 6.34E-02 | 0.0 | 9.09E-01 | -0.1 | 6.31E-01 | -0.3 | 1.73E-01 | 0.2 | 5.06E-01 | Peroxisome biogenesis protein 7 |
| AT2G37040 | -1.3 | 5.94E-07 | -0.6 | 5.39E-04 | -0.8 | 1.53E-04 | 0.0 | 9.69E-01 | 0.3 | 4.41E-02 | -0.3 | 9.28E-02 | Phenylalanine ammonia-lyase 1 |

| | | | | | | | | | | | | | |
|---|---|---|---|---|---|---|---|---|---|---|---|---|---|
| AT3G10340 | -1.0 | 3.47E-07 | 0.0 | 5.56E-01 | -1.0 | 1.09E-05 | 0.4 | 4.77E-03 | 0.4 | 2.19E-02 | 0.1 | 8.14E-01 | Phenylalanine ammonia-lyase 4 |
| AT3G53260 | -0.3 | 4.73E-03 | -0.1 | 2.71E-01 | -0.2 | 5.91E-02 | 0.5 | 1.89E-03 | 0.1 | 7.84E-01 | 0.5 | 2.94E-02 | Phenylalanine ammonia-lyase 2 |
| AT5G04230 | 0.5 | 4.82E-05 | 0.7 | 1.77E-05 | -0.1 | 1.41E-01 | 0.5 | 9.33E-03 | 0.3 | 1.48E-01 | 0.2 | 4.40E-01 | Phenylalanine ammonia-lyase |
| AT5G01650 | 0.7 | 1.73E-05 | 0.6 | 5.67E-05 | 0.1 | 6.20E-01 | 0.2 | 3.04E-01 | -0.1 | 8.03E-01 | 0.3 | 3.12E-01 | Tautomerase |
| AT5G57170 | 0.8 | 1.77E-05 | 1.1 | 4.99E-06 | -0.3 | 4.35E-02 | 0.0 | 8.71E-01 | 0.0 | 8.70E-01 | -0.1 | 7.76E-01 | Tautomerase |
| AT5G13800 | -1.4 | 7.75E-08 | -1.3 | 2.38E-07 | 0.0 | 8.62E-01 | -0.4 | 5.76E-03 | -0.1 | 5.18E-01 | -0.3 | 1.41E-01 | Pheophytinase |
| AT2G17280 | -1.3 | 3.02E-05 | -0.7 | 1.85E-03 | -0.6 | 1.66E-02 | 0.0 | 9.64E-01 | 0.4 | 3.13E-01 | -0.3 | 2.17E-01 | Phosphoglycerate mutase-like protein |
| AT3G59770 | -0.4 | 5.35E-04 | -0.4 | 3.08E-04 | 0.1 | 5.03E-01 | -0.2 | 2.18E-01 | -0.2 | 2.84E-01 | 0.0 | 1.00E+00 | SacI homology domain-containing protein |
| AT3G51830 | -0.3 | 5.71E-03 | -0.2 | 3.03E-02 | -0.1 | 4.65E-01 | -0.4 | 1.15E-01 | 0.0 | 9.42E-01 | -0.3 | 2.39E-01 | SAC domain-containing protein 8 |
| AT3G05170 | -0.3 | 3.29E-02 | -0.1 | 6.00E-01 | -0.2 | 1.49E-01 | 0.0 | 8.93E-01 | 0.2 | 6.73E-01 | -0.1 | 7.66E-01 | Phosphoglycerate mutase-like protein AT74 |
| AT3G55960 | -0.1 | 7.25E-02 | -0.1 | 5.09E-02 | 0.0 | 8.26E-01 | 0.2 | 1.92E-01 | -0.2 | 2.17E-01 | 0.3 | 4.81E-02 | hypothetical protein |
| AT4G15030 | -0.1 | 4.79E-01 | 0.0 | 6.93E-01 | 0.0 | 8.24E-01 | -0.1 | 6.31E-01 | -0.1 | 7.19E-01 | 0.0 | 9.84E-01 | Folate-sensitive fragile site protein |
| AT5G47760 | 0.0 | 8.29E-01 | -0.1 | 4.14E-01 | 0.0 | 6.06E-01 | -0.1 | 7.49E-01 | -0.1 | 5.94E-01 | 0.1 | 8.43E-01 | 2-phosphoglycolate phosphatase 2 |
| AT1G09932 | 0.2 | 3.95E-01 | -0.6 | 3.07E-02 | 0.8 | 1.84E-02 | 0.9 | 1.85E-02 | -0.1 | 9.08E-01 | 0.9 | 3.27E-02 | Phosphoglycerate mutase family protein |
| AT3G50520 | 0.3 | 6.69E-02 | 0.1 | 7.36E-01 | 0.2 | 1.98E-01 | -0.1 | 6.32E-01 | -0.1 | 6.86E-01 | 0.0 | 9.98E-01 | Phosphoglycerate mutase-like protein 4 |
| AT5G16480 | 0.3 | 9.48E-02 | 0.2 | 3.36E-01 | 0.1 | 5.61E-01 | 0.0 | 9.16E-01 | -0.5 | 6.77E-02 | 0.5 | 1.13E-01 | Phosphotyrosine protein phosphatases superfamily protein |
| AT5G45700 | 0.4 | 1.65E-03 | 0.7 | 9.87E-05 | -0.3 | 4.89E-02 | 0.2 | 5.52E-01 | 0.2 | 6.03E-01 | 0.0 | 9.92E-01 | hypothetical protein |
| AT5G36700 | 1.4 | 8.42E-07 | 1.1 | 7.79E-06 | 0.2 | 9.16E-02 | 0.3 | 2.09E-01 | 0.4 | 1.38E-01 | -0.1 | 8.13E-01 | Phosphoglycolate phosphatase 1A, chloroplastic |
| AT4G03960 | 0.1 | 5.33E-01 | 0.2 | 1.16E-01 | -0.1 | 3.96E-01 | 0.2 | 4.37E-01 | 0.1 | 8.35E-01 | 0.1 | 7.31E-01 | Probable tyrosine-protein phosphatase DSP4 |
| AT1G05000 | 0.2 | 1.58E-01 | 0.5 | 2.32E-03 | -0.3 | 5.32E-02 | 1.4 | 3.88E-05 | 0.6 | 6.13E-02 | 0.8 | 3.66E-03 | Phosphotyrosine protein phosphatases superfamily protein |
| AT5G22090 | -0.7 | 9.49E-06 | -0.5 | 2.09E-04 | -0.2 | 4.28E-02 | -0.1 | 7.27E-01 | 0.1 | 7.23E-01 | -0.1 | 5.81E-01 | Protein FAF-like, chloroplastic |
| AT1G63010 | -0.3 | 9.81E-04 | -0.2 | 1.09E-02 | -0.1 | 2.37E-01 | 0.0 | 8.18E-01 | 0.1 | 4.67E-01 | -0.2 | 4.32E-01 | Major Facilitator Superfamily with SPX domain-containing protein |
| AT4G26770 | -2.4 | 2.66E-06 | -0.7 | 1.77E-03 | -1.7 | 3.09E-04 | -2.2 | 2.32E-03 | -0.2 | 7.43E-01 | | | Phosphatidate cytidylyltransferase 3 |
| AT4G22340 | -1.1 | 1.40E-06 | -0.7 | 3.25E-05 | -0.3 | 1.46E-02 | 0.0 | 8.63E-01 | 0.2 | 2.85E-01 | -0.2 | 4.53E-01 | Phosphatidate cytidylyltransferase |
| AT1G62430 | -0.5 | 7.18E-05 | -0.4 | 1.12E-03 | -0.2 | 1.13E-01 | -0.5 | 2.25E-03 | 0.0 | 8.65E-01 | -0.4 | 2.63E-02 | Phosphatidate cytidylyltransferase 1 |
| AT2G45150 | 0.0 | 8.65E-01 | 0.1 | 5.80E-01 | 0.0 | 7.54E-01 | 0.1 | 7.38E-01 | 0.3 | 2.99E-01 | -0.2 | 5.77E-01 | Phosphatidate cytidylyltransferase 4, chloroplastic |
| AT3G47630 | 0.2 | 1.47E-01 | 0.0 | 7.74E-01 | 0.1 | 3.45E-01 | -0.1 | 7.02E-01 | -0.3 | 3.48E-01 | 0.2 | 6.13E-01 | Phosphatidate cytidylyltransferase, mitochondrial |
| AT5G42870 | -0.3 | 2.43E-03 | -0.4 | 8.61E-04 | 0.1 | 4.18E-01 | -0.1 | 6.03E-01 | 0.0 | 9.73E-01 | -0.1 | 6.80E-01 | phosphatidate phosphatase |
| AT3G09560 | -0.3 | 3.06E-03 | -0.3 | 2.20E-03 | 0.0 | 7.37E-01 | 0.2 | 2.66E-01 | 0.2 | 5.30E-01 | 0.1 | 8.33E-01 | Phosphatidate phosphatase PAH1 |
| AT3G29075 | -0.1 | 3.47E-01 | -0.1 | 6.48E-01 | 0.0 | 7.00E-01 | 0.0 | 9.98E-01 | -0.1 | 8.56E-01 | 0.1 | 8.80E-01 | At3g29075 |
| AT3G06960 | 0.2 | 8.65E-03 | 0.0 | 9.67E-01 | 0.2 | 2.29E-02 | -0.2 | 2.74E-01 | -0.2 | 4.52E-01 | 0.0 | 9.05E-01 | Pigment defective 320 |
| AT2G44640 | 0.8 | 6.78E-06 | 0.9 | 6.59E-06 | -0.1 | 3.63E-01 | 0.2 | 2.55E-01 | -0.1 | 7.13E-01 | 0.3 | 1.50E-01 | Expressed protein |
| AT5G63060 | 0.4 | 2.70E-02 | 0.4 | 6.70E-02 | 0.1 | 7.50E-01 | 0.2 | 6.89E-01 | 0.4 | 3.53E-01 | -0.2 | 6.81E-01 | Sec14p-like phosphatidylinositol transfer family protein |
| AT3G19420 | -0.3 | 1.19E-03 | -0.3 | 2.63E-04 | 0.1 | 1.90E-01 | -0.2 | 4.83E-02 | -0.2 | 2.62E-01 | -0.1 | 6.74E-01 | Phosphatidylinositol 3,4,5-trisphosphate 3-phosphatase and protein-tyrosine-phosphatase PTEN2A |
| AT1G11440 | 0.3 | 4.18E-03 | 0.4 | 3.36E-03 | 0.0 | 8.05E-01 | 0.1 | 8.24E-01 | 0.2 | 4.35E-01 | -0.2 | 6.78E-01 | At1g11440 |
| AT3G03530 | -3.2 | 1.55E-07 | -1.8 | 3.03E-06 | -1.4 | 3.93E-04 | 0.0 | 9.79E-01 | 0.7 | 1.38E-01 | -0.7 | 1.53E-01 | Non-specific phospholipase C4 |
| AT3G58830 | 0.3 | 1.14E-03 | 0.4 | 4.63E-04 | -0.1 | 4.33E-01 | 0.0 | 8.59E-01 | -0.1 | 7.10E-01 | 0.1 | 6.31E-01 | Uncharacterized protein |
| AT5G65480 | -0.5 | 3.34E-04 | -0.3 | 1.94E-02 | -0.2 | 4.49E-02 | -0.2 | 3.06E-01 | -0.2 | 5.85E-01 | -0.1 | 8.50E-01 | AT5g65480 |
| AT3G48195 | -0.5 | 1.96E-05 | -0.4 | 1.75E-04 | -0.1 | 1.94E-01 | -0.1 | 3.98E-01 | -0.1 | 4.65E-01 | 0.0 | 9.97E-01 | Uncharacterized protein T24C20_80 |
| AT5G07120 | -0.4 | 7.73E-04 | -0.1 | 1.20E-01 | -0.3 | 2.59E-02 | -0.1 | 3.68E-01 | -0.1 | 6.95E-01 | -0.1 | 7.99E-01 | Sorting nexin 2B |
| AT2G15900 | -0.4 | 1.60E-03 | -0.4 | 2.54E-03 | 0.0 | 9.44E-01 | 0.0 | 9.45E-01 | -0.2 | 6.36E-01 | 0.2 | 7.33E-01 | Phox domain-containing protein |
| AT2G25350 | -0.4 | 1.28E-02 | -0.4 | 7.07E-03 | 0.1 | 6.93E-01 | -0.5 | 7.85E-03 | -0.1 | 5.06E-01 | -0.4 | 9.88E-02 | Phox domain-containing protein |
| AT1G15240 | -0.4 | 4.01E-04 | -0.4 | 3.32E-04 | 0.0 | 6.82E-01 | -0.3 | 2.26E-01 | -0.2 | 4.20E-01 | -0.1 | 8.81E-01 | At1g15240 domain-containing protein) |
| AT3G15920 | -0.3 | 3.75E-03 | -0.3 | 2.60E-03 | 0.0 | 7.39E-01 | 0.0 | 8.23E-01 | 0.1 | 7.49E-01 | -0.1 | 6.60E-01 | PX domain-containing protein EREX |
| AT5G06140 | 0.0 | 7.82E-01 | 0.1 | 6.73E-02 | -0.1 | 1.39E-01 | -0.3 | 1.60E-02 | -0.3 | 4.11E-02 | 0.0 | 9.28E-01 | Sorting nexin 1 |
| AT5G58440 | -0.1 | 9.99E-02 | -0.2 | 1.25E-02 | 0.1 | 2.73E-01 | 0.0 | 9.32E-01 | 0.2 | 3.94E-01 | -0.2 | 4.18E-01 | Sorting nexin 2A |
| AT2G38410 | -0.7 | 2.05E-05 | -0.6 | 1.23E-04 | -0.1 | 3.04E-01 | -0.2 | 1.31E-01 | -0.2 | 2.62E-01 | 0.0 | 8.94E-01 | Uncharacterized protein |
| AT5G63640 | -0.6 | 4.87E-06 | -0.3 | 5.55E-04 | -0.3 | 4.28E-03 | -0.3 | 4.53E-02 | -0.3 | 9.07E-02 | 0.0 | 9.31E-01 | TOM1-like protein 5 |
| AT1G21380 | -0.5 | 1.01E-04 | -0.3 | 1.58E-03 | -0.1 | 1.22E-01 | -0.1 | 8.16E-01 | 0.0 | 9.72E-01 | -0.1 | 8.58E-01 | TOM1-like protein 3 |
| AT4G32760 | -0.4 | 1.27E-03 | -0.3 | 4.91E-03 | -0.1 | 5.68E-01 | -0.2 | 3.98E-02 | -0.4 | 5.60E-03 | 0.1 | 4.06E-01 | ENTH |
| AT1G06210 | -0.3 | 2.57E-03 | -0.3 | 5.53E-03 | 0.0 | 7.99E-01 | 0.1 | 5.02E-01 | 0.2 | 3.67E-01 | -0.1 | 8.40E-01 | TOM1-like protein 2 |
| AT1G76970 | -0.3 | 3.94E-02 | -0.7 | 8.22E-04 | 0.4 | 3.99E-02 | 0.2 | 2.49E-01 | -0.2 | 3.92E-01 | 0.4 | 1.12E-01 | Target of Myb protein 1 |
| AT5G16880 | -0.2 | 1.07E-02 | -0.3 | 1.11E-03 | 0.1 | 1.48E-01 | -0.2 | 5.86E-02 | -0.3 | 1.80E-02 | 0.1 | 6.30E-01 | Target of Myb protein 1 |
| AT3G27325 | -0.4 | 1.25E-04 | -0.3 | 1.14E-03 | -0.1 | 2.35E-01 | -0.2 | 1.79E-01 | -0.1 | 7.67E-01 | -0.1 | 5.39E-01 | GPI inositol-deacylase |
| AT3G45100 | 0.2 | 3.60E-02 | 0.2 | 4.04E-02 | 0.0 | 9.90E-01 | -0.2 | 1.41E-01 | -0.3 | 6.74E-02 | 0.1 | 7.46E-01 | Phosphatidylinositol N-acetylglucosaminyltransferase subunit A |
| AT3G50110 | -0.2 | 2.81E-02 | 0.1 | 3.82E-01 | -0.3 | 1.70E-02 | -0.1 | 8.97E-01 | 0.1 | 9.08E-01 | -0.1 | 8.40E-01 | Phosphatidylinositol 3,4,5-trisphosphate 3-phosphatase and protein-tyrosine-phosphatase PTEN2B |
| AT3G55940 | -2.5 | 2.95E-07 | -0.9 | 1.02E-04 | -1.6 | 7.19E-05 | -1.1 | 1.30E-02 | 0.1 | 9.01E-01 | -1.2 | 3.07E-02 | Phosphoinositide phospholipase C 7 |
| AT5G58690 | -1.7 | 1.72E-07 | -1.3 | 1.66E-06 | -0.4 | 1.17E-02 | -0.2 | 5.50E-01 | 0.1 | 7.12E-01 | -0.3 | 3.80E-01 | Phosphoinositide phospholipase C |
| AT3G47220 | -1.5 | 5.54E-08 | -0.4 | 6.15E-04 | -1.1 | 1.02E-05 | -0.6 | 6.85E-03 | 0.0 | 9.49E-01 | -0.5 | 2.32E-02 | Phosphoinositide phospholipase C 9 |
| AT5G58700 | -1.1 | 1.14E-06 | -0.6 | 5.80E-05 | -0.4 | 3.82E-03 | 0.0 | 8.75E-01 | 0.2 | 4.58E-01 | -0.2 | 4.33E-01 | Phosphoinositide phospholipase C |
| AT3G08510 | 0.0 | 9.25E-01 | 0.0 | 8.89E-01 | 0.0 | 9.63E-01 | -0.2 | 1.29E-01 | -0.2 | 6.44E-02 | 0.1 | 7.58E-01 | Phosphoinositide phospholipase C |
| AT3G47290 | 1.2 | 3.28E-07 | 0.7 | 5.35E-05 | 0.6 | 3.53E-04 | 0.3 | 2.79E-02 | 0.1 | 3.85E-01 | 0.2 | 3.73E-01 | Phosphoinositide phospholipase C |
| AT4G36640 | -0.7 | 3.98E-06 | -0.5 | 5.15E-05 | -0.2 | 6.28E-02 | -0.3 | 9.90E-02 | 0.3 | 1.41E-01 | -0.5 | 2.17E-02 | AT4G36640 protein |
| AT1G75170 | -0.5 | 2.44E-03 | -0.3 | 4.53E-02 | -0.2 | 1.78E-01 | 0.1 | 7.40E-01 | 0.0 | 9.34E-01 | 0.1 | 7.65E-01 | Sec14p-like phosphatidylinositol transfer family protein |
| AT1G22180 | -0.4 | 1.50E-03 | -0.4 | 3.65E-03 | 0.0 | 7.60E-01 | 0.3 | 2.02E-01 | 0.0 | 9.81E-01 | 0.3 | 2.71E-01 | Sec14p-like phosphatidylinositol transfer family protein |
| AT4G34660 | -0.4 | 1.01E-04 | -0.1 | 2.18E-01 | -0.3 | 1.95E-03 | -0.1 | 3.64E-01 | 0.1 | 6.90E-01 | -0.2 | 3.00E-01 | SH3 domain-containing protein 2 |
| AT3G10550 | -0.2 | 1.14E-02 | 0.0 | 7.60E-01 | -0.2 | 4.30E-02 | -0.5 | 5.86E-03 | 0.0 | 9.71E-01 | -0.5 | 1.49E-02 | Phosphatidylinositol-3-phosphatase myotubularin-1 |
| AT5G04540 | 0.1 | 3.66E-02 | 0.0 | 7.19E-01 | 0.1 | 1.21E-01 | 0.2 | 2.80E-01 | -0.2 | 4.83E-01 | 0.4 | 1.04E-01 | Phosphatidylinositol-3-phosphatase myotubularin-2 |
| AT1G17340 | -0.8 | 3.24E-06 | -0.4 | 4.42E-04 | -0.4 | 2.53E-03 | -0.2 | 2.72E-01 | -0.1 | 5.39E-01 | -0.1 | 8.32E-01 | SAC domain-containing protein |
| AT3G14205 | -0.4 | 9.55E-05 | -0.5 | 3.02E-05 | 0.1 | 1.33E-01 | 0.0 | 7.24E-01 | 0.1 | 2.85E-01 | -0.2 | 2.46E-01 | Phosphoinositide phosphatase family protein |
| AT3G43220 | -0.3 | 4.87E-03 | -0.3 | 4.05E-03 | 0.0 | 8.27E-01 | -0.1 | 5.28E-01 | -0.1 | 8.52E-01 | -0.1 | 8.13E-01 | Phosphoinositide phosphatase family protein |
| AT1G22620 | 0.0 | 7.98E-01 | -0.1 | 5.28E-01 | 0.1 | 4.28E-01 | -0.1 | 5.86E-01 | -0.3 | 2.09E-01 | 0.2 | 5.21E-01 | Phosphatidylinositol-3-phosphatase SAC1 |
| AT3G19190 | -0.7 | 3.77E-05 | -0.6 | 1.01E-04 | 0.0 | 7.55E-01 | 0.0 | 9.40E-01 | -0.1 | 8.59E-01 | 0.0 | 9.31E-01 | Autophagy-related protein 2 |
| AT2G29700 | -0.1 | 5.16E-02 | 0.0 | 5.83E-01 | -0.1 | 1.90E-01 | -0.3 | 1.01E-01 | -0.3 | 1.02E-01 | 0.0 | 9.06E-01 | Pleckstrin homology domain-containing protein 1 |
| AT5G04920 | 0.0 | 8.08E-01 | -0.1 | 1.68E-01 | 0.1 | 2.93E-01 | 0.0 | 8.50E-01 | 0.0 | 9.67E-01 | 0.0 | 8.69E-01 | Vacuolar protein sorting-associated protein 36 |
| AT5G13150 | -1.7 | 5.78E-07 | -0.3 | 3.93E-02 | -1.4 | 2.87E-05 | -0.7 | 3.89E-02 | 0.2 | 5.77E-01 | -0.8 | 1.05E-02 | Exocyst complex component EXO70C1 |
| AT3G55150 | -1.4 | 2.40E-07 | -0.4 | 8.80E-04 | -1.0 | 3.53E-05 | 0.1 | 6.05E-01 | 0.3 | 2.42E-01 | -0.2 | 5.94E-01 | Exocyst complex component EXO70H1 |
| AT5G13990 | -1.3 | 7.97E-07 | -0.1 | 1.68E-01 | -1.2 | 2.95E-05 | -0.8 | 8.41E-02 | -0.1 | 8.16E-01 | -0.6 | 2.74E-01 | Exocyst complex component EXO70C2 |
| AT1G72470 | -0.8 | 1.06E-05 | -0.4 | 1.82E-03 | -0.4 | 4.80E-03 | -0.4 | 7.13E-03 | 0.0 | 9.75E-01 | -0.4 | 3.45E-02 | Exocyst subunit Exo70 family protein |
| AT2G28650 | -0.7 | 3.24E-03 | -0.7 | 6.08E-03 | 0.0 | 9.01E-01 | 0.0 | 9.91E-01 | -0.1 | 8.25E-01 | 0.1 | 8.50E-01 | Exocyst subunit Exo70 family protein |
| AT1G54090 | -0.6 | 2.31E-06 | -0.6 | 1.18E-05 | -0.1 | 3.12E-01 | -0.3 | 6.11E-02 | -0.1 | 5.46E-01 | -0.2 | 3.90E-01 | Exocyst subunit Exo70 family protein |
| AT5G52350 | -0.5 | 1.50E-03 | 0.0 | 7.30E-01 | -0.4 | 8.74E-03 | 0.2 | 5.43E-01 | 0.2 | 3.90E-01 | -0.1 | 8.04E-01 | Exocyst complex component EXO70A3 |
| AT5G03540 | -0.4 | 6.18E-05 | -0.2 | 1.99E-03 | -0.1 | 4.88E-02 | -0.2 | 1.06E-01 | -0.2 | 1.08E-01 | 0.0 | 9.27E-01 | Exocyst subunit Exo70 family protein |

| | | | | | | | | | | | | | |
|---|---|---|---|---|---|---|---|---|---|---|---|---|---|
| AT5G58430 | -0.4 | 7.62E-05 | -0.4 | 3.79E-05 | 0.1 | 2.50E-01 | -0.4 | 8.99E-03 | -0.4 | 1.40E-02 | 0.0 | 9.70E-01 | Exocyst subunit Exo70 family protein |
| AT1G47560 | -0.4 | 2.86E-03 | -0.1 | 5.96E-01 | -0.3 | 2.04E-02 | -0.1 | 7.71E-01 | 0.2 | 5.36E-01 | -0.3 | 4.00E-01 | Exocyst complex component SEC3B |
| AT5G50380 | -0.3 | 2.54E-04 | -0.6 | 1.03E-05 | 0.2 | 5.04E-03 | -0.2 | 1.51E-01 | -0.2 | 2.26E-01 | 0.0 | 9.82E-01 | Exocyst subunit Exo70 family protein |
| AT5G61010 | -0.2 | 5.97E-03 | -0.6 | 2.56E-05 | 0.4 | 1.62E-03 | 0.4 | 1.13E-02 | 0.2 | 2.16E-01 | 0.2 | 4.02E-01 | Exocyst complex component EXO70E2 |
| AT3G14090 | -0.1 | 1.12E-01 | -0.3 | 6.27E-03 | 0.2 | 1.40E-01 | 0.0 | 9.31E-01 | -0.1 | 5.14E-01 | 0.1 | 6.32E-01 | Exocyst subunit Exo70 family protein |
| AT3G29400 | 0.1 | 4.19E-01 | -0.2 | 4.97E-02 | 0.3 | 2.28E-02 | 0.1 | 5.99E-01 | 0.1 | 7.45E-01 | 0.0 | 9.52E-01 | Exocyst subunit Exo70 family protein |
| AT3G09520 | 0.4 | 1.26E-02 | 0.0 | 7.97E-01 | 0.5 | 2.41E-02 | -0.2 | 6.83E-01 | -0.6 | 2.30E-01 | 0.4 | 5.30E-01 | Exocyst subunit Exo70 family protein |
| AT1G07000 | -0.5 | 5.35E-04 | -0.5 | 3.12E-04 | 0.1 | 5.20E-01 | 0.1 | 7.66E-01 | -0.3 | 6.92E-02 | 0.4 | 8.76E-02 | Exocyst complex component EXO70B2 |
| AT5G53280 | 0.7 | 7.63E-06 | 0.2 | 9.64E-03 | 0.4 | 1.07E-03 | 0.2 | 3.06E-01 | 0.3 | 1.51E-01 | -0.1 | 6.67E-01 | Plastid division protein PDV1 |
| AT3G51460 | -0.7 | 5.28E-06 | -0.2 | 3.57E-02 | -0.5 | 2.24E-04 | -0.3 | 1.02E-01 | 0.1 | 6.94E-01 | -0.4 | 1.11E-01 | Phosphoinositide phosphatase SAC7 |
| AT1G80860 | 0.0 | 6.88E-01 | 0.3 | 1.68E-02 | -0.2 | 5.98E-02 | -0.1 | 5.43E-01 | -0.2 | 3.35E-01 | 0.1 | 7.83E-01 | Phosphatidyl-N-methylethanolamine N-methyltransferase |
| AT4G25970 | -0.7 | 4.09E-06 | -0.5 | 6.05E-05 | -0.2 | 5.32E-02 | -0.5 | 1.05E-04 | -0.3 | 5.32E-03 | -0.2 | 3.03E-01 | Phosphatidylserine decarboxylase proenzyme 2 |
| AT4G16700 | 0.8 | 1.12E-03 | 0.4 | 9.28E-02 | 0.5 | 6.60E-02 | 0.5 | 3.49E-01 | 0.2 | 8.42E-01 | | | phosphatidylserine decarboxylase |
| AT4G21990 | -0.7 | 1.00E-05 | -0.4 | 6.19E-04 | -0.3 | 1.06E-02 | 1.8 | 8.02E-08 | 0.8 | 8.84E-04 | 1.0 | 7.98E-04 | APS reductase 3 |
| AT1G62180 | 0.6 | 4.93E-05 | 0.5 | 8.63E-04 | 0.2 | 1.10E-01 | 0.3 | 5.02E-01 | 0.0 | 9.57E-01 | 0.3 | 5.50E-01 | 5'adenylylphosphosulfate reductase 2 |
| AT3G14940 | -1.8 | 7.40E-07 | -0.8 | 4.44E-04 | -1.0 | 2.21E-04 | 0.7 | 2.66E-03 | 0.8 | 9.76E-04 | -0.2 | 5.29E-01 | Phosphoenolpyruvate carboxylase 3 |
| AT1G53310 | -0.5 | 5.94E-05 | -0.3 | 4.37E-03 | -0.2 | 2.22E-02 | 0.1 | 5.51E-01 | 0.3 | 7.32E-02 | -0.2 | 3.57E-01 | Phosphoenolpyruvate carboxylase 1 |
| AT2G42600 | 0.0 | 7.33E-01 | 0.0 | 9.34E-01 | 0.0 | 8.36E-01 | -0.7 | 3.22E-03 | -0.5 | 5.70E-02 | -0.2 | 4.95E-01 | Phosphoenolpyruvate carboxylase 2 |
| AT1G73600 | -0.7 | 6.32E-04 | 0.2 | 3.33E-01 | -0.9 | 6.17E-04 | 0.1 | 6.76E-01 | 0.6 | 1.94E-03 | -0.6 | 1.44E-02 | Phosphoethanolamine N-methyltransferase, putative |
| AT1G48600 | 0.0 | 9.02E-01 | 0.5 | 3.39E-03 | -0.5 | 4.75E-03 | -0.2 | 5.84E-02 | 0.0 | 8.03E-01 | -0.3 | 1.20E-01 | PMEAMT |
| AT1G01500 | 0.2 | 2.41E-03 | 0.1 | 1.13E-01 | 0.1 | 7.87E-02 | -0.2 | 3.12E-01 | -0.2 | 3.80E-01 | 0.0 | 9.99E-01 | Uncharacterized protein At1g01500 |
| AT1G75180 | 1.1 | 1.09E-07 | 0.8 | 1.57E-06 | 0.3 | 4.80E-03 | -0.4 | 4.74E-02 | -0.8 | 2.39E-03 | 0.4 | 1.61E-01 | AT1G75180 protein |
| AT5G36790 | 1.2 | 5.41E-07 | 1.1 | 2.54E-06 | 0.1 | 3.38E-01 | 0.4 | 1.58E-01 | 0.4 | 1.12E-01 | -0.1 | 8.38E-01 | Phosphoglycolate phosphatase 1B, chloroplastic |
| AT4G18550 | -1.3 | 3.85E-07 | -1.2 | 1.75E-06 | -0.2 | 2.68E-01 | -0.9 | 3.04E-02 | -0.1 | 8.42E-01 | -0.8 | 1.05E-01 | Phospholipase A1 |
| AT2G42690 | 0.8 | 1.52E-05 | 0.7 | 6.23E-05 | 0.1 | 5.13E-01 | -0.1 | 4.88E-01 | -0.2 | 1.66E-01 | 0.1 | 5.74E-01 | Phospholipase A1-IIdelta |
| AT1G06250 | | | | | | | 0.6 | 1.64E-01 | 0.2 | 8.21E-01 | 0.5 | 3.72E-01 | Phospholipase A1 |
| AT1G30370 | -1.6 | 7.04E-07 | -0.7 | 1.66E-04 | -0.9 | 3.39E-04 | 1.4 | 1.80E-04 | 0.2 | 6.06E-01 | 1.1 | 3.00E-03 | Alpha |
| AT4G29070 | 0.8 | 5.06E-05 | 0.4 | 1.02E-02 | 0.4 | 9.99E-03 | -0.1 | 4.00E-01 | 0.1 | 5.72E-01 | -0.2 | 2.28E-01 | Uncharacterized protein AT4g29070 |
| AT4G37060 | 1.8 | 4.49E-07 | 2.4 | 1.88E-07 | -0.6 | 8.45E-04 | 0.6 | 1.28E-02 | 0.7 | 8.95E-03 | -0.1 | 7.55E-01 | Patatin |
| AT3G48610 | 0.6 | 2.10E-05 | 0.2 | 4.70E-02 | 0.4 | 9.89E-04 | 0.1 | 6.42E-01 | 0.3 | 6.43E-02 | -0.2 | 2.65E-01 | Non-specific phospholipase C6 |
| AT2G26870 | 1.5 | 5.81E-06 | 1.0 | 2.83E-04 | 0.5 | 1.56E-02 | 0.1 | 7.99E-01 | 0.0 | 9.89E-01 | 0.1 | 8.75E-01 | Non-specific phospholipase C2 |
| AT2G04940 | -0.1 | 1.05E-01 | 0.0 | 9.61E-01 | -0.1 | 1.70E-01 | -0.2 | 1.38E-01 | -0.2 | 2.23E-01 | 0.0 | 9.58E-01 | Phospholipid scramblase |
| AT1G70820 | 0.8 | 3.68E-04 | 0.6 | 1.82E-03 | 0.1 | 4.66E-01 | -0.2 | 2.26E-01 | 0.3 | 2.25E-01 | -0.5 | 3.38E-02 | phosphoglucomutase |
| AT4G18270 | -0.3 | 2.42E-02 | -0.4 | 2.13E-02 | 0.0 | 8.89E-01 | -0.1 | 6.86E-01 | 0.5 | 6.41E-02 | -0.6 | 5.60E-02 | Phospho-N-acetylmuramoyl-pentapeptide-transferase |
| AT1G12350 | 0.1 | 1.79E-01 | -0.1 | 4.04E-01 | 0.2 | 7.13E-02 | -0.1 | 5.88E-01 | -0.2 | 2.87E-01 | 0.1 | 6.89E-01 | Phosphopantothenate--cysteine ligase 1 |
| AT5G58160 | -0.5 | 9.18E-05 | -0.3 | 2.33E-03 | -0.2 | 8.22E-02 | 0.0 | 9.34E-01 | 0.1 | 8.25E-01 | 0.0 | 9.03E-01 | Formin-like protein 13 |
| AT4G18140 | -0.2 | 1.28E-02 | -0.1 | 1.54E-01 | -0.1 | 2.67E-01 | -0.2 | 2.11E-01 | -0.3 | 1.43E-01 | 0.1 | 8.33E-01 | SCP1-like small phosphatase 4b |
| AT2G25050 | 0.1 | 2.44E-01 | 0.1 | 2.37E-01 | 0.0 | 9.60E-01 | 0.2 | 6.18E-01 | 0.3 | 4.10E-01 | -0.1 | 7.46E-01 | Formin-like protein 18 |
| AT5G45610 | 0.5 | 1.30E-02 | 0.3 | 9.63E-02 | 0.2 | 3.98E-01 | 0.1 | 8.76E-01 | 0.2 | 6.45E-01 | -0.1 | 7.52E-01 | Protein SENSITIVE TO UV 2 |
| AT1G22280 | -0.4 | 2.24E-04 | -0.5 | 6.01E-05 | 0.1 | 1.41E-01 | -0.1 | 6.96E-01 | -0.2 | 3.71E-01 | 0.1 | 6.86E-01 | Phytochrome-associated protein phosphatase type 2C |
| AT5G11860 | 0.1 | 1.00E-01 | 0.0 | 8.07E-01 | 0.2 | 1.03E-01 | 0.0 | 9.25E-01 | 0.0 | 9.30E-01 | 0.0 | 8.99E-01 | At5g11860 |
| AT2G40670 | -0.4 | 7.20E-02 | -0.5 | 5.21E-02 | 0.1 | 8.21E-01 | -0.3 | 5.73E-01 | 0.6 | 1.44E-01 | -0.9 | 6.35E-02 | Response regulator 16 |
| AT1G19050 | -0.1 | 4.37E-01 | 0.0 | 9.18E-01 | -0.1 | 5.94E-01 | 0.6 | 4.36E-02 | 0.4 | 2.37E-01 | 0.2 | 6.51E-01 | Two-component response regulator ARR7 |
| AT5G62920 | -0.1 | 6.62E-01 | -0.3 | 6.29E-02 | 0.2 | 1.87E-01 | 0.1 | 8.92E-01 | 0.3 | 4.38E-01 | -0.3 | 5.66E-01 | Two-component response regulator ARR6 |
| AT3G57040 | 0.9 | 2.71E-04 | 0.5 | 8.72E-03 | 0.3 | 9.10E-02 | -0.2 | 4.26E-01 | -0.4 | 7.09E-02 | 0.3 | 3.74E-01 | Two-component response regulator ARR9 |
| AT2G01830 | -0.9 | 7.51E-06 | -0.4 | 1.27E-03 | -0.5 | 3.64E-03 | 0.0 | 8.36E-01 | 0.3 | 1.25E-01 | -0.3 | 2.87E-01 | CHASE domain containing histidine kinase protein |
| AT1G27320 | -0.7 | 4.33E-05 | -0.8 | 3.61E-05 | 0.1 | 4.41E-01 | -0.2 | 3.13E-01 | -0.2 | 4.40E-01 | 0.0 | 9.65E-01 | hypothetical protein |
| AT1G09570 | -0.2 | 5.61E-02 | -0.1 | 1.35E-01 | 0.0 | 7.14E-01 | -0.4 | 5.04E-03 | -0.3 | 2.75E-02 | -0.1 | 7.64E-01 | Phytochrome |
| AT4G18130 | 0.0 | 8.19E-01 | -0.2 | 6.76E-02 | 0.1 | 1.35E-01 | 0.0 | 9.69E-01 | 0.1 | 5.89E-01 | -0.1 | 6.25E-01 | Phytochrome E |
| AT2G18790 | 0.6 | 9.75E-06 | 0.2 | 6.89E-03 | 0.4 | 1.32E-03 | 0.2 | 3.01E-01 | 0.2 | 3.60E-01 | 0.0 | 9.90E-01 | Phytochrome |
| AT1G29410 | 0.0 | 8.11E-01 | 0.2 | 1.63E-01 | -0.2 | 3.30E-01 | -0.3 | 2.65E-01 | 0.1 | 8.71E-01 | -0.4 | 1.96E-01 | phosphoribosylanthranilate isomerase |
| AT5G05590 | 0.1 | 6.32E-01 | 0.3 | 7.82E-02 | -0.2 | 2.56E-01 | 0.3 | 4.60E-01 | 0.4 | 3.01E-01 | -0.1 | 8.02E-01 | phosphoribosylanthranilate isomerase |
| AT1G07780 | 0.2 | 3.73E-01 | 0.3 | 3.42E-01 | 0.0 | 9.42E-01 | 0.5 | 3.09E-02 | 0.1 | 7.15E-01 | 0.4 | 6.58E-02 | phosphoribosylanthranilate isomerase |
| AT1G31220 | 0.3 | 9.79E-03 | 0.2 | 5.47E-02 | 0.1 | 4.78E-01 | 0.3 | 1.34E-01 | 0.0 | 9.60E-01 | 0.3 | 1.84E-01 | Phosphoribosylglycinamide formyltransferase, chloroplastic |
| AT4G34930 | -1.6 | 7.45E-06 | -0.4 | 2.05E-02 | -1.2 | 4.76E-04 | 0.0 | 1.00E+00 | -0.1 | 8.47E-01 | 0.1 | 8.65E-01 | At4g34930 |
| AT4G34920 | -0.3 | 1.59E-03 | -0.1 | 3.55E-01 | -0.2 | 1.57E-02 | 0.0 | 7.71E-01 | 0.2 | 2.06E-01 | -0.1 | 4.23E-01 | AT4g34920 |
| AT1G49740 | -0.2 | 2.93E-02 | -0.2 | 2.64E-02 | 0.0 | 9.11E-01 | 0.1 | 3.83E-01 | 0.3 | 1.35E-01 | -0.1 | 6.18E-01 | hypothetical protein |
| AT5G67130 | 0.0 | 7.00E-01 | 0.0 | 8.60E-01 | 0.0 | 6.25E-01 | -0.1 | 6.38E-01 | -0.2 | 3.01E-01 | 0.1 | 6.70E-01 | PI-PLC X domain-containing protein At5g67130 |
| AT4G38690 | 0.4 | 3.58E-03 | 0.6 | 4.91E-04 | -0.2 | 1.73E-01 | -0.6 | 1.25E-03 | -0.5 | 6.40E-03 | -0.1 | 7.92E-01 | hypothetical protein |
| AT3G25585 | -0.8 | 7.64E-07 | -0.4 | 2.36E-04 | -0.4 | 3.60E-04 | -0.1 | 5.27E-01 | 0.1 | 4.33E-01 | -0.2 | 1.53E-01 | Aminoalcoholphosphotransferase |
| AT2G29525 | -0.2 | 1.91E-01 | -0.2 | 1.91E-01 | 0.0 | 9.68E-01 | 0.1 | 8.19E-01 | 0.4 | 4.64E-01 | -0.3 | 6.70E-01 | Inositol phosphorylceramide synthase 3 |
| AT1G13560 | 0.0 | 8.16E-01 | -0.1 | 1.55E-01 | 0.1 | 2.62E-01 | -0.1 | 5.36E-01 | 0.0 | 8.89E-01 | -0.1 | 7.85E-01 | Aminoalcoholphosphotransferase 1 |
| AT2G18915 | -0.4 | 7.52E-04 | -0.4 | 7.71E-04 | 0.0 | 8.33E-01 | -0.6 | 2.02E-03 | -0.5 | 1.18E-02 | -0.1 | 7.24E-01 | LKP2 |
| AT1G68050 | -0.3 | 9.72E-02 | 0.1 | 4.74E-01 | -0.4 | 5.70E-02 | -0.2 | 4.23E-01 | 0.1 | 8.41E-01 | -0.3 | 3.44E-01 | Adagio protein 3 |
| AT2G02710 | 1.3 | 2.63E-06 | 1.1 | 2.21E-05 | 0.2 | 1.21E-01 | -0.7 | 4.33E-02 | -1.4 | 1.46E-03 | 0.7 | 1.03E-01 | PAS |
| AT5G35840 | -0.2 | 3.75E-02 | -0.3 | 9.40E-03 | 0.1 | 4.45E-01 | 0.0 | 9.14E-01 | 0.0 | 9.58E-01 | 0.0 | 9.77E-01 | Phytochrome C |
| AT5G07840 | -0.2 | 6.51E-03 | -0.2 | 1.16E-02 | 0.0 | 8.72E-01 | -0.4 | 3.35E-02 | -0.2 | 4.37E-01 | -0.2 | 3.40E-01 | Phytochrome-interacting ankyrin-repeat protein 1 |
| AT5G61230 | -0.2 | 3.95E-02 | -0.1 | 1.87E-01 | -0.1 | 4.67E-01 | -0.2 | 3.73E-02 | -0.3 | 4.11E-02 | 0.0 | 9.25E-01 | Phytochrome-interacting ankyrin-repeat protein 2 |
| AT4G14210 | 0.3 | 6.85E-03 | 0.1 | 2.87E-01 | 0.2 | 7.73E-02 | -0.1 | 7.43E-01 | 0.1 | 8.31E-01 | -0.1 | 6.76E-01 | 15-cis-phytoene desaturase, chloroplastic |
| AT4G13660 | -1.7 | 5.73E-08 | -0.9 | 5.33E-06 | -0.8 | 6.43E-05 | 0.0 | 9.36E-01 | 0.1 | 5.06E-01 | -0.2 | 3.67E-01 | Pinoresinol reductase 2 -pinoresinol reductase) |
| AT1G32100 | -1.0 | 3.46E-06 | 0.0 | 9.86E-01 | -1.0 | 4.09E-05 | 0.0 | 9.72E-01 | 0.3 | 2.20E-01 | -0.2 | 3.44E-01 | Pinoresinol reductase 1 -pinoresinol reductase) -pinoresinol reductase) |
| AT4G34540 | 0.0 | 6.44E-01 | 0.1 | 4.44E-01 | 0.0 | 7.97E-01 | -0.3 | 6.69E-02 | -0.2 | 2.17E-01 | -0.1 | 7.64E-01 | Probable pinoresinol-lariciresinol reductase 3 |
| AT4G00060 | -0.4 | 2.68E-04 | -0.5 | 2.06E-04 | 0.1 | 6.03E-01 | -0.1 | 4.27E-01 | 0.1 | 7.43E-01 | -0.2 | 3.20E-01 | AT4g00060 |
| AT2G31865 | -0.3 | 7.69E-02 | -0.9 | 3.46E-04 | 0.6 | 8.79E-03 | -0.1 | 8.22E-01 | -0.4 | 1.43E-01 | 0.3 | 3.27E-01 | poly glycohydrolase |
| AT2G31870 | 0.1 | 1.78E-01 | -0.1 | 4.21E-01 | 0.2 | 8.28E-02 | 0.0 | 9.11E-01 | -0.1 | 8.21E-01 | 0.1 | 9.23E-01 | poly glycohydrolase |
| AT1G31410 | 0.7 | 6.30E-05 | 0.4 | 3.23E-03 | 0.3 | 4.04E-02 | 0.1 | 6.37E-01 | 0.2 | 4.43E-01 | -0.1 | 8.15E-01 | At1g31410 |
| AT3G13620 | -1.1 | 2.31E-06 | -0.8 | 3.41E-05 | -0.3 | 4.09E-02 | -0.4 | 8.43E-02 | 0.0 | 9.34E-01 | -0.4 | 2.51E-01 | Probable polyamine transporter At3g13620 |
| AT3G19553 | -0.7 | 2.09E-05 | -0.6 | 1.39E-04 | -0.1 | 2.59E-01 | -0.4 | 9.70E-03 | -0.2 | 2.00E-01 | -0.2 | 4.10E-01 | Probable polyamine transporter At3g19553 |
| AT1G31830 | -0.7 | 4.15E-06 | -0.3 | 1.04E-03 | -0.3 | 1.79E-03 | -0.3 | 2.26E-02 | -0.1 | 5.06E-01 | -0.2 | 2.89E-01 | Probable polyamine transporter At1g31830 |
| AT1G31820 | -0.7 | 1.30E-03 | -1.0 | 2.00E-04 | 0.3 | 1.28E-01 | -0.4 | 1.50E-01 | -0.4 | 2.72E-01 | -0.1 | 9.05E-01 | Probable polyamine transporter At1g31820 |
| AT5G05630 | 0.2 | 1.64E-01 | 0.4 | 2.79E-02 | -0.2 | 3.88E-01 | 0.1 | 8.18E-01 | 0.3 | 4.97E-01 | -0.2 | 7.21E-01 | Polyamine transporter RMV1 |
| AT1G05660 | -1.7 | 2.32E-06 | -0.1 | 3.90E-01 | -1.6 | 4.75E-05 | -1.0 | 2.84E-02 | 0.1 | 8.65E-01 | -1.1 | 6.75E-02 | Polygalacturonase |
| AT3G59850 | -1.4 | 2.95E-04 | 0.0 | 9.76E-01 | -1.4 | 1.64E-03 | 0.3 | 5.25E-01 | 0.7 | 1.60E-01 | -0.4 | 5.07E-01 | hypothetical protein |

| | | | | | | | | | | | | | |
|---|---|---|---|---|---|---|---|---|---|---|---|---|---|
| AT3G07970 | -1.4 | 6.57E-06 | -0.8 | 2.19E-04 | -0.5 | 1.61E-02 | 0.6 | 2.73E-01 | 0.5 | 4.38E-01 | 0.1 | 8.94E-01 | Polygalacturonase QRT2 |
| AT1G48100 | -0.9 | 5.12E-02 | -1.1 | 3.15E-02 | 0.2 | 7.18E-01 | | | | | | | Polygalacturonase |
| AT4G33440 | -0.7 | 9.02E-05 | -0.2 | 1.37E-01 | -0.5 | 2.90E-03 | -0.1 | 6.80E-01 | -0.1 | 6.76E-01 | 0.0 | 9.69E-01 | hypothetical protein |
| AT1G56710 | -0.5 | 8.47E-02 | -0.4 | 1.48E-01 | -0.1 | 8.52E-01 | -0.3 | 4.23E-01 | 0.0 | 9.65E-01 | -0.4 | 4.18E-01 | hypothetical protein |
| AT3G42950 | -0.4 | 3.29E-04 | -0.1 | 4.77E-01 | -0.4 | 3.15E-03 | 0.0 | 9.15E-01 | 0.2 | 2.28E-01 | -0.2 | 3.98E-01 | Polygalacturonase |
| AT1G19170 | -0.3 | 3.01E-03 | 0.0 | 8.23E-01 | -0.3 | 1.25E-02 | 0.0 | 8.53E-01 | 0.0 | 9.77E-01 | 0.0 | 9.25E-01 | hypothetical protein |
| AT1G10640 | 0.1 | 5.56E-01 | -0.3 | 1.79E-01 | 0.4 | 1.05E-01 | -0.6 | 6.64E-03 | -0.1 | 6.31E-01 | -0.5 | 7.99E-02 | At1g10640 |
| AT4G23500 | 0.1 | 3.09E-01 | 0.3 | 9.76E-02 | -0.1 | 5.28E-01 | -0.2 | 9.56E-02 | 0.2 | 2.94E-01 | -0.4 | 4.57E-02 | Polygalacturonase |
| AT1G23460 | 0.2 | 1.73E-01 | 0.5 | 2.03E-03 | -0.3 | 3.87E-02 | -0.1 | 5.40E-01 | -0.1 | 6.73E-01 | 0.0 | 9.57E-01 | Polygalacturonase At1g80170 |
| AT3G57785 | 0.3 | 1.26E-02 | 0.4 | 6.00E-03 | -0.1 | 6.62E-01 | 0.0 | 9.33E-01 | 0.0 | 9.61E-01 | 0.1 | 9.21E-01 | Uncharacterized protein F15B8.20 |
| AT3G57790 | 0.3 | 6.97E-04 | 0.2 | 5.06E-02 | 0.2 | 5.26E-02 | -0.2 | 8.06E-02 | -0.1 | 4.63E-01 | -0.1 | 5.61E-01 | Uncharacterized protein F15B8.20 |
| AT3G61490 | 0.5 | 1.02E-03 | 0.3 | 1.20E-02 | 0.1 | 2.69E-01 | -0.1 | 5.44E-01 | -0.3 | 2.12E-01 | 0.2 | 6.04E-01 | hypothetical protein |
| AT1G60590 | 0.7 | 4.33E-02 | 0.2 | 5.37E-01 | 0.5 | 2.22E-01 | -0.7 | 1.22E-02 | 0.0 | 9.65E-01 | -0.7 | 5.34E-02 | At1g60590 |
| AT3G62110 | 1.0 | 5.24E-06 | 1.0 | 9.24E-06 | 0.0 | 8.21E-01 | 0.8 | 9.57E-06 | 0.2 | 1.60E-01 | 0.6 | 3.80E-03 | hypothetical protein |
| AT1G06780 | -1.5 | 1.09E-07 | -0.8 | 1.24E-05 | -0.7 | 1.28E-04 | 0.0 | 9.79E-01 | 0.2 | 5.46E-01 | -0.2 | 5.83E-01 | Hexosyltransferase |
| AT5G54690 | -1.4 | 4.56E-06 | -0.3 | 2.05E-02 | -1.1 | 2.94E-04 | -0.2 | 5.93E-01 | -0.1 | 8.90E-01 | -0.2 | 8.07E-01 | Probable galacturonosyltransferase 12 |
| AT3G62660 | -1.1 | 1.45E-07 | -0.6 | 1.61E-05 | -0.5 | 1.77E-04 | -0.7 | 2.78E-05 | -0.5 | 2.20E-03 | -0.2 | 2.00E-01 | Probable galacturonosyltransferase-like 7 |
| AT3G50760 | -1.0 | 6.63E-04 | -1.1 | 5.50E-04 | 0.1 | 7.06E-01 | -0.2 | 6.76E-01 | 0.3 | 4.25E-01 | -0.5 | 9.21E-02 | Probable galacturonosyltransferase-like 2 |
| AT1G19300 | -0.9 | 3.86E-06 | -0.3 | 1.03E-02 | -0.6 | 3.53E-04 | -0.4 | 4.23E-03 | -0.2 | 2.30E-01 | -0.2 | 1.75E-01 | Probable galacturonosyltransferase-like 1 |
| AT1G24170 | -0.7 | 1.77E-04 | -0.3 | 3.09E-02 | -0.4 | 1.36E-02 | -0.4 | 3.84E-04 | -0.3 | 1.07E-02 | -0.1 | 4.08E-01 | Probable galacturonosyltransferase-like 8 |
| AT5G15470 | -0.7 | 6.44E-06 | -0.4 | 7.88E-04 | -0.3 | 4.39E-03 | -0.2 | 2.54E-01 | 0.2 | 1.00E-01 | -0.4 | 2.75E-02 | Probable galacturonosyltransferase 14 |
| AT3G58790 | -0.6 | 2.64E-04 | -0.6 | 2.65E-04 | 0.0 | 7.44E-01 | -0.5 | 2.77E-01 | -0.2 | 7.67E-01 | -0.3 | 6.23E-01 | Hexosyltransferase |
| AT3G02350 | -0.2 | 1.72E-03 | -0.1 | 5.60E-02 | -0.1 | 9.94E-02 | 0.0 | 8.89E-01 | 0.1 | 5.12E-01 | -0.1 | 5.36E-01 | Probable galacturonosyltransferase 9 |
| AT3G28340 | -0.2 | 2.69E-01 | -0.4 | 6.41E-02 | 0.2 | 4.57E-01 | -0.1 | 7.31E-01 | 0.2 | 5.33E-01 | -0.3 | 3.45E-01 | Probable galacturonosyltransferase-like 10 |
| AT1G13250 | -0.2 | 9.95E-02 | -0.1 | 4.40E-01 | -0.1 | 4.40E-01 | -0.3 | 2.26E-02 | 0.1 | 6.36E-01 | -0.4 | 2.37E-02 | Probable galacturonosyltransferase-like 3 |
| AT2G20810 | -0.2 | 4.06E-02 | -0.1 | 1.03E-01 | 0.0 | 7.17E-01 | -0.4 | 3.40E-01 | -0.2 | 6.95E-01 | -0.2 | 7.65E-01 | Hexosyltransferase |
| AT4G38270 | -0.1 | 5.25E-02 | -0.3 | 2.44E-03 | 0.1 | 1.08E-01 | 0.2 | 4.65E-01 | 0.1 | 7.33E-01 | 0.1 | 8.55E-01 | Hexosyltransferase |
| AT3G61130 | -0.1 | 3.91E-01 | -0.1 | 2.64E-01 | 0.0 | 7.91E-01 | -0.2 | 2.49E-01 | 0.1 | 4.71E-01 | -0.3 | 1.30E-01 | Polygalacturonate 4-alpha-galacturonosyltransferase |
| AT3G01040 | -0.1 | 5.03E-01 | -0.1 | 1.78E-01 | 0.1 | 5.33E-01 | 0.1 | 7.10E-01 | 0.2 | 5.01E-01 | -0.1 | 7.99E-01 | Hexosyltransferase |
| AT2G38650 | 0.0 | 8.22E-01 | 0.1 | 2.06E-01 | -0.1 | 3.40E-01 | 0.2 | 1.79E-01 | 0.2 | 1.80E-01 | 0.0 | 9.28E-01 | Hexosyltransferase |
| AT2G30575 | 0.0 | 8.37E-01 | 0.0 | 7.17E-01 | 0.0 | 8.96E-01 | 0.0 | 9.41E-01 | 0.3 | 3.92E-01 | -0.3 | 4.39E-01 | Probable galacturonosyltransferase 5 |
| AT5G47780 | 0.3 | 5.61E-03 | 0.1 | 1.04E-01 | 0.1 | 1.71E-01 | -0.1 | 4.67E-01 | -0.1 | 4.85E-01 | 0.0 | 9.70E-01 | Probable galacturonosyltransferase 4 |
| AT4G02130 | 0.4 | 2.29E-02 | 0.4 | 2.05E-02 | 0.0 | 9.09E-01 | -0.6 | 1.92E-05 | -0.6 | 1.16E-04 | 0.0 | 9.26E-01 | Probable galacturonosyltransferase-like 6 |
| AT1G02720 | 0.4 | 1.20E-02 | 0.6 | 5.36E-04 | -0.3 | 7.68E-02 | -0.3 | 5.33E-02 | -0.1 | 6.50E-01 | -0.2 | 3.01E-01 | Probable galacturonosyltransferase-like 5 |
| AT1G70090 | 1.1 | 4.06E-05 | 0.4 | 2.53E-02 | 0.7 | 3.35E-03 | -0.6 | 1.44E-03 | -0.9 | 1.76E-04 | 0.3 | 1.76E-01 | Probable galacturonosyltransferase-like 9 |
| AT2G17570 | -0.4 | 2.77E-03 | -0.2 | 9.64E-02 | -0.2 | 1.14E-01 | -0.3 | 6.40E-02 | -0.3 | 1.44E-01 | 0.0 | 8.96E-01 | Dehydrodolichyl diphosphate synthase 6 |
| AT2G47220 | -0.1 | 4.46E-01 | 0.1 | 7.20E-01 | -0.2 | 3.54E-01 | -0.3 | 4.12E-01 | -0.5 | 1.41E-01 | 0.2 | 5.20E-01 | Uncharacterized protein |
| AT3G03710 | 0.6 | 9.47E-05 | 0.6 | 2.55E-04 | 0.0 | 7.34E-01 | 0.4 | 2.81E-02 | 0.1 | 7.73E-01 | 0.3 | 1.60E-01 | polyribonucleotide nucleotidyltransferase |
| AT1G10380 | -1.9 | 2.04E-08 | -1.1 | 7.82E-07 | -0.8 | 3.57E-05 | -0.1 | 3.99E-01 | 0.1 | 8.14E-01 | -0.2 | 4.01E-01 | hypothetical protein |
| AT2G46495 | -1.4 | 8.95E-06 | -0.8 | 3.93E-04 | -0.6 | 1.22E-02 | 1.7 | 2.23E-05 | 1.0 | 5.71E-03 | 0.7 | 7.62E-03 | RING-type E3 ubiquitin transferase |
| AT5G53110 | -0.9 | 9.81E-06 | -1.0 | 1.07E-05 | 0.1 | 4.57E-01 | 0.0 | 9.62E-01 | -0.2 | 3.55E-01 | 0.2 | 4.64E-01 | RING-type E3 ubiquitin transferase |
| AT1G05850 | -0.6 | 1.88E-05 | -0.2 | 1.84E-02 | -0.4 | 1.55E-03 | -0.4 | 2.51E-03 | -0.1 | 5.09E-01 | -0.3 | 8.35E-02 | Chitinase-like protein 1 |
| AT3G46550 | 0.1 | 9.90E-02 | 0.1 | 2.87E-01 | 0.0 | 6.10E-01 | 0.0 | 8.81E-01 | 0.1 | 5.96E-01 | -0.1 | 7.54E-01 | Fasciclin-like arabinogalactan protein 4 |
| AT1G16090 | 0.7 | 1.04E-03 | 0.1 | 4.26E-01 | 0.5 | 1.51E-02 | -0.2 | 6.55E-01 | -0.4 | 3.55E-01 | 0.2 | 6.85E-01 | Wall associated kinase-like 7 |
| AT4G14130 | -1.1 | 3.07E-06 | -0.4 | 1.75E-03 | -0.7 | 7.71E-04 | -0.8 | 2.00E-02 | -0.9 | 2.48E-02 | 0.0 | 9.72E-01 | Xyloglucan endotransglucosylase |
| AT3G44990 | -1.1 | 5.08E-05 | -0.5 | 1.05E-02 | -0.6 | 7.59E-03 | 0.2 | 2.97E-01 | 1.0 | 5.47E-05 | -0.8 | 1.25E-03 | Xyloglucan endotransglucosylase |
| AT2G29070 | -0.8 | 5.99E-06 | -0.6 | 7.73E-05 | -0.2 | 7.74E-02 | -0.2 | 5.26E-01 | 0.1 | 8.17E-01 | -0.3 | 4.21E-01 | Ubiquitin fusion degradation UFD1 family protein |
| AT2G21270 | -0.4 | 6.29E-04 | -0.2 | 4.27E-02 | -0.2 | 4.42E-02 | -0.1 | 3.12E-01 | 0.0 | 7.91E-01 | -0.1 | 6.83E-01 | AT2G21270 protein |
| AT2G12550 | -0.1 | 4.95E-01 | -0.3 | 8.17E-03 | 0.2 | 4.62E-02 | 0.0 | 9.94E-01 | 0.0 | 9.90E-01 | 0.0 | 9.99E-01 | Uncharacterized protein At2g12550 |
| AT3G08780 | 0.0 | 6.99E-01 | 0.1 | 1.29E-01 | -0.2 | 1.02E-01 | -0.3 | 6.86E-02 | -0.4 | 5.70E-02 | 0.0 | 8.63E-01 | AT3G08780 protein |
| AT3G02540 | 0.0 | 9.05E-01 | 0.1 | 2.64E-01 | -0.1 | 2.52E-01 | -0.4 | 6.57E-03 | -0.3 | 9.55E-02 | -0.1 | 5.11E-01 | Rad23 UV excision repair protein family |
| AT4G02970 | 1.6 | 6.00E-06 | 0.8 | 2.18E-03 | 0.8 | 1.99E-03 | 0.1 | 8.30E-01 | -0.3 | 3.91E-01 | 0.3 | 3.84E-01 | Ubiquitin domain-containing protein 7SL RNA1 |
| AT1G50400 | | | | | | | 2.797632317 | 6.68E-04 | 1.347087404 | 1.31E-01 | | | Probable mitochondrial import receptor subunit TOM40-2 |
| AT5G57490 | 0.0 | 7.90E-01 | 0.4 | 1.49E-03 | -0.3 | 4.28E-03 | 0.4 | 1.50E-03 | 0.2 | 6.22E-02 | 0.2 | 3.12E-01 | Mitochondrial outer membrane protein porin 4 |
| AT5G15090 | 0.1 | 4.79E-01 | 0.3 | 5.58E-02 | -0.2 | 2.12E-01 | 0.5 | 1.23E-02 | 0.1 | 5.62E-01 | 0.3 | 1.15E-01 | Mitochondrial outer membrane protein porin 3 |
| AT3G01280 | 0.4 | 3.45E-03 | 0.4 | 1.80E-03 | -0.1 | 5.64E-01 | 0.4 | 1.26E-03 | 0.3 | 4.27E-02 | 0.1 | 3.74E-01 | Mitochondrial outer membrane protein porin 1 |
| AT1G20816 | 1.0 | 2.08E-04 | 0.6 | 1.30E-02 | 0.4 | 5.57E-02 | 0.2 | 5.79E-01 | 0.2 | 7.78E-01 | 0.1 | 9.11E-01 | Outer envelope pore protein 21A, chloroplastic |
| AT1G76405 | 1.0 | 1.62E-05 | 0.9 | 6.43E-05 | 0.1 | 5.36E-01 | 0.2 | 2.00E-01 | -0.1 | 7.38E-01 | 0.3 | 2.02E-01 | Outer envelope pore protein 21B, chloroplastic |
| AT5G42960 | 0.1 | 4.46E-01 | 0.1 | 1.81E-01 | 0.0 | 5.94E-01 | 0.4 | 7.61E-03 | 0.0 | 9.99E-01 | 0.4 | 3.29E-02 | Outer envelope pore protein 24B, chloroplastic |
| AT1G45170 | 0.4 | 7.41E-04 | 0.5 | 4.57E-04 | -0.1 | 5.97E-01 | 0.1 | 8.74E-01 | -0.2 | 6.39E-01 | 0.2 | 5.74E-01 | hypothetical protein |
| AT1G69740 | 1.1 | 3.65E-07 | 0.9 | 4.10E-06 | 0.2 | 2.89E-02 | 0.1 | 3.01E-01 | -0.1 | 3.38E-01 | 0.3 | 1.03E-01 | Delta-aminolevulinic acid dehydratase 1, chloroplastic |
| AT4G13420 | -3.1 | 1.46E-07 | -2.8 | 4.72E-07 | -0.3 | 2.49E-01 | 1.2 | 4.30E-02 | 3.2 | 4.38E-05 | -2.0 | 4.95E-04 | Potassium transporter |
| AT1G70300 | -0.6 | 1.19E-04 | 0.0 | 6.63E-01 | -0.5 | 8.64E-04 | 0.4 | 3.03E-02 | 0.3 | 7.03E-02 | 0.0 | 9.11E-01 | Potassium transporter 6 |
| AT5G09400 | -0.4 | 4.65E-04 | -0.3 | 1.32E-03 | 0.0 | 6.82E-01 | -0.1 | 6.63E-01 | 0.0 | 9.49E-01 | -0.1 | 6.56E-01 | Potassium transporter |
| AT1G31120 | -0.2 | 1.49E-02 | -0.3 | 7.42E-03 | 0.0 | 6.64E-01 | -0.1 | 7.53E-01 | -0.1 | 8.03E-01 | 0.0 | 9.97E-01 | Potassium transporter |
| AT2G35060 | -0.2 | 3.25E-02 | -0.5 | 8.71E-04 | 0.2 | 4.67E-02 | -0.3 | 1.41E-01 | -0.3 | 2.58E-01 | 0.0 | 9.26E-01 | Potassium transporter |
| AT2G30070 | -0.2 | 3.59E-02 | -0.2 | 7.34E-02 | 0.0 | 7.95E-01 | -0.3 | 2.88E-02 | -0.2 | 2.34E-01 | -0.1 | 5.94E-01 | Potassium transporter |
| AT4G33530 | -0.1 | 1.08E-01 | 0.0 | 7.67E-01 | -0.1 | 2.37E-01 | -0.3 | 3.63E-02 | -0.2 | 3.31E-01 | -0.1 | 5.37E-01 | Potassium transporter |
| AT3G02050 | -0.1 | 1.85E-01 | -0.3 | 1.37E-02 | 0.1 | 1.76E-01 | -0.1 | 7.65E-01 | 0.1 | 7.72E-01 | -0.2 | 6.14E-01 | Potassium transporter 4 |
| AT4G19960 | -0.1 | 1.89E-01 | -0.3 | 1.32E-02 | 0.1 | 1.76E-01 | -0.1 | 5.74E-01 | 0.2 | 3.66E-01 | -0.4 | 2.14E-01 | Potassium transporter |
| AT4G23640 | -0.1 | 1.96E-01 | 0.0 | 5.49E-01 | 0.0 | 5.54E-01 | 0.0 | 9.89E-01 | 0.0 | 8.20E-01 | 0.0 | 8.66E-01 | Potassium transporter |
| AT2G40540 | 0.0 | 7.55E-01 | 0.1 | 2.69E-01 | -0.1 | 4.69E-01 | -0.4 | 2.24E-02 | -0.2 | 2.95E-01 | -0.2 | 4.52E-01 | Potassium transporter 2 |
| AT5G14880 | 0.3 | 2.69E-02 | 0.1 | 4.46E-01 | 0.2 | 1.56E-01 | 0.0 | 8.49E-01 | 0.0 | 8.39E-01 | 0.0 | 9.75E-01 | Potassium transporter 8 |
| AT1G60160 | 0.5 | 8.43E-03 | 0.2 | 2.85E-01 | 0.3 | 1.09E-01 | -0.4 | 9.90E-03 | -0.2 | 2.09E-01 | -0.2 | 3.36E-01 | Putative potassium transporter 12 |
| AT1G01790 | 0.9 | 1.87E-05 | 0.5 | 2.35E-03 | 0.4 | 7.75E-03 | 0.1 | 7.61E-01 | 0.2 | 3.32E-01 | -0.2 | 5.95E-01 | K efflux antiporter 1, chloroplastic |
| AT4G10310 | -1.4 | 3.09E-04 | -1.6 | 3.20E-04 | 0.2 | 6.89E-01 | -0.9 | 3.26E-02 | -0.6 | 1.66E-01 | -0.3 | 6.40E-01 | Sodium transporter HKT1 |
| AT5G11800 | -0.5 | 8.11E-05 | -0.2 | 5.28E-02 | -0.3 | 4.68E-03 | -0.2 | 4.23E-01 | -0.1 | 7.76E-01 | -0.1 | 7.35E-01 | K efflux antiporter 6 |
| AT4G00630 | -0.4 | 1.81E-03 | -0.4 | 1.88E-03 | 0.0 | 8.86E-01 | 0.0 | 9.53E-01 | 0.1 | 7.12E-01 | -0.1 | 7.24E-01 | K+ efflux antiporter 2 |
| AT5G51710 | -0.1 | 1.82E-01 | -0.3 | 1.18E-02 | 0.2 | 1.68E-01 | -0.4 | 1.49E-02 | 0.0 | 8.52E-01 | -0.4 | 7.05E-02 | K+ efflux antiporter 5 |
| AT2G19600 | 0.2 | 4.88E-02 | 0.3 | 6.79E-03 | -0.1 | 3.01E-01 | 0.1 | 4.85E-01 | 0.1 | 6.35E-01 | 0.0 | 9.53E-01 | K efflux antiporter 4 |
| AT5G55470 | 0.5 | 1.87E-02 | -0.1 | 5.43E-01 | 0.6 | 2.20E-02 | -0.7 | 7.33E-02 | -0.3 | 5.07E-01 | -0.4 | 4.65E-01 | Sodium |
| AT2G01980 | -0.7 | 2.43E-05 | -0.7 | 7.61E-05 | -0.1 | 6.78E-01 | -0.1 | 6.19E-01 | 0.1 | 5.10E-01 | -0.2 | 2.90E-01 | Sodium |
| AT5G23080 | -0.1 | 1.48E-01 | -0.1 | 1.54E-01 | 0.0 | 9.82E-01 | -0.3 | 2.14E-01 | -0.3 | 2.50E-01 | 0.0 | 9.80E-01 | G patch domain-containing protein TGH |

| | | | | | | | | | | | | | |
|---|---|---|---|---|---|---|---|---|---|---|---|---|---|
| AT1G77180 | 0.0 | 9.61E-01 | -0.1 | 6.88E-02 | 0.1 | 9.44E-02 | -0.2 | 7.61E-02 | -0.2 | 4.57E-02 | 0.1 | 8.00E-01 | SNW |
| AT2G02150 | 0.1 | 4.47E-01 | -0.1 | 5.92E-01 | 0.2 | 2.89E-01 | 0.6 | 9.30E-02 | 0.3 | 4.00E-01 | 0.2 | 5.71E-01 | Putative pentatricopeptide repeat-containing protein At2g02150 |
| AT5G15860 | -1.1 | 2.92E-07 | -0.9 | 2.18E-06 | -0.2 | 5.81E-02 | 0.1 | 4.58E-01 | 0.2 | 1.21E-01 | -0.1 | 5.12E-01 | Isoprenylcysteine alpha-carbonyl methylesterase ICME |
| AT1G26120 | -0.1 | 2.90E-02 | -0.3 | 1.35E-03 | 0.1 | 9.57E-02 | -0.5 | 4.81E-03 | -0.2 | 4.03E-01 | -0.4 | 9.01E-02 | Probable isoprenylcysteine alpha-carbonyl methylesterase ICMEL1 |
| AT2G18950 | -0.5 | 3.41E-04 | -0.6 | 1.85E-04 | 0.1 | 4.41E-01 | 0.3 | 7.65E-02 | 0.4 | 4.56E-02 | -0.1 | 7.93E-01 | Homogentisate phytyltransferase 1, chloroplastic |
| AT5G38710 | 0.5 | 8.44E-03 | 0.5 | 2.39E-02 | 0.1 | 7.04E-01 | 1.3 | 5.28E-03 | 0.4 | 4.59E-01 | 0.9 | 9.33E-02 | Proline dehydrogenase |
| AT2G16405 | -0.1 | 1.99E-01 | -0.2 | 1.52E-02 | 0.1 | 1.92E-01 | -0.3 | 1.65E-01 | -0.3 | 1.35E-01 | 0.0 | 8.81E-01 | At2g16405 |
| AT2G42430 | -0.8 | 2.23E-04 | -0.4 | 1.24E-02 | -0.4 | 5.47E-02 | 0.6 | 3.25E-01 | -0.1 | 9.20E-01 | | | LOB domain-containing protein 16 |
| AT5G42150 | 0.0 | 7.96E-01 | 0.0 | 9.36E-01 | 0.0 | 8.93E-01 | 0.9 | 2.42E-05 | 0.5 | 1.26E-02 | 0.4 | 2.81E-02 | Prostaglandin E synthase 2 |
| AT5G08335 | 0.2 | 7.09E-02 | -0.1 | 5.56E-01 | 0.2 | 5.27E-02 | 0.1 | 7.26E-01 | 0.0 | 9.96E-01 | 0.1 | 7.97E-01 | Protein-S-isoprenylcysteine O-methyltransferase B |
| AT2G28160 | -1.3 | 1.12E-05 | -0.8 | 4.76E-04 | -0.5 | 1.69E-02 | -0.9 | 2.93E-03 | -0.6 | 3.72E-02 | -0.3 | 1.49E-01 | FER-like regulator of iron uptake |
| AT4G29930 | -1.3 | 2.02E-05 | -1.3 | 3.83E-05 | 0.0 | 9.67E-01 | 0.0 | 9.65E-01 | 1.0 | 1.07E-02 | -1.0 | 2.50E-02 | BHLH domain-containing protein |
| AT5G50010 | -1.1 | 7.63E-06 | -0.2 | 7.34E-02 | -0.9 | 2.90E-04 | -0.8 | 9.66E-03 | -0.1 | 7.36E-01 | -0.7 | 3.48E-02 | BHLH domain-containing protein |
| AT4G30980 | -1.1 | 3.54E-05 | -0.3 | 5.76E-02 | -0.8 | 1.69E-03 | 0.2 | 4.85E-01 | 0.3 | 2.57E-01 | -0.1 | 7.00E-01 | LJRHL1-like 2 |
| AT1G05805 | -0.1 | 7.49E-02 | 0.0 | 8.65E-01 | -0.1 | 1.44E-01 | -0.2 | 1.55E-01 | -0.2 | 2.72E-01 | 0.0 | 9.28E-01 | BHLH domain-containing protein |
| AT1G79740 | -0.1 | 3.40E-01 | -0.2 | 3.35E-02 | 0.1 | 2.23E-01 | -0.3 | 2.95E-01 | -0.1 | 7.07E-01 | -0.1 | 7.30E-01 | F20B17.17 |
| AT2G24260 | 0.0 | 9.74E-01 | 0.0 | 6.70E-01 | 0.0 | 7.33E-01 | 0.4 | 2.30E-02 | 0.2 | 2.64E-01 | 0.2 | 3.90E-01 | LJRHL1-like 1 |
| AT3G13030 | 0.2 | 9.57E-02 | 0.0 | 7.85E-01 | 0.1 | 2.36E-01 | -0.1 | 7.74E-01 | 0.1 | 7.50E-01 | -0.2 | 4.85E-01 | HAT transposon superfamily protein |
| AT5G61270 | 0.2 | 2.63E-02 | 0.1 | 2.52E-01 | 0.1 | 2.84E-01 | 0.0 | 9.20E-01 | 0.3 | 4.67E-01 | -0.3 | 3.86E-01 | PIF7 |
| AT5G46690 | 0.3 | 1.36E-01 | 0.4 | 4.45E-02 | -0.1 | 5.66E-01 | -0.5 | 3.10E-02 | -0.3 | 3.18E-01 | -0.2 | 3.47E-01 | Beta HLH protein 71 |
| AT5G15160 | 0.5 | 9.06E-02 | 0.1 | 7.58E-01 | 0.4 | 2.59E-01 | 0.2 | 6.56E-01 | 0.1 | 8.46E-01 | 0.1 | 9.04E-01 | BNQ2 |
| AT1G26945 | 0.7 | 2.02E-03 | 0.8 | 9.55E-04 | -0.1 | 5.36E-01 | 0.4 | 1.61E-01 | 0.6 | 4.17E-02 | -0.2 | 4.36E-01 | Transcription factor PRE6 |
| AT2G46810 | 1.4 | 2.77E-07 | 0.9 | 1.24E-05 | 0.5 | 1.81E-03 | -0.4 | 1.42E-01 | -0.4 | 1.65E-01 | 0.0 | 9.87E-01 | hypothetical protein |
| AT1G72530 | | | | | | | 0.2 | 6.72E-01 | 0.5 | 3.85E-01 | -0.2 | 6.96E-01 | Plastid developmental protein DAG |
| AT3G47710 | | | | | | | -1.5 | 3.10E-02 | -1.1 | 1.21E-01 | -0.4 | 5.79E-01 | hypothetical protein |
| AT1G32580 | 0.1 | 3.45E-01 | 0.1 | 4.39E-01 | 0.0 | 9.14E-01 | 0.2 | 2.36E-01 | 0.0 | 9.75E-01 | 0.2 | 3.46E-01 | Multiple organellar RNA editing factor 5, chloroplastic |
| AT1G64440 | -0.9 | 2.94E-06 | -0.2 | 3.69E-02 | -0.7 | 1.29E-04 | -0.4 | 9.25E-03 | 0.2 | 1.98E-01 | -0.6 | 2.89E-03 | UDP-glucose 4-epimerase 4 |
| AT1G60850 | 0.1 | 3.31E-01 | 0.0 | 9.10E-01 | 0.1 | 3.63E-01 | 0.7 | 1.50E-03 | 0.4 | 6.17E-02 | 0.3 | 2.21E-01 | DNA-directed RNA polymerase family protein |
| AT1G47870 | -0.5 | 9.10E-04 | -0.3 | 9.50E-03 | -0.1 | 2.69E-01 | 0.3 | 1.48E-01 | 0.2 | 2.56E-01 | 0.0 | 9.30E-01 | E2FC |
| AT3G54960 | -0.8 | 2.43E-05 | 0.1 | 5.58E-01 | -0.9 | 8.87E-05 | 0.1 | 5.08E-01 | 0.4 | 4.60E-02 | -0.3 | 2.76E-01 | Protein disulfide isomerase-like 1-3 |
| AT1G21750 | -0.4 | 3.90E-03 | 0.0 | 7.60E-01 | -0.4 | 1.40E-02 | 0.2 | 1.35E-01 | 0.2 | 2.31E-01 | 0.0 | 9.66E-01 | Protein disulfide-isomerase |
| AT5G60640 | -0.3 | 6.12E-04 | -0.1 | 2.65E-01 | -0.2 | 8.28E-03 | 0.1 | 4.06E-01 | 0.2 | 2.18E-01 | -0.1 | 7.50E-01 | Protein disulfide-isomerase |
| AT2G47470 | -0.3 | 2.55E-02 | 0.1 | 5.66E-01 | -0.3 | 1.67E-02 | 0.2 | 3.29E-02 | 0.2 | 1.48E-01 | 0.1 | 7.88E-01 | Thioredoxin family protein |
| AT1G07960 | -0.2 | 2.42E-02 | -0.1 | 5.34E-01 | -0.2 | 1.23E-01 | 0.3 | 1.13E-01 | 0.2 | 3.29E-01 | 0.1 | 7.85E-01 | Protein disulfide-isomerase 5-1 |
| AT3G16110 | -0.2 | 2.06E-02 | 0.0 | 5.69E-01 | -0.1 | 1.00E-01 | 0.0 | 9.38E-01 | 0.0 | 8.55E-01 | -0.1 | 8.25E-01 | Protein disulfide isomerase-like 1-6 |
| AT1G52260 | -0.1 | 5.20E-01 | 0.0 | 7.36E-01 | -0.1 | 4.13E-01 | 0.0 | 9.12E-01 | 0.0 | 9.55E-01 | 0.0 | 9.04E-01 | Protein disulfide isomerase-like 1-5 |
| AT1G77510 | 0.0 | 8.19E-01 | -0.1 | 8.09E-01 | 0.0 | 9.82E-01 | 0.9 | 2.89E-07 | 0.6 | 1.76E-04 | 0.4 | 2.30E-02 | Protein disulfide isomerase-like 1-2 |
| AT3G20560 | 0.0 | 6.09E-01 | 0.2 | 1.65E-02 | -0.2 | 6.83E-02 | 0.1 | 7.32E-01 | 0.0 | 9.65E-01 | 0.1 | 7.81E-01 | Protein disulfide-isomerase 5-3 |
| AT1G35620 | 0.2 | 2.74E-02 | 0.3 | 2.17E-03 | -0.1 | 1.48E-01 | -0.1 | 2.14E-01 | -0.2 | 1.53E-01 | 0.0 | 8.48E-01 | Protein disulfide-isomerase 5-2 |
| AT1G04980 | 0.2 | 4.19E-01 | 0.3 | 1.27E-01 | -0.2 | 5.05E-01 | 0.4 | 3.08E-03 | 0.3 | 4.76E-02 | 0.1 | 5.20E-01 | PDIL2-2 |
| AT4G27080 | 0.4 | 1.10E-03 | 0.4 | 9.58E-04 | 0.0 | 7.76E-01 | 0.1 | 2.90E-01 | 0.2 | 2.10E-01 | 0.0 | 8.45E-01 | PDI-like 5-4 |
| AT1G11530 | 1.1 | 6.69E-03 | 0.7 | 6.43E-02 | 0.4 | 3.32E-01 | -0.1 | 6.91E-01 | -0.4 | 2.73E-01 | 0.2 | 6.02E-01 | Thioredoxin-like protein CXXS1 |
| AT2G32920 | 0.1 | 1.76E-01 | 0.3 | 6.78E-03 | -0.2 | 9.32E-02 | 0.4 | 1.66E-02 | 0.3 | 1.01E-01 | 0.1 | 7.00E-01 | Protein disulfide-isomerase 2-3 |
| AT5G01750 | 0.4 | 3.09E-04 | 0.6 | 6.38E-05 | -0.2 | 9.68E-02 | -0.3 | 3.27E-02 | -0.3 | 1.83E-02 | 0.1 | 7.62E-01 | Protein LURP-one-related 15 |
| AT2G06520 | 1.1 | 2.29E-06 | 1.2 | 2.91E-06 | -0.1 | 3.43E-01 | -0.1 | 4.03E-01 | -0.1 | 5.72E-01 | 0.0 | 9.44E-01 | Expressed protein |
| AT2G45180 | 1.5 | 1.02E-05 | 1.7 | 1.19E-05 | -0.2 | 4.74E-01 | 0.1 | 7.95E-01 | -0.7 | 8.90E-03 | 0.8 | 1.44E-02 | At2g45180 |
| AT3G48890 | 0.3 | 3.96E-03 | 0.1 | 4.02E-01 | 0.2 | 3.28E-02 | 0.2 | 1.52E-01 | 0.1 | 6.87E-01 | 0.1 | 5.85E-01 | Membrane steroid-binding protein 2 |
| AT5G40280 | 0.2 | 7.61E-02 | -0.2 | 7.61E-02 | 0.3 | 8.15E-03 | -0.2 | 1.62E-01 | -0.3 | 1.06E-01 | 0.1 | 8.07E-01 | Protein farnesyltransferase subunit beta |
| AT5G07960 | 0.2 | 1.82E-01 | 0.2 | 1.54E-01 | 0.0 | 9.09E-01 | 0.0 | 9.07E-01 | -0.1 | 8.74E-01 | 0.1 | 8.35E-01 | Protein Asterix |
| AT4G29520 | 0.3 | 1.05E-02 | 0.3 | 1.46E-02 | 0.0 | 9.56E-01 | 0.4 | 7.31E-02 | 0.4 | 1.42E-01 | 0.0 | 9.33E-01 | Nucleophosmin |
| AT5G23040 | 0.7 | 7.69E-06 | 0.6 | 4.70E-05 | 0.1 | 2.83E-01 | 0.0 | 8.30E-01 | -0.1 | 3.68E-01 | 0.1 | 5.82E-01 | Protein CHAPERONE-LIKE PROTEIN OF POR1, chloroplastic |
| AT5G19855 | 0.8 | 8.23E-04 | 0.4 | 5.12E-02 | 0.4 | 5.21E-02 | 0.1 | 7.64E-01 | 0.0 | 9.78E-01 | 0.1 | 7.94E-01 | Chaperonin-like RbcX protein |
| AT1G65260 | 0.8 | 3.27E-06 | 0.5 | 2.94E-04 | 0.3 | 4.11E-03 | -0.1 | 4.41E-01 | -0.1 | 5.85E-01 | 0.0 | 9.62E-01 | Plastid transcriptionally active 4 |
| AT4G04330 | 1.2 | 2.02E-04 | 1.6 | 2.63E-05 | -0.5 | 4.37E-02 | 0.4 | 4.65E-02 | 0.1 | 8.33E-01 | 0.3 | 2.19E-01 | Chaperonin-like RBCX protein 1, chloroplastic |
| AT3G62190 | 0.5 | 6.44E-03 | 0.0 | 9.57E-01 | 0.5 | 1.92E-02 | 0.0 | 8.86E-01 | -0.1 | 5.88E-01 | 0.1 | 7.24E-01 | AT3g62190 |
| AT1G56260 | | | | | | | 0.4 | 2.20E-01 | 0.1 | 8.57E-01 | 0.3 | 4.64E-01 | CST complex subunit TEN1 |
| AT3G22480 | 0.5 | 6.00E-05 | 0.4 | 7.31E-04 | 0.1 | 1.68E-01 | 0.1 | 6.57E-01 | -0.1 | 5.75E-01 | 0.2 | 4.04E-01 | Prefoldin subunit 2 |
| AT5G23090 | -0.5 | 2.08E-04 | -0.4 | 8.06E-04 | -0.1 | 5.54E-01 | -0.4 | 5.19E-02 | -0.3 | 1.32E-01 | -0.1 | 8.54E-01 | Nuclear factor Y, subunit B13 |
| AT1G17440 | -0.4 | 6.47E-04 | -0.5 | 2.08E-04 | 0.1 | 2.82E-01 | 0.3 | 1.15E-01 | 0.2 | 3.28E-01 | 0.1 | 7.86E-01 | Transcription initiation factor TFIID subunit 12 domain-containing protein |
| AT5G43130 | -0.3 | 4.71E-04 | -0.2 | 2.51E-03 | -0.1 | 4.15E-01 | -0.2 | 7.72E-02 | -0.1 | 7.36E-01 | -0.2 | 3.76E-01 | TBP-associated factor 4 |
| AT1G07980 | -0.1 | 2.50E-01 | 0.1 | 5.27E-01 | -0.2 | 1.35E-01 | -0.1 | 4.59E-01 | 0.0 | 9.31E-01 | -0.1 | 5.27E-01 | Nuclear factor Y, subunit C10 |
| AT4G32551 | -0.1 | 1.35E-01 | -0.2 | 1.96E-02 | 0.1 | 3.05E-01 | -0.1 | 7.42E-01 | 0.0 | 9.60E-01 | -0.1 | 7.84E-01 | Transcriptional corepressor LEUNIG |
| AT5G08190 | -0.1 | 3.89E-01 | -0.3 | 2.79E-02 | 0.2 | 1.74E-01 | 0.1 | 4.77E-01 | 0.2 | 4.72E-01 | 0.0 | 9.59E-01 | DR1-like protein |
| AT4G34340 | -0.1 | 5.76E-01 | 0.0 | 9.57E-01 | -0.1 | 7.04E-01 | 0.1 | 7.63E-01 | 0.0 | 9.53E-01 | 0.1 | 7.85E-01 | TBP-associated factor 8 |
| AT5G65540 | 0.0 | 8.84E-01 | -0.1 | 1.91E-01 | 0.1 | 3.12E-01 | -0.5 | 2.00E-01 | -0.3 | 5.21E-01 | -0.2 | 6.67E-01 | Transcription initiation factor TFIID subunit |
| AT5G15570 | 0.1 | 3.23E-01 | -0.1 | 4.71E-01 | 0.1 | 1.54E-01 | 0.1 | 8.19E-01 | 0.0 | 9.03E-01 | 0.1 | 7.84E-01 | Bromodomain associated domain-containing protein |
| AT1G02680 | 0.1 | 3.65E-01 | 0.1 | 6.07E-01 | 0.0 | 7.79E-01 | -0.3 | 1.69E-01 | -0.4 | 1.91E-01 | 0.0 | 9.77E-01 | Transcription initiation factor TFIID subunit 13 |
| AT3G02160 | 0.1 | 4.15E-01 | 0.1 | 5.15E-01 | 0.0 | 9.31E-01 | -0.1 | 8.22E-01 | -0.1 | 9.09E-01 | 0.0 | 9.57E-01 | hypothetical protein |
| AT1G78190 | 0.4 | 1.62E-02 | 0.1 | 7.24E-01 | 0.3 | 7.90E-02 | 0.5 | 1.64E-01 | 0.1 | 8.35E-01 | 0.4 | 2.95E-01 | Multifunctional methyltransferase subunit TRM112 homolog B |
| AT5G38140 | 0.4 | 6.29E-03 | 0.3 | 3.40E-02 | 0.1 | 4.92E-01 | 0.3 | 4.05E-01 | 0.2 | 6.25E-01 | 0.1 | 8.86E-01 | NF-YC12 |
| AT1G22270 | 0.8 | 1.47E-05 | 0.6 | 2.09E-04 | 0.2 | 9.53E-02 | 0.1 | 5.93E-01 | -0.2 | 3.95E-01 | 0.4 | 2.10E-01 | Multifunctional methyltransferase subunit TRM112 homolog A |
| AT4G26455 | -0.2 | 6.02E-03 | -0.1 | 9.10E-02 | -0.1 | 2.14E-01 | 0.0 | 9.72E-01 | -0.1 | 7.98E-01 | 0.1 | 7.80E-01 | WPP domain-interacting protein 1 |
| AT2G46260 | -0.2 | 8.50E-02 | -0.1 | 1.90E-01 | 0.0 | 7.22E-01 | -0.3 | 1.34E-01 | -0.3 | 2.05E-01 | 0.0 | 9.81E-01 | BTB |
| AT5G56210 | 0.2 | 1.75E-02 | 0.1 | 2.84E-01 | 0.1 | 2.03E-01 | -0.2 | 4.74E-01 | 0.0 | 9.27E-01 | -0.1 | 6.84E-01 | WPP domain-interacting protein 2 |
| AT5G42750 | 0.6 | 5.22E-04 | 0.1 | 3.76E-01 | 0.5 | 8.04E-03 | -0.3 | 2.53E-02 | -0.3 | 8.89E-02 | -0.1 | 8.19E-01 | BRI1 kinase inhibitor 1 |
| AT1G04950 | -0.1 | 5.10E-01 | 0.0 | 9.21E-01 | 0.0 | 6.55E-01 | -0.1 | 3.87E-01 | -0.1 | 3.75E-01 | 0.0 | 9.44E-01 | TATA box binding protein associated factor histone-like fold domain-containing protein |
| AT1G54360 | 0.2 | 1.73E-02 | 0.0 | 8.90E-01 | 0.2 | 4.35E-02 | 0.0 | 9.61E-01 | -0.2 | 5.92E-01 | 0.2 | 6.72E-01 | TBP-ASSOCIATED FACTOR 6B |
| AT1G54140 | -0.1 | 4.99E-01 | 0.0 | 7.91E-01 | -0.1 | 4.27E-01 | 0.1 | 8.50E-01 | 0.0 | 9.73E-01 | 0.0 | 9.24E-01 | Transcription initiation factor TFIID subunit 9 |
| AT4G20280 | 0.0 | 6.23E-01 | 0.0 | 9.79E-01 | 0.0 | 6.66E-01 | -0.2 | 1.78E-01 | -0.2 | 3.41E-01 | 0.0 | 8.98E-01 | Transcription initiation factor TFIID subunit 11 |
| AT5G62740 | -0.5 | 1.87E-04 | -0.3 | 6.22E-03 | -0.2 | 6.54E-02 | -0.2 | 9.01E-02 | -0.3 | 7.99E-02 | 0.0 | 8.96E-01 | Hypersensitive-induced response protein 1 |
| AT1G69840 | -0.2 | 1.29E-01 | -0.4 | 3.47E-02 | 0.1 | 4.81E-01 | 0.0 | 7.75E-01 | -0.4 | 1.32E-02 | 0.4 | 2.45E-02 | Hypersensitive-induced response protein 2 |
| AT5G43560 | -0.2 | 8.14E-03 | -0.3 | 1.10E-03 | 0.1 | 2.02E-01 | 0.0 | 9.40E-01 | 0.0 | 9.82E-01 | 0.0 | 9.78E-01 | TNF receptor-associated factor homolog 1a |
| AT4G12060 | 0.4 | 8.91E-04 | 0.6 | 2.18E-04 | -0.1 | 2.29E-01 | 0.4 | 4.43E-03 | 0.3 | 4.58E-02 | 0.1 | 5.94E-01 | ATP-dependent Clp protease ATP-binding subunit CLPT2, chloroplastic |

| | | | | | | | | | | | | | |
|---|---|---|---|---|---|---|---|---|---|---|---|---|---|
| AT4G15630 | 1.1 | 7.08E-07 | 0.9 | 6.65E-06 | 0.2 | 6.88E-02 | -0.2 | 3.68E-01 | -0.1 | 5.94E-01 | 0.0 | 9.04E-01 | CASP-like protein 1E1 |
| AT5G06200 | -2.2 | 7.56E-07 | 0.0 | 7.92E-01 | -2.1 | 1.89E-05 | -1.1 | 1.39E-03 | -0.1 | 6.92E-01 | -1.0 | 1.81E-02 | Casparian strip membrane protein 4 |
| AT2G27370 | -2.1 | 3.24E-08 | -0.3 | 2.36E-03 | -1.7 | 6.99E-06 | -0.6 | 1.51E-02 | -0.1 | 7.87E-01 | -0.5 | 9.58E-02 | Casparian strip membrane protein 3 |
| AT3G11550 | -1.9 | 3.77E-07 | -0.2 | 1.83E-01 | -1.7 | 1.47E-05 | -0.9 | 9.71E-04 | 0.1 | 6.66E-01 | -1.0 | 2.84E-03 | Casparian strip membrane protein 2 |
| AT2G36100 | -1.8 | 1.15E-07 | -0.2 | 9.45E-02 | -1.6 | 8.63E-06 | -0.7 | 4.64E-03 | 0.1 | 7.92E-01 | -0.8 | 7.46E-03 | Casparian strip membrane protein 1 |
| AT4G26750 | -0.6 | 3.62E-05 | -0.5 | 4.89E-04 | -0.2 | 1.16E-01 | -0.3 | 7.18E-02 | -0.2 | 2.73E-01 | -0.1 | 7.43E-01 | Protein HOMOLOG OF MAMMALIAN LYST-INTERACTING PROTEIN 5 |
| AT3G13290 | -0.4 | 6.85E-04 | -0.4 | 6.87E-04 | 0.0 | 8.12E-01 | 0.1 | 8.31E-01 | 0.1 | 8.24E-01 | 0.0 | 9.77E-01 | Varicose-related protein |
| AT3G58840 | -0.4 | 7.68E-03 | -0.1 | 4.25E-01 | -0.3 | 5.84E-02 | 0.1 | 7.77E-01 | 0.1 | 8.01E-01 | 0.0 | 9.83E-01 | Peroxisomal and mitochondrial division factor 1 |
| AT4G24840 | -0.3 | 6.53E-04 | -0.1 | 3.11E-01 | -0.3 | 8.17E-03 | 0.0 | 9.93E-01 | 0.0 | 8.76E-01 | 0.0 | 9.04E-01 | Conserved oligomeric Golgi complex subunit 2 |
| AT1G73430 | -0.3 | 7.20E-04 | -0.2 | 3.33E-02 | -0.2 | 6.78E-02 | 0.0 | 9.89E-01 | 0.0 | 8.86E-01 | -0.1 | 8.99E-01 | Conserved oligomeric Golgi complex subunit 3 |
| AT2G18840 | -0.3 | 7.36E-03 | 0.1 | 5.54E-01 | -0.3 | 7.05E-03 | -0.1 | 7.42E-01 | 0.0 | 8.90E-01 | -0.1 | 7.02E-01 | Protein YIP4a |
| AT1G72630 | 0.0 | 9.05E-01 | 0.1 | 5.39E-01 | -0.1 | 6.74E-01 | -0.4 | 8.03E-02 | -0.2 | 4.07E-01 | -0.2 | 5.61E-01 | Protein ELF4-LIKE 2 |
| AT1G67930 | 0.0 | 7.73E-01 | -0.1 | 1.22E-01 | 0.1 | 1.03E-01 | -0.1 | 6.32E-01 | 0.0 | 9.23E-01 | -0.1 | 6.56E-01 | Conserved oligomeric Golgi complex subunit 5 |
| AT2G40080 | 0.0 | 8.49E-01 | 1.1 | 8.01E-05 | -1.0 | 5.05E-04 | -0.2 | 4.06E-01 | 0.2 | 3.41E-01 | -0.4 | 8.36E-02 | Protein EARLY FLOWERING 4 |
| AT3G49250 | 0.1 | 5.16E-01 | 0.2 | 7.70E-02 | -0.1 | 3.04E-01 | 0.2 | 4.74E-01 | 0.1 | 8.63E-01 | 0.1 | 6.98E-01 | Protein DEFECTIVE IN MERISTEM SILENCING 3 |
| AT1G17455 | 0.1 | 6.03E-01 | 0.2 | 1.53E-01 | -0.1 | 4.23E-01 | -0.2 | 4.90E-01 | -0.1 | 7.14E-01 | -0.1 | 8.93E-01 | Protein ELF4-LIKE 4 |
| AT1G26540 | 1.6 | 1.38E-05 | 0.7 | 6.39E-03 | 0.8 | 3.98E-03 | 0.1 | 5.68E-01 | -0.2 | 3.80E-01 | 0.4 | 2.03E-01 | DUF724 domain-containing protein 3 |
| AT2G06255 | | | | | | | -0.5 | 3.23E-01 | -0.5 | 3.74E-01 | 0.0 | 9.97E-01 | Protein ELF4-LIKE 3 |
| AT5G67600 | 0.1 | 1.38E-01 | 0.2 | 4.46E-02 | -0.1 | 5.49E-01 | -0.2 | 4.62E-01 | -0.2 | 4.67E-01 | 0.0 | 9.57E-01 | Protein CYSTEINE-RICH TRANSMEMBRANE MODULE 13 |
| AT3G01680 | 0.2 | 4.45E-02 | 0.2 | 4.47E-02 | 0.0 | 9.59E-01 | -0.1 | 7.55E-01 | 0.0 | 9.29E-01 | -0.1 | 7.61E-01 | Protein SIEVE ELEMENT OCCLUSION B |
| AT4G26670 | 0.7 | 4.71E-05 | 0.9 | 7.33E-06 | -0.3 | 2.47E-02 | 0.1 | 4.28E-01 | -0.2 | 1.65E-01 | 0.3 | 8.72E-02 | Chloroplastic import inner membrane translocase subunit TIM22-2 |
| AT4G19003 | 0.3 | 8.59E-03 | 0.3 | 2.03E-02 | 0.0 | 7.65E-01 | 0.0 | 9.10E-01 | -0.2 | 3.54E-01 | 0.2 | 4.99E-01 | Vacuolar protein sorting-associated protein 25 |
| AT2G42870 | 1.1 | 6.94E-05 | 1.3 | 3.98E-05 | -0.2 | 3.51E-01 | -0.4 | 1.62E-01 | -0.5 | 7.34E-02 | 0.2 | 7.21E-01 | Transcription factor PAR1 |
| AT4G00810 | 0.9 | 9.62E-05 | 1.0 | 8.30E-05 | -0.1 | 5.19E-01 | 0.3 | 1.14E-01 | 0.1 | 4.86E-01 | 0.1 | 6.58E-01 | Large ribosomal subunit protein P1z |
| AT5G47700 | 1.0 | 1.48E-05 | 1.0 | 3.15E-05 | 0.0 | 9.91E-01 | 0.3 | 1.15E-01 | 0.1 | 7.65E-01 | 0.2 | 4.21E-01 | Large ribosomal subunit protein P1y |
| AT4G00300 | 0.4 | 3.09E-05 | 0.4 | 2.31E-04 | 0.1 | 2.53E-01 | 0.0 | 8.30E-01 | -0.2 | 1.08E-01 | 0.2 | 2.87E-01 | Receptor-like kinase |
| AT2G39190 | 0.7 | 3.66E-05 | 0.3 | 1.59E-02 | 0.4 | 3.85E-03 | 0.0 | 9.70E-01 | 0.0 | 9.24E-01 | 0.0 | 9.58E-01 | At2g39190 |
| AT3G63120 | -1.1 | 8.50E-07 | -0.5 | 3.67E-04 | -0.6 | 3.09E-04 | -0.6 | 6.03E-05 | -0.2 | 1.08E-01 | -0.4 | 2.40E-02 | Cyclin-U3-1 |
| AT2G44740 | -0.6 | 2.01E-03 | 0.4 | 1.50E-02 | -0.9 | 3.98E-04 | -0.9 | 3.64E-04 | -0.7 | 4.50E-03 | -0.2 | 4.49E-01 | Cyclin-U4-1 |
| AT3G60550 | -0.4 | 2.26E-02 | 0.2 | 3.12E-01 | -0.6 | 1.32E-02 | 0.3 | 4.44E-01 | 0.0 | 9.92E-01 | 0.3 | 5.64E-01 | Cyclin-U2-2 |
| AT2G32680 | 0.2 | 6.94E-01 | -0.8 | 1.12E-01 | 1.0 | 8.96E-02 | -0.4 | 4.42E-01 | -1.2 | 5.99E-02 | 0.8 | 3.29E-01 | Receptor like protein 23 |
| AT3G21870 | 2.6 | 1.13E-06 | 1.6 | 1.48E-04 | 1.0 | 1.15E-03 | -0.1 | 8.91E-01 | -0.5 | 1.45E-01 | 0.5 | 2.41E-01 | Cyclin-U1-1 |
| AT4G12970 | 2.4 | 1.18E-05 | 1.6 | 6.45E-04 | 0.8 | 1.23E-02 | -0.1 | 8.11E-01 | -0.3 | 4.98E-01 | 0.2 | 7.49E-01 | EPIDERMAL PATTERNING FACTOR-like protein 9 |
| AT5G02420 | -1.4 | 3.05E-04 | -0.6 | 2.34E-02 | -0.8 | 3.13E-02 | -0.9 | 1.16E-02 | 0.0 | 9.55E-01 | -0.9 | 2.09E-02 | Cyclin-dependent protein kinase inhibitor SMR3 |
| AT1G10690 | -1.2 | 1.05E-06 | -0.9 | 1.14E-05 | -0.3 | 5.11E-02 | 0.0 | 9.40E-01 | 0.2 | 5.24E-01 | -0.2 | 5.27E-01 | Cyclin-dependent protein kinase inhibitor SMR8 |
| AT2G28870 | -1.2 | 4.18E-04 | -0.2 | 2.47E-01 | -0.9 | 7.81E-03 | -0.2 | 6.10E-01 | 0.1 | 8.23E-01 | -0.3 | 4.95E-01 | Cyclin-dependent protein kinase inhibitor SMR10 |
| AT5G04470 | -1.0 | 2.07E-04 | 0.2 | 3.20E-01 | -1.1 | 4.32E-04 | -0.7 | 4.79E-02 | -0.1 | 9.14E-01 | -0.7 | 1.16E-01 | Uncharacterized protein |
| AT3G10525 | -0.6 | 2.09E-02 | -0.2 | 2.80E-01 | -0.3 | 2.33E-01 | -0.3 | 3.49E-01 | 0.1 | 8.78E-01 | -0.4 | 3.63E-01 | Cyclin-dependent protein kinase inhibitor SMR1 |
| AT2G28330 | -0.5 | 7.28E-03 | 0.0 | 7.49E-01 | -0.4 | 3.52E-02 | 0.0 | 9.68E-01 | -0.2 | 6.62E-01 | 0.2 | 7.51E-01 | Cyclin-dependent protein kinase inhibitor SMR11 |
| AT1G08180 | 0.1 | 6.82E-01 | 0.0 | 8.72E-01 | 0.1 | 6.46E-01 | -0.2 | 3.99E-01 | -0.1 | 8.49E-01 | -0.2 | 7.00E-01 | Cyclin-dependent protein kinase inhibitor SMR2 |
| AT5G02220 | 2.6 | 2.99E-05 | 0.7 | 1.44E-01 | 1.9 | 1.19E-03 | 0.4 | 3.16E-01 | 0.1 | 8.76E-01 | 0.3 | 5.61E-01 | Cyclin-dependent protein kinase inhibitor SMR4 |
| AT1G51355 | | | | | | | -0.3 | 3.85E-01 | 0.0 | 9.32E-01 | -0.4 | 3.47E-01 | Cyclin-dependent protein kinase inhibitor SMR9 |
| AT3G27630 | | | | | | | 0.2 | 6.79E-01 | -0.4 | 4.64E-01 | 0.5 | 2.21E-01 | Cyclin-dependent protein kinase inhibitor SMR7 |
| AT5G40460 | | | | | | | 0.3 | 6.30E-01 | 0.2 | 7.98E-01 | 0.1 | 9.19E-01 | Cyclin-dependent protein kinase inhibitor SMR6 |
| AT5G59360 | | | | | | | -0.4 | 4.55E-01 | 0.1 | 8.49E-01 | | | Cyclin-dependent protein kinase inhibitor SMR14 |
| AT2G44680 | -0.3 | 3.51E-03 | -0.3 | 1.02E-02 | 0.0 | 6.76E-01 | -0.3 | 3.84E-02 | -0.3 | 6.24E-02 | 0.0 | 9.97E-01 | Casein kinase II subunit beta |
| AT3G60250 | 0.1 | 3.34E-01 | 0.0 | 7.52E-01 | 0.1 | 2.59E-01 | -0.3 | 2.28E-02 | -0.4 | 1.87E-02 | 0.1 | 8.47E-01 | Casein kinase II subunit beta |
| AT3G29770 | -1.0 | 2.26E-06 | -0.5 | 6.65E-04 | -0.6 | 1.07E-03 | -0.2 | 4.38E-01 | 0.0 | 9.49E-01 | -0.2 | 6.28E-01 | Methylesterase |
| AT4G10050 | -0.4 | 1.94E-04 | -0.4 | 2.50E-04 | 0.0 | 8.82E-01 | -0.3 | 1.75E-01 | 0.0 | 8.66E-01 | -0.3 | 2.44E-01 | Protein phosphatase methylesterase 1 |
| AT5G10300 | -0.3 | 2.49E-02 | 0.1 | 6.47E-01 | -0.3 | 2.69E-02 | 0.1 | 7.08E-01 | 0.2 | 4.65E-01 | -0.1 | 7.90E-01 | Methylesterase |
| AT2G23610 | 0.1 | 5.37E-01 | 0.2 | 2.96E-02 | -0.2 | 1.12E-01 | -0.5 | 3.42E-03 | 0.0 | 9.88E-01 | -0.5 | 1.81E-02 | Methylesterase |
| AT3G50440 | 0.5 | 8.12E-04 | 0.7 | 1.29E-04 | -0.2 | 1.30E-01 | -0.5 | 9.49E-03 | -0.1 | 7.02E-01 | -0.4 | 1.13E-01 | Methylesterase |
| AT2G23600 | 0.8 | 6.84E-06 | 0.9 | 8.98E-06 | -0.1 | 5.02E-01 | 0.2 | 4.09E-01 | -0.3 | 1.14E-01 | 0.5 | 4.22E-02 | Methylesterase |
| AT2G26810 | 0.3 | 1.79E-02 | 0.1 | 6.22E-01 | 0.2 | 9.47E-02 | 0.1 | 7.92E-01 | 0.0 | 9.19E-01 | 0.0 | 9.21E-01 | Methyltransferase family protein |
| AT5G53920 | 0.2 | 1.33E-01 | 0.0 | 9.32E-01 | 0.2 | 2.53E-01 | 0.5 | 1.54E-01 | 0.3 | 5.67E-01 | 0.3 | 5.28E-01 | Ribosomal protein L11 methyltransferase-like protein |
| AT3G11540 | -0.4 | 6.66E-04 | -0.4 | 9.70E-04 | 0.0 | 1.00E+00 | -0.2 | 3.10E-01 | -0.3 | 5.63E-02 | 0.2 | 4.54E-01 | protein O-GlcNAc transferase |
| AT3G04240 | 0.0 | 6.86E-01 | -0.4 | 1.61E-03 | 0.4 | 1.85E-03 | 0.2 | 3.22E-01 | -0.1 | 5.60E-01 | 0.3 | 1.95E-01 | protein O-GlcNAc transferase |
| AT2G31305 | 0.0 | 8.84E-01 | -0.1 | 3.33E-01 | 0.1 | 4.85E-01 | -0.1 | 8.05E-01 | -0.1 | 7.21E-01 | 0.1 | 9.15E-01 | Protein phosphatase 1 regulatory subunit INH3 |
| AT3G06670 | 0.0 | 6.37E-01 | -0.2 | 2.83E-02 | 0.1 | 8.57E-02 | -0.1 | 3.64E-01 | -0.1 | 6.25E-01 | 0.0 | 8.73E-01 | At3g06670 |
| AT2G28360 | -0.4 | 8.44E-05 | -0.2 | 4.44E-03 | -0.2 | 3.55E-02 | -0.1 | 5.55E-01 | 0.1 | 7.35E-01 | -0.2 | 4.53E-01 | Uncharacterized protein At2g28360 |
| AT3G45190 | -0.3 | 9.26E-05 | -0.4 | 1.01E-04 | 0.0 | 6.91E-01 | -0.2 | 6.72E-02 | -0.1 | 3.41E-01 | -0.1 | 6.54E-01 | SIT4 phosphatase-associated family protein |
| AT1G07990 | -0.4 | 6.37E-05 | -0.4 | 1.58E-04 | 0.0 | 8.03E-01 | -0.3 | 4.87E-02 | -0.2 | 8.74E-02 | 0.0 | 9.75E-01 | At1g07990 |
| AT1G30470 | -0.1 | 7.52E-02 | -0.2 | 2.63E-02 | 0.0 | 5.66E-01 | 0.0 | 7.65E-01 | -0.1 | 5.18E-01 | 0.1 | 8.05E-01 | SIT4 phosphatase-associated family protein |
| AT5G19680 | -0.2 | 5.92E-03 | -0.1 | 3.88E-01 | -0.2 | 5.36E-02 | 0.0 | 9.84E-01 | 0.0 | 9.57E-01 | 0.0 | 9.49E-01 | Protein phosphatase 1 regulatory inhibitor subunit PPP1R7 homolog |
| AT5G52200 | -0.2 | 1.07E-01 | -0.1 | 3.58E-01 | -0.1 | 5.45E-01 | 0.0 | 8.08E-01 | 0.1 | 5.16E-01 | -0.2 | 4.27E-01 | Phosphoprotein phosphatase inhibitor |
| AT4G32870 | 0.3 | 1.43E-02 | 0.7 | 1.76E-04 | -0.4 | 1.61E-02 | -0.8 | 4.68E-03 | -0.4 | 1.94E-01 | -0.4 | 1.53E-01 | At4g32870 |
| AT5G53160 | 0.6 | 1.26E-04 | 0.3 | 5.91E-03 | 0.2 | 4.41E-02 | -0.3 | 1.43E-01 | -0.7 | 2.84E-03 | 0.4 | 1.16E-01 | Regulatory components of ABA receptor 3 |
| AT1G48870 | -0.7 | 1.53E-04 | -0.9 | 5.37E-05 | 0.2 | 1.72E-01 | | | | | | | Transducin |
| AT1G17720 | -0.3 | 1.97E-03 | -0.2 | 1.36E-02 | -0.1 | 3.72E-01 | 0.0 | 9.53E-01 | 0.0 | 8.12E-01 | -0.1 | 8.05E-01 | Serine |
| AT3G54930 | -0.2 | 1.26E-01 | 0.0 | 7.74E-01 | -0.2 | 3.08E-01 | -0.1 | 7.14E-01 | 0.1 | 8.14E-01 | -0.2 | 5.96E-01 | Serine |
| AT3G26020 | -0.2 | 1.01E-02 | -0.3 | 2.01E-03 | 0.1 | 3.16E-01 | -0.1 | 7.36E-01 | 0.0 | 9.90E-01 | -0.1 | 8.16E-01 | Serine |
| AT1G51690 | -0.2 | 1.49E-02 | -0.2 | 7.03E-03 | 0.0 | 6.28E-01 | -0.3 | 1.86E-02 | -0.2 | 1.51E-01 | -0.1 | 6.06E-01 | Serine |
| AT4G15415 | -0.1 | 4.53E-02 | -0.2 | 1.25E-02 | 0.1 | 4.84E-01 | 0.0 | 9.45E-01 | 0.0 | 8.73E-01 | 0.0 | 8.39E-01 | Serine |
| AT1G13460 | -0.1 | 1.75E-01 | -0.1 | 3.33E-01 | 0.0 | 7.65E-01 | 0.3 | 1.90E-02 | 0.2 | 3.39E-01 | 0.2 | 3.33E-01 | Serine |
| AT3G09880 | 0.0 | 6.05E-01 | -0.1 | 1.88E-01 | 0.1 | 4.69E-01 | -0.3 | 1.04E-01 | -0.5 | 9.50E-03 | 0.2 | 3.42E-01 | Serine |
| AT5G25510 | 0.0 | 6.59E-01 | 0.1 | 3.67E-01 | -0.1 | 2.34E-01 | -0.2 | 5.60E-02 | 0.0 | 9.56E-01 | -0.2 | 1.32E-01 | Serine |
| AT5G03470 | 0.1 | 3.78E-01 | 0.0 | 6.79E-01 | 0.0 | 7.27E-01 | -0.5 | 2.41E-03 | -0.6 | 7.23E-04 | 0.1 | 4.97E-01 | Serine |
| AT3G21650 | 0.1 | 1.32E-01 | -0.1 | 5.95E-01 | 0.2 | 9.17E-02 | -0.2 | 2.29E-01 | -0.5 | 3.01E-02 | 0.2 | 3.87E-01 | Serine |
| AT3G26030 | 0.3 | 9.99E-04 | 0.1 | 3.39E-01 | 0.3 | 1.08E-02 | -0.1 | 6.55E-01 | -0.2 | 8.76E-02 | 0.2 | 3.14E-01 | Serine |
| AT5G17070 | 0.4 | 7.93E-04 | 0.3 | 3.57E-03 | 0.1 | 5.23E-01 | 0.0 | 8.48E-01 | 0.1 | 7.55E-01 | -0.1 | 6.79E-01 | Uncharacterized protein F2K13_220 |
| AT1G25490 | -0.3 | 9.40E-04 | -0.1 | 1.20E-01 | -0.2 | 2.66E-02 | -0.2 | 3.49E-01 | -0.1 | 4.62E-01 | 0.0 | 9.77E-01 | Serine |
| AT4G09000 | -0.4 | 4.28E-04 | 0.0 | 8.98E-01 | -0.3 | 1.63E-03 | -0.2 | 3.62E-02 | -0.2 | 7.20E-02 | 0.0 | 9.73E-01 | General regulatory factor 1 |
| AT1G10095 | -0.3 | 1.77E-02 | -0.2 | 5.70E-02 | -0.1 | 6.63E-01 | -0.6 | 2.58E-02 | -0.6 | 3.12E-02 | 0.0 | 9.66E-01 | Protein prenyltransferase alpha subunit-related |

| | | | | | | | | | | | | | |
|---|---|---|---|---|---|---|---|---|---|---|---|---|---|
| AT3G59380 | 0.1 | 6.29E-02 | 0.1 | 4.61E-01 | 0.1 | 3.00E-01 | -0.3 | 7.19E-03 | -0.2 | 1.17E-01 | -0.1 | 4.52E-01 | Protein farnesyltransferase |
| AT1G32400 | -0.1 | 2.14E-01 | -0.1 | 6.99E-02 | 0.0 | 5.21E-01 | -0.2 | 2.57E-02 | -0.2 | 8.26E-02 | 0.0 | 8.66E-01 | Tobamovirus multiplication protein 2A |
| AT4G16444 | 0.0 | 8.89E-01 | 0.0 | 7.49E-01 | 0.0 | 8.81E-01 | -0.2 | 4.39E-01 | -0.1 | 6.94E-01 | -0.1 | 8.34E-01 | Protein GET1 |
| AT1G22882 | 0.0 | 8.51E-01 | -0.1 | 2.05E-01 | 0.2 | 1.96E-01 | -0.3 | 9.26E-02 | -0.3 | 6.24E-02 | 0.1 | 7.94E-01 | SUN domain-containing protein 3 |
| AT1G71360 | 0.2 | 3.65E-02 | 0.2 | 5.38E-02 | 0.0 | 9.16E-01 | 0.0 | 9.47E-01 | -0.1 | 4.49E-01 | 0.1 | 4.59E-01 | Galactose-binding protein |
| AT4G23950 | 0.2 | 3.36E-01 | 0.1 | 7.80E-01 | 0.1 | 6.07E-01 | 0.0 | 9.91E-01 | -0.2 | 6.39E-01 | 0.2 | 6.93E-01 | SUN domain-containing protein 5 |
| AT3G54170 | 0.2 | 5.67E-02 | -0.1 | 5.58E-01 | 0.3 | 4.13E-02 | -0.1 | 6.41E-01 | 0.1 | 8.12E-01 | -0.2 | 5.57E-01 | FKBP12-interacting protein of 37 kDa |
| AT3G19180 | 0.4 | 1.80E-03 | 0.3 | 2.89E-02 | 0.2 | 2.14E-01 | -0.3 | 2.74E-01 | -0.1 | 7.60E-01 | -0.2 | 5.27E-01 | Plastid division protein CDP1, chloroplastic |
| AT4G15545 | 0.7 | 1.44E-05 | 0.4 | 1.94E-03 | 0.3 | 6.41E-03 | 0.0 | 8.33E-01 | 0.0 | 9.86E-01 | 0.0 | 9.00E-01 | Uncharacterized protein At4g15545 |
| AT4G12470 | 1.1 | 2.50E-02 | -0.2 | 7.06E-01 | 1.3 | 2.26E-02 | 0.0 | 9.91E-01 | -2.3 | 1.58E-02 | 2.2 | 3.99E-02 | pEARLI1-like lipid transfer protein 1 |
| AT5G24650 | 0.3 | 9.60E-04 | 0.3 | 5.52E-03 | 0.1 | 4.14E-01 | 0.3 | 4.28E-02 | 0.2 | 1.14E-01 | 0.0 | 9.05E-01 | Chloroplastic import inner membrane translocase subunit HP30-2 |
| AT3G49560 | 0.6 | 8.95E-05 | 0.5 | 6.47E-04 | 0.1 | 2.86E-01 | 0.5 | 1.68E-03 | 0.1 | 3.96E-01 | 0.3 | 6.88E-02 | Chloroplastic import inner membrane translocase subunit HP30-1 |
| AT5G04990 | -0.2 | 2.60E-03 | -0.2 | 1.77E-02 | -0.1 | 3.74E-01 | -0.1 | 7.72E-01 | 0.1 | 5.18E-01 | -0.2 | 4.42E-01 | SUN domain-containing protein 1 |
| AT3G10730 | 0.5 | 7.10E-05 | 0.2 | 4.17E-02 | 0.3 | 4.73E-03 | 0.1 | 6.47E-01 | -0.1 | 7.05E-01 | 0.3 | 4.45E-01 | SUN domain-containing protein 2 |
| AT2G48120 | 0.9 | 7.78E-06 | 0.6 | 1.66E-04 | 0.2 | 4.98E-02 | 0.2 | 2.71E-01 | 0.0 | 9.76E-01 | 0.2 | 3.96E-01 | Protein PALE CRESS, chloroplastic |
| AT5G13680 | 0.0 | 9.93E-01 | -0.2 | 5.42E-02 | 0.2 | 7.86E-02 | 0.3 | 1.77E-01 | 0.0 | 9.71E-01 | 0.2 | 3.01E-01 | Elongator complex protein 1 |
| AT1G15130 | -0.4 | 6.47E-04 | -0.4 | 1.40E-03 | 0.0 | 8.26E-01 | -0.2 | 4.80E-01 | -0.1 | 8.21E-01 | -0.1 | 8.18E-01 | Vacuolar-sorting protein BRO1 |
| AT2G19310 | -0.4 | 1.43E-04 | -0.5 | 9.91E-05 | 0.1 | 4.87E-01 | 0.1 | 4.96E-01 | 0.1 | 6.22E-01 | 0.0 | 9.69E-01 | 18.5 kDa class IV heat shock protein |
| AT5G59720 | 0.0 | 8.11E-01 | 0.2 | 1.12E-01 | -0.3 | 1.33E-01 | | | | | | | 18.1 kDa class I heat shock protein |
| AT1G54050 | 0.3 | 4.04E-02 | 0.3 | 6.14E-02 | 0.0 | 9.26E-01 | -0.4 | 3.07E-01 | 0.1 | 8.01E-01 | -0.6 | 2.13E-01 | 17.4 kDa class III heat shock protein |
| AT2G29500 | 0.5 | 3.57E-02 | 0.0 | 9.21E-01 | 0.5 | 1.00E-01 | 1.3 | 1.35E-01 | 0.8 | 4.79E-01 | | | 17.6 kDa class I heat shock protein 2 |
| AT4G21870 | 1.8 | 2.69E-05 | 1.8 | 4.29E-05 | 0.0 | 8.71E-01 | 1.4 | 8.97E-04 | -0.2 | 7.97E-01 | 1.5 | 3.33E-03 | 15.4 kDa class V heat shock protein |
| AT1G59860 | | | | | | | 0.5 | 4.28E-01 | 0.6 | 4.25E-01 | | | 17.6 kDa class I heat shock protein 1 |
| AT5G12030 | | | | | | | -0.52007736 | 7.92E-01 | -2.186192833 | 3.05E-01 | | | 17.7 kDa class II heat shock protein |
| AT4G17270 | -0.2 | 5.81E-03 | -0.3 | 2.81E-03 | 0.0 | 6.07E-01 | -0.1 | 4.83E-01 | -0.1 | 6.90E-01 | 0.0 | 9.16E-01 | Putative MO25-like protein At4g17270 |
| AT5G18940 | -0.2 | 3.03E-02 | 0.0 | 8.25E-01 | -0.2 | 3.83E-02 | 0.0 | 8.49E-01 | 0.0 | 8.22E-01 | 0.0 | 9.67E-01 | Mo25 family protein |
| AT3G17420 | -1.4 | 4.62E-08 | -0.8 | 2.51E-06 | -0.6 | 9.04E-05 | 0.2 | 3.60E-01 | 0.3 | 1.42E-01 | -0.1 | 5.57E-01 | Serine |
| AT2G17930 | -0.6 | 9.36E-06 | -0.6 | 1.34E-05 | 0.0 | 7.18E-01 | -0.2 | 4.29E-01 | -0.1 | 5.35E-01 | 0.0 | 9.80E-01 | Phosphatidylinositol 3-and 4-kinase family protein with FAT domain-containing protein |
| AT4G36080 | -0.5 | 9.57E-05 | -0.5 | 2.02E-04 | 0.0 | 8.82E-01 | 0.0 | 8.36E-01 | -0.1 | 6.88E-01 | 0.1 | 6.12E-01 | Phosphotransferases |
| AT1G12970 | -0.3 | 2.72E-03 | -0.2 | 3.48E-02 | -0.1 | 2.55E-01 | -0.2 | 1.62E-01 | -0.3 | 4.50E-02 | 0.1 | 6.02E-01 | Plant intracellular Ras-group-related LRR protein 3 |
| AT5G27930 | -0.9 | 7.70E-07 | -0.5 | 8.12E-05 | -0.4 | 1.06E-03 | -0.7 | 1.63E-03 | -0.1 | 7.35E-01 | -0.6 | 2.52E-02 | Protein phosphatase 2C family protein |
| AT2G20630 | -0.8 | 2.03E-06 | -0.4 | 9.08E-04 | -0.5 | 6.36E-04 | 0.0 | 9.94E-01 | 0.2 | 2.12E-01 | -0.2 | 3.12E-01 | PPM-type phosphatase domain-containing protein |
| AT4G28400 | -0.7 | 1.30E-05 | -0.6 | 1.47E-04 | -0.2 | 1.10E-01 | 0.2 | 7.34E-02 | -0.1 | 6.21E-01 | 0.3 | 7.65E-02 | Protein phosphatase 2C family protein |
| AT3G16800 | -0.7 | 1.07E-04 | -0.2 | 8.47E-02 | -0.5 | 4.03E-03 | -0.8 | 4.52E-03 | -0.5 | 5.15E-02 | -0.3 | 4.17E-01 | Protein phosphatase 2C family protein |
| AT5G66080 | -0.7 | 1.01E-04 | -0.1 | 3.84E-01 | -0.6 | 1.59E-03 | -0.8 | 1.95E-03 | -0.2 | 5.45E-01 | -0.6 | 4.92E-02 | Protein phosphatase 2C family protein |
| AT4G32950 | -0.7 | 1.49E-04 | 0.2 | 8.70E-02 | -0.9 | 1.16E-04 | 0.0 | 9.68E-01 | 1.1 | 8.10E-05 | -1.1 | 3.62E-04 | Protein phosphatase 2C family protein |
| AT5G26010 | -0.4 | 7.81E-04 | 0.1 | 4.65E-01 | -0.5 | 1.41E-03 | 0.2 | 2.57E-01 | 0.0 | 9.00E-01 | 0.2 | 4.60E-01 | hypothetical protein |
| AT5G36250 | -0.3 | 4.33E-03 | -0.2 | 2.85E-02 | -0.1 | 3.95E-01 | -0.4 | 5.17E-02 | -0.2 | 2.45E-01 | -0.1 | 6.77E-01 | Protein phosphatase 2C family protein |
| AT2G46920 | -0.3 | 1.34E-03 | -0.4 | 1.95E-04 | 0.1 | 1.30E-01 | 0.0 | 9.23E-01 | 0.2 | 4.45E-01 | -0.1 | 5.78E-01 | AT2G46920 protein |
| AT5G06750 | -0.1 | 9.57E-02 | -0.2 | 1.01E-02 | 0.1 | 2.47E-01 | 0.1 | 5.74E-01 | 0.0 | 9.62E-01 | 0.1 | 6.80E-01 | Protein phosphatase 2C family protein |
| AT1G34750 | -0.1 | 2.19E-01 | -0.4 | 1.08E-03 | 0.3 | 1.21E-02 | 0.3 | 5.84E-02 | 0.0 | 8.70E-01 | 0.3 | 1.94E-01 | Protein phosphatase 2C family protein |
| AT1G78200 | 0.0 | 7.84E-01 | -0.3 | 3.38E-03 | 0.3 | 1.06E-02 | -0.2 | 1.13E-01 | -0.1 | 4.38E-01 | -0.1 | 6.89E-01 | Protein phosphatase 2C family protein |
| AT1G79630 | 0.0 | 9.63E-01 | -0.1 | 1.91E-01 | 0.1 | 2.35E-01 | -0.1 | 7.76E-01 | 0.0 | 8.90E-01 | -0.1 | 7.32E-01 | Protein phosphatase 2C family protein |
| AT4G33920 | 0.0 | 8.13E-01 | 0.0 | 9.19E-01 | 0.0 | 9.19E-01 | 0.3 | 1.15E-01 | 0.1 | 7.99E-01 | 0.2 | 3.32E-01 | Protein phosphatase 2C family protein |
| AT3G16560 | 1.4 | 3.65E-07 | 1.3 | 1.95E-06 | 0.1 | 2.00E-01 | -0.7 | 2.44E-04 | -0.4 | 3.40E-02 | -0.3 | 1.21E-01 | Protein phosphatase 2C family protein |
| AT3G09400 | | | | | | | 2.431900623 | 3.51E-04 | 1.49905968 | 3.01E-02 | 0.9 | 2.59E-02 | Pol-like 3 |
| AT5G59220 | | | | | | | -2.2 | 5.88E-02 | -0.3 | 7.80E-01 | -1.9 | 2.56E-01 | PP2C protein |
| AT5G55160 | -0.1 | 2.02E-01 | 0.1 | 2.48E-01 | -0.2 | 3.98E-02 | 0.0 | 9.01E-01 | -0.1 | 6.17E-01 | 0.1 | 7.69E-01 | Small ubiquitin-related modifier |
| AT5G42300 | 0.2 | 1.29E-01 | -0.1 | 6.31E-01 | 0.3 | 7.74E-02 | 0.2 | 1.95E-01 | 0.1 | 8.27E-01 | 0.2 | 5.17E-01 | Ubiquitin-like protein 5 |
| AT2G45695 | 0.7 | 5.74E-05 | 0.3 | 3.07E-02 | 0.4 | 5.97E-03 | -0.1 | 8.03E-01 | -0.2 | 6.25E-01 | 0.1 | 8.32E-01 | Ubiquitin-related modifier 1 homolog 1 |
| AT4G26840 | 0.4 | 4.46E-03 | 0.3 | 2.96E-02 | 0.1 | 3.92E-01 | 0.3 | 1.41E-01 | 0.1 | 6.75E-01 | 0.2 | 5.56E-01 | Small ubiquitin-related modifier 1 |
| AT2G32765 | 0.9 | 1.46E-04 | 0.6 | 4.53E-03 | 0.3 | 9.94E-02 | 0.3 | 3.69E-01 | 0.1 | 8.77E-01 | 0.2 | 6.08E-01 | Small ubiquitin-related modifier 5 |
| AT1G72750 | 0.5 | 1.32E-03 | 0.4 | 7.85E-03 | 0.1 | 4.10E-01 | 0.3 | 2.77E-01 | -0.2 | 6.06E-01 | 0.4 | 1.43E-01 | Mitochondrial import inner membrane translocase subunit TIM23-2 |
| AT5G50460 | 0.5 | 1.46E-03 | 0.5 | 2.48E-03 | 0.0 | 9.05E-01 | 0.8 | 3.23E-03 | 0.4 | 9.97E-02 | 0.3 | 3.74E-01 | Protein transport protein Sec61 subunit gamma |
| AT3G20000 | 0.5 | 1.00E-03 | 0.5 | 2.07E-03 | 0.0 | 8.42E-01 | 0.4 | 2.98E-03 | 0.2 | 2.24E-01 | 0.2 | 1.74E-01 | Translocase of the outer mitochondrial membrane 40 |
| AT3G48570 | 0.6 | 1.31E-03 | 0.2 | 1.59E-01 | 0.4 | 3.88E-02 | 0.2 | 4.15E-01 | 0.0 | 9.97E-01 | 0.2 | 5.15E-01 | Protein transport protein Sec61 subunit gamma-3 |
| AT5G01590 | 1.0 | 3.42E-07 | 0.9 | 2.39E-06 | 0.1 | 9.32E-02 | 0.2 | 6.19E-02 | 0.1 | 3.78E-01 | 0.1 | 5.74E-01 | Protein TIC 56, chloroplastic |
| AT5G55510 | 1.0 | 3.23E-05 | 1.0 | 8.59E-05 | 0.0 | 7.91E-01 | 0.3 | 7.74E-02 | -0.1 | 8.35E-01 | 0.4 | 1.14E-01 | Mitochondrial import inner membrane translocase subunit TIM22-3 |
| AT4G14870 | 1.7 | 6.49E-08 | 1.4 | 4.42E-07 | 0.2 | 2.48E-02 | 0.1 | 4.91E-01 | -0.1 | 5.30E-01 | 0.3 | 2.88E-01 | Preprotein translocase subunit SECE1 |
| AT3G04800 | | | | | | | 1.352116754 | 1.24E-01 | 0.775801708 | 4.93E-01 | | | Mitochondrial import inner membrane translocase subunit TIM23-3 |
| AT2G18710 | 0.6 | 2.92E-05 | 0.6 | 8.20E-05 | 0.0 | 7.41E-01 | 0.1 | 4.75E-01 | 0.0 | 8.19E-01 | 0.2 | 3.99E-01 | Preprotein translocase subunit SCY1, chloroplastic |
| AT1G71860 | -0.5 | 4.29E-05 | -0.4 | 2.03E-04 | -0.1 | 4.17E-01 | 0.0 | 9.55E-01 | 0.0 | 8.96E-01 | 0.0 | 9.49E-01 | Protein tyrosine phosphatase 1 |
| AT3G62010 | -0.3 | 3.53E-02 | -0.1 | 2.64E-01 | -0.1 | 3.23E-01 | 0.0 | 9.98E-01 | 0.1 | 8.41E-01 | -0.1 | 8.64E-01 | Metal ion-binding protein |
| AT3G02800 | -0.9 | 1.27E-04 | -0.6 | 2.41E-03 | -0.3 | 1.14E-01 | 1.3 | 3.05E-03 | 0.6 | 2.26E-01 | 0.7 | 9.86E-02 | Tyrosine-protein phosphatase domain-containing protein |
| AT5G23720 | -0.5 | 4.22E-05 | -0.3 | 7.14E-04 | -0.1 | 1.01E-01 | 0.0 | 9.56E-01 | 0.2 | 2.35E-01 | -0.2 | 3.04E-01 | Dual specificity protein phosphatase family protein |
| AT3G23610 | -0.4 | 3.29E-03 | -0.3 | 2.06E-02 | -0.1 | 4.45E-01 | -0.5 | 1.04E-01 | -0.3 | 4.22E-01 | -0.2 | 6.35E-01 | Uncharacterized protein |
| AT3G06110 | 0.4 | 3.75E-04 | 0.2 | 3.12E-02 | 0.2 | 4.11E-02 | -0.1 | 5.93E-01 | -0.2 | 2.59E-01 | 0.1 | 6.66E-01 | MKP2 |
| AT3G06930 | 0.1 | 6.47E-01 | 0.0 | 8.70E-01 | 0.0 | 8.25E-01 | 0.4 | 1.13E-02 | 0.3 | 1.61E-01 | 0.2 | 4.47E-01 | PRMT4B |
| AT3G12270 | 0.2 | 1.91E-01 | 0.2 | 1.23E-01 | 0.0 | 8.06E-01 | 0.5 | 4.06E-02 | 0.2 | 5.47E-01 | 0.3 | 3.43E-01 | PRMT3 |
| AT3G20020 | 0.3 | 2.09E-02 | 0.0 | 8.75E-01 | 0.2 | 6.27E-02 | 0.2 | 4.19E-01 | 0.0 | 9.13E-01 | 0.2 | 4.58E-01 | Protein arginine methyltransferase 6 |
| AT2G19670 | 0.5 | 1.84E-02 | 0.5 | 9.79E-03 | -0.1 | 7.11E-01 | 0.4 | 8.06E-02 | 0.3 | 2.88E-01 | 0.1 | 6.88E-01 | Probable protein arginine N-methyltransferase 1.2 |
| AT5G65860 | 0.5 | 7.33E-04 | 0.3 | 1.63E-02 | 0.2 | 1.48E-01 | 0.5 | 1.97E-02 | 0.3 | 1.31E-01 | 0.1 | 6.33E-01 | Protein arginine N-methyltransferase 2 |
| AT4G29510 | 0.3 | 7.72E-03 | 0.2 | 1.37E-01 | 0.2 | 1.80E-01 | 0.3 | 6.00E-02 | 0.1 | 4.48E-01 | 0.1 | 4.96E-01 | Protein arginine N-methyltransferase 1.1 |
| AT4G16570 | 0.3 | 8.01E-03 | 0.1 | 3.42E-01 | 0.2 | 1.02E-01 | -0.2 | 4.05E-01 | -0.2 | 4.63E-01 | 0.0 | 9.98E-01 | Protein arginine N-methyltransferase 1.6 |
| AT3G28700 | -0.1 | 3.05E-01 | -0.1 | 6.27E-01 | -0.1 | 6.82E-01 | 0.1 | 6.55E-01 | 0.2 | 5.18E-01 | -0.1 | 8.73E-01 | Protein arginine methyltransferase NDUFAF7 |
| AT1G04900 | -0.1 | 4.53E-01 | 0.1 | 3.00E-01 | -0.2 | 1.40E-01 | 0.5 | 6.63E-02 | 0.3 | 3.35E-01 | 0.2 | 5.97E-01 | Protein arginine methyltransferase NDUFAF7 |
| AT4G21560 | -0.3 | 1.32E-03 | -0.3 | 1.91E-03 | 0.0 | 9.80E-01 | -0.4 | 1.93E-02 | -0.3 | 7.49E-02 | -0.1 | 8.07E-01 | Vacuolar protein sorting-associated protein 28 homolog 1 |
| AT2G20360 | 0.0 | 7.54E-01 | 0.1 | 5.65E-02 | -0.2 | 4.40E-02 | 0.0 | 7.82E-01 | 0.1 | 6.92E-01 | 0.0 | 9.14E-01 | NADH dehydrogenase 1 alpha subcomplex subunit 9, mitochondrial |
| AT1G06515 | 0.5 | 5.16E-04 | 0.4 | 2.26E-03 | 0.1 | 5.31E-01 | 0.0 | 9.10E-01 | -0.1 | 7.00E-01 | 0.2 | 6.69E-01 | Transmembrane protein, putative |
| AT1G61000 | 0.6 | 1.27E-04 | 0.4 | 2.40E-03 | 0.2 | 1.35E-01 | 0.7 | 8.78E-04 | 0.6 | 5.34E-03 | 0.1 | 7.20E-01 | Kinetochore protein NUF2 homolog |
| AT5G40770 | 0.7 | 2.02E-04 | 0.4 | 9.44E-03 | 0.3 | 4.93E-02 | 0.5 | 2.14E-03 | 0.1 | 5.90E-01 | 0.4 | 4.42E-02 | Prohibitin-3, mitochondrial |
| AT1G32220 | 0.9 | 1.52E-05 | 0.8 | 5.14E-05 | 0.1 | 6.29E-01 | 0.1 | 7.41E-01 | 0.2 | 2.26E-01 | -0.1 | 4.66E-01 | Uncharacterized protein At1g32220, chloroplastic |
| AT3G60800 | -0.5 | 1.12E-04 | -0.4 | 1.27E-03 | -0.1 | 1.82E-01 | 0.0 | 9.55E-01 | 0.1 | 6.20E-01 | -0.1 | 7.24E-01 | Probable protein S-acyltransferase 14 |

| | | | | | | | | | | | | | |
|---|---|---|---|---|---|---|---|---|---|---|---|---|---|
| AT3G56930 | -0.9 | 1.85E-06 | -0.3 | 1.04E-02 | -0.7 | 1.44E-04 | -0.4 | 2.60E-02 | -0.2 | 3.41E-01 | -0.2 | 3.04E-01 | S-acyltransferase |
| AT3G48760 | -0.6 | 3.81E-06 | -0.5 | 4.97E-05 | -0.1 | 6.20E-02 | -0.3 | 2.09E-03 | -0.3 | 2.86E-02 | -0.1 | 5.70E-01 | Probable protein S-acyltransferase 5 |
| AT3G18620 | -0.6 | 3.72E-05 | -0.5 | 2.90E-04 | -0.1 | 2.58E-01 | -0.1 | 6.94E-01 | 0.3 | 3.66E-01 | -0.4 | 2.10E-01 | Protein S-acyltransferase 11 |
| AT4G15080 | -0.5 | 1.68E-05 | -0.4 | 7.64E-05 | -0.1 | 4.25E-01 | -0.2 | 4.09E-01 | -0.1 | 8.58E-01 | -0.1 | 7.01E-01 | Probable protein S-acyltransferase 19 |
| AT4G00840 | -0.4 | 5.04E-04 | -0.4 | 1.89E-03 | -0.1 | 5.84E-01 | -0.3 | 3.13E-02 | -0.5 | 1.22E-02 | 0.1 | 6.92E-01 | S-acyltransferase |
| AT5G50020 | -0.4 | 5.11E-04 | -0.2 | 1.59E-02 | -0.2 | 9.42E-02 | -0.1 | 7.09E-01 | 0.1 | 6.55E-01 | -0.2 | 4.40E-01 | S-acyltransferase |
| AT3G09320 | -0.4 | 2.07E-04 | -0.3 | 3.10E-03 | -0.1 | 1.59E-01 | -0.2 | 1.62E-01 | -0.3 | 9.73E-02 | 0.1 | 8.08E-01 | Probable protein S-acyltransferase 16 |
| AT3G04970 | -0.3 | 2.20E-03 | -0.3 | 6.57E-03 | 0.0 | 6.82E-01 | -0.2 | 2.55E-01 | 0.0 | 9.54E-01 | -0.2 | 2.97E-01 | S-acyltransferase |
| AT3G51390 | -0.3 | 6.48E-03 | -0.1 | 3.97E-01 | -0.2 | 6.30E-02 | 0.0 | 9.29E-01 | 0.1 | 7.64E-01 | -0.1 | 7.19E-01 | S-acyltransferase |
| AT5G41060 | -0.3 | 4.65E-03 | 0.0 | 9.64E-01 | -0.3 | 1.22E-02 | -0.2 | 2.96E-01 | -0.1 | 7.25E-01 | -0.1 | 6.76E-01 | S-acyltransferase |
| AT4G22750 | -0.2 | 2.18E-02 | 0.0 | 9.51E-01 | -0.1 | 4.40E-02 | -0.2 | 1.53E-01 | -0.3 | 3.57E-02 | 0.1 | 5.40E-01 | Probable protein S-acyltransferase 13 |
| AT3G26935 | -0.1 | 5.50E-02 | -0.1 | 4.13E-01 | -0.1 | 2.97E-01 | -0.1 | 5.10E-01 | 0.0 | 8.09E-01 | 0.0 | 8.48E-01 | Probable protein S-acyltransferase 7 |
| AT2G14255 | 0.0 | 7.45E-01 | -0.1 | 9.54E-02 | 0.1 | 2.15E-01 | 0.0 | 9.85E-01 | -0.2 | 5.82E-01 | 0.2 | 6.11E-01 | S-acyltransferase |
| AT3G22180 | 0.0 | 9.68E-01 | -0.1 | 2.29E-01 | 0.1 | 3.15E-01 | 0.1 | 7.64E-01 | 0.1 | 8.42E-01 | 0.0 | 9.77E-01 | Probable protein S-acyltransferase 20 |
| AT5G04270 | 0.2 | 4.40E-02 | 0.1 | 2.91E-01 | 0.1 | 3.94E-01 | -0.2 | 4.10E-01 | -0.2 | 4.59E-01 | 0.0 | 9.96E-01 | S-acyltransferase |
| AT1G69420 | 0.3 | 9.66E-04 | 0.3 | 3.58E-03 | 0.0 | 5.69E-01 | -0.2 | 8.88E-02 | -0.2 | 1.44E-01 | 0.0 | 9.81E-01 | Probable protein S-acyltransferase 22 |
| AT4G24630 | 0.3 | 9.60E-04 | 0.1 | 3.20E-01 | 0.2 | 1.44E-02 | -0.3 | 9.19E-02 | -0.2 | 3.18E-01 | -0.1 | 7.58E-01 | Protein S-acyltransferase 8 |
| AT4G01730 | 0.6 | 8.30E-05 | 0.6 | 9.29E-05 | 0.0 | 7.32E-01 | 0.1 | 6.56E-01 | 0.4 | 8.78E-02 | -0.3 | 2.41E-01 | Protein S-acyltransferase 18 |
| AT5G42850 | 0.2 | 5.10E-02 | 0.0 | 9.83E-01 | 0.3 | 8.08E-02 | 0.0 | 9.32E-01 | -0.1 | 8.48E-01 | 0.0 | 9.28E-01 | Thioredoxin-like protein Clot |
| AT5G15802 | 1.2 | 2.91E-05 | 0.9 | 4.96E-04 | 0.3 | 1.10E-01 | 0.2 | 6.37E-01 | -0.2 | 6.90E-01 | 0.3 | 4.52E-01 | Chaperone |
| AT5G23395 | 0.5 | 2.66E-03 | 0.4 | 4.41E-03 | 0.0 | 9.21E-01 | 0.3 | 6.36E-02 | -0.1 | 4.92E-01 | 0.4 | 3.10E-02 | Mitochondrial intermembrane space import and assembly protein 40 homolog |
| AT1G52590 | 0.7 | 9.58E-04 | 0.2 | 1.72E-01 | 0.5 | 2.67E-02 | 0.3 | 1.07E-01 | 0.2 | 2.53E-01 | 0.1 | 8.68E-01 | DCC family protein At1g52590, chloroplastic |
| AT4G03520 | 0.7 | 1.89E-06 | 0.6 | 1.07E-05 | 0.1 | 2.88E-01 | 0.0 | 8.40E-01 | 0.1 | 6.62E-01 | -0.1 | 5.74E-01 | Thioredoxin superfamily protein |
| AT5G50100 | 0.8 | 1.03E-03 | 0.2 | 4.07E-01 | 0.6 | 1.15E-02 | 0.1 | 7.94E-01 | 0.1 | 6.92E-01 | -0.1 | 9.07E-01 | Uncharacterized protein At5g50100, chloroplastic |
| AT1G24095 | 1.0 | 1.27E-04 | 0.1 | 5.90E-01 | 0.9 | 1.95E-03 | 1.4 | 4.12E-04 | 0.6 | 1.10E-01 | 0.8 | 1.16E-02 | Thiol-disulfide oxidoreductase DCC |
| AT5G62100 | -0.3 | 2.18E-01 | -0.5 | 5.90E-02 | 0.2 | 4.89E-01 | -0.4 | 2.20E-01 | -0.2 | 5.18E-01 | -0.1 | 7.17E-01 | BCL-2-associated athanogene 2 |
| AT4G11260 | 0.1 | 1.02E-01 | 0.1 | 4.80E-01 | 0.1 | 4.02E-01 | 0.1 | 3.24E-01 | 0.0 | 9.11E-01 | 0.2 | 4.49E-01 | Protein SGT1 homolog B |
| AT4G23570 | 0.6 | 2.29E-04 | 0.3 | 2.13E-02 | 0.3 | 2.69E-02 | -0.1 | 8.04E-01 | -0.5 | 2.50E-02 | 0.4 | 6.04E-02 | Protein SGT1 homolog A |
| AT3G07370 | 0.2 | 3.43E-02 | -0.1 | 5.50E-01 | 0.3 | 2.62E-02 | -0.1 | 5.68E-01 | -0.3 | 7.72E-02 | 0.2 | 3.47E-01 | E3 ubiquitin-protein ligase CHIP |
| AT2G20550 | -0.6 | 2.04E-04 | -0.4 | 3.04E-03 | -0.2 | 1.76E-01 | -1.1 | 1.67E-03 | -0.6 | 4.65E-02 | -0.5 | 2.44E-01 | At2g20550 |
| AT4G28480 | -0.2 | 6.41E-03 | -0.4 | 6.42E-04 | 0.1 | 1.32E-01 | 0.2 | 1.86E-01 | 0.3 | 8.12E-02 | -0.1 | 7.14E-01 | DNAJ heat shock family protein |
| AT5G01390 | 0.0 | 6.14E-01 | 0.0 | 5.93E-01 | 0.0 | 9.67E-01 | -0.1 | 7.71E-01 | 0.0 | 9.26E-01 | -0.1 | 7.60E-01 | DNAJ heat shock family protein |
| AT5G27280 | -0.5 | 5.80E-04 | -0.4 | 3.23E-03 | -0.1 | 4.30E-01 | -0.8 | 9.72E-04 | -0.2 | 4.93E-01 | -0.6 | 2.39E-02 | A_TM021B04.14 protein |
| AT3G54826 | 0.3 | 6.63E-03 | 0.1 | 2.15E-01 | 0.2 | 1.15E-01 | 0.1 | 5.68E-01 | -0.1 | 8.19E-01 | 0.2 | 5.20E-01 | Zim17-type zinc finger protein |
| AT5G50240 | -1.0 | 6.54E-05 | -1.2 | 3.28E-05 | 0.3 | 1.91E-01 | 0.0 | 9.70E-01 | 0.4 | 2.10E-01 | -0.4 | 2.61E-01 | Protein-L-isoaspartate O-methyltransferase |
| AT3G48330 | -0.4 | 1.05E-04 | -0.3 | 1.12E-03 | -0.1 | 2.00E-01 | -0.6 | 1.03E-03 | -0.2 | 1.41E-01 | -0.3 | 1.19E-01 | Protein-L-isoaspartate O-methyltransferase |
| AT1G66680 | -0.1 | 3.26E-01 | 0.1 | 4.04E-01 | -0.2 | 1.13E-01 | -0.2 | 1.05E-01 | -0.3 | 5.99E-02 | 0.1 | 7.93E-01 | Protein-lysine N-methyltransferase At1g66680 |
| AT1G01920 | 0.1 | 5.33E-01 | 0.0 | 7.97E-01 | 0.1 | 4.59E-01 | 0.3 | 1.48E-01 | 0.0 | 9.50E-01 | 0.3 | 2.46E-01 | SET domain-containing protein |
| AT5G14260 | 0.8 | 2.03E-05 | 0.5 | 1.74E-03 | 0.4 | 1.20E-02 | 0.0 | 9.69E-01 | 0.0 | 8.59E-01 | 0.0 | 9.12E-01 | Rubisco methyltransferase family protein |
| AT3G55080 | 0.9 | 3.82E-05 | 0.2 | 9.64E-02 | 0.6 | 1.82E-03 | -0.2 | 3.67E-01 | -0.2 | 3.14E-01 | 0.0 | 8.96E-01 | SET domain-containing protein |
| AT3G56570 | | | | | | | 0.5 | 1.86E-01 | 0.1 | 8.08E-01 | 0.3 | 4.07E-01 | N-lysine methyltransferase |
| AT3G51770 | -0.7 | 1.82E-05 | -0.3 | 6.11E-03 | -0.4 | 3.17E-03 | -0.3 | 5.27E-02 | 0.0 | 9.96E-01 | -0.3 | 1.00E-01 | Tetratricopeptide repeat -containing protein |
| AT1G10030 | -0.3 | 2.42E-02 | 0.4 | 5.37E-03 | -0.6 | 6.47E-04 | 0.0 | 8.85E-01 | 0.2 | 2.74E-01 | -0.2 | 4.61E-01 | Ergosterol biosynthetic protein 28 |
| AT3G05270 | -0.1 | 1.07E-01 | -0.1 | 4.05E-02 | 0.0 | 6.04E-01 | -0.1 | 4.55E-01 | 0.1 | 4.68E-01 | -0.2 | 2.40E-01 | Filament-like plant protein 3 |
| AT5G59140 | 0.0 | 6.54E-01 | 0.0 | 8.92E-01 | 0.0 | 6.16E-01 | 0.0 | 9.65E-01 | 0.0 | 9.03E-01 | 0.0 | 8.96E-01 | Elongin-C |
| AT2G31440 | 0.0 | 6.10E-01 | 0.2 | 2.00E-02 | -0.2 | 7.42E-02 | -0.1 | 7.29E-01 | -0.1 | 4.09E-01 | 0.1 | 7.26E-01 | Gamma-secretase subunit APH1-like |
| AT3G27640 | 0.5 | 1.26E-02 | 0.4 | 2.73E-02 | 0.0 | 8.27E-01 | 0.6 | 1.75E-01 | 0.5 | 2.95E-01 | 0.1 | 9.14E-01 | Transducin |
| AT3G58110 | -0.3 | 3.87E-04 | -0.2 | 3.38E-03 | -0.1 | 2.68E-01 | -0.1 | 3.48E-01 | -0.1 | 7.66E-01 | -0.1 | 7.36E-01 | Uncharacterized protein F9D24.20 |
| AT4G10790 | -0.2 | 2.54E-02 | -0.2 | 3.09E-03 | 0.1 | 2.22E-01 | -0.3 | 9.95E-02 | -0.2 | 4.92E-01 | -0.2 | 5.88E-01 | Plant UBX domain-containing protein 10 |
| AT1G08480 | 0.2 | 3.71E-02 | 0.3 | 7.35E-03 | -0.1 | 3.54E-01 | 0.1 | 4.66E-01 | 0.1 | 7.92E-01 | 0.1 | 8.10E-01 | Succinate dehydrogenase subunit 6, mitochondrial |
| AT3G21200 | 0.9 | 3.85E-07 | 0.7 | 5.58E-06 | 0.2 | 1.52E-02 | 0.3 | 2.63E-02 | 0.2 | 2.44E-01 | 0.1 | 4.63E-01 | Glutamyl-tRNA reductase-binding protein, chloroplastic |
| AT3G11820 | -0.3 | 1.03E-02 | -0.4 | 1.23E-03 | 0.1 | 1.74E-01 | 0.1 | 2.43E-01 | -0.2 | 1.61E-01 | 0.3 | 3.63E-02 | Syntaxin-121 |
| AT2G44420 | 0.2 | 1.23E-01 | -0.1 | 4.96E-01 | 0.4 | 7.13E-02 | -0.1 | 8.04E-01 | -0.1 | 7.99E-01 | 0.0 | 9.68E-01 | Protein N-terminal asparagine amidohydrolase family protein |
| AT2G41760 | 0.0 | 7.49E-01 | 0.0 | 7.09E-01 | 0.0 | 9.57E-01 | -0.2 | 3.35E-01 | -0.1 | 6.59E-01 | -0.1 | 8.06E-01 | Protein N-terminal glutamine amidohydrolase |
| AT1G04070 | 0.4 | 1.71E-02 | 0.1 | 6.26E-01 | 0.3 | 8.65E-02 | 0.5 | 9.76E-02 | 0.4 | 2.54E-01 | 0.1 | 8.31E-01 | Mitochondrial import receptor subunit TOM9-1 |
| AT3G27080 | 0.4 | 2.90E-02 | 0.4 | 5.19E-02 | 0.0 | 8.46E-01 | 0.5 | 1.50E-02 | 0.2 | 3.23E-01 | 0.3 | 2.96E-01 | Mitochondrial import receptor subunit TOM20-3 |
| AT3G27070 | 0.4 | 1.77E-02 | 0.1 | 6.65E-01 | 0.4 | 9.27E-02 | 1.1 | 2.34E-03 | 0.4 | 3.70E-01 | 0.8 | 1.08E-02 | Translocase outer membrane 20-1 |
| AT5G43970 | 0.8 | 1.34E-05 | 0.8 | 1.89E-05 | 0.0 | 7.51E-01 | 0.3 | 9.46E-02 | 0.2 | 5.27E-01 | 0.2 | 5.61E-01 | Mitochondrial import receptor subunit TOM9-2 |
| AT3G46740 | 0.9 | 1.33E-06 | 0.8 | 6.72E-06 | 0.1 | 3.16E-01 | 0.5 | 1.48E-04 | 0.1 | 5.42E-01 | 0.4 | 1.22E-02 | Protein TOC75-3, chloroplastic |
| AT1G03630 | 1.5 | 3.73E-07 | 1.4 | 1.18E-06 | 0.0 | 7.65E-01 | 0.8 | 7.63E-06 | 0.6 | 5.43E-04 | 0.2 | 2.25E-01 | NADPH-protochlorophyllide oxidoreductase |
| AT2G44520 | -0.1 | 1.04E-01 | 0.0 | 5.77E-01 | -0.1 | 3.40E-01 | 0.2 | 8.83E-02 | 0.2 | 3.52E-01 | 0.1 | 6.87E-01 | Protoheme IX farnesyltransferase, mitochondrial |
| AT5G52440 | 1.5 | 4.25E-08 | 1.2 | 2.97E-07 | 0.2 | 1.66E-02 | 0.3 | 2.75E-02 | 0.0 | 8.02E-01 | 0.2 | 1.73E-01 | Sec-independent protein translocase protein TATB, chloroplastic |
| AT4G26210 | 0.0 | 9.00E-01 | 0.4 | 8.70E-03 | -0.4 | 2.17E-02 | 0.2 | 1.76E-01 | 0.0 | 9.13E-01 | 0.2 | 3.80E-01 | Mitochondrial ATP synthase subunit G protein |
| AT3G52300 | 0.1 | 1.77E-01 | 0.3 | 6.60E-03 | -0.2 | 8.44E-02 | 0.1 | 3.36E-01 | 0.1 | 5.85E-01 | 0.0 | 8.81E-01 | ATP synthase subunit d, mitochondrial |
| AT4G29480 | 0.3 | 8.58E-03 | 0.5 | 7.95E-04 | -0.2 | 1.23E-01 | 0.3 | 8.66E-02 | 0.2 | 2.51E-01 | 0.1 | 8.37E-01 | Mitochondrial ATP synthase subunit G protein |
| AT2G19680 | 0.5 | 8.61E-05 | 0.6 | 9.29E-05 | 0.0 | 7.11E-01 | 0.4 | 5.09E-02 | 0.3 | 1.21E-01 | 0.0 | 9.01E-01 | Mitochondrial ATP synthase subunit G protein |
| AT5G13450 | 0.0 | 8.22E-01 | 0.3 | 3.81E-02 | -0.3 | 3.55E-02 | 0.2 | 4.03E-02 | 0.2 | 4.99E-02 | 0.0 | 9.42E-01 | AT5G13450 protein |
| AT2G33040 | 0.2 | 1.81E-02 | 0.3 | 2.63E-03 | -0.1 | 2.35E-01 | 0.3 | 1.88E-02 | 0.2 | 5.67E-02 | 0.0 | 9.07E-01 | ATP synthase subunit gamma |
| AT4G00895 | 0.6 | 1.15E-05 | 0.4 | 3.22E-04 | 0.2 | 3.30E-02 | 0.0 | 8.24E-01 | 0.1 | 7.58E-01 | 0.0 | 9.27E-01 | ATPase, F1 complex, OSCP |
| AT1G51650 | 0.8 | 1.23E-04 | 0.6 | 1.83E-03 | 0.2 | 1.42E-01 | 0.3 | 2.86E-01 | 0.2 | 5.76E-01 | 0.1 | 8.41E-01 | ATP synthase subunit epsilon, mitochondrial |
| AT3G01390 | 0.4 | 1.12E-04 | 0.5 | 4.91E-05 | -0.1 | 2.19E-01 | 0.1 | 8.38E-01 | 0.0 | 9.20E-01 | 0.1 | 8.26E-01 | V-type proton ATPase subunit G1 -ATPase subunit G isoform 1) |
| AT5G47030 | -0.1 | 5.26E-01 | 0.2 | 1.10E-01 | -0.2 | 4.94E-02 | 0.2 | 1.52E-01 | 0.2 | 3.35E-01 | 0.0 | 8.84E-01 | ATP synthase subunit delta', mitochondrial |
| AT2G28520 | -0.4 | 5.17E-05 | -0.2 | 8.23E-03 | -0.2 | 1.02E-02 | -0.1 | 8.42E-01 | 0.1 | 5.72E-01 | -0.2 | 4.66E-01 | V-type proton ATPase subunit a |
| AT4G23710 | -0.4 | 4.84E-04 | 0.2 | 7.18E-02 | -0.6 | 2.15E-04 | -0.2 | 2.57E-01 | 0.0 | 9.38E-01 | -0.2 | 4.18E-01 | V-type proton ATPase subunit G2 -ATPase subunit G isoform 2) |
| AT1G64200 | -0.4 | 1.22E-03 | 0.2 | 8.93E-02 | -0.6 | 5.18E-04 | -0.1 | 6.48E-01 | 0.0 | 9.21E-01 | -0.1 | 8.30E-01 | Vacuolar H+-ATPase subunit E isoform 3 |
| AT4G32530 | -0.4 | 4.15E-04 | 0.0 | 9.86E-01 | -0.4 | 1.30E-03 | -0.2 | 3.11E-01 | -0.1 | 7.61E-01 | -0.1 | 7.20E-01 | ATPase, F0 |
| AT1G75630 | -0.2 | 3.88E-02 | 0.1 | 4.47E-01 | -0.3 | 1.76E-02 | -0.3 | 9.87E-02 | -0.1 | 5.43E-01 | -0.1 | 5.44E-01 | V-type proton ATPase proteolipid subunit |
| AT1G19910 | -0.2 | 8.78E-02 | 0.0 | 7.68E-01 | -0.2 | 7.51E-02 | -0.4 | 1.75E-02 | -0.3 | 1.70E-01 | -0.1 | 5.02E-01 | V-type proton ATPase proteolipid subunit |
| AT1G12840 | -0.2 | 5.06E-02 | 0.1 | 2.57E-01 | -0.2 | 1.22E-02 | -0.1 | 4.21E-01 | 0.0 | 9.65E-01 | -0.1 | 6.52E-01 | V-type proton ATPase subunit C -ATPase subunit C) |
| AT1G78900 | -0.1 | 1.63E-01 | 0.1 | 3.50E-01 | -0.2 | 4.68E-02 | -0.1 | 2.84E-01 | 0.0 | 8.18E-01 | -0.1 | 6.62E-01 | Vacuolar-type H+-ATPase subunit A |
| AT3G58730 | -0.1 | 1.94E-01 | 0.1 | 9.02E-02 | -0.2 | 1.46E-02 | -0.1 | 3.60E-01 | -0.1 | 6.95E-01 | -0.1 | 8.14E-01 | V-type proton ATPase subunit D -ATPase subunit D) |
| AT2G25610 | -0.1 | 2.15E-01 | 0.2 | 1.66E-02 | -0.3 | 4.48E-03 | -0.1 | 7.23E-01 | 0.0 | 8.89E-01 | -0.1 | 6.33E-01 | ATPase, F0 |
| AT3G42050 | -0.1 | 2.62E-01 | 0.1 | 3.72E-01 | -0.2 | 7.88E-02 | -0.1 | 4.41E-01 | -0.1 | 7.16E-01 | 0.0 | 8.83E-01 | V-type proton ATPase subunit H |

| | | | | | | | | | | | | | |
|---|---|---|---|---|---|---|---|---|---|---|---|---|---|
| AT4G38920 | -0.1 | 3.90E-01 | 0.3 | 7.45E-03 | -0.3 | 3.59E-03 | -0.2 | 9.39E-02 | 0.0 | 9.84E-01 | -0.2 | 2.40E-01 | V-type proton ATPase proteolipid subunit |
| AT5G55290 | -0.1 | 4.24E-01 | 0.1 | 1.13E-01 | -0.2 | 4.43E-02 | -0.1 | 7.08E-01 | 0.0 | 9.59E-01 | -0.1 | 8.45E-01 | V-type proton ATPase subunit e1 -ATPase subunit e isoform 1) |
| AT3G28715 | 0.0 | 9.19E-01 | 0.2 | 1.96E-02 | -0.2 | 2.42E-02 | -0.2 | 8.19E-02 | -0.1 | 2.79E-01 | -0.1 | 7.88E-01 | V-type proton ATPase subunit |
| AT3G28710 | 0.0 | 9.82E-01 | 0.2 | 1.74E-02 | -0.2 | 2.58E-02 | -0.2 | 5.64E-02 | -0.2 | 1.05E-01 | 0.0 | 9.73E-01 | V-type proton ATPase subunit d1 -ATPase subunit d isoform 1) |
| AT4G11150 | 0.1 | 5.62E-01 | 0.4 | 3.16E-03 | -0.3 | 1.23E-02 | -0.1 | 7.69E-01 | -0.1 | 7.82E-01 | 0.0 | 9.80E-01 | hypothetical protein |
| AT4G34720 | 0.1 | 3.89E-01 | 0.2 | 8.68E-03 | -0.2 | 4.94E-02 | -0.3 | 1.21E-01 | -0.1 | 6.59E-01 | -0.2 | 5.13E-01 | V-type proton ATPase proteolipid subunit |
| AT4G26710 | 0.1 | 7.85E-02 | 0.3 | 2.77E-03 | -0.1 | 7.60E-02 | -0.1 | 5.72E-01 | 0.0 | 9.97E-01 | -0.1 | 7.24E-01 | V-type proton ATPase subunit e2 -ATPase subunit e isoform 2) |
| AT4G02620 | 0.2 | 1.11E-01 | 0.3 | 2.11E-02 | -0.1 | 3.86E-01 | 0.0 | 8.80E-01 | 0.0 | 8.70E-01 | 0.0 | 9.80E-01 | V-type proton ATPase subunit F -ATPase subunit F) |
| AT2G16510 | 0.3 | 2.82E-03 | 0.4 | 1.06E-03 | -0.1 | 4.14E-01 | -0.1 | 7.67E-01 | -0.1 | 6.69E-01 | 0.0 | 9.01E-01 | V-type proton ATPase subunit c5 -ATPase subunit c isoform 5) |
| AT1G78910 | -0.1 | 7.06E-01 | -0.2 | 2.51E-01 | 0.1 | 5.11E-01 | 0.0 | 9.86E-01 | 0.0 | 9.90E-01 | 0.0 | 9.99E-01 | Pseudouridine synthase family protein |
| AT1G76120 | 0.1 | 2.50E-01 | 0.0 | 9.74E-01 | 0.1 | 3.32E-01 | 0.5 | 2.47E-02 | 0.5 | 2.55E-02 | 0.0 | 9.18E-01 | At1g76120 |
| AT1G34150 | 0.2 | 4.61E-02 | -0.1 | 4.54E-01 | 0.3 | 3.01E-02 | -0.2 | 3.84E-01 | -0.3 | 4.04E-01 | 0.0 | 9.61E-01 | Pseudouridine synthase I TruA alpha |
| AT1G20370 | 0.2 | 4.38E-02 | 0.2 | 9.86E-02 | 0.0 | 7.58E-01 | 0.2 | 2.35E-01 | 0.0 | 9.19E-01 | 0.2 | 4.13E-01 | Pseudouridine synthase I TruA alpha |
| AT3G04820 | 0.2 | 5.16E-02 | 0.1 | 6.52E-01 | 0.2 | 1.90E-01 | 0.7 | 3.65E-03 | 0.6 | 1.55E-02 | 0.1 | 8.05E-01 | TRUD domain-containing protein |
| AT3G52260 | 0.2 | 1.46E-02 | 0.0 | 9.26E-01 | 0.2 | 4.39E-02 | -0.3 | 1.98E-01 | -0.1 | 8.68E-01 | -0.3 | 3.92E-01 | Pseudouridine synthase family protein |
| AT3G57150 | 0.3 | 6.48E-02 | 0.4 | 2.98E-02 | -0.1 | 6.44E-01 | 0.5 | 1.61E-02 | 0.2 | 5.07E-01 | 0.3 | 2.16E-01 | H |
| AT5G14460 | 0.4 | 1.24E-03 | 0.3 | 9.01E-03 | 0.1 | 3.70E-01 | 0.0 | 9.91E-01 | -0.1 | 6.66E-01 | 0.1 | 7.04E-01 | tRNA pseudouridine synthase |
| AT5G51140 | 0.4 | 1.55E-04 | 0.1 | 1.13E-01 | 0.3 | 5.58E-03 | 0.0 | 8.01E-01 | 0.0 | 8.35E-01 | 0.0 | 9.98E-01 | Pseudouridine synthase |
| AT3G19440 | 0.5 | 4.82E-03 | 0.3 | 6.98E-02 | 0.2 | 2.58E-01 | 0.4 | 2.45E-01 | 0.1 | 7.55E-01 | 0.2 | 6.24E-01 | Pseudouridine synthase family protein |
| AT4G21770 | 0.6 | 4.80E-04 | 0.5 | 3.11E-03 | 0.1 | 4.05E-01 | 0.1 | 7.58E-01 | 0.1 | 8.10E-01 | 0.0 | 9.95E-01 | Pseudouridine synthase family protein |
| AT1G76050 | 0.6 | 3.82E-06 | 0.2 | 4.80E-03 | 0.4 | 6.81E-04 | 0.0 | 9.76E-01 | 0.1 | 7.21E-01 | -0.1 | 7.48E-01 | RNA pseudouridine synthase 2, chloroplastic |
| AT2G39140 | 0.7 | 7.08E-06 | 0.6 | 9.09E-05 | 0.2 | 1.02E-01 | 0.5 | 8.98E-03 | 0.3 | 1.95E-01 | 0.2 | 3.12E-01 | Pseudouridine synthase family protein |
| AT2G30320 | 0.8 | 7.63E-05 | 0.5 | 1.69E-03 | 0.2 | 1.02E-01 | 0.4 | 4.43E-02 | 0.0 | 9.18E-01 | 0.3 | 8.96E-02 | Putative tRNA pseudouridine synthase |
| AT1G20410 | 0.3 | 3.71E-03 | 0.1 | 3.14E-01 | 0.2 | 5.21E-02 | 0.2 | 2.78E-01 | 0.2 | 3.69E-01 | 0.0 | 9.76E-01 | tRNA pseudouridine synthase |
| AT5G35400 | 0.3 | 6.89E-03 | 0.2 | 5.31E-02 | 0.1 | 3.93E-01 | 0.0 | 8.74E-01 | 0.0 | 9.21E-01 | 0.1 | 8.17E-01 | tRNA pseudouridine synthase |
| AT3G10960 | -0.3 | 8.05E-03 | -0.1 | 1.46E-01 | -0.1 | 1.89E-01 | 0.1 | 4.51E-01 | 0.1 | 5.21E-01 | 0.0 | 9.99E-01 | Adenine |
| AT1G30840 | -1.7 | 8.70E-07 | -1.2 | 1.46E-05 | -0.5 | 1.34E-02 | -0.1 | 5.25E-01 | -0.1 | 6.81E-01 | 0.0 | 9.41E-01 | Probable purine permease 4 |
| AT1G28230 | -1.7 | 5.66E-06 | -1.0 | 1.84E-04 | -0.6 | 1.52E-02 | -0.5 | 7.46E-02 | 0.6 | 2.74E-02 | -1.0 | 1.65E-03 | Purine permease 1 |
| AT1G47603 | -1.2 | 3.52E-04 | 0.3 | 1.89E-01 | -1.5 | 5.52E-04 | -1.5 | 6.12E-03 | 0.2 | 6.73E-01 | -1.7 | 7.91E-03 | Putative purine permease 19 |
| AT5G41160 | -1.2 | 1.13E-03 | -0.4 | 1.15E-01 | -0.8 | 3.77E-02 | | | | | | | Probable purine permease |
| AT1G47590 | -1.1 | 8.15E-04 | -0.1 | 5.58E-01 | -1.0 | 7.35E-03 | 0.2 | 7.79E-01 | 0.3 | 6.17E-01 | | | Putative purine permease 20 |
| AT1G57980 | -0.8 | 3.97E-05 | -0.4 | 3.90E-03 | -0.4 | 1.54E-02 | -0.5 | 2.85E-02 | -0.5 | 5.68E-02 | 0.0 | 9.33E-01 | Probable purine permease 23 |
| AT4G18205 | -0.7 | 1.17E-05 | -0.2 | 8.93E-02 | -0.5 | 3.26E-04 | 0.2 | 1.03E-01 | -0.3 | 1.08E-01 | 0.5 | 1.19E-02 | Probable purine permease 22 |
| AT1G44750 | -0.5 | 5.57E-05 | -0.4 | 1.13E-03 | -0.2 | 8.18E-02 | -0.2 | 9.39E-02 | 0.0 | 9.66E-01 | -0.2 | 2.71E-01 | Probable purine permease |
| AT4G18197 | -0.5 | 1.66E-03 | -0.3 | 3.48E-02 | -0.2 | 1.63E-01 | 0.2 | 4.66E-01 | 0.0 | 9.27E-01 | 0.2 | 6.83E-01 | Probable purine permease 7 |
| AT1G19770 | -0.5 | 1.80E-04 | -0.4 | 1.27E-03 | -0.1 | 3.14E-01 | -0.3 | 1.98E-02 | -0.4 | 1.34E-02 | 0.1 | 7.99E-01 | Purine permease 14 |
| AT2G24220 | -0.3 | 2.63E-02 | -0.1 | 3.17E-01 | -0.1 | 2.54E-01 | -0.1 | 8.13E-01 | 0.2 | 6.11E-01 | -0.2 | 5.15E-01 | Probable purine permease |
| AT1G57990 | -0.2 | 1.85E-01 | 0.0 | 9.51E-01 | -0.2 | 2.14E-01 | -0.3 | 1.05E-01 | -0.7 | 5.03E-03 | 0.4 | 1.83E-01 | Probable purine permease 18 |
| AT4G18210 | 0.3 | 3.84E-02 | -0.2 | 2.35E-01 | 0.5 | 1.59E-02 | | | | | | | Probable purine permease 10 |
| AT4G08700 | 0.3 | 1.12E-02 | 0.1 | 2.89E-01 | 0.2 | 1.46E-01 | -0.3 | 1.48E-01 | -0.2 | 4.96E-01 | -0.1 | 7.06E-01 | Probable purine permease |
| AT2G33750 | | | | | | | 0.2 | 5.91E-01 | -0.4 | 5.08E-01 | 0.6 | 1.98E-01 | Probable purine permease |
| AT2G32080 | 0.4 | 1.24E-04 | 0.2 | 5.41E-03 | 0.2 | 4.69E-02 | -0.2 | 5.97E-02 | -0.2 | 1.87E-01 | 0.0 | 8.39E-01 | hypothetical protein |
| AT5G37850 | 0.5 | 2.69E-05 | 0.4 | 2.56E-04 | 0.1 | 1.80E-01 | 0.1 | 3.85E-01 | -0.2 | 2.61E-01 | 0.3 | 8.44E-02 | pyridoxal kinase |
| AT3G48780 | -0.2 | 1.80E-02 | -0.2 | 2.11E-02 | 0.0 | 1.00E+00 | -0.2 | 1.98E-01 | -0.2 | 2.69E-01 | 0.0 | 9.96E-01 | Long chain base biosynthesis protein 2b |
| AT5G23670 | 0.1 | 1.19E-01 | 0.2 | 3.66E-02 | -0.1 | 5.26E-01 | -0.1 | 4.83E-01 | 0.1 | 7.50E-01 | -0.2 | 4.07E-01 | Long chain base biosynthesis protein 2a |
| AT1G43710 | -0.2 | 2.05E-02 | -0.1 | 3.29E-01 | -0.1 | 1.72E-01 | -0.2 | 9.61E-02 | -0.2 | 1.44E-01 | 0.0 | 9.98E-01 | Serine decarboxylase |
| AT1G72810 | 0.0 | 8.50E-01 | 0.1 | 2.17E-01 | -0.1 | 3.56E-01 | -0.2 | 1.89E-01 | 0.1 | 4.50E-01 | -0.3 | 1.03E-01 | Threonine synthase 2, chloroplastic |
| AT4G29840 | 0.3 | 2.55E-03 | 0.4 | 1.84E-03 | 0.0 | 7.05E-01 | 0.0 | 7.57E-01 | 0.0 | 8.08E-01 | 0.0 | 9.98E-01 | Threonine synthase 1, chloroplastic |
| AT2G38400 | -0.4 | 3.90E-03 | 0.1 | 6.04E-01 | -0.5 | 4.07E-03 | -1.3 | 6.81E-07 | -0.7 | 5.58E-04 | -0.6 | 1.35E-02 | Alanine:glyoxylate aminotransferase 3 |
| AT4G39660 | -0.3 | 1.07E-02 | 0.2 | 2.41E-02 | -0.5 | 7.30E-04 | -0.4 | 7.79E-03 | -0.2 | 3.02E-01 | -0.2 | 2.58E-01 | Alanine:glyoxylate aminotransferase 2 |
| AT3G08860 | 0.2 | 2.12E-01 | -1.7 | 1.96E-04 | 1.9 | 2.57E-04 | | | | | | | PYRIMIDINE 4 |
| AT4G23600 | 1.0 | 9.18E-03 | 0.7 | 6.78E-02 | 0.3 | 3.73E-01 | 0.4 | 2.81E-01 | 1.1 | 1.31E-02 | -0.6 | 2.12E-01 | Tyrosine transaminase family protein |
| AT5G41760 | -0.1 | 4.88E-01 | 0.0 | 7.73E-01 | -0.1 | 4.04E-01 | 0.1 | 5.89E-01 | 0.1 | 7.88E-01 | 0.0 | 9.17E-01 | CMP-sialic acid transporter 1 |
| AT2G43240 | -0.7 | 2.51E-05 | -0.3 | 5.22E-03 | -0.4 | 6.75E-03 | 0.0 | 9.97E-01 | 0.3 | 3.81E-01 | -0.3 | 3.33E-01 | CMP-sialic acid transporter 2 |
| AT3G59360 | -0.4 | 8.30E-05 | -0.1 | 8.38E-02 | -0.3 | 3.06E-03 | -0.1 | 6.64E-01 | -0.2 | 4.07E-01 | 0.1 | 7.51E-01 | CMP-sialic acid transporter 3 |
| AT4G35335 | -0.3 | 4.59E-03 | 0.0 | 8.18E-01 | -0.2 | 1.79E-02 | -0.3 | 3.34E-02 | -0.1 | 6.90E-01 | -0.3 | 1.62E-01 | CMP-sialic acid transporter 4 |
| AT1G56700 | -0.2 | 2.05E-02 | 0.2 | 3.82E-02 | -0.3 | 1.51E-03 | -0.3 | 7.59E-02 | -0.2 | 4.81E-01 | -0.2 | 5.28E-01 | Peptidase C15, pyroglutamyl peptidase I-like protein |
| AT1G73540 | -0.1 | 6.75E-01 | -0.6 | 8.78E-03 | 0.5 | 3.37E-02 | -0.3 | 3.88E-01 | -0.1 | 9.11E-01 | -0.2 | 6.38E-01 | Nudix hydrolase domain-containing protein |
| AT2G11890 | 0.2 | 1.06E-02 | 0.2 | 1.54E-02 | 0.0 | 9.36E-01 | 0.2 | 9.70E-02 | 0.1 | 5.70E-01 | 0.1 | 5.66E-01 | hypothetical protein |
| AT3G10620 | 0.2 | 5.87E-02 | 0.3 | 6.44E-02 | 0.0 | 9.88E-01 | 0.0 | 9.00E-01 | 0.0 | 9.74E-01 | 0.0 | 9.55E-01 | AT3G10620 protein |
| AT5G14800 | 0.4 | 7.27E-04 | 0.5 | 3.11E-04 | -0.1 | 3.79E-01 | 0.2 | 1.14E-01 | 0.1 | 4.89E-01 | 0.1 | 6.42E-01 | Pyrroline-5-carboxylate reductase |
| AT1G30120 | -0.2 | 2.64E-02 | -0.1 | 2.01E-01 | -0.1 | 3.36E-01 | -0.2 | 1.42E-01 | 0.0 | 8.32E-01 | -0.1 | 4.57E-01 | Pyruvate dehydrogenase E1 component subunit beta-2, chloroplastic |
| AT5G50850 | 0.0 | 9.19E-01 | 0.3 | 1.54E-02 | -0.3 | 1.91E-02 | 0.0 | 8.63E-01 | -0.1 | 4.89E-01 | 0.1 | 4.76E-01 | Pyruvate dehydrogenase E1 component subunit beta-1, mitochondrial |
| AT1G01090 | 0.1 | 3.49E-01 | 0.2 | 2.05E-02 | -0.2 | 1.24E-01 | 0.0 | 8.40E-01 | 0.0 | 8.03E-01 | 0.0 | 9.57E-01 | Pyruvate dehydrogenase E1 component subunit alpha-3, chloroplastic |
| AT1G59900 | 0.4 | 5.48E-05 | 0.2 | 3.76E-03 | 0.2 | 2.31E-02 | -0.3 | 2.64E-03 | -0.3 | 7.42E-03 | 0.0 | 9.61E-01 | Pyruvate dehydrogenase E1 component subunit alpha |
| AT2G34590 | 0.2 | 2.31E-03 | 0.3 | 1.04E-03 | -0.1 | 4.77E-01 | -0.2 | 1.08E-01 | -0.2 | 7.91E-02 | 0.0 | 8.38E-01 | Pyruvate dehydrogenase E1 component subunit beta-3, chloroplastic |
| AT2G26900 | 0.4 | 9.32E-04 | 0.3 | 2.52E-03 | 0.0 | 7.34E-01 | 0.4 | 2.13E-03 | 0.3 | 2.99E-02 | 0.1 | 5.78E-01 | Sodium |
| AT4G15260 | 0.7 | 3.39E-05 | 0.0 | 6.92E-01 | 0.7 | 4.10E-04 | 0.1 | 5.57E-01 | -0.6 | 5.69E-03 | 0.7 | 6.29E-03 | At4g15260 |
| AT5G38010 | -1.6 | 5.30E-07 | -0.7 | 1.01E-04 | -0.8 | 3.57E-04 | -0.4 | 5.48E-02 | 0.2 | 3.76E-01 | -0.7 | 1.17E-02 | UDP-glycosyltransferase 76E9 |
| AT3G46720 | -1.4 | 3.08E-06 | -0.7 | 3.24E-04 | -0.7 | 2.54E-03 | -0.3 | 2.26E-01 | 0.6 | 2.14E-02 | -0.9 | 6.13E-03 | UDP-glycosyltransferase 76E5 |
| AT3G55700 | -1.2 | 3.95E-05 | -0.5 | 6.74E-03 | -0.7 | 8.64E-03 | 1.0 | 1.51E-03 | 0.4 | 1.69E-01 | 0.5 | 7.15E-02 | UDP-glycosyltransferase 76F1 |
| AT3G46700 | -0.8 | 1.48E-04 | -0.8 | 3.00E-04 | 0.0 | 8.96E-01 | -0.1 | 9.43E-01 | 0.8 | 2.18E-01 | -0.8 | 2.44E-01 | UDP-glycosyltransferase 76E3 |
| AT2G26480 | -0.7 | 1.27E-04 | -1.1 | 9.08E-06 | 0.5 | 8.05E-03 | 0.8 | 4.17E-04 | 1.1 | 1.37E-04 | -0.2 | 3.28E-01 | UDP-glycosyltransferase 76D1 |
| AT5G38040 | 0.0 | 9.02E-01 | -0.4 | 5.85E-03 | 0.4 | 1.72E-02 | -0.2 | 5.57E-01 | -0.1 | 7.75E-01 | -0.1 | 8.84E-01 | UDP-glycosyltransferase 76E7 |
| AT5G05900 | 0.1 | 7.46E-01 | 0.5 | 3.10E-02 | -0.4 | 1.04E-01 | 0.2 | 5.52E-01 | 0.8 | 1.58E-02 | -0.6 | 1.00E-01 | UDP-glycosyltransferase 76C3 |
| AT4G15550 | 0.4 | 3.90E-03 | 0.1 | 6.05E-01 | 0.4 | 2.34E-02 | -0.1 | 6.60E-01 | 0.1 | 6.99E-01 | -0.2 | 4.46E-01 | UDP-glycosyltransferase 75D1 |
| AT2G31790 | 0.9 | 2.46E-05 | 0.6 | 8.46E-04 | 0.3 | 3.48E-02 | 0.0 | 9.39E-01 | 0.4 | 2.28E-02 | -0.4 | 3.74E-02 | UDP-glycosyltransferase 74C1 |
| AT3G46690 | 1.1 | 2.98E-05 | 0.9 | 2.90E-04 | 0.2 | 2.01E-01 | 0.2 | 5.09E-01 | -0.7 | 2.96E-02 | 0.9 | 1.62E-02 | UDP-glycosyltransferase 76E4 |
| AT3G55710 | 1.2 | 5.53E-04 | 1.0 | 4.79E-03 | 0.3 | 3.57E-01 | -0.4 | 1.37E-01 | -0.3 | 2.53E-01 | -0.1 | 9.00E-01 | UDP-glycosyltransferase 76F2 |
| AT1G05675 | 1.5 | 8.42E-05 | -0.2 | 4.80E-01 | 1.7 | 4.32E-04 | 0.0 | 9.29E-01 | -1.2 | 2.73E-02 | 1.2 | 2.79E-02 | UDP-glycosyltransferase 74E1 |
| AT4G34138 | 1.7 | 2.45E-05 | 1.3 | 4.03E-04 | 0.4 | 1.05E-01 | -0.4 | 3.11E-01 | -1.0 | 4.92E-02 | 0.5 | 4.02E-01 | UDP-glycosyltransferase 73B1 |
| AT2G29710 | | | | | | | 0.2 | 6.88E-01 | 0.4 | 4.42E-01 | -0.2 | 7.68E-01 | UDP-glycosyltransferase 71D2 |
| AT3G21560 | -0.9 | 3.10E-04 | -0.7 | 2.56E-03 | -0.2 | 2.66E-01 | -0.1 | 6.96E-01 | 0.5 | 1.22E-03 | -0.6 | 3.50E-03 | UDP-glycosyltransferase 84A2 |

| | | | | | | | | | | | | | |
|---|---|---|---|---|---|---|---|---|---|---|---|---|---|
| AT1G24100 | 0.3 | 9.91E-04 | 0.3 | 1.08E-03 | 0.0 | 8.53E-01 | -0.1 | 5.96E-01 | 0.2 | 2.64E-01 | -0.4 | 1.67E-01 | UDP-glycosyltransferase 74B1 |
| AT1G30530 | -0.7 | 1.84E-03 | -0.3 | 1.05E-01 | -0.4 | 5.90E-02 | 0.1 | 4.34E-01 | 0.6 | 1.76E-03 | -0.5 | 2.33E-02 | Flavonol-3-O-rhamnosyltransferase |
| AT3G16520 | 0.6 | 7.08E-05 | 0.5 | 1.09E-03 | 0.2 | 1.26E-01 | -0.2 | 5.28E-02 | -0.1 | 6.76E-01 | -0.2 | 3.46E-01 | UDP-glycosyltransferase 88A1 |
| AT2G29740 | -1.3 | 1.88E-06 | -0.3 | 2.67E-02 | -1.0 | 9.60E-05 | -1.0 | 1.51E-03 | -0.5 | 1.12E-01 | -0.6 | 1.56E-01 | UDP-glycosyltransferase 71C2 |
| AT4G23890 | 0.5 | 5.42E-04 | 0.4 | 3.19E-03 | 0.1 | 4.13E-01 | -0.3 | 2.49E-01 | -0.2 | 5.90E-01 | -0.1 | 7.37E-01 | NADH-quinone oxidoreductase subunit S, chloroplastic H dehydrogenase subunit S) |
| AT4G09350 | 0.7 | 2.31E-04 | 0.2 | 1.21E-01 | 0.5 | 7.84E-03 | 0.5 | 1.50E-02 | 0.5 | 2.11E-02 | 0.0 | 9.71E-01 | NADH-quinone oxidoreductase subunit T, chloroplastic H dehydrogenase subunit T) |
| AT5G21430 | 1.7 | 4.44E-07 | 1.1 | 2.52E-05 | 0.7 | 1.40E-03 | 0.1 | 2.73E-01 | 0.1 | 3.88E-01 | 0.0 | 9.76E-01 | NADH-quinone oxidoreductase subunit U, chloroplastic H dehydrogenase subunit U) |
| AT3G59920 | -0.6 | 1.66E-05 | -0.3 | 8.42E-03 | -0.4 | 2.25E-03 | 0.0 | 9.74E-01 | 0.1 | 3.55E-01 | -0.1 | 5.14E-01 | Guanosine nucleotide diphosphate dissociation inhibitor |
| AT4G24490 | -0.2 | 6.97E-03 | -0.1 | 8.37E-02 | -0.1 | 2.62E-01 | -0.1 | 4.25E-01 | 0.0 | 9.37E-01 | -0.1 | 6.03E-01 | Geranylgeranyl transferase type-2 subunit alpha 1 |
| AT1G15410 | -1.4 | 7.01E-07 | -1.2 | 4.69E-06 | -0.2 | 1.12E-01 | -0.1 | 7.33E-01 | 0.1 | 5.67E-01 | -0.2 | 4.40E-01 | hypothetical protein |
| AT1G68795 | -1.3 | 2.01E-05 | -0.7 | 1.13E-03 | -0.6 | 1.52E-02 | -0.2 | 6.96E-01 | 0.2 | 7.59E-01 | -0.4 | 4.36E-01 | CLAVATA3 |
| AT1G25425 | -0.4 | 8.15E-02 | -0.4 | 6.85E-02 | 0.0 | 8.92E-01 | -0.7 | 3.14E-02 | 0.1 | 8.11E-01 | -0.8 | 5.14E-02 | CLAVATA3 |
| AT3G24770 | 0.2 | 2.48E-02 | 0.1 | 1.31E-01 | 0.1 | 4.77E-01 | 0.3 | 2.26E-01 | 0.2 | 3.90E-01 | 0.0 | 9.18E-01 | CLAVATA3 |
| AT4G13195 | 0.4 | 1.06E-02 | 0.4 | 9.30E-03 | 0.0 | 8.63E-01 | 0.1 | 8.70E-01 | 0.6 | 1.40E-01 | -0.5 | 2.20E-01 | CLAVATA3 |
| AT1G06225 | 1.9 | 1.92E-05 | 3.0 | 9.57E-07 | -1.1 | 3.67E-04 | | | | | | | CLAVATA3 |
| AT1G73965 | | | | | | | 0.824288785 | 3.97E-01 | 0.97055631 | 3.68E-01 | | | CLAVATA3 |
| AT2G34925 | | | | | | | -0.1 | 8.03E-01 | 0.0 | 9.92E-01 | -0.1 | 8.51E-01 | CLAVATA3 |
| AT4G18510 | | | | | | | -0.9 | 6.96E-02 | -0.3 | 6.48E-01 | -0.7 | 3.76E-01 | CLAVATA3 |
| AT1G63245 | | | | | | | 1.1 | 4.65E-02 | 0.4 | 5.41E-01 | 0.6 | 3.22E-01 | CLAVATA3 |
| AT4G37000 | 0.0 | 7.98E-01 | -0.1 | 2.10E-01 | 0.2 | 1.67E-01 | 0.1 | 5.40E-01 | 0.3 | 1.77E-01 | -0.2 | 5.28E-01 | Red chlorophyll catabolite reductase, chloroplastic |
| AT2G20690 | 1.4 | 1.08E-07 | 1.0 | 2.17E-06 | 0.4 | 2.04E-03 | 0.4 | 5.72E-03 | 0.1 | 5.76E-01 | 0.3 | 9.22E-02 | At2g20690 |
| AT3G13740 | 1.1 | 2.22E-06 | 1.0 | 1.41E-05 | 0.1 | 2.17E-01 | 0.2 | 2.61E-01 | 0.0 | 9.03E-01 | 0.3 | 3.32E-01 | Ribonuclease III family protein |
| AT1G55140 | 1.2 | 1.92E-07 | 1.0 | 1.29E-06 | 0.2 | 8.20E-02 | 0.2 | 2.15E-01 | 0.1 | 6.07E-01 | 0.1 | 7.13E-01 | At1g55140 |
| AT4G15417 | -1.1 | 6.28E-06 | -0.5 | 1.80E-03 | -0.6 | 2.23E-03 | 0.3 | 4.20E-01 | 0.6 | 1.46E-01 | -0.3 | 5.39E-01 | RNAse II-like 1 |
| AT2G43190 | 0.2 | 2.14E-01 | -0.1 | 4.24E-01 | 0.3 | 1.02E-01 | 0.0 | 9.96E-01 | -0.2 | 6.73E-01 | 0.2 | 7.07E-01 | Ribonuclease P protein subunit p29 |
| AT1G04635 | | | | | | | 0.6 | 3.51E-01 | 0.5 | 5.20E-01 | 0.1 | 9.00E-01 | Probable ribonuclease P |
| AT1G14220 | -0.2 | 1.08E-01 | 0.6 | 2.73E-04 | -0.9 | 2.70E-04 | -0.4 | 3.42E-01 | 0.0 | 9.86E-01 | -0.4 | 3.22E-01 | hypothetical protein |
| AT2G39780 | 0.1 | 7.67E-02 | 0.1 | 1.04E-01 | 0.0 | 9.28E-01 | 0.0 | 7.64E-01 | 0.1 | 5.19E-01 | -0.1 | 7.87E-01 | Vacuolar ribonuclease |
| AT1G14210 | 0.4 | 2.55E-03 | 0.8 | 6.59E-05 | -0.4 | 1.40E-02 | 0.3 | 3.89E-02 | 0.2 | 1.22E-01 | 0.0 | 8.32E-01 | Uncharacterized protein |
| AT1G26820 | -0.8 | 1.80E-05 | -0.1 | 3.45E-01 | -0.7 | 3.18E-04 | -0.1 | 8.62E-01 | 0.1 | 7.96E-01 | -0.1 | 7.10E-01 | Ribonuclease 3 |
| AT2G02990 | | | | | | | 0.7 | 8.59E-01 | 3.9 | 1.86E-01 | -3.2 | 3.71E-01 | Ribonuclease 1 |
| AT2G41020 | 0.1 | 3.66E-01 | -0.2 | 3.58E-02 | 0.3 | 1.77E-02 | -0.3 | 2.88E-01 | 0.0 | 9.00E-01 | -0.3 | 3.53E-01 | Uncharacterized protein At2g41020 |
| AT2G40360 | 0.3 | 1.98E-02 | 0.2 | 5.89E-02 | 0.1 | 6.54E-01 | 0.6 | 1.31E-03 | 0.2 | 3.11E-01 | 0.4 | 9.07E-02 | Ribosome biogenesis protein BOP1 homolog |
| AT5G15550 | 0.3 | 1.64E-02 | 0.2 | 1.94E-01 | 0.2 | 2.57E-01 | 0.4 | 3.25E-02 | -0.1 | 8.19E-01 | 0.4 | 7.41E-02 | Ribosome biogenesis protein WDR12 homolog |
| AT1G71100 | 0.3 | 6.07E-02 | 0.1 | 7.30E-01 | 0.2 | 1.72E-01 | 0.4 | 2.44E-02 | 0.0 | 9.30E-01 | 0.4 | 9.43E-02 | Probable ribose-5-phosphate isomerase 1 |
| AT2G01290 | 0.9 | 6.86E-07 | 0.7 | 1.29E-05 | 0.2 | 1.82E-02 | 0.4 | 2.87E-01 | 0.3 | 4.94E-01 | 0.1 | 8.85E-01 | Probable ribose-5-phosphate isomerase 2 |
| AT3G04790 | 1.7 | 1.89E-08 | 1.3 | 1.62E-07 | 0.3 | 1.76E-03 | 0.6 | 1.49E-04 | 0.4 | 5.23E-03 | 0.2 | 3.49E-01 | Probable ribose-5-phosphate isomerase 3, chloroplastic |
| AT5G44520 | 1.2 | 4.34E-07 | 0.8 | 1.39E-05 | 0.4 | 4.15E-03 | -0.1 | 5.12E-01 | -0.2 | 2.53E-01 | 0.1 | 7.04E-01 | ribose-5-phosphate isomerase |
| AT3G01800 | 0.1 | 6.44E-01 | -0.1 | 7.04E-01 | 0.1 | 4.75E-01 | 0.5 | 2.07E-02 | 0.4 | 1.13E-01 | 0.1 | 6.42E-01 | Ribosome-recycling factor, chloroplastic |
| AT1G67620 | 0.4 | 6.99E-03 | 0.2 | 2.29E-01 | 0.2 | 1.24E-01 | 0.5 | 1.45E-01 | 0.3 | 3.45E-01 | 0.1 | 8.24E-01 | Protein Iojap-related, mitochondrial |
| AT3G12930 | 1.8 | 1.52E-07 | 1.5 | 1.25E-06 | 0.3 | 2.92E-02 | 0.4 | 1.85E-02 | 0.0 | 8.91E-01 | 0.4 | 1.34E-02 | Protein Iojap, chloroplastic |
| AT3G55620 | 0.9 | 2.66E-05 | 0.3 | 2.68E-02 | 0.6 | 1.77E-03 | 0.9 | 6.51E-06 | 0.3 | 6.24E-02 | 0.6 | 4.09E-03 | Eukaryotic translation initiation factor 6-2 |
| AT5G49930 | -0.3 | 4.43E-04 | -0.4 | 9.50E-05 | 0.1 | 1.45E-01 | 0.0 | 9.22E-01 | 0.0 | 9.49E-01 | 0.0 | 8.89E-01 | Gb\|AAF56406.1 family protein) |
| AT5G58410 | -0.2 | 1.85E-02 | -0.2 | 1.08E-02 | 0.0 | 7.30E-01 | -0.1 | 6.91E-01 | -0.2 | 6.17E-01 | 0.0 | 9.23E-01 | E3 ubiquitin-protein ligase listerin |
| AT4G34730 | 1.3 | 1.98E-07 | 0.8 | 1.28E-05 | 0.5 | 5.44E-04 | 0.4 | 7.62E-03 | 0.1 | 5.73E-01 | 0.3 | 1.23E-01 | Probable ribosome-binding factor A, chloroplastic |
| AT1G65540 | 0.0 | 5.04E-01 | -0.1 | 2.63E-01 | 0.1 | 1.15E-01 | 0.2 | 2.22E-01 | 0.1 | 6.22E-01 | 0.1 | 7.27E-01 | F5I14.7 protein |
| AT1G22730 | 0.2 | 1.38E-02 | 0.0 | 7.63E-01 | 0.2 | 4.65E-02 | -0.1 | 4.09E-01 | -0.3 | 5.35E-02 | 0.2 | 3.72E-01 | MA3 DOMAIN-CONTAINING TRANSLATION REGULATORY FACTOR 2 |
| AT3G48390 | 1.1 | 2.14E-04 | 1.0 | 1.03E-03 | 0.2 | 5.12E-01 | -0.9 | 1.06E-02 | -1.4 | 1.52E-03 | 0.5 | 2.42E-01 | MA3 DOMAIN-CONTAINING TRANSLATION REGULATORY FACTOR 4 |
| AT1G26630 | 0.3 | 8.68E-03 | 0.4 | 3.05E-03 | -0.1 | 4.67E-01 | 0.3 | 2.31E-02 | 0.1 | 3.70E-01 | 0.2 | 4.11E-01 | Eukaryotic translation initiation factor 5A |
| AT1G13950 | -0.7 | 1.51E-05 | -0.1 | 3.47E-01 | -0.6 | 2.63E-04 | 0.2 | 5.10E-01 | 0.2 | 4.77E-01 | 0.0 | 9.25E-01 | Eukaryotic translation initiation factor 5A-1 |
| AT1G69410 | -0.2 | 1.43E-02 | 0.0 | 7.71E-01 | -0.2 | 4.24E-02 | -0.1 | 6.31E-01 | 0.0 | 9.26E-01 | -0.1 | 8.37E-01 | Eukaryotic translation initiation factor 5A-3 |
| AT5G05470 | 0.5 | 3.09E-04 | 0.3 | 1.28E-02 | 0.2 | 7.43E-02 | 0.3 | 5.23E-02 | 0.2 | 2.42E-01 | 0.1 | 6.85E-01 | Eukaryotic translation initiation factor 2 subunit alpha |
| AT5G38410 | 2.4 | 4.25E-08 | 2.2 | 1.90E-07 | 0.2 | 2.57E-01 | 0.2 | 2.38E-01 | 0.2 | 1.85E-01 | -0.1 | 8.50E-01 | Ribulose bisphosphate carboxylase small subunit, chloroplastic |
| AT1G29550 | 0.2 | 7.86E-02 | 0.1 | 6.83E-01 | 0.2 | 2.57E-01 | 0.2 | 3.04E-01 | 0.1 | 7.61E-01 | 0.1 | 6.96E-01 | Eukaryotic translation initiation factor 4E-3 |
| AT5G18110 | 0.3 | 2.05E-03 | 0.0 | 8.58E-01 | 0.3 | 6.71E-03 | -0.2 | 3.45E-02 | -0.3 | 7.44E-03 | 0.1 | 5.40E-01 | Eukaryotic translation initiation factor NCBP |
| AT4G18040 | 0.5 | 1.25E-04 | 0.3 | 6.57E-03 | 0.2 | 3.94E-02 | 0.4 | 6.17E-03 | 0.2 | 3.13E-01 | 0.2 | 2.34E-01 | Eukaryotic translation initiation factor 4E-1 |
| AT5G35620 | 0.6 | 1.44E-04 | 0.6 | 1.42E-04 | -0.1 | 6.70E-01 | 0.3 | 8.06E-02 | 0.0 | 9.70E-01 | 0.3 | 1.85E-01 | Eukaryotic translation initiation factor isoform 4E 4E) 4F 25 kDa subunit) 4F p28 subunit) |
| AT1G29590 | | | | | | | 0.6 | 2.29E-01 | 0.7 | 1.69E-01 | | | Eukaryotic translation initiation factor 4E-2 |
| AT3G54770 | -1.6 | 5.72E-07 | -0.1 | 4.98E-01 | -1.5 | 1.80E-05 | -0.6 | 1.03E-02 | 0.2 | 3.33E-01 | -0.8 | 5.89E-03 | RNA-binding family protein |
| AT5G12280 | -1.5 | 6.46E-06 | -1.0 | 1.22E-04 | -0.5 | 3.42E-02 | | | | | | | Ubiquitin-like domain-containing protein |
| AT1G72800 | -1.4 | 1.24E-05 | -1.0 | 1.01E-04 | -0.3 | 1.77E-01 | -0.5 | 1.62E-01 | -0.3 | 4.45E-01 | -0.2 | 7.20E-01 | RRM domain-containing protein |
| AT5G46870 | -1.3 | 1.18E-06 | -0.5 | 7.10E-04 | -0.8 | 3.26E-04 | -0.2 | 5.11E-01 | 0.3 | 3.21E-01 | -0.5 | 1.14E-01 | At5g46870 family protein) |
| AT2G42890 | -1.3 | 4.63E-07 | -1.1 | 3.41E-06 | -0.2 | 7.80E-02 | -0.4 | 3.40E-02 | -0.3 | 2.01E-01 | -0.1 | 6.29E-01 | hypothetical protein |
| AT1G20880 | -1.1 | 6.26E-06 | -0.9 | 4.46E-05 | -0.2 | 1.94E-01 | -0.4 | 3.58E-02 | -0.2 | 2.82E-01 | -0.2 | 4.86E-01 | hypothetical protein |
| AT4G26990 | -1.0 | 3.66E-05 | -0.6 | 1.33E-03 | -0.4 | 3.91E-02 | 0.3 | 3.91E-01 | 0.1 | 8.99E-01 | 0.2 | 6.23E-01 | Ataxin 2 SM domain-containing protein |
| AT1G54130 | -0.9 | 1.39E-05 | -1.0 | 1.03E-05 | 0.1 | 2.23E-01 | -0.1 | 4.39E-01 | 0.0 | 8.82E-01 | -0.2 | 4.94E-01 | RSH3 |
| AT3G63450 | -0.8 | 2.42E-04 | -0.4 | 2.45E-02 | -0.4 | 2.93E-02 | -0.6 | 1.11E-02 | 0.0 | 9.66E-01 | -0.6 | 3.30E-02 | At3g63450 family protein) |
| AT2G46780 | -0.8 | 4.54E-05 | -0.2 | 1.01E-01 | -0.6 | 1.66E-03 | -0.5 | 1.00E-01 | -0.3 | 4.26E-01 | -0.2 | 6.27E-01 | RRM domain-containing protein |
| AT1G51520 | -0.7 | 1.81E-03 | -0.6 | 4.36E-03 | -0.1 | 8.02E-01 | -0.3 | 4.96E-01 | 0.4 | 3.50E-01 | -0.8 | 1.53E-01 | RNA-binding family protein |
| AT3G08620 | -0.7 | 9.09E-05 | -0.5 | 7.47E-04 | -0.2 | 2.66E-01 | 0.1 | 6.80E-01 | 0.0 | 9.68E-01 | 0.1 | 7.49E-01 | RNA-binding KH domain-containing protein |
| AT1G33470 | -0.7 | 1.59E-05 | -0.2 | 1.65E-02 | -0.4 | 1.25E-03 | -0.1 | 4.66E-01 | 0.0 | 8.99E-01 | -0.1 | 4.75E-01 | RNA-binding protein; 68390-68829 |
| AT1G33520 | -0.6 | 3.27E-06 | -0.3 | 2.22E-04 | -0.2 | 5.09E-03 | -0.3 | 5.73E-03 | -0.3 | 1.34E-02 | 0.0 | 9.71E-01 | Protein MOS2 |
| AT1G22830 | -0.6 | 1.57E-03 | -0.3 | 6.07E-02 | -0.3 | 1.07E-01 | 0.2 | 6.35E-01 | 0.1 | 8.38E-01 | 0.1 | 8.73E-01 | Pentatricopeptide repeat-containing protein At1g22830 |
| AT5G16840 | -0.6 | 1.09E-04 | -0.3 | 1.27E-02 | -0.3 | 1.67E-02 | -0.3 | 4.16E-02 | -0.2 | 2.28E-01 | -0.1 | 6.56E-01 | Binding partner of ACD11 1 |
| AT3G21100 | -0.6 | 1.21E-04 | -0.3 | 4.26E-03 | -0.2 | 6.39E-02 | -0.1 | 6.37E-01 | 0.0 | 9.60E-01 | -0.1 | 7.76E-01 | RNA-binding family protein |
| AT1G09140 | -0.5 | 1.63E-05 | -0.4 | 1.17E-04 | -0.1 | 2.25E-01 | -0.1 | 2.49E-01 | 0.0 | 8.14E-01 | -0.2 | 2.74E-01 | T12M4.19 protein |
| AT5G28390 | -0.5 | 8.55E-03 | -0.5 | 9.78E-03 | 0.0 | 9.29E-01 | | | | | | | RNA-binding family protein |
| AT1G14640 | -0.5 | 2.58E-02 | -1.0 | 1.13E-03 | 0.5 | 7.69E-02 | 1.1 | 1.28E-01 | 0.8 | 3.65E-01 | 0.3 | 7.30E-01 | SURP motif domain-containing protein |
| AT5G55550 | -0.5 | 3.90E-04 | -0.2 | 2.16E-02 | -0.2 | 5.47E-02 | -0.2 | 1.01E-01 | -0.1 | 7.95E-01 | -0.2 | 3.80E-01 | RNA-binding protein-like |
| AT4G18120 | -0.4 | 5.45E-03 | -0.7 | 5.57E-04 | 0.2 | 1.27E-01 | -0.2 | 5.36E-01 | -0.2 | 6.39E-01 | 0.0 | 9.77E-01 | Protein MEI2-like 3 |
| AT3G13224 | -0.4 | 1.30E-04 | -0.5 | 7.85E-05 | 0.1 | 4.02E-01 | -0.3 | 3.63E-02 | -0.2 | 3.94E-01 | -0.2 | 4.24E-01 | RNA-binding family protein |
| AT5G44500 | -0.4 | 1.19E-02 | 0.0 | 8.02E-01 | -0.4 | 2.05E-02 | 0.3 | 2.32E-01 | 0.2 | 5.02E-01 | 0.1 | 7.80E-01 | Small nuclear ribonucleoprotein-associated protein |
| AT3G14010 | -0.3 | 2.38E-04 | -0.3 | 6.09E-04 | 0.0 | 7.42E-01 | -0.1 | 3.56E-01 | -0.2 | 2.89E-01 | 0.0 | 8.78E-01 | CTC-interacting domain 4 |

| | | | | | | | | | | | | | |
|---|---|---|---|---|---|---|---|---|---|---|---|---|---|
| AT1G03457 | -0.3 | 2.66E-03 | 0.0 | 9.10E-01 | -0.3 | 9.88E-03 | -0.7 | 6.10E-04 | -0.4 | 4.89E-02 | -0.3 | 1.71E-01 | BRN2 |
| AT3G52120 | -0.3 | 8.33E-04 | -0.4 | 3.96E-04 | 0.1 | 4.60E-01 | -0.3 | 8.84E-02 | 0.0 | 9.93E-01 | -0.3 | 1.49E-01 | SWAP |
| AT2G31290 | -0.3 | 2.12E-02 | -0.3 | 5.21E-02 | 0.0 | 7.57E-01 | -0.1 | 7.21E-01 | -0.1 | 8.33E-01 | 0.0 | 9.46E-01 | PORR domain-containing protein |
| AT3G04750 | -0.3 | 1.46E-02 | -0.2 | 1.10E-01 | -0.1 | 3.87E-01 | 0.0 | 9.22E-01 | -0.1 | 8.58E-01 | 0.1 | 7.79E-01 | Pentatricopeptide repeat-containing protein At3g04750, mitochondrial |
| AT4G27000 | -0.3 | 1.42E-03 | -0.2 | 1.38E-02 | -0.1 | 2.83E-01 | -0.3 | 1.10E-02 | -0.3 | 1.14E-02 | 0.0 | 8.69E-01 | RRM domain-containing protein |
| AT4G31010 | -0.2 | 3.69E-02 | -0.1 | 1.78E-01 | -0.1 | 4.97E-01 | -0.1 | 7.03E-01 | -0.1 | 7.18E-01 | 0.0 | 9.81E-01 | RNA-binding CRS1 |
| AT4G26650 | -0.2 | 6.04E-03 | 0.0 | 6.84E-01 | -0.2 | 2.43E-02 | -0.2 | 2.33E-01 | 0.0 | 9.71E-01 | -0.1 | 4.52E-01 | HnRNP-like protein |
| AT3G56510 | -0.2 | 5.40E-02 | -0.1 | 2.63E-01 | -0.1 | 4.61E-01 | -0.1 | 6.42E-01 | 0.0 | 9.82E-01 | -0.1 | 7.76E-01 | At3g56510 family protein) |
| AT1G26900 | -0.2 | 6.46E-02 | 0.0 | 9.72E-01 | -0.2 | 1.15E-01 | 0.1 | 7.54E-01 | 0.2 | 6.83E-01 | -0.1 | 9.18E-01 | Pentatricopeptide repeat-containing protein At1g26900, mitochondrial |
| AT4G10610 | -0.2 | 3.93E-03 | 0.0 | 8.71E-01 | -0.2 | 1.12E-02 | -0.1 | 4.90E-01 | -0.2 | 1.31E-01 | 0.1 | 5.12E-01 | CTC-interacting domain 12 |
| AT1G14340 | -0.2 | 1.26E-01 | -0.4 | 1.02E-02 | 0.2 | 2.02E-01 | -0.2 | 2.21E-01 | -0.1 | 6.57E-01 | -0.1 | 6.50E-01 | hypothetical protein |
| AT4G31200 | -0.2 | 1.53E-02 | -0.2 | 1.21E-02 | 0.0 | 8.41E-01 | -0.2 | 5.88E-01 | 0.0 | 9.89E-01 | -0.2 | 7.08E-01 | hypothetical protein |
| AT1G62670 | -0.2 | 1.77E-01 | -0.1 | 7.24E-01 | -0.2 | 4.24E-01 | 0.2 | 6.64E-01 | -0.1 | 8.49E-01 | 0.2 | 5.70E-01 | Pentatricopeptide repeat-containing protein At1g62670, mitochondrial |
| AT4G39260 | -0.2 | 1.74E-01 | 0.1 | 4.17E-01 | -0.3 | 6.07E-02 | 0.2 | 2.19E-01 | 0.1 | 8.49E-01 | 0.2 | 4.90E-01 | Glycine-rich RNA-binding protein 8 splice variant 6 |
| AT3G07810 | -0.2 | 4.30E-02 | 0.0 | 8.84E-01 | -0.2 | 5.39E-02 | 0.1 | 3.09E-01 | 0.2 | 3.23E-01 | 0.0 | 9.48E-01 | hypothetical protein |
| AT4G25500 | -0.2 | 9.02E-03 | 0.0 | 6.75E-01 | -0.2 | 3.46E-02 | -0.5 | 1.72E-03 | -0.5 | 3.52E-03 | 0.0 | 9.80E-01 | RRM domain-containing protein |
| AT1G80930 | -0.2 | 3.32E-02 | -0.2 | 1.83E-02 | 0.0 | 7.08E-01 | 0.0 | 7.85E-01 | -0.1 | 1.71E-01 | 0.1 | 4.41E-01 | At1g80930 |
| AT1G11400 | -0.2 | 5.26E-02 | -0.2 | 3.07E-02 | 0.0 | 7.59E-01 | -0.1 | 7.13E-01 | -0.2 | 2.58E-01 | 0.1 | 5.59E-01 | Partner of Y14 and mago |
| AT1G55310 | -0.2 | 5.40E-02 | -0.2 | 3.52E-02 | 0.0 | 8.01E-01 | -0.1 | 2.12E-01 | -0.2 | 7.84E-02 | 0.1 | 6.74E-01 | SC35-like splicing factor 33 |
| AT5G61930 | -0.1 | 1.44E-01 | -0.1 | 2.66E-01 | 0.0 | 8.02E-01 | -0.1 | 6.54E-01 | -0.1 | 8.02E-01 | 0.0 | 9.41E-01 | APO protein 3, mitochondrial |
| AT1G09230 | -0.1 | 1.15E-01 | 0.0 | 7.91E-01 | -0.1 | 2.67E-01 | -0.4 | 9.05E-02 | -0.2 | 5.91E-01 | -0.3 | 4.14E-01 | RNA-binding family protein |
| AT4G36960 | -0.1 | 6.69E-02 | -0.1 | 2.94E-01 | -0.1 | 4.75E-01 | 0.0 | 9.44E-01 | 0.1 | 6.17E-01 | -0.1 | 6.79E-01 | RRM domain-containing protein |
| AT3G55460 | -0.1 | 7.34E-02 | 0.0 | 9.18E-01 | -0.1 | 1.26E-01 | -0.3 | 4.31E-02 | -0.3 | 3.10E-02 | 0.1 | 8.23E-01 | SCL30 |
| AT4G13070 | -0.1 | 2.61E-01 | -0.1 | 3.60E-01 | 0.0 | 9.00E-01 | -0.1 | 8.30E-01 | 0.1 | 7.60E-01 | -0.2 | 6.57E-01 | Uncharacterized protein At4g13070 |
| AT5G25060 | -0.1 | 6.01E-02 | -0.2 | 1.40E-02 | 0.1 | 4.25E-01 | -0.2 | 2.88E-01 | -0.2 | 4.11E-01 | 0.0 | 9.62E-01 | Protein RRC1 |
| AT1G17640 | -0.1 | 3.35E-01 | -0.2 | 1.97E-01 | 0.1 | 7.60E-01 | -0.6 | 1.40E-02 | -0.4 | 1.18E-01 | -0.2 | 4.93E-01 | RNA-binding family protein |
| AT5G45790 | -0.1 | 5.58E-01 | -0.3 | 2.57E-01 | 0.1 | 6.33E-01 | -0.1 | 7.34E-01 | 0.0 | 9.35E-01 | -0.1 | 8.65E-01 | Tyrosine-specific protein phosphatase-like protein |
| AT1G60900 | -0.1 | 1.78E-01 | -0.3 | 4.97E-03 | 0.2 | 6.81E-02 | 0.1 | 5.42E-01 | 0.1 | 5.75E-01 | 0.0 | 9.75E-01 | Splicing factor U2af large subunit ) |
| AT5G64390 | -0.1 | 2.50E-01 | -0.3 | 9.93E-03 | 0.2 | 1.06E-01 | 0.1 | 8.49E-01 | 0.1 | 8.29E-01 | 0.0 | 9.67E-01 | RNA-binding KH domain-containing protein |
| AT4G16470 | -0.1 | 3.00E-01 | -0.3 | 1.57E-02 | 0.2 | 1.45E-01 | 0.0 | 8.81E-01 | 0.1 | 7.67E-01 | -0.2 | 6.85E-01 | Pentatricopeptide repeat-containing protein At4g16470 |
| AT2G19430 | -0.1 | 2.69E-01 | -0.2 | 8.73E-02 | 0.1 | 5.46E-01 | -0.1 | 7.62E-01 | -0.2 | 3.52E-01 | 0.1 | 6.03E-01 | THO complex subunit 6 |
| AT1G14650 | -0.1 | 2.69E-01 | -0.1 | 9.81E-02 | 0.0 | 5.64E-01 | 0.0 | 8.96E-01 | -0.3 | 1.07E-01 | 0.2 | 2.00E-01 | Probable splicing factor 3A subunit 1 |
| AT5G37720 | -0.1 | 3.60E-01 | -0.1 | 4.26E-02 | 0.1 | 2.44E-01 | 0.0 | 9.54E-01 | -0.2 | 1.34E-01 | 0.2 | 1.78E-01 | ALWAYS EARLY 4 |
| AT1G31600 | 0.0 | 6.38E-01 | -0.2 | 4.93E-02 | 0.2 | 1.56E-01 | 0.1 | 5.79E-01 | 0.1 | 8.40E-01 | 0.1 | 8.40E-01 | RNA-binding family protein |
| AT5G52040 | 0.0 | 5.95E-01 | -0.1 | 7.99E-02 | 0.1 | 2.25E-01 | -0.2 | 1.81E-01 | -0.2 | 1.87E-01 | 0.0 | 9.38E-01 | hypothetical protein |
| AT4G34110 | 0.0 | 7.21E-01 | 0.0 | 7.24E-01 | -0.1 | 5.15E-01 | 0.0 | 7.71E-01 | 0.0 | 9.75E-01 | 0.1 | 8.38E-01 | Poly-binding protein |
| AT1G53650 | 0.0 | 7.54E-01 | 0.0 | 9.79E-01 | 0.0 | 8.25E-01 | 0.1 | 8.43E-01 | 0.3 | 3.28E-01 | -0.2 | 5.15E-01 | hypothetical protein |
| AT5G08510 | 0.0 | 8.29E-01 | -0.1 | 4.64E-01 | 0.1 | 6.68E-01 | -0.1 | 6.07E-01 | 0.0 | 9.66E-01 | -0.1 | 6.66E-01 | Pentatricopeptide repeat-containing protein At5g08510 |
| AT2G42240 | 0.0 | 8.19E-01 | -0.2 | 1.70E-01 | 0.1 | 3.24E-01 | 0.0 | 9.93E-01 | -0.1 | 7.79E-01 | 0.1 | 8.13E-01 | Uncharacterized protein At2g42245 |
| AT1G60170 | 0.0 | 7.00E-01 | -0.1 | 7.86E-02 | 0.1 | 1.82E-01 | 0.0 | 8.27E-01 | -0.2 | 2.51E-01 | 0.1 | 4.93E-01 | U4 |
| AT3G01150 | 0.0 | 7.62E-01 | -0.1 | 4.34E-01 | 0.0 | 6.76E-01 | -0.2 | 1.18E-01 | -0.2 | 2.79E-01 | -0.1 | 8.34E-01 | Polypyrimidine tract-binding protein 1 |
| AT2G33435 | 0.0 | 7.99E-01 | -0.1 | 1.55E-01 | 0.1 | 2.90E-01 | 0.3 | 5.17E-02 | 0.2 | 3.91E-01 | 0.1 | 4.63E-01 | hypothetical protein |
| AT3G13200 | 0.0 | 8.33E-01 | 0.0 | 6.71E-01 | 0.0 | 5.57E-01 | -0.2 | 9.89E-02 | -0.3 | 5.46E-02 | 0.1 | 7.58E-01 | Cwf15 |
| AT2G21090 | 0.0 | 9.37E-01 | 0.2 | 3.23E-01 | -0.2 | 3.69E-01 | 0.3 | 3.48E-01 | -0.1 | 8.69E-01 | 0.3 | 3.88E-01 | Pentatricopeptide repeat-containing protein At2g21090 |
| AT2G43370 | 0.0 | 9.13E-01 | -0.2 | 4.68E-02 | 0.2 | 9.72E-02 | -0.5 | 2.66E-02 | -0.3 | 1.80E-01 | -0.2 | 5.87E-01 | U11 |
| AT1G12300 | 0.0 | 9.75E-01 | 0.0 | 9.99E-01 | 0.0 | 9.81E-01 | -0.5 | 2.04E-01 | -0.1 | 7.92E-01 | -0.4 | 4.89E-01 | Pentatricopeptide repeat-containing protein At1g12300, mitochondrial |
| AT5G66010 | 0.0 | 9.75E-01 | 0.0 | 7.21E-01 | 0.0 | 7.81E-01 | -0.1 | 5.71E-01 | -0.3 | 1.51E-01 | 0.2 | 3.90E-01 | RNA-binding family protein |
| AT1G77170 | 0.0 | 9.48E-01 | -0.2 | 2.20E-01 | 0.2 | 2.71E-01 | 0.1 | 5.76E-01 | -0.2 | 4.39E-01 | 0.3 | 1.65E-01 | Pentatricopeptide repeat-containing protein At1g77170, mitochondrial |
| AT4G35785 | 0.0 | 8.66E-01 | 0.0 | 7.87E-01 | 0.0 | 9.28E-01 | -0.1 | 3.93E-01 | -0.1 | 4.60E-01 | 0.0 | 9.98E-01 | RNA-binding family protein |
| AT1G17370 | 0.0 | 8.74E-01 | 0.0 | 6.47E-01 | 0.0 | 5.69E-01 | -0.1 | 2.98E-01 | 0.1 | 7.58E-01 | -0.2 | 3.13E-01 | Oligouridylate binding protein 1B |
| AT1G13190 | 0.0 | 8.26E-01 | -0.1 | 8.58E-02 | 0.1 | 8.18E-02 | -0.3 | 2.76E-02 | -0.3 | 3.83E-02 | 0.0 | 9.71E-01 | hypothetical protein |
| AT2G43650 | 0.0 | 8.42E-01 | 0.0 | 8.13E-01 | 0.0 | 9.69E-01 | 0.6 | 6.24E-04 | 0.3 | 1.03E-01 | 0.3 | 1.03E-01 | Protein THALLO |
| AT2G33410 | 0.0 | 8.22E-01 | 0.0 | 9.87E-01 | 0.0 | 8.37E-01 | -0.1 | 6.82E-01 | 0.0 | 8.07E-01 | 0.0 | 9.61E-01 | RRM domain-containing protein |
| AT5G02770 | 0.0 | 7.20E-01 | -0.1 | 2.78E-01 | 0.1 | 1.85E-01 | 0.1 | 4.29E-01 | 0.0 | 9.39E-01 | 0.1 | 6.86E-01 | Protein MODIFIER OF SNC1 11 |
| AT5G23590 | 0.0 | 7.34E-01 | 0.0 | 8.75E-01 | 0.0 | 8.96E-01 | -0.1 | 4.20E-01 | -0.2 | 2.45E-01 | 0.1 | 7.67E-01 | hypothetical protein |
| AT5G46190 | 0.0 | 6.99E-01 | -0.1 | 4.57E-01 | 0.1 | 3.17E-01 | 0.0 | 9.93E-01 | 0.0 | 8.75E-01 | 0.0 | 8.93E-01 | RNA-binding KH domain-containing protein |
| AT5G59950 | 0.0 | 6.72E-01 | 0.1 | 4.26E-01 | 0.0 | 7.30E-01 | -0.2 | 2.60E-01 | -0.3 | 4.88E-02 | 0.2 | 4.34E-01 | RNA-binding family protein |
| AT1G09150 | 0.0 | 6.18E-01 | 0.2 | 3.40E-02 | -0.1 | 1.13E-01 | 0.0 | 7.92E-01 | 0.1 | 2.90E-01 | -0.1 | 5.40E-01 | hypothetical protein |
| AT5G19960 | 0.0 | 6.35E-01 | -0.1 | 4.80E-01 | 0.1 | 3.13E-01 | -0.3 | 1.74E-01 | -0.3 | 1.43E-01 | 0.1 | 8.96E-01 | RRM domain-containing protein |
| AT1G60200 | 0.0 | 5.05E-01 | -0.1 | 1.21E-01 | 0.2 | 5.22E-02 | -0.1 | 2.91E-01 | -0.2 | 1.35E-01 | 0.1 | 7.35E-01 | Splicing factor PWI domain-containing protein |
| AT3G59810 | 0.0 | 6.74E-01 | 0.1 | 4.06E-01 | 0.0 | 7.22E-01 | 0.0 | 8.45E-01 | 0.1 | 7.13E-01 | -0.1 | 6.05E-01 | Sm-like protein LSM6A |
| AT1G09290 | 0.0 | 7.12E-01 | 0.0 | 7.45E-01 | 0.0 | 9.85E-01 | 0.0 | 9.69E-01 | -0.2 | 4.59E-01 | 0.2 | 5.62E-01 | Uncharacterized protein At1g09290 |
| AT5G12410 | 0.1 | 5.32E-01 | 0.2 | 3.72E-02 | -0.2 | 1.49E-01 | 0.1 | 4.61E-01 | 0.0 | 9.91E-01 | 0.1 | 5.99E-01 | THUMP domain-containing protein |
| AT5G61960 | 0.1 | 5.04E-01 | -0.2 | 6.46E-02 | 0.3 | 2.98E-02 | 0.3 | 5.70E-02 | 0.1 | 5.93E-01 | 0.2 | 3.97E-01 | RRM domain-containing protein |
| AT5G54920 | 0.1 | 4.22E-01 | 0.1 | 4.62E-01 | 0.0 | 9.81E-01 | 0.2 | 5.72E-01 | 0.4 | 1.93E-01 | -0.2 | 5.18E-01 | hypothetical protein |
| AT5G10350 | 0.1 | 4.44E-01 | 0.2 | 2.85E-02 | -0.1 | 1.41E-01 | -0.2 | 1.70E-01 | -0.1 | 4.44E-01 | -0.1 | 8.04E-01 | RNA-binding family protein |
| AT3G13570 | 0.1 | 5.10E-01 | 0.1 | 5.83E-01 | 0.0 | 9.47E-01 | 0.1 | 6.83E-01 | 0.0 | 8.36E-01 | 0.1 | 5.90E-01 | SCL30A |
| AT3G04610 | 0.1 | 2.63E-01 | -0.1 | 5.67E-02 | 0.2 | 1.46E-02 | 0.0 | 9.17E-01 | -0.2 | 4.58E-01 | 0.2 | 6.08E-01 | RNA-binding KH domain-containing protein |
| AT1G16610 | 0.1 | 4.12E-01 | 0.0 | 9.03E-01 | 0.1 | 5.53E-01 | 0.1 | 3.88E-01 | -0.1 | 5.09E-01 | 0.2 | 2.40E-01 | Arginine |
| AT4G24880 | 0.1 | 4.26E-01 | 0.0 | 8.91E-01 | 0.1 | 6.07E-01 | -0.1 | 8.34E-01 | -0.1 | 7.49E-01 | 0.0 | 9.14E-01 | Snurportin-1 |
| AT2G21660 | 0.1 | 5.93E-01 | 0.5 | 1.00E-02 | -0.4 | 3.48E-02 | -0.4 | 3.04E-02 | -0.3 | 9.92E-02 | -0.1 | 8.65E-01 | Cold, circadian rhythm, and rna binding 2 |
| AT1G30480 | 0.1 | 2.76E-01 | -0.1 | 4.10E-01 | 0.2 | 1.09E-01 | 0.0 | 8.16E-01 | -0.1 | 4.78E-01 | 0.1 | 7.04E-01 | DNA-damage-repair |
| AT5G09880 | 0.1 | 2.81E-01 | -0.1 | 2.70E-01 | 0.2 | 6.14E-02 | 0.1 | 8.01E-01 | 0.0 | 8.66E-01 | 0.1 | 7.49E-01 | Splicing factor-like protein |
| AT5G19020 | 0.1 | 5.65E-01 | 0.0 | 8.61E-01 | 0.1 | 5.42E-01 | 0.0 | 9.72E-01 | 0.2 | 5.75E-01 | -0.2 | 6.54E-01 | Pentatricopeptide repeat-containing protein At5g19020, mitochondrial |
| AT4G14300 | 0.1 | 1.73E-01 | 0.2 | 4.53E-03 | -0.2 | 6.26E-02 | -0.1 | 3.72E-01 | -0.2 | 2.72E-01 | 0.0 | 8.55E-01 | RRM domain-containing protein |
| AT4G00420 | 0.1 | 2.91E-01 | 0.2 | 1.16E-01 | -0.1 | 6.19E-01 | 0.1 | 8.23E-01 | -0.1 | 6.15E-01 | 0.2 | 4.88E-01 | Double-stranded RNA-binding domain -containing protein |
| AT3G25150 | 0.1 | 1.74E-01 | 0.1 | 5.63E-02 | 0.0 | 5.39E-01 | -0.2 | 1.22E-01 | -0.2 | 7.46E-02 | 0.1 | 7.88E-01 | RNA-binding protein-like |
| AT2G02570 | 0.1 | 1.71E-01 | 0.0 | 6.87E-01 | 0.1 | 3.99E-01 | -0.1 | 5.28E-01 | 0.0 | 9.49E-01 | -0.1 | 6.39E-01 | Nucleic acid binding |
| AT5G54890 | 0.1 | 3.52E-01 | -0.1 | 2.35E-01 | 0.2 | 8.68E-02 | -0.1 | 8.15E-01 | 0.2 | 4.07E-01 | -0.3 | 3.46E-01 | CRS2-associated factor 2, mitochondrial |
| AT3G27550 | 0.1 | 4.82E-01 | 0.0 | 8.33E-01 | 0.1 | 4.54E-01 | 0.8 | 2.62E-03 | 0.5 | 9.31E-02 | 0.3 | 2.40E-01 | RNA-binding CRS1 |
| AT4G16280 | 0.1 | 3.04E-01 | -0.2 | 1.06E-01 | 0.3 | 3.22E-02 | -0.1 | 4.82E-01 | -0.1 | 6.12E-01 | 0.0 | 9.64E-01 | Flowering time control protein FCA |
| AT1G07350 | 0.1 | 7.85E-02 | -0.3 | 7.77E-04 | 0.4 | 2.93E-04 | -0.1 | 3.95E-01 | 0.0 | 9.96E-01 | -0.1 | 5.31E-01 | RRM domain-containing protein |
| AT1G63810 | 0.1 | 2.54E-01 | 0.0 | 7.07E-01 | 0.1 | 5.40E-01 | 0.9 | 7.65E-04 | 0.4 | 1.53E-01 | 0.5 | 6.71E-02 | Nucleolar protein |

| | | | | | | | | | | | | | |
|---|---|---|---|---|---|---|---|---|---|---|---|---|---|
| AT4G12640 | 0.1 | 1.28E-01 | -0.1 | 2.51E-01 | 0.2 | 3.27E-02 | -0.1 | 6.80E-01 | 0.1 | 6.68E-01 | -0.3 | 4.59E-01 | Uncharacterized protein AT4g12630 |
| AT3G52140 | 0.1 | 1.26E-01 | -0.1 | 4.28E-01 | 0.2 | 4.63E-02 | 0.4 | 3.05E-03 | 0.1 | 4.94E-01 | 0.3 | 9.07E-02 | Clustered mitochondria protein homolog |
| AT3G01160 | 0.1 | 2.33E-01 | 0.1 | 2.42E-01 | 0.0 | 9.89E-01 | 0.5 | 5.09E-03 | 0.2 | 3.02E-01 | 0.3 | 1.82E-01 | Pre-rRNA-processing ESF1-like protein |
| AT5G55100 | 0.1 | 8.97E-02 | -0.1 | 1.05E-01 | 0.3 | 1.02E-02 | -0.2 | 4.17E-01 | -0.4 | 4.47E-02 | 0.3 | 2.15E-01 | SURP motif domain-containing protein |
| AT4G10110 | 0.1 | 1.48E-01 | 0.1 | 1.37E-01 | 0.0 | 9.43E-01 | 0.1 | 4.38E-01 | 0.1 | 5.43E-01 | 0.0 | 9.80E-01 | Uncharacterized protein AT4g10110 |
| AT1G52730 | 0.1 | 1.03E-01 | 0.1 | 1.46E-01 | 0.0 | 9.04E-01 | 0.1 | 2.98E-01 | 0.2 | 9.31E-02 | -0.1 | 5.89E-01 | Transducin |
| AT2G25355 | 0.1 | 4.41E-01 | 0.1 | 6.70E-01 | 0.1 | 8.10E-01 | 0.4 | 1.39E-01 | 0.1 | 6.95E-01 | 0.2 | 4.32E-01 | Ribosomal RNA-processing protein 40 |
| AT4G36690 | 0.1 | 8.99E-02 | -0.1 | 4.79E-01 | 0.2 | 3.92E-02 | 0.0 | 9.40E-01 | -0.1 | 6.95E-01 | 0.1 | 8.15E-01 | Splicing factor like protein |
| AT4G19610 | 0.1 | 9.94E-02 | 0.0 | 8.75E-01 | 0.2 | 1.21E-01 | 0.1 | 2.91E-01 | 0.1 | 7.46E-01 | 0.1 | 7.11E-01 | Nucleotide |
| AT5G48040 | 0.1 | 3.20E-01 | 0.0 | 8.41E-01 | 0.1 | 5.41E-01 | 0.1 | 7.41E-01 | -0.1 | 8.99E-01 | 0.2 | 6.28E-01 | hypothetical protein |
| AT2G21440 | 0.1 | 5.93E-02 | 0.1 | 4.77E-01 | 0.1 | 2.80E-01 | 0.4 | 1.64E-02 | 0.1 | 4.50E-01 | 0.2 | 2.46E-01 | Expressed protein family protein) |
| AT2G35030 | 0.1 | 3.47E-01 | 0.1 | 5.28E-01 | 0.0 | 8.40E-01 | 0.3 | 4.20E-01 | 0.3 | 3.77E-01 | -0.1 | 9.18E-01 | Pentatricopeptide repeat-containing protein At2g35030, mitochondrial |
| AT3G21740 | 0.2 | 1.38E-01 | 0.2 | 1.12E-01 | 0.0 | 8.93E-01 | -0.3 | 2.28E-01 | -0.2 | 4.62E-01 | -0.1 | 8.37E-01 | APO domain-containing protein |
| AT3G25440 | 0.2 | 1.96E-01 | 0.1 | 3.11E-01 | 0.0 | 8.55E-01 | -0.3 | 1.84E-01 | -0.3 | 2.65E-01 | 0.0 | 9.66E-01 | hypothetical protein |
| AT1G03530 | 0.2 | 1.75E-01 | -0.1 | 6.37E-01 | 0.2 | 1.26E-01 | 0.3 | 2.63E-01 | -0.1 | 8.43E-01 | 0.4 | 2.71E-01 | H |
| AT2G03870 | 0.2 | 5.46E-02 | 0.3 | 1.05E-02 | -0.1 | 3.79E-01 | 0.0 | 9.69E-01 | -0.1 | 6.38E-01 | 0.1 | 7.02E-01 | Sm-like protein LSM7 |
| AT1G15470 | 0.2 | 1.08E-01 | 0.3 | 1.33E-02 | -0.1 | 2.90E-01 | 0.1 | 7.84E-01 | 0.3 | 5.41E-02 | -0.3 | 1.61E-01 | Transducin |
| AT1G43190 | 0.2 | 1.36E-02 | 0.2 | 1.01E-02 | 0.0 | 8.06E-01 | 0.1 | 4.56E-01 | 0.0 | 9.08E-01 | 0.1 | 7.32E-01 | Polypyrimidine tract-binding protein 3 |
| AT5G43790 | 0.2 | 5.39E-02 | 0.1 | 3.22E-01 | 0.1 | 4.17E-01 | 0.2 | 3.32E-01 | 0.3 | 3.18E-01 | 0.0 | 9.30E-01 | Pentatricopeptide repeat-containing protein At5g43790 |
| AT2G30260 | 0.2 | 6.84E-02 | 0.3 | 1.30E-02 | -0.1 | 3.78E-01 | -0.1 | 3.90E-01 | -0.2 | 3.36E-01 | 0.0 | 9.04E-01 | U2 small nuclear ribonucleoprotein B |
| AT4G26480 | 0.2 | 1.87E-02 | 0.1 | 7.68E-02 | 0.1 | 5.37E-01 | 0.0 | 8.59E-01 | -0.2 | 1.32E-01 | 0.2 | 2.88E-01 | RNA-binding KH domain-containing protein |
| AT1G74230 | 0.2 | 3.12E-02 | 0.2 | 6.70E-02 | 0.0 | 7.79E-01 | 0.1 | 4.89E-01 | 0.0 | 9.89E-01 | 0.1 | 6.51E-01 | RRM domain-containing protein |
| AT4G33495 | 0.2 | 5.31E-02 | -0.2 | 1.34E-01 | 0.4 | 1.10E-02 | 0.4 | 8.99E-02 | -0.1 | 8.47E-01 | 0.4 | 9.56E-02 | Protein ROOT PRIMORDIUM DEFECTIVE 1 |
| AT5G12190 | 0.2 | 1.67E-01 | 0.0 | 8.16E-01 | 0.2 | 1.70E-01 | 0.4 | 4.19E-02 | 0.2 | 2.69E-01 | 0.1 | 5.71E-01 | Splicing factor 3B subunit 6-like protein |
| AT3G04500 | 0.2 | 2.24E-02 | 0.2 | 6.72E-02 | 0.0 | 6.58E-01 | 0.1 | 7.08E-01 | 0.1 | 8.35E-01 | 0.0 | 9.58E-01 | hypothetical protein |
| AT4G04370 | 0.2 | 2.33E-01 | 0.0 | 8.82E-01 | 0.2 | 4.15E-01 | 0.0 | 9.57E-01 | 0.2 | 7.86E-01 | -0.2 | 7.30E-01 | Pentatricopeptide repeat-containing protein At4g04370 |
| AT3G20890 | 0.2 | 6.74E-02 | 0.1 | 6.56E-01 | 0.2 | 2.33E-01 | 0.1 | 4.46E-01 | -0.1 | 5.97E-01 | 0.2 | 3.03E-01 | RRM domain-containing protein |
| AT2G28480 | 0.2 | 6.20E-02 | 0.0 | 8.75E-01 | 0.2 | 9.11E-02 | 0.3 | 2.64E-01 | 0.2 | 5.80E-01 | 0.1 | 7.79E-01 | hypothetical protein |
| AT5G18440 | 0.2 | 1.99E-02 | 0.1 | 4.00E-01 | 0.1 | 1.62E-01 | 0.4 | 3.07E-02 | 0.3 | 1.12E-01 | 0.1 | 8.00E-01 | GATA zinc finger protein |
| AT5G10800 | 0.2 | 2.20E-01 | 0.2 | 2.26E-01 | 0.0 | 9.73E-01 | -0.6 | 2.08E-01 | -0.5 | 3.89E-01 | -0.2 | 8.54E-01 | RNA recognition motif -containing protein |
| AT4G02430 | 0.2 | 3.66E-02 | -0.1 | 2.24E-01 | 0.4 | 1.14E-02 | -0.2 | 2.19E-01 | -0.1 | 6.66E-01 | -0.1 | 5.85E-01 | RNA-binding family protein |
| AT5G21970 | 0.2 | 3.80E-02 | 0.0 | 8.29E-01 | 0.3 | 5.90E-02 | 0.3 | 7.74E-02 | 0.0 | 8.80E-01 | 0.2 | 2.57E-01 | hypothetical protein |
| AT2G43410 | 0.2 | 5.62E-02 | -0.1 | 3.90E-01 | 0.3 | 2.41E-02 | 0.0 | 9.32E-01 | -0.1 | 8.41E-01 | 0.1 | 7.93E-01 | RNA binding protein |
| AT5G53060 | 0.2 | 1.78E-02 | 0.1 | 5.80E-01 | 0.2 | 9.74E-02 | 0.1 | 7.34E-01 | 0.1 | 7.86E-01 | 0.0 | 9.98E-01 | RNA-binding KH domain-containing protein |
| AT1G02140 | 0.2 | 4.17E-03 | 0.2 | 5.86E-03 | 0.0 | 9.85E-01 | 0.2 | 2.46E-01 | 0.1 | 7.59E-01 | 0.1 | 6.54E-01 | Protein mago nashi homolog |
| AT5G08420 | 0.2 | 6.10E-03 | 0.2 | 1.38E-02 | 0.0 | 7.67E-01 | 0.2 | 8.81E-02 | 0.2 | 2.68E-01 | 0.0 | 8.04E-01 | KRR1 small subunit processome component |
| AT1G53645 | 0.2 | 5.40E-02 | 0.2 | 6.19E-02 | 0.0 | 9.92E-01 | 0.6 | 3.93E-04 | 0.3 | 8.20E-02 | 0.3 | 1.16E-01 | T3F20.1 protein |
| AT2G27330 | 0.2 | 8.99E-02 | 0.3 | 2.42E-02 | -0.1 | 5.15E-01 | 0.1 | 6.52E-01 | 0.1 | 8.31E-01 | 0.0 | 9.17E-01 | RRM domain-containing protein |
| AT4G20440 | 0.2 | 9.78E-02 | 0.3 | 6.97E-02 | 0.0 | 8.30E-01 | 0.2 | 6.47E-02 | 0.1 | 6.96E-01 | 0.2 | 3.48E-01 | Small nuclear ribonucleoprotein-associated protein |
| AT1G06380 | 0.2 | 1.35E-01 | 0.2 | 1.82E-01 | 0.0 | 9.22E-01 | 0.5 | 4.77E-02 | 0.0 | 9.36E-01 | 0.5 | 1.00E-01 | Ribosomal protein L1p |
| AT1G49600 | 0.3 | 3.94E-02 | 0.4 | 4.84E-03 | -0.2 | 2.32E-01 | 0.2 | 2.85E-02 | -0.1 | 1.84E-01 | 0.3 | 1.65E-02 | RNA-binding protein 47A |
| AT3G48670 | 0.3 | 4.81E-03 | 0.2 | 5.68E-02 | 0.1 | 2.66E-01 | 0.0 | 9.52E-01 | 0.1 | 7.09E-01 | -0.1 | 7.09E-01 | Protein INVOLVED IN DE NOVO 2 |
| AT5G32450 | 0.3 | 4.31E-03 | 0.5 | 7.88E-05 | -0.2 | 1.16E-02 | -0.1 | 3.38E-01 | -0.4 | 1.92E-02 | 0.3 | 2.38E-01 | RRM domain-containing protein |
| AT5G39350 | 0.3 | 6.85E-02 | 0.1 | 6.93E-01 | 0.2 | 2.39E-01 | 0.5 | 1.97E-01 | 0.1 | 9.11E-01 | 0.4 | 3.57E-01 | Pentatricopeptide repeat-containing protein At5g39350 |
| AT3G11500 | 0.3 | 5.50E-03 | 0.4 | 4.01E-04 | -0.2 | 1.00E-01 | 0.2 | 3.40E-01 | 0.2 | 5.30E-01 | 0.0 | 9.15E-01 | Small nuclear ribonucleoprotein G |
| AT2G34400 | 0.3 | 1.00E-01 | 0.0 | 8.17E-01 | 0.2 | 2.60E-01 | 0.4 | 1.54E-01 | 0.1 | 7.87E-01 | 0.3 | 4.06E-01 | Pentatricopeptide repeat-containing protein At2g34400 |
| AT4G18375 | 0.3 | 8.39E-02 | 0.1 | 5.61E-01 | 0.2 | 3.51E-01 | 0.3 | 4.45E-01 | 0.2 | 7.03E-01 | 0.1 | 8.41E-01 | RNA-binding KH domain-containing protein |
| AT3G20930 | 0.3 | 1.14E-02 | 0.5 | 4.63E-04 | -0.2 | 6.64E-02 | -0.1 | 7.93E-01 | 0.0 | 9.06E-01 | 0.0 | 9.34E-01 | RNA-binding family protein |
| AT5G53180 | 0.3 | 3.09E-03 | 0.1 | 2.87E-01 | 0.2 | 4.17E-02 | 0.1 | 6.65E-01 | 0.1 | 8.33E-01 | 0.0 | 9.31E-01 | Polypyrimidine tract-binding RNA transport protein-like |
| AT5G23880 | 0.3 | 6.31E-04 | 0.0 | 8.53E-01 | 0.3 | 3.27E-03 | 0.3 | 9.95E-02 | 0.3 | 2.74E-01 | 0.1 | 8.19E-01 | Cleavage and polyadenylation specificity factor subunit 2 |
| AT3G58660 | 0.3 | 8.16E-02 | 0.3 | 5.37E-02 | 0.0 | 8.00E-01 | 0.4 | 2.52E-02 | 0.1 | 5.46E-01 | 0.3 | 2.18E-01 | Ribosomal protein L1p |
| AT4G13850 | 0.3 | 5.38E-02 | 0.3 | 3.27E-02 | -0.1 | 7.53E-01 | 0.6 | 2.22E-04 | 0.3 | 5.27E-02 | 0.3 | 8.16E-02 | Glycine-rich RNA-binding protein 2 |
| AT3G66652 | 0.3 | 6.36E-03 | -0.1 | 4.20E-01 | 0.4 | 6.42E-03 | 0.2 | 4.88E-01 | 0.3 | 2.03E-01 | -0.2 | 6.45E-01 | FIP1-like protein ) polymerase-like 3) |
| AT5G02530 | 0.3 | 1.62E-03 | 0.3 | 3.07E-03 | 0.0 | 8.76E-01 | 0.1 | 5.50E-01 | -0.1 | 3.12E-01 | 0.2 | 1.97E-01 | RRM domain-containing protein |
| AT2G03640 | 0.3 | 9.82E-04 | 0.2 | 2.38E-02 | 0.1 | 1.22E-01 | -0.3 | 3.05E-02 | -0.5 | 6.06E-04 | 0.3 | 9.17E-02 | Nuclear transport factor 2 family protein with RNA binding domain-containing protein |
| AT5G56130 | 0.3 | 2.07E-03 | 0.1 | 1.63E-01 | 0.2 | 5.73E-02 | 0.2 | 3.47E-01 | 0.3 | 2.17E-01 | -0.1 | 8.15E-01 | THO complex subunit 3 |
| AT1G48920 | 0.3 | 3.89E-02 | 0.2 | 2.17E-01 | 0.1 | 4.17E-01 | 0.5 | 8.68E-04 | 0.2 | 2.13E-01 | 0.3 | 9.34E-02 | F27J15.29 |
| AT3G20430 | 0.3 | 9.32E-03 | 0.1 | 2.11E-01 | 0.2 | 1.65E-01 | 0.3 | 3.34E-01 | 0.0 | 9.75E-01 | 0.2 | 4.76E-01 | Phosphorylated adapter RNA export protein |
| AT5G46840 | 0.3 | 3.31E-04 | 0.4 | 2.54E-04 | 0.0 | 6.35E-01 | -0.1 | 3.57E-01 | -0.2 | 1.42E-01 | 0.1 | 6.87E-01 | At5g46840 family protein) |
| AT5G04430 | 0.3 | 6.12E-04 | 0.5 | 5.53E-05 | -0.2 | 3.57E-02 | 0.0 | 8.31E-01 | 0.0 | 9.87E-01 | 0.0 | 8.88E-01 | hypothetical protein |
| AT3G26540 | 0.3 | 1.33E-01 | -0.2 | 5.20E-01 | 0.5 | 8.92E-02 | | | | | | | Pentatricopeptide repeat-containing protein At3g26540 |
| AT1G21190 | 0.4 | 1.32E-02 | 0.1 | 3.70E-01 | 0.2 | 1.22E-01 | 0.3 | 2.88E-01 | 0.1 | 8.80E-01 | 0.2 | 5.74E-01 | Sm-like protein LSM3A |
| AT1G03360 | 0.4 | 2.29E-02 | 0.3 | 3.23E-02 | 0.0 | 9.36E-01 | 0.4 | 8.18E-02 | 0.1 | 7.72E-01 | 0.3 | 3.22E-01 | Exosome complex component RRP4 homolog |
| AT1G74600 | 0.4 | 2.14E-02 | 0.2 | 3.05E-01 | 0.2 | 2.42E-01 | 0.7 | 5.39E-03 | 0.2 | 5.23E-01 | 0.5 | 7.74E-02 | Pentatricopeptide repeat-containing protein At1g74600, chloroplastic |
| AT1G72320 | 0.4 | 5.53E-04 | 0.2 | 5.42E-02 | 0.2 | 3.66E-02 | 0.4 | 5.36E-03 | 0.1 | 3.51E-01 | 0.2 | 1.42E-01 | Pumilio homolog 23 |
| AT3G51320 | 0.4 | 5.94E-03 | 0.3 | 3.10E-02 | 0.1 | 5.08E-01 | 0.2 | 5.15E-01 | 0.1 | 6.95E-01 | 0.0 | 9.09E-01 | Pentatricopeptide repeat-containing protein At3g51320 |
| AT3G16230 | 0.4 | 1.88E-03 | 0.1 | 2.23E-01 | 0.3 | 3.62E-02 | 0.0 | 8.85E-01 | 0.1 | 5.89E-01 | -0.2 | 5.39E-01 | Uncharacterized protein At3g16300 |
| AT3G02950 | 0.4 | 2.64E-04 | 0.1 | 1.92E-01 | 0.3 | 6.05E-03 | 0.2 | 2.73E-01 | 0.2 | 3.07E-01 | 0.0 | 9.69E-01 | THO complex subunit 7B |
| AT4G13860 | 0.4 | 1.00E-02 | 0.3 | 3.30E-02 | 0.1 | 6.49E-01 | -1.1 | 2.60E-02 | -0.2 | 7.70E-01 | -0.9 | 1.62E-01 | RRM domain-containing protein |
| AT5G04600 | 0.4 | 1.49E-03 | 0.4 | 2.31E-03 | 0.0 | 9.48E-01 | 0.1 | 6.24E-01 | -0.1 | 5.24E-01 | 0.2 | 3.18E-01 | hypothetical protein |
| AT3G09040 | 0.4 | 1.38E-02 | 0.2 | 1.63E-01 | 0.2 | 3.08E-01 | 0.7 | 1.04E-01 | 0.2 | 7.02E-01 | 0.5 | 4.12E-01 | Pentatricopeptide repeat superfamily protein |
| AT4G15770 | 0.4 | 6.66E-03 | 0.5 | 4.85E-03 | 0.0 | 7.90E-01 | 0.6 | 3.89E-03 | 0.2 | 4.33E-01 | 0.4 | 1.07E-01 | At4g15770 |
| AT5G17930 | 0.4 | 4.40E-03 | 0.2 | 1.19E-01 | 0.2 | 1.40E-01 | 0.3 | 5.15E-02 | 0.1 | 3.94E-01 | 0.1 | 5.29E-01 | MIF4G domain-containing protein |
| AT1G49760 | 0.4 | 3.18E-04 | 0.4 | 3.82E-04 | 0.0 | 8.25E-01 | 0.3 | 3.98E-02 | 0.0 | 8.63E-01 | 0.2 | 2.13E-01 | Polyadenylate-binding protein |
| AT1G51580 | 0.4 | 6.46E-04 | 0.2 | 3.66E-02 | 0.2 | 6.61E-02 | -0.4 | 1.92E-01 | -0.5 | 1.48E-01 | 0.1 | 8.85E-01 | hypothetical protein |
| AT1G20580 | 0.4 | 2.53E-03 | 0.4 | 2.89E-03 | 0.0 | 9.39E-01 | 0.2 | 6.97E-02 | 0.0 | 9.58E-01 | 0.2 | 1.89E-01 | Small nuclear ribonucleoprotein Sm D3 |
| AT2G37310 | 0.4 | 2.86E-02 | 0.3 | 1.57E-01 | 0.2 | 4.84E-01 | 0.6 | 2.16E-02 | 0.4 | 1.41E-01 | 0.2 | 5.96E-01 | Pentatricopeptide repeat-containing protein At2g37310 |
| AT3G49430 | 0.5 | 4.73E-04 | 0.3 | 6.30E-03 | 0.1 | 1.91E-01 | 0.2 | 3.43E-02 | 0.0 | 8.77E-01 | 0.2 | 1.83E-01 | SRp34a |
| AT1G73490 | 0.5 | 6.99E-04 | 0.3 | 1.73E-02 | 0.2 | 1.17E-01 | 0.4 | 7.72E-03 | 0.1 | 4.29E-01 | 0.2 | 1.99E-01 | AT1G73490 protein family protein) |
| AT1G76860 | 0.5 | 7.18E-05 | 0.4 | 6.28E-04 | 0.1 | 2.36E-01 | 0.0 | 9.30E-01 | -0.2 | 3.23E-01 | 0.3 | 3.74E-01 | Sm-like protein LSM3B |
| AT4G22380 | 0.5 | 2.44E-03 | 0.2 | 7.48E-02 | 0.2 | 1.29E-01 | 0.7 | 1.00E-02 | 0.4 | 1.77E-01 | 0.3 | 3.37E-01 | Ribonucleoprotein |
| AT2G33680 | 0.5 | 9.01E-03 | 0.1 | 7.22E-01 | 0.4 | 5.01E-02 | 0.5 | 1.28E-01 | 0.5 | 1.63E-01 | 0.0 | 9.92E-01 | Tetratricopeptide repeat -like superfamily protein |
| AT1G13730 | 0.5 | 7.66E-04 | 0.5 | 6.29E-04 | 0.0 | 7.30E-01 | 0.2 | 1.45E-01 | 0.1 | 3.99E-01 | 0.1 | 8.09E-01 | hypothetical protein |

| | | | | | | | | | | | | | |
|---|---|---|---|---|---|---|---|---|---|---|---|---|---|
| AT3G13230 | 0.5 | 8.91E-03 | 0.5 | 1.44E-02 | 0.0 | 8.98E-01 | 0.6 | 4.48E-03 | 0.2 | 5.57E-01 | 0.5 | 5.70E-02 | hypothetical protein |
| AT4G17720 | 0.5 | 2.12E-04 | 0.2 | 2.80E-02 | 0.3 | 2.03E-02 | 0.2 | 1.45E-01 | -0.1 | 5.50E-01 | 0.3 | 8.56E-02 | Putative RRM-containing protein family protein) |
| AT4G23540 | 0.5 | 1.81E-04 | -0.1 | 3.90E-01 | 0.6 | 4.58E-04 | 0.0 | 9.85E-01 | 0.1 | 9.31E-01 | -0.1 | 9.57E-01 | ARM repeat superfamily protein |
| AT2G36730 | 0.5 | 3.93E-03 | 0.4 | 2.42E-02 | 0.1 | 4.76E-01 | 0.2 | 4.68E-01 | 0.2 | 6.46E-01 | 0.0 | 9.29E-01 | Pentatricopeptide repeat-containing protein At2g36730 |
| AT1G47500 | 0.5 | 1.48E-04 | 0.2 | 2.77E-02 | 0.3 | 1.44E-02 | 0.3 | 6.29E-02 | 0.2 | 2.97E-01 | 0.1 | 6.76E-01 | RNA-binding protein 47C |
| AT1G05750 | 0.6 | 5.18E-03 | 0.2 | 2.78E-01 | 0.4 | 9.52E-02 | 0.2 | 6.44E-01 | -0.1 | 7.13E-01 | 0.3 | 3.73E-01 | Pentatricopeptide repeat-containing protein At1g05750, chloroplastic |
| AT1G65700 | 0.6 | 4.43E-04 | 0.4 | 7.81E-03 | 0.2 | 1.71E-01 | 0.4 | 2.22E-01 | 0.1 | 7.60E-01 | 0.2 | 5.74E-01 | U6 snRNA-associated Sm-like protein LSm8 |
| AT3G14730 | 0.6 | 1.13E-03 | 0.3 | 7.44E-02 | 0.3 | 7.82E-02 | -0.2 | 5.38E-01 | -0.1 | 8.84E-01 | -0.1 | 7.58E-01 | Pentatricopeptide repeat-containing protein At3g14730 |
| AT2G18740 | 0.6 | 5.65E-05 | 0.4 | 1.99E-03 | 0.2 | 5.01E-02 | 0.6 | 7.89E-03 | 0.3 | 3.23E-01 | 0.4 | 2.40E-01 | Small nuclear ribonucleoprotein E |
| AT1G50270 | 0.6 | 2.26E-03 | 0.4 | 2.76E-02 | 0.2 | 3.05E-01 | -0.2 | 6.13E-01 | 0.2 | 6.37E-01 | -0.4 | 3.94E-01 | Pentatricopeptide repeat-containing protein At1g50270 |
| AT3G07590 | 0.6 | 6.36E-04 | 0.3 | 3.64E-02 | 0.3 | 6.90E-02 | 0.5 | 6.33E-02 | -0.2 | 6.81E-01 | 0.7 | 6.34E-02 | Small nuclear ribonucleoprotein SmD1a |
| AT2G23350 | 0.6 | 4.31E-05 | 0.5 | 2.75E-04 | 0.1 | 3.29E-01 | 0.5 | 3.94E-03 | 0.3 | 1.28E-01 | 0.2 | 3.68E-01 | Putative poly binding protein |
| AT4G20030 | 0.6 | 1.47E-04 | 0.3 | 1.99E-02 | 0.3 | 2.48E-02 | -0.1 | 8.16E-01 | -0.1 | 9.04E-01 | 0.0 | 9.64E-01 | At4g20030 family protein) |
| AT3G08010 | 0.6 | 4.07E-05 | 0.5 | 4.56E-04 | 0.2 | 1.55E-01 | 0.1 | 5.80E-01 | 0.1 | 6.61E-01 | 0.0 | 9.98E-01 | Protein TAB2 homolog, chloroplastic |
| AT1G09190 | 0.6 | 2.34E-03 | 0.4 | 6.22E-02 | 0.3 | 1.70E-01 | 0.8 | 1.57E-02 | 0.6 | 9.32E-02 | 0.2 | 6.63E-01 | Pentatricopeptide repeat-containing protein At1g09190 |
| AT4G32175 | 0.7 | 6.07E-04 | 0.4 | 1.08E-02 | 0.2 | 1.93E-01 | 0.4 | 1.28E-01 | 0.1 | 7.91E-01 | 0.3 | 3.55E-01 | Ribosomal RNA-processing protein 40 |
| AT3G58520 | 0.7 | 1.11E-03 | 0.2 | 3.50E-01 | 0.5 | 2.09E-02 | 0.4 | 3.20E-01 | 0.0 | 9.68E-01 | 0.4 | 4.13E-01 | Uncharacterized protein F14P22.110 |
| AT1G03330 | 0.7 | 2.78E-05 | 0.5 | 4.14E-04 | 0.2 | 1.12E-01 | 0.2 | 3.53E-01 | 0.0 | 9.71E-01 | 0.2 | 5.61E-01 | Sm-like protein LSM2 |
| AT2G37510 | 0.7 | 1.36E-03 | 0.6 | 3.04E-03 | 0.0 | 8.26E-01 | 0.7 | 1.42E-01 | 0.3 | 6.13E-01 | 0.4 | 3.62E-01 | RNA-binding family protein |
| AT2G47610 | 0.7 | 6.72E-04 | 0.7 | 8.40E-04 | 0.0 | 8.91E-01 | 0.4 | 1.07E-02 | 0.2 | 2.30E-01 | 0.2 | 3.79E-01 | 60S ribosomal protein L7a |
| AT2G46610 | 0.7 | 1.77E-04 | 0.2 | 2.61E-01 | 0.5 | 4.66E-03 | 0.0 | 9.88E-01 | 0.0 | 9.90E-01 | 0.0 | 9.90E-01 | RNA-binding family protein |
| AT3G62870 | 0.7 | 2.04E-04 | 0.7 | 4.81E-04 | 0.0 | 8.35E-01 | 0.4 | 3.25E-03 | 0.2 | 3.33E-01 | 0.3 | 1.65E-01 | 60S ribosomal protein L7a |
| AT5G48870 | 0.7 | 1.87E-04 | 0.4 | 8.61E-03 | 0.3 | 6.12E-02 | 0.7 | 4.43E-03 | 0.4 | 1.45E-01 | 0.3 | 2.89E-01 | Sm-like protein LSM5 |
| AT3G09160 | 0.7 | 6.32E-02 | 0.1 | 7.56E-01 | 0.6 | 2.00E-01 | | | | | | | MZB10.19 protein |
| AT1G06960 | 0.7 | 1.62E-05 | 0.5 | 4.77E-04 | 0.2 | 4.60E-02 | 0.0 | 8.03E-01 | -0.1 | 6.72E-01 | 0.0 | 8.84E-01 | RNA-binding family protein |
| AT5G04810 | 0.7 | 5.90E-05 | 0.6 | 5.87E-04 | 0.2 | 1.93E-01 | -0.1 | 7.46E-01 | 0.0 | 9.22E-01 | 0.0 | 8.88E-01 | Pentatricopeptide repeat-containing protein |
| AT1G53120 | 0.8 | 1.52E-05 | 0.2 | 3.72E-02 | 0.5 | 1.17E-03 | -0.5 | 2.59E-03 | -0.4 | 4.81E-02 | -0.2 | 4.31E-01 | RNA-binding S4 domain-containing protein |
| AT5G20160 | 0.8 | 3.17E-04 | 0.4 | 2.45E-02 | 0.4 | 3.48E-02 | 0.7 | 6.07E-04 | 0.3 | 1.90E-01 | 0.4 | 6.98E-02 | Ribonucleoprotein |
| AT3G46020 | 0.8 | 1.14E-03 | 0.7 | 2.72E-03 | 0.1 | 7.94E-01 | 0.0 | 9.91E-01 | -0.2 | 7.62E-01 | 0.2 | 7.88E-01 | RRM domain-containing protein |
| AT5G57930 | 0.8 | 1.24E-05 | 0.6 | 1.22E-04 | 0.2 | 1.52E-01 | 0.0 | 7.87E-01 | 0.0 | 9.02E-01 | -0.1 | 7.61E-01 | Emb1629 |
| AT2G14285 | 0.8 | 2.05E-05 | 0.4 | 5.08E-03 | 0.4 | 6.48E-03 | -0.1 | 7.83E-01 | 0.0 | 9.24E-01 | 0.0 | 9.22E-01 | Sm protein F |
| AT3G63370 | 0.8 | 1.79E-03 | 0.7 | 6.93E-03 | 0.1 | 6.29E-01 | 0.0 | 9.57E-01 | -0.2 | 7.67E-01 | 0.2 | 8.31E-01 | Tetratricopeptide repeat -like superfamily protein |
| AT5G27300 | 0.8 | 2.05E-03 | 0.5 | 2.93E-02 | 0.3 | 2.44E-01 | 0.0 | 8.58E-01 | -0.1 | 8.15E-01 | 0.0 | 9.53E-01 | RNA-binding protein 48 |
| AT2G23930 | 0.8 | 1.16E-04 | 0.7 | 3.67E-04 | 0.1 | 6.59E-01 | 0.4 | 6.47E-02 | 0.1 | 6.14E-01 | 0.2 | 4.16E-01 | Small nuclear ribonucleoprotein G |
| AT2G31890 | 0.9 | 1.49E-05 | 0.8 | 4.15E-05 | 0.0 | 7.78E-01 | 0.1 | 4.74E-01 | -0.1 | 7.36E-01 | 0.2 | 3.81E-01 | RAP domain-containing protein, chloroplastic |
| AT4G30220 | 0.9 | 9.80E-05 | 0.6 | 1.78E-03 | 0.3 | 1.13E-01 | 0.6 | 1.82E-02 | 0.1 | 7.40E-01 | 0.5 | 1.33E-01 | Sm protein F |
| AT4G30330 | 0.9 | 1.57E-04 | 0.9 | 3.02E-04 | 0.0 | 9.27E-01 | 0.3 | 3.14E-01 | 0.1 | 8.11E-01 | 0.2 | 6.40E-01 | Small nuclear ribonucleoprotein E |
| AT2G42920 | 0.9 | 1.88E-04 | 0.2 | 2.84E-01 | 0.7 | 5.06E-03 | -0.1 | 9.06E-01 | -0.3 | 8.17E-01 | 0.1 | 9.19E-01 | Pentatricopeptide repeat-containing protein At2g42920, chloroplastic |
| AT3G62840 | 1.0 | 1.47E-06 | 0.7 | 2.62E-05 | 0.3 | 2.35E-02 | 0.3 | 3.07E-02 | 0.0 | 9.66E-01 | 0.3 | 1.15E-01 | Small nuclear ribonucleoprotein Sm D2 |
| AT4G12600 | 1.0 | 1.11E-04 | 0.9 | 4.72E-04 | 0.1 | 4.98E-01 | 0.7 | 2.27E-03 | 0.3 | 2.86E-01 | 0.5 | 8.35E-02 | Ribonucleoprotein |
| AT1G76300 | 1.1 | 4.15E-06 | 0.5 | 2.06E-03 | 0.6 | 1.27E-03 | 0.5 | 1.77E-03 | 0.1 | 7.67E-01 | 0.5 | 2.23E-02 | Small nuclear ribonucleoprotein SmD3a |
| AT3G61620 | 1.2 | 2.02E-05 | 0.4 | 3.81E-02 | 0.8 | 1.47E-03 | 0.3 | 1.24E-01 | 0.0 | 8.92E-01 | 0.3 | 1.56E-01 | Exosome complex component RRP41 homolog |
| AT3G48500 | 1.2 | 8.40E-07 | 1.0 | 7.34E-06 | 0.2 | 8.46E-02 | 0.6 | 7.00E-04 | -0.1 | 8.25E-01 | 0.6 | 2.00E-03 | Protein PLASTID TRANSCRIPTIONALLY ACTIVE 10 |
| AT5G18810 | 1.5 | 2.19E-06 | 0.7 | 1.66E-03 | 0.8 | 8.07E-04 | 0.3 | 3.48E-01 | 0.4 | 3.01E-01 | -0.1 | 8.94E-01 | SC35-like splicing factor 28 |
| AT4G09040 | 1.7 | 6.09E-08 | 1.4 | 5.42E-07 | 0.3 | 7.33E-03 | 0.4 | 7.80E-04 | -0.1 | 3.48E-01 | 0.5 | 1.54E-03 | Uncharacterized protein AT4g09050 |
| AT3G08000 | 1.9 | 6.28E-05 | 0.9 | 1.59E-02 | 1.0 | 9.05E-03 | 0.7 | 1.72E-01 | 0.3 | 6.90E-01 | 0.4 | 4.86E-01 | RRM domain-containing protein |
| AT1G03510 | | | | | | | -0.1 | 8.85E-01 | -0.7 | 1.73E-01 | 0.6 | 2.81E-01 | Putative pentatricopeptide repeat-containing protein At1g03510 |
| AT1G06140 | | | | | | | 0.6 | 2.14E-01 | 0.5 | 2.85E-01 | 0.0 | 9.85E-01 | Pentatricopeptide repeat-containing protein At1g06140, mitochondrial |
| AT1G06440 | | | | | | | 0.9 | 1.51E-02 | 0.0 | 9.48E-01 | 1.0 | 2.12E-02 | hypothetical protein |
| AT1G09220 | | | | | | | 0.1 | 8.84E-01 | 0.0 | 9.88E-01 | 0.1 | 9.02E-01 | Uncharacterized protein |
| AT1G13410 | | | | | | | 0.6 | 7.27E-02 | 0.4 | 3.40E-01 | 0.2 | 6.23E-01 | T6J4.15 protein |
| AT1G17630 | | | | | | | 0.2 | 7.72E-01 | -0.3 | 5.73E-01 | 0.5 | 2.68E-01 | Putative pentatricopeptide repeat-containing protein At1g17630 |
| AT1G31430 | | | | | | | 0.6 | 1.23E-01 | 0.2 | 6.38E-01 | 0.3 | 4.52E-01 | Pentatricopeptide repeat-containing protein At1g31430 |
| AT1G31790 | | | | | | | 0.4 | 4.31E-01 | 0.3 | 6.57E-01 | 0.1 | 8.90E-01 | Pentatricopeptide repeat-containing protein At1g31790 |
| AT1G32415 | | | | | | | 0.6 | 1.16E-01 | 0.5 | 2.11E-01 | 0.1 | 9.21E-01 | Pentatricopeptide repeat-containing protein At1g32415, mitochondrial |
| AT2G13600 | | | | | | | 0.1 | 8.36E-01 | 0.4 | 3.97E-01 | -0.3 | 6.05E-01 | Pentatricopeptide repeat-containing protein At2g13600 |
| AT2G22410 | | | | | | | 0.4 | 2.56E-01 | 0.0 | 9.53E-01 | 0.4 | 3.04E-01 | Pentatricopeptide repeat-containing protein At2g22410, mitochondrial |
| AT2G36980 | | | | | | | 0.5 | 2.71E-01 | 0.5 | 3.36E-01 | 0.0 | 9.98E-01 | Tetratricopeptide repeat -like superfamily protein |
| AT2G37320 | | | | | | | 0.1 | 8.68E-01 | -0.3 | 3.61E-01 | 0.4 | 3.20E-01 | Pentatricopeptide repeat-containing protein At2g37320 |
| AT2G39120 | | | | | | | 0.2 | 7.20E-01 | 0.2 | 8.03E-01 | 0.0 | 9.79E-01 | Protein WHAT'S THIS FACTOR 9, mitochondrial |
| AT2G40720 | | | | | | | 0.1 | 8.45E-01 | -0.4 | 4.81E-01 | 0.5 | 4.06E-01 | Pentatricopeptide repeat-containing protein At2g40720 |
| AT3G13880 | | | | | | | 0.3 | 3.53E-01 | 0.1 | 9.19E-01 | 0.3 | 5.59E-01 | Pentatricopeptide repeat-containing protein At3g13880 |
| AT3G16610 | | | | | | | 1.2 | 1.29E-02 | 0.7 | 2.08E-01 | | | Pentatricopeptide repeat-containing protein At3g16610 |
| AT3G18610 | | | | | | | 1.8 | 5.96E-03 | 0.4 | 6.26E-01 | | | Nucleolin like 2 |
| AT3G18970 | | | | | | | 0.3 | 5.32E-01 | 0.3 | 5.74E-01 | 0.0 | 9.89E-01 | Pentatricopeptide repeat-containing protein At3g18970 |
| AT3G23370 | | | | | | | 0.5 | 2.30E-01 | 0.4 | 4.82E-01 | | | RNA-binding family protein |
| AT3G28660 | | | | | | | 0.6 | 1.77E-01 | 0.7 | 1.48E-01 | -0.1 | 8.88E-01 | Pentatricopeptide repeat-containing protein At3g28660 |
| AT3G53360 | | | | | | | -0.4 | 5.96E-01 | -0.1 | 9.22E-01 | | | Pentatricopeptide repeat-containing protein At3g53360, mitochondrial |
| AT3G58590 | | | | | | | 0.3 | 5.41E-01 | 0.3 | 5.65E-01 | | | Pentatricopeptide repeat-containing protein At3g58590 |
| AT4G14170 | | | | | | | 0.5 | 1.54E-01 | 0.2 | 6.24E-01 | 0.3 | 5.62E-01 | Pentatricopeptide repeat superfamily protein |
| AT4G24320 | | | | | | | -0.4 | 3.19E-01 | -0.8 | 7.56E-02 | 0.4 | 5.20E-01 | PORR domain-containing protein |
| AT4G32430 | | | | | | | 0.6 | 1.09E-01 | 0.4 | 3.22E-01 | 0.2 | 7.71E-01 | Pentatricopeptide repeat superfamily protein |
| AT4G39530 | | | | | | | 0.1 | 8.23E-01 | 0.1 | 7.99E-01 | 0.0 | 9.62E-01 | Pentatricopeptide repeat-containing protein At4g39530 |
| AT4G39952 | | | | | | | 0.3 | 4.62E-01 | 0.1 | 8.75E-01 | | | Pentatricopeptide repeat-containing protein At4g39952, mitochondrial |
| AT5G27110 | | | | | | | 1.1 | 1.61E-02 | 0.5 | 3.52E-01 | 0.6 | 2.14E-01 | Pentatricopeptide repeat-containing protein At5g27110 |
| AT5G37570 | | | | | | | 0.8 | 3.58E-02 | 0.4 | 3.19E-01 | 0.3 | 3.70E-01 | Putative pentatricopeptide repeat-containing protein At5g37570 |
| AT5G56310 | | | | | | | 0.8 | 2.33E-02 | 0.2 | 7.18E-01 | 0.7 | 1.54E-01 | Pentatricopeptide repeat-containing protein At5g56310 |
| AT5G61800 | | | | | | | 0.3 | 4.88E-01 | -0.3 | 5.95E-01 | 0.7 | 2.62E-01 | Pentatricopeptide repeat-containing protein At5g61800 |
| AT5G66500 | | | | | | | 0.5 | 2.15E-01 | 0.0 | 9.55E-01 | 0.4 | 2.58E-01 | Pentatricopeptide repeat-containing protein At5g66500, mitochondrial |
| AT1G19120 | 0.0 | 7.69E-01 | -0.1 | 5.69E-01 | 0.0 | 8.16E-01 | -0.1 | 7.07E-01 | -0.1 | 5.97E-01 | 0.1 | 8.98E-01 | Sm-like protein LSM1A |
| AT1G69440 | -1.4 | 3.52E-06 | -0.9 | 7.76E-05 | -0.5 | 2.45E-02 | -0.2 | 6.66E-01 | 0.3 | 3.43E-01 | -0.5 | 1.53E-01 | Protein argonaute 7 |
| AT4G15520 | 0.2 | 1.36E-01 | 0.2 | 1.01E-01 | 0.0 | 8.61E-01 | -0.1 | 7.60E-01 | -0.2 | 2.64E-01 | 0.2 | 5.20E-01 | OBP33pep like protein |
| AT4G38020 | 0.2 | 1.80E-01 | 0.3 | 6.91E-02 | -0.1 | 6.28E-01 | 0.0 | 9.01E-01 | 0.0 | 9.47E-01 | 0.0 | 9.84E-01 | tRNA |

| | | | | | | | | | | | | | |
|---|---|---|---|---|---|---|---|---|---|---|---|---|---|
| AT5G26880 | 0.7 | 8.07E-06 | 0.2 | 1.27E-02 | 0.4 | 8.63E-04 | 0.2 | 1.78E-01 | 0.1 | 5.13E-01 | 0.1 | 7.10E-01 | AGL26 |
| AT2G17510 | 0.0 | 9.03E-01 | -0.2 | 3.57E-02 | 0.2 | 6.62E-02 | 0.0 | 9.01E-01 | -0.1 | 5.91E-01 | 0.1 | 7.31E-01 | Ribonuclease II family protein |
| AT1G44910 | 0.0 | 5.11E-01 | -0.1 | 5.44E-02 | 0.1 | 2.11E-01 | -0.1 | 6.49E-01 | -0.1 | 5.25E-01 | 0.0 | 8.82E-01 | Pre-mRNA-processing protein 40A |
| AT2G19930 | -1.0 | 3.09E-05 | -0.8 | 1.97E-04 | -0.2 | 3.09E-01 | -0.3 | 4.47E-01 | 0.1 | 8.56E-01 | -0.3 | 4.13E-01 | RNA-dependent RNA polymerase |
| AT1G14790 | -0.4 | 6.33E-03 | -0.5 | 2.44E-03 | 0.1 | 5.19E-01 | 0.0 | 8.63E-01 | 0.0 | 9.10E-01 | 0.0 | 9.93E-01 | RNA-dependent RNA polymerase 1 |
| AT2G19910 | 0.6 | 3.92E-03 | 0.3 | 9.05E-02 | 0.3 | 1.93E-01 | 0.1 | 8.78E-01 | -0.3 | 7.22E-01 | | | RNA-dependent RNA polymerase |
| AT5G66360 | 0.0 | 6.55E-01 | 0.0 | 7.12E-01 | 0.0 | 9.67E-01 | -0.3 | 1.52E-01 | -0.1 | 5.62E-01 | -0.1 | 6.28E-01 | rRNA adenine N-methyltransferase |
| AT2G47420 | 0.5 | 2.01E-04 | 0.4 | 1.15E-03 | 0.1 | 3.95E-01 | 0.1 | 4.86E-01 | -0.1 | 2.57E-01 | 0.2 | 1.34E-01 | Ribosomal RNA small subunit methyltransferase -N |
| AT1G01860 | 0.7 | 2.93E-03 | 0.3 | 1.10E-01 | 0.3 | 1.23E-01 | 0.3 | 5.31E-01 | 0.2 | 7.52E-01 | 0.1 | 8.74E-01 | rRNA adenine N-methyltransferase |
| AT5G55920 | 0.4 | 4.00E-02 | 0.4 | 3.21E-02 | 0.0 | 8.73E-01 | 0.9 | 7.59E-06 | 0.5 | 6.54E-03 | 0.4 | 1.61E-02 | 25S rRNA )-methyltransferase NOP2A |
| AT1G52930 | 0.4 | 3.11E-03 | 0.4 | 5.02E-03 | 0.0 | 9.20E-01 | 0.6 | 4.68E-03 | 0.4 | 9.23E-02 | 0.2 | 3.92E-01 | Ribosome biogenesis protein BRX1 homolog 2 |
| AT5G58420 | 0.5 | 4.61E-03 | 0.6 | 1.89E-03 | -0.1 | 4.83E-01 | 0.4 | 1.50E-03 | 0.3 | 1.92E-02 | 0.1 | 6.21E-01 | Small ribosomal subunit protein eS4x |
| AT4G26600 | 0.5 | 6.79E-03 | 0.4 | 2.80E-02 | 0.1 | 5.70E-01 | 0.9 | 4.13E-04 | 0.2 | 4.29E-01 | 0.7 | 1.02E-02 | S-adenosyl-L-methionine-dependent methyltransferases superfamily protein |
| AT4G23920 | -0.8 | 9.94E-05 | -0.6 | 8.07E-04 | -0.2 | 2.60E-01 | 0.1 | 7.97E-01 | 0.3 | 1.45E-01 | -0.3 | 2.85E-01 | UDP-glucose 4-epimerase 2 |
| AT2G37020 | 0.7 | 1.41E-04 | 0.6 | 7.18E-04 | 0.1 | 4.53E-01 | 0.5 | 1.36E-02 | 0.3 | 2.06E-01 | 0.2 | 3.98E-01 | Translin-like protein |
| AT3G27750 | 0.7 | 3.18E-04 | 1.0 | 4.68E-05 | -0.3 | 8.22E-02 | 0.2 | 5.75E-01 | -0.2 | 6.45E-01 | 0.3 | 3.53E-01 | Protein THYLAKOID ASSEMBLY 8, chloroplastic |
| AT5G23535 | 0.2 | 1.94E-01 | 0.4 | 2.18E-02 | -0.2 | 2.65E-01 | 0.5 | 4.29E-03 | 0.3 | 1.06E-01 | 0.2 | 2.92E-01 | hypothetical protein |
| AT3G49080 | 0.3 | 1.28E-01 | 0.5 | 6.53E-03 | -0.3 | 1.31E-01 | 0.5 | 2.62E-03 | 0.3 | 9.29E-02 | 0.2 | 2.74E-01 | Small ribosomal subunit protein uS9m |
| AT5G22440 | 0.3 | 1.58E-01 | 0.7 | 8.06E-03 | -0.4 | 1.16E-01 | 0.2 | 2.50E-01 | 0.2 | 3.24E-01 | 0.0 | 9.97E-01 | Large ribosomal subunit protein uL1x |
| AT1G64880 | 0.5 | 5.92E-04 | 0.4 | 2.05E-03 | 0.1 | 5.97E-01 | 0.5 | 2.48E-04 | 0.3 | 2.71E-02 | 0.2 | 2.09E-01 | S5 DRBM domain-containing protein |
| AT5G09510 | 0.5 | 7.60E-03 | 0.6 | 3.07E-03 | -0.1 | 5.13E-01 | 0.5 | 3.36E-04 | 0.2 | 1.33E-01 | 0.3 | 8.56E-02 | Ribosomal protein S19 family protein |
| AT3G04230 | 0.6 | 7.55E-04 | 0.6 | 1.27E-03 | 0.0 | 9.36E-01 | 0.5 | 6.32E-02 | 0.3 | 3.75E-01 | 0.2 | 5.36E-01 | Small ribosomal subunit protein uS9y |
| AT4G27090 | 0.6 | 2.40E-04 | 0.6 | 2.10E-04 | -0.1 | 6.23E-01 | 0.1 | 4.42E-01 | 0.0 | 8.88E-01 | 0.2 | 4.95E-01 | Ribosomal protein L14e domain-containing protein |
| AT1G80750 | 0.6 | 4.50E-04 | 0.5 | 2.38E-03 | 0.1 | 4.29E-01 | 0.6 | 8.72E-04 | 0.2 | 1.89E-01 | 0.4 | 6.56E-02 | Large ribosomal subunit protein uL30z |
| AT1G08360 | 0.6 | 6.90E-04 | 0.7 | 5.79E-04 | -0.1 | 6.78E-01 | 0.4 | 8.38E-03 | 0.1 | 3.83E-01 | 0.2 | 2.16E-01 | Large ribosomal subunit protein uL1z |
| AT5G62300 | 0.6 | 8.39E-04 | 0.7 | 1.11E-03 | 0.0 | 9.30E-01 | 0.1 | 4.24E-01 | 0.0 | 9.55E-01 | 0.1 | 6.45E-01 | 40S ribosomal protein S20 |
| AT1G04270 | 0.7 | 1.53E-03 | 0.7 | 1.68E-03 | 0.0 | 8.67E-01 | 0.2 | 1.13E-01 | 0.1 | 5.73E-01 | 0.1 | 5.74E-01 | Cytosolic ribosomal protein S15 |
| AT5G23900 | 0.7 | 4.86E-04 | 0.7 | 7.85E-04 | 0.0 | 9.97E-01 | 0.1 | 7.90E-01 | -0.1 | 4.58E-01 | 0.2 | 3.80E-01 | Large ribosomal subunit protein eL13x |
| AT2G27530 | 0.7 | 1.01E-04 | 0.7 | 1.97E-04 | 0.0 | 9.63E-01 | 0.4 | 2.58E-02 | 0.1 | 5.63E-01 | 0.3 | 3.04E-01 | Large ribosomal subunit protein uL1y |
| AT1G61580 | 0.9 | 2.14E-05 | 0.5 | 2.00E-03 | 0.4 | 1.30E-02 | 1.0 | 6.52E-07 | 0.7 | 3.26E-04 | 0.4 | 2.18E-02 | Large ribosomal subunit protein uL3y |
| AT5G67510 | 1.0 | 5.78E-06 | 0.6 | 6.07E-04 | 0.5 | 4.80E-03 | 0.4 | 4.39E-02 | 0.1 | 6.85E-01 | 0.3 | 2.92E-01 | Large ribosomal subunit protein uL24y |
| AT1G22780 | 1.1 | 7.93E-06 | 1.0 | 2.45E-05 | 0.1 | 7.14E-01 | 0.3 | 1.13E-01 | 0.1 | 7.44E-01 | 0.2 | 4.61E-01 | 40S ribosomal protein S18 |
| AT5G54600 | 1.9 | 1.04E-07 | 1.6 | 7.87E-07 | 0.3 | 3.30E-02 | 0.2 | 4.55E-02 | 0.1 | 5.41E-01 | 0.1 | 3.72E-01 | Translation protein SH3-like family protein |
| AT5G14320 | 2.0 | 4.10E-08 | 1.6 | 3.37E-07 | 0.4 | 8.23E-03 | 0.1 | 5.69E-01 | -0.1 | 6.94E-01 | 0.1 | 3.96E-01 | Ribosomal protein S13 |
| AT4G33865 | 1.5 | 1.70E-06 | 1.0 | 4.58E-05 | 0.5 | 1.32E-02 | 0.5 | 3.89E-02 | 0.2 | 4.63E-01 | 0.3 | 4.43E-01 | 40S ribosomal protein S29 |
| AT3G60240 | -0.1 | 4.02E-02 | -0.3 | 5.19E-04 | 0.2 | 1.89E-02 | 0.1 | 4.21E-01 | 0.2 | 1.85E-01 | -0.1 | 6.93E-01 | Eukaryotic translation initiation factor 4G |
| AT5G54940 | -0.1 | 6.19E-01 | -0.2 | 2.75E-01 | 0.1 | 5.71E-01 | 0.7 | 2.90E-03 | 0.2 | 5.62E-01 | 0.5 | 8.41E-02 | AT5g54940 |
| AT5G35680 | 0.0 | 9.32E-01 | 0.0 | 9.97E-01 | 0.0 | 9.42E-01 | 0.4 | 7.10E-02 | 0.2 | 4.80E-01 | 0.2 | 5.29E-01 | Eukaryotic translation initiation factor 4C |
| AT2G40290 | 0.4 | 2.48E-03 | 0.4 | 1.97E-03 | 0.0 | 7.52E-01 | 0.2 | 8.80E-02 | 0.0 | 9.75E-01 | 0.2 | 2.49E-01 | At2g40290 |
| AT3G11400 | 0.7 | 1.44E-05 | 0.6 | 1.81E-04 | 0.2 | 1.12E-01 | 0.2 | 2.05E-02 | 0.1 | 6.90E-01 | 0.2 | 1.95E-01 | Eukaryotic translation initiation factor 3 subunit G |
| AT5G06000 | 0.8 | 2.60E-04 | 0.7 | 1.28E-03 | 0.1 | 4.86E-01 | 0.2 | 5.03E-01 | 0.2 | 6.09E-01 | 0.0 | 9.74E-01 | Eukaryotic translation initiation factor 3 subunit G |
| AT5G25780 | 0.2 | 7.95E-02 | 0.2 | 1.85E-01 | 0.1 | 7.15E-01 | 0.3 | 7.85E-03 | 0.0 | 7.90E-01 | 0.3 | 6.19E-02 | Eukaryotic translation initiation factor 3 subunit B |
| AT5G27640 | 0.2 | 1.38E-02 | 0.1 | 1.77E-01 | 0.1 | 2.28E-01 | 0.3 | 1.75E-02 | 0.0 | 8.10E-01 | 0.2 | 1.37E-01 | TRANSLATION INITIATION FACTOR 3 SUBUNIT 9-like protein |
| AT4G17610 | 0.8 | 1.16E-05 | 0.6 | 1.26E-04 | 0.2 | 1.53E-01 | 0.4 | 3.71E-02 | 0.3 | 2.90E-01 | 0.2 | 5.42E-01 | tRNA |
| AT3G06950 | -0.3 | 2.88E-02 | 0.2 | 1.41E-01 | -0.4 | 7.83E-03 | 0.0 | 9.42E-01 | -0.3 | 4.49E-01 | 0.3 | 4.41E-01 | tRNA pseudouridine synthase |
| AT1G20430 | 0.7 | 1.24E-03 | 0.5 | 7.92E-03 | 0.2 | 4.18E-01 | 0.2 | 3.85E-01 | 0.0 | 9.30E-01 | 0.2 | 6.14E-01 | tRNA pseudouridine synthase |
| AT2G16940 | 0.1 | 7.13E-02 | 0.0 | 4.93E-01 | 0.1 | 3.00E-01 | -0.1 | 5.06E-01 | -0.1 | 3.44E-01 | 0.0 | 8.11E-01 | Putative splicing factor |
| AT1G80670 | -0.5 | 4.46E-04 | -0.2 | 5.43E-02 | -0.3 | 2.69E-02 | -0.2 | 1.41E-01 | -0.1 | 3.50E-01 | 0.0 | 8.37E-01 | Protein RAE1 |
| AT3G45630 | 0.1 | 4.78E-01 | -0.1 | 2.22E-01 | 0.2 | 9.65E-02 | -0.1 | 6.63E-01 | 0.0 | 9.08E-01 | -0.1 | 6.52E-01 | RNA binding family protein |
| AT5G60170 | 0.1 | 1.06E-01 | -0.1 | 1.46E-01 | 0.2 | 1.41E-02 | 0.2 | 3.82E-01 | 0.2 | 3.90E-01 | 0.0 | 9.55E-01 | RNA binding family protein |
| AT1G12780 | 0.6 | 1.92E-03 | 0.4 | 2.34E-02 | 0.2 | 2.47E-01 | -0.4 | 2.29E-02 | -0.7 | 1.67E-03 | 0.3 | 2.39E-01 | Bifunctional UDP-glucose 4-epimerase and UDP-xylose 4-epimerase 1 |
| AT4G10960 | -1.8 | 1.59E-06 | -1.4 | 1.26E-05 | -0.4 | 6.81E-02 | -1.0 | 3.10E-03 | -0.4 | 1.84E-01 | -0.6 | 1.91E-01 | UDP-glucose 4-epimerase 5 |
| AT1G64810 | -1.2 | 8.81E-07 | -1.1 | 3.37E-06 | -0.1 | 4.67E-01 | -0.4 | 4.33E-02 | -0.3 | 1.19E-01 | -0.1 | 8.37E-01 | APO RNA-binding protein |
| AT3G47530 | -0.8 | 2.21E-05 | -0.8 | 4.94E-05 | 0.0 | 9.29E-01 | 0.4 | 1.63E-01 | 0.1 | 7.64E-01 | 0.3 | 4.60E-01 | Pentatricopeptide repeat-containing protein At3g47530 |
| AT5G52630 | -0.6 | 3.21E-03 | -0.2 | 1.88E-01 | -0.4 | 7.93E-02 | -0.2 | 5.01E-01 | -0.6 | 1.01E-01 | 0.4 | 3.62E-01 | Putative pentatricopeptide repeat-containing protein At5g52630 |
| AT3G03580 | -0.4 | 1.96E-02 | -0.3 | 6.59E-02 | -0.1 | 6.32E-01 | 0.2 | 5.53E-01 | 0.0 | 9.40E-01 | 0.2 | 7.21E-01 | Pentatricopeptide repeat-containing protein At3g03580 |
| AT5G50390 | -0.4 | 1.54E-03 | 0.1 | 5.99E-01 | -0.5 | 3.00E-03 | 0.0 | 8.81E-01 | 0.0 | 9.02E-01 | 0.1 | 8.37E-01 | Pentatricopeptide repeat-containing protein At5g50390, chloroplastic |
| AT4G32450 | -0.4 | 1.46E-02 | -0.4 | 2.03E-02 | 0.0 | 9.73E-01 | -0.3 | 4.91E-01 | 0.2 | 7.58E-01 | -0.5 | 3.48E-01 | Pentatricopeptide repeat-containing protein At4g32450, mitochondrial |
| AT3G46790 | -0.1 | 1.30E-01 | -0.2 | 5.50E-02 | 0.0 | 6.48E-01 | 0.3 | 2.69E-01 | 0.2 | 4.34E-01 | 0.0 | 9.13E-01 | Pentatricopeptide repeat-containing protein CRR2, chloroplastic |
| AT5G50990 | -0.1 | 6.25E-01 | 0.2 | 2.45E-01 | -0.3 | 1.76E-01 | 0.1 | 8.83E-01 | 0.2 | 5.54E-01 | -0.2 | 7.24E-01 | Pentatricopeptide repeat-containing protein At5g50990 |
| AT2G25580 | 0.0 | 9.04E-01 | 0.0 | 9.48E-01 | 0.0 | 9.70E-01 | 0.3 | 1.00E-01 | 0.1 | 6.71E-01 | 0.2 | 4.49E-01 | Pentatricopeptide repeat-containing protein At2g25580 |
| AT1G16480 | 0.1 | 5.24E-01 | 0.0 | 8.09E-01 | 0.1 | 4.80E-01 | 0.5 | 2.02E-01 | 0.4 | 4.07E-01 | 0.1 | 8.48E-01 | Tetratricopeptide repeat -like superfamily protein |
| AT4G33990 | 0.1 | 5.26E-01 | 0.1 | 7.17E-01 | 0.0 | 8.59E-01 | 0.2 | 6.33E-01 | 0.1 | 9.12E-01 | 0.1 | 8.24E-01 | Tetratricopeptide repeat -like superfamily protein |
| AT5G06160 | 0.1 | 1.11E-01 | 0.0 | 5.00E-01 | 0.2 | 5.56E-02 | 0.1 | 3.15E-01 | 0.0 | 9.81E-01 | 0.1 | 4.46E-01 | AT5G06160 protein |
| AT1G67210 | 0.1 | 2.57E-01 | -0.1 | 2.16E-01 | 0.2 | 6.08E-02 | 0.0 | 9.39E-01 | 0.1 | 9.16E-01 | -0.1 | 8.72E-01 | Proline-rich spliceosome-associated family protein |
| AT2G41080 | 0.1 | 2.49E-01 | 0.1 | 4.96E-01 | 0.0 | 7.32E-01 | 0.0 | 9.24E-01 | 0.1 | 7.11E-01 | -0.2 | 6.96E-01 | Pentatricopeptide repeat-containing protein At2g41080 |
| AT1G23860 | 0.2 | 7.26E-02 | 0.1 | 4.96E-01 | 0.1 | 3.13E-01 | -0.5 | 3.75E-03 | -0.5 | 5.00E-03 | 0.0 | 9.10E-01 | Uncharacterized protein |
| AT1G60650 | 0.2 | 4.99E-02 | 0.0 | 8.33E-01 | 0.2 | 1.24E-01 | 0.1 | 4.60E-01 | 0.2 | 4.41E-01 | 0.0 | 9.42E-01 | RNA-binding family protein with retrovirus zinc finger-like domain-containing protein |
| AT5G38600 | 0.2 | 2.13E-02 | 0.0 | 8.22E-01 | 0.2 | 6.05E-02 | -0.3 | 1.32E-01 | -0.2 | 3.06E-01 | -0.1 | 8.52E-01 | Proline-rich spliceosome-associated family protein |
| AT4G14050 | 0.2 | 3.17E-01 | 0.0 | 9.75E-01 | 0.2 | 4.46E-01 | 0.3 | 6.29E-01 | 0.3 | 6.95E-01 | 0.0 | 9.94E-01 | Pentatricopeptide repeat superfamily protein |
| AT5G03800 | 0.2 | 1.62E-01 | 0.3 | 4.38E-02 | -0.1 | 5.07E-01 | 0.2 | 4.94E-01 | 0.1 | 8.95E-01 | 0.2 | 6.29E-01 | Pentatricopeptide repeat-containing protein At5g03800 |
| AT1G15510 | 0.2 | 1.33E-01 | 0.1 | 5.41E-01 | 0.1 | 4.77E-01 | 0.6 | 1.87E-02 | 0.2 | 6.60E-01 | 0.5 | 1.16E-01 | Pentatricopeptide repeat-containing protein At1g15510, chloroplastic |
| AT1G34160 | 0.2 | 1.92E-01 | 0.3 | 6.76E-02 | -0.1 | 5.91E-01 | 0.2 | 5.96E-01 | -0.3 | 4.29E-01 | 0.4 | 1.94E-01 | Tetratricopeptide repeat -like superfamily protein |
| AT3G23330 | 0.2 | 1.95E-01 | 0.1 | 5.09E-01 | 0.1 | 6.38E-01 | 0.3 | 3.09E-01 | 0.4 | 3.35E-01 | 0.0 | 9.71E-01 | Tetratricopeptide repeat -like superfamily protein |
| AT5G44230 | 0.2 | 1.04E-01 | 0.2 | 1.52E-01 | 0.0 | 9.10E-01 | 0.2 | 4.59E-01 | 0.2 | 5.68E-01 | 0.0 | 9.69E-01 | Pentatricopeptide repeat superfamily protein |
| AT2G37340 | 0.2 | 8.96E-03 | 0.2 | 2.23E-02 | 0.0 | 7.33E-01 | -0.4 | 1.10E-02 | -0.5 | 5.25E-03 | 0.1 | 6.79E-01 | Serine |
| AT3G53500 | 0.3 | 2.77E-03 | 0.2 | 2.83E-02 | 0.1 | 2.81E-01 | 0.0 | 7.73E-01 | -0.2 | 1.33E-01 | 0.1 | 3.73E-01 | RSZ32 |
| AT3G24000 | 0.3 | 9.45E-02 | 0.2 | 2.28E-01 | 0.1 | 7.31E-01 | 0.1 | 6.50E-01 | 0.0 | 9.60E-01 | 0.2 | 6.76E-01 | Pentatricopeptide repeat-containing protein At3g24000, mitochondrial |
| AT1G29710 | 0.3 | 6.70E-02 | 0.2 | 2.75E-01 | 0.1 | 5.43E-01 | 0.1 | 7.69E-01 | 0.1 | 8.17E-01 | 0.0 | 9.96E-01 | Pentatricopeptide repeat-containing protein At1g29710, mitochondrial |
| AT3G26782 | 0.3 | 3.33E-02 | 0.0 | 9.29E-01 | 0.3 | 9.10E-02 | 0.1 | 8.61E-01 | 0.0 | 9.98E-01 | 0.1 | 8.97E-01 | Pentatricopeptide repeat-containing protein At3g26782, mitochondrial |
| AT5G65570 | 0.3 | 3.59E-02 | 0.1 | 4.31E-01 | 0.2 | 2.58E-01 | -0.3 | 2.82E-01 | -0.3 | 3.85E-01 | 0.0 | 9.60E-01 | Pentatricopeptide repeat-containing protein At5g65570 |
| AT4G02750 | 0.3 | 1.04E-01 | 0.2 | 2.68E-01 | 0.1 | 7.00E-01 | 0.3 | 4.09E-01 | 0.1 | 8.75E-01 | 0.2 | 6.18E-01 | Pentatricopeptide repeat-containing protein At4g02750 |
| AT5G04280 | 0.4 | 1.83E-04 | 0.0 | 8.12E-01 | 0.4 | 5.48E-04 | 0.2 | 1.30E-01 | 0.2 | 2.60E-01 | 0.0 | 9.19E-01 | AT5G04280 protein family protein with retrovirus zinc finger-like domain-containing protein) |

| | | | | | | | | | | | | | |
|---|---|---|---|---|---|---|---|---|---|---|---|---|---|
| AT4G13650 | 0.5 | 8.66E-03 | 0.5 | 1.88E-02 | 0.0 | 8.36E-01 | 0.4 | 2.81E-01 | 0.2 | 7.20E-01 | 0.2 | 6.74E-01 | Pentatricopeptide repeat-containing protein At4g13650 |
| AT2G24590 | 0.7 | 1.87E-05 | 0.3 | 3.57E-03 | 0.3 | 5.39E-03 | 0.4 | 5.83E-03 | 0.2 | 1.14E-01 | 0.1 | 4.57E-01 | Uncharacterized protein |
| AT4G37380 | 0.7 | 9.11E-05 | 0.4 | 6.15E-03 | 0.3 | 4.13E-02 | 0.2 | 4.32E-01 | 0.2 | 5.20E-01 | 0.0 | 9.87E-01 | Pentatricopeptide repeat-containing protein ELI1, chloroplastic |
| AT4G18750 | 0.9 | 3.10E-04 | 0.5 | 1.56E-02 | 0.4 | 7.30E-02 | 1.3 | 1.60E-02 | 0.8 | 2.39E-01 | 0.6 | 3.62E-01 | Pentatricopeptide repeat-containing protein DOT4, chloroplastic |
| AT4G14820 | 1.1 | 7.39E-06 | 0.5 | 5.06E-03 | 0.6 | 2.19E-03 | 0.4 | 1.39E-01 | -0.2 | 6.23E-01 | 0.5 | 1.01E-01 | Pentatricopeptide repeat-containing protein At4g14820 |
| AT5G16860 | 1.5 | 6.65E-05 | 0.7 | 1.53E-02 | 0.8 | 1.47E-02 | 0.5 | 1.64E-01 | 0.1 | 8.71E-01 | 0.5 | 3.82E-01 | Pentatricopeptide repeat-containing protein At5g16860 |
| AT2G02980 | 1.5 | 1.65E-06 | 1.1 | 3.78E-05 | 0.4 | 2.34E-02 | 0.6 | 5.56E-02 | 0.2 | 6.94E-01 | 0.4 | 2.86E-01 | Pentatricopeptide repeat-containing protein At2g02980, chloroplastic |
| AT1G08070 | | | | | | | -0.5 | 3.31E-01 | -0.1 | 8.71E-01 | -0.4 | 5.88E-01 | Tetratricopeptide repeat -like superfamily protein |
| AT1G20230 | | | | | | | 0.0 | 9.94E-01 | 0.0 | 9.88E-01 | | | Pentatricopeptide repeat-containing protein At1g20230 |
| AT1G25360 | | | | | | | 0.6 | 1.95E-01 | 0.5 | 2.83E-01 | 0.0 | 9.63E-01 | Pentatricopeptide repeat-containing protein At1g25360 |
| AT1G56690 | | | | | | | 1.0 | 1.40E-02 | 0.2 | 6.69E-01 | 0.7 | 9.54E-02 | Pentatricopeptide repeat-containing protein At1g56690, mitochondrial |
| AT2G01510 | | | | | | | 0.6 | 1.69E-01 | 0.0 | 9.90E-01 | 0.5 | 3.00E-01 | Pentatricopeptide repeat-containing protein At2g01510, mitochondrial |
| AT3G08820 | | | | | | | 0.8 | 4.64E-02 | 0.4 | 4.88E-01 | 0.5 | 3.20E-01 | Putative pentatricopeptide repeat-containing protein At3g08820 |
| AT3G15130 | | | | | | | 0.1 | 8.42E-01 | 0.1 | 9.28E-01 | 0.1 | 9.58E-01 | Putative pentatricopeptide repeat-containing protein At3g15130 |
| AT3G49710 | | | | | | | 0.8 | 7.47E-02 | 0.3 | 6.03E-01 | | | Pentatricopeptide repeat-containing protein At3g49710 |
| AT3G62890 | | | | | | | 0.4 | 4.28E-01 | 0.0 | 9.82E-01 | 0.5 | 4.69E-01 | Pentatricopeptide repeat-containing protein At3g62890 |
| AT4G16835 | | | | | | | 1.1 | 5.77E-02 | 0.9 | 1.50E-01 | 0.2 | 8.32E-01 | Pentatricopeptide repeat-containing protein At4g16835, mitochondrial |
| AT4G30700 | | | | | | | 0.4 | 1.72E-01 | 0.2 | 7.18E-01 | 0.3 | 4.68E-01 | Pentatricopeptide repeat-containing protein At4g30700 |
| AT4G33170 | | | | | | | 0.4 | 6.82E-01 | 0.6 | 5.48E-01 | -0.2 | 8.74E-01 | Pentatricopeptide repeat-containing protein At4g33170 |
| AT4G35130 | | | | | | | 0.6 | 1.53E-01 | 0.2 | 7.20E-01 | 0.4 | 3.99E-01 | Pentatricopeptide repeat-containing protein At4g35130, chloroplastic |
| AT4G37170 | | | | | | | 0.8 | 4.34E-02 | 0.4 | 3.50E-01 | 0.4 | 4.54E-01 | Pentatricopeptide repeat-containing protein At4g37170 |
| AT5G04780 | | | | | | | 0.5 | 8.67E-02 | 0.3 | 3.18E-01 | 0.2 | 6.98E-01 | CRAL-TRIO domain-containing protein |
| AT5G40405 | | | | | | | 0.6 | 1.06E-01 | 0.4 | 4.54E-01 | 0.3 | 6.16E-01 | Putative pentatricopeptide repeat-containing protein At5g40405 |
| AT5G52850 | | | | | | | 0.9 | 1.90E-02 | 0.4 | 4.41E-01 | 0.5 | 2.40E-01 | Pentatricopeptide repeat-containing protein At5g52850, chloroplastic |
| AT5G44200 | 0.2 | 2.34E-02 | 0.2 | 2.56E-02 | 0.0 | 9.81E-01 | 0.0 | 9.10E-01 | -0.2 | 2.42E-01 | 0.2 | 4.32E-01 | Nuclear cap-binding protein subunit 2 |
| AT3G14080 | 0.5 | 1.80E-03 | 0.3 | 1.86E-02 | 0.1 | 3.00E-01 | 0.3 | 1.05E-01 | 0.2 | 4.15E-01 | 0.1 | 6.63E-01 | U6 snRNA-associated Sm-like protein LSm1 |
| AT4G17100 | -0.2 | 1.02E-01 | -0.1 | 4.03E-01 | -0.1 | 4.98E-01 | -0.1 | 8.67E-01 | 0.3 | 7.56E-01 | -0.4 | 6.67E-01 | Poly-specific endoribonuclease-B protein |
| AT5G22100 | 0.5 | 9.72E-04 | 0.2 | 9.58E-02 | 0.3 | 4.58E-02 | 1.0 | 1.87E-04 | 0.3 | 2.58E-01 | 0.7 | 1.17E-02 | Probable RNA 3'-terminal phosphate cyclase-like protein |
| AT2G32940 | 0.7 | 3.45E-04 | 0.0 | 8.47E-01 | 0.7 | 3.24E-03 | | | | | | | Protein argonaute 6 |
| AT2G30800 | -0.3 | 7.11E-04 | -0.3 | 3.69E-03 | -0.1 | 4.57E-01 | -0.2 | 3.69E-01 | -0.2 | 3.88E-01 | 0.0 | 9.66E-01 | RNA helicase |
| AT2G01130 | -0.1 | 1.73E-01 | -0.4 | 1.61E-03 | 0.2 | 2.50E-02 | -0.1 | 7.75E-01 | -0.1 | 7.76E-01 | 0.0 | 9.73E-01 | RNA helicase |
| AT3G26560 | 0.0 | 6.35E-01 | -0.1 | 1.54E-01 | 0.1 | 3.64E-01 | 0.1 | 4.54E-01 | 0.0 | 9.80E-01 | 0.1 | 5.96E-01 | RNA helicase |
| AT1G54650 | -0.2 | 2.25E-01 | -0.1 | 3.08E-01 | 0.0 | 9.07E-01 | -0.3 | 2.00E-01 | -0.2 | 5.69E-01 | -0.1 | 7.06E-01 | tRNA N-methylcytidine methyltransferase |
| AT3G27180 | 0.8 | 6.66E-05 | 0.8 | 8.35E-05 | 0.0 | 7.82E-01 | 0.6 | 1.08E-02 | 0.0 | 9.21E-01 | 0.7 | 3.87E-02 | hypothetical protein |
| AT2G15560 | -0.2 | 2.27E-03 | -0.3 | 1.69E-03 | 0.0 | 7.42E-01 | -0.1 | 5.93E-01 | 0.0 | 8.99E-01 | -0.1 | 8.23E-01 | Uncharacterized protein At2g15560 |
| AT4G20480 | 0.1 | 1.03E-01 | -0.2 | 7.25E-03 | 0.3 | 1.46E-03 | 0.0 | 9.35E-01 | 0.0 | 8.35E-01 | 0.0 | 9.19E-01 | Uncharacterized protein AT4g20480 |
| AT4G12240 | 0.2 | 1.07E-01 | 0.0 | 8.35E-01 | 0.3 | 1.40E-01 | 0.3 | 4.34E-01 | 0.1 | 8.21E-01 | 0.2 | 7.46E-01 | C2H2-type domain-containing protein |
| AT3G62460 | 0.6 | 2.66E-03 | 0.2 | 2.67E-01 | 0.4 | 5.52E-02 | 0.3 | 3.99E-01 | -0.6 | 2.44E-01 | 0.9 | 7.46E-02 | NYN domain-containing protein |
| AT3G61100 | 0.9 | 1.05E-04 | 0.7 | 5.48E-04 | 0.1 | 4.43E-01 | 0.5 | 3.31E-02 | -0.1 | 8.01E-01 | 0.6 | 6.26E-02 | NYN domain-containing protein |
| AT3G62050 | | | | | | | -0.2 | 8.12E-01 | 0.1 | 8.72E-01 | -0.3 | 7.30E-01 | NYN domain-containing protein |
| AT3G14290 | 0.3 | 2.66E-02 | 0.3 | 7.73E-03 | -0.1 | 4.78E-01 | 0.0 | 8.00E-01 | 0.0 | 7.86E-01 | 0.1 | 6.54E-01 | Proteasome subunit alpha type-5-B |
| AT3G19840 | -0.1 | 2.06E-01 | -0.3 | 2.92E-03 | 0.2 | 4.07E-02 | -0.2 | 2.99E-01 | -0.1 | 6.68E-01 | -0.1 | 7.58E-01 | Pre-mRNA-processing protein 40C |
| AT5G64170 | 0.0 | 9.22E-01 | -0.5 | 2.91E-03 | 0.5 | 4.34E-03 | | | | | | | Protein LNK1 |
| AT2G34750 | 0.4 | 6.35E-04 | 0.1 | 2.69E-01 | 0.3 | 8.68E-03 | -0.2 | 8.95E-02 | -0.2 | 1.00E-01 | 0.0 | 9.39E-01 | RNA polymerase I specific transcription initiation factor RRN3 protein |
| AT1G30590 | 0.1 | 4.25E-01 | 0.0 | 8.74E-01 | 0.1 | 4.05E-01 | 0.1 | 3.93E-01 | 0.2 | 2.08E-01 | -0.1 | 7.32E-01 | RNA polymerase I specific transcription initiation factor RRN3 protein |
| AT5G14960 | 0.1 | 3.02E-01 | -0.4 | 1.38E-02 | 0.5 | 7.37E-03 | 0.2 | 7.11E-01 | 0.2 | 6.45E-01 | -0.1 | 9.22E-01 | DP-E2F-like 2 |
| AT3G61800 | -0.2 | 5.33E-02 | -0.2 | 7.09E-02 | 0.0 | 9.49E-01 | -0.1 | 7.67E-01 | 0.1 | 6.94E-01 | -0.2 | 5.53E-01 | UV-stimulated scaffold protein A homolog |
| AT3G22590 | 0.0 | 9.73E-01 | 0.0 | 8.34E-01 | 0.0 | 8.84E-01 | -0.1 | 5.44E-01 | -0.1 | 6.90E-01 | 0.0 | 9.46E-01 | Protein CDC73 homolog |
| AT5G65180 | 0.1 | 4.36E-01 | -0.1 | 6.19E-01 | 0.1 | 2.82E-01 | 0.1 | 7.60E-01 | 0.2 | 5.90E-01 | -0.1 | 8.26E-01 | ENTH |
| AT4G24440 | 0.4 | 1.50E-03 | 0.3 | 5.56E-03 | 0.1 | 5.83E-01 | 0.1 | 4.56E-01 | -0.2 | 4.83E-01 | 0.3 | 2.25E-01 | Transcription initiation factor IIA subunit 2 |
| AT3G09360 | -0.1 | 2.48E-01 | -0.1 | 8.46E-02 | 0.1 | 5.70E-01 | 0.1 | 5.06E-01 | 0.0 | 9.15E-01 | 0.1 | 7.27E-01 | Cyclin |
| AT2G45100 | 0.8 | 1.14E-05 | 0.6 | 2.19E-04 | 0.2 | 7.14E-02 | 0.0 | 9.16E-01 | 0.2 | 5.18E-01 | -0.2 | 5.21E-01 | Cyclin |
| AT1G30550 | -0.8 | 1.34E-03 | 0.0 | 8.20E-01 | -0.8 | 7.06E-03 | -0.8 | 2.74E-02 | -0.1 | 8.05E-01 | -0.7 | 1.23E-01 | Trimethylguanosine synthase |
| AT3G23910 | -0.3 | 1.16E-02 | -0.4 | 3.28E-03 | 0.1 | 4.34E-01 | -0.1 | 5.69E-01 | 0.0 | 9.86E-01 | -0.1 | 6.66E-01 | Reverse transcriptase-like protein |
| AT5G04050 | 0.0 | 7.91E-01 | -0.1 | 6.79E-01 | 0.1 | 5.73E-01 | -0.8 | 1.51E-01 | -0.2 | 7.32E-01 | -0.6 | 4.40E-01 | RNA-directed DNA polymerase |
| AT3G24255 | 0.5 | 2.69E-03 | 0.0 | 7.64E-01 | 0.5 | 1.61E-02 | -0.1 | 8.51E-01 | 0.0 | 9.71E-01 | -0.1 | 8.38E-01 | RNA-directed DNA polymerase -related family protein |
| AT5G57280 | 0.3 | 6.51E-02 | 0.2 | 1.14E-01 | 0.0 | 8.39E-01 | 0.5 | 1.51E-02 | 0.1 | 6.17E-01 | 0.3 | 1.41E-01 | 18S rRNA )-methyltransferase RID2 |
| AT3G57000 | 0.5 | 1.31E-03 | 0.2 | 7.01E-02 | 0.2 | 7.09E-02 | 0.4 | 1.43E-02 | 0.2 | 1.94E-01 | 0.1 | 4.47E-01 | At3g57000 |
| AT5G13830 | | | | | | | 1.1 | 1.08E-02 | 0.4 | 4.41E-01 | | | FtsJ |
| AT1G55805 | 0.9 | 1.16E-06 | 0.7 | 1.19E-05 | 0.2 | 7.44E-02 | -0.1 | 7.41E-01 | -0.2 | 4.13E-01 | 0.1 | 6.91E-01 | F20N2.18 |
| AT1G50000 | 1.1 | 5.27E-06 | 0.7 | 6.43E-04 | 0.5 | 6.51E-03 | 0.1 | 6.13E-01 | 0.0 | 9.63E-01 | 0.1 | 7.50E-01 | 16S rRNA -N)-methyltransferase |
| AT3G15460 | 0.2 | 2.35E-01 | 0.1 | 4.29E-01 | 0.0 | 7.71E-01 | 0.7 | 7.11E-03 | 0.3 | 2.62E-01 | 0.4 | 1.62E-01 | Brix domain-containing protein |
| AT5G61770 | 0.5 | 5.66E-03 | 0.3 | 6.06E-02 | 0.2 | 2.93E-01 | 0.5 | 1.22E-02 | 0.2 | 4.82E-01 | 0.3 | 1.14E-01 | PETER PAN-like protein |
| AT3G23620 | 0.7 | 1.17E-04 | 0.4 | 6.65E-03 | 0.3 | 3.88E-02 | 0.8 | 3.07E-05 | 0.3 | 1.51E-01 | 0.6 | 8.29E-03 | Ribosome production factor 2 homolog |
| AT1G70200 | 1.0 | 6.72E-07 | 1.0 | 1.80E-06 | 0.0 | 8.89E-01 | 0.2 | 2.57E-01 | 0.0 | 9.91E-01 | 0.2 | 4.12E-01 | RNA-binding family protein |
| AT5G15750 | 0.5 | 2.48E-03 | 0.4 | 1.68E-02 | 0.1 | 4.05E-01 | 0.7 | 1.89E-03 | 0.2 | 3.90E-01 | 0.5 | 5.48E-02 | Alpha-L RNA-binding motif |
| AT3G18880 | 0.0 | 8.12E-01 | 0.1 | 5.69E-01 | -0.1 | 5.07E-01 | -0.4 | 3.42E-01 | 0.0 | 9.53E-01 | -0.5 | 3.79E-01 | Small ribosomal subunit protein uS17c |
| AT1G49400 | 0.2 | 1.16E-01 | 0.3 | 1.20E-02 | -0.1 | 2.52E-01 | 0.3 | 9.87E-02 | 0.2 | 2.27E-01 | 0.0 | 8.66E-01 | Small ribosomal subunit protein uS17c |
| AT1G31817 | 0.3 | 1.31E-02 | 0.3 | 1.62E-02 | 0.0 | 9.87E-01 | 0.4 | 2.15E-02 | 0.1 | 5.59E-01 | 0.2 | 1.99E-01 | Small ribosomal subunit protein uS11m |
| AT1G77750 | 0.5 | 1.36E-02 | 0.6 | 3.19E-03 | -0.2 | 3.98E-01 | 0.2 | 2.64E-01 | 0.0 | 8.76E-01 | 0.2 | 5.43E-01 | Small ribosomal subunit protein uS13m |
| AT2G18400 | 0.5 | 1.80E-03 | 0.7 | 2.62E-04 | -0.2 | 1.60E-01 | 0.3 | 1.18E-01 | 0.2 | 3.90E-01 | 0.1 | 7.49E-01 | hypothetical protein |
| AT4G10450 | 0.5 | 1.35E-02 | 0.6 | 8.23E-03 | -0.1 | 7.29E-01 | 0.4 | 5.10E-02 | 0.2 | 3.60E-01 | 0.2 | 5.20E-01 | Ribosomal protein L6 family |
| AT5G15200 | 0.7 | 6.00E-04 | 0.7 | 1.20E-03 | 0.0 | 8.86E-01 | 0.2 | 2.51E-01 | 0.0 | 1.00E+00 | 0.2 | 4.13E-01 | Ribosomal protein S4 |
| AT1G16740 | 0.9 | 8.38E-05 | 0.5 | 5.01E-03 | 0.4 | 3.53E-02 | 0.5 | 5.81E-03 | 0.3 | 1.45E-01 | 0.2 | 3.48E-01 | 50S ribosomal protein L20 |
| AT2G17360 | 0.9 | 6.75E-05 | 0.8 | 2.61E-04 | 0.1 | 5.69E-01 | 0.3 | 1.52E-02 | 0.1 | 5.82E-01 | 0.2 | 2.08E-01 | Ribosomal protein S4 family protein |
| AT5G07090 | 0.9 | 2.47E-05 | 0.7 | 1.72E-04 | 0.2 | 2.84E-01 | 0.5 | 3.62E-04 | 0.2 | 9.18E-02 | 0.2 | 1.33E-01 | 40S ribosomal protein S4 |
| AT5G17870 | 1.8 | 2.37E-08 | 1.4 | 2.56E-07 | 0.4 | 2.04E-03 | 0.2 | 4.86E-01 | 0.0 | 9.66E-01 | 0.2 | 6.10E-01 | Plastid-specific 50S ribosomal protein 6 |
| AT5G13510 | 1.8 | 3.30E-08 | 1.5 | 2.40E-07 | 0.3 | 9.77E-03 | 0.3 | 1.79E-01 | 0.1 | 7.72E-01 | 0.2 | 4.99E-01 | Large ribosomal subunit protein uL10c |
| AT1G75350 | 2.0 | 1.10E-07 | 1.7 | 9.12E-07 | 0.4 | 2.50E-02 | 0.1 | 5.95E-01 | -0.1 | 8.16E-01 | 0.2 | 5.57E-01 | Large ribosomal subunit protein bL31c |
| AT3G54210 | 2.2 | 1.59E-08 | 1.7 | 1.07E-07 | 0.5 | 1.33E-03 | 0.2 | 9.53E-02 | 0.0 | 9.77E-01 | 0.2 | 2.25E-01 | Large ribosomal subunit protein bL17c |
| AT1G29070 | 2.4 | 2.75E-08 | 2.1 | 1.84E-07 | 0.3 | 2.90E-02 | 0.1 | 5.13E-01 | -0.1 | 4.29E-01 | 0.3 | 2.19E-01 | Large ribosomal subunit protein bL34c |
| AT4G30930 | 0.3 | 1.72E-02 | 0.5 | 3.67E-03 | -0.1 | 3.81E-01 | 0.4 | 2.49E-02 | 0.1 | 5.61E-01 | 0.3 | 2.70E-01 | Large ribosomal subunit protein bL21m |
| AT1G64510 | 2.2 | 2.19E-08 | 1.9 | 1.08E-07 | 0.2 | 5.24E-02 | 0.1 | 4.54E-01 | -0.1 | 7.36E-01 | 0.2 | 4.02E-01 | Translation elongation factor EF1B |

| | | | | | | | | | | | | | |
|---|---|---|---|---|---|---|---|---|---|---|---|---|---|
| AT1G17690 | 0.1 | 3.04E-01 | 0.1 | 5.12E-01 | 0.0 | 7.88E-01 | 0.4 | 2.73E-02 | 0.1 | 5.61E-01 | 0.3 | 1.58E-01 | Protein NUCLEOLAR FACTOR 1 |
| AT5G50110 | 0.6 | 1.04E-04 | 0.3 | 8.58E-03 | 0.3 | 3.42E-02 | 0.2 | 4.05E-01 | 0.1 | 8.76E-01 | 0.2 | 5.80E-01 | Glucose-inhibited division protein B-like protein |
| AT4G01560 | 0.2 | 1.86E-01 | 0.3 | 2.51E-02 | -0.2 | 3.04E-01 | 0.6 | 3.85E-04 | 0.1 | 4.70E-01 | 0.4 | 1.43E-02 | AT4g01560 |
| AT5G43940 | -0.1 | 3.53E-01 | 0.0 | 5.73E-01 | 0.0 | 7.83E-01 | -0.1 | 3.93E-01 | -0.1 | 5.70E-01 | 0.0 | 9.37E-01 | S-glutathione dehydrogenase |
| AT1G30070 | 0.3 | 4.95E-02 | 0.1 | 6.67E-01 | 0.2 | 1.83E-01 | 0.3 | 2.88E-01 | 0.3 | 3.07E-01 | 0.0 | 9.62E-01 | Calcyclin-binding protein |
| AT4G33150 | -0.9 | 3.39E-05 | -0.8 | 1.63E-04 | -0.1 | 4.21E-01 | -0.7 | 8.34E-03 | -0.5 | 6.45E-02 | -0.2 | 5.70E-01 | Lysine-ketoglutarate reductase |
| AT4G39460 | 0.9 | 1.93E-06 | 0.9 | 3.95E-06 | 0.0 | 8.88E-01 | 0.1 | 2.70E-01 | 0.0 | 9.25E-01 | 0.2 | 4.00E-01 | S-adenosylmethionine carrier 1, chloroplastic |
| AT5G42130 | 1.0 | 1.68E-07 | 0.7 | 3.96E-06 | 0.3 | 2.52E-03 | 0.1 | 4.56E-01 | 0.1 | 5.90E-01 | 0.0 | 9.69E-01 | Protein MITOFERRINLIKE 1, chloroplastic |
| AT4G00740 | -0.3 | 2.19E-03 | -0.1 | 2.73E-01 | -0.2 | 2.80E-02 | 0.0 | 8.82E-01 | 0.2 | 2.63E-01 | -0.2 | 4.52E-01 | Probable pectin methyltransferase QUA3 |
| AT2G32170 | -0.2 | 9.55E-03 | -0.1 | 1.96E-01 | -0.1 | 1.62E-01 | 0.1 | 6.17E-01 | 0.1 | 4.08E-01 | -0.1 | 8.04E-01 | S-adenosyl-L-methionine-dependent methyltransferases superfamily protein |
| AT2G32160 | 0.1 | 4.62E-01 | 0.0 | 8.84E-01 | 0.1 | 4.57E-01 | 0.6 | 2.34E-02 | 0.3 | 2.77E-01 | 0.3 | 4.15E-01 | S-adenosyl-L-methionine-dependent methyltransferases superfamily protein |
| AT3G59870 | 0.2 | 4.55E-02 | 0.3 | 2.74E-02 | 0.0 | 7.69E-01 | 0.1 | 5.94E-01 | 0.2 | 3.36E-01 | -0.1 | 7.36E-01 | Uncharacterized protein F24G16.140 |
| AT2G43910 | 0.7 | 6.36E-06 | 0.6 | 3.40E-05 | 0.1 | 3.44E-01 | 0.6 | 3.57E-04 | 0.4 | 1.86E-02 | 0.2 | 3.07E-01 | HARMLESS TO OZONE LAYER 1 |
| AT2G43920 | 2.1 | 6.39E-08 | 1.0 | 3.75E-05 | 1.2 | 2.95E-05 | -0.5 | 5.57E-02 | -0.3 | 2.30E-01 | -0.1 | 6.87E-01 | HOL2 |
| AT3G22740 | 0.3 | 1.81E-01 | 0.7 | 9.47E-03 | -0.4 | 1.43E-01 | -1.3 | 1.26E-01 | 0.6 | 3.61E-01 | -1.9 | 6.94E-02 | Homocysteine S-methyltransferase 3 |
| AT3G48880 | -0.7 | 1.00E-05 | -0.5 | 2.44E-04 | -0.2 | 3.62E-02 | -0.4 | 8.62E-03 | -0.1 | 5.75E-01 | -0.3 | 1.26E-01 | F-box |
| AT1G78000 | 1.5 | 5.03E-07 | 1.3 | 2.71E-06 | 0.2 | 2.16E-01 | 1.3 | 3.25E-06 | 0.9 | 3.72E-04 | 0.4 | 1.32E-01 | Sulfate transporter 12 |
| AT5G10180 | 0.5 | 3.01E-04 | 0.9 | 2.34E-05 | -0.3 | 1.91E-02 | -0.3 | 3.73E-02 | 0.0 | 9.83E-01 | -0.3 | 1.36E-01 | Sulfate transporter 2.1 |
| AT3G15990 | -1.0 | 2.86E-06 | -0.9 | 1.09E-05 | -0.1 | 5.03E-01 | 1.2 | 9.21E-05 | 0.6 | 6.49E-02 | 0.7 | 1.14E-02 | Probable sulfate transporter 3.4 |
| AT5G13550 | -0.8 | 1.51E-05 | -0.7 | 7.88E-05 | -0.1 | 3.57E-01 | 0.2 | 3.62E-01 | 0.3 | 2.59E-01 | -0.1 | 8.46E-01 | Sulfate transporter 4.1, chloroplastic |
| AT5G19600 | | | | | | | 2.0 | 1.54E-07 | 0.4 | 1.51E-01 | 1.6 | 4.28E-05 | Probable sulfate transporter 3.5 |
| AT3G23800 | -2.1 | 1.24E-07 | -1.3 | 2.94E-06 | -0.9 | 6.31E-04 | -1.9 | 1.51E-05 | -0.6 | 2.42E-02 | -1.3 | 3.20E-03 | Selenium-binding protein 3 |
| AT4G14040 | 0.2 | 1.46E-01 | 0.2 | 1.60E-01 | 0.0 | 9.93E-01 | -0.2 | 3.14E-01 | 0.4 | 2.25E-02 | -0.5 | 1.52E-02 | Uncharacterized protein |
| AT4G14030 | 0.2 | 1.86E-03 | 0.1 | 3.18E-01 | 0.2 | 2.07E-02 | 0.0 | 7.08E-01 | -0.1 | 5.56E-01 | 0.1 | 4.51E-01 | Selenium-binding protein 1 |
| AT3G53310 | -1.1 | 2.81E-05 | -0.9 | 1.62E-04 | -0.2 | 3.80E-01 | 0.5 | 1.66E-01 | -0.4 | 4.81E-01 | 0.9 | 6.45E-02 | B3 domain-containing protein REM20 |
| AT3G12977 | -0.1 | 5.94E-01 | 0.0 | 7.27E-01 | 0.0 | 9.06E-01 | -0.4 | 1.91E-01 | 0.3 | 4.04E-01 | -0.7 | 1.87E-02 | NAC domain transcriptional regulator superfamily protein protein-like) |
| AT3G53270 | 0.0 | 6.90E-01 | -0.2 | 8.28E-03 | 0.3 | 9.93E-03 | 0.1 | 3.73E-01 | 0.1 | 6.77E-01 | 0.1 | 8.05E-01 | Small nuclear RNA activating complex , subunit SNAP43 protein |
| AT2G03780 | 0.1 | 3.31E-01 | 0.1 | 4.69E-01 | 0.0 | 8.70E-01 | 0.2 | 2.19E-01 | 0.1 | 8.56E-01 | 0.2 | 4.53E-01 | Translin family protein |
| AT5G45830 | 2.1 | 5.85E-06 | 2.0 | 2.18E-05 | 0.1 | 5.59E-01 | | | | | | | Delay of germination 1 |
| AT5G49300 | -0.3 | 6.02E-02 | -0.2 | 1.78E-01 | -0.1 | 6.58E-01 | 0.4 | 2.28E-01 | 0.1 | 7.97E-01 | 0.3 | 5.11E-01 | GATA-type domain-containing protein |
| AT3G24050 | 0.1 | 6.75E-02 | 0.0 | 5.54E-01 | 0.1 | 2.52E-01 | -0.2 | 1.42E-01 | -0.2 | 1.03E-01 | 0.0 | 8.42E-01 | GATA transcription factor 1 |
| AT4G16141 | 0.7 | 1.89E-03 | 0.8 | 1.11E-03 | -0.1 | 6.35E-01 | 0.2 | 4.75E-01 | -0.2 | 6.22E-01 | 0.3 | 3.34E-01 | GATA type zinc finger transcription factor family protein |
| AT4G14605 | 0.2 | 1.14E-01 | 0.0 | 7.83E-01 | 0.1 | 2.58E-01 | 0.0 | 9.48E-01 | 0.1 | 6.69E-01 | -0.1 | 6.52E-01 | Transcription termination factor MTERF5, chloroplastic |
| AT5G47880 | 0.1 | 2.10E-01 | -0.3 | 4.38E-03 | 0.4 | 1.49E-03 | 0.4 | 4.70E-03 | 0.2 | 2.98E-01 | 0.3 | 1.20E-01 | Eukaryotic peptide chain release factor subunit 1-1 |
| AT3G26618 | 0.3 | 5.12E-04 | 0.2 | 1.22E-02 | 0.1 | 1.19E-01 | 0.2 | 2.92E-01 | 0.0 | 9.60E-01 | 0.2 | 5.21E-01 | Eukaryotic peptide chain release factor subunit 1-3 |
| AT1G12920 | 0.8 | 4.59E-06 | 0.5 | 1.51E-04 | 0.2 | 1.80E-02 | 0.4 | 2.26E-03 | 0.1 | 7.37E-01 | 0.3 | 5.34E-02 | Eukaryotic peptide chain release factor subunit 1-2 |
| AT5G56760 | 0.1 | 1.46E-01 | 0.2 | 1.78E-02 | -0.1 | 2.55E-01 | -0.1 | 5.83E-01 | -0.1 | 5.47E-01 | 0.0 | 9.34E-01 | Serine acetyltransferase 5 |
| AT2G17640 | 0.5 | 6.41E-03 | 0.1 | 4.64E-01 | 0.4 | 5.71E-02 | -0.2 | 3.72E-01 | -0.1 | 6.90E-01 | -0.1 | 7.89E-01 | SAT-106 |
| AT1G55920 | 0.7 | 1.83E-05 | 0.3 | 1.04E-02 | 0.4 | 2.52E-03 | 0.3 | 7.51E-02 | -0.3 | 1.34E-01 | 0.7 | 1.66E-02 | Serine acetyltransferase 1, chloroplastic |
| AT4G35640 | | | | | | | 1.604648006 | 9.47E-03 | 1.24877928 | 6.55E-02 | | | Serine acetyltransferase 4 |
| AT3G13110 | -0.3 | 4.05E-03 | -0.1 | 7.38E-02 | -0.1 | 1.73E-01 | -0.1 | 7.00E-01 | 0.1 | 4.52E-01 | -0.2 | 3.07E-01 | Serine acetyltransferase 3, mitochondrial |
| AT1G61130 | -2.1 | 2.79E-05 | 0.7 | 2.24E-03 | -2.8 | 3.57E-05 | -0.8 | 2.20E-01 | 0.6 | 2.64E-01 | -1.4 | 4.48E-02 | Serine carboxypeptidase-like 32 |
| AT2G22920 | -1.5 | 4.61E-07 | -0.8 | 3.84E-05 | -0.7 | 7.79E-04 | -0.5 | 2.44E-02 | 0.1 | 6.27E-01 | -0.7 | 2.79E-02 | Serine carboxypeptidase-like 12 |
| AT3G63470 | -1.2 | 1.37E-05 | -0.1 | 6.68E-01 | -1.1 | 1.69E-04 | -0.2 | 6.46E-01 | 0.4 | 3.41E-01 | -0.6 | 2.11E-01 | Serine carboxypeptidase-like 40 |
| AT2G35770 | -1.0 | 1.54E-06 | -0.2 | 2.06E-02 | -0.8 | 9.98E-05 | 1.0 | 2.51E-02 | 0.6 | 2.45E-01 | 0.4 | 4.68E-01 | Serine carboxypeptidase-like 28 |
| AT2G23010 | -0.8 | 3.39E-02 | -0.1 | 8.43E-01 | -0.7 | 9.70E-02 | 0.3 | 1.67E-01 | 1.0 | 9.83E-04 | -0.6 | 2.13E-02 | Serine carboxypeptidase-like 9 |
| AT4G12910 | -0.8 | 4.38E-05 | -0.4 | 5.59E-03 | -0.4 | 1.11E-02 | -0.2 | 2.28E-01 | 0.1 | 7.00E-01 | -0.3 | 1.85E-01 | Carboxypeptidase |
| AT2G22970 | -0.8 | 2.26E-06 | 0.0 | 5.10E-01 | -0.8 | 2.60E-05 | -0.4 | 2.63E-02 | -0.1 | 6.46E-01 | -0.3 | 1.89E-01 | Serine carboxypeptidase-like 11 |
| AT1G73280 | -0.7 | 6.71E-04 | 0.0 | 8.23E-01 | -0.7 | 4.11E-03 | -0.7 | 1.33E-01 | 0.1 | 8.32E-01 | -0.8 | 1.37E-01 | Serine carboxypeptidase-like 3 |
| AT3G45010 | -0.7 | 2.35E-05 | -0.5 | 5.08E-04 | -0.2 | 5.52E-02 | 0.2 | 2.97E-01 | 0.3 | 1.41E-01 | -0.1 | 7.35E-01 | Serine carboxypeptidase-like 48 |
| AT3G10410 | -0.6 | 3.41E-05 | -0.4 | 1.42E-03 | -0.2 | 3.05E-02 | -0.2 | 9.79E-02 | 0.0 | 8.05E-01 | -0.1 | 3.85E-01 | Serine carboxypeptidase-like 49 |
| AT3G07990 | -0.6 | 6.04E-05 | -0.2 | 2.61E-02 | -0.4 | 5.59E-03 | -0.4 | 8.20E-02 | -0.1 | 5.89E-01 | -0.2 | 4.63E-01 | Serine carboxypeptidase-like 27 |
| AT1G73310 | -0.6 | 1.58E-03 | -0.5 | 6.78E-03 | -0.1 | 5.43E-01 | -0.8 | 3.14E-02 | 0.2 | 5.40E-01 | -1.0 | 6.38E-03 | Serine carboxypeptidase-like 4 |
| AT5G36180 | -0.5 | 1.69E-03 | 0.0 | 7.69E-01 | -0.6 | 4.03E-03 | -1.1 | 1.01E-02 | 0.6 | 7.10E-02 | -1.7 | 8.65E-04 | Serine carboxypeptidase-like 1 |
| AT2G27920 | -0.2 | 3.29E-02 | 0.0 | 8.63E-01 | -0.2 | 4.58E-02 | 0.4 | 1.69E-02 | 0.3 | 8.33E-02 | 0.1 | 7.46E-01 | Carboxypeptidase |
| AT4G30810 | -0.1 | 1.98E-01 | 0.2 | 7.64E-02 | -0.3 | 1.39E-02 | 0.0 | 6.94E-01 | 0.2 | 1.06E-01 | -0.1 | 3.25E-01 | Serine carboxypeptidase-like 29 |
| AT3G02110 | 0.2 | 2.86E-02 | 0.2 | 1.73E-02 | 0.0 | 7.60E-01 | -0.1 | 5.07E-01 | -0.1 | 4.57E-01 | 0.0 | 9.22E-01 | Serine carboxypeptidase-like 25 |
| AT5G08260 | 0.2 | 3.82E-02 | 0.3 | 1.75E-02 | -0.1 | 6.50E-01 | 0.0 | 7.85E-01 | 0.1 | 6.30E-01 | -0.1 | 4.50E-01 | Serine carboxypeptidase-like 35 |
| AT5G23210 | 0.3 | 9.10E-04 | 0.6 | 2.45E-05 | -0.3 | 7.72E-03 | -0.1 | 4.65E-01 | -0.4 | 5.07E-02 | 0.2 | 3.18E-01 | Carboxypeptidase |
| AT1G28110 | 0.4 | 1.36E-03 | 0.4 | 2.77E-03 | 0.0 | 8.49E-01 | 0.4 | 3.91E-02 | 0.2 | 4.72E-01 | 0.3 | 4.12E-01 | Carboxypeptidase |
| AT2G33530 | 0.6 | 7.99E-05 | 0.2 | 1.09E-01 | 0.5 | 2.47E-03 | 0.2 | 1.41E-01 | 0.2 | 2.24E-01 | 0.0 | 9.68E-01 | Serine carboxypeptidase-like 46 |
| AT2G35780 | 0.7 | 5.32E-06 | 0.8 | 8.25E-06 | 0.0 | 7.10E-01 | 0.1 | 7.84E-01 | 0.0 | 8.47E-01 | 0.0 | 9.88E-01 | Serine carboxypeptidase-like 26 |
| AT3G12220 | 0.8 | 1.51E-03 | 0.4 | 5.18E-02 | 0.4 | 1.35E-01 | 0.3 | 5.12E-01 | 0.0 | 9.70E-01 | 0.3 | 5.85E-01 | Serine carboxypeptidase-like 16 |
| AT5G42240 | 1.0 | 1.32E-06 | 0.6 | 5.47E-05 | 0.3 | 6.34E-03 | 0.1 | 6.31E-01 | -0.1 | 4.96E-01 | 0.2 | 2.30E-01 | Serine carboxypeptidase-like 42 |
| AT3G10450 | 1.0 | 3.82E-06 | 0.7 | 1.75E-04 | 0.3 | 1.49E-02 | 0.2 | 2.37E-01 | 0.3 | 2.87E-01 | 0.0 | 9.95E-01 | Serine carboxypeptidase-like 7 |
| AT4G30610 | 1.1 | 3.20E-06 | 1.1 | 5.65E-06 | 0.0 | 7.69E-01 | -0.1 | 7.74E-01 | -0.1 | 8.61E-01 | 0.0 | 9.62E-01 | Serine carboxypeptidase 24 |
| AT2G22990 | 1.8 | 3.41E-06 | 1.7 | 1.34E-05 | 0.1 | 5.34E-01 | -0.2 | 3.24E-01 | -0.5 | 5.07E-03 | 0.4 | 5.32E-02 | Sinapoylglucose 1 |
| AT2G22980 | 1.9 | 7.93E-06 | 2.3 | 4.02E-06 | -0.4 | 7.33E-02 | -0.5 | 2.36E-02 | -0.7 | 5.07E-03 | 0.2 | 5.16E-01 | Putative serine carboxypeptidase I |
| AT1G11080 | 2.6 | 7.44E-08 | 2.4 | 3.32E-07 | 0.2 | 1.67E-01 | 1.3 | 5.90E-06 | 0.6 | 2.39E-02 | 0.8 | 3.50E-03 | Carboxypeptidase |
| AT1G33540 | | | | | | | 0.1 | 9.36E-01 | 0.3 | 5.92E-01 | -0.3 | 7.20E-01 | Serine carboxypeptidase-like 18 |
| AT2G24010 | | | | | | | -2.1 | 4.37E-03 | -0.6 | 3.00E-01 | -1.4 | 1.20E-01 | Carboxypeptidase |
| AT5G25640 | -1.7 | 7.69E-06 | -0.4 | 1.58E-02 | -1.3 | 5.61E-04 | -0.2 | 6.07E-01 | 0.7 | 4.78E-02 | -0.9 | 2.32E-02 | Peptidase S54 rhomboid domain-containing protein |
| AT4G23070 | -1.6 | 4.87E-05 | -0.1 | 7.01E-01 | -1.5 | 4.86E-04 | -1.4 | 7.47E-03 | -0.3 | 5.78E-01 | | | RHOMBOID-like protein 7 |
| AT3G14067 | -1.5 | 2.92E-07 | -0.9 | 2.01E-05 | -0.6 | 7.63E-04 | -0.3 | 8.22E-02 | 0.2 | 3.90E-01 | -0.5 | 5.78E-02 | Subtilisin-like protease SBT1.4 |
| AT1G32950 | -1.5 | 1.48E-04 | -0.7 | 1.21E-02 | -0.8 | 2.50E-02 | | | | | | | Subtilase family protein |
| AT5G67090 | -1.2 | 2.08E-05 | -0.5 | 4.44E-03 | -0.7 | 5.32E-03 | -0.4 | 2.02E-01 | -0.3 | 4.42E-01 | -0.1 | 8.02E-01 | Subtilisin-like protease SBT1.9 |
| AT4G10510 | -1.1 | 1.30E-06 | -0.8 | 1.69E-05 | -0.3 | 3.59E-02 | 0.8 | 8.38E-02 | 0.6 | 3.21E-01 | 0.3 | 6.74E-01 | Subtilase family protein |
| AT5G07250 | -1.0 | 1.21E-06 | -0.8 | 2.08E-05 | -0.3 | 2.05E-02 | -0.5 | 2.65E-03 | -0.2 | 3.03E-01 | -0.3 | 8.85E-02 | RHOMBOID-like protein 3 |
| AT4G10550 | -0.7 | 9.31E-05 | -0.3 | 1.17E-02 | -0.4 | 1.54E-02 | 0.2 | 2.63E-01 | 0.3 | 6.75E-02 | -0.1 | 5.02E-01 | Subtilisin-like protease SBT3.6 |
| AT1G63120 | -0.7 | 4.33E-06 | -0.2 | 4.66E-03 | -0.4 | 6.59E-04 | -0.5 | 3.14E-03 | -0.4 | 1.48E-02 | -0.1 | 7.91E-01 | RHOMBOID-like protein 2 |
| AT3G53780 | -0.5 | 1.19E-04 | -0.5 | 2.01E-04 | 0.0 | 9.93E-01 | 0.1 | 6.76E-01 | 0.1 | 6.23E-01 | 0.0 | 9.23E-01 | RHOMBOID-like protein |
| AT3G59520 | -0.5 | 6.74E-04 | -0.2 | 1.28E-01 | -0.3 | 2.31E-02 | -0.3 | 1.78E-01 | -0.1 | 7.14E-01 | -0.2 | 4.57E-01 | RHOMBOID-like protein 13 |
| AT1G06870 | -0.5 | 2.78E-04 | -0.4 | 2.38E-03 | -0.1 | 2.75E-01 | 0.2 | 2.68E-01 | 0.2 | 2.39E-01 | 0.0 | 9.05E-01 | Probable thylakoidal processing peptidase 2, chloroplastic |

| | | | | | | | | | | | | | |
|---|---|---|---|---|---|---|---|---|---|---|---|---|---|
| AT4G14570 | -0.5 | 1.70E-04 | -0.4 | 1.02E-03 | -0.1 | 3.52E-01 | -0.2 | 2.25E-01 | 0.0 | 9.79E-01 | -0.2 | 3.72E-01 | Acylamino-acid-releasing enzyme |
| AT2G05920 | -0.5 | 5.75E-05 | -0.3 | 5.95E-03 | -0.2 | 1.58E-02 | -0.1 | 3.28E-01 | 0.1 | 5.41E-01 | -0.3 | 2.25E-01 | Subtilisin-like protease SBT1.8 |
| AT3G56740 | -0.3 | 2.28E-03 | -0.2 | 3.99E-02 | -0.1 | 1.95E-01 | 0.1 | 4.76E-01 | 0.3 | 1.70E-01 | -0.1 | 5.74E-01 | Rhomboid-like protein 20 |
| AT1G69020 | -0.3 | 2.45E-02 | -0.4 | 9.30E-03 | 0.1 | 5.66E-01 | 0.1 | 8.88E-01 | 0.4 | 4.57E-01 | -0.3 | 6.32E-01 | Prolyl endopeptidase |
| AT4G00230 | -0.3 | 6.69E-03 | 0.2 | 2.16E-02 | -0.5 | 6.93E-04 | -0.1 | 7.65E-01 | -0.1 | 6.02E-01 | 0.1 | 8.44E-01 | Xylem serine peptidase 1 |
| AT3G03380 | -0.3 | 2.84E-03 | -0.3 | 6.29E-03 | 0.0 | 7.99E-01 | -0.2 | 3.12E-01 | -0.1 | 5.88E-01 | -0.1 | 8.59E-01 | DegP protease 7 |
| AT2G39850 | -0.3 | 5.03E-02 | -0.1 | 5.63E-01 | -0.2 | 2.00E-01 | 0.2 | 1.69E-01 | 0.4 | 2.05E-02 | -0.2 | 3.64E-01 | Subtilisin-like protease SBT4.1 |
| AT5G19660 | -0.2 | 2.33E-02 | -0.2 | 1.09E-02 | 0.0 | 6.52E-01 | -0.3 | 2.00E-01 | -0.2 | 4.13E-01 | -0.1 | 8.39E-01 | AT5G19660 protein |
| AT4G20430 | -0.2 | 1.03E-01 | 0.0 | 6.83E-01 | -0.2 | 8.90E-02 | -0.5 | 2.97E-01 | -0.1 | 8.08E-01 | -0.3 | 5.57E-01 | Subtilisin-like protease SBT2.2 |
| AT1G12750 | -0.2 | 1.99E-01 | 0.0 | 9.69E-01 | -0.2 | 2.85E-01 | -0.6 | 7.12E-04 | -0.2 | 1.69E-01 | -0.4 | 8.38E-02 | RHOMBOID-like protein |
| AT1G30600 | -0.1 | 2.03E-01 | -0.1 | 3.86E-01 | 0.0 | 7.62E-01 | 0.0 | 9.86E-01 | 0.1 | 6.73E-01 | -0.1 | 7.24E-01 | Subtilisin-like protease SBT2.1 |
| AT1G52600 | -0.1 | 2.56E-01 | 0.1 | 5.77E-01 | -0.2 | 1.45E-01 | 0.2 | 3.30E-01 | 0.1 | 6.17E-01 | 0.1 | 8.43E-01 | signal peptidase I |
| AT2G04160 | 0.0 | 5.92E-01 | 0.2 | 2.53E-02 | -0.2 | 1.80E-02 | 0.0 | 9.43E-01 | 0.3 | 1.47E-01 | -0.2 | 2.87E-01 | Subtilisin-like protease SBT5.3 |
| AT5G59130 | 0.0 | 8.64E-01 | -0.2 | 1.44E-01 | 0.2 | 2.38E-01 | 0.3 | 5.14E-01 | 0.2 | 6.42E-01 | 0.0 | 9.62E-01 | Subtilase family protein |
| AT3G58460 | 0.0 | 8.53E-01 | 0.0 | 4.96E-01 | 0.0 | 6.64E-01 | 0.2 | 2.45E-01 | 0.1 | 6.33E-01 | 0.1 | 7.51E-01 | RHOMBOID-like protein 15 |
| AT1G50380 | 0.1 | 2.96E-01 | -0.1 | 7.64E-02 | 0.2 | 2.20E-02 | 0.3 | 2.55E-02 | 0.1 | 5.53E-01 | 0.2 | 2.68E-01 | Prolyl endopeptidase |
| AT5G45650 | 0.1 | 4.32E-01 | 0.2 | 3.30E-02 | -0.2 | 1.66E-01 | -0.4 | 5.24E-02 | 0.6 | 5.01E-03 | -1.0 | 7.94E-04 | Subtilisin-like protease SBT5.6 |
| AT2G31140 | 0.2 | 1.03E-01 | 0.3 | 1.50E-02 | -0.1 | 3.27E-01 | 0.0 | 8.39E-01 | -0.2 | 4.17E-01 | 0.2 | 3.93E-01 | Peptidase S24 |
| AT5G67360 | 0.2 | 2.43E-02 | 0.0 | 7.15E-01 | 0.2 | 2.22E-02 | -0.4 | 6.41E-04 | -0.2 | 8.03E-02 | -0.2 | 2.01E-01 | Subtilisin-like protease SBT1.7 |
| AT5G44530 | 0.2 | 2.51E-01 | 0.6 | 2.09E-03 | -0.4 | 2.39E-02 | 0.0 | 9.73E-01 | 0.1 | 7.05E-01 | -0.1 | 7.20E-01 | Subtilase family protein |
| AT1G18600 | 0.2 | 3.20E-02 | 0.1 | 4.76E-01 | 0.1 | 1.90E-01 | 0.2 | 2.96E-01 | 0.2 | 4.06E-01 | 0.0 | 9.65E-01 | RHOMBOID-like protein 12, mitochondrial |
| AT1G76140 | 0.2 | 1.15E-02 | 0.2 | 4.97E-02 | 0.1 | 5.23E-01 | 0.2 | 6.96E-01 | 0.1 | 8.01E-01 | 0.0 | 9.71E-01 | Prolyl endopeptidase |
| AT2G29050 | 0.2 | 4.29E-02 | 0.3 | 1.69E-02 | -0.1 | 6.20E-01 | -0.1 | 3.61E-01 | -0.1 | 5.75E-01 | 0.0 | 9.02E-01 | RHOMBOID-like protein 1 |
| AT1G06200 | 0.3 | 1.90E-03 | 0.2 | 5.05E-03 | 0.0 | 7.12E-01 | 0.0 | 8.07E-01 | -0.2 | 3.10E-01 | 0.1 | 4.65E-01 | Peptidase S24 |
| AT3G15710 | 0.3 | 2.53E-03 | 0.3 | 6.26E-03 | 0.0 | 7.44E-01 | 0.0 | 9.87E-01 | 0.0 | 8.57E-01 | 0.0 | 9.00E-01 | signal peptidase I |
| AT4G30020 | 0.3 | 2.58E-03 | 0.3 | 1.86E-03 | 0.0 | 7.34E-01 | 0.0 | 8.79E-01 | 0.3 | 7.58E-02 | -0.3 | 1.05E-01 | Subtilisin-like protease SBT2.6 |
| AT4G10530 | 0.4 | 6.67E-02 | 1.1 | 1.48E-04 | -0.7 | 4.99E-03 | 2.4 | 1.14E-06 | 0.9 | 3.08E-02 | 1.5 | 3.99E-04 | Subtilisin-like protease SBT3.10 |
| AT3G14240 | 0.4 | 3.69E-03 | 0.5 | 1.82E-03 | -0.1 | 5.52E-01 | 0.0 | 1.00E+00 | -0.1 | 4.38E-01 | 0.1 | 4.85E-01 | AT3g14240 |
| AT5G36950 | 0.4 | 3.04E-03 | 0.4 | 3.16E-03 | 0.0 | 8.97E-01 | 0.2 | 4.35E-01 | 0.4 | 8.46E-02 | -0.2 | 4.04E-01 | Protease Do-like 10, mitochondrial |
| AT5G51750 | 0.4 | 1.74E-03 | 0.3 | 1.55E-02 | 0.1 | 3.06E-01 | 0.1 | 5.31E-01 | 0.0 | 9.26E-01 | 0.1 | 7.61E-01 | Subtilisin-like protease SBT1.3 |
| AT3G08980 | 0.4 | 5.28E-02 | 0.3 | 1.56E-01 | 0.1 | 6.73E-01 | 0.7 | 5.24E-02 | 0.3 | 4.80E-01 | 0.4 | 4.01E-01 | Mitochondrial inner membrane protease subunit 2 |
| AT4G21326 | 0.4 | 4.18E-03 | 0.0 | 9.02E-01 | 0.5 | 1.15E-02 | 0.0 | 9.77E-01 | 0.3 | 4.55E-01 | -0.3 | 5.45E-01 | Subtilase 3.12 |
| AT4G10540 | 0.5 | 3.34E-03 | -0.4 | 1.24E-02 | 0.9 | 4.63E-04 | 1.0 | 6.63E-04 | 0.4 | 1.16E-01 | 0.5 | 6.08E-02 | Subtilisin-like protease SBT3.8 |
| AT1G20160 | 0.5 | 1.15E-04 | 0.4 | 4.63E-04 | 0.1 | 5.57E-01 | 0.6 | 6.76E-02 | 0.2 | 6.60E-01 | 0.4 | 3.32E-01 | Uncharacterized protein |
| AT5G59090 | 0.5 | 1.46E-03 | 1.0 | 3.94E-05 | -0.4 | 1.19E-02 | -0.9 | 1.19E-03 | -0.2 | 4.27E-01 | -0.6 | 5.57E-02 | Subtilase 4.12 |
| AT1G53530 | 0.6 | 1.79E-04 | 0.5 | 2.04E-03 | 0.2 | 2.41E-01 | 0.4 | 6.40E-02 | 0.1 | 6.57E-01 | 0.3 | 3.62E-01 | Peptidase S24 |
| AT1G32940 | 0.6 | 4.56E-03 | -0.2 | 3.21E-01 | 0.8 | 4.18E-03 | 1.1 | 6.10E-04 | -0.1 | 8.47E-01 | 1.1 | 3.60E-03 | Subtilase family protein |
| AT1G25290 | 0.7 | 4.46E-04 | 0.6 | 2.58E-03 | 0.1 | 4.33E-01 | -0.1 | 4.51E-01 | -0.2 | 3.45E-01 | 0.1 | 8.54E-01 | RHOMBOID-like protein 10, chloroplastic |
| AT3G24590 | 0.8 | 2.56E-05 | 0.8 | 6.83E-05 | 0.0 | 7.81E-01 | 0.4 | 4.06E-02 | 0.2 | 3.12E-01 | 0.2 | 5.63E-01 | Plastidic type i signal peptidase 1 |
| AT5G27660 | 0.8 | 4.16E-04 | 0.5 | 2.06E-02 | 0.4 | 7.95E-02 | 0.4 | 2.19E-01 | -0.1 | 9.17E-01 | 0.5 | 1.82E-01 | Protease Do-like 14 |
| AT5G39830 | 1.1 | 5.06E-06 | 0.8 | 1.03E-04 | 0.3 | 3.55E-02 | 0.4 | 6.12E-03 | 0.3 | 2.79E-02 | 0.1 | 8.10E-01 | Trypsin family protein with PDZ domain-containing protein |
| AT5G38510 | 1.2 | 2.32E-05 | 0.8 | 6.18E-04 | 0.4 | 4.76E-02 | 0.3 | 1.39E-02 | 0.1 | 4.68E-01 | 0.2 | 2.25E-01 | RHOMBOID-like protein 9, chloroplastic |
| AT1G04110 | 1.2 | 2.55E-05 | 1.5 | 1.20E-05 | -0.3 | 1.33E-01 | 0.4 | 4.02E-01 | -0.2 | 7.80E-01 | 0.6 | 3.38E-01 | Subtilisin-like protease SBT1.2 |
| AT4G21650 | 1.4 | 1.69E-06 | 0.7 | 3.24E-04 | 0.6 | 1.64E-03 | 0.0 | 9.05E-01 | 0.1 | 6.91E-01 | -0.2 | 6.36E-01 | Subtilisin-like protease SBT3.13 |
| AT2G47940 | 1.6 | 1.04E-07 | 1.0 | 3.60E-06 | 0.5 | 6.52E-04 | 0.1 | 8.80E-01 | 0.1 | 7.79E-01 | -0.1 | 9.07E-01 | DEGP protease 2 |
| AT1G01900 | | | | | | | -0.3 | 5.99E-01 | 0.4 | 5.72E-01 | -0.7 | 3.04E-01 | Subtilisin-like protease SBT1.1 |
| AT5G66960 | | | | | | | 0.4 | 3.51E-01 | 0.4 | 4.10E-01 | | | Prolyl endopeptidase |
| AT4G34980 | -0.6 | 9.41E-05 | -0.3 | 9.00E-03 | -0.3 | 2.03E-02 | -0.1 | 1.69E-01 | 0.1 | 2.40E-01 | -0.3 | 6.56E-02 | Subtilisin-like protease SBT1.6 |
| AT2G19170 | -0.3 | 3.82E-03 | 0.1 | 2.13E-01 | -0.4 | 2.24E-03 | -0.2 | 2.81E-01 | 0.0 | 9.57E-01 | -0.2 | 3.86E-01 | Subtilisin-like protease SBT2.5 |
| AT5G40200 | 0.2 | 6.31E-02 | 0.1 | 5.03E-01 | 0.1 | 2.73E-01 | 0.1 | 3.80E-01 | 0.1 | 4.68E-01 | 0.0 | 9.94E-01 | Protease Do-like 9 |
| AT2G25240 | -1.5 | 2.76E-06 | 0.0 | 7.36E-01 | -1.6 | 3.04E-05 | -1.0 | 4.35E-04 | 0.0 | 9.27E-01 | -1.1 | 2.67E-03 | Serpin-Z10 |
| AT1G47710 | -0.8 | 2.35E-05 | -0.5 | 1.06E-03 | -0.3 | 2.30E-02 | 0.1 | 7.41E-01 | 0.0 | 9.81E-01 | 0.1 | 7.84E-01 | Serine protease inhibitor family protein |
| AT2G26390 | 0.2 | 1.28E-01 | 0.6 | 3.93E-04 | -0.4 | 1.02E-02 | 0.0 | 9.27E-01 | 0.2 | 5.14E-01 | -0.2 | 5.11E-01 | Serpin-Z3 |
| AT3G61980 | 0.5 | 6.12E-04 | 0.0 | 9.58E-01 | 0.5 | 3.12E-03 | -0.1 | 7.57E-01 | -0.3 | 3.16E-01 | 0.2 | 5.72E-01 | Serine protease inhibitor, Kazal-type family protein |
| AT5G43580 | 1.4 | 2.20E-02 | 0.5 | 4.53E-01 | 0.9 | 1.53E-01 | 1.1 | 1.36E-02 | 0.6 | 2.53E-01 | 0.5 | 2.83E-01 | Uncharacterized protein At5g43580 |
| AT1G73325 | | | | | | | 2.140412209 | 1.78E-01 | 2.246716487 | 2.07E-01 | -0.1 | 9.63E-01 | Kunitz trypsin inhibitor 3 |
| AT2G43510 | 0.8 | 2.88E-02 | 0.0 | 9.14E-01 | 0.8 | 8.15E-02 | -0.1 | 8.76E-01 | -0.6 | 4.63E-01 | 0.5 | 5.96E-01 | Defensin-like protein 195 |
| AT2G43520 | 1.9 | 1.95E-04 | 1.1 | 1.26E-02 | 0.8 | 4.52E-02 | 0.1 | 8.83E-01 | -0.3 | 5.70E-01 | 0.4 | 5.05E-01 | Defensin-like protein 193 |
| AT3G57680 | -1.6 | 1.82E-05 | -1.5 | 4.46E-05 | -0.1 | 8.28E-01 | -0.4 | 2.81E-01 | 0.0 | 9.97E-01 | | | Peptidase S41 family protein |
| AT5G65760 | -0.3 | 6.32E-04 | -0.2 | 2.54E-02 | -0.1 | 7.57E-02 | -0.3 | 8.15E-02 | -0.2 | 3.78E-01 | -0.1 | 6.08E-01 | AT5g65760 |
| AT5G20520 | 0.0 | 8.05E-01 | 0.1 | 3.00E-01 | -0.1 | 4.75E-01 | -0.3 | 2.69E-02 | -0.4 | 8.75E-03 | 0.1 | 6.15E-01 | Peptidase S9 prolyl oligopeptidase catalytic domain-containing protein |
| AT2G47390 | 0.0 | 7.06E-01 | 0.0 | 9.79E-01 | 0.0 | 7.67E-01 | 0.1 | 7.20E-01 | 0.3 | 3.24E-01 | -0.2 | 6.27E-01 | Peptidase S9 prolyl oligopeptidase catalytic domain-containing protein |
| AT5G36210 | 0.3 | 4.23E-04 | 0.0 | 5.28E-01 | 0.4 | 7.05E-04 | 0.1 | 6.13E-01 | 0.1 | 6.87E-01 | 0.0 | 9.98E-01 | Alpha |
| AT5G46390 | 0.5 | 4.36E-04 | 0.3 | 2.61E-02 | 0.3 | 5.86E-02 | 0.1 | 4.05E-01 | 0.1 | 4.58E-01 | 0.0 | 9.92E-01 | AT5g46390 |
| AT2G24280 | 0.7 | 6.34E-05 | 0.2 | 8.91E-02 | 0.5 | 2.12E-03 | 0.0 | 9.43E-01 | 0.2 | 3.89E-01 | -0.2 | 4.46E-01 | Lysosomal Pro-X carboxypeptidase |
| AT4G17740 | 1.0 | 1.20E-06 | 0.9 | 6.27E-06 | 0.1 | 2.82E-01 | 0.0 | 7.71E-01 | 0.1 | 3.96E-01 | -0.1 | 6.46E-01 | PDZ domain-containing protein |
| AT5G65000 | 0.1 | 4.59E-01 | -0.1 | 4.02E-01 | 0.1 | 1.84E-01 | -0.2 | 1.87E-01 | -0.2 | 3.60E-01 | 0.0 | 8.98E-01 | UDP-N-acetylglucosamine transporter ROCK1 |
| AT3G48820 | -0.1 | 3.99E-01 | 0.0 | 5.79E-01 | 0.0 | 8.41E-01 | 0.0 | 9.49E-01 | 0.1 | 4.81E-01 | -0.2 | 5.20E-01 | Sialyltransferase-like protein 2 |
| AT1G08660 | 0.0 | 5.27E-01 | 0.2 | 3.44E-02 | -0.1 | 1.44E-01 | -0.2 | 1.54E-01 | -0.2 | 2.63E-01 | 0.0 | 9.36E-01 | MALE GAMETOPHYTE DEFECTIVE 2 |
| AT4G30850 | -1.2 | 3.72E-07 | -0.5 | 2.01E-04 | -0.7 | 1.25E-04 | 0.0 | 8.21E-01 | -0.1 | 5.83E-01 | 0.1 | 8.04E-01 | Heptahelical transmembrane protein 2 |
| AT4G38320 | -0.2 | 7.19E-02 | -0.1 | 1.01E-01 | 0.0 | 9.25E-01 | -0.2 | 4.05E-01 | 0.3 | 2.43E-01 | -0.5 | 9.15E-02 | Heptahelical transmembrane protein 5 |
| AT5G20270 | -0.1 | 1.90E-01 | -0.4 | 3.04E-03 | 0.3 | 4.22E-02 | -0.2 | 4.19E-01 | 0.0 | 8.89E-01 | -0.2 | 4.39E-01 | Heptahelical transmembrane protein 1 |
| AT1G33600 | 1.0 | 3.13E-06 | 0.9 | 1.48E-05 | 0.1 | 3.70E-01 | 0.5 | 8.36E-04 | -0.5 | 9.76E-04 | 1.0 | 1.09E-04 | Leucine-rich repeat family protein |
| AT5G62810 | 0.1 | 7.38E-02 | -0.1 | 3.82E-01 | 0.2 | 2.53E-02 | -0.3 | 1.03E-01 | 0.0 | 9.66E-01 | -0.3 | 1.82E-01 | Peroxisomal membrane protein PEX14 |
| AT5G60370 | 0.0 | 6.41E-01 | -0.2 | 7.91E-02 | 0.1 | 2.35E-01 | -0.2 | 2.96E-01 | -0.1 | 6.54E-01 | -0.1 | 7.14E-01 | Exonuclease V-like protein |
| AT4G20010 | 0.1 | 1.96E-01 | 0.0 | 9.16E-01 | 0.1 | 3.12E-01 | 0.2 | 2.22E-01 | 0.0 | 8.47E-01 | 0.1 | 4.96E-01 | Protein OSB2, chloroplastic |
| AT5G44785 | 0.1 | 1.56E-01 | 0.4 | 2.80E-03 | -0.2 | 5.47E-02 | 0.3 | 1.11E-01 | 0.2 | 4.85E-01 | 0.1 | 5.72E-01 | Organellar single-stranded DNA binding protein 3 |
| AT1G31010 | 0.2 | 7.49E-02 | -0.2 | 1.13E-01 | 0.5 | 1.33E-02 | 0.4 | 1.79E-01 | 0.0 | 9.28E-01 | 0.4 | 2.03E-01 | Protein OSB4, chloroplastic |
| AT4G03090 | 0.3 | 3.12E-03 | 0.2 | 9.97E-02 | 0.2 | 1.28E-01 | 0.0 | 9.56E-01 | -0.1 | 8.58E-01 | 0.1 | 8.48E-01 | Sequence-specific DNA binding transcription factor ATNDX |
| AT3G49000 | 0.4 | 3.87E-04 | 0.2 | 3.12E-02 | 0.2 | 4.61E-02 | 0.1 | 4.16E-01 | 0.1 | 6.50E-01 | 0.0 | 8.79E-01 | DNA-directed RNA polymerase III subunit RPC3 |
| AT4G11060 | 0.6 | 5.37E-04 | 0.5 | 2.43E-03 | 0.1 | 5.22E-01 | 0.4 | 9.73E-02 | 0.1 | 7.34E-01 | 0.3 | 3.56E-01 | AT4G11060 protein |
| AT1G47720 | 0.6 | 6.12E-04 | 0.2 | 2.50E-01 | 0.4 | 1.44E-02 | 0.3 | 4.02E-01 | 0.1 | 7.20E-01 | 0.1 | 7.90E-01 | Protein OSB1, mitochondrial |
| AT4G14520 | -0.1 | 6.47E-01 | -0.2 | 1.98E-01 | 0.1 | 4.69E-01 | -0.1 | 5.44E-01 | 0.1 | 6.39E-01 | -0.3 | 3.05E-01 | DNA-directed RNA polymerase subunit 7-like protein |

| | | | | | | | | | | | | | |
|---|---|---|---|---|---|---|---|---|---|---|---|---|---|
| AT4G14660 | 0.9 | 2.18E-04 | 0.5 | 1.16E-02 | 0.4 | 6.27E-02 | 0.3 | 4.22E-01 | 0.0 | 9.75E-01 | 0.3 | 5.39E-01 | DNA-directed RNA polymerase V subunit 7 |
| AT3G22900 | | | | | | | 0.4 | 2.63E-01 | 0.0 | 9.54E-01 | 0.4 | 3.42E-01 | DNA-directed RNA polymerase IV subunit 7 |
| AT5G59180 | 0.2 | 4.81E-02 | 0.2 | 6.21E-02 | 0.0 | 9.58E-01 | 0.6 | 2.62E-02 | 0.3 | 3.69E-01 | 0.3 | 3.72E-01 | DNA-directed RNA polymerase II subunit 7 |
| AT2G24490 | 0.4 | 1.38E-02 | 0.4 | 1.52E-02 | 0.0 | 9.66E-01 | 0.2 | 2.78E-01 | 0.2 | 3.76E-01 | 0.0 | 9.69E-01 | Replication protein A 32 kDa subunit A |
| AT4G23910 | -0.4 | 2.21E-04 | -0.4 | 9.85E-05 | 0.1 | 3.20E-01 | 0.0 | 8.78E-01 | 0.0 | 8.82E-01 | 0.0 | 9.92E-01 | At4g23910 |
| AT2G39805 | -0.4 | 2.46E-04 | -0.2 | 3.35E-02 | -0.3 | 2.26E-02 | 0.0 | 9.27E-01 | -0.3 | 1.89E-01 | 0.3 | 3.31E-01 | Protein YIP |
| AT2G46180 | -0.3 | 7.11E-03 | -0.2 | 1.07E-02 | 0.0 | 9.34E-01 | -0.3 | 1.06E-01 | 0.0 | 8.82E-01 | -0.3 | 3.08E-01 | Golgin candidate 4 |
| AT3G05280 | -0.2 | 2.47E-02 | 0.0 | 7.55E-01 | -0.2 | 7.35E-02 | 0.0 | 7.43E-01 | -0.1 | 3.70E-01 | 0.1 | 6.87E-01 | Protein YIP |
| AT5G62600 | -0.2 | 1.07E-02 | -0.1 | 6.46E-02 | -0.1 | 4.33E-01 | -0.1 | 6.54E-01 | -0.1 | 4.93E-01 | 0.0 | 8.53E-01 | Transportin MOS14 |
| AT5G18410 | -0.2 | 1.42E-02 | 0.0 | 8.55E-01 | -0.2 | 3.74E-02 | -0.1 | 5.71E-01 | 0.0 | 9.17E-01 | -0.1 | 5.69E-01 | Transcription activator |
| AT1G26170 | -0.1 | 4.42E-02 | -0.1 | 1.05E-01 | 0.0 | 7.29E-01 | 0.1 | 4.91E-01 | 0.3 | 1.76E-01 | -0.1 | 5.57E-01 | ARM repeat superfamily protein |
| AT3G17340 | -0.1 | 7.18E-02 | -0.1 | 3.91E-01 | -0.1 | 3.93E-01 | 0.0 | 8.98E-01 | 0.0 | 9.48E-01 | 0.0 | 9.76E-01 | ARM repeat superfamily protein |
| AT4G27640 | -0.1 | 9.41E-02 | -0.2 | 1.66E-02 | 0.1 | 3.55E-01 | 0.2 | 2.48E-01 | 0.0 | 9.91E-01 | 0.2 | 3.61E-01 | Importin N-terminal domain-containing protein |
| AT1G27310 | -0.1 | 3.32E-01 | 0.3 | 3.55E-02 | -0.4 | 1.11E-02 | -0.1 | 6.70E-01 | -0.2 | 2.81E-01 | 0.1 | 6.31E-01 | Nuclear transport factor 2A |
| AT2G31660 | -0.1 | 2.03E-01 | -0.2 | 5.81E-02 | 0.1 | 4.92E-01 | 0.3 | 4.15E-02 | 0.1 | 5.51E-01 | 0.2 | 3.57E-01 | Importin beta-like SAD2 |
| AT3G59020 | -0.1 | 4.56E-01 | -0.2 | 1.10E-01 | 0.1 | 3.92E-01 | 0.1 | 5.89E-01 | 0.0 | 9.78E-01 | 0.2 | 6.78E-01 | ARM repeat superfamily protein |
| AT5G27490 | 0.0 | 8.25E-01 | 0.2 | 1.48E-01 | -0.2 | 1.62E-01 | -0.4 | 3.16E-01 | -0.1 | 8.40E-01 | -0.3 | 6.01E-01 | Protein YIP |
| AT2G34570 | 0.5 | 6.05E-03 | 0.4 | 9.81E-03 | 0.0 | 9.07E-01 | 0.8 | 1.21E-03 | 0.6 | 1.98E-02 | 0.2 | 4.16E-01 | PIN domain-like family protein |
| AT3G60770 | 0.8 | 7.63E-05 | 0.7 | 3.17E-04 | 0.1 | 5.33E-01 | 0.4 | 7.92E-03 | 0.2 | 1.93E-01 | 0.2 | 3.88E-01 | Small ribosomal subunit protein uS15z |
| AT3G18760 | 0.3 | 1.03E-01 | 0.5 | 2.39E-02 | -0.2 | 4.59E-01 | 0.6 | 9.61E-04 | 0.4 | 3.06E-02 | 0.2 | 2.87E-01 | Translation elongation factor EF1B |
| AT3G17170 | 1.5 | 1.22E-06 | 1.4 | 6.19E-06 | 0.2 | 2.83E-01 | 0.3 | 1.59E-01 | -0.2 | 4.84E-01 | 0.5 | 1.02E-01 | Protein REGULATOR OF FATTY ACID COMPOSITION 3, chloroplastic |
| AT2G05830 | -0.2 | 1.31E-02 | 0.0 | 5.85E-01 | -0.1 | 5.99E-02 | -0.2 | 2.37E-01 | -0.1 | 7.79E-01 | -0.1 | 5.92E-01 | Methylthioribose-1-phosphate isomerase |
| AT1G79590 | -0.5 | 2.67E-05 | -0.2 | 5.61E-03 | -0.2 | 5.71E-03 | -0.3 | 6.85E-02 | -0.3 | 1.09E-01 | 0.0 | 9.87E-01 | Syntaxin of plants 52 |
| AT3G17440 | -0.5 | 5.24E-05 | 0.0 | 9.32E-01 | -0.5 | 2.83E-04 | -0.4 | 7.48E-03 | -0.4 | 1.24E-02 | 0.0 | 9.80E-01 | Plant snare 13 |
| AT1G15880 | -0.4 | 3.40E-04 | -0.1 | 1.70E-01 | -0.3 | 7.35E-03 | -0.2 | 3.55E-02 | -0.3 | 2.68E-02 | 0.0 | 8.39E-01 | Golgi SNAP receptor complex member 1-1 |
| AT1G16240 | -0.3 | 1.98E-04 | -0.2 | 1.00E-02 | -0.2 | 4.39E-02 | -0.5 | 6.40E-04 | -0.5 | 1.67E-03 | 0.0 | 9.96E-01 | Syntaxin of plants 51 |
| AT4G02195 | -0.2 | 1.59E-01 | -0.4 | 7.86E-03 | 0.2 | 1.36E-01 | -0.5 | 4.77E-02 | -0.1 | 6.58E-01 | -0.4 | 2.31E-01 | t-SNARE coiled-coil homology domain-containing protein |
| AT1G28490 | -0.2 | 6.53E-02 | 0.0 | 8.59E-01 | -0.1 | 1.38E-01 | -0.1 | 4.15E-01 | -0.1 | 5.76E-01 | 0.0 | 9.41E-01 | Syntaxin of plants 61 |
| AT3G24350 | -0.1 | 1.83E-01 | 0.0 | 5.99E-01 | 0.0 | 4.95E-01 | 0.0 | 9.42E-01 | 0.1 | 7.48E-01 | -0.1 | 7.52E-01 | Syntaxin of plants 32 |
| AT1G11890 | 0.0 | 6.13E-01 | 0.2 | 1.37E-01 | -0.2 | 8.13E-02 | 0.0 | 8.20E-01 | 0.0 | 7.91E-01 | 0.0 | 9.57E-01 | Uncharacterized protein |
| AT4G14455 | 0.0 | 8.15E-01 | 0.0 | 7.60E-01 | -0.1 | 6.47E-01 | 0.2 | 4.69E-01 | 0.3 | 2.29E-01 | -0.1 | 6.90E-01 | Bet1-like SNARE 1-2 |
| AT4G17730 | 0.0 | 8.35E-01 | 0.0 | 7.80E-01 | 0.0 | 6.63E-01 | -0.3 | 2.90E-02 | -0.3 | 3.52E-02 | 0.0 | 9.40E-01 | AT4G17730 protein |
| AT5G61210 | 0.0 | 9.66E-01 | -0.1 | 3.21E-01 | 0.1 | 3.36E-01 | 0.0 | 7.79E-01 | -0.2 | 2.35E-01 | 0.2 | 2.32E-01 | SNAP25 homologous protein SNAP33 |
| AT3G24315 | 0.0 | 9.21E-01 | -0.1 | 2.42E-01 | 0.1 | 2.71E-01 | -0.3 | 7.38E-02 | -0.4 | 3.70E-02 | 0.1 | 7.44E-01 | Sec20 family protein |
| AT5G26980 | 0.1 | 4.68E-01 | -0.3 | 7.14E-02 | 0.4 | 4.13E-02 | -0.1 | 6.07E-01 | 0.0 | 9.88E-01 | -0.1 | 7.33E-01 | AT5G26980 protein |
| AT3G29100 | 0.1 | 3.82E-01 | 0.2 | 2.28E-01 | 0.0 | 7.61E-01 | -0.6 | 2.49E-02 | -0.2 | 5.33E-01 | -0.4 | 2.44E-01 | At3g29100 |
| AT2G36900 | 0.1 | 1.53E-01 | 0.1 | 1.64E-01 | 0.0 | 9.92E-01 | -0.4 | 3.77E-02 | -0.6 | 2.84E-03 | 0.3 | 3.03E-01 | Membrin |
| AT1G48240 | 0.3 | 2.67E-03 | 0.2 | 3.92E-02 | 0.1 | 2.29E-01 | -0.5 | 3.89E-03 | -0.2 | 1.57E-01 | -0.2 | 2.42E-01 | Plant snare 12 |
| AT1G54110 | 0.7 | 7.04E-04 | 0.2 | 1.58E-01 | 0.5 | 2.52E-02 | 0.1 | 7.56E-01 | 0.2 | 7.35E-01 | 0.0 | 9.59E-01 | Membrane fusion protein Use1 |
| AT4G03330 | -1.3 | 7.89E-06 | -0.1 | 5.95E-01 | -1.2 | 1.12E-04 | -0.7 | 7.42E-02 | 0.2 | 6.81E-01 | -0.9 | 5.14E-02 | Syntaxin-123 |
| AT5G46860 | -0.5 | 1.70E-04 | -0.1 | 2.10E-01 | -0.4 | 2.90E-03 | 0.0 | 8.74E-01 | -0.1 | 4.63E-01 | 0.1 | 6.51E-01 | Syntaxin-22 |
| AT5G05760 | -0.4 | 2.73E-03 | -0.3 | 2.53E-02 | -0.1 | 3.26E-01 | -0.1 | 5.65E-01 | -0.1 | 8.65E-01 | -0.1 | 7.98E-01 | Syntaxin-31 |
| AT3G52400 | -0.4 | 1.04E-02 | -0.2 | 9.86E-02 | -0.2 | 2.95E-01 | 0.2 | 9.99E-02 | -0.3 | 6.29E-02 | 0.6 | 4.55E-03 | Syntaxin-122 |
| AT5G39510 | -0.3 | 4.97E-04 | -0.2 | 2.22E-02 | -0.1 | 6.23E-02 | -0.1 | 6.06E-01 | 0.1 | 7.56E-01 | -0.1 | 4.95E-01 | Vesicle transport v-SNARE 11 |
| AT5G16830 | -0.2 | 4.70E-02 | -0.2 | 9.57E-02 | 0.0 | 7.87E-01 | 0.0 | 9.06E-01 | 0.0 | 9.34E-01 | -0.1 | 8.68E-01 | Syntaxin-21 |
| AT3G09740 | -0.1 | 1.46E-01 | 0.1 | 2.07E-01 | -0.2 | 2.41E-02 | -0.1 | 2.87E-01 | -0.1 | 3.96E-01 | 0.0 | 9.77E-01 | Syntaxin-71 |
| AT5G39630 | 0.0 | 9.90E-01 | 1.2 | 2.54E-05 | -1.2 | 1.53E-04 | -1.8 | 2.64E-04 | -0.3 | 3.83E-01 | -1.5 | 1.05E-02 | Vesicle transport v-SNARE family protein |
| AT3G05710 | 0.2 | 5.22E-02 | 0.0 | 9.44E-01 | 0.2 | 1.05E-01 | 0.0 | 8.43E-01 | 0.0 | 9.60E-01 | 0.0 | 9.31E-01 | Syntaxin-43 |
| AT5G50440 | 0.2 | 6.57E-02 | 0.1 | 5.53E-01 | 0.1 | 2.93E-01 | 0.0 | 9.36E-01 | 0.2 | 6.04E-01 | -0.1 | 7.30E-01 | Membrin-12 |
| AT1G08560 | 0.4 | 1.81E-02 | 0.7 | 7.05E-04 | -0.3 | 7.19E-02 | 0.0 | 9.23E-01 | 0.0 | 9.30E-01 | 0.0 | 9.95E-01 | Syntaxin-related protein KNOLLE |
| AT2G35190 | 0.6 | 9.74E-05 | 0.4 | 3.01E-03 | 0.2 | 6.42E-02 | -0.1 | 4.36E-01 | 0.2 | 2.64E-01 | -0.3 | 1.26E-01 | Novel plant SNARE 11 |
| AT5G58060 | -0.6 | 6.04E-06 | -0.4 | 4.51E-04 | -0.3 | 6.51E-03 | -0.3 | 1.36E-01 | 0.0 | 8.49E-01 | -0.2 | 3.98E-01 | SNARE-like superfamily protein |
| AT1G79070 | -0.2 | 4.94E-02 | -0.1 | 6.59E-01 | -0.2 | 1.90E-01 | -0.4 | 7.66E-02 | -0.4 | 1.12E-01 | 0.0 | 9.80E-01 | Biogenesis of lysosome-related organelles complex 1 subunit 7 |
| AT3G07100 | -0.4 | 5.21E-05 | -0.3 | 4.14E-04 | -0.1 | 2.48E-01 | -0.1 | 5.48E-01 | 0.0 | 8.29E-01 | 0.0 | 8.54E-01 | Protein transport protein SEC24 A |
| AT4G32640 | -0.2 | 1.89E-02 | -0.3 | 6.58E-03 | 0.1 | 5.24E-01 | 0.1 | 3.39E-01 | 0.1 | 6.28E-01 | 0.0 | 8.56E-01 | Protein transport protein SEC24 C |
| AT2G27460 | -0.1 | 2.30E-01 | 0.0 | 6.46E-01 | 0.0 | 5.48E-01 | 0.2 | 4.50E-01 | 0.1 | 8.68E-01 | 0.1 | 7.24E-01 | Protein transport protein SEC23 D |
| AT4G28200 | -0.1 | 5.44E-01 | -0.1 | 4.27E-01 | 0.0 | 8.64E-01 | 0.5 | 2.49E-03 | 0.3 | 1.09E-01 | 0.2 | 2.57E-01 | At4g28200 |
| AT1G56110 | 0.3 | 5.68E-02 | 0.3 | 9.41E-02 | 0.0 | 8.69E-01 | 0.7 | 9.79E-04 | 0.4 | 6.36E-02 | 0.3 | 2.12E-01 | NOP56 |
| AT2G20490 | 0.4 | 3.13E-02 | 0.5 | 7.23E-03 | -0.1 | 4.21E-01 | 0.4 | 4.76E-02 | 0.2 | 4.63E-01 | 0.2 | 3.88E-01 | Nucleolar protein 10 |
| AT4G05410 | 0.5 | 3.88E-03 | 0.2 | 1.78E-01 | 0.3 | 8.78E-02 | 0.7 | 2.44E-04 | 0.3 | 8.17E-02 | 0.4 | 4.48E-02 | U3 snoRNP-associated protein-like YAO |
| AT1G31660 | 0.5 | 3.76E-03 | 0.3 | 6.62E-02 | 0.2 | 1.93E-01 | 0.7 | 2.53E-04 | 0.3 | 1.15E-01 | 0.4 | 2.53E-02 | Bystin |
| AT3G12860 | 0.8 | 1.92E-03 | 0.6 | 1.49E-02 | 0.2 | 3.68E-01 | 0.6 | 2.47E-02 | 0.7 | 2.07E-02 | -0.1 | 8.38E-01 | Nop domain-containing protein |
| AT2G43550 | 1.5 | 3.49E-05 | 1.7 | 2.64E-05 | -0.2 | 3.83E-01 | 0.6 | 1.70E-02 | 0.9 | 2.68E-03 | -0.3 | 4.01E-01 | Knottin scorpion toxin-like domain-containing protein |
| AT2G43530 | 2.0 | 9.58E-04 | 1.0 | 6.68E-02 | 1.0 | 4.72E-02 | 0.6 | 3.37E-01 | 0.8 | 2.09E-01 | -0.2 | 7.99E-01 | Defensin-like protein 194 |
| AT2G43535 | 2.0 | 1.58E-05 | 1.6 | 1.32E-04 | 0.3 | 1.82E-01 | 0.1 | 7.65E-01 | -0.1 | 9.16E-01 | 0.2 | 7.33E-01 | Defensin-like protein 196 |
| AT3G06370 | -1.1 | 2.28E-06 | -0.4 | 2.50E-03 | -0.7 | 4.13E-04 | 0.2 | 3.42E-01 | 0.0 | 9.74E-01 | 0.2 | 4.61E-01 | Sodium hydrogen exchanger 4 |
| AT1G79610 | -0.9 | 6.68E-07 | -0.8 | 3.37E-06 | -0.1 | 2.61E-01 | 0.1 | 7.85E-01 | 0.4 | 1.40E-01 | -0.3 | 2.69E-01 | Cation |
| AT3G05030 | -0.7 | 9.38E-04 | -0.7 | 8.02E-04 | 0.1 | 7.46E-01 | -0.3 | 1.54E-01 | 0.6 | 4.01E-03 | -0.9 | 1.03E-03 | Sodium hydrogen exchanger 2 |
| AT1G14660 | -0.1 | 6.97E-01 | -0.3 | 6.96E-02 | 0.2 | 1.93E-01 | -0.3 | 3.53E-01 | -0.3 | 3.98E-01 | 0.0 | 9.94E-01 | Na+ |
| AT1G54370 | 0.1 | 4.96E-01 | -0.1 | 2.06E-01 | 0.2 | 9.53E-02 | -0.1 | 6.97E-01 | 0.0 | 9.80E-01 | -0.1 | 7.56E-01 | Cation |
| AT3G56450 | -0.1 | 5.22E-01 | -0.3 | 1.21E-01 | 0.2 | 4.03E-01 | -0.4 | 3.29E-01 | -0.5 | 2.44E-01 | 0.1 | 8.50E-01 | Alpha-soluble NSF attachment protein 1 |
| AT1G15460 | -3.5 | 5.56E-08 | -2.2 | 3.84E-07 | -1.2 | 6.80E-04 | | | | | | | HCO3-transporter family |
| AT1G74810 | -2.7 | 4.89E-04 | -3.0 | 6.34E-04 | 0.3 | 7.56E-01 | -0.4 | 4.57E-01 | -0.2 | 7.67E-01 | -0.2 | 7.56E-01 | HCO3-transporter family |
| AT3G62270 | -1.2 | 2.42E-07 | -0.6 | 7.21E-05 | -0.7 | 1.33E-04 | 0.2 | 4.17E-01 | 0.3 | 1.87E-01 | -0.1 | 5.55E-01 | Bicarbonate transporter-like transmembrane domain-containing protein |
| AT2G47160 | 0.0 | 7.09E-01 | -0.1 | 3.35E-01 | 0.1 | 6.01E-01 | -0.5 | 1.41E-02 | -0.3 | 2.01E-01 | -0.2 | 3.43E-01 | HCO3-transporter family |
| AT3G06450 | -0.2 | 8.45E-03 | -0.1 | 1.58E-01 | -0.1 | 1.69E-01 | -0.1 | 5.32E-01 | 0.0 | 9.07E-01 | -0.1 | 7.87E-01 | Probable boron transporter 3 |
| AT5G13750 | -0.6 | 7.61E-05 | -0.5 | 3.04E-04 | -0.1 | 5.25E-01 | 0.4 | 6.09E-03 | 0.4 | 9.49E-03 | 0.0 | 9.75E-01 | Protein ZINC INDUCED FACILITATOR-LIKE 1 |
| AT1G70310 | 0.3 | 2.16E-03 | 0.3 | 1.90E-03 | 0.0 | 8.07E-01 | 0.0 | 7.83E-01 | 0.0 | 8.08E-01 | 0.0 | 9.90E-01 | Spermidine synthase 2 |
| AT5G53120 | -1.4 | 1.34E-06 | -1.3 | 4.49E-06 | -0.1 | 5.89E-01 | -0.2 | 1.13E-01 | 0.2 | 3.62E-01 | -0.4 | 3.75E-02 | Spermidine synthase 3 |
| AT5G19530 | 0.3 | 1.29E-02 | 0.5 | 6.61E-04 | -0.2 | 7.57E-02 | -0.1 | 4.10E-01 | -0.4 | 5.69E-03 | 0.3 | 8.13E-02 | Thermospermine synthase ACAULIS5 |
| AT1G26200 | -1.2 | 8.42E-07 | -0.5 | 7.13E-04 | -0.8 | 2.14E-04 | -0.6 | 8.93E-03 | -0.4 | 9.85E-02 | -0.2 | 5.05E-01 | hypothetical protein |
| AT3G19260 | -0.7 | 1.92E-04 | -0.3 | 2.70E-02 | -0.4 | 1.99E-02 | -0.2 | 9.22E-02 | -0.2 | 2.59E-01 | -0.1 | 8.17E-01 | Ceramide synthase LOH2 |
| AT1G13580 | -0.6 | 1.06E-04 | -0.3 | 4.82E-03 | -0.2 | 4.80E-02 | 0.0 | 8.61E-01 | 0.3 | 1.80E-01 | -0.2 | 3.40E-01 | Ceramide synthase 1 LOH3 |

| | | | | | | | | | | | | | |
|---|---|---|---|---|---|---|---|---|---|---|---|---|---|
| AT3G25540 | 0.0 | 7.42E-01 | 0.1 | 8.35E-02 | -0.2 | 7.10E-02 | 0.0 | 8.10E-01 | 0.1 | 7.26E-01 | 0.0 | 9.14E-01 | TRAM, LAG1 and CLN8 lipid-sensing domain containing protein |
| AT4G22550 | 0.0 | 7.12E-01 | 0.3 | 3.76E-02 | -0.3 | 4.11E-02 | -0.4 | 1.31E-01 | -0.3 | 3.15E-01 | -0.1 | 8.37E-01 | Probable lipid phosphate phosphatase beta |
| AT3G55760 | -1.5 | 1.08E-06 | -0.9 | 4.65E-05 | -0.5 | 3.58E-03 | -0.2 | 3.30E-01 | 0.3 | 3.01E-01 | -0.5 | 8.95E-02 | Protein LIKE EARLY STARVATION, chloroplastic |
| AT5G26570 | -0.1 | 5.79E-01 | 0.0 | 7.84E-01 | -0.1 | 4.55E-01 | -0.2 | 3.84E-01 | 0.1 | 8.19E-01 | -0.2 | 3.88E-01 | Chloroplastidic phosphoglucan, water dikinase |
| AT5G01260 | 0.3 | 2.52E-02 | -0.1 | 3.66E-01 | 0.5 | 1.35E-02 | -0.1 | 6.46E-01 | 0.3 | 8.65E-02 | -0.4 | 5.55E-02 | Carbohydrate-binding-like fold |
| AT2G24940 | -0.6 | 9.26E-05 | -0.3 | 2.76E-02 | -0.4 | 7.37E-03 | -0.4 | 1.55E-01 | -0.2 | 6.15E-01 | -0.2 | 6.00E-01 | Probable steroid-binding protein 3 |
| AT5G13710 | -0.1 | 1.56E-01 | 0.2 | 8.55E-02 | -0.3 | 1.15E-02 | -0.3 | 1.59E-02 | -0.4 | 3.71E-03 | 0.1 | 5.32E-01 | Cycloartenol-C-24-methyltransferase |
| AT3G07020 | -0.7 | 3.39E-06 | -0.4 | 3.56E-04 | -0.3 | 3.29E-03 | -0.5 | 2.72E-03 | -0.2 | 1.30E-01 | -0.2 | 2.68E-01 | AT3G07020 protein |
| AT1G43620 | -0.3 | 2.69E-03 | -0.4 | 5.67E-04 | 0.1 | 2.50E-01 | 0.0 | 1.00E+00 | 0.2 | 4.12E-01 | -0.2 | 4.75E-01 | UDP-Glycosyltransferase superfamily protein |
| AT5G23840 | -0.7 | 7.57E-03 | 0.4 | 8.26E-02 | -1.1 | 2.19E-03 | -0.9 | 7.80E-04 | 0.3 | 2.27E-01 | -1.2 | 4.15E-04 | MD-2-related lipid recognition domain-containing protein |
| AT3G11780 | -0.4 | 1.77E-04 | -0.2 | 6.67E-03 | -0.2 | 5.79E-02 | -0.1 | 7.36E-01 | -0.1 | 5.25E-01 | 0.1 | 7.78E-01 | MD-2-related lipid recognition domain-containing protein |
| AT5G06480 | -0.4 | 8.68E-03 | -0.4 | 6.35E-03 | 0.0 | 7.90E-01 | -0.5 | 1.82E-01 | 0.3 | 5.44E-01 | -0.8 | 1.08E-01 | MD-2-related lipid-recognition domain-containing protein |
| AT3G44100 | 0.0 | 5.43E-01 | 0.0 | 7.10E-01 | 0.0 | 8.64E-01 | 0.2 | 2.92E-01 | 0.2 | 4.00E-01 | 0.0 | 9.78E-01 | At3g44100 |
| AT1G45015 | 0.1 | 3.88E-01 | 0.5 | 7.42E-03 | -0.4 | 6.90E-02 | -0.9 | 1.37E-02 | -0.1 | 8.57E-01 | -0.8 | 8.24E-02 | MD-2-related lipid recognition domain-containing protein |
| AT5G23820 | 1.3 | 2.67E-07 | 1.5 | 2.95E-07 | -0.2 | 5.30E-02 | 0.5 | 1.02E-01 | 0.5 | 1.32E-01 | 0.0 | 9.71E-01 | MD-2-related lipid-recognition protein 3 |
| AT3G09300 | -0.6 | 1.44E-05 | -0.1 | 1.17E-01 | -0.5 | 3.45E-04 | -0.3 | 5.62E-02 | -0.1 | 8.08E-01 | -0.2 | 2.60E-01 | Oxysterol-binding protein-related protein 3B |
| AT2G31020 | -0.6 | 5.39E-05 | 0.0 | 7.36E-01 | -0.6 | 2.51E-04 | 0.0 | 9.86E-01 | 0.3 | 2.71E-01 | -0.3 | 3.63E-01 | OSBP-related protein 1A |
| AT5G59420 | -0.3 | 2.72E-03 | -0.3 | 1.08E-03 | 0.1 | 4.42E-01 | -0.2 | 5.30E-02 | -0.2 | 6.44E-02 | 0.0 | 9.44E-01 | Oxysterol-binding protein-related protein 3C |
| AT5G22020 | -0.6 | 1.68E-03 | -0.7 | 2.03E-03 | 0.0 | 9.46E-01 | 0.0 | 8.45E-01 | 0.4 | 1.26E-02 | -0.5 | 2.63E-02 | Strictosidine synthase conserved region domain-containing protein |
| AT3G51430 | -0.3 | 9.74E-03 | -0.7 | 1.93E-04 | 0.4 | 2.11E-02 | 0.0 | 9.41E-01 | -0.1 | 6.83E-01 | 0.1 | 6.91E-01 | Calcium-dependent phosphotriesterase superfamily protein |
| AT5G06630 | -1.8 | 8.20E-06 | -0.2 | 4.21E-01 | -1.6 | 1.05E-04 | -2.0 | 9.44E-07 | -0.2 | 5.77E-01 | -1.8 | 2.22E-04 | Gb\|AAD23015.1 |
| AT4G08400 | -1.8 | 8.58E-06 | -0.4 | 8.71E-02 | -1.4 | 2.22E-04 | -1.9 | 1.22E-05 | 0.1 | 6.88E-01 | -2.0 | 2.29E-04 | Extensin-like protein |
| AT4G08410 | -1.7 | 1.15E-05 | -0.3 | 2.46E-01 | -1.5 | 1.78E-04 | -2.0 | 2.95E-06 | 0.1 | 6.86E-01 | -2.1 | 9.97E-05 | Extensin-like protein |
| AT1G23720 | -1.6 | 2.01E-06 | -0.3 | 1.42E-01 | -1.4 | 5.25E-05 | 0.1 | 8.93E-01 | 0.4 | 2.42E-01 | -0.4 | 3.61E-01 | F5O8.27 protein |
| AT5G35190 | -1.6 | 7.43E-06 | -0.2 | 2.60E-01 | -1.4 | 1.30E-04 | -0.6 | 8.74E-02 | 0.5 | 1.45E-01 | -1.2 | 1.61E-02 | Proline-rich extensin-like family protein |
| AT2G24980 | -1.4 | 2.43E-05 | 0.1 | 6.89E-01 | -1.5 | 1.01E-04 | -2.3 | 6.05E-07 | -0.1 | 8.81E-01 | -2.3 | 8.25E-05 | Proline-rich extensin-like family protein |
| AT5G06640 | -1.3 | 4.23E-05 | 0.4 | 7.21E-02 | -1.7 | 5.41E-05 | -2.6 | 7.89E-07 | -0.2 | 5.43E-01 | -2.4 | 1.91E-04 | Proline-rich extensin-like family protein |
| AT3G28550 | -1.3 | 1.21E-05 | -0.1 | 5.90E-01 | -1.2 | 1.25E-04 | -0.7 | 1.91E-02 | 0.2 | 5.72E-01 | -0.9 | 2.27E-02 | Extensin domain-containing protein |
| AT3G54590 | -1.3 | 6.75E-05 | 0.3 | 2.02E-01 | -1.6 | 1.05E-04 | -2.2 | 8.27E-06 | 0.0 | 9.81E-01 | -2.2 | 4.14E-04 | Extensin domain-containing protein |
| AT2G43150 | -1.2 | 1.19E-06 | -0.1 | 3.29E-01 | -1.1 | 2.86E-05 | -0.1 | 5.48E-01 | 0.1 | 6.74E-01 | -0.2 | 3.09E-01 | Proline-rich extensin-like family protein |
| AT3G54580 | -1.1 | 1.05E-04 | 0.2 | 2.52E-01 | -1.3 | 1.56E-04 | -1.1 | 1.57E-03 | 0.0 | 9.55E-01 | -1.1 | 9.89E-03 | Proline-rich extensin-like family protein |
| AT4G13390 | -0.7 | 1.08E-02 | -0.4 | 1.08E-01 | -0.3 | 3.09E-01 | -0.5 | 3.47E-01 | 0.7 | 1.36E-01 | -1.2 | 4.62E-02 | Extensin-like protein |
| AT1G26240 | -0.2 | 2.78E-01 | 0.0 | 8.88E-01 | -0.2 | 4.13E-01 | 3.2 | 9.77E-10 | 1.7 | 4.36E-06 | 1.6 | 1.95E-05 | Proline-rich extensin-like family protein |
| AT1G21310 | 0.2 | 1.40E-01 | 0.7 | 3.22E-04 | -0.5 | 4.07E-03 | 0.2 | 3.60E-01 | 0.2 | 2.95E-01 | -0.1 | 8.55E-01 | Extensin-3 |
| AT2G15880 | 0.2 | 4.90E-01 | 0.6 | 8.96E-02 | -0.3 | 3.77E-01 | -0.9 | 2.61E-02 | -1.7 | 1.91E-03 | 0.8 | 2.07E-01 | Pollen-specific leucine-rich repeat extensin-like protein 3 |
| AT4G18670 | 0.5 | 1.77E-03 | 0.5 | 2.96E-03 | 0.0 | 9.20E-01 | 0.1 | 5.76E-01 | 0.1 | 7.89E-01 | 0.0 | 9.09E-01 | Leucine-rich repeat extensin-like protein 5 |
| AT1G26770 | 1.0 | 1.25E-04 | 1.0 | 1.73E-04 | 0.0 | 8.78E-01 | 0.2 | 3.05E-01 | 0.5 | 3.95E-02 | -0.3 | 2.71E-01 | Expansin |
| AT1G76930 | 1.0 | 1.30E-02 | 0.3 | 4.13E-01 | 0.7 | 9.27E-02 | -0.2 | 5.94E-01 | -1.0 | 1.98E-02 | 0.8 | 9.41E-02 | Extensin-1 |
| AT3G60830 | 0.0 | 7.03E-01 | 0.1 | 2.07E-01 | -0.1 | 1.55E-01 | 0.0 | 9.15E-01 | -0.2 | 2.60E-01 | 0.2 | 4.27E-01 | Actin-related protein 7 |
| AT3G12380 | 0.0 | 8.69E-01 | -0.2 | 1.12E-01 | 0.2 | 1.25E-01 | -0.1 | 4.35E-01 | -0.1 | 7.39E-01 | -0.1 | 8.34E-01 | Actin-related protein 5 |
| AT5G05680 | -0.6 | 1.29E-04 | -0.4 | 3.20E-03 | -0.2 | 9.31E-02 | 0.0 | 8.75E-01 | -0.2 | 4.80E-01 | 0.3 | 3.84E-01 | MOS7 |
| AT2G45000 | -0.4 | 8.78E-03 | -0.1 | 3.74E-01 | -0.3 | 8.54E-02 | 0.0 | 8.93E-01 | -0.2 | 2.43E-01 | 0.2 | 3.90E-01 | NUP62 |
| AT1G59660 | -0.3 | 4.91E-03 | -0.5 | 8.64E-04 | 0.1 | 2.58E-01 | 0.1 | 6.01E-01 | 0.0 | 9.69E-01 | 0.1 | 6.98E-01 | Nucleoporin autopeptidase |
| AT1G55540 | -0.3 | 1.51E-03 | -0.3 | 2.96E-03 | 0.0 | 8.59E-01 | -0.3 | 2.45E-01 | 0.0 | 9.81E-01 | -0.3 | 2.31E-01 | Nuclear pore complex protein NUP214 |
| AT4G32910 | -0.3 | 5.73E-04 | -0.3 | 4.04E-04 | 0.0 | 6.27E-01 | -0.3 | 7.04E-02 | -0.3 | 1.10E-01 | 0.0 | 9.88E-01 | Nuclear pore complex protein NUP85 |
| AT1G79280 | -0.1 | 9.24E-02 | -0.2 | 6.44E-03 | 0.1 | 1.62E-01 | 0.3 | 3.10E-02 | 0.1 | 6.48E-01 | 0.2 | 2.29E-01 | Nuclear-pore anchor |
| AT4G38760 | -0.1 | 3.21E-01 | -0.3 | 7.12E-03 | 0.2 | 5.74E-02 | 0.0 | 9.68E-01 | 0.1 | 7.70E-01 | -0.1 | 8.38E-01 | Nucleoporin |
| AT1G14850 | 0.1 | 2.17E-01 | 0.0 | 7.88E-01 | 0.1 | 1.90E-01 | 0.2 | 2.80E-01 | 0.0 | 9.59E-01 | 0.2 | 4.07E-01 | Nuclear pore complex protein NUP155 |
| AT2G05120 | 0.2 | 6.36E-02 | 0.0 | 7.75E-01 | 0.2 | 7.13E-02 | 0.4 | 7.45E-03 | 0.1 | 4.28E-01 | 0.3 | 1.59E-01 | Nucleoporin, Nup133 |
| AT2G41620 | 0.3 | 3.77E-04 | 0.0 | 4.36E-01 | 0.3 | 5.75E-04 | -0.3 | 1.33E-01 | -0.2 | 3.46E-01 | -0.1 | 8.16E-01 | Nuclear pore complex protein NUP93A |
| AT3G14120 | 0.3 | 2.49E-03 | 0.0 | 6.53E-01 | 0.4 | 4.22E-03 | 0.3 | 1.05E-01 | -0.1 | 5.76E-01 | 0.4 | 6.94E-02 | Nuclear pore complex protein |
| AT3G57350 | 0.5 | 6.86E-05 | 0.3 | 4.72E-03 | 0.2 | 3.17E-02 | 0.3 | 1.07E-01 | 0.1 | 7.81E-01 | 0.2 | 3.72E-01 | Nuclear pore complex protein NUP93B |
| AT1G24310 | 0.7 | 1.03E-04 | 0.3 | 2.02E-02 | 0.4 | 1.41E-02 | 0.5 | 1.08E-02 | 0.1 | 7.86E-01 | 0.4 | 7.57E-02 | Nuclear pore complex protein NUP54 |
| AT5G39800 | -0.2 | 1.84E-01 | 0.1 | 4.87E-01 | -0.3 | 1.02E-01 | 0.2 | 6.58E-01 | 0.1 | 7.88E-01 | 0.0 | 9.53E-01 | Uncharacterized protein |
| AT1G24240 | -0.1 | 2.14E-01 | 0.2 | 8.08E-02 | -0.3 | 2.12E-02 | -0.1 | 7.60E-01 | 0.0 | 9.66E-01 | -0.1 | 7.96E-01 | At1g24240 |
| AT1G15810 | -0.1 | 2.59E-01 | 0.0 | 7.41E-01 | -0.1 | 5.10E-01 | 0.3 | 2.85E-02 | 0.4 | 1.46E-02 | -0.1 | 7.41E-01 | Small ribosomal subunit protein uS15c |
| AT4G21460 | -0.1 | 5.95E-01 | 0.1 | 2.24E-01 | -0.2 | 1.35E-01 | 0.1 | 7.29E-01 | 0.0 | 8.93E-01 | 0.0 | 9.25E-01 | Ribosomal protein S24 |
| AT3G18240 | 0.0 | 8.18E-01 | 0.2 | 1.32E-01 | -0.2 | 1.30E-01 | 0.3 | 7.89E-02 | 0.1 | 6.96E-01 | 0.2 | 3.76E-01 | Ribosomal protein S24 |
| AT5G55140 | 0.0 | 8.54E-01 | 0.1 | 2.72E-01 | -0.1 | 2.64E-01 | 0.0 | 8.96E-01 | -0.3 | 2.25E-01 | 0.2 | 4.03E-01 | Large ribosomal subunit protein uL30m |
| AT5G64650 | 0.1 | 3.83E-01 | 0.2 | 1.96E-01 | -0.1 | 7.04E-01 | 0.2 | 2.61E-01 | 0.1 | 8.20E-01 | 0.2 | 5.94E-01 | Putative 50S ribosomal protein L17 |
| AT5G11750 | 0.1 | 2.90E-01 | 0.3 | 3.78E-02 | -0.2 | 3.01E-01 | 0.0 | 8.99E-01 | 0.0 | 8.92E-01 | 0.0 | 9.78E-01 | Ribosomal protein L19 family protein |
| AT1G14620 | 0.2 | 8.96E-02 | 0.4 | 1.26E-02 | -0.2 | 3.22E-01 | 0.4 | 8.37E-03 | 0.0 | 8.64E-01 | 0.4 | 5.95E-02 | Decoy |
| AT4G34555 | 0.2 | 1.85E-01 | 0.4 | 2.67E-02 | -0.2 | 3.22E-01 | 0.3 | 1.49E-01 | 0.2 | 2.98E-01 | 0.0 | 9.05E-01 | Small ribosomal subunit protein eS25x |
| AT5G15220 | 0.2 | 1.18E-01 | 0.5 | 6.67E-03 | -0.3 | 1.52E-01 | 0.0 | 8.29E-01 | 0.1 | 8.35E-01 | 0.0 | 9.81E-01 | Ribosomal protein L27-like |
| AT2G16930 | 0.3 | 4.11E-03 | 0.2 | 1.51E-02 | 0.0 | 5.94E-01 | -0.3 | 5.66E-02 | -0.3 | 5.85E-02 | 0.0 | 9.19E-01 | hypothetical protein |
| AT3G45020 | 0.3 | 1.98E-01 | 0.0 | 9.29E-01 | 0.2 | 3.37E-01 | -0.2 | 5.96E-01 | -0.3 | 4.64E-01 | 0.1 | 8.71E-01 | Ribosomal L18p |
| AT5G52370 | 0.3 | 6.97E-03 | 0.3 | 7.56E-03 | 0.0 | 9.50E-01 | -0.2 | 2.07E-01 | -0.2 | 5.05E-01 | -0.1 | 7.99E-01 | 28S ribosomal S34 protein |
| AT3G02190 | 0.3 | 6.97E-02 | 0.4 | 4.65E-02 | 0.0 | 8.19E-01 | 0.5 | 8.38E-02 | 0.4 | 2.06E-01 | 0.1 | 8.74E-01 | Large ribosomal subunit protein eL39y |
| AT5G20180 | 0.4 | 8.72E-03 | 0.2 | 2.05E-01 | 0.2 | 1.49E-01 | 0.3 | 2.71E-01 | 0.2 | 4.14E-01 | 0.0 | 9.33E-01 | Ribosomal protein |
| AT1G52370 | 0.4 | 3.95E-03 | 0.1 | 2.91E-01 | 0.3 | 5.87E-02 | 0.1 | 5.22E-01 | 0.1 | 8.14E-01 | 0.1 | 8.39E-01 | Large ribosomal subunit protein uL22c |
| AT5G09770 | 0.4 | 6.46E-04 | 0.4 | 1.87E-03 | 0.0 | 6.90E-01 | 0.0 | 7.49E-01 | 0.0 | 9.24E-01 | 0.0 | 9.11E-01 | 50S ribosomal protein L17 |
| AT5G58990 | 0.4 | 6.88E-03 | 0.4 | 6.91E-03 | 0.0 | 9.21E-01 | 0.2 | 2.82E-01 | 0.2 | 4.42E-01 | 0.0 | 9.25E-01 | 28S ribosomal S34 protein |
| AT1G80620 | 0.4 | 3.48E-03 | 0.3 | 3.06E-02 | 0.1 | 3.31E-01 | 0.2 | 2.10E-01 | 0.1 | 7.91E-01 | 0.1 | 5.50E-01 | Small ribosomal subunit protein uS15c |
| AT3G03600 | 0.4 | 1.79E-03 | 0.4 | 3.85E-03 | 0.0 | 8.07E-01 | 0.4 | 5.94E-02 | 0.1 | 7.86E-01 | 0.3 | 1.85E-01 | Small ribosomal subunit protein uS2m |
| AT2G19720 | 0.4 | 1.74E-02 | 0.4 | 1.71E-02 | 0.0 | 9.28E-01 | 0.6 | 6.81E-02 | 0.7 | 3.33E-02 | -0.1 | 7.29E-01 | Small ribosomal subunit protein uS8mz |
| AT5G45590 | 0.4 | 1.41E-02 | 0.6 | 4.23E-03 | -0.1 | 4.82E-01 | 0.5 | 2.29E-02 | 0.4 | 8.32E-02 | 0.1 | 8.15E-01 | Uncharacterized protein |
| AT3G52580 | 0.4 | 8.36E-03 | 0.6 | 1.82E-03 | -0.2 | 3.15E-01 | 0.2 | 3.58E-01 | 0.1 | 6.04E-01 | 0.0 | 8.83E-01 | 40S ribosomal protein S14-3 |
| AT3G06040 | 0.4 | 2.40E-03 | 0.5 | 2.25E-03 | 0.0 | 8.53E-01 | 0.1 | 3.04E-01 | -0.1 | 7.95E-01 | 0.2 | 2.30E-01 | Ribosomal protein L7 |
| AT3G04770 | 0.5 | 3.04E-02 | 0.4 | 6.77E-02 | 0.1 | 7.70E-01 | 0.6 | 2.76E-02 | 0.2 | 5.80E-01 | 0.4 | 1.69E-01 | Small ribosomal subunit protein uS2 |
| AT1G07830 | 0.5 | 6.72E-04 | 0.5 | 1.01E-03 | 0.0 | 9.86E-01 | 0.3 | 1.25E-01 | -0.1 | 6.57E-01 | 0.4 | 7.51E-02 | Large ribosomal subunit protein uL29m |
| AT2G44065 | 0.5 | 1.87E-03 | 0.5 | 2.79E-03 | 0.0 | 9.57E-01 | 0.0 | 9.74E-01 | -0.1 | 7.84E-01 | 0.1 | 8.11E-01 | Ribosomal protein L2 |
| AT2G34480 | 0.5 | 9.11E-03 | 0.6 | 3.83E-03 | -0.1 | 5.37E-01 | 0.3 | 7.81E-02 | 0.0 | 9.13E-01 | 0.2 | 2.71E-01 | Ribosomal protein L18ae |
| AT3G09630 | 0.5 | 3.33E-03 | 0.5 | 5.05E-03 | 0.0 | 9.67E-01 | 0.3 | 1.37E-02 | 0.2 | 1.02E-01 | 0.1 | 6.92E-01 | Ribosomal protein L4 |

| | | | | | | | | | | | | | |
|---|---|---|---|---|---|---|---|---|---|---|---|---|---|
| AT4G11630 | 0.5 | 4.09E-04 | 0.3 | 1.63E-02 | 0.2 | 7.89E-02 | 0.5 | 6.90E-03 | 0.1 | 7.88E-01 | 0.4 | 7.16E-02 | Ribosomal protein L19 family protein |
| AT1G25260 | 0.5 | 4.00E-04 | 0.5 | 6.41E-04 | 0.0 | 9.71E-01 | 0.4 | 3.76E-02 | 0.0 | 9.27E-01 | 0.4 | 8.67E-02 | Ribosome assembly factor mrt4 |
| AT4G28360 | 0.5 | 1.38E-04 | 0.5 | 1.97E-04 | 0.0 | 9.23E-01 | 0.1 | 4.05E-01 | 0.1 | 5.27E-01 | 0.0 | 9.69E-01 | Large ribosomal subunit protein uL22c |
| AT3G44590 | 0.5 | 1.30E-02 | 0.9 | 8.24E-04 | -0.3 | 1.04E-01 | 0.4 | 2.08E-01 | 0.2 | 5.50E-01 | 0.1 | 7.24E-01 | Uncharacterized protein |
| AT3G58700 | 0.5 | 1.94E-03 | 0.7 | 4.64E-04 | -0.2 | 2.57E-01 | 0.3 | 1.65E-02 | 0.1 | 5.88E-01 | 0.2 | 1.67E-01 | 60S ribosomal protein L11 |
| AT3G26360 | 0.5 | 3.04E-03 | 0.5 | 5.97E-03 | 0.0 | 8.47E-01 | 0.4 | 6.86E-02 | 0.1 | 7.68E-01 | 0.3 | 3.11E-01 | Ribosomal protein S21 family protein |
| AT1G47278 | 0.5 | 4.69E-03 | 0.3 | 5.25E-02 | 0.2 | 2.96E-01 | 0.4 | 2.60E-01 | 0.1 | 8.52E-01 | 0.3 | 5.46E-01 | IMS import disulfide relay-system CHCH-CHCH-like Cx9C domain-containing protein |
| AT4G35490 | 0.5 | 2.43E-03 | 0.4 | 1.27E-02 | 0.1 | 4.61E-01 | 0.1 | 5.49E-01 | -0.1 | 8.39E-01 | 0.2 | 4.47E-01 | hypothetical protein |
| AT5G59240 | 0.6 | 1.90E-02 | 0.6 | 1.31E-02 | -0.1 | 8.12E-01 | 0.6 | 2.19E-01 | 0.9 | 6.18E-02 | -0.4 | 5.14E-01 | Small ribosomal subunit protein eS8y |
| AT4G36420 | 0.6 | 4.13E-04 | 0.3 | 2.74E-02 | 0.3 | 5.00E-02 | 0.5 | 5.22E-03 | 0.3 | 1.58E-01 | 0.2 | 2.82E-01 | hypothetical protein |
| AT1G26910 | 0.6 | 9.89E-04 | 0.7 | 3.17E-04 | -0.1 | 3.34E-01 | 0.4 | 3.91E-02 | 0.3 | 1.66E-01 | 0.1 | 7.29E-01 | Ribosomal protein L16p |
| AT3G49010 | 0.6 | 3.25E-03 | 0.7 | 2.04E-03 | -0.1 | 6.45E-01 | 0.5 | 5.94E-04 | 0.3 | 5.42E-02 | 0.2 | 2.17E-01 | Breast basic conserved 1 |
| AT3G06700 | 0.6 | 2.19E-03 | 0.8 | 6.50E-04 | -0.2 | 3.10E-01 | 0.5 | 7.33E-03 | 0.4 | 8.66E-02 | 0.2 | 5.90E-01 | Large ribosomal subunit protein eL29z |
| AT5G27820 | 0.6 | 4.92E-03 | 0.7 | 2.26E-03 | -0.1 | 6.08E-01 | 0.5 | 2.50E-02 | 0.2 | 4.68E-01 | 0.3 | 2.86E-01 | hypothetical protein |
| AT4G29430 | 0.6 | 2.26E-03 | 0.1 | 3.97E-01 | 0.4 | 2.81E-02 | 0.5 | 7.64E-02 | 0.4 | 2.93E-01 | 0.2 | 6.83E-01 | Small ribosomal subunit protein uS8my |
| AT1G18540 | 0.6 | 2.52E-03 | 0.7 | 1.82E-03 | -0.1 | 6.93E-01 | 0.4 | 6.66E-03 | 0.3 | 1.33E-01 | 0.2 | 4.38E-01 | AT1G18540 protein |
| AT5G03850 | 0.6 | 2.05E-03 | 0.7 | 8.59E-04 | -0.1 | 4.35E-01 | 0.3 | 9.38E-02 | 0.1 | 5.48E-01 | 0.2 | 5.14E-01 | 40S ribosomal protein S28 |
| AT1G09590 | 0.6 | 2.85E-03 | 0.7 | 2.12E-03 | -0.1 | 7.14E-01 | 0.5 | 1.01E-03 | 0.4 | 7.99E-03 | 0.1 | 7.35E-01 | Large ribosomal subunit protein eL21z |
| AT5G27330 | 0.6 | 2.53E-03 | 0.5 | 1.25E-02 | 0.1 | 4.84E-01 | 0.5 | 1.64E-03 | 0.5 | 3.83E-03 | 0.0 | 9.95E-01 | hypothetical protein |
| AT1G72370 | 0.6 | 2.34E-03 | 0.7 | 1.37E-03 | -0.1 | 5.94E-01 | 0.4 | 1.97E-02 | 0.1 | 4.40E-01 | 0.2 | 2.98E-01 | Small ribosomal subunit protein uS2 |
| AT1G43170 | 0.6 | 7.50E-04 | 0.6 | 1.16E-03 | 0.0 | 1.00E+00 | 0.3 | 2.37E-02 | 0.1 | 3.60E-01 | 0.1 | 4.49E-01 | Uncharacterized protein At1g43170 |
| AT5G18790 | 0.6 | 1.43E-03 | 0.8 | 6.71E-04 | -0.1 | 5.32E-01 | 0.2 | 3.99E-01 | 0.0 | 9.64E-01 | 0.3 | 5.07E-01 | 50S ribosomal protein L33, chloroplastic |
| AT4G37660 | 0.7 | 3.98E-04 | 0.6 | 1.23E-03 | 0.1 | 6.77E-01 | 0.1 | 7.64E-01 | 0.0 | 9.85E-01 | 0.1 | 8.15E-01 | hypothetical protein |
| AT4G15000 | 0.7 | 3.84E-04 | 0.7 | 4.06E-04 | 0.0 | 7.65E-01 | 0.4 | 4.65E-02 | 0.2 | 2.73E-01 | 0.1 | 6.53E-01 | hypothetical protein |
| AT3G59650 | 0.7 | 2.22E-04 | 0.7 | 2.19E-04 | 0.0 | 7.50E-01 | 0.3 | 1.42E-01 | 0.2 | 5.02E-01 | 0.1 | 6.59E-01 | Large ribosomal subunit protein mL43 |
| AT1G07070 | 0.7 | 9.82E-04 | 0.9 | 1.78E-04 | -0.2 | 1.71E-01 | 0.4 | 1.32E-01 | 0.1 | 7.21E-01 | 0.2 | 4.60E-01 | hypothetical protein |
| AT1G66580 | 0.7 | 6.85E-04 | 0.6 | 3.82E-03 | 0.1 | 4.35E-01 | 0.3 | 7.26E-02 | 0.1 | 8.26E-01 | 0.3 | 2.98E-01 | AT1G66580 protein |
| AT5G40080 | 0.7 | 2.66E-03 | 0.4 | 4.20E-02 | 0.3 | 2.33E-01 | 0.3 | 3.13E-01 | 0.2 | 6.73E-01 | 0.1 | 7.62E-01 | At5g40080 |
| AT3G55170 | 0.7 | 1.15E-04 | 0.6 | 4.49E-04 | 0.1 | 5.47E-01 | 0.4 | 3.29E-02 | 0.2 | 3.68E-01 | 0.2 | 4.60E-01 | Ribosomal L29 family protein |
| AT2G32060 | 0.7 | 1.85E-04 | 0.8 | 1.65E-04 | -0.1 | 6.05E-01 | 0.5 | 1.02E-02 | 0.2 | 2.21E-01 | 0.2 | 3.80E-01 | Small ribosomal subunit protein eS12y |
| AT2G34520 | 0.7 | 1.63E-05 | 0.4 | 1.25E-03 | 0.3 | 1.33E-02 | 0.2 | 3.79E-01 | 0.0 | 9.83E-01 | 0.2 | 5.29E-01 | Mitochondrial ribosomal protein S14 |
| AT5G02870 | 0.7 | 1.40E-04 | 0.6 | 9.43E-04 | 0.1 | 3.39E-01 | 0.4 | 2.23E-03 | 0.2 | 1.08E-01 | 0.2 | 3.06E-01 | 60S ribosomal protein-like |
| AT5G56710 | 0.7 | 7.76E-04 | 0.8 | 8.44E-04 | 0.0 | 8.26E-01 | 0.4 | 1.41E-02 | 0.2 | 2.90E-01 | 0.2 | 3.80E-01 | Ribosomal protein L31e family protein |
| AT5G56940 | 0.7 | 7.89E-05 | 0.5 | 1.22E-03 | 0.2 | 1.37E-01 | 0.4 | 1.80E-02 | 0.1 | 7.61E-01 | 0.3 | 1.47E-01 | Small ribosomal subunit protein bS16m |
| AT5G18380 | 0.7 | 1.92E-05 | 0.7 | 9.77E-05 | 0.1 | 4.10E-01 | 0.2 | 4.49E-01 | 0.0 | 9.18E-01 | 0.2 | 5.27E-01 | Ribosomal protein S5 domain 2-like superfamily protein |
| AT5G10360 | 0.7 | 1.88E-04 | 0.7 | 5.71E-04 | 0.1 | 6.98E-01 | 0.1 | 4.96E-01 | 0.0 | 8.20E-01 | 0.1 | 4.85E-01 | Ribosomal protein S6e |
| AT1G15930 | 0.8 | 3.59E-04 | 0.8 | 2.99E-04 | -0.1 | 6.27E-01 | 0.1 | 3.54E-01 | -0.1 | 7.33E-01 | 0.2 | 3.04E-01 | Small ribosomal subunit protein eS12z |
| AT1G26880 | 0.8 | 4.46E-04 | 0.7 | 1.03E-03 | 0.0 | 8.19E-01 | 0.5 | 1.03E-02 | 0.3 | 1.68E-01 | 0.2 | 4.66E-01 | Ribosomal protein L34e superfamily protein |
| AT2G27710 | 0.8 | 1.83E-04 | 0.8 | 2.97E-04 | 0.0 | 9.61E-01 | 0.5 | 2.06E-02 | 0.2 | 4.29E-01 | 0.3 | 3.04E-01 | 60S acidic ribosomal protein family |
| AT3G46040 | 0.8 | 4.30E-05 | 0.7 | 1.57E-04 | 0.1 | 5.93E-01 | 0.2 | 4.09E-01 | -0.1 | 8.39E-01 | 0.2 | 4.40E-01 | Small ribosomal subunit protein uS8x |
| AT5G15760 | 0.8 | 5.17E-05 | 0.5 | 1.74E-03 | 0.3 | 6.28E-02 | -0.2 | 5.90E-01 | -0.1 | 8.53E-01 | -0.1 | 8.52E-01 | Small ribosomal subunit protein cS23y |
| AT1G14320 | 0.8 | 9.13E-05 | 0.7 | 6.28E-04 | 0.2 | 3.22E-01 | 0.4 | 1.54E-03 | 0.2 | 3.22E-01 | 0.3 | 1.02E-01 | Ribosomal protein L16p |
| AT5G27700 | 0.8 | 1.88E-06 | 0.8 | 8.25E-06 | 0.1 | 4.46E-01 | 0.2 | 4.17E-01 | 0.0 | 9.79E-01 | 0.2 | 6.14E-01 | Small ribosomal subunit protein eS21y |
| AT3G56340 | 0.9 | 2.73E-04 | 0.9 | 3.32E-04 | 0.0 | 8.20E-01 | 0.2 | 1.79E-01 | 0.1 | 8.05E-01 | 0.2 | 5.15E-01 | 40S ribosomal protein S26 |
| AT4G14320 | 0.9 | 1.00E-04 | 0.8 | 2.26E-04 | 0.0 | 8.75E-01 | 0.4 | 5.21E-02 | 0.2 | 4.28E-01 | 0.2 | 5.23E-01 | Zinc-binding ribosomal protein family protein |
| AT1G09690 | 0.9 | 1.13E-04 | 0.8 | 3.65E-04 | 0.1 | 6.71E-01 | 0.4 | 9.38E-04 | 0.2 | 1.06E-01 | 0.2 | 2.00E-01 | Large ribosomal subunit protein eL21z |
| AT1G01100 | 0.9 | 2.83E-05 | 0.9 | 5.15E-05 | 0.0 | 9.21E-01 | 0.3 | 3.02E-01 | 0.0 | 9.87E-01 | 0.3 | 4.58E-01 | 60S acidic ribosomal protein family |
| AT4G31700 | 0.9 | 8.52E-05 | 0.8 | 3.70E-04 | 0.1 | 5.14E-01 | 0.5 | 1.43E-03 | 0.3 | 4.81E-02 | 0.2 | 3.62E-01 | Ribosomal protein S6 |
| AT4G25890 | 0.9 | 1.98E-05 | 0.8 | 5.77E-05 | 0.1 | 7.02E-01 | 0.5 | 1.99E-02 | 0.3 | 2.97E-01 | 0.2 | 4.44E-01 | Large ribosomal subunit protein P3z |
| AT1G07770 | 0.9 | 8.45E-06 | 0.9 | 1.71E-05 | 0.0 | 9.37E-01 | 0.0 | 9.27E-01 | -0.2 | 3.67E-01 | 0.3 | 4.01E-01 | Small ribosomal subunit protein uS8z |
| AT5G57290 | 0.9 | 4.58E-06 | 0.7 | 3.99E-05 | 0.2 | 1.49E-01 | 0.4 | 5.72E-02 | 0.2 | 4.12E-01 | 0.2 | 5.55E-01 | 60S acidic ribosomal protein family |
| AT3G13882 | 0.9 | 4.23E-06 | 0.6 | 2.22E-04 | 0.3 | 9.38E-03 | 0.1 | 4.82E-01 | -0.1 | 7.34E-01 | 0.2 | 3.86E-01 | Ribosomal protein L34 |
| AT2G32220 | 0.9 | 2.14E-04 | 0.4 | 3.83E-02 | 0.5 | 2.25E-02 | 0.8 | 5.27E-03 | 0.4 | 2.28E-01 | 0.4 | 1.88E-01 | Large ribosomal subunit protein eL27z |
| AT3G06680 | 0.9 | 1.76E-04 | 0.7 | 1.38E-03 | 0.2 | 2.89E-01 | 0.7 | 5.15E-03 | 0.5 | 6.13E-02 | 0.2 | 5.66E-01 | 60S ribosomal protein L29 |
| AT3G04400 | 0.9 | 6.03E-06 | 0.7 | 1.47E-04 | 0.3 | 3.27E-02 | 0.3 | 2.29E-01 | 0.0 | 9.99E-01 | 0.3 | 4.12E-01 | Ribosomal protein L14p |
| AT5G59850 | 1.0 | 4.10E-05 | 0.8 | 2.95E-04 | 0.2 | 2.79E-01 | 0.7 | 5.29E-03 | 0.4 | 1.13E-01 | 0.3 | 4.25E-01 | Small ribosomal subunit protein uS8z |
| AT1G67430 | 1.0 | 1.55E-05 | 0.8 | 1.07E-04 | 0.2 | 2.64E-01 | 0.4 | 2.12E-03 | 0.2 | 2.67E-01 | 0.3 | 1.68E-01 | Ribosomal protein L22p |
| AT1G04480 | 1.0 | 2.00E-04 | 0.8 | 1.52E-03 | 0.2 | 3.06E-01 | 0.3 | 7.27E-02 | 0.0 | 9.71E-01 | 0.3 | 1.32E-01 | F19P19.5 protein |
| AT3G23390 | 1.0 | 3.63E-05 | 0.8 | 2.40E-04 | 0.2 | 3.04E-01 | 0.6 | 1.00E-02 | 0.3 | 1.38E-01 | 0.2 | 5.15E-01 | Large ribosomal subunit protein eL42z |
| AT1G17560 | 1.0 | 4.13E-05 | 0.6 | 4.62E-03 | 0.4 | 1.90E-02 | 0.5 | 4.00E-02 | 0.3 | 2.11E-01 | 0.1 | 6.23E-01 | Ribosomal protein L14p |
| AT2G37600 | 1.0 | 9.25E-06 | 0.7 | 3.10E-04 | 0.3 | 2.58E-02 | 0.6 | 4.88E-02 | 0.1 | 9.09E-01 | 0.5 | 1.95E-01 | 60S ribosomal protein L36 |
| AT2G43460 | 1.0 | 8.57E-06 | 0.9 | 5.19E-05 | 0.1 | 2.95E-01 | 0.3 | 1.24E-01 | 0.1 | 6.95E-01 | 0.2 | 5.00E-01 | 60S ribosomal protein L38 |
| AT3G59540 | 1.1 | 9.42E-06 | 0.9 | 6.08E-05 | 0.2 | 2.71E-01 | 0.4 | 3.63E-02 | 0.1 | 6.72E-01 | 0.3 | 2.89E-01 | Large ribosomal subunit protein eL38z |
| AT3G06320 | 1.1 | 1.09E-03 | 0.6 | 4.44E-02 | 0.5 | 1.01E-01 | 0.9 | 2.26E-02 | 0.5 | 2.41E-01 | 0.4 | 4.68E-01 | 50S ribosomal protein L33, chloroplastic |
| AT3G53740 | 1.1 | 6.85E-06 | 0.9 | 4.36E-05 | 0.2 | 2.65E-01 | 0.4 | 9.45E-02 | 0.1 | 6.92E-01 | 0.3 | 4.48E-01 | Large ribosomal subunit protein eL36y |
| AT1G57660 | 1.2 | 3.84E-06 | 0.9 | 4.89E-05 | 0.3 | 7.27E-02 | 0.4 | 2.50E-02 | 0.1 | 5.33E-01 | 0.3 | 3.11E-01 | Large ribosomal subunit protein eL21x |
| AT2G27720 | 1.2 | 1.56E-06 | 0.9 | 2.09E-05 | 0.3 | 4.60E-02 | 0.2 | 3.28E-01 | -0.1 | 7.59E-01 | 0.3 | 2.99E-01 | 60S acidic ribosomal protein family |
| AT1G57860 | 1.2 | 7.17E-07 | 0.9 | 8.60E-06 | 0.3 | 3.51E-02 | 0.5 | 4.29E-03 | 0.1 | 5.22E-01 | 0.4 | 1.07E-01 | C-JID domain-containing protein |
| AT3G53890 | 1.2 | 2.02E-06 | 0.8 | 4.99E-05 | 0.4 | 1.72E-02 | 0.6 | 5.86E-03 | 0.4 | 1.02E-01 | 0.2 | 4.88E-01 | 40S ribosomal protein S21 |
| AT2G33370 | 1.3 | 1.91E-06 | 0.9 | 5.41E-05 | 0.4 | 1.32E-02 | 0.3 | 1.01E-01 | 0.1 | 8.76E-01 | 0.3 | 3.19E-01 | Ribosomal protein L14p |
| AT3G56020 | 1.4 | 3.63E-05 | 0.9 | 1.00E-03 | 0.5 | 5.09E-02 | 0.5 | 9.01E-02 | 0.2 | 5.58E-01 | 0.3 | 5.39E-01 | Large ribosomal subunit protein eL41z |
| AT3G08520 | 1.6 | 4.38E-06 | 1.1 | 1.59E-04 | 0.5 | 1.48E-02 | 0.5 | 1.60E-01 | 0.2 | 7.04E-01 | 0.3 | 5.66E-01 | Large ribosomal subunit protein eL41z |
| AT1G56045 | 1.6 | 3.30E-05 | 1.0 | 1.08E-03 | 0.6 | 3.94E-02 | 0.5 | 1.29E-01 | 0.2 | 6.89E-01 | 0.3 | 5.36E-01 | Large ribosomal subunit protein eL41z |
| AT3G11120 | 1.6 | 3.12E-06 | 1.1 | 1.23E-04 | 0.6 | 1.02E-02 | 0.6 | 7.09E-02 | 0.4 | 3.40E-01 | 0.2 | 6.77E-01 | Large ribosomal subunit protein eL41z |
| AT2G40205 | 1.7 | 2.20E-05 | 0.8 | 9.97E-03 | 0.9 | 4.18E-03 | 0.5 | 4.05E-01 | 0.5 | 5.17E-01 | 0.0 | 9.66E-01 | Large ribosomal subunit protein eL41z |
| AT5G65220 | 2.0 | 6.01E-07 | 1.4 | 1.34E-05 | 0.6 | 9.16E-03 | 0.1 | 7.06E-01 | -0.1 | 7.19E-01 | 0.2 | 5.59E-01 | Large ribosomal subunit protein uL29c |
| AT1G68590 | 2.4 | 1.44E-08 | 1.9 | 8.06E-08 | 0.5 | 7.01E-04 | 0.3 | 5.71E-02 | 0.2 | 1.48E-01 | 0.0 | 8.89E-01 | Ribosomal protein PSRP-3 |
| AT1G07320 | 2.5 | 1.44E-08 | 2.0 | 8.06E-08 | 0.5 | 8.24E-04 | 0.3 | 3.22E-02 | 0.0 | 9.76E-01 | 0.3 | 1.07E-01 | Ribosomal protein L4 |
| AT2G03130 | | | | | | | 0.990922021 | 6.18E-01 | -0.347654943 | 9.12E-01 | | | Ribosomal protein L7 |
| AT3G17626 | | | | | | | 0.0 | 9.51E-01 | -0.1 | 8.56E-01 | 0.2 | 8.38E-01 | hypothetical protein |
| AT3G43980 | 0.9 | 4.17E-05 | 0.7 | 5.16E-04 | 0.2 | 1.55E-01 | 0.5 | 5.14E-02 | 0.2 | 4.96E-01 | 0.3 | 4.41E-01 | Small ribosomal subunit protein uS14z |
| AT3G44010 | 1.0 | 2.70E-05 | 0.8 | 3.72E-04 | 0.3 | 1.16E-01 | 0.5 | 6.89E-02 | 0.2 | 5.40E-01 | 0.3 | 4.98E-01 | Small ribosomal subunit protein uS14z |
| AT3G51890 | -0.6 | 7.35E-04 | -0.7 | 3.83E-04 | 0.1 | 4.88E-01 | -0.1 | 7.34E-01 | -0.1 | 6.07E-01 | 0.1 | 8.82E-01 | Clathrin light chain |

| | | | | | | | | | | | | | |
|---|---|---|---|---|---|---|---|---|---|---|---|---|---|
| AT4G31480 | -0.3 | 6.53E-04 | -0.1 | 1.03E-01 | -0.2 | 2.12E-02 | 0.0 | 7.80E-01 | 0.0 | 9.40E-01 | -0.1 | 8.06E-01 | Coatomer subunit beta |
| AT1G62020 | -0.3 | 2.16E-03 | -0.2 | 1.41E-02 | -0.1 | 3.92E-01 | 0.0 | 9.31E-01 | 0.2 | 5.52E-01 | -0.2 | 6.78E-01 | Uncharacterized protein At1g62020 |
| AT4G34450 | -0.3 | 1.31E-03 | -0.1 | 1.81E-01 | -0.2 | 2.59E-02 | 0.0 | 7.96E-01 | 0.0 | 8.28E-01 | 0.0 | 9.97E-01 | Coatomer subunit gamma |
| AT2G34840 | -0.2 | 2.79E-02 | 0.1 | 3.71E-01 | -0.3 | 1.53E-02 | 0.0 | 9.72E-01 | 0.3 | 9.29E-02 | -0.3 | 1.37E-01 | Coatomer epsilon subunit |
| AT2G30050 | -0.2 | 7.96E-02 | 0.2 | 9.61E-02 | -0.3 | 7.46E-03 | -0.2 | 1.83E-01 | -0.1 | 5.50E-01 | -0.1 | 7.20E-01 | Protein transport protein SEC13 homolog B |
| AT1G29150 | -0.2 | 7.23E-02 | -0.2 | 1.13E-01 | 0.0 | 8.85E-01 | 0.1 | 2.02E-01 | 0.1 | 2.99E-01 | 0.0 | 9.81E-01 | 26S proteasome non-ATPase regulatory subunit 11 homolog |
| AT1G52360 | -0.2 | 1.24E-02 | -0.1 | 3.72E-01 | -0.1 | 9.88E-02 | 0.2 | 1.26E-01 | 0.2 | 2.64E-01 | 0.0 | 9.14E-01 | Coatomer subunit beta' |
| AT1G30630 | -0.1 | 2.45E-01 | 0.0 | 6.55E-01 | -0.1 | 1.49E-01 | 0.1 | 4.15E-01 | 0.2 | 9.31E-02 | -0.1 | 4.90E-01 | Coatomer subunit epsilon-1 |
| AT3G15980 | -0.1 | 2.60E-01 | 0.0 | 7.94E-01 | -0.1 | 4.55E-01 | 0.2 | 7.47E-02 | 0.1 | 3.76E-01 | 0.1 | 6.65E-01 | Coatomer subunit beta' |
| AT1G79990 | -0.1 | 2.31E-01 | 0.0 | 9.95E-01 | -0.1 | 2.87E-01 | 0.0 | 9.93E-01 | 0.1 | 7.35E-01 | -0.1 | 7.95E-01 | Coatomer subunit beta' |
| AT2G21390 | 0.0 | 5.03E-01 | -0.1 | 4.09E-01 | 0.0 | 8.73E-01 | 0.1 | 7.99E-01 | 0.0 | 9.76E-01 | 0.1 | 8.37E-01 | Coatomer subunit alpha-2 |
| AT3G01340 | 0.0 | 5.25E-01 | -0.1 | 1.67E-01 | 0.1 | 4.67E-01 | -0.2 | 1.52E-01 | -0.3 | 4.80E-02 | 0.1 | 6.02E-01 | Protein transport protein SEC13 homolog A |
| AT4G31490 | 0.0 | 5.17E-01 | 0.1 | 1.44E-01 | -0.1 | 4.25E-01 | 0.2 | 1.48E-01 | 0.1 | 5.86E-01 | 0.1 | 6.47E-01 | Coatomer subunit beta-2 |
| AT1G64350 | 0.2 | 4.45E-02 | 0.0 | 9.86E-01 | 0.2 | 7.69E-02 | 0.0 | 8.36E-01 | 0.1 | 6.95E-01 | -0.1 | 6.12E-01 | Protein SEH1 |
| AT5G51040 | -0.3 | 2.99E-03 | -0.1 | 1.81E-01 | -0.2 | 5.67E-02 | 0.1 | 3.20E-01 | 0.3 | 2.38E-02 | -0.2 | 2.30E-01 | Succinate dehydrogenase assembly factor |
| AT5G01340 | 0.1 | 3.63E-01 | 0.3 | 2.46E-02 | -0.2 | 1.83E-01 | 0.3 | 1.30E-01 | 0.1 | 8.65E-01 | 0.3 | 2.80E-01 | Mitochondrial succinate-fumarate transporter 1 |
| AT4G02280 | -2.6 | 2.92E-07 | -2.5 | 9.21E-07 | -0.2 | 5.24E-01 | -1.1 | 1.81E-02 | -0.6 | 2.30E-01 | -0.6 | 3.68E-01 | Sucrose synthase 3 |
| AT5G20830 | -0.9 | 7.19E-07 | -0.6 | 1.65E-05 | -0.3 | 7.90E-03 | -0.5 | 2.28E-03 | -0.1 | 4.23E-01 | -0.3 | 8.82E-02 | Sucrose synthase 1 |
| AT5G37180 | -0.9 | 3.20E-05 | -0.7 | 2.36E-04 | -0.2 | 2.54E-01 | -0.1 | 7.42E-01 | 0.3 | 5.09E-01 | -0.4 | 3.46E-01 | Sucrose synthase |
| AT1G73370 | -0.5 | 8.02E-04 | -0.3 | 9.16E-03 | -0.1 | 2.50E-01 | 0.3 | 2.32E-01 | 0.3 | 2.64E-01 | 0.0 | 9.75E-01 | Sucrose synthase |
| AT3G43190 | -0.1 | 4.53E-01 | 0.2 | 3.40E-01 | -0.3 | 1.63E-01 | -0.2 | 5.85E-01 | -0.2 | 6.68E-01 | 0.0 | 9.80E-01 | Sucrose synthase 4 |
| AT5G11110 | -2.5 | 2.91E-08 | -1.7 | 3.88E-07 | -0.8 | 3.75E-04 | -0.4 | 8.32E-02 | -0.3 | 3.52E-01 | -0.2 | 6.30E-01 | Probable sucrose-phosphate synthase 2 |
| AT4G10120 | 0.6 | 4.07E-04 | 0.3 | 6.53E-02 | 0.4 | 1.93E-02 | 0.7 | 8.78E-04 | 0.7 | 5.27E-03 | 0.1 | 7.76E-01 | Sucrose-phosphate synthase |
| AT1G09960 | -1.2 | 2.10E-06 | -0.6 | 1.81E-04 | -0.5 | 3.10E-03 | -0.2 | 2.25E-01 | 0.0 | 9.76E-01 | -0.2 | 4.28E-01 | Sucrose transport protein SUC4 |
| AT5G23660 | 0.1 | 2.99E-01 | 0.8 | 5.25E-05 | -0.7 | 4.71E-04 | 0.3 | 8.49E-02 | 0.2 | 3.24E-01 | 0.1 | 7.31E-01 | Bidirectional sugar transporter SWEET12 |
| AT3G48740 | 0.6 | 1.39E-04 | 0.6 | 2.15E-04 | 0.0 | 9.17E-01 | 0.1 | 3.18E-01 | -0.1 | 4.69E-01 | 0.2 | 1.59E-01 | Bidirectional sugar transporter SWEET11 |
| AT1G22710 | -0.7 | 5.14E-06 | -0.5 | 9.87E-05 | -0.2 | 4.23E-02 | -0.3 | 6.75E-02 | -0.2 | 3.15E-01 | -0.1 | 6.74E-01 | Sucrose transport protein SUC2 |
| AT1G71880 | 0.8 | 1.14E-05 | 0.6 | 2.51E-04 | 0.2 | 4.70E-02 | 0.4 | 4.07E-03 | 0.1 | 5.35E-01 | 0.3 | 1.04E-01 | Sucrose transport protein SUC1 |
| AT5G20280 | -0.7 | 9.29E-06 | -1.0 | 2.67E-06 | 0.3 | 1.72E-02 | -0.7 | 4.29E-04 | -0.1 | 7.68E-01 | -0.6 | 1.17E-02 | Sucrose-phosphate synthase 1 |
| AT3G05155 | -1.9 | 4.71E-06 | -0.5 | 9.71E-03 | -1.4 | 3.36E-04 | -1.0 | 2.38E-03 | -0.5 | 7.03E-02 | -0.5 | 2.20E-01 | Major facilitator superfamily profile domain-containing protein |
| AT4G04750 | -1.5 | 1.23E-04 | -0.9 | 2.37E-03 | -0.5 | 9.53E-02 | -0.2 | 4.00E-01 | 0.7 | 8.84E-03 | -0.9 | 5.81E-03 | Major facilitator superfamily protein |
| AT3G05150 | -1.1 | 9.66E-06 | 0.1 | 2.88E-01 | -1.3 | 5.41E-05 | -0.8 | 2.67E-02 | 0.2 | 6.03E-01 | -1.0 | 2.70E-02 | Major facilitator superfamily protein |
| AT1G08920 | -0.9 | 4.57E-06 | -0.5 | 2.95E-04 | -0.4 | 7.16E-03 | 0.1 | 4.69E-01 | 0.3 | 1.79E-01 | -0.1 | 6.05E-01 | ERD six-like 1 |
| AT3G05165 | -0.8 | 2.33E-06 | -0.4 | 3.74E-04 | -0.4 | 1.61E-03 | -0.3 | 5.27E-02 | 0.1 | 6.17E-01 | -0.4 | 4.48E-02 | Major facilitator superfamily protein |
| AT5G27350 | 0.0 | 8.74E-01 | 0.1 | 7.34E-01 | 0.0 | 8.74E-01 | -0.3 | 5.21E-02 | -0.3 | 8.42E-02 | 0.0 | 9.97E-01 | Major facilitator superfamily profile domain-containing protein |
| AT1G08900 | 0.1 | 1.83E-01 | 0.2 | 1.41E-01 | 0.0 | 8.73E-01 | -0.1 | 7.30E-01 | -0.2 | 3.04E-01 | 0.1 | 6.25E-01 | AT1G08900 protein |
| AT3G05160 | 0.2 | 6.65E-02 | 0.5 | 4.19E-03 | -0.2 | 1.53E-01 | 0.0 | 9.37E-01 | -0.1 | 5.02E-01 | 0.1 | 5.49E-01 | Major facilitator superfamily protein |
| AT1G54730 | 0.3 | 7.95E-03 | 0.3 | 8.51E-03 | 0.0 | 9.44E-01 | 0.1 | 8.04E-01 | 0.2 | 5.91E-01 | -0.1 | 7.94E-01 | Major facilitator superfamily protein |
| AT1G19450 | 0.3 | 9.01E-04 | 0.1 | 1.62E-01 | 0.2 | 1.96E-02 | -0.5 | 1.41E-02 | 0.2 | 2.62E-01 | -0.7 | 5.96E-03 | hypothetical protein |
| AT5G27360 | 0.5 | 2.19E-03 | 0.2 | 1.26E-01 | 0.3 | 8.74E-02 | -1.0 | 3.20E-02 | -1.1 | 2.92E-02 | 0.1 | 9.34E-01 | Major facilitator superfamily protein |
| AT3G14770 | 0.7 | 1.24E-04 | 0.5 | 2.48E-03 | 0.2 | 1.26E-01 | 0.0 | 9.36E-01 | -0.3 | 3.35E-01 | 0.3 | 3.97E-01 | Bidirectional sugar transporter SWEET2 |
| AT2G48020 | 1.1 | 3.75E-06 | 0.9 | 3.60E-05 | 0.2 | 1.07E-01 | 0.2 | 1.95E-01 | -0.2 | 3.78E-01 | 0.4 | 8.76E-02 | Major facilitator superfamily protein |
| AT3G20460 | -2.0 | 2.52E-06 | -0.5 | 1.52E-02 | -1.5 | 1.44E-04 | -2.4 | 8.99E-05 | -0.1 | 8.07E-01 | -2.3 | 2.81E-03 | Putative sugar transporter ERD6-like 13 |
| AT1G08930 | -0.1 | 2.23E-01 | 0.1 | 4.47E-01 | -0.2 | 8.32E-02 | 0.3 | 1.18E-01 | -0.6 | 2.97E-03 | 0.9 | 1.56E-03 | Sugar transporter ERD6 |
| AT1G75220 | 0.3 | 3.34E-03 | 0.7 | 1.59E-05 | -0.4 | 1.36E-03 | -0.3 | 4.60E-02 | -0.5 | 2.69E-03 | 0.2 | 2.03E-01 | Sugar transporter ERD6-like 6 |
| AT5G18840 | 2.0 | 1.56E-06 | 1.7 | 9.38E-06 | 0.2 | 2.12E-01 | 1.3 | 4.20E-05 | 0.8 | 5.65E-03 | 0.5 | 6.54E-02 | Sugar transporter ERD6-like 16 |
| AT3G45080 | -2.3 | 1.14E-06 | -0.5 | 3.72E-03 | -1.8 | 8.53E-05 | -2.0 | 1.16E-02 | 0.2 | 7.78E-01 | -2.2 | 1.83E-02 | Cytosolic sulfotransferase 6 |
| AT5G43690 | -2.0 | 1.74E-07 | -1.0 | 1.09E-05 | -1.0 | 2.33E-04 | 0.2 | 5.59E-01 | 0.4 | 3.29E-01 | -0.2 | 7.56E-01 | Cytosolic sulfotransferase 1 |
| AT1G28170 | -1.4 | 1.80E-05 | 0.3 | 2.53E-02 | -1.7 | 4.27E-05 | -0.8 | 8.24E-02 | 0.5 | 2.24E-01 | -1.3 | 2.08E-02 | Cytosolic sulfotransferase 7 |
| AT1G13430 | -1.2 | 7.59E-04 | -0.1 | 5.99E-01 | -1.1 | 6.11E-03 | 0.2 | 4.67E-01 | 0.3 | 3.53E-01 | -0.1 | 8.28E-01 | Cytosolic sulfotransferase 9 |
| AT3G50620 | -1.0 | 4.63E-05 | -0.4 | 1.10E-02 | -0.6 | 7.25E-03 | 0.0 | 9.77E-01 | 0.4 | 7.47E-02 | -0.4 | 6.31E-02 | Sulfotransferase |
| AT1G13420 | -0.5 | 5.87E-03 | 0.7 | 3.64E-04 | -1.3 | 8.84E-05 | -1.4 | 1.85E-04 | 0.0 | 9.08E-01 | -1.4 | 2.74E-03 | Cytosolic sulfotransferase 8 |
| AT5G07000 | -0.4 | 8.76E-03 | 0.0 | 9.08E-01 | -0.4 | 2.76E-02 | 0.7 | 6.66E-03 | 0.2 | 5.04E-01 | 0.5 | 8.94E-02 | Cytosolic sulfotransferase 14 |
| AT1G08030 | 0.0 | 6.94E-01 | -0.1 | 1.93E-01 | 0.1 | 4.17E-01 | -0.2 | 1.59E-01 | -0.3 | 1.22E-01 | 0.1 | 8.55E-01 | TPST |
| AT2G03750 | 1.1 | 1.49E-06 | 1.0 | 7.65E-06 | 0.1 | 3.00E-01 | 0.3 | 2.54E-01 | 0.1 | 7.60E-01 | 0.2 | 6.44E-01 | Sulfotransferase |
| AT4G33030 | 0.5 | 1.23E-04 | 0.4 | 5.54E-04 | 0.1 | 4.75E-01 | 0.4 | 2.42E-03 | 0.2 | 1.64E-01 | 0.2 | 2.17E-01 | UDP-sulfoquinovose synthase, chloroplastic |
| AT1G53580 | -1.2 | 2.14E-07 | -0.7 | 1.77E-05 | -0.5 | 3.43E-04 | -0.5 | 5.14E-04 | -0.2 | 2.54E-01 | -0.4 | 5.55E-02 | Glyoxalase II 3 |
| AT4G24940 | 0.2 | 1.99E-02 | 0.1 | 2.96E-01 | 0.1 | 2.03E-01 | 0.2 | 1.49E-01 | 0.2 | 2.62E-01 | 0.0 | 9.34E-01 | SUMO-activating enzyme subunit 1A |
| AT5G50580 | | | | | | | 0.3 | 8.46E-01 | 0.9 | 5.09E-01 | -0.6 | 7.22E-01 | SUMO-activating enzyme subunit 1B-1 |
| AT5G60410 | -0.1 | 2.23E-01 | -0.3 | 1.50E-03 | 0.2 | 1.51E-02 | 0.2 | 2.53E-01 | -0.1 | 7.11E-01 | 0.2 | 2.49E-01 | hypothetical protein |
| AT3G15150 | 0.2 | 2.64E-01 | 0.2 | 1.23E-01 | -0.1 | 6.85E-01 | 0.2 | 5.31E-01 | 0.2 | 6.53E-01 | 0.0 | 9.66E-01 | MMS21 |
| AT1G09730 | -0.4 | 2.15E-04 | -0.3 | 2.33E-03 | -0.1 | 2.06E-01 | -0.1 | 4.19E-01 | -0.1 | 6.38E-01 | 0.0 | 9.07E-01 | Cysteine proteinases superfamily protein |
| AT4G33620 | 0.4 | 7.38E-04 | 0.1 | 5.39E-01 | 0.4 | 7.35E-03 | 0.0 | 8.76E-01 | -0.1 | 5.70E-01 | 0.1 | 7.44E-01 | Probable ubiquitin-like-specific protease 2A |
| AT5G56780 | 0.0 | 7.23E-01 | 0.2 | 4.94E-02 | -0.3 | 5.51E-02 | 0.2 | 5.18E-01 | -0.3 | 2.76E-01 | 0.5 | 1.41E-01 | Protein EFFECTOR OF TRANSCRIPTION 2 |
| AT3G60070 | -1.0 | 1.37E-06 | -0.5 | 1.71E-04 | -0.4 | 1.62E-03 | -0.4 | 5.10E-02 | -0.1 | 7.79E-01 | -0.3 | 1.98E-01 | hypothetical protein |
| AT2G44280 | -0.9 | 3.79E-06 | -1.1 | 2.81E-06 | 0.2 | 1.13E-01 | -0.9 | 1.48E-03 | -0.1 | 7.55E-01 | -0.8 | 1.61E-02 | Major facilitator superfamily protein |
| AT1G23090 | 0.7 | 5.00E-04 | 0.7 | 4.45E-04 | -0.1 | 7.47E-01 | 0.0 | 9.89E-01 | -0.3 | 9.43E-02 | 0.3 | 1.59E-01 | Sulfate transporter 91 |
| AT5G45380 | -2.9 | 1.97E-08 | -1.7 | 1.08E-06 | -1.2 | 3.29E-05 | -2.2 | 1.25E-08 | 0.1 | 8.16E-01 | -2.3 | 7.22E-06 | Urea-proton symporter DUR3 |
| AT1G12360 | -0.3 | 3.64E-04 | -0.3 | 6.72E-04 | 0.0 | 9.20E-01 | -0.1 | 5.58E-01 | -0.2 | 1.89E-01 | 0.1 | 5.99E-01 | SNARE-interacting protein KEULE |
| AT4G19490 | -0.2 | 6.26E-03 | -0.2 | 3.25E-02 | -0.1 | 4.60E-01 | 0.0 | 9.96E-01 | 0.1 | 7.48E-01 | -0.1 | 8.03E-01 | Vacuolar protein sorting-associated protein 54, chloroplastic |
| AT2G17980 | -0.2 | 9.03E-03 | -0.1 | 5.84E-02 | -0.1 | 3.96E-01 | 0.0 | 9.41E-01 | 0.0 | 7.72E-01 | 0.0 | 8.70E-01 | SEC1 family transport protein SLY1 |
| AT1G71270 | -0.2 | 5.27E-02 | 0.0 | 9.48E-01 | -0.2 | 7.61E-02 | 0.2 | 1.91E-01 | 0.2 | 3.57E-01 | 0.0 | 8.83E-01 | Vacuolar protein sorting-associated protein 52 A |
| AT5G50840 | -0.1 | 1.12E-01 | -0.1 | 8.23E-02 | 0.0 | 8.50E-01 | 0.3 | 1.56E-01 | 0.3 | 2.78E-01 | 0.0 | 9.22E-01 | Alpha-taxilin-like protein |
| AT4G12120 | -0.1 | 5.12E-01 | -0.2 | 1.68E-02 | 0.2 | 8.38E-02 | 0.2 | 4.13E-01 | 0.2 | 5.08E-01 | 0.0 | 9.80E-01 | Protein transport Sec1b |
| AT4G36650 | -0.1 | 8.71E-02 | -0.2 | 9.43E-03 | 0.1 | 2.53E-01 | -0.1 | 5.21E-01 | -0.1 | 6.98E-01 | 0.0 | 9.14E-01 | Plant-specific TFIIB-related protein |
| AT2G04032 | -0.7 | 7.30E-02 | 0.0 | 9.36E-01 | -0.6 | 1.50E-01 | -0.6 | 1.08E-01 | -0.2 | 6.05E-01 | -0.4 | 2.39E-01 | ZIP7 |
| AT1G28260 | -0.2 | 8.50E-02 | -0.5 | 4.35E-03 | 0.3 | 1.26E-01 | -0.5 | 4.56E-02 | -0.5 | 9.45E-02 | 0.0 | 9.23E-01 | Nonsense-mediated mRNA decay factor SMG7-like |
| AT3G46590 | -0.3 | 3.87E-02 | -0.5 | 2.39E-03 | 0.2 | 1.22E-01 | -0.3 | 3.15E-02 | 0.1 | 7.92E-01 | -0.4 | 3.85E-02 | Uncharacterized protein |
| AT4G13280 | -0.5 | 3.89E-02 | -0.4 | 7.01E-02 | -0.1 | 8.37E-01 | -0.4 | 4.31E-01 | 0.0 | 9.80E-01 | -0.4 | 4.44E-01 | Terpene synthase N-terminal domain-containing protein |
| AT3G16990 | -0.1 | 3.89E-01 | -0.3 | 1.96E-02 | 0.2 | 1.25E-01 | -0.1 | 6.64E-01 | 0.0 | 9.45E-01 | -0.1 | 6.32E-01 | Bifunctional TENA-E protein |
| AT5G32470 | -0.2 | 3.97E-02 | 0.0 | 9.60E-01 | -0.2 | 6.74E-02 | -0.1 | 6.58E-01 | 0.0 | 8.94E-01 | -0.1 | 8.50E-01 | Bifunctional TH2 protein, mitochondrial |
| AT2G43940 | -0.6 | 7.99E-06 | -0.4 | 3.20E-04 | -0.2 | 1.90E-02 | 0.0 | 8.93E-01 | -0.1 | 6.63E-01 | 0.1 | 8.11E-01 | Probable thiol methyltransferase 2 |
| AT1G65980 | -0.2 | 1.13E-02 | 0.0 | 9.43E-01 | -0.2 | 2.39E-02 | -0.1 | 3.53E-01 | 0.0 | 9.44E-01 | -0.1 | 4.74E-01 | glutaredoxin-dependent peroxiredoxin |

| | | | | | | | | | | | | | |
|---|---|---|---|---|---|---|---|---|---|---|---|---|---|
| AT3G06050 | 0.2 | 3.46E-02 | 0.4 | 1.70E-03 | -0.2 | 8.94E-02 | 0.1 | 4.58E-01 | 0.1 | 7.58E-01 | 0.0 | 8.38E-01 | glutaredoxin-dependent peroxiredoxin |
| AT3G52960 | 0.8 | 2.43E-05 | 0.9 | 1.42E-05 | -0.2 | 1.89E-01 | 0.3 | 5.69E-02 | 0.3 | 9.80E-02 | 0.0 | 9.92E-01 | Peroxiredoxin-2E, chloroplastic |
| AT1G60740 | | | | | | | 1.809218806 | 1.98E-02 | 0.088778433 | 9.53E-01 | | | Peroxiredoxin-2D |
| AT1G21350 | 0.9 | 9.44E-06 | 0.8 | 5.56E-05 | 0.1 | 3.13E-01 | 0.3 | 4.52E-02 | 0.4 | 2.30E-02 | -0.1 | 7.25E-01 | thioredoxin-dependent peroxiredoxin |
| AT5G06290 | 1.9 | 4.70E-08 | 1.6 | 2.77E-07 | 0.2 | 4.63E-02 | 0.5 | 8.53E-04 | 0.4 | 1.04E-02 | 0.1 | 6.21E-01 | thioredoxin-dependent peroxiredoxin |
| AT4G31240 | -1.0 | 1.76E-05 | -1.1 | 2.42E-05 | 0.1 | 6.63E-01 | 0.1 | 7.30E-01 | -0.1 | 8.57E-01 | 0.2 | 6.04E-01 | Nucleoredoxin 3 |
| AT1G60420 | -0.6 | 7.09E-06 | -0.4 | 1.64E-04 | -0.2 | 3.33E-02 | -0.3 | 2.62E-02 | -0.1 | 6.71E-01 | -0.3 | 1.91E-01 | Probable nucleoredoxin 1 |
| AT2G17420 | 0.1 | 9.07E-02 | 0.2 | 3.32E-02 | 0.0 | 5.61E-01 | 0.2 | 9.65E-02 | 0.2 | 1.21E-01 | 0.0 | 9.62E-01 | Thioredoxin reductase |
| AT4G35460 | 0.2 | 8.71E-03 | 0.0 | 7.18E-01 | 0.2 | 3.61E-02 | 0.0 | 8.16E-01 | -0.1 | 7.91E-01 | 0.0 | 9.61E-01 | Thioredoxin reductase 1, mitochondrial |
| AT3G47300 | 0.7 | 3.25E-06 | 0.2 | 2.24E-02 | 0.5 | 2.62E-04 | -0.2 | 2.10E-01 | -0.4 | 5.81E-02 | 0.2 | 5.67E-01 | SelT-like protein |
| AT5G66170 | -2.7 | 3.17E-07 | -2.2 | 1.97E-06 | -0.5 | 5.70E-02 | -1.1 | 1.05E-03 | -0.1 | 8.65E-01 | -1.1 | 1.41E-02 | Sulfurtransferase 18 |
| AT1G16460 | -0.1 | 7.71E-02 | 0.0 | 6.66E-01 | -0.1 | 2.26E-01 | 0.0 | 9.02E-01 | -0.1 | 6.01E-01 | 0.1 | 6.04E-01 | Rhodanese homologue 2 |
| AT1G79230 | 0.0 | 7.80E-01 | 0.1 | 3.81E-01 | -0.1 | 5.82E-01 | 0.1 | 5.38E-01 | 0.2 | 2.25E-01 | -0.1 | 6.56E-01 | Sulfurtransferase |
| AT4G00590 | 0.0 | 8.40E-01 | 0.0 | 7.19E-01 | 0.0 | 8.96E-01 | -0.3 | 2.13E-01 | -0.4 | 1.15E-01 | 0.1 | 7.51E-01 | N-terminal nucleophile aminohydrolases superfamily protein |
| AT5G40580 | 0.1 | 3.39E-01 | 0.0 | 8.16E-01 | 0.1 | 5.43E-01 | 0.1 | 2.80E-01 | 0.1 | 6.11E-01 | 0.1 | 7.88E-01 | proteasome endopeptidase complex |
| AT3G27430 | 0.1 | 6.38E-02 | 0.2 | 1.98E-02 | -0.1 | 5.02E-01 | 0.4 | 1.50E-03 | 0.3 | 1.53E-02 | 0.1 | 6.84E-01 | proteasome endopeptidase complex |
| AT1G13490 | -0.1 | 3.48E-01 | 0.6 | 8.99E-04 | -0.7 | 1.30E-03 | | | | | | | F13B4.1 protein |
| AT1G66100 | 1.3 | 2.13E-04 | 2.3 | 8.56E-06 | -1.0 | 3.40E-03 | 0.5 | 1.11E-01 | 0.2 | 7.33E-01 | 0.4 | 4.49E-01 | Probable thionin-2.4 |
| AT5G36910 | 2.2 | 3.09E-06 | 1.9 | 1.83E-05 | 0.3 | 2.17E-01 | 0.9 | 9.20E-03 | 0.4 | 3.08E-01 | 0.5 | 1.43E-01 | Thionin-2.2 |
| AT5G13420 | -0.6 | 7.89E-05 | 0.0 | 8.38E-01 | -0.6 | 4.23E-04 | 0.3 | 9.82E-03 | 0.3 | 3.14E-02 | 0.0 | 9.07E-01 | transaldolase |
| AT1G12230 | 0.6 | 9.55E-06 | 0.6 | 2.56E-05 | 0.0 | 8.09E-01 | -0.2 | 2.62E-02 | -0.1 | 4.10E-01 | -0.1 | 3.80E-01 | transaldolase |
| AT5G27410 | -0.3 | 5.44E-03 | -0.4 | 6.69E-03 | 0.0 | 9.82E-01 | -0.1 | 5.28E-01 | 0.0 | 8.71E-01 | -0.1 | 7.52E-01 | D-aminoacid aminotransferase-like PLP-dependent enzymes superfamily protein |
| AT5G65780 | 0.0 | 6.64E-01 | 0.0 | 9.48E-01 | 0.0 | 6.65E-01 | 0.0 | 9.81E-01 | 0.1 | 3.75E-01 | -0.1 | 4.95E-01 | Branched-chain amino acid aminotransferase 5 |
| AT1G17930 | 0.1 | 4.99E-01 | -0.2 | 5.39E-02 | 0.3 | 3.36E-02 | -0.4 | 6.92E-02 | -0.3 | 2.46E-01 | -0.1 | 7.28E-01 | Protein MAINTENANCE OF MERISTEMS |
| AT2G25010 | 0.1 | 3.14E-01 | -0.3 | 3.11E-03 | 0.3 | 2.12E-03 | 0.1 | 6.90E-01 | 0.0 | 9.82E-01 | 0.1 | 7.70E-01 | Protein MAIN-LIKE 1 |
| AT2G04865 | 0.2 | 6.77E-03 | 0.0 | 7.88E-01 | 0.3 | 1.04E-02 | 0.0 | 8.61E-01 | -0.1 | 5.62E-01 | 0.1 | 7.27E-01 | Protein MAIN-LIKE 2 |
| AT5G57850 | 0.3 | 7.49E-04 | 0.3 | 1.40E-03 | 0.0 | 8.85E-01 | -0.1 | 4.12E-01 | -0.2 | 1.36E-01 | 0.1 | 5.46E-01 | D-aminoacid aminotransferase-like PLP-dependent enzymes superfamily protein |
| AT3G54970 | 0.7 | 7.84E-06 | 0.5 | 3.92E-04 | 0.3 | 2.00E-02 | -0.1 | 7.01E-01 | -0.1 | 8.36E-01 | 0.0 | 9.40E-01 | D-aminoacid aminotransferase-like PLP-dependent enzymes superfamily protein |
| AT5G04390 | -1.6 | 7.69E-06 | -0.7 | 8.38E-04 | -0.8 | 4.69E-03 | | | | | | | C2H2-type zinc finger family protein |
| AT5G67420 | -0.7 | 2.97E-02 | -0.9 | 1.16E-02 | 0.2 | 5.68E-01 | -0.2 | 1.57E-01 | -1.0 | 2.25E-05 | 0.8 | 1.12E-03 | LOB domain-containing protein 37 |
| AT1G31320 | -0.6 | 1.88E-04 | -0.6 | 3.23E-04 | 0.0 | 9.60E-01 | -0.2 | 6.06E-01 | 0.5 | 1.49E-01 | -0.7 | 7.60E-02 | LOB domain-containing protein 4 |
| AT1G16530 | -0.6 | 2.14E-02 | 0.0 | 9.36E-01 | -0.6 | 4.48E-02 | -0.1 | 8.36E-01 | 0.5 | 2.06E-01 | -0.6 | 2.21E-01 | LOB domain-containing protein 3 |
| AT4G17695 | -0.4 | 4.25E-03 | -0.6 | 4.03E-04 | 0.2 | 1.21E-01 | 0.1 | 8.04E-01 | 0.0 | 9.39E-01 | 0.1 | 9.23E-01 | Myb-like domain-containing protein |
| AT1G32240 | -0.4 | 1.18E-02 | -0.3 | 2.74E-02 | 0.0 | 7.65E-01 | 0.1 | 7.09E-01 | 0.1 | 6.48E-01 | 0.0 | 8.89E-01 | Homeodomain-like superfamily protein |
| AT1G44810 | -0.3 | 1.39E-03 | -0.2 | 3.48E-02 | -0.1 | 1.26E-01 | 0.0 | 9.27E-01 | -0.1 | 7.69E-01 | 0.1 | 8.69E-01 | Probable transcription factor At1g44810 |
| AT5G28040 | -0.3 | 7.48E-04 | -0.2 | 4.53E-03 | -0.1 | 3.95E-01 | -0.2 | 1.31E-01 | -0.1 | 5.88E-01 | -0.1 | 6.06E-01 | Probable transcription factor At5g28040 |
| AT2G25650 | -0.3 | 2.96E-02 | -0.4 | 7.75E-03 | 0.1 | 4.56E-01 | 0.5 | 3.31E-01 | 0.6 | 2.84E-01 | -0.1 | 9.01E-01 | GLABROUS1 enhancer-binding protein-like 1 |
| AT2G47900 | -0.2 | 3.69E-03 | -0.1 | 4.66E-02 | -0.1 | 2.36E-01 | -0.3 | 1.35E-02 | -0.2 | 1.70E-01 | -0.1 | 5.16E-01 | AT2G47900 protein |
| AT1G58220 | -0.1 | 7.12E-01 | -0.7 | 3.18E-02 | 0.6 | 9.89E-02 | | | | | | | At1g58220 |
| AT1G61730 | -0.1 | 4.23E-01 | 0.0 | 7.66E-01 | 0.0 | 6.85E-01 | 0.1 | 7.00E-01 | 0.0 | 9.54E-01 | 0.1 | 8.47E-01 | Probable transcription factor At1g61730 |
| AT1G76900 | 0.0 | 4.70E-01 | -0.1 | 1.82E-01 | 0.0 | 5.55E-01 | -0.1 | 7.00E-01 | 0.0 | 9.52E-01 | -0.1 | 8.33E-01 | Tubby-like F-box protein 1 |
| AT2G18280 | 0.0 | 9.58E-01 | 0.1 | 5.10E-01 | -0.1 | 5.13E-01 | -0.6 | 5.49E-04 | -0.5 | 5.25E-03 | -0.1 | 6.52E-01 | Tubby-like F-box protein 2 |
| AT1G12910 | 0.1 | 3.35E-01 | 0.1 | 3.61E-01 | 0.0 | 9.92E-01 | -0.4 | 3.95E-03 | -0.4 | 1.07E-02 | 0.0 | 9.33E-01 | WD repeat-containing protein LWD1 |
| AT4G25210 | 0.1 | 1.53E-01 | 0.1 | 3.65E-01 | 0.0 | 6.63E-01 | 0.1 | 5.17E-01 | 0.0 | 9.09E-01 | 0.1 | 5.31E-01 | GLABROUS1 enhancer-binding protein-like |
| AT4G00238 | 0.1 | 9.22E-02 | 0.1 | 4.51E-01 | 0.1 | 4.21E-01 | 0.0 | 8.85E-01 | -0.1 | 7.85E-01 | 0.1 | 7.19E-01 | Transcription factor STKL1 |
| AT5G63090 | 0.8 | 1.23E-04 | 0.4 | 2.24E-02 | 0.4 | 2.30E-02 | | | | | | | Protein LATERAL ORGAN BOUNDARIES |
| AT4G33925 | | | | | | | -0.2 | 6.66E-01 | -0.1 | 8.54E-01 | -0.1 | 8.96E-01 | At4g33925 |
| AT3G57480 | -0.4 | 5.35E-04 | -0.2 | 4.87E-02 | -0.2 | 4.13E-02 | -0.5 | 2.19E-02 | -0.6 | 8.75E-03 | 0.1 | 6.79E-01 | Zinc finger AN1 and C2H2 domain-containing stress-associated protein 13 |
| AT5G11680 | -0.5 | 2.43E-05 | -0.3 | 3.83E-03 | -0.3 | 6.98E-03 | -0.1 | 4.83E-01 | -0.1 | 7.08E-01 | 0.0 | 9.04E-01 | hypothetical protein |
| AT2G14850 | -0.2 | 2.42E-02 | -0.2 | 3.06E-02 | 0.0 | 9.73E-01 | 0.0 | 9.35E-01 | -0.2 | 3.74E-01 | 0.3 | 4.18E-01 | Transcriptional regulator of RNA polII, SAGA, subunit |
| AT3G21350 | -0.2 | 7.77E-02 | -0.1 | 4.33E-01 | -0.1 | 3.95E-01 | -0.2 | 3.04E-01 | -0.2 | 2.03E-01 | 0.1 | 7.93E-01 | Mediator of RNA polymerase II transcription subunit 6 |
| AT4G00850 | 0.0 | 8.58E-01 | -0.4 | 1.87E-02 | 0.4 | 4.92E-02 | 0.8 | 1.62E-02 | 0.6 | 5.93E-02 | 0.1 | 7.86E-01 | GRF1-interacting factor 3 |
| AT3G27100 | 0.1 | 4.91E-01 | 0.1 | 4.40E-01 | 0.0 | 9.33E-01 | -0.1 | 6.82E-01 | -0.1 | 7.77E-01 | 0.0 | 9.76E-01 | Transcription |
| AT1G01160 | 0.1 | 2.50E-01 | 0.0 | 7.42E-01 | 0.1 | 5.09E-01 | -0.1 | 5.84E-01 | -0.1 | 6.39E-01 | 0.0 | 9.98E-01 | GRF1-interacting factor 2 |
| AT2G14045 | 0.8 | 4.17E-04 | 0.3 | 1.39E-01 | 0.5 | 1.36E-02 | 0.1 | 5.01E-01 | 0.2 | 4.96E-01 | 0.0 | 9.56E-01 | c-Myc-binding protein |
| AT4G16420 | -0.4 | 5.52E-05 | -0.4 | 6.92E-05 | 0.0 | 7.67E-01 | -0.2 | 1.77E-01 | -0.2 | 3.17E-01 | 0.0 | 9.18E-01 | Transcriptional adapter |
| AT3G07740 | -0.2 | 1.71E-02 | -0.2 | 3.58E-02 | 0.0 | 8.07E-01 | -0.2 | 2.25E-01 | -0.2 | 3.31E-01 | 0.0 | 9.58E-01 | Transcriptional adapter |
| AT1G19330 | -0.7 | 5.27E-05 | -0.6 | 3.97E-04 | -0.1 | 2.66E-01 | -0.6 | 1.01E-03 | -0.1 | 4.40E-01 | -0.4 | 4.48E-02 | Histone deacetylase complex subunit |
| AT1G75060 | -0.6 | 7.04E-03 | -0.3 | 8.03E-02 | -0.2 | 2.95E-01 | -0.7 | 6.04E-02 | 0.0 | 9.94E-01 | -0.7 | 1.29E-01 | Histone deacetylase complex subunit |
| AT4G00450 | -0.5 | 2.90E-04 | -0.5 | 2.15E-04 | 0.1 | 5.94E-01 | -0.1 | 7.49E-01 | -0.1 | 8.02E-01 | 0.0 | 9.96E-01 | Mediator of RNA polymerase II transcription subunit 12 |
| AT1G55325 | -0.3 | 6.91E-03 | -0.5 | 3.22E-04 | 0.2 | 5.49E-02 | -0.2 | 4.77E-01 | 0.0 | 9.38E-01 | -0.1 | 7.03E-01 | Mediator of RNA polymerase II transcription subunit 13 |
| AT1G72390 | -0.2 | 6.96E-02 | -0.4 | 2.81E-03 | 0.2 | 9.38E-02 | 0.1 | 7.68E-01 | -0.1 | 8.83E-01 | 0.1 | 7.21E-01 | Protein PHYTOCHROME-DEPENDENT LATE-FLOWERING |
| AT5G19480 | -0.1 | 1.45E-01 | -0.1 | 2.77E-01 | 0.0 | 7.83E-01 | 0.2 | 3.07E-01 | 0.1 | 8.08E-01 | 0.2 | 6.51E-01 | Probable mediator of RNA polymerase II transcription subunit 19b |
| AT3G04740 | -0.1 | 1.06E-01 | -0.2 | 9.86E-02 | 0.0 | 9.42E-01 | -0.2 | 4.34E-01 | -0.4 | 8.25E-02 | 0.2 | 3.86E-01 | Mediator of RNA polymerase II transcription subunit 14 |
| AT2G28230 | -0.1 | 2.81E-01 | 0.0 | 7.55E-01 | -0.1 | 2.43E-01 | -0.1 | 7.41E-01 | 0.0 | 9.24E-01 | -0.1 | 8.97E-01 | Mediator of RNA polymerase II transcription subunit 20 |
| AT5G02850 | -0.1 | 2.61E-01 | -0.3 | 8.70E-03 | 0.2 | 9.38E-02 | 0.1 | 6.73E-01 | -0.2 | 4.24E-01 | 0.3 | 2.56E-01 | Mediator of RNA polymerase II transcription subunit 4 |
| AT1G26665 | -0.1 | 2.87E-01 | 0.1 | 4.80E-01 | -0.1 | 1.36E-01 | -0.4 | 3.29E-02 | -0.3 | 1.61E-01 | -0.1 | 7.31E-01 | Mediator of RNA polymerase II transcription subunit 10 |
| AT1G11760 | -0.1 | 4.52E-01 | -0.1 | 5.23E-01 | 0.0 | 9.51E-01 | -0.2 | 3.11E-01 | -0.2 | 3.74E-01 | 0.0 | 9.97E-01 | Mediator of RNA polymerase II transcription subunit 32 |
| AT1G07950 | -0.1 | 3.64E-01 | 0.1 | 2.88E-01 | -0.2 | 1.04E-01 | 0.2 | 2.46E-01 | 0.0 | 9.97E-01 | 0.2 | 4.03E-01 | Surfeit locus protein 5 subunit 22 of Mediator complex |
| AT5G19910 | -0.1 | 4.18E-01 | 0.0 | 7.09E-01 | -0.1 | 3.18E-01 | -0.2 | 3.53E-01 | 0.1 | 8.43E-01 | -0.3 | 3.53E-01 | Mediator of RNA polymerase II transcription subunit 31 |
| AT5G12230 | 0.0 | 6.86E-01 | -0.1 | 2.52E-01 | 0.0 | 5.11E-01 | -0.2 | 7.82E-02 | -0.3 | 4.62E-02 | 0.1 | 8.08E-01 | Mediator of RNA polymerase II transcription subunit 19a |
| AT5G20170 | 0.0 | 8.80E-01 | -0.2 | 4.52E-02 | 0.1 | 9.22E-02 | 0.1 | 6.66E-01 | -0.1 | 7.64E-01 | 0.2 | 4.93E-01 | Mediator of RNA polymerase II transcription subunit 17 |
| AT5G37190 | 0.1 | 5.44E-01 | 0.0 | 7.84E-01 | 0.0 | 8.09E-01 | 0.1 | 8.66E-01 | 0.2 | 6.15E-01 | -0.1 | 7.85E-01 | COP1-interacting protein 4 |
| AT3G01435 | 0.1 | 4.69E-01 | 0.1 | 5.30E-01 | 0.0 | 9.62E-01 | 0.3 | 2.47E-01 | 0.0 | 9.81E-01 | 0.3 | 4.33E-01 | Mediator of RNA polymerase II transcription subunit 11 |
| AT2G22370 | 0.1 | 3.99E-01 | 0.2 | 9.05E-02 | -0.1 | 4.29E-01 | 0.0 | 8.93E-01 | -0.2 | 6.49E-01 | 0.1 | 7.61E-01 | Mediator of RNA polymerase II transcription subunit 18 |
| AT1G26660 | 0.2 | 1.59E-01 | 0.0 | 9.81E-01 | 0.2 | 2.56E-01 | -0.1 | 8.08E-01 | -0.1 | 8.73E-01 | 0.0 | 9.72E-01 | Prefoldin chaperone subunit family protein |
| AT4G04780 | 0.2 | 3.19E-02 | 0.3 | 1.80E-02 | 0.0 | 7.50E-01 | 0.0 | 8.51E-01 | -0.2 | 3.29E-01 | 0.2 | 5.46E-01 | Mediator of RNA polymerase II transcription subunit 21 |
| AT5G03220 | 0.3 | 2.51E-02 | 0.2 | 1.14E-01 | 0.1 | 5.35E-01 | 0.1 | 8.36E-01 | -0.1 | 7.49E-01 | 0.2 | 6.08E-01 | Mediator of RNA polymerase II transcription subunit 7a |
| AT1G16430 | 0.6 | 1.20E-02 | 0.2 | 4.93E-01 | 0.4 | 9.66E-02 | 0.5 | 9.03E-02 | 0.5 | 1.41E-01 | 0.0 | 9.71E-01 | Mediator of RNA polymerase II transcription subunit 22a |
| AT5G41910 | 0.6 | 4.76E-04 | 0.1 | 3.79E-01 | 0.5 | 8.04E-03 | -0.1 | 8.65E-01 | 0.0 | 9.34E-01 | 0.0 | 9.72E-01 | Mediator of RNA polymerase II transcription subunit 10a |
| AT1G10450 | -0.9 | 9.53E-07 | -0.5 | 1.02E-04 | -0.4 | 1.43E-03 | 0.1 | 5.15E-01 | -0.1 | 6.96E-01 | 0.2 | 3.08E-01 | Paired amphipathic helix Sin3-like protein |
| AT1G24190 | -0.7 | 7.80E-06 | -0.6 | 4.02E-05 | -0.1 | 3.20E-01 | -0.1 | 6.34E-01 | -0.1 | 6.95E-01 | 0.0 | 9.98E-01 | SIN3-like 3 |
| AT1G09040 | -0.6 | 1.17E-03 | -0.1 | 3.69E-01 | -0.5 | 1.67E-02 | 0.0 | 9.59E-01 | -0.4 | 3.25E-01 | 0.3 | 4.26E-01 | Arginine-glutamic acid dipeptide repeat protein |

| | | | | | | | | | | | | | |
|---|---|---|---|---|---|---|---|---|---|---|---|---|---|
| AT4G40050 | -0.6 | 1.41E-05 | -0.4 | 2.91E-04 | -0.2 | 4.98E-02 | -0.3 | 1.85E-02 | -0.1 | 3.09E-01 | -0.2 | 4.11E-01 | Signal transducer, putative |
| AT1G09050 | -0.5 | 3.04E-03 | -0.3 | 2.60E-02 | -0.2 | 3.71E-01 | 0.4 | 2.76E-01 | 0.1 | 9.02E-01 | 0.3 | 4.76E-01 | Arginine-glutamic acid dipeptide repeat protein |
| AT5G15020 | -0.5 | 1.16E-03 | -0.6 | 2.85E-04 | 0.2 | 2.35E-01 | -0.2 | 3.79E-01 | 0.1 | 8.69E-01 | -0.2 | 4.06E-01 | SIN3-like 2 |
| AT1G70060 | -0.4 | 4.80E-04 | -0.4 | 1.01E-03 | 0.0 | 8.33E-01 | -0.1 | 6.97E-01 | -0.1 | 4.46E-01 | 0.1 | 7.85E-01 | Paired amphipathic helix protein Sin3-like 4 |
| AT2G47820 | -0.3 | 6.68E-03 | -0.3 | 1.87E-02 | 0.0 | 7.14E-01 | -0.3 | 3.73E-01 | 0.0 | 9.89E-01 | -0.2 | 4.75E-01 | Arginine-glutamic acid dipeptide repeat protein |
| AT3G01320 | -0.3 | 1.59E-02 | -0.5 | 5.91E-04 | 0.3 | 5.68E-02 | -0.1 | 5.09E-01 | 0.0 | 9.34E-01 | -0.1 | 6.99E-01 | SIN3-like 1 |
| AT5G15025 | -0.3 | 1.93E-01 | 0.2 | 4.37E-01 | -0.4 | 9.92E-02 | 0.0 | 9.75E-01 | 0.2 | 7.32E-01 | -0.2 | 7.74E-01 | Uncharacterized protein F2G14_150 |
| AT3G03570 | -0.3 | 1.73E-03 | -0.2 | 3.47E-03 | 0.0 | 8.40E-01 | -0.3 | 5.98E-02 | -0.3 | 1.88E-01 | -0.1 | 7.87E-01 | At3g03570 |
| AT1G59890 | -0.2 | 1.62E-02 | -0.3 | 6.97E-03 | 0.0 | 6.07E-01 | 0.2 | 3.04E-01 | 0.1 | 5.54E-01 | 0.1 | 8.62E-01 | SIN3-like 5 |
| AT2G32700 | -0.2 | 8.69E-02 | -0.3 | 4.59E-03 | 0.2 | 1.15E-01 | 0.0 | 9.83E-01 | 0.0 | 9.66E-01 | 0.0 | 9.64E-01 | MUM1 |
| AT5G37380 | 0.1 | 1.79E-01 | 0.1 | 2.26E-01 | 0.0 | 9.40E-01 | -0.1 | 3.34E-01 | -0.3 | 5.69E-02 | 0.2 | 4.29E-01 | AT5g37380 |
| AT4G15802 | 0.4 | 5.67E-04 | 0.6 | 8.26E-05 | -0.2 | 8.99E-02 | 0.1 | 5.41E-01 | 0.1 | 7.00E-01 | 0.0 | 9.55E-01 | HSBP |
| AT1G71080 | 0.0 | 9.45E-01 | -0.2 | 3.18E-02 | 0.2 | 5.19E-02 | -0.2 | 6.56E-02 | -0.2 | 1.11E-01 | 0.0 | 9.80E-01 | At1g71080 |
| AT5G38050 | 0.4 | 2.53E-03 | 0.0 | 7.04E-01 | 0.5 | 6.13E-03 | 0.0 | 9.37E-01 | 0.0 | 9.18E-01 | 0.0 | 9.76E-01 | Gb\|AAF26010.1 |
| AT3G20160 | -1.7 | 1.23E-05 | -0.2 | 1.53E-01 | -1.4 | 3.04E-04 | -0.8 | 2.05E-01 | -0.1 | 8.73E-01 | -0.7 | 4.49E-01 | Terpenoid synthases superfamily protein |
| AT1G61050 | -1.6 | 2.24E-07 | -1.1 | 3.25E-06 | -0.5 | 5.22E-03 | -0.8 | 6.58E-02 | 0.5 | 1.38E-01 | -1.3 | 8.97E-03 | Alpha 1,4-glycosyltransferase family protein |
| AT5G01250 | -1.4 | 4.84E-07 | -1.0 | 5.74E-06 | -0.4 | 2.19E-02 | -0.3 | 3.77E-01 | -0.3 | 4.36E-01 | 0.0 | 9.97E-01 | Alpha 1,4-glycosyltransferase family protein |
| AT2G28840 | -1.3 | 2.52E-07 | -0.9 | 4.63E-06 | -0.3 | 6.50E-03 | 0.0 | 9.51E-01 | -0.1 | 4.46E-01 | 0.1 | 6.11E-01 | RING-type domain-containing protein |
| AT1G01570 | -1.2 | 3.32E-05 | -0.5 | 7.29E-03 | -0.7 | 7.50E-03 | 0.9 | 5.83E-03 | 0.2 | 6.55E-01 | 0.7 | 2.40E-02 | Glycosyltransferase ) |
| AT2G44500 | -1.2 | 3.65E-06 | -0.4 | 5.65E-03 | -0.8 | 3.66E-04 | -0.3 | 3.35E-02 | -0.3 | 1.10E-01 | -0.1 | 8.34E-01 | O-fucosyltransferase family protein |
| AT1G20550 | -1.1 | 3.36E-06 | -0.3 | 5.10E-03 | -0.7 | 4.48E-04 | -0.3 | 3.80E-01 | 0.1 | 7.76E-01 | -0.4 | 3.27E-01 | O-fucosyltransferase family protein |
| AT2G35710 | -0.9 | 5.07E-05 | -0.3 | 2.48E-02 | -0.6 | 4.58E-03 | 0.0 | 9.85E-01 | 0.0 | 9.05E-01 | 0.0 | 9.11E-01 | Nucleotide-diphospho-sugar transferases superfamily protein |
| AT5G23850 | -0.8 | 3.37E-05 | -1.2 | 6.90E-06 | 0.4 | 2.59E-02 | -0.1 | 4.96E-01 | -0.2 | 4.28E-01 | 0.0 | 8.99E-01 | At5g23850 ) |
| AT4G15270 | -0.7 | 8.53E-06 | -0.7 | 2.38E-05 | 0.0 | 7.54E-01 | -1.0 | 7.62E-03 | -0.1 | 8.49E-01 | -0.9 | 5.34E-02 | At4g15270 |
| AT5G46340 | -0.7 | 3.04E-05 | -0.5 | 4.25E-04 | -0.2 | 1.12E-01 | -0.2 | 4.39E-01 | 0.0 | 9.80E-01 | -0.2 | 5.57E-01 | O-acetyltransferase family protein |
| AT5G60700 | -0.7 | 7.56E-06 | -0.3 | 1.69E-03 | -0.3 | 3.17E-03 | -0.4 | 2.83E-02 | -0.4 | 9.29E-02 | -0.1 | 8.14E-01 | Glycosyltransferase family protein 2 |
| AT1G63420 | -0.5 | 2.09E-03 | -0.1 | 5.13E-01 | -0.4 | 1.58E-02 | -0.1 | 6.97E-01 | 0.0 | 9.92E-01 | -0.1 | 7.68E-01 | Glycosyltransferase ) |
| AT3G03810 | -0.5 | 4.10E-04 | -0.4 | 1.67E-03 | -0.1 | 5.49E-01 | 0.0 | 9.30E-01 | 0.1 | 8.13E-01 | -0.1 | 7.68E-01 | O-fucosyltransferase family protein |
| AT2G35910 | -0.4 | 1.27E-02 | -0.5 | 6.65E-03 | 0.1 | 6.78E-01 | 0.1 | 7.71E-01 | -0.1 | 8.08E-01 | 0.2 | 5.16E-01 | hypothetical protein |
| AT5G15740 | -0.4 | 4.50E-03 | -0.3 | 2.92E-02 | -0.1 | 4.01E-01 | -0.3 | 3.14E-02 | -0.1 | 7.76E-01 | -0.3 | 1.73E-01 | O-fucosyltransferase family protein |
| AT2G01480 | -0.3 | 1.21E-03 | -0.3 | 4.50E-03 | 0.0 | 5.89E-01 | -0.1 | 5.26E-01 | -0.1 | 5.17E-01 | 0.0 | 9.61E-01 | O-fucosyltransferase family protein |
| AT1G52630 | -0.3 | 1.25E-02 | -0.1 | 1.65E-01 | -0.1 | 2.49E-01 | 0.0 | 8.66E-01 | 0.2 | 5.05E-01 | -0.2 | 4.75E-01 | O-fucosyltransferase family protein |
| AT4G08810 | -0.2 | 2.57E-03 | -0.1 | 1.17E-01 | -0.1 | 7.48E-02 | -0.2 | 3.16E-02 | -0.2 | 8.24E-02 | 0.0 | 9.12E-01 | O-fucosyltransferase family protein |
| AT3G06550 | -0.2 | 2.22E-02 | -0.3 | 8.73E-03 | 0.1 | 5.69E-01 | -0.1 | 5.44E-01 | 0.2 | 1.47E-01 | -0.3 | 1.00E-01 | Cas1p 10 TM acyl transferase domain-containing protein |
| AT2G39630 | -0.1 | 4.29E-02 | -0.1 | 5.11E-02 | 0.0 | 9.85E-01 | 0.0 | 8.47E-01 | 0.0 | 9.65E-01 | 0.0 | 9.27E-01 | Nucleotide-diphospho-sugar transferases superfamily protein |
| AT5G20680 | -0.1 | 1.36E-01 | -0.2 | 1.05E-01 | 0.0 | 8.76E-01 | 0.3 | 1.17E-01 | 0.2 | 3.34E-01 | 0.1 | 7.79E-01 | Trichome birefringence-like N-terminal domain-containing protein |
| AT3G27610 | -0.1 | 3.35E-01 | -0.3 | 6.91E-03 | 0.2 | 6.10E-02 | -0.1 | 6.26E-01 | -0.1 | 6.45E-01 | 0.0 | 9.76E-01 | Nucleotidylyl transferase superfamily protein |
| AT3G04830 | 0.0 | 8.50E-01 | 0.0 | 9.66E-01 | 0.0 | 8.43E-01 | 0.0 | 9.23E-01 | 0.0 | 8.86E-01 | 0.0 | 9.65E-01 | ER membrane protein complex subunit 2 |
| AT3G26370 | 0.0 | 7.94E-01 | 0.0 | 6.98E-01 | 0.0 | 9.10E-01 | 0.1 | 4.15E-01 | 0.1 | 6.01E-01 | 0.0 | 9.25E-01 | O-fucosyltransferase family protein |
| AT3G48980 | 0.0 | 7.23E-01 | 0.1 | 3.88E-01 | -0.1 | 6.70E-01 | -0.2 | 5.55E-01 | -0.3 | 3.74E-01 | 0.1 | 7.99E-01 | Glycosyltransferase ) |
| AT1G74390 | 0.1 | 3.91E-01 | 0.2 | 1.91E-01 | -0.1 | 6.85E-01 | -0.5 | 1.34E-01 | -0.2 | 6.17E-01 | -0.3 | 5.51E-01 | Polynucleotidyl transferase, ribonuclease H-like superfamily protein |
| AT1G29890 | 0.1 | 2.20E-01 | 0.1 | 1.69E-01 | 0.0 | 8.70E-01 | 0.0 | 8.93E-01 | 0.2 | 4.36E-01 | -0.1 | 6.25E-01 | O-acetyltransferase family protein |
| AT1G23820 | 0.2 | 8.47E-02 | 0.2 | 1.56E-02 | -0.1 | 3.52E-01 | 0.0 | 7.51E-01 | 0.1 | 3.00E-01 | -0.1 | 6.02E-01 | Spermidine synthase |
| AT1G14970 | 0.2 | 1.29E-01 | 0.1 | 5.28E-01 | 0.1 | 4.80E-01 | 0.1 | 7.34E-01 | -0.3 | 5.37E-01 | 0.4 | 4.05E-01 | O-fucosyltransferase family protein |
| AT1G07220 | 0.2 | 1.56E-02 | -0.2 | 1.03E-01 | 0.4 | 3.66E-03 | 0.4 | 7.79E-02 | 0.2 | 3.92E-01 | 0.2 | 6.02E-01 | F10K1.7 protein ) |
| AT4G12130 | 0.3 | 6.15E-03 | 0.1 | 5.88E-01 | 0.2 | 4.14E-02 | 0.4 | 3.79E-02 | 0.2 | 3.96E-01 | 0.2 | 3.98E-01 | Putative transferase At4g12130, mitochondrial |
| AT1G28710 | 0.5 | 4.04E-03 | 0.1 | 3.28E-01 | 0.3 | 5.51E-02 | 0.1 | 6.94E-01 | 0.0 | 9.55E-01 | 0.1 | 8.17E-01 | F1K23.9 |
| AT4G22570 | 0.7 | 8.97E-06 | 0.6 | 4.50E-05 | 0.1 | 3.69E-01 | 0.1 | 7.96E-01 | 0.0 | 8.73E-01 | 0.0 | 9.76E-01 | Adenine phosphoribosyl transferase 3 |
| AT1G60990 | 0.8 | 1.32E-05 | 0.5 | 4.49E-04 | 0.3 | 3.51E-02 | 0.1 | 8.34E-01 | 0.4 | 3.49E-01 | -0.3 | 5.05E-01 | Aminomethyltransferase folate-binding domain-containing protein |
| AT1G78510 | 0.8 | 2.28E-06 | 0.3 | 3.53E-03 | 0.5 | 3.10E-04 | 0.1 | 4.52E-01 | 0.1 | 6.11E-01 | 0.0 | 9.42E-01 | Uncharacterized protein |
| AT4G18230 | 0.9 | 3.78E-06 | 0.5 | 3.94E-04 | 0.4 | 5.35E-03 | -0.2 | 3.37E-01 | -0.3 | 3.18E-01 | 0.0 | 9.11E-01 | Glycosyltransferase |
| AT5G42830 | 0.9 | 2.26E-05 | 0.5 | 1.92E-03 | 0.4 | 1.75E-02 | 1.8 | 2.90E-06 | 0.3 | 4.14E-01 | 1.5 | 1.62E-04 | HXXXD-type acyl-transferase family protein |
| AT1G12244 | 1.0 | 2.21E-04 | 0.9 | 5.52E-04 | 0.0 | 8.31E-01 | 0.2 | 4.95E-01 | 0.3 | 3.97E-01 | -0.1 | 8.62E-01 | Polynucleotidyl transferase, ribonuclease H-like superfamily protein |
| AT3G50270 | 1.0 | 3.12E-05 | 1.1 | 3.87E-05 | -0.1 | 7.17E-01 | 0.1 | 6.00E-01 | 0.2 | 5.18E-01 | 0.0 | 9.04E-01 | Anthranilate N-hydroxycinnamoyl |
| AT4G13410 | 1.2 | 8.87E-05 | 1.2 | 1.05E-04 | -0.1 | 7.38E-01 | 1.1 | 8.32E-03 | 1.2 | 1.31E-02 | 0.0 | 9.80E-01 | Nucleotide-diphospho-sugar transferases superfamily protein |
| AT5G47610 | 1.6 | 3.96E-05 | 1.3 | 4.11E-04 | 0.3 | 2.01E-01 | -0.5 | 5.49E-02 | -0.4 | 1.61E-01 | -0.1 | 8.07E-01 | RING-type E3 ubiquitin transferase |
| AT1G29200 | | | | | | | -1.4 | 1.51E-02 | -0.2 | 7.60E-01 | -1.2 | 7.35E-02 | O-fucosyltransferase family protein |
| AT4G17905 | | | | | | | 0.3 | 5.06E-01 | 0.0 | 9.91E-01 | 0.3 | 5.89E-01 | RING-type E3 ubiquitin transferase |
| AT5G51920 | | | | | | | 0.8 | 1.61E-02 | 0.8 | 3.52E-02 | 0.0 | 9.23E-01 | Pyridoxal phosphate -dependent transferases superfamily protein |
| AT3G07300 | 0.3 | 5.45E-04 | 0.2 | 3.87E-02 | 0.2 | 4.76E-02 | 0.1 | 7.30E-01 | -0.2 | 3.82E-01 | 0.2 | 3.02E-01 | AT3G07300 protein |
| AT4G09100 | -0.4 | 1.17E-01 | 0.2 | 4.76E-01 | -0.6 | 7.08E-02 | 1.6 | 5.53E-03 | 0.3 | 7.17E-01 | 1.3 | 2.33E-02 | RING-H2 finger protein ATL39 |
| AT1G66660 | -0.3 | 1.02E-02 | -0.3 | 1.38E-02 | 0.0 | 9.69E-01 | -0.3 | 2.65E-01 | -0.2 | 4.10E-01 | 0.0 | 9.23E-01 | RING-type E3 ubiquitin transferase |
| AT5G53360 | 0.0 | 9.64E-01 | 0.0 | 9.63E-01 | 0.0 | 9.43E-01 | -0.3 | 1.80E-01 | -0.2 | 3.72E-01 | -0.1 | 8.64E-01 | RING-type E3 ubiquitin transferase |
| AT2G26350 | 0.1 | 3.37E-01 | -0.3 | 8.69E-03 | 0.3 | 5.23E-03 | -0.3 | 2.18E-01 | -0.1 | 6.83E-01 | -0.2 | 6.28E-01 | Peroxin 10 |
| AT1G79810 | 0.4 | 1.63E-04 | -0.1 | 1.50E-01 | 0.6 | 2.34E-04 | 0.1 | 8.36E-01 | 0.1 | 8.49E-01 | 0.0 | 9.92E-01 | RING-type E3 ubiquitin transferase |
| AT1G66620 | 0.8 | 2.35E-05 | 0.6 | 5.01E-04 | 0.2 | 8.75E-02 | 0.7 | 3.42E-02 | 0.3 | 5.08E-01 | 0.4 | 1.70E-01 | RING-type E3 ubiquitin transferase |
| AT5G58780 | -2.7 | 1.14E-06 | -1.7 | 1.61E-05 | -1.0 | 6.17E-03 | -0.4 | 4.15E-01 | 0.6 | 1.69E-01 | | | Alkyl transferase |
| AT4G17190 | -0.2 | 3.47E-02 | 0.2 | 5.63E-02 | -0.5 | 3.41E-03 | -0.1 | 6.20E-01 | 0.1 | 7.07E-01 | -0.2 | 4.31E-01 | Farnesyl diphosphate synthase 2 |
| AT1G47400 | 4.1 | 4.82E-05 | 2.8 | 1.46E-03 | 1.2 | 1.52E-02 | 2.7 | 5.21E-06 | 0.8 | 1.23E-01 | 1.9 | 9.12E-04 | Uncharacterized protein |
| AT5G43150 | -1.2 | 1.20E-06 | -0.8 | 2.85E-05 | -0.3 | 1.03E-02 | -0.2 | 4.40E-01 | 0.2 | 4.80E-01 | -0.4 | 2.00E-01 | At5g43150 |
| AT1G44770 | -1.0 | 4.77E-07 | -0.6 | 1.99E-05 | -0.3 | 2.23E-03 | -0.6 | 3.55E-04 | -0.3 | 5.54E-02 | -0.3 | 1.49E-01 | At1g44770 |
| AT2G42780 | -0.1 | 8.11E-02 | -0.2 | 2.14E-02 | 0.1 | 4.81E-01 | -0.1 | 4.66E-01 | -0.1 | 5.15E-01 | 0.0 | 9.93E-01 | Transcription elongation factor B polypeptide |
| AT1G32130 | -0.1 | 8.29E-02 | -0.2 | 2.32E-02 | 0.1 | 4.74E-01 | -0.1 | 4.58E-01 | 0.0 | 8.95E-01 | -0.1 | 5.29E-01 | Transcription elongation factor family protein |
| AT5G05140 | -0.1 | 2.86E-01 | -0.1 | 3.37E-01 | 0.0 | 9.56E-01 | 0.1 | 5.29E-01 | -0.2 | 2.77E-01 | 0.3 | 1.72E-01 | Probable mediator of RNA polymerase II transcription subunit 26b |
| AT5G25520 | -0.1 | 3.25E-01 | -0.3 | 4.79E-03 | 0.2 | 3.90E-02 | 0.0 | 9.65E-01 | 0.0 | 9.21E-01 | -0.1 | 9.07E-01 | Putative PHD finger protein |
| AT4G26310 | -0.1 | 4.81E-01 | 0.0 | 9.37E-01 | 0.0 | 6.17E-01 | -0.1 | 4.21E-01 | -0.1 | 7.41E-01 | -0.1 | 8.19E-01 | Elongation factor P family protein |
| AT3G05500 | 0.0 | 5.63E-01 | -0.2 | 1.30E-02 | 0.2 | 4.87E-02 | 0.2 | 2.94E-01 | 0.2 | 3.85E-01 | 0.0 | 9.92E-01 | Rubber elongation factor protein |
| AT5G09850 | 0.0 | 8.22E-01 | 0.0 | 7.94E-01 | 0.0 | 9.68E-01 | -0.1 | 5.95E-01 | 0.1 | 4.58E-01 | -0.2 | 3.19E-01 | Transcription elongation factor family protein |
| AT2G18110 | 0.1 | 2.76E-01 | 0.3 | 5.59E-03 | -0.2 | 4.90E-02 | 0.1 | 6.48E-01 | -0.1 | 5.44E-01 | 0.2 | 2.59E-01 | Translation elongation factor EF1B beta |
| AT4G11120 | 0.1 | 1.47E-01 | 0.1 | 1.77E-01 | 0.0 | 9.61E-01 | 0.4 | 1.99E-02 | 0.4 | 3.29E-02 | 0.0 | 9.97E-01 | Elongation factor Ts, mitochondrial |
| AT2G25640 | 0.4 | 7.67E-03 | 0.4 | 9.12E-03 | 0.0 | 9.80E-01 | 0.1 | 7.41E-01 | 0.1 | 8.09E-01 | 0.0 | 9.84E-01 | Putative transcription elongation factor S-II |
| AT1G30230 | 0.5 | 1.17E-04 | 0.5 | 4.29E-04 | 0.1 | 6.01E-01 | 0.1 | 6.22E-01 | -0.2 | 3.65E-01 | 0.3 | 2.44E-01 | Translation elongation factor EF1B |
| AT3G08740 | 2.7 | 1.41E-08 | 2.1 | 8.06E-08 | 0.6 | 1.35E-04 | 0.5 | 2.22E-03 | -0.1 | 6.88E-01 | 0.5 | 6.29E-03 | Putative elongation factor P ; 66839-65711 |

| | | | | | | | | | | | | | |
|---|---|---|---|---|---|---|---|---|---|---|---|---|---|
| AT2G38560 | 0.1 | 1.57E-01 | 0.2 | 4.06E-02 | -0.1 | 4.85E-01 | -0.4 | 1.31E-02 | -0.3 | 6.10E-02 | -0.1 | 7.86E-01 | Transcription elongation factor TFIIS |
| AT1G34360 | -0.1 | 1.73E-01 | 0.0 | 8.58E-01 | -0.2 | 1.93E-01 | 0.2 | 4.01E-01 | -0.2 | 6.53E-01 | 0.4 | 2.70E-01 | Translation initiation factor 3 family protein |
| AT2G36885 | 0.0 | 8.57E-01 | 0.3 | 1.10E-01 | -0.3 | 1.21E-01 | -1.6 | 7.62E-05 | 0.9 | 2.13E-03 | -2.4 | 1.01E-05 | Translation initiation factor |
| AT1G71350 | 0.0 | 9.43E-01 | -0.2 | 4.33E-02 | 0.2 | 6.39E-02 | -0.2 | 2.72E-01 | -0.3 | 6.65E-02 | 0.1 | 5.07E-01 | Eukaryotic translation initiation factor SUI1 family protein |
| AT5G11900 | 0.0 | 6.96E-01 | 0.2 | 7.10E-02 | -0.1 | 1.80E-01 | 0.3 | 4.01E-02 | 0.2 | 1.66E-01 | 0.1 | 7.67E-01 | Translation machinery-associated protein 22 |
| AT1G66070 | 0.1 | 4.28E-01 | 0.3 | 3.98E-02 | -0.2 | 2.04E-01 | 0.5 | 7.78E-03 | 0.3 | 1.77E-01 | 0.2 | 3.25E-01 | Eukaryotic translation initiation factor 3 subunit J |
| AT1G13020 | 0.2 | 2.91E-02 | 0.0 | 8.48E-01 | 0.2 | 6.74E-02 | 0.1 | 6.80E-01 | 0.1 | 8.41E-01 | 0.0 | 9.35E-01 | Eukaryotic translation initiation factor 4B2 |
| AT1G72340 | 0.2 | 4.56E-02 | 0.3 | 5.97E-03 | -0.1 | 2.77E-01 | 0.1 | 5.50E-01 | -0.2 | 3.46E-01 | 0.2 | 1.89E-01 | eIF-2B GDP-GTP exchange factor subunit alpha |
| AT2G45730 | 0.4 | 1.16E-03 | 0.3 | 3.48E-03 | 0.0 | 6.79E-01 | 0.2 | 2.43E-01 | 0.0 | 8.52E-01 | 0.2 | 2.53E-01 | tRNA -N)-methyltransferase non-catalytic subunit TRM6 |
| AT2G46290 | 0.4 | 2.51E-03 | 0.3 | 3.65E-02 | 0.1 | 2.46E-01 | 0.1 | 6.35E-01 | 0.2 | 5.98E-01 | 0.0 | 9.37E-01 | Eukaryotic translation initiation factor 3 subunit I |
| AT3G02200 | 0.5 | 2.51E-05 | 0.4 | 3.43E-04 | 0.1 | 1.12E-01 | 0.1 | 3.94E-01 | -0.1 | 7.87E-01 | 0.2 | 3.55E-01 | Eukaryotic translation initiation factor 3 subunit M |
| AT5G15610 | 0.5 | 1.22E-04 | 0.5 | 3.61E-04 | 0.0 | 6.80E-01 | 0.2 | 6.82E-02 | 0.1 | 4.24E-01 | 0.1 | 5.57E-01 | Eukaryotic translation initiation factor 3 subunit M |
| AT2G40780 | 0.5 | 6.53E-03 | 0.3 | 8.80E-02 | 0.2 | 2.69E-01 | 0.5 | 6.43E-02 | -0.1 | 8.73E-01 | 0.6 | 5.55E-02 | Nucleic acid-binding, OB-fold-like protein |
| AT3G43540 | 0.7 | 2.37E-04 | 0.3 | 1.80E-02 | 0.3 | 3.58E-02 | 0.1 | 7.82E-01 | 0.2 | 4.82E-01 | -0.1 | 7.25E-01 | Initiation factor 4F subunit |
| AT4G39630 | 0.9 | 5.70E-04 | 0.7 | 6.50E-03 | 0.2 | 2.78E-01 | 0.5 | 6.46E-02 | 0.2 | 5.07E-01 | 0.3 | 4.38E-01 | Translation initiation factor |
| AT5G37475 | 0.9 | 2.02E-06 | 0.6 | 9.20E-05 | 0.3 | 8.00E-03 | 0.3 | 1.73E-01 | -0.2 | 6.27E-01 | 0.5 | 1.25E-01 | Uncharacterized protein |
| AT4G30690 | 1.0 | 2.18E-05 | 1.0 | 7.51E-05 | 0.1 | 6.02E-01 | -0.3 | 3.13E-02 | -0.4 | 1.83E-02 | 0.1 | 7.78E-01 | Translation initiation factor 3 protein |
| AT4G30680 | 1.3 | 1.23E-06 | 1.2 | 9.67E-06 | 0.2 | 1.47E-01 | -0.2 | 3.86E-01 | -0.5 | 8.62E-02 | 0.3 | 4.45E-01 | Initiation factor eIF-4 gamma, MA3 |
| AT4G35250 | 1.5 | 1.26E-07 | 1.0 | 4.55E-06 | 0.5 | 8.10E-04 | 0.1 | 5.07E-01 | 0.3 | 6.94E-02 | -0.2 | 3.16E-01 | Protein HIGH CHLOROPHYLL FLUORESCENCE PHENOTYPE 244, chloroplastic |
| AT2G24060 | 1.9 | 2.25E-08 | 1.6 | 1.72E-07 | 0.3 | 9.18E-03 | 0.4 | 1.71E-03 | 0.3 | 8.22E-02 | 0.2 | 2.47E-01 | Translation initiation factor IF-3 |
| AT1G53880 | | | | | | | -1.682094766 | 1.16E-01 | -1.04370771 | 3.45E-01 | -0.6 | 7.46E-01 | eIF-2B GDP-GTP exchange factor subunit alpha |
| AT5G01940 | | | | | | | 0.2 | 6.09E-01 | 0.0 | 9.66E-01 | 0.2 | 7.44E-01 | Eukaryotic translation initiation factor 2B family protein |
| AT1G62850 | 0.5 | 5.94E-05 | 0.1 | 4.91E-01 | 0.5 | 8.23E-04 | -0.2 | 4.94E-01 | 0.0 | 9.86E-01 | -0.2 | 6.17E-01 | Class I peptide chain release factor |
| AT5G36170 | 0.8 | 4.08E-06 | 0.5 | 2.24E-04 | 0.3 | 8.39E-03 | 0.0 | 8.49E-01 | 0.1 | 7.32E-01 | -0.1 | 6.35E-01 | High chlorophyll fluorescent 109 |
| AT3G62910 | 1.4 | 1.59E-06 | 1.2 | 1.21E-05 | 0.2 | 1.40E-01 | 0.3 | 9.39E-02 | 0.2 | 3.15E-01 | 0.1 | 7.74E-01 | Peptide chain release factor APG3, chloroplastic |
| AT1G56350 | 0.3 | 5.82E-03 | 0.0 | 9.63E-01 | 0.3 | 1.73E-02 | 0.2 | 4.89E-01 | 0.2 | 4.86E-01 | 0.0 | 9.58E-01 | Peptide chain release factor PrfB2, chloroplastic |
| AT2G30890 | | | | | | | -0.2 | 5.91E-01 | 0.1 | 8.47E-01 | -0.3 | 4.99E-01 | Cytochrome b561 domain-containing protein |
| AT3G18710 | -0.7 | 4.59E-05 | -0.3 | 5.95E-03 | -0.4 | 1.36E-02 | -0.2 | 2.06E-01 | 0.0 | 9.44E-01 | -0.2 | 3.80E-01 | U-box domain-containing protein 29 |
| AT5G65200 | -0.9 | 5.97E-06 | -1.0 | 8.27E-06 | 0.1 | 5.80E-01 | -0.5 | 1.20E-02 | -0.3 | 1.60E-01 | -0.2 | 4.39E-01 | U-box domain-containing protein 38 |
| AT1G20780 | -0.1 | 9.60E-02 | -0.1 | 3.42E-01 | -0.1 | 5.25E-01 | 0.1 | 2.90E-01 | -0.1 | 4.01E-01 | 0.3 | 1.36E-01 | U-box domain-containing protein 44 |
| AT3G46510 | 0.0 | 8.39E-01 | -0.1 | 9.45E-02 | 0.1 | 1.61E-01 | 0.1 | 6.23E-01 | -0.1 | 5.59E-01 | 0.2 | 3.93E-01 | U-box domain-containing protein 13 |
| AT1G48270 | -0.1 | 3.74E-01 | -0.1 | 5.52E-01 | 0.0 | 8.45E-01 | -0.2 | 5.85E-01 | 0.0 | 9.76E-01 | -0.2 | 6.66E-01 | G-protein coupled receptors family 2 profile 2 domain-containing protein |
| AT1G60050 | -3.5 | 5.89E-07 | -1.2 | 8.17E-05 | -2.3 | 1.11E-04 | -0.1 | 8.16E-01 | -1.3 | 9.91E-03 | 1.2 | 1.98E-02 | WAT1-related protein At1g60050 |
| AT5G40210 | -2.3 | 1.35E-06 | -1.3 | 3.49E-05 | -1.0 | 4.46E-03 | | | | | | | UMAMIT42 |
| AT2G38100 | -2.3 | 1.35E-05 | -0.5 | 2.84E-02 | -1.8 | 6.21E-04 |  |  |  |  |  |  | Proton-dependent oligopeptide transport family protein |
| AT1G49960 | -2.2 | 1.18E-07 | -1.0 | 1.61E-05 | -1.2 | 7.31E-05 | 1.5 | 1.86E-02 | 1.4 | 5.05E-02 | 0.1 | 8.51E-01 | Nucleobase-ascorbate transporter 4 |
| AT2G34350 | -2.1 | 1.54E-06 | -1.0 | 1.01E-04 | -1.1 | 1.48E-03 | | | | | | | Nodulin-like |
| AT1G08890 | -1.7 | 7.93E-08 | -1.0 | 3.19E-06 | -0.7 | 2.72E-04 | -0.8 | 1.55E-03 | -0.4 | 6.15E-02 | -0.3 | 2.58E-01 | AT1G08890 protein |
| AT2G16980 | -1.7 | 1.69E-07 | -0.7 | 6.16E-05 | -1.0 | 7.42E-05 | -1.2 | 5.02E-04 | 0.0 | 9.68E-01 | -1.2 | 5.21E-03 | Major facilitator superfamily protein |
| AT5G11570 | -1.6 | 7.66E-07 | -0.7 | 1.06E-04 | -0.9 | 6.25E-04 | | | | | | | Protein NRT1 |
| AT5G49990 | -1.6 | 2.78E-07 | -1.3 | 2.10E-06 | -0.3 | 4.89E-02 | -0.2 | 3.61E-01 | -0.1 | 6.27E-01 | -0.1 | 8.65E-01 | Nucleobase-ascorbate transporter 5 |
| AT1G21890 | -1.6 | 1.44E-06 | 0.3 | 1.27E-02 | -1.9 | 1.09E-05 | | | | | | | WAT1-related protein |
| AT5G40240 | -1.5 | 8.79E-08 | -1.2 | 1.03E-06 | -0.4 | 4.57E-03 | 0.0 | 9.67E-01 | 0.2 | 4.30E-01 | -0.2 | 5.00E-01 | WAT1-related protein |
| AT5G42210 | -1.4 | 5.41E-06 | -1.2 | 3.65E-05 | -0.3 | 2.18E-01 | -0.5 | 1.55E-01 | -0.4 | 2.47E-01 | -0.1 | 9.28E-01 | Major facilitator superfamily profile domain-containing protein |
| AT5G40230 | -1.4 | 6.86E-07 | -0.9 | 1.61E-05 | -0.5 | 6.23E-03 | -0.7 | 7.31E-02 | -0.1 | 8.58E-01 | -0.6 | 1.41E-01 | WAT1-related protein |
| AT2G16990 | -1.4 | 1.62E-05 | -1.0 | 1.71E-04 | -0.4 | 1.13E-01 | -0.4 | 2.87E-01 | 0.4 | 2.95E-01 | -0.8 | 3.98E-02 | Major facilitator superfamily protein |
| AT4G35300 | -1.4 | 3.28E-07 | -1.1 | 2.88E-06 | -0.2 | 3.81E-02 | -0.2 | 4.35E-01 | 0.0 | 9.69E-01 | -0.2 | 6.25E-01 | Tonoplast monosaccharide transporter2 |
| AT3G28100 | -1.3 | 1.53E-06 | -1.1 | 7.93E-06 | -0.2 | 2.77E-01 | -0.2 | 2.54E-01 | 0.0 | 8.78E-01 | -0.2 | 4.58E-01 | WAT1-related protein |
| AT1G11450 | -1.3 | 4.28E-06 | -1.1 | 2.09E-05 | -0.2 | 3.51E-01 | 0.1 | 4.95E-01 | 0.1 | 8.35E-01 | 0.1 | 7.84E-01 | WAT1-related protein |
| AT2G03590 | -1.2 | 1.10E-07 | -0.6 | 1.28E-05 | -0.6 | 1.43E-04 | -0.2 | 5.13E-01 | -0.1 | 7.89E-01 | -0.1 | 8.33E-01 | UPS1 |
| AT4G25220 | -1.2 | 8.32E-04 | 0.0 | 9.16E-01 | -1.2 | 2.96E-03 | -2.1 | 8.65E-03 | -0.6 | 3.53E-01 | -1.5 | 1.34E-01 | Putative glycerol-3-phosphate transporter 2 |
| AT1G67300 | -1.1 | 6.36E-06 | -1.0 | 2.95E-05 | -0.1 | 3.84E-01 | -0.5 | 1.17E-02 | -0.1 | 7.35E-01 | -0.4 | 1.08E-01 | Major facilitator superfamily protein |
| AT3G45650 | -1.1 | 2.21E-06 | -0.7 | 1.30E-04 | -0.4 | 4.73E-03 | -0.1 | 6.84E-01 | 0.4 | 7.09E-02 | -0.6 | 7.22E-02 | Nitrate excretion transporter1 |
| AT1G11460 | -1.1 | 6.53E-06 | -0.3 | 2.81E-02 | -0.8 | 4.33E-04 | -0.4 | 4.09E-01 | 0.1 | 8.43E-01 | -0.5 | 2.72E-01 | WAT1-related protein At1g11460 |
| AT1G33440 | -1.1 | 3.17E-04 | -0.3 | 8.87E-02 | -0.7 | 1.26E-02 | -0.6 | 1.31E-02 | -0.1 | 6.73E-01 | -0.5 | 1.06E-01 | Protein NRT1 |
| AT3G28050 | -1.0 | 1.90E-07 | -0.8 | 2.15E-06 | -0.2 | 1.27E-02 | -0.9 | 7.01E-06 | 0.0 | 9.99E-01 | -0.9 | 4.04E-04 | WAT1-related protein At3g28050 |
| AT3G18200 | -0.9 | 5.20E-05 | 0.1 | 5.01E-01 | -1.0 | 1.99E-04 | 0.2 | 5.93E-01 | 0.6 | 5.04E-02 | -0.4 | 1.89E-01 | WAT1-related protein |
| AT3G56620 | -0.9 | 1.72E-04 | -0.2 | 1.33E-01 | -0.7 | 5.62E-03 | -0.2 | 3.04E-01 | 0.3 | 2.81E-01 | -0.5 | 5.14E-02 | WAT1-related protein At3g56620 |
| AT2G34355 | -0.9 | 9.48E-05 | -0.6 | 1.19E-03 | -0.3 | 1.59E-01 | -0.2 | 4.07E-01 | 0.0 | 9.09E-01 | -0.1 | 6.48E-01 | Major facilitator superfamily protein |
| AT4G30420 | -0.8 | 5.05E-04 | 0.0 | 9.86E-01 | -0.8 | 2.40E-03 | 0.8 | 1.41E-01 | 0.7 | 2.23E-01 | 0.1 | 9.46E-01 | WAT1-related protein |
| AT2G26510 | -0.8 | 2.13E-06 | -0.4 | 3.48E-04 | -0.4 | 1.47E-03 | -0.2 | 2.01E-01 | -0.1 | 6.61E-01 | -0.1 | 6.66E-01 | Xanthine |
| AT5G65687 | -0.8 | 2.85E-05 | -0.4 | 3.42E-03 | -0.4 | 1.16E-02 | -0.4 | 6.49E-02 | -0.1 | 5.92E-01 | -0.3 | 2.83E-01 | Probable sphingolipid transporter spinster homolog 1 |
| AT4G38050 | -0.7 | 4.61E-05 | -0.5 | 1.53E-03 | -0.3 | 5.19E-02 | 0.0 | 9.25E-01 | 0.2 | 6.68E-01 | -0.1 | 7.29E-01 | Nucleobase-ascorbate transporter 11 |
| AT2G16970 | -0.7 | 1.59E-05 | -0.3 | 6.96E-03 | -0.4 | 3.11E-03 | -0.5 | 7.33E-02 | -0.4 | 2.46E-01 | -0.2 | 7.24E-01 | Major facilitator superfamily protein |
| AT1G68570 | -0.7 | 2.64E-04 | -0.8 | 2.46E-04 | 0.1 | 7.11E-01 | 0.0 | 8.94E-01 | 0.3 | 1.03E-01 | -0.3 | 2.16E-01 | Major facilitator superfamily protein |
| AT1G26440 | -0.7 | 2.02E-05 | -0.7 | 6.97E-05 | -0.1 | 5.90E-01 | -0.5 | 4.44E-03 | -0.2 | 2.94E-01 | -0.3 | 1.59E-01 | Ureide permease 5 |
| AT1G25500 | -0.7 | 7.85E-05 | -0.4 | 2.05E-03 | -0.3 | 7.57E-02 | -0.4 | 5.65E-02 | -0.1 | 7.50E-01 | -0.3 | 2.86E-01 | Choline transporter-like protein |
| AT1G62200 | -0.7 | 1.18E-05 | -0.6 | 9.36E-05 | -0.1 | 1.90E-01 | -0.2 | 2.32E-01 | -0.2 | 2.02E-01 | 0.0 | 9.08E-01 | Major facilitator superfamily protein |
| AT3G43790 | -0.7 | 2.14E-04 | -0.3 | 4.69E-02 | -0.4 | 1.63E-02 | -0.4 | 1.02E-01 | 0.1 | 8.05E-01 | -0.5 | 6.59E-02 | Zinc induced facilitator-like 2 |
| AT4G15540 | -0.6 | 7.84E-05 | -0.5 | 7.98E-04 | -0.1 | 1.86E-01 | 0.2 | 3.83E-01 | 0.3 | 1.40E-01 | -0.1 | 5.70E-01 | WAT1-related protein |
| AT4G36790 | -0.6 | 1.80E-05 | -0.1 | 3.60E-01 | -0.5 | 2.55E-04 | -0.4 | 7.19E-03 | -0.3 | 9.42E-02 | -0.2 | 5.22E-01 | Major facilitator superfamily protein |
| AT2G01170 | -0.6 | 2.88E-05 | -0.4 | 4.38E-04 | -0.2 | 9.51E-02 | -0.4 | 5.41E-02 | -0.1 | 8.57E-01 | -0.3 | 1.98E-01 | At2g01170 |
| AT2G20780 | -0.6 | 1.65E-05 | -0.3 | 1.28E-03 | -0.2 | 1.09E-02 | 0.0 | 9.64E-01 | 0.0 | 8.40E-01 | 0.0 | 9.07E-01 | Major facilitator superfamily protein |
| AT5G64500 | -0.5 | 3.63E-05 | -0.3 | 1.32E-03 | -0.2 | 3.53E-02 | -0.2 | 1.01E-01 | -0.1 | 4.33E-01 | -0.1 | 6.68E-01 | Probable sphingolipid transporter spinster homolog 2 |
| AT4G01450 | -0.5 | 2.17E-05 | -0.4 | 1.87E-04 | -0.1 | 1.84E-01 | 0.0 | 9.75E-01 | 0.1 | 3.49E-01 | -0.1 | 4.63E-01 | WAT1-related protein |
| AT5G45370 | -0.5 | 1.38E-04 | -0.2 | 1.90E-02 | -0.3 | 2.12E-02 | -0.1 | 8.26E-01 | -0.1 | 8.90E-01 | 0.0 | 9.78E-01 | WAT1-related protein |
| AT1G60030 | -0.5 | 4.13E-03 | -0.5 | 7.30E-03 | 0.0 | 9.17E-01 | 0.1 | 9.03E-01 | 0.4 | 5.18E-01 | | | Nucleobase-ascorbate transporter 7 |
| AT4G21120 | -0.5 | 6.62E-02 | -0.7 | 1.49E-02 | 0.3 | 4.31E-01 | -1.2 | 2.19E-02 | -1.3 | 1.98E-02 | 0.1 | 9.04E-01 | Amino acid transporter 1 |
| AT5G64700 | -0.5 | 2.54E-03 | -0.3 | 2.23E-02 | -0.1 | 3.44E-01 | -0.7 | 2.53E-02 | -0.3 | 3.03E-01 | -0.3 | 4.19E-01 | WAT1-related protein |
| AT4G01440 | -0.5 | 2.44E-04 | -0.4 | 1.67E-03 | -0.1 | 3.50E-01 | 0.1 | 7.33E-01 | 0.2 | 3.68E-01 | -0.1 | 6.80E-01 | WAT1-related protein |
| AT2G13100 | -0.4 | 5.25E-05 | -0.2 | 4.73E-03 | -0.2 | 1.76E-02 | -0.3 | 1.09E-01 | -0.3 | 2.30E-01 | 0.0 | 8.91E-01 | Major facilitator superfamily protein |
| AT5G17010 | -0.4 | 4.31E-03 | -0.6 | 5.25E-04 | 0.2 | 1.51E-01 | 0.1 | 3.87E-01 | 0.4 | 3.15E-02 | -0.2 | 2.81E-01 | Major facilitator superfamily protein |
| AT4G19185 | -0.4 | 1.45E-03 | -0.1 | 1.85E-01 | -0.3 | 3.22E-02 | -0.2 | 1.98E-01 | 0.0 | 8.76E-01 | -0.2 | 4.56E-01 | WAT1-related protein |

| | | | | | | | | | | | | | |
|---|---|---|---|---|---|---|---|---|---|---|---|---|---|
| AT1G75500 | -0.4 | 5.36E-04 | -0.2 | 8.83E-02 | -0.2 | 2.00E-02 | 0.0 | 9.62E-01 | 0.0 | 9.68E-01 | 0.0 | 9.97E-01 | Protein WALLS ARE THIN 1 |
| AT4G08290 | -0.4 | 1.67E-02 | -0.5 | 1.03E-02 | 0.1 | 7.45E-01 | -1.1 | 4.96E-06 | -0.6 | 1.79E-03 | -0.5 | 3.49E-02 | WAT1-related protein At4g08290 |
| AT1G70260 | -0.4 | 9.27E-02 | 0.8 | 1.85E-03 | -1.2 | 9.48E-04 | -1.4 | 9.50E-02 | 0.6 | 3.79E-01 | | | WAT1-related protein At1g70260 |
| AT4G38640 | -0.4 | 1.72E-04 | -0.2 | 8.31E-03 | -0.2 | 4.49E-02 | -0.3 | 2.26E-02 | -0.2 | 1.38E-01 | -0.1 | 6.79E-01 | Choline transporter-like protein |
| AT2G37460 | -0.3 | 2.06E-04 | -0.3 | 4.51E-04 | 0.0 | 8.23E-01 | 0.0 | 9.29E-01 | 0.3 | 6.45E-02 | -0.4 | 6.86E-02 | WAT1-related protein |
| AT1G79820 | -0.3 | 2.28E-03 | -0.3 | 7.64E-03 | -0.1 | 6.38E-01 | -0.1 | 5.17E-01 | 0.1 | 6.38E-01 | -0.3 | 3.60E-01 | Major facilitator superfamily protein |
| AT1G05030 | -0.3 | 9.74E-04 | -0.1 | 2.08E-01 | -0.2 | 2.09E-02 | 0.0 | 9.98E-01 | 0.3 | 2.56E-01 | -0.3 | 3.40E-01 | Major facilitator superfamily protein |
| AT5G46050 | -0.3 | 2.28E-01 | -0.6 | 2.81E-02 | 0.3 | 2.84E-01 | -0.6 | 1.44E-01 | -1.1 | 2.57E-02 | 0.5 | 4.76E-01 | Peptide transporter 3 |
| AT3G03720 | -0.3 | 2.94E-03 | -0.4 | 6.12E-04 | 0.1 | 2.52E-01 | -0.2 | 2.18E-01 | -0.1 | 8.47E-01 | -0.2 | 4.63E-01 | Cationic amino acid transporter C-terminal domain-containing protein |
| AT3G07080 | -0.3 | 1.98E-02 | -0.4 | 3.95E-03 | 0.1 | 3.64E-01 | 0.2 | 5.52E-01 | 0.0 | 9.08E-01 | 0.1 | 7.38E-01 | hypothetical protein |
| AT3G28070 | -0.2 | 6.05E-02 | -0.2 | 1.76E-01 | -0.1 | 6.44E-01 | 0.5 | 2.20E-02 | 0.8 | 1.22E-03 | -0.3 | 1.58E-01 | WAT1-related protein |
| AT3G30340 | -0.2 | 2.28E-01 | -0.1 | 7.82E-01 | -0.2 | 4.68E-01 | 0.1 | 8.45E-01 | 0.5 | 1.55E-01 | -0.4 | 2.65E-01 | WAT1-related protein |
| AT1G20860 | -0.2 | 4.19E-01 | 0.4 | 1.10E-01 | -0.6 | 6.31E-02 | | | | | | | Phosphate transporter 18 |
| AT3G28130 | -0.2 | 2.31E-01 | 0.0 | 9.01E-01 | -0.2 | 3.73E-01 | -0.4 | 4.72E-02 | 0.1 | 8.56E-01 | -0.5 | 7.47E-02 | WAT1-related protein |
| AT2G43330 | -0.2 | 3.51E-02 | -0.2 | 6.29E-02 | 0.0 | 8.38E-01 | -0.6 | 5.95E-03 | -0.3 | 1.12E-01 | -0.2 | 3.77E-01 | INT1 |
| AT3G28080 | -0.2 | 2.03E-01 | -0.4 | 2.32E-02 | 0.2 | 2.63E-01 | 0.1 | 6.27E-01 | 0.5 | 4.11E-02 | -0.3 | 7.97E-02 | Nodulin MtN21 |
| AT1G43650 | -0.2 | 1.55E-01 | -0.2 | 2.16E-01 | 0.0 | 9.12E-01 | -0.6 | 7.82E-03 | -0.3 | 2.49E-01 | -0.4 | 2.27E-01 | WAT1-related protein |
| AT3G59320 | -0.2 | 1.47E-01 | -0.1 | 3.51E-01 | -0.1 | 6.82E-01 | -0.1 | 5.33E-01 | 0.2 | 3.70E-01 | -0.4 | 1.92E-01 | AT3G59320 protein ) |
| AT4G08300 | -0.1 | 2.63E-01 | 0.3 | 6.98E-03 | -0.5 | 2.45E-03 | -0.7 | 1.37E-05 | -0.3 | 1.07E-02 | -0.3 | 3.44E-02 | WAT1-related protein At4g08300 |
| AT1G69850 | -0.1 | 1.54E-01 | -0.2 | 6.92E-02 | 0.0 | 6.45E-01 | 0.0 | 9.32E-01 | -0.2 | 1.38E-01 | 0.2 | 2.62E-01 | Protein NRT1 |
| AT5G10190 | -0.1 | 1.55E-01 | -0.2 | 9.87E-03 | 0.1 | 1.62E-01 | 0.0 | 8.59E-01 | -0.1 | 4.44E-01 | 0.2 | 4.26E-01 | Major facilitator superfamily protein |
| AT3G46980 | -0.1 | 5.25E-01 | -0.1 | 2.64E-01 | 0.1 | 6.63E-01 | -0.1 | 5.22E-01 | 0.1 | 6.53E-01 | -0.2 | 2.92E-01 | Phosphate transporter 43 |
| AT5G62890 | 0.0 | 9.70E-01 | 0.0 | 6.74E-01 | 0.0 | 6.73E-01 | -0.1 | 5.77E-01 | -0.1 | 7.80E-01 | 0.0 | 9.18E-01 | AT5G62890 protein |
| AT1G05940 | 0.0 | 9.10E-01 | -0.1 | 1.04E-01 | 0.1 | 1.14E-01 | 0.0 | 8.97E-01 | 0.1 | 7.48E-01 | -0.1 | 7.03E-01 | Cationic amino acid transporter 9 |
| AT2G05760 | 0.0 | 7.92E-01 | 0.1 | 2.54E-01 | -0.1 | 4.61E-01 | 0.0 | 9.34E-01 | -0.1 | 8.22E-01 | 0.2 | 7.52E-01 | Nucleobase-ascorbate transporter 1 |
| AT1G17530 | 0.0 | 6.62E-01 | 0.0 | 8.73E-01 | 0.0 | 8.33E-01 | -0.7 | 2.42E-04 | -0.6 | 1.89E-03 | -0.1 | 7.19E-01 | Translocase of inner mitochondrial membrane 23 |
| AT5G04770 | 0.0 | 6.67E-01 | 0.6 | 2.15E-04 | -0.6 | 1.46E-03 | -0.1 | 8.04E-01 | -0.1 | 7.21E-01 | 0.0 | 9.14E-01 | Cationic amino acid transporter 6 |
| AT5G13760 | 0.1 | 3.97E-01 | 0.1 | 2.96E-01 | 0.0 | 8.50E-01 | -0.3 | 5.03E-02 | -0.2 | 2.10E-01 | -0.1 | 7.00E-01 | Choline transporter-like protein |
| AT4G22990 | 0.1 | 3.76E-01 | 0.0 | 7.74E-01 | 0.1 | 6.51E-01 | -0.5 | 1.48E-02 | -0.1 | 8.05E-01 | -0.4 | 8.40E-02 | Major Facilitator Superfamily with SPX domain-containing protein |
| AT5G03555 | 0.2 | 3.77E-02 | 0.0 | 5.90E-01 | 0.2 | 2.81E-02 | 0.1 | 5.18E-01 | 0.1 | 5.95E-01 | 0.0 | 9.97E-01 | Uncharacterized protein |
| AT2G34190 | 0.2 | 7.17E-02 | 0.5 | 6.22E-04 | -0.3 | 2.32E-02 | 0.0 | 9.00E-01 | 0.1 | 7.03E-01 | -0.1 | 8.27E-01 | Nucleobase-ascorbate transporter 2 |
| AT2G39210 | 0.2 | 8.77E-02 | -0.4 | 1.77E-02 | 0.6 | 2.20E-03 | 0.1 | 6.65E-01 | -0.1 | 7.08E-01 | 0.3 | 4.97E-01 | At2g39210 |
| AT2G02020 | 0.3 | 1.06E-02 | 0.4 | 1.83E-03 | -0.1 | 2.98E-01 | -0.1 | 7.66E-01 | 0.0 | 9.34E-01 | -0.1 | 8.78E-01 | Major facilitator superfamily protein |
| AT3G59310 | 0.4 | 2.66E-02 | 0.1 | 4.35E-01 | 0.2 | 1.99E-01 | -0.2 | 5.99E-01 | 0.0 | 9.34E-01 | -0.1 | 7.65E-01 | Solute carrier family 35 protein |
| AT3G47960 | 0.4 | 2.89E-04 | 0.3 | 5.80E-03 | 0.1 | 1.23E-01 | 0.8 | 1.31E-02 | 0.9 | 1.49E-02 | -0.1 | 9.12E-01 | Uncharacterized protein |
| AT1G74780 | 0.4 | 1.33E-03 | 0.3 | 6.47E-03 | 0.1 | 5.16E-01 | 0.6 | 2.42E-03 | 0.2 | 4.42E-01 | 0.4 | 5.69E-02 | Nodulin-like domain-containing protein |
| AT3G47420 | 0.5 | 4.52E-04 | 0.1 | 3.82E-01 | 0.4 | 5.88E-03 | 1.2 | 1.17E-05 | 0.7 | 3.86E-03 | 0.5 | 2.94E-02 | Putative glycerol-3-phosphate transporter 1 |
| AT5G13740 | 0.6 | 1.06E-03 | 0.3 | 5.33E-02 | 0.3 | 6.70E-02 | 1.7 | 7.74E-08 | 0.1 | 6.20E-01 | 1.5 | 3.73E-05 | Zinc induced facilitator 1 |
| AT2G27810 | 0.7 | 9.74E-06 | 0.3 | 1.69E-03 | 0.3 | 4.54E-03 | 0.1 | 7.54E-01 | 0.0 | 9.23E-01 | 0.1 | 7.34E-01 | Nucleobase-ascorbate transporter 12 |
| AT5G13400 | 0.8 | 1.42E-06 | 0.3 | 1.84E-03 | 0.5 | 3.01E-04 | -0.2 | 4.59E-01 | -0.1 | 8.21E-01 | -0.1 | 7.79E-01 | Protein NRT1 |
| AT2G26690 | 1.1 | 7.75E-07 | 0.9 | 9.27E-06 | 0.2 | 3.81E-02 | -0.4 | 9.32E-02 | -0.2 | 3.86E-01 | -0.1 | 6.66E-01 | Major facilitator superfamily protein |
| AT2G37450 | 1.7 | 1.29E-06 | 1.2 | 3.33E-05 | 0.5 | 1.29E-02 | 0.2 | 2.20E-01 | 0.1 | 5.81E-01 | 0.1 | 6.98E-01 | WAT1-related protein |
| AT3G09450 | 1.9 | 3.59E-05 | 0.9 | 1.19E-02 | 1.0 | 6.62E-03 | | | | | | | Uncharacterized protein |
| AT4G01430 | 2.4 | 1.48E-07 | 1.8 | 2.33E-06 | 0.5 | 3.90E-03 | 0.8 | 6.43E-02 | -0.5 | 4.24E-01 | 1.3 | 1.40E-02 | WAT1-related protein |
| AT4G28040 | 5.3 | 1.50E-04 | 4.5 | 7.84E-04 | 0.8 | 8.40E-02 | 0.7 | 5.97E-01 | 0.6 | 7.13E-01 | | | WAT1-related protein At4g28040 |
| AT3G01350 | | | | | | | -0.3 | 7.95E-01 | -0.6 | 5.89E-01 | 0.4 | 8.29E-01 | Protein NRT1 |
| AT3G53210 | | | | | | | -0.8 | 4.29E-03 | -0.4 | 1.00E-01 | -0.4 | 2.05E-01 | WAT1-related protein |
| AT3G59330 | | | | | | | -0.7 | 1.84E-01 | -0.3 | 6.23E-01 | -0.4 | 6.06E-01 | At3g59330 |
| AT1G35910 | 0.0 | 8.56E-01 | -0.4 | 9.14E-03 | 0.4 | 2.73E-02 | 0.2 | 5.44E-01 | 0.5 | 1.95E-01 | -0.2 | 5.32E-01 | Probable trehalose-phosphate phosphatase D |
| AT5G10100 | -1.4 | 4.70E-07 | -0.5 | 2.93E-04 | -0.9 | 1.48E-04 | -0.8 | 1.37E-02 | 0.3 | 4.04E-01 | -1.1 | 7.88E-03 | Trehalose 6-phosphate phosphatase |
| AT5G65140 | -1.3 | 3.24E-06 | -1.1 | 1.32E-05 | -0.1 | 4.18E-01 | -0.2 | 5.43E-01 | 0.5 | 5.19E-02 | -0.7 | 2.21E-02 | Trehalose 6-phosphate phosphatase |
| AT4G39770 | -0.9 | 1.39E-05 | -0.7 | 1.30E-04 | -0.2 | 1.62E-01 | -0.5 | 7.64E-02 | -0.5 | 8.39E-02 | 0.0 | 9.57E-01 | Trehalose 6-phosphate phosphatase |
| AT2G22190 | -0.6 | 1.51E-04 | -0.5 | 8.87E-04 | -0.1 | 3.65E-01 | -0.3 | 1.64E-01 | 0.2 | 5.34E-01 | -0.5 | 4.29E-02 | Trehalose 6-phosphate phosphatase |
| AT4G12430 | -0.1 | 7.03E-01 | -0.1 | 7.80E-01 | 0.0 | 9.53E-01 | 0.0 | 9.73E-01 | 0.3 | 4.55E-01 | -0.3 | 4.93E-01 | Probable trehalose-phosphate phosphatase F |
| AT4G22590 | 0.2 | 1.80E-01 | 0.4 | 1.03E-02 | -0.2 | 1.44E-01 | 0.7 | 2.84E-04 | 0.6 | 1.36E-03 | 0.0 | 8.64E-01 | Probable trehalose-phosphate phosphatase G |
| AT5G51460 | 0.8 | 1.23E-04 | 0.6 | 1.01E-03 | 0.2 | 2.91E-01 | -0.2 | 2.69E-01 | -0.8 | 4.91E-03 | 0.5 | 4.33E-02 | Trehalose 6-phosphate phosphatase |
| AT1G78090 | 0.8 | 1.64E-04 | 0.5 | 1.11E-02 | 0.4 | 4.53E-02 | 0.9 | 5.09E-04 | 0.1 | 7.83E-01 | 0.8 | 5.81E-03 | Trehalose 6-phosphate phosphatase |
| AT1G56630 | -1.2 | 3.41E-06 | 0.1 | 5.35E-01 | -1.3 | 3.46E-05 | -0.3 | 4.11E-01 | 0.1 | 8.01E-01 | -0.4 | 2.54E-01 | F25P12.93 protein |
| AT2G15230 | -0.6 | 2.43E-05 | -0.6 | 2.89E-05 | 0.0 | 6.58E-01 | -0.4 | 1.45E-02 | -0.1 | 5.81E-01 | -0.3 | 1.66E-01 | Triacylglycerol lipase 1 |
| AT3G57140 | -0.1 | 5.28E-01 | 0.1 | 1.91E-01 | -0.2 | 1.11E-01 | -0.6 | 3.32E-03 | -0.4 | 3.45E-02 | -0.2 | 5.05E-01 | Triacylglycerol lipase SDP1L |
| AT1G45201 | 0.0 | 8.14E-01 | 0.3 | 5.21E-02 | -0.2 | 9.90E-02 | 0.2 | 3.40E-01 | 0.7 | 5.03E-03 | -0.5 | 9.50E-02 | At1g45200 |
| AT5G04040 | 0.2 | 5.27E-02 | 0.1 | 1.84E-01 | 0.1 | 5.65E-01 | -0.3 | 1.62E-02 | -0.4 | 8.95E-03 | 0.1 | 7.56E-01 | Triacylglycerol lipase SDP1 |
| AT2G21170 | 1.2 | 1.09E-07 | 1.0 | 1.18E-06 | 0.2 | 1.22E-02 | 0.2 | 1.90E-01 | 0.1 | 4.26E-01 | 0.1 | 8.49E-01 | Triosephosphate isomerase |
| AT5G14600 | 0.3 | 2.27E-02 | 0.1 | 2.50E-01 | 0.2 | 2.83E-01 | 0.5 | 2.52E-02 | 0.4 | 7.73E-02 | 0.1 | 8.31E-01 | tRNA -N)-methyltransferase |
| AT5G01230 | 0.4 | 1.83E-03 | 0.2 | 3.62E-02 | 0.2 | 1.88E-01 | -0.1 | 8.68E-01 | 0.1 | 6.45E-01 | -0.2 | 5.40E-01 | Putative tRNA |
| AT4G27340 | 0.5 | 3.29E-04 | 0.2 | 3.42E-02 | 0.3 | 3.61E-02 | 0.7 | 6.18E-03 | 0.3 | 2.35E-01 | 0.4 | 2.37E-01 | tRNA -N1)-methyltransferase |
| AT3G56120 | 0.2 | 3.91E-02 | 0.1 | 5.16E-01 | 0.2 | 2.04E-01 | 0.2 | 2.28E-01 | 0.2 | 5.15E-01 | 0.1 | 7.96E-01 | tRNA -N1)-methyltransferase 1 |
| AT5G47680 | 0.3 | 4.35E-03 | 0.3 | 5.29E-03 | 0.0 | 9.81E-01 | 0.0 | 8.43E-01 | -0.2 | 1.70E-01 | 0.2 | 2.13E-01 | tRNA -N)-methyltransferase |
| AT3G02320 | -0.1 | 3.92E-01 | -0.1 | 2.34E-01 | 0.0 | 7.57E-01 | 0.3 | 2.50E-01 | 0.2 | 4.62E-01 | 0.1 | 8.69E-01 | tRNA -N)-dimethyltransferase 2 -N) methyltransferase 2) G26)dimethyltransferase 2) |
| AT5G15810 | 0.0 | 7.19E-01 | 0.1 | 1.28E-01 | -0.2 | 1.02E-01 | 0.3 | 7.02E-02 | 0.3 | 1.93E-01 | 0.1 | 8.37E-01 | tRNA -N)-dimethyltransferase 1 -N) methyltransferase 1) G26)dimethyltransferase 1) |
| AT3G56330 | 1.5 | 6.57E-07 | 1.3 | 4.71E-06 | 0.2 | 1.28E-01 | 1.3 | 3.57E-03 | 0.7 | 1.49E-01 | 0.6 | 1.63E-01 | tRNA -N)-dimethyltransferase |
| AT5G17660 | 0.4 | 2.22E-02 | 0.3 | 4.52E-02 | 0.0 | 8.17E-01 | 0.2 | 5.42E-01 | 0.2 | 5.67E-01 | 0.0 | 9.75E-01 | Gb\|AAB08476.1 |
| AT5G24840 | 0.4 | 1.51E-03 | 0.4 | 4.00E-03 | 0.0 | 7.34E-01 | 0.2 | 3.81E-01 | 0.1 | 7.86E-01 | 0.1 | 7.47E-01 | tRNA -)-methyltransferase -N)-methyltransferase) -methyltransferase) |
| AT5G15390 | 0.5 | 8.61E-05 | 0.5 | 1.16E-04 | 0.0 | 8.43E-01 | -0.1 | 5.23E-01 | -0.3 | 1.09E-01 | 0.2 | 4.47E-01 | rRNA methylase-like protein family protein) |
| AT2G45330 | -0.5 | 9.43E-06 | -0.2 | 1.33E-02 | -0.4 | 8.86E-04 | -0.4 | 1.10E-02 | -0.2 | 1.62E-01 | -0.2 | 4.40E-01 | 2'-phosphotransferase |
| AT4G04670 | 0.2 | 5.66E-02 | 0.0 | 9.93E-01 | 0.2 | 1.09E-01 | 0.3 | 3.97E-01 | 0.1 | 8.15E-01 | 0.2 | 6.91E-01 | tRNA wybutosine-synthesizing protein 2 |
| AT3G59980 | 1.8 | 3.55E-08 | 1.4 | 4.03E-07 | 0.4 | 1.96E-03 | 0.6 | 1.93E-04 | 0.1 | 5.18E-01 | 0.5 | 6.17E-03 | hypothetical protein |
| AT3G63510 | 1.3 | 2.23E-07 | 1.1 | 1.92E-06 | 0.2 | 4.73E-02 | 0.5 | 2.60E-02 | 0.2 | 5.42E-01 | 0.3 | 2.26E-01 | FMN-linked oxidoreductases superfamily protein |
| AT1G03687 | 0.6 | 8.42E-05 | 0.4 | 4.24E-03 | 0.2 | 4.80E-02 | 0.0 | 9.31E-01 | 0.0 | 9.77E-01 | 0.0 | 9.30E-01 | tRNA-uridine aminocarboxypropyltransferase |
| AT5G54880 | 0.8 | 2.02E-04 | 0.7 | 1.43E-03 | 0.2 | 3.61E-01 | 0.8 | 2.13E-02 | 0.4 | 3.12E-01 | 0.4 | 3.07E-01 | tRNA-uridine aminocarboxypropyltransferase |
| AT4G27070 | 0.2 | 2.22E-02 | 0.2 | 4.27E-02 | 0.0 | 8.29E-01 | 0.3 | 1.69E-01 | 0.2 | 4.74E-01 | 0.1 | 7.34E-01 | Tryptophan synthase beta chain 2, chloroplastic |
| AT5G54810 | 1.0 | 1.66E-06 | 0.9 | 1.07E-05 | 0.1 | 2.23E-01 | 0.3 | 1.28E-01 | -0.2 | 2.04E-01 | 0.5 | 1.63E-02 | tryptophan synthase |
| AT5G49510 | 0.1 | 1.56E-01 | 0.2 | 6.55E-02 | -0.1 | 6.43E-01 | 0.2 | 2.97E-01 | 0.2 | 5.08E-01 | 0.0 | 8.99E-01 | Prefoldin subunit 3 |

| | | | | | | | | | | | | | |
|---|---|---|---|---|---|---|---|---|---|---|---|---|---|
| AT4G39920 | 0.3 | 1.45E-02 | 0.3 | 3.15E-02 | 0.0 | 7.91E-01 | 0.1 | 5.47E-01 | 0.0 | 9.24E-01 | 0.2 | 5.75E-01 | Tubulin-folding cofactor C |
| AT2G47580 | 0.3 | 1.55E-02 | 0.3 | 2.15E-02 | 0.0 | 9.50E-01 | 0.3 | 2.43E-01 | 0.1 | 8.50E-01 | 0.2 | 4.05E-01 | U1 small nuclear ribonucleoprotein A |
| AT1G13030 | 0.2 | 1.28E-01 | 0.1 | 6.65E-01 | 0.1 | 3.73E-01 | 0.4 | 8.68E-02 | 0.3 | 3.52E-01 | 0.2 | 6.44E-01 | Coilin |
| AT3G11960 | -0.4 | 3.08E-04 | -0.4 | 3.13E-04 | 0.0 | 7.76E-01 | -0.1 | 4.31E-01 | -0.1 | 7.72E-01 | -0.1 | 7.87E-01 | Cleavage and polyadenylation specificity factor A subunit protein |
| AT3G55200 | -0.3 | 6.55E-02 | -0.4 | 2.58E-02 | 0.1 | 6.04E-01 | -0.7 | 5.91E-01 | -0.4 | 7.92E-01 | -0.3 | 8.94E-01 | Spliceosome-associated protein 130 A |
| AT1G09760 | 0.3 | 4.30E-04 | 0.3 | 6.70E-04 | 0.0 | 9.78E-01 | 0.2 | 1.91E-01 | 0.0 | 9.27E-01 | 0.1 | 3.85E-01 | U2 small nuclear ribonucleoprotein A' |
| AT5G16750 | 0.2 | 5.96E-02 | 0.1 | 2.97E-01 | 0.1 | 4.50E-01 | 0.6 | 2.95E-04 | 0.3 | 1.42E-01 | 0.4 | 3.44E-02 | Protein TORMOZ EMBRYO DEFECTIVE |
| AT2G41500 | 0.1 | 3.51E-01 | -0.1 | 1.32E-01 | 0.2 | 4.70E-02 | -0.2 | 3.86E-01 | -0.2 | 3.68E-01 | 0.0 | 9.35E-01 | U4 |
| AT4G24270 | 0.1 | 2.02E-01 | 0.0 | 8.17E-01 | 0.1 | 1.95E-01 | 0.3 | 7.22E-02 | 0.1 | 7.23E-01 | 0.2 | 3.12E-01 | EMBRYO DEFECTIVE 140 |
| AT5G27720 | 0.1 | 6.45E-02 | 0.1 | 4.75E-01 | 0.1 | 2.89E-01 | 0.1 | 7.03E-01 | -0.1 | 7.99E-01 | 0.1 | 5.85E-01 | Sm-like protein LSM4 |
| AT4G17650 | 0.1 | 1.39E-01 | 0.1 | 5.41E-01 | 0.1 | 4.68E-01 | -0.2 | 1.49E-01 | -0.2 | 2.09E-01 | 0.0 | 9.89E-01 | hypothetical protein |
| AT5G13860 | -0.6 | 4.18E-04 | -0.3 | 1.77E-02 | -0.3 | 7.15E-02 | -0.2 | 4.47E-01 | -0.2 | 4.07E-01 | 0.0 | 9.16E-01 | Protein ELC-like |
| AT5G53330 | -0.5 | 1.83E-04 | -0.3 | 1.44E-02 | -0.2 | 3.00E-02 | 0.0 | 9.49E-01 | 0.1 | 7.27E-01 | -0.1 | 8.14E-01 | hypothetical protein |
| AT5G25540 | -0.4 | 1.13E-03 | -0.1 | 3.25E-01 | -0.3 | 1.31E-02 | 0.0 | 8.62E-01 | 0.1 | 7.00E-01 | 0.0 | 8.53E-01 | AT5G25540 protein |
| AT3G23605 | -0.3 | 3.36E-02 | -0.5 | 9.41E-03 | 0.1 | 4.91E-01 | 0.1 | 8.48E-01 | 0.1 | 8.37E-01 | 0.0 | 9.76E-01 | Plant UBX domain-containing protein 12 |
| AT5G32440 | -0.2 | 7.09E-03 | -0.2 | 3.56E-02 | -0.1 | 4.71E-01 | -0.2 | 3.76E-02 | -0.2 | 7.06E-02 | 0.0 | 9.76E-01 | Ubiquitin system component Cue protein |
| AT1G27752 | -0.1 | 1.52E-01 | -0.2 | 2.56E-02 | 0.1 | 3.52E-01 | -0.1 | 5.37E-01 | -0.2 | 3.82E-01 | 0.1 | 8.28E-01 | Ubiquitin system component Cue protein |
| AT3G18860 | -0.1 | 9.53E-02 | -0.1 | 8.02E-02 | 0.0 | 9.03E-01 | 0.1 | 4.84E-01 | 0.0 | 9.94E-01 | 0.1 | 6.36E-01 | AT3g18860 |
| AT3G05090 | 0.0 | 6.87E-01 | -0.2 | 1.98E-02 | 0.2 | 1.72E-02 | -0.1 | 5.88E-01 | 0.0 | 8.84E-01 | -0.1 | 8.27E-01 | AT3g05090 |
| AT1G80040 | 0.1 | 1.58E-01 | 0.0 | 5.73E-01 | 0.1 | 8.40E-02 | 0.1 | 4.88E-01 | 0.1 | 7.00E-01 | 0.0 | 9.14E-01 | Ubiquitin system component Cue |
| AT4G22150 | 0.1 | 2.10E-01 | 0.1 | 6.89E-02 | -0.1 | 5.50E-01 | 0.0 | 8.62E-01 | -0.1 | 7.54E-01 | 0.0 | 9.06E-01 | Plant UBX domain-containing protein 3 |
| AT4G23040 | 0.1 | 1.82E-01 | -0.1 | 4.25E-01 | 0.2 | 7.93E-02 | -0.1 | 5.28E-01 | -0.3 | 1.68E-01 | 0.2 | 5.40E-01 | Plant UBX domain-containing protein 13 |
| AT4G00752 | 0.1 | 9.81E-02 | 0.0 | 7.92E-01 | 0.1 | 2.35E-01 | -0.6 | 2.22E-02 | -0.4 | 1.73E-01 | -0.2 | 5.40E-01 | Plant UBX domain-containing protein 9 |
| AT3G19590 | 1.3 | 3.77E-07 | 0.8 | 2.42E-05 | 0.5 | 1.49E-03 | 0.2 | 2.79E-01 | 0.2 | 2.78E-01 | 0.0 | 9.40E-01 | Mitotic checkpoint protein BUB3.1 |
| AT1G69400 | 1.9 | 3.35E-06 | 1.2 | 3.18E-04 | 0.7 | 4.67E-03 | 0.0 | 9.63E-01 | -0.1 | 8.54E-01 | 0.1 | 8.93E-01 | Mitotic checkpoint protein BUB3.3 |
| AT1G49910 | 2.8 | 7.52E-07 | 1.4 | 3.50E-04 | 1.4 | 2.67E-04 | 0.4 | 6.28E-02 | 0.3 | 2.50E-01 | 0.1 | 7.11E-01 | Mitotic checkpoint protein BUB3.2 |
| AT3G63000 | -0.4 | 8.01E-05 | -0.4 | 9.17E-05 | 0.0 | 7.43E-01 | -0.5 | 1.35E-03 | -0.3 | 4.20E-02 | -0.2 | 3.55E-01 | NPL4-like protein 1 |
| AT2G47970 | -0.4 | 4.80E-04 | -0.4 | 8.37E-04 | 0.0 | 9.36E-01 | -0.2 | 2.08E-01 | -0.1 | 5.49E-01 | -0.1 | 7.71E-01 | NPL4-like protein 2 |
| AT5G51450 | -0.3 | 2.72E-03 | -0.3 | 6.90E-03 | 0.0 | 7.50E-01 | -0.1 | 7.97E-01 | 0.0 | 8.97E-01 | 0.0 | 9.59E-01 | RPM1 interacting protein 3 |
| AT1G65430 | 0.0 | 9.26E-01 | -0.1 | 1.39E-01 | 0.1 | 1.56E-01 | 0.0 | 9.43E-01 | 0.1 | 5.75E-01 | -0.1 | 7.20E-01 | Probable E3 ubiquitin-protein ligase ARI8 |
| AT1G32340 | -0.5 | 5.12E-05 | -0.5 | 7.10E-05 | 0.0 | 8.17E-01 | -0.2 | 3.86E-01 | -0.3 | 3.00E-01 | 0.1 | 8.73E-01 | F5D14.10 protein |
| AT3G47180 | | | | | | | 0.3 | 6.94E-01 | 0.2 | 8.62E-01 | 0.1 | 9.10E-01 | At3g47180 |
| AT5G27920 | 0.2 | 1.28E-01 | 0.4 | 6.97E-03 | -0.2 | 1.39E-01 | -0.2 | 2.37E-01 | -0.4 | 2.83E-02 | 0.2 | 3.37E-01 | At5g27920 |
| AT4G10180 | -0.1 | 4.57E-01 | -0.2 | 2.80E-02 | 0.1 | 1.37E-01 | -0.1 | 6.98E-01 | -0.1 | 8.04E-01 | 0.0 | 9.69E-01 | Light-mediated development protein DET1 |
| AT5G09800 | -2.4 | 2.85E-04 | -2.3 | 5.94E-04 | -0.2 | 8.04E-01 | 1.0 | 1.22E-03 | 0.3 | 3.83E-01 | 0.7 | 3.10E-02 | U-box domain-containing protein 28 |
| AT5G40140 | -0.9 | 1.34E-05 | -0.5 | 1.48E-03 | -0.4 | 8.00E-03 | 0.1 | 8.19E-01 | 0.2 | 5.87E-01 | -0.1 | 8.08E-01 | U-box domain-containing protein 40 |
| AT5G64660 | -0.7 | 1.02E-04 | -0.7 | 2.03E-04 | 0.0 | 9.01E-01 | 0.3 | 1.38E-01 | 0.1 | 6.76E-01 | 0.2 | 4.55E-01 | U-box domain-containing protein 27 |
| AT1G72175 | -0.5 | 2.08E-04 | -0.2 | 5.37E-02 | -0.3 | 1.30E-02 | -0.3 | 2.26E-01 | 0.0 | 9.81E-01 | -0.3 | 1.76E-01 | RING-type E3 ubiquitin transferase |
| AT1G70320 | -0.3 | 2.67E-03 | -0.6 | 6.95E-05 | 0.3 | 1.20E-02 | -0.1 | 8.54E-01 | 0.0 | 9.90E-01 | -0.1 | 9.01E-01 | E3 ubiquitin-protein ligase UPL2 |
| AT3G49810 | -0.3 | 2.74E-02 | 0.1 | 5.32E-01 | -0.3 | 2.12E-02 | -0.1 | 5.57E-01 | 0.0 | 8.98E-01 | -0.1 | 7.61E-01 | U-box domain-containing protein 30 |
| AT2G23140 | -0.2 | 4.16E-03 | -0.2 | 5.07E-03 | 0.0 | 9.79E-01 | -0.4 | 5.31E-02 | -0.3 | 1.49E-01 | -0.1 | 8.27E-01 | RING-type E3 ubiquitin transferase |
| AT3G53090 | -0.2 | 5.75E-02 | -0.4 | 2.25E-03 | 0.2 | 9.41E-02 | 0.1 | 8.04E-01 | -0.2 | 5.75E-01 | 0.3 | 3.81E-01 | E3 ubiquitin-protein ligase UPL7 |
| AT4G24204 | -0.2 | 3.42E-01 | 1.4 | 6.75E-06 | -1.6 | 3.89E-05 | -4.6 | 1.58E-04 | -0.7 | 1.51E-01 | -3.9 | 6.72E-03 | RING |
| AT5G66070 | -0.1 | 5.02E-01 | -0.4 | 1.01E-02 | 0.3 | 5.37E-02 | 0.1 | 7.31E-01 | -0.1 | 6.91E-01 | 0.2 | 4.53E-01 | RING |
| AT3G60300 | -0.1 | 2.48E-01 | -0.1 | 1.49E-01 | 0.0 | 7.61E-01 | -0.4 | 2.35E-03 | -0.5 | 1.99E-03 | 0.1 | 7.75E-01 | Uncharacterized protein At3g60300 |
| AT5G01160 | 0.0 | 6.80E-01 | -0.1 | 4.94E-01 | 0.0 | 8.11E-01 | -0.3 | 1.91E-02 | -0.4 | 7.54E-03 | 0.1 | 6.71E-01 | C2H2-type domain-containing protein |
| AT3G19380 | 0.0 | 6.94E-01 | -0.1 | 7.57E-02 | 0.1 | 2.01E-01 | -0.3 | 7.58E-02 | -0.3 | 5.72E-02 | 0.0 | 8.52E-01 | U-box domain-containing protein 25 |
| AT2G33340 | 0.1 | 1.82E-01 | 0.0 | 9.01E-01 | 0.1 | 1.87E-01 | -0.2 | 1.55E-01 | -0.2 | 1.88E-01 | 0.0 | 9.71E-01 | Pre-mRNA-processing factor 19 |
| AT1G49780 | 0.1 | 4.29E-01 | -0.3 | 3.89E-02 | 0.4 | 1.97E-02 | -0.6 | 3.58E-03 | -0.2 | 3.50E-01 | -0.4 | 1.31E-01 | U-box domain-containing protein 26 |
| AT2G15580 | 0.1 | 1.25E-01 | 0.0 | 9.70E-01 | 0.1 | 1.81E-01 | -0.4 | 1.81E-02 | -0.2 | 1.87E-01 | -0.2 | 5.64E-01 | AT2G15580 protein |
| AT2G24330 | 0.2 | 1.46E-02 | 0.1 | 3.58E-01 | 0.1 | 1.35E-01 | 0.1 | 5.64E-01 | -0.2 | 4.40E-01 | 0.3 | 2.28E-01 | Uncharacterized protein At2g24330 |
| AT2G32950 | 0.2 | 6.72E-02 | -0.2 | 6.29E-02 | 0.4 | 5.31E-03 | 0.0 | 9.17E-01 | 0.0 | 9.62E-01 | 0.0 | 8.89E-01 | Transducin |
| AT1G04510 | 0.3 | 4.76E-03 | 0.2 | 2.41E-02 | 0.1 | 4.72E-01 | 0.2 | 2.75E-01 | 0.1 | 8.17E-01 | 0.1 | 6.25E-01 | Pre-mRNA-processing factor 19 |
| AT5G65920 | 0.3 | 1.66E-02 | 0.1 | 5.05E-01 | 0.2 | 1.12E-01 | -0.4 | 7.31E-03 | -0.2 | 2.81E-01 | -0.3 | 1.74E-01 | U-box domain-containing protein 31 |
| AT3G11840 | 0.4 | 1.10E-02 | -0.2 | 2.29E-01 | 0.5 | 5.63E-03 | 0.9 | 9.07E-04 | 0.4 | 1.60E-01 | 0.5 | 6.40E-02 | U-box domain-containing protein |
| AT1G66160 | 1.5 | 7.69E-04 | 0.9 | 2.18E-02 | 0.6 | 1.24E-01 | 0.1 | 7.85E-01 | -0.7 | 1.41E-01 | 0.8 | 1.47E-01 | U-box domain-containing protein |
| AT5G37490 | 3.6 | 6.49E-05 | 0.6 | 4.12E-01 | 3.0 | 9.16E-04 | 4.3 | 2.11E-05 | 1.5 | 1.35E-01 | 2.8 | 4.16E-04 | U-box domain-containing protein 21 |
| AT3G02840 | | | | | | | 2.200094161 | 1.62E-03 | 0.347065545 | 7.35E-01 | 1.9 | 1.59E-02 | U-box domain-containing protein |
| AT2G35930 | -0.6 | 1.09E-02 | -0.5 | 3.12E-02 | -0.1 | 6.68E-01 | 0.6 | 5.66E-02 | 0.1 | 8.21E-01 | 0.5 | 1.26E-01 | E3 ubiquitin-protein ligase PUB23 |
| AT1G10560 | -0.5 | 1.19E-02 | 0.2 | 1.88E-01 | -0.7 | 5.16E-03 | -0.7 | 1.39E-01 | 0.4 | 3.43E-01 | -1.1 | 3.42E-02 | U-box domain-containing protein 18 |
| AT1G55860 | -0.4 | 1.93E-03 | -0.7 | 2.28E-04 | 0.2 | 1.05E-01 | -0.1 | 8.78E-01 | 0.2 | 7.84E-01 | -0.2 | 7.17E-01 | E3 ubiquitin-protein ligase UPL1 |
| AT3G52450 | -0.2 | 9.65E-02 | 0.2 | 2.07E-01 | -0.4 | 2.26E-02 | 0.2 | 2.53E-01 | 0.1 | 8.49E-01 | 0.2 | 4.13E-01 | E3 ubiquitin-protein ligase PUB22 |
| AT5G18650 | 0.1 | 2.18E-01 | 0.0 | 7.95E-01 | 0.1 | 4.00E-01 | -0.3 | 5.66E-02 | -0.5 | 5.74E-03 | 0.2 | 2.87E-01 | E3 ubiquitin-protein ligase MIEL1 |
| AT5G15400 | -0.3 | 5.65E-04 | -0.4 | 1.43E-04 | 0.1 | 1.89E-01 | 0.1 | 5.96E-01 | 0.2 | 4.07E-01 | -0.1 | 8.08E-01 | Probable ubiquitin conjugation factor E4 |
| AT3G58040 | -0.5 | 2.92E-05 | -0.4 | 2.47E-04 | -0.1 | 2.07E-01 | -0.1 | 3.92E-01 | -0.1 | 7.16E-01 | -0.1 | 8.27E-01 | E3 ubiquitin-protein ligase SINAT2 |
| AT5G02310 | -0.4 | 5.07E-04 | -0.5 | 1.55E-04 | 0.1 | 2.36E-01 | -0.1 | 7.01E-01 | -0.1 | 7.07E-01 | 0.0 | 9.72E-01 | E3 ubiquitin-protein ligase |
| AT3G61460 | -0.3 | 7.70E-03 | 0.0 | 8.19E-01 | -0.3 | 2.25E-02 | 0.0 | 9.53E-01 | -0.3 | 1.80E-01 | 0.3 | 2.46E-01 | Brassinosteroid-responsive RING protein 1 |
| AT3G61790 | -0.3 | 7.78E-04 | -0.2 | 1.55E-02 | -0.1 | 1.44E-01 | -0.4 | 6.18E-03 | -0.3 | 1.87E-02 | 0.0 | 8.75E-01 | E3 ubiquitin-protein ligase SINAT3 |
| AT5G11620 | -0.1 | 5.81E-01 | -0.1 | 5.77E-01 | 0.0 | 9.78E-01 | 0.4 | 6.17E-01 | 0.2 | 8.68E-01 | 0.2 | 8.45E-01 | SWIM-type domain-containing protein |
| AT4G17410 | 0.1 | 4.07E-01 | -0.1 | 2.04E-01 | 0.2 | 8.22E-02 | -0.2 | 2.85E-01 | -0.1 | 6.73E-01 | -0.1 | 7.33E-01 | DWNN domain, a CCHC-type zinc finger |
| AT4G27880 | 0.1 | 2.80E-01 | 0.0 | 9.50E-01 | 0.1 | 3.33E-01 | -0.5 | 4.30E-03 | -0.4 | 3.38E-02 | -0.1 | 6.46E-01 | E3 ubiquitin-protein ligase SINAT4 |
| AT4G23860 | 0.1 | 6.59E-02 | 0.0 | 8.29E-01 | 0.1 | 1.52E-01 | 0.0 | 8.92E-01 | -0.1 | 4.92E-01 | 0.1 | 4.46E-01 | Uncharacterized protein AT4g23860 |
| AT5G37890 | 0.4 | 2.69E-03 | 0.2 | 8.93E-02 | 0.2 | 1.25E-01 | -0.1 | 6.74E-01 | 0.1 | 8.01E-01 | -0.2 | 5.63E-01 | E3 ubiquitin-protein ligase SINA-like 7 |
| AT2G41980 | 0.9 | 7.86E-07 | 0.5 | 8.59E-05 | 0.4 | 1.53E-03 | 0.3 | 6.17E-02 | 0.0 | 9.18E-01 | 0.3 | 2.07E-01 | Putative E3 ubiquitin-protein ligase SINAT1 |
| AT1G26830 | -0.2 | 1.98E-03 | -0.3 | 1.68E-03 | 0.0 | 7.81E-01 | -0.1 | 6.58E-01 | 0.0 | 9.80E-01 | -0.1 | 8.04E-01 | Cullin-3A |
| AT1G69670 | -0.2 | 9.93E-03 | -0.3 | 1.47E-03 | 0.1 | 2.41E-01 | 0.1 | 3.98E-01 | 0.1 | 8.19E-01 | 0.1 | 7.40E-01 | Cullin-3B |
| AT5G46210 | -0.2 | 9.85E-03 | -0.2 | 5.20E-03 | 0.0 | 6.43E-01 | -0.1 | 5.04E-01 | -0.1 | 6.59E-01 | 0.0 | 9.57E-01 | Cullin-4 |
| AT4G02570 | -0.1 | 5.87E-02 | -0.3 | 2.59E-03 | 0.1 | 8.69E-02 | 0.0 | 7.00E-01 | -0.1 | 7.06E-01 | 0.1 | 5.54E-01 | Cullin-1 |
| AT4G08455 | -0.1 | 2.18E-01 | -0.1 | 2.77E-01 | 0.0 | 9.37E-01 | 0.0 | 9.27E-01 | -0.2 | 6.47E-01 | 0.1 | 7.32E-01 | BTB |
| AT2G04660 | 0.2 | 5.31E-02 | -0.1 | 1.38E-01 | 0.3 | 1.08E-02 | 0.4 | 1.03E-01 | 0.1 | 7.48E-01 | 0.3 | 3.45E-01 | Anaphase-promoting complex subunit 2 |
| AT2G32410 | -0.2 | 5.50E-02 | -0.1 | 4.87E-01 | -0.1 | 2.85E-01 | 0.5 | 6.97E-02 | 0.4 | 1.71E-01 | 0.1 | 8.33E-01 | AXR1-like protein |
| AT5G37530 | 0.5 | 3.42E-04 | 0.1 | 3.41E-01 | 0.4 | 6.67E-03 | 0.6 | 1.46E-02 | 0.2 | 4.36E-01 | 0.4 | 1.96E-01 | THIF-type NAD |
| AT5G50680 | | | | | | | 0.091355241 | 9.55E-01 | -0.116661791 | 9.58E-01 | | | Ubiquitin activating enzyme |

| | | | | | | | | | | | | | |
|---|---|---|---|---|---|---|---|---|---|---|---|---|---|
| AT5G61500 | -0.6 | 6.72E-06 | -0.5 | 3.32E-05 | -0.1 | 3.44E-01 | -0.2 | 5.40E-02 | 0.1 | 7.40E-01 | -0.3 | 9.15E-02 | Autophagocytosis protein-like |
| AT3G07525 | 0.5 | 1.45E-02 | 0.1 | 5.00E-01 | 0.3 | 1.11E-01 | 0.1 | 7.81E-01 | 0.0 | 9.52E-01 | 0.1 | 9.05E-01 | ATG10 |
| AT1G60190 | -1.9 | 2.43E-03 | -1.7 | 5.75E-03 | -0.3 | 7.32E-01 | -0.7 | 4.72E-01 | -0.2 | 8.63E-01 | -0.5 | 7.56E-01 | U-box domain-containing protein 19 |
| AT1G01660 | -1.3 | 1.29E-05 | 0.0 | 8.14E-01 | -1.3 | 1.35E-04 | -1.1 | 3.05E-02 | -0.1 | 9.15E-01 | -1.1 | 9.02E-02 | RING |
| AT5G62560 | -1.1 | 4.18E-07 | -0.7 | 2.33E-05 | -0.4 | 1.20E-03 | -0.4 | 1.79E-02 | -0.3 | 7.57E-02 | -0.1 | 7.61E-01 | U-box domain-containing protein 41 |
| AT4G23450 | -1.1 | 5.67E-06 | -0.9 | 2.76E-05 | -0.1 | 3.64E-01 | 0.0 | 9.51E-01 | 0.0 | 9.41E-01 | -0.1 | 9.01E-01 | RING |
| AT4G21350 | -0.8 | 4.92E-06 | -0.6 | 4.65E-05 | -0.2 | 1.24E-01 | -0.6 | 1.71E-04 | -0.4 | 1.22E-02 | -0.2 | 1.88E-01 | U-box domain-containing protein 8 |
| AT3G47820 | -0.8 | 9.75E-07 | -0.6 | 8.17E-06 | -0.1 | 8.55E-02 | -0.3 | 2.12E-02 | -0.1 | 3.05E-01 | -0.1 | 4.34E-01 | U-box domain-containing protein 39 |
| AT5G19970 | -0.7 | 9.48E-03 | -0.4 | 8.73E-02 | -0.3 | 3.18E-01 | -0.9 | 3.27E-03 | -0.2 | 4.59E-01 | -0.6 | 5.32E-02 | GRAS family transcription factor family protein |
| AT3G55070 | -0.7 | 3.44E-06 | -0.4 | 2.79E-04 | -0.3 | 4.74E-03 | -0.3 | 8.06E-02 | -0.1 | 4.51E-01 | -0.1 | 5.55E-01 | LisH |
| AT5G05230 | -0.6 | 1.36E-05 | -0.6 | 3.44E-05 | 0.0 | 8.08E-01 | -0.2 | 3.30E-01 | 0.0 | 9.80E-01 | -0.2 | 4.76E-01 | U-box domain-containing protein 62 |
| AT1G67530 | -0.6 | 7.33E-05 | -0.5 | 5.23E-04 | -0.1 | 2.98E-01 | -0.4 | 5.12E-02 | -0.2 | 3.32E-01 | -0.2 | 5.23E-01 | RING-type E3 ubiquitin transferase |
| AT5G01830 | -0.6 | 1.41E-04 | 0.0 | 8.51E-01 | -0.5 | 8.02E-04 | -0.2 | 2.14E-01 | -0.1 | 5.79E-01 | -0.1 | 7.11E-01 | U-box domain-containing protein 16 |
| AT1G76390 | -0.5 | 2.85E-03 | -0.5 | 4.08E-03 | 0.0 | 9.71E-01 | -0.5 | 1.72E-01 | -0.2 | 7.09E-01 | -0.3 | 5.14E-01 | U-box domain-containing protein 43 |
| AT5G43190 | -0.5 | 4.87E-04 | -0.4 | 3.36E-03 | -0.1 | 3.47E-01 | -0.5 | 2.68E-03 | -0.7 | 8.34E-04 | 0.2 | 5.27E-01 | F-box |
| AT5G15710 | -0.5 | 3.97E-04 | -0.3 | 3.31E-03 | -0.1 | 3.05E-01 | -0.1 | 6.60E-01 | -0.2 | 5.75E-01 | 0.0 | 9.07E-01 | F-box |
| AT5G67250 | -0.4 | 5.90E-05 | -0.3 | 2.32E-03 | -0.2 | 3.90E-02 | -0.5 | 1.64E-03 | -0.4 | 1.19E-02 | -0.1 | 7.50E-01 | F-box protein SKIP2 |
| AT3G54790 | -0.4 | 9.93E-04 | -0.2 | 1.56E-02 | -0.1 | 1.85E-01 | -0.1 | 3.87E-01 | -0.1 | 6.34E-01 | 0.0 | 8.85E-01 | RING-type E3 ubiquitin transferase |
| AT3G54850 | -0.3 | 9.94E-04 | -0.5 | 1.01E-04 | 0.2 | 6.92E-02 | -0.2 | 1.54E-01 | -0.4 | 1.99E-02 | 0.2 | 4.31E-01 | AT3G54850 protein |
| AT3G24760 | -0.3 | 2.56E-03 | -0.4 | 6.24E-04 | 0.1 | 2.92E-01 | -0.4 | 3.49E-02 | 0.0 | 8.75E-01 | -0.3 | 1.49E-01 | F-box |
| AT1G47056 | -0.3 | 4.04E-03 | -0.3 | 6.17E-03 | 0.0 | 9.41E-01 | -0.4 | 1.02E-01 | -0.3 | 1.59E-01 | 0.0 | 9.75E-01 | F-box protein At1g47056 |
| AT3G50080 | -0.3 | 1.79E-03 | -0.1 | 5.35E-02 | -0.1 | 1.23E-01 | -0.2 | 1.66E-01 | -0.2 | 2.14E-01 | 0.0 | 9.98E-01 | F-box |
| AT3G05545 | -0.2 | 1.66E-02 | -0.2 | 1.27E-02 | 0.0 | 8.19E-01 | 0.1 | 3.90E-01 | 0.1 | 4.68E-01 | 0.0 | 9.99E-01 | hypothetical protein |
| AT3G63530 | -0.2 | 9.27E-02 | 0.0 | 8.03E-01 | -0.2 | 2.27E-01 | -0.2 | 3.79E-01 | 0.0 | 9.68E-01 | -0.2 | 4.95E-01 | RING-type domain-containing protein |
| AT2G28830 | -0.2 | 1.44E-01 | -0.2 | 1.44E-01 | 0.0 | 9.61E-01 | 0.3 | 3.98E-01 | -0.1 | 8.43E-01 | 0.3 | 3.46E-01 | U-box domain-containing protein 12 |
| AT1G27910 | -0.2 | 1.44E-01 | -0.3 | 3.52E-02 | 0.1 | 4.61E-01 | 0.0 | 1.00E+00 | 0.1 | 8.89E-01 | -0.1 | 9.08E-01 | RING-type E3 ubiquitin transferase |
| AT4G36550 | -0.1 | 1.51E-01 | -0.3 | 1.62E-02 | 0.1 | 2.58E-01 | 0.0 | 9.45E-01 | 0.2 | 5.34E-01 | -0.2 | 5.50E-01 | U-box domain-containing protein 5 |
| AT1G29340 | -0.1 | 8.48E-02 | -0.3 | 6.05E-03 | 0.1 | 1.53E-01 | -0.2 | 1.54E-01 | -0.2 | 8.67E-02 | 0.1 | 7.71E-01 | U-box domain-containing protein 17 |
| AT3G47990 | -0.1 | 1.34E-01 | -0.2 | 1.90E-02 | 0.1 | 3.07E-01 | 0.0 | 9.53E-01 | 0.1 | 7.11E-01 | -0.1 | 8.07E-01 | SUGAR-INSENSITIVE 3 |
| AT3G48070 | -0.1 | 2.69E-01 | -0.1 | 1.59E-01 | 0.0 | 7.62E-01 | -0.2 | 2.42E-01 | -0.2 | 3.58E-01 | 0.0 | 9.59E-01 | RING |
| AT4G02440 | 0.0 | 7.54E-01 | -0.1 | 5.36E-01 | 0.0 | 7.91E-01 | -0.7 | 2.79E-05 | -0.8 | 4.33E-05 | 0.1 | 7.00E-01 | Phytochrome A-associated F-box protein |
| AT2G40640 | 0.0 | 9.70E-01 | 0.1 | 5.22E-01 | -0.1 | 5.74E-01 | -0.4 | 4.00E-01 | -0.2 | 7.18E-01 | -0.2 | 7.89E-01 | RING |
| AT1G24330 | 0.0 | 8.13E-01 | 0.0 | 9.07E-01 | 0.1 | 7.77E-01 | | | | | | | U-box domain-containing protein 6 |
| AT5G67340 | 0.1 | 3.79E-01 | -0.5 | 8.21E-03 | 0.7 | 5.35E-03 | 0.4 | 2.22E-02 | 0.1 | 8.47E-01 | 0.4 | 9.76E-02 | U-box domain-containing protein 2 |
| AT5G03450 | 0.1 | 1.53E-01 | 0.1 | 5.43E-01 | 0.1 | 5.09E-01 | -0.2 | 4.52E-01 | -0.2 | 5.26E-01 | 0.0 | 9.97E-01 | hypothetical protein |
| AT2G47700 | 0.2 | 5.52E-02 | 0.1 | 5.72E-01 | 0.1 | 2.11E-01 | -0.6 | 1.69E-02 | -0.6 | 2.06E-02 | 0.0 | 9.28E-01 | hypothetical protein |
| AT1G01670 | 0.2 | 7.80E-02 | 0.0 | 8.81E-01 | 0.3 | 1.21E-01 | | | | | | | RING |
| AT5G62910 | 0.3 | 7.18E-03 | 0.0 | 6.76E-01 | 0.2 | 3.60E-02 | -0.5 | 1.51E-03 | -0.6 | 2.00E-03 | 0.0 | 9.11E-01 | RING-type domain-containing protein |
| AT2G28540 | 0.3 | 1.93E-03 | -0.1 | 5.22E-01 | 0.4 | 2.44E-03 | -0.1 | 4.82E-01 | -0.2 | 2.24E-01 | 0.1 | 6.93E-01 | Uncharacterized protein At2g28540 |
| AT1G49990 | 0.3 | 3.97E-02 | 0.5 | 1.27E-02 | -0.1 | 5.40E-01 | -0.1 | 7.81E-01 | 0.0 | 9.55E-01 | -0.1 | 7.99E-01 | F-box protein At1g49990 |
| AT1G23030 | 0.4 | 8.45E-03 | 0.3 | 9.13E-02 | 0.2 | 2.71E-01 | -0.3 | 3.83E-02 | -0.3 | 6.33E-02 | 0.0 | 9.89E-01 | U-box domain-containing protein 11 |
| AT4G07400 | 0.5 | 8.53E-03 | 0.5 | 1.20E-02 | 0.0 | 9.66E-01 | 0.0 | 9.78E-01 | -0.3 | 2.74E-01 | 0.3 | 3.36E-01 | Putative F-box |
| AT1G15165 | 0.6 | 1.69E-02 | 0.2 | 4.73E-01 | 0.4 | 1.40E-01 | 0.0 | 9.53E-01 | -0.4 | 5.93E-01 | 0.5 | 6.02E-01 | RING |
| AT1G71020 | 0.6 | 4.16E-05 | 0.5 | 1.75E-04 | 0.1 | 5.02E-01 | -0.4 | 4.20E-03 | -0.5 | 3.09E-03 | 0.1 | 7.76E-01 | RING-type E3 ubiquitin transferase |
| AT1G02610 | 0.7 | 3.00E-03 | 0.5 | 2.96E-02 | 0.2 | 3.33E-01 | -0.4 | 2.67E-01 | -1.0 | 2.25E-02 | 0.6 | 2.77E-01 | RING |
| AT1G21410 | 0.8 | 3.88E-04 | 0.1 | 4.20E-01 | 0.7 | 5.71E-03 | -0.6 | 1.27E-02 | -0.4 | 1.10E-01 | -0.2 | 5.52E-01 | F-box protein SKP2A |
| AT1G01680 | 2.0 | 5.06E-04 | -0.5 | 4.07E-01 | 2.5 | 2.45E-03 | 0.9 | 4.68E-02 | 0.4 | 4.58E-01 | 0.5 | 3.89E-01 | U-box domain-containing protein 54 |
| AT1G14260 | -0.9 | 1.13E-02 | -0.2 | 4.42E-01 | -0.7 | 9.11E-02 | 0.5 | 3.63E-02 | 0.4 | 1.48E-01 | 0.1 | 7.19E-01 | At1g14260 |
| AT4G28370 | -0.6 | 2.45E-05 | -0.4 | 1.04E-03 | -0.2 | 2.90E-02 | -0.1 | 8.14E-01 | 0.2 | 5.23E-01 | -0.3 | 4.70E-01 | Transmembrane E3 ubiquitin-protein ligase FLY1 |
| AT2G02960 | -0.6 | 2.83E-04 | -0.2 | 6.12E-02 | -0.4 | 1.39E-02 | -0.4 | 3.00E-02 | 0.0 | 9.75E-01 | -0.4 | 7.74E-02 | RING |
| AT2G31510 | -0.4 | 1.99E-03 | -0.5 | 3.09E-04 | 0.1 | 1.53E-01 | -0.4 | 1.60E-02 | -0.1 | 5.05E-01 | -0.2 | 1.78E-01 | RBR-type E3 ubiquitin transferase |
| AT3G06330 | -0.3 | 1.77E-03 | -0.1 | 2.37E-01 | -0.2 | 2.86E-02 | -0.5 | 1.76E-03 | -0.5 | 9.50E-03 | -0.1 | 7.58E-01 | AT3g06330 |
| AT4G34370 | -0.3 | 2.22E-03 | -0.4 | 3.67E-04 | 0.1 | 1.83E-01 | -0.2 | 3.22E-01 | -0.3 | 1.50E-01 | 0.1 | 7.13E-01 | RING |
| AT3G47550 | -0.3 | 1.03E-03 | -0.3 | 6.20E-04 | 0.0 | 5.90E-01 | -0.2 | 5.44E-02 | -0.2 | 1.01E-01 | 0.0 | 9.57E-01 | RING |
| AT5G63760 | -0.2 | 1.00E-01 | -0.5 | 6.13E-04 | 0.3 | 1.44E-02 | 0.3 | 6.03E-02 | 0.1 | 6.98E-01 | 0.2 | 2.56E-01 | RING |
| AT2G16090 | -0.1 | 4.07E-01 | 0.0 | 5.33E-01 | 0.0 | 8.97E-01 | 0.1 | 7.08E-01 | 0.0 | 9.06E-01 | 0.1 | 9.01E-01 | RBR-type E3 ubiquitin transferase |
| AT1G05890 | 0.0 | 5.80E-01 | -0.2 | 4.30E-02 | 0.2 | 2.53E-02 | 0.0 | 8.66E-01 | -0.1 | 4.86E-01 | 0.2 | 4.52E-01 | RBR-type E3 ubiquitin transferase |
| AT4G02075 | 0.6 | 2.12E-02 | 0.0 | 9.96E-01 | 0.6 | 5.32E-02 | 0.3 | 4.25E-01 | 0.2 | 7.20E-01 | 0.1 | 7.84E-01 | At4g02075 |
| AT1G05880 | 2.3 | 2.43E-07 | 1.4 | 2.76E-05 | 0.9 | 3.53E-04 | 2.6 | 1.53E-07 | 1.1 | 4.22E-03 | 1.5 | 6.27E-05 | RING |
| AT2G34850 | -1.4 | 2.81E-04 | -1.6 | 3.57E-04 | 0.1 | 7.56E-01 | | | | | | | Putative UDP-arabinose 4-epimerase 2 |
| AT4G20460 | -0.8 | 3.97E-05 | 0.0 | 9.22E-01 | -0.9 | 2.31E-04 | 0.0 | 9.67E-01 | 0.2 | 4.15E-01 | -0.2 | 4.49E-01 | Probable UDP-arabinose 4-epimerase 3 |
| AT5G44480 | -1.3 | 3.80E-08 | -0.6 | 9.52E-06 | -0.7 | 2.60E-05 | 0.4 | 2.26E-02 | 0.2 | 1.93E-01 | 0.1 | 4.38E-01 | NAD-binding Rossmann-fold superfamily protein |
| AT1G30620 | -0.6 | 2.40E-04 | -0.5 | 1.74E-03 | -0.1 | 3.23E-01 | -0.1 | 7.49E-01 | 0.0 | 8.81E-01 | 0.0 | 9.35E-01 | NAD-binding Rossmann-fold superfamily protein |
| AT1G63180 | 1.3 | 1.38E-05 | 1.2 | 4.55E-05 | 0.1 | 6.52E-01 | -0.5 | 3.79E-02 | -0.7 | 1.96E-02 | 0.1 | 7.35E-01 | UDP-glucose 4-epimerase |
| AT2G37330 | -0.6 | 4.25E-05 | -0.2 | 2.26E-02 | -0.4 | 4.36E-03 | -0.5 | 2.20E-02 | -0.2 | 5.31E-01 | -0.4 | 2.42E-01 | Protein ALUMINUM SENSITIVE 3 |
| AT1G71220 | -0.2 | 9.43E-03 | -0.2 | 1.23E-02 | 0.0 | 9.84E-01 | 0.1 | 6.15E-01 | 0.3 | 2.87E-01 | -0.1 | 6.81E-01 | EMS-MUTAGENIZED BRI1 SUPPRESSOR 1 |
| AT1G64920 | -0.8 | 1.20E-03 | 0.1 | 4.56E-01 | -1.0 | 2.12E-03 | -0.4 | 5.41E-01 | 0.7 | 1.63E-01 | -1.1 | 4.38E-02 | UDP-glycosyltransferase 79B11 |
| AT1G10400 | -0.8 | 2.20E-03 | -0.1 | 6.78E-01 | -0.7 | 1.32E-02 | 0.0 | 9.77E-01 | 0.3 | 2.62E-01 | -0.3 | 2.77E-01 | UDP-glycosyltransferase 90A2 |
| AT4G09500 | -0.6 | 1.98E-03 | -0.6 | 4.99E-03 | -0.1 | 7.50E-01 | -0.6 | 3.85E-02 | 0.2 | 5.62E-01 | -0.7 | 3.22E-02 | UDP-glycosyltransferase 79B7 |
| AT5G12890 | -0.6 | 8.86E-04 | -0.6 | 1.40E-03 | 0.0 | 9.60E-01 | 0.0 | 9.04E-01 | -0.1 | 7.17E-01 | 0.1 | 8.45E-01 | UDP-glycosyltransferase 92A1 |
| AT2G36970 | -0.5 | 3.09E-03 | -0.3 | 2.35E-02 | -0.1 | 3.54E-01 | 0.2 | 1.57E-01 | -0.2 | 2.69E-01 | 0.5 | 4.95E-02 | UDP-glycosyltransferase 86A1 |
| AT1G50580 | -0.5 | 4.18E-03 | 0.3 | 2.36E-02 | -0.8 | 7.21E-04 | 0.1 | 8.43E-01 | 0.0 | 9.77E-01 | 0.1 | 9.07E-01 | UDP-glycosyltransferase 79B5 |
| AT1G64910 | -0.3 | 2.16E-01 | 0.4 | 7.20E-02 | -0.6 | 2.21E-02 | 0.1 | 7.64E-01 | 0.7 | 4.47E-02 | -0.6 | 8.71E-02 | UDP-glycosyltransferase 79B10 |
| AT2G28080 | -0.3 | 2.29E-03 | 0.1 | 1.31E-01 | -0.4 | 9.61E-04 | -0.1 | 7.32E-01 | 0.0 | 9.08E-01 | 0.0 | 9.06E-01 | UDP-glycosyltransferase 86A2 |
| AT2G30140 | 0.4 | 7.08E-02 | -0.5 | 3.60E-02 | 0.9 | 4.22E-03 | 0.0 | 9.64E-01 | -0.5 | 7.22E-02 | 0.5 | 9.51E-02 | UDP-glycosyltransferase 87A2 |
| AT2G36780 | 1.0 | 3.34E-03 | 0.3 | 3.78E-01 | 0.7 | 5.19E-02 | -0.4 | 5.78E-01 | -1.6 | 6.22E-02 | 1.2 | 1.63E-01 | UDP-glycosyltransferase 73C3 |
| AT2G22590 | | | | | | | -0.3 | 8.86E-01 | 2.9 | 1.65E-02 | | | UDP-glycosyltransferase 91A1 |
| AT2G36770 | | | | | | | 0.0 | 9.98E-01 | -1.4 | 3.00E-02 | 1.4 | 5.48E-02 | UDP-glycosyltransferase 73C4 |
| AT3G53160 | | | | | | | 0.939668234 | 2.37E-01 | 0.425877526 | 7.02E-01 | 0.5 | 6.29E-01 | UDP-glycosyltransferase 73C7 |
| AT5G53990 | | | | | | | 1.6 | 4.07E-03 | 0.9 | 1.16E-01 | 0.6 | 1.21E-01 | UDP-glycosyltransferase 79B9 |
| AT4G27560 | -1.3 | 2.38E-05 | -0.8 | 8.19E-04 | -0.5 | 2.84E-02 | 0.1 | 6.33E-01 | 0.8 | 1.61E-03 | -0.7 | 1.28E-02 | UDP-glycosyltransferase 79B2 |
| AT5G03490 | -0.7 | 8.58E-06 | -0.7 | 1.24E-05 | 0.0 | 6.81E-01 | -0.3 | 1.80E-01 | -0.5 | 4.49E-02 | 0.2 | 4.57E-01 | UDP-glycosyltransferase 89A2 |
| AT3G23820 | -0.8 | 7.76E-06 | -0.5 | 4.09E-04 | -0.3 | 1.28E-02 | -0.5 | 2.13E-04 | 0.0 | 9.03E-01 | -0.5 | 4.12E-03 | UDP-glucuronate 4-epimerase 6 |

| | | | | | | | | | | | | | |
|---|---|---|---|---|---|---|---|---|---|---|---|---|---|
| AT4G12250 | -0.7 | 8.94E-06 | -0.6 | 7.29E-05 | -0.1 | 1.83E-01 | -0.1 | 7.10E-01 | -0.3 | 6.78E-02 | 0.2 | 2.54E-01 | UDP-glucuronate 4-epimerase 5 |
| AT1G02000 | -0.5 | 1.92E-04 | -0.4 | 9.37E-04 | -0.1 | 4.59E-01 | -0.6 | 1.79E-03 | -0.2 | 2.14E-01 | -0.3 | 1.09E-01 | UDP-glucuronate 4-epimerase 2 |
| AT4G30440 | -0.4 | 3.66E-03 | -0.4 | 4.39E-03 | 0.0 | 9.42E-01 | -0.7 | 1.43E-05 | -0.2 | 6.96E-02 | -0.4 | 1.34E-02 | UDP-glucuronate 4-epimerase 1 |
| AT3G29630 | -1.9 | 2.18E-06 | -0.7 | 8.36E-04 | -1.2 | 5.06E-04 | 0.0 | 8.88E-01 | 0.4 | 1.02E-01 | -0.4 | 1.50E-01 | UDP-Glycosyltransferase superfamily protein |
| AT4G01070 | -1.3 | 3.42E-07 | -1.0 | 7.31E-06 | -0.4 | 5.59E-03 | -0.5 | 2.27E-03 | 0.4 | 1.84E-02 | -0.8 | 6.13E-04 | UDP-Glycosyltransferase superfamily protein |
| AT2G15480 | -1.2 | 3.72E-06 | -0.8 | 8.74E-05 | -0.4 | 2.24E-02 | 0.6 | 2.41E-03 | 0.2 | 2.33E-01 | 0.3 | 1.22E-01 | Glycosyltransferase |
| AT2G16890 | -1.0 | 9.58E-04 | -1.2 | 5.75E-04 | 0.2 | 5.43E-01 | 0.3 | 1.99E-01 | 0.8 | 2.16E-03 | -0.5 | 4.61E-02 | Glycosyltransferase |
| AT1G01420 | -0.7 | 2.45E-04 | -0.8 | 1.92E-04 | 0.1 | 5.88E-01 | 0.4 | 1.39E-02 | 0.3 | 6.49E-02 | 0.1 | 7.49E-01 | Glycosyltransferase |
| AT4G34135 | -0.5 | 2.84E-02 | -1.0 | 1.07E-03 | 0.5 | 6.84E-02 | 0.0 | 8.57E-01 | -0.2 | 6.42E-02 | 0.2 | 1.73E-01 | UDP-glucosyltransferase 73B2 |
| AT2G18570 | -0.2 | 1.62E-02 | 0.0 | 8.04E-01 | -0.2 | 5.10E-02 | 0.1 | 5.78E-01 | 0.0 | 8.63E-01 | 0.1 | 8.40E-01 | Putative flavonol 3-O-glucosyltransferase |
| AT5G24750 | 0.2 | 1.18E-01 | 0.1 | 6.45E-01 | 0.1 | 3.75E-01 | -0.3 | 2.41E-01 | -0.2 | 4.95E-01 | -0.1 | 7.96E-01 | Uncharacterized protein |
| AT3G21790 | 0.2 | 2.16E-02 | 0.1 | 1.70E-01 | 0.1 | 3.54E-01 | -0.2 | 2.23E-01 | -0.4 | 1.17E-02 | 0.2 | 2.32E-01 | Glycosyltransferase |
| AT3G53150 | 0.2 | 4.00E-01 | -0.5 | 1.25E-01 | 0.7 | 6.16E-02 | 1.8 | 2.23E-06 | 1.6 | 5.41E-05 | 0.2 | 3.22E-01 | UDP-glycosyltransferase 73D1 |
| AT1G22360 | 0.4 | 8.57E-03 | 0.6 | 1.62E-03 | -0.2 | 2.80E-01 | 0.0 | 9.74E-01 | -0.4 | 2.41E-02 | 0.4 | 3.62E-02 | Glycosyltransferase |
| AT1G01390 | 0.6 | 7.48E-03 | 0.4 | 3.33E-02 | 0.1 | 5.55E-01 | 1.1 | 5.35E-03 | 0.8 | 5.66E-02 | 0.3 | 4.92E-01 | Glycosyltransferase |
| AT2G18560 | 0.7 | 1.46E-05 | 0.7 | 3.21E-05 | 0.0 | 9.32E-01 | -0.1 | 5.02E-01 | -0.5 | 3.29E-02 | 0.3 | 1.49E-01 | Glycosyltransferase |
| AT2G30150 | 1.1 | 2.92E-07 | 0.6 | 4.33E-05 | 0.5 | 4.30E-04 | -0.3 | 1.35E-01 | 0.0 | 9.07E-01 | -0.2 | 3.53E-01 | UDP-Glycosyltransferase superfamily protein |
| AT5G17040 | | | | | | | -0.4 | 5.80E-01 | 0.6 | 3.43E-01 | | | UDP-Glycosyltransferase superfamily protein |
| AT2G35020 | -0.1 | 6.82E-02 | -0.1 | 3.04E-01 | -0.1 | 4.77E-01 | -0.3 | 7.80E-02 | -0.2 | 3.24E-01 | -0.1 | 6.92E-01 | UDP-N-acetylglucosamine diphosphorylase 2 |
| AT3G56040 | 0.9 | 8.48E-05 | 0.7 | 8.24E-04 | 0.2 | 2.06E-01 | -0.4 | 5.61E-02 | 0.0 | 9.68E-01 | -0.4 | 1.02E-01 | UTP--glucose-1-phosphate uridylyltransferase 3, chloroplastic |
| AT4G01750 | -0.6 | 1.93E-03 | -0.1 | 4.53E-01 | -0.5 | 1.64E-02 | 0.1 | 6.74E-01 | 0.0 | 9.19E-01 | 0.1 | 8.52E-01 | UDP-D-xylose:L-fucose alpha-1,3-D-xylosyltransferase |
| AT5G33290 | -1.4 | 2.02E-07 | -1.1 | 2.33E-06 | -0.3 | 1.14E-02 | 0.1 | 6.23E-01 | 0.1 | 7.31E-01 | 0.0 | 9.77E-01 | Xylogalacturonan beta-1,3-xylosyltransferase |
| AT1G27530 | 0.1 | 2.12E-01 | 0.2 | 7.56E-02 | -0.1 | 5.58E-01 | 0.0 | 8.97E-01 | 0.0 | 8.78E-01 | 0.0 | 9.77E-01 | Ubiquitin-fold modifier-conjugating enzyme 1 |
| AT1G59725 | -1.4 | 1.44E-05 | -0.2 | 2.53E-01 | -1.3 | 3.09E-04 | -1.8 | 1.71E-03 | 0.1 | 9.19E-01 | | | J domain-containing protein |
| AT2G20560 | -1.3 | 9.46E-07 | -1.2 | 3.94E-06 | -0.1 | 3.92E-01 | -0.3 | 5.85E-01 | 0.3 | 5.06E-01 | -0.6 | 2.67E-01 | J domain-containing protein |
| AT1G44160 | -1.1 | 1.67E-06 | -0.9 | 1.41E-05 | -0.2 | 9.46E-02 | -1.3 | 4.77E-04 | -0.1 | 7.91E-01 | -1.2 | 1.41E-03 | At1g44160 |
| AT3G47940 | -0.6 | 3.04E-05 | -0.3 | 4.29E-03 | -0.3 | 1.09E-02 | -0.6 | 1.19E-02 | -0.4 | 8.39E-02 | -0.2 | 5.99E-01 | hypothetical protein |
| AT3G08910 | -0.2 | 1.57E-02 | 0.1 | 2.13E-01 | -0.3 | 5.84E-03 | -0.3 | 8.84E-02 | 0.0 | 9.38E-01 | -0.4 | 1.51E-01 | J domain-containing protein |
| AT1G10350 | -0.2 | 2.87E-02 | 0.0 | 8.06E-01 | -0.2 | 3.69E-02 | -0.1 | 7.69E-01 | -0.1 | 7.79E-01 | 0.0 | 9.81E-01 | J domain-containing protein |
| AT5G18140 | -0.1 | 4.56E-01 | -0.3 | 8.59E-03 | 0.2 | 4.93E-02 | -0.5 | 6.67E-03 | -0.4 | 4.50E-02 | -0.1 | 6.18E-01 | Chaperone DnaJ-domain superfamily protein |
| AT5G53400 | 0.0 | 9.26E-01 | -0.1 | 6.14E-01 | 0.0 | 7.17E-01 | 0.3 | 5.75E-02 | 0.3 | 1.69E-01 | 0.1 | 8.36E-01 | Protein BOBBER 1 |
| AT5G13850 | 0.3 | 2.86E-02 | 0.4 | 3.45E-03 | -0.2 | 2.18E-01 | 0.1 | 3.53E-01 | 0.0 | 9.24E-01 | 0.1 | 6.28E-01 | Nascent polypeptide-associated complex subunit alpha-like protein 3 |
| AT1G72420 | 0.3 | 1.11E-02 | 0.2 | 1.90E-01 | 0.2 | 2.23E-01 | 0.3 | 2.00E-01 | -0.3 | 3.21E-01 | 0.6 | 5.75E-02 | NADH:ubiquinone oxidoreductase intermediate-associated protein 30 |
| AT1G49245 | 0.4 | 2.65E-02 | 0.4 | 2.72E-02 | 0.0 | 9.51E-01 | 0.3 | 3.70E-01 | 0.0 | 9.99E-01 | 0.3 | 3.12E-01 | Prefoldin chaperone subunit family protein |
| AT1G33040 | 0.4 | 3.89E-04 | 0.5 | 1.82E-04 | -0.1 | 3.83E-01 | 0.2 | 7.22E-02 | 0.1 | 3.45E-01 | 0.1 | 6.91E-01 | Nascent polypeptide-associated complex subunit alpha-like protein 5 |
| AT1G20770 | 0.5 | 1.90E-04 | 0.3 | 3.76E-03 | 0.2 | 1.26E-01 | 0.3 | 4.33E-02 | 0.1 | 3.55E-01 | 0.1 | 5.22E-01 | Coiled-coil protein |
| AT3G12390 | 0.5 | 1.87E-03 | 0.5 | 2.26E-03 | 0.0 | 9.23E-01 | 0.2 | 4.17E-02 | 0.0 | 7.17E-01 | 0.1 | 3.17E-01 | Nascent polypeptide-associated complex subunit alpha-like protein 1 |
| AT4G18810 | 0.6 | 1.51E-04 | -0.1 | 5.75E-01 | 0.6 | 3.43E-04 | 0.3 | 2.51E-01 | 0.3 | 2.42E-01 | 0.0 | 9.22E-01 | NAD-binding Rossmann-fold superfamily protein |
| AT1G55820 | 0.6 | 5.50E-04 | 0.2 | 1.79E-01 | 0.4 | 1.61E-02 | 0.1 | 7.55E-01 | 0.0 | 9.76E-01 | 0.1 | 8.58E-01 | GBF-interacting protein 1-like |
| AT5G23290 | 0.6 | 8.85E-04 | 0.5 | 3.56E-03 | 0.1 | 5.61E-01 | 0.5 | 7.80E-02 | 0.1 | 7.18E-01 | 0.3 | 3.78E-01 | Prefoldin subunit 5 |
| AT2G07340 | 0.7 | 6.07E-05 | 0.6 | 2.15E-04 | 0.1 | 6.03E-01 | 0.3 | 1.53E-01 | 0.0 | 9.27E-01 | 0.4 | 2.56E-01 | PFD1 |
| AT3G18630 | | | | | | | 0.0 | 9.96E-01 | 0.0 | 9.90E-01 | 0.0 | 9.96E-01 | Uracil-DNA glycosylase, mitochondrial |
| AT2G26230 | -0.6 | 5.72E-05 | -0.4 | 1.25E-03 | -0.2 | 7.31E-02 | 0.0 | 9.88E-01 | 0.1 | 4.44E-01 | -0.1 | 5.26E-01 | Uricase |
| AT2G36830 | -0.1 | 3.11E-01 | 0.7 | 1.13E-04 | -0.8 | 1.25E-04 | -0.8 | 1.03E-05 | -0.4 | 1.86E-02 | -0.4 | 2.27E-02 | Aquaporin TIP1-1 |
| AT4G17050 | 0.2 | 1.04E-02 | 0.1 | 1.37E-01 | 0.1 | 2.27E-01 | -0.1 | 5.85E-01 | 0.0 | 9.76E-01 | -0.1 | 7.04E-01 | UGLYAH |
| AT2G35810 | -0.2 | 2.99E-02 | 0.0 | 8.02E-01 | -0.2 | 3.44E-02 | -0.1 | 5.94E-01 | 0.1 | 7.72E-01 | -0.2 | 3.48E-01 | Expressed protein |
| AT2G35830 | 0.9 | 2.05E-05 | 0.4 | 7.53E-03 | 0.5 | 3.46E-03 | 0.0 | 9.73E-01 | 0.0 | 9.27E-01 | 0.0 | 9.65E-01 | AT2G35830 protein |
| AT2G35820 | 1.8 | 3.28E-07 | 0.8 | 1.94E-04 | 0.9 | 1.53E-04 | 0.1 | 6.28E-01 | -0.1 | 6.73E-01 | 0.2 | 4.42E-01 | Ureidoglycolate hydrolase |
| AT1G31070 | -0.8 | 4.58E-06 | -0.4 | 1.35E-03 | -0.4 | 1.58E-03 | 0.0 | 9.43E-01 | 0.2 | 1.66E-01 | -0.2 | 2.49E-01 | At1g31070 |
| AT2G40490 | 2.0 | 2.25E-08 | 1.6 | 2.01E-07 | 0.4 | 3.88E-03 | 0.4 | 4.04E-03 | 0.1 | 6.90E-01 | 0.4 | 5.05E-02 | Uroporphyrinogen decarboxylase 2, chloroplastic |
| AT3G14930 | 2.0 | 1.97E-08 | 1.5 | 2.38E-07 | 0.5 | 5.75E-04 | 0.1 | 4.90E-01 | -0.1 | 3.58E-01 | 0.2 | 1.98E-01 | Uroporphyrinogen decarboxylase 1, chloroplastic |
| AT2G26540 | 1.0 | 1.11E-06 | 0.9 | 6.78E-06 | 0.1 | 2.09E-01 | 0.2 | 2.45E-01 | 0.3 | 1.03E-01 | -0.1 | 6.52E-01 | Uroporphyrinogen-III synthase |
| AT3G03250 | -0.9 | 2.39E-05 | -0.5 | 2.82E-03 | -0.4 | 9.25E-03 | 0.0 | 8.38E-01 | 0.4 | 1.70E-02 | -0.4 | 3.93E-02 | UTP--glucose-1-phosphate uridylyltransferase |
| AT5G17310 | 0.7 | 1.91E-06 | 0.8 | 2.77E-06 | -0.1 | 5.17E-01 | 0.4 | 2.61E-02 | 0.3 | 1.23E-01 | 0.1 | 7.42E-01 | UTP--glucose-1-phosphate uridylyltransferase |
| AT2G46720 | -0.2 | 2.44E-01 | 0.3 | 1.04E-01 | -0.5 | 3.22E-02 | 0.1 | 7.67E-01 | 0.1 | 9.00E-01 | 0.1 | 9.35E-01 | 3-ketoacyl-CoA synthase |
| AT1G19440 | -0.1 | 2.67E-01 | 0.2 | 7.66E-02 | -0.3 | 2.02E-02 | -0.3 | 9.13E-02 | -0.2 | 2.68E-01 | -0.1 | 8.07E-01 | 3-ketoacyl-CoA synthase 4 |
| AT1G07720 | 0.2 | 2.66E-01 | -0.4 | 5.00E-02 | 0.5 | 1.26E-02 | -0.3 | 6.65E-02 | -0.1 | 3.91E-01 | -0.1 | 5.62E-01 | 3-ketoacyl-CoA synthase |
| AT2G26640 | 0.2 | 5.48E-02 | 0.5 | 7.60E-04 | -0.3 | 2.95E-02 | -0.1 | 4.24E-01 | -0.2 | 3.06E-01 | 0.0 | 8.28E-01 | 3-ketoacyl-CoA synthase 11 |
| AT1G68530 | 0.3 | 1.01E-02 | 0.2 | 2.67E-02 | 0.0 | 7.22E-01 | 0.0 | 8.66E-01 | 0.0 | 8.63E-01 | 0.0 | 9.83E-01 | 3-ketoacyl-CoA synthase |
| AT2G16280 | 0.6 | 3.75E-05 | 0.6 | 1.16E-04 | 0.0 | 6.99E-01 | -0.1 | 3.72E-01 | -0.2 | 9.31E-02 | 0.1 | 5.25E-01 | 3-ketoacyl-CoA synthase 9 |
| AT2G26250 | 0.7 | 2.65E-05 | 0.3 | 2.88E-03 | 0.3 | 1.09E-02 | -0.1 | 2.87E-01 | -0.2 | 9.03E-02 | 0.1 | 6.25E-01 | 3-ketoacyl-CoA synthase 10 |
| AT5G04530 | 0.7 | 3.33E-04 | 0.6 | 2.07E-03 | 0.1 | 3.92E-01 | -0.3 | 9.53E-02 | -0.3 | 1.14E-01 | 0.0 | 9.55E-01 | 3-ketoacyl-CoA synthase 19 |
| AT4G34250 | 0.8 | 1.48E-04 | 0.7 | 7.53E-04 | 0.1 | 4.74E-01 | -0.2 | 4.86E-01 | -0.2 | 4.17E-01 | 0.0 | 9.06E-01 | 3-ketoacyl-CoA synthase 16 |
| AT2G15090 | 0.9 | 7.26E-05 | 0.8 | 2.47E-04 | 0.1 | 5.98E-01 | -0.3 | 4.71E-02 | -0.6 | 4.88E-03 | 0.2 | 2.86E-01 | 3-ketoacyl-CoA synthase 8 |
| AT4G34510 | 1.5 | 7.19E-06 | 0.7 | 5.55E-03 | 0.9 | 1.91E-03 | 0.6 | 6.09E-02 | 0.5 | 1.42E-01 | 0.1 | 8.63E-01 | 3-ketoacyl-CoA synthase 17 |
| AT2G28630 | 2.2 | 8.24E-07 | 1.6 | 1.77E-05 | 0.6 | 9.97E-03 | 0.0 | 9.29E-01 | -1.3 | 4.00E-03 | 1.3 | 9.21E-03 | 3-ketoacyl-CoA synthase 12 |
| AT1G08550 | 0.9 | 1.22E-05 | 1.1 | 1.12E-05 | -0.1 | 3.54E-01 | -0.2 | 8.49E-02 | -0.2 | 7.56E-02 | 0.0 | 8.74E-01 | NPQ1 |
| AT2G45870 | -0.4 | 7.30E-02 | -0.2 | 2.81E-01 | -0.2 | 5.51E-01 | 0.3 | 4.00E-01 | 0.1 | 8.82E-01 | 0.3 | 5.62E-01 | Bestrophin-like protein |
| AT1G55620 | -0.2 | 6.59E-02 | -0.2 | 1.42E-02 | 0.1 | 4.19E-01 | -0.1 | 4.24E-01 | 0.0 | 9.26E-01 | -0.1 | 6.06E-01 | Chloride channel protein CLC-f |
| AT5G26240 | -0.1 | 1.61E-01 | -0.4 | 6.49E-04 | 0.3 | 9.13E-03 | -0.2 | 2.40E-01 | -0.1 | 7.25E-01 | -0.1 | 6.56E-01 | Putative CLC-d chloride channel protein |
| AT3G61320 | 0.0 | 7.46E-01 | 0.0 | 6.25E-01 | 0.0 | 8.88E-01 | 0.2 | 3.69E-01 | 0.1 | 7.17E-01 | 0.1 | 7.88E-01 | Bestrophin-like protein |
| AT4G35440 | 0.9 | 6.29E-05 | 0.7 | 4.55E-04 | 0.2 | 3.21E-01 | -0.4 | 3.30E-01 | 0.1 | 9.21E-01 | -0.5 | 2.15E-01 | Chloride channel protein |
| AT5G40890 | 1.4 | 2.09E-06 | 1.0 | 4.94E-05 | 0.4 | 1.79E-02 | -0.1 | 4.98E-01 | -0.5 | 6.15E-03 | 0.4 | 4.82E-02 | Chloride channel protein |
| AT1G62262 | -2.5 | 2.62E-07 | -1.3 | 7.89E-06 | -1.2 | 5.88E-04 | | | | | | | S-type anion channel SLAH4 |
| AT3G49920 | -0.6 | 3.21E-03 | -0.2 | 1.97E-01 | -0.4 | 7.39E-02 | -0.2 | 6.31E-01 | -0.2 | 7.05E-01 | 0.0 | 9.84E-01 | Voltage dependent anion channel 5 |
| AT5G67500 | -0.1 | 1.80E-01 | 0.1 | 3.24E-01 | -0.2 | 4.65E-02 | 0.3 | 3.62E-02 | 0.2 | 7.92E-02 | 0.0 | 9.55E-01 | Voltage dependent anion channel 2 |
| AT5G24030 | 0.8 | 8.36E-05 | 0.2 | 6.62E-02 | 0.5 | 3.95E-03 | -0.4 | 2.07E-01 | -0.1 | 8.33E-01 | -0.3 | 4.52E-01 | SLAC1 homologue 3 |
| AT1G62280 | 8.2 | 1.58E-06 | 7.3 | 9.95E-06 | 0.9 | 2.22E-02 | | | | | | | S-type anion channel SLAH1 |
| AT1G04690 | -0.4 | 7.74E-04 | -0.3 | 6.70E-03 | -0.1 | 2.97E-01 | -0.1 | 1.64E-01 | -0.1 | 4.67E-01 | 0.0 | 7.84E-01 | Probable voltage-gated potassium channel subunit beta channel subunit beta) |
| AT2G24610 | -1.4 | 1.94E-06 | -0.1 | 4.35E-01 | -1.3 | 4.04E-05 | -0.6 | 3.55E-02 | 0.2 | 4.84E-01 | -0.8 | 2.97E-02 | Cyclic nucleotide-gated channel 14 |
| AT4G32650 | -0.8 | 5.97E-06 | -0.6 | 1.38E-04 | -0.3 | 3.05E-02 | -0.5 | 2.25E-02 | -0.1 | 5.19E-01 | -0.3 | 1.85E-01 | Potassium channel |
| AT5G37500 | -0.8 | 3.86E-04 | -0.8 | 6.45E-04 | 0.0 | 9.19E-01 | 1.1 | 2.22E-03 | 0.4 | 4.02E-01 | 0.8 | 1.43E-02 | Potassium channel |
| AT2G26650 | -0.7 | 5.03E-05 | -0.6 | 2.78E-04 | -0.1 | 3.49E-01 | 0.0 | 9.17E-01 | -0.1 | 3.60E-01 | 0.1 | 5.09E-01 | Potassium channel |

| | | | | | | | | | | | | | |
|---|---|---|---|---|---|---|---|---|---|---|---|---|---|
| AT4G22200 | 0.3 | 1.33E-01 | 0.4 | 9.23E-02 | -0.1 | 8.30E-01 | -0.3 | 2.62E-01 | 0.6 | 1.36E-02 | -0.9 | 3.50E-03 | Potassium channel |
| AT4G18290 | 0.4 | 1.41E-02 | 0.3 | 2.87E-02 | 0.0 | 8.43E-01 | 0.2 | 4.14E-01 | 0.0 | 9.20E-01 | 0.2 | 5.99E-01 | Potassium channel |
| AT3G02850 | 0.7 | 5.87E-03 | -0.2 | 4.64E-01 | 0.9 | 8.26E-03 | -0.7 | 8.41E-02 | -0.3 | 5.35E-01 | -0.4 | 4.39E-01 | Potassium channel |
| AT5G60660 | -1.8 | 3.68E-07 | -0.1 | 6.82E-01 | -1.7 | 1.09E-05 | -2.1 | 1.01E-07 | -0.8 | 1.02E-03 | -1.3 | 5.53E-04 | Probable aquaporin PIP2-4 |
| AT2G25810 | -1.3 | 4.33E-06 | 0.0 | 9.40E-01 | -1.3 | 3.95E-05 | -0.2 | 3.82E-01 | 0.3 | 2.83E-01 | -0.5 | 1.00E-01 | Aquaporin TIP4-1 |
| AT4G23400 | -0.6 | 1.26E-03 | 0.1 | 5.77E-01 | -0.7 | 1.64E-03 | -0.9 | 3.97E-04 | -0.7 | 6.82E-03 | -0.2 | 5.41E-01 | Probable aquaporin PIP1-5 |
| AT3G04090 | -0.4 | 8.57E-05 | -0.3 | 9.28E-04 | -0.1 | 1.90E-01 | -0.1 | 2.59E-01 | 0.0 | 9.88E-01 | -0.1 | 4.32E-01 | Aquaporin SIP1-1 |
| AT5G18290 | -0.3 | 3.49E-02 | 0.1 | 2.96E-01 | -0.4 | 1.59E-02 | 0.4 | 4.15E-02 | -0.1 | 8.51E-01 | 0.5 | 7.05E-02 | Probable aquaporin SIP1-2 |
| AT3G61430 | -0.1 | 1.10E-01 | 0.3 | 3.81E-03 | -0.4 | 8.64E-04 | -1.0 | 1.52E-06 | -0.4 | 9.99E-03 | -0.6 | 6.81E-03 | Aquaporin PIP1-1 |
| AT4G17340 | -0.1 | 3.78E-01 | 0.5 | 8.15E-04 | -0.6 | 6.01E-04 | -1.2 | 5.43E-07 | -0.3 | 1.07E-01 | -0.9 | 2.77E-04 | Probable aquaporin TIP2-2 |
| AT2G16850 | 0.6 | 4.67E-04 | 0.8 | 1.21E-04 | -0.2 | 2.08E-01 | -0.1 | 4.25E-01 | 0.0 | 9.61E-01 | -0.1 | 6.50E-01 | Probable aquaporin PIP2-8 |
| AT2G37180 | 0.7 | 2.13E-03 | 0.9 | 1.07E-03 | -0.1 | 5.61E-01 | -0.8 | 1.93E-02 | -0.1 | 9.12E-01 | -0.7 | 8.47E-02 | Aquaporin PIP2-3 |
| AT2G32150 | -0.6 | 1.08E-04 | -0.3 | 1.62E-02 | -0.3 | 1.36E-02 | -1.0 | 3.25E-06 | -0.6 | 5.58E-04 | -0.3 | 6.42E-02 | At2g32150 |
| AT5G64570 | 1.4 | 1.45E-04 | 0.8 | 8.83E-03 | 0.6 | 4.11E-02 | 0.0 | 9.96E-01 | -1.0 | 1.31E-01 | 1.0 | 1.45E-01 | Beta-D-xylosidase 4 |
| AT5G49360 | 1.5 | 4.77E-04 | 1.3 | 2.03E-03 | 0.2 | 5.09E-01 | -0.6 | 8.90E-02 | -1.7 | 3.10E-04 | 1.2 | 2.97E-03 | Xylosidase |
| AT2G01275 | -1.3 | 1.20E-04 | -0.3 | 7.26E-02 | -0.9 | 5.16E-03 | | | | | | | RING-CH-type domain-containing protein |
| AT1G35330 | -1.2 | 1.87E-05 | -0.4 | 2.46E-02 | -0.9 | 1.49E-03 | -0.3 | 4.13E-01 | 0.3 | 4.80E-01 | -0.6 | 9.02E-02 | RING |
| AT2G47890 | -1.2 | 4.66E-07 | -0.8 | 1.00E-05 | -0.3 | 6.37E-03 | 0.0 | 9.91E-01 | -0.1 | 7.76E-01 | 0.1 | 7.84E-01 | B-box type zinc finger protein with CCT domain-containing protein |
| AT4G32670 | -1.0 | 4.41E-06 | -0.7 | 1.02E-04 | -0.3 | 2.84E-02 | 0.4 | 2.01E-01 | 0.2 | 5.59E-01 | 0.2 | 6.82E-01 | RING-CH-type domain-containing protein |
| AT3G11950 | -1.0 | 2.16E-06 | -0.2 | 1.65E-02 | -0.7 | 1.58E-04 | 0.1 | 7.99E-01 | 0.3 | 2.58E-01 | -0.2 | 3.62E-01 | F26K24.24 protein |
| AT5G60580 | -1.0 | 4.01E-07 | -0.8 | 2.59E-06 | -0.1 | 1.18E-01 | -0.4 | 1.22E-02 | -0.2 | 1.44E-01 | -0.1 | 5.10E-01 | hypothetical protein |
| AT3G26760 | -0.8 | 5.16E-06 | -0.1 | 2.25E-01 | -0.7 | 1.29E-04 | -0.2 | 2.88E-01 | -0.3 | 1.89E-01 | 0.1 | 8.31E-01 | Alcohol dehydrogenase-like protein -binding Rossmann-fold superfamily protein) |
| AT5G59000 | -0.8 | 7.44E-05 | -0.4 | 4.07E-03 | -0.3 | 3.43E-02 | -0.3 | 1.18E-01 | -0.2 | 3.04E-01 | -0.1 | 8.05E-01 | At5g59000 |
| AT2G34200 | -0.6 | 2.00E-03 | -0.2 | 1.78E-01 | -0.4 | 4.86E-02 | 0.0 | 9.06E-01 | -0.2 | 4.55E-01 | 0.3 | 3.42E-01 | Uncharacterized protein At2g34200 |
| AT5G08750 | -0.6 | 1.46E-05 | -0.7 | 1.09E-05 | 0.1 | 2.67E-01 | -0.3 | 2.51E-01 | 0.0 | 9.91E-01 | -0.3 | 3.58E-01 | RING |
| AT2G26000 | -0.5 | 3.60E-04 | -0.5 | 5.69E-04 | 0.0 | 9.88E-01 | 0.0 | 9.73E-01 | -0.2 | 4.59E-01 | 0.2 | 4.75E-01 | Zinc finger family protein |
| AT3G02260 | -0.4 | 2.95E-04 | -0.5 | 1.98E-04 | 0.1 | 4.99E-01 | 0.0 | 8.78E-01 | -0.1 | 6.73E-01 | 0.1 | 6.54E-01 | Auxin transport protein |
| AT1G76320 | -0.4 | 2.82E-04 | -0.3 | 9.75E-04 | 0.0 | 6.09E-01 | 0.1 | 7.63E-01 | 0.2 | 5.68E-01 | -0.1 | 8.37E-01 | Protein FAR1-RELATED SEQUENCE |
| AT2G42160 | -0.4 | 9.66E-04 | -0.4 | 1.08E-03 | 0.0 | 8.88E-01 | -0.4 | 4.27E-02 | -0.3 | 1.28E-01 | -0.1 | 8.19E-01 | Zinc finger domain-containing protein |
| AT4G14160 | -0.4 | 3.05E-04 | -0.3 | 2.50E-03 | -0.1 | 2.86E-01 | -0.2 | 1.53E-01 | -0.1 | 4.83E-01 | -0.1 | 7.22E-01 | Protein transport protein SEC23 |
| AT1G05520 | -0.4 | 3.58E-04 | -0.2 | 1.16E-02 | -0.2 | 7.72E-02 | 0.2 | 4.68E-01 | 0.2 | 4.79E-01 | 0.0 | 9.62E-01 | Protein transport protein SEC23 |
| AT1G80010 | -0.4 | 9.56E-04 | -0.5 | 2.91E-04 | 0.1 | 2.95E-01 | -0.2 | 1.28E-01 | -0.1 | 5.93E-01 | -0.1 | 5.61E-01 | Protein FAR1-RELATED SEQUENCE |
| AT1G17210 | -0.3 | 2.17E-03 | -0.4 | 1.59E-03 | 0.0 | 7.15E-01 | -0.3 | 2.85E-02 | -0.3 | 7.16E-02 | 0.0 | 9.22E-01 | F20D23.9 protein |
| AT1G52520 | -0.3 | 6.68E-03 | -0.3 | 1.04E-02 | 0.0 | 9.28E-01 | -0.3 | 3.65E-01 | -0.1 | 7.79E-01 | -0.2 | 6.68E-01 | Protein FAR1-RELATED SEQUENCE 6 |
| AT5G13190 | -0.3 | 2.99E-02 | -0.2 | 1.12E-01 | -0.1 | 5.58E-01 | -0.1 | 7.65E-01 | -0.3 | 1.52E-01 | 0.3 | 2.94E-01 | GSH-induced LITAF domain protein |
| AT2G22120 | -0.3 | 1.15E-03 | -0.1 | 1.14E-01 | -0.2 | 4.08E-02 | -0.2 | 1.31E-01 | -0.1 | 7.10E-01 | -0.1 | 4.77E-01 | RING |
| AT2G27110 | -0.3 | 3.52E-03 | -0.3 | 3.93E-03 | 0.0 | 9.36E-01 | -0.2 | 5.30E-01 | -0.1 | 6.98E-01 | 0.0 | 9.12E-01 | Protein FAR1-RELATED SEQUENCE |
| AT1G06740 | -0.3 | 5.97E-02 | -0.5 | 5.93E-03 | 0.2 | 2.31E-01 | | | | | | | hypothetical protein |
| AT3G62900 | -0.3 | 7.40E-03 | -0.4 | 1.27E-03 | 0.1 | 2.58E-01 | 0.2 | 2.97E-01 | 0.0 | 9.02E-01 | 0.2 | 5.45E-01 | CW-type Zinc Finger |
| AT2G33500 | -0.2 | 3.09E-02 | -0.2 | 7.24E-02 | 0.0 | 7.55E-01 | -0.4 | 3.11E-02 | -0.3 | 1.44E-01 | -0.1 | 7.29E-01 | Zinc finger protein CONSTANS-LIKE 14 |
| AT2G21630 | -0.2 | 1.96E-02 | -0.1 | 1.15E-01 | -0.1 | 4.46E-01 | 0.2 | 3.56E-01 | -0.1 | 7.38E-01 | 0.3 | 2.91E-01 | Protein transport protein SEC23 |
| AT4G15730 | -0.2 | 1.17E-01 | -0.5 | 4.61E-03 | 0.3 | 1.09E-01 | 0.5 | 1.70E-01 | 0.5 | 2.83E-01 | 0.1 | 9.22E-01 | At4g15730 |
| AT3G09760 | -0.2 | 8.62E-03 | -0.2 | 8.07E-03 | 0.0 | 8.95E-01 | 0.0 | 8.50E-01 | 0.0 | 9.15E-01 | 0.0 | 9.78E-01 | At3g09760 |
| AT5G45290 | -0.2 | 1.66E-02 | -0.2 | 9.51E-03 | 0.0 | 7.21E-01 | -0.2 | 3.43E-01 | -0.1 | 7.47E-01 | -0.1 | 7.42E-01 | RING |
| AT3G44340 | -0.2 | 2.37E-02 | -0.1 | 4.63E-02 | 0.0 | 8.08E-01 | 0.1 | 6.36E-01 | 0.0 | 9.75E-01 | 0.1 | 7.35E-01 | Protein transport protein Sec24-like CEF |
| AT3G05850 | -0.2 | 1.05E-01 | -0.2 | 2.13E-02 | 0.1 | 4.07E-01 | -0.4 | 7.13E-02 | -0.1 | 5.95E-01 | -0.2 | 3.69E-01 | hypothetical protein |
| AT3G04460 | -0.1 | 2.74E-01 | -0.1 | 4.80E-01 | 0.0 | 7.81E-01 | -0.1 | 8.42E-01 | -0.2 | 5.02E-01 | 0.1 | 6.58E-01 | Peroxin-12 |
| AT5G03180 | -0.1 | 5.51E-01 | -0.2 | 2.82E-01 | 0.1 | 6.72E-01 | -0.2 | 5.70E-01 | 0.0 | 9.68E-01 | -0.2 | 6.50E-01 | Uncharacterized protein F15A17_210 |
| AT3G47610 | -0.1 | 2.71E-01 | -0.1 | 2.55E-01 | 0.0 | 9.49E-01 | -0.4 | 5.67E-03 | -0.3 | 3.12E-02 | -0.1 | 7.36E-01 | hypothetical protein |
| AT1G11020 | -0.1 | 3.32E-01 | -0.1 | 1.83E-01 | 0.0 | 7.32E-01 | -0.3 | 8.78E-02 | -0.2 | 3.07E-01 | -0.1 | 6.82E-01 | hypothetical protein |
| AT5G25560 | 0.0 | 7.44E-01 | -0.3 | 6.47E-03 | 0.3 | 1.73E-02 | -0.2 | 4.13E-01 | 0.0 | 9.56E-01 | -0.1 | 6.21E-01 | CHY-type |
| AT5G18960 | 0.0 | 8.24E-01 | -0.1 | 3.27E-01 | 0.1 | 5.07E-01 | -0.3 | 3.08E-01 | -0.2 | 4.83E-01 | 0.0 | 9.16E-01 | Protein FAR1-RELATED SEQUENCE |
| AT1G50440 | 0.0 | 8.68E-01 | 0.0 | 7.88E-01 | 0.0 | 9.29E-01 | 0.0 | 8.18E-01 | 0.0 | 9.33E-01 | 0.0 | 9.45E-01 | RING |
| AT3G23660 | 0.0 | 8.95E-01 | 0.0 | 9.36E-01 | 0.0 | 9.71E-01 | 0.2 | 3.18E-01 | 0.3 | 2.86E-01 | 0.0 | 9.14E-01 | Protein transport protein SEC23 |
| AT5G37340 | 0.0 | 9.56E-01 | -0.1 | 5.18E-01 | 0.1 | 6.26E-01 | 0.3 | 2.86E-01 | 0.2 | 6.49E-01 | 0.1 | 7.40E-01 | ZPR1 zinc-finger domain protein |
| AT5G18760 | 0.0 | 9.42E-01 | -0.1 | 1.71E-01 | 0.1 | 2.42E-01 | -0.2 | 3.51E-01 | -0.2 | 3.54E-01 | 0.0 | 9.57E-01 | At5g18760 |
| AT2G41835 | 0.0 | 6.55E-01 | -0.2 | 6.17E-02 | 0.3 | 5.40E-02 | 0.2 | 3.29E-01 | -0.1 | 7.69E-01 | 0.3 | 2.46E-01 | Zinc finger AN1 and C2H2 domain-containing stress-associated protein 11 |
| AT3G06250 | 0.1 | 3.14E-01 | 0.0 | 7.59E-01 | 0.1 | 2.61E-01 | 0.1 | 7.13E-01 | 0.0 | 9.00E-01 | 0.1 | 6.96E-01 | Protein FAR1-RELATED SEQUENCE 7 |
| AT2G19260 | 0.1 | 4.09E-01 | -0.1 | 5.73E-01 | 0.1 | 2.50E-01 | -0.2 | 6.00E-01 | -0.5 | 1.73E-01 | 0.3 | 5.08E-01 | RING |
| AT5G20570 | 0.1 | 2.37E-01 | 0.1 | 8.67E-02 | -0.1 | 5.68E-01 | 0.0 | 8.42E-01 | -0.1 | 3.53E-01 | 0.1 | 5.80E-01 | RING-box 1 |
| AT5G62460 | 0.1 | 1.85E-01 | 0.1 | 4.96E-01 | 0.0 | 5.91E-01 | -0.2 | 2.62E-01 | -0.1 | 6.30E-01 | -0.1 | 7.63E-01 | AT5g62460 |
| AT1G06040 | 0.1 | 1.37E-01 | 0.2 | 2.99E-02 | -0.1 | 3.95E-01 | 0.0 | 9.73E-01 | -0.1 | 7.34E-01 | 0.1 | 7.36E-01 | B-box zinc finger family protein |
| AT3G62970 | 0.1 | 1.32E-01 | 0.0 | 6.00E-01 | 0.2 | 9.23E-02 | -0.3 | 9.70E-02 | -0.2 | 3.64E-01 | -0.1 | 6.34E-01 | At3g62970 |
| AT4G32600 | 0.1 | 5.40E-02 | 0.0 | 9.85E-01 | 0.1 | 8.01E-02 | 0.1 | 3.93E-01 | 0.1 | 3.91E-01 | 0.0 | 9.42E-01 | hypothetical protein |
| AT2G14825 | 0.1 | 1.51E-01 | 0.2 | 1.29E-01 | 0.0 | 9.13E-01 | -0.2 | 5.68E-01 | -0.4 | 1.96E-01 | 0.2 | 5.79E-01 | UBP-type domain-containing protein |
| AT4G19990 | 0.1 | 7.97E-02 | -0.2 | 3.12E-02 | 0.3 | 4.59E-03 | 0.0 | 9.72E-01 | 0.1 | 8.14E-01 | -0.1 | 8.54E-01 | Protein FAR1-RELATED SEQUENCE |
| AT4G13970 | 0.1 | 5.50E-02 | -0.1 | 2.65E-01 | 0.2 | 1.79E-02 | -0.2 | 3.08E-01 | 0.0 | 9.17E-01 | -0.2 | 3.76E-01 | Zinc ion binding protein |
| AT1G74770 | 0.2 | 8.22E-02 | -0.2 | 3.87E-02 | 0.4 | 4.30E-03 | 1.1 | 5.53E-04 | -0.2 | 6.39E-01 | 1.3 | 2.27E-03 | Zinc ion binding protein |
| AT5G28530 | 0.2 | 5.00E-02 | 0.2 | 5.58E-02 | 0.0 | 9.92E-01 | -0.3 | 1.81E-01 | -0.2 | 4.96E-01 | -0.1 | 7.20E-01 | Protein FAR1-RELATED SEQUENCE |
| AT2G32250 | 0.2 | 1.67E-02 | -0.2 | 4.12E-02 | 0.4 | 1.89E-03 | 0.1 | 7.87E-01 | 0.0 | 9.55E-01 | 0.0 | 9.10E-01 | Protein FAR1-RELATED SEQUENCE |
| AT4G38170 | 0.2 | 4.41E-02 | 0.0 | 6.78E-01 | 0.2 | 4.77E-02 | 0.3 | 2.88E-01 | -0.1 | 8.90E-01 | 0.4 | 2.99E-01 | Protein FAR1-RELATED SEQUENCE |
| AT1G80400 | 0.2 | 1.04E-02 | 0.0 | 5.70E-01 | 0.2 | 6.09E-02 | -0.3 | 4.52E-02 | -0.2 | 4.38E-01 | -0.2 | 4.39E-01 | hypothetical protein |
| AT5G05830 | 0.2 | 4.73E-02 | 0.2 | 1.43E-01 | 0.1 | 6.50E-01 | -0.2 | 3.68E-01 | -0.2 | 3.55E-01 | 0.0 | 9.36E-01 | RING-CH-type domain-containing protein |
| AT3G28210 | 0.3 | 1.42E-01 | -0.9 | 1.74E-03 | 1.2 | 1.11E-03 | 1.6 | 1.06E-03 | 1.8 | 7.19E-04 | -0.2 | 5.00E-01 | Zinc finger AN1 domain-containing stress-associated protein 12 |
| AT1G05055 | 0.3 | 2.23E-02 | 0.2 | 4.58E-02 | 0.0 | 8.08E-01 | 0.1 | 8.13E-01 | 0.1 | 6.54E-01 | -0.1 | 8.65E-01 | Putative transcription factor |
| AT1G10240 | 0.3 | 6.50E-03 | 0.1 | 3.06E-01 | 0.2 | 9.11E-02 | -0.1 | 6.47E-01 | -0.3 | 4.41E-01 | 0.1 | 7.86E-01 | Protein FAR1-RELATED SEQUENCE 11 |
| AT4G27585 | 0.3 | 1.48E-02 | 0.1 | 6.70E-01 | 0.3 | 6.19E-02 | 1.3 | 8.45E-07 | 0.3 | 1.72E-01 | 1.0 | 2.49E-04 | SPFH |
| AT4G32930 | 0.3 | 1.92E-03 | 0.3 | 4.37E-03 | 0.0 | 7.83E-01 | 0.0 | 9.39E-01 | -0.1 | 5.67E-01 | 0.1 | 6.87E-01 | Uncharacterized protein |
| AT2G24790 | 0.3 | 1.39E-03 | 0.1 | 1.29E-01 | 0.2 | 3.79E-02 | -0.4 | 3.16E-02 | -0.5 | 7.83E-03 | 0.2 | 5.57E-01 | CONSTANS-like 3 |
| AT5G22480 | 0.4 | 5.35E-05 | 0.0 | 7.53E-01 | 0.4 | 4.54E-04 | -0.2 | 3.73E-01 | -0.2 | 4.31E-01 | 0.0 | 9.97E-01 | Zinc finger ZPR1-type domain-containing protein |
| AT5G63670 | 0.5 | 1.33E-04 | 0.4 | 5.03E-04 | 0.1 | 5.66E-01 | 0.1 | 5.64E-01 | 0.0 | 9.02E-01 | 0.1 | 8.05E-01 | SPT4 homolog 2 |
| AT2G37950 | 0.6 | 2.62E-03 | 0.9 | 2.01E-04 | -0.3 | 8.78E-02 | -0.4 | 1.21E-01 | -0.9 | 2.53E-03 | 0.5 | 8.87E-02 | RING-CH-type domain-containing protein |
| AT5G24930 | 0.7 | 2.63E-05 | 0.5 | 5.59E-04 | 0.2 | 6.83E-02 | -0.3 | 7.16E-03 | -0.2 | 1.77E-01 | -0.1 | 4.03E-01 | Zinc finger protein CONSTANS-LIKE 4 |

| | | | | | | | | | | | | | |
|---|---|---|---|---|---|---|---|---|---|---|---|---|---|
| AT1G47580 | 1.1 | 4.63E-05 | 0.7 | 2.78E-03 | 0.4 | 3.15E-02 | 0.3 | 1.20E-01 | 0.1 | 7.04E-01 | 0.2 | 3.89E-01 | Pentatricopeptide repeat-containing protein DWY1, chloroplastic |
| AT1G68730 | 1.1 | 3.05E-06 | 0.8 | 9.73E-05 | 0.3 | 1.91E-02 | 0.6 | 1.20E-02 | 0.3 | 2.59E-01 | 0.3 | 2.17E-01 | DNL-type domain-containing protein |
| AT1G68520 | 1.3 | 1.15E-04 | 1.1 | 4.14E-04 | 0.1 | 5.96E-01 | -0.1 | 3.97E-01 | -0.4 | 2.12E-02 | 0.2 | 1.59E-01 | Uncharacterized protein |
| AT5G63780 | 1.7 | 1.48E-07 | 1.3 | 2.16E-06 | 0.4 | 5.49E-03 | 0.3 | 1.58E-01 | 0.2 | 5.00E-01 | 0.1 | 7.14E-01 | At5g63780 |
| AT1G68190 | 2.1 | 5.94E-07 | 2.0 | 2.62E-06 | 0.2 | 3.52E-01 | -0.5 | 7.16E-02 | -0.4 | 1.69E-01 | -0.1 | 8.53E-01 | B box-type domain-containing protein |
| AT1G49920 | 2.7 | 6.86E-07 | 1.9 | 2.64E-05 | 0.8 | 2.81E-03 | 0.7 | 1.28E-01 | 0.6 | 3.03E-01 | 0.2 | 8.25E-01 | SWIM-type domain-containing protein |
| AT5G08565 | | | | | | | 0.5 | 2.08E-01 | 0.9 | 1.89E-02 | -0.4 | 2.40E-01 | Transcription elongation factor SPT4 homolog |
| AT4G33020 | -0.6 | 1.24E-03 | 0.1 | 4.30E-01 | -0.7 | 2.11E-03 | -0.4 | 3.66E-01 | -0.1 | 8.47E-01 | -0.3 | 6.13E-01 | ZIP metal ion transporter family |
| AT2G30080 | -0.4 | 2.82E-04 | -0.1 | 2.04E-01 | -0.3 | 6.65E-03 | -0.2 | 3.07E-01 | -0.2 | 4.13E-01 | 0.0 | 9.57E-01 | Zinc transporter 6, chloroplastic |
| AT3G12750 | -0.4 | 1.08E-03 | -0.3 | 7.49E-03 | -0.1 | 3.82E-01 | 0.0 | 9.92E-01 | 0.2 | 3.32E-01 | -0.2 | 3.38E-01 | Zinc transporter 1 |
| AT3G12100 | -0.3 | 1.19E-03 | -0.2 | 5.78E-03 | -0.1 | 5.01E-01 | -0.2 | 4.15E-01 | -0.1 | 6.04E-01 | 0.0 | 9.08E-01 | Cation efflux family protein |
| AT2G04620 | -0.3 | 3.33E-02 | -0.4 | 6.45E-03 | 0.1 | 3.71E-01 | -0.2 | 6.77E-01 | 0.0 | 9.62E-01 | -0.3 | 6.80E-01 | Uncharacterized protein |
| AT2G32270 | -0.2 | 1.49E-02 | 0.4 | 7.94E-04 | -0.6 | 1.69E-04 | 1.8 | 8.37E-07 | 1.0 | 2.17E-03 | 0.9 | 2.97E-03 | Zinc transporter 3 |
| AT1G60960 | -0.1 | 1.80E-01 | 0.2 | 1.05E-02 | -0.3 | 3.38E-03 | 0.6 | 4.35E-03 | 0.8 | 2.26E-03 | -0.1 | 6.67E-01 | Iron regulated transporter 3 |
| AT1G10970 | -0.1 | 4.76E-01 | 0.2 | 9.65E-02 | -0.2 | 5.73E-02 | 0.3 | 3.17E-01 | 0.2 | 5.22E-01 | 0.1 | 8.74E-01 | hypothetical protein |
| AT1G05300 | 0.4 | 9.44E-03 | 0.2 | 1.86E-01 | 0.2 | 1.81E-01 | 0.4 | 9.41E-02 | 0.1 | 8.62E-01 | 0.3 | 1.74E-01 | Zinc transporter 5 |
| AT1G55910 | 0.9 | 4.79E-06 | 0.2 | 8.99E-02 | 0.7 | 1.70E-04 | 1.0 | 2.41E-06 | 0.5 | 2.39E-03 | 0.4 | 1.61E-02 | Zinc transporter 11 |
| AT4G19680 | | | | | | | 1.221755263 | 8.20E-03 | -0.681852423 | 2.86E-01 | 1.9 | 3.25E-03 | Fe transport protein 2 transport protein 2) |
| AT2G25625 | -5.0 | 9.76E-06 | -6.6 | 3.26E-05 | 1.6 | 1.64E-01 | | | | | | | Uncharacterized protein |
| AT1G80130 | -4.2 | 3.65E-07 | -3.3 | 2.86E-06 | -1.0 | 1.65E-02 | 0.3 | 6.52E-01 | 1.0 | 1.36E-01 | -0.6 | 3.31E-01 | hypothetical protein |
| AT5G03210 | -4.0 | 6.08E-03 | -3.8 | 7.38E-03 | -0.2 | 9.24E-01 | -0.9 | 5.69E-01 | 0.7 | 6.62E-01 | | | Arabidopsis thaliana genomic DNA, chromosome 5, P1 clone:MOK16 |
| AT2G41810 | -3.9 | 1.83E-06 | 0.9 | 1.47E-04 | -4.8 | 1.09E-05 | -3.2 | 1.94E-06 | 0.1 | 8.19E-01 | -3.3 | 1.09E-04 | hypothetical protein |
| AT5G16740 | -3.9 | 4.41E-06 | -5.2 | 9.83E-06 | 1.3 | 1.03E-01 | | | | | | | hypothetical protein |
| AT5G11930 | -3.8 | 2.98E-07 | -2.7 | 1.42E-06 | -1.1 | 1.14E-02 | -0.4 | 3.88E-01 | 0.7 | 7.58E-02 | -1.1 | 1.61E-02 | Glutaredoxin-C10 |
| AT5G15500 | -3.8 | 1.09E-05 | -3.3 | 2.66E-05 | -0.5 | 4.38E-01 | -1.0 | 1.52E-01 | 0.2 | 7.81E-01 | -1.2 | 1.81E-01 | Ankyrin repeat family protein |
| AT5G25390 | -3.7 | 3.72E-05 | -2.6 | 1.51E-04 | -1.1 | 1.28E-01 | | | | | | | SHN3 |
| AT1G29395 | -3.7 | 4.05E-07 | -1.3 | 2.44E-04 | -2.3 | 6.88E-05 | -1.7 | 4.97E-07 | -0.2 | 4.24E-01 | -1.5 | 1.44E-04 | Uncharacterized protein |
| AT4G00670 | -3.4 | 6.53E-05 | -2.3 | 3.50E-04 | -1.1 | 1.47E-01 | -0.7 | 1.91E-01 | -0.2 | 7.99E-01 | | | hypothetical protein |
| AT5G26070 | -3.1 | 6.28E-06 | -0.6 | 1.00E-02 | -2.5 | 2.70E-04 | -4.9 | 4.12E-04 | -0.9 | 1.30E-01 | -4.0 | 1.51E-02 | Transmembrane protein |
| AT4G33905 | -3.0 | 1.36E-04 | -2.8 | 2.82E-04 | -0.2 | 7.98E-01 | -0.2 | 8.85E-01 | 0.2 | 8.94E-01 | | | hypothetical protein |
| AT5G24860 | -3.0 | 3.72E-07 | -2.3 | 1.94E-06 | -0.7 | 4.49E-02 | | | | | | | Flowering promoting factor 1 |
| AT1G55760 | -3.0 | 1.44E-08 | -2.4 | 8.06E-08 | -0.6 | 1.49E-03 | -0.6 | 1.11E-01 | 0.4 | 2.29E-01 | -1.0 | 2.38E-02 | BTB |
| AT4G26050 | -3.0 | 5.33E-07 | -1.3 | 3.21E-05 | -1.7 | 2.70E-04 | -2.2 | 3.84E-04 | -0.1 | 8.04E-01 | -2.1 | 6.00E-03 | Plant intracellular Ras-group-related LRR protein 8 |
| AT1G64590 | -2.9 | 1.44E-08 | -2.1 | 1.07E-07 | -0.8 | 8.04E-05 | -2.2 | 4.57E-09 | -0.1 | 6.96E-01 | -2.1 | 1.12E-06 | NAD-binding Rossmann-fold superfamily protein |
| AT2G02610 | -2.9 | 4.06E-08 | -1.3 | 1.34E-06 | -1.6 | 3.66E-05 | -0.6 | 7.34E-02 | 0.1 | 8.01E-01 | -0.7 | 7.05E-02 | Cysteine |
| AT2G47880 | -2.9 | 4.80E-05 | -1.1 | 3.66E-03 | -1.8 | 5.57E-03 | -0.9 | 1.07E-02 | -0.6 | 1.08E-01 | -0.4 | 4.69E-01 | Glutaredoxin domain-containing protein |
| AT5G23750 | -2.9 | 3.81E-08 | -1.3 | 3.69E-06 | -1.5 | 2.81E-05 | -1.1 | 6.63E-04 | -0.2 | 5.22E-01 | -0.9 | 2.43E-02 | Remorin family protein |
| AT5G52310 | -2.8 | 2.54E-05 | -2.6 | 8.08E-05 | -0.2 | 6.24E-01 | -1.2 | 6.34E-02 | 0.1 | 8.78E-01 | -1.3 | 1.23E-01 | AT5G52310 protein |
| AT5G47635 | -2.8 | 2.07E-06 | -1.1 | 2.52E-04 | -1.7 | 5.63E-04 | | | | | | | hypothetical protein |
| AT5G59050 | -2.8 | 1.59E-08 | -1.7 | 2.75E-07 | -1.1 | 3.35E-05 | -0.3 | 1.25E-01 | 0.4 | 3.10E-02 | -0.7 | 6.29E-03 | G patch domain protein |
| AT4G33040 | -2.7 | 2.19E-08 | -2.0 | 1.96E-07 | -0.7 | 8.94E-04 | -0.3 | 1.32E-01 | 0.6 | 2.08E-03 | -0.9 | 6.21E-04 | Glutaredoxin-C6 |
| AT5G39520 | -2.7 | 7.67E-05 | -3.5 | 7.94E-05 | 0.8 | 2.74E-01 | -2.0 | 2.47E-01 | -2.4 | 2.30E-01 | 0.4 | 8.88E-01 | Uncharacterized protein At5g39520 |
| AT5G67210 | -2.7 | 4.90E-08 | -0.8 | 8.56E-05 | -1.9 | 1.09E-05 | -0.7 | 2.35E-03 | 0.4 | 8.95E-02 | -1.1 | 7.95E-04 | Protein IRX15-LIKE |
| AT4G33610 | -2.6 | 7.44E-08 | -1.1 | 3.15E-05 | -1.6 | 2.69E-05 | 1.1 | 4.22E-05 | 0.7 | 6.39E-03 | 0.4 | 7.21E-02 | Glycine-rich protein |
| AT5G09520 | -2.6 | 1.06E-07 | -1.1 | 1.98E-05 | -1.6 | 4.37E-05 | 1.0 | 1.05E-03 | 0.7 | 1.89E-02 | 0.3 | 3.35E-01 | Protein PELPK2 |
| AT1G78990 | -2.6 | 2.04E-08 | -1.4 | 7.29E-07 | -1.2 | 2.95E-05 | 0.8 | 5.01E-03 | 1.4 | 1.46E-04 | -0.6 | 5.14E-02 | YUP8H12R.39 protein |
| AT1G55990 | -2.5 | 9.51E-08 | -1.1 | 1.55E-05 | -1.4 | 5.00E-05 | 1.4 | 4.22E-05 | 0.9 | 3.23E-03 | 0.4 | 9.17E-02 | Glycine-rich protein |
| AT1G67360 | -2.5 | 1.18E-07 | -1.7 | 3.52E-06 | -0.8 | 7.50E-04 | -0.5 | 3.29E-02 | 0.4 | 8.37E-02 | -0.9 | 1.83E-03 | REF |
| AT4G37700 | -2.5 | 2.34E-06 | -1.2 | 1.41E-04 | -1.4 | 1.65E-03 | -2.4 | 1.10E-03 | 0.1 | 9.17E-01 | -2.5 | 7.26E-03 | Uncharacterized protein |
| AT1G10385 | -2.5 | 2.05E-06 | -0.5 | 1.42E-02 | -2.0 | 1.01E-04 | | | | | | | Exocyst complex component EXO84A |
| AT5G09480 | -2.5 | 1.62E-07 | -1.0 | 9.50E-05 | -1.4 | 4.37E-05 | 1.2 | 7.74E-06 | 1.2 | 4.38E-05 | 0.0 | 9.81E-01 | Uncharacterized protein |
| AT4G20390 | -2.5 | 1.89E-08 | -1.0 | 6.65E-06 | -1.5 | 8.88E-06 | 1.4 | 8.52E-07 | 1.3 | 6.49E-06 | 0.0 | 8.85E-01 | CASP-like protein 1B2 |
| AT3G62960 | -2.5 | 3.42E-07 | -1.7 | 3.11E-06 | -0.7 | 8.96E-03 | -0.3 | 3.53E-01 | 0.0 | 9.76E-01 | -0.3 | 4.94E-01 | Glutaredoxin-C14 |
| AT4G38080 | -2.4 | 3.80E-08 | -1.0 | 4.70E-05 | -1.5 | 1.20E-05 | 1.3 | 1.66E-06 | 1.2 | 4.26E-05 | 0.1 | 5.59E-01 | hypothetical protein |
| AT2G22510 | -2.4 | 5.56E-08 | -1.1 | 1.84E-05 | -1.4 | 2.60E-05 | 1.5 | 1.41E-05 | 1.1 | 5.58E-04 | 0.3 | 2.57E-01 | Hydroxyproline-rich glycoprotein family protein |
| AT3G04370 | -2.4 | 7.88E-07 | -0.7 | 6.43E-04 | -1.7 | 1.02E-04 | 2.2 | 7.01E-05 | 1.6 | 3.23E-03 | 0.6 | 1.17E-01 | Plasmodesmata-located protein 4 |
| AT2G22470 | -2.4 | 2.35E-07 | -0.9 | 4.14E-04 | -1.5 | 3.70E-05 | -0.7 | 6.25E-03 | 0.1 | 8.49E-01 | -0.8 | 7.96E-03 | Classical arabinogalactan protein 2 |
| AT5G24880 | -2.4 | 3.52E-08 | -0.8 | 3.14E-05 | -1.6 | 1.09E-05 | -1.9 | 4.90E-06 | -0.1 | 6.80E-01 | -1.7 | 1.84E-04 | Chromo domain cec-like protein |
| AT5G42510 | -2.4 | 3.15E-07 | -0.4 | 1.01E-02 | -1.9 | 2.39E-05 | -1.2 | 2.33E-04 | 0.0 | 8.92E-01 | -1.3 | 1.71E-03 | Dirigent protein 1 |
| AT4G01140 | -2.4 | 8.05E-07 | -0.4 | 1.17E-02 | -1.9 | 4.16E-05 | -1.3 | 1.32E-03 | -0.2 | 5.76E-01 | -1.1 | 1.17E-02 | At4g01140 ) |
| AT3G12540 | -2.3 | 5.72E-07 | -0.6 | 1.30E-03 | -1.7 | 6.06E-05 | -1.0 | 1.57E-02 | 0.4 | 2.98E-01 | -1.4 | 4.04E-03 | hypothetical protein |
| AT3G05890 | -2.3 | 2.02E-07 | -0.3 | 5.42E-02 | -2.0 | 1.09E-05 | -0.7 | 7.12E-03 | 0.1 | 8.03E-01 | -0.8 | 7.52E-03 | Hydrophobic protein RCI2B |
| AT3G22620 | -2.3 | 2.75E-08 | -1.0 | 1.39E-05 | -1.4 | 1.09E-05 | 0.7 | 3.53E-03 | 0.3 | 1.45E-01 | 0.3 | 2.66E-01 | Uncharacterized protein |
| AT1G03940 | -2.3 | 7.67E-04 | -2.2 | 1.47E-03 | -0.2 | 8.22E-01 | 0.4 | 2.43E-01 | 0.6 | 1.47E-01 | -0.1 | 7.82E-01 | HXXXD-type acyl-transferase family protein |
| AT1G80120 | -2.3 | 2.20E-07 | -1.1 | 2.52E-05 | -1.3 | 1.39E-04 | -0.4 | 9.68E-02 | 0.3 | 3.21E-01 | -0.7 | 9.89E-03 | Protein LURP-one-related 5 |
| AT4G02830 | -2.3 | 2.77E-07 | -1.0 | 2.43E-05 | -1.3 | 1.96E-04 | 0.6 | 1.42E-01 | 0.4 | 4.07E-01 | 0.2 | 6.20E-01 | Uncharacterized protein AT4g02830 |
| AT1G02570 | -2.3 | 1.27E-05 | -0.8 | 4.77E-03 | -1.5 | 1.50E-03 | -2.1 | 1.91E-03 | -0.5 | 3.46E-01 | -1.6 | 3.42E-02 | Transmembrane protein |
| AT5G36870 | -2.3 | 5.48E-07 | -1.7 | 5.16E-06 | -0.6 | 2.20E-02 | | | | | | | Similarity to callose synthase catalytic subunit |
| AT3G63360 | -2.3 | 3.39E-06 | -0.6 | 5.87E-03 | -1.7 | 2.57E-04 | | | | | | | Defensin-like protein 11 |
| AT5G50800 | -2.3 | 6.62E-07 | -1.7 | 8.69E-06 | -0.6 | 1.57E-02 | -0.6 | 6.50E-02 | 0.5 | 9.29E-02 | -1.1 | 8.41E-03 | SWEET13 |
| AT4G07815 | -2.3 | 6.26E-03 | -2.5 | 6.34E-03 | 0.2 | 8.37E-01 | -0.4 | 7.32E-01 | 0.8 | 4.83E-01 | | | NA |
| AT2G39050 | -2.2 | 5.61E-08 | -1.4 | 1.54E-06 | -0.8 | 2.40E-04 | -1.1 | 5.47E-05 | -0.3 | 2.11E-01 | -0.8 | 2.00E-03 | Hydroxyproline-rich glycoprotein family protein |
| AT5G49270 | -2.2 | 4.84E-07 | -0.4 | 2.05E-02 | -1.9 | 2.77E-05 | -1.6 | 1.04E-04 | -0.1 | 6.73E-01 | -1.4 | 3.98E-03 | COBRA-like protein 9 |
| AT5G19260 | -2.2 | 1.20E-06 | -1.3 | 4.85E-05 | -0.9 | 2.67E-03 | 0.0 | 8.70E-01 | 0.4 | 1.48E-01 | -0.3 | 1.81E-01 | Protein FANTASTIC FOUR 3 |
| AT3G28007 | -2.2 | 1.88E-03 | -1.9 | 4.65E-03 | -0.3 | 6.89E-01 | -1.2 | 2.51E-03 | -0.2 | 5.68E-01 | -1.0 | 1.78E-02 | SWEET4 |
| AT3G51642 | -2.2 | 1.05E-06 | -0.9 | 2.15E-04 | -1.3 | 4.13E-04 | 0.9 | 4.38E-02 | 0.6 | 2.70E-01 | 0.3 | 5.37E-01 | Uncharacterized protein |
| AT5G63560 | -2.2 | 5.72E-08 | -1.6 | 8.67E-07 | -0.6 | 9.00E-04 | 0.0 | 8.71E-01 | 0.0 | 8.79E-01 | -0.1 | 7.63E-01 | FACT |
| AT1G69260 | -2.2 | 4.83E-05 | -1.4 | 4.90E-04 | -0.8 | 8.37E-02 | -1.1 | 3.94E-03 | -0.6 | 8.54E-02 | -0.5 | 3.17E-01 | Ninja-family protein AFP1 |
| AT4G37220 | -2.2 | 5.30E-05 | -0.1 | 6.67E-01 | -2.1 | 4.49E-04 | -3.6 | 2.04E-04 | -2.0 | 2.52E-03 | -1.7 | 1.71E-01 | Cold-regulated 413 plasma membrane protein 4 |
| AT4G24580 | -2.2 | 1.64E-07 | -0.7 | 1.83E-04 | -1.5 | 3.23E-05 | -0.7 | 1.07E-01 | 0.3 | 4.57E-01 | -1.0 | 6.45E-02 | Rho GTPase activation protein with PH domain-containing protein |
| AT2G33790 | -2.2 | 4.78E-05 | -0.2 | 4.20E-01 | -1.9 | 4.87E-04 | -1.3 | 2.55E-02 | 0.3 | 5.98E-01 | -1.6 | 2.82E-02 | ATAGP30 |
| AT4G03540 | -2.2 | 8.81E-06 | -2.0 | 2.51E-05 | -0.1 | 7.11E-01 | 2.0 | 3.45E-05 | 1.4 | 2.23E-03 | 0.6 | 3.54E-02 | CASP-like protein 1C1 |
| AT5G43230 | -2.2 | 2.91E-06 | -0.5 | 1.33E-02 | -1.7 | 1.59E-04 | -1.6 | 1.50E-02 | 0.4 | 3.89E-01 | -2.0 | 1.41E-02 | Gb\|AAF03435.1 |

| | | | | | | | | | | | | | |
|---|---|---|---|---|---|---|---|---|---|---|---|---|---|
| AT2G47530 | -2.2 | 1.03E-05 | -0.7 | 4.54E-03 | -1.5 | 1.40E-03 | -0.8 | 3.29E-01 | -0.4 | 6.50E-01 | | | Pollen Ole e 1 allergen and extensin family protein |
| AT3G58550 | -2.2 | 1.66E-07 | -1.6 | 2.39E-06 | -0.6 | 4.02E-03 | -0.4 | 1.66E-01 | 0.2 | 5.63E-01 | -0.6 | 8.14E-02 | Non-specific lipid transfer protein GPI-anchored 22 |
| AT1G56320 | -2.2 | 2.54E-07 | -1.0 | 3.09E-05 | -1.2 | 1.69E-04 | 1.6 | 3.51E-05 | 0.9 | 1.31E-02 | 0.7 | 2.03E-02 | Uncharacterized protein |
| AT1G58037 | -2.2 | 5.00E-06 | -0.2 | 2.12E-01 | -1.9 | 1.10E-04 | | | | | | | Cysteine |
| AT5G55110 | -2.2 | 1.81E-05 | -0.6 | 1.34E-02 | -1.6 | 1.22E-03 | -0.3 | 4.91E-01 | 0.5 | 1.35E-01 | -0.8 | 5.32E-02 | Stigma-specific STIG1-like protein 2 |
| AT5G24313 | -2.1 | 3.79E-06 | -0.2 | 1.99E-01 | -1.9 | 8.50E-05 | -2.0 | 1.16E-03 | -0.3 | 5.78E-01 | -1.8 | 1.75E-02 | hypothetical protein |
| AT5G23030 | -2.1 | 8.64E-06 | -1.0 | 8.40E-04 | -1.2 | 4.03E-03 | | | | | | | Tetraspanin-12 |
| AT3G55500 | -2.1 | 1.27E-02 | -1.1 | 8.76E-02 | -1.0 | 2.82E-01 | -1.0 | 1.84E-04 | 0.2 | 3.33E-01 | -1.2 | 3.92E-04 | Expansin-A16 |
| AT1G63530 | -2.1 | 1.75E-07 | -1.1 | 7.39E-06 | -1.0 | 3.11E-04 | 0.4 | 4.15E-01 | 0.1 | 9.05E-01 | 0.3 | 6.16E-01 | F2K11.11 |
| AT3G21680 | -2.1 | 8.42E-07 | -0.6 | 1.25E-03 | -1.5 | 1.10E-04 | -1.1 | 8.69E-03 | 0.2 | 6.72E-01 | -1.3 | 7.22E-03 | Uncharacterized protein |
| AT5G18130 | -2.1 | 7.44E-08 | -1.3 | 3.53E-06 | -0.8 | 1.96E-04 | -0.2 | 2.68E-01 | -0.1 | 7.52E-01 | -0.1 | 6.93E-01 | Transmembrane protein |
| AT4G21250 | -2.1 | 1.02E-07 | -0.9 | 4.31E-05 | -1.2 | 3.57E-05 | -0.1 | 5.82E-01 | -0.4 | 1.49E-01 | 0.2 | 4.93E-01 | Sulfite exporter TauE |
| AT1G55240 | -2.1 | 1.13E-06 | -0.9 | 1.82E-04 | -1.2 | 5.48E-04 | -0.6 | 1.09E-01 | -0.2 | 5.79E-01 | -0.4 | 4.84E-01 | Uncharacterized protein F7A10.18 |
| AT3G48020 | -2.1 | 1.74E-07 | -1.6 | 1.18E-06 | -0.4 | 2.75E-02 | -0.5 | 1.42E-01 | -0.7 | 7.41E-02 | 0.2 | 7.15E-01 | hypothetical protein |
| AT5G09530 | -2.1 | 3.80E-08 | -1.0 | 9.53E-06 | -1.1 | 2.60E-05 | 1.7 | 4.17E-08 | 1.1 | 4.28E-05 | 0.7 | 1.92E-03 | Protein PELPK1 |
| AT1G49320 | -2.1 | 5.61E-08 | -1.1 | 5.99E-06 | -0.9 | 5.97E-05 | -1.5 | 3.21E-06 | -0.6 | 8.32E-03 | -0.9 | 3.42E-03 | BURP domain protein USPL1 |
| AT4G25560 | -2.1 | 1.66E-07 | -1.0 | 9.95E-06 | -1.0 | 2.30E-04 | -0.4 | 3.68E-01 | 0.3 | 5.61E-01 | -0.7 | 2.02E-01 | Myb domain protein 18 |
| AT5G19810 | -2.1 | 9.95E-05 | 0.3 | 1.62E-01 | -2.4 | 2.15E-04 | -2.4 | 2.24E-05 | -0.4 | 1.39E-01 | -1.9 | 1.48E-03 | Proline-rich extensin-like family protein |
| AT4G15610 | -2.1 | 1.02E-07 | -1.5 | 1.92E-06 | -0.6 | 1.72E-03 | -0.5 | 4.40E-02 | -0.7 | 2.24E-02 | 0.2 | 7.19E-01 | CASP-like protein |
| AT1G76960 | -2.0 | 9.91E-08 | -1.9 | 3.99E-07 | -0.2 | 2.34E-01 | -0.2 | 7.85E-01 | -0.3 | 7.10E-01 | 0.1 | 9.19E-01 | F22K20.6 protein |
| AT3G50130 | -2.0 | 1.33E-06 | -0.3 | 6.41E-02 | -1.8 | 4.65E-05 | -1.8 | 8.91E-04 | -0.3 | 4.07E-01 | -1.5 | 2.44E-02 | Transmembrane protein, |
| AT3G62680 | -2.0 | 1.05E-06 | -0.4 | 2.41E-02 | -1.6 | 4.65E-05 | -2.4 | 3.06E-07 | -0.2 | 4.34E-01 | -2.2 | 7.98E-05 | Proline-rich protein 3 |
| AT3G00880 | -2.0 | 1.82E-05 | -0.8 | 3.51E-03 | -1.2 | 3.52E-03 | 1.0 | 4.05E-02 | 0.4 | 5.50E-01 | 0.6 | 1.27E-01 | NA |
| AT5G64550 | -2.0 | 3.03E-07 | -1.4 | 6.97E-06 | -0.6 | 3.00E-03 | -0.3 | 2.72E-01 | 1.2 | 2.38E-04 | -1.5 | 3.65E-04 | Uncharacterized protein At5g64550 |
| AT2G19200 | -2.0 | 7.22E-07 | -0.9 | 1.00E-04 | -1.1 | 4.52E-04 | 1.3 | 3.05E-03 | 1.3 | 5.00E-03 | 0.0 | 9.70E-01 | UPF0725 protein At2g19200 |
| AT1G30850 | -2.0 | 1.04E-05 | -0.3 | 1.32E-01 | -1.7 | 2.44E-04 | -2.0 | 1.82E-04 | -0.4 | 3.34E-01 | -1.7 | 5.46E-03 | hypothetical protein |
| AT5G42500 | -2.0 | 2.37E-07 | -0.4 | 4.42E-03 | -1.6 | 2.37E-05 | -0.6 | 6.85E-03 | 0.2 | 4.41E-01 | -0.8 | 4.08E-03 | Dirigent protein 2 |
| AT1G03700 | -2.0 | 8.19E-07 | -1.1 | 3.50E-05 | -0.9 | 1.81E-03 | 0.6 | 7.58E-02 | 0.4 | 3.76E-01 | 0.3 | 5.48E-01 | CASP-like protein 1C2 |
| AT5G52260 | -2.0 | 3.68E-06 | -1.0 | 2.48E-04 | -1.0 | 3.02E-03 | -1.4 | 2.60E-02 | -0.2 | 8.05E-01 | -1.2 | 7.27E-02 | Uncharacterized protein |
| AT1G64700 | -2.0 | 2.68E-05 | -1.7 | 1.09E-04 | -0.3 | 4.12E-01 | -0.2 | 3.82E-01 | 0.2 | 5.07E-01 | -0.4 | 1.99E-01 | At1g64700 |
| AT5G35110 | -2.0 | 3.97E-06 | -0.8 | 6.48E-04 | -1.2 | 1.56E-03 | 0.4 | 4.24E-01 | 0.1 | 8.97E-01 | | | hypothetical protein |
| AT4G22666 | -2.0 | 3.10E-07 | -0.1 | 3.98E-01 | -1.9 | 1.09E-05 | -1.6 | 6.91E-05 | -0.3 | 4.14E-01 | -1.4 | 2.01E-03 | Bifunctional inhibitor |
| AT2G37670 | -2.0 | 2.53E-06 | -0.5 | 5.16E-03 | -1.5 | 2.29E-04 | | | | | | | Putative WD-40 repeat protein |
| AT4G14280 | -2.0 | 7.75E-06 | -0.2 | 3.49E-01 | -1.8 | 1.31E-04 | -2.2 | 5.38E-04 | 0.3 | 5.32E-01 | -2.4 | 2.74E-03 | ARM repeat superfamily protein |
| AT3G48940 | -2.0 | 1.58E-05 | -0.1 | 5.50E-01 | -1.9 | 1.86E-04 | -2.4 | 1.67E-05 | -0.7 | 2.03E-02 | -1.7 | 6.08E-03 | Remorin |
| AT2G21100 | -2.0 | 1.36E-07 | -0.9 | 1.79E-05 | -1.1 | 9.04E-05 | 1.8 | 2.35E-07 | 1.1 | 2.03E-04 | 0.7 | 2.81E-03 | Dirigent protein 23 |
| AT3G01760 | -2.0 | 2.93E-05 | -0.4 | 7.33E-02 | -1.6 | 9.12E-04 | -0.3 | 6.29E-01 | 0.3 | 6.92E-01 | | | Amino acid transporter transmembrane domain-containing protein |
| AT1G54540 | -2.0 | 7.72E-07 | -0.9 | 7.97E-05 | -1.0 | 5.98E-04 | 1.3 | 7.14E-04 | 0.7 | 5.02E-02 | 0.5 | 9.58E-02 | Late embryogenesis abundant protein LEA-2 subgroup domain-containing protein |
| AT1G54575 | -2.0 | 1.48E-07 | -2.1 | 3.13E-07 | 0.1 | 5.35E-01 | -0.2 | 6.28E-01 | 0.3 | 4.48E-01 | -0.5 | 2.95E-01 | Uncharacterized protein |
| AT1G27380 | -2.0 | 1.36E-05 | -1.0 | 7.51E-04 | -1.0 | 9.01E-03 | | | | | | | ROP-interactive CRIB motif-containing protein 2 |
| AT5G15960 | -2.0 | 1.42E-02 | -2.8 | 6.49E-03 | 0.8 | 4.75E-01 | -1.9 | 7.06E-02 | 1.3 | 5.79E-02 | -3.2 | 1.61E-02 | Stress-induced protein KIN1 |
| AT1G30750 | -1.9 | 3.45E-07 | -0.4 | 5.24E-03 | -1.5 | 2.86E-05 | -0.1 | 7.15E-01 | 0.4 | 1.49E-01 | -0.5 | 8.21E-02 | T5I8.20 protein |
| AT1G12040 | -1.9 | 4.01E-06 | -0.4 | 4.49E-02 | -1.5 | 1.40E-04 | -1.9 | 8.95E-06 | -0.1 | 7.90E-01 | -1.8 | 4.23E-04 | Leucine-rich repeat-containing N-terminal plant-type domain-containing protein |
| AT3G24927 | -1.9 | 1.48E-07 | -1.2 | 4.38E-06 | -0.7 | 7.81E-04 | -0.9 | 2.87E-04 | -0.6 | 8.95E-03 | -0.3 | 3.27E-01 | NA |
| AT3G03280 | -1.9 | 2.99E-05 | -0.7 | 6.82E-03 | -1.2 | 4.57E-03 | -1.8 | 6.55E-03 | -0.3 | 6.52E-01 | -1.6 | 6.94E-02 | Uncharacterized protein |
| AT1G24430 | -1.9 | 2.62E-07 | -1.2 | 5.51E-06 | -0.7 | 1.80E-03 | 0.4 | 3.36E-01 | 0.0 | 9.85E-01 | 0.4 | 4.08E-01 | F21J9.9 |
| AT2G13910 | -1.9 | 1.91E-06 | -0.9 | 2.56E-04 | -1.1 | 1.16E-03 | -1.1 | 2.60E-02 | -0.2 | 6.35E-01 | -0.8 | 1.44E-01 | NA |
| AT5G26080 | -1.9 | 1.34E-05 | 0.0 | 8.70E-01 | -2.0 | 9.77E-05 | -3.6 | 1.84E-04 | -0.2 | 6.01E-01 | -3.4 | 3.22E-03 | Proline-rich extensin-like family protein |
| AT1G24485 | -1.9 | 3.51E-07 | -0.3 | 1.72E-02 | -1.6 | 2.60E-05 | -2.5 | 5.86E-05 | 0.1 | 8.84E-01 | -2.5 | 1.17E-03 | ER protein carbohydrate-binding protein |
| AT2G34910 | -1.9 | 2.63E-05 | -0.2 | 2.61E-01 | -1.7 | 4.48E-04 | | | | | | | At2g34910 |
| AT2G39430 | -1.9 | 3.97E-07 | -0.4 | 9.01E-03 | -1.5 | 2.86E-05 | -0.4 | 1.65E-01 | 0.1 | 8.21E-01 | -0.5 | 1.28E-01 | Dirigent protein 9 |
| AT1G62440 | -1.9 | 5.72E-06 | -0.9 | 3.89E-04 | -1.0 | 4.02E-03 | -0.4 | 5.53E-01 | 0.4 | 4.72E-01 | -0.8 | 2.39E-01 | Leucine-rich repeat extensin-like protein 2 |
| AT1G75590 | -1.9 | 2.64E-04 | -1.3 | 2.15E-03 | -0.6 | 1.85E-01 | | | | | | | F10A5.20 |
| AT5G46900 | -1.9 | 3.24E-06 | 0.2 | 3.56E-01 | -2.1 | 2.69E-05 | -2.2 | 1.17E-05 | -0.8 | 1.94E-02 | -1.4 | 5.20E-03 | hypothetical protein |
| AT3G12720 | -1.9 | 2.04E-07 | -0.9 | 1.58E-05 | -1.0 | 2.32E-04 | | | | | | | MYB67 |
| AT2G48140 | -1.9 | 2.74E-07 | -0.8 | 1.75E-04 | -1.1 | 7.29E-05 | 0.5 | 3.56E-02 | 0.5 | 4.11E-02 | 0.0 | 9.28E-01 | Bifunctional inhibitor |
| AT4G12510 | -1.9 | 3.19E-06 | 0.0 | 9.39E-01 | -1.9 | 3.76E-05 | -1.7 | 4.02E-04 | -0.9 | 2.11E-02 | -0.8 | 7.21E-02 | hypothetical protein |
| AT5G44550 | -1.9 | 1.16E-07 | -0.7 | 3.27E-04 | -1.2 | 2.46E-05 | 1.7 | 2.68E-08 | 1.3 | 2.61E-06 | 0.4 | 5.32E-02 | CASP-like protein 1B1 |
| AT4G20190 | -1.9 | 2.62E-04 | -0.3 | 2.69E-01 | -1.6 | 4.13E-03 | | | | | | | hypothetical protein |
| AT2G43440 | -1.9 | 1.49E-04 | -1.2 | 1.86E-03 | -0.7 | 1.39E-01 | 0.0 | 9.82E-01 | -0.2 | 7.99E-01 | 0.2 | 8.08E-01 | F-box protein At2g43440 |
| AT4G14650 | -1.9 | 1.53E-07 | -0.8 | 3.49E-05 | -1.1 | 7.61E-05 | -0.4 | 1.77E-01 | 0.5 | 1.05E-01 | -1.0 | 1.12E-02 | At4g14650 |
| AT4G12520 | -1.9 | 2.91E-06 | 0.1 | 6.71E-01 | -2.0 | 2.86E-05 | -1.3 | 7.64E-04 | -0.4 | 3.10E-01 | -0.9 | 1.93E-02 | hypothetical protein |
| AT1G73410 | -1.9 | 1.90E-06 | -0.9 | 1.74E-04 | -1.0 | 1.57E-03 | 1.8 | 2.59E-03 | 1.5 | 1.58E-02 | 0.3 | 6.44E-01 | Myb domain protein 54 |
| AT5G46890 | -1.9 | 1.89E-06 | 0.3 | 9.72E-02 | -2.2 | 1.45E-05 | -2.5 | 8.45E-07 | -1.3 | 3.26E-04 | -1.2 | 1.34E-02 | Bifunctional inhibitor |
| AT3G26040 | -1.9 | 3.12E-05 | -0.1 | 6.85E-01 | -1.8 | 3.20E-04 | | | | | | | Acetyltranferase-like protein |
| AT5G25020 | -1.9 | 3.96E-06 | -0.4 | 2.99E-02 | -1.5 | 1.78E-04 | -0.8 | 9.20E-02 | -0.1 | 8.88E-01 | -0.7 | 1.95E-01 | Protein ENHANCED DISEASE RESISTANCE 2 C-terminal domain-containing protein |
| AT5G24655 | -1.9 | 5.08E-05 | -1.7 | 1.54E-04 | -0.2 | 6.73E-01 | -0.5 | 3.03E-01 | 0.1 | 9.08E-01 | -0.5 | 2.19E-01 | Protein RESPONSE TO LOW SULFUR 4 |
| AT5G05500 | -1.8 | 2.39E-06 | -0.2 | 3.36E-01 | -1.7 | 4.27E-05 | -1.3 | 4.01E-04 | 0.0 | 9.97E-01 | -1.3 | 2.81E-03 | At5g05500 |
| AT1G18290 | -1.8 | 1.28E-05 | -1.0 | 5.37E-04 | -0.9 | 1.29E-02 | -1.1 | 3.28E-02 | -0.4 | 4.72E-01 | -0.7 | 2.57E-01 | DUF4228 domain protein |
| AT5G54370 | -1.8 | 4.01E-06 | -0.1 | 6.15E-01 | -1.7 | 5.00E-05 | -0.5 | 1.38E-01 | 0.3 | 5.61E-01 | -0.8 | 3.44E-02 | At5g54370 protein-like protein) |
| AT3G20850 | -1.8 | 6.94E-04 | 0.4 | 1.17E-01 | -2.3 | 6.90E-04 | -4.1 | 7.38E-05 | -0.3 | 4.63E-01 | -3.8 | 1.43E-03 | Proline-rich extensin-like family protein |
| AT3G43110 | -1.8 | 2.49E-07 | -0.6 | 3.19E-04 | -1.3 | 4.59E-05 | 0.0 | 9.67E-01 | 0.4 | 6.46E-01 | | | At3g43110 |
| AT1G58170 | -1.8 | 7.22E-07 | -1.3 | 6.90E-06 | -0.5 | 2.99E-02 | 0.8 | 4.53E-03 | 0.8 | 5.64E-03 | 0.0 | 8.87E-01 | Dirigent protein 19 |
| AT2G35860 | -1.8 | 1.10E-05 | -0.5 | 1.54E-02 | -1.3 | 7.32E-04 | -0.7 | 2.22E-01 | 0.4 | 4.37E-01 | -1.1 | 8.76E-02 | Fasciclin-like arabinogalactan protein 16 |
| AT3G44510 | -1.8 | 1.58E-07 | -0.4 | 8.12E-04 | -1.4 | 2.47E-05 | -0.8 | 4.56E-02 | 0.1 | 9.09E-01 | -0.8 | 7.05E-02 | At3g44510 |
| AT2G43670 | -1.8 | 2.64E-05 | -0.8 | 2.24E-03 | -1.0 | 8.74E-03 | 1.5 | 4.56E-02 | 1.3 | 1.23E-01 | 0.2 | 8.37E-01 | Putative beta-1,3-glucanase, C terminal |
| AT3G19430 | -1.8 | 6.25E-06 | -0.3 | 9.65E-02 | -1.5 | 1.59E-04 | -0.6 | 1.73E-01 | 0.3 | 4.72E-01 | -0.9 | 6.86E-02 | Late embryogenesis abundant hydroxyproline-rich glycoprotein family |
| AT2G01580 | -1.8 | 3.44E-06 | -0.7 | 9.55E-04 | -1.1 | 1.09E-03 | | | | | | | hypothetical protein |
| AT5G47950 | -1.8 | 1.71E-06 | -0.7 | 5.93E-04 | -1.1 | 4.69E-04 | -2.4 | 1.10E-06 | -0.3 | 2.08E-01 | -2.1 | 2.05E-04 | Acetyl-CoA:benzylalcohol acetyltranferase-like protein |
| AT2G43390 | -1.8 | 1.76E-04 | -1.0 | 3.37E-03 | -0.8 | 8.28E-02 | 1.0 | 7.97E-03 | 0.4 | 3.21E-01 | 0.6 | 1.26E-01 | Uncharacterized protein |
| AT4G36010 | -1.8 | 3.26E-05 | -1.8 | 5.86E-05 | 0.0 | 9.46E-01 | 0.5 | 6.62E-03 | 0.2 | 2.86E-01 | 0.3 | 2.12E-01 | Thaumatin-like protein 1 |
| AT3G19320 | -1.8 | 2.84E-05 | -0.2 | 2.46E-01 | -1.6 | 5.55E-04 | -1.0 | 1.91E-01 | 0.5 | 5.55E-01 | | | Leucine-rich repeat family protein |
| AT3G46240 | -1.8 | 4.33E-06 | 0.0 | 9.21E-01 | -1.8 | 5.25E-05 | -3.1 | 2.48E-04 | -0.3 | 5.01E-01 | -2.8 | 7.37E-03 | Uncharacterized protein F12M12_210 |

| | | | | | | | | | | | | | |
|---|---|---|---|---|---|---|---|---|---|---|---|---|---|
| AT1G78070 | -1.8 | 2.78E-06 | -1.6 | 1.34E-05 | -0.2 | 2.96E-01 | -1.9 | 4.38E-06 | -0.2 | 4.28E-01 | -1.7 | 6.88E-04 | hypothetical protein |
| AT5G54050 | -1.8 | 1.02E-05 | -0.7 | 2.17E-03 | -1.1 | 2.69E-03 | | | | | | | Cysteine |
| AT1G66440 | -1.8 | 5.45E-06 | -1.0 | 2.70E-04 | -0.8 | 6.65E-03 | -0.6 | 2.92E-01 | 0.5 | 2.72E-01 | -1.1 | 1.43E-02 | Cysteine |
| AT4G36610 | -1.8 | 7.44E-08 | -1.0 | 4.19E-06 | -0.8 | 1.54E-04 | 1.5 | 2.38E-06 | 1.1 | 3.50E-04 | 0.4 | 2.25E-02 | hypothetical protein |
| AT2G33460 | -1.8 | 5.50E-06 | 0.0 | 9.26E-01 | -1.8 | 6.82E-05 | | | | | | | CRIB domain-containing protein RIC1 |
| AT3G55230 | -1.8 | 5.24E-08 | -0.3 | 1.20E-02 | -1.5 | 6.99E-06 | -0.2 | 3.49E-01 | 0.1 | 7.28E-01 | -0.3 | 2.24E-01 | Dirigent protein 24 |
| AT5G03360 | -1.8 | 8.57E-06 | -0.8 | 1.29E-03 | -1.0 | 3.28E-03 | | | | | | | Uncharacterized protein F12E4_100 |
| AT4G31470 | -1.8 | 1.34E-06 | -0.2 | 2.17E-01 | -1.6 | 3.65E-05 | -0.6 | 9.16E-02 | 0.1 | 8.35E-01 | -0.7 | 8.21E-02 | hypothetical protein |
| AT3G29780 | -1.8 | 2.18E-05 | -1.0 | 6.35E-04 | -0.8 | 2.83E-02 | -0.2 | 6.70E-01 | 0.6 | 2.30E-01 | -0.9 | 6.41E-02 | hypothetical protein |
| AT1G50630 | -1.8 | 7.93E-08 | -1.1 | 2.57E-06 | -0.6 | 3.93E-04 | -0.7 | 2.30E-04 | 0.0 | 8.36E-01 | -0.7 | 4.00E-03 | Extracellular ligand-gated ion channel protein |
| AT2G28315 | -1.8 | 1.72E-07 | -0.7 | 3.52E-05 | -1.0 | 9.47E-05 | -0.4 | 3.47E-01 | 0.1 | 7.96E-01 | -0.5 | 2.91E-01 | Nucleotide |
| AT4G33120 | -1.8 | 2.23E-05 | 0.2 | 3.56E-01 | -1.9 | 1.07E-04 | -2.8 | 1.51E-05 | -1.1 | 2.84E-03 | -1.6 | 6.01E-03 | Uncharacterized protein |
| AT2G45360 | -1.8 | 3.22E-06 | -0.7 | 9.84E-04 | -1.1 | 9.23E-04 | -1.5 | 6.94E-04 | -0.8 | 3.07E-02 | -0.7 | 2.00E-01 | Ankyrin repeat |
| AT1G11210 | -1.8 | 4.95E-05 | -1.3 | 4.38E-04 | -0.5 | 1.47E-01 | -0.1 | 6.37E-01 | 0.0 | 9.61E-01 | -0.1 | 7.27E-01 | DUF4408 domain-containing protein |
| AT4G25790 | -1.7 | 7.43E-06 | -0.1 | 5.04E-01 | -1.6 | 8.82E-05 | -2.1 | 2.13E-06 | -0.2 | 4.17E-01 | -1.9 | 3.10E-04 | hypothetical protein |
| AT5G19800 | -1.7 | 1.27E-05 | 0.2 | 2.23E-01 | -2.0 | 5.25E-05 | -2.7 | 1.61E-05 | -0.2 | 4.53E-01 | -2.4 | 1.20E-03 | Uncharacterized protein |
| AT5G02360 | -1.7 | 1.82E-06 | -0.7 | 4.08E-04 | -1.0 | 7.78E-04 | -0.8 | 1.03E-01 | -0.2 | 7.96E-01 | -0.7 | 2.99E-01 | DC1 domain-containing protein |
| AT1G02575 | -1.7 | 1.79E-05 | 0.0 | 7.87E-01 | -1.8 | 1.26E-04 | -3.5 | 6.48E-05 | -0.6 | 1.39E-01 | -2.9 | 2.97E-03 | Transmembrane protein |
| AT1G03070 | -1.7 | 1.83E-04 | -0.3 | 1.92E-01 | -1.4 | 3.50E-03 | -0.5 | 1.87E-01 | 0.1 | 7.81E-01 | -0.7 | 1.95E-01 | Protein LIFEGUARD 4 |
| AT1G13970 | -1.7 | 4.44E-07 | -0.6 | 6.13E-04 | -1.2 | 8.07E-05 | -0.2 | 4.71E-01 | 0.4 | 2.00E-01 | -0.7 | 7.82E-02 | Beta-hexosaminidase |
| AT5G65300 | -1.7 | 2.80E-05 | -1.3 | 1.64E-04 | -0.4 | 2.35E-01 | 0.5 | 2.92E-01 | -0.1 | 9.28E-01 | 0.6 | 3.61E-01 | Uncharacterized protein |
| AT4G22270 | -1.7 | 4.10E-08 | -1.1 | 7.82E-07 | -0.6 | 3.66E-04 | -0.6 | 2.71E-03 | 0.0 | 9.15E-01 | -0.6 | 2.44E-02 | Transmembrane protein, putative |
| AT5G19875 | -1.7 | 1.53E-06 | -1.8 | 3.15E-06 | 0.0 | 8.38E-01 | -0.2 | 3.83E-01 | -0.1 | 7.95E-01 | -0.1 | 6.84E-01 | hypothetical protein |
| AT1G04570 | -1.7 | 1.07E-03 | -1.3 | 5.02E-03 | -0.4 | 4.60E-01 | | | | | | | Major facilitator superfamily protein |
| AT5G42710 | -1.7 | 5.37E-05 | -0.8 | 5.37E-03 | -1.0 | 1.24E-02 | | | | | | | DUF3741 domain-containing protein |
| AT5G54020 | -1.7 | 5.13E-07 | -0.7 | 1.03E-04 | -1.0 | 2.72E-04 | -0.6 | 2.37E-01 | 0.1 | 9.12E-01 | -0.6 | 3.05E-01 | CHP-rich zinc finger protein-like |
| AT5G42785 | -1.7 | 1.22E-05 | 0.0 | 8.59E-01 | -1.7 | 1.25E-04 | -0.9 | 2.63E-02 | 0.1 | 8.28E-01 | -1.0 | 3.74E-02 | Transmembrane protein |
| AT5G65030 | -1.7 | 5.09E-05 | -1.5 | 1.80E-04 | -0.2 | 5.61E-01 | | | | | | | hypothetical protein |
| AT2G02630 | -1.7 | 2.48E-06 | -0.5 | 3.69E-03 | -1.2 | 2.76E-04 | -0.4 | 3.71E-01 | 0.4 | 4.14E-01 | -0.8 | 1.18E-01 | Cysteine |
| AT4G01110 | -1.7 | 8.78E-06 | -0.4 | 4.26E-02 | -1.4 | 3.53E-04 | -1.2 | 9.19E-04 | -0.6 | 5.99E-02 | -0.6 | 8.56E-02 | Uncharacterized protein AT4g01110 |
| AT2G36090 | -1.7 | 7.71E-08 | -1.2 | 1.25E-06 | -0.5 | 1.58E-03 | 0.0 | 8.40E-01 | 0.2 | 3.32E-01 | -0.2 | 4.66E-01 | Probable F-box protein At2g36090 |
| AT4G34580 | -1.7 | 9.89E-07 | -0.2 | 1.76E-01 | -1.5 | 2.86E-05 | -1.7 | 1.03E-05 | -0.3 | 2.75E-01 | -1.4 | 1.29E-03 | Phosphatidylinositol |
| AT3G52180 | -1.7 | 1.20E-05 | -1.5 | 5.86E-05 | -0.2 | 3.37E-01 | -0.2 | 1.46E-01 | 0.4 | 1.61E-02 | -0.6 | 2.41E-03 | Uncharacterized protein At3g52180 |
| AT5G61340 | -1.7 | 7.44E-08 | -0.7 | 2.75E-05 | -1.0 | 3.23E-05 | -0.1 | 6.58E-01 | 0.4 | 2.36E-02 | -0.4 | 2.04E-02 | Transmembrane protein |
| AT3G25930 | -1.7 | 2.19E-08 | -0.6 | 4.04E-05 | -1.2 | 8.39E-06 | -1.3 | 1.56E-04 | -0.7 | 2.50E-02 | -0.6 | 5.48E-02 | Uncharacterized protein |
| AT4G26320 | -1.7 | 1.28E-06 | -0.1 | 7.03E-01 | -1.7 | 2.69E-05 | -0.8 | 1.64E-02 | 0.1 | 8.07E-01 | -0.9 | 9.52E-03 | Arabinogalactan protein 13 |
| AT5G06990 | -1.7 | 7.97E-06 | -0.1 | 5.35E-01 | -1.6 | 1.12E-04 | -2.1 | 2.18E-03 | -0.1 | 9.05E-01 | -2.1 | 1.09E-02 | hypothetical protein |
| AT2G43120 | -1.7 | 8.06E-08 | -0.8 | 1.55E-05 | -0.9 | 5.44E-05 | 0.2 | 2.46E-01 | 0.4 | 1.09E-02 | -0.2 | 1.99E-01 | RmlC-like cupins superfamily protein |
| AT4G04990 | -1.7 | 3.49E-05 | -2.7 | 1.34E-05 | 1.0 | 2.24E-02 | 0.1 | 7.16E-01 | 0.4 | 8.04E-02 | -0.3 | 2.49E-01 | AT4g04990 protein ) |
| AT5G10430 | -1.7 | 8.28E-06 | -0.2 | 2.72E-01 | -1.5 | 1.66E-04 | -0.8 | 1.59E-01 | -0.1 | 8.95E-01 | -0.7 | 2.91E-01 | Classical arabinogalactan protein 4 |
| AT4G26580 | -1.7 | 6.30E-06 | -0.8 | 7.97E-04 | -0.9 | 3.30E-03 | 0.1 | 9.06E-01 | 0.2 | 7.67E-01 | | | Uncharacterized protein AT4g26580 |
| AT4G28085 | -1.7 | 7.12E-06 | -0.6 | 2.72E-03 | -1.1 | 1.32E-03 | 1.1 | 8.65E-04 | 0.3 | 5.03E-01 | 0.8 | 9.08E-03 | Transmembrane protein |
| AT2G27180 | -1.7 | 1.29E-04 | -2.0 | 1.04E-04 | 0.3 | 4.38E-01 | | | | | | | Transmembrane protein |
| AT5G49080 | -1.7 | 1.21E-05 | -0.2 | 2.89E-01 | -1.5 | 1.77E-04 | -2.3 | 5.37E-05 | 0.2 | 7.08E-01 | -2.5 | 5.38E-04 | NA |
| AT3G13277 | -1.7 | 1.91E-06 | -1.0 | 5.35E-05 | -0.7 | 7.96E-03 | -0.6 | 7.80E-02 | -0.1 | 8.62E-01 | -0.5 | 2.21E-01 | NA |
| AT5G61290 | -1.7 | 6.61E-05 | -1.2 | 7.11E-04 | -0.5 | 1.44E-01 | -0.6 | 1.62E-02 | 0.1 | 6.88E-01 | -0.7 | 2.16E-02 | Uncharacterized protein |
| AT3G01730 | -1.7 | 6.95E-05 | 0.2 | 3.55E-01 | -1.8 | 2.20E-04 | -2.8 | 3.93E-04 | 0.0 | 1.00E+00 | -2.8 | 3.78E-03 | F28J7.6 protein |
| AT1G43020 | -1.7 | 4.51E-06 | -0.2 | 2.19E-01 | -1.5 | 1.07E-04 | -0.3 | 5.32E-01 | -0.2 | 6.47E-01 | 0.0 | 9.61E-01 | DUF547 domain-containing protein |
| AT2G47360 | -1.7 | 1.20E-06 | -0.3 | 2.77E-02 | -1.4 | 6.01E-05 | -0.7 | 2.61E-02 | -0.1 | 8.47E-01 | -0.6 | 5.70E-02 | hypothetical protein |
| AT5G17390 | -1.7 | 1.00E-06 | -0.2 | 1.61E-01 | -1.5 | 3.35E-05 | -0.8 | 4.98E-02 | 0.0 | 9.88E-01 | -0.8 | 1.13E-01 | hypothetical protein |
| AT5G54040 | -1.7 | 2.92E-07 | -1.2 | 5.01E-06 | -0.5 | 5.56E-03 | -0.1 | 5.12E-01 | 0.3 | 1.06E-01 | -0.5 | 3.30E-02 | CHP-rich zinc finger protein-like |
| AT1G63820 | -1.7 | 3.92E-05 | -1.0 | 1.08E-03 | -0.7 | 3.97E-02 | -0.1 | 7.69E-01 | 0.4 | 4.04E-01 | -0.5 | 3.04E-01 | Uncharacterized protein T12P18.16 |
| AT3G01930 | -1.7 | 1.56E-07 | -1.1 | 3.69E-06 | -0.5 | 1.51E-03 | 0.0 | 9.80E-01 | 0.3 | 2.74E-01 | -0.3 | 3.51E-01 | AT.I.24-7 protein |
| AT1G10480 | -1.7 | 1.89E-07 | -1.3 | 1.24E-06 | -0.3 | 4.38E-02 | 1.3 | 6.40E-03 | 1.0 | 5.34E-02 | 0.3 | 5.42E-01 | hypothetical protein |
| AT3G59370 | -1.7 | 3.71E-07 | 0.2 | 1.80E-01 | -1.8 | 8.63E-06 | -1.3 | 2.87E-04 | -0.4 | 2.73E-01 | -1.0 | 1.10E-02 | hypothetical protein |
| AT3G27150 | -1.6 | 1.24E-07 | -0.8 | 2.68E-05 | -0.9 | 7.49E-05 | -0.3 | 1.20E-01 | 0.0 | 8.72E-01 | -0.3 | 1.78E-01 | F-box |
| AT3G50220 | -1.6 | 2.10E-06 | 0.0 | 9.85E-01 | -1.6 | 3.35E-05 | -0.2 | 5.36E-01 | -0.1 | 8.46E-01 | -0.1 | 7.85E-01 | Protein IRREGULAR XYLEM 15 |
| AT4G01060 | -1.6 | 6.04E-03 | -1.6 | 9.04E-03 | 0.0 | 9.53E-01 | 1.7 | 7.51E-02 | 1.8 | 8.64E-02 | | | CAPRICE-like MYB3 |
| AT4G37235 | -1.6 | 3.61E-05 | -0.1 | 4.36E-01 | -1.5 | 4.83E-04 | -1.3 | 1.55E-02 | -0.3 | 6.01E-01 | -1.0 | 3.09E-02 | CASP-like protein 5C1 |
| AT2G47540 | -1.6 | 1.88E-06 | 0.0 | 9.16E-01 | -1.6 | 2.99E-05 | -0.2 | 5.25E-01 | 0.1 | 7.84E-01 | -0.3 | 3.14E-01 | Pollen Ole e 1 allergen and extensin family protein |
| AT1G50590 | -1.6 | 2.52E-06 | -0.7 | 4.85E-04 | -0.9 | 1.19E-03 | -0.2 | 4.43E-01 | -0.2 | 5.89E-01 | 0.0 | 9.45E-01 | RmlC-like cupins superfamily protein |
| AT4G34950 | -1.6 | 3.89E-04 | -1.5 | 1.07E-03 | -0.2 | 6.76E-01 | -0.8 | 2.38E-06 | -0.4 | 5.01E-03 | -0.4 | 1.43E-02 | Nodulin-like domain-containing protein |
| AT5G40860 | -1.6 | 7.96E-05 | -0.1 | 7.62E-01 | -1.6 | 6.55E-04 | | | | | | | Transmembrane protein |
| AT5G57770 | -1.6 | 7.64E-07 | -0.8 | 6.83E-05 | -0.8 | 9.61E-04 | -0.6 | 2.95E-01 | 0.3 | 6.73E-01 | -0.9 | 2.07E-01 | Auxin canalization protein |
| AT1G62980 | -1.6 | 8.97E-06 | 0.0 | 9.32E-01 | -1.6 | 6.71E-05 | -1.8 | 4.91E-05 | 0.1 | 7.67E-01 | -2.0 | 4.87E-04 | Expansin-A18 |
| AT1G24420 | -1.6 | 3.51E-06 | 0.0 | 8.51E-01 | -1.6 | 4.93E-05 | -0.2 | 6.93E-01 | 0.6 | 8.26E-02 | -0.8 | 6.90E-02 | F21J9.8 |
| AT1G63930 | -1.6 | 1.78E-06 | -0.2 | 8.47E-02 | -1.4 | 6.25E-05 | -0.4 | 1.42E-01 | 0.4 | 1.43E-01 | -0.8 | 1.74E-02 | Protein ROH1A |
| AT5G13250 | -1.6 | 6.92E-06 | -0.5 | 5.55E-03 | -1.1 | 9.58E-04 | -0.1 | 8.74E-01 | 0.7 | 2.14E-01 | -0.8 | 2.25E-01 | Uncharacterized protein T31B5_70 |
| AT3G47210 | -1.6 | 1.88E-07 | -0.6 | 1.88E-04 | -1.0 | 4.57E-05 | -0.3 | 1.79E-01 | 0.1 | 7.48E-01 | -0.4 | 8.76E-02 | AT3g47210 |
| AT1G59740 | -1.6 | 1.50E-06 | -1.5 | 5.51E-06 | -0.1 | 5.31E-01 | -0.2 | 3.82E-01 | 0.0 | 9.83E-01 | -0.2 | 5.15E-01 | Putative nitrate transporter NTL1 |
| AT1G12560 | -1.6 | 4.43E-06 | 0.0 | 9.28E-01 | -1.6 | 4.37E-05 | -1.3 | 3.09E-03 | -0.2 | 6.90E-01 | -1.1 | 1.94E-02 | Expansin-A7 |
| AT3G56230 | -1.6 | 1.01E-06 | -0.2 | 1.52E-01 | -1.4 | 3.35E-05 | -0.7 | 3.61E-02 | -0.1 | 7.33E-01 | -0.6 | 1.08E-01 | BTB |
| AT2G28440 | -1.6 | 9.51E-06 | -0.3 | 7.82E-02 | -1.3 | 3.01E-04 | -0.5 | 2.93E-01 | 0.3 | 6.35E-01 | -0.8 | 2.05E-01 | En |
| AT4G16146 | -1.6 | 1.16E-03 | -0.8 | 3.71E-02 | -0.9 | 8.00E-02 | 0.1 | 7.27E-01 | 0.5 | 2.02E-01 | -0.3 | 4.83E-01 | cAMP-regulated phosphoprotein 19-related protein |
| AT4G20260 | -1.6 | 8.29E-08 | -1.0 | 5.79E-06 | -0.6 | 2.03E-04 | -0.7 | 2.95E-04 | -0.5 | 8.83E-03 | -0.2 | 3.99E-01 | Plasma-membrane associated cation-binding protein 1 |
| AT4G33730 | -1.6 | 3.86E-05 | 0.1 | 5.94E-01 | -1.7 | 1.84E-04 | -1.9 | 1.04E-03 | -0.4 | 4.23E-01 | -1.6 | 2.75E-02 | hypothetical protein |
| AT5G17460 | -1.6 | 2.05E-06 | -1.3 | 1.41E-05 | -0.3 | 1.36E-01 | -0.6 | 9.90E-02 | -0.1 | 8.68E-01 | -0.5 | 2.73E-01 | Uncharacterized protein |
| AT5G57625 | -1.6 | 4.63E-06 | 0.1 | 6.67E-01 | -1.7 | 3.35E-05 | -2.2 | 4.78E-07 | -0.2 | 4.58E-01 | -2.0 | 1.09E-04 | hypothetical protein |
| AT5G45220 | -1.6 | 1.81E-07 | -0.6 | 8.97E-05 | -1.0 | 6.82E-05 | -0.8 | 1.44E-03 | -0.1 | 5.80E-01 | -0.7 | 2.33E-02 | Disease resistance protein family |
| AT2G30395 | -1.6 | 2.34E-06 | -0.2 | 8.40E-02 | -1.4 | 8.50E-05 | -0.9 | 8.09E-02 | 0.1 | 8.39E-01 | -1.0 | 6.90E-02 | Transcription repressor OFP17 |
| AT4G13680 | -1.6 | 6.76E-06 | -0.4 | 1.92E-02 | -1.2 | 4.71E-04 | | | | | | | Uncharacterized protein AT4g13680 |
| AT2G38500 | -1.6 | 2.57E-06 | -0.4 | 1.42E-02 | -1.2 | 1.69E-04 | -0.4 | 1.46E-01 | 0.0 | 9.36E-01 | -0.4 | 2.33E-01 | Uncharacterized protein At2g38500 |
| AT5G01280 | -1.6 | 4.22E-05 | -0.2 | 2.10E-01 | -1.3 | 8.81E-04 | -0.8 | 1.60E-01 | 0.0 | 9.76E-01 | -0.8 | 2.23E-01 | GPI-anchored protein |

| | | | | | | | | | | | | | |
|---|---|---|---|---|---|---|---|---|---|---|---|---|---|
| AT1G61930 | -1.6 | 7.43E-06 | -1.3 | 4.04E-05 | -0.3 | 2.39E-01 | -0.1 | 7.68E-01 | 0.4 | 1.56E-01 | -0.5 | 5.64E-02 | Protein S40-6 |
| AT2G38180 | -1.6 | 5.94E-07 | -0.8 | 1.10E-04 | -0.8 | 3.95E-04 | -0.1 | 5.71E-01 | 0.1 | 5.40E-01 | -0.2 | 3.15E-01 | Uncharacterized protein |
| AT2G37700 | -1.6 | 1.91E-06 | -0.2 | 6.27E-02 | -1.3 | 7.42E-05 | -0.4 | 1.13E-01 | 0.3 | 1.89E-01 | -0.7 | 1.57E-02 | Fatty acid hydroxylase superfamily |
| AT2G34610 | -1.6 | 3.73E-05 | -2.2 | 1.51E-05 | 0.7 | 6.44E-02 | 1.2 | 2.42E-02 | 1.8 | 3.99E-03 | -0.5 | 2.67E-01 | Cotton fiber protein |
| AT5G67620 | -1.6 | 9.39E-06 | -0.4 | 1.19E-02 | -1.1 | 7.60E-04 | -0.3 | 4.85E-01 | 0.6 | 8.27E-02 | -0.8 | 2.44E-02 | DUF4228 domain protein |
| AT4G18630 | -1.6 | 3.20E-06 | -0.7 | 4.35E-04 | -0.8 | 1.94E-03 | -0.4 | 1.62E-01 | 0.1 | 7.12E-01 | -0.5 | 1.03E-01 | Uncharacterized protein At4g18630 |
| AT1G52080 | -1.6 | 3.26E-07 | -1.0 | 6.11E-06 | -0.5 | 4.07E-03 | -0.4 | 2.63E-01 | 0.2 | 4.93E-01 | -0.6 | 8.84E-02 | Actin binding protein family |
| AT3G49845 | -1.6 | 2.29E-06 | 0.0 | 9.76E-01 | -1.5 | 2.99E-05 | 0.2 | 4.25E-01 | 0.2 | 5.39E-01 | 0.0 | 9.71E-01 | Cysteine-rich TM module stress tolerance protein |
| AT4G03500 | -1.5 | 3.68E-06 | -0.5 | 2.59E-03 | -1.0 | 6.40E-04 | -1.3 | 6.79E-03 | -0.1 | 9.27E-01 | -1.2 | 2.15E-02 | Ankyrin repeat family protein |
| AT4G32780 | -1.5 | 7.18E-07 | -1.0 | 1.41E-05 | -0.5 | 8.68E-03 | 0.1 | 8.01E-01 | 0.2 | 4.83E-01 | -0.1 | 7.19E-01 | At4g32785 |
| AT2G12400 | -1.5 | 2.33E-07 | -0.6 | 1.51E-04 | -0.9 | 6.66E-05 | -0.6 | 3.45E-03 | 0.1 | 7.99E-01 | -0.7 | 3.76E-03 | Uncharacterized protein At2g12400 |
| AT3G48920 | -1.5 | 6.87E-07 | -0.6 | 2.31E-04 | -0.9 | 2.70E-04 | 1.4 | 9.24E-04 | 1.2 | 6.11E-03 | 0.2 | 6.35E-01 | Uncharacterized protein |
| AT4G40090 | -1.5 | 1.27E-05 | 0.1 | 6.00E-01 | -1.6 | 6.05E-05 | -2.5 | 1.24E-06 | -0.3 | 3.79E-01 | -2.2 | 1.36E-04 | Classical arabinogalactan protein 3 |
| AT2G15440 | -1.5 | 1.23E-06 | -0.7 | 1.44E-04 | -0.8 | 1.11E-03 | -0.2 | 4.14E-01 | 0.4 | 1.34E-01 | -0.6 | 4.55E-02 | Uncharacterized protein |
| AT3G48450 | -1.5 | 1.42E-06 | -0.8 | 7.94E-05 | -0.7 | 2.97E-03 | 2.0 | 6.82E-05 | 0.9 | 7.46E-02 | 1.1 | 3.46E-03 | hypothetical protein |
| AT3G09925 | -1.5 | 3.68E-06 | 0.1 | 5.90E-01 | -1.6 | 3.29E-05 | -1.3 | 6.82E-04 | 0.0 | 9.16E-01 | -1.4 | 3.94E-03 | Pollen Ole e 1 allergen and extensin family protein |
| AT2G02300 | -1.5 | 9.11E-05 | -0.9 | 2.07E-03 | -0.6 | 6.73E-02 | | | | | | | PP2-B6 |
| AT4G06536 | -1.5 | 6.14E-06 | -0.5 | 5.09E-03 | -1.0 | 8.64E-04 | -0.8 | 2.15E-02 | -0.5 | 2.05E-01 | -0.4 | 3.62E-01 | At4g06536 domain-containing protein) |
| AT5G06610 | -1.5 | 1.38E-07 | -0.6 | 6.35E-05 | -0.9 | 5.23E-05 | -0.2 | 1.04E-01 | -0.1 | 7.21E-01 | -0.2 | 3.87E-01 | Gb\|AAD30234.1 ) |
| AT4G25190 | -1.5 | 1.43E-04 | -0.7 | 1.10E-02 | -0.8 | 2.62E-02 | | | | | | | QWRF7 |
| AT3G52370 | -1.5 | 1.16E-06 | -0.6 | 7.92E-04 | -0.9 | 2.47E-04 | -0.1 | 7.83E-01 | 0.5 | 5.41E-03 | -0.6 | 9.12E-03 | FASCICLIN-like arabinogalactan protein 15 |
| AT1G75620 | -1.5 | 8.09E-06 | -0.4 | 2.27E-02 | -1.1 | 4.86E-04 | -0.9 | 2.16E-02 | -0.2 | 6.76E-01 | -0.7 | 1.74E-01 | F10A5.18 |
| AT2G47860 | -1.5 | 5.41E-08 | -1.0 | 1.42E-06 | -0.5 | 3.28E-04 | -0.5 | 2.44E-02 | 0.0 | 9.45E-01 | -0.5 | 4.80E-02 | Phototropic-responsive NPH3 family protein |
| AT4G13580 | -1.5 | 1.90E-06 | 0.1 | 2.91E-01 | -1.6 | 2.33E-05 | -0.6 | 2.52E-02 | -0.1 | 7.89E-01 | -0.5 | 8.53E-02 | Dirigent protein 18 |
| AT1G13830 | -1.5 | 9.63E-06 | -0.6 | 3.70E-03 | -0.9 | 1.79E-03 | -0.4 | 1.99E-01 | 0.4 | 1.88E-01 | -0.7 | 1.71E-02 | Carbohydrate-binding X8 domain superfamily protein |
| AT3G00260 | -1.5 | 1.82E-04 | 0.2 | 3.47E-01 | -1.7 | 4.32E-04 | -1.2 | 1.06E-02 | 0.1 | 8.64E-01 | -1.3 | 1.05E-02 | NA |
| AT5G16490 | -1.5 | 1.01E-04 | -0.9 | 2.08E-03 | -0.6 | 8.86E-02 | 0.0 | 9.37E-01 | 0.2 | 7.11E-01 | | | CRIB domain-containing protein RIC4 |
| AT2G17710 | -1.5 | 3.17E-08 | -1.0 | 8.67E-07 | -0.5 | 1.42E-04 | -1.1 | 8.72E-06 | -0.4 | 4.30E-02 | -0.7 | 7.35E-03 | Uncharacterized protein |
| AT4G02270 | -1.5 | 6.93E-06 | 0.1 | 6.85E-01 | -1.6 | 4.25E-05 | -2.0 | 2.47E-06 | -0.2 | 4.67E-01 | -1.8 | 1.08E-04 | Protein SEED AND ROOT HAIR PROTECTIVE PROTEIN |
| AT1G16850 | -1.5 | 7.69E-04 | -0.9 | 9.67E-03 | -0.6 | 1.71E-01 | -0.7 | 1.18E-01 | 0.0 | 9.41E-01 | -0.7 | 1.56E-01 | At1g16880 |
| AT2G29620 | -1.5 | 4.40E-06 | -0.4 | 6.11E-03 | -1.0 | 5.10E-04 | -1.2 | 6.95E-02 | 0.0 | 9.58E-01 | -1.3 | 1.24E-01 | Uncharacterized protein At2g29620 |
| AT5G40470 | -1.5 | 1.53E-05 | -1.1 | 1.54E-04 | -0.4 | 1.23E-01 | -0.3 | 6.05E-02 | -0.4 | 4.29E-02 | 0.1 | 8.42E-01 | AT5g40470 |
| AT1G21000 | -1.5 | 1.51E-07 | -1.2 | 1.25E-06 | -0.3 | 2.05E-02 | -0.4 | 2.68E-02 | -0.7 | 1.22E-03 | 0.3 | 1.95E-01 | F9H16.1 protein |
| AT2G16760 | -1.5 | 3.93E-07 | -0.4 | 3.47E-03 | -1.1 | 3.78E-05 | 0.7 | 2.49E-03 | 0.6 | 1.63E-02 | 0.1 | 5.83E-01 | SMP-30 |
| AT4G17030 | -1.5 | 3.63E-08 | -1.6 | 9.13E-08 | 0.1 | 2.54E-01 | 0.3 | 5.16E-02 | 0.3 | 1.31E-01 | 0.0 | 8.92E-01 | Expansin-like B1 |
| AT3G03870 | -1.5 | 9.43E-07 | -1.2 | 6.65E-06 | -0.2 | 1.06E-01 | -0.6 | 2.61E-04 | -0.4 | 8.50E-03 | -0.2 | 3.80E-01 | Transmembrane protein |
| AT2G16660 | -1.5 | 3.71E-04 | -0.9 | 6.39E-03 | -0.5 | 1.24E-01 | -0.9 | 1.90E-03 | -0.6 | 2.29E-02 | -0.2 | 5.01E-01 | hypothetical protein |
| AT4G17480 | -1.5 | 1.82E-05 | -0.7 | 2.19E-03 | -0.8 | 7.74E-03 | 0.0 | 9.81E-01 | -0.1 | 8.47E-01 | 0.2 | 8.37E-01 | Thioesterase like protein |
| AT1G45545 | -1.5 | 3.40E-05 | -0.1 | 5.06E-01 | -1.4 | 4.63E-04 | | | | | | | WEAK CHLOROPLAST MOVEMENT UNDER BLUE LIGHT-like protein |
| AT1G56550 | -1.5 | 1.06E-07 | -0.9 | 3.96E-06 | -0.6 | 4.69E-04 | 0.0 | 9.21E-01 | 0.3 | 2.27E-01 | -0.3 | 2.35E-01 | At1g56550 |
| AT2G33770 | -1.5 | 1.34E-06 | -1.3 | 7.39E-06 | -0.2 | 2.82E-01 | -0.3 | 1.43E-01 | 0.0 | 9.85E-01 | -0.3 | 3.27E-01 | AT2G33770 protein |
| AT1G03550 | -1.5 | 4.31E-06 | -0.5 | 4.64E-03 | -1.0 | 5.80E-04 | -0.4 | 3.89E-01 | -0.4 | 4.47E-01 | 0.0 | 9.96E-01 | Secretory carrier-associated membrane protein |
| AT5G01640 | -1.5 | 2.17E-05 | -1.1 | 2.32E-04 | -0.4 | 1.18E-01 | 0.1 | 8.75E-01 | 0.5 | 9.66E-02 | -0.4 | 2.24E-01 | PRA1 family protein B5 |
| AT3G11385 | -1.5 | 3.62E-05 | -0.1 | 5.59E-01 | -1.4 | 4.52E-04 | -0.2 | 7.34E-01 | 0.5 | 3.53E-01 | -0.7 | 1.68E-01 | Cysteine |
| AT3G55910 | -1.5 | 1.72E-05 | -1.4 | 4.13E-05 | 0.0 | 8.76E-01 | | | | | | | hypothetical protein |
| AT2G18360 | -1.4 | 3.97E-07 | -1.1 | 4.72E-06 | -0.4 | 1.99E-02 | 0.5 | 1.13E-01 | 0.3 | 4.49E-01 | 0.2 | 4.90E-01 | Alpha |
| AT4G15620 | -1.4 | 3.02E-04 | -0.6 | 4.38E-02 | -0.9 | 1.92E-02 | -0.4 | 2.95E-02 | 0.3 | 1.68E-01 | -0.7 | 8.89E-03 | CASP-like protein |
| AT2G45400 | -1.4 | 7.19E-08 | -0.7 | 1.66E-05 | -0.8 | 4.16E-05 | -0.7 | 1.51E-03 | -0.2 | 4.08E-01 | -0.5 | 5.32E-02 | NAD-binding Rossmann-fold superfamily protein |
| AT4G01630 | -1.4 | 7.93E-06 | 0.3 | 4.30E-02 | -1.7 | 2.95E-05 | -1.9 | 1.70E-05 | 0.2 | 4.29E-01 | -2.2 | 2.32E-04 | Putative expansin-A17 |
| AT2G05632 | -1.4 | 5.07E-06 | -0.8 | 1.96E-04 | -0.6 | 1.10E-02 | -0.7 | 1.09E-03 | -0.2 | 4.31E-01 | -0.5 | 2.89E-02 | Uncharacterized protein |
| AT2G22930 | -1.4 | 6.74E-06 | -0.4 | 1.37E-02 | -1.0 | 5.24E-04 | -1.5 | 4.47E-03 | 0.2 | 5.94E-01 | -1.7 | 6.53E-03 | UDP-Glycosyltransferase superfamily protein |
| AT1G62870 | -1.4 | 1.59E-06 | -1.0 | 2.14E-05 | -0.4 | 3.24E-02 | -0.5 | 2.69E-02 | 0.2 | 4.02E-01 | -0.7 | 9.35E-03 | F16P17.2 protein |
| AT2G35150 | -1.4 | 6.78E-06 | -0.9 | 1.51E-04 | -0.5 | 2.78E-02 | -0.2 | 6.07E-01 | 0.4 | 3.15E-01 | -0.6 | 1.53E-01 | Protein EXORDIUM-like 7 |
| AT3G56260 | -1.4 | 7.24E-04 | -1.0 | 4.87E-03 | -0.4 | 3.30E-01 | 0.6 | 2.23E-01 | 1.1 | 4.49E-02 | -0.4 | 4.52E-01 | Uncharacterized protein F18O21_220 |
| AT1G35250 | -1.4 | 1.20E-06 | -0.8 | 6.05E-05 | -0.6 | 3.03E-03 | 0.5 | 2.04E-01 | 0.5 | 3.21E-01 | 0.1 | 9.32E-01 | Thioesterase superfamily protein |
| AT1G50060 | -1.4 | 1.71E-04 | 0.0 | 8.92E-01 | -1.4 | 1.01E-03 | 0.2 | 6.44E-01 | 0.6 | 1.01E-01 | -0.4 | 3.04E-01 | SCP domain-containing protein |
| AT2G26410 | -1.4 | 6.15E-07 | -0.5 | 9.31E-04 | -0.9 | 1.12E-04 | 0.0 | 9.62E-01 | 0.3 | 3.75E-01 | -0.3 | 2.78E-01 | Putative SF16 protein |
| AT2G24400 | -1.4 | 3.03E-05 | 0.0 | 9.90E-01 | -1.4 | 2.44E-04 | -0.6 | 8.30E-02 | 0.0 | 9.24E-01 | -0.6 | 1.86E-01 | SAUR-like auxin-responsive protein family |
| AT5G49680 | -1.4 | 3.02E-06 | -0.7 | 4.92E-04 | -0.7 | 1.80E-03 | -0.3 | 2.04E-01 | 0.1 | 7.72E-01 | -0.4 | 1.73E-01 | Protein KINKY POLLEN |
| AT5G15630 | -1.4 | 5.81E-07 | -0.3 | 1.82E-02 | -1.2 | 3.70E-05 | -0.2 | 5.81E-01 | 0.0 | 9.71E-01 | -0.2 | 7.06E-01 | COBRA-like extracellular glycosyl-phosphatidyl inositol-anchored protein family |
| AT4G31320 | -1.4 | 1.58E-05 | -0.1 | 3.93E-01 | -1.3 | 2.29E-04 | -0.3 | 2.37E-01 | 0.3 | 2.16E-01 | -0.5 | 2.66E-02 | SAUR-like auxin-responsive protein family |
| AT1G11655 | -1.4 | 5.93E-04 | 0.0 | 8.55E-01 | -1.5 | 2.21E-03 | -3.7 | 1.19E-04 | -0.7 | 1.03E-01 | -3.0 | 9.24E-03 | F25C20.20 |
| AT4G15740 | -1.4 | 8.34E-06 | -0.6 | 2.16E-03 | -0.8 | 2.39E-03 | -0.7 | 2.01E-02 | 0.1 | 8.73E-01 | -0.7 | 4.57E-02 | Calcium-dependent lipid-binding family protein |
| AT2G17840 | -1.4 | 1.51E-07 | -0.9 | 8.61E-06 | -0.5 | 4.18E-04 | -0.3 | 9.77E-03 | -0.1 | 5.70E-01 | -0.2 | 1.43E-01 | Senescence |
| AT4G31910 | -1.4 | 1.51E-06 | -0.7 | 2.80E-04 | -0.7 | 8.90E-04 | -0.3 | 8.53E-02 | -0.1 | 7.57E-01 | -0.2 | 3.74E-01 | PIZ |
| AT5G40960 | -1.4 | 1.07E-05 | -0.8 | 3.59E-04 | -0.6 | 1.88E-02 | -0.2 | 4.06E-01 | 0.2 | 5.38E-01 | -0.5 | 1.36E-01 | Transmembrane protein, |
| AT1G67920 | -1.4 | 9.00E-05 | -1.4 | 1.70E-04 | 0.0 | 9.87E-01 | 0.3 | 4.35E-01 | -0.5 | 3.10E-01 | 0.8 | 1.30E-01 | Uncharacterized protein |
| AT3G23470 | -1.4 | 4.37E-06 | 0.0 | 9.79E-01 | -1.4 | 4.65E-05 | -0.4 | 2.76E-01 | 0.4 | 1.66E-01 | -0.8 | 2.00E-03 | At3g23470 |
| AT5G40020 | -1.4 | 2.37E-06 | -0.2 | 1.37E-01 | -1.2 | 7.58E-05 | 0.3 | 3.79E-01 | 0.3 | 5.06E-01 | 0.0 | 9.48E-01 | Thaumatin-like protein |
| AT5G44350 | -1.4 | 8.51E-05 | -1.0 | 7.06E-04 | -0.4 | 2.34E-01 | 0.1 | 7.18E-01 | 0.4 | 2.72E-01 | -0.3 | 3.54E-01 | Ethylene-regulated nuclear protein ERT2-like protein |
| AT1G17090 | -1.4 | 7.50E-06 | -0.5 | 7.37E-03 | -1.0 | 8.81E-04 | 0.1 | 8.47E-01 | 0.4 | 2.31E-01 | -0.3 | 3.04E-01 | Transmembrane protein |
| AT5G17860 | -1.4 | 6.27E-05 | -1.9 | 2.80E-05 | 0.5 | 1.40E-01 | 0.4 | 4.92E-01 | -0.1 | 8.79E-01 | 0.5 | 4.15E-01 | Calcium exchanger 7 |
| AT5G53250 | -1.4 | 4.26E-05 | 0.2 | 2.18E-01 | -1.6 | 9.47E-05 | -1.4 | 5.03E-05 | -0.2 | 5.99E-01 | -1.2 | 1.87E-03 | Arabinogalactan protein 22 |
| AT5G41040 | -1.4 | 1.44E-07 | -0.8 | 7.57E-06 | -0.6 | 3.67E-04 | 0.9 | 1.41E-04 | 0.6 | 9.29E-03 | 0.3 | 1.58E-01 | HXXXD-type acyl-transferase family protein |
| AT2G41800 | -1.4 | 1.46E-04 | -0.1 | 5.72E-01 | -1.3 | 1.18E-03 | -0.3 | 1.50E-01 | 0.3 | 1.77E-01 | -0.7 | 1.43E-02 | At2g41800 |
| AT4G27400 | -1.4 | 2.79E-04 | 0.0 | 9.34E-01 | -1.4 | 1.47E-03 | -0.3 | 6.08E-01 | 0.0 | 9.55E-01 | -0.2 | 7.42E-01 | Late embryogenesis abundant protein-like protein |
| AT1G48500 | -1.4 | 8.56E-07 | -0.8 | 4.13E-05 | -0.6 | 2.94E-03 | -0.5 | 1.28E-01 | 0.2 | 5.14E-01 | -0.8 | 8.03E-02 | Protein TIFY |
| AT5G21950 | -1.4 | 4.43E-07 | -1.0 | 6.58E-06 | -0.4 | 1.22E-02 | -0.8 | 8.64E-03 | -0.3 | 3.89E-01 | -0.6 | 1.05E-01 | Uncharacterized protein At5g21950 |
| AT3G29680 | -1.4 | 5.51E-07 | -0.7 | 5.73E-05 | -0.7 | 7.33E-04 | 0.1 | 8.15E-01 | 0.2 | 4.03E-01 | -0.2 | 6.20E-01 | Uncharacterized protein |
| AT3G00650 | -1.4 | 1.45E-03 | -0.5 | 1.04E-01 | -0.9 | 5.29E-02 | -0.2 | 6.32E-01 | -0.5 | 3.50E-01 | 0.3 | 7.27E-01 | NA |
| AT3G44940 | -1.4 | 1.40E-06 | -0.8 | 7.02E-05 | -0.6 | 3.67E-03 | -0.1 | 7.25E-01 | 0.3 | 3.15E-01 | -0.4 | 1.12E-01 | Uncharacterized protein F14D17_10 |
| AT1G74660 | -1.4 | 3.95E-07 | -0.6 | 8.58E-05 | -0.8 | 2.58E-04 | -0.2 | 5.16E-01 | -0.2 | 7.24E-01 | -0.1 | 8.93E-01 | Mini zinc finger 1 |
| AT1G30250 | -1.4 | 1.88E-02 | 0.1 | 9.15E-01 | -1.4 | 3.63E-02 | -1.8 | 1.66E-06 | -0.7 | 7.66E-03 | -1.1 | 4.03E-03 | Uncharacterized protein |

| | | | | | | | | | | | | | |
|---|---|---|---|---|---|---|---|---|---|---|---|---|---|
| AT1G28680 | -1.4 | 1.80E-06 | -0.4 | 6.65E-03 | -1.0 | 1.65E-04 | -0.1 | 8.29E-01 | 0.2 | 6.78E-01 | -0.3 | 5.16E-01 | F1K23.12 |
| AT5G56160 | -1.4 | 1.33E-06 | -0.8 | 5.18E-05 | -0.6 | 5.10E-03 | 0.1 | 8.73E-01 | 0.2 | 5.96E-01 | -0.2 | 7.59E-01 | Sec14p-like phosphatidylinositol transfer family protein |
| AT4G17215 | -1.4 | 4.44E-07 | -0.5 | 5.99E-04 | -0.9 | 9.04E-05 | 1.5 | 1.11E-06 | 0.9 | 7.40E-04 | 0.6 | 7.71E-03 | Pollen Ole e 1 allergen and extensin family protein |
| AT1G25370 | -1.4 | 2.87E-07 | -1.0 | 4.26E-06 | -0.4 | 7.67E-03 | -0.1 | 5.81E-01 | -0.2 | 3.58E-01 | 0.1 | 7.90E-01 | Uncharacterized protein F4F7.24 |
| AT2G15830 | -1.4 | 7.71E-07 | -1.1 | 7.36E-06 | -0.3 | 5.06E-02 | -0.6 | 7.90E-03 | -0.2 | 4.47E-01 | -0.4 | 9.05E-02 | Transmembrane protein |
| AT5G64900 | -1.4 | 2.71E-06 | -1.0 | 3.09E-05 | -0.4 | 5.77E-02 | 0.2 | 5.93E-01 | 0.0 | 9.80E-01 | 0.2 | 6.88E-01 | Precursor of elicitor peptide 1 |
| AT1G55390 | -1.4 | 4.67E-05 | -0.2 | 1.56E-01 | -1.1 | 1.26E-03 | -0.6 | 1.82E-01 | 0.3 | 4.43E-01 | -0.9 | 8.47E-02 | Cysteine |
| AT3G18450 | -1.4 | 4.10E-06 | 0.4 | 1.64E-02 | -1.7 | 1.47E-05 | -1.1 | 3.14E-04 | 0.0 | 9.74E-01 | -1.0 | 4.03E-03 | Protein PLANT CADMIUM RESISTANCE 5 |
| AT4G23496 | -1.4 | 1.63E-04 | -0.6 | 1.50E-02 | -0.8 | 2.39E-02 | -0.5 | 2.66E-01 | 0.2 | 7.41E-01 | -0.7 | 2.17E-01 | Protein SPIRAL1-like 5 |
| AT5G51850 | -1.4 | 9.09E-06 | -0.6 | 1.32E-03 | -0.7 | 4.44E-03 | -0.2 | 7.09E-01 | 0.4 | 4.37E-01 | -0.7 | 2.69E-01 | DUF3741 domain-containing protein |
| AT5G20470 | -1.4 | 3.13E-06 | -0.8 | 1.94E-04 | -0.6 | 5.01E-03 | -0.9 | 2.31E-02 | -0.1 | 7.47E-01 | -0.7 | 7.10E-02 | Myosin |
| AT5G56540 | -1.4 | 6.57E-07 | 0.2 | 1.39E-01 | -1.5 | 1.02E-05 | -0.7 | 2.62E-02 | 0.1 | 8.51E-01 | -0.8 | 1.36E-02 | Arabinogalactan protein 14 |
| AT4G22640 | -1.4 | 1.71E-05 | 0.4 | 2.94E-02 | -1.7 | 3.35E-05 | 0.5 | 3.10E-02 | 0.6 | 1.56E-02 | -0.1 | 7.06E-01 | Non-specific lipid transfer protein GPI-anchored 33 |
| AT5G49350 | -1.4 | 3.58E-06 | -0.6 | 8.68E-04 | -0.8 | 1.37E-03 | 2.4 | 1.47E-05 | 1.9 | 6.83E-04 | 0.6 | 1.03E-01 | At5g49350 |
| AT1G71740 | -1.4 | 7.29E-06 | 0.0 | 9.31E-01 | -1.3 | 8.22E-05 | -0.7 | 1.01E-01 | -0.2 | 6.83E-01 | -0.5 | 3.43E-01 | hypothetical protein |
| AT3G49750 | -1.4 | 3.30E-06 | -0.5 | 3.48E-03 | -0.9 | 4.96E-04 | -0.2 | 3.89E-01 | 0.0 | 9.23E-01 | -0.3 | 4.18E-01 | Receptor like protein |
| AT5G65390 | -1.4 | 4.45E-06 | -0.2 | 1.83E-01 | -1.2 | 9.04E-05 | -1.0 | 3.31E-04 | -0.5 | 5.64E-02 | -0.5 | 9.72E-02 | Classical arabinogalactan protein 7 |
| AT3G56500 | -1.4 | 2.55E-04 | -0.3 | 2.01E-01 | -1.1 | 5.66E-03 | | | | | | | hypothetical protein |
| AT3G52790 | -1.3 | 1.38E-04 | -0.3 | 1.09E-01 | -1.0 | 4.11E-03 | 1.8 | 5.41E-05 | 1.3 | 2.42E-03 | 0.5 | 8.22E-02 | hypothetical protein |
| AT5G43400 | -1.3 | 6.57E-07 | -0.3 | 7.06E-03 | -1.0 | 5.03E-05 | -0.3 | 2.70E-01 | 0.2 | 5.01E-01 | -0.5 | 1.31E-01 | AT5g43400 |
| AT2G04780 | -1.3 | 2.02E-06 | -0.4 | 1.30E-02 | -0.9 | 1.26E-04 | -0.3 | 5.59E-02 | 0.1 | 4.59E-01 | -0.4 | 4.61E-02 | Fasciclin-like arabinogalactan protein 7 |
| AT3G11370 | -1.3 | 4.34E-05 | -0.2 | 2.00E-01 | -1.1 | 9.82E-04 | -1.4 | 3.31E-02 | 0.3 | 6.02E-01 | -1.7 | 3.38E-02 | DC1 domain-containing protein |
| AT4G02090 | -1.3 | 3.94E-06 | 0.0 | 9.96E-01 | -1.3 | 4.76E-05 | -0.5 | 4.77E-02 | -0.1 | 8.02E-01 | -0.5 | 1.26E-01 | Uncharacterized protein |
| AT2G45720 | -1.3 | 7.71E-08 | -0.8 | 3.69E-06 | -0.5 | 2.52E-04 | -0.7 | 5.54E-03 | -0.2 | 5.44E-01 | -0.5 | 9.93E-02 | hypothetical protein |
| AT2G33205 | -1.3 | 1.82E-05 | -1.1 | 1.05E-04 | -0.2 | 2.99E-01 | | | | | | | Serinc-domain containing serine and sphingolipid biosynthesis protein |
| AT1G17620 | -1.3 | 5.55E-07 | -0.6 | 1.98E-04 | -0.7 | 2.42E-04 | -0.5 | 3.80E-03 | -0.1 | 8.09E-01 | -0.5 | 2.83E-02 | hypothetical protein |
| AT3G21710 | -1.3 | 3.88E-07 | -0.8 | 1.40E-05 | -0.5 | 2.12E-03 | -0.3 | 3.35E-01 | 0.2 | 4.70E-01 | -0.5 | 6.29E-02 | Vascular-related unknown protein 1 |
| AT5G42860 | -1.3 | 9.26E-07 | -0.5 | 9.06E-04 | -0.8 | 1.88E-04 | 0.0 | 8.99E-01 | 0.5 | 1.69E-02 | -0.5 | 3.04E-02 | Emb\|CAB53482.1 |
| AT1G17030 | -1.3 | 4.27E-05 | -0.7 | 2.08E-03 | -0.6 | 2.46E-02 | | | | | | | F20D23.27 protein |
| AT5G09440 | -1.3 | 3.91E-07 | -0.9 | 1.16E-05 | -0.4 | 3.01E-03 | -0.7 | 1.66E-03 | -0.6 | 5.45E-03 | 0.0 | 9.21E-01 | Protein EXORDIUM-like 4 |
| AT1G20990 | -1.3 | 7.80E-05 | -1.5 | 7.97E-05 | 0.2 | 5.73E-01 | | | | | | | F9H16.2 protein |
| AT5G43030 | -1.3 | 1.05E-05 | -0.1 | 6.83E-01 | -1.3 | 1.40E-04 | -0.5 | 1.96E-01 | 0.3 | 4.59E-01 | -0.9 | 8.55E-02 | CHP-rich zinc finger protein-like |
| AT3G50840 | -1.3 | 6.85E-06 | -0.7 | 4.45E-04 | -0.6 | 8.12E-03 | -0.4 | 5.36E-02 | -0.2 | 3.90E-01 | -0.2 | 4.94E-01 | Phototropic-responsive NPH3 family protein |
| AT5G11790 | -1.3 | 4.45E-07 | -1.0 | 9.31E-06 | -0.4 | 7.21E-03 | -0.4 | 1.68E-03 | 0.3 | 2.14E-02 | -0.7 | 7.58E-04 | N-MYC downregulated-like 2 |
| AT5G19580 | -1.3 | 1.41E-04 | -0.8 | 2.60E-03 | -0.5 | 1.12E-01 | 0.1 | 7.83E-01 | 0.0 | 9.79E-01 | 0.2 | 8.19E-01 | Uncharacterized protein |
| AT4G18425 | -1.3 | 2.36E-04 | -0.7 | 8.57E-03 | -0.6 | 6.33E-02 | -0.1 | 9.29E-01 | 0.6 | 3.78E-01 | | | Protein DMP4 |
| AT3G27210 | -1.3 | 6.57E-07 | -0.8 | 2.44E-05 | -0.5 | 3.24E-03 | 0.0 | 9.01E-01 | 0.1 | 6.87E-01 | -0.1 | 8.08E-01 | Uncharacterized protein At3g27210 |
| AT4G24130 | -1.3 | 3.08E-07 | -0.6 | 6.43E-05 | -0.7 | 2.24E-04 | 0.8 | 3.41E-04 | 0.5 | 2.11E-02 | 0.3 | 1.13E-01 | hypothetical protein |
| AT4G06534 | -1.3 | 2.87E-05 | -0.8 | 7.83E-04 | -0.5 | 4.46E-02 | -0.1 | 9.12E-01 | 0.1 | 8.43E-01 | -0.1 | 7.84E-01 | Uncharacterized protein |
| AT5G37550 | -1.3 | 3.57E-05 | -1.0 | 2.96E-04 | -0.3 | 2.01E-01 | 0.0 | 9.84E-01 | 0.9 | 4.01E-02 | -0.9 | 2.49E-02 | At5g37550 |
| AT3G26460 | -1.3 | 1.02E-05 | -0.7 | 1.06E-03 | -0.6 | 6.00E-03 | 0.4 | 3.32E-01 | 1.0 | 2.43E-02 | -0.6 | 2.39E-01 | Bet v I |
| AT5G13200 | -1.3 | 4.76E-06 | -1.0 | 3.78E-05 | -0.3 | 1.45E-01 | -0.1 | 8.72E-01 | 0.0 | 9.75E-01 | 0.0 | 9.34E-01 | GRAM domain family protein |
| AT1G71910 | -1.3 | 9.43E-06 | -1.1 | 4.43E-05 | -0.2 | 3.98E-01 | 0.0 | 9.86E-01 | 0.1 | 8.20E-01 | -0.1 | 8.17E-01 | hypothetical protein |
| AT1G16225 | -1.3 | 7.22E-05 | -0.9 | 8.96E-04 | -0.4 | 1.42E-01 | -0.5 | 3.45E-01 | 0.0 | 9.81E-01 | -0.5 | 3.50E-01 | Target SNARE coiled-coil domain protein |
| AT1G64890 | -1.3 | 1.06E-06 | -0.8 | 3.09E-05 | -0.4 | 7.21E-03 | -0.4 | 3.32E-02 | 0.1 | 8.35E-01 | -0.5 | 4.60E-02 | Probable folate-biopterin transporter 7 |
| AT1G22065 | -1.3 | 1.26E-06 | -1.1 | 7.83E-06 | -0.2 | 1.69E-01 | 0.1 | 9.07E-01 | 0.1 | 7.72E-01 | -0.1 | 8.58E-01 | hypothetical protein |
| AT3G21600 | -1.3 | 9.29E-07 | -0.8 | 2.72E-05 | -0.4 | 6.48E-03 | 0.0 | 9.01E-01 | -0.2 | 4.40E-01 | 0.2 | 5.65E-01 | Senescence |
| AT5G47590 | -1.3 | 6.73E-05 | -1.1 | 3.00E-04 | -0.2 | 4.45E-01 | -1.0 | 7.33E-02 | -1.7 | 1.12E-02 | 0.8 | 3.64E-01 | Heat shock protein HSP20 |
| AT5G63270 | -1.3 | 3.34E-04 | 0.3 | 1.76E-01 | -1.5 | 4.52E-04 | -1.4 | 3.67E-02 | 0.3 | 6.78E-01 | -1.7 | 4.67E-02 | hypothetical protein |
| AT5G66595 | -1.3 | 7.62E-03 | 0.4 | 1.82E-01 | -1.7 | 4.09E-03 | -1.2 | 1.71E-01 | 0.5 | 5.66E-01 | | | hypothetical protein |
| AT1G79910 | -1.3 | 4.20E-05 | -0.3 | 1.23E-01 | -1.0 | 1.29E-03 | -0.4 | 1.29E-01 | -0.1 | 8.82E-01 | -0.4 | 2.87E-01 | Regulator of Vps4 activity in the MVB pathway protein |
| AT4G30650 | -1.3 | 1.35E-04 | -0.4 | 9.52E-02 | -0.9 | 3.95E-03 | -0.2 | 4.68E-01 | 0.5 | 1.02E-01 | -0.7 | 5.55E-02 | UPF0057 membrane protein At4g30650 |
| AT5G39970 | -1.3 | 5.22E-06 | -0.5 | 3.16E-03 | -0.8 | 1.02E-03 | -0.2 | 5.20E-01 | 0.1 | 8.00E-01 | -0.3 | 4.53E-01 | Catalytics |
| AT4G22214 | -1.3 | 5.11E-04 | 0.0 | 8.26E-01 | -1.2 | 2.84E-03 | -1.4 | 9.62E-03 | -0.9 | 6.85E-02 | -0.5 | 4.14E-01 | Defensin-like protein 99 |
| AT5G06300 | -1.3 | 9.58E-08 | -1.0 | 1.06E-06 | -0.3 | 5.93E-03 | 0.1 | 5.42E-01 | 0.3 | 2.38E-01 | -0.2 | 4.55E-01 | Lysine decarboxylase-like protein |
| AT3G03440 | -1.3 | 6.44E-07 | -0.9 | 7.89E-06 | -0.3 | 2.61E-02 | 0.6 | 2.38E-02 | 0.5 | 8.54E-02 | 0.1 | 7.90E-01 | ARM repeat superfamily protein |
| AT5G23950 | -1.3 | 4.41E-06 | -1.1 | 2.11E-05 | -0.2 | 3.79E-01 | -0.1 | 9.31E-01 | 0.1 | 9.04E-01 | | | Calcium-dependent lipid-binding family protein |
| AT2G30340 | -1.3 | 4.79E-05 | -0.3 | 9.30E-02 | -1.0 | 1.59E-03 | -0.3 | 2.96E-01 | 0.3 | 4.16E-01 | -0.6 | 1.33E-01 | LOB domain-containing protein 13 |
| AT4G00880 | -1.3 | 1.30E-03 | -0.4 | 1.31E-01 | -0.8 | 4.02E-02 | -0.7 | 1.35E-01 | -0.1 | 8.10E-01 | -0.6 | 3.59E-01 | AT4g00880 protein |
| AT2G17300 | -1.3 | 5.23E-06 | -0.5 | 2.94E-03 | -0.8 | 1.11E-03 | -1.0 | 5.05E-04 | -0.1 | 7.94E-01 | -0.9 | 4.40E-03 | Uncharacterized protein |
| AT5G11290 | -1.3 | 6.25E-06 | -1.1 | 2.88E-05 | -0.2 | 4.01E-01 | | | | | | | At5g11290 ) |
| AT1G22220 | -1.3 | 9.96E-07 | -0.6 | 1.66E-04 | -0.6 | 8.33E-04 | -1.3 | 1.90E-05 | -0.3 | 1.10E-01 | -0.9 | 1.61E-03 | F-box protein AUF2 |
| AT3G59120 | -1.3 | 8.12E-05 | -0.2 | 1.66E-01 | -1.0 | 2.26E-03 | | | | | | | DC1 domain-containing protein |
| AT1G54217 | -1.3 | 4.97E-06 | -0.4 | 7.55E-03 | -0.9 | 5.48E-04 | -0.1 | 8.81E-01 | 0.2 | 6.19E-01 | -0.2 | 4.77E-01 | hypothetical protein |
| AT1G29195 | -1.2 | 5.99E-05 | -0.5 | 1.16E-02 | -0.7 | 9.54E-03 | -0.1 | 6.95E-01 | -0.1 | 8.87E-01 | -0.1 | 8.84E-01 | hypothetical protein |
| AT1G79180 | -1.2 | 4.30E-06 | -0.7 | 1.77E-04 | -0.5 | 1.23E-02 | 0.2 | 7.23E-01 | 0.5 | 2.93E-01 | -0.3 | 5.55E-01 | Myb domain protein 63 |
| AT3G22840 | -1.2 | 1.92E-04 | -1.4 | 1.53E-04 | 0.2 | 5.61E-01 | 0.4 | 6.19E-01 | 1.6 | 3.63E-02 | | | Early light-induced protein 1, chloroplastic |
| AT3G50640 | -1.2 | 1.94E-06 | 0.1 | 4.86E-01 | -1.3 | 2.69E-05 | -0.3 | 3.71E-01 | 0.3 | 5.06E-01 | -0.6 | 9.22E-02 | hypothetical protein |
| AT5G26660 | -1.2 | 4.45E-07 | -0.3 | 1.01E-02 | -1.0 | 3.54E-05 | 0.0 | 9.73E-01 | 0.4 | 2.01E-01 | -0.4 | 3.08E-01 | Myb domain protein 86 |
| AT3G50350 | -1.2 | 8.68E-07 | -0.6 | 2.18E-04 | -0.6 | 4.58E-04 | 0.7 | 2.83E-04 | 0.6 | 1.57E-03 | 0.1 | 8.10E-01 | Membrane insertase, putative |
| AT3G19340 | -1.2 | 7.71E-08 | -0.7 | 4.39E-06 | -0.5 | 1.87E-04 | -0.3 | 1.59E-02 | 0.0 | 8.58E-01 | -0.3 | 1.12E-01 | Uncharacterized protein At3g19340 |
| AT1G27620 | -1.2 | 1.27E-05 | -0.6 | 1.58E-03 | -0.7 | 6.03E-03 | 0.4 | 2.29E-01 | 0.5 | 1.17E-01 | -0.1 | 6.88E-01 | Putative hypersensitivity-related protein |
| AT3G05660 | -1.2 | 1.35E-04 | -1.0 | 5.82E-04 | -0.2 | 4.45E-01 | 0.6 | 1.91E-01 | 0.0 | 9.98E-01 | 0.6 | 2.91E-01 | RLP33 |
| AT2G01150 | -1.2 | 1.36E-05 | -0.8 | 2.53E-04 | -0.4 | 5.26E-02 | -0.3 | 1.50E-01 | 0.1 | 6.11E-01 | -0.4 | 4.80E-02 | RING-H2 finger protein 2B |
| AT4G28330 | -1.2 | 2.14E-05 | -1.2 | 4.97E-05 | 0.0 | 8.85E-01 | -0.6 | 1.91E-01 | -0.2 | 6.28E-01 | -0.3 | 5.79E-01 | Uncharacterized protein AT4g28330 |
| AT4G29700 | -1.2 | 4.44E-07 | -0.4 | 1.12E-03 | -0.8 | 6.47E-05 | -0.3 | 3.65E-01 | 0.6 | 6.21E-02 | -0.8 | 3.10E-02 | Uncharacterized protein |
| AT3G06020 | -1.2 | 6.96E-06 | -0.5 | 1.92E-03 | -0.7 | 2.16E-03 | 0.2 | 5.85E-01 | 0.1 | 7.92E-01 | 0.1 | 8.95E-01 | Protein FANTASTIC FOUR 4 |
| AT4G03820 | -1.2 | 4.90E-06 | -0.9 | 5.76E-05 | -0.3 | 7.63E-02 | -0.3 | 3.27E-01 | -0.1 | 8.49E-01 | -0.2 | 5.79E-01 | Transmembrane protein, putative |
| AT5G23830 | -1.2 | 4.59E-07 | -0.3 | 6.06E-03 | -0.9 | 3.57E-05 | -0.4 | 6.84E-02 | 0.3 | 2.03E-01 | -0.7 | 1.34E-02 | hypothetical protein |
| AT1G61080 | -1.2 | 3.90E-05 | -0.6 | 6.88E-03 | -0.7 | 7.86E-03 | 0.1 | 8.16E-01 | 0.0 | 9.30E-01 | 0.0 | 9.29E-01 | Hydroxyproline-rich glycoprotein family protein |
| AT1G03495 | -1.2 | 2.70E-03 | -1.5 | 1.49E-03 | 0.3 | 5.68E-01 | 0.8 | 3.56E-03 | 0.5 | 1.03E-01 | 0.3 | 2.12E-01 | Uncharacterized protein At1g03495 |
| AT1G18750 | -1.2 | 4.84E-04 | -0.5 | 3.32E-02 | -0.7 | 4.80E-02 | -0.4 | 6.49E-01 | 0.2 | 8.14E-01 | | | AGAMOUS-like 65 |
| AT1G49450 | -1.2 | 2.31E-07 | -1.1 | 1.08E-06 | -0.1 | 1.88E-01 | -0.4 | 3.87E-02 | 0.6 | 8.81E-03 | -1.0 | 8.65E-04 | hypothetical protein |

| | | | | | | | | | | | | | |
|---|---|---|---|---|---|---|---|---|---|---|---|---|---|
| AT3G22410 | -1.2 | 1.87E-05 | -1.1 | 5.96E-05 | -0.1 | 7.40E-01 | 0.2 | 5.79E-01 | 0.2 | 5.03E-01 | -0.1 | 9.11E-01 | Sec14p-like phosphatidylinositol transfer family protein |
| AT3G12830 | -1.2 | 2.56E-05 | -0.5 | 8.44E-03 | -0.7 | 3.91E-03 | -0.1 | 7.02E-01 | -0.3 | 2.04E-01 | 0.2 | 4.64E-01 | Auxin-responsive protein SAUR72 |
| AT5G16360 | -1.2 | 7.99E-07 | -1.0 | 5.32E-06 | -0.2 | 1.39E-01 | -0.5 | 8.82E-02 | 0.1 | 8.96E-01 | -0.5 | 8.25E-02 | LRAT domain-containing protein |
| AT3G19240 | -1.2 | 7.79E-08 | -0.8 | 2.29E-06 | -0.4 | 5.70E-04 | -0.2 | 1.71E-01 | -0.1 | 5.95E-01 | -0.1 | 5.94E-01 | Vacuolar import |
| AT2G23530 | -1.2 | 2.32E-05 | -0.7 | 7.73E-04 | -0.5 | 3.33E-02 | -0.4 | 3.99E-01 | 0.0 | 9.47E-01 | -0.4 | 4.21E-01 | Zinc-finger domain-containing protein |
| AT5G20885 | -1.2 | 7.72E-07 | -0.6 | 2.09E-04 | -0.6 | 3.77E-04 | -0.1 | 7.66E-01 | -0.1 | 7.82E-01 | 0.0 | 9.81E-01 | RING-type domain-containing protein |
| AT1G25530 | -1.2 | 2.68E-07 | -0.7 | 1.67E-05 | -0.5 | 7.14E-04 | 0.2 | 2.51E-01 | 0.3 | 5.69E-02 | -0.1 | 5.05E-01 | AT1G25530 protein |
| AT5G37210 | -1.2 | 7.02E-06 | -0.4 | 1.27E-02 | -0.9 | 7.20E-04 | -0.4 | 3.93E-01 | 0.0 | 9.86E-01 | -0.4 | 5.14E-01 | Cysteine |
| AT4G28250 | -1.2 | 6.23E-05 | -0.6 | 9.81E-03 | -0.6 | 1.03E-02 | -0.4 | 3.16E-03 | 0.4 | 1.11E-02 | -0.8 | 3.88E-04 | Putative beta-expansin |
| AT3G61270 | -1.2 | 1.34E-06 | -0.3 | 1.53E-02 | -0.9 | 8.64E-05 | -0.3 | 9.90E-02 | 0.2 | 3.85E-01 | -0.5 | 3.94E-02 | Glycosyl transferase CAP10 domain-containing protein |
| AT4G16400 | -1.2 | 1.00E-05 | -0.3 | 2.32E-02 | -0.9 | 7.05E-04 | 0.2 | 6.74E-01 | 0.4 | 2.10E-01 | -0.3 | 5.22E-01 | Transmembrane protein |
| AT5G14150 | -1.2 | 1.67E-05 | 0.0 | 7.52E-01 | -1.2 | 1.14E-04 | -0.4 | 1.47E-01 | 0.0 | 9.56E-01 | -0.5 | 2.34E-01 | DUF642 domain-containing protein |
| AT2G03260 | -1.2 | 9.07E-06 | -0.4 | 9.35E-03 | -0.8 | 1.03E-03 | 0.0 | 8.46E-01 | 0.1 | 8.12E-01 | -0.1 | 7.27E-01 | EXS family protein |
| AT2G24170 | -1.2 | 1.25E-06 | -0.4 | 2.15E-03 | -0.8 | 1.77E-04 | 0.1 | 7.27E-01 | 0.4 | 7.80E-02 | -0.3 | 2.63E-01 | Transmembrane 9 superfamily member |
| AT5G66820 | -1.2 | 1.44E-07 | -0.9 | 1.91E-06 | -0.3 | 5.96E-03 | -0.4 | 5.05E-02 | 0.0 | 8.78E-01 | -0.3 | 1.11E-01 | Transmembrane protein |
| AT5G09978 | -1.2 | 1.10E-04 | -0.3 | 8.85E-02 | -0.9 | 3.93E-03 | -1.0 | 4.93E-03 | 0.1 | 9.13E-01 | -1.1 | 5.24E-03 | Elicitor peptide 7 |
| AT5G60720 | -1.2 | 1.55E-05 | -0.4 | 1.03E-02 | -0.8 | 2.19E-03 | -0.1 | 8.15E-01 | 0.2 | 7.13E-01 | -0.3 | 5.87E-01 | AT5g60720 |
| AT5G26300 | -1.2 | 5.40E-05 | 0.1 | 6.48E-01 | -1.3 | 2.03E-04 | -0.1 | 7.60E-01 | 0.0 | 9.61E-01 | -0.1 | 7.46E-01 | TRAF-like family protein |
| AT4G15390 | -1.2 | 8.02E-07 | -0.3 | 8.97E-03 | -0.9 | 5.91E-05 | -1.2 | 3.73E-06 | -0.2 | 3.23E-01 | -1.0 | 5.73E-04 | hypothetical protein |
| AT4G35985 | -1.2 | 1.52E-07 | -0.8 | 4.55E-06 | -0.4 | 1.25E-03 | -0.6 | 2.57E-02 | 0.3 | 2.79E-01 | -0.9 | 6.00E-03 | Senescence |
| AT5G63130 | -1.2 | 6.96E-04 | -1.1 | 1.61E-03 | -0.1 | 7.46E-01 | 0.4 | 2.80E-01 | 0.4 | 4.03E-01 | 0.0 | 9.41E-01 | hypothetical protein |
| AT1G69970 | -1.2 | 4.28E-04 | -0.9 | 2.44E-03 | -0.3 | 4.13E-01 | -0.4 | 1.24E-01 | -0.1 | 7.57E-01 | -0.3 | 4.01E-01 | CLAVATA3 |
| AT2G35585 | -1.2 | 4.33E-06 | -0.6 | 2.93E-04 | -0.5 | 5.83E-03 | 0.0 | 9.63E-01 | 0.2 | 7.13E-01 | -0.2 | 6.83E-01 | Cystic fibrosis transmembrane conductance regulator |
| AT5G05350 | -1.2 | 2.10E-05 | -0.5 | 6.78E-03 | -0.7 | 4.33E-03 | | | | | | | Gb\|AAF01523.1 |
| AT2G03090 | -1.2 | 4.28E-05 | -0.3 | 1.38E-01 | -0.9 | 9.45E-04 | 0.2 | 9.18E-02 | 0.6 | 6.26E-04 | -0.3 | 3.27E-02 | Expansin-A15 |
| AT3G26440 | -1.2 | 1.32E-07 | -0.9 | 1.17E-06 | -0.2 | 1.67E-02 | 0.2 | 3.00E-01 | 0.2 | 4.82E-01 | 0.0 | 9.14E-01 | Transmembrane protein, putative |
| AT2G16580 | -1.2 | 1.31E-03 | -0.3 | 2.60E-01 | -0.9 | 2.09E-02 | -0.7 | 5.93E-02 | 0.1 | 7.86E-01 | -0.8 | 7.33E-02 | Protein SMALL AUXIN UP-REGULATED RNA 8 |
| AT3G47200 | -1.2 | 1.05E-06 | -0.5 | 5.04E-04 | -0.7 | 3.70E-04 | 0.2 | 4.59E-01 | 0.1 | 8.03E-01 | 0.1 | 7.01E-01 | UPF0481 protein At3g47200 |
| AT5G01790 | -1.2 | 1.49E-05 | -0.5 | 3.47E-03 | -0.6 | 3.81E-03 | -0.3 | 2.70E-01 | 0.6 | 1.99E-02 | -0.9 | 3.49E-03 | At5g01790 |
| AT4G27435 | -1.2 | 1.12E-06 | -0.2 | 3.23E-02 | -1.0 | 6.15E-05 | -0.5 | 2.33E-02 | -0.2 | 4.84E-01 | -0.4 | 2.25E-01 | hypothetical protein |
| AT5G44690 | -1.2 | 1.90E-04 | -0.6 | 1.07E-02 | -0.6 | 4.72E-02 | | | | | | | RING finger PFF0165c-like protein |
| AT2G32140 | -1.2 | 9.67E-04 | -1.4 | 6.41E-04 | 0.2 | 6.02E-01 | 0.2 | 7.23E-01 | 0.2 | 8.16E-01 | | | Transmembrane receptor |
| AT5G06930 | -1.2 | 7.75E-07 | -0.3 | 3.71E-03 | -0.9 | 8.89E-05 | -0.3 | 5.03E-01 | 0.1 | 7.99E-01 | -0.4 | 4.55E-01 | Nucleolar-like protein |
| AT5G62900 | -1.2 | 7.15E-06 | -0.8 | 1.23E-04 | -0.4 | 4.78E-02 | -0.5 | 7.47E-03 | -0.5 | 9.23E-03 | 0.0 | 9.26E-01 | hypothetical protein |
| AT1G76820 | -1.2 | 4.57E-04 | -0.7 | 6.66E-03 | -0.4 | 1.71E-01 | 0.0 | 9.73E-01 | -0.1 | 8.01E-01 | 0.1 | 8.51E-01 | NA |
| AT3G49070 | -1.2 | 8.13E-06 | -0.4 | 3.60E-03 | -0.7 | 1.96E-03 | | | | | | | Transmembrane protein, putative |
| AT1G55280 | -1.2 | 3.11E-05 | -0.8 | 6.00E-04 | -0.4 | 7.71E-02 | -0.6 | 7.10E-02 | -0.3 | 4.64E-01 | -0.3 | 5.10E-01 | Lipase |
| AT5G65980 | -1.2 | 8.45E-05 | -0.3 | 8.46E-02 | -0.8 | 2.62E-03 | -0.2 | 4.19E-01 | -0.1 | 7.41E-01 | -0.1 | 8.22E-01 | Auxin efflux carrier family protein |
| AT4G20160 | -1.2 | 8.26E-06 | 0.2 | 1.32E-01 | -1.3 | 3.97E-05 | -0.5 | 2.96E-01 | 0.2 | 7.62E-01 | -0.6 | 2.19E-01 | Glu-rich protein |
| AT1G76952 | -1.2 | 6.63E-04 | -0.6 | 1.90E-02 | -0.5 | 9.98E-02 | 0.3 | 4.05E-01 | 0.3 | 5.20E-01 | 0.0 | 9.65E-01 | Inflorescence deficient in abscission -like 5 |
| AT4G35560 | -1.2 | 1.60E-07 | -0.8 | 2.90E-06 | -0.3 | 3.61E-03 | -0.4 | 1.23E-01 | 0.3 | 3.18E-01 | -0.6 | 2.13E-02 | Uncharacterized protein At4g35560 |
| AT4G24960 | -1.2 | 1.27E-05 | -0.8 | 2.72E-04 | -0.4 | 4.22E-02 | -0.2 | 4.26E-01 | 0.3 | 9.53E-02 | -0.5 | 3.95E-02 | HVA22-like protein |
| AT3G09790 | -1.2 | 3.03E-05 | -0.9 | 2.09E-04 | -0.2 | 2.86E-01 | -0.8 | 1.83E-02 | -0.2 | 5.17E-01 | -0.6 | 1.87E-01 | Ubiquitin-like domain-containing protein |
| AT1G62370 | -1.2 | 1.57E-06 | -0.9 | 1.89E-05 | -0.3 | 4.14E-02 | -0.8 | 3.91E-03 | 0.2 | 4.76E-01 | -1.0 | 5.23E-03 | F2401.10 |
| AT4G12360 | -1.2 | 5.86E-04 | 0.8 | 2.77E-03 | -2.0 | 9.04E-05 | -2.0 | 9.21E-05 | -0.1 | 7.98E-01 | -1.9 | 2.97E-03 | Non-specific lipid transfer protein GPI-anchored 24 |
| AT1G53790 | -1.2 | 1.02E-05 | -0.8 | 1.65E-04 | -0.4 | 6.83E-02 | -0.6 | 3.04E-01 | -0.1 | 9.26E-01 | -0.5 | 4.66E-01 | hypothetical protein |
| AT1G68600 | -1.2 | 4.36E-05 | 0.6 | 1.91E-03 | -1.8 | 2.69E-05 | -1.3 | 5.61E-07 | 0.4 | 1.33E-02 | -1.7 | 1.01E-05 | Aluminum-activated malate transporter 5 |
| AT4G23680 | -1.2 | 1.07E-05 | -1.9 | 1.38E-06 | 0.7 | 2.07E-03 | 0.2 | 5.64E-01 | 0.6 | 1.26E-01 | -0.4 | 4.49E-01 | Bet v I |
| AT1G11700 | -1.1 | 9.84E-07 | -0.8 | 2.41E-05 | -0.4 | 8.32E-03 | -0.1 | 3.81E-01 | 0.5 | 5.70E-03 | -0.6 | 6.38E-03 | hypothetical protein |
| AT1G56020 | -1.1 | 3.38E-06 | -0.6 | 3.52E-04 | -0.5 | 3.42E-03 | -0.1 | 5.55E-01 | 0.0 | 9.83E-01 | -0.2 | 5.89E-01 | At1g56020 |
| AT1G27420 | -1.1 | 1.01E-05 | -0.5 | 3.37E-03 | -0.7 | 2.54E-03 | -0.9 | 1.68E-02 | -0.5 | 1.88E-01 | -0.4 | 3.46E-01 | Putative F-box |
| AT2G46140 | -1.1 | 9.72E-06 | 0.0 | 7.55E-01 | -1.1 | 8.75E-05 | -0.2 | 4.39E-01 | 0.2 | 3.98E-01 | -0.4 | 9.25E-02 | Desiccation-related protein At2g46140 |
| AT3G44326 | -1.1 | 2.88E-04 | -0.8 | 2.53E-03 | -0.3 | 2.44E-01 | -0.3 | 6.32E-01 | 0.3 | 6.07E-01 | -0.6 | 3.56E-01 | F-box protein At3g44326 |
| AT5G54240 | -1.1 | 5.49E-06 | 0.1 | 6.02E-01 | -1.2 | 5.17E-05 | -0.2 | 6.33E-01 | 0.1 | 8.10E-01 | -0.2 | 5.17E-01 | hypothetical protein |
| AT3G61380 | -1.1 | 2.33E-06 | -0.4 | 2.15E-03 | -0.7 | 4.23E-04 | -0.7 | 4.09E-04 | 0.0 | 9.43E-01 | -0.7 | 4.72E-03 | Phosphatidylinositol N-acetyglucosaminlytransferase subunit P-like protein |
| AT1G79760 | -1.1 | 4.98E-06 | 0.0 | 9.84E-01 | -1.1 | 6.05E-05 | -0.5 | 4.07E-02 | -0.2 | 5.83E-01 | -0.3 | 2.04E-01 | Downstream target of AGL15-4 |
| AT4G26950 | -1.1 | 6.88E-03 | -0.8 | 2.91E-02 | -0.3 | 5.25E-01 | 0.4 | 3.96E-01 | 0.9 | 4.55E-02 | -0.5 | 2.78E-01 | Senescence regulator |
| AT3G53950 | -1.1 | 2.42E-07 | -0.7 | 9.11E-06 | -0.4 | 1.42E-03 | -0.4 | 2.99E-03 | -0.3 | 5.07E-02 | -0.1 | 4.52E-01 | hypothetical protein |
| AT5G65160 | -1.1 | 3.81E-06 | -0.2 | 1.73E-01 | -1.0 | 1.01E-04 | -0.3 | 2.74E-01 | 0.2 | 5.50E-01 | -0.4 | 8.09E-02 | At5g65160 -containing protein) |
| AT5G02090 | -1.1 | 2.54E-04 | -0.7 | 4.03E-03 | -0.4 | 1.41E-01 | -0.7 | 4.82E-02 | -0.2 | 6.15E-01 | -0.5 | 3.19E-01 | Uncharacterized protein |
| AT1G23040 | -1.1 | 2.76E-06 | -0.5 | 8.46E-04 | -0.6 | 1.02E-03 | -0.2 | 4.61E-01 | -0.1 | 8.42E-01 | -0.1 | 7.28E-01 | hypothetical protein |
| AT1G65985 | -1.1 | 6.53E-07 | -0.5 | 3.94E-04 | -0.7 | 2.00E-04 | 0.0 | 8.81E-01 | 0.5 | 1.11E-02 | -0.4 | 3.56E-02 | At1g65985 ) |
| AT5G03170 | -1.1 | 1.58E-06 | -0.2 | 8.79E-02 | -1.0 | 5.83E-05 | -0.2 | 3.35E-01 | 0.0 | 8.56E-01 | -0.2 | 3.46E-01 | Fasciclin-like arabinogalactan protein 11 |
| AT3G54000 | -1.1 | 1.57E-06 | -0.6 | 9.47E-05 | -0.5 | 3.82E-03 | -0.5 | 4.46E-02 | -0.3 | 3.20E-01 | -0.2 | 5.14E-01 | Uncharacterized protein F5K20_300 |
| AT5G14330 | -1.1 | 3.52E-05 | 0.6 | 6.03E-03 | -1.7 | 2.69E-05 | -1.1 | 4.10E-04 | -0.1 | 8.62E-01 | -1.1 | 2.67E-03 | hypothetical protein |
| AT4G39720 | -1.1 | 1.51E-06 | -0.8 | 3.47E-05 | -0.4 | 1.41E-02 | 0.4 | 2.09E-01 | 0.3 | 4.09E-01 | 0.1 | 8.56E-01 | VQ domain-containing protein |
| AT3G24210 | -1.1 | 2.75E-05 | 0.1 | 5.96E-01 | -1.2 | 1.44E-04 | -0.6 | 3.97E-02 | 0.4 | 1.50E-01 | -1.0 | 9.43E-03 | Uncharacterized protein |
| AT2G17070 | -1.1 | 6.36E-03 | 0.3 | 3.27E-01 | -1.4 | 5.54E-03 | -0.4 | 4.62E-01 | -0.1 | 8.95E-01 | | | Uncharacterized protein |
| AT4G03490 | -1.1 | 8.62E-05 | -0.2 | 2.03E-01 | -0.9 | 2.16E-03 | | | | | | | Ankyrin repeat family protein |
| AT5G27030 | -1.1 | 1.95E-07 | -0.9 | 2.10E-06 | -0.2 | 1.51E-02 | 0.3 | 3.44E-01 | 0.4 | 2.32E-01 | -0.1 | 8.16E-01 | TOPLESS-related 3 |
| AT4G03610 | -1.1 | 4.51E-06 | -0.7 | 1.23E-04 | -0.4 | 2.25E-02 | | | | | | | Metallo-hydrolase |
| AT3G27997 | -1.1 | 6.27E-06 | -0.8 | 9.14E-05 | -0.3 | 7.29E-02 | -0.3 | 1.85E-01 | -0.1 | 8.20E-01 | -0.3 | 4.40E-01 | NA |
| AT1G29240 | -1.1 | 2.80E-05 | -0.7 | 6.35E-04 | -0.4 | 5.72E-02 | 0.2 | 5.20E-01 | 0.1 | 6.53E-01 | 0.0 | 9.64E-01 | hypothetical protein |
| AT5G60520 | -1.1 | 4.05E-05 | 0.0 | 8.52E-01 | -1.2 | 2.21E-04 | -0.2 | 3.72E-01 | 0.2 | 3.62E-01 | -0.5 | 9.72E-02 | hypothetical protein |
| AT1G54950 | -1.1 | 6.77E-07 | -0.1 | 2.91E-01 | -1.0 | 2.50E-05 | -0.2 | 3.76E-01 | 0.2 | 5.46E-01 | -0.4 | 6.53E-02 | Transmembrane protein |
| AT5G52990 | -1.1 | 3.27E-05 | -0.1 | 4.11E-01 | -1.0 | 4.76E-04 | -0.7 | 1.51E-02 | -0.1 | 6.53E-01 | -0.5 | 1.35E-01 | Phytolongin Phyl2.2 |
| AT3G52460 | -1.1 | 2.46E-05 | 0.0 | 8.22E-01 | -1.1 | 1.59E-04 | -0.2 | 4.65E-01 | 0.0 | 9.08E-01 | -0.2 | 6.76E-01 | Late embryogenesis abundant protein LEA-2 subgroup domain-containing protein |
| AT1G70990 | -1.1 | 1.39E-05 | 0.0 | 7.20E-01 | -1.1 | 1.59E-04 | -0.2 | 3.99E-01 | 0.0 | 9.20E-01 | -0.2 | 5.48E-01 | At1g70990 |
| AT3G44820 | -1.1 | 1.14E-06 | -0.9 | 1.22E-05 | -0.3 | 5.27E-02 | -0.8 | 1.32E-03 | -0.5 | 3.83E-02 | -0.3 | 3.05E-01 | BTB |
| AT5G53420 | -1.1 | 3.24E-05 | -1.4 | 1.21E-05 | 0.3 | 8.64E-02 | -0.1 | 6.18E-01 | 0.5 | 4.65E-03 | -0.6 | 7.34E-03 | CCT domain-containing protein |
| AT3G27884 | -1.1 | 2.86E-05 | -1.0 | 8.95E-05 | -0.1 | 6.38E-01 | | | | | | | NA |
| AT1G23340 | -1.1 | 1.33E-05 | -0.4 | 1.01E-02 | -0.7 | 1.72E-03 | -0.1 | 7.44E-01 | 0.2 | 4.68E-01 | -0.3 | 3.64E-01 | At1g23340 ) |
| AT5G40510 | -1.1 | 5.38E-07 | 0.0 | 8.38E-01 | -1.1 | 1.26E-05 | -0.3 | 8.19E-02 | 0.4 | 1.17E-02 | -0.7 | 9.75E-04 | Sucrose cleavage protein-like |

| | | | | | | | | | | | | | |
|---|---|---|---|---|---|---|---|---|---|---|---|---|---|
| AT5G53590 | -1.1 | 6.01E-06 | -0.7 | 1.65E-04 | -0.4 | 2.35E-02 | -0.2 | 1.64E-01 | 0.3 | 9.31E-02 | -0.6 | 5.95E-03 | At5g53590 |
| AT1G30860 | -1.1 | 3.40E-04 | -0.6 | 8.58E-03 | -0.5 | 1.12E-01 | | | | | | | At1g30860 |
| AT4G11210 | -1.1 | 1.65E-03 | 0.4 | 4.94E-02 | -1.5 | 8.00E-04 | -1.9 | 7.73E-04 | -1.4 | 4.32E-03 | -0.5 | 5.43E-01 | Dirigent protein 14 |
| AT1G70690 | -1.1 | 2.72E-05 | -0.8 | 3.65E-04 | -0.3 | 1.07E-01 | 0.4 | 1.52E-02 | 0.1 | 6.42E-01 | 0.3 | 6.50E-02 | Plasmodesmata-located protein 5 |
| AT3G62570 | -1.1 | 9.36E-06 | 0.0 | 9.63E-01 | -1.1 | 9.47E-05 | -0.5 | 7.78E-03 | -0.1 | 7.99E-01 | -0.5 | 6.07E-02 | hypothetical protein |
| AT3G01513 | -1.1 | 2.23E-05 | -0.2 | 1.89E-01 | -0.9 | 5.89E-04 | -0.2 | 6.88E-01 | -0.3 | 5.76E-01 | 0.1 | 8.94E-01 | Uncharacterized protein |
| AT5G60530 | -1.1 | 1.33E-04 | 0.1 | 7.66E-01 | -1.2 | 3.85E-04 | -0.2 | 5.34E-01 | 0.3 | 3.75E-01 | -0.5 | 9.66E-02 | Late embryogenesis abundant protein-like protein |
| AT4G30320 | -1.1 | 2.50E-04 | 0.5 | 9.23E-03 | -1.6 | 1.07E-04 | -2.5 | 1.66E-05 | -0.4 | 2.16E-01 | -2.1 | 1.73E-03 | hypothetical protein |
| AT1G52910 | -1.1 | 6.94E-06 | -0.2 | 8.57E-02 | -0.9 | 2.45E-04 | -0.6 | 3.35E-02 | 0.1 | 6.14E-01 | -0.7 | 1.49E-02 | Fiber |
| AT2G41830 | -1.1 | 2.60E-06 | -0.5 | 8.65E-04 | -0.6 | 1.05E-03 | -0.4 | 1.70E-01 | -0.2 | 6.54E-01 | -0.3 | 5.84E-01 | ARM repeat superfamily protein |
| AT4G39180 | -1.1 | 1.90E-05 | -0.9 | 1.47E-04 | -0.2 | 2.22E-01 | -0.4 | 1.64E-01 | 0.0 | 9.13E-01 | -0.4 | 2.10E-01 | Sec14p-like phosphatidylinositol transfer family protein |
| AT3G60110 | -1.1 | 1.15E-06 | -0.9 | 1.04E-05 | -0.2 | 7.53E-02 | -0.1 | 8.33E-01 | 0.0 | 9.94E-01 | -0.1 | 8.77E-01 | Bromo domain-containing protein |
| AT1G80820 | -1.1 | 3.02E-06 | -1.0 | 1.42E-05 | -0.1 | 3.26E-01 | 2.6 | 6.05E-07 | 1.4 | 1.71E-03 | 1.2 | 4.99E-04 | Cinnamoyl coa reductase |
| AT5G16380 | -1.1 | 3.32E-06 | -0.6 | 2.87E-04 | -0.5 | 4.04E-03 | 0.0 | 8.93E-01 | 0.0 | 9.70E-01 | 0.0 | 8.81E-01 | Gb\|AAF02163.1 |
| AT4G14060 | -1.1 | 3.03E-06 | -1.2 | 4.71E-06 | 0.1 | 6.27E-01 | -0.9 | 5.22E-02 | -0.3 | 5.70E-01 | -0.6 | 3.52E-01 | AT4g14060 |
| AT4G35320 | -1.1 | 4.99E-05 | -0.8 | 7.71E-04 | -0.3 | 1.01E-01 | -0.8 | 2.42E-04 | 0.1 | 8.37E-01 | -0.9 | 2.43E-03 | Uncharacterized protein AT4g35320 |
| AT1G07710 | -1.1 | 1.04E-04 | -0.6 | 5.53E-03 | -0.5 | 4.05E-02 | | | | | | | Ankyrin repeat family protein |
| AT5G21050 | -1.1 | 1.25E-04 | 0.1 | 7.03E-01 | -1.1 | 5.63E-04 | -0.3 | 3.43E-01 | 0.4 | 2.79E-01 | -0.7 | 9.11E-02 | Hyccin |
| AT2G40113 | -1.1 | 1.85E-05 | 0.0 | 9.98E-01 | -1.1 | 1.59E-04 | -0.1 | 7.64E-01 | 0.0 | 9.46E-01 | -0.2 | 7.56E-01 | Pollen Ole e 1 allergen and extensin family protein |
| AT4G10720 | -1.1 | 1.55E-04 | -1.0 | 4.11E-04 | -0.1 | 7.49E-01 | 0.6 | 2.19E-01 | 0.1 | 9.41E-01 | 0.5 | 3.84E-01 | Ankyrin repeat family protein |
| AT3G06035 | -1.1 | 1.26E-05 | -0.4 | 1.58E-02 | -0.7 | 9.47E-04 | 0.1 | 7.84E-01 | 0.6 | 4.01E-03 | -0.5 | 2.58E-02 | Uncharacterized GPI-anchored protein At3g06035 |
| AT1G75160 | -1.1 | 9.65E-04 | 0.1 | 5.63E-01 | -1.2 | 2.20E-03 | | | | | | | Uncharacterized protein F22H5.11 |
| AT2G22000 | -1.1 | 1.10E-05 | -0.6 | 5.03E-04 | -0.4 | 1.70E-02 | 0.3 | 2.64E-01 | 0.2 | 4.31E-01 | 0.1 | 9.08E-01 | Elicitor peptide 6 |
| AT5G24990 | -1.1 | 4.81E-05 | -0.7 | 7.37E-04 | -0.3 | 1.12E-01 | 0.0 | 9.72E-01 | 0.5 | 7.63E-02 | -0.5 | 1.14E-01 | Enhanced disease resistance-like protein |
| AT5G11970 | -1.1 | 2.82E-04 | 0.0 | 8.93E-01 | -1.0 | 1.77E-03 | -0.2 | 6.44E-01 | -0.2 | 7.67E-01 | 0.0 | 9.53E-01 | hypothetical protein |
| AT3G42180 | -1.1 | 1.31E-05 | -0.6 | 4.78E-04 | -0.4 | 2.71E-02 | 1.2 | 2.39E-02 | 0.8 | 1.65E-01 | 0.4 | 4.08E-01 | Exostosin family protein |
| AT1G79030 | -1.1 | 3.08E-07 | -0.9 | 2.52E-06 | -0.2 | 5.12E-02 | -0.5 | 2.11E-03 | 0.0 | 9.98E-01 | -0.5 | 1.06E-02 | hypothetical protein |
| AT1G22470 | -1.1 | 4.61E-04 | -0.4 | 5.53E-02 | -0.7 | 2.89E-02 | -0.4 | 3.19E-01 | 0.0 | 9.65E-01 | -0.5 | 3.09E-01 | F12K8.18 protein |
| AT1G77400 | -1.1 | 6.25E-04 | -0.6 | 1.86E-02 | -0.5 | 9.42E-02 | 0.1 | 8.01E-01 | -0.2 | 5.92E-01 | 0.2 | 5.03E-01 | Uncharacterized protein |
| AT2G45820 | -1.1 | 2.25E-07 | -0.3 | 1.10E-03 | -0.7 | 3.12E-05 | -0.6 | 1.46E-03 | -0.4 | 3.87E-02 | -0.2 | 4.02E-01 | Remorin |
| AT4G01925 | -1.1 | 3.47E-05 | -0.2 | 1.87E-01 | -0.9 | 9.38E-04 | -0.7 | 1.59E-01 | -0.4 | 4.37E-01 | -0.3 | 6.77E-01 | Cysteine |
| AT5G39090 | -1.1 | 1.12E-05 | -1.1 | 1.65E-05 | 0.1 | 7.10E-01 | 0.4 | 4.17E-01 | 0.0 | 9.52E-01 | 0.3 | 5.50E-01 | Uncharacterized protein |
| AT5G56980 | -1.1 | 1.57E-06 | -0.8 | 3.03E-05 | -0.3 | 1.92E-02 | 0.2 | 5.85E-01 | 0.3 | 4.36E-01 | -0.1 | 8.40E-01 | Pathogen-associated molecular patterns-induced protein A70 |
| AT1G13920 | -1.1 | 6.84E-05 | -0.7 | 1.85E-03 | -0.4 | 6.20E-02 | -0.5 | 1.20E-01 | -0.2 | 5.90E-01 | -0.3 | 5.27E-01 | F7A19.1 protein |
| AT5G66600 | -1.1 | 4.26E-06 | -0.3 | 1.00E-02 | -0.7 | 4.16E-04 | -0.6 | 4.65E-02 | 0.0 | 9.76E-01 | -0.6 | 1.09E-01 | Uncharacterized protein |
| AT2G25737 | -1.1 | 3.67E-07 | -0.6 | 2.86E-05 | -0.5 | 8.17E-04 | 0.2 | 2.63E-01 | 0.2 | 2.80E-01 | 0.0 | 9.59E-01 | Sulfite exporter TauE |
| AT3G18777 | -1.1 | 3.69E-04 | -0.2 | 2.41E-01 | -0.8 | 7.31E-03 | | | | | | | NA |
| AT4G00440 | -1.1 | 1.16E-06 | -0.7 | 3.83E-05 | -0.4 | 6.17E-03 | -0.2 | 3.88E-01 | 0.2 | 4.60E-01 | -0.4 | 1.63E-01 | GPI-anchored adhesin-like protein, putative |
| AT2G39435 | -1.1 | 2.13E-05 | -0.7 | 3.81E-04 | -0.3 | 7.33E-02 | -0.4 | 1.02E-01 | 0.1 | 7.49E-01 | -0.5 | 8.69E-02 | Uncharacterized protein At2g39440 |
| AT3G00720 | -1.1 | 5.87E-05 | -0.7 | 7.74E-04 | -0.3 | 1.45E-01 | 0.1 | 8.05E-01 | 0.0 | 9.92E-01 | 0.1 | 8.37E-01 | NA |
| AT4G39140 | -1.1 | 1.83E-07 | -1.1 | 3.88E-07 | 0.0 | 6.16E-01 | -0.4 | 5.22E-02 | -0.3 | 2.06E-01 | -0.1 | 7.00E-01 | hypothetical protein |
| AT3G50300 | -1.1 | 2.10E-05 | -0.5 | 3.66E-03 | -0.6 | 6.25E-03 | -0.6 | 1.24E-02 | 0.4 | 1.04E-01 | -1.0 | 2.97E-03 | HXXXD-type acyl-transferase family protein |
| AT4G38060 | -1.1 | 5.20E-06 | -0.9 | 2.38E-05 | -0.1 | 3.74E-01 | -0.2 | 4.49E-01 | 0.0 | 9.24E-01 | -0.2 | 4.64E-01 | Putative protei |
| AT3G24750 | -1.1 | 7.76E-05 | 0.3 | 3.11E-02 | -1.4 | 1.00E-04 | -0.6 | 1.26E-01 | 0.2 | 6.61E-01 | -0.8 | 5.83E-02 | At3g24750 |
| AT5G03510 | -1.0 | 3.34E-06 | -0.3 | 6.65E-03 | -0.7 | 3.86E-04 | -0.5 | 1.53E-01 | -0.2 | 6.20E-01 | -0.3 | 4.62E-01 | C2H2-type domain-containing protein |
| AT3G58000 | -1.0 | 1.03E-03 | 0.6 | 6.80E-03 | -1.7 | 2.19E-04 | -1.5 | 3.02E-03 | -0.1 | 8.19E-01 | -1.4 | 8.43E-03 | VQ motif-containing protein 25 |
| AT3G50150 | -1.0 | 4.28E-04 | 0.1 | 5.65E-01 | -1.2 | 1.21E-03 | | | | | | | Transmembrane protein, putative |
| AT5G60490 | -1.0 | 7.38E-06 | -0.2 | 1.54E-01 | -0.9 | 2.03E-04 | -0.4 | 9.99E-02 | -0.2 | 4.96E-01 | -0.2 | 4.65E-01 | FLA12 |
| AT3G17860 | -1.0 | 1.06E-06 | -0.4 | 1.55E-03 | -0.7 | 1.77E-04 | -0.2 | 4.59E-01 | 0.1 | 6.21E-01 | -0.3 | 3.23E-01 | Protein TIFY |
| AT3G10250 | -1.0 | 4.13E-07 | -0.8 | 5.47E-06 | -0.3 | 1.78E-02 | -0.4 | 1.11E-02 | -0.3 | 6.15E-02 | -0.1 | 7.22E-01 | Histidine-tRNA ligase |
| AT2G17080 | -1.0 | 2.11E-04 | -0.3 | 9.94E-02 | -0.8 | 8.15E-03 | 0.1 | 9.04E-01 | 0.4 | 3.62E-01 | -0.3 | 4.54E-01 | Uncharacterized protein |
| AT4G15563 | -1.0 | 3.90E-06 | -0.8 | 4.16E-05 | -0.2 | 9.17E-02 | -0.1 | 7.40E-01 | 0.0 | 9.65E-01 | -0.1 | 8.37E-01 | At4g15563 |
| AT3G53350 | -1.0 | 9.51E-07 | -0.8 | 1.07E-05 | -0.2 | 4.38E-02 | -0.3 | 2.78E-01 | 0.1 | 7.04E-01 | -0.4 | 1.74E-01 | ROP interactive partner 4 |
| AT1G60390 | -1.0 | 6.97E-07 | -0.6 | 3.37E-05 | -0.4 | 2.50E-03 | 0.0 | 9.15E-01 | 0.3 | 2.70E-01 | -0.2 | 3.92E-01 | Polygalacturonase 1 beta-like protein 2 |
| AT2G36290 | -1.0 | 1.16E-06 | -0.5 | 4.35E-04 | -0.6 | 4.21E-04 | 0.1 | 5.52E-01 | 0.4 | 4.44E-02 | -0.3 | 1.05E-01 | Alpha |
| AT1G44050 | -1.0 | 1.24E-05 | -0.4 | 8.87E-03 | -0.7 | 1.64E-03 | -0.7 | 1.34E-02 | 0.1 | 6.82E-01 | -0.8 | 2.16E-02 | Cysteine |
| AT4G30670 | -1.0 | 2.40E-05 | 0.2 | 1.74E-01 | -1.2 | 5.25E-05 | -0.9 | 1.26E-03 | -0.1 | 6.73E-01 | -0.7 | 1.14E-02 | Uncharacterized protein AT4g30670 |
| AT5G56747 | -1.0 | 8.04E-06 | -0.5 | 1.63E-03 | -0.6 | 3.33E-03 | -0.1 | 5.54E-01 | -0.1 | 7.18E-01 | 0.0 | 9.36E-01 | NA |
| AT1G06930 | -1.0 | 2.33E-04 | 0.3 | 6.09E-02 | -1.4 | 2.35E-04 | -0.4 | 2.06E-01 | 0.5 | 1.60E-01 | -0.9 | 2.75E-02 | hypothetical protein |
| AT4G34380 | -1.0 | 1.90E-03 | -0.2 | 3.27E-01 | -0.8 | 2.60E-02 | 0.0 | 9.80E-01 | 0.3 | 5.25E-01 | -0.4 | 3.70E-01 | Uncharacterized protein |
| AT4G17230 | -1.0 | 1.39E-06 | -1.0 | 3.09E-06 | 0.0 | 9.56E-01 | -0.3 | 6.90E-02 | -0.1 | 5.81E-01 | -0.2 | 4.32E-01 | SCARECROW-like 13 |
| AT4G33550 | -1.0 | 2.93E-02 | -0.9 | 4.69E-02 | -0.1 | 8.80E-01 | -0.4 | 5.08E-01 | 0.2 | 7.80E-01 | | | At4g33550 |
| AT1G30990 | -1.0 | 2.37E-03 | -0.4 | 1.19E-01 | -0.6 | 7.49E-02 | -0.1 | 8.26E-01 | 0.6 | 1.77E-01 | -0.7 | 1.35E-01 | hypothetical protein |
| AT5G64510 | -1.0 | 1.70E-02 | -0.6 | 1.28E-01 | -0.5 | 3.56E-01 | | | | | | | Tunicamycin induced protein |
| AT1G07620 | -1.0 | 1.72E-05 | -1.1 | 1.85E-05 | 0.1 | 5.46E-01 | 0.7 | 3.20E-03 | 0.4 | 1.28E-01 | 0.3 | 2.52E-01 | GTP-binding protein Obg |
| AT5G55970 | -1.0 | 1.05E-05 | -0.7 | 1.63E-04 | -0.3 | 7.14E-02 | 0.3 | 1.48E-01 | 0.0 | 9.59E-01 | 0.3 | 2.52E-01 | Similarity to unknown protein |
| AT4G30240 | -1.0 | 3.75E-07 | -0.7 | 7.33E-06 | -0.3 | 7.24E-03 | -0.3 | 4.45E-02 | -0.1 | 5.53E-01 | -0.2 | 3.34E-01 | Uncharacterized protein AT4g30240 |
| AT2G28710 | -1.0 | 9.20E-04 | -0.6 | 1.14E-02 | -0.4 | 2.17E-01 | -0.3 | 3.31E-01 | -0.6 | 1.43E-01 | 0.2 | 6.90E-01 | hypothetical protein |
| AT1G68340 | -1.0 | 2.68E-04 | -0.6 | 9.37E-03 | -0.5 | 6.76E-02 | 0.2 | 5.26E-01 | 0.2 | 6.83E-01 | 0.0 | 9.42E-01 | Uncharacterized protein |
| AT1G29300 | -1.0 | 9.10E-07 | -0.8 | 1.07E-05 | -0.2 | 4.00E-02 | 0.0 | 8.26E-01 | 0.4 | 5.56E-02 | -0.4 | 6.33E-02 | UNE1 |
| AT3G27270 | -1.0 | 7.17E-06 | -0.4 | 2.42E-03 | -0.6 | 2.04E-03 | 1.4 | 1.82E-04 | 0.9 | 1.23E-02 | 0.5 | 5.76E-02 | TLC domain-containing protein |
| AT1G10040 | -1.0 | 4.09E-06 | -0.7 | 5.57E-05 | -0.3 | 6.35E-02 | -0.2 | 6.48E-01 | -0.1 | 8.34E-01 | -0.1 | 9.10E-01 | Alpha |
| AT1G22930 | -1.0 | 8.07E-06 | -1.1 | 1.10E-05 | 0.1 | 5.63E-01 | -0.4 | 2.41E-03 | -0.3 | 4.35E-02 | -0.1 | 4.98E-01 | T-complex protein 11 |
| AT3G05990 | -1.0 | 2.94E-06 | -0.5 | 1.09E-03 | -0.6 | 8.81E-04 | -0.4 | 1.74E-02 | 0.0 | 9.46E-01 | -0.4 | 6.83E-02 | At3g05990 family protein) |
| AT5G47180 | -1.0 | 7.19E-07 | -0.6 | 6.39E-05 | -0.4 | 1.25E-03 | -0.5 | 3.74E-03 | -0.3 | 1.03E-01 | -0.2 | 3.20E-01 | Vesicle-associated protein 2-1 |
| AT1G64360 | -1.0 | 2.62E-04 | -0.8 | 1.22E-03 | -0.2 | 4.57E-01 | -0.3 | 1.82E-01 | -0.1 | 7.79E-01 | -0.2 | 5.12E-01 | hypothetical protein |
| AT1G61840 | -1.0 | 1.29E-04 | -0.3 | 9.04E-02 | -0.7 | 5.63E-03 | 0.2 | 6.79E-01 | 0.9 | 1.69E-02 | -0.7 | 6.16E-02 | Cysteine |
| AT5G40730 | -1.0 | 1.47E-04 | 0.5 | 1.27E-02 | -1.5 | 6.46E-05 | -1.5 | 3.66E-05 | -0.3 | 3.01E-01 | -1.2 | 2.32E-03 | Arabinogalactan protein 24 |
| AT2G44010 | -1.0 | 6.53E-06 | -0.5 | 5.85E-04 | -0.5 | 6.29E-03 | 0.4 | 1.03E-01 | 0.5 | 6.90E-02 | -0.1 | 8.13E-01 | hypothetical protein |
| AT3G19970 | -1.0 | 5.71E-06 | -0.9 | 2.35E-05 | -0.1 | 4.67E-01 | -0.3 | 3.38E-01 | -0.1 | 8.33E-01 | -0.2 | 6.31E-01 | hypothetical protein |
| AT1G09925 | -1.0 | 7.92E-04 | -1.0 | 1.09E-03 | 0.0 | 9.57E-01 | -0.5 | 2.81E-01 | 0.1 | 8.77E-01 | -0.5 | 3.09E-01 | NA |
| AT3G21550 | -1.0 | 4.16E-06 | -0.1 | 5.29E-01 | -0.9 | 7.12E-05 | 0.0 | 9.75E-01 | 0.0 | 9.27E-01 | 0.0 | 8.99E-01 | Protein DMP2 |
| AT4G04020 | -1.0 | 2.13E-04 | -1.0 | 4.57E-04 | 0.0 | 8.84E-01 | 0.5 | 4.31E-02 | 0.8 | 3.60E-03 | -0.3 | 2.99E-01 | Probable plastid-lipid-associated protein 1, chloroplastic |

| | | | | | | | | | | | | | |
|---|---|---|---|---|---|---|---|---|---|---|---|---|---|
| AT1G03010 | -1.0 | 3.92E-05 | -0.2 | 8.45E-02 | -0.8 | 1.70E-03 | 0.0 | 9.88E-01 | 0.0 | 9.40E-01 | 0.0 | 9.50E-01 | Phototropic-responsive NPH3 family protein |
| AT2G27310 | -1.0 | 2.46E-06 | -0.6 | 1.24E-04 | -0.4 | 6.35E-03 | -0.1 | 6.82E-01 | -0.1 | 5.89E-01 | 0.0 | 9.06E-01 | F-box protein At2g27310 |
| AT5G49710 | -1.0 | 4.24E-07 | -0.9 | 1.41E-06 | 0.0 | 6.13E-01 | -0.5 | 7.72E-03 | -0.2 | 2.26E-01 | -0.3 | 2.81E-01 | hypothetical protein |
| AT4G23020 | -1.0 | 9.39E-04 | -0.8 | 3.54E-03 | -0.2 | 5.70E-01 | -0.2 | 6.43E-01 | 0.1 | 7.18E-01 | -0.3 | 3.92E-01 | DUF4378 domain-containing protein |
| AT5G15190 | -1.0 | 3.85E-03 | -0.9 | 8.46E-03 | -0.1 | 7.79E-01 | 0.2 | 6.53E-01 | -0.6 | 3.42E-01 | 0.8 | 1.69E-01 | Uncharacterized protein F8M21_80 |
| AT3G07730 | -1.0 | 3.22E-04 | -1.0 | 5.22E-04 | 0.0 | 9.75E-01 | -0.1 | 9.03E-01 | 0.1 | 9.15E-01 | -0.1 | 8.48E-01 | F17A17.7 protein |
| AT1G67105 | -1.0 | 3.58E-03 | -1.0 | 4.88E-03 | 0.0 | 9.88E-01 | -1.3 | 2.85E-03 | -0.5 | 2.08E-01 | -0.8 | 1.43E-01 | NA |
| AT5G15265 | -1.0 | 3.27E-04 | 0.4 | 1.80E-02 | -1.4 | 1.71E-04 | -0.3 | 3.44E-01 | 0.0 | 9.65E-01 | -0.3 | 4.45E-01 | hypothetical protein |
| AT5G42010 | -1.0 | 3.79E-06 | -0.8 | 4.33E-05 | -0.2 | 7.89E-02 | 0.2 | 4.05E-01 | 0.1 | 5.38E-01 | 0.0 | 9.57E-01 | Transducin |
| AT1G07680 | -1.0 | 4.11E-03 | 0.7 | 1.30E-02 | -1.7 | 5.68E-04 | -3.2 | 6.60E-06 | 0.0 | 9.32E-01 | -3.2 | 1.97E-04 | Transmembrane protein |
| AT1G11592 | -1.0 | 6.32E-05 | -0.5 | 4.08E-03 | -0.5 | 2.84E-02 | -0.7 | 1.92E-01 | -0.2 | 7.55E-01 | -0.5 | 4.71E-01 | NA |
| AT2G02820 | -1.0 | 5.00E-06 | -0.6 | 1.57E-04 | -0.3 | 1.87E-02 | 0.1 | 4.49E-01 | 0.1 | 4.35E-01 | 0.0 | 9.37E-01 | Myb domain protein 88 |
| AT3G63310 | -1.0 | 1.33E-06 | -0.7 | 2.45E-05 | -0.3 | 1.77E-02 | -0.2 | 7.15E-02 | -0.3 | 2.41E-02 | 0.1 | 6.31E-01 | Protein LIFEGUARD 2 |
| AT3G13965 | -1.0 | 3.33E-06 | -0.3 | 1.78E-02 | -0.7 | 2.35E-04 | -0.4 | 4.34E-02 | -0.1 | 8.19E-01 | -0.4 | 1.54E-01 | NA |
| AT1G63220 | -1.0 | 2.89E-05 | 0.1 | 6.11E-01 | -1.1 | 1.09E-04 | -0.3 | 1.07E-01 | 0.2 | 4.41E-01 | -0.5 | 3.81E-02 | Calcium-dependent lipid-binding family protein |
| AT2G42060 | -1.0 | 5.52E-07 | -0.3 | 3.11E-03 | -0.7 | 6.25E-05 | 0.2 | 3.67E-01 | 0.4 | 7.12E-02 | -0.2 | 4.12E-01 | Phorbol-ester |
| AT5G59020 | -1.0 | 1.01E-05 | -0.6 | 6.20E-04 | -0.4 | 1.31E-02 | 0.0 | 9.55E-01 | 0.3 | 5.22E-01 | -0.2 | 5.09E-01 | Gb\|AAC95187.1 ) |
| AT3G22260 | -1.0 | 1.22E-06 | -0.7 | 2.95E-05 | -0.3 | 1.14E-02 | 0.0 | 9.26E-01 | -0.1 | 5.75E-01 | 0.2 | 5.77E-01 | hypothetical protein |
| AT5G17910 | -1.0 | 1.59E-07 | -0.5 | 1.93E-05 | -0.5 | 1.75E-04 | -0.2 | 1.20E-01 | 0.0 | 9.02E-01 | -0.2 | 2.94E-01 | Cardiomyopathy-associated protein |
| AT1G13500 | -1.0 | 9.29E-06 | -0.2 | 1.43E-01 | -0.8 | 2.62E-04 | -0.7 | 4.40E-02 | 0.3 | 3.23E-01 | -1.1 | 1.90E-02 | F13B4.2 protein |
| AT4G05590 | -1.0 | 4.59E-04 | -1.1 | 3.86E-04 | 0.1 | 6.84E-01 | 0.0 | 9.10E-01 | 0.2 | 6.59E-01 | -0.1 | 7.62E-01 | Mitochondrial pyruvate carrier |
| AT3G60090 | -1.0 | 6.51E-04 | 0.0 | 9.75E-01 | -1.0 | 3.23E-03 | -0.4 | 4.09E-01 | 0.6 | 2.05E-01 | -1.1 | 5.15E-02 | Uncharacterized protein T2O9.70 |
| AT2G05910 | -1.0 | 3.57E-05 | -0.8 | 2.02E-04 | -0.2 | 3.55E-01 | 0.0 | 9.48E-01 | 0.5 | 2.94E-01 | -0.5 | 3.71E-01 | Protein LURP-one-related 6 |
| AT4G24160 | -1.0 | 2.31E-06 | -0.7 | 3.84E-05 | -0.3 | 3.15E-02 | 0.2 | 3.88E-01 | 0.0 | 9.65E-01 | 0.2 | 4.68E-01 | Alpha |
| AT3G46845 | -1.0 | 1.00E-04 | -0.4 | 1.19E-02 | -0.5 | 1.71E-02 | -0.1 | 8.97E-01 | 0.0 | 9.56E-01 | -0.1 | 8.75E-01 | Uncharacterized protein |
| AT1G17240 | -1.0 | 6.61E-05 | -0.4 | 1.03E-02 | -0.5 | 1.41E-02 | 0.5 | 1.53E-01 | 0.1 | 8.35E-01 | 0.4 | 3.80E-01 | Receptor like protein 2 |
| AT5G42050 | -1.0 | 4.00E-06 | -1.2 | 2.81E-06 | 0.2 | 9.92E-02 | 0.2 | 1.48E-01 | 0.1 | 6.47E-01 | 0.1 | 5.72E-01 | DCD domain-containing protein NRP |
| AT5G40780 | -1.0 | 1.31E-05 | -0.8 | 8.35E-05 | -0.1 | 2.88E-01 | -0.3 | 6.86E-02 | 0.0 | 9.50E-01 | -0.3 | 2.28E-01 | Lysine histidine transporter 1 |
| AT5G26320 | -1.0 | 1.26E-04 | 0.4 | 1.45E-02 | -1.4 | 8.00E-05 | -1.3 | 1.51E-04 | -0.4 | 9.96E-02 | -0.9 | 1.87E-02 | MATH domain-containing protein |
| AT5G53750 | -1.0 | 2.95E-04 | -0.5 | 1.24E-02 | -0.5 | 6.32E-02 | -0.4 | 1.05E-01 | 0.3 | 1.94E-01 | -0.7 | 1.87E-02 | CBS domain-containing protein |
| AT2G37370 | -1.0 | 9.70E-05 | -0.3 | 7.07E-02 | -0.7 | 5.15E-03 | | | | | | | Centrosomal protein of 135 kDa-like protein |
| AT5G10965 | -1.0 | 2.35E-04 | -0.8 | 8.95E-04 | -0.1 | 5.90E-01 | | | | | | | NA |
| AT5G07322 | -1.0 | 4.71E-03 | -0.4 | 1.43E-01 | -0.6 | 1.14E-01 | -0.1 | 7.44E-01 | 0.1 | 7.75E-01 | -0.2 | 5.54E-01 | NA |
| AT5G49280 | -1.0 | 6.50E-06 | -0.5 | 7.89E-04 | -0.5 | 4.58E-03 | -0.4 | 8.26E-02 | 0.3 | 2.03E-01 | -0.7 | 2.22E-02 | Uncharacterized protein |
| AT5G62330 | -1.0 | 5.87E-03 | 0.2 | 4.49E-01 | -1.2 | 6.66E-03 | 0.2 | 8.14E-01 | 1.0 | 7.29E-02 | -0.8 | 1.69E-01 | Uncharacterized protein |
| AT1G23760 | -1.0 | 2.05E-04 | 0.1 | 5.51E-01 | -1.1 | 5.99E-04 | -0.6 | 1.51E-01 | 0.0 | 9.99E-01 | -0.6 | 2.60E-01 | Polygalacturonase 1 beta-like protein 1 |
| AT5G22870 | -1.0 | 2.76E-04 | -0.3 | 1.10E-01 | -0.7 | 1.09E-02 | -0.1 | 9.00E-01 | -0.3 | 5.85E-01 | 0.3 | 7.28E-01 | Gb\|AAF18257.1 hydroxyproline-rich glycoprotein family) |
| AT5G48930 | -1.0 | 7.68E-07 | -0.3 | 2.03E-03 | -0.6 | 1.02E-04 | -0.3 | 2.75E-02 | -0.1 | 5.48E-01 | -0.2 | 3.27E-01 | AT5G48930 protein |
| AT5G02640 | -1.0 | 6.84E-04 | -0.7 | 4.09E-03 | -0.2 | 4.02E-01 | -0.9 | 2.69E-02 | -0.1 | 9.04E-01 | -0.8 | 1.08E-01 | Uncharacterized protein T22P11_230 |
| AT4G39320 | -1.0 | 1.48E-03 | -0.6 | 1.48E-02 | -0.3 | 2.86E-01 | -0.1 | 8.76E-01 | 0.1 | 8.89E-01 | -0.2 | 7.93E-01 | AT4g39320 |
| AT5G10820 | -1.0 | 6.60E-06 | -0.3 | 9.49E-03 | -0.6 | 7.08E-04 | -0.4 | 1.31E-02 | 0.0 | 8.10E-01 | -0.3 | 1.02E-01 | Probable folate-biopterin transporter 6 |
| AT5G24310 | -1.0 | 1.34E-05 | -0.4 | 2.45E-03 | -0.5 | 5.04E-03 | -0.2 | 5.34E-01 | -0.1 | 8.57E-01 | -0.1 | 7.87E-01 | ABL interactor-like protein 3 |
| AT3G57570 | -1.0 | 1.01E-06 | -0.8 | 9.56E-06 | -0.2 | 6.92E-02 | 0.0 | 8.66E-01 | 0.0 | 9.19E-01 | 0.0 | 9.82E-01 | ARM repeat superfamily protein |
| AT2G05260 | -1.0 | 2.30E-06 | -0.4 | 7.45E-04 | -0.5 | 9.61E-04 | -0.3 | 1.39E-01 | 0.1 | 7.14E-01 | -0.4 | 1.01E-01 | Alpha |
| AT2G27740 | -1.0 | 2.92E-05 | -0.1 | 3.73E-01 | -0.8 | 5.18E-04 | -0.3 | 4.34E-01 | 0.0 | 9.73E-01 | -0.3 | 4.74E-01 | RAB6-interacting golgin |
| AT2G29125 | -1.0 | 1.66E-03 | -0.5 | 4.09E-02 | -0.5 | 1.42E-01 | -0.4 | 2.53E-01 | 0.0 | 9.42E-01 | -0.4 | 2.70E-01 | Small polypeptide DEVIL 13 |
| AT1G21200 | -1.0 | 2.14E-07 | -0.6 | 1.42E-05 | -0.4 | 5.63E-04 | -0.4 | 1.57E-02 | -0.2 | 4.02E-01 | -0.3 | 2.40E-01 | Myb |
| AT5G16730 | -1.0 | 3.03E-07 | -0.6 | 1.96E-05 | -0.4 | 8.10E-04 | -0.6 | 1.02E-04 | -0.2 | 2.55E-01 | -0.4 | 1.41E-02 | WEB family protein At5g16730, chloroplastic |
| AT5G11990 | -1.0 | 6.33E-05 | -0.5 | 2.52E-03 | -0.4 | 4.72E-02 | -0.1 | 8.47E-01 | 0.2 | 7.66E-01 | -0.3 | 6.44E-01 | Proline-rich family protein |
| AT1G23330 | -1.0 | 1.02E-06 | -0.7 | 1.51E-05 | -0.2 | 2.49E-02 | 0.4 | 1.16E-01 | 0.7 | 1.01E-02 | -0.3 | 2.59E-01 | At1g23330 |
| AT4G13110 | -1.0 | 1.34E-04 | -1.1 | 9.79E-05 | 0.2 | 4.81E-01 | -0.3 | 3.57E-01 | 0.3 | 2.44E-01 | -0.6 | 5.68E-02 | Uncharacterized protein At4g13110 |
| AT2G40480 | -0.9 | 1.30E-05 | -0.5 | 1.42E-03 | -0.5 | 7.12E-03 | -0.2 | 1.61E-01 | 0.2 | 1.48E-01 | -0.5 | 1.34E-02 | WEB family protein At2g40480 |
| AT3G46450 | -0.9 | 5.22E-07 | -1.1 | 4.64E-07 | 0.2 | 4.35E-02 | -0.6 | 2.42E-03 | -0.2 | 4.43E-01 | -0.5 | 4.51E-02 | SEC14 cytosolic factor family protein |
| AT4G18580 | -0.9 | 8.32E-06 | -0.6 | 2.14E-04 | -0.3 | 3.07E-02 | 0.0 | 9.44E-01 | 0.0 | 9.70E-01 | 0.0 | 9.92E-01 | Uncharacterized protein |
| AT3G05880 | -0.9 | 4.59E-06 | -0.4 | 4.74E-03 | -0.6 | 5.73E-04 | -0.6 | 2.05E-02 | -0.2 | 4.44E-01 | -0.3 | 2.39E-01 | Hydrophobic protein RCI2A |
| AT1G21390 | -0.9 | 1.01E-05 | -1.1 | 7.23E-06 | 0.2 | 2.04E-01 | 0.3 | 1.38E-01 | 0.1 | 8.62E-01 | 0.2 | 2.79E-01 | Transmembrane protein |
| AT4G32070 | -0.9 | 2.45E-07 | -0.6 | 1.31E-05 | -0.4 | 7.43E-04 | 0.0 | 7.85E-01 | 0.0 | 8.59E-01 | 0.1 | 6.80E-01 | Protein PHOX4 |
| AT5G15600 | -0.9 | 3.02E-05 | 0.3 | 9.81E-03 | -1.3 | 3.70E-05 | -1.5 | 6.81E-04 | -0.2 | 6.17E-01 | -1.4 | 1.05E-02 | Protein SPIRAL1-like 4 |
| AT2G35760 | -0.9 | 2.65E-06 | -0.5 | 3.92E-04 | -0.5 | 1.92E-03 | -0.3 | 1.26E-01 | 0.1 | 6.73E-01 | -0.4 | 9.24E-02 | CASP-like protein |
| AT1G76260 | -0.9 | 1.35E-06 | -0.3 | 6.86E-03 | -0.7 | 1.36E-04 | -0.3 | 3.21E-01 | -0.1 | 8.76E-01 | -0.2 | 5.05E-01 | WD repeat-containing protein DWA2 |
| AT1G80490 | -0.9 | 1.68E-07 | -0.7 | 4.08E-06 | -0.3 | 1.96E-03 | -0.1 | 7.14E-01 | 0.1 | 7.10E-01 | -0.2 | 5.00E-01 | Topless-related protein 1 |
| AT2G27080 | -0.9 | 9.08E-04 | -0.3 | 1.03E-01 | -0.6 | 3.24E-02 | -0.2 | 4.07E-01 | -0.1 | 7.48E-01 | -0.1 | 7.84E-01 | NDR1 |
| AT4G17350 | -0.9 | 3.98E-06 | -0.5 | 6.22E-04 | -0.5 | 2.91E-03 | -0.5 | 4.31E-02 | -0.1 | 7.13E-01 | -0.4 | 2.08E-01 | At4g17350 ) |
| AT3G05937 | -0.9 | 4.15E-05 | -0.8 | 1.44E-04 | -0.1 | 5.96E-01 | 0.5 | 1.16E-01 | 0.0 | 9.88E-01 | 0.5 | 2.12E-01 | Uncharacterized protein |
| AT1G19900 | -0.9 | 3.53E-04 | 0.3 | 4.71E-02 | -1.3 | 2.30E-04 | -0.6 | 1.15E-01 | 0.1 | 8.01E-01 | -0.7 | 9.11E-02 | F6F9.4 protein |
| AT2G36410 | -0.9 | 5.38E-07 | -0.4 | 4.35E-04 | -0.5 | 1.36E-04 | -0.4 | 2.31E-02 | -0.2 | 1.70E-01 | -0.1 | 6.06E-01 | RAB6-interacting golgin |
| AT5G42690 | -0.9 | 1.60E-06 | -0.5 | 2.71E-04 | -0.5 | 1.39E-03 | -0.1 | 7.32E-01 | 0.1 | 7.78E-01 | -0.2 | 5.74E-01 | Expressed protein |
| AT5G41400 | -0.9 | 1.97E-06 | -0.7 | 2.33E-05 | -0.2 | 5.87E-02 | 0.6 | 2.58E-02 | 0.8 | 7.85E-03 | -0.2 | 4.85E-01 | hypothetical protein |
| AT2G28690 | -0.9 | 1.04E-03 | -1.1 | 6.59E-04 | 0.1 | 6.27E-01 | 0.7 | 1.42E-01 | 0.6 | 2.73E-01 | 0.1 | 8.80E-01 | Uncharacterized protein At2g28690 |
| AT2G45830 | -0.9 | 2.70E-06 | -0.5 | 1.51E-04 | -0.4 | 6.48E-03 | 0.1 | 6.76E-01 | 0.0 | 8.84E-01 | 0.0 | 8.92E-01 | At2g45830 |
| AT5G23160 | -0.9 | 8.32E-04 | 0.3 | 1.09E-01 | -1.2 | 6.86E-04 | -1.5 | 4.63E-03 | -0.3 | 6.01E-01 | -1.2 | 6.68E-02 | Uncharacterized protein At5g23160 |
| AT4G22758 | -0.9 | 7.13E-06 | -0.3 | 8.28E-03 | -0.6 | 9.41E-04 | 0.5 | 1.42E-01 | 0.4 | 2.95E-01 | 0.1 | 8.54E-01 | PPR containing protein |
| AT1G17300 | -0.9 | 3.99E-04 | 0.4 | 2.59E-02 | -1.3 | 2.21E-04 | -0.2 | 6.44E-01 | 0.0 | 9.64E-01 | -0.2 | 7.18E-01 | Uncharacterized protein |
| AT4G14290 | -0.9 | 3.63E-06 | -0.5 | 3.40E-04 | -0.4 | 4.28E-03 | -0.1 | 5.53E-01 | 0.2 | 3.14E-01 | -0.4 | 1.71E-01 | Alpha |
| AT5G22550 | -0.9 | 6.44E-04 | -0.7 | 5.12E-03 | -0.2 | 3.44E-01 | | | | | | | Emb\|CAB72473.1 ) |
| AT2G44600 | -0.9 | 1.85E-05 | -0.5 | 7.58E-04 | -0.4 | 2.59E-02 | 0.0 | 8.94E-01 | 0.1 | 8.51E-01 | -0.1 | 7.52E-01 | hypothetical protein |
| AT2G40470 | -0.9 | 3.93E-03 | 0.0 | 9.80E-01 | -0.9 | 1.27E-02 | 0.1 | 8.70E-01 | 0.5 | 3.94E-01 | | | LOB domain-containing protein 15 |
| AT5G46600 | -0.9 | 2.85E-05 | -0.3 | 3.96E-02 | -0.7 | 1.82E-03 | 0.5 | 1.92E-01 | 0.7 | 1.41E-01 | -0.1 | 8.45E-01 | Aluminum-activated malate transporter 13 |
| AT1G19870 | -0.9 | 3.52E-07 | -0.5 | 3.85E-05 | -0.4 | 4.68E-04 | -0.2 | 2.36E-01 | -0.2 | 2.28E-01 | 0.0 | 9.24E-01 | IQ-domain 32 |
| AT3G02290 | -0.9 | 1.61E-06 | -0.6 | 4.80E-05 | -0.3 | 1.12E-02 | -0.3 | 6.78E-02 | -0.1 | 6.34E-01 | -0.2 | 2.71E-01 | RING |
| AT2G46535 | -0.9 | 2.59E-05 | -0.5 | 1.25E-03 | -0.4 | 2.51E-02 | -0.1 | 6.74E-01 | 0.4 | 2.27E-02 | -0.5 | 2.82E-02 | Uncharacterized protein |
| AT5G24760 | -0.9 | 2.39E-06 | -0.4 | 1.10E-03 | -0.5 | 7.33E-04 | 0.0 | 9.85E-01 | 0.2 | 3.83E-01 | -0.2 | 4.21E-01 | GroES-like zinc-binding dehydrogenase family protein |
| AT5G54700 | -0.9 | 6.36E-05 | -1.0 | 6.99E-05 | 0.1 | 6.02E-01 | 0.2 | 5.53E-01 | 0.6 | 7.93E-02 | -0.4 | 2.38E-01 | Ankyrin repeat family protein |

| | | | | | | | | | | | | | |
|---|---|---|---|---|---|---|---|---|---|---|---|---|---|
| AT1G08320 | -0.9 | 3.35E-06 | -0.6 | 1.36E-04 | -0.3 | 1.02E-02 | -0.4 | 3.99E-02 | -0.1 | 8.21E-01 | -0.3 | 1.84E-01 | T27G7.3 |
| AT1G73950 | -0.9 | 1.13E-06 | -0.7 | 1.90E-05 | -0.2 | 2.22E-02 | -0.2 | 2.14E-01 | 0.0 | 8.77E-01 | -0.2 | 4.66E-01 | Transmembrane Fragile-X-F-associated protein |
| AT4G25070 | -0.9 | 8.74E-06 | -0.3 | 2.68E-02 | -0.7 | 5.24E-04 | -0.3 | 4.42E-01 | -0.1 | 8.91E-01 | -0.2 | 6.79E-01 | Uncharacterized protein AT4g25060 |
| AT3G14280 | -0.9 | 1.26E-05 | -0.4 | 2.50E-03 | -0.5 | 3.67E-03 | -1.3 | 1.41E-04 | 0.1 | 7.86E-01 | -1.4 | 1.36E-04 | AT3G14280 protein |
| AT1G55420 | -0.9 | 5.85E-05 | -0.4 | 1.12E-02 | -0.5 | 1.05E-02 | -0.1 | 6.42E-01 | 0.3 | 2.58E-01 | -0.5 | 8.84E-02 | Cysteine |
| AT5G43880 | -0.9 | 1.81E-06 | -0.5 | 1.76E-04 | -0.4 | 2.19E-03 | -0.2 | 1.55E-01 | 0.1 | 7.22E-01 | -0.3 | 1.61E-01 | Methyl-coenzyme M reductase II subunit gamma, putative |
| AT4G32350 | -0.9 | 5.71E-06 | -0.3 | 8.30E-03 | -0.6 | 6.82E-04 | -0.4 | 4.11E-02 | -0.2 | 2.99E-01 | -0.2 | 5.57E-01 | Uncharacterized protein AT4g32350 |
| AT1G63720 | -0.9 | 1.15E-05 | -0.7 | 1.06E-04 | -0.2 | 1.51E-01 | 0.6 | 2.32E-02 | -0.1 | 7.57E-01 | 0.8 | 2.78E-02 | Uncharacterized protein F24D7.9 |
| AT5G48170 | -0.9 | 5.78E-04 | -0.5 | 1.26E-02 | -0.4 | 1.41E-01 | 0.0 | 9.98E-01 | -0.2 | 5.86E-01 | 0.2 | 6.49E-01 | F-box protein SNE |
| AT3G25640 | -0.9 | 1.61E-05 | -0.6 | 3.23E-04 | -0.3 | 5.42E-02 | -0.1 | 7.30E-01 | 0.2 | 6.09E-01 | -0.3 | 4.03E-01 | Protein MIZU-KUSSEI 1 |
| AT1G44608 | -0.9 | 7.50E-04 | 0.2 | 1.46E-01 | -1.1 | 7.49E-04 | -0.9 | 4.12E-02 | 0.2 | 6.81E-01 | -1.1 | 4.90E-02 | Uncharacterized protein |
| AT3G27960 | -0.9 | 1.65E-06 | -0.4 | 4.98E-04 | -0.5 | 7.83E-04 | -0.6 | 2.13E-04 | -0.1 | 4.10E-01 | -0.5 | 1.84E-02 | Tetratricopeptide repeat -like superfamily protein |
| AT3G55420 | -0.9 | 3.97E-06 | -0.4 | 1.70E-03 | -0.5 | 1.19E-03 | -0.2 | 2.54E-01 | 0.1 | 7.57E-01 | -0.3 | 1.81E-01 | Uncharacterized protein |
| AT3G20340 | -0.9 | 3.57E-04 | -0.4 | 3.62E-02 | -0.5 | 2.98E-02 | -1.2 | 3.01E-03 | -1.0 | 1.22E-02 | -0.2 | 7.41E-01 | hypothetical protein |
| AT3G13520 | -0.9 | 4.86E-05 | -0.2 | 1.06E-01 | -0.7 | 1.32E-03 | -1.0 | 8.09E-04 | -1.0 | 2.15E-03 | 0.0 | 9.98E-01 | Arabinogalactan protein 12 |
| AT1G67365 | -0.9 | 1.07E-03 | -0.9 | 2.04E-03 | 0.0 | 9.00E-01 | -0.7 | 1.02E-01 | -0.5 | 2.66E-01 | -0.2 | 8.02E-01 | NA |
| AT2G04170 | -0.9 | 2.02E-06 | 0.0 | 8.33E-01 | -0.9 | 2.95E-05 | 0.3 | 1.81E-02 | 0.6 | 1.33E-03 | -0.2 | 2.40E-01 | TRAF-like family protein |
| AT4G28380 | -0.9 | 2.46E-04 | -0.7 | 2.09E-03 | -0.2 | 2.79E-01 | 0.0 | 9.51E-01 | 0.2 | 6.11E-01 | | | hypothetical protein |
| AT2G12462 | -0.9 | 2.65E-05 | -0.2 | 6.58E-02 | -0.7 | 1.11E-03 | -0.2 | 4.67E-01 | 0.2 | 5.20E-01 | -0.4 | 1.28E-01 | Sterile alpha motif domain protein |
| AT5G65280 | -0.9 | 1.61E-05 | -0.6 | 4.35E-04 | -0.3 | 4.16E-02 | -1.1 | 9.00E-03 | -0.2 | 6.73E-01 | -0.9 | 1.02E-01 | GCR2-like 1 |
| AT5G63595 | -0.9 | 5.01E-03 | -0.5 | 6.49E-02 | -0.4 | 2.45E-01 | | | | | | | FLS4 |
| AT5G28610 | -0.9 | 1.82E-02 | -0.4 | 2.67E-01 | -0.5 | 2.18E-01 | 1.2 | 8.67E-04 | 0.3 | 5.89E-01 | 1.0 | 1.98E-02 | At5g28610 |
| AT4G23630 | -0.9 | 8.09E-06 | -0.6 | 1.54E-04 | -0.2 | 5.22E-02 | -0.3 | 6.70E-02 | 0.0 | 9.65E-01 | -0.2 | 1.43E-01 | Reticulon-like protein B1 |
| AT4G02810 | -0.9 | 4.30E-04 | 0.5 | 7.55E-03 | -1.4 | 1.48E-04 | 1.0 | 1.81E-02 | 0.9 | 5.33E-02 | 0.1 | 8.29E-01 | Protein FANTASTIC FOUR 1 |
| AT3G16860 | -0.9 | 1.59E-05 | -1.0 | 1.39E-05 | 0.1 | 3.63E-01 | 0.2 | 3.29E-01 | 0.1 | 8.17E-01 | 0.2 | 6.31E-01 | COBRA-like protein 8 |
| AT2G36220 | -0.9 | 2.33E-04 | -0.7 | 1.16E-03 | -0.2 | 4.77E-01 | | | | | | | At2g36220 |
| AT3G50280 | -0.9 | 8.34E-05 | -0.6 | 1.66E-03 | -0.3 | 9.65E-02 | -0.2 | 6.21E-01 | 0.7 | 1.04E-01 | -0.9 | 5.34E-02 | HXXXD-type acyl-transferase family protein |
| AT3G23200 | -0.9 | 3.39E-06 | -0.5 | 1.69E-04 | -0.4 | 8.59E-03 | -0.7 | 1.58E-03 | -0.4 | 8.35E-02 | -0.3 | 1.93E-01 | CASP-like protein |
| AT1G49310 | -0.9 | 3.69E-04 | -0.5 | 2.30E-02 | -0.4 | 4.04E-02 | -0.5 | 7.24E-02 | 0.0 | 9.17E-01 | -0.6 | 7.95E-02 | hypothetical protein |
| AT2G38465 | -0.9 | 1.96E-03 | -0.6 | 1.59E-02 | -0.3 | 3.42E-01 | -0.5 | 1.58E-01 | -0.3 | 4.83E-01 | -0.2 | 6.95E-01 | hypothetical protein |
| AT2G14520 | -0.9 | 5.67E-05 | -0.5 | 3.08E-03 | -0.4 | 3.40E-02 | -0.1 | 7.18E-01 | 0.3 | 2.95E-01 | -0.5 | 1.94E-01 | CBS domain protein |
| AT3G06390 | -0.9 | 3.43E-05 | 0.1 | 5.65E-01 | -0.9 | 1.25E-04 | -0.2 | 5.43E-01 | -0.1 | 8.42E-01 | -0.1 | 8.37E-01 | CASP-like protein 1D2 |
| AT1G08160 | -0.9 | 3.76E-06 | -0.3 | 2.78E-03 | -0.5 | 6.95E-04 | -0.1 | 8.02E-01 | 0.1 | 6.36E-01 | -0.2 | 5.31E-01 | Uncharacterized protein At1g08160 |
| AT2G15760 | -0.9 | 3.56E-04 | -0.3 | 7.99E-02 | -0.6 | 1.68E-02 | 0.5 | 5.82E-02 | -0.1 | 8.88E-01 | 0.5 | 6.51E-02 | hypothetical protein |
| AT3G53820 | -0.9 | 8.18E-05 | -0.7 | 4.09E-04 | -0.1 | 4.59E-01 | | | | | | | C2H2-type domain-containing protein |
| AT4G11780 | -0.9 | 3.66E-03 | 0.2 | 4.08E-01 | -1.1 | 4.34E-03 | -1.4 | 5.54E-04 | -0.4 | 1.47E-01 | -0.9 | 4.26E-02 | GAR2-like protein |
| AT1G13510 | -0.9 | 1.32E-05 | -0.4 | 2.70E-03 | -0.5 | 4.78E-03 | 0.0 | 9.37E-01 | 1.1 | 2.06E-03 | -1.0 | 3.63E-03 | F13B4.2 protein |
| AT5G13810 | -0.9 | 1.19E-06 | -0.6 | 2.11E-05 | -0.2 | 1.80E-02 | -0.3 | 7.52E-02 | 0.0 | 9.92E-01 | -0.3 | 1.43E-01 | hypothetical protein |
| AT1G14530 | -0.9 | 5.35E-05 | -0.4 | 1.11E-02 | -0.5 | 9.29E-03 | -0.2 | 3.05E-01 | -0.1 | 8.67E-01 | -0.2 | 5.68E-01 | Tobamovirus multiplication-like protein |
| AT1G70860 | -0.9 | 1.66E-04 | -0.5 | 5.10E-03 | -0.4 | 7.08E-02 | -1.5 | 1.61E-02 | 0.1 | 9.20E-01 | -1.6 | 4.85E-03 | Major latex-like protein |
| AT3G27510 | -0.9 | 2.77E-05 | -0.3 | 1.04E-02 | -0.5 | 4.84E-03 | -0.6 | 1.26E-01 | -0.1 | 7.75E-01 | -0.4 | 4.19E-01 | Cysteine |
| AT3G19370 | -0.9 | 3.95E-07 | -0.6 | 1.60E-05 | -0.3 | 1.93E-03 | -0.2 | 3.41E-01 | 0.3 | 9.31E-02 | -0.4 | 2.38E-02 | Filament-like plant protein 7 |
| AT1G13340 | -0.9 | 3.48E-03 | -1.1 | 1.69E-03 | 0.2 | 5.47E-01 | | | | | | | Regulator of Vps4 activity in the MVB pathway protein |
| AT1G22050 | -0.9 | 3.77E-05 | -0.3 | 2.26E-02 | -0.6 | 3.61E-03 | -0.1 | 6.16E-01 | 0.2 | 6.00E-01 | -0.3 | 2.57E-01 | Membrane-anchored ubiquitin-fold protein 6 |
| AT1G13390 | -0.9 | 1.63E-05 | -0.7 | 1.80E-04 | -0.2 | 1.28E-01 | -0.1 | 6.08E-01 | -0.2 | 5.03E-01 | 0.1 | 8.93E-01 | hypothetical protein |
| AT1G31470 | -0.9 | 1.60E-04 | -0.7 | 8.19E-04 | -0.2 | 4.37E-01 | -0.1 | 9.16E-01 | 0.4 | 7.06E-01 | -0.5 | 6.49E-01 | Protein NUCLEAR FUSION DEFECTIVE 4 |
| AT4G31130 | -0.9 | 7.50E-07 | -0.3 | 8.06E-04 | -0.5 | 1.53E-04 | -0.3 | 1.24E-01 | 0.0 | 9.81E-01 | -0.3 | 2.43E-01 | hypothetical protein |
| AT1G16510 | -0.9 | 3.84E-05 | -0.6 | 6.38E-04 | -0.3 | 9.52E-02 | -0.1 | 7.48E-01 | -0.3 | 2.85E-01 | 0.2 | 5.21E-01 | Auxin-responsive protein SAUR41 |
| AT2G20562 | -0.9 | 7.09E-04 | 0.1 | 6.57E-01 | -0.9 | 1.82E-03 | 0.2 | 6.84E-01 | -0.1 | 8.77E-01 | 0.3 | 5.98E-01 | Signaling peptide TAXIMIN 2 |
| AT1G56660 | -0.9 | 2.94E-04 | 0.0 | 9.05E-01 | -0.9 | 8.78E-04 | -0.1 | 8.04E-01 | 0.2 | 5.67E-01 | -0.3 | 4.85E-01 | Uncharacterized protein |
| AT4G30710 | -0.9 | 1.77E-05 | -0.5 | 9.19E-04 | -0.4 | 2.02E-02 | -0.2 | 3.33E-01 | 0.1 | 7.92E-01 | -0.3 | 2.78E-01 | QWRF motif protein |
| AT3G01400 | -0.9 | 1.63E-06 | -0.6 | 2.99E-05 | -0.2 | 2.25E-02 | -0.3 | 2.36E-02 | -0.2 | 2.27E-01 | -0.1 | 4.96E-01 | hypothetical protein |
| AT1G10810 | -0.9 | 4.03E-05 | 0.1 | 1.84E-01 | -1.0 | 1.15E-04 | -0.3 | 3.73E-01 | 0.2 | 7.33E-01 | -0.5 | 2.56E-01 | NAD-linked oxidoreductase superfamily protein |
| AT4G28150 | -0.9 | 1.13E-04 | -0.4 | 1.15E-02 | -0.5 | 2.41E-02 | 0.2 | 3.06E-01 | -0.3 | 2.39E-01 | 0.6 | 4.36E-02 | Uncharacterized protein AT4g28150 |
| AT1G52155 | -0.9 | 7.27E-05 | -0.4 | 1.18E-02 | -0.5 | 1.35E-02 | 0.0 | 9.91E-01 | 0.4 | 8.23E-02 | -0.4 | 1.32E-01 | Transmembrane protein |
| AT5G56050 | -0.9 | 2.21E-04 | -0.8 | 7.05E-04 | -0.1 | 6.76E-01 | 0.9 | 4.84E-02 | 0.3 | 6.03E-01 | 0.6 | 1.92E-01 | Late embryogenesis abundant protein LEA-2 subgroup domain-containing protein |
| AT3G42658 | -0.9 | 2.67E-02 | -0.2 | 6.08E-01 | -0.7 | 1.22E-01 | -1.1 | 5.84E-03 | -0.7 | 5.57E-02 | -0.4 | 4.12E-01 | NA |
| AT2G14820 | -0.9 | 4.47E-03 | 0.2 | 4.06E-01 | -1.0 | 4.82E-03 | 0.1 | 7.94E-01 | -0.1 | 8.74E-01 | 0.2 | 7.40E-01 | BTB |
| AT1G64065 | -0.9 | 5.53E-03 | -0.6 | 3.31E-02 | -0.3 | 4.37E-01 | 0.1 | 6.80E-01 | -0.5 | 1.22E-01 | 0.7 | 5.29E-02 | Late embryogenesis abundant protein At1g64065 |
| AT5G39480 | -0.9 | 2.26E-03 | -0.1 | 5.28E-01 | -0.7 | 2.00E-02 | 0.6 | 3.34E-01 | 1.1 | 9.73E-02 | -0.5 | 4.57E-01 | Putative F-box protein At5g39480 |
| AT1G15625 | -0.9 | 1.19E-04 | 0.1 | 3.31E-01 | -1.0 | 2.74E-04 | 1.3 | 3.78E-04 | 1.1 | 4.25E-03 | 0.2 | 4.86E-01 | E3 ubiquitin-protein ligase |
| AT2G26355 | -0.9 | 2.04E-04 | -1.0 | 1.20E-04 | 0.2 | 4.17E-01 | -0.9 | 6.90E-04 | -0.5 | 3.02E-02 | -0.4 | 2.56E-01 | NA |
| AT5G54790 | -0.9 | 5.53E-03 | 0.0 | 8.69E-01 | -0.9 | 1.33E-02 | -1.3 | 1.04E-01 | -0.6 | 5.12E-01 | | | CTD small phosphatase-like protein |
| AT4G32020 | -0.9 | 1.81E-06 | -0.6 | 8.01E-05 | -0.3 | 6.89E-03 | -0.1 | 3.65E-01 | -0.1 | 5.63E-01 | 0.0 | 9.14E-01 | AT4g32020 |
| AT3G50340 | -0.9 | 2.22E-05 | -0.3 | 9.70E-03 | -0.5 | 3.25E-03 | 0.0 | 9.96E-01 | 0.2 | 2.65E-01 | -0.2 | 3.27E-01 | Knr4 |
| AT1G01225 | -0.9 | 2.55E-04 | -0.5 | 9.99E-03 | -0.4 | 6.71E-02 | 0.3 | 3.63E-01 | 0.3 | 3.36E-01 | 0.0 | 9.24E-01 | LRAT domain-containing protein |
| AT4G12450 | -0.9 | 2.47E-05 | -0.2 | 9.23E-02 | -0.7 | 9.61E-04 | -0.5 | 1.08E-01 | -0.1 | 8.21E-01 | -0.4 | 2.56E-01 | At4g12450 family protein) |
| AT5G62820 | -0.8 | 2.40E-05 | -0.8 | 9.87E-05 | -0.1 | 5.57E-01 | -0.5 | 2.97E-02 | -0.3 | 3.54E-01 | -0.3 | 3.82E-01 | CASP-like protein 4A2 |
| AT2G29670 | -0.8 | 2.75E-05 | -0.6 | 5.41E-04 | -0.2 | 7.35E-02 | 0.0 | 9.27E-01 | 0.0 | 9.71E-01 | 0.0 | 9.27E-01 | hypothetical protein |
| AT2G21020 | -0.8 | 8.58E-06 | -0.1 | 2.17E-01 | -0.7 | 1.81E-04 | -0.3 | 1.60E-01 | 0.2 | 3.36E-01 | -0.5 | 1.61E-02 | NA |
| AT3G54680 | -0.8 | 2.26E-06 | -0.5 | 8.68E-05 | -0.3 | 8.98E-03 | -0.4 | 1.36E-02 | -0.2 | 3.13E-01 | -0.2 | 3.33E-01 | Uncharacterized protein T5N23_40 |
| AT1G77855 | -0.8 | 1.68E-04 | -0.4 | 2.19E-02 | -0.5 | 2.31E-02 | 0.0 | 9.17E-01 | 0.0 | 9.46E-01 | 0.1 | 8.91E-01 | BPS1-like protein |
| AT5G60050 | -0.8 | 1.17E-05 | -0.5 | 8.45E-04 | -0.4 | 1.16E-02 | -0.3 | 1.42E-01 | -0.3 | 2.97E-01 | -0.1 | 8.60E-01 | BTB |
| AT1G79420 | -0.8 | 2.93E-06 | -0.6 | 6.26E-05 | -0.3 | 2.72E-02 | -0.2 | 4.37E-01 | -0.2 | 4.76E-01 | 0.0 | 9.88E-01 | hypothetical protein |
| AT1G75310 | -0.8 | 6.79E-07 | -0.5 | 3.88E-05 | -0.3 | 2.10E-03 | -0.4 | 1.09E-01 | -0.3 | 3.70E-01 | -0.1 | 7.29E-01 | Auxin-like 1 protein |
| AT4G17760 | -0.8 | 1.85E-04 | -0.6 | 1.41E-03 | -0.2 | 3.23E-01 | 0.5 | 2.07E-01 | 0.7 | 1.35E-01 | -0.1 | 8.07E-01 | PCNA domain-containing protein |
| AT1G61260 | -0.8 | 1.66E-05 | -0.9 | 2.59E-05 | 0.0 | 8.44E-01 | -0.6 | 1.16E-03 | 0.0 | 9.18E-01 | -0.6 | 9.19E-03 | F11P17.2 protein |
| AT4G24310 | -0.8 | 1.23E-05 | -0.1 | 5.43E-01 | -0.8 | 1.57E-04 | -0.1 | 8.44E-01 | 0.2 | 4.40E-01 | -0.3 | 2.59E-01 | Protein DMP3 |
| AT5G19380 | -0.8 | 3.35E-05 | -0.6 | 4.51E-04 | -0.2 | 1.23E-01 | -0.1 | 6.89E-01 | 0.2 | 4.46E-01 | -0.3 | 2.67E-01 | CRT -like transporter 1 |
| AT2G38360 | -0.8 | 1.86E-04 | -0.2 | 1.31E-01 | -0.6 | 4.74E-03 | -0.1 | 7.37E-01 | 0.3 | 2.11E-01 | -0.3 | 1.10E-01 | PRA1 family protein |
| AT5G18640 | -0.8 | 3.20E-06 | -0.6 | 8.52E-05 | -0.3 | 1.81E-02 | -0.7 | 5.96E-04 | -0.4 | 5.48E-02 | -0.3 | 1.59E-01 | Alpha |
| AT3G25950 | -0.8 | 3.07E-03 | -0.1 | 5.93E-01 | -0.7 | 2.28E-02 | -0.7 | 3.77E-02 | -0.5 | 1.81E-01 | -0.2 | 6.58E-01 | TLC domain-containing protein |
| AT1G23060 | -0.8 | 1.70E-04 | -0.1 | 7.08E-01 | -0.8 | 1.47E-03 | -0.3 | 3.04E-01 | -0.1 | 8.05E-01 | -0.2 | 5.93E-01 | T26J12.16 protein |

| | | | | | | | | | | | | | |
|---|---|---|---|---|---|---|---|---|---|---|---|---|---|
| AT4G14550 | -0.8 | 2.48E-05 | 0.0 | 7.58E-01 | -0.8 | 2.37E-04 | 0.0 | 9.93E-01 | 0.1 | 6.90E-01 | -0.1 | 7.22E-01 | Auxin-responsive protein |
| AT5G07650 | -0.8 | 1.66E-04 | -0.5 | 5.39E-03 | -0.3 | 7.94E-02 | -0.7 | 1.70E-01 | 0.2 | 7.21E-01 | -0.9 | 1.45E-01 | Formin-like protein |
| AT2G42760 | -0.8 | 1.04E-05 | -0.3 | 1.44E-02 | -0.6 | 1.00E-03 | 0.1 | 4.69E-01 | 0.4 | 2.17E-02 | -0.3 | 1.62E-01 | DUF1685 family protein |
| AT3G50160 | -0.8 | 1.33E-03 | 0.0 | 8.63E-01 | -0.8 | 6.66E-03 | 0.3 | 5.46E-01 | 0.6 | 1.51E-01 | -0.3 | 4.62E-01 | Transmembrane protein, putative |
| AT5G57070 | -0.8 | 1.83E-06 | -0.4 | 3.86E-04 | -0.4 | 1.09E-03 | -0.2 | 9.48E-02 | 0.0 | 9.22E-01 | -0.3 | 1.01E-01 | Gb\|AAF21150.1 |
| AT5G14020 | -0.8 | 4.64E-04 | 0.3 | 3.71E-02 | -1.2 | 2.70E-04 | -0.3 | 4.40E-01 | 0.1 | 8.97E-01 | -0.4 | 4.30E-01 | Endosomal targeting BRO1-like domain-containing protein |
| AT1G16500 | -0.8 | 1.03E-04 | -0.6 | 1.33E-03 | -0.2 | 1.54E-01 | 0.0 | 7.87E-01 | -0.1 | 8.02E-01 | 0.0 | 9.88E-01 | Uncharacterized protein |
| AT5G18970 | -0.8 | 2.38E-05 | -0.4 | 2.97E-03 | -0.4 | 9.46E-03 | 0.2 | 3.54E-01 | 0.3 | 2.70E-01 | -0.1 | 8.66E-01 | hypothetical protein |
| AT1G08820 | -0.8 | 5.55E-07 | -0.6 | 8.25E-06 | -0.2 | 1.64E-02 | -0.1 | 4.40E-01 | 0.0 | 9.69E-01 | -0.1 | 6.33E-01 | MSP domain-containing protein |
| AT1G16860 | -0.8 | 8.26E-07 | -0.3 | 6.42E-04 | -0.5 | 1.99E-04 | -0.4 | 4.05E-03 | -0.1 | 6.94E-01 | -0.4 | 3.22E-02 | Uncharacterized membrane protein At1g16860 |
| AT2G30933 | -0.8 | 4.34E-04 | -0.3 | 8.88E-02 | -0.5 | 2.02E-02 | 0.1 | 8.32E-01 | 0.4 | 2.25E-01 | -0.4 | 3.81E-01 | X8 domain-containing protein |
| AT1G14780 | -0.8 | 1.99E-05 | -0.8 | 4.13E-05 | 0.0 | 9.70E-01 | -0.2 | 5.15E-01 | 0.3 | 3.44E-01 | -0.5 | 1.56E-01 | MACPF domain-containing protein At1g14780 |
| AT3G54290 | -0.8 | 1.77E-05 | -0.6 | 2.46E-04 | -0.2 | 9.38E-02 | -0.2 | 1.69E-01 | 0.1 | 8.32E-01 | -0.3 | 2.33E-01 | Uncharacterized protein F24B22.250 |
| AT1G77600 | -0.8 | 1.00E-05 | -0.7 | 6.51E-05 | -0.1 | 2.56E-01 | -0.2 | 3.85E-01 | 0.2 | 4.00E-01 | -0.4 | 9.58E-02 | ARM repeat superfamily protein |
| AT4G13530 | -0.8 | 3.65E-07 | -0.6 | 8.68E-06 | -0.2 | 4.75E-03 | -0.7 | 3.35E-04 | -0.4 | 2.36E-02 | -0.3 | 2.02E-01 | Transmembrane protein |
| AT1G76980 | -0.8 | 9.96E-05 | -0.9 | 8.15E-05 | 0.1 | 5.30E-01 | 0.3 | 3.46E-01 | 0.4 | 1.78E-01 | -0.1 | 7.19E-01 | Patatin-like phospholipase domain protein |
| AT3G58350 | -0.8 | 2.02E-04 | -0.7 | 1.04E-03 | -0.1 | 4.53E-01 | -0.3 | 4.44E-01 | 0.0 | 9.78E-01 | -0.3 | 5.01E-01 | MATH domain-containing protein |
| AT1G66450 | -0.8 | 1.20E-03 | -0.3 | 9.92E-02 | -0.5 | 5.19E-02 | -0.3 | 6.69E-01 | 0.1 | 9.19E-01 | -0.4 | 6.32E-01 | Cysteine |
| AT5G55860 | -0.8 | 3.65E-07 | -0.6 | 6.46E-06 | -0.2 | 7.90E-03 | -0.1 | 4.93E-01 | -0.1 | 6.65E-01 | 0.0 | 9.39E-01 | WEB family protein At5g55860 |
| AT4G11211 | -0.8 | 5.69E-04 | -0.4 | 1.96E-02 | -0.4 | 8.92E-02 | -0.1 | 7.09E-01 | 0.2 | 5.57E-01 | -0.3 | 4.25E-01 | Uncharacterized protein |
| AT1G28765 | -0.8 | 3.78E-04 | -0.8 | 5.17E-04 | 0.0 | 9.04E-01 | -0.1 | 8.83E-01 | 0.1 | 8.23E-01 | -0.2 | 7.50E-01 | Uncharacterized protein |
| AT1G04540 | -0.8 | 1.43E-05 | -0.2 | 6.36E-02 | -0.6 | 7.14E-04 | -0.4 | 2.63E-01 | 0.2 | 7.05E-01 | -0.6 | 2.00E-01 | T1G11.21 protein |
| AT3G62780 | -0.8 | 7.64E-05 | 0.0 | 7.97E-01 | -0.8 | 5.77E-04 | -0.4 | 1.76E-01 | 0.1 | 8.10E-01 | -0.5 | 1.61E-01 | At3g62780 family protein) |
| AT5G55960 | -0.8 | 2.93E-06 | -0.5 | 9.65E-05 | -0.3 | 1.20E-02 | -0.3 | 1.50E-02 | 0.0 | 9.92E-01 | -0.3 | 8.42E-02 | Transmembrane protein 245 |
| AT1G02860 | -0.8 | 6.11E-07 | -0.6 | 8.56E-06 | -0.2 | 2.35E-02 | 0.2 | 3.58E-01 | -0.1 | 6.82E-01 | 0.3 | 2.75E-01 | RING-type E3 ubiquitin transferase |
| AT5G46230 | -0.8 | 8.16E-05 | -0.2 | 1.52E-01 | -0.6 | 2.25E-03 | 0.0 | 9.69E-01 | 0.2 | 3.94E-01 | -0.2 | 4.12E-01 | hypothetical protein |
| AT3G13600 | -0.8 | 5.91E-04 | -0.2 | 3.27E-01 | -0.7 | 8.47E-03 | 0.9 | 6.60E-03 | 0.3 | 5.09E-01 | 0.7 | 6.93E-02 | IQ domain-containing protein IQM2 |
| AT5G07110 | -0.8 | 7.06E-05 | -0.2 | 1.77E-01 | -0.6 | 1.83E-03 | -0.5 | 6.75E-02 | -0.1 | 8.43E-01 | -0.5 | 2.48E-01 | PRA1 family protein B6 |
| AT2G15320 | -0.8 | 1.78E-06 | -0.7 | 1.50E-05 | -0.2 | 1.01E-01 | -0.2 | 4.89E-02 | 0.0 | 8.76E-01 | -0.2 | 2.03E-01 | Leucine-rich repeat family protein |
| AT1G75400 | -0.8 | 1.49E-06 | -0.6 | 2.12E-05 | -0.2 | 3.31E-02 | -0.4 | 1.64E-02 | 0.0 | 9.91E-01 | -0.4 | 4.57E-02 | F1B16.7 protein |
| AT3G23175 | -0.8 | 5.10E-06 | 0.2 | 5.10E-02 | -1.0 | 2.10E-05 | 0.3 | 2.10E-01 | 0.5 | 4.58E-02 | -0.2 | 3.23E-01 | HR-like lesion-inducing protein |
| AT3G16650 | -0.8 | 4.88E-04 | -0.4 | 2.48E-02 | -0.4 | 6.62E-02 | 1.2 | 1.53E-02 | 0.8 | 1.23E-01 | | | Protein pleiotropic regulator PRL2 |
| AT2G36840 | -0.8 | 1.09E-05 | -0.8 | 2.19E-05 | 0.0 | 1.00E+00 | -0.4 | 2.61E-02 | -0.1 | 7.28E-01 | -0.4 | 1.83E-01 | ACT domain-containing protein ACR10 |
| AT4G36120 | -0.8 | 4.91E-05 | -0.5 | 1.32E-03 | -0.3 | 6.39E-02 | 0.0 | 9.17E-01 | 0.0 | 9.80E-01 | 0.0 | 9.60E-01 | Filament-like plant protein 5 |
| AT1G65510 | -0.8 | 2.48E-04 | -0.9 | 1.78E-04 | 0.1 | 5.74E-01 | -0.3 | 2.51E-01 | -0.1 | 8.64E-01 | -0.2 | 4.70E-01 | Transmembrane protein |
| AT2G19580 | -0.8 | 4.55E-04 | -0.1 | 4.05E-01 | -0.7 | 6.05E-03 | 0.0 | 9.44E-01 | 0.1 | 9.04E-01 | -0.1 | 8.61E-01 | Tetraspanin-2 |
| AT1G28480 | -0.8 | 1.26E-02 | -1.0 | 6.47E-03 | 0.2 | 6.64E-01 | 1.4 | 1.02E-02 | 1.2 | 4.44E-02 | 0.2 | 6.66E-01 | Glutaredoxin-C9 |
| AT2G20750 | -0.8 | 2.74E-04 | 0.4 | 1.07E-02 | -1.2 | 8.64E-05 | -0.7 | 2.60E-03 | 0.0 | 9.80E-01 | -0.7 | 1.12E-02 | Expansin B1 |
| AT5G66870 | -0.8 | 8.30E-05 | -0.2 | 6.85E-02 | -0.6 | 4.11E-03 | -0.1 | 8.49E-01 | 0.3 | 3.23E-01 | -0.4 | 1.59E-01 | LOB domain-containing protein 36 |
| AT5G67140 | -0.8 | 4.70E-05 | -0.9 | 5.32E-05 | 0.1 | 6.66E-01 | 0.1 | 8.56E-01 | 0.4 | 3.55E-01 | -0.3 | 5.21E-01 | Uncharacterized protein |
| AT3G07360 | -0.8 | 1.17E-06 | -0.8 | 2.01E-06 | 0.0 | 5.69E-01 | -0.8 | 2.97E-05 | -0.4 | 1.70E-02 | -0.4 | 3.99E-02 | PUB9 |
| AT2G39530 | -0.8 | 2.18E-04 | 0.2 | 2.48E-01 | -1.0 | 2.58E-04 | 0.0 | 9.40E-01 | 0.5 | 8.75E-02 | -0.5 | 6.02E-02 | CASP-like protein 4D1 |
| AT3G15300 | -0.8 | 6.19E-05 | -0.6 | 9.54E-04 | -0.2 | 1.20E-01 | 0.1 | 6.15E-01 | 0.0 | 9.66E-01 | 0.1 | 6.67E-01 | VQ motif-containing protein 19 |
| AT1G23750 | -0.8 | 9.94E-05 | 0.3 | 4.79E-02 | -1.1 | 9.94E-05 | -0.3 | 2.26E-01 | 0.1 | 7.35E-01 | -0.4 | 1.85E-01 | Nucleic acid-binding, OB-fold-like protein |
| AT5G43980 | -0.8 | 2.24E-03 | -0.9 | 2.26E-03 | 0.1 | 8.44E-01 | -0.3 | 6.65E-01 | 0.2 | 8.35E-01 | -0.4 | 5.72E-01 | Plasmodesmata-located protein 1 |
| AT3G29810 | -0.8 | 1.71E-03 | -0.2 | 3.76E-01 | -0.6 | 2.09E-02 | -0.7 | 2.57E-01 | -0.7 | 3.03E-01 | 0.0 | 9.91E-01 | COBRA-like protein 2 |
| AT2G41330 | -0.8 | 2.90E-04 | -0.2 | 1.10E-01 | -0.6 | 1.10E-02 | -0.7 | 4.80E-03 | -0.3 | 1.88E-01 | -0.4 | 2.24E-01 | Glutaredoxin domain-containing protein |
| AT1G78120 | -0.8 | 1.52E-05 | -0.2 | 3.92E-02 | -0.6 | 8.48E-04 | -0.2 | 3.42E-01 | 0.3 | 2.31E-01 | -0.6 | 3.87E-02 | Tetratricopeptide repeat -like superfamily protein |
| AT2G20515 | -0.8 | 1.12E-03 | 0.3 | 1.42E-01 | -1.0 | 7.77E-04 | -0.3 | 1.80E-01 | -0.1 | 8.57E-01 | -0.3 | 4.31E-01 | At2g20515 |
| AT5G62210 | -0.8 | 1.22E-02 | -0.6 | 5.39E-02 | -0.2 | 5.27E-01 | -0.1 | 9.01E-01 | 0.6 | 1.12E-01 | -0.7 | 1.01E-01 | At5g62210 |
| AT2G35600 | -0.8 | 5.00E-03 | -0.5 | 3.53E-02 | -0.2 | 4.23E-01 | | | | | | | Protein Brevis radix-like 1 |
| AT1G07900 | -0.8 | 1.29E-01 | -1.9 | 1.14E-02 | 1.1 | 1.61E-01 | 0.4 | 4.73E-01 | 0.0 | 9.78E-01 | 0.4 | 5.02E-01 | LOB domain-containing protein 1 |
| AT5G55950 | -0.8 | 9.14E-03 | -0.3 | 3.10E-01 | -0.5 | 1.13E-01 | -0.2 | 7.69E-01 | 0.4 | 5.58E-01 | -0.5 | 4.22E-01 | Nucleotide |
| AT3G60690 | -0.8 | 2.22E-04 | -0.4 | 2.13E-02 | -0.4 | 2.72E-02 | -0.4 | 1.34E-01 | -0.4 | 9.84E-02 | 0.1 | 8.62E-01 | AT3g60690 |
| AT3G24840 | -0.8 | 2.47E-05 | -0.4 | 3.14E-03 | -0.4 | 9.89E-03 | 0.0 | 9.41E-01 | 0.4 | 1.75E-01 | -0.4 | 1.70E-01 | Sec14p-like phosphatidylinositol transfer family protein |
| AT5G05840 | -0.8 | 1.47E-04 | -0.2 | 1.87E-01 | -0.6 | 3.63E-03 | -1.6 | 8.29E-04 | 0.0 | 9.56E-01 | -1.6 | 9.66E-03 | Uncharacterized protein |
| AT5G20900 | -0.8 | 1.06E-06 | -0.6 | 1.03E-05 | -0.2 | 6.19E-02 | -0.1 | 5.05E-01 | 0.1 | 5.70E-01 | -0.3 | 3.01E-01 | Protein TIFY 3B |
| AT5G48920 | -0.8 | 7.22E-05 | 0.0 | 8.59E-01 | -0.8 | 3.94E-04 | 0.3 | 1.97E-01 | 0.3 | 2.53E-01 | 0.0 | 9.99E-01 | Protein TRACHEARY ELEMENT DIFFERENTIATION-RELATED 7 |
| AT5G19050 | -0.8 | 1.54E-06 | -0.4 | 2.15E-04 | -0.4 | 1.46E-03 | -0.3 | 6.27E-02 | -0.1 | 5.68E-01 | -0.2 | 3.63E-01 | Uncharacterized protein |
| AT1G08760 | -0.8 | 1.47E-06 | -0.4 | 1.41E-04 | -0.3 | 2.25E-03 | -0.2 | 1.95E-01 | 0.0 | 8.35E-01 | -0.2 | 4.93E-01 | Uncharacterized protein |
| AT3G16510 | -0.8 | 3.88E-05 | -0.5 | 1.06E-03 | -0.3 | 5.75E-02 | 0.1 | 8.16E-01 | 0.1 | 8.88E-01 | 0.0 | 9.77E-01 | hypothetical protein |
| AT2G01660 | -0.8 | 5.69E-06 | -0.4 | 1.51E-03 | -0.4 | 2.25E-03 | -0.3 | 6.47E-02 | -0.1 | 6.10E-01 | -0.2 | 4.06E-01 | PDLP6 |
| AT4G32860 | -0.8 | 7.39E-05 | -0.4 | 4.01E-03 | -0.4 | 3.68E-02 | -0.1 | 7.81E-01 | 0.1 | 8.25E-01 | -0.2 | 5.69E-01 | Avr9 |
| AT3G13720 | -0.8 | 1.36E-06 | -0.2 | 6.65E-03 | -0.5 | 1.10E-04 | -0.4 | 6.82E-02 | -0.3 | 1.51E-01 | 0.0 | 9.20E-01 | PRA1 family protein F3 |
| AT5G54870 | -0.8 | 1.12E-06 | -0.7 | 4.95E-06 | -0.1 | 3.36E-01 | -0.3 | 5.06E-02 | 0.0 | 9.87E-01 | -0.3 | 1.31E-01 | Inositol-1,4,5-trisphosphate 5-phosphatase |
| AT4G02100 | -0.8 | 1.57E-05 | -0.2 | 4.00E-02 | -0.6 | 8.00E-04 | 0.0 | 9.65E-01 | -0.1 | 6.54E-01 | 0.1 | 7.21E-01 | J domain-containing protein |
| AT1G28400 | -0.8 | 1.10E-05 | -0.3 | 4.49E-03 | -0.4 | 2.05E-03 | 0.3 | 4.89E-02 | 0.4 | 1.65E-02 | -0.1 | 5.80E-01 | F3M18.16 |
| AT5G66930 | -0.8 | 4.77E-06 | -0.6 | 6.58E-05 | -0.2 | 6.56E-02 | -0.6 | 3.12E-03 | -0.1 | 5.66E-01 | -0.4 | 7.24E-02 | AT5G66930 protein |
| AT3G18950 | -0.8 | 1.15E-04 | -0.7 | 3.59E-04 | -0.1 | 6.77E-01 | 0.0 | 9.64E-01 | 0.2 | 3.23E-01 | -0.3 | 3.85E-01 | En |
| AT1G29520 | -0.8 | 6.53E-06 | -0.2 | 4.71E-02 | -0.6 | 2.68E-04 | 0.1 | 2.88E-01 | 0.0 | 9.95E-01 | 0.1 | 4.41E-01 | AWPM-19-like family protein |
| AT3G25130 | -0.8 | 2.77E-05 | -0.6 | 3.77E-04 | -0.2 | 1.27E-01 | 0.0 | 9.04E-01 | 0.4 | 1.08E-01 | -0.4 | 1.07E-01 | Uncharacterized protein |
| AT5G12880 | -0.8 | 1.65E-04 | 0.1 | 3.66E-01 | -0.9 | 3.77E-04 | 0.8 | 1.23E-03 | 0.5 | 3.23E-02 | 0.3 | 2.24E-01 | hypothetical protein |
| AT1G73850 | -0.8 | 2.63E-06 | -0.6 | 3.10E-05 | -0.2 | 6.88E-02 | -0.1 | 6.04E-01 | -0.1 | 6.17E-01 | 0.0 | 9.78E-01 | At1g73850 |
| AT4G33130 | -0.8 | 1.72E-03 | -0.2 | 3.70E-01 | -0.6 | 2.16E-02 | -0.4 | 1.21E-01 | -0.1 | 8.35E-01 | -0.3 | 3.46E-01 | Uncharacterized protein At4g33130 |
| AT5G46910 | -0.8 | 4.76E-06 | -0.2 | 1.20E-02 | -0.5 | 4.47E-04 | 0.0 | 8.21E-01 | 0.1 | 8.36E-01 | -0.1 | 7.11E-01 | Lysine-specific demethylase JMJ706-like |
| AT5G24230 | -0.8 | 9.53E-05 | -0.5 | 1.85E-03 | -0.3 | 1.04E-01 | 0.0 | 8.85E-01 | 0.3 | 3.34E-01 | -0.3 | 3.25E-01 | Fungal lipase-like domain-containing protein |
| AT1G15740 | -0.8 | 3.26E-04 | -0.8 | 5.55E-04 | 0.0 | 9.62E-01 | -0.8 | 2.66E-05 | -0.7 | 2.64E-04 | -0.1 | 7.35E-01 | Uncharacterized protein At1g15740 |
| AT1G70900 | -0.8 | 1.52E-04 | -0.6 | 1.14E-03 | -0.2 | 2.94E-01 | -0.2 | 2.36E-01 | 0.0 | 9.01E-01 | -0.3 | 2.91E-01 | Uncharacterized protein |
| AT3G48185 | -0.8 | 6.16E-05 | -0.3 | 3.68E-02 | -0.5 | 4.80E-03 | 0.0 | 9.61E-01 | 0.0 | 9.75E-01 | 0.0 | 9.97E-01 | Transmembrane protein |
| AT4G35840 | -0.8 | 1.05E-04 | -0.2 | 8.90E-02 | -0.6 | 4.23E-03 | 0.1 | 9.05E-01 | 0.3 | 5.38E-01 | -0.2 | 6.49E-01 | RING |
| AT1G29390 | -0.8 | 3.02E-05 | -0.4 | 2.34E-03 | -0.3 | 1.47E-02 | -0.5 | 1.36E-03 | 0.0 | 9.21E-01 | -0.5 | 1.17E-02 | Uncharacterized protein |
| AT1G03380 | -0.8 | 4.82E-05 | -0.7 | 1.50E-04 | -0.1 | 6.49E-01 | -0.2 | 7.31E-02 | -0.1 | 6.61E-01 | -0.2 | 3.90E-01 | Autophagy-related protein 18g |
| AT1G49030 | -0.8 | 2.13E-04 | 0.5 | 3.83E-03 | -1.2 | 5.96E-05 | -0.4 | 2.56E-02 | 0.3 | 7.73E-02 | -0.7 | 4.08E-03 | Protein PLANT CADMIUM RESISTANCE 6 |

| | | | | | | | | | | | | | |
|---|---|---|---|---|---|---|---|---|---|---|---|---|---|
| AT2G25490 | -0.8 | 4.37E-05 | -0.6 | 3.88E-04 | -0.2 | 2.12E-01 | 0.1 | 5.02E-01 | 0.1 | 6.83E-01 | 0.0 | 9.44E-01 | EIN3-binding F-box protein 1 |
| AT1G15760 | -0.8 | 2.45E-03 | -0.2 | 3.48E-01 | -0.6 | 3.29E-02 | 0.4 | 1.60E-01 | 0.2 | 5.69E-01 | 0.2 | 5.86E-01 | At1g15760 domain-containing protein) |
| AT1G32410 | -0.8 | 5.36E-06 | -0.5 | 1.88E-04 | -0.3 | 1.61E-02 | -0.4 | 2.40E-03 | -0.3 | 4.73E-02 | -0.1 | 4.62E-01 | Vacuolar protein sorting-associated protein 55 homolog |
| AT5G01610 | -0.8 | 1.66E-04 | -0.1 | 4.39E-01 | -0.7 | 1.83E-03 | -0.6 | 2.42E-02 | -0.2 | 5.83E-01 | -0.4 | 1.01E-01 | Uncharacterized protein At5g01610 |
| AT3G11320 | -0.8 | 2.68E-05 | -0.2 | 5.42E-02 | -0.5 | 1.12E-03 | -0.2 | 2.49E-01 | 0.1 | 8.34E-01 | -0.3 | 3.12E-01 | Sugar phosphate transporter domain-containing protein |
| AT3G01860 | -0.8 | 1.18E-05 | -0.6 | 1.67E-04 | -0.2 | 7.69E-02 | -0.1 | 4.60E-01 | -0.2 | 3.04E-01 | 0.1 | 8.06E-01 | F28J7.19 protein |
| AT1G54710 | -0.8 | 2.24E-06 | -0.4 | 1.59E-04 | -0.3 | 3.99E-03 | -0.2 | 1.83E-01 | 0.1 | 3.39E-01 | -0.3 | 7.26E-02 | BCAS3 domain-containing protein |
| AT5G14230 | -0.8 | 4.34E-05 | -0.7 | 1.69E-04 | -0.1 | 5.46E-01 | 1.0 | 8.94E-02 | 0.8 | 2.46E-01 | 0.2 | 7.99E-01 | Ankyrin |
| AT3G05980 | -0.8 | 9.68E-04 | 0.3 | 6.96E-02 | -1.0 | 5.44E-04 | -0.2 | 2.39E-01 | 0.1 | 8.26E-01 | -0.3 | 2.16E-01 | hypothetical protein |
| AT4G19390 | -0.8 | 2.95E-04 | -0.6 | 1.94E-03 | -0.2 | 3.46E-01 | -0.4 | 1.25E-02 | -0.6 | 2.79E-03 | 0.2 | 4.84E-01 | AT4g19390 |
| AT4G33160 | -0.8 | 1.33E-04 | -0.4 | 5.11E-03 | -0.3 | 6.16E-02 | -0.4 | 2.27E-01 | -0.3 | 4.41E-01 | -0.1 | 8.15E-01 | F-box family protein |
| AT5G51670 | -0.8 | 2.68E-05 | -0.5 | 7.05E-04 | -0.3 | 5.41E-02 | -0.2 | 2.82E-01 | -0.1 | 6.72E-01 | -0.1 | 7.11E-01 | hypothetical protein |
| AT5G42570 | -0.8 | 1.99E-05 | -0.4 | 2.30E-03 | -0.4 | 7.85E-03 | -0.1 | 4.75E-01 | 0.0 | 9.91E-01 | -0.1 | 5.46E-01 | Endoplasmic reticulum transmembrane protein |
| AT5G64430 | -0.8 | 2.71E-04 | -0.8 | 2.80E-04 | 0.0 | 7.76E-01 | -0.2 | 3.26E-01 | -0.3 | 3.15E-01 | 0.0 | 9.30E-01 | PB1 domain-containing protein |
| AT2G23050 | -0.8 | 8.91E-03 | 0.2 | 4.71E-01 | -0.9 | 8.81E-03 | 0.0 | 9.55E-01 | 0.2 | 5.40E-01 | -0.2 | 5.90E-01 | Phototropic-responsive NPH3 family protein |
| AT1G31750 | -0.8 | 5.48E-04 | -0.2 | 1.54E-01 | -0.5 | 1.72E-02 | 0.0 | 9.26E-01 | 0.6 | 1.62E-01 | -0.5 | 2.17E-01 | At1g31750 |
| AT2G39705 | -0.8 | 1.16E-04 | -0.5 | 1.40E-03 | -0.2 | 1.73E-01 | -0.3 | 3.21E-01 | -0.1 | 7.48E-01 | -0.2 | 7.30E-01 | Small polypeptide DEVIL 11 |
| AT5G46060 | -0.8 | 1.33E-03 | 0.0 | 9.91E-01 | -0.8 | 5.28E-03 | -0.6 | 5.47E-02 | -0.3 | 3.37E-01 | -0.3 | 5.64E-01 | Expressed protein |
| AT4G12730 | -0.8 | 1.59E-05 | -0.2 | 1.32E-01 | -0.6 | 3.67E-04 | -0.5 | 2.79E-03 | -0.2 | 2.78E-01 | -0.3 | 1.35E-01 | Fasciclin-like arabinogalactan protein 2 |
| AT3G54990 | -0.8 | 5.16E-04 | -0.4 | 1.84E-02 | -0.3 | 9.45E-02 | -0.2 | 7.80E-01 | 0.0 | 9.66E-01 | -0.2 | 8.70E-01 | SMZ |
| AT5G56230 | -0.8 | 8.44E-04 | -0.6 | 6.14E-03 | -0.2 | 3.72E-01 | -0.2 | 6.84E-01 | 0.0 | 9.82E-01 | -0.2 | 7.90E-01 | PRA1 family protein |
| AT1G78310 | -0.7 | 5.15E-06 | -0.5 | 8.24E-05 | -0.2 | 5.61E-02 | -0.3 | 2.77E-01 | -0.3 | 2.66E-01 | 0.0 | 9.29E-01 | VQ motif-containing protein 9 |
| AT4G00355 | -0.7 | 1.16E-06 | -0.5 | 4.17E-05 | -0.3 | 5.71E-03 | -0.7 | 1.92E-05 | -0.4 | 7.57E-03 | -0.3 | 7.46E-02 | ATG8-interacting protein 2 |
| AT2G37035 | -0.7 | 1.92E-06 | -0.4 | 1.27E-04 | -0.3 | 4.24E-03 | -0.5 | 3.87E-03 | -0.4 | 3.37E-02 | -0.1 | 5.95E-01 | Transmembrane protein |
| AT5G52280 | -0.7 | 1.21E-05 | -0.2 | 3.04E-02 | -0.5 | 7.03E-04 | -0.3 | 9.18E-02 | 0.1 | 6.56E-01 | -0.4 | 9.97E-02 | At5g52280 |
| AT5G42146 | -0.7 | 1.76E-04 | -0.1 | 2.96E-01 | -0.6 | 3.21E-03 | -0.2 | 3.24E-01 | -0.1 | 8.65E-01 | -0.2 | 6.13E-01 | Transmembrane protein |
| AT1G22230 | -0.7 | 2.21E-05 | -0.2 | 2.77E-02 | -0.5 | 1.74E-03 | -0.6 | 3.40E-02 | 0.0 | 9.66E-01 | -0.6 | 9.13E-02 | Uncharacterized protein |
| AT5G06330 | -0.7 | 2.15E-04 | -0.5 | 3.27E-03 | -0.2 | 1.58E-01 | -0.3 | 3.43E-01 | 0.0 | 9.91E-01 | -0.3 | 4.45E-01 | hypothetical protein |
| AT1G67800 | -0.7 | 1.39E-05 | -1.0 | 4.34E-06 | 0.2 | 3.56E-02 | 0.7 | 4.59E-04 | 0.2 | 2.27E-01 | 0.4 | 5.47E-02 | Copine family |
| AT4G06744 | -0.7 | 5.94E-06 | -0.3 | 1.05E-02 | -0.5 | 4.90E-04 | 0.0 | 9.51E-01 | 0.2 | 2.17E-01 | -0.2 | 2.56E-01 | Uncharacterized protein At4g06744 |
| AT3G01750 | -0.7 | 1.14E-04 | -0.5 | 1.79E-03 | -0.2 | 1.39E-01 | -0.4 | 1.59E-02 | -0.3 | 1.73E-01 | -0.2 | 5.11E-01 | Ankyrin repeat family protein |
| AT5G02060 | -0.7 | 7.22E-04 | -0.3 | 9.39E-02 | -0.5 | 3.38E-02 | 0.5 | 2.12E-01 | 0.1 | 8.43E-01 | 0.4 | 3.82E-01 | CASP-like protein 5B1 |
| AT1G03290 | -0.7 | 2.48E-05 | -0.9 | 1.29E-05 | 0.2 | 1.68E-01 | 0.1 | 6.10E-01 | 0.0 | 9.24E-01 | 0.1 | 8.23E-01 | ELKS |
| AT3G29180 | -0.7 | 5.16E-06 | -0.2 | 1.23E-02 | -0.5 | 4.12E-04 | -0.4 | 6.52E-03 | -0.1 | 6.35E-01 | -0.3 | 9.50E-02 | AT3G29180 protein ) |
| AT4G23870 | -0.7 | 2.59E-05 | 0.0 | 9.62E-01 | -0.7 | 1.76E-04 | -1.1 | 1.86E-03 | -0.5 | 1.50E-01 | -0.6 | 1.26E-01 | Uncharacterized protein |
| AT4G10360 | -0.7 | 1.52E-06 | -0.5 | 5.41E-05 | -0.3 | 6.88E-03 | 0.0 | 8.40E-01 | 0.1 | 4.20E-01 | -0.2 | 2.99E-01 | AT4G10360 protein lipid-sensing domain containing protein) |
| AT3G06170 | -0.7 | 1.18E-06 | -0.7 | 5.04E-06 | -0.1 | 3.99E-01 | -0.5 | 7.62E-05 | -0.3 | 2.40E-02 | -0.2 | 9.79E-02 | F28L1.11 protein |
| AT3G12090 | -0.7 | 2.62E-06 | -0.3 | 4.56E-03 | -0.5 | 3.47E-04 | 0.3 | 8.60E-02 | 0.3 | 2.32E-01 | 0.1 | 8.21E-01 | Tetraspanin-6 |
| AT4G02715 | -0.7 | 8.09E-06 | -0.5 | 1.91E-04 | -0.2 | 3.25E-02 | -0.4 | 5.53E-03 | -0.1 | 5.05E-01 | -0.3 | 9.10E-02 | At4g02715 |
| AT4G27900 | -0.7 | 5.05E-05 | -0.6 | 5.55E-04 | -0.2 | 1.59E-01 | 0.0 | 8.99E-01 | 0.0 | 8.46E-01 | -0.1 | 7.86E-01 | CCT domain-containing protein |
| AT3G09032 | -0.7 | 2.40E-03 | -0.1 | 5.88E-01 | -0.6 | 1.77E-02 | -0.3 | 3.84E-01 | -0.4 | 2.57E-01 | 0.1 | 8.13E-01 | Josephin-like protein |
| AT1G69610 | -0.7 | 5.27E-03 | -0.9 | 3.59E-03 | 0.1 | 7.09E-01 | -0.1 | 8.70E-01 | 0.3 | 2.81E-01 | -0.4 | 2.76E-01 | Uncharacterized protein F24J1.24 ) |
| AT5G02260 | -0.7 | 1.33E-03 | 0.2 | 1.61E-01 | -1.0 | 8.02E-04 | 0.1 | 8.39E-01 | 0.2 | 5.01E-01 | -0.1 | 6.85E-01 | Expansin-A9 |
| AT2G24040 | -0.7 | 4.22E-04 | -0.3 | 5.72E-02 | -0.4 | 2.59E-02 | -0.2 | 6.00E-01 | -0.3 | 4.26E-01 | 0.1 | 8.37E-01 | Low temperature and salt responsive protein family |
| AT5G48370 | -0.7 | 3.21E-06 | -0.5 | 5.43E-05 | -0.2 | 3.99E-02 | -0.3 | 1.02E-01 | 0.0 | 9.68E-01 | -0.3 | 2.25E-01 | hypothetical protein |
| AT4G14990 | -0.7 | 6.12E-06 | -0.5 | 1.85E-04 | -0.2 | 2.01E-02 | -0.3 | 7.72E-03 | 0.0 | 9.49E-01 | -0.3 | 5.11E-02 | Topoisomerase II-associated protein PAT1 |
| AT1G16930 | -0.7 | 4.93E-06 | -0.4 | 7.98E-04 | -0.4 | 3.35E-03 | -0.2 | 3.52E-01 | 0.0 | 9.15E-01 | -0.2 | 4.02E-01 | F-box |
| AT1G53400 | -0.7 | 2.34E-05 | -0.3 | 4.89E-03 | -0.4 | 6.00E-03 | -0.5 | 8.84E-02 | -0.3 | 2.62E-01 | -0.1 | 8.10E-01 | hypothetical protein |
| AT2G01070 | -0.7 | 3.12E-06 | -0.2 | 9.38E-03 | -0.5 | 2.79E-04 | -0.1 | 6.78E-01 | 0.1 | 7.16E-01 | -0.2 | 4.79E-01 | Lung seven transmembrane receptor family protein |
| AT5G52740 | -0.7 | 1.34E-01 | -1.4 | 2.56E-02 | 0.7 | 3.55E-01 | | | | | | | Copper transport protein family |
| AT2G04110 | -0.7 | 3.90E-05 | -0.5 | 1.16E-03 | -0.3 | 5.24E-02 | -0.2 | 4.45E-01 | 0.1 | 7.76E-01 | -0.2 | 3.89E-01 | NA |
| AT1G20030 | -0.7 | 2.50E-03 | 0.0 | 7.95E-01 | -0.8 | 5.57E-03 | 0.1 | 6.01E-01 | 0.1 | 7.38E-01 | 0.0 | 9.50E-01 | At1g20030 |
| AT2G38640 | -0.7 | 8.52E-05 | -0.3 | 1.37E-02 | -0.4 | 1.34E-02 | -0.2 | 4.37E-01 | -0.1 | 7.91E-01 | -0.1 | 7.58E-01 | Protein LURP-one-related 8 |
| AT3G52470 | -0.7 | 1.52E-06 | -0.2 | 5.62E-03 | -0.5 | 1.38E-04 | -0.1 | 6.84E-01 | 0.3 | 7.03E-02 | -0.3 | 6.82E-02 | hypothetical protein |
| AT1G70210 | -0.7 | 1.07E-04 | -0.3 | 3.26E-02 | -0.4 | 8.03E-03 | -0.1 | 3.82E-01 | 0.0 | 9.88E-01 | -0.1 | 4.90E-01 | CYCD1 |
| AT5G17520 | -0.7 | 1.25E-03 | -0.5 | 1.85E-02 | -0.3 | 2.06E-01 | -0.1 | 3.71E-01 | 0.1 | 4.15E-01 | -0.3 | 1.63E-01 | Root cap 1 |
| AT3G07195 | -0.7 | 5.32E-05 | -0.3 | 1.46E-02 | -0.4 | 5.98E-03 | -0.1 | 3.61E-01 | 0.0 | 9.36E-01 | -0.2 | 4.59E-01 | RPM1-interacting protein 4 family protein |
| AT2G15970 | -0.7 | 4.34E-05 | -0.1 | 2.76E-01 | -0.6 | 6.34E-04 | -0.7 | 1.10E-03 | -0.1 | 6.20E-01 | -0.6 | 2.83E-02 | Cold regulated 413 plasma membrane 1 |
| AT3G58710 | -0.7 | 2.06E-05 | -0.1 | 1.88E-01 | -0.6 | 4.66E-04 | -0.6 | 9.57E-03 | -0.2 | 5.02E-01 | -0.4 | 1.11E-01 | WRKY transcription factor 69 |
| AT5G10130 | -0.7 | 7.40E-03 | 0.3 | 2.53E-01 | -1.0 | 3.75E-03 | 0.0 | 9.38E-01 | 0.3 | 2.00E-01 | -0.4 | 2.40E-01 | Protein DOWNSTREAM OF FLC |
| AT4G10840 | -0.7 | 2.67E-06 | -0.3 | 5.79E-03 | -0.5 | 2.71E-04 | -0.4 | 2.66E-03 | -0.2 | 1.79E-01 | -0.2 | 2.48E-01 | Protein KINESIN LIGHT CHAIN-RELATED 1 |
| AT5G40690 | -0.7 | 6.57E-06 | -0.2 | 2.64E-02 | -0.5 | 3.39E-04 | 1.2 | 1.04E-04 | 0.3 | 2.51E-01 | 0.9 | 4.72E-03 | At5g40690 |
| AT3G23190 | -0.7 | 3.43E-05 | 0.2 | 4.46E-02 | -0.9 | 4.37E-05 | 0.2 | 2.97E-01 | 0.5 | 1.96E-02 | -0.3 | 9.62E-02 | Uncharacterized protein At3g23190 |
| AT5G38220 | -0.7 | 1.24E-05 | -0.4 | 8.28E-04 | -0.3 | 1.27E-02 | -0.2 | 1.61E-01 | 0.2 | 3.60E-01 | -0.4 | 6.62E-02 | Alpha |
| AT3G59210 | -0.7 | 9.76E-06 | -0.6 | 4.30E-05 | -0.1 | 4.19E-01 | -0.1 | 4.52E-01 | 0.1 | 3.83E-01 | -0.3 | 1.33E-01 | F-box |
| AT1G09070 | -0.7 | 4.76E-05 | -0.8 | 3.16E-05 | 0.1 | 3.09E-01 | -0.2 | 3.41E-02 | -0.4 | 2.18E-03 | 0.2 | 2.57E-01 | Protein SRC2 homolog |
| AT3G52920 | -0.7 | 4.06E-05 | -0.4 | 3.43E-03 | -0.3 | 1.51E-02 | -0.3 | 1.21E-01 | -0.1 | 7.87E-01 | -0.2 | 4.23E-01 | RAB6-interacting golgin |
| AT3G14920 | -0.7 | 4.94E-06 | -0.6 | 5.13E-05 | -0.2 | 9.83E-02 | 0.3 | 1.94E-01 | 0.4 | 8.24E-02 | -0.1 | 7.04E-01 | Peptide-N4-asparagine amidase A protein |
| AT5G02550 | -0.7 | 4.95E-04 | 0.0 | 7.55E-01 | -0.7 | 3.24E-03 | 0.1 | 7.60E-01 | 0.2 | 4.36E-01 | -0.1 | 6.36E-01 | hypothetical protein |
| AT5G10946 | -0.7 | 2.76E-02 | -0.8 | 2.24E-02 | 0.1 | 8.50E-01 | -0.4 | 1.68E-01 | 0.0 | 9.92E-01 | -0.4 | 3.03E-01 | Transmembrane protein |
| AT3G06420 | -0.7 | 1.74E-04 | 0.1 | 2.29E-01 | -0.9 | 2.55E-04 | -0.1 | 8.45E-01 | 0.0 | 9.53E-01 | -0.1 | 8.28E-01 | Autophagy-related protein 8h |
| AT3G03020 | -0.7 | 4.15E-05 | -0.6 | 1.54E-04 | -0.1 | 5.56E-01 | -0.5 | 3.50E-02 | -0.3 | 2.36E-01 | -0.2 | 5.95E-01 | Ovule protein |
| AT2G43680 | -0.7 | 6.51E-05 | -0.6 | 5.15E-04 | -0.1 | 2.68E-01 | -0.1 | 5.85E-01 | -0.2 | 4.11E-01 | 0.1 | 8.24E-01 | IQD14 |
| AT1G04030 | -0.7 | 1.12E-04 | -0.6 | 4.73E-04 | -0.1 | 5.14E-01 | -0.5 | 2.78E-01 | 0.3 | 5.64E-01 | -0.7 | 1.77E-01 | F21M11.3 protein |
| AT2G21500 | -0.7 | 1.83E-05 | -0.6 | 1.05E-04 | -0.1 | 3.31E-01 | 0.1 | 6.08E-01 | -0.2 | 4.02E-01 | 0.3 | 2.39E-01 | Uncharacterized protein At2g21500 |
| AT1G21760 | -0.7 | 1.33E-06 | -0.7 | 2.33E-06 | 0.0 | 6.65E-01 | -0.6 | 1.39E-04 | -0.3 | 1.48E-02 | -0.3 | 1.74E-01 | F-box protein |
| AT2G32240 | -0.7 | 1.66E-06 | -0.4 | 1.47E-04 | -0.3 | 2.60E-03 | 0.5 | 1.32E-03 | 0.2 | 1.31E-01 | 0.3 | 1.93E-01 | At2g32240 |
| AT2G18100 | -0.7 | 9.06E-05 | -0.4 | 2.68E-03 | -0.3 | 7.66E-02 | -0.9 | 2.00E-02 | -0.4 | 2.46E-01 | -0.4 | 4.21E-01 | Transmembrane |
| AT1G35160 | -0.7 | 4.38E-06 | -0.2 | 2.31E-02 | -0.5 | 2.34E-04 | 0.0 | 9.65E-01 | 0.1 | 5.38E-01 | -0.1 | 5.50E-01 | At1g35160 |
| AT1G69295 | -0.7 | 3.26E-05 | -0.3 | 8.37E-03 | -0.4 | 5.04E-03 | -0.2 | 1.31E-01 | 0.0 | 8.46E-01 | -0.2 | 4.10E-01 | X8 domain-containing protein |
| AT1G11740 | -0.7 | 2.40E-03 | 0.0 | 9.77E-01 | -0.7 | 8.47E-03 | -0.9 | 9.43E-04 | -0.8 | 3.56E-03 | -0.1 | 8.21E-01 | At1g11740 |
| AT4G12000 | -0.7 | 1.14E-04 | -0.6 | 7.23E-04 | -0.1 | 3.47E-01 | -0.4 | 9.23E-02 | -0.1 | 6.67E-01 | -0.3 | 3.14E-01 | hypothetical protein |
| AT1G68450 | -0.7 | 8.41E-04 | -0.7 | 1.17E-03 | 0.0 | 9.64E-01 | 0.8 | 2.00E-02 | 0.2 | 7.63E-01 | 0.7 | 3.32E-02 | VQ motif-containing protein 8, chloroplastic |

| | | | | | | | | | | | | | |
|---|---|---|---|---|---|---|---|---|---|---|---|---|---|
| AT2G45540 | -0.7 | 4.08E-05 | -0.5 | 6.00E-04 | -0.2 | 1.07E-01 | -0.1 | 8.00E-01 | 0.1 | 9.37E-01 | -0.2 | 7.99E-01 | BEACH domain-containing protein C2 |
| AT5G20880 | -0.7 | 1.36E-03 | -0.4 | 2.08E-02 | -0.3 | 2.00E-01 | 0.1 | 8.45E-01 | 0.0 | 8.96E-01 | 0.0 | 9.86E-01 | NA |
| AT2G23700 | -0.7 | 2.51E-05 | -0.4 | 1.48E-03 | -0.3 | 2.23E-02 | -0.2 | 3.74E-01 | 0.1 | 7.35E-01 | -0.2 | 2.65E-01 | Uncharacterized protein At2g23700 |
| AT2G30930 | -0.7 | 5.16E-04 | 0.2 | 2.51E-01 | -0.9 | 4.32E-04 | -0.8 | 6.93E-04 | -0.5 | 2.83E-02 | -0.3 | 2.73E-01 | Expressed protein |
| AT5G28430 | -0.7 | 1.69E-02 | -0.6 | 3.94E-02 | -0.1 | 7.65E-01 | | | | | | | NA |
| AT5G43950 | -0.7 | 1.69E-04 | -0.3 | 2.04E-02 | -0.4 | 2.33E-02 | -0.4 | 3.05E-02 | 0.0 | 9.18E-01 | -0.4 | 1.04E-01 | Uncharacterized protein At5g43950 ) |
| AT3G50910 | -0.7 | 1.44E-05 | -0.8 | 1.36E-05 | 0.1 | 4.26E-01 | 0.2 | 1.75E-01 | 0.3 | 1.35E-01 | 0.0 | 8.56E-01 | Uncharacterized protein F18B3.190 |
| AT1G54120 | -0.7 | 1.60E-03 | -0.6 | 7.64E-03 | -0.2 | 5.02E-01 | 0.1 | 8.81E-01 | -0.3 | 4.11E-01 | 0.4 | 2.49E-01 | hypothetical protein |
| AT1G08230 | -0.7 | 3.62E-04 | -0.5 | 5.16E-03 | -0.2 | 1.81E-01 | -0.2 | 3.23E-01 | 0.1 | 7.46E-01 | -0.2 | 3.04E-01 | Transmembrane amino acid transporter family protein |
| AT3G61900 | -0.7 | 2.01E-03 | -0.4 | 5.20E-02 | -0.3 | 1.45E-01 | | | | | | | SAUR-like auxin-responsive protein family |
| AT3G55390 | -0.7 | 1.93E-03 | -0.3 | 7.12E-02 | -0.4 | 1.07E-01 | -0.3 | 2.81E-01 | -0.2 | 5.89E-01 | -0.1 | 7.66E-01 | CASP-like protein 4C1 |
| AT1G63580 | -0.7 | 1.41E-04 | -0.4 | 6.99E-03 | -0.3 | 5.00E-02 | 1.1 | 6.06E-03 | 0.8 | 7.84E-02 | 0.3 | 3.89E-01 | Probable cysteine-rich repeat secretory protein 6 |
| AT5G16550 | -0.7 | 4.91E-05 | -0.3 | 1.34E-02 | -0.4 | 5.78E-03 | -0.6 | 4.02E-04 | -0.4 | 9.94E-03 | -0.2 | 3.63E-01 | Voltage-dependent L-type calcium channel subunit |
| AT2G15292 | -0.7 | 1.86E-05 | -0.1 | 2.89E-01 | -0.6 | 2.91E-04 | -0.1 | 7.09E-01 | -0.1 | 6.72E-01 | 0.0 | 9.43E-01 | NA |
| AT2G37930 | -0.7 | 4.19E-05 | -0.4 | 1.35E-03 | -0.3 | 5.33E-02 | -0.3 | 1.89E-01 | -0.2 | 4.30E-01 | -0.1 | 7.82E-01 | At2g37930 |
| AT1G65000 | -0.7 | 6.51E-05 | -0.3 | 7.10E-03 | -0.4 | 1.77E-02 | -0.4 | 1.14E-01 | 0.0 | 9.80E-01 | -0.4 | 2.36E-01 | Uncharacterized protein |
| AT2G19710 | -0.7 | 2.12E-05 | -0.6 | 9.44E-05 | -0.1 | 4.61E-01 | 0.1 | 7.30E-01 | -0.1 | 8.40E-01 | 0.1 | 6.41E-01 | Uncharacterized protein At2g19710 |
| AT5G42880 | -0.7 | 7.27E-05 | -0.1 | 2.09E-01 | -0.6 | 1.63E-03 | -0.4 | 1.92E-01 | 0.1 | 8.43E-01 | -0.5 | 1.95E-01 | Protein WEAK CHLOROPLAST MOVEMENT UNDER BLUE LIGHT-like 3 |
| AT2G03810 | -0.7 | 4.21E-04 | -0.5 | 5.25E-03 | -0.2 | 2.11E-01 | 0.2 | 2.42E-01 | 0.2 | 2.62E-01 | 0.0 | 9.59E-01 | hypothetical protein |
| AT5G23460 | -0.7 | 3.82E-05 | -0.2 | 3.45E-02 | -0.5 | 2.88E-03 | -0.1 | 8.62E-01 | -0.3 | 3.60E-01 | 0.3 | 5.72E-01 | Uncharacterized protein |
| AT5G45275 | -0.7 | 2.78E-04 | -0.5 | 1.94E-03 | -0.2 | 3.45E-01 | -0.1 | 6.83E-01 | 0.3 | 4.02E-01 | -0.4 | 2.16E-01 | Major facilitator superfamily protein |
| AT3G58720 | -0.7 | 1.35E-04 | -0.4 | 7.97E-03 | -0.3 | 4.08E-02 | 0.2 | 5.57E-01 | 0.0 | 9.81E-01 | 0.2 | 6.54E-01 | RING |
| AT5G19930 | -0.7 | 2.63E-05 | -0.2 | 2.43E-02 | -0.5 | 2.02E-03 | -0.3 | 6.47E-02 | -0.1 | 6.76E-01 | -0.2 | 3.19E-01 | Protein PGR |
| AT4G20880 | -0.7 | 2.51E-05 | -0.4 | 7.67E-04 | -0.2 | 4.43E-02 | -0.5 | 1.97E-02 | -0.2 | 3.70E-01 | -0.3 | 3.29E-01 | Ethylene-regulated transcript 2 ) |
| AT1G80240 | -0.7 | 1.84E-03 | -0.4 | 2.45E-02 | -0.3 | 2.23E-01 | 0.9 | 4.02E-03 | 0.3 | 4.28E-01 | 0.6 | 3.80E-02 | hypothetical protein |
| AT5G58000 | -0.7 | 2.91E-05 | -0.4 | 2.39E-03 | -0.3 | 1.64E-02 | -0.4 | 2.09E-01 | 0.2 | 4.48E-01 | -0.6 | 9.34E-02 | Reticulon-like protein |
| AT5G25050 | -0.7 | 9.41E-05 | -0.6 | 4.97E-04 | -0.1 | 4.26E-01 | -0.3 | 5.34E-01 | -0.4 | 4.31E-01 | 0.1 | 8.86E-01 | Probable folate-biopterin transporter 2 |
| AT4G32150 | -0.7 | 2.58E-05 | -0.4 | 3.26E-03 | -0.3 | 8.95E-03 | -0.5 | 5.82E-05 | -0.3 | 1.20E-02 | -0.2 | 1.35E-01 | Vesicle-associated membrane protein 711 |
| AT2G15220 | -0.7 | 8.65E-05 | -0.5 | 6.22E-04 | -0.1 | 2.96E-01 | 1.7 | 2.94E-05 | 1.2 | 3.21E-03 | 0.6 | 8.97E-02 | Plant basic secretory protein family protein |
| AT3G02550 | -0.7 | 4.88E-03 | -0.4 | 8.56E-02 | -0.3 | 1.84E-01 | 0.3 | 2.78E-01 | -0.3 | 3.28E-01 | 0.6 | 5.16E-02 | LOB domain-containing protein 41 |
| AT1G32690 | -0.7 | 2.87E-03 | -0.3 | 1.13E-01 | -0.4 | 9.76E-02 | -0.1 | 7.28E-01 | 0.0 | 9.71E-01 | -0.1 | 8.14E-01 | Uncharacterized protein |
| AT5G16530 | -0.7 | 3.02E-04 | -0.3 | 5.04E-02 | -0.4 | 2.31E-02 | 0.0 | 9.40E-01 | 0.2 | 7.67E-01 | -0.2 | 7.16E-01 | Auxin efflux carrier component |
| AT1G75850 | -0.7 | 2.56E-05 | -0.6 | 1.42E-04 | -0.1 | 3.43E-01 | -0.3 | 1.65E-01 | -0.1 | 5.74E-01 | -0.1 | 6.55E-01 | VPS35 homolog B |
| AT5G02230 | -0.7 | 2.41E-05 | -0.4 | 1.48E-03 | -0.3 | 1.83E-02 | -0.8 | 5.81E-05 | 0.2 | 2.84E-01 | -0.9 | 2.85E-04 | Haloacid dehalogenase-like hydrolase superfamily protein |
| AT5G09540 | -0.7 | 5.50E-04 | -0.5 | 3.39E-03 | -0.2 | 4.19E-01 | -0.5 | 1.68E-01 | -0.1 | 8.52E-01 | -0.4 | 3.78E-01 | At5g09540 |
| AT2G41660 | -0.7 | 1.71E-03 | -0.3 | 7.57E-02 | -0.4 | 8.20E-02 | 0.1 | 8.07E-01 | 0.2 | 4.13E-01 | -0.1 | 5.02E-01 | Protein MIZU-KUSSEI 1 |
| AT1G33230 | -0.7 | 3.53E-06 | -0.4 | 5.13E-04 | -0.3 | 2.43E-03 | -0.4 | 7.88E-03 | -0.1 | 4.51E-01 | -0.3 | 1.69E-01 | TMPIT-like protein |
| AT4G27350 | -0.7 | 1.45E-04 | 0.1 | 6.15E-01 | -0.7 | 4.86E-04 | -0.5 | 4.98E-02 | -0.2 | 4.87E-01 | -0.3 | 3.81E-01 | Uncharacterized protein |
| AT1G12380 | -0.7 | 1.28E-04 | -0.6 | 4.40E-04 | -0.1 | 6.19E-01 | -0.3 | 3.15E-01 | 0.2 | 5.43E-01 | -0.5 | 1.93E-01 | F5O11.10 |
| AT4G32140 | -0.7 | 2.53E-05 | -0.2 | 6.37E-02 | -0.5 | 1.07E-03 | 0.0 | 7.76E-01 | -0.1 | 4.80E-01 | 0.1 | 7.46E-01 | EamA domain-containing protein |
| AT2G37780 | -0.7 | 3.08E-03 | 0.4 | 2.36E-02 | -1.1 | 6.58E-04 | -1.6 | 1.66E-03 | -0.1 | 7.64E-01 | -1.5 | 1.83E-02 | DC1 domain-containing protein |
| AT2G28410 | -0.7 | 3.48E-04 | -0.1 | 3.85E-01 | -0.6 | 4.56E-03 | -0.1 | 6.07E-01 | 0.1 | 8.21E-01 | -0.2 | 5.44E-01 | hypothetical protein |
| AT1G07310 | -0.7 | 7.44E-06 | -0.5 | 6.99E-05 | -0.1 | 1.31E-01 | -0.5 | 7.30E-05 | -0.3 | 1.70E-02 | -0.2 | 1.20E-01 | Calcium-dependent lipid-binding family protein |
| AT2G30550 | -0.7 | 8.26E-04 | -0.7 | 1.14E-03 | 0.0 | 9.88E-01 | 0.4 | 4.98E-02 | 0.3 | 1.21E-01 | 0.0 | 9.08E-01 | Alpha |
| AT1G72510 | -0.7 | 2.70E-06 | -0.2 | 9.40E-03 | -0.5 | 2.09E-04 | -0.3 | 1.23E-01 | -0.1 | 7.49E-01 | -0.2 | 4.33E-01 | hypothetical protein |
| AT1G76620 | -0.7 | 5.65E-03 | -0.5 | 3.65E-02 | -0.2 | 4.13E-01 | -0.7 | 1.66E-03 | 0.0 | 9.03E-01 | -0.7 | 2.97E-03 | DUF547 domain-containing protein |
| AT2G31260 | -0.7 | 2.28E-06 | -0.6 | 1.75E-05 | -0.1 | 1.37E-01 | -0.1 | 6.32E-01 | 0.1 | 8.49E-01 | -0.2 | 5.65E-01 | Autophagy-related protein 9 |
| AT3G09470 | -0.7 | 1.11E-05 | -0.3 | 1.75E-03 | -0.3 | 4.49E-03 | -0.2 | 1.24E-01 | -0.2 | 1.23E-01 | 0.0 | 9.24E-01 | UNC93-like protein 3 |
| AT5G23410 | -0.7 | 6.31E-04 | -0.3 | 6.26E-02 | -0.4 | 3.93E-02 | -0.5 | 1.76E-01 | -0.2 | 6.55E-01 | -0.3 | 5.32E-01 | NA |
| AT5G43403 | -0.7 | 2.31E-03 | -0.6 | 7.02E-03 | -0.1 | 6.86E-01 | -0.4 | 2.10E-01 | 0.0 | 9.77E-01 | -0.4 | 3.03E-01 | NA |
| AT5G50900 | -0.7 | 3.64E-06 | -0.3 | 7.32E-04 | -0.3 | 1.85E-03 | -0.5 | 1.51E-03 | -0.3 | 3.41E-02 | -0.2 | 3.53E-01 | ARM repeat superfamily protein |
| AT1G22770 | -0.7 | 1.97E-05 | -0.5 | 2.33E-04 | -0.2 | 1.25E-01 | | | | | | | Protein GIGANTEA |
| AT4G09830 | -0.7 | 9.05E-06 | -0.6 | 5.86E-05 | -0.1 | 2.46E-01 | -0.5 | 1.22E-03 | -0.5 | 6.51E-03 | -0.1 | 8.04E-01 | Holocarboxylase synthetase |
| AT2G19350 | -0.7 | 2.17E-05 | -0.5 | 1.77E-04 | -0.1 | 2.18E-01 | -0.5 | 1.05E-01 | -0.3 | 4.40E-01 | -0.2 | 6.41E-01 | Transmembrane protein 230 |
| AT4G38930 | -0.7 | 5.41E-06 | -0.3 | 1.89E-03 | -0.4 | 1.70E-03 | -0.3 | 1.12E-01 | 0.0 | 8.79E-01 | -0.3 | 1.59E-01 | Ubiquitin fusion degradation UFD1 family protein |
| AT4G24990 | -0.7 | 1.52E-05 | -0.5 | 1.50E-04 | -0.1 | 1.39E-01 | -0.3 | 1.26E-01 | -0.2 | 2.33E-01 | 0.0 | 9.38E-01 | Membrane-anchored ubiquitin-fold protein |
| AT5G17210 | -0.7 | 1.28E-04 | -0.4 | 3.46E-03 | -0.3 | 8.38E-02 | 0.4 | 2.11E-01 | 0.5 | 1.86E-01 | -0.1 | 9.05E-01 | Transmembrane protein, putative |
| AT5G05380 | -0.7 | 1.93E-04 | -0.3 | 1.14E-02 | -0.3 | 4.56E-02 | -0.3 | 1.29E-01 | -0.1 | 7.58E-01 | -0.2 | 4.09E-01 | PRA1 family protein |
| AT4G15160 | -0.7 | 2.27E-03 | -0.2 | 2.59E-01 | -0.5 | 3.21E-02 | -0.3 | 1.13E-01 | -0.5 | 2.50E-02 | 0.2 | 5.44E-01 | Bifunctional inhibitor |
| AT4G03320 | -0.7 | 7.31E-04 | -1.0 | 9.29E-05 | 0.3 | 7.97E-02 | 1.0 | 1.92E-03 | 0.3 | 3.32E-01 | 0.6 | 3.04E-02 | Protein TIC 20 |
| AT1G11175 | -0.7 | 9.00E-05 | -0.4 | 3.66E-03 | -0.3 | 5.04E-02 | -0.9 | 4.42E-04 | -0.8 | 2.65E-03 | -0.1 | 7.27E-01 | NA |
| AT3G09490 | -0.7 | 1.45E-05 | -0.3 | 3.38E-03 | -0.4 | 4.67E-03 | 0.3 | 2.70E-01 | 0.3 | 1.67E-01 | -0.1 | 7.98E-01 | Uncharacterized protein |
| AT4G01200 | -0.7 | 2.40E-04 | -0.1 | 4.12E-01 | -0.6 | 3.54E-03 | 0.0 | 9.27E-01 | -0.1 | 7.36E-01 | 0.1 | 8.48E-01 | A_IG002N01.24 protein |
| AT1G77770 | -0.7 | 5.95E-06 | -0.5 | 6.56E-05 | -0.2 | 1.02E-01 | -0.4 | 7.03E-02 | 0.0 | 9.35E-01 | -0.4 | 1.29E-01 | At1g77770 ) |
| AT5G57840 | -0.7 | 4.22E-05 | -0.4 | 2.24E-03 | -0.3 | 3.04E-02 | 0.1 | 6.93E-01 | 0.4 | 2.10E-01 | -0.3 | 4.76E-01 | Uncharacterized protein |
| AT1G09176 | -0.7 | 1.15E-03 | 0.0 | 7.59E-01 | -0.6 | 7.39E-03 | -0.8 | 2.19E-01 | 0.1 | 8.61E-01 | -0.9 | 2.82E-01 | Transmembrane protein |
| AT5G24390 | -0.7 | 1.67E-03 | -0.3 | 5.68E-02 | -0.3 | 1.23E-01 | -0.5 | 1.22E-01 | 0.0 | 9.26E-01 | -0.5 | 1.85E-01 | Rab-GAP TBC domain-containing protein |
| AT1G04770 | -0.7 | 3.91E-05 | -0.6 | 1.66E-04 | -0.1 | 4.95E-01 | 1.7 | 4.45E-06 | 0.8 | 1.19E-02 | 0.9 | 2.47E-03 | Protein SULFUR DEFICIENCY-INDUCED 2 |
| AT5G65170 | -0.7 | 1.03E-03 | -0.9 | 2.53E-04 | 0.3 | 1.97E-01 | 0.1 | 7.52E-01 | 0.3 | 5.98E-01 | -0.1 | 8.33E-01 | VQ motif-containing protein |
| AT1G48090 | -0.7 | 3.01E-05 | -0.7 | 4.97E-05 | 0.0 | 9.19E-01 | -0.2 | 3.65E-01 | -0.2 | 5.50E-01 | -0.1 | 9.20E-01 | Calcium-dependent lipid-binding family protein |
| AT3G44220 | -0.7 | 4.27E-03 | 0.1 | 6.96E-01 | -0.7 | 8.18E-03 | -0.4 | 2.13E-01 | -0.2 | 6.65E-01 | -0.2 | 5.84E-01 | hypothetical protein |
| AT3G51090 | -0.7 | 3.71E-04 | -0.3 | 1.97E-02 | -0.3 | 6.00E-02 | 0.1 | 7.09E-01 | 0.5 | 9.66E-02 | -0.4 | 2.84E-01 | Coiled-coil 90B-like protein |
| AT1G17460 | -0.7 | 2.27E-03 | -0.8 | 1.01E-03 | 0.2 | 5.10E-01 | 0.4 | 2.26E-01 | 0.1 | 8.30E-01 | 0.3 | 5.20E-01 | MYB transcription factor |
| AT3G11600 | -0.7 | 4.52E-04 | -0.1 | 3.24E-01 | -0.5 | 6.63E-03 | -0.4 | 5.27E-02 | -0.4 | 4.39E-02 | 0.1 | 8.78E-01 | Protein GL2-INTERACTING REPRESSOR 2 |
| AT5G47445 | -0.7 | 9.41E-05 | -0.4 | 2.48E-03 | -0.2 | 8.64E-02 | 0.0 | 9.45E-01 | 0.4 | 3.38E-01 | -0.4 | 2.16E-01 | NA |
| AT2G15730 | -0.7 | 5.02E-04 | -0.5 | 5.43E-03 | -0.2 | 2.52E-01 | -0.2 | 4.50E-01 | -0.1 | 8.36E-01 | -0.1 | 7.46E-01 | P-loop containing nucleoside triphosphate hydrolases superfamily protein |
| AT4G29440 | -0.7 | 1.10E-04 | -0.5 | 6.06E-04 | -0.1 | 4.08E-01 | 0.0 | 9.82E-01 | 0.1 | 9.02E-01 | -0.1 | 9.05E-01 | Uncharacterized protein AT4g29440 |
| AT5G66490 | -0.7 | 7.12E-04 | -0.4 | 1.54E-02 | -0.3 | 1.39E-01 | -0.6 | 2.65E-02 | -0.6 | 4.97E-02 | 0.0 | 9.61E-01 | Uncharacterized protein At5g66490 |
| AT5G16320 | -0.7 | 8.71E-05 | -0.6 | 3.34E-04 | -0.1 | 6.00E-01 | -0.3 | 6.27E-01 | 0.0 | 9.67E-01 | -0.2 | 7.72E-01 | FRIGIDA-like protein 1 |
| AT1G24460 | -0.7 | 1.09E-06 | -0.6 | 4.89E-06 | -0.1 | 3.31E-01 | -0.1 | 5.90E-01 | 0.0 | 8.77E-01 | -0.2 | 5.99E-01 | TGN-related, localized SYP41 interacting protein |
| AT1G56090 | -0.7 | 9.21E-05 | -0.3 | 1.16E-02 | -0.4 | 1.80E-02 | -0.6 | 4.08E-03 | -0.6 | 9.23E-03 | 0.0 | 9.37E-01 | Tetratricopeptide repeat -like superfamily protein |
| AT3G04020 | -0.7 | 5.06E-03 | -0.5 | 2.69E-02 | -0.2 | 4.69E-01 | 0.3 | 2.68E-01 | 0.2 | 4.77E-01 | 0.1 | 8.70E-01 | T11I18.13 protein |
| AT1G68650 | -0.7 | 2.60E-04 | -0.2 | 9.85E-02 | -0.4 | 9.56E-03 | -0.4 | 3.72E-02 | 0.2 | 3.73E-01 | -0.6 | 8.13E-03 | GDT1 family protein |

| | | | | | | | | | | | | | |
|---|---|---|---|---|---|---|---|---|---|---|---|---|---|
| AT1G15350 | -0.7 | 4.67E-06 | -0.2 | 4.10E-02 | -0.5 | 2.20E-04 | -0.3 | 9.83E-02 | -0.3 | 1.37E-01 | 0.0 | 9.98E-01 | DUF4050 domain-containing protein |
| AT4G01600 | -0.7 | 3.92E-04 | -0.4 | 6.33E-03 | -0.2 | 1.64E-01 | -0.2 | 3.40E-01 | 0.2 | 3.47E-01 | -0.4 | 1.11E-01 | GRAM domain family protein |
| AT4G28240 | -0.7 | 6.43E-05 | -0.2 | 1.32E-01 | -0.5 | 1.51E-03 | -0.3 | 7.03E-02 | -0.4 | 3.12E-02 | 0.1 | 7.19E-01 | Wound-responsive family protein |
| AT2G42540 | -0.7 | 1.93E-01 | -0.2 | 6.69E-01 | -0.4 | 4.56E-01 | -1.2 | 6.83E-04 | 1.1 | 3.90E-03 | -2.3 | 7.98E-05 | Cold-regulated 15a |
| AT2G38480 | -0.7 | 4.77E-04 | -0.2 | 1.09E-01 | -0.4 | 1.85E-02 | -1.1 | 5.24E-03 | -0.2 | 5.36E-01 | -0.9 | 3.55E-02 | CASP-like protein 4B1 |
| AT2G35610 | -0.7 | 7.56E-06 | -0.2 | 4.41E-02 | -0.5 | 2.93E-04 | 0.1 | 6.21E-01 | 0.2 | 1.73E-01 | -0.1 | 4.71E-01 | Nucleotide-diphospho-sugar transferase domain-containing protein |
| AT1G79450 | -0.7 | 2.63E-04 | -0.2 | 5.44E-02 | -0.4 | 1.67E-02 | 0.7 | 2.59E-02 | 0.3 | 4.89E-01 | 0.4 | 2.28E-01 | ALA-interacting subunit 5 |
| AT3G13650 | -0.7 | 9.63E-05 | -0.3 | 2.66E-02 | -0.4 | 9.42E-03 | 0.7 | 2.71E-02 | 0.7 | 3.63E-02 | 0.0 | 9.48E-01 | Dirigent protein 7 |
| AT5G42280 | -0.7 | 1.16E-04 | -0.2 | 1.21E-01 | -0.5 | 4.11E-03 | -0.2 | 4.59E-01 | 0.1 | 8.47E-01 | -0.3 | 4.20E-01 | Uncharacterized protein At5g42280 |
| AT4G02845 | -0.7 | 1.04E-03 | -0.3 | 3.90E-02 | -0.3 | 9.65E-02 | -0.4 | 4.10E-02 | -0.2 | 3.52E-01 | -0.2 | 5.01E-01 | NA |
| AT1G58420 | -0.7 | 2.39E-02 | -1.2 | 2.02E-03 | 0.5 | 1.51E-01 | 0.1 | 8.88E-01 | 0.3 | 3.76E-01 | -0.3 | 4.86E-01 | Uncharacterized protein |
| AT2G03505 | -0.7 | 4.26E-03 | 0.3 | 8.88E-02 | -0.9 | 1.77E-03 | 0.7 | 6.93E-03 | 0.5 | 8.23E-02 | 0.2 | 5.06E-01 | X8 domain-containing protein |
| AT2G27430 | -0.7 | 6.99E-04 | 0.0 | 7.52E-01 | -0.7 | 2.16E-03 | -0.2 | 5.63E-01 | 0.1 | 8.75E-01 | -0.3 | 5.22E-01 | Uncharacterized protein At2g27430 |
| AT3G43440 | -0.7 | 1.03E-05 | -0.5 | 1.44E-04 | -0.2 | 8.82E-02 | 0.0 | 9.35E-01 | 0.2 | 4.28E-01 | -0.2 | 4.51E-01 | Protein TIFY |
| AT4G02815 | -0.6 | 7.61E-04 | -0.3 | 4.41E-02 | -0.3 | 6.80E-02 | 0.2 | 6.90E-01 | 0.4 | 4.38E-01 | -0.2 | 7.72E-01 | NA |
| AT5G11390 | -0.6 | 6.30E-06 | -0.4 | 1.75E-04 | -0.2 | 2.78E-02 | -0.2 | 1.01E-01 | -0.3 | 1.31E-01 | 0.0 | 9.93E-01 | WPP domain-interacting tail-anchored protein 1 |
| AT1G62790 | -0.6 | 2.52E-04 | -0.2 | 6.85E-02 | -0.4 | 1.05E-02 | -0.2 | 4.33E-01 | 0.1 | 8.49E-01 | -0.2 | 4.52E-01 | Bifunctional inhibitor |
| AT5G17350 | -0.6 | 6.23E-03 | -1.0 | 8.99E-04 | 0.4 | 1.80E-01 | 0.4 | 3.30E-01 | -0.1 | 8.40E-01 | 0.5 | 2.88E-01 | hypothetical protein |
| AT1G69900 | -0.6 | 4.92E-04 | -0.2 | 7.66E-02 | -0.4 | 2.22E-02 | 0.0 | 9.91E-01 | -0.1 | 8.32E-01 | 0.1 | 8.55E-01 | DUF569 domain-containing protein |
| AT2G19830 | -0.6 | 2.16E-05 | -0.4 | 1.14E-03 | -0.3 | 1.94E-02 | 0.0 | 9.23E-01 | -0.1 | 8.21E-01 | 0.0 | 9.15E-01 | SNF7 family protein |
| AT1G04960 | -0.6 | 3.61E-06 | -0.4 | 9.16E-05 | -0.2 | 2.20E-02 | -0.1 | 4.11E-01 | -0.1 | 7.05E-01 | -0.1 | 8.23E-01 | Golgin family A protein |
| AT5G07760 | -0.6 | 1.77E-05 | -0.2 | 3.91E-02 | -0.5 | 1.05E-03 | 0.0 | 9.01E-01 | 0.1 | 8.14E-01 | -0.1 | 7.47E-01 | Formin-like protein |
| AT3G27670 | -0.6 | 2.47E-05 | -0.6 | 8.59E-05 | -0.1 | 6.17E-01 | -0.3 | 2.02E-01 | -0.3 | 3.59E-01 | -0.1 | 9.05E-01 | Protein RST1 |
| AT1G11540 | -0.6 | 9.09E-05 | -0.1 | 3.79E-01 | -0.6 | 1.07E-03 | 0.1 | 5.06E-01 | 0.3 | 1.10E-01 | -0.2 | 3.60E-01 | Sulfite exporter TauE |
| AT4G14900 | -0.6 | 5.37E-06 | -0.6 | 1.75E-05 | 0.0 | 6.36E-01 | -0.3 | 1.78E-01 | -0.2 | 4.84E-01 | -0.1 | 7.58E-01 | FRIGIDA-like protein 4b |
| AT3G61750 | -0.6 | 3.66E-05 | -0.4 | 2.64E-03 | -0.3 | 2.06E-02 | -0.4 | 2.31E-01 | 0.2 | 6.11E-01 | -0.6 | 9.22E-02 | Cytochrome b561 |
| AT1G04240 | -0.6 | 2.68E-04 | -0.4 | 3.61E-03 | -0.2 | 1.86E-01 | -0.1 | 8.01E-01 | -0.1 | 8.37E-01 | 0.0 | 1.00E+00 | Auxin-responsive protein |
| AT1G52330 | -0.6 | 1.88E-04 | -0.4 | 4.46E-03 | -0.2 | 1.11E-01 | -0.1 | 8.01E-01 | 0.1 | 7.92E-01 | -0.3 | 5.60E-01 | Expressed protein hydroxyproline-rich glycoprotein family) |
| AT2G39300 | -0.6 | 4.15E-05 | -0.3 | 5.58E-03 | -0.3 | 1.24E-02 | -0.2 | 2.73E-01 | 0.0 | 9.02E-01 | -0.3 | 2.74E-01 | Uncharacterized protein At2g39300 |
| AT4G05010 | -0.6 | 8.41E-03 | -0.5 | 2.48E-02 | -0.1 | 6.91E-01 | 0.1 | 8.66E-01 | 0.2 | 5.85E-01 | -0.1 | 7.20E-01 | F-box protein At4g05010 |
| AT5G38895 | -0.6 | 1.47E-03 | -0.6 | 2.24E-03 | 0.0 | 9.65E-01 | -0.4 | 7.10E-02 | -0.4 | 1.21E-01 | 0.0 | 9.65E-01 | RING-type domain-containing protein |
| AT3G55720 | -0.6 | 4.01E-02 | -0.3 | 2.41E-01 | -0.3 | 4.15E-01 | 0.1 | 8.12E-01 | 0.7 | 1.31E-01 | -0.6 | 2.40E-01 | hypothetical protein |
| AT3G13772 | -0.6 | 1.74E-05 | -0.5 | 2.44E-04 | -0.2 | 8.74E-02 | -0.1 | 8.70E-01 | 0.0 | 9.19E-01 | -0.1 | 8.37E-01 | Transmembrane 9 superfamily member 7 |
| AT1G18470 | -0.6 | 5.71E-06 | -0.6 | 1.89E-05 | 0.0 | 6.18E-01 | -0.5 | 1.48E-03 | -0.2 | 2.87E-01 | -0.3 | 1.02E-01 | Transmembrane Fragile-X-F-associated protein |
| AT4G17140 | -0.6 | 2.69E-06 | -0.6 | 1.17E-05 | -0.1 | 4.06E-01 | -0.4 | 7.61E-03 | -0.2 | 1.40E-01 | -0.2 | 4.62E-01 | Uncharacterized protein |
| AT3G21295 | -0.6 | 9.39E-06 | -0.6 | 3.60E-05 | -0.1 | 5.19E-01 | -0.2 | 5.44E-01 | 0.0 | 9.90E-01 | -0.2 | 5.92E-01 | PWWP domain-containing protein |
| AT1G17080 | -0.6 | 3.82E-05 | -0.3 | 3.34E-03 | -0.3 | 1.47E-02 | -0.3 | 2.77E-02 | -0.2 | 2.93E-01 | -0.1 | 4.60E-01 | 60S ribosomal protein L18a-like protein |
| AT2G45245 | -0.6 | 2.13E-02 | -0.6 | 3.50E-02 | 0.0 | 9.19E-01 | 0.0 | 9.81E-01 | 0.3 | 6.51E-01 | -0.3 | 6.96E-01 | NA |
| AT5G51160 | -0.6 | 2.26E-04 | -0.4 | 4.40E-03 | -0.2 | 1.39E-01 | | | | | | | Ankyrin repeat family protein |
| AT1G32700 | -0.6 | 3.70E-04 | -0.5 | 1.99E-03 | -0.1 | 4.08E-01 | -0.4 | 4.61E-03 | -0.4 | 1.68E-02 | 0.0 | 8.81E-01 | F6N18.8 |
| AT1G52347 | -0.6 | 4.43E-03 | -0.7 | 4.08E-03 | 0.0 | 8.47E-01 | -0.4 | 2.32E-01 | -0.1 | 8.05E-01 | -0.3 | 5.24E-01 | NA |
| AT5G26770 | -0.6 | 6.10E-04 | -0.7 | 5.12E-04 | 0.1 | 7.23E-01 | -0.2 | 5.05E-01 | -0.3 | 2.55E-01 | 0.1 | 7.05E-01 | Myosin heavy chain, cardiac protein |
| AT3G03180 | -0.6 | 7.68E-05 | -0.1 | 3.91E-01 | -0.5 | 1.18E-03 | -0.5 | 3.65E-02 | -0.3 | 3.28E-01 | -0.2 | 4.53E-01 | Got1 |
| AT1G77020 | -0.6 | 6.33E-04 | 0.0 | 9.49E-01 | -0.6 | 2.65E-03 | 0.0 | 9.15E-01 | 0.2 | 5.34E-01 | -0.1 | 6.64E-01 | DNAJ heat shock N-terminal domain-containing protein |
| AT1G80610 | -0.6 | 9.76E-05 | -0.4 | 1.33E-03 | -0.2 | 1.55E-01 | 0.1 | 7.60E-01 | 0.0 | 9.40E-01 | 0.1 | 7.83E-01 | At1g80610 |
| AT2G19660 | -0.6 | 1.18E-03 | 0.1 | 3.70E-01 | -0.8 | 1.82E-03 | -0.5 | 2.26E-01 | -0.2 | 7.20E-01 | -0.3 | 5.69E-01 | At2g19660 |
| AT4G00500 | -0.6 | 3.24E-05 | -0.5 | 1.55E-04 | -0.1 | 4.38E-01 | 0.2 | 4.90E-01 | 0.2 | 4.49E-01 | 0.0 | 9.30E-01 | hypothetical protein |
| AT5G26731 | -0.6 | 2.88E-02 | -0.7 | 2.89E-02 | 0.0 | 9.46E-01 | 0.3 | 3.20E-01 | 0.2 | 7.23E-01 | 0.2 | 7.00E-01 | Uncharacterized protein |
| AT2G01735 | -0.6 | 1.30E-05 | -0.6 | 2.50E-05 | 0.0 | 9.68E-01 | -0.6 | 1.01E-03 | -0.3 | 7.36E-02 | -0.3 | 1.48E-01 | RING-finger protein for embryogenesi |
| AT4G38400 | -0.6 | 8.00E-03 | -0.6 | 1.57E-02 | -0.1 | 8.28E-01 | -0.9 | 4.44E-03 | -0.5 | 7.68E-02 | -0.4 | 4.12E-01 | Expansin-like A2 |
| AT5G23110 | -0.6 | 6.07E-05 | -0.6 | 9.85E-05 | 0.0 | 9.43E-01 | -0.2 | 4.79E-01 | 0.0 | 9.58E-01 | -0.2 | 5.79E-01 | Zinc finger, C3HC4 type family protein |
| AT5G39785 | -0.6 | 1.30E-04 | -0.7 | 1.60E-04 | 0.0 | 8.16E-01 | 0.5 | 1.06E-01 | 0.2 | 5.21E-01 | 0.2 | 5.39E-01 | Gb\|AAF22924.1 |
| AT5G55170 | -0.6 | 1.85E-04 | -0.2 | 1.67E-01 | -0.5 | 5.06E-03 | 0.0 | 9.41E-01 | -0.1 | 8.59E-01 | 0.1 | 9.34E-01 | Small ubiquitin-like modifier 3 |
| AT1G76250 | -0.6 | 6.27E-04 | 0.1 | 2.22E-01 | -0.8 | 8.02E-04 | 0.2 | 5.82E-01 | 0.2 | 4.33E-01 | -0.1 | 8.37E-01 | T23E18.18 |
| AT2G37650 | -0.6 | 1.32E-04 | -0.5 | 8.99E-04 | -0.1 | 3.26E-01 | -0.4 | 8.41E-02 | -0.1 | 7.54E-01 | -0.3 | 2.58E-01 | GRAS family transcription factor |
| AT1G15950 | -0.6 | 2.26E-05 | -0.9 | 4.62E-06 | 0.3 | 1.96E-02 | -0.3 | 5.71E-01 | 0.1 | 9.19E-01 | -0.4 | 6.09E-01 | Cinnamoyl coa reductase 1 |
| AT5G08770 | -0.6 | 1.44E-04 | -0.4 | 4.56E-03 | -0.3 | 7.53E-02 | -0.3 | 1.68E-01 | -0.5 | 3.08E-02 | 0.2 | 4.36E-01 | Uncharacterized protein |
| AT1G01240 | -0.6 | 1.42E-04 | -0.3 | 1.26E-02 | -0.3 | 2.55E-02 | -0.9 | 1.15E-04 | -0.7 | 1.02E-03 | -0.1 | 6.53E-01 | hypothetical protein |
| AT2G20010 | -0.6 | 2.53E-06 | -0.5 | 3.39E-05 | -0.2 | 4.93E-02 | 0.0 | 8.07E-01 | 0.1 | 7.79E-01 | 0.0 | 9.55E-01 | Gls protein |
| AT4G11300 | -0.6 | 7.17E-05 | -0.6 | 2.62E-04 | -0.1 | 5.66E-01 | -0.4 | 5.28E-02 | -0.4 | 6.45E-02 | 0.0 | 9.55E-01 | ROH1, |
| AT4G18760 | -0.6 | 5.16E-05 | 0.0 | 8.21E-01 | -0.6 | 4.13E-04 | -0.5 | 1.30E-02 | -0.4 | 6.43E-02 | -0.1 | 7.11E-01 | Receptor-like protein 51 |
| AT4G14740 | -0.6 | 1.77E-04 | -0.4 | 5.73E-03 | -0.2 | 6.94E-02 | -0.2 | 8.75E-02 | 0.0 | 9.41E-01 | -0.2 | 1.73E-01 | Auxin canalization protein |
| AT3G28370 | -0.6 | 5.78E-03 | -0.2 | 2.48E-01 | -0.4 | 9.39E-02 | -0.2 | 5.47E-01 | -0.2 | 6.05E-01 | 0.0 | 9.98E-01 | At3g28370 |
| AT5G55850 | -0.6 | 8.48E-06 | -0.4 | 3.11E-04 | -0.2 | 1.91E-02 | -0.1 | 8.03E-01 | -0.1 | 6.87E-01 | 0.1 | 8.96E-01 | RPM1-interacting protein 4 family protein |
| AT4G02760 | -0.6 | 3.55E-05 | -0.5 | 4.52E-04 | -0.2 | 1.40E-01 | -0.1 | 6.91E-01 | 0.1 | 8.66E-01 | -0.2 | 6.51E-01 | RNI-like superfamily protein |
| AT5G66250 | -0.6 | 2.66E-06 | -0.5 | 2.82E-05 | -0.1 | 7.55E-02 | -0.4 | 5.83E-03 | -0.2 | 2.24E-01 | -0.2 | 2.43E-01 | Kinectin-like protein |
| AT2G05590 | -0.6 | 6.23E-05 | -0.4 | 8.68E-04 | -0.2 | 1.42E-01 | -0.3 | 3.00E-02 | -0.2 | 1.54E-01 | -0.1 | 6.67E-01 | Expressed protein |
| AT2G47460 | -0.6 | 1.39E-02 | -0.2 | 4.37E-01 | -0.5 | 1.10E-01 | -0.1 | 8.94E-01 | 1.0 | 4.39E-02 | -1.1 | 4.00E-02 | PFG1 |
| AT5G06390 | -0.6 | 7.10E-06 | -0.1 | 6.70E-02 | -0.5 | 2.26E-04 | -0.3 | 2.20E-02 | 0.0 | 8.25E-01 | -0.4 | 6.47E-02 | Fasciclin-like arabinogalactan protein 17 |
| AT5G24660 | -0.6 | 6.86E-03 | -0.7 | 3.29E-03 | 0.1 | 6.19E-01 | 2.2 | 1.51E-04 | 1.5 | 1.02E-02 | 0.7 | 7.38E-02 | Protein RESPONSE TO LOW SULFUR 2 |
| AT2G34730 | -0.6 | 3.25E-06 | -0.4 | 1.04E-04 | -0.2 | 1.56E-02 | 0.1 | 4.40E-01 | 0.0 | 9.73E-01 | 0.1 | 6.21E-01 | Myosin heavy chain-like protein |
| AT1G01640 | -0.6 | 6.38E-05 | -0.3 | 1.33E-02 | -0.3 | 1.01E-02 | -0.2 | 3.73E-01 | -0.1 | 7.83E-01 | -0.1 | 7.12E-01 | BTB |
| AT3G45600 | -0.6 | 9.99E-05 | -0.3 | 1.25E-02 | -0.3 | 1.51E-02 | -0.3 | 1.08E-01 | 0.0 | 9.98E-01 | -0.3 | 2.52E-01 | Tetraspanin3 |
| AT4G18070 | -0.6 | 4.07E-06 | -0.3 | 1.07E-03 | -0.3 | 1.51E-03 | -0.1 | 2.68E-01 | 0.1 | 5.88E-01 | -0.2 | 1.57E-01 | hypothetical protein |
| AT2G44340 | -0.6 | 2.56E-05 | -0.1 | 1.29E-01 | -0.5 | 8.14E-04 | 0.6 | 9.22E-02 | 0.6 | 1.45E-01 | 0.0 | 9.48E-01 | VQ motif-containing protein 18 |
| AT5G04235 | -0.6 | 1.93E-04 | -0.6 | 4.03E-04 | 0.0 | 8.88E-01 | 0.3 | 4.14E-01 | 0.3 | 5.02E-01 | 0.0 | 9.83E-01 | NA |
| AT3G60640 | -0.6 | 1.46E-03 | -0.7 | 6.81E-04 | 0.1 | 5.11E-01 | -0.1 | 6.96E-01 | -0.1 | 8.80E-01 | -0.1 | 9.08E-01 | Autophagy-related protein 8g |
| AT3G20380 | -0.6 | 1.27E-05 | -0.1 | 3.11E-01 | -0.5 | 1.93E-04 | 0.4 | 1.08E-02 | 0.4 | 1.45E-02 | 0.0 | 9.42E-01 | TRAF-like family protein |
| AT5G23903 | -0.6 | 1.69E-03 | 0.8 | 1.39E-04 | -1.4 | 4.16E-05 | -1.4 | 1.89E-04 | 0.3 | 2.24E-01 | -1.8 | 4.74E-04 | Transmembrane protein |
| AT1G03340 | -0.6 | 3.73E-04 | -0.5 | 1.82E-03 | -0.1 | 4.80E-01 | 0.1 | 8.23E-01 | 0.0 | 9.90E-01 | 0.1 | 8.45E-01 | hypothetical protein |
| AT1G31780 | -0.6 | 1.77E-05 | -0.3 | 7.94E-03 | -0.4 | 2.58E-03 | -0.3 | 2.65E-01 | -0.1 | 7.36E-01 | -0.2 | 6.49E-01 | Conserved oligomeric Golgi complex subunit 6 |
| AT5G46410 | -0.6 | 7.84E-06 | -0.4 | 1.75E-04 | -0.2 | 3.85E-02 | -0.4 | 3.21E-03 | -0.4 | 2.42E-03 | 0.1 | 7.77E-01 | Similarity to |
| AT4G23880 | -0.6 | 1.33E-03 | -0.5 | 4.27E-03 | -0.1 | 6.39E-01 | -0.1 | 7.68E-01 | -0.2 | 5.28E-01 | 0.1 | 7.85E-01 | Uncharacterized protein |

| | | | | | | | | | | | | | |
|---|---|---|---|---|---|---|---|---|---|---|---|---|---|
| AT1G76020 | -0.6 | 1.22E-04 | -0.3 | 1.46E-02 | -0.3 | 1.86E-02 | -0.2 | 2.38E-01 | 0.3 | 1.19E-01 | -0.5 | 2.98E-02 | T4O12.23 |
| AT3G14172 | -0.6 | 1.08E-04 | -0.4 | 1.34E-03 | -0.2 | 1.61E-01 | 0.2 | 5.59E-01 | 0.1 | 7.42E-01 | 0.1 | 9.28E-01 | GPI-anchored adhesin-like protein |
| AT1G12710 | -0.6 | 2.54E-03 | -0.3 | 8.45E-02 | -0.3 | 1.13E-01 | -0.6 | 1.04E-02 | -0.3 | 2.62E-01 | -0.3 | 2.08E-01 | Phloem protein 2-A12 |
| AT5G23340 | -0.6 | 8.75E-06 | -0.4 | 1.47E-04 | -0.2 | 6.08E-02 | -0.7 | 6.82E-05 | -0.5 | 1.79E-03 | -0.2 | 4.34E-01 | hypothetical protein |
| AT3G61260 | -0.6 | 6.03E-06 | 0.0 | 9.32E-01 | -0.6 | 5.23E-05 | -0.9 | 1.95E-05 | -0.8 | 1.94E-04 | -0.1 | 7.90E-01 | Uncharacterized protein At3g61260 |
| AT3G60680 | -0.6 | 6.54E-05 | -0.3 | 6.03E-03 | -0.3 | 2.21E-02 | 0.2 | 5.85E-01 | 0.2 | 4.79E-01 | -0.1 | 8.78E-01 | Uncharacterized protein At3g60680 |
| AT1G68820 | -0.6 | 1.89E-05 | -0.4 | 5.30E-04 | -0.2 | 4.01E-02 | 0.1 | 4.75E-01 | 0.1 | 6.91E-01 | 0.0 | 9.04E-01 | Transmembrane Fragile-X-F-associated protein |
| AT1G23710 | -0.6 | 5.23E-04 | -0.8 | 1.91E-04 | 0.2 | 3.08E-01 | 0.2 | 2.08E-01 | -0.2 | 2.60E-01 | 0.4 | 4.65E-02 | At1g23710 |
| AT2G01190 | -0.6 | 2.15E-04 | -0.5 | 1.69E-03 | -0.1 | 2.72E-01 | -0.1 | 6.30E-01 | -0.5 | 1.84E-02 | 0.4 | 9.19E-02 | Expressed protein |
| AT3G25870 | -0.6 | 1.01E-03 | -0.5 | 7.21E-03 | -0.2 | 3.84E-01 | -0.3 | 3.36E-01 | -0.1 | 7.82E-01 | -0.2 | 6.97E-01 | Uncharacterized protein At3g25870 |
| AT5G20190 | -0.6 | 4.21E-03 | -0.7 | 3.32E-03 | 0.1 | 7.63E-01 | -0.4 | 5.86E-02 | 0.3 | 1.63E-01 | -0.7 | 5.72E-03 | Uncharacterized protein |
| AT2G31130 | -0.6 | 8.93E-05 | -0.6 | 1.65E-04 | 0.0 | 9.58E-01 | -0.2 | 3.38E-01 | 0.0 | 9.58E-01 | -0.2 | 4.77E-01 | Uncharacterized protein |
| AT3G61560 | -0.6 | 8.61E-05 | 0.0 | 6.14E-01 | -0.7 | 2.72E-04 | -0.1 | 6.86E-01 | 0.1 | 6.33E-01 | -0.3 | 3.53E-01 | Reticulon-like protein |
| AT1G56080 | -0.6 | 1.03E-03 | -0.1 | 6.48E-01 | -0.5 | 7.52E-03 | 0.1 | 8.12E-01 | 0.2 | 5.58E-01 | -0.1 | 7.61E-01 | hypothetical protein |
| AT2G38970 | -0.6 | 3.17E-04 | -0.2 | 9.97E-02 | -0.4 | 1.23E-02 | 0.1 | 7.48E-01 | 0.4 | 7.32E-02 | -0.3 | 2.31E-01 | Putative retroelement pol polyprotein |
| AT4G20300 | -0.6 | 8.93E-06 | -0.3 | 2.30E-03 | -0.3 | 2.49E-03 | -0.2 | 1.85E-01 | -0.1 | 4.76E-01 | -0.1 | 7.63E-01 | Serine |
| AT4G15120 | -0.6 | 1.62E-03 | -0.5 | 9.76E-03 | -0.2 | 4.33E-01 | 1.2 | 1.36E-04 | 0.5 | 1.01E-01 | 0.7 | 6.19E-03 | VQ domain-containing protein |
| AT2G20840 | -0.6 | 8.17E-05 | 0.0 | 6.87E-01 | -0.6 | 7.33E-04 | -0.2 | 4.70E-01 | 0.0 | 9.47E-01 | -0.2 | 5.16E-01 | Secretory carrier-associated membrane protein |
| AT5G02580 | -0.6 | 5.98E-02 | -0.7 | 5.11E-02 | 0.0 | 9.17E-01 | -0.2 | 8.67E-01 | 0.6 | 6.11E-01 | | | Uncharacterized protein T22P11_170 |
| AT5G41350 | -0.6 | 3.86E-06 | -0.6 | 7.71E-06 | 0.0 | 8.98E-01 | -0.2 | 2.70E-01 | 0.0 | 9.96E-01 | -0.2 | 4.25E-01 | RING-type domain-containing protein |
| AT1G76460 | -0.6 | 2.15E-04 | -0.3 | 1.55E-02 | -0.3 | 3.51E-02 | -0.4 | 6.29E-02 | -0.2 | 3.46E-01 | -0.2 | 6.32E-01 | AT1G76460 protein |
| AT1G28200 | -0.6 | 4.02E-06 | -0.4 | 7.94E-05 | -0.2 | 3.14E-02 | -0.3 | 3.04E-02 | -0.1 | 5.05E-01 | -0.2 | 3.49E-01 | GEM-like protein 1 |
| AT2G36390 | -0.6 | 4.45E-04 | -0.6 | 7.73E-04 | 0.0 | 9.29E-01 | 0.0 | 8.85E-01 | 0.2 | 4.03E-01 | -0.2 | 4.25E-01 | SBE2.1 |
| AT5G58750 | -0.6 | 3.33E-03 | 0.6 | 1.77E-03 | -1.2 | 1.81E-04 | -0.1 | 8.63E-01 | 0.4 | 2.84E-01 | -0.5 | 1.26E-01 | NAD-binding Rossmann-fold superfamily protein |
| AT1G52320 | -0.6 | 1.92E-05 | -0.6 | 4.90E-05 | 0.0 | 8.07E-01 | -0.1 | 5.88E-01 | -0.2 | 4.78E-01 | 0.1 | 8.86E-01 | BZIP protein, putative; 48652-45869 |
| AT3G26580 | -0.6 | 4.80E-06 | -0.5 | 5.01E-05 | -0.1 | 1.06E-01 | -0.5 | 3.94E-04 | -0.3 | 3.06E-02 | -0.2 | 2.49E-01 | Orf03 protein -like superfamily protein) |
| AT3G63430 | -0.6 | 6.90E-03 | -0.1 | 7.75E-01 | -0.5 | 2.91E-02 | -0.3 | 1.43E-01 | 0.0 | 9.55E-01 | -0.3 | 2.85E-01 | At3g63430 |
| AT1G23140 | -0.6 | 7.09E-03 | -0.4 | 4.04E-02 | -0.2 | 4.61E-01 | 0.4 | 1.61E-01 | 0.3 | 4.94E-01 | 0.2 | 7.20E-01 | C2 domain-containing protein |
| AT1G77700 | -0.6 | 4.13E-04 | -0.3 | 4.49E-02 | -0.3 | 3.48E-02 | 0.1 | 7.19E-01 | 0.3 | 2.95E-01 | -0.2 | 5.62E-01 | Pathogenesis-related thaumatin superfamily protein |
| AT1G67060 | -0.6 | 1.72E-05 | -0.5 | 1.00E-04 | -0.1 | 3.18E-01 | 0.1 | 6.90E-01 | -0.1 | 6.52E-01 | 0.2 | 4.55E-01 | hypothetical protein |
| AT5G63905 | -0.6 | 2.18E-04 | -0.4 | 5.12E-03 | -0.2 | 1.05E-01 | -0.1 | 8.29E-01 | 0.1 | 7.47E-01 | 0.1 | 9.18E-01 | Transmembrane protein |
| AT4G28770 | -0.6 | 1.48E-05 | -0.1 | 7.83E-02 | -0.5 | 4.50E-04 | -0.3 | 6.59E-02 | -0.2 | 3.31E-01 | -0.1 | 6.69E-01 | Tetraspanin family protein |
| AT5G41590 | -0.6 | 1.57E-03 | 0.0 | 8.23E-01 | -0.6 | 4.66E-03 | | | | | | | Uncharacterized protein |
| AT1G21670 | -0.6 | 7.32E-05 | -0.2 | 6.51E-02 | -0.4 | 3.02E-03 | -0.3 | 3.81E-02 | -0.3 | 5.07E-02 | 0.0 | 9.69E-01 | hypothetical protein |
| AT1G79660 | -0.6 | 4.37E-05 | -0.3 | 2.44E-03 | -0.3 | 2.88E-02 | -0.4 | 2.00E-02 | -0.3 | 1.08E-01 | -0.1 | 6.80E-01 | Uncharacterized protein |
| AT2G38120 | -0.6 | 1.05E-04 | -0.1 | 3.39E-01 | -0.5 | 1.23E-03 | -0.4 | 5.62E-03 | -0.1 | 5.35E-01 | -0.3 | 9.33E-02 | Transmembrane amino acid transporter family protein |
| AT4G12070 | -0.6 | 1.01E-05 | -0.5 | 7.60E-05 | -0.1 | 1.95E-01 | 0.1 | 5.29E-01 | -0.1 | 6.22E-01 | 0.2 | 3.46E-01 | Uncharacterized protein At4g12070 |
| AT5G40450 | -0.6 | 2.25E-04 | -0.2 | 4.94E-02 | -0.3 | 1.29E-02 | 0.1 | 6.38E-01 | -0.2 | 4.90E-01 | 0.3 | 3.37E-01 | Uncharacterized protein At5g40450 |
| AT4G01910 | -0.6 | 5.15E-04 | -0.4 | 6.31E-03 | -0.2 | 2.26E-01 | 0.1 | 8.38E-01 | 0.3 | 2.95E-01 | -0.2 | 4.28E-01 | Cysteine |
| AT3G51110 | -0.6 | 5.62E-05 | -0.4 | 1.11E-03 | -0.2 | 9.55E-02 | -0.2 | 2.32E-01 | -0.3 | 1.49E-01 | 0.1 | 8.20E-01 | Tetratricopeptide repeat -like superfamily protein |
| AT4G31890 | -0.6 | 6.76E-05 | -0.4 | 6.65E-04 | -0.1 | 2.14E-01 | -0.2 | 2.97E-01 | 0.2 | 4.87E-01 | -0.4 | 1.39E-01 | ARM repeat superfamily protein |
| AT4G06676 | -0.6 | 3.81E-05 | -0.5 | 2.85E-04 | -0.1 | 2.67E-01 | -0.1 | 7.36E-01 | 0.0 | 9.40E-01 | -0.1 | 7.53E-01 | Etoposide-induced protein |
| AT2G46000 | -0.6 | 6.15E-04 | 0.1 | 3.46E-01 | -0.7 | 9.65E-04 | -0.1 | 8.14E-01 | 0.2 | 4.38E-01 | -0.3 | 3.05E-01 | hypothetical protein |
| AT5G22460 | -0.6 | 4.22E-02 | -0.4 | 2.01E-01 | -0.2 | 5.06E-01 | 0.6 | 3.00E-01 | 1.1 | 8.95E-02 | -0.4 | 5.32E-01 | Alpha |
| AT1G17380 | -0.6 | 3.05E-02 | -0.8 | 1.05E-02 | 0.2 | 5.21E-01 | -0.1 | 9.04E-01 | 0.3 | 7.78E-01 | -0.4 | 7.06E-01 | Protein TIFY |
| AT2G39518 | -0.6 | 6.71E-03 | -0.2 | 1.92E-01 | -0.3 | 1.27E-01 | -0.4 | 2.40E-01 | -0.3 | 3.64E-01 | -0.1 | 8.89E-01 | CASP-like protein |
| AT2G33310 | -0.6 | 1.08E-05 | -0.5 | 1.19E-04 | -0.1 | 1.14E-01 | -0.2 | 2.97E-01 | -0.3 | 1.08E-01 | 0.1 | 6.32E-01 | Auxin-responsive protein |
| AT3G09810 | -0.6 | 1.07E-04 | 0.1 | 4.34E-01 | -0.7 | 2.19E-04 | 0.3 | 1.76E-02 | 0.1 | 3.88E-01 | 0.2 | 3.11E-01 | Putative dehydrogenase |
| AT4G09750 | -0.6 | 1.18E-03 | -0.9 | 1.33E-04 | 0.3 | 8.06E-02 | 0.7 | 3.07E-02 | 0.7 | 2.88E-02 | -0.1 | 8.73E-01 | NAD-binding Rossmann-fold superfamily protein |
| AT1G48760 | -0.6 | 7.13E-06 | -0.3 | 4.51E-04 | -0.2 | 1.04E-02 | -0.2 | 3.51E-01 | -0.1 | 6.63E-01 | -0.1 | 8.21E-01 | AP-3 complex subunit delta |
| AT2G25950 | -0.6 | 1.19E-03 | -0.3 | 7.20E-02 | -0.3 | 6.27E-02 | -0.1 | 5.75E-01 | 0.0 | 9.52E-01 | -0.1 | 6.32E-01 | PITH domain protein |
| AT4G14750 | -0.6 | 7.26E-05 | 0.0 | 6.60E-01 | -0.6 | 2.92E-04 | -0.4 | 2.63E-02 | -0.1 | 4.97E-01 | -0.3 | 2.91E-01 | DUF4005 domain-containing protein |
| AT5G48320 | -0.6 | 1.43E-04 | -0.4 | 3.53E-03 | -0.2 | 9.65E-02 | 0.2 | 4.07E-01 | 0.3 | 3.37E-01 | -0.1 | 8.89E-01 | Cysteine |
| AT3G17710 | -0.6 | 4.42E-04 | -0.3 | 2.67E-02 | -0.3 | 5.75E-02 | -0.2 | 6.41E-01 | -0.2 | 7.34E-01 | 0.0 | 9.75E-01 | F-box protein At3g17710 |
| AT3G23160 | -0.6 | 2.01E-05 | -0.3 | 1.08E-03 | -0.2 | 2.22E-02 | 0.2 | 2.50E-01 | 0.2 | 1.26E-01 | -0.1 | 7.46E-01 | Uncharacterized protein |
| AT2G14890 | -0.6 | 9.33E-04 | 0.2 | 1.47E-01 | -0.8 | 4.62E-04 | -0.3 | 1.46E-01 | 0.1 | 7.76E-01 | -0.3 | 1.73E-01 | Arabinogalactan protein 9 |
| AT1G04780 | -0.6 | 5.45E-06 | -0.7 | 3.47E-06 | 0.1 | 1.15E-01 | -0.2 | 3.51E-01 | -0.2 | 1.78E-01 | 0.1 | 7.47E-01 | F13M7.23 protein |
| AT4G14305 | -0.6 | 4.83E-03 | -0.3 | 5.82E-02 | -0.2 | 2.69E-01 | -0.7 | 2.08E-01 | -0.2 | 7.76E-01 | -0.5 | 4.38E-01 | Peroxisomal membrane 22 kDa family protein |
| AT1G64690 | -0.6 | 7.14E-04 | -0.1 | 4.78E-01 | -0.5 | 7.65E-03 | -0.2 | 5.04E-01 | 0.2 | 4.48E-01 | -0.4 | 1.95E-01 | Branchless trichome |
| AT2G40810 | -0.6 | 1.77E-04 | -0.4 | 1.76E-03 | -0.2 | 2.32E-01 | -0.5 | 1.85E-02 | -0.2 | 5.47E-01 | -0.4 | 1.85E-01 | Autophagy-like protein |
| AT5G56020 | -0.6 | 6.10E-05 | -0.2 | 2.17E-02 | -0.4 | 6.69E-03 | -0.5 | 5.99E-03 | -0.4 | 3.47E-02 | -0.1 | 7.05E-01 | Vesicle transport protein |
| AT3G52520 | -0.6 | 5.03E-03 | 0.1 | 5.93E-01 | -0.7 | 7.75E-03 | 0.2 | 5.36E-01 | 0.1 | 7.35E-01 | 0.1 | 9.15E-01 | At3g52520 |
| AT5G39570 | -0.6 | 9.35E-06 | -0.3 | 7.06E-04 | -0.2 | 9.29E-03 | -0.2 | 4.89E-02 | 0.0 | 8.25E-01 | -0.2 | 2.42E-01 | Transmembrane protein |
| AT1G31130 | -0.6 | 5.21E-05 | -0.5 | 2.32E-04 | -0.1 | 4.73E-01 | 0.0 | 9.90E-01 | 0.0 | 8.65E-01 | 0.0 | 9.00E-01 | Polyadenylate-binding protein 1-B-binding protein |
| AT5G23100 | -0.6 | 2.40E-05 | -0.3 | 3.54E-03 | -0.3 | 9.23E-03 | -0.2 | 1.88E-01 | -0.1 | 5.67E-01 | -0.1 | 7.18E-01 | At5g23100 |
| AT3G49490 | -0.6 | 6.72E-06 | -0.5 | 2.71E-05 | -0.1 | 4.66E-01 | -0.3 | 9.70E-03 | -0.2 | 6.33E-02 | -0.1 | 7.09E-01 | Uncharacterized protein T9C5.90 |
| AT1G71240 | -0.6 | 5.64E-04 | -0.7 | 2.15E-04 | 0.1 | 3.28E-01 | -0.2 | 5.49E-01 | 0.2 | 5.89E-01 | -0.3 | 2.82E-01 | Uncharacterized protein At1g71240 |
| AT5G08200 | -0.6 | 4.10E-05 | -0.3 | 2.03E-03 | -0.2 | 3.20E-02 | -0.1 | 7.27E-01 | 0.1 | 6.53E-01 | -0.2 | 4.84E-01 | Uncharacterized protein T22D6_140 |
| AT3G19470 | -0.6 | 6.40E-04 | -0.5 | 2.85E-03 | -0.1 | 5.50E-01 | -0.9 | 1.91E-02 | -0.6 | 9.98E-02 | -0.3 | 6.26E-01 | F-box protein At3g19470 |
| AT3G62770 | -0.6 | 2.71E-05 | -0.4 | 5.80E-04 | -0.2 | 6.01E-02 | -0.2 | 2.15E-01 | -0.1 | 4.00E-01 | 0.0 | 8.87E-01 | Transducin |
| AT3G27460 | -0.6 | 1.32E-04 | -0.5 | 4.03E-04 | -0.1 | 6.74E-01 | -0.3 | 1.82E-01 | -0.2 | 4.83E-01 | -0.1 | 7.53E-01 | SGF29 tudor-like domain-containing protein |
| AT1G13530 | -0.6 | 8.70E-04 | -0.1 | 4.60E-01 | -0.5 | 9.55E-03 | 0.3 | 2.74E-01 | 0.4 | 2.73E-01 | 0.0 | 9.31E-01 | hypothetical protein |
| AT4G11670 | -0.6 | 2.43E-04 | -0.7 | 1.26E-04 | 0.1 | 3.87E-01 | -0.2 | 7.52E-01 | 0.0 | 9.80E-01 | -0.2 | 7.88E-01 | Uncharacterized protein At4g11670 |
| AT4G37890 | -0.6 | 1.05E-03 | -0.5 | 2.62E-03 | -0.1 | 7.60E-01 | 0.0 | 9.00E-01 | -0.3 | 3.07E-01 | 0.4 | 3.14E-01 | Zinc finger family protein |
| AT3G50810 | -0.6 | 1.00E-03 | -0.5 | 2.39E-03 | -0.1 | 7.80E-01 | -0.3 | 2.10E-01 | -0.3 | 2.58E-01 | 0.0 | 9.97E-01 | CASP-like protein |
| AT5G57090 | -0.6 | 4.11E-03 | 0.3 | 1.10E-01 | -0.8 | 1.26E-03 | 0.0 | 8.98E-01 | 0.4 | 1.40E-01 | -0.3 | 2.49E-01 | Auxin efflux carrier component |
| AT2G45260 | -0.6 | 1.03E-05 | -0.3 | 7.05E-04 | -0.2 | 1.08E-02 | -0.1 | 5.46E-01 | -0.1 | 5.90E-01 | 0.0 | 9.90E-01 | Myosin-4 protein |
| AT5G47940 | -0.6 | 1.36E-05 | -0.4 | 1.33E-04 | -0.1 | 1.58E-01 | 0.0 | 8.55E-01 | 0.2 | 3.98E-01 | -0.1 | 5.91E-01 | 40S ribosomal protein S27 |
| AT3G57070 | -0.6 | 5.63E-05 | -0.3 | 2.89E-03 | -0.2 | 3.04E-02 | -0.2 | 3.08E-01 | 0.0 | 9.47E-01 | -0.2 | 4.70E-01 | Uncharacterized protein At3g57070 |
| AT1G06473 | -0.6 | 6.89E-03 | 0.0 | 8.30E-01 | -0.6 | 1.56E-02 | -1.1 | 1.03E-02 | -0.4 | 2.68E-01 | -0.6 | 2.52E-01 | NA |
| AT2G47600 | -0.6 | 1.06E-04 | -0.3 | 3.53E-03 | -0.2 | 6.22E-02 | -0.8 | 2.42E-04 | -0.2 | 2.73E-01 | -0.6 | 2.24E-02 | Magnesium |
| AT3G07760 | -0.6 | 1.39E-05 | -0.2 | 6.79E-03 | -0.3 | 2.12E-03 | -0.4 | 1.69E-02 | -0.3 | 9.29E-02 | -0.1 | 7.18E-01 | hypothetical protein |
| AT1G70518 | -0.6 | 7.03E-04 | -0.4 | 4.04E-03 | -0.1 | 4.30E-01 | -0.5 | 3.26E-02 | -0.4 | 8.62E-02 | -0.1 | 8.53E-01 | NA |

| | | | | | | | | | | | | | |
|---|---|---|---|---|---|---|---|---|---|---|---|---|---|
| AT3G50430 | -0.6 | 1.83E-04 | -0.5 | 9.19E-04 | -0.1 | 4.69E-01 | -0.2 | 6.88E-01 | -0.1 | 8.73E-01 | -0.1 | 8.96E-01 | Golgin |
| AT4G26120 | -0.6 | 3.65E-03 | -1.0 | 2.02E-04 | 0.5 | 4.92E-02 | 0.7 | 1.51E-02 | 0.2 | 6.05E-01 | 0.5 | 9.85E-02 | Ankyrin repeat family protein |
| AT3G54780 | -0.6 | 1.38E-03 | -0.5 | 3.07E-03 | 0.0 | 8.06E-01 | 0.2 | 6.36E-01 | 0.5 | 2.73E-01 | -0.3 | 5.37E-01 | Zinc finger family protein |
| AT1G53380 | -0.6 | 3.30E-04 | -0.2 | 9.68E-02 | -0.4 | 1.20E-02 | 0.0 | 8.05E-01 | -0.1 | 5.46E-01 | 0.1 | 7.79E-01 | DUF641 domain-containing protein |
| AT4G29400 | -0.6 | 1.00E-02 | -0.3 | 1.79E-01 | -0.3 | 1.95E-01 | 0.5 | 8.58E-02 | 0.2 | 5.94E-01 | 0.3 | 4.31E-01 | Uncharacterized protein |
| AT1G09812 | -0.6 | 1.09E-02 | -0.7 | 6.16E-03 | 0.1 | 6.93E-01 | 0.1 | 8.82E-01 | 0.5 | 2.86E-01 | -0.4 | 4.14E-01 | Transmembrane protein |
| AT1G47565 | -0.6 | 8.21E-04 | -0.6 | 6.23E-04 | 0.1 | 6.61E-01 | -0.4 | 1.62E-01 | -0.1 | 7.34E-01 | -0.3 | 4.99E-01 | NA |
| AT5G01990 | -0.6 | 1.66E-04 | -0.5 | 5.66E-04 | -0.1 | 6.32E-01 | -0.6 | 1.77E-01 | -0.1 | 8.57E-01 | -0.5 | 4.35E-01 | Protein PIN-LIKES 6 |
| AT1G10410 | -0.6 | 1.84E-05 | -0.5 | 7.38E-05 | -0.1 | 4.88E-01 | 0.2 | 2.69E-01 | 0.0 | 9.67E-01 | 0.2 | 3.82E-01 | hypothetical protein |
| AT3G50780 | -0.6 | 8.69E-05 | -0.3 | 9.45E-03 | -0.3 | 1.95E-02 | -0.7 | 3.88E-05 | -0.2 | 2.20E-01 | -0.5 | 8.38E-03 | BTB |
| AT1G80980 | -0.6 | 1.01E-04 | -0.5 | 2.97E-04 | 0.0 | 7.14E-01 | -0.3 | 2.92E-01 | -0.2 | 4.21E-01 | 0.0 | 9.22E-01 | At1g80980 |
| AT1G62045 | -0.6 | 3.42E-03 | 0.1 | 6.96E-01 | -0.6 | 6.81E-03 | -0.1 | 9.22E-01 | 0.2 | 6.95E-01 | -0.3 | 6.78E-01 | Uncharacterized protein |
| AT5G42730 | -0.6 | 3.06E-04 | -0.4 | 6.16E-03 | -0.2 | 1.39E-01 | -0.6 | 1.20E-02 | -0.4 | 1.29E-01 | -0.3 | 4.78E-01 | NA |
| AT2G34920 | -0.6 | 2.10E-03 | -0.1 | 3.28E-01 | -0.4 | 3.01E-02 | -0.1 | 8.67E-01 | 0.4 | 2.64E-01 | -0.5 | 1.85E-01 | RING |
| AT1G05960 | -0.6 | 2.14E-05 | -0.2 | 9.58E-03 | -0.3 | 2.67E-03 | -0.1 | 7.94E-01 | 0.0 | 9.28E-01 | 0.0 | 9.34E-01 | At1g05960 |
| AT1G56280 | -0.6 | 2.53E-04 | -0.5 | 1.06E-03 | -0.1 | 5.35E-01 | -0.6 | 4.12E-04 | -0.4 | 1.41E-02 | -0.2 | 4.11E-01 | Drought-induced 19 |
| AT5G41620 | -0.6 | 1.04E-03 | -0.3 | 2.24E-02 | -0.2 | 1.52E-01 | -0.3 | 1.98E-01 | -0.1 | 7.95E-01 | -0.2 | 5.11E-01 | Intracellular protein transporter USO1-like protein |
| AT1G23970 | -0.6 | 2.35E-04 | -0.5 | 5.40E-04 | 0.0 | 8.17E-01 | 0.0 | 8.97E-01 | 0.0 | 9.45E-01 | -0.1 | 8.52E-01 | Uncharacterized protein |
| AT5G44005 | -0.6 | 7.26E-02 | -0.4 | 2.39E-01 | -0.2 | 5.89E-01 | 0.0 | 9.25E-01 | 0.1 | 8.33E-01 | -0.1 | 7.92E-01 | hypothetical protein |
| AT3G09030 | -0.6 | 4.45E-05 | -0.4 | 4.33E-04 | -0.1 | 1.92E-01 | -0.4 | 6.09E-03 | -0.3 | 8.65E-02 | -0.1 | 4.96E-01 | Protein ENDOPLASMIC RETICULUM-ARRESTED PEN3 |
| AT1G29970 | -0.6 | 3.12E-05 | -0.4 | 7.56E-04 | -0.2 | 5.49E-02 | -0.5 | 5.24E-04 | -0.5 | 1.01E-03 | 0.0 | 9.40E-01 | 60S ribosomal protein L18A-1 |
| AT3G28850 | -0.6 | 1.41E-04 | -0.5 | 8.44E-04 | -0.1 | 3.84E-01 | 0.7 | 3.25E-03 | 0.4 | 7.25E-02 | 0.2 | 3.35E-01 | Uncharacterized protein At3g28850 |
| AT5G15820 | -0.6 | 8.89E-04 | -0.3 | 3.70E-02 | -0.3 | 8.99E-02 | 0.0 | 9.87E-01 | 0.1 | 8.43E-01 | -0.1 | 8.73E-01 | RING-type domain-containing protein |
| AT4G35140 | -0.6 | 9.64E-05 | -0.5 | 6.39E-04 | -0.1 | 3.41E-01 | -0.3 | 8.38E-02 | -0.2 | 2.15E-01 | -0.1 | 8.24E-01 | DDB1- and CUL4-associated factor 8 |
| AT1G61240 | -0.6 | 1.23E-04 | -0.3 | 1.66E-02 | -0.3 | 1.95E-02 | -0.1 | 7.29E-01 | -0.1 | 6.94E-01 | 0.0 | 9.53E-01 | At1g61240 ) |
| AT1G01448 | -0.6 | 2.34E-04 | -0.4 | 3.95E-03 | -0.2 | 1.62E-01 | -0.5 | 5.87E-02 | -0.6 | 3.78E-02 | 0.1 | 7.99E-01 | NA |
| AT5G19060 | -0.6 | 7.35E-03 | -0.2 | 1.71E-01 | -0.3 | 1.72E-01 | 0.1 | 7.73E-01 | -0.4 | 4.08E-01 | 0.5 | 3.01E-01 | Uncharacterized protein |
| AT4G31410 | -0.6 | 6.02E-06 | -0.5 | 1.71E-05 | 0.0 | 7.47E-01 | -0.5 | 1.28E-02 | -0.5 | 1.83E-02 | 0.0 | 9.83E-01 | Uncharacterized protein AT4g31410 |
| AT5G14285 | -0.6 | 4.04E-05 | -0.4 | 3.03E-04 | -0.1 | 2.69E-01 | -0.3 | 2.04E-01 | -0.2 | 4.79E-01 | -0.1 | 7.48E-01 | TLC domain-containing protein At5g14285 |
| AT2G04800 | -0.6 | 4.22E-03 | -0.3 | 8.89E-02 | -0.3 | 1.69E-01 | -0.3 | 3.42E-01 | 0.0 | 9.40E-01 | -0.3 | 5.24E-01 | At2g04800 |
| AT1G22030 | -0.6 | 6.59E-05 | 0.0 | 5.83E-01 | -0.5 | 6.94E-04 | 0.3 | 6.74E-02 | 0.1 | 7.60E-01 | 0.2 | 2.40E-01 | BPS1-like protein |
| AT2G02370 | -0.6 | 3.34E-05 | -0.6 | 2.51E-05 | 0.1 | 3.69E-01 | -0.1 | 6.62E-01 | -0.2 | 3.71E-01 | 0.1 | 7.24E-01 | At2g02370 |
| AT4G01670 | -0.6 | 2.42E-03 | -0.2 | 2.48E-01 | -0.4 | 4.69E-02 | -0.2 | 6.10E-01 | -0.3 | 4.19E-01 | 0.1 | 7.99E-01 | Uncharacterized protein AT4g01670 |
| AT1G74510 | -0.6 | 1.62E-05 | -0.1 | 8.74E-02 | -0.4 | 4.99E-04 | -0.3 | 3.87E-02 | 0.0 | 9.82E-01 | -0.3 | 1.03E-01 | F-box |
| AT5G18690 | -0.6 | 1.66E-03 | -0.2 | 8.17E-02 | -0.3 | 7.89E-02 | 0.0 | 9.52E-01 | 0.1 | 6.83E-01 | -0.1 | 7.75E-01 | Classical arabinogalactan protein 25 |
| AT1G08190 | -0.6 | 2.94E-05 | -0.4 | 8.46E-04 | -0.2 | 4.66E-02 | -0.2 | 4.49E-02 | -0.1 | 6.27E-01 | -0.2 | 3.63E-01 | Vacuolar protein sorting-associated protein 41 homolog |
| AT4G16770 | -0.6 | 4.19E-05 | -0.3 | 1.76E-03 | -0.2 | 3.72E-02 | 0.0 | 8.60E-01 | 0.3 | 2.75E-01 | -0.3 | 2.58E-01 | Gibberellin oxidase-like protein |
| AT5G01170 | -0.6 | 1.75E-05 | -0.3 | 7.49E-04 | -0.2 | 2.64E-02 | 0.0 | 9.60E-01 | 0.2 | 4.28E-01 | -0.2 | 5.55E-01 | Protein OCTOPUS-like |
| AT4G29420 | -0.6 | 2.03E-04 | -0.4 | 4.09E-03 | -0.2 | 1.28E-01 | -0.5 | 1.60E-02 | -0.5 | 5.18E-02 | -0.1 | 8.07E-01 | F-box |
| AT1G49850 | -0.6 | 1.63E-05 | -0.3 | 9.44E-04 | -0.2 | 1.79E-02 | -0.3 | 7.29E-02 | -0.2 | 2.94E-01 | -0.1 | 7.27E-01 | Uncharacterized protein |
| AT4G36210 | -0.6 | 1.51E-05 | -0.8 | 2.59E-06 | 0.2 | 7.55E-03 | 0.0 | 8.39E-01 | -0.1 | 6.92E-01 | 0.0 | 8.83E-01 | Transmembrane |
| AT3G58620 | -0.5 | 5.53E-04 | -0.1 | 2.22E-01 | -0.4 | 1.05E-02 | -0.7 | 9.08E-04 | -0.4 | 3.04E-02 | -0.3 | 3.26E-01 | Tetratricopetide-repeat thioredoxin-like 4 |
| AT4G03070 | -0.5 | 9.97E-04 | -0.4 | 1.19E-02 | -0.2 | 2.48E-01 | -0.2 | 4.59E-01 | 0.1 | 7.87E-01 | -0.3 | 3.08E-01 | Oxidoreductase like protein |
| AT3G50830 | -0.5 | 1.77E-05 | -0.4 | 1.79E-04 | -0.1 | 1.49E-01 | -0.6 | 4.77E-04 | -0.3 | 8.72E-02 | -0.3 | 1.30E-01 | Cold-regulated 413 plasma membrane protein 2 |
| AT2G34380 | -0.5 | 9.21E-05 | -0.4 | 6.28E-04 | -0.1 | 3.33E-01 | -0.2 | 2.88E-01 | -0.2 | 3.85E-01 | 0.0 | 9.69E-01 | Seipin-3 |
| AT1G10720 | -0.5 | 7.59E-06 | -0.5 | 3.21E-05 | -0.1 | 4.36E-01 | -0.1 | 4.51E-01 | -0.1 | 7.77E-01 | -0.1 | 8.11E-01 | BSD domain-containing protein |
| AT3G26470 | -0.5 | 1.22E-04 | -0.9 | 1.14E-05 | 0.3 | 1.48E-02 | 1.5 | 1.07E-05 | 1.0 | 1.89E-03 | 0.5 | 2.78E-02 | RPW8 domain-containing protein |
| AT2G20740 | -0.5 | 3.12E-05 | -0.2 | 1.00E-02 | -0.3 | 4.58E-03 | -0.6 | 2.59E-04 | -0.5 | 3.89E-03 | -0.1 | 5.40E-01 | Tetraspanin-19 |
| AT5G64190 | -0.5 | 2.78E-01 | -0.6 | 2.82E-01 | 0.0 | 9.61E-01 | -0.8 | 3.06E-01 | -3.1 | 2.20E-02 | 2.3 | 1.72E-01 | Gb\|AAD25673.1 |
| AT2G18680 | -0.5 | 2.45E-02 | -0.6 | 2.26E-02 | 0.0 | 9.14E-01 | 0.5 | 1.76E-01 | 0.1 | 8.91E-01 | 0.4 | 3.51E-01 | Uncharacterized protein At2g18680 |
| AT5G60920 | -0.5 | 6.03E-05 | -0.2 | 3.90E-02 | -0.3 | 3.51E-03 | -0.2 | 5.71E-02 | 0.0 | 9.69E-01 | -0.3 | 1.32E-01 | COBRA-like protein |
| AT2G22170 | -0.5 | 1.02E-03 | 0.5 | 2.42E-03 | -1.1 | 6.96E-05 | -0.3 | 1.59E-01 | 0.1 | 7.67E-01 | -0.4 | 1.30E-01 | PLAT domain-containing protein 2 |
| AT3G14830 | -0.5 | 8.03E-06 | -0.5 | 2.62E-05 | 0.0 | 6.18E-01 | -0.2 | 2.43E-01 | -0.2 | 2.94E-01 | 0.0 | 9.96E-01 | Uncharacterized protein |
| AT3G24150 | -0.5 | 1.01E-04 | -0.4 | 2.02E-03 | -0.2 | 1.09E-01 | 0.1 | 6.22E-01 | 0.4 | 9.84E-02 | -0.3 | 3.51E-01 | Uncharacterized protein |
| AT2G37750 | -0.5 | 9.72E-04 | 0.1 | 5.02E-01 | -0.6 | 1.26E-03 | -0.1 | 6.59E-01 | 0.0 | 9.26E-01 | -0.1 | 8.46E-01 | hypothetical protein |
| AT2G41000 | -0.5 | 6.85E-04 | -0.6 | 3.67E-04 | 0.1 | 4.85E-01 | 0.1 | 7.85E-01 | 0.2 | 5.80E-01 | -0.1 | 7.85E-01 | Chaperone DnaJ-domain superfamily protein |
| AT5G66330 | -0.5 | 1.17E-03 | 0.4 | 6.08E-03 | -0.9 | 1.93E-04 | -0.7 | 1.71E-02 | -0.3 | 3.92E-01 | -0.4 | 2.32E-01 | hypothetical protein |
| AT3G19150 | -0.5 | 1.78E-03 | -1.1 | 5.01E-05 | 0.5 | 1.23E-02 | -0.3 | 2.02E-01 | 0.2 | 3.52E-01 | -0.5 | 7.57E-02 | KIP-related protein 6 |
| AT5G15330 | -0.5 | 1.33E-03 | -0.3 | 6.32E-02 | -0.3 | 8.15E-02 | -0.6 | 1.29E-02 | -0.5 | 4.02E-02 | -0.1 | 8.34E-01 | SPX domain-containing protein 4 |
| AT2G44770 | -0.5 | 2.30E-05 | -0.3 | 4.49E-03 | -0.3 | 6.00E-03 | -0.1 | 4.97E-01 | 0.0 | 9.91E-01 | -0.1 | 6.26E-01 | hypothetical protein |
| AT3G01520 | -0.5 | 3.26E-05 | -0.1 | 1.80E-01 | -0.4 | 6.23E-04 | -0.6 | 3.58E-04 | -0.4 | 1.69E-02 | -0.2 | 3.03E-01 | UspA domain-containing protein |
| AT1G74860 | -0.5 | 5.05E-05 | -0.5 | 1.03E-04 | 0.0 | 9.35E-01 | -0.2 | 2.32E-01 | 0.1 | 6.51E-01 | -0.3 | 1.81E-01 | At1g74860 |
| AT4G13830 | -0.5 | 9.89E-04 | -0.3 | 3.15E-02 | -0.2 | 9.36E-02 | -0.2 | 3.28E-02 | -0.4 | 4.65E-03 | 0.1 | 4.21E-01 | DNAJ-like 20 |
| AT2G41150 | -0.5 | 3.19E-03 | 0.0 | 9.39E-01 | -0.5 | 1.19E-02 | -0.4 | 2.93E-01 | 0.4 | 3.39E-01 | -0.8 | 7.84E-02 | Uncharacterized protein At2g41150 |
| AT3G26740 | -0.5 | 1.91E-03 | 0.1 | 6.65E-01 | -0.6 | 2.60E-03 | -0.5 | 2.22E-03 | -0.3 | 1.27E-01 | -0.3 | 2.70E-01 | Light-regulated protein 1, chloroplastic |
| AT5G10730 | -0.5 | 9.64E-04 | -0.2 | 7.21E-02 | -0.3 | 4.63E-02 | -0.3 | 7.03E-02 | 0.1 | 4.31E-01 | -0.4 | 2.38E-02 | NAD-binding domain-containing protein |
| AT1G35115 | -0.5 | 8.45E-05 | -0.4 | 7.18E-04 | -0.1 | 2.40E-01 | 0.3 | 2.67E-01 | 0.3 | 4.73E-01 | 0.1 | 8.86E-01 | NA |
| AT3G57780 | -0.5 | 7.15E-05 | -0.4 | 5.92E-04 | -0.1 | 2.53E-01 | -0.2 | 5.36E-01 | 0.2 | 4.12E-01 | -0.4 | 1.99E-01 | Nucleolar-like protein |
| AT4G33565 | -0.5 | 2.95E-05 | -0.4 | 3.43E-04 | -0.1 | 1.44E-01 | -0.2 | 1.57E-01 | -0.5 | 5.29E-03 | 0.3 | 1.94E-01 | RING-type E3 ubiquitin transferase |
| AT4G11450 | -0.5 | 1.01E-03 | -0.2 | 8.79E-02 | -0.3 | 4.80E-02 | -0.1 | 8.26E-01 | 0.0 | 9.90E-01 | -0.1 | 8.64E-01 | Bromo-adjacent homology domain-containing protein |
| AT5G56530 | -0.5 | 4.12E-05 | -0.2 | 2.29E-02 | -0.3 | 3.84E-03 | 0.1 | 6.67E-01 | 0.1 | 5.45E-01 | 0.0 | 8.87E-01 | Carboxyl-terminal peptidase |
| AT3G25800 | -0.5 | 5.01E-06 | -0.4 | 5.13E-05 | -0.1 | 1.04E-01 | -0.1 | 2.49E-01 | -0.1 | 5.76E-01 | 0.0 | 8.07E-01 | Protein phosphatase 2A subunit A2 |
| AT4G36840 | -0.5 | 2.25E-04 | -0.4 | 1.21E-03 | -0.1 | 4.34E-01 | -0.3 | 8.58E-02 | 0.1 | 6.65E-01 | -0.4 | 4.86E-02 | Galactose oxidase |
| AT1G55590 | -0.5 | 1.41E-05 | -0.5 | 3.26E-05 | 0.0 | 8.85E-01 | -0.4 | 5.49E-02 | -0.4 | 8.74E-02 | 0.0 | 9.75E-01 | F-box domain-containing protein |
| AT5G04860 | -0.5 | 3.58E-05 | -0.4 | 2.74E-04 | -0.1 | 2.37E-01 | -0.2 | 2.50E-01 | -0.1 | 6.08E-01 | -0.1 | 7.19E-01 | Splicing factor 3A subunit |
| AT5G15537 | -0.5 | 3.34E-02 | -1.1 | 1.52E-03 | 0.6 | 8.86E-02 | 0.1 | 9.18E-01 | -0.3 | 6.87E-01 | 0.4 | 5.99E-01 | Transmembrane protein |
| AT3G18260 | -0.5 | 6.66E-03 | -0.4 | 2.19E-02 | -0.1 | 6.40E-01 | 0.1 | 7.61E-01 | 0.3 | 5.96E-01 | -0.1 | 8.44E-01 | Reticulon-like protein B9 |
| AT4G14020 | -0.5 | 2.41E-02 | -1.0 | 1.65E-03 | 0.4 | 1.25E-01 | 0.0 | 9.51E-01 | 0.2 | 4.27E-01 | -0.2 | 4.80E-01 | At4g14020 family protein) |
| AT1G01440 | -0.5 | 4.16E-05 | -0.4 | 2.24E-04 | -0.1 | 3.73E-01 | -0.2 | 2.83E-01 | -0.1 | 5.06E-01 | -0.1 | 8.82E-01 | hypothetical protein |
| AT1G77500 | -0.5 | 5.21E-05 | -0.4 | 3.27E-04 | -0.1 | 3.25E-01 | 0.3 | 1.30E-01 | 0.2 | 4.17E-01 | 0.1 | 7.10E-01 | Uncharacterized protein |
| AT1G10160 | -0.5 | 2.74E-04 | -0.4 | 3.84E-03 | -0.2 | 1.89E-01 | -0.6 | 3.66E-02 | -0.4 | 2.75E-01 | -0.3 | 4.57E-01 | NA |
| AT5G42420 | -0.5 | 7.84E-05 | 0.0 | 8.72E-01 | -0.5 | 4.32E-04 | -0.1 | 4.55E-01 | 0.0 | 9.98E-01 | -0.1 | 6.18E-01 | Nucleotide-sugar transporter family protein |
| AT5G18980 | -0.5 | 1.27E-04 | -0.5 | 4.90E-04 | -0.1 | 5.50E-01 | -0.6 | 4.07E-03 | -0.4 | 3.36E-02 | -0.1 | 5.71E-01 | hypothetical protein |

| | | | | | | | | | | | | | |
|---|---|---|---|---|---|---|---|---|---|---|---|---|---|
| AT1G01470 | -0.5 | 2.47E-03 | -0.3 | 3.44E-02 | -0.2 | 2.11E-01 | -0.4 | 8.84E-02 | -0.1 | 7.35E-01 | -0.3 | 3.49E-01 | Probable desiccation-related protein LEA14 |
| AT2G30710 | -0.5 | 2.94E-05 | -0.3 | 8.41E-04 | -0.2 | 4.69E-02 | -0.2 | 1.60E-01 | -0.1 | 6.03E-01 | -0.1 | 6.42E-01 | hypothetical protein |
| AT3G04930 | -0.5 | 2.78E-05 | -0.4 | 2.02E-04 | -0.1 | 2.38E-01 | -0.6 | 1.09E-04 | -0.6 | 2.38E-04 | 0.0 | 9.08E-01 | Probable transcription factor At3g04930 |
| AT5G48610 | -0.5 | 8.69E-05 | -0.2 | 1.50E-02 | -0.3 | 1.38E-02 | -0.2 | 4.15E-01 | 0.1 | 8.54E-01 | -0.2 | 4.11E-01 | Gb\|AAF20218.1 |
| AT3G27390 | -0.5 | 1.16E-05 | -0.3 | 5.92E-04 | -0.2 | 1.46E-02 | -0.3 | 1.45E-02 | -0.1 | 6.92E-01 | -0.3 | 1.02E-01 | Uncharacterized membrane protein At3g27390 |
| AT2G25280 | -0.5 | 4.18E-05 | -0.3 | 2.91E-03 | -0.2 | 2.23E-02 | -0.1 | 7.71E-01 | -0.1 | 7.98E-01 | 0.0 | 9.97E-01 | AT2G25280 protein |
| AT4G25760 | -0.5 | 1.35E-03 | 0.3 | 1.03E-02 | -0.9 | 2.50E-04 | 1.1 | 6.58E-04 | 0.5 | 1.49E-01 | 0.6 | 4.29E-02 | Protein GLUTAMINE DUMPER 2 |
| AT4G24590 | -0.5 | 5.05E-04 | -0.7 | 1.82E-04 | 0.1 | 3.05E-01 | -0.5 | 5.83E-03 | -0.3 | 1.08E-01 | -0.2 | 4.29E-01 | hypothetical protein |
| AT3G49350 | -0.5 | 1.08E-05 | -0.4 | 2.62E-04 | -0.2 | 3.77E-02 | 0.0 | 7.76E-01 | 0.0 | 9.52E-01 | -0.1 | 7.74E-01 | Rab-GAP TBC domain-containing protein |
| AT3G11420 | -0.5 | 6.58E-05 | -0.2 | 1.07E-02 | -0.3 | 1.07E-02 | 0.0 | 7.78E-01 | 0.1 | 5.57E-01 | -0.2 | 4.58E-01 | Uncharacterized protein |
| AT1G47860 | -0.5 | 2.17E-04 | -0.1 | 2.18E-01 | -0.4 | 3.94E-03 | -0.2 | 6.65E-02 | -0.1 | 3.54E-01 | -0.1 | 6.06E-01 | NA |
| AT5G19900 | -0.5 | 9.98E-06 | -0.4 | 1.27E-04 | -0.1 | 8.82E-02 | 0.1 | 7.55E-01 | 0.1 | 5.75E-01 | 0.0 | 8.48E-01 | PRLI-interacting factor A |
| AT5G44160 | -0.5 | 1.63E-04 | -0.2 | 8.48E-02 | -0.4 | 7.50E-03 | -0.3 | 4.88E-01 | 0.0 | 9.56E-01 | -0.2 | 6.56E-01 | C2H2-like zinc finger protein |
| AT1G05060 | -0.5 | 6.09E-05 | -0.8 | 6.90E-06 | 0.3 | 1.30E-02 | 2.0 | 2.08E-07 | 0.1 | 6.57E-01 | 1.9 | 5.28E-06 | hypothetical protein |
| AT5G06265 | -0.5 | 1.48E-04 | -0.2 | 3.74E-02 | -0.3 | 1.24E-02 | 0.1 | 6.85E-01 | 0.3 | 2.64E-01 | -0.2 | 6.00E-01 | Hyaluronan mediated motility receptor-like protein |
| AT2G17990 | -0.5 | 5.65E-06 | -0.4 | 5.85E-05 | -0.1 | 1.02E-01 | -0.4 | 7.48E-04 | -0.3 | 3.23E-02 | -0.2 | 3.32E-01 | At2g17990 |
| AT3G13435 | -0.5 | 6.35E-02 | -0.6 | 5.40E-02 | 0.1 | 8.94E-01 | -0.1 | 9.06E-01 | 0.4 | 4.26E-01 | -0.4 | 3.51E-01 | Transmembrane protein |
| AT2G20920 | -0.5 | 1.83E-05 | -0.4 | 3.14E-04 | -0.1 | 7.42E-02 | 0.6 | 1.15E-03 | -0.3 | 5.90E-02 | 0.9 | 4.44E-04 | hypothetical protein |
| AT1G58270 | -0.5 | 5.05E-05 | 0.2 | 4.32E-02 | -0.7 | 4.90E-05 | -0.4 | 8.67E-03 | 0.1 | 6.69E-01 | -0.4 | 1.65E-02 | Expressed protein |
| AT5G47120 | -0.5 | 6.59E-05 | -0.4 | 6.88E-04 | -0.1 | 1.93E-01 | 0.3 | 5.63E-02 | 0.3 | 1.00E-01 | 0.0 | 9.81E-01 | Bax inhibitor 1 |
| AT5G10660 | -0.5 | 1.29E-02 | 0.2 | 3.46E-01 | -0.7 | 9.50E-03 | | | | | | | Calmodulin-binding protein-like protein |
| AT2G43250 | -0.5 | 1.03E-04 | -0.4 | 1.23E-03 | -0.1 | 1.84E-01 | -0.3 | 2.36E-01 | -0.1 | 7.49E-01 | -0.2 | 5.32E-01 | hypothetical protein |
| AT5G62865 | -0.5 | 1.57E-03 | -0.1 | 6.22E-01 | -0.5 | 9.33E-03 | -0.9 | 4.25E-05 | -0.7 | 8.29E-04 | -0.2 | 4.44E-01 | hypothetical protein |
| AT2G25760 | -0.5 | 3.77E-05 | -0.6 | 3.28E-05 | 0.1 | 4.92E-01 | -0.1 | 7.69E-01 | -0.1 | 7.84E-01 | 0.0 | 9.84E-01 | Uncharacterized protein At2g25750 |
| AT1G67410 | -0.5 | 3.83E-04 | -0.2 | 5.44E-02 | -0.3 | 2.59E-02 | -0.6 | 2.73E-03 | -0.2 | 3.29E-01 | -0.4 | 8.76E-02 | F1N21.23 |
| AT1G35260 | -0.5 | 6.56E-03 | -0.5 | 9.18E-03 | 0.0 | 9.56E-01 | 0.9 | 1.31E-03 | 0.5 | 7.56E-02 | 0.4 | 2.05E-01 | MLP-like protein 165 |
| AT3G15880 | -0.5 | 4.05E-05 | -0.5 | 7.34E-05 | 0.0 | 9.66E-01 | 0.1 | 7.67E-01 | 0.1 | 7.59E-01 | 0.0 | 9.66E-01 | WUS-interacting protein 2 |
| AT3G48860 | -0.5 | 2.91E-05 | -0.4 | 5.29E-04 | -0.2 | 8.00E-02 | -0.3 | 2.69E-02 | -0.3 | 3.55E-02 | 0.0 | 9.64E-01 | Coiled-coil protein |
| AT1G61760 | -0.5 | 2.87E-03 | -0.5 | 4.13E-03 | 0.0 | 9.98E-01 | 0.5 | 4.29E-01 | 0.9 | 1.09E-01 | -0.5 | 3.80E-01 | Late embryogenesis abundant hydroxyproline-rich glycoprotein family |
| AT4G15470 | -0.5 | 4.33E-05 | -0.3 | 7.00E-03 | -0.3 | 9.26E-03 | -0.1 | 3.49E-01 | 0.0 | 9.56E-01 | -0.1 | 5.73E-01 | BI1-like protein |
| AT2G38710 | -0.5 | 3.19E-05 | -0.4 | 4.42E-04 | -0.1 | 1.09E-01 | 0.0 | 9.06E-01 | 0.2 | 4.59E-01 | -0.2 | 4.54E-01 | Uncharacterized protein At2g38710 |
| AT5G06470 | -0.5 | 9.36E-03 | 0.2 | 1.87E-01 | -0.7 | 4.75E-03 | -0.7 | 6.11E-02 | -0.4 | 3.14E-01 | -0.3 | 6.23E-01 | Glutaredoxin domain-containing protein |
| AT1G31460 | -0.5 | 3.01E-03 | -0.3 | 7.95E-02 | -0.3 | 1.51E-01 | 0.1 | 7.53E-01 | 0.0 | 9.89E-01 | 0.1 | 8.36E-01 | At1g31460 |
| AT3G15780 | -0.5 | 1.45E-03 | -0.5 | 3.30E-03 | 0.0 | 7.98E-01 | -0.4 | 2.28E-02 | -0.3 | 1.83E-01 | -0.2 | 5.50E-01 | hypothetical protein |
| AT1G68230 | -0.5 | 2.22E-02 | -0.5 | 2.21E-02 | 0.0 | 9.35E-01 | 0.1 | 8.03E-01 | 0.0 | 9.50E-01 | 0.1 | 8.06E-01 | Reticulon-like protein B14 |
| AT3G15040 | -0.5 | 3.38E-04 | -0.3 | 6.29E-03 | -0.2 | 1.39E-01 | -0.5 | 4.77E-03 | -0.2 | 2.24E-01 | -0.3 | 2.43E-01 | hypothetical protein |
| AT1G70550 | -0.5 | 2.63E-03 | -0.3 | 4.72E-02 | -0.2 | 1.89E-01 | 0.0 | 8.36E-01 | 0.2 | 3.28E-01 | -0.2 | 3.16E-01 | Uncharacterized protein F24J13.12 |
| AT1G07280 | -0.5 | 1.36E-03 | -0.3 | 1.86E-02 | -0.2 | 2.09E-01 | -0.2 | 1.31E-01 | -0.2 | 3.57E-01 | -0.1 | 8.21E-01 | At1g07280 |
| AT5G35160 | -0.5 | 4.26E-05 | -0.2 | 1.14E-02 | -0.3 | 5.72E-03 | 0.0 | 1.00E+00 | 0.1 | 7.39E-01 | -0.1 | 7.88E-01 | Transmembrane 9 superfamily member |
| AT2G39890 | -0.5 | 1.27E-04 | -0.7 | 2.66E-05 | 0.2 | 7.51E-02 | -0.5 | 2.23E-02 | -0.1 | 8.49E-01 | -0.4 | 1.25E-01 | Proline transporter 1 |
| AT3G59150 | -0.5 | 2.31E-03 | -0.4 | 1.44E-02 | -0.1 | 4.31E-01 | -0.2 | 4.44E-01 | -0.3 | 3.95E-01 | 0.0 | 9.11E-01 | F-box domain-containing protein |
| AT5G57900 | -0.5 | 4.88E-05 | -0.5 | 9.44E-05 | 0.0 | 9.60E-01 | -0.5 | 5.69E-03 | -0.3 | 1.03E-01 | -0.2 | 3.91E-01 | F-box protein SKIP1 |
| AT3G26100 | -0.5 | 5.02E-04 | -0.6 | 2.13E-04 | 0.1 | 3.54E-01 | -0.6 | 3.88E-04 | -0.2 | 2.30E-01 | -0.4 | 2.12E-02 | Regulator of chromosome condensation family protein |
| AT2G20520 | -0.5 | 1.30E-01 | 1.1 | 2.08E-03 | -1.6 | 1.51E-03 | -4.7 | 5.02E-05 | -0.3 | 5.13E-01 | -4.5 | 1.47E-03 | Fasciclin-like arabinogalactan protein 6 |
| AT4G30750 | -0.5 | 7.29E-05 | -0.4 | 2.90E-04 | -0.1 | 5.17E-01 | -0.2 | 1.17E-01 | -0.2 | 9.90E-02 | 0.0 | 8.95E-01 | AT4g30750 |
| AT1G65690 | -0.5 | 2.79E-04 | -0.6 | 1.35E-04 | 0.1 | 3.70E-01 | 1.7 | 2.71E-06 | 0.7 | 2.81E-02 | 1.1 | 9.00E-04 | NDR1 |
| AT1G15670 | -0.5 | 5.78E-03 | -0.5 | 6.81E-03 | 0.0 | 9.75E-01 | -0.5 | 2.27E-02 | -1.0 | 1.02E-03 | 0.4 | 1.97E-01 | F-box |
| AT1G01180 | -0.5 | 7.61E-05 | -0.1 | 1.15E-01 | -0.4 | 2.43E-03 | -0.3 | 2.60E-02 | -0.5 | 1.58E-03 | 0.2 | 2.29E-01 | T25K16.17 |
| AT5G18661 | -0.5 | 7.42E-02 | -0.1 | 6.03E-01 | -0.4 | 2.76E-01 | -0.8 | 8.55E-02 | 0.0 | 9.88E-01 | -0.8 | 1.14E-01 | Transmembrane protein |
| AT3G60590 | -0.5 | 1.34E-04 | -0.5 | 2.35E-04 | 0.0 | 9.78E-01 | -0.5 | 1.50E-02 | -0.2 | 2.21E-01 | -0.2 | 3.68E-01 | Cytochrome P450 family protein |
| AT5G26260 | -0.5 | 1.60E-02 | 0.4 | 6.63E-02 | -0.9 | 2.19E-03 | -0.9 | 5.81E-05 | -0.2 | 2.42E-01 | -0.7 | 6.43E-03 | F9D12.8 protein |
| AT3G25670 | -0.5 | 4.76E-02 | 0.5 | 3.17E-02 | -1.0 | 4.11E-03 | -0.3 | 3.56E-01 | -0.1 | 8.92E-01 | -0.3 | 5.53E-01 | Uncharacterized protein |
| AT4G24380 | -0.5 | 1.28E-02 | -0.6 | 6.65E-03 | 0.1 | 6.48E-01 | -0.3 | 5.49E-01 | 0.1 | 8.08E-01 | -0.4 | 4.93E-01 | Uncharacterized protein AT4g24380 |
| AT3G51130 | -0.5 | 2.25E-05 | -0.5 | 8.45E-05 | 0.0 | 5.33E-01 | -0.4 | 3.20E-03 | -0.1 | 7.03E-01 | -0.3 | 5.73E-02 | PHAF1 protein At3g51130 |
| AT3G48410 | -0.5 | 1.10E-03 | -0.1 | 5.41E-01 | -0.4 | 9.29E-03 | -0.5 | 1.19E-02 | -0.1 | 6.40E-01 | -0.4 | 1.11E-01 | Alpha |
| AT1G68220 | -0.5 | 6.24E-05 | -0.2 | 2.60E-02 | -0.3 | 5.32E-03 | -0.4 | 1.42E-02 | -0.2 | 4.45E-01 | -0.3 | 2.60E-01 | hypothetical protein |
| AT1G06890 | -0.5 | 1.97E-05 | -0.2 | 4.36E-03 | -0.3 | 4.54E-03 | -0.2 | 9.33E-02 | -0.1 | 5.49E-01 | -0.1 | 5.63E-01 | Sugar phosphate transporter domain-containing protein |
| AT4G17970 | -0.5 | 4.72E-03 | -0.1 | 5.14E-01 | -0.4 | 3.67E-02 | 0.0 | 8.99E-01 | 0.4 | 1.48E-01 | -0.4 | 2.68E-01 | Uncharacterized protein At4g17970 |
| AT3G53670 | -0.5 | 1.18E-04 | -0.2 | 7.91E-02 | -0.3 | 4.19E-03 | -0.1 | 4.91E-01 | -0.1 | 7.48E-01 | 0.0 | 8.81E-01 | hypothetical protein |
| AT2G20950 | -0.5 | 4.42E-04 | -0.4 | 2.65E-03 | -0.1 | 3.77E-01 | 0.0 | 8.28E-01 | -0.1 | 7.78E-01 | 0.1 | 6.91E-01 | Phospholipase-like protein family protein |
| AT5G56220 | -0.5 | 5.62E-04 | -0.2 | 1.37E-01 | -0.3 | 1.64E-02 | -0.4 | 1.32E-01 | 0.0 | 9.72E-01 | -0.4 | 2.63E-01 | TIR domain-containing protein |
| AT1G50600 | -0.5 | 1.52E-05 | -0.4 | 1.37E-04 | -0.1 | 1.63E-01 | -0.2 | 8.02E-02 | -0.2 | 2.47E-01 | -0.1 | 8.15E-01 | AT1G50600 protein |
| AT4G09890 | -0.5 | 4.25E-02 | -0.6 | 1.90E-02 | 0.1 | 6.44E-01 | -0.2 | 4.62E-01 | 0.0 | 9.28E-01 | -0.2 | 7.01E-01 | hypothetical protein |
| AT5G59210 | -0.5 | 5.05E-05 | -0.3 | 4.35E-03 | -0.2 | 1.85E-02 | -0.2 | 2.86E-01 | 0.2 | 4.09E-01 | -0.3 | 1.12E-01 | Myosin heavy chain-like protein |
| AT4G39870 | -0.5 | 9.87E-05 | -0.4 | 3.87E-04 | -0.1 | 5.50E-01 | -0.2 | 1.96E-01 | -0.3 | 9.26E-02 | 0.1 | 7.27E-01 | TLD-domain containing nucleolar protein |
| AT5G23370 | -0.5 | 1.12E-02 | 0.4 | 3.56E-02 | -0.9 | 1.93E-03 | | | | | | | Putative GEM-like protein 8 |
| AT3G45830 | -0.5 | 1.49E-05 | -0.6 | 1.45E-05 | 0.1 | 4.60E-01 | -0.3 | 2.19E-01 | 0.0 | 8.85E-01 | -0.2 | 4.91E-01 | DEUBAD domain-containing protein |
| AT1G27850 | -0.5 | 3.60E-05 | -0.5 | 9.44E-05 | 0.0 | 7.68E-01 | -0.1 | 4.77E-01 | 0.0 | 8.55E-01 | -0.1 | 7.66E-01 | hypothetical protein |
| AT1G18200 | -0.5 | 2.04E-03 | -0.4 | 7.63E-03 | -0.1 | 5.92E-01 | -0.1 | 8.45E-01 | -0.1 | 8.44E-01 | 0.0 | 9.82E-01 | RABA6b |
| AT1G63840 | -0.5 | 6.88E-05 | -0.1 | 3.17E-01 | -0.4 | 1.03E-03 | -0.1 | 5.27E-01 | 0.0 | 8.90E-01 | -0.1 | 7.53E-01 | RING-type domain-containing protein |
| AT5G59470 | -0.5 | 5.76E-03 | -0.4 | 3.22E-02 | -0.1 | 4.81E-01 | 0.0 | 9.57E-01 | 0.0 | 9.34E-01 | -0.1 | 9.04E-01 | Mannose-P-dolichol utilization defect 1 protein homolog 1 |
| AT4G29790 | -0.5 | 2.74E-04 | -0.5 | 2.80E-04 | 0.0 | 7.66E-01 | -0.1 | 3.42E-01 | -0.2 | 3.25E-01 | 0.0 | 9.37E-01 | hypothetical protein |
| AT2G21340 | -0.5 | 4.66E-04 | -0.5 | 7.71E-04 | 0.0 | 9.48E-01 | -0.2 | 4.63E-01 | 0.1 | 7.91E-01 | -0.3 | 3.72E-01 | MATE efflux family protein |
| AT1G53035 | -0.5 | 1.37E-03 | 0.2 | 9.19E-02 | -0.7 | 7.33E-04 | 0.2 | 5.27E-01 | 0.4 | 2.27E-01 | -0.2 | 6.13E-01 | Transmembrane protein |
| AT3G49590 | -0.5 | 9.97E-05 | -0.4 | 3.99E-04 | -0.1 | 5.20E-01 | -0.7 | 6.77E-03 | -0.3 | 2.02E-01 | -0.3 | 2.41E-01 | Autophagy-related protein 13 |
| AT2G17110 | -0.5 | 6.07E-05 | -0.4 | 3.35E-04 | -0.1 | 3.86E-01 | -0.1 | 6.91E-01 | -0.2 | 3.06E-01 | 0.1 | 6.05E-01 | AT4g35240 |
| AT5G14750 | -0.5 | 2.25E-02 | 0.0 | 8.00E-01 | -0.5 | 3.55E-02 | 0.1 | 7.77E-01 | 0.5 | 8.04E-02 | -0.4 | 2.23E-01 | Myb domain protein 66 |
| AT4G00335 | -0.5 | 9.71E-05 | -0.4 | 3.22E-04 | 0.0 | 6.32E-01 | -0.1 | 8.05E-01 | -0.1 | 6.67E-01 | 0.1 | 8.73E-01 | RHB1A |
| AT3G04485 | -0.5 | 1.29E-02 | -0.2 | 3.17E-01 | -0.3 | 1.53E-01 | -0.7 | 2.90E-02 | -0.3 | 3.26E-01 | -0.4 | 4.00E-01 | NA |
| AT2G43320 | -0.5 | 8.35E-05 | -0.5 | 1.66E-04 | 0.0 | 9.33E-01 | 0.4 | 5.89E-02 | 0.1 | 5.50E-01 | 0.2 | 2.99E-01 | hypothetical protein |
| AT1G71400 | -0.5 | 2.13E-04 | -0.2 | 4.37E-02 | -0.3 | 1.61E-02 | 0.9 | 3.19E-03 | 0.6 | 7.07E-02 | 0.3 | 3.40E-01 | Receptor-like protein 12 |
| AT1G14160 | -0.5 | 9.02E-04 | 0.1 | 4.89E-01 | -0.6 | 1.72E-03 | 0.4 | 1.52E-01 | 0.5 | 8.69E-02 | -0.1 | 7.45E-01 | CASP-like protein |
| AT3G10260 | -0.5 | 1.66E-05 | -0.2 | 3.00E-02 | -0.3 | 9.33E-04 | -0.1 | 4.40E-01 | -0.2 | 4.51E-01 | 0.0 | 9.61E-01 | Reticulon-like protein B8 |

| | | | | | | | | | | | | | |
|---|---|---|---|---|---|---|---|---|---|---|---|---|---|
| AT4G00910 | -0.5 | 4.74E-02 | -0.1 | 7.24E-01 | -0.4 | 1.65E-01 | | | | | | | Aluminum activated malate transporter family protein |
| AT1G58120 | -0.5 | 1.03E-01 | 0.2 | 5.23E-01 | -0.7 | 7.47E-02 | | | | | | | Uncharacterized protein T18I24.4 |
| AT5G57340 | -0.5 | 3.31E-03 | -0.6 | 1.41E-03 | 0.1 | 5.23E-01 | -0.4 | 1.81E-01 | -0.4 | 2.99E-01 | -0.1 | 9.16E-01 | Ras guanine nucleotide exchange factor Q-like protein |
| AT3G47350 | -0.5 | 1.29E-04 | 0.0 | 8.82E-01 | -0.5 | 4.32E-04 | 0.0 | 9.43E-01 | 0.0 | 9.88E-01 | 0.0 | 9.76E-01 | Hydroxysteroid dehydrogenase 2 |
| AT1G02260 | -0.5 | 3.27E-05 | -0.6 | 3.01E-05 | 0.1 | 5.05E-01 | -0.2 | 3.67E-01 | -0.1 | 8.30E-01 | -0.1 | 6.93E-01 | Citrate transporter-like domain-containing protein |
| AT1G12080 | -0.5 | 1.34E-03 | 0.7 | 2.78E-04 | -1.2 | 3.23E-05 | -0.5 | 4.52E-02 | -0.1 | 7.47E-01 | -0.4 | 1.81E-01 | hypothetical protein |
| AT3G01516 | -0.5 | 1.58E-02 | 0.2 | 2.37E-01 | -0.7 | 7.89E-03 | 0.1 | 9.19E-01 | 0.4 | 3.17E-01 | -0.4 | 4.14E-01 | At3g01516 |
| AT3G46385 | -0.5 | 1.10E-02 | -0.3 | 5.63E-02 | -0.1 | 4.98E-01 | -0.2 | 3.74E-01 | 0.0 | 9.36E-01 | -0.3 | 4.66E-01 | NA |
| AT2G39130 | -0.5 | 1.30E-04 | -0.1 | 5.44E-01 | -0.4 | 1.10E-03 | 0.0 | 9.72E-01 | 0.3 | 1.51E-01 | -0.3 | 2.04E-01 | Transmembrane amino acid transporter family protein |
| AT3G06210 | -0.5 | 3.07E-04 | -0.5 | 5.75E-04 | 0.0 | 9.21E-01 | -0.8 | 1.39E-03 | -0.8 | 2.53E-03 | 0.0 | 9.98E-01 | hypothetical protein |
| AT2G46490 | -0.5 | 5.68E-04 | 0.0 | 8.89E-01 | -0.5 | 2.09E-03 | -0.6 | 2.30E-04 | -0.4 | 2.28E-02 | -0.3 | 2.00E-01 | Uncharacterized protein |
| AT1G26300 | -0.5 | 5.60E-04 | -0.4 | 3.07E-03 | -0.1 | 4.36E-01 | -0.2 | 1.75E-01 | -0.2 | 4.16E-01 | -0.1 | 7.95E-01 | BSD domain-containing protein |
| AT4G00800 | -0.5 | 3.63E-04 | -0.6 | 1.66E-04 | 0.1 | 3.65E-01 | -0.1 | 7.48E-01 | 0.1 | 7.83E-01 | -0.2 | 6.22E-01 | AT4g00790 protein |
| AT5G28630 | -0.5 | 2.53E-04 | 0.5 | 2.92E-04 | -1.0 | 2.69E-05 | 0.3 | 2.95E-01 | 0.0 | 9.27E-01 | 0.2 | 5.27E-01 | Uncharacterized protein |
| AT1G07300 | -0.5 | 5.24E-03 | -0.1 | 3.02E-01 | -0.3 | 7.19E-02 | -0.5 | 1.37E-01 | 0.0 | 9.48E-01 | -0.4 | 2.68E-01 | Josephin-like protein |
| AT5G53830 | -0.5 | 2.61E-03 | -0.4 | 9.74E-03 | -0.1 | 5.89E-01 | 0.5 | 2.88E-02 | 0.1 | 6.85E-01 | 0.4 | 1.28E-01 | VQ motif-containing protein 33 |
| AT3G11810 | -0.5 | 4.77E-05 | -0.5 | 1.05E-04 | 0.0 | 8.64E-01 | -0.4 | 3.81E-02 | -0.3 | 2.09E-01 | -0.1 | 6.93E-01 | F26K24.10 protein |
| AT1G59520 | -0.5 | 1.49E-04 | -0.4 | 7.71E-04 | -0.1 | 4.32E-01 | -0.1 | 6.70E-01 | 0.0 | 9.40E-01 | -0.1 | 7.08E-01 | At1g59520 |
| AT1G26930 | -0.5 | 5.36E-04 | -0.5 | 8.00E-04 | 0.0 | 9.95E-01 | -0.5 | 9.17E-03 | -0.2 | 1.87E-01 | -0.2 | 3.71E-01 | Galactose oxidase |
| AT3G10640 | -0.5 | 6.52E-04 | -0.3 | 1.88E-02 | -0.2 | 9.99E-02 | 0.2 | 3.05E-01 | 0.0 | 9.24E-01 | 0.1 | 5.48E-01 | SNF7 family protein |
| AT3G06890 | -0.5 | 1.83E-03 | -0.4 | 5.25E-03 | -0.1 | 7.13E-01 | -0.7 | 2.00E-02 | -0.4 | 1.86E-01 | -0.3 | 4.66E-01 | hypothetical protein |
| AT5G04890 | -0.5 | 3.19E-04 | -0.3 | 4.48E-03 | -0.2 | 1.86E-01 | -0.1 | 7.00E-01 | -0.2 | 4.55E-01 | 0.1 | 7.90E-01 | Protein RESTRICTED TEV MOVEMENT 2 |
| AT4G00930 | -0.5 | 1.11E-03 | -0.4 | 4.22E-03 | -0.1 | 5.80E-01 | 0.3 | 2.55E-01 | 0.1 | 7.49E-01 | 0.2 | 5.49E-01 | COP1-interacting protein 4.1 |
| AT2G29510 | -0.5 | 3.83E-04 | -0.5 | 5.63E-04 | 0.0 | 9.88E-01 | -0.2 | 1.62E-01 | -0.1 | 7.36E-01 | -0.2 | 5.30E-01 | At2g29510 |
| AT5G19025 | -0.5 | 9.89E-04 | -0.2 | 7.08E-02 | -0.3 | 5.84E-02 | -0.3 | 1.98E-01 | -0.3 | 2.46E-01 | 0.0 | 9.99E-01 | Ribosomal protein L34e superfamily protein |
| AT3G59570 | -0.5 | 1.46E-04 | -0.3 | 4.37E-03 | -0.2 | 8.19E-02 | 0.4 | 8.84E-02 | 0.5 | 6.55E-02 | -0.1 | 8.48E-01 | Uncharacterized protein At3g59570 |
| AT1G47900 | -0.5 | 2.28E-04 | -0.2 | 2.53E-02 | -0.3 | 2.32E-02 | 0.2 | 1.75E-01 | 0.3 | 4.14E-02 | -0.1 | 5.45E-01 | Filament-like protein |
| AT5G02320 | -0.5 | 2.53E-03 | -0.5 | 3.81E-03 | 0.0 | 9.77E-01 | -0.1 | 7.59E-01 | -0.4 | 2.20E-01 | 0.3 | 4.29E-01 | Myb domain protein 3r-5 |
| AT5G35732 | -0.5 | 3.13E-03 | -0.3 | 5.02E-02 | -0.2 | 2.08E-01 | -0.1 | 6.31E-01 | -0.3 | 2.95E-01 | 0.2 | 6.80E-01 | At5g35732 |
| AT1G23900 | -0.5 | 1.75E-05 | -0.4 | 1.03E-04 | -0.1 | 2.92E-01 | 0.0 | 8.41E-01 | 0.0 | 9.44E-01 | 0.1 | 8.41E-01 | AP-1 complex subunit gamma |
| AT1G27300 | -0.5 | 3.90E-04 | -0.3 | 8.45E-03 | -0.2 | 1.28E-01 | -0.7 | 8.44E-04 | -0.7 | 1.67E-03 | 0.0 | 9.84E-01 | F17L21.9 |
| AT5G65630 | -0.5 | 1.14E-03 | -0.3 | 1.13E-02 | -0.1 | 2.67E-01 | -0.5 | 9.32E-05 | 0.0 | 9.47E-01 | -0.5 | 3.52E-03 | Transcription factor GTE7 |
| AT4G19645 | -0.5 | 6.52E-03 | -0.4 | 2.06E-02 | -0.1 | 6.83E-01 | 0.4 | 3.10E-01 | 0.0 | 9.99E-01 | 0.4 | 4.38E-01 | TLC domain-containing protein |
| AT1G28050 | -0.5 | 6.77E-05 | -0.3 | 2.05E-03 | -0.2 | 6.10E-02 | -0.4 | 6.39E-03 | -0.1 | 4.32E-01 | -0.3 | 1.63E-01 | Uncharacterized protein At1g28050 |
| AT1G22790 | -0.5 | 7.19E-05 | -0.2 | 1.58E-02 | -0.3 | 9.28E-03 | -0.3 | 2.81E-02 | 0.0 | 9.03E-01 | -0.3 | 1.16E-01 | Low affinity potassium transport system protein |
| AT1G11170 | -0.5 | 1.37E-02 | -0.5 | 1.09E-02 | 0.1 | 8.27E-01 | -0.3 | 4.83E-01 | -0.1 | 9.09E-01 | -0.2 | 7.01E-01 | At1g11170 |
| AT5G23575 | -0.5 | 1.29E-05 | -0.1 | 9.66E-02 | -0.4 | 3.45E-04 | 0.1 | 7.12E-01 | 0.2 | 3.33E-01 | -0.1 | 6.70E-01 | Uncharacterized protein At5g23575 |
| AT2G25250 | -0.5 | 2.21E-04 | -0.2 | 8.47E-02 | -0.3 | 8.04E-03 | -0.3 | 2.92E-02 | -0.1 | 7.34E-01 | -0.3 | 2.15E-01 | At2g25250 |
| AT4G20720 | -0.5 | 9.80E-04 | -0.3 | 3.12E-02 | -0.2 | 1.19E-01 | -0.4 | 2.92E-01 | 0.1 | 8.42E-01 | -0.6 | 2.70E-01 | Uncharacterized protein AT4g20720 |
| AT5G55896 | -0.5 | 1.17E-04 | -0.5 | 1.77E-04 | 0.0 | 9.46E-01 | -0.1 | 5.02E-01 | 0.0 | 9.76E-01 | -0.1 | 6.31E-01 | NA |
| AT5G64970 | -0.5 | 1.63E-03 | -0.3 | 2.64E-02 | -0.2 | 2.11E-01 | -0.1 | 8.94E-01 | 0.3 | 5.31E-01 | -0.3 | 4.68E-01 | Mitochondrial substrate carrier family protein |
| AT1G23530 | -0.5 | 7.10E-03 | -0.4 | 2.89E-02 | -0.1 | 5.67E-01 | 0.0 | 9.09E-01 | 0.1 | 7.44E-01 | -0.1 | 7.16E-01 | hypothetical protein |
| AT5G10750 | -0.5 | 4.20E-04 | -0.4 | 2.78E-03 | -0.1 | 3.50E-01 | -0.7 | 2.47E-04 | -0.3 | 1.09E-01 | -0.4 | 3.45E-02 | Protein ENHANCED DISEASE RESISTANCE 2 C-terminal domain-containing protein |
| AT3G18310 | -0.5 | 2.68E-04 | -0.4 | 9.73E-04 | -0.1 | 5.95E-01 | 0.2 | 3.87E-01 | 0.2 | 4.95E-01 | 0.0 | 9.71E-01 | Uncharacterized protein |
| AT3G54860 | -0.5 | 2.16E-04 | -0.3 | 8.07E-03 | -0.2 | 6.61E-02 | -0.3 | 6.68E-02 | -0.2 | 2.13E-01 | -0.1 | 7.88E-01 | Sec1 |
| AT3G55470 | -0.5 | 6.77E-05 | -0.4 | 4.71E-04 | -0.1 | 2.87E-01 | 0.5 | 2.62E-02 | 0.0 | 9.55E-01 | 0.5 | 7.19E-02 | Calcium-dependent lipid-binding family protein |
| AT2G23960 | -0.5 | 3.01E-03 | -0.2 | 1.21E-01 | -0.3 | 1.06E-01 | 0.2 | 4.49E-01 | 0.4 | 2.44E-01 | -0.1 | 6.93E-01 | Class I glutamine amidotransferase-like superfamily protein |
| AT5G03660 | -0.5 | 1.38E-04 | -0.4 | 1.44E-03 | -0.1 | 1.96E-01 | -0.1 | 5.25E-01 | 0.0 | 9.13E-01 | -0.1 | 7.54E-01 | RAB6-interacting golgin |
| AT4G18530 | -0.5 | 2.40E-03 | -0.5 | 1.92E-03 | 0.0 | 7.73E-01 | -0.2 | 3.88E-01 | 0.1 | 7.87E-01 | -0.3 | 3.23E-01 | Lysine ketoglutarate reductase trans-splicing-like protein, putative |
| AT1G14380 | -0.5 | 1.75E-04 | -0.2 | 1.82E-02 | -0.2 | 2.29E-02 | -0.2 | 2.36E-01 | 0.0 | 9.41E-01 | -0.2 | 3.62E-01 | IQ-domain 28 |
| AT3G14810 | -0.5 | 2.77E-02 | -0.5 | 1.98E-02 | 0.1 | 8.16E-01 | -0.3 | 1.88E-01 | 0.1 | 8.43E-01 | -0.4 | 2.42E-01 | Mechanosensitive ion channel protein |
| AT3G15430 | -0.5 | 1.06E-04 | -0.3 | 7.09E-03 | -0.2 | 2.77E-02 | -0.1 | 4.29E-01 | 0.0 | 9.79E-01 | -0.1 | 5.54E-01 | Regulator of chromosome condensation family protein |
| AT2G31090 | -0.5 | 2.67E-04 | -0.2 | 3.51E-02 | -0.3 | 2.21E-02 | -0.3 | 1.45E-01 | -0.4 | 9.85E-02 | 0.1 | 8.26E-01 | Signaling peptide TAXIMIN 1 |
| AT4G14390 | -0.5 | 4.98E-02 | -0.2 | 3.34E-01 | -0.3 | 3.77E-01 | 0.4 | 2.13E-01 | 0.9 | 1.03E-02 | -0.5 | 1.75E-01 | Ankyrin repeat family protein |
| AT1G27110 | -0.5 | 7.91E-03 | 0.0 | 8.18E-01 | -0.4 | 3.28E-02 | -0.2 | 6.81E-01 | 0.2 | 6.48E-01 | -0.4 | 4.21E-01 | Tetratricopeptide repeat protein 38 |
| AT1G80570 | -0.5 | 8.69E-04 | -0.5 | 7.33E-04 | 0.0 | 7.42E-01 | -0.5 | 9.23E-02 | -0.5 | 1.30E-01 | 0.0 | 9.80E-01 | F-box |
| AT4G12230 | -0.5 | 3.49E-05 | -0.3 | 2.30E-03 | -0.2 | 1.86E-02 | -0.1 | 2.12E-01 | -0.1 | 5.18E-01 | -0.1 | 7.88E-01 | Maspardin |
| AT4G17900 | -0.5 | 1.32E-04 | -0.3 | 2.67E-03 | -0.2 | 9.74E-02 | -0.1 | 4.46E-01 | -0.2 | 1.78E-01 | 0.1 | 5.94E-01 | PLATZ transcription factor family protein |
| AT4G28900 | -0.5 | 2.24E-02 | 0.1 | 5.46E-01 | -0.6 | 2.32E-02 | -1.2 | 8.37E-03 | -1.0 | 2.64E-02 | -0.2 | 8.20E-01 | NA |
| AT4G22240 | -0.5 | 1.45E-03 | -0.7 | 1.75E-04 | 0.2 | 8.65E-02 | 0.1 | 6.72E-01 | 0.3 | 7.41E-02 | -0.2 | 2.88E-01 | Probable plastid-lipid-associated protein 2, chloroplastic |
| AT3G51760 | -0.5 | 2.73E-02 | 0.1 | 7.76E-01 | -0.5 | 4.01E-02 | -0.2 | 5.59E-01 | 0.0 | 9.76E-01 | -0.3 | 6.33E-01 | Uncharacterized protein |
| AT1G02400 | -0.5 | 2.68E-03 | -0.4 | 1.42E-02 | -0.1 | 4.75E-01 | 0.3 | 1.99E-01 | 0.2 | 4.83E-01 | 0.1 | 7.30E-01 | Gibberellin 2-oxidase 6 |
| AT1G75260 | -0.5 | 1.41E-02 | -0.2 | 2.30E-01 | -0.3 | 2.18E-01 | | | | | | | Uncharacterized protein |
| AT5G24320 | -0.5 | 3.82E-04 | -0.6 | 1.53E-04 | 0.1 | 3.12E-01 | 0.0 | 8.60E-01 | 0.1 | 7.75E-01 | -0.1 | 6.83E-01 | Transducin |
| AT3G13300 | -0.5 | 4.18E-05 | -0.4 | 1.86E-04 | -0.1 | 4.77E-01 | -0.1 | 5.94E-01 | -0.1 | 7.52E-01 | 0.0 | 9.49E-01 | Transducin |
| AT1G11360 | -0.5 | 1.24E-05 | -0.2 | 3.17E-03 | -0.3 | 3.21E-03 | -0.3 | 1.32E-02 | -0.2 | 1.77E-01 | -0.1 | 4.71E-01 | At1g11360 |
| AT3G28430 | -0.5 | 3.18E-04 | -0.5 | 2.79E-04 | 0.0 | 6.86E-01 | -0.4 | 6.82E-02 | 0.0 | 9.90E-01 | -0.4 | 1.80E-01 | Protein TRANSPARENT TESTA 9 |
| AT2G17790 | -0.5 | 3.20E-05 | -0.3 | 6.92E-04 | -0.1 | 6.27E-02 | 0.0 | 9.53E-01 | 0.1 | 4.55E-01 | -0.2 | 5.26E-01 | Vacuolar protein sorting-associated protein 35A |
| AT1G74950 | -0.5 | 5.11E-05 | -0.4 | 4.46E-04 | -0.1 | 2.16E-01 | -0.4 | 2.95E-02 | -0.1 | 7.01E-01 | -0.3 | 2.26E-01 | Protein TIFY |
| AT1G19110 | -0.5 | 2.68E-05 | -0.2 | 6.00E-03 | -0.2 | 5.28E-03 | -0.1 | 7.57E-01 | -0.1 | 5.07E-01 | 0.1 | 7.96E-01 | VWFA domain-containing protein |
| AT1G22640 | -0.5 | 2.11E-04 | -0.2 | 3.74E-02 | -0.3 | 1.59E-02 | -0.8 | 7.60E-04 | -0.2 | 2.97E-01 | -0.6 | 4.58E-02 | Myb domain protein 3 |
| AT5G46620 | -0.5 | 4.88E-04 | -0.3 | 1.21E-02 | -0.2 | 1.18E-01 | -0.1 | 5.95E-01 | 0.1 | 7.20E-01 | -0.2 | 4.45E-01 | Uncharacterized protein |
| AT2G28400 | -0.5 | 2.43E-02 | -0.5 | 2.60E-02 | 0.0 | 9.81E-01 | 0.3 | 2.52E-01 | -0.3 | 3.65E-01 | 0.7 | 4.77E-02 | Protein S40-4 |
| AT1G74160 | -0.5 | 7.58E-05 | -0.3 | 1.55E-03 | -0.2 | 8.72E-02 | -0.2 | 2.40E-01 | 0.1 | 6.95E-01 | -0.3 | 2.31E-01 | LONGIFOLIA protein |
| AT3G23180 | -0.5 | 1.48E-04 | 0.1 | 7.50E-02 | -0.6 | 1.63E-04 | 0.0 | 8.76E-01 | 0.0 | 9.08E-01 | 0.0 | 9.98E-01 | HR-like lesion-inducing protein-like protein |
| AT2G21180 | -0.5 | 5.86E-04 | -0.3 | 5.47E-03 | -0.1 | 2.88E-01 | -0.2 | 3.24E-01 | -0.3 | 3.07E-01 | 0.0 | 9.27E-01 | hypothetical protein |
| AT2G01180 | -0.5 | 6.75E-04 | -0.3 | 6.15E-03 | -0.1 | 2.85E-01 | 0.2 | 3.57E-01 | 0.2 | 3.51E-01 | 0.0 | 9.42E-01 | Phosphatidic acid phosphatase 1 |
| AT1G59650 | -0.5 | 1.88E-04 | -0.2 | 1.63E-02 | -0.2 | 3.19E-02 | -0.2 | 2.42E-01 | 0.0 | 9.92E-01 | -0.2 | 3.64E-01 | CW14 |
| AT5G02040 | -0.5 | 1.26E-04 | -0.3 | 8.06E-03 | -0.2 | 3.39E-02 | -0.1 | 3.98E-01 | -0.1 | 5.43E-01 | 0.0 | 9.47E-01 | PRA1 family protein A1 |
| AT1G43790 | -0.5 | 1.43E-03 | 0.2 | 8.02E-02 | -0.7 | 6.81E-04 | 0.1 | 6.40E-01 | -0.1 | 7.55E-01 | 0.3 | 4.56E-01 | Protein TRACHEARY ELEMENT DIFFERENTIATION-RELATED 6 |
| AT4G23885 | -0.5 | 2.23E-04 | -0.2 | 8.93E-02 | -0.3 | 8.86E-03 | 0.1 | 5.07E-01 | 0.2 | 2.96E-01 | -0.1 | 7.55E-01 | Uncharacterized protein |
| AT1G67860 | -0.5 | 1.90E-03 | -0.3 | 4.00E-02 | -0.2 | 1.44E-01 | -0.5 | 1.99E-03 | -0.1 | 3.80E-01 | -0.3 | 7.40E-02 | hypothetical protein |
| AT4G16857 | -0.5 | 2.34E-02 | 0.3 | 5.66E-02 | -0.8 | 3.75E-03 | -0.7 | 9.90E-02 | -1.1 | 3.14E-02 | 0.4 | 5.58E-01 | Disease resistance protein family protein |

| | | | | | | | | | | | | | |
|---|---|---|---|---|---|---|---|---|---|---|---|---|---|
| AT1G19650 | -0.5 | 2.03E-03 | -0.4 | 4.67E-03 | 0.0 | 7.89E-01 | 0.1 | 7.18E-01 | 0.2 | 5.63E-01 | -0.1 | 8.58E-01 | Sec14p-like phosphatidylinositol transfer family protein |
| AT1G55680 | -0.5 | 5.37E-05 | -0.4 | 1.63E-04 | 0.0 | 7.10E-01 | -0.3 | 1.38E-02 | -0.2 | 2.76E-01 | -0.2 | 3.89E-01 | hypothetical protein |
| AT5G08240 | -0.5 | 2.84E-03 | -0.4 | 7.45E-03 | -0.1 | 7.30E-01 | 0.5 | 2.99E-02 | -0.3 | 2.62E-01 | 0.7 | 1.27E-02 | Transmembrane protein |
| AT1G23230 | -0.5 | 4.00E-05 | -0.6 | 9.99E-06 | 0.2 | 4.93E-02 | -0.1 | 7.38E-01 | 0.2 | 5.96E-01 | -0.3 | 4.49E-01 | Mediator of RNA polymerase II transcription subunit |
| AT3G61480 | -0.5 | 4.32E-04 | -0.3 | 5.07E-03 | -0.1 | 2.17E-01 | -0.2 | 3.70E-01 | 0.0 | 9.21E-01 | -0.2 | 6.01E-01 | Quinoprotein amine dehydrogenase, beta chain-like RIC1-like guanyl-nucleotide exchange factor |
| AT5G11840 | -0.5 | 7.11E-05 | -0.5 | 6.05E-05 | 0.1 | 5.37E-01 | -0.2 | 1.33E-01 | -0.1 | 7.79E-01 | -0.1 | 4.40E-01 | YCF36, |
| AT2G27830 | -0.5 | 7.68E-02 | -0.5 | 7.33E-02 | 0.0 | 9.45E-01 | -0.4 | 5.30E-02 | -1.2 | 8.26E-05 | 0.8 | 8.23E-03 | Uncharacterized protein |
| AT4G27430 | -0.5 | 2.14E-04 | -0.2 | 1.94E-02 | -0.2 | 2.66E-02 | -0.3 | 6.62E-03 | -0.1 | 4.36E-01 | -0.2 | 1.88E-01 | COP1-interacting protein 7 |
| AT3G27880 | -0.5 | 2.35E-04 | -0.2 | 3.68E-02 | -0.3 | 2.09E-02 | 0.4 | 1.36E-01 | 0.0 | 9.23E-01 | 0.3 | 1.64E-01 | Uncharacterized protein |
| AT3G33530 | -0.5 | 1.00E-04 | -0.4 | 3.21E-04 | 0.0 | 6.29E-01 | 0.0 | 8.43E-01 | 0.0 | 9.45E-01 | -0.1 | 8.40E-01 | Transducin family protein |
| AT4G16070 | -0.5 | 1.90E-04 | -0.2 | 1.59E-02 | -0.2 | 3.36E-02 | -0.1 | 5.27E-01 | 0.1 | 6.91E-01 | -0.2 | 3.78E-01 | Lipase class 3 family protein |
| AT5G23955 | -0.5 | 3.99E-03 | -0.5 | 5.01E-03 | 0.0 | 9.67E-01 | -0.5 | 3.95E-02 | -0.3 | 3.25E-01 | -0.2 | 4.86E-01 | NA |
| AT4G19950 | -0.5 | 9.27E-04 | -0.1 | 3.06E-01 | -0.4 | 1.55E-02 | -0.2 | 5.77E-01 | -0.1 | 8.43E-01 | -0.1 | 8.28E-01 | Uncharacterized protein At4g19950 |
| AT5G11600 | -0.5 | 2.96E-04 | -0.5 | 4.03E-04 | 0.0 | 9.25E-01 | -0.3 | 1.53E-01 | -0.1 | 7.19E-01 | -0.2 | 5.04E-01 | Uncharacterized protein |
| AT1G24560 | -0.5 | 3.03E-05 | -0.3 | 7.29E-04 | -0.1 | 5.50E-02 | -0.2 | 7.28E-02 | -0.2 | 2.14E-01 | 0.0 | 8.34E-01 | Paramyosin |
| AT1G54040 | -0.5 | 9.97E-02 | 0.3 | 2.38E-01 | -0.8 | 2.67E-02 | -0.6 | 2.00E-02 | 0.2 | 5.70E-01 | -0.7 | 1.23E-02 | Epithiospecifier protein |
| AT5G19520 | -0.5 | 8.50E-03 | 0.3 | 3.69E-02 | -0.8 | 1.13E-03 | -0.1 | 7.27E-01 | 0.4 | 1.51E-01 | -0.5 | 1.30E-01 | Mechanosensitive ion channel protein |
| AT1G56460 | -0.5 | 2.54E-04 | -0.6 | 4.99E-05 | 0.2 | 9.65E-02 | -0.2 | 2.80E-01 | -0.2 | 2.49E-01 | 0.0 | 9.05E-01 | HIT zinc finger and PAPA-1-like domain-containing protein |
| AT5G64090 | -0.5 | 2.46E-05 | -0.3 | 1.16E-03 | -0.2 | 2.48E-02 | -0.2 | 1.38E-01 | -0.2 | 9.66E-02 | 0.0 | 8.43E-01 | At5g64090 |
| AT5G14680 | -0.5 | 5.76E-05 | -0.4 | 3.18E-04 | -0.1 | 3.72E-01 | -0.7 | 2.61E-04 | -0.3 | 1.13E-01 | -0.4 | 5.40E-02 | hypothetical protein |
| AT1G78780 | -0.5 | 9.27E-03 | -0.9 | 2.94E-04 | 0.5 | 3.77E-02 | -0.4 | 2.75E-01 | -0.6 | 1.94E-01 | 0.1 | 8.39E-01 | Uncharacterized protein At1g78780 |
| AT3G11330 | -0.5 | 4.29E-04 | -0.1 | 4.29E-01 | -0.4 | 3.89E-03 | 0.2 | 2.73E-01 | 0.2 | 3.53E-01 | 0.0 | 9.98E-01 | Plant intracellular Ras-group-related LRR protein 9 |
| AT2G28140 | -0.5 | 2.64E-02 | 0.0 | 8.53E-01 | -0.4 | 8.45E-02 | 0.1 | 8.96E-01 | 0.5 | 1.94E-01 | -0.5 | 3.22E-01 | Enabled-like protein |
| AT1G74650 | -0.5 | 7.36E-03 | -1.3 | 3.79E-05 | 0.8 | 2.52E-03 | -0.2 | 6.77E-01 | -0.2 | 6.88E-01 | 0.0 | 9.78E-01 | hypothetical protein |
| AT5G12260 | -0.5 | 1.16E-04 | -0.2 | 2.63E-02 | -0.3 | 1.25E-02 | -0.3 | 5.46E-02 | -0.2 | 3.59E-01 | -0.2 | 5.64E-01 | Uncharacterized protein |
| AT3G08630 | -0.5 | 1.48E-03 | -0.1 | 5.72E-01 | -0.4 | 9.29E-03 | -0.2 | 1.84E-01 | 0.1 | 8.20E-01 | -0.3 | 1.56E-01 | Protein RETICULATA-RELATED 2, chloroplastic |
| AT3G17920 | -0.5 | 2.00E-03 | -0.4 | 4.60E-03 | 0.0 | 7.90E-01 | -0.3 | 2.64E-01 | -0.2 | 4.61E-01 | -0.1 | 8.85E-01 | Outer arm dynein light chain 1 protein |
| AT3G59490 | -0.5 | 3.52E-03 | -0.3 | 1.98E-02 | -0.1 | 4.63E-01 | -0.1 | 6.36E-01 | -0.2 | 4.71E-01 | 0.1 | 8.35E-01 | At3g59490 |
| AT2G15042 | -0.5 | 1.76E-03 | -0.2 | 7.24E-02 | -0.2 | 9.28E-02 | 0.4 | 4.16E-01 | -0.1 | 8.33E-01 | 0.5 | 3.59E-01 | Receptor-like protein 18 |
| AT2G48060 | -0.5 | 2.46E-04 | -0.3 | 5.05E-03 | -0.2 | 1.11E-01 | -0.3 | 8.99E-02 | -0.1 | 7.13E-01 | -0.2 | 3.97E-01 | Piezo-type mechanosensitive ion channel component |
| AT3G27260 | -0.5 | 1.83E-05 | -0.5 | 1.03E-05 | 0.1 | 1.64E-01 | -0.2 | 3.09E-01 | -0.1 | 7.78E-01 | -0.1 | 7.00E-01 | AT3g27260 |
| AT2G37110 | -0.5 | 1.55E-04 | -0.3 | 3.29E-03 | -0.1 | 1.02E-01 | -0.1 | 4.77E-01 | -0.1 | 5.01E-01 | 0.0 | 9.69E-01 | hypothetical protein |
| AT2G45200 | -0.5 | 9.53E-04 | 0.0 | 8.18E-01 | -0.4 | 4.46E-03 | -0.4 | 1.34E-03 | -0.3 | 1.23E-02 | -0.1 | 6.39E-01 | hypothetical protein |
| AT3G02930 | -0.5 | 2.00E-03 | -0.3 | 1.60E-02 | -0.1 | 3.52E-01 | -0.2 | 3.49E-01 | 0.0 | 9.02E-01 | -0.1 | 5.94E-01 | Weak chloroplast movement under blue light protein |
| AT3G27990 | -0.5 | 8.69E-04 | -0.5 | 8.21E-04 | 0.0 | 7.95E-01 | -0.3 | 1.68E-01 | -0.2 | 4.24E-01 | -0.1 | 7.79E-01 | NA |
| AT5G13890 | -0.5 | 7.14E-04 | -0.2 | 7.79E-02 | -0.3 | 3.22E-02 | 0.0 | 8.03E-01 | -0.1 | 7.13E-01 | 0.0 | 9.19E-01 | hypothetical protein |
| AT5G41330 | -0.5 | 5.18E-04 | -0.5 | 3.82E-04 | 0.1 | 6.32E-01 | -0.3 | 4.02E-01 | -0.1 | 7.66E-01 | -0.2 | 7.84E-01 | BTB |
| AT1G16180 | -0.5 | 4.98E-05 | -0.3 | 6.22E-04 | -0.1 | 1.52E-01 | -0.1 | 2.24E-01 | -0.1 | 3.21E-01 | 0.0 | 9.83E-01 | Serinc-domain containing serine and sphingolipid biosynthesis protein |
| AT2G13650 | -0.5 | 5.35E-04 | -0.2 | 2.18E-02 | -0.2 | 7.82E-02 | -0.1 | 5.67E-01 | 0.0 | 9.05E-01 | -0.2 | 5.76E-01 | Golgi nucleotide sugar transporter 1 |
| AT1G71940 | -0.5 | 2.30E-03 | -0.1 | 5.37E-01 | -0.4 | 1.84E-02 | -0.1 | 7.18E-01 | 0.1 | 6.88E-01 | -0.2 | 4.59E-01 | SNARE associated Golgi protein family |
| AT1G35470 | -0.5 | 5.14E-05 | -0.3 | 3.04E-03 | -0.2 | 2.79E-02 | -0.1 | 4.33E-01 | -0.1 | 8.01E-01 | -0.1 | 7.88E-01 | Ran-binding protein M homolog |
| AT4G35510 | -0.4 | 2.17E-04 | -0.3 | 4.54E-03 | -0.2 | 1.20E-01 | -0.1 | 4.59E-01 | 0.0 | 8.82E-01 | -0.2 | 4.87E-01 | Uncharacterized protein At4g35510 |
| AT1G76070 | -0.4 | 3.09E-04 | -0.4 | 5.40E-04 | 0.0 | 9.51E-01 | 0.0 | 9.42E-01 | -0.1 | 7.03E-01 | 0.1 | 7.71E-01 | Uncharacterized protein At1g76070 |
| AT5G57130 | -0.4 | 7.84E-05 | -0.4 | 1.81E-04 | 0.0 | 8.49E-01 | -0.2 | 2.34E-01 | -0.1 | 7.80E-01 | -0.1 | 5.69E-01 | Protein SMAX1-LIKE 5 |
| AT2G36810 | -0.4 | 1.21E-04 | -0.4 | 3.94E-04 | 0.0 | 6.23E-01 | -0.4 | 2.85E-02 | -0.2 | 1.95E-01 | -0.1 | 6.34E-01 | ARM repeat superfamily protein |
| AT1G25682 | -0.4 | 1.51E-04 | -0.3 | 2.25E-03 | -0.1 | 1.47E-01 | -0.1 | 3.18E-01 | -0.2 | 3.73E-01 | 0.0 | 9.98E-01 | Uncharacterized protein |
| AT1G56610 | -0.4 | 9.59E-03 | -0.7 | 9.35E-04 | 0.3 | 1.46E-01 | -0.2 | 4.10E-01 | 0.1 | 8.27E-01 | -0.3 | 4.13E-01 | Protein with RNI-like |
| AT4G32440 | -0.4 | 1.15E-03 | -0.5 | 1.33E-03 | 0.0 | 9.07E-01 | -0.1 | 5.88E-01 | -0.3 | 1.72E-01 | 0.2 | 4.71E-01 | ENT domain-containing protein |
| AT3G26990 | -0.4 | 9.74E-04 | -0.3 | 5.84E-03 | -0.1 | 4.23E-01 | -0.1 | 6.32E-01 | -0.1 | 7.17E-01 | 0.0 | 9.82E-01 | CID domain-containing protein |
| AT1G28620 | -0.4 | 9.20E-04 | -0.4 | 2.88E-03 | -0.1 | 6.72E-01 | -0.7 | 1.46E-02 | -0.1 | 6.91E-01 | -0.6 | 1.23E-01 | NA |
| AT2G17550 | -0.4 | 9.86E-04 | 0.0 | 8.13E-01 | -0.4 | 5.06E-03 | -0.6 | 1.92E-03 | -0.4 | 1.53E-02 | -0.1 | 6.76E-01 | RB1-inducible coiled-coil protein |
| AT4G22740 | -0.4 | 1.10E-04 | -0.6 | 3.33E-05 | 0.1 | 1.37E-01 | -0.2 | 2.24E-01 | 0.0 | 8.19E-01 | -0.1 | 5.54E-01 | Glycine-rich protein |
| AT1G61667 | -0.4 | 5.47E-03 | 0.0 | 8.27E-01 | -0.5 | 1.10E-02 | -0.3 | 1.80E-01 | -0.3 | 2.05E-01 | 0.0 | 9.78E-01 | hypothetical protein |
| AT1G30300 | -0.4 | 1.69E-03 | -0.2 | 7.09E-02 | -0.2 | 9.19E-02 | 0.0 | 9.90E-01 | 0.0 | 9.15E-01 | 0.0 | 9.14E-01 | Uncharacterized protein F12P21.8 |
| AT3G12400 | -0.4 | 9.82E-05 | -0.4 | 2.26E-04 | 0.0 | 8.09E-01 | -0.5 | 7.17E-05 | -0.5 | 7.93E-04 | -0.1 | 6.86E-01 | Ubiquitin-conjugating enzyme |
| AT2G02480 | -0.4 | 1.75E-04 | -0.1 | 2.96E-01 | -0.4 | 2.91E-03 | 0.0 | 9.17E-01 | 0.2 | 3.99E-01 | -0.2 | 4.80E-01 | STICHEL |
| AT1G78110 | -0.4 | 4.25E-04 | -0.4 | 2.53E-03 | -0.1 | 4.08E-01 | -0.7 | 3.94E-04 | -0.5 | 9.60E-03 | -0.2 | 3.81E-01 | At1g78110 |
| AT1G55690 | -0.4 | 5.26E-04 | -0.3 | 1.40E-02 | -0.2 | 1.10E-01 | 0.3 | 4.73E-02 | 0.3 | 3.69E-02 | 0.0 | 8.53E-01 | Sec14p-like phosphatidylinositol transfer family protein |
| AT4G28550 | -0.4 | 2.10E-03 | -0.2 | 1.01E-01 | -0.2 | 7.71E-02 | -0.3 | 7.18E-02 | 0.1 | 8.32E-01 | -0.4 | 1.07E-01 | Rab-GAP TBC domain-containing protein |
| AT4G35750 | -0.4 | 4.74E-03 | -0.2 | 2.01E-01 | -0.3 | 7.93E-02 | -0.1 | 7.84E-01 | -0.4 | 8.03E-02 | 0.3 | 2.59E-01 | hypothetical protein |
| AT5G25770 | -0.4 | 4.86E-05 | -0.1 | 2.96E-01 | -0.4 | 7.39E-04 | -0.6 | 2.22E-03 | -0.2 | 3.29E-01 | -0.4 | 1.09E-01 | Alpha |
| AT1G30890 | -0.4 | 6.43E-04 | 0.0 | 8.52E-01 | -0.5 | 1.84E-03 | -0.1 | 5.50E-01 | 0.0 | 9.38E-01 | -0.2 | 5.11E-01 | hypothetical protein |
| AT3G11080 | -0.4 | 1.39E-02 | -0.6 | 2.74E-03 | 0.2 | 3.43E-01 | 0.3 | 4.37E-01 | 0.1 | 7.63E-01 | 0.1 | 8.13E-01 | Receptor-like protein 35 |
| AT4G23510 | -0.4 | 9.22E-04 | -0.3 | 5.24E-03 | -0.1 | 4.61E-01 | 0.4 | 2.56E-01 | 0.3 | 4.19E-01 | 0.1 | 8.98E-01 | Disease resistance protein family |
| AT3G17712 | -0.4 | 1.91E-02 | -0.6 | 5.98E-03 | 0.2 | 4.88E-01 | | | | | | | Protein NRDE2 homolog |
| AT1G36940 | -0.4 | 2.34E-02 | 0.0 | 8.06E-01 | -0.4 | 8.19E-02 | -0.7 | 1.60E-02 | -0.9 | 1.10E-02 | 0.1 | 8.33E-01 | Myotubularin-like protein |
| AT1G47550 | -0.4 | 5.69E-05 | -0.2 | 5.62E-03 | -0.2 | 1.54E-02 | -0.2 | 1.91E-01 | -0.1 | 4.89E-01 | -0.1 | 7.94E-01 | Exocyst complex component sec3A |
| AT4G13400 | -0.4 | 8.63E-04 | -0.4 | 4.32E-03 | -0.1 | 4.98E-01 | -0.4 | 2.85E-01 | -0.1 | 8.64E-01 | -0.3 | 5.05E-01 | Uncharacterized protein At4g13400 |
| AT1G03060 | -0.4 | 6.39E-04 | -0.4 | 1.12E-03 | 0.0 | 9.10E-01 | -0.1 | 4.20E-01 | 0.0 | 9.15E-01 | -0.1 | 6.79E-01 | BEACH-DOMAIN HOMOLOG A1 |
| AT4G08330 | -0.4 | 1.72E-03 | -0.3 | 2.41E-02 | -0.2 | 2.27E-01 | -0.3 | 3.01E-01 | -0.4 | 1.84E-01 | 0.1 | 7.87E-01 | Uncharacterized protein |
| AT1G54320 | -0.4 | 1.55E-04 | -0.1 | 7.55E-02 | -0.3 | 5.82E-03 | 0.1 | 7.34E-01 | 0.1 | 6.82E-01 | 0.0 | 9.38E-01 | ALA-interacting subunit 3 |
| AT2G36300 | -0.4 | 2.91E-04 | -0.2 | 3.46E-02 | -0.2 | 2.35E-02 | -0.1 | 4.80E-01 | 0.0 | 8.47E-01 | -0.2 | 4.47E-01 | Protein YIP |
| AT1G20970 | -0.4 | 8.01E-05 | 0.0 | 5.72E-01 | -0.5 | 2.06E-04 | -0.3 | 6.43E-03 | -0.2 | 4.27E-02 | -0.1 | 7.39E-01 | Uncharacterized protein At1g20970 |
| AT1G08040 | -0.4 | 1.37E-04 | -0.3 | 1.22E-03 | -0.1 | 2.48E-01 | -0.4 | 5.25E-03 | -0.1 | 5.01E-01 | -0.3 | 1.06E-01 | Putative storage protein ) |
| AT3G01670 | -0.4 | 1.47E-04 | -0.3 | 1.23E-03 | -0.1 | 2.48E-01 | 0.0 | 8.79E-01 | 0.1 | 6.65E-01 | -0.1 | 8.07E-01 | Sieve element occlusion protein |
| AT4G26060 | -0.4 | 5.69E-04 | -0.2 | 2.06E-02 | -0.2 | 8.13E-02 | 0.1 | 6.18E-01 | -0.1 | 6.83E-01 | 0.3 | 3.75E-01 | At4g26060 |
| AT2G37300 | -0.4 | 1.38E-01 | -0.4 | 2.13E-01 | -0.1 | 8.80E-01 | -0.1 | 7.39E-01 | 0.1 | 8.63E-01 | -0.2 | 6.39E-01 | At2g37300 |
| AT3G47680 | -0.4 | 2.13E-03 | -0.4 | 3.90E-03 | 0.0 | 8.82E-01 | -0.1 | 8.08E-01 | -0.1 | 7.80E-01 | 0.0 | 9.46E-01 | No apical meristem-associated C-terminal domain-containing protein |
| AT1G67230 | -0.4 | 5.41E-05 | -0.1 | 1.08E-01 | -0.3 | 1.48E-03 | 0.1 | 3.85E-01 | 0.0 | 9.10E-01 | 0.1 | 6.68E-01 | Protein CROWDED NUCLEI 1 |
| AT1G69680 | -0.4 | 2.94E-03 | -0.3 | 2.63E-02 | -0.1 | 3.32E-01 | -0.3 | 2.83E-01 | 0.0 | 9.80E-01 | -0.4 | 3.17E-01 | Uncharacterized protein |
| AT4G35420 | -0.4 | 3.37E-03 | -0.2 | 1.41E-01 | -0.3 | 1.00E-01 | 0.4 | 1.28E-01 | 0.5 | 4.96E-02 | -0.1 | 6.54E-01 | TKPR1 |
| AT3G12740 | -0.4 | 7.58E-05 | -0.6 | 2.66E-05 | 0.1 | 1.34E-01 | 0.0 | 7.34E-01 | -0.2 | 5.85E-02 | 0.2 | 1.81E-01 | ALA-interacting subunit 1 |
| AT3G49800 | -0.4 | 6.80E-05 | -0.4 | 1.75E-04 | 0.0 | 7.64E-01 | -0.4 | 2.80E-03 | -0.2 | 6.49E-02 | -0.1 | 4.21E-01 | AT3g49800 |

| | | | | | | | | | | | | | |
|---|---|---|---|---|---|---|---|---|---|---|---|---|---|
| AT4G27320 | -0.4 | 1.24E-04 | -0.1 | 2.13E-01 | -0.3 | 2.10E-03 | -0.3 | 1.55E-02 | 0.0 | 8.04E-01 | -0.2 | 1.34E-01 | UspA domain-containing protein |
| AT5G60460 | -0.4 | 8.25E-04 | 0.1 | 1.87E-01 | -0.6 | 5.99E-04 | -0.1 | 5.53E-01 | 0.0 | 9.66E-01 | -0.1 | 6.72E-01 | Protein transport protein Sec61 subunit beta |
| AT5G64230 | -0.4 | 2.59E-02 | -0.6 | 6.07E-03 | 0.2 | 4.10E-01 | 0.0 | 9.81E-01 | 0.1 | 7.67E-01 | -0.1 | 7.93E-01 | Uncharacterized protein At5g64230 |
| AT5G24350 | -0.4 | 1.93E-03 | -0.5 | 1.04E-03 | 0.1 | 5.77E-01 | 0.1 | 8.96E-01 | -0.1 | 8.99E-01 | 0.1 | 8.36E-01 | Neuroblastoma-amplified sequence protein |
| AT5G28330 | -0.4 | 1.67E-03 | -0.5 | 1.23E-03 | 0.1 | 6.98E-01 | -0.4 | 3.73E-01 | -0.2 | 6.24E-01 | -0.1 | 8.27E-01 | NA |
| AT3G28455 | -0.4 | 3.38E-03 | -0.3 | 2.84E-02 | -0.1 | 3.77E-01 | 0.1 | 7.94E-01 | 0.3 | 4.62E-01 | -0.2 | 7.13E-01 | CLAVATA3 |
| AT3G60720 | -0.4 | 2.76E-03 | -0.3 | 2.96E-02 | -0.2 | 3.04E-01 | 0.3 | 3.45E-01 | -0.1 | 8.28E-01 | 0.4 | 3.47E-01 | Plasmodesmata-located protein 8 |
| AT1G16250 | -0.4 | 9.35E-04 | -0.4 | 3.49E-03 | -0.1 | 5.86E-01 | -0.9 | 7.70E-05 | -0.8 | 8.23E-04 | -0.2 | 5.88E-01 | F-box domain-containing protein |
| AT5G40640 | -0.4 | 3.75E-04 | -0.3 | 4.84E-03 | -0.1 | 2.08E-01 | 0.2 | 3.56E-01 | 0.5 | 6.99E-02 | -0.3 | 3.93E-01 | Transmembrane protein |
| AT2G26260 | -0.4 | 4.38E-03 | 0.0 | 8.06E-01 | -0.5 | 8.56E-03 | -0.2 | 3.82E-01 | 0.2 | 3.86E-01 | -0.4 | 1.49E-01 | Reticulon-like protein |
| AT2G37195 | -0.4 | 1.08E-02 | 0.0 | 8.40E-01 | -0.5 | 2.26E-02 | -0.2 | 6.31E-01 | 0.1 | 8.61E-01 | -0.3 | 5.67E-01 | Acyl-CoA-binding domain protein |
| AT2G30060 | -0.4 | 2.14E-04 | 0.0 | 7.24E-01 | -0.4 | 1.19E-03 | 0.1 | 5.16E-01 | 0.2 | 2.73E-01 | -0.1 | 7.18E-01 | Ran-binding protein 1 homolog b |
| AT5G08560 | -0.4 | 2.29E-05 | -0.4 | 8.13E-05 | 0.0 | 5.73E-01 | -0.1 | 2.73E-01 | 0.0 | 9.98E-01 | -0.1 | 4.73E-01 | WD repeat-containing protein 26 homolog |
| AT4G31020 | -0.4 | 4.96E-02 | 0.2 | 4.26E-01 | -0.6 | 3.22E-02 | 0.7 | 9.78E-02 | 0.4 | 4.15E-01 | 0.3 | 5.66E-01 | hypothetical protein |
| AT5G05850 | -0.4 | 4.43E-04 | -0.1 | 1.60E-01 | -0.3 | 1.27E-02 | -0.3 | 1.52E-01 | -0.3 | 3.03E-01 | -0.1 | 8.57E-01 | Plant intracellular Ras-group-related LRR protein 1 |
| AT1G76780 | -0.4 | 6.70E-03 | -0.3 | 4.82E-02 | -0.1 | 3.97E-01 | 0.1 | 6.66E-01 | 0.3 | 3.22E-01 | -0.2 | 5.87E-01 | HSP20-like chaperones superfamily protein |
| AT2G29970 | -0.4 | 2.60E-04 | -0.1 | 1.12E-01 | -0.3 | 7.29E-03 | -0.4 | 2.56E-03 | 0.0 | 8.09E-01 | -0.3 | 3.78E-02 | Protein SMAX1-LIKE 7 |
| AT1G15490 | -0.4 | 4.50E-04 | -0.1 | 1.30E-01 | -0.3 | 1.47E-02 | -0.2 | 4.80E-01 | 0.0 | 9.78E-01 | -0.2 | 5.89E-01 | hypothetical protein |
| AT5G41600 | -0.4 | 3.72E-04 | -0.2 | 3.66E-02 | -0.2 | 2.80E-02 | -0.3 | 4.07E-02 | 0.1 | 7.39E-01 | -0.4 | 3.47E-02 | Reticulon-like protein B4 |
| AT1G16840 | -0.4 | 6.65E-05 | -0.3 | 1.03E-03 | -0.1 | 1.12E-01 | -0.3 | 8.66E-02 | -0.1 | 6.38E-01 | -0.2 | 4.41E-01 | At1g16840 |
| AT1G73130 | -0.4 | 2.17E-03 | -0.4 | 2.64E-03 | 0.0 | 9.59E-01 | 0.1 | 5.72E-01 | -0.2 | 5.59E-01 | 0.3 | 3.03E-01 | F3N23.33 protein |
| AT4G14950 | -0.4 | 1.67E-04 | -0.2 | 2.42E-02 | -0.2 | 1.89E-02 | -0.1 | 4.69E-01 | 0.2 | 1.52E-01 | -0.4 | 6.32E-02 | SNARE associated Golgi protein family |
| AT5G58930 | -0.4 | 1.62E-04 | -0.4 | 2.02E-04 | 0.0 | 8.58E-01 | -0.3 | 1.92E-02 | -0.2 | 7.41E-02 | 0.0 | 8.23E-01 | Emb\|CAB61945.1 |
| AT1G27070 | -0.4 | 4.83E-04 | -0.3 | 4.95E-03 | -0.1 | 2.62E-01 | -0.1 | 8.00E-01 | -0.1 | 7.36E-01 | 0.0 | 9.27E-01 | hypothetical protein |
| AT1G35660 | -0.4 | 2.37E-03 | -0.5 | 2.36E-03 | 0.0 | 8.64E-01 | -0.1 | 6.59E-01 | -0.1 | 7.98E-01 | 0.0 | 9.45E-01 | Erythroid differentiation factor-like protein |
| AT5G18120 | -0.4 | 3.12E-04 | -0.1 | 2.69E-01 | -0.3 | 4.80E-03 | -0.2 | 1.48E-01 | 0.0 | 8.81E-01 | -0.2 | 3.76E-01 | APR-like 7 |
| AT2G24945 | -0.4 | 9.06E-04 | -0.2 | 2.56E-02 | -0.2 | 1.17E-01 | -0.3 | 3.34E-01 | 0.0 | 9.71E-01 | -0.2 | 4.56E-01 | Transmembrane protein |
| AT2G12490 | -0.4 | 1.25E-02 | -0.4 | 2.73E-02 | -0.1 | 8.06E-01 | -0.2 | 4.77E-01 | -0.1 | 7.24E-01 | -0.1 | 8.67E-01 | NA |
| AT2G26780 | -0.4 | 1.83E-04 | -0.5 | 8.70E-05 | 0.1 | 3.23E-01 | 0.0 | 9.50E-01 | 0.0 | 9.68E-01 | 0.0 | 9.97E-01 | At2g26780 |
| AT3G01810 | -0.4 | 2.35E-04 | -0.2 | 6.23E-02 | -0.3 | 1.08E-02 | -0.5 | 1.69E-02 | -0.1 | 5.70E-01 | -0.4 | 1.94E-01 | EEIG1 |
| AT5G66985 | -0.4 | 9.66E-03 | 0.2 | 2.50E-01 | -0.6 | 4.83E-03 | 0.9 | 4.99E-03 | 0.6 | 6.40E-02 | 0.3 | 2.94E-01 | Uncharacterized protein |
| AT1G01550 | -0.4 | 5.66E-05 | -0.4 | 2.30E-04 | 0.0 | 5.41E-01 | 0.1 | 5.80E-01 | -0.1 | 3.57E-01 | 0.2 | 2.50E-01 | Protein BPS1, chloroplastic |
| AT2G41310 | -0.4 | 5.16E-04 | -0.1 | 3.37E-01 | -0.3 | 6.80E-03 | -0.3 | 2.76E-02 | 0.2 | 1.64E-01 | -0.6 | 7.34E-03 | Response regulatory domain-containing protein |
| AT1G58230 | -0.4 | 2.86E-04 | -0.4 | 1.06E-03 | -0.1 | 5.63E-01 | -0.2 | 2.22E-01 | -0.1 | 5.38E-01 | -0.1 | 7.52E-01 | Uncharacterized protein F16M22.7 |
| AT4G31570 | -0.4 | 2.81E-04 | -0.4 | 3.06E-04 | 0.0 | 8.09E-01 | 0.0 | 9.86E-01 | 0.1 | 8.79E-01 | -0.1 | 8.84E-01 | Nucleoporin |
| AT2G20815 | -0.4 | 1.56E-02 | -0.2 | 2.00E-01 | -0.2 | 2.60E-01 | | | | | | | QWRF motif protein |
| AT3G47590 | -0.4 | 5.00E-04 | -0.4 | 2.26E-03 | -0.1 | 4.99E-01 | -0.3 | 2.07E-01 | -0.1 | 8.56E-01 | -0.2 | 4.63E-01 | Serine aminopeptidase S33 domain-containing protein |
| AT3G57590 | -0.4 | 1.17E-02 | -0.2 | 1.82E-01 | -0.2 | 2.28E-01 | -0.5 | 5.54E-02 | -0.3 | 1.96E-01 | -0.1 | 7.49E-01 | F-box and associated interaction domains-containing protein |
| AT3G09570 | -0.4 | 4.20E-04 | -0.1 | 3.20E-01 | -0.3 | 4.97E-03 | -0.2 | 9.37E-02 | -0.2 | 2.10E-01 | 0.0 | 8.93E-01 | Lung seven transmembrane receptor family protein |
| AT1G48510 | -0.4 | 1.82E-02 | 0.5 | 5.66E-03 | -0.9 | 8.37E-04 | 0.0 | 8.98E-01 | 0.5 | 7.52E-02 | -0.6 | 4.57E-02 | SURF1-like protein |
| AT2G28370 | -0.4 | 1.95E-04 | -0.3 | 4.45E-03 | -0.1 | 9.46E-02 | -0.3 | 3.09E-02 | -0.2 | 1.71E-01 | -0.1 | 7.14E-01 | CASP-like protein 5A2 |
| AT5G14540 | -0.4 | 8.16E-04 | -0.3 | 3.67E-03 | -0.1 | 4.91E-01 | -0.1 | 3.41E-01 | -0.1 | 4.52E-01 | 0.0 | 9.73E-01 | AT5G14540 protein ) |
| AT3G18480 | -0.4 | 6.03E-05 | -0.3 | 6.86E-04 | -0.1 | 1.64E-01 | 0.0 | 9.39E-01 | 0.1 | 5.11E-01 | -0.2 | 5.33E-01 | Protein CASP |
| AT1G12810 | -0.4 | 3.04E-04 | -0.3 | 4.13E-03 | -0.1 | 1.81E-01 | -0.3 | 4.65E-02 | -0.1 | 7.17E-01 | -0.2 | 2.43E-01 | Proline-rich family protein |
| AT5G46530 | -0.4 | 7.41E-04 | -0.1 | 2.24E-01 | -0.3 | 1.62E-02 | 0.0 | 9.01E-01 | -0.2 | 6.15E-01 | 0.2 | 5.74E-01 | AWPM-19-like family protein |
| AT5G21280 | -0.4 | 8.04E-03 | 0.1 | 6.21E-01 | -0.5 | 1.22E-02 | 0.2 | 7.09E-01 | 0.2 | 7.92E-01 | 0.0 | 9.72E-01 | Hydroxyproline-rich glycoprotein family protein |
| AT5G05480 | -0.4 | 2.07E-03 | -0.1 | 1.80E-01 | -0.3 | 4.97E-02 | -0.6 | 1.05E-02 | -0.3 | 2.24E-01 | -0.3 | 2.89E-01 | AT5g05480 |
| AT4G27595 | -0.4 | 2.56E-03 | -0.3 | 1.91E-02 | -0.1 | 3.79E-01 | 0.1 | 6.97E-01 | 0.5 | 3.32E-02 | -0.4 | 1.56E-01 | WEB family protein |
| AT4G30630 | -0.4 | 3.69E-03 | -0.5 | 3.10E-03 | 0.0 | 8.21E-01 | -0.1 | 6.38E-01 | 0.0 | 9.29E-01 | -0.1 | 8.15E-01 | hypothetical protein |
| AT4G09580 | -0.4 | 2.93E-04 | -0.3 | 3.99E-03 | -0.1 | 1.84E-01 | -0.4 | 8.84E-02 | -0.2 | 5.42E-01 | -0.2 | 5.11E-01 | Uncharacterized membrane protein At4g09580 |
| AT3G05220 | -0.4 | 1.16E-03 | -0.2 | 6.70E-02 | -0.2 | 5.75E-02 | -0.3 | 6.66E-03 | -0.2 | 1.04E-01 | -0.1 | 5.33E-01 | Heavy metal transport |
| AT3G20300 | -0.4 | 2.40E-03 | -0.5 | 8.15E-04 | 0.1 | 4.00E-01 | -0.2 | 2.87E-01 | -0.3 | 1.88E-01 | 0.1 | 8.02E-01 | Uncharacterized protein At3g20300 |
| AT1G08770 | -0.4 | 8.64E-05 | -0.3 | 5.16E-04 | -0.1 | 3.63E-01 | -0.3 | 1.56E-01 | -0.4 | 8.56E-02 | 0.1 | 7.71E-01 | PRA1 family protein E |
| AT1G34050 | -0.4 | 2.18E-02 | -1.1 | 2.31E-04 | 0.7 | 1.29E-02 | 2.3 | 4.43E-05 | 0.7 | 1.95E-01 | 1.6 | 4.88E-04 | Ankyrin repeat family protein |
| AT4G02740 | -0.4 | 1.12E-04 | -0.3 | 6.58E-04 | -0.1 | 3.80E-01 | -0.3 | 6.35E-02 | -0.3 | 6.66E-02 | 0.0 | 9.32E-01 | F-box |
| AT4G11440 | -0.4 | 8.04E-03 | -0.4 | 2.00E-02 | -0.1 | 7.49E-01 | 0.4 | 3.96E-01 | 0.1 | 8.73E-01 | 0.3 | 6.38E-01 | Uncharacterized protein AT4g11440 |
| AT3G18230 | -0.4 | 2.71E-03 | -0.4 | 4.91E-03 | 0.0 | 8.75E-01 | -0.2 | 5.47E-01 | -0.2 | 6.10E-01 | 0.0 | 9.99E-01 | PB1 domain-containing protein |
| AT2G05630 | -0.4 | 2.49E-03 | -0.4 | 2.92E-03 | 0.0 | 9.52E-01 | -0.1 | 7.40E-01 | 0.1 | 7.92E-01 | -0.1 | 6.13E-01 | Autophagy-related protein |
| AT5G60570 | -0.4 | 2.26E-04 | -0.2 | 5.17E-02 | -0.3 | 1.21E-02 | -0.3 | 5.98E-02 | -0.1 | 4.81E-01 | -0.1 | 4.92E-01 | F-box |
| AT5G18780 | -0.4 | 5.71E-04 | -0.6 | 1.23E-04 | 0.2 | 1.65E-01 | -0.5 | 5.66E-02 | -0.1 | 8.58E-01 | -0.4 | 1.67E-01 | F-box |
| AT1G34370 | -0.4 | 1.11E-04 | -0.4 | 2.29E-04 | 0.0 | 9.00E-01 | -0.3 | 4.18E-02 | -0.4 | 2.16E-02 | 0.1 | 7.40E-01 | AT1G34370 protein |
| AT2G32900 | -0.4 | 8.30E-05 | -0.2 | 4.60E-03 | -0.2 | 3.67E-02 | -0.3 | 1.91E-01 | -0.1 | 6.45E-01 | -0.1 | 6.56E-01 | Centromere |
| AT2G47960 | -0.4 | 3.17E-04 | -0.2 | 4.40E-02 | -0.2 | 2.04E-02 | -0.3 | 1.67E-02 | -0.2 | 2.60E-01 | -0.1 | 4.34E-01 | At2g47960 |
| AT5G49610 | -0.4 | 1.14E-03 | -0.2 | 4.77E-02 | -0.2 | 9.36E-02 | -0.1 | 5.94E-01 | -0.3 | 2.37E-01 | 0.2 | 6.08E-01 | F-box protein At5g49610 |
| AT3G03680 | -0.4 | 6.45E-04 | -0.1 | 3.49E-01 | -0.3 | 8.29E-03 | -0.1 | 7.21E-01 | 0.1 | 7.98E-01 | -0.2 | 6.09E-01 | hypothetical protein |
| AT1G24070 | -0.4 | 1.35E-01 | -0.6 | 5.84E-02 | 0.2 | 6.35E-01 | 0.5 | 3.59E-01 | 1.0 | 5.60E-02 | -0.5 | 3.82E-01 | Cellulose synthase-like A10 |
| AT2G35330 | -0.4 | 5.21E-05 | -0.2 | 1.88E-02 | -0.2 | 5.05E-03 | -0.2 | 2.73E-01 | -0.1 | 6.69E-01 | -0.1 | 7.31E-01 | RING |
| AT1G22060 | -0.4 | 7.84E-04 | -0.3 | 1.04E-02 | -0.1 | 2.00E-01 | -0.1 | 4.82E-01 | 0.1 | 8.56E-01 | -0.2 | 4.71E-01 | Sporulation-specific protein |
| AT1G64180 | -0.4 | 1.45E-03 | -0.4 | 4.20E-03 | -0.1 | 7.00E-01 | -0.2 | 4.76E-01 | -0.1 | 6.13E-01 | 0.0 | 9.47E-01 | At1g64180 |
| AT3G06780 | -0.4 | 1.72E-03 | -0.4 | 1.49E-03 | 0.0 | 8.00E-01 | -0.7 | 4.70E-03 | -0.6 | 1.53E-02 | -0.1 | 8.56E-01 | F3E22.8 protein |
| AT5G54660 | -0.4 | 2.08E-02 | -0.3 | 4.80E-02 | -0.1 | 7.56E-01 | -0.1 | 4.79E-01 | -0.3 | 1.03E-01 | 0.2 | 4.81E-01 | 21.7 kDa class VI heat shock protein |
| AT1G74580 | -0.4 | 6.98E-02 | -0.4 | 1.01E-01 | 0.0 | 9.28E-01 | | | | | | | Putative pentatricopeptide repeat-containing protein At1g74580 |
| AT1G80060 | -0.4 | 8.58E-03 | -0.1 | 3.79E-01 | -0.3 | 9.24E-02 | -0.1 | 7.75E-01 | 0.0 | 9.33E-01 | -0.1 | 7.31E-01 | hypothetical protein |
| AT2G16595 | -0.4 | 7.81E-04 | -0.4 | 2.76E-03 | -0.1 | 6.08E-01 | -0.4 | 8.26E-02 | -0.2 | 3.45E-01 | -0.2 | 6.56E-01 | Translocon-associated protein subunit alpha |
| AT3G27470 | -0.4 | 1.76E-04 | -0.1 | 7.69E-02 | -0.3 | 7.74E-03 | 0.1 | 5.06E-01 | 0.3 | 1.47E-01 | -0.1 | 5.09E-01 | Lysine ketoglutarate reductase trans-splicing protein ) |
| AT2G26890 | -0.4 | 1.84E-04 | -0.4 | 3.63E-04 | 0.0 | 9.02E-01 | -0.1 | 6.31E-01 | 0.0 | 9.72E-01 | -0.1 | 7.23E-01 | DnaJ homolog subfamily C GRV2 |
| AT1G27290 | -0.4 | 1.30E-03 | -0.2 | 7.73E-02 | -0.2 | 5.54E-02 | -0.6 | 3.52E-03 | -0.6 | 3.60E-03 | 0.1 | 8.30E-01 | F17L21.8 |
| AT5G02140 | -0.4 | 1.21E-01 | -0.2 | 3.71E-01 | -0.2 | 6.04E-01 | | | | | | | Pathogenesis-related thaumatin superfamily protein |
| AT5G40840 | -0.4 | 5.51E-03 | -0.3 | 3.79E-02 | -0.1 | 4.13E-01 | 0.3 | 3.38E-01 | 0.2 | 6.73E-01 | 0.1 | 7.52E-01 | Rad21 |
| AT1G04850 | -0.4 | 9.45E-05 | -0.2 | 2.70E-02 | -0.2 | 7.49E-03 | 0.0 | 8.86E-01 | 0.0 | 9.38E-01 | 0.0 | 9.80E-01 | Ubiquitin-associated |
| AT3G02340 | -0.4 | 1.03E-03 | -0.3 | 6.16E-03 | -0.1 | 4.10E-01 | -0.6 | 3.59E-04 | -0.4 | 1.01E-02 | -0.2 | 3.51E-01 | RING-type domain-containing protein |
| AT4G15450 | -0.4 | 9.15E-03 | -0.3 | 3.77E-02 | -0.1 | 5.53E-01 | -0.3 | 9.65E-02 | 0.3 | 1.10E-01 | -0.7 | 1.38E-02 | At4g15450 |
| AT1G12850 | -0.4 | 4.17E-04 | -0.2 | 1.25E-02 | -0.2 | 8.47E-02 | -0.3 | 2.68E-02 | 0.0 | 9.06E-01 | -0.3 | 1.43E-01 | F13K23.10 protein |

| | | | | | | | | | | | | | |
|---|---|---|---|---|---|---|---|---|---|---|---|---|---|
| AT1G31440 | -0.4 | 2.76E-04 | -0.4 | 2.45E-04 | 0.0 | 6.92E-01 | -0.3 | 2.78E-02 | -0.1 | 6.85E-01 | -0.2 | 2.29E-01 | SH3 domain-containing protein 1 |
| AT4G30350 | -0.4 | 1.76E-03 | -0.3 | 1.67E-02 | -0.1 | 2.83E-01 | -0.6 | 3.11E-05 | -0.2 | 2.20E-01 | -0.4 | 8.95E-03 | Protein SMAX1-LIKE 2 |
| AT3G14595 | -0.4 | 4.78E-04 | -0.2 | 2.07E-02 | -0.2 | 6.30E-02 | -0.4 | 9.28E-04 | -0.2 | 1.16E-01 | -0.2 | 1.26E-01 | 60S ribosomal protein L18a-like protein |
| AT1G29060 | -0.4 | 6.32E-04 | -0.2 | 1.78E-02 | -0.2 | 1.15E-01 | -0.1 | 7.76E-01 | 0.0 | 9.58E-01 | -0.1 | 7.96E-01 | Target SNARE coiled-coil domain protein |
| AT1G34220 | -0.4 | 4.88E-04 | -0.3 | 4.12E-03 | -0.1 | 2.82E-01 | -0.4 | 4.33E-02 | -0.3 | 1.73E-01 | -0.1 | 7.41E-01 | F23M19.10 |
| AT3G21480 | -0.4 | 1.60E-03 | -0.3 | 6.08E-03 | -0.1 | 5.90E-01 | 0.3 | 3.65E-01 | 0.3 | 3.46E-01 | 0.0 | 9.31E-01 | At3g21480 protein |
| AT1G50570 | -0.4 | 2.38E-04 | -0.4 | 7.05E-04 | 0.0 | 6.67E-01 | -0.1 | 3.06E-01 | -0.1 | 5.95E-01 | -0.1 | 8.39E-01 | At1g50570 family protein) |
| AT3G12200 | -0.4 | 2.38E-04 | -0.5 | 8.97E-05 | 0.1 | 2.56E-01 | -0.1 | 6.60E-01 | 0.1 | 8.05E-01 | -0.1 | 5.92E-01 | Uncharacterized protein At3g12200 |
| AT4G17310 | -0.4 | 3.32E-03 | -0.2 | 5.82E-02 | -0.2 | 2.07E-01 | -0.1 | 7.20E-01 | -0.3 | 2.17E-01 | 0.2 | 4.79E-01 | Protein NUCLEAR FUSION DEFECTIVE 6, chloroplastic |
| AT1G50970 | -0.4 | 4.43E-02 | -0.5 | 1.83E-02 | 0.1 | 6.12E-01 | -0.2 | 5.70E-01 | 0.1 | 8.41E-01 | -0.3 | 3.88E-01 | Membrane trafficking VPS53 family protein |
| AT5G16610 | -0.4 | 3.42E-03 | -0.2 | 1.17E-01 | -0.2 | 1.15E-01 | -0.2 | 5.20E-01 | -0.3 | 2.96E-01 | 0.1 | 7.18E-01 | Uncharacterized protein |
| AT2G37480 | -0.4 | 2.87E-04 | 0.0 | 9.00E-01 | -0.4 | 1.16E-03 | -0.2 | 1.91E-01 | -0.1 | 3.51E-01 | 0.0 | 9.19E-01 | Uncharacterized protein At2g37490 |
| AT5G66480 | -0.4 | 4.78E-03 | -1.0 | 4.80E-05 | 0.6 | 5.39E-03 | 0.6 | 9.82E-02 | -0.6 | 2.39E-01 | 1.2 | 1.74E-02 | Uncharacterized protein |
| AT3G62580 | -0.4 | 9.05E-04 | -0.1 | 2.27E-01 | -0.3 | 1.50E-02 | -0.4 | 5.95E-02 | 0.0 | 9.28E-01 | -0.4 | 1.94E-01 | DUF4149 domain-containing protein |
| AT3G05650 | -0.4 | 2.81E-02 | -0.8 | 1.53E-03 | 0.4 | 1.10E-01 | 0.3 | 6.10E-01 | 0.1 | 9.36E-01 | 0.3 | 7.49E-01 | Receptor-like protein 32 |
| AT2G15040 | -0.4 | 1.82E-01 | -0.7 | 3.93E-02 | 0.3 | 4.10E-01 | | | | | | | Receptor-like protein 18 |
| AT5G04460 | -0.4 | 7.38E-04 | -0.5 | 4.31E-04 | 0.1 | 5.44E-01 | 0.0 | 9.31E-01 | 0.0 | 9.15E-01 | -0.1 | 8.69E-01 | RING |
| AT5G44417 | -0.4 | 7.22E-02 | 1.0 | 3.01E-04 | -1.4 | 3.00E-04 | | | | | | | NA |
| AT5G47690 | -0.4 | 5.72E-05 | -0.3 | 4.63E-04 | -0.1 | 2.50E-01 | -0.1 | 4.02E-01 | -0.1 | 5.48E-01 | 0.0 | 9.61E-01 | Binding protein |
| AT4G07390 | -0.4 | 1.09E-03 | -0.2 | 5.96E-02 | -0.2 | 5.99E-02 | 0.0 | 8.10E-01 | 0.0 | 8.98E-01 | -0.1 | 7.62E-01 | Mannose-P-dolichol utilization defect 1 protein homolog 2 |
| AT5G56750 | -0.4 | 6.53E-04 | -0.3 | 7.88E-03 | -0.1 | 2.31E-01 | -0.7 | 2.30E-04 | -0.7 | 5.01E-04 | 0.0 | 9.51E-01 | N-MYC downregulated-like 1 |
| AT3G17740 | -0.4 | 6.71E-04 | -0.3 | 4.73E-03 | -0.1 | 3.71E-01 | -0.4 | 5.31E-02 | -0.1 | 7.83E-01 | -0.3 | 2.31E-01 | At3g17740 |
| AT5G25100 | -0.4 | 3.28E-04 | -0.2 | 3.96E-02 | -0.2 | 2.35E-02 | -0.1 | 3.76E-01 | 0.0 | 8.44E-01 | -0.2 | 4.23E-01 | Transmembrane 9 superfamily member |
| AT3G50590 | -0.4 | 1.20E-04 | -0.4 | 5.19E-04 | -0.1 | 5.01E-01 | 0.0 | 9.63E-01 | 0.0 | 9.68E-01 | 0.0 | 9.49E-01 | Uncharacterized protein At3g50600 |
| AT2G18240 | -0.4 | 1.06E-03 | -0.4 | 2.99E-03 | 0.0 | 6.97E-01 | 0.0 | 9.04E-01 | 0.1 | 7.03E-01 | -0.1 | 8.23E-01 | Protein RER1D |
| AT5G01970 | -0.4 | 2.00E-04 | -0.2 | 5.97E-03 | -0.2 | 8.33E-02 | -0.5 | 1.70E-03 | -0.3 | 3.90E-02 | -0.2 | 3.56E-01 | hypothetical protein |
| AT2G20320 | -0.4 | 6.39E-04 | -0.6 | 1.24E-04 | 0.2 | 1.45E-01 | 0.5 | 2.03E-02 | 0.3 | 2.13E-01 | 0.2 | 4.73E-01 | Uncharacterized protein At2g20320 |
| AT1G13990 | -0.4 | 1.09E-02 | -0.8 | 2.02E-04 | 0.4 | 1.92E-02 | -0.5 | 4.94E-02 | -0.5 | 9.13E-02 | 0.0 | 9.56E-01 | Plant |
| AT5G23810 | -0.4 | 1.20E-03 | -0.2 | 2.09E-02 | -0.2 | 1.92E-01 | -0.3 | 3.62E-01 | 0.2 | 5.62E-01 | -0.6 | 1.88E-01 | Amino acid permease 7 |
| AT1G71696 | -0.4 | 1.93E-03 | -0.1 | 5.82E-01 | -0.3 | 1.42E-02 | -0.3 | 1.82E-01 | 0.0 | 9.81E-01 | -0.3 | 3.03E-01 | SOL1 protein |
| AT5G01220 | -0.4 | 1.65E-03 | 0.0 | 6.47E-01 | -0.4 | 8.11E-03 | 0.1 | 4.81E-01 | 0.1 | 5.74E-01 | 0.0 | 9.96E-01 | Sulfoquinovosyldiacylglycerol 2 |
| AT4G26180 | -0.4 | 2.20E-03 | -0.5 | 1.55E-03 | 0.1 | 7.11E-01 | -0.1 | 8.50E-01 | -0.2 | 5.65E-01 | 0.1 | 7.24E-01 | Mitochondrial substrate carrier family protein |
| AT3G15480 | -0.4 | 3.91E-04 | -0.1 | 1.13E-01 | -0.3 | 1.12E-02 | -0.4 | 5.28E-03 | -0.2 | 9.18E-02 | -0.1 | 4.68E-01 | AT3g15480 |
| AT5G57010 | -0.4 | 1.09E-01 | -0.8 | 7.97E-03 | 0.4 | 1.86E-01 | 0.7 | 1.17E-01 | 0.1 | 8.73E-01 | 0.6 | 2.94E-01 | Uncharacterized protein At5g57010 |
| AT3G17120 | -0.4 | 1.16E-02 | -0.2 | 1.60E-01 | -0.2 | 2.32E-01 | 0.0 | 9.57E-01 | 0.4 | 3.30E-02 | -0.4 | 7.65E-02 | hypothetical protein |
| AT4G25770 | -0.4 | 4.86E-03 | -0.5 | 1.52E-03 | 0.1 | 4.20E-01 | -0.4 | 2.43E-02 | -0.3 | 1.63E-01 | -0.1 | 6.33E-01 | Alpha |
| AT1G61600 | -0.4 | 2.22E-03 | -0.3 | 1.85E-02 | -0.1 | 3.50E-01 | -0.1 | 8.73E-01 | 0.2 | 5.27E-01 | -0.3 | 4.85E-01 | DUF1262 family protein |
| AT3G11660 | -0.4 | 1.12E-02 | -0.2 | 2.05E-01 | -0.2 | 1.65E-01 | 0.0 | 7.68E-01 | 0.1 | 5.26E-01 | -0.2 | 4.53E-01 | NDR1 |
| AT5G54855 | -0.4 | 7.23E-03 | -0.3 | 1.86E-02 | -0.1 | 7.27E-01 | -0.2 | 1.73E-01 | -0.1 | 5.82E-01 | -0.1 | 6.34E-01 | AT1g27100 |
| AT3G48430 | -0.4 | 3.06E-04 | -0.3 | 2.49E-03 | -0.1 | 2.89E-01 | 0.0 | 8.99E-01 | 0.1 | 7.31E-01 | -0.1 | 8.50E-01 | Relative of early flowering 6 |
| AT2G45140 | -0.4 | 8.05E-05 | -0.2 | 1.19E-02 | -0.2 | 1.23E-02 | -0.1 | 6.18E-01 | 0.0 | 9.60E-01 | -0.1 | 7.10E-01 | Plant VAP homolog 12 |
| AT5G44280 | -0.4 | 2.14E-03 | 0.0 | 7.90E-01 | -0.4 | 1.08E-02 | -0.3 | 1.01E-01 | 0.0 | 8.97E-01 | -0.3 | 2.71E-01 | RING 1A |
| AT4G28300 | -0.4 | 3.81E-03 | -0.3 | 1.17E-02 | -0.1 | 6.50E-01 | -0.3 | 2.96E-02 | -0.4 | 3.23E-02 | 0.0 | 9.20E-01 | Formin-like protein |
| AT5G58920 | -0.4 | 3.47E-03 | -0.1 | 4.69E-01 | -0.3 | 2.88E-02 | -0.2 | 3.24E-01 | -0.4 | 1.15E-01 | 0.2 | 6.39E-01 | hypothetical protein |
| AT1G30040 | -0.4 | 9.50E-02 | -0.8 | 6.26E-03 | 0.4 | 1.64E-01 | 0.2 | 4.98E-01 | 0.0 | 9.77E-01 | 0.1 | 6.23E-01 | GA2OX2 |
| AT1G26670 | -0.4 | 1.61E-03 | -0.3 | 1.01E-02 | -0.1 | 3.89E-01 | -0.5 | 6.15E-04 | -0.4 | 1.01E-02 | -0.1 | 5.60E-01 | At1g26670 |
| AT1G04200 | -0.4 | 2.68E-04 | -0.3 | 1.38E-03 | -0.1 | 4.23E-01 | -0.1 | 3.58E-01 | 0.1 | 7.17E-01 | -0.2 | 2.98E-01 | Dymeclin |
| AT4G36648 | -0.4 | 3.34E-02 | -0.6 | 5.10E-03 | 0.2 | 2.82E-01 | 0.2 | 3.53E-01 | 0.0 | 9.68E-01 | 0.2 | 5.08E-01 | NA |
| AT5G04730 | -0.4 | 1.55E-02 | 0.7 | 6.40E-04 | -1.0 | 1.59E-04 | -1.6 | 2.38E-06 | -1.2 | 6.54E-05 | -0.4 | 2.85E-01 | Uncharacterized protein T1E3_90 |
| AT4G25610 | -0.4 | 4.40E-04 | -0.2 | 2.45E-02 | -0.2 | 6.06E-02 | -0.1 | 6.00E-01 | -0.1 | 7.21E-01 | 0.0 | 9.59E-01 | C2H2-like zinc finger protein |
| AT1G72820 | -0.4 | 3.97E-04 | -0.2 | 2.96E-02 | -0.2 | 3.91E-02 | -0.6 | 1.28E-03 | -0.7 | 1.22E-03 | 0.1 | 8.00E-01 | Solute carrier family 25 member 44 |
| AT1G76955 | -0.4 | 2.56E-02 | -0.3 | 1.15E-01 | -0.1 | 5.22E-01 | 0.8 | 8.29E-02 | 0.7 | 1.98E-01 | 0.1 | 8.45E-01 | Transmembrane protein |
| AT5G44820 | -0.4 | 3.25E-03 | -1.1 | 1.01E-05 | 0.7 | 7.16E-04 | 1.5 | 1.95E-04 | 0.3 | 4.55E-01 | 1.2 | 5.58E-03 | Nucleotide-diphospho-sugar transferase domain-containing protein |
| AT5G23150 | -0.4 | 6.65E-04 | -0.5 | 1.98E-04 | 0.1 | 2.59E-01 | 0.0 | 8.87E-01 | -0.1 | 5.37E-01 | 0.1 | 7.21E-01 | ENHANCER OF AG-4 protein 2 |
| AT2G46560 | -0.4 | 4.19E-05 | -0.3 | 3.56E-04 | -0.1 | 2.11E-01 | -0.1 | 3.55E-01 | 0.0 | 9.08E-01 | -0.1 | 6.34E-01 | Transducin family protein |
| AT1G74280 | -0.4 | 2.06E-03 | -0.1 | 4.01E-01 | -0.3 | 2.30E-02 | -0.2 | 4.91E-01 | -0.2 | 4.11E-01 | 0.1 | 8.96E-01 | Alpha |
| AT1G75800 | -0.4 | 1.26E-04 | -0.1 | 1.10E-01 | -0.3 | 3.56E-03 | -0.6 | 1.15E-04 | -0.7 | 1.38E-04 | 0.1 | 6.76E-01 | AT1G75800 protein |
| AT1G25280 | -0.4 | 1.46E-04 | -0.3 | 7.01E-04 | -0.1 | 4.38E-01 | 0.0 | 7.44E-01 | 0.0 | 9.23E-01 | 0.0 | 9.10E-01 | Tubby-like F-box protein |
| AT1G32127 | -0.4 | 3.27E-03 | 0.0 | 8.86E-01 | -0.4 | 1.31E-02 | 0.0 | 9.32E-01 | 0.1 | 6.41E-01 | -0.1 | 7.65E-01 | Uncharacterized protein F3C3.9 |
| AT1G80520 | -0.4 | 4.20E-02 | -0.1 | 4.69E-01 | -0.3 | 2.44E-01 | 0.0 | 9.72E-01 | 0.3 | 4.87E-01 | -0.2 | 5.80E-01 | SAM domain-containing protein |
| AT5G25010 | -0.4 | 8.60E-03 | 0.0 | 8.09E-01 | -0.4 | 3.42E-02 | -0.3 | 2.72E-01 | 0.1 | 7.19E-01 | -0.4 | 2.37E-01 | Enhanced disease resistance-like protein |
| AT5G47860 | -0.4 | 6.66E-04 | -0.8 | 1.20E-05 | 0.4 | 2.84E-03 | -0.1 | 6.95E-01 | -0.1 | 7.36E-01 | 0.0 | 9.93E-01 | AT5g47860 |
| AT3G02430 | -0.4 | 4.80E-02 | 0.5 | 8.09E-03 | -0.9 | 1.82E-03 | -1.7 | 6.30E-03 | -0.5 | 3.24E-01 | -1.1 | 1.01E-01 | Protein DMP5 |
| AT2G30100 | -0.4 | 1.29E-02 | -0.1 | 3.25E-01 | -0.3 | 1.29E-01 | -0.1 | 4.35E-01 | 0.1 | 5.09E-01 | -0.3 | 2.31E-01 | Pentatricopeptide repeat-containing protein At2g30100, chloroplastic |
| AT1G29760 | -0.4 | 6.95E-04 | -0.4 | 1.02E-03 | 0.0 | 9.99E-01 | -0.4 | 1.35E-02 | -0.4 | 2.16E-02 | 0.0 | 9.94E-01 | Seipin-2 |
| AT1G22350 | -0.4 | 5.39E-03 | -0.4 | 7.29E-03 | 0.0 | 9.86E-01 | 0.1 | 7.11E-01 | 0.2 | 5.16E-01 | -0.1 | 8.14E-01 | NA |
| AT4G34460 | -0.4 | 9.87E-05 | -0.5 | 4.89E-05 | 0.1 | 2.57E-01 | -0.2 | 9.38E-02 | -0.3 | 1.94E-02 | 0.1 | 5.39E-01 | At4g34460 |
| AT3G19020 | -0.4 | 1.22E-03 | -0.3 | 3.42E-03 | 0.0 | 7.08E-01 | 0.4 | 2.22E-01 | 0.1 | 9.19E-01 | 0.3 | 3.95E-01 | Leucine-rich repeat-containing N-terminal plant-type domain-containing protein |
| AT4G27020 | -0.4 | 6.93E-04 | -0.6 | 4.80E-05 | 0.2 | 3.06E-02 | -0.2 | 3.54E-01 | -0.1 | 8.21E-01 | -0.1 | 6.93E-01 | AT4g27020 |
| AT3G16830 | -0.4 | 2.12E-04 | -0.4 | 2.59E-04 | 0.0 | 8.56E-01 | 0.1 | 4.88E-01 | -0.1 | 5.74E-01 | 0.3 | 2.79E-01 | Topless-related protein 2 |
| AT5G62150 | -0.4 | 1.90E-01 | -0.4 | 1.97E-01 | 0.0 | 9.81E-01 | 1.4 | 2.72E-03 | 0.5 | 3.90E-01 | 0.9 | 5.51E-02 | hypothetical protein |
| AT3G21610 | -0.4 | 1.05E-03 | -0.2 | 1.91E-02 | -0.1 | 1.67E-01 | -0.3 | 5.12E-02 | -0.2 | 2.55E-01 | -0.1 | 6.83E-01 | Acid phosphatase |
| AT5G53530 | -0.4 | 1.15E-04 | -0.2 | 1.05E-02 | -0.2 | 2.34E-02 | 0.1 | 5.96E-01 | 0.0 | 8.33E-01 | 0.0 | 8.80E-01 | Vacuolar protein sorting-associated protein 26A |
| AT1G12320 | -0.4 | 1.76E-03 | -0.2 | 2.94E-02 | -0.2 | 2.01E-01 | -0.2 | 3.55E-01 | -0.2 | 4.51E-01 | 0.0 | 9.78E-01 | Ankyrin repeat |
| AT4G08690 | -0.4 | 5.19E-04 | -0.1 | 4.36E-01 | -0.3 | 5.07E-03 | -0.3 | 5.66E-02 | 0.1 | 5.70E-01 | -0.4 | 4.03E-02 | hypothetical protein |
| AT2G33360 | -0.4 | 6.48E-03 | -0.4 | 4.88E-03 | 0.0 | 7.88E-01 | -0.2 | 4.61E-01 | 0.0 | 9.94E-01 | -0.2 | 6.08E-01 | Uncharacterized protein At2g33360 |
| AT5G12470 | -0.4 | 2.71E-04 | 0.0 | 6.43E-01 | -0.3 | 1.66E-03 | -0.1 | 5.07E-01 | 0.1 | 5.74E-01 | -0.2 | 3.30E-01 | Protein RETICULATA-RELATED 4, chloroplastic |
| AT3G51980 | -0.4 | 9.59E-04 | -0.2 | 1.66E-02 | -0.1 | 1.69E-01 | 0.1 | 7.37E-01 | 0.1 | 4.70E-01 | -0.1 | 7.53E-01 | Nucleotide exchange factor Fes1 domain-containing protein |
| AT1G66480 | -0.4 | 2.49E-03 | -0.4 | 2.13E-03 | 0.0 | 8.06E-01 | 0.5 | 9.58E-02 | 0.1 | 8.25E-01 | 0.4 | 3.32E-01 | Uncharacterized protein At1g66480 |
| AT2G24550 | -0.4 | 3.26E-02 | 0.0 | 7.85E-01 | -0.4 | 3.39E-02 | -0.5 | 4.76E-02 | -0.4 | 9.42E-02 | 0.0 | 9.54E-01 | Protein OXIDATIVE STRESS 3 LIKE 6 |
| AT5G65910 | -0.4 | 1.14E-03 | -0.4 | 2.48E-03 | 0.0 | 7.93E-01 | -0.4 | 2.85E-03 | -0.3 | 3.23E-02 | -0.1 | 5.82E-01 | BSD domain-containing protein |
| AT4G13180 | -0.4 | 5.94E-02 | -0.2 | 4.03E-01 | -0.2 | 3.30E-01 | 0.5 | 1.48E-02 | 0.1 | 6.22E-01 | 0.4 | 1.24E-01 | Uncharacterized protein |
| AT5G35935 | -0.4 | 7.16E-02 | -0.4 | 6.65E-02 | 0.0 | 9.44E-01 | -0.3 | 4.43E-01 | -0.1 | 8.17E-01 | -0.2 | 7.47E-01 | NA |

| | | | | | | | | | | | | | |
|---|---|---|---|---|---|---|---|---|---|---|---|---|---|
| AT1G62880 | -0.4 | 1.77E-03 | -0.3 | 4.65E-03 | 0.0 | 7.38E-01 | -0.4 | 1.62E-01 | -0.1 | 8.00E-01 | -0.3 | 4.83E-01 | Cornichon family protein |
| AT1G59510 | -0.4 | 6.85E-03 | -0.3 | 2.63E-02 | -0.1 | 6.09E-01 | 0.0 | 9.12E-01 | 0.0 | 9.79E-01 | -0.1 | 9.18E-01 | Uncharacterized protein T4M14.9 |
| AT4G32750 | -0.4 | 5.68E-04 | -0.4 | 6.80E-04 | 0.0 | 9.00E-01 | -0.4 | 3.99E-02 | -0.4 | 6.42E-02 | 0.0 | 9.82E-01 | hypothetical protein |
| AT3G59090 | -0.4 | 3.63E-04 | -0.2 | 2.25E-02 | -0.2 | 4.42E-02 | -0.2 | 1.76E-01 | -0.1 | 5.79E-01 | -0.1 | 6.76E-01 | AT3G59090 protein |
| AT1G06460 | -0.4 | 9.29E-03 | -0.1 | 5.63E-01 | -0.3 | 4.54E-02 | -0.4 | 9.76E-03 | -0.1 | 4.20E-01 | -0.2 | 1.61E-01 | F12K11.19 |
| AT1G79985 | -0.4 | 6.33E-04 | -0.2 | 5.33E-02 | -0.2 | 4.02E-02 | -0.3 | 4.75E-02 | -0.3 | 6.72E-02 | 0.0 | 9.91E-01 | Protein SYS1 homolog |
| AT1G56230 | -0.4 | 2.67E-04 | -0.3 | 3.92E-03 | -0.1 | 1.65E-01 | -0.2 | 1.03E-01 | 0.0 | 9.72E-01 | -0.2 | 2.33E-01 | Enolase |
| AT4G21810 | -0.4 | 2.02E-03 | -0.2 | 1.26E-01 | -0.2 | 5.55E-02 | -0.4 | 1.17E-03 | -0.4 | 2.79E-03 | 0.0 | 9.92E-01 | Derlin |
| AT2G40070 | -0.4 | 4.41E-04 | -0.1 | 8.18E-02 | -0.2 | 1.89E-02 | 0.0 | 8.21E-01 | 0.1 | 5.10E-01 | -0.2 | 4.49E-01 | At2g40070 |
| AT2G46630 | -0.4 | 4.06E-03 | 0.0 | 9.80E-01 | -0.4 | 1.10E-02 | -0.2 | 4.09E-01 | -0.1 | 8.11E-01 | -0.1 | 7.38E-01 | Uncharacterized protein |
| AT1G18190 | -0.4 | 6.16E-04 | -0.3 | 9.08E-03 | -0.1 | 1.95E-01 | 0.2 | 5.06E-01 | 0.0 | 9.24E-01 | 0.1 | 7.36E-01 | Golgin candidate 2 |
| AT5G65290 | -0.4 | 2.68E-04 | -0.3 | 3.03E-03 | -0.1 | 2.20E-01 | 0.0 | 9.07E-01 | 0.2 | 4.75E-01 | -0.2 | 6.24E-01 | LMBR1-like membrane protein |
| AT3G01780 | -0.4 | 9.16E-05 | -0.3 | 4.04E-04 | -0.1 | 4.64E-01 | -0.1 | 5.32E-01 | 0.0 | 9.87E-01 | -0.1 | 6.80E-01 | ARM repeat superfamily protein |
| AT4G23430 | -0.4 | 1.72E-04 | -0.4 | 3.77E-04 | 0.0 | 8.28E-01 | -0.4 | 5.18E-02 | 0.1 | 5.28E-01 | -0.5 | 4.71E-02 | NAD-binding Rossmann-fold superfamily protein |
| AT3G20450 | -0.4 | 1.56E-02 | 0.0 | 8.56E-01 | -0.4 | 5.51E-02 | 0.7 | 1.16E-01 | 0.5 | 3.44E-01 | 0.2 | 6.62E-01 | Endoplasmic reticulum transmembrane protein |
| AT5G51570 | -0.4 | 1.21E-03 | -0.1 | 4.77E-01 | -0.3 | 9.34E-03 | 0.1 | 6.74E-01 | -0.2 | 2.02E-01 | 0.2 | 1.51E-01 | Hypersensitive-induced response protein 4 |
| AT3G03790 | -0.4 | 1.37E-03 | -0.6 | 1.30E-04 | 0.2 | 6.88E-02 | -0.1 | 8.23E-01 | 0.2 | 4.98E-01 | -0.2 | 4.61E-01 | Ankyrin repeat family protein |
| AT4G27840 | -0.4 | 4.21E-03 | -0.1 | 5.45E-01 | -0.3 | 3.05E-02 | -0.4 | 2.87E-02 | -0.2 | 4.14E-01 | -0.2 | 3.62E-01 | Phytolongin Phyl2.1 |
| AT1G08350 | -0.4 | 1.52E-03 | 0.0 | 6.48E-01 | -0.4 | 3.03E-03 | -0.1 | 7.17E-01 | 0.1 | 6.55E-01 | -0.2 | 4.51E-01 | Transmembrane 9 superfamily member |
| AT4G32820 | -0.4 | 1.22E-03 | -0.4 | 1.35E-03 | 0.0 | 8.87E-01 | 0.0 | 9.48E-01 | 0.1 | 8.58E-01 | -0.1 | 8.23E-01 | Tetratricopeptide repeat -like superfamily protein |
| AT3G07790 | -0.4 | 5.21E-05 | -0.4 | 9.57E-05 | 0.0 | 9.78E-01 | -0.3 | 1.47E-02 | -0.2 | 9.09E-02 | -0.1 | 6.99E-01 | hypothetical protein |
| AT3G19660 | -0.4 | 1.78E-01 | -0.8 | 1.61E-02 | 0.5 | 2.15E-01 | 0.0 | 9.55E-01 | -0.1 | 8.44E-01 | 0.2 | 8.33E-01 | Uncharacterized protein |
| AT2G02880 | -0.4 | 1.68E-02 | 0.0 | 8.95E-01 | -0.4 | 5.01E-02 | 0.2 | 3.94E-01 | -0.4 | 1.20E-01 | 0.5 | 3.78E-02 | hypothetical protein |
| AT5G02890 | -0.4 | 6.82E-02 | 0.2 | 4.45E-01 | -0.5 | 3.29E-02 | 0.1 | 7.40E-01 | 0.3 | 2.04E-01 | -0.2 | 3.81E-01 | HXXXD-type acyl-transferase family protein |
| AT4G25515 | -0.4 | 4.83E-03 | -0.2 | 7.41E-02 | -0.2 | 2.31E-01 | -0.1 | 8.26E-01 | 0.1 | 7.12E-01 | -0.2 | 5.72E-01 | Probable transcriptional regulator SLK3 |
| AT3G03610 | -0.4 | 1.39E-03 | -0.2 | 7.72E-02 | -0.2 | 7.04E-02 | -0.2 | 4.21E-01 | -0.2 | 4.96E-01 | 0.0 | 9.92E-01 | T12J13.11 protein |
| AT3G59680 | -0.4 | 2.83E-03 | 0.0 | 6.83E-01 | -0.3 | 1.69E-02 | 0.1 | 8.10E-01 | 0.0 | 9.24E-01 | 0.0 | 9.40E-01 | At3g59680 |
| AT2G40475 | -0.4 | 1.28E-01 | -0.1 | 7.46E-01 | -0.3 | 2.99E-01 | -0.1 | 7.27E-01 | 0.3 | 1.26E-01 | -0.4 | 8.30E-02 | Uncharacterized protein |
| AT1G62860 | -0.4 | 2.02E-02 | -0.3 | 3.46E-02 | 0.0 | 8.76E-01 | -0.3 | 3.53E-01 | 0.0 | 9.85E-01 | -0.3 | 4.51E-01 | NA |
| AT1G21660 | -0.4 | 1.51E-04 | -0.2 | 6.31E-03 | -0.2 | 4.90E-02 | -0.3 | 1.80E-02 | -0.1 | 7.98E-01 | -0.3 | 1.05E-01 | AT1G21660 protein |
| AT1G59590 | -0.4 | 1.36E-02 | -0.6 | 1.46E-03 | 0.2 | 1.88E-01 | 0.5 | 5.78E-02 | -0.5 | 1.16E-01 | 0.9 | 8.17E-03 | At1g59590 |
| AT1G79060 | -0.4 | 1.59E-02 | 0.0 | 9.06E-01 | -0.4 | 4.60E-02 | -0.2 | 5.20E-01 | 0.1 | 7.91E-01 | -0.3 | 3.36E-01 | Uncharacterized protein |
| AT1G26650 | -0.4 | 1.52E-03 | -0.2 | 7.93E-02 | -0.2 | 6.91E-02 | -0.1 | 6.89E-01 | 0.1 | 7.55E-01 | -0.2 | 5.55E-01 | Uncharacterized protein At1g26650 |
| AT2G41550 | -0.4 | 2.65E-02 | -0.4 | 3.12E-02 | 0.0 | 9.75E-01 | -1.1 | 4.83E-02 | -0.5 | 3.49E-01 | -0.6 | 4.16E-01 | Rho termination factor N-terminal domain-containing protein |
| AT4G30510 | -0.4 | 2.51E-03 | -0.1 | 4.11E-01 | -0.3 | 2.83E-02 | -0.2 | 4.12E-01 | -0.2 | 4.14E-01 | 0.0 | 9.66E-01 | Autophagy 18 B-like protein |
| AT1G49470 | -0.4 | 2.42E-03 | 0.0 | 6.71E-01 | -0.4 | 3.95E-03 | -0.1 | 5.63E-01 | -0.1 | 4.32E-01 | 0.0 | 8.53E-01 | At1g49470 ) |
| AT1G49360 | -0.4 | 1.00E-03 | -0.5 | 3.56E-04 | 0.1 | 3.68E-01 | -0.3 | 1.91E-01 | 0.0 | 9.66E-01 | -0.3 | 3.36E-01 | F-box family protein |
| AT3G51540 | -0.4 | 6.23E-03 | 0.2 | 5.42E-02 | -0.6 | 1.36E-03 | 0.4 | 2.04E-01 | 0.6 | 5.17E-02 | -0.2 | 5.22E-01 | Mucin-5AC-like protein |
| AT3G61350 | -0.4 | 2.65E-03 | -0.1 | 1.75E-01 | -0.2 | 6.75E-02 | -0.3 | 1.54E-01 | -0.5 | 7.80E-02 | 0.1 | 7.10E-01 | F-box |
| AT3G63460 | -0.4 | 4.58E-04 | -0.3 | 1.64E-03 | 0.0 | 6.02E-01 | 0.1 | 4.20E-01 | 0.2 | 2.83E-01 | -0.1 | 8.21E-01 | Transducin family protein |
| AT2G41312 | -0.4 | 8.84E-02 | -0.1 | 6.00E-01 | -0.3 | 3.26E-01 | 0.4 | 2.50E-01 | 0.5 | 1.69E-01 | -0.1 | 8.20E-01 | NA |
| AT4G30640 | -0.4 | 1.66E-02 | 0.1 | 5.28E-01 | -0.5 | 1.72E-02 | -0.7 | 9.62E-02 | -0.4 | 3.44E-01 | -0.3 | 6.57E-01 | Putative F-box |
| AT3G16500 | -0.4 | 6.47E-04 | -0.2 | 1.16E-02 | -0.1 | 1.54E-01 | -0.1 | 7.47E-01 | -0.2 | 3.29E-01 | 0.1 | 6.17E-01 | Auxin-responsive protein |
| AT3G18770 | -0.4 | 5.94E-03 | -0.7 | 2.30E-04 | 0.3 | 4.27E-02 | -0.8 | 1.56E-03 | -0.2 | 4.11E-01 | -0.6 | 4.76E-02 | Autophagy-related protein 13b |
| AT5G52410 | -0.4 | 2.49E-02 | -0.4 | 2.05E-02 | 0.0 | 8.73E-01 | 0.1 | 7.44E-01 | 0.6 | 5.64E-02 | -0.4 | 1.09E-01 | Oxidoreductase |
| AT2G32580 | -0.4 | 1.55E-03 | -0.2 | 3.35E-02 | -0.2 | 1.38E-01 | -0.2 | 2.77E-01 | 0.0 | 8.16E-01 | -0.1 | 5.62E-01 | Expressed protein ) |
| AT1G30835 | -0.4 | 1.68E-04 | -0.3 | 1.28E-03 | -0.1 | 2.94E-01 | -0.1 | 5.94E-01 | 0.0 | 9.49E-01 | -0.1 | 5.77E-01 | NA |
| AT5G24810 | -0.4 | 1.53E-04 | -0.6 | 1.20E-05 | 0.2 | 1.06E-02 | 0.0 | 8.57E-01 | 0.1 | 6.66E-01 | -0.1 | 5.85E-01 | ABC1 family protein |
| AT5G08270 | -0.4 | 5.95E-03 | -0.1 | 6.23E-01 | -0.3 | 3.81E-02 | -0.1 | 8.38E-01 | 0.4 | 2.10E-01 | -0.5 | 1.27E-01 | hypothetical protein |
| AT2G20495 | -0.4 | 1.76E-03 | -0.4 | 1.52E-03 | 0.0 | 7.96E-01 | -0.4 | 6.23E-02 | -0.1 | 7.82E-01 | -0.3 | 2.45E-01 | At2g20495 |
| AT3G51940 | -0.4 | 3.32E-03 | -0.2 | 6.13E-02 | -0.2 | 1.96E-01 | -0.3 | 3.97E-01 | -0.1 | 7.65E-01 | -0.1 | 7.24E-01 | Uncharacterized protein F4F15.50 |
| AT3G52900 | -0.4 | 1.39E-02 | 0.2 | 1.74E-01 | -0.5 | 5.04E-03 | 0.2 | 3.10E-01 | 0.3 | 1.55E-01 | -0.1 | 6.83E-01 | RAB6-interacting golgin |
| AT5G11700 | -0.4 | 7.33E-04 | -0.3 | 1.53E-03 | 0.0 | 8.42E-01 | -0.1 | 5.63E-01 | 0.0 | 9.07E-01 | -0.1 | 7.87E-01 | AT5g11700 |
| AT5G11040 | -0.4 | 1.85E-04 | -0.3 | 1.25E-03 | -0.1 | 3.17E-01 | -0.4 | 1.74E-02 | -0.3 | 5.77E-02 | -0.1 | 8.40E-01 | Trafficking protein particle complex II-specific subunit 120 homolog |
| AT5G22360 | -0.4 | 1.62E-04 | -0.2 | 7.18E-03 | -0.2 | 4.85E-02 | -0.2 | 5.49E-02 | -0.3 | 3.95E-02 | 0.0 | 8.37E-01 | Vesicle-associated membrane protein 714 |
| AT1G69360 | -0.4 | 1.28E-03 | -0.2 | 4.94E-02 | -0.2 | 9.67E-02 | -0.6 | 2.73E-02 | -0.2 | 4.42E-01 | -0.4 | 2.63E-01 | Uncharacterized protein F10D13.5 |
| AT1G06060 | -0.4 | 1.04E-03 | -0.1 | 4.64E-01 | -0.3 | 9.57E-03 | -0.4 | 2.72E-02 | -0.3 | 1.15E-01 | -0.1 | 7.38E-01 | LisH and RanBPM domains containing protein |
| AT5G35730 | -0.4 | 1.85E-04 | -0.1 | 3.88E-01 | -0.3 | 2.08E-03 | -0.2 | 3.22E-01 | 0.0 | 9.68E-01 | -0.2 | 4.41E-01 | EXS family protein |
| AT4G00380 | -0.4 | 1.04E-02 | -0.3 | 2.68E-02 | -0.1 | 7.55E-01 | 0.0 | 9.29E-01 | -0.1 | 8.14E-01 | 0.1 | 7.45E-01 | XH |
| AT5G44860 | -0.4 | 2.16E-04 | -0.2 | 6.88E-03 | -0.1 | 7.55E-02 | -0.2 | 1.02E-01 | -0.2 | 3.42E-01 | -0.1 | 7.58E-01 | hypothetical protein |
| AT5G50970 | -0.4 | 5.28E-04 | -0.4 | 2.99E-04 | 0.1 | 5.16E-01 | -0.2 | 5.78E-01 | 0.3 | 4.90E-01 | -0.5 | 2.74E-01 | Uncharacterized protein |
| AT5G58005 | -0.4 | 1.52E-03 | -0.2 | 2.10E-02 | -0.1 | 2.20E-01 | -0.1 | 6.96E-01 | -0.1 | 8.14E-01 | 0.0 | 9.40E-01 | Cytochrome c oxidase, subunit Vib family protein |
| AT4G39390 | -0.4 | 4.42E-03 | -0.1 | 3.14E-01 | -0.3 | 5.20E-02 | 0.1 | 6.82E-01 | 0.2 | 2.40E-01 | -0.1 | 5.62E-01 | Nucleotide sugar transporter-KT 1 |
| AT3G23540 | -0.4 | 1.15E-04 | -0.4 | 2.39E-04 | 0.0 | 8.70E-01 | 0.1 | 6.37E-01 | 0.0 | 8.73E-01 | 0.2 | 5.86E-01 | Alpha |
| AT2G22795 | -0.4 | 1.28E-03 | 0.1 | 1.86E-01 | -0.5 | 6.65E-04 | 0.0 | 8.37E-01 | 0.3 | 1.93E-01 | -0.2 | 3.82E-01 | Transmembrane protein |
| AT3G12550 | -0.4 | 3.65E-03 | -0.1 | 1.62E-01 | -0.2 | 9.33E-02 | 0.0 | 9.31E-01 | 0.2 | 4.11E-01 | -0.3 | 3.83E-01 | XH |
| AT5G03995 | -0.4 | 2.59E-01 | -0.4 | 2.39E-01 | 0.0 | 9.33E-01 | 0.1 | 7.51E-01 | 0.3 | 4.53E-01 | -0.2 | 7.46E-01 | Uncharacterized protein |
| AT5G23400 | -0.4 | 9.72E-03 | 0.0 | 8.61E-01 | -0.4 | 1.95E-02 | 0.0 | 9.86E-01 | 0.3 | 2.69E-01 | -0.3 | 3.59E-01 | hypothetical protein |
| AT3G63290 | -0.4 | 3.79E-03 | -0.1 | 6.09E-01 | -0.3 | 2.63E-02 | -0.2 | 4.47E-01 | 0.0 | 9.81E-01 | -0.2 | 5.15E-01 | Uncharacterized protein F16M2_140 |
| AT5G48800 | -0.4 | 1.02E-02 | -0.4 | 4.79E-03 | 0.1 | 6.19E-01 | 0.1 | 7.98E-01 | 0.1 | 7.75E-01 | 0.0 | 9.59E-01 | BTB |
| AT5G41810 | -0.4 | 4.68E-03 | -0.1 | 5.67E-01 | -0.3 | 2.76E-02 | -0.2 | 2.90E-01 | 0.0 | 9.77E-01 | -0.2 | 4.73E-01 | AT5g41810 |
| AT3G09230 | -0.4 | 4.99E-03 | -0.1 | 4.57E-01 | -0.3 | 4.43E-02 | -0.3 | 4.41E-01 | 0.2 | 5.09E-01 | -0.5 | 1.83E-01 | MYB1 |
| AT3G22380 | -0.4 | 1.69E-03 | -0.4 | 7.24E-04 | 0.1 | 4.27E-01 | -0.2 | 1.74E-01 | -0.2 | 1.71E-01 | 0.0 | 9.22E-01 | Time for coffee |
| AT4G09630 | -0.4 | 1.26E-03 | -0.2 | 6.84E-02 | -0.2 | 6.64E-02 | -0.2 | 6.98E-01 | -0.1 | 8.64E-01 | -0.1 | 9.15E-01 | Transmembrane protein |
| AT2G38000 | -0.4 | 3.39E-04 | -0.2 | 3.60E-02 | -0.2 | 3.07E-02 | -0.4 | 2.11E-03 | -0.6 | 4.92E-04 | 0.2 | 4.31E-01 | AT2G38000 protein |
| AT2G46690 | -0.4 | 1.20E-03 | 0.4 | 1.72E-03 | -0.7 | 6.82E-05 | 0.1 | 7.16E-01 | 0.2 | 4.67E-01 | -0.1 | 7.68E-01 | Auxin-responsive protein SAUR32 |
| AT1G70850 | -0.4 | 1.70E-02 | 0.2 | 1.58E-01 | -0.6 | 3.59E-03 | -1.6 | 3.20E-08 | -0.5 | 5.03E-03 | -1.1 | 1.83E-04 | MLP-like protein 34 |
| AT2G33060 | -0.4 | 9.22E-03 | -0.4 | 1.12E-02 | 0.0 | 9.89E-01 | 0.1 | 7.16E-01 | -0.2 | 4.26E-01 | 0.3 | 3.09E-01 | Receptor like protein 27 |
| AT1G24300 | -0.4 | 1.54E-04 | -0.3 | 4.46E-04 | 0.0 | 6.82E-01 | 0.0 | 9.37E-01 | 0.1 | 7.52E-01 | -0.1 | 8.48E-01 | F3I6.24 protein |
| AT3G09370 | -0.4 | 2.01E-03 | -0.3 | 6.97E-03 | -0.1 | 6.21E-01 | -0.1 | 6.62E-01 | 0.0 | 8.57E-01 | 0.0 | 8.98E-01 | Myb domain protein 3r-3 |
| AT2G44200 | -0.4 | 3.45E-04 | -0.3 | 2.77E-03 | -0.1 | 2.95E-01 | 0.0 | 9.12E-01 | 0.1 | 7.90E-01 | -0.1 | 7.14E-01 | At2g44200 |
| AT5G61530 | -0.4 | 1.60E-04 | -0.4 | 6.74E-05 | 0.1 | 2.54E-01 | -0.2 | 2.07E-01 | -0.2 | 1.89E-01 | 0.0 | 9.07E-01 | Small G protein family protein |
| AT5G24710 | -0.4 | 3.16E-03 | -0.4 | 2.07E-03 | 0.0 | 6.94E-01 | -0.1 | 5.94E-01 | -0.1 | 8.28E-01 | -0.1 | 8.87E-01 | Transducin |

| | | | | | | | | | | | | | |
|---|---|---|---|---|---|---|---|---|---|---|---|---|---|
| AT3G57950 | -0.4 | 4.59E-02 | -0.3 | 1.03E-01 | -0.1 | 7.61E-01 | 0.0 | 9.32E-01 | 0.1 | 8.49E-01 | -0.1 | 9.22E-01 | Cotton fiber protein |
| AT1G17940 | -0.4 | 1.14E-02 | -0.1 | 2.56E-01 | -0.2 | 1.54E-01 | -0.3 | 1.82E-01 | 0.0 | 9.93E-01 | -0.3 | 3.33E-01 | Uncharacterized protein At1g17940 |
| AT1G69252 | -0.4 | 1.59E-02 | -0.4 | 9.92E-03 | 0.1 | 7.51E-01 | -0.8 | 8.09E-05 | -0.5 | 9.35E-03 | -0.3 | 1.87E-01 | NA |
| AT1G17680 | -0.4 | 7.95E-04 | -0.5 | 2.70E-04 | 0.1 | 3.22E-01 | -0.3 | 1.43E-01 | -0.1 | 7.65E-01 | -0.2 | 4.06E-01 | hypothetical protein |
| AT3G54300 | -0.4 | 1.46E-04 | -0.2 | 7.49E-03 | -0.2 | 4.09E-02 | -0.1 | 4.25E-01 | -0.1 | 5.09E-01 | 0.0 | 9.92E-01 | Vesicle-associated membrane protein 727 |
| AT3G14410 | -0.4 | 4.71E-04 | 0.0 | 7.21E-01 | -0.3 | 3.08E-03 | -0.3 | 6.43E-02 | 0.0 | 8.54E-01 | -0.2 | 2.49E-01 | Sugar phosphate transporter domain-containing protein |
| AT5G08630 | -0.4 | 3.12E-04 | -0.3 | 1.00E-03 | 0.0 | 6.46E-01 | 0.1 | 8.12E-01 | 0.0 | 9.78E-01 | 0.1 | 8.43E-01 | DDT domain-containing protein |
| AT3G15358 | -0.4 | 1.62E-02 | -0.3 | 7.35E-02 | -0.1 | 5.44E-01 | -0.3 | 3.79E-01 | -0.3 | 4.75E-01 | 0.0 | 9.75E-01 | Transmembrane protein |
| AT1G70340 | -0.4 | 1.56E-02 | 0.0 | 8.46E-01 | -0.3 | 5.49E-02 | -0.1 | 8.27E-01 | 0.2 | 7.28E-01 | -0.3 | 6.04E-01 | hypothetical protein |
| AT1G65010 | -0.4 | 7.49E-04 | 0.0 | 8.51E-01 | -0.3 | 2.83E-03 | -0.2 | 2.58E-01 | 0.1 | 5.16E-01 | -0.3 | 1.70E-01 | Putative WEB family protein At1g65010, chloroplastic |
| AT5G63490 | -0.4 | 1.15E-03 | -0.3 | 1.98E-03 | 0.0 | 9.12E-01 | 0.3 | 1.19E-01 | 0.0 | 9.46E-01 | 0.3 | 1.22E-01 | CBS domain-containing protein CBSCBSPB1 |
| AT2G18630 | -0.4 | 4.23E-03 | 0.0 | 9.32E-01 | -0.3 | 1.26E-02 | -0.2 | 1.93E-01 | -0.1 | 4.49E-01 | -0.1 | 8.26E-01 | Transmembrane protein, putative |
| AT4G10810 | -0.4 | 2.26E-03 | -0.1 | 1.68E-01 | -0.2 | 4.72E-02 | -0.2 | 3.00E-01 | -0.2 | 4.11E-01 | 0.0 | 9.71E-01 | Uncharacterized protein At4g10810 |
| AT1G46554 | -0.4 | 6.58E-02 | -0.4 | 3.76E-02 | 0.1 | 7.49E-01 | -0.2 | 5.13E-01 | 0.2 | 5.64E-01 | -0.4 | 3.01E-01 | NA |
| AT4G30500 | -0.4 | 5.58E-02 | 0.1 | 7.24E-01 | -0.4 | 6.36E-02 | -0.1 | 6.94E-01 | -0.1 | 6.81E-01 | 0.0 | 9.63E-01 | At4g30500 ) |
| AT1G61250 | -0.4 | 2.20E-03 | -0.2 | 4.14E-02 | -0.1 | 1.65E-01 | -0.2 | 1.29E-01 | -0.3 | 5.68E-02 | 0.1 | 6.91E-01 | Secretory carrier-associated membrane protein |
| AT5G15260 | -0.4 | 2.81E-03 | -0.3 | 8.49E-03 | -0.1 | 6.74E-01 | -0.5 | 2.06E-02 | -0.2 | 3.83E-01 | -0.3 | 3.37E-01 | Uncharacterized protein F8M21_150 |
| AT4G00110 | -0.4 | 6.97E-03 | 0.1 | 4.56E-01 | -0.4 | 6.35E-03 | -0.1 | 7.39E-01 | -0.1 | 8.31E-01 | 0.0 | 9.65E-01 | Putative nucleotide sugar epimerase |
| AT5G47480 | -0.4 | 7.41E-04 | -0.5 | 1.23E-04 | 0.1 | 1.15E-01 | 0.2 | 6.44E-01 | 0.1 | 7.91E-01 | 0.0 | 9.50E-01 | Protein transport protein sec16 |
| AT4G25710 | -0.4 | 1.14E-03 | -0.1 | 1.17E-01 | -0.2 | 3.92E-02 | 0.0 | 8.16E-01 | -0.1 | 6.84E-01 | 0.1 | 5.81E-01 | Galactose oxidase |
| AT3G30380 | -0.4 | 3.32E-04 | -0.2 | 9.37E-03 | -0.1 | 1.00E-01 | -0.3 | 2.17E-01 | 0.0 | 9.38E-01 | -0.3 | 2.50E-01 | Alpha |
| AT1G06230 | -0.4 | 4.72E-04 | -0.4 | 4.71E-04 | 0.0 | 7.88E-01 | -0.2 | 7.31E-02 | -0.2 | 1.06E-01 | 0.0 | 9.96E-01 | Transcription factor GTE4 |
| AT5G02910 | -0.4 | 1.38E-02 | -0.3 | 1.73E-02 | 0.0 | 9.88E-01 | 0.1 | 8.38E-01 | 0.0 | 9.15E-01 | 0.0 | 9.66E-01 | F-box |
| AT1G55380 | -0.4 | 3.03E-02 | 0.3 | 4.69E-02 | -0.6 | 3.87E-03 | -0.3 | 2.08E-01 | 0.0 | 9.60E-01 | -0.3 | 3.93E-01 | Cysteine |
| AT4G21215 | -0.4 | 7.62E-03 | -0.4 | 6.43E-03 | 0.0 | 8.47E-01 | -0.4 | 1.88E-02 | 0.0 | 8.62E-01 | -0.5 | 2.80E-02 | Transmembrane protein |
| AT5G64310 | -0.4 | 3.94E-02 | 0.1 | 4.11E-01 | -0.5 | 1.85E-02 | 0.0 | 8.39E-01 | 0.1 | 6.92E-01 | -0.2 | 5.95E-01 | Classical arabinogalactan protein 1 |
| AT5G07740 | -0.4 | 1.32E-03 | -0.4 | 9.43E-04 | 0.0 | 6.90E-01 | -0.1 | 3.59E-01 | -0.1 | 5.35E-01 | 0.0 | 9.30E-01 | Formin-like protein |
| AT1G36380 | -0.4 | 3.80E-02 | -0.2 | 1.32E-01 | -0.1 | 6.16E-01 | -0.1 | 7.82E-01 | 0.0 | 9.53E-01 | -0.1 | 8.85E-01 | Transmembrane protein |
| AT3G04560 | -0.4 | 1.79E-03 | -0.2 | 1.91E-02 | -0.1 | 2.82E-01 | -0.3 | 5.42E-02 | -0.2 | 1.45E-01 | 0.0 | 8.48E-01 | F7O18.3 protein |
| AT3G26090 | -0.4 | 1.15E-03 | -0.4 | 1.22E-03 | 0.0 | 8.70E-01 | 0.2 | 4.64E-01 | 0.0 | 8.76E-01 | 0.1 | 7.39E-01 | REGULATOR OF G-PROTEIN SIGNALING 1 |
| AT3G47700 | -0.4 | 4.11E-04 | -0.2 | 1.51E-02 | -0.2 | 8.43E-02 | -0.2 | 4.11E-01 | 0.2 | 2.50E-01 | -0.4 | 1.03E-01 | RINT1-like protein MAG2 |
| AT4G32970 | -0.4 | 8.08E-03 | -0.2 | 4.98E-02 | -0.1 | 4.43E-01 | -0.1 | 6.26E-01 | -0.2 | 6.12E-01 | 0.0 | 9.59E-01 | BRISC and BRCA1-A complex member 1 |
| AT5G15790 | -0.4 | 1.18E-02 | -0.2 | 2.09E-01 | -0.2 | 1.95E-01 | -0.1 | 6.64E-01 | 0.3 | 2.76E-01 | -0.4 | 1.19E-01 | At5g15790 |
| AT2G17787 | -0.3 | 2.04E-03 | -0.3 | 7.84E-03 | -0.1 | 5.98E-01 | -0.2 | 3.57E-01 | -0.4 | 1.39E-01 | 0.2 | 6.55E-01 | At2g17787 |
| AT1G23130 | -0.3 | 5.84E-02 | -0.3 | 1.20E-01 | -0.1 | 7.84E-01 | -0.1 | 7.61E-01 | 0.0 | 9.13E-01 | 0.0 | 9.29E-01 | Bet v I |
| AT2G35230 | -0.3 | 1.29E-03 | -0.5 | 1.63E-04 | 0.2 | 1.06E-01 | 0.1 | 8.71E-01 | 0.0 | 9.47E-01 | 0.1 | 8.17E-01 | VQ motif-containing protein |
| AT2G22122 | -0.3 | 2.75E-01 | -0.3 | 3.39E-01 | 0.0 | 9.46E-01 | -0.2 | 5.65E-01 | -0.5 | 1.43E-01 | 0.3 | 4.61E-01 | Uncharacterized protein |
| AT1G18940 | -0.3 | 2.54E-02 | 0.2 | 2.22E-01 | -0.5 | 9.29E-03 | -0.5 | 8.15E-02 | 0.1 | 7.00E-01 | -0.6 | 8.63E-02 | Nodulin-like domain-containing protein |
| AT1G54210 | -0.3 | 9.39E-03 | -0.4 | 3.57E-03 | 0.1 | 5.42E-01 | -0.1 | 7.61E-01 | -0.2 | 5.81E-01 | 0.1 | 8.39E-01 | Ubiquitin-like superfamily protein |
| AT5G07940 | -0.3 | 4.34E-04 | -0.4 | 1.59E-04 | 0.1 | 2.96E-01 | 0.0 | 8.88E-01 | 0.1 | 8.06E-01 | 0.0 | 9.26E-01 | Dentin sialophosphoprotein-like protein |
| AT2G10950 | -0.3 | 1.86E-04 | -0.4 | 1.91E-04 | 0.0 | 7.41E-01 | -0.1 | 4.39E-01 | -0.1 | 8.06E-01 | -0.1 | 7.56E-01 | hypothetical protein |
| AT1G27540 | -0.3 | 6.08E-04 | -0.1 | 1.58E-01 | -0.2 | 1.72E-02 | -0.2 | 2.88E-01 | -0.1 | 7.13E-01 | -0.1 | 6.73E-01 | F-box protein |
| AT2G19620 | -0.3 | 4.95E-03 | 0.0 | 8.06E-01 | -0.4 | 1.02E-02 | -0.5 | 3.40E-02 | -0.1 | 6.55E-01 | -0.4 | 1.90E-01 | N-MYC downregulated-like 3 |
| AT5G66640 | -0.3 | 2.10E-02 | -0.5 | 4.10E-03 | 0.2 | 3.63E-01 | -0.5 | 2.40E-01 | -0.5 | 2.73E-01 | 0.0 | 9.85E-01 | DA1-related protein 3 |
| AT3G14870 | -0.3 | 3.38E-02 | -0.4 | 2.05E-02 | 0.1 | 7.56E-01 | -0.3 | 2.33E-01 | -0.1 | 5.91E-01 | -0.1 | 7.02E-01 | Uncharacterized protein At3g14870 |
| AT4G14980 | -0.3 | 3.71E-02 | 0.3 | 4.09E-02 | -0.7 | 3.93E-03 | -0.2 | 5.03E-01 | 0.1 | 7.78E-01 | -0.4 | 3.80E-01 | At4g14980 |
| AT5G12050 | -0.3 | 7.63E-02 | 0.6 | 7.66E-03 | -0.9 | 1.78E-03 | -0.1 | 6.93E-01 | 0.0 | 9.13E-01 | -0.1 | 8.67E-01 | Protein BIG GRAIN 1-like D |
| AT2G23810 | -0.3 | 6.09E-03 | -0.4 | 2.81E-03 | 0.1 | 5.57E-01 | 0.0 | 7.96E-01 | -0.3 | 1.51E-01 | 0.2 | 3.17E-01 | Tetraspanin-8 |
| AT1G02960 | -0.3 | 1.76E-03 | -0.2 | 1.69E-02 | -0.1 | 3.08E-01 | -0.5 | 6.36E-03 | -0.1 | 5.36E-01 | -0.4 | 7.77E-02 | Kinetochore protein |
| AT4G24290 | -0.3 | 1.32E-04 | -0.4 | 1.02E-04 | 0.0 | 5.23E-01 | 0.0 | 9.06E-01 | 0.0 | 9.75E-01 | 0.0 | 9.65E-01 | MACPF domain-containing protein At4g24290 |
| AT1G79710 | -0.3 | 1.45E-03 | -0.4 | 4.33E-04 | 0.1 | 3.29E-01 | -0.2 | 5.50E-01 | -0.1 | 9.20E-01 | -0.2 | 6.91E-01 | Major facilitator superfamily protein |
| AT2G07360 | -0.3 | 2.34E-04 | -0.2 | 4.74E-03 | -0.1 | 1.13E-01 | -0.1 | 3.97E-01 | -0.2 | 1.62E-01 | 0.1 | 6.74E-01 | SH3 domain-containing protein |
| AT5G52880 | -0.3 | 1.63E-03 | -0.2 | 8.65E-02 | -0.2 | 7.73E-02 | -0.2 | 4.72E-01 | 0.2 | 3.66E-01 | -0.4 | 1.82E-01 | AT5G52880 protein |
| AT1G35220 | -0.3 | 2.43E-03 | -0.3 | 3.18E-03 | 0.0 | 9.88E-01 | -0.2 | 3.77E-01 | 0.0 | 8.86E-01 | -0.2 | 6.45E-01 | At1g35220 |
| AT1G05360 | -0.3 | 9.57E-03 | -0.2 | 1.31E-01 | -0.2 | 2.41E-01 | -0.1 | 6.20E-01 | 0.2 | 3.39E-01 | -0.3 | 1.81E-01 | AT1G05360 protein |
| AT1G12930 | -0.3 | 1.40E-03 | -0.3 | 3.72E-03 | 0.0 | 7.27E-01 | -0.1 | 6.32E-01 | -0.2 | 5.11E-01 | 0.1 | 8.73E-01 | Exportin-1 |
| AT1G54200 | -0.3 | 3.94E-03 | -0.1 | 3.10E-01 | -0.2 | 5.65E-02 | 0.2 | 5.55E-01 | 0.2 | 6.01E-01 | 0.0 | 9.92E-01 | Protein BIG GRAIN 1-like B |
| AT5G18520 | -0.3 | 3.97E-04 | -0.1 | 1.13E-01 | -0.2 | 1.15E-02 | 0.0 | 7.89E-01 | 0.1 | 4.49E-01 | -0.1 | 7.11E-01 | hypothetical protein |
| AT2G27140 | -0.3 | 2.56E-03 | -0.2 | 2.03E-02 | -0.1 | 3.61E-01 | 0.0 | 9.46E-01 | -0.1 | 7.86E-01 | 0.1 | 7.56E-01 | At2g27140 |
| AT5G20540 | -0.3 | 1.75E-03 | -0.2 | 1.95E-02 | -0.1 | 2.75E-01 | 0.1 | 3.77E-01 | 0.1 | 4.36E-01 | 0.0 | 9.96E-01 | BRXL4 |
| AT3G04350 | -0.3 | 1.38E-03 | -0.2 | 3.43E-02 | -0.1 | 1.30E-01 | -0.4 | 4.57E-03 | -0.4 | 1.53E-02 | 0.0 | 8.87E-01 | Vacuolar sorting-associated protein |
| AT5G62770 | -0.3 | 7.91E-03 | -0.1 | 3.02E-01 | -0.2 | 1.00E-01 | 0.8 | 1.20E-03 | 0.3 | 2.20E-01 | 0.5 | 8.36E-02 | Uncharacterized protein |
| AT2G43160 | -0.3 | 6.66E-03 | -0.4 | 4.61E-03 | 0.0 | 7.29E-01 | 0.0 | 9.21E-01 | -0.3 | 4.17E-01 | 0.2 | 5.82E-01 | ENTH domain-containing protein |
| AT5G65770 | -0.3 | 2.36E-03 | -0.4 | 1.32E-03 | 0.1 | 6.10E-01 | 0.1 | 8.52E-01 | 0.1 | 8.19E-01 | 0.0 | 9.58E-01 | Protein CROWDED NUCLEI 4 |
| AT4G10710 | -0.3 | 1.18E-04 | -0.4 | 1.11E-04 | 0.0 | 6.33E-01 | -0.1 | 1.66E-01 | -0.2 | 1.67E-01 | 0.0 | 9.31E-01 | FACT complex subunit |
| AT4G17940 | -0.3 | 2.79E-03 | 0.0 | 7.02E-01 | -0.3 | 1.34E-02 | 0.1 | 5.20E-01 | -0.1 | 6.45E-01 | 0.2 | 3.18E-01 | Tetratricopeptide repeat -like superfamily protein |
| AT1G76590 | -0.3 | 5.11E-02 | -0.5 | 1.73E-02 | 0.1 | 5.61E-01 | -0.8 | 7.08E-03 | -0.9 | 3.57E-03 | 0.2 | 7.21E-01 | PLATZ transcription factor family protein |
| AT3G01560 | -0.3 | 1.32E-03 | -0.4 | 9.77E-04 | 0.0 | 7.09E-01 | -0.1 | 4.05E-01 | -0.4 | 4.72E-02 | 0.2 | 3.44E-01 | F4P13.11 protein |
| AT1G14560 | -0.3 | 8.97E-04 | -0.2 | 1.53E-02 | -0.1 | 1.79E-01 | -0.2 | 4.11E-01 | -0.1 | 7.57E-01 | -0.1 | 7.99E-01 | Mitochondrial substrate carrier family protein |
| AT5G16210 | -0.3 | 7.97E-04 | -0.4 | 3.69E-04 | 0.1 | 4.44E-01 | -0.2 | 3.68E-01 | 0.0 | 9.32E-01 | -0.3 | 4.55E-01 | AT5g16210 |
| AT4G08980 | -0.3 | 3.83E-04 | -0.3 | 7.50E-04 | 0.0 | 8.81E-01 | -0.3 | 6.26E-02 | -0.2 | 3.36E-01 | -0.1 | 6.74E-01 | F-box protein FBW2 |
| AT3G29034 | -0.3 | 6.03E-02 | 0.0 | 7.92E-01 | -0.4 | 6.32E-02 | 0.1 | 5.73E-01 | 0.1 | 7.67E-01 | 0.0 | 9.12E-01 | Transmembrane protein |
| AT1G76185 | -0.3 | 5.00E-03 | -0.2 | 3.39E-02 | -0.1 | 4.14E-01 | -0.1 | 7.74E-01 | 0.1 | 7.55E-01 | -0.3 | 5.97E-01 | hypothetical protein |
| AT1G68790 | -0.3 | 2.64E-04 | -0.3 | 1.56E-03 | -0.1 | 3.67E-01 | 0.1 | 5.97E-01 | 0.3 | 2.53E-01 | -0.2 | 6.21E-01 | Protein CROWDED NUCLEI 3 |
| AT3G60850 | -0.3 | 1.64E-03 | -0.3 | 2.36E-03 | 0.0 | 9.88E-01 | -0.3 | 1.31E-01 | -0.4 | 6.11E-02 | 0.1 | 7.25E-01 | At3g60850 |
| AT3G62860 | -0.3 | 9.17E-03 | 0.0 | 8.00E-01 | -0.3 | 3.71E-02 | -0.5 | 1.36E-02 | -0.4 | 4.49E-02 | -0.1 | 8.25E-01 | AT3g62860 |
| AT3G54200 | -0.3 | 1.05E-02 | -0.2 | 6.02E-02 | -0.1 | 4.43E-01 | -0.4 | 1.16E-01 | -0.3 | 3.81E-01 | -0.2 | 7.13E-01 | Late embryogenesis abundant hydroxyproline-rich glycoprotein family |
| AT3G04605 | -0.3 | 3.13E-04 | -0.3 | 1.31E-03 | -0.1 | 5.17E-01 | -0.1 | 7.73E-01 | 0.0 | 8.92E-01 | 0.0 | 9.42E-01 | NA |
| AT2G39725 | -0.3 | 4.93E-02 | -0.2 | 1.89E-01 | -0.1 | 5.69E-01 | -0.3 | 4.31E-01 | -0.2 | 7.17E-01 | -0.1 | 8.38E-01 | hypothetical protein |
| AT3G54366 | -0.3 | 2.51E-03 | 0.1 | 4.72E-01 | -0.4 | 2.49E-03 | -0.1 | 6.32E-01 | -0.1 | 6.59E-01 | 0.0 | 9.78E-01 | NA |
| AT1G47340 | -0.3 | 1.22E-02 | -0.2 | 1.73E-01 | -0.2 | 2.41E-01 | 0.0 | 9.68E-01 | 0.1 | 8.02E-01 | -0.1 | 8.48E-01 | F-box and associated interaction domains-containing protein |
| AT3G09020 | -0.3 | 8.04E-03 | 0.0 | 8.17E-01 | -0.3 | 2.94E-02 | 0.1 | 8.36E-01 | -0.4 | 1.16E-01 | 0.5 | 1.40E-01 | hypothetical protein |
| AT1G62970 | -0.3 | 4.14E-03 | -0.2 | 1.01E-01 | -0.2 | 1.53E-01 | -0.2 | 3.35E-01 | -0.3 | 2.25E-01 | 0.1 | 8.27E-01 | F16P17.12 protein |

| | | | | | | | | | | | | | |
|---|---|---|---|---|---|---|---|---|---|---|---|---|---|
| AT4G04900 | -0.3 | 1.11E-01 | -0.3 | 1.98E-01 | -0.1 | 8.23E-01 | 0.1 | 9.33E-01 | 0.2 | 8.01E-01 | -0.1 | 8.73E-01 | CRIB domain-containing protein RIC10 |
| AT1G28520 | -0.3 | 8.42E-04 | -0.5 | 1.87E-04 | 0.1 | 1.90E-01 | -0.3 | 2.54E-02 | -0.4 | 1.74E-02 | 0.1 | 8.24E-01 | Vascular plant one zinc finger protein |
| AT3G11880 | -0.3 | 1.73E-03 | -0.4 | 5.69E-04 | 0.1 | 3.68E-01 | 0.0 | 7.61E-01 | 0.0 | 9.30E-01 | 0.0 | 9.08E-01 | F26K24.17 protein |
| AT5G20610 | -0.3 | 4.78E-04 | -0.3 | 3.06E-03 | -0.1 | 3.57E-01 | -0.1 | 5.12E-01 | -0.2 | 1.49E-01 | 0.1 | 5.37E-01 | Protein PLASTID MOVEMENT IMPAIRED 1-RELATED 1 |
| AT1G07323 | -0.3 | 3.11E-02 | 0.2 | 1.78E-01 | -0.5 | 8.80E-03 | 0.2 | 2.99E-01 | 0.0 | 9.04E-01 | 0.2 | 5.59E-01 | NA |
| AT2G27660 | -0.3 | 1.73E-02 | -0.2 | 1.16E-01 | -0.1 | 3.77E-01 | 0.5 | 4.47E-03 | 0.4 | 5.01E-02 | 0.1 | 5.72E-01 | Cysteine |
| AT4G01395 | -0.3 | 2.64E-04 | -0.1 | 7.45E-02 | -0.2 | 1.05E-02 | -0.2 | 1.36E-01 | -0.1 | 5.92E-01 | -0.1 | 6.23E-01 | A_IG002N01.1 protein |
| AT4G16957 | -0.3 | 2.02E-03 | -0.4 | 6.78E-04 | 0.1 | 3.89E-01 | 0.3 | 1.84E-01 | 0.2 | 4.61E-01 | 0.1 | 7.80E-01 | SUPPRESSOR OF, CONSTITUTIVE protein |
| AT1G09980 | -0.3 | 9.91E-04 | -0.3 | 5.38E-03 | -0.1 | 4.36E-01 | 0.0 | 9.31E-01 | 0.2 | 3.32E-01 | -0.2 | 3.40E-01 | Serine esterase family protein |
| AT1G70420 | -0.3 | 1.48E-01 | 0.0 | 9.79E-01 | -0.3 | 2.28E-01 | 0.8 | 1.17E-03 | 0.1 | 8.73E-01 | 0.8 | 3.76E-03 | Uncharacterized protein |
| AT1G70950 | -0.3 | 2.81E-02 | 0.0 | 8.96E-01 | -0.3 | 7.60E-02 | 0.1 | 6.42E-01 | 0.1 | 6.62E-01 | 0.0 | 9.80E-01 | Protein WVD2-like 7 |
| AT3G04780 | -0.3 | 2.97E-03 | 0.0 | 7.94E-01 | -0.3 | 1.14E-02 | 0.1 | 7.49E-01 | 0.0 | 9.98E-01 | 0.1 | 8.25E-01 | PITH domain-containing protein At3g04780 |
| AT1G75730 | -0.3 | 3.39E-03 | -0.3 | 6.97E-03 | 0.0 | 8.26E-01 | -0.2 | 2.81E-01 | -0.1 | 6.73E-01 | -0.1 | 7.31E-01 | At1g75730 |
| AT2G04305 | -0.3 | 6.16E-04 | -0.1 | 5.40E-02 | -0.2 | 3.94E-02 | -0.4 | 2.86E-02 | -0.3 | 7.79E-02 | 0.0 | 8.69E-01 | Uncharacterized protein At2g04305 |
| AT1G23950 | -0.3 | 4.75E-04 | -0.3 | 1.48E-03 | 0.0 | 6.38E-01 | 0.0 | 9.27E-01 | 0.0 | 9.64E-01 | 0.0 | 9.12E-01 | UPF0725 protein At1g23950 |
| AT2G01505 | -0.3 | 5.25E-02 | 0.0 | 7.68E-01 | -0.4 | 6.36E-02 | -0.2 | 3.61E-01 | 0.1 | 7.54E-01 | -0.3 | 2.77E-01 | CLAVATA3 |
| AT4G12710 | -0.3 | 3.21E-03 | -0.2 | 1.10E-01 | -0.2 | 1.15E-01 | -0.4 | 2.84E-02 | -0.5 | 6.39E-03 | 0.2 | 4.71E-01 | ARM repeat superfamily protein |
| AT1G02380 | -0.3 | 1.24E-01 | 0.3 | 1.51E-01 | -0.6 | 2.57E-02 | -0.6 | 1.61E-01 | -0.3 | 5.61E-01 | -0.3 | 5.99E-01 | At1g02380 |
| AT5G25240 | -0.3 | 1.75E-01 | -0.1 | 8.10E-01 | -0.3 | 3.66E-01 | -0.9 | 1.71E-02 | -1.0 | 1.63E-02 | 0.1 | 9.16E-01 | Stress induced protein |
| AT2G26920 | -0.3 | 4.01E-03 | -0.5 | 7.88E-04 | 0.1 | 2.58E-01 | -0.2 | 5.12E-01 | 0.0 | 9.56E-01 | -0.2 | 7.04E-01 | hypothetical protein |
| AT3G60415 | -0.3 | 2.04E-01 | -1.0 | 4.04E-03 | 0.7 | 5.14E-02 | -0.1 | 5.47E-01 | -0.3 | 2.96E-01 | 0.1 | 7.15E-01 | Phosphoglycerate mutase family protein |
| AT1G23890 | -0.3 | 1.74E-03 | -0.2 | 4.12E-02 | -0.1 | 1.51E-01 | -0.2 | 2.47E-01 | 0.0 | 9.50E-01 | -0.2 | 4.28E-01 | At1g23890 |
| AT3G52240 | -0.3 | 3.18E-04 | -0.4 | 7.39E-05 | 0.1 | 1.40E-01 | -0.2 | 4.77E-02 | -0.5 | 4.92E-04 | 0.3 | 6.34E-02 | Uncharacterized protein |
| AT5G16780 | -0.3 | 1.88E-03 | -0.1 | 1.54E-01 | -0.2 | 4.23E-02 | -0.1 | 6.20E-01 | -0.2 | 1.30E-01 | 0.2 | 4.21E-01 | SART-1 family protein DOT2 |
| AT3G11690 | -0.3 | 5.65E-04 | 0.0 | 8.68E-01 | -0.3 | 1.59E-03 | -0.2 | 2.76E-01 | -0.3 | 2.23E-01 | 0.1 | 8.59E-01 | hypothetical protein |
| AT4G32160 | -0.3 | 9.23E-04 | -0.3 | 1.55E-03 | 0.0 | 9.26E-01 | -0.1 | 6.96E-01 | -0.1 | 6.72E-01 | 0.0 | 9.51E-01 | Uncharacterized protein F10M6.200 |
| AT5G16220 | -0.3 | 1.17E-03 | -0.2 | 8.58E-03 | -0.1 | 3.54E-01 | -0.2 | 4.67E-01 | -0.2 | 5.03E-01 | 0.0 | 9.81E-01 | hypothetical protein |
| AT4G28260 | -0.3 | 1.08E-03 | -0.2 | 4.54E-02 | -0.2 | 7.34E-02 | -0.6 | 4.29E-04 | -0.4 | 8.89E-03 | -0.1 | 4.83E-01 | hypothetical protein |
| AT1G11060 | -0.3 | 4.54E-04 | -0.4 | 4.63E-04 | 0.0 | 7.98E-01 | -0.2 | 3.55E-01 | -0.2 | 2.36E-01 | 0.1 | 7.62E-01 | Wings apart-like protein C-terminal domain-containing protein |
| AT1G72690 | -0.3 | 2.51E-01 | -0.3 | 3.30E-01 | 0.0 | 9.22E-01 | -0.5 | 3.20E-01 | 0.1 | 8.04E-01 | -0.6 | 2.87E-01 | hypothetical protein |
| AT3G62450 | -0.3 | 2.24E-03 | -0.2 | 3.70E-02 | -0.1 | 2.00E-01 | -0.1 | 6.96E-01 | -0.2 | 2.08E-01 | 0.2 | 4.98E-01 | hypothetical protein |
| AT4G27040 | -0.3 | 4.06E-04 | -0.3 | 1.76E-03 | -0.1 | 5.04E-01 | -0.4 | 1.20E-02 | -0.2 | 3.21E-01 | -0.2 | 2.44E-01 | Vacuolar protein sorting-associated protein 22 homolog 1 |
| AT5G06910 | -0.3 | 4.83E-03 | -0.2 | 6.05E-02 | -0.1 | 2.60E-01 | 0.2 | 3.34E-01 | 0.1 | 6.00E-01 | 0.1 | 8.42E-01 | Chaperone protein dnaJ 6 |
| AT1G45120 | -0.3 | 4.10E-02 | -0.2 | 1.75E-01 | -0.1 | 5.39E-01 | -0.1 | 9.10E-01 | 0.2 | 7.95E-01 | -0.2 | 7.39E-01 | NA |
| AT4G08540 | -0.3 | 9.66E-04 | -0.6 | 3.70E-05 | 0.3 | 1.47E-02 | -0.1 | 7.22E-01 | 0.1 | 7.13E-01 | -0.2 | 4.27E-01 | AT4G08540 protein |
| AT3G58980 | -0.3 | 2.65E-02 | 0.1 | 5.56E-01 | -0.4 | 2.48E-02 | -0.3 | 3.68E-01 | 0.1 | 9.17E-01 | -0.4 | 3.85E-01 | F-box domain-containing protein |
| AT2G17705 | -0.3 | 2.04E-03 | -0.2 | 4.04E-02 | -0.1 | 1.74E-01 | 0.2 | 4.56E-01 | -0.2 | 3.67E-01 | 0.4 | 1.36E-01 | MsrB domain-containing protein |
| AT4G39420 | -0.3 | 1.78E-03 | -0.3 | 2.51E-03 | 0.0 | 9.81E-01 | -0.2 | 2.03E-01 | 0.0 | 8.94E-01 | -0.2 | 4.03E-01 | Uncharacterized protein AT4g39420 |
| AT2G39700 | -0.3 | 2.43E-03 | -0.1 | 2.17E-01 | -0.2 | 3.91E-02 | 0.0 | 7.91E-01 | 0.3 | 1.21E-02 | -0.3 | 7.19E-02 | Expansin-A4 |
| AT4G32050 | -0.3 | 1.22E-03 | -0.2 | 5.03E-02 | -0.2 | 9.30E-02 | -0.2 | 2.81E-01 | -0.3 | 1.11E-01 | 0.1 | 6.51E-01 | Uncharacterized protein AT4g32050 |
| AT5G54520 | -0.3 | 5.72E-03 | -0.1 | 2.73E-01 | -0.2 | 9.05E-02 | -0.2 | 4.80E-01 | -0.2 | 6.08E-01 | 0.0 | 9.53E-01 | Uncharacterized protein At5g54520 |
| AT3G52490 | -0.3 | 9.91E-03 | -0.3 | 1.25E-02 | 0.0 | 9.90E-01 | -0.2 | 4.99E-01 | -0.1 | 9.03E-01 | -0.1 | 7.11E-01 | Protein SMAX1-LIKE 3 |
| AT4G27990 | -0.3 | 1.12E-03 | -0.4 | 5.30E-04 | 0.1 | 4.77E-01 | -0.3 | 1.59E-02 | -0.1 | 3.94E-01 | -0.2 | 3.18E-01 | YlmG homolog protein 1-2, chloroplastic |
| AT2G45380 | -0.3 | 3.83E-03 | -0.4 | 2.17E-03 | 0.1 | 6.49E-01 | -0.4 | 4.20E-03 | -0.2 | 1.88E-01 | -0.2 | 2.21E-01 | Myeloid leukemia factor |
| AT5G24450 | -0.3 | 1.63E-03 | -0.4 | 5.16E-04 | 0.1 | 3.47E-01 | -0.3 | 1.39E-02 | -0.2 | 1.32E-01 | -0.1 | 5.64E-01 | Uncharacterized protein At5g24450 |
| AT3G14560 | -0.3 | 1.47E-01 | -0.6 | 2.60E-02 | 0.3 | 3.66E-01 | 0.2 | 6.26E-01 | 0.5 | 1.59E-01 | -0.3 | 3.59E-01 | Uncharacterized protein |
| AT2G01400 | -0.3 | 3.17E-02 | -0.3 | 6.22E-02 | 0.0 | 8.37E-01 | -0.2 | 5.29E-01 | -0.2 | 5.91E-01 | 0.0 | 9.97E-01 | Uncharacterized protein |
| AT5G27650 | -0.3 | 1.75E-03 | -0.3 | 6.95E-03 | -0.1 | 5.58E-01 | -0.2 | 4.24E-01 | -0.1 | 8.74E-01 | -0.1 | 6.89E-01 | PWWP domain-containing protein 1 |
| AT4G02680 | -0.3 | 7.38E-04 | -0.3 | 1.23E-03 | 0.0 | 9.39E-01 | 0.0 | 8.92E-01 | 0.1 | 7.51E-01 | 0.0 | 8.86E-01 | ETO1-like protein 1 |
| AT4G24550 | -0.3 | 2.82E-04 | -0.2 | 1.73E-02 | -0.2 | 4.05E-02 | -0.2 | 5.46E-02 | -0.2 | 2.43E-01 | -0.1 | 7.40E-01 | Clathrin adaptor complexes medium subunit family protein |
| AT3G27530 | -0.3 | 2.95E-04 | -0.2 | 2.83E-03 | -0.1 | 2.36E-01 | -0.3 | 1.91E-02 | -0.2 | 1.69E-01 | -0.1 | 5.78E-01 | Golgin candidate 6 |
| AT5G41950 | -0.3 | 8.90E-04 | -0.2 | 5.68E-02 | -0.2 | 4.89E-02 | -0.2 | 1.85E-01 | -0.1 | 4.00E-01 | 0.0 | 8.60E-01 | Protein HLB1 |
| AT2G25730 | -0.3 | 1.03E-03 | -0.2 | 2.18E-02 | -0.1 | 1.44E-01 | 0.1 | 7.09E-01 | 0.0 | 8.46E-01 | 0.0 | 9.46E-01 | Uncharacterized protein At2g25730 |
| AT1G56590 | -0.3 | 3.74E-03 | 0.0 | 9.85E-01 | -0.3 | 9.91E-03 | -0.2 | 3.42E-01 | -0.1 | 7.61E-01 | -0.1 | 7.20E-01 | Clathrin-associated protein, putative |
| AT1G27690 | -0.3 | 6.54E-03 | -0.2 | 3.21E-02 | -0.1 | 5.11E-01 | -0.3 | 1.20E-01 | 0.0 | 9.37E-01 | -0.2 | 2.85E-01 | hypothetical protein |
| AT5G40340 | -0.3 | 6.23E-03 | -0.1 | 1.99E-01 | -0.2 | 1.02E-01 | -0.3 | 1.21E-01 | -0.3 | 2.43E-01 | 0.0 | 9.21E-01 | Tudor |
| AT1G14570 | -0.3 | 2.30E-04 | -0.2 | 9.45E-03 | -0.1 | 5.57E-02 | -0.2 | 1.75E-01 | -0.2 | 3.21E-01 | 0.0 | 9.12E-01 | UBX domain-containing protein |
| AT4G34100 | -0.3 | 1.21E-03 | -0.3 | 1.22E-03 | 0.0 | 8.17E-01 | -0.4 | 9.19E-04 | -0.3 | 1.11E-02 | -0.1 | 6.24E-01 | Uncharacterized protein At4g34100 |
| AT5G58550 | -0.3 | 8.57E-02 | -0.2 | 2.25E-01 | -0.1 | 6.81E-01 | -0.1 | 8.30E-01 | -0.3 | 5.76E-01 | 0.2 | 7.84E-01 | ETO1-like protein 2 |
| AT3G16740 | -0.3 | 2.23E-03 | -0.3 | 5.37E-03 | 0.0 | 7.65E-01 | -0.2 | 1.47E-01 | -0.3 | 1.03E-01 | 0.1 | 8.45E-01 | F-box |
| AT3G09085 | -0.3 | 2.21E-02 | -0.2 | 9.75E-02 | -0.1 | 5.46E-01 | -0.5 | 2.51E-01 | -0.5 | 3.20E-01 | 0.0 | 9.82E-01 | hypothetical protein |
| AT5G11810 | -0.3 | 1.76E-03 | 0.0 | 7.04E-01 | -0.3 | 8.79E-03 | 0.0 | 9.26E-01 | 0.1 | 6.40E-01 | -0.1 | 7.76E-01 | Uncharacterized protein |
| AT3G16690 | -0.3 | 3.38E-03 | 0.6 | 8.16E-05 | -0.9 | 3.33E-05 | -0.8 | 8.78E-04 | -0.5 | 3.69E-02 | -0.3 | 2.08E-01 | Bidirectional sugar transporter SWEET |
| AT1G22300 | -0.3 | 1.12E-03 | 0.0 | 7.09E-01 | -0.3 | 1.86E-03 | 0.2 | 1.99E-01 | 0.2 | 3.49E-01 | 0.0 | 9.28E-01 | General regulatory factor 10 |
| AT3G59910 | -0.3 | 3.25E-04 | -0.5 | 1.76E-05 | 0.2 | 1.11E-02 | -0.2 | 3.07E-01 | 0.1 | 7.72E-01 | -0.2 | 3.00E-01 | Ankyrin repeat protein SKIP35 |
| AT1G76850 | -0.3 | 1.38E-03 | -0.3 | 2.09E-03 | 0.0 | 9.57E-01 | -0.2 | 4.77E-01 | -0.1 | 8.49E-01 | -0.1 | 7.63E-01 | Exocyst complex component SEC5 |
| AT1G58235 | -0.3 | 1.96E-03 | -0.1 | 2.49E-01 | -0.2 | 3.06E-02 | -0.4 | 4.89E-02 | -0.3 | 2.41E-01 | -0.1 | 6.71E-01 | At1g58243 |
| AT1G76920 | -0.3 | 3.02E-04 | -0.3 | 6.25E-04 | 0.0 | 8.49E-01 | -0.5 | 9.36E-03 | -0.5 | 1.90E-02 | 0.0 | 9.78E-01 | SKP1-interacting partner 15 |
| AT1G01830 | -0.3 | 1.32E-03 | -0.3 | 6.74E-03 | -0.1 | 4.53E-01 | -0.5 | 2.27E-03 | -0.2 | 3.42E-01 | -0.3 | 7.24E-02 | T1N6.25 protein |
| AT5G38820 | -0.3 | 7.45E-02 | -0.9 | 1.06E-03 | 0.6 | 3.28E-02 | -1.3 | 2.37E-02 | -1.4 | 2.38E-02 | 0.1 | 9.19E-01 | hypothetical protein |
| AT1G27150 | -0.3 | 2.85E-03 | -0.2 | 4.37E-02 | -0.1 | 2.09E-01 | -0.2 | 1.62E-01 | -0.2 | 1.03E-01 | 0.1 | 8.09E-01 | Tetratricopeptide repeat protein 38 |
| AT2G33120 | -0.3 | 4.09E-04 | -0.2 | 2.19E-02 | -0.1 | 5.04E-02 | 0.1 | 4.41E-01 | -0.1 | 5.68E-01 | 0.2 | 3.13E-01 | Synaptobrevin-related protein 1 |
| AT4G03410 | -0.3 | 2.61E-03 | -0.2 | 1.22E-02 | -0.1 | 5.17E-01 | 0.0 | 9.61E-01 | 0.1 | 8.37E-01 | -0.1 | 8.88E-01 | Uncharacterized protein AT4g03410 |
| AT4G39190 | -0.3 | 6.79E-02 | -0.5 | 1.01E-02 | 0.2 | 3.14E-01 | 0.0 | 9.92E-01 | 0.3 | 1.79E-01 | -0.3 | 2.17E-01 | Nucleolar-like protein |
| AT2G39940 | -0.3 | 3.85E-04 | -0.2 | 2.56E-03 | -0.1 | 3.38E-01 | -0.3 | 2.72E-02 | -0.3 | 1.53E-02 | 0.1 | 7.47E-01 | Coronatine-insensitive protein 1 |
| AT1G71110 | -0.3 | 2.17E-03 | -0.2 | 1.19E-02 | -0.1 | 4.72E-01 | 0.1 | 5.50E-01 | 0.2 | 3.71E-01 | -0.1 | 7.89E-01 | Transmembrane protein |
| AT1G03950 | -0.3 | 5.10E-02 | -0.3 | 1.11E-01 | -0.1 | 7.59E-01 | -0.2 | 4.43E-01 | 0.0 | 8.95E-01 | -0.2 | 4.41E-01 | Vacuolar protein sorting-associated protein 2 homolog 3 |
| AT1G49430 | -0.3 | 1.74E-02 | -0.3 | 1.56E-02 | 0.0 | 9.00E-01 | 0.7 | 5.12E-04 | 0.5 | 1.81E-02 | 0.2 | 3.60E-01 | Long-chain acyl-CoA synthetase 2 |
| AT4G29160 | -0.3 | 4.77E-04 | -0.3 | 1.45E-03 | 0.0 | 6.74E-01 | -0.2 | 5.61E-02 | 0.0 | 9.61E-01 | -0.2 | 1.80E-01 | Vacuolar protein sorting-associated protein 32 homolog 2 |
| AT3G14060 | -0.3 | 5.04E-02 | -0.7 | 1.57E-03 | 0.4 | 7.50E-02 | -0.7 | 5.50E-02 | -0.7 | 8.04E-02 | 0.0 | 9.83E-01 | hypothetical protein |
| AT4G33210 | -0.3 | 1.46E-03 | -0.4 | 3.92E-04 | 0.1 | 2.79E-01 | -0.2 | 2.40E-01 | -0.2 | 2.57E-01 | 0.0 | 9.57E-01 | F-box |
| AT3G50460 | -0.3 | 5.86E-02 | 0.0 | 9.42E-01 | -0.3 | 1.25E-01 | 0.4 | 2.22E-01 | 0.4 | 2.41E-01 | 0.0 | 9.57E-01 | RPW8-like protein 2 |
| AT1G21080 | -0.3 | 4.71E-04 | -0.4 | 2.21E-04 | 0.1 | 4.08E-01 | -0.3 | 4.92E-02 | -0.1 | 7.19E-01 | -0.2 | 2.63E-01 | At1g21080 |

| | | | | | | | | | | | | | |
|---|---|---|---|---|---|---|---|---|---|---|---|---|---|
| AT3G14910 | -0.3 | 9.84E-03 | -0.2 | 3.66E-02 | -0.1 | 5.91E-01 | -0.1 | 6.47E-01 | -0.1 | 7.55E-01 | 0.0 | 9.71E-01 | Rab3-GAP regulatory subunit N-terminal domain-containing protein |
| AT1G19180 | -0.3 | 2.79E-02 | -0.5 | 5.32E-03 | 0.1 | 3.50E-01 | 0.2 | 3.84E-01 | 0.3 | 3.80E-01 | 0.0 | 9.49E-01 | Protein TIFY |
| AT4G13750 | -0.3 | 3.79E-03 | -0.4 | 6.59E-04 | 0.1 | 2.37E-01 | 0.7 | 4.60E-02 | 0.5 | 1.92E-01 | 0.2 | 6.91E-01 | Protein NO VEIN |
| AT5G38380 | -0.3 | 1.09E-03 | 0.0 | 5.40E-01 | -0.3 | 8.85E-03 | 0.0 | 1.00E+00 | -0.3 | 1.01E-01 | 0.3 | 1.67E-01 | Uncharacterized protein At5g38380 |
| AT4G19900 | -0.3 | 3.87E-03 | -0.3 | 1.27E-02 | -0.1 | 6.46E-01 | -0.3 | 1.55E-01 | 0.0 | 9.81E-01 | -0.3 | 2.55E-01 | Alpha 1,4-glycosyltransferase family protein |
| AT1G17780 | -0.3 | 2.76E-03 | -0.4 | 1.23E-03 | 0.1 | 5.18E-01 | -0.1 | 3.97E-01 | -0.4 | 1.85E-02 | 0.3 | 1.98E-01 | Transmembrane protein |
| AT3G48120 | -0.3 | 3.61E-02 | -0.4 | 1.34E-02 | 0.1 | 6.04E-01 | -0.2 | 2.14E-01 | -0.4 | 8.48E-02 | 0.1 | 6.57E-01 | Uncharacterized protein T17F15.10 |
| AT1G67140 | -0.3 | 3.42E-04 | -0.2 | 3.94E-03 | -0.1 | 2.10E-01 | -0.2 | 4.22E-01 | 0.1 | 7.78E-01 | -0.2 | 3.87E-01 | Protein SWEETIE |
| AT5G58740 | -0.3 | 1.38E-03 | -0.2 | 2.68E-02 | -0.1 | 1.50E-01 | -0.2 | 7.93E-02 | 0.0 | 9.47E-01 | -0.2 | 2.28E-01 | NudC domain-containing protein 2 |
| AT5G20165 | -0.3 | 6.81E-03 | -0.1 | 4.85E-01 | -0.2 | 4.69E-02 | -0.2 | 3.60E-01 | -0.2 | 3.75E-01 | 0.0 | 9.62E-01 | Protein kish |
| AT2G41520 | -0.3 | 7.45E-04 | -0.3 | 3.16E-03 | -0.1 | 5.33E-01 | -0.1 | 7.31E-01 | 0.0 | 9.58E-01 | -0.1 | 7.66E-01 | Uncharacterized protein At2g41520 |
| AT3G61710 | -0.3 | 2.48E-03 | -0.4 | 1.38E-03 | 0.1 | 6.11E-01 | -0.2 | 3.89E-01 | 0.1 | 6.90E-01 | -0.3 | 2.46E-01 | AUTOPHAGY 6 |
| AT1G69060 | -0.3 | 2.17E-03 | -0.5 | 2.89E-04 | 0.1 | 1.45E-01 | -0.7 | 1.53E-02 | -0.4 | 1.23E-01 | -0.2 | 5.76E-01 | F4N2.3 |
| AT1G68140 | -0.3 | 5.05E-04 | -0.2 | 6.16E-03 | -0.1 | 2.02E-01 | -0.2 | 1.26E-01 | -0.2 | 3.74E-01 | -0.1 | 7.97E-01 | Zinc finger |
| AT5G15930 | -0.3 | 3.75E-02 | 0.0 | 9.10E-01 | -0.3 | 8.78E-02 | -0.2 | 2.68E-01 | -0.1 | 6.38E-01 | -0.1 | 7.25E-01 | At5g15930 |
| AT4G33740 | -0.3 | 6.76E-04 | -0.1 | 2.06E-01 | -0.2 | 1.18E-02 | -0.1 | 3.08E-01 | -0.1 | 5.83E-01 | 0.0 | 8.28E-01 | Myb-like protein X |
| AT5G16200 | -0.3 | 8.06E-02 | -0.3 | 1.39E-01 | 0.0 | 8.49E-01 | -0.6 | 1.26E-01 | -0.1 | 8.94E-01 | -0.5 | 3.14E-01 | 50S ribosomal protein-like protein |
| AT5G40550 | -0.3 | 6.07E-03 | -0.1 | 3.55E-01 | -0.2 | 6.95E-02 | -0.1 | 8.15E-01 | 0.2 | 3.64E-01 | -0.3 | 3.04E-01 | SGF29 tudor-like domain-containing protein |
| AT1G11710 | -0.3 | 3.58E-02 | -0.3 | 2.60E-02 | 0.0 | 8.35E-01 | 0.1 | 8.94E-01 | -0.1 | 9.30E-01 | 0.1 | 8.53E-01 | Pentatricopeptide repeat superfamily protein |
| AT1G80910 | -0.3 | 3.09E-03 | -0.3 | 3.50E-03 | 0.0 | 9.42E-01 | 0.2 | 2.32E-01 | 0.2 | 2.53E-01 | 0.0 | 9.56E-01 | At1g80910 |
| AT1G73380 | -0.3 | 1.42E-03 | -0.4 | 8.14E-04 | 0.1 | 5.92E-01 | -0.7 | 6.77E-04 | -0.6 | 4.72E-03 | -0.1 | 6.59E-01 | Uncharacterized protein At1g73380 |
| AT5G61240 | -0.3 | 3.97E-03 | -0.1 | 1.62E-01 | -0.2 | 8.25E-02 | 0.2 | 1.89E-01 | 0.3 | 4.90E-02 | -0.1 | 5.63E-01 | Leucine-rich repeat family protein |
| AT4G01090 | -0.3 | 1.12E-03 | -0.4 | 1.55E-04 | 0.1 | 1.11E-01 | 0.0 | 9.01E-01 | 0.0 | 9.92E-01 | 0.0 | 9.28E-01 | AT4G01090 protein |
| AT1G24120 | -0.3 | 1.54E-03 | -0.1 | 2.84E-01 | -0.2 | 2.12E-02 | -0.4 | 2.00E-02 | 0.0 | 9.45E-01 | -0.3 | 7.91E-02 | Chaperone protein dnaJ 16 |
| AT4G10690 | -0.3 | 2.51E-02 | -0.1 | 2.36E-01 | -0.2 | 3.10E-01 | 0.2 | 4.76E-01 | 0.1 | 5.86E-01 | 0.0 | 9.76E-01 | NA |
| AT4G36720 | -0.3 | 3.11E-03 | -0.6 | 1.29E-04 | 0.2 | 3.52E-02 | -0.3 | 2.39E-01 | -0.2 | 5.05E-01 | -0.1 | 8.07E-01 | HVA22-like protein k |
| AT1G48780 | -0.3 | 9.00E-03 | 0.0 | 8.38E-01 | -0.3 | 3.54E-02 | 0.0 | 9.94E-01 | -0.1 | 9.23E-01 | 0.1 | 9.29E-01 | Uncharacterized protein |
| AT1G08400 | -0.3 | 1.07E-02 | -0.3 | 9.58E-03 | 0.0 | 8.79E-01 | 0.1 | 6.31E-01 | 0.1 | 7.51E-01 | 0.0 | 9.58E-01 | RINT1-like protein MAG2L |
| AT2G17030 | -0.3 | 1.88E-03 | -0.2 | 1.48E-02 | -0.1 | 3.60E-01 | 0.0 | 8.13E-01 | -0.1 | 8.20E-01 | 0.0 | 9.87E-01 | F-box protein SKIP23 |
| AT2G16900 | -0.3 | 4.47E-03 | -0.5 | 2.29E-04 | 0.2 | 5.54E-02 | -0.1 | 6.33E-01 | -0.2 | 1.66E-01 | 0.2 | 5.05E-01 | Expressed protein family protein) |
| AT3G03150 | -0.3 | 2.55E-03 | 0.2 | 7.69E-02 | -0.5 | 6.04E-04 | -0.4 | 6.34E-03 | -0.3 | 8.49E-02 | -0.1 | 5.58E-01 | T17B22.16 protein |
| AT5G46090 | -0.3 | 5.94E-03 | 0.0 | 8.23E-01 | -0.3 | 1.27E-02 | 0.3 | 2.75E-01 | 0.1 | 8.89E-01 | 0.2 | 4.60E-01 | Protein DMP6 |
| AT1G50120 | -0.3 | 1.54E-03 | -0.2 | 3.64E-02 | -0.1 | 1.40E-01 | 0.1 | 5.03E-01 | 0.2 | 3.33E-01 | -0.1 | 7.79E-01 | Immunoglobulin |
| AT2G35110 | -0.3 | 2.56E-03 | -0.3 | 2.87E-03 | 0.0 | 9.36E-01 | 0.1 | 8.82E-01 | 0.2 | 7.15E-01 | -0.1 | 8.53E-01 | Uncharacterized protein At2g35110 |
| AT2G19180 | -0.3 | 1.58E-03 | -0.3 | 2.00E-03 | 0.0 | 9.68E-01 | -0.1 | 4.00E-01 | -0.2 | 3.32E-01 | 0.0 | 8.94E-01 | hypothetical protein |
| AT5G45500 | -0.3 | 3.49E-03 | 0.0 | 7.54E-01 | -0.3 | 1.27E-02 | -0.1 | 2.61E-01 | 0.0 | 9.72E-01 | -0.1 | 4.74E-01 | RNI-like superfamily protein |
| AT2G41170 | -0.3 | 2.36E-02 | -0.3 | 2.98E-02 | 0.0 | 9.74E-01 | 0.2 | 4.07E-01 | 0.0 | 8.89E-01 | 0.2 | 3.96E-01 | F-box family protein |
| AT1G52140 | -0.3 | 6.63E-03 | -0.1 | 1.33E-01 | -0.2 | 1.78E-01 | 0.0 | 9.97E-01 | 0.0 | 9.67E-01 | 0.0 | 9.76E-01 | Avr9 |
| AT5G47800 | -0.3 | 5.67E-03 | -0.2 | 2.83E-02 | -0.1 | 5.02E-01 | 0.3 | 2.06E-01 | 0.2 | 4.32E-01 | 0.1 | 8.38E-01 | Phototropic-responsive NPH3 family protein |
| AT2G19390 | -0.3 | 9.52E-04 | -0.3 | 1.20E-03 | 0.0 | 9.39E-01 | 0.1 | 7.76E-01 | 0.1 | 6.46E-01 | 0.0 | 8.64E-01 | Expressed protein |
| AT3G13275 | -0.3 | 2.01E-02 | -0.1 | 5.32E-01 | -0.2 | 1.20E-01 | 0.0 | 9.19E-01 | 0.2 | 5.91E-01 | -0.2 | 5.88E-01 | Transmembrane protein |
| AT3G10210 | -0.3 | 2.13E-02 | -0.5 | 1.84E-03 | 0.2 | 1.70E-01 | -0.5 | 9.94E-02 | 0.1 | 8.49E-01 | -0.5 | 6.88E-02 | hypothetical protein |
| AT2G19560 | -0.3 | 6.60E-03 | -0.2 | 2.96E-02 | -0.1 | 5.32E-01 | -0.5 | 1.90E-02 | -0.5 | 3.15E-02 | 0.0 | 9.91E-01 | Proteasome family protein |
| AT4G05631 | -0.3 | 6.74E-02 | -0.3 | 8.70E-02 | 0.0 | 9.75E-01 | -0.2 | 6.95E-01 | 0.0 | 9.72E-01 | -0.2 | 7.23E-01 | Uncharacterized protein |
| AT1G23260 | -0.3 | 4.21E-03 | -0.1 | 1.48E-01 | -0.2 | 9.74E-02 | -0.2 | 1.87E-01 | -0.2 | 2.47E-01 | 0.0 | 9.98E-01 | MMS ZWEI homologue 1 |
| AT2G06530 | -0.3 | 4.46E-04 | -0.2 | 4.58E-03 | -0.1 | 2.41E-01 | 0.0 | 9.81E-01 | 0.0 | 9.27E-01 | 0.0 | 9.57E-01 | SNF7 family protein |
| AT3G02700 | -0.3 | 2.71E-03 | -0.2 | 2.67E-02 | -0.1 | 2.90E-01 | -0.2 | 1.35E-01 | -0.2 | 2.95E-01 | 0.0 | 8.70E-01 | At3g02700 |
| AT5G28442 | -0.3 | 7.11E-02 | -0.1 | 4.98E-01 | -0.2 | 3.47E-01 | -0.4 | 3.51E-01 | -0.5 | 2.95E-01 | 0.1 | 8.69E-01 | Quinoprotein amine dehydrogenase, beta chain-like RIC1-like guanyl-nucleotide exchange factor |
| AT1G79970 | -0.3 | 6.68E-03 | 0.1 | 1.55E-01 | -0.4 | 2.04E-03 | -0.2 | 1.92E-01 | -0.1 | 3.98E-01 | 0.0 | 8.69E-01 | F18B13.5 protein |
| AT1G69870 | -0.3 | 5.07E-02 | -0.7 | 9.45E-04 | 0.4 | 3.56E-02 | -0.1 | 2.79E-01 | 0.0 | 9.63E-01 | -0.2 | 4.12E-01 | At1g69870 |
| AT3G13980 | -0.3 | 1.01E-02 | 0.2 | 1.32E-01 | -0.5 | 3.16E-03 | -0.2 | 4.05E-01 | -0.1 | 5.63E-01 | 0.0 | 9.40E-01 | Protein BIG GRAIN 1-like A |
| AT3G26500 | -0.3 | 4.22E-02 | -0.2 | 9.35E-02 | -0.1 | 7.55E-01 | 0.3 | 8.22E-02 | -0.1 | 7.86E-01 | 0.4 | 1.07E-01 | Plant intracellular Ras-group-related LRR protein 2 |
| AT5G19400 | -0.3 | 7.35E-04 | -0.5 | 6.72E-05 | 0.2 | 4.16E-02 | -0.1 | 3.78E-01 | 0.0 | 9.23E-01 | -0.1 | 4.95E-01 | Telomerase activating protein Est1 |
| AT2G21600 | -0.3 | 3.08E-03 | -0.1 | 3.30E-01 | -0.2 | 3.20E-02 | -0.3 | 2.85E-02 | -0.2 | 1.90E-01 | -0.1 | 6.72E-01 | Protein RER1B |
| AT5G07690 | -0.3 | 2.04E-01 | 0.0 | 9.01E-01 | -0.3 | 3.34E-01 | -0.9 | 1.00E-01 | 0.6 | 2.30E-01 | -1.5 | 2.93E-02 | Uncharacterized protein |
| AT4G10140 | -0.3 | 5.15E-03 | 0.1 | 4.69E-01 | -0.4 | 5.11E-03 | 0.0 | 8.31E-01 | 0.3 | 2.02E-01 | -0.2 | 3.89E-01 | hypothetical protein |
| AT2G05755 | -0.3 | 1.41E-02 | 0.0 | 6.82E-01 | -0.3 | 6.46E-02 | 0.1 | 5.95E-01 | 0.2 | 5.44E-01 | 0.0 | 9.23E-01 | At2g05755 |
| AT4G24760 | -0.3 | 8.48E-03 | -0.1 | 2.54E-01 | -0.2 | 1.17E-01 | -0.2 | 1.38E-01 | 0.1 | 8.01E-01 | -0.3 | 1.78E-01 | Uncharacterized protein At4g24760 |
| AT5G66580 | -0.3 | 4.00E-02 | 0.4 | 5.38E-03 | -0.7 | 1.15E-03 | 0.0 | 9.75E-01 | 0.3 | 4.52E-01 | -0.3 | 5.16E-01 | hypothetical protein |
| AT5G04850 | -0.3 | 8.61E-04 | -0.3 | 8.34E-04 | 0.0 | 8.17E-01 | 0.1 | 5.00E-01 | 0.0 | 8.19E-01 | 0.1 | 8.25E-01 | SNF7 family protein |
| AT4G14096 | -0.3 | 1.06E-01 | -0.2 | 2.98E-01 | -0.1 | 6.40E-01 | -0.2 | 4.21E-01 | 0.0 | 9.19E-01 | -0.1 | 6.07E-01 | F-box |
| AT1G34418 | -0.3 | 1.58E-03 | -0.4 | 6.93E-04 | 0.1 | 4.79E-01 | 0.2 | 2.55E-01 | 0.2 | 4.81E-01 | 0.1 | 8.72E-01 | NA |
| AT3G62160 | -0.3 | 4.81E-02 | 0.5 | 1.84E-03 | -0.8 | 6.79E-04 | -0.3 | 1.78E-01 | 0.0 | 9.19E-01 | -0.4 | 2.43E-01 | HXXXD-type acyl-transferase family protein |
| AT3G10500 | -0.3 | 8.34E-03 | -0.6 | 2.57E-04 | 0.3 | 3.48E-02 | 0.5 | 5.41E-03 | 0.2 | 2.78E-01 | 0.3 | 2.11E-01 | NAC domain containing protein 53 |
| AT5G57250 | -0.3 | 2.12E-02 | -0.3 | 2.38E-02 | 0.0 | 9.67E-01 | -0.1 | 9.14E-01 | 0.1 | 9.13E-01 | -0.3 | 8.23E-01 | Pentatricopeptide repeat-containing protein At5g57250, mitochondrial |
| AT1G51160 | -0.3 | 2.05E-03 | 0.0 | 9.28E-01 | -0.3 | 4.67E-03 | -0.1 | 3.77E-01 | 0.0 | 9.59E-01 | -0.1 | 5.17E-01 | Trafficking protein particle complex subunit |
| AT2G32280 | -0.3 | 3.32E-02 | 0.3 | 2.77E-02 | -0.6 | 2.84E-03 | 0.0 | 9.03E-01 | 0.1 | 8.17E-01 | -0.1 | 7.01E-01 | Protein VASCULATURE COMPLEXITY AND CONNECTIVITY |
| AT2G36420 | -0.3 | 4.55E-03 | -0.1 | 1.81E-01 | -0.2 | 8.77E-02 | -0.2 | 1.61E-01 | -0.3 | 7.23E-02 | 0.1 | 6.99E-01 | hypothetical protein |
| AT5G06280 | -0.3 | 1.56E-03 | -0.2 | 1.31E-02 | -0.1 | 3.35E-01 | -0.4 | 2.04E-01 | 0.2 | 6.03E-01 | -0.5 | 1.58E-01 | Uncharacterized protein At5g06280; MHF15.20 |
| AT5G50230 | -0.3 | 4.23E-03 | -0.3 | 1.02E-02 | 0.0 | 7.50E-01 | -0.4 | 3.03E-03 | -0.3 | 6.49E-02 | -0.2 | 3.70E-01 | Autophagy-related protein 16 domain-containing protein |
| AT5G52550 | -0.3 | 9.21E-04 | 0.0 | 5.07E-01 | -0.2 | 7.16E-03 | -0.1 | 7.36E-01 | -0.1 | 6.41E-01 | 0.0 | 9.05E-01 | AT5g52550 |
| AT5G04290 | -0.3 | 8.25E-03 | -0.4 | 2.48E-03 | 0.1 | 4.41E-01 | 0.0 | 9.15E-01 | -0.2 | 5.46E-01 | 0.1 | 7.17E-01 | Kow domain-containing transcription factor 1 |
| AT5G15680 | -0.3 | 3.39E-03 | -0.4 | 1.20E-03 | 0.1 | 4.41E-01 | 0.0 | 8.90E-01 | -0.1 | 8.43E-01 | 0.0 | 9.57E-01 | Uncharacterized protein At5g15680 |
| AT5G06370 | -0.3 | 1.88E-03 | -0.2 | 1.55E-02 | -0.1 | 3.29E-01 | -0.6 | 3.27E-04 | -0.4 | 2.32E-02 | -0.3 | 2.26E-01 | Protein LEAD-SENSITIVE 1 |
| AT1G48330 | -0.3 | 1.57E-01 | 0.5 | 1.56E-02 | -0.8 | 5.23E-03 | -0.2 | 5.54E-01 | 0.4 | 1.46E-01 | -0.6 | 8.76E-02 | hypothetical protein |
| AT5G45620 | -0.3 | 5.03E-03 | -0.1 | 3.82E-01 | -0.2 | 4.31E-02 | 0.1 | 5.12E-01 | 0.0 | 8.52E-01 | 0.0 | 8.15E-01 | Proteasome component domain protein |
| AT2G35470 | -0.3 | 8.04E-03 | 0.0 | 6.31E-01 | -0.2 | 3.92E-02 | -0.4 | 2.94E-02 | -0.2 | 3.09E-01 | -0.2 | 4.74E-01 | Transmembrane protein |
| AT3G24740 | -0.3 | 1.54E-02 | -0.4 | 2.56E-03 | 0.1 | 2.88E-01 | -0.4 | 6.23E-03 | -0.2 | 1.63E-01 | -0.2 | 3.33E-01 | hypothetical protein |
| AT5G01430 | -0.3 | 6.91E-03 | 0.0 | 8.21E-01 | -0.3 | 1.26E-02 | 0.2 | 4.04E-01 | 0.1 | 8.28E-01 | 0.1 | 7.24E-01 | At3g49420 |
| AT5G65660 | -0.3 | 6.31E-03 | 0.1 | 5.50E-01 | -0.3 | 5.34E-03 | -0.5 | 1.09E-03 | -0.7 | 4.22E-04 | 0.1 | 4.79E-01 | Hydroxyproline-rich glycoprotein family protein |
| AT1G10820 | -0.3 | 5.11E-03 | -0.1 | 1.57E-01 | -0.2 | 1.19E-01 | -0.1 | 5.01E-01 | -0.1 | 5.81E-01 | 0.0 | 9.95E-01 | At1g10820 |
| AT1G74450 | -0.3 | 2.06E-02 | -0.3 | 1.07E-02 | 0.1 | 6.94E-01 | 0.6 | 1.76E-03 | -0.2 | 3.41E-01 | 0.8 | 1.01E-03 | Protein ROH1D |
| AT5G56320 | -0.3 | 4.40E-02 | 0.4 | 9.95E-03 | -0.7 | 1.54E-03 | -0.2 | 4.64E-01 | 0.0 | 9.58E-01 | -0.2 | 5.95E-01 | Expansin |

| | | | | | | | | | | | | | |
|---|---|---|---|---|---|---|---|---|---|---|---|---|---|
| AT2G27900 | -0.3 | 7.97E-04 | -0.2 | 5.88E-03 | -0.1 | 3.29E-01 | -0.1 | 7.01E-01 | -0.1 | 6.84E-01 | 0.0 | 9.62E-01 | Uncharacterized protein At2g27910 |
| AT2G25560 | -0.3 | 2.37E-03 | -0.2 | 1.86E-02 | -0.1 | 3.66E-01 | -0.1 | 8.25E-01 | 0.0 | 8.91E-01 | 0.0 | 9.78E-01 | J domain-containing protein |
| AT2G39980 | -0.3 | 9.16E-03 | -0.6 | 1.84E-04 | 0.3 | 2.05E-02 | -0.2 | 2.99E-01 | -0.5 | 7.93E-02 | 0.2 | 5.68E-01 | hypothetical protein |
| AT3G19895 | -0.3 | 3.39E-03 | -0.5 | 1.95E-04 | 0.2 | 5.64E-02 | -0.1 | 7.43E-01 | 0.1 | 8.55E-01 | -0.1 | 6.39E-01 | At3g19895 |
| AT3G61570 | -0.3 | 3.44E-02 | -0.6 | 1.75E-03 | 0.3 | 1.13E-01 | -0.4 | 3.44E-01 | -0.1 | 8.56E-01 | -0.3 | 5.82E-01 | GRIP-related ARF-binding domain-containing protein 1 |
| AT5G40190 | -0.3 | 1.87E-02 | 0.0 | 9.25E-01 | -0.3 | 4.56E-02 | -0.2 | 2.82E-01 | -0.2 | 4.89E-01 | -0.1 | 8.84E-01 | RNA ligase |
| AT5G57815 | -0.3 | 5.47E-02 | -0.2 | 1.35E-01 | -0.1 | 7.08E-01 | -0.2 | 4.57E-01 | -0.2 | 5.38E-01 | 0.0 | 9.94E-01 | Cytochrome c oxidase subunit 6b-2 |
| AT5G62090 | -0.3 | 3.64E-03 | -0.4 | 8.38E-04 | 0.1 | 2.98E-01 | -0.2 | 2.51E-01 | -0.1 | 5.94E-01 | -0.1 | 7.49E-01 | Probable transcriptional regulator SLK2 |
| AT4G25670 | -0.3 | 4.87E-03 | -0.2 | 4.89E-02 | -0.1 | 2.90E-01 | -0.1 | 7.09E-01 | 0.0 | 9.25E-01 | -0.1 | 8.79E-01 | Uncharacterized protein AT4g25670 |
| AT5G61450 | -0.3 | 3.26E-03 | -0.3 | 6.91E-03 | 0.0 | 8.10E-01 | 0.0 | 9.25E-01 | 0.1 | 5.82E-01 | -0.1 | 7.24E-01 | P-loop containing nucleoside triphosphate hydrolases superfamily protein |
| AT3G58900 | -0.3 | 1.44E-02 | -0.3 | 2.23E-02 | 0.0 | 9.21E-01 | -0.4 | 5.98E-02 | -0.1 | 7.52E-01 | -0.3 | 2.56E-01 | F-box |
| AT1G47250 | -0.3 | 3.09E-02 | -0.1 | 3.20E-01 | -0.2 | 2.80E-01 | 0.1 | 8.35E-01 | 0.3 | 5.08E-01 | -0.2 | 7.20E-01 | Proteasome subunit alpha type-1-B |
| AT5G28350 | -0.3 | 1.51E-03 | -0.3 | 2.30E-03 | 0.0 | 9.59E-01 | -0.2 | 2.83E-01 | 0.0 | 9.32E-01 | -0.3 | 3.53E-01 | Quinoprotein amine dehydrogenase, beta chain-like RIC1-like guanyl-nucleotide exchange factor |
| AT3G06340 | -0.3 | 1.39E-03 | -0.3 | 6.72E-04 | 0.1 | 5.06E-01 | 0.0 | 8.81E-01 | -0.1 | 7.17E-01 | 0.2 | 6.25E-01 | DNAJ heat shock N-terminal domain-containing protein |
| AT4G21700 | -0.3 | 2.97E-02 | -0.3 | 1.53E-02 | 0.1 | 6.97E-01 | 0.0 | 9.52E-01 | 0.0 | 9.72E-01 | 0.0 | 9.94E-01 | DUF2921 family protein |
| AT3G03220 | -0.3 | 6.45E-03 | 0.0 | 7.27E-01 | -0.3 | 9.54E-03 | 0.0 | 9.03E-01 | 0.2 | 1.23E-01 | -0.2 | 1.86E-01 | Expansin-A13 |
| AT1G17145 | -0.3 | 8.86E-04 | -0.3 | 2.15E-03 | 0.0 | 7.69E-01 | -0.5 | 3.58E-03 | -0.3 | 4.97E-02 | -0.2 | 5.19E-01 | RING |
| AT5G49290 | -0.3 | 1.50E-01 | -0.3 | 1.47E-01 | 0.0 | 9.50E-01 | -0.4 | 3.69E-01 | 0.1 | 8.57E-01 | -0.5 | 3.30E-01 | Receptor like protein 56 |
| AT5G08230 | -0.3 | 1.26E-01 | -0.5 | 3.06E-02 | 0.2 | 4.59E-01 | -0.1 | 7.66E-01 | -0.1 | 8.11E-01 | 0.0 | 9.97E-01 | Protein HUA2-LIKE 1 |
| AT2G25964 | -0.3 | 1.12E-01 | -0.3 | 1.71E-01 | 0.0 | 8.93E-01 | -0.4 | 1.77E-01 | -0.3 | 4.38E-01 | -0.1 | 7.87E-01 | Uncharacterized protein |
| AT5G58970 | -0.3 | 6.56E-03 | -0.1 | 2.45E-01 | -0.2 | 1.06E-01 | -0.2 | 5.50E-01 | 0.1 | 8.08E-01 | -0.3 | 4.74E-01 | UCP2 |
| AT1G34320 | -0.3 | 6.74E-03 | -0.3 | 4.08E-03 | 0.0 | 7.06E-01 | -0.1 | 3.73E-01 | -0.1 | 7.84E-01 | -0.1 | 7.26E-01 | Protein PSK SIMULATOR 1 |
| AT1G57600 | -0.3 | 2.56E-03 | -0.1 | 3.20E-01 | -0.2 | 3.54E-02 | -0.1 | 8.19E-01 | 0.2 | 3.33E-01 | -0.3 | 2.52E-01 | Membrane-bound O-acyltransferase C24H6.01c |
| AT5G65950 | -0.3 | 1.00E-03 | -0.2 | 5.24E-03 | -0.1 | 4.39E-01 | -0.1 | 7.53E-01 | -0.1 | 7.91E-01 | 0.0 | 9.97E-01 | Trafficking protein particle complex subunit 11 domain-containing protein |
| AT5G64780 | -0.3 | 1.10E-01 | -0.2 | 1.81E-01 | 0.0 | 8.64E-01 | 0.1 | 8.19E-01 | 0.2 | 4.35E-01 | -0.2 | 6.75E-01 | hypothetical protein |
| AT2G24250 | -0.3 | 3.26E-02 | -0.1 | 3.83E-01 | -0.2 | 2.59E-01 | -0.4 | 2.41E-01 | -0.2 | 6.03E-01 | -0.2 | 7.31E-01 | F-box |
| AT2G26110 | -0.3 | 1.68E-03 | 0.1 | 4.37E-01 | -0.3 | 1.91E-03 | -0.1 | 6.49E-01 | 0.0 | 9.28E-01 | -0.1 | 8.38E-01 | At2g26110 ) |
| AT4G16520 | -0.3 | 1.02E-02 | -0.2 | 9.12E-02 | -0.1 | 3.11E-01 | -0.5 | 8.72E-04 | -0.5 | 3.96E-03 | 0.0 | 8.82E-01 | Autophagy-related protein |
| AT3G06240 | -0.3 | 1.38E-03 | -0.2 | 3.53E-02 | -0.1 | 1.35E-01 | -0.3 | 8.93E-02 | -0.2 | 2.33E-01 | -0.1 | 8.07E-01 | F-box |
| AT4G10330 | -0.3 | 7.58E-03 | -0.2 | 5.04E-02 | -0.1 | 4.17E-01 | -0.2 | 4.93E-01 | -0.2 | 6.94E-01 | -0.1 | 9.07E-01 | hypothetical protein |
| AT2G46640 | -0.3 | 5.61E-02 | 0.0 | 8.67E-01 | -0.3 | 1.37E-01 | 0.2 | 5.72E-01 | 0.3 | 3.65E-01 | -0.1 | 7.64E-01 | Protein TILLER ANGLE CONTROL 1 |
| AT3G46280 | -0.3 | 1.52E-01 | 0.0 | 8.74E-01 | -0.2 | 2.55E-01 | -0.2 | 5.15E-01 | 0.0 | 9.47E-01 | -0.2 | 5.30E-01 | Uncharacterized protein F12M12_250 |
| AT4G04920 | -0.3 | 1.84E-03 | -0.4 | 3.99E-04 | 0.1 | 2.35E-01 | 0.0 | 8.12E-01 | 0.0 | 8.10E-01 | 0.1 | 7.17E-01 | Sensitive to freezing 6 |
| AT1G13360 | -0.3 | 4.42E-02 | 0.0 | 8.45E-01 | -0.3 | 9.58E-02 | -0.2 | 4.73E-01 | -0.2 | 5.67E-01 | 0.0 | 9.92E-01 | Uncharacterized protein |
| AT5G53030 | -0.3 | 6.72E-02 | -0.6 | 4.45E-03 | 0.3 | 1.59E-01 | 0.4 | 1.99E-01 | 0.1 | 8.40E-01 | 0.3 | 4.57E-01 | Uncharacterized protein |
| AT1G27430 | -0.3 | 3.50E-03 | -0.4 | 8.80E-04 | 0.1 | 3.22E-01 | -0.2 | 2.74E-01 | -0.1 | 4.65E-01 | 0.0 | 8.98E-01 | GYF domain-containing protein |
| AT4G32270 | -0.3 | 4.24E-03 | 0.0 | 7.99E-01 | -0.3 | 8.16E-03 | 0.0 | 8.07E-01 | 0.1 | 4.50E-01 | -0.2 | 3.81E-01 | hypothetical protein |
| AT5G39250 | -0.3 | 2.33E-02 | -0.2 | 1.54E-01 | -0.1 | 4.08E-01 | 0.0 | 9.42E-01 | 0.1 | 7.79E-01 | -0.1 | 7.63E-01 | F-box protein At5g39250 |
| AT2G18210 | -0.3 | 2.39E-01 | 0.0 | 9.96E-01 | -0.3 | 3.41E-01 | 1.0 | 6.07E-02 | 0.8 | 1.87E-01 | 0.2 | 7.70E-01 | At2g18210 |
| AT5G08450 | -0.3 | 4.07E-03 | -0.2 | 7.57E-02 | -0.1 | 1.71E-01 | 0.1 | 3.15E-01 | 0.2 | 1.46E-01 | -0.1 | 7.16E-01 | Uncharacterized protein At5g08450 |
| AT3G56190 | -0.3 | 7.71E-04 | -0.1 | 1.91E-01 | -0.2 | 1.44E-02 | -0.1 | 1.57E-01 | -0.1 | 4.59E-01 | -0.1 | 7.79E-01 | Alpha-soluble NSF attachment protein 2 |
| AT5G37660 | -0.3 | 2.17E-02 | 0.0 | 8.16E-01 | -0.3 | 6.90E-02 | 0.0 | 9.20E-01 | 0.0 | 9.38E-01 | -0.1 | 8.64E-01 | Plasmodesmata-located protein 7 |
| AT3G15180 | -0.3 | 7.79E-03 | -0.6 | 1.64E-04 | 0.3 | 2.06E-02 | 0.3 | 1.38E-01 | 0.2 | 3.48E-01 | 0.1 | 7.99E-01 | ARM repeat superfamily protein |
| AT3G51620 | -0.3 | 1.13E-03 | -0.5 | 4.27E-05 | 0.2 | 1.60E-02 | 0.1 | 6.47E-01 | 0.1 | 6.09E-01 | 0.0 | 9.46E-01 | Uncharacterized protein F26O13.260 |
| AT3G15410 | -0.3 | 6.61E-03 | -0.1 | 1.99E-01 | -0.2 | 1.11E-01 | -0.2 | 1.32E-01 | -0.1 | 6.49E-01 | -0.1 | 5.12E-01 | Leucine-rich repeat family protein |
| AT3G03855 | -0.3 | 8.70E-02 | -0.7 | 2.27E-03 | 0.4 | 6.51E-02 | -0.1 | 9.19E-01 | 0.0 | 9.66E-01 | -0.1 | 9.08E-01 | NA |
| AT1G78300 | -0.3 | 1.02E-02 | 0.1 | 1.60E-01 | -0.4 | 2.46E-03 | -0.2 | 1.03E-01 | -0.1 | 7.51E-01 | -0.1 | 4.52E-01 | 14-3-3-like protein GF14 omega |
| AT2G27790 | -0.3 | 8.29E-02 | -0.5 | 1.06E-02 | 0.2 | 2.87E-01 | -0.2 | 5.10E-01 | -0.1 | 7.35E-01 | -0.1 | 8.93E-01 | RNA-binding family protein |
| AT5G52580 | -0.3 | 1.24E-03 | -0.3 | 8.25E-04 | 0.0 | 6.49E-01 | -0.4 | 2.81E-02 | -0.2 | 1.95E-01 | -0.1 | 5.69E-01 | RabGAP |
| AT3G08930 | -0.3 | 2.40E-03 | 0.0 | 9.06E-01 | -0.3 | 4.78E-03 | -0.1 | 3.13E-01 | -0.1 | 4.98E-01 | 0.0 | 9.23E-01 | LIMR family protein At3g08930 |
| AT2G26570 | -0.3 | 1.82E-03 | -0.3 | 4.14E-03 | 0.0 | 7.87E-01 | -0.2 | 3.28E-01 | -0.1 | 8.49E-01 | -0.1 | 6.69E-01 | Protein WEAK CHLOROPLAST MOVEMENT UNDER BLUE LIGHT 1 |
| AT5G47580 | -0.3 | 3.75E-03 | -0.3 | 1.75E-03 | 0.1 | 5.64E-01 | -0.1 | 6.43E-01 | -0.1 | 6.90E-01 | 0.0 | 9.97E-01 | ASG7 |
| AT5G17190 | -0.3 | 8.40E-03 | 0.0 | 9.84E-01 | -0.3 | 1.69E-02 | 0.0 | 8.96E-01 | 0.0 | 9.59E-01 | 0.0 | 9.67E-01 | Endoplasmic reticulum transmembrane protein |
| AT1G50500 | -0.3 | 3.09E-03 | -0.2 | 3.30E-02 | -0.1 | 2.67E-01 | -0.1 | 6.02E-01 | 0.0 | 8.98E-01 | -0.1 | 8.03E-01 | Membrane trafficking VPS53 family protein |
| AT3G15095 | -0.3 | 1.33E-02 | -0.1 | 5.02E-01 | -0.2 | 7.29E-02 | -0.4 | 7.76E-03 | -0.2 | 1.96E-01 | -0.2 | 3.62E-01 | Chloroplast protein HCF243 |
| AT1G33415 | -0.3 | 1.26E-01 | -0.2 | 2.42E-01 | -0.1 | 8.00E-01 | -0.2 | 6.67E-01 | 0.3 | 4.67E-01 | -0.5 | 2.88E-01 | NA |
| AT2G35540 | -0.3 | 2.17E-02 | -0.3 | 3.82E-02 | 0.0 | 8.61E-01 | -0.4 | 6.21E-02 | -0.3 | 2.23E-01 | -0.1 | 6.85E-01 | J domain-containing protein |
| AT5G43930 | -0.3 | 3.40E-02 | -0.6 | 1.02E-03 | 0.3 | 5.98E-02 | -0.4 | 1.31E-01 | -0.2 | 5.19E-01 | -0.2 | 6.38E-01 | AT5G43930 protein |
| AT5G58040 | -0.3 | 7.28E-04 | -0.3 | 8.53E-04 | 0.0 | 9.05E-01 | -0.4 | 9.25E-03 | -0.4 | 6.82E-03 | 0.1 | 8.02E-01 | FIP1-like protein |
| AT1G64610 | -0.3 | 3.28E-02 | -0.8 | 1.72E-04 | 0.5 | 7.66E-03 | 0.1 | 7.37E-01 | -0.3 | 3.20E-01 | 0.4 | 2.22E-01 | Transducin |
| AT1G47640 | -0.3 | 3.91E-02 | 0.1 | 4.12E-01 | -0.4 | 1.71E-02 | -0.2 | 5.05E-02 | -0.1 | 2.53E-01 | -0.1 | 6.91E-01 | BOS complex subunit TMEM147 |
| AT4G37390 | -0.3 | 1.02E-02 | -0.1 | 5.26E-01 | -0.2 | 6.78E-02 | 1.3 | 1.40E-04 | 0.1 | 7.36E-01 | 1.1 | 3.43E-04 | hypothetical protein |
| AT3G03160 | -0.3 | 2.03E-02 | 0.2 | 1.13E-01 | -0.5 | 3.24E-03 | -0.2 | 1.61E-01 | -0.2 | 2.58E-01 | 0.0 | 9.67E-01 | Endoplasmic reticulum transmembrane protein |
| AT1G45150 | -0.3 | 1.22E-03 | -0.4 | 2.40E-04 | 0.1 | 1.92E-01 | -0.4 | 3.30E-02 | -0.2 | 4.29E-01 | -0.2 | 3.73E-01 | Alpha-1,6-mannosyl-glycoprotein 2-beta-N-acetylglucosaminyltransferase |
| AT5G41770 | -0.3 | 1.83E-03 | -0.2 | 2.35E-02 | -0.1 | 2.20E-01 | 0.1 | 3.47E-01 | 0.1 | 5.55E-01 | 0.0 | 9.12E-01 | Crooked neck protein, putative |
| AT5G35753 | -0.3 | 7.56E-03 | -0.1 | 1.58E-01 | -0.1 | 1.76E-01 | 0.0 | 9.52E-01 | 0.1 | 7.78E-01 | -0.1 | 8.54E-01 | DnaJ heat shock amino-terminal domain protein |
| AT5G50430 | -0.3 | 3.37E-03 | -0.3 | 6.53E-03 | 0.0 | 8.42E-01 | -0.1 | 7.23E-01 | 0.1 | 5.97E-01 | -0.2 | 4.23E-01 | UBC33 |
| AT3G20590 | -0.3 | 1.75E-01 | 0.3 | 1.78E-01 | -0.5 | 3.94E-02 | 0.7 | 5.60E-02 | 0.7 | 9.59E-02 | 0.0 | 9.62E-01 | Late embryogenesis abundant hydroxyproline-rich glycoprotein family |
| AT5G64260 | -0.3 | 2.06E-02 | -0.1 | 5.11E-01 | -0.2 | 1.04E-01 | -0.5 | 8.14E-04 | -0.9 | 4.28E-05 | 0.3 | 8.33E-02 | Protein EXORDIUM-like 2 |
| AT4G12770 | -0.3 | 1.29E-03 | -0.3 | 1.85E-03 | 0.0 | 9.79E-01 | 0.0 | 7.78E-01 | 0.1 | 6.80E-01 | 0.0 | 9.06E-01 | Chaperone DnaJ-domain superfamily protein |
| AT4G31880 | -0.3 | 1.04E-03 | -0.2 | 6.41E-03 | -0.1 | 3.89E-01 | -0.1 | 4.35E-01 | 0.0 | 7.92E-01 | -0.1 | 8.15E-01 | Transcriptional regulator |
| AT3G60900 | -0.3 | 1.16E-01 | 0.0 | 8.06E-01 | -0.2 | 2.67E-01 | 0.1 | 6.93E-01 | 0.3 | 2.18E-01 | -0.2 | 5.20E-01 | Fasciclin-like arabinogalactan protein 10 |
| AT4G10170 | -0.3 | 4.05E-03 | -0.1 | 1.08E-01 | -0.1 | 1.41E-01 | -0.4 | 2.09E-02 | -0.2 | 2.28E-01 | -0.2 | 4.63E-01 | Phytolongin Phyl1.1 |
| AT3G56880 | -0.3 | 4.45E-02 | -0.6 | 1.34E-03 | 0.3 | 5.71E-02 | -0.2 | 1.85E-01 | -0.2 | 8.74E-02 | 0.1 | 7.10E-01 | hypothetical protein |
| AT5G51640 | -0.3 | 5.03E-02 | -0.7 | 1.08E-03 | 0.4 | 4.63E-02 | 0.0 | 9.92E-01 | 0.4 | 6.66E-01 | -0.3 | 7.21E-01 | YLS7 |
| AT2G11240 | -0.3 | 1.36E-03 | -0.2 | 7.80E-03 | -0.1 | 4.28E-01 | -0.1 | 3.47E-01 | -0.1 | 5.86E-01 | 0.0 | 8.74E-01 | NA |
| AT4G17790 | -0.3 | 1.54E-03 | -0.1 | 4.18E-01 | -0.2 | 1.46E-02 | -0.5 | 8.67E-04 | -0.3 | 4.80E-02 | -0.2 | 2.40E-01 | SNARE associated Golgi protein family |
| AT2G14680 | -0.3 | 3.26E-03 | -0.3 | 1.14E-03 | 0.1 | 4.45E-01 | 0.1 | 3.67E-01 | 0.0 | 9.99E-01 | 0.1 | 5.27E-01 | Myosin heavy chain-like protein |
| AT1G06800 | -0.3 | 3.47E-02 | -0.3 | 3.62E-02 | 0.0 | 9.48E-01 | -0.1 | 9.03E-01 | -0.2 | 8.21E-01 | 0.1 | 9.28E-01 | Alpha |
| AT1G30440 | -0.3 | 3.72E-03 | -0.1 | 3.24E-01 | -0.2 | 3.88E-02 | -0.3 | 1.01E-01 | -0.1 | 6.88E-01 | -0.2 | 4.50E-01 | BTB |
| AT1G28340 | -0.3 | 2.10E-03 | -0.1 | 7.37E-02 | -0.1 | 9.39E-02 | -0.2 | 4.50E-01 | 0.2 | 3.44E-01 | -0.4 | 1.66E-01 | Receptor-like protein 4 |
| AT3G56270 | -0.3 | 2.57E-02 | -0.1 | 2.46E-01 | -0.1 | 3.13E-01 | 0.1 | 8.90E-01 | 0.4 | 1.80E-01 | -0.4 | 2.74E-01 | WEB family protein At3g56270 |
| AT3G17380 | -0.3 | 5.15E-02 | -0.4 | 2.13E-02 | 0.1 | 6.23E-01 | 0.1 | 7.96E-01 | 0.3 | 6.01E-01 | -0.2 | 8.28E-01 | MATH domain-containing protein |

| | | | | | | | | | | | | | |
|---|---|---|---|---|---|---|---|---|---|---|---|---|---|
| AT1G64080 | -0.3 | 7.30E-02 | -0.5 | 6.72E-03 | 0.2 | 2.09E-01 | -0.1 | 6.79E-01 | 0.1 | 6.95E-01 | -0.2 | 4.30E-01 | Probable membrane-associated kinase regulator 2 |
| AT5G56520 | -0.3 | 2.15E-01 | -0.5 | 7.03E-02 | 0.2 | 5.45E-01 | -0.1 | 7.27E-01 | 0.0 | 9.25E-01 | -0.1 | 7.05E-01 | Uncharacterized protein |
| AT2G19950 | -0.3 | 8.22E-04 | -0.1 | 3.53E-02 | -0.1 | 7.32E-02 | -0.1 | 6.82E-01 | 0.0 | 9.90E-01 | -0.1 | 7.73E-01 | GC1 |
| AT5G19000 | -0.3 | 1.39E-03 | -0.3 | 1.96E-03 | 0.0 | 9.84E-01 | -0.2 | 4.10E-02 | -0.3 | 4.54E-02 | 0.0 | 9.28E-01 | BPM1 |
| AT1G08465 | -0.3 | 9.68E-03 | 0.0 | 8.08E-01 | -0.2 | 3.51E-02 | -0.3 | 1.50E-01 | -0.3 | 1.43E-01 | 0.0 | 9.20E-01 | YAB2 |
| AT5G53930 | -0.3 | 9.00E-03 | -0.2 | 8.31E-02 | -0.1 | 3.27E-01 | -0.1 | 7.61E-01 | -0.1 | 7.46E-01 | 0.0 | 9.61E-01 | Uncharacterized protein |
| AT2G48110 | -0.3 | 9.18E-03 | -0.3 | 3.13E-03 | 0.1 | 4.94E-01 | -0.1 | 7.83E-01 | 0.0 | 9.55E-01 | -0.1 | 7.75E-01 | Mediator of RNA polymerase II transcription subunit 33B |
| AT1G29380 | -0.3 | 1.57E-02 | -0.3 | 1.75E-02 | 0.0 | 9.72E-01 | 0.1 | 7.30E-01 | 0.3 | 3.96E-01 | -0.2 | 6.84E-01 | Beta-1,3 glucanase, putative; 26636-27432 |
| AT2G30880 | -0.3 | 6.08E-03 | -0.2 | 1.75E-02 | 0.0 | 6.78E-01 | -0.2 | 3.10E-01 | 0.1 | 6.79E-01 | -0.3 | 2.04E-01 | At2g30880 |
| AT2G46225 | -0.3 | 2.90E-03 | -0.1 | 3.49E-01 | -0.2 | 3.22E-02 | -0.1 | 5.17E-01 | 0.1 | 7.93E-01 | -0.1 | 4.47E-01 | ABI-1-like 1 |
| AT4G13250 | -0.3 | 1.45E-03 | -0.3 | 2.27E-03 | 0.0 | 9.59E-01 | -1.8 | 1.07E-05 | 0.0 | 9.16E-01 | -1.8 | 5.05E-05 | NAD-binding Rossmann-fold superfamily protein |
| AT1G80020 | -0.3 | 1.52E-03 | -0.5 | 7.79E-05 | 0.2 | 2.93E-02 | 0.0 | 8.88E-01 | 0.0 | 9.65E-01 | 0.0 | 9.62E-01 | NA |
| AT1G17790 | -0.3 | 1.73E-02 | -0.2 | 4.51E-02 | 0.0 | 7.22E-01 | -0.2 | 5.41E-01 | 0.0 | 9.00E-01 | -0.2 | 4.96E-01 | Transcription factor GTE5, chloroplastic |
| AT3G58600 | -0.3 | 1.24E-03 | -0.1 | 1.07E-01 | -0.2 | 4.40E-02 | 0.0 | 9.81E-01 | -0.1 | 4.55E-01 | 0.1 | 5.48E-01 | NECAP PHear domain-containing protein |
| AT5G20090 | -0.3 | 5.66E-03 | 0.1 | 4.62E-01 | -0.3 | 4.04E-03 | 0.0 | 8.59E-01 | 0.0 | 8.89E-01 | 0.0 | 1.00E+00 | Mitochondrial pyruvate carrier |
| AT4G02350 | -0.3 | 1.01E-03 | -0.3 | 8.40E-04 | 0.0 | 7.47E-01 | -0.3 | 3.05E-02 | -0.2 | 1.35E-01 | -0.1 | 7.69E-01 | Exocyst complex component SEC15B |
| AT2G01690 | -0.3 | 4.41E-02 | -0.4 | 1.50E-02 | 0.1 | 5.48E-01 | -0.5 | 4.33E-01 | 0.2 | 8.27E-01 | -0.7 | 4.06E-01 | ARM repeat superfamily protein |
| AT5G56420 | -0.3 | 5.81E-02 | -0.3 | 8.22E-02 | 0.0 | 9.30E-01 | -0.2 | 2.29E-01 | -0.2 | 2.81E-01 | 0.0 | 9.98E-01 | F-box |
| AT5G48412 | -0.3 | 1.80E-02 | 0.1 | 4.09E-01 | -0.4 | 8.70E-03 | -0.6 | 6.99E-04 | -0.4 | 1.69E-02 | -0.2 | 4.34E-01 | NA |
| AT5G54290 | -0.3 | 9.38E-03 | 0.0 | 9.51E-01 | -0.3 | 2.20E-02 | 0.0 | 8.67E-01 | 0.0 | 8.76E-01 | 0.0 | 9.92E-01 | Cytochrome c biogenesis protein family |
| AT1G04555 | -0.3 | 4.13E-02 | -0.4 | 1.14E-02 | 0.1 | 4.93E-01 | 0.2 | 5.24E-01 | 0.0 | 9.60E-01 | 0.3 | 6.16E-01 | Transmembrane protein |
| AT5G62960 | -0.3 | 6.57E-02 | -0.3 | 6.28E-02 | 0.0 | 9.33E-01 | -0.1 | 6.32E-01 | 0.1 | 8.21E-01 | -0.2 | 5.64E-01 | UDP-N-acetylglucosamine-N-acetylmuramyl-pyrophosphoryl-undecaprenol N-acetylglucosamine protein |
| AT2G35520 | -0.3 | 9.40E-02 | 0.3 | 3.71E-02 | -0.6 | 5.92E-03 | 0.0 | 9.11E-01 | 0.1 | 7.59E-01 | -0.1 | 8.64E-01 | Dolichyl-diphosphooligosaccharide--protein glycosyltransferase subunit DAD1 |
| AT2G15860 | -0.3 | 1.69E-03 | -0.2 | 2.69E-02 | -0.1 | 1.84E-01 | 0.0 | 8.59E-01 | 0.0 | 9.87E-01 | 0.0 | 9.06E-01 | BAT2 domain protein |
| AT1G06110 | -0.3 | 2.23E-03 | -0.2 | 1.34E-02 | -0.1 | 4.14E-01 | -0.3 | 4.27E-02 | -0.2 | 1.49E-01 | -0.1 | 8.12E-01 | SKP1 |
| AT1G48230 | -0.3 | 7.22E-03 | 0.0 | 8.48E-01 | -0.3 | 1.12E-02 | -0.1 | 4.21E-01 | 0.0 | 8.68E-01 | -0.1 | 4.72E-01 | Nodulin MtN21 |
| AT1G20560 | -0.3 | 3.24E-03 | -0.2 | 1.15E-02 | 0.0 | 5.94E-01 | -0.4 | 1.13E-03 | -0.1 | 7.38E-01 | -0.4 | 1.87E-02 | Acyl activating enzyme 1 |
| AT5G14970 | -0.3 | 4.67E-02 | -0.1 | 6.26E-01 | -0.2 | 1.58E-01 | 0.0 | 7.89E-01 | 0.1 | 2.62E-01 | -0.1 | 5.12E-01 | hypothetical protein |
| AT1G04230 | -0.3 | 4.50E-03 | -0.3 | 1.48E-03 | 0.1 | 4.45E-01 | -0.2 | 3.82E-01 | -0.1 | 6.09E-01 | -0.1 | 8.83E-01 | rRNA-processing protein EFG1 |
| AT5G10950 | -0.3 | 4.93E-03 | -0.1 | 2.49E-01 | -0.2 | 7.18E-02 | -0.2 | 2.86E-01 | -0.2 | 2.95E-01 | 0.0 | 9.50E-01 | Tudor |
| AT1G69340 | -0.3 | 1.05E-03 | -0.4 | 2.32E-04 | 0.1 | 2.03E-01 | -0.2 | 7.20E-02 | -0.1 | 2.53E-01 | -0.1 | 7.81E-01 | At1g69340 |
| AT1G33050 | -0.3 | 8.51E-02 | -0.4 | 2.79E-02 | 0.1 | 5.43E-01 | -0.4 | 4.99E-03 | -0.5 | 1.70E-03 | 0.1 | 5.59E-01 | Uncharacterized protein At1g33050 |
| AT4G15070 | -0.3 | 5.11E-02 | -0.3 | 6.32E-02 | 0.0 | 9.73E-01 | 0.2 | 4.84E-01 | 0.3 | 4.75E-01 | 0.0 | 9.44E-01 | Cysteine |
| AT3G60600 | -0.3 | 2.33E-03 | 0.0 | 9.82E-01 | -0.3 | 5.67E-03 | -0.4 | 1.15E-03 | -0.3 | 2.29E-02 | -0.1 | 5.14E-01 | Vesicle-associated protein 1-1 |
| AT2G18780 | -0.3 | 1.10E-01 | -0.2 | 2.65E-01 | -0.1 | 7.16E-01 | 0.2 | 6.52E-01 | 0.0 | 9.97E-01 | | | F-box protein At2g18780 |
| AT1G24148 | -0.3 | 2.15E-01 | -0.5 | 3.01E-02 | 0.3 | 3.05E-01 | 0.6 | 1.36E-02 | 0.6 | 3.69E-02 | 0.1 | 8.89E-01 | Uncharacterized protein |
| AT3G08890 | -0.3 | 4.25E-02 | -0.1 | 4.53E-01 | -0.2 | 2.49E-01 | -0.6 | 1.71E-03 | 0.0 | 9.17E-01 | -0.5 | 1.58E-02 | DUF538 family protein |
| AT3G11397 | -0.3 | 6.44E-04 | -0.3 | 1.07E-03 | 0.0 | 9.42E-01 | 0.1 | 6.96E-01 | -0.1 | 6.43E-01 | 0.1 | 4.90E-01 | PRA1 family protein A3 |
| AT5G50150 | -0.3 | 3.48E-03 | -0.1 | 9.55E-02 | -0.1 | 1.33E-01 | 0.0 | 9.38E-01 | 0.1 | 6.29E-01 | -0.1 | 7.22E-01 | AT5g50150 |
| AT5G03970 | -0.3 | 1.10E-02 | -0.2 | 4.99E-02 | -0.1 | 5.38E-01 | 0.1 | 6.27E-01 | -0.3 | 3.20E-01 | 0.4 | 1.30E-01 | F-box associated ubiquitination effector family protein |
| AT2G43210 | -0.3 | 3.08E-03 | -0.2 | 7.16E-03 | 0.0 | 7.68E-01 | -0.3 | 3.65E-02 | -0.4 | 1.42E-02 | 0.1 | 6.83E-01 | Plant UBX domain-containing protein 11 |
| AT1G68160 | -0.3 | 5.51E-03 | -0.1 | 5.14E-01 | -0.2 | 4.07E-02 | 0.1 | 6.44E-01 | 0.0 | 9.38E-01 | 0.1 | 6.92E-01 | Uncharacterized protein |
| AT1G21170 | -0.3 | 3.46E-03 | -0.3 | 5.66E-03 | 0.0 | 9.13E-01 | -0.1 | 7.69E-01 | 0.0 | 9.05E-01 | 0.0 | 9.33E-01 | Exocyst complex component SEC5 |
| AT5G37740 | -0.3 | 9.51E-03 | -0.2 | 5.25E-02 | -0.1 | 4.51E-01 | 0.8 | 2.33E-04 | 0.2 | 4.72E-01 | 0.6 | 6.48E-03 | Calcium-dependent lipid-binding family protein |
| AT5G61820 | -0.3 | 4.60E-02 | -0.8 | 1.09E-04 | 0.6 | 2.69E-03 | -0.4 | 5.08E-02 | -0.2 | 3.10E-01 | -0.1 | 6.30E-01 | Stress up-regulated Nod 19 protein |
| AT5G05010 | -0.3 | 2.03E-03 | 0.0 | 7.66E-01 | -0.2 | 7.62E-03 | -0.1 | 1.80E-01 | 0.0 | 9.66E-01 | -0.1 | 3.49E-01 | Coatomer subunit delta |
| AT1G16750 | -0.3 | 5.95E-02 | -0.1 | 6.20E-01 | -0.2 | 2.27E-01 | -0.3 | 1.53E-01 | 0.1 | 8.40E-01 | -0.4 | 2.13E-01 | At1g16750 |
| AT3G61670 | -0.3 | 7.17E-03 | -0.3 | 6.99E-03 | 0.0 | 9.06E-01 | -0.1 | 5.07E-01 | -0.1 | 8.06E-01 | -0.1 | 8.35E-01 | Probable zinc-ribbon domain-containing protein |
| AT2G44850 | -0.3 | 8.80E-02 | -0.3 | 6.62E-02 | 0.0 | 8.42E-01 | -0.5 | 1.05E-01 | -0.3 | 3.11E-01 | -0.1 | 7.39E-01 | Uncharacterized protein At2g44850 |
| AT1G09645 | -0.3 | 5.42E-02 | -0.3 | 3.71E-02 | 0.0 | 8.20E-01 | 0.0 | 8.92E-01 | -0.2 | 3.73E-01 | 0.2 | 5.58E-01 | hypothetical protein |
| AT3G54620 | -0.3 | 1.67E-03 | -0.3 | 3.05E-03 | 0.0 | 8.76E-01 | -0.1 | 4.20E-01 | -0.2 | 1.06E-01 | 0.1 | 5.35E-01 | BZO2H4 |
| AT5G39360 | -0.3 | 2.40E-03 | -0.2 | 9.01E-03 | 0.0 | 5.95E-01 | -0.1 | 5.65E-01 | 0.1 | 7.51E-01 | -0.2 | 4.38E-01 | EID1-like F-box protein 2 |
| AT1G27000 | -0.3 | 4.01E-03 | -0.3 | 4.25E-03 | 0.0 | 9.01E-01 | 0.0 | 7.50E-01 | -0.1 | 5.15E-01 | 0.2 | 4.12E-01 | At1g27000 |
| AT1G18740 | -0.3 | 1.24E-02 | -0.3 | 1.71E-02 | 0.0 | 9.48E-01 | -0.4 | 6.67E-02 | -0.4 | 7.06E-02 | 0.0 | 9.23E-01 | Protein BYPASS1-LIKE |
| AT5G63880 | -0.3 | 4.20E-02 | -0.2 | 7.90E-02 | 0.0 | 8.22E-01 | 0.1 | 8.58E-01 | 0.0 | 9.51E-01 | 0.0 | 9.52E-01 | SNF7 family protein |
| AT2G32910 | -0.3 | 9.79E-04 | -0.2 | 2.69E-03 | 0.0 | 6.97E-01 | -0.1 | 6.19E-01 | 0.2 | 4.04E-01 | -0.3 | 2.02E-01 | DCD domain protein |
| AT3G63300 | -0.3 | 3.61E-02 | -0.1 | 5.91E-01 | -0.2 | 1.62E-01 | -0.4 | 5.60E-02 | -0.2 | 4.44E-01 | -0.2 | 4.71E-01 | FORKED 1 |
| AT1G13670 | -0.3 | 3.06E-02 | 0.4 | 5.40E-03 | -0.6 | 6.94E-04 | -0.5 | 3.04E-02 | -0.1 | 6.95E-01 | -0.4 | 1.78E-01 | Uncharacterized protein |
| AT1G17280 | -0.3 | 1.71E-03 | -0.2 | 9.34E-03 | -0.1 | 4.35E-01 | -0.1 | 6.69E-01 | -0.1 | 7.92E-01 | 0.0 | 9.63E-01 | Ubiquitin-conjugating enzyme 34 |
| AT5G28960 | -0.3 | 1.57E-01 | 0.6 | 3.04E-03 | -0.9 | 1.93E-03 | -0.1 | 7.68E-01 | 0.3 | 5.76E-01 | -0.4 | 4.38E-01 | Alpha--fucosyltransferase |
| AT5G18950 | -0.3 | 1.84E-02 | -0.2 | 3.99E-02 | 0.0 | 7.92E-01 | 0.2 | 4.52E-01 | 0.1 | 8.05E-01 | 0.1 | 7.80E-01 | Pentatricopeptide repeat-containing protein At5g18950 |
| AT5G17270 | -0.3 | 1.69E-02 | -0.3 | 1.54E-02 | 0.0 | 9.08E-01 | 0.5 | 2.76E-02 | 0.0 | 9.66E-01 | 0.5 | 1.08E-01 | Protein prenylyltransferase superfamily protein |
| AT3G11100 | -0.3 | 8.05E-03 | -0.3 | 1.10E-02 | 0.0 | 9.61E-01 | -0.4 | 3.10E-02 | -0.4 | 1.50E-02 | 0.1 | 7.23E-01 | MADF domain-containing protein |
| AT5G22930 | -0.3 | 4.65E-02 | -0.2 | 1.76E-01 | -0.1 | 5.73E-01 | 0.3 | 4.36E-01 | 0.7 | 6.55E-02 | -0.4 | 3.65E-01 | Enabled-like protein |
| AT5G03150 | -0.3 | 3.56E-02 | 0.0 | 9.72E-01 | -0.3 | 7.14E-02 | -0.3 | 2.29E-01 | -0.2 | 5.74E-01 | -0.1 | 7.38E-01 | hypothetical protein |
| AT2G17190 | -0.3 | 2.79E-03 | -0.3 | 1.36E-03 | 0.0 | 5.32E-01 | 0.1 | 6.06E-01 | 0.1 | 4.19E-01 | -0.1 | 8.18E-01 | UBA domain-containing protein |
| AT1G22040 | -0.3 | 2.62E-03 | -0.3 | 3.09E-03 | 0.0 | 9.48E-01 | 0.0 | 9.48E-01 | 0.1 | 7.95E-01 | 0.0 | 8.66E-01 | F-box |
| AT4G24200 | -0.3 | 4.05E-03 | -0.4 | 2.86E-04 | 0.2 | 7.83E-02 | 0.1 | 8.22E-01 | -0.1 | 6.77E-01 | 0.2 | 5.41E-01 | Uncharacterized protein AT4g24200 |
| AT5G46700 | -0.3 | 2.41E-02 | 0.1 | 5.23E-01 | -0.3 | 1.79E-02 | 0.1 | 5.51E-01 | 0.1 | 6.47E-01 | 0.0 | 9.85E-01 | Protein TORNADO 2 |
| AT2G46060 | -0.3 | 7.42E-03 | -0.3 | 4.24E-03 | 0.0 | 6.87E-01 | -0.1 | 8.76E-01 | -0.2 | 5.99E-01 | 0.1 | 7.23E-01 | At2g46060 |
| AT2G40380 | -0.3 | 3.85E-03 | -0.2 | 2.98E-02 | -0.1 | 3.45E-01 | -0.1 | 7.84E-01 | 0.1 | 8.20E-01 | -0.1 | 6.77E-01 | PRA1 family protein B2 |
| AT5G06180 | -0.3 | 4.74E-02 | -0.2 | 1.36E-01 | -0.1 | 6.74E-01 | -0.5 | 3.96E-02 | -0.8 | 9.22E-03 | 0.2 | 5.23E-01 | Fission ELM1-like protein |
| AT3G50010 | -0.3 | 6.21E-02 | 0.0 | 8.83E-01 | -0.2 | 1.40E-01 | 0.0 | 9.44E-01 | 0.0 | 9.02E-01 | 0.0 | 9.60E-01 | Cysteine |
| AT1G31935 | -0.3 | 1.78E-01 | 0.0 | 9.86E-01 | -0.3 | 2.72E-01 | 0.5 | 4.63E-02 | 0.4 | 1.65E-01 | 0.1 | 7.43E-01 | NA |
| AT1G19980 | -0.3 | 8.83E-02 | -0.1 | 3.53E-01 | -0.1 | 5.20E-01 | 0.2 | 5.06E-01 | 0.2 | 6.34E-01 | 0.0 | 9.61E-01 | Cytomatrix protein-like protein |
| AT3G59900 | -0.3 | 4.57E-02 | -0.5 | 3.20E-03 | 0.2 | 1.64E-01 | 1.1 | 1.65E-02 | 0.5 | 4.16E-01 | 0.6 | 4.26E-02 | ARGOS-like protein |
| AT5G55730 | -0.3 | 1.39E-02 | 0.2 | 1.19E-01 | -0.4 | 2.83E-03 | -0.1 | 5.45E-01 | 0.2 | 4.83E-01 | -0.3 | 2.92E-01 | Fasciclin-like arabinogalactan protein 1 |
| AT1G05035 | -0.3 | 4.01E-01 | 0.4 | 1.57E-01 | -0.7 | 7.93E-02 | 1.3 | 1.02E-01 | 0.3 | 8.28E-01 | 1.0 | 2.32E-01 | Uncharacterized protein |
| AT1G05785 | -0.3 | 1.22E-02 | -0.1 | 1.58E-01 | -0.1 | 2.53E-01 | 0.2 | 5.72E-01 | -0.1 | 8.43E-01 | 0.2 | 4.59E-01 | hypothetical protein |
| AT4G33780 | -0.3 | 2.18E-03 | -0.2 | 5.94E-03 | 0.0 | 7.06E-01 | -0.1 | 7.35E-01 | 0.0 | 8.36E-01 | -0.1 | 6.63E-01 | ATP phosphoribosyltransferase regulatory subunit |
| AT2G36895 | -0.3 | 6.73E-02 | -0.2 | 2.64E-01 | -0.1 | 5.10E-01 | -1.1 | 7.91E-05 | 0.1 | 6.48E-01 | -1.2 | 3.97E-04 | D-tagatose-1,6-bisphosphate aldolase subunit |
| AT5G24290 | -0.3 | 1.00E-01 | 0.2 | 2.64E-01 | -0.4 | 3.01E-02 | -0.1 | 6.78E-01 | 0.2 | 4.87E-01 | -0.4 | 3.05E-01 | MEB2 |
| AT5G08500 | -0.3 | 7.44E-03 | -0.2 | 3.11E-02 | -0.1 | 5.50E-01 | 0.0 | 9.48E-01 | -0.1 | 5.20E-01 | 0.1 | 6.61E-01 | Cleft lip and palate associated transmembrane protein-like |

| | | | | | | | | | | | | | |
|---|---|---|---|---|---|---|---|---|---|---|---|---|---|
| AT3G17900 | -0.3 | 2.56E-03 | -0.1 | 5.25E-02 | -0.1 | 1.55E-01 | -0.1 | 7.04E-01 | -0.1 | 6.04E-01 | 0.0 | 9.04E-01 | AT3g17900 |
| AT1G01630 | -0.3 | 1.08E-02 | 0.0 | 5.93E-01 | -0.2 | 4.86E-02 | 0.0 | 9.23E-01 | 0.1 | 4.83E-01 | -0.1 | 6.19E-01 | CRAL-TRIO domain-containing protein |
| AT3G63070 | -0.3 | 6.12E-03 | -0.5 | 2.72E-04 | 0.2 | 4.92E-02 | 0.1 | 7.99E-01 | 0.0 | 9.66E-01 | 0.1 | 8.95E-01 | Protein HUA2-LIKE 3 |
| AT2G26360 | -0.3 | 5.34E-02 | 0.1 | 3.70E-01 | -0.4 | 2.72E-02 | 0.0 | 9.93E-01 | 0.2 | 6.17E-01 | -0.2 | 5.28E-01 | Mitochondrial substrate carrier family protein |
| AT5G10200 | -0.3 | 1.04E-02 | -0.1 | 2.56E-01 | -0.2 | 1.45E-01 | 0.0 | 9.93E-01 | 0.0 | 9.52E-01 | 0.0 | 9.44E-01 | Uncharacterized protein T31P16_190 |
| AT5G12290 | -0.3 | 2.00E-03 | -0.2 | 4.14E-03 | 0.0 | 8.20E-01 | 0.0 | 9.29E-01 | 0.0 | 9.22E-01 | -0.1 | 8.64E-01 | Dgd1 suppressor 1 |
| AT1G64405 | -0.3 | 1.07E-01 | -0.4 | 2.92E-02 | 0.1 | 5.02E-01 | -0.1 | 9.17E-01 | 0.3 | 6.08E-01 | -0.4 | 5.79E-01 | hypothetical protein |
| AT2G30110 | -0.3 | 2.79E-03 | -0.3 | 2.67E-03 | 0.0 | 8.29E-01 | 0.1 | 4.87E-01 | 0.1 | 4.80E-01 | 0.0 | 9.48E-01 | Putative ubiquitin activating enzyme |
| AT1G64270 | -0.3 | 1.46E-01 | -0.5 | 1.90E-02 | 0.2 | 2.91E-01 | -0.4 | 4.39E-01 | -0.3 | 6.08E-01 | -0.1 | 9.14E-01 | NA |
| AT1G21780 | -0.3 | 1.03E-02 | -0.3 | 8.63E-03 | 0.0 | 8.58E-01 | -0.4 | 7.81E-03 | -0.4 | 1.67E-02 | 0.0 | 9.75E-01 | BTB |
| AT2G27550 | -0.3 | 2.58E-01 | -1.2 | 1.11E-03 | 1.0 | 1.22E-02 | 0.5 | 1.26E-01 | 0.4 | 3.36E-01 | 0.1 | 7.49E-01 | Protein CENTRORADIALIS-like |
| AT4G12590 | -0.3 | 1.42E-02 | 0.0 | 7.75E-01 | -0.3 | 1.63E-02 | -0.1 | 3.42E-01 | -0.1 | 6.23E-01 | -0.1 | 8.48E-01 | ER membrane protein complex subunit 3 |
| AT1G03350 | -0.3 | 1.50E-03 | -0.2 | 2.68E-03 | 0.0 | 8.92E-01 | 0.0 | 9.44E-01 | 0.0 | 7.98E-01 | 0.0 | 8.72E-01 | BSD domain-containing protein |
| AT1G03280 | -0.3 | 1.19E-03 | -0.3 | 2.65E-04 | 0.1 | 2.16E-01 | 0.0 | 8.83E-01 | 0.0 | 9.40E-01 | 0.0 | 8.54E-01 | TFIIE-alpha 1 |
| AT4G23440 | -0.3 | 1.22E-02 | -0.2 | 1.07E-01 | -0.1 | 3.39E-01 | -0.4 | 8.94E-02 | -0.1 | 6.02E-01 | -0.2 | 4.35E-01 | Disease resistance protein |
| AT3G56640 | -0.3 | 2.03E-02 | -0.1 | 5.63E-01 | -0.2 | 1.10E-01 | 0.1 | 8.06E-01 | 0.0 | 9.47E-01 | 0.1 | 8.13E-01 | Exocyst complex component SEC15A |
| AT5G13390 | -0.3 | 9.72E-03 | -0.3 | 6.37E-03 | 0.0 | 7.53E-01 | -0.1 | 6.37E-01 | 0.0 | 9.48E-01 | -0.1 | 8.15E-01 | AT5g13390 |
| AT1G70780 | -0.3 | 4.14E-02 | 0.0 | 8.48E-01 | -0.2 | 9.03E-02 | -0.9 | 1.41E-05 | -0.6 | 1.89E-03 | -0.3 | 1.61E-01 | AT1G70780 protein |
| AT4G08510 | -0.3 | 2.31E-02 | -0.3 | 7.72E-03 | 0.1 | 5.34E-01 | 0.2 | 4.70E-01 | 0.1 | 7.03E-01 | 0.1 | 8.78E-01 | At4g08510 |
| AT4G17330 | -0.3 | 1.57E-03 | -0.5 | 2.85E-05 | 0.2 | 4.97E-03 | -0.1 | 5.98E-01 | 0.0 | 9.41E-01 | -0.1 | 6.74E-01 | Uncharacterized protein AT4g17320 |
| AT5G04240 | -0.3 | 5.31E-03 | -0.2 | 1.58E-02 | 0.0 | 6.68E-01 | -0.2 | 3.65E-01 | -0.1 | 8.78E-01 | -0.2 | 6.54E-01 | ELF6 |
| AT2G38630 | -0.3 | 1.81E-02 | -0.2 | 6.66E-02 | -0.1 | 5.96E-01 | 0.0 | 9.11E-01 | 0.3 | 7.55E-02 | -0.3 | 1.65E-01 | Uncharacterized protein At2g38630 |
| AT4G39170 | -0.3 | 7.72E-03 | -0.2 | 2.99E-02 | -0.1 | 5.57E-01 | -0.2 | 1.59E-01 | 0.0 | 9.24E-01 | -0.2 | 3.89E-01 | Sec14p-like phosphatidylinositol transfer family protein |
| AT5G07730 | -0.3 | 8.38E-02 | -0.4 | 2.60E-02 | 0.1 | 5.50E-01 | 0.0 | 9.77E-01 | 0.1 | 7.68E-01 | -0.1 | 8.16E-01 | Transmembrane protein |
| AT4G11190 | -0.3 | 2.75E-01 | 0.4 | 8.05E-02 | -0.6 | 3.06E-02 | -0.2 | 5.50E-01 | 0.7 | 5.81E-02 | -0.9 | 2.98E-02 | Dirigent protein 13 |
| AT1G13940 | -0.3 | 2.40E-02 | -0.2 | 3.10E-02 | 0.0 | 9.70E-01 | -0.1 | 7.72E-01 | -0.2 | 6.24E-01 | 0.1 | 8.72E-01 | F16A14.15 ) |
| AT5G37930 | -0.3 | 9.46E-03 | -0.4 | 1.82E-03 | 0.1 | 3.14E-01 | -0.5 | 5.32E-03 | -0.3 | 1.24E-01 | -0.2 | 3.33E-01 | Uncharacterized protein |
| AT1G63830 | -0.2 | 2.05E-02 | -0.3 | 1.73E-02 | 0.0 | 8.72E-01 | -0.3 | 1.61E-02 | -0.4 | 2.83E-03 | 0.1 | 4.53E-01 | PLAC8 family protein |
| AT3G59000 | -0.2 | 1.93E-02 | -0.2 | 1.30E-01 | -0.1 | 4.13E-01 | -0.5 | 2.26E-02 | -0.3 | 1.67E-01 | -0.2 | 5.64E-01 | F-box protein At3g59000 |
| AT4G38360 | -0.2 | 1.34E-03 | -0.3 | 1.64E-03 | 0.0 | 9.50E-01 | -0.2 | 2.15E-01 | -0.2 | 2.27E-01 | 0.0 | 9.54E-01 | Protein LAZ1 |
| AT1G77000 | -0.2 | 1.19E-01 | -0.4 | 3.72E-02 | 0.1 | 5.41E-01 | -0.6 | 2.79E-02 | -0.1 | 7.99E-01 | -0.5 | 1.58E-01 | RNI-like superfamily protein |
| AT3G06290 | -0.2 | 3.06E-03 | -0.4 | 3.62E-04 | 0.1 | 1.33E-01 | -0.3 | 1.83E-01 | -0.2 | 4.33E-01 | -0.1 | 8.00E-01 | SAC3 |
| AT4G01290 | -0.2 | 1.44E-02 | -0.4 | 9.68E-04 | 0.2 | 9.84E-02 | 0.0 | 9.25E-01 | 0.1 | 8.91E-01 | 0.0 | 9.66E-01 | Uncharacterized protein At4g01300 |
| AT3G26910 | -0.2 | 1.62E-02 | -0.6 | 8.99E-05 | 0.4 | 5.43E-03 | -0.3 | 4.21E-01 | -0.4 | 3.82E-01 | 0.1 | 9.15E-01 | Hydroxyproline-rich glycoprotein family protein |
| AT3G55020 | -0.2 | 1.12E-03 | -0.3 | 6.71E-04 | 0.0 | 5.73E-01 | 0.0 | 9.99E-01 | 0.1 | 8.09E-01 | -0.1 | 8.40E-01 | AT3g55020 |
| AT1G73970 | -0.2 | 2.96E-02 | -0.4 | 2.40E-03 | 0.2 | 1.69E-01 | 0.0 | 9.71E-01 | 0.2 | 3.92E-01 | -0.2 | 4.52E-01 | Obscurin-like protein |
| AT5G16280 | -0.2 | 8.72E-03 | -0.2 | 1.03E-02 | 0.0 | 9.88E-01 | 0.1 | 5.67E-01 | 0.2 | 4.79E-01 | -0.1 | 8.82E-01 | Tetratricopeptide repeat -like superfamily protein |
| AT5G12940 | -0.2 | 1.59E-01 | 0.4 | 3.53E-02 | -0.6 | 7.12E-03 | 0.0 | 9.74E-01 | 0.4 | 1.25E-01 | -0.4 | 1.38E-01 | Leucine-rich repeat-containing N-terminal plant-type domain-containing protein |
| AT5G64880 | -0.2 | 1.38E-02 | 0.1 | 5.08E-01 | -0.3 | 1.21E-02 | -0.3 | 1.07E-01 | -0.4 | 7.00E-02 | 0.1 | 8.26E-01 | hypothetical protein |
| AT3G07280 | -0.2 | 2.55E-03 | 0.0 | 5.57E-01 | -0.2 | 1.73E-02 | 0.0 | 9.11E-01 | -0.1 | 6.36E-01 | 0.1 | 6.34E-01 | NA |
| AT4G19920 | -0.2 | 2.57E-01 | -0.1 | 7.80E-01 | -0.2 | 5.10E-01 | 0.7 | 7.85E-03 | 0.4 | 1.52E-01 | 0.3 | 3.23E-01 | TIR domain-containing protein |
| AT2G02148 | -0.2 | 1.97E-02 | -0.2 | 1.26E-01 | -0.1 | 4.27E-01 | -0.1 | 5.90E-01 | -0.1 | 6.73E-01 | 0.0 | 9.82E-01 | PPR containing protein |
| AT3G51930 | -0.2 | 3.25E-02 | 0.0 | 7.60E-01 | -0.3 | 4.43E-02 | 0.2 | 6.51E-01 | 0.1 | 8.44E-01 | 0.1 | 8.99E-01 | hypothetical protein |
| AT2G30640 | -0.2 | 6.10E-02 | -0.5 | 4.89E-03 | 0.2 | 1.90E-01 | 0.4 | 3.50E-01 | 0.0 | 9.69E-01 | 0.4 | 5.33E-01 | NA |
| AT1G04340 | -0.2 | 5.91E-03 | 0.1 | 1.48E-01 | -0.4 | 2.12E-03 | 0.1 | 5.02E-01 | 0.1 | 7.14E-01 | 0.0 | 9.13E-01 | hypothetical protein |
| AT1G02290 | -0.2 | 2.27E-02 | -0.2 | 1.41E-01 | -0.1 | 4.28E-01 | 0.1 | 7.91E-01 | 0.0 | 9.86E-01 | 0.1 | 8.62E-01 | DEUBAD domain-containing protein |
| AT5G16520 | -0.2 | 1.43E-02 | -0.3 | 1.51E-02 | 0.0 | 9.47E-01 | -0.3 | 1.65E-01 | 0.0 | 9.66E-01 | -0.3 | 2.57E-01 | At5g16520 |
| AT1G64572 | -0.2 | 2.66E-02 | -0.1 | 2.99E-01 | -0.1 | 2.63E-01 | -0.2 | 3.26E-01 | -0.2 | 5.63E-01 | -0.1 | 8.68E-01 | NA |
| AT5G64930 | -0.2 | 3.01E-03 | -0.3 | 1.12E-03 | 0.1 | 4.59E-01 | -0.3 | 2.52E-02 | -0.3 | 8.53E-02 | -0.1 | 8.22E-01 | Protein CPR-5 |
| AT4G00231 | -0.2 | 4.10E-02 | -0.1 | 2.37E-01 | -0.1 | 4.27E-01 | -0.2 | 3.29E-01 | -0.3 | 1.75E-01 | 0.1 | 7.43E-01 | MEE50 |
| AT3G22270 | -0.2 | 2.82E-03 | -0.2 | 6.33E-03 | 0.0 | 7.80E-01 | -0.1 | 4.95E-01 | 0.0 | 9.92E-01 | -0.1 | 6.73E-01 | hypothetical protein |
| AT4G36630 | -0.2 | 9.56E-03 | -0.2 | 2.03E-02 | 0.0 | 7.86E-01 | 0.0 | 9.57E-01 | 0.1 | 4.97E-01 | -0.1 | 5.58E-01 | Vacuolar sorting protein 39 |
| AT5G55040 | -0.2 | 6.29E-03 | -0.2 | 1.03E-02 | 0.0 | 8.96E-01 | 0.0 | 8.65E-01 | 0.0 | 9.18E-01 | 0.0 | 9.87E-01 | Gb\|AAF16663.1 |
| AT1G27760 | -0.2 | 1.27E-02 | -0.3 | 3.37E-03 | 0.1 | 4.03E-01 | 0.4 | 6.85E-03 | 0.4 | 1.36E-02 | 0.0 | 9.93E-01 | At1g27760 |
| AT5G45350 | -0.2 | 9.04E-03 | -0.2 | 3.61E-02 | -0.1 | 5.60E-01 | 0.0 | 9.90E-01 | 0.1 | 6.76E-01 | -0.1 | 7.34E-01 | hypothetical protein |
| AT4G14905 | -0.2 | 8.37E-03 | -0.1 | 1.39E-01 | -0.1 | 2.12E-01 | -0.3 | 9.76E-02 | -0.2 | 3.42E-01 | -0.1 | 7.14E-01 | F-box |
| AT3G12280 | -0.2 | 5.60E-03 | -0.3 | 5.25E-03 | 0.0 | 8.86E-01 | 0.3 | 1.38E-01 | 0.2 | 3.34E-01 | 0.1 | 8.39E-01 | Retinoblastoma-related 1 |
| AT3G13225 | -0.2 | 1.13E-02 | -0.3 | 1.90E-03 | 0.1 | 2.78E-01 | 0.0 | 8.71E-01 | 0.0 | 9.19E-01 | 0.1 | 8.48E-01 | WW domain-containing protein |
| AT3G05680 | -0.2 | 7.41E-02 | -0.4 | 9.86E-03 | 0.2 | 2.83E-01 | 0.0 | 9.95E-01 | 0.0 | 9.67E-01 | 0.0 | 9.78E-01 | Virilizer N-terminal domain-containing protein |
| AT4G12650 | -0.2 | 5.80E-03 | -0.1 | 4.51E-01 | -0.2 | 4.00E-02 | 0.0 | 7.39E-01 | 0.0 | 7.80E-01 | -0.1 | 6.36E-01 | Transmembrane 9 superfamily member 12 |
| AT2G01080 | -0.2 | 1.01E-02 | -0.1 | 2.55E-01 | -0.1 | 1.38E-01 | -0.2 | 1.92E-01 | 0.0 | 8.61E-01 | -0.2 | 4.68E-01 | Late embryogenesis abundant protein LEA-2 subgroup domain-containing protein |
| AT1G24160 | -0.2 | 3.17E-03 | -0.2 | 1.18E-02 | 0.0 | 5.73E-01 | -0.2 | 6.53E-02 | -0.3 | 2.15E-02 | 0.1 | 6.56E-01 | Uncharacterized protein |
| AT3G25495 | -0.2 | 8.46E-02 | -0.6 | 2.51E-03 | 0.3 | 7.68E-02 | 0.1 | 6.80E-01 | 0.4 | 2.01E-01 | -0.3 | 3.82E-01 | NA |
| AT2G13130 | -0.2 | 2.10E-01 | -0.3 | 1.06E-01 | 0.1 | 7.03E-01 | 0.1 | 8.26E-01 | 0.2 | 7.42E-01 | -0.1 | 9.16E-01 | NA |
| AT5G47490 | -0.2 | 1.45E-02 | -0.2 | 9.78E-02 | -0.1 | 4.02E-01 | -0.1 | 6.67E-01 | 0.0 | 9.65E-01 | -0.2 | 7.32E-01 | Protein transport protein sec16 |
| AT2G31585 | -0.2 | 2.73E-02 | 0.1 | 6.11E-01 | -0.3 | 2.50E-02 | -0.4 | 1.23E-01 | -0.4 | 9.29E-02 | 0.1 | 8.50E-01 | NA |
| AT5G07890 | -0.2 | 2.26E-02 | -0.1 | 2.20E-01 | -0.1 | 2.99E-01 | -0.4 | 1.73E-02 | -0.2 | 3.05E-01 | -0.2 | 3.77E-01 | Myosin heavy chain-like protein |
| AT3G03030 | -0.2 | 1.20E-01 | 0.2 | 2.47E-01 | -0.4 | 3.60E-02 | -0.1 | 8.22E-01 | 0.0 | 9.97E-01 | -0.1 | 8.69E-01 | F-box |
| AT1G27840 | -0.2 | 2.97E-03 | -0.2 | 1.06E-02 | 0.0 | 6.14E-01 | -0.4 | 3.51E-02 | -0.1 | 4.80E-01 | -0.2 | 3.81E-01 | Transducin |
| AT5G51430 | -0.2 | 1.65E-03 | -0.3 | 8.82E-04 | 0.0 | 5.61E-01 | -0.2 | 2.12E-01 | -0.1 | 7.16E-01 | -0.1 | 6.08E-01 | Conserved oligomeric Golgi complex subunit 7 |
| AT1G62040 | -0.2 | 2.40E-01 | -0.4 | 5.30E-02 | 0.2 | 4.19E-01 | -0.3 | 2.09E-01 | -0.3 | 3.53E-01 | 0.0 | 9.23E-01 | Autophagy-related protein |
| AT5G40970 | -0.2 | 6.65E-02 | -0.1 | 2.88E-01 | -0.1 | 5.02E-01 | 0.1 | 7.37E-01 | 0.1 | 6.95E-01 | 0.0 | 9.44E-01 | Transmembrane protein |
| AT4G16870 | -0.2 | 1.97E-01 | 0.3 | 1.67E-01 | -0.5 | 3.43E-02 | 0.1 | 8.49E-01 | -0.2 | 6.87E-01 | 0.3 | 5.20E-01 | NA |
| AT1G05420 | -0.2 | 3.07E-01 | 0.5 | 4.33E-02 | -0.7 | 2.23E-02 | -0.7 | 2.49E-01 | -0.6 | 3.78E-01 | | | Transcription repressor |
| AT3G60260 | -0.2 | 1.28E-02 | 0.1 | 4.60E-01 | -0.3 | 9.96E-03 | -0.4 | 2.66E-02 | -0.4 | 4.20E-02 | 0.0 | 9.92E-01 | hypothetical protein |
| AT2G03530 | -0.2 | 6.18E-02 | -0.2 | 1.01E-01 | 0.0 | 8.61E-01 | 0.2 | 3.49E-01 | 0.2 | 4.51E-01 | 0.0 | 9.75E-01 | Uncharacterized protein At2g03540 |
| AT5G65840 | -0.2 | 7.88E-02 | -0.2 | 2.39E-01 | -0.1 | 5.98E-01 | -0.2 | 2.36E-01 | 0.4 | 1.31E-02 | -0.6 | 5.67E-03 | Thioredoxin superfamily protein |
| AT1G15800 | -0.2 | 1.97E-01 | -0.1 | 6.70E-01 | -0.2 | 4.90E-01 | -0.6 | 7.48E-03 | -0.7 | 4.47E-03 | 0.1 | 7.75E-01 | Uncharacterized protein |
| AT3G51040 | -0.2 | 6.76E-03 | 0.0 | 6.43E-01 | -0.2 | 3.38E-02 | -0.5 | 4.81E-03 | -0.4 | 1.93E-02 | -0.1 | 8.07E-01 | Protein RTE1-HOMOLOG |
| AT4G39820 | -0.2 | 1.67E-02 | 0.0 | 8.00E-01 | -0.2 | 4.92E-02 | 0.1 | 2.65E-01 | -0.1 | 5.35E-01 | 0.2 | 1.58E-01 | At4g39820 -like superfamily protein) |
| AT5G06830 | -0.2 | 3.64E-03 | -0.1 | 3.20E-01 | -0.2 | 4.32E-02 | 0.0 | 9.61E-01 | 0.0 | 9.21E-01 | 0.0 | 9.08E-01 | CDK5RAP3-like protein |
| AT3G48380 | -0.2 | 8.85E-03 | -0.1 | 3.86E-01 | -0.2 | 7.81E-02 | -0.5 | 4.12E-03 | -0.4 | 1.53E-02 | -0.1 | 8.55E-01 | Peptidase C78, ubiquitin fold modifier-specific peptidase 1 |
| AT3G22470 | -0.2 | 1.35E-01 | -0.2 | 2.73E-01 | -0.1 | 7.82E-01 | 0.0 | 9.35E-01 | 0.1 | 8.54E-01 | -0.1 | 8.09E-01 | Pentatricopeptide repeat superfamily protein |

| | | | | | | | | | | | | | |
|---|---|---|---|---|---|---|---|---|---|---|---|---|---|
| AT5G41850 | -0.2 | 1.56E-01 | -0.2 | 2.88E-01 | -0.1 | 7.96E-01 | 0.1 | 6.56E-01 | 0.2 | 5.79E-01 | -0.1 | 9.14E-01 | KANL3 |
| AT4G31080 | -0.2 | 3.08E-03 | -0.1 | 6.70E-02 | -0.1 | 1.46E-01 | -0.1 | 2.54E-01 | -0.2 | 4.34E-02 | 0.1 | 4.70E-01 | Uncharacterized protein AT4g31090 |
| AT3G56820 | -0.2 | 1.16E-02 | -0.1 | 1.65E-01 | -0.1 | 2.38E-01 | -0.4 | 5.84E-02 | -0.3 | 3.14E-01 | -0.2 | 6.02E-01 | hypothetical protein |
| AT1G44960 | -0.2 | 2.14E-02 | -0.2 | 6.49E-02 | -0.1 | 6.63E-01 | 0.2 | 2.72E-01 | 0.2 | 3.21E-01 | 0.0 | 9.90E-01 | F27F5.5 |
| AT1G74520 | -0.2 | 7.93E-02 | -0.1 | 6.49E-01 | -0.2 | 2.43E-01 | -0.5 | 1.65E-02 | -0.3 | 2.51E-01 | -0.2 | 4.44E-01 | HVA22-like protein a |
| AT5G13590 | -0.2 | 4.75E-02 | -0.4 | 7.46E-03 | 0.1 | 3.13E-01 | 0.1 | 6.97E-01 | -0.1 | 6.63E-01 | 0.2 | 5.08E-01 | Uncharacterized protein |
| AT1G72650 | -0.2 | 3.05E-02 | -0.4 | 4.67E-03 | 0.1 | 2.98E-01 | -0.4 | 7.98E-03 | -0.2 | 2.84E-01 | -0.2 | 2.70E-01 | At1g72650 |
| AT1G48540 | -0.2 | 9.69E-02 | -0.2 | 1.23E-01 | 0.0 | 9.51E-01 | -0.2 | 5.30E-01 | 0.0 | 9.69E-01 | -0.2 | 5.91E-01 | Outer arm dynein light chain 1 protein |
| AT5G46780 | -0.2 | 4.89E-02 | -0.2 | 8.20E-02 | 0.0 | 8.56E-01 | 0.0 | 9.62E-01 | -0.3 | 1.71E-01 | 0.3 | 2.04E-01 | VQ domain-containing protein |
| AT1G23780 | -0.2 | 1.93E-03 | -0.3 | 3.85E-04 | 0.1 | 2.17E-01 | -0.1 | 5.81E-01 | -0.3 | 1.34E-01 | 0.2 | 3.79E-01 | F-box protein SKIP22 |
| AT4G14200 | -0.2 | 1.98E-02 | 0.0 | 9.24E-01 | -0.2 | 5.14E-02 | -0.1 | 5.31E-01 | 0.1 | 7.66E-01 | -0.2 | 4.00E-01 | Uncharacterized protein |
| AT5G28030 | -0.2 | 2.96E-01 | -0.2 | 3.59E-01 | 0.0 | 9.49E-01 | -0.4 | 3.59E-01 | 0.0 | 9.62E-01 | -0.4 | 5.36E-01 | L-cysteine desulfhydrase 1 |
| AT3G58800 | -0.2 | 9.06E-02 | 0.1 | 7.05E-01 | -0.3 | 9.03E-02 | -0.4 | 1.65E-01 | -0.1 | 7.98E-01 | -0.3 | 4.51E-01 | hypothetical protein |
| AT4G15240 | -0.2 | 1.03E-02 | -0.2 | 2.65E-02 | 0.0 | 7.33E-01 | 0.0 | 9.52E-01 | 0.0 | 9.41E-01 | 0.0 | 9.04E-01 | Uncharacterized protein |
| AT5G14790 | -0.2 | 2.91E-02 | 0.2 | 1.13E-01 | -0.4 | 5.09E-03 | 0.0 | 8.02E-01 | 0.3 | 5.88E-02 | -0.3 | 6.69E-02 | TOG domain-containing protein |
| AT1G70360 | -0.2 | 1.26E-01 | -0.2 | 2.17E-01 | 0.0 | 8.40E-01 | -0.2 | 4.24E-01 | -0.3 | 3.29E-01 | 0.1 | 8.73E-01 | F-box family protein |
| AT5G25070 | -0.2 | 3.51E-03 | -0.1 | 8.39E-02 | -0.1 | 1.45E-01 | 0.1 | 6.03E-01 | 0.1 | 7.79E-01 | 0.0 | 9.23E-01 | Neurofilament light protein |
| AT3G07870 | -0.2 | 1.02E-02 | -0.2 | 2.12E-02 | 0.0 | 8.08E-01 | -0.5 | 3.93E-04 | -0.8 | 7.11E-06 | 0.4 | 2.89E-02 | F-box protein At3g07870 |
| AT1G73150 | -0.2 | 7.47E-02 | -0.1 | 2.88E-01 | -0.1 | 5.41E-01 | 0.0 | 1.00E+00 | 0.0 | 9.60E-01 | 0.0 | 9.66E-01 | Global transcription factor group E3 |
| AT3G02140 | -0.2 | 1.13E-01 | 0.6 | 1.18E-03 | -0.8 | 5.58E-04 | -0.1 | 7.73E-01 | -0.1 | 8.06E-01 | 0.0 | 9.97E-01 | Ninja-family protein AFP4 |
| AT3G63500 | -0.2 | 9.90E-03 | -0.4 | 4.63E-04 | 0.2 | 5.82E-02 | 0.0 | 9.29E-01 | -0.1 | 6.71E-01 | 0.1 | 7.79E-01 | Fibronectin type III domain protein |
| AT1G63240 | -0.2 | 1.35E-01 | -0.2 | 2.24E-01 | 0.0 | 8.45E-01 | -0.3 | 1.16E-01 | -0.2 | 3.46E-01 | -0.1 | 7.64E-01 | MBD domain-containing protein |
| AT2G13370 | -0.2 | 4.62E-03 | -0.3 | 1.51E-03 | 0.1 | 4.35E-01 | 0.0 | 8.93E-01 | -0.1 | 8.61E-01 | 0.1 | 7.90E-01 | Chromodomain-helicase-DNA-binding protein 1-like C-terminal domain-containing protein |
| AT5G14390 | -0.2 | 2.86E-02 | -0.1 | 2.02E-01 | -0.1 | 3.52E-01 | -0.5 | 3.31E-04 | -0.3 | 1.34E-02 | -0.2 | 3.50E-01 | Serine aminopeptidase S33 domain-containing protein |
| AT1G27500 | -0.2 | 6.12E-02 | -0.1 | 6.30E-01 | -0.2 | 2.27E-01 | -0.2 | 3.48E-01 | -0.2 | 3.68E-01 | 0.0 | 9.75E-01 | Tetratricopeptide repeat -like superfamily protein |
| AT4G36500 | -0.2 | 2.42E-01 | -0.1 | 4.61E-01 | -0.1 | 7.34E-01 | 0.2 | 3.68E-01 | -0.1 | 7.31E-01 | 0.4 | 2.85E-01 | hypothetical protein |
| AT2G36500 | -0.2 | 1.32E-01 | -0.1 | 4.24E-01 | -0.1 | 5.78E-01 | -0.1 | 7.81E-01 | 0.1 | 9.19E-01 | -0.2 | 7.23E-01 | CBS domain-containing protein CBSCBSPB2 |
| AT3G25010 | -0.2 | 7.27E-01 | -1.2 | 1.47E-01 | 0.9 | 3.13E-01 | -0.1 | 9.47E-01 | -0.9 | 2.24E-01 | 0.9 | 2.83E-01 | Receptor like protein 41 |
| AT1G65270 | -0.2 | 4.93E-03 | 0.0 | 7.33E-01 | -0.2 | 1.81E-02 | 0.1 | 2.47E-01 | 0.2 | 1.05E-01 | -0.1 | 6.96E-01 | ER membrane protein complex subunit 10 |
| AT4G04620 | -0.2 | 2.98E-02 | -0.7 | 9.14E-05 | 0.5 | 3.84E-03 | 0.2 | 3.96E-01 | -0.3 | 3.22E-01 | 0.5 | 8.13E-02 | Autophagy-related protein 8b |
| AT3G50370 | -0.2 | 3.80E-02 | -0.4 | 1.51E-03 | 0.2 | 7.15E-02 | 0.0 | 8.85E-01 | -0.1 | 7.40E-01 | 0.1 | 6.87E-01 | Uncharacterized protein At3g50370 |
| AT4G03440 | -0.2 | 7.55E-02 | 0.0 | 9.85E-01 | -0.2 | 1.35E-01 | 0.1 | 7.44E-01 | 0.4 | 2.02E-01 | -0.3 | 4.23E-01 | Ankyrin repeat family protein |
| AT3G11720 | -0.2 | 1.70E-02 | -0.2 | 3.22E-02 | 0.0 | 8.40E-01 | 0.0 | 9.89E-01 | 0.2 | 5.52E-01 | -0.2 | 6.29E-01 | At3g11720 |
| AT5G23490 | -0.2 | 1.19E-02 | -0.4 | 1.47E-03 | 0.1 | 2.15E-01 | 0.1 | 8.06E-01 | 0.1 | 7.76E-01 | 0.0 | 9.55E-01 | Gb\|AAF01580.1 |
| AT2G41475 | -0.2 | 5.52E-03 | 0.2 | 2.06E-02 | -0.4 | 4.91E-04 | 0.0 | 8.62E-01 | 0.1 | 4.89E-01 | -0.1 | 6.59E-01 | Uncharacterized protein At2g41470 |
| AT4G33480 | -0.2 | 5.15E-02 | -0.1 | 3.27E-01 | -0.1 | 3.68E-01 | -0.3 | 2.77E-02 | -0.3 | 3.95E-02 | 0.0 | 9.77E-01 | Uncharacterized protein At4g33480 |
| AT3G51580 | -0.2 | 2.69E-03 | -0.2 | 4.60E-03 | 0.0 | 8.96E-01 | -0.1 | 4.31E-01 | 0.0 | 9.99E-01 | -0.1 | 6.03E-01 | Transmembrane protein |
| AT5G10580 | -0.2 | 7.69E-02 | -0.8 | 2.08E-04 | 0.5 | 4.79E-03 | 0.6 | 2.77E-02 | 0.2 | 4.81E-01 | 0.4 | 2.75E-01 | Uncharacterized protein |
| AT3G14000 | -0.2 | 2.47E-02 | 0.0 | 8.92E-01 | -0.2 | 6.60E-02 | -0.4 | 3.57E-02 | -0.1 | 6.97E-01 | -0.3 | 1.70E-01 | Protein Brevis radix-like 2 |
| AT1G66510 | -0.2 | 1.06E-01 | -0.2 | 2.83E-01 | -0.1 | 6.65E-01 | 0.2 | 4.53E-01 | 0.0 | 9.66E-01 | 0.2 | 5.95E-01 | AAR2 protein family |
| AT1G11905 | -0.2 | 1.83E-02 | -0.2 | 2.57E-02 | 0.0 | 9.45E-01 | -0.1 | 6.00E-01 | -0.2 | 4.24E-01 | 0.1 | 8.15E-01 | Endoplasmic reticulum transmembrane protein |
| AT5G16410 | -0.2 | 1.77E-01 | 0.2 | 3.49E-01 | -0.4 | 7.40E-02 | 0.3 | 4.23E-01 | 0.0 | 9.39E-01 | 0.2 | 6.20E-01 | At5g16410 |
| AT1G62300 | -0.2 | 7.32E-02 | -0.4 | 8.28E-03 | 0.2 | 2.50E-01 | 0.3 | 3.77E-01 | -0.4 | 2.39E-01 | 0.7 | 4.71E-02 | Truncated WRKY6 protein |
| AT2G38070 | -0.2 | 2.07E-01 | 0.0 | 9.07E-01 | -0.2 | 3.62E-01 | 0.3 | 3.79E-01 | 0.0 | 9.89E-01 | 0.2 | 5.38E-01 | LOW protein: UPF0503-like protein, putative |
| AT5G28010 | -0.2 | 1.57E-01 | 0.1 | 3.62E-01 | -0.4 | 6.71E-02 | -1.0 | 2.28E-02 | -0.2 | 6.08E-01 | -0.7 | 1.54E-01 | Polyketide cyclase |
| AT5G42660 | -0.2 | 1.15E-02 | -0.1 | 3.51E-01 | -0.2 | 1.15E-01 | -0.1 | 5.67E-01 | 0.1 | 6.69E-01 | -0.2 | 3.75E-01 | DNA-directed RNA polymerase subunit beta |
| AT4G14590 | -0.2 | 1.02E-01 | -0.1 | 3.12E-01 | -0.1 | 6.18E-01 | 0.0 | 9.23E-01 | -0.6 | 2.13E-01 | 0.6 | 2.01E-01 | Embryo defective 2739 |
| AT4G16790 | -0.2 | 1.26E-02 | -0.3 | 4.45E-03 | 0.1 | 5.18E-01 | 0.1 | 5.17E-01 | 0.2 | 3.74E-01 | -0.1 | 8.42E-01 | AT4g16790 |
| AT2G01970 | -0.2 | 7.37E-03 | -0.1 | 1.17E-01 | -0.1 | 1.94E-01 | 0.0 | 9.21E-01 | 0.1 | 5.04E-01 | -0.1 | 6.88E-01 | Transmembrane 9 superfamily member 3 |
| AT1G31814 | -0.2 | 4.51E-02 | 0.0 | 7.72E-01 | -0.3 | 5.72E-02 | 0.4 | 1.49E-01 | 0.1 | 6.94E-01 | 0.2 | 3.97E-01 | Inactive FRIGIDA-like protein 2 |
| AT4G13270 | -0.2 | 2.27E-03 | -0.1 | 3.52E-02 | -0.1 | 2.00E-01 | -0.1 | 4.05E-01 | -0.1 | 4.82E-01 | 0.0 | 9.97E-01 | At4g13270 hydroxyproline-rich glycoprotein family) |
| AT3G50140 | -0.2 | 1.79E-01 | 0.2 | 3.33E-01 | -0.4 | 6.91E-02 | 0.0 | 9.85E-01 | -0.1 | 8.69E-01 | 0.1 | 8.64E-01 | Transmembrane protein, |
| AT4G24060 | -0.2 | 1.07E-02 | 0.0 | 8.89E-01 | -0.2 | 1.88E-02 | 0.0 | 9.23E-01 | -0.1 | 7.51E-01 | 0.0 | 8.61E-01 | Uncharacterized protein |
| AT5G26720 | -0.2 | 2.50E-01 | -0.3 | 1.71E-01 | 0.1 | 8.20E-01 | -0.4 | 4.30E-01 | -0.2 | 7.77E-01 | -0.2 | 7.79E-01 | hypothetical protein |
| AT4G39240 | -0.2 | 8.94E-03 | -0.1 | 7.87E-02 | -0.1 | 3.25E-01 | 0.2 | 2.27E-01 | 0.2 | 2.76E-01 | 0.0 | 9.81E-01 | F-box |
| AT3G52760 | -0.2 | 2.11E-02 | 0.1 | 1.51E-01 | -0.4 | 5.17E-03 | -0.3 | 1.72E-01 | -0.4 | 1.25E-01 | 0.1 | 8.42E-01 | Protein YIP |
| AT4G00650 | -0.2 | 5.96E-02 | -0.4 | 8.01E-03 | 0.2 | 2.99E-01 | -0.1 | 7.65E-01 | -0.2 | 7.42E-01 | 0.0 | 9.63E-01 | FRIGIDA-like protein |
| AT2G18500 | -0.2 | 7.26E-02 | -0.4 | 7.06E-03 | 0.2 | 2.26E-01 | 0.1 | 6.48E-01 | 0.3 | 3.27E-01 | -0.2 | 7.01E-01 | Transcription repressor OFP7 |
| AT1G76660 | -0.2 | 9.65E-03 | -0.2 | 2.53E-02 | 0.0 | 7.14E-01 | 0.0 | 7.76E-01 | -0.1 | 7.24E-01 | 0.0 | 9.40E-01 | Hydroxyproline-rich glycoprotein family protein |
| AT1G76700 | -0.2 | 5.18E-03 | -0.5 | 7.59E-05 | 0.2 | 1.02E-02 | -0.3 | 3.85E-02 | -0.4 | 5.39E-02 | 0.0 | 9.81E-01 | Chaperone protein dnaJ 10 |
| AT1G55475 | -0.2 | 1.78E-01 | -0.4 | 2.63E-02 | 0.2 | 3.28E-01 | -0.1 | 6.97E-01 | 0.0 | 9.47E-01 | -0.1 | 8.37E-01 | hypothetical protein |
| AT5G23390 | -0.2 | 5.50E-03 | -0.2 | 2.31E-02 | 0.0 | 5.43E-01 | -0.2 | 2.33E-01 | -0.1 | 6.88E-01 | -0.1 | 5.96E-01 | AT5g23390 |
| AT5G22450 | -0.2 | 1.16E-02 | -0.5 | 1.80E-04 | 0.2 | 1.56E-02 | -0.2 | 3.40E-01 | -0.2 | 4.07E-01 | 0.0 | 1.00E+00 | Spectrin beta chain, brain |
| AT4G23460 | -0.2 | 9.69E-03 | -0.2 | 1.13E-02 | 0.0 | 9.78E-01 | 0.2 | 2.27E-01 | 0.0 | 9.76E-01 | 0.2 | 4.42E-01 | Adaptin family protein |
| AT1G17510 | -0.2 | 3.18E-02 | -0.1 | 4.95E-01 | -0.2 | 1.82E-01 | 0.0 | 8.84E-01 | 0.0 | 9.18E-01 | 0.0 | 9.94E-01 | hypothetical protein |
| AT2G30230 | -0.2 | 2.05E-01 | -0.4 | 3.64E-02 | 0.2 | 3.73E-01 | -0.4 | 5.00E-02 | 0.1 | 8.50E-01 | -0.5 | 7.14E-02 | hypothetical protein |
| AT2G27090 | -0.2 | 8.45E-03 | -0.3 | 2.72E-03 | 0.1 | 4.67E-01 | 0.1 | 5.48E-01 | -0.1 | 7.19E-01 | 0.2 | 3.89E-01 | At2g27090 ) |
| AT3G18160 | -0.2 | 1.33E-02 | -0.2 | 7.24E-02 | -0.1 | 4.67E-01 | 0.3 | 1.15E-01 | 0.3 | 1.59E-01 | 0.0 | 9.98E-01 | Peroxin 3-1 |
| AT4G31820 | -0.2 | 2.85E-02 | 0.1 | 1.97E-01 | -0.3 | 7.10E-03 | -0.4 | 8.57E-03 | -0.4 | 1.58E-02 | 0.0 | 9.97E-01 | Phototropic-responsive NPH3 family protein |
| AT4G32620 | -0.2 | 3.36E-03 | -0.3 | 5.49E-04 | 0.1 | 2.00E-01 | -0.1 | 7.84E-01 | 0.0 | 8.45E-01 | 0.0 | 9.90E-01 | Enhancer of polycomb-like protein |
| AT2G42450 | -0.2 | 6.65E-02 | -0.3 | 3.52E-02 | 0.1 | 7.29E-01 | -0.3 | 2.25E-01 | -0.2 | 3.72E-01 | 0.0 | 9.11E-01 | Alpha |
| AT4G16060 | -0.2 | 3.65E-02 | -0.1 | 3.17E-01 | -0.1 | 3.05E-01 | -0.4 | 3.48E-03 | 0.0 | 9.40E-01 | -0.4 | 2.11E-02 | Uncharacterized protein |
| AT1G34760 | -0.2 | 5.69E-02 | 0.1 | 5.33E-01 | -0.3 | 3.91E-02 | -0.2 | 1.65E-01 | -0.7 | 5.07E-04 | 0.5 | 1.87E-02 | 14-3-3 domain-containing protein |
| AT2G36330 | -0.2 | 2.40E-02 | -0.1 | 3.40E-01 | -0.1 | 2.08E-01 | 0.0 | 9.94E-01 | 0.1 | 8.52E-01 | -0.1 | 8.77E-01 | CASP-like protein |
| AT1G03820 | -0.2 | 6.73E-02 | -0.2 | 8.20E-02 | 0.0 | 9.72E-01 | 0.2 | 2.45E-01 | 0.4 | 5.68E-02 | -0.2 | 5.22E-01 | hypothetical protein |
| AT1G14090 | -0.2 | 9.02E-02 | -0.3 | 2.15E-02 | 0.1 | 4.58E-01 | 0.5 | 2.08E-01 | 0.1 | 8.61E-01 | 0.4 | 4.41E-01 | NA |
| AT2G14860 | -0.2 | 2.46E-01 | -0.1 | 7.95E-01 | -0.2 | 4.74E-01 | 0.0 | 9.88E-01 | 0.0 | 9.90E-01 | 0.0 | 9.88E-01 | 22 kDa peroxisomal membrane protein family protein) |
| AT3G59760 | -0.2 | 3.62E-02 | 0.2 | 1.20E-01 | -0.4 | 5.02E-03 | 0.1 | 2.66E-01 | 0.2 | 1.21E-01 | -0.1 | 7.21E-01 | Cysteine synthase |
| AT5G01350 | -0.2 | 4.06E-02 | -0.2 | 7.46E-02 | 0.0 | 8.20E-01 | -0.1 | 7.83E-01 | -0.2 | 5.92E-01 | 0.1 | 8.44E-01 | Uncharacterized protein T10O8_60 |
| AT5G53090 | -0.2 | 3.72E-02 | 0.1 | 5.48E-01 | -0.3 | 3.11E-02 | 0.0 | 8.79E-01 | 0.3 | 1.92E-01 | -0.4 | 2.01E-01 | Protochlorophyllide reductase; oxidoreductase required for shoot apex development |
| AT1G29690 | -0.2 | 1.26E-02 | -0.3 | 2.56E-03 | 0.1 | 3.37E-01 | 0.0 | 7.91E-01 | -0.1 | 5.67E-01 | 0.1 | 8.26E-01 | CAD1 |
| AT5G13720 | -0.2 | 1.80E-02 | -0.2 | 5.04E-02 | 0.0 | 6.76E-01 | -0.2 | 5.18E-02 | 0.0 | 9.21E-01 | -0.2 | 1.94E-01 | Uncharacterized protein At5g13720 ) |
| AT2G21080 | -0.2 | 1.26E-01 | 0.2 | 7.89E-02 | -0.5 | 1.41E-02 | 0.1 | 5.96E-01 | -0.2 | 3.97E-01 | 0.3 | 2.28E-01 | Uncharacterized protein At2g21080 |

| | | | | | | | | | | | | | |
|---|---|---|---|---|---|---|---|---|---|---|---|---|---|
| AT2G10931 | -0.2 | 1.67E-01 | -0.1 | 7.39E-01 | -0.2 | 3.91E-01 | -0.2 | 5.06E-01 | -0.1 | 7.14E-01 | -0.1 | 9.07E-01 | Uncharacterized protein |
| AT1G80530 | -0.2 | 7.00E-03 | -0.1 | 6.31E-02 | -0.1 | 3.08E-01 | 0.3 | 1.22E-02 | 0.1 | 4.80E-01 | 0.2 | 2.01E-01 | Major facilitator superfamily protein |
| AT1G11270 | -0.2 | 1.49E-01 | -0.3 | 1.02E-01 | 0.0 | 8.14E-01 | 0.1 | 7.32E-01 | 0.0 | 9.32E-01 | 0.1 | 8.74E-01 | F-box and associated interaction domains-containing protein |
| AT5G54440 | -0.2 | 1.01E-02 | -0.2 | 2.29E-02 | 0.0 | 7.56E-01 | -0.3 | 1.54E-01 | -0.1 | 6.90E-01 | -0.2 | 5.67E-01 | CLUB |
| AT3G11010 | -0.2 | 5.40E-01 | -0.7 | 6.68E-02 | 0.5 | 2.31E-01 | 1.4 | 2.11E-04 | 0.7 | 6.21E-02 | 0.7 | 4.18E-02 | Receptor like protein 34 |
| AT3G58170 | -0.2 | 4.42E-02 | -0.1 | 2.02E-01 | -0.1 | 4.82E-01 | -0.1 | 5.61E-01 | -0.1 | 7.17E-01 | 0.0 | 9.44E-01 | AT3G58170 protein |
| AT5G28220 | -0.2 | 5.67E-02 | -0.1 | 2.38E-01 | -0.1 | 5.14E-01 | 0.2 | 3.77E-01 | 0.1 | 6.34E-01 | 0.1 | 8.52E-01 | ER membrane protein complex subunit 2 |
| AT1G54920 | -0.2 | 5.58E-03 | -0.2 | 3.64E-02 | -0.1 | 3.91E-01 | -0.2 | 2.65E-01 | 0.0 | 8.69E-01 | -0.2 | 5.44E-01 | At1g54920 |
| AT1G73210 | -0.2 | 6.65E-02 | 0.0 | 7.45E-01 | -0.2 | 2.02E-01 | 0.0 | 9.39E-01 | 0.2 | 6.73E-01 | -0.2 | 6.66E-01 | Uncharacterized protein |
| AT3G56440 | -0.2 | 6.65E-02 | -0.2 | 6.14E-02 | 0.0 | 9.28E-01 | 0.0 | 8.92E-01 | 0.3 | 2.96E-01 | -0.4 | 3.24E-01 | Autophagy 18 D-like protein |
| AT4G17150 | -0.2 | 9.57E-03 | -0.1 | 1.49E-01 | -0.1 | 2.06E-01 | -0.2 | 2.18E-01 | -0.1 | 7.05E-01 | -0.1 | 6.24E-01 | Alpha |
| AT3G26950 | -0.2 | 8.34E-03 | -0.1 | 1.96E-01 | -0.1 | 1.38E-01 | -0.3 | 1.22E-01 | 0.1 | 7.91E-01 | -0.3 | 1.47E-01 | Uncharacterized protein At3g26950 |
| AT5G57950 | -0.2 | 6.07E-02 | -0.2 | 1.20E-01 | 0.0 | 7.97E-01 | -0.2 | 3.06E-01 | -0.3 | 1.94E-01 | 0.1 | 8.16E-01 | 26S proteasome regulatory subunit |
| AT2G46150 | -0.2 | 8.66E-02 | 0.0 | 6.94E-01 | -0.2 | 2.56E-01 | -0.3 | 7.96E-02 | -0.1 | 7.99E-01 | -0.3 | 2.14E-01 | Late embryogenesis abundant protein LEA-2 subgroup domain-containing protein |
| AT3G12350 | -0.2 | 2.02E-02 | -0.3 | 1.77E-03 | 0.1 | 1.65E-01 | -0.2 | 3.25E-01 | -0.1 | 4.12E-01 | 0.0 | 9.81E-01 | AT3G12350 protein |
| AT3G50845 | -0.2 | 2.48E-01 | -0.1 | 7.93E-01 | -0.2 | 4.79E-01 | -0.3 | 2.85E-01 | -0.5 | 1.33E-01 | 0.2 | 6.73E-01 | Uncharacterized protein |
| AT4G00780 | -0.2 | 2.52E-02 | 0.0 | 8.16E-01 | -0.2 | 6.28E-02 | 0.0 | 9.35E-01 | 0.1 | 6.20E-01 | -0.1 | 7.61E-01 | MATH domain-containing protein |
| AT5G57860 | -0.2 | 1.89E-02 | -0.3 | 1.98E-03 | 0.1 | 2.04E-01 | 0.0 | 9.22E-01 | -0.1 | 7.67E-01 | 0.2 | 7.26E-01 | Ubiquitin-like domain-containing protein |
| AT2G37570 | -0.2 | 1.04E-02 | -0.5 | 9.44E-05 | 0.3 | 7.12E-03 | -0.1 | 2.96E-01 | -0.2 | 1.69E-01 | 0.1 | 7.78E-01 | HSP20-like chaperones superfamily protein |
| AT2G47350 | -0.2 | 1.39E-02 | -0.2 | 1.37E-02 | 0.0 | 9.25E-01 | -0.4 | 2.06E-02 | -0.2 | 3.21E-01 | -0.2 | 3.74E-01 | HIT zinc finger and PAPA-1-like domain-containing protein |
| AT3G63445 | -0.2 | 1.49E-01 | -0.2 | 2.76E-01 | 0.0 | 8.07E-01 | 0.0 | 9.84E-01 | 0.0 | 9.75E-01 | 0.0 | 9.63E-01 | NA |
| AT5G59980 | -0.2 | 1.55E-01 | -0.3 | 4.07E-02 | 0.1 | 4.96E-01 | 0.2 | 5.01E-01 | -0.1 | 9.04E-01 | 0.3 | 5.01E-01 | Polymerase |
| AT5G41280 | -0.2 | 2.69E-01 | -0.8 | 3.05E-03 | 0.6 | 3.59E-02 | 1.8 | 6.21E-04 | 2.2 | 2.67E-04 | -0.4 | 2.49E-01 | Cysteine-rich repeat secretory protein 57 |
| AT1G21610 | -0.2 | 2.25E-02 | -0.2 | 2.41E-02 | 0.0 | 9.68E-01 | -0.2 | 3.69E-01 | -0.1 | 5.95E-01 | -0.1 | 8.59E-01 | Wound-responsive family protein |
| AT3G21290 | -0.2 | 2.30E-02 | -0.2 | 4.35E-02 | 0.0 | 8.34E-01 | -0.2 | 3.04E-01 | -0.2 | 4.72E-01 | 0.0 | 9.24E-01 | hypothetical protein |
| AT1G61030 | -0.2 | 3.95E-02 | -0.4 | 3.34E-03 | 0.2 | 1.89E-01 | -0.3 | 2.10E-01 | -0.2 | 5.30E-01 | -0.1 | 7.52E-01 | Wings apart-like protein 2 |
| AT5G35320 | -0.2 | 2.83E-02 | -0.1 | 1.19E-01 | -0.1 | 5.43E-01 | 0.0 | 9.13E-01 | 0.0 | 9.48E-01 | 0.0 | 9.86E-01 | Uncharacterized protein |
| AT3G22520 | -0.2 | 1.02E-02 | -0.2 | 4.35E-02 | -0.1 | 5.29E-01 | 0.1 | 5.16E-01 | 0.1 | 7.38E-01 | 0.0 | 9.11E-01 | Spindle assembly abnormal protein |
| AT1G65320 | -0.2 | 2.55E-02 | -0.1 | 2.84E-01 | -0.1 | 2.51E-01 | -0.2 | 9.08E-02 | 0.0 | 7.95E-01 | -0.2 | 3.68E-01 | CBS domain-containing protein CBSX6 |
| AT5G18680 | -0.2 | 1.42E-02 | 0.0 | 6.71E-01 | -0.2 | 5.32E-02 | -0.3 | 7.06E-03 | -0.3 | 4.97E-02 | -0.1 | 6.98E-01 | Tubby like protein 11 |
| AT5G23520 | -0.2 | 2.06E-02 | -0.4 | 9.31E-04 | 0.2 | 8.22E-02 | -0.2 | 2.08E-01 | -0.1 | 6.71E-01 | -0.1 | 6.33E-01 | Smr domain-containing protein |
| AT4G24210 | -0.2 | 3.31E-02 | 0.0 | 9.27E-01 | -0.2 | 5.35E-02 | -0.3 | 2.62E-01 | -0.1 | 7.80E-01 | -0.2 | 5.99E-01 | F-box protein GID2 |
| AT1G68490 | -0.2 | 1.59E-02 | -0.1 | 4.44E-01 | -0.1 | 1.00E-01 | -0.2 | 4.46E-01 | -0.1 | 8.46E-01 | -0.1 | 7.61E-01 | Translocase subunit seca |
| AT1G04390 | -0.2 | 1.11E-01 | -0.4 | 1.30E-02 | 0.2 | 2.72E-01 | -0.5 | 8.21E-02 | -0.2 | 4.44E-01 | -0.2 | 5.70E-01 | Uncharacterized protein |
| AT1G54660 | -0.2 | 4.04E-02 | -0.1 | 2.05E-01 | -0.1 | 4.67E-01 | -1.0 | 3.84E-04 | 0.1 | 8.59E-01 | -1.0 | 1.87E-03 | NA |
| AT1G15400 | -0.2 | 6.22E-02 | -0.1 | 1.82E-01 | -0.1 | 6.24E-01 | -0.3 | 9.63E-02 | -0.4 | 3.11E-02 | 0.1 | 6.34E-01 | hypothetical protein |
| AT5G03553 | -0.2 | 4.93E-01 | 0.3 | 2.27E-01 | -0.6 | 1.28E-01 | -0.6 | 9.39E-02 | -0.1 | 8.25E-01 | -0.5 | 2.65E-01 | Uncharacterized protein |
| AT3G06940 | -0.2 | 3.03E-02 | -0.1 | 3.04E-01 | -0.1 | 2.76E-01 | -0.1 | 7.59E-01 | 0.0 | 9.99E-01 | -0.1 | 8.37E-01 | NA |
| AT1G19370 | -0.2 | 8.57E-03 | 0.2 | 2.33E-02 | -0.4 | 8.29E-04 | -0.2 | 2.08E-01 | 0.0 | 9.87E-01 | -0.2 | 3.15E-01 | Uncharacterized protein |
| AT2G04410 | -0.2 | 6.13E-03 | 0.0 | 8.72E-01 | -0.2 | 9.57E-03 | -0.2 | 3.69E-01 | -0.2 | 2.75E-01 | 0.1 | 8.52E-01 | hypothetical protein |
| AT2G38020 | -0.2 | 4.16E-03 | -0.2 | 4.14E-03 | 0.0 | 9.01E-01 | 0.0 | 9.02E-01 | 0.1 | 7.04E-01 | -0.1 | 8.28E-01 | Uncharacterized protein At2g38020 |
| AT3G06365 | -0.2 | 5.85E-01 | -0.2 | 7.07E-01 | -0.1 | 9.10E-01 | 0.4 | 7.81E-01 | 0.0 | 9.90E-01 | 0.4 | 8.53E-01 | NA |
| AT2G25920 | -0.2 | 2.26E-02 | -0.1 | 1.93E-01 | -0.1 | 3.36E-01 | -0.3 | 8.82E-02 | 0.0 | 9.04E-01 | -0.3 | 1.94E-01 | Pollen-specific LRR extensin-like protein |
| AT4G28600 | -0.2 | 1.87E-02 | -0.4 | 4.31E-04 | 0.2 | 3.34E-02 | -0.1 | 6.51E-01 | -0.2 | 6.47E-01 | 0.0 | 9.62E-01 | No pollen germination related 2 |
| AT5G58590 | -0.2 | 4.68E-02 | 0.0 | 8.24E-01 | -0.2 | 1.09E-01 | 0.1 | 7.81E-01 | 0.0 | 9.88E-01 | 0.1 | 8.69E-01 | Ran-binding protein 1 homolog c |
| AT1G60670 | -0.2 | 1.10E-02 | -0.2 | 7.69E-03 | 0.0 | 7.88E-01 | -0.1 | 5.18E-01 | -0.1 | 6.61E-01 | 0.0 | 9.51E-01 | Uncharacterized protein At1g60670 |
| AT4G09340 | -0.2 | 1.84E-02 | -0.3 | 1.65E-03 | 0.1 | 1.66E-01 | 0.1 | 6.91E-01 | 0.1 | 4.87E-01 | -0.1 | 8.26E-01 | Uncharacterized protein AT4g09330 |
| AT2G17200 | -0.2 | 1.19E-02 | -0.4 | 5.70E-04 | 0.2 | 6.00E-02 | -0.2 | 1.05E-01 | -0.2 | 2.31E-01 | 0.0 | 9.09E-01 | DSK2b |
| AT5G48385 | -0.2 | 7.12E-03 | -0.3 | 1.55E-03 | 0.1 | 2.94E-01 | 0.0 | 8.88E-01 | -0.2 | 4.75E-01 | 0.1 | 6.67E-01 | FRIGIDA-like protein 3 |
| AT5G65495 | -0.2 | 5.40E-02 | 0.0 | 8.15E-01 | -0.2 | 1.42E-01 | 0.1 | 7.29E-01 | 0.0 | 9.35E-01 | 0.1 | 7.27E-01 | Protein SKIP34 |
| AT3G03740 | -0.2 | 9.44E-03 | -0.3 | 2.55E-03 | 0.1 | 4.12E-01 | -0.1 | 5.39E-01 | 0.0 | 9.41E-01 | -0.1 | 5.83E-01 | BPM4 |
| AT1G01990 | -0.2 | 9.56E-03 | -0.2 | 1.69E-02 | 0.0 | 8.67E-01 | -0.1 | 7.73E-01 | -0.3 | 1.04E-01 | 0.2 | 2.72E-01 | Transmembrane protein |
| AT5G10745 | -0.2 | 4.04E-02 | 0.0 | 7.78E-01 | -0.2 | 4.74E-02 | 0.0 | 8.37E-01 | -0.2 | 3.88E-01 | 0.1 | 5.98E-01 | hypothetical protein |
| AT2G47630 | -0.2 | 1.18E-01 | 0.0 | 8.60E-01 | -0.2 | 1.30E-01 | 0.1 | 4.35E-01 | 0.2 | 2.83E-01 | -0.1 | 8.15E-01 | Serine aminopeptidase S33 domain-containing protein |
| AT1G03365 | -0.2 | 9.10E-03 | -0.1 | 7.69E-02 | -0.1 | 3.50E-01 | -0.2 | 3.29E-01 | -0.1 | 5.02E-01 | 0.0 | 9.12E-01 | RING-type domain-containing protein |
| AT1G05320 | -0.2 | 1.56E-02 | 0.1 | 1.58E-01 | -0.3 | 3.98E-03 | 0.1 | 3.54E-01 | 0.2 | 1.20E-01 | -0.1 | 6.12E-01 | Uncharacterized protein At1g05320 |
| AT1G68070 | -0.2 | 8.08E-03 | -0.3 | 1.57E-03 | 0.1 | 2.99E-01 | -0.4 | 4.22E-03 | -0.3 | 4.29E-02 | -0.1 | 5.53E-01 | RING-type domain-containing protein |
| AT3G27570 | -0.2 | 9.97E-03 | -0.1 | 6.18E-02 | -0.1 | 4.10E-01 | 0.2 | 4.92E-02 | 0.1 | 4.38E-01 | 0.1 | 4.73E-01 | Sucrose cleavage protein-like |
| AT3G16760 | -0.2 | 1.66E-02 | -0.1 | 3.32E-01 | -0.1 | 1.45E-01 | 0.0 | 8.22E-01 | 0.1 | 5.51E-01 | -0.1 | 7.49E-01 | Tetratricopeptide repeat -like superfamily protein |
| AT1G02990 | -0.2 | 5.11E-02 | -0.4 | 2.48E-03 | 0.2 | 1.10E-01 | 0.0 | 9.44E-01 | 0.1 | 7.64E-01 | -0.1 | 7.46E-01 | Uncharacterized protein At1g02990 |
| AT1G16680 | -0.2 | 4.37E-02 | -0.2 | 4.75E-02 | 0.0 | 9.82E-01 | -0.2 | 2.03E-01 | 0.0 | 9.69E-01 | -0.2 | 4.00E-01 | F19K19.3 protein |
| AT5G03500 | -0.2 | 4.18E-02 | -0.1 | 1.74E-01 | -0.1 | 5.35E-01 | 0.0 | 9.82E-01 | -0.2 | 5.22E-01 | 0.1 | 5.72E-01 | Med7b |
| AT1G13320 | -0.2 | 4.28E-03 | 0.0 | 6.17E-01 | -0.2 | 2.05E-02 | -0.1 | 3.92E-01 | -0.1 | 7.10E-01 | 0.0 | 8.39E-01 | Protein phosphatase 2A subunit A3 |
| AT3G13360 | -0.2 | 1.02E-02 | -0.1 | 7.24E-02 | -0.1 | 3.74E-01 | -0.4 | 7.74E-03 | -0.2 | 2.17E-01 | -0.2 | 3.39E-01 | WPP domain-interacting protein 3 |
| AT1G66190 | -0.2 | 3.69E-02 | 0.0 | 6.75E-01 | -0.2 | 1.39E-01 | 0.0 | 9.77E-01 | -0.1 | 6.92E-01 | 0.1 | 6.77E-01 | Uncharacterized protein |
| AT4G25030 | -0.2 | 2.99E-02 | 0.1 | 5.45E-01 | -0.3 | 1.80E-02 | -0.6 | 1.76E-04 | -0.4 | 1.47E-02 | -0.2 | 1.94E-01 | AT4G25030 protein |
| AT5G13560 | -0.2 | 1.19E-02 | -0.2 | 4.03E-02 | 0.0 | 6.07E-01 | 0.0 | 9.44E-01 | -0.1 | 7.37E-01 | 0.1 | 7.43E-01 | At5g13560 |
| AT3G62630 | -0.2 | 1.03E-01 | -0.1 | 4.88E-01 | -0.1 | 4.10E-01 | -0.2 | 1.64E-01 | -0.1 | 4.57E-01 | -0.1 | 7.57E-01 | Stress response NST1-like protein |
| AT1G80810 | -0.2 | 4.02E-02 | -0.1 | 4.08E-01 | -0.1 | 2.59E-01 | -0.3 | 1.29E-01 | -0.3 | 1.68E-01 | 0.0 | 9.97E-01 | Tudor |
| AT1G12760 | -0.2 | 7.47E-03 | -0.3 | 3.68E-04 | 0.1 | 5.34E-02 | -0.2 | 1.63E-02 | -0.2 | 1.35E-01 | -0.1 | 6.51E-01 | AT1G12760 protein family protein) |
| AT4G27860 | -0.2 | 3.63E-02 | 0.1 | 4.12E-01 | -0.3 | 1.76E-02 | 0.1 | 6.82E-01 | 0.2 | 4.28E-01 | -0.1 | 7.64E-01 | Membrane protein of ER body-like protein |
| AT4G27690 | -0.2 | 6.98E-02 | -0.1 | 4.67E-01 | -0.1 | 3.38E-01 | -0.1 | 6.72E-01 | 0.1 | 7.69E-01 | -0.2 | 4.38E-01 | Vacuolar protein sorting 26B |
| AT5G64030 | -0.2 | 9.13E-03 | 0.0 | 7.01E-01 | -0.2 | 3.35E-02 | 0.3 | 5.50E-02 | 0.1 | 6.70E-01 | 0.2 | 3.47E-01 | Putative ankyrin |
| AT5G04420 | -0.2 | 8.59E-03 | -0.1 | 1.59E-01 | -0.1 | 1.73E-01 | -0.2 | 3.41E-01 | -0.1 | 6.52E-01 | -0.1 | 8.28E-01 | Galactose oxidase |
| AT2G41780 | -0.2 | 1.37E-01 | 0.2 | 2.05E-01 | -0.4 | 3.17E-02 | -0.1 | 6.84E-01 | 0.0 | 9.94E-01 | -0.1 | 7.37E-01 | hypothetical protein |
| AT1G69890 | -0.2 | 1.28E-01 | 0.0 | 7.05E-01 | -0.2 | 1.09E-01 | -0.4 | 2.13E-02 | -0.7 | 1.58E-03 | 0.3 | 2.76E-01 | hypothetical protein |
| AT5G42350 | -0.2 | 1.03E-01 | -0.3 | 5.32E-02 | 0.1 | 7.17E-01 | -0.4 | 1.71E-01 | -0.5 | 9.18E-02 | 0.1 | 7.86E-01 | F-box |
| AT3G06180 | -0.2 | 6.71E-02 | -0.1 | 2.36E-01 | -0.1 | 5.77E-01 | -0.5 | 4.63E-03 | -0.3 | 1.24E-01 | -0.2 | 3.44E-01 | Ribosomal protein L34e superfamily protein |
| AT3G08670 | -0.2 | 4.91E-02 | 0.0 | 6.79E-01 | -0.2 | 4.16E-02 | -0.2 | 1.61E-01 | 0.1 | 8.08E-01 | -0.3 | 1.73E-01 | At3g08670 |
| AT2G35900 | -0.2 | 4.18E-02 | -0.2 | 6.92E-02 | 0.0 | 8.67E-01 | 0.0 | 8.55E-01 | -0.2 | 1.44E-01 | 0.2 | 1.72E-01 | hypothetical protein |
| AT2G41160 | -0.2 | 1.82E-02 | -0.2 | 5.16E-02 | 0.0 | 6.79E-01 | -0.2 | 2.19E-01 | 0.1 | 5.42E-01 | -0.3 | 1.16E-01 | Ubiquitin-associated protein |
| AT1G61670 | -0.2 | 1.37E-02 | -0.2 | 9.25E-03 | 0.0 | 7.78E-01 | 0.1 | 5.90E-01 | 0.0 | 8.56E-01 | 0.0 | 8.59E-01 | Lung seven transmembrane receptor family protein |
| AT4G12780 | -0.2 | 6.81E-03 | -0.3 | 1.71E-03 | 0.1 | 3.59E-01 | 0.0 | 8.16E-01 | 0.0 | 8.68E-01 | 0.1 | 7.57E-01 | Chaperone DnaJ-domain superfamily protein |

| | | | | | | | | | | | | | |
|---|---|---|---|---|---|---|---|---|---|---|---|---|---|
| AT1G52200 | -0.2 | 1.53E-01 | -0.1 | 4.66E-01 | -0.1 | 5.45E-01 | 0.4 | 2.94E-02 | -0.3 | 1.03E-01 | 0.7 | 8.87E-03 | PLAC8 family protein |
| AT1G25570 | -0.2 | 1.60E-02 | -0.1 | 1.33E-01 | -0.1 | 3.29E-01 | -0.1 | 4.27E-01 | 0.0 | 8.76E-01 | -0.2 | 4.58E-01 | Di-glucose binding protein with Leucine-rich repeat domain-containing protein |
| AT5G19590 | -0.2 | 1.11E-02 | -0.1 | 1.54E-01 | -0.1 | 2.16E-01 | 0.1 | 6.23E-01 | 0.1 | 7.17E-01 | 0.0 | 9.88E-01 | hypothetical protein |
| AT4G25520 | -0.2 | 3.80E-02 | -0.4 | 1.86E-03 | 0.2 | 1.04E-01 | 0.0 | 9.80E-01 | 0.1 | 8.25E-01 | -0.1 | 8.02E-01 | Probable transcriptional regulator SLK1 |
| AT3G02420 | -0.2 | 3.17E-02 | -0.1 | 4.07E-01 | -0.1 | 1.98E-01 | 0.0 | 9.82E-01 | 0.1 | 5.45E-01 | -0.1 | 5.99E-01 | AT3g02420 |
| AT1G04140 | -0.2 | 1.44E-02 | -0.3 | 4.48E-03 | 0.1 | 4.53E-01 | -0.1 | 3.81E-01 | 0.0 | 9.59E-01 | -0.1 | 6.25E-01 | Transducin family protein |
| AT2G44760 | -0.2 | 2.57E-02 | 0.0 | 8.12E-01 | -0.2 | 7.24E-02 | 0.1 | 2.98E-01 | 0.1 | 4.64E-01 | 0.0 | 9.28E-01 | Dihydroorotate dehydrogenase |
| AT5G04070 | -0.2 | 1.09E-01 | -0.2 | 6.12E-02 | 0.1 | 7.48E-01 | 0.2 | 4.51E-01 | 0.4 | 1.88E-01 | -0.2 | 6.61E-01 | At5g04070 -binding Rossmann-fold superfamily protein) |
| AT5G35980 | -0.2 | 4.94E-02 | -0.4 | 1.72E-03 | 0.2 | 6.57E-02 | 0.0 | 9.95E-01 | 0.0 | 9.89E-01 | 0.0 | 9.90E-01 | Uncharacterized protein |
| AT5G60100 | -0.2 | 2.31E-01 | 0.0 | 9.03E-01 | -0.2 | 2.80E-01 | 0.1 | 7.30E-01 | 0.3 | 3.08E-01 | -0.2 | 6.07E-01 | Pseudo-response regulator 3 |
| AT4G27050 | -0.2 | 3.60E-02 | -0.2 | 6.04E-02 | 0.0 | 8.69E-01 | -0.1 | 6.65E-01 | 0.0 | 9.46E-01 | -0.1 | 6.95E-01 | F-box protein At4g27050 |
| AT2G23360 | -0.2 | 7.11E-02 | -0.1 | 4.45E-01 | -0.1 | 3.63E-01 | -0.1 | 7.07E-01 | 0.1 | 5.91E-01 | -0.2 | 4.03E-01 | Filament-like plant protein 7 |
| AT4G11270 | -0.2 | 3.64E-02 | -0.2 | 1.41E-02 | 0.1 | 5.95E-01 | 0.1 | 4.75E-01 | 0.1 | 6.48E-01 | 0.0 | 9.34E-01 | Transducin |
| AT4G38660 | -0.2 | 7.11E-02 | 0.1 | 5.57E-01 | -0.3 | 4.16E-02 | -0.1 | 4.13E-01 | -0.1 | 8.06E-01 | -0.1 | 7.67E-01 | Pathogenesis-related thaumatin superfamily protein |
| AT1G64570 | -0.2 | 6.73E-02 | -0.5 | 1.41E-03 | 0.3 | 4.96E-02 | -0.3 | 5.89E-01 | 0.1 | 8.59E-01 | -0.5 | 5.47E-01 | F1N19.14 |
| AT1G63310 | -0.2 | 1.03E-01 | -0.1 | 4.84E-01 | -0.1 | 4.28E-01 | -0.1 | 7.42E-01 | 0.2 | 4.67E-01 | -0.3 | 3.61E-01 | hypothetical protein |
| AT1G60070 | -0.2 | 7.28E-03 | 0.0 | 6.71E-01 | -0.2 | 7.46E-03 | 0.1 | 6.23E-01 | 0.2 | 1.13E-01 | -0.2 | 4.05E-01 | AP-1 complex subunit gamma |
| AT3G07890 | -0.2 | 1.07E-02 | 0.0 | 5.78E-01 | -0.2 | 9.27E-03 | -0.1 | 3.14E-01 | 0.0 | 9.88E-01 | -0.1 | 5.23E-01 | Putative GTPase activator protein |
| AT5G42470 | -0.2 | 1.97E-02 | -0.3 | 2.50E-03 | 0.1 | 2.40E-01 | -0.2 | 1.58E-01 | -0.2 | 2.18E-01 | 0.0 | 9.88E-01 | BRISC and BRCA1-A complex member 2 |
| AT4G18380 | -0.2 | 3.23E-02 | 0.0 | 9.35E-01 | -0.2 | 6.81E-02 | -0.2 | 1.05E-01 | 0.0 | 8.97E-01 | -0.3 | 1.87E-01 | F-box protein At4g18380 |
| AT5G55610 | -0.2 | 5.01E-02 | 0.1 | 4.91E-01 | -0.3 | 2.57E-02 | 0.1 | 3.36E-01 | 0.2 | 1.03E-01 | -0.1 | 5.72E-01 | Isopentenyl-diphosphate delta-isomerase |
| AT5G63500 | -0.2 | 4.22E-02 | 0.2 | 7.77E-02 | -0.4 | 5.62E-03 | -0.6 | 4.90E-03 | -0.4 | 6.67E-02 | -0.2 | 4.87E-01 | hypothetical protein |
| AT5G60820 | -0.2 | 5.52E-02 | -0.3 | 1.72E-02 | 0.1 | 5.42E-01 | -0.2 | 2.75E-01 | -0.2 | 4.60E-01 | -0.1 | 8.95E-01 | hypothetical protein |
| AT4G39860 | -0.2 | 9.05E-02 | 0.2 | 5.29E-02 | -0.4 | 6.21E-03 | 0.1 | 5.37E-01 | 0.2 | 2.86E-01 | -0.1 | 7.29E-01 | DUF4057 domain-containing protein |
| AT2G30105 | -0.2 | 7.11E-02 | -0.2 | 1.45E-01 | 0.0 | 7.81E-01 | 0.0 | 8.67E-01 | 0.0 | 9.63E-01 | -0.1 | 8.73E-01 | LRR repeats and ubiquitin-like domain-containing protein At2g30105 |
| AT2G19280 | -0.2 | 1.29E-01 | -0.1 | 4.15E-01 | -0.1 | 5.78E-01 | 0.2 | 6.31E-01 | -0.1 | 8.72E-01 | 0.2 | 5.87E-01 | Pentatricopeptide repeat-containing protein At2g19280 |
| AT4G21790 | -0.2 | 1.56E-02 | 0.0 | 9.21E-01 | -0.2 | 2.34E-02 | -0.3 | 1.03E-02 | -0.2 | 2.57E-01 | -0.2 | 3.24E-01 | Tobamovirus multiplication protein 1 |
| AT1G50950 | -0.2 | 2.29E-01 | -0.3 | 8.47E-02 | 0.1 | 5.96E-01 | 0.1 | 6.74E-01 | -0.2 | 6.36E-01 | 0.3 | 4.27E-01 | Protein disulfide-isomerase 5-like protein |
| AT5G10060 | -0.2 | 2.09E-02 | -0.2 | 2.05E-02 | 0.0 | 9.37E-01 | -0.2 | 4.07E-01 | 0.0 | 9.80E-01 | -0.2 | 4.99E-01 | CID domain-containing protein |
| AT3G20800 | -0.2 | 2.99E-02 | -0.1 | 1.58E-01 | -0.1 | 4.53E-01 | -0.3 | 6.75E-02 | -0.1 | 4.68E-01 | -0.1 | 5.16E-01 | RCD1 |
| AT3G22970 | -0.2 | 9.02E-02 | 0.0 | 7.76E-01 | -0.2 | 2.00E-01 | -0.3 | 1.11E-02 | -0.1 | 4.14E-01 | -0.2 | 2.58E-01 | At3g22970 |
| AT2G45690 | -0.2 | 1.50E-01 | -0.3 | 2.68E-02 | 0.1 | 3.81E-01 | 0.0 | 9.18E-01 | -0.2 | 5.24E-01 | 0.2 | 6.47E-01 | Peroxisome biogenesis protein 16 |
| AT3G06920 | -0.2 | 1.49E-01 | -0.3 | 7.45E-02 | 0.1 | 7.10E-01 | 0.6 | 3.97E-02 | 0.0 | 9.81E-01 | 0.6 | 1.12E-01 | Tetratricopeptide repeat -like superfamily protein |
| AT1G10750 | -0.2 | 2.52E-01 | 0.0 | 8.27E-01 | -0.2 | 2.63E-01 | 0.0 | 9.09E-01 | 0.2 | 5.21E-01 | -0.2 | 6.69E-01 | Carboxyl-terminal peptidase, putative |
| AT5G41110 | -0.2 | 8.35E-02 | -0.1 | 4.32E-01 | -0.1 | 4.20E-01 | -0.2 | 6.07E-01 | 0.2 | 5.87E-01 | -0.4 | 2.61E-01 | Uncharacterized protein At5g41110 |
| AT4G27390 | -0.2 | 3.62E-02 | -0.2 | 1.99E-02 | 0.0 | 7.29E-01 | -0.1 | 7.93E-01 | 0.1 | 5.03E-01 | -0.2 | 4.02E-01 | Transmembrane protein |
| AT1G12390 | -0.2 | 2.08E-02 | -0.1 | 1.59E-01 | -0.1 | 3.66E-01 | 0.0 | 8.72E-01 | -0.1 | 6.85E-01 | 0.1 | 8.07E-01 | Protein cornichon homolog 4 |
| AT3G18215 | -0.2 | 7.96E-02 | 0.0 | 9.76E-01 | -0.2 | 1.32E-01 | -0.4 | 2.15E-03 | -0.3 | 1.19E-02 | -0.1 | 7.72E-01 | hypothetical protein |
| AT3G58640 | -0.2 | 7.74E-03 | -0.3 | 1.30E-03 | 0.1 | 2.52E-01 | 0.0 | 8.10E-01 | 0.0 | 9.58E-01 | 0.1 | 8.28E-01 | AT3g58640 |
| AT2G20130 | -0.2 | 6.42E-02 | 0.0 | 7.07E-01 | -0.2 | 6.10E-02 | -0.2 | 2.41E-01 | -0.1 | 8.15E-01 | -0.2 | 5.55E-01 | Protein LIKE COV 1 |
| AT5G53020 | -0.2 | 1.21E-01 | 0.0 | 9.18E-01 | -0.2 | 2.21E-01 | 0.0 | 8.83E-01 | 0.2 | 3.16E-01 | -0.3 | 3.27E-01 | Emb\|CAB82814.1 |
| AT4G37680 | -0.2 | 3.61E-02 | -0.2 | 1.14E-02 | 0.1 | 5.23E-01 | -0.2 | 2.21E-01 | 0.0 | 9.80E-01 | -0.2 | 3.65E-01 | Heptahelical protein 4 |
| AT2G36370 | -0.2 | 1.05E-01 | -0.3 | 1.39E-02 | 0.1 | 3.03E-01 | -0.1 | 7.81E-01 | 0.0 | 9.95E-01 | -0.1 | 8.27E-01 | BTB |
| AT3G04860 | -0.2 | 2.48E-01 | 0.2 | 1.80E-01 | -0.4 | 5.18E-02 | -0.7 | 7.24E-03 | -0.5 | 3.55E-02 | -0.1 | 7.10E-01 | hypothetical protein |
| AT4G16440 | -0.2 | 6.65E-02 | -0.3 | 6.63E-03 | 0.1 | 2.33E-01 | -0.1 | 5.60E-01 | 0.0 | 9.77E-01 | -0.1 | 6.33E-01 | NAR1 |
| AT5G53800 | -0.2 | 1.00E-01 | 0.0 | 7.14E-01 | -0.1 | 2.57E-01 | -0.1 | 7.03E-01 | -0.2 | 4.03E-01 | 0.1 | 7.34E-01 | AT5g53800 |
| AT4G38120 | -0.2 | 4.42E-02 | -0.3 | 1.36E-02 | 0.1 | 5.22E-01 | 0.1 | 7.97E-01 | 0.1 | 8.00E-01 | 0.0 | 9.78E-01 | Uncharacterized protein AT4g38120 |
| AT4G04470 | -0.2 | 2.69E-01 | -0.5 | 1.53E-02 | 0.3 | 1.47E-01 | 0.3 | 2.69E-01 | 0.5 | 2.63E-02 | -0.3 | 3.20E-01 | Peroxisomal membrane protein PMP22 |
| AT5G58530 | -0.2 | 9.88E-02 | 0.2 | 7.76E-02 | -0.4 | 1.05E-02 | -0.3 | 5.88E-02 | -0.2 | 2.24E-01 | -0.1 | 7.25E-01 | Glutaredoxin domain-containing protein |
| AT5G07980 | -0.2 | 6.13E-02 | -0.2 | 3.53E-02 | 0.0 | 7.55E-01 | 0.0 | 9.41E-01 | 0.1 | 7.89E-01 | -0.1 | 8.20E-01 | Dentin sialophosphoprotein-like protein |
| AT1G10950 | -0.2 | 1.36E-02 | 0.0 | 7.82E-01 | -0.2 | 3.93E-02 | -0.1 | 2.15E-01 | -0.1 | 3.96E-01 | 0.0 | 9.12E-01 | Transmembrane 9 superfamily member |
| AT2G42040 | -0.2 | 2.26E-01 | 0.2 | 1.92E-01 | -0.4 | 4.46E-02 | -0.4 | 3.83E-02 | -0.5 | 3.21E-02 | 0.1 | 8.69E-01 | WRC domain-containing protein |
| AT5G23550 | -0.2 | 3.57E-02 | 0.1 | 4.41E-01 | -0.2 | 2.23E-02 | -0.1 | 6.66E-01 | -0.2 | 4.05E-01 | 0.1 | 7.63E-01 | Vesicle transport protein |
| AT1G54180 | -0.2 | 1.58E-01 | -0.1 | 4.78E-01 | -0.1 | 5.82E-01 | 0.2 | 5.73E-01 | 0.2 | 6.68E-01 | 0.0 | 9.79E-01 | BRX-LIKE3 |
| AT2G27950 | -0.2 | 1.26E-02 | -0.3 | 6.89E-04 | 0.1 | 8.25E-02 | -0.2 | 3.34E-01 | 0.0 | 9.58E-01 | -0.2 | 4.66E-01 | Uncharacterized protein At2g27950 |
| AT3G47810 | -0.2 | 3.28E-02 | -0.2 | 2.92E-02 | 0.0 | 9.01E-01 | -0.1 | 4.29E-01 | 0.0 | 9.26E-01 | -0.1 | 5.08E-01 | Vacuolar protein sorting-associated protein 29 |
| AT2G44060 | -0.2 | 5.28E-02 | 0.1 | 1.84E-01 | -0.3 | 9.25E-03 | -0.2 | 2.02E-01 | 0.0 | 9.17E-01 | -0.1 | 4.24E-01 | Similar to late embryogenesis abundant proteins |
| AT1G07470 | -0.2 | 1.33E-02 | -0.1 | 1.37E-01 | -0.1 | 2.90E-01 | -0.2 | 9.83E-02 | -0.1 | 4.85E-01 | -0.1 | 5.84E-01 | Uncharacterized protein |
| AT1G30130 | -0.2 | 4.36E-02 | 0.1 | 5.32E-01 | -0.2 | 2.62E-02 | -0.4 | 9.10E-03 | -0.2 | 3.28E-01 | -0.2 | 2.42E-01 | AT1G30130 protein |
| AT5G16300 | -0.2 | 7.55E-03 | -0.2 | 3.08E-03 | 0.0 | 5.50E-01 | 0.1 | 8.03E-01 | 0.1 | 6.28E-01 | -0.1 | 8.51E-01 | Conserved oligomeric Golgi complex subunit 1 |
| AT2G24440 | -0.2 | 9.24E-02 | 0.2 | 3.13E-02 | -0.4 | 5.78E-03 | 0.2 | 5.26E-01 | 0.1 | 7.60E-01 | 0.1 | 8.83E-01 | Uncharacterized protein At2g24440 |
| AT1G19840 | -0.2 | 2.26E-01 | 0.1 | 5.09E-01 | -0.3 | 1.30E-01 | -0.3 | 4.66E-01 | 0.1 | 9.08E-01 | -0.3 | 4.60E-01 | Uncharacterized protein |
| AT3G52748 | -0.2 | 3.83E-01 | -0.3 | 1.61E-01 | 0.1 | 6.23E-01 | 1.0 | 5.29E-03 | 0.9 | 2.39E-02 | 0.1 | 6.99E-01 | NA |
| AT2G47410 | -0.2 | 1.09E-01 | -0.3 | 9.30E-03 | 0.2 | 1.99E-01 | 0.2 | 5.17E-01 | 0.1 | 7.18E-01 | 0.0 | 9.07E-01 | WD40 domain-containing protein |
| AT1G24265 | -0.2 | 2.51E-01 | -0.1 | 4.76E-01 | -0.1 | 7.59E-01 | 0.0 | 9.29E-01 | 0.0 | 9.54E-01 | 0.0 | 9.94E-01 | BZIP transcription factor, putative |
| AT5G10710 | -0.2 | 3.82E-02 | -0.1 | 1.69E-01 | -0.1 | 5.20E-01 | -0.1 | 5.93E-01 | -0.2 | 3.63E-01 | 0.1 | 7.49E-01 | Uncharacterized protein AT5g10710 |
| AT1G04750 | -0.2 | 3.32E-02 | -0.1 | 1.21E-01 | -0.1 | 5.68E-01 | 0.2 | 2.49E-01 | 0.0 | 9.09E-01 | 0.1 | 5.32E-01 | Vesicle-associated membrane protein 721 |
| AT3G07950 | -0.2 | 4.64E-02 | 0.1 | 4.87E-01 | -0.2 | 2.48E-02 | 0.1 | 4.58E-01 | 0.0 | 8.82E-01 | 0.1 | 7.39E-01 | Rhomboid-like protein 19 |
| AT3G16330 | -0.2 | 3.05E-01 | 0.0 | 8.32E-01 | -0.2 | 3.10E-01 | 0.6 | 4.88E-02 | 0.3 | 4.77E-01 | 0.3 | 2.60E-01 | hypothetical protein |
| AT2G17350 | -0.2 | 2.44E-02 | -0.2 | 2.06E-02 | 0.0 | 8.85E-01 | -0.2 | 5.39E-01 | -0.1 | 6.35E-01 | 0.0 | 9.80E-01 | Beta-mannosyltransferase-like protein |
| AT5G44910 | -0.2 | 1.84E-01 | -0.3 | 2.65E-02 | 0.2 | 3.21E-01 | 0.2 | 5.03E-01 | 0.0 | 9.43E-01 | 0.2 | 5.82E-01 | Similarity to disease resistance protein domain family protein) |
| AT5G54300 | -0.2 | 1.72E-01 | -0.4 | 8.51E-03 | 0.3 | 1.36E-01 | -0.5 | 8.15E-02 | -0.5 | 1.15E-01 | 0.0 | 9.90E-01 | DUF4408 domain-containing protein |
| AT1G55675 | -0.2 | 3.20E-02 | -0.1 | 1.31E-01 | -0.1 | 5.46E-01 | -0.6 | 2.73E-03 | -0.4 | 7.49E-02 | -0.2 | 3.03E-01 | Transmembrane protein |
| AT4G12090 | -0.2 | 2.40E-01 | 0.2 | 1.05E-01 | -0.4 | 3.32E-02 | 1.0 | 2.03E-02 | 0.3 | 5.42E-01 | 0.6 | 1.58E-01 | Protein cornichon homolog 5 |
| AT3G05940 | -0.2 | 5.03E-02 | -0.2 | 5.68E-02 | 0.0 | 9.94E-01 | -0.1 | 7.02E-01 | 0.0 | 9.19E-01 | -0.1 | 6.91E-01 | hypothetical protein |
| AT4G35940 | -0.2 | 5.57E-02 | -0.1 | 1.16E-01 | 0.0 | 7.80E-01 | 0.0 | 9.46E-01 | -0.3 | 1.20E-01 | 0.3 | 2.44E-01 | Myb-like protein X |
| AT5G06970 | -0.2 | 1.90E-02 | 0.0 | 8.85E-01 | -0.2 | 2.58E-02 | -0.1 | 4.74E-01 | 0.1 | 8.23E-01 | -0.2 | 4.36E-01 | Protein unc-13 homolog |
| AT3G19540 | -0.2 | 1.52E-02 | -0.1 | 5.27E-02 | 0.0 | 5.97E-01 | -0.2 | 1.74E-01 | -0.1 | 5.59E-01 | -0.1 | 6.72E-01 | AT3g19540 |
| AT5G49830 | -0.2 | 1.08E-02 | -0.1 | 8.85E-02 | -0.1 | 3.30E-01 | -0.1 | 3.61E-01 | -0.3 | 1.22E-02 | 0.2 | 1.89E-01 | Exocyst complex component 84B |
| AT4G34200 | -0.2 | 1.50E-01 | 0.1 | 3.13E-01 | -0.3 | 3.79E-02 | 0.2 | 2.18E-01 | 0.1 | 5.19E-01 | 0.1 | 7.89E-01 | Phosphoglycerate dehydrogenase-like protein |
| AT1G33250 | -0.2 | 5.90E-02 | 0.0 | 7.73E-01 | -0.2 | 5.87E-02 | -0.2 | 1.04E-01 | -0.4 | 7.57E-03 | 0.2 | 3.01E-01 | Uncharacterized protein |
| AT5G45360 | -0.2 | 4.06E-02 | -0.2 | 4.36E-02 | 0.0 | 9.81E-01 | -0.5 | 1.45E-03 | -0.5 | 2.92E-03 | 0.0 | 9.82E-01 | F-box protein SKIP31 |
| AT1G60430 | -0.2 | 1.43E-01 | -0.2 | 2.12E-01 | 0.0 | 8.89E-01 | -0.6 | 3.29E-03 | -0.2 | 3.00E-01 | -0.4 | 9.58E-02 | Actin-related protein 2 |

| | | | | | | | | | | | | | |
|---|---|---|---|---|---|---|---|---|---|---|---|---|---|
| AT2G20210 | -0.2 | 1.11E-01 | 0.0 | 8.10E-01 | -0.1 | 2.66E-01 | -0.6 | 1.03E-01 | -0.4 | 3.44E-01 | -0.2 | 6.32E-01 | Uncharacterized protein At2g20220 |
| AT5G08290 | -0.2 | 1.98E-02 | -0.1 | 4.33E-01 | -0.1 | 1.23E-01 | -0.1 | 4.42E-01 | -0.1 | 4.49E-01 | 0.0 | 9.56E-01 | mRNA splicing factor, thioredoxin-like U5 snRNP |
| AT5G50170 | -0.2 | 1.12E-01 | -0.2 | 1.50E-01 | 0.0 | 9.40E-01 | -0.1 | 8.06E-01 | -0.1 | 8.05E-01 | 0.0 | 9.75E-01 | C2 and GRAM domain-containing protein At5g50170 |
| AT3G07390 | -0.2 | 1.16E-01 | 0.1 | 4.65E-01 | -0.3 | 4.71E-02 | 0.0 | 9.93E-01 | 0.1 | 4.06E-01 | -0.1 | 5.12E-01 | Auxin-induced in root cultures-like protein |
| AT1G31175 | -0.2 | 1.33E-01 | -0.3 | 3.16E-02 | 0.1 | 4.62E-01 | -0.4 | 9.34E-02 | -0.3 | 2.68E-01 | -0.1 | 7.89E-01 | Uncharacterized protein |
| AT2G03330 | -0.2 | 1.73E-01 | -0.1 | 5.22E-01 | -0.1 | 5.65E-01 | -0.4 | 1.41E-02 | -0.3 | 6.69E-02 | -0.1 | 7.33E-01 | Transmembrane protein |
| AT1G07140 | -0.2 | 7.90E-02 | 0.1 | 2.59E-01 | -0.3 | 1.78E-02 | 0.2 | 7.68E-02 | 0.0 | 8.21E-01 | 0.2 | 3.41E-01 | Ran-binding protein 1 homolog a |
| AT1G63410 | -0.2 | 2.75E-01 | -0.1 | 6.33E-01 | -0.1 | 6.45E-01 | 0.0 | 9.53E-01 | 0.3 | 6.08E-01 | -0.2 | 6.78E-01 | hypothetical protein |
| AT2G19090 | -0.2 | 1.56E-01 | -0.1 | 5.99E-01 | -0.1 | 4.74E-01 | 0.0 | 9.31E-01 | -0.2 | 7.21E-01 | 0.2 | 7.14E-01 | Uncharacterized protein |
| AT3G56710 | -0.2 | 2.02E-01 | -0.9 | 1.33E-04 | 0.7 | 1.52E-03 | 1.4 | 1.89E-05 | 0.2 | 6.19E-01 | 1.2 | 5.20E-04 | Sigma factor binding protein 1, chloroplastic |
| AT3G54910 | -0.2 | 1.21E-01 | -0.1 | 3.10E-01 | -0.1 | 6.76E-01 | -0.2 | 4.22E-01 | 0.0 | 9.62E-01 | -0.2 | 5.12E-01 | Uncharacterized protein F28P10.110 |
| AT4G26620 | -0.2 | 2.20E-02 | -0.1 | 8.52E-02 | 0.0 | 5.61E-01 | -0.1 | 2.88E-01 | -0.2 | 7.56E-02 | 0.1 | 5.78E-01 | hypothetical protein |
| AT1G05640 | -0.2 | 2.08E-01 | -0.2 | 1.76E-01 | 0.0 | 9.07E-01 | -0.3 | 4.48E-01 | 0.0 | 9.45E-01 | -0.2 | 6.49E-01 | PGG domain-containing protein |
| AT3G12590 | -0.2 | 2.86E-02 | 0.0 | 9.10E-01 | -0.2 | 4.33E-02 | -0.2 | 1.37E-01 | -0.1 | 4.25E-01 | -0.1 | 7.43E-01 | ARM repeat superfamily protein |
| AT3G10980 | -0.2 | 1.62E-02 | 0.0 | 4.80E-01 | -0.1 | 9.62E-02 | -0.1 | 3.98E-01 | 0.0 | 9.27E-01 | -0.1 | 4.51E-01 | hypothetical protein |
| AT1G53300 | -0.2 | 1.09E-01 | -0.3 | 2.13E-02 | 0.1 | 3.83E-01 | -0.2 | 2.13E-01 | 0.0 | 9.79E-01 | -0.3 | 3.53E-01 | TPR repeat-containing thioredoxin TTL1 |
| AT5G11030 | -0.2 | 7.60E-02 | -0.3 | 5.62E-03 | 0.2 | 1.80E-01 | 0.1 | 7.87E-01 | 0.0 | 9.44E-01 | 0.0 | 9.12E-01 | Aberrant root formation protein |
| AT1G12330 | -0.2 | 9.15E-02 | -0.1 | 5.87E-01 | -0.1 | 3.15E-01 | -0.2 | 1.18E-01 | -0.1 | 4.25E-01 | -0.1 | 7.11E-01 | Uncharacterized protein At1g12330 |
| AT5G58375 | -0.2 | 1.64E-02 | 0.1 | 4.10E-02 | -0.3 | 1.45E-03 | -0.1 | 5.83E-01 | -0.3 | 2.41E-01 | 0.2 | 6.38E-01 | Methyltransferase-related protein |
| AT5G23050 | -0.2 | 1.53E-01 | -0.2 | 2.20E-01 | 0.0 | 8.98E-01 | -0.2 | 2.40E-01 | -0.2 | 4.21E-01 | 0.0 | 8.89E-01 | Acyl-activating enzyme 17 |
| AT5G08080 | -0.2 | 3.57E-02 | 0.0 | 8.94E-01 | -0.2 | 4.73E-02 | -0.2 | 1.24E-01 | -0.1 | 6.11E-01 | -0.1 | 5.57E-01 | Syntaxin of plants 132 |
| AT3G07565 | -0.2 | 1.95E-02 | -0.1 | 6.99E-02 | 0.0 | 5.96E-01 | -0.2 | 2.46E-01 | -0.2 | 2.84E-01 | 0.0 | 9.75E-01 | Histone H2A deubiquitinase |
| AT3G18295 | -0.2 | 1.03E-01 | -0.1 | 5.56E-01 | -0.1 | 3.80E-01 | 0.1 | 7.63E-01 | 0.2 | 5.53E-01 | -0.1 | 7.61E-01 | hypothetical protein |
| AT5G54730 | -0.2 | 8.70E-02 | -0.4 | 2.30E-03 | 0.2 | 6.54E-02 | -0.3 | 3.44E-02 | -0.2 | 2.90E-01 | -0.1 | 5.40E-01 | Autophagy 18 F-like protein |
| AT4G03430 | -0.2 | 1.88E-02 | -0.4 | 1.95E-04 | 0.2 | 1.11E-02 | -0.1 | 7.37E-01 | -0.1 | 7.40E-01 | 0.0 | 9.74E-01 | Protein STABILIZED1 |
| AT1G71780 | -0.2 | 1.18E-01 | 0.2 | 1.20E-01 | -0.3 | 1.44E-02 | 0.2 | 2.29E-01 | 0.1 | 3.68E-01 | 0.0 | 9.37E-01 | F14O23.16 protein |
| AT5G21990 | -0.2 | 1.16E-02 | -0.1 | 1.06E-01 | -0.1 | 3.03E-01 | -0.1 | 5.21E-01 | 0.0 | 7.67E-01 | 0.0 | 8.96E-01 | Outer envelope protein 61 |
| AT4G30470 | -0.2 | 2.74E-01 | -0.6 | 2.44E-03 | 0.5 | 2.44E-02 | -0.5 | 1.80E-02 | -0.3 | 2.00E-01 | -0.2 | 4.76E-01 | NAD-dependent epimerase |
| AT3G54380 | -0.2 | 7.92E-02 | -0.1 | 4.88E-01 | -0.1 | 3.57E-01 | -0.1 | 5.16E-01 | -0.3 | 3.08E-01 | 0.1 | 7.62E-01 | SAC3 |
| AT1G69220 | -0.2 | 3.39E-02 | -0.2 | 5.91E-02 | 0.0 | 8.46E-01 | -0.2 | 3.61E-01 | -0.4 | 7.06E-02 | 0.2 | 4.60E-01 | SIK1 |
| AT1G80680 | -0.2 | 2.88E-02 | -0.3 | 1.27E-03 | 0.1 | 8.54E-02 | 0.2 | 2.35E-01 | 0.2 | 3.81E-01 | 0.0 | 9.34E-01 | Nucleoporin PRECOZ like protein |
| AT3G17950 | -0.2 | 6.15E-02 | -0.2 | 3.12E-02 | 0.0 | 7.13E-01 | -0.6 | 7.30E-03 | -0.2 | 3.68E-01 | -0.4 | 1.94E-01 | Transmembrane protein |
| AT4G16850 | -0.2 | 6.92E-02 | -0.1 | 4.65E-01 | -0.1 | 3.42E-01 | -0.1 | 5.78E-01 | 0.0 | 9.38E-01 | -0.1 | 5.71E-01 | Transmembrane protein |
| AT4G32680 | -0.2 | 2.35E-02 | -0.1 | 6.42E-02 | 0.0 | 6.90E-01 | -0.5 | 2.24E-03 | -0.5 | 3.54E-03 | 0.0 | 9.68E-01 | Uncharacterized protein AT4g32680 |
| AT3G05945 | -0.2 | 1.26E-01 | 0.3 | 2.43E-02 | -0.4 | 5.25E-03 | -0.4 | 1.94E-01 | -0.4 | 2.16E-01 | 0.0 | 9.61E-01 | NA |
| AT4G38630 | -0.2 | 6.75E-02 | -0.1 | 5.01E-01 | -0.1 | 2.82E-01 | 0.1 | 5.57E-01 | 0.1 | 4.19E-01 | 0.0 | 8.49E-01 | Multiubiquitin chain binding protein |
| AT1G43130 | -0.2 | 2.13E-02 | -0.2 | 1.49E-02 | 0.0 | 8.06E-01 | -0.3 | 7.03E-02 | -0.2 | 2.47E-01 | -0.1 | 7.36E-01 | Like COV 2 |
| AT1G10730 | -0.2 | 7.58E-02 | -0.2 | 2.37E-02 | 0.1 | 5.46E-01 | -0.1 | 6.37E-01 | -0.1 | 8.59E-01 | -0.1 | 8.76E-01 | AP-1 complex subunit mu-1 |
| AT4G05000 | -0.2 | 5.12E-02 | -0.1 | 3.35E-01 | -0.1 | 3.70E-01 | -0.4 | 4.13E-02 | -0.2 | 4.07E-01 | -0.2 | 3.93E-01 | Vacuolar protein sorting-associated protein 28 homolog |
| AT1G80790 | -0.2 | 8.43E-02 | 0.0 | 9.20E-01 | -0.2 | 1.46E-01 | 0.0 | 7.58E-01 | 0.0 | 9.66E-01 | 0.0 | 8.75E-01 | XH |
| AT5G01010 | -0.2 | 1.37E-02 | -0.2 | 7.01E-03 | 0.0 | 6.46E-01 | 0.1 | 3.05E-01 | -0.1 | 7.18E-01 | 0.2 | 3.04E-01 | Retinal-binding protein |
| AT1G11390 | -0.2 | 2.92E-02 | 0.0 | 6.72E-01 | -0.1 | 1.12E-01 | -0.1 | 6.88E-01 | -0.1 | 8.44E-01 | 0.0 | 9.14E-01 | T23J18.5 |
| AT5G27710 | -0.2 | 9.78E-02 | 0.0 | 8.30E-01 | -0.2 | 1.14E-01 | 0.0 | 9.85E-01 | 0.1 | 8.20E-01 | 0.0 | 8.62E-01 | T-box protein |
| AT5G37710 | -0.2 | 1.06E-01 | 0.1 | 1.64E-01 | -0.3 | 2.09E-02 | -0.3 | 4.33E-02 | -0.5 | 6.67E-03 | 0.2 | 4.40E-01 | Calmodulin-binding heat-shock protein |
| AT4G20410 | -0.2 | 3.58E-02 | -0.1 | 2.46E-01 | -0.1 | 3.51E-01 | -0.2 | 2.48E-01 | -0.1 | 3.77E-01 | 0.0 | 9.52E-01 | Gamma-soluble NSF attachment protein |
| AT4G34310 | -0.2 | 1.11E-01 | -0.4 | 2.86E-03 | 0.2 | 6.92E-02 | -0.3 | 3.37E-01 | 0.0 | 9.18E-01 | -0.2 | 4.57E-01 | Alpha |
| AT5G66440 | -0.2 | 9.31E-02 | -0.1 | 4.30E-01 | -0.1 | 4.51E-01 | 0.1 | 4.75E-01 | 0.0 | 9.79E-01 | 0.1 | 6.49E-01 | hypothetical protein |
| AT4G36980 | -0.2 | 1.18E-01 | -0.4 | 3.04E-03 | 0.2 | 5.68E-02 | -0.3 | 9.58E-02 | 0.0 | 8.79E-01 | -0.3 | 2.09E-01 | CLK4-associating serine |
| AT5G13310 | -0.2 | 3.62E-02 | -0.1 | 8.99E-02 | 0.0 | 7.27E-01 | 0.0 | 9.52E-01 | -0.1 | 8.32E-01 | 0.0 | 9.05E-01 | U5 small nuclear ribonucleoprotein TSSC4 |
| AT1G12620 | -0.2 | 1.23E-01 | 0.0 | 9.22E-01 | -0.2 | 2.33E-01 | 0.0 | 9.41E-01 | 0.3 | 4.07E-01 | -0.3 | 3.90E-01 | Pentatricopeptide repeat-containing protein At1g12620 |
| AT2G31820 | -0.2 | 9.66E-02 | -0.4 | 1.98E-03 | 0.2 | 4.82E-02 | -0.2 | 1.96E-01 | -0.1 | 7.93E-01 | -0.2 | 5.32E-01 | PGG domain-containing protein |
| AT3G19780 | -0.2 | 1.26E-01 | -0.2 | 5.77E-02 | 0.1 | 6.81E-01 | -0.2 | 6.07E-01 | 0.2 | 5.73E-01 | -0.4 | 3.29E-01 | Thioredoxin domain-containing protein |
| AT2G26660 | -0.2 | 1.53E-01 | -0.1 | 3.87E-01 | -0.1 | 6.53E-01 | 0.0 | 9.50E-01 | -0.1 | 8.02E-01 | 0.1 | 8.07E-01 | SPX domain-containing protein 2 |
| AT2G21520 | -0.2 | 1.70E-02 | -0.2 | 2.23E-02 | 0.0 | 9.79E-01 | 0.1 | 5.04E-01 | -0.1 | 6.92E-01 | 0.2 | 4.18E-01 | Sec14p-like phosphatidylinositol transfer family protein |
| AT3G51250 | -0.2 | 3.10E-02 | -0.2 | 5.02E-03 | 0.1 | 3.05E-01 | -0.2 | 3.39E-01 | 0.2 | 2.79E-01 | -0.4 | 9.92E-02 | Senescence |
| AT5G35430 | -0.2 | 2.85E-02 | -0.1 | 8.75E-02 | 0.0 | 6.38E-01 | 0.1 | 5.73E-01 | 0.0 | 9.21E-01 | 0.1 | 6.34E-01 | Gb\|AAD55417.1 |
| AT2G17130 | -0.2 | 3.59E-02 | 0.1 | 2.59E-01 | -0.3 | 9.50E-03 | 0.1 | 3.95E-01 | 0.1 | 7.36E-01 | 0.0 | 8.22E-01 | hypothetical protein |
| AT5G56795 | -0.2 | 6.05E-01 | 0.4 | 1.95E-01 | -0.6 | 1.40E-01 | -0.4 | 2.99E-01 | -0.5 | 2.65E-01 | 0.1 | 9.11E-01 | hypothetical protein |
| AT1G08940 | -0.2 | 1.91E-01 | -0.2 | 1.33E-01 | 0.0 | 8.35E-01 | 0.4 | 4.02E-02 | -0.2 | 4.24E-01 | 0.6 | 1.62E-02 | Phosphoglycerate mutase family protein |
| AT1G27970 | -0.2 | 1.33E-01 | 0.1 | 2.75E-01 | -0.3 | 3.00E-02 | 0.1 | 5.17E-01 | 0.0 | 8.47E-01 | 0.1 | 4.96E-01 | Nuclear transport factor 2, putative |
| AT1G10890 | -0.2 | 4.26E-02 | -0.3 | 2.15E-03 | 0.1 | 1.06E-01 | -0.2 | 2.15E-01 | -0.2 | 3.03E-01 | 0.0 | 9.82E-01 | Uncharacterized protein At1g10890 |
| AT4G25690 | -0.2 | 6.19E-02 | 0.0 | 9.77E-01 | -0.2 | 1.07E-01 | -0.2 | 3.74E-01 | -0.5 | 1.22E-02 | 0.3 | 1.06E-01 | hypothetical protein |
| AT1G77320 | -0.2 | 2.01E-01 | -0.2 | 1.04E-01 | 0.1 | 7.19E-01 | 0.3 | 5.20E-01 | 0.4 | 3.83E-01 | -0.1 | 8.29E-01 | Transcription coactivator |
| AT1G21370 | -0.2 | 4.47E-02 | 0.0 | 7.94E-01 | -0.2 | 5.27E-02 | 0.1 | 5.24E-01 | 0.1 | 7.99E-01 | 0.1 | 8.45E-01 | hypothetical protein |
| AT2G31340 | -0.2 | 9.10E-02 | -0.1 | 1.35E-01 | 0.0 | 8.99E-01 | 0.5 | 4.31E-02 | 0.1 | 8.65E-01 | 0.4 | 1.70E-01 | Embryo defective 1381 |
| AT5G24520 | -0.2 | 3.39E-02 | -0.1 | 8.77E-02 | 0.0 | 6.91E-01 | -0.3 | 1.17E-02 | -0.4 | 4.56E-03 | 0.1 | 6.32E-01 | Mutant protein of TTG1 |
| AT2G40090 | -0.2 | 1.19E-01 | 0.0 | 7.12E-01 | -0.2 | 1.09E-01 | -0.1 | 5.49E-01 | -0.1 | 8.52E-01 | -0.1 | 8.30E-01 | ABC transporter like protein |
| AT4G37445 | -0.2 | 2.55E-01 | 0.1 | 6.54E-01 | -0.2 | 1.96E-01 | 0.3 | 3.96E-01 | 0.6 | 1.59E-01 | -0.2 | 6.25E-01 | Calcium ion-binding protein |
| AT3G58010 | -0.2 | 8.85E-02 | 0.1 | 3.07E-01 | -0.3 | 2.79E-02 | -0.3 | 1.18E-02 | -0.2 | 1.15E-01 | -0.1 | 5.40E-01 | Probable plastid-lipid-associated protein 9, chloroplastic |
| AT4G04692 | -0.2 | 8.95E-02 | 0.0 | 9.59E-01 | -0.2 | 1.43E-01 | -0.2 | 4.35E-01 | -0.2 | 4.83E-01 | 0.0 | 9.83E-01 | NA |
| AT5G11000 | -0.2 | 7.74E-02 | 0.2 | 9.49E-02 | -0.3 | 7.65E-03 | 0.0 | 9.05E-01 | -0.1 | 5.74E-01 | 0.1 | 7.34E-01 | DUF868 family protein |
| AT4G22290 | -0.2 | 4.79E-02 | -0.1 | 3.32E-01 | -0.1 | 3.42E-01 | -0.3 | 1.68E-02 | -0.1 | 7.86E-01 | -0.3 | 1.13E-01 | hypothetical protein |
| AT2G36960 | -0.2 | 3.71E-02 | -0.4 | 6.72E-04 | 0.2 | 3.57E-02 | -0.4 | 4.53E-02 | 0.0 | 8.73E-01 | -0.3 | 1.13E-01 | TSL-kinase interacting protein 1 |
| AT2G15240 | -0.2 | 3.83E-02 | 0.0 | 8.39E-01 | -0.1 | 9.09E-02 | -0.2 | 2.25E-01 | -0.2 | 2.73E-01 | 0.0 | 9.86E-01 | At2g15240 |
| AT3G59660 | -0.2 | 3.69E-02 | -0.2 | 5.68E-03 | 0.1 | 2.98E-01 | 0.1 | 7.05E-01 | 0.0 | 9.30E-01 | 0.1 | 7.23E-01 | C2 domain-containing protein |
| AT2G41420 | -0.2 | 3.63E-02 | -0.1 | 3.32E-01 | -0.1 | 2.68E-01 | -0.3 | 1.99E-01 | -0.2 | 3.51E-01 | 0.0 | 9.26E-01 | Cysteine-rich and transmembrane domain-containing protein WIH2 |
| AT5G09620 | -0.2 | 4.09E-02 | -0.4 | 1.59E-04 | 0.3 | 4.44E-03 | -0.2 | 4.44E-01 | 0.0 | 9.28E-01 | -0.2 | 5.48E-01 | Gb\|AAD14519.1 |
| AT3G09000 | -0.2 | 1.42E-01 | 0.0 | 9.49E-01 | -0.2 | 2.18E-01 | -0.3 | 6.54E-02 | -0.2 | 1.42E-01 | 0.0 | 9.08E-01 | AT3g09000 |
| AT4G03030 | -0.2 | 5.38E-02 | -0.5 | 2.02E-04 | 0.3 | 5.22E-03 | -0.6 | 2.42E-04 | -0.6 | 4.54E-04 | 0.0 | 8.80E-01 | F-box |
| AT1G55160 | -0.2 | 2.47E-02 | 0.0 | 5.58E-01 | -0.2 | 1.77E-02 | -0.2 | 1.66E-01 | -0.2 | 3.06E-01 | 0.0 | 9.19E-01 | WAS |
| AT1G67150 | -0.2 | 4.15E-01 | 0.3 | 8.84E-02 | -0.5 | 4.71E-02 | 0.2 | 7.37E-01 | 0.7 | 1.17E-01 | -0.5 | 3.07E-01 | F5A8.6 protein ) |
| AT3G14430 | -0.2 | 2.37E-01 | -0.2 | 1.13E-01 | 0.1 | 6.84E-01 | -0.3 | 3.79E-01 | -0.2 | 5.91E-01 | -0.1 | 8.99E-01 | hypothetical protein |
| AT2G44230 | -0.2 | 2.27E-01 | -0.2 | 1.25E-01 | 0.1 | 7.34E-01 | -0.4 | 2.00E-02 | -0.2 | 4.14E-01 | -0.2 | 3.29E-01 | Vacuolar protein sorting-associated protein 62 |

| | | | | | | | | | | | | | |
|---|---|---|---|---|---|---|---|---|---|---|---|---|---|
| AT5G22730 | -0.2 | 1.09E-01 | 0.1 | 5.53E-01 | -0.2 | 7.22E-02 | -0.2 | 4.40E-01 | -0.2 | 4.34E-01 | 0.0 | 9.55E-01 | F-box domain-containing protein |
| AT1G05790 | -0.2 | 1.31E-01 | -0.3 | 9.63E-03 | 0.2 | 1.82E-01 | -0.2 | 3.73E-01 | -0.2 | 4.80E-01 | 0.0 | 9.62E-01 | Lipase class 3 family protein |
| AT1G34350 | -0.2 | 1.59E-01 | 0.2 | 1.08E-01 | -0.3 | 2.05E-02 | 0.1 | 5.82E-01 | 0.1 | 8.05E-01 | 0.1 | 8.83E-01 | Transmembrane protein 18 |
| AT1G36980 | -0.2 | 4.75E-02 | 0.0 | 7.45E-01 | -0.1 | 1.25E-01 | -0.4 | 5.34E-03 | -0.4 | 2.52E-02 | -0.1 | 8.16E-01 | hypothetical protein |
| AT4G17440 | -0.2 | 2.03E-01 | -0.1 | 3.48E-01 | 0.0 | 8.21E-01 | 0.1 | 8.51E-01 | -0.1 | 8.39E-01 | 0.2 | 7.40E-01 | At4g17440 ) |
| AT4G33985 | -0.2 | 4.98E-01 | -0.3 | 1.81E-01 | 0.2 | 5.49E-01 | 0.1 | 7.48E-01 | 0.0 | 9.96E-01 | 0.1 | 7.89E-01 | hypothetical protein |
| AT5G12370 | -0.2 | 3.61E-02 | -0.2 | 1.35E-02 | 0.0 | 5.57E-01 | -0.2 | 1.26E-01 | -0.1 | 4.19E-01 | -0.1 | 7.32E-01 | Exocyst complex component sec10 |
| AT2G01650 | -0.2 | 6.43E-02 | -0.1 | 1.45E-01 | 0.0 | 7.31E-01 | 0.1 | 3.98E-01 | -0.2 | 1.77E-01 | 0.3 | 8.55E-02 | Plant UBX domain-containing protein 2 |
| AT4G09770 | -0.2 | 2.28E-01 | -0.2 | 9.08E-02 | 0.1 | 6.30E-01 | 0.1 | 8.70E-01 | 0.5 | 1.45E-01 | -0.4 | 2.99E-01 | TRAF-like family protein |
| AT4G32030 | -0.2 | 6.42E-02 | 0.0 | 7.32E-01 | -0.1 | 1.70E-01 | -0.3 | 9.18E-02 | -0.4 | 3.62E-02 | 0.1 | 6.93E-01 | Uncharacterized protein |
| AT2G20960 | -0.2 | 8.08E-02 | -0.4 | 5.60E-04 | 0.3 | 1.21E-02 | 0.0 | 7.68E-01 | -0.3 | 2.84E-02 | 0.3 | 4.12E-02 | PEARLI 4 protein family protein) |
| AT4G08580 | -0.2 | 5.31E-02 | -0.1 | 2.13E-01 | -0.1 | 5.31E-01 | 0.0 | 8.97E-01 | -0.2 | 3.71E-01 | 0.1 | 5.07E-01 | hypothetical protein |
| AT1G18260 | -0.2 | 6.38E-02 | -0.2 | 4.46E-02 | 0.0 | 8.24E-01 | -0.1 | 4.35E-01 | -0.1 | 5.96E-01 | 0.0 | 9.50E-01 | ERAD-associated E3 ubiquitin-protein ligase component HRD3A |
| AT5G19390 | -0.2 | 9.57E-02 | -0.1 | 2.19E-01 | 0.0 | 7.06E-01 | 0.0 | 9.34E-01 | 0.1 | 8.98E-01 | -0.1 | 8.67E-01 | Rho GTPase activation protein with PH domain-containing protein |
| AT1G77140 | -0.2 | 4.10E-02 | -0.2 | 3.38E-02 | 0.0 | 8.86E-01 | 0.0 | 7.81E-01 | 0.2 | 3.76E-01 | -0.1 | 6.35E-01 | Vacuolar protein sorting-associated protein 45 homolog |
| AT4G16380 | -0.2 | 1.35E-01 | 0.0 | 9.21E-01 | -0.2 | 1.50E-01 | 0.0 | 9.63E-01 | -0.1 | 5.03E-01 | 0.1 | 6.56E-01 | Heavy metal transport |
| AT3G20720 | -0.2 | 6.58E-02 | -0.2 | 4.14E-02 | 0.0 | 7.90E-01 | 0.2 | 2.72E-01 | 0.2 | 4.54E-01 | 0.0 | 9.05E-01 | Uncharacterized protein |
| AT2G02910 | -0.2 | 2.53E-01 | -0.1 | 3.21E-01 | 0.0 | 9.33E-01 | 0.1 | 6.03E-01 | 0.1 | 6.80E-01 | 0.0 | 9.94E-01 | Uncharacterized protein At2g02910 |
| AT1G72480 | -0.2 | 2.92E-02 | -0.1 | 3.25E-01 | -0.1 | 2.38E-01 | 0.2 | 1.79E-01 | 0.2 | 1.01E-01 | -0.1 | 7.63E-01 | Lung seven transmembrane receptor family protein |
| AT3G45230 | -0.2 | 2.91E-01 | 0.2 | 1.66E-01 | -0.4 | 4.79E-02 | -0.1 | 5.74E-01 | 0.0 | 8.97E-01 | -0.1 | 5.87E-01 | Hydroxyproline-rich glycoprotein family protein |
| AT2G33390 | -0.2 | 2.40E-01 | -0.1 | 3.99E-01 | 0.0 | 8.16E-01 | -0.1 | 7.65E-01 | -0.1 | 5.79E-01 | 0.1 | 8.42E-01 | Uncharacterized protein |
| AT5G03100 | -0.2 | 2.36E-01 | 0.0 | 7.63E-01 | -0.2 | 2.23E-01 | -0.1 | 8.95E-01 | 0.0 | 9.76E-01 | 0.0 | 9.48E-01 | F-box protein At5g03100 |
| AT1G10140 | -0.2 | 1.00E-01 | 0.0 | 6.11E-01 | -0.2 | 6.97E-02 | 0.1 | 7.69E-01 | -0.4 | 1.97E-01 | 0.5 | 1.80E-01 | hypothetical protein |
| AT1G58250 | -0.2 | 5.12E-02 | -0.3 | 3.68E-03 | 0.1 | 1.57E-01 | -0.1 | 7.48E-01 | -0.1 | 8.17E-01 | 0.0 | 9.83E-01 | HYPERSENSITIVE TO PI STARVATION 4 |
| AT3G01450 | -0.2 | 7.89E-02 | 0.1 | 5.50E-01 | -0.2 | 5.71E-02 | -0.1 | 7.69E-01 | -0.1 | 6.94E-01 | 0.0 | 9.28E-01 | TOG domain-containing protein |
| AT4G33890 | -0.2 | 6.22E-02 | -0.2 | 4.31E-02 | 0.0 | 8.26E-01 | 0.0 | 9.92E-01 | -0.1 | 8.01E-01 | 0.1 | 8.39E-01 | Transcriptional regulator of RNA polII, SAGA, subunit |
| AT2G41370 | -0.2 | 1.34E-01 | -0.1 | 2.81E-01 | 0.0 | 7.49E-01 | -0.2 | 5.02E-01 | -0.2 | 5.34E-01 | 0.0 | 9.86E-01 | BTB domain-containing protein |
| AT1G32260 | -0.2 | 1.14E-01 | -0.1 | 4.50E-01 | -0.1 | 4.89E-01 | -0.1 | 6.89E-01 | 0.0 | 8.59E-01 | 0.0 | 9.19E-01 | Envelope glycoprotein |
| AT4G27500 | -0.2 | 2.29E-02 | -0.1 | 1.51E-01 | -0.1 | 3.83E-01 | -0.1 | 6.55E-01 | 0.0 | 8.65E-01 | 0.0 | 9.04E-01 | Uncharacterized protein At4g27500 |
| AT2G41960 | -0.2 | 1.74E-02 | -0.2 | 1.62E-03 | 0.1 | 1.67E-01 | 0.2 | 2.22E-01 | 0.1 | 6.91E-01 | 0.1 | 6.62E-01 | At2g41960 |
| AT5G20360 | -0.2 | 8.50E-02 | -0.4 | 2.19E-03 | 0.2 | 6.67E-02 | 0.1 | 7.38E-01 | 0.0 | 9.04E-01 | 0.0 | 9.08E-01 | Octicosapeptide |
| AT2G45320 | -0.2 | 1.58E-01 | -0.2 | 9.33E-02 | 0.0 | 7.69E-01 | -0.5 | 1.25E-01 | -0.4 | 3.30E-01 | -0.2 | 7.62E-01 | Uncharacterized protein At2g45320 |
| AT3G12360 | -0.2 | 8.28E-02 | -0.3 | 4.36E-03 | 0.2 | 1.15E-01 | -0.3 | 6.65E-02 | -0.4 | 3.68E-02 | 0.1 | 7.54E-01 | Ankyrin repeat family protein |
| AT1G75140 | -0.2 | 1.09E-01 | 0.1 | 5.29E-01 | -0.2 | 6.32E-02 | 0.0 | 9.97E-01 | 0.2 | 2.97E-01 | -0.2 | 3.76E-01 | Uncharacterized membrane protein At1g75140 |
| AT3G63180 | -0.2 | 2.02E-01 | -0.2 | 2.07E-01 | 0.0 | 9.81E-01 | 0.0 | 9.19E-01 | -0.1 | 7.72E-01 | 0.1 | 8.84E-01 | TIC-like protein |
| AT2G27285 | -0.2 | 5.60E-02 | -0.1 | 2.25E-01 | -0.1 | 5.21E-01 | 0.0 | 9.17E-01 | 0.0 | 9.27E-01 | 0.0 | 8.70E-01 | Nuclear speckle splicing regulatory protein 1 N-terminal domain-containing protein |
| AT3G22240 | -0.1 | 6.71E-01 | 0.5 | 1.34E-01 | -0.7 | 9.40E-02 | 0.5 | 3.15E-01 | 0.3 | 5.75E-01 | 0.1 | 5.79E-01 | Protein CYSTEINE-RICH TRANSMEMBRANE MODULE 9 |
| AT1G09460 | -0.1 | 2.63E-01 | 0.3 | 5.05E-02 | -0.4 | 1.75E-02 | -0.1 | 6.73E-01 | -0.5 | 8.22E-02 | 0.4 | 1.70E-01 | F14J9.12 protein |
| AT1G79830 | -0.1 | 2.76E-02 | -0.1 | 1.34E-01 | -0.1 | 4.75E-01 | 0.0 | 8.22E-01 | -0.1 | 5.46E-01 | 0.1 | 7.90E-01 | Golgin Putative 5 |
| AT4G19140 | -0.1 | 5.34E-02 | -0.1 | 9.29E-02 | 0.0 | 8.47E-01 | -0.2 | 4.68E-01 | -0.1 | 5.90E-01 | 0.0 | 9.71E-01 | At4g19140 |
| AT1G04300 | -0.1 | 8.40E-02 | -0.4 | 1.04E-03 | 0.2 | 2.52E-02 | -0.1 | 4.26E-01 | -0.3 | 1.91E-01 | 0.1 | 6.67E-01 | TRAF-like superfamily protein |
| AT1G43245 | -0.1 | 1.52E-01 | -0.3 | 1.97E-02 | 0.1 | 3.09E-01 | -0.2 | 3.53E-01 | -0.3 | 3.64E-01 | 0.0 | 9.62E-01 | SET domain-containing protein |
| AT2G30870 | -0.1 | 2.00E-01 | 0.5 | 2.81E-03 | -0.6 | 1.03E-03 | 0.5 | 3.63E-02 | 0.2 | 3.55E-01 | 0.2 | 4.74E-01 | GST N-terminal domain-containing protein |
| AT3G43955 | -0.1 | 2.59E-01 | -0.1 | 6.31E-01 | -0.1 | 6.15E-01 | 0.3 | 2.35E-01 | 0.3 | 1.70E-01 | -0.1 | 8.26E-01 | NA |
| AT1G69510 | -0.1 | 6.99E-02 | 0.1 | 4.05E-01 | -0.2 | 2.56E-02 | -0.3 | 8.07E-03 | 0.0 | 9.22E-01 | -0.3 | 6.82E-02 | cAMP-regulated phosphoprotein 19-related protein |
| AT5G24180 | -0.1 | 5.16E-01 | -0.1 | 7.42E-01 | -0.1 | 8.31E-01 | | | | | | | Gb\|AAD29063.1 |
| AT1G26620 | -0.1 | 9.58E-02 | -0.2 | 3.60E-02 | 0.1 | 6.04E-01 | -0.1 | 7.59E-01 | -0.1 | 6.84E-01 | 0.0 | 9.18E-01 | T-box transcription factor, putative |
| AT5G35995 | -0.1 | 6.68E-02 | -0.1 | 2.48E-01 | -0.1 | 5.56E-01 | -0.2 | 3.28E-01 | -0.1 | 7.01E-01 | -0.1 | 7.41E-01 | F-box |
| AT1G15940 | -0.1 | 5.12E-02 | -0.3 | 5.10E-03 | 0.1 | 2.15E-01 | 0.1 | 5.37E-01 | 0.2 | 3.95E-01 | -0.1 | 8.43E-01 | Sister chromatid cohesion protein PDS5 homolog E |
| AT3G55490 | -0.1 | 2.28E-01 | 0.1 | 2.85E-01 | -0.3 | 7.09E-02 | -0.4 | 1.39E-01 | -0.1 | 7.34E-01 | -0.3 | 4.49E-01 | At1g19080 |
| AT3G60200 | -0.1 | 2.47E-01 | 0.0 | 7.69E-01 | -0.1 | 4.82E-01 | -0.1 | 7.91E-01 | -0.2 | 5.58E-01 | 0.1 | 8.06E-01 | hypothetical protein |
| AT3G12570 | -0.1 | 3.71E-02 | -0.3 | 2.52E-03 | 0.1 | 1.36E-01 | -0.2 | 2.62E-01 | -0.2 | 2.97E-01 | 0.0 | 9.71E-01 | FYD |
| AT5G58720 | -0.1 | 4.43E-02 | -0.3 | 2.37E-03 | 0.1 | 1.12E-01 | -0.5 | 4.12E-04 | -0.4 | 2.73E-03 | -0.1 | 7.46E-01 | Smr domain-containing protein |
| AT3G60350 | -0.1 | 9.17E-02 | -0.3 | 8.36E-03 | 0.1 | 2.14E-01 | 0.0 | 8.62E-01 | -0.1 | 7.95E-01 | 0.0 | 9.33E-01 | Protein ARABIDILLO 2 |
| AT1G69230 | -0.1 | 4.82E-01 | 0.0 | 9.34E-01 | -0.2 | 5.23E-01 | 0.0 | 9.81E-01 | 0.2 | 4.72E-01 | -0.2 | 5.70E-01 | Protein SPIRAL1-like 2 |
| AT3G05675 | -0.1 | 1.21E-01 | 0.1 | 3.33E-01 | -0.2 | 4.59E-02 | -0.1 | 8.05E-01 | -0.1 | 7.67E-01 | 0.0 | 9.39E-01 | BTB |
| AT5G26740 | -0.1 | 5.84E-02 | 0.2 | 1.88E-02 | -0.3 | 1.98E-03 | -0.3 | 3.18E-02 | -0.3 | 1.12E-01 | -0.1 | 8.03E-01 | Organic solute transporter ostalpha protein |
| AT3G29120 | -0.1 | 2.80E-01 | -0.2 | 1.59E-01 | 0.1 | 7.51E-01 | 0.0 | 9.92E-01 | 0.1 | 7.56E-01 | -0.1 | 7.91E-01 | NA |
| AT2G18440 | -0.1 | 2.91E-01 | -0.1 | 3.67E-01 | 0.0 | 9.30E-01 | -0.3 | 1.19E-01 | -0.3 | 1.55E-01 | 0.0 | 9.80E-01 | NA |
| AT5G15580 | -0.1 | 9.61E-02 | 0.2 | 9.05E-02 | -0.3 | 9.54E-03 | -0.2 | 2.02E-01 | -0.1 | 4.37E-01 | -0.1 | 8.40E-01 | Protein LONGIFOLIA 1 |
| AT5G28913 | -0.1 | 1.79E-01 | 0.2 | 1.10E-01 | -0.3 | 2.29E-02 | -0.4 | 5.27E-02 | 0.0 | 9.62E-01 | -0.4 | 1.19E-01 | NA |
| AT3G12260 | -0.1 | 8.18E-02 | 0.2 | 7.84E-02 | -0.3 | 6.45E-03 | 0.0 | 8.94E-01 | 0.1 | 7.79E-01 | 0.0 | 8.84E-01 | NADH dehydrogenase 1 alpha subcomplex subunit 6 |
| AT2G42140 | -0.1 | 4.71E-01 | 0.3 | 2.05E-01 | -0.4 | 1.11E-01 | -0.5 | 3.41E-01 | 0.0 | 9.61E-01 | | | VQ motif-containing protein 17 |
| AT5G24740 | -0.1 | 6.20E-02 | -0.4 | 2.37E-04 | 0.3 | 6.61E-03 | 0.0 | 8.61E-01 | -0.2 | 4.78E-01 | 0.2 | 4.68E-01 | Vacuolar protein sorting-associated protein, putative |
| AT5G53440 | -0.1 | 1.05E-01 | -0.4 | 2.22E-03 | 0.2 | 5.17E-02 | -0.1 | 6.53E-01 | -0.2 | 5.73E-01 | 0.1 | 9.11E-01 | LOW protein: zinc finger CCCH domain protein |
| AT3G12010 | -0.1 | 4.82E-02 | -0.2 | 8.02E-03 | 0.1 | 3.30E-01 | -0.1 | 6.88E-01 | 0.1 | 6.04E-01 | -0.2 | 4.47E-01 | C18orf8 |
| AT1G64990 | -0.1 | 4.42E-02 | -0.2 | 9.20E-03 | 0.1 | 3.94E-01 | 0.0 | 7.47E-01 | 0.1 | 6.63E-01 | 0.0 | 9.11E-01 | hypothetical protein |
| AT3G12180 | -0.1 | 1.50E-01 | 0.1 | 3.77E-01 | -0.2 | 6.06E-02 | 0.1 | 7.16E-01 | 0.0 | 8.77E-01 | 0.0 | 8.99E-01 | Protein cornichon homolog 1 |
| AT1G03370 | -0.1 | 5.62E-02 | -0.3 | 9.87E-04 | 0.2 | 3.42E-02 | 0.1 | 4.32E-01 | 0.1 | 6.55E-01 | 0.0 | 9.13E-01 | C2 calcium |
| AT3G49601 | -0.1 | 6.87E-02 | -0.1 | 8.90E-02 | 0.0 | 9.52E-01 | -0.2 | 1.54E-01 | -0.2 | 1.99E-01 | 0.0 | 9.86E-01 | Pre-mRNA-splicing factor |
| AT2G25690 | -0.1 | 3.97E-01 | -0.2 | 2.65E-01 | 0.1 | 8.06E-01 | 0.3 | 1.56E-01 | 0.3 | 3.56E-01 | 0.1 | 8.22E-01 | DUF581 family protein, putative |
| AT1G56440 | -0.1 | 1.35E-01 | -0.4 | 1.75E-03 | 0.3 | 3.70E-02 | 0.1 | 8.22E-01 | -0.1 | 6.65E-01 | 0.2 | 5.28E-01 | Tetratricopeptide repeat -like superfamily protein |
| AT1G51540 | -0.1 | 1.63E-01 | -0.1 | 3.35E-01 | 0.0 | 7.44E-01 | -0.3 | 4.45E-01 | -0.1 | 8.19E-01 | -0.2 | 7.70E-01 | Uncharacterized protein F5D21.24 |
| AT4G02030 | -0.1 | 4.61E-02 | -0.1 | 1.03E-01 | 0.0 | 7.51E-01 | 0.1 | 8.05E-01 | -0.1 | 8.56E-01 | 0.1 | 7.12E-01 | Vacuolar protein sorting-associated protein 51 homolog |
| AT3G24100 | -0.1 | 6.85E-02 | 0.1 | 3.04E-01 | -0.2 | 2.12E-02 | 0.0 | 8.62E-01 | 0.0 | 9.32E-01 | 0.1 | 8.39E-01 | Uncharacterized protein AT3g24100 |
| AT4G02370 | -0.1 | 5.70E-02 | 0.0 | 6.90E-01 | -0.2 | 4.46E-02 | 0.1 | 6.04E-01 | -0.1 | 8.47E-01 | 0.2 | 5.57E-01 | AT4g02370 protein |
| AT1G55270 | -0.1 | 1.83E-01 | -0.2 | 4.90E-02 | 0.1 | 4.89E-01 | 0.0 | 9.58E-01 | 0.2 | 7.33E-01 | -0.2 | 7.21E-01 | Galactose oxidase |
| AT4G38940 | -0.1 | 9.84E-02 | -0.2 | 9.20E-02 | 0.0 | 9.40E-01 | 0.1 | 7.83E-01 | -0.3 | 1.91E-01 | 0.4 | 1.39E-01 | F-box |
| AT1G60995 | -0.1 | 4.48E-02 | -0.2 | 1.18E-02 | 0.1 | 4.70E-01 | -0.1 | 4.79E-01 | -0.1 | 6.94E-01 | 0.0 | 9.01E-01 | Uncharacterized protein |
| AT5G10850 | -0.1 | 5.63E-01 | 0.4 | 1.42E-01 | -0.5 | 9.95E-02 | 0.7 | 1.06E-01 | 0.7 | 1.45E-01 | 0.0 | 1.00E+00 | NA |
| AT2G02080 | -0.1 | 1.21E-01 | -0.2 | 2.14E-02 | 0.1 | 3.65E-01 | -0.4 | 1.15E-02 | -0.4 | 7.79E-03 | 0.1 | 7.93E-01 | C2H2-type domain-containing protein |
| AT1G01230 | -0.1 | 8.13E-02 | 0.1 | 2.76E-01 | -0.2 | 2.14E-02 | -0.2 | 7.97E-02 | -0.1 | 4.45E-01 | -0.1 | 6.01E-01 | F6F3.4 protein |
| AT1G50730 | -0.1 | 2.88E-01 | -0.4 | 9.35E-03 | 0.3 | 8.82E-02 | -0.1 | 8.16E-01 | 0.0 | 9.76E-01 | -0.1 | 8.28E-01 | Uncharacterized protein At1g50730 |

| | | | | | | | | | | | | |
|---|---|---|---|---|---|---|---|---|---|---|---|---|
| AT2G25660 | -0.1 | 9.04E-02 | -0.1 | 1.10E-01 | 0.0 | 9.66E-01 | 0.0 | 9.78E-01 | 0.1 | 8.56E-01 | -0.1 | 8.95E-01 | Embryo defective 2410 |
| AT2G00680 | -0.1 | 3.35E-01 | 0.0 | 7.68E-01 | -0.2 | 3.08E-01 | 0.0 | 8.97E-01 | 0.0 | 9.63E-01 | -0.1 | 8.95E-01 | NA |
| AT5G41140 | -0.1 | 3.09E-01 | -0.4 | 2.51E-02 | 0.2 | 1.81E-01 | -0.1 | 7.49E-01 | 0.2 | 2.81E-01 | -0.3 | 2.41E-01 | Myosin heavy chain-related protein |
| AT5G01240 | -0.1 | 3.46E-01 | 0.2 | 2.10E-01 | -0.3 | 6.07E-02 | -0.4 | 3.16E-03 | -0.2 | 1.07E-01 | -0.2 | 3.61E-01 | Like AUXIN RESISTANT 1 |
| AT3G11590 | -0.1 | 6.12E-02 | -0.1 | 7.37E-02 | 0.0 | 9.74E-01 | -0.1 | 6.52E-01 | -0.1 | 8.23E-01 | 0.0 | 9.26E-01 | At3g11590 |
| AT5G47020 | -0.1 | 1.96E-01 | -0.2 | 6.02E-02 | 0.1 | 5.22E-01 | -0.4 | 1.22E-02 | -0.3 | 9.34E-02 | -0.1 | 6.53E-01 | MraZ |
| AT2G43080 | -0.1 | 6.05E-02 | -0.1 | 2.73E-01 | -0.1 | 4.77E-01 | 0.0 | 8.46E-01 | -0.1 | 7.76E-01 | 0.0 | 9.34E-01 | P4H isoform 1 |
| AT5G22310 | -0.1 | 1.60E-01 | 0.1 | 2.98E-01 | -0.2 | 4.99E-02 | -0.3 | 1.54E-01 | -0.5 | 4.90E-02 | 0.2 | 6.28E-01 | AT5g22310 |
| AT5G19340 | -0.1 | 2.63E-01 | 0.5 | 1.10E-03 | -0.7 | 9.97E-04 | -0.4 | 2.75E-02 | -0.2 | 2.64E-01 | -0.2 | 5.05E-01 | At5g19340 |
| AT4G17010 | -0.1 | 3.35E-01 | 0.1 | 3.62E-01 | -0.3 | 1.20E-01 | -0.1 | 6.10E-01 | -0.1 | 5.76E-01 | 0.0 | 9.47E-01 | AT4g17010 |
| AT4G03200 | -0.1 | 1.36E-01 | -0.3 | 4.20E-03 | 0.2 | 7.48E-02 | -0.1 | 5.56E-01 | -0.1 | 6.55E-01 | 0.0 | 9.81E-01 | At4g03200 |
| AT5G05987 | -0.1 | 1.17E-01 | -0.1 | 2.98E-01 | 0.0 | 6.74E-01 | -0.3 | 8.99E-02 | -0.3 | 1.11E-01 | 0.0 | 9.78E-01 | PRA1 family protein A2 |
| AT2G20330 | -0.1 | 4.13E-02 | -0.3 | 2.10E-03 | 0.1 | 1.07E-01 | -0.1 | 6.17E-01 | -0.1 | 6.94E-01 | 0.0 | 9.92E-01 | At2g20330 |
| AT3G56110 | -0.1 | 1.48E-01 | 0.1 | 5.97E-01 | -0.2 | 8.78E-02 | -0.1 | 5.50E-01 | -0.1 | 4.84E-01 | 0.0 | 9.14E-01 | PRA1 family protein B1 |
| AT1G72790 | -0.1 | 2.37E-01 | -0.1 | 6.83E-01 | -0.1 | 5.35E-01 | -0.3 | 1.38E-01 | -0.2 | 3.68E-01 | -0.1 | 7.96E-01 | Hydroxyproline-rich glycoprotein family protein |
| AT1G06630 | -0.1 | 2.28E-01 | -0.2 | 1.16E-01 | 0.1 | 7.10E-01 | -0.2 | 1.31E-01 | -0.3 | 5.34E-02 | 0.1 | 6.91E-01 | F-box |
| AT1G62930 | -0.1 | 3.36E-01 | -0.1 | 6.56E-01 | -0.1 | 7.08E-01 | 0.1 | 8.36E-01 | 0.0 | 9.92E-01 | 0.1 | 8.85E-01 | Pentatricopeptide repeat-containing protein At1g62930, chloroplastic |
| AT2G30960 | -0.1 | 4.09E-01 | 0.2 | 3.40E-01 | -0.3 | 1.50E-01 | -0.4 | 4.89E-02 | -0.1 | 5.58E-01 | -0.3 | 3.42E-01 | At3g04090 |
| AT1G36070 | -0.1 | 1.28E-01 | -0.3 | 6.18E-03 | 0.2 | 1.28E-01 | -0.9 | 3.85E-04 | -0.5 | 1.78E-02 | -0.4 | 1.85E-01 | At1g36070 |
| AT3G50860 | -0.1 | 1.12E-01 | -0.1 | 4.46E-01 | -0.1 | 4.89E-01 | 0.0 | 8.94E-01 | 0.1 | 5.75E-01 | -0.1 | 7.21E-01 | AP complex subunit sigma |
| AT2G33320 | -0.1 | 1.13E-01 | -0.2 | 4.73E-02 | 0.0 | 6.43E-01 | -0.3 | 9.19E-02 | -0.1 | 5.82E-01 | -0.2 | 4.19E-01 | C2 domain-containing protein |
| AT4G12700 | -0.1 | 1.73E-01 | 0.0 | 7.88E-01 | -0.2 | 1.67E-01 | 0.0 | 8.46E-01 | 0.1 | 6.23E-01 | -0.1 | 5.49E-01 | At4g12700 |
| AT2G02350 | -0.1 | 3.51E-01 | -0.1 | 4.61E-01 | 0.0 | 9.10E-01 | 0.2 | 5.87E-01 | 0.2 | 6.63E-01 | 0.0 | 9.92E-01 | F-box protein SKIP3 |
| AT4G16442 | -0.1 | 2.43E-01 | 0.0 | 7.42E-01 | -0.1 | 4.89E-01 | -0.4 | 6.85E-02 | -0.2 | 6.06E-01 | -0.3 | 3.79E-01 | CASP-like protein 2B1 |
| AT3G05000 | -0.1 | 8.96E-02 | 0.1 | 7.80E-02 | -0.3 | 8.41E-03 | -0.1 | 6.59E-01 | -0.1 | 7.08E-01 | 0.0 | 9.96E-01 | Transport protein particle component |
| AT5G18040 | -0.1 | 1.73E-01 | -0.4 | 3.39E-03 | 0.3 | 5.88E-02 | -0.1 | 7.57E-01 | -0.3 | 4.31E-01 | 0.2 | 7.13E-01 | Target of trans acting-siR480 |
| AT1G14480 | -0.1 | 2.62E-01 | -0.5 | 2.69E-03 | 0.4 | 3.30E-02 | 0.3 | 4.91E-01 | 0.0 | 9.57E-01 | 0.2 | 5.61E-01 | Ankyrin repeat family protein |
| AT2G46080 | -0.1 | 3.22E-02 | -0.2 | 2.42E-03 | 0.1 | 1.52E-01 | 0.0 | 9.36E-01 | -0.1 | 7.77E-01 | 0.0 | 8.78E-01 | hypothetical protein |
| AT5G24170 | -0.1 | 1.64E-01 | 0.2 | 9.74E-02 | -0.3 | 1.96E-02 | 0.1 | 7.13E-01 | 0.3 | 2.41E-01 | -0.2 | 5.29E-01 | Vesicle transport protein |
| AT4G19450 | -0.1 | 4.93E-02 | -0.1 | 4.85E-02 | 0.0 | 9.53E-01 | 0.1 | 5.12E-01 | -0.1 | 4.95E-01 | 0.2 | 2.69E-01 | Uncharacterized protein AT4g19450 |
| AT1G16520 | -0.1 | 1.02E-01 | -0.1 | 4.68E-01 | -0.1 | 4.29E-01 | 0.0 | 9.70E-01 | 0.0 | 9.33E-01 | 0.0 | 9.71E-01 | hypothetical protein |
| AT4G34150 | -0.1 | 3.39E-01 | -0.5 | 5.05E-03 | 0.4 | 3.37E-02 | 0.1 | 5.82E-01 | -0.1 | 2.67E-01 | 0.2 | 1.76E-01 | hypothetical protein |
| AT5G40170 | -0.1 | 2.50E-01 | -0.1 | 4.67E-01 | 0.0 | 7.41E-01 | 0.4 | 4.70E-03 | -0.1 | 8.01E-01 | 0.5 | 1.71E-02 | Receptor-like protein 54 |
| AT5G19130 | -0.1 | 6.01E-02 | -0.1 | 4.60E-01 | -0.1 | 3.00E-01 | -0.2 | 2.62E-01 | -0.2 | 1.77E-01 | 0.1 | 8.39E-01 | GPI transamidase component family protein |
| AT5G03670 | -0.1 | 3.36E-01 | 0.4 | 5.59E-03 | -0.6 | 5.02E-03 | 0.3 | 5.75E-01 | 0.4 | 4.47E-01 | -0.1 | 8.62E-01 | Histone-lysine N-methyltransferase SETD1B-like protein |
| AT1G13245 | -0.1 | 5.10E-01 | -0.2 | 4.09E-01 | 0.0 | 8.76E-01 | 0.0 | 9.20E-01 | -0.7 | 7.72E-02 | 0.7 | 1.87E-01 | Small polypeptide DEVIL 4 |
| AT5G10450 | -0.1 | 7.55E-02 | 0.0 | 9.76E-01 | -0.1 | 1.03E-01 | -0.1 | 3.82E-01 | -0.2 | 1.21E-01 | 0.1 | 5.92E-01 | G-box regulating factor 6 |
| AT3G09690 | -0.1 | 1.43E-01 | 0.0 | 6.31E-01 | -0.1 | 4.09E-01 | 0.0 | 9.98E-01 | 0.2 | 4.55E-01 | -0.2 | 5.26E-01 | Alpha |
| AT3G20360 | -0.1 | 4.99E-01 | 1.2 | 3.71E-05 | -1.4 | 1.65E-04 | 0.0 | 9.82E-01 | 0.2 | 5.65E-01 | -0.2 | 6.10E-01 | MATH domain-containing protein |
| AT3G08570 | -0.1 | 4.97E-01 | -0.3 | 2.25E-01 | 0.1 | 6.35E-01 | 0.2 | 6.38E-01 | 0.2 | 7.16E-01 | 0.0 | 9.84E-01 | BTB |
| AT5G58340 | -0.1 | 2.38E-01 | -0.1 | 3.05E-01 | 0.0 | 9.30E-01 | -0.3 | 2.66E-01 | -0.3 | 3.80E-01 | 0.0 | 9.40E-01 | hypothetical protein |
| AT2G36060 | -0.1 | 6.07E-02 | 0.0 | 6.65E-01 | -0.1 | 1.82E-01 | 0.0 | 9.48E-01 | -0.1 | 8.02E-01 | 0.0 | 8.86E-01 | UEV1C |
| AT5G49980 | -0.1 | 1.04E-01 | 0.0 | 7.79E-01 | -0.1 | 2.25E-01 | -0.3 | 7.78E-03 | -0.2 | 6.60E-02 | -0.1 | 6.83E-01 | hypothetical protein |
| AT5G35690 | -0.1 | 9.67E-02 | -0.4 | 3.76E-04 | 0.3 | 7.16E-03 | 0.2 | 1.50E-01 | 0.0 | 9.91E-01 | 0.2 | 3.25E-01 | Uncharacterized protein At5g35690 |
| AT2G32295 | -0.1 | 5.67E-02 | -0.1 | 7.72E-02 | 0.0 | 9.38E-01 | 0.1 | 6.13E-01 | 0.4 | 1.24E-01 | -0.2 | 4.30E-01 | EXS domain-containing protein |
| AT4G29860 | -0.1 | 1.91E-01 | -0.2 | 6.53E-02 | 0.1 | 5.81E-01 | -0.1 | 6.38E-01 | -0.2 | 5.02E-01 | 0.1 | 8.72E-01 | Protein DECREASED SIZE EXCLUSION LIMIT 1 |
| AT3G51050 | -0.1 | 7.35E-02 | -0.1 | 7.67E-02 | 0.0 | 9.77E-01 | 0.1 | 5.26E-01 | 0.0 | 9.92E-01 | 0.1 | 6.91E-01 | FG-GAP repeat-containing protein |
| AT1G31880 | -0.1 | 4.24E-01 | -0.4 | 2.86E-02 | 0.3 | 1.67E-01 | -0.5 | 3.09E-01 | -0.3 | 6.86E-01 | -0.3 | 7.47E-01 | BRX domain-containing protein |
| AT5G49430 | -0.1 | 9.79E-02 | -0.1 | 3.22E-01 | -0.1 | 5.74E-01 | 0.0 | 9.91E-01 | -0.1 | 8.92E-01 | 0.0 | 9.18E-01 | WD40 |
| AT1G70620 | -0.1 | 1.43E-01 | -0.2 | 3.48E-02 | 0.1 | 4.41E-01 | -0.1 | 7.54E-01 | -0.1 | 5.31E-01 | 0.1 | 8.20E-01 | Uncharacterized protein F24J13.19 |
| AT3G15470 | -0.1 | 8.54E-02 | -0.2 | 1.21E-02 | 0.1 | 3.04E-01 | -0.3 | 3.08E-02 | -0.2 | 4.00E-01 | -0.2 | 3.75E-01 | Transducin |
| AT3G02300 | -0.1 | 1.29E-01 | -0.3 | 2.36E-03 | 0.2 | 4.54E-02 | -0.2 | 2.51E-01 | 0.0 | 8.46E-01 | -0.2 | 2.85E-01 | Uncharacterized protein F14P3.5 |
| AT2G28310 | -0.1 | 7.87E-02 | 0.1 | 4.72E-01 | -0.2 | 3.88E-02 | -0.2 | 2.28E-01 | 0.0 | 9.24E-01 | -0.2 | 3.37E-01 | hypothetical protein |
| AT4G03010 | -0.1 | 3.97E-01 | 0.0 | 8.51E-01 | -0.2 | 4.01E-01 | 0.1 | 8.56E-01 | 0.3 | 3.44E-01 | -0.2 | 4.82E-01 | Leucine-rich repeat family protein |
| AT1G54890 | -0.1 | 5.25E-01 | 0.7 | 4.07E-03 | -0.8 | 4.48E-03 | 0.1 | 5.83E-01 | 0.5 | 8.81E-02 | -0.3 | 2.52E-01 | Late embryogenesis abundant protein-like protein |
| AT5G03230 | -0.1 | 3.39E-01 | 0.0 | 7.67E-01 | -0.2 | 2.95E-01 | -0.6 | 1.25E-02 | -0.9 | 3.02E-03 | 0.2 | 4.70E-01 | Arabidopsis thaliana genomic DNA, chromosome 5, P1 clone:MOK16 |
| AT5G65670 | -0.1 | 5.98E-02 | 0.0 | 7.70E-01 | -0.1 | 1.44E-01 | 0.1 | 2.65E-01 | 0.0 | 9.18E-01 | 0.1 | 5.08E-01 | Auxin-responsive protein |
| AT1G10865 | -0.1 | 1.63E-01 | -0.2 | 9.44E-02 | 0.0 | 7.64E-01 | -0.1 | 6.13E-01 | -0.4 | 7.46E-02 | 0.3 | 2.52E-01 | Cytochrome C oxidase assembly factor |
| AT2G34655 | -0.1 | 5.28E-01 | -0.3 | 1.79E-01 | 0.2 | 5.31E-01 | -0.3 | 5.84E-01 | -0.2 | 8.02E-01 | -0.1 | 8.84E-01 | At2g34655 |
| AT1G65150 | -0.1 | 3.74E-01 | -0.2 | 1.61E-01 | 0.1 | 6.32E-01 | 0.3 | 2.93E-01 | 0.4 | 2.71E-01 | -0.1 | 9.20E-01 | MATH domain-containing protein |
| AT1G79940 | -0.1 | 1.31E-01 | 0.0 | 9.98E-01 | -0.1 | 1.72E-01 | 0.1 | 2.00E-01 | 0.0 | 8.85E-01 | 0.1 | 4.85E-01 | Putative DnaJ protein |
| AT4G01410 | -0.1 | 3.05E-01 | 0.1 | 5.51E-01 | -0.2 | 1.73E-01 | -0.4 | 2.55E-02 | -0.2 | 2.67E-01 | -0.2 | 4.47E-01 | At4g01410 hydroxyproline-rich glycoprotein family) |
| AT5G52530 | -0.1 | 5.69E-02 | -0.3 | 1.68E-03 | 0.1 | 6.00E-02 | 0.2 | 2.56E-01 | 0.0 | 9.72E-01 | 0.2 | 4.01E-01 | Dentin sialophosphoprotein-like protein |
| AT1G79245 | -0.1 | 3.06E-01 | -0.5 | 3.66E-03 | 0.3 | 2.69E-02 | 0.0 | 9.20E-01 | 0.0 | 8.58E-01 | 0.0 | 9.45E-01 | NA |
| AT2G13420 | -0.1 | 4.35E-01 | 0.2 | 1.51E-01 | -0.4 | 8.12E-02 | 0.5 | 9.25E-02 | 0.2 | 6.41E-01 | 0.3 | 4.14E-01 | Pentatricopeptide repeat-containing protein At2g13420, mitochondrial |
| AT5G11740 | -0.1 | 3.47E-01 | 0.1 | 4.28E-01 | -0.2 | 1.23E-01 | -0.3 | 1.26E-01 | -0.3 | 2.67E-01 | 0.0 | 9.01E-01 | Arabinogalactan protein 15 |
| AT3G05230 | -0.1 | 2.11E-01 | 0.2 | 1.03E-01 | -0.3 | 2.40E-02 | 0.4 | 2.22E-02 | 0.6 | 7.92E-03 | -0.1 | 6.23E-01 | Signal peptidase complex subunit 3 |
| AT3G07210 | -0.1 | 1.82E-01 | 0.0 | 9.35E-01 | -0.1 | 3.01E-01 | -0.4 | 1.02E-01 | 0.1 | 8.42E-01 | -0.5 | 1.61E-01 | hypothetical protein |
| AT2G13550 | -0.1 | 3.37E-01 | 0.1 | 4.98E-01 | -0.2 | 1.85E-01 | -0.4 | 2.10E-01 | -0.3 | 4.46E-01 | -0.1 | 7.96E-01 | Uncharacterized protein At2g13550 |
| AT3G18280 | -0.1 | 2.69E-01 | 0.1 | 3.53E-01 | -0.2 | 8.24E-02 | 0.2 | 5.17E-01 | 0.0 | 9.21E-01 | 0.2 | 5.49E-01 | hypothetical protein |
| AT4G22360 | -0.1 | 8.95E-02 | -0.1 | 3.50E-01 | -0.1 | 5.01E-01 | -0.4 | 2.82E-02 | -0.4 | 4.27E-02 | 0.0 | 9.91E-01 | SWIB complex BAF60b domain-containing protein |
| AT2G21188 | -0.1 | 2.96E-01 | -0.3 | 5.13E-02 | 0.1 | 3.66E-01 | -0.2 | 6.60E-01 | -0.5 | 2.63E-01 | 0.3 | 5.65E-01 | NA |
| AT1G53000 | -0.1 | 1.41E-01 | 0.0 | 9.74E-01 | -0.1 | 1.85E-01 | -0.2 | 8.11E-02 | -0.1 | 7.02E-01 | -0.1 | 3.88E-01 | At1g53000 |
| AT2G30990 | -0.1 | 9.13E-02 | 0.1 | 3.20E-01 | -0.2 | 2.89E-02 | 0.0 | 9.31E-01 | -0.2 | 4.38E-01 | 0.2 | 5.33E-01 | Uncharacterized protein |
| AT1G63550 | -0.1 | 3.48E-01 | 0.8 | 6.54E-05 | -1.0 | 1.65E-04 | 0.1 | 8.36E-01 | -0.5 | 9.31E-02 | 0.5 | 7.87E-02 | Cysteine-rich repeat secretory protein 9 |
| AT3G23590 | -0.1 | 7.35E-02 | -0.1 | 3.67E-01 | -0.1 | 4.19E-01 | -0.2 | 3.69E-01 | 0.0 | 8.46E-01 | -0.1 | 6.79E-01 | Mediator of RNA polymerase II transcription subunit 33A |
| AT2G29910 | -0.1 | 2.24E-01 | -0.3 | 1.48E-02 | 0.2 | 1.72E-01 | 0.0 | 9.88E-01 | 0.1 | 8.05E-01 | -0.1 | 8.39E-01 | F-box |
| AT5G22700 | -0.1 | 3.45E-01 | -0.2 | 1.16E-01 | 0.1 | 5.46E-01 | -0.2 | 4.00E-01 | 0.0 | 9.90E-01 | -0.2 | 4.68E-01 | LOW protein: F-box |
| AT5G14270 | -0.1 | 6.87E-02 | -0.1 | 5.80E-02 | 0.0 | 9.01E-01 | 0.0 | 9.01E-01 | -0.2 | 3.16E-01 | 0.1 | 5.14E-01 | Bromodomain and extraterminal domain protein 9 |
| AT3G48115 | -0.1 | 2.38E-01 | -0.1 | 2.87E-01 | 0.0 | 9.51E-01 | -0.7 | 1.87E-03 | -0.5 | 1.98E-02 | -0.2 | 5.70E-01 | NA |
| AT5G12900 | -0.1 | 2.44E-01 | 0.0 | 6.61E-01 | -0.2 | 1.76E-01 | 0.0 | 9.12E-01 | 0.0 | 8.78E-01 | -0.1 | 8.36E-01 | IRK-interacting protein |
| AT4G24680 | -0.1 | 1.19E-01 | -0.3 | 8.11E-03 | 0.1 | 1.59E-01 | -0.1 | 5.93E-01 | -0.1 | 5.70E-01 | 0.0 | 9.44E-01 | Modifier of snc1 |
| AT2G28320 | -0.1 | 1.50E-01 | -0.4 | 5.79E-04 | 0.3 | 8.77E-03 | -0.1 | 7.36E-01 | 0.0 | 9.08E-01 | -0.1 | 7.04E-01 | At2g28320 |

| | | | | | | | | | | | | | |
|---|---|---|---|---|---|---|---|---|---|---|---|---|---|
| AT1G22860 | -0.1 | 7.38E-02 | -0.2 | 6.22E-03 | 0.1 | 2.01E-01 | -0.3 | 3.42E-01 | -0.2 | 6.37E-01 | -0.1 | 8.04E-01 | Vacuolar sorting protein 3 |
| AT1G55210 | -0.1 | 1.58E-01 | -0.1 | 2.57E-01 | 0.0 | 8.37E-01 | 0.3 | 1.06E-01 | 0.5 | 1.87E-02 | -0.2 | 4.34E-01 | Dirigent protein 20 |
| AT5G62280 | -0.1 | 6.31E-01 | 0.7 | 1.66E-02 | -0.8 | 1.60E-02 | 0.0 | 9.45E-01 | 0.0 | 9.76E-01 | 0.0 | 9.86E-01 | hypothetical protein |
| AT3G07580 | -0.1 | 3.90E-01 | -0.1 | 5.81E-01 | 0.0 | 8.34E-01 | 0.1 | 8.24E-01 | 0.0 | 9.36E-01 | 0.0 | 9.43E-01 | At3g07580 |
| AT5G46150 | -0.1 | 1.06E-01 | -0.2 | 3.98E-02 | 0.0 | 6.05E-01 | -0.2 | 2.88E-01 | -0.2 | 4.16E-01 | 0.0 | 9.56E-01 | ALA-interacting subunit |
| AT4G30996 | -0.1 | 1.37E-01 | 0.0 | 6.05E-01 | -0.2 | 8.58E-02 | -0.3 | 9.38E-02 | -0.2 | 3.99E-01 | -0.1 | 6.49E-01 | Ubiquitin-associated protein |
| AT1G16960 | -0.1 | 7.83E-01 | -0.3 | 5.08E-01 | 0.2 | 7.44E-01 | 0.3 | 3.48E-01 | 0.3 | 3.68E-01 | 0.0 | 9.70E-01 | DC-UbP |
| AT4G03635 | -0.1 | 3.47E-01 | -0.1 | 7.09E-01 | -0.1 | 6.76E-01 | -0.3 | 1.53E-01 | -0.3 | 3.24E-01 | -0.1 | 8.47E-01 | F-box |
| AT3G22430 | -0.1 | 1.51E-01 | -0.2 | 2.92E-02 | 0.1 | 4.04E-01 | -0.2 | 2.52E-01 | -0.2 | 4.69E-01 | -0.1 | 8.60E-01 | AT3g22430 |
| AT1G77890 | -0.1 | 2.23E-01 | -0.3 | 8.54E-03 | 0.2 | 1.10E-01 | 0.3 | 2.04E-01 | 0.3 | 2.55E-01 | 0.0 | 9.97E-01 | F28K19.11 |
| AT3G49550 | -0.1 | 5.03E-01 | -0.1 | 4.85E-01 | 0.0 | 9.64E-01 | -0.4 | 2.46E-01 | 0.1 | 8.63E-01 | -0.4 | 2.65E-01 | Uncharacterized protein |
| AT1G13520 | -0.1 | 6.01E-01 | -0.9 | 7.03E-03 | 0.7 | 3.38E-02 | 1.9 | 3.44E-06 | 0.7 | 4.17E-02 | 1.2 | 3.79E-04 | F13B4.3 protein |
| AT4G34120 | -0.1 | 2.30E-01 | -0.2 | 1.23E-01 | 0.0 | 7.10E-01 | 0.1 | 6.91E-01 | -0.2 | 3.14E-01 | 0.2 | 1.96E-01 | CBS domain-containing protein CBSX2, chloroplastic |
| AT3G60450 | -0.1 | 2.46E-01 | 0.0 | 9.66E-01 | -0.1 | 2.90E-01 | 0.6 | 3.20E-03 | 0.4 | 4.10E-02 | 0.1 | 5.12E-01 | hypothetical protein |
| AT1G03910 | -0.1 | 7.30E-02 | -0.2 | 3.54E-02 | 0.0 | 6.89E-01 | -0.1 | 4.95E-01 | -0.2 | 1.96E-01 | 0.1 | 6.60E-01 | Splicing factor Cactin |
| AT5G66290 | -0.1 | 4.45E-01 | -0.2 | 1.61E-01 | 0.1 | 5.61E-01 | -0.4 | 9.94E-02 | -0.5 | 6.89E-02 | 0.1 | 8.39E-01 | BLOC-1-related complex subunit 5 |
| AT5G04700 | -0.1 | 4.13E-01 | -0.1 | 3.53E-01 | 0.0 | 9.08E-01 | | | | | | | PGG domain-containing protein |
| AT1G55360 | -0.1 | 1.31E-01 | 0.1 | 3.93E-01 | -0.2 | 4.30E-02 | 0.5 | 2.12E-03 | 0.0 | 9.43E-01 | 0.5 | 1.32E-02 | Carboxyl-terminal peptidase |
| AT2G24240 | -0.1 | 4.87E-01 | -0.1 | 7.69E-01 | -0.1 | 7.68E-01 | -0.2 | 4.02E-01 | -0.3 | 2.83E-01 | 0.1 | 8.39E-01 | BTB |
| AT4G16490 | -0.1 | 9.17E-02 | -0.2 | 5.02E-03 | 0.1 | 1.16E-01 | -0.3 | 3.16E-02 | -0.1 | 5.74E-01 | -0.2 | 3.18E-01 | hypothetical protein |
| AT1G63670 | -0.1 | 2.24E-01 | -0.4 | 2.51E-03 | 0.3 | 3.24E-02 | -0.2 | 3.20E-01 | 0.0 | 9.21E-01 | -0.2 | 3.27E-01 | Uncharacterized protein F24D7.14 |
| AT5G17650 | -0.1 | 1.50E-01 | 0.0 | 7.47E-01 | -0.1 | 3.53E-01 | 0.2 | 3.81E-01 | 0.2 | 4.31E-01 | 0.0 | 9.93E-01 | hypothetical protein |
| AT4G31440 | -0.1 | 3.14E-01 | -0.3 | 4.05E-02 | 0.2 | 2.92E-01 | -0.8 | 1.52E-01 | -0.2 | 7.02E-01 | -0.5 | 5.17E-01 | Transcriptional regulator of RNA polII, SAGA, subunit |
| AT1G65720 | -0.1 | 8.99E-02 | -0.1 | 3.29E-01 | 0.0 | 5.19E-01 | -0.1 | 4.92E-01 | -0.1 | 6.73E-01 | 0.0 | 9.34E-01 | hypothetical protein |
| AT5G07610 | -0.1 | 2.31E-01 | -0.2 | 8.35E-02 | 0.1 | 5.92E-01 | 0.2 | 5.51E-01 | -0.1 | 6.75E-01 | 0.3 | 3.97E-01 | F-box protein At5g07610 |
| AT5G11430 | -0.1 | 1.27E-01 | 0.0 | 8.89E-01 | -0.1 | 1.52E-01 | -0.2 | 1.96E-01 | -0.1 | 8.41E-01 | -0.2 | 4.77E-01 | Uncharacterized protein F15N18_20 |
| AT5G44060 | -0.1 | 2.71E-01 | -0.1 | 5.80E-01 | -0.1 | 6.70E-01 | 0.1 | 7.28E-01 | 0.2 | 4.07E-01 | -0.1 | 6.80E-01 | Embryo sac development arrest protein |
| AT4G34070 | -0.1 | 1.53E-01 | 0.0 | 5.59E-01 | -0.2 | 8.77E-02 | -0.1 | 3.45E-01 | -0.1 | 4.28E-01 | 0.0 | 9.94E-01 | Uncharacterized protein |
| AT5G48657 | -0.1 | 3.43E-01 | 0.0 | 7.99E-01 | -0.1 | 5.71E-01 | 0.2 | 1.55E-01 | -0.1 | 6.68E-01 | 0.3 | 1.24E-01 | Defense protein-like protein |
| AT1G02120 | -0.1 | 7.13E-02 | -0.3 | 5.33E-04 | 0.2 | 1.23E-02 | 0.0 | 9.39E-01 | -0.1 | 5.64E-01 | 0.1 | 5.90E-01 | GRAM domain family protein |
| AT2G17020 | -0.1 | 8.51E-02 | 0.0 | 8.16E-01 | -0.1 | 1.95E-01 | -0.5 | 2.91E-04 | -0.5 | 1.89E-03 | -0.1 | 7.70E-01 | F-box |
| AT4G08520 | -0.1 | 2.36E-01 | 0.3 | 2.00E-02 | -0.4 | 5.39E-03 | 0.2 | 2.53E-01 | 0.3 | 1.75E-01 | -0.1 | 8.32E-01 | Coatomer subunit zeta-3 |
| AT3G13970 | -0.1 | 4.24E-01 | 0.1 | 5.79E-01 | -0.2 | 2.69E-01 | -0.4 | 2.64E-01 | -0.1 | 8.62E-01 | -0.3 | 4.66E-01 | Ubiquitin-like protein ATG12 |
| AT2G14878 | -0.1 | 1.46E-01 | -0.1 | 3.22E-01 | 0.0 | 7.03E-01 | -0.5 | 5.28E-03 | -0.5 | 8.49E-03 | 0.0 | 9.51E-01 | NA |
| AT5G27460 | -0.1 | 2.55E-01 | -0.1 | 3.40E-01 | 0.0 | 9.10E-01 | -0.3 | 9.56E-02 | -0.3 | 1.41E-01 | 0.0 | 9.80E-01 | Pentatricopeptide repeat-containing protein At5g27460 |
| AT3G20170 | -0.1 | 2.15E-01 | -0.2 | 1.10E-01 | 0.0 | 7.15E-01 | -0.2 | 2.59E-01 | -0.1 | 5.95E-01 | -0.1 | 7.86E-01 | ARM repeat superfamily protein |
| AT4G34600 | -0.1 | 6.02E-01 | -0.2 | 4.95E-01 | 0.0 | 8.85E-01 | 0.1 | 8.26E-01 | -0.1 | 8.18E-01 | 0.2 | 7.09E-01 | Transmembrane protein |
| AT5G21020 | -0.1 | 5.16E-01 | -0.3 | 7.05E-02 | 0.2 | 2.38E-01 | -0.7 | 9.36E-03 | -0.5 | 8.04E-02 | -0.2 | 6.39E-01 | Transmembrane protein |
| AT3G12140 | -0.1 | 8.23E-02 | -0.1 | 4.32E-02 | 0.0 | 7.24E-01 | -0.1 | 2.34E-01 | -0.1 | 4.30E-01 | 0.0 | 8.85E-01 | EML1 |
| AT1G02816 | -0.1 | 2.91E-01 | -0.1 | 2.15E-01 | 0.0 | 8.57E-01 | -0.2 | 4.42E-01 | -0.2 | 3.88E-01 | 0.0 | 9.08E-01 | Uncharacterized protein |
| AT4G00310 | -0.1 | 4.61E-01 | 0.1 | 3.45E-01 | -0.3 | 1.67E-01 | -0.2 | 5.92E-01 | 0.3 | 4.33E-01 | -0.4 | 2.56E-01 | AT4G00310 protein |
| AT5G11550 | -0.1 | 2.86E-01 | 0.1 | 4.99E-01 | -0.2 | 1.34E-01 | -0.7 | 3.08E-05 | -0.2 | 1.47E-01 | -0.5 | 1.01E-02 | ARM repeat superfamily protein |
| AT2G34585 | -0.1 | 1.55E-01 | 0.1 | 3.41E-01 | -0.2 | 4.34E-02 | -0.2 | 2.85E-01 | -0.2 | 3.44E-01 | 0.0 | 9.87E-01 | Outer envelope membrane protein 7 |
| AT4G05040 | -0.1 | 1.95E-01 | 0.0 | 6.41E-01 | -0.1 | 5.05E-01 | 0.2 | 5.19E-01 | 0.2 | 4.72E-01 | 0.0 | 9.18E-01 | AT4g05040 protein |
| AT5G17290 | -0.1 | 2.96E-01 | -0.2 | 6.08E-02 | 0.1 | 4.01E-01 | -0.1 | 8.46E-01 | -0.2 | 4.55E-01 | 0.2 | 6.81E-01 | Autophagy protein 5 |
| AT5G56240 | -0.1 | 2.12E-01 | -0.4 | 1.75E-03 | 0.3 | 2.05E-02 | 0.1 | 7.96E-01 | 0.0 | 9.24E-01 | 0.1 | 7.49E-01 | Hapless 8 |
| AT1G23080 | -0.1 | 3.37E-01 | 0.2 | 1.03E-01 | -0.3 | 2.81E-02 | -0.3 | 4.20E-02 | 0.0 | 9.92E-01 | -0.3 | 1.60E-01 | Auxin efflux carrier component |
| AT2G20120 | -0.1 | 1.89E-01 | 0.1 | 2.00E-01 | -0.2 | 3.46E-02 | -0.4 | 5.07E-03 | -0.5 | 1.66E-03 | 0.1 | 5.11E-01 | Protein CONTINUOUS VASCULAR RING 1 |
| AT4G31430 | -0.1 | 8.90E-02 | -0.1 | 6.42E-02 | 0.0 | 8.43E-01 | -0.2 | 3.97E-02 | -0.2 | 6.20E-02 | 0.0 | 9.98E-01 | Protein KAKU4 |
| AT5G03110 | -0.1 | 2.93E-01 | -0.2 | 1.09E-01 | 0.1 | 5.95E-01 | -0.2 | 2.97E-01 | -0.2 | 4.40E-01 | 0.0 | 9.28E-01 | Arabidopsis thaliana genomic DNA, chromosome 5, P1 clone:MOK16 |
| AT5G13610 | -0.1 | 2.55E-01 | -0.1 | 3.60E-01 | 0.0 | 8.85E-01 | 0.2 | 3.30E-01 | 0.3 | 1.91E-01 | -0.1 | 7.80E-01 | Protein RETARDED ROOT GROWTH-LIKE |
| AT1G67080 | -0.1 | 3.24E-01 | 0.1 | 5.02E-01 | -0.2 | 1.56E-01 | 0.1 | 7.25E-01 | 0.1 | 5.90E-01 | -0.1 | 8.74E-01 | Abscisic acid -deficient 4 |
| AT3G11440 | -0.1 | 4.52E-01 | -0.2 | 2.64E-01 | 0.1 | 7.44E-01 | -0.1 | 8.58E-01 | -0.1 | 9.23E-01 | 0.0 | 9.71E-01 | Myb domain protein 65 |
| AT2G33590 | -0.1 | 1.70E-01 | -0.2 | 3.21E-02 | 0.1 | 3.74E-01 | 0.0 | 7.67E-01 | 0.0 | 9.92E-01 | 0.1 | 8.22E-01 | CRL1 |
| AT5G22950 | -0.1 | 1.41E-01 | -0.1 | 2.44E-01 | 0.0 | 8.11E-01 | 0.0 | 9.46E-01 | 0.1 | 3.23E-01 | -0.2 | 3.84E-01 | Vacuolar protein sorting-associated protein 24 homolog 1 |
| AT5G47730 | -0.1 | 9.34E-02 | -0.1 | 5.56E-02 | 0.0 | 7.73E-01 | 0.1 | 3.53E-01 | 0.2 | 2.76E-01 | 0.0 | 8.45E-01 | AT5G47730 protein |
| AT1G58280 | -0.1 | 2.97E-01 | 0.0 | 8.74E-01 | -0.1 | 4.70E-01 | 0.0 | 8.80E-01 | 0.4 | 9.29E-02 | -0.4 | 2.14E-01 | Phosphoglycerate mutase family protein |
| AT4G34280 | -0.1 | 2.58E-01 | -0.2 | 6.99E-02 | 0.1 | 4.85E-01 | -0.5 | 5.34E-02 | 0.0 | 9.93E-01 | -0.5 | 1.03E-01 | Protein DWD HYPERSENSITIVE TO UV-B 1 |
| AT1G56423 | -0.1 | 1.30E-01 | 0.0 | 9.02E-01 | -0.1 | 2.33E-01 | 0.2 | 2.37E-01 | 0.0 | 9.75E-01 | 0.2 | 3.72E-01 | Uncharacterized protein |
| AT5G53080 | -0.1 | 5.23E-01 | 0.0 | 9.63E-01 | -0.1 | 6.46E-01 | -0.3 | 3.70E-01 | -0.1 | 9.01E-01 | -0.2 | 6.32E-01 | Uncharacterized protein |
| AT1G64980 | -0.1 | 2.11E-01 | -0.3 | 6.57E-03 | 0.2 | 8.00E-02 | 1.5 | 7.43E-07 | 0.2 | 4.75E-01 | 1.3 | 1.58E-04 | Protein CDI |
| AT3G24160 | -0.1 | 1.24E-01 | 0.1 | 1.11E-01 | -0.2 | 1.18E-02 | 0.0 | 9.31E-01 | 0.1 | 5.25E-01 | -0.1 | 7.04E-01 | AT3g24160 |
| AT1G75810 | -0.1 | 1.48E-01 | -0.2 | 4.35E-02 | 0.1 | 5.21E-01 | -0.5 | 6.66E-02 | -0.5 | 9.11E-02 | 0.0 | 9.97E-01 | Small hydrophobic protein |
| AT5G08010 | -0.1 | 3.42E-01 | 0.0 | 9.46E-01 | -0.1 | 4.75E-01 | -0.2 | 3.21E-01 | 0.0 | 9.51E-01 | -0.2 | 4.66E-01 | Uncharacterized protein F13G24.210 |
| AT4G33625 | -0.1 | 1.46E-01 | 0.1 | 1.77E-01 | -0.2 | 2.15E-02 | -0.6 | 1.50E-04 | -0.3 | 4.01E-02 | -0.3 | 1.01E-01 | Vacuole protein |
| AT1G79090 | -0.1 | 1.55E-01 | -0.3 | 4.28E-03 | 0.2 | 6.67E-02 | 0.0 | 7.87E-01 | 0.1 | 7.12E-01 | 0.0 | 9.22E-01 | Uncharacterized protein |
| AT2G37200 | -0.1 | 2.61E-01 | -0.2 | 9.51E-02 | 0.1 | 5.81E-01 | -0.4 | 1.79E-03 | -0.3 | 3.40E-02 | -0.1 | 4.50E-01 | CASP-like protein |
| AT3G07180 | -0.1 | 8.83E-02 | -0.1 | 9.72E-02 | 0.0 | 1.00E+00 | -0.1 | 7.01E-01 | 0.0 | 9.82E-01 | -0.1 | 8.12E-01 | GPI transamidase component PIG-S-like protein |
| AT3G51650 | -0.1 | 1.35E-01 | -0.1 | 1.49E-01 | 0.0 | 9.99E-01 | 0.3 | 9.84E-02 | 0.2 | 4.62E-01 | 0.2 | 5.41E-01 | Uncharacterized protein T18N14.30 |
| AT5G58100 | -0.1 | 8.28E-02 | -0.1 | 7.71E-02 | 0.0 | 9.37E-01 | 0.2 | 2.08E-01 | 0.1 | 5.54E-01 | 0.1 | 7.63E-01 | Transmembrane protein |
| AT4G35110 | -0.1 | 3.33E-01 | 0.0 | 9.91E-01 | -0.1 | 4.15E-01 | -0.6 | 6.73E-03 | -0.5 | 3.79E-02 | -0.1 | 6.88E-01 | Phospholipase-like protein family protein |
| AT5G51220 | -0.1 | 2.87E-01 | -0.2 | 3.69E-02 | 0.1 | 2.86E-01 | 0.4 | 2.81E-02 | 0.3 | 2.23E-01 | 0.2 | 5.56E-01 | Ubiquinol-cytochrome c chaperone domain-containing protein |
| AT4G01920 | -0.1 | 4.23E-01 | -0.2 | 2.66E-01 | 0.0 | 7.81E-01 | 0.4 | 2.09E-01 | 0.3 | 3.56E-01 | 0.1 | 9.11E-01 | AT4g01920 |
| AT2G47710 | -0.1 | 1.82E-01 | -0.2 | 1.28E-02 | 0.1 | 1.55E-01 | -0.3 | 2.65E-03 | -0.3 | 1.41E-02 | 0.0 | 8.16E-01 | hypothetical protein |
| AT4G00860 | -0.1 | 3.33E-01 | 0.3 | 4.21E-02 | -0.4 | 1.27E-02 | 0.1 | 6.80E-01 | 0.1 | 5.03E-01 | -0.1 | 8.26E-01 | AT0ZI1 protein ) |
| AT5G54530 | -0.1 | 4.71E-01 | 0.5 | 2.73E-03 | -0.6 | 3.43E-03 | -0.1 | 7.28E-01 | 0.1 | 8.36E-01 | -0.2 | 5.95E-01 | Gb\|AAF02153.1 |
| AT3G57090 | -0.1 | 1.92E-01 | -0.1 | 1.90E-01 | 0.0 | 9.66E-01 | 0.2 | 1.01E-01 | 0.0 | 8.66E-01 | 0.3 | 1.13E-01 | Mitochondrial fission 1 protein A |
| AT4G33540 | -0.1 | 3.89E-01 | -0.5 | 4.13E-03 | 0.3 | 2.84E-02 | -0.4 | 8.00E-03 | -0.4 | 1.40E-02 | 0.0 | 1.00E+00 | Metallo-beta-lactamase domain-containing protein |
| AT5G12430 | -0.1 | 2.52E-01 | -0.2 | 2.25E-02 | 0.1 | 2.05E-01 | 0.1 | 7.00E-01 | 0.1 | 6.81E-01 | 0.0 | 9.53E-01 | Heat shock protein DnaJ with tetratricopeptide repeat-containing protein |
| AT1G28240 | -0.1 | 1.52E-01 | -0.3 | 1.42E-03 | 0.2 | 2.30E-02 | 0.0 | 9.45E-01 | 0.2 | 5.09E-01 | -0.1 | 6.42E-01 | Uncharacterized protein |
| AT3G52030 | -0.1 | 3.70E-01 | -0.2 | 7.90E-02 | 0.1 | 4.05E-01 | -0.2 | 3.78E-01 | 0.1 | 8.59E-01 | -0.3 | 3.82E-01 | F-box domain-containing protein |
| AT2G25320 | -0.1 | 2.87E-01 | -0.3 | 6.11E-03 | 0.2 | 5.68E-02 | 0.0 | 9.91E-01 | 0.1 | 7.11E-01 | -0.1 | 7.68E-01 | CGS1 mRNA stability 1 |
| AT2G35155 | -0.1 | 2.48E-01 | -0.1 | 5.70E-01 | -0.1 | 6.45E-01 | -0.1 | 5.79E-01 | 0.0 | 9.76E-01 | -0.1 | 6.77E-01 | Trypsin family protein |

| | | | | | | | | | | | | | |
|---|---|---|---|---|---|---|---|---|---|---|---|---|---|
| AT2G32380 | -0.1 | 4.81E-01 | 0.5 | 5.92E-03 | -0.6 | 4.73E-03 | 0.0 | 9.21E-01 | 0.0 | 9.83E-01 | 0.0 | 9.62E-01 | EXPERA domain-containing protein |
| AT5G63200 | -0.1 | 1.84E-01 | -0.1 | 1.57E-01 | 0.0 | 9.12E-01 | -0.2 | 2.24E-01 | -0.2 | 3.07E-01 | 0.0 | 9.80E-01 | Uncharacterized protein |
| AT4G01210 | -0.1 | 1.08E-01 | -0.1 | 2.88E-01 | 0.0 | 6.35E-01 | -0.2 | 2.73E-01 | -0.1 | 3.82E-01 | 0.0 | 9.68E-01 | Uncharacterized protein At4g01210 |
| AT3G05345 | -0.1 | 5.86E-01 | -0.2 | 2.33E-01 | 0.1 | 5.62E-01 | 0.0 | 8.65E-01 | 0.2 | 5.04E-01 | -0.2 | 4.31E-01 | J domain-containing protein |
| AT4G38440 | -0.1 | 2.14E-01 | -0.4 | 5.67E-04 | 0.3 | 7.24E-03 | -0.1 | 4.78E-01 | -0.3 | 1.24E-01 | 0.2 | 5.04E-01 | Transcriptional elongation regulator MINIYO |
| AT5G55900 | -0.1 | 5.96E-01 | -0.2 | 3.18E-01 | 0.1 | 6.83E-01 | -0.7 | 8.52E-03 | -0.5 | 8.73E-02 | -0.3 | 4.15E-01 | At5g55900 |
| AT3G62140 | -0.1 | 2.02E-01 | -0.2 | 4.60E-02 | 0.1 | 4.44E-01 | -0.1 | 7.61E-01 | 0.0 | 9.30E-01 | -0.1 | 7.58E-01 | FAM192A |
| AT1G67400 | -0.1 | 4.30E-01 | 0.1 | 3.14E-01 | -0.2 | 1.46E-01 | 0.0 | 8.94E-01 | -0.2 | 5.59E-01 | 0.1 | 6.93E-01 | ELMO |
| AT3G08600 | -0.1 | 3.16E-01 | 0.3 | 2.47E-02 | -0.4 | 1.04E-02 | -0.6 | 1.02E-04 | -0.4 | 9.02E-03 | -0.3 | 1.53E-01 | AT3g08600 |
| AT4G22250 | -0.1 | 4.80E-01 | 0.1 | 3.10E-01 | -0.2 | 1.56E-01 | -0.1 | 6.04E-01 | 0.1 | 7.89E-01 | -0.2 | 4.91E-01 | RING |
| AT1G73710 | -0.1 | 1.85E-01 | -0.3 | 7.25E-03 | 0.2 | 1.08E-01 | 0.2 | 3.09E-01 | 0.1 | 6.34E-01 | 0.1 | 7.87E-01 | Pentatricopeptide repeat-containing protein At1g73710 |
| AT5G56190 | -0.1 | 2.20E-01 | -0.3 | 4.18E-03 | 0.2 | 5.09E-02 | 0.0 | 8.62E-01 | 0.1 | 7.61E-01 | 0.0 | 9.12E-01 | Uncharacterized protein |
| AT1G50660 | -0.1 | 1.59E-01 | -0.1 | 1.73E-01 | 0.0 | 1.00E+00 | -0.1 | 6.97E-01 | 0.1 | 7.22E-01 | -0.1 | 5.32E-01 | Uncharacterized protein |
| AT5G58960 | -0.1 | 2.25E-01 | -0.1 | 1.49E-01 | 0.0 | 8.09E-01 | -0.3 | 5.36E-02 | -0.3 | 7.88E-02 | 0.0 | 9.97E-01 | hypothetical protein |
| AT4G06634 | -0.1 | 2.20E-01 | -0.1 | 1.51E-01 | 0.0 | 8.26E-01 | 0.1 | 7.77E-01 | 0.3 | 1.35E-01 | -0.2 | 3.11E-01 | Zinc finger family protein |
| AT2G47010 | -0.1 | 5.30E-01 | -0.3 | 8.82E-02 | 0.2 | 3.16E-01 | -0.8 | 3.70E-02 | 0.0 | 9.97E-01 | -0.8 | 1.04E-01 | Expressed protein |
| AT1G32920 | -0.1 | 7.23E-01 | -0.8 | 1.85E-02 | 0.7 | 5.46E-02 | 0.6 | 1.45E-02 | -0.4 | 1.87E-01 | 1.0 | 4.09E-03 | At1g32920 |
| AT3G14170 | -0.1 | 5.61E-01 | 0.2 | 2.45E-01 | -0.3 | 1.53E-01 | 0.2 | 5.54E-01 | 0.6 | 9.45E-02 | -0.4 | 3.19E-01 | At3g14170 |
| AT4G38670 | -0.1 | 3.75E-01 | -0.1 | 2.21E-01 | 0.0 | 7.56E-01 | -0.3 | 2.41E-01 | -0.3 | 2.76E-01 | 0.0 | 9.80E-01 | Putative thaumatin-like protein |
| AT2G20790 | -0.1 | 2.34E-01 | -0.1 | 2.94E-01 | 0.0 | 9.36E-01 | 0.1 | 5.41E-01 | 0.0 | 9.48E-01 | 0.1 | 7.47E-01 | Clathrin adaptor complexes medium subunit family protein |
| AT2G42820 | -0.1 | 5.03E-01 | -0.2 | 2.57E-01 | 0.1 | 6.83E-01 | 0.3 | 5.28E-01 | 0.0 | 9.58E-01 | 0.3 | 6.59E-01 | HVA22-like protein f |
| AT2G16460 | -0.1 | 3.60E-01 | 0.1 | 4.17E-01 | -0.2 | 1.46E-01 | 0.2 | 3.67E-01 | 0.4 | 4.44E-02 | -0.2 | 3.25E-01 | Coiled-coil 90B-like protein |
| AT4G08930 | -0.1 | 3.10E-01 | -0.2 | 1.07E-01 | 0.1 | 5.48E-01 | -0.3 | 9.94E-02 | -0.2 | 3.01E-01 | -0.1 | 7.86E-01 | 5'-adenylylsulfate reductase-like 6 |
| AT5G37070 | -0.1 | 3.40E-01 | -0.4 | 4.76E-03 | 0.3 | 4.19E-02 | 0.1 | 7.96E-01 | -0.1 | 7.16E-01 | 0.2 | 5.90E-01 | hypothetical protein |
| AT2G25590 | -0.1 | 4.69E-01 | -0.4 | 1.64E-02 | 0.3 | 8.69E-02 | -0.3 | 1.60E-01 | 0.0 | 9.02E-01 | -0.2 | 3.24E-01 | Plant Tudor-like protein |
| AT3G56590 | -0.1 | 1.93E-01 | -0.1 | 3.53E-01 | 0.0 | 7.68E-01 | 0.3 | 2.33E-01 | 0.5 | 9.67E-02 | -0.2 | 6.55E-01 | Hydroxyproline-rich glycoprotein family protein |
| AT4G36140 | -0.1 | 4.09E-01 | -0.2 | 9.05E-02 | 0.1 | 4.00E-01 | -0.1 | 7.06E-01 | -0.2 | 5.72E-01 | 0.1 | 8.80E-01 | Putative disease resistance protein |
| AT3G22440 | -0.1 | 3.26E-01 | -0.2 | 2.88E-02 | 0.1 | 1.81E-01 | -0.2 | 4.05E-01 | -0.2 | 6.01E-01 | 0.0 | 9.21E-01 | FRIGIDA-like protein 4a |
| AT2G42230 | -0.1 | 2.28E-01 | 0.0 | 7.52E-01 | -0.1 | 4.52E-01 | 0.0 | 9.23E-01 | 0.1 | 5.39E-01 | -0.1 | 6.74E-01 | TBCC domain-containing protein 1 |
| AT4G04790 | -0.1 | 4.84E-01 | -0.3 | 5.39E-02 | 0.2 | 2.42E-01 | 0.4 | 3.61E-01 | 0.5 | 3.23E-01 | -0.1 | 9.11E-01 | Tetratricopeptide repeat -like superfamily protein |
| AT1G14670 | -0.1 | 2.45E-01 | 0.0 | 8.59E-01 | -0.1 | 3.88E-01 | 0.0 | 9.34E-01 | 0.0 | 9.98E-01 | 0.0 | 9.62E-01 | Transmembrane 9 superfamily member 2 |
| AT5G49000 | -0.1 | 4.26E-01 | 0.0 | 9.74E-01 | -0.1 | 5.04E-01 | 0.1 | 8.30E-01 | -0.1 | 8.78E-01 | 0.1 | 7.49E-01 | F-box |
| AT5G11890 | -0.1 | 2.57E-01 | 0.0 | 6.96E-01 | -0.1 | 5.33E-01 | -0.2 | 1.17E-01 | -0.2 | 2.61E-01 | 0.0 | 8.89E-01 | EMB3135 |
| AT4G03400 | -0.1 | 4.81E-01 | 0.0 | 7.75E-01 | -0.1 | 7.39E-01 | -0.1 | 7.43E-01 | -0.7 | 7.84E-03 | 0.6 | 3.56E-02 | Putative GH3-like protein |
| AT5G15640 | -0.1 | 3.57E-01 | -0.1 | 4.21E-01 | 0.0 | 9.50E-01 | 0.0 | 9.88E-01 | 0.1 | 8.01E-01 | -0.1 | 8.24E-01 | Mitochondrial substrate carrier family protein |
| AT4G31160 | -0.1 | 2.10E-01 | -0.2 | 1.35E-02 | 0.1 | 1.59E-01 | 0.0 | 8.44E-01 | -0.1 | 5.79E-01 | 0.2 | 5.39E-01 | DDB1- and CUL4-associated factor homolog 1 |
| AT4G15475 | -0.1 | 2.48E-01 | -0.2 | 2.33E-02 | 0.1 | 2.25E-01 | -0.4 | 9.02E-02 | -0.1 | 6.93E-01 | -0.3 | 3.65E-01 | F-box domain-containing protein |
| AT5G56170 | -0.1 | 2.34E-01 | 0.1 | 1.75E-01 | -0.2 | 3.31E-02 | -0.2 | 2.66E-01 | -0.3 | 1.91E-01 | 0.1 | 8.36E-01 | GPI-anchored protein LLG1 |
| AT1G27100 | -0.1 | 3.40E-01 | -0.3 | 7.90E-03 | 0.2 | 5.33E-02 | 0.0 | 8.86E-01 | -0.5 | 1.18E-03 | 0.5 | 4.97E-03 | T7N9.16 |
| AT3G20260 | -0.1 | 2.45E-01 | 0.0 | 6.07E-01 | -0.1 | 1.61E-01 | -0.3 | 2.17E-01 | 0.1 | 8.17E-01 | -0.3 | 2.21E-01 | hypothetical protein |
| AT2G28120 | -0.1 | 5.28E-01 | 0.3 | 2.82E-02 | -0.4 | 2.40E-02 | -1.2 | 3.93E-04 | -1.4 | 2.64E-04 | 0.2 | 6.55E-01 | Nodulin-like domain-containing protein |
| AT1G15270 | -0.1 | 4.07E-01 | 0.0 | 7.75E-01 | -0.1 | 3.12E-01 | 0.3 | 1.54E-01 | 0.2 | 5.13E-01 | 0.1 | 7.17E-01 | hypothetical protein |
| AT1G03730 | -0.1 | 2.96E-01 | -0.1 | 1.38E-01 | 0.0 | 6.70E-01 | 0.3 | 3.53E-01 | 0.1 | 8.14E-01 | 0.2 | 6.81E-01 | F21B7.33 |
| AT1G71970 | -0.1 | 4.60E-01 | 0.0 | 9.34E-01 | -0.1 | 4.77E-01 | 0.1 | 8.09E-01 | -0.3 | 2.39E-01 | 0.4 | 2.04E-01 | hypothetical protein |
| AT3G43510 | -0.1 | 4.11E-01 | -0.4 | 4.82E-03 | 0.3 | 3.55E-02 | 0.1 | 7.39E-01 | -0.1 | 8.59E-01 | 0.1 | 6.84E-01 | NA |
| AT1G72500 | -0.1 | 2.59E-01 | 0.0 | 7.49E-01 | -0.1 | 4.91E-01 | -0.1 | 8.21E-01 | 0.2 | 5.59E-01 | -0.3 | 3.98E-01 | Inter alpha-trypsin inhibitor, heavy chain-like protein |
| AT1G26690 | -0.1 | 3.07E-01 | -0.1 | 3.22E-01 | 0.0 | 9.97E-01 | 0.3 | 1.26E-01 | 0.2 | 3.73E-01 | 0.1 | 7.71E-01 | hypothetical protein |
| AT5G19540 | -0.1 | 2.51E-01 | -0.1 | 1.91E-01 | 0.0 | 8.53E-01 | -0.1 | 5.27E-01 | 0.1 | 7.37E-01 | -0.1 | 3.64E-01 | Thermosome subunit gamma |
| AT5G13340 | -0.1 | 3.70E-01 | -0.1 | 5.97E-01 | 0.0 | 7.93E-01 | -0.2 | 4.69E-01 | -0.2 | 5.92E-01 | 0.0 | 9.62E-01 | hypothetical protein |
| AT5G50720 | -0.1 | 6.30E-01 | 0.0 | 8.50E-01 | -0.1 | 8.33E-01 | -0.2 | 4.12E-01 | -0.2 | 4.76E-01 | 0.0 | 9.95E-01 | HVA22-like protein e |
| AT5G09860 | -0.1 | 1.76E-01 | 0.0 | 6.46E-01 | -0.1 | 4.44E-01 | -0.1 | 4.54E-01 | -0.2 | 1.27E-01 | 0.1 | 5.18E-01 | Nuclear matrix protein-like protein |
| AT5G25590 | -0.1 | 2.71E-01 | 0.1 | 2.70E-01 | -0.2 | 7.21E-02 | -0.3 | 6.21E-02 | -0.2 | 3.71E-01 | -0.1 | 5.21E-01 | Uncharacterized protein |
| AT4G11740 | -0.1 | 3.06E-01 | -0.4 | 6.95E-04 | 0.4 | 5.58E-03 | 0.0 | 9.68E-01 | 0.0 | 9.52E-01 | 0.0 | 9.41E-01 | Ubiquitin-like superfamily protein |
| AT1G61170 | -0.1 | 4.07E-01 | 0.1 | 2.99E-01 | -0.2 | 1.23E-01 | -0.6 | 2.07E-03 | -0.4 | 2.66E-02 | -0.2 | 4.54E-01 | At1g61170 |
| AT5G63280 | -0.1 | 2.54E-01 | -0.2 | 6.60E-02 | 0.1 | 4.76E-01 | 0.0 | 9.21E-01 | 0.0 | 9.15E-01 | 0.1 | 8.59E-01 | C2H2-type domain-containing protein |
| AT5G04670 | -0.1 | 2.33E-01 | -0.3 | 5.78E-03 | 0.2 | 6.76E-02 | 0.1 | 7.23E-01 | 0.1 | 6.70E-01 | 0.0 | 9.41E-01 | Enhancer of polycomb-like protein |
| AT5G41940 | -0.1 | 1.59E-01 | -0.1 | 1.59E-01 | 0.0 | 9.71E-01 | -0.3 | 1.71E-02 | -0.2 | 2.72E-01 | -0.2 | 4.22E-01 | Transmembrane protein |
| AT1G13910 | -0.1 | 5.24E-01 | -0.4 | 2.87E-02 | 0.3 | 1.33E-01 | 0.0 | 9.29E-01 | 0.2 | 4.66E-01 | -0.3 | 4.40E-01 | F16A14.12 |
| AT2G22475 | -0.1 | 2.72E-01 | 0.0 | 8.68E-01 | -0.1 | 4.33E-01 | 0.0 | 8.32E-01 | 0.2 | 2.61E-01 | -0.2 | 4.59E-01 | GLABRA2 expression modulator |
| AT1G10270 | -0.1 | 3.69E-01 | -0.2 | 3.21E-02 | 0.2 | 2.07E-01 | 0.9 | 5.09E-03 | 0.3 | 4.37E-01 | 0.6 | 9.05E-02 | Pentatricopeptide repeat-containing protein At1g10270 |
| AT3G13510 | -0.1 | 2.75E-01 | 0.0 | 9.79E-01 | -0.1 | 3.30E-01 | -0.1 | 6.37E-01 | 0.0 | 9.57E-01 | -0.1 | 7.23E-01 | hypothetical protein |
| AT1G16550 | -0.1 | 4.43E-01 | 0.0 | 9.95E-01 | -0.1 | 5.46E-01 | -0.2 | 4.88E-01 | 0.2 | 5.75E-01 | -0.5 | 2.13E-01 | NA |
| AT4G01650 | -0.1 | 2.79E-01 | 0.0 | 6.41E-01 | -0.1 | 1.91E-01 | -0.2 | 4.71E-01 | 0.0 | 9.85E-01 | -0.2 | 6.38E-01 | Polyketide cyclase |
| AT4G29960 | -0.1 | 1.95E-01 | 0.0 | 5.67E-01 | -0.1 | 1.11E-01 | -0.1 | 6.01E-01 | -0.1 | 4.66E-01 | 0.0 | 8.78E-01 | Transmembrane protein |
| AT4G35470 | -0.1 | 3.94E-01 | 0.1 | 4.90E-01 | -0.2 | 1.81E-01 | -0.7 | 5.15E-03 | -0.5 | 7.10E-02 | -0.2 | 4.62E-01 | Plant intracellular ras group-related LRR 4 |
| AT1G26580 | -0.1 | 2.40E-01 | -0.2 | 9.91E-03 | 0.1 | 9.79E-02 | -0.1 | 4.34E-01 | -0.1 | 7.70E-01 | -0.1 | 8.15E-01 | At1g26580 |
| AT3G24500 | -0.1 | 5.49E-01 | -0.4 | 3.51E-02 | 0.3 | 1.50E-01 | -0.3 | 4.65E-01 | -0.3 | 5.88E-01 | 0.0 | 9.55E-01 | Multiprotein bridging factor 1C |
| AT5G24620 | -0.1 | 3.09E-01 | -0.3 | 8.88E-03 | 0.2 | 7.41E-02 | -0.4 | 5.61E-01 | 0.0 | 9.83E-01 | -0.3 | 6.95E-01 | Thaumatin-like protein |
| AT5G17900 | -0.1 | 2.04E-01 | -0.1 | 1.35E-01 | 0.0 | 8.09E-01 | -0.1 | 4.05E-01 | -0.1 | 3.99E-01 | 0.0 | 9.50E-01 | Microfibril-associated protein-like |
| AT3G19990 | -0.1 | 1.91E-01 | -0.2 | 2.28E-02 | 0.1 | 2.76E-01 | -0.3 | 4.33E-02 | -0.1 | 6.23E-01 | -0.2 | 2.73E-01 | E3 ubiquitin-protein ligase |
| AT4G16180 | -0.1 | 1.48E-01 | -0.1 | 1.10E-01 | 0.0 | 8.60E-01 | -0.1 | 7.16E-01 | -0.2 | 2.32E-01 | 0.1 | 4.41E-01 | Transmembrane protein |
| AT3G15400 | -0.1 | 5.60E-01 | 0.0 | 9.81E-01 | -0.1 | 6.66E-01 | 0.0 | 9.21E-01 | -0.1 | 9.23E-01 | 0.0 | 9.93E-01 | ATA20 protein |
| AT3G16200 | -0.1 | 2.42E-01 | 0.2 | 1.39E-02 | -0.3 | 5.24E-03 | -0.2 | 3.91E-01 | 0.0 | 9.87E-01 | -0.2 | 4.55E-01 | DNA-directed RNA polymerase subunit beta |
| AT4G37900 | -0.1 | 2.81E-01 | -0.1 | 1.19E-01 | 0.0 | 6.38E-01 | 0.5 | 2.53E-02 | 0.3 | 1.51E-01 | 0.1 | 6.49E-01 | Glycine-rich domain-containing protein 2 |
| AT3G56210 | -0.1 | 3.24E-01 | 0.0 | 9.77E-01 | -0.1 | 4.26E-01 | 0.2 | 4.57E-01 | 0.0 | 9.97E-01 | 0.2 | 6.24E-01 | ARM repeat superfamily protein |
| AT5G17580 | -0.1 | 5.24E-01 | -0.1 | 4.20E-01 | 0.0 | 8.74E-01 | -0.1 | 7.05E-01 | 0.0 | 9.99E-01 | -0.1 | 7.42E-01 | Uncharacterized protein |
| AT1G33410 | -0.1 | 2.15E-01 | -0.3 | 1.12E-03 | 0.2 | 1.36E-02 | 0.2 | 1.23E-01 | 0.0 | 9.40E-01 | 0.2 | 2.37E-01 | SAR1 |
| AT1G34550 | -0.1 | 2.69E-01 | -0.1 | 1.18E-01 | 0.0 | 6.44E-01 | 0.0 | 9.56E-01 | 0.1 | 7.18E-01 | -0.1 | 7.93E-01 | F12K21.12 |
| AT2G39340 | -0.1 | 4.61E-01 | -0.3 | 1.54E-02 | 0.2 | 7.78E-02 | -0.1 | 5.30E-01 | 0.0 | 9.92E-01 | -0.1 | 6.98E-01 | SAC3 family protein A |
| AT5G03990 | -0.1 | 4.30E-01 | -0.1 | 2.91E-01 | 0.0 | 8.11E-01 | 0.3 | 3.75E-01 | 0.0 | 9.24E-01 | 0.2 | 5.23E-01 | Uncharacterized protein F8F6_200 |
| AT2G38470 | -0.1 | 6.15E-01 | -0.9 | 4.18E-04 | 0.8 | 1.59E-03 | 0.8 | 1.27E-04 | -0.2 | 2.64E-01 | 1.0 | 4.09E-04 | Truncated WRKY33 protein |
| AT3G53940 | -0.1 | 4.77E-01 | -0.3 | 5.47E-02 | 0.2 | 2.53E-01 | 0.9 | 1.26E-02 | 0.6 | 1.11E-01 | 0.3 | 5.11E-01 | hypothetical protein |
| AT4G33700 | -0.1 | 3.16E-01 | 0.0 | 8.68E-01 | -0.1 | 4.92E-01 | -0.3 | 4.47E-02 | 0.2 | 1.99E-01 | -0.5 | 1.63E-02 | AT4G33700 protein |

| | | | | | | | | | | | | | |
|---|---|---|---|---|---|---|---|---|---|---|---|---|---|
| AT5G66030 | -0.1 | 2.30E-01 | -0.1 | 5.42E-02 | 0.1 | 4.28E-01 | -0.3 | 8.98E-02 | -0.3 | 1.95E-01 | 0.0 | 9.03E-01 | Golgi-localized GRIP domain-containing protein |
| AT1G02160 | -0.1 | 3.77E-01 | 0.1 | 1.81E-01 | -0.2 | 6.81E-02 | -0.1 | 6.31E-01 | -0.2 | 4.28E-01 | 0.1 | 7.88E-01 | Cox19 family protein |
| AT1G12530 | -0.1 | 3.62E-01 | 0.1 | 3.41E-01 | -0.2 | 1.19E-01 | -0.1 | 8.02E-01 | 0.2 | 4.80E-01 | -0.2 | 3.79E-01 | T12C24.7 |
| AT2G23840 | -0.1 | 6.28E-01 | -0.4 | 5.61E-02 | 0.3 | 1.68E-01 | -0.1 | 7.82E-01 | 0.0 | 9.27E-01 | -0.1 | 7.67E-01 | Uncharacterized protein At2g23840 |
| AT1G27595 | -0.1 | 3.76E-01 | -0.2 | 6.18E-02 | 0.1 | 3.22E-01 | 0.0 | 9.74E-01 | -0.1 | 8.44E-01 | 0.1 | 8.72E-01 | At1g27590 |
| AT5G56360 | -0.1 | 3.14E-01 | 0.2 | 1.55E-02 | -0.3 | 5.65E-03 | 0.2 | 1.43E-01 | 0.1 | 3.99E-01 | 0.1 | 8.12E-01 | AT5g56360 |
| AT1G71820 | -0.1 | 2.04E-01 | -0.1 | 1.60E-01 | 0.0 | 8.69E-01 | 0.0 | 7.56E-01 | -0.1 | 6.95E-01 | 0.0 | 9.39E-01 | SEC6 |
| AT4G23840 | -0.1 | 3.65E-01 | -0.2 | 7.80E-02 | 0.1 | 4.10E-01 | -0.1 | 5.95E-01 | -0.4 | 8.74E-02 | 0.3 | 3.25E-01 | Uncharacterized protein AT4g23840 |
| AT5G64730 | -0.1 | 5.12E-01 | 0.0 | 7.80E-01 | 0.0 | 7.81E-01 | 0.0 | 9.03E-01 | -0.1 | 5.68E-01 | 0.1 | 7.29E-01 | Similarity to unknown protein |
| AT5G04680 | -0.1 | 5.75E-01 | 0.0 | 8.96E-01 | -0.1 | 7.50E-01 | -0.2 | 6.33E-01 | 0.1 | 8.18E-01 | -0.3 | 4.88E-01 | Ankyrin repeat family protein |
| AT2G04690 | -0.1 | 4.74E-01 | 0.0 | 8.26E-01 | -0.1 | 6.94E-01 | -0.4 | 1.81E-02 | -0.3 | 6.96E-02 | -0.1 | 8.29E-01 | Uncharacterized protein At2g04690 |
| AT5G07970 | -0.1 | 2.30E-01 | -0.2 | 2.80E-02 | 0.1 | 2.73E-01 | 0.0 | 8.94E-01 | -0.1 | 6.06E-01 | 0.2 | 5.85E-01 | Dentin sialophosphoprotein-like protein |
| AT1G77590 | -0.1 | 2.87E-01 | -0.1 | 3.75E-01 | 0.0 | 9.10E-01 | 0.1 | 7.71E-01 | 0.3 | 6.24E-02 | -0.3 | 1.55E-01 | Long chain acyl-CoA synthetase 9 |
| AT1G05860 | -0.1 | 4.67E-01 | 0.0 | 8.33E-01 | -0.1 | 6.94E-01 | -0.2 | 3.21E-01 | -0.2 | 3.00E-01 | 0.0 | 9.23E-01 | KAT8 regulatory NSL complex subunit 2 |
| AT1G55535 | -0.1 | 3.16E-01 | 0.1 | 1.63E-01 | -0.2 | 5.33E-02 | 0.2 | 4.13E-01 | 0.0 | 8.77E-01 | 0.2 | 4.40E-01 | At1g55535 |
| AT3G54190 | -0.1 | 1.64E-01 | -0.1 | 2.79E-02 | 0.1 | 3.43E-01 | -0.2 | 1.70E-01 | -0.2 | 1.17E-01 | 0.1 | 8.31E-01 | hypothetical protein |
| AT1G77580 | -0.1 | 5.70E-01 | 0.1 | 4.95E-01 | -0.2 | 2.81E-01 | 0.0 | 9.96E-01 | 0.2 | 3.85E-01 | -0.2 | 4.20E-01 | Filament-like protein |
| AT1G31940 | -0.1 | 2.75E-01 | -0.2 | 1.35E-02 | 0.1 | 1.28E-01 | -0.1 | 6.90E-01 | 0.0 | 9.46E-01 | -0.1 | 8.42E-01 | Uncharacterized protein F5M6.6 |
| AT3G23980 | -0.1 | 2.12E-01 | -0.1 | 4.10E-02 | 0.1 | 3.82E-01 | -0.1 | 4.83E-01 | -0.1 | 5.33E-01 | 0.0 | 9.87E-01 | Protein BLISTER |
| AT4G26240 | -0.1 | 3.80E-01 | 0.0 | 6.11E-01 | -0.1 | 2.42E-01 | -0.1 | 5.68E-01 | -0.3 | 1.56E-01 | 0.2 | 5.07E-01 | hypothetical protein |
| AT4G37480 | -0.1 | 5.86E-01 | -0.1 | 4.70E-01 | 0.0 | 8.76E-01 | -0.3 | 3.40E-01 | -0.3 | 3.72E-01 | 0.0 | 9.74E-01 | hypothetical protein |
| AT2G16050 | -0.1 | 4.60E-01 | 0.1 | 5.72E-01 | -0.1 | 2.79E-01 | 0.2 | 4.44E-01 | 0.3 | 1.08E-01 | -0.2 | 4.98E-01 | Uncharacterized protein At2g16050 |
| AT1G61770 | -0.1 | 4.75E-01 | 0.3 | 1.85E-02 | -0.4 | 1.03E-02 | 0.0 | 9.00E-01 | 0.0 | 9.12E-01 | 0.0 | 9.93E-01 | Chaperone protein dnaJ 50 |
| AT5G18800 | -0.1 | 3.49E-01 | 0.2 | 1.46E-02 | -0.3 | 5.57E-03 | -0.1 | 2.72E-01 | -0.2 | 3.38E-01 | 0.0 | 9.96E-01 | NADH dehydrogenase 1 alpha subcomplex subunit 8-B |
| AT3G03320 | -0.1 | 5.32E-01 | 0.2 | 6.95E-02 | -0.3 | 4.72E-02 | 0.3 | 2.46E-01 | 0.5 | 4.20E-02 | -0.2 | 4.60E-01 | ASCH domain-containing protein |
| AT5G12300 | -0.1 | 6.92E-01 | -0.1 | 6.64E-01 | 0.0 | 9.62E-01 | -0.3 | 5.52E-01 | -0.1 | 8.08E-01 | -0.1 | 8.60E-01 | hypothetical protein |
| AT3G07440 | -0.1 | 5.18E-01 | 0.1 | 4.04E-01 | -0.2 | 2.15E-01 | -0.1 | 5.23E-01 | -0.1 | 7.79E-01 | -0.1 | 8.70E-01 | Uncharacterized protein |
| AT2G42400 | -0.1 | 3.21E-01 | -0.2 | 6.21E-02 | 0.1 | 3.78E-01 | -0.3 | 1.78E-01 | -0.3 | 1.90E-01 | 0.0 | 9.48E-01 | Transcription factor VOZ2 |
| AT3G19030 | -0.1 | 6.27E-01 | 0.4 | 1.23E-02 | -0.5 | 1.32E-02 | -0.4 | 1.88E-01 | -1.1 | 2.13E-03 | 0.7 | 8.35E-02 | hypothetical protein |
| AT1G70160 | -0.1 | 2.98E-01 | -0.2 | 5.86E-02 | 0.1 | 3.52E-01 | -0.2 | 1.31E-01 | -0.2 | 6.71E-02 | 0.1 | 7.66E-01 | hypothetical protein |
| AT1G80960 | -0.1 | 5.89E-01 | -0.1 | 4.00E-01 | 0.0 | 7.95E-01 | -0.4 | 3.12E-01 | -0.3 | 4.48E-01 | -0.1 | 9.29E-01 | F-box and Leucine Rich Repeat domains containing protein |
| AT1G68980 | -0.1 | 4.23E-01 | -0.1 | 4.87E-01 | 0.0 | 9.60E-01 | -0.1 | 7.10E-01 | -0.1 | 8.70E-01 | 0.0 | 9.10E-01 | Pentatricopeptide repeat-containing protein At1g68980, mitochondrial |
| AT1G73030 | -0.1 | 2.62E-01 | -0.2 | 1.07E-02 | 0.1 | 1.01E-01 | -0.3 | 3.06E-01 | -0.3 | 3.17E-01 | 0.0 | 9.51E-01 | ESCRT-related protein CHMP1B |
| AT4G00760 | -0.1 | 4.73E-01 | -0.2 | 1.40E-01 | 0.1 | 4.89E-01 | -0.3 | 3.50E-01 | -0.1 | 8.17E-01 | -0.2 | 6.54E-01 | Response regulatory domain-containing protein |
| AT3G60310 | -0.1 | 4.95E-01 | 0.0 | 9.90E-01 | -0.1 | 5.79E-01 | -0.5 | 1.15E-01 | -0.5 | 1.03E-01 | 0.1 | 9.12E-01 | Acyl-CoA synthetase family protein |
| AT3G15610 | -0.1 | 2.49E-01 | 0.1 | 1.26E-01 | -0.2 | 3.00E-02 | 0.0 | 9.23E-01 | 0.0 | 9.13E-01 | 0.0 | 9.82E-01 | Uncharacterized protein |
| AT4G08230 | -0.1 | 3.90E-01 | -0.1 | 4.19E-01 | 0.0 | 9.88E-01 | 0.4 | 3.77E-02 | 0.4 | 7.72E-02 | 0.0 | 9.53E-01 | Glycine-rich protein |
| AT3G10380 | -0.1 | 3.44E-01 | -0.2 | 3.92E-02 | 0.1 | 2.29E-01 | 0.0 | 8.81E-01 | 0.1 | 7.67E-01 | 0.0 | 8.91E-01 | Exocyst complex component Sec8 |
| AT4G16143 | -0.1 | 2.96E-01 | -0.2 | 9.03E-03 | 0.2 | 6.89E-02 | 0.1 | 2.02E-01 | 0.0 | 9.39E-01 | 0.1 | 4.71E-01 | Importin subunit alpha |
| AT1G13570 | -0.1 | 3.35E-01 | 0.0 | 6.51E-01 | -0.1 | 2.29E-01 | -0.4 | 2.42E-02 | -0.3 | 9.42E-02 | -0.1 | 7.58E-01 | F-box |
| AT5G04260 | -0.1 | 4.56E-01 | -0.2 | 1.61E-01 | 0.1 | 5.53E-01 | -0.2 | 4.71E-01 | 0.1 | 8.56E-01 | -0.2 | 4.55E-01 | WCRKC2 |
| AT1G51370 | -0.1 | 4.89E-01 | 0.0 | 7.88E-01 | 0.0 | 7.59E-01 | 0.1 | 7.21E-01 | -0.2 | 5.34E-01 | 0.3 | 3.66E-01 | F-box |
| AT3G01170 | -0.1 | 4.02E-01 | -0.3 | 1.42E-02 | 0.2 | 9.10E-02 | -0.2 | 9.86E-02 | 0.0 | 9.75E-01 | -0.2 | 1.99E-01 | T4P13.14 protein |
| AT4G30130 | -0.1 | 6.70E-01 | 0.0 | 8.37E-01 | -0.1 | 6.08E-01 | -0.1 | 8.39E-01 | -0.2 | 7.44E-01 | 0.1 | 8.89E-01 | hypothetical protein |
| AT5G66590 | -0.1 | 6.62E-01 | 0.7 | 1.11E-03 | -0.8 | 2.16E-03 | -0.2 | 4.34E-01 | 0.0 | 9.66E-01 | -0.2 | 5.40E-01 | hypothetical protein |
| AT4G39560 | -0.1 | 3.57E-01 | 0.0 | 7.29E-01 | 0.0 | 6.64E-01 | -0.3 | 1.31E-01 | -0.2 | 4.00E-01 | -0.1 | 7.47E-01 | F-box |
| AT4G19006 | -0.1 | 5.36E-01 | 0.0 | 9.36E-01 | -0.1 | 5.39E-01 | 0.3 | 3.50E-02 | 0.2 | 1.98E-01 | 0.1 | 6.93E-01 | Proteasome component domain protein |
| AT3G11700 | -0.1 | 3.28E-01 | 0.0 | 8.00E-01 | -0.1 | 2.66E-01 | -0.4 | 5.91E-03 | -0.5 | 5.01E-03 | 0.1 | 7.97E-01 | Fasciclin-like arabinogalactan protein 18 |
| AT3G28720 | -0.1 | 3.47E-01 | 0.0 | 6.69E-01 | 0.0 | 6.86E-01 | 0.1 | 5.20E-01 | 0.1 | 8.64E-01 | 0.1 | 8.11E-01 | hypothetical protein |
| AT5G50310 | -0.1 | 3.85E-01 | 0.1 | 1.16E-01 | -0.2 | 4.19E-02 | 0.2 | 8.64E-02 | 0.0 | 9.56E-01 | 0.2 | 2.07E-01 | DUF4110 domain-containing protein |
| AT3G49050 | -0.1 | 4.01E-01 | -0.2 | 1.06E-01 | 0.1 | 4.60E-01 | 0.1 | 7.52E-01 | 0.2 | 4.17E-01 | -0.1 | 7.21E-01 | Alpha |
| AT3G58100 | -0.1 | 4.94E-01 | 0.4 | 3.90E-03 | -0.4 | 4.71E-03 | 0.1 | 6.49E-01 | 0.4 | 2.14E-01 | -0.2 | 5.15E-01 | Plasmodesmata callose-binding protein 5 |
| AT3G17910 | -0.1 | 4.28E-01 | -0.1 | 1.54E-01 | 0.1 | 5.61E-01 | 0.5 | 1.13E-02 | 0.2 | 3.28E-01 | 0.3 | 1.77E-01 | Surfeit locus protein 1 |
| AT1G53280 | -0.1 | 3.92E-01 | -0.3 | 1.18E-02 | 0.2 | 6.90E-02 | -0.1 | 4.18E-01 | 0.0 | 9.56E-01 | -0.1 | 5.32E-01 | Class I glutamine amidotransferase-like superfamily protein |
| AT4G01960 | -0.1 | 3.08E-01 | -0.1 | 1.54E-01 | 0.0 | 6.95E-01 | -0.3 | 1.46E-02 | -0.4 | 9.28E-03 | 0.1 | 7.99E-01 | hypothetical protein |
| AT3G47675 | -0.1 | 6.08E-01 | -0.1 | 3.34E-01 | 0.1 | 6.97E-01 | -0.7 | 1.20E-02 | -0.2 | 4.12E-01 | -0.5 | 2.04E-01 | Ternary complex factor MIP1 leucine-zipper domain-containing protein |
| AT4G13345 | -0.1 | 5.53E-01 | 0.1 | 6.77E-01 | -0.1 | 3.96E-01 | 0.2 | 4.27E-01 | 0.1 | 8.62E-01 | 0.1 | 7.01E-01 | Serinc-domain containing serine and sphingolipid biosynthesis protein |
| AT2G23310 | -0.1 | 3.20E-01 | 0.1 | 6.19E-02 | -0.2 | 1.85E-02 | -0.1 | 3.26E-01 | -0.3 | 2.35E-02 | 0.2 | 2.36E-01 | Protein RER1 |
| AT5G01400 | -0.1 | 3.37E-01 | -0.3 | 4.01E-03 | 0.2 | 3.32E-02 | 0.2 | 3.08E-01 | 0.1 | 6.91E-01 | 0.1 | 7.49E-01 | HEAT repeat-containing protein |
| AT5G38840 | -0.1 | 3.26E-01 | -0.2 | 5.09E-02 | 0.1 | 3.20E-01 | 0.3 | 1.29E-01 | 0.2 | 3.30E-01 | 0.1 | 8.33E-01 | FHA domain-containing protein |
| AT3G56720 | -0.1 | 2.32E-01 | 0.0 | 5.47E-01 | 0.0 | 6.26E-01 | -0.2 | 2.19E-01 | -0.1 | 3.47E-01 | 0.0 | 9.46E-01 | Pre-mRNA-splicing factor |
| AT1G56612 | -0.1 | 5.90E-01 | -0.1 | 6.74E-01 | 0.0 | 9.50E-01 | -0.4 | 2.86E-01 | -0.5 | 1.77E-01 | 0.1 | 7.36E-01 | NA |
| AT2G46940 | -0.1 | 7.15E-01 | -0.4 | 6.71E-02 | 0.3 | 1.87E-01 | -0.6 | 6.29E-02 | -0.3 | 3.70E-01 | -0.3 | 4.47E-01 | Fold protein |
| AT3G22810 | -0.1 | 5.55E-01 | -0.1 | 4.23E-01 | 0.0 | 8.51E-01 | -0.1 | 7.82E-01 | 0.0 | 9.07E-01 | 0.0 | 9.41E-01 | Auxin canalization protein |
| AT5G20580 | -0.1 | 3.88E-01 | -0.2 | 5.11E-02 | 0.1 | 2.80E-01 | 0.0 | 9.05E-01 | 0.0 | 9.76E-01 | 0.0 | 9.11E-01 | TMEM192 family protein |
| AT1G64280 | -0.1 | 3.22E-01 | -0.3 | 1.15E-03 | 0.3 | 8.63E-03 | 0.0 | 8.20E-01 | 0.0 | 9.92E-01 | 0.0 | 8.84E-01 | Regulatory protein NPR1 |
| AT3G61190 | -0.1 | 8.35E-01 | -0.5 | 1.75E-01 | 0.4 | 3.32E-01 | 1.6 | 1.13E-03 | 0.1 | 9.32E-01 | 1.5 | 7.50E-03 | C2 domain-containing protein |
| AT2G01450 | -0.1 | 3.71E-01 | -0.2 | 1.45E-02 | 0.2 | 8.72E-02 | -0.5 | 5.15E-03 | -0.2 | 1.83E-01 | -0.2 | 2.44E-01 | Mitogen-activated protein kinase |
| AT5G01720 | -0.1 | 5.13E-01 | -0.1 | 3.55E-01 | 0.0 | 8.13E-01 | 0.1 | 5.77E-01 | -0.2 | 3.91E-01 | 0.4 | 1.64E-01 | RNI-like superfamily protein |
| AT5G14240 | -0.1 | 3.74E-01 | 0.1 | 1.45E-01 | -0.2 | 4.76E-02 | -0.3 | 2.30E-02 | -0.4 | 8.60E-03 | 0.1 | 6.43E-01 | hypothetical protein |
| AT2G44900 | -0.1 | 3.91E-01 | -0.2 | 3.80E-02 | 0.1 | 2.17E-01 | -0.2 | 3.00E-01 | -0.3 | 2.19E-01 | 0.1 | 8.49E-01 | Protein ARABIDILLO 1 |
| AT1G13780 | -0.1 | 6.10E-01 | 0.0 | 7.38E-01 | -0.1 | 4.85E-01 | -0.2 | 4.23E-01 | -0.3 | 3.33E-01 | 0.1 | 8.64E-01 | F-box |
| AT3G52950 | -0.1 | 3.35E-01 | -0.2 | 3.09E-02 | 0.1 | 2.12E-01 | 0.2 | 1.80E-01 | 0.1 | 4.81E-01 | 0.1 | 7.71E-01 | CBS domain-containing protein CBSCBSPB3 |
| AT1G71480 | -0.1 | 5.63E-01 | -0.1 | 6.18E-01 | 0.0 | 9.72E-01 | 0.1 | 7.57E-01 | -0.3 | 5.99E-01 | 0.4 | 4.21E-01 | Nuclear transport factor 2 domain-containing protein |
| AT5G23610 | -0.1 | 2.51E-01 | -0.2 | 4.95E-03 | 0.1 | 5.21E-02 | -0.1 | 2.45E-01 | -0.3 | 6.34E-02 | 0.1 | 5.64E-01 | Gb\|AAD12714.1 |
| AT2G19780 | -0.1 | 5.85E-01 | 0.1 | 3.58E-01 | -0.2 | 2.21E-01 | -0.4 | 1.13E-01 | 0.1 | 8.54E-01 | -0.5 | 1.27E-01 | Leucine-rich repeat-containing N-terminal plant-type domain-containing protein |
| AT3G20550 | -0.1 | 3.75E-01 | 0.0 | 7.69E-01 | 0.0 | 6.36E-01 | -0.3 | 3.56E-02 | -0.3 | 4.39E-02 | 0.0 | 9.59E-01 | DDL |
| AT3G22290 | -0.1 | 3.43E-01 | 0.0 | 5.45E-01 | -0.1 | 1.74E-01 | 0.0 | 8.88E-01 | -0.1 | 8.04E-01 | 0.0 | 9.26E-01 | Endoplasmic reticulum vesicle transporter protein |
| AT5G19070 | -0.1 | 3.52E-01 | -0.2 | 8.00E-03 | 0.2 | 6.53E-02 | -0.1 | 7.30E-01 | 0.1 | 7.70E-01 | -0.2 | 5.91E-01 | SNARE associated Golgi protein family |
| AT1G15230 | -0.1 | 4.07E-01 | 0.0 | 7.50E-01 | -0.1 | 3.15E-01 | -0.4 | 5.07E-02 | -0.2 | 3.33E-01 | -0.1 | 5.90E-01 | Uncharacterized protein |
| AT1G30260 | -0.1 | 7.85E-01 | -0.3 | 2.41E-01 | 0.2 | 4.34E-01 | -0.2 | 2.65E-01 | -0.1 | 6.37E-01 | -0.1 | 7.47E-01 | hypothetical protein |
| AT1G12450 | -0.1 | 4.84E-01 | -0.1 | 4.56E-01 | 0.0 | 9.52E-01 | -0.7 | 1.01E-02 | -0.2 | 5.56E-01 | -0.6 | 1.32E-01 | SNARE associated Golgi protein family |
| AT1G13060 | -0.1 | 5.27E-01 | 0.1 | 3.65E-01 | -0.2 | 1.63E-01 | 0.0 | 9.35E-01 | 0.0 | 8.54E-01 | 0.0 | 8.41E-01 | 20S proteasome beta subunit E1 |

| | | | | | | | | | | | | | |
|---|---|---|---|---|---|---|---|---|---|---|---|---|---|
| AT5G65490 | -0.1 | 3.98E-01 | 0.0 | 6.02E-01 | 0.0 | 8.19E-01 | 0.0 | 8.49E-01 | -0.2 | 2.60E-01 | 0.2 | 2.71E-01 | Protein ecdysoneless homolog |
| AT3G22800 | -0.1 | 5.77E-01 | 0.1 | 2.77E-01 | -0.2 | 1.51E-01 | 0.7 | 6.99E-04 | 0.3 | 1.63E-01 | 0.4 | 8.00E-02 | Leucine-rich repeat-containing N-terminal plant-type domain-containing protein |
| AT5G03030 | -0.1 | 5.94E-01 | -0.1 | 3.56E-01 | 0.1 | 7.32E-01 | 0.6 | 4.64E-03 | 0.2 | 4.52E-01 | 0.4 | 3.62E-02 | Chaperone DnaJ-domain superfamily protein |
| AT2G21560 | -0.1 | 4.37E-01 | -0.1 | 1.81E-01 | 0.1 | 6.00E-01 | -0.5 | 1.33E-03 | -0.3 | 7.04E-02 | -0.2 | 2.31E-01 | Uncharacterized protein |
| AT3G07560 | -0.1 | 3.51E-01 | -0.2 | 7.23E-03 | 0.2 | 4.64E-02 | -0.2 | 3.73E-02 | -0.3 | 7.55E-03 | 0.1 | 5.13E-01 | Peroxisomal membrane protein 13 |
| AT5G55000 | -0.1 | 5.29E-01 | -0.1 | 3.57E-01 | 0.0 | 7.98E-01 | -0.3 | 4.68E-02 | -0.3 | 8.23E-02 | 0.0 | 9.65E-01 | Potassium channel tetramerization domain-containing protein |
| AT1G80800 | -0.1 | 4.54E-01 | -0.2 | 3.99E-02 | 0.1 | 1.98E-01 | -0.7 | 1.42E-02 | -0.3 | 2.19E-01 | -0.3 | 4.15E-01 | NA |
| AT4G35240 | -0.1 | 4.19E-01 | -0.2 | 2.27E-02 | 0.1 | 1.29E-01 | -0.4 | 1.45E-02 | -0.1 | 3.90E-01 | -0.2 | 2.70E-01 | DNA-directed RNA polymerase subunit beta, putative |
| AT5G04830 | -0.1 | 6.36E-01 | -0.5 | 5.06E-03 | 0.4 | 1.76E-02 | 0.2 | 3.69E-01 | 0.4 | 1.68E-01 | -0.1 | 7.12E-01 | hypothetical protein |
| AT1G17730 | -0.1 | 4.13E-01 | -0.1 | 1.31E-01 | 0.1 | 4.98E-01 | -0.1 | 6.84E-01 | 0.1 | 4.46E-01 | -0.2 | 2.99E-01 | VPS46.1 |
| AT4G12680 | -0.1 | 5.76E-01 | -0.1 | 2.56E-01 | 0.1 | 6.17E-01 | -0.3 | 2.25E-01 | 0.0 | 8.94E-01 | -0.2 | 4.42E-01 | Transmembrane protein |
| AT5G57870 | -0.1 | 3.84E-01 | 0.0 | 6.17E-01 | 0.0 | 7.80E-01 | 0.1 | 4.31E-01 | -0.1 | 5.92E-01 | 0.2 | 3.20E-01 | AT5g57870 |
| AT4G29910 | -0.1 | 4.61E-01 | 0.0 | 6.77E-01 | 0.0 | 8.25E-01 | 0.0 | 8.33E-01 | 0.0 | 9.52E-01 | 0.1 | 8.34E-01 | Origin recognition complex protein 5 |
| AT5G24680 | -0.1 | 6.33E-01 | -0.1 | 3.31E-01 | 0.1 | 6.70E-01 | -0.3 | 3.66E-01 | -0.3 | 3.42E-01 | 0.0 | 9.22E-01 | Zinc finger with UFM1-specific peptidase domain protein |
| AT3G51720 | -0.1 | 6.23E-01 | 0.3 | 5.76E-02 | -0.3 | 4.31E-02 | 0.2 | 2.09E-01 | 0.4 | 3.27E-02 | -0.2 | 4.20E-01 | WEB family protein At3g51720 |
| AT3G60380 | -0.1 | 4.34E-01 | 0.3 | 5.19E-03 | -0.3 | 3.63E-03 | -0.1 | 4.42E-01 | -0.1 | 7.34E-01 | -0.1 | 8.07E-01 | Cotton fiber protein |
| AT1G73400 | -0.1 | 6.53E-01 | -0.2 | 1.69E-01 | 0.1 | 4.11E-01 | -0.2 | 6.56E-01 | 0.0 | 9.60E-01 | -0.1 | 7.91E-01 | Pentatricopeptide repeat-containing protein At1g73400, mitochondrial |
| AT5G25350 | -0.1 | 5.21E-01 | -0.4 | 3.21E-03 | 0.3 | 1.35E-02 | -0.1 | 8.36E-01 | -0.4 | 2.06E-01 | 0.3 | 3.91E-01 | EIN3-binding F-box protein 2 |
| AT4G20060 | -0.1 | 6.61E-01 | -0.4 | 3.39E-02 | 0.3 | 1.16E-01 | -0.1 | 8.42E-01 | 0.0 | 9.84E-01 | -0.1 | 9.12E-01 | Uncharacterized protein |
| AT1G03140 | -0.1 | 3.26E-01 | 0.0 | 5.67E-01 | 0.0 | 7.49E-01 | -0.5 | 5.18E-04 | -0.3 | 1.25E-02 | -0.1 | 4.44E-01 | Pre-mRNA-splicing factor 18 |
| AT5G63460 | -0.1 | 4.66E-01 | 0.0 | 8.52E-01 | -0.1 | 4.37E-01 | -0.1 | 6.09E-01 | -0.1 | 7.77E-01 | 0.0 | 9.26E-01 | DET1- and DDB1-associated protein 1 |
| AT1G70895 | -0.1 | 6.48E-01 | -0.1 | 5.99E-01 | 0.0 | 9.44E-01 | -0.2 | 4.36E-01 | 0.0 | 9.27E-01 | -0.2 | 4.50E-01 | CLAVATA3 |
| AT2G46900 | -0.1 | 3.50E-01 | -0.1 | 3.23E-01 | 0.0 | 9.44E-01 | -0.3 | 1.44E-02 | -0.4 | 1.31E-02 | 0.0 | 8.66E-01 | Transcription factor 25 |
| AT1G57765 | -0.1 | 6.43E-01 | -0.1 | 5.10E-01 | 0.0 | 8.64E-01 | 0.0 | 9.15E-01 | -0.2 | 5.89E-01 | 0.1 | 7.34E-01 | Uncharacterized protein T8L23.22 |
| AT5G57460 | -0.1 | 3.01E-01 | 0.0 | 9.02E-01 | -0.1 | 3.00E-01 | -0.2 | 1.54E-01 | 0.0 | 9.66E-01 | -0.2 | 3.52E-01 | At5g57460 |
| AT1G47200 | -0.1 | 4.95E-01 | 0.2 | 7.85E-02 | -0.2 | 3.51E-02 | 0.1 | 6.38E-01 | 0.1 | 8.05E-01 | 0.0 | 9.32E-01 | WPP domain-containing protein 2 |
| AT3G63220 | -0.1 | 4.46E-01 | 0.0 | 8.21E-01 | -0.1 | 4.04E-01 | -0.4 | 4.59E-02 | -0.6 | 9.25E-03 | 0.2 | 4.87E-01 | hypothetical protein |
| AT5G13020 | -0.1 | 3.12E-01 | -0.3 | 5.52E-04 | 0.2 | 4.03E-03 | 0.0 | 9.84E-01 | 0.0 | 9.19E-01 | 0.0 | 9.31E-01 | Emsy N Terminus |
| AT3G13780 | -0.1 | 5.78E-01 | 0.0 | 9.59E-01 | -0.1 | 6.95E-01 | 0.0 | 8.71E-01 | -0.1 | 7.76E-01 | 0.0 | 9.12E-01 | SMAD |
| AT3G07550 | -0.1 | 5.44E-01 | -0.2 | 6.26E-02 | 0.1 | 2.41E-01 | -0.7 | 4.30E-03 | -0.4 | 8.26E-02 | -0.3 | 3.83E-01 | RNI-like superfamily protein |
| AT5G54780 | -0.1 | 4.43E-01 | 0.1 | 1.49E-01 | -0.2 | 6.64E-02 | 0.0 | 8.68E-01 | 0.1 | 8.17E-01 | 0.0 | 9.34E-01 | Similarity to GTPase activating protein |
| AT5G17690 | -0.1 | 4.33E-01 | -0.2 | 2.16E-02 | 0.1 | 1.24E-01 | -0.2 | 2.12E-01 | -0.2 | 1.53E-01 | 0.0 | 8.51E-01 | Like heterochromatin protein |
| AT3G55530 | -0.1 | 5.48E-01 | -0.2 | 7.80E-02 | 0.1 | 2.74E-01 | 0.0 | 8.71E-01 | 0.0 | 8.86E-01 | -0.1 | 7.87E-01 | RING-type domain-containing protein |
| AT4G15840 | -0.1 | 3.30E-01 | -0.2 | 1.61E-02 | 0.1 | 1.21E-01 | -0.2 | 2.46E-01 | -0.1 | 7.01E-01 | -0.1 | 6.41E-01 | BTB |
| AT5G42890 | -0.1 | 6.39E-01 | 0.1 | 3.78E-01 | -0.2 | 2.36E-01 | 0.0 | 9.71E-01 | 0.1 | 7.59E-01 | -0.1 | 8.25E-01 | Sterol carrier protein 2 |
| AT5G01460 | -0.1 | 4.54E-01 | 0.1 | 3.14E-01 | -0.1 | 1.22E-01 | -0.2 | 2.47E-01 | 0.0 | 9.65E-01 | -0.2 | 3.50E-01 | LIMR family protein At5g01460 |
| AT4G30390 | -0.1 | 5.66E-01 | -0.1 | 2.02E-01 | 0.1 | 5.31E-01 | 0.0 | 9.62E-01 | -0.1 | 7.75E-01 | 0.1 | 7.84E-01 | Uncharacterized protein |
| AT2G35290 | -0.1 | 8.22E-01 | -0.3 | 2.82E-01 | 0.2 | 4.70E-01 | | | | | | | Uncharacterized protein At2g35290 |
| AT2G15960 | -0.1 | 8.79E-01 | 0.3 | 3.51E-01 | -0.4 | 3.49E-01 | -0.7 | 3.86E-02 | -1.3 | 1.91E-03 | 0.6 | 2.09E-01 | Uncharacterized protein |
| AT2G30530 | -0.1 | 3.60E-01 | -0.2 | 6.12E-03 | 0.2 | 4.62E-02 | -0.2 | 2.18E-01 | -0.2 | 2.69E-01 | 0.0 | 9.95E-01 | hypothetical protein |
| AT1G30090 | -0.1 | 3.67E-01 | -0.3 | 6.41E-04 | 0.3 | 4.59E-03 | -0.1 | 5.57E-01 | 0.0 | 9.22E-01 | -0.1 | 5.70E-01 | F-box |
| AT1G66240 | -0.1 | 4.82E-01 | 0.1 | 1.96E-01 | -0.2 | 7.75E-02 | 0.0 | 9.60E-01 | 0.0 | 9.23E-01 | 0.0 | 9.68E-01 | Homolog of anti-oxidant 1 |
| AT1G18950 | -0.1 | 3.75E-01 | -0.1 | 1.54E-01 | 0.0 | 6.04E-01 | 0.0 | 9.27E-01 | 0.0 | 8.61E-01 | 0.0 | 9.41E-01 | DDT domain superfamily |
| AT5G54200 | -0.1 | 5.30E-01 | 0.0 | 8.71E-01 | -0.1 | 5.03E-01 | -0.5 | 1.27E-01 | -0.2 | 6.47E-01 | -0.3 | 5.26E-01 | Transducin |
| AT4G04180 | -0.1 | 6.96E-01 | -0.3 | 9.36E-02 | 0.2 | 2.55E-01 | 1.0 | 1.76E-02 | 0.3 | 5.63E-01 | 0.7 | 1.67E-01 | Putative vesicle transfer ATPase |
| AT4G12990 | -0.1 | 6.51E-01 | -0.1 | 5.31E-01 | 0.0 | 8.81E-01 | -0.4 | 1.83E-01 | -0.1 | 6.84E-01 | -0.2 | 4.04E-01 | At4g12990 |
| AT4G21570 | -0.1 | 5.97E-01 | 0.0 | 9.73E-01 | -0.1 | 6.77E-01 | 0.1 | 5.75E-01 | 0.2 | 2.66E-01 | -0.1 | 6.83E-01 | Organic solute transporter ostalpha protein |
| AT4G36230 | -0.1 | 7.62E-01 | 0.4 | 4.50E-02 | -0.4 | 5.89E-02 | -0.1 | 8.16E-01 | 0.2 | 5.71E-01 | -0.3 | 4.67E-01 | Glycine-rich protein |
| AT2G31600 | -0.1 | 5.86E-01 | -0.1 | 3.21E-01 | 0.1 | 7.04E-01 | 0.1 | 6.76E-01 | 0.1 | 8.58E-01 | 0.1 | 9.08E-01 | KAT8 regulatory NSL complex subunit 2 |
| AT4G21740 | -0.1 | 5.35E-01 | -0.2 | 4.16E-02 | 0.1 | 1.72E-01 | 0.1 | 8.36E-01 | 0.2 | 4.43E-01 | -0.2 | 6.75E-01 | Transmembrane protein |
| AT2G18876 | -0.1 | 6.28E-01 | -0.1 | 4.69E-01 | 0.0 | 8.33E-01 | -0.3 | 9.17E-02 | -0.1 | 8.01E-01 | -0.2 | 2.49E-01 | Afadin |
| AT4G16447 | -0.1 | 7.53E-01 | 0.3 | 1.02E-01 | -0.3 | 1.05E-01 | -0.2 | 3.41E-01 | 0.2 | 3.83E-01 | -0.5 | 7.97E-02 | Uncharacterized protein |
| AT2G26990 | -0.1 | 4.24E-01 | 0.0 | 5.35E-01 | -0.1 | 2.07E-01 | 0.1 | 4.41E-01 | -0.1 | 7.12E-01 | 0.2 | 3.60E-01 | Putative PCI domain protein |
| AT3G46270 | -0.1 | 6.50E-01 | 0.1 | 3.62E-01 | -0.2 | 2.45E-01 | 0.3 | 3.02E-01 | 0.5 | 4.40E-02 | -0.3 | 3.68E-01 | Uncharacterized protein F12M12_240 |
| AT1G20460 | -0.1 | 6.06E-01 | 0.1 | 5.75E-01 | -0.1 | 3.68E-01 | 0.3 | 2.86E-01 | -0.1 | 8.46E-01 | 0.3 | 2.08E-01 | NADH-ubiquinone oxidoreductase chain |
| AT4G05060 | -0.1 | 5.37E-01 | 0.2 | 2.06E-02 | -0.3 | 1.63E-02 | -0.5 | 3.03E-02 | -0.4 | 7.73E-02 | -0.1 | 8.84E-01 | Vesicle-associated protein 4-3 |
| AT2G01940 | -0.1 | 5.91E-01 | 0.2 | 8.21E-02 | -0.2 | 6.34E-02 | -0.1 | 7.74E-01 | -0.1 | 8.57E-01 | 0.0 | 9.69E-01 | SGR5 |
| AT3G07140 | -0.1 | 5.05E-01 | 0.0 | 6.81E-01 | -0.1 | 3.46E-01 | 0.1 | 4.91E-01 | 0.2 | 3.47E-01 | -0.1 | 8.34E-01 | GPI transamidase component Gpi16 subunit family protein |
| AT1G15370 | -0.1 | 4.34E-01 | 0.2 | 2.83E-02 | -0.2 | 1.39E-02 | -0.2 | 1.66E-01 | -0.1 | 4.44E-01 | -0.1 | 7.88E-01 | Coatomer subunit zeta |
| AT3G54363 | -0.1 | 7.33E-01 | 0.4 | 1.08E-02 | -0.5 | 1.79E-02 | -0.1 | 7.13E-01 | -0.4 | 3.73E-01 | 0.2 | 6.39E-01 | Uncharacterized protein |
| AT1G19020 | -0.1 | 8.03E-01 | -0.7 | 1.16E-02 | 0.6 | 3.39E-02 | 1.2 | 1.37E-03 | -0.2 | 7.55E-01 | 1.4 | 4.03E-03 | CDP-diacylglycerol-glycerol-3-phosphate 3-phosphatidyltransferase |
| AT1G56300 | -0.1 | 7.64E-01 | 0.2 | 1.93E-01 | -0.3 | 1.55E-01 | -0.5 | 6.34E-02 | -0.6 | 3.24E-02 | 0.1 | 7.70E-01 | DnaJ protein, putative |
| AT5G62200 | -0.1 | 5.70E-01 | 0.0 | 6.82E-01 | -0.1 | 4.03E-01 | 0.0 | 8.94E-01 | -0.1 | 3.47E-01 | 0.2 | 3.82E-01 | Embryo-specific protein ATS3B |
| AT3G53860 | -0.1 | 7.07E-01 | 0.1 | 5.75E-01 | -0.1 | 4.40E-01 | 0.0 | 8.94E-01 | 0.2 | 6.36E-01 | -0.2 | 5.85E-01 | KAT8 regulatory NSL complex subunit 2 |
| AT4G05150 | -0.1 | 6.01E-01 | 0.0 | 9.98E-01 | -0.1 | 6.49E-01 | -0.7 | 3.91E-04 | -0.6 | 4.02E-03 | -0.1 | 6.52E-01 | Uncharacterized protein AT4g05150 |
| AT3G53120 | -0.1 | 6.34E-01 | -0.2 | 8.29E-02 | 0.2 | 2.43E-01 | 0.1 | 4.89E-01 | 0.1 | 5.82E-01 | 0.0 | 9.88E-01 | Vacuolar protein-sorting-associated protein 37 homolog 1 |
| AT4G38260 | -0.1 | 5.39E-01 | -0.2 | 1.54E-02 | 0.2 | 6.87E-02 | -0.1 | 7.69E-01 | -0.1 | 7.10E-01 | 0.0 | 9.28E-01 | Uncharacterized protein At4g38260 |
| AT3G44430 | -0.1 | 5.73E-01 | -0.1 | 3.97E-01 | 0.0 | 8.02E-01 | 0.3 | 6.29E-02 | 0.1 | 7.00E-01 | 0.2 | 2.99E-01 | Transmembrane protein |
| AT3G08065 | -0.1 | 7.94E-01 | -0.3 | 1.78E-01 | 0.2 | 3.44E-01 | -0.2 | 6.04E-01 | -0.1 | 7.70E-01 | -0.1 | 9.22E-01 | NA |
| AT3G13340 | -0.1 | 3.89E-01 | -0.1 | 1.60E-01 | 0.0 | 5.89E-01 | -0.4 | 3.52E-03 | -0.3 | 2.85E-02 | -0.1 | 6.28E-01 | Transducin |
| AT4G38480 | -0.1 | 7.55E-01 | 0.0 | 9.14E-01 | -0.1 | 7.36E-01 | 0.0 | 9.79E-01 | 0.1 | 8.49E-01 | 0.0 | 8.94E-01 | Transducin |
| AT3G08800 | -0.1 | 6.28E-01 | -0.1 | 3.05E-01 | 0.1 | 6.45E-01 | -0.2 | 4.21E-01 | 0.1 | 7.91E-01 | -0.3 | 3.41E-01 | ARM repeat superfamily protein |
| AT5G28830 | -0.1 | 6.62E-01 | -0.5 | 2.26E-03 | 0.4 | 9.50E-03 | -0.3 | 1.24E-01 | -0.5 | 1.91E-02 | 0.2 | 4.45E-01 | Calcium-binding EF hand family protein |
| AT1G54060 | -0.1 | 4.44E-01 | -0.1 | 1.34E-01 | 0.1 | 4.80E-01 | -0.2 | 1.62E-01 | -0.2 | 4.58E-01 | -0.1 | 7.71E-01 | Myb |
| AT2G25800 | -0.1 | 5.72E-01 | -0.1 | 3.18E-01 | 0.0 | 6.94E-01 | -0.6 | 1.34E-02 | -0.4 | 8.78E-02 | -0.2 | 6.51E-01 | hypothetical protein |
| AT1G65020 | 0.0 | 5.62E-01 | -0.1 | 3.97E-01 | 0.0 | 8.11E-01 | 0.1 | 4.85E-01 | -0.1 | 7.90E-01 | 0.2 | 4.30E-01 | NERD domain-containing protein |
| AT5G59350 | 0.0 | 7.91E-01 | -0.2 | 4.14E-01 | 0.1 | 6.37E-01 | -0.4 | 9.62E-03 | 0.1 | 4.52E-01 | -0.6 | 9.11E-03 | Transmembrane protein |
| AT2G13970 | 0.0 | 6.33E-01 | -0.3 | 1.83E-02 | 0.2 | 7.24E-02 | -0.5 | 2.85E-01 | -0.4 | 4.32E-01 | -0.1 | 9.19E-01 | NA |
| AT5G38300 | 0.0 | 7.64E-01 | 0.2 | 3.17E-01 | -0.2 | 2.78E-01 | 0.2 | 5.13E-01 | 0.3 | 3.56E-01 | -0.1 | 8.14E-01 | Uncharacterized protein |
| AT5G20150 | 0.0 | 7.51E-01 | 0.2 | 1.93E-01 | -0.3 | 1.67E-01 | 1.2 | 1.93E-04 | 0.6 | 4.07E-02 | 0.6 | 6.64E-02 | SPX domain-containing protein 1 |
| AT3G62500 | 0.0 | 7.80E-01 | 0.2 | 2.38E-01 | -0.3 | 2.24E-01 | -0.1 | 8.22E-01 | 0.0 | 9.87E-01 | -0.1 | 8.86E-01 | F-box protein RMF |
| AT5G34853 | 0.0 | 5.70E-01 | -0.3 | 5.49E-03 | 0.3 | 2.47E-02 | -0.2 | 5.09E-01 | -0.2 | 5.20E-01 | 0.0 | 9.70E-01 | NA |
| AT3G09070 | 0.0 | 6.46E-01 | -0.2 | 6.14E-02 | 0.2 | 1.84E-01 | 0.0 | 8.36E-01 | 0.1 | 6.51E-01 | -0.2 | 5.50E-01 | Protein OCTOPUS |

| | | | | | | | | | | | | | |
|---|---|---|---|---|---|---|---|---|---|---|---|---|---|
| AT4G27652 | 0.0 | 7.26E-01 | -0.1 | 5.49E-01 | 0.0 | 8.36E-01 | 0.6 | 2.11E-01 | 0.2 | 7.98E-01 | 0.4 | 3.77E-01 | hypothetical protein |
| AT3G56320 | 0.0 | 6.97E-01 | -0.3 | 2.38E-02 | 0.3 | 8.18E-02 | 0.0 | 9.05E-01 | 0.0 | 9.41E-01 | 0.0 | 9.86E-01 | Polymerase nucleotidyl transferase domain-containing protein |
| AT1G56290 | 0.0 | 6.28E-01 | -0.2 | 9.57E-02 | 0.1 | 2.82E-01 | -0.1 | 7.94E-01 | 0.0 | 9.74E-01 | -0.1 | 8.19E-01 | Uncharacterized protein F14G9.10 |
| AT2G02730 | 0.0 | 6.63E-01 | -0.1 | 5.33E-01 | 0.0 | 8.71E-01 | -0.1 | 8.12E-01 | -0.3 | 2.40E-01 | 0.2 | 4.52E-01 | hypothetical protein |
| AT3G59220 | 0.0 | 7.00E-01 | -0.3 | 5.42E-02 | 0.2 | 1.53E-01 | 0.5 | 2.20E-01 | 1.1 | 7.83E-03 | -0.6 | 6.58E-02 | Pirin-1 |
| AT1G33480 | 0.0 | 7.79E-01 | 0.3 | 9.67E-02 | -0.3 | 1.04E-01 | -0.5 | 5.75E-02 | 0.0 | 9.32E-01 | -0.5 | 1.95E-01 | RING-type domain-containing protein |
| AT4G37095 | 0.0 | 8.14E-01 | 0.2 | 2.82E-01 | -0.3 | 2.73E-01 | 0.2 | 5.31E-01 | 0.2 | 6.45E-01 | 0.0 | 9.71E-01 | Uncharacterized protein |
| AT1G32760 | 0.0 | 7.07E-01 | 0.2 | 7.66E-02 | -0.3 | 7.60E-02 | -0.5 | 2.84E-02 | -0.2 | 3.16E-01 | -0.2 | 3.64E-01 | At1g32760 |
| AT1G23960 | 0.0 | 6.76E-01 | -0.2 | 7.85E-02 | 0.2 | 2.03E-01 | -0.3 | 8.64E-02 | -0.3 | 6.90E-02 | 0.1 | 8.67E-01 | UPF0725 protein At1g23960 |
| AT5G14730 | 0.0 | 6.28E-01 | -0.4 | 1.81E-03 | 0.4 | 9.36E-03 | 1.8 | 5.72E-06 | -0.1 | 7.70E-01 | 1.9 | 2.88E-05 | At5g14730 |
| AT3G53390 | 0.0 | 4.73E-01 | -0.2 | 2.27E-02 | 0.1 | 1.15E-01 | -0.1 | 8.54E-01 | 0.2 | 5.36E-01 | -0.2 | 4.80E-01 | Transducin |
| AT4G14230 | 0.0 | 4.48E-01 | -0.1 | 1.21E-01 | 0.1 | 4.37E-01 | -0.2 | 1.53E-01 | -0.1 | 6.19E-01 | -0.1 | 6.16E-01 | DUF21 domain-containing protein At4g14230 |
| AT1G68330 | 0.0 | 7.35E-01 | 0.0 | 8.53E-01 | 0.0 | 9.22E-01 | -0.1 | 7.34E-01 | -0.2 | 6.05E-01 | 0.1 | 8.88E-01 | hypothetical protein |
| AT5G24500 | 0.0 | 6.26E-01 | 0.0 | 8.75E-01 | 0.0 | 8.01E-01 | 0.0 | 9.46E-01 | 0.0 | 9.66E-01 | 0.0 | 9.31E-01 | Uncharacterized protein |
| AT5G24190 | 0.0 | 8.21E-01 | 0.7 | 3.40E-03 | -0.7 | 8.39E-03 | -1.6 | 1.96E-02 | 0.0 | 9.56E-01 | -1.7 | 4.47E-02 | Gb\|AAD29063.1 |
| AT5G64330 | 0.0 | 4.50E-01 | 0.0 | 9.45E-01 | 0.0 | 5.58E-01 | -0.3 | 5.24E-02 | -0.2 | 2.01E-01 | -0.1 | 7.57E-01 | Phototropic-responsive NPH3 family protein |
| AT5G48330 | 0.0 | 6.41E-01 | -0.1 | 5.19E-01 | 0.0 | 8.75E-01 | 0.1 | 5.35E-01 | 0.3 | 8.68E-02 | -0.2 | 3.64E-01 | Regulator of chromosome condensation like family protein) |
| AT5G24220 | 0.0 | 8.32E-01 | -0.3 | 1.59E-01 | 0.3 | 3.00E-01 | -1.3 | 1.12E-04 | 0.4 | 6.96E-02 | -1.7 | 1.97E-04 | Gb\|AAD29063.1 |
| AT3G12940 | 0.0 | 5.50E-01 | -0.2 | 5.06E-02 | 0.1 | 1.99E-01 | -0.5 | 3.79E-02 | -0.4 | 1.06E-01 | -0.1 | 8.30E-01 | 2-oxoglutarate and Fe-dependent oxygenase superfamily protein |
| AT5G25360 | 0.0 | 6.23E-01 | 0.1 | 4.47E-01 | -0.1 | 2.65E-01 | -0.1 | 8.24E-01 | 0.1 | 8.37E-01 | -0.1 | 7.35E-01 | DUF4050 domain-containing protein |
| AT1G67870 | 0.0 | 6.52E-01 | 0.2 | 1.59E-01 | -0.2 | 9.45E-02 | 0.0 | 9.26E-01 | 0.0 | 8.47E-01 | -0.1 | 8.27E-01 | Uncharacterized protein T23K23.28 |
| AT5G14170 | 0.0 | 4.81E-01 | -0.1 | 3.42E-01 | 0.0 | 8.17E-01 | -0.3 | 3.99E-03 | -0.4 | 6.20E-03 | 0.0 | 9.51E-01 | SWI |
| AT1G78730 | 0.0 | 7.42E-01 | 0.0 | 8.09E-01 | -0.1 | 6.44E-01 | 0.3 | 4.05E-01 | 0.4 | 2.65E-01 | -0.1 | 8.07E-01 | Putative FBD-associated F-box protein At1g78730 |
| AT2G20310 | 0.0 | 6.01E-01 | -0.2 | 6.20E-02 | 0.1 | 2.12E-01 | -0.2 | 3.77E-01 | -0.2 | 5.82E-01 | -0.1 | 8.95E-01 | RPM1 interacting protein 13 |
| AT5G03830 | 0.0 | 7.35E-01 | -0.1 | 4.96E-01 | 0.0 | 7.75E-01 | -0.2 | 3.50E-01 | -0.1 | 7.46E-01 | -0.1 | 7.36E-01 | CDK inhibitor P21 binding protein |
| AT5G22820 | 0.0 | 6.74E-01 | -0.1 | 2.25E-01 | 0.1 | 4.91E-01 | -0.1 | 7.66E-01 | -0.2 | 6.35E-01 | 0.1 | 8.73E-01 | ARM repeat superfamily protein |
| AT3G57965 | 0.0 | 7.80E-01 | 0.0 | 8.22E-01 | -0.1 | 6.82E-01 | 0.0 | 9.39E-01 | 0.1 | 8.44E-01 | -0.1 | 7.93E-01 | NA |
| AT3G01360 | 0.0 | 6.87E-01 | 0.1 | 3.35E-01 | -0.1 | 2.45E-01 | -0.1 | 7.81E-01 | -0.1 | 8.46E-01 | 0.0 | 9.81E-01 | At3g01360 ) |
| AT4G29330 | 0.0 | 6.39E-01 | 0.0 | 7.45E-01 | 0.0 | 9.25E-01 | 0.1 | 5.75E-01 | 0.1 | 5.64E-01 | 0.0 | 9.49E-01 | Derlin |
| AT2G27260 | 0.0 | 4.98E-01 | -0.2 | 2.10E-02 | 0.1 | 1.02E-01 | -0.2 | 3.18E-01 | -0.2 | 3.57E-01 | 0.0 | 9.86E-01 | hypothetical protein |
| AT1G80450 | 0.0 | 6.40E-01 | 0.2 | 2.02E-02 | -0.3 | 2.13E-02 | 0.4 | 5.93E-02 | -0.1 | 8.68E-01 | 0.5 | 8.04E-02 | VQ motif-containing protein 11 |
| AT3G02520 | 0.0 | 6.85E-01 | 0.0 | 9.87E-01 | 0.0 | 7.41E-01 | 0.0 | 9.50E-01 | 0.0 | 9.19E-01 | 0.0 | 8.89E-01 | 14-3-3-like protein GF14 nu |
| AT1G51740 | 0.0 | 6.68E-01 | 0.1 | 4.52E-01 | -0.1 | 3.15E-01 | -0.3 | 1.00E-01 | -0.2 | 2.73E-01 | -0.1 | 8.09E-01 | Syntaxin of plants 81 |
| AT5G25150 | 0.0 | 6.18E-01 | -0.1 | 1.29E-01 | 0.1 | 3.49E-01 | 0.0 | 8.78E-01 | 0.1 | 8.47E-01 | -0.1 | 7.42E-01 | TBP-associated factor 5 |
| AT5G54750 | 0.0 | 6.35E-01 | 0.2 | 2.51E-02 | -0.3 | 1.99E-02 | -0.1 | 6.17E-01 | -0.1 | 7.94E-01 | 0.0 | 9.27E-01 | Trafficking protein particle complex subunit |
| AT5G22000 | 0.0 | 5.42E-01 | -0.3 | 3.38E-03 | 0.2 | 1.38E-02 | -0.1 | 3.06E-01 | -0.1 | 3.99E-01 | 0.0 | 9.90E-01 | RING-H2 group F2A |
| AT1G47410 | 0.0 | 7.32E-01 | 0.4 | 5.18E-03 | -0.4 | 8.92E-03 | 0.2 | 4.56E-01 | 0.2 | 5.51E-01 | 0.0 | 9.79E-01 | At1g47410 |
| AT2G12461 | 0.0 | 7.91E-01 | -0.1 | 4.38E-01 | 0.1 | 6.69E-01 | 0.0 | 9.58E-01 | 0.0 | 9.30E-01 | 0.0 | 9.71E-01 | Uncharacterized protein |
| AT4G11390 | 0.0 | 8.31E-01 | 0.9 | 2.26E-04 | -1.0 | 1.04E-03 | 0.1 | 8.39E-01 | 0.2 | 7.11E-01 | -0.1 | 8.88E-01 | Cysteine |
| AT1G47310 | 0.0 | 5.38E-01 | -0.2 | 1.49E-02 | 0.2 | 6.71E-02 | -0.2 | 1.48E-01 | -0.2 | 2.95E-01 | 0.0 | 8.89E-01 | Uncharacterized protein |
| AT3G21430 | 0.0 | 6.05E-01 | -0.3 | 6.48E-03 | 0.2 | 2.69E-02 | -0.1 | 6.68E-01 | -0.1 | 6.78E-01 | 0.0 | 9.76E-01 | DNA binding protein |
| AT5G54850 | 0.0 | 7.18E-01 | 0.1 | 6.02E-01 | -0.1 | 4.58E-01 | -0.2 | 4.84E-01 | -0.3 | 2.19E-01 | 0.1 | 6.82E-01 | hypothetical protein |
| AT5G51380 | 0.0 | 6.99E-01 | 0.3 | 1.26E-02 | -0.3 | 1.68E-02 | -0.3 | 1.91E-01 | -0.5 | 2.99E-02 | 0.2 | 4.01E-01 | F-box protein At5g51380 |
| AT3G02555 | 0.0 | 6.94E-01 | 0.1 | 2.01E-01 | -0.2 | 1.63E-01 | 0.3 | 2.86E-01 | 0.0 | 9.25E-01 | 0.3 | 3.22E-01 | hypothetical protein |
| AT5G18630 | 0.0 | 6.55E-01 | 0.0 | 8.42E-01 | 0.0 | 8.53E-01 | -0.7 | 7.21E-04 | -0.7 | 1.38E-03 | 0.0 | 9.70E-01 | AT5g18630 |
| AT4G31600 | 0.0 | 7.03E-01 | 0.2 | 9.84E-02 | -0.2 | 8.41E-02 | 0.4 | 3.59E-02 | 0.3 | 1.71E-01 | 0.1 | 6.43E-01 | UDP-N-acetylglucosamine transporter family |
| AT3G02070 | 0.0 | 6.06E-01 | -0.2 | 4.74E-02 | 0.1 | 1.63E-01 | -0.5 | 3.67E-03 | -0.5 | 5.07E-03 | 0.0 | 9.49E-01 | OTU domain-containing protein |
| AT3G15970 | 0.0 | 6.23E-01 | -0.1 | 4.61E-01 | 0.0 | 8.27E-01 | -0.1 | 4.37E-01 | 0.0 | 9.55E-01 | -0.2 | 5.29E-01 | Nuclear pore complex protein NUP50B |
| AT5G13260 | 0.0 | 6.18E-01 | 0.1 | 5.32E-01 | -0.1 | 3.12E-01 | -0.1 | 6.31E-01 | -0.2 | 2.92E-01 | 0.1 | 6.59E-01 | Coiled-coil domain-containing protein SCD2 |
| AT1G61150 | 0.0 | 6.09E-01 | -0.1 | 4.91E-01 | 0.0 | 8.71E-01 | 0.0 | 7.74E-01 | -0.1 | 3.49E-01 | 0.1 | 6.14E-01 | LisH and RanBPM domains containing protein |
| AT5G25940 | 0.0 | 7.76E-01 | 0.1 | 4.54E-01 | -0.1 | 3.67E-01 | 0.0 | 8.45E-01 | 0.0 | 8.46E-01 | -0.1 | 7.43E-01 | Nodulin-like protein |
| AT5G61150 | 0.0 | 5.59E-01 | -0.2 | 3.11E-02 | 0.1 | 1.12E-01 | 0.0 | 8.62E-01 | -0.1 | 8.05E-01 | 0.0 | 9.39E-01 | Leo1-like family protein |
| AT2G06005 | 0.0 | 6.83E-01 | -0.1 | 6.03E-01 | 0.0 | 9.20E-01 | -0.3 | 3.72E-01 | 0.2 | 6.41E-01 | -0.4 | 2.58E-01 | At2g06010 |
| AT1G21740 | 0.0 | 6.97E-01 | 0.1 | 5.35E-01 | -0.1 | 3.92E-01 | 0.2 | 4.93E-01 | 0.2 | 5.64E-01 | 0.0 | 9.99E-01 | DUF630 family protein, putative |
| AT3G58050 | 0.0 | 5.90E-01 | -0.2 | 6.34E-03 | 0.2 | 2.61E-02 | -0.1 | 6.56E-01 | -0.2 | 5.17E-01 | 0.1 | 8.55E-01 | Uncharacterized protein |
| AT1G03560 | 0.0 | 7.36E-01 | 0.1 | 4.73E-01 | -0.1 | 3.70E-01 | 0.0 | 9.27E-01 | -0.1 | 8.35E-01 | 0.0 | 9.11E-01 | Pentatricopeptide repeat-containing protein At1g03560, mitochondrial |
| AT2G02560 | 0.0 | 6.12E-01 | -0.2 | 4.34E-02 | 0.1 | 1.34E-01 | 0.1 | 7.60E-01 | 0.1 | 8.08E-01 | 0.0 | 1.00E+00 | Cullin-associated NEDD8-dissociated protein 1 |
| AT2G43340 | 0.0 | 8.71E-01 | 0.0 | 9.44E-01 | -0.1 | 8.48E-01 | -0.5 | 2.93E-02 | 0.0 | 9.77E-01 | -0.5 | 6.41E-02 | Uncharacterized protein |
| AT4G33690 | 0.0 | 6.87E-01 | -0.2 | 1.24E-01 | 0.1 | 3.13E-01 | -0.3 | 2.00E-01 | -0.3 | 2.79E-01 | 0.0 | 9.70E-01 | hypothetical protein |
| AT5G06440 | 0.0 | 7.29E-01 | -0.2 | 5.66E-02 | 0.2 | 1.41E-01 | 0.0 | 8.26E-01 | 0.0 | 8.67E-01 | -0.1 | 7.65E-01 | Polyketide cyclase |
| AT2G01720 | 0.0 | 6.40E-01 | 0.1 | 3.10E-01 | -0.1 | 1.77E-01 | 0.0 | 9.81E-01 | 0.1 | 4.92E-01 | -0.1 | 5.94E-01 | Dolichyl-diphosphooligosaccharide--protein glycosyltransferase subunit 1B |
| AT1G77350 | 0.0 | 6.37E-01 | 0.1 | 9.49E-02 | -0.2 | 6.61E-02 | 0.1 | 5.80E-01 | 0.1 | 4.96E-01 | 0.0 | 8.97E-01 | Dolichyl-diphosphooligosaccharide--protein glycosyltransferase subunit KCP2 |
| AT2G40316 | 0.0 | 7.21E-01 | 0.2 | 4.32E-02 | -0.3 | 4.10E-02 | -0.1 | 7.41E-01 | 0.1 | 6.95E-01 | -0.2 | 4.82E-01 | Autophagy-like protein |
| AT5G37310 | 0.0 | 6.62E-01 | 0.0 | 8.28E-01 | 0.0 | 8.71E-01 | 0.2 | 1.60E-01 | 0.1 | 4.21E-01 | 0.1 | 8.01E-01 | Transmembrane 9 superfamily member 4 |
| AT3G51230 | 0.0 | 8.76E-01 | 0.4 | 5.46E-02 | -0.5 | 8.30E-02 | 0.4 | 2.42E-01 | 0.0 | 9.43E-01 | 0.4 | 2.99E-01 | Uncharacterized protein |
| AT2G40600 | 0.0 | 7.11E-01 | -0.1 | 3.77E-01 | 0.1 | 6.47E-01 | 0.1 | 5.72E-01 | -0.1 | 7.05E-01 | 0.1 | 4.65E-01 | Expressed protein |
| AT2G38260 | 0.0 | 6.94E-01 | -0.1 | 4.21E-01 | 0.0 | 7.23E-01 | -0.1 | 6.33E-01 | 0.0 | 9.79E-01 | -0.1 | 6.51E-01 | NA |
| AT3G23255 | 0.0 | 7.80E-01 | 0.1 | 6.82E-01 | -0.1 | 5.66E-01 | -0.3 | 1.13E-01 | -0.2 | 2.95E-01 | -0.1 | 7.96E-01 | Uncharacterized protein |
| AT5G09900 | 0.0 | 6.63E-01 | 0.1 | 4.94E-01 | -0.1 | 3.04E-01 | 0.1 | 2.43E-01 | 0.0 | 8.37E-01 | 0.2 | 3.33E-01 | 26S proteasome regulatory subunit, putative |
| AT5G20120 | 0.0 | 7.65E-01 | -0.1 | 5.68E-01 | 0.0 | 8.20E-01 | -0.4 | 2.28E-02 | -0.5 | 3.36E-02 | 0.0 | 9.95E-01 | At5g20120 |
| AT5G42560 | 0.0 | 5.87E-01 | -0.1 | 1.46E-01 | 0.1 | 3.90E-01 | -0.2 | 8.58E-02 | -0.1 | 3.17E-01 | -0.1 | 7.23E-01 | HVA22-like protein |
| AT3G10680 | 0.0 | 8.45E-01 | -0.2 | 1.85E-01 | 0.2 | 3.34E-01 | -0.1 | 9.21E-01 | -0.6 | 2.53E-01 | 0.5 | 3.53E-01 | SHSP domain-containing protein |
| AT5G53730 | 0.0 | 8.20E-01 | 0.2 | 2.65E-01 | -0.2 | 2.61E-01 | -0.5 | 1.26E-02 | -0.2 | 2.47E-01 | -0.3 | 3.19E-01 | NDR1 |
| AT5G45410 | 0.0 | 6.49E-01 | 0.0 | 7.69E-01 | 0.0 | 9.11E-01 | -0.8 | 1.14E-03 | 0.0 | 9.98E-01 | -0.8 | 5.33E-03 | AT5G45410 protein |
| AT5G65925 | 0.0 | 8.76E-01 | 0.0 | 9.69E-01 | 0.0 | 8.76E-01 | 0.0 | 9.23E-01 | -0.3 | 2.27E-01 | 0.3 | 2.71E-01 | Uncharacterized protein |
| AT4G19200 | 0.0 | 6.84E-01 | 0.0 | 9.47E-01 | 0.0 | 7.80E-01 | -0.2 | 2.20E-01 | -0.1 | 6.63E-01 | -0.1 | 6.98E-01 | hypothetical protein |
| AT4G21720 | 0.0 | 7.51E-01 | 0.1 | 1.76E-01 | -0.2 | 1.61E-01 | 0.2 | 3.50E-01 | 0.0 | 8.92E-01 | 0.3 | 3.27E-01 | hypothetical protein |
| AT5G64870 | 0.0 | 9.08E-01 | -0.2 | 5.19E-01 | 0.2 | 6.67E-01 | -0.1 | 8.81E-01 | -0.1 | 9.27E-01 | 0.0 | 9.81E-01 | Flotillin-like protein 3 |
| AT3G21080 | 0.0 | 9.30E-01 | -0.3 | 4.53E-01 | 0.3 | 5.81E-01 | 0.3 | 5.37E-01 | -1.5 | 4.51E-02 | 1.9 | 2.66E-02 | Uncharacterized protein |
| AT1G62680 | 0.0 | 7.89E-01 | 0.0 | 7.97E-01 | -0.1 | 6.68E-01 | 0.0 | 9.71E-01 | 0.0 | 9.95E-01 | 0.0 | 9.85E-01 | Pentatricopeptide repeat-containing protein At1g62680, mitochondrial |
| AT4G36040 | 0.0 | 8.27E-01 | 0.0 | 9.67E-01 | 0.0 | 8.22E-01 | -0.5 | 1.84E-01 | -1.7 | 1.19E-03 | 1.2 | 4.20E-02 | J11 |
| AT4G10430 | 0.0 | 6.61E-01 | -0.3 | 3.48E-03 | 0.2 | 1.32E-02 | 0.0 | 8.84E-01 | 0.2 | 2.64E-01 | -0.2 | 3.93E-01 | TMPIT-like protein |
| AT4G09150 | 0.0 | 7.49E-01 | -0.2 | 1.00E-01 | 0.1 | 2.07E-01 | -0.2 | 2.90E-01 | -0.1 | 8.04E-01 | -0.1 | 6.51E-01 | T-complex protein 11 |

| | | | | | | | | | | | | | |
|---|---|---|---|---|---|---|---|---|---|---|---|---|---|
| AT5G17370 | 0.0 | 7.97E-01 | -0.2 | 9.08E-02 | 0.2 | 2.06E-01 | -0.1 | 8.73E-01 | -0.2 | 7.33E-01 | 0.1 | 8.64E-01 | Transducin |
| AT3G13810 | 0.0 | 7.84E-01 | -0.3 | 1.44E-02 | 0.3 | 3.94E-02 | -0.2 | 3.68E-01 | 0.0 | 9.19E-01 | -0.2 | 4.26E-01 | Indeterminate-domain 11 |
| AT2G30700 | 0.0 | 6.72E-01 | -0.1 | 1.23E-01 | 0.1 | 2.97E-01 | 0.0 | 8.74E-01 | 0.0 | 9.62E-01 | 0.0 | 9.55E-01 | GPI-anchored protein |
| AT3G09550 | 0.0 | 8.74E-01 | -0.2 | 4.22E-01 | 0.1 | 5.85E-01 | 0.1 | 8.94E-01 | 0.3 | 4.57E-01 | -0.2 | 5.94E-01 | At3g09550 |
| AT1G56450 | 0.0 | 7.49E-01 | 0.1 | 4.71E-01 | -0.1 | 3.36E-01 | 0.3 | 2.85E-02 | 0.1 | 3.68E-01 | 0.1 | 4.31E-01 | Proteasome subunit beta type-4 |
| AT3G07400 | 0.0 | 7.78E-01 | -0.2 | 1.52E-01 | 0.1 | 3.02E-01 | -0.4 | 1.16E-01 | -0.3 | 3.20E-01 | -0.1 | 7.85E-01 | F21O3.11 protein |
| AT2G42700 | 0.0 | 6.72E-01 | -0.2 | 1.56E-02 | 0.2 | 5.54E-02 | 0.1 | 8.00E-01 | 0.0 | 9.66E-01 | 0.0 | 9.04E-01 | Vesicle docking protein |
| AT5G27970 | 0.0 | 6.43E-01 | -0.1 | 3.39E-01 | 0.0 | 6.59E-01 | 0.0 | 9.41E-01 | 0.1 | 5.90E-01 | -0.2 | 5.53E-01 | ARM repeat superfamily protein |
| AT5G10490 | 0.0 | 6.32E-01 | -0.2 | 3.72E-03 | 0.2 | 1.40E-02 | 0.1 | 5.97E-01 | 0.2 | 4.10E-01 | -0.1 | 8.19E-01 | AT5g10490 |
| AT2G33490 | 0.0 | 7.43E-01 | -0.3 | 5.94E-03 | 0.3 | 1.99E-02 | 0.3 | 1.03E-01 | 0.2 | 3.59E-01 | 0.1 | 7.27E-01 | Hydroxyproline-rich glycoprotein family protein |
| AT3G03070 | 0.0 | 6.63E-01 | 0.2 | 2.67E-02 | -0.2 | 1.91E-02 | 0.1 | 6.98E-01 | 0.1 | 6.60E-01 | 0.0 | 9.40E-01 | NADH dehydrogenase iron-sulfur protein 6, mitochondrial |
| AT5G42000 | 0.0 | 7.53E-01 | 0.0 | 6.23E-01 | 0.0 | 8.79E-01 | 0.1 | 7.68E-01 | 0.0 | 9.79E-01 | 0.1 | 7.62E-01 | At5g42000 |
| AT1G67190 | 0.0 | 7.56E-01 | 0.0 | 6.74E-01 | 0.0 | 9.22E-01 | -0.3 | 1.11E-01 | -0.2 | 2.36E-01 | 0.0 | 8.87E-01 | F-box |
| AT2G14910 | 0.0 | 7.79E-01 | -0.2 | 6.00E-02 | 0.2 | 1.20E-01 | -0.4 | 3.76E-03 | -0.3 | 2.28E-02 | -0.1 | 7.51E-01 | MAR-binding filament-like protein |
| AT3G02710 | 0.0 | 7.36E-01 | 0.0 | 9.28E-01 | 0.0 | 8.47E-01 | 0.0 | 8.52E-01 | 0.1 | 5.69E-01 | -0.1 | 7.74E-01 | ARM repeat superfamily protein |
| AT3G17780 | 0.0 | 8.25E-01 | 0.5 | 1.33E-03 | -0.6 | 2.26E-03 | 0.0 | 7.81E-01 | -0.1 | 6.73E-01 | 0.1 | 5.82E-01 | Endoplasmic reticulum transmembrane protein |
| AT4G38500 | 0.0 | 6.83E-01 | -0.1 | 3.14E-01 | 0.0 | 5.91E-01 | -0.2 | 3.03E-01 | -0.1 | 4.20E-01 | 0.0 | 9.64E-01 | AT4g38500 |
| AT4G08110 | 0.0 | 7.79E-01 | -0.1 | 2.50E-01 | 0.1 | 4.44E-01 | -0.3 | 2.15E-01 | -0.4 | 9.29E-02 | 0.1 | 6.97E-01 | NA |
| AT5G46400 | 0.0 | 7.97E-01 | -0.3 | 1.99E-02 | 0.3 | 5.65E-02 | 0.0 | 9.53E-01 | 0.0 | 9.48E-01 | 0.0 | 9.88E-01 | Tetratricopeptide repeat -like superfamily protein |
| AT1G48960 | 0.0 | 8.55E-01 | 0.1 | 5.48E-01 | -0.1 | 5.15E-01 | 0.1 | 7.54E-01 | -0.2 | 5.61E-01 | 0.3 | 4.44E-01 | F27J15.25 |
| AT5G43310 | 0.0 | 7.95E-01 | -0.2 | 1.54E-01 | 0.1 | 2.88E-01 | 0.0 | 8.64E-01 | 0.0 | 9.59E-01 | 0.0 | 9.46E-01 | Similarity to COP1-interacting protein 7 |
| AT1G07020 | 0.0 | 9.11E-01 | -0.2 | 5.12E-01 | 0.1 | 6.56E-01 | -0.1 | 8.38E-01 | 0.2 | 5.76E-01 | -0.3 | 4.75E-01 | F10K1.26 protein |
| AT3G57450 | 0.0 | 8.77E-01 | -0.3 | 1.65E-01 | 0.2 | 2.69E-01 | 0.4 | 2.05E-01 | -0.2 | 6.75E-01 | 0.6 | 1.62E-01 | hypothetical protein |
| AT5G67350 | 0.0 | 8.10E-01 | -0.4 | 3.75E-03 | 0.4 | 1.28E-02 | -0.1 | 4.42E-01 | 0.0 | 9.72E-01 | -0.1 | 6.17E-01 | Uncharacterized protein |
| AT4G15420 | 0.0 | 7.77E-01 | -0.3 | 7.53E-03 | 0.3 | 2.24E-02 | 0.0 | 9.56E-01 | -0.1 | 6.27E-01 | 0.1 | 7.46E-01 | Ubiquitin fusion degradation UFD1 family protein |
| AT1G34630 | 0.0 | 7.97E-01 | -0.4 | 2.41E-03 | 0.4 | 8.70E-03 | -0.6 | 2.44E-02 | -0.7 | 1.97E-02 | 0.1 | 8.81E-01 | At1g34630 |
| AT5G12340 | 0.0 | 8.89E-01 | 0.2 | 2.44E-01 | -0.2 | 2.78E-01 | 1.8 | 5.39E-06 | 1.2 | 1.77E-03 | 0.7 | 1.66E-02 | DUF4228 domain protein |
| AT5G64180 | 0.0 | 7.57E-01 | 0.1 | 2.66E-01 | -0.1 | 2.14E-01 | -0.2 | 3.57E-01 | -0.4 | 1.49E-01 | 0.2 | 6.91E-01 | Tropomyosin |
| AT1G66345 | 0.0 | 8.69E-01 | 0.0 | 8.92E-01 | 0.0 | 9.88E-01 | 0.1 | 8.04E-01 | 0.2 | 4.81E-01 | -0.1 | 7.30E-01 | Pentatricopeptide repeat-containing protein At1g66345, mitochondrial |
| AT2G46200 | 0.0 | 8.04E-01 | -0.2 | 1.61E-01 | 0.1 | 3.16E-01 | -0.2 | 4.65E-01 | -0.2 | 6.60E-01 | -0.1 | 9.22E-01 | U11 |
| AT4G17250 | 0.0 | 7.51E-01 | -0.4 | 5.35E-04 | 0.4 | 2.18E-03 | -0.1 | 5.40E-01 | -0.6 | 1.92E-02 | 0.5 | 1.13E-01 | AT4g17250 |
| AT3G19460 | 0.0 | 7.87E-01 | 0.0 | 6.11E-01 | -0.1 | 4.98E-01 | -0.1 | 7.55E-01 | -0.2 | 3.26E-01 | 0.1 | 6.12E-01 | Reticulon-like protein |
| AT5G18460 | 0.0 | 8.03E-01 | 0.2 | 1.04E-01 | -0.2 | 1.00E-01 | -0.2 | 2.08E-01 | 0.0 | 8.58E-01 | -0.2 | 2.33E-01 | Neprosin activation peptide domain-containing protein |
| AT5G01370 | 0.0 | 8.83E-01 | 0.2 | 1.44E-01 | -0.3 | 1.72E-01 | 0.1 | 6.62E-01 | -0.1 | 8.88E-01 | 0.2 | 6.35E-01 | ALC-interacting protein 1 |
| AT1G13810 | 0.0 | 8.52E-01 | -0.2 | 2.42E-01 | 0.1 | 4.03E-01 | 0.4 | 8.46E-02 | 0.1 | 7.50E-01 | 0.3 | 2.52E-01 | hypothetical protein |
| AT2G25930 | 0.0 | 7.51E-01 | 0.0 | 8.40E-01 | 0.0 | 6.51E-01 | -0.2 | 2.25E-01 | -0.3 | 1.51E-01 | 0.1 | 8.22E-01 | PYK20 |
| AT1G30755 | 0.0 | 7.74E-01 | -0.4 | 8.23E-04 | 0.4 | 3.16E-03 | 0.0 | 9.11E-01 | -0.3 | 2.23E-01 | 0.3 | 2.07E-01 | T5I8.20 protein |
| AT1G68430 | 0.0 | 8.74E-01 | 0.1 | 3.41E-01 | -0.2 | 3.56E-01 | 0.1 | 8.49E-01 | 0.1 | 7.67E-01 | -0.1 | 9.11E-01 | hypothetical protein |
| AT1G49840 | 0.0 | 8.25E-01 | 0.1 | 6.01E-01 | -0.1 | 5.23E-01 | -0.6 | 1.67E-03 | -0.2 | 3.70E-01 | -0.4 | 5.95E-02 | Uncharacterized protein |
| AT1G68300 | 0.0 | 7.36E-01 | 0.1 | 1.19E-01 | -0.1 | 8.90E-02 | -0.2 | 1.65E-01 | -0.2 | 4.63E-01 | -0.1 | 7.61E-01 | hypothetical protein |
| AT3G60410 | 0.0 | 8.29E-01 | -0.2 | 4.46E-02 | 0.2 | 9.49E-02 | 0.1 | 2.68E-01 | 0.0 | 9.54E-01 | 0.2 | 3.80E-01 | hypothetical protein |
| AT4G13100 | 0.0 | 7.48E-01 | 0.0 | 9.24E-01 | 0.0 | 8.63E-01 | -0.2 | 1.16E-01 | 0.0 | 9.88E-01 | -0.3 | 2.56E-01 | Uncharacterized protein At4g13100 |
| AT4G26980 | 0.0 | 8.04E-01 | -0.2 | 2.58E-02 | 0.2 | 7.09E-02 | 0.0 | 9.44E-01 | 0.0 | 9.21E-01 | 0.0 | 8.90E-01 | At4g26980 |
| AT5G63830 | 0.0 | 8.01E-01 | -0.1 | 2.49E-01 | 0.1 | 4.38E-01 | 0.2 | 3.08E-01 | 0.1 | 6.81E-01 | 0.1 | 7.41E-01 | At5g63830 |
| AT3G05570 | 0.0 | 8.55E-01 | 0.1 | 5.98E-01 | -0.1 | 5.46E-01 | 0.1 | 8.50E-01 | 0.0 | 9.53E-01 | 0.0 | 9.44E-01 | hypothetical protein |
| AT3G57890 | 0.0 | 7.42E-01 | -0.1 | 2.53E-01 | 0.1 | 4.45E-01 | 0.0 | 9.91E-01 | 0.1 | 6.07E-01 | -0.1 | 6.68E-01 | TBCC domain-containing protein 1 |
| AT2G11000 | 0.0 | 7.93E-01 | -0.2 | 1.17E-02 | 0.2 | 3.31E-02 | 0.1 | 7.52E-01 | 0.0 | 9.17E-01 | 0.1 | 7.17E-01 | MAK10 homologue |
| AT2G45900 | 0.0 | 9.28E-01 | -0.1 | 6.44E-01 | 0.1 | 7.59E-01 | -0.1 | 8.93E-01 | 0.2 | 6.83E-01 | -0.3 | 6.12E-01 | hypothetical protein |
| AT3G21580 | 0.0 | 8.46E-01 | 0.1 | 3.97E-01 | -0.1 | 3.70E-01 | 0.4 | 3.05E-02 | 0.6 | 9.35E-03 | -0.1 | 5.99E-01 | Protein ABCI12, chloroplastic |
| AT5G02970 | 0.0 | 8.36E-01 | -0.1 | 5.55E-01 | 0.0 | 7.45E-01 | -0.4 | 1.50E-02 | -0.3 | 7.08E-02 | -0.1 | 7.65E-01 | AT5g02970 |
| AT4G02210 | 0.0 | 8.56E-01 | -0.1 | 6.00E-01 | 0.0 | 7.80E-01 | -0.6 | 5.01E-02 | -0.4 | 2.62E-01 | -0.3 | 6.11E-01 | Uncharacterized protein AT4g02210 |
| AT3G26680 | 0.0 | 8.49E-01 | -0.3 | 1.29E-02 | 0.3 | 3.61E-02 | -0.1 | 7.37E-01 | -0.1 | 7.91E-01 | 0.0 | 9.98E-01 | DNA repair metallo-beta-lactamase family protein |
| AT2G26070 | 0.0 | 8.28E-01 | 0.1 | 1.71E-01 | -0.1 | 1.69E-01 | -0.1 | 3.45E-01 | -0.1 | 6.23E-01 | 0.0 | 8.47E-01 | Protein REVERSION-TO-ETHYLENE SENSITIVITY1 |
| AT1G73390 | 0.0 | 9.08E-01 | -0.3 | 7.98E-02 | 0.3 | 1.28E-01 | -0.1 | 6.64E-01 | 0.2 | 2.74E-01 | -0.3 | 1.87E-01 | At1g73390 |
| AT2G03680 | 0.0 | 8.19E-01 | 0.1 | 2.70E-01 | -0.1 | 2.17E-01 | 0.0 | 9.13E-01 | 0.1 | 5.99E-01 | -0.1 | 5.87E-01 | Protein SPIRAL1 |
| AT3G10915 | 0.0 | 8.12E-01 | 0.0 | 7.35E-01 | 0.0 | 9.29E-01 | 0.0 | 9.95E-01 | -0.1 | 6.63E-01 | 0.1 | 7.27E-01 | Reticulon-like protein |
| AT3G06190 | 0.0 | 7.99E-01 | -0.1 | 1.20E-01 | 0.1 | 2.27E-01 | -0.4 | 2.77E-03 | -0.5 | 2.48E-03 | 0.1 | 8.14E-01 | BTB-POZ and MATH domain 2 |
| AT1G07705 | 0.0 | 8.11E-01 | 0.0 | 9.23E-01 | 0.0 | 7.78E-01 | 0.2 | 4.11E-01 | 0.1 | 8.04E-01 | 0.1 | 7.62E-01 | NOT2 |
| AT2G26190 | 0.0 | 8.73E-01 | -0.3 | 3.83E-02 | 0.2 | 6.90E-02 | 0.3 | 7.02E-02 | -0.2 | 4.04E-01 | 0.4 | 3.90E-02 | Calmodulin-binding family protein |
| AT5G18750 | 0.0 | 8.59E-01 | -0.3 | 2.91E-02 | 0.2 | 6.98E-02 | 0.2 | 2.74E-01 | -0.1 | 5.98E-01 | 0.4 | 1.92E-01 | J domain-containing protein |
| AT5G35560 | 0.0 | 8.48E-01 | -0.2 | 2.63E-02 | 0.2 | 5.52E-02 | -0.1 | 6.09E-01 | 0.0 | 9.76E-01 | -0.1 | 7.12E-01 | DENN domain-containing protein |
| AT2G32760 | 0.0 | 8.25E-01 | -0.1 | 2.77E-01 | 0.1 | 4.54E-01 | 0.1 | 7.83E-01 | 0.2 | 2.91E-01 | -0.2 | 4.80E-01 | UV radiation resistance-associated protein |
| AT4G18570 | 0.0 | 8.51E-01 | 0.0 | 9.38E-01 | 0.0 | 9.35E-01 | -0.3 | 9.55E-02 | -0.1 | 5.80E-01 | -0.2 | 5.21E-01 | Tetratricopeptide repeat -like superfamily protein |
| AT4G02200 | 0.0 | 9.12E-01 | -0.3 | 5.01E-02 | 0.3 | 1.02E-01 | -0.3 | 3.08E-01 | 0.2 | 5.31E-01 | -0.6 | 1.80E-01 | Drought-responsive family protein |
| AT4G22230 | 0.0 | 9.12E-01 | 0.7 | 3.41E-04 | -0.7 | 1.35E-03 | -1.0 | 1.81E-02 | 0.1 | 8.21E-01 | -1.1 | 3.59E-02 | Defensin-like protein 96 |
| AT5G65650 | 0.0 | 8.58E-01 | 0.3 | 1.01E-02 | -0.3 | 1.59E-02 | 0.0 | 9.68E-01 | 0.1 | 8.46E-01 | -0.1 | 8.37E-01 | Emb\|CAB16809.1 |
| AT1G08125 | 0.0 | 8.18E-01 | 0.1 | 4.63E-01 | -0.1 | 4.04E-01 | -0.1 | 6.79E-01 | 0.0 | 9.23E-01 | 0.0 | 8.51E-01 | Uncharacterized protein At1g08125 |
| AT1G76570 | 0.0 | 9.29E-01 | -0.4 | 4.77E-02 | 0.4 | 8.77E-02 | -0.2 | 5.52E-01 | 0.1 | 6.66E-01 | -0.3 | 3.88E-01 | Transmembrane protein |
| AT3G28270 | 0.0 | 9.34E-01 | 0.0 | 8.19E-01 | 0.0 | 9.01E-01 | 0.6 | 8.29E-02 | 1.5 | 6.86E-04 | -0.9 | 4.49E-02 | UPF0496 protein At3g28270 |
| AT1G53345 | 0.0 | 9.10E-01 | 0.1 | 5.71E-01 | -0.1 | 5.74E-01 | 0.3 | 3.90E-01 | 0.2 | 5.96E-01 | 0.1 | 8.96E-01 | DHHA1 domain protein |
| AT5G26990 | 0.0 | 9.07E-01 | 0.0 | 9.94E-01 | 0.0 | 9.23E-01 | -0.1 | 8.43E-01 | -0.2 | 5.79E-01 | 0.1 | 7.93E-01 | Protein DEHYDRATION-INDUCED 19 homolog 6 |
| AT2G18245 | 0.0 | 8.58E-01 | 0.1 | 1.49E-01 | -0.1 | 1.62E-01 | 0.1 | 7.54E-01 | 0.1 | 6.53E-01 | 0.0 | 9.05E-01 | Alpha |
| AT3G57400 | 0.0 | 8.46E-01 | -0.2 | 7.75E-03 | 0.2 | 1.75E-02 | -0.3 | 9.95E-03 | -0.3 | 3.24E-02 | 0.0 | 8.90E-01 | Transmembrane protein |
| AT2G02870 | 0.0 | 8.51E-01 | 0.2 | 1.71E-02 | -0.2 | 2.05E-02 | -0.2 | 3.77E-02 | -0.1 | 6.72E-01 | -0.2 | 2.49E-01 | F-box |
| AT1G28040 | 0.0 | 9.28E-01 | 0.5 | 3.01E-03 | -0.6 | 8.05E-03 | 0.1 | 8.71E-01 | 0.1 | 7.69E-01 | -0.1 | 9.07E-01 | RING-type E3 ubiquitin transferase |
| AT4G28230 | 0.0 | 9.11E-01 | 0.3 | 4.02E-02 | -0.3 | 6.26E-02 | 0.3 | 2.29E-01 | 0.3 | 3.73E-01 | 0.0 | 9.29E-01 | hypothetical protein |
| AT5G52430 | 0.0 | 8.81E-01 | 0.0 | 7.85E-01 | 0.0 | 9.17E-01 | -0.5 | 3.06E-03 | -0.2 | 3.43E-01 | -0.3 | 1.04E-01 | At5g52430 |
| AT5G24890 | 0.0 | 9.12E-01 | -0.1 | 2.98E-01 | 0.1 | 4.07E-01 | -0.4 | 5.65E-02 | -0.7 | 9.06E-03 | 0.3 | 4.54E-01 | Protein OXIDATIVE STRESS 3 LIKE 4 |
| AT5G50120 | 0.0 | 9.12E-01 | 0.2 | 8.63E-02 | -0.2 | 1.19E-01 | 0.1 | 7.99E-01 | -0.1 | 8.18E-01 | 0.2 | 6.73E-01 | Transducin |
| AT4G30260 | 0.0 | 8.81E-01 | 0.1 | 1.37E-01 | -0.1 | 1.36E-01 | -0.2 | 3.12E-01 | -0.1 | 6.68E-01 | -0.1 | 7.85E-01 | Protein YIP |
| AT1G04550 | 0.0 | 8.74E-01 | -0.1 | 4.16E-01 | 0.1 | 5.65E-01 | 0.1 | 5.48E-01 | 0.0 | 9.28E-01 | 0.1 | 7.81E-01 | Auxin-responsive protein |
| AT1G65900 | 0.0 | 9.53E-01 | 0.2 | 4.30E-01 | -0.2 | 4.73E-01 | -0.2 | 4.31E-01 | 0.2 | 3.48E-01 | -0.4 | 1.51E-01 | At1g65900 |
| AT2G41430 | 0.0 | 8.96E-01 | 0.1 | 2.72E-01 | -0.1 | 2.53E-01 | -0.1 | 5.92E-01 | -0.4 | 4.44E-02 | 0.3 | 2.55E-01 | Protein EARLY RESPONSIVE TO DEHYDRATION 15 -binding protein-interacting protein 1) |

| | | | | | | | | | | | | | |
|---|---|---|---|---|---|---|---|---|---|---|---|---|---|
| AT5G28150 | 0.0 | 8.93E-01 | -0.1 | 4.00E-01 | 0.1 | 5.42E-01 | 0.2 | 3.77E-01 | 0.2 | 4.96E-01 | 0.0 | 9.63E-01 | hypothetical protein |
| AT1G60780 | 0.0 | 8.29E-01 | 0.0 | 6.04E-01 | 0.0 | 7.85E-01 | 0.1 | 3.63E-01 | 0.2 | 2.63E-01 | 0.0 | 8.37E-01 | Putative Clathrin Coat Assembly protein |
| AT4G19440 | 0.0 | 9.28E-01 | -0.1 | 3.41E-01 | 0.1 | 4.67E-01 | 0.2 | 4.60E-01 | 0.2 | 5.57E-01 | 0.0 | 9.80E-01 | Pentatricopeptide repeat-containing protein At4g19440, chloroplastic |
| AT1G04000 | 0.0 | 8.68E-01 | 0.0 | 6.27E-01 | 0.0 | 5.74E-01 | 0.1 | 5.55E-01 | 0.1 | 8.08E-01 | 0.0 | 8.73E-01 | Uncharacterized protein |
| AT2G19080 | 0.0 | 8.71E-01 | 0.1 | 4.87E-01 | -0.1 | 4.47E-01 | 0.0 | 8.18E-01 | -0.1 | 8.10E-01 | 0.1 | 7.00E-01 | Metaxin-like protein |
| AT1G78810 | 0.0 | 9.06E-01 | 0.1 | 4.93E-01 | -0.1 | 4.99E-01 | 0.0 | 9.12E-01 | -0.1 | 6.63E-01 | 0.1 | 7.70E-01 | Uncharacterized protein At1g78810 |
| AT1G29980 | 0.0 | 9.21E-01 | 0.1 | 6.24E-01 | -0.1 | 6.07E-01 | 0.1 | 5.75E-01 | 0.3 | 5.02E-02 | -0.2 | 2.44E-01 | Uncharacterized protein T1P2.9 |
| AT4G37630 | 0.0 | 9.24E-01 | 0.2 | 1.62E-01 | -0.2 | 2.08E-01 | 0.3 | 3.79E-01 | 0.3 | 4.72E-01 | 0.0 | 9.80E-01 | Cyclin d51 |
| AT4G39220 | 0.0 | 8.95E-01 | 0.2 | 8.80E-02 | -0.2 | 9.83E-02 | -0.2 | 2.29E-01 | -0.1 | 3.84E-01 | 0.0 | 9.24E-01 | Protein RER1A |
| AT3G29390 | 0.0 | 8.95E-01 | -0.2 | 3.13E-02 | 0.2 | 5.87E-02 | 0.1 | 6.21E-01 | 0.2 | 5.09E-01 | -0.1 | 8.51E-01 | RIK |
| AT4G38495 | 0.0 | 9.31E-01 | 0.1 | 6.57E-01 | -0.1 | 6.63E-01 | -0.1 | 7.62E-01 | -0.1 | 7.45E-01 | 0.0 | 9.59E-01 | Protein EIN6 ENHANCER |
| AT1G53110 | 0.0 | 9.34E-01 | -0.1 | 7.14E-01 | 0.0 | 8.15E-01 | 0.3 | 2.39E-01 | 0.4 | 1.91E-01 | -0.1 | 8.62E-01 | Proton pump-interactor 4 |
| AT4G16450 | 0.0 | 9.22E-01 | 0.4 | 3.58E-03 | -0.4 | 5.22E-03 | 0.0 | 9.72E-01 | 0.1 | 8.56E-01 | 0.0 | 9.06E-01 | At4g16450 |
| AT1G19080 | 0.0 | 9.45E-01 | -0.1 | 4.18E-01 | 0.1 | 5.36E-01 | -0.1 | 8.57E-01 | 0.0 | 9.18E-01 | 0.0 | 9.71E-01 | At1g19080 |
| AT5G54110 | 0.0 | 8.83E-01 | 0.1 | 4.32E-01 | -0.1 | 3.98E-01 | -0.1 | 6.24E-01 | -0.3 | 6.18E-02 | 0.2 | 2.77E-01 | Membrane-associated mannitol-induced |
| AT1G09920 | 0.0 | 8.85E-01 | -0.1 | 2.26E-01 | 0.1 | 3.36E-01 | 0.0 | 9.23E-01 | 0.0 | 9.76E-01 | 0.0 | 9.71E-01 | TRAF-type domain-containing protein |
| AT1G03270 | 0.0 | 9.62E-01 | -0.1 | 5.72E-01 | 0.1 | 6.71E-01 | -0.1 | 8.64E-01 | 0.5 | 3.40E-01 | -0.5 | 3.27E-01 | Putative DUF21 domain-containing protein At1g03270 |
| AT3G52560 | 0.0 | 8.92E-01 | -0.1 | 2.94E-01 | 0.1 | 3.96E-01 | -0.1 | 4.23E-01 | -0.1 | 4.76E-01 | 0.0 | 9.87E-01 | Ubiquitin E2 variant 1D-4 |
| AT1G21090 | 0.0 | 9.45E-01 | 0.2 | 8.13E-02 | -0.3 | 1.17E-01 | -0.1 | 8.26E-01 | -0.1 | 7.34E-01 | 0.0 | 9.23E-01 | Cupredoxin superfamily protein |
| AT2G32880 | 0.0 | 9.55E-01 | 0.5 | 5.92E-03 | -0.5 | 1.33E-02 | -0.1 | 6.41E-01 | -0.3 | 7.85E-02 | 0.2 | 2.74E-01 | TRAF-like family protein |
| AT1G63110 | 0.0 | 8.81E-01 | 0.2 | 1.38E-02 | -0.2 | 1.94E-02 | -0.2 | 6.65E-02 | -0.3 | 7.06E-02 | 0.0 | 9.32E-01 | GPI transamidase subunit PIG-U |
| AT2G22790 | 0.0 | 9.48E-01 | -0.2 | 2.40E-01 | 0.2 | 3.49E-01 | -0.6 | 6.19E-02 | -0.5 | 1.10E-01 | 0.0 | 9.31E-01 | Uncharacterized protein At2g22790 |
| AT1G67850 | 0.0 | 9.32E-01 | -0.4 | 2.42E-03 | 0.4 | 6.33E-03 | 0.0 | 9.38E-01 | -0.1 | 7.53E-01 | 0.2 | 7.19E-01 | hypothetical protein |
| AT5G66675 | 0.0 | 9.23E-01 | -0.3 | 1.21E-02 | 0.3 | 2.49E-02 | 0.0 | 8.59E-01 | -0.2 | 3.35E-01 | 0.1 | 4.62E-01 | Uncharacterized protein |
| AT4G36515 | 0.0 | 9.39E-01 | 0.0 | 7.63E-01 | 0.0 | 8.54E-01 | 0.0 | 9.33E-01 | 0.1 | 7.09E-01 | -0.2 | 6.74E-01 | Uncharacterized protein |
| AT5G10940 | 0.0 | 9.10E-01 | -0.1 | 1.27E-01 | 0.1 | 1.99E-01 | -0.4 | 2.09E-02 | -0.2 | 3.33E-01 | -0.2 | 3.80E-01 | hypothetical protein |
| AT5G48020 | 0.0 | 9.19E-01 | -0.1 | 3.28E-01 | 0.1 | 4.25E-01 | -0.2 | 1.25E-01 | 0.1 | 7.58E-01 | -0.3 | 1.73E-01 | 2-oxoglutarate and Fe-dependent oxygenase superfamily protein |
| AT2G02860 | 0.0 | 9.26E-01 | -0.2 | 4.12E-02 | 0.2 | 7.41E-02 | 0.1 | 6.94E-01 | 0.1 | 6.18E-01 | 0.0 | 9.16E-01 | Sucrose transporter 2 |
| AT2G15695 | 0.0 | 9.24E-01 | -0.2 | 5.35E-02 | 0.2 | 9.22E-02 | -0.4 | 1.23E-02 | -0.3 | 5.54E-02 | -0.1 | 7.58E-01 | At2g15695 |
| AT3G45443 | 0.0 | 9.51E-01 | 0.1 | 2.77E-01 | -0.1 | 3.31E-01 | -0.2 | 4.56E-01 | -0.2 | 3.75E-01 | 0.1 | 8.94E-01 | AT3g45441 |
| AT3G01740 | 0.0 | 9.20E-01 | 0.1 | 3.84E-01 | -0.1 | 3.85E-01 | -0.1 | 5.37E-01 | -0.2 | 3.90E-01 | 0.1 | 8.33E-01 | Large ribosomal subunit protein mL54 |
| AT3G21175 | 0.0 | 9.16E-01 | -0.1 | 4.23E-01 | 0.0 | 5.46E-01 | -0.4 | 1.44E-02 | -0.3 | 6.79E-02 | -0.1 | 7.61E-01 | ZIM-like 1 |
| AT2G06210 | 0.0 | 9.12E-01 | -0.1 | 1.53E-01 | 0.1 | 2.28E-01 | 0.1 | 5.29E-01 | 0.2 | 2.50E-01 | -0.1 | 6.66E-01 | Protein CTR9 homolog |
| AT5G09310 | 0.0 | 9.49E-01 | 0.0 | 6.80E-01 | -0.1 | 6.77E-01 | -0.3 | 9.54E-02 | -0.4 | 3.95E-02 | 0.1 | 6.83E-01 | Probable gamma-secretase subunit PEN-2 |
| AT5G48120 | 0.0 | 9.30E-01 | -0.1 | 3.48E-01 | 0.1 | 4.59E-01 | 0.2 | 5.07E-01 | -0.1 | 8.75E-01 | 0.2 | 4.91E-01 | MMS19 nucleotide excision repair protein homolog |
| AT5G11560 | 0.0 | 9.27E-01 | 0.0 | 9.65E-01 | 0.0 | 9.72E-01 | 0.3 | 9.78E-02 | 0.2 | 2.82E-01 | 0.1 | 8.36E-01 | ER membrane protein complex subunit 1 |
| AT1G65280 | 0.0 | 9.27E-01 | 0.0 | 7.36E-01 | 0.0 | 8.31E-01 | -0.2 | 9.32E-02 | -0.2 | 1.17E-01 | 0.0 | 9.76E-01 | DNAJ heat shock N-terminal domain-containing protein |
| AT2G46550 | 0.0 | 9.60E-01 | 0.0 | 7.97E-01 | 0.0 | 8.55E-01 | -0.5 | 1.73E-03 | -0.4 | 5.85E-03 | 0.0 | 9.19E-01 | Transmembrane protein |
| AT2G24592 | 0.0 | 9.57E-01 | -0.2 | 1.64E-01 | 0.2 | 2.44E-01 | 0.1 | 7.97E-01 | 0.2 | 4.31E-01 | -0.2 | 7.04E-01 | NA |
| AT1G27670 | 0.0 | 9.76E-01 | 0.1 | 7.07E-01 | -0.1 | 7.42E-01 | 0.2 | 6.51E-01 | 0.1 | 7.58E-01 | 0.0 | 9.65E-01 | hypothetical protein |
| AT2G26460 | 0.0 | 9.29E-01 | -0.1 | 1.26E-01 | 0.1 | 1.93E-01 | 0.2 | 2.09E-01 | 0.1 | 4.86E-01 | 0.1 | 8.11E-01 | Suppressor of mec-8 and unc-52 protein homolog 2 |
| AT1G14300 | 0.0 | 9.65E-01 | 0.1 | 2.74E-01 | -0.1 | 3.28E-01 | 0.3 | 6.29E-02 | 0.1 | 8.01E-01 | 0.3 | 2.25E-01 | Protein HGH1 homolog |
| AT5G51510 | 0.0 | 9.48E-01 | -0.1 | 4.14E-01 | 0.1 | 5.08E-01 | -0.1 | 3.74E-01 | -0.2 | 1.76E-01 | 0.1 | 7.14E-01 | Jagunal-like protein |
| AT1G49040 | 0.0 | 9.27E-01 | 0.0 | 9.52E-01 | 0.0 | 9.83E-01 | 0.3 | 4.51E-02 | 0.1 | 5.22E-01 | 0.2 | 3.91E-01 | Stomatal cytokinesis defective |
| AT1G06470 | 0.0 | 9.47E-01 | 0.1 | 3.28E-01 | -0.1 | 3.54E-01 | 0.0 | 8.83E-01 | 0.2 | 2.55E-01 | -0.2 | 4.23E-01 | Sugar phosphate transporter domain-containing protein |
| AT5G06240 | 0.0 | 9.60E-01 | 0.0 | 9.60E-01 | 0.0 | 9.36E-01 | -0.2 | 3.11E-01 | -0.3 | 2.81E-01 | 0.0 | 9.12E-01 | Emb2735 |
| AT1G65390 | 0.0 | 9.80E-01 | 0.4 | 8.59E-02 | -0.4 | 1.23E-01 | 0.1 | 7.58E-01 | -0.8 | 4.69E-02 | 1.0 | 4.55E-02 | Phloem protein 2 A5 |
| AT3G21060 | 0.0 | 9.51E-01 | -0.1 | 3.06E-01 | 0.1 | 4.05E-01 | 0.0 | 8.49E-01 | 0.2 | 5.20E-01 | -0.1 | 7.28E-01 | Protein RBL |
| AT1G11220 | 0.0 | 9.78E-01 | 0.1 | 6.74E-01 | -0.1 | 7.12E-01 | -0.1 | 8.51E-01 | 0.3 | 4.95E-01 | -0.4 | 3.33E-01 | Cotton fiber, putative |
| AT4G14720 | 0.0 | 9.58E-01 | 0.0 | 6.94E-01 | 0.0 | 7.04E-01 | 0.1 | 6.28E-01 | -0.2 | 2.47E-01 | 0.2 | 1.82E-01 | Protein TIFY |
| AT5G14250 | 0.0 | 9.51E-01 | 0.0 | 9.82E-01 | 0.0 | 9.76E-01 | 0.4 | 1.13E-02 | 0.1 | 4.08E-01 | 0.2 | 2.11E-01 | Proteasome component domain protein |
| AT2G33560 | 0.0 | 9.75E-01 | 0.3 | 2.53E-02 | -0.3 | 4.90E-02 | 0.3 | 1.95E-01 | 0.3 | 2.57E-01 | 0.0 | 9.96E-01 | BUBR1 |
| AT1G08570 | 0.0 | 9.64E-01 | 0.1 | 3.40E-01 | -0.1 | 3.61E-01 | -0.4 | 7.45E-03 | -0.5 | 3.77E-03 | 0.1 | 6.58E-01 | Atypical CYS HIS rich thioredoxin 4 |
| AT1G69800 | 0.0 | 9.62E-01 | 0.0 | 7.48E-01 | 0.0 | 7.56E-01 | -0.4 | 6.30E-02 | -0.3 | 1.52E-01 | -0.1 | 8.68E-01 | Uncharacterized protein At1g69800; T17F3.17 |
| AT4G24900 | 0.0 | 9.68E-01 | -0.1 | 5.01E-01 | 0.1 | 5.95E-01 | -0.2 | 3.54E-01 | -0.1 | 7.67E-01 | -0.1 | 6.80E-01 | Coiled coil protein |
| AT5G51840 | 0.0 | 9.74E-01 | 0.2 | 1.35E-01 | -0.2 | 1.76E-01 | -0.1 | 5.63E-01 | 0.1 | 6.84E-01 | -0.2 | 3.81E-01 | RAB6-interacting golgin |
| AT1G76630 | 0.0 | 9.67E-01 | -0.3 | 7.45E-03 | 0.3 | 1.60E-02 | -0.1 | 6.24E-01 | 0.0 | 9.27E-01 | -0.1 | 8.02E-01 | Tetratricopeptide repeat -like superfamily protein |
| AT4G29240 | 0.0 | 9.80E-01 | 0.3 | 1.78E-02 | -0.3 | 3.77E-02 | 0.1 | 8.39E-01 | 0.2 | 7.18E-01 | -0.1 | 8.79E-01 | hypothetical protein |
| AT2G42730 | 0.0 | 9.85E-01 | -0.3 | 1.86E-01 | 0.3 | 2.63E-01 | 0.1 | 8.19E-01 | 0.6 | 2.81E-01 | -0.4 | 4.02E-01 | F-box |
| AT5G42790 | 0.0 | 9.75E-01 | 0.1 | 3.40E-01 | -0.1 | 3.59E-01 | 0.1 | 7.31E-01 | -0.1 | 8.26E-01 | 0.1 | 6.38E-01 | Proteasome subunit alpha type-1-A |
| AT3G22990 | 0.0 | 9.69E-01 | -0.1 | 1.74E-01 | 0.1 | 2.32E-01 | -0.1 | 4.78E-01 | 0.0 | 9.04E-01 | -0.1 | 7.39E-01 | Armadillo repeat-containing protein LFR |
| AT1G24706 | 0.0 | 9.63E-01 | -0.2 | 2.28E-02 | 0.2 | 3.80E-02 | 0.0 | 9.95E-01 | -0.1 | 6.94E-01 | 0.1 | 7.47E-01 | THO complex subunit 2 |
| AT3G07660 | 0.0 | 9.67E-01 | 0.1 | 3.31E-01 | -0.1 | 3.56E-01 | 0.1 | 7.52E-01 | 0.0 | 8.35E-01 | 0.1 | 6.73E-01 | hypothetical protein |
| AT2G18860 | 0.0 | 9.80E-01 | 0.0 | 9.15E-01 | 0.0 | 9.11E-01 | 0.0 | 9.60E-01 | -0.1 | 5.27E-01 | 0.1 | 5.85E-01 | At2g18860 |
| AT1G79260 | 0.0 | 9.86E-01 | 0.3 | 5.23E-02 | -0.3 | 8.73E-02 | 0.0 | 9.82E-01 | 0.1 | 7.23E-01 | -0.1 | 7.82E-01 | Nitrobindin heme-binding domain protein |
| AT3G17300 | 0.0 | 9.84E-01 | 0.2 | 5.23E-02 | -0.2 | 8.38E-02 | 0.2 | 5.11E-01 | 0.1 | 8.37E-01 | 0.1 | 8.27E-01 | COMPLEX 1 LYR-like protein |
| AT1G14990 | 0.0 | 9.87E-01 | 0.1 | 5.02E-01 | -0.1 | 5.62E-01 | 0.3 | 1.77E-01 | 0.2 | 2.89E-01 | 0.0 | 9.45E-01 | T15D22.3 |
| AT3G47850 | 0.0 | 9.82E-01 | -0.2 | 5.33E-02 | 0.2 | 8.74E-02 | 0.0 | 9.27E-01 | -0.2 | 2.66E-01 | 0.2 | 3.72E-01 | AT3g47850 |
| AT4G23530 | 0.0 | 9.77E-01 | 0.0 | 7.59E-01 | 0.0 | 8.16E-01 | -0.3 | 1.46E-01 | -0.3 | 1.94E-01 | 0.0 | 9.97E-01 | hypothetical protein |
| AT4G39550 | 0.0 | 9.87E-01 | 0.0 | 7.84E-01 | 0.0 | 8.14E-01 | 0.0 | 8.82E-01 | 0.0 | 9.11E-01 | 0.1 | 8.35E-01 | F-box |
| AT5G08660 | 0.0 | 9.81E-01 | -0.2 | 1.46E-02 | 0.2 | 2.66E-02 | 0.0 | 9.74E-01 | -0.2 | 2.42E-01 | 0.2 | 3.38E-01 | D-lactate dehydrogenase |
| AT2G26530 | 0.0 | 9.94E-01 | -0.1 | 5.70E-01 | 0.1 | 6.34E-01 | 0.2 | 4.35E-01 | -0.2 | 5.93E-01 | 0.4 | 1.95E-01 | AR781 |
| AT3G08960 | 0.0 | 9.84E-01 | 0.0 | 5.62E-01 | 0.0 | 6.24E-01 | 0.1 | 5.55E-01 | -0.1 | 8.22E-01 | 0.2 | 5.22E-01 | At3g08955 |
| AT2G26060 | 0.0 | 9.89E-01 | 0.0 | 7.04E-01 | 0.0 | 7.50E-01 | 0.0 | 9.47E-01 | -0.1 | 5.86E-01 | 0.1 | 7.27E-01 | Protein CIA1 |
| AT3G44530 | 0.0 | 9.93E-01 | -0.2 | 1.54E-01 | 0.2 | 2.15E-01 | 0.1 | 7.32E-01 | -0.1 | 8.17E-01 | 0.2 | 5.38E-01 | Protein HIRA |
| AT5G46630 | 0.0 | 9.89E-01 | 0.0 | 5.48E-01 | 0.0 | 5.90E-01 | 0.0 | 9.46E-01 | -0.1 | 4.80E-01 | 0.1 | 6.48E-01 | Clathrin adaptor complexes medium subunit family protein |
| AT5G61990 | 0.0 | 9.99E-01 | 0.1 | 2.85E-01 | -0.1 | 3.70E-01 | 0.2 | 5.01E-01 | 0.1 | 7.23E-01 | 0.1 | 8.85E-01 | Pentatricopeptide repeat-containing protein At5g61990, mitochondrial |
| AT2G22670 | 0.0 | 9.96E-01 | 0.2 | 1.97E-02 | -0.2 | 2.81E-02 | -0.3 | 4.45E-02 | -0.1 | 5.73E-01 | -0.2 | 4.00E-01 | Auxin-responsive protein |
| AT2G24530 | 0.0 | 9.96E-01 | -0.2 | 5.54E-02 | 0.2 | 9.49E-02 | 0.2 | 4.06E-01 | 0.0 | 1.00E+00 | 0.2 | 5.27E-01 | Uncharacterized protein At2g24530 |
| AT3G56750 | 0.0 | 9.93E-01 | 0.2 | 4.66E-02 | -0.2 | 7.84E-02 | 0.1 | 5.16E-01 | 0.1 | 7.21E-01 | 0.0 | 9.14E-01 | AT3g56750 |
| AT4G07990 | 0.0 | 9.92E-01 | 0.0 | 7.62E-01 | 0.0 | 7.88E-01 | 0.2 | 1.13E-01 | 0.2 | 2.13E-01 | 0.0 | 9.28E-01 | J domain-containing protein |
| AT3G55600 | 0.0 | 9.92E-01 | -0.1 | 2.60E-01 | 0.1 | 3.32E-01 | -0.3 | 1.04E-01 | -0.4 | 5.39E-02 | 0.1 | 7.61E-01 | Uncharacterized protein F1I16_10 |
| AT5G13970 | 0.0 | 9.88E-01 | -0.1 | 4.01E-01 | 0.1 | 4.61E-01 | -0.2 | 5.10E-01 | -0.1 | 6.60E-01 | 0.0 | 9.42E-01 | U5 small nuclear ribonucleoprotein TSSC4 |

| | | | | | | | | | | | | | |
|---|---|---|---|---|---|---|---|---|---|---|---|---|---|
| AT3G04470 | 0.0 | 9.80E-01 | 0.0 | 5.98E-01 | 0.0 | 6.18E-01 | -0.3 | 1.41E-01 | -0.3 | 1.47E-01 | 0.0 | 9.37E-01 | Uncharacterized protein At3g04470 |
| AT1G31300 | 0.0 | 9.82E-01 | 0.0 | 9.39E-01 | 0.0 | 9.61E-01 | -0.4 | 5.83E-03 | -0.2 | 1.92E-01 | -0.2 | 3.27E-01 | TLC domain-containing protein |
| AT1G76060 | 0.0 | 9.85E-01 | 0.0 | 7.30E-01 | 0.0 | 7.60E-01 | 0.2 | 4.86E-01 | 0.0 | 9.44E-01 | 0.1 | 6.74E-01 | T4O12.26 |
| AT5G09450 | 0.0 | 9.78E-01 | 0.0 | 9.48E-01 | 0.0 | 9.72E-01 | 0.2 | 1.64E-01 | 0.0 | 8.37E-01 | 0.2 | 4.44E-01 | Pentatricopeptide repeat-containing protein At5g09450, mitochondrial |
| AT1G63130 | 0.0 | 9.84E-01 | 0.0 | 8.62E-01 | 0.0 | 8.76E-01 | 0.0 | 8.99E-01 | 0.1 | 8.29E-01 | 0.0 | 9.29E-01 | Pentatricopeptide repeat-containing protein At1g63130, mitochondrial |
| AT5G57700 | 0.0 | 9.66E-01 | 0.2 | 4.18E-03 | -0.2 | 1.03E-02 | 0.0 | 8.78E-01 | 0.2 | 4.87E-01 | -0.1 | 6.72E-01 | BNR |
| AT1G78895 | 0.0 | 9.78E-01 | 0.0 | 7.15E-01 | 0.0 | 7.82E-01 | -0.3 | 1.38E-01 | -0.2 | 3.55E-01 | -0.1 | 8.18E-01 | Reticulon domain-containing protein |
| AT4G34630 | 0.0 | 9.72E-01 | -0.3 | 1.14E-02 | 0.3 | 1.78E-02 | 0.0 | 9.77E-01 | 0.3 | 1.50E-01 | -0.3 | 2.15E-01 | Uncharacterized protein |
| AT1G04260 | 0.0 | 9.73E-01 | -0.1 | 5.79E-01 | 0.1 | 6.10E-01 | -0.1 | 4.41E-01 | -0.3 | 2.09E-01 | 0.1 | 7.14E-01 | PRA1 family protein D |
| AT1G13380 | 0.0 | 9.64E-01 | 0.2 | 3.25E-02 | -0.2 | 5.75E-02 | -0.4 | 2.75E-03 | -0.5 | 3.98E-03 | 0.0 | 9.24E-01 | Uncharacterized protein |
| AT1G64840 | 0.0 | 9.59E-01 | -0.1 | 1.48E-01 | 0.1 | 1.87E-01 | 0.0 | 9.27E-01 | -0.1 | 5.89E-01 | 0.2 | 6.04E-01 | F-box |
| AT3G52480 | 0.0 | 9.78E-01 | 0.3 | 5.33E-02 | -0.3 | 9.87E-02 | -0.5 | 6.07E-02 | -0.9 | 5.07E-03 | 0.4 | 3.23E-01 | hypothetical protein |
| AT1G55130 | 0.0 | 9.56E-01 | 0.2 | 2.65E-02 | -0.2 | 5.25E-02 | 0.1 | 5.46E-01 | 0.1 | 8.71E-01 | 0.1 | 8.06E-01 | Transmembrane 9 superfamily member 6 |
| AT5G63320 | 0.0 | 9.67E-01 | -0.2 | 4.17E-02 | 0.2 | 5.96E-02 | 0.3 | 1.39E-01 | 0.3 | 2.28E-01 | 0.0 | 9.43E-01 | Transcription factor GTE10 |
| AT1G55840 | 0.0 | 9.56E-01 | 0.0 | 5.97E-01 | 0.0 | 5.96E-01 | -0.1 | 4.29E-01 | -0.1 | 4.96E-01 | 0.0 | 9.97E-01 | Sec14p-like phosphatidylinositol transfer family protein |
| AT3G42052 | 0.0 | 9.68E-01 | -0.1 | 5.59E-01 | 0.1 | 5.93E-01 | -0.1 | 6.74E-01 | 0.0 | 8.91E-01 | -0.2 | 6.42E-01 | NA |
| AT5G12170 | 0.0 | 9.56E-01 | -0.1 | 3.18E-01 | 0.1 | 3.43E-01 | 0.1 | 6.09E-01 | 0.1 | 6.45E-01 | 0.0 | 9.80E-01 | CRT -like transporter 3 |
| AT4G03000 | 0.0 | 9.51E-01 | -0.1 | 3.52E-01 | 0.1 | 3.78E-01 | -0.3 | 8.74E-02 | -0.1 | 4.35E-01 | -0.1 | 5.83E-01 | hypothetical protein |
| AT1G61100 | 0.0 | 9.65E-01 | 0.0 | 8.76E-01 | 0.0 | 8.59E-01 | -0.6 | 1.64E-03 | -0.6 | 3.02E-03 | 0.0 | 9.65E-01 | CSL1 ) |
| AT4G14145 | 0.0 | 9.66E-01 | 0.2 | 1.78E-01 | -0.2 | 2.61E-01 | -0.4 | 1.22E-02 | -0.4 | 4.61E-02 | -0.1 | 8.07E-01 | At4g14145 |
| AT5G54980 | 0.0 | 9.68E-01 | 0.2 | 2.05E-01 | -0.2 | 2.91E-01 | -0.1 | 4.22E-01 | -0.1 | 6.73E-01 | -0.1 | 8.76E-01 | CASP-like protein 2D1 |
| AT5G42360 | 0.0 | 9.50E-01 | -0.1 | 1.01E-01 | 0.1 | 1.39E-01 | -0.1 | 6.74E-01 | 0.0 | 9.32E-01 | -0.2 | 7.12E-01 | F-box |
| AT5G03160 | 0.0 | 9.56E-01 | 0.1 | 2.97E-01 | -0.1 | 3.75E-01 | 0.4 | 3.43E-02 | 0.5 | 4.30E-03 | -0.2 | 3.63E-01 | DnaJ protein P58IPK homolog |
| AT4G23410 | 0.0 | 9.73E-01 | 0.0 | 9.64E-01 | 0.0 | 9.50E-01 | 0.1 | 8.50E-01 | 0.0 | 9.38E-01 | 0.1 | 8.13E-01 | TET5 |
| AT4G18060 | 0.0 | 9.43E-01 | 0.1 | 1.75E-01 | -0.1 | 2.46E-01 | -0.1 | 5.61E-01 | 0.0 | 9.81E-01 | -0.1 | 6.15E-01 | SH3 domain-containing protein 3 |
| AT2G33850 | 0.0 | 9.75E-01 | 0.4 | 3.48E-02 | -0.4 | 5.37E-02 | 0.1 | 7.53E-01 | 0.2 | 3.92E-01 | -0.2 | 6.43E-01 | hypothetical protein |
| AT4G05460 | 0.0 | 9.41E-01 | -0.1 | 2.78E-01 | 0.1 | 3.15E-01 | -0.2 | 3.26E-01 | -0.1 | 5.02E-01 | 0.0 | 9.11E-01 | F-box domain-containing protein |
| AT5G53570 | 0.0 | 9.30E-01 | -0.2 | 4.05E-02 | 0.2 | 5.23E-02 | 0.1 | 3.80E-01 | -0.1 | 5.59E-01 | 0.2 | 1.90E-01 | Ypt |
| AT3G51640 | 0.0 | 9.44E-01 | 0.0 | 7.37E-01 | 0.0 | 7.25E-01 | 0.2 | 3.53E-01 | 0.1 | 7.51E-01 | 0.1 | 7.32E-01 | Uncharacterized protein T18N14.20 |
| AT5G64320 | 0.0 | 9.58E-01 | 0.0 | 8.09E-01 | 0.0 | 8.11E-01 | 0.0 | 9.68E-01 | 0.1 | 8.25E-01 | -0.1 | 8.76E-01 | Pentatricopeptide repeat-containing protein At5g64320, mitochondrial |
| AT2G20590 | 0.0 | 9.69E-01 | 0.5 | 1.53E-02 | -0.4 | 3.83E-02 | 0.2 | 6.55E-01 | 0.4 | 3.55E-01 | -0.2 | 6.32E-01 | Reticulon-like protein |
| AT3G58530 | 0.0 | 9.55E-01 | 0.0 | 9.42E-01 | 0.0 | 9.86E-01 | -0.1 | 4.34E-01 | -0.3 | 1.53E-01 | 0.1 | 5.93E-01 | F-box protein At3g58530 |
| AT2G18735 | 0.0 | 9.53E-01 | -0.1 | 6.67E-01 | 0.1 | 6.84E-01 | -0.1 | 7.61E-01 | -0.2 | 4.88E-01 | 0.1 | 7.68E-01 | NA |
| AT5G63220 | 0.0 | 9.34E-01 | 0.1 | 1.16E-01 | -0.1 | 1.86E-01 | 0.3 | 1.04E-01 | 0.2 | 3.91E-01 | 0.1 | 6.30E-01 | Golgi-to-ER traffic-like protein |
| AT4G32470 | 0.0 | 9.51E-01 | 0.2 | 6.67E-02 | -0.2 | 9.35E-02 | 0.1 | 6.82E-01 | 0.1 | 6.37E-01 | 0.0 | 9.31E-01 | Cytochrome b-c1 complex subunit 7 |
| AT4G37450 | 0.0 | 9.51E-01 | 0.1 | 2.60E-01 | -0.1 | 3.38E-01 | -0.1 | 5.35E-01 | 0.1 | 5.23E-01 | -0.3 | 2.57E-01 | Arabinogalactan protein 18 |
| AT4G11410 | 0.0 | 9.48E-01 | -0.1 | 2.51E-01 | 0.1 | 2.88E-01 | 0.0 | 9.55E-01 | 0.2 | 2.02E-01 | -0.2 | 3.19E-01 | Uncharacterized protein At4g11410 |
| AT1G57620 | 0.0 | 9.33E-01 | 0.2 | 1.58E-01 | -0.1 | 2.40E-01 | 0.0 | 9.01E-01 | 0.0 | 8.54E-01 | 0.0 | 9.50E-01 | GOLD domain-containing protein |
| AT1G16630 | 0.0 | 9.45E-01 | 0.4 | 1.26E-02 | -0.4 | 3.08E-02 | 0.3 | 2.18E-01 | 0.3 | 2.41E-01 | 0.0 | 9.64E-01 | At1g16630 |
| AT5G03460 | 0.0 | 9.05E-01 | 0.1 | 1.56E-01 | -0.1 | 2.28E-01 | 0.0 | 8.96E-01 | -0.2 | 5.56E-01 | 0.1 | 7.27E-01 | Transmembrane protein |
| AT4G17420 | 0.0 | 8.95E-01 | -0.1 | 3.93E-01 | 0.1 | 3.92E-01 | -0.2 | 3.10E-01 | 0.1 | 7.37E-01 | -0.3 | 2.53E-01 | Altered inheritance of mitochondria protein 24, mitochondrial |
| AT4G28050 | 0.0 | 9.24E-01 | 0.4 | 6.17E-03 | -0.4 | 1.52E-02 | -0.1 | 6.48E-01 | 0.0 | 9.40E-01 | -0.1 | 6.86E-01 | Tetraspanin-7 |
| AT2G40800 | 0.0 | 9.00E-01 | 0.1 | 1.77E-01 | -0.1 | 2.69E-01 | 0.0 | 9.26E-01 | 0.2 | 3.49E-01 | -0.2 | 3.27E-01 | Import inner membrane translocase subunit |
| AT1G50020 | 0.0 | 9.20E-01 | -0.1 | 2.21E-01 | 0.2 | 2.31E-01 | 0.0 | 9.22E-01 | 0.3 | 2.60E-02 | -0.4 | 5.55E-02 | F2J10.10 protein |
| AT5G17610 | 0.0 | 8.96E-01 | 0.1 | 1.23E-01 | -0.1 | 2.14E-01 | -0.1 | 8.31E-01 | -0.1 | 7.00E-01 | 0.1 | 8.92E-01 | hypothetical protein |
| AT3G01345 | 0.0 | 8.81E-01 | 0.1 | 4.71E-01 | 0.0 | 6.00E-01 | 0.2 | 1.72E-01 | 0.0 | 8.71E-01 | 0.1 | 4.31E-01 | Expressed protein |
| AT4G08035 | 0.0 | 9.24E-01 | -0.1 | 3.85E-01 | 0.1 | 4.00E-01 | -0.1 | 7.14E-01 | 0.0 | 9.87E-01 | -0.1 | 7.99E-01 | NA |
| AT4G28060 | 0.0 | 8.91E-01 | 0.2 | 1.40E-02 | -0.2 | 2.74E-02 | -0.1 | 8.18E-01 | 0.1 | 7.98E-01 | -0.1 | 7.05E-01 | Cytochrome c oxidase subunit 6b-3 |
| AT2G31560 | 0.0 | 9.52E-01 | -0.1 | 6.32E-01 | 0.1 | 6.42E-01 | -0.7 | 3.09E-03 | 0.1 | 8.28E-01 | -0.8 | 1.10E-02 | hypothetical protein |
| AT5G01980 | 0.0 | 8.85E-01 | -0.2 | 8.57E-02 | 0.2 | 9.65E-02 | -0.4 | 6.95E-03 | -0.5 | 4.82E-03 | 0.1 | 7.65E-01 | hypothetical protein |
| AT5G38590 | 0.0 | 9.13E-01 | 0.0 | 7.97E-01 | 0.0 | 7.61E-01 | -0.5 | 8.49E-02 | -0.1 | 8.23E-01 | -0.4 | 3.06E-01 | F-box |
| AT1G73720 | 0.0 | 8.74E-01 | 0.0 | 7.98E-01 | 0.0 | 9.33E-01 | -0.1 | 4.27E-01 | -0.1 | 5.45E-01 | 0.0 | 9.72E-01 | Suppressor of mec-8 and unc-52 protein homolog 1 |
| AT5G66200 | 0.0 | 8.99E-01 | -0.2 | 7.04E-02 | 0.2 | 7.29E-02 | -0.1 | 7.60E-01 | -0.2 | 3.69E-01 | 0.1 | 6.62E-01 | Armadillo repeat only 2 |
| AT2G44390 | 0.0 | 9.43E-01 | 0.6 | 5.09E-03 | -0.5 | 1.71E-02 | -0.3 | 3.54E-01 | 0.6 | 4.72E-02 | -1.0 | 2.27E-02 | Cysteine |
| AT2G04740 | 0.0 | 8.69E-01 | -0.2 | 4.01E-02 | 0.2 | 5.39E-02 | 0.1 | 6.55E-01 | -0.2 | 4.38E-01 | 0.3 | 1.77E-01 | Ankyrin repeat family protein |
| AT5G22080 | 0.0 | 8.53E-01 | 0.0 | 9.25E-01 | 0.0 | 9.46E-01 | 0.0 | 8.69E-01 | -0.1 | 4.79E-01 | 0.1 | 6.96E-01 | Chaperone DnaJ-domain superfamily protein |
| AT4G40065 | 0.0 | 9.16E-01 | 0.1 | 4.22E-01 | -0.1 | 5.66E-01 | -0.2 | 7.06E-01 | -0.5 | 2.51E-01 | 0.3 | 5.57E-01 | NA |
| AT2G20230 | 0.0 | 8.64E-01 | 0.2 | 3.54E-02 | -0.2 | 6.31E-02 | -0.3 | 2.58E-03 | -0.3 | 1.04E-02 | 0.0 | 8.82E-01 | Tetraspanin-18 |
| AT1G78100 | 0.0 | 9.08E-01 | -0.1 | 6.11E-01 | 0.1 | 5.65E-01 | -0.3 | 2.42E-02 | 0.1 | 5.79E-01 | -0.4 | 2.13E-02 | F-box protein AUF1 |
| AT3G16910 | 0.0 | 8.43E-01 | 0.0 | 7.15E-01 | 0.0 | 6.05E-01 | 0.1 | 5.57E-01 | 0.1 | 4.80E-01 | 0.0 | 8.99E-01 | ACN1 |
| AT5G18250 | 0.0 | 9.24E-01 | -0.1 | 6.06E-01 | 0.1 | 6.10E-01 | 0.6 | 5.23E-02 | 0.4 | 2.24E-01 | 0.2 | 6.54E-01 | Transmembrane protein |
| AT5G21040 | 0.0 | 8.47E-01 | -0.1 | 3.97E-01 | 0.1 | 3.51E-01 | -0.1 | 2.67E-01 | -0.1 | 7.49E-01 | -0.1 | 6.84E-01 | F-box |
| AT4G26550 | 0.0 | 9.07E-01 | 0.2 | 6.81E-02 | -0.2 | 1.30E-01 | -0.1 | 6.62E-01 | -0.3 | 1.54E-01 | 0.2 | 3.74E-01 | Vesicle transport protein |
| AT1G63300 | 0.0 | 8.83E-01 | 0.3 | 9.33E-03 | -0.3 | 2.64E-02 | -0.2 | 3.56E-01 | 0.0 | 9.83E-01 | -0.2 | 5.04E-01 | Myosin heavy chain-related protein |
| AT3G26780 | 0.0 | 8.46E-01 | 0.1 | 9.66E-02 | -0.1 | 1.70E-01 | 0.1 | 5.28E-01 | 0.1 | 3.79E-01 | 0.0 | 8.41E-01 | Uncharacterized protein At3g26780 |
| AT3G44205 | 0.0 | 8.93E-01 | -0.3 | 1.64E-02 | 0.4 | 2.90E-02 | | | | | | | NA |
| AT3G55640 | 0.0 | 8.65E-01 | -0.2 | 8.85E-02 | 0.2 | 1.08E-01 | 0.0 | 9.27E-01 | 0.0 | 9.66E-01 | 0.0 | 9.85E-01 | Ca-dependent solute carrier-like protein |
| AT3G48030 | 0.0 | 8.96E-01 | 0.3 | 1.58E-02 | -0.3 | 3.37E-02 | -0.2 | 4.11E-01 | -0.1 | 6.20E-01 | 0.0 | 9.11E-01 | Hypoxia-responsive family protein |
| AT2G23470 | 0.0 | 8.94E-01 | -0.3 | 2.00E-02 | 0.4 | 3.42E-02 | 0.4 | 4.02E-01 | 0.2 | 7.52E-01 | | | RUS4 |
| AT1G56000 | 0.0 | 9.09E-01 | 0.1 | 5.66E-01 | -0.1 | 7.06E-01 | 0.1 | 5.01E-01 | -0.1 | 7.10E-01 | 0.3 | 3.68E-01 | AT1G56000 protein -binding oxidoreductase family protein) |
| AT1G14330 | 0.0 | 8.83E-01 | 0.1 | 3.04E-01 | -0.1 | 4.45E-01 | -0.3 | 2.53E-02 | -0.3 | 7.46E-02 | 0.0 | 8.63E-01 | hypothetical protein |
| AT5G26960 | 0.0 | 8.61E-01 | 0.0 | 6.83E-01 | 0.0 | 8.41E-01 | -0.3 | 1.86E-02 | -0.2 | 2.37E-01 | -0.1 | 4.57E-01 | F-box |
| AT5G02130 | 0.0 | 8.64E-01 | -0.2 | 1.05E-01 | 0.2 | 1.25E-01 | 0.3 | 2.90E-01 | 0.1 | 8.09E-01 | 0.2 | 6.30E-01 | Tetratricopeptide repeat -like superfamily protein |
| AT4G02995 | 0.0 | 8.44E-01 | 0.0 | 8.33E-01 | 0.0 | 7.32E-01 | 0.0 | 9.81E-01 | -0.1 | 7.09E-01 | 0.1 | 7.80E-01 | NA |
| AT2G34680 | 0.0 | 8.49E-01 | 0.0 | 8.97E-01 | 0.0 | 9.67E-01 | -0.3 | 1.20E-01 | -0.1 | 6.04E-01 | -0.2 | 5.54E-01 | Outer arm dynein light chain 1 protein |
| AT3G04710 | 0.0 | 8.68E-01 | 0.0 | 7.68E-01 | 0.0 | 6.94E-01 | 0.5 | 1.17E-02 | 0.4 | 5.07E-02 | 0.1 | 7.56E-01 | Ankyrin repeat family protein |
| AT5G11980 | 0.0 | 8.02E-01 | 0.1 | 9.65E-02 | -0.1 | 1.92E-01 | 0.0 | 8.94E-01 | 0.1 | 6.25E-01 | -0.1 | 7.94E-01 | Conserved oligomeric Golgi complex subunit 8 |
| AT2G47620 | 0.0 | 8.24E-01 | 0.1 | 1.23E-01 | -0.1 | 2.45E-01 | 0.0 | 8.95E-01 | -0.1 | 6.09E-01 | 0.1 | 7.51E-01 | SWITCH |
| AT5G24105 | 0.0 | 9.08E-01 | 0.7 | 6.42E-04 | -0.7 | 2.76E-03 | 0.0 | 9.39E-01 | 0.0 | 9.73E-01 | 0.0 | 9.83E-01 | Arabinogalactan protein 41 |
| AT5G20660 | 0.0 | 7.85E-01 | 0.0 | 6.95E-01 | 0.0 | 9.12E-01 | -0.1 | 6.33E-01 | 0.0 | 9.73E-01 | -0.1 | 7.30E-01 | Uncharacterized protein |
| AT1G23880 | 0.0 | 8.88E-01 | 0.0 | 8.07E-01 | 0.0 | 9.32E-01 | -0.1 | 7.90E-01 | 0.0 | 9.67E-01 | -0.1 | 8.07E-01 | At1g23880 |
| AT4G30940 | 0.0 | 8.93E-01 | 0.0 | 7.39E-01 | 0.0 | 8.69E-01 | -0.1 | 8.21E-01 | 0.0 | 9.88E-01 | -0.1 | 8.45E-01 | BTB |
| AT4G12500 | 0.0 | 9.83E-01 | -0.3 | 7.21E-01 | 0.4 | 7.45E-01 | -1.2 | 1.56E-01 | -3.5 | 2.29E-02 | 2.3 | 1.47E-01 | pEARLI1-like lipid transfer protein 3 |

| | | | | | | | | | | | | | |
|---|---|---|---|---|---|---|---|---|---|---|---|---|---|
| AT1G71950 | 0.0 | 8.50E-01 | -0.2 | 1.32E-01 | 0.2 | 1.33E-01 | 0.0 | 9.34E-01 | -0.1 | 5.00E-01 | 0.1 | 5.24E-01 | Inhibitor I9 domain-containing protein |
| AT1G49540 | 0.0 | 8.29E-01 | -0.1 | 4.33E-01 | 0.1 | 3.95E-01 | 0.3 | 3.24E-01 | -0.2 | 5.76E-01 | 0.5 | 1.67E-01 | Elongator complex protein 2 |
| AT2G19270 | 0.0 | 8.20E-01 | 0.1 | 4.46E-01 | 0.0 | 6.43E-01 | 0.0 | 8.73E-01 | -0.2 | 1.89E-01 | 0.1 | 3.93E-01 | Uncharacterized protein At2g19270 |
| AT2G01680 | 0.0 | 8.19E-01 | -0.1 | 1.43E-01 | 0.2 | 1.24E-01 | -0.1 | 4.52E-01 | 0.0 | 8.32E-01 | -0.2 | 4.59E-01 | Ankyrin repeat-containing protein At2g01680 |
| AT3G18560 | 0.0 | 8.71E-01 | 0.1 | 2.78E-01 | -0.1 | 4.16E-01 | 0.3 | 2.33E-01 | -0.3 | 3.17E-01 | 0.6 | 3.39E-02 | Transmembrane protein |
| AT2G18410 | 0.0 | 8.44E-01 | 0.0 | 6.86E-01 | 0.1 | 6.12E-01 | 0.1 | 4.09E-01 | -0.1 | 8.41E-01 | 0.2 | 3.79E-01 | Elongator complex protein 5 |
| AT5G22790 | 0.0 | 8.67E-01 | 0.2 | 1.32E-01 | -0.2 | 2.39E-01 | 0.3 | 1.24E-01 | 0.1 | 7.87E-01 | 0.2 | 4.16E-01 | Protein RETICULATA-RELATED 1, chloroplastic |
| AT4G33180 | 0.0 | 7.93E-01 | -0.1 | 1.99E-01 | 0.1 | 1.76E-01 | 0.0 | 9.92E-01 | 0.0 | 9.06E-01 | 0.0 | 9.27E-01 | Uncharacterized protein AT4g33190 |
| AT2G13690 | 0.0 | 7.87E-01 | 0.1 | 4.70E-01 | 0.0 | 7.06E-01 | 0.0 | 8.97E-01 | 0.0 | 9.52E-01 | 0.0 | 9.76E-01 | BTB |
| AT2G20410 | 0.0 | 8.51E-01 | -0.1 | 2.12E-01 | 0.2 | 2.17E-01 | 0.2 | 4.99E-01 | 0.2 | 3.60E-01 | -0.1 | 8.45E-01 | Expressed protein |
| AT3G53440 | 0.0 | 8.87E-01 | 0.1 | 4.91E-01 | -0.1 | 6.56E-01 | -0.2 | 5.49E-01 | 0.0 | 9.74E-01 | -0.2 | 6.47E-01 | hypothetical protein |
| AT2G38580 | 0.0 | 8.26E-01 | 0.1 | 5.95E-01 | 0.0 | 7.94E-01 | 0.2 | 3.06E-01 | 0.3 | 6.70E-02 | -0.2 | 4.66E-01 | Mitochondrial ATP synthase D chain-related protein |
| AT5G14830 | 0.0 | 8.72E-01 | 0.2 | 1.04E-01 | -0.2 | 2.12E-01 | 0.0 | 8.98E-01 | 0.1 | 8.26E-01 | -0.1 | 7.41E-01 | NA |
| AT2G11880 | 0.0 | 8.61E-01 | -0.1 | 2.65E-01 | 0.2 | 2.76E-01 | 0.1 | 7.44E-01 | 0.2 | 6.71E-01 | -0.1 | 9.22E-01 | NA |
| AT1G09330 | 0.0 | 7.56E-01 | 0.2 | 4.03E-02 | -0.1 | 9.97E-02 | 0.1 | 7.41E-01 | 0.0 | 8.66E-01 | 0.0 | 9.42E-01 | Golgi apparatus membrane protein TVP23 |
| AT1G78170 | 0.0 | 9.30E-01 | 0.1 | 5.67E-01 | -0.1 | 6.83E-01 | -0.2 | 4.19E-01 | -0.4 | 9.55E-02 | 0.2 | 4.60E-01 | Uncharacterized protein |
| AT4G15010 | 0.0 | 8.15E-01 | -0.3 | 6.53E-03 | 0.4 | 9.77E-03 | 0.0 | 8.87E-01 | -0.1 | 7.04E-01 | 0.1 | 8.47E-01 | AT4G15010 protein |
| AT1G11420 | 0.0 | 8.74E-01 | 0.0 | 8.43E-01 | 0.1 | 7.73E-01 | 0.5 | 2.87E-01 | 0.3 | 6.19E-01 | 0.2 | 7.21E-01 | DUF724 domain-containing protein 2 |
| AT5G07910 | 0.0 | 7.28E-01 | 0.0 | 9.32E-01 | 0.0 | 8.42E-01 | 0.0 | 9.68E-01 | -0.1 | 7.50E-01 | 0.1 | 7.55E-01 | Uncharacterized protein F13G24.110 |
| AT5G57100 | 0.0 | 7.84E-01 | 0.1 | 1.37E-01 | -0.1 | 2.91E-01 | -0.2 | 2.24E-01 | -0.1 | 5.17E-01 | -0.1 | 7.55E-01 | Nucleotide |
| AT1G26920 | 0.0 | 7.69E-01 | 0.3 | 2.49E-03 | -0.3 | 7.66E-03 | -0.8 | 5.67E-04 | -0.7 | 3.36E-03 | -0.1 | 7.79E-01 | hypothetical protein |
| AT4G04614 | 0.0 | 7.83E-01 | 0.1 | 2.80E-01 | -0.1 | 4.85E-01 | -0.2 | 1.35E-01 | -0.2 | 2.06E-01 | 0.0 | 9.74E-01 | hypothetical protein |
| AT2G01990 | 0.0 | 8.81E-01 | 0.0 | 8.48E-01 | 0.0 | 9.67E-01 | -0.1 | 7.87E-01 | 0.1 | 8.20E-01 | -0.2 | 6.29E-01 | Uncharacterized protein At2g01990 |
| AT5G18540 | 0.0 | 8.48E-01 | 0.1 | 5.59E-01 | -0.1 | 7.51E-01 | -0.2 | 5.57E-01 | -0.3 | 5.50E-01 | 0.0 | 9.52E-01 | E3 ubiquitin-protein ligase |
| AT1G64370 | 0.0 | 8.18E-01 | 0.1 | 6.48E-01 | 0.0 | 8.36E-01 | 0.6 | 3.57E-04 | 0.5 | 4.07E-03 | 0.1 | 6.76E-01 | Filaggrin-2-like |
| AT3G01770 | 0.0 | 7.81E-01 | 0.0 | 6.44E-01 | 0.1 | 5.08E-01 | 0.0 | 9.34E-01 | -0.1 | 4.31E-01 | 0.1 | 5.81E-01 | Bromodomain and extraterminal domain protein 10 |
| AT5G65430 | 0.0 | 7.65E-01 | 0.2 | 6.38E-02 | -0.1 | 1.30E-01 | -0.2 | 3.08E-01 | -0.2 | 2.06E-01 | 0.1 | 8.21E-01 | General regulatory factor 8 |
| AT3G05700 | 0.0 | 7.86E-01 | -0.1 | 2.86E-01 | 0.1 | 2.52E-01 | -0.1 | 6.71E-01 | 0.3 | 2.50E-01 | -0.4 | 1.50E-01 | Drought-responsive family protein |
| AT5G44170 | 0.0 | 8.28E-01 | 0.0 | 9.68E-01 | 0.0 | 8.96E-01 | 0.0 | 9.94E-01 | -0.2 | 5.61E-01 | 0.2 | 6.49E-01 | S-adenosyl-L-methionine-dependent methyltransferases superfamily protein |
| AT5G46730 | 0.0 | 8.99E-01 | -0.5 | 2.25E-02 | 0.6 | 3.17E-02 | -0.5 | 4.14E-03 | -0.3 | 1.04E-01 | -0.2 | 3.66E-01 | Glycine-rich protein |
| AT1G75710 | 0.0 | 7.89E-01 | 0.0 | 7.49E-01 | 0.0 | 9.57E-01 | -0.3 | 3.77E-02 | 0.0 | 8.97E-01 | -0.3 | 9.61E-02 | C2H2-type domain-containing protein |
| AT4G37440 | 0.0 | 7.51E-01 | -0.2 | 2.83E-02 | 0.2 | 2.74E-02 | -0.2 | 1.43E-01 | -0.3 | 3.95E-02 | 0.1 | 6.02E-01 | Uncharacterized protein |
| AT5G50315 | 0.0 | 8.71E-01 | -0.1 | 6.53E-01 | 0.1 | 6.16E-01 | 0.8 | 7.31E-02 | 0.4 | 4.37E-01 | 0.4 | 4.98E-01 | NA |
| AT1G66900 | 0.0 | 7.24E-01 | 0.0 | 7.64E-01 | 0.0 | 9.78E-01 | -0.3 | 2.61E-02 | 0.0 | 9.70E-01 | -0.3 | 1.01E-01 | hypothetical protein |
| AT1G08300 | 0.0 | 8.56E-01 | -0.1 | 4.80E-01 | 0.1 | 4.61E-01 | 0.1 | 7.61E-01 | -0.2 | 7.17E-01 | 0.4 | 4.80E-01 | No vein-like protein |
| AT5G15880 | 0.0 | 7.72E-01 | 0.0 | 7.14E-01 | 0.0 | 9.43E-01 | -0.2 | 1.65E-01 | -0.2 | 3.17E-01 | 0.0 | 8.99E-01 | At5g15880 |
| AT5G58640 | 0.0 | 7.75E-01 | 0.1 | 4.35E-01 | 0.0 | 6.68E-01 | 0.1 | 6.16E-01 | 0.0 | 8.14E-01 | 0.1 | 5.39E-01 | Similarity to unknown protein |
| AT2G45920 | 0.0 | 7.69E-01 | -0.8 | 4.37E-05 | 0.8 | 1.45E-04 | 1.0 | 9.43E-06 | 1.0 | 8.26E-05 | 0.1 | 7.59E-01 | U-box domain-containing protein |
| AT1G53040 | 0.0 | 7.63E-01 | 0.1 | 1.51E-01 | -0.1 | 3.06E-01 | 0.1 | 3.53E-01 | 0.2 | 1.89E-01 | -0.1 | 7.41E-01 | tRNA cytidine acetyltransferase, |
| AT3G11290 | 0.0 | 7.71E-01 | 0.0 | 6.66E-01 | 0.1 | 5.30E-01 | -0.3 | 5.01E-02 | -0.5 | 4.50E-03 | 0.2 | 2.94E-01 | At3g11290 |
| AT3G16190 | 0.0 | 8.64E-01 | -0.3 | 1.26E-01 | 0.3 | 1.35E-01 | -0.5 | 2.73E-02 | -0.2 | 3.33E-01 | -0.2 | 4.44E-01 | Probable inactive nicotinamidase At3g16190 |
| AT4G01570 | 0.0 | 7.23E-01 | 0.0 | 6.04E-01 | 0.1 | 4.57E-01 | 0.0 | 9.79E-01 | -0.1 | 7.52E-01 | 0.1 | 7.84E-01 | Pentatricopeptide repeat-containing protein At4g01570 |
| AT5G03050 | 0.0 | 8.73E-01 | -0.3 | 2.12E-01 | 0.3 | 2.30E-01 | -0.3 | 1.98E-01 | -0.4 | 1.14E-01 | 0.1 | 7.93E-01 | At5g03050 |
| AT1G14580 | 0.0 | 7.30E-01 | 0.1 | 3.53E-01 | -0.1 | 6.04E-01 | 0.1 | 6.58E-01 | 0.1 | 6.38E-01 | 0.0 | 9.54E-01 | C2H2-like zinc finger protein |
| AT2G29020 | 0.0 | 7.05E-01 | 0.2 | 1.62E-02 | -0.2 | 4.72E-02 | -0.2 | 1.35E-01 | -0.1 | 5.38E-01 | -0.1 | 6.46E-01 | hypothetical protein |
| AT4G24050 | 0.0 | 7.97E-01 | 0.2 | 7.61E-02 | -0.2 | 1.79E-01 | -0.6 | 8.09E-03 | -0.5 | 3.79E-02 | -0.1 | 7.24E-01 | AT4g24050 |
| AT3G52870 | 0.0 | 7.61E-01 | 0.0 | 6.83E-01 | 0.0 | 9.25E-01 | 0.2 | 7.46E-02 | -0.1 | 2.78E-01 | 0.4 | 3.39E-02 | IQ domain-containing protein IQM3 |
| AT3G02180 | 0.0 | 6.95E-01 | 0.3 | 6.27E-03 | -0.2 | 1.90E-02 | -0.2 | 8.84E-02 | -0.3 | 5.90E-02 | 0.1 | 8.20E-01 | SPIRAL1-like3 |
| AT5G59560 | 0.0 | 7.07E-01 | 0.1 | 4.82E-01 | 0.0 | 7.84E-01 | 0.1 | 7.69E-01 | 0.1 | 6.58E-01 | 0.0 | 8.85E-01 | Protein SENSITIVITY TO RED LIGHT REDUCED 1 |
| AT3G54830 | 0.0 | 8.25E-01 | 0.2 | 1.83E-01 | -0.2 | 3.43E-01 | -1.2 | 1.28E-04 | -1.2 | 2.22E-04 | 0.0 | 9.26E-01 | Amino acid transporter transmembrane domain-containing protein |
| AT5G39450 | 0.0 | 7.19E-01 | -0.2 | 2.45E-02 | 0.3 | 2.72E-02 | 0.2 | 3.32E-01 | 0.2 | 4.43E-01 | 0.0 | 9.67E-01 | hypothetical protein |
| AT1G47270 | 0.0 | 6.41E-01 | 0.0 | 6.69E-01 | 0.1 | 4.41E-01 | -0.4 | 7.88E-03 | -0.2 | 2.37E-01 | -0.2 | 3.16E-01 | Tubby like protein 6 |
| AT5G19151 | 0.0 | 8.75E-01 | 0.0 | 9.67E-01 | 0.0 | 9.36E-01 | -0.2 | 6.17E-01 | 0.1 | 8.02E-01 | -0.3 | 4.95E-01 | Transmembrane protein |
| AT3G02600 | 0.0 | 6.21E-01 | 0.2 | 3.64E-02 | -0.1 | 1.15E-01 | 0.0 | 8.86E-01 | -0.2 | 3.18E-01 | 0.2 | 3.48E-01 | Lipid phosphate phosphatase 3 |
| AT4G32920 | 0.0 | 6.45E-01 | -0.2 | 1.08E-02 | 0.3 | 1.14E-02 | 0.4 | 7.53E-02 | 0.4 | 1.22E-01 | 0.0 | 9.71E-01 | AT4g32920 |
| AT4G21450 | 0.0 | 5.53E-01 | -0.1 | 1.57E-01 | 0.1 | 7.47E-02 | -0.2 | 4.57E-02 | -0.3 | 1.62E-02 | 0.1 | 6.67E-01 | PapD-like superfamily protein |
| AT3G14750 | 0.0 | 7.31E-01 | 0.0 | 9.50E-01 | 0.0 | 7.46E-01 | -0.2 | 3.19E-01 | -0.3 | 2.54E-01 | 0.1 | 8.79E-01 | Protein FLX-like 1 |
| AT5G41020 | 0.0 | 7.24E-01 | -0.1 | 3.98E-01 | 0.1 | 2.94E-01 | 0.2 | 1.98E-01 | 0.0 | 9.58E-01 | 0.2 | 3.66E-01 | Gb\|AAC33480.1 |
| AT1G77110 | 0.0 | 8.48E-01 | 0.2 | 2.02E-01 | -0.2 | 3.63E-01 | 0.8 | 8.13E-02 | 0.4 | 4.73E-01 | 0.4 | 5.05E-01 | Auxin efflux carrier component |
| AT3G60500 | 0.0 | 6.98E-01 | 0.1 | 2.26E-01 | -0.1 | 4.66E-01 | 0.1 | 3.77E-01 | 0.0 | 9.70E-01 | 0.2 | 5.04E-01 | 3'-5'-exoribonuclease family protein |
| AT1G64520 | 0.0 | 7.43E-01 | 0.2 | 9.01E-02 | -0.2 | 1.85E-01 | 0.0 | 9.37E-01 | 0.0 | 9.66E-01 | 0.0 | 9.32E-01 | 26S proteasome non-ATPase regulatory subunit 8 homolog A |
| AT5G21090 | 0.0 | 7.02E-01 | 0.0 | 8.13E-01 | 0.1 | 5.74E-01 | 0.2 | 1.28E-01 | 0.0 | 8.38E-01 | 0.3 | 1.95E-01 | Leucine-rich repeat protein 1 |
| AT4G18593 | 0.0 | 7.53E-01 | -0.1 | 5.46E-01 | 0.1 | 4.30E-01 | 0.1 | 8.65E-01 | 0.0 | 9.61E-01 | 0.0 | 9.48E-01 | Probable inactive dual specificity protein phosphatase-like At4g18593 |
| AT1G28380 | 0.0 | 6.33E-01 | -0.4 | 2.14E-04 | 0.5 | 3.21E-04 | 0.0 | 9.10E-01 | -0.2 | 3.52E-01 | 0.2 | 5.13E-01 | MACPF domain-containing protein NSL1 |
| AT1G69460 | 0.0 | 5.33E-01 | 0.2 | 1.89E-02 | -0.1 | 7.46E-02 | 0.0 | 8.41E-01 | 0.0 | 9.65E-01 | 0.0 | 9.34E-01 | AT1G69460 protein |
| AT5G03420 | 0.0 | 6.60E-01 | -0.2 | 5.10E-02 | 0.2 | 4.05E-02 | -0.1 | 6.40E-01 | 0.0 | 9.41E-01 | -0.1 | 8.00E-01 | Protein PTST homolog 3, chloroplastic |
| AT5G18230 | 0.0 | 6.08E-01 | -0.1 | 2.72E-01 | 0.1 | 1.44E-01 | 0.0 | 8.98E-01 | 0.0 | 9.32E-01 | -0.1 | 8.72E-01 | Transcription regulator NOT2 |
| AT3G05430 | 0.0 | 7.72E-01 | 0.1 | 3.24E-01 | -0.1 | 5.64E-01 | 0.2 | 6.06E-01 | 0.1 | 8.98E-01 | 0.1 | 8.26E-01 | Tudor |
| AT2G32100 | 0.0 | 7.91E-01 | 0.0 | 9.80E-01 | 0.0 | 8.10E-01 | -0.3 | 1.77E-01 | -0.9 | 2.33E-03 | 0.6 | 5.03E-02 | Transcription repressor OFP16 |
| AT2G36470 | 0.0 | 7.07E-01 | -0.1 | 2.21E-01 | 0.2 | 1.71E-01 | 0.0 | 8.06E-01 | -0.1 | 7.65E-01 | 0.1 | 6.20E-01 | DUF868 family protein, putative |
| AT3G07680 | 0.0 | 6.92E-01 | 0.2 | 2.42E-02 | -0.2 | 6.63E-02 | 0.0 | 8.04E-01 | 0.0 | 8.50E-01 | 0.0 | 9.98E-01 | Transmembrane emp24 domain-containing protein p24beta2 |
| AT3G42630 | 0.0 | 7.31E-01 | 0.0 | 8.12E-01 | 0.0 | 9.47E-01 | -0.3 | 3.72E-02 | -0.3 | 9.66E-02 | 0.0 | 8.67E-01 | Pentatricopeptide repeat-containing protein At3g42630 |
| AT1G69980 | 0.0 | 6.77E-01 | 0.2 | 9.90E-02 | -0.1 | 2.60E-01 | 0.0 | 9.60E-01 | -0.2 | 3.96E-01 | 0.2 | 4.53E-01 | Structural polyprotein |
| AT4G01525 | 0.0 | 7.97E-01 | 0.2 | 1.81E-01 | -0.2 | 3.51E-01 | -0.1 | 7.02E-01 | 0.0 | 9.38E-01 | -0.1 | 8.42E-01 | NA |
| AT3G29170 | 0.0 | 8.05E-01 | -0.2 | 1.34E-01 | 0.3 | 1.35E-01 | 0.0 | 9.28E-01 | -0.2 | 6.39E-01 | 0.2 | 7.50E-01 | Transmembrane protein 230 |
| AT3G10650 | 0.0 | 6.21E-01 | -0.1 | 1.60E-01 | 0.2 | 9.27E-02 | 0.1 | 4.70E-01 | 0.0 | 8.75E-01 | 0.1 | 4.99E-01 | Nuclear pore complex protein NUP1 |
| AT5G06780 | 0.0 | 6.52E-01 | 0.0 | 8.19E-01 | 0.0 | 8.75E-01 | -0.1 | 4.69E-01 | -0.3 | 1.15E-01 | 0.2 | 4.98E-01 | Protein EMSY-LIKE 2 |
| AT3G18535 | 0.0 | 6.87E-01 | 0.0 | 9.18E-01 | 0.0 | 8.19E-01 | -0.6 | 1.50E-02 | -0.6 | 3.71E-02 | -0.1 | 8.89E-01 | Transmembrane protein |
| AT2G26200 | 0.0 | 6.61E-01 | 0.2 | 2.57E-02 | -0.2 | 8.89E-02 | 0.1 | 5.63E-01 | 0.0 | 8.58E-01 | 0.1 | 5.28E-01 | At2g26200 |
| AT1G76240 | 0.0 | 6.05E-01 | 0.4 | 2.83E-04 | -0.4 | 1.39E-03 | -0.3 | 3.11E-01 | 0.1 | 8.45E-01 | -0.3 | 2.79E-01 | T23E18.17 |
| AT4G10400 | 0.0 | 7.79E-01 | 0.1 | 4.71E-01 | -0.1 | 7.18E-01 | -0.1 | 8.40E-01 | -0.1 | 8.65E-01 | 0.0 | 1.00E+00 | FBD-associated F-box protein At4g10400 |
| AT1G05720 | 0.0 | 6.98E-01 | 0.1 | 1.97E-01 | -0.1 | 4.35E-01 | 0.1 | 7.54E-01 | 0.0 | 9.26E-01 | 0.1 | 7.20E-01 | F3F20.17 protein |
| AT4G35080 | 0.0 | 6.04E-01 | 0.2 | 4.05E-02 | -0.1 | 1.32E-01 | 0.1 | 6.30E-01 | -0.1 | 7.45E-01 | 0.1 | 5.29E-01 | High-affinity nickel-transport family protein |

| | | | | | | | | | | | | | |
|---|---|---|---|---|---|---|---|---|---|---|---|---|---|
| AT4G14620 | 0.0 | 7.59E-01 | 0.1 | 5.26E-01 | 0.0 | 7.82E-01 | 0.0 | 8.50E-01 | 0.2 | 4.19E-01 | -0.2 | 3.72E-01 | DUF506 family protein |
| AT2G21540 | 0.0 | 7.20E-01 | 0.5 | 1.30E-03 | -0.4 | 5.66E-03 | -0.5 | 1.07E-03 | -0.2 | 2.37E-01 | -0.3 | 6.50E-02 | CRAL-TRIO domain-containing protein |
| AT3G50380 | 0.0 | 5.33E-01 | -0.2 | 1.62E-02 | 0.2 | 1.16E-02 | 0.0 | 9.86E-01 | 0.1 | 7.03E-01 | -0.1 | 7.71E-01 | Vacuolar protein sorting-associated protein, putative |
| AT2G02790 | 0.0 | 5.85E-01 | -0.1 | 1.91E-01 | 0.1 | 1.14E-01 | 0.3 | 7.67E-02 | 0.2 | 2.64E-01 | 0.1 | 7.65E-01 | IQ-domain 29 |
| AT3G14200 | 0.0 | 6.31E-01 | -0.2 | 6.52E-02 | 0.2 | 4.40E-02 | -0.1 | 7.20E-01 | 0.2 | 4.70E-01 | -0.4 | 2.74E-01 | hypothetical protein |
| AT4G11550 | 0.0 | 7.79E-01 | 0.0 | 9.57E-01 | 0.0 | 7.94E-01 | -0.1 | 7.25E-01 | 0.1 | 7.83E-01 | -0.2 | 5.81E-01 | Cysteine |
| AT5G44150 | 0.0 | 7.34E-01 | 0.0 | 8.79E-01 | 0.1 | 6.87E-01 | 0.0 | 9.75E-01 | 0.0 | 9.52E-01 | 0.0 | 9.75E-01 | Protein ECERIFERUM 16 |
| AT2G23680 | 0.0 | 8.51E-01 | -0.2 | 4.17E-01 | 0.2 | 4.01E-01 | 0.5 | 9.80E-02 | -0.1 | 7.66E-01 | 0.6 | 7.13E-02 | Uncharacterized protein |
| AT5G15460 | 0.0 | 7.42E-01 | 0.1 | 4.85E-01 | 0.0 | 7.56E-01 | 0.2 | 3.13E-01 | -0.1 | 8.77E-01 | 0.3 | 3.58E-01 | Membrane-anchored ubiquitin-fold protein |
| AT3G03860 | 0.0 | 5.65E-01 | 0.1 | 4.19E-01 | 0.0 | 8.32E-01 | -0.1 | 5.71E-01 | -0.1 | 5.18E-01 | 0.0 | 9.27E-01 | 5'-adenylylsulfate reductase-like 5 |
| AT3G25030 | 0.0 | 7.56E-01 | -0.1 | 5.66E-01 | 0.1 | 4.56E-01 | 0.1 | 7.99E-01 | 0.3 | 2.93E-01 | -0.2 | 5.33E-01 | RING-type domain-containing protein |
| AT4G21150 | 0.0 | 5.52E-01 | 0.1 | 1.13E-01 | -0.1 | 3.29E-01 | 0.2 | 5.19E-02 | 0.2 | 2.14E-01 | 0.1 | 7.81E-01 | Dolichyl-diphosphooligosaccharide--protein glycosyltransferase subunit 2 |
| AT3G06570 | 0.0 | 6.49E-01 | 0.0 | 7.30E-01 | 0.1 | 4.98E-01 | -0.6 | 4.66E-03 | -0.5 | 1.93E-02 | -0.1 | 7.84E-01 | F-box |
| AT3G61640 | 0.0 | 6.95E-01 | 0.4 | 1.52E-03 | -0.4 | 6.64E-03 | -0.3 | 1.91E-01 | -0.1 | 8.21E-01 | -0.2 | 5.02E-01 | Arabinogalactan protein 20 |
| AT1G36050 | 0.0 | 5.39E-01 | 0.1 | 3.11E-01 | 0.0 | 7.17E-01 | -0.1 | 5.88E-01 | -0.2 | 2.18E-01 | 0.1 | 5.99E-01 | Endoplasmic reticulum vesicle transporter protein |
| AT4G22160 | 0.0 | 6.98E-01 | 0.1 | 2.70E-01 | -0.1 | 5.41E-01 | -0.2 | 5.00E-01 | -0.1 | 6.95E-01 | -0.1 | 9.14E-01 | Expressed protein |
| AT1G15060 | 0.0 | 5.82E-01 | -0.3 | 4.56E-03 | 0.3 | 4.30E-03 | 0.0 | 9.27E-01 | 0.0 | 9.71E-01 | 0.0 | 9.12E-01 | T15D22.11 protein |
| AT5G09960 | 0.0 | 6.16E-01 | 0.2 | 9.14E-02 | -0.1 | 2.77E-01 | -0.1 | 4.36E-01 | -0.1 | 5.43E-01 | 0.0 | 9.78E-01 | Sorbin |
| AT3G19670 | 0.0 | 5.00E-01 | -0.2 | 8.22E-03 | 0.3 | 5.94E-03 | 0.0 | 8.71E-01 | -0.1 | 5.01E-01 | 0.1 | 6.76E-01 | Pre-mRNA-processing protein 40B |
| AT2G03690 | 0.0 | 6.12E-01 | -0.1 | 4.89E-01 | 0.1 | 3.07E-01 | -0.4 | 2.41E-02 | -0.1 | 5.67E-01 | -0.3 | 2.42E-01 | Ubiquinone biosynthesis protein COQ4 homolog, mitochondrial |
| AT1G75370 | 0.0 | 6.41E-01 | -0.2 | 6.16E-02 | 0.2 | 4.08E-02 | -0.2 | 3.63E-01 | -0.1 | 5.21E-01 | 0.0 | 9.48E-01 | Sec14p-like phosphatidylinositol transfer family protein |
| AT3G02490 | 0.0 | 6.85E-01 | 0.1 | 4.60E-01 | 0.0 | 7.81E-01 | 0.3 | 8.85E-02 | 0.0 | 9.94E-01 | 0.3 | 1.97E-01 | Pentatricopeptide repeat-containing protein At3g02490, mitochondrial |
| AT3G59280 | 0.0 | 6.54E-01 | 0.0 | 6.71E-01 | 0.0 | 1.00E+00 | 0.3 | 1.61E-01 | 0.2 | 4.44E-01 | 0.1 | 7.87E-01 | Protein Transporter, Pam16 |
| AT1G69950 | 0.0 | 5.88E-01 | -0.1 | 3.49E-01 | 0.1 | 2.16E-01 | 0.2 | 4.65E-01 | 0.1 | 6.82E-01 | 0.1 | 9.01E-01 | NA |
| AT5G49530 | 0.0 | 5.06E-01 | 0.0 | 5.94E-01 | 0.1 | 3.02E-01 | 0.2 | 2.01E-01 | 0.1 | 5.24E-01 | 0.1 | 7.42E-01 | SIN-like family protein |
| AT2G35720 | 0.0 | 7.27E-01 | -0.1 | 5.40E-01 | 0.1 | 4.07E-01 | 0.5 | 3.96E-03 | 0.1 | 5.26E-01 | 0.4 | 6.44E-02 | Chaperone protein dnaJ 13 |
| AT5G49540 | 0.0 | 5.80E-01 | 0.2 | 1.92E-02 | -0.2 | 7.48E-02 | 0.1 | 7.01E-01 | 0.1 | 8.01E-01 | 0.0 | 9.78E-01 | ER membrane protein complex subunit 6 |
| AT3G51500 | 0.0 | 8.36E-01 | -0.2 | 4.35E-01 | 0.2 | 3.96E-01 | 0.0 | 9.51E-01 | -0.2 | 4.66E-01 | 0.2 | 5.99E-01 | Uncharacterized protein |
| AT4G14220 | 0.0 | 6.07E-01 | -0.1 | 1.51E-01 | 0.2 | 9.08E-02 | 0.4 | 9.77E-03 | 0.3 | 1.28E-01 | 0.2 | 4.93E-01 | AT4g14220 |
| AT3G54480 | 0.0 | 5.52E-01 | -0.1 | 7.24E-02 | 0.2 | 4.13E-02 | -0.1 | 4.41E-01 | -0.1 | 5.70E-01 | 0.0 | 9.66E-01 | SKP1 |
| AT2G32970 | 0.0 | 6.92E-01 | 0.0 | 9.60E-01 | 0.0 | 7.88E-01 | -0.2 | 1.98E-01 | -0.1 | 8.47E-01 | -0.2 | 4.71E-01 | Expressed protein |
| AT1G05870 | 0.0 | 6.97E-01 | -0.1 | 5.41E-01 | 0.1 | 3.82E-01 | -0.4 | 4.25E-02 | 0.1 | 8.25E-01 | -0.4 | 6.82E-02 | Uncharacterized protein At1g05870 |
| AT5G20635 | 0.0 | 7.29E-01 | 0.3 | 4.06E-02 | -0.2 | 1.31E-01 | -0.4 | 7.74E-02 | -0.2 | 3.64E-01 | -0.2 | 6.46E-01 | Guanine nucleotide-binding protein subunit gamma |
| AT1G35350 | 0.0 | 6.55E-01 | -0.1 | 5.84E-01 | 0.1 | 3.81E-01 | 0.1 | 8.16E-01 | -0.4 | 5.09E-02 | 0.5 | 5.28E-02 | EXS family protein |
| AT1G67960 | 0.0 | 5.48E-01 | -0.1 | 5.18E-01 | 0.1 | 2.92E-01 | 0.1 | 6.82E-01 | 0.2 | 5.89E-01 | -0.1 | 9.04E-01 | Protein POLLEN DEFECTIVE IN GUIDANCE 1 |
| AT1G66810 | 0.0 | 8.22E-01 | 0.2 | 3.62E-01 | -0.1 | 5.71E-01 | 0.2 | 4.04E-01 | 0.4 | 1.88E-01 | -0.2 | 6.58E-01 | Zinc finger C-x8-C-x5-C-x3-H type family protein |
| AT3G09670 | 0.0 | 5.66E-01 | 0.0 | 9.02E-01 | 0.1 | 5.63E-01 | 0.0 | 8.82E-01 | -0.3 | 2.71E-01 | 0.2 | 4.36E-01 | PWWP domain-containing protein 2 |
| AT4G00530 | 0.0 | 7.59E-01 | 0.0 | 7.66E-01 | 0.0 | 9.94E-01 | -0.1 | 8.35E-01 | 0.0 | 9.83E-01 | -0.1 | 9.06E-01 | Uncharacterized protein |
| AT3G52860 | 0.0 | 6.72E-01 | 0.0 | 9.64E-01 | 0.1 | 7.12E-01 | -0.1 | 8.24E-01 | -0.4 | 1.06E-01 | 0.3 | 2.46E-01 | Mediator of RNA polymerase II transcription subunit 28 |
| AT4G23420 | 0.0 | 7.42E-01 | 0.0 | 9.53E-01 | 0.1 | 7.63E-01 | -0.4 | 4.32E-01 | 0.5 | 3.15E-01 | -0.8 | 1.12E-01 | NAD-binding Rossmann-fold superfamily protein |
| AT5G10840 | 0.0 | 4.64E-01 | -0.1 | 9.45E-02 | 0.2 | 3.93E-02 | 0.1 | 9.26E-01 | 0.2 | 8.19E-01 | -0.1 | 9.12E-01 | Transmembrane 9 superfamily member 8 |
| AT3G05100 | 0.0 | 4.57E-01 | 0.2 | 3.33E-02 | -0.1 | 1.66E-01 | -0.2 | 1.69E-01 | -0.1 | 6.06E-01 | -0.1 | 6.44E-01 | S-adenosyl-L-methionine-dependent methyltransferases superfamily protein |
| AT5G59900 | 0.0 | 7.62E-01 | 0.0 | 9.57E-01 | 0.0 | 8.57E-01 | -0.2 | 6.27E-01 | 0.2 | 6.73E-01 | -0.5 | 3.59E-01 | Putative pentatricopeptide repeat-containing protein At5g59900 |
| AT5G40480 | 0.0 | 5.24E-01 | -0.1 | 1.99E-01 | 0.1 | 9.80E-02 | 0.6 | 2.45E-02 | 0.2 | 6.40E-01 | 0.5 | 2.20E-01 | Nuclear pore complex protein GP210 |
| AT1G65040 | 0.0 | 5.86E-01 | -0.2 | 8.54E-02 | 0.2 | 5.51E-02 | 0.4 | 6.17E-02 | 0.4 | 6.18E-02 | 0.0 | 9.11E-01 | RING |
| AT5G61040 | 0.0 | 6.01E-01 | 0.1 | 4.47E-01 | 0.0 | 8.40E-01 | -0.4 | 2.31E-01 | -0.2 | 5.59E-01 | -0.2 | 7.23E-01 | Uncharacterized protein |
| AT2G12646 | 0.0 | 7.89E-01 | 0.4 | 1.94E-02 | -0.4 | 6.51E-02 | -0.1 | 7.82E-01 | 0.2 | 5.10E-01 | -0.3 | 4.18E-01 | PLATZ transcription factor family protein |
| AT5G21070 | 0.0 | 5.39E-01 | 0.0 | 8.47E-01 | 0.0 | 7.42E-01 | -0.1 | 6.53E-01 | 0.0 | 9.34E-01 | -0.1 | 6.91E-01 | hypothetical protein |
| AT4G35530 | 0.0 | 6.17E-01 | 0.0 | 9.77E-01 | 0.1 | 6.74E-01 | 0.0 | 8.27E-01 | 0.1 | 6.24E-01 | -0.1 | 8.31E-01 | hypothetical protein |
| AT1G07390 | 0.0 | 7.18E-01 | -0.3 | 3.67E-02 | 0.4 | 4.13E-02 | 1.0 | 1.83E-01 | 0.2 | 8.32E-01 | 0.7 | 4.33E-01 | Receptor like protein 1 |
| AT3G13480 | 0.0 | 5.22E-01 | -0.2 | 9.57E-03 | 0.3 | 8.13E-03 | -0.1 | 6.03E-01 | 0.0 | 9.48E-01 | -0.1 | 7.81E-01 | Uncharacterized protein |
| AT4G31720 | 0.0 | 5.53E-01 | 0.2 | 6.26E-02 | -0.1 | 2.14E-01 | -0.3 | 7.91E-03 | -0.3 | 4.31E-02 | -0.1 | 7.79E-01 | Transcription initiation factor TFIID subunit 10 |
| AT1G77660 | 0.0 | 7.29E-01 | 0.0 | 8.76E-01 | 0.1 | 6.79E-01 | 0.0 | 8.88E-01 | -0.1 | 8.62E-01 | 0.1 | 7.94E-01 | hypothetical protein |
| AT3G28160 | 0.0 | 8.22E-01 | 0.0 | 9.21E-01 | 0.1 | 7.90E-01 | 0.2 | 6.51E-01 | 0.0 | 9.96E-01 | 0.2 | 7.22E-01 | NA |
| AT2G32870 | 0.1 | 7.50E-01 | 0.0 | 8.00E-01 | 0.0 | 9.71E-01 | 0.0 | 8.46E-01 | 0.0 | 9.66E-01 | 0.0 | 9.27E-01 | Uncharacterized protein At2g32870 |
| AT1G02090 | 0.1 | 6.30E-01 | 0.0 | 6.97E-01 | 0.0 | 9.56E-01 | 0.0 | 8.53E-01 | -0.1 | 7.32E-01 | 0.1 | 6.51E-01 | FUS5 |
| AT1G51270 | 0.1 | 6.32E-01 | -0.1 | 2.11E-01 | 0.2 | 1.27E-01 | -0.3 | 1.72E-01 | -0.3 | 1.34E-01 | 0.1 | 8.14E-01 | Vesicle-associated protein 1-4 |
| AT1G06500 | 0.1 | 6.59E-01 | 0.2 | 1.98E-01 | -0.1 | 4.66E-01 | 0.0 | 9.47E-01 | -0.1 | 8.90E-01 | 0.0 | 9.54E-01 | AT1G06500 protein |
| AT1G06720 | 0.1 | 5.67E-01 | -0.1 | 4.92E-01 | 0.1 | 2.69E-01 | 0.6 | 7.27E-04 | 0.3 | 4.78E-02 | 0.2 | 1.96E-01 | Uncharacterized protein F4H5.19 |
| AT4G24610 | 0.1 | 5.79E-01 | -0.3 | 7.93E-03 | 0.4 | 7.63E-03 | -0.3 | 5.36E-01 | 0.3 | 5.90E-01 | -0.5 | 3.21E-01 | Pesticidal crystal cry8Ba protein |
| AT5G37370 | 0.1 | 4.53E-01 | -0.1 | 1.21E-01 | 0.2 | 4.84E-02 | -0.2 | 9.75E-02 | -0.3 | 2.38E-02 | 0.1 | 5.42E-01 | Pre-mRNA-splicing factor 38 |
| AT4G12030 | 0.1 | 6.53E-01 | 0.4 | 4.16E-03 | -0.4 | 1.45E-02 | -0.5 | 4.76E-02 | 0.7 | 1.53E-02 | -1.2 | 2.66E-03 | Bile acid transporter 5 |
| AT5G10780 | 0.1 | 4.58E-01 | 0.1 | 4.59E-01 | 0.0 | 9.76E-01 | -0.1 | 6.69E-01 | -0.1 | 4.70E-01 | 0.1 | 8.32E-01 | ER membrane protein complex subunit 4 |
| AT5G13100 | 0.1 | 5.72E-01 | 0.4 | 2.15E-03 | -0.3 | 1.01E-02 | -0.3 | 2.76E-02 | -0.4 | 6.71E-03 | 0.1 | 5.49E-01 | hypothetical protein |
| AT1G29370 | 0.1 | 5.00E-01 | 0.0 | 7.68E-01 | 0.0 | 7.75E-01 | 0.1 | 7.46E-01 | -0.2 | 1.85E-01 | 0.3 | 1.35E-01 | RNA polymerase II degradation factor-like protein |
| AT1G49890 | 0.1 | 6.28E-01 | 0.2 | 1.05E-01 | -0.1 | 2.96E-01 | 0.0 | 9.23E-01 | -0.1 | 5.97E-01 | 0.1 | 6.25E-01 | QWRF motif-containing protein 2 |
| AT4G17060 | 0.1 | 6.02E-01 | 0.1 | 2.59E-01 | -0.1 | 5.89E-01 | -0.1 | 2.43E-01 | -0.1 | 3.90E-01 | 0.0 | 9.26E-01 | hypothetical protein |
| AT1G32610 | 0.1 | 7.75E-01 | 0.2 | 2.27E-01 | -0.2 | 4.41E-01 | 0.5 | 4.35E-01 | 0.3 | 7.04E-01 | 0.2 | 8.58E-01 | Hydroxyproline-rich glycoprotein family protein |
| AT3G29130 | 0.1 | 7.15E-01 | 0.0 | 9.18E-01 | 0.0 | 8.49E-01 | 0.1 | 6.91E-01 | 0.0 | 9.57E-01 | 0.1 | 8.38E-01 | KxDL motif protein |
| AT4G02485 | 0.1 | 7.46E-01 | 0.0 | 9.00E-01 | 0.0 | 8.92E-01 | -0.2 | 5.84E-01 | 0.2 | 6.92E-01 | -0.4 | 3.50E-01 | 2-oxoglutarate and Fe-dependent oxygenase superfamily protein |
| AT1G21790 | 0.1 | 6.25E-01 | -0.1 | 2.67E-01 | 0.2 | 1.74E-01 | -0.7 | 1.95E-03 | 0.1 | 7.68E-01 | -0.7 | 6.10E-03 | F8K7.23 lipid-sensing domain containing protein) |
| AT3G51610 | 0.1 | 3.99E-01 | 0.0 | 5.60E-01 | 0.0 | 8.54E-01 | -0.1 | 3.69E-01 | -0.1 | 4.42E-01 | 0.0 | 9.97E-01 | Uncharacterized protein F26O13.250 |
| AT1G09870 | 0.1 | 4.52E-01 | 0.0 | 9.90E-01 | 0.1 | 5.16E-01 | -0.1 | 7.66E-01 | 0.0 | 8.68E-01 | -0.1 | 7.11E-01 | At1g09870-1 |
| AT4G10925 | 0.1 | 5.33E-01 | 0.0 | 9.06E-01 | 0.0 | 7.00E-01 | -0.3 | 1.69E-01 | -0.7 | 4.65E-03 | 0.4 | 1.14E-01 | F-box protein SKIP8 |
| AT5G11150 | 0.1 | 5.43E-01 | -0.1 | 4.33E-01 | 0.1 | 2.23E-01 | 0.0 | 8.84E-01 | 0.0 | 8.77E-01 | 0.0 | 9.81E-01 | Vesicle-associated membrane protein 713 |
| AT2G03875 | 0.1 | 9.05E-01 | 0.0 | 9.80E-01 | 0.1 | 9.13E-01 | 0.5 | 5.99E-01 | -0.1 | 9.75E-01 | 0.6 | 6.69E-01 | NA |
| AT5G12390 | 0.1 | 5.58E-01 | 0.0 | 6.39E-01 | 0.1 | 3.68E-01 | -0.1 | 4.86E-01 | -0.2 | 2.44E-01 | 0.1 | 7.13E-01 | Mitochondrial fission 1 protein B |
| AT1G78770 | 0.1 | 6.32E-01 | 0.1 | 2.66E-01 | -0.1 | 5.90E-01 | 0.4 | 2.07E-01 | 0.2 | 4.99E-01 | 0.1 | 7.76E-01 | Anaphase-promoting complex subunit 6 |
| AT2G38160 | 0.1 | 7.48E-01 | 0.2 | 3.90E-01 | -0.1 | 6.57E-01 | 0.3 | 4.60E-01 | 0.4 | 2.42E-01 | -0.2 | 7.21E-01 | Uncharacterized protein At2g38160 |
| AT3G18690 | 0.1 | 6.23E-01 | -0.1 | 3.79E-01 | 0.2 | 2.50E-01 | 0.3 | 1.92E-01 | -0.4 | 1.42E-01 | 0.6 | 2.23E-02 | Protein MKS1 |
| AT3G03100 | 0.1 | 4.18E-01 | 0.1 | 1.84E-01 | 0.0 | 6.06E-01 | -0.1 | 4.53E-01 | -0.1 | 7.59E-01 | -0.1 | 8.41E-01 | NADH dehydrogenase 1 alpha subcomplex subunit 12 |
| AT1G28060 | 0.1 | 4.52E-01 | 0.0 | 8.18E-01 | 0.1 | 3.83E-01 | 0.2 | 2.92E-01 | 0.1 | 6.27E-01 | 0.1 | 7.99E-01 | Protein RDM16 |
| AT5G46170 | 0.1 | 4.48E-01 | 0.0 | 7.12E-01 | 0.0 | 7.68E-01 | 0.1 | 7.42E-01 | 0.1 | 6.38E-01 | 0.0 | 9.03E-01 | F-box protein At5g46170 |

| | | | | | | | | | | | | | |
|---|---|---|---|---|---|---|---|---|---|---|---|---|---|
| AT4G38280 | 0.1 | 6.08E-01 | 0.2 | 6.62E-02 | -0.2 | 2.30E-01 | -0.2 | 6.07E-01 | 0.1 | 8.13E-01 | -0.3 | 5.22E-01 | hypothetical protein |
| AT1G15260 | 0.1 | 6.50E-01 | 0.5 | 1.69E-03 | -0.5 | 7.32E-03 | 0.0 | 9.04E-01 | 0.0 | 9.18E-01 | 0.0 | 9.95E-01 | F9L1.20 protein |
| AT3G56900 | 0.1 | 5.89E-01 | 0.0 | 7.41E-01 | 0.1 | 4.64E-01 | 0.2 | 6.00E-01 | 0.1 | 9.09E-01 | 0.1 | 8.15E-01 | Aladin |
| AT1G45229 | 0.1 | 7.54E-01 | -0.2 | 4.09E-01 | 0.2 | 3.36E-01 | 0.3 | 3.35E-01 | 0.2 | 5.34E-01 | 0.1 | 8.90E-01 | Transmembrane protein |
| AT5G14370 | 0.1 | 4.25E-01 | 0.0 | 6.52E-01 | 0.0 | 7.97E-01 | -0.3 | 1.55E-02 | -0.2 | 1.55E-01 | -0.1 | 5.81E-01 | CCT domain-containing protein |
| AT3G13990 | 0.1 | 4.98E-01 | -0.2 | 6.55E-02 | 0.2 | 3.37E-02 | 0.4 | 6.89E-02 | 0.2 | 3.75E-01 | 0.2 | 5.59E-01 | GBF-interacting protein 1 N-terminal domain-containing protein |
| AT5G27730 | 0.1 | 5.97E-01 | 0.2 | 1.44E-01 | -0.1 | 3.92E-01 | -0.4 | 3.66E-02 | -0.3 | 1.26E-01 | -0.1 | 8.27E-01 | Heparan-alpha-glucosaminide N-acetyltransferase catalytic domain-containing protein |
| AT5G19950 | 0.1 | 4.54E-01 | 0.0 | 7.04E-01 | 0.0 | 7.93E-01 | 0.1 | 6.43E-01 | -0.2 | 5.47E-01 | 0.3 | 3.53E-01 | Tudor domain protein |
| AT5G41685 | 0.1 | 5.85E-01 | 0.1 | 3.50E-01 | 0.0 | 7.39E-01 | 0.3 | 2.08E-01 | 0.1 | 7.55E-01 | 0.2 | 5.23E-01 | Mitochondrial import receptor subunit TOM7-1 |
| AT2G05840 | 0.1 | 5.91E-01 | 0.0 | 8.08E-01 | 0.1 | 4.78E-01 | 0.3 | 6.90E-03 | 0.1 | 4.78E-01 | 0.2 | 1.83E-01 | Proteasome subunit alpha type |
| AT2G48030 | 0.1 | 5.96E-01 | 0.2 | 9.26E-02 | -0.1 | 2.76E-01 | 0.2 | 4.00E-01 | -0.1 | 5.36E-01 | 0.3 | 2.09E-01 | DNAse I-like superfamily protein |
| AT5G13690 | 0.1 | 5.08E-01 | 0.1 | 1.50E-01 | -0.1 | 4.79E-01 | -0.4 | 1.02E-01 | -0.1 | 7.27E-01 | -0.3 | 4.25E-01 | Alpha-N-acetylglucosaminidase |
| AT2G18850 | 0.1 | 4.92E-01 | -0.2 | 5.41E-02 | 0.3 | 3.46E-02 | 0.0 | 9.04E-01 | -0.3 | 2.74E-01 | 0.3 | 4.34E-01 | Uncharacterized protein At2g18850 |
| AT5G23200 | 0.1 | 5.62E-01 | 0.2 | 1.17E-01 | -0.1 | 3.70E-01 | 0.1 | 7.85E-01 | 0.1 | 6.58E-01 | 0.0 | 8.74E-01 | Uncharacterized protein |
| AT5G58450 | 0.1 | 3.15E-01 | 0.0 | 5.03E-01 | 0.1 | 1.47E-01 | 0.3 | 3.05E-01 | 0.4 | 2.79E-01 | -0.1 | 9.14E-01 | Tetratricopeptide repeat -like superfamily protein |
| AT1G31870 | 0.1 | 3.74E-01 | -0.1 | 3.03E-01 | 0.1 | 1.04E-01 | 0.1 | 3.10E-01 | 0.0 | 9.50E-01 | 0.1 | 4.51E-01 | AT1G31870 protein |
| AT3G03590 | 0.1 | 6.27E-01 | 0.0 | 7.85E-01 | 0.1 | 5.30E-01 | -0.2 | 3.54E-01 | -0.2 | 4.31E-01 | 0.0 | 9.86E-01 | DM2 domain-containing protein |
| AT5G43870 | 0.1 | 5.91E-01 | 0.2 | 2.01E-01 | -0.1 | 5.04E-01 | -0.4 | 2.32E-02 | -0.4 | 5.97E-02 | 0.0 | 8.79E-01 | At5g43870 ) |
| AT1G05340 | 0.1 | 5.52E-01 | 0.0 | 7.62E-01 | 0.1 | 4.45E-01 | -0.1 | 7.99E-01 | -0.7 | 6.70E-02 | 0.6 | 1.60E-01 | Protein CYSTEINE-RICH TRANSMEMBRANE MODULE 1 |
| AT2G36360 | 0.1 | 4.41E-01 | -0.2 | 3.87E-02 | 0.3 | 2.09E-02 | -0.1 | 7.27E-01 | 0.1 | 8.99E-01 | -0.2 | 7.17E-01 | AT2G36360 protein |
| AT3G48660 | 0.1 | 7.69E-01 | 0.3 | 1.51E-01 | -0.2 | 3.39E-01 | 0.6 | 6.27E-02 | 0.5 | 1.53E-01 | 0.1 | 8.57E-01 | Transmembrane protein |
| AT5G46850 | 0.1 | 5.91E-01 | 0.3 | 9.92E-03 | -0.3 | 5.18E-02 | 0.1 | 8.40E-01 | 0.0 | 9.32E-01 | 0.1 | 8.07E-01 | Phosphatidylinositol-glycan biosynthesis class X-like protein |
| AT1G48200 | 0.1 | 6.28E-01 | 0.2 | 8.85E-02 | -0.2 | 2.73E-01 | 0.3 | 1.35E-01 | 0.4 | 4.17E-02 | -0.1 | 5.81E-01 | F21D18.7 |
| AT1G11185 | 0.1 | 7.49E-01 | 0.4 | 6.86E-02 | -0.3 | 1.99E-01 | 0.9 | 6.09E-03 | 0.0 | 9.87E-01 | 0.9 | 2.94E-02 | NA |
| AT2G23330 | 0.1 | 5.67E-01 | -0.3 | 1.55E-02 | 0.4 | 1.56E-02 | 0.0 | 9.95E-01 | -0.3 | 4.32E-01 | 0.3 | 4.28E-01 | NA |
| AT1G24267 | 0.1 | 5.02E-01 | -0.1 | 5.33E-01 | 0.1 | 2.53E-01 | 0.1 | 5.40E-01 | -0.1 | 6.66E-01 | 0.2 | 3.77E-01 | At1g24268 ) |
| AT5G40700 | 0.1 | 4.72E-01 | 0.2 | 9.91E-02 | -0.1 | 3.98E-01 | 0.0 | 9.06E-01 | 0.1 | 6.87E-01 | -0.1 | 8.26E-01 | Dbj\|BAA95710.1 |
| AT5G28740 | 0.1 | 3.37E-01 | 0.1 | 2.12E-01 | 0.0 | 7.76E-01 | 0.2 | 2.22E-01 | 0.0 | 9.79E-01 | 0.2 | 4.19E-01 | Tetratricopeptide repeat -like superfamily protein |
| AT5G54540 | 0.1 | 3.78E-01 | 0.1 | 3.02E-01 | 0.0 | 8.68E-01 | -0.4 | 3.07E-02 | -0.4 | 2.10E-02 | 0.1 | 8.04E-01 | Uncharacterized protein |
| AT1G70590 | 0.1 | 5.70E-01 | -0.2 | 1.83E-01 | 0.2 | 1.07E-01 | -0.5 | 1.66E-02 | -0.3 | 2.14E-01 | -0.2 | 4.51E-01 | F-box protein At1g70590 |
| AT4G14600 | 0.1 | 4.27E-01 | 0.1 | 1.16E-01 | -0.1 | 4.73E-01 | 0.0 | 9.65E-01 | -0.1 | 5.76E-01 | 0.1 | 6.78E-01 | Bet1-like protein At4g14600 |
| AT3G50690 | 0.1 | 3.55E-01 | -0.1 | 3.43E-01 | 0.1 | 1.02E-01 | 0.0 | 7.48E-01 | -0.2 | 1.88E-01 | 0.1 | 4.64E-01 | Acidic leucine-rich nuclear phosphoprotein 32-related protein |
| AT4G37590 | 0.1 | 4.18E-01 | 0.2 | 4.81E-02 | -0.1 | 2.35E-01 | -0.2 | 1.07E-01 | -0.1 | 4.83E-01 | -0.1 | 6.50E-01 | BTB |
| AT3G42150 | 0.1 | 3.60E-01 | 0.1 | 1.82E-01 | 0.0 | 6.94E-01 | -0.4 | 4.37E-02 | 0.0 | 9.60E-01 | -0.4 | 9.21E-02 | Transmembrane protein |
| AT1G14450 | 0.1 | 6.95E-01 | 0.2 | 2.96E-01 | -0.1 | 5.88E-01 | 0.1 | 7.49E-01 | 0.2 | 6.47E-01 | -0.1 | 8.96E-01 | NADH dehydrogenase s |
| AT4G27610 | 0.1 | 3.97E-01 | 0.1 | 1.25E-01 | -0.1 | 5.18E-01 | -0.5 | 9.32E-04 | -0.1 | 7.31E-01 | -0.5 | 2.04E-02 | hypothetical protein |
| AT1G53180 | 0.1 | 4.18E-01 | 0.3 | 2.06E-03 | -0.3 | 1.64E-02 | 0.3 | 1.08E-01 | 0.3 | 2.27E-01 | 0.0 | 8.91E-01 | F12M16.8 |
| AT1G27700 | 0.1 | 5.91E-01 | 0.1 | 4.57E-01 | 0.0 | 8.57E-01 | -0.2 | 2.23E-01 | -0.2 | 2.80E-01 | 0.0 | 9.98E-01 | At1g27700 |
| AT1G28250 | 0.1 | 6.09E-01 | -0.1 | 6.59E-01 | 0.1 | 4.12E-01 | -0.1 | 5.21E-01 | -0.1 | 5.38E-01 | 0.0 | 9.73E-01 | Transmembrane protein |
| AT5G16505 | 0.1 | 3.13E-01 | -0.1 | 2.89E-01 | 0.1 | 7.99E-02 | -0.4 | 8.25E-04 | -0.5 | 7.74E-04 | 0.1 | 7.82E-01 | NA |
| AT3G06760 | 0.1 | 4.65E-01 | -0.1 | 1.80E-01 | 0.2 | 8.19E-02 | -0.2 | 2.31E-01 | -0.1 | 5.35E-01 | -0.1 | 7.89E-01 | Drought-responsive family protein |
| AT3G13410 | 0.1 | 4.27E-01 | 0.3 | 7.84E-03 | -0.2 | 4.02E-02 | -0.1 | 3.55E-01 | 0.0 | 9.33E-01 | -0.1 | 6.14E-01 | V-type proton ATPase subunit S1 |
| AT2G37890 | 0.1 | 6.80E-01 | 0.0 | 9.98E-01 | 0.1 | 7.51E-01 | 0.6 | 2.23E-02 | 0.2 | 6.11E-01 | 0.4 | 1.72E-01 | Mitochondrial substrate carrier family protein |
| AT1G22750 | 0.1 | 2.56E-01 | -0.1 | 9.31E-02 | 0.2 | 2.04E-02 | 0.2 | 1.44E-01 | 0.2 | 2.69E-01 | 0.0 | 9.24E-01 | Transmembrane protein |
| AT4G24500 | 0.1 | 5.30E-01 | 0.1 | 4.42E-01 | 0.0 | 8.99E-01 | 0.0 | 9.24E-01 | -0.1 | 7.05E-01 | 0.1 | 6.29E-01 | hypothetical protein |
| AT3G12970 | 0.1 | 7.35E-01 | 0.6 | 6.28E-03 | -0.6 | 2.79E-02 | -0.3 | 4.34E-01 | 0.1 | 9.07E-01 | -0.3 | 4.40E-01 | hypothetical protein |
| AT5G17160 | 0.1 | 6.16E-01 | 0.4 | 2.27E-02 | -0.3 | 8.58E-02 | 0.1 | 5.57E-01 | 0.3 | 1.22E-01 | -0.2 | 4.27E-01 | Uncharacterized protein F2K13_310 |
| AT1G23120 | 0.1 | 5.31E-01 | 0.2 | 4.56E-02 | -0.2 | 2.04E-01 | -0.6 | 6.66E-02 | -0.3 | 4.98E-01 | -0.4 | 4.55E-01 | Polyketide cyclase |
| AT5G07870 | 0.1 | 4.65E-01 | 0.1 | 2.92E-01 | 0.0 | 7.76E-01 | 0.3 | 2.85E-01 | 0.0 | 9.16E-01 | 0.2 | 4.99E-01 | HXXXD-type acyl-transferase family protein |
| AT5G55210 | 0.1 | 5.12E-01 | 0.1 | 1.82E-01 | -0.1 | 5.38E-01 | 0.1 | 8.20E-01 | -0.2 | 4.43E-01 | 0.2 | 4.21E-01 | At5g55210 |
| AT1G55170 | 0.1 | 3.86E-01 | -0.1 | 4.70E-01 | 0.1 | 1.87E-01 | 0.3 | 1.56E-01 | 0.1 | 6.44E-01 | 0.2 | 5.61E-01 | Protein FLX-like 3 |
| AT5G02620 | 0.1 | 3.60E-01 | -0.3 | 2.81E-03 | 0.4 | 1.89E-03 | 0.0 | 9.75E-01 | 0.2 | 3.20E-01 | -0.2 | 4.50E-01 | Ankyrin-like1 |
| AT5G24830 | 0.1 | 5.01E-01 | 0.1 | 2.96E-01 | 0.0 | 7.48E-01 | 0.5 | 1.66E-01 | 0.3 | 5.18E-01 | 0.2 | 6.25E-01 | Pentatricopeptide repeat-containing protein At5g24830 |
| AT5G09670 | 0.1 | 2.70E-01 | 0.0 | 6.55E-01 | 0.1 | 1.66E-01 | 0.0 | 8.27E-01 | 0.0 | 8.76E-01 | 0.1 | 7.82E-01 | At5g09670 |
| AT5G43920 | 0.1 | 3.04E-01 | 0.1 | 1.00E-01 | -0.1 | 5.47E-01 | -0.3 | 1.01E-01 | -0.2 | 4.00E-01 | -0.1 | 6.46E-01 | WD repeat-containing protein WDS homolog |
| AT5G55940 | 0.1 | 3.16E-01 | 0.2 | 1.86E-02 | -0.1 | 1.40E-01 | -0.1 | 6.37E-01 | 0.0 | 8.21E-01 | 0.0 | 9.20E-01 | MPN domain-containing protein |
| AT2G30460 | 0.1 | 4.15E-01 | 0.2 | 2.33E-02 | -0.2 | 1.44E-01 | 0.1 | 6.37E-01 | 0.3 | 1.75E-01 | -0.2 | 4.96E-01 | Nucleotide |
| AT4G28910 | 0.1 | 2.42E-01 | 0.0 | 9.64E-01 | 0.1 | 3.21E-01 | 0.1 | 5.71E-01 | 0.1 | 5.43E-01 | 0.0 | 9.40E-01 | AFP homolog 2 |
| AT4G24460 | 0.1 | 3.74E-01 | 0.1 | 2.50E-01 | 0.0 | 8.06E-01 | 0.1 | 6.00E-01 | 0.3 | 2.23E-01 | -0.2 | 5.79E-01 | CRT -like transporter 2 |
| AT1G10180 | 0.1 | 2.78E-01 | -0.1 | 2.07E-01 | 0.2 | 5.23E-02 | 0.0 | 8.77E-01 | 0.2 | 5.07E-01 | -0.1 | 6.82E-01 | Exocyst complex component EXO84C |
| AT3G06150 | 0.1 | 3.45E-01 | 0.1 | 5.24E-01 | 0.0 | 8.23E-01 | -0.3 | 1.27E-02 | -0.3 | 2.00E-02 | 0.0 | 9.81E-01 | hypothetical protein |
| AT5G64010 | 0.1 | 4.36E-01 | -0.1 | 3.44E-01 | 0.2 | 1.60E-01 | 0.2 | 5.05E-01 | 0.3 | 3.39E-01 | -0.1 | 8.13E-01 | U2 small nuclear ribonucleoprotein auxiliary factor-like protein |
| AT1G58350 | 0.1 | 3.18E-01 | 0.1 | 1.18E-01 | 0.0 | 5.85E-01 | 0.1 | 6.63E-01 | 0.1 | 5.96E-01 | 0.0 | 9.14E-01 | Serine esterase family protein |
| AT1G18700 | 0.1 | 4.40E-01 | -0.1 | 3.68E-01 | 0.2 | 1.52E-01 | -0.1 | 3.15E-01 | -0.2 | 1.86E-01 | 0.1 | 7.72E-01 | DNAJ heat shock N-terminal domain-containing protein |
| AT4G33640 | 0.1 | 3.81E-01 | 0.4 | 1.50E-03 | -0.3 | 1.25E-02 | -0.1 | 5.41E-01 | -0.3 | 1.36E-01 | 0.2 | 4.93E-01 | Costars family protein |
| AT5G20030 | 0.1 | 6.16E-01 | 0.0 | 9.49E-01 | 0.1 | 7.47E-01 | 0.0 | 9.28E-01 | -0.3 | 3.26E-01 | 0.3 | 4.31E-01 | ENT domain-containing protein |
| AT5G49320 | 0.1 | 5.74E-01 | 0.3 | 2.27E-02 | -0.3 | 1.13E-01 | | | | | | | At5g49320 ) |
| AT4G29030 | 0.1 | 6.04E-01 | 0.6 | 2.70E-03 | -0.5 | 1.09E-02 | 0.5 | 1.89E-02 | 0.1 | 6.39E-01 | 0.3 | 1.76E-01 | Glycine-rich protein |
| AT3G14075 | 0.1 | 3.20E-01 | -0.1 | 3.10E-01 | 0.2 | 9.44E-02 | -0.1 | 7.58E-01 | 0.0 | 8.68E-01 | 0.0 | 9.51E-01 | AT3g14070 |
| AT3G30214 | 0.1 | 6.65E-01 | 0.2 | 2.02E-01 | -0.2 | 4.75E-01 | 0.4 | 1.55E-01 | 0.3 | 3.47E-01 | 0.1 | 8.10E-01 | NA |
| AT1G32210 | 0.1 | 4.06E-01 | 0.4 | 2.15E-03 | -0.3 | 1.39E-02 | 0.1 | 4.61E-01 | 0.0 | 8.63E-01 | 0.2 | 4.88E-01 | Dolichyl-diphosphooligosaccharide--protein glycosyltransferase subunit DAD1 |
| AT1G77885 | 0.1 | 5.63E-01 | 0.1 | 6.90E-01 | 0.0 | 9.06E-01 | 0.1 | 8.42E-01 | -0.2 | 5.48E-01 | 0.3 | 4.42E-01 | Uncharacterized protein |
| AT3G18900 | 0.1 | 6.01E-01 | -0.1 | 4.62E-01 | 0.2 | 2.99E-01 | 0.0 | 9.18E-01 | 0.0 | 9.81E-01 | 0.0 | 9.62E-01 | Ternary complex factor MIP1 leucine-zipper protein |
| AT3G58490 | 0.1 | 3.46E-01 | 0.0 | 9.05E-01 | 0.1 | 4.86E-01 | -0.2 | 2.94E-01 | -0.3 | 4.22E-02 | 0.2 | 3.89E-01 | Phosphatidic acid phosphatase family protein |
| AT5G62170 | 0.1 | 5.72E-01 | 0.3 | 2.30E-02 | -0.3 | 1.07E-01 | 0.1 | 7.17E-01 | 0.3 | 4.86E-01 | -0.1 | 7.94E-01 | LOW protein: M-phase inducer phosphatase-like protein |
| AT2G38370 | 0.1 | 3.80E-01 | -0.1 | 1.81E-01 | 0.2 | 7.06E-02 | -0.7 | 1.57E-03 | -0.4 | 5.54E-02 | -0.3 | 2.08E-01 | Weak chloroplast movement under blue light protein |
| AT5G64830 | 0.1 | 3.54E-01 | 0.0 | 9.01E-01 | 0.1 | 3.78E-01 | 0.0 | 8.87E-01 | 0.1 | 7.46E-01 | 0.0 | 8.86E-01 | AT5G64830 protein |
| AT5G40300 | 0.1 | 3.50E-01 | -0.1 | 1.85E-01 | 0.2 | 6.42E-02 | 0.1 | 4.18E-01 | 0.1 | 5.83E-01 | 0.0 | 9.36E-01 | CASP-like protein 4A1 |
| AT1G51350 | 0.1 | 3.18E-01 | 0.0 | 7.29E-01 | 0.1 | 2.49E-01 | 0.1 | 6.08E-01 | 0.1 | 8.42E-01 | 0.0 | 8.86E-01 | Uncharacterized protein |
| AT3G15115 | 0.1 | 4.80E-01 | 0.1 | 3.05E-01 | 0.0 | 7.78E-01 | -0.7 | 2.25E-02 | -0.1 | 7.25E-01 | -0.6 | 1.32E-01 | hypothetical protein |
| AT5G18490 | 0.1 | 4.44E-01 | 0.0 | 6.96E-01 | 0.0 | 7.80E-01 | -0.1 | 5.31E-01 | -0.4 | 7.91E-03 | 0.3 | 7.31E-02 | At5g18490 ) |
| AT5G63010 | 0.1 | 4.90E-01 | -0.1 | 3.90E-01 | 0.2 | 2.00E-01 | -0.3 | 3.24E-01 | -0.4 | 2.36E-01 | 0.1 | 8.54E-01 | Uncharacterized protein At5g63010 |
| AT3G51100 | 0.1 | 3.13E-01 | 0.2 | 4.60E-02 | -0.1 | 3.07E-01 | 0.1 | 3.18E-01 | 0.0 | 9.25E-01 | 0.1 | 5.27E-01 | Altered inheritance of mitochondria protein |

| | | | | | | | | | | | | | |
|---|---|---|---|---|---|---|---|---|---|---|---|---|---|
| AT5G52980 | 0.1 | 4.38E-01 | 0.2 | 1.13E-01 | -0.1 | 4.53E-01 | 0.2 | 2.09E-01 | 0.2 | 2.67E-01 | 0.0 | 1.00E+00 | At5g52980 |
| AT3G14260 | 0.1 | 7.13E-01 | 1.1 | 1.99E-04 | -1.0 | 1.68E-03 | -0.2 | 5.23E-01 | -0.6 | 1.31E-01 | 0.3 | 3.46E-01 | hypothetical protein |
| AT3G27220 | 0.1 | 5.83E-01 | 0.0 | 9.25E-01 | 0.1 | 7.31E-01 | 0.4 | 1.72E-01 | 0.4 | 2.15E-01 | 0.0 | 9.96E-01 | Kelch repeat-containing protein At3g27220 |
| AT2G44080 | 0.1 | 5.73E-01 | -0.3 | 7.72E-02 | 0.4 | 4.85E-02 | 1.5 | 3.88E-05 | 0.4 | 2.69E-01 | 1.1 | 3.18E-03 | ARGOS-like protein |
| AT5G41100 | 0.1 | 3.32E-01 | -0.1 | 2.12E-01 | 0.2 | 6.95E-02 | 0.1 | 8.12E-01 | -0.3 | 3.74E-01 | 0.3 | 3.09E-01 | Hydroxyproline-rich glycoprotein family protein |
| AT1G26760 | 0.1 | 7.41E-01 | 0.5 | 4.27E-02 | -0.4 | 1.33E-01 | 0.3 | 4.98E-01 | 0.2 | 6.44E-01 | 0.0 | 9.22E-01 | T24P13.14 |
| AT2G39760 | 0.1 | 2.10E-01 | 0.0 | 5.61E-01 | 0.1 | 1.14E-01 | 0.0 | 8.31E-01 | 0.0 | 9.32E-01 | 0.0 | 9.55E-01 | BTB |
| AT3G59640 | 0.1 | 2.92E-01 | -0.1 | 3.43E-01 | 0.2 | 1.01E-01 | 0.3 | 1.02E-01 | -0.1 | 7.80E-01 | 0.4 | 9.76E-02 | Glycine-rich protein |
| AT4G13150 | 0.1 | 3.14E-01 | -0.1 | 3.77E-01 | 0.2 | 1.13E-01 | 0.0 | 9.77E-01 | 0.0 | 9.30E-01 | 0.0 | 9.46E-01 | Uncharacterized protein AT4g13140 |
| AT4G14965 | 0.1 | 3.35E-01 | -0.1 | 4.78E-01 | 0.1 | 1.53E-01 | 0.0 | 9.20E-01 | 0.0 | 9.19E-01 | -0.1 | 8.70E-01 | Membrane-associated progesterone binding protein 4 |
| AT5G50410 | 0.1 | 3.85E-01 | -0.1 | 5.48E-01 | 0.1 | 2.17E-01 | 0.2 | 2.90E-01 | 0.2 | 3.68E-01 | 0.0 | 9.91E-01 | Uncharacterized protein |
| AT3G27030 | 0.1 | 4.03E-01 | 0.3 | 4.43E-03 | -0.3 | 3.45E-02 | 0.0 | 9.99E-01 | -0.1 | 8.76E-01 | 0.1 | 9.05E-01 | Transmembrane protein |
| AT5G03905 | 0.1 | 3.50E-01 | -0.2 | 7.05E-02 | 0.3 | 2.83E-02 | -0.2 | 3.79E-01 | 0.0 | 9.02E-01 | -0.3 | 3.76E-01 | Iron-sulfur cluster biosynthesis family protein |
| AT4G36440 | 0.1 | 2.00E-01 | 0.1 | 8.77E-02 | 0.0 | 6.55E-01 | -0.1 | 2.87E-01 | 0.0 | 7.83E-01 | -0.1 | 6.59E-01 | G-protein coupled receptor |
| AT1G14010 | 0.1 | 4.08E-01 | 0.3 | 2.06E-02 | -0.2 | 1.25E-01 | 0.2 | 2.48E-01 | 0.0 | 8.22E-01 | 0.1 | 5.60E-01 | Transmembrane emp24 domain-containing protein p24delta7 |
| AT3G15351 | 0.1 | 5.49E-01 | -0.2 | 1.43E-01 | 0.3 | 8.92E-02 | -0.2 | 5.34E-01 | -0.4 | 3.12E-01 | 0.2 | 7.71E-01 | p53 and DNA damage-regulated protein 1 |
| AT5G03330 | 0.1 | 2.53E-01 | 0.1 | 3.84E-01 | 0.0 | 8.52E-01 | 0.3 | 1.82E-01 | -0.2 | 4.42E-01 | 0.4 | 3.12E-02 | Cysteine proteinases superfamily protein |
| AT1G33780 | 0.1 | 4.00E-01 | -0.2 | 1.21E-01 | 0.3 | 5.32E-02 | -0.3 | 3.11E-01 | -0.1 | 8.40E-01 | -0.2 | 5.79E-01 | Electron transporter, putative |
| AT4G04630 | 0.1 | 4.75E-01 | 0.2 | 2.19E-01 | -0.1 | 6.54E-01 | -0.9 | 1.93E-04 | -0.5 | 1.16E-02 | -0.3 | 2.11E-01 | Uncharacterized protein AT4g04630 |
| AT4G11680 | 0.1 | 2.38E-01 | 0.0 | 8.99E-01 | 0.1 | 2.69E-01 | -0.1 | 4.56E-01 | 0.1 | 5.73E-01 | -0.2 | 2.13E-01 | Zinc finger, C3HC4 type family protein |
| AT4G00290 | 0.1 | 3.21E-01 | 0.0 | 6.03E-01 | 0.0 | 7.19E-01 | 0.6 | 1.21E-04 | 0.3 | 7.49E-02 | 0.3 | 4.44E-02 | Mechanosensitive ion channel domain-containing protein |
| AT2G46915 | 0.1 | 5.34E-01 | -0.3 | 5.60E-02 | 0.4 | 3.08E-02 | 0.1 | 7.92E-01 | 0.3 | 2.69E-01 | -0.2 | 4.46E-01 | DUF3754 family protein, putative |
| AT4G01740 | 0.1 | 3.74E-01 | -0.2 | 8.31E-02 | 0.3 | 3.78E-02 | -0.1 | 6.59E-01 | 0.0 | 8.84E-01 | -0.1 | 8.70E-01 | Cysteine |
| AT5G43460 | 0.1 | 2.06E-01 | 0.2 | 1.38E-02 | -0.1 | 1.49E-01 | -0.3 | 9.80E-02 | -0.4 | 1.09E-01 | 0.0 | 9.35E-01 | AT5g43460 |
| AT2G32980 | 0.1 | 2.37E-01 | 0.1 | 9.09E-02 | 0.0 | 6.07E-01 | -0.2 | 1.31E-01 | -0.2 | 1.87E-01 | 0.0 | 9.93E-01 | AUGMIN subunit 2 |
| AT1G12650 | 0.1 | 5.65E-01 | 0.3 | 9.84E-02 | -0.2 | 3.15E-01 | 0.1 | 3.77E-01 | -0.1 | 8.13E-01 | 0.2 | 3.73E-01 | rRNA biogenesis protein RRP36 |
| AT4G39570 | 0.1 | 3.05E-01 | 0.1 | 2.29E-01 | 0.0 | 8.65E-01 | -0.1 | 6.97E-01 | 0.0 | 8.77E-01 | 0.0 | 9.05E-01 | F-box |
| AT1G02370 | 0.1 | 4.91E-01 | 0.0 | 8.93E-01 | 0.1 | 4.96E-01 | 0.6 | 9.73E-03 | 0.2 | 4.11E-01 | 0.4 | 1.26E-01 | Pentatricopeptide repeat-containing protein At1g02370, mitochondrial |
| AT5G67390 | 0.1 | 6.44E-01 | 0.0 | 8.71E-01 | 0.1 | 8.25E-01 | -0.5 | 6.24E-02 | -0.4 | 1.56E-01 | -0.1 | 8.30E-01 | Uncharacterized protein |
| AT4G15930 | 0.1 | 2.63E-01 | 0.0 | 6.71E-01 | 0.1 | 5.67E-01 | -0.1 | 5.57E-01 | 0.0 | 8.74E-01 | -0.1 | 8.19E-01 | Dynein light chain |
| AT2G33845 | 0.1 | 4.44E-01 | 0.2 | 5.00E-02 | -0.2 | 2.43E-01 | 0.4 | 1.58E-01 | 0.2 | 5.72E-01 | 0.2 | 6.11E-01 | Nucleic acid-binding, OB-fold-like protein |
| AT5G19840 | 0.1 | 4.85E-01 | -0.2 | 2.73E-01 | 0.2 | 1.44E-01 | 0.1 | 7.51E-01 | 0.1 | 7.92E-01 | 0.0 | 1.00E+00 | 2-oxoglutarate and Fe-dependent oxygenase superfamily protein |
| AT2G26210 | 0.1 | 2.70E-01 | 0.0 | 8.87E-01 | 0.1 | 2.77E-01 | 0.1 | 7.36E-01 | -0.1 | 3.89E-01 | 0.2 | 3.39E-01 | Ankyrin repeat family protein |
| AT1G64050 | 0.1 | 3.56E-01 | -0.3 | 7.99E-03 | 0.4 | 4.32E-03 | 0.2 | 4.89E-01 | -0.1 | 6.55E-01 | 0.3 | 2.91E-01 | F22C12.19 |
| AT1G51745 | 0.1 | 3.25E-01 | 0.2 | 4.09E-02 | -0.1 | 2.86E-01 | -0.2 | 9.00E-02 | -0.3 | 3.59E-02 | 0.1 | 7.12E-01 | Tudor |
| AT2G32340 | 0.1 | 5.98E-01 | -0.2 | 2.96E-01 | 0.3 | 1.97E-01 | -0.4 | 3.92E-01 | 0.1 | 8.03E-01 | -0.5 | 3.70E-01 | TraB family protein |
| AT2G21160 | 0.1 | 3.66E-01 | 0.4 | 5.01E-03 | -0.3 | 3.07E-02 | 0.3 | 3.00E-02 | 0.2 | 7.51E-02 | 0.0 | 9.29E-01 | Translocon-associated protein subunit alpha |
| AT1G63100 | 0.1 | 4.01E-01 | 0.5 | 8.38E-04 | -0.4 | 7.46E-03 | 0.2 | 5.25E-01 | 0.4 | 1.37E-01 | -0.2 | 3.70E-01 | GRAS family transcription factor |
| AT1G01800 | 0.1 | 1.99E-01 | 0.0 | 8.63E-01 | 0.1 | 3.27E-01 | -0.2 | 3.06E-01 | -0.1 | 5.07E-01 | 0.0 | 9.05E-01 | T1N6.22 protein |
| AT5G01030 | 0.1 | 2.63E-01 | -0.2 | 6.66E-02 | 0.3 | 1.98E-02 | 0.2 | 3.57E-01 | 0.3 | 2.06E-01 | -0.1 | 7.53E-01 | Uncharacterized protein |
| AT1G12120 | 0.1 | 2.05E-01 | 0.0 | 9.64E-01 | 0.1 | 2.41E-01 | -0.1 | 3.44E-01 | 0.0 | 9.32E-01 | -0.1 | 6.08E-01 | hypothetical protein |
| AT2G43280 | 0.1 | 4.80E-01 | -0.1 | 6.62E-01 | 0.1 | 3.39E-01 | -0.1 | 6.59E-01 | -0.2 | 4.96E-01 | 0.1 | 8.56E-01 | Far-red impaired responsive family protein |
| AT5G40520 | 0.1 | 3.57E-01 | -0.1 | 5.97E-01 | 0.1 | 2.26E-01 | 0.3 | 1.91E-01 | 0.2 | 4.99E-01 | 0.1 | 7.58E-01 | Uncharacterized protein |
| AT1G25375 | 0.1 | 2.66E-01 | -0.2 | 3.74E-02 | 0.3 | 1.32E-02 | -0.2 | 4.85E-01 | -0.1 | 8.10E-01 | -0.1 | 7.99E-01 | Uncharacterized protein F4F7.24 |
| AT1G67050 | 0.1 | 3.93E-01 | 0.3 | 1.19E-02 | -0.2 | 8.78E-02 | -0.2 | 3.23E-01 | -0.3 | 3.13E-01 | 0.0 | 9.35E-01 | Uncharacterized protein |
| AT4G23895 | 0.1 | 3.65E-01 | 0.1 | 3.45E-01 | 0.0 | 9.54E-01 | -0.3 | 1.46E-01 | -0.2 | 4.23E-01 | -0.1 | 7.31E-01 | Uncharacterized protein At4g23895 |
| AT3G57580 | 0.1 | 5.75E-01 | 0.3 | 1.23E-01 | -0.2 | 3.92E-01 | 0.1 | 6.37E-01 | -0.2 | 5.75E-01 | 0.3 | 3.25E-01 | F-box protein At3g57580 |
| AT3G62790 | 0.1 | 3.32E-01 | 0.3 | 1.53E-02 | -0.2 | 1.15E-01 | 0.0 | 9.59E-01 | 0.1 | 8.32E-01 | -0.1 | 8.22E-01 | NADH dehydrogenase iron-sulfur protein 5-B |
| AT2G33110 | 0.1 | 4.28E-01 | -0.1 | 3.88E-01 | 0.2 | 1.74E-01 | -0.1 | 7.36E-01 | -0.3 | 2.74E-01 | 0.2 | 5.20E-01 | Vesicle-associated membrane protein 723 |
| AT4G32960 | 0.1 | 2.39E-01 | 0.0 | 7.85E-01 | 0.1 | 2.18E-01 | -0.2 | 2.65E-01 | -0.1 | 6.94E-01 | -0.1 | 7.13E-01 | BRISC and BRCA1-A complex member 1 |
| AT2G39580 | 0.1 | 4.04E-01 | -0.2 | 1.32E-01 | 0.3 | 4.93E-02 | -0.1 | 5.37E-01 | -0.1 | 5.39E-01 | 0.0 | 9.61E-01 | Zinc finger C3H1 domain protein |
| AT2G21150 | 0.1 | 2.69E-01 | 0.1 | 5.05E-01 | 0.0 | 7.37E-01 | 0.0 | 9.13E-01 | 0.1 | 7.76E-01 | 0.0 | 8.62E-01 | Protein XAP5 CIRCADIAN TIMEKEEPER |
| AT1G25250 | 0.1 | 5.53E-01 | 0.1 | 7.06E-01 | 0.0 | 8.91E-01 | -0.6 | 1.11E-01 | 0.3 | 3.41E-01 | -0.9 | 3.11E-02 | IDD16 |
| AT4G24690 | 0.1 | 2.59E-01 | -0.2 | 1.03E-02 | 0.3 | 3.26E-03 | -0.2 | 2.58E-02 | -0.3 | 1.51E-02 | 0.1 | 7.74E-01 | Uncharacterized protein At4g24690 |
| AT3G03130 | 0.1 | 5.70E-01 | 0.5 | 7.53E-03 | -0.4 | 3.95E-02 | 0.3 | 1.11E-01 | 0.5 | 2.25E-02 | -0.2 | 5.26E-01 | LisH domain-like protein |
| AT3G62920 | 0.1 | 7.12E-01 | 0.3 | 2.56E-01 | -0.2 | 5.27E-01 | 0.2 | 5.17E-01 | -0.2 | 6.27E-01 | 0.5 | 2.16E-01 | hypothetical protein |
| AT1G20890 | 0.1 | 3.43E-01 | -0.1 | 5.90E-01 | 0.1 | 2.18E-01 | 0.0 | 9.63E-01 | -0.3 | 3.31E-01 | 0.3 | 4.28E-01 | Caveolin-1 protein |
| AT4G12560 | 0.1 | 3.71E-01 | 0.0 | 8.89E-01 | 0.1 | 3.66E-01 | -0.3 | 3.25E-02 | -0.2 | 1.63E-01 | -0.1 | 7.30E-01 | F-box domain-containing protein |
| AT5G56380 | 0.1 | 4.06E-01 | 0.1 | 3.96E-01 | 0.0 | 9.68E-01 | 0.1 | 6.26E-01 | 0.1 | 6.68E-01 | 0.0 | 9.92E-01 | FBD-associated F-box protein At5g56380 |
| AT3G19520 | 0.1 | 2.30E-01 | 0.0 | 6.67E-01 | 0.1 | 1.58E-01 | -0.3 | 6.11E-02 | -0.3 | 5.72E-02 | 0.0 | 9.04E-01 | Uncharacterized protein |
| AT2G29210 | 0.1 | 1.64E-01 | -0.1 | 4.01E-01 | 0.2 | 5.65E-02 | 0.0 | 7.25E-01 | 0.0 | 8.05E-01 | 0.0 | 9.80E-01 | At2g29210 |
| AT4G22310 | 0.1 | 2.00E-01 | 0.2 | 1.78E-02 | -0.1 | 1.92E-01 | 0.3 | 2.67E-02 | 0.3 | 6.10E-02 | 0.0 | 9.53E-01 | Mitochondrial pyruvate carrier 4 |
| AT5G48660 | 0.1 | 4.09E-01 | 0.0 | 7.41E-01 | 0.1 | 7.19E-01 | 0.4 | 7.88E-02 | 0.3 | 3.28E-01 | 0.1 | 6.51E-01 | Endoplasmic reticulum transmembrane protein |
| AT5G04550 | 0.1 | 2.13E-01 | -0.1 | 9.87E-02 | 0.2 | 1.74E-02 | -0.1 | 3.74E-01 | -0.2 | 2.47E-01 | 0.1 | 8.21E-01 | hypothetical protein |
| AT5G60870 | 0.1 | 4.96E-01 | 0.1 | 6.34E-01 | 0.0 | 9.01E-01 | 0.0 | 9.13E-01 | -0.1 | 8.56E-01 | 0.0 | 9.47E-01 | Regulator of chromosome condensation family protein |
| AT1G09915 | 0.1 | 3.89E-01 | 0.2 | 5.86E-02 | -0.1 | 3.30E-01 | 0.0 | 9.28E-01 | -0.2 | 5.76E-01 | 0.2 | 5.55E-01 | NA |
| AT3G59800 | 0.1 | 3.18E-01 | 0.1 | 3.10E-01 | 0.0 | 9.67E-01 | -0.2 | 4.26E-01 | -0.2 | 3.63E-01 | 0.0 | 8.94E-01 | At3g59800 |
| AT4G21890 | 0.1 | 4.57E-01 | -0.2 | 1.27E-01 | 0.3 | 6.36E-02 | 0.2 | 6.72E-01 | 0.2 | 6.87E-01 | 0.0 | 9.76E-01 | Uncharacterized protein |
| AT4G11790 | 0.1 | 1.63E-01 | 0.0 | 4.80E-01 | 0.1 | 7.51E-02 | 0.1 | 5.38E-01 | 0.2 | 2.68E-01 | -0.1 | 7.17E-01 | AT4g11790 |
| AT1G51965 | 0.1 | 2.71E-01 | 0.0 | 6.21E-01 | 0.1 | 1.83E-01 | 0.0 | 9.66E-01 | 0.0 | 1.00E+00 | 0.0 | 9.80E-01 | ABA Overly-Sensitive 5 |
| AT2G35736 | 0.1 | 3.91E-01 | 0.0 | 7.95E-01 | 0.1 | 6.60E-01 | 0.6 | 4.08E-02 | 0.4 | 2.15E-01 | 0.2 | 6.28E-01 | Uncharacterized protein |
| AT5G20840 | 0.1 | 2.68E-01 | 0.0 | 7.52E-01 | 0.1 | 2.06E-01 | 0.1 | 4.66E-01 | -0.1 | 5.28E-01 | 0.2 | 2.71E-01 | Uncharacterized protein |
| AT4G29870 | 0.1 | 3.02E-01 | 0.2 | 3.13E-02 | -0.1 | 2.35E-01 | -0.1 | 6.45E-01 | -0.1 | 6.63E-01 | 0.0 | 9.76E-01 | Oligosaccharyltransferase complex |
| AT1G34315 | 0.1 | 3.58E-01 | 0.0 | 8.37E-01 | 0.1 | 3.56E-01 | -0.2 | 4.22E-01 | -0.3 | 3.57E-01 | 0.1 | 9.01E-01 | Transmembrane protein |
| AT5G16190 | 0.1 | 5.13E-01 | 0.2 | 1.12E-01 | -0.1 | 3.97E-01 | 0.3 | 1.93E-01 | 0.4 | 1.22E-01 | -0.1 | 8.15E-01 | Cellulose synthase like A11 |
| AT3G17800 | 0.1 | 2.17E-01 | -0.1 | 1.13E-01 | 0.2 | 2.03E-02 | -0.2 | 1.78E-01 | -0.3 | 1.44E-01 | 0.0 | 8.56E-01 | UV-B-induced protein At3g17800, chloroplastic |
| AT5G41300 | 0.1 | 5.80E-01 | 0.3 | 8.22E-02 | -0.2 | 2.96E-01 | 0.3 | 4.53E-01 | 0.8 | 1.94E-02 | -0.5 | 9.23E-02 | Cysteine-rich repeat secretory protein 59 |
| AT4G13550 | 0.1 | 4.59E-01 | -0.3 | 2.27E-02 | 0.4 | 1.28E-02 | 0.0 | 9.40E-01 | 0.1 | 7.79E-01 | -0.1 | 7.51E-01 | Uncharacterized protein At4g13550 |
| AT1G01730 | 0.1 | 1.67E-01 | -0.3 | 2.64E-03 | 0.4 | 1.05E-03 | -0.2 | 1.10E-01 | -0.2 | 2.07E-01 | 0.0 | 9.29E-01 | T1N6.14 protein |
| AT5G57060 | 0.1 | 4.83E-01 | -0.1 | 4.48E-01 | 0.2 | 2.19E-01 | 0.1 | 7.40E-01 | -0.1 | 8.14E-01 | 0.2 | 5.96E-01 | 60S ribosomal L18a-like protein |
| AT3G61590 | 0.1 | 3.47E-01 | 0.3 | 9.74E-03 | -0.2 | 8.51E-02 | -0.1 | 6.83E-01 | 0.1 | 8.19E-01 | -0.2 | 5.60E-01 | Galactose oxidase |
| AT5G05190 | 0.1 | 5.00E-01 | -0.4 | 2.28E-02 | 0.5 | 1.65E-02 | 0.7 | 1.68E-02 | -0.1 | 7.59E-01 | 0.8 | 2.68E-02 | Protein ENHANCED DISEASE RESISTANCE 4 |
| AT1G48770 | 0.1 | 5.24E-01 | 0.0 | 9.11E-01 | 0.1 | 5.45E-01 | 0.0 | 9.06E-01 | -0.4 | 3.28E-01 | 0.3 | 4.13E-01 | DUF1639 family protein |

| | | | | | | | | | | | | | |
|---|---|---|---|---|---|---|---|---|---|---|---|---|---|
| AT4G00170 | 0.1 | 3.09E-01 | -0.1 | 5.13E-01 | 0.2 | 1.53E-01 | 0.6 | 3.01E-03 | 0.1 | 5.61E-01 | 0.4 | 3.97E-02 | MSP domain-containing protein |
| AT1G61065 | 0.1 | 4.43E-01 | 0.1 | 3.67E-01 | 0.0 | 8.94E-01 | -0.5 | 1.09E-01 | -0.1 | 7.37E-01 | -0.3 | 4.06E-01 | 1,3-beta-glucan synthase component |
| AT5G48500 | 0.1 | 4.08E-01 | 0.3 | 2.63E-02 | -0.2 | 1.65E-01 | 0.1 | 5.66E-01 | 0.0 | 8.98E-01 | 0.2 | 6.04E-01 | Gb\|AAD28645.1 |
| AT1G21010 | 0.1 | 3.55E-01 | 0.1 | 4.17E-01 | 0.0 | 9.56E-01 | 0.1 | 6.59E-01 | -0.1 | 8.46E-01 | 0.2 | 5.87E-01 | hypothetical protein |
| AT3G59500 | 0.1 | 3.07E-01 | 0.1 | 3.58E-01 | 0.0 | 9.58E-01 | -0.1 | 4.46E-01 | -0.2 | 3.82E-01 | 0.0 | 9.05E-01 | Uncharacterized protein T16L24.50 |
| AT2G42320 | 0.1 | 3.36E-01 | 0.0 | 8.08E-01 | 0.1 | 5.51E-01 | -0.2 | 3.00E-01 | 0.0 | 9.87E-01 | -0.2 | 4.47E-01 | Nucleolar protein gar2-like protein |
| AT1G61340 | 0.1 | 6.87E-01 | -0.4 | 1.12E-01 | 0.5 | 1.01E-01 | 1.0 | 3.99E-02 | 0.4 | 5.36E-01 | 0.6 | 3.12E-01 | FBS1 |
| AT1G32050 | 0.1 | 1.46E-01 | 0.1 | 6.54E-02 | 0.0 | 6.40E-01 | 0.0 | 9.63E-01 | -0.1 | 5.99E-01 | 0.1 | 6.42E-01 | Secretory carrier-associated membrane protein 4 |
| AT5G64160 | 0.1 | 3.47E-01 | 0.1 | 2.20E-01 | 0.0 | 7.93E-01 | -0.2 | 4.78E-01 | -0.3 | 2.81E-01 | 0.1 | 7.49E-01 | hypothetical protein |
| AT2G01340 | 0.1 | 5.81E-01 | 0.5 | 8.62E-03 | -0.4 | 4.78E-02 | 0.4 | 5.18E-02 | -0.2 | 4.12E-01 | 0.6 | 2.44E-02 | At2g01340 |
| AT3G55005 | 0.1 | 1.29E-01 | 0.1 | 4.07E-02 | 0.0 | 5.47E-01 | 0.0 | 8.10E-01 | 0.0 | 8.59E-01 | 0.1 | 7.06E-01 | Protein TONNEAU 1b |
| AT2G32190 | 0.1 | 6.67E-01 | -0.6 | 4.92E-02 | 0.6 | 4.90E-02 | -0.2 | 5.80E-01 | -0.3 | 3.74E-01 | 0.1 | 7.81E-01 | Cysteine-rich |
| AT5G49960 | 0.1 | 2.96E-01 | -0.1 | 3.42E-01 | 0.2 | 1.03E-01 | -0.3 | 1.52E-01 | 0.0 | 8.62E-01 | -0.3 | 1.62E-01 | Ion channel protein |
| AT2G40630 | 0.1 | 3.73E-01 | -0.1 | 3.04E-01 | 0.2 | 1.15E-01 | 0.1 | 7.21E-01 | -0.2 | 3.74E-01 | 0.3 | 2.72E-01 | At2g40630 |
| AT1G12400 | 0.1 | 3.76E-01 | 0.2 | 1.78E-01 | -0.1 | 6.78E-01 | 0.4 | 2.27E-01 | 0.4 | 2.35E-01 | 0.0 | 9.48E-01 | General transcription and DNA repair factor IIH subunit TFB5 |
| AT4G19870 | 0.1 | 2.49E-01 | 0.2 | 1.51E-02 | -0.1 | 1.72E-01 | -0.1 | 5.55E-01 | -0.1 | 5.96E-01 | 0.0 | 9.97E-01 | F-box |
| AT1G23560 | 0.1 | 4.68E-01 | 0.4 | 1.68E-02 | -0.3 | 1.08E-01 | 0.3 | 3.96E-01 | 0.0 | 9.85E-01 | 0.3 | 5.25E-01 | F28C11.19 |
| AT5G57015 | 0.1 | 1.61E-01 | 0.0 | 8.54E-01 | 0.1 | 2.97E-01 | -0.2 | 2.59E-01 | 0.0 | 9.01E-01 | -0.2 | 5.25E-01 | Ckl12 |
| AT2G45520 | 0.1 | 2.00E-01 | 0.1 | 2.56E-01 | 0.0 | 9.32E-01 | 0.2 | 3.81E-01 | 0.1 | 5.37E-01 | 0.0 | 9.48E-01 | Uncharacterized protein |
| AT2G37540 | 0.1 | 1.91E-01 | 0.0 | 5.90E-01 | 0.1 | 5.26E-01 | -0.2 | 2.10E-01 | -0.4 | 4.71E-02 | 0.2 | 4.44E-01 | NAD-binding Rossmann-fold superfamily protein |
| AT5G59960 | 0.1 | 2.89E-01 | 0.2 | 1.03E-01 | -0.1 | 5.67E-01 | 0.1 | 5.92E-01 | 0.0 | 8.82E-01 | 0.1 | 6.01E-01 | K-stimulated pyrophosphate-energized sodium pump protein |
| AT5G48965 | 0.1 | 2.26E-01 | -0.2 | 9.98E-02 | 0.3 | 2.68E-02 | -0.1 | 7.83E-01 | -0.2 | 5.64E-01 | 0.1 | 7.85E-01 | NA |
| AT2G47930 | 0.1 | 5.37E-01 | 0.5 | 4.81E-03 | -0.4 | 3.30E-02 | -0.7 | 7.31E-02 | -0.7 | 8.34E-02 | 0.0 | 9.60E-01 | Classical arabinogalactan protein 26 |
| AT1G13000 | 0.1 | 1.46E-01 | -0.1 | 2.25E-01 | 0.2 | 3.07E-02 | 0.1 | 4.30E-01 | 0.0 | 9.38E-01 | 0.1 | 6.40E-01 | Transmembrane protein, |
| AT5G52240 | 0.1 | 3.15E-01 | 0.1 | 5.80E-01 | 0.0 | 7.21E-01 | 0.0 | 9.64E-01 | -0.1 | 6.83E-01 | 0.1 | 7.88E-01 | Membrane steroid binding protein 1 |
| AT3G45750 | 0.1 | 3.93E-01 | -0.1 | 6.95E-01 | 0.2 | 2.99E-01 | -0.1 | 6.20E-01 | -0.1 | 6.82E-01 | 0.0 | 9.95E-01 | Nucleotidyltransferase family protein |
| AT5G06590 | 0.1 | 5.48E-01 | 0.2 | 3.31E-01 | -0.1 | 7.58E-01 | 0.4 | 2.79E-01 | 0.3 | 5.35E-01 | 0.1 | 8.16E-01 | Spindle and kinetochore-associated protein 3 |
| AT5G09390 | 0.1 | 1.65E-01 | 0.0 | 9.72E-01 | 0.1 | 2.19E-01 | 0.0 | 8.03E-01 | -0.1 | 6.59E-01 | 0.0 | 8.68E-01 | CD2-binding protein-like protein |
| AT4G08220 | 0.1 | 4.01E-01 | 0.1 | 2.68E-01 | 0.0 | 8.09E-01 | -0.1 | 6.43E-01 | 0.0 | 8.81E-01 | -0.1 | 8.73E-01 | NA |
| AT5G16790 | 0.1 | 5.37E-01 | 0.1 | 4.16E-01 | 0.0 | 8.63E-01 | 0.5 | 2.88E-01 | 0.2 | 7.95E-01 | 0.3 | 5.56E-01 | THO complex subunit 7A |
| AT1G69070 | 0.1 | 2.95E-01 | 0.0 | 9.80E-01 | 0.1 | 3.62E-01 | 0.7 | 2.93E-02 | 0.3 | 4.63E-01 | 0.4 | 3.06E-01 | Nucleolar-like protein |
| AT3G45900 | 0.1 | 2.23E-01 | 0.0 | 6.42E-01 | 0.1 | 1.50E-01 | -0.4 | 2.63E-02 | -0.2 | 1.81E-01 | -0.1 | 6.00E-01 | Ribonuclease P protein subunit P38-like protein |
| AT3G27350 | 0.1 | 4.09E-01 | -0.2 | 1.85E-01 | 0.3 | 7.00E-02 | -0.3 | 1.22E-01 | -0.3 | 3.14E-01 | -0.1 | 8.23E-01 | Transcriptional regulator ATRX-like protein |
| AT5G26160 | 0.1 | 3.40E-01 | -0.1 | 3.12E-01 | 0.2 | 1.12E-01 | 0.0 | 9.17E-01 | 0.0 | 9.84E-01 | -0.1 | 9.26E-01 | Light-independent protochlorophyllide reductase subunit |
| AT3G15840 | 0.1 | 2.84E-01 | 0.2 | 1.31E-01 | -0.1 | 6.37E-01 | -0.2 | 2.59E-01 | 0.1 | 5.89E-01 | -0.2 | 1.76E-01 | Post-illumination chlorophyll fluorescence increase |
| AT1G62050 | 0.1 | 2.72E-01 | -0.2 | 1.22E-01 | 0.3 | 3.54E-02 | -0.2 | 5.32E-01 | 0.3 | 3.53E-01 | -0.5 | 1.62E-01 | Ankyrin repeat family protein |
| AT5G64760 | 0.1 | 2.56E-01 | 0.0 | 9.42E-01 | 0.1 | 3.10E-01 | 0.3 | 8.33E-02 | 0.2 | 4.81E-01 | 0.2 | 4.65E-01 | Regulatory particle non-ATPase subunit 5B |
| AT1G53025 | 0.1 | 3.52E-01 | 0.0 | 9.68E-01 | 0.1 | 4.33E-01 | 0.1 | 6.59E-01 | 0.4 | 2.55E-01 | -0.2 | 5.49E-01 | Ubiquitin-conjugating enzyme family protein |
| AT5G67150 | 0.1 | 3.73E-01 | 0.2 | 6.61E-02 | -0.1 | 3.51E-01 | -0.1 | 5.94E-01 | 0.5 | 3.10E-02 | -0.6 | 1.52E-02 | HXXXD-type acyl-transferase family protein |
| AT3G18350 | 0.1 | 1.92E-01 | -0.1 | 8.73E-02 | 0.3 | 1.91E-02 | -0.3 | 1.55E-01 | -0.1 | 6.43E-01 | -0.2 | 5.67E-01 | Uncharacterized protein |
| AT4G16140 | 0.1 | 2.44E-01 | 0.3 | 1.02E-02 | -0.2 | 1.18E-01 | 0.0 | 8.36E-01 | 0.2 | 3.67E-01 | -0.2 | 5.00E-01 | Uncharacterized protein |
| AT2G25420 | 0.1 | 5.52E-01 | -0.1 | 6.14E-01 | 0.2 | 3.56E-01 | 0.1 | 7.16E-01 | 0.1 | 8.16E-01 | 0.0 | 9.61E-01 | Transducin family protein |
| AT3G06455 | 0.1 | 3.07E-01 | 0.2 | 1.60E-01 | 0.0 | 7.19E-01 | -0.4 | 7.35E-02 | -0.6 | 2.22E-02 | 0.2 | 6.25E-01 | SDE2-like C-terminal domain-containing protein |
| AT1G76310 | 0.1 | 4.28E-01 | 0.4 | 1.39E-02 | -0.3 | 9.99E-02 | 0.1 | 6.85E-01 | 0.2 | 4.41E-01 | -0.1 | 7.58E-01 | CYCLIN B24 |
| AT1G22140 | 0.1 | 2.67E-01 | 0.0 | 6.34E-01 | 0.1 | 6.10E-01 | 0.3 | 2.59E-01 | 0.2 | 6.55E-01 | 0.1 | 7.39E-01 | Zinc finger CCCH domain protein |
| AT2G46790 | 0.1 | 5.13E-01 | -0.3 | 1.16E-01 | 0.4 | 5.98E-02 | | | | | | | Uncharacterized protein At2g46670 |
| AT3G60820 | 0.1 | 2.24E-01 | 0.0 | 6.57E-01 | 0.1 | 5.11E-01 | 0.1 | 4.44E-01 | 0.1 | 6.70E-01 | 0.0 | 9.15E-01 | Proteasome subunit beta |
| AT5G05920 | 0.1 | 3.39E-01 | 0.0 | 9.02E-01 | 0.1 | 5.02E-01 | 0.4 | 6.47E-02 | 0.2 | 3.36E-01 | 0.1 | 6.20E-01 | Deoxyhypusine synthase |
| AT2G42080 | 0.1 | 2.09E-01 | 0.1 | 3.16E-01 | 0.0 | 8.68E-01 | 0.1 | 8.37E-01 | -0.1 | 8.63E-01 | 0.1 | 7.21E-01 | Chaperone DnaJ-domain superfamily protein |
| AT3G26920 | 0.1 | 2.48E-01 | -0.1 | 3.31E-01 | 0.2 | 8.50E-02 | -0.1 | 5.12E-01 | -0.1 | 7.10E-01 | 0.0 | 9.16E-01 | FBD |
| AT4G08460 | 0.1 | 2.48E-01 | 0.1 | 2.28E-01 | 0.0 | 9.42E-01 | -0.2 | 1.17E-01 | -0.2 | 1.21E-01 | 0.0 | 9.30E-01 | hypothetical protein |
| AT1G57540 | 0.1 | 3.80E-01 | 0.1 | 5.09E-01 | 0.0 | 8.88E-01 | 0.0 | 8.61E-01 | -0.2 | 4.89E-01 | 0.2 | 4.07E-01 | hypothetical protein |
| AT1G10570 | 0.1 | 2.77E-01 | -0.2 | 9.28E-02 | 0.3 | 2.77E-02 | -0.2 | 3.91E-01 | 0.1 | 8.49E-01 | -0.2 | 3.88E-01 | ULP1C |
| AT1G70480 | 0.1 | 1.05E-01 | -0.2 | 6.66E-03 | 0.3 | 1.61E-03 | -0.3 | 4.73E-02 | -0.2 | 1.86E-01 | -0.1 | 7.49E-01 | OBP32pep, putative |
| AT3G05727 | 0.1 | 5.38E-01 | 0.4 | 5.11E-02 | -0.3 | 1.73E-01 | 0.0 | 9.98E-01 | -0.4 | 2.60E-01 | 0.4 | 3.14E-01 | Defensin-like protein 204 |
| AT2G27020 | 0.1 | 3.63E-01 | 0.3 | 4.82E-02 | -0.2 | 2.51E-01 | 0.2 | 1.27E-01 | 0.1 | 5.87E-01 | 0.1 | 5.88E-01 | 20S proteasome alpha subunit G1 |
| AT4G24820 | 0.1 | 2.52E-01 | 0.0 | 7.38E-01 | 0.1 | 4.88E-01 | 0.2 | 7.99E-02 | 0.0 | 8.09E-01 | 0.2 | 3.19E-01 | 26S proteasome non-ATPase regulatory subunit 6 homolog |
| AT1G22330 | 0.1 | 4.19E-01 | 0.8 | 1.19E-04 | -0.7 | 9.73E-04 | -0.4 | 1.06E-01 | -0.6 | 6.24E-02 | 0.1 | 7.96E-01 | Uncharacterized protein |
| AT2G40935 | 0.1 | 3.58E-01 | 0.1 | 5.26E-01 | 0.0 | 8.51E-01 | -0.2 | 5.24E-01 | 0.0 | 9.60E-01 | -0.2 | 6.32E-01 | Protein PLANT CADMIUM RESISTANCE 10 |
| AT1G59730 | 0.1 | 6.87E-01 | 0.7 | 1.24E-02 | -0.6 | 5.26E-02 | 1.0 | 6.29E-02 | 0.8 | 1.98E-01 | 0.2 | 6.86E-01 | Thioredoxin H7 |
| AT5G36230 | 0.1 | 1.83E-01 | 0.2 | 1.67E-02 | -0.1 | 1.94E-01 | 0.0 | 7.97E-01 | -0.2 | 1.35E-01 | 0.1 | 3.56E-01 | ARM repeat superfamily protein |
| AT1G77310 | 0.1 | 3.78E-01 | 0.2 | 1.49E-01 | -0.1 | 6.08E-01 | 0.1 | 8.81E-01 | 0.0 | 9.97E-01 | 0.1 | 9.12E-01 | Wound-responsive family protein |
| AT5G55530 | 0.1 | 1.90E-01 | 0.0 | 6.02E-01 | 0.1 | 5.04E-01 | -0.4 | 3.16E-03 | -0.3 | 2.43E-02 | -0.1 | 7.02E-01 | C2 domain-containing protein |
| AT4G26130 | 0.1 | 1.47E-01 | 0.1 | 8.62E-02 | 0.0 | 7.45E-01 | 0.1 | 4.13E-01 | 0.1 | 6.48E-01 | 0.0 | 9.05E-01 | DUF4408 domain-containing protein |
| AT5G15440 | 0.1 | 2.32E-01 | 0.1 | 1.71E-01 | 0.0 | 8.53E-01 | -0.2 | 4.20E-01 | -0.2 | 4.13E-01 | 0.0 | 9.44E-01 | EID1-like F-box protein 1 |
| AT5G13480 | 0.1 | 2.59E-01 | 0.0 | 9.58E-01 | 0.1 | 3.20E-01 | 0.2 | 6.21E-01 | 0.2 | 6.76E-01 | 0.0 | 9.99E-01 | Transducin |
| AT2G17540 | 0.1 | 2.46E-01 | 0.0 | 6.79E-01 | 0.1 | 5.46E-01 | -0.2 | 9.75E-02 | -0.3 | 8.19E-02 | 0.0 | 8.87E-01 | Uncharacterized protein At2g17540 |
| AT4G00585 | 0.1 | 2.20E-01 | 0.4 | 4.97E-04 | -0.3 | 5.11E-03 | 0.0 | 8.04E-01 | 0.0 | 8.71E-01 | 0.0 | 9.81E-01 | Transmembrane protein |
| AT4G26450 | 0.1 | 1.62E-01 | -0.2 | 2.31E-02 | 0.3 | 4.51E-03 | 0.0 | 8.12E-01 | 0.1 | 7.00E-01 | -0.1 | 5.90E-01 | Uncharacterized protein At4g26450 |
| AT4G32560 | 0.1 | 3.78E-01 | -0.2 | 1.70E-01 | 0.3 | 6.80E-02 | -0.1 | 8.04E-01 | 0.2 | 5.05E-01 | -0.2 | 4.03E-01 | AT4G32560 protein |
| AT5G18310 | 0.1 | 3.06E-01 | -0.2 | 6.61E-02 | 0.3 | 1.93E-02 | 0.3 | 2.00E-01 | 0.2 | 5.22E-01 | 0.1 | 7.80E-01 | Ubiquitin hydrolase |
| AT1G64090 | 0.1 | 1.94E-01 | 0.2 | 4.03E-02 | -0.1 | 3.82E-01 | 0.1 | 6.44E-01 | 0.0 | 9.89E-01 | 0.1 | 7.56E-01 | Reticulon-like protein |
| AT1G80700 | 0.1 | 4.04E-01 | 0.0 | 7.93E-01 | 0.1 | 6.65E-01 | -0.1 | 8.18E-01 | -0.1 | 6.14E-01 | 0.1 | 8.33E-01 | F23A5.5 |
| AT2G34310 | 0.1 | 2.39E-01 | 0.1 | 2.56E-01 | 0.0 | 1.00E+00 | -0.1 | 6.74E-01 | -0.1 | 7.28E-01 | 0.0 | 9.99E-01 | Uncharacterized protein |
| AT5G62640 | 0.1 | 1.79E-01 | 0.0 | 5.58E-01 | 0.1 | 5.19E-01 | 0.1 | 5.44E-01 | 0.0 | 8.90E-01 | 0.2 | 5.68E-01 | Proline-rich family protein |
| AT1G22885 | 0.1 | 4.00E-01 | 0.4 | 8.41E-03 | -0.3 | 6.10E-02 | 0.3 | 2.07E-01 | 0.0 | 9.81E-01 | 0.3 | 2.81E-01 | Transmembrane protein |
| AT4G17080 | 0.1 | 1.88E-01 | 0.1 | 2.85E-01 | 0.0 | 8.59E-01 | 0.2 | 1.37E-01 | -0.2 | 4.58E-01 | 0.4 | 7.89E-02 | Histone H3 K4-specific methyltransferase SET7 |
| AT1G21560 | 0.1 | 3.07E-01 | 0.1 | 3.86E-01 | 0.0 | 9.30E-01 | 0.0 | 9.10E-01 | 0.1 | 6.69E-01 | -0.1 | 7.83E-01 | Uncharacterized protein At1g21560 |
| AT3G52730 | 0.1 | 3.07E-01 | 0.4 | 5.48E-03 | -0.3 | 4.12E-02 | 0.2 | 1.65E-01 | 0.1 | 4.94E-01 | 0.1 | 7.36E-01 | Complex III subunit 9 |
| AT3G03880 | 0.1 | 3.81E-01 | -0.1 | 3.19E-01 | 0.2 | 1.24E-01 | 0.0 | 9.66E-01 | -0.2 | 4.63E-01 | 0.2 | 4.79E-01 | hypothetical protein |
| AT4G13540 | 0.1 | 4.06E-01 | 0.2 | 1.81E-01 | -0.1 | 6.50E-01 | -0.2 | 5.93E-01 | -0.1 | 7.63E-01 | -0.1 | 9.23E-01 | Golgin family A protein |
| AT1G42430 | 0.1 | 2.02E-01 | 0.2 | 7.60E-02 | -0.1 | 6.04E-01 | 0.2 | 3.18E-01 | 0.2 | 5.99E-01 | 0.1 | 8.07E-01 | Inactive purple acid phosphatase-like protein |
| AT3G13310 | 0.1 | 3.16E-01 | 0.3 | 9.91E-03 | -0.2 | 7.74E-02 | 0.4 | 7.95E-02 | -0.3 | 2.86E-01 | 0.6 | 1.34E-02 | At3g13310 |

| | | | | | | | | | | | | | |
|---|---|---|---|---|---|---|---|---|---|---|---|---|---|
| AT4G03600 | 0.1 | 4.01E-01 | 0.0 | 8.55E-01 | 0.1 | 4.09E-01 | | | | | | | Pyrroline-5-carboxylate reductase |
| AT2G31060 | 0.1 | 2.83E-01 | -0.2 | 1.54E-01 | 0.3 | 3.84E-02 | 0.7 | 9.64E-06 | 0.2 | 1.46E-01 | 0.5 | 3.89E-03 | Putative GTP-binding protein |
| AT1G49520 | 0.1 | 4.27E-01 | 0.0 | 7.62E-01 | 0.2 | 3.70E-01 | 0.0 | 9.62E-01 | 0.1 | 8.76E-01 | -0.1 | 9.08E-01 | SWIB complex BAF60b domain-containing protein |
| AT1G77370 | 0.1 | 2.61E-01 | 0.0 | 6.66E-01 | 0.2 | 1.75E-01 | -0.1 | 8.00E-01 | -0.1 | 8.01E-01 | 0.0 | 9.77E-01 | Glutaredoxin family protein |
| AT3G58220 | 0.1 | 2.24E-01 | 0.2 | 3.95E-02 | -0.1 | 3.75E-01 | 0.1 | 7.42E-01 | -0.1 | 7.14E-01 | 0.1 | 5.40E-01 | TRAF-like family protein |
| AT3G62550 | 0.1 | 6.87E-01 | 0.9 | 5.03E-03 | -0.8 | 1.73E-02 | 0.4 | 3.83E-02 | 0.0 | 9.36E-01 | 0.3 | 1.76E-01 | hypothetical protein |
| AT1G59710 | 0.1 | 2.79E-01 | 0.0 | 8.72E-01 | 0.1 | 2.78E-01 | -0.2 | 1.21E-01 | -0.2 | 2.09E-01 | 0.0 | 9.49E-01 | DUF569 domain-containing protein |
| AT3G26000 | 0.1 | 2.44E-01 | -0.1 | 5.35E-01 | 0.2 | 1.21E-01 | -0.1 | 5.50E-01 | -0.1 | 3.67E-01 | 0.1 | 8.15E-01 | hypothetical protein |
| AT2G40410 | 0.1 | 1.17E-01 | 0.2 | 4.29E-02 | 0.0 | 5.96E-01 | -0.2 | 7.65E-02 | -0.4 | 1.96E-02 | 0.1 | 5.15E-01 | Ca-dependent nuclease family protein |
| AT3G18140 | 0.1 | 2.11E-01 | -0.1 | 5.03E-01 | 0.2 | 9.72E-02 | -0.2 | 1.69E-01 | -0.1 | 6.50E-01 | -0.1 | 6.32E-01 | Target of rapamycin complex subunit LST8 |
| AT3G11530 | 0.1 | 1.96E-01 | 0.1 | 1.79E-01 | 0.0 | 9.40E-01 | -0.1 | 7.03E-01 | -0.2 | 4.44E-01 | 0.1 | 7.79E-01 | Uncharacterized protein F24K9.21 |
| AT2G17670 | 0.1 | 3.23E-01 | 0.0 | 8.39E-01 | 0.1 | 5.35E-01 | 0.5 | 6.09E-02 | 0.1 | 8.04E-01 | 0.4 | 1.84E-01 | Tetratricopeptide repeat -like superfamily protein |
| AT4G00090 | 0.1 | 1.68E-01 | 0.2 | 6.65E-02 | 0.0 | 6.24E-01 | 0.1 | 5.84E-01 | -0.1 | 6.67E-01 | 0.2 | 4.10E-01 | AT4g00090 |
| AT4G14548 | 0.1 | 5.68E-01 | 0.2 | 4.34E-01 | 0.0 | 8.56E-01 | 0.1 | 8.62E-01 | 0.3 | 5.44E-01 | -0.2 | 7.44E-01 | NA |
| AT3G11964 | 0.1 | 2.93E-01 | 0.1 | 6.27E-01 | 0.1 | 6.48E-01 | 0.6 | 8.36E-04 | 0.4 | 3.85E-02 | 0.2 | 2.56E-01 | Uncharacterized protein |
| AT2G43720 | 0.1 | 6.09E-01 | -0.4 | 9.20E-02 | 0.6 | 7.21E-02 | 0.6 | 1.29E-01 | -0.1 | 9.19E-01 | 0.7 | 1.17E-01 | FAM136A-like protein |
| AT1G28100 | 0.1 | 4.38E-01 | 0.2 | 1.68E-01 | -0.1 | 5.91E-01 | -0.1 | 7.41E-01 | -0.2 | 6.18E-01 | 0.1 | 8.75E-01 | Protein NEOXANTHIN-DEFICIENT 1 |
| AT5G12120 | 0.1 | 2.04E-01 | -0.1 | 2.59E-01 | 0.2 | 4.93E-02 | -0.6 | 4.49E-01 | -0.4 | 6.56E-01 | -0.2 | 8.97E-01 | hypothetical protein |
| AT2G46100 | 0.1 | 3.54E-01 | -0.3 | 5.91E-02 | 0.4 | 1.96E-02 | -0.2 | 2.65E-01 | 0.1 | 4.66E-01 | -0.3 | 1.25E-01 | Nuclear transport factor 2 family protein |
| AT4G37460 | 0.1 | 1.03E-01 | -0.2 | 2.05E-02 | 0.3 | 2.89E-03 | -0.2 | 1.40E-01 | -0.1 | 5.38E-01 | -0.1 | 6.67E-01 | Tetratricopeptide repeat -like superfamily protein |
| AT1G77440 | 0.1 | 3.36E-01 | 0.1 | 2.57E-01 | 0.0 | 8.60E-01 | 0.1 | 5.31E-01 | 0.1 | 8.15E-01 | 0.1 | 8.39E-01 | Proteasome subunit beta |
| AT3G13437 | 0.1 | 6.26E-01 | -0.2 | 4.87E-01 | 0.3 | 3.30E-01 | 0.4 | 2.58E-01 | 0.0 | 9.66E-01 | 0.4 | 2.99E-01 | Transmembrane protein |
| AT3G62300 | 0.1 | 1.89E-01 | 0.1 | 2.37E-01 | 0.0 | 9.43E-01 | 0.0 | 9.36E-01 | 0.0 | 9.31E-01 | 0.0 | 8.90E-01 | Agenet domain-containing protein |
| AT3G49645 | 0.1 | 3.58E-01 | -0.1 | 7.00E-01 | 0.2 | 2.78E-01 | 0.0 | 9.54E-01 | 0.0 | 9.28E-01 | 0.1 | 9.14E-01 | Uncharacterized protein |
| AT1G01130 | 0.1 | 4.15E-01 | 0.2 | 1.35E-01 | -0.1 | 5.42E-01 | 0.0 | 9.22E-01 | -0.1 | 9.22E-01 | 0.1 | 8.66E-01 | CBL-interacting Serine |
| AT5G21010 | 0.1 | 1.33E-01 | 0.0 | 9.65E-01 | 0.1 | 1.84E-01 | -0.2 | 2.46E-01 | -0.1 | 7.84E-01 | -0.2 | 6.12E-01 | BTB |
| AT1G09840 | 0.1 | 7.04E-02 | 0.1 | 1.74E-01 | 0.0 | 6.91E-01 | 0.1 | 5.30E-01 | 0.1 | 6.84E-01 | 0.0 | 9.53E-01 | AT1G09840 protein |
| AT1G18850 | 0.1 | 4.24E-01 | 0.3 | 3.38E-02 | -0.2 | 1.92E-01 | 0.5 | 1.06E-02 | 0.1 | 6.85E-01 | 0.4 | 5.71E-02 | Uncharacterized protein |
| AT2G38185 | 0.1 | 5.59E-01 | -0.1 | 5.26E-01 | 0.3 | 3.15E-01 | 0.0 | 9.99E-01 | 0.1 | 9.56E-01 | -0.1 | 9.63E-01 | At2g38190 |
| AT2G19530 | 0.1 | 5.62E-01 | 0.2 | 3.49E-01 | -0.1 | 7.65E-01 | 0.3 | 2.88E-01 | 0.2 | 6.00E-01 | 0.1 | 7.79E-01 | Transmembrane protein |
| AT3G57062 | 0.1 | 1.05E-01 | 0.2 | 7.12E-03 | -0.1 | 1.77E-01 | 0.0 | 8.96E-01 | 0.0 | 9.74E-01 | 0.0 | 9.55E-01 | hypothetical protein |
| AT1G69030 | 0.1 | 2.94E-01 | 0.3 | 3.16E-02 | -0.1 | 2.54E-01 | 0.2 | 3.45E-01 | 0.2 | 4.47E-01 | 0.0 | 9.80E-01 | BSD domain-containing protein |
| AT3G53240 | 0.1 | 1.76E-01 | 0.0 | 6.76E-01 | 0.2 | 1.38E-01 | -0.3 | 8.54E-02 | -0.2 | 2.43E-01 | -0.1 | 8.17E-01 | Receptor like protein 45 |
| AT2G20142 | 0.1 | 6.17E-01 | -0.8 | 1.54E-02 | 0.9 | 1.66E-02 | 0.1 | 8.07E-01 | -0.1 | 9.14E-01 | 0.2 | 7.77E-01 | Toll-Interleukin-Resistance domain family protein |
| AT1G70180 | 0.1 | 1.47E-01 | 0.0 | 5.84E-01 | 0.2 | 9.32E-02 | -0.3 | 9.30E-02 | -0.3 | 2.06E-01 | -0.1 | 8.85E-01 | Sterile alpha motif domain-containing protein |
| AT2G03865 | 0.1 | 7.99E-01 | 0.1 | 8.25E-01 | 0.0 | 9.88E-01 | 0.2 | 9.18E-01 | 0.1 | 9.71E-01 | 0.1 | 9.66E-01 | NA |
| AT2G37080 | 0.1 | 2.20E-01 | 0.1 | 2.11E-01 | 0.0 | 9.58E-01 | 0.0 | 9.47E-01 | 0.0 | 9.17E-01 | 0.0 | 9.70E-01 | ROP interactive partner 3 |
| AT3G56810 | 0.1 | 2.45E-01 | 0.0 | 9.81E-01 | 0.1 | 3.25E-01 | -0.1 | 7.31E-01 | 0.0 | 8.36E-01 | 0.0 | 9.62E-01 | Uncharacterized protein |
| AT1G24575 | 0.1 | 4.25E-01 | -0.2 | 2.85E-01 | 0.3 | 1.18E-01 | 0.7 | 3.73E-02 | 0.4 | 3.54E-01 | 0.3 | 4.54E-01 | hypothetical protein |
| AT3G58930 | 0.1 | 2.54E-01 | 0.0 | 9.64E-01 | 0.1 | 3.77E-01 | 0.2 | 5.22E-01 | -0.1 | 6.82E-01 | 0.3 | 3.19E-01 | F-box |
| AT5G65520 | 0.1 | 1.44E-01 | 0.0 | 9.70E-01 | 0.1 | 2.06E-01 | -0.1 | 5.79E-01 | -0.3 | 3.00E-01 | 0.1 | 7.07E-01 | hypothetical protein |
| AT3G44330 | 0.1 | 1.95E-01 | 0.3 | 7.88E-03 | -0.2 | 9.45E-02 | 0.0 | 7.73E-01 | 0.0 | 8.71E-01 | 0.0 | 9.65E-01 | Nicalin |
| AT5G48310 | 0.1 | 3.63E-01 | 0.2 | 8.50E-02 | -0.1 | 4.42E-01 | 0.3 | 3.30E-01 | 0.2 | 6.04E-01 | 0.1 | 8.37E-01 | Portal protein |
| AT4G30150 | 0.1 | 1.31E-01 | -0.2 | 2.17E-02 | 0.3 | 4.85E-03 | 0.8 | 3.89E-03 | 0.4 | 2.79E-01 | 0.5 | 1.18E-01 | Urb2 |
| AT1G35430 | 0.1 | 2.78E-01 | 0.1 | 3.20E-01 | 0.0 | 9.69E-01 | -0.6 | 3.77E-02 | -0.4 | 2.17E-01 | -0.2 | 6.17E-01 | Uncharacterized protein F12A4.3 |
| AT1G74750 | 0.1 | 1.94E-01 | -0.1 | 2.21E-01 | 0.2 | 4.76E-02 | 0.4 | 4.73E-01 | 0.2 | 7.88E-01 | 0.2 | 8.11E-01 | Pentatricopeptide repeat-containing protein At1g74750 |
| AT3G11890 | 0.1 | 1.06E-01 | 0.0 | 8.17E-01 | 0.1 | 1.16E-01 | -0.2 | 2.25E-01 | -0.1 | 8.25E-01 | -0.2 | 5.20E-01 | F26K24.18 protein domain-containing protein) |
| AT5G14580 | 0.1 | 3.57E-01 | 0.1 | 2.98E-01 | 0.0 | 9.07E-01 | 0.9 | 2.47E-03 | 0.6 | 6.74E-02 | 0.3 | 3.01E-01 | Putative polynucleotide phosphorylase |
| AT2G27730 | 0.1 | 2.93E-01 | 0.4 | 8.39E-03 | -0.3 | 6.53E-02 | 0.1 | 6.77E-01 | 0.1 | 6.14E-01 | 0.0 | 9.20E-01 | hypothetical protein |
| AT1G15200 | 0.1 | 1.44E-01 | 0.0 | 8.52E-01 | 0.1 | 2.58E-01 | -0.1 | 3.87E-01 | -0.1 | 4.66E-01 | 0.0 | 9.97E-01 | Protein-protein interaction regulator family protein |
| AT1G76280 | 0.1 | 2.63E-01 | 0.1 | 2.80E-01 | 0.0 | 9.96E-01 | 0.0 | 8.34E-01 | -0.3 | 2.20E-01 | 0.3 | 2.16E-01 | Tetratricopeptide repeat -like superfamily protein |
| AT5G67610 | 0.1 | 2.15E-01 | 0.1 | 4.48E-01 | 0.0 | 7.15E-01 | -0.1 | 7.47E-01 | 0.0 | 9.23E-01 | -0.1 | 6.79E-01 | Transmembrane protein |
| AT5G08440 | 0.1 | 3.20E-01 | -0.1 | 3.63E-01 | 0.2 | 1.14E-01 | -0.2 | 4.66E-01 | 0.0 | 8.79E-01 | -0.2 | 4.88E-01 | Uncharacterized protein F8L15.170 |
| AT3G52420 | 0.1 | 1.70E-01 | 0.0 | 9.81E-01 | 0.1 | 2.40E-01 | 0.0 | 9.55E-01 | 0.0 | 8.87E-01 | 0.0 | 9.24E-01 | Outer envelope membrane protein 7 |
| AT3G27906 | 0.1 | 1.46E-01 | 0.1 | 9.56E-02 | 0.0 | 8.16E-01 | -0.2 | 3.38E-01 | -0.1 | 7.31E-01 | -0.1 | 7.57E-01 | At3g27906 |
| AT4G21660 | 0.1 | 7.70E-02 | 0.1 | 3.78E-01 | 0.1 | 4.14E-01 | -0.1 | 2.20E-01 | -0.2 | 6.71E-02 | 0.1 | 6.24E-01 | At4g21660 family protein) |
| AT2G20820 | 0.1 | 1.82E-01 | 0.4 | 3.04E-03 | -0.2 | 3.76E-02 | 0.2 | 2.40E-01 | 0.2 | 4.54E-01 | 0.0 | 8.87E-01 | Expressed protein |
| AT1G04945 | 0.1 | 3.96E-01 | -0.1 | 4.40E-01 | 0.2 | 1.81E-01 | 0.6 | 4.40E-02 | 0.1 | 7.73E-01 | 0.5 | 1.07E-01 | HIT-type Zinc finger family protein |
| AT1G06980 | 0.1 | 6.07E-01 | 0.8 | 5.40E-03 | -0.6 | 2.89E-02 | -0.9 | 3.38E-02 | -0.3 | 4.92E-01 | -0.6 | 2.28E-01 | hypothetical protein |
| AT3G52360 | 0.1 | 1.73E-01 | 0.5 | 2.21E-04 | -0.4 | 3.07E-03 | 0.4 | 5.92E-02 | 0.4 | 1.03E-01 | 0.0 | 9.80E-01 | hypothetical protein |
| AT5G59710 | 0.1 | 9.11E-02 | -0.2 | 3.73E-02 | 0.3 | 4.66E-03 | 0.0 | 9.17E-01 | 0.1 | 7.66E-01 | -0.1 | 8.70E-01 | VIRE2 interacting protein 2 |
| AT3G01430 | 0.1 | 3.22E-01 | 0.0 | 7.41E-01 | 0.2 | 2.76E-01 | -0.2 | 4.65E-01 | -0.4 | 1.10E-01 | 0.2 | 4.82E-01 | hypothetical protein |
| AT4G27870 | 0.1 | 1.54E-01 | 0.3 | 1.14E-02 | -0.1 | 1.70E-01 | -0.3 | 3.89E-03 | -0.2 | 8.74E-02 | -0.1 | 4.27E-01 | Vacuolar iron transporter family protein |
| AT1G13180 | 0.1 | 1.33E-01 | 0.0 | 6.13E-01 | 0.1 | 4.09E-01 | -0.2 | 2.64E-01 | 0.1 | 5.45E-01 | -0.3 | 1.36E-01 | DIS1 |
| AT3G55060 | 0.1 | 1.06E-01 | 0.1 | 5.00E-01 | 0.1 | 4.34E-01 | 0.2 | 3.88E-01 | 0.1 | 7.62E-01 | 0.1 | 7.60E-01 | CAP-gly domain linker |
| AT5G37480 | 0.1 | 2.02E-01 | 0.1 | 3.68E-01 | 0.0 | 7.84E-01 | 0.4 | 2.56E-02 | 0.1 | 7.26E-01 | 0.3 | 1.62E-01 | Transmembrane protein |
| AT1G79200 | 0.1 | 3.02E-01 | 0.3 | 2.58E-02 | -0.2 | 2.17E-01 | 0.1 | 8.36E-01 | 0.0 | 8.80E-01 | 0.0 | 9.92E-01 | Style cell-cycle inhibitor 1 |
| AT1G52870 | 0.1 | 2.14E-01 | 0.0 | 9.50E-01 | 0.1 | 2.41E-01 | 0.0 | 7.72E-01 | 0.2 | 2.06E-01 | -0.1 | 3.85E-01 | Peroxisomal membrane 22 kDa family protein |
| AT1G03260 | 0.1 | 1.89E-01 | 0.2 | 5.84E-02 | -0.1 | 5.46E-01 | -0.2 | 3.97E-01 | -0.3 | 2.73E-01 | 0.1 | 8.01E-01 | SNARE associated Golgi protein family |
| AT1G26740 | 0.1 | 3.33E-01 | 0.3 | 4.10E-02 | -0.2 | 2.79E-01 | 0.3 | 1.02E-01 | 0.0 | 9.31E-01 | 0.3 | 2.57E-01 | T24P13.12 |
| AT4G29660 | 0.1 | 2.71E-01 | -0.1 | 4.09E-01 | 0.2 | 1.15E-01 | 0.2 | 4.52E-01 | 0.0 | 9.83E-01 | 0.2 | 5.33E-01 | Uncharacterized protein At4g29660 |
| AT2G35605 | 0.1 | 4.33E-01 | 0.5 | 9.59E-03 | -0.4 | 6.34E-02 | 0.3 | 1.42E-01 | 0.1 | 6.15E-01 | 0.2 | 5.84E-01 | hypothetical protein |
| AT3G23740 | 0.1 | 2.58E-01 | 0.1 | 5.02E-01 | 0.0 | 7.47E-01 | 0.5 | 1.11E-01 | 0.5 | 1.51E-01 | 0.0 | 9.97E-01 | Uncharacterized protein |
| AT4G14800 | 0.1 | 1.75E-01 | 0.0 | 6.44E-01 | 0.1 | 4.38E-01 | 0.1 | 2.61E-01 | 0.1 | 7.27E-01 | 0.1 | 6.97E-01 | Proteasome subunit beta |
| AT3G22110 | 0.1 | 1.00E-01 | 0.2 | 5.52E-02 | 0.0 | 7.20E-01 | 0.1 | 3.35E-01 | 0.1 | 7.20E-01 | 0.1 | 7.96E-01 | Proteasome subunit alpha type-4-A |
| AT1G30320 | 0.1 | 1.63E-01 | 0.0 | 7.48E-01 | 0.2 | 1.54E-01 | -0.6 | 1.42E-03 | -0.2 | 1.44E-01 | -0.3 | 1.51E-01 | Remorin family protein |
| AT2G04900 | 0.1 | 1.04E-01 | 0.1 | 4.16E-01 | 0.1 | 4.74E-01 | 0.0 | 8.03E-01 | -0.2 | 4.12E-01 | 0.1 | 6.37E-01 | Transmembrane protein |
| AT2G34780 | 0.1 | 8.18E-02 | -0.1 | 1.69E-01 | 0.2 | 1.63E-02 | 0.2 | 2.97E-01 | 0.2 | 3.88E-01 | 0.0 | 9.80E-01 | Uncharacterized protein At2g34780 |
| AT5G53220 | 0.1 | 2.96E-01 | -0.2 | 1.60E-01 | 0.3 | 5.40E-02 | -0.1 | 7.82E-01 | 0.0 | 9.87E-01 | -0.1 | 8.50E-01 | Uncharacterized protein |
| AT3G44798 | 0.1 | 5.84E-01 | 0.4 | 1.44E-01 | -0.2 | 4.23E-01 | -0.3 | 1.82E-01 | -0.5 | 4.15E-02 | 0.2 | 5.27E-01 | NA |
| AT4G16695 | 0.1 | 2.36E-01 | 0.3 | 1.29E-02 | -0.2 | 1.53E-01 | 0.2 | 2.48E-01 | 0.0 | 9.89E-01 | 0.2 | 3.44E-01 | Transmembrane protein |
| AT3G17020 | 0.1 | 3.08E-01 | 0.1 | 6.83E-01 | 0.1 | 6.14E-01 | -0.4 | 6.59E-03 | -0.3 | 9.85E-02 | -0.1 | 5.02E-01 | hypothetical protein |
| AT2G30120 | 0.1 | 3.11E-01 | 0.0 | 8.06E-01 | 0.2 | 2.96E-01 | -0.1 | 8.58E-01 | 0.0 | 9.71E-01 | -0.1 | 8.64E-01 | Protein FLC EXPRESSOR |

| | | | | | | | | | | | | | |
|---|---|---|---|---|---|---|---|---|---|---|---|---|---|
| AT1G13160 | 0.1 | 1.68E-01 | 0.1 | 1.70E-01 | 0.0 | 9.75E-01 | 0.3 | 2.37E-01 | 0.3 | 3.01E-01 | 0.0 | 9.99E-01 | Protein SDA1 |
| AT5G13070 | 0.1 | 1.06E-01 | 0.2 | 2.17E-02 | -0.1 | 4.16E-01 | 0.3 | 6.50E-02 | 0.1 | 5.61E-01 | 0.2 | 4.30E-01 | hypothetical protein |
| AT1G78040 | 0.1 | 2.39E-01 | 0.3 | 3.78E-02 | -0.1 | 3.05E-01 | -0.3 | 7.56E-02 | -0.2 | 4.31E-01 | -0.1 | 6.00E-01 | Pollen Ole e 1 allergen and extensin family protein |
| AT1G79020 | 0.1 | 1.59E-01 | -0.2 | 1.29E-01 | 0.3 | 2.29E-02 | -0.1 | 5.17E-01 | 0.1 | 8.19E-01 | -0.2 | 4.57E-01 | Enhancer of polycomb-like protein |
| AT5G37590 | 0.1 | 2.61E-01 | -0.1 | 3.53E-01 | 0.2 | 9.71E-02 | 0.2 | 4.98E-01 | 0.2 | 5.18E-01 | 0.0 | 9.77E-01 | Similarity to Krm protein |
| AT4G20090 | 0.1 | 3.79E-01 | -0.1 | 6.00E-01 | 0.2 | 2.52E-01 | 0.3 | 3.13E-01 | 0.0 | 9.56E-01 | 0.2 | 4.91E-01 | Pentatricopeptide repeat-containing protein At4g20090 |
| AT2G45980 | 0.1 | 1.06E-01 | 0.0 | 8.19E-01 | 0.2 | 1.02E-01 | -0.3 | 6.09E-02 | -0.5 | 8.04E-03 | 0.2 | 3.79E-01 | ATG8-interacting protein 1 |
| AT4G16100 | 0.1 | 1.68E-01 | -0.1 | 4.24E-01 | 0.2 | 7.88E-02 | 0.1 | 6.96E-01 | 0.0 | 8.74E-01 | 0.1 | 6.40E-01 | At4g16100 ) |
| AT4G22850 | 0.1 | 2.07E-01 | 0.2 | 3.39E-02 | -0.1 | 3.62E-01 | 0.3 | 1.28E-01 | 0.4 | 1.01E-01 | -0.1 | 8.75E-01 | SNARE associated Golgi protein family |
| AT1G79190 | 0.1 | 1.18E-01 | -0.2 | 1.74E-02 | 0.4 | 3.99E-03 | 0.2 | 2.65E-01 | -0.1 | 8.49E-01 | 0.3 | 2.81E-01 | ARM repeat superfamily protein |
| AT3G20920 | 0.1 | 1.17E-01 | 0.2 | 2.05E-02 | -0.1 | 3.61E-01 | -0.1 | 6.52E-01 | 0.0 | 7.74E-01 | 0.0 | 9.64E-01 | Translocation protein SEC62 |
| AT2G47230 | 0.1 | 1.95E-01 | 0.1 | 6.01E-01 | 0.1 | 5.49E-01 | 0.0 | 9.68E-01 | 0.0 | 9.27E-01 | 0.1 | 8.84E-01 | Agenet domain-containing protein |
| AT2G14090 | 0.1 | 4.02E-01 | 0.1 | 5.57E-01 | 0.0 | 8.79E-01 | 0.4 | 1.84E-01 | 0.1 | 9.12E-01 | 0.4 | 3.23E-01 | NA |
| AT2G18690 | 0.1 | 5.30E-01 | 0.0 | 9.95E-01 | 0.1 | 5.80E-01 | 0.9 | 2.38E-04 | 0.0 | 9.38E-01 | 0.9 | 3.50E-03 | At2g18690 |
| AT1G78560 | 0.1 | 1.03E-01 | 0.1 | 1.70E-01 | 0.0 | 8.46E-01 | 0.0 | 9.83E-01 | 0.2 | 2.59E-01 | -0.2 | 3.86E-01 | Probable sodium |
| AT1G56420 | 0.1 | 2.84E-01 | 0.0 | 8.55E-01 | 0.2 | 3.07E-01 | 0.1 | 8.42E-01 | 0.3 | 4.68E-01 | -0.2 | 6.77E-01 | hypothetical protein |
| AT5G49880 | 0.1 | 9.65E-02 | -0.2 | 4.98E-02 | 0.3 | 5.89E-03 | 0.1 | 3.81E-01 | 0.2 | 3.17E-01 | 0.0 | 8.72E-01 | MAD1 |
| AT3G15160 | 0.1 | 7.84E-02 | 0.2 | 3.52E-02 | 0.0 | 6.68E-01 | -0.2 | 1.26E-01 | -0.1 | 3.69E-01 | -0.1 | 7.67E-01 | AP-5 complex subunit zeta-1 |
| AT1G02110 | 0.1 | 6.49E-02 | 0.2 | 1.09E-02 | -0.1 | 3.47E-01 | -0.3 | 1.47E-01 | -0.1 | 5.75E-01 | -0.2 | 6.05E-01 | hypothetical protein |
| AT1G04190 | 0.1 | 1.40E-01 | 0.3 | 4.25E-03 | -0.2 | 8.22E-02 | 0.4 | 3.47E-03 | 0.3 | 8.16E-02 | 0.2 | 3.96E-01 | TPR3 |
| AT1G72410 | 0.1 | 7.08E-02 | 0.3 | 4.51E-03 | -0.1 | 1.52E-01 | -0.2 | 3.37E-01 | -0.2 | 5.02E-01 | 0.0 | 9.39E-01 | COP1-interacting protein-like protein |
| AT3G24820 | 0.1 | 5.99E-01 | -0.1 | 8.02E-01 | 0.2 | 5.20E-01 | -0.3 | 2.77E-01 | -0.4 | 1.37E-01 | 0.1 | 7.31E-01 | BSD domain-containing protein |
| AT1G70505 | 0.1 | 4.79E-01 | 0.0 | 9.13E-01 | 0.2 | 5.04E-01 | -0.2 | 6.54E-01 | -0.1 | 8.95E-01 | -0.1 | 7.91E-01 | Transmembrane protein |
| AT5G48340 | 0.1 | 1.36E-01 | 0.0 | 8.20E-01 | 0.2 | 1.49E-01 | 0.1 | 7.84E-01 | 0.0 | 9.87E-01 | 0.1 | 8.66E-01 | Uncharacterized protein At5g48340 |
| AT1G32730 | 0.1 | 1.86E-01 | 0.4 | 3.25E-03 | -0.2 | 5.88E-02 | -0.2 | 5.77E-01 | 0.0 | 9.59E-01 | -0.2 | 7.40E-01 | Electron carrier |
| AT1G06130 | 0.1 | 3.70E-02 | -0.1 | 4.24E-02 | 0.3 | 2.70E-03 | -0.1 | 4.63E-01 | -0.2 | 1.25E-01 | 0.1 | 5.39E-01 | GLX2-4 |
| AT3G60520 | 0.1 | 4.10E-01 | 0.1 | 4.56E-01 | 0.0 | 9.74E-01 | -0.2 | 4.90E-01 | -0.4 | 2.44E-01 | 0.2 | 6.90E-01 | Zinc ion-binding protein |
| AT4G04940 | 0.1 | 2.77E-01 | 0.3 | 4.19E-02 | -0.1 | 3.35E-01 | 0.6 | 1.17E-02 | 0.5 | 6.72E-02 | 0.1 | 6.76E-01 | Transducin family protein |
| AT5G45090 | 0.1 | 3.78E-01 | 0.3 | 4.21E-02 | -0.2 | 2.73E-01 | 1.2 | 1.36E-03 | 0.4 | 3.10E-01 | 0.7 | 2.43E-02 | Phloem protein 2-A7 |
| AT3G19900 | 0.1 | 2.96E-01 | 0.0 | 9.15E-01 | 0.2 | 3.17E-01 | -0.1 | 7.56E-01 | -0.2 | 2.80E-01 | 0.1 | 4.30E-01 | AT3G19900 protein |
| AT1G04590 | 0.1 | 1.98E-01 | 0.4 | 5.61E-03 | -0.2 | 8.50E-02 | 0.3 | 9.90E-02 | 0.2 | 3.23E-01 | 0.1 | 7.21E-01 | PPR containing-like protein |
| AT5G23680 | 0.1 | 2.63E-01 | 0.1 | 4.74E-01 | 0.0 | 7.73E-01 | -0.3 | 2.62E-01 | -0.1 | 6.95E-01 | -0.2 | 6.79E-01 | SAM domain-containing protein |
| AT5G62270 | 0.1 | 2.69E-01 | 0.2 | 7.06E-02 | -0.1 | 4.68E-01 | 0.5 | 1.14E-02 | 0.2 | 3.03E-01 | 0.2 | 2.36E-01 | GCD1 |
| AT4G08240 | 0.1 | 2.18E-01 | 0.2 | 1.69E-01 | 0.0 | 8.76E-01 | -0.4 | 7.46E-02 | -0.2 | 5.25E-01 | -0.2 | 4.32E-01 | hypothetical protein |
| AT5G00420 | 0.1 | 3.99E-01 | 0.1 | 7.46E-01 | 0.1 | 7.11E-01 | 0.1 | 8.99E-01 | 0.2 | 8.21E-01 | -0.1 | 9.12E-01 | NA |
| AT1G19950 | 0.1 | 2.84E-01 | 0.3 | 2.67E-02 | -0.2 | 2.37E-01 | -0.1 | 6.59E-01 | 0.3 | 1.92E-01 | -0.3 | 1.05E-01 | HVA22-like protein h |
| AT3G62650 | 0.1 | 1.93E-01 | 0.2 | 3.76E-02 | -0.1 | 3.77E-01 | -0.3 | 9.25E-02 | -0.4 | 5.90E-02 | 0.1 | 8.09E-01 | Uncharacterized protein At3g62650; F26K9_80 |
| AT4G13120 | 0.1 | 2.25E-01 | 0.3 | 3.66E-02 | -0.1 | 3.76E-01 | -0.1 | 7.25E-01 | 0.2 | 5.48E-01 | -0.4 | 3.89E-01 | NA |
| AT1G11120 | 0.1 | 5.47E-01 | 0.5 | 4.83E-02 | -0.3 | 2.05E-01 | -0.3 | 3.65E-01 | 0.1 | 8.84E-01 | -0.3 | 3.74E-01 | CTTNBP 2 amino-terminal-like protein |
| AT3G23030 | 0.1 | 1.86E-01 | 0.0 | 9.85E-01 | 0.1 | 2.36E-01 | -0.1 | 7.40E-01 | -0.3 | 1.81E-01 | 0.2 | 4.19E-01 | Auxin-responsive protein |
| AT3G44690 | 0.1 | 4.06E-01 | -0.1 | 4.55E-01 | 0.3 | 2.01E-01 | 0.4 | 4.21E-01 | 0.1 | 9.13E-01 | 0.3 | 6.63E-01 | FIP1-like protein |
| AT1G16810 | 0.1 | 7.22E-02 | 0.1 | 4.08E-01 | 0.1 | 3.69E-01 | -0.2 | 1.51E-01 | -0.3 | 9.60E-02 | 0.1 | 8.15E-01 | hypothetical protein |
| AT5G19860 | 0.1 | 7.64E-02 | 0.3 | 3.01E-03 | -0.2 | 8.36E-02 | -0.1 | 4.22E-01 | 0.0 | 8.01E-01 | -0.1 | 7.99E-01 | Uncharacterized protein |
| AT3G48180 | 0.1 | 8.90E-02 | 0.5 | 1.41E-04 | -0.3 | 3.60E-03 | 0.3 | 1.92E-01 | 0.2 | 3.89E-01 | 0.1 | 8.49E-01 | hypothetical protein |
| AT1G03870 | 0.1 | 3.57E-01 | 0.6 | 2.41E-03 | -0.5 | 1.53E-02 | -0.4 | 1.90E-02 | -0.5 | 2.63E-02 | 0.0 | 9.48E-01 | FAS1 domain-containing protein |
| AT1G05270 | 0.1 | 6.65E-02 | 0.1 | 1.50E-01 | 0.0 | 7.33E-01 | -0.2 | 1.56E-01 | -0.2 | 2.62E-01 | 0.0 | 9.42E-01 | hypothetical protein |
| AT5G55620 | 0.1 | 3.13E-01 | 0.1 | 5.62E-01 | 0.1 | 7.37E-01 | -0.1 | 7.82E-01 | 0.1 | 7.80E-01 | -0.2 | 6.42E-01 | Gb\|AAF04428.1 |
| AT5G47400 | 0.1 | 1.23E-01 | -0.1 | 4.93E-01 | 0.2 | 7.09E-02 | -0.2 | 3.62E-01 | -0.2 | 3.41E-01 | 0.0 | 9.28E-01 | Uncharacterized protein |
| AT5G47090 | 0.1 | 1.03E-01 | 0.1 | 4.84E-01 | 0.1 | 4.33E-01 | 0.0 | 9.00E-01 | -0.2 | 3.07E-01 | 0.2 | 3.41E-01 | AT5g47090 |
| AT2G40230 | 0.1 | 2.27E-01 | 0.2 | 9.24E-02 | -0.1 | 6.33E-01 | 0.1 | 8.29E-01 | 0.3 | 1.66E-01 | -0.3 | 3.19E-01 | hypothetical protein |
| AT3G19870 | 0.1 | 9.06E-02 | 0.0 | 9.09E-01 | 0.1 | 1.63E-01 | -0.1 | 4.89E-01 | -0.1 | 6.21E-01 | 0.0 | 9.61E-01 | AP-5 complex subunit beta-1 |
| AT5G18390 | 0.1 | 1.34E-01 | 0.0 | 6.55E-01 | 0.1 | 3.87E-01 | -0.3 | 1.38E-01 | -0.3 | 1.98E-01 | 0.0 | 9.80E-01 | Pentatricopeptide repeat-containing protein At5g18390, mitochondrial |
| AT2G11910 | 0.1 | 2.38E-01 | 0.3 | 1.57E-02 | -0.2 | 1.45E-01 | 0.1 | 6.01E-01 | -0.1 | 8.60E-01 | 0.2 | 5.86E-01 | Uncharacterized protein |
| AT3G53540 | 0.1 | 2.34E-01 | -0.1 | 6.48E-01 | 0.2 | 1.54E-01 | 0.0 | 9.97E-01 | 0.0 | 9.51E-01 | 0.0 | 9.57E-01 | AT3g53540 |
| AT4G34412 | 0.1 | 3.47E-01 | 0.0 | 8.48E-01 | 0.1 | 5.66E-01 | -0.2 | 4.64E-01 | -0.1 | 6.96E-01 | -0.1 | 8.59E-01 | hypothetical protein |
| AT5G54585 | 0.1 | 7.11E-01 | 0.3 | 4.68E-01 | -0.1 | 7.65E-01 | | | | | | | Uncharacterized protein |
| AT3G56990 | 0.1 | 2.23E-01 | 0.0 | 7.70E-01 | 0.1 | 4.44E-01 | 1.1 | 5.14E-06 | 0.4 | 3.48E-02 | 0.7 | 4.72E-03 | AT3g56990 |
| AT1G19800 | 0.1 | 8.43E-02 | 0.1 | 4.71E-01 | 0.1 | 3.75E-01 | 0.2 | 3.51E-01 | 0.1 | 7.70E-01 | 0.1 | 7.44E-01 | AT1G19800 protein |
| AT1G10470 | 0.1 | 8.84E-02 | 0.0 | 5.55E-01 | 0.1 | 3.10E-01 | 0.3 | 3.18E-01 | 0.3 | 3.97E-01 | 0.0 | 9.99E-01 | Response regulator 4 |
| AT5G11280 | 0.1 | 6.69E-02 | 0.3 | 1.22E-03 | -0.2 | 4.13E-02 | -0.2 | 1.31E-01 | -0.1 | 5.80E-01 | -0.1 | 5.22E-01 | hypothetical protein |
| AT3G12587 | 0.1 | 1.26E-01 | 0.2 | 2.77E-02 | -0.1 | 4.51E-01 | 0.2 | 3.00E-01 | 0.1 | 6.95E-01 | 0.1 | 7.22E-01 | Uncharacterized protein T2E22.10 |
| AT2G09795 | 0.1 | 3.89E-01 | 0.1 | 4.31E-01 | 0.0 | 9.79E-01 | 0.1 | 7.00E-01 | 0.1 | 8.01E-01 | 0.0 | 9.67E-01 | NA |
| AT5G64080 | 0.1 | 4.58E-01 | 0.3 | 1.78E-01 | -0.1 | 5.88E-01 | 0.5 | 9.72E-03 | 0.4 | 5.47E-02 | 0.1 | 6.63E-01 | XYP1 |
| AT1G10020 | 0.1 | 3.18E-01 | 0.2 | 1.48E-01 | -0.1 | 6.61E-01 | -0.9 | 3.88E-05 | -1.0 | 4.33E-05 | 0.1 | 6.06E-01 | hypothetical protein |
| AT2G37860 | 0.1 | 1.74E-01 | 0.4 | 2.48E-03 | -0.3 | 3.82E-02 | 0.3 | 1.61E-02 | 0.3 | 8.25E-02 | 0.1 | 7.41E-01 | Protein RETICULATA, chloroplastic |
| AT3G25545 | 0.1 | 2.07E-01 | 0.0 | 9.77E-01 | 0.1 | 2.73E-01 | 0.3 | 6.68E-02 | 0.2 | 4.06E-01 | 0.2 | 4.64E-01 | hypothetical protein |
| AT3G10820 | 0.1 | 2.71E-01 | -0.3 | 4.95E-02 | 0.4 | 1.82E-02 | -0.2 | 5.79E-01 | 0.0 | 9.97E-01 | -0.2 | 6.90E-01 | T7M13.9 protein |
| AT3G58650 | 0.1 | 1.62E-01 | 0.5 | 8.52E-04 | -0.3 | 1.75E-02 | 0.1 | 7.00E-01 | 0.2 | 4.93E-01 | -0.1 | 8.05E-01 | GPI-anchored adhesin-like protein |
| AT1G07840 | 0.1 | 2.05E-01 | 0.1 | 3.54E-01 | 0.0 | 8.04E-01 | 0.3 | 9.92E-02 | 0.1 | 5.79E-01 | 0.2 | 4.03E-01 | F24B9.6 |
| AT3G26115 | 0.1 | 3.23E-01 | 0.2 | 2.27E-01 | 0.0 | 8.40E-01 | 0.3 | 4.17E-01 | 0.4 | 4.02E-01 | 0.0 | 9.41E-01 | Pyridoxal-5'-phosphate-dependent enzyme family protein |
| AT1G27410 | 0.1 | 2.39E-01 | -0.1 | 6.43E-01 | 0.2 | 1.85E-01 | -0.7 | 1.02E-01 | -0.3 | 5.37E-01 | -0.4 | 4.73E-01 | DNA repair metallo-beta-lactamase family protein |
| AT2G02400 | 0.1 | 6.55E-02 | 0.0 | 7.39E-01 | 0.1 | 1.91E-01 | -0.4 | 2.63E-02 | -0.2 | 2.89E-01 | -0.2 | 3.47E-01 | NAD-dependent epimerase |
| AT1G09310 | 0.1 | 1.89E-01 | 0.3 | 1.77E-02 | -0.2 | 2.00E-01 | 0.4 | 4.08E-03 | 0.5 | 2.94E-03 | -0.1 | 7.58E-01 | T31J12.3 protein |
| AT4G32610 | 0.1 | 3.57E-02 | 0.2 | 1.46E-02 | 0.0 | 6.18E-01 | 0.1 | 4.77E-01 | 0.1 | 4.86E-01 | 0.0 | 9.62E-01 | Uncharacterized protein AT4g32610 |
| AT4G18975 | 0.1 | 1.06E-01 | 0.0 | 9.71E-01 | 0.1 | 1.57E-01 | 0.3 | 1.60E-01 | 0.2 | 3.10E-01 | 0.0 | 8.93E-01 | Pentatricopeptide repeat-containing protein At4g18975, chloroplastic |
| AT1G68680 | 0.1 | 9.75E-02 | 0.2 | 2.06E-02 | -0.1 | 4.27E-01 | 0.0 | 8.19E-01 | -0.1 | 5.49E-01 | 0.2 | 4.99E-01 | hypothetical protein |
| AT2G41200 | 0.1 | 2.77E-01 | 0.0 | 7.68E-01 | 0.2 | 2.57E-01 | -0.4 | 2.04E-01 | -0.3 | 3.20E-01 | 0.0 | 9.40E-01 | hypothetical protein |
| AT1G58190 | 0.1 | 3.67E-01 | 0.2 | 1.45E-01 | -0.1 | 6.18E-01 | 0.1 | 8.37E-01 | -0.3 | 4.60E-01 | 0.4 | 3.91E-01 | Uncharacterized protein At1g58190 |
| AT5G49820 | 0.1 | 3.64E-02 | 0.2 | 8.46E-03 | -0.1 | 4.23E-01 | 0.0 | 9.93E-01 | -0.2 | 2.26E-01 | 0.2 | 3.36E-01 | Protein root UVB sensitive 6 |
| AT1G02330 | 0.1 | 8.01E-02 | 0.1 | 2.95E-01 | 0.1 | 5.33E-01 | 0.0 | 8.16E-01 | 0.1 | 6.87E-01 | 0.0 | 8.83E-01 | Protein COP1 SUPPRESSOR 2 |
| AT1G73840 | 0.1 | 1.09E-01 | 0.1 | 4.62E-01 | 0.1 | 4.49E-01 | 0.1 | 8.04E-01 | -0.1 | 8.10E-01 | 0.1 | 6.80E-01 | Hydroxyproline-rich glycoprotein family protein |
| AT3G48250 | 0.1 | 2.34E-01 | 0.1 | 3.87E-01 | 0.0 | 8.27E-01 | 0.6 | 2.22E-02 | 0.4 | 1.20E-01 | 0.1 | 6.51E-01 | Pentatricopeptide repeat-containing protein At3g48250, chloroplastic |
| AT5G02590 | 0.1 | 3.47E-01 | -0.1 | 7.22E-01 | 0.2 | 2.85E-01 | 0.3 | 4.29E-01 | 0.0 | 9.50E-01 | 0.3 | 5.82E-01 | Uncharacterized protein |

| | | | | | | | | | | | | | |
|---|---|---|---|---|---|---|---|---|---|---|---|---|---|
| AT2G39795 | 0.1 | 2.54E-01 | 0.2 | 8.73E-02 | -0.1 | 5.73E-01 | 0.7 | 7.06E-03 | 0.4 | 1.38E-01 | 0.3 | 3.01E-01 | Uncharacterized protein At2g39795, mitochondrial |
| AT5G44710 | 0.1 | 1.59E-01 | 0.3 | 5.35E-03 | -0.2 | 9.73E-02 | 0.1 | 7.42E-01 | 0.2 | 6.11E-01 | -0.1 | 8.65E-01 | Small ribosomal subunit protein mS33 |
| AT1G80280 | 0.1 | 1.07E-01 | 0.5 | 2.54E-04 | -0.3 | 5.13E-03 | -0.2 | 3.39E-01 | 0.0 | 9.35E-01 | -0.2 | 4.57E-01 | Uncharacterized protein At1g80280 |
| AT5G41270 | 0.1 | 1.39E-01 | 0.1 | 3.79E-01 | 0.1 | 6.43E-01 | -0.5 | 4.40E-02 | -0.6 | 2.24E-02 | 0.1 | 7.45E-01 | Uncharacterized protein |
| AT1G52825 | 0.1 | 4.00E-01 | 0.2 | 3.13E-01 | 0.0 | 8.74E-01 | 0.4 | 4.58E-01 | 0.4 | 3.83E-01 | -0.1 | 8.89E-01 | Uncharacterized protein |
| AT1G11630 | 0.1 | 1.39E-01 | 0.0 | 8.03E-01 | 0.2 | 1.51E-01 | 0.0 | 8.53E-01 | -0.2 | 5.38E-01 | 0.2 | 4.76E-01 | Pentatricopeptide repeat-containing protein At1g11630, mitochondrial |
| AT3G23170 | 0.1 | 5.85E-01 | -0.2 | 3.98E-01 | 0.4 | 2.35E-01 | -1.3 | 5.14E-04 | -0.3 | 3.05E-01 | -0.9 | 2.84E-02 | At3g23170 |
| AT2G20240 | 0.1 | 2.19E-01 | 0.3 | 2.80E-02 | -0.1 | 3.15E-01 | -0.1 | 6.18E-01 | -0.1 | 7.80E-01 | 0.0 | 9.23E-01 | DUF3741 domain-containing protein |
| AT5G43260 | 0.1 | 2.76E-01 | 0.0 | 9.60E-01 | 0.1 | 3.75E-01 | 0.0 | 8.67E-01 | 0.0 | 8.73E-01 | 0.0 | 9.93E-01 | Chaperone protein dnaJ-like protein |
| AT1G20300 | 0.1 | 1.47E-01 | 0.2 | 9.17E-02 | 0.0 | 7.98E-01 | 0.4 | 6.00E-02 | 0.3 | 2.30E-01 | 0.1 | 6.93E-01 | Pentatricopeptide repeat-containing protein At1g20300, mitochondrial |
| AT4G10000 | 0.1 | 7.39E-02 | 0.0 | 6.44E-01 | 0.2 | 6.16E-02 | 0.0 | 8.49E-01 | 0.3 | 2.32E-01 | -0.3 | 2.40E-01 | Thioredoxin family protein |
| AT5G08060 | 0.1 | 1.31E-01 | 0.3 | 2.06E-02 | -0.1 | 3.10E-01 | 0.1 | 7.69E-01 | 0.0 | 9.54E-01 | 0.0 | 9.02E-01 | Furry |
| AT4G22320 | 0.2 | 2.04E-01 | 0.1 | 2.27E-01 | 0.0 | 9.85E-01 | 0.1 | 6.48E-01 | 0.3 | 1.64E-01 | -0.2 | 4.55E-01 | Golgin family A protein |
| AT5G18590 | 0.2 | 1.11E-01 | 0.0 | 6.17E-01 | 0.2 | 7.30E-02 | -0.3 | 3.28E-01 | -0.2 | 6.42E-01 | -0.1 | 8.15E-01 | Galactose oxidase |
| AT5G66680 | 0.2 | 6.24E-02 | 0.2 | 1.91E-02 | -0.1 | 4.97E-01 | 0.2 | 5.42E-02 | 0.1 | 4.19E-01 | 0.1 | 5.51E-01 | Dolichyl-diphosphooligosaccharide--protein glycosyltransferase 48 kDa subunit |
| AT4G17616 | 0.2 | 1.59E-01 | -0.2 | 9.23E-02 | 0.4 | 1.86E-02 | 0.1 | 8.09E-01 | 0.0 | 8.91E-01 | 0.1 | 7.44E-01 | Pentatricopeptide repeat-containing protein At4g17616 |
| AT1G20540 | 0.2 | 1.67E-01 | 0.2 | 7.16E-02 | -0.1 | 6.58E-01 | 0.0 | 9.74E-01 | 0.0 | 9.95E-01 | 0.0 | 9.88E-01 | Transducin |
| AT5G01740 | 0.2 | 4.42E-01 | -0.7 | 1.26E-02 | 0.8 | 1.00E-02 | 0.3 | 3.53E-01 | -0.4 | 3.64E-01 | 0.7 | 1.16E-01 | Wound-induced protein 1 |
| AT1G71865 | 0.2 | 3.19E-01 | 0.0 | 8.58E-01 | 0.1 | 5.19E-01 | -0.3 | 1.45E-01 | -0.3 | 2.35E-01 | 0.0 | 9.44E-01 | hypothetical protein |
| AT3G14800 | 0.2 | 8.48E-02 | -0.1 | 3.11E-01 | 0.2 | 3.01E-02 | 0.0 | 9.57E-01 | 0.0 | 9.27E-01 | 0.0 | 9.07E-01 | NA |
| AT1G09580 | 0.2 | 2.44E-01 | 0.5 | 2.07E-03 | -0.4 | 2.70E-02 | -0.2 | 2.66E-01 | 0.2 | 2.22E-01 | -0.4 | 6.02E-02 | Transmembrane emp24 domain-containing protein p24delta3 |
| AT1G72416 | 0.2 | 5.21E-01 | 0.6 | 2.52E-02 | -0.4 | 1.19E-01 | 0.7 | 2.40E-02 | 0.2 | 5.64E-01 | 0.4 | 2.28E-01 | Chaperone DnaJ-domain superfamily protein |
| AT1G48440 | 0.2 | 2.48E-02 | 0.2 | 2.77E-03 | -0.1 | 2.09E-01 | -0.1 | 6.05E-01 | 0.1 | 7.14E-01 | -0.2 | 4.59E-01 | Endoplasmic reticulum transmembrane protein |
| AT4G09810 | 0.2 | 2.41E-01 | -0.1 | 6.88E-01 | 0.2 | 2.01E-01 | 0.0 | 9.30E-01 | -0.1 | 8.05E-01 | 0.1 | 8.92E-01 | Nucleotide-sugar transporter family protein |
| AT2G36340 | 0.2 | 1.29E-01 | -0.1 | 2.65E-01 | 0.3 | 3.89E-02 | 0.1 | 6.06E-01 | -0.1 | 8.39E-01 | 0.2 | 5.50E-01 | GLABROUS1 enhancer-binding protein-like 3 |
| AT3G11220 | 0.2 | 1.10E-01 | 0.2 | 8.85E-02 | 0.0 | 8.93E-01 | 0.3 | 2.43E-01 | 0.0 | 9.03E-01 | 0.3 | 2.36E-01 | Elongator complex protein 4 |
| AT4G24180 | 0.2 | 3.27E-01 | 0.9 | 8.59E-05 | -0.8 | 1.15E-03 | | | | | | | THAUMATIN-LIKE PROTEIN 1 |
| AT4G34140 | 0.2 | 2.13E-01 | -0.2 | 1.08E-01 | 0.4 | 2.46E-02 | -0.4 | 1.00E-01 | 0.0 | 8.76E-01 | -0.3 | 2.66E-01 | Uncharacterized protein AT4g34140 |
| AT5G49015 | 0.2 | 5.27E-01 | -0.2 | 3.98E-01 | 0.4 | 2.18E-01 | -0.3 | 4.60E-01 | -0.4 | 4.12E-01 | 0.1 | 9.16E-01 | hypothetical protein |
| AT2G17250 | 0.2 | 2.38E-01 | 0.2 | 8.62E-02 | -0.1 | 5.88E-01 | 0.4 | 2.38E-02 | 0.3 | 1.23E-01 | 0.1 | 6.67E-01 | Protein NUCLEOLAR COMPLEX ASSOCIATED 4 |
| AT3G29776 | 0.2 | 2.72E-01 | -0.2 | 1.88E-01 | 0.4 | 5.95E-02 | -0.4 | 3.85E-01 | -0.5 | 2.90E-01 | 0.1 | 8.47E-01 | NA |
| AT2G26860 | 0.2 | 1.57E-01 | 0.1 | 6.38E-01 | 0.1 | 4.49E-01 | 0.3 | 2.91E-01 | 0.2 | 4.96E-01 | 0.1 | 8.78E-01 | FBD-associated F-box protein At2g26860 |
| AT5G62550 | 0.2 | 6.87E-02 | 0.1 | 9.59E-02 | 0.0 | 9.26E-01 | 0.0 | 8.40E-01 | 0.3 | 1.11E-01 | -0.3 | 2.23E-01 | Microtubule-associated futsch-like protein |
| AT1G03760 | 0.2 | 1.31E-01 | -0.2 | 1.67E-01 | 0.3 | 2.71E-02 | 0.1 | 6.15E-01 | -0.3 | 3.58E-01 | 0.4 | 2.03E-01 | F11M21.31 protein |
| AT1G08710 | 0.2 | 9.91E-02 | 0.0 | 9.93E-01 | 0.2 | 1.65E-01 | 0.0 | 8.51E-01 | -0.1 | 7.62E-01 | 0.0 | 9.18E-01 | hypothetical protein |
| AT2G02180 | 0.2 | 3.81E-02 | 0.0 | 5.65E-01 | 0.1 | 1.55E-01 | -0.2 | 2.51E-01 | -0.3 | 7.52E-02 | 0.1 | 6.06E-01 | Tobamovirus multiplication protein 3 |
| AT5G53500 | 0.2 | 9.86E-02 | 0.3 | 1.47E-02 | -0.1 | 3.20E-01 | -0.5 | 2.74E-02 | -0.4 | 8.62E-02 | -0.1 | 8.35E-01 | Uncharacterized protein |
| AT1G64385 | 0.2 | 5.26E-02 | 0.0 | 7.88E-01 | 0.1 | 1.36E-01 | 0.0 | 7.34E-01 | -0.1 | 6.07E-01 | 0.0 | 8.87E-01 | At1g64385 |
| AT5G55790 | 0.2 | 6.20E-02 | 0.0 | 8.19E-01 | 0.1 | 1.54E-01 | 0.0 | 9.41E-01 | 0.1 | 8.00E-01 | -0.1 | 8.70E-01 | At5g55790 |
| AT3G13674 | 0.2 | 4.74E-01 | 0.5 | 2.25E-02 | -0.4 | 1.38E-01 | 0.5 | 7.66E-02 | 0.1 | 8.38E-01 | 0.4 | 2.02E-01 | Uncharacterized protein |
| AT1G76400 | 0.2 | 4.34E-02 | 0.2 | 9.64E-03 | -0.1 | 4.11E-01 | 0.2 | 1.51E-01 | 0.1 | 5.15E-01 | 0.1 | 6.98E-01 | Dolichyl-diphosphooligosaccharide--protein glycosyltransferase subunit 1 |
| AT5G11500 | 0.2 | 7.16E-02 | 0.1 | 1.52E-01 | 0.0 | 7.63E-01 | 0.1 | 4.45E-01 | 0.1 | 5.83E-01 | 0.0 | 9.71E-01 | Coiled-coil protein |
| AT3G04140 | 0.2 | 1.32E-01 | 0.0 | 9.32E-01 | 0.1 | 2.15E-01 | -0.1 | 7.94E-01 | 0.2 | 5.40E-01 | -0.2 | 4.03E-01 | Ankyrin repeat family protein |
| AT1G33400 | 0.2 | 6.65E-02 | 0.0 | 7.93E-01 | 0.2 | 7.79E-02 | -0.3 | 5.65E-02 | 0.0 | 9.23E-01 | -0.3 | 1.69E-01 | Tetratricopeptide repeat -like superfamily protein |
| AT2G45250 | 0.2 | 8.09E-02 | 0.2 | 5.17E-02 | 0.0 | 8.01E-01 | -0.1 | 6.05E-01 | -0.1 | 5.72E-01 | 0.0 | 9.40E-01 | Integral membrane hemolysin-III-like protein |
| AT3G12990 | 0.2 | 6.60E-02 | -0.1 | 5.29E-01 | 0.2 | 4.78E-02 | -0.1 | 6.69E-01 | -0.1 | 8.38E-01 | 0.0 | 9.17E-01 | Ribonuclease PH45A |
| AT4G28820 | 0.2 | 3.76E-01 | 0.4 | 4.20E-02 | -0.2 | 2.76E-01 | 0.1 | 8.43E-01 | -0.1 | 8.98E-01 | 0.2 | 7.80E-01 | At4g28820 |
| AT2G41440 | 0.2 | 1.60E-01 | -0.1 | 2.09E-01 | 0.3 | 3.62E-02 | -0.2 | 3.12E-01 | -0.1 | 6.45E-01 | -0.1 | 7.52E-01 | Agamous-like MADS-box protein |
| AT3G20620 | 0.2 | 8.75E-02 | 0.1 | 3.09E-01 | 0.1 | 5.63E-01 | 0.0 | 9.70E-01 | 0.1 | 5.44E-01 | -0.1 | 5.11E-01 | F-box associated domain-containing protein |
| AT3G11402 | 0.2 | 1.81E-01 | -0.3 | 4.68E-02 | 0.4 | 1.28E-02 | 0.4 | 9.84E-02 | 0.2 | 6.17E-01 | 0.3 | 4.27E-01 | Cysteine |
| AT5G18420 | 0.2 | 2.77E-02 | 0.0 | 7.04E-01 | 0.1 | 9.99E-02 | 0.1 | 7.47E-01 | 0.0 | 9.55E-01 | 0.1 | 7.81E-01 | CCR4-NOT transcription complex subunit 11 |
| AT5G46020 | 0.2 | 5.28E-02 | 0.2 | 9.34E-03 | -0.1 | 3.25E-01 | 0.0 | 9.32E-01 | -0.1 | 5.99E-01 | 0.1 | 6.25E-01 | 28 kDa heat |
| AT5G07770 | 0.2 | 2.56E-01 | -0.2 | 2.59E-01 | 0.3 | 5.87E-02 | 0.1 | 7.28E-01 | 0.0 | 8.79E-01 | 0.0 | 9.28E-01 | Formin-like protein |
| AT2G28430 | 0.2 | 8.37E-02 | 0.3 | 8.33E-03 | -0.1 | 2.39E-01 | 0.1 | 6.95E-01 | 0.0 | 9.58E-01 | 0.1 | 8.39E-01 | Uncharacterized protein |
| AT2G39445 | 0.2 | 3.59E-01 | 0.0 | 8.62E-01 | 0.2 | 3.77E-01 | 0.0 | 9.24E-01 | -0.1 | 9.09E-01 | 0.1 | 8.62E-01 | hypothetical protein |
| AT4G19160 | 0.2 | 1.26E-01 | 0.2 | 9.21E-02 | 0.0 | 8.34E-01 | -0.3 | 9.79E-03 | -0.6 | 5.59E-04 | 0.2 | 1.34E-01 | Transglutaminase family protein |
| AT3G15550 | 0.2 | 1.18E-01 | 0.3 | 1.38E-02 | -0.1 | 2.90E-01 | -0.2 | 7.39E-01 | 0.2 | 7.65E-01 | -0.4 | 5.79E-01 | MJK13.21 protein |
| AT5G48970 | 0.2 | 1.34E-01 | 0.1 | 6.44E-01 | 0.1 | 3.88E-01 | 0.3 | 5.05E-02 | -0.2 | 3.73E-01 | 0.5 | 1.86E-02 | Mitochondrial substrate carrier family protein |
| AT5G23130 | 0.2 | 1.84E-01 | 0.2 | 8.56E-02 | -0.1 | 6.91E-01 | -0.2 | 4.82E-01 | -0.4 | 1.74E-01 | 0.2 | 5.69E-01 | hypothetical protein |
| AT1G27340 | 0.2 | 2.73E-02 | 0.1 | 1.40E-01 | 0.1 | 4.58E-01 | -0.1 | 7.20E-01 | 0.1 | 6.38E-01 | -0.1 | 4.81E-01 | F-box only protein 6 |
| AT5G03440 | 0.2 | 2.44E-01 | 0.1 | 4.66E-01 | 0.1 | 7.55E-01 | -0.3 | 1.95E-01 | -0.5 | 1.03E-01 | 0.1 | 7.66E-01 | hypothetical protein |
| AT1G07480 | 0.2 | 3.33E-02 | 0.0 | 9.47E-01 | 0.2 | 7.30E-02 | -0.4 | 9.24E-03 | -0.4 | 2.80E-02 | 0.0 | 8.50E-01 | Uncharacterized protein |
| AT1G05205 | 0.2 | 3.19E-01 | 0.5 | 5.90E-03 | -0.4 | 6.27E-02 | 0.1 | 8.98E-01 | 0.2 | 7.23E-01 | -0.1 | 8.39E-01 | Hexokinase-1 protein |
| AT1G30845 | 0.2 | 3.39E-01 | 0.3 | 9.28E-02 | -0.1 | 5.05E-01 | 0.0 | 9.16E-01 | -0.2 | 7.12E-01 | 0.1 | 8.28E-01 | T17H7.16 |
| AT5G05800 | 0.2 | 8.37E-02 | 0.0 | 9.49E-01 | 0.2 | 1.50E-01 | -0.3 | 1.41E-02 | -0.4 | 5.69E-03 | 0.1 | 6.68E-01 | L10-interacting MYB domain-containing protein |
| AT5G12240 | 0.2 | 5.68E-02 | 0.3 | 1.67E-03 | -0.2 | 7.11E-02 | -0.1 | 6.28E-01 | -0.3 | 3.21E-01 | 0.1 | 7.01E-01 | Octanoyltransferase |
| AT5G45460 | 0.2 | 1.71E-01 | 0.3 | 2.02E-02 | -0.1 | 2.98E-01 | -0.1 | 8.26E-01 | -0.2 | 7.53E-01 | 0.1 | 9.24E-01 | Transmembrane protein |
| AT1G64140 | 0.2 | 7.04E-02 | -0.1 | 1.90E-01 | 0.3 | 1.20E-02 | -0.1 | 2.98E-01 | 0.0 | 9.28E-01 | -0.2 | 3.70E-01 | WRKY transcription factor 19 |
| AT3G45970 | 0.2 | 5.73E-01 | -0.2 | 4.49E-01 | 0.4 | 2.68E-01 | -1.2 | 1.42E-04 | -1.0 | 5.46E-04 | -0.1 | 8.24E-01 | Uncharacterized protein |
| AT5G21900 | 0.2 | 1.27E-01 | -0.4 | 6.96E-03 | 0.5 | 2.48E-03 | -0.1 | 8.55E-01 | -0.1 | 8.25E-01 | 0.0 | 9.66E-01 | RNI-like superfamily protein |
| AT1G21065 | 0.2 | 8.71E-02 | 0.3 | 7.01E-03 | -0.1 | 1.76E-01 | -0.2 | 2.46E-01 | 0.1 | 7.99E-01 | -0.2 | 2.49E-01 | Secondary thiamine-phosphate synthase enzyme |
| AT2G42710 | 0.2 | 2.47E-01 | 0.3 | 6.86E-02 | -0.1 | 4.95E-01 | 0.5 | 2.99E-03 | 0.4 | 5.79E-02 | 0.2 | 3.59E-01 | Uncharacterized protein At2g42710 |
| AT2G01100 | 0.2 | 5.71E-02 | 0.0 | 9.59E-01 | 0.2 | 1.01E-01 | -0.5 | 4.61E-03 | -0.5 | 5.23E-03 | 0.0 | 8.79E-01 | Uncharacterized protein |
| AT2G27770 | 0.2 | 1.77E-01 | 0.2 | 6.92E-02 | -0.1 | 6.29E-01 | 0.0 | 9.38E-01 | -0.1 | 7.38E-01 | 0.1 | 8.34E-01 | DUF868 family protein |
| AT4G16840 | 0.2 | 2.67E-01 | 0.1 | 6.21E-01 | 0.1 | 6.44E-01 | 0.0 | 9.44E-01 | -0.4 | 1.36E-01 | 0.5 | 1.70E-01 | Transmembrane protein |
| AT4G38560 | 0.2 | 5.24E-01 | -1.4 | 2.35E-03 | 1.5 | 3.24E-03 | -0.6 | 4.46E-01 | -1.0 | 2.32E-01 | | | hypothetical protein |
| AT2G06000 | 0.2 | 6.00E-02 | 0.1 | 2.82E-01 | 0.1 | 4.79E-01 | 0.2 | 3.42E-01 | 0.1 | 7.69E-01 | 0.1 | 7.10E-01 | Pentatricopeptide repeat-containing protein At2g06000 |
| AT5G03890 | 0.2 | 3.39E-01 | 0.1 | 5.68E-01 | 0.1 | 7.92E-01 | 0.0 | 9.90E-01 | 0.1 | 8.49E-01 | -0.2 | 8.70E-01 | DUF4228 domain protein |
| AT3G18910 | 0.2 | 9.06E-02 | 0.3 | 1.14E-02 | -0.1 | 3.09E-01 | -0.2 | 3.38E-01 | -0.4 | 1.17E-01 | 0.2 | 6.09E-01 | F-box protein ETP2 |
| AT4G23470 | 0.2 | 1.59E-01 | 0.1 | 2.75E-01 | 0.0 | 8.16E-01 | 0.1 | 2.64E-01 | -0.1 | 5.71E-01 | 0.2 | 1.90E-01 | PLAC8 family protein |
| AT3G55690 | 0.2 | 2.91E-01 | 0.1 | 6.96E-01 | 0.1 | 6.20E-01 | 0.5 | 2.84E-02 | 0.2 | 4.63E-01 | 0.3 | 2.58E-01 | At3g55690 |
| AT2G03470 | 0.2 | 3.79E-02 | -0.2 | 3.09E-02 | 0.3 | 2.19E-03 | -0.2 | 1.49E-01 | -0.3 | 7.41E-02 | 0.1 | 7.50E-01 | ELM2 domain-containing protein |
| AT3G59430 | 0.2 | 1.28E-01 | -0.1 | 4.12E-01 | 0.3 | 6.33E-02 | 0.1 | 8.47E-01 | 0.2 | 6.62E-01 | -0.1 | 8.38E-01 | At3g59430 |

| | | | | | | | | | | | | | |
|---|---|---|---|---|---|---|---|---|---|---|---|---|---|
| AT1G51560 | 0.2 | 2.85E-01 | 0.0 | 9.88E-01 | 0.2 | 3.80E-01 | -0.1 | 5.37E-01 | 0.0 | 9.34E-01 | -0.1 | 7.42E-01 | Pyridoxamine 5'-phosphate oxidase putative domain-containing protein |
| AT1G48635 | 0.2 | 2.75E-02 | -0.1 | 4.95E-01 | 0.2 | 1.96E-02 | 0.1 | 4.50E-01 | -0.1 | 8.32E-01 | 0.2 | 3.44E-01 | Peroxin 3 |
| AT5G04820 | 0.2 | 1.56E-01 | 0.5 | 1.16E-03 | -0.3 | 2.31E-02 | -0.1 | 5.17E-01 | -0.2 | 2.87E-01 | 0.1 | 7.31E-01 | Transcription repressor OFP13 |
| AT5G05520 | 0.2 | 7.73E-02 | 0.1 | 2.88E-01 | 0.1 | 5.28E-01 | 0.1 | 5.68E-01 | 0.0 | 9.34E-01 | 0.1 | 7.88E-01 | Outer membrane OMP85 family protein |
| AT4G01390 | 0.2 | 1.44E-01 | 0.2 | 6.33E-02 | -0.1 | 6.53E-01 | 0.6 | 8.26E-04 | 0.6 | 1.94E-03 | 0.0 | 9.96E-01 | Uncharacterized protein AT4g01390 |
| AT2G26590 | 0.2 | 2.63E-02 | 0.2 | 2.36E-02 | 0.0 | 9.11E-01 | 0.1 | 4.67E-01 | 0.0 | 9.79E-01 | 0.1 | 6.43E-01 | Uncharacterized protein At2g26590 |
| AT5G51400 | 0.2 | 2.09E-02 | 0.2 | 5.21E-03 | -0.1 | 4.10E-01 | -0.1 | 5.83E-01 | -0.2 | 2.73E-01 | 0.1 | 6.95E-01 | Gb\|AAB68038.1 |
| AT4G14713 | 0.2 | 1.61E-01 | 0.3 | 2.59E-02 | -0.1 | 3.68E-01 | 0.1 | 6.11E-01 | -0.1 | 5.76E-01 | 0.3 | 3.44E-01 | Protein TIFY |
| AT2G41050 | 0.2 | 1.55E-01 | 0.2 | 1.52E-01 | 0.0 | 9.60E-01 | 0.0 | 9.72E-01 | 0.2 | 7.48E-01 | -0.2 | 7.41E-01 | PQ-loop repeat family protein |
| AT1G16210 | 0.2 | 3.79E-02 | -0.1 | 3.47E-01 | 0.2 | 1.31E-02 | -0.1 | 2.46E-01 | -0.1 | 3.06E-01 | 0.0 | 9.94E-01 | F3O9.2 protein |
| AT3G16660 | 0.2 | 4.46E-01 | 0.0 | 9.18E-01 | 0.1 | 5.75E-01 | 0.0 | 9.89E-01 | -0.4 | 1.13E-01 | 0.4 | 6.40E-02 | Pollen Ole e 1 allergen and extensin family protein |
| AT4G11000 | 0.2 | 4.69E-01 | -0.4 | 1.39E-01 | 0.6 | 7.59E-02 | -0.4 | 2.05E-01 | -0.9 | 1.80E-02 | 0.5 | 2.45E-01 | Uncharacterized protein AT4g11000 |
| AT5G53620 | 0.2 | 2.22E-02 | 0.1 | 7.97E-02 | 0.0 | 5.79E-01 | 0.2 | 2.97E-01 | 0.1 | 6.03E-01 | 0.1 | 8.15E-01 | Uncharacterized protein |
| AT3G22845 | 0.2 | 6.42E-02 | 0.1 | 1.84E-01 | 0.0 | 6.33E-01 | 0.3 | 2.96E-02 | 0.2 | 1.39E-01 | 0.1 | 7.49E-01 | Emp24 |
| AT2G04378 | 0.2 | 1.64E-01 | 0.3 | 3.24E-02 | -0.1 | 4.20E-01 | 0.1 | 6.18E-01 | 0.0 | 9.66E-01 | 0.2 | 6.68E-01 | Beta-galactosidase related protein |
| AT1G35210 | 0.2 | 5.24E-01 | -0.8 | 2.96E-02 | 0.9 | 2.46E-02 | 0.9 | 6.40E-02 | 0.3 | 6.92E-01 | 0.6 | 2.87E-01 | At1g35210 |
| AT5G64400 | 0.2 | 1.62E-01 | 0.1 | 2.87E-01 | 0.0 | 8.04E-01 | 0.0 | 8.17E-01 | 0.1 | 4.01E-01 | -0.1 | 6.28E-01 | CHCH domain protein |
| AT1G22200 | 0.2 | 6.06E-02 | 0.0 | 7.79E-01 | 0.1 | 1.52E-01 | 0.3 | 9.97E-03 | 0.0 | 7.91E-01 | 0.3 | 9.32E-02 | Endoplasmic reticulum vesicle transporter protein |
| AT5G15010 | 0.2 | 1.86E-01 | 0.2 | 9.57E-02 | -0.1 | 7.27E-01 | 0.2 | 5.80E-01 | 0.0 | 9.40E-01 | 0.2 | 7.71E-01 | Pentatricopeptide repeat-containing protein At5g15010, mitochondrial |
| AT2G19250 | 0.2 | 3.34E-01 | -0.4 | 9.11E-02 | 0.5 | 3.71E-02 | -0.1 | 7.18E-01 | 0.0 | 9.25E-01 | -0.2 | 7.00E-01 | NA |
| AT4G39366 | 0.2 | 2.83E-01 | 0.3 | 6.22E-02 | -0.1 | 4.46E-01 | 0.3 | 4.09E-01 | -0.1 | 8.39E-01 | 0.3 | 3.43E-01 | NA |
| AT3G45890 | 0.2 | 1.02E-01 | -0.1 | 5.93E-01 | 0.2 | 8.12E-02 | 0.2 | 7.82E-01 | -0.2 | 8.55E-01 | 0.4 | 6.87E-01 | Protein root UVB sensitive 1, chloroplastic |
| AT2G37805 | 0.2 | 2.10E-01 | 0.8 | 1.60E-04 | -0.6 | 2.38E-03 | 2.2 | 1.53E-07 | 0.7 | 2.58E-02 | 1.6 | 3.74E-05 | Cysteine |
| AT1G42960 | 0.2 | 2.95E-02 | 0.3 | 4.82E-03 | -0.1 | 2.85E-01 | 0.1 | 6.00E-01 | 0.2 | 3.17E-01 | -0.1 | 7.21E-01 | Inner membrane localized protein |
| AT1G52618 | 0.2 | 2.03E-01 | 0.0 | 8.91E-01 | 0.2 | 3.71E-01 | -0.1 | 7.29E-01 | -0.1 | 8.40E-01 | 0.0 | 9.57E-01 | Uncharacterized protein |
| AT1G34065 | 0.2 | 2.17E-01 | 0.3 | 3.76E-02 | -0.1 | 3.86E-01 | -0.1 | 7.92E-01 | -0.2 | 5.92E-01 | 0.1 | 8.28E-01 | Allinase like protein |
| AT4G21520 | 0.2 | 1.48E-01 | 0.2 | 1.88E-01 | 0.0 | 9.44E-01 | 0.1 | 7.12E-01 | -0.1 | 8.59E-01 | 0.2 | 6.35E-01 | Transducin |
| AT1G19680 | 0.2 | 3.13E-02 | 0.0 | 7.55E-01 | 0.2 | 3.66E-02 | -0.1 | 6.64E-01 | 0.0 | 9.26E-01 | -0.1 | 8.35E-01 | RING-type domain-containing protein |
| AT1G35320 | 0.2 | 2.28E-01 | 0.2 | 2.94E-01 | 0.0 | 9.27E-01 | 0.5 | 1.52E-01 | 0.4 | 2.68E-01 | 0.1 | 9.08E-01 | Transmembrane protein |
| AT3G58820 | 0.2 | 3.31E-01 | 0.3 | 1.54E-01 | -0.1 | 6.71E-01 | -0.1 | 8.45E-01 | -0.1 | 7.78E-01 | 0.1 | 9.35E-01 | Putative F-box protein At3g58820 |
| AT5G27750 | 0.2 | 1.27E-01 | 0.1 | 3.92E-01 | 0.1 | 5.96E-01 | -0.1 | 7.82E-01 | -0.1 | 7.89E-01 | 0.0 | 9.80E-01 | F-box |
| AT1G59980 | 0.2 | 2.87E-01 | 0.2 | 3.36E-01 | 0.0 | 9.66E-01 | -0.4 | 1.39E-01 | -0.2 | 5.19E-01 | -0.2 | 6.23E-01 | ARG1-like 2 |
| AT4G15140 | 0.2 | 4.15E-01 | 0.3 | 1.45E-01 | -0.2 | 5.56E-01 | 0.5 | 1.04E-01 | 0.1 | 8.10E-01 | 0.4 | 3.43E-01 | Uncharacterized protein At4g15140 |
| AT5G61910 | 0.2 | 6.92E-02 | 0.0 | 8.71E-01 | 0.2 | 1.55E-01 | 0.5 | 6.18E-02 | 0.2 | 6.32E-01 | 0.3 | 3.43E-01 | AT5g61910 |
| AT3G25590 | 0.2 | 2.28E-01 | -0.2 | 1.31E-01 | 0.4 | 3.51E-02 | -0.1 | 6.84E-01 | 0.2 | 2.23E-01 | -0.3 | 1.47E-01 | At3g25590 |
| AT5G02280 | 0.2 | 3.26E-01 | 0.2 | 2.65E-01 | 0.0 | 8.94E-01 | 0.2 | 5.93E-01 | 0.4 | 2.23E-01 | -0.2 | 5.67E-01 | Trafficking protein particle complex subunit |
| AT3G27520 | 0.2 | 1.87E-01 | 0.4 | 1.29E-02 | -0.2 | 1.99E-01 | -0.4 | 1.50E-01 | -0.3 | 3.32E-01 | -0.1 | 8.40E-01 | Uncharacterized protein |
| AT5G16250 | 0.2 | 5.37E-02 | 0.5 | 3.27E-04 | -0.3 | 1.02E-02 | -0.3 | 9.80E-02 | -0.1 | 4.75E-01 | -0.1 | 6.36E-01 | hypothetical protein |
| AT2G44440 | 0.2 | 4.89E-02 | 0.2 | 6.59E-02 | 0.0 | 9.45E-01 | 0.0 | 9.57E-01 | -0.2 | 4.40E-01 | 0.2 | 4.68E-01 | Protein EMSY-LIKE 4 |
| AT2G06010 | 0.2 | 5.50E-02 | 0.1 | 4.94E-01 | 0.1 | 2.76E-01 | 0.0 | 9.52E-01 | -0.1 | 6.20E-01 | 0.1 | 7.41E-01 | OBP3-responsive protein 4 |
| AT1G30880 | 0.2 | 5.08E-02 | 0.3 | 5.39E-03 | -0.1 | 2.08E-01 | -0.3 | 1.92E-01 | -0.4 | 8.03E-02 | 0.1 | 6.73E-01 | Uncharacterized protein |
| AT5G26290 | 0.2 | 1.80E-01 | 0.2 | 1.33E-01 | 0.0 | 8.63E-01 | -0.1 | 9.11E-01 | 0.7 | 9.37E-02 | -0.8 | 1.17E-01 | TRAF-like family protein |
| AT5G19330 | 0.2 | 4.45E-02 | 0.0 | 9.15E-01 | 0.2 | 5.75E-02 | -0.2 | 9.95E-02 | -0.2 | 7.33E-02 | 0.0 | 8.45E-01 | ARM repeat protein interacting with ABF2 |
| AT4G37030 | 0.2 | 1.59E-01 | 0.1 | 4.22E-01 | 0.1 | 6.54E-01 | 2.7 | 4.12E-07 | 0.9 | 3.40E-02 | 1.8 | 1.85E-05 | Membrane protein |
| AT3G06530 | 0.2 | 6.56E-02 | 0.1 | 2.12E-01 | 0.1 | 5.95E-01 | 0.8 | 4.65E-04 | 0.4 | 5.24E-02 | 0.3 | 1.57E-01 | At3g06530 |
| AT1G21720 | 0.2 | 6.06E-02 | 0.2 | 1.11E-01 | 0.0 | 8.17E-01 | 0.1 | 3.81E-01 | 0.1 | 7.89E-01 | 0.1 | 7.47E-01 | Proteasome subunit beta type-3-A |
| AT3G05830 | 0.2 | 5.35E-02 | 0.0 | 7.39E-01 | 0.2 | 6.02E-02 | 0.2 | 2.90E-01 | 0.2 | 5.34E-01 | 0.1 | 8.21E-01 | Spindle pole body component-like protein |
| AT5G18850 | 0.2 | 2.15E-01 | 0.2 | 1.53E-01 | 0.0 | 8.44E-01 | 0.1 | 6.20E-01 | -0.3 | 3.05E-01 | 0.5 | 1.89E-01 | Low-density receptor-like protein |
| AT3G60930 | 0.2 | 2.58E-01 | 0.0 | 7.73E-01 | 0.1 | 5.14E-01 | -1.0 | 8.69E-04 | -0.9 | 3.58E-03 | -0.1 | 8.15E-01 | Uncharacterized protein At3g60930, chloroplastic |
| AT2G17730 | 0.2 | 7.62E-02 | 0.1 | 1.52E-01 | 0.0 | 7.94E-01 | 0.0 | 8.86E-01 | -0.4 | 3.46E-02 | 0.4 | 6.62E-02 | NEP-interacting protein 2 |
| AT5G02650 | 0.2 | 2.03E-01 | -0.1 | 4.99E-01 | 0.3 | 1.15E-01 | 0.0 | 9.12E-01 | 0.1 | 7.14E-01 | -0.2 | 6.23E-01 | Uncharacterized protein |
| AT3G52610 | 0.2 | 9.36E-02 | -0.1 | 5.96E-01 | 0.2 | 7.09E-02 | -0.3 | 2.33E-01 | -0.3 | 3.53E-01 | 0.0 | 9.36E-01 | Uncharacterized protein F3C22_10 |
| AT1G72020 | 0.2 | 7.01E-02 | 0.4 | 3.28E-03 | -0.2 | 9.57E-02 | 0.1 | 6.95E-01 | 0.1 | 7.19E-01 | 0.0 | 9.78E-01 | TonB-dependent heme receptor A |
| AT4G27120 | 0.2 | 1.49E-02 | 0.0 | 9.03E-01 | 0.2 | 2.49E-02 | -0.1 | 7.19E-01 | -0.1 | 7.91E-01 | 0.0 | 9.85E-01 | DDRGK domain protein |
| AT5G65440 | 0.2 | 7.63E-03 | 0.0 | 8.40E-01 | 0.2 | 2.41E-02 | -0.1 | 6.76E-01 | -0.1 | 4.47E-01 | 0.1 | 8.02E-01 | Transmembrane protein |
| AT1G28410 | 0.2 | 2.11E-01 | 0.2 | 2.76E-01 | 0.0 | 9.28E-01 | -0.1 | 7.09E-01 | 0.1 | 7.30E-01 | -0.2 | 4.90E-01 | Myosin heavy chain-like protein |
| AT1G74180 | 0.2 | 1.74E-01 | -0.1 | 6.14E-01 | 0.3 | 1.30E-01 | 0.1 | 7.93E-01 | 0.3 | 5.26E-01 | -0.1 | 7.56E-01 | Receptor like protein 14 |
| AT5G09420 | 0.2 | 1.21E-01 | 0.1 | 5.61E-01 | 0.1 | 4.25E-01 | 1.6 | 2.67E-05 | 0.5 | 2.03E-01 | 1.1 | 9.78E-04 | Outer envelope protein 64, mitochondrial |
| AT1G01930 | 0.2 | 2.89E-02 | 0.0 | 8.84E-01 | 0.2 | 4.74E-02 | -0.1 | 6.77E-01 | 0.0 | 9.57E-01 | -0.1 | 7.04E-01 | Uncharacterized protein At1g01930 |
| AT5G45110 | 0.2 | 8.21E-02 | -0.2 | 5.87E-02 | 0.4 | 6.11E-03 | 0.7 | 4.09E-04 | 0.4 | 6.35E-02 | 0.4 | 9.19E-02 | NPR1-like protein 3 |
| AT4G38090 | 0.2 | 1.58E-01 | -0.1 | 4.24E-01 | 0.3 | 7.24E-02 | -0.5 | 8.31E-03 | -0.3 | 2.00E-01 | -0.3 | 3.47E-01 | Uncharacterized protein AT4g38090 |
| AT1G49140 | 0.2 | 6.60E-02 | 0.3 | 4.10E-03 | -0.2 | 1.29E-01 | -0.2 | 2.24E-01 | -0.1 | 6.69E-01 | -0.1 | 6.60E-01 | NADH dehydrogenase 1 beta subcomplex subunit 10-A |
| AT4G37820 | 0.2 | 2.91E-02 | 0.2 | 5.48E-02 | 0.0 | 8.17E-01 | 0.1 | 4.55E-01 | 0.2 | 3.50E-01 | 0.0 | 8.51E-01 | Transmembrane protein |
| AT3G26612 | 0.2 | 4.12E-01 | 0.0 | 9.74E-01 | 0.2 | 5.41E-01 | -0.6 | 1.08E-01 | -0.2 | 5.60E-01 | -0.3 | 4.79E-01 | NA |
| AT5G16950 | 0.2 | 1.41E-01 | 0.1 | 5.27E-01 | 0.1 | 4.97E-01 | 0.3 | 1.92E-01 | 0.0 | 9.70E-01 | 0.3 | 2.14E-01 | Uncharacterized protein |
| AT4G30900 | 0.2 | 2.99E-02 | 0.0 | 8.64E-01 | 0.2 | 7.90E-02 | -0.2 | 2.82E-01 | 0.0 | 9.32E-01 | -0.1 | 4.76E-01 | Uncharacterized protein AT4g30900 |
| AT1G55080 | 0.2 | 1.21E-01 | 0.1 | 6.35E-01 | 0.1 | 3.74E-01 | 0.0 | 9.77E-01 | 0.0 | 8.89E-01 | 0.1 | 8.97E-01 | Mediator of RNA polymerase II transcription subunit 9 |
| AT1G75980 | 0.2 | 1.55E-01 | 0.0 | 9.34E-01 | 0.2 | 2.12E-01 | 0.1 | 6.63E-01 | 0.1 | 8.31E-01 | 0.0 | 9.24E-01 | Single hybrid motif superfamily protein |
| AT2G43540 | 0.2 | 1.89E-02 | -0.1 | 1.32E-01 | 0.3 | 4.41E-03 | -0.6 | 1.95E-02 | -0.4 | 1.35E-01 | -0.2 | 6.37E-01 | hypothetical protein |
| AT4G21110 | 0.2 | 3.86E-02 | 0.2 | 7.30E-02 | 0.0 | 8.12E-01 | -0.2 | 3.23E-01 | -0.3 | 3.45E-01 | 0.0 | 9.61E-01 | hypothetical protein |
| AT2G29060 | 0.2 | 1.40E-01 | 0.2 | 9.90E-02 | 0.0 | 8.38E-01 | -0.7 | 1.60E-02 | -0.4 | 1.23E-01 | -0.2 | 4.97E-01 | GRAS family transcription factor |
| AT5G54145 | 0.2 | 1.32E-01 | 0.1 | 2.94E-01 | 0.1 | 7.23E-01 | -0.1 | 7.36E-01 | 0.0 | 9.21E-01 | -0.1 | 8.96E-01 | Transmembrane protein |
| AT5G47390 | 0.2 | 4.17E-02 | 0.0 | 8.95E-01 | 0.2 | 5.63E-02 | -0.2 | 1.35E-01 | -0.2 | 1.73E-01 | 0.0 | 9.89E-01 | Uncharacterized protein |
| AT1G06050 | 0.2 | 6.80E-02 | 0.1 | 5.98E-01 | 0.1 | 2.68E-01 | 0.1 | 7.63E-01 | -0.2 | 6.78E-01 | 0.3 | 5.12E-01 | T21E18.10 protein |
| AT2G01021 | 0.2 | 7.66E-01 | 0.0 | 9.52E-01 | 0.2 | 7.88E-01 | 0.2 | 9.21E-01 | 0.1 | 9.59E-01 | 0.1 | 9.81E-01 | Uncharacterized protein |
| AT4G06701 | 0.2 | 1.61E-01 | 0.1 | 5.04E-01 | 0.1 | 5.60E-01 | -0.2 | 3.49E-01 | -0.2 | 5.03E-01 | 0.0 | 9.28E-01 | NA |
| AT3G27650 | 0.2 | 1.85E-01 | 0.1 | 3.34E-01 | 0.0 | 8.08E-01 | 0.1 | 8.75E-01 | 0.2 | 7.57E-01 | -0.1 | 8.85E-01 | hypothetical protein |
| AT1G19400 | 0.2 | 6.80E-02 | 0.1 | 5.15E-01 | 0.1 | 2.76E-01 | -0.3 | 3.08E-02 | -0.6 | 5.55E-04 | 0.3 | 8.75E-02 | F18O14.16 |
| AT5G44040 | 0.2 | 1.38E-01 | 0.1 | 2.42E-01 | 0.0 | 8.23E-01 | 0.0 | 9.92E-01 | 0.0 | 9.70E-01 | 0.0 | 9.70E-01 | At5g44040 |
| AT1G16870 | 0.2 | 4.85E-02 | 0.2 | 5.49E-02 | 0.0 | 9.99E-01 | 0.4 | 5.52E-03 | 0.2 | 1.66E-01 | 0.2 | 3.56E-01 | Small ribosomal subunit protein mS29 |
| AT4G39690 | 0.2 | 1.81E-02 | 0.2 | 1.64E-02 | 0.0 | 9.07E-01 | 0.2 | 2.34E-01 | 0.2 | 2.64E-01 | 0.0 | 9.67E-01 | MICOS complex subunit MIC60 |
| AT1G18900 | 0.2 | 4.42E-02 | -0.2 | 4.22E-02 | 0.4 | 3.50E-03 | 0.5 | 2.36E-02 | 0.0 | 9.67E-01 | 0.4 | 6.35E-02 | Pentatricopeptide repeat superfamily protein |
| AT4G32360 | 0.2 | 6.52E-02 | 0.0 | 7.46E-01 | 0.2 | 7.24E-02 | -0.1 | 7.34E-01 | 0.1 | 6.43E-01 | -0.2 | 4.40E-01 | AT4g32360 |

| | | | | | | | | | | | | | |
|---|---|---|---|---|---|---|---|---|---|---|---|---|---|
| AT4G02400 | 0.2 | 6.48E-02 | 0.1 | 2.82E-01 | 0.1 | 4.85E-01 | 0.6 | 1.05E-02 | 0.4 | 6.33E-02 | 0.1 | 7.19E-01 | U3 ribonucleoprotein family protein |
| AT5G27270 | 0.2 | 1.26E-01 | -0.1 | 2.73E-01 | 0.3 | 3.70E-02 | -0.1 | 7.30E-01 | 0.1 | 8.56E-01 | -0.2 | 6.02E-01 | Tetratricopeptide repeat -like superfamily protein |
| AT2G44210 | 0.2 | 3.62E-02 | -0.1 | 2.30E-01 | 0.3 | 8.38E-03 | 0.0 | 9.43E-01 | 0.0 | 7.71E-01 | -0.1 | 7.79E-01 | AT2G44210 protein ) |
| AT1G80150 | 0.2 | 1.95E-01 | 0.1 | 3.35E-01 | 0.0 | 8.19E-01 | 0.3 | 2.87E-01 | 0.1 | 8.34E-01 | 0.2 | 5.61E-01 | Tetratricopeptide repeat -like superfamily protein |
| AT3G54040 | 0.2 | 2.02E-01 | 0.1 | 6.61E-01 | 0.1 | 4.71E-01 | 1.2 | 1.66E-06 | 0.8 | 2.94E-04 | 0.4 | 2.58E-02 | PAR1 protein |
| AT2G37410 | 0.2 | 2.63E-02 | 0.0 | 9.74E-01 | 0.2 | 4.61E-02 | -0.2 | 2.11E-01 | -0.3 | 1.98E-02 | 0.2 | 3.36E-01 | Mitochondrial import inner membrane translocase subunit TIM17-2 |
| AT4G28690 | 0.2 | 1.08E-01 | 0.0 | 9.93E-01 | 0.2 | 1.91E-01 | 0.8 | 9.90E-02 | 0.2 | 7.90E-01 | 0.6 | 3.04E-01 | RPM1 interacting protein 13 |
| AT4G07507 | 0.2 | 3.56E-01 | -0.2 | 5.07E-01 | 0.3 | 1.96E-01 | 0.1 | 7.47E-01 | 0.1 | 7.81E-01 | 0.0 | 9.92E-01 | NA |
| AT4G14240 | 0.2 | 3.13E-02 | 0.0 | 6.08E-01 | 0.2 | 2.38E-02 | -0.2 | 6.20E-02 | 0.1 | 6.48E-01 | -0.3 | 6.74E-02 | CBS domain protein with a domain protein |
| AT3G18165 | 0.2 | 3.45E-02 | 0.2 | 7.18E-02 | 0.0 | 7.77E-01 | 0.2 | 1.02E-01 | 0.1 | 6.35E-01 | 0.1 | 5.06E-01 | Pre-mRNA-splicing factor SPF27 homolog |
| AT1G53850 | 0.2 | 3.91E-02 | 0.0 | 8.55E-01 | 0.2 | 8.75E-02 | 0.4 | 9.35E-03 | 0.2 | 1.12E-01 | 0.1 | 5.07E-01 | Proteasome subunit alpha type |
| AT3G07190 | 0.2 | 5.12E-02 | 0.3 | 1.12E-02 | -0.1 | 4.34E-01 | 0.0 | 9.31E-01 | -0.1 | 5.94E-01 | 0.1 | 7.27E-01 | Endoplasmic reticulum transmembrane protein |
| AT4G28100 | 0.2 | 2.55E-02 | 0.3 | 5.56E-03 | -0.1 | 4.00E-01 | -0.2 | 1.66E-01 | -0.1 | 7.31E-01 | -0.1 | 4.86E-01 | SPARK domain-containing protein |
| AT4G02425 | 0.2 | 1.38E-01 | 0.1 | 4.42E-01 | 0.1 | 5.74E-01 | 0.0 | 9.95E-01 | 0.2 | 5.49E-01 | -0.2 | 6.25E-01 | LYR motif protein |
| AT4G29110 | 0.2 | 2.78E-01 | 0.0 | 9.02E-01 | 0.2 | 4.18E-01 | 0.5 | 3.75E-02 | 0.0 | 9.13E-01 | 0.5 | 7.08E-02 | hypothetical protein |
| AT1G13880 | 0.2 | 4.83E-02 | -0.3 | 9.07E-03 | 0.5 | 1.21E-03 | -0.3 | 4.97E-02 | -0.2 | 2.42E-01 | -0.1 | 6.93E-01 | hypothetical protein |
| AT1G30290 | 0.2 | 3.26E-02 | 0.0 | 7.76E-01 | 0.2 | 1.10E-01 | -0.1 | 7.36E-01 | -0.1 | 7.60E-01 | 0.0 | 9.88E-01 | Tetratricopeptide repeat -like superfamily protein |
| AT1G80410 | 0.2 | 1.06E-02 | 0.1 | 5.58E-02 | 0.1 | 4.59E-01 | 0.3 | 5.21E-02 | 0.1 | 7.53E-01 | 0.2 | 2.48E-01 | Tetratricopeptide repeat -containing protein |
| AT2G35790 | 0.2 | 2.46E-01 | 0.4 | 3.76E-02 | -0.2 | 3.25E-01 | 0.4 | 2.19E-03 | 0.0 | 9.34E-01 | 0.5 | 5.72E-03 | hypothetical protein |
| AT3G22104 | 0.2 | 1.35E-01 | 0.0 | 7.41E-01 | 0.1 | 3.26E-01 | 0.0 | 9.59E-01 | 0.0 | 9.39E-01 | 0.0 | 9.14E-01 | Phototropic-responsive NPH3 family protein |
| AT1G80630 | 0.2 | 1.04E-01 | 0.2 | 7.56E-02 | 0.0 | 8.53E-01 | -0.1 | 7.50E-01 | -0.2 | 5.96E-01 | 0.1 | 8.61E-01 | hypothetical protein |
| AT3G58090 | 0.2 | 8.59E-02 | 0.1 | 3.73E-01 | 0.1 | 4.95E-01 | -0.3 | 1.00E-01 | -0.3 | 2.08E-01 | -0.1 | 8.92E-01 | Dirigent protein 17 |
| AT3G26922 | 0.2 | 1.86E-01 | 0.1 | 5.91E-01 | 0.1 | 5.47E-01 | 0.1 | 8.99E-01 | 0.1 | 7.48E-01 | -0.1 | 8.58E-01 | F-box |
| AT3G20350 | 0.2 | 1.24E-01 | 0.1 | 3.00E-01 | 0.1 | 6.96E-01 | -0.2 | 3.79E-01 | -0.2 | 3.98E-01 | 0.0 | 9.67E-01 | Uncharacterized protein |
| AT1G69587 | 0.2 | 2.24E-01 | 0.3 | 5.48E-02 | -0.1 | 4.76E-01 | -0.6 | 2.79E-02 | -0.3 | 2.02E-01 | -0.2 | 4.85E-01 | NA |
| AT3G61360 | 0.2 | 3.23E-01 | 0.4 | 6.18E-02 | -0.2 | 3.93E-01 | 0.0 | 9.95E-01 | 0.3 | 4.39E-01 | -0.3 | 5.14E-01 | Pentatricopeptide repeat-containing protein At3g61360 |
| AT4G13810 | 0.2 | 1.03E-01 | -0.3 | 3.26E-02 | 0.5 | 5.20E-03 | 0.2 | 3.88E-01 | 0.2 | 3.79E-01 | 0.0 | 9.42E-01 | Receptor like protein |
| AT3G26640 | 0.2 | 5.94E-02 | 0.1 | 4.07E-01 | 0.1 | 3.49E-01 | -0.1 | 5.80E-01 | -0.3 | 9.47E-02 | 0.2 | 3.50E-01 | WD repeat-containing protein LWD2 |
| AT1G80500 | 0.2 | 1.13E-01 | 0.2 | 1.09E-01 | 0.0 | 9.56E-01 | 0.2 | 3.19E-01 | 0.1 | 5.56E-01 | 0.0 | 8.84E-01 | At1g80500 |
| AT5G40400 | 0.2 | 1.90E-01 | 0.2 | 2.69E-01 | 0.0 | 9.11E-01 | 0.2 | 4.03E-01 | 0.0 | 9.34E-01 | 0.3 | 4.67E-01 | Pentatricopeptide repeat superfamily protein |
| AT4G26965 | 0.2 | 6.18E-02 | 0.1 | 3.11E-01 | 0.1 | 4.58E-01 | 0.0 | 8.91E-01 | 0.1 | 8.49E-01 | 0.0 | 9.60E-01 | NADH dehydrogenase 1 alpha subcomplex subunit 12 |
| AT5G56910 | 0.2 | 7.86E-02 | 0.0 | 9.55E-01 | 0.2 | 1.47E-01 | -0.1 | 7.20E-01 | -0.1 | 5.60E-01 | 0.0 | 8.64E-01 | Cystatin |
| AT1G73350 | 0.2 | 5.66E-02 | 0.0 | 6.76E-01 | 0.2 | 5.35E-02 | -0.1 | 5.53E-01 | 0.0 | 9.17E-01 | -0.1 | 7.02E-01 | Uncharacterized protein At1g73350 |
| AT2G27385 | 0.2 | 3.19E-01 | 0.6 | 8.41E-03 | -0.4 | 6.06E-02 | -0.9 | 5.98E-05 | -0.7 | 1.57E-03 | -0.2 | 4.66E-01 | Pollen Ole e 1 allergen and extensin family protein |
| AT2G06040 | 0.2 | 6.81E-02 | 0.1 | 4.46E-01 | 0.1 | 3.71E-01 | 0.2 | 3.67E-01 | -0.2 | 5.83E-01 | 0.4 | 1.80E-01 | Uncharacterized protein At2g06040 |
| AT2G42650 | 0.2 | 1.77E-01 | 0.0 | 9.46E-01 | 0.2 | 2.96E-01 | 0.2 | 3.94E-01 | 0.1 | 8.28E-01 | 0.1 | 7.13E-01 | hypothetical protein |
| AT3G54650 | 0.2 | 1.87E-02 | 0.4 | 7.51E-04 | -0.2 | 7.16E-02 | 0.1 | 4.73E-01 | 0.0 | 9.27E-01 | 0.1 | 5.57E-01 | F-box |
| AT4G07825 | 0.2 | 4.96E-02 | 0.2 | 7.39E-02 | 0.0 | 8.99E-01 | 0.0 | 8.60E-01 | 0.0 | 9.30E-01 | 0.0 | 9.77E-01 | At4g07825 |
| AT4G03260 | 0.2 | 3.57E-02 | -0.3 | 3.43E-03 | 0.5 | 4.10E-04 | -0.1 | 5.31E-01 | 0.0 | 9.24E-01 | -0.2 | 5.70E-01 | Uncharacterized protein AT4g03260 |
| AT5G63440 | 0.2 | 4.60E-02 | 0.2 | 8.16E-02 | 0.0 | 8.51E-01 | 0.2 | 3.71E-01 | 0.2 | 3.35E-01 | 0.0 | 9.20E-01 | hypothetical protein |
| AT5G44660 | 0.2 | 7.43E-02 | 0.0 | 8.74E-01 | 0.2 | 1.62E-01 | -0.2 | 3.79E-01 | 0.1 | 5.79E-01 | -0.3 | 1.99E-01 | Uncharacterized protein |
| AT5G04080 | 0.2 | 5.66E-01 | 0.2 | 6.38E-01 | 0.0 | 9.55E-01 | -0.5 | 8.17E-02 | -0.3 | 4.63E-01 | -0.3 | 4.64E-01 | Cysteine-rich TM module stress tolerance protein |
| AT1G32370 | 0.2 | 2.88E-02 | 0.2 | 6.66E-02 | 0.0 | 7.62E-01 | -0.2 | 5.06E-01 | -0.4 | 2.08E-01 | 0.2 | 6.64E-01 | Tobamovirus multiplication protein 2B |
| AT2G16070 | 0.2 | 1.15E-02 | -0.1 | 2.98E-01 | 0.3 | 5.32E-03 | -0.2 | 1.65E-01 | -0.2 | 1.72E-01 | 0.0 | 9.50E-01 | PDV2 |
| AT5G58020 | 0.2 | 7.96E-03 | 0.0 | 4.65E-01 | 0.2 | 5.39E-03 | -0.1 | 2.84E-01 | -0.1 | 2.46E-01 | 0.0 | 9.01E-01 | Replication termination factor 2 |
| AT1G48550 | 0.2 | 9.79E-02 | 0.1 | 2.67E-01 | 0.1 | 6.61E-01 | -0.1 | 6.34E-01 | -0.1 | 8.10E-01 | 0.0 | 9.18E-01 | Vacuolar protein sorting-associated protein 26 |
| AT5G55125 | 0.2 | 1.86E-01 | 0.5 | 7.93E-03 | -0.3 | 1.28E-01 | 0.5 | 2.27E-02 | 0.2 | 2.74E-01 | 0.2 | 4.54E-01 | Ribosomal protein L31 |
| AT2G37100 | 0.2 | 1.86E-01 | 0.4 | 2.41E-02 | -0.2 | 3.12E-01 | -0.3 | 1.10E-01 | 0.0 | 9.38E-01 | -0.3 | 1.84E-01 | hypothetical protein |
| AT4G38770 | 0.2 | 1.44E-01 | 0.6 | 1.94E-03 | -0.4 | 2.91E-02 | -0.1 | 3.84E-01 | 0.0 | 9.25E-01 | -0.1 | 6.25E-01 | Proline-rich protein 4 |
| AT2G29690 | 0.2 | 1.27E-02 | 0.1 | 1.75E-01 | 0.1 | 2.28E-01 | 0.2 | 3.53E-01 | 0.2 | 3.47E-01 | 0.0 | 9.33E-01 | Anthranilate synthase 2 |
| AT1G09740 | 0.2 | 1.34E-02 | 0.3 | 1.50E-03 | -0.1 | 1.89E-01 | -0.2 | 2.86E-01 | -0.2 | 4.17E-01 | 0.0 | 9.56E-01 | hypothetical protein |
| AT3G21465 | 0.2 | 1.08E-02 | 0.2 | 3.64E-02 | 0.0 | 6.39E-01 | 0.4 | 7.29E-02 | 0.2 | 4.59E-01 | 0.2 | 5.27E-01 | Adenylyl cyclase |
| AT3G57980 | 0.2 | 6.17E-02 | 0.0 | 9.93E-01 | 0.2 | 1.20E-01 | 0.1 | 8.40E-01 | -0.3 | 4.02E-01 | 0.4 | 3.69E-01 | Uncharacterized protein T10K17.190 |
| AT5G06270 | 0.2 | 1.14E-01 | 0.4 | 3.57E-03 | -0.2 | 8.81E-02 | 0.1 | 8.06E-01 | -0.1 | 8.01E-01 | 0.1 | 6.70E-01 | Protein GL2-INTERACTING REPRESSOR 1 |
| AT5G19570 | 0.2 | 9.69E-02 | 0.5 | 1.01E-03 | -0.3 | 2.92E-02 | -0.2 | 3.83E-01 | -0.1 | 7.05E-01 | -0.1 | 8.16E-01 | Transmembrane protein |
| AT1G24050 | 0.2 | 4.53E-02 | 0.3 | 9.59E-03 | -0.1 | 3.87E-01 | 0.1 | 6.37E-01 | 0.0 | 9.90E-01 | 0.1 | 7.68E-01 | AT1G24050 protein |
| AT2G24600 | 0.2 | 3.58E-01 | -0.4 | 8.95E-02 | 0.6 | 3.18E-02 | 0.4 | 3.97E-02 | -0.1 | 7.70E-01 | 0.5 | 6.56E-02 | Ankyrin repeat family protein |
| AT1G73620 | 0.2 | 2.83E-01 | 0.5 | 2.45E-02 | -0.3 | 2.16E-01 | 0.2 | 5.88E-01 | 0.5 | 5.51E-02 | -0.4 | 1.99E-01 | Thaumatin-like protein |
| AT5G24735 | 0.2 | 3.80E-01 | -0.2 | 4.98E-01 | 0.4 | 1.76E-01 | -0.4 | 1.58E-01 | -0.4 | 1.44E-01 | 0.1 | 9.06E-01 | NA |
| AT4G14830 | 0.2 | 1.63E-01 | 0.2 | 1.71E-01 | 0.0 | 9.84E-01 | -0.2 | 5.28E-01 | -0.2 | 5.01E-01 | 0.0 | 9.40E-01 | hypothetical protein |
| AT4G18905 | 0.2 | 4.56E-02 | 0.0 | 8.37E-01 | 0.2 | 5.61E-02 | 0.5 | 1.65E-03 | 0.2 | 1.77E-01 | 0.3 | 1.53E-01 | AT4g18900 |
| AT1G52670 | 0.2 | 4.38E-02 | 0.5 | 4.63E-04 | -0.3 | 1.90E-02 | 0.1 | 6.15E-01 | 0.0 | 9.54E-01 | 0.1 | 7.11E-01 | F6D8.11 protein |
| AT1G49000 | 0.2 | 3.42E-01 | -0.5 | 4.45E-02 | 0.7 | 2.05E-02 | 0.4 | 1.02E-01 | -0.1 | 6.94E-01 | 0.5 | 4.40E-02 | At1g49000 |
| AT4G28190 | 0.2 | 4.94E-02 | 0.4 | 1.24E-03 | -0.2 | 6.47E-02 | -0.8 | 3.57E-04 | -0.6 | 5.65E-03 | -0.2 | 3.12E-01 | Developmental regulator, ULTRAPETALA |
| AT2G10410 | 0.2 | 3.03E-02 | 0.2 | 8.78E-02 | 0.0 | 6.66E-01 | -0.3 | 2.19E-02 | -0.3 | 3.80E-02 | 0.0 | 9.98E-01 | NA |
| AT5G22875 | 0.2 | 4.97E-02 | 0.1 | 2.97E-01 | 0.1 | 4.06E-01 | 0.1 | 7.81E-01 | 0.3 | 3.17E-01 | -0.2 | 5.68E-01 | hypothetical protein |
| AT1G61790 | 0.2 | 6.86E-03 | 0.4 | 2.11E-04 | -0.2 | 2.79E-02 | 0.1 | 5.46E-01 | 0.0 | 9.86E-01 | 0.1 | 6.71E-01 | Probable dolichyl-diphosphooligosaccharide--protein glycosyltransferase subunit 3B |
| AT1G79160 | 0.2 | 1.67E-01 | 0.2 | 3.12E-01 | 0.0 | 7.83E-01 | -0.1 | 6.84E-01 | -0.5 | 1.36E-02 | 0.4 | 9.35E-02 | Uncharacterized protein |
| AT1G17430 | 0.2 | 4.39E-02 | 0.2 | 4.79E-02 | 0.0 | 9.83E-01 | -0.4 | 4.34E-02 | 0.1 | 8.32E-01 | -0.4 | 5.78E-02 | F28G4.9 protein |
| AT1G28280 | 0.2 | 3.65E-02 | 0.1 | 1.63E-01 | 0.1 | 4.91E-01 | 0.3 | 4.71E-02 | -0.1 | 3.70E-01 | 0.4 | 2.39E-02 | VQ motif-containing protein |
| AT1G61215 | 0.2 | 1.76E-01 | 0.0 | 8.57E-01 | 0.2 | 3.54E-01 | -0.3 | 5.06E-01 | 0.2 | 6.18E-01 | -0.5 | 2.59E-01 | At1g61215 |
| AT3G59390 | 0.2 | 1.19E-01 | 0.0 | 9.94E-01 | 0.2 | 1.96E-01 | 0.1 | 8.08E-01 | 0.0 | 9.49E-01 | 0.1 | 7.85E-01 | SREBP regulating gene protein |
| AT5G40920 | 0.2 | 5.64E-02 | -0.2 | 4.74E-02 | 0.4 | 4.93E-03 | 0.3 | 3.03E-01 | 0.2 | 5.46E-01 | 0.1 | 8.50E-01 | NA |
| AT5G48520 | 0.2 | 1.37E-02 | 0.0 | 7.82E-01 | 0.2 | 1.99E-02 | 0.0 | 8.57E-01 | 0.0 | 9.88E-01 | 0.0 | 8.87E-01 | HAUS augmin-like complex subunit |
| AT1G28290 | 0.2 | 1.20E-01 | 0.7 | 2.80E-04 | -0.5 | 3.96E-03 | 0.0 | 8.85E-01 | 0.2 | 3.73E-01 | -0.2 | 5.59E-01 | AGP31 |
| AT5G02740 | 0.2 | 1.88E-02 | 0.2 | 2.13E-02 | 0.0 | 9.86E-01 | 0.3 | 5.20E-02 | 0.3 | 2.52E-02 | -0.1 | 7.36E-01 | Ribosomal protein S24e family protein |
| AT3G61600 | 0.2 | 1.03E-02 | 0.0 | 6.68E-01 | 0.2 | 3.99E-02 | -0.3 | 3.56E-02 | -0.3 | 5.07E-02 | 0.0 | 9.74E-01 | hypothetical protein |
| AT5G14440 | 0.2 | 4.01E-02 | 0.3 | 7.93E-03 | -0.1 | 3.83E-01 | 0.1 | 5.75E-01 | -0.1 | 7.33E-01 | 0.3 | 4.45E-01 | Surfeit locus protein 2 |
| AT5G54930 | 0.2 | 1.10E-01 | 0.1 | 4.04E-01 | 0.1 | 5.41E-01 | 0.0 | 9.51E-01 | 0.0 | 9.55E-01 | 0.0 | 9.95E-01 | hypothetical protein |
| AT1G08910 | 0.2 | 2.19E-02 | -0.2 | 3.14E-02 | 0.4 | 1.87E-03 | 0.0 | 9.97E-01 | -0.1 | 7.95E-01 | 0.1 | 8.16E-01 | Uncharacterized protein |
| AT5G04520 | 0.2 | 5.66E-02 | 0.0 | 8.03E-01 | 0.2 | 7.33E-02 | -0.3 | 9.04E-02 | -0.2 | 2.42E-01 | -0.1 | 8.17E-01 | hypothetical protein |
| AT5G50335 | 0.2 | 5.90E-01 | 0.0 | 9.19E-01 | 0.3 | 6.07E-01 | -0.9 | 1.28E-01 | 0.2 | 8.02E-01 | -1.1 | 1.58E-01 | Transmembrane protein |
| AT3G57170 | 0.2 | 1.32E-02 | 0.0 | 9.64E-01 | 0.2 | 2.81E-02 | -0.4 | 8.46E-02 | -0.3 | 2.09E-01 | -0.1 | 8.32E-01 | Uncharacterized protein At3g57170 |

| | | | | | | | | | | | | | |
|---|---|---|---|---|---|---|---|---|---|---|---|---|---|
| AT5G54090 | 0.2 | 1.93E-01 | 0.0 | 8.52E-01 | 0.2 | 3.77E-01 | 0.1 | 8.88E-01 | 0.5 | 4.61E-01 | -0.4 | 6.04E-01 | DNA mismatch repair protein MutS2-like |
| AT2G24160 | 0.2 | 2.50E-01 | -0.4 | 7.31E-02 | 0.6 | 2.25E-02 | 0.1 | 6.73E-01 | 0.0 | 9.77E-01 | 0.1 | 7.14E-01 | NA |
| AT3G58270 | 0.2 | 3.00E-01 | -0.9 | 2.90E-03 | 1.1 | 2.34E-03 | -0.4 | 2.36E-01 | -0.6 | 1.26E-01 | 0.2 | 7.50E-01 | MATH domain and coiled-coil domain-containing protein At3g58270 |
| AT3G01710 | 0.2 | 1.59E-01 | 0.6 | 2.07E-03 | -0.4 | 4.61E-02 | -0.1 | 8.45E-01 | 0.2 | 4.99E-01 | -0.3 | 4.33E-01 | TPX2 protein family |
| AT1G56150 | 0.2 | 2.27E-01 | 0.3 | 1.43E-01 | -0.1 | 8.01E-01 | 0.1 | 5.72E-01 | -0.4 | 1.76E-01 | 0.5 | 7.21E-02 | Auxin-responsive protein SAUR71 |
| AT5G20200 | 0.2 | 1.70E-02 | 0.1 | 1.24E-01 | 0.1 | 3.71E-01 | 0.2 | 1.84E-01 | 0.0 | 9.65E-01 | 0.3 | 1.91E-01 | Nucleoporin-like protein |
| AT2G18910 | 0.2 | 2.03E-02 | 0.4 | 4.34E-04 | -0.2 | 3.61E-02 | 0.1 | 3.62E-01 | 0.0 | 8.82E-01 | 0.1 | 6.67E-01 | Hydroxyproline-rich glycoprotein family protein |
| AT1G23480 | 0.2 | 6.76E-02 | 0.5 | 4.52E-04 | -0.3 | 1.49E-02 | 0.0 | 8.46E-01 | -0.2 | 3.30E-01 | 0.2 | 2.88E-01 | hypothetical protein |
| AT4G31510 | 0.2 | 6.52E-02 | 0.0 | 9.93E-01 | 0.2 | 1.16E-01 | -0.4 | 1.21E-01 | -0.5 | 4.81E-02 | 0.2 | 6.76E-01 | Protein OXIDATIVE STRESS 3 LIKE 5 |
| AT5G63150 | 0.2 | 1.41E-01 | 0.2 | 1.48E-01 | 0.0 | 9.88E-01 | 0.1 | 5.66E-01 | 0.1 | 7.17E-01 | 0.0 | 9.48E-01 | hypothetical protein |
| AT1G22900 | 0.2 | 1.38E-01 | 0.3 | 3.28E-02 | -0.1 | 4.77E-01 | -0.2 | 4.37E-01 | -0.4 | 3.16E-01 | 0.1 | 8.16E-01 | Dirigent protein 11 |
| AT1G06910 | 0.2 | 9.08E-02 | 0.0 | 8.43E-01 | 0.2 | 2.18E-01 | -0.2 | 6.88E-01 | 0.0 | 9.76E-01 | -0.2 | 7.83E-01 | TRFL7 |
| AT5G07790 | 0.2 | 9.28E-02 | 0.0 | 9.17E-01 | 0.2 | 1.38E-01 | -0.2 | 3.30E-01 | 0.0 | 9.14E-01 | -0.3 | 3.61E-01 | Gb\|AAF67766.1 |
| AT1G67250 | 0.2 | 6.64E-02 | 0.3 | 8.05E-03 | -0.1 | 2.65E-01 | 0.3 | 1.06E-01 | 0.1 | 5.30E-01 | 0.1 | 5.16E-01 | hypothetical protein |
| AT3G15810 | 0.2 | 5.96E-02 | -0.1 | 2.28E-01 | 0.3 | 1.30E-02 | 0.2 | 4.55E-01 | 0.0 | 9.59E-01 | 0.1 | 6.74E-01 | Protein LURP-one-related 12 |
| AT1G47420 | 0.2 | 8.36E-03 | 0.3 | 4.34E-03 | 0.0 | 6.15E-01 | 0.0 | 7.96E-01 | 0.0 | 9.86E-01 | 0.0 | 8.89E-01 | Succinate dehydrogenase subunit 5, mitochondrial |
| AT2G31710 | 0.2 | 1.57E-01 | 0.3 | 5.37E-02 | -0.1 | 5.86E-01 | 0.4 | 1.70E-01 | 0.1 | 7.37E-01 | 0.2 | 4.77E-01 | Vacuolar ATPase assembly integral membrane protein VMA21 homolog |
| AT5G66810 | 0.2 | 2.97E-02 | -0.1 | 4.94E-01 | 0.3 | 2.01E-02 | -0.2 | 2.77E-01 | -0.1 | 6.76E-01 | -0.1 | 6.71E-01 | Gb\|AAB71479.1\| |
| AT1G68540 | 0.2 | 3.11E-01 | 0.3 | 2.35E-01 | 0.0 | 8.60E-01 | 0.0 | 9.47E-01 | 0.2 | 5.45E-01 | -0.2 | 5.99E-01 | NAD-binding Rossmann-fold superfamily protein |
| AT4G15570 | 0.2 | 3.76E-02 | 0.0 | 7.25E-01 | 0.2 | 1.33E-01 | 0.1 | 6.22E-01 | 0.0 | 8.89E-01 | 0.2 | 6.23E-01 | SEN1 like protein |
| AT1G28140 | 0.2 | 1.69E-02 | 0.1 | 1.38E-01 | 0.1 | 3.28E-01 | -0.1 | 7.82E-01 | 0.1 | 8.19E-01 | -0.1 | 6.74E-01 | F3H9.20 protein |
| AT5G38690 | 0.2 | 7.31E-03 | 0.0 | 6.95E-01 | 0.2 | 3.57E-02 | 0.2 | 1.81E-01 | 0.4 | 2.75E-02 | -0.2 | 4.23E-01 | AT5g38690 |
| AT5G47890 | 0.2 | 1.52E-01 | 0.2 | 2.42E-01 | 0.0 | 8.47E-01 | -0.1 | 6.31E-01 | -0.2 | 5.58E-01 | 0.1 | 9.14E-01 | NADH dehydrogenase 1 alpha subcomplex subunit 2 |
| AT2G03070 | 0.2 | 3.51E-02 | 0.0 | 8.07E-01 | 0.2 | 1.02E-01 | -0.1 | 8.07E-01 | -0.1 | 8.27E-01 | 0.0 | 9.94E-01 | Mediator of RNA polymerase II transcription subunit 8 |
| AT1G28190 | 0.2 | 2.80E-01 | -0.6 | 1.82E-02 | 0.8 | 8.58E-03 | 0.1 | 7.69E-01 | -0.3 | 5.73E-01 | 0.4 | 4.77E-01 | hypothetical protein |
| AT3G03890 | 0.2 | 5.47E-03 | -0.1 | 5.00E-02 | 0.4 | 8.91E-04 | -0.1 | 6.43E-01 | 0.0 | 8.50E-01 | -0.1 | 6.25E-01 | F20H23.6 protein |
| AT2G36680 | 0.2 | 1.18E-01 | 0.0 | 7.34E-01 | 0.3 | 1.09E-01 | -0.1 | 4.76E-01 | -0.1 | 8.33E-01 | -0.1 | 7.86E-01 | Vacuolar protein-sorting-associated protein 37 homolog 2 |
| AT2G25110 | 0.2 | 6.03E-02 | 0.2 | 5.77E-02 | 0.0 | 9.42E-01 | 0.2 | 3.04E-02 | 0.3 | 2.67E-02 | 0.0 | 8.64E-01 | Stromal cell-derived factor 2-like protein |
| AT5G23940 | 0.2 | 1.07E-01 | 0.2 | 1.80E-01 | 0.0 | 8.35E-01 | 0.4 | 9.79E-03 | 0.2 | 1.64E-01 | 0.2 | 3.38E-01 | PEL3 |
| AT3G09180 | 0.2 | 6.26E-03 | 0.0 | 5.41E-01 | 0.2 | 4.13E-02 | 0.0 | 8.18E-01 | 0.1 | 7.15E-01 | 0.0 | 9.11E-01 | Mediator of RNA polymerase II transcription subunit |
| AT3G06125 | 0.2 | 7.49E-03 | 0.3 | 2.64E-03 | -0.1 | 4.98E-01 | 0.2 | 5.02E-01 | 0.0 | 9.36E-01 | 0.2 | 5.50E-01 | NA |
| AT4G27620 | 0.2 | 3.63E-02 | 0.1 | 3.71E-01 | 0.1 | 2.72E-01 | -0.2 | 1.77E-01 | -0.4 | 2.72E-02 | 0.2 | 4.30E-01 | Intracellular protein transporter |
| AT3G10970 | 0.2 | 9.10E-03 | 0.2 | 2.12E-02 | 0.0 | 7.55E-01 | -0.1 | 5.43E-01 | 0.0 | 9.25E-01 | -0.1 | 7.70E-01 | Uncharacterized protein F9F8.21 |
| AT3G60980 | 0.2 | 6.60E-02 | 0.0 | 8.25E-01 | 0.2 | 7.99E-02 | 0.1 | 7.73E-01 | -0.1 | 7.18E-01 | 0.2 | 5.71E-01 | Pentatricopeptide repeat-containing protein At3g60980, mitochondrial |
| AT5G45660 | 0.2 | 6.39E-02 | 0.4 | 3.52E-03 | -0.2 | 1.53E-01 | -0.1 | 4.68E-01 | -0.2 | 3.07E-01 | 0.1 | 7.79E-01 | Uncharacterized protein |
| AT3G21865 | 0.2 | 2.81E-02 | 0.1 | 4.38E-01 | 0.1 | 1.66E-01 | 0.0 | 9.74E-01 | -0.1 | 3.53E-01 | 0.1 | 4.94E-01 | Peroxisome biogenesis protein 22 |
| AT3G18790 | 0.2 | 2.61E-02 | 0.2 | 1.89E-02 | 0.0 | 8.29E-01 | 0.1 | 7.40E-01 | -0.1 | 5.56E-01 | 0.2 | 4.40E-01 | AT3G18790 protein |
| AT1G22970 | 0.2 | 2.45E-02 | 0.0 | 9.77E-01 | 0.2 | 5.09E-02 | 0.2 | 2.09E-01 | 0.1 | 5.99E-01 | 0.1 | 6.75E-01 | Cyclin-D1-binding protein |
| AT1G52720 | 0.2 | 1.65E-01 | 0.5 | 7.38E-03 | -0.3 | 1.19E-01 | 0.1 | 5.58E-01 | 0.2 | 4.87E-01 | -0.1 | 9.01E-01 | hypothetical protein |
| AT1G21326 | 0.2 | 2.04E-01 | 0.3 | 9.82E-02 | -0.1 | 6.98E-01 | 0.9 | 1.41E-02 | -0.3 | 5.40E-01 | 1.2 | 1.37E-02 | VQ domain-containing protein |
| AT2G45450 | 0.2 | 1.55E-01 | 0.1 | 4.25E-01 | 0.1 | 6.35E-01 | 0.1 | 8.54E-01 | -0.1 | 7.51E-01 | 0.2 | 6.73E-01 | hypothetical protein |
| AT2G31240 | 0.2 | 4.39E-02 | 0.0 | 6.58E-01 | 0.3 | 4.18E-02 | 0.5 | 3.89E-03 | 0.1 | 8.24E-01 | 0.4 | 2.94E-02 | Putative kinesin light chain |
| AT5G14030 | 0.2 | 9.68E-02 | 0.5 | 4.83E-03 | -0.2 | 1.07E-01 | 0.2 | 2.49E-01 | 0.1 | 4.63E-01 | 0.0 | 8.95E-01 | Translocon-associated protein subunit beta |
| AT5G19230 | 0.2 | 4.10E-02 | 0.6 | 1.05E-04 | -0.4 | 3.59E-03 | 0.1 | 5.97E-01 | -0.4 | 1.71E-01 | 0.5 | 1.25E-01 | Uncharacterized GPI-anchored protein At5g19230 |
| AT3G51280 | 0.2 | 2.14E-01 | 0.5 | 1.94E-02 | -0.2 | 2.32E-01 | 0.3 | 1.25E-01 | 0.2 | 2.73E-01 | 0.0 | 8.83E-01 | Protein POLLENLESS 3-LIKE 2 |
| AT1G77220 | 0.2 | 3.35E-02 | 0.1 | 5.27E-01 | 0.2 | 1.73E-01 | 0.0 | 8.10E-01 | -0.1 | 7.95E-01 | 0.0 | 9.71E-01 | Protein LAZ1 homolog 1 |
| AT4G21105 | 0.2 | 2.39E-02 | 0.3 | 1.08E-02 | -0.1 | 6.09E-01 | 0.2 | 3.31E-01 | 0.1 | 6.95E-01 | 0.1 | 7.94E-01 | Cytochrome-c oxidase |
| AT4G00755 | 0.2 | 8.71E-03 | 0.2 | 8.32E-03 | 0.0 | 9.05E-01 | -0.3 | 3.49E-02 | -0.3 | 1.07E-01 | -0.1 | 8.33E-01 | F-box protein At4g00755 |
| AT2G45470 | 0.2 | 3.75E-02 | 0.7 | 9.53E-05 | -0.4 | 2.45E-03 | -0.2 | 1.73E-01 | -0.2 | 3.44E-01 | 0.0 | 9.07E-01 | Fasciclin-like arabinogalactan protein 8 |
| AT2G44970 | 0.2 | 1.02E-02 | 0.1 | 2.14E-01 | 0.1 | 1.53E-01 | 0.0 | 9.89E-01 | -0.1 | 7.29E-01 | 0.1 | 7.92E-01 | Alpha |
| AT1G48950 | 0.2 | 1.24E-02 | 0.2 | 1.27E-02 | 0.0 | 9.37E-01 | -0.1 | 6.28E-01 | -0.2 | 2.02E-01 | 0.1 | 5.10E-01 | Uncharacterized protein At1g48950 |
| AT1G65032 | 0.2 | 1.91E-01 | 0.3 | 1.43E-01 | 0.0 | 8.65E-01 | 0.1 | 8.86E-01 | -0.2 | 5.76E-01 | 0.3 | 4.96E-01 | COMPLEX 1 LYR-like protein |
| AT3G10040 | 0.2 | 9.50E-02 | 0.5 | 3.80E-03 | -0.2 | 1.22E-01 | 1.1 | 2.48E-03 | 0.3 | 5.45E-01 | 0.9 | 4.71E-02 | Uncharacterized protein T22K18.13 |
| AT5G57040 | 0.2 | 4.13E-02 | 0.2 | 9.12E-02 | 0.0 | 7.57E-01 | -0.2 | 1.91E-01 | 0.1 | 5.58E-01 | -0.3 | 1.46E-01 | VOC domain-containing protein |
| AT1G73570 | 0.2 | 6.60E-02 | 0.3 | 4.25E-02 | 0.0 | 8.02E-01 | -0.3 | 6.06E-01 | 0.2 | 8.03E-01 | -0.4 | 4.82E-01 | HCP-like superfamily protein |
| AT1G16170 | 0.2 | 5.36E-02 | 0.3 | 1.68E-02 | -0.1 | 5.51E-01 | -0.7 | 7.51E-04 | -1.0 | 1.76E-04 | 0.3 | 3.59E-01 | T24D18.25 protein |
| AT5G27830 | 0.2 | 1.83E-02 | 0.0 | 8.25E-01 | 0.2 | 6.03E-02 | 0.1 | 5.39E-01 | -0.2 | 4.08E-01 | 0.3 | 2.32E-01 | Folate receptor family protein |
| AT3G24420 | 0.2 | 4.67E-01 | 0.5 | 9.30E-02 | -0.3 | 3.48E-01 | -0.9 | 3.97E-04 | -0.4 | 4.52E-02 | -0.4 | 1.41E-01 | hypothetical protein |
| AT5G02480 | 0.2 | 9.44E-03 | 0.0 | 6.58E-01 | 0.2 | 3.75E-02 | 0.2 | 3.89E-02 | 0.0 | 9.22E-01 | 0.2 | 1.27E-01 | Uncharacterized protein T22P11_70 |
| AT3G21400 | 0.2 | 1.63E-01 | 0.3 | 4.30E-02 | -0.1 | 5.10E-01 | -0.1 | 7.30E-01 | 0.0 | 8.88E-01 | -0.1 | 6.76E-01 | Dynein beta chain, ciliary protein |
| AT1G07135 | 0.2 | 4.62E-01 | -0.3 | 3.90E-01 | 0.5 | 1.78E-01 | -0.1 | 7.34E-01 | -0.8 | 8.70E-03 | 0.7 | 3.43E-02 | At1g07135 |
| AT1G55750 | 0.2 | 6.95E-02 | -0.1 | 2.93E-01 | 0.4 | 2.49E-02 | 0.1 | 5.59E-01 | 0.1 | 8.60E-01 | 0.1 | 8.20E-01 | BSD domain |
| AT5G67570 | 0.2 | 5.25E-02 | 0.2 | 9.43E-02 | 0.0 | 8.41E-01 | -0.3 | 1.66E-01 | -0.6 | 1.34E-02 | 0.3 | 2.90E-01 | Uncharacterized protein At5g67570 |
| AT1G74170 | 0.2 | 4.25E-02 | 0.1 | 4.03E-01 | 0.1 | 2.88E-01 | 0.2 | 4.61E-01 | 0.1 | 8.27E-01 | 0.1 | 7.48E-01 | Receptor like protein 13 |
| AT2G45460 | 0.2 | 3.47E-02 | -0.2 | 1.21E-01 | 0.4 | 7.42E-03 | 0.0 | 9.26E-01 | 0.2 | 6.09E-01 | -0.2 | 5.40E-01 | Uncharacterized protein At2g45460 |
| AT1G02180 | 0.2 | 1.03E-01 | 0.5 | 4.49E-03 | -0.2 | 1.28E-01 | -0.4 | 2.00E-02 | 0.0 | 9.02E-01 | -0.5 | 4.29E-02 | Ferredoxin-like protein |
| AT3G02220 | 0.2 | 3.95E-02 | 0.3 | 1.86E-02 | -0.1 | 6.73E-01 | 0.2 | 2.81E-01 | -0.1 | 7.31E-01 | 0.2 | 2.21E-01 | Small acidic-like protein |
| AT5G49580 | 0.2 | 1.04E-02 | -0.1 | 4.46E-01 | 0.3 | 7.52E-03 | -0.1 | 6.55E-01 | 0.1 | 7.14E-01 | -0.1 | 5.10E-01 | Chaperone DnaJ-domain superfamily protein |
| AT1G08050 | 0.2 | 1.67E-01 | 0.0 | 8.39E-01 | 0.3 | 1.62E-01 | 0.6 | 5.65E-03 | 0.1 | 6.25E-01 | 0.4 | 7.60E-02 | Zinc finger family protein |
| AT4G23330 | 0.2 | 8.11E-02 | 0.0 | 7.87E-01 | 0.3 | 9.22E-02 | -0.1 | 6.76E-01 | -0.3 | 1.67E-01 | 0.2 | 4.57E-01 | Uncharacterized protein AT4g23330 |
| AT5G03545 | 0.2 | 4.10E-02 | 0.0 | 7.53E-01 | 0.2 | 1.29E-01 | 0.2 | 6.84E-01 | 0.7 | 1.17E-01 | -0.5 | 2.52E-01 | At5g03545 |
| AT5G60680 | 0.2 | 1.27E-01 | 0.3 | 9.52E-02 | 0.0 | 8.63E-01 | -0.4 | 3.32E-02 | -1.0 | 5.98E-04 | 0.5 | 7.90E-02 | Senescence regulator |
| AT5G51150 | 0.2 | 1.25E-02 | 0.2 | 1.21E-02 | 0.0 | 9.20E-01 | -0.2 | 2.68E-01 | -0.3 | 1.64E-01 | 0.1 | 7.50E-01 | Gb\|AAD46013.1 |
| AT5G37130 | 0.2 | 9.97E-03 | 0.0 | 9.22E-01 | 0.2 | 1.88E-02 | 0.0 | 9.99E-01 | -0.2 | 3.22E-01 | 0.2 | 4.27E-01 | Uncharacterized protein At5g37130 |
| AT5G40600 | 0.2 | 1.50E-01 | 0.2 | 1.60E-01 | 0.0 | 9.87E-01 | 0.0 | 9.51E-01 | 0.2 | 6.86E-01 | -0.2 | 6.12E-01 | Bromodomain testis-specific protein |
| AT4G22490 | 0.2 | 5.60E-02 | 0.4 | 3.26E-03 | -0.2 | 1.26E-01 | 0.4 | 1.21E-01 | -0.3 | 2.37E-01 | 0.7 | 1.77E-02 | hypothetical protein |
| AT3G26890 | 0.2 | 8.93E-02 | 0.1 | 2.72E-01 | 0.1 | 5.95E-01 | -0.4 | 7.28E-03 | -0.4 | 2.37E-02 | 0.0 | 8.77E-01 | hypothetical protein |
| AT4G32605 | 0.2 | 1.47E-01 | 0.2 | 1.84E-01 | 0.0 | 9.49E-01 | 0.4 | 2.24E-02 | 0.2 | 3.50E-01 | 0.2 | 3.19E-01 | Uncharacterized protein AT4g32610 |
| AT1G52380 | 0.2 | 1.49E-02 | 0.1 | 9.70E-02 | 0.1 | 4.02E-01 | 0.1 | 3.40E-01 | -0.1 | 7.05E-01 | 0.2 | 2.71E-01 | Nuclear pore complex protein NUP50A |
| AT5G09260 | 0.2 | 1.42E-01 | -0.1 | 6.11E-01 | 0.3 | 1.03E-01 | 0.3 | 1.28E-01 | 0.0 | 9.75E-01 | 0.3 | 2.24E-01 | hypothetical protein |
| AT4G14840 | 0.2 | 1.31E-01 | 0.0 | 9.01E-01 | 0.2 | 2.71E-01 | -0.3 | 5.37E-01 | -0.1 | 9.19E-01 | -0.2 | 7.49E-01 | At4g14840 |
| AT4G02380 | 0.2 | 3.07E-01 | -0.5 | 5.49E-02 | 0.7 | 1.57E-02 | 0.2 | 5.83E-01 | -1.7 | 1.58E-03 | 1.9 | 3.56E-03 | Senescence-associated gene 21 |
| AT1G03620 | 0.2 | 1.93E-01 | 0.3 | 1.40E-01 | 0.0 | 8.45E-01 | -0.8 | 1.21E-01 | 0.0 | 9.79E-01 | -0.8 | 2.68E-01 | hypothetical protein |

| | | | | | | | | | | | | | |
|---|---|---|---|---|---|---|---|---|---|---|---|---|---|
| AT1G25420 | 0.2 | 2.44E-02 | -0.1 | 2.61E-01 | 0.3 | 8.81E-03 | -0.2 | 1.49E-01 | -0.1 | 5.25E-01 | -0.1 | 6.78E-01 | At1g25420 |
| AT2G31530 | 0.2 | 2.93E-02 | 0.1 | 3.08E-01 | 0.1 | 2.79E-01 | 0.1 | 5.63E-01 | 0.2 | 3.04E-01 | -0.1 | 7.19E-01 | SCY2 |
| AT5G25270 | 0.2 | 5.19E-03 | 0.0 | 7.24E-01 | 0.2 | 2.06E-02 | -0.3 | 1.30E-01 | -0.4 | 6.35E-02 | 0.1 | 6.97E-01 | Ubiquitin-like domain-containing protein |
| AT2G15080 | 0.2 | 1.77E-01 | -0.1 | 7.84E-01 | 0.3 | 1.56E-01 | 0.5 | 5.74E-02 | -0.6 | 9.07E-02 | 1.2 | 5.99E-03 | Receptor-like protein 19 |
| AT1G63090 | 0.2 | 2.42E-02 | -0.1 | 1.88E-01 | 0.4 | 6.87E-03 | 0.2 | 3.73E-01 | -0.1 | 8.05E-01 | 0.3 | 3.46E-01 | PP2-A11 |
| AT3G06380 | 0.2 | 3.71E-02 | 0.2 | 7.34E-02 | 0.0 | 7.96E-01 | -0.3 | 6.76E-02 | -0.4 | 8.98E-03 | 0.2 | 4.25E-01 | Tubby-like protein 9 |
| AT1G67040 | 0.2 | 1.56E-01 | 0.1 | 6.90E-01 | 0.2 | 4.17E-01 | -0.1 | 8.44E-01 | 0.0 | 8.90E-01 | -0.1 | 7.66E-01 | DnaA initiator-associating protein |
| AT1G70790 | 0.2 | 5.18E-02 | 0.2 | 1.83E-01 | 0.1 | 5.95E-01 | -0.2 | 3.26E-01 | 0.0 | 8.83E-01 | -0.2 | 5.66E-01 | Calcium-dependent lipid-binding family protein |
| AT4G01935 | 0.2 | 3.61E-02 | 0.2 | 1.31E-01 | 0.1 | 6.06E-01 | -0.4 | 1.25E-01 | -0.1 | 7.67E-01 | -0.3 | 3.90E-01 | Insulin-induced protein |
| AT2G03310 | 0.2 | 2.15E-02 | 0.4 | 1.77E-03 | -0.1 | 1.64E-01 | 0.1 | 4.31E-01 | 0.0 | 9.44E-01 | 0.1 | 6.35E-01 | Uncharacterized protein At2g03310 |
| AT4G28760 | 0.2 | 3.43E-03 | 0.0 | 5.04E-01 | 0.3 | 3.64E-03 | -0.2 | 4.40E-01 | -0.4 | 9.45E-02 | 0.2 | 4.54E-01 | Methyl-coenzyme M reductase II subunit gamma, putative |
| AT3G02510 | 0.2 | 6.42E-02 | 0.3 | 1.83E-02 | -0.1 | 5.15E-01 | 0.2 | 4.41E-01 | 0.3 | 2.59E-01 | -0.1 | 7.81E-01 | At3g02510 |
| AT1G29220 | 0.2 | 1.20E-02 | 0.0 | 8.35E-01 | 0.3 | 2.09E-02 | -0.1 | 6.79E-01 | -0.1 | 7.54E-01 | 0.0 | 9.86E-01 | Transcriptional regulator family protein |
| AT5G45330 | 0.2 | 5.33E-02 | 0.2 | 1.17E-01 | 0.0 | 7.73E-01 | 0.1 | 7.97E-01 | 0.4 | 3.49E-01 | -0.3 | 5.64E-01 | Decapping 5-like protein |
| AT3G24515 | 0.2 | 1.65E-01 | 0.3 | 7.94E-02 | -0.1 | 7.05E-01 | -0.1 | 6.52E-01 | 0.0 | 9.32E-01 | -0.1 | 7.76E-01 | Ubiquitin-conjugating enzyme 37 |
| AT1G02475 | 0.2 | 2.62E-02 | -0.1 | 5.58E-01 | 0.3 | 1.88E-02 | 0.2 | 2.01E-01 | 0.3 | 7.09E-02 | -0.1 | 6.43E-01 | Polyketide cyclase |
| AT1G64790 | 0.2 | 8.00E-03 | 0.1 | 1.52E-01 | 0.1 | 1.65E-01 | 0.4 | 4.75E-02 | 0.2 | 3.03E-01 | 0.1 | 6.25E-01 | ILITYHIA |
| AT4G26410 | 0.2 | 2.63E-02 | 0.3 | 7.94E-03 | -0.1 | 4.93E-01 | 0.0 | 9.41E-01 | 0.0 | 9.97E-01 | 0.0 | 9.61E-01 | RGS1-HXK1-interacting protein 1 |
| AT4G14190 | 0.2 | 1.60E-01 | -0.2 | 2.60E-01 | 0.4 | 4.89E-02 | 0.2 | 6.47E-01 | 0.5 | 1.16E-01 | -0.4 | 3.64E-01 | Pentatricopeptide repeat-containing protein At4g14190, chloroplastic |
| AT3G57930 | 0.2 | 5.33E-02 | 0.3 | 1.23E-02 | -0.1 | 4.47E-01 | -0.2 | 4.09E-01 | -0.4 | 1.61E-01 | 0.2 | 6.50E-01 | hypothetical protein |
| AT5G61330 | 0.2 | 2.57E-02 | 0.2 | 3.41E-02 | 0.0 | 9.55E-01 | -0.1 | 4.90E-01 | -0.3 | 3.10E-02 | 0.2 | 1.93E-01 | Uncharacterized protein |
| AT1G19520 | 0.2 | 3.20E-02 | 0.3 | 1.42E-02 | -0.1 | 6.45E-01 | 0.1 | 4.02E-01 | 0.0 | 9.75E-01 | 0.1 | 5.43E-01 | Pentacotripeptide-repeat region of PRORP domain-containing protein |
| AT1G04080 | 0.2 | 2.18E-03 | 0.0 | 5.24E-01 | 0.3 | 2.24E-03 | 0.2 | 4.95E-01 | 0.1 | 7.67E-01 | 0.1 | 8.69E-01 | Tetratricopeptide repeat -like superfamily protein |
| AT2G42580 | 0.2 | 5.30E-02 | 0.3 | 3.15E-02 | 0.0 | 7.64E-01 | -0.6 | 2.60E-03 | -0.7 | 2.11E-03 | 0.1 | 7.46E-01 | Inactive TPR repeat-containing thioredoxin TTL3 |
| AT5G59410 | 0.2 | 2.13E-01 | 0.3 | 1.92E-01 | 0.0 | 9.42E-01 | -0.2 | 5.18E-01 | -0.2 | 5.76E-01 | 0.0 | 1.00E+00 | Gb\|AAF17201.1 |
| AT3G52640 | 0.2 | 4.76E-03 | 0.1 | 1.19E-01 | 0.1 | 1.36E-01 | 0.3 | 1.42E-01 | 0.2 | 2.57E-01 | 0.0 | 9.37E-01 | Nicastrin |
| AT3G03040 | 0.2 | 1.86E-02 | 0.0 | 7.09E-01 | 0.3 | 2.39E-02 | -0.2 | 2.33E-01 | -0.4 | 6.56E-02 | 0.1 | 5.89E-01 | F-box protein At3g03040 |
| AT1G15430 | 0.2 | 5.92E-02 | 0.0 | 9.28E-01 | 0.2 | 9.58E-02 | -0.2 | 1.80E-01 | -0.5 | 1.15E-02 | 0.3 | 2.53E-01 | F9L1.39 |
| AT5G51370 | 0.2 | 6.14E-02 | 0.0 | 7.91E-01 | 0.3 | 8.37E-02 | -0.1 | 7.25E-01 | 0.0 | 9.25E-01 | -0.1 | 7.21E-01 | F-box protein At5g51370 |
| AT1G73940 | 0.2 | 6.55E-02 | 0.5 | 2.26E-03 | -0.3 | 8.13E-02 | 0.3 | 5.54E-02 | 0.1 | 6.38E-01 | 0.2 | 3.14E-01 | hypothetical protein |
| AT3G04950 | 0.2 | 1.79E-01 | 0.5 | 1.06E-02 | -0.3 | 1.82E-01 | 0.7 | 1.73E-02 | 0.4 | 3.29E-01 | 0.4 | 2.88E-01 | Uncharacterized protein |
| AT1G28540 | 0.2 | 9.17E-02 | 0.0 | 9.05E-01 | 0.3 | 1.31E-01 | 0.1 | 8.88E-01 | 0.0 | 9.88E-01 | 0.0 | 9.32E-01 | hypothetical protein |
| AT1G06010 | 0.2 | 9.05E-02 | 0.5 | 2.14E-03 | -0.3 | 6.69E-02 | 0.0 | 9.79E-01 | -0.2 | 4.63E-01 | 0.2 | 5.35E-01 | Basic leucine zipper |
| AT4G20340 | 0.2 | 3.53E-02 | 0.1 | 4.37E-01 | 0.2 | 2.29E-01 | 0.3 | 1.24E-01 | 0.1 | 6.07E-01 | 0.2 | 5.03E-01 | Transcription factor TFIIE, alpha subunit |
| AT2G24290 | 0.2 | 2.36E-02 | 0.4 | 2.09E-03 | -0.1 | 1.83E-01 | -0.2 | 3.49E-01 | -0.1 | 5.36E-01 | 0.0 | 9.05E-01 | Expressed protein ) |
| AT3G58020 | 0.2 | 2.28E-02 | 0.1 | 1.40E-01 | 0.1 | 4.42E-01 | 0.2 | 3.32E-01 | 0.0 | 9.74E-01 | 0.2 | 4.27E-01 | Chaperone DnaJ-domain superfamily protein |
| AT1G01540 | 0.2 | 2.27E-03 | 0.1 | 3.96E-01 | 0.2 | 2.00E-02 | -0.6 | 2.77E-04 | -0.5 | 2.07E-03 | -0.1 | 7.72E-01 | Uncharacterized protein At1g01540 |
| AT3G14190 | 0.2 | 5.44E-02 | 0.5 | 1.38E-03 | -0.3 | 7.03E-02 | 0.0 | 8.89E-01 | 0.4 | 1.04E-01 | -0.4 | 1.89E-01 | Protein PATRONUS 1 |
| AT2G19990 | 0.2 | 2.93E-01 | -0.1 | 7.73E-01 | 0.3 | 2.80E-01 | -0.7 | 1.41E-01 | 0.3 | 4.76E-01 | -1.0 | 4.07E-02 | PR-1-like protein |
| AT3G61370 | 0.2 | 7.93E-02 | 0.0 | 7.59E-01 | 0.2 | 2.29E-01 | 0.3 | 3.34E-01 | 0.1 | 7.44E-01 | 0.2 | 7.27E-01 | Uncharacterized protein |
| AT4G39150 | 0.2 | 2.05E-02 | 0.2 | 1.24E-01 | 0.1 | 4.09E-01 | 0.1 | 7.74E-01 | 0.1 | 7.53E-01 | 0.0 | 9.58E-01 | hypothetical protein |
| AT3G14780 | 0.2 | 3.18E-02 | 0.4 | 1.69E-03 | -0.2 | 1.26E-01 | 0.4 | 2.51E-01 | 0.3 | 5.37E-01 | 0.1 | 7.63E-01 | Uncharacterized protein |
| AT1G55000 | 0.2 | 5.17E-03 | 0.2 | 3.09E-02 | 0.1 | 4.33E-01 | -0.4 | 2.55E-03 | -0.4 | 7.55E-03 | 0.0 | 9.29E-01 | F-box protein At1g55000 |
| AT2G16380 | 0.2 | 3.91E-02 | 0.7 | 1.54E-04 | -0.4 | 5.60E-03 | 0.0 | 9.77E-01 | -0.1 | 4.08E-01 | 0.1 | 5.37E-01 | Sec14p-like phosphatidylinositol transfer family protein |
| AT1G74088 | 0.2 | 8.77E-02 | 0.2 | 2.49E-01 | 0.1 | 6.67E-01 | 0.2 | 5.21E-01 | 0.2 | 7.24E-01 | 0.1 | 8.79E-01 | hypothetical protein |
| AT5G60940 | 0.2 | 1.52E-02 | 0.1 | 2.20E-01 | 0.1 | 2.29E-01 | -0.1 | 7.91E-01 | -0.1 | 6.84E-01 | 0.0 | 9.05E-01 | Cleavage stimulation factor 50 kDa subunit |
| AT5G42970 | 0.2 | 1.08E-02 | 0.2 | 3.96E-02 | 0.1 | 5.94E-01 | 0.2 | 9.39E-02 | 0.0 | 9.70E-01 | 0.2 | 1.91E-01 | FUS8 |
| AT2G23940 | 0.2 | 1.79E-02 | 0.3 | 1.36E-02 | 0.0 | 8.35E-01 | 0.1 | 7.87E-01 | -0.1 | 8.04E-01 | 0.1 | 6.54E-01 | Expressed protein ) |
| AT3G17160 | 0.2 | 1.71E-02 | 0.3 | 9.46E-03 | 0.0 | 7.12E-01 | -0.1 | 6.05E-01 | -0.2 | 3.80E-01 | 0.1 | 7.92E-01 | Uncharacterized protein |
| AT1G63850 | 0.2 | 1.08E-02 | 0.3 | 9.44E-03 | 0.0 | 8.78E-01 | -0.1 | 5.82E-01 | 0.0 | 8.32E-01 | 0.0 | 8.49E-01 | BTB |
| AT5G53150 | 0.2 | 3.33E-03 | 0.0 | 5.47E-01 | 0.2 | 2.42E-02 | 0.2 | 3.47E-01 | 0.1 | 8.17E-01 | 0.1 | 6.68E-01 | DnaJ heat shock amino-terminal domain protein |
| AT1G30200 | 0.2 | 4.91E-03 | 0.2 | 1.23E-02 | 0.0 | 7.31E-01 | -0.5 | 2.78E-04 | -0.6 | 5.88E-04 | 0.0 | 9.49E-01 | F-box protein At1g30200 |
| AT3G12300 | 0.2 | 1.04E-02 | 0.1 | 4.70E-01 | 0.2 | 7.93E-02 | -0.1 | 8.39E-01 | -0.2 | 5.00E-01 | 0.1 | 7.17E-01 | Uncharacterized protein F28J15.10 |
| AT5G22350 | 0.2 | 5.29E-03 | 0.1 | 1.41E-01 | 0.1 | 1.30E-01 | 0.4 | 6.71E-04 | 0.0 | 8.49E-01 | 0.5 | 6.92E-03 | Mitochondrial fission protein ELM1 |
| AT4G37830 | 0.2 | 1.37E-02 | 0.4 | 1.75E-03 | -0.1 | 1.94E-01 | 0.0 | 8.84E-01 | 0.0 | 9.41E-01 | 0.0 | 9.80E-01 | Cytochrome c oxidase-like protein |
| AT5G05950 | 0.2 | 1.91E-02 | 0.1 | 2.69E-01 | 0.1 | 2.27E-01 | -0.4 | 3.77E-02 | -0.4 | 4.27E-02 | 0.0 | 9.54E-01 | hypothetical protein |
| AT2G20000 | 0.2 | 5.90E-03 | 0.1 | 1.98E-01 | 0.1 | 1.11E-01 | -0.1 | 4.46E-01 | 0.0 | 9.38E-01 | -0.2 | 5.07E-01 | CDC27 family protein |
| AT2G47370 | 0.2 | 2.73E-02 | 0.1 | 3.12E-01 | 0.1 | 2.67E-01 | 0.0 | 9.46E-01 | 0.0 | 9.60E-01 | 0.0 | 9.24E-01 | Calcium-dependent phosphotriesterase superfamily protein |
| AT4G17960 | 0.2 | 4.66E-02 | 0.1 | 3.97E-01 | 0.1 | 3.15E-01 | 0.0 | 9.24E-01 | 0.1 | 7.98E-01 | -0.1 | 7.79E-01 | hypothetical protein |
| AT5G62290 | 0.2 | 3.57E-02 | 0.1 | 2.18E-01 | 0.1 | 4.21E-01 | 0.3 | 1.77E-01 | 0.0 | 8.89E-01 | 0.2 | 4.31E-01 | Chloride conductance regulatory protein ICln ) |
| AT3G53830 | 0.2 | 6.40E-03 | -0.3 | 6.26E-03 | 0.5 | 3.35E-04 | | | | | | | Regulator of chromosome condensation family protein |
| AT2G37160 | 0.2 | 8.93E-03 | 0.2 | 8.17E-02 | 0.1 | 3.34E-01 | 0.2 | 3.47E-01 | 0.1 | 8.69E-01 | 0.2 | 5.85E-01 | Uncharacterized protein At2g37160 |
| AT2G47790 | 0.2 | 1.31E-02 | 0.1 | 4.29E-01 | 0.2 | 1.05E-01 | 0.2 | 4.06E-01 | 0.1 | 7.14E-01 | 0.1 | 8.39E-01 | WD repeat-containing protein GTS1 |
| AT2G42485 | 0.2 | 1.32E-01 | 0.1 | 7.27E-01 | 0.2 | 3.54E-01 | 0.5 | 1.55E-01 | 0.3 | 4.84E-01 | 0.2 | 6.56E-01 | NA |
| AT5G48680 | 0.2 | 4.40E-02 | 0.0 | 9.72E-01 | 0.2 | 8.99E-02 | 0.1 | 4.43E-01 | 0.0 | 9.33E-01 | 0.2 | 5.15E-01 | hypothetical protein |
| AT4G31360 | 0.2 | 8.01E-02 | 0.3 | 2.80E-02 | -0.1 | 5.95E-01 | 0.1 | 6.37E-01 | 0.1 | 7.79E-01 | 0.0 | 9.48E-01 | Selenium binding protein |
| AT3G15690 | 0.2 | 1.55E-02 | 0.2 | 8.51E-02 | 0.1 | 4.47E-01 | 0.0 | 9.94E-01 | 0.0 | 8.36E-01 | 0.0 | 8.66E-01 | Putative acetyl-CoA carboxylase biotin-containing subunit |
| AT5G25210 | 0.2 | 4.73E-02 | 0.3 | 1.47E-02 | -0.1 | 5.46E-01 | -0.3 | 1.68E-01 | -0.4 | 1.14E-01 | 0.1 | 8.42E-01 | Uncharacterized protein |
| AT2G39870 | 0.2 | 2.65E-02 | 0.2 | 4.05E-02 | 0.0 | 9.00E-01 | 0.3 | 1.34E-01 | -0.2 | 2.42E-01 | 0.5 | 3.08E-02 | At2g39870 |
| AT4G07410 | 0.2 | 1.10E-02 | 0.1 | 4.37E-01 | 0.2 | 7.58E-02 | 0.3 | 4.28E-02 | 0.0 | 8.38E-01 | 0.2 | 1.98E-01 | Transducin family protein |
| AT3G55000 | 0.2 | 7.42E-03 | 0.1 | 1.14E-01 | 0.1 | 2.17E-01 | 0.1 | 7.61E-01 | 0.0 | 8.32E-01 | 0.0 | 9.86E-01 | Protein TONNEAU 1a |
| AT1G12020 | 0.2 | 3.73E-02 | 0.5 | 1.26E-03 | -0.2 | 8.09E-02 | 0.2 | 3.29E-01 | -0.2 | 3.96E-01 | 0.4 | 1.35E-01 | hypothetical protein |
| AT1G05410 | 0.2 | 6.76E-03 | 0.1 | 2.04E-01 | 0.1 | 1.30E-01 | -0.1 | 7.40E-01 | -0.1 | 6.97E-01 | 0.0 | 9.46E-01 | AT1G05410 protein ) |
| AT1G28715 | 0.2 | 1.73E-01 | 0.0 | 9.26E-01 | 0.3 | 2.41E-01 | 0.0 | 9.69E-01 | 0.1 | 8.65E-01 | -0.1 | 8.29E-01 | NA |
| AT4G09060 | 0.2 | 6.97E-02 | 0.7 | 2.71E-04 | -0.5 | 1.06E-02 | -0.1 | 6.98E-01 | 0.0 | 9.53E-01 | -0.1 | 7.23E-01 | Uncharacterized protein |
| AT3G27930 | 0.3 | 4.43E-03 | 0.0 | 5.74E-01 | 0.2 | 3.05E-02 | 0.1 | 6.36E-01 | 0.0 | 8.82E-01 | 0.0 | 8.63E-01 | Beta-galactosidase |
| AT2G02170 | 0.3 | 5.40E-02 | 0.2 | 2.38E-01 | 0.1 | 5.08E-01 | 0.0 | 9.61E-01 | -0.1 | 6.75E-01 | 0.1 | 7.63E-01 | At2g02170 |
| AT3G47250 | 0.3 | 6.56E-03 | -0.1 | 2.02E-01 | 0.4 | 2.23E-03 | 0.1 | 6.28E-01 | -0.3 | 3.63E-02 | 0.4 | 2.93E-02 | Transmembrane protein, putative |
| AT4G21880 | 0.3 | 1.04E-01 | 0.3 | 5.03E-02 | -0.1 | 7.01E-01 | 0.5 | 1.78E-01 | 0.3 | 5.02E-01 | 0.2 | 6.83E-01 | Pentacotripeptide-repeat region of PRORP domain-containing protein |
| AT2G19540 | 0.3 | 1.81E-02 | -0.1 | 2.74E-01 | 0.4 | 6.11E-03 | 0.6 | 3.38E-04 | 0.2 | 3.81E-01 | 0.5 | 2.71E-02 | Protein HEAT STRESS TOLERANT DWD 1 |
| AT4G15900 | 0.3 | 2.97E-03 | 0.0 | 7.73E-01 | 0.2 | 1.07E-02 | 0.0 | 9.08E-01 | -0.1 | 6.29E-01 | 0.1 | 7.89E-01 | Protein pleiotropic regulatory locus 1 |
| AT5G55840 | 0.3 | 6.80E-02 | 0.0 | 9.18E-01 | 0.3 | 1.15E-01 | -0.2 | 5.92E-01 | -0.2 | 7.36E-01 | -0.1 | 9.46E-01 | PPR superfamily protein |
| AT5G44250 | 0.3 | 7.80E-03 | 0.2 | 2.69E-02 | 0.0 | 6.26E-01 | -0.2 | 9.87E-02 | -0.2 | 3.49E-01 | -0.1 | 7.37E-01 | At5g44250 |

| | | | | | | | | | | | | | |
|---|---|---|---|---|---|---|---|---|---|---|---|---|---|
| AT5G16040 | 0.3 | 7.66E-03 | 0.3 | 8.13E-03 | 0.0 | 9.43E-01 | 0.3 | 1.08E-01 | 0.2 | 4.96E-01 | 0.1 | 5.76E-01 | Regulator of chromosome condensation family protein |
| AT3G19790 | 0.3 | 4.29E-02 | 0.2 | 1.37E-01 | 0.1 | 6.37E-01 | 0.0 | 9.32E-01 | -0.2 | 6.88E-01 | 0.1 | 7.99E-01 | Uncharacterized protein |
| AT4G17000 | 0.3 | 7.46E-02 | 0.3 | 2.67E-02 | -0.1 | 6.02E-01 | 0.2 | 5.12E-01 | 0.2 | 5.67E-01 | 0.0 | 9.97E-01 | Neurofilament heavy protein |
| AT3G29575 | 0.3 | 4.58E-02 | 0.1 | 6.98E-01 | 0.2 | 1.65E-01 | -0.5 | 4.87E-02 | 0.0 | 9.04E-01 | -0.5 | 1.51E-01 | Ninja-family protein AFP3 |
| AT2G28570 | 0.3 | 2.81E-01 | -0.7 | 1.98E-02 | 1.0 | 8.46E-03 | 0.3 | 4.20E-01 | -0.1 | 8.56E-01 | 0.4 | 4.46E-01 | hypothetical protein |
| AT1G34010 | 0.3 | 2.22E-02 | 0.5 | 4.53E-04 | -0.3 | 3.89E-02 | -0.3 | 8.60E-02 | -0.4 | 7.86E-02 | 0.0 | 8.96E-01 | F12G12.17 protein |
| AT5G60710 | 0.3 | 2.58E-03 | 0.3 | 1.49E-03 | 0.0 | 6.29E-01 | -0.2 | 4.92E-02 | -0.4 | 2.84E-03 | 0.2 | 2.42E-01 | hypothetical protein |
| AT1G62520 | 0.3 | 8.89E-02 | 0.5 | 3.62E-03 | -0.3 | 1.19E-01 | -0.6 | 3.39E-02 | -0.2 | 4.73E-01 | -0.4 | 2.02E-01 | Sulfated surface-like glycoprotein |
| AT3G22850 | 0.3 | 1.19E-02 | 0.6 | 5.60E-05 | -0.4 | 3.24E-03 | 0.4 | 9.26E-02 | 0.1 | 8.08E-01 | 0.3 | 2.71E-01 | hypothetical protein |
| AT1G62422 | 0.3 | 5.11E-02 | -0.1 | 2.81E-01 | 0.4 | 1.90E-02 | 0.0 | 9.74E-01 | -0.5 | 7.12E-02 | 0.5 | 1.14E-01 | Uncharacterized protein |
| AT5G63660 | 0.3 | 2.96E-01 | 0.6 | 2.69E-02 | -0.3 | 2.48E-01 | 0.5 | 1.31E-01 | 0.6 | 8.24E-02 | -0.1 | 8.02E-01 | Defensin-like protein 6 |
| AT2G37680 | 0.3 | 3.89E-02 | 0.1 | 6.66E-01 | 0.2 | 1.53E-01 | 0.1 | 5.94E-01 | 0.0 | 9.90E-01 | 0.1 | 6.87E-01 | Glucose-induced degradation protein 4 homolog |
| AT3G17668 | 0.3 | 5.52E-02 | 0.3 | 4.83E-02 | 0.0 | 9.05E-01 | -0.2 | 3.00E-01 | -0.3 | 2.08E-01 | 0.1 | 8.45E-01 | ENA |
| AT5G39600 | 0.3 | 2.79E-02 | 0.3 | 3.16E-02 | 0.0 | 9.91E-01 | 0.0 | 9.91E-01 | -0.1 | 8.01E-01 | 0.1 | 8.48E-01 | Large ribosomal subunit protein mL53 |
| AT5G39940 | 0.3 | 8.25E-02 | 0.1 | 6.82E-01 | 0.2 | 2.70E-01 | 0.4 | 2.12E-01 | 0.5 | 8.53E-02 | -0.2 | 6.93E-01 | FAD |
| AT4G05070 | 0.3 | 1.54E-01 | 0.4 | 6.05E-02 | -0.1 | 6.02E-01 | -0.9 | 1.23E-03 | -1.4 | 9.05E-05 | 0.5 | 1.41E-01 | Wound-responsive family protein |
| AT2G44360 | 0.3 | 1.55E-01 | 0.0 | 9.64E-01 | 0.2 | 2.47E-01 | 0.0 | 9.37E-01 | -0.1 | 7.91E-01 | 0.1 | 8.75E-01 | Ecotropic viral integration site protein |
| AT1G79490 | 0.3 | 1.76E-02 | 0.0 | 9.26E-01 | 0.2 | 5.11E-02 | 0.4 | 1.36E-01 | 0.0 | 9.62E-01 | 0.5 | 2.17E-01 | Pentatricopeptide repeat-containing protein At1g79490, mitochondrial |
| AT2G17440 | 0.3 | 5.65E-03 | 0.3 | 1.40E-03 | -0.1 | 3.64E-01 | -0.2 | 3.47E-01 | -0.3 | 2.86E-01 | 0.1 | 8.81E-01 | Plant intracellular Ras-group-related LRR protein 5 |
| AT5G66380 | 0.3 | 3.02E-02 | 0.1 | 5.42E-01 | 0.2 | 1.66E-01 | 0.2 | 4.45E-01 | 0.1 | 6.90E-01 | 0.1 | 8.64E-01 | Folate transporter 1, chloroplastic |
| AT4G23670 | 0.3 | 6.25E-02 | 0.4 | 7.17E-03 | -0.2 | 2.27E-01 | 0.2 | 1.34E-01 | 0.6 | 2.52E-03 | -0.3 | 1.05E-01 | Bet v I |
| AT1G67950 | 0.3 | 1.78E-02 | 0.0 | 6.76E-01 | 0.2 | 7.81E-02 | 0.1 | 7.41E-01 | 0.1 | 7.71E-01 | 0.0 | 9.90E-01 | Uncharacterized protein T23K23.20 |
| AT5G43140 | 0.3 | 6.23E-02 | -0.1 | 5.89E-01 | 0.3 | 5.00E-02 | 0.1 | 7.23E-01 | 0.4 | 2.39E-01 | -0.3 | 5.28E-01 | Similarity to 22 kDa peroxisomal membrane protein |
| AT1G54380 | 0.3 | 3.48E-02 | 0.1 | 5.60E-01 | 0.2 | 1.69E-01 | 0.2 | 3.22E-01 | 0.1 | 5.30E-01 | 0.0 | 8.94E-01 | Spliceosome protein-like protein |
| AT1G51360 | 0.3 | 1.45E-02 | 0.4 | 3.31E-03 | -0.1 | 3.98E-01 | -0.3 | 1.25E-01 | -0.3 | 1.77E-01 | 0.0 | 9.89E-01 | Stress-response A |
| AT3G12950 | 0.3 | 8.33E-03 | 0.1 | 4.26E-01 | 0.2 | 6.54E-02 | 0.2 | 2.57E-01 | 0.2 | 3.71E-01 | 0.0 | 9.63E-01 | Trypsin family protein |
| AT3G29010 | 0.3 | 2.13E-01 | 0.0 | 9.35E-01 | 0.2 | 3.55E-01 | 0.0 | 9.17E-01 | -0.2 | 6.76E-01 | 0.2 | 6.25E-01 | BPL |
| AT4G28430 | 0.3 | 1.57E-01 | 0.6 | 8.91E-03 | -0.3 | 1.67E-01 | 0.2 | 6.60E-01 | 0.3 | 5.82E-01 | -0.1 | 9.08E-01 | Reticulon-like protein B18 |
| AT5G46550 | 0.3 | 9.31E-03 | 0.1 | 5.28E-01 | 0.2 | 6.28E-02 | 0.1 | 7.03E-01 | -0.1 | 6.76E-01 | 0.2 | 5.04E-01 | Transcription factor GTE12 |
| AT5G54860 | 0.3 | 8.37E-02 | 0.0 | 7.78E-01 | 0.2 | 2.11E-01 | 0.7 | 3.03E-03 | 0.6 | 3.12E-02 | 0.2 | 5.90E-01 | Probable folate-biopterin transporter 4 |
| AT2G22080 | 0.3 | 1.04E-02 | 0.6 | 1.01E-04 | -0.3 | 7.91E-03 | 0.3 | 6.68E-02 | 0.1 | 8.13E-01 | 0.2 | 2.81E-01 | Transmembrane protein |
| AT2G43235 | 0.3 | 3.23E-02 | -0.3 | 1.97E-02 | 0.6 | 2.02E-03 | 0.5 | 7.67E-02 | 0.1 | 7.34E-01 | 0.4 | 2.02E-01 | Phosphoribosylformylglycinamidine synthase |
| AT3G16750 | 0.3 | 4.63E-02 | 0.4 | 8.08E-03 | -0.1 | 3.73E-01 | -0.3 | 4.03E-01 | -0.3 | 4.80E-01 | 0.0 | 9.92E-01 | Uncharacterized protein |
| AT1G63860 | 0.3 | 2.73E-02 | -0.1 | 2.21E-01 | 0.4 | 9.45E-03 | 0.1 | 8.18E-01 | -0.3 | 3.48E-01 | 0.3 | 3.07E-01 | Uncharacterized protein |
| AT1G18610 | 0.3 | 1.43E-02 | 0.2 | 1.04E-01 | 0.1 | 4.08E-01 | -0.3 | 1.72E-01 | 0.0 | 9.03E-01 | -0.4 | 2.40E-01 | Galactose oxidase |
| AT4G10080 | 0.3 | 1.98E-03 | 0.2 | 5.88E-03 | 0.0 | 6.83E-01 | -0.3 | 2.76E-02 | -0.6 | 1.18E-03 | 0.3 | 1.94E-01 | Uncharacterized protein AT4g10080 |
| AT3G62620 | 0.3 | 1.68E-02 | 0.1 | 1.73E-01 | 0.1 | 3.17E-01 | 0.0 | 8.53E-01 | 0.1 | 6.74E-01 | -0.1 | 5.72E-01 | At3g62620 |
| AT4G33250 | 0.3 | 1.24E-02 | 0.3 | 4.95E-03 | -0.1 | 5.31E-01 | -0.1 | 2.59E-01 | -0.2 | 1.90E-01 | 0.0 | 8.50E-01 | AT4g33250 |
| AT5G16420 | 0.3 | 6.28E-02 | 0.0 | 9.35E-01 | 0.2 | 1.44E-01 | 0.1 | 8.15E-01 | 0.0 | 9.78E-01 | 0.1 | 8.99E-01 | Pentatricopeptide repeat-containing protein At5g16420, mitochondrial |
| AT1G76200 | 0.3 | 4.41E-03 | 0.5 | 2.18E-04 | -0.2 | 4.14E-02 | -0.1 | 4.77E-01 | 0.0 | 9.76E-01 | -0.1 | 6.61E-01 | NADH dehydrogenase 1 beta subcomplex subunit 2 |
| AT5G08180 | 0.3 | 1.41E-01 | 0.4 | 2.50E-02 | -0.2 | 3.48E-01 | 0.3 | 1.26E-01 | 0.3 | 2.49E-01 | 0.0 | 9.20E-01 | Ribosomal protein L7Ae |
| AT5G62440 | 0.3 | 1.04E-01 | 0.2 | 2.07E-01 | 0.1 | 7.69E-01 | 0.5 | 8.33E-03 | 0.3 | 9.30E-02 | 0.1 | 4.93E-01 | Protein EMBRYO DEFECTIVE 514 |
| AT3G27770 | 0.3 | 1.15E-02 | 0.3 | 9.71E-03 | 0.0 | 8.65E-01 | -0.4 | 5.60E-02 | -0.6 | 1.88E-02 | 0.2 | 5.11E-01 | Transmembrane protein |
| AT5G19500 | 0.3 | 2.53E-02 | 0.0 | 7.68E-01 | 0.3 | 3.30E-02 | 0.0 | 9.43E-01 | -0.1 | 7.70E-01 | 0.1 | 7.49E-01 | At5g19500 |
| AT1G19690 | 0.3 | 2.13E-02 | 0.1 | 2.41E-01 | 0.1 | 2.78E-01 | 0.1 | 4.74E-01 | 0.2 | 3.73E-01 | -0.1 | 8.57E-01 | F6F9.26 protein |
| AT5G03345 | 0.3 | 1.97E-02 | 0.3 | 5.52E-03 | -0.1 | 4.73E-01 | 0.1 | 8.14E-01 | 0.0 | 9.49E-01 | 0.0 | 9.28E-01 | Membrane magnesium transporter-like protein |
| AT5G42460 | 0.3 | 4.19E-02 | 0.1 | 6.22E-01 | 0.2 | 1.81E-01 | 0.0 | 9.72E-01 | -0.7 | 6.12E-02 | 0.7 | 2.71E-02 | F-box protein At5g42460 |
| AT5G23350 | 0.3 | 1.92E-01 | 0.9 | 8.44E-04 | -0.6 | 1.79E-02 | 0.8 | 1.12E-01 | 0.1 | 9.47E-01 | 0.7 | 1.80E-01 | GRAM domain-containing protein |
| AT1G47970 | 0.3 | 2.31E-03 | 0.1 | 9.70E-02 | 0.1 | 8.13E-02 | -0.1 | 6.01E-01 | -0.2 | 2.63E-01 | 0.1 | 6.72E-01 | At1g47970 |
| AT3G01130 | 0.3 | 4.08E-03 | 0.3 | 1.08E-03 | -0.1 | 3.46E-01 | 0.3 | 1.25E-01 | 0.1 | 7.21E-01 | 0.2 | 4.95E-01 | ATP synthase E chain |
| AT1G14200 | 0.3 | 3.83E-02 | -0.2 | 8.48E-02 | 0.5 | 5.19E-03 | -0.2 | 4.97E-01 | -0.5 | 1.21E-01 | 0.3 | 5.03E-01 | RING-type domain-containing protein |
| AT1G77930 | 0.3 | 2.99E-02 | -0.3 | 1.41E-02 | 0.6 | 1.04E-03 | -0.3 | 1.11E-01 | -0.1 | 4.89E-01 | -0.1 | 6.19E-01 | Chaperone DnaJ-domain superfamily protein |
| AT3G50900 | 0.3 | 1.48E-01 | 0.2 | 2.02E-01 | 0.0 | 9.12E-01 | 0.5 | 2.84E-02 | 0.3 | 3.88E-01 | 0.3 | 3.29E-01 | At3g50900 |
| AT1G24350 | 0.3 | 1.18E-02 | 0.0 | 9.51E-01 | 0.3 | 3.21E-02 | -0.2 | 3.83E-01 | -0.2 | 2.79E-01 | 0.1 | 8.07E-01 | Acid phosphatase |
| AT4G31930 | 0.3 | 4.63E-02 | 0.2 | 2.07E-01 | 0.1 | 5.31E-01 | -0.1 | 7.17E-01 | 0.0 | 9.27E-01 | -0.1 | 8.65E-01 | Mitochondrial glycoprotein family protein |
| AT1G57680 | 0.3 | 3.41E-02 | 0.2 | 7.47E-02 | 0.0 | 7.68E-01 | -0.3 | 1.41E-02 | -0.6 | 5.07E-04 | 0.3 | 1.37E-01 | Uncharacterized protein |
| AT5G23730 | 0.3 | 5.91E-03 | 0.1 | 3.78E-01 | 0.2 | 6.42E-02 | -0.3 | 2.61E-01 | -0.1 | 7.91E-01 | -0.2 | 5.36E-01 | WD repeat-containing protein RUP2 |
| AT4G34750 | 0.3 | 3.70E-02 | 0.5 | 2.67E-03 | -0.2 | 1.73E-01 | -0.4 | 6.53E-02 | -0.4 | 9.92E-02 | 0.0 | 9.86E-01 | hypothetical protein |
| AT4G21100 | 0.3 | 5.13E-03 | 0.2 | 2.53E-02 | 0.1 | 4.99E-01 | 0.2 | 1.31E-01 | 0.1 | 8.01E-01 | 0.2 | 4.20E-01 | Cleavage |
| AT2G34357 | 0.3 | 2.86E-02 | 0.0 | 8.51E-01 | 0.3 | 3.66E-02 | 0.7 | 8.19E-03 | 0.4 | 1.80E-01 | 0.3 | 3.53E-01 | Ribosomal RNA-processing protein 12-like conserved domain-containing protein |
| AT3G24480 | 0.3 | 2.89E-02 | 0.1 | 3.70E-01 | 0.2 | 2.02E-01 | 0.2 | 3.37E-01 | 0.2 | 4.96E-01 | 0.0 | 9.41E-01 | Leucine-rich repeat-containing N-terminal plant-type domain-containing protein |
| AT2G40815 | 0.3 | 5.02E-02 | 0.2 | 1.08E-01 | 0.0 | 7.90E-01 | 0.1 | 7.63E-01 | 0.1 | 7.36E-01 | 0.0 | 9.54E-01 | C2 domain-containing protein |
| AT5G50350 | 0.3 | 3.64E-03 | -0.2 | 9.39E-03 | 0.5 | 3.09E-04 | -0.1 | 5.79E-01 | 0.0 | 8.36E-01 | -0.1 | 5.43E-01 | Uncharacterized protein At5g50350 |
| AT1G33490 | 0.3 | 7.68E-03 | -0.1 | 5.59E-01 | 0.3 | 8.32E-03 | 0.3 | 4.02E-02 | 0.3 | 1.27E-01 | 0.1 | 8.13E-01 | DUF2062 domain-containing protein |
| AT2G43260 | 0.3 | 3.90E-03 | 0.2 | 1.13E-02 | 0.0 | 7.03E-01 | 0.2 | 2.75E-01 | 0.0 | 9.76E-01 | 0.2 | 4.10E-01 | F-box and associated interaction domains-containing protein |
| AT4G37020 | 0.3 | 1.80E-02 | 0.1 | 4.48E-01 | 0.2 | 1.35E-01 | -0.1 | 6.79E-01 | -0.1 | 7.58E-01 | 0.0 | 9.83E-01 | Uncharacterized protein AT4g37020 |
| AT1G05540 | 0.3 | 2.60E-01 | 0.6 | 2.91E-02 | -0.3 | 2.67E-01 | 0.2 | 5.41E-01 | 0.5 | 1.34E-01 | -0.3 | 4.68E-01 | Uncharacterized protein |
| AT1G71060 | 0.3 | 4.80E-02 | 0.1 | 5.04E-01 | 0.2 | 2.64E-01 | 0.1 | 7.07E-01 | 0.0 | 9.68E-01 | 0.1 | 7.87E-01 | Pentatricopeptide repeat-containing protein At1g71060, mitochondrial |
| AT1G06710 | 0.3 | 2.22E-01 | -0.3 | 2.10E-01 | 0.6 | 5.55E-02 | | | | | | | Tetratricopeptide repeat -like superfamily protein |
| AT3G12650 | 0.3 | 1.27E-01 | 0.3 | 8.60E-02 | 0.0 | 8.40E-01 | -0.2 | 5.21E-01 | -0.4 | 1.69E-01 | 0.2 | 4.95E-01 | Transmembrane protein |
| AT5G62650 | 0.3 | 6.61E-03 | -0.1 | 2.40E-01 | 0.4 | 3.33E-03 | 0.0 | 7.82E-01 | -0.1 | 4.52E-01 | 0.2 | 4.01E-01 | Tic22-like family protein |
| AT3G51310 | 0.3 | 4.03E-03 | 0.1 | 3.07E-01 | 0.2 | 4.56E-02 | 0.0 | 8.33E-01 | 0.0 | 9.83E-01 | 0.0 | 9.11E-01 | Vacuolar sorting protein 35 homolog |
| AT3G25680 | 0.3 | 7.01E-03 | 0.2 | 4.66E-02 | 0.1 | 4.24E-01 | 0.0 | 9.17E-01 | -0.1 | 7.41E-01 | 0.1 | 7.22E-01 | Uncharacterized protein T5M7.13 |
| AT1G15120 | 0.3 | 9.36E-02 | 0.4 | 2.82E-02 | -0.1 | 5.09E-01 | 0.4 | 8.90E-02 | 0.2 | 6.38E-01 | 0.3 | 4.69E-01 | Ubiquinol-cytochrome C reductase hinge domain-containing protein |
| AT2G47840 | 0.3 | 3.21E-02 | 0.3 | 2.19E-02 | 0.0 | 8.09E-01 | -0.1 | 5.44E-01 | 0.0 | 9.28E-01 | -0.1 | 7.53E-01 | Protein TIC 20-II, chloroplastic |
| AT4G10030 | 0.3 | 1.05E-02 | 0.1 | 4.65E-01 | 0.2 | 7.87E-02 | -0.4 | 1.01E-01 | -0.2 | 5.67E-01 | -0.2 | 4.24E-01 | hypothetical protein |
| AT4G22560 | 0.3 | 6.16E-02 | 0.4 | 2.33E-02 | -0.1 | 6.20E-01 | 0.1 | 6.70E-01 | 0.2 | 6.04E-01 | 0.0 | 9.24E-01 | hypothetical protein |
| AT5G57120 | 0.3 | 7.00E-02 | 0.4 | 1.20E-02 | -0.2 | 3.37E-01 | 0.3 | 1.41E-01 | 0.1 | 7.95E-01 | 0.2 | 4.78E-01 | AT5g57120 |
| AT5G40590 | 0.3 | 1.89E-01 | 0.0 | 9.90E-01 | 0.3 | 2.80E-01 | 1.9 | 4.91E-05 | 1.1 | 9.34E-03 | 0.8 | 1.17E-02 | DC1 domain-containing protein |
| AT1G27470 | 0.3 | 5.71E-03 | 0.1 | 1.53E-01 | 0.1 | 1.38E-01 | 0.6 | 9.37E-05 | 0.3 | 7.58E-02 | 0.4 | 3.07E-02 | Transducin family protein |
| AT4G34560 | 0.3 | 1.68E-01 | -0.1 | 7.77E-01 | 0.3 | 1.64E-01 | 0.1 | 5.00E-01 | 0.0 | 8.78E-01 | 0.2 | 5.23E-01 | DUF4408 domain-containing protein |
| AT3G60480 | 0.3 | 1.10E-02 | 0.2 | 2.90E-02 | 0.0 | 7.28E-01 | -0.4 | 4.33E-02 | -0.1 | 5.90E-01 | -0.3 | 3.32E-01 | Uncharacterized protein T8B10_140 |
| AT1G15440 | 0.3 | 4.13E-02 | 0.1 | 6.09E-01 | 0.2 | 1.65E-01 | 0.5 | 4.72E-02 | 0.1 | 8.71E-01 | 0.4 | 1.90E-01 | Periodic tryptophan protein 2 |

| | | | | | | | | | | | | | |
|---|---|---|---|---|---|---|---|---|---|---|---|---|---|
| AT4G00030 | 0.3 | 4.81E-03 | 0.0 | 7.80E-01 | 0.3 | 8.39E-03 | 0.0 | 9.37E-01 | 0.1 | 8.46E-01 | 0.0 | 9.11E-01 | Probable plastid-lipid-associated protein 11, chloroplastic |
| AT2G37990 | 0.3 | 1.40E-02 | 0.3 | 1.11E-02 | 0.0 | 8.49E-01 | 0.6 | 2.33E-03 | 0.4 | 5.63E-02 | 0.2 | 3.93E-01 | Ribosome biogenesis regulatory protein homolog |
| AT3G03560 | 0.3 | 1.27E-02 | -0.1 | 1.91E-01 | 0.4 | 4.23E-03 | 0.5 | 1.34E-03 | 0.4 | 1.55E-02 | 0.1 | 5.49E-01 | Uncharacterized protein |
| AT4G36660 | 0.3 | 1.26E-02 | 0.2 | 4.21E-02 | 0.1 | 6.28E-01 | -0.2 | 3.91E-01 | 0.0 | 9.80E-01 | -0.2 | 5.07E-01 | At4g36660 ) |
| AT3G58850 | 0.3 | 9.46E-02 | 0.6 | 2.86E-03 | -0.3 | 9.40E-02 | -0.3 | 2.76E-01 | 0.0 | 9.81E-01 | -0.3 | 3.73E-01 | Transcription factor PAR2 |
| AT4G19370 | 0.3 | 5.59E-02 | -0.4 | 2.85E-02 | 0.6 | 3.95E-03 | 0.0 | 9.63E-01 | -0.9 | 1.86E-02 | 1.0 | 2.67E-02 | Transmembrane protein |
| AT5G11690 | 0.3 | 4.90E-03 | 0.4 | 6.19E-04 | -0.1 | 1.88E-01 | 0.0 | 8.68E-01 | -0.1 | 8.09E-01 | 0.0 | 9.42E-01 | Mitochondrial import inner membrane translocase subunit TIM17-3 |
| AT1G44780 | 0.3 | 1.02E-02 | 0.2 | 4.72E-02 | 0.1 | 5.41E-01 | 0.1 | 6.17E-01 | 0.2 | 6.32E-01 | 0.0 | 9.76E-01 | T12C22.5 protein |
| AT2G48160 | 0.3 | 1.42E-03 | -0.1 | 3.15E-01 | 0.3 | 1.14E-03 | 0.1 | 6.64E-01 | 0.1 | 6.54E-01 | 0.0 | 9.57E-01 | Tudor |
| AT1G19580 | 0.3 | 1.05E-03 | 0.3 | 1.85E-03 | 0.0 | 8.97E-01 | 0.1 | 2.40E-01 | 0.1 | 6.61E-01 | 0.1 | 7.08E-01 | Gamma carbonic anhydrase 1 |
| AT5G06350 | 0.3 | 5.81E-03 | 0.1 | 5.35E-01 | 0.2 | 4.19E-02 | 0.4 | 3.05E-02 | 0.3 | 1.66E-01 | 0.1 | 6.72E-01 | Uncharacterized protein |
| AT2G46170 | 0.3 | 1.00E-02 | 0.4 | 2.97E-03 | -0.1 | 4.25E-01 | 0.2 | 2.66E-01 | 0.3 | 1.44E-01 | -0.1 | 7.14E-01 | Reticulon-like protein |
| AT5G56990 | 0.3 | 8.71E-02 | 0.1 | 5.68E-01 | 0.2 | 3.56E-01 | -0.4 | 1.87E-01 | -0.1 | 8.77E-01 | -0.3 | 3.86E-01 | Proteinase inhibitor I25, cystatin, motif protein |
| AT5G51180 | 0.3 | 1.73E-02 | 0.1 | 2.44E-01 | 0.1 | 2.28E-01 | -0.1 | 4.29E-01 | -0.3 | 1.28E-01 | 0.1 | 4.76E-01 | DUF676 domain-containing protein |
| AT2G16800 | 0.3 | 1.10E-03 | 0.3 | 9.58E-04 | 0.0 | 7.83E-01 | -0.3 | 8.47E-03 | -0.3 | 9.23E-03 | 0.0 | 9.02E-01 | Chloroplast protein FOR GROWTH AND FERTILITY 2 |
| AT5G06360 | 0.3 | 9.52E-03 | 0.3 | 6.88E-03 | 0.0 | 7.76E-01 | 0.2 | 2.46E-01 | -0.1 | 6.58E-01 | 0.2 | 2.20E-01 | Ribosome biogenesis protein NSA2 homolog |
| AT3G59470 | 0.3 | 8.22E-03 | 0.1 | 3.89E-01 | 0.2 | 8.41E-02 | -0.3 | 2.32E-01 | 0.0 | 8.88E-01 | -0.2 | 4.90E-01 | Uncharacterized protein T16L24.20 |
| AT2G14220 | 0.3 | 5.13E-02 | 0.2 | 1.16E-01 | 0.0 | 7.78E-01 | 0.1 | 7.80E-01 | 0.0 | 9.76E-01 | 0.1 | 8.70E-01 | NA |
| AT3G57500 | 0.3 | 1.33E-01 | 0.5 | 2.17E-02 | -0.2 | 3.86E-01 | 0.3 | 5.49E-01 | 0.4 | 4.53E-01 | -0.1 | 8.89E-01 | Fission regulator-like protein |
| AT1G23850 | 0.3 | 3.13E-02 | 0.4 | 5.88E-03 | -0.1 | 3.86E-01 | 0.1 | 6.26E-01 | -0.1 | 6.79E-01 | 0.3 | 3.79E-01 | Transmembrane protein |
| AT5G30490 | 0.3 | 3.85E-03 | 0.0 | 5.81E-01 | 0.2 | 2.67E-02 | 0.1 | 5.30E-01 | 0.1 | 8.00E-01 | 0.1 | 8.50E-01 | hypothetical protein |
| AT3G61780 | 0.3 | 4.37E-03 | 0.1 | 3.40E-01 | 0.2 | 5.50E-02 | 0.1 | 5.74E-01 | 0.0 | 9.29E-01 | 0.1 | 7.64E-01 | Embryo defective 1703 |
| AT1G76940 | 0.3 | 2.66E-03 | 0.0 | 6.99E-01 | 0.2 | 1.56E-02 | -0.2 | 3.71E-01 | -0.3 | 2.65E-01 | 0.1 | 8.14E-01 | RNA-binding family protein |
| AT2G27775 | 0.3 | 5.10E-02 | 0.5 | 3.18E-03 | -0.2 | 1.68E-01 | 1.0 | 2.61E-04 | 0.6 | 1.68E-02 | 0.4 | 1.89E-01 | Uncharacterized protein |
| AT5G07670 | 0.3 | 6.82E-03 | 0.2 | 3.30E-02 | 0.1 | 5.17E-01 | 0.0 | 9.39E-01 | -0.1 | 7.95E-01 | 0.1 | 7.47E-01 | F-box protein At5g07670 |
| AT3G26600 | 0.3 | 1.23E-02 | -0.4 | 5.43E-03 | 0.6 | 3.93E-04 | 0.3 | 1.82E-01 | 0.1 | 7.33E-01 | 0.2 | 5.50E-01 | hypothetical protein |
| AT2G19650 | 0.3 | 4.50E-03 | 0.2 | 3.98E-02 | 0.1 | 3.43E-01 | 0.5 | 3.22E-02 | 0.3 | 2.99E-01 | 0.2 | 4.76E-01 | Cysteine |
| AT4G27750 | 0.3 | 9.62E-03 | 0.0 | 9.03E-01 | 0.3 | 2.96E-02 | -0.1 | 5.49E-01 | -0.1 | 6.44E-01 | 0.0 | 9.83E-01 | ISI1 |
| AT3G11760 | 0.3 | 9.66E-03 | 0.5 | 4.12E-04 | -0.2 | 6.82E-02 | -0.4 | 2.56E-02 | -0.6 | 8.69E-03 | 0.1 | 6.17E-01 | Uncharacterized protein At3g11760 |
| AT5G08000 | 0.3 | 2.03E-02 | 0.3 | 1.12E-02 | 0.0 | 7.21E-01 | 0.1 | 4.73E-01 | 0.0 | 9.78E-01 | 0.1 | 6.54E-01 | Glucan endo-1,3-beta-glucosidase-like protein 3 |
| AT3G62880 | 0.3 | 8.08E-03 | 0.0 | 6.97E-01 | 0.3 | 1.28E-02 | 0.0 | 8.93E-01 | 0.0 | 9.47E-01 | -0.1 | 8.71E-01 | Mitochondrial import inner membrane translocase subunit Tim17 |
| AT4G30990 | 0.3 | 1.93E-02 | 0.0 | 7.80E-01 | 0.2 | 6.17E-02 | 0.5 | 1.26E-03 | 0.3 | 8.61E-02 | 0.2 | 1.87E-01 | ARM repeat superfamily protein |
| AT4G34270 | 0.3 | 6.46E-04 | 0.1 | 9.65E-02 | 0.2 | 2.55E-02 | 0.0 | 9.11E-01 | 0.0 | 8.77E-01 | 0.1 | 8.31E-01 | TIP41-like protein |
| AT1G67590 | 0.3 | 1.88E-02 | 0.6 | 1.94E-04 | -0.3 | 1.49E-02 | -0.2 | 1.52E-01 | -0.2 | 2.59E-01 | 0.0 | 9.44E-01 | hypothetical protein |
| AT1G17350 | 0.3 | 1.95E-02 | 0.4 | 3.89E-03 | -0.1 | 3.79E-01 | -0.1 | 5.94E-01 | -0.2 | 6.45E-01 | 0.0 | 9.92E-01 | NADH:ubiquinone oxidoreductase intermediate-associated protein 30 |
| AT3G57990 | 0.3 | 5.17E-04 | 0.0 | 6.41E-01 | 0.3 | 3.00E-03 | -0.1 | 6.09E-01 | 0.0 | 8.81E-01 | 0.0 | 8.62E-01 | Uncharacterized protein |
| AT3G55605 | 0.3 | 1.17E-01 | 0.2 | 2.28E-01 | 0.1 | 7.80E-01 | 0.5 | 5.20E-02 | 0.1 | 6.62E-01 | 0.3 | 2.71E-01 | Mitochondrial glycoprotein family protein |
| AT3G28670 | 0.3 | 1.94E-03 | 0.1 | 2.11E-01 | 0.2 | 4.11E-02 | 0.1 | 6.08E-01 | -0.1 | 8.04E-01 | 0.2 | 4.47E-01 | hypothetical protein |
| AT4G33500 | 0.3 | 2.65E-02 | 0.2 | 1.62E-01 | 0.1 | 4.11E-01 | 0.4 | 3.76E-02 | 0.4 | 5.23E-02 | 0.0 | 9.75E-01 | AT4G33500 protein |
| AT2G19970 | 0.3 | 8.69E-02 | -0.1 | 4.36E-01 | 0.4 | 4.96E-02 | -0.6 | 1.56E-01 | 0.3 | 5.50E-01 | -0.9 | 8.30E-02 | hypothetical protein |
| AT3G21540 | 0.3 | 1.05E-02 | 0.0 | 8.79E-01 | 0.3 | 3.33E-02 | 0.9 | 4.30E-03 | 0.3 | 4.57E-01 | 0.6 | 9.04E-02 | Small-subunit processome Utp12 domain-containing protein |
| AT3G45210 | 0.3 | 3.96E-02 | 0.5 | 2.06E-03 | -0.2 | 1.21E-01 | 0.0 | 1.00E+00 | -0.1 | 7.93E-01 | 0.1 | 7.96E-01 | AT3g45210 |
| AT5G11240 | 0.3 | 4.89E-02 | 0.1 | 3.37E-01 | 0.1 | 3.58E-01 | 0.8 | 1.23E-04 | 0.5 | 1.57E-02 | 0.3 | 8.76E-02 | Uncharacterized protein At5g11240 |
| AT2G34050 | 0.3 | 2.88E-02 | 0.3 | 3.22E-02 | 0.0 | 9.90E-01 | 0.3 | 2.04E-01 | 0.4 | 1.44E-01 | -0.1 | 8.45E-01 | ATP synthase mitochondrial F1 complex assembly factor 1 |
| AT4G21170 | 0.3 | 5.04E-02 | 0.1 | 5.35E-01 | 0.2 | 2.65E-01 | 0.5 | 7.83E-02 | 0.3 | 4.72E-01 | 0.3 | 4.51E-01 | Pentatricopeptide repeat-containing protein At4g21170 |
| AT5G13980 | 0.3 | 1.91E-03 | 0.5 | 1.48E-04 | -0.2 | 5.37E-02 | -0.1 | 2.34E-01 | -0.1 | 2.85E-01 | 0.0 | 9.83E-01 | Alpha-mannosidase |
| AT4G14420 | 0.3 | 2.51E-03 | 0.4 | 2.60E-04 | -0.1 | 1.04E-01 | 0.4 | 1.87E-02 | 0.1 | 6.24E-01 | 0.3 | 2.14E-01 | hypothetical protein |
| AT4G14270 | 0.3 | 1.40E-01 | 0.1 | 7.03E-01 | 0.2 | 3.42E-01 | -0.7 | 2.48E-03 | -0.4 | 7.41E-02 | -0.3 | 3.74E-01 | Polyadenylate-binding protein-interacting protein 2 -binding protein-interacting protein 2) |
| AT2G04230 | 0.3 | 5.19E-02 | 0.1 | 5.79E-01 | 0.2 | 2.25E-01 | 0.1 | 8.16E-01 | 0.0 | 9.04E-01 | 0.0 | 9.66E-01 | F-box |
| AT2G12460 | 0.3 | 5.67E-02 | -0.2 | 3.19E-01 | 0.5 | 2.76E-02 | 0.3 | 4.44E-01 | 0.6 | 1.97E-01 | -0.3 | 6.80E-01 | NA |
| AT5G25790 | 0.3 | 4.30E-02 | 0.0 | 8.84E-01 | 0.3 | 1.20E-01 | -0.3 | 5.85E-01 | 0.2 | 7.31E-01 | -0.4 | 4.44E-01 | Tesmin |
| AT5G56250 | 0.3 | 9.83E-03 | -0.1 | 1.98E-01 | 0.4 | 3.32E-03 | 0.1 | 4.68E-01 | 0.0 | 9.15E-01 | 0.1 | 7.16E-01 | Hapless 8 |
| AT4G39730 | 0.3 | 1.58E-02 | 0.5 | 1.25E-03 | -0.2 | 1.19E-01 | 0.2 | 2.48E-01 | 0.4 | 4.80E-02 | -0.2 | 4.04E-01 | PLAT domain-containing protein 1 |
| AT5G43822 | 0.3 | 4.29E-02 | -0.2 | 1.59E-01 | 0.5 | 1.21E-02 | -0.1 | 7.99E-01 | -0.2 | 4.57E-01 | 0.2 | 6.89E-01 | Uncharacterized protein |
| AT1G11200 | 0.3 | 1.22E-02 | 0.2 | 7.34E-02 | 0.1 | 4.40E-01 | 0.2 | 2.65E-01 | 0.1 | 7.83E-01 | 0.1 | 5.76E-01 | Organic solute transporter ostalpha protein |
| AT1G05210 | 0.3 | 3.22E-02 | 0.6 | 1.08E-03 | -0.3 | 8.14E-02 | -0.3 | 2.24E-01 | 0.1 | 7.39E-01 | -0.4 | 2.12E-01 | hypothetical protein |
| AT4G05400 | 0.3 | 2.16E-02 | 0.5 | 1.33E-03 | -0.2 | 1.27E-01 | 0.4 | 2.62E-02 | 0.4 | 6.44E-02 | 0.0 | 9.08E-01 | hypothetical protein |
| AT5G14710 | 0.3 | 1.13E-01 | 0.1 | 6.54E-01 | 0.2 | 3.52E-01 | 0.1 | 8.14E-01 | 0.0 | 9.85E-01 | 0.1 | 8.52E-01 | hypothetical protein |
| AT4G25225 | 0.3 | 1.09E-01 | 0.2 | 3.28E-01 | 0.1 | 6.24E-01 | -0.3 | 3.71E-01 | -0.2 | 5.16E-01 | 0.0 | 9.31E-01 | Transmembrane protein |
| AT5G45600 | 0.3 | 8.63E-03 | 0.2 | 5.67E-02 | 0.1 | 4.26E-01 | 0.3 | 8.38E-02 | 0.2 | 4.41E-01 | 0.1 | 5.63E-01 | YAF9a |
| AT2G44090 | 0.3 | 9.79E-04 | 0.0 | 7.13E-01 | 0.3 | 4.79E-03 | 0.0 | 9.59E-01 | 0.0 | 9.77E-01 | 0.0 | 9.96E-01 | hypothetical protein |
| AT1G61900 | 0.3 | 4.90E-03 | 0.3 | 9.95E-03 | 0.0 | 8.20E-01 | -0.4 | 4.44E-03 | -0.4 | 4.02E-03 | 0.0 | 8.42E-01 | SPARK domain-containing protein |
| AT5G24060 | 0.3 | 8.97E-03 | 0.2 | 5.04E-02 | 0.1 | 4.64E-01 | -0.1 | 7.43E-01 | 0.0 | 9.88E-01 | -0.1 | 8.34E-01 | Emb\|CAB61996.1 superfamily protein) |
| AT3G51290 | 0.3 | 1.52E-02 | 0.4 | 5.26E-03 | -0.1 | 5.40E-01 | -0.4 | 1.28E-01 | -0.2 | 4.51E-01 | -0.2 | 6.21E-01 | Pyridoxal-phosphate-dependent serine hydroxymethyltransferase, putative |
| AT5G00980 | 0.3 | 1.28E-01 | 0.2 | 3.69E-01 | 0.1 | 6.41E-01 | -0.2 | 5.02E-01 | -0.2 | 6.46E-01 | 0.0 | 9.41E-01 | NA |
| AT1G17360 | 0.3 | 8.05E-03 | -0.1 | 1.57E-01 | 0.4 | 2.45E-03 | 0.1 | 8.62E-01 | 0.1 | 8.79E-01 | 0.0 | 9.97E-01 | F28G4.18 protein |
| AT3G13950 | 0.3 | 4.08E-01 | -0.2 | 5.72E-01 | 0.5 | 2.65E-01 | 0.8 | 2.27E-01 | -0.9 | 3.41E-01 | 1.6 | 8.26E-02 | At3g13950 |
| AT1G49870 | 0.3 | 2.42E-02 | 0.5 | 2.07E-03 | -0.2 | 1.96E-01 | 0.2 | 5.34E-01 | 0.2 | 6.57E-01 | 0.0 | 9.66E-01 | Myosin-2 heavy chain-like protein |
| AT3G26240 | 0.3 | 3.88E-02 | 0.0 | 9.35E-01 | 0.3 | 7.33E-02 | -0.3 | 4.88E-01 | -0.5 | 2.67E-01 | 0.2 | 7.47E-01 | Cysteine |
| AT5G47420 | 0.3 | 7.89E-03 | 0.1 | 3.06E-01 | 0.2 | 9.49E-02 | 0.1 | 6.89E-01 | 0.2 | 4.33E-01 | -0.1 | 7.20E-01 | Altered inheritance of mitochondria protein 24, mitochondrial |
| AT4G26660 | 0.3 | 1.87E-02 | 0.5 | 7.68E-04 | -0.2 | 8.20E-02 | 0.2 | 5.52E-01 | 0.2 | 5.99E-01 | 0.0 | 9.93E-01 | At4g26660 |
| AT3G56870 | 0.3 | 3.71E-02 | 0.3 | 3.33E-02 | 0.0 | 9.03E-01 | 0.2 | 6.90E-01 | 0.1 | 9.02E-01 | | | Uncharacterized protein T8M16_200 |
| AT3G61420 | 0.3 | 4.41E-02 | -0.1 | 4.66E-01 | 0.4 | 2.56E-02 | -0.2 | 2.75E-01 | -0.2 | 3.49E-01 | 0.0 | 9.92E-01 | BSD domain |
| AT2G32040 | 0.3 | 2.01E-03 | 0.1 | 2.18E-01 | 0.2 | 3.80E-02 | -0.1 | 6.17E-01 | -0.1 | 8.01E-01 | 0.0 | 9.21E-01 | Major facilitator superfamily protein |
| AT4G22190 | 0.3 | 3.99E-03 | 0.2 | 2.21E-02 | 0.1 | 4.43E-01 | -0.5 | 1.64E-03 | -0.5 | 7.03E-03 | -0.1 | 8.60E-01 | Serine |
| AT1G72440 | 0.3 | 2.08E-02 | 0.2 | 1.20E-01 | 0.1 | 4.27E-01 | 0.5 | 6.09E-03 | 0.3 | 1.75E-01 | 0.2 | 2.71E-01 | Protein SLOW WALKER 2 |
| AT1G65680 | 0.3 | 4.08E-01 | 0.3 | 3.58E-01 | 0.0 | 9.23E-01 | | | | | | | Putative expansin-B2 |
| AT4G15820 | 0.3 | 2.46E-02 | 0.1 | 3.79E-01 | 0.2 | 2.21E-01 | -0.2 | 4.15E-01 | -0.2 | 6.16E-01 | -0.1 | 9.06E-01 | ABC subfamily C protein |
| AT3G07460 | 0.3 | 2.03E-02 | 0.4 | 5.91E-03 | -0.1 | 4.43E-01 | -0.4 | 9.14E-03 | -0.6 | 2.91E-03 | 0.1 | 5.44E-01 | F21O3.17 protein |
| AT5G19920 | 0.3 | 2.99E-02 | 0.0 | 7.67E-01 | 0.3 | 1.05E-01 | 0.3 | 4.37E-01 | 0.4 | 2.86E-01 | -0.1 | 8.07E-01 | Uncharacterized protein |
| AT5G58490 | 0.3 | 1.09E-03 | 0.3 | 3.92E-03 | 0.0 | 5.86E-01 | 0.0 | 9.31E-01 | 0.0 | 8.30E-01 | 0.0 | 8.20E-01 | NAD-dependent epimerase |
| AT5G20600 | 0.3 | 1.52E-02 | 0.4 | 4.76E-03 | -0.1 | 4.91E-01 | 0.0 | 8.70E-01 | -0.3 | 4.07E-02 | 0.3 | 1.20E-01 | AT5g20600 |
| AT5G20110 | 0.3 | 7.18E-03 | -0.1 | 2.10E-01 | 0.4 | 3.87E-03 | -0.6 | 4.43E-02 | -0.1 | 7.51E-01 | -0.5 | 1.77E-01 | hypothetical protein |

| | | | | | | | | | | | | | |
|---|---|---|---|---|---|---|---|---|---|---|---|---|---|
| AT4G13615 | 0.3 | 1.57E-02 | 0.4 | 3.44E-03 | -0.1 | 3.34E-01 | 0.0 | 9.12E-01 | -0.1 | 8.11E-01 | 0.0 | 9.10E-01 | At4g13612 |
| AT2G22270 | 0.3 | 2.77E-03 | 0.1 | 1.01E-01 | 0.2 | 1.15E-01 | 0.0 | 9.88E-01 | 0.0 | 9.82E-01 | 0.0 | 9.97E-01 | Hematological |
| AT1G03780 | 0.3 | 1.07E-02 | 0.5 | 1.18E-03 | -0.2 | 2.01E-01 | -0.1 | 6.87E-01 | 0.2 | 5.53E-01 | -0.3 | 4.07E-01 | hypothetical protein |
| AT5G38640 | 0.3 | 4.45E-03 | 0.1 | 2.80E-01 | 0.2 | 6.82E-02 | 0.5 | 3.69E-01 | 0.2 | 8.28E-01 | 0.3 | 6.73E-01 | hypothetical protein |
| AT3G29375 | 0.3 | 4.29E-03 | 0.3 | 4.37E-03 | 0.0 | 9.03E-01 | 0.0 | 9.64E-01 | 0.2 | 4.16E-01 | -0.2 | 4.31E-01 | XH domain-containing protein |
| AT1G76440 | 0.3 | 1.94E-02 | 0.2 | 1.21E-01 | 0.1 | 4.37E-01 | 0.1 | 7.81E-01 | -0.2 | 5.97E-01 | 0.3 | 3.46E-01 | hypothetical protein |
| AT1G36320 | 0.3 | 3.56E-03 | 0.3 | 3.33E-03 | 0.0 | 8.59E-01 | 0.2 | 7.27E-02 | 0.0 | 8.99E-01 | 0.2 | 1.08E-01 | Uncharacterized protein |
| AT3G12370 | 0.3 | 1.13E-02 | 0.3 | 1.55E-02 | 0.0 | 9.60E-01 | 0.2 | 3.24E-01 | 0.1 | 7.25E-01 | 0.1 | 6.74E-01 | hypothetical protein |
| AT3G11560 | 0.3 | 7.45E-02 | 0.1 | 7.55E-01 | 0.2 | 1.86E-01 | -0.1 | 8.90E-01 | 0.0 | 9.68E-01 | -0.1 | 9.59E-01 | LETM1-like protein |
| AT4G10090 | 0.3 | 1.40E-02 | 0.2 | 6.76E-02 | 0.1 | 5.31E-01 | 0.0 | 9.96E-01 | -0.2 | 3.70E-01 | 0.2 | 3.54E-01 | ELP6 |
| AT5G25280 | 0.3 | 1.22E-02 | 0.2 | 1.19E-01 | 0.1 | 2.92E-01 | -0.3 | 2.60E-02 | -0.6 | 8.40E-04 | 0.3 | 1.07E-01 | hypothetical protein |
| AT1G75030 | 0.3 | 1.38E-01 | 1.0 | 4.40E-04 | -0.7 | 9.85E-03 | 0.5 | 2.67E-01 | 0.4 | 3.85E-01 | 0.0 | 9.48E-01 | TLP-3 |
| AT2G36430 | 0.3 | 1.11E-02 | 0.3 | 9.77E-03 | 0.0 | 8.80E-01 | -0.1 | 3.35E-01 | -0.3 | 7.37E-02 | 0.1 | 5.23E-01 | hypothetical protein |
| AT5G03870 | 0.3 | 3.00E-02 | 0.5 | 2.18E-03 | -0.2 | 1.75E-01 | 0.1 | 6.22E-01 | 0.0 | 9.81E-01 | 0.1 | 7.19E-01 | Emb\|CAB85507.1 |
| AT1G19130 | 0.3 | 1.01E-02 | 0.3 | 2.21E-02 | 0.0 | 7.84E-01 | 0.2 | 3.08E-01 | -0.1 | 7.55E-01 | 0.2 | 3.04E-01 | hypothetical protein |
| AT4G19865 | 0.3 | 3.10E-02 | 0.2 | 1.99E-01 | 0.1 | 4.40E-01 | 0.0 | 9.06E-01 | -0.1 | 7.57E-01 | 0.2 | 7.03E-01 | F-box |
| AT5G05640 | 0.3 | 2.41E-02 | 0.4 | 5.53E-03 | -0.1 | 4.28E-01 | -0.2 | 7.32E-01 | -0.2 | 7.10E-01 | 0.0 | 9.62E-01 | At5g05640 |
| AT1G08220 | 0.3 | 5.42E-02 | 0.1 | 5.40E-01 | 0.2 | 2.58E-01 | 0.4 | 8.60E-02 | 0.1 | 8.28E-01 | 0.4 | 2.74E-01 | ATPase complex subunit |
| AT1G09765 | 0.3 | 9.41E-02 | 0.4 | 5.01E-02 | -0.1 | 7.37E-01 | -0.4 | 2.35E-01 | -0.2 | 5.18E-01 | -0.1 | 7.31E-01 | NA |
| AT1G70280 | 0.3 | 6.45E-03 | 0.4 | 1.37E-03 | -0.1 | 3.21E-01 | -0.2 | 2.65E-01 | 0.0 | 9.42E-01 | -0.2 | 3.60E-01 | NHL domain-containing protein |
| AT5G57685 | 0.3 | 6.54E-02 | 0.4 | 2.13E-02 | -0.1 | 5.38E-01 | -0.1 | 5.94E-01 | 0.1 | 6.53E-01 | -0.2 | 3.20E-01 | Protein GLUTAMINE DUMPER 3 |
| AT5G58210 | 0.3 | 3.82E-02 | -0.2 | 1.45E-01 | 0.5 | 8.71E-03 | 0.5 | 5.69E-03 | -0.3 | 1.48E-01 | 0.8 | 1.73E-03 | AT5G58210 protein |
| AT4G34700 | 0.3 | 1.00E-02 | 0.4 | 2.33E-03 | -0.1 | 3.30E-01 | 0.1 | 5.49E-01 | 0.1 | 6.52E-01 | 0.0 | 9.89E-01 | NADH dehydrogenase 1 beta subcomplex subunit 9 |
| AT1G42480 | 0.3 | 1.86E-02 | 0.4 | 6.11E-03 | -0.1 | 5.20E-01 | 0.2 | 2.11E-01 | 0.3 | 1.17E-01 | -0.1 | 7.61E-01 | TLR4 regulator |
| AT3G46600 | 0.3 | 3.98E-02 | -0.4 | 1.54E-02 | 0.7 | 1.30E-03 | 0.3 | 9.18E-02 | -0.3 | 8.68E-02 | 0.6 | 9.62E-03 | GRAS family transcription factor |
| AT2G26030 | 0.3 | 6.49E-02 | -0.1 | 4.89E-01 | 0.4 | 4.34E-02 | -0.4 | 1.28E-01 | -0.1 | 8.21E-01 | -0.3 | 4.03E-01 | F-box |
| AT4G33945 | 0.3 | 2.47E-03 | 0.2 | 1.03E-02 | 0.1 | 5.51E-01 | 0.0 | 8.45E-01 | 0.0 | 9.99E-01 | 0.0 | 9.05E-01 | Armadillo repeat-containing protein 6 |
| AT1G55300 | 0.3 | 5.10E-03 | 0.4 | 1.85E-03 | -0.1 | 5.10E-01 | -0.2 | 3.46E-01 | -0.4 | 1.60E-01 | 0.1 | 7.11E-01 | TAFII55 protein conserved region domain-containing protein |
| AT5G22560 | 0.3 | 1.22E-01 | 0.4 | 8.09E-02 | -0.1 | 8.22E-01 | -0.2 | 7.50E-01 | 0.3 | 6.17E-01 | -0.5 | 3.71E-01 | Emb\|CAB72473.1 ) |
| AT3G49990 | 0.3 | 4.07E-03 | 0.2 | 5.01E-02 | 0.1 | 2.44E-01 | 0.2 | 7.94E-02 | 0.0 | 9.55E-01 | 0.2 | 2.40E-01 | At3g49990 |
| AT5G13240 | 0.3 | 3.13E-03 | 0.3 | 3.06E-03 | 0.0 | 8.87E-01 | -0.1 | 6.64E-01 | -0.1 | 5.74E-01 | 0.0 | 9.07E-01 | Repressor of RNA polymerase III transcription |
| AT2G47690 | 0.3 | 2.08E-03 | 0.4 | 4.48E-04 | -0.1 | 2.44E-01 | -0.1 | 4.87E-01 | -0.2 | 3.60E-01 | 0.1 | 8.16E-01 | NADH-ubiquinone oxidoreductase-like protein |
| AT1G53200 | 0.3 | 9.31E-03 | 0.1 | 2.34E-01 | 0.2 | 1.57E-01 | 0.3 | 9.54E-02 | -0.3 | 1.59E-01 | 0.7 | 1.61E-02 | TAF RNA polymerase I subunit A |
| AT4G01897 | 0.3 | 3.83E-01 | -0.4 | 3.70E-01 | 0.7 | 1.50E-01 | 0.1 | 7.52E-01 | 0.3 | 5.36E-01 | -0.1 | 8.17E-01 | At4g01897 |
| AT1G62030 | 0.3 | 4.85E-02 | 0.2 | 1.55E-01 | 0.1 | 6.47E-01 | 0.0 | 9.41E-01 | 0.1 | 8.88E-01 | -0.1 | 8.33E-01 | Cysteine |
| AT5G01770 | 0.3 | 1.19E-02 | 0.1 | 3.36E-01 | 0.2 | 1.35E-01 | 0.7 | 6.32E-02 | 0.7 | 6.64E-02 | -0.1 | 9.31E-01 | Regulatory-associated protein of TOR 2 |
| AT4G29730 | 0.3 | 1.34E-02 | 0.2 | 8.46E-02 | 0.1 | 4.47E-01 | 0.1 | 8.24E-01 | 0.2 | 6.11E-01 | -0.1 | 8.23E-01 | Nucleosome |
| AT5G58130 | 0.3 | 4.17E-02 | -0.3 | 9.87E-02 | 0.6 | 7.49E-03 | 0.0 | 9.74E-01 | 0.2 | 8.49E-01 | -0.2 | 8.50E-01 | Uncharacterized protein |
| AT5G38720 | 0.3 | 5.07E-02 | 0.2 | 1.32E-01 | 0.1 | 6.97E-01 | 0.6 | 1.35E-02 | 0.3 | 3.95E-01 | 0.4 | 2.04E-01 | Ribosomal RNA-processing 7 protein |
| AT1G14060 | 0.3 | 7.58E-02 | 0.2 | 1.60E-01 | 0.1 | 7.66E-01 | 0.8 | 2.40E-03 | 0.2 | 3.92E-01 | 0.5 | 4.59E-02 | F16A14.27 |
| AT2G40430 | 0.3 | 1.10E-02 | 0.4 | 4.34E-03 | -0.1 | 5.48E-01 | 0.5 | 1.76E-03 | 0.3 | 5.04E-02 | 0.2 | 3.30E-01 | Ribosome biogenesis protein NOP53 |
| AT4G22540 | 0.3 | 5.97E-03 | 0.2 | 2.12E-02 | 0.1 | 5.94E-01 | -0.1 | 4.62E-01 | -0.2 | 4.58E-01 | 0.0 | 9.48E-01 | Uncharacterized protein At4g22540 |
| AT2G39170 | 0.3 | 7.56E-03 | 0.1 | 3.18E-01 | 0.2 | 8.84E-02 | -0.2 | 2.41E-01 | -0.1 | 4.66E-01 | 0.0 | 8.51E-01 | hypothetical protein |
| AT3G25070 | 0.3 | 8.60E-03 | 0.2 | 1.16E-01 | 0.1 | 2.28E-01 | 0.3 | 8.71E-02 | 0.0 | 9.52E-01 | 0.3 | 1.73E-01 | RPM1 interacting protein 4 |
| AT1G21900 | 0.3 | 2.65E-03 | 0.2 | 5.83E-02 | 0.1 | 1.49E-01 | 0.3 | 1.52E-01 | 0.0 | 8.72E-01 | 0.2 | 3.77E-01 | Transmembrane emp24 domain-containing protein p24delta5 |
| AT2G40650 | 0.3 | 5.55E-03 | 0.2 | 9.98E-02 | 0.1 | 1.82E-01 | 0.1 | 3.20E-01 | -0.2 | 3.82E-01 | 0.3 | 1.46E-01 | Pre-mRNA-splicing factor 38 |
| AT4G33630 | 0.3 | 1.97E-03 | -0.1 | 2.76E-01 | 0.4 | 1.26E-03 | -0.5 | 4.69E-03 | -0.2 | 2.25E-01 | -0.3 | 1.21E-01 | AT4G33630 protein |
| AT3G01690 | 0.3 | 1.83E-02 | 0.4 | 9.48E-03 | -0.1 | 6.50E-01 | -0.3 | 2.28E-02 | -0.4 | 1.69E-02 | 0.1 | 8.14E-01 | hypothetical protein |
| AT4G23930 | 0.3 | 8.60E-02 | 0.2 | 3.18E-01 | 0.1 | 5.73E-01 | 0.2 | 5.12E-01 | 0.2 | 4.98E-01 | 0.0 | 9.53E-01 | Late embryogenesis abundant hydroxyproline-rich glycoprotein family |
| AT1G77145 | 0.3 | 2.06E-02 | 0.0 | 9.15E-01 | 0.3 | 4.38E-02 | -1.1 | 1.92E-03 | -0.2 | 4.83E-01 | -0.9 | 3.28E-02 | T14N5.3 protein ) |
| AT3G15357 | 0.3 | 1.41E-01 | 0.7 | 4.60E-03 | -0.4 | 1.05E-01 | 0.1 | 7.22E-01 | 0.2 | 5.04E-01 | -0.1 | 7.49E-01 | hypothetical protein |
| AT2G16860 | 0.3 | 4.79E-04 | 0.2 | 1.14E-02 | 0.1 | 1.12E-01 | -0.1 | 7.43E-01 | 0.0 | 9.79E-01 | -0.1 | 8.11E-01 | Pre-mRNA-splicing factor SYF2 |
| AT5G27860 | 0.3 | 1.04E-01 | 0.0 | 8.65E-01 | 0.3 | 1.22E-01 | -0.1 | 4.91E-01 | -0.3 | 1.02E-01 | 0.2 | 4.54E-01 | Uncharacterized protein |
| AT5G47360 | 0.3 | 2.69E-02 | 0.1 | 4.11E-01 | 0.2 | 2.17E-01 | -0.1 | 8.51E-01 | -0.1 | 6.84E-01 | 0.1 | 8.56E-01 | Pentatricopeptide repeat-containing protein At5g47360 |
| AT5G47830 | 0.3 | 1.04E-02 | 0.1 | 1.90E-01 | 0.2 | 2.01E-01 | -0.5 | 2.41E-02 | -0.6 | 1.53E-02 | 0.1 | 8.15E-01 | Uncharacterized protein At5g47830 |
| AT3G04310 | 0.3 | 3.68E-03 | 0.2 | 8.38E-02 | 0.1 | 1.75E-01 | -0.1 | 7.58E-01 | -0.2 | 3.94E-01 | 0.1 | 6.90E-01 | T6K12.7 protein |
| AT1G29785 | 0.3 | 1.55E-02 | 0.5 | 8.61E-04 | -0.2 | 1.14E-01 | 0.2 | 5.59E-01 | 0.0 | 9.54E-01 | 0.2 | 7.04E-01 | NA |
| AT2G39950 | 0.3 | 3.57E-03 | 0.1 | 1.69E-01 | 0.2 | 7.71E-02 | 0.3 | 3.07E-02 | 0.1 | 5.75E-01 | 0.2 | 2.52E-01 | Uncharacterized protein At2g39950 |
| AT1G09800 | 0.3 | 5.38E-02 | 0.3 | 4.35E-02 | 0.0 | 8.79E-01 | 0.6 | 8.91E-02 | 0.5 | 2.15E-01 | 0.1 | 8.39E-01 | F21M12.18 protein |
| AT2G26280 | 0.3 | 1.88E-03 | 0.0 | 8.53E-01 | 0.3 | 3.61E-03 | 0.1 | 6.41E-01 | 0.0 | 7.76E-01 | 0.1 | 5.77E-01 | CTC-interacting domain 7 |
| AT3G16220 | 0.3 | 2.49E-03 | 0.0 | 6.20E-01 | 0.3 | 1.48E-02 | 0.0 | 8.13E-01 | -0.1 | 7.38E-01 | 0.1 | 6.15E-01 | hypothetical protein |
| AT3G25410 | 0.3 | 3.10E-03 | 0.1 | 1.72E-01 | 0.2 | 6.99E-02 | 0.0 | 9.77E-01 | 0.0 | 9.55E-01 | 0.0 | 9.82E-01 | Probable sodium |
| AT3G44280 | 0.3 | 5.69E-02 | 0.2 | 2.56E-01 | 0.1 | 5.00E-01 | 0.2 | 6.18E-01 | 0.1 | 8.19E-01 | 0.1 | 9.10E-01 | hypothetical protein |
| AT3G18940 | 0.3 | 5.93E-03 | 0.4 | 2.92E-03 | -0.1 | 6.26E-01 | 0.0 | 8.77E-01 | -0.2 | 3.50E-01 | 0.2 | 3.32E-01 | Proteasome assembly chaperone 2 |
| AT1G04760 | 0.3 | 9.61E-02 | 0.3 | 1.46E-01 | 0.0 | 9.15E-01 | -0.2 | 5.84E-01 | 0.2 | 5.89E-01 | -0.4 | 2.52E-01 | hypothetical protein |
| AT1G45207 | 0.3 | 1.19E-03 | 0.3 | 1.40E-03 | 0.0 | 9.21E-01 | -0.4 | 9.80E-03 | -0.3 | 3.40E-02 | 0.0 | 8.44E-01 | Remorin family protein |
| AT4G31115 | 0.3 | 5.79E-02 | 0.1 | 6.34E-01 | 0.2 | 2.19E-01 | 0.2 | 2.32E-01 | 0.1 | 7.44E-01 | 0.1 | 6.20E-01 | AT4g31110 |
| AT1G78890 | 0.3 | 5.36E-04 | 0.3 | 1.74E-03 | 0.0 | 6.37E-01 | 0.0 | 9.39E-01 | 0.0 | 1.00E+00 | 0.0 | 9.62E-01 | Uncharacterized protein |
| AT4G25620 | 0.3 | 1.39E-02 | 0.1 | 3.85E-01 | 0.2 | 1.19E-01 | -0.4 | 9.65E-03 | -0.6 | 1.99E-03 | 0.2 | 4.47E-01 | At4g25620 |
| AT4G09805 | 0.3 | 4.65E-02 | 0.4 | 1.90E-02 | -0.1 | 6.46E-01 | 0.4 | 1.73E-01 | 0.2 | 4.51E-01 | 0.1 | 7.68E-01 | NA |
| AT3G61180 | 0.3 | 1.80E-03 | 0.1 | 2.17E-01 | 0.2 | 3.15E-02 | -0.1 | 7.34E-01 | -0.3 | 1.05E-01 | 0.2 | 2.61E-01 | RING-type domain-containing protein |
| AT5G40370 | 0.3 | 4.12E-03 | 0.3 | 1.60E-02 | 0.1 | 5.74E-01 | 0.3 | 1.68E-01 | 0.2 | 5.06E-01 | 0.1 | 7.50E-01 | Glutaredoxin family protein |
| AT1G68945 | 0.3 | 1.43E-02 | 0.2 | 5.46E-02 | 0.1 | 5.95E-01 | -0.2 | 2.91E-01 | -0.2 | 4.35E-01 | 0.0 | 9.34E-01 | Uncharacterized protein |
| AT3G48800 | 0.3 | 4.29E-03 | 0.3 | 5.24E-03 | 0.0 | 9.75E-01 | 0.1 | 5.79E-01 | -0.2 | 4.03E-01 | 0.3 | 2.26E-01 | hypothetical protein |
| AT2G19790 | 0.3 | 6.29E-04 | 0.4 | 4.32E-04 | 0.0 | 6.24E-01 | -0.3 | 9.89E-02 | -0.2 | 1.98E-01 | 0.0 | 9.26E-01 | hypothetical protein |
| AT5G15320 | 0.3 | 1.83E-02 | 0.7 | 2.60E-04 | -0.4 | 1.77E-02 | 0.2 | 2.52E-01 | 0.1 | 6.84E-01 | 0.1 | 7.21E-01 | ATP synthase E chain |
| AT1G28330 | 0.3 | 7.08E-02 | 0.7 | 2.03E-03 | -0.4 | 5.66E-02 | -0.9 | 4.32E-04 | -1.4 | 5.41E-05 | 0.4 | 1.57E-01 | Uncharacterized protein |
| AT3G18460 | 0.3 | 3.08E-01 | 1.8 | 8.54E-05 | -1.4 | 1.14E-03 | -3.6 | 2.87E-04 | -0.4 | 4.92E-01 | -3.3 | 4.39E-03 | Protein PLANT CADMIUM RESISTANCE 4 |
| AT2G18465 | 0.3 | 3.95E-03 | 0.0 | 8.59E-01 | 0.3 | 1.67E-02 | 0.1 | 5.44E-01 | 0.2 | 4.42E-01 | -0.1 | 8.56E-01 | J domain-containing protein |
| AT1G05070 | 0.3 | 3.10E-02 | 0.1 | 5.71E-01 | 0.2 | 1.48E-01 | -0.1 | 7.54E-01 | -0.2 | 3.48E-01 | 0.1 | 6.54E-01 | At1g05070 |
| AT5G53045 | 0.3 | 1.63E-02 | 0.4 | 1.04E-02 | 0.0 | 7.78E-01 | 0.5 | 2.86E-02 | 0.3 | 2.93E-01 | 0.2 | 4.51E-01 | At5g53043 |
| AT5G02810 | 0.3 | 7.87E-03 | 0.1 | 4.21E-01 | 0.2 | 5.74E-02 | 0.6 | 2.10E-01 | -0.5 | 3.95E-01 | 1.1 | 6.51E-02 | Pseudo-response regulator 7 |
| AT3G43690 | 0.3 | 4.61E-03 | 0.1 | 5.95E-01 | 0.3 | 2.84E-02 | 0.0 | 9.07E-01 | 0.1 | 3.67E-01 | -0.2 | 3.90E-01 | NA |

| | | | | | | | | | | | | | |
|---|---|---|---|---|---|---|---|---|---|---|---|---|---|
| AT1G04790 | 0.3 | 2.76E-04 | 0.0 | 8.37E-01 | 0.3 | 1.03E-03 | 0.0 | 9.06E-01 | 0.0 | 9.16E-01 | 0.1 | 8.47E-01 | RING-type domain-containing protein |
| AT2G18900 | 0.3 | 3.04E-02 | 0.2 | 1.90E-01 | 0.1 | 4.09E-01 | 0.8 | 8.03E-04 | 0.2 | 4.50E-01 | 0.6 | 2.41E-02 | Uncharacterized protein |
| AT3G61140 | 0.3 | 3.75E-04 | 0.1 | 8.12E-02 | 0.2 | 1.55E-02 | 0.4 | 8.82E-03 | 0.3 | 6.63E-02 | 0.1 | 6.73E-01 | COP9 signalosome complex subunit 1 |
| AT5G61300 | 0.3 | 4.08E-02 | 0.3 | 6.52E-02 | 0.0 | 8.94E-01 | -0.6 | 2.03E-01 | 0.0 | 9.79E-01 | -0.6 | 2.15E-01 | Uncharacterized protein |
| AT4G17370 | 0.3 | 1.28E-02 | -0.1 | 3.37E-01 | 0.4 | 7.05E-03 | -0.2 | 3.58E-01 | -0.1 | 5.72E-01 | 0.0 | 8.99E-01 | Oxidoreductase family protein |
| AT5G14840 | 0.3 | 1.74E-02 | 0.1 | 2.85E-01 | 0.2 | 2.07E-01 | -0.2 | 1.08E-01 | -0.3 | 4.34E-02 | 0.1 | 7.02E-01 | NA |
| AT1G16470 | 0.3 | 2.84E-03 | 0.1 | 1.69E-01 | 0.2 | 6.00E-02 | 0.5 | 6.53E-03 | 0.2 | 2.44E-01 | 0.2 | 2.73E-01 | Proteasome subunit alpha type-2-A |
| AT5G47455 | 0.3 | 9.88E-03 | 0.3 | 3.64E-02 | 0.1 | 6.08E-01 | 0.4 | 2.37E-01 | 0.3 | 4.21E-01 | 0.1 | 8.75E-01 | Uncharacterized protein At5g47455 |
| AT1G07700 | 0.3 | 2.19E-04 | 0.2 | 4.16E-03 | 0.1 | 1.20E-01 | 0.3 | 2.86E-02 | 0.1 | 5.58E-01 | 0.2 | 2.58E-01 | Thioredoxin domain-containing protein |
| AT1G05430 | 0.3 | 3.96E-02 | -0.1 | 6.51E-01 | 0.4 | 3.67E-02 | 0.1 | 6.42E-01 | 0.1 | 4.66E-01 | -0.1 | 8.19E-01 | Transmembrane protein |
| AT2G01818 | 0.3 | 8.42E-02 | 0.2 | 2.08E-01 | 0.1 | 7.23E-01 | -0.3 | 5.20E-01 | -0.4 | 4.20E-01 | 0.1 | 8.81E-01 | PLATZ transcription factor family protein |
| AT1G22403 | 0.3 | 4.37E-02 | 0.2 | 3.47E-01 | 0.2 | 3.50E-01 | -0.4 | 4.14E-02 | -0.2 | 4.36E-01 | -0.3 | 4.07E-01 | NA |
| AT5G23360 | 0.3 | 5.23E-02 | 0.5 | 1.01E-02 | -0.2 | 3.98E-01 | 0.1 | 8.78E-01 | 0.1 | 8.59E-01 | 0.0 | 9.73E-01 | GEM-like protein 7 |
| AT3G07230 | 0.3 | 1.50E-02 | 0.4 | 8.23E-03 | -0.1 | 6.89E-01 | 0.3 | 9.61E-02 | 0.2 | 3.80E-01 | 0.1 | 6.93E-01 | Wound-induced basic protein |
| AT5G27395 | 0.3 | 1.38E-02 | 0.1 | 2.40E-01 | 0.2 | 1.97E-01 | 0.6 | 2.40E-03 | 0.1 | 5.95E-01 | 0.5 | 3.98E-02 | Mitochondrial inner membrane translocase complex, subunit Tim44-related protein |
| AT1G65220 | 0.3 | 4.91E-04 | 0.3 | 1.74E-03 | 0.0 | 6.04E-01 | 0.0 | 8.73E-01 | 0.0 | 7.79E-01 | 0.1 | 7.31E-01 | W2 domain-containing protein |
| AT5G58030 | 0.3 | 4.53E-03 | 0.2 | 3.96E-02 | 0.1 | 3.30E-01 | 0.1 | 4.92E-01 | 0.0 | 9.75E-01 | 0.1 | 6.30E-01 | Trafficking protein particle complex subunit |
| AT2G31040 | 0.3 | 4.59E-03 | 0.4 | 2.61E-03 | -0.1 | 6.28E-01 | -0.1 | 6.36E-01 | 0.2 | 2.84E-01 | -0.3 | 1.73E-01 | Protein CONSERVED ONLY IN THE GREEN LINEAGE 160, chloroplastic |
| AT5G09225 | 0.3 | 1.25E-02 | 0.1 | 5.20E-01 | 0.2 | 8.54E-02 | -0.3 | 2.15E-01 | -0.3 | 2.83E-01 | 0.0 | 9.92E-01 | hypothetical protein |
| AT2G33220 | 0.3 | 3.77E-03 | 0.3 | 3.96E-03 | 0.0 | 8.90E-01 | -0.1 | 6.89E-01 | 0.0 | 9.14E-01 | -0.1 | 8.80E-01 | NADH dehydrogenase 1 alpha subcomplex subunit 13 |
| AT5G52540 | 0.3 | 3.63E-03 | 0.2 | 1.21E-01 | 0.2 | 1.02E-01 | 0.0 | 9.31E-01 | 0.0 | 8.03E-01 | 0.1 | 7.95E-01 | hypothetical protein |
| AT3G16890 | 0.3 | 1.78E-02 | 0.4 | 9.97E-03 | -0.1 | 7.19E-01 | 0.2 | 5.47E-01 | 0.1 | 8.99E-01 | 0.1 | 7.81E-01 | Pentatricopeptide domain protein 40 |
| AT3G55480 | 0.3 | 5.11E-03 | 0.0 | 9.63E-01 | 0.3 | 1.40E-02 | 0.0 | 9.83E-01 | -0.1 | 8.65E-01 | 0.1 | 9.00E-01 | AP-3 complex subunit beta |
| AT1G60080 | 0.3 | 2.57E-02 | 0.2 | 8.98E-02 | 0.1 | 6.14E-01 | 0.5 | 2.60E-02 | 0.0 | 9.66E-01 | 0.5 | 9.11E-02 | 3'-5'-exoribonuclease family protein |
| AT3G47833 | 0.3 | 4.05E-02 | 0.2 | 1.98E-01 | 0.1 | 4.70E-01 | 0.2 | 4.37E-01 | 0.1 | 8.54E-01 | 0.2 | 7.35E-01 | Succinate dehydrogenase subunit 7A, mitochondrial |
| AT3G27090 | 0.3 | 9.57E-04 | 0.2 | 3.15E-02 | 0.1 | 9.39E-02 | 0.2 | 1.79E-01 | 0.0 | 9.79E-01 | 0.2 | 3.88E-01 | DCD domain-containing protein |
| AT5G65830 | 0.3 | 4.34E-03 | 0.2 | 2.39E-02 | 0.1 | 4.79E-01 | 0.3 | 1.83E-01 | 0.3 | 3.03E-01 | 0.0 | 9.34E-01 | Receptor-like protein 57 |
| AT2G43780 | 0.3 | 3.18E-02 | 0.4 | 1.12E-02 | -0.1 | 5.71E-01 | 0.5 | 2.15E-02 | 0.2 | 4.58E-01 | 0.3 | 1.62E-01 | Cytochrome c oxidase assembly factor 3 mitochondrial coiled-coil domain-containing protein |
| AT3G28970 | 0.3 | 1.45E-02 | 0.1 | 5.50E-01 | 0.3 | 8.84E-02 | -0.1 | 8.12E-01 | -0.4 | 6.43E-02 | 0.4 | 1.64E-01 | Defective in cullin neddylation protein |
| AT3G25020 | 0.3 | 2.71E-01 | -0.1 | 7.71E-01 | 0.4 | 2.45E-01 | 0.7 | 4.09E-03 | 0.1 | 8.46E-01 | 0.6 | 2.66E-02 | Receptor-like protein 42 |
| AT4G24275 | 0.3 | 9.43E-02 | 0.4 | 3.45E-02 | -0.1 | 6.21E-01 | 0.1 | 6.95E-01 | -0.1 | 8.26E-01 | 0.2 | 5.85E-01 | Uncharacterized protein |
| AT3G20370 | 0.3 | 1.20E-03 | 0.6 | 2.32E-05 | -0.3 | 5.16E-03 | 0.1 | 5.67E-01 | 0.5 | 7.60E-03 | -0.4 | 3.80E-02 | MATH domain-containing protein |
| AT1G18650 | 0.3 | 1.20E-02 | 0.6 | 6.82E-04 | -0.2 | 9.69E-02 | -0.4 | 6.14E-02 | -0.2 | 4.97E-01 | -0.2 | 4.64E-01 | Plasmodesmata callose-binding protein 3 |
| AT1G73240 | 0.3 | 2.91E-03 | 0.2 | 8.56E-02 | 0.2 | 1.35E-01 | 0.1 | 7.35E-01 | 0.1 | 8.76E-01 | 0.0 | 9.31E-01 | Nucleoporin protein Ndc1-Nup |
| AT5G60210 | 0.3 | 6.39E-03 | 0.2 | 7.69E-02 | 0.1 | 2.69E-01 | 0.0 | 7.18E-01 | 0.1 | 3.76E-01 | -0.2 | 2.93E-01 | ROP interactive partner 5 |
| AT1G76520 | 0.3 | 4.34E-04 | 0.1 | 2.58E-01 | 0.3 | 6.11E-03 | 0.2 | 1.28E-01 | -0.3 | 1.11E-01 | 0.5 | 7.35E-03 | Protein PIN-LIKES 3 |
| AT2G05075 | 0.3 | 1.44E-01 | -0.1 | 6.84E-01 | 0.4 | 1.33E-01 | -0.2 | 6.38E-01 | -0.3 | 4.00E-01 | 0.2 | 7.80E-01 | NA |
| AT2G31670 | 0.3 | 1.44E-03 | 0.3 | 2.05E-03 | 0.0 | 9.95E-01 | 0.0 | 9.83E-01 | 0.1 | 5.56E-01 | -0.1 | 6.62E-01 | Stress-response A |
| AT5G27430 | 0.3 | 1.55E-03 | 0.4 | 5.25E-04 | -0.1 | 3.85E-01 | 0.1 | 6.56E-01 | -0.1 | 7.53E-01 | 0.1 | 5.26E-01 | Signal peptidase complex subunit 3B |
| AT4G10800 | 0.3 | 3.33E-02 | 0.4 | 2.80E-02 | 0.0 | 8.92E-01 | 0.1 | 6.64E-01 | 0.1 | 7.58E-01 | 0.0 | 9.80E-01 | BTB |
| AT3G24900 | 0.3 | 6.01E-01 | -0.9 | 2.75E-01 | 1.3 | 2.01E-01 | | | | | | | Receptor like protein 39 |
| AT1G11380 | 0.3 | 7.33E-03 | 0.6 | 1.32E-04 | -0.3 | 1.96E-02 | -0.5 | 8.36E-03 | -0.2 | 1.78E-01 | -0.2 | 3.28E-01 | hypothetical protein |
| AT3G09455 | 0.3 | 1.15E-01 | 0.3 | 1.52E-01 | 0.0 | 9.38E-01 | 0.7 | 4.79E-02 | 0.5 | 2.18E-01 | 0.2 | 6.97E-01 | Dolichyl-diphosphooligosaccharide--protein glycosyltransferase subunit 4C |
| AT5G40660 | 0.3 | 1.67E-02 | 0.1 | 3.68E-01 | 0.2 | 1.48E-01 | 0.1 | 4.79E-01 | -0.1 | 8.19E-01 | 0.2 | 4.46E-01 | ATP synthase mitochondrial F1 complex assembly factor 2 |
| AT2G31190 | 0.3 | 2.16E-02 | 0.3 | 6.01E-02 | 0.1 | 7.08E-01 | -0.4 | 9.22E-02 | -0.2 | 5.70E-01 | -0.2 | 4.40E-01 | Protein root UVB sensitive 2, chloroplastic |
| AT3G54600 | 0.3 | 1.07E-02 | 0.4 | 5.85E-03 | -0.1 | 6.54E-01 | -0.3 | 1.59E-01 | 0.6 | 7.63E-03 | -0.9 | 3.63E-03 | DJ-1 protein homolog F |
| AT5G53486 | 0.3 | 4.08E-02 | 0.6 | 1.74E-03 | -0.3 | 1.11E-01 | -0.2 | 3.68E-01 | 0.1 | 8.40E-01 | -0.3 | 2.53E-01 | Transmembrane protein |
| AT2G30330 | 0.3 | 4.27E-02 | 0.2 | 2.81E-01 | 0.2 | 3.91E-01 | 0.0 | 9.24E-01 | -0.1 | 8.63E-01 | 0.0 | 9.46E-01 | Biogenesis of lysosome-related organelles complex 1 subunit 1 |
| AT1G26730 | 0.3 | 3.14E-02 | 0.4 | 1.59E-02 | -0.1 | 6.96E-01 | -0.2 | 6.98E-01 | 0.3 | 3.76E-01 | -0.5 | 2.77E-01 | EXS family protein |
| AT5G11580 | 0.3 | 2.51E-04 | 0.2 | 2.74E-03 | 0.1 | 2.09E-01 | -0.1 | 2.64E-01 | -0.3 | 2.22E-02 | 0.2 | 3.12E-01 | Regulator of chromosome condensation family protein |
| AT2G02510 | 0.3 | 1.90E-02 | 0.3 | 2.30E-02 | 0.0 | 9.88E-01 | 0.1 | 5.88E-01 | 0.0 | 9.76E-01 | 0.1 | 7.44E-01 | NADH dehydrogenase 1 beta subcomplex subunit 3-A |
| AT5G11910 | 0.3 | 5.21E-04 | 0.1 | 9.08E-02 | 0.2 | 2.45E-02 | -0.1 | 6.07E-01 | -0.1 | 7.07E-01 | 0.0 | 9.80E-01 | Alpha |
| AT5G38150 | 0.3 | 5.79E-03 | 0.1 | 1.98E-01 | 0.2 | 1.16E-01 | 0.3 | 3.72E-01 | 0.3 | 3.84E-01 | 0.0 | 9.60E-01 | Protein PLASTID MOVEMENT IMPAIRED 15 |
| AT1G54095 | 0.3 | 8.54E-02 | -0.1 | 6.23E-01 | 0.4 | 7.68E-02 | 1.1 | 9.68E-03 | 1.2 | 1.01E-02 | -0.1 | 8.46E-01 | F15I1.18 |
| AT3G12530 | 0.3 | 4.52E-02 | 0.4 | 1.44E-02 | -0.1 | 5.51E-01 | 0.3 | 1.80E-01 | -0.3 | 2.43E-01 | 0.7 | 3.44E-02 | DNA replication complex GINS protein PSF2 |
| AT2G47020 | 0.3 | 7.88E-03 | 0.2 | 4.23E-02 | 0.1 | 4.99E-01 | -0.1 | 8.52E-01 | -0.2 | 5.50E-01 | 0.2 | 7.01E-01 | Peptide chain release factor 1 |
| AT5G24610 | 0.3 | 1.52E-03 | 0.2 | 6.55E-02 | 0.2 | 8.18E-02 | -0.3 | 8.41E-02 | -0.4 | 8.04E-02 | 0.0 | 9.09E-01 | hypothetical protein |
| AT1G18810 | 0.3 | 3.17E-02 | 0.3 | 3.52E-02 | 0.0 | 9.79E-01 | 0.1 | 6.50E-01 | 0.0 | 8.96E-01 | 0.2 | 6.26E-01 | Protein PHYTOCHROME KINASE SUBSTRATE 3 |
| AT2G17830 | 0.3 | 1.23E-01 | 0.5 | 3.97E-02 | -0.1 | 5.76E-01 | 0.0 | 9.54E-01 | 0.0 | 9.78E-01 | 0.0 | 9.46E-01 | F-box protein At2g17830 |
| AT4G27540 | 0.3 | 2.23E-03 | 0.3 | 8.56E-03 | 0.1 | 5.96E-01 | 0.2 | 4.71E-01 | -0.3 | 3.71E-01 | 0.4 | 1.67E-01 | PRA1 family protein |
| AT1G71840 | 0.3 | 1.40E-02 | 0.0 | 9.80E-01 | 0.3 | 2.92E-02 | 0.3 | 2.16E-02 | 0.0 | 8.62E-01 | 0.3 | 8.21E-02 | Anaphase-promoting complex subunit 4-like WD40 domain-containing protein |
| AT2G43770 | 0.3 | 5.23E-04 | 0.2 | 4.11E-02 | 0.2 | 4.08E-02 | 0.2 | 1.51E-01 | 0.0 | 8.21E-01 | 0.2 | 1.64E-01 | hypothetical protein |
| AT1G14590 | 0.3 | 7.40E-03 | 0.3 | 8.48E-03 | 0.0 | 9.61E-01 | 0.4 | 1.37E-02 | 0.0 | 8.40E-01 | 0.4 | 2.75E-02 | At1g14590 |
| AT4G16120 | 0.3 | 1.29E-02 | 0.1 | 2.94E-01 | 0.2 | 1.45E-01 | -0.2 | 3.42E-01 | -0.1 | 5.90E-01 | -0.1 | 8.65E-01 | COBRA-like protein 7 |
| AT1G70370 | 0.3 | 2.68E-03 | 0.4 | 9.02E-04 | -0.1 | 3.70E-01 | -0.2 | 8.78E-02 | 0.1 | 6.65E-01 | -0.3 | 1.03E-01 | Polygalacturonase 1 beta-like protein 3 |
| AT5G49210 | 0.3 | 3.94E-03 | 0.5 | 6.43E-04 | -0.1 | 2.27E-01 | 0.1 | 5.17E-01 | 0.0 | 9.01E-01 | 0.2 | 5.49E-01 | hypothetical protein |
| AT1G08530 | 0.3 | 9.57E-03 | 0.1 | 4.03E-01 | 0.2 | 9.39E-02 | 0.3 | 2.48E-01 | 0.4 | 1.12E-01 | -0.1 | 7.09E-01 | F22O13.1 |
| AT3G62080 | 0.3 | 1.20E-02 | -0.1 | 2.68E-01 | 0.5 | 6.72E-03 | -0.2 | 4.76E-01 | -0.3 | 2.37E-01 | 0.1 | 7.10E-01 | SNF7 family protein |
| AT2G47380 | 0.3 | 6.31E-03 | 0.4 | 2.29E-03 | -0.1 | 5.08E-01 | 0.0 | 9.96E-01 | 0.0 | 1.00E+00 | 0.0 | 9.97E-01 | Probable cytochrome c oxidase subunit 5C-1 |
| AT2G34260 | 0.3 | 2.61E-02 | 0.2 | 2.98E-01 | 0.2 | 2.66E-01 | 0.7 | 3.21E-03 | 0.1 | 7.45E-01 | 0.6 | 2.54E-02 | WD repeat-containing protein 55 |
| AT2G23120 | 0.3 | 1.12E-01 | 0.1 | 5.78E-01 | 0.2 | 3.54E-01 | -0.2 | 4.69E-01 | 0.0 | 9.27E-01 | -0.3 | 5.27E-01 | Uncharacterized protein |
| AT4G16510 | 0.3 | 2.87E-02 | 0.2 | 2.19E-01 | 0.2 | 3.80E-01 | 0.0 | 9.29E-01 | 0.0 | 9.65E-01 | 0.0 | 9.80E-01 | Uncharacterized protein AT4g16510 |
| AT1G27090 | 0.3 | 7.77E-04 | 0.2 | 2.06E-02 | 0.1 | 1.11E-01 | -0.2 | 2.38E-02 | -0.4 | 3.55E-03 | 0.1 | 4.34E-01 | hypothetical protein |
| AT1G30240 | 0.3 | 2.37E-03 | 0.2 | 1.06E-01 | 0.2 | 9.88E-02 | 0.5 | 2.22E-02 | 0.0 | 9.71E-01 | 0.5 | 7.63E-02 | Uncharacterized protein At1g30240 |
| AT2G41945 | 0.3 | 1.86E-03 | 0.1 | 3.70E-01 | 0.3 | 2.41E-02 | -0.1 | 6.23E-01 | -0.1 | 6.80E-01 | 0.0 | 9.98E-01 | At2g41945 |
| AT3G19910 | 0.3 | 7.90E-04 | 0.2 | 1.55E-02 | 0.1 | 1.43E-01 | -0.3 | 1.66E-02 | -0.2 | 6.55E-02 | 0.0 | 8.06E-01 | RING-type domain-containing protein |
| AT1G54520 | 0.3 | 1.14E-03 | 0.1 | 1.91E-01 | 0.2 | 2.14E-02 | 0.1 | 4.97E-01 | 0.0 | 9.21E-01 | 0.1 | 5.72E-01 | Myelin-associated oligodendrocyte basic protein |
| AT2G23370 | 0.3 | 1.59E-03 | 0.2 | 1.62E-02 | 0.1 | 3.03E-01 | 0.0 | 9.02E-01 | 0.1 | 4.85E-01 | -0.1 | 6.59E-01 | Cyclin delta-3 |
| AT2G02590 | 0.3 | 1.68E-02 | 0.5 | 1.94E-03 | -0.2 | 2.35E-01 | 0.3 | 3.09E-01 | 0.1 | 7.57E-01 | 0.2 | 5.14E-01 | Small multi-drug export protein |
| AT3G24630 | 0.3 | 4.92E-02 | 0.1 | 4.46E-01 | 0.2 | 3.07E-01 | -0.5 | 2.26E-01 | -0.3 | 5.67E-01 | -0.2 | 6.99E-01 | DUF3741 domain-containing protein |
| AT1G60010 | 0.3 | 1.60E-02 | 0.2 | 2.26E-01 | 0.2 | 2.17E-01 | -0.4 | 4.03E-02 | -0.4 | 4.06E-02 | 0.0 | 9.05E-01 | Uncharacterized protein |
| AT3G12870 | 0.3 | 1.02E-02 | 0.7 | 1.00E-04 | -0.4 | 1.17E-02 | -0.1 | 7.74E-01 | 0.0 | 9.25E-01 | -0.1 | 9.13E-01 | Transmembrane protein |
| AT3G21140 | 0.3 | 1.91E-03 | 0.1 | 1.38E-01 | 0.2 | 5.90E-02 | 0.4 | 1.12E-02 | 0.0 | 8.33E-01 | 0.4 | 3.39E-02 | hypothetical protein |

| | | | | | | | | | | | | |
|---|---|---|---|---|---|---|---|---|---|---|---|---|
| AT1G16690 | 0.3 | 1.00E-03 | 0.2 | 1.06E-02 | 0.1 | 2.88E-01 | 0.0 | 9.47E-01 | 0.0 | 9.65E-01 | 0.0 | 9.98E-01 | Enhancer of polycomb-like protein |
| AT4G35905 | 0.3 | 3.34E-03 | 0.3 | 4.25E-03 | 0.0 | 9.88E-01 | 0.0 | 9.77E-01 | 0.0 | 9.41E-01 | 0.0 | 9.68E-01 | hypothetical protein |
| AT4G21140 | 0.3 | 6.02E-02 | 0.4 | 3.82E-02 | -0.1 | 8.06E-01 | 0.6 | 2.18E-02 | 0.3 | 2.53E-01 | 0.2 | 4.33E-01 | Copper ion binding protein |
| AT5G11310 | 0.3 | 1.82E-02 | 0.0 | 8.84E-01 | 0.3 | 6.04E-02 | 0.2 | 4.26E-01 | -0.2 | 5.67E-01 | 0.4 | 2.08E-01 | Pentatricopeptide repeat-containing protein At5g11310, mitochondrial |
| AT3G10630 | 0.3 | 2.68E-03 | 0.2 | 2.46E-02 | 0.1 | 3.36E-01 | 0.0 | 9.64E-01 | -0.3 | 1.86E-01 | 0.3 | 2.54E-01 | Uncharacterized protein F13M14.8 |
| AT4G32280 | 0.3 | 5.28E-01 | 1.4 | 1.32E-02 | -1.1 | 7.85E-02 | 0.0 | 9.40E-01 | 0.9 | 5.30E-02 | -0.8 | 1.20E-01 | Auxin-responsive protein |
| AT5G67440 | 0.3 | 1.18E-03 | 0.4 | 1.07E-03 | 0.0 | 8.05E-01 | -0.1 | 7.16E-01 | -0.5 | 2.05E-02 | 0.4 | 4.29E-02 | Phototropic-responsive NPH3 family protein |
| AT3G59670 | 0.3 | 1.63E-02 | 0.2 | 1.38E-01 | 0.1 | 3.47E-01 | 0.5 | 4.30E-03 | 0.4 | 2.76E-02 | 0.1 | 7.00E-01 | At3g59670 |
| AT1G24145 | 0.3 | 1.26E-01 | -0.1 | 6.35E-01 | 0.5 | 1.01E-01 | 0.8 | 2.47E-02 | 0.0 | 9.80E-01 | 0.8 | 4.40E-02 | At1g24145 |
| AT2G25720 | 0.3 | 7.99E-03 | 0.2 | 6.86E-02 | 0.1 | 3.56E-01 | 0.3 | 2.01E-01 | 0.2 | 6.34E-01 | 0.2 | 6.56E-01 | Uncharacterized protein |
| AT1G65620 | 0.3 | 5.76E-02 | 0.4 | 6.30E-02 | 0.0 | 9.75E-01 | 0.0 | 9.27E-01 | 0.1 | 8.05E-01 | -0.1 | 8.73E-01 | Protein ASYMMETRIC LEAVES 2 |
| AT3G05810 | 0.3 | 1.71E-02 | 0.4 | 8.46E-03 | -0.1 | 6.84E-01 | 0.2 | 5.36E-01 | 0.0 | 9.16E-01 | 0.1 | 7.41E-01 | IGR protein motif domain-containing protein |
| AT1G74240 | 0.3 | 6.44E-02 | 0.2 | 3.24E-01 | 0.2 | 4.69E-01 | 1.0 | 1.71E-03 | 0.7 | 5.55E-02 | 0.4 | 2.42E-01 | Mitochondrial substrate carrier family protein |
| AT5G35590 | 0.4 | 2.48E-03 | 0.3 | 1.57E-02 | 0.1 | 4.08E-01 | 0.2 | 2.69E-02 | 0.2 | 1.09E-01 | 0.0 | 8.16E-01 | Proteasome subunit alpha type-6-A |
| AT5G04880 | 0.4 | 4.37E-02 | 0.0 | 8.21E-01 | 0.3 | 1.35E-01 | -0.5 | 6.97E-02 | -0.5 | 9.34E-02 | 0.0 | 9.95E-01 | NA |
| AT5G64630 | 0.4 | 1.04E-02 | 0.4 | 5.74E-03 | -0.1 | 6.98E-01 | 0.3 | 4.50E-01 | 0.1 | 7.67E-01 | 0.1 | 8.21E-01 | Transducin |
| AT1G31350 | 0.4 | 1.95E-02 | 0.1 | 4.78E-01 | 0.2 | 1.26E-01 | -0.6 | 5.01E-04 | -0.6 | 1.81E-03 | 0.0 | 9.40E-01 | KAR-UP F-box 1 |
| AT2G25200 | 0.4 | 1.17E-01 | 0.2 | 4.11E-01 | 0.2 | 5.51E-01 | -0.8 | 2.27E-03 | -1.7 | 2.08E-05 | 0.9 | 1.44E-02 | At2g25200 |
| AT5G48990 | 0.4 | 2.02E-03 | 0.3 | 4.35E-03 | 0.0 | 8.28E-01 | -0.2 | 4.19E-01 | -0.2 | 4.09E-01 | 0.0 | 9.56E-01 | F-box |
| AT2G46250 | 0.4 | 2.12E-02 | 0.4 | 2.24E-02 | 0.0 | 9.58E-01 | 0.0 | 9.11E-01 | 0.0 | 9.44E-01 | 0.0 | 8.85E-01 | Myosin heavy chain-like protein |
| AT5G56180 | 0.4 | 9.78E-04 | 0.1 | 5.49E-01 | 0.3 | 8.09E-03 | -0.3 | 2.77E-01 | -0.5 | 8.72E-02 | 0.2 | 5.63E-01 | Actin-related protein 8 |
| AT5G52120 | 0.4 | 9.51E-02 | 0.4 | 5.61E-02 | -0.1 | 7.73E-01 | 0.2 | 3.48E-01 | 0.0 | 9.90E-01 | 0.2 | 4.70E-01 | F-box domain-containing protein |
| AT5G07950 | 0.4 | 2.41E-02 | -0.2 | 1.81E-01 | 0.6 | 8.68E-03 | 0.0 | 9.35E-01 | 0.0 | 9.79E-01 | 0.0 | 9.22E-01 | Uncharacterized protein |
| AT1G02870 | 0.4 | 8.03E-03 | 0.4 | 3.53E-03 | -0.1 | 5.98E-01 | 0.3 | 4.19E-02 | 0.0 | 8.26E-01 | 0.3 | 7.72E-02 | Nucleolar protein 16 |
| AT4G40045 | 0.4 | 1.13E-03 | 0.2 | 2.64E-02 | 0.1 | 1.45E-01 | -0.1 | 7.82E-01 | -0.1 | 5.06E-01 | 0.1 | 7.59E-01 | Transmembrane protein |
| AT3G04640 | 0.4 | 4.50E-02 | -0.2 | 3.21E-01 | 0.5 | 2.09E-02 | 0.4 | 5.77E-02 | 0.0 | 9.89E-01 | 0.5 | 1.27E-01 | At3g04640 |
| AT1G72450 | 0.4 | 1.14E-03 | 0.0 | 6.00E-01 | 0.3 | 6.68E-03 | 0.2 | 5.52E-01 | 0.1 | 9.09E-01 | 0.1 | 7.89E-01 | Protein TIFY |
| AT3G52110 | 0.4 | 1.30E-02 | 0.4 | 5.37E-03 | -0.1 | 5.99E-01 | 0.2 | 2.98E-01 | -0.1 | 7.86E-01 | 0.3 | 2.94E-01 | At3g52110 |
| AT4G05440 | 0.4 | 1.36E-02 | 0.2 | 1.68E-01 | 0.2 | 2.71E-01 | 0.7 | 6.90E-03 | 0.2 | 5.87E-01 | 0.5 | 7.40E-02 | Cell division cycle protein 123 homolog |
| AT5G65120 | 0.4 | 6.29E-03 | 0.1 | 3.70E-01 | 0.3 | 7.28E-02 | 0.2 | 3.89E-01 | 0.2 | 4.16E-01 | 0.0 | 9.76E-01 | hypothetical protein |
| AT2G45010 | 0.4 | 1.97E-03 | 0.3 | 1.49E-02 | 0.1 | 3.58E-01 | 0.4 | 2.81E-03 | 0.2 | 3.07E-01 | 0.3 | 1.43E-01 | PLAC8 family protein |
| AT5G22555 | 0.4 | 8.76E-02 | 0.8 | 1.49E-03 | -0.5 | 4.90E-02 | 1.0 | 1.01E-03 | 1.3 | 2.11E-04 | -0.3 | 2.54E-01 | Transmembrane protein |
| AT3G23440 | 0.4 | 2.73E-02 | -0.1 | 7.15E-01 | 0.4 | 4.09E-02 | -0.4 | 5.57E-01 | -0.1 | 9.32E-01 | -0.3 | 6.90E-01 | Embryo sac development arrest 6 |
| AT2G44510 | 0.4 | 2.57E-02 | 0.2 | 1.99E-01 | 0.2 | 3.75E-01 | 0.6 | 3.75E-02 | 0.1 | 8.21E-01 | 0.5 | 9.24E-02 | Protein BCCIP homolog |
| AT4G34881 | 0.4 | 3.34E-02 | 0.1 | 6.10E-01 | 0.3 | 1.55E-01 | -0.4 | 6.87E-02 | -0.3 | 2.66E-01 | -0.1 | 7.26E-01 | Transmembrane protein |
| AT5G14690 | 0.4 | 2.82E-02 | 0.8 | 2.21E-04 | -0.5 | 1.52E-02 | -0.3 | 2.51E-01 | 0.2 | 5.55E-01 | -0.5 | 1.18E-01 | Transmembrane protein |
| AT5G28626 | 0.4 | 6.82E-03 | 0.4 | 5.10E-03 | 0.0 | 7.99E-01 | -0.3 | 1.28E-01 | -0.4 | 3.93E-02 | 0.1 | 5.85E-01 | NA |
| AT1G05780 | 0.4 | 3.13E-02 | 0.2 | 2.02E-01 | 0.1 | 4.25E-01 | 0.1 | 6.36E-01 | -0.1 | 7.08E-01 | 0.3 | 4.52E-01 | Vacuolar ATPase assembly integral membrane protein VMA21 homolog |
| AT5G14050 | 0.4 | 9.71E-03 | 0.1 | 2.95E-01 | 0.2 | 1.08E-01 | 0.3 | 1.38E-01 | 0.1 | 6.62E-01 | 0.2 | 5.00E-01 | U3 small nucleolar RNA-associated protein 18 homolog |
| AT2G22490 | 0.4 | 1.00E-03 | 0.3 | 8.61E-03 | 0.1 | 3.33E-01 | 0.2 | 4.51E-01 | 0.0 | 9.18E-01 | 0.1 | 6.66E-01 | Cyclin D21 |
| AT5G64480 | 0.4 | 2.56E-03 | 0.1 | 2.04E-01 | 0.2 | 5.65E-02 | 0.2 | 2.44E-01 | 0.1 | 7.47E-01 | 0.1 | 6.35E-01 | Uncharacterized protein |
| AT5G52250 | 0.4 | 1.03E-02 | 0.6 | 5.79E-04 | -0.2 | 1.05E-01 | -0.4 | 1.96E-01 | -1.1 | 3.86E-03 | 0.7 | 9.61E-02 | WD repeat-containing protein RUP1 |
| AT3G13175 | 0.4 | 1.26E-02 | 0.3 | 2.62E-02 | 0.0 | 8.20E-01 | -0.3 | 2.62E-01 | -0.1 | 7.67E-01 | -0.2 | 6.07E-01 | At3g13171 |
| AT3G20070 | 0.4 | 1.29E-03 | 0.2 | 1.43E-02 | 0.1 | 2.81E-01 | -0.4 | 3.57E-02 | -0.4 | 6.72E-02 | 0.0 | 9.40E-01 | Titan9 |
| AT5G13470 | 0.4 | 3.87E-03 | 0.1 | 4.20E-01 | 0.3 | 4.31E-02 | 0.0 | 8.92E-01 | 0.1 | 6.50E-01 | -0.1 | 7.84E-01 | Uncharacterized protein |
| AT5G13320 | 0.4 | 3.03E-01 | -0.8 | 9.33E-02 | 1.1 | 3.80E-02 | 1.1 | 2.34E-03 | 0.2 | 6.71E-01 | 0.9 | 2.01E-02 | Auxin-responsive GH3 family protein |
| AT3G17670 | 0.4 | 1.24E-02 | 0.5 | 2.97E-03 | -0.1 | 4.04E-01 | -0.1 | 7.53E-01 | 0.2 | 3.51E-01 | -0.2 | 2.48E-01 | Tetratricopeptide repeat -containing protein |
| AT5G07630 | 0.4 | 3.32E-03 | 0.2 | 1.22E-01 | 0.2 | 1.10E-01 | 0.1 | 7.57E-01 | 0.1 | 5.20E-01 | -0.1 | 7.97E-01 | Protein RFT1 homolog |
| AT1G52550 | 0.4 | 1.42E-03 | 0.0 | 9.04E-01 | 0.4 | 5.16E-03 | -0.1 | 6.88E-01 | 0.1 | 8.46E-01 | -0.2 | 5.85E-01 | Transmembrane protein |
| AT5G04910 | 0.4 | 1.73E-03 | 0.0 | 8.31E-01 | 0.3 | 8.48E-03 | -0.2 | 2.79E-01 | -0.4 | 7.13E-02 | 0.2 | 5.27E-01 | DNA repair REX1-B protein |
| AT3G45620 | 0.4 | 1.79E-03 | -0.1 | 4.40E-01 | 0.4 | 1.85E-03 | 0.3 | 5.11E-02 | 0.1 | 5.01E-01 | 0.2 | 4.12E-01 | Transducin |
| AT3G27420 | 0.4 | 2.00E-03 | 0.2 | 2.83E-02 | 0.1 | 2.44E-01 | -0.4 | 1.05E-01 | -0.2 | 5.19E-01 | -0.2 | 5.56E-01 | NET domain-containing protein |
| AT3G12820 | 0.4 | 4.18E-02 | -0.5 | 2.86E-02 | 0.8 | 3.73E-03 | -0.8 | 9.04E-02 | -1.4 | 1.91E-02 | 0.5 | 4.39E-01 | Myb domain protein 10 |
| AT1G31910 | 0.4 | 7.16E-04 | 0.0 | 5.80E-01 | 0.3 | 4.37E-03 | 0.3 | 1.21E-02 | 0.2 | 1.06E-01 | 0.1 | 6.54E-01 | Uncharacterized protein At1g31910 |
| AT3G45740 | 0.4 | 1.44E-03 | 0.2 | 5.07E-02 | 0.2 | 1.16E-01 | 0.1 | 8.07E-01 | 0.0 | 9.50E-01 | 0.0 | 9.10E-01 | Hydrolase family protein |
| AT3G29030 | 0.4 | 5.79E-02 | 1.0 | 3.70E-04 | -0.6 | 1.23E-02 | -0.1 | 4.61E-01 | 0.1 | 6.80E-01 | -0.3 | 2.99E-01 | Expansin |
| AT5G04440 | 0.4 | 1.68E-04 | 0.1 | 5.00E-02 | 0.2 | 9.33E-03 | 0.1 | 5.40E-01 | 0.0 | 7.49E-01 | 0.0 | 9.22E-01 | RAP release 2, galactose-binding-like domain protein, putative |
| AT1G26750 | 0.4 | 5.53E-03 | 0.4 | 6.58E-03 | 0.0 | 9.76E-01 | 0.6 | 3.12E-03 | 0.4 | 7.04E-02 | 0.2 | 3.64E-01 | T24P13.13 |
| AT3G56830 | 0.4 | 1.67E-02 | 0.4 | 1.55E-02 | 0.0 | 8.93E-01 | -0.3 | 4.11E-01 | -0.2 | 5.34E-01 | 0.0 | 9.58E-01 | At3g56830 ) |
| AT3G11230 | 0.4 | 6.65E-02 | 0.1 | 7.19E-01 | 0.3 | 2.10E-01 | -0.1 | 8.13E-01 | -0.1 | 7.99E-01 | 0.0 | 9.70E-01 | AT3G11230 protein |
| AT3G61415 | 0.4 | 4.49E-04 | 0.4 | 3.77E-04 | 0.0 | 7.04E-01 | -0.1 | 7.17E-01 | -0.2 | 4.54E-01 | 0.1 | 7.71E-01 | SK21 |
| AT1G12845 | 0.4 | 6.89E-02 | 0.4 | 4.60E-02 | -0.1 | 8.21E-01 | -0.9 | 1.07E-03 | -0.2 | 3.08E-01 | -0.6 | 3.91E-02 | F13K23.10 protein |
| AT3G60810 | 0.4 | 6.46E-03 | 0.3 | 1.99E-02 | 0.1 | 6.78E-01 | -0.2 | 4.45E-01 | 0.0 | 9.50E-01 | -0.2 | 6.33E-01 | DUF1499 family protein |
| AT4G12840 | 0.4 | 1.24E-02 | 0.2 | 1.74E-01 | 0.2 | 2.58E-01 | 0.3 | 4.74E-01 | 0.2 | 7.19E-01 | 0.1 | 8.48E-01 | GTPase Der |
| AT4G14385 | 0.4 | 2.62E-03 | 0.3 | 1.27E-02 | 0.1 | 5.14E-01 | -0.2 | 2.85E-01 | -0.3 | 3.70E-02 | 0.2 | 3.58E-01 | Histone acetyltransferase subunit NuA4-domain protein |
| AT1G12700 | 0.4 | 1.14E-02 | 0.2 | 1.33E-01 | 0.2 | 3.07E-01 | 0.4 | 3.26E-01 | 0.2 | 6.40E-01 | 0.1 | 7.78E-01 | RNA processing FACTOR |
| AT5G05710 | 0.4 | 1.69E-03 | 0.2 | 3.35E-02 | 0.1 | 1.84E-01 | -0.2 | 4.35E-01 | -0.5 | 2.44E-02 | 0.3 | 2.25E-01 | PH domain-containing protein |
| AT1G72050 | 0.4 | 1.31E-03 | 0.2 | 2.24E-02 | 0.1 | 1.84E-01 | 0.3 | 5.66E-02 | 0.1 | 5.05E-01 | 0.1 | 4.21E-01 | Transcription factor IIIA |
| AT4G38062 | 0.4 | 2.52E-02 | 0.6 | 2.23E-03 | -0.2 | 1.97E-01 | 0.1 | 6.62E-01 | 0.1 | 7.30E-01 | 0.0 | 9.97E-01 | Uncharacterized protein At4g38062 |
| AT3G01980 | 0.4 | 1.71E-03 | 0.4 | 3.04E-03 | 0.0 | 8.96E-01 | -0.1 | 7.46E-01 | 0.1 | 7.67E-01 | -0.1 | 5.74E-01 | F28J7.36 protein -binding Rossmann-fold superfamily protein) |
| AT3G24080 | 0.4 | 1.02E-02 | 0.3 | 4.41E-02 | 0.1 | 5.31E-01 | 0.5 | 1.92E-03 | 0.2 | 1.70E-01 | 0.3 | 1.85E-01 | Kri1-like C-terminal domain-containing protein |
| AT2G25270 | 0.4 | 6.44E-03 | 0.2 | 9.65E-02 | 0.2 | 2.48E-01 | -0.1 | 8.70E-01 | 0.0 | 9.93E-01 | -0.1 | 9.03E-01 | Transmembrane protein |
| AT3G07640 | 0.4 | 4.83E-04 | 0.3 | 2.07E-03 | 0.1 | 5.38E-01 | 0.0 | 9.97E-01 | 0.1 | 7.44E-01 | -0.1 | 7.89E-01 | MLP3.9 protein |
| AT3G52220 | 0.4 | 5.84E-04 | 0.1 | 3.63E-01 | 0.3 | 6.18E-03 | 0.1 | 7.42E-01 | 0.1 | 5.20E-01 | -0.1 | 8.10E-01 | hypothetical protein |
| AT1G25400 | 0.4 | 8.49E-02 | -0.1 | 8.12E-01 | 0.4 | 1.01E-01 | -0.3 | 1.21E-01 | -1.0 | 5.84E-04 | 0.7 | 3.97E-02 | Transmembrane protein |
| AT4G13590 | 0.4 | 2.35E-04 | 0.2 | 2.89E-02 | 0.2 | 2.16E-02 | -0.1 | 6.26E-01 | 0.1 | 7.32E-01 | -0.1 | 4.39E-01 | GDT1 family protein |
| AT1G75550 | 0.4 | 4.04E-02 | 0.6 | 6.48E-03 | -0.2 | 3.40E-01 | 0.0 | 9.07E-01 | 0.3 | 4.94E-01 | -0.3 | 4.21E-01 | Glycine-rich protein |
| AT1G65950 | 0.4 | 3.12E-03 | 0.2 | 1.21E-01 | 0.2 | 1.10E-01 | -0.1 | 7.79E-01 | 0.2 | 5.10E-01 | -0.3 | 4.20E-01 | ABC1 atypical kinase-like domain-containing protein |
| AT5G63135 | 0.4 | 1.09E-01 | -0.1 | 8.43E-01 | 0.4 | 1.40E-01 | 0.3 | 3.53E-01 | 0.2 | 7.14E-01 | 0.1 | 7.82E-01 | Uncharacterized protein |
| AT1G53450 | 0.4 | 2.75E-03 | 0.1 | 1.72E-01 | 0.2 | 7.81E-02 | -0.1 | 6.84E-01 | -0.1 | 7.55E-01 | 0.0 | 9.90E-01 | At1g53450 |
| AT5G28020 | 0.4 | 5.39E-04 | 0.5 | 1.82E-04 | -0.1 | 2.71E-01 | -0.4 | 1.17E-02 | -0.2 | 3.45E-01 | -0.2 | 2.97E-01 | Cysteine synthase D2 |
| AT3G19570 | 0.4 | 4.29E-03 | 0.1 | 3.03E-01 | 0.3 | 6.52E-02 | -0.2 | 6.28E-01 | -0.1 | 8.43E-01 | -0.1 | 8.96E-01 | SNOWY COTYLEDON protein |
| AT1G34355 | 0.4 | 2.05E-02 | 0.5 | 5.89E-03 | -0.1 | 4.92E-01 | 0.2 | 5.49E-01 | 0.1 | 7.92E-01 | 0.1 | 8.62E-01 | Forkhead-associated domain-containing protein |

| | | | | | | | | | | | | | |
|---|---|---|---|---|---|---|---|---|---|---|---|---|---|
| AT3G15000 | 0.4 | 4.55E-03 | 0.2 | 5.02E-02 | 0.1 | 2.69E-01 | 0.5 | 4.60E-04 | 0.2 | 1.67E-01 | 0.3 | 8.85E-02 | RIP1 |
| AT1G13770 | 0.4 | 2.88E-03 | 0.1 | 2.95E-01 | 0.3 | 4.65E-02 | -0.1 | 5.43E-01 | -0.1 | 7.67E-01 | -0.1 | 8.97E-01 | Protein root UVB sensitive 3 |
| AT5G10810 | 0.4 | 5.24E-04 | 0.3 | 6.33E-03 | 0.1 | 2.37E-01 | 0.2 | 4.05E-01 | 0.0 | 9.24E-01 | 0.2 | 4.55E-01 | Enhancer of rudimentary homolog |
| AT2G47490 | 0.4 | 4.77E-05 | 0.1 | 2.86E-01 | 0.3 | 7.43E-04 | 0.1 | 5.47E-01 | 0.4 | 5.84E-02 | -0.3 | 2.72E-01 | NDT1 |
| AT1G24996 | 0.4 | 3.32E-01 | -0.5 | 3.16E-01 | 0.8 | 1.14E-01 | 0.8 | 4.06E-01 | 0.3 | 8.36E-01 | 0.5 | 5.94E-01 | F5A9.10 |
| AT2G02050 | 0.4 | 4.72E-03 | 0.4 | 4.08E-03 | 0.0 | 8.28E-01 | 0.2 | 4.18E-01 | 0.1 | 6.92E-01 | 0.1 | 8.70E-01 | NADH dehydrogenase 1 beta subcomplex subunit 7 |
| AT1G73920 | 0.4 | 2.24E-03 | 0.2 | 5.53E-02 | 0.2 | 1.32E-01 | -0.1 | 4.93E-01 | -0.4 | 2.38E-02 | 0.3 | 1.74E-01 | At1g73920 |
| AT5G49220 | 0.4 | 9.65E-04 | 0.2 | 6.16E-02 | 0.2 | 6.06E-02 | 0.2 | 2.64E-01 | 0.2 | 1.56E-01 | -0.1 | 7.92E-01 | hypothetical protein |
| AT2G06950 | 0.4 | 2.79E-02 | 0.0 | 9.97E-01 | 0.4 | 5.24E-02 | 0.1 | 7.40E-01 | -0.2 | 3.33E-01 | 0.3 | 2.47E-01 | NA |
| AT4G07455 | 0.4 | 1.46E-01 | 0.1 | 7.61E-01 | 0.3 | 3.56E-01 | 0.1 | 8.89E-01 | -0.1 | 9.24E-01 | 0.1 | 8.52E-01 | Uncharacterized protein |
| AT1G58225 | 0.4 | 1.09E-01 | -0.3 | 3.33E-01 | 0.6 | 4.90E-02 | -0.4 | 3.37E-01 | -0.7 | 1.51E-01 | 0.3 | 7.02E-01 | Transmembrane protein |
| AT1G06240 | 0.4 | 5.25E-03 | 0.2 | 7.27E-02 | 0.2 | 2.51E-01 | -0.2 | 1.46E-01 | -0.1 | 4.44E-01 | -0.1 | 7.32E-01 | DUF455 family protein |
| AT3G60360 | 0.4 | 2.70E-02 | 0.4 | 3.47E-02 | 0.0 | 9.66E-01 | 0.5 | 7.15E-03 | 0.1 | 6.89E-01 | 0.4 | 8.76E-02 | Probable U3 small nucleolar RNA-associated protein 11 |
| AT2G38450 | 0.4 | 2.46E-02 | 0.1 | 5.18E-01 | 0.3 | 1.45E-01 | 0.2 | 4.56E-01 | -0.1 | 8.81E-01 | 0.3 | 4.43E-01 | At2g38450 |
| AT1G60560 | 0.4 | 2.11E-03 | 0.2 | 7.36E-02 | 0.2 | 1.21E-01 | -0.1 | 6.65E-01 | -0.1 | 6.27E-01 | 0.0 | 9.32E-01 | SWIM zinc finger family protein |
| AT3G51530 | 0.4 | 3.21E-03 | 0.3 | 3.21E-02 | 0.1 | 3.33E-01 | 0.0 | 9.05E-01 | 0.0 | 9.77E-01 | 0.0 | 9.57E-01 | F-box |
| AT1G74530 | 0.4 | 1.17E-03 | 0.2 | 8.19E-02 | 0.2 | 6.00E-02 | 0.0 | 9.79E-01 | 0.1 | 7.61E-01 | -0.1 | 7.71E-01 | At1g74530 |
| AT3G49320 | 0.4 | 4.01E-02 | 0.3 | 1.52E-01 | 0.1 | 5.90E-01 | 0.9 | 5.95E-03 | 0.1 | 9.15E-01 | 0.9 | 1.58E-02 | Metal-dependent protein hydrolase |
| AT5G64680 | 0.4 | 1.09E-02 | 0.0 | 8.92E-01 | 0.4 | 2.37E-02 | 0.2 | 4.23E-01 | 0.0 | 9.80E-01 | 0.2 | 5.40E-01 | hypothetical protein |
| AT1G03110 | 0.4 | 2.02E-03 | 0.4 | 3.21E-03 | 0.0 | 9.42E-01 | 0.3 | 8.31E-02 | 0.1 | 6.35E-01 | 0.2 | 4.10E-01 | tRNA -)-methyltransferase non-catalytic subunit |
| AT1G78460 | 0.4 | 9.36E-03 | 0.7 | 4.27E-04 | -0.3 | 6.33E-02 | 0.3 | 1.63E-01 | -0.8 | 9.73E-03 | 1.1 | 2.41E-03 | At1g78460 |
| AT3G46110 | 0.4 | 3.33E-02 | 0.1 | 5.79E-01 | 0.3 | 1.70E-01 | 0.6 | 2.78E-02 | 0.1 | 9.01E-01 | 0.5 | 8.47E-02 | UPSTREAM OF FLC-like protein |
| AT4G34360 | 0.4 | 8.48E-02 | 0.5 | 3.20E-02 | -0.1 | 6.24E-01 | 0.6 | 7.55E-02 | 0.1 | 8.14E-01 | 0.5 | 2.37E-01 | Uncharacterized protein AT4g34360 |
| AT2G01420 | 0.4 | 5.10E-03 | 0.6 | 5.88E-04 | -0.2 | 1.45E-01 | -0.2 | 5.29E-02 | -0.2 | 6.18E-02 | 0.0 | 9.34E-01 | Auxin efflux carrier component |
| AT1G78030 | 0.4 | 5.92E-02 | 0.6 | 1.09E-02 | -0.2 | 4.10E-01 | 0.3 | 6.08E-01 | 0.4 | 5.01E-01 | -0.1 | 8.94E-01 | Uncharacterized protein |
| AT3G47600 | 0.4 | 7.31E-03 | 0.1 | 5.61E-01 | 0.3 | 5.11E-02 | -0.1 | 6.68E-01 | 0.1 | 8.56E-01 | -0.1 | 6.42E-01 | Myb domain protein 94 |
| AT2G43465 | 0.4 | 1.35E-01 | -0.1 | 7.30E-01 | 0.5 | 1.40E-01 | -0.1 | 7.92E-01 | -0.8 | 1.03E-01 | 0.7 | 2.45E-01 | RNA-binding ASCH domain protein |
| AT3G19650 | 0.4 | 1.65E-03 | 0.3 | 2.17E-02 | 0.1 | 2.45E-01 | 0.2 | 3.66E-01 | 0.1 | 6.14E-01 | 0.0 | 8.75E-01 | At3g19650 |
| AT1G69380 | 0.4 | 8.53E-03 | 0.2 | 1.38E-01 | 0.2 | 2.28E-01 | 0.4 | 1.16E-01 | 0.2 | 5.93E-01 | 0.2 | 4.99E-01 | Protein RETARDED ROOT GROWTH, mitochondrial |
| AT5G64850 | 0.4 | 1.73E-02 | 0.4 | 2.38E-02 | 0.0 | 9.48E-01 | 0.4 | 2.08E-01 | 0.3 | 4.19E-01 | 0.1 | 8.55E-01 | Emb\|CAB61744.1 |
| AT5G46500 | 0.4 | 2.00E-02 | -0.1 | 4.68E-01 | 0.5 | 1.89E-02 | -0.7 | 1.26E-01 | -0.8 | 7.41E-02 | 0.2 | 7.47E-01 | Protein VARIATION IN COMPOUND TRIGGERED ROOT growth protein |
| AT3G05410 | 0.4 | 4.22E-03 | 0.4 | 2.70E-03 | 0.0 | 7.12E-01 | 0.0 | 8.76E-01 | 0.0 | 9.47E-01 | 0.1 | 8.56E-01 | Photosystem II reaction center PsbP family protein |
| AT5G60030 | 0.4 | 4.59E-02 | 0.4 | 6.14E-02 | 0.0 | 9.44E-01 | 0.2 | 4.54E-01 | -0.1 | 8.49E-01 | 0.2 | 4.52E-01 | Uncharacterized protein |
| AT3G07750 | 0.4 | 1.82E-02 | 0.2 | 1.77E-01 | 0.2 | 3.15E-01 | 0.1 | 6.32E-01 | -0.1 | 6.28E-01 | 0.3 | 3.44E-01 | 3'-5'-exoribonuclease family protein |
| AT5G59790 | 0.4 | 2.61E-03 | 0.0 | 9.47E-01 | 0.4 | 8.21E-03 | -0.1 | 5.75E-01 | -0.2 | 2.83E-01 | 0.1 | 7.02E-01 | UPSTREAM OF FLC protein |
| AT1G49032 | 0.4 | 6.14E-02 | 0.3 | 1.07E-01 | 0.0 | 8.54E-01 | -0.3 | 3.20E-01 | -0.7 | 1.11E-02 | 0.5 | 1.79E-01 | Transmembrane protein |
| AT5G52030 | 0.4 | 3.26E-04 | 0.2 | 9.06E-03 | 0.1 | 1.04E-01 | -0.1 | 7.60E-01 | 0.1 | 4.96E-01 | -0.2 | 4.01E-01 | TraB family protein |
| AT5G02050 | 0.4 | 2.50E-02 | 0.4 | 1.73E-02 | 0.0 | 8.13E-01 | 0.7 | 3.91E-04 | 0.4 | 3.95E-02 | 0.3 | 5.70E-02 | hypothetical protein |
| AT1G27390 | 0.4 | 6.57E-03 | 0.3 | 2.29E-02 | 0.1 | 6.00E-01 | 0.6 | 8.65E-04 | 0.3 | 7.58E-02 | 0.3 | 1.97E-01 | Translocase outer membrane 20-2 |
| AT1G16020 | 0.4 | 9.43E-04 | 0.3 | 4.01E-03 | 0.1 | 5.43E-01 | 0.3 | 1.45E-01 | 0.2 | 4.46E-01 | 0.1 | 7.28E-01 | Uncharacterized protein At1g16020 ) |
| AT2G40765 | 0.4 | 1.87E-02 | 0.5 | 4.50E-03 | -0.1 | 4.10E-01 | 0.1 | 8.10E-01 | 0.0 | 9.13E-01 | 0.0 | 9.59E-01 | Cytochrome b-c1 complex subunit 10, mitochondrial |
| AT3G08943 | 0.4 | 6.32E-04 | 0.2 | 1.46E-02 | 0.1 | 1.20E-01 | 0.4 | 4.30E-03 | -0.1 | 7.58E-01 | 0.4 | 1.87E-02 | NA |
| AT3G01210 | 0.4 | 2.13E-02 | 0.0 | 8.69E-01 | 0.4 | 6.24E-02 | -0.2 | 4.55E-01 | 0.2 | 3.41E-01 | -0.4 | 1.16E-01 | At3g01210 |
| AT5G17840 | 0.4 | 3.75E-04 | 0.3 | 1.31E-03 | 0.0 | 6.03E-01 | 0.1 | 7.93E-01 | 0.0 | 9.98E-01 | 0.1 | 8.62E-01 | hypothetical protein |
| AT5G49010 | 0.4 | 3.52E-03 | 0.4 | 4.79E-03 | 0.0 | 9.93E-01 | 0.5 | 4.15E-02 | 0.1 | 7.13E-01 | 0.4 | 2.12E-01 | DNA replication complex GINS protein SLD5 |
| AT2G37975 | 0.4 | 1.03E-02 | 0.4 | 9.92E-03 | 0.0 | 9.16E-01 | 0.0 | 9.02E-01 | 0.0 | 9.66E-01 | 0.0 | 9.62E-01 | Yos1-like protein |
| AT5G12980 | 0.4 | 5.32E-03 | 0.4 | 3.81E-03 | 0.0 | 7.73E-01 | -0.2 | 3.48E-01 | 0.0 | 9.17E-01 | -0.2 | 5.63E-01 | Cell differentiation, Rcd1-like protein |
| AT2G31840 | 0.4 | 2.30E-03 | 0.5 | 7.83E-04 | -0.1 | 4.13E-01 | 0.4 | 1.41E-02 | -0.1 | 7.75E-01 | 0.5 | 3.28E-02 | Uncharacterized protein |
| AT1G11150 | 0.4 | 1.55E-03 | 0.3 | 1.46E-02 | 0.1 | 3.38E-01 | -0.2 | 4.53E-01 | 0.2 | 3.17E-01 | -0.4 | 1.50E-01 | NA |
| AT1G58590 | 0.4 | 2.85E-01 | -0.1 | 7.66E-01 | 0.5 | 2.59E-01 | 0.6 | 5.89E-02 | 0.2 | 6.32E-01 | 0.4 | 3.17E-01 | NA |
| AT3G60320 | 0.4 | 1.76E-03 | 0.2 | 2.61E-02 | 0.1 | 2.00E-01 | -0.4 | 4.22E-02 | -0.6 | 6.24E-03 | 0.2 | 3.90E-01 | Nitrate regulatory gene2 protein |
| AT3G55160 | 0.4 | 2.49E-03 | -0.1 | 2.19E-01 | 0.5 | 1.89E-03 | 0.1 | 8.67E-01 | 0.1 | 8.47E-01 | 0.0 | 9.72E-01 | hypothetical protein |
| AT3G56140 | 0.4 | 8.99E-03 | 0.1 | 6.52E-01 | 0.3 | 3.82E-02 | -0.2 | 3.25E-01 | -0.3 | 2.05E-01 | 0.1 | 7.82E-01 | DUF399 family protein, putative |
| AT3G59880 | 0.4 | 2.82E-02 | 0.1 | 6.84E-01 | 0.3 | 1.22E-01 | 0.3 | 3.55E-01 | -0.4 | 4.38E-01 | 0.7 | 1.16E-01 | Uncharacterized protein F24G16.150 |
| AT1G59600 | 0.4 | 2.21E-04 | 0.2 | 4.17E-02 | 0.2 | 1.92E-02 | 0.3 | 1.20E-01 | 0.1 | 8.43E-01 | 0.2 | 2.99E-01 | ZCW7 |
| AT1G29120 | 0.4 | 6.36E-04 | 0.1 | 5.23E-01 | 0.3 | 5.11E-03 | 0.1 | 6.79E-01 | 0.1 | 6.71E-01 | 0.0 | 9.65E-01 | F28N24.17 protein |
| AT1G32520 | 0.4 | 5.16E-02 | 0.1 | 5.28E-01 | 0.3 | 2.45E-01 | -0.5 | 7.61E-02 | 0.1 | 6.32E-01 | -0.6 | 7.89E-02 | TLDc domain protein |
| AT2G46330 | 0.4 | 2.81E-03 | 0.6 | 3.33E-04 | -0.2 | 1.13E-01 | -0.4 | 9.84E-02 | -1.0 | 7.31E-04 | 0.6 | 5.22E-02 | Arabinogalactan protein 16 |
| AT5G61880 | 0.4 | 1.06E-02 | 0.3 | 7.23E-02 | 0.1 | 4.21E-01 | 0.1 | 8.04E-01 | 0.3 | 3.10E-01 | -0.2 | 4.86E-01 | Protein Transporter, Pam16 |
| AT3G06433 | 0.4 | 4.17E-02 | 0.1 | 6.00E-01 | 0.3 | 2.00E-01 | 0.1 | 8.82E-01 | 0.0 | 9.51E-01 | 0.1 | 8.75E-01 | NA |
| AT5G08710 | 0.4 | 1.30E-02 | 0.1 | 7.46E-01 | 0.3 | 5.46E-02 | 0.1 | 8.38E-01 | 0.0 | 9.82E-01 | 0.1 | 9.09E-01 | Regulator of chromosome condensation family protein |
| AT3G14860 | 0.4 | 5.42E-04 | 0.2 | 1.63E-02 | 0.2 | 9.99E-02 | 0.2 | 1.98E-01 | 0.0 | 8.95E-01 | 0.2 | 2.91E-01 | NHL domain-containing protein |
| AT1G27385 | 0.4 | 6.14E-04 | 0.3 | 5.01E-03 | 0.1 | 3.22E-01 | 0.0 | 9.37E-01 | 0.0 | 9.31E-01 | 0.0 | 8.96E-01 | At1g27385 |
| AT3G08880 | 0.4 | 6.67E-03 | 0.3 | 3.58E-02 | 0.1 | 5.01E-01 | -0.6 | 6.68E-02 | -0.3 | 4.91E-01 | -0.3 | 4.83E-01 | Kinetochore protein SPC24 homolog |
| AT2G45640 | 0.4 | 1.58E-03 | 0.2 | 7.72E-02 | 0.2 | 7.29E-02 | 0.1 | 7.38E-01 | 0.0 | 8.82E-01 | 0.1 | 7.13E-01 | SIN3 associated polypeptide P18 |
| AT4G08685 | 0.4 | 1.28E-03 | 0.4 | 2.53E-03 | 0.0 | 8.45E-01 | 0.1 | 6.65E-01 | 0.2 | 3.39E-01 | -0.1 | 6.75E-01 | SAH7 |
| AT1G70100 | 0.4 | 4.04E-04 | 0.3 | 7.95E-03 | 0.1 | 1.47E-01 | -0.2 | 3.11E-01 | -0.2 | 5.43E-01 | -0.1 | 8.54E-01 | Neurofilament heavy protein |
| AT1G80245 | 0.4 | 4.51E-03 | 0.1 | 4.41E-01 | 0.3 | 4.79E-02 | 0.4 | 2.25E-01 | -0.1 | 8.87E-01 | 0.4 | 2.14E-01 | At1g80245 component) |
| AT1G27510 | 0.4 | 8.93E-05 | 0.2 | 9.26E-03 | 0.2 | 1.91E-02 | -0.2 | 7.67E-02 | -0.3 | 1.57E-02 | 0.1 | 5.05E-01 | Protein EXECUTER 2, chloroplastic |
| AT5G06430 | 0.4 | 1.59E-03 | 0.1 | 4.39E-01 | 0.3 | 2.05E-02 | 0.3 | 1.98E-01 | 0.1 | 8.36E-01 | 0.3 | 4.86E-01 | Thioredoxin domain-containing protein |
| AT3G49840 | 0.4 | 1.76E-02 | 0.1 | 4.57E-01 | 0.3 | 1.43E-01 | 0.2 | 6.47E-01 | 0.2 | 6.95E-01 | 0.0 | 9.98E-01 | Transmembrane protein |
| AT4G21705 | 0.4 | 1.42E-03 | 0.1 | 6.21E-01 | 0.3 | 1.16E-02 | 0.3 | 7.76E-02 | 0.3 | 1.62E-01 | 0.0 | 9.09E-01 | Pentatricopeptide repeat-containing protein At4g21705, mitochondrial |
| AT2G21320 | 0.4 | 4.82E-02 | -0.4 | 6.91E-02 | 0.8 | 5.48E-03 | | | | | | | |
| | | | | | | | | | | | | | hypothetical protein |
| AT5G26910 | 0.4 | 4.18E-04 | 0.2 | 3.28E-02 | 0.2 | 4.36E-02 | -0.2 | 2.25E-01 | -0.1 | 5.79E-01 | -0.1 | 7.55E-01 | At5g26910 |
| AT1G13690 | 0.4 | 2.61E-03 | 0.4 | 4.58E-03 | 0.0 | 8.92E-01 | 0.1 | 6.87E-01 | -0.1 | 7.59E-01 | 0.2 | 4.85E-01 | AtE1 |
| AT5G67260 | 0.4 | 1.21E-03 | 0.4 | 8.99E-04 | 0.0 | 7.09E-01 | -0.2 | 6.14E-02 | -0.2 | 1.04E-01 | 0.0 | 9.82E-01 | CYCLIN D32 |
| AT4G11980 | 0.4 | 8.77E-03 | 0.4 | 1.65E-02 | 0.0 | 8.58E-01 | 0.2 | 3.64E-01 | 0.1 | 6.99E-01 | 0.1 | 7.94E-01 | NUDX14 |
| AT3G25580 | 0.4 | 1.09E-02 | 0.2 | 1.24E-01 | 0.2 | 2.87E-01 | 0.1 | 6.48E-01 | 0.2 | 5.10E-01 | -0.1 | 8.67E-01 | Thioredoxin domain-containing protein |
| AT1G49490 | 0.4 | 2.69E-02 | 0.9 | 3.23E-04 | -0.5 | 2.46E-02 | -0.4 | 1.30E-01 | -0.4 | 1.61E-01 | 0.0 | 9.93E-01 | Leucine-rich repeat-containing N-terminal plant-type domain-containing protein |
| AT4G18280 | 0.4 | 2.61E-03 | 0.1 | 2.14E-01 | 0.3 | 5.53E-02 | -0.6 | 1.78E-02 | -0.3 | 3.28E-01 | -0.3 | 3.36E-01 | Uncharacterized protein |
| AT1G71430 | 0.4 | 3.28E-02 | 0.1 | 5.49E-01 | 0.3 | 1.79E-01 | -0.1 | 8.61E-01 | -0.3 | 4.26E-01 | 0.2 | 5.95E-01 | Uncharacterized protein |
| AT4G39900 | 0.4 | 1.65E-03 | 0.3 | 5.67E-03 | 0.1 | 6.17E-01 | -0.4 | 3.06E-03 | -0.3 | 2.71E-02 | -0.1 | 6.25E-01 | Transmembrane protein |
| AT5G52820 | 0.4 | 1.97E-04 | 0.2 | 2.05E-02 | 0.2 | 2.57E-02 | 0.3 | 2.51E-02 | 0.1 | 8.08E-01 | 0.3 | 1.84E-01 | Notchless protein homolog |

| | | | | | | | | | | | | | |
|---|---|---|---|---|---|---|---|---|---|---|---|---|---|
| AT3G23290 | 0.4 | 7.31E-04 | 0.2 | 5.33E-02 | 0.2 | 5.37E-02 | 0.3 | 2.22E-01 | 0.3 | 1.97E-01 | 0.0 | 9.05E-01 | hypothetical protein |
| AT2G23690 | 0.4 | 4.05E-02 | 0.8 | 1.04E-03 | -0.4 | 6.75E-02 | -0.7 | 5.39E-02 | -0.6 | 1.11E-01 | -0.1 | 8.96E-01 | HTH-type transcriptional regulator |
| AT3G01050 | 0.4 | 3.81E-02 | 0.0 | 8.02E-01 | 0.3 | 1.11E-01 | 0.1 | 8.25E-01 | 0.0 | 9.44E-01 | 0.1 | 8.27E-01 | Membrane-anchored ubiquitin-fold protein 1 |
| AT2G28150 | 0.4 | 1.49E-03 | 0.3 | 7.44E-03 | 0.1 | 5.02E-01 | -0.4 | 7.47E-03 | -0.3 | 1.87E-02 | 0.0 | 9.34E-01 | UPSTREAM OF FLC protein |
| AT5G07590 | 0.4 | 3.20E-04 | 0.4 | 9.94E-04 | 0.0 | 6.68E-01 | 0.1 | 4.37E-01 | -0.2 | 2.73E-01 | 0.3 | 1.31E-01 | WD repeat-containing protein WRAP73 |
| AT1G13220 | 0.4 | 3.87E-04 | 0.2 | 6.13E-02 | 0.2 | 2.35E-02 | 0.4 | 4.17E-02 | 0.3 | 1.43E-01 | 0.1 | 7.89E-01 | Nuclear matrix constituent protein-like protein |
| AT5G66052 | 0.4 | 2.82E-03 | 0.2 | 7.83E-02 | 0.2 | 1.57E-01 | -0.4 | 7.64E-02 | -0.4 | 1.00E-01 | 0.0 | 9.77E-01 | Transmembrane protein |
| AT5G52840 | 0.4 | 4.64E-03 | 0.4 | 3.36E-03 | 0.0 | 7.30E-01 | 0.0 | 7.01E-01 | 0.1 | 7.58E-01 | -0.1 | 5.54E-01 | NADH-ubiquinone oxidoreductase-like protein |
| AT3G44750 | 0.4 | 9.28E-02 | 0.6 | 3.21E-02 | -0.2 | 5.65E-01 | 0.8 | 8.29E-05 | 0.4 | 3.49E-02 | 0.4 | 4.89E-02 | Histone deacetylase 3 |
| AT2G42310 | 0.4 | 6.57E-04 | 0.3 | 4.13E-03 | 0.1 | 3.87E-01 | -0.3 | 1.69E-01 | -0.2 | 4.29E-01 | -0.1 | 8.18E-01 | Uncharacterized protein |
| AT3G02885 | 0.4 | 5.50E-02 | 0.8 | 1.48E-03 | -0.4 | 7.98E-02 | -1.3 | 7.74E-03 | 0.4 | 2.72E-01 | -1.8 | 4.03E-03 | GAST1 protein homolog 5 |
| AT5G65400 | 0.4 | 8.46E-03 | -0.2 | 1.25E-01 | 0.6 | 3.22E-03 | -0.4 | 1.14E-01 | -0.3 | 1.67E-01 | 0.0 | 9.70E-01 | Similarity to unknown protein |
| AT1G67170 | 0.4 | 6.81E-04 | 0.2 | 2.27E-02 | 0.2 | 1.07E-01 | 0.4 | 3.65E-02 | 0.1 | 8.42E-01 | 0.3 | 1.75E-01 | Protein FLX-like 2 |
| AT1G14140 | 0.4 | 1.07E-03 | 0.4 | 2.92E-03 | 0.0 | 7.22E-01 | -0.1 | 4.19E-01 | -0.2 | 2.25E-01 | 0.1 | 7.52E-01 | Putative mitochondrial uncoupling protein |
| AT3G23250 | 0.4 | 2.75E-01 | -0.6 | 1.88E-01 | 1.0 | 6.60E-02 | 2.5 | 6.70E-06 | 1.1 | 2.67E-02 | 1.4 | 1.11E-03 | Myb domain protein 15 |
| AT3G54120 | 0.4 | 1.39E-02 | 0.1 | 4.28E-01 | 0.3 | 1.30E-01 | 0.2 | 5.13E-01 | 0.1 | 7.46E-01 | 0.1 | 8.88E-01 | Reticulon-like protein B12 |
| AT3G49940 | 0.4 | 1.52E-01 | 0.4 | 1.76E-01 | 0.0 | 9.82E-01 | 1.5 | 1.64E-06 | 0.0 | 9.44E-01 | 1.4 | 9.44E-05 | LOB domain-containing protein 38 |
| AT3G06790 | 0.4 | 6.37E-04 | 0.3 | 3.18E-03 | 0.1 | 4.80E-01 | 0.2 | 1.49E-01 | 0.1 | 7.09E-01 | 0.2 | 5.24E-01 | Multiple organellar RNA editing factor 3, mitochondrial |
| AT2G21140 | 0.4 | 4.01E-02 | 0.5 | 1.33E-02 | -0.1 | 5.44E-01 | 0.2 | 3.15E-01 | 0.2 | 5.79E-01 | 0.1 | 8.39E-01 | Proline-rich protein 2 |
| AT1G73320 | 0.4 | 2.44E-02 | 0.1 | 6.17E-01 | 0.3 | 1.28E-01 | 0.0 | 9.85E-01 | 0.0 | 9.35E-01 | 0.1 | 9.28E-01 | At1g73320 |
| AT2G37400 | 0.4 | 7.97E-04 | 0.2 | 6.16E-02 | 0.2 | 4.99E-02 | 0.4 | 3.53E-02 | 0.1 | 7.92E-01 | 0.4 | 1.71E-01 | Uncharacterized protein |
| AT2G39440 | 0.4 | 1.57E-02 | 0.2 | 2.20E-01 | 0.2 | 2.45E-01 | 0.0 | 9.12E-01 | 0.1 | 8.73E-01 | 0.0 | 9.51E-01 | Uncharacterized protein At2g39440 |
| AT1G80720 | 0.4 | 2.88E-02 | 0.2 | 2.23E-01 | 0.2 | 3.73E-01 | 0.7 | 8.64E-03 | 0.0 | 9.92E-01 | 0.7 | 1.72E-02 | At1g80720 |
| AT2G42975 | 0.4 | 8.03E-03 | 0.0 | 8.91E-01 | 0.4 | 1.70E-02 | -0.1 | 6.42E-01 | 0.4 | 4.03E-02 | -0.5 | 3.45E-02 | hypothetical protein |
| AT4G28740 | 0.4 | 4.21E-04 | 0.0 | 5.66E-01 | 0.5 | 7.33E-04 | 0.3 | 6.27E-02 | 0.2 | 2.11E-01 | 0.1 | 8.07E-01 | Uncharacterized protein AT4g28740 |
| AT1G54770 | 0.4 | 1.11E-02 | 0.4 | 1.90E-02 | 0.0 | 8.79E-01 | 0.2 | 6.04E-01 | -0.1 | 7.68E-01 | 0.3 | 4.77E-01 | hypothetical protein |
| AT3G49420 | 0.4 | 6.12E-03 | 0.1 | 4.44E-01 | 0.3 | 6.41E-02 | 0.3 | 2.89E-01 | 0.0 | 9.53E-01 | 0.3 | 3.29E-01 | At3g49420 |
| AT3G62940 | 0.4 | 1.59E-03 | 0.3 | 1.06E-02 | 0.1 | 3.85E-01 | 0.2 | 1.73E-01 | 0.1 | 4.68E-01 | 0.1 | 7.61E-01 | OTU domain-containing protein |
| AT5G02540 | 0.4 | 4.24E-02 | 0.9 | 6.46E-04 | -0.5 | 3.47E-02 | -0.1 | 8.01E-01 | -0.1 | 7.46E-01 | 0.0 | 9.26E-01 | NAD-binding Rossmann-fold superfamily protein |
| AT5G49400 | 0.4 | 3.75E-03 | 0.2 | 5.91E-02 | 0.2 | 2.16E-01 | -0.2 | 2.53E-01 | -0.4 | 4.38E-02 | 0.2 | 4.76E-01 | Similarity to unknown protein |
| AT5G11760 | 0.4 | 6.99E-04 | 0.3 | 7.56E-03 | 0.1 | 2.50E-01 | 0.0 | 9.39E-01 | -0.2 | 2.25E-01 | 0.2 | 2.44E-01 | hypothetical protein |
| AT5G61670 | 0.4 | 1.29E-03 | 0.5 | 3.48E-04 | -0.1 | 2.64E-01 | -0.2 | 2.24E-01 | 0.3 | 6.49E-02 | -0.4 | 2.19E-02 | Protein ORANGE, chloroplastic |
| AT1G10657 | 0.4 | 8.64E-03 | 0.8 | 9.91E-05 | -0.4 | 1.11E-02 | 0.1 | 7.68E-01 | 0.5 | 7.90E-02 | -0.4 | 2.12E-01 | Transmembrane protein |
| AT3G61280 | 0.4 | 9.43E-03 | -0.2 | 2.55E-01 | 0.6 | 4.67E-03 | 0.5 | 1.58E-01 | -0.4 | 3.96E-01 | 0.8 | 8.13E-02 | Uncharacterized protein T20K12.180 |
| AT5G20450 | 0.4 | 7.69E-02 | 0.2 | 3.95E-01 | 0.2 | 4.49E-01 | -0.2 | 6.89E-01 | 0.1 | 9.07E-01 | -0.2 | 6.65E-01 | At5g20450 |
| AT1G13740 | 0.4 | 1.89E-02 | 0.7 | 1.44E-03 | -0.3 | 1.65E-01 | 0.5 | 1.31E-01 | 0.1 | 8.65E-01 | 0.4 | 2.06E-01 | Ninja-family protein AFP2 |
| AT5G15230 | 0.4 | 1.24E-02 | 0.7 | 7.90E-04 | -0.3 | 9.26E-02 | -0.1 | 4.90E-01 | 0.0 | 9.65E-01 | -0.1 | 6.06E-01 | GAST1 protein homolog 4 |
| AT4G39300 | 0.4 | 5.53E-03 | 0.5 | 3.11E-03 | -0.1 | 6.80E-01 | 0.6 | 1.60E-02 | 0.3 | 2.43E-01 | 0.3 | 3.26E-01 | Protein NUCLEAR FUSION DEFECTIVE 6, chloroplastic |
| AT1G45010 | 0.4 | 3.69E-02 | 0.0 | 8.37E-01 | 0.4 | 1.01E-01 | 0.1 | 6.79E-01 | 0.3 | 2.98E-01 | -0.2 | 5.79E-01 | TRAM, LAG1 and CLN8 lipid-sensing domain containing protein |
| AT2G35360 | 0.4 | 1.52E-02 | 0.1 | 4.30E-01 | 0.3 | 1.28E-01 | -0.1 | 6.73E-01 | -0.2 | 5.67E-01 | 0.1 | 8.89E-01 | Uncharacterized protein At2g35360 |
| AT2G08765 | 0.4 | 2.20E-02 | 0.5 | 7.18E-03 | -0.1 | 5.46E-01 | 0.3 | 4.84E-01 | -0.1 | 8.46E-01 | 0.4 | 4.53E-01 | NA |
| AT4G04745 | 0.4 | 7.46E-02 | 0.1 | 5.64E-01 | 0.3 | 3.02E-01 | 0.7 | 3.07E-02 | 0.4 | 2.80E-01 | 0.3 | 4.56E-01 | At4g04745 |
| AT4G35730 | 0.4 | 1.60E-03 | 0.5 | 8.10E-04 | -0.1 | 5.70E-01 | -0.1 | 8.22E-01 | -0.2 | 6.69E-01 | 0.1 | 8.48E-01 | Uncharacterized protein AT4g35730 |
| AT1G01020 | 0.4 | 2.51E-02 | 0.2 | 3.16E-01 | 0.2 | 2.48E-01 | 0.2 | 4.57E-01 | 0.0 | 9.76E-01 | 0.2 | 4.38E-01 | Protein ARV |
| AT5G57230 | 0.4 | 2.47E-03 | 0.4 | 2.83E-03 | 0.0 | 9.39E-01 | 0.2 | 3.52E-01 | 0.0 | 9.62E-01 | 0.3 | 4.68E-01 | Thioredoxin superfamily protein |
| AT2G35490 | 0.4 | 1.54E-04 | 0.4 | 1.47E-04 | 0.0 | 6.30E-01 | 0.0 | 9.36E-01 | 0.1 | 7.28E-01 | -0.1 | 7.10E-01 | Probable plastid-lipid-associated protein 3, chloroplastic |
| AT1G61350 | 0.4 | 5.73E-03 | 0.2 | 2.04E-01 | 0.2 | 1.13E-01 | 0.1 | 3.78E-01 | 0.0 | 8.45E-01 | 0.2 | 4.00E-01 | ARM repeat superfamily protein |
| AT4G12490 | 0.4 | 6.77E-01 | -0.3 | 7.89E-01 | 0.7 | 5.72E-01 | -2.8 | 3.99E-02 | -6.4 | 5.56E-02 | 3.6 | 3.51E-01 | pEARLI1-like lipid transfer protein 2 |
| AT5G22270 | 0.4 | 1.40E-02 | -0.2 | 1.86E-01 | 0.6 | 5.35E-03 | 0.0 | 9.50E-01 | -0.2 | 5.02E-01 | 0.2 | 5.59E-01 | Protein salt-induced and EIN3 |
| AT1G48745 | 0.4 | 2.52E-02 | 0.0 | 9.15E-01 | 0.4 | 5.25E-02 | 0.0 | 9.99E-01 | -0.4 | 2.11E-01 | 0.4 | 2.87E-01 | Uncharacterized protein |
| AT4G13330 | 0.4 | 5.94E-02 | 0.0 | 9.42E-01 | 0.4 | 1.10E-01 | 0.4 | 3.37E-01 | 0.4 | 4.03E-01 | 0.0 | 9.97E-01 | hypothetical protein |
| AT3G22160 | 0.4 | 2.05E-03 | -0.3 | 1.82E-02 | 0.7 | 4.20E-04 | 0.7 | 1.03E-02 | -0.1 | 7.70E-01 | 0.8 | 8.98E-03 | VQ motif-containing protein 22 |
| AT3G51260 | 0.4 | 9.20E-04 | 0.1 | 3.79E-01 | 0.3 | 8.63E-03 | 0.3 | 7.20E-03 | 0.3 | 4.82E-02 | 0.1 | 7.41E-01 | Proteasome subunit alpha type |
| AT1G07170 | 0.4 | 1.34E-03 | 0.3 | 5.59E-03 | 0.1 | 5.61E-01 | 0.0 | 8.97E-01 | -0.3 | 2.73E-01 | 0.3 | 3.11E-01 | PHD finger-like domain-containing protein 5B |
| AT2G19460 | 0.4 | 2.58E-02 | 0.2 | 2.64E-01 | 0.2 | 2.99E-01 | 0.1 | 7.83E-01 | -0.3 | 4.06E-01 | 0.4 | 2.88E-01 | DUF3511 domain protein |
| AT1G33240 | 0.4 | 4.18E-04 | 0.3 | 6.93E-03 | 0.1 | 1.54E-01 | -0.5 | 8.76E-03 | -0.5 | 1.12E-02 | 0.0 | 9.25E-01 | GT-2-like 1 |
| AT1G07030 | 0.4 | 1.02E-04 | 0.3 | 2.13E-03 | 0.1 | 1.03E-01 | 0.4 | 1.65E-02 | 0.0 | 8.63E-01 | 0.5 | 1.57E-02 | F10K1.26 protein |
| AT5G48530 | 0.4 | 2.39E-02 | 0.1 | 4.59E-01 | 0.3 | 1.72E-01 | -0.3 | 2.49E-01 | -0.3 | 3.74E-01 | 0.0 | 9.33E-01 | Uncharacterized protein |
| AT4G33720 | 0.4 | 1.12E-02 | 0.6 | 1.41E-03 | -0.2 | 1.83E-01 | -2.7 | 6.25E-09 | -1.1 | 2.93E-04 | -1.7 | 1.58E-06 | hypothetical protein |
| AT1G19880 | 0.4 | 1.59E-04 | 0.2 | 1.55E-02 | 0.2 | 2.36E-02 | -0.1 | 5.19E-01 | -0.2 | 2.19E-01 | 0.1 | 6.51E-01 | Regulator of chromosome condensation family protein |
| AT2G47300 | 0.4 | 1.08E-02 | 0.1 | 5.07E-01 | 0.3 | 9.11E-02 | 0.8 | 5.93E-02 | 0.2 | 7.50E-01 | 0.6 | 2.29E-01 | Ribonuclease Ps |
| AT5G42330 | 0.4 | 6.01E-03 | 0.6 | 1.23E-03 | -0.1 | 3.34E-01 | 0.5 | 9.18E-02 | 0.5 | 1.68E-01 | 0.0 | 9.29E-01 | At5g42330 |
| AT5G45000 | 0.4 | 4.77E-03 | 0.1 | 4.98E-01 | 0.3 | 4.49E-02 | 0.4 | 1.48E-01 | 0.3 | 4.42E-01 | 0.2 | 6.73E-01 | Disease resistance protein family |
| AT2G34170 | 0.4 | 3.27E-03 | 0.8 | 7.69E-05 | -0.4 | 1.71E-02 | -0.7 | 1.21E-03 | -0.3 | 1.72E-01 | -0.4 | 7.54E-02 | Uncharacterized protein |
| AT5G55540 | 0.4 | 4.71E-03 | 0.4 | 7.85E-03 | 0.0 | 9.11E-01 | -0.1 | 5.38E-01 | -0.2 | 3.77E-01 | 0.1 | 8.08E-01 | Tornado 1 |
| AT1G19600 | 0.4 | 4.88E-05 | 0.5 | 2.62E-05 | -0.1 | 2.68E-01 | -0.1 | 5.02E-01 | -0.2 | 2.73E-01 | 0.1 | 7.30E-01 | hypothetical protein |
| AT5G17550 | 0.4 | 1.75E-03 | 0.0 | 6.74E-01 | 0.4 | 1.06E-02 | 0.2 | 3.01E-01 | -0.1 | 7.69E-01 | 0.3 | 2.75E-01 | Peroxin 19-2 |
| AT2G44525 | 0.4 | 2.82E-04 | 0.3 | 1.48E-03 | 0.1 | 4.43E-01 | 0.2 | 2.78E-01 | 0.0 | 9.27E-01 | 0.2 | 3.47E-01 | NADH dehydrogenase 1 alpha subcomplex assembly factor 3 |
| AT1G61640 | 0.4 | 5.87E-03 | 0.2 | 1.65E-01 | 0.2 | 1.43E-01 | 0.9 | 4.32E-04 | 0.3 | 2.67E-01 | 0.6 | 2.93E-02 | T25B24.1 protein |
| AT4G33380 | 0.4 | 5.86E-05 | 0.3 | 5.45E-04 | 0.1 | 2.06E-01 | 0.0 | 8.11E-01 | 0.0 | 9.82E-01 | 0.0 | 9.10E-01 | Uncharacterized protein AT4g33380 |
| AT2G34900 | 0.4 | 2.83E-03 | 0.1 | 3.03E-01 | 0.3 | 4.16E-02 | 0.3 | 2.48E-02 | 0.1 | 7.52E-01 | 0.3 | 1.55E-01 | Transcription factor GTE6 |
| AT4G37080 | 0.4 | 4.36E-04 | 0.1 | 3.05E-01 | 0.3 | 6.71E-03 | -0.4 | 2.29E-02 | -0.1 | 7.72E-01 | -0.3 | 1.78E-01 | AT4g37080 |
| AT1G51080 | 0.4 | 3.47E-03 | 0.4 | 3.21E-03 | 0.0 | 8.62E-01 | 0.5 | 5.44E-02 | 0.3 | 2.25E-01 | 0.1 | 7.20E-01 | Golgin family A protein |
| AT4G03115 | 0.4 | 3.36E-03 | 0.1 | 6.55E-01 | 0.4 | 2.48E-02 | -0.2 | 6.56E-01 | 0.0 | 1.00E+00 | -0.2 | 7.44E-01 | Mitochondrial substrate carrier family protein |
| AT2G20390 | 0.4 | 2.02E-03 | 0.3 | 7.96E-03 | 0.1 | 5.82E-01 | 0.4 | 5.03E-02 | 0.0 | 9.05E-01 | 0.4 | 1.09E-01 | Cytochrome oxidase complex assembly protein |
| AT5G25840 | 0.4 | 2.14E-02 | 0.2 | 1.65E-01 | 0.2 | 3.64E-01 | 0.2 | 4.76E-01 | -0.1 | 7.56E-01 | 0.2 | 4.25E-01 | hypothetical protein |
| AT2G41350 | 0.4 | 2.13E-04 | 0.2 | 5.06E-02 | 0.3 | 1.53E-02 | 0.2 | 3.33E-01 | 0.2 | 4.55E-01 | 0.0 | 9.63E-01 | EMB2819 |
| AT3G17930 | 0.4 | 2.00E-04 | 0.1 | 1.22E-01 | 0.3 | 5.30E-03 | 0.1 | 5.80E-01 | 0.2 | 1.34E-01 | -0.2 | 4.71E-01 | Transmembrane protein |
| AT4G01860 | 0.4 | 2.45E-04 | 0.2 | 7.01E-02 | 0.3 | 1.32E-02 | 0.3 | 2.66E-02 | 0.1 | 5.02E-01 | 0.2 | 3.25E-01 | At4g01860 |
| AT5G28640 | 0.4 | 8.45E-03 | 0.5 | 4.65E-03 | -0.1 | 6.88E-01 | -0.4 | 6.70E-03 | -0.2 | 2.53E-01 | -0.2 | 1.93E-01 | SSXT family protein |
| AT5G17240 | 0.4 | 4.60E-03 | 0.3 | 2.00E-02 | 0.1 | 5.76E-01 | 0.7 | 1.75E-02 | 0.4 | 3.10E-01 | 0.4 | 2.63E-01 | SET domain group 40 |
| AT1G52905 | 0.4 | 1.36E-02 | 0.1 | 4.23E-01 | 0.3 | 1.19E-01 | 0.8 | 7.30E-03 | 0.5 | 1.17E-01 | 0.3 | 4.16E-01 | Uncharacterized protein |
| AT1G49410 | 0.4 | 1.50E-02 | 0.6 | 3.84E-03 | -0.1 | 4.05E-01 | 0.6 | 1.11E-02 | 0.4 | 9.82E-02 | 0.2 | 5.87E-01 | Mitochondrial import receptor subunit TOM6 homolog |

| | | | | | | | | | | | | | |
|---|---|---|---|---|---|---|---|---|---|---|---|---|---|
| AT1G04800 | 0.4 | 3.73E-02 | 0.7 | 4.55E-03 | -0.2 | 2.62E-01 | 0.1 | 6.50E-01 | 0.0 | 8.98E-01 | 0.1 | 6.32E-01 | Uncharacterized protein |
| AT5G51200 | 0.4 | 2.06E-04 | 0.1 | 8.73E-02 | 0.3 | 7.47E-03 | 0.1 | 4.27E-01 | 0.1 | 5.03E-01 | 0.0 | 9.99E-01 | Nuclear pore complex protein |
| AT4G09970 | 0.4 | 3.10E-02 | 0.1 | 6.27E-01 | 0.3 | 1.52E-01 | 0.3 | 2.90E-01 | 0.2 | 5.40E-01 | 0.1 | 8.40E-01 | Transmembrane protein |
| AT5G13950 | 0.4 | 1.34E-03 | 0.2 | 7.90E-02 | 0.2 | 5.94E-02 | -0.3 | 2.16E-01 | -0.5 | 4.65E-02 | 0.2 | 4.63E-01 | DEUBAD domain-containing protein |
| AT3G11070 | 0.4 | 2.07E-03 | 0.5 | 1.74E-03 | 0.0 | 7.99E-01 | 0.0 | 9.43E-01 | 0.0 | 9.21E-01 | 0.0 | 8.99E-01 | Outer membrane OMP85 family protein |
| AT2G01740 | 0.4 | 1.92E-02 | 0.1 | 7.38E-01 | 0.4 | 8.70E-02 | 0.7 | 6.84E-02 | 0.2 | 7.14E-01 | 0.5 | 2.86E-01 | Pentatricopeptide repeat-containing protein At2g01740 |
| AT3G49890 | 0.4 | 1.03E-02 | 0.3 | 6.84E-02 | 0.1 | 4.38E-01 | 0.8 | 4.07E-03 | 0.4 | 1.73E-01 | 0.4 | 1.32E-01 | At3g49890 |
| AT1G79770 | 0.4 | 1.34E-01 | 0.6 | 4.30E-02 | -0.2 | 5.80E-01 | | | | | | | hypothetical protein |
| AT5G14090 | 0.4 | 6.26E-03 | 0.2 | 1.05E-01 | 0.2 | 2.32E-01 | -0.1 | 8.19E-01 | -0.4 | 1.43E-01 | 0.3 | 2.11E-01 | LAZY1 |
| AT4G29020 | 0.4 | 1.84E-03 | 0.4 | 2.92E-03 | 0.0 | 9.55E-01 | -0.5 | 1.91E-02 | -0.3 | 3.31E-01 | -0.3 | 3.51E-01 | Glycine-rich protein |
| AT3G26850 | 0.4 | 2.28E-03 | 0.1 | 3.34E-01 | 0.3 | 3.52E-02 | 0.7 | 5.73E-03 | 0.5 | 5.48E-02 | 0.2 | 5.39E-01 | hypothetical protein |
| AT5G02670 | 0.4 | 5.38E-03 | 0.0 | 9.49E-01 | 0.4 | 1.94E-02 | 0.1 | 8.88E-01 | 0.0 | 9.31E-01 | 0.1 | 8.45E-01 | Uncharacterized protein T22P11_260 |
| AT2G38570 | 0.4 | 9.57E-03 | 0.4 | 2.41E-02 | 0.1 | 7.65E-01 | 0.2 | 6.08E-01 | 0.1 | 7.92E-01 | 0.1 | 9.14E-01 | PRC-barrel domain-containing protein |
| AT5G55520 | 0.4 | 4.67E-03 | 0.7 | 2.32E-04 | -0.3 | 6.39E-02 | -0.1 | 6.03E-01 | 0.1 | 6.35E-01 | -0.3 | 3.56E-01 | Uncharacterized protein |
| AT2G01620 | 0.4 | 8.15E-04 | 0.2 | 6.87E-02 | 0.2 | 4.81E-02 | -0.4 | 1.63E-03 | -0.3 | 1.04E-02 | -0.1 | 7.40E-01 | F-box protein SKIP28 |
| AT5G23920 | 0.4 | 4.84E-05 | 0.4 | 9.33E-05 | 0.0 | 9.66E-01 | 0.2 | 2.43E-01 | 0.1 | 5.06E-01 | 0.1 | 8.46E-01 | Transmembrane protein |
| AT4G39740 | 0.4 | 4.73E-04 | 0.0 | 8.48E-01 | 0.4 | 3.16E-03 | 1.1 | 2.93E-04 | 0.1 | 6.90E-01 | 0.9 | 1.25E-03 | Uncharacterized protein At4g39740 |
| AT2G20724 | 0.4 | 1.07E-01 | 0.2 | 5.51E-01 | 0.3 | 3.97E-01 | -0.4 | 5.77E-02 | -0.4 | 8.33E-02 | 0.0 | 9.95E-01 | NA |
| AT1G68780 | 0.4 | 1.49E-02 | 0.8 | 6.86E-04 | -0.3 | 8.66E-02 | 0.1 | 4.22E-01 | -0.2 | 4.63E-01 | 0.3 | 2.12E-01 | Piriformospora indica-insensitive protein 2 |
| AT4G32240 | 0.4 | 9.51E-04 | 0.2 | 3.93E-02 | 0.2 | 8.55E-02 | 0.2 | 1.80E-01 | 0.1 | 4.58E-01 | 0.1 | 7.90E-01 | Uncharacterized protein AT4g32240 |
| AT2G21870 | 0.4 | 1.31E-04 | 0.4 | 4.71E-04 | 0.0 | 6.12E-01 | 0.3 | 4.04E-02 | 0.1 | 4.36E-01 | 0.2 | 4.45E-01 | MALE GAMETOPHYTE DEFECTIVE 1 |
| AT3G32930 | 0.4 | 1.99E-04 | 0.3 | 5.71E-03 | 0.2 | 7.88E-02 | 0.0 | 9.51E-01 | 0.0 | 8.84E-01 | 0.0 | 8.58E-01 | 6,7-dimethyl-8-ribityllumazine synthase |
| AT5G59610 | 0.4 | 5.26E-02 | 0.1 | 6.83E-01 | 0.3 | 1.89E-01 | 0.4 | 1.86E-01 | 0.5 | 1.74E-01 | -0.1 | 9.22E-01 | At5g59610 |
| AT5G12150 | 0.4 | 2.27E-04 | 0.2 | 8.31E-02 | 0.3 | 8.24E-03 | 0.1 | 6.84E-01 | 0.1 | 7.06E-01 | 0.0 | 9.79E-01 | Rho GTPase activation protein with PH domain-containing protein |
| AT3G07530 | 0.4 | 5.87E-03 | -0.2 | 3.14E-01 | 0.6 | 5.21E-03 | 0.1 | 7.74E-01 | 0.0 | 9.54E-01 | 0.1 | 8.85E-01 | F21O3.24 protein |
| AT5G10460 | 0.4 | 5.22E-05 | 0.4 | 3.89E-04 | 0.1 | 2.92E-01 | -0.5 | 2.61E-02 | -0.3 | 2.11E-01 | -0.2 | 5.66E-01 | Uncharacterized protein F12B17_190 |
| AT2G33255 | 0.4 | 5.91E-04 | 0.2 | 9.57E-02 | 0.3 | 2.64E-02 | 0.0 | 8.25E-01 | 0.1 | 6.73E-01 | -0.1 | 8.70E-01 | Haloacid dehalogenase-like hydrolase superfamily protein |
| AT5G16110 | 0.4 | 4.98E-04 | 0.6 | 1.03E-04 | -0.2 | 1.24E-01 | -0.4 | 2.98E-03 | -0.7 | 1.78E-04 | 0.3 | 1.53E-01 | Uncharacterized protein T21H19_30 |
| AT5G05930 | 0.4 | 1.50E-03 | 0.0 | 8.76E-01 | 0.4 | 6.83E-03 | -0.1 | 4.69E-01 | -0.3 | 1.27E-01 | 0.2 | 5.41E-01 | Guanylyl cyclase 1 |
| AT5G61130 | 0.4 | 5.29E-03 | 0.5 | 2.08E-03 | -0.1 | 5.16E-01 | 0.3 | 5.33E-02 | 0.1 | 4.28E-01 | 0.1 | 5.15E-01 | PDCB1 |
| AT3G19680 | 0.4 | 1.59E-01 | -0.2 | 5.04E-01 | 0.7 | 7.12E-02 | -1.0 | 5.04E-04 | -1.0 | 1.09E-03 | 0.0 | 9.86E-01 | Uncharacterized protein At3g19680 |
| AT3G50800 | 0.4 | 5.40E-02 | -0.1 | 6.28E-01 | 0.6 | 5.69E-02 | 0.0 | 9.98E-01 | 0.2 | 8.04E-01 | -0.2 | 8.38E-01 | Uncharacterized protein |
| AT5G10110 | 0.4 | 3.33E-03 | 0.3 | 1.77E-02 | 0.1 | 4.90E-01 | -0.1 | 4.27E-01 | -0.1 | 5.49E-01 | 0.0 | 9.65E-01 | DNA-directed RNA polymerase subunit beta |
| AT1G60640 | 0.4 | 5.63E-03 | 0.2 | 9.92E-02 | 0.2 | 2.02E-01 | 0.8 | 1.32E-04 | 0.3 | 9.05E-02 | 0.5 | 2.27E-02 | Stress response protein |
| AT4G32400 | 0.4 | 9.32E-04 | 0.3 | 1.28E-02 | 0.1 | 2.10E-01 | 0.2 | 6.35E-02 | 0.0 | 8.47E-01 | 0.2 | 2.48E-01 | Adenine nucleotide transporter BT1, chloroplastic |
| AT3G15310 | 0.4 | 1.20E-01 | 0.5 | 8.41E-02 | -0.1 | 8.29E-01 | | | | | | | NA |
| AT1G07650 | 0.4 | 2.96E-04 | 0.2 | 4.15E-02 | 0.3 | 2.06E-02 | -0.1 | 7.39E-01 | -0.2 | 1.48E-01 | 0.2 | 3.42E-01 | Uncharacterized protein At1g07655 |
| AT1G26180 | 0.4 | 3.01E-04 | 0.4 | 1.74E-03 | 0.1 | 4.22E-01 | -0.5 | 3.07E-02 | -0.4 | 1.19E-01 | -0.1 | 7.19E-01 | Membrane protein |
| AT1G02750 | 0.4 | 7.16E-02 | -0.2 | 4.65E-01 | 0.6 | 4.47E-02 | 0.3 | 2.82E-01 | -0.3 | 3.43E-01 | 0.6 | 1.00E-01 | Drought-responsive family protein |
| AT1G79210 | 0.4 | 1.90E-03 | 0.1 | 4.55E-01 | 0.4 | 1.58E-02 | 0.1 | 3.22E-01 | 0.0 | 9.66E-01 | 0.1 | 5.40E-01 | Proteasome subunit alpha type-2-B |
| AT3G02170 | 0.4 | 1.34E-02 | 0.3 | 6.91E-02 | 0.1 | 4.70E-01 | -0.4 | 2.85E-01 | -0.2 | 7.51E-01 | -0.2 | 6.83E-01 | Longifolia2 |
| AT3G07170 | 0.4 | 9.01E-05 | 0.3 | 1.79E-03 | 0.1 | 1.01E-01 | 0.2 | 1.01E-01 | 0.2 | 3.64E-01 | 0.1 | 7.19E-01 | AT3g07170 |
| AT1G12730 | 0.4 | 3.22E-03 | 0.3 | 1.86E-02 | 0.1 | 4.83E-01 | 0.0 | 8.74E-01 | -0.2 | 3.53E-01 | 0.2 | 5.39E-01 | GPI transamidase subunit PIG-U |
| AT5G47660 | 0.4 | 4.14E-04 | 0.2 | 2.41E-02 | 0.2 | 6.13E-02 | -0.3 | 5.91E-02 | -0.4 | 2.36E-02 | 0.1 | 6.86E-01 | Uncharacterized protein At5g47660 |
| AT4G27657 | 0.4 | 7.16E-02 | -0.1 | 6.26E-01 | 0.6 | 6.49E-02 | 0.3 | 5.83E-01 | 0.0 | 9.95E-01 | 0.3 | 6.80E-01 | hypothetical protein |
| AT3G58975 | 0.4 | 3.34E-02 | 0.3 | 1.31E-01 | 0.1 | 5.98E-01 | -0.5 | 7.57E-02 | -0.3 | 3.45E-01 | -0.2 | 6.30E-01 | FBD domain-containing protein At3g58975 |
| AT1G35340 | 0.4 | 7.32E-05 | 0.4 | 1.86E-04 | 0.0 | 7.71E-01 | 0.0 | 8.79E-01 | 0.2 | 4.54E-01 | -0.2 | 4.27E-01 | Uncharacterized protein T9I1.11 |
| AT2G34000 | 0.4 | 1.49E-02 | 1.1 | 5.47E-05 | -0.6 | 4.80E-03 | 0.2 | 6.30E-01 | 0.0 | 9.87E-01 | 0.2 | 6.33E-01 | hypothetical protein |
| AT1G10580 | 0.4 | 1.57E-04 | 0.1 | 8.19E-02 | 0.3 | 5.88E-03 | 0.4 | 5.73E-03 | 0.1 | 4.63E-01 | 0.2 | 1.55E-01 | T10O24.21 |
| AT4G30340 | 0.4 | 3.18E-04 | 0.1 | 2.20E-01 | 0.3 | 6.48E-03 | 0.2 | 1.51E-01 | -0.1 | 6.86E-01 | 0.3 | 1.27E-01 | Uncharacterized protein At4g30340 |
| AT3G63390 | 0.4 | 1.64E-04 | 0.1 | 2.55E-01 | 0.4 | 3.56E-03 | 0.0 | 9.90E-01 | -0.1 | 6.00E-01 | 0.1 | 6.38E-01 | Uncharacterized protein MAA21_20 |
| AT5G41970 | 0.4 | 1.62E-03 | 0.5 | 1.92E-03 | 0.0 | 9.45E-01 | 0.3 | 3.92E-02 | 0.0 | 9.37E-01 | 0.3 | 6.73E-02 | GAMM1 protein-like |
| AT5G53880 | 0.4 | 2.35E-03 | 0.5 | 1.89E-03 | 0.0 | 7.85E-01 | 0.1 | 7.78E-01 | -0.2 | 5.88E-01 | 0.2 | 5.00E-01 | Uncharacterized protein |
| AT2G33820 | 0.4 | 1.79E-03 | 0.4 | 1.03E-02 | 0.1 | 4.68E-01 | 0.0 | 9.19E-01 | -0.1 | 6.67E-01 | 0.1 | 7.66E-01 | Mitochondrial substrate carrier family protein |
| AT1G28530 | 0.4 | 1.28E-02 | 0.2 | 2.43E-01 | 0.2 | 2.02E-01 | 0.3 | 3.06E-01 | 0.1 | 9.16E-01 | 0.3 | 4.52E-01 | Deneddylase |
| AT5G49230 | 0.4 | 6.05E-04 | 0.2 | 6.17E-02 | 0.2 | 3.99E-02 | -0.2 | 2.93E-01 | -0.1 | 7.30E-01 | -0.1 | 6.99E-01 | Uncharacterized protein |
| AT2G04340 | 0.4 | 4.88E-03 | 0.1 | 6.81E-01 | 0.4 | 2.83E-02 | -0.1 | 8.51E-01 | -0.3 | 2.83E-01 | 0.2 | 4.95E-01 | At2g04340 |
| AT5G50760 | 0.4 | 4.20E-03 | 0.3 | 5.04E-02 | 0.2 | 2.79E-01 | 2.1 | 1.35E-05 | 0.6 | 2.02E-01 | 1.5 | 6.49E-04 | SAUR-like auxin-responsive protein family |
| AT3G10530 | 0.4 | 8.21E-03 | 0.3 | 1.07E-01 | 0.2 | 2.58E-01 | 0.8 | 7.30E-05 | 0.5 | 6.30E-03 | 0.3 | 1.29E-01 | F18K10.11 protein |
| AT1G67570 | 0.5 | 6.56E-04 | 0.0 | 8.48E-01 | 0.5 | 2.52E-03 | -0.1 | 6.84E-01 | 0.1 | 7.76E-01 | -0.2 | 5.34E-01 | Zinc finger CONSTANS-like protein |
| AT1G64960 | 0.5 | 1.92E-03 | 0.0 | 7.66E-01 | 0.4 | 1.08E-02 | 0.1 | 6.97E-01 | 0.2 | 3.89E-01 | -0.1 | 7.04E-01 | ARM repeat superfamily protein |
| AT1G05950 | 0.5 | 1.58E-02 | 0.2 | 2.95E-01 | 0.3 | 2.03E-01 | 0.1 | 7.42E-01 | 0.0 | 9.75E-01 | 0.1 | 8.21E-01 | T21E18.2 protein |
| AT1G35180 | 0.5 | 8.38E-02 | 1.0 | 2.17E-03 | -0.5 | 7.20E-02 | 0.2 | 5.02E-01 | 0.3 | 4.29E-01 | -0.1 | 9.01E-01 | Uncharacterized protein T32G9.28 |
| AT5G25260 | 0.5 | 1.40E-01 | 0.4 | 1.85E-01 | 0.0 | 9.29E-01 | 1.3 | 2.07E-04 | -0.1 | 8.78E-01 | 1.4 | 3.23E-04 | Flotillin-like protein 2 |
| AT2G25670 | 0.5 | 1.47E-03 | 0.4 | 6.11E-03 | 0.1 | 5.48E-01 | 0.2 | 4.93E-02 | 0.0 | 7.72E-01 | 0.2 | 2.76E-01 | hypothetical protein |
| AT3G05480 | 0.5 | 3.83E-03 | 0.3 | 4.90E-02 | 0.2 | 3.00E-01 | 0.5 | 1.76E-01 | 0.6 | 9.31E-02 | -0.1 | 7.36E-01 | Cell cycle checkpoint control protein family |
| AT1G10340 | 0.5 | 1.13E-01 | 0.2 | 5.13E-01 | 0.3 | 4.42E-01 | 0.3 | 3.23E-01 | -0.3 | 4.04E-01 | 0.5 | 1.37E-01 | Ankyrin repeat family protein |
| AT5G38360 | 0.5 | 2.76E-03 | 0.4 | 1.50E-02 | 0.1 | 4.84E-01 | 0.1 | 7.80E-01 | 0.0 | 9.44E-01 | 0.1 | 7.98E-01 | At5g38360 |
| AT1G23220 | 0.5 | 2.93E-03 | 0.2 | 1.07E-01 | 0.2 | 1.13E-01 | 0.3 | 1.93E-01 | 0.1 | 8.72E-01 | 0.2 | 4.66E-01 | Dynein light chain |
| AT1G31258 | 0.5 | 9.48E-02 | -0.2 | 4.99E-01 | 0.7 | 6.40E-02 | 0.0 | 9.57E-01 | 0.0 | 9.82E-01 | 0.0 | 9.57E-01 | NA |
| AT4G00560 | 0.5 | 1.39E-04 | 0.4 | 3.17E-04 | 0.0 | 8.35E-01 | -0.3 | 1.06E-02 | -0.2 | 1.73E-01 | -0.1 | 4.43E-01 | NAD-binding Rossmann-fold superfamily protein |
| AT5G13060 | 0.5 | 3.77E-03 | 0.4 | 8.32E-03 | 0.0 | 8.17E-01 | 0.4 | 2.19E-01 | 0.5 | 1.63E-01 | -0.1 | 8.45E-01 | hypothetical protein |
| AT2G25605 | 0.5 | 4.41E-04 | 0.2 | 8.89E-02 | 0.3 | 2.08E-02 | -0.3 | 1.25E-01 | -0.2 | 4.30E-01 | -0.1 | 7.08E-01 | DNA-directed RNA polymerase subunit beta |
| AT5G18770 | 0.5 | 4.48E-04 | 0.3 | 1.13E-02 | 0.2 | 1.32E-01 | 0.0 | 9.85E-01 | 0.1 | 8.20E-01 | -0.1 | 8.30E-01 | F-box |
| AT3G49180 | 0.5 | 1.83E-03 | 0.2 | 8.71E-02 | 0.2 | 9.49E-02 | 0.4 | 1.08E-01 | 0.1 | 8.52E-01 | 0.3 | 2.52E-01 | Protein ROOT INITIATION DEFECTIVE 3 |
| AT5G42060 | 0.5 | 8.83E-03 | 0.0 | 9.80E-01 | 0.5 | 2.43E-02 | 0.6 | 4.25E-02 | 0.5 | 1.45E-01 | 0.1 | 7.59E-01 | Mediator-associated protein 3 |
| AT5G58790 | 0.5 | 2.72E-02 | 0.0 | 8.90E-01 | 0.4 | 8.30E-02 | 0.0 | 9.32E-01 | 0.3 | 4.76E-01 | -0.3 | 5.08E-01 | Uncharacterized protein |
| AT5G65250 | 0.5 | 2.00E-04 | 0.2 | 1.65E-02 | 0.2 | 3.60E-02 | -0.1 | 7.47E-01 | -0.3 | 1.24E-01 | 0.2 | 3.07E-01 | Transmembrane protein |
| AT1G67148 | 0.5 | 1.23E-01 | -0.2 | 6.28E-01 | 0.6 | 1.01E-01 | 2.1 | 1.46E-04 | 1.2 | 2.58E-02 | 0.9 | 4.20E-02 | Uncharacterized protein |
| AT5G45760 | 0.5 | 2.71E-02 | 0.0 | 9.81E-01 | 0.5 | 6.12E-02 | -0.1 | 8.12E-01 | -0.2 | 6.13E-01 | 0.1 | 8.28E-01 | Transducin |
| AT5G41960 | 0.5 | 2.86E-02 | -0.3 | 2.12E-01 | 0.7 | 1.04E-02 | 0.1 | 7.92E-01 | -0.1 | 8.37E-01 | 0.2 | 7.11E-01 | At5g41960 |
| AT3G47965 | 0.5 | 7.18E-02 | 0.2 | 3.41E-01 | 0.2 | 4.77E-01 | -0.1 | 7.60E-01 | -0.3 | 4.75E-01 | 0.2 | 7.05E-01 | AT3g47961 |
| AT1G29320 | 0.5 | 9.37E-03 | 0.2 | 1.30E-01 | 0.2 | 2.45E-01 | 0.6 | 1.03E-02 | 0.3 | 2.36E-01 | 0.3 | 1.78E-01 | Transducin |

| | | | | | | | | | | | | | |
|---|---|---|---|---|---|---|---|---|---|---|---|---|---|
| AT2G27590 | 0.5 | 4.03E-04 | 0.3 | 8.90E-03 | 0.2 | 1.39E-01 | 0.2 | 2.12E-01 | 0.0 | 9.94E-01 | 0.2 | 3.42E-01 | S-adenosyl-L-methionine-dependent methyltransferases superfamily protein |
| AT3G18410 | 0.5 | 2.80E-03 | 0.7 | 4.15E-04 | -0.2 | 1.51E-01 | 0.1 | 6.28E-01 | 0.0 | 8.54E-01 | 0.0 | 8.94E-01 | NADH dehydrogenase 1 beta subcomplex subunit 10-B |
| AT3G55510 | 0.5 | 1.40E-03 | 0.1 | 2.45E-01 | 0.3 | 2.72E-02 | 0.8 | 2.29E-04 | 0.4 | 7.28E-02 | 0.4 | 2.68E-02 | Noc2p family |
| AT5G49550 | 0.5 | 3.43E-02 | 0.1 | 7.43E-01 | 0.4 | 1.26E-01 | -0.3 | 2.67E-01 | -0.2 | 5.37E-01 | -0.1 | 8.19E-01 | Biogenesis of lysosome-related organelles complex 1 subunit 2 |
| AT2G16365 | 0.5 | 1.47E-04 | 0.3 | 7.18E-03 | 0.2 | 4.38E-02 | -0.1 | 4.55E-01 | -0.1 | 5.57E-01 | 0.0 | 9.80E-01 | F-box protein At2g16365 |
| AT2G01260 | 0.5 | 1.71E-04 | 0.3 | 6.18E-03 | 0.2 | 6.60E-02 | -0.1 | 3.85E-01 | 0.0 | 8.38E-01 | -0.2 | 3.47E-01 | DUF789 family protein |
| AT5G51620 | 0.5 | 6.86E-03 | -0.1 | 7.27E-01 | 0.5 | 1.20E-02 | 0.1 | 7.84E-01 | 0.0 | 9.47E-01 | 0.1 | 7.86E-01 | Uncharacterized protein |
| AT5G13090 | 0.5 | 1.50E-03 | 0.6 | 4.51E-04 | -0.1 | 3.39E-01 | -0.3 | 1.43E-01 | -0.5 | 4.07E-02 | 0.2 | 5.84E-01 | Uncharacterized protein T19L5_50 |
| AT3G02640 | 0.5 | 1.93E-03 | 0.8 | 7.29E-05 | -0.3 | 2.46E-02 | 0.2 | 2.07E-01 | 0.3 | 1.51E-01 | -0.1 | 8.45E-01 | hypothetical protein |
| AT3G52540 | 0.5 | 1.57E-02 | 0.5 | 1.54E-02 | 0.0 | 9.11E-01 | -0.1 | 8.78E-01 | -0.1 | 8.02E-01 | 0.1 | 9.20E-01 | Transcription repressor OFP18 |
| AT2G34670 | 0.5 | 1.48E-03 | 0.4 | 2.72E-03 | 0.0 | 8.96E-01 | 0.2 | 3.61E-01 | 0.1 | 6.92E-01 | 0.1 | 7.98E-01 | Benzoyl-CoA reductase subunit C, putative |
| AT3G48140 | 0.5 | 6.78E-05 | 0.4 | 2.89E-04 | 0.1 | 5.21E-01 | 0.1 | 5.86E-01 | 0.0 | 9.46E-01 | 0.1 | 6.67E-01 | hypothetical protein |
| AT1G64670 | 0.5 | 6.74E-04 | 0.3 | 1.21E-02 | 0.2 | 1.73E-01 | 0.2 | 5.75E-01 | 0.0 | 9.38E-01 | 0.1 | 7.56E-01 | Alpha |
| AT3G18320 | 0.5 | 2.93E-03 | 0.2 | 1.20E-01 | 0.2 | 1.19E-01 | 0.2 | 6.84E-01 | 0.0 | 9.76E-01 | 0.2 | 7.29E-01 | F-box associated domain-containing protein |
| AT3G13190 | 0.5 | 8.03E-04 | 0.2 | 1.01E-01 | 0.3 | 3.77E-02 | 0.0 | 9.26E-01 | -0.1 | 6.20E-01 | 0.2 | 5.88E-01 | WEB family protein At3g13190 |
| AT2G20930 | 0.5 | 1.16E-04 | 0.4 | 3.98E-04 | 0.0 | 6.13E-01 | 0.0 | 9.44E-01 | 0.0 | 9.60E-01 | 0.0 | 9.28E-01 | AT2G20930 protein |
| AT2G39090 | 0.5 | 1.34E-04 | 0.2 | 1.70E-02 | 0.2 | 2.30E-02 | -0.3 | 8.12E-02 | -0.1 | 4.80E-01 | -0.2 | 5.38E-01 | AtAPC7 |
| AT4G03020 | 0.5 | 4.56E-05 | 0.1 | 3.11E-01 | 0.4 | 6.04E-04 | 0.1 | 1.88E-01 | -0.1 | 4.82E-01 | 0.2 | 9.61E-02 | LEC14B homolog |
| AT3G29644 | 0.5 | 2.63E-03 | 0.4 | 5.11E-03 | 0.0 | 8.81E-01 | -0.2 | 4.45E-01 | -0.3 | 2.48E-01 | 0.1 | 7.27E-01 | NA |
| AT4G13520 | 0.5 | 1.30E-02 | 0.2 | 1.61E-01 | 0.2 | 2.41E-01 | 0.1 | 7.82E-01 | -0.1 | 7.75E-01 | 0.2 | 6.52E-01 | Small acidic protein 1 |
| AT5G01510 | 0.5 | 4.39E-03 | 0.1 | 3.64E-01 | 0.3 | 5.68E-02 | -0.1 | 8.42E-01 | 0.2 | 4.96E-01 | -0.3 | 4.58E-01 | Uncharacterized protein At5g01510 |
| AT3G51010 | 0.5 | 3.52E-04 | 0.6 | 8.59E-05 | -0.1 | 1.65E-01 | 0.2 | 3.50E-01 | -0.1 | 7.68E-01 | 0.2 | 3.31E-01 | hypothetical protein |
| AT3G04980 | 0.5 | 2.35E-03 | 0.2 | 1.25E-01 | 0.3 | 9.17E-02 | 0.2 | 5.28E-01 | 0.1 | 8.17E-01 | 0.1 | 8.30E-01 | DNAJ heat shock N-terminal domain-containing protein |
| AT1G69700 | 0.5 | 4.18E-03 | 0.5 | 2.60E-03 | -0.1 | 7.04E-01 | 0.0 | 8.95E-01 | -0.1 | 7.92E-01 | 0.1 | 9.11E-01 | HVA22-like protein c |
| AT5G11630 | 0.5 | 6.74E-03 | 0.4 | 2.02E-02 | 0.1 | 6.78E-01 | 0.3 | 1.82E-01 | 0.0 | 8.92E-01 | 0.2 | 3.64E-01 | Protein NONRESPONDING TO OXYLIPINS 2, mitochondrial |
| AT1G78430 | 0.5 | 1.24E-03 | 0.7 | 1.74E-04 | -0.2 | 1.44E-01 | 0.0 | 9.48E-01 | 0.2 | 5.84E-01 | -0.2 | 6.07E-01 | Interactor of constitutive active ROPs 4 |
| AT3G20390 | 0.5 | 1.55E-04 | 0.6 | 4.42E-05 | -0.1 | 1.21E-01 | 0.1 | 3.08E-01 | 0.2 | 1.01E-01 | -0.1 | 6.23E-01 | Endoribonuclease L-PSP family protein |
| AT2G33585 | 0.5 | 9.02E-04 | 0.3 | 1.50E-02 | 0.2 | 2.05E-01 | 0.8 | 7.79E-02 | 0.3 | 6.09E-01 | 0.5 | 3.40E-01 | Transmembrane protein |
| AT2G23390 | 0.5 | 3.91E-04 | 0.1 | 1.52E-01 | 0.3 | 9.28E-03 | 0.2 | 1.95E-01 | 0.1 | 4.90E-01 | 0.1 | 8.02E-01 | Acyl-CoA |
| AT5G63520 | 0.5 | 1.25E-03 | 0.2 | 1.52E-01 | 0.3 | 4.09E-02 | -0.2 | 4.02E-01 | -0.2 | 3.37E-01 | 0.1 | 8.94E-01 | F-box |
| AT3G49260 | 0.5 | 1.28E-02 | 0.6 | 3.13E-03 | -0.2 | 3.91E-01 | -0.3 | 1.08E-01 | -0.1 | 5.70E-01 | -0.2 | 5.50E-01 | IQ-domain 21 |
| AT3G22630 | 0.5 | 9.42E-04 | 0.3 | 8.38E-03 | 0.1 | 2.97E-01 | 0.3 | 3.38E-02 | 0.1 | 8.19E-01 | 0.3 | 1.61E-01 | Proteasome subunit beta type-2-A |
| AT1G54850 | 0.5 | 1.82E-03 | 0.2 | 7.53E-02 | 0.2 | 1.06E-01 | 0.1 | 6.96E-01 | 0.0 | 9.48E-01 | 0.1 | 8.40E-01 | Alpha-crystallin domain-containing protein 22.3 |
| AT3G05070 | 0.5 | 2.45E-04 | 0.4 | 1.05E-03 | 0.1 | 5.11E-01 | 0.4 | 2.67E-02 | 0.2 | 2.30E-01 | 0.1 | 5.79E-01 | Coiled-coil domain-containing protein 12 |
| AT3G47080 | 0.5 | 1.15E-03 | 0.2 | 6.08E-02 | 0.2 | 7.74E-02 | -0.3 | 1.28E-01 | 0.0 | 8.71E-01 | -0.2 | 3.70E-01 | hypothetical protein |
| AT5G55640 | 0.5 | 1.14E-03 | 0.3 | 1.11E-02 | 0.1 | 3.17E-01 | 0.1 | 7.13E-01 | 0.2 | 5.01E-01 | -0.1 | 8.20E-01 | hypothetical protein |
| AT2G17230 | 0.5 | 1.49E-01 | 0.2 | 5.65E-01 | 0.3 | 4.49E-01 | -0.6 | 6.98E-03 | -0.7 | 3.36E-03 | 0.1 | 6.74E-01 | Protein EXORDIUM-like 5 |
| AT2G30270 | 0.5 | 1.09E-04 | 0.3 | 4.74E-03 | 0.2 | 5.57E-02 | -0.1 | 7.89E-01 | -0.3 | 1.89E-01 | 0.2 | 4.12E-01 | Protein LURP-one-related 7 |
| AT4G35980 | 0.5 | 1.26E-04 | 0.4 | 1.46E-03 | 0.1 | 2.10E-01 | 0.1 | 7.40E-01 | -0.1 | 8.58E-01 | 0.1 | 6.74E-01 | Uncharacterized protein |
| AT3G10860 | 0.5 | 7.33E-04 | 0.4 | 5.51E-03 | 0.1 | 3.38E-01 | 0.2 | 3.13E-01 | 0.2 | 5.50E-01 | 0.1 | 8.94E-01 | Cytochrome b-c1 complex subunit 8-1, mitochondrial |
| AT4G39470 | 0.5 | 2.73E-03 | 0.4 | 1.87E-02 | 0.1 | 4.19E-01 | 0.6 | 7.63E-03 | 0.1 | 6.84E-01 | 0.5 | 6.12E-02 | Uncharacterized protein AT4g39470 |
| AT2G20940 | 0.5 | 6.81E-03 | 0.4 | 1.67E-02 | 0.1 | 7.50E-01 | 0.9 | 4.63E-03 | 0.3 | 3.35E-01 | 0.5 | 1.19E-01 | DUF1279 domain-containing protein |
| AT5G67640 | 0.5 | 3.04E-02 | 0.0 | 9.68E-01 | 0.5 | 7.94E-02 | 0.4 | 2.01E-01 | -0.1 | 7.50E-01 | 0.5 | 1.71E-01 | Uncharacterized protein |
| AT3G30122 | 0.5 | 5.37E-02 | 0.2 | 5.37E-01 | 0.3 | 2.67E-01 | 0.8 | 7.09E-02 | 0.3 | 5.99E-01 | 0.5 | 3.77E-01 | NA |
| AT2G30000 | 0.5 | 3.11E-02 | 0.3 | 1.22E-01 | 0.1 | 5.76E-01 | 0.3 | 3.99E-01 | -0.1 | 9.15E-01 | 0.3 | 4.18E-01 | PHD finger-like domain-containing protein 5A |
| AT4G38980 | 0.5 | 6.69E-04 | 0.2 | 1.45E-01 | 0.3 | 2.33E-02 | -0.1 | 6.59E-01 | -0.3 | 3.94E-01 | 0.1 | 7.65E-01 | Uncharacterized protein At4g38980 |
| AT4G20020 | 0.5 | 1.47E-03 | 0.3 | 2.31E-02 | 0.2 | 1.96E-01 | 0.6 | 7.21E-04 | 0.2 | 2.59E-01 | 0.4 | 5.37E-02 | Putative DAG protein |
| AT4G26960 | 0.5 | 1.32E-02 | 0.7 | 1.79E-03 | -0.2 | 2.65E-01 | 0.5 | 1.92E-01 | 0.5 | 2.40E-01 | 0.0 | 9.96E-01 | hypothetical protein |
| AT5G03700 | 0.5 | 6.14E-04 | 0.3 | 1.42E-02 | 0.2 | 1.39E-01 | 0.2 | 4.60E-01 | -0.2 | 5.01E-01 | 0.3 | 1.64E-01 | PAN domain-containing protein At5g03700 |
| AT3G15395 | 0.5 | 7.46E-03 | 0.3 | 1.35E-01 | 0.2 | 2.12E-01 | 0.0 | 9.66E-01 | 0.0 | 9.26E-01 | 0.1 | 9.14E-01 | AT3G15395 protein |
| AT2G32560 | 0.5 | 1.12E-04 | 0.2 | 2.25E-02 | 0.3 | 1.14E-02 | 0.2 | 2.90E-01 | 0.4 | 2.87E-02 | -0.2 | 2.43E-01 | Uncharacterized protein At2g32570 |
| AT5G25250 | 0.5 | 2.22E-02 | -0.1 | 5.41E-01 | 0.6 | 1.58E-02 | 0.7 | 1.91E-03 | -0.2 | 3.55E-01 | 0.9 | 2.67E-03 | Flotillin-like |
| AT1G67900 | 0.5 | 1.04E-03 | 0.4 | 8.17E-03 | 0.1 | 3.28E-01 | 0.3 | 2.00E-01 | 0.2 | 4.90E-01 | 0.1 | 7.56E-01 | BTB |
| AT1G07970 | 0.5 | 6.24E-05 | 0.1 | 1.03E-01 | 0.3 | 2.65E-03 | 0.0 | 9.53E-01 | -0.2 | 4.12E-01 | 0.1 | 5.29E-01 | Cytochrome B561, amino-terminal protein |
| AT3G52680 | 0.5 | 2.82E-04 | 0.3 | 6.02E-03 | 0.2 | 1.50E-01 | 0.3 | 3.46E-01 | 0.2 | 5.92E-01 | 0.1 | 8.59E-01 | F-box |
| AT1G50055 | 0.5 | 9.05E-03 | -0.1 | 7.81E-01 | 0.5 | 1.96E-02 | 0.6 | 2.95E-02 | 0.2 | 6.76E-01 | 0.4 | 1.87E-01 | NA |
| AT2G37820 | 0.5 | 2.71E-02 | 0.8 | 1.36E-03 | -0.4 | 1.16E-01 | | | | | | | hypothetical protein |
| AT3G19080 | 0.5 | 7.45E-04 | 0.5 | 1.25E-03 | 0.0 | 9.44E-01 | 0.1 | 7.55E-01 | 0.1 | 7.50E-01 | 0.0 | 9.66E-01 | SWIB complex BAF60b domain-containing protein |
| AT3G48650 | 0.5 | 7.74E-02 | -0.2 | 5.04E-01 | 0.7 | 5.19E-02 | 0.2 | 7.81E-01 | -1.0 | 1.30E-01 | 1.1 | 1.12E-01 | UPF0496 protein At3g48650 |
| AT5G54100 | 0.5 | 6.01E-03 | 0.2 | 3.37E-01 | 0.3 | 6.71E-02 | 1.6 | 1.71E-07 | 0.1 | 5.32E-01 | 1.4 | 2.94E-05 | At5g54100 |
| AT1G65030 | 0.5 | 3.81E-03 | 0.2 | 1.15E-01 | 0.2 | 1.32E-01 | 0.6 | 1.26E-03 | 0.3 | 1.08E-01 | 0.3 | 1.63E-01 | hypothetical protein |
| AT1G15420 | 0.5 | 3.61E-03 | 0.3 | 4.92E-02 | 0.2 | 2.44E-01 | 0.4 | 2.79E-02 | 0.0 | 9.37E-01 | 0.4 | 4.03E-02 | WD repeat protein |
| AT2G15020 | 0.5 | 3.46E-02 | 0.4 | 6.09E-02 | 0.1 | 8.43E-01 | | | | | | | Uncharacterized protein |
| AT5G25630 | 0.5 | 1.13E-03 | 0.1 | 2.17E-01 | 0.3 | 1.88E-02 | -0.4 | 2.99E-02 | -0.2 | 4.12E-01 | -0.2 | 3.80E-01 | Tetratricopeptide repeat -like superfamily protein |
| AT5G61200 | 0.5 | 4.22E-03 | 0.4 | 1.20E-02 | 0.1 | 7.35E-01 | 0.0 | 9.35E-01 | 0.0 | 9.77E-01 | 0.0 | 9.75E-01 | Myosin heavy chain-like protein |
| AT1G23170 | 0.5 | 2.31E-04 | 0.4 | 1.74E-03 | 0.1 | 3.31E-01 | 0.1 | 7.83E-01 | 0.1 | 5.65E-01 | -0.1 | 8.21E-01 | Transmembrane protein |
| AT5G52420 | 0.5 | 2.61E-05 | 0.3 | 7.84E-04 | 0.2 | 4.54E-02 | 0.0 | 9.73E-01 | 0.0 | 9.69E-01 | 0.0 | 9.49E-01 | Transmembrane protein |
| AT4G15640 | 0.5 | 4.38E-03 | 0.3 | 3.36E-02 | 0.1 | 3.79E-01 | 0.7 | 5.60E-04 | 0.4 | 3.40E-02 | 0.3 | 2.00E-01 | Uncharacterized protein AT4g15640 |
| AT1G34770 | 0.5 | 2.31E-03 | 0.2 | 1.37E-01 | 0.3 | 8.69E-02 | 0.5 | 8.21E-02 | 0.3 | 3.55E-01 | 0.2 | 6.44E-01 | AT1G34770 protein |
| AT1G04985 | 0.5 | 1.33E-04 | 0.3 | 7.48E-03 | 0.2 | 4.07E-02 | 0.2 | 1.54E-01 | -0.1 | 7.64E-01 | 0.3 | 1.90E-01 | Triacylglycerol lipase-like protein |
| AT5G37010 | 0.5 | 1.21E-03 | 0.4 | 3.62E-03 | 0.1 | 6.83E-01 | -0.2 | 1.30E-01 | 0.0 | 8.64E-01 | -0.2 | 3.42E-01 | Uncharacterized protein |
| AT5G35740 | 0.5 | 1.77E-03 | 0.7 | 2.67E-04 | -0.2 | 1.69E-01 | 0.1 | 8.16E-01 | 0.1 | 8.08E-01 | 0.0 | 9.75E-01 | X8 domain-containing protein |
| AT2G39740 | 0.5 | 3.60E-04 | 0.2 | 3.71E-02 | 0.3 | 3.44E-02 | 0.1 | 6.33E-01 | 0.1 | 8.62E-01 | 0.1 | 8.74E-01 | Uncharacterized protein |
| AT5G47570 | 0.5 | 1.07E-03 | 0.5 | 1.16E-03 | 0.0 | 8.47E-01 | -0.2 | 3.39E-01 | 0.0 | 8.83E-01 | -0.1 | 6.14E-01 | NADH dehydrogenase 1 beta subcomplex subunit 8, mitochondrial |
| AT5G15310 | 0.5 | 9.02E-04 | 0.4 | 6.51E-03 | 0.1 | 3.81E-01 | -0.2 | 3.49E-01 | 0.0 | 8.90E-01 | -0.3 | 4.12E-01 | Myb domain protein 16 |
| AT3G47510 | 0.5 | 7.07E-02 | 0.1 | 7.25E-01 | 0.4 | 2.29E-01 | 0.1 | 7.99E-01 | 0.0 | 9.89E-01 | 0.1 | 8.33E-01 | hypothetical protein |
| AT4G38150 | 0.5 | 4.50E-03 | 0.1 | 5.09E-01 | 0.4 | 3.92E-02 | 0.4 | 1.05E-01 | -0.1 | 7.46E-01 | 0.5 | 7.99E-02 | Pentatricopeptide repeat-containing protein At4g38150 |
| AT2G41190 | 0.5 | 4.98E-02 | -0.2 | 5.56E-01 | 0.7 | 4.46E-02 | -0.5 | 4.84E-01 | 0.0 | 9.98E-01 | -0.5 | 6.16E-01 | Transmembrane amino acid transporter family protein |
| AT5G24470 | 0.5 | 2.09E-03 | 0.7 | 4.10E-04 | -0.2 | 2.28E-01 | -0.2 | 5.06E-01 | -0.5 | 1.20E-01 | 0.3 | 4.50E-01 | Two-component response regulator-like APRR5 |
| AT5G61440 | 0.5 | 1.32E-02 | 1.1 | 7.07E-05 | -0.6 | 5.80E-03 | -0.1 | 7.22E-01 | -0.7 | 4.65E-03 | 0.6 | 1.75E-02 | Atypical CYS HIS rich thioredoxin 5 |
| AT4G01590 | 0.5 | 1.06E-03 | 0.4 | 2.60E-03 | 0.0 | 7.52E-01 | 0.4 | 2.53E-02 | 0.2 | 5.18E-01 | 0.3 | 3.08E-01 | DNA-directed RNA polymerase III subunit |
| AT4G00525 | 0.5 | 1.55E-03 | 0.2 | 2.04E-01 | 0.3 | 4.56E-02 | -0.3 | 2.58E-01 | -0.3 | 3.59E-01 | 0.0 | 9.55E-01 | Uncharacterized protein |
| AT5G05360 | 0.5 | 5.35E-04 | 0.4 | 5.59E-03 | 0.1 | 2.56E-01 | 0.0 | 9.40E-01 | -0.1 | 5.50E-01 | 0.1 | 6.32E-01 | Uncharacterized protein |

| | | | | | | | | | | | | | |
|---|---|---|---|---|---|---|---|---|---|---|---|---|---|
| AT1G67865 | 0.5 | 2.51E-01 | 0.1 | 7.94E-01 | 0.4 | 4.64E-01 | 0.0 | 9.08E-01 | 0.0 | 9.59E-01 | 0.0 | 9.80E-01 | At1g67862 |
| AT1G34780 | 0.5 | 1.44E-03 | 0.2 | 7.54E-02 | 0.3 | 7.59E-02 | -0.2 | 3.30E-01 | -0.3 | 1.33E-01 | 0.1 | 6.56E-01 | hypothetical protein |
| AT5G52910 | 0.5 | 8.11E-03 | 0.3 | 9.86E-02 | 0.2 | 2.95E-01 | 0.6 | 1.64E-01 | 0.1 | 9.32E-01 | 0.6 | 3.49E-01 | Timeless family protein |
| AT4G32570 | 0.5 | 1.02E-03 | 0.3 | 2.71E-02 | 0.2 | 1.29E-01 | -0.5 | 1.89E-01 | -0.3 | 4.48E-01 | -0.2 | 7.52E-01 | Protein TIFY |
| AT4G02820 | 0.5 | 3.03E-04 | 0.2 | 3.09E-02 | 0.3 | 3.49E-02 | 0.2 | 1.90E-01 | 0.2 | 2.90E-01 | 0.0 | 9.57E-01 | Pentatricopeptide repeat-containing protein At4g02820, mitochondrial |
| AT2G39080 | 0.5 | 1.78E-04 | 0.2 | 3.09E-02 | 0.3 | 1.73E-02 | 0.1 | 3.87E-01 | -0.1 | 4.28E-01 | 0.3 | 1.48E-01 | Uncharacterized protein At2g39080 |
| AT5G26850 | 0.5 | 1.74E-04 | 0.3 | 1.29E-02 | 0.2 | 3.14E-02 | -0.3 | 7.76E-02 | -0.2 | 3.85E-01 | -0.1 | 6.54E-01 | Uncharacterized protein At5g26850 |
| AT2G03010 | 0.5 | 6.49E-04 | 0.2 | 7.44E-02 | 0.3 | 4.55E-02 | 0.0 | 9.97E-01 | 0.0 | 9.32E-01 | 0.0 | 9.46E-01 | DUF577 domain-containing protein |
| AT2G03440 | 0.5 | 1.79E-03 | 0.4 | 7.32E-03 | 0.1 | 5.58E-01 | 0.1 | 7.85E-01 | -0.2 | 4.09E-01 | 0.2 | 3.87E-01 | Nodulin-related protein 1 |
| AT1G27350 | 0.5 | 4.12E-02 | 0.3 | 1.49E-01 | 0.1 | 5.75E-01 | 0.5 | 5.17E-02 | 0.4 | 1.22E-01 | 0.1 | 9.14E-01 | At1g27330 |
| AT4G38932 | 0.5 | 2.15E-03 | 0.4 | 9.43E-03 | 0.1 | 5.47E-01 | -0.6 | 2.33E-02 | -0.5 | 7.50E-02 | -0.1 | 8.46E-01 | NA |
| AT1G29500 | 0.5 | 2.01E-02 | 0.7 | 3.14E-03 | -0.2 | 3.09E-01 | 0.1 | 8.29E-01 | -0.1 | 8.49E-01 | 0.2 | 7.14E-01 | Auxin-responsive protein SAUR66 |
| AT3G44380 | 0.5 | 1.11E-04 | 0.3 | 2.28E-03 | 0.2 | 1.19E-01 | 0.3 | 2.18E-01 | 0.0 | 9.02E-01 | 0.2 | 4.66E-01 | Water stress and hypersensitive response domain-containing protein |
| AT4G13235 | 0.5 | 2.31E-02 | 0.2 | 3.44E-01 | 0.3 | 2.23E-01 | 0.2 | 5.27E-01 | 0.2 | 6.20E-01 | 0.0 | 9.80E-01 | hypothetical protein |
| AT3G01890 | 0.5 | 9.44E-05 | 0.0 | 7.71E-01 | 0.5 | 9.53E-04 | 0.1 | 8.35E-01 | 0.0 | 9.91E-01 | 0.1 | 8.93E-01 | F1C9.33 protein |
| AT4G28450 | 0.5 | 6.26E-04 | 0.2 | 1.30E-01 | 0.3 | 2.10E-02 | 0.7 | 1.95E-04 | 0.1 | 5.86E-01 | 0.6 | 7.30E-03 | DDB1- and CUL4-associated factor 13 |
| AT1G62250 | 0.5 | 1.12E-02 | 0.1 | 5.26E-01 | 0.4 | 6.55E-02 | -0.1 | 4.94E-01 | 0.0 | 8.51E-01 | -0.2 | 4.28E-01 | F19K23.17 protein |
| AT5G25910 | 0.5 | 3.77E-02 | -0.1 | 5.72E-01 | 0.6 | 4.13E-02 | 0.4 | 4.47E-01 | -0.3 | 7.59E-01 | 0.7 | 3.58E-01 | Receptor-like protein 52 |
| AT1G28510 | 0.5 | 6.52E-04 | 0.5 | 7.75E-04 | 0.0 | 9.04E-01 | 0.2 | 9.92E-02 | 0.0 | 9.15E-01 | 0.2 | 2.96E-01 | hypothetical protein |
| AT2G13975 | 0.5 | 1.08E-01 | -0.3 | 4.17E-01 | 0.8 | 5.81E-02 | 0.1 | 8.76E-01 | -0.2 | 7.75E-01 | 0.2 | 6.91E-01 | NA |
| AT5G50130 | 0.5 | 5.45E-03 | -0.2 | 2.90E-01 | 0.7 | 4.69E-03 | 0.4 | 2.89E-01 | 0.3 | 4.04E-01 | 0.0 | 9.47E-01 | NAD-binding Rossmann-fold superfamily protein |
| AT2G31400 | 0.5 | 1.09E-04 | 0.2 | 1.93E-02 | 0.3 | 1.24E-02 | 0.4 | 9.57E-02 | 0.3 | 3.13E-01 | 0.1 | 7.85E-01 | S uncoupled 1 |
| AT3G57860 | 0.5 | 1.41E-03 | 0.4 | 4.24E-03 | 0.1 | 7.13E-01 | -0.2 | 3.14E-01 | -0.1 | 7.87E-01 | -0.1 | 6.81E-01 | Protein GIGAS CELL1 |
| AT4G30010 | 0.5 | 1.37E-03 | 0.3 | 1.73E-02 | 0.2 | 2.27E-01 | 0.1 | 4.27E-01 | 0.1 | 5.98E-01 | 0.0 | 9.48E-01 | hypothetical protein |
| AT1G33820 | 0.5 | 1.89E-01 | 1.8 | 3.13E-04 | -1.3 | 4.79E-03 | | | | | | | Uncharacterized protein |
| AT5G23890 | 0.5 | 5.83E-06 | 0.2 | 2.72E-03 | 0.3 | 1.30E-03 | -0.1 | 5.49E-01 | -0.1 | 5.72E-01 | 0.0 | 9.75E-01 | GPI-anchored adhesin-like protein |
| AT2G16270 | 0.5 | 2.95E-03 | 0.6 | 8.95E-04 | -0.1 | 4.01E-01 | 0.1 | 6.59E-01 | 0.3 | 2.26E-01 | -0.2 | 5.01E-01 | Transmembrane protein |
| AT5G43720 | 0.5 | 1.17E-04 | 0.3 | 4.91E-03 | 0.2 | 5.88E-02 | 0.1 | 7.62E-01 | -0.1 | 6.67E-01 | 0.1 | 5.16E-01 | rRNA-processing protein EFG1 |
| AT4G27490 | 0.5 | 3.00E-03 | 0.3 | 8.88E-02 | 0.2 | 1.39E-01 | 0.4 | 1.04E-02 | 0.1 | 5.36E-01 | 0.3 | 1.51E-01 | 3'-5'-exoribonuclease family protein |
| AT4G38370 | 0.5 | 1.11E-03 | 0.4 | 3.77E-03 | 0.1 | 6.43E-01 | -0.2 | 3.56E-01 | -0.2 | 3.47E-01 | 0.0 | 9.52E-01 | Phosphoglycerate mutase family protein |
| AT1G14280 | 0.5 | 4.26E-04 | 0.5 | 1.37E-03 | 0.1 | 6.30E-01 | -0.3 | 1.54E-01 | -0.5 | 2.39E-02 | 0.2 | 4.23E-01 | Protein PHYTOCHROME KINASE SUBSTRATE 2 |
| AT3G28890 | 0.5 | 1.25E-01 | -0.3 | 4.31E-01 | 0.8 | 7.78E-02 | | | | | | | Receptor-like protein 43 |
| AT2G15280 | 0.5 | 2.48E-03 | 0.5 | 6.82E-03 | 0.1 | 7.25E-01 | 0.1 | 5.60E-01 | 0.0 | 9.87E-01 | 0.1 | 6.82E-01 | Reticulon-like protein B10 |
| AT1G12090 | 0.5 | 2.89E-04 | 0.8 | 3.02E-05 | -0.3 | 3.01E-02 | -0.8 | 3.27E-04 | -0.4 | 2.81E-02 | -0.3 | 1.91E-01 | hypothetical protein |
| AT1G61795 | 0.5 | 4.29E-02 | 0.7 | 1.09E-02 | -0.2 | 4.71E-01 | -0.6 | 1.49E-01 | 0.0 | 9.63E-01 | -0.5 | 1.90E-01 | CRIB domain-containing protein |
| AT2G41600 | 0.5 | 2.03E-04 | 0.5 | 7.15E-04 | 0.1 | 6.16E-01 | 0.2 | 2.23E-01 | -0.1 | 4.84E-01 | 0.4 | 6.51E-02 | Mitochondrial glycoprotein family protein |
| AT2G44820 | 0.5 | 1.41E-03 | 0.2 | 9.33E-02 | 0.3 | 6.26E-02 | 0.1 | 4.75E-01 | -0.1 | 6.22E-01 | 0.2 | 3.22E-01 | Axoneme-associated protein MST101 protein |
| AT2G43110 | 0.5 | 5.59E-03 | 0.5 | 8.31E-03 | 0.0 | 9.42E-01 | 0.4 | 9.75E-02 | 0.4 | 1.60E-01 | 0.0 | 9.64E-01 | U3 containing 90S pre-ribosomal complex subunit |
| AT1G15830 | 0.5 | 1.40E-02 | 1.1 | 1.01E-04 | -0.6 | 9.58E-03 | | | | | | | Uncharacterized protein |
| AT1G10990 | 0.5 | 2.40E-02 | 0.6 | 1.29E-02 | -0.1 | 7.28E-01 | 0.7 | 1.10E-03 | 0.1 | 7.95E-01 | 0.7 | 4.25E-03 | Transmembrane protein |
| AT5G09995 | 0.5 | 4.37E-04 | 0.5 | 1.28E-03 | 0.0 | 7.09E-01 | 0.2 | 3.92E-01 | 0.2 | 6.46E-01 | 0.1 | 8.62E-01 | Transmembrane protein |
| AT1G18460 | 0.5 | 5.59E-04 | 0.2 | 1.31E-01 | 0.3 | 1.74E-02 | -0.2 | 3.96E-01 | -0.2 | 5.64E-01 | 0.0 | 9.14E-01 | F15H18.6 |
| AT1G16825 | 0.5 | 1.58E-03 | 0.3 | 4.99E-02 | 0.2 | 1.40E-01 | -0.1 | 6.65E-01 | -0.1 | 7.92E-01 | 0.0 | 9.57E-01 | hypothetical protein |
| AT1G19990 | 0.5 | 1.04E-03 | 0.6 | 8.18E-04 | 0.0 | 7.31E-01 | 0.1 | 8.45E-01 | 0.0 | 9.84E-01 | 0.1 | 8.76E-01 | T20H2.23 protein |
| AT5G15910 | 0.5 | 8.86E-04 | 0.2 | 1.48E-01 | 0.3 | 2.39E-02 | 0.4 | 1.42E-01 | 0.5 | 8.24E-02 | -0.1 | 7.76E-01 | Uncharacterized protein F1N13_50 |
| AT2G04795 | 0.5 | 2.64E-03 | 0.2 | 1.59E-01 | 0.3 | 7.94E-02 | 0.0 | 9.34E-01 | -1.0 | 3.90E-02 | 0.9 | 8.16E-02 | AT2g04793 |
| AT2G20585 | 0.5 | 2.81E-03 | 0.6 | 1.98E-03 | -0.1 | 7.41E-01 | 0.8 | 4.87E-04 | 0.2 | 3.16E-01 | 0.5 | 2.12E-02 | Nuclear fusion defective 6 |
| AT2G32840 | 0.5 | 6.76E-05 | 0.1 | 3.10E-01 | 0.4 | 1.39E-03 | -0.1 | 4.97E-01 | 0.2 | 4.38E-01 | -0.3 | 2.20E-01 | Proline-rich family protein |
| AT5G06790 | 0.5 | 2.31E-02 | 0.5 | 5.03E-02 | 0.1 | 8.06E-01 | 0.4 | 1.71E-01 | 0.5 | 1.84E-01 | 0.0 | 9.55E-01 | Cotton fiber protein |
| AT3G06355 | 0.5 | 2.95E-04 | -0.1 | 5.76E-01 | 0.6 | 7.41E-04 | -0.3 | 5.67E-01 | -0.5 | 3.77E-01 | 0.2 | 8.01E-01 | NA |
| AT5G49560 | 0.5 | 3.07E-03 | 0.5 | 8.32E-03 | 0.1 | 7.45E-01 | 0.4 | 1.51E-01 | 0.2 | 6.53E-01 | 0.2 | 5.11E-01 | Methyltransferase family protein |
| AT1G66590 | 0.5 | 1.49E-02 | 0.4 | 4.56E-02 | 0.1 | 6.85E-01 | 0.8 | 2.80E-02 | -0.1 | 9.19E-01 | 0.8 | 5.32E-02 | Cytochrome c oxidase 19-1 |
| AT3G09890 | 0.5 | 3.41E-04 | 0.4 | 1.57E-03 | 0.1 | 5.03E-01 | 0.2 | 3.36E-01 | 0.0 | 9.67E-01 | 0.2 | 5.15E-01 | Ankyrin repeat family protein |
| AT1G66890 | 0.5 | 9.14E-03 | 0.3 | 7.51E-02 | 0.2 | 3.68E-01 | 0.0 | 9.52E-01 | -0.1 | 8.33E-01 | 0.1 | 8.15E-01 | hypothetical protein |
| AT3G09700 | 0.5 | 9.88E-03 | 0.2 | 3.28E-01 | 0.3 | 1.28E-01 | 0.7 | 4.08E-02 | 0.6 | 1.42E-01 | 0.1 | 7.79E-01 | J domain-containing protein |
| AT3G05370 | 0.5 | 5.91E-04 | 0.5 | 1.55E-03 | 0.0 | 7.56E-01 | 0.5 | 9.23E-02 | 0.1 | 8.40E-01 | 0.4 | 2.10E-01 | hypothetical protein |
| AT4G27010 | 0.5 | 1.32E-03 | 0.1 | 5.13E-01 | 0.4 | 1.36E-02 | 0.6 | 6.48E-02 | 0.5 | 2.50E-01 | 0.2 | 7.16E-01 | Ribosome 60S biogenesis amino-terminal protein |
| AT2G46280 | 0.5 | 8.96E-04 | 0.5 | 2.34E-03 | 0.0 | 7.50E-01 | 0.3 | 2.74E-02 | 0.0 | 9.88E-01 | 0.3 | 9.58E-02 | TGF-beta receptor interacting protein 1 |
| AT5G66140 | 0.5 | 6.34E-04 | 0.2 | 1.12E-01 | 0.3 | 1.93E-02 | 0.2 | 2.13E-01 | 0.1 | 6.32E-01 | 0.1 | 6.84E-01 | Proteasome subunit alpha type-7-B |
| AT3G06610 | 0.5 | 2.87E-03 | 0.6 | 1.84E-03 | -0.1 | 6.91E-01 | 0.2 | 3.51E-01 | -0.1 | 8.29E-01 | 0.3 | 3.73E-01 | Nascent polypeptide-associated complex subunit alpha-like UBA domain-containing protein |
| AT3G11650 | 0.5 | 7.04E-04 | 0.3 | 1.90E-02 | 0.2 | 1.30E-01 | 0.2 | 3.28E-01 | 0.1 | 6.11E-01 | 0.1 | 8.45E-01 | NDR1 |
| AT4G11640 | 0.5 | 9.69E-04 | 0.3 | 1.72E-02 | 0.2 | 2.13E-01 | 0.2 | 5.73E-01 | 0.0 | 9.28E-01 | 0.1 | 7.50E-01 | Uncharacterized protein At4g11640 |
| AT4G19070 | 0.5 | 2.24E-03 | 0.3 | 9.73E-02 | 0.3 | 1.09E-01 | 0.0 | 9.29E-01 | -0.1 | 8.68E-01 | 0.1 | 8.33E-01 | Cadmium-induced protein |
| AT3G52155 | 0.5 | 1.86E-04 | 0.3 | 4.70E-03 | 0.2 | 1.05E-01 | 0.0 | 8.07E-01 | 0.0 | 8.73E-01 | 0.0 | 9.81E-01 | Phosphoglycerate mutase family protein |
| AT1G80440 | 0.5 | 1.94E-01 | 0.3 | 4.98E-01 | 0.2 | 6.03E-01 | -0.9 | 3.04E-04 | -2.2 | 5.32E-07 | 1.3 | 1.17E-03 | F-box |
| AT2G46830 | 0.5 | 4.96E-03 | -0.3 | 5.36E-02 | 0.9 | 8.02E-04 | | | | | | | CCA1 |
| AT4G21530 | 0.5 | 2.30E-04 | 0.2 | 4.75E-02 | 0.3 | 2.09E-02 | 0.1 | 5.42E-01 | -0.1 | 6.06E-01 | 0.3 | 3.38E-01 | Anaphase-promoting complex subunit 4 |
| AT3G08610 | 0.5 | 7.07E-05 | 0.5 | 3.12E-04 | 0.1 | 4.95E-01 | 0.2 | 4.40E-01 | 0.0 | 9.15E-01 | 0.1 | 7.10E-01 | NADH dehydrogenase 1 alpha subcomplex subunit 1 |
| AT3G18510 | 0.5 | 2.42E-02 | 0.6 | 1.27E-02 | -0.1 | 7.29E-01 | 0.5 | 1.74E-01 | 0.3 | 5.65E-01 | 0.2 | 6.38E-01 | ATP-dependent helicase |
| AT1G19730 | 0.5 | 6.00E-03 | 0.3 | 1.11E-01 | 0.2 | 2.05E-01 | 0.2 | 5.14E-01 | 0.3 | 4.78E-01 | -0.1 | 9.33E-01 | hypothetical protein |
| AT5G42030 | 0.5 | 8.57E-05 | 0.2 | 1.42E-02 | 0.3 | 1.24E-02 | -0.3 | 1.47E-02 | 0.0 | 9.81E-01 | -0.3 | 6.89E-02 | ABL interactor-like protein 4 |
| AT3G07310 | 0.5 | 1.58E-03 | 0.5 | 2.64E-03 | 0.0 | 9.25E-01 | -0.6 | 9.47E-04 | -0.3 | 7.84E-02 | -0.3 | 1.33E-01 | Uncharacterized protein |
| AT1G64430 | 0.5 | 1.63E-04 | 0.1 | 1.85E-01 | 0.4 | 3.61E-03 | -0.1 | 6.19E-01 | -0.1 | 7.92E-01 | 0.0 | 9.08E-01 | Pentatricopeptide repeat superfamily protein |
| AT2G03430 | 0.5 | 3.61E-04 | 0.2 | 6.23E-02 | 0.3 | 2.43E-02 | 0.3 | 6.04E-02 | 0.1 | 5.80E-01 | 0.2 | 3.76E-01 | hypothetical protein |
| AT2G35650 | 0.5 | 1.10E-04 | 0.5 | 2.27E-04 | 0.0 | 8.89E-01 | -0.4 | 2.86E-02 | -0.2 | 3.09E-01 | -0.2 | 4.61E-01 | Cellulose synthase like |
| AT2G45170 | 0.5 | 3.10E-02 | 0.3 | 2.81E-01 | 0.3 | 2.95E-01 | 0.6 | 9.70E-03 | -0.4 | 1.35E-01 | 0.9 | 3.69E-03 | Autophagy-related protein 8e |
| AT1G01770 | 0.5 | 6.29E-04 | 0.3 | 4.12E-02 | 0.3 | 6.13E-02 | -0.4 | 4.13E-02 | -0.4 | 9.43E-02 | 0.0 | 9.11E-01 | Propionyl-CoA carboxylase |
| AT1G07590 | 0.5 | 9.57E-02 | 0.1 | 8.73E-01 | 0.5 | 1.73E-01 | -0.3 | 4.07E-01 | -0.4 | 3.50E-01 | 0.1 | 8.90E-01 | Pentatricopeptide repeat-containing protein At1g07590, mitochondrial |
| AT2G01640 | 0.5 | 2.62E-03 | 0.4 | 1.80E-02 | 0.1 | 4.15E-01 | 0.4 | 2.20E-01 | 0.2 | 7.17E-01 | 0.3 | 6.23E-01 | Ribosome biogenesis protein slx9-like |
| AT1G02010 | 0.5 | 1.36E-03 | 0.0 | 8.08E-01 | 0.6 | 4.16E-03 | -0.2 | 5.82E-01 | 0.0 | 9.40E-01 | -0.2 | 7.30E-01 | Secretory 1A |
| AT2G44870 | 0.5 | 6.23E-06 | 0.4 | 9.25E-05 | 0.1 | 6.53E-02 | 0.4 | 2.02E-03 | 0.2 | 1.32E-01 | 0.2 | 2.51E-01 | Replicase polyprotein 1ab protein |
| AT2G35170 | 0.5 | 2.81E-04 | 0.3 | 6.58E-03 | 0.2 | 1.27E-01 | 0.0 | 9.65E-01 | 0.1 | 6.56E-01 | -0.1 | 7.49E-01 | Histone H3 K4-specific methyltransferase SET7 |
| AT1G74640 | 0.5 | 7.47E-06 | 0.4 | 8.87E-05 | 0.1 | 9.84E-02 | 0.3 | 7.51E-02 | 0.1 | 5.06E-01 | 0.1 | 5.14E-01 | Uncharacterized protein F1M20.32 |

| | | | | | | | | | | | | | |
|---|---|---|---|---|---|---|---|---|---|---|---|---|---|
| AT3G23050 | 0.5 | 8.61E-05 | 0.7 | 2.80E-05 | -0.2 | 1.19E-01 | -0.3 | 8.39E-03 | -0.4 | 4.79E-03 | 0.1 | 7.34E-01 | Auxin-responsive protein |
| AT4G22000 | 0.5 | 1.82E-04 | 0.4 | 1.75E-03 | 0.1 | 2.43E-01 | 0.3 | 2.30E-01 | 0.1 | 6.92E-01 | 0.2 | 6.80E-01 | hypothetical protein |
| AT5G44578 | 0.5 | 7.40E-02 | -0.2 | 5.79E-01 | 0.7 | 6.28E-02 | 0.0 | 9.39E-01 | 0.3 | 4.41E-01 | -0.3 | 4.62E-01 | Transmembrane protein |
| AT2G22425 | 0.5 | 6.75E-05 | 0.4 | 7.38E-04 | 0.1 | 1.79E-01 | -0.1 | 7.84E-01 | -0.2 | 4.36E-01 | 0.1 | 7.20E-01 | Signal peptidase complex subunit 1 |
| AT5G66540 | 0.5 | 1.95E-03 | 0.3 | 3.41E-02 | 0.2 | 1.93E-01 | 0.9 | 5.41E-05 | 0.5 | 1.20E-02 | 0.4 | 5.34E-02 | U3 small nucleolar ribonucleoprotein protein MPP10 |
| AT1G60870 | 0.5 | 3.60E-03 | 0.3 | 7.38E-02 | 0.2 | 1.64E-01 | -0.1 | 7.96E-01 | -0.3 | 1.93E-01 | 0.3 | 4.43E-01 | Maternal effect embryo arrest 9 |
| AT1G78620 | 0.5 | 8.58E-06 | 0.2 | 7.21E-03 | 0.3 | 1.15E-03 | 0.4 | 3.60E-03 | 0.3 | 5.66E-02 | 0.1 | 4.27E-01 | Protein VTE6, chloroplastic |
| AT2G01110 | 0.5 | 2.91E-05 | 0.4 | 2.35E-04 | 0.1 | 2.31E-01 | -0.2 | 7.78E-02 | -0.1 | 7.22E-01 | -0.2 | 3.52E-01 | AT2G01110 protein |
| AT3G17890 | 0.5 | 1.08E-02 | 0.4 | 5.82E-02 | 0.1 | 5.09E-01 | 0.0 | 9.52E-01 | -0.3 | 5.63E-01 | 0.2 | 6.80E-01 | Uncharacterized protein At3g17890 |
| AT2G35795 | 0.5 | 8.23E-05 | 0.4 | 1.40E-03 | 0.2 | 1.31E-01 | 0.2 | 2.70E-01 | 0.0 | 8.92E-01 | 0.2 | 5.18E-01 | J domain-containing protein |
| AT1G77122 | 0.5 | 1.29E-03 | 0.4 | 1.13E-02 | 0.1 | 3.39E-01 | -0.3 | 2.68E-01 | -0.1 | 7.76E-01 | -0.2 | 5.94E-01 | At1g77122 |
| AT1G11240 | 0.5 | 1.52E-03 | 0.5 | 4.16E-03 | 0.1 | 7.09E-01 | 0.1 | 3.83E-01 | -0.1 | 7.23E-01 | 0.2 | 2.75E-01 | Ribosomal RNA-processing protein 17 |
| AT4G17085 | 0.5 | 7.09E-03 | 0.5 | 2.05E-02 | 0.1 | 7.05E-01 | -0.2 | 6.05E-01 | -0.3 | 4.05E-01 | 0.1 | 8.07E-01 | Membrane lipoprotein |
| AT2G39910 | 0.5 | 3.20E-04 | 0.2 | 1.01E-01 | 0.3 | 1.56E-02 | -0.2 | 3.85E-01 | -0.2 | 3.94E-01 | 0.0 | 9.63E-01 | ARM repeat superfamily protein |
| AT3G07060 | 0.5 | 5.37E-04 | 0.2 | 9.29E-02 | 0.3 | 2.81E-02 | 0.4 | 4.99E-02 | 0.1 | 6.09E-01 | 0.3 | 3.36E-01 | NHL domain-containing protein |
| AT3G07500 | 0.5 | 2.77E-03 | 0.0 | 8.71E-01 | 0.5 | 1.50E-02 | 0.1 | 7.49E-01 | -0.6 | 1.20E-01 | 0.7 | 8.94E-02 | Far-red impaired responsive family protein |
| AT1G14180 | 0.5 | 4.07E-04 | 0.6 | 4.99E-04 | 0.0 | 8.85E-01 | -0.1 | 7.67E-01 | 0.1 | 6.63E-01 | -0.2 | 4.68E-01 | F7A19.26 protein |
| AT1G70890 | 0.5 | 3.51E-05 | 0.8 | 4.90E-06 | -0.3 | 1.14E-02 | 0.4 | 1.18E-01 | 0.2 | 4.95E-01 | 0.2 | 6.34E-01 | MLP-like protein 43 |
| AT5G57370 | 0.5 | 2.77E-04 | 0.4 | 1.49E-03 | 0.1 | 4.28E-01 | 0.1 | 5.53E-01 | -0.1 | 4.83E-01 | 0.2 | 3.00E-01 | U4 |
| AT1G29840 | 0.5 | 4.72E-03 | 0.5 | 1.54E-02 | 0.1 | 6.82E-01 | -0.3 | 4.11E-01 | 0.1 | 8.75E-01 | -0.4 | 4.21E-01 | At1g29840 |
| AT3G07568 | 0.5 | 4.83E-03 | 0.7 | 1.23E-03 | -0.2 | 3.82E-01 | 0.3 | 1.80E-01 | 0.1 | 8.59E-01 | 0.3 | 2.57E-01 | Uncharacterized protein |
| AT5G30495 | 0.5 | 1.79E-03 | 0.1 | 4.40E-01 | 0.4 | 2.12E-02 | 0.3 | 3.02E-01 | 0.2 | 7.07E-01 | 0.2 | 7.37E-01 | hypothetical protein |
| AT2G05995 | 0.5 | 3.32E-02 | 1.4 | 9.56E-05 | -0.9 | 4.80E-03 | 0.6 | 1.45E-01 | 0.1 | 8.39E-01 | 0.5 | 3.55E-01 | NA |
| AT1G04650 | 0.5 | 4.16E-03 | 0.3 | 8.23E-02 | 0.2 | 2.19E-01 | | | | | | | Holliday junction resolvase |
| AT2G41650 | 0.5 | 4.10E-04 | 0.4 | 2.64E-03 | 0.1 | 3.66E-01 | 0.6 | 4.45E-02 | 0.3 | 4.07E-01 | 0.3 | 4.77E-01 | hypothetical protein |
| AT3G02210 | 0.5 | 3.74E-04 | 0.4 | 3.45E-03 | 0.1 | 3.19E-01 | -0.2 | 5.99E-01 | -0.4 | 2.34E-01 | 0.2 | 4.66E-01 | COBRA-like protein 1 |
| AT5G03455 | 0.5 | 1.88E-03 | 0.1 | 3.11E-01 | 0.4 | 2.68E-02 | 0.0 | 9.02E-01 | 0.1 | 7.85E-01 | 0.0 | 9.04E-01 | Rhodanese |
| AT3G41768 | 0.5 | 2.17E-01 | -0.1 | 8.35E-01 | 0.6 | 1.96E-01 | 0.6 | 5.80E-01 | 0.2 | 9.14E-01 | 0.4 | 7.92E-01 | NA |
| AT1G21680 | 0.5 | 1.43E-03 | 0.4 | 8.56E-03 | 0.1 | 4.16E-01 | -0.1 | 6.54E-01 | -0.6 | 1.47E-02 | 0.5 | 5.13E-02 | DPP6 N-terminal domain-like protein |
| AT2G21580 | 0.5 | 3.11E-03 | 0.7 | 9.55E-04 | -0.2 | 3.47E-01 | 0.3 | 9.25E-02 | 0.2 | 4.89E-01 | 0.1 | 5.76E-01 | 40S ribosomal protein S25 |
| AT5G06220 | 0.5 | 1.30E-03 | 0.0 | 7.49E-01 | 0.5 | 5.89E-03 | 0.1 | 9.22E-01 | 0.0 | 9.92E-01 | 0.0 | 9.53E-01 | LETM1-like protein |
| AT2G37920 | 0.5 | 1.15E-03 | 0.4 | 1.55E-02 | 0.2 | 2.24E-01 | 0.0 | 8.85E-01 | -0.1 | 8.29E-01 | 0.1 | 7.49E-01 | Uncharacterized protein At2g37920 |
| AT5G52810 | 0.5 | 2.27E-02 | -0.2 | 3.63E-01 | 0.8 | 1.14E-02 | 1.0 | 1.49E-04 | 0.6 | 1.01E-02 | 0.3 | 1.36E-01 | AT5G52810 protein |
| AT3G26510 | 0.5 | 2.91E-02 | 0.3 | 2.60E-01 | 0.3 | 3.24E-01 | -0.1 | 7.18E-01 | -0.6 | 1.48E-02 | 0.5 | 6.96E-02 | PB1 domain-containing protein |
| AT1G35830 | 0.5 | 2.38E-03 | 0.4 | 1.71E-02 | 0.1 | 4.43E-01 | -0.2 | 7.60E-01 | 0.2 | 7.90E-01 | -0.3 | 6.13E-01 | hypothetical protein |
| AT1G48170 | 0.5 | 2.01E-03 | 0.3 | 7.44E-02 | 0.3 | 1.21E-01 | 0.3 | 3.82E-01 | -0.1 | 9.21E-01 | 0.4 | 4.33E-01 | Proteasome assembly chaperone 4 |
| AT3G15390 | 0.5 | 1.98E-03 | 0.2 | 1.39E-01 | 0.3 | 6.83E-02 | 0.2 | 5.41E-01 | 0.0 | 9.54E-01 | 0.2 | 6.04E-01 | DUF1771 domain-containing protein |
| AT5G57780 | 0.5 | 1.32E-02 | 0.8 | 1.77E-03 | -0.3 | 2.51E-01 | -0.7 | 5.33E-03 | -0.7 | 7.24E-03 | 0.0 | 9.71E-01 | At5g57780 |
| AT4G24920 | 0.5 | 9.96E-03 | 0.4 | 7.04E-02 | 0.2 | 3.75E-01 | 0.5 | 5.95E-02 | 0.2 | 5.47E-01 | 0.3 | 4.53E-01 | SecE |
| AT2G29680 | 0.5 | 2.63E-03 | 0.4 | 3.69E-02 | 0.2 | 2.68E-01 | 0.0 | 9.93E-01 | 0.1 | 8.16E-01 | -0.1 | 8.47E-01 | Cell division control protein 6 homolog |
| AT3G58940 | 0.5 | 8.90E-04 | 0.6 | 8.93E-04 | 0.0 | 8.29E-01 | -0.1 | 8.75E-01 | -0.2 | 6.15E-01 | 0.1 | 7.70E-01 | F-box |
| AT3G09800 | 0.5 | 8.78E-04 | 0.6 | 1.14E-03 | 0.0 | 9.48E-01 | 0.1 | 8.04E-01 | 0.0 | 9.59E-01 | 0.1 | 8.15E-01 | Coatomer subunit zeta |
| AT5G48240 | 0.5 | 2.88E-03 | 0.4 | 1.90E-02 | 0.1 | 4.19E-01 | 0.4 | 2.34E-02 | 0.0 | 9.96E-01 | 0.4 | 6.74E-02 | Rrp15p protein |
| AT3G26980 | 0.5 | 3.45E-04 | 0.2 | 1.29E-01 | 0.4 | 1.32E-02 | 0.3 | 2.05E-01 | 0.1 | 8.51E-01 | 0.2 | 4.51E-01 | Membrane-anchored ubiquitin-fold protein 4 |
| AT1G71730 | 0.5 | 2.63E-04 | 0.4 | 3.67E-03 | 0.2 | 1.73E-01 | -0.1 | 4.05E-01 | -0.3 | 1.42E-01 | 0.1 | 6.23E-01 | Embryogenesis-like protein |
| AT3G56430 | 0.6 | 3.27E-04 | 0.4 | 4.08E-03 | 0.2 | 2.20E-01 | 0.2 | 3.07E-01 | 0.0 | 9.18E-01 | 0.1 | 5.72E-01 | At3g56430 |
| AT3G06660 | 0.6 | 6.50E-04 | 0.3 | 1.54E-02 | 0.2 | 1.50E-01 | -0.1 | 5.49E-01 | -0.1 | 6.10E-01 | 0.0 | 1.00E+00 | PAPA-1-like family protein |
| AT1G72240 | 0.6 | 1.57E-01 | -0.5 | 2.83E-01 | 1.1 | 5.65E-02 | -1.1 | 5.29E-02 | -0.2 | 7.95E-01 | | | Uncharacterized protein |
| AT3G26395 | 0.6 | 1.47E-02 | 0.4 | 5.30E-02 | 0.1 | 6.17E-01 | 0.1 | 7.86E-01 | -0.1 | 7.00E-01 | 0.2 | 5.90E-01 | Uncharacterized protein |
| AT4G24370 | 0.6 | 1.01E-02 | 0.3 | 1.95E-01 | 0.3 | 2.12E-01 | 0.0 | 8.81E-01 | -0.4 | 6.34E-02 | 0.5 | 9.58E-02 | Uncharacterized protein At4g24370 |
| AT1G23280 | 0.6 | 3.25E-03 | 0.3 | 4.45E-02 | 0.2 | 2.40E-01 | 0.6 | 4.14E-03 | 0.3 | 2.25E-01 | 0.3 | 1.46E-01 | Protein MAK16 homolog |
| AT5G46280 | 0.6 | 1.64E-04 | 0.2 | 3.15E-02 | 0.3 | 1.70E-02 | 0.2 | 3.35E-01 | 0.2 | 4.33E-01 | 0.0 | 9.78E-01 | At5g46280 |
| AT4G21190 | 0.6 | 1.09E-03 | 0.5 | 2.76E-03 | 0.0 | 7.64E-01 | 0.0 | 8.75E-01 | 0.0 | 9.29E-01 | 0.1 | 8.52E-01 | Pentatricopeptide repeat-containing protein At4g21190 |
| AT5G47380 | 0.6 | 4.05E-03 | -0.1 | 4.19E-01 | 0.7 | 4.65E-03 | 0.0 | 8.84E-01 | 0.2 | 5.32E-01 | -0.1 | 7.11E-01 | Uncharacterized protein At5g47380 |
| AT1G31335 | 0.6 | 1.02E-04 | 0.7 | 4.38E-05 | -0.1 | 2.62E-01 | -0.4 | 2.62E-01 | 0.0 | 9.74E-01 | -0.4 | 4.38E-01 | hypothetical protein |
| AT3G10440 | 0.6 | 1.64E-04 | 0.1 | 5.05E-01 | 0.5 | 2.12E-03 | -0.3 | 1.76E-01 | -0.1 | 5.52E-01 | -0.1 | 6.94E-01 | Shugoshin C terminus |
| AT1G21460 | 0.6 | 6.30E-03 | 0.2 | 1.85E-01 | 0.3 | 1.32E-01 | -0.5 | 1.03E-02 | -0.1 | 6.35E-01 | -0.4 | 1.07E-01 | Nodulin MtN3 family protein |
| AT3G53630 | 0.6 | 3.78E-04 | 0.4 | 2.52E-03 | 0.1 | 3.56E-01 | 0.5 | 2.58E-03 | 0.0 | 8.35E-01 | 0.5 | 8.36E-03 | Uncharacterized protein F4P12_330 |
| AT4G22670 | 0.6 | 4.04E-04 | 0.5 | 2.18E-03 | 0.1 | 4.37E-01 | 0.2 | 1.13E-01 | -0.1 | 4.61E-01 | 0.3 | 7.68E-02 | TPR11 |
| AT3G46870 | 0.6 | 3.83E-04 | 0.2 | 5.31E-02 | 0.3 | 3.09E-02 | 0.8 | 2.27E-03 | 0.6 | 2.80E-02 | 0.2 | 5.14E-01 | Pentatricopeptide repeat superfamily protein |
| AT1G03860 | 0.6 | 9.79E-04 | 0.3 | 3.49E-02 | 0.3 | 8.64E-02 | 0.4 | 2.60E-03 | 0.1 | 3.68E-01 | 0.3 | 1.32E-01 | Prohibitin |
| AT5G59460 | 0.6 | 7.39E-04 | 0.3 | 2.60E-02 | 0.2 | 1.06E-01 | 0.1 | 7.82E-01 | -0.1 | 7.38E-01 | 0.2 | 5.80E-01 | At5g59460 ) |
| AT3G62400 | 0.6 | 7.93E-03 | 0.4 | 4.63E-02 | 0.2 | 4.34E-01 | 0.3 | 1.79E-01 | 0.1 | 7.87E-01 | 0.2 | 4.54E-01 | Cytochrome c oxidase subunit 5C-2 |
| AT2G03350 | 0.6 | 2.79E-04 | 0.5 | 1.73E-03 | 0.1 | 3.88E-01 | 0.0 | 8.78E-01 | -0.2 | 1.52E-01 | 0.2 | 3.07E-01 | hypothetical protein |
| AT3G46430 | 0.6 | 6.45E-04 | 0.4 | 6.11E-03 | 0.2 | 2.67E-01 | 0.4 | 2.91E-02 | 0.3 | 1.09E-01 | 0.1 | 8.26E-01 | Uncharacterized protein |
| AT3G24570 | 0.6 | 2.04E-05 | 0.5 | 9.29E-05 | 0.1 | 4.82E-01 | -0.1 | 7.38E-01 | -0.3 | 1.66E-01 | 0.2 | 4.06E-01 | Peroxisomal membrane 22 kDa family protein |
| AT1G55340 | 0.6 | 4.88E-05 | 0.5 | 2.59E-04 | 0.1 | 3.99E-01 | 0.1 | 3.91E-01 | -0.1 | 4.98E-01 | 0.2 | 1.50E-01 | Uncharacterized protein |
| AT4G37190 | 0.6 | 3.85E-05 | 0.4 | 5.53E-04 | 0.1 | 1.40E-01 | 0.5 | 1.67E-02 | -0.1 | 7.55E-01 | 0.6 | 1.83E-02 | At4g37190 |
| AT2G24860 | 0.6 | 1.17E-04 | 0.4 | 1.61E-03 | 0.2 | 1.57E-01 | 0.0 | 8.97E-01 | -0.1 | 8.01E-01 | 0.0 | 9.18E-01 | At2g24860 |
| AT3G49290 | 0.6 | 1.50E-03 | 0.4 | 1.37E-02 | 0.2 | 3.31E-01 | -0.1 | 8.48E-01 | 0.2 | 4.08E-01 | -0.3 | 3.54E-01 | Protein ABIL2 |
| AT1G67350 | 0.6 | 1.74E-03 | 0.4 | 1.65E-02 | 0.2 | 2.88E-01 | 0.2 | 4.80E-01 | 0.0 | 9.64E-01 | 0.2 | 5.95E-01 | At1g67350 |
| AT2G36630 | 0.6 | 1.77E-04 | -0.2 | 4.04E-02 | 0.8 | 1.00E-04 | -1.0 | 6.05E-03 | -0.5 | 1.40E-01 | -0.5 | 2.48E-01 | Sulfite exporter TauE |
| AT4G14110 | 0.6 | 7.39E-05 | 0.4 | 2.53E-03 | 0.2 | 6.04E-02 | 0.2 | 4.65E-01 | -0.1 | 8.03E-01 | 0.3 | 4.33E-01 | COP9 signalosome complex subunit 8 |
| AT2G47640 | 0.6 | 3.50E-04 | 0.4 | 5.56E-03 | 0.2 | 1.67E-01 | 0.2 | 3.85E-01 | 0.0 | 8.91E-01 | 0.2 | 4.32E-01 | Small nuclear ribonucleoprotein Sm D2 |
| AT2G38780 | 0.6 | 2.19E-03 | 0.3 | 8.76E-02 | 0.3 | 1.03E-01 | 0.0 | 9.47E-01 | 0.2 | 4.18E-01 | -0.2 | 4.52E-01 | At2g38780 |
| AT5G50810 | 0.6 | 4.87E-03 | 0.6 | 6.31E-03 | 0.0 | 9.90E-01 | 0.4 | 1.43E-01 | 0.2 | 4.62E-01 | 0.1 | 6.95E-01 | Mitochondrial import inner membrane translocase subunit |
| AT2G38980 | 0.6 | 1.25E-03 | 0.3 | 6.50E-02 | 0.3 | 8.19E-02 | -0.4 | 8.21E-02 | -0.3 | 3.07E-01 | -0.1 | 6.85E-01 | NA |
| AT5G01910 | 0.6 | 1.19E-03 | 0.5 | 3.50E-03 | 0.1 | 7.32E-01 | 0.3 | 4.30E-01 | 0.3 | 5.42E-01 | 0.0 | 9.70E-01 | Myelin transcription factor |
| AT3G22540 | 0.6 | 7.38E-02 | 0.6 | 5.98E-02 | -0.1 | 8.64E-01 | -0.2 | 7.84E-01 | 0.3 | 6.96E-01 | -0.4 | 4.13E-01 | hypothetical protein |
| AT3G02900 | 0.6 | 1.73E-04 | 0.6 | 1.95E-04 | 0.0 | 8.01E-01 | 0.0 | 8.23E-01 | 0.1 | 7.92E-01 | -0.1 | 6.95E-01 | At3g02900 |
| AT1G19010 | 0.6 | 1.26E-04 | 0.3 | 6.68E-03 | 0.2 | 5.53E-02 | 0.0 | 9.66E-01 | 0.0 | 9.79E-01 | 0.0 | 9.56E-01 | Uncharacterized protein |
| AT4G00026 | 0.6 | 7.57E-04 | 0.4 | 9.15E-03 | 0.2 | 2.41E-01 | 0.5 | 4.49E-03 | 0.1 | 4.94E-01 | 0.4 | 9.22E-02 | Probable mitochondrial import inner membrane translocase subunit TIM21 |
| AT5G25580 | 0.6 | 1.43E-03 | 0.5 | 3.71E-03 | 0.0 | 7.77E-01 | 0.0 | 9.95E-01 | -0.1 | 7.81E-01 | 0.1 | 8.07E-01 | DDT domain-containing protein DDR4 |

| | | | | | | | | | | | | | |
|---|---|---|---|---|---|---|---|---|---|---|---|---|---|
| AT2G14560 | 0.6 | 1.96E-01 | 0.1 | 8.43E-01 | 0.5 | 3.35E-01 | 2.9 | 1.87E-07 | 1.5 | 4.92E-04 | 1.4 | 4.17E-03 | LURP-one-like protein |
| AT4G25740 | 0.6 | 1.36E-03 | 0.8 | 3.11E-04 | -0.2 | 2.03E-01 | 0.2 | 1.76E-01 | 0.1 | 8.38E-01 | 0.2 | 4.51E-01 | RNA binding Plectin |
| AT3G62810 | 0.6 | 1.35E-04 | 0.4 | 1.46E-03 | 0.1 | 2.04E-01 | 0.4 | 7.44E-02 | 0.2 | 4.35E-01 | 0.2 | 5.37E-01 | hypothetical protein |
| AT2G43945 | 0.6 | 1.47E-04 | 0.3 | 9.53E-03 | 0.3 | 3.37E-02 | -0.1 | 5.63E-01 | 0.0 | 9.39E-01 | -0.1 | 7.40E-01 | Uncharacterized protein At2g43940 |
| AT3G15620 | 0.6 | 2.13E-02 | 0.4 | 7.81E-02 | 0.1 | 6.38E-01 | -0.2 | 5.29E-01 | -0.7 | 6.60E-02 | 0.5 | 2.94E-01 | Photolyase |
| AT5G03900 | 0.6 | 7.07E-05 | 0.1 | 2.02E-01 | 0.5 | 1.37E-03 | 0.1 | 4.83E-01 | 0.2 | 3.68E-01 | -0.1 | 8.41E-01 | Uncharacterized protein At5g03900, chloroplastic |
| AT1G17880 | 0.6 | 7.63E-04 | 0.7 | 3.61E-04 | -0.1 | 4.06E-01 | 0.3 | 7.22E-02 | 0.1 | 8.04E-01 | 0.2 | 3.34E-01 | Nascent polypeptide-associated complex subunit beta |
| AT5G54720 | 0.6 | 2.40E-02 | -0.3 | 2.46E-01 | 0.9 | 1.16E-02 | 1.6 | 2.00E-02 | 0.6 | 5.02E-01 | 1.0 | 1.62E-01 | Uncharacterized protein |
| AT5G45520 | 0.6 | 3.91E-03 | 0.4 | 2.04E-02 | 0.1 | 5.25E-01 | 0.2 | 6.64E-01 | 0.4 | 2.20E-01 | -0.3 | 5.11E-01 | Leucine-rich repeat family protein |
| AT1G78915 | 0.6 | 4.07E-03 | 0.5 | 8.05E-03 | 0.0 | 8.53E-01 | 0.3 | 1.83E-01 | 0.1 | 6.09E-01 | 0.1 | 6.53E-01 | F9K20.3 |
| AT1G57790 | 0.6 | 2.53E-03 | 0.1 | 4.87E-01 | 0.5 | 2.74E-02 | -0.3 | 1.72E-01 | -0.1 | 7.54E-01 | -0.2 | 4.57E-01 | F-box |
| AT5G56120 | 0.6 | 1.05E-03 | 0.7 | 3.67E-04 | -0.1 | 3.75E-01 | 0.2 | 3.26E-01 | 0.0 | 9.65E-01 | 0.2 | 5.31E-01 | Uncharacterized protein |
| AT3G52905 | 0.6 | 4.23E-04 | 0.2 | 6.76E-02 | 0.3 | 2.94E-02 | 0.1 | 6.77E-01 | 0.1 | 7.35E-01 | 0.0 | 9.99E-01 | YqgF |
| AT4G19660 | 0.6 | 2.75E-04 | 0.2 | 5.15E-02 | 0.3 | 1.89E-02 | 0.9 | 1.18E-04 | 0.6 | 3.82E-03 | 0.2 | 3.56E-01 | NPR1-like protein 4 |
| AT2G48090 | 0.6 | 2.44E-04 | -0.3 | 2.73E-02 | 0.9 | 1.74E-04 | 0.5 | 1.90E-01 | 0.4 | 4.13E-01 | 0.1 | 7.97E-01 | Uncharacterized protein |
| AT4G10470 | 0.6 | 1.88E-04 | 0.4 | 1.76E-03 | 0.1 | 2.79E-01 | -0.1 | 8.29E-01 | -0.2 | 4.33E-01 | 0.1 | 6.30E-01 | Uncharacterized protein |
| AT2G01090 | 0.6 | 7.42E-05 | 0.5 | 5.55E-04 | 0.1 | 3.10E-01 | 0.1 | 6.83E-01 | 0.1 | 8.54E-01 | 0.0 | 9.15E-01 | Cytochrome b-c1 complex subunit 6-2, mitochondrial |
| AT3G07215 | 0.6 | 1.30E-04 | 0.3 | 1.59E-02 | 0.3 | 2.70E-02 | -0.1 | 6.03E-01 | -0.1 | 5.89E-01 | 0.0 | 9.59E-01 | NA |
| AT2G23090 | 0.6 | 1.27E-02 | 0.1 | 5.84E-01 | 0.5 | 6.03E-02 | 0.2 | 5.29E-01 | 0.0 | 9.76E-01 | 0.2 | 6.54E-01 | Uncharacterized protein At2g23090 |
| AT1G48970 | 0.6 | 6.46E-05 | 0.3 | 1.12E-02 | 0.3 | 1.23E-02 | 0.4 | 1.91E-02 | 0.1 | 5.28E-01 | 0.3 | 2.52E-01 | F27J15.24 |
| AT2G46540 | 0.6 | 4.72E-03 | 0.5 | 9.55E-03 | 0.0 | 8.19E-01 | 0.2 | 4.42E-01 | 0.1 | 8.17E-01 | 0.1 | 7.84E-01 | Fiber protein Fb11 |
| AT2G47440 | 0.6 | 2.16E-02 | 0.2 | 3.38E-01 | 0.4 | 1.76E-01 | -0.8 | 1.16E-04 | -0.6 | 2.54E-03 | -0.2 | 3.99E-01 | Tetratricopeptide repeat -like superfamily protein |
| AT3G57157 | 0.6 | 8.89E-02 | 0.3 | 3.83E-01 | 0.3 | 5.17E-01 | 1.2 | 5.92E-04 | -0.6 | 1.37E-01 | 1.8 | 2.64E-04 | NA |
| AT5G61310 | 0.6 | 5.50E-04 | 0.4 | 5.57E-03 | 0.2 | 2.79E-01 | 0.5 | 1.25E-01 | 0.3 | 4.04E-01 | 0.2 | 7.32E-01 | Probable cytochrome c oxidase subunit 5C-3 |
| AT5G20640 | 0.6 | 2.06E-03 | 0.7 | 5.98E-04 | -0.2 | 3.65E-01 | -0.2 | 5.55E-01 | 0.3 | 5.27E-01 | -0.5 | 2.63E-01 | Protein LURP-one-related 16 |
| AT3G18420 | 0.6 | 4.10E-04 | 0.6 | 7.24E-04 | 0.0 | 9.30E-01 | 0.6 | 3.46E-03 | 0.2 | 4.02E-01 | 0.4 | 9.86E-02 | protein SLOW GREEN 1, chloroplastic |
| AT5G49100 | 0.6 | 1.78E-03 | 0.5 | 1.11E-02 | 0.1 | 4.30E-01 | -0.6 | 1.89E-03 | -0.3 | 1.08E-01 | -0.3 | 2.11E-01 | Vitellogenin-like protein |
| AT3G26490 | 0.6 | 3.14E-03 | 0.4 | 2.37E-02 | 0.2 | 4.17E-01 | -0.2 | 6.53E-01 | 0.1 | 8.27E-01 | -0.3 | 5.47E-01 | Phototropic-responsive NPH3 family protein |
| AT4G22212 | 0.6 | 6.19E-04 | 1.3 | 6.65E-06 | -0.7 | 1.01E-03 | -0.5 | 1.49E-03 | 0.1 | 4.15E-01 | -0.6 | 2.13E-03 | Defensin-like protein 98 |
| AT5G60950 | 0.6 | 4.39E-05 | 0.7 | 3.02E-05 | -0.1 | 3.84E-01 | 0.3 | 1.74E-01 | 0.0 | 8.90E-01 | 0.2 | 3.98E-01 | COBRA-like protein 5 |
| AT1G31810 | 0.6 | 8.49E-05 | 0.1 | 2.48E-01 | 0.5 | 1.51E-03 | 0.3 | 1.33E-01 | 0.2 | 2.39E-01 | 0.0 | 9.27E-01 | Formin-like protein |
| AT5G43745 | 0.6 | 2.19E-03 | 0.4 | 3.20E-02 | 0.2 | 2.31E-01 | 0.3 | 2.66E-01 | 0.7 | 2.65E-02 | -0.4 | 2.52E-01 | Putative ion channel POLLUX-like 2 |
| AT1G62480 | 0.6 | 4.73E-05 | 0.8 | 1.07E-05 | -0.2 | 3.61E-02 | -0.1 | 6.83E-01 | -0.3 | 2.96E-01 | 0.2 | 6.25E-01 | Cytosolic calcium-binding protein 1 -binding protein 1) |
| AT4G21510 | 0.6 | 1.26E-02 | 0.3 | 2.11E-01 | 0.3 | 2.40E-01 | 0.1 | 6.88E-01 | -0.5 | 1.87E-01 | 0.6 | 1.33E-01 | F-box domain-containing protein |
| AT3G47836 | 0.6 | 4.31E-04 | 0.4 | 6.47E-03 | 0.2 | 1.82E-01 | 0.2 | 3.45E-01 | 0.1 | 7.19E-01 | 0.1 | 7.77E-01 | Transmembrane protein |
| AT5G03880 | 0.6 | 9.00E-05 | 0.5 | 3.60E-04 | 0.1 | 5.56E-01 | -0.2 | 1.41E-01 | 0.0 | 9.34E-01 | -0.2 | 3.40E-01 | Thioredoxin family protein |
| AT1G51130 | 0.6 | 4.57E-04 | 0.1 | 2.63E-01 | 0.4 | 1.05E-02 | 0.1 | 6.56E-01 | -0.1 | 8.60E-01 | 0.2 | 5.11E-01 | Non-structural maintenance of chromosomes element 4 homolog A |
| AT5G05240 | 0.6 | 8.26E-05 | 0.5 | 7.07E-04 | 0.1 | 2.76E-01 | 0.5 | 6.09E-02 | 0.3 | 3.53E-01 | 0.2 | 5.57E-01 | Cation-transporting ATPase |
| AT1G72170 | 0.6 | 8.11E-05 | 0.6 | 1.58E-04 | 0.0 | 9.36E-01 | 0.4 | 3.14E-02 | 0.3 | 1.04E-01 | 0.1 | 8.33E-01 | MICOS complex subunit Mic10 |
| AT3G13226 | 0.6 | 1.92E-05 | 0.4 | 7.20E-04 | 0.2 | 3.74E-02 | 0.2 | 3.66E-01 | 0.0 | 9.80E-01 | 0.2 | 5.05E-01 | Regulatory protein RecX family protein |
| AT1G04520 | 0.6 | 1.65E-04 | 0.5 | 5.16E-04 | 0.1 | 6.87E-01 | 0.2 | 3.41E-01 | 0.2 | 4.29E-01 | 0.0 | 9.82E-01 | Gnk2-homologous domain-containing protein |
| AT5G27990 | 0.6 | 1.49E-05 | 0.5 | 1.16E-04 | 0.1 | 2.28E-01 | 0.1 | 7.34E-01 | -0.3 | 1.66E-01 | 0.3 | 1.64E-01 | Pre-rRNA-processing protein TSR2 |
| AT3G06310 | 0.6 | 2.74E-05 | 0.6 | 5.13E-05 | 0.0 | 1.00E+00 | 0.4 | 2.86E-02 | 0.2 | 3.46E-01 | 0.2 | 4.33E-01 | Cox19-like CHCH family protein |
| AT4G21192 | 0.6 | 4.64E-02 | 0.3 | 3.13E-01 | 0.3 | 3.86E-01 | 0.3 | 5.06E-01 | 0.2 | 7.43E-01 | 0.1 | 8.89E-01 | COX assembly mitochondrial protein |
| AT1G77710 | 0.6 | 1.07E-04 | 0.4 | 2.48E-03 | 0.2 | 8.45E-02 | 0.0 | 9.94E-01 | -0.1 | 7.31E-01 | 0.1 | 7.91E-01 | Ubiquitin-fold modifier 1 |
| AT1G76600 | 0.6 | 8.93E-03 | -0.4 | 8.36E-02 | 1.0 | 2.66E-03 | 0.7 | 2.15E-02 | -0.4 | 3.41E-01 | 1.1 | 8.97E-03 | hypothetical protein |
| AT4G29830 | 0.6 | 1.75E-04 | 0.3 | 1.18E-02 | 0.3 | 4.24E-02 | 0.2 | 4.34E-01 | 0.0 | 8.79E-01 | 0.1 | 6.72E-01 | VIP3 |
| AT3G50470 | 0.6 | 6.16E-03 | 0.1 | 5.48E-01 | 0.5 | 4.76E-02 | -0.3 | 4.53E-01 | -0.5 | 1.93E-01 | 0.2 | 6.76E-01 | RPW8-like protein 3 |
| AT1G20390 | 0.6 | 3.74E-04 | -0.2 | 1.50E-01 | 0.8 | 5.34E-04 | -0.8 | 1.02E-01 | -0.5 | 3.55E-01 | -0.3 | 6.64E-01 | NA |
| AT3G10985 | 0.6 | 2.02E-05 | 0.7 | 1.77E-05 | -0.1 | 3.68E-01 | -0.2 | 3.19E-01 | -0.4 | 1.49E-01 | 0.1 | 7.24E-01 | Senescence associated gene 20 |
| AT1G53760 | 0.6 | 6.67E-05 | 0.4 | 2.28E-03 | 0.2 | 6.39E-02 | 0.2 | 4.21E-01 | -0.1 | 6.40E-01 | 0.3 | 3.05E-01 | K+-H+ exchange-like protein |
| AT1G24530 | 0.6 | 7.71E-03 | 0.3 | 2.17E-01 | 0.3 | 1.23E-01 | -0.2 | 5.41E-01 | -0.8 | 8.03E-03 | 0.7 | 5.23E-02 | F21J9.19 |
| AT1G14870 | 0.6 | 1.86E-04 | 0.6 | 1.85E-04 | -0.1 | 6.87E-01 | 0.8 | 2.04E-03 | 0.2 | 6.18E-01 | 0.7 | 5.29E-02 | PLANT CADMIUM RESISTANCE 2 |
| AT4G20150 | 0.6 | 1.29E-04 | 0.4 | 2.03E-03 | 0.2 | 1.36E-01 | 0.3 | 3.81E-01 | 0.1 | 7.54E-01 | 0.1 | 7.90E-01 | Uncharacterized protein AT4g20150 |
| AT4G23760 | 0.6 | 1.90E-04 | 0.6 | 1.97E-04 | 0.0 | 7.63E-01 | 0.1 | 8.57E-01 | -0.1 | 6.56E-01 | 0.2 | 4.96E-01 | hypothetical protein |
| AT2G37240 | 0.6 | 3.57E-04 | 0.0 | 7.19E-01 | 0.6 | 1.95E-03 | 0.5 | 2.51E-03 | 0.6 | 6.54E-04 | -0.1 | 4.39E-01 | Thioredoxin-like protein AAED1, chloroplastic |
| AT3G05936 | 0.6 | 2.82E-03 | 0.5 | 1.60E-02 | 0.1 | 5.19E-01 | -0.7 | 9.10E-02 | -0.1 | 7.72E-01 | -0.5 | 3.46E-01 | Uncharacterized protein |
| AT5G54710 | 0.6 | 1.69E-03 | 0.1 | 5.62E-01 | 0.5 | 1.04E-02 | 0.5 | 4.48E-03 | -0.2 | 4.31E-01 | 0.6 | 8.58E-03 | Similarity to ankyrin-like protein |
| AT1G74300 | 0.6 | 6.62E-05 | 0.2 | 3.29E-02 | 0.4 | 6.63E-03 | 0.4 | 8.09E-02 | 0.4 | 1.87E-01 | 0.1 | 8.41E-01 | Alpha |
| AT5G24200 | 0.6 | 3.51E-01 | -0.3 | 6.99E-01 | 0.9 | 3.18E-01 | | | | | | | Alpha |
| AT1G68310 | 0.6 | 4.23E-05 | 0.4 | 8.37E-04 | 0.2 | 9.21E-02 | 0.1 | 7.39E-01 | -0.1 | 7.38E-01 | 0.2 | 5.58E-01 | MIP18 family-like domain-containing protein |
| AT5G44560 | 0.6 | 1.52E-03 | 0.4 | 1.99E-02 | 0.2 | 2.75E-01 | 0.0 | 1.00E+00 | -0.1 | 7.68E-01 | 0.1 | 7.93E-01 | SNF7 family protein |
| AT1G55205 | 0.6 | 2.21E-03 | 0.5 | 6.65E-03 | 0.1 | 7.05E-01 | 0.5 | 1.96E-01 | 0.5 | 2.69E-01 | 0.0 | 9.81E-01 | Uncharacterized protein |
| AT5G43680 | 0.6 | 6.57E-05 | 0.3 | 6.11E-03 | 0.3 | 2.59E-02 | 0.4 | 1.01E-01 | 0.0 | 9.53E-01 | 0.4 | 1.72E-01 | Sec-independent protein translocase protein TatB |
| AT2G42620 | 0.6 | 8.47E-05 | 0.4 | 1.46E-03 | 0.2 | 1.20E-01 | -0.2 | 4.52E-01 | -0.4 | 8.27E-02 | 0.2 | 3.70E-01 | F-box protein MAX2 |
| AT3G19508 | 0.6 | 2.58E-02 | 0.7 | 1.79E-02 | -0.1 | 8.41E-01 | 0.9 | 1.68E-02 | 0.5 | 2.73E-01 | 0.4 | 3.44E-01 | LYR motif-containing protein At3g19508 |
| AT2G45950 | 0.6 | 3.37E-05 | 0.4 | 1.55E-03 | 0.2 | 3.68E-02 | 0.1 | 7.32E-01 | -0.2 | 5.06E-01 | 0.2 | 4.12E-01 | AT2G45950 protein |
| AT1G51730 | 0.6 | 5.02E-06 | 0.4 | 9.33E-05 | 0.2 | 4.36E-02 | 0.0 | 8.45E-01 | -0.2 | 1.91E-01 | 0.2 | 2.21E-01 | hypothetical protein |
| AT1G72430 | 0.6 | 5.89E-04 | 0.4 | 1.68E-02 | 0.2 | 9.91E-02 | 0.0 | 9.77E-01 | -0.6 | 5.07E-03 | 0.6 | 2.13E-02 | Auxin-responsive protein SAUR78 |
| AT1G64490 | 0.6 | 3.40E-04 | 0.1 | 5.77E-01 | 0.5 | 3.73E-03 | 0.5 | 7.16E-02 | 0.2 | 6.71E-01 | 0.3 | 3.79E-01 | hypothetical protein |
| AT1G56345 | 0.6 | 2.19E-04 | 0.4 | 9.75E-03 | 0.2 | 7.82E-02 | 0.4 | 1.87E-01 | 0.2 | 5.22E-01 | 0.1 | 7.02E-01 | Pseudouridine synthase family protein |
| AT4G21865 | 0.6 | 1.53E-02 | 0.2 | 3.50E-01 | 0.4 | 1.65E-01 | 0.5 | 1.04E-01 | 0.2 | 7.22E-01 | 0.4 | 3.65E-01 | Uncharacterized protein AT4g21860 |
| AT5G01630 | 0.6 | 9.43E-04 | 0.5 | 8.46E-03 | 0.2 | 3.46E-01 | 0.6 | 1.03E-01 | 0.4 | 3.53E-01 | 0.2 | 6.99E-01 | BRCA2B |
| AT2G46735 | 0.6 | 6.58E-05 | 0.0 | 7.92E-01 | 0.6 | 6.04E-04 | 0.0 | 9.12E-01 | -0.1 | 8.51E-01 | 0.0 | 9.40E-01 | Uncharacterized protein |
| AT5G50890 | 0.6 | 6.15E-04 | 0.4 | 9.22E-03 | 0.2 | 2.22E-01 | 0.7 | 1.41E-02 | 0.5 | 9.07E-02 | 0.2 | 6.15E-01 | Alpha |
| AT5G52750 | 0.6 | 4.92E-03 | -0.1 | 6.45E-01 | 0.7 | 6.76E-03 | 0.7 | 1.18E-02 | -0.3 | 3.36E-01 | 1.0 | 4.56E-03 | Heavy metal transport |
| AT1G17490 | 0.6 | 5.20E-03 | 0.2 | 4.14E-01 | 0.4 | 5.37E-02 | 0.4 | 2.41E-01 | 0.3 | 4.11E-01 | 0.1 | 9.08E-01 | hypothetical protein |
| AT1G42550 | 0.6 | 6.32E-05 | 0.2 | 5.61E-02 | 0.4 | 2.82E-03 | -0.2 | 2.56E-01 | -0.4 | 1.60E-02 | 0.2 | 1.92E-01 | Protein PLASTID MOVEMENT IMPAIRED 1 |
| AT3G43740 | 0.6 | 2.38E-05 | 0.3 | 1.14E-02 | 0.4 | 3.83E-03 | 0.4 | 1.52E-02 | 0.0 | 8.77E-01 | 0.4 | 8.49E-02 | Leucine-rich repeat protein 2 |
| AT1G04930 | 0.6 | 8.64E-05 | 0.3 | 1.10E-02 | 0.3 | 2.30E-02 | -0.2 | 5.13E-01 | -0.1 | 6.94E-01 | 0.0 | 9.10E-01 | Uncharacterized protein At1g04930 |
| AT1G68585 | 0.6 | 1.39E-02 | 0.2 | 3.38E-01 | 0.4 | 1.58E-01 | 0.0 | 9.24E-01 | -0.4 | 2.40E-01 | 0.4 | 2.46E-01 | hypothetical protein |
| AT3G13677 | 0.6 | 1.43E-03 | 0.4 | 3.68E-02 | 0.3 | 1.57E-01 | 0.1 | 7.15E-01 | 0.0 | 9.82E-01 | 0.1 | 8.09E-01 | Uncharacterized protein |
| AT4G04925 | 0.6 | 5.35E-05 | 0.4 | 1.25E-03 | 0.2 | 8.57E-02 | -0.4 | 7.86E-03 | -0.4 | 1.91E-02 | 0.0 | 9.33E-01 | At4g04925 |

| | | | | | | | | | | | | | |
|---|---|---|---|---|---|---|---|---|---|---|---|---|---|
| AT3G05750 | 0.6 | 3.08E-05 | 0.3 | 3.12E-03 | 0.3 | 1.73E-02 | 0.0 | 8.26E-01 | 0.1 | 5.72E-01 | -0.1 | 7.57E-01 | F10A16.3 protein |
| AT3G59300 | 0.6 | 2.56E-04 | 0.3 | 2.08E-02 | 0.3 | 3.58E-02 | 0.1 | 6.54E-01 | 0.2 | 4.70E-01 | -0.1 | 8.28E-01 | Uncharacterized protein F25L23_160 |
| AT3G18800 | 0.6 | 1.15E-03 | 0.6 | 1.28E-03 | 0.0 | 8.88E-01 | -0.3 | 1.84E-01 | -0.6 | 2.87E-02 | 0.3 | 4.47E-01 | hypothetical protein |
| AT5G23420 | 0.6 | 2.15E-04 | 0.5 | 9.85E-04 | 0.1 | 4.82E-01 | 0.5 | 3.45E-02 | 0.2 | 4.69E-01 | 0.3 | 3.27E-01 | High-mobility group box 6 |
| AT3G49900 | 0.6 | 5.15E-03 | 0.5 | 3.09E-02 | 0.1 | 5.04E-01 | 0.1 | 8.26E-01 | 0.2 | 6.30E-01 | -0.1 | 8.16E-01 | Phototropic-responsive NPH3 family protein |
| AT1G06660 | 0.6 | 8.42E-03 | 0.7 | 5.73E-03 | -0.1 | 7.79E-01 | -0.4 | 3.01E-01 | -0.1 | 7.51E-01 | -0.2 | 6.64E-01 | Protein JASON |
| AT5G16660 | 0.6 | 1.66E-04 | 0.3 | 1.14E-02 | 0.3 | 3.35E-02 | 0.0 | 9.95E-01 | -0.2 | 5.99E-01 | 0.2 | 6.76E-01 | Uncharacterized protein F5E19_10 |
| AT4G31560 | 0.6 | 2.00E-05 | 0.4 | 7.12E-04 | 0.2 | 3.31E-02 | -0.2 | 2.98E-01 | -0.3 | 1.83E-01 | 0.1 | 8.02E-01 | HCF153 |
| AT2G21790 | 0.6 | 8.39E-06 | 0.3 | 9.23E-04 | 0.3 | 5.78E-03 | 0.6 | 6.82E-05 | 0.2 | 8.56E-02 | 0.4 | 2.94E-02 | At2g21790 |
| AT2G45210 | 0.6 | 7.03E-04 | 0.4 | 9.65E-03 | 0.2 | 2.30E-01 | -0.4 | 3.43E-01 | -0.9 | 5.35E-02 | 0.5 | 3.03E-01 | Auxin-responsive protein SAUR36 |
| AT1G59840 | 0.6 | 6.29E-03 | 0.8 | 2.30E-03 | -0.1 | 5.01E-01 | 0.1 | 7.27E-01 | 0.1 | 7.59E-01 | 0.0 | 9.90E-01 | Uncharacterized protein At1g59840 |
| AT1G64750 | 0.6 | 5.92E-03 | 0.4 | 8.97E-02 | 0.3 | 2.09E-01 | 0.2 | 5.02E-01 | 0.0 | 9.70E-01 | 0.2 | 6.97E-01 | 26S proteasome complex subunit SEM1 |
| AT1G23790 | 0.6 | 1.03E-03 | 0.7 | 1.02E-03 | 0.0 | 8.15E-01 | 0.0 | 9.17E-01 | -0.1 | 9.18E-01 | 0.1 | 8.50E-01 | Uncharacterized protein |
| AT1G17140 | 0.6 | 3.01E-04 | 0.6 | 4.09E-04 | 0.0 | 9.40E-01 | -0.3 | 3.35E-02 | -0.1 | 5.04E-01 | -0.2 | 3.71E-01 | Interactor of constitutive active ROPs 1 |
| AT4G14342 | 0.6 | 2.29E-05 | 0.6 | 3.38E-05 | 0.0 | 8.34E-01 | 0.3 | 2.06E-01 | 0.0 | 9.95E-01 | 0.3 | 3.79E-01 | Splicing factor 3B subunit 5 |
| AT2G33600 | 0.6 | 4.86E-06 | 0.6 | 2.35E-05 | 0.1 | 3.77E-01 | 0.4 | 1.20E-02 | 0.0 | 9.28E-01 | 0.4 | 6.78E-02 | CRL2 |
| AT4G33100 | 0.6 | 2.06E-03 | 0.2 | 3.03E-01 | 0.4 | 3.84E-02 | 0.1 | 7.67E-01 | -0.2 | 7.03E-01 | 0.3 | 5.52E-01 | Uncharacterized protein At4g33100 |
| AT4G28510 | 0.6 | 1.38E-04 | 0.3 | 2.43E-02 | 0.3 | 1.48E-02 | 1.1 | 1.11E-06 | 0.4 | 1.13E-02 | 0.6 | 1.23E-03 | Prohibitin-1, mitochondrial |
| AT1G67195 | 0.6 | 6.80E-02 | 0.1 | 7.15E-01 | 0.5 | 2.27E-01 | -0.3 | 4.07E-01 | -0.5 | 1.93E-01 | 0.2 | 7.19E-01 | NA |
| AT2G44860 | 0.6 | 1.96E-03 | 0.5 | 8.39E-03 | 0.1 | 5.24E-01 | 0.6 | 2.87E-03 | 0.2 | 4.93E-01 | 0.4 | 7.18E-02 | hypothetical protein |
| AT1G23180 | 0.6 | 6.01E-06 | 0.3 | 1.55E-03 | 0.3 | 2.63E-03 | 0.4 | 8.75E-02 | 0.5 | 9.83E-02 | 0.0 | 9.51E-01 | ARM repeat superfamily protein |
| AT1G03610 | 0.6 | 8.98E-05 | 0.5 | 4.17E-04 | 0.1 | 4.65E-01 | -0.7 | 1.28E-03 | -0.6 | 3.71E-03 | 0.0 | 9.22E-01 | Plant |
| AT3G24460 | 0.6 | 1.52E-03 | 0.0 | 9.13E-01 | 0.6 | 7.42E-03 | -0.2 | 6.23E-01 | -0.1 | 8.10E-01 | -0.1 | 8.62E-01 | Serinc-domain containing serine and sphingolipid biosynthesis protein |
| AT5G39790 | 0.6 | 5.73E-04 | 0.7 | 6.41E-04 | 0.0 | 8.54E-01 | 0.6 | 5.65E-02 | 0.4 | 2.72E-01 | 0.2 | 6.30E-01 | AMP-activated protein kinase glycogen-binding domain-containing protein |
| AT1G26450 | 0.6 | 2.23E-04 | 0.6 | 6.85E-04 | 0.1 | 6.90E-01 | -0.1 | 7.44E-01 | -0.1 | 7.95E-01 | 0.0 | 9.92E-01 | X8 domain-containing protein |
| AT1G25220 | 0.6 | 1.71E-03 | 0.6 | 4.92E-03 | 0.1 | 6.84E-01 | 0.4 | 6.88E-02 | 0.1 | 8.27E-01 | 0.4 | 2.99E-01 | Anthranilate synthase beta subunit 1 |
| AT4G40042 | 0.6 | 8.05E-03 | 0.3 | 1.45E-01 | 0.3 | 2.08E-01 | 0.4 | 1.98E-01 | 0.2 | 6.92E-01 | 0.2 | 6.07E-01 | At4g40042 ) |
| AT1G03180 | 0.6 | 3.43E-04 | 0.2 | 2.23E-01 | 0.5 | 1.02E-02 | -0.6 | 4.31E-02 | -0.2 | 6.27E-01 | -0.5 | 1.89E-01 | F15k9.21 |
| AT2G01250 | 0.6 | 4.12E-04 | 0.7 | 4.35E-04 | 0.0 | 7.71E-01 | 0.4 | 1.60E-03 | 0.2 | 8.18E-02 | 0.2 | 3.31E-01 | AT2G01250 protein |
| AT2G46390 | 0.6 | 3.99E-03 | 0.3 | 1.85E-01 | 0.4 | 8.11E-02 | 0.2 | 5.13E-01 | 0.1 | 8.64E-01 | 0.1 | 8.06E-01 | Succinate dehydrogenase subunit 8, mitochondrial |
| AT2G18990 | 0.6 | 2.18E-05 | 0.7 | 3.20E-05 | 0.0 | 8.09E-01 | 0.5 | 9.49E-03 | 0.3 | 1.71E-01 | 0.2 | 3.69E-01 | Thioredoxin domain-containing protein 9 homolog |
| AT4G10480 | 0.6 | 1.05E-03 | 0.8 | 5.65E-04 | -0.1 | 4.92E-01 | 0.4 | 1.44E-02 | 0.2 | 3.11E-01 | 0.2 | 3.63E-01 | Nascent polypeptide-associated complex , alpha subunit family protein |
| AT5G26622 | 0.6 | 3.58E-05 | 0.3 | 1.23E-02 | 0.4 | 6.06E-03 | -0.2 | 3.68E-01 | 0.1 | 8.33E-01 | -0.2 | 3.15E-01 | Beta-galactosidase related protein |
| AT2G26340 | 0.6 | 7.25E-05 | 0.6 | 3.33E-04 | 0.1 | 4.56E-01 | -0.2 | 1.14E-01 | -0.2 | 3.07E-01 | 0.0 | 8.15E-01 | Uncharacterized protein |
| AT1G02820 | 0.6 | 1.33E-03 | 0.5 | 1.53E-02 | 0.2 | 2.80E-01 | | | | | | | Late embryogenis abundant protein 2 |
| AT4G35930 | 0.6 | 1.77E-04 | 0.3 | 2.07E-02 | 0.3 | 2.80E-02 | -0.1 | 8.30E-01 | -0.2 | 4.55E-01 | 0.1 | 6.69E-01 | F-box family protein |
| AT3G60440 | 0.6 | 1.67E-05 | 0.6 | 6.19E-05 | 0.1 | 5.66E-01 | 0.4 | 1.01E-02 | 0.4 | 2.85E-02 | 0.0 | 9.11E-01 | At3g60440 |
| AT1G22520 | 0.6 | 3.34E-05 | 0.6 | 7.92E-05 | 0.0 | 8.45E-01 | 0.3 | 1.64E-01 | 0.3 | 2.85E-01 | 0.0 | 9.34E-01 | MICOS complex subunit Mic10-like protein |
| AT4G28020 | 0.7 | 3.59E-04 | 0.4 | 2.18E-02 | 0.3 | 6.68E-02 | -0.1 | 7.74E-01 | 0.1 | 8.67E-01 | -0.2 | 6.74E-01 | Uncharacterized protein AT4g28020 |
| AT3G53020 | 0.7 | 6.29E-04 | 0.6 | 1.57E-03 | 0.0 | 7.79E-01 | 0.4 | 1.00E-02 | 0.2 | 2.99E-01 | 0.2 | 3.08E-01 | hypothetical protein |
| AT5G25040 | 0.7 | 1.28E-05 | 0.2 | 5.65E-02 | 0.5 | 6.31E-04 | 0.2 | 3.28E-01 | 0.2 | 2.55E-01 | -0.1 | 8.62E-01 | Major facilitator superfamily protein |
| AT5G40930 | 0.7 | 5.35E-04 | 0.2 | 1.17E-01 | 0.4 | 2.00E-02 | 0.5 | 1.04E-02 | 0.4 | 6.72E-02 | 0.1 | 6.79E-01 | Translocase of outer membrane 20-4 |
| AT1G29450 | 0.7 | 9.53E-03 | 0.9 | 1.50E-03 | -0.3 | 2.69E-01 | -0.5 | 2.52E-01 | -0.2 | 6.18E-01 | -0.2 | 7.14E-01 | Auxin-responsive protein SAUR64 |
| AT1G68440 | 0.7 | 4.87E-05 | 0.9 | 1.20E-05 | -0.2 | 4.65E-02 | -0.5 | 4.34E-02 | -0.7 | 9.62E-03 | 0.2 | 5.19E-01 | Transmembrane protein |
| AT1G68660 | 0.7 | 8.77E-03 | 0.1 | 6.70E-01 | 0.6 | 3.76E-02 | 0.2 | 2.91E-01 | 0.3 | 2.20E-01 | -0.1 | 8.56E-01 | hypothetical protein |
| AT5G23570 | 0.7 | 6.29E-06 | 0.4 | 5.45E-04 | 0.3 | 8.40E-03 | 0.2 | 4.83E-01 | 0.0 | 8.87E-01 | 0.1 | 7.50E-01 | XS domain-containing protein |
| AT1G29700 | 0.7 | 5.50E-05 | 0.2 | 1.54E-01 | 0.5 | 1.32E-03 | 0.0 | 9.35E-01 | 0.0 | 9.13E-01 | 0.0 | 9.73E-01 | hypothetical protein |
| AT1G61780 | 0.7 | 2.36E-03 | 0.3 | 1.59E-01 | 0.4 | 6.11E-02 | 0.1 | 5.67E-01 | -0.2 | 5.42E-01 | 0.3 | 3.36E-01 | Cysteine-rich PDZ-binding protein |
| AT5G43520 | 0.7 | 1.10E-05 | 0.8 | 3.78E-06 | -0.2 | 4.82E-02 | 0.1 | 5.05E-01 | 0.0 | 9.76E-01 | 0.1 | 6.38E-01 | DC1 domain-containing protein |
| AT5G24460 | 0.7 | 1.12E-05 | 0.6 | 3.39E-05 | 0.0 | 6.70E-01 | -0.1 | 3.21E-01 | -0.4 | 1.18E-02 | 0.3 | 1.34E-01 | Uncharacterized protein |
| AT1G29530 | 0.7 | 2.50E-05 | 0.4 | 2.10E-03 | 0.3 | 1.75E-02 | -0.3 | 1.13E-01 | 0.2 | 2.82E-01 | -0.5 | 3.59E-02 | Uncharacterized protein |
| AT2G32690 | 0.7 | 4.27E-04 | 1.0 | 3.70E-05 | -0.4 | 2.71E-02 | -0.2 | 1.27E-01 | -0.2 | 2.64E-01 | 0.0 | 9.06E-01 | Glycine-rich protein 23 |
| AT2G27840 | 0.7 | 1.30E-03 | 0.5 | 7.12E-03 | 0.1 | 4.53E-01 | 1.1 | 5.91E-05 | 0.8 | 3.97E-03 | 0.3 | 1.87E-01 | Histone deacetylase-related |
| AT2G16630 | 0.7 | 1.80E-04 | 0.9 | 2.87E-05 | -0.3 | 6.79E-02 | 0.4 | 3.51E-01 | 0.7 | 1.31E-01 | -0.3 | 6.30E-01 | Uncharacterized protein At2g16630 |
| AT1G21050 | 0.7 | 1.99E-04 | 0.4 | 9.30E-03 | 0.3 | 5.70E-02 | 0.4 | 3.79E-02 | -0.3 | 1.16E-01 | 0.7 | 6.02E-03 | Protein MIZU-KUSSEI 1 |
| AT2G33250 | 0.7 | 9.05E-06 | 0.4 | 7.37E-04 | 0.3 | 9.85E-03 | 0.0 | 9.47E-01 | 0.2 | 3.93E-01 | -0.2 | 4.89E-01 | At2g33250 |
| AT5G48335 | 0.7 | 3.63E-05 | 0.4 | 1.23E-03 | 0.2 | 5.03E-02 | -0.2 | 2.77E-01 | -0.2 | 2.14E-01 | 0.0 | 8.83E-01 | Uncharacterized protein |
| AT1G14820 | 0.7 | 3.75E-05 | 0.4 | 2.64E-03 | 0.3 | 2.42E-02 | 0.0 | 9.41E-01 | -0.1 | 7.61E-01 | 0.1 | 7.47E-01 | Putative phosphatidylinositol |
| AT5G66090 | 0.7 | 7.69E-05 | 0.6 | 3.23E-04 | 0.1 | 5.00E-01 | 0.0 | 9.39E-01 | -0.2 | 2.25E-01 | 0.2 | 3.63E-01 | hypothetical protein |
| AT1G09470 | 0.7 | 4.80E-04 | 0.5 | 3.11E-03 | 0.1 | 4.27E-01 | 0.2 | 4.06E-01 | 0.1 | 6.55E-01 | 0.1 | 8.64E-01 | Uncharacterized protein |
| AT5G63040 | 0.7 | 1.84E-05 | 0.5 | 2.90E-04 | 0.2 | 1.04E-01 | 0.3 | 3.23E-01 | 0.3 | 3.73E-01 | 0.0 | 9.90E-01 | Emb\|CAB81845.1 |
| AT1G67480 | 0.7 | 3.86E-06 | 0.6 | 1.62E-05 | 0.1 | 4.75E-01 | -0.1 | 6.77E-01 | 0.0 | 9.81E-01 | 0.0 | 8.16E-01 | F-box |
| AT2G15000 | 0.7 | 1.58E-03 | 0.7 | 2.46E-03 | 0.0 | 9.50E-01 | 0.7 | 2.19E-03 | 0.3 | 3.09E-01 | 0.5 | 7.90E-02 | At2g15000 |
| AT3G07470 | 0.7 | 2.63E-04 | 0.6 | 1.15E-03 | 0.1 | 5.05E-01 | -0.1 | 4.74E-01 | -0.5 | 1.89E-02 | 0.4 | 1.73E-01 | F21O3.18 protein |
| AT2G37970 | 0.7 | 1.72E-04 | 0.0 | 7.47E-01 | 0.7 | 7.13E-04 | 0.5 | 1.31E-01 | 0.1 | 7.79E-01 | 0.3 | 3.87E-01 | Uncharacterized protein At2g37970 |
| AT2G25735 | 0.7 | 1.81E-03 | 0.5 | 2.08E-02 | 0.2 | 2.78E-01 | 0.3 | 3.93E-01 | -0.6 | 1.42E-01 | 0.9 | 6.23E-02 | Uncharacterized protein |
| AT5G14105 | 0.7 | 2.30E-04 | 0.5 | 3.56E-03 | 0.2 | 1.73E-01 | 0.6 | 1.45E-02 | 0.1 | 8.78E-01 | 0.5 | 5.62E-02 | Transmembrane protein |
| AT1G32160 | 0.7 | 2.08E-05 | 0.2 | 3.26E-02 | 0.5 | 1.17E-03 | 0.1 | 2.62E-01 | 0.1 | 7.01E-01 | 0.1 | 7.09E-01 | At1g32160 |
| AT4G14400 | 0.7 | 8.35E-03 | 0.1 | 7.30E-01 | 0.6 | 2.90E-02 | 1.6 | 1.20E-06 | 0.7 | 5.34E-03 | 0.9 | 7.35E-03 | Protein ACCELERATED CELL DEATH 6 |
| AT1G16916 | 0.7 | 2.22E-02 | 0.3 | 3.42E-01 | 0.4 | 2.21E-01 | 0.2 | 6.48E-01 | -0.2 | 6.34E-01 | 0.4 | 4.33E-01 | Transmembrane protein |
| AT3G52740 | 0.7 | 9.20E-03 | 0.2 | 4.85E-01 | 0.5 | 7.03E-02 | -0.1 | 7.23E-01 | 0.4 | 2.53E-01 | -0.5 | 1.60E-01 | Protein BIC1 |
| AT5G17640 | 0.7 | 3.34E-05 | 0.4 | 1.22E-03 | 0.2 | 4.13E-02 | -0.3 | 8.98E-02 | -0.3 | 1.96E-01 | 0.0 | 8.45E-01 | At5g17640 ) |
| AT3G09200 | 0.7 | 4.99E-04 | 0.6 | 1.60E-03 | 0.1 | 6.57E-01 | 0.2 | 1.01E-01 | 0.1 | 7.67E-01 | 0.2 | 4.37E-01 | 60S acidic ribosomal protein P0 |
| AT4G24972 | 0.7 | 6.37E-04 | 0.7 | 8.34E-04 | 0.0 | 9.42E-01 | -1.0 | 7.91E-04 | -0.7 | 9.40E-03 | -0.3 | 4.26E-01 | At4g24973 |
| AT1G58025 | 0.7 | 6.29E-06 | 0.1 | 1.03E-01 | 0.5 | 2.21E-04 | 0.1 | 4.22E-01 | 0.1 | 5.28E-01 | 0.0 | 9.79E-01 | Uncharacterized protein T15M6.22 |
| AT1G64580 | 0.7 | 4.57E-04 | 0.5 | 7.52E-03 | 0.2 | 2.01E-01 | 0.3 | 2.81E-01 | 0.0 | 9.32E-01 | 0.3 | 4.40E-01 | Pentatricopeptide repeat-containing protein At1g64580 |
| AT3G09250 | 0.7 | 5.11E-06 | 0.4 | 2.45E-04 | 0.2 | 1.60E-02 | 0.5 | 2.73E-02 | 0.2 | 3.86E-01 | 0.3 | 3.55E-01 | Nuclear transport factor 2 family protein |
| AT1G04630 | 0.7 | 1.25E-05 | 0.6 | 7.97E-05 | 0.1 | 2.65E-01 | -0.1 | 6.80E-01 | 0.0 | 8.63E-01 | 0.0 | 9.07E-01 | NADH dehydrogenase 1 alpha subcomplex subunit 13-A |
| AT3G23010 | 0.7 | 7.78E-03 | 0.1 | 5.71E-01 | 0.5 | 6.40E-02 | 1.3 | 3.06E-03 | 0.3 | 5.57E-01 | 1.0 | 1.70E-02 | Receptor like protein |
| AT5G08400 | 0.7 | 8.97E-04 | 0.5 | 5.85E-03 | 0.1 | 4.25E-01 | 0.7 | 6.04E-02 | 0.4 | 3.94E-01 | 0.3 | 5.33E-01 | Uncharacterized protein F8L15_130 |
| AT5G41520 | 0.7 | 4.97E-04 | 0.7 | 9.77E-04 | 0.0 | 9.02E-01 | 0.3 | 1.43E-01 | 0.0 | 9.07E-01 | 0.2 | 3.39E-01 | RNA binding Plectin |
| AT3G01290 | 0.7 | 1.17E-03 | 0.4 | 1.85E-02 | 0.2 | 1.85E-01 | 0.2 | 2.08E-01 | -0.6 | 8.32E-03 | 0.8 | 4.25E-03 | Hypersensitive-induced response protein 3 |
| AT2G25570 | 0.7 | 2.75E-04 | 0.4 | 8.36E-03 | 0.3 | 1.01E-01 | 0.3 | 1.35E-01 | 0.3 | 2.13E-01 | 0.0 | 9.61E-01 | Binding protein |

| | | | | | | | | | | | | |
|---|---|---|---|---|---|---|---|---|---|---|---|---|
| AT4G34265 | 0.7 | 3.46E-04 | 0.5 | 2.54E-03 | 0.2 | 3.15E-01 | 0.2 | 3.20E-01 | 0.0 | 8.91E-01 | 0.1 | 5.92E-01 | hypothetical protein |
| AT3G08660 | 0.7 | 1.14E-03 | 0.5 | 1.07E-02 | 0.2 | 3.45E-01 | 0.3 | 5.07E-01 | 0.2 | 7.71E-01 | 0.1 | 8.58E-01 | Phototropic-responsive NPH3 family protein |
| AT3G00850 | 0.7 | 2.67E-02 | -0.1 | 7.58E-01 | 0.8 | 4.59E-02 | 0.2 | 7.81E-01 | -0.7 | 4.57E-01 | 0.9 | 2.83E-01 | NA |
| AT5G67370 | 0.7 | 4.73E-04 | 0.2 | 2.35E-01 | 0.5 | 7.45E-03 | 2.6 | 1.08E-08 | 0.3 | 3.39E-01 | 2.3 | 1.01E-05 | CGLD27 |
| AT4G33350 | 0.7 | 4.35E-05 | 0.5 | 4.70E-04 | 0.2 | 1.73E-01 | 0.3 | 5.66E-02 | 0.1 | 4.73E-01 | 0.2 | 5.01E-01 | Protein TIC 22, chloroplastic |
| AT4G39235 | 0.7 | 2.51E-03 | 0.5 | 1.42E-02 | 0.1 | 4.91E-01 | 0.5 | 2.33E-01 | -0.1 | 8.14E-01 | 0.6 | 2.04E-01 | At4g39235 |
| AT1G52855 | 0.7 | 2.96E-02 | -0.9 | 3.75E-02 | 1.5 | 3.80E-03 | | | | | | | Uncharacterized protein |
| AT5G44730 | 0.7 | 4.76E-05 | 0.6 | 1.81E-04 | 0.1 | 5.74E-01 | 0.5 | 2.60E-02 | 0.2 | 5.22E-01 | 0.3 | 2.30E-01 | hypothetical protein |
| AT1G48840 | 0.7 | 9.50E-05 | 0.5 | 7.76E-04 | 0.1 | 2.93E-01 | -0.2 | 4.96E-01 | -0.1 | 7.21E-01 | -0.1 | 8.75E-01 | hypothetical protein |
| AT1G12370 | 0.7 | 1.02E-05 | 0.2 | 7.06E-02 | 0.5 | 5.21E-04 | 0.2 | 4.22E-01 | 0.5 | 6.88E-02 | -0.3 | 3.72E-01 | Photolyase 1 |
| AT5G38890 | 0.7 | 1.70E-03 | 0.4 | 4.80E-02 | 0.3 | 1.40E-01 | 0.7 | 2.02E-03 | 0.1 | 6.82E-01 | 0.6 | 3.14E-02 | Uncharacterized protein At5g38885 |
| AT5G02020 | 0.7 | 1.32E-02 | -0.1 | 7.10E-01 | 0.8 | 2.45E-02 | -1.2 | 2.32E-03 | -1.5 | 1.51E-03 | 0.3 | 6.88E-01 | AT5G02020 protein |
| AT3G61700 | 0.7 | 2.36E-05 | 0.3 | 5.49E-03 | 0.4 | 6.49E-03 | 0.1 | 6.49E-01 | 0.2 | 3.09E-01 | -0.1 | 6.27E-01 | hypothetical protein |
| AT1G09794 | 0.7 | 1.47E-02 | 0.2 | 5.33E-01 | 0.5 | 1.03E-01 | 0.6 | 7.79E-02 | 0.3 | 5.07E-01 | 0.3 | 4.95E-01 | Cx9C motif-containing protein 4 |
| AT2G46380 | 0.7 | 1.15E-04 | 0.3 | 1.91E-02 | 0.4 | 1.90E-02 | -0.1 | 5.62E-01 | 0.2 | 3.36E-01 | -0.4 | 1.83E-01 | Extra-large G-like protein, putative |
| AT5G52760 | 0.7 | 1.30E-01 | -0.5 | 3.22E-01 | 1.2 | 5.96E-02 | 0.2 | 7.32E-01 | -1.0 | 1.84E-01 | 1.2 | 1.19E-01 | Copper transport protein family |
| AT1G50575 | 0.7 | 2.34E-04 | 0.7 | 2.18E-04 | -0.1 | 7.09E-01 | -0.1 | 6.73E-01 | 0.0 | 9.02E-01 | -0.1 | 8.56E-01 | cytokinin riboside 5'-monophosphate phosphoribohydrolase |
| AT4G20325 | 0.7 | 1.02E-03 | 0.4 | 3.07E-02 | 0.3 | 1.37E-01 | 0.1 | 8.36E-01 | 0.0 | 9.43E-01 | 0.1 | 8.21E-01 | Ribonuclease H2 subunit B |
| AT4G14615 | 0.7 | 4.65E-04 | 0.5 | 6.14E-03 | 0.2 | 2.09E-01 | 0.3 | 2.82E-01 | 0.1 | 7.87E-01 | 0.2 | 6.66E-01 | Uncharacterized protein |
| AT5G06130 | 0.7 | 8.90E-07 | 0.4 | 8.00E-05 | 0.3 | 1.63E-03 | 0.0 | 9.68E-01 | 0.1 | 3.17E-01 | -0.2 | 3.42E-01 | Chaperone protein dnaJ-like protein |
| AT5G35520 | 0.7 | 1.73E-04 | 0.3 | 2.85E-02 | 0.4 | 2.10E-02 | 0.3 | 1.76E-01 | 0.2 | 4.90E-01 | 0.1 | 7.35E-01 | Protein MIS12 homolog |
| AT1G69760 | 0.7 | 1.03E-04 | 0.4 | 7.64E-03 | 0.3 | 3.14E-02 | 0.3 | 1.84E-01 | 0.0 | 9.80E-01 | 0.3 | 3.66E-01 | Uncharacterized protein |
| AT5G49440 | 0.7 | 3.38E-05 | 0.7 | 6.93E-05 | 0.0 | 9.97E-01 | 0.0 | 6.84E-01 | -0.2 | 1.21E-01 | 0.1 | 4.08E-01 | AT5g49440 |
| AT2G26270 | 0.7 | 2.75E-04 | 0.4 | 9.53E-03 | 0.3 | 9.80E-02 | 0.5 | 1.48E-01 | 0.2 | 6.90E-01 | 0.3 | 4.42E-01 | BRCT domain-containing protein |
| AT1G20830 | 0.7 | 1.59E-05 | 0.6 | 1.51E-04 | 0.1 | 1.88E-01 | 0.4 | 1.62E-02 | 0.4 | 4.61E-02 | 0.0 | 8.78E-01 | Multiple chloroplast division site 1 |
| AT1G62500 | 0.7 | 1.56E-03 | 0.5 | 1.87E-02 | 0.2 | 2.61E-01 | 0.2 | 1.80E-01 | -0.3 | 1.11E-01 | 0.5 | 5.38E-03 | Bifunctional inhibitor |
| AT5G60840 | 0.7 | 6.02E-05 | 0.7 | 7.72E-05 | 0.0 | 7.94E-01 | 0.2 | 6.23E-01 | 0.1 | 8.28E-01 | 0.1 | 8.97E-01 | Uncharacterized protein |
| AT1G10910 | 0.7 | 4.88E-05 | 0.6 | 3.96E-04 | 0.1 | 2.62E-01 | 0.1 | 5.34E-01 | 0.0 | 9.90E-01 | 0.1 | 6.20E-01 | Pentatricopeptide repeat superfamily protein |
| AT5G42480 | 0.7 | 9.13E-06 | 0.4 | 1.17E-03 | 0.3 | 5.62E-03 | 0.1 | 7.20E-01 | 0.0 | 9.40E-01 | 0.1 | 8.64E-01 | Chaperone DnaJ-domain superfamily protein |
| AT4G13900 | 0.7 | 6.67E-03 | -0.1 | 7.60E-01 | 0.8 | 1.22E-02 | 1.3 | 4.53E-03 | 0.4 | 5.42E-01 | 0.9 | 5.62E-02 | Receptor-like protein 49 |
| AT2G38860 | 0.7 | 5.84E-04 | 0.5 | 6.70E-03 | 0.2 | 2.28E-01 | 0.5 | 4.94E-02 | -0.3 | 2.56E-01 | 0.8 | 1.98E-02 | Class I glutamine amidotransferase-like superfamily protein |
| AT1G03440 | 0.7 | 7.40E-05 | 0.4 | 2.78E-03 | 0.3 | 5.46E-02 | -0.2 | 2.65E-01 | -0.2 | 3.10E-01 | 0.0 | 9.85E-01 | F21B7.6 |
| AT5G42680 | 0.7 | 1.11E-04 | 0.3 | 1.78E-02 | 0.4 | 2.12E-02 | -0.4 | 3.50E-02 | -0.5 | 1.87E-02 | 0.1 | 7.47E-01 | At5g42680 |
| AT5G61220 | 0.7 | 3.46E-03 | 0.3 | 1.11E-01 | 0.4 | 1.28E-01 | 0.5 | 4.39E-02 | 0.3 | 2.75E-01 | 0.2 | 5.92E-01 | hypothetical protein |
| AT2G31410 | 0.7 | 2.50E-04 | 0.6 | 1.29E-03 | 0.1 | 4.31E-01 | 0.5 | 1.72E-02 | 0.1 | 6.08E-01 | 0.3 | 2.03E-01 | Coiled-coil domain-containing protein 86 |
| AT3G00590 | 0.7 | 2.39E-03 | 0.5 | 2.67E-02 | 0.2 | 3.15E-01 | -0.3 | 4.83E-01 | 0.0 | 9.92E-01 | -0.3 | 5.86E-01 | NA |
| AT4G30460 | 0.7 | 2.04E-04 | 0.6 | 6.51E-04 | 0.1 | 6.55E-01 | -0.5 | 8.25E-02 | -0.2 | 4.91E-01 | -0.2 | 5.05E-01 | Glycine-rich protein |
| AT2G32660 | 0.7 | 2.87E-03 | 0.6 | 1.27E-02 | 0.1 | 5.68E-01 | | | | | | | Receptor like protein 22 |
| AT2G45070 | 0.7 | 1.75E-04 | 0.4 | 4.93E-03 | 0.3 | 7.88E-02 | 0.4 | 4.07E-02 | 0.2 | 3.28E-01 | 0.2 | 5.70E-01 | Protein transport protein Sec61 subunit beta |
| AT5G50140 | 0.7 | 1.10E-03 | 0.4 | 4.35E-02 | 0.3 | 1.19E-01 | 0.3 | 1.98E-01 | 0.2 | 6.15E-01 | 0.2 | 5.87E-01 | Ankyrin repeat family protein |
| AT5G22320 | 0.7 | 1.29E-04 | 0.3 | 2.68E-02 | 0.4 | 1.50E-02 | 0.5 | 1.50E-02 | 0.2 | 3.02E-01 | 0.3 | 3.41E-01 | Leucine-rich repeat family protein |
| AT1G65710 | 0.7 | 1.28E-04 | 0.7 | 2.62E-04 | 0.0 | 9.22E-01 | 0.3 | 2.72E-01 | 0.1 | 7.78E-01 | 0.2 | 6.38E-01 | Uncharacterized protein At1g65710 |
| AT5G23510 | 0.7 | 6.89E-05 | 0.4 | 2.30E-03 | 0.2 | 6.46E-02 | 0.5 | 2.64E-02 | 0.1 | 7.78E-01 | 0.4 | 1.41E-01 | At5g23510 |
| AT3G15950 | 0.7 | 8.80E-06 | 0.9 | 5.15E-06 | -0.2 | 1.07E-01 | 0.2 | 2.16E-01 | 0.3 | 5.05E-02 | -0.1 | 4.85E-01 | DNA topoisomerase-like protein |
| AT5G06620 | 0.7 | 3.38E-03 | 0.5 | 3.15E-02 | 0.2 | 3.63E-01 | 0.4 | 2.22E-01 | 0.2 | 6.52E-01 | 0.2 | 6.49E-01 | SET domain protein 38 |
| AT2G17695 | 0.7 | 1.40E-04 | 0.5 | 1.21E-03 | 0.2 | 2.56E-01 | 0.1 | 6.86E-01 | 0.3 | 1.73E-01 | -0.2 | 4.27E-01 | Outer envelope protein |
| AT5G26270 | 0.7 | 1.21E-03 | 0.3 | 1.08E-01 | 0.4 | 5.78E-02 | -0.1 | 8.95E-01 | -0.1 | 8.61E-01 | 0.0 | 9.65E-01 | Transmembrane protein |
| AT1G04530 | 0.7 | 3.45E-05 | 0.4 | 1.38E-03 | 0.3 | 3.38E-02 | 0.2 | 2.65E-01 | -0.1 | 7.53E-01 | 0.3 | 2.39E-01 | Tetratricopeptide repeat -like superfamily protein |
| AT4G11521 | 0.7 | 9.41E-03 | 0.3 | 2.21E-01 | 0.4 | 1.65E-01 | 0.5 | 6.80E-02 | -0.4 | 2.64E-01 | 0.9 | 2.04E-02 | Putative cysteine-rich receptor-like protein kinase At4g11521 |
| AT3G26770 | 0.7 | 9.56E-05 | 0.1 | 4.82E-01 | 0.6 | 1.45E-03 | -0.3 | 2.04E-01 | -0.4 | 1.16E-01 | 0.1 | 7.89E-01 | Alcohol dehydrogenase-like protein -binding Rossmann-fold superfamily protein) |
| AT3G24506 | 0.7 | 1.44E-05 | 0.6 | 5.60E-05 | 0.1 | 5.43E-01 | -0.2 | 3.34E-01 | -0.3 | 2.66E-01 | 0.1 | 8.85E-01 | Uncharacterized protein At3g24506 |
| AT5G08320 | 0.7 | 4.07E-04 | 0.5 | 5.24E-03 | 0.2 | 2.27E-01 | -0.3 | 4.02E-01 | -0.4 | 4.09E-01 | 0.0 | 9.62E-01 | E2F-associated phosphoprotein |
| AT5G43540 | 0.7 | 1.09E-02 | 0.6 | 3.51E-02 | 0.1 | 6.88E-01 | -1.0 | 1.05E-02 | -0.8 | 3.37E-02 | -0.2 | 7.32E-01 | C2H2-type domain-containing protein |
| AT1G78210 | 0.7 | 2.99E-04 | 0.4 | 1.74E-02 | 0.3 | 5.12E-02 | 0.0 | 9.51E-01 | -0.1 | 7.95E-01 | 0.1 | 8.67E-01 | Uncharacterized protein T11I11.15 |
| AT5G03780 | 0.7 | 4.56E-05 | -0.1 | 2.76E-01 | 0.8 | 1.80E-04 | -0.2 | 6.69E-01 | -0.3 | 5.97E-01 | 0.1 | 9.16E-01 | MYB transcription factor |
| AT1G72920 | 0.7 | 1.66E-02 | -0.3 | 2.75E-01 | 1.1 | 1.01E-02 | 1.0 | 2.68E-02 | 0.2 | 7.83E-01 | 0.8 | 9.97E-02 | Similar to part of disease resistance protein domain family protein) |
| AT4G32295 | 0.7 | 8.30E-05 | 0.8 | 7.39E-05 | -0.1 | 5.90E-01 | -0.5 | 1.02E-01 | -0.5 | 1.66E-01 | 0.0 | 9.42E-01 | Histone acetyltransferase |
| AT1G62730 | 0.7 | 2.22E-04 | 0.4 | 1.60E-02 | 0.3 | 5.01E-02 | 0.8 | 1.47E-02 | 0.1 | 8.12E-01 | 0.7 | 8.96E-02 | Terpenoid synthases superfamily protein |
| AT1G80920 | 0.7 | 1.07E-02 | 0.8 | 8.83E-03 | -0.1 | 8.28E-01 | -0.6 | 7.53E-02 | -1.0 | 1.22E-02 | 0.4 | 4.57E-01 | Chaperone protein dnaJ 8, chloroplastic |
| AT5G11070 | 0.7 | 5.40E-06 | 0.5 | 1.89E-04 | 0.2 | 2.25E-02 | -0.2 | 3.99E-01 | -0.4 | 7.41E-02 | 0.2 | 4.62E-01 | Uncharacterized protein |
| AT2G36590 | 0.7 | 7.91E-03 | 0.5 | 6.07E-02 | 0.2 | 4.14E-01 | -0.7 | 2.42E-01 | 0.5 | 3.33E-01 | -1.2 | 8.96E-02 | Proline transporter 3 |
| AT1G58200 | 0.7 | 4.86E-05 | 0.5 | 1.03E-03 | 0.2 | 8.19E-02 | -0.2 | 2.64E-01 | 0.0 | 9.83E-01 | -0.2 | 4.59E-01 | MSL3 |
| AT1G79390 | 0.7 | 1.37E-04 | 0.3 | 3.20E-02 | 0.4 | 1.41E-02 | 0.4 | 4.58E-02 | 0.2 | 4.23E-01 | 0.2 | 4.41E-01 | hypothetical protein |
| AT3G07910 | 0.7 | 3.53E-05 | 0.6 | 1.76E-04 | 0.1 | 4.04E-01 | 0.6 | 1.69E-02 | 0.2 | 5.42E-01 | 0.4 | 1.89E-01 | hypothetical protein |
| AT1G65230 | 0.7 | 4.57E-05 | 0.7 | 1.14E-04 | 0.0 | 8.07E-01 | 0.0 | 8.13E-01 | 0.1 | 5.06E-01 | -0.2 | 4.39E-01 | T23K8.14 protein |
| AT3G27280 | 0.7 | 6.84E-05 | 0.5 | 2.27E-03 | 0.3 | 5.37E-02 | 0.8 | 2.11E-05 | 0.2 | 2.13E-01 | 0.6 | 2.41E-03 | Prohibitin-4, mitochondrial |
| AT1G34640 | 0.7 | 9.15E-03 | 0.3 | 3.28E-01 | 0.5 | 1.21E-01 | 0.2 | 6.48E-01 | 0.1 | 7.70E-01 | 0.0 | 9.61E-01 | At1g34640 |
| AT3G17970 | 0.7 | 4.49E-06 | 0.6 | 3.11E-05 | 0.1 | 2.04E-01 | 0.4 | 4.49E-03 | 0.0 | 8.34E-01 | 0.5 | 6.66E-03 | Translocon at the outer membrane of chloroplasts 64-III |
| AT4G36970 | 0.7 | 2.69E-05 | 0.3 | 2.71E-02 | 0.5 | 2.41E-03 | -0.4 | 3.58E-03 | -0.3 | 4.65E-02 | -0.1 | 5.39E-01 | hypothetical protein |
| AT4G16000 | 0.7 | 2.70E-04 | 0.2 | 1.36E-01 | 0.5 | 1.22E-02 | 0.0 | 9.10E-01 | 0.3 | 3.54E-01 | -0.4 | 3.73E-01 | Uncharacterized protein AT4g16000 |
| AT2G30520 | 0.7 | 3.68E-05 | 0.6 | 2.57E-04 | 0.1 | 2.94E-01 | -0.2 | 1.91E-01 | -0.6 | 2.02E-03 | 0.4 | 7.46E-02 | Phototropic-responsive NPH3 family protein |
| AT2G46320 | 0.7 | 6.29E-06 | 0.2 | 5.18E-02 | 0.5 | 3.53E-04 | -0.2 | 4.23E-01 | -0.1 | 8.17E-01 | -0.1 | 7.34E-01 | Uncharacterized protein At2g46320 |
| AT3G48100 | 0.7 | 2.02E-04 | 0.5 | 2.34E-03 | 0.2 | 2.28E-01 | -0.2 | 6.43E-01 | -0.1 | 8.01E-01 | 0.0 | 9.41E-01 | RR5 |
| AT5G24215 | 0.7 | 2.73E-02 | 0.9 | 1.16E-02 | -0.2 | 6.14E-01 | 0.9 | 1.08E-02 | 0.5 | 1.50E-01 | 0.3 | 4.27E-01 | Uncharacterized protein |
| AT1G65845 | 0.7 | 1.67E-04 | 0.5 | 3.11E-03 | 0.2 | 1.17E-01 | 0.0 | 9.98E-01 | -0.5 | 6.34E-02 | 0.5 | 9.68E-02 | At1g65844 |
| AT2G14580 | 0.7 | 6.98E-03 | 0.1 | 7.52E-01 | 0.6 | 3.83E-02 | 0.9 | 3.48E-02 | 0.0 | 9.55E-01 | 1.0 | 3.59E-02 | Pathogenesis-related protein 1 |
| AT3G13845 | 0.7 | 2.56E-04 | 0.5 | 5.18E-03 | 0.2 | 1.43E-01 | 0.2 | 3.40E-01 | 0.1 | 6.81E-01 | 0.1 | 7.97E-01 | hypothetical protein |
| AT2G31150 | 0.7 | 4.70E-05 | 0.6 | 2.37E-04 | 0.1 | 4.48E-01 | -0.1 | 6.07E-01 | -0.2 | 3.92E-01 | 0.1 | 7.80E-01 | ATP binding |
| AT5G37630 | 0.7 | 1.34E-05 | 0.4 | 9.86E-04 | 0.3 | 1.60E-02 | 0.2 | 4.60E-01 | 0.3 | 3.83E-01 | -0.1 | 8.83E-01 | ARM repeat superfamily protein |
| AT2G20530 | 0.7 | 3.96E-05 | 0.5 | 8.16E-04 | 0.2 | 7.48E-02 | 0.5 | 6.57E-04 | 0.1 | 4.31E-01 | 0.4 | 3.53E-02 | Prohibitin-6, mitochondrial |
| AT5G08040 | 0.7 | 4.56E-05 | 0.7 | 7.64E-05 | 0.0 | 9.55E-01 | 0.4 | 7.62E-02 | 0.3 | 3.35E-01 | 0.2 | 7.02E-01 | Mitochondrial import receptor subunit TOM5 homolog |
| AT4G00180 | 0.7 | 3.25E-05 | 0.5 | 4.63E-04 | 0.2 | 1.16E-01 | -0.1 | 6.68E-01 | -0.2 | 1.60E-01 | 0.1 | 4.73E-01 | AT4G00180 protein |
| AT1G67785 | 0.7 | 2.73E-03 | 0.4 | 5.21E-02 | 0.3 | 1.69E-01 | 0.3 | 2.24E-01 | 0.2 | 5.37E-01 | 0.1 | 7.97E-01 | hypothetical protein |

| | | | | | | | | | | | |
|---|---|---|---|---|---|---|---|---|---|---|---|
| AT5G12920 | 0.7 | 2.19E-03 | 0.4 | 4.73E-02 | 0.3 | 1.92E-01 | 0.3 | 5.57E-01 | 0.2 | 7.56E-01 | | | Uncharacterized protein T24H18_100 |
| AT5G48720 | 0.7 | 9.17E-06 | 0.0 | 6.50E-01 | 0.7 | 1.53E-04 | 0.0 | 9.27E-01 | 0.0 | 8.76E-01 | 0.1 | 8.44E-01 | Protein XRI1 |
| AT4G01915 | 0.7 | 1.09E-03 | 0.3 | 1.31E-01 | 0.4 | 4.33E-02 | 0.0 | 1.00E+00 | 0.1 | 8.25E-01 | -0.1 | 8.62E-01 | Uncharacterized protein |
| AT2G04495 | 0.7 | 4.08E-02 | -0.4 | 3.02E-01 | 1.2 | 2.44E-02 | 0.4 | 4.63E-01 | -0.5 | 4.14E-01 | 0.9 | 1.67E-01 | At2g04495 |
| AT1G48460 | 0.7 | 8.07E-05 | 0.6 | 4.72E-04 | 0.1 | 3.94E-01 | 0.7 | 8.71E-04 | 0.2 | 3.29E-01 | 0.5 | 4.89E-02 | tRNA-processing ribonuclease BN |
| AT2G32210 | 0.7 | 3.61E-02 | 0.0 | 9.52E-01 | 0.8 | 7.38E-02 | -0.5 | 2.46E-01 | -0.5 | 2.48E-01 | 0.0 | 9.56E-01 | Cysteine-rich |
| AT4G37300 | 0.7 | 1.20E-05 | 0.7 | 4.85E-05 | 0.1 | 5.15E-01 | -0.8 | 1.01E-04 | -0.8 | 4.18E-04 | 0.0 | 9.75E-01 | hypothetical protein |
| AT1G28150 | 0.7 | 6.00E-05 | 0.5 | 1.00E-03 | 0.2 | 1.09E-01 | -0.1 | 4.09E-01 | -0.1 | 6.04E-01 | 0.0 | 9.23E-01 | UPF0426 protein At1g28150, chloroplastic |
| AT2G00840 | 0.7 | 1.21E-02 | 1.0 | 2.26E-03 | -0.3 | 3.42E-01 | 1.0 | 7.68E-02 | 0.6 | 3.82E-01 | 0.4 | 4.98E-01 | NA |
| AT1G16320 | 0.7 | 3.29E-05 | 0.1 | 2.47E-01 | 0.6 | 5.74E-04 | -0.2 | 2.13E-01 | 0.0 | 9.19E-01 | -0.2 | 2.56E-01 | Uncharacterized protein |
| AT3G43700 | 0.7 | 1.94E-05 | 0.3 | 2.58E-02 | 0.5 | 1.92E-03 | -0.1 | 4.11E-01 | -0.1 | 5.76E-01 | 0.0 | 9.36E-01 | BTB-POZ and MATH domain 6 |
| AT1G69750 | 0.7 | 6.32E-04 | 0.4 | 3.18E-02 | 0.3 | 7.57E-02 | 0.3 | 3.11E-01 | 0.1 | 8.99E-01 | 0.2 | 4.82E-01 | Uncharacterized protein T12I7.4 |
| AT5G14350 | 0.7 | 9.82E-04 | 0.6 | 3.67E-03 | 0.1 | 6.44E-01 | 0.4 | 2.05E-01 | 0.4 | 2.81E-01 | 0.0 | 9.81E-01 | Pentatricopeptide repeat-containing protein |
| AT5G52070 | 0.7 | 1.17E-03 | -0.3 | 2.15E-01 | 1.0 | 1.25E-03 | 0.2 | 2.85E-01 | 0.1 | 7.94E-01 | 0.1 | 6.43E-01 | Agenet domain-containing protein |
| AT1G73470 | 0.7 | 2.50E-05 | 0.6 | 1.66E-04 | 0.1 | 2.99E-01 | 0.1 | 7.85E-01 | 0.1 | 8.28E-01 | 0.0 | 9.97E-01 | Uncharacterized protein |
| AT3G61930 | 0.7 | 7.41E-04 | -0.2 | 3.68E-01 | 0.9 | 1.70E-03 | -2.3 | 2.03E-03 | -2.8 | 1.77E-03 | 0.5 | 6.18E-01 | hypothetical protein |
| AT2G46450 | 0.7 | 3.45E-06 | 0.2 | 1.32E-02 | 0.5 | 2.44E-04 | 0.9 | 1.37E-04 | 0.5 | 1.85E-02 | 0.4 | 1.10E-01 | Putative cyclic nucleotide-regulated ion channel protein |
| AT1G47820 | 0.7 | 7.37E-04 | 0.5 | 7.70E-03 | 0.2 | 2.98E-01 | 0.1 | 9.08E-01 | 0.0 | 9.76E-01 | 0.0 | 9.63E-01 | hypothetical protein |
| AT3G48550 | 0.7 | 2.24E-03 | 0.6 | 1.06E-02 | 0.1 | 5.47E-01 | -0.1 | 9.23E-01 | -0.1 | 8.57E-01 | 0.1 | 9.43E-01 | hypothetical protein |
| AT4G03180 | 0.7 | 4.25E-05 | 0.4 | 4.71E-03 | 0.3 | 1.77E-02 | 0.0 | 8.99E-01 | -0.1 | 6.55E-01 | 0.2 | 6.30E-01 | rRNA-processing protein |
| AT3G10110 | 0.7 | 7.43E-03 | 0.7 | 1.04E-02 | 0.0 | 9.50E-01 | 1.0 | 3.81E-03 | 0.2 | 5.76E-01 | 0.8 | 3.42E-02 | Mitochondrial import inner membrane translocase subunit Tim17 |
| AT4G18372 | 0.7 | 7.12E-03 | 0.1 | 8.34E-01 | 0.7 | 3.35E-02 | 0.3 | 4.42E-01 | 0.0 | 9.47E-01 | 0.3 | 4.97E-01 | At4g18372 |
| AT5G62140 | 0.7 | 6.03E-05 | 0.4 | 6.19E-03 | 0.4 | 1.79E-02 | -0.2 | 4.03E-01 | 0.2 | 4.14E-01 | -0.3 | 1.75E-01 | Uncharacterized protein |
| AT2G42840 | 0.7 | 5.57E-05 | 0.6 | 6.40E-04 | 0.2 | 1.70E-01 | -0.2 | 2.23E-01 | -0.3 | 1.35E-01 | 0.1 | 7.79E-01 | Protodermal factor 1 |
| AT3G13550 | 0.7 | 1.18E-04 | 0.4 | 6.74E-03 | 0.3 | 4.72E-02 | -0.1 | 8.46E-01 | -0.3 | 2.53E-01 | 0.2 | 4.27E-01 | AT3G13550 protein |
| AT2G28950 | 0.7 | 3.51E-05 | 0.9 | 2.62E-05 | -0.1 | 3.47E-01 | -0.1 | 4.26E-01 | 0.1 | 5.53E-01 | -0.2 | 2.70E-01 | Expansin-A6 |
| AT5G22210 | 0.7 | 9.26E-03 | 0.2 | 5.10E-01 | 0.6 | 6.99E-02 | 0.0 | 9.98E-01 | 0.0 | 9.87E-01 | 0.0 | 9.86E-01 | hypothetical protein |
| AT3G59765 | 0.8 | 9.35E-04 | 0.8 | 1.30E-03 | 0.0 | 9.76E-01 | 0.6 | 1.88E-01 | 0.7 | 2.10E-01 | 0.0 | 9.63E-01 | NA |
| AT1G32120 | 0.8 | 1.75E-04 | 0.7 | 4.68E-04 | 0.0 | 7.86E-01 | -0.2 | 7.26E-01 | 0.1 | 9.24E-01 | -0.2 | 6.98E-01 | Serine |
| AT5G02440 | 0.8 | 5.75E-04 | 0.4 | 2.78E-02 | 0.3 | 8.90E-02 | 0.0 | 8.88E-01 | -0.4 | 2.28E-01 | 0.4 | 1.66E-01 | Uncharacterized protein |
| AT1G27060 | 0.8 | 1.37E-04 | 0.2 | 2.29E-01 | 0.6 | 4.19E-03 | 0.0 | 9.13E-01 | -0.2 | 6.59E-01 | 0.2 | 5.48E-01 | hypothetical protein |
| AT1G06590 | 0.8 | 4.65E-05 | 0.2 | 1.88E-01 | 0.6 | 1.38E-03 | -0.1 | 5.43E-01 | -0.2 | 4.29E-01 | 0.1 | 8.81E-01 | Anaphase-promoting complex subunit 5 |
| AT1G25277 | 0.8 | 5.30E-03 | 0.2 | 4.77E-01 | 0.6 | 5.84E-02 | 0.2 | 5.98E-01 | 0.4 | 2.73E-01 | -0.2 | 6.71E-01 | Uncharacterized protein |
| AT1G50290 | 0.8 | 2.23E-04 | 0.3 | 4.57E-02 | 0.4 | 2.25E-02 | -0.2 | 6.53E-01 | -0.2 | 5.35E-01 | 0.1 | 8.72E-01 | At1g50290 |
| AT3G25717 | 0.8 | 1.09E-02 | 0.7 | 2.62E-02 | 0.1 | 7.52E-01 | 0.3 | 3.58E-01 | 0.2 | 4.59E-01 | 0.0 | 9.80E-01 | Small polypeptide DEVIL 6 |
| AT4G08950 | 0.8 | 2.00E-02 | 0.0 | 9.86E-01 | 0.8 | 4.24E-02 | -0.9 | 1.95E-04 | -1.2 | 9.05E-05 | 0.2 | 3.84E-01 | EXO |
| AT4G39200 | 0.8 | 4.18E-04 | 0.7 | 8.94E-04 | 0.0 | 8.63E-01 | 0.0 | 8.29E-01 | -0.1 | 6.17E-01 | 0.2 | 5.52E-01 | 40S ribosomal protein S25 |
| AT2G46230 | 0.8 | 5.95E-06 | 0.2 | 1.60E-02 | 0.5 | 5.15E-04 | 0.3 | 6.97E-02 | 0.1 | 5.03E-01 | 0.2 | 5.15E-01 | PIN domain-like family protein |
| AT4G15780 | 0.8 | 7.53E-05 | 0.5 | 1.69E-03 | 0.2 | 8.47E-02 | -0.1 | 6.08E-01 | -0.3 | 7.50E-02 | 0.2 | 2.97E-01 | Vesicle-associated membrane protein 724 |
| AT3G56130 | 0.8 | 4.66E-06 | 0.6 | 3.65E-05 | 0.1 | 1.66E-01 | 0.2 | 1.73E-01 | 0.1 | 5.24E-01 | 0.1 | 7.39E-01 | Biotin |
| AT5G12130 | 0.8 | 2.55E-06 | 0.5 | 9.87E-05 | 0.3 | 1.12E-02 | 0.3 | 1.22E-01 | 0.3 | 1.86E-01 | 0.0 | 9.73E-01 | Integral membrane TerC family protein |
| AT4G33760 | 0.8 | 7.17E-06 | 0.8 | 1.34E-05 | 0.0 | 9.27E-01 | 0.3 | 2.34E-02 | 0.1 | 6.59E-01 | 0.3 | 1.73E-01 | OB domain-containing protein |
| AT3G53850 | 0.8 | 9.56E-05 | 0.7 | 3.97E-04 | 0.1 | 5.38E-01 | 0.0 | 9.25E-01 | -0.2 | 4.03E-01 | 0.2 | 4.13E-01 | CASP-like protein |
| AT4G10280 | 0.8 | 4.86E-03 | 0.1 | 7.73E-01 | 0.7 | 2.58E-02 | 0.5 | 8.60E-02 | 0.2 | 5.32E-01 | 0.3 | 4.84E-01 | hypothetical protein |
| AT2G41120 | 0.8 | 1.04E-05 | 0.1 | 4.41E-01 | 0.7 | 2.27E-04 | -0.1 | 7.84E-01 | 0.1 | 9.07E-01 | -0.2 | 7.14E-01 | DUF309 domain-containing protein |
| AT2G28790 | 0.8 | 6.40E-04 | 0.8 | 4.95E-04 | -0.1 | 6.79E-01 | 0.2 | 1.65E-01 | 0.3 | 1.52E-01 | 0.0 | 9.06E-01 | At2g28790 |
| AT3G56360 | 0.8 | 4.80E-06 | 0.5 | 1.10E-04 | 0.2 | 3.14E-02 | 1.0 | 6.05E-07 | 0.2 | 2.36E-01 | 0.8 | 3.42E-04 | Uncharacterized protein |
| AT1G08580 | 0.8 | 1.11E-04 | 0.5 | 2.16E-03 | 0.2 | 1.10E-01 | 0.8 | 1.13E-03 | 0.5 | 6.04E-02 | 0.3 | 2.44E-01 | hypothetical protein |
| AT3G02120 | 0.8 | 6.29E-04 | 0.9 | 2.63E-04 | -0.2 | 4.03E-01 | 0.2 | 2.47E-01 | 0.1 | 6.21E-01 | 0.1 | 7.44E-01 | Uncharacterized protein |
| AT3G52230 | 0.8 | 1.84E-06 | 0.4 | 1.94E-04 | 0.3 | 2.31E-03 | 0.1 | 7.10E-01 | 0.0 | 9.12E-01 | 0.1 | 7.32E-01 | AT3g52230 |
| AT2G25460 | 0.8 | 1.78E-04 | 0.6 | 1.45E-03 | 0.2 | 3.20E-01 | 1.1 | 1.54E-03 | 0.3 | 4.73E-01 | 0.8 | 1.23E-02 | EEIG1 |
| AT1G15030 | 0.8 | 8.67E-06 | 0.4 | 1.92E-03 | 0.4 | 3.74E-03 | -0.2 | 2.01E-01 | -0.3 | 1.45E-01 | 0.1 | 8.33E-01 | T15D22.8 |
| AT4G33660 | 0.8 | 4.59E-05 | 0.9 | 3.42E-05 | -0.1 | 4.23E-01 | -0.1 | 7.32E-01 | 0.2 | 6.50E-01 | -0.3 | 4.91E-01 | Protein CYSTEINE-RICH TRANSMEMBRANE MODULE 11 |
| AT1G60730 | 0.8 | 1.75E-04 | -0.1 | 4.28E-01 | 0.9 | 4.36E-04 | 0.8 | 2.13E-04 | -0.2 | 4.75E-01 | 1.0 | 1.28E-03 | NAD-linked oxidoreductase superfamily protein |
| AT5G63000 | 0.8 | 2.80E-05 | 0.4 | 4.97E-03 | 0.4 | 9.86E-03 | 0.3 | 1.49E-01 | 0.1 | 8.36E-01 | 0.3 | 3.70E-01 | hypothetical protein |
| AT5G47540 | 0.8 | 4.53E-05 | 0.5 | 9.99E-04 | 0.2 | 8.12E-02 | -0.1 | 6.90E-01 | -0.1 | 5.16E-01 | 0.1 | 8.32E-01 | MO25-like protein At5g47540 |
| AT3G26070 | 0.8 | 2.56E-06 | 0.5 | 1.29E-04 | 0.3 | 7.14E-03 | 0.0 | 9.38E-01 | 0.1 | 2.79E-01 | -0.1 | 4.57E-01 | Probable plastid-lipid-associated protein 4, chloroplastic |
| AT1G14630 | 0.8 | 4.51E-04 | 0.3 | 8.88E-02 | 0.5 | 2.72E-02 | 0.2 | 7.20E-01 | 0.3 | 4.54E-01 | -0.2 | 7.47E-01 | XRI1-like protein |
| AT5G06980 | 0.8 | 3.82E-05 | 0.4 | 5.08E-03 | 0.4 | 1.06E-02 | | | | | | | Protein LNK4 |
| AT2G35980 | 0.8 | 2.27E-02 | 0.2 | 6.65E-01 | 0.6 | 1.18E-01 | 1.2 | 1.00E-02 | -1.2 | 8.08E-02 | 2.4 | 2.46E-03 | NDR1 |
| AT4G29735 | 0.8 | 9.78E-04 | 0.5 | 1.72E-02 | 0.3 | 1.78E-01 | 0.2 | 6.11E-01 | 0.0 | 9.13E-01 | 0.1 | 8.30E-01 | Dolichyl-diphosphooligosaccharide-protein glycosyltransferase subunit OST5 |
| AT4G03460 | 0.8 | 2.58E-02 | 0.2 | 6.45E-01 | 0.6 | 1.29E-01 | 0.8 | 2.68E-02 | -0.1 | 8.72E-01 | 0.9 | 3.28E-02 | Ankyrin repeat family protein |
| AT1G21830 | 0.8 | 2.12E-05 | 0.4 | 2.70E-03 | 0.3 | 1.23E-02 | -0.4 | 4.39E-02 | -0.5 | 1.79E-02 | 0.1 | 6.72E-01 | F8K7.25 protein |
| AT3G26165 | 0.8 | 1.90E-04 | 0.6 | 1.52E-03 | 0.2 | 3.15E-01 | 0.1 | 6.22E-01 | -0.4 | 1.12E-01 | 0.5 | 6.53E-02 | NA |
| AT1G69935 | 0.8 | 3.74E-06 | 0.2 | 2.61E-02 | 0.6 | 2.61E-04 | 0.0 | 9.29E-01 | 0.0 | 9.56E-01 | 0.0 | 9.12E-01 | Short hypocotyl in white light1 |
| AT1G53560 | 0.8 | 4.38E-06 | 0.5 | 1.46E-04 | 0.2 | 2.37E-02 | -0.2 | 5.49E-01 | -0.3 | 3.28E-01 | 0.1 | 7.48E-01 | At1g53560 |
| AT5G66000 | 0.8 | 1.23E-04 | 0.6 | 9.99E-04 | 0.1 | 3.27E-01 | 0.2 | 5.22E-01 | 0.2 | 6.68E-01 | 0.0 | 9.46E-01 | Uncharacterized protein At5g66000 |
| AT5G63480 | 0.8 | 5.11E-05 | 0.6 | 5.20E-04 | 0.2 | 2.27E-01 | -0.2 | 5.18E-01 | 0.0 | 9.13E-01 | -0.1 | 7.38E-01 | Mediator of RNA polymerase II transcription subunit |
| AT1G21260 | 0.8 | 9.59E-05 | 0.4 | 7.79E-03 | 0.3 | 3.66E-02 | 0.3 | 4.22E-01 | 0.1 | 8.66E-01 | 0.2 | 7.02E-01 | NA |
| AT5G10320 | 0.8 | 2.20E-04 | 0.1 | 4.65E-01 | 0.7 | 3.46E-03 | 0.2 | 3.56E-01 | 0.2 | 5.07E-01 | 0.0 | 9.35E-01 | ATP synthase subunit B |
| AT5G63100 | 0.8 | 8.66E-05 | 0.6 | 7.56E-04 | 0.2 | 2.76E-01 | 0.2 | 3.86E-01 | 0.1 | 7.91E-01 | 0.1 | 6.60E-01 | S-adenosyl-L-methionine-dependent methyltransferases superfamily protein |
| AT4G03420 | 0.8 | 8.21E-06 | 0.5 | 3.41E-04 | 0.3 | 1.99E-02 | 0.2 | 6.19E-01 | -0.7 | 7.35E-02 | 0.9 | 6.11E-02 | Uncharacterized protein AT4g03420 |
| AT3G12210 | 0.8 | 1.42E-05 | 0.5 | 4.54E-04 | 0.2 | 4.02E-02 | -0.2 | 3.87E-01 | -0.2 | 4.20E-01 | 0.0 | 9.81E-01 | DNA binding protein |
| AT3G51660 | 0.8 | 5.72E-03 | 0.1 | 6.49E-01 | 0.7 | 3.43E-02 | 0.6 | 1.26E-01 | 0.3 | 4.80E-01 | 0.2 | 6.24E-01 | LS1-like protein |
| AT4G08280 | 0.8 | 3.73E-06 | 0.4 | 3.86E-04 | 0.3 | 4.34E-03 | 0.0 | 8.63E-01 | 0.0 | 8.72E-01 | 0.0 | 9.94E-01 | Glutaredoxin-like protein |
| AT2G27400 | 0.8 | 7.84E-05 | 0.4 | 2.06E-02 | 0.4 | 1.28E-02 | 0.6 | 2.28E-02 | 0.7 | 1.58E-02 | -0.1 | 8.10E-01 | NA |
| AT5G53650 | 0.8 | 1.15E-04 | 0.5 | 4.22E-03 | 0.3 | 6.62E-02 | 0.2 | 5.15E-01 | -0.1 | 8.05E-01 | 0.3 | 4.68E-01 | Transmembrane protein |
| AT1G17850 | 0.8 | 9.02E-04 | 0.7 | 3.10E-03 | 0.1 | 6.47E-01 | 0.3 | 4.63E-01 | 0.4 | 3.35E-01 | -0.1 | 8.31E-01 | Rhodanese |
| AT5G04320 | 0.8 | 4.15E-05 | 0.4 | 9.84E-03 | 0.4 | 8.68E-03 | 0.3 | 5.20E-02 | 0.2 | 3.09E-01 | 0.1 | 5.51E-01 | Shugoshin C terminus |
| AT1G45230 | 0.8 | 9.63E-06 | 0.7 | 6.72E-05 | 0.1 | 2.33E-01 | -0.1 | 6.04E-01 | -0.1 | 5.80E-01 | 0.0 | 9.44E-01 | DCL protein |
| AT2G39675 | 0.8 | 2.55E-04 | 0.2 | 1.84E-01 | 0.6 | 8.54E-03 | 0.6 | 1.88E-01 | 0.5 | 2.75E-01 | 0.0 | 9.61E-01 | NA |
| AT1G73770 | 0.8 | 3.18E-04 | 0.5 | 7.03E-03 | 0.3 | 1.48E-01 | 0.4 | 1.94E-01 | 0.2 | 6.16E-01 | 0.2 | 5.61E-01 | Coiled-coil protein |
| AT3G09600 | 0.8 | 7.22E-04 | 0.0 | 9.34E-01 | 0.8 | 2.90E-03 | -0.2 | 7.94E-01 | -0.4 | 6.17E-01 | | | Uncharacterized protein F11F8_20 |

| | | | | | | | | | | | | | |
|---|---|---|---|---|---|---|---|---|---|---|---|---|---|
| AT5G52650 | 0.8 | 2.44E-04 | 0.8 | 2.91E-04 | 0.0 | 8.04E-01 | 0.4 | 4.12E-02 | 0.1 | 7.98E-01 | 0.3 | 2.21E-01 | AT5G52650 protein |
| AT5G18080 | 0.8 | 4.51E-02 | 1.1 | 9.46E-03 | -0.3 | 4.09E-01 | 0.0 | 9.46E-01 | 0.1 | 8.01E-01 | | | Auxin-responsive protein SAUR24 |
| AT5G19750 | 0.8 | 2.65E-05 | 0.6 | 3.55E-04 | 0.2 | 1.21E-01 | 0.6 | 5.73E-04 | 0.1 | 6.34E-01 | 0.5 | 7.59E-03 | Peroxisomal membrane 22 kDa family protein |
| AT1G17200 | 0.8 | 6.23E-05 | 0.9 | 5.09E-05 | -0.1 | 5.08E-01 | 0.0 | 9.10E-01 | -0.2 | 3.55E-01 | 0.2 | 3.42E-01 | CASP-like protein |
| AT4G38490 | 0.8 | 3.56E-06 | 0.5 | 1.09E-04 | 0.2 | 2.31E-02 | -0.2 | 4.26E-01 | -0.2 | 5.15E-01 | 0.0 | 9.80E-01 | Transmembrane protein |
| AT5G19240 | 0.8 | 1.55E-03 | 0.5 | 2.53E-02 | 0.3 | 1.82E-01 | 0.8 | 3.97E-04 | 0.2 | 5.41E-01 | 0.7 | 6.13E-03 | Uncharacterized GPI-anchored protein At5g19240 |
| AT1G08847 | 0.8 | 3.33E-04 | 0.3 | 9.32E-02 | 0.5 | 1.88E-02 | 0.7 | 3.11E-02 | 0.4 | 2.73E-01 | 0.3 | 4.65E-01 | NA |
| AT2G02130 | 0.8 | 9.18E-04 | 0.8 | 1.09E-03 | 0.0 | 8.91E-01 | 0.2 | 3.61E-01 | 0.2 | 3.18E-01 | 0.0 | 9.02E-01 | Defensin-like protein 1 |
| AT4G28310 | 0.8 | 1.27E-04 | 0.6 | 1.86E-03 | 0.2 | 1.69E-01 | 0.5 | 4.63E-02 | 0.3 | 3.32E-01 | 0.2 | 5.40E-01 | Uncharacterized protein AT4g28310 |
| AT5G24690 | 0.8 | 1.84E-06 | 0.5 | 1.79E-04 | 0.3 | 2.52E-03 | 0.0 | 8.73E-01 | -0.2 | 3.03E-01 | 0.1 | 4.59E-01 | Gb\|AAD25674.1 |
| AT1G16730 | 0.8 | 2.71E-04 | -0.1 | 4.90E-01 | 0.9 | 7.55E-04 | | | | | | | F17F16.6 protein |
| AT5G02502 | 0.8 | 4.32E-03 | 0.6 | 1.89E-02 | 0.2 | 5.46E-01 | 0.3 | 2.39E-01 | 0.0 | 9.68E-01 | 0.3 | 3.92E-01 | Dolichyl-diphosphooligosaccharide--protein glycosyltransferase subunit 4B |
| AT3G07350 | 0.8 | 1.77E-01 | -0.4 | 6.22E-01 | 1.2 | 1.69E-01 | -0.2 | 6.31E-01 | -1.4 | 7.21E-03 | 1.2 | 3.79E-02 | Uncharacterized protein |
| AT5G19250 | 0.8 | 4.10E-05 | 0.9 | 4.00E-05 | -0.1 | 5.11E-01 | 0.5 | 4.73E-03 | 0.1 | 6.67E-01 | 0.4 | 7.05E-02 | Uncharacterized GPI-anchored protein At5g19250 |
| AT1G77270 | 0.8 | 2.70E-04 | 0.6 | 3.19E-03 | 0.2 | 2.66E-01 | 0.0 | 9.93E-01 | 0.5 | 2.02E-01 | -0.5 | 2.87E-01 | Uncharacterized protein At1g77270 |
| AT3G13000 | 0.8 | 1.10E-05 | 0.8 | 2.21E-05 | 0.0 | 9.82E-01 | -0.8 | 2.96E-03 | -0.7 | 1.18E-02 | -0.1 | 8.12E-01 | At3g13000 |
| AT1G71810 | 0.8 | 2.00E-05 | 0.6 | 2.98E-04 | 0.2 | 1.07E-01 | 0.4 | 3.60E-02 | 0.3 | 9.44E-02 | 0.0 | 8.78E-01 | Uncharacterized protein |
| AT3G47348 | 0.8 | 3.01E-04 | 0.3 | 5.83E-02 | 0.5 | 2.53E-02 | 0.2 | 4.14E-01 | 0.0 | 9.54E-01 | 0.3 | 4.84E-01 | NA |
| AT5G01660 | 0.8 | 3.23E-04 | 0.4 | 5.17E-02 | 0.4 | 3.17E-02 | 0.8 | 9.95E-02 | 0.6 | 2.33E-01 | 0.1 | 8.37E-01 | Influenza virus NS1A-binding protein |
| AT1G44890 | 0.8 | 3.41E-03 | 0.1 | 6.26E-01 | 0.7 | 2.96E-02 | 0.2 | 5.92E-01 | 0.1 | 8.56E-01 | 0.1 | 8.15E-01 | T12C22.17 protein |
| AT3G12150 | 0.8 | 1.22E-05 | 0.8 | 2.51E-05 | 0.0 | 9.88E-01 | -0.1 | 4.63E-01 | -0.2 | 2.58E-01 | 0.1 | 7.36E-01 | Protein ABHD18 |
| AT1G11320 | 0.8 | 7.08E-05 | 0.2 | 1.38E-01 | 0.6 | 3.21E-03 | 0.8 | 3.14E-02 | 0.1 | 8.03E-01 | 0.7 | 1.30E-01 | Uncharacterized protein |
| AT5G65207 | 0.8 | 3.87E-02 | 0.4 | 2.77E-01 | 0.4 | 3.68E-01 | -1.0 | 3.45E-03 | -0.9 | 9.87E-03 | -0.1 | 8.89E-01 | Uncharacterized protein |
| AT1G66330 | 0.8 | 2.17E-04 | 0.4 | 2.31E-02 | 0.4 | 2.37E-02 | 0.1 | 7.18E-01 | 0.0 | 9.80E-01 | 0.1 | 8.27E-01 | Senescence-associated protein AAF, chloroplastic |
| AT3G15750 | 0.8 | 1.13E-03 | 0.7 | 5.68E-03 | 0.1 | 4.98E-01 | 0.3 | 5.93E-01 | 0.0 | 9.86E-01 | 0.3 | 6.67E-01 | Essential protein Yae1, N-terminal |
| AT5G57785 | 0.8 | 2.93E-03 | 0.7 | 7.83E-03 | 0.1 | 7.63E-01 | -0.4 | 1.28E-01 | -0.4 | 1.83E-01 | 0.0 | 9.82E-01 | Uncharacterized protein |
| AT3G12320 | 0.8 | 3.31E-05 | 0.4 | 1.03E-02 | 0.4 | 4.98E-03 | | | | | | | Protein LNK3 |
| AT3G44450 | 0.8 | 1.80E-03 | 0.4 | 5.62E-02 | 0.4 | 1.11E-01 | 0.8 | 2.07E-02 | 0.9 | 1.22E-02 | -0.1 | 7.19E-01 | Protein BIC2 |
| AT3G13910 | 0.8 | 8.51E-03 | 0.3 | 3.49E-01 | 0.5 | 9.65E-02 | 0.5 | 2.30E-01 | 0.2 | 7.52E-01 | 0.3 | 6.25E-01 | hypothetical protein |
| AT2G29530 | 0.8 | 9.45E-04 | 0.5 | 1.53E-02 | 0.3 | 2.05E-01 | 0.5 | 2.10E-02 | 0.3 | 2.24E-01 | 0.2 | 4.77E-01 | Mitochondrial import inner membrane translocase subunit |
| AT4G20760 | 0.8 | 3.97E-06 | 0.7 | 1.91E-05 | 0.1 | 3.68E-01 | 0.2 | 2.51E-01 | 0.2 | 4.02E-01 | 0.0 | 9.33E-01 | AT4g20760 |
| AT4G03150 | 0.8 | 2.25E-04 | 0.9 | 2.02E-04 | -0.1 | 6.82E-01 | -0.2 | 5.34E-01 | -0.1 | 6.88E-01 | 0.0 | 9.40E-01 | Uncharacterized protein |
| AT1G27330 | 0.8 | 3.15E-04 | 0.4 | 2.37E-02 | 0.4 | 3.97E-02 | 0.6 | 1.68E-02 | 0.4 | 1.92E-01 | 0.2 | 4.98E-01 | F17L21.12 |
| AT4G12480 | 0.8 | 2.81E-01 | -0.4 | 6.35E-01 | 1.2 | 1.74E-01 | -0.8 | 2.87E-01 | -3.7 | 1.85E-02 | 2.9 | 1.17E-01 | Lipid transfer protein EARLI 1 |
| AT1G08640 | 0.8 | 3.97E-06 | 0.8 | 1.39E-05 | 0.1 | 5.68E-01 | 0.7 | 7.79E-04 | 0.2 | 4.44E-01 | 0.5 | 3.10E-02 | Protein CHLOROPLAST J-LIKE DOMAIN 1, chloroplastic |
| AT1G25275 | 0.8 | 4.10E-05 | 0.8 | 1.09E-04 | 0.0 | 7.98E-01 | -0.3 | 1.71E-01 | -0.7 | 1.04E-02 | 0.4 | 2.44E-01 | Thionin-like protein |
| AT4G15790 | 0.8 | 4.95E-05 | 0.5 | 1.34E-03 | 0.3 | 7.13E-02 | -0.1 | 7.57E-01 | 0.0 | 9.78E-01 | -0.1 | 7.83E-01 | Uveal autoantigen with coiled-coil |
| AT2G31490 | 0.8 | 4.99E-04 | 0.6 | 4.91E-03 | 0.2 | 2.52E-01 | 0.3 | 1.20E-01 | 0.1 | 7.55E-01 | 0.2 | 4.62E-01 | AT2G31490 protein |
| AT4G31730 | 0.8 | 9.60E-04 | 0.7 | 4.65E-03 | 0.1 | 5.01E-01 | 1.8 | 1.54E-04 | 0.5 | 3.51E-01 | 1.3 | 4.28E-03 | Protein GLUTAMINE DUMPER 1 |
| AT3G02690 | 0.8 | 1.80E-05 | 0.4 | 5.98E-03 | 0.4 | 3.49E-03 | -0.2 | 1.38E-01 | -0.2 | 3.30E-01 | -0.1 | 8.40E-01 | WAT1-related protein At3g02690, chloroplastic |
| AT5G66840 | 0.8 | 3.37E-03 | 0.5 | 4.86E-02 | 0.3 | 2.86E-01 | 0.3 | 5.11E-01 | -0.1 | 9.38E-01 | 0.4 | 5.18E-01 | SAP domain-containing protein |
| AT3G46630 | 0.8 | 2.11E-06 | 0.8 | 6.68E-06 | 0.0 | 6.76E-01 | 0.2 | 3.96E-01 | 0.0 | 9.02E-01 | 0.3 | 4.48E-01 | Uncharacterized protein |
| AT1G70700 | 0.8 | 2.48E-04 | 0.6 | 2.07E-03 | 0.2 | 2.74E-01 | -0.4 | 3.76E-01 | -0.1 | 9.02E-01 | -0.3 | 6.58E-01 | Protein TIFY |
| AT5G37360 | 0.8 | 1.43E-05 | 0.6 | 1.82E-04 | 0.2 | 1.08E-01 | -0.1 | 7.47E-01 | 0.1 | 4.59E-01 | -0.2 | 3.22E-01 | Uncharacterized protein |
| AT2G43090 | 0.8 | 1.06E-05 | 0.5 | 3.21E-04 | 0.3 | 2.98E-02 | 0.4 | 2.66E-03 | 0.2 | 1.84E-01 | 0.2 | 2.07E-01 | Aconitase |
| AT1G70830 | 0.8 | 2.55E-05 | 0.9 | 1.90E-05 | -0.1 | 3.10E-01 | 0.0 | 9.01E-01 | 0.0 | 9.79E-01 | 0.0 | 9.11E-01 | MLP-like protein 28 |
| AT5G12320 | 0.8 | 8.17E-05 | 0.4 | 8.09E-03 | 0.4 | 2.50E-02 | 0.4 | 1.46E-01 | 0.3 | 3.43E-01 | 0.1 | 8.37E-01 | AT5G12320 protein |
| AT2G21290 | 0.8 | 3.58E-05 | 0.5 | 1.28E-03 | 0.3 | 4.19E-02 | 0.1 | 6.53E-01 | -0.1 | 5.74E-01 | 0.2 | 4.08E-01 | Small ribosomal subunit protein bTHXm |
| AT1G27435 | 0.8 | 1.34E-05 | 0.8 | 4.17E-05 | 0.1 | 6.66E-01 | 0.5 | 1.03E-02 | 0.2 | 3.22E-01 | 0.3 | 2.71E-01 | Uncharacterized protein |
| AT5G59080 | 0.8 | 4.87E-03 | 0.7 | 1.69E-02 | 0.1 | 6.35E-01 | -0.1 | 7.33E-01 | -0.3 | 4.11E-01 | 0.2 | 7.03E-01 | Uncharacterized protein |
| AT3G21390 | 0.8 | 2.73E-05 | 0.5 | 2.89E-03 | 0.4 | 1.44E-02 | -0.1 | 7.95E-01 | 0.3 | 1.32E-01 | -0.4 | 8.97E-02 | Mitochondrial thiamine diphosphate carrier 2 |
| AT4G26810 | 0.8 | 3.23E-04 | 0.4 | 2.17E-02 | 0.4 | 6.11E-02 | 0.3 | 1.92E-01 | -0.1 | 8.85E-01 | 0.3 | 2.21E-01 | hypothetical protein |
| AT1G26470 | 0.8 | 4.65E-04 | 0.5 | 1.15E-02 | 0.3 | 1.27E-01 | 0.3 | 2.56E-01 | -0.1 | 6.72E-01 | 0.4 | 1.91E-01 | Chromatin modification-like protein |
| AT5G39080 | 0.8 | 4.77E-05 | 0.6 | 4.51E-04 | 0.2 | 2.00E-01 | 0.1 | 6.52E-01 | -0.5 | 4.51E-02 | 0.6 | 7.46E-03 | Uncharacterized protein |
| AT5G64130 | 0.8 | 4.94E-06 | 0.7 | 4.69E-05 | 0.2 | 1.23E-01 | 0.0 | 9.93E-01 | -0.1 | 5.94E-01 | 0.1 | 6.80E-01 | cAMP-regulated phosphoprotein 19-related protein |
| AT3G47980 | 0.8 | 4.36E-06 | 1.2 | 6.71E-07 | -0.4 | 2.52E-03 | 0.3 | 1.30E-01 | 0.0 | 8.49E-01 | 0.2 | 3.51E-01 | Uncharacterized protein T17F15.160 |
| AT5G23380 | 0.8 | 1.05E-04 | 0.6 | 2.10E-03 | 0.2 | 1.28E-01 | -0.4 | 1.22E-01 | -0.4 | 1.64E-01 | 0.0 | 9.97E-01 | Gb\|AAD11584.1 |
| AT3G29280 | 0.8 | 4.55E-05 | 0.8 | 1.40E-04 | 0.1 | 7.09E-01 | 0.4 | 1.42E-01 | 0.3 | 3.96E-01 | 0.1 | 7.59E-01 | Uncharacterized protein |
| AT1G57630 | 0.8 | 5.15E-02 | -0.4 | 4.23E-01 | 1.2 | 4.17E-02 | 0.7 | 1.38E-01 | -0.5 | 4.50E-01 | 1.2 | 6.79E-02 | Disease resistance protein RPP1-WsB, putative domain family protein) |
| AT3G15110 | 0.8 | 9.37E-06 | 0.7 | 6.02E-05 | 0.1 | 2.55E-01 | 0.2 | 2.49E-01 | 0.3 | 5.93E-02 | -0.1 | 4.55E-01 | Transmembrane protein |
| AT1G33840 | 0.8 | 6.68E-05 | 1.1 | 1.20E-05 | -0.3 | 4.09E-02 | 0.3 | 2.03E-01 | 0.5 | 3.71E-02 | -0.2 | 3.26E-01 | Protein LURP-one-related 15 |
| AT3G06778 | 0.8 | 2.05E-03 | 0.3 | 1.84E-01 | 0.5 | 6.03E-02 | -0.3 | 4.98E-01 | -0.1 | 8.28E-01 | -0.2 | 7.37E-01 | Uncharacterized protein |
| AT1G52770 | 0.8 | 2.14E-02 | 0.5 | 1.94E-01 | 0.4 | 3.46E-01 | -0.1 | 7.09E-01 | 0.3 | 2.64E-01 | -0.5 | 1.27E-01 | At1g52770 |
| AT4G36910 | 0.8 | 9.65E-06 | 0.4 | 2.26E-03 | 0.4 | 4.48E-03 | 0.1 | 5.04E-01 | 0.1 | 4.00E-01 | 0.0 | 8.57E-01 | hypothetical protein |
| AT1G16080 | 0.8 | 8.58E-06 | 0.8 | 1.62E-05 | 0.0 | 9.10E-01 | 0.2 | 1.36E-01 | 0.3 | 1.21E-01 | 0.0 | 8.89E-01 | AT1G16080 protein |
| AT2G14660 | 0.8 | 1.09E-04 | 1.1 | 2.26E-05 | -0.3 | 8.37E-02 | 0.4 | 2.07E-01 | 0.1 | 9.01E-01 | 0.4 | 4.52E-01 | At2g14660 |
| AT3G20240 | 0.8 | 8.07E-05 | 0.5 | 3.01E-03 | 0.3 | 5.98E-02 | 0.8 | 9.32E-04 | 0.4 | 7.10E-02 | 0.4 | 1.23E-01 | Probable mitochondrial adenine nucleotide transporter BTL1 |
| AT4G31530 | 0.8 | 5.90E-05 | 0.6 | 8.64E-04 | 0.2 | 1.23E-01 | 0.1 | 3.68E-01 | 0.0 | 7.60E-01 | 0.1 | 7.79E-01 | NAD-binding Rossmann-fold superfamily protein |
| AT1G65470 | 0.8 | 8.58E-06 | 0.6 | 1.15E-04 | 0.2 | 1.01E-01 | 0.6 | 4.32E-04 | 0.2 | 1.69E-01 | 0.4 | 4.67E-02 | F5I14.1 protein |
| AT5G51960 | 0.8 | 3.97E-05 | 0.7 | 1.82E-04 | 0.1 | 4.75E-01 | 0.4 | 1.90E-01 | 0.2 | 6.59E-01 | 0.2 | 6.30E-01 | hypothetical protein |
| AT1G53625 | 0.8 | 2.28E-02 | -0.2 | 6.24E-01 | 1.0 | 2.92E-02 | 2.6 | 3.44E-06 | 0.8 | 9.44E-02 | 1.8 | 3.65E-04 | Uncharacterized protein |
| AT5G48830 | 0.8 | 2.26E-06 | 0.7 | 1.42E-05 | 0.1 | 2.21E-01 | 0.2 | 2.46E-01 | 0.0 | 8.62E-01 | 0.2 | 4.50E-01 | Phosphoglycolate phosphatase |
| AT1G23390 | 0.8 | 2.47E-03 | 0.9 | 3.33E-03 | 0.0 | 9.73E-01 | -0.4 | 8.41E-02 | -1.5 | 7.63E-05 | 1.1 | 6.64E-03 | F-box |
| AT1G75580 | 0.8 | 2.04E-04 | 0.9 | 1.98E-04 | -0.1 | 7.01E-01 | -0.2 | 4.83E-01 | -0.1 | 7.62E-01 | -0.1 | 8.46E-01 | Protein SMALL AUXIN UP-REGULATED RNA 51 |
| AT4G27380 | 0.8 | 1.23E-04 | 0.6 | 2.03E-03 | 0.2 | 1.44E-01 | 0.6 | 2.02E-02 | 0.1 | 8.87E-01 | 0.5 | 8.16E-02 | Uncharacterized protein |
| AT5G24000 | 0.8 | 1.88E-04 | 0.6 | 2.69E-03 | 0.2 | 2.01E-01 | 0.2 | 4.64E-01 | 0.1 | 8.72E-01 | 0.1 | 7.20E-01 | Uncharacterized protein |
| AT5G19620 | 0.8 | 4.85E-07 | 0.4 | 1.43E-04 | 0.4 | 3.35E-04 | 0.2 | 1.59E-01 | 0.2 | 3.08E-01 | 0.0 | 8.81E-01 | TOC75-V |
| AT1G16000 | 0.8 | 4.41E-03 | 0.5 | 6.60E-02 | 0.3 | 2.35E-01 | 0.4 | 1.87E-02 | 0.2 | 3.36E-01 | 0.2 | 3.18E-01 | GAG1At protein |
| AT1G02910 | 0.8 | 1.59E-06 | 0.5 | 1.43E-04 | 0.4 | 2.55E-03 | 0.0 | 8.56E-01 | -0.1 | 4.52E-01 | 0.1 | 6.62E-01 | Tetratricopeptide repeat -containing protein |
| AT1G78230 | 0.8 | 2.01E-05 | 0.6 | 6.92E-04 | 0.3 | 4.33E-02 | -0.8 | 1.22E-02 | -0.5 | 1.04E-01 | -0.3 | 5.64E-01 | F3F9.22 |
| AT5G16060 | 0.8 | 9.37E-05 | 0.5 | 3.48E-03 | 0.3 | 6.33E-02 | 0.6 | 1.31E-02 | 0.3 | 2.83E-01 | 0.3 | 3.01E-01 | COX assembly mitochondrial protein |
| AT2G44120 | 0.8 | 1.44E-05 | 0.8 | 3.54E-05 | 0.0 | 8.86E-01 | 0.5 | 1.43E-03 | 0.1 | 5.11E-01 | 0.4 | 6.00E-02 | At2g44120 |
| AT3G49790 | 0.8 | 2.41E-03 | 0.9 | 1.94E-03 | -0.1 | 7.84E-01 | -0.2 | 2.48E-01 | -1.1 | 1.61E-04 | 0.9 | 5.96E-03 | hypothetical protein |

| | | | | | | | | | | | | | |
|---|---|---|---|---|---|---|---|---|---|---|---|---|---|
| AT5G03285 | 0.8 | 2.63E-05 | 0.2 | 1.41E-01 | 0.6 | 1.23E-03 | 0.2 | 8.01E-01 | 0.5 | 3.57E-01 | -0.4 | 5.87E-01 | NA |
| AT1G34570 | 0.8 | 1.37E-04 | 0.2 | 1.82E-01 | 0.6 | 4.00E-03 | 0.0 | 9.09E-01 | 0.1 | 7.83E-01 | -0.1 | 9.03E-01 | At1g34570 |
| AT4G29310 | 0.8 | 4.42E-06 | 0.3 | 1.15E-02 | 0.6 | 4.86E-04 | -0.7 | 1.32E-03 | -0.6 | 8.63E-03 | -0.1 | 7.20E-01 | DUF1005 family protein |
| AT5G40710 | 0.8 | 2.14E-04 | 0.3 | 5.92E-02 | 0.5 | 1.60E-02 | 0.3 | 2.78E-01 | 0.1 | 7.22E-01 | 0.1 | 6.79E-01 | C2H2-type domain-containing protein |
| AT1G51405 | 0.8 | 1.01E-03 | 0.6 | 1.06E-02 | 0.2 | 3.05E-01 | 0.1 | 8.07E-01 | 0.5 | 3.47E-01 | -0.3 | 5.62E-01 | hypothetical protein |
| AT4G25170 | 0.8 | 1.38E-05 | 0.5 | 1.10E-03 | 0.4 | 1.03E-02 | -0.4 | 7.14E-03 | -0.5 | 2.18E-03 | 0.1 | 4.42E-01 | Uncharacterized protein AT4g25170 |
| AT5G20935 | 0.9 | 2.11E-06 | 0.4 | 7.83E-04 | 0.4 | 1.23E-03 | 0.3 | 2.78E-01 | 0.2 | 6.67E-01 | 0.1 | 6.60E-01 | DUF3148 family protein |
| AT5G05370 | 0.9 | 1.09E-03 | 0.5 | 2.28E-02 | 0.3 | 1.52E-01 | 0.4 | 1.03E-01 | 0.1 | 8.05E-01 | 0.3 | 3.57E-01 | Cytochrome b-c1 complex subunit 8-2, mitochondrial |
| AT5G24980 | 0.9 | 5.04E-03 | 0.3 | 2.39E-01 | 0.5 | 8.99E-02 | 0.5 | 1.26E-01 | 0.0 | 9.41E-01 | 0.4 | 2.76E-01 | hypothetical protein |
| AT5G05060 | 0.9 | 3.58E-06 | 0.4 | 8.07E-04 | 0.4 | 2.26E-03 | 0.1 | 7.58E-01 | -0.2 | 4.08E-01 | 0.3 | 3.44E-01 | Cystatin |
| AT1G28395 | 0.9 | 7.32E-05 | 0.7 | 6.79E-04 | 0.2 | 2.36E-01 | 0.7 | 3.24E-04 | 0.3 | 9.27E-02 | 0.4 | 6.60E-02 | Protein NUCLEAR FUSION DEFECTIVE 6, chloroplastic |
| AT5G55710 | 0.9 | 4.58E-07 | 0.6 | 1.23E-05 | 0.3 | 5.24E-03 | 0.1 | 4.15E-01 | 0.1 | 5.22E-01 | 0.0 | 9.81E-01 | Protein TIC 20-v, chloroplastic |
| AT3G17030 | 0.9 | 2.28E-05 | 0.4 | 8.06E-03 | 0.5 | 5.91E-03 | 0.3 | 2.80E-01 | 0.2 | 6.83E-01 | 0.2 | 6.85E-01 | hypothetical protein |
| AT5G17230 | 0.9 | 1.20E-05 | 0.8 | 3.78E-05 | 0.0 | 7.06E-01 | 0.1 | 2.92E-01 | 0.3 | 3.49E-02 | -0.1 | 3.64E-01 | Phytoene synthase |
| AT1G31580 | 0.9 | 2.14E-03 | 0.9 | 3.12E-03 | 0.0 | 9.92E-01 | 0.4 | 9.63E-03 | 0.1 | 6.04E-01 | 0.3 | 1.30E-01 | Protein ECS1 |
| AT1G29418 | 0.9 | 7.63E-05 | 0.6 | 1.26E-03 | 0.2 | 1.42E-01 | 0.4 | 2.25E-01 | -0.2 | 7.09E-01 | 0.5 | 1.49E-01 | Transmembrane protein |
| AT5G65880 | 0.9 | 1.08E-04 | 0.7 | 8.40E-04 | 0.2 | 3.00E-01 | 0.8 | 3.60E-03 | 0.4 | 2.57E-01 | 0.5 | 1.35E-01 | At5g65880 |
| AT3G57060 | 0.9 | 5.74E-05 | 0.6 | 1.90E-03 | 0.3 | 5.97E-02 | -0.2 | 5.31E-01 | -0.1 | 7.21E-01 | -0.1 | 9.20E-01 | Condensin-1 complex subunit CAP-D2 |
| AT1G70350 | 0.9 | 5.27E-05 | 0.5 | 3.13E-03 | 0.3 | 3.45E-02 | 0.6 | 5.61E-02 | 0.1 | 7.46E-01 | 0.4 | 2.21E-01 | F17O7.11 |
| AT1G75770 | 0.9 | 2.07E-03 | 0.5 | 6.91E-02 | 0.4 | 1.40E-01 | -0.8 | 1.87E-02 | -1.1 | 5.71E-03 | 0.3 | 5.35E-01 | Uncharacterized protein |
| AT1G10780 | 0.9 | 1.57E-05 | 0.7 | 1.12E-04 | 0.1 | 2.76E-01 | 0.0 | 9.37E-01 | 0.1 | 6.95E-01 | -0.1 | 7.85E-01 | F-box |
| AT1G49510 | 0.9 | 5.08E-06 | 0.9 | 8.78E-06 | 0.0 | 7.94E-01 | 0.3 | 4.94E-02 | 0.2 | 2.66E-01 | 0.1 | 6.56E-01 | At1g49510 |
| AT2G01010 | 0.9 | 6.07E-02 | 0.1 | 9.18E-01 | 0.8 | 1.55E-01 | 0.0 | 9.76E-01 | 0.0 | 9.80E-01 | 0.0 | 9.98E-01 | NA |
| AT3G62040 | 0.9 | 3.34E-06 | 1.2 | 8.42E-07 | -0.4 | 4.74E-03 | 0.8 | 8.68E-04 | 0.6 | 1.70E-02 | 0.2 | 3.73E-01 | At3g62040 |
| AT2G40760 | 0.9 | 2.60E-06 | 0.7 | 2.06E-05 | 0.1 | 1.59E-01 | 0.0 | 8.56E-01 | -0.2 | 4.04E-01 | 0.1 | 5.59E-01 | Rhodanese-like domain-containing protein 7 |
| AT2G39650 | 0.9 | 6.43E-04 | 0.1 | 5.69E-01 | 0.7 | 6.87E-03 | 1.5 | 5.39E-06 | 0.4 | 1.38E-01 | 1.1 | 5.52E-04 | At2g39650 ) |
| AT4G29950 | 0.9 | 1.19E-05 | 0.4 | 3.79E-03 | 0.5 | 2.94E-03 | 0.0 | 9.62E-01 | -0.5 | 1.28E-02 | 0.5 | 4.26E-02 | Microtubule-associated protein |
| AT5G14140 | 0.9 | 1.45E-05 | 0.6 | 2.35E-04 | 0.2 | 9.27E-02 | 0.3 | 3.34E-01 | 0.2 | 5.56E-01 | 0.1 | 8.67E-01 | C2H2-type domain-containing protein |
| AT3G23710 | 0.9 | 4.95E-06 | 0.6 | 1.72E-04 | 0.3 | 2.32E-02 | 0.2 | 4.34E-01 | 0.0 | 9.88E-01 | 0.2 | 5.47E-01 | Tic22-like family protein |
| AT1G52342 | 0.9 | 3.31E-03 | 0.6 | 3.42E-02 | 0.3 | 3.10E-01 | 0.6 | 1.57E-02 | 0.1 | 8.40E-01 | 0.5 | 1.02E-01 | Uncharacterized protein |
| AT3G53470 | 0.9 | 1.05E-05 | 0.4 | 2.70E-03 | 0.5 | 3.29E-03 | -0.3 | 1.66E-01 | -0.2 | 4.23E-01 | -0.1 | 7.75E-01 | 2,3-bisphosphoglycerate-independent phosphoglycerate mutase |
| AT3G19810 | 0.9 | 6.68E-06 | 0.6 | 1.69E-04 | 0.3 | 3.66E-02 | 0.2 | 1.73E-01 | 0.0 | 8.73E-01 | 0.3 | 2.42E-01 | Large ribosomal RNA subunit accumulation protein YCED homolog 1, chloroplastic |
| AT5G52010 | 0.9 | 7.50E-06 | 0.9 | 1.17E-05 | 0.0 | 7.40E-01 | 0.2 | 2.65E-01 | 0.1 | 6.24E-01 | 0.1 | 7.60E-01 | C2H2-type domain-containing protein |
| AT4G32340 | 0.9 | 3.70E-03 | 1.1 | 1.04E-03 | -0.2 | 3.73E-01 | -0.2 | 4.66E-01 | -0.4 | 8.67E-02 | 0.3 | 4.14E-01 | Tetratricopeptide repeat -like superfamily protein |
| AT3G16140 | 0.9 | 3.29E-06 | 0.8 | 9.54E-06 | 0.0 | 8.31E-01 | -0.1 | 5.07E-01 | 0.0 | 8.66E-01 | -0.1 | 7.90E-01 | Photosystem I reaction center subunit VI-1, chloroplastic |
| AT1G03850 | 0.9 | 5.85E-04 | 0.8 | 1.42E-03 | 0.1 | 7.65E-01 | -0.2 | 6.40E-01 | -0.3 | 4.63E-01 | 0.1 | 8.34E-01 | Glutaredoxin family protein |
| AT1G15290 | 0.9 | 3.12E-06 | 0.6 | 1.01E-04 | 0.3 | 1.45E-02 | -0.1 | 5.62E-01 | 0.0 | 9.71E-01 | -0.1 | 7.48E-01 | Tetratricopeptide repeat -like superfamily protein |
| AT1G10660 | 0.9 | 8.57E-07 | 0.3 | 2.81E-03 | 0.6 | 1.35E-04 | -0.2 | 3.65E-01 | 0.0 | 8.49E-01 | -0.1 | 6.07E-01 | hypothetical protein |
| AT4G12800 | 0.9 | 7.15E-06 | 0.9 | 1.30E-05 | 0.0 | 8.21E-01 | 0.0 | 8.41E-01 | 0.0 | 9.92E-01 | 0.0 | 9.09E-01 | PSI subunit V |
| AT2G31725 | 0.9 | 6.03E-04 | 0.4 | 8.53E-02 | 0.5 | 3.30E-02 | 0.5 | 6.60E-02 | 0.4 | 2.39E-01 | 0.1 | 7.52E-01 | hypothetical protein |
| AT5G38980 | 0.9 | 1.79E-05 | 0.7 | 1.74E-04 | 0.2 | 1.70E-01 | 0.3 | 6.59E-02 | 0.3 | 8.35E-02 | 0.0 | 9.51E-01 | Transmembrane protein |
| AT1G21060 | 0.9 | 1.17E-05 | 0.3 | 1.10E-02 | 0.5 | 1.93E-03 | -0.3 | 1.57E-01 | -0.2 | 5.19E-01 | -0.1 | 6.44E-01 | T22I11.12 protein |
| AT3G25690 | 0.9 | 1.51E-04 | 0.4 | 2.28E-02 | 0.5 | 1.59E-02 | 0.1 | 6.50E-01 | 0.1 | 6.93E-01 | 0.0 | 9.87E-01 | Hydroxyproline-rich glycoprotein family protein |
| AT5G13770 | 0.9 | 1.00E-04 | 0.5 | 3.99E-03 | 0.3 | 4.76E-02 | 0.0 | 9.24E-01 | 0.2 | 3.38E-01 | -0.2 | 3.57E-01 | Uncharacterized protein At5g13770 |
| AT2G39681 | 0.9 | 9.63E-03 | 0.1 | 7.14E-01 | 0.8 | 4.96E-02 | 0.4 | 2.22E-01 | 0.1 | 8.82E-01 | 0.3 | 4.24E-01 | NA |
| AT1G23840 | 0.9 | 2.30E-03 | 0.3 | 2.24E-01 | 0.6 | 5.12E-02 | 0.0 | 9.99E-01 | -0.7 | 6.72E-02 | 0.7 | 1.05E-01 | Transmembrane protein |
| AT2G39570 | 0.9 | 6.23E-06 | 1.0 | 6.82E-06 | -0.1 | 3.71E-01 | 0.2 | 2.08E-01 | -0.4 | 4.85E-02 | 0.6 | 1.08E-02 | ACT domain-containing protein ACR9 |
| AT5G09270 | 0.9 | 6.43E-05 | 0.4 | 2.26E-02 | 0.5 | 9.93E-03 | 0.5 | 8.54E-02 | 0.1 | 8.47E-01 | 0.4 | 2.44E-01 | hypothetical protein |
| AT4G10630 | 0.9 | 6.85E-05 | 0.7 | 5.94E-04 | 0.2 | 2.67E-01 | 0.0 | 8.53E-01 | 0.2 | 4.47E-01 | -0.1 | 6.61E-01 | F3H7.9 protein |
| AT4G25370 | 0.9 | 4.94E-07 | 0.5 | 2.87E-05 | 0.3 | 1.66E-03 | 0.1 | 3.81E-01 | 0.0 | 9.46E-01 | 0.1 | 4.75E-01 | ATP-dependent Clp protease ATP-binding subunit CLPT1, chloroplastic |
| AT4G13670 | 0.9 | 1.92E-05 | 0.6 | 3.23E-04 | 0.2 | 7.83E-02 | 0.0 | 7.58E-01 | -0.1 | 4.28E-01 | 0.1 | 7.08E-01 | Plastid transcriptionally active 5 |
| AT3G44740 | 0.9 | 1.08E-03 | 0.2 | 3.23E-01 | 0.7 | 1.99E-02 | 0.2 | 4.89E-01 | 0.2 | 6.49E-01 | 0.0 | 9.40E-01 | Class II aaRS and biotin synthetases superfamily protein |
| AT1G35612 | 0.9 | 1.53E-03 | 0.9 | 1.64E-03 | 0.0 | 8.82E-01 | -0.1 | 7.43E-01 | -0.6 | 4.13E-02 | 0.5 | 1.27E-01 | NA |
| AT1G33610 | 0.9 | 6.63E-04 | 0.8 | 2.33E-03 | 0.1 | 6.06E-01 | 0.0 | 8.42E-01 | -0.5 | 4.87E-03 | 0.5 | 1.11E-02 | Leucine-rich repeat-containing N-terminal plant-type domain-containing protein |
| AT2G18290 | 0.9 | 3.89E-05 | 0.9 | 5.18E-05 | 0.0 | 7.78E-01 | 0.0 | 8.97E-01 | -0.4 | 2.08E-01 | 0.3 | 3.79E-01 | Anaphase-promoting complex subunit 10 |
| AT2G05540 | 0.9 | 2.89E-03 | 0.8 | 1.04E-02 | 0.1 | 6.24E-01 | -1.0 | 4.77E-03 | -1.9 | 7.72E-05 | 0.9 | 2.97E-03 | Glycine-rich protein |
| AT2G20875 | 0.9 | 1.66E-04 | 0.6 | 4.44E-03 | 0.3 | 1.20E-01 | -0.4 | 4.14E-01 | -0.1 | 8.15E-01 | -0.2 | 7.20E-01 | Protein EPIDERMAL PATTERNING FACTOR 1 |
| AT3G07860 | 0.9 | 1.02E-04 | 0.3 | 5.92E-02 | 0.6 | 8.05E-03 | 0.6 | 1.25E-02 | 0.1 | 7.81E-01 | 0.5 | 8.67E-02 | Ubiquitin-like superfamily protein |
| AT2G44580 | 0.9 | 1.60E-04 | 0.7 | 1.59E-03 | 0.2 | 2.77E-01 | 0.1 | 6.58E-01 | -0.2 | 6.83E-01 | 0.3 | 4.11E-01 | Zinc ion binding protein |
| AT1G69210 | 0.9 | 4.40E-04 | 0.5 | 1.54E-02 | 0.4 | 1.04E-01 | 0.5 | 1.34E-01 | -0.1 | 9.19E-01 | 0.5 | 1.67E-01 | Uncharacterized protein family UPF0090 |
| AT1G06190 | 0.9 | 1.40E-06 | 0.7 | 1.94E-05 | 0.2 | 3.57E-02 | 0.1 | 4.26E-01 | -0.2 | 3.00E-01 | 0.3 | 1.25E-01 | Rho termination factor |
| AT1G13470 | 0.9 | 1.78E-02 | -0.4 | 2.73E-01 | 1.3 | 9.25E-03 | -0.2 | 8.49E-01 | -1.2 | 2.38E-01 | 1.0 | 4.01E-01 | Uncharacterized protein At1g13470 |
| AT4G29905 | 0.9 | 2.93E-04 | 0.7 | 3.32E-03 | 0.2 | 2.34E-01 | -1.5 | 8.92E-05 | -1.6 | 2.01E-04 | 0.1 | 9.15E-01 | hypothetical protein |
| AT1G16330 | 0.9 | 1.74E-05 | 0.5 | 2.29E-03 | 0.4 | 1.22E-02 | 0.2 | 5.96E-01 | 0.2 | 7.05E-01 | 0.0 | 9.66E-01 | CYCB3 |
| AT3G51140 | 0.9 | 5.88E-06 | 0.8 | 2.76E-05 | 0.1 | 3.71E-01 | 0.3 | 4.15E-02 | 0.2 | 3.26E-01 | 0.1 | 5.50E-01 | DnaJ |
| AT3G18980 | 0.9 | 1.18E-06 | 0.7 | 1.50E-05 | 0.2 | 4.11E-02 | 0.0 | 8.70E-01 | -0.3 | 8.07E-02 | 0.3 | 2.22E-01 | F-box protein ETP1 |
| AT2G14460 | 0.9 | 4.32E-04 | 0.4 | 4.45E-02 | 0.5 | 3.95E-02 | 0.7 | 1.49E-02 | 0.5 | 1.87E-01 | 0.3 | 4.55E-01 | Uncharacterized protein |
| AT3G26050 | 0.9 | 1.50E-04 | 0.8 | 7.91E-04 | 0.1 | 4.77E-01 | 0.8 | 6.01E-02 | 0.5 | 3.16E-01 | 0.3 | 6.05E-01 | TPX2 protein family |
| AT5G13140 | 0.9 | 7.33E-07 | 0.9 | 2.51E-06 | 0.0 | 5.98E-01 | -0.3 | 8.48E-02 | -0.2 | 1.92E-01 | 0.0 | 8.98E-01 | Pollen Ole e 1 allergen and extensin family protein |
| AT1G34000 | 0.9 | 5.98E-06 | 0.7 | 7.65E-05 | 0.2 | 8.00E-02 | 0.0 | 8.88E-01 | 0.1 | 7.78E-01 | 0.0 | 9.06E-01 | Light-harvesting complex-like protein OHP2, chloroplastic |
| AT4G28030 | 0.9 | 7.33E-07 | 0.7 | 1.03E-05 | 0.2 | 2.64E-02 | 0.0 | 7.97E-01 | 0.1 | 7.72E-01 | 0.0 | 9.53E-01 | AT4G28030 protein |
| AT5G10010 | 0.9 | 5.40E-06 | 0.6 | 1.28E-04 | 0.3 | 2.99E-02 | 0.4 | 2.67E-03 | 0.1 | 4.63E-01 | 0.3 | 6.42E-02 | HIT4 |
| AT2G42260 | 0.9 | 5.93E-05 | 0.7 | 1.18E-03 | 0.3 | 1.21E-01 | 0.3 | 2.82E-01 | 0.3 | 3.39E-01 | 0.0 | 9.98E-01 | Protein POLYCHOME |
| AT3G15354 | 0.9 | 2.44E-04 | 0.1 | 6.87E-01 | 0.8 | 1.72E-03 | 0.6 | 5.88E-02 | 0.6 | 5.61E-02 | -0.1 | 9.07E-01 | SPA1-related 3 |
| AT1G73120 | 0.9 | 8.20E-04 | 0.0 | 9.34E-01 | 0.9 | 4.09E-03 | -0.3 | 3.82E-01 | -3.7 | 2.14E-04 | 3.4 | 1.36E-03 | Uncharacterized protein |
| AT1G73177 | 0.9 | 4.88E-03 | 0.5 | 8.52E-02 | 0.4 | 2.24E-01 | 0.4 | 3.79E-01 | 0.5 | 2.90E-01 | -0.1 | 8.47E-01 | Bonsai |
| AT5G16620 | 0.9 | 1.58E-06 | 0.9 | 3.96E-06 | 0.0 | 9.32E-01 | 0.4 | 7.05E-04 | 0.1 | 5.77E-01 | 0.3 | 2.69E-02 | hypothetical protein |
| AT3G25805 | 0.9 | 1.54E-04 | 0.6 | 5.12E-03 | 0.3 | 8.08E-02 | 0.1 | 6.87E-01 | 0.0 | 9.68E-01 | 0.1 | 7.51E-01 | Transmembrane protein |
| AT1G15885 | 0.9 | 4.91E-04 | 0.6 | 1.61E-02 | 0.4 | 1.18E-01 | 0.0 | 9.11E-01 | -0.2 | 6.01E-01 | 0.3 | 5.29E-01 | Uncharacterized protein |
| AT3G28910 | 0.9 | 5.41E-06 | 0.4 | 4.83E-03 | 0.5 | 1.13E-03 | -0.2 | 2.58E-01 | -0.3 | 8.05E-02 | 0.1 | 5.81E-01 | Myb domain protein 30 |
| AT1G10740 | 0.9 | 1.31E-06 | 0.7 | 2.29E-05 | 0.2 | 2.46E-02 | -0.1 | 7.90E-01 | -0.2 | 3.71E-01 | 0.1 | 5.84E-01 | Alpha |
| AT3G46560 | 0.9 | 2.53E-04 | 0.5 | 1.62E-02 | 0.4 | 4.51E-02 | 0.6 | 3.18E-02 | 0.3 | 4.00E-01 | 0.3 | 4.23E-01 | Mitochondrial import inner membrane translocase subunit |
| AT3G52170 | 0.9 | 5.42E-06 | 0.6 | 3.34E-04 | 0.3 | 1.16E-02 | 0.6 | 2.94E-03 | 0.2 | 4.19E-01 | 0.4 | 8.55E-02 | Uncharacterized protein F4F15.280 |

| | | | | | | | | | | | | | |
|---|---|---|---|---|---|---|---|---|---|---|---|---|---|
| AT2G45190 | 0.9 | 2.71E-05 | 0.9 | 6.93E-05 | 0.0 | 8.19E-01 | 0.0 | 9.52E-01 | -0.2 | 4.83E-01 | 0.2 | 5.09E-01 | Uncharacterized protein At2g45190 |
| AT1G80190 | 0.9 | 5.83E-06 | 0.7 | 1.55E-04 | 0.3 | 4.05E-02 | 0.0 | 9.27E-01 | -0.5 | 2.69E-02 | 0.5 | 6.60E-02 | hypothetical protein |
| AT5G42670 | 0.9 | 4.41E-06 | 0.4 | 3.93E-03 | 0.5 | 1.03E-03 | 0.3 | 2.27E-01 | 0.1 | 8.13E-01 | 0.2 | 5.13E-01 | Agenet domain-containing protein |
| AT1G79040 | 0.9 | 5.10E-06 | 0.8 | 3.13E-05 | 0.1 | 2.70E-01 | -0.2 | 8.56E-02 | -0.2 | 1.93E-01 | 0.0 | 9.10E-01 | Photosystem II 10 kDa polypeptide, chloroplastic |
| AT4G34020 | 0.9 | 1.52E-06 | 0.6 | 6.74E-05 | 0.3 | 7.96E-03 | -0.1 | 5.55E-01 | -0.3 | 1.43E-01 | 0.2 | 4.53E-01 | DJ-1 |
| AT4G24175 | 0.9 | 5.44E-04 | 0.9 | 1.44E-03 | 0.1 | 7.73E-01 | 0.3 | 1.65E-01 | 0.2 | 4.09E-01 | 0.1 | 8.00E-01 | Kinesin-like protein |
| AT3G60620 | 0.9 | 1.09E-06 | 0.6 | 3.83E-05 | 0.3 | 7.49E-03 | 0.3 | 3.57E-02 | 0.3 | 1.03E-01 | 0.0 | 8.64E-01 | Cytidinediphosphate diacylglycerol synthase 5 |
| AT1G68185 | 0.9 | 1.06E-06 | 0.5 | 1.66E-04 | 0.4 | 1.47E-03 | 0.3 | 1.46E-01 | 0.0 | 9.44E-01 | 0.3 | 2.21E-01 | Ubiquitin-like domain-containing protein |
| AT4G28210 | 0.9 | 1.43E-05 | 0.9 | 2.66E-05 | 0.0 | 9.67E-01 | 0.4 | 1.21E-02 | 0.2 | 1.79E-01 | 0.2 | 4.29E-01 | hypothetical protein |
| AT5G63050 | 0.9 | 1.22E-06 | 0.7 | 2.58E-05 | 0.3 | 1.70E-02 | 0.1 | 3.29E-01 | 0.0 | 9.32E-01 | 0.1 | 5.19E-01 | EMB2759 |
| AT3G20820 | 0.9 | 1.93E-05 | 0.8 | 8.59E-05 | 0.1 | 4.76E-01 | -0.3 | 5.49E-02 | -0.1 | 5.31E-01 | -0.2 | 4.42E-01 | Leucine-rich repeat-containing N-terminal plant-type domain-containing protein |
| AT5G37290 | 0.9 | 2.12E-04 | 0.6 | 4.81E-03 | 0.3 | 1.33E-01 | 0.4 | 7.27E-02 | 0.1 | 6.78E-01 | 0.3 | 3.68E-01 | ARM repeat superfamily protein |
| AT2G35960 | 0.9 | 1.63E-05 | 0.6 | 4.75E-04 | 0.3 | 3.58E-02 | 0.2 | 2.16E-01 | 0.3 | 3.96E-02 | -0.1 | 4.93E-01 | NDR1 |
| AT1G08390 | 0.9 | 9.89E-04 | 1.0 | 7.38E-04 | -0.1 | 6.49E-01 | | | | | | | RecQ-mediated instability-like protein |
| AT5G35525 | 0.9 | 4.66E-04 | 1.3 | 7.95E-05 | -0.3 | 1.10E-01 | -0.3 | 2.58E-01 | 0.1 | 7.33E-01 | -0.4 | 1.67E-01 | Protein PLANT CADMIUM RESISTANCE 3 |
| AT5G51550 | 0.9 | 2.51E-04 | 0.9 | 6.72E-04 | 0.1 | 7.57E-01 | -0.1 | 3.40E-01 | -0.6 | 8.29E-04 | 0.5 | 1.87E-02 | Protein EXORDIUM-like 3 |
| AT2G33400 | 0.9 | 4.55E-06 | 0.8 | 2.10E-05 | 0.1 | 4.23E-01 | -0.2 | 3.32E-01 | 0.1 | 5.73E-01 | -0.3 | 2.08E-01 | Uncharacterized protein At2g33400 |
| AT1G19240 | 0.9 | 2.34E-04 | 0.3 | 1.22E-01 | 0.6 | 9.67E-03 | 0.5 | 9.85E-02 | 0.2 | 7.18E-01 | 0.4 | 3.79E-01 | T29M8.11 |
| AT2G40920 | 0.9 | 7.66E-06 | 0.8 | 4.69E-05 | 0.1 | 2.96E-01 | 0.0 | 9.76E-01 | -0.3 | 7.65E-02 | 0.4 | 1.24E-01 | F-box and associated interaction domains-containing protein |
| AT2G42190 | 0.9 | 2.18E-05 | 0.7 | 2.57E-04 | 0.2 | 1.27E-01 | -0.2 | 4.05E-01 | -0.3 | 1.61E-01 | 0.2 | 6.67E-01 | Expressed protein |
| AT2G21195 | 0.9 | 6.71E-03 | 0.6 | 5.89E-02 | 0.3 | 3.63E-01 | -0.1 | 7.40E-01 | -0.2 | 7.20E-01 | 0.0 | 9.63E-01 | Uncharacterized protein |
| AT2G36620 | 0.9 | 2.84E-05 | 0.9 | 9.91E-05 | 0.1 | 6.39E-01 | 0.5 | 6.96E-03 | 0.3 | 1.87E-01 | 0.2 | 3.79E-01 | AT2G36620 protein |
| AT5G26800 | 0.9 | 1.27E-05 | 0.5 | 1.26E-03 | 0.4 | 1.08E-02 | 0.6 | 6.04E-03 | 0.2 | 4.10E-01 | 0.4 | 1.58E-01 | NA |
| AT4G13200 | 0.9 | 3.75E-05 | 0.6 | 1.50E-03 | 0.3 | 4.11E-02 | 0.2 | 5.19E-01 | -0.1 | 8.52E-01 | 0.2 | 5.05E-01 | Uncharacterized protein At4g13200, chloroplastic |
| AT1G03660 | 0.9 | 1.89E-04 | -0.1 | 7.07E-01 | 1.0 | 1.06E-03 | 0.7 | 1.66E-01 | -0.6 | 3.60E-01 | 1.3 | 2.54E-02 | Ankyrin-repeat containing protein |
| AT3G08640 | 0.9 | 6.48E-07 | 0.7 | 8.95E-06 | 0.2 | 2.48E-02 | 0.5 | 3.80E-04 | 0.2 | 2.36E-01 | 0.4 | 4.64E-02 | Protein RETICULATA-RELATED 3, chloroplastic |
| AT2G34460 | 0.9 | 6.44E-06 | 0.6 | 2.03E-04 | 0.3 | 2.18E-02 | 0.0 | 8.39E-01 | 0.0 | 9.53E-01 | 0.0 | 8.58E-01 | Uncharacterized protein At2g34460, chloroplastic |
| AT4G35450 | 0.9 | 1.10E-06 | 0.6 | 9.04E-05 | 0.4 | 2.13E-03 | -0.1 | 3.68E-01 | -0.2 | 2.03E-01 | 0.1 | 7.63E-01 | Ankyrin repeat-containing protein 2 |
| AT3G15900 | 0.9 | 1.14E-06 | 0.7 | 2.38E-05 | 0.2 | 2.05E-02 | -0.2 | 2.95E-01 | 0.0 | 9.87E-01 | -0.2 | 4.70E-01 | hypothetical protein |
| AT2G02950 | 0.9 | 3.21E-05 | 0.8 | 2.07E-04 | 0.1 | 3.21E-01 | -0.1 | 5.06E-01 | -0.4 | 2.34E-02 | 0.3 | 1.92E-01 | Protein PHYTOCHROME KINASE SUBSTRATE 1 |
| AT1G19380 | 0.9 | 1.02E-04 | 0.5 | 9.23E-03 | 0.4 | 2.78E-02 | 0.0 | 9.98E-01 | -0.3 | 3.20E-01 | 0.3 | 3.85E-01 | Sugar transporter |
| AT3G12685 | 0.9 | 2.91E-04 | 0.9 | 6.67E-04 | 0.0 | 8.25E-01 | -0.1 | 8.32E-01 | 0.1 | 6.69E-01 | -0.2 | 4.93E-01 | Acid phosphatase |
| AT2G02410 | 0.9 | 4.97E-05 | 0.6 | 1.61E-03 | 0.3 | 6.05E-02 | -0.1 | 5.28E-01 | -0.2 | 5.03E-01 | 0.0 | 9.40E-01 | Uncharacterized protein At2g02410 |
| AT4G21610 | 0.9 | 2.19E-05 | 0.6 | 1.07E-03 | 0.3 | 3.38E-02 | 0.1 | 7.37E-01 | -0.1 | 7.85E-01 | 0.2 | 4.89E-01 | Lsd one like 2 |
| AT1G69160 | 0.9 | 9.01E-05 | 0.8 | 3.23E-04 | 0.1 | 5.75E-01 | 0.0 | 8.93E-01 | -0.5 | 4.53E-02 | 0.4 | 6.91E-02 | Protein BIG GRAIN 1-like E |
| AT5G48590 | 0.9 | 1.30E-06 | 0.7 | 2.13E-05 | 0.2 | 2.69E-02 | 0.0 | 7.92E-01 | -0.3 | 3.85E-02 | 0.2 | 1.44E-01 | Uncharacterized protein |
| AT5G09820 | 0.9 | 3.10E-04 | 0.5 | 2.66E-02 | 0.5 | 4.31E-02 | -0.1 | 6.55E-01 | -0.1 | 8.27E-01 | 0.0 | 9.30E-01 | Plastid-lipid associated protein PAP |
| AT1G49170 | 0.9 | 2.35E-04 | 0.7 | 3.51E-03 | 0.3 | 1.98E-01 | -0.1 | 7.08E-01 | -0.3 | 1.98E-01 | 0.2 | 4.31E-01 | F27J15.6 |
| AT2G36050 | 0.9 | 5.36E-04 | 1.0 | 3.99E-04 | -0.1 | 6.58E-01 | -0.6 | 3.86E-03 | -0.8 | 1.38E-03 | 0.2 | 5.55E-01 | Transcription repressor OFP15 |
| AT2G42130 | 0.9 | 1.99E-05 | 0.9 | 4.58E-05 | 0.0 | 8.69E-01 | 0.4 | 4.07E-03 | 0.2 | 1.35E-01 | 0.2 | 3.23E-01 | Plastid-lipid associated protein PAP |
| AT3G54880 | 1.0 | 2.10E-03 | 0.6 | 4.48E-02 | 0.4 | 1.68E-01 | 0.0 | 9.53E-01 | -0.4 | 1.43E-01 | 0.4 | 2.23E-01 | hypothetical protein |
| AT2G39920 | 1.0 | 1.72E-03 | 1.9 | 1.26E-05 | -1.0 | 1.88E-03 | -0.5 | 1.22E-01 | 0.1 | 8.37E-01 | -0.5 | 1.10E-01 | hypothetical protein |
| AT1G80180 | 1.0 | 1.48E-04 | 0.6 | 3.77E-03 | 0.3 | 8.30E-02 | -0.2 | 2.34E-01 | -0.7 | 4.76E-04 | 0.5 | 2.20E-02 | MAPK kinase substrate protein At1g80180 |
| AT3G12145 | 1.0 | 3.63E-05 | 0.7 | 7.44E-04 | 0.3 | 9.33E-02 | 0.3 | 5.12E-01 | 0.6 | 1.34E-01 | -0.3 | 5.15E-01 | Leucine-rich repeat protein FLOR 1 |
| AT4G25990 | 1.0 | 2.32E-03 | 1.6 | 7.89E-05 | -0.6 | 2.60E-02 | 0.1 | 6.91E-01 | -0.5 | 1.69E-01 | 0.7 | 9.84E-02 | CCT motif family protein |
| AT4G12690 | 1.0 | 1.09E-05 | 1.0 | 1.85E-05 | 0.0 | 8.65E-01 | -0.4 | 4.77E-02 | -0.9 | 1.24E-03 | 0.5 | 8.69E-02 | hypothetical protein |
| AT3G43430 | 1.0 | 1.85E-04 | 1.4 | 1.44E-05 | -0.5 | 1.84E-02 | 0.3 | 3.13E-01 | 0.0 | 9.65E-01 | 0.4 | 3.48E-01 | RING-type domain-containing protein |
| AT4G04410 | 1.0 | 4.09E-04 | 1.1 | 3.28E-04 | -0.1 | 6.68E-01 | -0.6 | 1.47E-01 | -0.5 | 3.18E-01 | -0.1 | 8.54E-01 | NA |
| AT4G32460 | 1.0 | 2.17E-04 | 0.6 | 4.41E-03 | 0.3 | 1.17E-01 | -0.2 | 3.82E-01 | -0.1 | 7.41E-01 | -0.1 | 7.95E-01 | hypothetical protein |
| AT1G52315 | 1.0 | 3.50E-03 | 0.1 | 8.06E-01 | 0.9 | 2.14E-02 | -0.4 | 4.27E-01 | 0.0 | 9.86E-01 | | | Regulator of Vps4 activity in the MVB pathway protein |
| AT3G50550 | 1.0 | 2.54E-06 | 0.5 | 8.50E-04 | 0.5 | 1.26E-03 | 0.2 | 3.26E-01 | 0.2 | 6.25E-01 | 0.1 | 8.39E-01 | Uncharacterized protein At3g50550 |
| AT1G65370 | 1.0 | 2.60E-05 | 0.4 | 8.99E-03 | 0.5 | 5.32E-03 | 0.3 | 2.37E-01 | 0.0 | 9.16E-01 | 0.2 | 3.97E-01 | At1g65370 |
| AT2G21280 | 1.0 | 9.38E-06 | 0.9 | 2.45E-05 | 0.0 | 7.85E-01 | 0.3 | 6.65E-02 | 0.2 | 1.45E-01 | 0.0 | 9.18E-01 | NAD-binding Rossmann-fold superfamily protein |
| AT1G36622 | 1.0 | 7.80E-02 | 0.1 | 8.94E-01 | 0.9 | 1.74E-01 | 1.8 | 1.66E-03 | 0.3 | 7.06E-01 | 1.5 | 1.73E-02 | Transmembrane protein |
| AT5G66040 | 1.0 | 6.74E-06 | 0.7 | 1.31E-04 | 0.3 | 4.69E-02 | -0.4 | 2.03E-02 | -0.1 | 5.69E-01 | -0.2 | 2.48E-01 | Sulfurtransferase protein 16 |
| AT5G59613 | 1.0 | 1.24E-03 | 0.6 | 3.20E-02 | 0.4 | 1.19E-01 | 0.2 | 3.43E-01 | 0.2 | 4.28E-01 | 0.0 | 9.98E-01 | ATP synthase small subunit 6-B, mitochondrial |
| AT5G43500 | 1.0 | 1.94E-06 | 0.5 | 6.20E-04 | 0.5 | 1.18E-03 | 0.1 | 5.72E-01 | -0.1 | 7.99E-01 | 0.2 | 4.42E-01 | Actin-related protein 9 |
| AT1G09815 | 1.0 | 1.96E-06 | 0.4 | 1.34E-03 | 0.6 | 4.92E-04 | 0.3 | 4.08E-02 | 0.2 | 3.53E-01 | 0.1 | 4.79E-01 | hypothetical protein |
| AT2G27290 | 1.0 | 8.76E-05 | 0.6 | 2.52E-03 | 0.3 | 6.40E-02 | -0.1 | 5.47E-01 | -0.1 | 8.12E-01 | 0.0 | 8.52E-01 | DUF1279 domain-containing protein |
| AT1G73090 | 1.0 | 3.88E-07 | 0.6 | 2.08E-05 | 0.3 | 2.09E-03 | -0.1 | 8.18E-01 | -0.3 | 1.45E-01 | 0.3 | 2.26E-01 | WD repeat protein |
| AT1G22700 | 1.0 | 1.73E-07 | 0.8 | 2.07E-06 | 0.2 | 1.26E-02 | 0.1 | 5.27E-01 | -0.1 | 7.17E-01 | 0.2 | 4.20E-01 | Tetratricopeptide repeat -like superfamily protein |
| AT5G39590 | 1.0 | 2.53E-07 | 0.7 | 4.10E-06 | 0.2 | 7.57E-03 | -0.2 | 5.84E-02 | -0.4 | 2.39E-03 | 0.2 | 2.21E-01 | AT5G39590 protein |
| AT5G14410 | 1.0 | 7.07E-03 | 0.5 | 1.68E-01 | 0.5 | 1.71E-01 | 0.0 | 9.95E-01 | -0.3 | 5.58E-01 | 0.3 | 6.36E-01 | Uncharacterized protein |
| AT3G23325 | 1.0 | 4.13E-06 | 0.5 | 1.62E-03 | 0.5 | 1.78E-03 | 0.4 | 9.18E-02 | 0.2 | 5.75E-01 | 0.2 | 4.90E-01 | Uncharacterized protein At3g23325 |
| AT1G01320 | 1.0 | 4.27E-06 | 0.5 | 8.35E-04 | 0.5 | 2.23E-03 | 0.4 | 1.01E-01 | 0.1 | 7.86E-01 | 0.3 | 3.01E-01 | Tetratricopeptide repeat -like superfamily protein |
| AT4G32190 | 1.0 | 7.99E-07 | 0.7 | 2.59E-05 | 0.3 | 5.74E-03 | 0.0 | 9.00E-01 | 0.0 | 9.15E-01 | 0.0 | 9.97E-01 | Myosin heavy chain-related protein |
| AT2G27130 | 1.0 | 4.36E-06 | 0.8 | 3.91E-05 | 0.2 | 1.54E-01 | 0.0 | 9.87E-01 | 0.1 | 8.66E-01 | -0.1 | 8.79E-01 | Non-specific lipid transfer protein GPI-anchored 12 |
| AT1G06690 | 1.0 | 1.45E-06 | 0.6 | 1.02E-04 | 0.4 | 3.04E-03 | 0.2 | 1.53E-01 | 0.3 | 1.03E-01 | -0.1 | 8.12E-01 | NADP-dependent oxidoreductase domain-containing protein |
| AT1G80200 | 1.0 | 1.65E-04 | 0.6 | 8.85E-03 | 0.4 | 6.69E-02 | 0.9 | 3.40E-02 | 0.6 | 1.88E-01 | 0.3 | 6.11E-01 | Transmembrane protein |
| AT3G06600 | 1.0 | 3.77E-04 | 0.7 | 7.14E-03 | 0.3 | 1.80E-01 | 0.0 | 9.42E-01 | 0.9 | 4.75E-02 | -0.9 | 6.52E-02 | Protein SINE3 |
| AT3G15760 | 1.0 | 4.01E-04 | 0.7 | 4.82E-03 | 0.3 | 2.44E-01 | 0.0 | 9.56E-01 | -0.6 | 1.12E-01 | 0.6 | 1.54E-01 | Transmembrane protein |
| AT5G62575 | 1.0 | 3.93E-04 | 0.7 | 8.26E-03 | 0.3 | 1.43E-01 | 0.1 | 6.55E-01 | 0.1 | 8.44E-01 | 0.1 | 8.96E-01 | SDH7B |
| AT3G26080 | 1.0 | 1.26E-06 | 0.5 | 3.23E-04 | 0.5 | 1.06E-03 | 0.1 | 7.30E-01 | 0.0 | 9.60E-01 | 0.1 | 7.82E-01 | Plastid-lipid associated protein PAP |
| AT3G06840 | 1.0 | 7.27E-05 | 1.0 | 1.03E-04 | 0.0 | 8.17E-01 | -0.3 | 3.09E-01 | 0.0 | 9.96E-01 | -0.3 | 3.70E-01 | hypothetical protein |
| AT3G11090 | 1.0 | 2.72E-05 | 0.6 | 1.53E-03 | 0.4 | 2.75E-02 | 0.3 | 2.51E-01 | 0.1 | 5.89E-01 | 0.1 | 7.76E-01 | LOB domain-containing protein 21 |
| AT1G56220 | 1.0 | 1.85E-05 | 0.9 | 5.43E-05 | 0.0 | 7.59E-01 | -0.1 | 5.49E-01 | -0.7 | 1.75E-03 | 0.6 | 2.68E-02 | Dormancy |
| AT4G31040 | 1.0 | 2.06E-07 | 0.7 | 3.29E-06 | 0.2 | 6.58E-03 | -0.1 | 3.99E-01 | -0.1 | 6.76E-01 | -0.1 | 8.39E-01 | Chloroplast envelope membrane protein |
| AT1G10510 | 1.0 | 4.60E-07 | 0.6 | 2.51E-05 | 0.4 | 2.02E-03 | 0.6 | 1.74E-03 | 0.3 | 1.33E-01 | 0.3 | 1.17E-01 | Emb2004 |
| AT2G47485 | 1.0 | 3.64E-06 | 0.8 | 2.64E-05 | 0.1 | 2.03E-01 | -0.3 | 1.76E-01 | -0.3 | 2.12E-01 | 0.0 | 9.92E-01 | Uncharacterized protein |
| AT1G13480 | 1.0 | 1.52E-03 | 0.5 | 1.03E-01 | 0.5 | 6.99E-02 | 2.7 | 5.64E-08 | 0.8 | 1.40E-02 | 1.8 | 2.88E-05 | Uncharacterized protein |
| AT4G27450 | 1.0 | 1.81E-03 | 1.1 | 1.64E-03 | -0.1 | 8.20E-01 | -1.1 | 1.42E-02 | -1.8 | 1.68E-03 | 0.7 | 2.59E-01 | hypothetical protein |
| AT3G56290 | 1.0 | 1.09E-05 | 0.6 | 4.06E-04 | 0.3 | 2.29E-02 | 0.5 | 1.27E-02 | 0.3 | 1.45E-01 | 0.2 | 4.93E-01 | GATA-type transcription activator N-terminal domain-containing protein |
| AT5G52890 | 1.0 | 1.43E-05 | 0.6 | 1.56E-03 | 0.4 | 1.29E-02 | -0.1 | 8.76E-01 | 0.1 | 8.13E-01 | -0.1 | 7.35E-01 | AT hook motif-containing protein |

| | | | | | | | | | | | | | |
|---|---|---|---|---|---|---|---|---|---|---|---|---|---|
| AT3G53560 | 1.0 | 1.49E-06 | 0.8 | 1.34E-05 | 0.2 | 9.17E-02 | 1.0 | 3.44E-06 | 0.4 | 1.81E-02 | 0.6 | 2.97E-03 | Tetratricopeptide repeat -like superfamily protein |
| AT1G74850 | 1.0 | 3.08E-07 | 0.7 | 7.45E-06 | 0.3 | 4.66E-03 | 0.3 | 2.05E-01 | 0.1 | 7.50E-01 | 0.2 | 5.10E-01 | Pentatricopeptide repeat-containing protein At1g74850, chloroplastic |
| AT1G74458 | 1.0 | 3.26E-04 | 0.7 | 7.85E-03 | 0.3 | 1.38E-01 | 1.1 | 4.80E-02 | 0.6 | 3.35E-01 | 0.4 | 4.68E-01 | Transmembrane protein |
| AT3G61440 | 1.0 | 4.64E-07 | 0.6 | 2.05E-05 | 0.4 | 2.43E-03 | 0.2 | 8.29E-02 | -0.2 | 1.78E-01 | 0.4 | 2.25E-02 | Cysteine synthase C1 |
| AT1G31760 | 1.0 | 6.79E-04 | 0.7 | 1.43E-02 | 0.3 | 1.76E-01 | 0.2 | 5.19E-01 | 0.1 | 8.17E-01 | 0.1 | 8.31E-01 | DM2 domain-containing protein |
| AT1G18250 | 1.0 | 1.73E-05 | 1.0 | 2.76E-05 | 0.0 | 8.64E-01 | 0.4 | 8.92E-03 | 0.5 | 5.91E-03 | -0.1 | 7.72E-01 | Pathogenesis-related thaumatin superfamily protein |
| AT2G22260 | 1.0 | 2.01E-05 | 0.7 | 7.94E-04 | 0.3 | 4.39E-02 | 0.1 | 7.25E-01 | 0.3 | 5.64E-01 | -0.1 | 8.47E-01 | Oxidoreductase, 2OG-Fe oxygenase family protein |
| AT1G70985 | 1.0 | 2.76E-05 | 0.8 | 2.24E-04 | 0.2 | 2.59E-01 | 0.3 | 6.64E-01 | 0.3 | 7.19E-01 | 0.0 | 9.98E-01 | Hydroxyproline-rich glycoprotein family protein |
| AT4G17540 | 1.0 | 1.15E-04 | 0.8 | 9.85E-04 | 0.2 | 2.89E-01 | 0.6 | 5.26E-02 | 0.3 | 3.62E-01 | 0.2 | 4.96E-01 | Transmembrane protein |
| AT5G38865 | 1.0 | 9.90E-04 | 0.6 | 2.19E-02 | 0.4 | 1.88E-01 | -0.2 | 8.04E-01 | 0.0 | 9.68E-01 | -0.2 | 8.87E-01 | NA |
| AT3G12170 | 1.0 | 1.94E-04 | 0.7 | 3.48E-03 | 0.3 | 1.69E-01 | 0.4 | 4.01E-01 | 0.3 | 5.44E-01 | 0.1 | 9.44E-01 | At3g12170 |
| AT1G01970 | 1.0 | 1.33E-06 | 0.7 | 2.66E-05 | 0.3 | 2.12E-02 | 0.0 | 7.88E-01 | 0.1 | 6.38E-01 | 0.0 | 8.59E-01 | Pentatricopeptide repeat-containing protein At1g01970 |
| AT3G59780 | 1.0 | 8.64E-06 | 0.8 | 1.04E-04 | 0.2 | 1.04E-01 | 0.1 | 7.68E-01 | 0.3 | 4.94E-02 | -0.3 | 1.48E-01 | Rhodanese |
| AT5G17560 | 1.0 | 3.74E-05 | 0.5 | 9.40E-03 | 0.5 | 6.58E-03 | 0.4 | 7.47E-02 | 0.3 | 2.64E-01 | 0.1 | 7.79E-01 | BolA-like family protein |
| AT3G51510 | 1.0 | 2.21E-05 | 0.8 | 2.43E-04 | 0.2 | 1.42E-01 | -0.1 | 5.85E-01 | -0.1 | 6.79E-01 | 0.0 | 9.89E-01 | hypothetical protein |
| AT2G41950 | 1.0 | 1.22E-05 | 0.9 | 5.38E-05 | 0.1 | 4.67E-01 | 0.6 | 6.05E-03 | 0.1 | 5.89E-01 | 0.4 | 7.95E-02 | hypothetical protein |
| AT3G08375 | 1.0 | 2.83E-03 | 0.1 | 7.24E-01 | 0.9 | 1.83E-02 | | | | | | | NA |
| AT1G78450 | 1.0 | 7.45E-06 | 1.4 | 1.92E-06 | -0.3 | 1.54E-02 | 0.3 | 2.14E-01 | -0.2 | 5.42E-01 | 0.5 | 7.98E-02 | Uncharacterized protein At1g78450 |
| AT3G02410 | 1.0 | 2.16E-04 | 0.3 | 2.57E-01 | 0.8 | 6.48E-03 | -0.8 | 5.85E-02 | -0.6 | 1.72E-01 | -0.2 | 7.93E-01 | Alpha |
| AT2G39690 | 1.0 | 4.37E-05 | 0.1 | 4.87E-01 | 0.9 | 8.96E-04 | 0.0 | 9.86E-01 | -0.1 | 8.59E-01 | 0.1 | 8.37E-01 | DUF547 domain-containing protein |
| AT5G05990 | 1.0 | 1.43E-05 | 0.6 | 9.71E-04 | 0.4 | 1.85E-02 | 0.8 | 1.85E-02 | 0.4 | 3.68E-01 | 0.4 | 2.87E-01 | AT5g05990 |
| AT1G18773 | 1.0 | 7.85E-05 | 0.7 | 2.37E-03 | 0.3 | 8.80E-02 | -0.6 | 1.50E-02 | -0.6 | 4.20E-02 | -0.1 | 8.52E-01 | Acyl thioesterase-like protein |
| AT4G15910 | 1.0 | 3.72E-05 | 0.9 | 2.09E-04 | 0.1 | 3.63E-01 | -0.1 | 6.42E-01 | -0.2 | 5.58E-01 | 0.1 | 8.89E-01 | Late embryogenis abundant protein 41 |
| AT5G23120 | 1.0 | 5.24E-06 | 1.1 | 9.36E-06 | 0.0 | 7.44E-01 | 0.0 | 7.52E-01 | 0.0 | 9.90E-01 | 0.0 | 8.48E-01 | Photosystem II stability |
| AT5G25570 | 1.0 | 9.40E-05 | 0.5 | 2.02E-02 | 0.5 | 1.58E-02 | 0.1 | 7.64E-01 | 0.1 | 8.09E-01 | 0.0 | 9.97E-01 | Polyamine-modulated factor 1-binding protein |
| AT2G21385 | 1.0 | 2.51E-06 | 1.0 | 6.17E-06 | 0.0 | 9.16E-01 | 0.3 | 1.04E-02 | 0.0 | 8.12E-01 | 0.3 | 9.15E-02 | Uncharacterized protein |
| AT4G02920 | 1.0 | 8.76E-07 | 0.6 | 4.73E-05 | 0.4 | 3.13E-03 | 0.3 | 2.12E-02 | 0.0 | 9.78E-01 | 0.3 | 8.97E-02 | hypothetical protein |
| AT5G42765 | 1.0 | 2.05E-06 | 0.9 | 9.84E-06 | 0.1 | 3.62E-01 | 0.3 | 1.13E-01 | 0.2 | 3.88E-01 | 0.1 | 7.49E-01 | Uncharacterized protein |
| AT3G17640 | 1.0 | 2.82E-04 | 1.0 | 5.04E-04 | 0.0 | 9.60E-01 | 0.1 | 6.30E-01 | 0.3 | 2.97E-01 | -0.1 | 6.53E-01 | Leucine-rich repeat-containing N-terminal plant-type domain-containing protein |
| AT4G21180 | 1.0 | 9.97E-06 | 0.6 | 1.60E-03 | 0.5 | 6.35E-03 | 0.2 | 7.46E-01 | 0.2 | 8.05E-01 | 0.0 | 9.91E-01 | DnaJ |
| AT3G20490 | 1.0 | 1.47E-06 | 0.2 | 4.79E-02 | 0.8 | 8.64E-05 | -0.1 | 7.86E-01 | 0.0 | 9.75E-01 | -0.1 | 8.54E-01 | hypothetical protein |
| AT3G04260 | 1.0 | 3.29E-07 | 0.8 | 5.73E-06 | 0.3 | 8.55E-03 | 0.3 | 1.89E-01 | -0.2 | 5.77E-01 | 0.4 | 8.21E-02 | Plastid transcriptionally active 3 |
| AT2G25470 | 1.0 | 1.54E-02 | 0.1 | 8.95E-01 | 1.0 | 5.58E-02 | -0.5 | 5.24E-01 | -2.0 | 4.13E-02 | 1.5 | 1.94E-01 | Receptor like protein 21 |
| AT5G27760 | 1.0 | 2.77E-04 | 0.3 | 2.23E-01 | 0.8 | 6.26E-03 | 0.5 | 1.04E-01 | -0.1 | 8.00E-01 | 0.6 | 1.43E-01 | Hypoxia-responsive family protein |
| AT5G48540 | 1.0 | 9.42E-05 | 0.7 | 2.98E-03 | 0.4 | 6.51E-02 | 1.0 | 2.23E-02 | -0.5 | 4.02E-01 | 1.5 | 1.43E-02 | Cysteine-rich repeat secretory protein 55 |
| AT1G67780 | 1.0 | 1.76E-05 | 0.7 | 8.36E-04 | 0.4 | 3.37E-02 | 0.0 | 8.96E-01 | 0.0 | 9.41E-01 | 0.1 | 8.56E-01 | Zinc-finger domain of monoamine-oxidase A repressor R1 protein |
| AT1G06510 | 1.0 | 3.31E-06 | 0.6 | 4.46E-04 | 0.5 | 3.78E-03 | 0.2 | 2.36E-01 | 0.0 | 9.23E-01 | 0.2 | 4.39E-01 | F12K11.16 |
| AT2G27680 | 1.0 | 4.20E-07 | 0.8 | 6.84E-06 | 0.3 | 1.33E-02 | 0.1 | 7.50E-01 | 0.1 | 4.17E-01 | -0.1 | 6.95E-01 | NADP-dependent oxidoreductase domain-containing protein |
| AT1G17700 | 1.0 | 1.64E-05 | 0.8 | 1.86E-04 | 0.2 | 1.74E-01 | 0.2 | 6.36E-01 | -0.2 | 5.03E-01 | 0.4 | 2.52E-01 | PRA1 family protein F1 |
| AT2G45860 | 1.0 | 3.57E-04 | 0.4 | 7.97E-02 | 0.6 | 2.08E-02 | 0.7 | 5.63E-02 | 0.6 | 1.13E-01 | 0.1 | 9.29E-01 | hypothetical protein |
| AT2G43630 | 1.0 | 1.44E-06 | 0.7 | 4.19E-05 | 0.3 | 1.17E-02 | 0.0 | 8.44E-01 | -0.1 | 6.19E-01 | 0.1 | 8.23E-01 | Nucleusenvelope protein |
| AT5G57410 | 1.0 | 3.65E-06 | 0.5 | 1.23E-03 | 0.5 | 1.68E-03 | -0.1 | 3.53E-01 | -0.2 | 1.24E-01 | 0.1 | 6.19E-01 | Afadin |
| AT4G33560 | 1.0 | 1.68E-05 | 0.4 | 1.33E-02 | 0.6 | 2.93E-03 | -0.2 | 5.77E-01 | 0.0 | 9.89E-01 | -0.2 | 6.73E-01 | At4g33560 |
| AT3G25597 | 1.0 | 5.28E-04 | 0.4 | 1.01E-01 | 0.6 | 2.81E-02 | 1.3 | 7.30E-03 | 0.3 | 6.96E-01 | 1.0 | 3.77E-02 | Transmembrane protein |
| AT4G37920 | 1.0 | 2.00E-06 | 0.7 | 7.01E-05 | 0.3 | 1.30E-02 | 0.5 | 7.28E-03 | 0.2 | 3.01E-01 | 0.3 | 2.29E-01 | Uncharacterized protein At4g37920 |
| AT5G47900 | 1.0 | 8.44E-05 | 1.0 | 3.32E-04 | 0.1 | 6.17E-01 | 0.0 | 9.94E-01 | 0.0 | 9.41E-01 | 0.0 | 9.41E-01 | Heparan-alpha-glucosaminide N-acetyltransferase catalytic domain-containing protein |
| AT2G40390 | 1.0 | 8.31E-06 | 0.4 | 1.04E-02 | 0.6 | 1.56E-03 | 0.0 | 8.97E-01 | -0.1 | 7.98E-01 | 0.2 | 7.31E-01 | Uncharacterized protein At2g40390 |
| AT2G48070 | 1.0 | 2.64E-07 | 1.0 | 9.11E-07 | 0.1 | 5.05E-01 | 0.0 | 9.28E-01 | 0.0 | 8.99E-01 | 0.0 | 8.64E-01 | Resistance to phytophthora 1 |
| AT2G17033 | 1.0 | 1.26E-07 | 0.7 | 6.81E-06 | 0.4 | 5.25E-04 | 0.2 | 2.90E-01 | 0.0 | 9.53E-01 | 0.2 | 5.11E-01 | Pentatricopeptide repeat-containing protein |
| AT1G75460 | 1.0 | 1.42E-05 | 1.0 | 3.18E-05 | 0.0 | 9.59E-01 | 0.3 | 3.63E-02 | 0.4 | 2.36E-02 | -0.1 | 7.93E-01 | hypothetical protein |
| AT5G54970 | 1.0 | 1.47E-05 | 0.8 | 3.11E-04 | 0.3 | 6.30E-02 | 0.6 | 7.47E-03 | 0.3 | 1.56E-01 | 0.2 | 2.28E-01 | hypothetical protein |
| AT1G35230 | 1.0 | 1.26E-02 | 0.1 | 7.67E-01 | 0.9 | 5.23E-02 | 0.1 | 8.61E-01 | -0.6 | 2.06E-01 | 0.7 | 1.52E-01 | hypothetical protein |
| AT1G22850 | 1.1 | 4.44E-07 | 0.7 | 1.37E-05 | 0.3 | 3.88E-03 | 0.1 | 4.50E-01 | 0.1 | 7.79E-01 | 0.1 | 8.31E-01 | SNARE associated Golgi protein family |
| AT5G60400 | 1.1 | 8.84E-04 | 0.6 | 3.87E-02 | 0.5 | 8.47E-02 | 0.3 | 2.19E-01 | -0.1 | 8.08E-01 | 0.4 | 2.37E-01 | AT5g60400 |
| AT3G20600 | 1.1 | 7.22E-06 | 0.6 | 1.42E-03 | 0.5 | 4.23E-03 | 1.0 | 5.91E-04 | 0.4 | 2.10E-01 | 0.6 | 5.32E-02 | Protein NDR1 |
| AT5G06690 | 1.1 | 1.02E-04 | 1.4 | 2.29E-05 | -0.3 | 8.12E-02 | 0.0 | 8.78E-01 | -0.1 | 6.45E-01 | 0.2 | 6.25E-01 | Putative thioredoxin |
| AT5G27560 | 1.1 | 3.97E-07 | 0.9 | 3.85E-06 | 0.2 | 4.16E-02 | 0.2 | 1.38E-01 | 0.1 | 4.55E-01 | 0.1 | 7.43E-01 | DUF1995 domain protein, putative |
| AT3G56160 | 1.1 | 5.32E-05 | 0.6 | 3.82E-03 | 0.5 | 2.35E-02 | 0.0 | 9.89E-01 | 0.1 | 6.65E-01 | -0.1 | 6.99E-01 | Sodium Bile acid symporter family |
| AT5G43070 | 1.1 | 3.13E-07 | 0.9 | 2.86E-06 | 0.2 | 5.04E-02 | 0.3 | 1.62E-01 | 0.1 | 6.51E-01 | 0.2 | 6.07E-01 | WPP domain-containing protein 1 |
| AT1G20350 | 1.1 | 5.41E-03 | 0.0 | 9.68E-01 | 1.0 | 2.53E-02 | | | | | | | Mitochondrial import inner membrane translocase subunit TIM17-1 |
| AT1G19960 | 1.1 | 1.03E-03 | 0.4 | 9.80E-02 | 0.6 | 3.61E-02 | 1.3 | 3.39E-05 | 0.9 | 1.99E-03 | 0.4 | 2.13E-01 | Uncharacterized protein |
| AT4G33110 | 1.1 | 3.62E-06 | 0.8 | 7.23E-05 | 0.3 | 3.43E-02 | 0.1 | 5.93E-01 | 0.1 | 6.96E-01 | 0.0 | 9.71E-01 | Uncharacterized protein AT4G33110 |
| AT3G19800 | 1.1 | 5.41E-07 | 0.6 | 8.79E-05 | 0.5 | 6.04E-04 | -0.1 | 5.63E-01 | 0.2 | 3.76E-01 | -0.3 | 1.98E-01 | hypothetical protein |
| AT4G27660 | 1.1 | 2.75E-05 | 0.8 | 5.76E-04 | 0.3 | 9.88E-02 | 0.7 | 6.06E-02 | 0.4 | 4.20E-01 | 0.3 | 4.48E-01 | Uncharacterized protein |
| AT4G02725 | 1.1 | 1.38E-05 | 0.7 | 3.65E-04 | 0.3 | 4.25E-02 | -0.1 | 5.72E-01 | -0.2 | 5.06E-01 | 0.0 | 9.19E-01 | At4g02725 |
| AT1G27460 | 1.1 | 4.61E-06 | 1.0 | 2.40E-05 | 0.1 | 3.47E-01 | -0.2 | 3.31E-01 | -0.5 | 5.58E-02 | 0.2 | 4.32E-01 | No pollen germination related 1 |
| AT5G19940 | 1.1 | 1.60E-06 | 0.9 | 1.26E-05 | 0.2 | 1.45E-01 | -0.1 | 3.12E-01 | -0.3 | 5.19E-02 | 0.1 | 4.53E-01 | Plastid-lipid associated protein PAP |
| AT3G54750 | 1.1 | 1.10E-06 | 0.7 | 7.97E-05 | 0.4 | 2.88E-03 | 0.3 | 5.97E-02 | 0.1 | 4.43E-01 | 0.1 | 5.01E-01 | AT3G54750 protein |
| AT5G24570 | 1.1 | 2.48E-06 | 0.3 | 1.23E-02 | 0.7 | 2.72E-04 | -0.2 | 4.09E-01 | -0.2 | 4.78E-01 | 0.0 | 9.99E-01 | hypothetical protein |
| AT2G32640 | 1.1 | 3.58E-06 | 0.8 | 7.41E-05 | 0.3 | 3.79E-02 | 0.4 | 1.10E-01 | 0.3 | 3.35E-01 | 0.1 | 7.69E-01 | Lycopene beta |
| AT4G10300 | 1.1 | 9.15E-06 | 0.8 | 1.23E-04 | 0.3 | 8.39E-02 | 0.3 | 1.71E-01 | 0.2 | 5.27E-01 | 0.1 | 7.34E-01 | #NAME? |
| AT1G53490 | 1.1 | 1.59E-05 | 0.5 | 6.12E-03 | 0.6 | 4.48E-03 | 0.0 | 9.77E-01 | -0.2 | 6.83E-01 | 0.2 | 7.39E-01 | RING |
| AT3G05625 | 1.1 | 4.01E-06 | 1.0 | 1.19E-05 | 0.0 | 7.86E-01 | 0.4 | 5.29E-02 | 0.3 | 1.25E-01 | 0.0 | 9.04E-01 | hypothetical protein |
| AT2G46910 | 1.1 | 1.20E-05 | 0.6 | 1.43E-03 | 0.5 | 7.42E-03 | -0.2 | 3.34E-01 | 0.0 | 9.91E-01 | -0.2 | 4.42E-01 | Probable plastid-lipid-associated protein 10, chloroplastic |
| AT1G74790 | 1.1 | 6.27E-07 | 0.6 | 1.46E-04 | 0.5 | 4.45E-04 | -0.1 | 5.78E-01 | -0.5 | 2.28E-03 | 0.4 | 2.71E-02 | Glucose |
| AT5G59500 | 1.1 | 1.07E-06 | 0.8 | 1.90E-05 | 0.3 | 2.09E-02 | 0.5 | 2.87E-03 | 0.2 | 2.34E-01 | 0.3 | 1.37E-01 | Protein-S-isoprenylcysteine O-methyltransferase |
| AT4G18740 | 1.1 | 2.83E-04 | 0.6 | 2.07E-02 | 0.5 | 4.43E-02 | -0.4 | 6.09E-02 | -0.7 | 6.11E-03 | 0.3 | 2.68E-01 | Rho termination factor |
| AT3G27050 | 1.1 | 1.05E-05 | 0.8 | 2.40E-04 | 0.3 | 4.76E-02 | 0.1 | 7.24E-01 | -0.2 | 3.94E-01 | 0.3 | 3.04E-01 | Uncharacterized protein |
| AT1G35140 | 1.1 | 4.95E-02 | -0.4 | 5.50E-01 | 1.5 | 4.96E-02 | -1.5 | 1.17E-03 | -1.3 | 4.69E-03 | -0.2 | 7.49E-01 | Protein EXORDIUM-like 1 |
| AT4G13370 | 1.1 | 2.04E-06 | 0.5 | 7.12E-04 | 0.6 | 1.18E-03 | -0.1 | 6.43E-01 | -0.1 | 8.77E-01 | -0.1 | 8.60E-01 | Uncharacterized protein |
| AT4G15890 | 1.1 | 4.57E-05 | 0.6 | 3.56E-03 | 0.5 | 2.70E-02 | 0.2 | 3.74E-01 | 0.0 | 9.78E-01 | 0.2 | 4.13E-01 | Condensin-2 complex subunit |
| AT4G14746 | 1.1 | 7.07E-06 | 0.5 | 4.48E-03 | 0.6 | 1.62E-03 | 0.2 | 3.30E-01 | -0.2 | 5.70E-01 | 0.4 | 1.93E-01 | Neurogenic locus notch-like protein |
| AT5G64980 | 1.1 | 2.34E-05 | 0.6 | 3.75E-03 | 0.5 | 1.20E-02 | | | | | | | hypothetical protein |

| | | | | | | | | | | | | | |
|---|---|---|---|---|---|---|---|---|---|---|---|---|---|
| AT4G13220 | 1.1 | 1.72E-06 | 0.7 | 9.62E-05 | 0.4 | 5.76E-03 | 0.4 | 5.85E-02 | 0.1 | 6.43E-01 | 0.3 | 3.15E-01 | AT4G13220 protein |
| AT4G35180 | 1.1 | 1.42E-02 | -0.3 | 6.24E-01 | 1.4 | 2.51E-02 | 1.3 | 1.57E-02 | 0.4 | 5.94E-01 | 0.9 | 1.27E-01 | LYS |
| AT5G27890 | 1.1 | 2.04E-05 | 0.5 | 9.86E-03 | 0.6 | 4.81E-03 | 0.2 | 4.11E-01 | 0.1 | 5.77E-01 | 0.0 | 9.28E-01 | Uncharacterized protein |
| AT1G21520 | 1.1 | 6.74E-04 | -0.9 | 1.35E-02 | 2.0 | 2.23E-04 | -0.3 | 2.86E-01 | -0.4 | 2.73E-01 | 0.0 | 9.37E-01 | hypothetical protein |
| AT2G07042 | 1.1 | 2.13E-05 | 0.4 | 2.35E-02 | 0.7 | 2.93E-03 | -1.0 | 2.84E-02 | -0.8 | 8.17E-02 | -0.2 | 7.89E-01 | NA |
| AT3G48200 | 1.1 | 2.56E-06 | 0.6 | 3.06E-04 | 0.5 | 2.48E-03 | 0.2 | 3.23E-01 | 0.2 | 2.08E-01 | -0.1 | 8.05E-01 | Transmembrane protein |
| AT5G62840 | 1.1 | 9.97E-06 | 1.0 | 5.13E-05 | 0.1 | 3.85E-01 | 0.4 | 2.83E-02 | 0.2 | 2.37E-01 | 0.1 | 5.59E-01 | Phosphoglycerate mutase family protein |
| AT4G13495 | 1.1 | 5.22E-07 | 1.2 | 8.67E-07 | -0.1 | 3.86E-01 | -0.1 | 7.65E-01 | 0.1 | 7.13E-01 | -0.2 | 4.75E-01 | NA |
| AT4G29170 | 1.1 | 2.19E-05 | 0.5 | 6.65E-03 | 0.6 | 6.89E-03 | 0.4 | 1.74E-01 | 0.0 | 9.73E-01 | 0.3 | 3.37E-01 | Mnd1 family protein |
| AT5G12360 | 1.1 | 2.49E-05 | 0.7 | 2.45E-03 | 0.5 | 1.95E-02 | 0.1 | 8.48E-01 | 0.1 | 9.22E-01 | 0.0 | 9.70E-01 | Uncharacterized protein |
| AT3G20680 | 1.1 | 6.29E-06 | 0.9 | 9.14E-05 | 0.3 | 8.27E-02 | -0.3 | 1.21E-01 | -0.3 | 1.66E-01 | 0.0 | 9.96E-01 | hypothetical protein |
| AT1G01690 | 1.1 | 6.89E-07 | 0.4 | 1.22E-03 | 0.7 | 1.78E-04 | 0.1 | 7.16E-01 | 0.1 | 6.24E-01 | 0.0 | 9.09E-01 | Recombination initiation defects 3 |
| AT1G10682 | 1.1 | 6.52E-04 | 1.2 | 5.81E-04 | -0.1 | 7.39E-01 | 0.4 | 1.78E-01 | 0.3 | 3.21E-01 | 0.0 | 9.22E-01 | NA |
| AT4G28590 | 1.1 | 8.97E-06 | 0.6 | 2.41E-03 | 0.6 | 4.03E-03 | 0.3 | 1.67E-01 | 0.1 | 8.60E-01 | 0.3 | 3.63E-01 | Thioredoxin-like fold domain-containing protein MRL7, chloroplastic |
| AT2G30480 | 1.1 | 4.76E-05 | 0.2 | 2.37E-01 | 0.9 | 1.30E-03 | -0.2 | 5.51E-01 | 0.2 | 6.34E-01 | -0.4 | 2.74E-01 | Uncharacterized protein |
| AT5G03355 | 1.1 | 8.90E-05 | 0.5 | 1.72E-02 | 0.6 | 1.57E-02 | -0.1 | 7.07E-01 | -0.5 | 1.07E-01 | 0.4 | 2.74E-01 | Cysteine |
| AT1G65484 | 1.1 | 1.85E-03 | -0.3 | 4.57E-01 | 1.4 | 3.50E-03 | 0.8 | 8.27E-02 | 0.2 | 6.92E-01 | 0.5 | 3.51E-01 | Transmembrane protein |
| AT4G14310 | 1.1 | 1.52E-06 | 0.6 | 3.65E-04 | 0.5 | 1.17E-03 | 0.1 | 6.24E-01 | 0.2 | 3.71E-01 | -0.1 | 7.02E-01 | hypothetical protein |
| AT4G28730 | 1.1 | 2.25E-04 | 0.6 | 2.60E-02 | 0.6 | 3.01E-02 | 0.3 | 9.56E-02 | 0.1 | 6.51E-01 | 0.2 | 4.40E-01 | Glutaredoxin family protein |
| AT3G51750 | 1.1 | 1.27E-05 | 0.7 | 1.00E-03 | 0.4 | 1.50E-02 | 1.8 | 2.10E-03 | 1.2 | 5.99E-02 | 0.6 | 2.41E-01 | Uncharacterized protein |
| AT4G28410 | 1.1 | 8.90E-05 | 0.4 | 4.60E-02 | 0.7 | 8.65E-03 | 0.2 | 5.47E-01 | 0.4 | 2.42E-01 | -0.2 | 5.95E-01 | Uncharacterized protein AT4g28410 |
| AT1G63260 | 1.1 | 1.37E-05 | 0.7 | 7.70E-04 | 0.4 | 2.09E-02 | -0.2 | 3.08E-01 | -0.5 | 4.57E-02 | 0.3 | 3.58E-01 | Tetraspanin-10 |
| AT3G18050 | 1.1 | 7.75E-07 | 1.2 | 1.75E-06 | 0.0 | 8.31E-01 | -0.7 | 3.97E-04 | -0.7 | 1.46E-03 | 0.0 | 9.55E-01 | hypothetical protein |
| AT4G38100 | 1.1 | 4.83E-06 | 0.7 | 2.06E-04 | 0.4 | 1.55E-02 | 0.8 | 1.13E-03 | 0.4 | 1.04E-01 | 0.4 | 1.24E-01 | CURVATURE THYLAKOID 1D-like protein |
| AT3G04760 | 1.1 | 2.87E-07 | 0.8 | 6.75E-06 | 0.3 | 3.96E-03 | 0.4 | 6.85E-03 | 0.1 | 5.89E-01 | 0.3 | 1.09E-01 | Pentatricopeptide repeat superfamily protein |
| AT1G63610 | 1.1 | 9.75E-07 | 0.8 | 3.78E-05 | 0.4 | 5.41E-03 | 0.2 | 1.38E-01 | 0.2 | 3.66E-01 | 0.1 | 8.18E-01 | At1g63610 |
| AT3G44020 | 1.1 | 3.44E-05 | 0.7 | 2.75E-03 | 0.5 | 2.09E-02 | 0.3 | 1.80E-01 | 0.2 | 5.20E-01 | 0.1 | 7.42E-01 | Uncharacterized protein |
| AT1G17147 | 1.1 | 1.09E-03 | 0.3 | 2.59E-01 | 0.8 | 2.39E-02 | 0.9 | 6.55E-03 | -0.2 | 7.62E-01 | 1.1 | 3.75E-03 | VQ motif-containing protein 1 |
| AT4G25290 | 1.1 | 1.18E-05 | 0.5 | 4.14E-03 | 0.6 | 3.03E-03 | -0.4 | 8.79E-02 | 0.0 | 9.53E-01 | -0.4 | 1.87E-01 | DNA photolyase |
| AT5G40500 | 1.1 | 1.35E-03 | 0.3 | 2.63E-01 | 0.8 | 2.48E-02 | -0.2 | 7.19E-01 | 0.1 | 7.77E-01 | -0.3 | 5.46E-01 | Uncharacterized protein At5g40500 |
| AT3G45050 | 1.1 | 6.95E-07 | 1.1 | 2.29E-06 | 0.0 | 6.75E-01 | 0.6 | 1.26E-02 | 0.4 | 1.60E-01 | 0.2 | 4.77E-01 | At3g45050 |
| AT3G23400 | 1.1 | 3.72E-07 | 0.9 | 5.15E-06 | 0.3 | 1.81E-02 | 0.3 | 1.85E-02 | 0.2 | 6.92E-02 | 0.0 | 8.40E-01 | Plastid-lipid associated protein PAP |
| AT1G09415 | 1.1 | 8.35E-06 | 0.8 | 2.50E-04 | 0.3 | 3.44E-02 | 0.3 | 9.87E-02 | -0.1 | 7.18E-01 | 0.4 | 1.20E-01 | Uncharacterized protein |
| AT1G32500 | 1.1 | 1.75E-07 | 0.8 | 4.89E-06 | 0.4 | 1.88E-03 | 0.5 | 8.38E-04 | 0.3 | 6.45E-02 | 0.2 | 1.86E-01 | Protein ABCI7, chloroplastic |
| AT1G29430 | 1.1 | 6.22E-05 | 1.1 | 1.83E-04 | 0.1 | 7.23E-01 | 0.6 | 1.29E-01 | 0.3 | 4.94E-01 | 0.2 | 5.61E-01 | Auxin-responsive protein SAUR62 |
| AT3G49470 | 1.1 | 3.72E-07 | 1.0 | 2.36E-06 | 0.2 | 1.19E-01 | 0.0 | 9.81E-01 | -0.1 | 3.75E-01 | 0.1 | 4.53E-01 | Nascent polypeptide-associated complex subunit alpha-like protein 2 |
| AT3G15570 | 1.1 | 3.01E-05 | 0.8 | 8.41E-04 | 0.4 | 6.15E-02 | -0.9 | 7.09E-02 | -0.6 | 2.62E-01 | -0.3 | 6.76E-01 | Non-phototropic hypocotyl protein, putative |
| AT1G64500 | 1.2 | 1.89E-05 | 0.5 | 9.47E-03 | 0.7 | 2.49E-03 | 0.3 | 1.67E-01 | -0.2 | 3.74E-01 | 0.5 | 2.75E-02 | F1N19.7 |
| AT4G20000 | 1.2 | 1.22E-03 | -0.1 | 7.95E-01 | 1.2 | 4.82E-03 | 0.6 | 2.85E-02 | -0.2 | 5.69E-01 | 0.9 | 3.20E-02 | hypothetical protein |
| AT2G04360 | 1.2 | 2.27E-07 | 0.8 | 7.86E-06 | 0.4 | 1.91E-03 | 0.3 | 1.40E-02 | 0.1 | 6.18E-01 | 0.3 | 1.62E-01 | Transmembrane protein |
| AT2G30570 | 1.2 | 1.66E-06 | 1.3 | 2.31E-06 | -0.1 | 3.69E-01 | -0.2 | 2.15E-01 | -0.1 | 8.21E-01 | -0.1 | 5.48E-01 | PSII 6.1 kDa protein |
| AT1G74290 | 1.2 | 1.41E-05 | 0.4 | 1.93E-02 | 0.7 | 1.68E-03 | 0.7 | 1.34E-01 | 0.7 | 1.84E-01 | | | Alpha |
| AT1G78995 | 1.2 | 3.87E-06 | 0.9 | 6.85E-05 | 0.3 | 4.72E-02 | 0.3 | 5.05E-02 | 0.2 | 1.62E-01 | 0.1 | 8.23E-01 | YUP8H12R.39 protein |
| AT4G30840 | 1.2 | 2.83E-06 | 0.5 | 1.57E-03 | 0.6 | 1.06E-03 | 0.7 | 1.06E-03 | 0.1 | 6.97E-01 | 0.6 | 2.04E-02 | Nuclear pore complex protein NUP43 |
| AT5G16650 | 1.2 | 4.95E-05 | 0.7 | 4.12E-03 | 0.5 | 2.41E-02 | -0.1 | 7.99E-01 | -0.3 | 3.31E-01 | 0.2 | 5.90E-01 | J domain-containing protein |
| AT5G56550 | 1.2 | 5.22E-04 | 1.2 | 7.54E-04 | 0.0 | 9.86E-01 | -0.7 | 1.33E-02 | -1.4 | 5.46E-04 | 0.6 | 1.56E-01 | Protein OXIDATIVE STRESS 3 |
| AT5G43180 | 1.2 | 4.90E-07 | 1.1 | 1.65E-06 | 0.0 | 6.50E-01 | 0.1 | 6.92E-01 | -0.2 | 5.32E-01 | 0.2 | 3.23E-01 | DUF599 domain-containing protein |
| AT3G53800 | 1.2 | 2.55E-05 | 1.9 | 1.69E-06 | -0.7 | 1.77E-03 | -0.4 | 2.65E-02 | -0.4 | 2.68E-02 | 0.0 | 9.16E-01 | TOG domain-containing protein |
| AT1G75750 | 1.2 | 7.93E-04 | 0.5 | 6.68E-02 | 0.7 | 3.88E-02 | -0.8 | 1.05E-02 | -0.9 | 1.57E-02 | 0.0 | 9.49E-01 | GAST1 protein homolog 1 |
| AT5G43750 | 1.2 | 1.74E-05 | 1.1 | 6.17E-05 | 0.1 | 5.88E-01 | 0.2 | 2.82E-01 | 0.4 | 3.91E-02 | -0.2 | 3.37E-01 | Photosynthetic NDH subunit of subcomplex B 5, chloroplastic H dehydrogenase 18) |
| AT2G24150 | 1.2 | 3.05E-07 | 0.9 | 3.96E-06 | 0.3 | 1.55E-02 | -0.2 | 1.35E-01 | -0.1 | 3.59E-01 | -0.1 | 8.10E-01 | Heptahelical protein 3 |
| AT1G70880 | 1.2 | 7.97E-06 | 1.0 | 3.78E-05 | 0.1 | 3.94E-01 | -1.6 | 9.64E-06 | -0.2 | 4.24E-01 | -1.4 | 5.53E-04 | Major latex-like protein |
| AT5G52110 | 1.2 | 1.36E-06 | 0.7 | 1.33E-04 | 0.5 | 3.06E-03 | 0.3 | 6.72E-02 | 0.2 | 1.92E-01 | 0.1 | 8.30E-01 | Protein COFACTOR ASSEMBLY OF COMPLEX C SUBUNIT B CCB2, chloroplastic |
| AT2G02100 | 1.2 | 1.51E-06 | 1.1 | 5.39E-06 | 0.1 | 6.31E-01 | -0.1 | 6.07E-01 | -0.2 | 3.67E-01 | 0.1 | 7.77E-01 | Defensin-like protein 2 |
| AT1G62510 | 1.2 | 1.17E-03 | 1.7 | 1.06E-04 | -0.5 | 6.33E-02 | -1.0 | 3.16E-02 | -1.2 | 1.47E-02 | 0.3 | 7.21E-01 | At1g62510 |
| AT5G02950 | 1.2 | 1.49E-05 | 0.7 | 1.40E-03 | 0.5 | 1.54E-02 | -0.1 | 8.50E-01 | 0.0 | 9.69E-01 | -0.1 | 8.56E-01 | PWWP domain-containing protein 5 |
| AT5G44130 | 1.2 | 4.48E-05 | 1.1 | 1.57E-04 | 0.1 | 6.24E-01 | -0.3 | 5.99E-02 | -0.4 | 2.41E-02 | 0.1 | 6.86E-01 | Fasciclin-like arabinogalactan protein 13 |
| AT2G05510 | 1.2 | 1.80E-04 | 1.5 | 5.10E-05 | -0.3 | 1.59E-01 | 2.3 | 3.68E-07 | 1.4 | 2.64E-04 | 0.9 | 8.87E-03 | At2g05510 |
| AT3G26900 | 1.2 | 5.47E-07 | 0.8 | 1.42E-05 | 0.4 | 6.23E-03 | 0.1 | 3.07E-01 | -0.1 | 7.36E-01 | 0.2 | 3.09E-01 | Probable inactive shikimate kinase like 1, chloroplastic |
| AT3G29185 | 1.2 | 5.51E-07 | 1.0 | 4.76E-06 | 0.2 | 6.36E-02 | 0.1 | 5.57E-01 | 0.1 | 4.54E-01 | 0.0 | 8.89E-01 | Glutamate NMDA receptor subunit epsilon-1, putative |
| AT1G16720 | 1.2 | 1.36E-05 | 0.8 | 4.49E-04 | 0.4 | 2.97E-02 | 0.0 | 7.99E-01 | -0.2 | 1.67E-01 | 0.2 | 3.72E-01 | High chlorophyll fluorescence phenotype 173 |
| AT3G10020 | 1.2 | 1.67E-04 | 1.0 | 1.07E-03 | 0.2 | 3.45E-01 | -0.7 | 9.42E-03 | -1.0 | 1.53E-03 | 0.3 | 3.51E-01 | AT3g10020 |
| AT1G78410 | 1.2 | 1.09E-03 | -0.1 | 8.41E-01 | 1.3 | 4.78E-03 | 0.7 | 1.29E-01 | -0.5 | 4.28E-01 | 1.3 | 3.80E-02 | VQ motif-containing protein 10 |
| AT2G14900 | 1.2 | 1.91E-04 | 0.9 | 2.64E-03 | 0.3 | 1.90E-01 | 0.0 | 9.44E-01 | -0.3 | 4.13E-01 | 0.3 | 5.44E-01 | Gibberellin-regulated protein 7 |
| AT3G61770 | 1.2 | 7.19E-08 | 0.9 | 1.17E-06 | 0.3 | 1.96E-03 | 0.4 | 2.35E-02 | 0.3 | 2.10E-01 | 0.2 | 5.62E-01 | Uncharacterized protein F15G16.160 |
| AT2G46980 | 1.2 | 6.55E-06 | 0.4 | 1.33E-02 | 0.8 | 8.94E-04 | 0.0 | 9.29E-01 | 0.1 | 8.40E-01 | -0.1 | 7.96E-01 | Asynaptic protein |
| AT3G55240 | 1.2 | 1.84E-03 | 1.1 | 4.16E-03 | 0.1 | 8.27E-01 | | | | | | | Uncharacterized protein At3g55240 |
| AT3G55250 | 1.2 | 9.75E-06 | 0.9 | 1.88E-04 | 0.3 | 4.96E-02 | 0.1 | 3.77E-01 | 0.0 | 8.69E-01 | 0.2 | 4.48E-01 | Photosystem I assembly factor PSA3, chloroplastic |
| AT2G39560 | 1.2 | 2.05E-04 | 1.1 | 8.45E-04 | 0.1 | 5.75E-01 | -0.2 | 7.35E-01 | -0.1 | 8.25E-01 | 0.0 | 9.65E-01 | hypothetical protein |
| AT5G44582 | 1.2 | 9.04E-05 | 0.5 | 5.57E-02 | 0.7 | 7.69E-03 | -0.3 | 2.98E-01 | -0.4 | 2.77E-01 | 0.1 | 9.28E-01 | Uncharacterized protein |
| AT2G38230 | 1.2 | 1.04E-07 | 0.6 | 1.34E-05 | 0.6 | 1.27E-04 | -0.3 | 1.60E-02 | -0.3 | 5.40E-02 | 0.0 | 8.81E-01 | PdxS |
| AT5G53860 | 1.2 | 7.03E-08 | 1.0 | 6.40E-07 | 0.2 | 9.12E-03 | 0.0 | 8.44E-01 | -0.2 | 5.66E-02 | 0.2 | 1.76E-01 | AT5g53860 |
| AT5G67385 | 1.2 | 4.58E-07 | 0.8 | 1.78E-05 | 0.4 | 2.63E-03 | -0.3 | 7.38E-02 | -0.4 | 4.93E-02 | 0.1 | 7.79E-01 | Phototropic-responsive NPH3 family protein |
| AT3G09210 | 1.2 | 3.28E-07 | 1.0 | 2.57E-06 | 0.2 | 6.16E-02 | 0.3 | 1.80E-02 | 0.0 | 9.65E-01 | 0.3 | 8.36E-02 | F3L24.8 protein |
| AT1G21930 | 1.2 | 8.59E-04 | 0.6 | 7.77E-02 | 0.7 | 4.86E-02 | 0.2 | 5.57E-01 | -0.2 | 6.35E-01 | 0.3 | 3.53E-01 | hypothetical protein |
| AT1G55670 | 1.2 | 1.54E-06 | 1.5 | 1.07E-06 | -0.3 | 4.46E-02 | -0.1 | 7.83E-01 | -0.2 | 6.21E-01 | 0.1 | 8.53E-01 | Photosystem I reaction center subunit V, chloroplastic |
| AT1G33810 | 1.2 | 1.52E-07 | 1.1 | 8.03E-07 | 0.1 | 1.24E-01 | 0.4 | 4.31E-02 | 0.0 | 9.19E-01 | 0.3 | 1.80E-01 | Uncharacterized protein At1g33810 |
| AT1G66725 | 1.2 | 4.76E-04 | 0.5 | 6.27E-02 | 0.7 | 3.22E-02 | 0.8 | 2.36E-01 | 1.1 | 1.21E-01 | | | NA |
| AT2G38695 | 1.2 | 4.31E-06 | 0.8 | 2.11E-04 | 0.4 | 1.59E-02 | 0.2 | 3.88E-01 | -0.1 | 7.67E-01 | 0.3 | 2.59E-01 | Gag-Pol polyprotein |
| AT5G64460 | 1.2 | 1.42E-06 | 0.9 | 1.74E-05 | 0.3 | 4.53E-02 | 0.0 | 9.50E-01 | -0.2 | 1.47E-01 | 0.2 | 2.91E-01 | Phosphoglycerate mutase family protein |
| AT2G37640 | 1.2 | 1.03E-06 | 1.2 | 2.15E-06 | 0.0 | 8.46E-01 | -0.2 | 2.40E-01 | 0.0 | 8.89E-01 | -0.2 | 4.91E-01 | Expansin |
| AT2G42770 | 1.2 | 5.42E-08 | 0.8 | 1.85E-06 | 0.4 | 2.72E-04 | 0.1 | 6.13E-01 | 0.1 | 4.80E-01 | 0.0 | 8.65E-01 | Expressed protein family protein) |
| AT1G14960 | 1.2 | 6.56E-04 | 0.9 | 7.69E-03 | 0.3 | 2.70E-01 | -0.5 | 4.92E-01 | 1.5 | 1.93E-02 | -2.0 | 1.15E-02 | hypothetical protein |

| | | | | | | | | | | | | | |
|---|---|---|---|---|---|---|---|---|---|---|---|---|---|
| AT1G33590 | 1.2 | 7.61E-06 | 1.2 | 2.11E-05 | 0.0 | 8.09E-01 | 0.2 | 5.42E-02 | -0.1 | 4.90E-01 | 0.3 | 3.80E-02 | Leucine-rich repeat family protein |
| AT1G66840 | 1.2 | 7.14E-06 | 0.6 | 2.91E-03 | 0.6 | 2.30E-03 | 0.2 | 3.98E-01 | 0.2 | 5.08E-01 | 0.0 | 9.74E-01 | Protein PLASTID MOVEMENT IMPAIRED 2 |
| AT3G61870 | 1.2 | 3.48E-07 | 1.1 | 2.29E-06 | 0.2 | 1.24E-01 | -0.7 | 8.75E-05 | 0.5 | 3.32E-03 | -1.2 | 1.33E-05 | AT3g61870 |
| AT2G04235 | 1.2 | 1.18E-05 | 0.5 | 8.03E-03 | 0.7 | 2.31E-03 | 0.1 | 7.92E-01 | 0.4 | 1.48E-01 | -0.3 | 2.45E-01 | At2g04235 |
| AT2G04530 | 1.2 | 4.41E-06 | 1.2 | 1.27E-05 | 0.0 | 7.80E-01 | 0.7 | 3.56E-03 | 0.1 | 8.73E-01 | 0.7 | 1.57E-02 | Metallo-beta-lactamase domain-containing protein |
| AT4G30993 | 1.2 | 6.55E-07 | 0.7 | 7.86E-05 | 0.5 | 9.25E-04 | 0.1 | 5.47E-01 | 0.0 | 8.22E-01 | 0.1 | 4.94E-01 | Calcineurin-like metallo-phosphoesterase superfamily protein |
| AT1G61740 | 1.2 | 8.85E-07 | 0.7 | 7.09E-05 | 0.5 | 1.81E-03 | -0.6 | 6.79E-04 | -0.2 | 1.92E-01 | -0.4 | 7.64E-02 | Sulfite exporter TauE |
| AT3G06750 | 1.2 | 2.94E-07 | 1.0 | 2.82E-06 | 0.2 | 3.47E-02 | -0.3 | 9.76E-02 | -0.2 | 2.38E-01 | -0.1 | 8.41E-01 | Uncharacterized protein |
| AT1G56200 | 1.2 | 1.73E-05 | 0.8 | 7.09E-04 | 0.4 | 2.40E-02 | 0.0 | 9.81E-01 | -0.4 | 1.33E-01 | 0.4 | 2.49E-01 | At1g56200 |
| AT2G17880 | 1.2 | 3.01E-06 | 1.1 | 1.20E-05 | 0.1 | 5.23E-01 | -0.4 | 1.86E-01 | -1.1 | 3.36E-03 | 0.7 | 1.12E-01 | J domain-containing protein |
| AT3G48210 | 1.2 | 5.72E-07 | 0.5 | 1.41E-03 | 0.8 | 1.15E-04 | 0.3 | 6.08E-02 | 0.3 | 2.20E-02 | -0.1 | 6.66E-01 | Kinetochore protein SPC25 homolog |
| AT3G09860 | 1.2 | 4.67E-05 | 0.9 | 9.40E-04 | 0.4 | 8.08E-02 | 0.4 | 5.11E-02 | 0.1 | 6.14E-01 | 0.3 | 3.80E-01 | Uncharacterized protein |
| AT4G21620 | 1.2 | 1.18E-07 | 0.8 | 5.83E-06 | 0.5 | 4.46E-04 | -0.1 | 8.36E-01 | -0.4 | 1.56E-01 | 0.3 | 3.22E-01 | Glycine-rich protein |
| AT3G08030 | 1.2 | 2.37E-06 | 1.2 | 7.31E-06 | 0.0 | 7.57E-01 | 0.0 | 9.29E-01 | -0.1 | 4.04E-01 | 0.1 | 4.61E-01 | hypothetical protein |
| AT3G50685 | 1.2 | 1.69E-06 | 1.2 | 3.96E-06 | 0.0 | 9.42E-01 | -0.1 | 4.96E-01 | -0.1 | 5.66E-01 | 0.0 | 9.98E-01 | Uncharacterized protein T3A5.60 |
| AT3G49140 | 1.2 | 3.20E-06 | 0.7 | 2.27E-04 | 0.5 | 5.19E-03 | 0.0 | 9.93E-01 | 0.0 | 9.57E-01 | 0.0 | 9.53E-01 | Uncharacterized protein At3g49140 |
| AT2G39500 | 1.2 | 8.90E-05 | 0.6 | 1.70E-02 | 0.6 | 1.57E-02 | 0.6 | 1.25E-01 | 0.3 | 5.08E-01 | 0.3 | 6.29E-01 | Transmembrane protein |
| AT4G33400 | 1.2 | 3.42E-07 | 0.7 | 2.94E-05 | 0.5 | 7.15E-04 | 0.0 | 8.72E-01 | 0.0 | 8.82E-01 | 0.0 | 9.93E-01 | Dem-like protein |
| AT2G25830 | 1.2 | 3.57E-07 | 1.1 | 2.20E-06 | 0.2 | 1.28E-01 | 0.5 | 5.51E-04 | 0.5 | 1.41E-03 | 0.0 | 9.92E-01 | Probable transcriptional regulatory protein At2g25830 |
| AT4G23215 | 1.2 | 2.48E-03 | 0.0 | 9.29E-01 | 1.2 | 1.33E-02 | -0.4 | 6.58E-01 | -1.6 | 1.11E-01 | 1.2 | 3.37E-01 | NA |
| AT2G01870 | 1.2 | 3.71E-04 | 0.7 | 1.85E-02 | 0.5 | 7.80E-02 | 0.3 | 5.03E-01 | 0.1 | 9.03E-01 | 0.2 | 6.87E-01 | At2g01870 |
| AT2G01590 | 1.2 | 1.19E-06 | 1.0 | 1.08E-05 | 0.2 | 8.44E-02 | 0.0 | 8.72E-01 | 0.0 | 8.73E-01 | 0.0 | 9.88E-01 | Chlororespiratory reduction 3 |
| AT1G53800 | 1.2 | 8.66E-06 | 0.7 | 8.48E-04 | 0.5 | 8.72E-03 | 0.1 | 8.85E-01 | -0.2 | 5.35E-01 | 0.3 | 3.72E-01 | At1g53800 |
| AT4G16008 | 1.2 | 1.03E-06 | 1.2 | 2.79E-06 | 0.0 | 8.72E-01 | 0.7 | 2.59E-03 | 0.6 | 2.71E-02 | 0.2 | 5.47E-01 | Uncharacterized protein |
| AT1G26761 | 1.2 | 4.49E-07 | 1.1 | 3.11E-06 | 0.2 | 1.01E-01 | 0.2 | 2.15E-01 | 0.3 | 1.44E-01 | -0.1 | 7.85E-01 | T24P13.14 |
| AT1G68740 | 1.2 | 3.21E-05 | 0.7 | 3.66E-03 | 0.5 | 1.82E-02 | 1.2 | 4.06E-03 | 0.4 | 3.90E-01 | 0.8 | 8.89E-02 | EXS family protein |
| AT3G50480 | 1.2 | 1.04E-03 | 0.4 | 2.18E-01 | 0.9 | 1.71E-02 | -0.1 | 8.89E-01 | -1.2 | 1.45E-02 | 1.1 | 5.76E-02 | Homolog of RPW8 4 |
| AT1G73060 | 1.2 | 9.91E-08 | 0.9 | 1.65E-06 | 0.3 | 2.98E-03 | 0.1 | 4.96E-01 | 0.1 | 4.92E-01 | 0.0 | 9.51E-01 | Protein LPA3 |
| AT5G07860 | 1.3 | 2.40E-06 | 0.9 | 4.25E-05 | 0.3 | 4.03E-02 | 0.4 | 3.69E-03 | 0.2 | 2.84E-01 | 0.3 | 1.48E-01 | HXXXD-type acyl-transferase family protein |
| AT2G46730 | 1.3 | 6.79E-05 | 0.1 | 6.67E-01 | 1.1 | 1.02E-03 | | | | | | | NA |
| AT5G57180 | 1.3 | 7.72E-07 | 1.3 | 1.21E-06 | -0.1 | 4.66E-01 | -0.1 | 7.36E-01 | -0.2 | 4.31E-01 | 0.1 | 6.62E-01 | Chloroplast import apparatus 2 |
| AT2G20260 | 1.3 | 1.34E-07 | 1.2 | 5.25E-07 | 0.1 | 3.80E-01 | -0.2 | 1.89E-01 | -0.2 | 1.54E-01 | 0.0 | 8.74E-01 | Photosystem I reaction center subunit IV B, chloroplast |
| AT1G76530 | 1.3 | 1.56E-04 | 0.6 | 2.22E-02 | 0.6 | 3.01E-02 | | | | | | | Auxin efflux carrier family protein |
| AT3G00800 | 1.3 | 3.64E-03 | -1.4 | 2.45E-02 | 2.6 | 1.20E-03 | -1.6 | 1.18E-03 | -0.9 | 3.02E-02 | -0.7 | 2.78E-01 | NA |
| AT1G49380 | 1.3 | 3.82E-06 | 1.1 | 1.67E-05 | 0.1 | 4.18E-01 | 0.3 | 4.62E-02 | 0.1 | 6.62E-01 | 0.2 | 2.73E-01 | Cytochrome c biogenesis protein CCS1, chloroplastic |
| AT4G16880 | 1.3 | 2.14E-04 | 0.6 | 3.26E-02 | 0.7 | 2.09E-02 | 1.2 | 2.37E-02 | 1.2 | 3.69E-02 | 0.0 | 9.98E-01 | Disease resistance RPP5 like protein |
| AT2G34510 | 1.3 | 2.22E-06 | 1.1 | 1.44E-05 | 0.2 | 1.85E-01 | -0.4 | 2.29E-03 | -0.2 | 9.84E-02 | -0.2 | 1.94E-01 | Uncharacterized protein At2g34510 |
| AT1G06950 | 1.3 | 1.46E-07 | 0.9 | 4.00E-06 | 0.4 | 1.51E-03 | 0.8 | 2.01E-05 | 0.1 | 5.95E-01 | 0.7 | 1.94E-03 | Protein TIC110, chloroplastic |
| AT1G14430 | 1.3 | 6.99E-06 | 1.2 | 2.11E-05 | 0.1 | 7.28E-01 | 0.2 | 3.66E-01 | -0.2 | 4.05E-01 | 0.5 | 1.38E-01 | Glyoxal oxidase-related protein |
| AT2G20270 | 1.3 | 3.20E-07 | 0.7 | 3.20E-05 | 0.5 | 5.87E-04 | 0.0 | 9.73E-01 | -0.1 | 6.84E-01 | 0.1 | 7.67E-01 | Thioredoxin superfamily protein |
| AT5G48480 | 1.3 | 5.01E-06 | 1.2 | 1.27E-05 | 0.0 | 8.49E-01 | 0.5 | 2.90E-02 | 0.1 | 7.18E-01 | 0.4 | 1.99E-01 | Uncharacterized protein At5g48480 |
| AT3G09050 | 1.3 | 5.83E-06 | 1.1 | 5.51E-05 | 0.2 | 1.52E-01 | 0.0 | 9.46E-01 | 0.1 | 7.14E-01 | -0.1 | 8.15E-01 | 8-amino-7-oxononanoate synthase |
| AT3G56010 | 1.3 | 2.35E-07 | 0.8 | 8.31E-06 | 0.4 | 1.58E-03 | 0.2 | 1.25E-01 | 0.2 | 2.68E-01 | 0.0 | 9.05E-01 | Transmembrane protein |
| AT5G44600 | 1.3 | 2.59E-05 | 0.9 | 9.68E-04 | 0.4 | 4.29E-02 | 0.2 | 2.97E-01 | 0.3 | 2.40E-01 | -0.1 | 8.81E-01 | S-adenosyl-L-methionine-dependent methyltransferases superfamily protein |
| AT2G28130 | 1.3 | 2.08E-06 | 0.5 | 5.23E-03 | 0.8 | 3.35E-04 | -0.5 | 8.16E-03 | -0.1 | 6.54E-01 | -0.4 | 9.68E-02 | Actin protein 2 |
| AT4G28750 | 1.3 | 1.50E-06 | 1.4 | 2.33E-06 | -0.1 | 4.57E-01 | -0.2 | 4.65E-01 | -0.1 | 7.67E-01 | -0.1 | 8.52E-01 | Putative photosystem I subunit PSI-E protein |
| AT2G41720 | 1.3 | 6.45E-07 | 1.1 | 4.13E-06 | 0.2 | 1.48E-01 | 0.2 | 1.95E-01 | 0.0 | 8.70E-01 | 0.2 | 2.45E-01 | Tetratricopeptide repeat -like superfamily protein |
| AT1G65486 | 1.3 | 5.73E-06 | 0.3 | 5.61E-02 | 1.0 | 2.93E-04 | 1.0 | 2.12E-04 | 0.6 | 2.63E-02 | 0.4 | 1.14E-01 | Transmembrane protein |
| AT3G54500 | 1.3 | 8.01E-05 | 0.5 | 2.05E-02 | 0.7 | 7.81E-03 | | | | | | | Uncharacterized protein At3g54500 |
| AT5G42110 | 1.3 | 3.23E-05 | 0.8 | 1.97E-03 | 0.5 | 2.71E-02 | 0.6 | 4.81E-02 | 0.2 | 7.03E-01 | 0.4 | 2.52E-01 | Uncharacterized protein |
| AT1G04107 | 1.3 | 1.14E-06 | 1.2 | 3.96E-06 | 0.1 | 6.01E-01 | 0.6 | 3.11E-02 | 0.2 | 5.94E-01 | 0.4 | 2.21E-01 | NA |
| AT5G10150 | 1.3 | 3.57E-06 | 1.1 | 2.16E-05 | 0.2 | 2.64E-01 | -0.4 | 1.66E-02 | -0.7 | 5.07E-04 | 0.3 | 1.01E-01 | UPSTREAM OF FLC protein |
| AT3G29240 | 1.3 | 4.97E-05 | 0.6 | 1.17E-02 | 0.7 | 6.87E-03 | -0.1 | 6.36E-01 | -0.5 | 2.28E-03 | 0.4 | 2.08E-02 | hypothetical protein |
| AT5G43050 | 1.3 | 4.45E-07 | 0.9 | 1.68E-05 | 0.4 | 3.62E-03 | 0.4 | 6.40E-02 | 0.1 | 7.68E-01 | 0.3 | 2.44E-01 | At5g43050 ) |
| AT1G29465 | 1.3 | 4.77E-05 | 1.0 | 6.22E-04 | 0.3 | 1.67E-01 | 0.5 | 1.14E-01 | -0.3 | 4.11E-01 | 0.8 | 3.82E-02 | Transmembrane protein |
| AT4G34090 | 1.3 | 1.24E-05 | 1.1 | 7.40E-05 | 0.2 | 2.99E-01 | 0.0 | 9.53E-01 | 0.0 | 9.32E-01 | 0.0 | 9.77E-01 | Cyclin delta-3 |
| AT4G25830 | 1.3 | 7.84E-07 | 1.2 | 3.08E-06 | 0.1 | 4.94E-01 | -0.3 | 3.59E-02 | -0.2 | 2.70E-01 | -0.1 | 5.69E-01 | CASP-like protein 2C1 |
| AT5G54630 | 1.3 | 3.62E-06 | 1.1 | 3.38E-05 | 0.2 | 1.18E-01 | 0.3 | 8.50E-02 | -0.2 | 4.82E-01 | 0.5 | 3.77E-02 | hypothetical protein |
| AT4G03450 | 1.3 | 4.73E-03 | 0.4 | 3.19E-01 | 0.9 | 7.50E-02 | 0.8 | 3.53E-01 | -0.6 | 6.26E-01 | 1.4 | 1.59E-01 | Ankyrin repeat family protein |
| AT1G51110 | 1.3 | 1.19E-04 | 0.8 | 4.84E-03 | 0.5 | 4.86E-02 | 0.2 | 2.86E-01 | 0.3 | 1.49E-01 | -0.1 | 7.13E-01 | Uncharacterized protein |
| AT1G73885 | 1.3 | 1.42E-06 | 0.8 | 6.39E-05 | 0.5 | 5.64E-03 | 0.0 | 8.65E-01 | -0.1 | 5.54E-01 | 0.1 | 5.26E-01 | AT-rich interactive domain protein |
| AT1G14030 | 1.3 | 1.88E-06 | 1.2 | 1.19E-05 | 0.2 | 2.31E-01 | 0.3 | 2.09E-01 | 0.1 | 5.94E-01 | 0.1 | 7.16E-01 | Uncharacterized protein |
| AT3G45160 | 1.3 | 2.73E-06 | 1.2 | 9.98E-06 | 0.1 | 5.67E-01 | 0.2 | 4.96E-01 | -0.1 | 7.85E-01 | 0.3 | 4.52E-01 | hypothetical protein |
| AT2G35480 | 1.3 | 4.13E-06 | 0.4 | 2.43E-02 | 0.9 | 3.72E-04 | 1.0 | 1.75E-03 | 0.3 | 4.38E-01 | 0.7 | 4.80E-02 | Envelope glycoprotein |
| AT2G04039 | 1.3 | 3.63E-05 | 0.6 | 1.13E-02 | 0.7 | 5.34E-03 | 0.6 | 3.48E-03 | 0.5 | 1.54E-02 | 0.1 | 7.60E-01 | At2g04039 |
| AT3G14110 | 1.3 | 1.08E-07 | 1.1 | 9.11E-07 | 0.2 | 1.98E-02 | 0.1 | 4.25E-01 | -0.1 | 3.55E-01 | 0.2 | 1.63E-01 | Tetratricopeptide repeat -like superfamily protein |
| AT5G05510 | 1.3 | 1.09E-06 | 0.9 | 4.19E-05 | 0.4 | 8.24E-03 | -0.1 | 8.49E-01 | -0.1 | 8.55E-01 | 0.0 | 9.88E-01 | hypothetical protein |
| AT1G21600 | 1.3 | 1.75E-07 | 1.0 | 2.32E-06 | 0.3 | 9.42E-03 | 0.3 | 5.97E-02 | -0.1 | 4.29E-01 | 0.4 | 4.65E-02 | PLASTID TRANSCRIPTIONALLY ACTIVE protein 6, chloroplastic |
| AT4G24090 | 1.3 | 1.16E-07 | 0.9 | 5.15E-06 | 0.5 | 5.63E-04 | 0.1 | 6.90E-01 | 0.0 | 9.88E-01 | 0.1 | 8.07E-01 | At4g24090 |
| AT3G48150 | 1.3 | 8.09E-07 | 0.7 | 2.27E-04 | 0.6 | 6.45E-04 | 0.0 | 9.30E-01 | 0.1 | 7.50E-01 | -0.1 | 8.46E-01 | Anaphase-promoting complex subunit 8 |
| AT3G06880 | 1.3 | 1.19E-05 | 0.8 | 7.58E-04 | 0.5 | 1.59E-02 | -0.5 | 4.08E-02 | -0.2 | 5.90E-01 | -0.4 | 3.17E-01 | Transducin |
| AT2G33830 | 1.4 | 5.75E-05 | 1.4 | 1.12E-04 | 0.0 | 9.93E-01 | -1.0 | 3.82E-03 | -1.7 | 1.76E-04 | 0.7 | 9.30E-02 | Dormancy-associated protein homolog 1 |
| AT4G25780 | 1.4 | 3.53E-06 | 1.0 | 9.69E-05 | 0.4 | 2.70E-02 | -0.2 | 5.51E-01 | -0.4 | 2.71E-01 | 0.2 | 6.74E-01 | SCP domain-containing protein |
| AT2G17240 | 1.4 | 6.68E-07 | 1.2 | 2.91E-06 | 0.1 | 3.44E-01 | 0.5 | 1.67E-02 | 0.0 | 9.90E-01 | 0.5 | 8.74E-02 | Arginine |
| AT5G51545 | 1.4 | 2.43E-07 | 1.3 | 1.04E-06 | 0.1 | 3.07E-01 | 0.1 | 6.96E-01 | -0.1 | 4.76E-01 | 0.2 | 3.74E-01 | Protein LPA2 |
| AT2G34640 | 1.4 | 3.29E-07 | 1.1 | 3.45E-06 | 0.3 | 2.90E-02 | 0.8 | 3.60E-05 | 0.1 | 6.19E-01 | 0.7 | 1.85E-03 | Protein PLASTID TRANSCRIPTIONALLY ACTIVE 12, chloroplastic |
| AT2G28780 | 1.4 | 1.84E-05 | 1.6 | 9.62E-06 | -0.3 | 1.52E-01 | -0.4 | 8.22E-02 | -0.4 | 9.82E-02 | 0.0 | 9.73E-01 | Uncharacterized protein At2g28780 |
| AT1G53140 | 1.4 | 1.05E-06 | 0.9 | 6.11E-05 | 0.5 | 4.13E-03 | 0.0 | 9.17E-01 | 0.0 | 9.46E-01 | 0.0 | 9.92E-01 | F12M16.1 |
| AT1G30610 | 1.4 | 1.19E-06 | 0.8 | 1.50E-04 | 0.6 | 1.74E-03 | 0.3 | 2.02E-01 | 0.1 | 8.43E-01 | 0.2 | 4.84E-01 | Pentatricopeptide repeat-containing protein |
| AT1G44575 | 1.4 | 8.15E-07 | 1.3 | 3.85E-06 | 0.1 | 3.69E-01 | 0.4 | 6.93E-03 | 0.4 | 4.30E-02 | 0.1 | 7.44E-01 | Chlorophyll A-B binding family protein |
| AT1G51100 | 1.4 | 2.79E-06 | 1.2 | 1.93E-05 | 0.2 | 1.93E-01 | 0.6 | 2.49E-02 | 0.2 | 5.40E-01 | 0.4 | 2.42E-01 | Protein CHLORORESPIRATORY REDUCTION 41, chloroplastic |
| AT3G63490 | 1.4 | 1.08E-07 | 1.1 | 8.62E-07 | 0.2 | 3.01E-02 | 0.2 | 2.94E-01 | 0.1 | 4.58E-01 | 0.0 | 9.37E-01 | Ribosomal protein |
| AT5G35480 | 1.4 | 1.55E-03 | 0.7 | 6.78E-02 | 0.6 | 9.68E-02 | 0.0 | 9.77E-01 | -0.2 | 6.35E-01 | 0.2 | 7.15E-01 | Uncharacterized protein |

| | | | | | | | | | | | | | |
|---|---|---|---|---|---|---|---|---|---|---|---|---|---|
| AT1G22690 | 1.4 | 2.92E-05 | 1.3 | 8.40E-05 | 0.1 | 7.32E-01 | 0.9 | 5.16E-03 | -0.2 | 6.69E-01 | 1.1 | 6.80E-03 | Gibberellin-regulated protein 9 |
| AT3G50560 | 1.4 | 2.18E-05 | 0.8 | 2.43E-03 | 0.6 | 1.44E-02 | -0.2 | 5.84E-01 | -1.4 | 4.31E-03 | 1.2 | 2.00E-02 | AT3g50560 |
| AT5G28500 | 1.4 | 5.24E-08 | 1.0 | 1.12E-06 | 0.4 | 6.34E-04 | 0.4 | 7.99E-02 | 0.3 | 2.64E-01 | 0.1 | 8.06E-01 | Rubisco accumulation factor 1.1, chloroplastic |
| AT1G80650 | 1.4 | 1.23E-06 | 1.1 | 1.50E-05 | 0.3 | 5.42E-02 | 0.2 | 3.24E-01 | -0.2 | 3.33E-01 | 0.5 | 1.07E-01 | RNAse THREE-like protein 1 |
| AT5G27250 | 1.4 | 1.10E-06 | 0.5 | 8.07E-03 | 0.9 | 1.66E-04 | 0.0 | 8.84E-01 | 0.2 | 5.34E-01 | -0.2 | 4.84E-01 | NA |
| AT5G55820 | 1.4 | 4.33E-07 | 0.9 | 1.92E-05 | 0.5 | 2.67E-03 | -0.1 | 5.36E-01 | 0.0 | 8.83E-01 | -0.1 | 7.99E-01 | Inner centromere protein, ARK-binding region protein |
| AT2G36145 | 1.4 | 1.43E-07 | 1.1 | 1.93E-06 | 0.3 | 6.58E-03 | 0.3 | 2.19E-01 | 0.4 | 7.13E-02 | -0.2 | 5.87E-01 | hypothetical protein |
| AT4G01150 | 1.4 | 4.22E-07 | 1.3 | 1.43E-06 | 0.1 | 6.88E-01 | -0.1 | 7.22E-01 | 0.1 | 6.18E-01 | -0.1 | 4.51E-01 | CURVATURE THYLAKOID 1A-like protein |
| AT5G52900 | 1.4 | 1.02E-06 | 0.5 | 2.18E-03 | 0.9 | 2.15E-04 | 0.0 | 9.37E-01 | 0.2 | 6.34E-01 | -0.2 | 5.97E-01 | Probable membrane-associated kinase regulator 6 |
| AT4G02660 | 1.4 | 1.65E-07 | 0.5 | 7.46E-04 | 0.9 | 3.70E-05 | 0.4 | 2.53E-01 | 0.3 | 3.90E-01 | 0.0 | 9.32E-01 | Beige |
| AT1G12250 | 1.4 | 7.30E-06 | 1.0 | 1.95E-04 | 0.4 | 2.98E-02 | 0.2 | 2.11E-01 | 0.2 | 2.75E-01 | 0.0 | 1.00E+00 | Pentapeptide repeat-containing protein |
| AT1G18060 | 1.4 | 5.17E-07 | 1.2 | 4.34E-06 | 0.2 | 7.60E-02 | 0.0 | 9.22E-01 | 0.1 | 6.11E-01 | -0.1 | 5.92E-01 | Uncharacterized protein |
| AT3G25882 | 1.4 | 5.75E-05 | -0.2 | 5.09E-01 | 1.6 | 3.45E-04 | 1.9 | 4.80E-04 | 1.1 | 4.23E-02 | 0.8 | 7.84E-02 | Protein NIM1-INTERACTING 2 |
| AT5G48470 | 1.4 | 4.51E-06 | 1.3 | 1.88E-05 | 0.1 | 4.62E-01 | 1.3 | 1.07E-05 | 0.6 | 2.86E-02 | 0.7 | 1.05E-02 | Protein PEP-RELATED DEVELOPMENT ARRESTED 1, chloroplastic |
| AT5G42530 | 1.4 | 2.95E-04 | 1.1 | 1.83E-03 | 0.3 | 3.81E-01 | 0.4 | 1.96E-02 | 0.0 | 9.89E-01 | 0.4 | 9.02E-02 | hypothetical protein |
| AT2G30820 | 1.4 | 1.60E-05 | 0.7 | 5.99E-03 | 0.7 | 5.09E-03 | -0.2 | 7.41E-01 | 0.2 | 6.79E-01 | -0.4 | 4.58E-01 | Protein JASON |
| AT5G44565 | 1.4 | 4.61E-05 | 0.6 | 2.29E-02 | 0.8 | 6.01E-03 | 0.8 | 6.13E-02 | 0.4 | 4.67E-01 | 0.4 | 4.48E-01 | Transmembrane protein |
| AT2G38823 | 1.4 | 3.55E-05 | 0.2 | 4.55E-01 | 1.2 | 7.71E-04 | 3.0 | 1.45E-07 | 0.2 | 7.60E-01 | 2.8 | 1.01E-05 | Uncharacterized protein |
| AT1G52220 | 1.4 | 1.69E-06 | 1.2 | 1.06E-05 | 0.2 | 2.28E-01 | 0.3 | 1.44E-01 | 0.2 | 3.67E-01 | 0.1 | 8.27E-01 | CURVATURE THYLAKOID protein |
| AT5G20790 | 1.4 | 9.15E-05 | 1.4 | 1.65E-04 | 0.0 | 9.93E-01 | 1.6 | 2.44E-03 | 0.6 | 3.81E-01 | 1.1 | 6.20E-02 | Transmembrane protein |
| AT1G49750 | 1.4 | 3.38E-06 | 1.1 | 3.50E-05 | 0.3 | 1.00E-01 | 0.9 | 4.01E-07 | 0.7 | 3.38E-05 | 0.2 | 2.46E-01 | AT1G49750 protein family protein) |
| AT4G01050 | 1.4 | 4.89E-08 | 1.1 | 6.75E-07 | 0.4 | 1.82E-03 | 0.3 | 8.72E-02 | 0.1 | 5.32E-01 | 0.1 | 5.26E-01 | Thylakoid rhodanese-like protein |
| AT4G22830 | 1.4 | 3.26E-07 | 1.1 | 6.14E-06 | 0.4 | 8.61E-03 | 0.1 | 6.38E-01 | 0.1 | 8.81E-01 | 0.1 | 8.67E-01 | YCF49-like protein |
| AT5G04190 | 1.4 | 1.15E-04 | 1.1 | 1.53E-03 | 0.3 | 1.89E-01 | | | | | | | Protein PHYTOCHROME KINASE SUBSTRATE 4 |
| AT2G34860 | 1.4 | 1.41E-07 | 1.1 | 1.47E-06 | 0.3 | 1.41E-02 | 0.3 | 3.57E-02 | 0.2 | 1.77E-01 | 0.1 | 7.42E-01 | hypothetical protein |
| AT5G18030 | 1.4 | 7.32E-04 | 1.9 | 1.52E-04 | -0.4 | 1.45E-01 | -0.2 | 7.25E-01 | 0.2 | 7.48E-01 | -0.4 | 3.88E-01 | Auxin-responsive protein SAUR21 |
| AT3G15770 | 1.4 | 2.51E-07 | 1.0 | 8.77E-06 | 0.4 | 2.42E-03 | -0.5 | 1.12E-01 | -0.7 | 6.13E-02 | 0.2 | 7.79E-01 | Uncharacterized protein At3g15770 |
| AT5G14920 | 1.4 | 1.02E-04 | 1.2 | 6.78E-04 | 0.3 | 3.34E-01 | -0.2 | 2.98E-01 | -0.8 | 6.26E-04 | 0.6 | 2.07E-02 | Gibberellin-regulated family protein |
| AT5G23060 | 1.4 | 5.24E-07 | 1.0 | 1.31E-05 | 0.4 | 6.88E-03 | 0.2 | 1.84E-01 | 0.3 | 1.19E-02 | -0.2 | 2.34E-01 | Calcium sensing receptor |
| AT1G05135 | 1.4 | 7.34E-05 | 1.6 | 6.74E-05 | -0.2 | 5.23E-01 | 0.1 | 4.89E-01 | -0.5 | 1.37E-02 | 0.6 | 1.74E-02 | NA |
| AT4G23170 | 1.4 | 3.56E-06 | 0.7 | 1.23E-03 | 0.8 | 1.23E-03 | 1.2 | 6.87E-07 | 0.6 | 2.41E-03 | 0.6 | 3.49E-03 | Putative cysteine-rich receptor-like protein kinase 9 |
| AT5G11780 | 1.4 | 8.57E-06 | 0.4 | 5.06E-02 | 1.0 | 7.12E-04 | -0.2 | 7.48E-01 | 0.5 | 3.13E-01 | -0.6 | 2.30E-01 | Uncharacterized protein |
| AT5G24280 | 1.4 | 2.19E-04 | -0.7 | 8.08E-02 | 2.1 | 4.44E-04 | 0.2 | 7.40E-01 | 0.1 | 8.59E-01 | 0.1 | 9.45E-01 | Gamma-irradiation and mitomycin c induced 1 |
| AT1G05385 | 1.4 | 1.34E-06 | 1.3 | 6.65E-06 | 0.1 | 3.20E-01 | 0.2 | 2.30E-01 | 0.2 | 3.74E-01 | 0.0 | 9.24E-01 | Photosystem II 11 kDa protein-like protein |
| AT3G23120 | 1.4 | 3.14E-04 | 0.0 | 9.07E-01 | 1.5 | 2.19E-03 | 2.3 | 6.79E-05 | 1.5 | 7.42E-03 | 0.8 | 3.59E-02 | Receptor-like protein 38 |
| AT5G11420 | 1.4 | 8.66E-07 | 1.5 | 2.02E-06 | 0.0 | 8.54E-01 | -0.3 | 3.49E-02 | -0.5 | 7.88E-03 | 0.1 | 4.97E-01 | hypothetical protein |
| AT1G55260 | 1.4 | 1.64E-05 | 1.1 | 2.09E-04 | 0.3 | 1.06E-01 | 0.2 | 3.59E-01 | 0.3 | 3.37E-01 | 0.0 | 9.21E-01 | Bifunctional inhibitor |
| AT1G67910 | 1.4 | 2.01E-05 | 1.4 | 4.07E-05 | 0.0 | 9.92E-01 | -0.2 | 5.11E-01 | -0.5 | 4.54E-02 | 0.4 | 2.07E-01 | Serine-rich protein-like protein |
| AT5G08600 | 1.4 | 3.64E-06 | 0.8 | 5.80E-04 | 0.6 | 3.80E-03 | 0.3 | 1.40E-01 | 0.2 | 5.46E-01 | 0.2 | 6.19E-01 | U3 ribonucleoprotein family protein |
| AT1G68560 | 1.4 | 5.91E-08 | 1.4 | 2.16E-07 | 0.1 | 5.81E-01 | 0.3 | 1.42E-02 | 0.0 | 9.85E-01 | 0.3 | 8.54E-02 | Alpha-xylosidase |
| AT1G78180 | 1.4 | 8.24E-07 | 0.9 | 4.65E-05 | 0.5 | 3.44E-03 | 0.2 | 4.47E-01 | -0.1 | 8.58E-01 | 0.2 | 4.54E-01 | Probable mitochondrial adenine nucleotide transporter BTL2 |
| AT5G02710 | 1.4 | 2.63E-06 | 0.9 | 1.72E-04 | 0.5 | 6.67E-03 | 0.0 | 8.62E-01 | -0.2 | 5.22E-01 | 0.2 | 4.20E-01 | At5g02710 |
| AT2G20890 | 1.4 | 4.72E-07 | 1.1 | 7.93E-06 | 0.4 | 1.48E-02 | 0.3 | 6.40E-03 | 0.2 | 9.58E-02 | 0.1 | 5.24E-01 | Protein THYLAKOID FORMATION 1, chloroplastic |
| AT3G14760 | 1.5 | 1.55E-04 | 1.0 | 3.90E-03 | 0.5 | 1.13E-01 | -0.8 | 7.12E-03 | -0.7 | 1.53E-02 | -0.1 | 9.25E-01 | hypothetical protein |
| AT5G53490 | 1.5 | 9.65E-08 | 1.5 | 2.35E-07 | -0.1 | 5.79E-01 | 0.1 | 5.68E-01 | 0.0 | 8.66E-01 | 0.1 | 5.73E-01 | Tetratricopeptide repeat -like superfamily protein |
| AT3G16320 | 1.5 | 5.97E-06 | 0.5 | 1.59E-02 | 0.9 | 7.18E-04 | 0.3 | 2.79E-01 | 0.0 | 9.65E-01 | 0.3 | 4.19E-01 | Tetratricopeptide repeat -like superfamily protein |
| AT2G40010 | 1.5 | 5.62E-05 | 0.7 | 1.83E-02 | 0.8 | 9.86E-03 | 0.9 | 8.46E-02 | 0.6 | 3.36E-01 | 0.3 | 6.80E-01 | 60S acidic ribosomal protein P0 |
| AT5G14120 | 1.5 | 1.83E-06 | 1.5 | 3.95E-06 | 0.0 | 8.37E-01 | -0.5 | 8.55E-02 | -1.3 | 7.46E-04 | 0.8 | 1.07E-02 | hypothetical protein |
| AT4G34550 | 1.5 | 7.94E-05 | 0.7 | 2.85E-02 | 0.8 | 9.60E-03 | | | | | | | hypothetical protein |
| AT3G62530 | 1.5 | 5.61E-08 | 1.3 | 3.32E-07 | 0.2 | 5.22E-02 | 0.7 | 2.78E-05 | 0.3 | 7.34E-02 | 0.5 | 1.98E-02 | Uncharacterized protein T12C14_230 |
| AT3G23880 | 1.5 | 2.49E-05 | 1.1 | 3.88E-04 | 0.4 | 1.13E-01 | -0.3 | 8.27E-02 | -1.1 | 6.59E-05 | 0.8 | 4.13E-03 | F-box |
| AT3G47070 | 1.5 | 1.41E-06 | 1.2 | 1.34E-05 | 0.3 | 8.78E-02 | -0.4 | 8.16E-02 | 0.4 | 8.95E-02 | -0.8 | 1.00E-02 | Thylakoid soluble phosphoprotein |
| AT5G40160 | 1.5 | 2.23E-07 | 0.9 | 1.44E-05 | 0.6 | 7.45E-04 | 0.3 | 7.15E-02 | 0.0 | 9.65E-01 | 0.3 | 1.03E-01 | Ankyrin repeat family protein |
| AT4G14100 | 1.5 | 7.25E-07 | 1.0 | 3.42E-05 | 0.5 | 3.80E-03 | 0.2 | 2.10E-01 | 0.1 | 5.05E-01 | 0.1 | 7.71E-01 | Uncharacterized protein |
| AT3G04550 | 1.5 | 1.25E-07 | 1.3 | 6.67E-07 | 0.2 | 1.22E-01 | 0.5 | 2.47E-03 | 0.2 | 2.42E-01 | 0.3 | 1.67E-01 | Rubisco accumulation factor 1.2, chloroplastic |
| AT2G47450 | 1.5 | 3.02E-07 | 1.1 | 6.78E-06 | 0.4 | 5.20E-03 | -0.1 | 4.95E-01 | -0.3 | 9.76E-03 | 0.3 | 9.33E-02 | Chromo domain-containing protein |
| AT4G37110 | 1.5 | 2.59E-06 | 1.0 | 1.08E-04 | 0.5 | 1.13E-02 | 0.1 | 7.63E-01 | 0.0 | 9.23E-01 | 0.0 | 9.11E-01 | Zinc-finger domain of monoamine-oxidase A repressor R1 |
| AT5G22390 | 1.5 | 8.05E-06 | 1.4 | 3.40E-05 | 0.1 | 4.69E-01 | 0.2 | 3.80E-01 | 0.0 | 9.55E-01 | 0.2 | 4.55E-01 | FAF domain-containing protein |
| AT5G56850 | 1.5 | 2.09E-06 | 1.0 | 5.70E-05 | 0.4 | 1.54E-02 | 0.1 | 7.82E-01 | 0.2 | 1.68E-01 | -0.2 | 2.68E-01 | At5g56850 |
| AT4G12917 | 1.5 | 1.08E-04 | 0.8 | 9.98E-03 | 0.7 | 2.89E-02 | 0.6 | 2.75E-02 | 0.3 | 2.97E-01 | 0.3 | 4.49E-01 | NA |
| AT2G14880 | 1.5 | 1.48E-06 | 1.2 | 1.29E-05 | 0.3 | 9.37E-02 | 0.6 | 4.17E-03 | 0.1 | 7.31E-01 | 0.5 | 6.84E-02 | DM2 domain-containing protein |
| AT4G01270 | 1.5 | 3.62E-06 | 1.0 | 2.65E-04 | 0.5 | 8.26E-03 | 0.8 | 9.66E-02 | 0.6 | 3.09E-01 | 0.2 | 7.32E-01 | RING-type domain-containing protein |
| AT5G41050 | 1.5 | 3.07E-05 | 0.9 | 1.76E-03 | 0.6 | 2.57E-02 | 0.3 | 2.07E-01 | 0.1 | 7.48E-01 | 0.2 | 5.81E-01 | hypothetical protein |
| AT2G30600 | 1.5 | 5.69E-05 | 1.2 | 5.39E-04 | 0.3 | 1.97E-01 | -0.1 | 8.36E-01 | -0.8 | 2.00E-01 | 0.7 | 3.42E-01 | BTB |
| AT3G17185 | 1.5 | 4.25E-05 | 1.2 | 4.68E-04 | 0.3 | 1.89E-01 | 0.2 | 5.58E-01 | 0.2 | 5.34E-01 | 0.0 | 9.47E-01 | NA |
| AT4G38840 | 1.5 | 5.98E-06 | 1.3 | 2.91E-05 | 0.2 | 3.57E-01 | 0.4 | 1.44E-01 | 0.0 | 9.43E-01 | 0.3 | 3.30E-01 | Uncharacterized protein |
| AT1G74730 | 1.5 | 2.04E-07 | 1.3 | 1.56E-06 | 0.2 | 5.75E-02 | 0.1 | 7.95E-01 | 0.2 | 3.02E-01 | -0.2 | 5.43E-01 | Transmembrane protein, putative |
| AT5G22640 | 1.5 | 3.84E-08 | 1.2 | 4.25E-07 | 0.3 | 2.53E-03 | 0.3 | 1.29E-02 | -0.1 | 5.49E-01 | 0.4 | 1.52E-02 | MORN repeat-containing protein |
| AT4G27700 | 1.5 | 1.25E-07 | 1.3 | 7.96E-07 | 0.2 | 6.97E-02 | 0.3 | 1.94E-02 | 0.2 | 3.35E-01 | 0.2 | 3.66E-01 | Rhodanese-like domain-containing protein 14, chloroplastic |
| AT4G11100 | 1.5 | 1.09E-07 | 1.2 | 1.67E-06 | 0.4 | 5.06E-03 | -0.1 | 5.22E-01 | -0.3 | 4.60E-02 | 0.2 | 2.80E-01 | Gelsolin protein |
| AT5G64040 | 1.5 | 9.76E-05 | 1.5 | 2.15E-04 | 0.0 | 8.93E-01 | -0.8 | 1.84E-04 | 0.0 | 9.01E-01 | -0.7 | 2.99E-03 | Photosystem I reaction center subunit PSI-N, chloroplast, putative |
| AT4G38860 | 1.5 | 4.82E-05 | 1.4 | 1.87E-04 | 0.1 | 5.44E-01 | -0.6 | 1.25E-02 | -0.4 | 1.11E-01 | -0.2 | 5.27E-01 | Protein SMALL AUXIN UP-REGULATED RNA 16 |
| AT3G05730 | 1.6 | 1.24E-05 | 1.3 | 1.00E-04 | 0.3 | 2.04E-01 | -0.1 | 5.75E-01 | -0.2 | 3.47E-01 | 0.1 | 7.71E-01 | hypothetical protein |
| AT3G12965 | 1.6 | 1.36E-05 | 1.0 | 6.61E-04 | 0.6 | 1.91E-02 | 0.6 | 5.49E-02 | 0.4 | 3.40E-01 | 0.3 | 6.05E-01 | NA |
| AT5G02940 | 1.6 | 1.62E-05 | 1.1 | 4.89E-04 | 0.5 | 3.51E-02 | 0.6 | 2.03E-02 | 0.7 | 7.83E-03 | -0.2 | 6.21E-01 | Ion channel POLLUX-like protein, putative |
| AT1G14345 | 1.6 | 5.59E-08 | 1.1 | 8.96E-07 | 0.4 | 1.27E-03 | 0.1 | 6.51E-01 | 0.1 | 7.55E-01 | 0.0 | 9.81E-01 | hypothetical protein |
| AT5G02430 | 1.6 | 1.21E-05 | 0.8 | 3.13E-03 | 0.8 | 4.52E-03 | -0.1 | 7.79E-01 | 0.4 | 2.01E-01 | -0.5 | 1.73E-01 | Transducin |
| AT5G07020 | 1.6 | 5.56E-08 | 1.3 | 4.77E-07 | 0.3 | 9.42E-03 | 0.3 | 7.53E-02 | 0.3 | 5.13E-02 | -0.1 | 8.00E-01 | Protein MAINTENANCE OF PSII UNDER HIGH LIGHT 1 |
| AT4G34760 | 1.6 | 3.67E-06 | 1.2 | 7.35E-05 | 0.4 | 3.82E-02 | -0.6 | 1.68E-02 | -0.5 | 5.54E-02 | -0.1 | 8.48E-01 | Auxin-responsive protein SAUR50 |
| AT3G45730 | 1.6 | 5.40E-05 | 0.2 | 4.64E-01 | 1.4 | 7.30E-04 | 0.2 | 3.20E-01 | -0.3 | 2.14E-01 | 0.5 | 7.90E-02 | Uncharacterized protein |
| AT1G33615 | 1.6 | 5.34E-05 | 0.6 | 5.42E-02 | 1.0 | 3.98E-03 | -0.1 | 7.67E-01 | 0.1 | 7.76E-01 | -0.2 | 6.14E-01 | NA |
| AT1G55480 | 1.6 | 1.69E-06 | 1.1 | 3.15E-05 | 0.4 | 2.47E-02 | 0.4 | 4.59E-04 | 0.4 | 3.98E-03 | 0.1 | 7.04E-01 | Protein MET1, chloroplastic |
| AT5G26170 | 1.6 | 3.24E-05 | 0.0 | 8.87E-01 | 1.5 | 5.50E-04 | 0.5 | 2.69E-01 | 0.0 | 9.90E-01 | 0.5 | 3.58E-01 | WRKY50 |

| | | | | | | | | | | | | | |
|---|---|---|---|---|---|---|---|---|---|---|---|---|---|
| AT1G14680 | 1.6 | 3.69E-06 | 0.8 | 2.08E-03 | 0.8 | 1.45E-03 | -0.2 | 6.01E-01 | -0.6 | 7.62E-02 | 0.4 | 2.92E-01 | Uncharacterized protein |
| AT2G33330 | 1.6 | 3.18E-06 | 1.5 | 1.19E-05 | 0.1 | 5.44E-01 | 0.2 | 3.93E-01 | -0.1 | 7.94E-01 | 0.2 | 3.58E-01 | Gnk2-homologous domain-containing protein |
| AT5G52780 | 1.6 | 1.80E-07 | 1.3 | 1.66E-06 | 0.3 | 2.62E-02 | 0.6 | 3.63E-03 | 0.3 | 1.62E-01 | 0.3 | 2.78E-01 | Uncharacterized protein PAM68-like |
| AT1G23150 | 1.6 | 3.42E-07 | 1.6 | 6.81E-07 | -0.1 | 6.46E-01 | 0.8 | 2.45E-02 | 0.0 | 9.42E-01 | 0.7 | 3.88E-02 | Senescence-associated protein |
| AT5G38870 | 1.6 | 2.74E-06 | 1.7 | 3.09E-06 | -0.2 | 2.99E-01 | -0.6 | 5.14E-02 | -0.1 | 8.40E-01 | -0.5 | 1.95E-01 | NA |
| AT1G06680 | 1.6 | 1.58E-07 | 1.5 | 5.71E-07 | 0.1 | 5.05E-01 | 0.2 | 1.86E-01 | 0.1 | 4.30E-01 | 0.1 | 8.40E-01 | AT1G06680 protein |
| AT3G25480 | 1.6 | 7.38E-08 | 1.1 | 1.68E-06 | 0.5 | 1.10E-03 | 0.4 | 4.17E-02 | 0.3 | 1.44E-01 | 0.1 | 8.05E-01 | Rhodanese-like domain-containing protein 4A, chloroplastic |
| AT2G17780 | 1.6 | 9.75E-07 | 1.2 | 1.90E-05 | 0.4 | 1.82E-02 | -0.1 | 8.37E-01 | -0.2 | 3.37E-01 | 0.2 | 4.93E-01 | PLAC8 family protein |
| AT4G24700 | 1.6 | 3.82E-05 | 0.8 | 1.02E-02 | 0.8 | 8.04E-03 | 0.7 | 8.24E-02 | 0.3 | 5.31E-01 | 0.4 | 3.97E-01 | Uncharacterized protein |
| AT2G47910 | 1.6 | 3.66E-08 | 1.1 | 1.07E-06 | 0.5 | 2.06E-04 | 0.1 | 6.52E-01 | -0.1 | 4.85E-01 | 0.2 | 3.05E-01 | Chlororespiratory reduction 6 |
| AT4G08380 | 1.6 | 1.78E-06 | 1.3 | 1.59E-05 | 0.3 | 1.01E-01 | 0.3 | 1.39E-01 | 0.2 | 4.89E-01 | 0.1 | 7.16E-01 | Extensin-like protein |
| AT5G46710 | 1.6 | 2.06E-05 | -0.1 | 6.24E-01 | 1.7 | 2.02E-04 | 0.4 | 2.94E-01 | -0.2 | 7.99E-01 | 0.6 | 1.72E-01 | PLATZ transcription factor family protein |
| AT3G47560 | 1.6 | 1.63E-06 | 1.2 | 2.80E-05 | 0.4 | 3.17E-02 | 0.1 | 8.59E-01 | -0.1 | 8.53E-01 | 0.1 | 7.44E-01 | Alpha |
| AT3G07430 | 1.6 | 1.68E-06 | 1.0 | 6.54E-05 | 0.6 | 7.21E-03 | 0.7 | 2.17E-04 | 0.0 | 9.66E-01 | 0.7 | 2.17E-03 | YlmG homolog protein 1-1, chloroplastic |
| AT5G36120 | 1.6 | 7.71E-08 | 1.3 | 6.13E-07 | 0.3 | 1.81E-02 | 0.7 | 6.74E-05 | 0.4 | 2.29E-02 | 0.3 | 7.45E-02 | Protein COFACTOR ASSEMBLY OF COMPLEX C SUBUNIT B CCB3, chloroplastic |
| AT1G30380 | 1.6 | 1.84E-07 | 1.6 | 5.99E-07 | 0.0 | 7.15E-01 | -0.4 | 7.48E-03 | -0.1 | 4.97E-01 | -0.3 | 1.62E-01 | PSI-K |
| AT2G24395 | 1.6 | 1.84E-05 | 1.0 | 1.47E-03 | 0.6 | 1.54E-02 | 0.1 | 8.66E-01 | 0.0 | 9.69E-01 | 0.1 | 8.78E-01 | Uncharacterized protein |
| AT4G38700 | 1.6 | 1.30E-05 | 1.0 | 1.11E-03 | 0.6 | 1.41E-02 | | | | | | | Dirigent protein |
| AT3G22060 | 1.6 | 1.20E-05 | 0.9 | 9.00E-04 | 0.7 | 9.51E-03 | -0.2 | 7.19E-01 | -1.4 | 8.22E-03 | 1.2 | 4.75E-02 | Cysteine-rich repeat secretory protein 38 |
| AT4G13572 | 1.6 | 3.07E-06 | 1.6 | 6.73E-06 | 0.0 | 9.28E-01 | 0.1 | 8.53E-01 | 0.1 | 8.82E-01 | 0.0 | 9.98E-01 | Uncharacterized protein |
| AT1G53640 | 1.6 | 1.11E-05 | 1.3 | 1.37E-04 | 0.3 | 1.11E-01 | -0.1 | 7.32E-01 | 0.1 | 8.37E-01 | -0.2 | 5.88E-01 | Transmembrane protein |
| AT1G18730 | 1.6 | 2.51E-05 | 1.1 | 5.92E-04 | 0.5 | 5.85E-02 | 0.2 | 1.03E-01 | 0.2 | 2.89E-01 | 0.1 | 8.04E-01 | NDH dependent flow 6 |
| AT1G66465 | 1.6 | 9.21E-04 | 0.5 | 2.67E-01 | 1.1 | 2.30E-02 | 0.0 | 9.73E-01 | -0.3 | 6.05E-01 | 0.2 | 6.84E-01 | Transmembrane protein |
| AT1G52230 | 1.6 | 8.74E-06 | 1.4 | 5.62E-05 | 0.2 | 2.73E-01 | 0.5 | 1.47E-03 | 0.4 | 5.70E-03 | 0.0 | 9.06E-01 | Photosystem I reaction center subunit VI-2, chloroplastic |
| AT1G11180 | 1.6 | 2.67E-06 | 1.0 | 1.66E-04 | 0.6 | 6.82E-03 | -0.3 | 4.02E-02 | -0.3 | 1.03E-01 | 0.0 | 8.94E-01 | Secretory carrier-associated membrane protein |
| AT3G10405 | 1.6 | 3.44E-07 | 1.1 | 1.18E-05 | 0.5 | 3.04E-03 | -0.1 | 7.84E-01 | -0.1 | 7.98E-01 | 0.0 | 9.77E-01 | hypothetical protein |
| AT5G19850 | 1.6 | 4.44E-07 | 0.7 | 5.42E-04 | 1.0 | 1.26E-04 | 0.9 | 2.67E-03 | 0.4 | 2.55E-01 | 0.5 | 8.16E-02 | AB hydrolase-1 domain-containing protein |
| AT1G11850 | 1.6 | 7.17E-06 | 1.3 | 7.25E-05 | 0.3 | 1.25E-01 | -0.2 | 2.89E-01 | -0.1 | 6.83E-01 | -0.1 | 7.38E-01 | Transmembrane protein |
| AT1G54740 | 1.6 | 7.96E-06 | 1.5 | 3.21E-05 | 0.1 | 4.96E-01 | -0.4 | 9.90E-02 | -1.3 | 1.39E-04 | 0.9 | 1.14E-02 | T22H22.16 protein |
| AT3G50570 | 1.7 | 3.47E-05 | 0.4 | 1.36E-01 | 1.2 | 1.63E-03 | -0.5 | 2.54E-01 | -0.1 | 8.99E-01 | -0.4 | 4.18E-01 | At3g50570 |
| AT3G16670 | 1.7 | 1.14E-03 | 1.5 | 3.40E-03 | 0.2 | 6.79E-01 | -0.1 | 5.96E-01 | -0.7 | 1.38E-02 | 0.5 | 4.92E-02 | hypothetical protein |
| AT1G49975 | 1.7 | 1.35E-06 | 1.4 | 1.08E-05 | 0.3 | 1.22E-01 | 0.4 | 8.28E-02 | 0.2 | 4.57E-01 | 0.2 | 5.81E-01 | Uncharacterized protein |
| AT3G09162 | 1.7 | 3.21E-05 | 1.1 | 1.76E-03 | 0.6 | 2.94E-02 | 0.8 | 7.17E-02 | 0.3 | 6.72E-01 | 0.5 | 2.82E-01 | At3g09162 |
| AT5G15240 | 1.7 | 3.90E-06 | 1.4 | 3.60E-05 | 0.3 | 1.43E-01 | 0.0 | 9.18E-01 | 0.3 | 4.78E-01 | -0.3 | 3.97E-01 | Transmembrane amino acid transporter family protein |
| AT1G16445 | 1.7 | 1.14E-06 | 1.4 | 1.37E-05 | 0.3 | 5.51E-02 | 0.2 | 3.52E-01 | -0.4 | 1.15E-01 | 0.7 | 4.53E-02 | S-adenosyl-L-methionine-dependent methyltransferases superfamily protein |
| AT1G29715 | 1.7 | 2.71E-05 | 0.2 | 3.93E-01 | 1.4 | 7.26E-04 | | | | | | | NA |
| AT5G24165 | 1.7 | 1.06E-04 | 1.2 | 1.87E-03 | 0.5 | 1.08E-01 | 0.4 | 1.77E-01 | 0.2 | 5.39E-01 | 0.2 | 7.42E-01 | hypothetical protein |
| AT2G24762 | 1.7 | 1.30E-06 | 1.9 | 1.48E-06 | -0.2 | 2.37E-01 | 1.2 | 4.57E-05 | 0.3 | 3.15E-01 | 0.9 | 6.07E-03 | Protein GLUTAMINE DUMPER 4 |
| AT3G04650 | 1.7 | 1.97E-08 | 1.1 | 6.49E-07 | 0.6 | 6.46E-05 | 0.8 | 4.60E-04 | 0.3 | 2.27E-01 | 0.5 | 3.98E-02 | AT3g04650 |
| AT3G21055 | 1.7 | 1.45E-07 | 1.9 | 2.38E-07 | -0.2 | 1.68E-01 | -0.2 | 7.66E-02 | -0.1 | 8.02E-01 | -0.2 | 3.23E-01 | Photosystem II subunit T |
| AT4G31805 | 1.7 | 2.48E-05 | 0.9 | 4.60E-03 | 0.8 | 9.62E-03 | 0.2 | 5.05E-01 | 0.2 | 7.20E-01 | 0.1 | 8.98E-01 | Protein POLAR LOCALIZATION DURING ASYMMETRIC DIVISION AND REDISTRIBUTION |
| AT2G01860 | 1.7 | 1.22E-06 | 0.9 | 2.31E-04 | 0.7 | 1.35E-03 | 0.5 | 1.42E-01 | -0.1 | 7.99E-01 | 0.6 | 1.37E-01 | Pentatricopeptide repeat-containing protein At2g01860 |
| AT2G20110 | 1.7 | 6.73E-06 | 0.7 | 6.58E-03 | 1.0 | 1.42E-03 | 0.1 | 8.19E-01 | -0.1 | 8.73E-01 | 0.2 | 7.58E-01 | Tesmin |
| AT2G29180 | 1.7 | 1.20E-05 | 1.1 | 8.52E-04 | 0.6 | 1.27E-02 | 0.4 | 5.86E-02 | 0.1 | 7.23E-01 | 0.3 | 2.17E-01 | Transmembrane protein |
| AT1G50040 | 1.7 | 1.82E-03 | 1.0 | 4.18E-02 | 0.7 | 1.59E-01 | -1.0 | 2.00E-02 | -1.7 | 2.76E-03 | 0.7 | 3.53E-01 | F2J10.8 protein ) |
| AT4G21445 | 1.7 | 2.82E-05 | 1.2 | 8.09E-04 | 0.5 | 5.02E-02 | 0.8 | 2.90E-03 | 0.5 | 6.59E-02 | 0.3 | 3.80E-01 | Receptor-interacting protein |
| AT3G18250 | 1.7 | 1.38E-02 | 0.5 | 4.53E-01 | 1.2 | 1.03E-01 | 1.0 | 7.11E-02 | -1.2 | 1.49E-01 | 2.1 | 1.01E-02 | Transmembrane protein |
| AT3G14900 | 1.7 | 1.83E-06 | 1.1 | 9.04E-05 | 0.6 | 6.58E-03 | 1.6 | 4.09E-06 | 0.5 | 1.18E-01 | 1.1 | 7.72E-04 | Uncharacterized protein At3g14900 |
| AT1G18620 | 1.7 | 1.96E-06 | 1.5 | 1.06E-05 | 0.2 | 2.75E-01 | -0.1 | 5.37E-01 | -0.5 | 6.11E-03 | 0.4 | 5.67E-02 | LONGIFOLIA protein |
| AT4G35770 | 1.7 | 1.44E-02 | 1.5 | 3.27E-02 | 0.2 | 7.63E-01 | -0.7 | 3.27E-01 | -2.4 | 6.82E-03 | 1.8 | 1.00E-02 | Rhodanese |
| AT4G19100 | 1.7 | 3.20E-06 | 1.6 | 1.46E-05 | 0.1 | 4.09E-01 | 0.1 | 7.89E-01 | 0.0 | 9.23E-01 | 0.0 | 9.24E-01 | PAM68-like protein |
| AT1G24147 | 1.7 | 1.00E-04 | 0.8 | 1.93E-02 | 0.9 | 1.39E-02 | 1.5 | 3.95E-03 | 0.4 | 5.21E-01 | 1.1 | 6.58E-02 | Transmembrane protein |
| AT1G72640 | 1.7 | 2.62E-06 | 1.2 | 7.94E-05 | 0.5 | 1.52E-02 | -0.1 | 6.44E-01 | 0.0 | 9.76E-01 | -0.1 | 7.76E-01 | At1g72640 -binding Rossmann-fold superfamily protein) |
| AT2G44920 | 1.7 | 8.20E-08 | 1.6 | 2.97E-07 | 0.1 | 4.46E-01 | 0.0 | 9.39E-01 | -0.2 | 3.74E-01 | 0.2 | 4.40E-01 | Thylakoid lumenal 15 kDa protein 1, chloroplastic |
| AT4G28025 | 1.7 | 4.96E-06 | 1.1 | 2.02E-04 | 0.6 | 1.28E-02 | 0.5 | 6.94E-03 | 0.2 | 4.50E-01 | 0.3 | 1.79E-01 | Uncharacterized protein AT4g28020 |
| AT2G37800 | 1.7 | 4.54E-05 | 2.6 | 2.97E-06 | -0.9 | 2.48E-03 | | | | | | | Phorbol-ester |
| AT5G44650 | 1.7 | 1.67E-08 | 1.2 | 2.42E-07 | 0.5 | 1.38E-04 | 0.5 | 2.82E-04 | 0.3 | 4.90E-02 | 0.2 | 1.14E-01 | Ycf3-interacting protein 1, chloroplastic |
| AT2G05520 | 1.7 | 5.94E-07 | 1.5 | 3.96E-06 | 0.2 | 1.44E-01 | -0.2 | 3.91E-01 | -0.5 | 2.05E-02 | 0.3 | 2.08E-01 | Glycine-rich protein 3 |
| AT1G15980 | 1.7 | 1.43E-07 | 1.3 | 1.97E-06 | 0.4 | 6.79E-03 | 0.3 | 6.33E-02 | 0.4 | 7.32E-02 | 0.0 | 9.31E-01 | Photosynthetic NDH subunit of subcomplex B 1, chloroplastic H DEHYDROGENASE SUBUNIT 48) |
| AT5G05180 | 1.7 | 6.03E-06 | 0.9 | 1.84E-03 | 0.9 | 2.35E-03 | -0.3 | 1.36E-02 | -0.6 | 4.92E-04 | 0.3 | 1.62E-01 | Myosin heavy chain, striated protein |
| AT2G42900 | 1.7 | 1.51E-06 | 1.7 | 5.16E-06 | 0.1 | 6.83E-01 | -0.2 | 3.56E-01 | -0.4 | 1.77E-01 | 0.1 | 7.33E-01 | Plant basic secretory protein family protein |
| AT1G32928 | 1.7 | 8.55E-05 | 1.8 | 1.31E-04 | 0.0 | 9.32E-01 | 1.2 | 2.11E-03 | -0.6 | 2.75E-01 | 1.8 | 2.47E-03 | Avr9 |
| AT5G18020 | 1.7 | 5.23E-04 | 2.1 | 2.18E-04 | -0.3 | 3.25E-01 | | | | | | | Auxin-responsive protein SAUR20 |
| AT4G23770 | 1.7 | 1.82E-05 | 1.1 | 1.26E-03 | 0.7 | 1.69E-02 | 0.4 | 4.37E-01 | -0.1 | 8.86E-01 | 0.5 | 4.40E-01 | hypothetical protein |
| AT2G10940 | 1.8 | 6.63E-07 | 1.7 | 2.30E-06 | 0.1 | 6.61E-01 | -0.9 | 3.11E-04 | -0.6 | 2.30E-02 | -0.4 | 1.59E-01 | At2g10940 |
| AT1G09340 | 1.8 | 5.59E-08 | 1.6 | 2.77E-07 | 0.2 | 1.15E-01 | 0.3 | 1.47E-02 | 0.3 | 6.28E-02 | 0.1 | 8.15E-01 | Chloroplast RNA binding protein |
| AT3G13210 | 1.8 | 2.48E-06 | 0.6 | 9.23E-03 | 1.1 | 3.87E-04 | | | | | | | Crooked neck protein, putative |
| AT5G06870 | 1.8 | 2.97E-06 | 1.3 | 5.53E-05 | 0.4 | 3.84E-02 | 0.0 | 9.43E-01 | -0.2 | 7.71E-01 | 0.1 | 8.58E-01 | Polygalacturonase inhibiting protein |
| AT2G44380 | 1.8 | 9.09E-07 | 1.7 | 3.23E-06 | 0.1 | 5.46E-01 | 0.4 | 9.68E-02 | 0.4 | 8.98E-02 | 0.0 | 8.73E-01 | hypothetical protein |
| AT2G35500 | 1.8 | 1.89E-08 | 1.5 | 1.12E-07 | 0.3 | 4.32E-03 | 0.6 | 1.26E-03 | 0.2 | 1.82E-01 | 0.3 | 1.03E-01 | Probable inactive shikimate kinase like 2, chloroplastic |
| AT5G52970 | 1.8 | 2.62E-07 | 1.9 | 4.74E-07 | -0.1 | 4.79E-01 | 0.2 | 2.86E-01 | 0.0 | 9.53E-01 | 0.1 | 4.93E-01 | Thylakoid lumenal 15.0 kDa protein 2, chloroplastic |
| AT2G01300 | 1.8 | 1.12E-04 | 0.9 | 2.02E-02 | 0.9 | 1.82E-02 | -0.3 | 5.78E-01 | -0.5 | 4.27E-01 | | | hypothetical protein |
| AT2G25510 | 1.8 | 2.48E-04 | 1.4 | 1.48E-03 | 0.3 | 3.91E-01 | 0.5 | 2.29E-02 | 0.0 | 9.22E-01 | 0.5 | 1.13E-01 | Transmembrane protein |
| AT3G08920 | 1.8 | 4.25E-08 | 1.6 | 2.01E-07 | 0.1 | 1.48E-01 | 0.1 | 7.49E-01 | -0.2 | 4.92E-01 | 0.2 | 3.71E-01 | Rhodanese-like domain-containing protein 10 |
| AT5G15780 | 1.8 | 4.25E-08 | 1.4 | 3.99E-07 | 0.4 | 5.17E-03 | -0.8 | 1.00E-03 | -0.4 | 8.33E-02 | -0.4 | 1.31E-01 | At5g15780 |
| AT3G23760 | 1.8 | 5.18E-06 | 1.5 | 5.35E-05 | 0.3 | 1.13E-01 | 0.2 | 4.27E-01 | -0.1 | 7.11E-01 | 0.3 | 3.09E-01 | hypothetical protein |
| AT5G36000 | 1.8 | 2.75E-05 | 0.7 | 2.46E-02 | 1.1 | 3.38E-03 | | | | | | | Probable F-box protein At5g36000 |
| AT5G57345 | 1.8 | 1.67E-07 | 1.6 | 8.76E-07 | 0.2 | 1.58E-01 | -0.3 | 2.95E-01 | 0.1 | 8.28E-01 | -0.4 | 2.80E-01 | At5g57345 |
| AT3G20760 | 1.8 | 1.05E-04 | 1.1 | 4.68E-03 | 0.7 | 4.61E-02 | 0.6 | 9.68E-02 | 0.2 | 6.63E-01 | 0.4 | 4.27E-01 | Non-structural maintenance of chromosomes element 4 homolog B |
| AT5G39210 | 1.8 | 2.97E-06 | 1.4 | 4.87E-05 | 0.4 | 4.15E-02 | 0.4 | 1.02E-01 | 0.2 | 6.45E-01 | 0.3 | 4.75E-01 | Protein CHLORORESPIRATORY REDUCTION 7, chloroplastic |
| AT4G34830 | 1.8 | 2.78E-07 | 1.1 | 1.32E-05 | 0.7 | 1.09E-03 | 0.4 | 8.06E-02 | 0.2 | 4.23E-01 | 0.2 | 5.43E-01 | Pentatricopeptide repeat-containing protein MRL1, chloroplastic |
| AT5G35490 | 1.8 | 5.82E-04 | 1.1 | 2.19E-02 | 0.7 | 8.93E-02 | | | | | | | Uncharacterized protein |
| AT1G10417 | 1.8 | 3.90E-05 | 0.5 | 1.50E-01 | 1.3 | 1.82E-03 | 0.4 | 3.21E-01 | 0.2 | 7.46E-01 | 0.2 | 6.51E-01 | T10O24.3 |

| | | | | | | | | | | | | | |
|---|---|---|---|---|---|---|---|---|---|---|---|---|---|
| AT4G24750 | 1.8 | 9.53E-08 | 1.4 | 1.03E-06 | 0.4 | 7.50E-03 | 0.3 | 2.15E-02 | -0.1 | 5.03E-01 | 0.4 | 2.89E-02 | Rhodanese-like domain-containing protein 11, chloroplastic |
| AT1G50900 | 1.8 | 2.25E-08 | 1.3 | 4.72E-07 | 0.5 | 2.41E-04 | 0.3 | 1.29E-01 | 0.1 | 8.06E-01 | 0.3 | 3.53E-01 | Protein LHCP TRANSLOCATION DEFECT |
| AT3G06145 | 1.8 | 2.43E-07 | 1.3 | 7.15E-06 | 0.5 | 2.49E-03 | 0.8 | 1.53E-03 | 0.2 | 4.72E-01 | 0.6 | 3.93E-02 | At3g06145 |
| AT2G05310 | 1.8 | 8.94E-05 | 0.9 | 1.12E-02 | 0.9 | 1.53E-02 | 0.1 | 8.19E-01 | 0.0 | 8.98E-01 | 0.1 | 7.93E-01 | Expressed protein |
| AT4G16240 | 1.8 | 1.44E-04 | 1.3 | 2.90E-03 | 0.5 | 1.15E-01 | 0.6 | 3.96E-01 | -0.6 | 5.59E-01 | 1.2 | 2.28E-01 | Glycine-rich protein homolog |
| AT2G33855 | 1.8 | 1.18E-05 | 0.9 | 3.67E-03 | 0.9 | 3.14E-03 | 0.1 | 7.87E-01 | 0.1 | 8.93E-01 | 0.0 | 9.62E-01 | hypothetical protein |
| AT5G52710 | 1.8 | 1.25E-05 | 0.7 | 1.58E-02 | 1.1 | 2.02E-03 | 2.8 | 1.84E-07 | 1.6 | 4.18E-04 | 1.2 | 1.83E-04 | Copper transport protein family |
| AT2G38210 | 1.8 | 2.67E-06 | 0.7 | 5.87E-03 | 1.1 | 3.80E-04 | -0.2 | 5.72E-01 | -0.3 | 4.04E-01 | 0.1 | 8.39E-01 | Pyridoxal 5'-phosphate synthase PDX1-like 4 |
| AT5G48790 | 1.8 | 1.48E-07 | 1.4 | 2.02E-06 | 0.4 | 6.15E-03 | 0.3 | 2.88E-02 | 0.1 | 7.13E-01 | 0.2 | 2.07E-01 | LOW PSII ACCUMULATION protein |
| AT2G41990 | 1.8 | 6.24E-06 | 2.2 | 2.82E-06 | -0.4 | 4.50E-02 | 0.4 | 2.08E-01 | -0.5 | 2.62E-01 | 0.9 | 4.08E-02 | Late embryogenesis abundant protein LEA-2 subgroup domain-containing protein |
| AT1G75190 | 1.8 | 7.98E-07 | 1.8 | 2.26E-06 | 0.0 | 8.58E-01 | -0.1 | 8.36E-01 | -0.6 | 7.94E-02 | 0.5 | 1.67E-01 | AT1G75190 protein |
| AT5G19190 | 1.8 | 4.63E-06 | 1.0 | 7.64E-04 | 0.8 | 3.43E-03 | -0.8 | 4.18E-03 | -1.3 | 5.98E-04 | 0.4 | 2.24E-01 | hypothetical protein |
| AT3G46780 | 1.8 | 7.71E-08 | 1.7 | 3.51E-07 | 0.1 | 2.52E-01 | -0.1 | 4.23E-01 | 0.1 | 8.05E-01 | -0.2 | 4.15E-01 | Protein PLASTID TRANSCRIPTIONALLY ACTIVE 16, chloroplastic |
| AT3G19850 | 1.8 | 1.30E-06 | 1.8 | 3.09E-06 | 0.0 | 9.99E-01 | -0.3 | 7.66E-02 | -0.4 | 2.25E-02 | 0.1 | 5.94E-01 | BTB |
| AT5G39530 | 1.8 | 2.12E-06 | 1.3 | 5.40E-05 | 0.6 | 1.50E-02 | -0.3 | 1.88E-01 | -0.2 | 4.44E-01 | -0.1 | 8.14E-01 | Uncharacterized protein |
| AT1G21525 | 1.8 | 6.96E-03 | -0.3 | 7.06E-01 | 2.2 | 2.25E-02 | -0.6 | 3.94E-01 | -0.8 | 3.45E-01 | | | NA |
| AT4G34290 | 1.8 | 1.74E-07 | 1.6 | 1.25E-06 | 0.3 | 4.63E-02 | 0.8 | 1.72E-04 | 0.3 | 1.99E-01 | 0.5 | 3.04E-02 | hypothetical protein |
| AT3G60540 | 1.8 | 1.02E-04 | 1.0 | 1.03E-02 | 0.8 | 3.11E-02 | 0.6 | 1.75E-01 | 0.2 | 7.58E-01 | 0.4 | 4.99E-01 | Protein transport protein Sec61 subunit beta |
| AT2G20250 | 1.8 | 4.01E-06 | 1.5 | 5.37E-05 | 0.4 | 8.27E-02 | | | | | | | Uncharacterized protein |
| AT5G26280 | 1.9 | 4.13E-08 | 1.8 | 1.62E-07 | 0.1 | 4.50E-01 | 0.0 | 9.31E-01 | 0.1 | 7.54E-01 | -0.1 | 8.53E-01 | AT5g26280 |
| AT1G22630 | 1.9 | 9.93E-05 | 1.1 | 6.00E-03 | 0.8 | 3.21E-02 | 0.5 | 1.23E-01 | 0.2 | 5.16E-01 | 0.2 | 6.54E-01 | Uncharacterized protein |
| AT1G65490 | 1.9 | 1.06E-05 | 0.4 | 1.07E-01 | 1.4 | 4.17E-04 | 0.3 | 3.87E-01 | 0.6 | 9.99E-02 | -0.3 | 5.10E-01 | Transmembrane protein |
| AT5G49170 | 1.9 | 1.64E-05 | 1.6 | 1.40E-04 | 0.3 | 2.18E-01 | 0.3 | 3.72E-01 | -0.1 | 9.17E-01 | 0.3 | 4.15E-01 | Uncharacterized protein |
| AT4G02110 | 1.9 | 3.34E-07 | 0.9 | 2.22E-04 | 1.0 | 1.47E-04 | 0.1 | 7.92E-01 | 0.1 | 7.62E-01 | 0.0 | 9.51E-01 | Transcription coactivator |
| AT2G38250 | 1.9 | 2.61E-05 | -0.5 | 2.08E-01 | 2.3 | 1.71E-04 | 0.7 | 1.33E-01 | -0.6 | 3.56E-01 | 1.4 | 1.88E-02 | Uncharacterized protein |
| AT2G36835 | 1.9 | 3.84E-08 | 1.2 | 1.48E-06 | 0.7 | 1.34E-04 | 0.6 | 1.32E-03 | 0.0 | 9.87E-01 | 0.6 | 1.60E-02 | hypothetical protein |
| AT3G12345 | 1.9 | 2.25E-08 | 1.5 | 2.42E-07 | 0.4 | 1.73E-03 | 0.1 | 3.81E-01 | -0.1 | 5.64E-01 | 0.2 | 2.19E-01 | hypothetical protein |
| AT5G24314 | 1.9 | 9.66E-06 | 1.4 | 2.01E-04 | 0.5 | 4.20E-02 | 0.5 | 2.76E-03 | 0.3 | 1.51E-01 | 0.3 | 2.48E-01 | Protein PLASTID TRANSCRIPTIONALLY ACTIVE 7 |
| AT5G42070 | 1.9 | 4.28E-07 | 1.3 | 1.21E-05 | 0.6 | 4.11E-03 | 0.3 | 2.48E-01 | 0.3 | 3.21E-01 | 0.0 | 9.94E-01 | Uncharacterized protein At5g42070 |
| AT4G08555 | 1.9 | 4.37E-04 | -0.8 | 1.78E-01 | 2.7 | 1.05E-03 | 2.6 | 6.72E-04 | -0.4 | 7.73E-01 | 2.9 | 1.65E-03 | Uncharacterized protein |
| AT5G28750 | 1.9 | 4.63E-07 | 1.3 | 1.45E-05 | 0.6 | 3.44E-03 | 0.6 | 1.23E-03 | 0.3 | 1.96E-01 | 0.4 | 1.23E-01 | Sec-independent protein translocase protein TATA, chloroplastic |
| AT1G72645 | 1.9 | 3.47E-04 | 0.6 | 1.58E-01 | 1.3 | 1.08E-02 | 0.0 | 9.40E-01 | -0.1 | 8.69E-01 | 0.1 | 8.40E-01 | Transmembrane protein |
| AT5G11510 | 1.9 | 1.20E-06 | 0.8 | 2.20E-03 | 1.2 | 2.28E-04 | 0.6 | 6.66E-03 | 0.5 | 3.40E-02 | 0.1 | 7.71E-01 | Myb domain protein 3r-4 |
| AT4G02530 | 1.9 | 3.08E-06 | 1.4 | 4.88E-05 | 0.5 | 3.86E-02 | 0.3 | 1.48E-02 | 0.2 | 7.34E-02 | 0.1 | 7.73E-01 | Chloroplast thylakoid lumen protein |
| AT1G04420 | 1.9 | 1.01E-07 | 1.4 | 1.67E-06 | 0.5 | 2.56E-03 | 0.2 | 2.31E-01 | 0.1 | 4.11E-01 | 0.0 | 9.05E-01 | NADP-dependent oxidoreductase domain-containing protein |
| AT1G21810 | 1.9 | 1.34E-06 | 1.2 | 1.09E-04 | 0.7 | 3.16E-03 | -0.3 | 6.80E-01 | 0.2 | 7.62E-01 | | | Filament-like protein |
| AT4G09215 | 1.9 | 8.22E-04 | -0.6 | 3.43E-01 | 2.5 | 2.44E-03 | 2.4 | 6.23E-04 | 0.4 | 6.46E-01 | 2.0 | 7.24E-04 | NA |
| AT2G35370 | 1.9 | 3.22E-05 | 1.3 | 8.54E-04 | 0.7 | 5.14E-02 | 0.4 | 4.67E-03 | 0.3 | 4.72E-02 | 0.1 | 6.33E-01 | Glycine cleavage system H protein 1, mitochondrial |
| AT1G21500 | 1.9 | 7.44E-08 | 1.9 | 2.38E-07 | 0.0 | 8.81E-01 | -0.3 | 1.11E-01 | -0.4 | 5.63E-02 | 0.1 | 7.43E-01 | AT1G21500 protein |
| AT1G32470 | 2.0 | 9.13E-08 | 1.6 | 6.42E-07 | 0.3 | 4.02E-02 | 0.3 | 9.71E-03 | 0.3 | 8.73E-02 | 0.1 | 6.32E-01 | Glycine cleavage system H protein 3, mitochondrial |
| AT3G22231 | 2.0 | 1.35E-02 | 0.9 | 1.92E-01 | 1.0 | 2.04E-01 | 2.3 | 9.98E-07 | 1.4 | 4.92E-04 | 0.9 | 3.98E-02 | Cysteine-rich and transmembrane domain-containing protein PCC1 |
| AT1G01170 | 2.0 | 3.54E-07 | 1.3 | 1.49E-05 | 0.6 | 1.92E-03 | 0.0 | 1.00E+00 | -0.4 | 1.24E-01 | 0.4 | 2.14E-01 | hypothetical protein |
| AT3G10930 | 2.0 | 1.88E-04 | -0.4 | 4.73E-01 | 2.3 | 1.03E-03 | 1.2 | 8.81E-03 | 0.1 | 9.39E-01 | 1.1 | 1.57E-02 | Uncharacterized protein At3g10930 |
| AT5G16030 | 2.0 | 2.00E-06 | 1.9 | 6.65E-06 | 0.1 | 6.61E-01 | -0.7 | 9.58E-04 | -0.7 | 2.06E-03 | 0.0 | 9.71E-01 | Mental retardation GTPase activating protein |
| AT3G48640 | 2.0 | 6.88E-05 | 0.4 | 3.52E-01 | 1.6 | 2.05E-03 | 0.6 | 2.93E-01 | -0.7 | 3.56E-01 | 1.4 | 7.76E-02 | Transmembrane protein |
| AT5G52960 | 2.0 | 9.61E-08 | 1.5 | 1.70E-06 | 0.5 | 2.36E-03 | 0.1 | 6.15E-01 | 0.0 | 9.74E-01 | 0.1 | 7.20E-01 | hypothetical protein |
| AT5G13210 | 2.0 | 4.83E-06 | -0.4 | 1.48E-01 | 2.3 | 3.16E-05 | 3.3 | 4.05E-09 | 0.6 | 7.06E-02 | 2.7 | 9.04E-07 | AT5g13210 |
| AT3G42806 | 2.0 | 8.09E-07 | 1.0 | 3.44E-04 | 1.0 | 4.10E-04 | 0.1 | 7.08E-01 | -0.3 | 4.07E-01 | 0.4 | 2.95E-01 | NA |
| AT5G25460 | 2.0 | 2.16E-07 | 1.9 | 8.39E-07 | 0.1 | 4.57E-01 | 0.0 | 9.55E-01 | -0.5 | 5.65E-03 | 0.6 | 1.88E-02 | Protein DUF642 L-GALACTONO-1,4-LACTONE-RESPONSIVE GENE 2 |
| AT3G21351 | 2.0 | 6.86E-04 | 1.1 | 3.10E-02 | 0.8 | 7.94E-02 | 0.1 | 8.20E-01 | -1.0 | 1.54E-01 | 1.2 | 9.29E-02 | Transmembrane protein |
| AT4G01883 | 2.0 | 1.34E-07 | 1.4 | 3.68E-06 | 0.6 | 1.32E-03 | 0.6 | 2.75E-03 | 0.4 | 4.60E-02 | 0.2 | 4.39E-01 | Coenzyme Q-binding protein COQ10 START domain-containing protein |
| AT4G37240 | 2.0 | 1.07E-05 | 1.4 | 3.99E-04 | 0.6 | 2.35E-02 | -0.5 | 3.91E-02 | -0.8 | 7.21E-03 | 0.3 | 4.75E-01 | Uncharacterized protein AT4g37240 |
| AT4G11650 | 2.0 | 9.99E-07 | 0.4 | 3.75E-02 | 1.6 | 5.44E-05 | 2.5 | 2.53E-07 | 1.5 | 2.24E-04 | 1.0 | 2.05E-03 | Osmotin-like protein OSM34 |
| AT2G32500 | 2.0 | 1.13E-05 | 1.4 | 3.16E-04 | 0.6 | 3.19E-02 | 0.3 | 3.16E-02 | 0.3 | 3.42E-02 | 0.0 | 9.22E-01 | Stress responsive alpha-beta barrel domain protein |
| AT5G51440 | 2.0 | 4.98E-05 | -0.9 | 7.41E-02 | 2.9 | 2.27E-04 | 3.8 | 3.44E-06 | 2.1 | 3.12E-03 | 1.6 | 3.64E-04 | 23.5 kDa heat shock protein, mitochondrial |
| AT4G02825 | 2.0 | 1.49E-03 | 1.6 | 1.09E-02 | 0.4 | 3.61E-01 | | | | | | | NA |
| AT1G71760 | 2.0 | 1.33E-05 | 1.0 | 5.82E-03 | 1.0 | 3.05E-03 | 0.4 | 4.31E-01 | 0.2 | 8.13E-01 | | | MRN complex-interacting protein |
| AT3G06070 | 2.0 | 2.87E-07 | 1.2 | 2.81E-05 | 0.8 | 5.20E-04 | 0.0 | 9.21E-01 | -0.5 | 5.86E-02 | 0.5 | 1.35E-01 | Uncharacterized protein F24F17.5 |
| AT1G47520 | 2.0 | 1.30E-05 | 1.9 | 4.07E-05 | 0.1 | 7.08E-01 | 0.3 | 3.05E-01 | -0.1 | 7.85E-01 | 0.5 | 2.91E-01 | NA |
| AT3G22235 | 2.0 | 2.30E-03 | 0.8 | 1.21E-01 | 1.2 | 6.34E-02 | 1.8 | 1.54E-05 | 0.9 | 9.07E-03 | 0.8 | 3.52E-02 | Protein CYSTEINE-RICH TRANSMEMBRANE MODULE 8 |
| AT4G30450 | 2.0 | 1.37E-07 | 1.6 | 1.86E-06 | 0.4 | 6.55E-03 | 0.0 | 8.84E-01 | -0.1 | 5.50E-01 | 0.1 | 7.34E-01 | Transmembrane protein |
| AT1G69530 | 2.0 | 1.88E-06 | 2.2 | 2.73E-06 | -0.2 | 3.97E-01 | 0.1 | 5.25E-01 | -0.3 | 4.65E-02 | 0.4 | 1.44E-02 | Expansin |
| AT1G02450 | 2.0 | 3.07E-03 | 0.1 | 8.73E-01 | 1.9 | 2.36E-02 | 1.5 | 1.95E-02 | 1.0 | 1.56E-01 | 0.5 | 4.57E-01 | hypothetical protein |
| AT5G22920 | 2.1 | 1.78E-06 | 2.1 | 3.09E-06 | -0.1 | 6.65E-01 | -0.1 | 7.05E-01 | -1.1 | 4.68E-04 | 1.0 | 4.03E-03 | PGPD14 protein |
| AT2G27402 | 2.1 | 6.93E-07 | 1.5 | 1.34E-05 | 0.5 | 1.13E-02 | 0.8 | 6.77E-04 | 0.2 | 3.86E-01 | 0.6 | 4.70E-02 | Uncharacterized protein |
| AT2G04790 | 2.1 | 1.27E-05 | 0.9 | 1.23E-02 | 1.2 | 1.65E-03 | 0.0 | 9.73E-01 | -0.1 | 7.64E-01 | 0.1 | 8.31E-01 | PTB domain engulfment adapter |
| AT1G49940 | 2.1 | 1.82E-05 | 1.1 | 5.08E-03 | 1.0 | 5.60E-03 | -0.1 | 8.43E-01 | 0.2 | 6.73E-01 | -0.3 | 5.21E-01 | Uncharacterized protein |
| AT3G15630 | 2.1 | 4.11E-07 | 2.4 | 4.26E-07 | -0.3 | 6.18E-02 | -0.2 | 5.32E-01 | -1.0 | 8.59E-03 | 0.8 | 9.19E-02 | Uncharacterized protein |
| AT4G04595 | 2.1 | 6.32E-06 | 1.3 | 7.13E-04 | 0.8 | 5.49E-03 | 0.2 | 6.44E-01 | -0.1 | 8.70E-01 | 0.3 | 5.59E-01 | NA |
| AT2G26500 | 2.1 | 8.89E-06 | 1.3 | 3.48E-04 | 0.8 | 1.94E-02 | -0.4 | 5.60E-02 | 0.1 | 6.45E-01 | -0.5 | 6.70E-02 | Cytochrome b6f complex subunit |
| AT4G18490 | 2.1 | 1.12E-05 | 1.0 | 6.94E-03 | 1.1 | 2.58E-03 | 0.1 | 7.83E-01 | 0.1 | 8.14E-01 | 0.0 | 9.97E-01 | Uncharacterized protein |
| AT1G22590 | 2.1 | 7.19E-08 | 1.4 | 2.71E-06 | 0.7 | 3.20E-04 | 0.3 | 1.80E-01 | 0.0 | 9.34E-01 | 0.3 | 3.34E-01 | Uncharacterized protein |
| AT3G56910 | 2.1 | 5.61E-08 | 1.8 | 4.10E-07 | 0.4 | 1.42E-02 | 0.1 | 6.40E-01 | -0.2 | 6.11E-01 | 0.3 | 3.85E-01 | Large ribosomal subunit protein cL37 |
| AT1G79075 | 2.1 | 1.36E-05 | 1.0 | 5.14E-03 | 1.1 | 2.58E-03 | 0.4 | 2.71E-01 | 0.2 | 6.81E-01 | 0.2 | 7.34E-01 | NA |
| AT1G15175 | 2.1 | 3.71E-06 | 2.1 | 9.69E-06 | 0.0 | 9.23E-01 | -0.4 | 1.42E-01 | -0.8 | 2.55E-02 | 0.3 | 3.68E-01 | NA |
| AT1G44000 | 2.1 | 5.24E-08 | 1.5 | 1.04E-06 | 0.7 | 6.31E-04 | 0.3 | 3.98E-02 | 0.2 | 1.20E-01 | 0.0 | 8.59E-01 | Staygreen protein domain-containing protein |
| AT1G32540 | 2.1 | 3.16E-06 | 2.3 | 3.76E-06 | -0.2 | 3.26E-01 | -1.0 | 3.42E-03 | -0.4 | 2.46E-01 | -0.6 | 1.33E-01 | Zinc finger LSD1-type domain-containing protein |
| AT4G13577 | 2.1 | 7.47E-06 | 2.5 | 4.77E-06 | -0.4 | 8.78E-02 | | | | | | | Uncharacterized protein |
| AT5G03390 | 2.2 | 8.18E-07 | 1.8 | 1.10E-05 | 0.4 | 3.50E-02 | 1.9 | 4.43E-05 | 0.8 | 8.43E-02 | 1.1 | 4.17E-03 | At5g03390 |
| AT5G12930 | 2.2 | 8.83E-07 | 0.9 | 1.38E-03 | 1.3 | 1.98E-04 | 0.4 | 1.92E-02 | 0.2 | 3.39E-01 | 0.2 | 2.89E-01 | Inactive rhomboid protein |
| AT5G21920 | 2.2 | 1.44E-08 | 1.7 | 9.13E-08 | 0.4 | 6.68E-04 | 0.1 | 7.67E-01 | -0.1 | 8.71E-01 | 0.1 | 6.99E-01 | YGGT family protein |
| AT5G56100 | 2.2 | 4.43E-06 | 1.8 | 3.76E-05 | 0.3 | 1.35E-01 | -0.2 | 5.80E-01 | -1.3 | 3.52E-03 | 1.1 | 2.18E-02 | Glycine-rich protein |
| AT3G18890 | 2.2 | 4.07E-08 | 1.8 | 3.32E-07 | 0.4 | 7.75E-03 | 1.0 | 9.22E-04 | 0.6 | 6.81E-02 | 0.5 | 1.86E-01 | Protein TIC 62, chloroplastic |

| | | | | | | | | | | | | | |
|---|---|---|---|---|---|---|---|---|---|---|---|---|---|
| AT1G50050 | 2.2 | 3.31E-06 | 2.1 | 9.38E-06 | 0.1 | 7.65E-01 | 0.3 | 4.22E-01 | -0.1 | 8.68E-01 | 0.4 | 3.62E-01 | CAP superfamily protein |
| AT4G17240 | 2.2 | 4.25E-07 | 1.3 | 5.73E-05 | 0.9 | 5.40E-04 | 0.0 | 9.49E-01 | -0.1 | 3.25E-01 | 0.2 | 3.78E-01 | Structural maintenance of chromosomes protein |
| AT4G24930 | 2.2 | 1.44E-08 | 1.6 | 1.12E-07 | 0.6 | 1.93E-04 | 0.3 | 3.44E-02 | 0.2 | 3.33E-01 | 0.2 | 5.05E-01 | Thylakoid lumenal 17.9 kDa protein, chloroplastic |
| AT1G67090 | 2.2 | 2.92E-08 | 1.8 | 2.26E-07 | 0.4 | 1.23E-02 | 0.2 | 1.01E-01 | 0.1 | 6.46E-01 | 0.1 | 5.14E-01 | Ribulose bisphosphate carboxylase small subunit |
| AT5G52930 | 2.2 | 3.98E-07 | 0.8 | 1.75E-03 | 1.4 | 7.37E-05 | 1.3 | 8.99E-02 | 0.9 | 3.26E-01 | | | Expressed protein |
| AT2G21210 | 2.2 | 1.08E-06 | 1.9 | 9.09E-06 | 0.3 | 8.46E-02 | -0.9 | 2.32E-04 | -1.0 | 2.62E-04 | 0.1 | 7.92E-01 | SAUR-like auxin-responsive protein family |
| AT2G14610 | 2.2 | 3.99E-02 | 0.8 | 4.74E-01 | 1.4 | 2.33E-01 | -1.5 | 4.39E-01 | -4.9 | 3.00E-01 | 3.5 | 4.54E-01 | Pathogenesis-related protein 1 |
| AT4G01870 | 2.2 | 1.09E-05 | 0.9 | 1.42E-02 | 1.4 | 1.14E-03 | -0.4 | 2.94E-01 | -1.7 | 1.38E-03 | 1.3 | 1.33E-02 | TolB protein-like protein |
| AT5G22340 | 2.3 | 2.37E-08 | 1.7 | 4.10E-07 | 0.6 | 3.54E-04 | 0.3 | 4.43E-02 | -0.1 | 5.88E-01 | 0.4 | 3.06E-02 | At5g22340 |
| AT3G05130 | 2.3 | 1.21E-07 | 1.5 | 5.78E-06 | 0.8 | 4.52E-04 | 0.4 | 1.75E-02 | 0.1 | 6.39E-01 | 0.3 | 1.13E-01 | Uncharacterized protein |
| AT2G20510 | 2.3 | 1.41E-05 | 1.4 | 1.73E-03 | 0.9 | 8.90E-03 | -0.1 | 9.37E-01 | 0.4 | 4.99E-01 | | | Translocase inner membrane subunit 44-1 |
| AT2G42220 | 2.3 | 1.59E-08 | 2.0 | 8.81E-08 | 0.3 | 2.24E-02 | 0.0 | 8.57E-01 | -0.1 | 3.37E-01 | 0.2 | 3.44E-01 | Rhodanese-like domain-containing protein 9, chloroplastic |
| AT1G20190 | 2.3 | 1.16E-06 | 2.0 | 6.19E-06 | 0.3 | 2.26E-01 | -0.5 | 6.96E-03 | -0.6 | 7.92E-03 | 0.0 | 9.20E-01 | Expansin |
| AT4G12540 | 2.3 | 3.00E-07 | 1.1 | 2.18E-04 | 1.2 | 1.34E-04 | 0.2 | 6.43E-01 | 0.5 | 1.01E-01 | -0.4 | 2.98E-01 | Uncharacterized protein AT4g12540 |
| AT5G24490 | 2.3 | 1.50E-07 | 2.1 | 6.75E-07 | 0.2 | 2.43E-01 | -0.3 | 2.28E-02 | -0.7 | 4.40E-04 | 0.3 | 5.76E-02 | Similarity to ribosomal protein 30S subunit |
| AT1G30520 | 2.3 | 3.15E-07 | 1.7 | 6.90E-06 | 0.6 | 4.67E-03 | -0.2 | 1.59E-01 | -0.2 | 2.43E-01 | 0.0 | 9.70E-01 | Acyl-activating enzyme 14 |
| AT3G26710 | 2.3 | 7.44E-08 | 1.7 | 1.24E-06 | 0.6 | 1.51E-03 | 0.5 | 6.96E-03 | 0.0 | 9.47E-01 | 0.5 | 4.13E-02 | Protein COFACTOR ASSEMBLY OF COMPLEX C SUBUNIT B CCB1, chloroplastic |
| AT5G62720 | 2.3 | 1.98E-07 | 2.4 | 4.03E-07 | -0.1 | 5.58E-01 | -0.5 | 5.16E-04 | -0.5 | 9.47E-04 | 0.0 | 9.05E-01 | Uncharacterized protein |
| AT3G20470 | 2.3 | 3.07E-05 | 2.1 | 1.01E-04 | 0.2 | 6.14E-01 | 0.7 | 2.17E-02 | 0.0 | 9.46E-01 | 0.7 | 6.96E-02 | Uncharacterized protein |
| AT4G06120 | 2.4 | 2.87E-06 | 3.1 | 9.11E-07 | -0.7 | 3.88E-03 | | | | | | | NA |
| AT3G60470 | 2.4 | 2.50E-05 | 1.3 | 4.20E-03 | 1.1 | 9.73E-03 | 1.5 | 1.61E-03 | 1.1 | 1.89E-02 | 0.3 | 3.79E-01 | Transmembrane protein, putative |
| AT2G33793 | 2.4 | 5.24E-08 | 1.5 | 3.52E-06 | 0.9 | 9.50E-05 | -0.1 | 6.59E-01 | -0.2 | 3.66E-01 | 0.1 | 7.37E-01 | Meiosis-specific protein ASY3-like coiled-coil domain-containing protein |
| AT5G54610 | 2.4 | 6.98E-05 | 1.1 | 2.05E-02 | 1.3 | 7.78E-03 | 2.1 | 1.72E-04 | 0.5 | 4.04E-01 | 1.6 | 5.49E-03 | Ankyrin repeat-containing protein BDA1 |
| AT1G65870 | 2.4 | 2.30E-06 | 1.7 | 1.03E-04 | 0.7 | 6.79E-03 | | | | | | | Dirigent protein 21 |
| AT1G03600 | 2.4 | 1.67E-08 | 2.1 | 9.13E-08 | 0.3 | 1.60E-02 | 0.1 | 5.32E-01 | 0.0 | 9.87E-01 | 0.1 | 7.06E-01 | Photosystem II repair protein PSB27-H1, chloroplastic |
| AT5G03120 | 2.4 | 1.27E-05 | 1.7 | 3.63E-04 | 0.7 | 2.89E-02 | -0.4 | 1.01E-01 | -0.6 | 3.95E-02 | 0.2 | 6.70E-01 | Transmembrane protein |
| AT2G46220 | 2.5 | 9.61E-08 | 2.2 | 5.37E-07 | 0.3 | 7.98E-02 | -0.3 | 6.87E-02 | -0.7 | 2.94E-04 | 0.4 | 1.52E-02 | NTF2-like domain-containing protein |
| AT2G38820 | 2.5 | 3.48E-07 | 2.2 | 2.22E-06 | 0.3 | 9.72E-02 | 0.1 | 7.25E-01 | -0.4 | 1.44E-01 | 0.5 | 1.14E-01 | DNA-directed RNA polymerase subunit beta-beta protein, putative |
| AT1G19530 | 2.5 | 3.02E-05 | 1.3 | 4.98E-03 | 1.1 | 6.65E-03 | 1.2 | 2.55E-02 | 0.1 | 9.05E-01 | 1.1 | 4.01E-02 | F18O14.29 |
| AT3G09430 | 2.5 | 9.07E-06 | 1.2 | 6.20E-03 | 1.3 | 1.90E-03 | -0.3 | 4.38E-01 | 0.0 | 9.88E-01 | -0.3 | 5.33E-01 | F3L24.32 protein |
| AT5G07380 | 2.5 | 9.07E-06 | 0.8 | 4.24E-02 | 1.7 | 7.29E-04 | -0.1 | 9.09E-01 | -0.1 | 8.59E-01 | 0.1 | 9.56E-01 | Uncharacterized protein |
| AT5G23700 | 2.5 | 3.07E-05 | 1.7 | 1.45E-03 | 0.8 | 2.74E-02 | -0.4 | 3.78E-01 | 0.0 | 9.71E-01 | -0.4 | 5.59E-01 | Coiled-coil protein |
| AT3G10880 | 2.5 | 5.94E-07 | 1.4 | 1.63E-04 | 1.1 | 4.23E-04 | 0.2 | 6.53E-01 | 0.2 | 5.73E-01 | -0.1 | 9.10E-01 | T7M13.4 protein |
| AT4G15990 | 2.5 | 4.96E-07 | 1.5 | 7.21E-05 | 1.0 | 5.18E-04 | 0.6 | 1.04E-01 | -0.1 | 8.40E-01 | 0.7 | 1.29E-01 | hypothetical protein |
| AT4G07820 | 2.6 | 1.10E-06 | 2.1 | 1.34E-05 | 0.5 | 3.16E-02 | -1.1 | 8.09E-05 | -1.1 | 2.31E-04 | 0.0 | 9.98E-01 | hypothetical protein |
| AT4G13500 | 2.6 | 4.60E-06 | 1.5 | 4.58E-04 | 1.1 | 3.71E-03 | 0.4 | 7.13E-02 | 0.2 | 4.77E-01 | 0.2 | 5.32E-01 | hypothetical protein |
| AT2G15128 | 2.6 | 2.33E-07 | 1.5 | 3.40E-05 | 1.1 | 2.34E-04 | 0.7 | 1.03E-02 | 0.0 | 9.32E-01 | 0.7 | 3.44E-02 | NA |
| AT2G20670 | 2.6 | 3.86E-08 | 2.3 | 2.22E-07 | 0.3 | 3.72E-02 | -0.2 | 5.29E-01 | -2.0 | 2.38E-04 | 1.7 | 4.39E-03 | hypothetical protein |
| AT1G33055 | 2.6 | 1.42E-05 | 3.1 | 8.54E-06 | -0.5 | 9.77E-02 | -0.3 | 6.38E-01 | -1.4 | 1.05E-01 | 1.0 | 2.77E-01 | Uncharacterized protein unannotated coding sequence from BAC F9L11 |
| AT2G36120 | 2.6 | 1.76E-07 | 2.5 | 6.00E-07 | 0.1 | 6.31E-01 | -0.2 | 1.08E-01 | -0.1 | 6.53E-01 | -0.1 | 5.28E-01 | Glycine-rich protein DOT1 |
| AT3G17050 | 2.7 | 2.09E-05 | 2.1 | 2.32E-04 | 0.5 | 1.14E-01 | 0.1 | 5.67E-01 | -0.2 | 4.02E-01 | 0.4 | 2.22E-01 | Glycine-rich cell wall structural protein |
| AT2G33180 | 2.7 | 2.25E-08 | 1.8 | 5.40E-07 | 0.8 | 1.08E-04 | 0.9 | 3.53E-04 | 0.5 | 2.93E-02 | 0.4 | 1.81E-01 | hypothetical protein |
| AT5G02160 | 2.7 | 6.32E-08 | 2.5 | 2.62E-07 | 0.2 | 2.76E-01 | -0.4 | 4.40E-02 | -0.5 | 1.50E-02 | 0.1 | 6.47E-01 | Uncharacterized protein T7H20_210 |
| AT3G59450 | 2.7 | 2.10E-05 | 1.0 | 3.15E-02 | 1.7 | 1.45E-03 | 0.0 | 9.57E-01 | 0.3 | 5.15E-01 | -0.3 | 5.40E-01 | Calcium-binding EF-hand family protein |
| AT3G25920 | 2.7 | 9.35E-09 | 2.1 | 5.54E-08 | 0.6 | 2.21E-04 | 0.2 | 5.51E-02 | 0.0 | 8.01E-01 | 0.2 | 2.10E-01 | 50S ribosomal protein L15, chloroplast |
| AT4G28660 | 2.7 | 2.75E-08 | 2.0 | 3.67E-07 | 0.7 | 7.08E-04 | 0.7 | 4.47E-04 | 0.4 | 5.63E-02 | 0.3 | 1.41E-01 | Photosystem II reaction center Psb28 protein |
| AT5G08050 | 2.7 | 6.86E-07 | 1.8 | 1.90E-05 | 0.9 | 4.73E-03 | 0.0 | 9.08E-01 | 0.3 | 1.87E-01 | -0.3 | 3.47E-01 | Uncharacterized protein F13G24.250 ) |
| AT5G55570 | 2.7 | 4.28E-06 | 1.2 | 4.16E-03 | 1.5 | 5.63E-04 | | | | | | | Transmembrane protein |
| AT4G13575 | 2.7 | 3.07E-06 | 2.6 | 1.01E-05 | 0.1 | 6.33E-01 | 0.0 | 9.95E-01 | -0.3 | 4.53E-01 | 0.3 | 4.80E-01 | Uncharacterized protein |
| AT4G32480 | 2.7 | 4.42E-07 | 2.7 | 1.08E-06 | 0.0 | 9.47E-01 | 0.3 | 8.55E-02 | -1.4 | 4.33E-05 | 1.8 | 9.97E-05 | Sugar phosphate exchanger, |
| AT1G76740 | 2.7 | 1.07E-06 | 1.2 | 1.08E-03 | 1.5 | 2.29E-04 | 0.2 | 5.52E-01 | 0.3 | 4.06E-01 | -0.1 | 8.49E-01 | Uncharacterized protein |
| AT3G22210 | 2.8 | 2.79E-04 | 1.1 | 8.35E-02 | 1.6 | 1.01E-02 | 0.7 | 2.64E-01 | 0.2 | 8.65E-01 | 0.5 | 5.19E-01 | hypothetical protein |
| AT1G50732 | 2.8 | 3.95E-07 | 1.9 | 2.04E-05 | 0.9 | 9.49E-04 | 0.2 | 2.23E-01 | 0.0 | 9.03E-01 | 0.3 | 3.23E-01 | Transmembrane protein |
| AT3G28220 | 2.8 | 1.18E-04 | 2.0 | 2.65E-03 | 0.9 | 5.19E-02 | 1.1 | 1.85E-02 | 1.1 | 3.95E-02 | 0.0 | 9.70E-01 | AT3g28220 |
| AT2G23130 | 2.8 | 8.09E-07 | 2.0 | 3.02E-05 | 0.9 | 3.06E-03 | -1.1 | 2.02E-03 | -1.1 | 2.48E-03 | 0.1 | 9.28E-01 | Arabinogalactan protein 17 |
| AT3G55646 | 2.8 | 2.60E-05 | 3.4 | 1.20E-05 | -0.6 | 6.17E-02 | | | | | | | hypothetical protein |
| AT3G62430 | 2.8 | 4.48E-07 | 1.8 | 3.78E-05 | 1.0 | 8.09E-04 | 0.2 | 6.94E-01 | 0.2 | 6.96E-01 | 0.0 | 9.70E-01 | F-box protein At3g62430 |
| AT5G44420 | 2.9 | 1.23E-03 | 1.6 | 5.11E-02 | 1.3 | 6.19E-02 | 1.0 | 2.99E-01 | -0.2 | 8.97E-01 | 1.2 | 3.53E-01 | Defensin-like protein 16 |
| AT5G52290 | 2.9 | 1.96E-05 | 1.6 | 3.16E-03 | 1.2 | 5.99E-03 | -0.2 | 7.99E-01 | -0.2 | 8.15E-01 | 0.0 | 9.88E-01 | Protein SHORTAGE IN CHIASMATA 1 |
| AT2G42340 | 2.9 | 1.09E-05 | 1.7 | 1.44E-03 | 1.2 | 5.75E-03 | -0.5 | 3.34E-01 | 0.4 | 3.67E-01 | -0.9 | 7.66E-02 | Uncharacterized protein At2g42340 |
| AT2G40610 | 2.9 | 2.46E-05 | 3.0 | 3.93E-05 | -0.1 | 8.47E-01 | -0.3 | 9.78E-02 | -0.1 | 4.74E-01 | -0.1 | 5.90E-01 | Expansin-A8 |
| AT3G00400 | 2.9 | 9.14E-07 | 3.4 | 8.46E-07 | -0.4 | 3.71E-02 | 0.5 | 2.86E-01 | 0.8 | 1.23E-01 | -0.3 | 6.19E-01 | NA |
| AT3G15450 | 3.0 | 1.52E-07 | 3.1 | 3.46E-07 | -0.1 | 6.09E-01 | -0.9 | 4.93E-02 | -2.3 | 2.76E-04 | 1.4 | 3.79E-02 | Aluminum induced protein with YGL and LRDR motifs |
| AT2G37025 | 3.0 | 2.18E-06 | 2.4 | 4.09E-05 | 0.7 | 1.67E-02 | -1.1 | 8.33E-03 | -1.4 | 3.08E-03 | 0.3 | 6.32E-01 | TRF-like 8 |
| AT5G44430 | 3.1 | 2.48E-03 | 2.5 | 1.30E-02 | 0.6 | 3.88E-01 | 0.2 | 8.62E-01 | -0.3 | 8.49E-01 | 0.6 | 7.49E-01 | Defensin-like protein 17 |
| AT1G31173 | 3.2 | 1.08E-05 | 2.4 | 2.56E-04 | 0.8 | 2.59E-02 | | | | | | | NA |
| AT3G54065 | 3.2 | 1.87E-06 | 1.8 | 4.69E-04 | 1.4 | 7.24E-04 | 1.2 | 1.01E-02 | 1.1 | 2.40E-02 | 0.1 | 8.97E-01 | Uncharacterized protein F24B22.30 |
| AT5G24205 | 3.2 | 8.97E-06 | 1.8 | 2.07E-03 | 1.4 | 2.50E-03 | 0.5 | 3.22E-01 | 0.3 | 6.68E-01 | 0.2 | 7.72E-01 | NA |
| AT2G21640 | 3.3 | 3.78E-06 | 0.2 | 6.63E-01 | 3.1 | 9.80E-05 | 3.8 | 1.30E-06 | 0.9 | 2.07E-01 | 3.0 | 4.20E-05 | Uncharacterized protein |
| AT2G41730 | 3.4 | 1.56E-06 | 0.7 | 8.89E-02 | 2.6 | 7.42E-05 | 2.8 | 7.74E-08 | -0.4 | 3.04E-01 | 3.2 | 9.04E-07 | Calcium-binding site protein |
| AT5G41761 | 3.4 | 2.52E-06 | 2.3 | 1.28E-04 | 1.1 | 2.95E-03 | 0.3 | 3.91E-01 | -0.7 | 1.31E-01 | 1.1 | 4.38E-02 | Uncharacterized protein |
| AT2G26010 | 3.6 | 9.70E-05 | 2.0 | 1.09E-02 | 1.6 | 8.68E-03 | 1.2 | 2.50E-01 | -0.7 | 6.90E-01 | 1.9 | 2.27E-01 | Defensin-like protein 14 |
| AT3G57440 | 3.6 | 3.89E-05 | 1.9 | 1.03E-02 | 1.7 | 5.11E-03 | -0.1 | 9.22E-01 | -0.2 | 7.67E-01 | | | Uncharacterized protein T8H10.40 |
| AT1G47395 | 3.6 | 3.77E-07 | 2.4 | 2.68E-05 | 1.2 | 3.43E-04 | 2.6 | 1.71E-06 | 0.2 | 6.92E-01 | 2.3 | 1.49E-04 | Uncharacterized protein |
| AT1G11070 | 3.7 | 3.78E-06 | 2.1 | 7.65E-04 | 1.6 | 9.27E-04 | | | | | | | Hydroxyproline-rich glycoprotein family protein |
| AT3G62950 | 3.7 | 7.50E-07 | 3.2 | 6.25E-06 | 0.5 | 2.95E-02 | 0.1 | 8.76E-01 | 0.0 | 9.84E-01 | 0.1 | 9.26E-01 | Glutaredoxin-C11 |
| AT5G24920 | 3.8 | 3.00E-07 | 4.2 | 4.03E-07 | -0.4 | 3.36E-02 | 0.0 | 9.84E-01 | -0.1 | 8.06E-01 | 0.1 | 8.47E-01 | Protein GLUTAMINE DUMPER 5 |
| AT4G37540 | 3.9 | 1.42E-07 | 3.4 | 7.82E-07 | 0.4 | 4.47E-02 | 0.2 | 4.41E-01 | -1.0 | 4.57E-03 | 1.2 | 3.06E-03 | LOB domain-containing protein 39 |
| AT5G05250 | 3.9 | 1.92E-08 | 3.3 | 1.12E-07 | 0.6 | 1.41E-03 | 2.4 | 6.25E-09 | 0.9 | 5.13E-04 | 1.5 | 8.05E-05 | AT5g05250 |
| AT1G13650 | 3.9 | 1.70E-05 | 3.9 | 3.33E-05 | 0.0 | 9.63E-01 | | | | | | | Uncharacterized protein |
| AT4G14690 | 3.9 | 4.29E-07 | 1.3 | 4.13E-03 | 2.6 | 2.99E-05 | | | | | | | Early light-induced protein 2, chloroplastic |
| AT1G61450 | 4.0 | 1.08E-06 | 2.3 | 1.78E-04 | 1.6 | 2.70E-04 | 0.1 | 5.96E-01 | 0.1 | 7.95E-01 | 0.0 | 9.01E-01 | hypothetical protein |
| AT2G02120 | 4.1 | 7.71E-08 | 2.1 | 2.62E-05 | 2.0 | 1.89E-05 | 1.3 | 3.88E-05 | 0.4 | 2.06E-01 | 0.9 | 4.77E-03 | Defensin-like protein 4 |

| | | | | | | | | | | | | | |
|---|---|---|---|---|---|---|---|---|---|---|---|---|---|
| AT2G40400 | 4.3 | 2.19E-08 | 2.9 | 4.65E-07 | 1.4 | 5.92E-05 | 1.0 | 9.92E-05 | 0.0 | 9.45E-01 | 1.0 | 9.21E-04 | DUF399 family protein, putative |
| AT2G05380 | 4.4 | 4.60E-07 | 1.7 | 3.77E-04 | 2.7 | 1.07E-04 | -0.6 | 9.10E-02 | -1.7 | 4.77E-04 | 1.1 | 1.24E-02 | AT2G05380 protein |
| AT2G34580 | 4.4 | 8.47E-07 | 2.6 | 1.47E-04 | 1.8 | 2.46E-04 | 0.6 | 3.55E-01 | 0.8 | 2.51E-01 | -0.2 | 7.85E-01 | Cytomegalovirus UL139 protein |
| AT1G49500 | 4.5 | 9.70E-07 | 3.5 | 1.64E-05 | 1.0 | 2.49E-03 | -0.5 | 1.09E-01 | -1.5 | 3.36E-04 | 1.0 | 1.89E-02 | Uncharacterized protein |
| AT4G12735 | 4.8 | 2.50E-05 | -0.9 | 4.33E-01 | 5.6 | 3.16E-04 | 3.5 | 1.99E-04 | -0.6 | 6.94E-01 | 4.1 | 2.08E-04 | Transmembrane protein |
| AT5G52940 | 4.8 | 7.44E-08 | 1.3 | 4.55E-03 | 3.5 | 1.02E-05 | 1.9 | 8.89E-02 | 0.6 | 7.12E-01 | | | DUF295 domain-containing protein |
| AT1G74670 | 4.8 | 2.37E-05 | 4.6 | 6.24E-05 | 0.2 | 6.79E-01 | -0.4 | 7.30E-02 | -1.0 | 1.78E-03 | 0.6 | 1.13E-01 | hypothetical protein |
| AT5G09570 | 4.9 | 3.13E-07 | -0.5 | 4.35E-01 | 5.4 | 1.80E-05 | 4.0 | 1.14E-03 | 0.1 | 9.50E-01 | 3.8 | 8.47E-04 | hypothetical protein |
| AT1G68238 | 5.4 | 2.28E-06 | 4.1 | 4.93E-05 | 1.3 | 2.15E-03 | -0.5 | 6.00E-02 | -0.8 | 1.18E-02 | 0.3 | 4.94E-01 | hypothetical protein |
| AT2G26020 | 5.6 | 8.15E-04 | 3.4 | 2.73E-02 | 2.1 | 3.88E-02 | | | | | | | Putative defensin-like protein 15 |
| AT2G05530 | 5.8 | 3.65E-07 | 4.8 | 3.37E-06 | 0.9 | 1.48E-03 | 1.5 | 1.67E-02 | 0.6 | 4.47E-01 | 0.9 | 2.06E-01 | Glycine-rich protein family |
| AT4G05380 | 5.9 | 7.95E-06 | -0.1 | 9.44E-01 | 6.0 | 9.04E-05 | | | | | | | At4g05370 |
| AT2G34317 | 6.0 | 4.33E-06 | 4.6 | 7.38E-05 | 1.3 | 3.24E-03 | -4.6 | 2.98E-05 | -1.8 | 1.80E-03 | -2.8 | 1.27E-02 | Avirulence induced family protein |
| AT5G24640 | 6.0 | 1.29E-05 | 1.7 | 1.04E-01 | 4.3 | 3.17E-04 | 5.8 | 6.24E-04 | 0.3 | 9.27E-01 | 5.5 | 2.72E-03 | Uncharacterized protein |
| AT2G05440 | 6.1 | 1.08E-07 | 5.1 | 7.41E-07 | 1.0 | 9.30E-03 | 2.8 | 4.40E-07 | 1.4 | 1.57E-03 | 1.4 | 4.60E-03 | GLYCINE RICH PROTEIN 9 |
| AT2G30766 | 7.0 | 2.25E-05 | 5.6 | 2.42E-04 | 1.4 | 4.85E-03 | 2.9 | 5.54E-05 | 0.7 | 3.64E-01 | 2.2 | 1.55E-03 | Uncharacterized protein |
| AT2G35570 | 7.1 | 6.42E-06 | 0.6 | 6.14E-01 | 6.5 | 1.07E-04 | | | | | | | NA |
| AT4G36850 | 7.2 | 2.77E-05 | 4.6 | 1.38E-03 | 2.6 | 4.88E-04 | 0.5 | 6.09E-01 | -2.4 | 2.06E-01 | 3.0 | 4.89E-02 | PQ-loop repeat family protein |
| AT5G26130 | 7.7 | 3.22E-06 | 6.7 | 1.86E-05 | 1.0 | 1.27E-02 | -1.7 | 1.13E-04 | -1.3 | 1.71E-03 | -0.4 | 4.53E-01 | T1N24.14 protein |
| AT1G13609 | 8.9 | 4.13E-05 | 6.6 | 7.68E-04 | 2.3 | 4.63E-03 | 3.8 | 8.37E-07 | 0.3 | 7.10E-01 | 3.5 | 1.14E-04 | Defensin-like family protein |
| AT2G14247 | 9.4 | 1.15E-05 | 8.0 | 8.54E-05 | 1.4 | 1.58E-02 | 3.8 | 1.32E-06 | 0.0 | 9.91E-01 | 3.8 | 1.09E-04 | hypothetical protein |
| AT2G00370 | 9.7 | 7.39E-05 | 1.9 | 3.81E-01 | 7.8 | 1.33E-03 | | | | | | | NA |
| AT1G01715 | | | | | | | 0.0 | 9.69E-01 | -0.1 | 8.19E-01 | 0.1 | 8.53E-01 | Uncharacterized protein |
| AT1G01725 | | | | | | | 0.1 | 8.95E-01 | -0.1 | 8.15E-01 | | | Transmembrane protein |
| AT1G02470 | | | | | | | -1.1 | 1.14E-01 | -1.2 | 9.96E-02 | 0.2 | 8.90E-01 | Coenzyme Q-binding protein COQ10 START domain-containing protein |
| AT1G02700 | | | | | | | 0.1 | 7.91E-01 | -0.7 | 1.54E-01 | 0.8 | 1.19E-01 | hypothetical protein |
| AT1G03580 | | | | | | | 0.2 | 7.97E-01 | -1.9 | 1.97E-02 | 2.1 | 2.69E-02 | NA |
| AT1G04330 | | | | | | | -0.9 | 1.49E-01 | 0.1 | 8.49E-01 | | | hypothetical protein |
| AT1G04425 | | | | | | | 0.1 | 8.55E-01 | 0.3 | 6.69E-01 | -0.2 | 8.25E-01 | NA |
| AT1G04660 | | | | | | | -0.726416107 | 1.12E-01 | -0.769682449 | 1.21E-01 | 0.0 | 9.61E-01 | Uncharacterized protein |
| AT1G05370 | | | | | | | 0.6 | 1.20E-01 | 0.3 | 5.83E-01 | 0.3 | 4.46E-01 | Sec14p-like phosphatidylinositol transfer family protein |
| AT1G05450 | | | | | | | 1.2 | 5.54E-03 | 0.2 | 7.77E-01 | 1.0 | 1.64E-02 | At1g05450 |
| AT1G05503 | | | | | | | 0.4 | 5.28E-01 | 0.5 | 4.76E-01 | | | NA |
| AT1G05575 | | | | | | | 0.6 | 1.90E-01 | 0.2 | 7.38E-01 | 0.4 | 5.06E-01 | hypothetical protein |
| AT1G05730 | | | | | | | 1.4 | 5.81E-03 | 0.6 | 3.00E-01 | | | FAM136A-like protein |
| AT1G06137 | | | | | | | 4.851307377 | 1.63E-03 | 2.229877772 | 2.16E-01 | 2.6 | 4.44E-02 | Transmembrane protein |
| AT1G06265 | | | | | | | -0.2 | 7.33E-01 | -0.2 | 8.05E-01 | | | NA |
| AT1G06370 | | | | | | | -0.3 | 4.57E-01 | -0.5 | 2.30E-01 | 0.2 | 7.05E-01 | NA |
| AT1G06580 | | | | | | | -0.1 | 8.16E-01 | -0.2 | 8.15E-01 | 0.0 | 9.75E-01 | Pentatricopeptide repeat superfamily protein |
| AT1G07050 | | | | | | | 0.1 | 8.76E-01 | 0.2 | 4.30E-01 | -0.2 | 6.30E-01 | F10K1.24 protein |
| AT1G07167 | | | | | | | 0.18849609 | 6.40E-01 | -0.045301825 | 9.39E-01 | 0.2 | 6.54E-01 | NA |
| AT1G07175 | | | | | | | -0.1 | 8.74E-01 | 0.2 | 7.52E-01 | | | Secreted protein |
| AT1G07367 | | | | | | | 3.364298051 | 3.97E-04 | 0.197872405 | 9.02E-01 | 3.2 | 1.80E-03 | NA |
| AT1G07373 | | | | | | | 3.790579684 | 3.32E-05 | 0.402157597 | 7.39E-01 | 3.4 | 1.64E-04 | NA |
| AT1G07690 | | | | | | | -0.4 | 4.05E-01 | -0.2 | 7.67E-01 | -0.2 | 7.51E-01 | receptor protein-tyrosine kinase |
| AT1G07795 | | | | | | | -1.2 | 3.45E-02 | -0.2 | 7.67E-01 | | | Forkhead box protein G1 |
| AT1G07902 | | | | | | | 0.1 | 7.82E-01 | 0.2 | 6.89E-01 | -0.1 | 9.07E-01 | Uncharacterized protein |
| AT1G07917 | | | | | | | 2.804753907 | 1.99E-04 | 1.888254874 | 1.24E-02 | 0.9 | 1.62E-02 | NA |
| AT1G08115 | | | | | | | -0.191847763 | 7.67E-01 | -1.152873528 | 1.06E-01 | 1.0 | 2.57E-01 | NA |
| AT1G08173 | | | | | | | 0.5 | 2.88E-01 | 0.8 | 1.34E-01 | | | NA |
| AT1G08283 | | | | | | | 0.2 | 7.43E-01 | 0.2 | 8.36E-01 | | | NA |
| AT1G08643 | | | | | | | 0.144726958 | 8.57E-01 | 0.492561275 | 5.17E-01 | | | NA |
| AT1G08763 | | | | | | | 2.525373486 | 5.65E-04 | 1.408098276 | 6.65E-02 | 1.1 | 8.38E-03 | NA |
| AT1G08765 | | | | | | | 0.2 | 6.97E-01 | 0.4 | 5.30E-01 | -0.1 | 8.27E-01 | NA |
| AT1G08903 | | | | | | | 0.0 | 9.73E-01 | 0.7 | 1.20E-01 | -0.7 | 2.02E-01 | NA |
| AT1G09155 | | | | | | | -0.2 | 6.85E-01 | -0.1 | 9.30E-01 | | | PP2-B15 |
| AT1G09467 | | | | | | | 0.1 | 8.22E-01 | -0.3 | 6.49E-01 | | | NA |
| AT1G09486 | | | | | | | 0.2 | 7.40E-01 | 0.4 | 4.92E-01 | -0.2 | 7.53E-01 | NA |
| AT1G09505 | | | | | | | 5.179989772 | 2.84E-04 | -0.449547132 | 8.59E-01 | 5.6 | 1.20E-03 | NA |
| AT1G09605 | | | | | | | 1.5 | 1.39E-02 | 0.5 | 5.86E-01 | 1.0 | 9.23E-02 | NA |
| AT1G09995 | | | | | | | 0.0 | 9.86E-01 | -0.2 | 6.83E-01 | 0.2 | 6.74E-01 | Uncharacterized protein |
| AT1G10220 | | | | | | | 0.8 | 1.27E-01 | -0.4 | 6.09E-01 | 1.2 | 3.44E-02 | ZCF37 |
| AT1G10790 | | | | | | | 0.6 | 3.30E-01 | 0.6 | 3.84E-01 | | | hypothetical protein |
| AT1G10875 | | | | | | | 5.566346012 | 2.44E-04 | -0.063149724 | 9.83E-01 | | | Core-2 |
| AT1G12030 | | | | | | | 3.822884179 | 2.17E-04 | -1.430298579 | 3.79E-01 | 5.3 | 1.28E-03 | Uncharacterized protein |
| AT1G12775 | | | | | | | 0.4 | 2.59E-01 | 0.6 | 1.08E-01 | -0.2 | 6.49E-01 | Pentatricopeptide repeat-containing protein At1g12775, mitochondrial |
| AT1G13145 | | | | | | | 0.0 | 9.78E-01 | 0.1 | 7.72E-01 | -0.1 | 7.64E-01 | NA |
| AT1G13540 | | | | | | | 1.2 | 4.39E-02 | 0.4 | 5.67E-01 | 0.7 | 2.79E-01 | F13B4.4 protein |
| AT1G13550 | | | | | | | 1.0 | 1.62E-02 | 0.4 | 4.56E-01 | | | F13B4.4 protein |
| AT1G13607 | | | | | | | 2.067065433 | 6.21E-03 | -1.002015848 | 3.97E-01 | 3.1 | 8.72E-03 | Defensin-like protein 286 |
| AT1G13608 | | | | | | | 3.190598127 | 4.01E-05 | 0.062066537 | 9.62E-01 | 3.1 | 5.26E-04 | Putative defensin-like protein 288 |
| AT1G13630 | | | | | | | 0.5 | 3.65E-01 | 0.4 | 5.95E-01 | | | Tetratricopeptide repeat -like superfamily protein |
| AT1G15010 | | | | | | | -0.807042062 | 2.91E-01 | -1.425359319 | 1.16E-01 | 0.6 | 6.71E-01 | T15D22.5 protein |
| AT1G15610 | | | | | | | 2.156702943 | 6.86E-03 | 2.633362286 | 2.84E-03 | -0.5 | 4.49E-01 | T16N11.11 protein |
| AT1G15630 | | | | | | | 0.836668165 | 1.33E-01 | 1.312445067 | 2.50E-02 | -0.5 | 2.74E-01 | T16N11.14 protein |
| AT1G15640 | | | | | | | 0.3 | 4.60E-01 | 1.2 | 7.45E-03 | -0.9 | 3.77E-02 | Transmembrane protein |
| AT1G15870 | | | | | | | 1.1 | 3.18E-02 | 0.2 | 7.60E-01 | 0.8 | 1.59E-01 | Mitochondrial glycoprotein family protein |
| AT1G15900 | | | | | | | -0.4 | 4.19E-01 | -0.5 | 2.94E-01 | 0.1 | 8.23E-01 | Transmembrane protein |
| AT1G16515 | | | | | | | -0.1 | 8.84E-01 | 0.2 | 8.52E-01 | | | At1g16515 |
| AT1G17285 | | | | | | | -0.019220537 | 9.77E-01 | 0.368850062 | 5.15E-01 | | | Transmembrane protein |
| AT1G17665 | | | | | | | 0.3 | 4.96E-02 | -0.2 | 2.95E-01 | 0.5 | 1.51E-02 | F1L3.36 |
| AT1G17830 | | | | | | | 0.1 | 8.08E-01 | 0.1 | 9.19E-01 | 0.1 | 9.49E-01 | At1g17830 |
| AT1G19025 | | | | | | | 0.3 | 3.82E-01 | 0.6 | 9.66E-02 | -0.3 | 4.98E-01 | DNA repair metallo-beta-lactamase domain-containing protein |

| | | | | | | | | | |
|---|---|---|---|---|---|---|---|---|---|
| AT1G19610 | | | -1.9 | 2.25E-01 | -3.4 | 1.42E-01 | 1.5 | 5.11E-01 | F18O14.36 |
| AT1G21530 | | | -0.3 | 5.43E-01 | 0.3 | 6.71E-01 | -0.6 | 3.89E-01 | Uncharacterized protein |
| AT1G24577 | | | -0.8 | 5.35E-02 | -0.1 | 8.63E-01 | -0.7 | 1.30E-01 | Uncharacterized protein |
| AT1G24580 | | | 3.3 | 1.73E-03 | -0.9 | 6.11E-01 | 4.2 | 5.24E-03 | At1g24580 |
| AT1G25422 | | | -1.6 | 1.12E-03 | -0.3 | 4.70E-01 | -1.3 | 2.52E-02 | Uncharacterized protein |
| AT1G27285 | | | 0.0 | 9.53E-01 | -0.4 | 4.36E-01 | 0.4 | 5.11E-01 | NA |
| AT1G29270 | | | -0.6 | 2.77E-01 | -0.4 | 5.12E-01 | -0.2 | 8.24E-01 | F28N24.5 protein |
| AT1G29510 | | | -0.1 | 8.25E-01 | -0.1 | 7.92E-01 | 0.0 | 9.61E-01 | Auxin-responsive protein SAUR67 |
| AT1G29640 | | | -0.1 | 9.35E-01 | 0.8 | 2.01E-01 | -0.8 | 1.11E-01 | hypothetical protein |
| AT1G31040 | | | -0.67669858 | 1.53E-01 | -0.303795033 | 5.79E-01 | -0.4 | 4.27E-01 | PLATZ transcription factor family protein |
| AT1G31310 | | | -0.784590006 | 2.60E-03 | -0.525061699 | 3.49E-02 | -0.3 | 3.72E-01 | At1g31310 |
| AT1G31355 | | | -0.4 | 4.93E-01 | -0.1 | 8.76E-01 | -0.3 | 7.53E-01 | NA |
| AT1G32630 | | | 0.1 | 8.82E-01 | -0.3 | 7.21E-01 | 0.4 | 6.49E-01 | hypothetical protein |
| AT1G33102 | | | 0.3 | 4.99E-01 | -0.1 | 9.32E-01 | 0.4 | 5.62E-01 | Uncharacterized protein |
| AT1G33475 | | | -0.1 | 7.61E-01 | -0.4 | 3.98E-01 | 0.2 | 6.09E-01 | SNARE-like superfamily protein |
| AT1G34245 | | | 0.6 | 1.67E-01 | 0.7 | 1.86E-01 | 0.0 | 9.57E-01 | Epidermal patterning factor-like protein |
| AT1G34460 | | | -0.1 | 7.75E-01 | 0.1 | 8.29E-01 | -0.2 | 6.44E-01 | CYCLIN B15 |
| AT1G35310 | | | 0.4 | 4.09E-01 | 0.1 | 8.97E-01 | | | Bet v I |
| AT1G43605 | | | 0.134320292 | 7.41E-01 | 0.172122407 | 7.08E-01 | 0.0 | 9.51E-01 | Uncharacterized protein |
| AT1G44010 | | | 1.0 | 2.60E-02 | 0.0 | 9.60E-01 | 0.9 | 3.11E-02 | Transmembrane protein |
| AT1G44020 | | | -0.4 | 3.57E-01 | 0.0 | 9.97E-01 | -0.4 | 4.58E-01 | Cysteine |
| AT1G44740 | | | -0.2 | 6.86E-01 | -0.2 | 8.01E-01 | 0.0 | 9.62E-01 | Uncharacterized protein |
| AT1G47130 | | | 4.02656029 | 3.27E-04 | 2.379801415 | 3.79E-02 | 1.6 | 2.69E-02 | Uncharacterized protein |
| AT1G47405 | | | 2.269268738 | 5.97E-02 | -2.176391647 | 2.14E-01 | | | NA |
| AT1G50790 | | | -0.3 | 5.81E-01 | 0.0 | 9.65E-01 | -0.3 | 7.04E-01 | Plant mobile domain protein family |
| AT1G52270 | | | 2.3 | 4.04E-04 | 1.5 | 2.27E-02 | 0.8 | 1.18E-01 | Uncharacterized protein F19K6.11 |
| AT1G52565 | | | -0.3 | 5.66E-01 | -0.3 | 5.46E-01 | 0.0 | 9.53E-01 | Uncharacterized protein At1g52565 |
| AT1G52690 | | | -3.764664195 | 1.64E-01 | -2.576024991 | 2.91E-01 | -1.2 | 8.37E-01 | Late embryogenesis abundant protein 7 |
| AT1G53130 | | | -0.2 | 7.19E-01 | 0.3 | 5.41E-01 | -0.5 | 3.29E-01 | Protein GRIM REAPER |
| AT1G53330 | | | 0.5 | 1.18E-01 | 0.2 | 6.63E-01 | 0.3 | 4.54E-01 | Pentatricopeptide repeat superfamily protein |
| AT1G53610 | | | 0.4 | 2.87E-01 | 0.4 | 2.60E-01 | -0.1 | 8.77E-01 | T3F20.7 protein |
| AT1G53620 | | | 2.409938247 | 3.53E-04 | 0.912318681 | 2.12E-01 | 1.5 | 2.47E-03 | T3F20.6 protein |
| AT1G53633 | | | 0.4 | 2.58E-01 | 0.0 | 9.73E-01 | 0.4 | 2.15E-01 | Uncharacterized protein |
| AT1G53635 | | | 0.1 | 7.94E-01 | -0.3 | 6.03E-01 | 0.4 | 4.11E-01 | T3F20.3 protein |
| AT1G54840 | | | 0.6 | 1.91E-01 | -0.2 | 7.12E-01 | 0.8 | 1.64E-01 | Increased DNA methylation 2 |
| AT1G55010 | | | -0.3 | 6.20E-01 | -0.5 | 4.62E-01 | 0.2 | 8.14E-01 | Plant defensin 1.5 |
| AT1G55230 | | | 3.165223899 | 8.62E-03 | 0.89701189 | 6.04E-01 | 2.3 | 8.41E-03 | hypothetical protein |
| AT1G59940 | | | -0.6 | 1.52E-01 | -0.9 | 6.42E-02 | 0.3 | 6.35E-01 | Response regulator 3 |
| AT1G61255 | | | 0.8 | 3.15E-01 | 0.3 | 7.68E-01 | | | Uncharacterized protein |
| AT1G61750 | | | -0.8 | 2.66E-02 | -0.9 | 2.76E-02 | 0.1 | 9.30E-01 | Cysteine-rich repeat secretory protein 1 |
| AT1G62350 | | | 1.0 | 2.06E-02 | 0.6 | 2.35E-01 | 0.4 | 4.16E-01 | Pentatricopeptide repeat superfamily protein |
| AT1G62420 | | | 2.357397172 | 3.69E-02 | -3.932921845 | 5.90E-02 | | | hypothetical protein |
| AT1G62920 | | | -0.1 | 9.10E-01 | 0.1 | 9.24E-01 | -0.1 | 8.59E-01 | Proteasome maturation factor |
| AT1G62981 | | | -0.8 | 9.78E-02 | -0.2 | 7.31E-01 | -0.6 | 3.85E-01 | F16P17.15 ) |
| AT1G63080 | | | -0.6 | 1.38E-01 | -0.2 | 6.87E-01 | -0.4 | 4.60E-01 | Pentatricopeptide repeat-containing protein At1g63080, mitochondrial |
| AT1G63150 | | | 0.3 | 5.64E-01 | 0.4 | 4.23E-01 | -0.1 | 8.45E-01 | Pentatricopeptide repeat-containing protein At1g63150 |
| AT1G63230 | | | 0.3 | 4.80E-01 | 0.3 | 5.04E-01 | 0.0 | 9.76E-01 | Uncharacterized protein F9N12.15 |
| AT1G63295 | | | 0.5 | 2.45E-01 | -0.2 | 7.76E-01 | 0.7 | 2.18E-01 | Remorin C-terminal domain-containing protein |
| AT1G63400 | | | 0.1 | 8.20E-01 | 0.5 | 3.18E-01 | -0.3 | 4.81E-01 | Pentatricopeptide repeat-containing protein At1g63400 |
| AT1G64160 | | | 4.5 | 2.41E-06 | 2.1 | 1.12E-02 | 2.5 | 1.08E-04 | Dirigent protein |
| AT1G64185 | | | 0.7 | 9.32E-02 | 0.1 | 8.44E-01 | 0.6 | 2.60E-01 | hypothetical protein |
| AT1G64220 | | | 0.9 | 5.66E-02 | 0.7 | 2.38E-01 | | | Mitochondrial import receptor subunit TOM7-1 |
| AT1G65480 | | | -0.52183443 | 2.57E-01 | -0.009930963 | 9.89E-01 | | | PEBP family protein |
| AT1G65481 | | | 1.9 | 6.37E-04 | 0.9 | 9.72E-02 | 0.9 | 1.80E-02 | Transmembrane protein |
| AT1G65485 | | | 2.29046326 | 1.95E-03 | 1.61746717 | 3.95E-02 | 0.7 | 1.27E-01 | NA |
| AT1G67035 | | | -0.1 | 8.62E-01 | -0.2 | 7.92E-01 | 0.1 | 9.28E-01 | F1O19.9 protein |
| AT1G67600 | | | 0.3 | 7.01E-01 | 0.4 | 5.57E-01 | | | hypothetical protein |
| AT1G68500 | | | -0.1 | 6.95E-01 | -0.1 | 7.26E-01 | 0.0 | 9.92E-01 | hypothetical protein |
| AT1G68725 | | | -0.3 | 3.73E-01 | -0.1 | 7.86E-01 | -0.2 | 7.26E-01 | Lysine-rich arabinogalactan protein 19 |
| AT1G69140 | -2.5 | 2.65E-02 | -1.396826734 | 2.95E-02 | 0.42020141 | 4.38E-01 | -1.8 | 3.59E-02 | NA |
| AT1G69150 | | | 0.9 | 2.44E-02 | 1.1 | 1.33E-02 | -0.2 | 7.24E-01 | DC1 domain-containing protein |
| AT1G69280 | | | 0.3 | 5.45E-01 | 0.4 | 3.91E-01 | | | Uncharacterized protein F23O10.14 |
| AT1G69290 | | | -0.6 | 5.43E-01 | 0.1 | 9.42E-01 | -0.6 | 5.64E-01 | Pentatricopeptide repeat-containing protein At1g69290 |
| AT1G70470 | | | 0.4 | 3.53E-01 | 0.5 | 3.51E-01 | -0.1 | 9.39E-01 | hypothetical protein |
| AT1G70885 | | | -1.7 | 7.74E-03 | -0.6 | 2.50E-01 | -1.0 | 1.59E-01 | NA |
| AT1G71015 | | | -0.8 | 9.56E-02 | 0.3 | 6.14E-01 | -1.1 | 7.21E-02 | At1g71015 |
| AT1G73780 | | | 0.1 | 8.96E-01 | 0.4 | 4.80E-01 | | | hypothetical protein |
| AT1G73890 | | | -0.5 | 2.65E-01 | -0.5 | 2.92E-01 | 0.0 | 9.80E-01 | Non-specific lipid transfer protein GPI-anchored 10 |
| AT1G74453 | | | 0.5 | 1.92E-01 | 0.0 | 9.69E-01 | 0.5 | 3.30E-01 | NA |
| AT1G74456 | | | 0.8 | 9.40E-02 | -0.1 | 9.08E-01 | 0.9 | 1.57E-01 | NA |
| AT1G75166 | | | 0.1 | 8.93E-01 | 0.4 | 6.96E-01 | -0.2 | 8.33E-01 | NA |
| AT1G75335 | | | 0.7 | 1.89E-01 | 0.2 | 7.79E-01 | 0.5 | 4.84E-01 | hypothetical protein |
| AT1G75360 | | | -0.3 | 2.88E-01 | -0.6 | 7.58E-02 | 0.3 | 5.08E-01 | Transmembrane protein |
| AT1G75720 | | | 0.6 | 2.65E-01 | 0.3 | 6.45E-01 | | | WEB family protein |
| AT1G76705 | | | -0.2 | 7.32E-01 | -0.2 | 7.96E-01 | | | Uncharacterized protein |
| AT1G76870 | | | -0.5 | 1.95E-01 | -0.1 | 8.73E-01 | -0.4 | 3.72E-01 | hypothetical protein |
| AT1G77131 | | | -0.2 | 8.13E-01 | -0.1 | 8.71E-01 | | | NA |
| AT1G77360 | | | -0.4 | 3.14E-01 | -0.6 | 1.21E-01 | 0.3 | 6.25E-01 | Pentatricopeptide repeat-containing protein At1g77360, mitochondrial |
| AT1G78865 | | | 0.5 | 2.09E-01 | 0.7 | 8.67E-02 | -0.2 | 6.77E-01 | NA |
| AT1G80880 | | | 0.3 | 4.85E-01 | 0.2 | 7.67E-01 | 0.1 | 8.37E-01 | Pentatricopeptide repeat-containing protein At1g80880, mitochondrial |
| AT1G80890 | | | 0.4 | 2.79E-01 | 0.0 | 9.80E-01 | 0.4 | 3.27E-01 | Uncharacterized protein |
| AT1G80970 | | | 0.2 | 8.01E-01 | 0.1 | 9.31E-01 | 0.1 | 9.30E-01 | XH domain-containing protein |
| AT2G00340 | | | -0.4 | 3.86E-01 | -0.5 | 3.55E-01 | 0.1 | 9.22E-01 | NA |

| | | | | | | | |
|---|---|---|---|---|---|---|---|
| AT2G00410 | -0.6 | 2.49E-01 | -0.7 | 2.22E-01 | 0.1 | 9.11E-01 | NA |
| AT2G00670 | -0.352541636 | 6.68E-01 | -0.963303955 | 2.94E-01 | 0.6 | 5.86E-01 | NA |
| AT2G00920 | 5.101882977 | 1.98E-04 | 0.045657137 | 9.86E-01 | 5.1 | 5.53E-04 | NA |
| AT2G01020 | -0.5 | 5.68E-01 | -0.6 | 5.49E-01 | 0.1 | 9.43E-01 | NA |
| AT2G01390 | -0.9 | 2.97E-02 | -0.5 | 2.58E-01 | -0.4 | 4.85E-01 | Pentatricopeptide repeat-containing protein At2g01390 |
| AT2G01913 | 0.1 | 7.71E-01 | -0.1 | 8.91E-01 | 0.2 | 7.42E-01 | Uncharacterized protein |
| AT2G04025 | 0.5 | 4.18E-01 | 1.0 | 1.11E-01 | -0.5 | 4.80E-01 | Root meristem growth factor |
| AT2G04190 | -0.7 | 1.97E-01 | -0.3 | 6.62E-01 | -0.4 | 5.78E-01 | TRAF-like family protein |
| AT2G04460 | 1.980228149 | 2.30E-02 | 2.476559461 | 8.95E-03 | | | NA |
| AT2G04955 | 0.8 | 5.29E-02 | 0.2 | 8.11E-01 | 0.7 | 1.85E-01 | NA |
| AT2G04965 | 1.1 | 1.02E-02 | 0.3 | 6.57E-01 | 0.8 | 1.02E-01 | NA |
| AT2G05518 | 0.0 | 9.94E-01 | 0.4 | 4.42E-01 | -0.4 | 5.11E-01 | NA |
| AT2G06995 | 0.2 | 5.78E-01 | 0.2 | 6.28E-01 | 0.0 | 9.97E-01 | NA |
| AT2G08145 | -0.3 | 5.31E-01 | -0.5 | 3.75E-01 | 0.2 | 8.28E-01 | NA |
| AT2G14095 | -0.6 | 3.18E-01 | 0.3 | 7.02E-01 | | | Expressed protein |
| AT2G14115 | -0.1 | 8.09E-01 | -0.3 | 3.30E-01 | 0.2 | 5.68E-01 | NA |
| AT2G16367 | 1.5 | 1.05E-02 | 2.0 | 2.71E-03 | -0.5 | 3.66E-01 | NA |
| AT2G17036 | 0.6 | 9.14E-02 | 0.3 | 5.48E-01 | 0.3 | 3.99E-01 | F-box domain-containing protein |
| AT2G17442 | 0.1 | 9.21E-01 | -0.1 | 9.19E-01 | 0.1 | 8.71E-01 | Uncharacterized protein |
| AT2G20362 | -0.6 | 3.22E-01 | -0.2 | 8.21E-01 | | | hypothetical protein |
| AT2G20480 | 0.328776069 | 3.61E-01 | 0.286637148 | 4.94E-01 | 0.0 | 9.48E-01 | hypothetical protein |
| AT2G20835 | 0.1 | 8.39E-01 | 0.1 | 8.81E-01 | 0.0 | 9.92E-01 | Uncharacterized protein |
| AT2G23040 | 0.3 | 5.09E-01 | 0.1 | 9.52E-01 | 0.3 | 6.75E-01 | NA |
| AT2G23270 | 4.1769507 | 6.15E-05 | 2.354311419 | 1.81E-02 | 1.8 | 2.94E-03 | Transmembrane protein |
| AT2G23830 | 1.6 | 3.21E-03 | 0.8 | 2.18E-01 | 0.9 | 3.90E-02 | Vesicle-associated protein 3-1 |
| AT2G25297 | 1.0 | 1.92E-02 | 0.1 | 9.23E-01 | 0.9 | 6.59E-02 | Transmembrane protein |
| AT2G26790 | 0.9 | 3.57E-02 | 0.6 | 2.05E-01 | | | Pentatricopeptide repeat-containing protein At2g26790, mitochondrial |
| AT2G28050 | 0.2 | 6.88E-01 | 0.5 | 1.95E-01 | -0.4 | 4.66E-01 | Pentatricopeptide repeat-containing protein At2g28050 |
| AT2G28085 | 0.163349246 | 7.36E-01 | -0.020355676 | 9.78E-01 | 0.2 | 7.52E-01 | SAUR-like auxin-responsive protein family |
| AT2G28270 | -0.2 | 6.54E-01 | -1.2 | 3.19E-02 | 1.0 | 1.31E-01 | hypothetical protein |
| AT2G28500 | -0.3 | 4.27E-01 | 0.3 | 5.73E-01 | -0.6 | 2.23E-01 | LOB domain-containing protein |
| AT2G30370 | 0.3 | 4.71E-01 | -0.2 | 6.44E-01 | 0.5 | 1.84E-01 | Epidermal patterning factor-like protein |
| AT2G30505 | -0.1 | 6.47E-01 | 0.0 | 9.82E-01 | -0.1 | 7.51E-01 | At2g30505 |
| AT2G30760 | 2.603355878 | 4.84E-03 | 0.347664803 | 8.19E-01 | | | Uncharacterized protein At2g30760 |
| AT2G30942 | -0.1 | 8.35E-01 | -0.4 | 5.44E-01 | | | Transmembrane protein, putative |
| AT2G30945 | -0.6 | 1.36E-01 | -0.8 | 6.45E-02 | 0.2 | 7.13E-01 | NA |
| AT2G31860 | 3.736121829 | 3.15E-05 | -0.240923563 | 8.62E-01 | 4.0 | 2.32E-04 | NA |
| AT2G32487 | 0.465359689 | 3.53E-01 | -0.094515088 | 9.08E-01 | 0.6 | 4.09E-01 | Secreted protein |
| AT2G32630 | 0.1 | 8.44E-01 | 0.4 | 3.08E-01 | -0.3 | 4.83E-01 | Pentatricopeptide repeat-containing protein At2g32630 |
| AT2G32650 | 0.3 | 5.02E-01 | -0.5 | 3.46E-01 | 0.9 | 1.63E-01 | At2g32650 |
| AT2G32795 | -0.1 | 7.68E-01 | -0.2 | 6.92E-01 | 0.1 | 9.28E-01 | NA |
| AT2G33735 | 0.4 | 4.67E-01 | 0.3 | 6.46E-01 | 0.1 | 9.25E-01 | J domain-containing protein |
| AT2G34340 | 0.1 | 8.78E-01 | 0.1 | 7.77E-01 | -0.1 | 9.08E-01 | Senescence regulator |
| AT2G36026 | -0.9 | 1.79E-02 | 0.1 | 8.63E-01 | -1.0 | 4.24E-02 | Transcription repressor |
| AT2G36295 | 0.5 | 2.37E-01 | 0.5 | 3.08E-01 | 0.0 | 9.90E-01 | At2g36295 |
| AT2G36307 | 0.1 | 8.82E-01 | 0.2 | 7.79E-01 | -0.1 | 9.01E-01 | Protein LITTLE ZIPPER 4 |
| AT2G36650 | -0.2 | 7.00E-01 | 0.2 | 7.67E-01 | -0.4 | 5.30E-01 | CHUP1-like protein |
| AT2G37530 | 0.4 | 2.73E-01 | 0.4 | 3.98E-01 | 0.0 | 9.47E-01 | Transmembrane protein |
| AT2G38430 | 0.6 | 1.21E-01 | 0.0 | 9.92E-01 | 0.6 | 1.76E-01 | Uncharacterized protein |
| AT2G38830 | 0.0 | 9.74E-01 | 0.0 | 9.23E-01 | 0.0 | 9.58E-01 | At2g38830 |
| AT2G40085 | -0.2 | 5.86E-01 | -0.3 | 4.13E-01 | 0.1 | 8.21E-01 | Uncharacterized protein |
| AT2G41082 | 0.2 | 6.14E-01 | -0.1 | 9.30E-01 | 0.3 | 6.08E-01 | Uncharacterized protein |
| AT2G41230 | 0.189154529 | 8.38E-01 | -1.080102618 | 3.07E-01 | 1.3 | 3.15E-01 | Protein ORGAN SIZE RELATED 1 |
| AT2G41905 | 0.0 | 9.64E-01 | 0.0 | 9.71E-01 | 0.0 | 1.00E+00 | hypothetical protein |
| AT2G42065 | 3.4 | 6.87E-03 | 0.3 | 8.97E-01 | | | J domain-containing protein |
| AT2G42110 | 0.3 | 3.30E-01 | 0.5 | 1.96E-01 | -0.1 | 7.61E-01 | At2g42110 |
| AT2G42800 | 0.6 | 1.11E-01 | 0.4 | 3.57E-01 | 0.2 | 6.90E-01 | Receptor like protein 29 |
| AT2G42860 | 0.3 | 5.91E-01 | -0.3 | 6.43E-01 | 0.6 | 3.76E-01 | Expressed protein |
| AT2G42865 | 0.4 | 1.39E-01 | 0.3 | 2.94E-01 | 0.1 | 8.69E-01 | Uncharacterized protein |
| AT2G43445 | -0.2 | 5.64E-01 | -0.4 | 4.07E-01 | 0.1 | 8.38E-01 | F-box |
| AT2G43990 | 0.0 | 9.98E-01 | 0.0 | 9.83E-01 | 0.0 | 9.86E-01 | GPI-anchored adhesin-like protein |
| AT2G44240 | -0.9 | 6.42E-01 | -3.3 | 2.51E-01 | 2.3 | 5.64E-01 | hypothetical protein |
| AT2G44578 | 3.0 | 1.22E-04 | 1.5 | 4.74E-02 | 1.4 | 5.32E-03 | Putative RING zinc finger protein |
| AT2G44700 | -1.1 | 2.16E-02 | -0.4 | 3.35E-01 | -0.6 | 3.24E-01 | F-box |
| AT2G44735 | 0.2 | 5.03E-01 | -0.3 | 3.39E-01 | 0.5 | 1.31E-01 | Transmembrane protein |
| AT2G45023 | -0.4 | 4.95E-01 | -0.1 | 9.15E-01 | -0.3 | 7.01E-01 | NA |
| AT2G45760 | 3.073139355 | 5.03E-05 | 1.058320045 | 1.67E-01 | 2.0 | 1.26E-03 | BON1-associated protein 2 |
| AT2G46410 | 0.0 | 9.29E-01 | -0.1 | 8.52E-01 | 0.1 | 9.35E-01 | CPC |
| AT2G47780 | -0.552421703 | 3.94E-01 | -0.26883177 | 7.35E-01 | -0.3 | 7.62E-01 | REF |
| AT3G00700 | 5.344471303 | 5.44E-04 | -0.591470558 | 8.26E-01 | | | NA |
| AT3G01175 | -0.7 | 1.53E-01 | -0.8 | 1.10E-01 | 0.2 | 8.39E-01 | T4P13.14 protein |
| AT3G01940 | 0.1 | 8.97E-01 | 0.0 | 9.95E-01 | 0.1 | 9.22E-01 | Transmembrane protein |
| AT3G01960 | 0.4 | 3.58E-01 | 0.4 | 2.86E-01 | -0.1 | 8.64E-01 | At3g01960 |
| AT3G02480 | 0.6 | 5.58E-01 | -0.8 | 5.92E-01 | 1.5 | 2.89E-01 | Stress-induced protein KIN2-like |
| AT3G02500 | -0.1 | 9.19E-01 | 0.4 | 4.14E-01 | | | hypothetical protein |
| AT3G03170 | -0.2 | 7.47E-01 | 0.4 | 3.96E-01 | -0.6 | 3.04E-01 | Protein OXIDATIVE STRESS 3 LIKE 3 |
| AT3G03341 | -2.403852754 | 2.85E-02 | -0.638129563 | 4.72E-01 | -1.8 | 2.49E-01 | hypothetical protein |
| AT3G03370 | -0.2 | 7.48E-01 | -0.2 | 8.00E-01 | 0.0 | 9.96E-01 | Uncharacterized protein |
| AT3G03850 | 1.2 | 1.78E-02 | 0.4 | 5.22E-01 | 0.8 | 1.16E-01 | Auxin-responsive protein SAUR26 |
| AT3G04960 | 0.1 | 8.26E-01 | 0.1 | 9.28E-01 | | | Trichohyalin, putative |
| AT3G05858 | -0.1 | 9.31E-01 | 0.4 | 4.96E-01 | | | hypothetical protein |
| AT3G06345 | -0.6 | 3.00E-01 | -0.4 | 4.92E-01 | -0.1 | 8.99E-01 | NA |
| AT3G06868 | -0.190323876 | 7.11E-01 | -0.257384431 | 6.59E-01 | 0.1 | 9.34E-01 | Vitellogenin-like protein |

| | | | | | | | |
|---|---|---|---|---|---|---|---|
| AT3G07425 | 0.620497893 | 2.52E-01 | 0.50983947 | 4.24E-01 | | | hypothetical protein |
| AT3G07600 | 0.7 | 1.76E-01 | 0.8 | 1.09E-01 | | | Heavy metal transport |
| AT3G07675 | 0.283348479 | 5.24E-01 | -0.225479601 | 7.00E-01 | 0.5 | 3.74E-01 | NA |
| AT3G09730 | 0.0 | 9.36E-01 | 0.2 | 6.86E-01 | -0.2 | 7.82E-01 | POLAR LOCALIZATION DURING ASYMMETRIC DIVISION AND protein |
| AT3G11745 | 0.0 | 9.62E-01 | 0.4 | 4.82E-01 | -0.4 | 6.06E-01 | Transmembrane protein |
| AT3G12955 | 0.0 | 9.62E-01 | -0.1 | 9.24E-01 | | | At3g12955 |
| AT3G13270 | -0.1 | 7.78E-01 | -0.7 | 1.42E-01 | 0.6 | 1.84E-01 | NA |
| AT3G13432 | 2.9 | 1.34E-04 | 1.7 | 2.40E-02 | 1.2 | 3.30E-02 | Transmembrane protein |
| AT3G14700 | 1.1 | 1.14E-01 | -1.2 | 2.84E-01 | | | hypothetical protein |
| AT3G15340 | 0.7 | 7.66E-02 | -0.6 | 2.94E-01 | 1.2 | 1.98E-02 | Uncharacterized protein |
| AT3G17580 | 0.2 | 6.48E-01 | 0.4 | 4.04E-01 | -0.2 | 7.21E-01 | hypothetical protein |
| AT3G19200 | 0.8 | 4.78E-02 | 0.8 | 7.06E-02 | 0.0 | 9.97E-01 | hypothetical protein |
| AT3G19330 | 0.8 | 1.20E-01 | 0.6 | 3.21E-01 | 0.2 | 7.75E-01 | UPF0496 protein At3g19330 |
| AT3G20810 | -0.6 | 3.09E-04 | -0.3 | 4.65E-02 | -0.3 | 1.54E-01 | 2-oxoglutarate and Fe-dependent oxygenase superfamily protein |
| AT3G21010 | -0.2 | 7.45E-01 | 0.0 | 9.88E-01 | -0.2 | 7.96E-01 | NA |
| AT3G21520 | 2.1 | 7.69E-04 | 1.1 | 7.78E-02 | 0.9 | 6.65E-02 | Protein DMP1 |
| AT3G21755 | 0.0 | 9.98E-01 | 0.4 | 3.08E-01 | -0.4 | 3.91E-01 | NA |
| AT3G22820 | -0.1 | 8.43E-01 | 0.0 | 9.69E-01 | -0.1 | 9.18E-01 | EPIDERMAL PATTERNING FACTOR-like protein 5 |
| AT3G23260 | -0.2 | 6.49E-01 | 0.2 | 6.77E-01 | | | Putative F-box protein At3g23260 |
| AT3G23930 | 0.6 | 7.99E-02 | 0.2 | 6.59E-01 | 0.4 | 3.13E-01 | Troponin T, skeletal protein |
| AT3G24020 | -0.5 | 4.86E-01 | 0.2 | 7.79E-01 | -0.7 | 3.72E-01 | Dirigent protein 16 |
| AT3G24535 | -0.4 | 5.45E-01 | -0.1 | 9.40E-01 | -0.3 | 7.38E-01 | Uncharacterized protein |
| AT3G24730 | -0.2 | 5.31E-01 | -0.2 | 6.06E-01 | 0.0 | 9.92E-01 | At3g24730 |
| AT3G24890 | 0.0 | 9.41E-01 | 0.0 | 9.81E-01 | 0.0 | 9.80E-01 | V-SNARE coiled-coil homology domain-containing protein |
| AT3G25655 | 1.4 | 8.90E-02 | 0.4 | 7.49E-01 | | | Protein IDA-LIKE 1 |
| AT3G25905 | 0.38660306 | 5.08E-01 | 0.58692311 | 3.44E-01 | | | CLAVATA3 |
| AT3G26135 | 0.125953186 | 8.69E-01 | 0.578074298 | 3.92E-01 | | | 2-oxoglutarate |
| AT3G26960 | 0.0 | 9.53E-01 | 1.0 | 1.06E-01 | | | Pollen Ole e 1 allergen and extensin family protein |
| AT3G27027 | 0.4 | 3.82E-01 | -0.2 | 6.76E-01 | | | GPI-anchored-like protein |
| AT3G27250 | -2.2 | 3.31E-03 | 0.0 | 9.76E-01 | -2.2 | 1.87E-03 | hypothetical protein |
| AT3G27290 | -0.4 | 5.32E-01 | 0.3 | 6.38E-01 | | | F-box protein At3g27290 |
| AT3G27660 | 0.2 | 7.71E-01 | 0.2 | 8.36E-01 | 0.0 | 9.86E-01 | Oleosin |
| AT3G27940 | 0.1 | 8.59E-01 | -0.3 | 5.46E-01 | 0.4 | 4.52E-01 | LOB domain-containing protein 26 |
| AT3G28290 | -0.077271406 | 9.08E-01 | 0.290987369 | 6.55E-01 | -0.4 | 6.35E-01 | UPF0496 protein At3g28290 |
| AT3G28420 | 0.846274644 | 2.25E-01 | 0.459412369 | 6.15E-01 | | | Transmembrane protein |
| AT3G29140 | -0.6 | 1.11E-01 | -0.5 | 2.43E-01 | -0.1 | 8.28E-01 | Uncharacterized protein |
| AT3G29330 | -0.5 | 1.56E-01 | -0.6 | 1.26E-01 | 0.1 | 8.88E-01 | Uncharacterized protein |
| AT3G29370 | -0.87520991 | 3.10E-02 | -0.934880252 | 3.28E-02 | 0.1 | 9.33E-01 | P1R3 |
| AT3G30350 | 0.6 | 8.34E-02 | 0.5 | 2.65E-01 | 0.2 | 7.65E-01 | Root meristem growth factor-like protein |
| AT3G42725 | 0.2 | 6.80E-01 | 0.6 | 2.99E-01 | -0.3 | 6.02E-01 | Membrane lipoprotein |
| AT3G42800 | -0.1 | 8.11E-01 | 0.0 | 9.97E-01 | -0.1 | 8.59E-01 | Protein BIG GRAIN 1-like C |
| AT3G44420 | 0.0 | 9.58E-01 | 0.1 | 9.23E-01 | -0.1 | 9.05E-01 | NA |
| AT3G44710 | -0.8 | 8.21E-02 | 0.1 | 9.00E-01 | -0.8 | 1.12E-01 | At3g44710 ) |
| AT3G44716 | 0.2 | 6.37E-01 | 0.2 | 7.09E-01 | 0.0 | 9.92E-01 | Uncharacterized protein |
| AT3G44960 | 0.4 | 5.23E-01 | 0.4 | 6.01E-01 | | | Shugoshin |
| AT3G45638 | -0.2 | 8.23E-01 | -0.3 | 6.88E-01 | 0.2 | 8.88E-01 | NA |
| AT3G46880 | 0.469671735 | 2.39E-01 | 0.327429566 | 4.94E-01 | 0.1 | 8.19E-01 | Uncharacterized protein T6H20.90 |
| AT3G48490 | 0.8 | 3.50E-02 | 0.5 | 2.35E-01 | 0.3 | 5.27E-01 | hypothetical protein |
| AT3G48600 | -0.6 | 1.11E-01 | -0.2 | 5.95E-01 | -0.3 | 4.77E-01 | SWIB complex BAF60b domain-containing protein |
| AT3G49570 | 3.5 | 5.50E-04 | 2.4 | 2.35E-02 | 1.1 | 2.08E-02 | Protein RESPONSE TO LOW SULFUR 3 |
| AT3G49580 | 2.904798503 | 1.60E-03 | 2.103787907 | 3.14E-02 | | | Response to low sulfur 1 |
| AT3G50030 | 2.661420305 | 3.77E-03 | 1.72495671 | 8.81E-02 | | | ARM-repeat |
| AT3G50040 | 0.6 | 1.80E-01 | 0.2 | 7.14E-01 | | | At3g50040 |
| AT3G50120 | -0.9 | 1.55E-02 | -0.3 | 5.27E-01 | -0.7 | 1.80E-01 | Transmembrane protein, |
| AT3G50825 | 0.9 | 7.68E-02 | 0.4 | 5.44E-01 | 0.5 | 3.08E-01 | NA |
| AT3G51220 | -0.1 | 8.93E-01 | 0.4 | 5.97E-01 | | | WEB family protein At3g51220 |
| AT3G52525 | 0.0 | 9.44E-01 | -0.3 | 5.15E-01 | 0.3 | 5.38E-01 | Transcription repressor OFP6 |
| AT3G52550 | -0.3 | 5.30E-01 | 0.1 | 8.55E-01 | -0.4 | 5.14E-01 | At3g52550 |
| AT3G52670 | 0.5 | 1.54E-01 | 0.5 | 2.75E-01 | 0.1 | 9.08E-01 | FBD-associated F-box protein At3g52670 |
| AT3G52770 | 0.255360461 | 5.87E-01 | 0.304086329 | 5.67E-01 | 0.0 | 9.49E-01 | Protein LITTLE ZIPPER 3 |
| AT3G53170 | 0.2 | 5.86E-01 | 0.2 | 6.23E-01 | 0.0 | 9.88E-01 | Pentatricopeptide repeat-containing protein At3g53170 |
| AT3G53220 | 0.9 | 5.89E-02 | 0.5 | 3.58E-01 | 0.4 | 5.57E-01 | Thioredoxin-like 3-3 |
| AT3G53232 | -0.718285687 | 2.05E-01 | -0.722785449 | 2.58E-01 | 0.0 | 9.98E-01 | Small polypeptide DEVIL 20 |
| AT3G53250 | -0.009443921 | 9.90E-01 | -0.946500521 | 1.53E-01 | 0.9 | 2.07E-01 | Uncharacterized protein T4D2.180 |
| AT3G54520 | 6.039183016 | 2.67E-05 | 1.511271449 | 3.32E-01 | 4.5 | 2.05E-04 | Uncharacterized protein |
| AT3G54530 | 9.448781276 | 3.11E-05 | 3.230183266 | 1.78E-01 | 6.2 | 2.86E-04 | Uncharacterized protein |
| AT3G55790 | 4.539398401 | 8.33E-05 | 3.760180925 | 1.53E-03 | 0.8 | 5.77E-02 | Transmembrane protein |
| AT3G56275 | -1.201437889 | 5.49E-02 | -0.447863606 | 4.95E-01 | -0.8 | 3.68E-01 | NA |
| AT3G56825 | -0.05728578 | 9.39E-01 | 0.043411645 | 9.65E-01 | | | NA |
| AT3G57160 | 0.1 | 8.33E-01 | 0.3 | 3.33E-01 | -0.3 | 4.83E-01 | Cysteine-rich TM module stress tolerance protein |
| AT3G57200 | 0.4 | 2.37E-01 | 0.0 | 9.62E-01 | 0.4 | 3.38E-01 | Glycosyltransferase family 92 protein |
| AT3G57360 | -0.2 | 7.10E-01 | 0.0 | 9.91E-01 | -0.2 | 7.17E-01 | hypothetical protein |
| AT3G58150 | 2.705337154 | 2.08E-02 | 0.543901918 | 7.80E-01 | | | Uncharacterized protein F9D24.60 |
| AT3G58200 | 0.1 | 8.96E-01 | -0.1 | 8.59E-01 | 0.2 | 7.61E-01 | MATH domain and coiled-coil domain-containing protein At3g58200 |
| AT3G58430 | 0.6 | 1.72E-01 | 0.2 | 6.99E-01 | 0.4 | 5.15E-01 | MATH domain-containing protein |
| AT3G58865 | -0.6 | 2.34E-01 | -0.2 | 7.73E-01 | -0.4 | 5.70E-01 | NA |
| AT3G59930 | -0.966955738 | 2.30E-01 | -1.368381783 | 1.41E-01 | 0.4 | 7.43E-01 | Defensin-like protein 206 |
| AT3G60150 | -0.1 | 8.74E-01 | 0.0 | 9.75E-01 | -0.1 | 8.74E-01 | NADH dehydrogenase 1 alpha subcomplex assembly factor 3 |
| AT3G61198 | -0.5 | 2.59E-01 | -0.9 | 6.27E-02 | 0.4 | 5.36E-01 | NA |
| AT3G61390 | 0.8 | 4.70E-02 | -0.5 | 3.71E-01 | 1.2 | 2.27E-02 | RING |
| AT3G61840 | 0.726232314 | 1.14E-01 | -0.11446388 | 8.80E-01 | 0.8 | 1.15E-01 | Uncharacterized protein F21F14.10 |
| AT3G61898 | 0.588179115 | 3.76E-01 | 0.747759349 | 3.00E-01 | -0.2 | 8.65E-01 | Transmembrane protein |
| AT3G62070 | -0.5 | 5.31E-02 | -0.6 | 4.06E-02 | 0.1 | 8.59E-01 | Fold protein |

| | | | | | | | |
|---|---|---|---|---|---|---|---|
| AT3G63050 | -0.4 | 5.73E-01 | 0.4 | 6.52E-01 | -0.8 | 3.60E-01 | Uncharacterized protein T20O10_150 |
| AT4G01510 | 0.1 | 9.19E-01 | 0.1 | 9.23E-01 | 0.0 | 9.93E-01 | Protein ARV |
| AT4G01790 | 0.8 | 5.84E-02 | 0.2 | 7.05E-01 | 0.6 | 2.42E-01 | hypothetical protein |
| AT4G01895 | 0.71230756 | 2.35E-01 | -0.158859943 | 8.70E-01 | 0.9 | 2.61E-01 | At4g01895 regulator protein NIMIN-1-like protein) |
| AT4G02005 | 0.0 | 9.51E-01 | 0.4 | 3.96E-01 | -0.4 | 4.56E-01 | NA |
| AT4G02360 | -0.1 | 9.28E-01 | 0.3 | 6.76E-01 | -0.4 | 6.66E-01 | T14P8.17 |
| AT4G03475 | -0.2 | 6.37E-01 | -0.2 | 5.76E-01 | 0.1 | 9.27E-01 | NA |
| AT4G03935 | -0.1 | 7.96E-01 | 0.0 | 9.60E-01 | -0.1 | 7.95E-01 | NA |
| AT4G03940 | -0.3 | 5.94E-01 | -0.7 | 3.39E-01 | 0.3 | 6.14E-01 | T24M8.7 protein |
| AT4G04190 | 0.2 | 5.18E-01 | -0.1 | 8.36E-01 | 0.3 | 3.87E-01 | Transmembrane protein |
| AT4G04360 | -0.3 | 6.89E-01 | 0.1 | 8.88E-01 | -0.4 | 6.37E-01 | Uncharacterized protein AT4g04360 |
| AT4G06195 | 2.4 | 1.03E-04 | -0.4 | 6.08E-01 | 2.8 | 2.85E-04 | NA |
| AT4G06880 | 0.5 | 2.88E-01 | 0.5 | 3.10E-01 | 0.0 | 9.63E-01 | NA |
| AT4G08991 | 0.0 | 9.44E-01 | 0.5 | 4.02E-01 | -0.4 | 5.22E-01 | NA |
| AT4G09200 | -0.3 | 9.31E-01 | 0.1 | 9.81E-01 | -0.4 | 9.25E-01 | At4g09200 domain-containing protein) |
| AT4G10860 | 1.797375194 | 6.30E-04 | -0.647800227 | 3.67E-01 | 2.4 | 1.80E-03 | F8M12.2 protein |
| AT4G11547 | -0.6 | 2.74E-01 | -0.1 | 8.78E-01 | | | Cysteine |
| AT4G11655 | 0.6 | 1.49E-01 | 0.6 | 1.52E-01 | 0.0 | 9.40E-01 | CASP-like protein |
| AT4G12545 | -0.8 | 9.20E-02 | 0.1 | 8.08E-01 | -0.9 | 1.22E-01 | Putative lipid-binding protein AIR1B |
| AT4G12850 | 0.0 | 9.39E-01 | -0.1 | 8.85E-01 | 0.0 | 9.53E-01 | Uncharacterized protein AT4g12850 |
| AT4G13560 | 0.0 | 9.31E-01 | 0.3 | 4.84E-01 | -0.3 | 5.39E-01 | Late embryogenesis abundant protein family protein |
| AT4G13690 | 0.6 | 1.74E-01 | 0.6 | 2.08E-01 | 0.0 | 9.83E-01 | RNA-binding protein |
| AT4G13820 | 0.1 | 8.76E-01 | -0.3 | 6.92E-01 | | | Disease resistance like protein family protein) |
| AT4G13880 | -0.9 | 3.25E-02 | -0.4 | 4.29E-01 | -0.6 | 3.31E-01 | Receptor-like protein 48 |
| AT4G14103 | -0.2 | 7.30E-01 | -0.1 | 9.13E-01 | -0.1 | 8.99E-01 | F-box |
| AT4G14380 | -0.5 | 1.23E-01 | -0.2 | 6.90E-01 | -0.4 | 4.54E-01 | Cotton fiber protein |
| AT4G14450 | 0.9 | 3.14E-02 | -0.8 | 1.77E-01 | 1.7 | 3.28E-03 | Uncharacterized protein |
| AT4G14695 | 0.0 | 9.83E-01 | 1.0 | 4.36E-02 | | | Mitochondrial pyruvate carrier |
| AT4G14860 | 0.0 | 9.37E-01 | 0.3 | 5.87E-01 | -0.3 | 5.87E-01 | Transcription repressor OFP11 |
| AT4G15150 | 0.8 | 3.11E-02 | 0.5 | 1.94E-01 | 0.2 | 5.62E-01 | Glycine-rich protein |
| AT4G15165 | 0.1 | 8.84E-01 | 0.0 | 9.99E-01 | | | Putative proteasome subunit alpha type-4-B |
| AT4G15700 | -0.2 | 8.63E-01 | -1.9 | 9.16E-02 | 1.7 | 6.90E-02 | hypothetical protein |
| AT4G16590 | 2.646839633 | 1.73E-01 | 3.940413149 | 5.48E-02 | -1.3 | 4.00E-01 | Uncharacterized protein |
| AT4G16610 | -0.1 | 8.26E-01 | 0.5 | 2.62E-01 | -0.6 | 1.67E-01 | hypothetical protein |
| AT4G18400 | -0.3 | 3.96E-01 | -0.4 | 3.53E-01 | 0.1 | 9.09E-01 | At4g18400 |
| AT4G19090 | 0.4 | 3.24E-01 | 0.2 | 6.59E-01 | 0.2 | 7.70E-01 | Transmembrane protein, putative |
| AT4G19550 | 0.6 | 2.66E-01 | 0.4 | 5.49E-01 | | | Zinc ion binding |
| AT4G19980 | 1.2 | 8.12E-02 | 1.0 | 1.77E-01 | 0.1 | 8.70E-01 | hypothetical protein |
| AT4G20350 | 0.4 | 3.65E-01 | 0.2 | 7.70E-01 | 0.2 | 7.39E-01 | Oxidoreductase |
| AT4G20370 | 1.3 | 1.10E-01 | 1.7 | 4.70E-02 | | | Protein of the phosphatidylethanolamine-binding protein family |
| AT4G20730 | 0.5 | 4.08E-01 | 0.5 | 4.49E-01 | | | NA |
| AT4G21200 | 1.7 | 8.22E-03 | 1.2 | 6.86E-02 | | | Gibberellin 2-oxidase 8 |
| AT4G21320 | 0.0 | 9.69E-01 | -0.1 | 8.08E-01 | 0.2 | 7.41E-01 | Protein HEAT-STRESS-ASSOCIATED 32 |
| AT4G21500 | -0.4 | 1.46E-01 | -0.4 | 1.93E-01 | 0.0 | 9.90E-01 | At4g21500 |
| AT4G21780 | -0.2 | 7.88E-01 | -0.2 | 7.85E-01 | 0.0 | 9.73E-01 | Uncharacterized protein |
| AT4G21930 | -0.5 | 6.81E-01 | 1.1 | 2.53E-01 | | | Uncharacterized protein AT4g21930 |
| AT4G22030 | 5.272193821 | 1.35E-05 | 2.562564333 | 2.19E-02 | 2.7 | 4.93E-04 | F-box protein with a domain protein |
| AT4G22475 | 0.2 | 6.52E-01 | -0.3 | 6.27E-01 | 0.6 | 4.27E-01 | Putative glycine-rich protein |
| AT4G23493 | -0.6 | 3.03E-01 | 0.6 | 2.82E-01 | -1.2 | 6.00E-02 | Uncharacterized protein AT4g23490 |
| AT4G24026 | -1.2 | 6.21E-02 | -0.4 | 5.26E-01 | -0.8 | 4.14E-01 | Transmembrane protein |
| AT4G24110 | -0.3 | 5.70E-01 | -0.6 | 3.90E-01 | 0.2 | 8.03E-01 | hypothetical protein |
| AT4G24910 | 0.3 | 5.46E-01 | 0.4 | 5.02E-01 | | | Glucuronoxylan 4-O-methyltransferase-like protein |
| AT4G25930 | 2.4 | 1.59E-01 | 1.2 | 6.03E-01 | | | DUF295 domain-containing protein |
| AT4G26055 | 0.6 | 3.43E-01 | 0.3 | 6.76E-01 | | | hypothetical protein |
| AT4G26255 | -0.3 | 6.93E-01 | -0.1 | 8.71E-01 | | | NA |
| AT4G27460 | -1.0 | 3.57E-02 | -1.4 | 9.56E-03 | 0.4 | 5.69E-01 | CBS domain-containing protein CBSX5 |
| AT4G28162 | -0.356382438 | 5.40E-01 | -0.591685902 | 3.62E-01 | | | NA |
| AT4G28460 | 3.3 | 4.19E-05 | 1.3 | 8.54E-02 | 1.9 | 8.47E-04 | PAMP-induced secreted peptide 1 |
| AT4G28703 | -1.1 | 9.58E-02 | -1.3 | 6.66E-02 | 0.3 | 8.29E-01 | hypothetical protein |
| AT4G29560 | 0.0 | 9.77E-01 | 0.3 | 6.53E-01 | -0.3 | 6.69E-01 | Fanconi Anaemia group E protein C-terminal domain-containing protein |
| AT4G30090 | 0.5 | 2.85E-01 | 0.4 | 3.92E-01 | 0.0 | 9.63E-01 | Golgin family A protein |
| AT4G30230 | 1.8 | 2.43E-04 | 0.9 | 6.20E-02 | 0.9 | 3.35E-02 | Uncharacterized protein AT4g30230 |
| AT4G30660 | -0.7 | 6.56E-02 | 0.1 | 8.90E-01 | -0.8 | 7.82E-02 | UPF0057 membrane protein At4g30660 |
| AT4G30845 | 0.1 | 8.09E-01 | -0.4 | 4.57E-01 | 0.5 | 3.77E-01 | hypothetical protein |
| AT4G31330 | 0.083362768 | 9.00E-01 | -0.000257209 | 1.00E+00 | | | DUF599 domain-containing protein |
| AT4G33960 | -1.4 | 5.15E-05 | -0.7 | 1.12E-02 | -0.7 | 6.65E-02 | Transmembrane protein |
| AT4G33980 | -0.4 | 5.76E-02 | -0.8 | 5.07E-03 | 0.3 | 3.42E-01 | Uncharacterized protein |
| AT4G34419 | 0.5 | 4.09E-01 | -0.5 | 5.32E-01 | 0.9 | 1.43E-01 | Uncharacterized protein AT4g34420 |
| AT4G34420 | 0.8 | 5.66E-02 | 0.8 | 7.98E-02 | 0.0 | 9.97E-01 | P-loop containing nucleoside triphosphate hydrolases superfamily protein |
| AT4G34770 | 0.4 | 4.37E-01 | -0.8 | 1.47E-01 | 1.2 | 6.35E-02 | hypothetical protein |
| AT4G34790 | -0.5 | 1.76E-01 | -0.9 | 2.80E-02 | 0.4 | 4.21E-01 | SAUR-like auxin-responsive protein family |
| AT4G34800 | 0.0 | 9.65E-01 | -0.3 | 5.90E-01 | 0.3 | 6.24E-01 | Uncharacterized protein |
| AT4G35720 | 0.3 | 5.56E-01 | 0.6 | 1.98E-01 | -0.3 | 4.95E-01 | hypothetical protein |
| AT4G36700 | 6.389930494 | 1.93E-03 | -0.590615591 | 8.89E-01 | 7.0 | 2.04E-03 | Vicilin-like seed storage protein At4g36700 |
| AT4G37290 | 2.919171066 | 3.32E-03 | 0.852679975 | 5.18E-01 | 2.1 | 1.72E-02 | PAMP-induced secreted peptide 2 |
| AT4G37295 | 0.3 | 5.94E-01 | -0.4 | 5.87E-01 | | | hypothetical protein |
| AT4G37483 | 0.0 | 9.35E-01 | -0.5 | 3.45E-01 | 0.5 | 3.80E-01 | Uncharacterized protein |
| AT4G37608 | 0.736986508 | 1.24E-01 | 0.547997747 | 3.30E-01 | | | hypothetical protein |
| AT4G37810 | -0.2 | 6.76E-01 | 0.1 | 8.57E-01 | -0.3 | 5.64E-01 | EPIDERMAL PATTERNING FACTOR-like protein 2 |
| AT4G38330 | 0.5 | 2.68E-01 | 0.2 | 7.20E-01 | 0.3 | 6.76E-01 | Uncharacterized protein |
| AT4G39580 | 0.4 | 1.67E-01 | -0.1 | 7.64E-01 | 0.6 | 1.63E-01 | F-box |
| AT4G39600 | 0.3 | 5.50E-01 | 0.0 | 9.60E-01 | 0.2 | 7.16E-01 | Galactose oxidase |
| AT4G39675 | -0.6 | 2.06E-01 | -2.0 | 3.08E-03 | 1.4 | 2.89E-02 | Uncharacterized protein |

| | | | | | | | |
|---|---|---|---|---|---|---|---|
| AT4G39925 | -0.5 | 4.02E-01 | -0.1 | 8.64E-01 | -0.3 | 6.98E-01 | Uncharacterized protein |
| AT5G00445 | -0.1 | 8.35E-01 | 0.2 | 7.01E-01 | -0.3 | 6.08E-01 | NA |
| AT5G00560 | 0.161969494 | 7.59E-01 | -0.807825428 | 1.55E-01 | | | NA |
| AT5G01175 | 0.0 | 9.55E-01 | 0.5 | 2.44E-01 | -0.4 | 3.03E-01 | NA |
| AT5G01445 | 0.3 | 5.81E-01 | -0.2 | 7.92E-01 | 0.5 | 4.48E-01 | Uncharacterized protein |
| AT5G02520 | 0.7 | 1.33E-01 | 0.4 | 4.63E-01 | 0.3 | 6.67E-01 | Dentin sialophosphoprotein-like protein |
| AT5G02565 | 1.251553704 | 1.42E-01 | 0.755899911 | 4.86E-01 | | | NA |
| AT5G02575 | 0.865434953 | 1.25E-01 | 0.731748952 | 2.63E-01 | | | NA |
| AT5G03250 | 0.3 | 4.99E-01 | 0.5 | 3.23E-01 | -0.2 | 7.97E-01 | Phototropic-responsive NPH3 family protein |
| AT5G05300 | 0.4 | 5.13E-01 | -0.4 | 6.59E-01 | 0.8 | 3.52E-01 | Gb\|AAF01528.1 |
| AT5G05435 | 0.1 | 8.08E-01 | -0.6 | 3.31E-01 | 0.7 | 2.90E-01 | NA |
| AT5G08150 | -0.3 | 6.04E-01 | -1.0 | 7.26E-02 | 0.8 | 2.60E-01 | Uncharacterized protein |
| AT5G08360 | -0.5 | 1.63E-01 | -0.4 | 2.30E-01 | 0.0 | 9.73E-01 | Uncharacterized protein |
| AT5G09580 | 0.6 | 4.92E-02 | 0.2 | 6.56E-01 | 0.4 | 2.47E-01 | Heat shock protein |
| AT5G09935 | -0.546057976 | 3.77E-01 | -0.55598196 | 4.31E-01 | 0.0 | 9.95E-01 | NA |
| AT5G09980 | 0.0 | 9.72E-01 | 0.2 | 7.48E-01 | -0.2 | 7.70E-01 | Elicitor peptide 4 |
| AT5G10210 | 0.392686657 | 3.79E-01 | -1.420654136 | 2.68E-02 | 1.8 | 1.87E-02 | hypothetical protein |
| AT5G10310 | 0.0 | 9.29E-01 | 0.0 | 9.74E-01 | 0.0 | 9.76E-01 | Epidermal patterning factor-like protein |
| AT5G10440 | 0.2 | 6.49E-01 | 0.2 | 7.64E-01 | 0.0 | 9.58E-01 | Cyclin d42 |
| AT5G11242 | 0.2 | 6.56E-01 | -0.1 | 9.12E-01 | 0.3 | 6.44E-01 | NA |
| AT5G13220 | -0.4 | 5.51E-01 | 0.0 | 9.86E-01 | -0.4 | 6.91E-01 | Protein TIFY |
| AT5G13880 | -0.2 | 7.24E-01 | 0.1 | 8.65E-01 | -0.3 | 6.37E-01 | Cotton fiber protein |
| AT5G14070 | 0.7 | 9.90E-02 | -0.1 | 8.64E-01 | 0.8 | 6.54E-02 | Glutaredoxin-C8 |
| AT5G14280 | -0.09006047 | 8.83E-01 | -0.127051167 | 8.56E-01 | 0.0 | 9.71E-01 | GLABROUS1 enhancer-binding protein-like 2 |
| AT5G14380 | -0.662230631 | 2.22E-01 | -0.379533859 | 5.38E-01 | -0.3 | 6.75E-01 | Classical arabinogalactan protein 6 |
| AT5G14890 | 0.0 | 9.98E-01 | 0.2 | 7.20E-01 | | | NHL domain-containing protein |
| AT5G16640 | 0.2 | 7.45E-01 | -0.1 | 8.39E-01 | 0.3 | 5.66E-01 | Pentatricopeptide repeat-containing protein At5g16640, mitochondrial |
| AT5G18060 | -0.221634479 | 6.20E-01 | 0.066223964 | 9.13E-01 | -0.3 | 5.99E-01 | Auxin-responsive protein SAUR23 |
| AT5G18065 | 0.7 | 7.17E-02 | 0.4 | 4.59E-01 | | | Protein HEAT-INDUCED TAS1 TARGET 3 |
| AT5G18245 | -0.6 | 2.36E-01 | -0.1 | 9.23E-01 | -0.5 | 4.60E-01 | NA |
| AT5G18920 | 0.3 | 6.53E-01 | 0.2 | 7.81E-01 | 0.1 | 9.52E-01 | CHCH domain-containing protein |
| AT5G19170 | -0.3 | 3.28E-01 | -0.3 | 5.09E-01 | -0.1 | 9.04E-01 | NEP-interacting protein, putative |
| AT5G19650 | 0.0 | 9.41E-01 | 0.5 | 3.30E-01 | -0.5 | 3.82E-01 | Transcription repressor OFP8 |
| AT5G20045 | -0.5 | 5.43E-01 | 0.5 | 5.95E-01 | -1.0 | 2.87E-01 | Transmembrane protein |
| AT5G20670 | -0.4 | 5.37E-01 | 0.6 | 3.24E-01 | -1.0 | 1.93E-01 | At5g20670 ) |
| AT5G20820 | 0.3 | 5.69E-01 | -0.2 | 6.90E-01 | 0.5 | 3.75E-01 | Auxin-responsive protein SAUR76 |
| AT5G22520 | 1.6 | 7.31E-02 | 0.0 | 9.90E-01 | 1.6 | 9.44E-02 | At5g22520 |
| AT5G22530 | 2.7 | 8.20E-04 | 0.7 | 4.38E-01 | 1.9 | 1.32E-02 | Uncharacterized protein At5g22530 |
| AT5G22540 | 0.4 | 6.07E-01 | -0.8 | 4.35E-01 | 1.2 | 2.74E-01 | At5g22540 ) |
| AT5G22608 | 0.2 | 7.13E-01 | -0.2 | 7.90E-01 | | | Uncharacterized protein |
| AT5G23800 | 0.2 | 6.92E-01 | 0.5 | 1.96E-01 | -0.4 | 4.39E-01 | Agenet domain-containing protein |
| AT5G24040 | 0.7 | 2.58E-01 | 0.5 | 4.66E-01 | | | F-box SKIP23-like protein |
| AT5G24600 | 1.510014658 | 3.43E-02 | 0.140609302 | 9.12E-01 | 1.4 | 8.67E-02 | DUF599 domain-containing protein |
| AT5G26200 | -1.8 | 5.67E-03 | -1.1 | 6.04E-02 | -0.7 | 3.56E-01 | AT5g26200 |
| AT5G26730 | -0.8 | 2.62E-02 | -0.5 | 1.86E-01 | -0.3 | 5.64E-01 | Fasciclin-like arabinogalactan family protein |
| AT5G26790 | -0.1 | 7.60E-01 | -0.2 | 6.67E-01 | 0.1 | 9.08E-01 | Transmembrane protein |
| AT5G37400 | 0.5 | 3.36E-01 | 0.1 | 9.21E-01 | | | Similarity to unknown protein |
| AT5G37410 | 0.2 | 7.71E-01 | -0.1 | 8.43E-01 | 0.3 | 6.14E-01 | Similarity to unknown protein |
| AT5G37485 | 1.1 | 7.54E-02 | 0.9 | 2.54E-01 | | | NA |
| AT5G37840 | 5.764659169 | 1.40E-03 | 3.525190264 | 7.10E-02 | 2.2 | 3.59E-02 | hypothetical protein |
| AT5G39240 | 0.329585488 | 4.84E-01 | 0.247170876 | 6.76E-01 | | | Transcription factor UPBEAT-like protein |
| AT5G39865 | -0.2 | 6.73E-01 | -0.1 | 8.40E-01 | -0.1 | 9.05E-01 | Uncharacterized protein At5g39865 |
| AT5G40630 | -0.7 | 3.73E-01 | 0.1 | 9.17E-01 | -0.8 | 4.23E-01 | Ubiquitin-like domain-containing protein |
| AT5G41170 | 0.5 | 2.10E-01 | 0.3 | 5.15E-01 |  |  | Pentatricopeptide repeat superfamily protein |
| AT5G41290 | 1.3 | 6.79E-04 | 0.8 | 3.24E-02 | 0.5 | 1.95E-01 | Cysteine-rich repeat secretory protein 58 |
| AT5G41471 | 0.747410874 | 3.96E-01 | 0.76658536 | 4.46E-01 | | | NA |
| AT5G42090 | -0.1 | 9.08E-01 | -0.6 | 2.50E-01 | 0.5 | 3.68E-01 | Dbj\|BAA84809.1 |
| AT5G42370 | 0.3 | 5.50E-01 | -0.2 | 7.60E-01 | | | Calcineurin-like metallo-phosphoesterase superfamily protein |
| AT5G42655 | 0.3 | 6.42E-01 | 0.2 | 8.52E-01 | 0.1 | 8.62E-01 | Dirigent protein |
| AT5G42900 | -0.592140129 | 1.80E-03 | -0.632732124 | 2.56E-03 | 0.0 | 9.09E-01 | COR27 |
| AT5G44010 | 0.2 | 5.81E-01 | -0.2 | 7.34E-01 | 0.3 | 4.23E-01 | Fanconi anemia group F protein |
| AT5G44569 | 0.618717475 | 1.84E-01 | -0.272068064 | 6.96E-01 | 0.9 | 1.19E-01 | NA |
| AT5G45115 | -0.025155639 | 9.71E-01 | -0.360548245 | 5.94E-01 | 0.3 | 6.90E-01 | Transcription termination factor family protein |
| AT5G45310 | -0.6 | 4.90E-02 | -0.5 | 9.84E-02 | -0.1 | 9.11E-01 | Uncharacterized protein |
| AT5G45530 | 0.0 | 9.55E-01 | 0.0 | 9.41E-01 | 0.0 | 9.84E-01 | Transmembrane protein, putative |
| AT5G47330 | 0.4 | 8.18E-01 | 1.7 | 2.29E-01 | -1.3 | 4.36E-01 | Uncharacterized protein |
| AT5G47510 | 0.4 | 2.29E-01 | -0.2 | 6.04E-01 | 0.6 | 1.07E-01 | Sec14p-like phosphatidylinositol transfer family protein |
| AT5G48175 | 1.7 | 2.92E-03 | 0.8 | 2.31E-01 | | | At5g48175 |
| AT5G48690 | 0.1 | 8.02E-01 | 0.1 | 8.50E-01 | 0.0 | 9.93E-01 | UBA domain-containing protein |
| AT5G48850 | 3.0 | 4.43E-05 | 1.5 | 3.38E-02 | 1.5 | 2.71E-03 | Protein SULFUR DEFICIENCY-INDUCED 1 |
| AT5G49330 | -0.2 | 7.67E-01 | 0.2 | 7.25E-01 | -0.4 | 4.84E-01 | PFG3 |
| AT5G49390 | 0.4 | 3.50E-01 | 0.6 | 2.09E-01 | | | Similarity to unknown protein |
| AT5G49410 | 0.2 | 6.36E-01 | 0.1 | 8.55E-01 | 0.1 | 8.81E-01 | At5g49410 |
| AT5G49645 | -0.2 | 5.83E-01 | -0.1 | 8.52E-01 | -0.1 | 8.39E-01 | Uncharacterized protein |
| AT5G50090 | -0.1 | 7.64E-01 | -0.2 | 7.21E-01 | | | D-ribose-binding periplasmic protein |
| AT5G50175 | -0.6 | 3.72E-01 | -0.2 | 7.99E-01 | | | Transmembrane protein |
| AT5G50360 | -3.22797136 | 1.89E-01 | -2.750291392 | 3.34E-01 | -0.4 | 9.45E-01 | Uncharacterized protein At5g50360 |
| AT5G50530 | -0.1 | 9.63E-01 | -0.2 | 9.00E-01 | 0.1 | 9.26E-01 | CBS domain-containing protein CBSCBSPB4 |
| AT5G51390 | 0.0 | 9.46E-01 | -0.3 | 7.21E-01 | 0.3 | 7.22E-01 | At5g51390 |
| AT5G52300 | -1.540498126 | 4.37E-01 | -1.738167885 | 4.51E-01 | 0.2 | 9.68E-01 | CAP160 protein |
| AT5G52545 | 0.2 | 4.60E-01 | 0.0 | 9.51E-01 | 0.3 | 5.32E-01 | hypothetical protein |
| AT5G52547 | 0.3 | 5.62E-01 | 0.1 | 9.40E-01 | 0.3 | 7.49E-01 | hypothetical protein |
| AT5G52670 | 3.209661189 | 2.13E-04 | 1.987030099 | 2.24E-02 | 1.2 | 3.26E-02 | HMA domain-containing protein |

| | | | | | | | |
|---|---|---|---|---|---|---|---|
| AT5G52680 | 5.554524645 | 6.96E-05 | 3.025489108 | 2.66E-02 | 2.5 | 2.97E-03 | Copper transport protein family |
| AT5G52700 | 2.083392981 | 3.49E-03 | 0.650955605 | 4.71E-01 | 1.4 | 5.62E-02 | Copper transport protein family |
| AT5G54064 | 0.0 | 9.74E-01 | 0.1 | 8.17E-01 | -0.1 | 8.64E-01 | NA |
| AT5G54450 | 1.9 | 1.99E-01 | 0.4 | 8.51E-01 | | | DUF295 domain-containing protein |
| AT5G54550 | 3.1 | 2.00E-02 | 0.5 | 8.28E-01 | | | DUF295 domain-containing protein |
| AT5G54560 | 3.6 | 2.24E-03 | 0.4 | 8.25E-01 | | | DUF295 domain-containing protein |
| AT5G55150 | 5.4 | 2.59E-04 | 0.2 | 9.53E-01 | 5.2 | 1.95E-03 | Putative F-box protein At5g55150 |
| AT5G55680 | -0.2 | 7.25E-01 | 0.5 | 2.64E-01 | -0.7 | 1.91E-01 | Glycine-rich protein |
| AT5G57080 | 0.7 | 4.91E-02 | 0.2 | 6.56E-01 | 0.5 | 2.53E-01 | hypothetical protein |
| AT5G57123 | -0.537654981 | 4.04E-01 | -0.147838047 | 8.61E-01 | | | Methionyl-tRNA synthetase |
| AT5G57790 | -0.7 | 3.03E-02 | -0.2 | 5.16E-01 | -0.4 | 2.52E-01 | At5g57790 |
| AT5G58570 | -1.0 | 1.45E-02 | -1.0 | 1.94E-02 | 0.0 | 9.92E-01 | Transmembrane protein |
| AT5G58680 | 0.3 | 5.68E-01 | -0.2 | 7.44E-01 | 0.5 | 3.84E-01 | ARM repeat superfamily protein |
| AT5G59990 | -0.7 | 1.36E-01 | -0.1 | 8.96E-01 | -0.6 | 3.14E-01 | CCT motif family protein |
| AT5G61360 | 0.0 | 9.32E-01 | -0.5 | 1.30E-01 | 0.5 | 2.43E-01 | Transmembrane protein |
| AT5G61865 | 0.0 | 9.35E-01 | -0.1 | 8.25E-01 | 0.2 | 7.97E-01 | Uncharacterized protein |
| AT5G62490 | 5.476090395 | 1.23E-03 | 2.064169066 | 3.21E-01 | 3.4 | 2.65E-03 | HVA22-like protein b |
| AT5G64667 | -0.1 | 7.39E-01 | -0.1 | 8.79E-01 | -0.1 | 9.12E-01 | Protein IDA-LIKE 2 |
| AT5G64690 | -0.1 | 8.32E-01 | 0.6 | 3.52E-01 | | | Uncharacterized protein |
| AT5G64890 | 2.090058908 | 1.09E-03 | 1.486346801 | 2.59E-02 | 0.6 | 2.75E-01 | Elicitor peptide 2 |
| AT5G64905 | 3.0 | 5.91E-04 | 1.4 | 1.47E-01 | 1.7 | 1.88E-02 | Elicitor peptide 3 |
| AT5G65420 | 0.0 | 9.86E-01 | 0.2 | 6.53E-01 | -0.2 | 7.13E-01 | CYCLIN D41 |
| AT5G65575 | -0.3 | 4.35E-01 | -0.1 | 8.17E-01 | -0.2 | 7.45E-01 | NA |
| AT5G65850 | 0.1 | 8.92E-01 | 0.1 | 9.12E-01 | | | F-box protein At5g65850 |
| AT5G66631 | 0.3 | 4.19E-01 | 0.2 | 6.08E-01 | | | Protein DA1-like domain-containing protein |
| AT5G66670 | 1.6 | 2.29E-02 | 1.0 | 2.47E-01 | 0.6 | 3.27E-01 | UPF0496 protein At5g66670 |
| AT5G67020 | 0.3 | 5.67E-01 | 0.2 | 8.14E-01 | | | At5g67020 |
| AT5G67245 | -0.3 | 5.63E-01 | -0.1 | 8.08E-01 | -0.1 | 8.56E-01 | Uncharacterized protein |

**Table S5 Functional enrichment analysis of genes differentially regulated by NUF versus urea**

**Up-regulated (NUF vs Urea at 7 d)**

| GO Biological Process | Normed Freq. | *P* -value |
|---|---|---|
| cell death | 3.47 | 4.6E-09 |
| abscission | 2.72 | 4.3E-02 |
| response to biotic stimulus | 2.6 | 3.5E-45 |
| response to external stimulus | 2.49 | 4.4E-47 |
| cellular homeostasis | 2.44 | 3.6E-05 |
| DNA metabolic process | 2.27 | 1.9E-07 |
| photosynthesis | 2.2 | 7.0E-04 |
| cell cycle | 2.18 | 5.5E-09 |
| response to stress | 2.16 | 3.3E-64 |
| response to abiotic stimulus | 2.16 | 5.3E-28 |
| response to light stimulus | 2.02 | 1.6E-17 |
| secondary metabolic process | 1.97 | 1.3E-07 |
| circadian rhythm | 1.93 | 1.6E-02 |
| response to chemical | 1.82 | 4.6E-32 |
| other biological processes | 1.75 | 2.4E-08 |
| response to endogenous stimulus | 1.69 | 5.0E-12 |
| signal transduction | 1.65 | 7.2E-11 |
| lipid metabolic process | 1.51 | 1.5E-04 |
| cell communication | 1.48 | 2.8E-03 |
| generation of precursor metabolites and energy | 1.32 | 4.4E-02 |
| other cellular processes | 1.2 | 1.7E-14 |
| cellular component organization | 1.16 | 6.9E-03 |
| other metabolic processes | 1.11 | 6.8E-05 |
| post-embryonic development | 1.11 | 2.5E-02 |
| biosynthetic process | 1.1 | 6.7E-03 |
| nucleobase-containing compound metabolic process | 1.01 | 3.5E-02 |
| reproduction | 1 | 4.2E-02 |
| catabolic process | 0.99 | 4.8E-02 |
| multicellular organism development | 0.97 | 3.4E-02 |
| anatomical structure development | 0.91 | 1.5E-02 |
| transport | 0.89 | 2.7E-02 |
| protein metabolic process | 0.7 | 3.6E-03 |
| carbohydrate metabolic process | 0.69 | 7.7E-03 |
| pollination | 0.59 | 3.8E-02 |
| flower development | 0.56 | 7.3E-03 |
| growth | 0.46 | 2.5E-06 |
| translation | 0.44 | 4.7E-06 |
| unknown biological processes | 0.42 | 7.1E-28 |
| cell differentiation | 0.37 | 2.8E-07 |
| cell growth | 0.34 | 1.4E-04 |
| **GO Molecular function** | | |
| motor activity | 5.51 | 2.1E-08 |
| signaling receptor activity | 1.82 | 1.2E-02 |
| nucleotide binding | 1.8 | 4.0E-07 |
| carbohydrate binding | 1.6 | 3.5E-02 |
| kinase activity | 1.56 | 4.8E-05 |
| other binding | 1.48 | 8.2E-09 |
| DNA binding | 1.35 | 5.7E-04 |
| DNA-binding transcription factor activity | 1.35 | 1.1E-03 |
| catalytic activity | 1.31 | 5.9E-07 |
| nucleic acid binding | 1.26 | 1.1E-02 |
| protein binding | 1.22 | 6.2E-05 |
| transferase activity | 1.2 | 2.0E-03 |
| transporter activity | 1.18 | 2.6E-02 |
| hydrolase activity | 1.15 | 1.2E-02 |
| unknown molecular functions | 0.72 | 5.0E-12 |
| RNA binding | 0.52 | 2.0E-07 |
| enzyme regulator activity | 0.33 | 6.4E-04 |
| lipid binding | 0.17 | 6.1E-04 |
| **GO Cellular component** | | |
| thylakoid | 2.47 | 4.1E-09 |
| cytoskeleton | 1.93 | 4.8E-04 |
| peroxisome | 1.78 | 4.0E-03 |
| plastid | 1.74 | 4.9E-10 |
| cell wall | 1.69 | 1.2E-03 |
| chloroplast | 1.55 | 2.1E-16 |
| endoplasmic reticulum | 1.41 | 2.1E-03 |
| other intracellular components | 1.26 | 2.8E-04 |
| plasma membrane | 1.17 | 6.2E-03 |
| cytoplasm | 1.15 | 1.8E-03 |
| other membranes | 1.1 | 2.0E-02 |
| extracellular region | 1.07 | 2.8E-02 |
| cytosol | 0.99 | 4.3E-02 |
| nucleus | 0.95 | 1.2E-02 |
| mitochondrion | 0.9 | 1.3E-02 |

**Up-regulated (NUF vs Urea at 12 h)**

| GO Biological Process | Normed Freq. | *P* -value |
|---|---|---|
| secondary metabolic process | 4.06 | 2.0E-30 |
| response to biotic stimulus | 3.44 | 2.6E-64 |
| response to external stimulus | 3.4 | 4.7E-74 |
| cell death | 2.72 | 2.1E-04 |
| response to abiotic stimulus | 2.66 | 2.0E-36 |
| response to stress | 2.64 | 1.5E-83 |
| other biological processes | 2.62 | 2.4E-19 |
| cellular homeostasis | 2.46 | 4.1E-04 |
| response to chemical | 2.37 | 1.7E-56 |
| response to endogenous stimulus | 2.28 | 1.3E-23 |
| cell communication | 1.96 | 3.1E-05 |
| signal transduction | 1.95 | 4.3E-14 |
| catabolic process | 1.23 | 1.1E-02 |
| other cellular processes | 1.21 | 7.8E-11 |
| response to light stimulus | 1.2 | 1.7E-02 |
| transport | 1.13 | 2.9E-02 |
| other metabolic processes | 1.09 | 2.5E-03 |
| biosynthetic process | 1.08 | 1.9E-02 |
| multicellular organism development | 1.04 | 3.6E-02 |
| nucleobase-containing compound metabolic process | 1.04 | 3.8E-02 |
| anatomical structure development | 0.96 | 3.5E-02 |
| post-embryonic development | 0.87 | 3.6E-02 |
| cellular component organization | 0.74 | 2.6E-03 |
| reproduction | 0.73 | 2.2E-03 |
| cell differentiation | 0.71 | 1.8E-02 |
| growth | 0.65 | 5.6E-03 |
| cell growth | 0.55 | 1.8E-02 |
| protein metabolic process | 0.5 | 1.5E-04 |
| flower development | 0.49 | 1.0E-02 |
| DNA metabolic process | 0.43 | 8.4E-03 |
| unknown biological processes | 0.31 | 4.0E-29 |
| translation | 0.15 | 5.2E-10 |
| cell cycle | 0.15 | 1.9E-06 |
| **GO Molecular function** | | |
| unknown molecular functions | 0.74 | 5.8E-08 |
| RNA binding | 0.52 | 1.1E-05 |
| other binding | 1.4 | 1.8E-05 |
| DNA-binding transcription factor activity | 1.54 | 1.4E-04 |
| DNA binding | 1.5 | 1.5E-04 |
| structural molecule activity | 0.32 | 3.4E-03 |
| nucleotide binding | 1.45 | 3.8E-03 |
| protein binding | 1.15 | 3.9E-03 |
| signaling receptor activity | 2.22 | 5.2E-03 |
| transferase activity | 1.19 | 7.1E-03 |
| transporter activity | 1.31 | 1.3E-02 |
| nucleic acid binding | 1.31 | 1.4E-02 |
| other molecular functions | 0.28 | 1.9E-02 |
| catalytic activity | 1.09 | 2.0E-02 |
| transcription regulator activity | 0.21 | 4.3E-02 |
| hydrolase activity | 1.03 | 4.8E-02 |
| **GO Cellular component** | | |
| cell wall | 2.69 | 4.1E-08 |
| extracellular region | 1.47 | 3.5E-06 |
| unknown cellular components | 0.6 | 9.9E-06 |
| plasma membrane | 1.42 | 3.1E-05 |
| cytoskeleton | 0.1 | 4.7E-04 |
| nucleoplasm | 0.17 | 6.7E-04 |
| chloroplast | 1.22 | 6.8E-04 |
| Golgi apparatus | 0.53 | 1.7E-03 |
| vacuole | 1.51 | 2.6E-03 |
| cytoplasm | 1.14 | 6.9E-03 |
| other membranes | 1.17 | 1.3E-02 |
| nucleus | 0.93 | 1.3E-02 |
| endosome | 0.38 | 1.3E-02 |
| plastid | 1.15 | 2.9E-02 |
| cytosol | 1.11 | 3.0E-02 |
| mitochondrion | 1.04 | 3.5E-02 |
| other cellular components | 0.87 | 3.7E-02 |
| ribosome | 0.54 | 4.0E-02 |
| endoplasmic reticulum | 1.19 | 4.1E-02 |
| other intracellular components | 0.95 | 4.2E-02 |

**Down-regulated (NUF vs Urea at 7 d)**

| GO Biological Process | Normed F | *P* -value |
|---|---|---|
| cell differentiation | 3.31 | 1.62E-72 |
| growth | 2.83 | 8.88E-58 |
| secondary metabolic process | 2.72 | 1.11E-30 |
| cell growth | 2.53 | 8.58E-19 |
| carbohydrate metabolic process | 2.21 | 8.89E-22 |
| anatomical structure development | 1.93 | 1.49E-70 |
| multicellular organism development | 1.75 | 2.41E-41 |
| lipid metabolic process | 1.66 | 8.90E-10 |
| pollination | 1.3 | 0.025 |
| response to endogenous stimulus | 1.21 | 4.06E-04 |
| response to chemical | 1.2 | 3.02E-06 |
| response to stress | 1.18 | 1.06E-05 |
| other biological processes | 1.17 | 8.29E-03 |
| transport | 1.16 | 2.90E-03 |
| response to external stimulus | 1.11 | 7.19E-03 |
| response to biotic stimulus | 1.1 | 0.011 |
| other cellular processes | 1.09 | 1.20E-06 |
| response to abiotic stimulus | 1.06 | 0.022 |
| biosynthetic process | 1.02 | 0.019 |
| other metabolic processes | 0.99 | 0.018 |
| signal transduction | 0.93 | 0.022 |
| cellular component organization | 0.9 | 0.01 |
| catabolic process | 0.88 | 0.012 |
| cell communication | 0.85 | 0.035 |
| response to light stimulus | 0.72 | 6.48E-05 |
| reproduction | 0.69 | 7.10E-07 |
| cell cycle | 0.69 | 3.82E-03 |
| cell death | 0.67 | 0.05 |
| post-embryonic development | 0.63 | 1.16E-07 |
| protein metabolic process | 0.61 | 4.29E-06 |
| generation of precursor metabolites and energy | 0.59 | 6.35E-03 |
| nucleobase-containing compound metabolic process | 0.58 | 8.63E-16 |
| flower development | 0.51 | 1.56E-04 |
| unknown biological processes | 0.43 | 1.15E-47 |
| photosynthesis | 0.39 | 3.25E-03 |
| embryo development | 0.34 | 1.55E-08 |
| regulation of gene expression, epigenetic | 0.33 | 0.042 |
| DNA metabolic process | 0.11 | 1.08E-11 |
| translation | 0.02 | 1.15E-35 |
| **GO Molecular function** | | |
| carbohydrate binding | 2.48 | 8.19E-07 |
| transporter activity | 1.69 | 2.57E-10 |
| transferase activity | 1.4 | 7.90E-11 |
| other binding | 1.3 | 3.14E-07 |
| kinase activity | 1.27 | 1.52E-03 |
| enzyme regulator activity | 1.26 | 0.022 |
| catalytic activity | 1.23 | 3.89E-07 |
| lipid binding | 1.23 | 0.047 |
| hydrolase activity | 1.18 | 1.17E-03 |
| DNA-binding transcription factor activity | 1.05 | 0.034 |
| nucleotide binding | 0.98 | 0.049 |
| nucleic acid binding | 0.9 | 0.033 |
| DNA binding | 0.83 | 5.75E-03 |
| protein binding | 0.79 | 8.93E-07 |
| unknown molecular functions | 0.75 | 2.25E-16 |
| structural molecule activity | 0.53 | 1.30E-03 |
| translation factor activity, RNA binding | 0.3 | 0.026 |
| nuclease activity | 0.27 | 1.77E-04 |
| other molecular functions | 0.22 | 4.41E-05 |
| chromatin binding | 0.22 | 5.78E-04 |
| RNA binding | 0.18 | 3.47E-38 |
| transcription regulator activity | 0.17 | 4.24E-04 |
| **GO Cellular component** | | |
| lysosome | 3.06 | 2.91E-03 |
| cell wall | 2.39 | 7.50E-14 |
| extracellular region | 1.73 | 5.76E-27 |
| plasma membrane | 1.66 | 8.55E-22 |
| Golgi apparatus | 1.43 | 3.67E-05 |
| other membranes | 1.38 | 1.19E-08 |
| cytoskeleton | 1.33 | 0.019 |
| endoplasmic reticulum | 1.18 | 0.013 |
| vacuole | 1.16 | 0.017 |
| cytoplasm | 0.88 | 1.08E-03 |
| other cellular components | 0.88 | 0.013 |
| nucleus | 0.8 | 5.96E-12 |

**Down-regulated (NUF vs Urea at 12 h)**

| GO Biological Process | Normed Freq. | *P* -value |
|---|---|---|
| secondary metabolic process | 3.28 | 1.14E-18 |
| cell growth | 2.55 | 5.06E-08 |
| cell differentiation | 2.4 | 1.37E-12 |
| growth | 2.37 | 3.94E-14 |
| carbohydrate metabolic process | 2.22 | 3.73E-09 |
| cellular homeostasis | 1.95 | 9.74E-03 |
| lipid metabolic process | 1.87 | 1.24E-06 |
| photosynthesis | 1.87 | 0.019 |
| response to abiotic stimulus | 1.72 | 2.22E-09 |
| anatomical structure development | 1.67 | 2.15E-15 |
| multicellular organism development | 1.65 | 6.40E-13 |
| response to chemical | 1.57 | 2.18E-12 |
| response to endogenous stimulus | 1.56 | 1.12E-06 |
| response to stress | 1.49 | 1.64E-10 |
| cell communication | 1.4 | 0.018 |
| response to light stimulus | 1.36 | 1.68E-03 |
| transport | 1.22 | 0.011 |
| response to external stimulus | 1.2 | 8.58E-03 |
| other cellular processes | 1.17 | 2.96E-08 |
| other metabolic processes | 1.09 | 2.38E-03 |
| response to biotic stimulus | 1.09 | 0.038 |
| biosynthetic process | 1.08 | 0.018 |
| catabolic process | 1.08 | 0.044 |
| cellular component organization | 0.96 | 0.048 |
| signal transduction | 0.88 | 0.034 |
| post-embryonic development | 0.82 | 0.021 |
| reproduction | 0.68 | 7.45E-04 |
| flower development | 0.63 | 0.038 |
| protein metabolic process | 0.57 | 1.05E-03 |
| nucleobase-containing compound metabolic | 0.44 | 1.83E-11 |
| pollination | 0.43 | 0.031 |
| embryo development | 0.36 | 7.74E-04 |
| unknown biological processes | 0.34 | 1.64E-25 |
| cell cycle | 0.31 | 2.34E-04 |
| translation | 0.03 | 1.23E-13 |
| **GO Molecular function** | | |
| carbohydrate binding | 3.14 | 4.21E-05 |
| transporter activity | 1.96 | 2.11E-07 |
| transferase activity | 1.46 | 3.57E-06 |
| other binding | 1.43 | 9.23E-06 |
| catalytic activity | 1.29 | 5.77E-05 |
| hydrolase activity | 1.26 | 4.60E-03 |
| kinase activity | 1.22 | 0.029 |
| protein binding | 0.88 | 0.013 |
| DNA binding | 0.83 | 0.033 |
| nucleic acid binding | 0.73 | 0.025 |
| unknown molecular functions | 0.64 | 2.33E-13 |
| RNA binding | 0.24 | 1.33E-12 |
| nuclease activity | 0.14 | 6.63E-03 |
| **GO Cellular component** | | |
| cell wall | 1.73 | 3.90E-03 |
| extracellular region | 1.6 | 2.63E-08 |
| other membranes | 1.46 | 2.09E-05 |
| plasma membrane | 1.35 | 3.10E-04 |
| vacuole | 1.21 | 0.041 |
| chloroplast | 1.15 | 6.63E-03 |
| plastid | 1.13 | 0.035 |
| cytoplasm | 0.97 | 0.038 |
| cytosol | 0.94 | 0.047 |
| other cellular components | 0.89 | 0.042 |
| nucleus | 0.71 | 5.16E-10 |
| other intracellular components | 0.68 | 2.31E-04 |
| unknown cellular components | 0.68 | 3.54E-04 |
| mitochondrion | 0.65 | 5.17E-06 |
| endosome | 0.39 | 0.017 |
| ribosome | 0.28 | 4.45E-03 |
| nucleoplasm | 0.27 | 3.52E-03 |
| nucleolus | 0.08 | 4.58E-05 |

| | | | | | |
|---|---|---|---|---|---|
| other cellular components | 0.76 | 4.1E-03 | chloroplast | 0.78 | 4.74E-07 |
| Golgi apparatus | 0.73 | 1.2E-02 | cytosol | 0.75 | 5.47E-05 |
| nucleoplasm | 0.68 | 4.7E-02 | endosome | 0.71 | 0.027 |
| unknown cellular components | 0.56 | 1.3E-08 | unknown cellular components | 0.69 | 4.76E-08 |
| endosome | 0.2 | 1.5E-04 | peroxisome | 0.67 | 0.028 |
| | | | mitochondrion | 0.57 | 9.06E-19 |
| | | | plastid | 0.52 | 2.88E-10 |
| | | | other intracellular components | 0.47 | 1.42E-21 |
| | | | thylakoid | 0.31 | 6.96E-07 |
| | | | nucleoplasm | 0.21 | 1.43E-07 |
| | | | nucleolus | 0.18 | 6.97E-09 |
| | | | ribosome | 0.18 | 5.76E-08 |

**Table S6 Enriched KEGG pathways in genes differentially regulated by NUF versus urea**

| Up-regulated (NUF vs Urea at 7 d) | | |
|---|---|---|
| **Enriched Pathway** | **Normed F** | ***P*-value** |
| Homologous recombination | 4.8 | 6.2E-05 |
| Plant-pathogen interaction | 2.2 | 3.6E-03 |
| Alanine, aspartate and glutamate metabolism | 3.7 | 4.8E-03 |
| Sesquiterpenoid and triterpenoid biosynthesis | 4.1 | 1.5E-02 |
| Glutathione metabolism | 2.1 | 3.7E-02 |
| Porphyrin metabolism | 2.6 | 4.0E-02 |
| Ribosome | 2.5 | 4.7E-02 |
| Base excision repair | 3.6 | 4.8E-02 |
| MAPK signaling pathway - plant | 1.8 | 5.0E-02 |
| ABC transporters | 2.7 | 5.9E-02 |
| Protein export | 3.9 | 9.0E-02 |
| Lipoic acid metabolism | 10.7 | 9.1E-02 |
| Carotenoid biosynthesis | 2.7 | 9.8E-02 |
| Fatty acid elongation | 2.7 | 9.8E-02 |

| Up-regulated (NUF vs Urea at 12 h) | | |
|---|---|---|
| **Enriched Pathway** | **Normed F** | ***P*-value** |
| Glutathione metabolism | 4.0 | 5.9E-05 |
| MAPK signaling pathway - plant | 3.3 | 1.0E-04 |
| Alanine, aspartate and glutamate metabolism | 5.2 | 7.9E-04 |
| Arginine biosynthesis | 5.8 | 4.2E-03 |
| Plant-pathogen interaction | 2.3 | 1.2E-02 |
| Carbon fixation in photosynthetic organisms | 3.0 | 4.0E-02 |
| Cysteine and methionine metabolism | 2.2 | 5.5E-02 |
| alpha-Linolenic acid metabolism | 3.3 | 6.2E-02 |
| ABC transporters | 2.9 | 8.6E-02 |
| Glycolysis / Gluconeogenesis | 2.3 | 9.4E-02 |
| Tyrosine metabolism | 3.8 | 9.5E-02 |
| Flavonoid biosynthesis | 3.8 | 9.5E-02 |
| Biosynthesis of amino acids | 1.7 | 9.7E-02 |

| Down-regulated (NUF vs Urea at 7 d) | | |
|---|---|---|
| **Enriched Pathway** | **Normed F** | ***P*-value** |
| Metabolic pathways | 1.4 | 2.6E-10 |
| Phenylpropanoid biosynthesis | 3.0 | 1.2E-05 |
| Nitrogen metabolism | 4.4 | 1.6E-05 |
| Biosynthesis of nucleotide sugars | 2.8 | 1.5E-04 |
| Amino sugar and nucleotide sugar metabolism | 2.3 | 1.4E-03 |
| Stilbenoid, diarylheptanoid and gingerol biosynthesis | 8.2 | 3.5E-03 |
| Pentose and glucuronate interconversions | 3.3 | 3.9E-03 |
| Flavonoid biosynthesis | 4.3 | 4.5E-03 |
| Biosynthesis of various secondary metabolites - part 2 | 13.7 | 5.3E-03 |
| Cutin, suberine and wax biosynthesis | 2.9 | 1.4E-02 |
| Biosynthesis of secondary metabolites | 1.2 | 1.7E-02 |
| Ubiquinone and other terpenoid-quinone biosynthesis | 3.0 | 2.3E-02 |
| Tryptophan metabolism | 2.2 | 2.4E-02 |
| Alanine, aspartate and glutamate metabolism | 2.3 | 3.9E-02 |
| Biosynthesis of various secondary metabolites - part 3 | 4.6 | 6.6E-02 |
| Steroid biosynthesis | 2.1 | 8.0E-02 |
| Sulfur metabolism | 2.1 | 8.9E-02 |
| Phenylalanine metabolism | 2.7 | 9.1E-02 |
| Carbon metabolism | 1.4 | 9.3E-02 |

| Down-regulated (NUF vs Urea at 12 h) | | |
|---|---|---|
| **Enriched Pathway** | **Normed F** | ***P*-value** |
| Glucosinolate biosynthesis | 10.8 | 5.7E-09 |
| 2-Oxocarboxylic acid metabolism | 5.0 | 1.9E-05 |
| Biosynthesis of secondary metabolites | 1.6 | 2.3E-04 |
| Nitrogen metabolism | 4.6 | 1.6E-03 |
| Flavone and flavonol biosynthesis | 13.0 | 8.3E-03 |
| Valine, leucine and isoleucine biosynthesis | 5.6 | 1.5E-02 |
| Metabolic pathways | 1.2 | 1.9E-02 |
| Flavonoid biosynthesis | 4.9 | 2.1E-02 |
| Monobactam biosynthesis | 6.5 | 3.5E-02 |
| Sulfur metabolism | 3.2 | 3.6E-02 |
| Ubiquinone and other terpenoid-quinone biosynthesis | 3.4 | 5.6E-02 |
| Starch and sucrose metabolism | 1.8 | 5.7E-02 |
| Porphyrin metabolism | 2.5 | 7.0E-02 |
| Peroxisome | 2.2 | 7.4E-02 |
| Selenocompound metabolism | 4.3 | 7.5E-02 |
| MAPK signaling pathway - plant | 1.7 | 8.7E-02 |
| Phenylpropanoid biosynthesis | 1.9 | 8.9E-02 |
| Cutin, suberine and wax biosynthesis | 2.8 | 9.0E-02 |
